# Mutated clones driving leukemic transformation are already detectable at the single cell level in CD34-positive cells in the chronic phase of primary myelofibrosis

**DOI:** 10.1101/2020.12.07.414177

**Authors:** Sandra Parenti, Sebastiano Rontauroli, Chiara Carretta, Selene Mallia, Elena Genovese, Chiara Chiereghin, Clelia Peano, Lara Tavernari, Elisa Bianchi, Sebastian Fantini, Stefano Sartini, Oriana Romano, Silvio Bicciato, Enrico Tagliafico, Matteo Della Porta, Rossella Manfredini

**Author notes:** Corresponding author: Rossella Manfredini, PhD, Centre for Regenerative Medicine “Stefano Ferrari”, University of Modena and Reggio Emilia, via Gottardi n.100, 41125 Modena, Italy. ‘These authors contributed equally to this work’ (Sandra Parenti, Sebastiano Rontauroli, Chiara Carretta, Selene Mallia). ‘These authors contributed equally to this work’ (Matteo Della Porta, Rossella Manfredini).

## Abstract

Disease progression of myeloproliferative neoplasms is the result of increased genomic complexity. Since the ability to predict disease evolution is crucial for clinical decision, we studied single cell genomics and transcriptomics of CD34-positive cells from a primary myelofibrosis (PMF) patient who progressed to acute myeloid Leukemia (AML) while receiving Ruxolitinib.

Single cell genomics allowed the reconstruction of clonal hierarchy and demonstrated that *TET2* was the first mutated gene while *FLT3* was the last one. Disease evolution was accompanied by increased clonal heterogeneity and mutational rate, but clones carrying *TP53* and *FLT3* mutations were already present in chronic phase.

Single cell transcriptomics unraveled repression of interferon signaling suggesting an immunosuppressive effect exerted by Ruxolitinib. Moreover, AML transformation was associated with a differentiative block and immune escape.

These results suggest that single cell analysis can unmask tumor heterogeneity and provide meaningful insights about PMF progression that might guide personalized therapy.

## INTRODUCTION

Clonal evolution, mediated by the serial acquisition of somatic mutations at the stem cell level, is the basis of Myeloproliferative Neoplasms (MPNs) such as Polycythemia Vera, Essential Thrombocythemia and Primary Myelofibrosis (PMF). PMF is a heterogeneous disorder characterized by bone marrow fibrosis, megakaryocyte hyperplasia and extramedullary hematopoiesis (EMH). PMF has the worst prognosis among MPNs also due to evolution to Acute Myeloid Leukemia (AML), which occurs in 15-20% of cases and is unresponsive to conventional therapy^1,2^.

Three driver mutations leading to constitutive activation of JAK/STAT pathway were identified in *JAK2*, *MPL* or *CALR* genes^3^. The complex molecular phenotype of these disorders is however characterized by other somatic mutations. Some of these, called “High Molecular Risk” (HMR) mutations (e.g. in *ASXL1* and *SRSF2*), are associated with a worse prognosis and leukemic transformation^4^. Moreover, other pathogenic variants affecting genes such as *TET2*, *TP53* and *FLT3* are related to preleukemic and leukemic conditions^5^.

Disease onset and evolution are the result of the sequential acquisition of somatic mutations in different subclones, giving to each clone phenotypic traits that influence their competition and disease progression. The temporal order in which these variants accumulate is crucial for the fate of the subclones and for disease evolution. Recent studies at single cell level shed light into intratumoral heterogeneity and identified therapy-resistant clones^6^. For instance, the acquisition of a *TET2* mutation preceding JAK2V617F confers a lower sensitivity to Ruxolitinib, which is nowadays the best available therapy^7^. Up to date, several genomic lesions with potential pathogenetic implications have been described, but the molecular mechanisms underlying progression to leukemia have not been defined yet. Several issues remain to be addressed: what are the molecular mechanisms leading to disease progression? What are the relationships between the clones maintaining the chronic phase and the ones driving the leukemic phase? Can a consistent pattern of clonal evolution be identified in MPN progression? Can specific signaling pathways activated during disease evolution be identified?

In order to answer these questions, here we show the single cell-based genomic profiling of CD34-positive (CD34+) cells from a patient with PMF at three different timepoints.

Moreover, we analyzed the single cell transcriptome of CD34+ cells from the same patient to identify signaling pathways abnormally activated during disease progression and/or leukemic transformation that could represent novel therapeutic targets.

## RESULTS

### Single cell analysis in CD34+ population reveals *TET2* and *FLT3* as the first and the last mutated genes

In order to reconstruct the clonal architecture of the stem cell compartment during MPN evolution, peripheral blood (PB) CD34+ cells were analyzed at three different stages of disease: at diagnosis (T1), during the accelerated phase (T2) and in AML phase (T3). The single cell tree based on the mutational profiles of 900 cells (300/sample) is shown in **Supplementary Fig. 1a**. The heatmap was built according to the presence/absence of the mutations. The tree reveals that just a small number of parental clones seem to generate clonal complexity. Fifty-two clones are shown in the phylogenetic tree during temporal evolution (**Supplementary Fig. 1b).**

The main branch arising from a TET2 p.Leu1248Pro (hereafter called TET2a) mutated cell is shown in **Fig. 1a** along with its mutational evolution, while **Fig. 1b** represents the clonal prevalence during phylogenesis. Two main findings emerge from this reconstruction: the expansion of the clone harboring the *TP53* mutation during disease progression (**Fig. 1a,b**) and the early identification of FLT3-mutant cells in chronic phase (**Fig. 1a, b).** However, *FLT3* mutation has not been detected by the bulk diagnostic NGS (**Table 1**). Interestingly, *FLT3* mutation represents also the last evolutionary event of the clone in which all mutations accumulate (**Fig. 1a, b**). This clone, although already present in low percentage in T1 (9%), expands in T2 (15%) and in T3 (23%).

**Figure 1.**
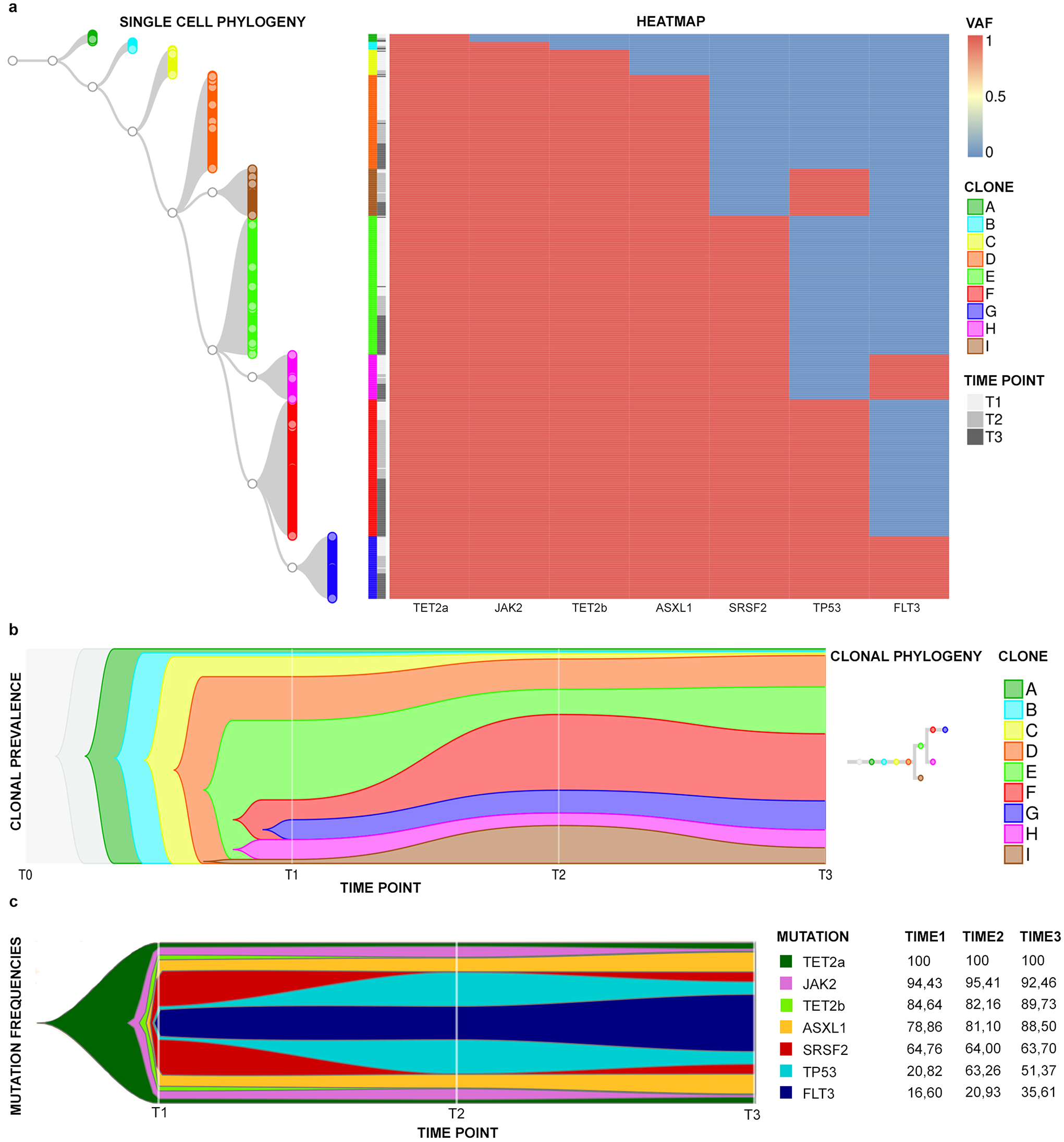
Single Cell Phylogeny and mutation acquisition order. Panel **a** shows the phylogenetic tree of the most represented clones identified in T1, T2 and T3. On the left is shown the tree starting from the founder cell carrying a single TET2a mutation (clone A). From the parental TET2a mutated cell, 8 clones originate (B-I) and each clone is represented by a specific colour; nodes are represented as white circles. On the right is the heatmap showing the mutational event (blue: wild-type, red: mutated) of the genes indicated at the bottom. Each cell in the phylogenetic tree corresponds to a row in the heatmap, identifying its mutational profile. In Panel **b**, the Fish Plot shows the abundance of 9 clones (A-I), identified in Panel **a**, and their temporal evolution. Each clone is represented by the same color used in Panel **a**. In particular, F clone in red and I clone in brown represent TP53-mutated clones, while G clone in blue and H clone in purple represent FLT3-mutated ones. In Panel **c**, the Fish Plot indicated the variation of mutation frequencies through time and the mutational acquisition order. Each variant is represented by a color. On the right the table summarizing the mutation frequencies in single cells during time is shown.

**Table 1.** Clinical data and mutational characterization of the patient in three different stages.

| CLINICAL-HEMATOLOGICAL DATA |  |  |  | Time point 1 | Time point 2 | Time point 3 |
| --- | --- | --- | --- | --- | --- | --- |
| White Blood Cells (x10<sup>9</sup>/l) |  |  |  | 31.03 | 105.2 | 151.18 |
| Neutrophils absolute count (x10<sup>9</sup>/l) |  |  |  | 27.8 | 95.6 | 118.5 |
| Peripheral blasts (%) |  |  |  | 2 | 2 | 20 |
| Peripheral CD34+ (%) |  |  |  | 1 | 1 | 15 |
| Leukoerythroblastosis |  |  |  | yes | yes | yes |
| Hemoglobin (g/l) |  |  |  | 9.9 | 9.4 | 8.4 |
| MCV (fl) |  |  |  | 87.7 | 92.6 | 91.3 |
| Platelets (x10<sup>9</sup>/l) |  |  |  | 79 | 84 | 41 |
| LDH |  |  |  | 1153 | 2630 | 1848 |
| Hepatosplenomegaly |  |  |  | yes | yes | yes |
| Transfusion dependency |  |  |  | no | yes | yes |
| Bone marrow cellularity (%) |  |  |  | 55 | 50 | NA |
| Fibrosis (EUMNET consensus grade) |  |  |  | 3 | 3 | Fibrotic substitution |
| Karyotype (ISCN) |  |  |  | 46,XY[25] | failed | 47,XY,+21[3]/46,XY,i(21)(q10)[2]/46,XY[25] |
| DIPSS risk group |  |  |  | high | high | / |
| MUTATIONAL STATE |  |  |  |  |  |  |
| GENE | TRANSCRIPT | VARIANT | PROTEIN MUTATION | VAF (%) | VAF (%) | VAF (%) |
| TET2 (a) | NM_001127208 | c.3743T>C | p.Leu1248Pro | 48.1 | 42.4 | 46.1 |
| JAK2 | NM_004972 | c.1849G>T | p.Val617Phe | 75.8 | 71.5 | 51.6 |
| TET2 (b) | NM_017628 | c.3409_3416delGGTAATGT | p.Gly1137Profs*5 | 37.7 | 40.3 | 47 |
| ASXL1 | NM_015338 | c.1934dupG | p.Gly646Trpfs*12 | 43.6 | 40.6 | 40.6 |
| SRSF2 | NM_003016 | c.284C>A | p.Pro95His | 46.1 | 45.6 | 45.3 |
| TP53 | NM_000546 | c.713G>C | p.Cys238Ser | 5.7 | 25.6 | 13.7 |
| FLT3 | NM_004119 | c.2503G>C | p.Asp835Tyr | / | 6.3 | 35.3 |

The study of the frequencies and associations of the variants shows, as depicted in the fishplot in **Fig. 1c,** that TET2a mutation is the most frequent, thus the first to be acquired, followed by the JAK2V617F and by the second TET2 mutation (p.Gly1137Profs*5, hereafter called TET2b) (**Table 1**). The frequency of *ASXL1* and *SRSF2* variants remains stable during disease progression, while the percentage of *TP53-*mutated cells increases from 20.8% in T1 to 63% in T2. Finally, the frequency of FLT3-mutated cells grows linearly from 16.6% in T1 to 25% in T2 up to 35.6% in T3, indicating probable support for AML onset.

### Clonal heterogeneity increases during disease progression

**Supplementary Fig. 2, 3 and 4** show the phylogenetic trees in each timepoint built considering the zygosity of the different mutations. As shown by this heatmaps, the clonal heterogeneity increases during disease progression, both in terms of mutation’ combinations and variants’ zygosity.

**Figure 2.**
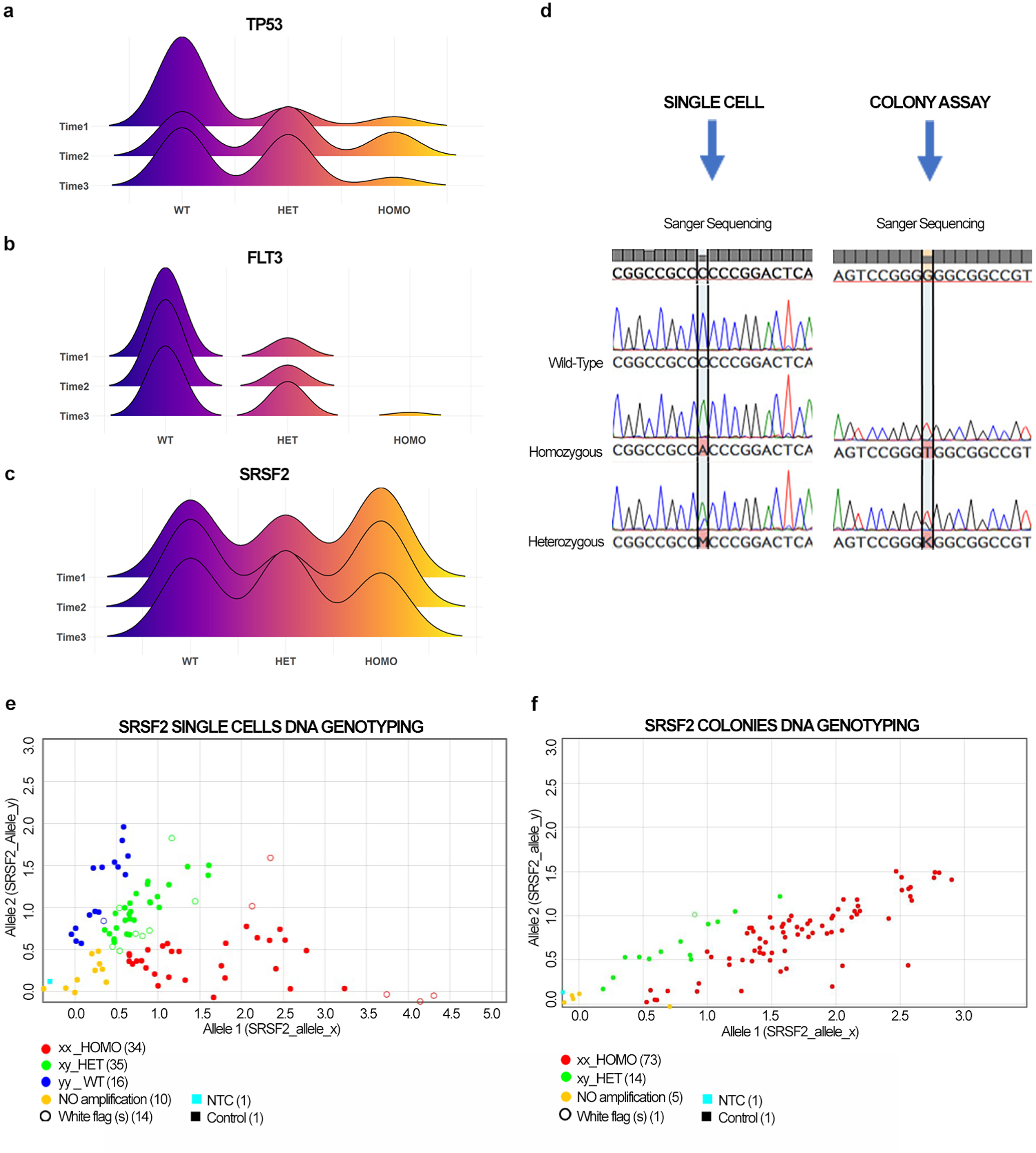
TP53, FLT3 and SRSF2 mutation evolution during time and detection of SRSF2 homozygous mutated cells. In Panel **a**, **b** and **c**, the Joy Plot shows the distribution of zygosity of TP53 (**a**), FLT3 (**b**) and SRSF2 (**c**) in T1, T2 and T3 respectively. Panel **d** shows the results of Sanger sequencing on single cells (SC) and colonies obtained from PB CD34+ cells. On the left are shown the electropherograms representing wild-type, heterozygous and homozygous SRSF2 gene variant in SC, while on the right there are electropherograms from colonies indicating heterozygous and homozygous mutational state. In Panels **e-f**, representative dot plots show genotyping results obtained from SC and colonies. Allele 1 (x) indicates SRSF2 mutated gene and allele 2 (y) refers to SRSF2 wild-type gene. The graph represents the distribution of amplified DNA according to a wild-type (y/y, blue dot), heterozygous (x/y, green dot) and homozygous (x/x, red dot) configuration. In Panel **e**, the dot plots highlights the presence of wild-type (blue), heterozygous (green) and homozygous (red) single cells. In Panel **f**, the dot plot shows heterozygous (green) and homozygous (red) colonies.

**Figure 3:**
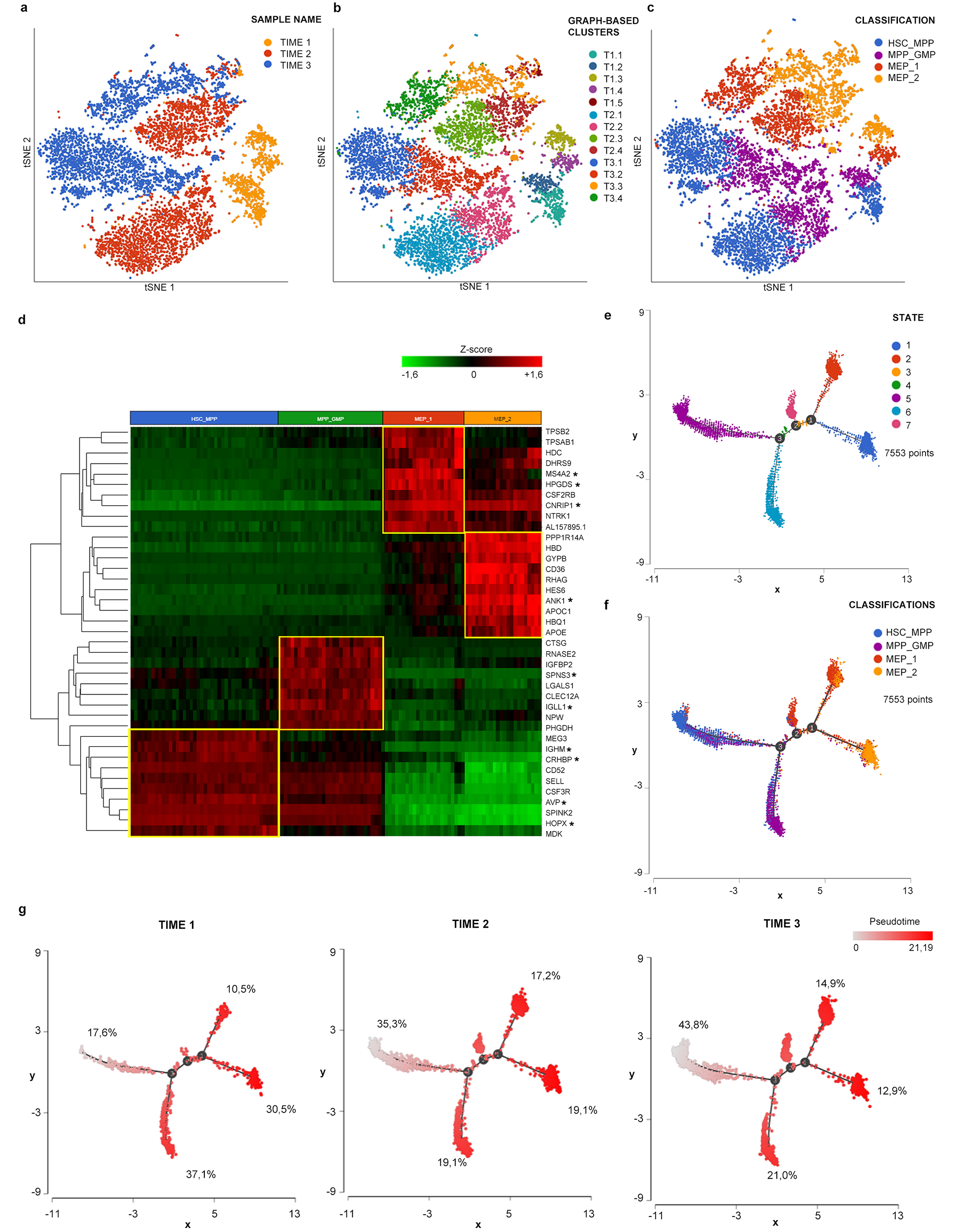
scRNA-seq clustering and trajectory analysis. **a, b, c:** tSNE projections of cells from all three samples in which 7553 cells were included. In panel **a** cells are distinguished according to the sample of origin. In panel **b** different colors indicate clusters identified by clustering analysis. Panel **c** represents the classification we applied to these clusters. We identified 4 cell groups namely HSC_MPP, MPP_GMP, MEP_1 and MEP_2. **d:** The heatmap was generated using the top 10 marker genes for each cell group. Marker genes shared by the three samples are highlighted by the star next to the gene symbol. Panel **e** represents the distribution of single cells colored according to the 7 states identified by trajectory analysis. As shown in panel **f, s**tate 5 represented the most primitive one enriched in cells belonging to HSC_MPP cluster. Starting from state 5 the first branch point corresponds to the appearance of a population enriched in MPP_GMP cells. State 7 originates from the second branch point. Two different MEP populations originated from the third branchpoint, corresponding to MEP_1 and MEP_2 clusters. In panel **g** trajectory cells are located according to the sample of origin. Cells are colored according to pseudotime. Percentages represent the frequency of cells belonging to the indicated sample included in that specific state.

**Figure 4:**
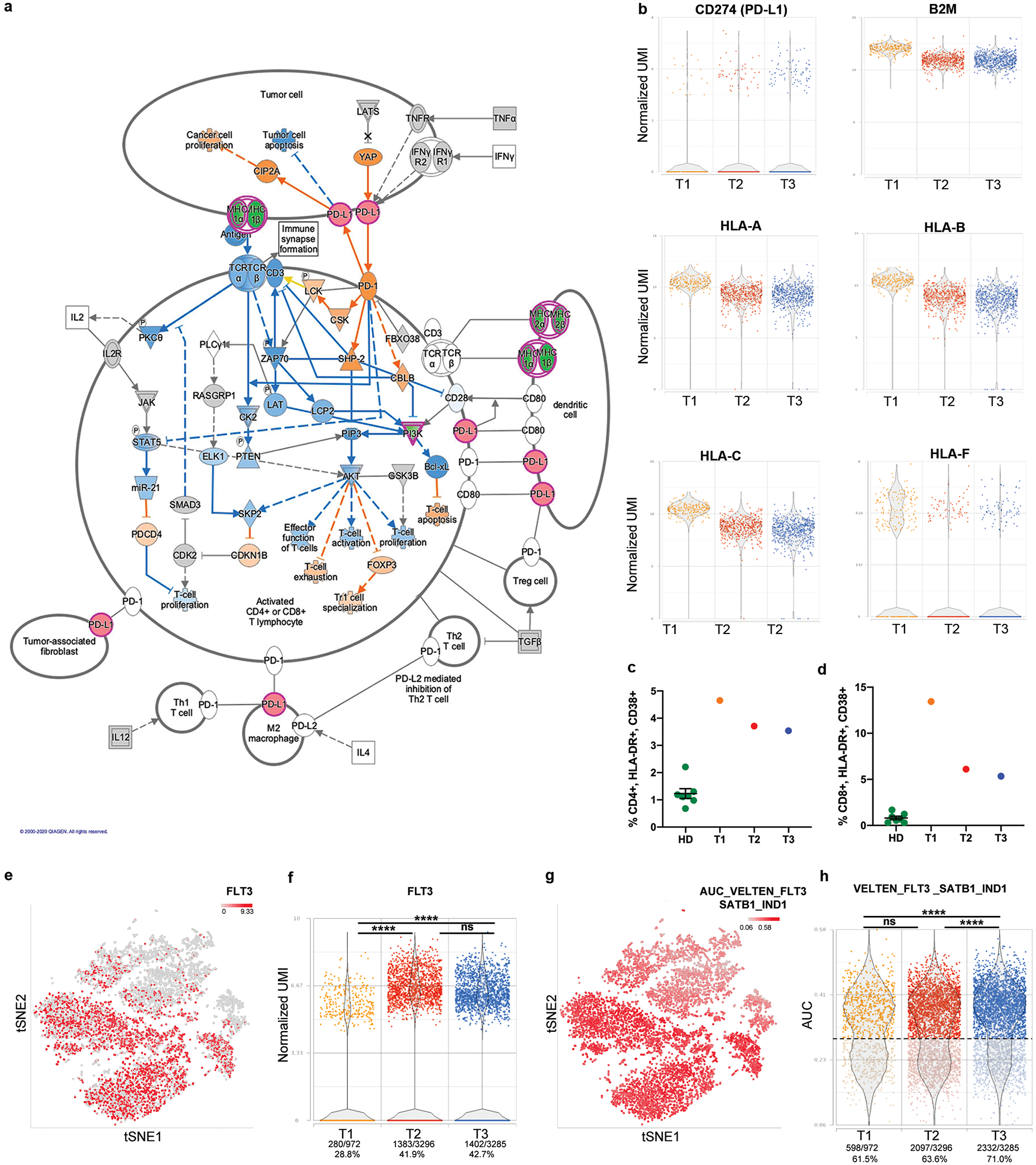
PDL-1 and FLT3 increased expression during disease progression: Panel **a** represents the canonical pathway “PD-1, PD-L1 cancer immunotherapy pathway” according to IPA. Green indicates downregulated genes, red indicates overexpressed ones in the comparison T3 vs T1 MEP_2 cluster. Blue genes are predicted inhibited, while orange indicates predicted activated ones. Panel **b** includes dot plots representing the expression of genes involved in PD-1/PD-L1 pathway in cells belonging to the MEP_2 cluster. Panel **c and d** shows the percentages of activated CD4+ and CD8+ T cells assessed in HD (n=7), T1, T2 and T3 samples respectively. T cells activation was evaluated by flow cytometry; the gating strategy used to identify activated subpopulations, defined as CD38+ HLA-DR+, is reported in Supplementary Fig. 10. Percentages are reported as mean ± standard error of the mean (s.e.m.). In panel **e** cells in tSNE plot are colored in red scale according to FLT3 expression. Dot plot in panel **f** represents the distribution of FLT3 expression values. Percentages represent the frequency of FLT3 expressing cells in each sample. In panel **g** cells in tSNE plot are colored based on AUC values for VELTEN_FLT3_SATB1_IND1 gene module^18^. Panel **h** shows the distribution of AUC values. Dots are colored based on sample name. Based on this distribution we identified cells were the gene signature is active with an AUC > 0.29. Two sided Fisher’s exact test was used to compare the proportion of cells expressing FLT3 or with active VELTEN_FLT3_SATB1_IND1 gene module in three time points (**** p≤0.0001).

Noteworthy is the modulation of *TP53* and *FLT3* variants’ zygosity. The joy plots in **Fig. 2a and b** show that, while the number of *TP53* and *FLT3* wild-type cells decreases during progression, the number of mutated cells increase, particularly those carrying heterozygous mutations. Particularly interesting is the presence of *FLT3* homozygous mutation only in the leukemic phase. The change in zygosity of all mutations during time is shown in **Supplementary Fig. 5 and Supplementary Table 2**.

### Single cell sequencing allows the characterization of a novel TP53 variant

In order to assess whether Copy Number Variations (CNVs) were present during disease progression, bulk NGS data were analyzed using the CNV algorithm implemented in SOPHIA DDM platform which revealed no detectable alterations.

*TP53* loss-of-function is a crucial event in the activation of mechanisms underlying tumor progression^8^. In order to identify CNVs in TP53 gene, we analyzed this region by MLPA in bulk CD34+ cells of T2 and T3, because of the significant expansion of the clones carrying TP53 p.Cys238Ser mutation in these timepoints. As shown in **Supplementary Fig. 6a-d**, this analysis did not highlight any gene rearrangement.

Finally, in order to identify any smaller CNVs affecting *TP53*, we sequenced all of its exons in single cells of each timepoint. Interestingly, Sanger sequencing identified a 454bp deletion between *TP53* exons 2 and 4 (**Supplementary Fig. 6e**). This new likely pathogenic variant was found in a small subpopulation of cells already harboring the p.Cys238Ser homozygous mutation and its frequency increases in T2 following the trend of p.Cys238Ser mutated cells (**Supplementary Fig. 6f)**.

### Single cell sequencing allows the characterization of SRSF2 homozygous mutation

**Table 1** shows that VAF of *SRSF2* P95H in the diagnostic NGS was 50%. However, P95H was detected as homozygous in 39.4%, heterozygous in 26.7%, and wild-type in 33.8% of T1 single cells (**Fig. 2c, d**). The zygosity of this variant does not change significantly during disease evolution (**Supplementary Table 2**), but P95H homozygosity is detectable only at single cell level.

To exclude a possible Allele Drop Out (ADO) effect and validate our results, we performed the same analysis on CD34+-derived colonies not subjected to WGA. Sanger Sequencing detected only heterozygous and homozygous colonies, as shown in **Fig. 2d**. To further validate these findings, we performed a SNP genotyping assay on both single cell WGAs and colonies’ DNA. The results confirmed the presence of 36% of wild-type, 28% of heterozygous and 36% of homozygous cells in T2 (**Fig. 2e, Supplementary Fig. 5**). We identified 16% of heterozygous and 84% of homozygous colonies in T2 (**Fig. 2f**). Surprisingly, no wild-type colonies are detected. This difference between single cell and colonies analysis suggests that the semisolid culture could introduce some bias due to the growth of selected clones.

### Leukemic transformation is characterized by a less differentiated phenotype

To investigate the molecular processes and transcriptional changes occurring during disease progression, we performed scRNA-seq of CD34+ cells isolated at same timepoints as those of genomic analysis. Clustering analysis led to the identification of 5 clusters in T1 and 4 in T2 and T3 (**Fig. 3b**).

As detailed in **Supplementary Discussion** we were able to classify these clusters in 4 cell populations shared by the three samples, namely HSC_MPP, MPP_GMP, MEP_1 and MEP_2 (**Fig. 3c and Supplementary Fig. 7**) according to the expression of genes related to specific lineages (**Fig. 3d, Supplementary Data File 1, 2 and Supplementary Table 3**) and to a more primitive or differentiated phenotype (**Supplementary Fig. 8**). To better characterize the differentiation status of cells in the three samples, we performed trajectory analysis that allowed the identification of 7 cell states (**Fig. 3e**). The distribution of cells belonging to different samples (**Fig. 3g**) shows that 43.8% of T3 cells are included in state 5, the most primitive one enriched in HSC_MPP cells, while only 17.6% of T1 cells are included in this state. On the contrary, T1 cells are present at a higher frequency in more differentiated states, including state 6 (myeloid biased progenitors, 37.1 %) and state 1 (MEPs, 30.5 %), thus confirming that CD34+ cells from secondary AML are characterized by a less differentiated phenotype compared to the same cells from the chronic phase of the disease.

### Gene expression changes during disease progression

Gene expression analysis, made by comparing the same cluster of each timepoint, highlighted the major transcriptional variation between T3 and T1 rather than in T2 vs T1 comparison (**Supplementary Table 4 and Supplementary Data 3**).

To better understand biological processes deregulated during disease progression we interrogated Differentially Expressed Genes (DEGs) lists by means of IPA^®^. Malignant cells put up several strategies in order to avoid immune system control^9^.One of the contributors to this mechanism is Ruxolitinib^10^ that induces a marked downregulation of Interferon (IFN) signaling. “Interferon signaling” canonical pathway is significantly inactivated according to IPA^®^ in T2 and T3 vs T1, increasing its significance in more differentiated clusters (**Supplementary Fig. 9 and Supplementary Discussion**). In particular, IFN-γ pathway blockade leads to the downregulation of type I and II HLA antigens, together with B2M protein (**Fig. 4b**), in peripheral blood mononuclear cells (PBMCs), thus causing the immune escape of AML malignant cells^11^. IFN pathway influences also apoptosis through BAK1 downregulation (**Supplementary Fig. 9c**). The predicted inactivation of several functional categories related to cell death together with the activation of cell quiescence might cooperate, favoring resistance to therapy and escape of leukemic cells from the immune surveillance (**Supplementary Discussion**).

PD-L1/PD-1 axis is involved in this process since it can inhibit T cell proliferation while favoring regulatory T cells apoptosis^12^. According to IPA^®^ analysis, this pathway was found significantly activated in HSC_MPP, MEP_1 and MEP_2 clusters of T2 and in MEP clusters of T3 (**Fig. 4a and Supplementary Data 4**), increasing its significance in more differentiated clusters (**Supplementary Data 4**). This observation is strengthened by the upregulation of PD-L1 expression in MEP clusters (MEP_2 in T2 vs. T1; MEP_1 and MEP_2 in T3 vs. T1) (**Fig. 4b and Supplementary Data 3**). Flow cytometry analysis of T cell activation in PBMCs from the three timepoints supported these results showing a global induction of CD4+ and CD8+ cell activation compared to healthy donors (**Fig. 4c, d**). Moreover, disease progression displays a progressive reduction of T cell activation (**Fig. 4c, d**), thus representing an acquired immune escape mechanism and confirming transcriptomic data.

Mobilization of Hematopoietic Stem Cells (HSCs) and extramedullary hematopoiesis (EMH) are distinctive traits in PMF^13^. In this patient we observed a complete fibrotic substitution at T3 associated with marked splenomegaly; these clinical features are supported by transcriptomic analysis which reveals the activation of molecular processes involved in EMH (**Supplementary Discussion**).

### Disease progression correlates with increased FLT3 expression

Single cell sequencing demonstrated a significant increase in the number of FLT3 mutated cells during disease progression. Since FLT3 tyrosine kinase domain (TKD) mutations, like p.Asp835Tyr, are associated with increased expression of FLT3 in AML^14^, we analyzed its transcriptional level in the three timepoints. FLT3 transcript is significantly upregulated in T2 HSC_MPP and MPP_GMP clusters compared to T1, while in T3 a significant increase is evident in MEP_1 and MEP_2 clusters (**Supplementary Data 3**). As shown in **Fig. 4e**, FLT3 expression is detected in more primitive clusters (HSC_MPP and MPP_GMP), while only a small fraction of MEP_1 and MEP_2 cells express this gene. Furthermore, a higher proportion of T2 and T3 cells shows detectable levels of FLT3 transcript if compared with T1 (**Fig. 4f**). By means of AUCell task in Partek^®^ Flow^®^ we assayed a gene module related to FLT3^15^ and revealed that its activation increases in HSC_MPP and MPP_GMP clusters, in agreement with FLT3 expression (**Fig. 4g**). The frequency of cells with higher AUC values is significantly increased in T3 sample compared with T1 and T2 (**Fig. 4h**). These results are therefore in line with genomic data showing the expansion of FLT3 mutated clones during disease progression.

## DISCUSSION

Clonal evolution is the result of the serial accumulation of genomic alterations that determine the fate of a neoplastic population, clinical history and response to therapy. PMF pathogenesis is characterized by the acquisition at the HSC level of somatic mutations in genes involved in JAK/STAT pathway, such as *JAK2*, *MPL* and *CALR*. Moreover, other genes implicated in epigenetic modification (*TET2*, *ASXL1*), splicing regulation (*SRSF2*), DNA repair (*TP53*) and cell proliferation (*FLT3*), are frequently mutated in MPNs.

It has been demonstrated since 2015 that the order of mutation acquisition in HSCs influences disease progression^7^. In order to describe the mutation acquisition order and characterize the clonal architecture of PMF in stem and progenitor cell compartment, we performed genomic single cell analysis on CD34+ cells from a PMF patient during Ruxolitinib treatment and disease progression.

Our results demonstrated that in this patient the p.Leu1248Pro substitution in *TET2* gene was the first mutational event, followed by the JAK2V617F mutation. Treatment with Ruxolitinib did not significantly change the clinical history of the patient we described, and did not prevent progression to leukemia, in agreement with the data from Ortmann et al, which demonstrated that “TET2-first patients” are less sensitive to Ruxolitinib^7^.

Clonal phylogeny reconstruction demonstrates that clones carrying *TP53* mutations expand during disease progression (T2 vs T1) and show a slight decrease in T3 vs T2, suggesting that *TP53* mutations could promote the accelerated phase (T2) of the disease, but was not sufficient to support leukemic transformation. Since a recent work by Bernard *et al.* demonstrated that *TP53* impairment determines CNVs which correlate with poor prognosis in hematological malignancies^8^, we deeply analyzed *TP53* mutational state and identified a novel deletion at single cell level. Although this variant affects a small cell subpopulation, it could support the genomic instability that characterizes the accelerated phase of the disease. Interestingly, this mutation was found in co-occurrence with the *TP53* p.Cys238Ser homozygous mutation. This mutational asset assigns our patient to the multi-hit state described in the paper by Bernard *et al.*, which is associated with lower overall survival and higher incidence of AML transformation^8^.

On the other hand, the linear expansion of *FLT3*-mutated cells from T1 to T3 seems to trigger AML onset. To date, *FLT3* mutations are a widely recognized AML-associated driver mutations. The FLT3-TKD mutation, carried by the patient, like the canonical FLT3-ITD, constitutively activates the receptor and promotes cell proliferation^16^.

Regarding the global mutational profile of single cells studied at the zygosity level, we observed a significant increase in the clonal heterogeneity during disease evolution, in agreement with recent data by Mylonas et al^17^. This causes the expansion of highly mutated clones in later stages of the disease, accounting for the worsening of the patient’ conditions. The study of mutation zygosity further confirmed that *TP53* and *FLT3* are responsible for disease evolution, indeed the VAFs of these mutations are the most modulated during disease progression. Noteworthy, the clone recapitulating all mutational events, in which *FLT3* mutation is the last event, is already detectable at low frequency in T1, when *FLT3* mutation is undetectable by the bulk diagnostic NGS. Furthermore, homozygous *FLT3* mutations are detected only in T3 by single cell analysis.

Spliceosome machinery gene mutations are usually mutually exclusive and heterozygous, suggesting that cells cannot tolerate significant malfunction of the normal splicing activity^18^. Although the presence of *SRSF2* homozygous mutation has not been yet well characterized in MPNs^19^, single cell genomic analysis has demonstrated the presence of a significant stem cell fraction carrying P95H mutation in homozygous status. Altogether these data demonstrate that single cell genomics represents a promising and powerful method to describe a real scenario of mutational zygosity state in complex genotypes and high heterogeneous contexts such as PMF.

As well as genomic analysis, single cell transcriptomics depicted the worsening of patient condition during disease progression. Gene enrichment analysis demonstrated the deregulation of several pathways and molecular mechanisms that might contribute to the clinical characteristics of the patient. Trajectory analysis clearly showed that CD34+ cells in T3 display a more primitive phenotype compared to those in T1 demonstrating that these cells underwent a progressive differentiative block resulting in the development of secondary AML. Interestingly, despite Ruxolitinib treatment, the marked splenomegaly observed during the chronic phase has only been partially reduced. This is consistent with the activation of pathways favoring EMH^13^, such as *CXCR4, MMP7*, and several chemoattractant cytokines (i.e. IL-8, PDGFB and PF4) that displayed increased activity in T3. The activation of EMH pathway is related to disease progression since the patient suffered from a grade 3 of bone marrow fibrosis at T1 and T2, which evolved in a complete fibrotic substitution with a severe bone marrow damage in T3.

One of the most relevant processes deregulated during disease progression is represented by immune escape. In particular, gene expression analysis showed that hematopoietic stem and progenitor cells activated several mechanisms that reduce their sensitivity to immune surveillance, such as PD-L1 increased expression, downregulation of molecules involved in antigen presentation and inactivation of IFN signaling. Our data suggest that Ruxolitinib exerts a direct inhibition of IFN signaling, mediated by JAK/STAT pathway. The immunosuppressive side of Ruxolitinib has already been identified in different studies, as it is able to reduce leukemic cells’ sensitivity to NK cells^20^. This observation was confirmed by the decrease of CD8+ and CD4+ cell activation during disease progression, which could be due to the anti-inflammatory effect of Ruxolitinib^20^ and to the interaction between T cells and leukemic CD34+ cells which activate PD-1/PD-L1 axis.

On the other hand, IFN-signaling inhibition was related to apoptosis inhibition. Together with the activation of molecular pathways favoring HSC quiescence in T3, this observation suggests that leukemic cells acquired phenotypic traits that protect them from cell death thus resisting to therapy.

The activation of PD-1/PD-L1 pathway was predicted by IPA^®^ due to the downregulation of class I and II HLA and B2M and to the increased expression of PD-L1 in T3 vs T1. A recent work described how a *TP53* mutation inhibits miR34 transcription, which in turn cannot inhibit the translation of its target PD-L1^21^. In agreement with these data, the patient we studied harbors a loss-of-function mutation in *TP53* (p.Cys238Ser), which contributes to PD-L1 upregulation.

It has been demonstrated that FLT3-p.Asp835Tyr is associated with FLT3 increased expression in AML samples^14^. Consistently, we observed an increase in FLT3 expressing cells during disease progression and higher frequency of cells with activated FLT3 gene module in T3. These results confirm single cell genomics findings and strengthen the hypothesis that FLT3 mutation plays a pivotal role in the induction of AML transformation. As a whole, in this work we characterized by genomic stem cell analysis the clonal architecture of the stem cell compartment of a PMF patient in different stages of disease evolution. This analysis allowed us to identify the first mutational hit, the increasing clonal heterogeneity during the disease progression and the presence of TP53 and FLT3-mutated clones also in the chronic phase of the disease, months before leukemic transformation.

In conclusion, the results described so far suggest that the single cell genomic study could provide information with possible predictive value of the evolution of the clinical history of the disease. It is difficult to speculate on the actual clinical applicability of such a complex and expensive study, but certainly our data serve as proof of principle that the possibility to identify these clones in early stages of disease could allow a better prognostic evaluation and could address to personalized therapeutic strategies. Finally, the single cell transcriptome analysis of the stem cell compartment in the same patient allowed us to identify several pathways deranged during disease evolution which could lead to the development of new targeted therapies.

## METHODS

### Ethics statement

A PMF patient (**Table 1**) who evolved to AML was studied at three timepoints: during chronic phase (Time1, T1), after 8 months of Ruxolitinib treatment (Time2, T2) and after 11 months of Ruxolitinib treatment at AML diagnosis (Time3, T3). The diagnosis was performed in agreement with World Health Organization (WHO) criteria updated in 2016^22^. The study was conducted in accordance with the Declaration of Helsinki and with ethical approval obtained from the local ethics committee of the Humanitas Research Hospital – Milan, Italy. (Approval date: 28 jan 2019; approval file # 2175). The patient provided written informed consent to take part in the study.

### Bulk Next Generation Sequencing (NGS) analysis

Targeted DNA-sequencing on genomic DNA extracted from whole peripheral blood of the patient was performed through Capture-based target enrichment kit - CE-IVD Myeloid Solution™ by Sophia Genetics. Sequencing was performed on the Illumina MiSeq instrument. Minimum required coverage was set to 1000X. After demultiplexing the FASTQ files were further processed using the Sequence Pilot software version 4.1.1 Build 510 (JSI Medical Systems, Ettenheim, Germany) for alignment and variant calling. Analysis parameters were set according to manufacturers’ default recommendation. Validity of the somatic mutations was checked against the publicly accessible COSMIC v69 database (http://cancer.sanger.ac.uk/cancergenome/projects/cosmic) and functional interpretation was performed using SIFT 1.03 (http://sift.jcvi.org), PolyPhen 2.0 (http://genetics.bwh.harvard.edu/pph2) and MutationTaster 1.0 algorithms (http://www.mutationtaster.org). Additionally, TP53 variants were verified using the IARC repository. Single nucleotide polymorphisms (SNP) were annotated according to the NCBI dbSNP (http://www.ncbi.nlm.nih.gov/snp; Build 137) and gnomAD (http://gnomad.broadinstitute.org; gnomAD r2.0.1) databases.

### Purification of CD34+ cells

Frozen peripheral blood mononuclear cells (PBMCs) were thawed following 10X Genomics®◻ “Sample Preparation Demostrated Protocol” (10X Genomics, Pleasanton, CA, USA). Immunomagnetic selection of CD34+ cell population was then performed by means of “CD34 Microbead kit, human” (Miltenyi Biotec, Bergisch Gladbach, Germany) following the protocol provided by the manufacturer. The sample was then split between genomic and transcriptomic analyses.

Moreover, a fraction of the CD34+ cell population was seeded in MethoCult™ GF H4434 (StemCell Technologies Inc.; Vancouver), as previously described^23^. Colonies were picked and genotyped for SRSF2 p.P95H mutation, as previously described^7^.

### CD34+ cell immunostaining, fixation and single cell sorting

Cells were stained with anti-human CD34 (AC136, APC, Miltenyi Biotec, Cat No 130-113-738, 1:50) and anti-human CD38 (IB6, PE, Miltenyi Biotec, Cat No 130-113-989, 1:50) antibodies. Immunostained cells were then fixed with Paraformaldehyde (PFA) 0.5% in PBS1X at 4°C for 15’ and washed with PBS1X before being resuspended in autoMACS Running Buffer (Miltenyi Biotec). Immunostained CD34+ cells were then subjected to single cell sorting by means of DEPArray™ Technology (Menarini Silicon Biosystems). Each single cell was resuspended in 1ul PBS1X.

### Whole Genome Amplification

Whole Genome Amplification (WGA) was performed on 300 single cells for each Time by means of SMARTer PicoPLEX Single Cell WGA kit (Takara) following manufacturer’s protocol as previously described^24^. For each WGA experiment, a genomic DNA sample and 1ul Low TE (Tris-EDTA) buffer (Thermo Fisher Scientific) were included as positive and negative control respectively. WGA product was then purified by means of AMPure XP (Beckman Coulter) immunomagnetic beads and eluted in 20ul Low TE buffer. Quality control on WGA yield and amplicons’ size was performed through Bioanalyzer High Sensitivity DNA Analysis (Agilent) on 1ul of 1:20 diluted WGAs.

### Mutation detection

PCR and sequencing analysis were performed by using BigDye® Direct Cycle Sequencing Kit (Applied Biosystem®) following the manufacturer’s protocol. Primers used for PCR reactions were synthesized (Supplementary Table 1) (Thermo Fisher Scientific and Integrated DNA Technologies). Sequencing products were purified by ethanol/EDTA precipitation. Sequencing was performed by capillary electrophoresis on 3130×l Genetic Analyzer (Applied Biosystems®).

SRSF2 p.P95H variant was also analyzed by means of SNP genotyping through the Custom TaqMan™ SNP Genotyping Assay (rs751713049_g_t, ANAAJWD, SNP, ThermoFisher Scientific) in picked colonies and in single cell WGAs.

### Clonal hierarchy reconstruction

The clonal tree reconstruction of the mutations’ acquisition order was performed through the CellScape R package (script retrieved from https://github.com/shahcompbio/cellscape/blob/511a8b6a2d7c6eb28fddb83f1b29c2c1092d5bae/R/cellscape.R)^25^. For the trees representing the presence/absence of the variants, Variant Allele Frequency (VAF)=0 was assigned to a wild-type status and VAF=1 was assigned to a mutant status in the mutational matrix. Clonal evolution analysis was performed through the TimeScape R package (script retrieved from https://github.com/shahcompbio/cellscape/blob/511a8b6a2d7c6eb28fddb83f1b29c2c1092d5bae/R/cellscape.R)^25^. Each clone was assigned to a letter and Timepoints were indicated as T1, T2 and T3. For the trees representing the zygosity of the mutations, VAF=0 was assigned to a wild-type status, VAF=0.5 was assigned to heterozygosity and VAF=1 was assigned to homozygosity. All analyses were performed using R (version 3.6.2) and R Studio (version 1.3.959) software.

### Genomic data analysis

The fishplot representing mutational frequencies was performed through the Fishplot R package (script retrieved from https://github.com/chrisamiller/fishplot)^26^. Mutational frequency for each variant was computed as the percentage of mutated cells (either heterozygous or homozygous) within the totality of the cells of each timepoint.

Joy Plots of the frequencies through time of TP53, FLT3 and SRSF2 mutations through time were performed by means of ggridges R package (retrieved from https://CRAN.R-project.org/package=ggridges; script retrieved from https://cran.r-project.org/web/packages/ggridges/vignettes/introduction.html).

The bubble plot of the frequencies of mutations through time was built through ggplot2 R package (script retrieved from https://www.r-graph-gallery.com/320-the-basis-of-bubble-plot.html)^27^. All analyses were performed using R (version 3.6.2) and R Studio (version 1.3.959) software.

### TP53 Multiplex Ligation-dependent Probe Amplification (MLPA)

DNA coming from CD34+ cells of the patient and Mononuclear cells of Healthy Donors was extracted by means of DNeasy Blood & Tissue Kit (Qiagen, Hilden, Germany). 50ng of DNA were analysed through P056 MLPA kits (MRC Holland, Amsterdam, Netherlands), following manufacturer’s instructions. At least 5 controls were included in each run. Fragment analysis was performed by capillary electrophoresis on 3130xl Genetic Analyzer device (Applied Biosystems). The software Coffalyser.net (MRC-Holland), was used to analyse all MLPA data as previously described^28^.

### Single-cell RNA sequencing (10X Chromium)

Unfixed CD34+ cells were subjected to single-cell RNA sequencing analysis. Single-cell suspensions were prepared and cells were resuspended in 0.5 ml PBS 1X plus 0.04% BSA and washed once by centrifugation at 450 rcf for 7min. After the wash cells were resuspended in 50 ul and counted with an automatic cell counter (Countess II, Thermo Fisher) to get a precise estimation of the total number of cells recovered and of cells concentration. Afterwards we loaded about 10.000 cells of each sample into one channel of the Chromium Chip B using the Single Cell reagent kit v3 (10X Genomic) for Gel bead Emulsion generation into the Chromium system. Following capture and lysis, cDNA was synthesized and amplified for 14 cycles following the manufacturer’s protocol. 50 ng of the amplified cDNA were then used for each sample to construct Illumina sequencing libraries. Sequencing was performed on the NextSeq550 Illumina sequencing platform using High Output Kit v2.5 chemistry and following 10xGenomics instruction for read generation, reaching at least 50.000 reads as mean reads per cell.

### Processing and analysis of single-cell RNA sequencing data

Raw base call (BCL) files generated by NextSeq 550 sequencer were demultiplexed using Cell Ranger software (version 3.1.0). The FASTQ files obtained were then processed using Partek^®^ Flow^®^ software (version 9.0). Reads were trimmed, aligned to the human reference genome hg38 (GRCh38) then deduplicated, to obtain one alignment per unique molecular identifier (UMI), choosing Ensemble Transcripts release 91 for the annotation. After filtering out barcodes associated to droplets containing no cells, aligned reads were quantified generating a single cell count matrix. Cells meeting the following quality control (QC) parameters were included in the analysis: total reads between 6.500 to 52.000; expressed genes between 1.000 and the maximum detected number; mitochondrial reads percentage less than 15%. Following this selection, we obtained 7.717 cells that passed QC filters: 1.043 fo rT1, 3.313 for T2 and 3.361 for T3. Next, features were filtered in order to include only genes expressed in more than 0.1% of cells and 15.296 genes were retained. UMI counts were then normalized following Partek^®^ Flow^®^ recommendations: for each UMI in each sample the number of raw reads was divided by the number of total mapped reads in that sample and multiplied by 1.000.000, obtaining a count per million value (CPM), the normalized expression value was log transformed (pseudocount=1).Starting from the normalized data node we performed clustering analysis for each sample separately by means of graph based clustering task in Partek® Flow® software which employs Louvain algorithm. Clustering analysis was done based on the first 100 principal components. To visualize single cells in a two-dimensional space we performed a t-distributed statistical neighbor embedding (tSNE) dimensional reduction using the 50 principal components for each sample separately and for the entire dataset.

By means of “compute biomarker” function we were able to identify marker genes for each identified cell group. In Partek® Flow® software this task performs an ANOVA test comparing each cluster to all the other cells in the dataset and returns a list of genes with FC>1.5 ranked according to ascending p-value. To define cluster identity in each sample we compared the generated lists of marker genes with lineage signatures derived from different hematopoietic datasets recently published^15,29,30^. Furthermore, we exported the normalized expression matrix from Partek® Flow® software and classified single cells by means of SingleR R package according to Blueprint Encode dataset. We took advantage of single cells classification to better define cluster identity in each sample as detailed in supplementary results. A cell group mainly composed of monocytes and lymphoid cells was identified in all samples and considered as contaminant cells, therefore it was excluded from analysis which were performed on the remaining 7.553 cells. To evaluate the activation state of specific gene modules within our dataset we used the AUCell task provided by Partek® Flow®. This function evaluates the activity of a gene signature in every single cell based on its gene expression profile and represents this activity by an AUC (“area under the curve”) score. By evaluating the distribution of AUC scores we were able to identify cells where gene modules results activated^31^. To identify differentially expressed genes we performed paired sample comparison within each cell cluster by means of ANOVA analysis and considered genes with FC>2 or FC<−2 and step-up p-value<0.05 as differentially expressed. Core analysis was performed by means of Ingenuity® Pathway Analysis (IPA®, Qiagen) software in order to identify canonical pathways, upstream regulators and biological processes deregulated in our dataset according to differential gene expression results. Differentiation trajectory reconstruction was performed by means of Partek® Flow® software which uses Monocle 2 algorithm. Analysis was performed considering only the 5.000 genes with the highest variance within our dataset.

### T cell immunophenotype by flow cytometry

In order to identify activated T cells up to 1 million of thawed PBMCs were washed twice in PBS and stained with the viability dye LIVE/DEAD Fixable Aqua Dead Cell Stain Kit (ThermoFisher, Cat no. L34966, 1:100). Then, cells were washed in PBS supplemented with 2mmM EDTA and 2% FBS, incubated with FcR Blocking Reagent (Miltenyi Biotec, Cat no. 130-059-901, 1:50) and subsequently stained at 4°C with antibodies against following human antigens: CD3 (REA613, APC-Vio770, Miltenyi Biotec, Cat no.130-113-698, 1:50), CD4 (A161A1, FITC, BioLegend, San Diego, CA, USA, Cat no. 357405, 1:40), CD8 (SK1, PerCP-Cy5.5, BioLegend, Cat no. 344709, 1:40), CD38 (HB-7, PE-Cy7, BioLegend, Cat no. 356608, 1:80) and HLA-DR (L243, APC, BioLegend, Cat no. 307609, 1:80). For cytometric analysis the FACS Canto II (Becton Dickinson, Franklin Lakes, New Jersey, USA) was used. Data were analysed by FlowJo (version 10.7.1).

### Data Availability

The data generated and/or analysed during the related study are described in the figshare metadata record: https://doi.org/10.6084/m9.figshare.13259285^32^. Single-cell RNA sequencing data is available via the NCBI Gene Expression Omnibus repository with accession: https://identifiers.org/geo:GSE153319^33^. These scRNA sequencing data underlie Figures 3–4 and Supplementary Figures 7-8, 10-11. Data supporting Figure 3 and Supplementary Figure 9 are contained in the file ‘T cells.fcs’, which is not openly available to protect patient privacy. Data requests should be made to the corresponding author. The patient clinical data are contained in the file ‘Patient clinical data.xlsx’, which is also not available to protect patient privacy. Data requests should be made to the corresponding author. All other data are shared openly as part of the metadata record^32^. Data supporting Figures 1–2 and Supplementary Figures 1–6 are contained in the .zip file ‘Parenti_et_al_supporting_data.zip’ and arranged in folders named according to the figures they underlie. Data supporting Supplementary Table 4 and Supplementary Data Files 4-6 are contained in the file ‘Supplementary Data File 3.xlsx’.

### Code Availability

Single cell genomics analysis pipelines and codes can be obtained from the original author’s open source R package and are indicated in Methods section. Analyses were performed using default standard parameters. All analyses were performed using R (version 3.6.2) and R Studio (version 1.3.959) software.

## Acknowledgements

This work was supported by Associazione Italiana per la Ricerca sul Cancro (AIRC), IG project number #19818; AIRC 5 per 1000 project #21267; Italian Ministry of Health (project #RF-2016-02362930); Ministry of Education, University and Research (project #2017WXR7ZT_005).

## Author contributions

These authors contributed equally as first authors: Sandra Parenti, Sebastiano Rontauroli, Chiara Carretta, Selene Mallia.

These authors contributed equally as last authors: Matteo Della Porta and Rossella Manfredini.

SP, CC and SM performed genomic single cell analysis; SR performed transcriptomic single cell analysis; EG performed Single cell and colony analysis; CCh selected and recruited patient; CP performed 10X Genomics and scRNA sequencing experiments; LT performed T cell activation analysis by flow cytometry; EB and SF performed CD34+ cell separation; SS and LT performed flow cytometry evaluation of stem cell isolation, OR, SB and ET performed scRNA sequencing analysis; MDP and RM designed the research and wrote the paper. These authors jointly supervised this work.

