## Supplementary Material for "Mutated clones driving leukemic transformation are already detectable at the single cell level in CD34-positive cells in the chronic phase of primary myelofibrosis"

***SUPPLEMENTARY DISCUSSION***

**ScRNA-seq clustering analysis**

By means of graph based clustering task on Partek® Flow® we defined cell clusters specific for each sample; in particular this analysis identified 5 clusters in T1 and 4 clusters in T2 and T3 **(Supplementary Fig. 7).** In order to define the identity of these clusters we classified cells according to their expression profile using Blueprint ENCODE dataset in SingleR package. Furthermore, we defined clusters’ marker genes by means of Partek® Flow® and we compared them with those identified by several authors in different subpopulations of hematopoietic stem/progenitor cells[1-3].

As shown in **Supplementary Fig. 7c,** T1 cluster 1 (T1.1) includes foremost hematopoietic stem cells (HSCs) and multipotent progenitors (MPPs) according to SingleR classification and is characterized by the expression of HSC markers such as *AVP*[4]*, HOPX*[5], *CRHBP*[6], *MEG3*[7] and *IGHM*[8] (**Supplementary Data 1 and Supplementary Fig. 7d**).

MPP cells are also included in cluster 2 (T1.2) which is composed primarily by granulocyte and monocyte progenitors (GMPs) (**Supplementary Figure 7c).** This cluster is characterized by the expression of genes associated to multilineage and granulocytic progenitors such as *SPINK2*, *SPINS3* and *CLEC12A*[9] (**Supplementary Data 1 and Supplementary Fig. 7d**).

According to SingleR classification, clusters 3 and 4 (T1.3 and T1.4) includes almost exclusively megakaryocyte-erythroid progenitors (MEPs) (**Supplementary Fig. 7c**). Marker gene analysis revealed that T1.4 cluster is characterized by the expression of *MS4A2*[10], reported as a marker gene of Eosinophil Basophil and Mastocyte (Eo/B/Mast) progenitors, but also by megakaryocytic and erythroid progenitors’ genes such as *CNRIP1*[11], *FCER1A*[12] and *HPGDS* (**Supplementary Data 1 and Supplementary Fig. 7d**).

Conversely, in T1.3 cluster, which includes not only MEP but also Erythrocytes according to SingleR (**Supplementary Figure 7c**), cells express erythroid progenitors’ specific genes such as *ANK1*[13,14] (**Supplementary Data 1 and Supplementary Fig. 7d**).

According to SingleR classification cluster 5 (T1.5) is composed almost exclusively by megakaryocytes (**Supplementary Figure 7c**) and is characterized by the expression of specific MK progenitor genes such as *GP9* and *ITGB3* (CD61) (**Supplementary Data 1**).
Finally, another cluster was identified; it is composed by different cell types according to SingleR classification, including monocytes (data not shown). Gene expression analysis revealed the expression of marker genes identified in different lymphoid and myeloid populations according to literature findings (data not shown). We considered these 71 cells as contaminants and excluded them from analysis.
As told before, clustering analysis identified 4 cell clusters in both T2 and T3 samples and the same strategy described for Time 1 was applied in order to define cluster identity in these specimens.
As you can see in **Supplementary Fig. 7** we detected high level similarities between clusters identified in the three samples in terms of cell composition (SingleR Blueprint classifications) and marker genes expression. Indeed, T2 cluster 1 (T2.1) and T3 cluster 1 (T3.1) include HSCs, MPPs, GMPs and a small fraction of MEPs like T1 cluster 1 (T1.1) and display common marker genes such as *AVP*, *HOPX* and *CRHBP* specific for hematopoietic stem cells as told before (**Supplementary Data 1 and Supplementary Fig. 7d,h and n**). Therefore, we classified cells belonging to these clusters as HSC_MPP.
T1 cluster 2 (T1.2) displayes a cellular composition highly similar to T2 cluster 2 (T2.2) and T3 cluster 2 (T3.2) including foremost GMPs (**Supplementary Fig. 7c, g and m).** Once again, these clusters display common marker genes associated to granulocytic or multilineage progenitors such as *IGLL1, CLEC12A* and *SPNS3* (**Supplementary Data 1 and Fig. Supplementary 7d, h and n**), therefore we decided to define these clusters as MPP_GMP.
Cells belonging to megakaryocytic or erythroid lineages (MEPs but also erythrocytes and megakaryocytes) were included in T2 and T3 clusters 3 and 4 (**Supplementary Fig. 7g and m)**. T2 cluster 3 (T2.3) and T3 cluster 4 (T3.4) are composed almost exclusively by MEPs such as T1 cluster 4 (T1.4) and shared the expression of genes reported as expressed in Eo/B/Mast progenitors but also MEPs and ERPs (i.e. *MS4A2*, *HPGDS* and *CNRIP1*) (**Supplementary Data 1 and Supplementary Fig. 7d, h and n**). This observation was consistent with recent findings demonstrating the common origin of MEPs and Eo/B/Mast progenitors[15], leading us to hypothesize that these clusters represent a more primitive state in megakaryocytic and erythroid lineage differentiation. For these reasons we grouped these clusters in MEP_1 category.

Conversely T2 cluster 4 (T2.4) and T3 cluster 3 (T3.3) include not only MEPs but also erythrocytes and megakaryocytes like T1 clusters 3 (T1.3) and 5 (T1.5) (**Supplementary Fig. 7c, g and m**), suggesting that these clusters represent late stages along this lineage. Among marker genes we identified the expression of erythroid (*ANK1* and *KLF1*) and megakaryocytic genes (*ITGA2B*) (**Supplementary Data 1 and Supplementary Fig. 7d, h and n).** These clusters were then included in a common MEP_2 group.

By means of AUCell task in Partek® Flow® we evaluated the activation state of gene signatures specific for hematopoietic and leukemic stem cells and progenitor cells identified by Eppert et al.[1]. As shown in **Supplementary Fig. 8** clusters belonging to HSC_MPP (T1.1, T2.1, T3.1) and MPP_GMP (T1.2, T2.2, T3.2) groups display increased activation of gene modules related to HSC phenotype (EPPERT_HSC_R and EPPERT_CE_HSC_LSC modules) (**Supplementary Fig. 8b and c**), while clusters we included in MEP_1 (T1.4, T2.3, T3.4) and MEP_2 (T1.3, T1.5, T2.4, T3.3) groups display activation of gene signatures specific for progenitor subpopulations (EPPERT_PROGENITOR) (**Supplementary Fig. 8d**). This result are therefore in agreement with our classification.

Finally, in T2 and T3 clusters 5 we identified 17 and 76 contaminating dispersed cells respectively, including myeloid and lymphoid cells (data not shown) that were removed from the analysis.

**Inhibition of interferon signaling characterizes disease progression**

By comparing clusters identified in the three samples we were able to identify cellular groups that share the same characteristics in T1, T2 and T3 named HSC_MPP, MPP_GMP, MEP_1 and MEP_2. Therefore, we compared gene expression between samples within the shared clusters. To better understand the molecular mechanisms responsible for changes undergoing during disease progression we studied lists of differentially expressed genes by means of IPA®.

According to IPA® analysis all cell clusters in T3 and T2 samples display a significant inactivation of interferon (IFN) signaling compared to T1 (**Supplementary Data 4 and Supplementary Fig. 9a**); this might be the effect of Ruxolitinib treatment which is able to inhibit this pathway through the downmodulation of JAK/STAT signaling. In all cell clusters belonging to T3, IFN pathway inactivation is enhanced by the upregulation of SOCS1 (**Supplementary Fig. 9a, b**), a known inhibitor of JAK/STAT signaling pathway. Furthermore, several Interferon Regulatory Factors (IRFs) were identified as inhibited upstream regulators by IPA® analysis in all clusters (**Supplementary Data 5**) due to the downregulation of IFN-induced genes such as: STAT1, IFITM1, MX1, OAS1, IFI6 and IFIT3 (**Supplementary Fig. 9b**). Among them, IRF1, IRF3 and IRF7, are key mediators of IFN-induced immune surveillance, involved in type I HLA molecules expression. As long as the disease progresses and acquires a more aggressive phenotype, malignant cells achieve the capability to resist against immune surveillance. Surprisingly, Ruxolitinib actively contributes to this process since it is able to reduce leukemic cells' sensitivity to NK cells[16,17]. Moreover, it was demonstrated that the inhibition of IFN-γ signaling causes the downregulation of type I and type II HLA and B2M[18]. This is consistent with the enhanced immune escaping of malignant cells that lose the ability to present antigens to cytotoxic T cells and therefore can't be eradicated[19]. Our data confirm the deregulation of this axis, which is altered in other stem cells from hematological malignancies such as chronic myeloid leukemia[20].

Moreover, the downregulation of IFN signaling may also be responsible for the less differentiated state of cells in T3 and T2 samples as demonstrated by trajectory analysis (**Fig. 3g**). IFN-signaling was found to be more downregulated in MEP clusters compared to HSC_MPP and MPP_GMP (**Supplementary Data 4**). It has been described that IFN-γ is able to induce myeloid differentiation[21], and its administration has already been proposed as a therapeutic strategy in AML in order to escape the differentiative block[22]. Our results suggest that IFN-γ signaling was strong during the PMF phase, accounting for the accumulation of differentiated myeloid cells. This signal is progressively lost during disease progression, partially due to Ruxolitinib effect, therefore allowing the generation of a less differentiated blast cell population. Again, IPA® analysis supported this hypothesis since "Maturation of blood cells" category was identified among the most significantly inhibited disease and functions in T3 MEP_2 cluster (**Supplementary Table 10**).

**Leukemic transformation is associated to reduced apoptosis and increased quiescence of HSPCs**

According to gene enrichment analysis several disease and functions categories related to "Cell death and survival" were predicted inactivated in all the comparisons. In particular, "Cell Death of hematopoietic cells" and "Cell Death of hematopoietic progenitor cells" turned out as two of the most inhibited functions in MPP_GMP cluster in T3 vs T1 (**Supplementary Data 6**). In this comparison, IRF1 is predicted by IPA® as inactivated upstream regulator due to the downregulation of several direct targets, including the proapoptotic gene BAK1 **(Supplementary Fig. 9c and Supplementary Data 5).** Bak1 protein increases apoptosis of mouse bone marrow-derived neutrophils in cell culture that is increased by Fasl protein[23].

We can therefore hypothesize that IFN-signaling inhibition we observed in our patient during the leukemic transformation, may favor immune escape mechanisms but can also inhibit apoptosis through the regulation of *BAK1*.

Moreover, we observed the activation of genes related to cell quiescence, a process involved in the resistance to therapy of the neoplastic clone. In particular, it has been recently demonstrated in a mouse model that Ruxolitinib treatment is more effective in proliferating cell populations rather than quiescent ones[24]. Our data shows that FOXO1 and FOXO3 are predicted by IPA® analysis as upstream regulators activated in T3 HSC_MPP and MPP_GMP cells (**Supplementary Data 5**) thus demonstrating that quiescence is activated also in leukemic stem cells. FOXO3 is also active in T3 MEP_1 cluster (**Supplementary Data 5).** These transcription factors have already been described as able to promote the maintenance of hematopoietic and leukemic stem cells[25]. FOXO transcriptional factors expression may also be related to *TP53* mutation. It has in fact been described that TP53-mutant is unable to promote FOXO1 and FOXO3 proteasome-mediated degradation in glioblastoma, therefore favoring the stemness of cancer cells[26].

Our results demonstrated the upregulation of several targets of the FOXO family proteins, such as *CDKN1A* (p21), *PIK3IP1*, *EGR1*, *KLF4* and *KLF7*. Moreover, we observed in T3 vs T1 an increased expression of other transcriptional factors such as *EGR1*[27], *EGR3*[28], *NR4A2*[29], *MAFF*[30] and *HES1*[31] (**Supplementary Data 3**), which co-operate to the quiescent state of HSCs and precocious myeloid-biased progenitor cells. In particular, EGR1 is shown also as active upstream regulator in all cell clusters (T3 vs. T1) and stimulates stem cell quiescence by transactivating *AREG*, *SOCS1* and *CDKN1A* (p21) expression (**Supplementary Data 5).**

**Extramedullary hematopoiesis (EMH) increases during disease progression**

IPA® analysis shows that extramedullary hematopoiesis (EMH), which establishes an alternative niche for HSCs because of the fibrotic state of bone marrow (BM)[32], increases during the progression of the disease. In this niche, HSCs are protected from chemical treatments and maintain a quiescent state in order to preserve the malignant population[33]. The activation of this mechanism is confirmed by the clinical features of the patient, who has a marked splenomegaly which has only been partially reduced by Ruxolitinib treatment.

The establishment of a new niche is facilitated by stem cells' mobilizing molecules produced by the BM, the expression of pro-invasive factors by HSCs and the presence of chemoattractant molecules in the microenvironment[34]. Our data demonstrates that the chemokine receptor *CXCR4* is significantly upregulated in T2 and T3 vs T1, especially in HSC_MPP and MPP_GMP clusters (**Supplementary Data 3 and Supplementary Fig. 11**). In T3, CXCR4 upregulation extends to the more mature clusters. This modulation is a sign for disease progression, since CXCR4 is downregulated in PMF circulating CD34+ cells in comparison with healthy donors[35], but its expression is upregulated in AML being indicated as a prognostic marker for a shorter relapse-free survival and overall survival[36].

Several targets involved in cell migration have been described as induced by CXCR4 signaling and are upregulated in our dataset. Among them CXCR4 itself, the pro-leukemogenic factors *CD69*, *ID1* and the pro-EMH factors *EGR1* and *IL8* can be found (**Supplementary Data 3**).

The invasion of ectopic sites by HSCs is promoted by several enzymes, such as metalloproteases, which enable extracellular matrix disruption[37]. *MMP7* is strongly upregulated in our data (HSC_MPP and MPP_GMP clusters in T3 vs. T1 comparison) (**Supplementary Data 3 and Supplementary Fig. 11**). One of the sources for MMP7 production is *PTGS2* (COX2)[38], an enzyme that is highly expressed in T3 HSPCs compared to T1. In particular, the production of prostaglandin E2 (PGE2) mediated by PTSG2 stimulates its receptors. In our dataset, *PTGS2* is strongly upregulated in all T3 vs T1 clusters, while in T2 vs T1 its upregulation is detected only in MEP_1 cluster **(Supplementary Data 3 and Supplementary Fig. 11**). Moreover, PGE2 and its receptors PTGER2 and 4 are active upstream regulators, according to IPA predictions, in MPP_GMP and MEP clusters of both T2 and T3 vs. T1 (**Supplementary Data 5).** PTGER2 and 4 promote through cAMP signaling pathway the transcription of several pro-EMH factors upregulated in our dataset such as *CXCR4*[39], *IL8*[40] and *EGR1*[41]. Of note, upregulation of *PTGS2* (COX2) may also enhance immune escape since it is able to impair NK and cytotoxic T cells function[42].

Together with the upregulation of *MMP7*, the infiltrative process is facilitated by the downregulation of *CDH1* in HSC_MPP and MEP_2 clusters (T3 vs. T1) and the increased expression of *RHOB* in HSC_MPP and MPP_GMP clusters of both T2 and T3 vs T1 (**Supplementary Data 3**). This protein in fact facilitates membrane blebbing and blebby ameboid migration of leukemic cells[43].

As previously said, EMH requires the cooperation between cells and microenvironment. Besides IL8, other chemoattractant cytokines have been identified by IPA analysis as activated upstream regulators in our dataset. Indeed, PDFGB and PF4 turned out among the strongest activated upstream regulators in all cell clusters in T3 sample (**Supplementary**

**Data 5)**. Among the pro-migratory targets induced by PDGFB we found *AREG, PTGS2, SGK1, SERPINE1, IER2* and *RHOB*.

**Leukemic transformation is associated to reduced differentiation**

IPA® analysis identified "Maturation of blood cells" among the most significantly inhibited disease and functions in T3 MEP_2 cluster (**Supplementary Data 6).** This is consistent with the less primed state of leukemic hematopoietic progenitors. In this category we found upregulated genes favoring the differentiative block, such as *CD69* and *KLF2* (**Supplementary Data 3**). On the other hand, we identified several downregulated genes which stimulate hematopoietic differentiation, such as *CDH1*, *DDX58* and *HOXA3* (**Supplementary Data 3).**

Moreover, the downregulation of IFN signaling may contribute to immune escape but may also be responsible for the less differentiated state of these cells. Indeed, as described in the main text, IFN-signaling inhibition was more evident in MEP clusters compared to the stem ones. IFN-γ has been described as able to induce myeloid differentiation and its administration has already been proposed as a therapeutic strategy in AML in order to escape the differentiative block[22].

### SUPPLEMENTARY FIGURES

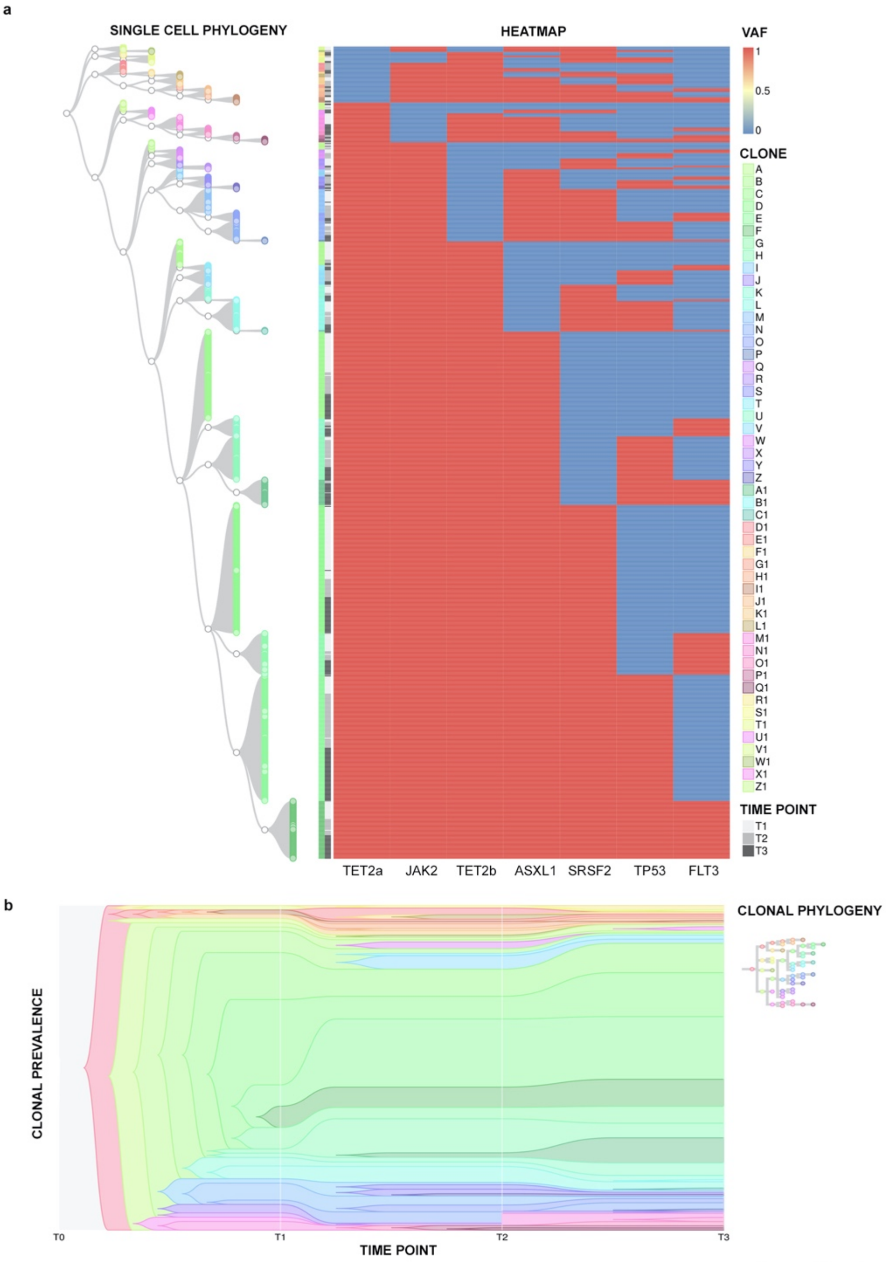


Figure 1 Suppl.

**Supplementary Fig. 1. Single cell phylogeny and clone temporal evolution.** Panel **a** shows the phylogenetic tree of 900 single cells analyzed in T1, T2 and T3. *Left part:* tree starting from a group of founder clones. Each clone (A-Z1) is represented by a color; nodes are represented as white circles. *Right part:* heatmap describing a mutational event (blue: wild-type, red: mutated) of the variants indicated in the bottom part of the graph. Each cell in the phylogenetic tree corresponds to a row in the heatmap, identifying its mutational profile. In Panel **b**, the Fishplot shows the abundance of the 52 clones (A-Z1), identified in Panel a, and their temporal evolution. Each clone is represented by the same color used in Panel a.

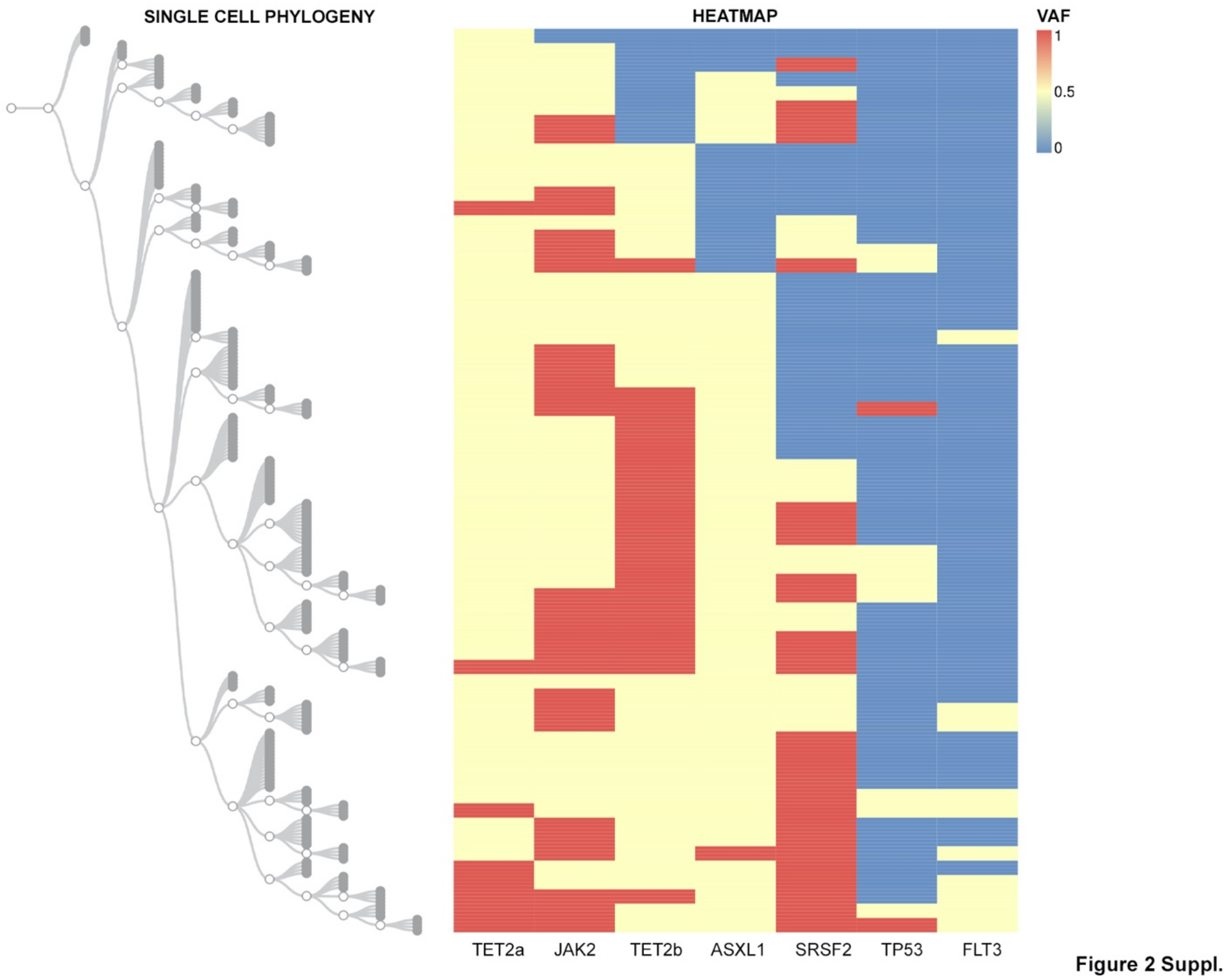


**Supplementary Fig. 2 Single cell phylogeny of Time 1.** Phylogenetic tree reconstruction built based on the zygosity of the different mutations in Time 1 and performed using CellScape R package. *Left part:* phylogenetic tree, black circles represents every single cell, the white circles represents the nodes of different clones. *Right part:* heatmap describing mutational event (blue: wild-type, yellow: heterozygous, red: homozygous) of the variants indicated in the bottom part of the graph. Each cell in the phylogenetic tree corresponds to a row in the heatmap, identifying its mutational profile.

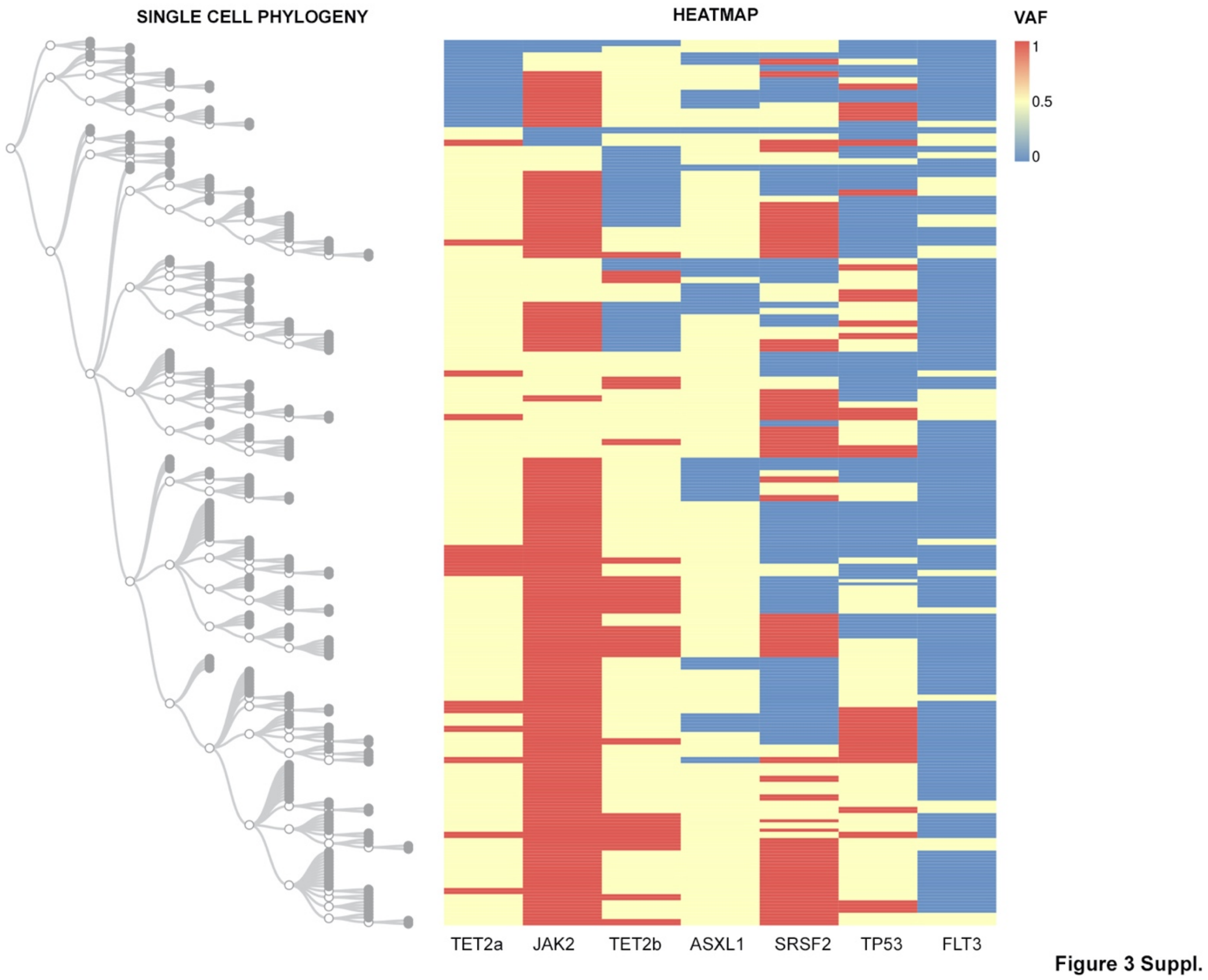


**Supplementary Fig. 3 Single cell phylogeny of Time 2.** Phylogenetic tree reconstruction built based on the zygosity of the different mutations in Time 2 and performed using CellScape R package. *Left part:* phylogenetic tree, black circles represents every single cell, the white circles represents the nodes of different clones. *Right part:* heatmap describing mutational event (blue: wild-type, yellow: heterozygous, red: homozygous) of the variants indicated in the bottom part of the graph. Each cell in the phylogenetic tree corresponds to a row in the heatmap, identifying its mutational profile.

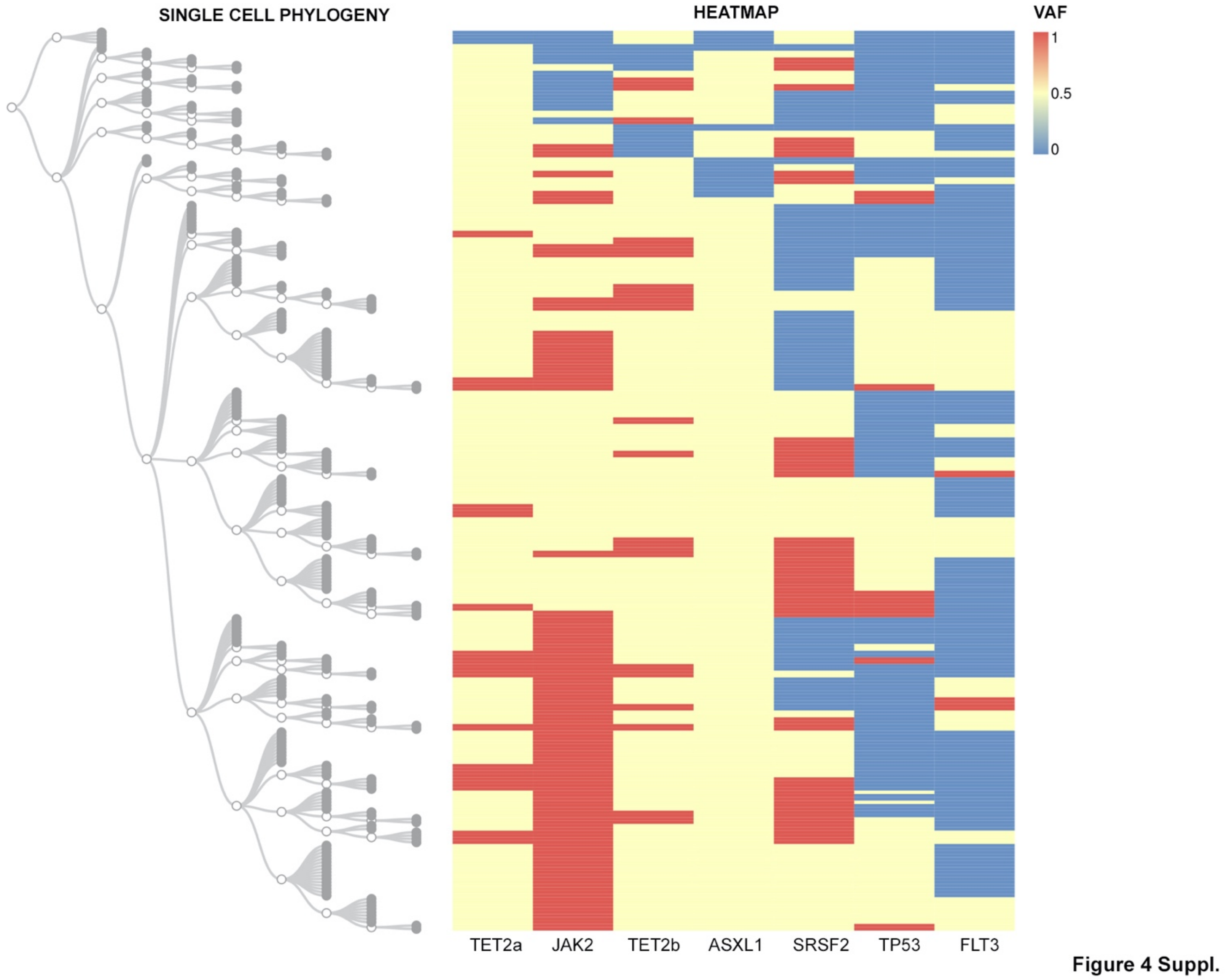


**Supplementary Fig. 4 Single cell phylogeny of Time 3.** Phylogenetic tree reconstruction built based on the zygosity of the different mutations in Time 3 and performed using CellScape R package. *Left part:* phylogenetic tree, black circles represents every single cell, the white circles represents the nodes of different clones. *Right part:* heatmap describing mutational event (blue: wild-type, yellow: heterozygous, red: homozygous) of the variants indicated in the bottom part of the graph. Each cell in the phylogenetic tree corresponds to a row in the heatmap, identifying its mutational profile.

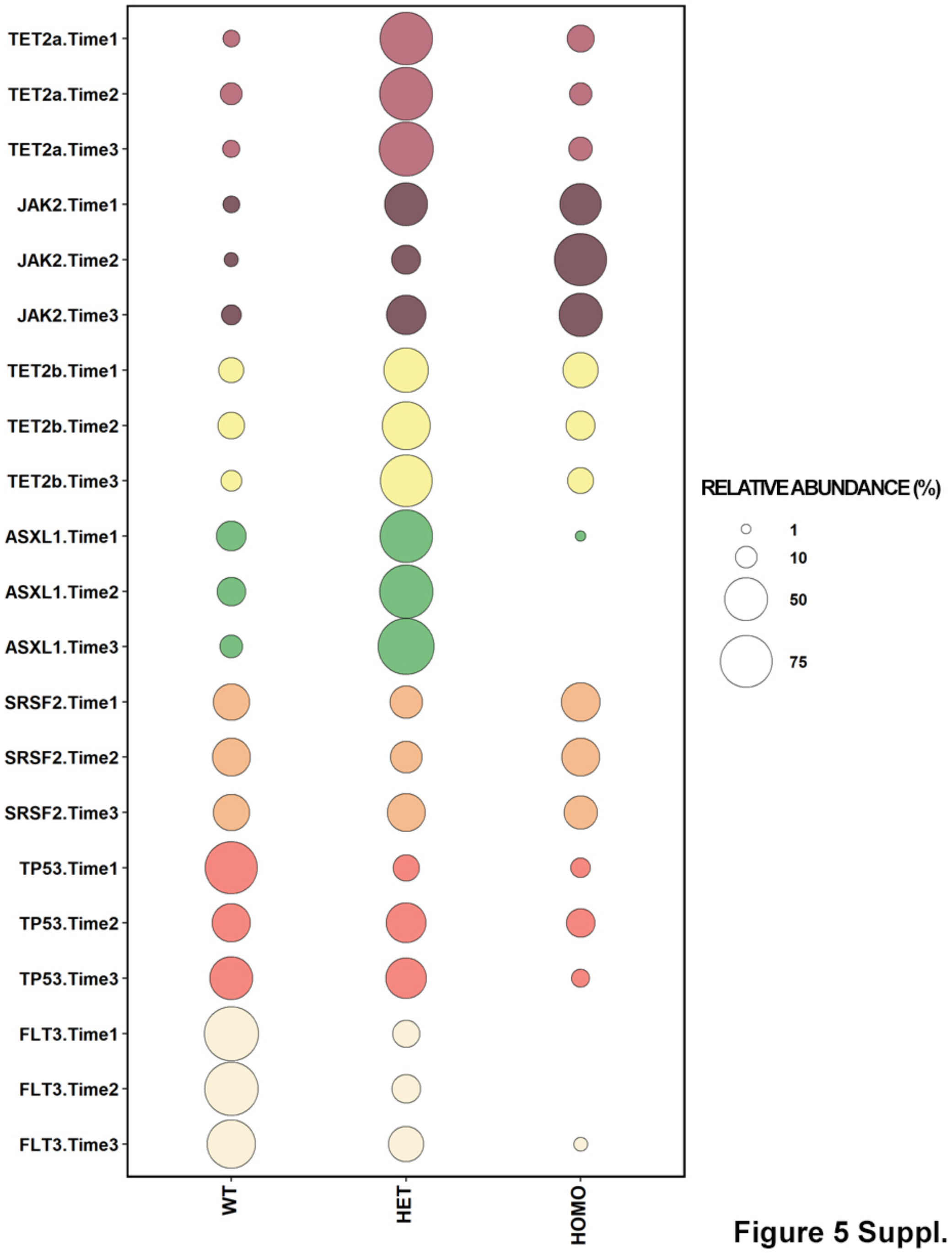


**Supplementary Figure 5**. **Zygosity distribution of all mutations in each time point.** Bubble plot of the frequencies of mutations through time performed by ggplot2 R package. On the y axis are represented all variants in Time 1, 2 and 3; on the x axis is shown the mutational status (WT: wild-type; HET: heterozygous; HOMO: homozygous). Each colour indicates a different variant. Circles' dimension indicates the relative abundance of the event.

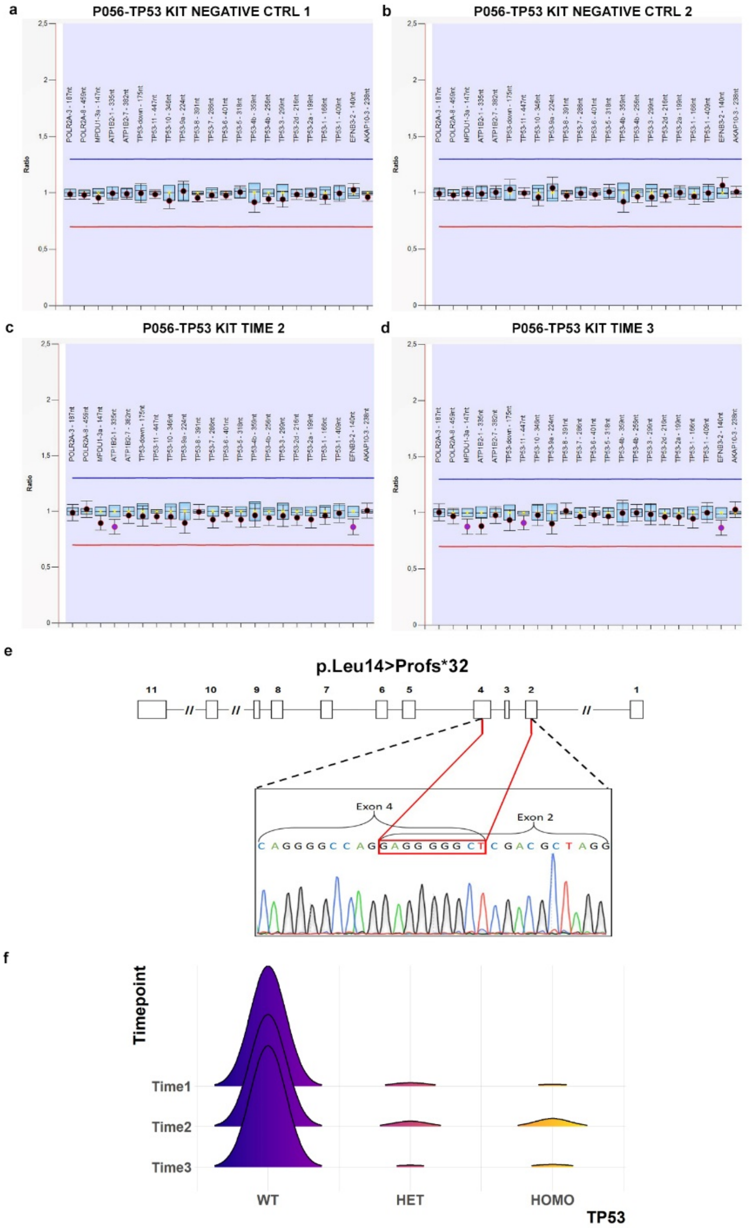


Figure 6 Suppl.

**Supplementary Fig. 6. Characterization of *TP53* deletion**. Panels **a-d** shows the results of Multiplex Ligation-dependent Probe Amplification (MLPA) analysis of *TP53* gene in bulk CD34+ cells. The data obtained from two healthy controls (Ctrl) (**a**, **b**), T2 (**c**) and T3 (**d**) do not show any CNVs. Panel **e** shows a deletion of 454bp in *TP53* gene (c.41_152del, p.Leu14>Profs*32) affecting a region between exons 2 and 4, detected by Sanger sequencing. In Panel **f** the Joy Plot shows the distribution of 454bp *TP53* deletion referred to all CD34+ cells analyzed in T1 (wt 96%, het 2,6%, homo 1,4%), T2 (wt 89,5%, het 4,2%, homo 6,3%) and T3 (wt 97,4%, het 0,65%, homo 1,95%).

Wt: wild-type; het: heterozygous; homo: homozygous.

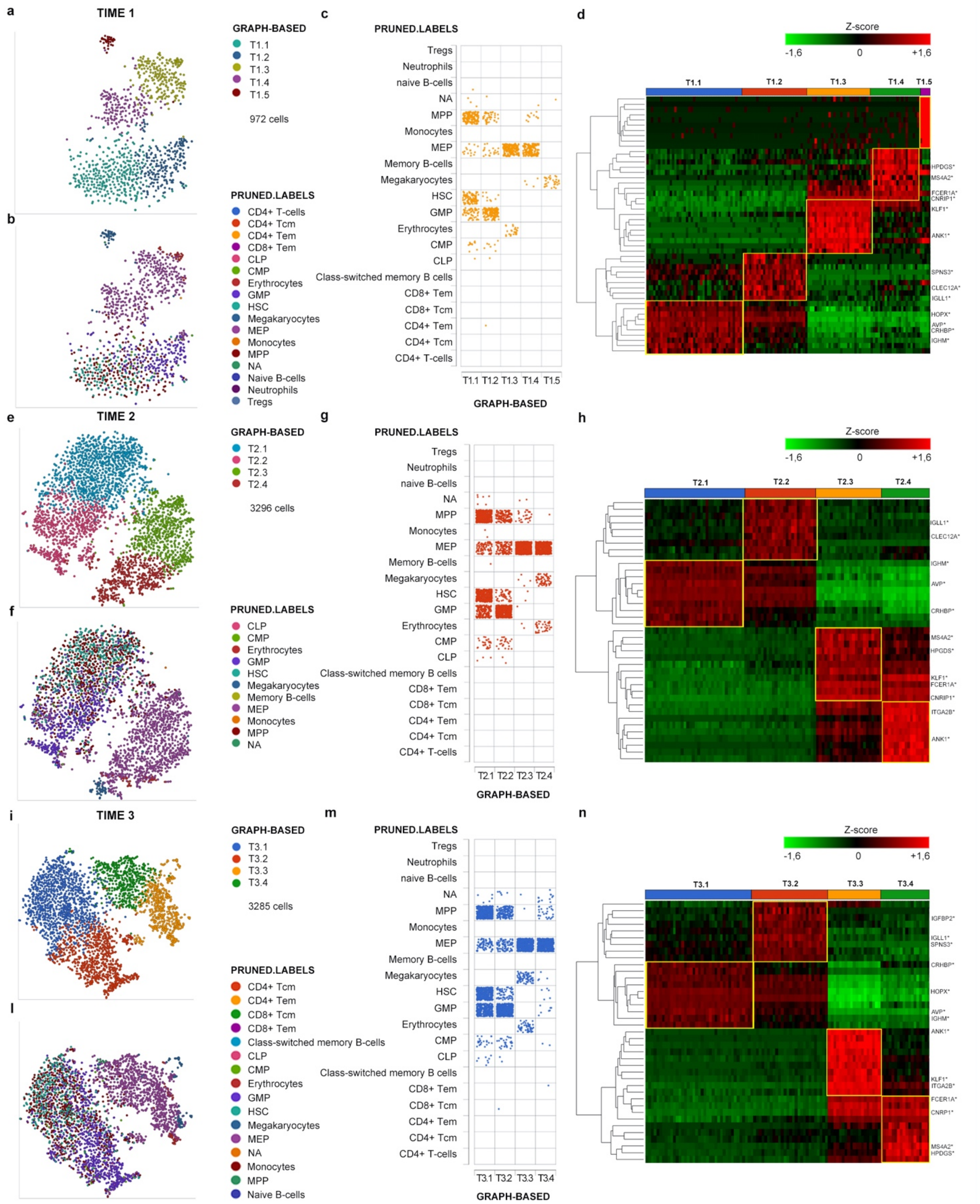


Figure 7 Suppl.

**Supplementary figure 7: a** tSNE projection representing clustering analysis result for T1 sample; in this sample 6 clusters were identified according to Partek Flow. Only 5 clusters are represented since one cluster was excluded due to the presence of contaminant cells. **b** Cells from T1 sample are colored in the tSNE projection according to the classification obtained using SingleR package. The distribution of cells classified using SingleR in the 5 clusters identified in T1 is represented in panel **c. d**: The heatmap represents the supervised hierarchical clustering of cells in T1 and was obtained using the top 10 marker genes for each identified cluster. The same images were generated for T2 (**e** to **h**) and T3 (**i** to **n**) in order to allow the comparison between samples. In panels **d, h** and **n** marker genes shared by the three samples are shown.

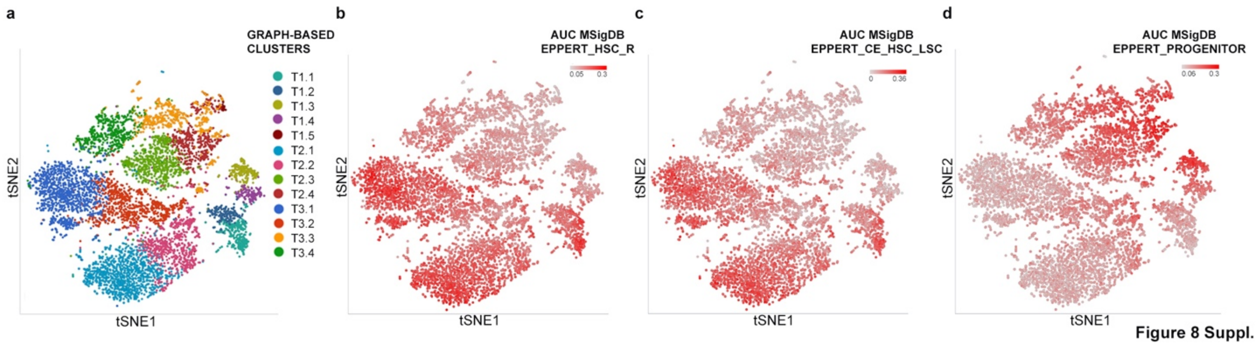


**Supplementary Figure 8:** tSNE projection of cells from all three samples. Data processing and analysis was performed by means of Partek® Flow® software; in these graphs 7553 cells were included. Panel **a** shows the different clusters identified by clustering analysis: 5 in T1 sample and 4 in both T2 and T3. In panels **b, c and d** cells are colored in red scale according to gene module activation, predicted using AUCell function in Partek® Flow® software. Gene signatures derived from Eppert et al. publication and retrieved from MSigDB was analyzed, in particular: **b** represents HSC related signature (EPPERT_HSC_R), **c** represents core enriched (CE) HSC-LSC genes (EPPERT_CE_HSC_LSC) (genes related to HSC features differentially expressed in LSC from AML samples) while panel **d** shows results relative to the progenitor cells' specific signature (EPPERT_PROGENITORS).

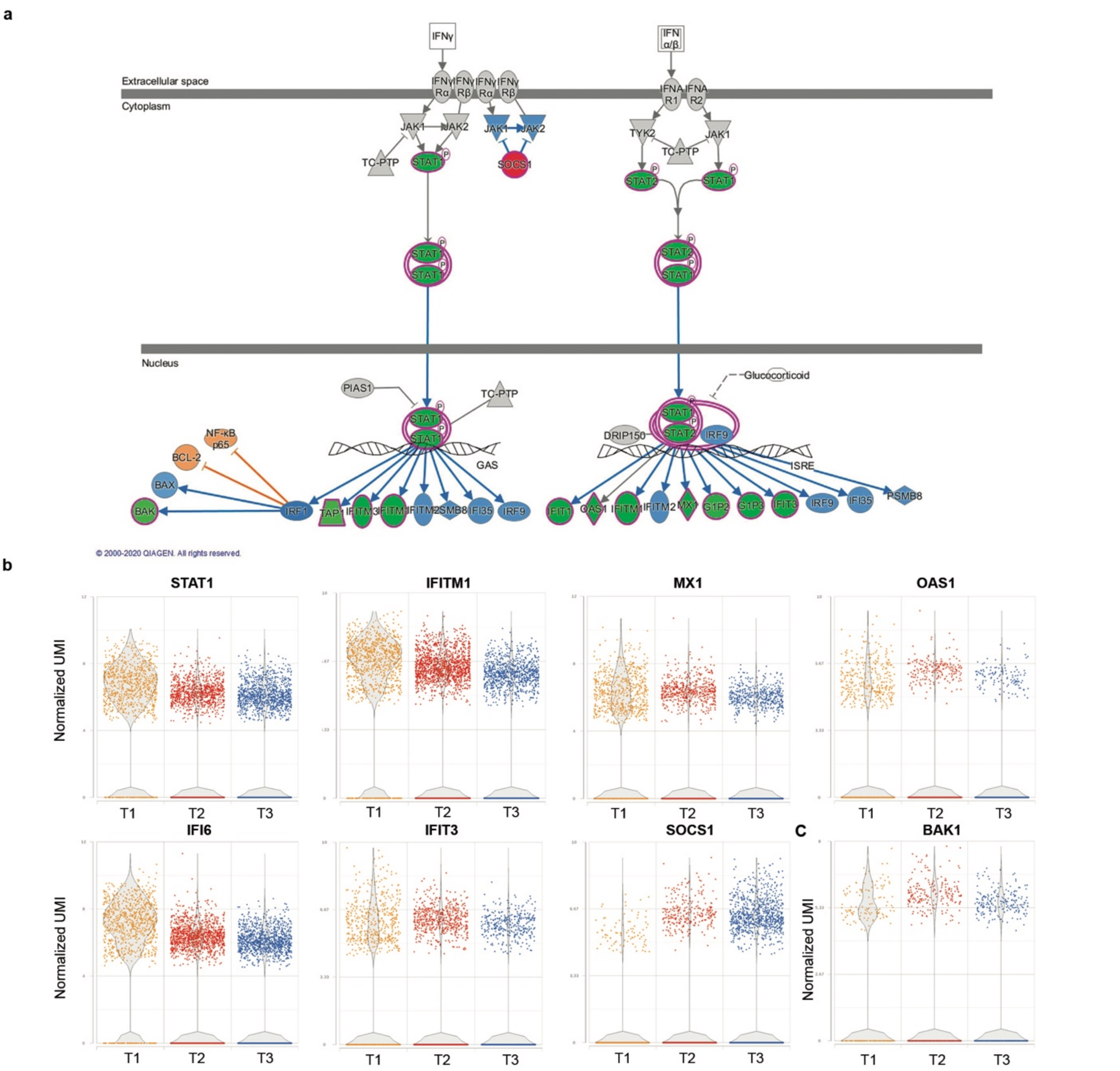


Figure 9 Suppl.

**Supplementary Figure 9: Interferon signaling pathway.** Panel **a** represents the canonical pathway "Interferon Signaling" as it is reported by IPA. Genes in green are downregulated while those in red are upregulated. Blue genes represent predicted inhibited upstream regulators, while orange genes are predicted activated ones. Figure shows differentially expressed genes in the comparison T3 vs T1 MPP_GMP cluster. Panel **b** includes dot plots generated from Partek® Flow® representing the expression of genes involved in IFN signaling according to IPA® in the three timepoints. Panel **c** shows the expression of BAK1 in cells belonging to MPP_GMP cluster in all the three samples.

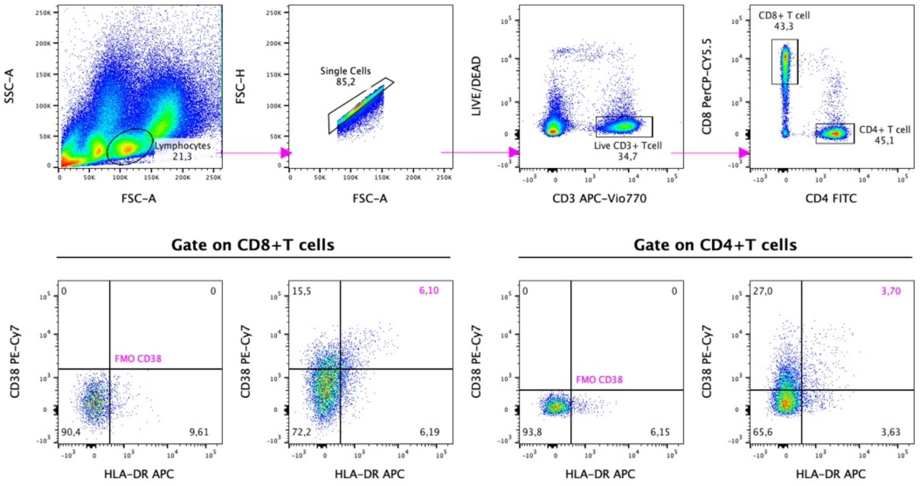


Figure 10 Suppl.

**Supplementary Figure 10. Gating strategy for the detection of activated CD4+ and CD8+ T cells .** Peripheral blood lymphocytes were gated based on SSC and FSC properties and doublets were excluded. Dead cells were excluded as negative to LIVE/DEAD dye. The expression of CD4 and CD8 were evaluated and the activation markers CD38 and HLA-DR were assessed within CD4+ and CD8+ T cells. FMO controls for CD38 and HLA-DR were used to sharply define the CD38+/HLA-DR+ population. The data refer to T2 and are representative of 10 independent experiments.

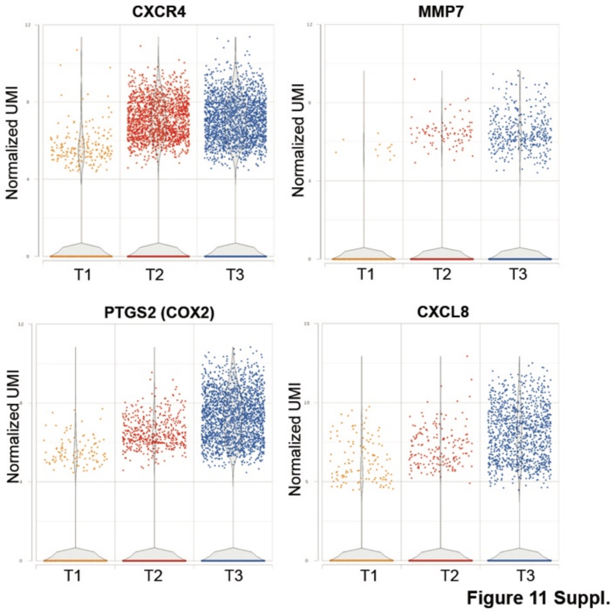


**Supplementary Figure 11:** Dot plots display the increased expression of genes related to extramedullary hematopoiesis in T3 sample compared to T1. Cells belonging to all the four clusters (HSC_MPP, MPP_GMP, MEP_1 and MEP_2) are included in this representation.

### SUPPLEMENTARY TABLES

**Supplementary Table 1: Primers used for PCR reactions in single cell sequencing**

| Gene | ID | Mutation (genomic coordinate hg38) | F | R |
|---|---|---|---|---|
| JAK2 | Hs00532916_CE | g.5073770 | TGAAGCAGCAAGTATGATGAGCAA | CTGACACCTAGCTGTGATCCTG |
| ASXL1 | custom | g.32434646 | CGGATCATCCCCACCACGGAGT | GGCGGCAGTAGTTGTGTTCGC |
| TET2 | Hs00732331_CE | g.105243718 | ATTGTGATTCTCATCCTGGTGTGG | AATGTAAAAGTGCACGCTGAACTC |
| TET2 | Hs00629106_CE | g.105237351_105237358 | ATTGGATACACCTGTCAAGACTCAATATG | CAAGACACAAGCATCGGTAACTTG |
| SRSF2 | Hs00170120_CE | g.76736877 | CGGCTGTGGTGTGAGTCCG | GCTGCCCGCCCAGTTGTTA |
| TP53 | custom | g.7674250 | AAGCCACAGGTTAAGAGGTCCC | AAAAGGCCTCCCCTGCTTGCC |
| FLT3 | Hs00785316_CE | g.28018505 | CCACAGTGAGTGCAGTTGTTTAC | AGAAGAAAGATTGCACTCCAGGATAATAC |

**Supplementary Table 2: % of TP53, FLT3 and SRSF2 mutated cells during disease progression**

| TP53 | WT | HET | HOMO |
|---|---|---|---|
| Time1 | 76 | 16 | 8 |
| Time2 | 38.02 | 42.25 | 19.71 |
| Time3 | 49.34 | 44.07 | 6.57 |

| FLT3 | WT | HET | HOMO |
|---|---|---|---|
| Time1 | 82.6 | 17.3 | 0 |
| Time2 | 80.4 | 19.5 | 0 |
| Time3 | 64.9 | 31.8 | 3.2 |

| SRSF2 | WT | HET | HOMO |
|---|---|---|---|
| Time1 | 33.8 | 26.7 | 39.4 |
| Time2 | 37.8 | 25 | 37.1 |
| Time3 | 34.4 | 37.6 | 27.9 |

**Supplementary Table 3: Frequency of cells classified according to SingleR using Blueprint ENCODE dataset in each cluster**

| | HSC_MPP | | MPP_GMP | | MEP_1 | | MEP_2 | |
|---|---|---|---|---|---|---|---|---|
| SingleR Blueprint Classification | number | frequency | number | frequency | number | frequency | number | frequency |
| Tregs | | 0.00% | | 0.00% | | 0.00% | | 0.00% |
| Neutrophils | | 0.00% | | 0.00% | | 0.00% | | 0.00% |
| Naive B-cells | 1 | 0.04% | | 0.00% | | 0.00% | | 0.00% |
| NA | 14 | 0.52% | 6 | 0.31% | 15 | 1.00% | 2 | 0.14% |
| MPP | 1089 | 40.07% | 332 | 17.26% | 65 | 4.35% | 2 | 0.14% |
| Monocytes | 2 | 0.07% | | 0.00% | | 0.00% | | 0.00% |
| MEP | 172 | 6.33% | 238 | 12.38% | 1381 | 92.50% | 1119 | 78.86% |
| Memory B-Cells | 1 | 0.04% | | 0.00% | | 0.00% | | 0.00% |
| Megakaryocytes | | 0.00% | | 0.00% | 12 | 0.80% | 179 | 12.61% |
| HSC | 839 | 30.87% | 119 | 6.19% | 7 | 0.47% | | 0.00% |
| GMP | 527 | 19.39% | 1161 | 60.37% | 8 | 0.54% | | 0.00% |
| Erythrocytes | | 0.00% | | 0.00% | 2 | 0.13% | 117 | 8.25% |
| CMP | 61 | 2.24% | 60 | 3.12% | 2 | 0.13% | | 0.00% |
| CLP | 12 | 0.44% | 5 | 0.26% | | 0.00% | | 0.00% |
| Class-switched memory B-cells | | 0.00% | | 0.00% | | 0.00% | | 0.00% |
| CD8+ Tem | | 0.00% | | 0.00% | 1 | 0.07% | | 0.00% |
| CD8+ Tcm | | 0.00% | 1 | 0.05% | | 0.00% | | 0.00% |
| CD4+ Tem | | 0.00% | 1 | 0.05% | | 0.00% | | 0.00% |
| CD4+ Tcm | | 0.00% | | 0.00% | | 0.00% | | 0.00% |
| CD4+ T-cells | | 0.00% | | 0.00% | | 0.00% | | 0.00% |
| Total | 2718 | | 1923 | | 1493 | | 1419 | |

**Supplementary Table 4: number of differentially expressed genes (DEGs) in all comparisons**

| | | DEGs \|FC\|>2 and FDR<0.05 | | |
|---|---|---|---|---|
| Comparison | Cluster | Tot | Up | Down |
| T3 vs T1 | HSC_MPP | 1091 | 340 | 751 |
| T2 vs T1 | HSC_MPP | 576 | 268 | 308 |
| T3 vs T1 | MPP_GMP | 854 | 266 | 588 |
| T2 vs T1 | MPP_GMP | 472 | 226 | 246 |
| T3 vs T1 | MEP_1 | 1186 | 264 | 922 |
| T2 vs T1 | MEP_1 | 672 | 243 | 429 |
| T3 vs T1 | MEP_2 | 819 | 283 | 536 |
| T2 vs T1 | MEP_2 | 520 | 234 | 286 |

### Supplementary Table 5: Lists of genes retrieved from literature and used for AUCell analysis in Partek Flow

| VELTEN_FLT3_ SATB1_IND1 | EPPERT_HSC_R | | EPPERT_CE_ HSC_LSC | EPPERT_PROGENITOR | |
|---|---|---|---|---|---|
| TMSB4X | ABCB1 | KDM2A | ABCB1 | ACO1 | MIPEP |
| ITM2C | ADGRG6 | KLF4 | ADGRG6 | ACSM3 | MRE11 |
| FLT3 | ALCAM | KMT2A | ALCAM | ADGRA3 | MRPL35 |
| LSP1 | ANK3 | KSR1 | BAALC | AHCYL1 | MSH3 |
| SPNS3 | ASXL1 | LONP2 | BCL11A | AKAP2 | MTHFD2 |
| PRAM1 | ATP8B4 | LPP | CACNB2 | ANK1 | MTMR2 |
| CORO1A | BAALC | MAFK | CRHBP | ANKRD27 | MTPAP |
| CAPG | BCL11A | MCTP1 | DAPK1 | ARMC8 | MTREX |
| SEPT1 | BEX1 | MECOM | DRAM1 | ASRGL1 | MYCN |
| HLA-A | BTBD11 | MEIS1 | ELK3 | BAMBI | NCBP1 |
| HLA-B | C2orf69 | MLLT3 | ERG | BMS1 | NDC1 |
| SLC2A5 | CACNB2 | MREG | FAM30A | BTK | NDC80 |
| VAMP5 | CALN1 | MSI2 | FLT3 | C5orf30 | NET1 |
| GSTP1 | CD109 | MSMO1 | FRMD4B | CALU | NOMO1 |
| SASH3 | CFH | MYO5C | GUCY1A1 | CCNB2 | ODC1 |
| SH3BGRL3 | COL5A1 | NPR3 | HLA-DRB4 | CCNJ | P2RY1 |
| SORL1 | CRHBP | OGT | HLF | CDK6 | PAICS |
| ITGAL | CRIM1 | OSER1 | HOXA5 | CDK7 | PCLAF |
| SATB1 | CYLD | PAM | HOXB2 | CENPU | PCTP |
| HLA-DQB1 | DACH1 | PAN3-AS1 | HOXB3 | CHEK1 | PDLIM1 |
| LCP1 | DAPK1 | PCNX1 | HTR1F | CHEK2 | PDZD8 |
| FNBP1 | DDX5 | PDE10A | INPP4B | CKAP5 | PEX5 |
| LAIR1 | DLK1 | PLSCR4 | KAT6A | CNRIP1 | PHF6 |
| CD53 | DNAJB9 | PNISR | MECOM | CSNK2A1 | PLIN2 |
| GAPT | DRAM1 | PNP | MEIS1 | CTNNBL1 | PLK4 |
| SDK2 | DST | PPP1R16B | MYO5C | CTPS1 | PMP22 |
| TPM4 | DUSP6 | PRKCH | PLSCR4 | DCAF7 | POLA1 |
| ARPC2 | EIF5 | PROM1 | PPP1R16B | DDAH1 | POLE2 |
| HLX | ELK3 | PTK2 | PRKCH | DLAT | PRKAR2B |
| RNASET2 | EPC1 | RBM7 | RBPMS | DLC1 | PRKDC |
| NEGR1 | ERG | RBPMS | RNF125 | DNAJC6 | PSMD1 |
| CD44 | FAM106A | RCSD1 | SLC25A36 | DTL | RACGAP1 |
| TNFSF8 | FAM169A | RGCC | SMARCA1 | DUT | RFC3 |
| BCL11A | FAM30A | RIMKLB | SOCS2 | EIF2AK1 | RFX7 |
| STK17B | FAM3C | RLIM | SPINK2 | EIF2S1 | RHAG |
| MAP1A | FGD5 | RNF125 | SPTBN1 | EIF5B | RMDN1 |
| LITAF | FLT3 | RPL31 | TCEAL9 | EREG | RRM1 |
| | FNBP1 | RPS20P22 | TFPI | FAM171A1 | RYR3 |
| | FOXO1 | RSL1D1 | TMEM38B | FANCI | SART3 |
| | FRMD4B | RUNX2 | YES1 | FBXO7 | SCD |
| | GCNT2 | SEL1L3 | ZNF165 | FECH | SLC27A2 |
| | GNL1 | SLC17A9 | | FKBP14 | SLF1 |
| | GUCY1A1 | SLC25A36 | | GINS2 | SMC1A |
| | HES1 | SMARCA1 | | HAUS6 | SORD |
| | HIST1H2BD | SMARCA2 | | HBS1L | SPTA1 |
| | HIST1H2BE | SOCS2 | | HDC | SSX2IP |
| | HIST1H2BG | SPINK2 | | HIRA | STAM |
| | HIST1H2BI | SPTBN1 | | HMGB1 | STEAP3 |
| | HIST2H2AA3 | ST3GAL6 | | HMGXB4 | SUPT16H |
| | HLA-DRB4 | TCEAL9 | | HNRNPR | TAL1 |
| | HLF | TCF12 | | HS2ST1 | TFRC |
| | HOPX | TFPI | | HSP90AA1 | TIPIN |
| | HOXA5 | TMEM107 | | IDE | TM9SF3 |
| | HOXB2 | TMEM200A | | IDH3A | TMEM97 |
| | HOXB3 | TMEM38B | | IPO5 | TMOD3 |
| | HTR1F | TPT1 | | KCNQ5 | TNIK |
| | INPP4B | WDR91 | | KIFAP3 | TPM1 |
| | INSIG1 | YES1 | | KNTC1 | TPR |
| | IPO11 | ZBTB4 | | LANCL2 | TRIP13 |
| | ITSN2 | ZDHHC21 | | LDHB | TRIT1 |
| | JUN | ZEB1 | | LHFPL2 | TXNRD1 |
| | KAT6A | ZNF165 | | LRPPRC | UBE2FP1 |
| | KBTBD8 | | | LRRFIP2 | UMPS |
| | | | | MARCH5 | WASF1 |
| | | | | METAP2 | WRN |
| | | | | METTL14 | XK |
| | | | | MEX3B | ZNF225 |
| | | | | MICAL2 | ZWINT |
| | | | | MINPP1 | |

### SUPPLEMENTARY DATA FILES LEGENDS

**Supplementary Data File 1:** The list of top-25 marker genes for each cluster in each sample ordered according to ascending p-value. It is also specified whether they are reported as marker genes for different hematopoietic stem/progenitor cells populations according to literature.

**Supplementary Data File 2:** The list of top-25 marker genes for each cell cluster identified by our classification ordered according to ascending p-value. It is also specified whether they are reported as marker genes for different hematopoietic stem/progenitor cells populations according to literature.

**Supplementary Data File 3**: The results of ANOVA analysis. For each identified cluster T3 and T2 sample was compared to T1. For each comparison Storey Q-value, false discovery rate step up p-value and fold change are reported. These data were used to perform core analysis by means of Ingenuity Pathway Analysis (IPA).

**Supplementary Data File 4**: The results of core analysis performed with Ingenuity Pathway Analysis (IPA). This sheet contains the list of canonical pathways predicted activated (z-score >2) or inhibited (z-score <2) in all comparisons.

**Supplementary Data File 5**: The results of core analysis performed with Ingenuity Pathway Analysis (IPA). This sheet contains the list of upstream regulators predicted activated (z-score >2) or inhibited (z-score <-2) in all comparisons.

**Supplementary Data File 6**: The results of core analysis performed with Ingenuity Pathway Analysis (IPA). This sheet contains the list of disease and function categories predicted activated (z-score >2) or inhibited (z-score <-2) in all comparisons.
