## Supplementary Data File 1 for "Mutated clones driving leukemic transformation are already detectable at the single cell level in CD34-positive cells in the chronic phase of primary myelofibrosis"

| Sample | Cluster | Rank | Gene | Cluster p-value | Cluster fold change | Hay et al. marker genes | Velten et al. marker genes | Eppert et al marker genes |
|---|---|---|---|---|---|---|---|---|
| Time 1 | Cluster 1 | 1 | AVP | 6.7863E-113 | 7.191645089 | HSC | Imm1 | #N/D |
| Time 1 | Cluster 1 | 2 | HOPX | 1.69147E-94 | 4.358444867 | HSC | Imm1 | EPPERT_HSC |
| Time 1 | Cluster 1 | 3 | SPINK2 | 2.26924E-93 | 2.85585882 | MultiLin | Imm1 | EPPERT_HSC |
| Time 1 | Cluster 1 | 4 | CRHBP | 1.08428E-88 | 4.208555882 | HSC | Imm1 | EPPERT_HSC |
| Time 1 | Cluster 1 | 5 | MEG3 | 1.05724E-80 | 5.71910098 | #N/D | #N/D | #N/D |
| Time 1 | Cluster 1 | 6 | MDK | 4.49699E-71 | 4.153704055 | HSC | Imm2 High Cycl | #N/D |
| Time 1 | Cluster 1 | 7 | AC026369.3 | 1.07277E-62 | 5.26159117 | #N/D | #N/D | #N/D |
| Time 1 | Cluster 1 | 8 | IGHM | 2.02358E-61 | 4.66522745 | #N/D | #N/D | #N/D |
| Time 1 | Cluster 1 | 9 | AJ009632.2 | 7.836E-57 | 7.927901461 | #N/D | #N/D | #N/D |
| Time 1 | Cluster 1 | 10 | NPR3 | 4.53703E-56 | 2.71122751 | HSC | STEMNET MK | EPPERT_HSC |
| Time 1 | Cluster 1 | 11 | ADGRG6 | 1.38643E-54 | 5.899175564 | #N/D | Imm1 | EPPERT_HSC |
| Time 1 | Cluster 1 | 12 | RBPMS | 2.22057E-53 | 4.923854128 | HSC | Imm1 | EPPERT_HSC |
| Time 1 | Cluster 1 | 13 | CSF3R | 9.54214E-53 | 2.543012196 | MultiLin | Imm1 | #N/D |
| Time 1 | Cluster 1 | 14 | BAALC | 7.59291E-51 | 3.142275115 | HSC | STEMNET B | EPPERT_HSC |
| Time 1 | Cluster 1 | 15 | VWF | 3.79597E-49 | 4.206750229 | MKP | #N/D | #N/D |
| Time 1 | Cluster 1 | 16 | DLK1 | 1.54468E-47 | 4.55955266 | HSC | #N/D | EPPERT_HSC |
| Time 1 | Cluster 1 | 17 | RASSF6 | 1.72194E-47 | 13.18455135 | HSC | STEMNET Ery | #N/D |
| Time 1 | Cluster 1 | 18 | AC011139.1 | 1.13373E-44 | 5.045847116 | #N/D | #N/D | #N/D |
| Time 1 | Cluster 1 | 19 | SELL | 2.85305E-43 | 1.999558103 | MultiLin | Imm1 | #N/D |
| Time 1 | Cluster 1 | 20 | HLA-DQA1 | 6.89947E-43 | 2.642315 | pre-B | #N/D | #N/D |
| Time 1 | Cluster 1 | 21 | HLA-DPA1 | 9.84313E-43 | 2.113953961 | pre-PC | #N/D | #N/D |
| Time 1 | Cluster 1 | 22 | PCDH9 | 2.27715E-42 | 6.783892702 | pre-B | #N/D | #N/D |
| Time 1 | Cluster 1 | 23 | CD74 | 1.14135E-41 | 2.002659703 | pre-PC | STEMNET Ery | #N/D |
| Time 1 | Cluster 1 | 24 | NRIP1 | 7.74478E-40 | 2.631918516 | HSC | #N/D | #N/D |
| Time 1 | Cluster 1 | 25 | NPDC1 | 2.97682E-39 | 3.110075773 | HSC | #N/D | #N/D |
| Time 1 | Cluster 2 | 1 | IGLL1 | 2.58216E-62 | 5.460078827 | pre-PC | STEMNET MK | #N/D |
| Time 1 | Cluster 2 | 2 | SPINK2 | 1.41372E-50 | 2.212249125 | MultiLin | Imm1 | EPPERT_HSC |
| Time 1 | Cluster 2 | 3 | MPO | 8.88673E-49 | 4.144011538 | Gran | STEMNET Gran | #N/D |
| Time 1 | Cluster 2 | 4 | SPNS3 | 2.48719E-47 | 2.848688653 | MultiLin | #N/D | #N/D |
| Time 1 | Cluster 2 | 5 | TSPOAP1 | 1.96095E-45 | 8.007286399 | #N/D | #N/D | #N/D |
| Time 1 | Cluster 2 | 6 | TRGC2 | 7.43182E-45 | 4.033437229 | #N/D | #N/D | #N/D |
| Time 1 | Cluster 2 | 7 | CLEC12A | 5.26748E-42 | 2.034856973 | Gran | #N/D | #N/D |
| Time 1 | Cluster 2 | 8 | TOP1MT | 2.64443E-41 | 2.528855659 | Gran | STEMNET Gran | #N/D |
| Time 1 | Cluster 2 | 9 | PRAM1 | 1.09768E-40 | 2.543634751 | MultiLin | STEMNET Gran | #N/D |
| Time 1 | Cluster 2 | 10 | RASGRP3 | 2.39017E-40 | 3.258980879 | MKP | STEMNET EBM | #N/D |
| Time 1 | Cluster 2 | 11 | CTSG | 8.98404E-40 | 6.613701828 | Gran | STEMNET Gran | #N/D |
| Time 1 | Cluster 2 | 12 | PHGDH | 2.04972E-36 | 2.647608122 | MultiLin | Imm2 High Cycl | #N/D |
| Time 1 | Cluster 2 | 13 | ATP8B4 | 4.49204E-36 | 2.135966235 | MultiLin | STEMNET Gran | EPPERT_HSC |
| Time 1 | Cluster 2 | 14 | IGFBP2 | 1.56054E-35 | 3.330906139 | Gran | #N/D | #N/D |
| Time 1 | Cluster 2 | 15 | RNASE2 | 4.66083E-35 | 3.777255034 | Gran | STEMNET Gran | #N/D |
| Time 1 | Cluster 2 | 16 | C1QTNF4 | 1.71412E-34 | 2.968024448 | MultiLin | STEMNET Gran | #N/D |
| Time 1 | Cluster 2 | 17 | RFLNB | 6.38236E-34 | 2.070555646 | #N/D | #N/D | #N/D |
| Time 1 | Cluster 2 | 18 | BAHCC1 | 1.63699E-33 | 3.487140589 | CLP | #N/D | #N/D |
| Time 1 | Cluster 2 | 19 | ADGRG3 | 1.64884E-33 | 3.782932207 | #N/D | #N/D | #N/D |
| Time 1 | Cluster 2 | 20 | FUT7 | 5.36977E-33 | 3.464077719 | LMPP | #N/D | #N/D |
| Time 1 | Cluster 2 | 21 | LGALS1 | 8.39508E-32 | 2.388146706 | MDP-2 | STEMNET MD | #N/D |
| Time 1 | Cluster 2 | 22 | MGST1 | 9.20234E-32 | 2.452887184 | Gran | STEMNET Gran | #N/D |
| Time 1 | Cluster 2 | 23 | TRGC1 | 1.06171E-31 | 3.575777028 | #N/D | #N/D | #N/D |
| Time 1 | Cluster 2 | 24 | KBTBD11 | 2.7688E-31 | 3.391815013 | Gran | STEMNET Gran | #N/D |
| Time 1 | Cluster 2 | 25 | NPW | 3.07558E-31 | 3.208909185 | Gran | #N/D | #N/D |
| Time 1 | Cluster 3 | 1 | ANK1 | 5.9879E-246 | 27.37217877 | ERP | STEMNET Ery | EPPERT_PROGENITOR |
| Time 1 | Cluster 3 | 2 | DHRS3 | 5.8406E-181 | 12.45654916 | ERP | #N/D | #N/D |
| Time 1 | Cluster 3 | 3 | APOC1 | 5.4164E-172 | 23.2322569 | ERP | STEMNET Ery | #N/D |
| Time 1 | Cluster 3 | 4 | KLF1 | 3.4946E-158 | 8.80199049 | ERP | STEMNET Ery | #N/D |
| Time 1 | Cluster 3 | 5 | TRIB2 | 1.435E-155 | 13.56653206 | ERP | #N/D | #N/D |
| Time 1 | Cluster 3 | 6 | GYPB | 1.0074E-152 | 44.25682423 | ERP-Early | #N/D | #N/D |
| Time 1 | Cluster 3 | 7 | MYL4 | 3.6495E-144 | 6.365493245 | ERP | #N/D | #N/D |
| Time 1 | Cluster 3 | 8 | STXBP6 | 3.6232E-126 | 6.163920908 | ERP-Early | #N/D | #N/D |
| Time 1 | Cluster 3 | 9 | HES6 | 7.7048E-117 | 20.13542614 | ERP | #N/D | #N/D |
| Time 1 | Cluster 3 | 10 | TST | 1.5672E-115 | 8.616842319 | ERP | #N/D | #N/D |
| Time 1 | Cluster 3 | 11 | PVT1 | 3.079E-114 | 8.893980024 | #N/D | #N/D | #N/D |
| Time 1 | Cluster 3 | 12 | PPP1R14A | 5.9496E-113 | 4.994153541 | ERP | #N/D | #N/D |
| Time 1 | Cluster 3 | 13 | AC055874.1 | 1.6192E-112 | 9.968354054 | #N/D | #N/D | #N/D |
| Time 1 | Cluster 3 | 14 | CDH1 | 1.9882E-110 | 24.77994013 | #N/D | STEMNET Ery | #N/D |
| Time 1 | Cluster 3 | 15 | GATA1 | 1.7846E-107 | 4.85060441 | ERP | #N/D | #N/D |
| Time 1 | Cluster 3 | 16 | GYPE | 2.1237E-107 | 12.49572469 | #N/D | #N/D | #N/D |
| Time 1 | Cluster 3 | 17 | KCNH2 | 2.0198E-106 | 6.234193549 | ERP | STEMNET Ery | #N/D |
| Time 1 | Cluster 3 | 18 | RNASE1 | 9.0325E-106 | 26.59905425 | ERP | #N/D | #N/D |
| Time 1 | Cluster 3 | 19 | PRKAR2B | 2.3421E-104 | 5.228835987 | ERP | #N/D | EPPERT_PROGENITOR |
| Time 1 | Cluster 3 | 20 | CNRIP1 | 4.0388E-103 | 3.488010253 | ERP | STEMNET Ery | EPPERT_PROGENITOR |
| Time 1 | Cluster 3 | 21 | HBQ1 | 1.2705E-99 | 11.9978495 | ERP | STEMNET Ery | #N/D |
| Time 1 | Cluster 3 | 22 | APOE | 4.1833E-98 | 22.44841146 | ERP | STEMNET Ery | #N/D |
| Time 1 | Cluster 3 | 23 | ITGA2B | 3.3228E-89 | 2.704170836 | MKP | #N/D | #N/D |
| Time 1 | Cluster 3 | 24 | ADD2 | 2.73292E-86 | 10.43940637 | ERP | STEMNET Ery | #N/D |
| Time 1 | Cluster 3 | 25 | HBD | 6.27385E-86 | 6.339720961 | ERP | #N/D | #N/D |
| Time 1 | Cluster 4 | 1 | HPGDS | 2.4431E-111 | 11.4044606 | Eo/B/Mast | STEMNET MK | #N/D |
| Time 1 | Cluster 4 | 2 | TPSB2 | 4.895E-92 | 21.46224925 | Eo/B/Mast | STEMNET EBM | #N/D |
| Time 1 | Cluster 4 | 3 | TPSAB1 | 6.56804E-88 | 19.12256265 | Eo/B/Mast | #N/D | #N/D |
| Time 1 | Cluster 4 | 4 | DHRS9 | 7.8351E-79 | 6.401579376 | #N/D | #N/D | #N/D |
| Time 1 | Cluster 4 | 5 | CNRIP1 | 2.12036E-74 | 4.284472539 | ERP | STEMNET Ery | EPPERT_PROGENITOR |
| Time 1 | Cluster 4 | 6 | AL157895.1 | 5.15229E-71 | 5.832962217 | #N/D | #N/D | #N/D |
| Time 1 | Cluster 4 | 7 | SLC45A3 | 1.65455E-60 | 5.268106437 | #N/D | STEMNET EBM | #N/D |
| Time 1 | Cluster 4 | 8 | MS4A2 | 3.01814E-59 | 35.10322942 | Eo/B/Mast | STEMNET EBM | #N/D |
| Time 1 | Cluster 4 | 9 | NTRK1 | 1.86581E-58 | 6.610055437 | #N/D | STEMNET EBM | #N/D |
| Time 1 | Cluster 4 | 10 | FCER1A | 6.19296E-58 | 4.547253416 | MEP | STEMNET MK | #N/D |
| Time 1 | Cluster 4 | 11 | GLOD5 | 4.38829E-54 | 6.907254396 | #N/D | #N/D | #N/D |
| Time 1 | Cluster 4 | 12 | HDC | 4.9707E-54 | 8.86091104 | Eo/B/Mast | STEMNET EBM | EPPERT_PROGENITOR |
| Time 1 | Cluster 4 | 13 | CSF2RB | 1.54112E-52 | 4.372658451 | ERP-Early | STEMNET EBM | #N/D |
| Time 1 | Cluster 4 | 14 | GRAP2 | 2.97662E-50 | 8.410394042 | pre-T | #N/D | #N/D |
| Time 1 | Cluster 4 | 15 | SIGLEC10 | 6.8991E-47 | 4.193156796 | Eo/B/Mast | #N/D | #N/D |
| Time 1 | Cluster 4 | 16 | NLK | 9.33842E-44 | 4.049737009 | MEP | #N/D | #N/D |
| Time 1 | Cluster 4 | 17 | IL1RL1 | 2.78555E-43 | 10.18834057 | #N/D | #N/D | #N/D |
| Time 1 | Cluster 4 | 18 | CPA3 | 4.0642E-41 | 3.349424197 | Gran | #N/D | #N/D |
| Time 1 | Cluster 4 | 19 | STXBP6 | 1.03747E-40 | 2.158114471 | ERP-Early | #N/D | #N/D |
| Time 1 | Cluster 4 | 20 | CPPED1 | 1.20509E-39 | 3.763859251 | MEP | #N/D | #N/D |
| Time 1 | Cluster 4 | 21 | ALOX5AP | 4.05088E-39 | 9.092192233 | Eo/B/Mast | STEMNET EBM | #N/D |
| Time 1 | Cluster 4 | 22 | PIEZO2 | 1.36162E-38 | 4.230577281 | #N/D | #N/D | #N/D |
| Time 1 | Cluster 4 | 23 | DEPTOR | 1.46519E-36 | 3.387292549 | MEP | STEMNET MK | #N/D |
| Time 1 | Cluster 4 | 24 | TIMP3 | 9.63105E-36 | 3.261739408 | MEP | STEMNET MK | #N/D |
| Time 1 | Cluster 4 | 25 | ANGPT2 | 2.74448E-35 | 6.178948011 | Eo/B/Mast | STEMNET EBM | #N/D |
| Time 1 | Cluster 6 | 1 | GP9 | 1.3704E-173 | 39.11878922 | MKP | #N/D | #N/D |
| Time 1 | Cluster 6 | 2 | TRPC6 | 2.3485E-154 | 26.87453707 | #N/D | #N/D | #N/D |
| Time 1 | Cluster 6 | 3 | KALRN | 3.5307E-153 | 57.31531198 | MKP | #N/D | #N/D |
| Time 1 | Cluster 6 | 4 | ITGB3 | 3.9018E-146 | 25.29234075 | MKP | #N/D | #N/D |
| Time 1 | Cluster 6 | 5 | PROS1 | 9.6115E-144 | 29.31681353 | MKP | #N/D | #N/D |
| Time 1 | Cluster 6 | 6 | MYEOV | 1.2221E-143 | 36.77337169 | #N/D | #N/D | #N/D |
| Time 1 | Cluster 6 | 7 | CCND1 | 3.5577E-125 | 35.62451332 | MKP | #N/D | #N/D |
| Time 1 | Cluster 6 | 8 | CR1L | 1.6919E-118 | 33.16267323 | #N/D | #N/D | #N/D |
| Time 1 | Cluster 6 | 9 | TUBB1 | 4.2748E-112 | 38.76140501 | MKP | #N/D | #N/D |
| Time 1 | Cluster 6 | 10 | SPX | 2.9987E-106 | 29.66309115 | #N/D | #N/D | #N/D |
| Time 1 | Cluster 6 | 11 | SELP | 3.973E-106 | 29.11692509 | MKP | #N/D | #N/D |
| Time 1 | Cluster 6 | 12 | LRRC32 | 1.7131E-101 | 28.03243582 | #N/D | #N/D | #N/D |
| Time 1 | Cluster 6 | 13 | SLC35D3 | 2.1375E-101 | 19.07917529 | MKP | #N/D | #N/D |
| Time 1 | Cluster 6 | 14 | VEGFC | 3.8134E-101 | 28.96171119 | #N/D | #N/D | #N/D |
| Time 1 | Cluster 6 | 15 | PBK | 5.0992E-101 | 30.06047716 | Lymphoid UNK | #N/D | #N/D |
| Time 1 | Cluster 6 | 16 | GP1BA | 8.5268E-99 | 26.21119084 | MKP | #N/D | #N/D |
| Time 1 | Cluster 6 | 17 | PPBP | 3.039E-98 | 20.69891447 | MKP | #N/D | #N/D |
| Time 1 | Cluster 6 | 18 | F2RL3 | 6.17774E-94 | 34.97371488 | pre-PC | #N/D | #N/D |
| Time 1 | Cluster 6 | 19 | DKK1 | 7.30971E-91 | 19.35084345 | #N/D | #N/D | #N/D |
| Time 1 | Cluster 6 | 20 | CLEC1B | 3.92704E-87 | 20.94628707 | MKP | #N/D | #N/D |
| Time 1 | Cluster 6 | 21 | CCDC68 | 1.29866E-86 | 17.68176498 | #N/D | #N/D | #N/D |
| Time 1 | Cluster 6 | 22 | CMTM5 | 2.82745E-86 | 30.63607713 | MKP | #N/D | #N/D |
| Time 1 | Cluster 6 | 23 | LTBP1 | 1.01902E-80 | 29.25289453 | MKP | STEMNET EBM | #N/D |
| Time 1 | Cluster 6 | 24 | GP6 | 3.81712E-80 | 20.11127125 | MKP | #N/D | #N/D |
| Time 1 | Cluster 6 | 25 | LY6G6F | 2.32719E-76 | 22.83664123 | MKP | #N/D | #N/D |
| Time 2 | Cluster 1 | 1 | AVP | 0 | 5.327850703 | HSC | Imm1 | #N/D |
| Time 2 | Cluster 1 | 2 | HOPX | 1.1253287856676E-310 | 3.686116583 | HSC | Imm1 | EPPERT_HSC |
| Time 2 | Cluster 1 | 3 | SPINK2 | 1.8478E-276 | 3.01373705 | MultiLin | Imm1 | EPPERT_HSC |
| Time 2 | Cluster 1 | 4 | IGHM | 1.2216E-247 | 6.395660523 | #N/D | #N/D | #N/D |
| Time 2 | Cluster 1 | 5 | CRHBP | 6.0214E-230 | 3.622961125 | HSC | Imm1 | EPPERT_HSC |
| Time 2 | Cluster 1 | 6 | CD52 | 3.6218E-201 | 3.064531417 | HSC | Imm1 | #N/D |
| Time 2 | Cluster 1 | 7 | MDK | 1.0382E-175 | 4.527356159 | HSC | Imm2 High Cycl | #N/D |
| Time 2 | Cluster 1 | 8 | SELL | 7.3524E-174 | 2.904237982 | MultiLin | Imm1 | #N/D |
| Time 2 | Cluster 1 | 9 | GIMAP7 | 5.1023E-160 | 6.805948875 | pre-T | #N/D | #N/D |
| Time 2 | Cluster 1 | 10 | CSF3R | 2.0137E-153 | 2.6506747 | MultiLin | Imm1 | #N/D |
| Time 2 | Cluster 1 | 11 | GIMAP1 | 4.6207E-149 | 5.408794281 | pre-T | #N/D | #N/D |
| Time 2 | Cluster 1 | 12 | HLA-DPA1 | 1.0509E-143 | 2.343840399 | pre-PC | #N/D | #N/D |
| Time 2 | Cluster 1 | 13 | MEG3 | 5.6143E-143 | 3.665914069 | #N/D | #N/D | #N/D |
| Time 2 | Cluster 1 | 14 | CD74 | 5.667E-139 | 2.323802557 | pre-PC | STEMNET Ery | #N/D |
| Time 2 | Cluster 1 | 15 | AJ009632.2 | 1.4035E-135 | 6.139714973 | #N/D | #N/D | #N/D |
| Time 2 | Cluster 1 | 16 | HLA-DQA1 | 2.5644E-130 | 2.699635283 | pre-B | #N/D | #N/D |
| Time 2 | Cluster 1 | 17 | AL031733.2 | 2.6551E-126 | 3.356223829 | #N/D | #N/D | #N/D |
| Time 2 | Cluster 1 | 18 | AC026369.3 | 3.5221E-120 | 5.930478688 | #N/D | #N/D | #N/D |
| Time 2 | Cluster 1 | 19 | ISG20 | 2.1352E-119 | 3.545192094 | pre-PC | #N/D | #N/D |
| Time 2 | Cluster 1 | 20 | HIF3A | 2.7592E-119 | 7.364073872 | #N/D | #N/D | #N/D |
| Time 2 | Cluster 1 | 21 | RCSD1 | 1.8551E-118 | 2.548267127 | pre-B | Imm1 | EPPERT_HSC |
| Time 2 | Cluster 1 | 22 | HLA-DPB1 | 2.4364E-116 | 2.162693359 | pre-PC | #N/D | #N/D |
| Time 2 | Cluster 1 | 23 | HLA-DRA | 3.6638E-116 | 2.13570606 | pre-PC | #N/D | #N/D |
| Time 2 | Cluster 1 | 24 | C9orf139 | 1.4983E-109 | 3.652324698 | MultiLin | Imm1 | #N/D |
| Time 2 | Cluster 1 | 25 | HLA-E | 4.1569E-109 | 1.920614288 | pre-B | Imm1 | #N/D |
| Time 2 | Cluster 2 | 1 | IGLL1 | 2.363E-186 | 7.574066884 | pre-PC | STEMNET MK | #N/D |
| Time 2 | Cluster 2 | 2 | RNASE2 | 7.5871E-139 | 9.734701986 | Gran | STEMNET Gran | #N/D |
| Time 2 | Cluster 2 | 3 | LGALS1 | 4.6519E-134 | 4.056776426 | MDP-2 | STEMNET MD | #N/D |
| Time 2 | Cluster 2 | 4 | IGFBP2 | 5.189E-134 | 4.812753998 | Gran | #N/D | #N/D |
| Time 2 | Cluster 2 | 5 | SPINK2 | 4.4028E-130 | 1.699923816 | MultiLin | Imm1 | EPPERT_HSC |
| Time 2 | Cluster 2 | 6 | CLEC12A | 1.2597E-124 | 4.926538366 | Gran | #N/D | #N/D |
| Time 2 | Cluster 2 | 7 | SPNS3 | 3.5431E-121 | 3.371748293 | MultiLin | #N/D | #N/D |
| Time 2 | Cluster 2 | 8 | TRGC2 | 3.3557E-119 | 4.132909562 | #N/D | #N/D | #N/D |
| Time 2 | Cluster 2 | 9 | NPW | 2.6548E-115 | 3.368969856 | Gran | #N/D | #N/D |
| Time 2 | Cluster 2 | 10 | TRGC1 | 1.6013E-113 | 4.20730258 | #N/D | #N/D | #N/D |
| Time 2 | Cluster 2 | 11 | C16orf74 | 6.9985E-107 | 6.142218141 | pre-B | STEMNET Gran | #N/D |
| Time 2 | Cluster 2 | 12 | RFLNB | 1.6161E-105 | 2.193532331 | #N/D | #N/D | #N/D |
| Time 2 | Cluster 2 | 13 | PHGDH | 2.6204E-104 | 2.355646527 | MultiLin | Imm2 High Cycl | #N/D |
| Time 2 | Cluster 2 | 14 | NKG7 | 3.6954E-101 | 2.854128733 | pre-T | STEMNET Gran | #N/D |
| Time 2 | Cluster 2 | 15 | CSF3R | 9.6056E-101 | 1.75735052 | MultiLin | Imm1 | #N/D |
| Time 2 | Cluster 2 | 16 | CTSG | 3.13864E-98 | 10.72265754 | Gran | STEMNET Gran | #N/D |
| Time 2 | Cluster 2 | 17 | RAB32 | 6.77756E-97 | 2.199076888 | Gran | STEMNET Gran | #N/D |
| Time 2 | Cluster 2 | 18 | CEBPD | 3.33647E-95 | 7.745253615 | Gran | STEMNET Gran | #N/D |
| Time 2 | Cluster 2 | 19 | F13A1 | 2.2069E-94 | 4.361316283 | Gran | STEMNET Gran | #N/D |
| Time 2 | Cluster 2 | 20 | TOP1MT | 2.85825E-93 | 2.39504998 | Gran | STEMNET Gran | #N/D |
| Time 2 | Cluster 2 | 21 | PRAM1 | 1.50838E-91 | 2.517569484 | MultiLin | STEMNET Gran | #N/D |
| Time 2 | Cluster 2 | 22 | MGST1 | 1.15123E-90 | 2.238824903 | Gran | STEMNET Gran | #N/D |
| Time 2 | Cluster 2 | 23 | RASGRP3 | 2.46493E-86 | 3.685827887 | MKP | STEMNET EBM | #N/D |
| Time 2 | Cluster 2 | 24 | C1orf186 | 6.56195E-85 | 1.92032792 | MultiLin | #N/D | #N/D |
| Time 2 | Cluster 2 | 25 | ENO1 | 1.11336E-84 | 1.576410676 | MultiLin | #N/D | #N/D |
| Time 2 | Cluster 3 | 1 | CNRIP1 | 0 | 5.878098528 | ERP | STEMNET Ery | EPPERT_PROGENITOR |
| Time 2 | Cluster 3 | 2 | FCER1A | 3.5831E-307 | 5.619312795 | MEP | STEMNET MK | #N/D |
| Time 2 | Cluster 3 | 3 | MS4A2 | 5.086E-272 | 12.50544636 | Eo/B/Mast | STEMNET EBM | #N/D |
| Time 2 | Cluster 3 | 4 | HPGDS | 5.3285E-262 | 10.67218365 | Eo/B/Mast | STEMNET MK | #N/D |
| Time 2 | Cluster 3 | 5 | CSF2RB | 3.2458E-242 | 5.635249522 | ERP-Early | STEMNET EBM | #N/D |
| Time 2 | Cluster 3 | 6 | STXBP6 | 2.6438E-213 | 3.919824101 | ERP-Early | #N/D | #N/D |
| Time 2 | Cluster 3 | 7 | AL157895.1 | 3.0169E-174 | 4.905632561 | #N/D | #N/D | #N/D |
| Time 2 | Cluster 3 | 8 | DHRS9 | 4.4175E-169 | 6.438275281 | #N/D | #N/D | #N/D |
| Time 2 | Cluster 3 | 9 | CTNNBL1 | 7.898E-161 | 3.395329563 | ERP-Early | STEMNET MK | EPPERT_PROGENITOR |
| Time 2 | Cluster 3 | 10 | HDC | 1.0522E-154 | 11.89322417 | Eo/B/Mast | STEMNET EBM | EPPERT_PROGENITOR |
| Time 2 | Cluster 3 | 11 | NTRK1 | 1.1578E-147 | 7.950265241 | #N/D | STEMNET EBM | #N/D |
| Time 2 | Cluster 3 | 12 | TPSAB1 | 1.3029E-139 | 13.06013368 | Eo/B/Mast | #N/D | #N/D |
| Time 2 | Cluster 3 | 13 | MINPP1 | 4.9349E-138 | 3.628815568 | ERP | STEMNET MK | EPPERT_PROGENITOR |
| Time 2 | Cluster 3 | 14 | MYL4 | 1.7178E-134 | 2.632167757 | ERP | #N/D | #N/D |
| Time 2 | Cluster 3 | 15 | CPPED1 | 1.5085E-133 | 3.763268482 | MEP | #N/D | #N/D |
| Time 2 | Cluster 3 | 16 | TPSB2 | 1.5112E-133 | 14.27409112 | Eo/B/Mast | STEMNET EBM | #N/D |
| Time 2 | Cluster 3 | 17 | TFR2 | 1.5124E-124 | 2.823873488 | ERP | STEMNET Ery | #N/D |
| Time 2 | Cluster 3 | 18 | [illegible] | 9.5259E-124 | 3.061494706 | #N/D | #N/D | #N/D |
| Time 2 | Cluster 3 | 19 | GATA1 | 1.5882E-111 | 2.974946106 | ERP | #N/D | #N/D |
| Time 2 | Cluster 3 | 20 | NLK | 3.3133E-109 | 4.006062184 | MEP | #N/D | #N/D |
| Time 2 | Cluster 3 | 21 | PKIG | 2.0569E-108 | 2.790024443 | ERP-Early | #N/D | #N/D |
| Time 2 | Cluster 3 | 22 | CPA3 | 1.4013E-103 | 2.499352984 | Gran | #N/D | #N/D |
| Time 2 | Cluster 3 | 23 | ITGA2B | 4.6996E-101 | 2.119938186 | MKP | #N/D | #N/D |
| Time 2 | Cluster 3 | 24 | KLF1 | 4.7578E-101 | 2.437800341 | ERP | STEMNET Ery | #N/D |
| Time 2 | Cluster 3 | 25 | FTH1P20 | 1.1779E-99 | 1.820098624 | #N/D | #N/D | #N/D |
| Time 2 | Cluster 4 | 1 | HBD | 0 | 18.81501 | ERP | #N/D | #N/D |
| Time 2 | Cluster 4 | 2 | FCGR2A | 4.9E-324 | 12.14502672 | MKP | #N/D | #N/D |
| Time 2 | Cluster 4 | 3 | ITGA2B | 2.89104328890767E-310 | 6.327253386 | MKP | #N/D | #N/D |
| Time 2 | Cluster 4 | 4 | ANK1 | 1.1519E-307 | 7.516729496 | ERP | STEMNET Ery | EPPERT_PROGENITOR |
| Time 2 | Cluster 4 | 5 | CD36 | 1.7576E-299 | 9.078345065 | ERP | STEMNET Ery | #N/D |
| Time 2 | Cluster 4 | 6 | RHAG | 9.7632E-284 | 11.6983136 | #N/D | STEMNET Ery | EPPERT_PROGENITOR |
| Time 2 | Cluster 4 | 7 | MYL4 | 3.4052E-279 | 5.41318492 | ERP | #N/D | #N/D |
| Time 2 | Cluster 4 | 8 | PRKAR2B | 1.5085E-270 | 5.849656824 | ERP | #N/D | EPPERT_PROGENITOR |
| Time 2 | Cluster 4 | 9 | DHRS3 | 1.6257E-257 | 4.450886679 | ERP | #N/D | #N/D |
| Time 2 | Cluster 4 | 10 | KLF1 | 8.152E-255 | 5.501708125 | ERP | STEMNET Ery | #N/D |
| Time 2 | Cluster 4 | 11 | PPP1R14A | 2.2714E-248 | 8.400156744 | ERP | #N/D | #N/D |
| Time 2 | Cluster 4 | 12 | APOC1 | 5.7889E-246 | 10.13169745 | ERP | STEMNET Ery | #N/D |
| Time 2 | Cluster 4 | 13 | GATA1 | 5.1802E-233 | 4.594967074 | ERP | #N/D | #N/D |
| Time 2 | Cluster 4 | 14 | HES6 | 7.2107E-218 | 7.38839973 | ERP | #N/D | #N/D |
| Time 2 | Cluster 4 | 15 | GYPB | 1.0751E-190 | 7.507883372 | ERP-Early | #N/D | #N/D |
| Time 2 | Cluster 4 | 16 | KCNH2 | 7.5603E-184 | 4.471742957 | ERP | STEMNET Ery | #N/D |
| Time 2 | Cluster 4 | 17 | STXBP6 | 1.248E-181 | 3.136555733 | ERP-Early | #N/D | #N/D |
| Time 2 | Cluster 4 | 18 | SPTA1 | 1.5165E-172 | 6.500720308 | ERP | STEMNET Ery | EPPERT_PROGENITOR |
| Time 2 | Cluster 4 | 19 | BLVRB | 4.2772E-167 | 6.711186873 | ERP | STEMNET Ery | #N/D |
| Time 2 | Cluster 4 | 20 | XK | 2.7792E-164 | 6.167343276 | ERP | #N/D | EPPERT_PROGENITOR |
| Time 2 | Cluster 4 | 21 | PCNA | 2.0908E-158 | 4.288755967 | pre-B cycling | Imm2 High Cycl | #N/D |
| Time 2 | Cluster 4 | 22 | CNRIP1 | 1.2935E-152 | 2.406492164 | ERP | STEMNET Ery | EPPERT_PROGENITOR |
| Time 2 | Cluster 4 | 23 | DLC1 | 7.2661E-149 | 4.511404123 | ERP | #N/D | EPPERT_PROGENITOR |
| Time 2 | Cluster 4 | 24 | SMIM1 | 1.3181E-146 | 3.6667545 | ERP | STEMNET Ery | #N/D |
| Time 2 | Cluster 4 | 25 | TYMS | 2.0158E-144 | 4.082605073 | pre-B cycling | Imm2 High Cycl | #N/D |
| Time 3 | Cluster 1 | 1 | HOPX | 1.0515E-307 | 3.858771555 | HSC | Imm1 | EPPERT_HSC |
| Time 3 | Cluster 1 | 2 | AVP | 5.9461E-282 | 4.972328473 | HSC | Imm1 | #N/D |
| Time 3 | Cluster 1 | 3 | SPINK2 | 3.4357E-248 | 2.527194622 | MultiLin | Imm1 | EPPERT_HSC |
| Time 3 | Cluster 1 | 4 | CD52 | 1.5616E-227 | 3.059236404 | HSC | Imm1 | #N/D |
| Time 3 | Cluster 1 | 5 | IGHM | 1.0221E-174 | 4.455355362 | #N/D | #N/D | #N/D |
| Time 3 | Cluster 1 | 6 | ID1 | 7.1255E-169 | 3.846408308 | HSC | #N/D | #N/D |
| Time 3 | Cluster 1 | 7 | CRHBP | 1.6308E-166 | 2.932191248 | HSC | Imm1 | EPPERT_HSC |
| Time 3 | Cluster 1 | 8 | CSF3R | 1.2478E-158 | 2.652296565 | MultiLin | Imm1 | #N/D |
| Time 3 | Cluster 1 | 9 | MDK | 7.3231E-155 | 4.290262039 | HSC | Imm2 High Cycl | #N/D |
| Time 3 | Cluster 1 | 10 | SELL | 1.5327E-151 | 2.519550298 | MultiLin | Imm1 | #N/D |
| Time 3 | Cluster 1 | 11 | HLA-DPA1 | 1.6573E-148 | 2.359338942 | pre-PC | #N/D | #N/D |
| Time 3 | Cluster 1 | 12 | CD74 | 6.0591E-145 | 2.283199992 | pre-PC | STEMNET Ery | #N/D |
| Time 3 | Cluster 1 | 13 | MEG3 | 6.5212E-135 | 3.594613048 | #N/D | #N/D | #N/D |
| Time 3 | Cluster 1 | 14 | HLA-DPB1 | 4.0977E-134 | 2.1969648 | pre-PC | #N/D | #N/D |
| Time 3 | Cluster 1 | 15 | HLA-DRA | 2.2362E-129 | 2.148218733 | pre-PC | #N/D | #N/D |
| Time 3 | Cluster 1 | 16 | HLA-DQA1 | 5.5468E-123 | 2.641748404 | pre-B | #N/D | #N/D |
| Time 3 | Cluster 1 | 17 | HLA-DRB6 | 3.6074E-115 | 2.073371148 | #N/D | #N/D | #N/D |
| Time 3 | Cluster 1 | 18 | BAALC | 7.0733E-107 | 2.746151865 | HSC | STEMNET B | EPPERT_HSC |
| Time 3 | Cluster 1 | 19 | MAL | 1.0401E-105 | 5.698698993 | pre-B | #N/D | #N/D |
| Time 3 | Cluster 1 | 20 | HLA-DRB1 | 2.6422E-104 | 2.060553889 | pre-PC | #N/D | #N/D |
| Time 3 | Cluster 1 | 21 | CD44 | 2.69443E-95 | 1.817209916 | HSC | #N/D | #N/D |
| Time 3 | Cluster 1 | 22 | NFKBIA | 9.41441E-95 | 2.367633846 | MultiLin | #N/D | #N/D |
| Time 3 | Cluster 1 | 23 | HIF3A | 1.66741E-94 | 4.838779662 | #N/D | #N/D | #N/D |
| Time 3 | Cluster 1 | 24 | GIMAP1 | 1.5359E-93 | 3.982927352 | pre-T | #N/D | #N/D |
| Time 3 | Cluster 1 | 25 | ID3 | 1.61684E-93 | 6.593600773 | Lymphoid UNK | #N/D | #N/D |
| Time 3 | Cluster 2 | 1 | IGLL1 | 4.5716E-214 | 6.647192029 | pre-PC | STEMNET MK | #N/D |
| Time 3 | Cluster 2 | 2 | IGFBP2 | 7.1817E-160 | 5.165920543 | Gran | #N/D | #N/D |
| Time 3 | Cluster 2 | 3 | SPNS3 | 3.4389E-141 | 2.786195949 | MultiLin | #N/D | #N/D |
| Time 3 | Cluster 2 | 4 | SPINK2 | 2.1422E-125 | 1.715456433 | MultiLin | Imm1 | EPPERT_HSC |
| Time 3 | Cluster 2 | 5 | PHGDH | 1.9184E-122 | 2.490081196 | MultiLin | Imm2 High Cycl | #N/D |
| Time 3 | Cluster 2 | 6 | NPW | 1.0306E-117 | 3.301935497 | Gran | #N/D | #N/D |
| Time 3 | Cluster 2 | 7 | CDT1 | 4.1665E-114 | 2.648955322 | pre-B cycling | #N/D | #N/D |
| Time 3 | Cluster 2 | 8 | CTSG | 1.6342E-113 | 4.821001762 | Gran | STEMNET Gran | #N/D |
| Time 3 | Cluster 2 | 9 | PKM | 1.5161E-111 | 2.104342863 | Lymphoid UNK | Imm2 High Cycl | #N/D |
| Time 3 | Cluster 2 | 10 | LGALS1 | 2.1622E-109 | 2.51719114 | MDP-2 | STEMNET MD | #N/D |
| Time 3 | Cluster 2 | 11 | TOP1MT | 7.941E-107 | 2.312374668 | Gran | STEMNET Gran | #N/D |
| Time 3 | Cluster 2 | 12 | CLEC12A | 4.3516E-105 | 3.284658279 | Gran | #N/D | #N/D |
| Time 3 | Cluster 2 | 13 | MCM4 | 1.2058E-103 | 2.710505417 | Lymphoid UNK | Imm2 High Cycl | #N/D |
| Time 3 | Cluster 2 | 14 | PCLAF | 1.7214E-103 | 2.25644638 | #N/D | #N/D | EPPERT_PROGENITOR |
| Time 3 | Cluster 2 | 15 | RFLNB | 6.8876E-103 | 2.076820318 | #N/D | #N/D | #N/D |
| Time 3 | Cluster 2 | 16 | RNASE2 | 4.3476E-99 | 3.496896841 | Gran | STEMNET Gran | #N/D |
| Time 3 | Cluster 2 | 17 | TYMS | 5.4694E-99 | 2.343285708 | pre-B cycling | Imm2 High Cycl | #N/D |
| Time 3 | Cluster 2 | 18 | TNFSF13B | 6.35445E-98 | 3.734524502 | MultiLin | STEMNET Gran | #N/D |
| Time 3 | Cluster 2 | 19 | CDCA7 | 2.50302E-95 | 2.22801206 | pre-B cycling | #N/D | #N/D |
| Time 3 | Cluster 2 | 20 | AC133555.4 | 5.19344E-88 | 1.95854497 | #N/D | #N/D | #N/D |
| Time 3 | Cluster 2 | 21 | ADA | 1.51034E-87 | 1.977533434 | CLP | #N/D | #N/D |
| Time 3 | Cluster 2 | 22 | PYGL | 4.86754E-87 | 2.154423992 | MultiLin | #N/D | #N/D |
| Time 3 | Cluster 2 | 23 | ANXA2 | 1.00888E-84 | 1.928415131 | pre-T | #N/D | #N/D |
| Time 3 | Cluster 2 | 24 | CITED4 | 2.99337E-84 | 2.982057429 | Gran | #N/D | #N/D |
| Time 3 | Cluster 2 | 25 | ENO1 | 2.25916E-83 | 1.644046614 | MultiLin | #N/D | #N/D |
| Time 3 | Cluster 3 | 1 | CD36 | 0 | 34.81445848 | ERP | STEMNET Ery | #N/D |
| Time 3 | Cluster 3 | 2 | APOC1 | 0 | 27.04966851 | ERP | STEMNET Ery | #N/D |
| Time 3 | Cluster 3 | 3 | RHAG | 0 | 20.81314749 | #N/D | STEMNET Ery | EPPERT_PROGENITOR |
| Time 3 | Cluster 3 | 4 | PPP1R14A | 0 | 16.90193239 | ERP | #N/D | #N/D |
| Time 3 | Cluster 3 | 5 | ANK1 | 0 | 14.06500927 | ERP | STEMNET Ery | EPPERT_PROGENITOR |
| Time 3 | Cluster 3 | 6 | PRKAR2B | 0 | 12.92743013 | ERP | #N/D | EPPERT_PROGENITOR |
| Time 3 | Cluster 3 | 7 | FCGR2A | 0 | 12.89180046 | MKP | #N/D | #N/D |
| Time 3 | Cluster 3 | 8 | HBD | 0 | 12.51592629 | ERP | #N/D | #N/D |
| Time 3 | Cluster 3 | 9 | KLF1 | 0 | 10.35013348 | ERP | STEMNET Ery | #N/D |
| Time 3 | Cluster 3 | 10 | ITGA2B | 0 | 9.890078881 | MKP | #N/D | #N/D |
| Time 3 | Cluster 3 | 11 | DHRS3 | 0 | 9.767708948 | ERP | #N/D | #N/D |
| Time 3 | Cluster 3 | 12 | GATA1 | 0 | 8.471174455 | ERP | #N/D | #N/D |
| Time 3 | Cluster 3 | 13 | MYL4 | 0 | 7.865626233 | ERP | #N/D | #N/D |
| Time 3 | Cluster 3 | 14 | GYPB | 1.16905E-318 | 18.36088266 | ERP-Early | #N/D | #N/D |
| Time 3 | Cluster 3 | 15 | STXBP6 | 2.21095158717463E-310 | 5.491466651 | ERP-Early | #N/D | #N/D |
| Time 3 | Cluster 3 | 16 | HES6 | 1.4745E-266 | 11.97362451 | ERP | #N/D | #N/D |
| Time 3 | Cluster 3 | 17 | CNRIP1 | 2.0979E-266 | 3.802385045 | ERP | STEMNET Ery | EPPERT_PROGENITOR |
| Time 3 | Cluster 3 | 18 | CA1 | 1.0129E-264 | 8.437444143 | ERP | STEMNET Ery | #N/D |
| Time 3 | Cluster 3 | 19 | AC055874.1 | 3.5766E-258 | 10.05999103 | #N/D | #N/D | #N/D |
| Time 3 | Cluster 3 | 20 | SMIM1 | 2.8957E-255 | 5.20179207 | ERP | STEMNET Ery | #N/D |
| Time 3 | Cluster 3 | 21 | SPTA1 | 1.6068E-243 | 8.092344852 | ERP | STEMNET Ery | EPPERT_PROGENITOR |
| Time 3 | Cluster 3 | 22 | RYR3 | 4.7996E-232 | 6.460103964 | #N/D | #N/D | EPPERT_PROGENITOR |
| Time 3 | Cluster 3 | 23 | FHL2 | 1.9412E-228 | 5.028577499 | ERP | #N/D | #N/D |
| Time 3 | Cluster 3 | 24 | XK | 2.2067E-226 | 8.939269901 | ERP | #N/D | EPPERT_PROGENITOR |
| Time 3 | Cluster 3 | 25 | BLVRB | 6.0704E-221 | 7.735491487 | ERP | STEMNET Ery | #N/D |
| Time 3 | Cluster 4 | 1 | MS4A2 | 2.324007614934E-312 | 20.85251848 | Eo/B/Mast | STEMNET EBM | #N/D |
| Time 3 | Cluster 4 | 2 | HPGDS | 6.0265E-306 | 12.65922169 | Eo/B/Mast | STEMNET MK | #N/D |
| Time 3 | Cluster 4 | 3 | CNRIP1 | 4.6666E-258 | 4.969716289 | ERP | STEMNET Ery | EPPERT_PROGENITOR |
| Time 3 | Cluster 4 | 4 | AL157895.1 | 9.7569E-209 | 5.924893315 | #N/D | #N/D | #N/D |
| Time 3 | Cluster 4 | 5 | FCER1A | 3.68E-198 | 4.95509633 | MEP | STEMNET MK | #N/D |
| Time 3 | Cluster 4 | 6 | TPSAB1 | 6.2824E-185 | 16.54506264 | Eo/B/Mast | #N/D | #N/D |
| Time 3 | Cluster 4 | 7 | TPSB2 | 1.1845E-172 | 15.82088572 | Eo/B/Mast | STEMNET EBM | #N/D |
| Time 3 | Cluster 4 | 8 | HDC | 3.6183E-137 | 15.36970951 | Eo/B/Mast | STEMNET EBM | EPPERT_PROGENITOR |
| Time 3 | Cluster 4 | 9 | STXBP6 | 1.2797E-135 | 3.140345068 | ERP-Early | #N/D | #N/D |
| Time 3 | Cluster 4 | 10 | CTNNBL1 | 2.0997E-127 | 3.68729007 | ERP-Early | STEMNET MK | EPPERT_PROGENITOR |
| Time 3 | Cluster 4 | 11 | NTRK1 | 3.6522E-124 | 8.941145098 | #N/D | STEMNET EBM | #N/D |
| Time 3 | Cluster 4 | 12 | CPPED1 | 4.43E-121 | 4.429575836 | MEP | #N/D | #N/D |
| Time 3 | Cluster 4 | 13 | CSF2RB | 1.6306E-120 | 4.725604878 | ERP-Early | STEMNET EBM | #N/D |
| Time 3 | Cluster 4 | 14 | DHRS9 | 1.2787E-117 | 6.506402122 | #N/D | #N/D | #N/D |
| Time 3 | Cluster 4 | 15 | CPA3 | 5.9835E-116 | 3.13242463 | Gran | #N/D | #N/D |
| Time 3 | Cluster 4 | 16 | IL1RL1 | 4.4663E-99 | 18.31212005 | #N/D | #N/D | #N/D |
| Time 3 | Cluster 4 | 17 | PNMT | 1.93208E-98 | 3.316137217 | ERP | STEMNET Ery | #N/D |
| Time 3 | Cluster 4 | 18 | FTH1P20 | 6.48447E-97 | 1.97006643 | #N/D | #N/D | #N/D |
| Time 3 | Cluster 4 | 19 | CSF1 | 3.94641E-95 | 6.653135132 | ERP | STEMNET Ery | #N/D |
| Time 3 | Cluster 4 | 20 | SLC40A1 | 2.31308E-83 | 2.790019121 | ERP-Early | STEMNET MK | #N/D |
| Time 3 | Cluster 4 | 21 | FREM1 | 5.0659E-79 | 3.352636549 | #N/D | STEMNET MK | #N/D |
| Time 3 | Cluster 4 | 22 | MALAT1 | 2.39001E-77 | 1.756975952 | #N/D | #N/D | #N/D |
| Time 3 | Cluster 4 | 23 | MYL4 | 1.79123E-76 | 2.109816396 | ERP | #N/D | #N/D |
| Time 3 | Cluster 4 | 24 | GRAP2 | 1.72736E-75 | 4.746672242 | pre-T | #N/D | #N/D |
| Time 3 | Cluster 4 | 25 | PKIG | 5.95787E-73 | 2.526458663 | ERP-Early | #N/D | #N/D |
