## Supplementary Data File 2 for "Mutated clones driving leukemic transformation are already detectable at the single cell level in CD34-positive cells in the chronic phase of primary myelofibrosis"

| Sample | Cluster | Rank | Gene | Cluster p-value | Cluster fold change | Hay et al. marker genes | Velten et al. marker genes | Eppert et al marker genes |
|---|---|---|---|---|---|---|---|---|
| All | HSC_MPP | 1 | IGHM | 0 | 5.398792994 | #N/D | #N/D | #N/D |
| All | HSC_MPP | 2 | AVP | 0 | 5.367551564 | HSC | Imm1 | #N/D |
| All | HSC_MPP | 3 | MDK | 0 | 4.332193626 | HSC | Imm2 High Cycl | #N/D |
| All | HSC_MPP | 4 | HOPX | 0 | 3.856227549 | HSC | Imm1 | EPPERT_HSC |
| All | HSC_MPP | 5 | MEG3 | 0 | 3.808108408 | #N/D | #N/D | #N/D |
| All | HSC_MPP | 6 | CRHBP | 0 | 3.392836952 | HSC | Imm1 | EPPERT_HSC |
| All | HSC_MPP | 7 | CD52 | 0 | 2.89673109 | HSC | Imm1 | #N/D |
| All | HSC_MPP | 8 | SPINK2 | 0 | 2.784589821 | MultiLin | Imm1 | EPPERT_HSC |
| All | HSC_MPP | 9 | CSF3R | 0 | 2.636435963 | MultiLin | Imm1 | #N/D |
| All | HSC_MPP | 10 | SELL | 0 | 2.588877425 | MultiLin | Imm1 | #N/D |
| All | HSC_MPP | 11 | HLA-DPA1 | 1.42471476566E-313 | 2.279977559 | pre-PC | #N/D | #N/D |
| All | HSC_MPP | 12 | CD74 | 1.6079E-298 | 2.216877814 | pre-PC | STEMNET Ery | #N/D |
| All | HSC_MPP | 13 | HLA-DQA1 | 2.9614E-275 | 2.623756296 | pre-B | #N/D | #N/D |
| All | HSC_MPP | 14 | HLA-DPB1 | 2.3213E-265 | 2.135719833 | pre-PC | #N/D | #N/D |
| All | HSC_MPP | 15 | HLA-DRA | 1.456E-260 | 2.097245077 | pre-PC | #N/D | #N/D |
| All | HSC_MPP | 16 | AJ009632.2 | 3.4119E-259 | 5.91069746 | #N/D | #N/D | #N/D |
| All | HSC_MPP | 17 | BAALC | 8.9566E-249 | 2.845278084 | HSC | STEMNET B | EPPERT_HSC |
| All | HSC_MPP | 18 | GIMAP1 | 3.5667E-242 | 4.368251053 | pre-T | #N/D | #N/D |
| All | HSC_MPP | 19 | AC026369.3 | 6.4602E-240 | 4.836496276 | #N/D | #N/D | #N/D |
| All | HSC_MPP | 20 | NPR3 | 8.1493E-237 | 2.416887387 | HSC | STEMNET MK | EPPERT_HSC |
| All | HSC_MPP | 21 | RCSD1 | 1.7365E-235 | 2.295790496 | pre-B | Imm1 | EPPERT_HSC |
| All | HSC_MPP | 22 | HLA-DRB6 | 2.7151E-215 | 1.987683887 | #N/D | #N/D | #N/D |
| All | HSC_MPP | 23 | CD44 | 1.1481E-214 | 1.868068995 | HSC | #N/D | #N/D |
| All | HSC_MPP | 24 | HLA-DRB1 | 2.6749E-212 | 2.035794183 | pre-PC | #N/D | #N/D |
| All | HSC_MPP | 25 | HLA-E | 2.8268E-210 | 1.760765711 | pre-B | Imm1 | #N/D |
| All | MEP_1 | 1 | MS4A2 | 0 | 16.54973009 | Eo/B/Mast | STEMNET EBM | #N/D |
| All | MEP_1 | 2 | TPSB2 | 0 | 15.9553614 | Eo/B/Mast | STEMNET EBM | #N/D |
| All | MEP_1 | 3 | TPSAB1 | 0 | 15.1389513 | Eo/B/Mast | #N/D | #N/D |
| All | MEP_1 | 4 | HDC | 0 | 11.76145059 | Eo/B/Mast | STEMNET EBM | EPPERT_PROGENITOR |
| All | MEP_1 | 5 | HPGDS | 0 | 11.61111948 | Eo/B/Mast | STEMNET MK | #N/D |
| All | MEP_1 | 6 | NTRK1 | 0 | 8.046538109 | #N/D | STEMNET EBM | #N/D |
| All | MEP_1 | 7 | DHRS9 | 0 | 6.773339639 | #N/D | #N/D | #N/D |
| All | MEP_1 | 8 | CNRIP1 | 0 | 5.452071145 | ERP | STEMNET Ery | EPPERT_PROGENITOR |
| All | MEP_1 | 9 | AL157895.1 | 0 | 5.416620972 | #N/D | #N/D | #N/D |
| All | MEP_1 | 10 | CSF2RB | 0 | 5.254155437 | ERP-Early | STEMNET EBM | #N/D |
| All | MEP_1 | 11 | FCER1A | 0 | 5.232798329 | MEP | STEMNET MK | #N/D |
| All | MEP_1 | 12 | STXBP6 | 0 | 3.470026429 | ERP-Early | #N/D | #N/D |
| All | MEP_1 | 13 | CTNNBL1 | 1.094948E-317 | 3.449562549 | ERP-Early | STEMNET MK | EPPERT_PROGENITOR |
| All | MEP_1 | 14 | CPPED1 | 2.8919E-287 | 4.023618292 | MEP | #N/D | #N/D |
| All | MEP_1 | 15 | CPA3 | 1.5657E-253 | 2.843001843 | Gran | #N/D | #N/D |
| All | MEP_1 | 16 | MINPP1 | 2.2517E-240 | 3.63171803 | ERP | STEMNET MK | EPPERT_PROGENITOR |
| All | MEP_1 | 17 | NLK | 4.1892E-213 | 3.938600994 | MEP | #N/D | #N/D |
| All | MEP_1 | 18 | GRAP2 | 5.5986E-211 | 6.624792016 | pre-T | #N/D | #N/D |
| All | MEP_1 | 19 | MIR4435-2HG | 9.3528E-209 | 2.721687978 | #N/D | #N/D | #N/D |
| All | MEP_1 | 20 | FTH1P20 | 1.1221E-205 | 1.816663196 | #N/D | #N/D | #N/D |
| All | MEP_1 | 21 | MYL4 | 9.0479E-205 | 2.155409594 | ERP | #N/D | #N/D |
| All | MEP_1 | 22 | FREM1 | 3.2267E-201 | 3.039327636 | #N/D | STEMNET MK | #N/D |
| All | MEP_1 | 23 | PNMT | 6.7613E-200 | 3.062944412 | ERP | STEMNET Ery | #N/D |
| All | MEP_1 | 24 | SLC40A1 | 5.6392E-192 | 2.560661552 | ERP-Early | STEMNET MK | #N/D |
| All | MEP_1 | 25 | TIMP3 | 1.9683E-189 | 2.652064491 | MEP | STEMNET MK | #N/D |
| All | MEP_2 | 1 | APOC1 | 0 | 16.19417752 | ERP | STEMNET Ery | #N/D |
| All | MEP_2 | 2 | HBD | 0 | 15.4782632 | ERP | #N/D | #N/D |
| All | MEP_2 | 3 | RHAG | 0 | 14.99985832 | #N/D | STEMNET Ery | EPPERT_PROGENITOR |
| All | MEP_2 | 4 | CD36 | 0 | 14.5945766 | ERP | STEMNET Ery | #N/D |
| All | MEP_2 | 5 | APOE | 0 | 13.15500212 | ERP | STEMNET Ery | #N/D |
| All | MEP_2 | 6 | GYPB | 0 | 12.88652156 | ERP-Early | #N/D | #N/D |
| All | MEP_2 | 7 | PPP1R14A | 0 | 11.82556455 | ERP | #N/D | #N/D |
| All | MEP_2 | 8 | ANK1 | 0 | 10.61963562 | ERP | STEMNET Ery | EPPERT_PROGENITOR |
| All | MEP_2 | 9 | HES6 | 0 | 9.345447677 | ERP | #N/D | #N/D |
| All | MEP_2 | 10 | HBQ1 | 0 | 9.277463636 | ERP | STEMNET Ery | #N/D |
| All | MEP_2 | 11 | ADD2 | 0 | 9.224508982 | ERP | STEMNET Ery | #N/D |
| All | MEP_2 | 12 | CA1 | 0 | 9.152764282 | ERP | STEMNET Ery | #N/D |
| All | MEP_2 | 13 | FCGR2A | 0 | 9.027939904 | MKP | #N/D | #N/D |
| All | MEP_2 | 14 | PRKAR2B | 0 | 8.424057265 | ERP | #N/D | EPPERT_PROGENITOR |
| All | MEP_2 | 15 | XK | 0 | 7.419360926 | ERP | #N/D | EPPERT_PROGENITOR |
| All | MEP_2 | 16 | ITGA2B | 0 | 7.362404572 | MKP | #N/D | #N/D |
| All | MEP_2 | 17 | SPTA1 | 0 | 7.224652001 | ERP | STEMNET Ery | EPPERT_PROGENITOR |
| All | MEP_2 | 18 | BLVRB | 0 | 7.060477878 | ERP | STEMNET Ery | #N/D |
| All | MEP_2 | 19 | AC055874.1 | 0 | 6.975525489 | #N/D | #N/D | #N/D |
| All | MEP_2 | 20 | MYL4 | 0 | 6.789306704 | ERP | #N/D | #N/D |
| All | MEP_2 | 21 | KLF1 | 0 | 6.78603344 | ERP | STEMNET Ery | #N/D |
| All | MEP_2 | 22 | DHRS3 | 0 | 6.523001397 | ERP | #N/D | #N/D |
| All | MEP_2 | 23 | GATA1 | 0 | 5.949238038 | ERP | #N/D | #N/D |
| All | MEP_2 | 24 | PVT1 | 0 | 5.820437813 | #N/D | #N/D | #N/D |
| All | MEP_2 | 25 | DLC1 | 0 | 5.517163116 | ERP | #N/D | EPPERT_PROGENITOR |
| All | MPP_GMP | 1 | IGLL1 | 0 | 6.613519343 | pre-PC | STEMNET MK | #N/D |
| All | MPP_GMP | 2 | IGFBP2 | 3.151E-320 | 4.751074935 | Gran | #N/D | #N/D |
| All | MPP_GMP | 3 | SPNS3 | 1.5414E-304 | 3.016711965 | MultiLin | #N/D | #N/D |
| All | MPP_GMP | 4 | SPINK2 | 4.5624E-303 | 1.766509967 | MultiLin | Imm1 | EPPERT_HSC |
| All | MPP_GMP | 5 | LGALS1 | 4.872E-260 | 2.909658167 | MDP-2 | STEMNET MD | #N/D |
| All | MPP_GMP | 6 | RNASE2 | 1.0972E-259 | 5.009899035 | Gran | STEMNET Gran | #N/D |
| All | MPP_GMP | 7 | CLEC12A | 2.5981E-259 | 3.407125173 | Gran | #N/D | #N/D |
| All | MPP_GMP | 8 | NPW | 2.8176E-259 | 3.323485914 | Gran | #N/D | #N/D |
| All | MPP_GMP | 9 | PHGDH | 2.1646E-258 | 2.452373598 | MultiLin | Imm2 High Cycl | #N/D |
| All | MPP_GMP | 10 | CTSG | 2.4828E-246 | 6.363000293 | Gran | STEMNET Gran | #N/D |
| All | MPP_GMP | 11 | RFLNB | 3.0298E-236 | 2.11325588 | #N/D | #N/D | #N/D |
| All | MPP_GMP | 12 | TOP1MT | 3.028E-235 | 2.377080679 | Gran | STEMNET Gran | #N/D |
| All | MPP_GMP | 13 | TRGC2 | 1.9535E-216 | 3.363268763 | #N/D | #N/D | #N/D |
| All | MPP_GMP | 14 | C16orf74 | 1.1011E-212 | 4.727008313 | pre-B | STEMNET Gran | #N/D |
| All | MPP_GMP | 15 | TRGC1 | 1.1097E-202 | 3.546220472 | #N/D | #N/D | #N/D |
| All | MPP_GMP | 16 | PKM | 2.9292E-198 | 1.867894796 | Lymphoid UNK | Imm2 High Cycl | #N/D |
| All | MPP_GMP | 17 | CSF3R | 7.3264E-196 | 1.597518457 | MultiLin | Imm1 | #N/D |
| All | MPP_GMP | 18 | RAB32 | 2.8051E-194 | 1.964805182 | Gran | STEMNET Gran | #N/D |
| All | MPP_GMP | 19 | SELL | 1.041E-191 | 1.50797394 | MultiLin | Imm1 | #N/D |
| All | MPP_GMP | 20 | MGST1 | 5.6647E-187 | 2.169912766 | Gran | STEMNET Gran | #N/D |
| All | MPP_GMP | 21 | ENO1 | 4.6728E-185 | 1.618697419 | MultiLin | #N/D | #N/D |
| All | MPP_GMP | 22 | CDCA7 | 7.0783E-185 | 2.090253166 | pre-B cycling | #N/D | #N/D |
| All | MPP_GMP | 23 | PRAM1 | 2.911E-184 | 2.186135809 | MultiLin | STEMNET Gran | #N/D |
| All | MPP_GMP | 24 | CDT1 | 6.7584E-184 | 2.347904158 | pre-B cycling | #N/D | #N/D |
| All | MPP_GMP | 25 | MPO | 4.0835E-179 | 4.458100508 | Gran | STEMNET Gran | #N/D |
