## Supplementary Data File 3 for "Mutated clones driving leukemic transformation are already detectable at the single cell level in CD34-positive cells in the chronic phase of primary myelofibrosis"

| Gene symbol | Q-value (T3 vs. T1 HSC MPP) | FDR step up (T3 vs. T1 HSC MPP) | Fold change (T3 vs. T1 HSC MPP) | Q-value (T2 vs. T1 HSC MPP) | FDR step up (T2 vs. T1 HSC MPP) | Fold change (T2 vs. T1 HSC MPP) | Q-value (T3 vs. T1 MPP GMP) | FDR step up (T3 vs. T1 MPP GMP) | Fold change (T3 vs. T1 MPP GMP) | Q-value (T2 vs. T1 MPP GMP) | FDR step up (T2 vs. T1 MPP GMP) | Fold change (T2 vs. T1 MPP GMP) | Q-value (T3 vs. T1 MEP 1) | FDR step up (T3 vs. T1 MEP 1) | Fold change (T3 vs. T1 MEP 1) | Q-value (T2 vs. T1 MEP 1) | FDR step up (T2 vs. T1 MEP 1) | Fold change (T2 vs. T1 MEP 1) | Q-value (T3 vs. T1 MEP 2) | FDR step up (T3 vs. T1 MEP 2) | Fold change (T3 vs. T1 MEP 2) | Q-value (T2 vs. T1 MEP 2) | FDR step up (T2 vs. T1 MEP 2) | Fold change (T2 vs. T1 MEP 2) |
|---|---|---|---|---|---|---|---|---|---|---|---|---|---|---|---|---|---|---|---|---|---|---|---|---|
| IFI44L | 0.00E+00 | 0.00E+00 | -2.49E+01 | 0.00E+00 | 0.00E+00 | -8.67E+00 | 1.11E-299 | 4.73E-299 | -4.30E+01 | 7.59E-280 | 2.90E-279 | -1.67E+01 | 7.72E-148 | 2.75E-147 | -4.18E+01 | 4.88E-152 | 1.38E-151 | -2.57E+01 | 1.39E-281 | 5.09E-281 | -6.48E+01 | 2.47E-261 | 6.80E-261 | -3.73E+01 |
| TSC22D3 | 4.14E-292 | 2.16E-291 | 6.32E+00 | 4.63E-298 | 2.00E-297 | 6.65E+00 | 9.35E-184 | 3.97E-183 | 5.34E+00 | 2.35E-200 | 8.98E-200 | 6.10E+00 | 1.00E-268 | 3.57E-268 | 1.22E+01 | 7.07E-239 | 1.99E-238 | 8.97E+00 | 0.00E+00 | 0.00E+00 | 1.33E+01 | 1.29E-280 | 3.55E-280 | 9.11E+00 |
| KLF2 | 2.84E-284 | 1.48E-283 | 5.79E+01 | 9.52E-89 | 4.12E-88 | 2.12E+01 | 1.25E-143 | 5.29E-143 | 4.04E+01 | 2.52E-29 | 9.61E-29 | 1.68E+01 | 3.56E-131 | 1.27E-130 | 6.63E+01 | 1.83E-20 | 5.15E-20 | 1.43E+01 | 3.58E-51 | 1.32E-50 | 1.60E+01 | 1.74E-01 | 4.77E-01 | 2.22E+00 |
| MT-RNR2 | 2.16E-207 | 1.13E-206 | 1.96E+00 | 1.71E-55 | 7.38E-55 | 1.40E+00 | 1.70E-120 | 7.22E-120 | 1.86E+00 | 1.22E-32 | 4.66E-32 | 1.37E+00 | 4.21E-119 | 1.50E-118 | 2.05E+00 | 1.38E-49 | 3.90E-49 | 1.55E+00 | 6.26E-133 | 2.30E-132 | 1.95E+00 | 1.72E-49 | 4.74E-49 | 1.48E+00 |
| CD69 | 2.25E-198 | 1.17E-197 | 1.52E+01 | 3.29E-29 | 1.42E-28 | 3.66E+00 | 1.15E-96 | 4.89E-96 | 1.00E+01 | 5.65E-25 | 2.16E-24 | 3.79E+00 | 2.59E-131 | 9.23E-131 | 1.56E+01 | 7.84E-42 | 2.21E-41 | 4.79E+00 | 2.77E-87 | 1.02E-86 | 8.18E+00 | 3.71E-05 | 1.02E-04 | 1.75E+00 |
| MT-ND1 | 1.76E-185 | 9.19E-185 | 2.32E+00 | 4.47E-95 | 1.93E-94 | 1.82E+00 | 1.10E-103 | 4.66E-103 | 2.11E+00 | 1.08E-52 | 4.14E-52 | 1.71E+00 | 4.32E-88 | 1.54E-87 | 2.25E+00 | 1.75E-56 | 4.94E-56 | 1.87E+00 | 1.61E-111 | 5.92E-111 | 2.24E+00 | 5.16E-80 | 1.42E-79 | 1.93E+00 |
| AREG | 7.54E-175 | 3.93E-174 | 2.18E+01 | 1.51E-47 | 6.52E-47 | 6.20E+00 | 7.88E-128 | 3.35E-127 | 3.28E+01 | 1.49E-39 | 5.68E-39 | 1.00E+01 | 1.60E-102 | 5.71E-102 | 7.64E+01 | 3.49E-30 | 9.83E-30 | 1.87E+01 | 1.40E-71 | 5.13E-71 | 3.09E+01 | 1.07E-14 | 2.94E-14 | 8.24E+00 |
| RPS7 | 4.09E-167 | 2.13E-166 | 1.99E+00 | 2.88E-86 | 1.24E-85 | 1.64E+00 | 3.78E-125 | 1.61E-124 | 2.05E+00 | 1.81E-63 | 6.91E-63 | 1.67E+00 | 5.61E-85 | 2.00E-84 | 1.98E+00 | 1.56E-48 | 4.41E-48 | 1.63E+00 | 2.72E-92 | 9.98E-92 | 1.94E+00 | 2.93E-50 | 8.05E-50 | 1.62E+00 |
| RPL13AP7 | 1.06E-166 | 5.53E-166 | 2.33E+00 | 1.17E-83 | 5.07E-83 | 1.83E+00 | 5.74E-130 | 2.44E-129 | 2.43E+00 | 1.27E-77 | 4.85E-77 | 1.99E+00 | 1.57E-90 | 5.59E-90 | 2.34E+00 | 3.24E-57 | 9.13E-57 | 1.91E+00 | 3.10E-93 | 1.14E-92 | 2.24E+00 | 5.06E-69 | 1.39E-68 | 1.90E+00 |
| RPL13P12 | 1.24E-158 | 6.48E-158 | 1.96E+00 | 5.77E-192 | 2.49E-191 | 2.12E+00 | 1.59E-116 | 6.75E-116 | 2.00E+00 | 3.46E-148 | 1.32E-147 | 2.21E+00 | 2.78E-84 | 9.91E-84 | 1.97E+00 | 9.23E-114 | 2.60E-113 | 2.16E+00 | 2.85E-87 | 1.05E-86 | 1.93E+00 | 1.21E-122 | 3.33E-122 | 2.10E+00 |
| ID1 | 1.15E-152 | 6.00E-152 | 1.24E+01 | 9.38E-03 | 4.05E-02 | 1.05E+00 | 1.86E-42 | 7.90E-42 | 9.60E+00 | 8.42E-04 | 3.22E-03 | 1.04E+00 | 7.12E-12 | 2.54E-11 | 1.28E+01 | 2.99E-01 | 8.42E-01 | 1.01E+00 | 6.35E-06 | 2.33E-05 | 3.82E+00 | 1.13E-03 | 3.10E-03 | -2.04E+00 |
| FP236383.3 | 1.27E-151 | 6.62E-151 | 1.92E+00 | 1.72E-37 | 7.42E-37 | 1.31E+00 | 8.60E-68 | 3.65E-67 | 1.68E+00 | 2.66E-13 | 1.02E-12 | 1.18E+00 | 1.25E-122 | 4.46E-122 | 2.51E+00 | 2.52E-22 | 7.10E-22 | 1.15E+00 | 1.01E-125 | 3.71E-125 | 2.12E+00 | 3.18E-21 | 8.74E-21 | 1.09E+00 |
| MT-CO3 | 1.49E-141 | 7.77E-141 | 1.91E+00 | 2.00E-69 | 8.64E-69 | 1.56E+00 | 5.81E-76 | 2.47E-75 | 1.78E+00 | 1.07E-39 | 4.09E-39 | 1.52E+00 | 1.71E-63 | 6.10E-63 | 1.84E+00 | 1.23E-56 | 3.46E-56 | 1.76E+00 | 1.41E-95 | 5.19E-95 | 1.93E+00 | 1.54E-73 | 4.22E-73 | 1.76E+00 |
| RPS3AP5 | 8.34E-132 | 4.35E-131 | -2.08E+00 | 6.91E-40 | 2.99E-39 | -1.48E+00 | 1.71E-89 | 7.25E-89 | -2.08E+00 | 9.31E-23 | 3.56E-22 | -1.43E+00 | 6.14E-64 | 2.19E-63 | -2.01E+00 | 8.59E-21 | 2.42E-20 | -1.46E+00 | 7.66E-128 | 2.81E-127 | -2.15E+00 | 1.18E-30 | 3.23E-30 | -1.48E+00 |
| HLA-C | 8.06E-131 | 4.20E-130 | -2.59E+00 | 4.00E-88 | 1.73E-87 | -2.14E+00 | 3.07E-62 | 1.30E-61 | -2.23E+00 | 3.89E-48 | 1.49E-47 | -2.05E+00 | 2.96E-104 | 1.05E-103 | -2.96E+00 | 3.21E-78 | 9.06E-78 | -2.59E+00 | 6.57E-165 | 2.41E-164 | -3.48E+00 | 9.51E-112 | 2.61E-111 | -2.98E+00 |
| KLF6 | 9.16E-125 | 4.77E-124 | 4.07E+00 | 3.89E-43 | 1.68E-42 | 2.20E+00 | 4.27E-42 | 1.82E-41 | 3.15E+00 | 6.74E-15 | 2.58E-14 | 1.99E+00 | 2.35E-58 | 8.36E-58 | 4.20E+00 | 1.09E-09 | 3.07E-09 | 1.85E+00 | 7.96E-101 | 2.92E-100 | 6.23E+00 | 1.39E-15 | 3.83E-15 | 2.36E+00 |
| MT-RNR1 | 1.32E-121 | 6.88E-121 | 1.83E+00 | 1.03E-09 | 4.46E-09 | -1.17E+00 | 3.82E-72 | 1.63E-71 | 1.76E+00 | 3.36E-09 | 1.28E-08 | -1.20E+00 | 1.51E-53 | 5.40E-53 | 1.78E+00 | 8.28E-02 | 2.33E-01 | -1.03E+00 | 1.60E-63 | 5.87E-63 | 1.73E+00 | 7.29E-03 | 2.00E-02 | -1.07E+00 |
| IFITM1 | 5.99E-119 | 3.12E-118 | -4.73E+00 | 9.11E-63 | 3.94E-62 | -2.27E+00 | 1.93E-104 | 8.21E-104 | -6.89E+00 | 7.62E-52 | 2.91E-51 | -2.99E+00 | 5.31E-60 | 1.89E-59 | -7.10E+00 | 3.23E-40 | 9.12E-40 | -3.30E+00 | 3.18E-79 | 1.17E-78 | -6.23E+00 | 4.20E-44 | 1.15E-43 | -3.38E+00 |
| FOS | 1.86E-118 | 9.69E-118 | 5.90E+00 | 1.70E-05 | 7.37E-05 | 1.61E+00 | 1.34E-67 | 5.69E-67 | 5.22E+00 | 1.84E-06 | 7.04E-06 | 1.67E+00 | 6.50E-57 | 2.32E-56 | 5.57E+00 | 2.51E-08 | 7.07E-08 | 1.72E+00 | 2.56E-64 | 9.39E-64 | 4.84E+00 | 4.47E-04 | 1.23E-03 | 1.39E+00 |
| RPL10P9 | 2.35E-117 | 1.23E-116 | 1.76E+00 | 1.41E-81 | 6.09E-81 | 1.60E+00 | 3.20E-79 | 1.36E-78 | 1.76E+00 | 1.93E-61 | 7.35E-61 | 1.65E+00 | 2.97E-48 | 1.06E-47 | 1.65E+00 | 2.58E-29 | 7.28E-29 | 1.46E+00 | 1.90E-52 | 6.98E-52 | 1.65E+00 | 4.75E-36 | 1.31E-35 | 1.46E+00 |
| RPS23 | 1.26E-112 | 6.55E-112 | -2.10E+00 | 1.26E-47 | 5.44E-47 | -1.61E+00 | 6.36E-68 | 2.70E-67 | -2.02E+00 | 5.17E-25 | 1.97E-24 | -1.52E+00 | 8.50E-54 | 3.03E-53 | -2.00E+00 | 6.91E-19 | 1.95E-18 | -1.48E+00 | 4.61E-95 | 1.69E-94 | -2.04E+00 | 1.15E-31 | 3.15E-31 | -1.54E+00 |
| ID3 | 5.15E-111 | 2.69E-110 | 3.05E+01 | 1.28E-01 | 5.52E-01 | -1.08E+00 | 2.78E-16 | 1.18E-15 | 1.39E+01 | 2.02E-01 | 7.70E-01 | 1.50E+00 | 5.50E-02 | 1.96E-01 | 4.59E+00 | 3.55E-01 | 1.00E+00 | 1.00E+00 | 7.20E-03 | 2.64E-02 | 5.42E+00 | 3.64E-01 | 1.00E+00 | 1.00E+00 |
| RPL10 | 1.48E-110 | 7.71E-110 | 1.88E+00 | 8.27E-187 | 3.57E-186 | 2.33E+00 | 5.81E-71 | 2.47E-70 | 1.87E+00 | 1.14E-134 | 4.37E-134 | 2.40E+00 | 1.49E-42 | 5.30E-42 | 1.74E+00 | 2.69E-81 | 7.59E-81 | 2.10E+00 | 3.88E-40 | 1.42E-39 | 1.62E+00 | 3.13E-96 | 8.60E-96 | 2.06E+00 |
| RPL9 | 2.44E-109 | 1.27E-108 | 2.16E+00 | 2.80E-38 | 1.21E-37 | 1.64E+00 | 1.18E-78 | 4.99E-78 | 2.18E+00 | 2.96E-32 | 1.13E-31 | 1.66E+00 | 3.99E-50 | 1.42E-49 | 2.10E+00 | 2.96E-17 | 8.35E-17 | 1.54E+00 | 2.77E-50 | 1.02E-49 | 1.95E+00 | 3.40E-22 | 9.34E-22 | 1.54E+00 |
| PTGS2 | 3.38E-107 | 1.76E-106 | 2.11E+01 | 6.22E-02 | 2.69E-01 | 2.08E+00 | 2.21E-59 | 9.39E-59 | 1.96E+01 | 9.52E-02 | 3.64E-01 | 2.28E+00 | 2.68E-66 | 9.55E-66 | 2.63E+01 | 1.62E-02 | 4.56E-02 | 2.99E+00 | 1.98E-53 | 7.28E-53 | 1.52E+01 | 7.84E-02 | 2.15E-01 | 2.15E+00 |
| B2M | 6.19E-107 | 3.22E-106 | -1.61E+00 | 1.08E-78 | 4.65E-78 | -1.50E+00 | 7.44E-49 | 3.16E-48 | -1.48E+00 | 3.40E-50 | 1.30E-49 | -1.49E+00 | 1.49E-115 | 5.32E-115 | -1.99E+00 | 7.97E-116 | 2.25E-115 | -1.98E+00 | 3.01E-181 | 1.11E-180 | -2.11E+00 | 5.31E-176 | 1.46E-175 | -2.15E+00 |
| RPL9P18 | 1.73E-104 | 9.02E-104 | 2.30E+00 | 1.82E-125 | 7.87E-125 | 2.58E+00 | 5.02E-64 | 2.13E-63 | 2.30E+00 | 8.90E-85 | 3.40E-84 | 2.67E+00 | 4.02E-57 | 1.43E-56 | 2.35E+00 | 4.05E-77 | 1.14E-76 | 2.68E+00 | 3.28E-43 | 1.21E-42 | 2.16E+00 | 1.39E-73 | 3.82E-73 | 2.53E+00 |
| STAT1 | 6.25E-104 | 3.26E-103 | -4.62E+00 | 2.78E-108 | 1.20E-107 | -4.70E+00 | 1.41E-71 | 5.98E-71 | -4.89E+00 | 1.51E-87 | 5.77E-87 | -5.56E+00 | 2.39E-42 | 8.52E-42 | -4.82E+00 | 3.63E-41 | 1.02E-40 | -4.26E+00 | 8.90E-67 | 3.27E-66 | -4.71E+00 | 1.53E-58 | 4.21E-58 | -4.03E+00 |
| CXCR4 | 1.89E-103 | 9.85E-103 | 9.97E+00 | 3.57E-95 | 1.54E-94 | 9.70E+00 | 8.85E-61 | 3.76E-60 | 7.18E+00 | 1.05E-43 | 4.00E-43 | 6.48E+00 | 3.83E-16 | 1.37E-15 | 4.75E+00 | 7.42E-03 | 2.09E-02 | 1.52E+00 | 3.26E-05 | 1.20E-04 | 2.84E+00 | 3.31E-02 | 9.10E-02 | 1.62E+00 |
| RPL9P28 | 2.51E-103 | 1.31E-102 | 2.14E+00 | 1.59E-33 | 6.88E-33 | 1.59E+00 | 8.83E-71 | 3.75E-70 | 2.15E+00 | 4.34E-30 | 1.66E-29 | 1.66E+00 | 1.53E-51 | 5.44E-51 | 2.09E+00 | 5.41E-20 | 1.53E-19 | 1.58E+00 | 3.14E-53 | 1.15E-52 | 2.01E+00 | 2.47E-23 | 6.78E-23 | 1.58E+00 |
| EGR1 | 8.59E-100 | 4.48E-99 | 8.45E+00 | 3.27E-04 | 1.41E-03 | 1.85E+00 | 1.40E-70 | 5.96E-70 | 9.64E+00 | 7.40E-11 | 2.83E-10 | 3.03E+00 | 4.03E-35 | 1.44E-34 | 5.66E+00 | 2.36E-06 | 6.66E-06 | 2.37E+00 | 7.89E-32 | 2.90E-31 | 7.47E+00 | 3.50E-11 | 9.61E-11 | 3.64E+00 |
| RPL7AP50 | 2.42E-98 | 1.26E-97 | -2.80E+00 | 1.66E-27 | 7.15E-27 | -1.72E+00 | 8.38E-40 | 3.56E-39 | -2.56E+00 | 2.06E-09 | 7.88E-09 | -1.52E+00 | 1.61E-47 | 5.74E-47 | -2.68E+00 | 2.08E-09 | 5.87E-09 | -1.56E+00 | 1.46E-56 | 5.36E-56 | -2.58E+00 | 1.51E-11 | 4.16E-11 | -1.50E+00 |
| MT-ATP6 | 6.24E-97 | 3.25E-96 | 1.85E+00 | 2.22E-124 | 9.60E-124 | 2.03E+00 | 6.29E-70 | 2.67E-69 | 1.89E+00 | 2.52E-99 | 9.63E-99 | 2.19E+00 | 1.09E-52 | 3.89E-52 | 1.89E+00 | 5.02E-91 | 1.42E-90 | 2.26E+00 | 3.05E-73 | 1.12E-72 | 1.95E+00 | 2.63E-130 | 7.23E-130 | 2.40E+00 |
| ZFP36 | 1.17E-94 | 6.09E-94 | 6.41E+00 | 3.90E-02 | 1.69E-01 | 1.74E+00 | 3.75E-44 | 1.59E-43 | 4.88E+00 | 6.28E-02 | 2.40E-01 | 1.67E+00 | 7.59E-38 | 2.70E-37 | 5.41E+00 | 1.60E-01 | 4.50E-01 | 1.43E+00 | 3.79E-21 | 1.39E-20 | 4.37E+00 | 4.73E-02 | 1.30E-01 | -1.09E+00 |
| FP671120.4 | 3.24E-92 | 1.69E-91 | 1.56E+00 | 1.08E-16 | 4.67E-16 | 1.16E+00 | 1.05E-33 | 4.44E-33 | 1.12E+00 | 4.97E-03 | 1.90E-02 | -1.34E+00 | 3.29E-77 | 1.17E-76 | 1.88E+00 | 1.14E-12 | 3.21E-12 | -1.20E+00 | 9.43E-79 | 3.46E-78 | 1.80E+00 | 9.07E-10 | 2.49E-09 | -1.15E+00 |
| RPS3A | 1.02E-91 | 5.30E-91 | 1.66E+00 | 3.49E-83 | 1.51E-82 | 1.63E+00 | 4.90E-69 | 2.08E-68 | 1.71E+00 | 4.72E-69 | 1.80E-68 | 1.72E+00 | 6.95E-58 | 2.48E-57 | 1.76E+00 | 3.24E-57 | 9.13E-57 | 1.73E+00 | 3.09E-47 | 1.13E-46 | 1.67E+00 | 3.89E-63 | 1.07E-62 | 1.68E+00 |
| AC006511.3 | 1.02E-91 | 5.30E-91 | -1.87E+00 | 4.95E-47 | 2.14E-46 | -1.56E+00 | 4.98E-42 | 2.12E-41 | -1.67E+00 | 6.22E-22 | 2.38E-21 | -1.44E+00 | 1.94E-32 | 6.93E-32 | -1.66E+00 | 2.40E-18 | 6.77E-18 | -1.45E+00 | 5.05E-59 | 1.85E-58 | -1.69E+00 | 2.47E-27 | 6.79E-27 | -1.47E+00 |
| ARL4C | 5.18E-91 | 2.70E-90 | 9.79E+00 | 7.94E-03 | 3.43E-02 | 1.59E+00 | 3.13E-72 | 1.33E-71 | 1.60E+01 | 1.70E-02 | 6.50E-02 | 2.57E+00 | 1.66E-12 | 5.93E-12 | 2.56E+01 | 2.43E-01 | 6.85E-01 | 2.38E+00 | 2.32E-03 | 8.51E-03 | 5.27E+00 | 3.54E-01 | 9.73E-01 | 1.21E+00 |
| AC064799.1 | 2.24E-89 | 1.17E-88 | 1.97E+00 | 3.68E-69 | 1.59E-68 | 1.84E+00 | 2.52E-76 | 1.07E-75 | 2.12E+00 | 1.00E-59 | 3.82E-59 | 1.96E+00 | 3.25E-49 | 1.16E-48 | 2.01E+00 | 8.46E-40 | 2.38E-39 | 1.85E+00 | 5.46E-56 | 2.00E-55 | 1.98E+00 | 5.40E-52 | 1.48E-51 | 1.86E+00 |
| IFI6 | 8.60E-88 | 4.48E-87 | -4.47E+00 | 1.09E-68 | 4.73E-68 | -3.15E+00 | 1.98E-83 | 8.39E-83 | -6.95E+00 | 4.95E-68 | 1.89E-67 | -5.06E+00 | 5.65E-79 | 2.01E-78 | -9.43E+00 | 9.87E-78 | 2.78E-77 | -7.21E+00 | 1.42E-126 | 5.23E-126 | -1.36E+01 | 2.68E-101 | 7.36E-101 | -9.72E+00 |
| RPL7P23 | 2.57E-87 | 1.34E-86 | 1.76E+00 | 2.71E-40 | 1.17E-39 | 1.49E+00 | 2.82E-64 | 1.20E-63 | 1.79E+00 | 2.29E-36 | 8.76E-36 | 1.56E+00 | 5.82E-42 | 2.08E-41 | 1.76E+00 | 1.11E-21 | 3.14E-21 | 1.49E+00 | 2.96E-33 | 1.09E-32 | 1.65E+00 | 1.06E-27 | 2.90E-27 | 1.48E+00 |
| RPL15 | 4.23E-87 | 2.20E-86 | 1.54E+00 | 1.14E-83 | 4.95E-83 | 1.53E+00 | 7.81E-61 | 3.32E-60 | 1.54E+00 | 2.20E-58 | 8.39E-58 | 1.54E+00 | 8.97E-56 | 3.20E-55 | 1.62E+00 | 7.49E-63 | 2.11E-62 | 1.63E+00 | 1.89E-59 | 6.95E-59 | 1.56E+00 | 1.46E-49 | 4.01E-49 | 1.49E+00 |
| EEF1A1P9 | 6.13E-87 | 3.19E-86 | -1.98E+00 | 9.23E-03 | 3.99E-02 | -1.06E+00 | 1.32E-40 | 5.63E-40 | -1.81E+00 | 2.13E-01 | 8.14E-01 | 1.03E+00 | 3.55E-28 | 1.27E-27 | -1.75E+00 | 3.34E-01 | 9.41E-01 | 1.03E+00 | 3.11E-53 | 1.14E-52 | -1.86E+00 | 2.76E-01 | 7.57E-01 | 1.01E+00 |
| AC020916.1 | 6.22E-86 | 3.24E-85 | 5.85E+00 | 1.70E-04 | 7.33E-04 | -1.03E+00 | 1.23E-61 | 5.22E-61 | 6.28E+00 | 2.20E-01 | 8.42E-01 | 1.36E+00 | 1.88E-66 | 6.69E-66 | 8.23E+00 | 2.70E-01 | 7.61E-01 | 1.49E+00 | 2.65E-66 | 9.71E-66 | 1.05E+01 | 2.36E-01 | 6.47E-01 | 1.35E+00 |
| FAUP1 | 7.34E-86 | 3.83E-85 | 1.86E+00 | 1.58E-42 | 6.82E-42 | 1.60E+00 | 7.54E-68 | 3.20E-67 | 1.96E+00 | 2.67E-40 | 1.02E-39 | 1.70E+00 | 9.93E-40 | 3.54E-39 | 1.82E+00 | 2.10E-23 | 5.92E-23 | 1.57E+00 | 3.20E-69 | 1.17E-68 | 1.98E+00 | 2.73E-45 | 7.49E-45 | 1.75E+00 |
| MX1 | 5.63E-84 | 2.94E-83 | -4.72E+00 | 4.15E-58 | 1.79E-57 | -2.71E+00 | 1.88E-91 | 8.00E-91 | -5.82E+00 | 2.13E-86 | 8.13E-86 | -4.76E+00 | 3.67E-62 | 1.31E-61 | -1.92E+01 | 3.54E-64 | 9.99E-64 | -1.07E+01 | 4.30E-61 | 1.58E-60 | -1.42E+01 | 2.75E-60 | 7.57E-60 | -1.63E+01 |
| CLU | 7.69E-84 | 4.01E-83 | -4.51E+00 | 2.78E-67 | 1.20E-66 | -3.12E+00 | 1.64E-52 | 6.97E-52 | -4.56E+00 | 2.06E-38 | 7.86E-38 | -2.78E+00 | 2.03E-26 | 7.22E-26 | -2.50E+00 | 2.13E-13 | 6.00E-13 | -1.70E+00 | 1.30E-07 | 4.78E-07 | -1.34E+00 | 8.61E-11 | 2.36E-10 | -1.50E+00 |
| AC113367.1 | 8.96E-83 | 4.67E-82 | 2.78E+00 | 1.46E-55 | 6.32E-55 | 2.42E+00 | 1.61E-40 | 6.84E-40 | 2.75E+00 | 5.78E-25 | 2.21E-24 | 2.35E+00 | 4.18E-33 | 1.49E-32 | 2.69E+00 | 3.57E-21 | 1.01E-20 | 2.21E+00 | 2.87E-38 | 1.05E-37 | 2.65E+00 | 7.54E-23 | 2.07E-22 | 2.19E+00 |
| AC004453.1 | 2.46E-82 | 1.28E-81 | 1.70E+00 | 4.86E-26 | 2.10E-25 | 1.34E+00 | 2.48E-60 | 1.06E-59 | 1.76E+00 | 4.00E-24 | 1.53E-23 | 1.44E+00 | 6.62E-45 | 2.36E-44 | 1.74E+00 | 6.60E-19 | 1.86E-18 | 1.41E+00 | 6.27E-40 | 2.30E-39 | 1.66E+00 | 1.03E-12 | 2.82E-12 | 1.33E+00 |
| PFN1P1 | 3.34E-82 | 1.74E-81 | -3.42E+00 | 9.64E-49 | 4.17E-48 | -1.93E+00 | 7.11E-33 | 3.02E-32 | -2.97E+00 | 3.49E-13 | 1.33E-12 | -1.80E+00 | 1.17E-49 | 4.16E-49 | -3.58E+00 | 1.14E-21 | 3.20E-21 | -2.02E+00 | 1.07E-44 | 3.93E-44 | -3.47E+00 | 2.53E-12 | 6.95E-12 | -1.96E+00 |
| BTF3P9 | 3.29E-79 | 1.71E-78 | -3.15E+00 | 4.81E-24 | 2.08E-23 | -1.64E+00 | 8.86E-41 | 3.77E-40 | -3.24E+00 | 1.02E-08 | 3.88E-08 | -1.52E+00 | 4.97E-46 | 1.77E-45 | -3.18E+00 | 5.77E-13 | 1.63E-12 | -1.53E+00 | 1.92E-44 | 7.05E-44 | -3.30E+00 | 1.66E-08 | 4.57E-08 | -1.61E+00 |
| AL139095.2 | 1.02E-77 | 5.31E-77 | 1.94E+00 | 7.21E-95 | 3.12E-94 | 2.09E+00 | 7.83E-65 | 3.33E-64 | 2.07E+00 | 4.18E-80 | 1.60E-79 | 2.27E+00 | 1.78E-56 | 6.36E-56 | 2.17E+00 | 1.05E-69 | 2.97E-69 | 2.31E+00 | 3.06E-56 | 1.12E-55 | 2.08E+00 | 6.50E-79 | 1.78E-78 | 2.30E+00 |
| RPS27AP11 | 1.90E-77 | 9.92E-77 | 2.10E+00 | 1.71E-50 | 7.40E-50 | 1.88E+00 | 5.49E-54 | 2.33E-53 | 2.15E+00 | 1.20E-39 | 4.57E-39 | 1.95E+00 | 1.39E-40 | 4.95E-40 | 2.04E+00 | 1.18E-30 | 3.33E-30 | 1.83E+00 | 9.99E-37 | 3.67E-36 | 2.01E+00 | 5.82E-31 | 1.60E-30 | 1.81E+00 |
| AC136632.1 | 3.38E-77 | 1.76E-76 | 2.07E+00 | 4.67E-68 | 2.02E-67 | 2.00E+00 | 5.08E-47 | 2.16E-46 | 2.03E+00 | 2.61E-41 | 9.98E-41 | 1.97E+00 | 1.64E-37 | 5.83E-37 | 2.05E+00 | 1.96E-41 | 5.54E-41 | 2.08E+00 | 2.30E-34 | 8.45E-34 | 2.07E+00 | 2.10E-41 | 5.76E-41 | 2.06E+00 |
| AC079250.1 | 4.13E-76 | 2.15E-75 | 1.85E+00 | 3.00E-25 | 1.29E-24 | 1.44E+00 | 1.97E-52 | 8.38E-52 | 1.86E+00 | 1.67E-20 | 6.37E-20 | 1.47E+00 | 2.14E-33 | 7.65E-33 | 1.78E+00 | 3.23E-19 | 9.12E-19 | 1.51E+00 | 3.53E-39 | 1.29E-38 | 1.84E+00 | 2.62E-22 | 7.19E-22 | 1.52E+00 |
| CDH7 | 3.21E-75 | 1.67E-74 | -6.41E+00 | 8.10E-71 | 3.50E-70 | -5.48E+00 | 8.36E-02 | 3.55E-01 | -1.96E+00 | 2.16E-01 | 8.24E-01 | -1.38E+00 | 2.80E-01 | 1.00E+00 | 1.00E+00 | 3.55E-01 | 1.00E+00 | 1.00E+00 | 1.82E-01 | 6.70E-01 | -1.17E+00 | 2.23E-01 | 6.14E-01 | -1.22E+00 |
| AC116366.1 | 5.79E-74 | 3.02E-73 | -2.38E+00 | 1.84E-149 | 7.94E-149 | -9.19E+00 | 2.66E-28 | 1.13E-27 | -2.84E+00 | 2.10E-65 | 8.01E-65 | -9.66E+00 | 1.74E-14 | 6.20E-14 | -3.23E+00 | 2.47E-18 | 6.97E-18 | -5.19E+00 | 3.82E-26 | 1.40E-25 | -6.00E+00 | 7.68E-29 | 2.11E-28 | -7.54E+00 |
| RPL29P30 | 2.87E-72 | 1.49E-71 | -2.86E+00 | 9.83E-38 | 4.25E-37 | -1.83E+00 | 2.87E-28 | 1.22E-27 | -2.72E+00 | 7.76E-14 | 2.96E-13 | -1.81E+00 | 2.82E-33 | 1.01E-32 | -2.77E+00 | 9.10E-16 | 2.57E-15 | -1.76E+00 | 2.36E-33 | 8.67E-33 | -2.81E+00 | 4.47E-12 | 1.23E-11 | -1.73E+00 |
| RPS27A | 4.16E-71 | 2.17E-70 | 1.54E+00 | 1.70E-53 | 7.34E-53 | 1.46E+00 | 1.26E-46 | 5.35E-46 | 1.52E+00 | 5.25E-39 | 2.00E-38 | 1.48E+00 | 1.10E-34 | 3.94E-34 | 1.52E+00 | 7.33E-32 | 2.07E-31 | 1.48E+00 | 1.08E-30 | 3.96E-30 | 1.49E+00 | 2.53E-31 | 6.94E-31 | 1.44E+00 |
| SESN1 | 8.94E-71 | 4.66E-70 | 5.91E+00 | 1.37E-25 | 5.94E-25 | 3.79E+00 | 1.01E-21 | 4.27E-21 | 3.63E+00 | 3.99E-04 | 1.52E-03 | 2.32E+00 | 5.73E-03 | 2.04E-02 | 2.62E+00 | 1.12E-01 | 3.16E-01 | 1.29E+00 | 8.05E-02 | 2.96E-01 | 1.57E+00 | 9.30E-02 | 2.55E-01 | 1.13E+00 |
| AL009174.1 | 1.10E-70 | 5.71E-70 | 1.61E+00 | 1.82E-76 | 7.86E-76 | 1.65E+00 | 8.17E-52 | 3.47E-51 | 1.63E+00 | 2.69E-61 | 1.03E-60 | 1.72E+00 | 1.20E-36 | 4.28E-36 | 1.62E+00 | 1.06E-40 | 2.98E-40 | 1.63E+00 | 2.45E-39 | 8.98E-39 | 1.62E+00 | 8.10E-44 | 2.23E-43 | 1.60E+00 |
| RPS26P47 | 4.94E-70 | 2.57E-69 | -2.99E+00 | 3.65E-69 | 1.58E-68 | -2.59E+00 | 1.31E-39 | 5.58E-39 | -2.65E+00 | 1.30E-41 | 4.97E-41 | -2.46E+00 | 9.15E-33 | 3.26E-32 | -2.90E+00 | 1.70E-30 | 4.80E-30 | -2.47E+00 | 4.28E-32 | 1.57E-31 | -2.64E+00 | 7.85E-31 | 2.16E-30 | -2.46E+00 |
| NEAT1 | 7.92E-70 | 4.13E-69 | 2.46E+00 | 6.27E-96 | 2.71E-95 | 3.00E+00 | 1.82E-24 | 7.72E-24 | 2.00E+00 | 2.41E-30 | 9.20E-30 | 2.26E+00 | 5.67E-28 | 2.02E-27 | 2.48E+00 | 2.47E-32 | 6.95E-32 | 2.54E+00 | 2.42E-23 | 8.89E-23 | 2.31E+00 | 4.02E-20 | 1.11E-19 | 2.17E+00 |
| RPL29P33 | 1.89E-68 | 9.83E-68 | -2.87E+00 | 9.67E-44 | 4.18E-43 | -1.86E+00 | 1.33E-24 | 5.64E-24 | -2.52E+00 | 9.20E-11 | 3.51E-10 | -1.58E+00 | 7.89E-37 | 2.81E-36 | -2.93E+00 | 1.12E-12 | 3.15E-12 | -1.71E+00 | 1.35E-36 | 4.97E-36 | -2.81E+00 | 1.30E-11 | 3.57E-11 | -1.69E+00 |
| RPL8P2 | 5.42E-68 | 2.83E-67 | -2.78E+00 | 3.04E-52 | 1.31E-51 | -1.99E+00 | 2.04E-30 | 8.68E-30 | -2.61E+00 | 3.11E-24 | 1.19E-23 | -1.91E+00 | 3.99E-26 | 1.42E-25 | -2.26E+00 | 9.28E-17 | 2.62E-16 | -1.70E+00 | 5.28E-34 | 1.94E-33 | -2.49E+00 | 5.90E-14 | 1.62E-13 | -1.71E+00 |
| RPS12 | 9.56E-68 | 4.98E-67 | 1.54E+00 | 2.46E-74 | 1.06E-73 | 1.58E+00 | 9.12E-56 | 3.88E-55 | 1.62E+00 | 9.47E-65 | 3.62E-64 | 1.69E+00 | 2.08E-36 | 7.42E-36 | 1.56E+00 | 1.73E-46 | 4.87E-46 | 1.62E+00 | 1.49E-32 | 5.48E-32 | 1.55E+00 | 8.62E-48 | 2.37E-47 | 1.62E+00 |
| FKBP5 | 1.12E-66 | 5.84E-66 | 3.68E+00 | 2.74E-33 | 1.18E-32 | 2.93E+00 | 1.52E-34 | 6.47E-34 | 3.06E+00 | 1.38E-25 | 5.28E-25 | 2.78E+00 | 1.24E-24 | 4.41E-24 | 3.28E+00 | 4.44E-18 | 1.25E-17 | 2.60E+00 | 1.27E-19 | 4.65E-19 | 2.50E+00 | 3.84E-25 | 1.05E-24 | 2.63E+00 |
| BTG1 | 2.27E-66 | 1.18E-65 | 2.71E+00 | 5.05E-64 | 2.18E-63 | 2.72E+00 | 2.97E-32 | 1.26E-31 | 2.56E+00 | 6.73E-23 | 2.57E-22 | 2.35E+00 | 8.31E-40 | 2.96E-39 | 3.12E+00 | 2.37E-36 | 6.69E-36 | 2.80E+00 | 2.23E-35 | 8.19E-35 | 2.59E+00 | 3.93E-24 | 1.08E-23 | 2.25E+00 |
| RPL19P16 | 2.27E-66 | 1.18E-65 | -1.83E+00 | 5.09E-68 | 2.20E-67 | -1.83E+00 | 7.86E-42 | 3.34E-41 | -1.84E+00 | 1.47E-35 | 5.62E-35 | -1.73E+00 | 2.52E-35 | 8.97E-35 | -1.82E+00 | 3.78E-26 | 1.07E-25 | -1.69E+00 | 1.94E-55 | 7.11E-55 | -1.83E+00 | 5.95E-37 | 1.63E-36 | -1.73E+00 |
| BTF3P5 | 3.46E-66 | 1.80E-65 | -2.63E+00 | 2.55E-26 | 1.10E-25 | -1.53E+00 | 6.61E-23 | 2.81E-22 | -2.44E+00 | 1.27E-06 | 4.83E-06 | -1.40E+00 | 4.48E-28 | 1.60E-27 | -2.56E+00 | 1.44E-07 | 4.05E-07 | -1.49E+00 | 9.54E-40 | 3.50E-39 | -2.78E+00 | 1.95E-09 | 5.34E-09 | -1.55E+00 |
| ADGRG6 | 8.53E-66 | 4.45E-65 | -2.95E+00 | 4.84E-39 | 2.09E-38 | -1.72E+00 | 9.21E-08 | 3.92E-07 | -1.69E+00 | 3.52E-06 | 1.35E-05 | -1.24E+00 | 1.57E-02 | 5.58E-02 | -1.68E+00 | 4.38E-02 | 1.23E-01 | -1.32E+00 | 1.81E-01 | 6.63E-01 | -1.11E+00 | 3.59E-01 | 9.87E-01 | 1.22E+00 |
| AC016739.1 | 1.50E-65 | 7.84E-65 | 1.57E+00 | 5.76E-57 | 2.49E-56 | 1.53E+00 | 4.15E-51 | 1.76E-50 | 1.61E+00 | 7.91E-55 | 3.02E-54 | 1.65E+00 | 1.97E-25 | 7.04E-25 | 1.49E+00 | 5.33E-34 | 1.50E-33 | 1.55E+00 | 3.11E-26 | 1.14E-25 | 1.51E+00 | 1.93E-37 | 5.30E-37 | 1.57E+00 |
| TXNIP | 1.52E-65 | 7.91E-65 | 2.87E+00 | 6.17E-103 | 2.67E-102 | 3.83E+00 | 1.06E-29 | 4.49E-29 | 2.57E+00 | 7.14E-58 | 2.73E-57 | 3.73E+00 | 2.36E-12 | 8.40E-12 | 2.44E+00 | 1.08E-20 | 3.04E-20 | 2.68E+00 | 3.40E-20 | 1.25E-19 | 2.50E+00 | 1.65E-14 | 4.52E-14 | 2.29E+00 |
| RPL12 | 2.83E-65 | 1.48E-64 | 1.61E+00 | 4.55E-39 | 1.97E-38 | 1.45E+00 | 1.07E-45 | 4.56E-45 | 1.62E+00 | 1.29E-36 | 4.94E-36 | 1.55E+00 | 2.48E-29 | 8.86E-29 | 1.58E+00 | 1.74E-25 | 4.91E-25 | 1.50E+00 | 1.92E-27 | 7.03E-27 | 1.54E+00 | 6.03E-27 | 1.66E-26 | 1.48E+00 |
| AC104619.1 | 1.39E-64 | 7.25E-64 | 2.04E+00 | 3.98E-55 | 1.72E-54 | 1.98E+00 | 2.82E-49 | 1.20E-48 | 2.18E+00 | 1.19E-43 | 4.55E-43 | 2.14E+00 | 7.68E-42 | 2.74E-41 | 2.24E+00 | 5.41E-46 | 1.53E-45 | 2.22E+00 | 5.16E-46 | 1.89E-45 | 2.10E+00 | 2.50E-45 | 6.86E-45 | 2.02E+00 |
| SLC40A1 | 1.55E-64 | 8.10E-64 | -2.90E+00 | 4.03E-13 | 1.74E-12 | -1.28E+00 | 1.09E-26 | 4.64E-26 | -2.60E+00 | 3.62E-04 | 1.38E-03 | -1.11E+00 | 1.20E-06 | 4.29E-06 | -1.51E+00 | 2.99E-01 | 8.43E-01 | 1.04E+00 | 5.98E-03 | 2.20E-02 | -1.17E+00 | 1.48E-01 | 4.08E-01 | 1.21E+00 |
| RCN1 | 2.05E-63 | 1.07E-62 | -2.86E+00 | 2.80E-29 | 1.21E-28 | -1.62E+00 | 7.06E-27 | 3.00E-26 | -2.21E+00 | 8.41E-13 | 3.21E-12 | -1.47E+00 | 1.09E-28 | 3.89E-28 | -2.97E+00 | 3.68E-14 | 1.04E-13 | -1.65E+00 | 6.37E-17 | 2.34E-16 | -1.86E+00 | 6.09E-05 | 1.67E-04 | -1.22E+00 |
| OAS1 | 3.98E-63 | 2.07E-62 | -5.08E+00 | 4.54E-37 | 1.96E-36 | -2.25E+00 | 9.39E-46 | 3.99E-45 | -5.64E+00 | 6.34E-43 | 2.42E-42 | -4.71E+00 | 4.72E-13 | 1.68E-12 | -1.73E+01 | 1.03E-11 | 2.89E-11 | -6.48E+00 | 7.49E-76 | 2.75E-75 | -2.12E+01 | 8.60E-74 | 2.36E-73 | -2.28E+01 |
| AL590139.1 | 4.87E-63 | 2.54E-62 | 2.48E+00 | 7.75E-28 | 3.35E-27 | 2.01E+00 | 7.43E-36 | 3.16E-35 | 2.24E+00 | 2.10E-23 | 8.03E-23 | 1.92E+00 | 1.35E-09 | 4.80E-09 | 1.79E+00 | 2.56E-05 | 7.21E-05 | 1.55E+00 | 1.84E-21 | 6.76E-21 | 2.09E+00 | 3.11E-13 | 8.54E-13 | 1.73E+00 |
| HSP90AA2P | 4.25E-62 | 2.21E-61 | -2.31E+00 | 6.21E-34 | 2.68E-33 | -1.66E+00 | 4.66E-14 | 1.98E-13 | -1.93E+00 | 7.19E-06 | 2.75E-05 | -1.38E+00 | 1.14E-30 | 4.08E-30 | -2.46E+00 | 9.96E-15 | 2.81E-14 | -1.74E+00 | 1.35E-16 | 4.95E-16 | -2.09E+00 | 1.26E-03 | 3.47E-03 | -1.30E+00 |
| AL591846.1 | 4.94E-62 | 2.58E-61 | -2.31E+00 | 1.69E-09 | 7.28E-09 | -1.17E+00 | 4.25E-25 | 1.80E-24 | -2.42E+00 | 1.84E-03 | 7.01E-03 | -1.24E+00 | 5.97E-29 | 2.13E-28 | -2.21E+00 | 5.98E-03 | 1.69E-02 | -1.14E+00 | 7.35E-30 | 2.70E-29 | -2.42E+00 | 8.37E-04 | 2.30E-03 | -1.25E+00 |
| AC113398.1 | 1.75E-61 | 9.14E-61 | 1.70E+00 | 4.42E-36 | 1.91E-35 | 1.53E+00 | 4.00E-44 | 1.70E-43 | 1.73E+00 | 6.35E-30 | 2.43E-29 | 1.59E+00 | 1.63E-23 | 5.82E-23 | 1.61E+00 | 1.16E-19 | 3.26E-19 | 1.53E+00 | 7.49E-28 | 2.75E-27 | 1.66E+00 | 6.68E-31 | 1.84E-30 | 1.61E+00 |
| TNFSF13B | 3.04E-61 | 1.59E-60 | -5.15E+00 | 2.00E-58 | 8.63E-58 | -4.18E+00 | 2.43E-48 | 1.03E-47 | -2.80E+00 | 1.48E-65 | 5.66E-65 | -3.02E+00 | 2.65E-06 | 9.45E-06 | -2.80E+00 | 7.16E-07 | 2.02E-06 | -3.94E+00 | 3.39E-15 | 1.24E-14 | -5.78E+00 | 1.30E-16 | 3.57E-16 | -9.99E+00 |
| RPS4XP16 | 1.20E-60 | 6.28E-60 | -2.55E+00 | 1.29E-12 | 5.57E-12 | -1.29E+00 | 2.98E-27 | 1.27E-26 | -2.35E+00 | 4.30E-04 | 1.64E-03 | -1.14E+00 | 7.26E-29 | 2.59E-28 | -2.33E+00 | 1.21E-04 | 3.41E-04 | -1.17E+00 | 1.89E-25 | 6.93E-25 | -2.15E+00 | 5.03E-04 | 1.38E-03 | -1.18E+00 |
| SAP30 | 2.88E-60 | 1.50E-59 | 6.38E+00 | 3.22E-07 | 1.39E-06 | 2.81E+00 | 1.00E-28 | 4.26E-28 | 4.29E+00 | 4.06E-04 | 1.55E-03 | 2.19E+00 | 5.65E-02 | 2.02E-01 | 1.83E+00 | 2.86E-02 | 8.07E-02 | 1.14E+00 | 2.66E-01 | 9.77E-01 | 1.33E+00 | 5.45E-02 | 1.50E-01 | 1.11E+00 |
| EGR3 | 3.03E-60 | 1.58E-59 | 3.42E+01 | 1.64E-01 | 7.09E-01 | 1.99E+00 | 3.30E-36 | 1.40E-35 | 2.68E+01 | 1.50E-01 | 5.73E-01 | 2.15E+00 | 2.82E-35 | 1.01E-34 | 2.35E+01 | 1.88E-01 | 5.31E-01 | 2.42E+00 | 4.49E-24 | 1.65E-23 | 2.20E+01 | 1.46E-01 | 4.02E-01 | 2.56E+00 |
| PCDH9 | 3.77E-60 | 1.97E-59 | -6.30E+00 | 2.18E-35 | 9.43E-35 | -2.88E+00 | 1.87E-02 | 7.93E-02 | -1.66E+00 | 1.76E-01 | 6.70E-01 | 1.17E+00 | 1.20E-01 | 4.28E-01 | -1.14E+00 | 3.54E-02 | 9.98E-02 | -1.14E+00 | 1.93E-01 | 7.09E-01 | 1.49E+00 | 2.33E-01 | 6.40E-01 | 1.65E+00 |
| EPSTI1 | 4.25E-60 | 2.21E-59 | -3.13E+00 | 1.32E-60 | 5.70E-60 | -2.71E+00 | 5.21E-47 | 2.21E-46 | -3.46E+00 | 9.22E-48 | 3.52E-47 | -3.22E+00 | 7.95E-30 | 2.83E-29 | -3.12E+00 | 1.02E-20 | 2.87E-20 | -2.18E+00 | 2.56E-28 | 9.41E-28 | -2.56E+00 | 3.66E-24 | 1.00E-23 | -2.52E+00 |
| EEF1DP1 | 1.86E-59 | 9.69E-59 | -2.51E+00 | 3.97E-54 | 1.72E-53 | -2.13E+00 | 6.23E-21 | 2.65E-20 | -2.16E+00 | 1.06E-21 | 4.06E-21 | -1.90E+00 | 4.89E-27 | 1.74E-26 | -2.37E+00 | 6.58E-19 | 1.86E-18 | -1.89E+00 | 1.45E-20 | 5.31E-20 | -2.04E+00 | 5.60E-14 | 1.54E-13 | -1.74E+00 |
| HLA-B | 4.82E-59 | 2.51E-58 | -1.93E+00 | 3.81E-40 | 1.64E-39 | -1.68E+00 | 4.28E-36 | 1.82E-35 | -1.85E+00 | 2.60E-27 | 9.92E-27 | -1.71E+00 | 1.12E-65 | 4.01E-65 | -2.38E+00 | 1.98E-52 | 5.59E-52 | -2.18E+00 | 4.02E-137 | 1.48E-136 | -3.07E+00 | 5.48E-89 | 1.51E-88 | -2.64E+00 |
| NPM1P24 | 6.78E-59 | 3.54E-58 | -2.51E+00 | 1.82E-21 | 7.88E-21 | -1.49E+00 | 2.81E-24 | 1.19E-23 | -2.45E+00 | 1.22E-09 | 4.66E-09 | -1.53E+00 | 2.90E-29 | 1.03E-28 | -2.51E+00 | 3.61E-07 | 1.02E-06 | -1.30E+00 | 3.13E-23 | 1.15E-22 | -2.37E+00 | 8.71E-05 | 2.39E-04 | -1.43E+00 |
| CACNB4 | 7.26E-59 | 3.78E-58 | -3.73E+00 | 4.15E-35 | 1.79E-34 | -1.91E+00 | 3.76E-20 | 1.60E-19 | -4.01E+00 | 3.16E-14 | 1.21E-13 | -2.01E+00 | 5.99E-02 | 2.14E-01 | -1.84E+00 | 2.33E-01 | 6.57E-01 | 1.16E+00 | 6.63E-02 | 2.43E-01 | -2.19E+00 | 1.91E-01 | 5.25E-01 | -1.41E+00 |
| HIF3A | 4.31E-58 | 2.24E-57 | 1.26E+01 | 8.42E-55 | 3.64E-54 | 1.41E+01 | 1.22E-07 | 5.19E-07 | 5.30E+00 | 1.69E-05 | 6.47E-05 | 5.05E+00 | 5.60E-03 | 1.99E-02 | 4.49E+00 | 2.91E-01 | 8.19E-01 | 1.76E+00 | 1.12E-01 | 4.12E-01 | 2.07E+00 | 3.64E-01 | 1.00E+00 | 1.23E+00 |
| PDZRN4 | 6.99E-58 | 3.64E-57 | -5.83E+00 | 4.84E-52 | 2.09E-51 | -4.07E+00 | 6.95E-03 | 2.95E-02 | -1.66E+00 | 2.82E-02 | 1.08E-01 | -1.15E+00 | 1.36E-03 | 4.85E-03 | -1.83E+00 | 1.78E-03 | 5.03E-03 | -1.90E+00 | 2.49E-01 | 9.15E-01 | 1.05E+00 | 2.94E-01 | 8.07E-01 | 1.19E+00 |
| EIF4A1P7 | 7.54E-58 | 3.93E-57 | -2.58E+00 | 7.61E-34 | 3.29E-33 | -1.77E+00 | 6.94E-26 | 2.95E-25 | -2.47E+00 | 3.69E-18 | 1.41E-17 | -1.93E+00 | 2.77E-31 | 9.87E-31 | -2.56E+00 | 8.56E-22 | 2.41E-21 | -1.92E+00 | 1.51E-20 | 5.53E-20 | -2.07E+00 | 1.92E-10 | 5.28E-10 | -1.61E+00 |
| RPS14 | 1.27E-57 | 6.61E-57 | 1.41E+00 | 2.05E-37 | 8.85E-37 | 1.32E+00 | 3.65E-38 | 1.55E-37 | 1.41E+00 | 3.17E-28 | 1.21E-27 | 1.35E+00 | 1.10E-25 | 3.94E-25 | 1.37E+00 | 2.54E-26 | 7.15E-26 | 1.36E+00 | 3.21E-26 | 1.18E-25 | 1.37E+00 | 2.70E-24 | 7.42E-24 | 1.32E+00 |
| CYSLTR1 | 1.27E-57 | 6.61E-57 | -3.52E+00 | 1.94E-35 | 8.36E-35 | -1.83E+00 | 5.04E-14 | 2.14E-13 | -1.64E+00 | 2.47E-11 | 9.45E-11 | -1.42E+00 | 2.10E-10 | 7.50E-10 | -2.29E+00 | 6.48E-09 | 1.83E-08 | -1.81E+00 | 5.34E-05 | 1.96E-04 | -2.79E+00 | 6.21E-04 | 1.70E-03 | -2.11E+00 |
| STAT2 | 1.37E-57 | 7.14E-57 | -4.08E+00 | 8.38E-37 | 3.62E-36 | -2.07E+00 | 2.03E-51 | 8.64E-51 | -4.68E+00 | 3.59E-28 | 1.37E-27 | -2.32E+00 | 2.20E-19 | 7.84E-19 | -2.64E+00 | 3.80E-15 | 1.07E-14 | -1.99E+00 | 1.84E-32 | 6.75E-32 | -3.41E+00 | 2.83E-17 | 7.77E-17 | -2.08E+00 |
| IMP3 | 1.12E-56 | 5.81E-56 | -2.98E+00 | 2.95E-18 | 1.27E-17 | -1.22E+00 | 2.82E-23 | 1.20E-22 | -2.04E+00 | 4.68E-04 | 1.79E-03 | -1.01E+00 | 5.78E-38 | 2.06E-37 | -3.13E+00 | 5.87E-09 | 1.66E-08 | -1.24E+00 | 3.45E-20 | 1.27E-19 | -1.95E+00 | 4.18E-02 | 1.15E-01 | -1.01E+00 |
| DTX3L | 5.46E-56 | 2.85E-55 | -2.90E+00 | 1.03E-35 | 4.46E-35 | -1.92E+00 | 1.17E-41 | 4.96E-41 | -3.04E+00 | 1.38E-30 | 5.29E-30 | -2.20E+00 | 3.31E-22 | 1.18E-21 | -3.07E+00 | 7.13E-13 | 2.01E-12 | -1.86E+00 | 7.24E-34 | 2.66E-33 | -3.15E+00 | 1.45E-22 | 3.98E-22 | -2.21E+00 |
| PARP9 | 1.04E-55 | 5.42E-55 | -3.20E+00 | 4.34E-43 | 1.88E-42 | -2.22E+00 | 4.44E-59 | 1.88E-58 | -4.17E+00 | 2.96E-48 | 1.13E-47 | -2.90E+00 | 1.30E-34 | 4.64E-34 | -3.94E+00 | 2.67E-29 | 7.52E-29 | -2.84E+00 | 5.59E-64 | 2.05E-63 | -4.61E+00 | 6.15E-45 | 1.69E-44 | -3.25E+00 |
| RPS18P9 | 1.84E-55 | 9.58E-55 | 2.25E+00 | 3.28E-65 | 1.42E-64 | 2.51E+00 | 5.37E-37 | 2.28E-36 | 2.25E+00 | 7.42E-48 | 2.83E-47 | 2.57E+00 | 2.30E-24 | 8.19E-24 | 2.27E+00 | 2.09E-40 | 5.90E-40 | 2.62E+00 | 1.84E-30 | 6.74E-30 | 2.28E+00 | 2.78E-42 | 7.63E-42 | 2.59E+00 |
| NFE2 | 9.40E-55 | 4.90E-54 | -2.65E+00 | 1.02E-25 | 4.42E-25 | -1.44E+00 | 4.03E-30 | 1.71E-29 | -2.63E+00 | 4.18E-07 | 1.59E-06 | -1.26E+00 | 1.37E-38 | 4.88E-38 | -3.25E+00 | 1.97E-07 | 5.55E-07 | -1.39E+00 | 2.10E-34 | 7.72E-34 | -2.40E+00 | 1.73E-09 | 4.74E-09 | -1.59E+00 |
| MAL | 1.08E-54 | 5.64E-54 | 1.36E+01 | 1.50E-59 | 6.48E-59 | 1.66E+01 | 1.28E-06 | 5.43E-06 | 7.06E+00 | 7.47E-06 | 2.85E-05 | 7.64E+00 | 1.19E-02 | 4.24E-02 | 7.83E+00 | 1.77E-03 | 4.99E-03 | 1.31E+01 | 1.07E-01 | 3.92E-01 | 2.63E+00 | 1.53E-01 | 4.20E-01 | 2.88E+00 |
| AC026403.1 | 1.85E-54 | 9.63E-54 | 1.67E+00 | 4.13E-55 | 1.79E-54 | 1.70E+00 | 1.16E-36 | 4.94E-36 | 1.68E+00 | 1.10E-41 | 4.22E-41 | 1.75E+00 | 1.68E-26 | 6.00E-26 | 1.67E+00 | 4.01E-32 | 1.13E-31 | 1.72E+00 | 9.29E-19 | 3.41E-18 | 1.58E+00 | 2.78E-32 | 7.65E-32 | 1.67E+00 |
| AC097658.1 | 6.10E-54 | 3.18E-53 | 1.77E+00 | 4.66E-131 | 2.01E-130 | 2.47E+00 | 8.10E-40 | 3.44E-39 | 1.86E+00 | 6.50E-89 | 2.48E-88 | 2.57E+00 | 3.89E-23 | 1.39E-22 | 1.73E+00 | 3.97E-73 | 1.12E-72 | 2.54E+00 | 5.10E-22 | 1.87E-21 | 1.74E+00 | 1.39E-73 | 3.82E-73 | 2.39E+00 |
| AC004224.1 | 7.75E-54 | 4.04E-53 | -2.44E+00 | 1.30E-29 | 5.60E-29 | -1.51E+00 | 6.20E-27 | 2.63E-26 | -2.41E+00 | 1.80E-14 | 6.86E-14 | -1.46E+00 | 2.82E-27 | 1.01E-26 | -2.42E+00 | 6.54E-12 | 1.84E-11 | -1.44E+00 | 1.51E-28 | 5.55E-28 | -2.50E+00 | 1.39E-09 | 3.82E-09 | -1.38E+00 |
| ATP2B1-AS1 | 8.84E-54 | 4.61E-53 | 4.40E+00 | 2.45E-12 | 1.06E-11 | 2.60E+00 | 1.94E-23 | 8.22E-23 | 3.74E+00 | 2.62E-08 | 1.00E-07 | 2.53E+00 | 6.75E-18 | 2.41E-17 | 4.98E+00 | 5.07E-13 | 1.43E-12 | 3.76E+00 | 4.70E-08 | 1.73E-07 | 2.72E+00 | 2.81E-04 | 7.71E-04 | 2.10E+00 |
| AC091429.1 | 1.63E-53 | 8.51E-53 | -2.10E+00 | 1.78E-04 | 7.71E-04 | -1.16E+00 | 1.52E-20 | 6.44E-20 | -1.88E+00 | 6.53E-02 | 2.50E-01 | -1.05E+00 | 3.35E-25 | 1.20E-24 | -1.95E+00 | 9.24E-02 | 2.61E-01 | -1.02E+00 | 4.70E-36 | 1.72E-35 | -2.17E+00 | 1.58E-02 | 4.34E-02 | -1.18E+00 |
| EIF3LP2 | 2.80E-53 | 1.46E-52 | -2.24E+00 | 2.25E-26 | 9.73E-26 | -1.49E+00 | 5.76E-17 | 2.45E-16 | -1.88E+00 | 1.80E-09 | 6.88E-09 | -1.37E+00 | 3.77E-27 | 1.35E-26 | -2.19E+00 | 1.89E-09 | 5.33E-09 | -1.37E+00 | 5.87E-17 | 2.16E-16 | -1.87E+00 | 3.02E-07 | 8.29E-07 | -1.40E+00 |
| HLA-A | 2.85E-53 | 1.49E-52 | -1.57E+00 | 4.97E-35 | 2.15E-34 | -1.43E+00 | 1.82E-33 | 7.75E-33 | -1.56E+00 | 9.51E-30 | 3.63E-29 | -1.53E+00 | 3.41E-60 | 1.22E-59 | -1.88E+00 | 3.96E-54 | 1.12E-53 | -1.82E+00 | 1.90E-115 | 6.96E-115 | -2.12E+00 | 9.81E-102 | 2.69E-101 | -2.17E+00 |
| ATP6V0E1 | 6.06E-53 | 3.16E-52 | -2.12E+00 | 2.75E-14 | 1.19E-13 | -1.16E+00 | 8.00E-19 | 3.40E-18 | -1.77E+00 | 1.71E-03 | 6.52E-03 | 1.04E+00 | 2.59E-19 | 9.24E-19 | -1.64E+00 | 5.06E-02 | 1.43E-01 | 1.06E+00 | 4.30E-04 | 1.58E-03 | -1.12E+00 | 3.60E-01 | 9.90E-01 | 1.14E+00 |
| GBP4 | 6.20E-53 | 3.23E-52 | -2.31E+00 | 5.67E-25 | 2.45E-24 | -1.42E+00 | 5.56E-28 | 2.36E-27 | -2.56E+00 | 3.99E-18 | 1.52E-17 | -1.71E+00 | 7.67E-08 | 2.74E-07 | -1.55E+00 | 1.25E-06 | 3.53E-06 | -1.48E+00 | 2.92E-27 | 1.07E-26 | -3.21E+00 | 1.20E-27 | 3.31E-27 | -3.61E+00 |
| MTND4P12 | 6.43E-53 | 3.35E-52 | -2.47E+00 | 7.05E-50 | 3.05E-49 | -2.05E+00 | 6.73E-24 | 2.86E-23 | -2.47E+00 | 2.55E-17 | 9.73E-17 | -1.93E+00 | 4.01E-25 | 1.43E-24 | -2.44E+00 | 1.30E-14 | 3.66E-14 | -1.84E+00 | 2.41E-16 | 8.84E-16 | -2.06E+00 | 2.52E-07 | 6.92E-07 | -1.60E+00 |
| AC068993.1 | 1.28E-52 | 6.70E-52 | -3.11E+00 | 1.37E-37 | 5.93E-37 | -1.97E+00 | 4.95E-29 | 2.10E-28 | -2.42E+00 | 4.26E-24 | 1.63E-23 | -1.77E+00 | 3.26E-21 | 1.16E-20 | -2.49E+00 | 4.68E-16 | 1.32E-15 | -1.79E+00 | 2.09E-09 | 7.69E-09 | -1.64E+00 | 7.30E-07 | 2.00E-06 | -1.46E+00 |
| RPSAP19 | 2.50E-52 | 1.30E-51 | -1.87E+00 | 1.69E-27 | 7.30E-27 | -1.53E+00 | 6.54E-30 | 2.78E-29 | -1.85E+00 | 8.53E-17 | 3.26E-16 | -1.56E+00 | 3.25E-26 | 1.16E-25 | -1.82E+00 | 4.36E-11 | 1.23E-10 | -1.42E+00 | 4.78E-40 | 1.75E-39 | -1.81E+00 | 3.16E-14 | 8.68E-14 | -1.46E+00 |
| SOCS3 | 3.06E-52 | 1.59E-51 | 6.67E+00 | 8.97E-03 | 3.87E-02 | 1.05E+00 | 2.14E-11 | 9.08E-11 | 3.74E+00 | 8.58E-05 | 3.28E-04 | -1.47E+00 | 7.57E-26 | 2.70E-25 | 2.20E+01 | 2.48E-01 | 7.00E-01 | 1.36E+00 | 1.07E-19 | 3.92E-19 | 1.15E+01 | 3.15E-01 | 8.64E-01 | 1.38E+00 |
| FTH1P2 | 3.44E-52 | 1.79E-51 | -2.30E+00 | 3.96E-31 | 1.71E-30 | -1.74E+00 | 1.42E-25 | 6.05E-25 | -2.19E+00 | 6.54E-13 | 2.50E-12 | -1.67E+00 | 2.08E-16 | 7.42E-16 | -2.40E+00 | 1.02E-08 | 2.88E-08 | -1.92E+00 | 2.29E-20 | 8.39E-20 | -2.34E+00 | 1.08E-07 | 2.96E-07 | -1.75E+00 |
| BTN3A2 | 7.80E-52 | 4.07E-51 | -2.51E+00 | 6.75E-31 | 2.92E-30 | -1.52E+00 | 5.01E-27 | 2.13E-26 | -2.39E+00 | 6.50E-14 | 2.48E-13 | -1.50E+00 | 4.36E-40 | 1.56E-39 | -3.24E+00 | 5.25E-22 | 1.48E-21 | -1.84E+00 | 2.49E-46 | 9.14E-46 | -3.38E+00 | 2.30E-27 | 6.32E-27 | -2.33E+00 |
| PABPC1P4 | 3.35E-51 | 1.75E-50 | -2.89E+00 | 5.19E-36 | 2.24E-35 | -1.83E+00 | 7.64E-50 | 3.25E-49 | -3.75E+00 | 3.93E-29 | 1.50E-28 | -2.09E+00 | 4.42E-22 | 1.58E-21 | -2.91E+00 | 1.93E-14 | 5.45E-14 | -1.85E+00 | 6.31E-32 | 2.31E-31 | -3.21E+00 | 4.06E-21 | 1.11E-20 | -2.11E+00 |
| PTAFR | 5.87E-51 | 3.06E-50 | -7.26E+00 | 8.76E-29 | 3.78E-28 | -2.12E+00 | 2.47E-27 | 1.05E-26 | -3.94E+00 | 7.76E-14 | 2.97E-13 | -1.63E+00 | 3.16E-24 | 1.13E-23 | -3.15E+00 | 1.22E-11 | 3.43E-11 | -1.53E+00 | 8.31E-18 | 3.05E-17 | -2.71E+00 | 5.87E-06 | 1.61E-05 | -1.32E+00 |
| HLA-H | 1.55E-50 | 8.09E-50 | -3.50E+00 | 2.73E-46 | 1.18E-45 | -2.61E+00 | 3.68E-34 | 1.57E-33 | -3.48E+00 | 4.05E-25 | 1.55E-24 | -2.26E+00 | 6.64E-25 | 2.37E-24 | -3.54E+00 | 1.19E-22 | 3.35E-22 | -2.64E+00 | 4.52E-48 | 1.66E-47 | -5.26E+00 | 1.53E-39 | 4.19E-39 | -3.65E+00 |
| AC067942.1 | 1.98E-50 | 1.03E-49 | -1.78E+00 | 1.65E-14 | 7.14E-14 | -1.34E+00 | 3.61E-24 | 1.53E-23 | -1.70E+00 | 2.76E-06 | 1.05E-05 | -1.25E+00 | 8.46E-21 | 3.02E-20 | -1.69E+00 | 2.24E-05 | 6.30E-05 | -1.25E+00 | 5.72E-41 | 2.10E-40 | -1.80E+00 | 3.09E-06 | 8.49E-06 | -1.27E+00 |
| RPS18 | 2.07E-50 | 1.08E-49 | 1.59E+00 | 9.10E-73 | 3.93E-72 | 1.76E+00 | 3.93E-39 | 1.67E-38 | 1.65E+00 | 1.15E-59 | 4.38E-59 | 1.87E+00 | 6.80E-27 | 2.42E-26 | 1.61E+00 | 3.91E-48 | 1.10E-47 | 1.86E+00 | 7.47E-23 | 2.74E-22 | 1.61E+00 | 1.12E-52 | 3.07E-52 | 1.82E+00 |
| AL122020.1 | 3.84E-50 | 2.00E-49 | -1.61E+00 | 3.81E-48 | 1.64E-47 | -1.59E+00 | 2.91E-30 | 1.24E-29 | -1.56E+00 | 4.50E-27 | 1.72E-26 | -1.53E+00 | 1.29E-24 | 4.60E-24 | -1.57E+00 | 1.74E-20 | 4.91E-20 | -1.50E+00 | 1.54E-45 | 5.67E-45 | -1.61E+00 | 1.94E-31 | 5.34E-31 | -1.58E+00 |
| EEF1B2 | 7.42E-50 | 3.87E-49 | -1.64E+00 | 8.22E-57 | 3.55E-56 | -1.71E+00 | 1.43E-14 | 6.07E-14 | -1.39E+00 | 4.74E-20 | 1.81E-19 | -1.49E+00 | 8.53E-14 | 3.04E-13 | -1.40E+00 | 1.57E-13 | 4.42E-13 | -1.41E+00 | 2.08E-22 | 7.63E-22 | -1.39E+00 | 1.26E-18 | 3.47E-18 | -1.44E+00 |
| UBE2L6 | 8.68E-50 | 4.52E-49 | -2.14E+00 | 1.21E-28 | 5.21E-28 | -1.56E+00 | 2.50E-16 | 1.06E-15 | -2.02E+00 | 3.54E-11 | 1.35E-10 | -1.61E+00 | 4.36E-17 | 1.56E-16 | -2.09E+00 | 1.55E-07 | 4.38E-07 | -1.54E+00 | 1.86E-14 | 6.84E-14 | -1.84E+00 | 2.15E-06 | 5.89E-06 | -1.41E+00 |
| HOPX | 1.94E-49 | 1.01E-48 | 2.43E+00 | 3.13E-47 | 1.35E-46 | 2.47E+00 | 1.76E-22 | 7.47E-22 | 2.30E+00 | 1.17E-28 | 4.48E-28 | 2.74E+00 | 1.67E-21 | 5.97E-21 | 7.16E+00 | 1.93E-07 | 5.45E-07 | 4.25E+00 | 1.44E-04 | 5.28E-04 | 6.10E+00 | 5.85E-03 | 1.61E-02 | 5.94E+00 |
| SPN | 2.97E-49 | 1.55E-48 | -2.01E+00 | 2.55E-08 | 1.10E-07 | -1.05E+00 | 1.47E-17 | 6.27E-17 | -1.84E+00 | 1.99E-03 | 7.59E-03 | -1.05E+00 | 9.33E-11 | 3.33E-10 | -1.53E+00 | 2.17E-01 | 6.12E-01 | 1.11E+00 | 7.84E-11 | 2.88E-10 | -1.69E+00 | 5.73E-02 | 1.57E-01 | -1.07E+00 |
| STK17B | 1.59E-48 | 8.29E-48 | 3.59E+00 | 2.25E-01 | 9.71E-01 | 1.49E+00 | 3.42E-18 | 1.45E-17 | 2.91E+00 | 8.58E-03 | 3.28E-02 | 1.14E+00 | 3.01E-31 | 1.07E-30 | 4.62E+00 | 2.45E-01 | 6.92E-01 | 1.31E+00 | 9.34E-03 | 3.43E-02 | 1.67E+00 | 9.85E-13 | 2.71E-12 | -1.63E+00 |
| RPL30 | 2.53E-48 | 1.32E-47 | -1.38E+00 | 3.94E-29 | 1.70E-28 | -1.28E+00 | 2.04E-23 | 8.65E-23 | -1.31E+00 | 9.51E-13 | 3.63E-12 | -1.21E+00 | 1.82E-16 | 6.47E-16 | -1.30E+00 | 3.49E-12 | 9.84E-12 | -1.24E+00 | 1.02E-37 | 3.74E-37 | -1.34E+00 | 5.70E-21 | 1.57E-20 | -1.27E+00 |
| AL031733.2 | 2.55E-48 | 1.33E-47 | 4.05E+00 | 3.03E-104 | 1.31E-103 | 7.67E+00 | 5.80E-16 | 2.47E-15 | 3.34E+00 | 2.90E-38 | 1.11E-37 | 6.05E+00 | 3.35E-07 | 1.19E-06 | 3.67E+00 | 1.05E-09 | 2.95E-09 | 3.84E+00 | 5.42E-05 | 1.99E-04 | 4.09E+00 | 1.83E-05 | 5.03E-05 | 4.49E+00 |
| HSPB1 | 3.01E-48 | 1.57E-47 | -2.45E+00 | 6.87E-19 | 2.97E-18 | -1.37E+00 | 5.84E-19 | 2.48E-18 | -2.07E+00 | 9.74E-17 | 3.72E-16 | -1.76E+00 | 4.79E-23 | 1.71E-22 | -2.53E+00 | 1.32E-16 | 3.72E-16 | -1.87E+00 | 8.74E-14 | 3.21E-13 | -1.52E+00 | 5.76E-13 | 1.58E-12 | -1.60E+00 |
| RPL14P1 | 3.24E-48 | 1.69E-47 | -1.61E+00 | 1.10E-46 | 4.76E-46 | -1.57E+00 | 1.33E-34 | 5.63E-34 | -1.65E+00 | 7.47E-27 | 2.85E-26 | -1.55E+00 | 2.86E-33 | 1.02E-32 | -1.74E+00 | 4.40E-20 | 1.24E-19 | -1.52E+00 | 7.73E-61 | 2.84E-60 | -1.73E+00 | 5.29E-24 | 1.45E-23 | -1.50E+00 |
| RPL3 | 3.25E-48 | 1.70E-47 | 1.70E+00 | 3.63E-123 | 1.57E-122 | 2.39E+00 | 5.05E-38 | 2.15E-37 | 1.79E+00 | 3.82E-95 | 1.46E-94 | 2.57E+00 | 5.60E-27 | 2.00E-26 | 1.74E+00 | 3.34E-76 | 9.41E-76 | 2.44E+00 | 9.90E-21 | 3.63E-20 | 1.69E+00 | 4.26E-78 | 1.17E-77 | 2.36E+00 |
| KLF4 | 7.60E-48 | 3.96E-47 | 5.41E+00 | 2.68E-03 | 1.16E-02 | -1.04E+00 | 3.95E-23 | 1.68E-22 | 4.27E+00 | 2.94E-04 | 1.12E-03 | -1.46E+00 | 8.36E-11 | 2.98E-10 | 5.99E+00 | 7.75E-02 | 2.19E-01 | -1.09E+00 | 1.41E-11 | 5.18E-11 | 6.06E+00 | 2.42E-01 | 6.65E-01 | 1.18E+00 |
| AGPS | 7.89E-48 | 4.11E-47 | 3.60E+00 | 4.40E-32 | 1.90E-31 | 3.58E+00 | 2.45E-19 | 1.04E-18 | 2.85E+00 | 4.76E-09 | 1.82E-08 | 2.46E+00 | 1.59E-01 | 5.66E-01 | 1.62E+00 | 2.96E-01 | 8.35E-01 | 1.34E+00 | 1.53E-02 | 5.63E-02 | 1.19E+00 | 4.01E-02 | 1.10E-01 | 1.18E+00 |
| HLA-F | 1.64E-47 | 8.55E-47 | -2.30E+00 | 4.07E-48 | 1.76E-47 | -2.07E+00 | 9.78E-25 | 4.16E-24 | -2.35E+00 | 1.15E-25 | 4.39E-25 | -2.20E+00 | 1.06E-38 | 3.78E-38 | -5.34E+00 | 1.75E-40 | 4.93E-40 | -4.68E+00 | 2.49E-51 | 9.13E-51 | -4.86E+00 | 9.17E-51 | 2.52E-50 | -5.18E+00 |
| TRIM69 | 1.65E-47 | 8.62E-47 | -3.70E+00 | 2.81E-44 | 1.21E-43 | -3.01E+00 | 4.20E-54 | 1.78E-53 | -3.72E+00 | 8.32E-59 | 3.18E-58 | -4.01E+00 | 4.50E-36 | 1.60E-35 | -6.91E+00 | 8.27E-31 | 2.33E-30 | -3.76E+00 | 3.22E-49 | 1.18E-48 | -4.59E+00 | 1.79E-47 | 4.92E-47 | -4.34E+00 |
| OAS2 | 1.98E-47 | 1.03E-46 | -3.45E+00 | 7.07E-28 | 3.05E-27 | -1.83E+00 | 1.29E-51 | 5.49E-51 | -4.23E+00 | 1.19E-42 | 4.55E-42 | -2.86E+00 | 1.64E-04 | 5.84E-04 | -6.88E+00 | 4.20E-04 | 1.19E-03 | -4.97E+00 | 5.14E-12 | 1.89E-11 | -8.11E+00 | 1.27E-10 | 3.48E-10 | -6.10E+00 |
| RPS4XP13 | 3.65E-47 | 1.90E-46 | -2.08E+00 | 6.59E-08 | 2.85E-07 | -1.23E+00 | 1.14E-18 | 4.83E-18 | -1.93E+00 | 3.84E-03 | 1.47E-02 | -1.15E+00 | 1.10E-21 | 3.93E-21 | -1.91E+00 | 5.67E-03 | 1.60E-02 | -1.14E+00 | 1.14E-25 | 4.19E-25 | -1.97E+00 | 6.30E-03 | 1.73E-02 | -1.19E+00 |
| IFITM3 | 4.77E-47 | 2.49E-46 | -2.56E+00 | 1.84E-19 | 7.97E-19 | -1.66E+00 | 2.45E-33 | 1.04E-32 | -3.26E+00 | 4.60E-16 | 1.76E-15 | -2.23E+00 | 1.23E-46 | 4.37E-46 | -3.49E+00 | 8.84E-30 | 2.49E-29 | -2.59E+00 | 1.76E-49 | 6.47E-49 | -3.50E+00 | 3.09E-28 | 8.50E-28 | -2.72E+00 |
| RPL6P27 | 2.73E-46 | 1.42E-45 | 1.44E+00 | 4.34E-26 | 1.87E-25 | 1.32E+00 | 5.43E-39 | 2.31E-38 | 1.49E+00 | 2.43E-25 | 9.30E-25 | 1.39E+00 | 5.34E-12 | 1.90E-11 | 1.30E+00 | 6.00E-11 | 1.69E-10 | 1.27E+00 | 3.72E-12 | 1.36E-11 | 1.33E+00 | 1.57E-13 | 4.32E-13 | 1.29E+00 |
| AC010343.1 | 4.72E-46 | 2.46E-45 | 1.77E+00 | 3.62E-15 | 1.56E-14 | 1.41E+00 | 6.66E-36 | 2.83E-35 | 1.83E+00 | 1.51E-17 | 5.76E-17 | 1.52E+00 | 2.41E-19 | 8.59E-19 | 1.72E+00 | 1.23E-12 | 3.46E-12 | 1.52E+00 | 1.06E-17 | 3.87E-17 | 1.69E+00 | 1.25E-10 | 3.43E-10 | 1.42E+00 |
| JUND | 8.72E-46 | 4.55E-45 | 2.23E+00 | 1.03E-22 | 4.46E-22 | -1.35E+00 | 4.73E-27 | 2.01E-26 | 2.18E+00 | 8.19E-06 | 3.13E-05 | -1.22E+00 | 4.36E-29 | 1.55E-28 | 2.70E+00 | 1.34E-03 | 3.79E-03 | -1.10E+00 | 1.06E-26 | 3.91E-26 | 2.37E+00 | 2.86E-03 | 7.86E-03 | -1.12E+00 |
| RPL12P19 | 2.60E-45 | 1.35E-44 | -2.16E+00 | 7.34E-18 | 3.17E-17 | -1.38E+00 | 7.27E-18 | 3.09E-17 | -2.01E+00 | 9.80E-05 | 3.74E-04 | -1.17E+00 | 8.49E-25 | 3.03E-24 | -2.08E+00 | 1.84E-04 | 5.18E-04 | -1.23E+00 | 2.30E-18 | 8.45E-18 | -1.96E+00 | 2.06E-03 | 5.65E-03 | -1.24E+00 |
| KLF7 | 4.78E-45 | 2.49E-44 | 5.31E+00 | 6.88E-06 | 2.97E-05 | 2.77E+00 | 1.64E-17 | 6.95E-17 | 3.42E+00 | 1.93E-02 | 7.39E-02 | 1.84E+00 | 5.56E-02 | 1.98E-01 | 2.26E+00 | 1.53E-01 | 4.32E-01 | 1.38E+00 | 1.10E-01 | 4.04E-01 | 1.69E+00 | 1.95E-01 | 5.36E-01 | 1.24E+00 |
| AL450063.1 | 1.95E-44 | 1.02E-43 | -2.31E+00 | 1.53E-25 | 6.59E-25 | -1.53E+00 | 4.05E-16 | 1.72E-15 | -1.78E+00 | 2.74E-12 | 1.05E-11 | -1.39E+00 | 2.44E-23 | 8.69E-23 | -2.20E+00 | 3.26E-15 | 9.19E-15 | -1.51E+00 | 1.98E-15 | 7.27E-15 | -1.73E+00 | 2.12E-07 | 5.81E-07 | -1.33E+00 |
| AL136454.1 | 2.36E-44 | 1.23E-43 | -1.95E+00 | 1.37E-28 | 5.93E-28 | -1.65E+00 | 6.32E-21 | 2.69E-20 | -2.04E+00 | 1.84E-12 | 7.02E-12 | -1.65E+00 | 2.06E-19 | 7.35E-19 | -1.93E+00 | 1.19E-14 | 3.36E-14 | -1.68E+00 | 1.10E-26 | 4.02E-26 | -2.01E+00 | 4.09E-12 | 1.12E-11 | -1.65E+00 |
| PIK3IP1 | 2.93E-44 | 1.52E-43 | 4.11E+00 | 1.24E-31 | 5.34E-31 | 3.81E+00 | 1.39E-14 | 5.92E-14 | 3.00E+00 | 6.67E-08 | 2.55E-07 | 2.84E+00 | 2.63E-07 | 9.36E-07 | 3.05E+00 | 1.56E-03 | 4.39E-03 | 2.34E+00 | 1.46E-08 | 5.37E-08 | 3.16E+00 | 6.71E-02 | 1.84E-01 | 1.85E+00 |
| ARF5 | 4.39E-44 | 2.29E-43 | -2.28E+00 | 2.35E-21 | 1.02E-20 | -1.37E+00 | 1.29E-20 | 5.49E-20 | -1.95E+00 | 2.92E-10 | 1.12E-09 | -1.35E+00 | 2.53E-17 | 9.03E-17 | -1.97E+00 | 2.62E-08 | 7.39E-08 | -1.23E+00 | 2.29E-20 | 8.39E-20 | -1.86E+00 | 9.21E-07 | 2.53E-06 | -1.20E+00 |
| NACA2 | 4.87E-44 | 2.54E-43 | 1.98E+00 | 8.46E-51 | 3.66E-50 | 2.11E+00 | 5.81E-23 | 2.47E-22 | 1.85E+00 | 2.38E-35 | 9.08E-35 | 2.14E+00 | 1.77E-13 | 6.29E-13 | 1.88E+00 | 1.15E-25 | 3.24E-25 | 2.13E+00 | 1.63E-17 | 5.97E-17 | 1.79E+00 | 8.49E-30 | 2.33E-29 | 2.00E+00 |
| AC095059.2 | 7.61E-44 | 3.97E-43 | -3.09E+00 | 4.15E-27 | 1.79E-26 | -1.68E+00 | 4.30E-30 | 1.83E-29 | -2.77E+00 | 5.91E-14 | 2.26E-13 | -1.38E+00 | 1.43E-24 | 5.11E-24 | -3.12E+00 | 6.96E-15 | 1.96E-14 | -1.75E+00 | 4.19E-25 | 1.54E-24 | -2.87E+00 | 5.97E-11 | 1.64E-10 | -1.55E+00 |
| GLUL | 1.06E-43 | 5.51E-43 | 2.37E+00 | 2.13E-39 | 9.21E-39 | 2.39E+00 | 1.60E-17 | 6.79E-17 | 1.78E+00 | 1.98E-20 | 7.55E-20 | 1.93E+00 | 5.77E-06 | 2.06E-05 | 1.60E+00 | 2.15E-06 | 6.07E-06 | 1.70E+00 | 1.44E-07 | 5.29E-07 | 1.46E+00 | 3.54E-12 | 9.72E-12 | 1.67E+00 |
| PNMT | 1.06E-43 | 5.51E-43 | 1.19E+01 | 1.52E-22 | 6.59E-22 | 9.15E+00 | 1.93E-08 | 8.20E-08 | 7.78E+00 | 8.52E-08 | 3.26E-07 | 7.95E+00 | 8.10E-49 | 2.89E-48 | 6.22E+00 | 1.15E-33 | 3.24E-33 | 4.57E+00 | 1.05E-43 | 3.85E-43 | 5.21E+00 | 5.26E-35 | 1.45E-34 | 4.68E+00 |
| ALDH9A1 | 1.39E-43 | 7.25E-43 | -2.70E+00 | 6.81E-20 | 2.94E-19 | -1.35E+00 | 5.29E-21 | 2.25E-20 | -1.92E+00 | 1.88E-15 | 7.18E-15 | -1.43E+00 | 7.56E-08 | 2.70E-07 | -2.36E+00 | 1.23E-05 | 3.47E-05 | -1.57E+00 | 2.06E-13 | 7.55E-13 | -2.13E+00 | 1.44E-06 | 3.94E-06 | -1.41E+00 |
| AC131235.1 | 2.55E-43 | 1.33E-42 | 1.59E+00 | 6.50E-59 | 2.81E-58 | 1.73E+00 | 2.87E-32 | 1.22E-31 | 1.63E+00 | 1.05E-42 | 4.03E-42 | 1.76E+00 | 1.19E-27 | 4.23E-27 | 1.72E+00 | 3.23E-43 | 9.10E-43 | 1.89E+00 | 5.16E-34 | 1.90E-33 | 1.75E+00 | 6.58E-58 | 1.81E-57 | 1.90E+00 |
| AC093249.4 | 2.62E-43 | 1.37E-42 | -2.08E+00 | 2.86E-36 | 1.24E-35 | -1.72E+00 | 2.95E-18 | 1.25E-17 | -2.06E+00 | 3.59E-14 | 1.37E-13 | -1.69E+00 | 6.29E-21 | 2.24E-20 | -2.02E+00 | 1.01E-19 | 2.86E-19 | -1.86E+00 | 6.67E-20 | 2.45E-19 | -2.15E+00 | 6.21E-13 | 1.71E-12 | -1.82E+00 |
| RPL6 | 4.17E-43 | 2.17E-42 | -1.49E+00 | 4.40E-23 | 1.90E-22 | -1.33E+00 | 9.00E-25 | 3.82E-24 | -1.46E+00 | 1.92E-10 | 7.32E-10 | -1.26E+00 | 5.98E-35 | 2.13E-34 | -1.66E+00 | 7.69E-15 | 2.17E-14 | -1.37E+00 | 2.25E-57 | 8.26E-57 | -1.68E+00 | 4.09E-20 | 1.12E-19 | -1.36E+00 |
| ATRX | 4.18E-43 | 2.18E-42 | -1.95E+00 | 1.46E-11 | 6.32E-11 | -1.17E+00 | 6.83E-15 | 2.90E-14 | -1.81E+00 | 4.88E-05 | 1.86E-04 | -1.20E+00 | 6.59E-18 | 2.35E-17 | -1.85E+00 | 2.53E-05 | 7.13E-05 | -1.22E+00 | 8.14E-12 | 2.99E-11 | -1.68E+00 | 8.26E-04 | 2.27E-03 | -1.31E+00 |
| JUNB | 4.90E-43 | 2.55E-42 | 4.63E+00 | 2.29E-21 | 9.91E-21 | -1.15E+00 | 2.08E-16 | 8.85E-16 | 3.15E+00 | 2.05E-15 | 7.85E-15 | -1.47E+00 | 6.69E-09 | 2.39E-08 | 2.72E+00 | 1.47E-09 | 4.13E-09 | -1.27E+00 | 2.08E-05 | 7.64E-05 | 2.73E+00 | 7.26E-20 | 1.99E-19 | -1.63E+00 |
| AC004130.1 | 5.17E-43 | 2.69E-42 | -3.37E+00 | 3.14E-33 | 1.35E-32 | -2.15E+00 | 1.09E-14 | 4.62E-14 | -2.96E+00 | 2.67E-16 | 1.02E-15 | -3.11E+00 | 1.62E-09 | 5.78E-09 | -3.88E+00 | 1.97E-10 | 5.55E-10 | -4.12E+00 | 5.35E-02 | 1.96E-01 | -1.36E+00 | 1.04E-01 | 2.86E-01 | -1.16E+00 |
| NAP1L1P1 | 7.46E-43 | 3.89E-42 | -2.03E+00 | 9.98E-50 | 4.31E-49 | -1.97E+00 | 9.03E-17 | 3.84E-16 | -2.01E+00 | 9.38E-19 | 3.58E-18 | -1.95E+00 | 1.51E-13 | 5.39E-13 | -2.04E+00 | 1.42E-14 | 3.99E-14 | -1.99E+00 | 8.07E-16 | 2.96E-15 | -1.70E+00 | 6.87E-18 | 1.89E-17 | -1.84E+00 |
| PRDX1P1 | 1.62E-42 | 8.45E-42 | -2.88E+00 | 2.73E-18 | 1.18E-17 | -1.42E+00 | 3.88E-11 | 1.65E-10 | -1.82E+00 | 7.88E-04 | 3.01E-03 | -1.10E+00 | 4.35E-10 | 1.55E-09 | -2.70E+00 | 6.95E-04 | 1.96E-03 | -1.19E+00 | 6.08E-11 | 2.23E-10 | -1.85E+00 | 1.24E-02 | 3.41E-02 | -1.03E+00 |
| HSP90AB3P | 1.83E-42 | 9.52E-42 | -2.12E+00 | 1.09E-34 | 4.70E-34 | -1.92E+00 | 4.24E-12 | 1.80E-11 | -1.82E+00 | 1.59E-10 | 6.08E-10 | -1.70E+00 | 6.75E-18 | 2.41E-17 | -1.90E+00 | 1.97E-14 | 5.55E-14 | -1.77E+00 | 1.42E-14 | 5.20E-14 | -1.67E+00 | 1.64E-07 | 4.51E-07 | -1.56E+00 |
| AC034207.1 | 3.41E-42 | 1.78E-41 | -2.16E+00 | 8.31E-23 | 3.59E-22 | -1.49E+00 | 1.13E-23 | 4.78E-23 | -2.25E+00 | 1.92E-12 | 7.32E-12 | -1.53E+00 | 6.18E-24 | 2.20E-23 | -2.09E+00 | 1.13E-09 | 3.19E-09 | -1.45E+00 | 1.66E-25 | 6.10E-25 | -2.25E+00 | 1.02E-08 | 2.81E-08 | -1.51E+00 |
| DUSP2 | 5.21E-42 | 2.71E-41 | 9.15E+00 | 7.24E-03 | 3.13E-02 | -1.16E+00 | 2.65E-12 | 1.12E-11 | 3.83E+00 | 1.19E-02 | 4.53E-02 | -1.35E+00 | 1.85E-01 | 6.59E-01 | 1.71E+00 | 5.93E-02 | 1.67E-01 | -1.32E+00 | 2.30E-01 | 8.45E-01 | 1.38E+00 | 1.28E-01 | 3.51E-01 | -1.19E+00 |
| AC007688.1 | 5.52E-42 | 2.88E-41 | -2.13E+00 | 4.60E-34 | 1.99E-33 | -1.75E+00 | 1.94E-15 | 8.22E-15 | -2.17E+00 | 4.25E-13 | 1.62E-12 | -1.80E+00 | 1.90E-18 | 6.78E-18 | -1.96E+00 | 5.57E-13 | 1.57E-12 | -1.68E+00 | 2.41E-22 | 8.84E-22 | -2.20E+00 | 1.50E-13 | 4.13E-13 | -1.83E+00 |
| STON2 | 6.37E-42 | 3.32E-41 | -2.98E+00 | 2.22E-13 | 9.58E-13 | -1.25E+00 | 2.10E-19 | 8.93E-19 | -2.97E+00 | 2.03E-04 | 7.74E-04 | -1.09E+00 | 7.47E-22 | 2.66E-21 | -1.86E+00 | 7.69E-08 | 2.17E-07 | -1.14E+00 | 1.63E-19 | 5.98E-19 | -1.84E+00 | 1.86E-08 | 5.10E-08 | -1.33E+00 |
| IRF7 | 1.32E-41 | 6.88E-41 | -3.64E+00 | 5.05E-32 | 2.18E-31 | -2.48E+00 | 6.64E-48 | 2.82E-47 | -4.33E+00 | 4.52E-36 | 1.73E-35 | -2.85E+00 | 2.16E-29 | 7.69E-29 | -3.88E+00 | 2.44E-25 | 6.89E-25 | -2.75E+00 | 1.75E-43 | 6.41E-43 | -4.57E+00 | 1.45E-41 | 3.98E-41 | -5.02E+00 |
| DUSP1 | 2.40E-41 | 1.25E-40 | 2.64E+00 | 3.11E-19 | 1.34E-18 | 2.22E+00 | 2.31E-37 | 9.81E-37 | 3.05E+00 | 7.36E-14 | 2.81E-13 | 2.10E+00 | 3.51E-35 | 1.25E-34 | 3.32E+00 | 1.43E-11 | 4.02E-11 | 1.88E+00 | 4.98E-37 | 1.83E-36 | 2.96E+00 | 1.24E-02 | 3.40E-02 | 1.33E+00 |
| MCL1 | 2.85E-41 | 1.48E-40 | 3.29E+00 | 4.01E-03 | 1.73E-02 | 1.32E+00 | 3.34E-13 | 1.42E-12 | 2.55E+00 | 2.35E-01 | 8.98E-01 | 1.45E+00 | 4.71E-10 | 1.68E-09 | 2.35E+00 | 6.42E-03 | 1.81E-02 | 1.05E+00 | 1.21E-01 | 4.43E-01 | 1.59E+00 | 7.37E-05 | 2.02E-04 | -1.03E+00 |
| SAMD9L | 2.97E-41 | 1.55E-40 | -2.77E+00 | 5.96E-26 | 2.57E-25 | -1.79E+00 | 3.22E-32 | 1.37E-31 | -3.35E+00 | 8.55E-29 | 3.27E-28 | -2.60E+00 | 3.81E-25 | 1.36E-24 | -3.51E+00 | 7.10E-22 | 2.00E-21 | -2.76E+00 | 1.14E-43 | 4.19E-43 | -3.63E+00 | 6.85E-42 | 1.88E-41 | -3.49E+00 |
| RPL34P18 | 3.52E-41 | 1.83E-40 | 1.55E+00 | 9.36E-04 | 4.04E-03 | 1.12E+00 | 1.84E-29 | 7.80E-29 | 1.60E+00 | 3.35E-05 | 1.28E-04 | 1.21E+00 | 7.04E-23 | 2.51E-22 | 1.60E+00 | 8.68E-03 | 2.45E-02 | 1.14E+00 | 3.32E-13 | 1.22E-12 | 1.48E+00 | 2.55E-01 | 7.01E-01 | 1.07E+00 |
| DUSP6 | 3.83E-41 | 2.00E-40 | 4.72E+00 | 1.02E-02 | 4.39E-02 | 1.91E+00 | 1.01E-29 | 4.29E-29 | 4.43E+00 | 5.01E-03 | 1.91E-02 | 2.02E+00 | 1.61E-08 | 5.75E-08 | 4.02E+00 | 1.89E-01 | 5.32E-01 | -1.03E+00 | 6.86E-03 | 2.52E-02 | 1.81E+00 | 3.49E-01 | 9.59E-01 | -1.02E+00 |
| AL121871.1 | 4.34E-41 | 2.26E-40 | 1.59E+00 | 1.76E-02 | 7.61E-02 | 1.09E+00 | 6.21E-39 | 2.64E-38 | 1.72E+00 | 1.20E-03 | 4.59E-03 | 1.15E+00 | 6.99E-24 | 2.49E-23 | 1.62E+00 | 3.03E-02 | 8.53E-02 | 1.10E+00 | 3.36E-23 | 1.23E-22 | 1.65E+00 | 1.38E-03 | 3.79E-03 | 1.16E+00 |
| YWHAG | 5.02E-41 | 2.62E-40 | -2.08E+00 | 4.88E-42 | 2.11E-41 | -1.88E+00 | 8.53E-11 | 3.62E-10 | -1.70E+00 | 2.18E-08 | 8.34E-08 | -1.37E+00 | 5.76E-17 | 2.05E-16 | -1.91E+00 | 1.07E-14 | 3.00E-14 | -1.71E+00 | 1.42E-17 | 5.23E-17 | -1.97E+00 | 4.90E-08 | 1.35E-07 | -1.37E+00 |
| RPL7P9 | 5.24E-41 | 2.73E-40 | -1.51E+00 | 7.46E-20 | 3.22E-19 | -1.31E+00 | 4.14E-28 | 1.76E-27 | -1.53E+00 | 1.45E-11 | 5.55E-11 | -1.30E+00 | 2.71E-25 | 9.65E-25 | -1.54E+00 | 4.90E-08 | 1.38E-07 | -1.27E+00 | 3.84E-57 | 1.41E-56 | -1.66E+00 | 5.39E-14 | 1.48E-13 | -1.30E+00 |
| RPL27A | 7.30E-41 | 3.80E-40 | 1.39E+00 | 1.06E-01 | 4.58E-01 | -1.02E+00 | 6.85E-22 | 2.91E-21 | 1.34E+00 | 2.10E-01 | 8.03E-01 | 1.02E+00 | 3.04E-20 | 1.09E-19 | 1.39E+00 | 2.99E-02 | 8.44E-02 | 1.07E+00 | 6.20E-07 | 2.28E-06 | 1.23E+00 | 3.24E-01 | 8.89E-01 | 1.04E+00 |
| BANF1 | 8.22E-41 | 4.28E-40 | -2.01E+00 | 1.21E-23 | 5.23E-23 | -1.35E+00 | 6.36E-15 | 2.70E-14 | -1.74E+00 | 3.50E-08 | 1.34E-07 | -1.25E+00 | 5.58E-22 | 1.99E-21 | -1.94E+00 | 3.50E-10 | 9.86E-10 | -1.28E+00 | 5.57E-17 | 2.04E-16 | -2.01E+00 | 1.74E-04 | 4.78E-04 | -1.25E+00 |
| HCP5 | 1.31E-40 | 6.83E-40 | -2.96E+00 | 2.48E-33 | 1.07E-32 | -2.21E+00 | 7.14E-41 | 3.04E-40 | -3.50E+00 | 3.68E-34 | 1.41E-33 | -2.45E+00 | 9.57E-24 | 3.41E-23 | -4.71E+00 | 2.52E-19 | 7.11E-19 | -2.90E+00 | 4.70E-73 | 1.72E-72 | -8.23E+00 | 1.89E-67 | 5.18E-67 | -7.05E+00 |
| EEF1A1P5 | 1.42E-40 | 7.40E-40 | -1.66E+00 | 4.93E-02 | 2.13E-01 | 1.16E+00 | 9.67E-13 | 4.11E-12 | -1.50E+00 | 1.17E-04 | 4.48E-04 | 1.32E+00 | 3.82E-16 | 1.36E-15 | -1.59E+00 | 3.48E-02 | 9.81E-02 | 1.24E+00 | 1.03E-20 | 3.76E-20 | -1.63E+00 | 3.48E-03 | 9.56E-03 | 1.24E+00 |
| RPL10AP2 | 1.42E-40 | 7.40E-40 | 1.96E+00 | 3.20E-17 | 1.38E-16 | 1.71E+00 | 3.49E-36 | 1.48E-35 | 2.25E+00 | 4.88E-24 | 1.87E-23 | 1.96E+00 | 3.73E-17 | 1.33E-16 | 2.10E+00 | 7.20E-12 | 2.03E-11 | 1.83E+00 | 2.20E-21 | 8.06E-21 | 2.05E+00 | 7.04E-17 | 1.93E-16 | 1.78E+00 |
| HSP90AB7P | 1.63E-40 | 8.51E-40 | -4.53E+00 | 1.74E-30 | 7.51E-30 | -2.56E+00 | 3.08E-39 | 1.31E-38 | -3.58E+00 | 9.91E-28 | 3.79E-27 | -2.16E+00 | 1.74E-20 | 6.22E-20 | -2.81E+00 | 2.14E-16 | 6.04E-16 | -2.01E+00 | 1.28E-27 | 4.69E-27 | -3.24E+00 | 1.81E-16 | 4.98E-16 | -1.84E+00 |
| RPS13 | 6.19E-40 | 3.22E-39 | 1.41E+00 | 9.20E-18 | 3.97E-17 | 1.26E+00 | 3.91E-32 | 1.66E-31 | 1.45E+00 | 1.74E-16 | 6.65E-16 | 1.30E+00 | 5.06E-21 | 1.80E-20 | 1.41E+00 | 1.62E-15 | 4.56E-15 | 1.32E+00 | 2.00E-19 | 7.33E-19 | 1.39E+00 | 6.49E-17 | 1.78E-16 | 1.30E+00 |
| RPL24P7 | 6.96E-40 | 3.63E-39 | -1.88E+00 | 1.24E-29 | 5.37E-29 | -1.48E+00 | 4.74E-13 | 2.02E-12 | -1.73E+00 | 1.44E-13 | 5.48E-13 | -1.47E+00 | 6.05E-14 | 2.16E-13 | -1.70E+00 | 8.53E-12 | 2.41E-11 | -1.40E+00 | 5.40E-13 | 1.98E-12 | -1.68E+00 | 1.57E-07 | 4.31E-07 | -1.33E+00 |
| H3F3BP1 | 9.43E-40 | 4.91E-39 | -2.33E+00 | 7.25E-42 | 3.13E-41 | -2.08E+00 | 2.29E-14 | 9.74E-14 | -1.79E+00 | 4.25E-16 | 1.62E-15 | -1.64E+00 | 1.94E-27 | 6.91E-27 | -2.57E+00 | 8.84E-25 | 2.49E-24 | -2.28E+00 | 7.23E-28 | 2.65E-27 | -2.09E+00 | 6.03E-24 | 1.66E-23 | -1.96E+00 |
| CHST12 | 1.09E-39 | 5.70E-39 | -1.91E+00 | 2.40E-08 | 1.04E-07 | -1.00E+00 | 1.51E-22 | 6.42E-22 | -2.25E+00 | 1.61E-03 | 6.14E-03 | -1.09E+00 | 5.94E-10 | 2.12E-09 | -1.49E+00 | 2.72E-02 | 7.67E-02 | 1.03E+00 | 5.47E-10 | 2.01E-09 | -1.35E+00 | 9.62E-05 | 2.64E-04 | -1.07E+00 |
| FAM117A | 1.20E-39 | 6.24E-39 | -2.42E+00 | 8.38E-07 | 3.62E-06 | -1.26E+00 | 2.93E-33 | 1.24E-32 | -3.08E+00 | 4.18E-06 | 1.60E-05 | -1.44E+00 | 8.82E-32 | 3.15E-31 | -2.99E+00 | 1.49E-06 | 4.19E-06 | -1.44E+00 | 5.93E-27 | 2.18E-26 | -2.92E+00 | 2.71E-10 | 7.45E-10 | -1.92E+00 |
| TP53INP1 | 1.59E-39 | 8.29E-39 | 6.08E+00 | 4.58E-29 | 1.98E-28 | 6.28E+00 | 9.93E-05 | 4.22E-04 | 2.74E+00 | 5.08E-04 | 1.94E-03 | 2.85E+00 | 1.87E-05 | 6.65E-05 | 2.49E+00 | 1.22E-02 | 3.43E-02 | 2.28E+00 | 3.71E-05 | 1.36E-04 | 2.37E+00 | 4.67E-02 | 1.28E-01 | 1.88E+00 |
| HNRNPF | 3.77E-39 | 1.96E-38 | -1.74E+00 | 1.74E-12 | 7.51E-12 | -1.12E+00 | 7.91E-08 | 3.36E-07 | -1.55E+00 | 2.99E-02 | 1.14E-01 | -1.08E+00 | 3.62E-31 | 1.29E-30 | -2.18E+00 | 3.34E-05 | 9.40E-05 | -1.16E+00 | 1.91E-10 | 7.02E-10 | -1.65E+00 | 3.35E-01 | 9.20E-01 | 1.04E+00 |
| MMP7 | 3.85E-39 | 2.01E-38 | 1.17E+01 | 2.18E-02 | 9.43E-02 | 2.73E+00 | 1.21E-12 | 5.15E-12 | 9.99E+00 | 4.20E-02 | 1.61E-01 | 2.81E+00 | 1.69E-01 | 6.01E-01 | 2.07E+00 | 3.33E-01 | 9.37E-01 | 1.16E+00 | 1.81E-01 | 6.66E-01 | 1.58E+00 | 3.63E-01 | 9.97E-01 | 1.02E+00 |
| MLEC | 1.19E-38 | 6.22E-38 | -2.16E+00 | 2.72E-20 | 1.18E-19 | -1.44E+00 | 1.47E-08 | 6.23E-08 | -1.45E+00 | 3.60E-03 | 1.38E-02 | -1.09E+00 | 1.01E-22 | 3.61E-22 | -2.01E+00 | 7.83E-12 | 2.21E-11 | -1.42E+00 | 1.57E-07 | 5.77E-07 | -1.41E+00 | 5.46E-02 | 1.50E-01 | -1.04E+00 |
| CCDC47 | 1.61E-38 | 8.37E-38 | -1.94E+00 | 4.48E-29 | 1.93E-28 | -1.50E+00 | 7.99E-15 | 3.40E-14 | -1.53E+00 | 1.44E-09 | 5.48E-09 | -1.15E+00 | 4.85E-10 | 1.73E-09 | -1.32E+00 | 1.29E-05 | 3.64E-05 | -1.07E+00 | 1.47E-11 | 5.39E-11 | -1.52E+00 | 3.08E-06 | 8.46E-06 | -1.18E+00 |
| NACA3P | 1.92E-38 | 9.99E-38 | 2.27E+00 | 1.68E-39 | 7.28E-39 | 2.39E+00 | 2.18E-28 | 9.27E-28 | 2.27E+00 | 7.43E-34 | 2.84E-33 | 2.51E+00 | 2.36E-15 | 8.43E-15 | 2.16E+00 | 1.53E-22 | 4.30E-22 | 2.38E+00 | 2.63E-19 | 9.64E-19 | 2.23E+00 | 6.57E-24 | 1.81E-23 | 2.34E+00 |
| IQGAP2 | 2.72E-38 | 1.42E-37 | -1.98E+00 | 1.01E-06 | 4.38E-06 | -1.17E+00 | 4.26E-09 | 1.81E-08 | -1.67E+00 | 3.34E-02 | 1.28E-01 | -1.09E+00 | 5.09E-29 | 1.81E-28 | -2.19E+00 | 4.01E-12 | 1.13E-11 | -1.58E+00 | 1.32E-13 | 4.85E-13 | -1.57E+00 | 1.45E-08 | 3.99E-08 | -1.46E+00 |
| GAPT | 8.47E-38 | 4.41E-37 | -2.32E+00 | 1.09E-26 | 4.69E-26 | -1.66E+00 | 4.24E-23 | 1.80E-22 | -1.98E+00 | 6.90E-17 | 2.64E-16 | -1.58E+00 | 5.50E-33 | 1.96E-32 | -2.86E+00 | 7.02E-33 | 1.98E-32 | -2.54E+00 | 7.01E-10 | 2.57E-09 | -2.73E+00 | 1.80E-06 | 4.94E-06 | -1.96E+00 |
| PRKG2 | 9.79E-38 | 5.10E-37 | -2.59E+00 | 3.35E-21 | 1.45E-20 | -1.48E+00 | 1.39E-13 | 5.91E-13 | -3.18E+00 | 3.01E-05 | 1.15E-04 | -1.44E+00 | 1.88E-33 | 6.69E-33 | -3.78E+00 | 1.16E-23 | 3.28E-23 | -2.08E+00 | 1.18E-01 | 4.32E-01 | -1.08E+00 | 2.18E-01 | 6.00E-01 | 1.48E+00 |
| HLA-DMB | 1.10E-37 | 5.71E-37 | -1.97E+00 | 1.82E-18 | 7.85E-18 | -1.27E+00 | 1.87E-15 | 7.93E-15 | -1.75E+00 | 1.97E-09 | 7.51E-09 | -1.27E+00 | 1.75E-07 | 6.24E-07 | -2.29E+00 | 9.37E-05 | 2.64E-04 | -1.46E+00 | 3.74E-08 | 1.37E-07 | -3.64E+00 | 1.26E-05 | 3.47E-05 | -2.37E+00 |
| RPL7A | 1.14E-37 | 5.93E-37 | 1.60E+00 | 9.44E-113 | 4.08E-112 | 2.33E+00 | 9.17E-29 | 3.89E-28 | 1.66E+00 | 5.57E-85 | 2.13E-84 | 2.42E+00 | 2.96E-18 | 1.05E-17 | 1.62E+00 | 5.03E-71 | 1.42E-70 | 2.42E+00 | 6.17E-22 | 2.27E-21 | 1.65E+00 | 3.59E-81 | 9.86E-81 | 2.42E+00 |
| LAG3 | 1.26E-37 | 6.54E-37 | -2.17E+00 | 2.61E-24 | 1.13E-23 | -1.42E+00 | 6.68E-14 | 2.84E-13 | -2.44E+00 | 1.69E-09 | 6.46E-09 | -1.65E+00 | 1.03E-01 | 3.67E-01 | -1.67E+00 | 1.73E-01 | 4.88E-01 | -1.39E+00 | 2.44E-01 | 8.95E-01 | 1.21E+00 | 3.38E-01 | 9.28E-01 | 1.16E+00 |
| GBP1 | 2.04E-37 | 1.06E-36 | -2.50E+00 | 3.00E-41 | 1.30E-40 | -2.44E+00 | 5.02E-37 | 2.13E-36 | -2.92E+00 | 7.27E-57 | 2.78E-56 | -3.72E+00 | 1.51E-02 | 5.38E-02 | -3.00E+00 | 2.66E-02 | 7.48E-02 | -2.55E+00 | 5.46E-33 | 2.01E-32 | -7.80E+00 | 1.76E-30 | 4.85E-30 | -7.71E+00 |
| RPL7AP30 | 2.26E-37 | 1.18E-36 | 1.72E+00 | 9.71E-52 | 4.19E-51 | 1.99E+00 | 8.92E-28 | 3.79E-27 | 1.83E+00 | 1.07E-39 | 4.10E-39 | 2.10E+00 | 6.61E-21 | 2.36E-20 | 1.86E+00 | 5.98E-37 | 1.69E-36 | 2.21E+00 | 2.16E-19 | 7.91E-19 | 1.83E+00 | 5.10E-41 | 1.40E-40 | 2.16E+00 |
| LUC7L3 | 4.56E-37 | 2.38E-36 | -1.55E+00 | 4.74E-06 | 2.05E-05 | 1.65E+00 | 4.19E-15 | 1.78E-14 | -1.63E+00 | 2.13E-02 | 8.13E-02 | 1.34E+00 | 1.37E-11 | 4.90E-11 | -1.27E+00 | 1.65E-02 | 4.65E-02 | 1.44E+00 | 3.46E-07 | 1.27E-06 | -1.35E+00 | 1.97E-01 | 5.42E-01 | 1.12E+00 |
| ZKSCAN1 | 7.44E-37 | 3.88E-36 | -1.99E+00 | 1.06E-07 | 4.57E-07 | -1.03E+00 | 7.32E-20 | 3.11E-19 | -2.06E+00 | 1.59E-06 | 6.07E-06 | -1.19E+00 | 2.37E-14 | 8.46E-14 | -1.70E+00 | 2.50E-06 | 7.06E-06 | -1.20E+00 | 3.00E-11 | 1.10E-10 | -1.56E+00 | 9.69E-05 | 2.66E-04 | -1.15E+00 |
| FABP5P7 | 9.54E-37 | 4.97E-36 | -2.75E+00 | 1.58E-27 | 6.84E-27 | -1.86E+00 | 2.79E-19 | 1.18E-18 | -1.93E+00 | 8.92E-12 | 3.41E-11 | -1.32E+00 | 2.24E-19 | 7.98E-19 | -2.38E+00 | 1.14E-12 | 3.21E-12 | -1.76E+00 | 2.05E-19 | 7.52E-19 | -1.95E+00 | 7.99E-05 | 2.20E-04 | -1.14E+00 |
| GPR27 | 1.47E-36 | 7.64E-36 | -3.81E+00 | 1.22E-31 | 5.28E-31 | -2.71E+00 | 2.51E-20 | 1.06E-19 | -2.46E+00 | 3.03E-15 | 1.16E-14 | -1.64E+00 | 2.92E-13 | 1.04E-12 | -2.97E+00 | 2.73E-15 | 7.68E-15 | -4.26E+00 | 3.86E-09 | 1.42E-08 | -3.93E+00 | 4.84E-07 | 1.33E-06 | -2.90E+00 |
| RNF24 | 1.55E-36 | 8.09E-36 | -2.18E+00 | 1.47E-19 | 6.35E-19 | -1.44E+00 | 4.60E-20 | 1.96E-19 | -2.18E+00 | 2.89E-07 | 1.10E-06 | -1.33E+00 | 2.52E-20 | 8.97E-20 | -2.03E+00 | 4.99E-08 | 1.41E-07 | -1.28E+00 | 1.31E-18 | 4.83E-18 | -2.01E+00 | 3.80E-09 | 1.04E-08 | -1.40E+00 |
| TOP1 | 1.65E-36 | 8.60E-36 | -1.90E+00 | 5.66E-18 | 2.45E-17 | -1.25E+00 | 2.05E-17 | 8.70E-17 | -1.75E+00 | 7.75E-11 | 2.96E-10 | -1.37E+00 | 6.98E-21 | 2.49E-20 | -2.03E+00 | 1.34E-04 | 3.78E-04 | -1.16E+00 | 2.89E-11 | 1.06E-10 | -1.70E+00 | 3.26E-04 | 8.97E-04 | -1.38E+00 |
| TMEM98 | 2.11E-36 | 1.10E-35 | -2.14E+00 | 7.48E-22 | 3.23E-21 | -1.40E+00 | 3.69E-05 | 1.57E-04 | -1.52E+00 | 6.33E-05 | 2.42E-04 | -1.24E+00 | 1.22E-15 | 4.36E-15 | -2.12E+00 | 1.00E-18 | 2.82E-18 | -2.50E+00 | 1.95E-06 | 7.16E-06 | -2.03E+00 | 2.08E-03 | 5.72E-03 | -1.30E+00 |
| RPL29P11 | 2.25E-36 | 1.17E-35 | -1.48E+00 | 3.51E-59 | 1.52E-58 | -1.64E+00 | 1.71E-20 | 7.29E-20 | -1.42E+00 | 1.14E-29 | 4.34E-29 | -1.53E+00 | 8.18E-16 | 2.92E-15 | -1.40E+00 | 8.67E-24 | 2.44E-23 | -1.52E+00 | 5.80E-25 | 2.13E-24 | -1.40E+00 | 2.17E-31 | 5.96E-31 | -1.54E+00 |
| RPL12P13 | 2.37E-36 | 1.23E-35 | -2.16E+00 | 5.22E-38 | 2.26E-37 | -2.04E+00 | 9.87E-20 | 4.20E-19 | -1.98E+00 | 3.49E-19 | 1.33E-18 | -1.66E+00 | 2.29E-11 | 8.17E-11 | -1.80E+00 | 5.55E-10 | 1.56E-09 | -1.49E+00 | 4.57E-26 | 1.68E-25 | -2.22E+00 | 8.03E-19 | 2.20E-18 | -1.87E+00 |
| [illegible] | [illegible] | [illegible] | [illegible] | [illegible] | [illegible] | [illegible] | [illegible] | [illegible] | [illegible] | [illegible] | [illegible] | [illegible] | [illegible] | [illegible] | [illegible] | [illegible] | [illegible] | [illegible] | [illegible] | [illegible] | [illegible] | [illegible] | [illegible] | [illegible] |

| | | | | | | | | | | | | | | | | | | | | | | | | |
|---|---|---|---|---|---|---|---|---|---|---|---|---|---|---|---|---|---|---|---|---|---|---|---|---|
| LYPLA1 | 1.20E-34 | 6.25E-34 | -1.72E+00 | 1.11E-11 | 4.80E-11 | -1.07E+00 | 5.73E-06 | 2.43E-05 | -1.31E+00 | 2.87E-04 | 1.10E-03 | -1.02E+00 | 7.43E-06 | 2.65E-05 | -1.17E+00 | 2.54E-02 | 7.17E-02 | 1.10E+00 | 2.77E-05 | 1.02E-04 | -1.31E+00 | 2.54E-01 | 6.98E-01 | 1.04E+00 |
| PARP1 | 1.28E-34 | 6.69E-34 | -1.76E+00 | 5.75E-11 | 2.49E-10 | -1.14E+00 | 2.41E-04 | 1.02E-03 | -1.28E+00 | 2.18E-01 | 8.34E-01 | 1.11E+00 | 2.78E-10 | 9.93E-10 | -1.36E+00 | 1.42E-03 | 3.99E-03 | -1.01E+00 | 1.48E-04 | 5.42E-04 | -1.14E+00 | 2.11E-01 | 5.80E-01 | 1.18E+00 |
| RPS18P12 | 1.31E-34 | 6.81E-34 | -1.88E+00 | 2.26E-13 | 9.77E-13 | -1.43E+00 | 1.14E-14 | 4.85E-14 | -1.79E+00 | 1.26E-05 | 4.83E-05 | -1.32E+00 | 4.06E-18 | 1.45E-17 | -1.78E+00 | 1.57E-05 | 4.42E-05 | -1.32E+00 | 1.33E-21 | 4.87E-21 | -1.84E+00 | 1.30E-05 | 3.57E-05 | -1.39E+00 |
| RPL41P2 | 1.38E-34 | 7.20E-34 | -1.38E+00 | 5.62E-177 | 2.43E-176 | -2.15E+00 | 5.56E-23 | 2.36E-22 | -1.39E+00 | 1.68E-130 | 6.44E-130 | -2.24E+00 | 1.27E-21 | 4.53E-21 | -1.43E+00 | 9.70E-96 | 2.73E-95 | -2.13E+00 | 5.84E-29 | 2.14E-28 | -1.39E+00 | 5.12E-118 | 1.41E-117 | -2.12E+00 |
| AL592293.2 | 2.31E-34 | 1.20E-33 | -1.76E+00 | 1.36E-25 | 5.88E-25 | -1.35E+00 | 1.05E-04 | 4.45E-04 | -1.16E+00 | 1.51E-06 | 5.78E-06 | -1.16E+00 | 4.47E-10 | 1.59E-09 | -1.46E+00 | 6.45E-09 | 1.82E-08 | -1.28E+00 | 1.82E-05 | 6.66E-05 | -1.24E+00 | 1.43E-04 | 3.92E-04 | -1.20E+00 |
| ZNF462 | 2.70E-34 | 1.41E-33 | -4.19E+00 | 4.33E-21 | 1.87E-20 | -1.98E+00 | 9.85E-31 | 4.18E-30 | -5.54E+00 | 6.03E-23 | 2.30E-22 | -2.69E+00 | 3.82E-08 | 1.36E-07 | -4.41E+00 | 3.87E-05 | 1.09E-04 | -1.80E+00 | 3.03E-24 | 1.11E-23 | -3.59E+00 | 2.52E-15 | 6.92E-15 | -2.22E+00 |
| EEF1A1P13 | 4.29E-34 | 2.24E-33 | -1.68E+00 | 1.68E-03 | 7.24E-03 | 1.04E+00 | 1.05E-08 | 4.45E-08 | -1.51E+00 | 2.56E-01 | 9.79E-01 | 1.10E+00 | 1.33E-13 | 4.74E-13 | -1.48E+00 | 2.80E-01 | 7.89E-01 | 1.15E+00 | 1.13E-11 | 4.15E-11 | -1.48E+00 | 1.45E-01 | 3.97E-01 | 1.16E+00 |
| TP53 | 6.40E-34 | 3.33E-33 | -2.45E+00 | 3.40E-18 | 1.47E-17 | -1.40E+00 | 2.31E-19 | 9.83E-19 | -1.87E+00 | 5.79E-09 | 2.21E-08 | -1.20E+00 | 1.93E-16 | 6.88E-16 | -2.26E+00 | 1.54E-04 | 4.33E-04 | -1.09E+00 | 6.84E-24 | 2.51E-23 | -1.96E+00 | 7.45E-07 | 2.05E-06 | -1.26E+00 |
| GCSAML | 6.84E-34 | 3.57E-33 | -2.60E+00 | 2.29E-21 | 9.91E-21 | -1.60E+00 | 2.16E-16 | 9.18E-16 | -2.80E+00 | 2.60E-09 | 9.93E-09 | -1.67E+00 | 6.05E-17 | 2.16E-16 | -2.14E+00 | 6.27E-08 | 1.77E-07 | -1.31E+00 | 1.44E-08 | 5.29E-08 | -1.31E+00 | 2.03E-08 | 5.57E-08 | -1.58E+00 |
| IFIT3 | 7.01E-34 | 3.65E-33 | -4.87E+00 | 1.83E-22 | 7.91E-22 | -2.53E+00 | 2.17E-37 | 9.20E-37 | -6.14E+00 | 4.30E-32 | 1.64E-31 | -4.43E+00 | 3.59E-27 | 1.28E-26 | -6.42E+00 | 4.22E-10 | 1.19E-09 | -1.95E+00 | 6.29E-84 | 2.31E-83 | -5.34E+00 | 1.37E-55 | 3.75E-55 | -3.48E+00 |
| EIF3LP3 | 7.18E-34 | 3.74E-33 | -2.74E+00 | 1.48E-22 | 6.39E-22 | -1.72E+00 | 5.85E-20 | 2.49E-19 | -2.15E+00 | 4.08E-15 | 1.56E-14 | -1.51E+00 | 3.40E-13 | 1.21E-12 | -2.51E+00 | 1.71E-09 | 4.81E-09 | -1.79E+00 | 4.37E-21 | 1.60E-20 | -2.17E+00 | 2.44E-13 | 6.70E-13 | -1.57E+00 |
| SETP14 | 8.39E-34 | 4.37E-33 | -2.02E+00 | 1.16E-36 | 5.03E-36 | -1.86E+00 | 2.36E-14 | 1.00E-13 | -1.66E+00 | 2.07E-18 | 7.92E-18 | -1.70E+00 | 2.36E-11 | 8.41E-11 | -1.45E+00 | 7.24E-16 | 2.04E-15 | -1.60E+00 | 7.44E-09 | 2.73E-08 | -1.40E+00 | 1.09E-09 | 2.99E-09 | -1.55E+00 |
| TM9SF2 | 8.67E-34 | 4.52E-33 | -2.21E+00 | 3.83E-20 | 1.65E-19 | -1.43E+00 | 1.18E-12 | 5.03E-12 | -1.48E+00 | 6.43E-08 | 2.46E-07 | -1.14E+00 | 3.20E-14 | 1.14E-13 | -1.81E+00 | 6.11E-09 | 1.72E-08 | -1.41E+00 | 3.53E-13 | 1.30E-12 | -1.48E+00 | 3.17E-07 | 8.71E-07 | -1.20E+00 |
| NCOR1 | 9.04E-34 | 4.71E-33 | -1.59E+00 | 5.52E-11 | 2.38E-10 | -1.03E+00 | 8.43E-08 | 3.58E-07 | -1.44E+00 | 1.83E-03 | 6.98E-03 | -1.04E+00 | 3.82E-16 | 1.36E-15 | -1.57E+00 | 4.97E-05 | 1.40E-04 | -1.16E+00 | 1.94E-05 | 7.13E-05 | -1.26E+00 | 9.88E-02 | 2.71E-01 | 1.03E+00 |
| RPS6 | 1.12E-33 | 5.82E-33 | 1.35E+00 | 5.77E-29 | 2.50E-28 | 1.33E+00 | 2.20E-27 | 9.33E-27 | 1.39E+00 | 9.79E-29 | 3.74E-28 | 1.41E+00 | 4.14E-18 | 1.48E-17 | 1.37E+00 | 5.91E-23 | 1.66E-22 | 1.41E+00 | 2.41E-08 | 8.84E-08 | 1.28E+00 | 6.78E-21 | 1.86E-20 | 1.34E+00 |
| TRIM22 | 1.17E-33 | 6.09E-33 | -1.99E+00 | 3.92E-19 | 1.69E-18 | -1.34E+00 | 1.85E-29 | 7.86E-29 | -2.28E+00 | 3.94E-18 | 1.50E-17 | -1.59E+00 | 1.65E-02 | 5.90E-02 | -1.37E+00 | 5.29E-02 | 1.49E-01 | -1.18E+00 | 2.27E-14 | 8.32E-14 | -3.81E+00 | 2.58E-13 | 7.10E-13 | -3.39E+00 |
| PSMB9 | 1.18E-33 | 6.13E-33 | -1.73E+00 | 1.12E-23 | 4.83E-23 | -1.41E+00 | 7.41E-10 | 3.15E-09 | -1.71E+00 | 4.29E-13 | 1.64E-12 | -1.70E+00 | 1.89E-20 | 6.74E-20 | -1.78E+00 | 3.82E-15 | 1.08E-14 | -1.34E+00 | 5.63E-25 | 2.07E-24 | -1.83E+00 | 2.24E-14 | 6.15E-14 | -1.30E+00 |
| RPSA | 1.42E-33 | 7.38E-33 | 1.74E+00 | 1.11E-53 | 4.80E-53 | 2.01E+00 | 3.85E-21 | 1.64E-20 | 1.71E+00 | 3.44E-33 | 1.32E-32 | 1.99E+00 | 5.67E-15 | 2.02E-14 | 1.71E+00 | 1.50E-30 | 4.22E-30 | 2.11E+00 | 4.92E-12 | 1.80E-11 | 1.60E+00 | 7.10E-32 | 1.95E-31 | 1.98E+00 |
| HLF | 1.42E-33 | 7.38E-33 | -2.66E+00 | 3.97E-13 | 1.72E-12 | -1.31E+00 | 1.81E-02 | 7.67E-02 | -1.32E+00 | 4.91E-02 | 1.88E-01 | -1.11E+00 | 2.25E-01 | 8.03E-01 | 1.49E+00 | 3.48E-01 | 9.80E-01 | 1.10E+00 | 2.55E-01 | 9.34E-01 | 1.14E+00 | 3.31E-01 | 9.10E-01 | 1.21E+00 |
| AL358472.1 | 1.75E-33 | 9.10E-33 | -1.84E+00 | 6.24E-46 | 2.69E-45 | -1.91E+00 | 9.02E-14 | 3.83E-13 | -1.69E+00 | 3.75E-25 | 1.43E-24 | -1.91E+00 | 1.60E-19 | 5.70E-19 | -1.95E+00 | 1.09E-20 | 3.07E-20 | -1.89E+00 | 1.63E-25 | 6.00E-25 | -2.11E+00 | 1.90E-22 | 5.21E-22 | -2.09E+00 |
| RPL12P6 | 1.84E-33 | 9.60E-33 | -1.97E+00 | 1.04E-39 | 4.50E-39 | -1.94E+00 | 5.55E-13 | 2.36E-12 | -1.94E+00 | 2.23E-13 | 8.53E-13 | -1.75E+00 | 5.32E-13 | 1.90E-12 | -1.73E+00 | 2.33E-12 | 6.56E-12 | -1.72E+00 | 2.56E-16 | 9.41E-16 | -1.99E+00 | 2.11E-11 | 5.80E-11 | -1.81E+00 |
| LRRC58 | 3.09E-33 | 1.61E-32 | -1.92E+00 | 4.86E-17 | 2.10E-16 | -1.25E+00 | 6.39E-13 | 2.72E-12 | -1.57E+00 | 8.40E-08 | 3.21E-07 | -1.11E+00 | 2.14E-05 | 7.63E-05 | -1.19E+00 | 1.06E-02 | 3.00E-02 | 1.01E+00 | 5.39E-15 | 1.98E-14 | -1.67E+00 | 1.97E-04 | 5.42E-04 | -1.06E+00 |
| HLTF | 4.86E-33 | 2.53E-32 | -2.30E+00 | 1.27E-13 | 5.49E-13 | -1.27E+00 | 3.21E-14 | 1.36E-13 | -1.71E+00 | 1.89E-06 | 7.21E-06 | -1.06E+00 | 3.29E-24 | 1.17E-23 | -2.01E+00 | 6.14E-09 | 1.73E-08 | -1.24E+00 | 1.18E-12 | 4.35E-12 | -1.68E+00 | 4.20E-03 | 1.15E-02 | -1.18E+00 |
| AL049830.1 | 7.82E-33 | 4.08E-32 | -1.90E+00 | 3.16E-24 | 1.37E-23 | -1.42E+00 | 7.12E-16 | 3.03E-15 | -1.85E+00 | 7.81E-15 | 2.98E-14 | -1.56E+00 | 1.36E-21 | 4.85E-21 | -1.84E+00 | 7.56E-14 | 2.13E-13 | -1.39E+00 | 2.08E-22 | 7.65E-22 | -2.09E+00 | 1.81E-09 | 4.97E-09 | -1.47E+00 |
| DUSP3 | 8.04E-33 | 4.19E-32 | -1.88E+00 | 4.93E-27 | 2.13E-26 | -1.45E+00 | 8.90E-33 | 3.78E-32 | -2.80E+00 | 5.99E-16 | 2.29E-15 | -1.44E+00 | 1.41E-02 | 5.04E-02 | -1.18E+00 | 1.30E-02 | 3.65E-02 | -1.14E+00 | 7.75E-03 | 2.85E-02 | -1.05E+00 | 1.20E-01 | 3.31E-01 | 1.09E+00 |
| SP110 | 1.01E-32 | 5.26E-32 | -2.08E+00 | 1.88E-24 | 8.12E-24 | -1.65E+00 | 3.48E-34 | 1.48E-33 | -2.60E+00 | 3.04E-24 | 1.16E-23 | -1.91E+00 | 1.01E-20 | 3.60E-20 | -2.43E+00 | 1.16E-15 | 3.28E-15 | -1.94E+00 | 1.61E-25 | 5.89E-25 | -2.52E+00 | 1.45E-22 | 3.98E-22 | -2.27E+00 |
| CPLX1 | 1.30E-32 | 6.77E-32 | -2.98E+00 | 4.09E-28 | 1.77E-27 | -2.49E+00 | 1.94E-09 | 8.24E-09 | -3.23E+00 | 1.79E-08 | 6.82E-08 | -2.99E+00 | 2.80E-01 | 1.00E+00 | -1.00E+00 | 3.55E-01 | 1.00E+00 | -1.00E+00 | 2.40E-01 | 8.79E-01 | 1.08E+00 | 3.22E-01 | 8.85E-01 | 1.08E+00 |
| ORAI3 | 1.65E-32 | 8.58E-32 | -1.91E+00 | 4.43E-15 | 1.91E-14 | -1.22E+00 | 5.55E-15 | 2.36E-14 | -1.79E+00 | 2.93E-08 | 1.12E-07 | -1.31E+00 | 1.75E-18 | 6.25E-18 | -2.35E+00 | 2.45E-13 | 6.91E-13 | -1.75E+00 | 2.01E-07 | 7.36E-07 | -1.45E+00 | 2.36E-05 | 6.47E-05 | -1.33E+00 |
| AC004057.1 | 2.24E-32 | 1.17E-31 | -1.84E+00 | 1.24E-70 | 5.35E-70 | -2.32E+00 | 1.76E-11 | 7.46E-11 | -1.71E+00 | 5.72E-28 | 2.19E-27 | -2.17E+00 | 5.37E-15 | 1.91E-14 | -1.79E+00 | 8.03E-33 | 2.26E-32 | -2.39E+00 | 4.18E-15 | 1.53E-14 | -1.62E+00 | 3.29E-25 | 9.05E-25 | -2.04E+00 |
| DYNLL1 | 2.26E-32 | 1.18E-31 | -1.75E+00 | 3.11E-04 | 1.34E-03 | -1.06E+00 | 8.06E-09 | 3.42E-08 | -1.60E+00 | 2.85E-02 | 1.09E-01 | -1.12E+00 | 2.77E-31 | 9.87E-31 | -2.25E+00 | 4.79E-05 | 1.35E-04 | -1.23E+00 | 2.09E-07 | 7.67E-07 | -1.42E+00 | 1.44E-01 | 3.97E-01 | -1.07E+00 |
| GRSF1 | 3.08E-32 | 1.60E-31 | -1.97E+00 | 1.79E-16 | 7.74E-16 | -1.29E+00 | 6.05E-10 | 2.57E-09 | -1.58E+00 | 1.69E-06 | 6.45E-06 | -1.18E+00 | 5.84E-14 | 2.08E-13 | -1.70E+00 | 8.37E-05 | 2.36E-04 | -1.05E+00 | 3.66E-09 | 1.34E-08 | -1.39E+00 | 5.60E-03 | 1.54E-02 | -1.04E+00 |
| SAMD9 | 3.76E-32 | 1.96E-31 | -2.13E+00 | 1.09E-19 | 4.70E-19 | -1.47E+00 | 2.35E-22 | 9.96E-22 | -2.38E+00 | 2.53E-14 | 9.65E-14 | -1.64E+00 | 3.87E-19 | 1.38E-18 | -2.17E+00 | 2.35E-10 | 6.63E-10 | -1.55E+00 | 1.97E-16 | 7.22E-16 | -1.94E+00 | 1.21E-11 | 3.32E-11 | -1.74E+00 |
| AC099340.1 | 4.26E-32 | 2.22E-31 | 3.02E+00 | 6.12E-27 | 2.64E-26 | 3.28E+00 | 8.73E-27 | 3.71E-26 | 3.07E+00 | 3.04E-24 | 1.16E-23 | 3.51E+00 | 5.48E-13 | 1.95E-12 | 2.96E+00 | 5.42E-26 | 1.53E-25 | 3.71E+00 | 2.73E-12 | 1.00E-11 | 2.49E+00 | 5.87E-24 | 1.61E-23 | 3.08E+00 |
| IRF2 | 4.74E-32 | 2.47E-31 | -2.16E+00 | 9.57E-16 | 4.14E-15 | -1.23E+00 | 2.57E-15 | 1.09E-14 | -2.05E+00 | 3.58E-06 | 1.37E-05 | -1.17E+00 | 1.87E-12 | 6.68E-12 | -2.49E+00 | 3.15E-06 | 8.88E-06 | -1.39E+00 | 5.75E-16 | 2.11E-15 | -1.99E+00 | 1.87E-06 | 5.14E-06 | -1.35E+00 |
| AC023232.1 | 6.08E-32 | 3.17E-31 | -1.95E+00 | 7.41E-25 | 3.20E-24 | -1.49E+00 | 2.76E-13 | 1.17E-12 | -1.71E+00 | 1.88E-13 | 7.17E-13 | -1.52E+00 | 1.29E-22 | 4.60E-22 | -2.05E+00 | 7.55E-21 | 2.13E-20 | -1.75E+00 | 6.73E-20 | 2.47E-19 | -1.99E+00 | 1.57E-14 | 4.30E-14 | -1.58E+00 |
| THRAP3 | 7.19E-32 | 3.75E-31 | -1.63E+00 | 1.77E-03 | 7.65E-03 | 1.06E+00 | 3.04E-05 | 1.29E-04 | -1.24E+00 | 1.24E-01 | 4.74E-01 | 1.11E+00 | 4.24E-11 | 1.51E-10 | -1.53E+00 | 5.34E-02 | 1.51E-01 | -1.01E+00 | 1.08E-04 | 3.98E-04 | -1.28E+00 | 1.86E-01 | 5.10E-01 | -1.01E+00 |
| PABPC3 | 9.92E-32 | 5.17E-31 | -1.91E+00 | 9.49E-29 | 4.10E-28 | -1.74E+00 | 1.61E-25 | 6.85E-25 | -2.48E+00 | 3.54E-17 | 1.35E-16 | -1.87E+00 | 1.54E-14 | 5.50E-14 | -1.89E+00 | 2.81E-20 | 7.92E-20 | -2.05E+00 | 2.96E-28 | 1.09E-27 | -2.56E+00 | 1.52E-16 | 4.17E-16 | -1.97E+00 |
| XBP1 | 1.16E-31 | 6.07E-31 | -1.84E+00 | 7.01E-07 | 3.03E-06 | -1.18E+00 | 5.85E-08 | 2.49E-07 | -1.76E+00 | 2.98E-02 | 1.14E-01 | -1.17E+00 | 3.94E-11 | 1.41E-10 | -1.39E+00 | 2.06E-06 | 5.81E-06 | -1.17E+00 | 9.92E-06 | 3.64E-05 | -1.21E+00 | 3.37E-01 | 9.26E-01 | 1.20E+00 |
| MTCO1P12 | 1.23E-31 | 6.43E-31 | -1.40E+00 | 4.37E-106 | 1.89E-105 | -1.92E+00 | 6.94E-26 | 2.95E-25 | -1.46E+00 | 1.16E-79 | 4.42E-79 | -1.97E+00 | 1.38E-25 | 4.92E-25 | -1.55E+00 | 3.97E-55 | 1.12E-54 | -1.89E+00 | 3.97E-36 | 1.46E-35 | -1.51E+00 | 4.18E-62 | 1.15E-61 | -1.85E+00 |
| EEF1A1P12 | 1.24E-31 | 6.44E-31 | -1.64E+00 | 3.11E-06 | 1.34E-05 | -1.05E+00 | 3.87E-10 | 1.64E-09 | -1.56E+00 | 1.58E-01 | 6.04E-01 | 1.04E+00 | 3.38E-14 | 1.20E-13 | -1.49E+00 | 9.95E-02 | 2.80E-01 | 1.03E+00 | 5.07E-13 | 1.86E-12 | -1.55E+00 | 1.45E-01 | 3.98E-01 | 1.03E+00 |
| RPS2P5 | 1.60E-31 | 8.34E-31 | -1.62E+00 | 3.31E-02 | 1.43E-01 | 1.13E+00 | 6.73E-13 | 2.86E-12 | -1.55E+00 | 8.77E-02 | 3.35E-01 | 1.12E+00 | 6.95E-20 | 2.48E-19 | -1.71E+00 | 5.96E-02 | 1.68E-01 | 1.13E+00 | 9.51E-21 | 3.49E-20 | -1.54E+00 | 3.42E-01 | 9.40E-01 | 1.11E+00 |
| AC007969.1 | 1.78E-31 | 9.28E-31 | 1.31E+00 | 3.84E-07 | 1.66E-06 | 1.13E+00 | 1.85E-24 | 7.87E-24 | 1.34E+00 | 3.08E-06 | 1.18E-05 | 1.15E+00 | 3.06E-12 | 1.09E-11 | 1.26E+00 | 2.92E-04 | 8.22E-04 | 1.13E+00 | 1.61E-14 | 5.90E-14 | 1.30E+00 | 1.62E-07 | 4.44E-07 | 1.16E+00 |
| ARPC3P2 | 1.82E-31 | 9.48E-31 | -1.95E+00 | 2.30E-28 | 9.93E-28 | -1.60E+00 | 1.04E-20 | 4.43E-20 | -1.92E+00 | 3.83E-21 | 1.46E-20 | -1.74E+00 | 9.91E-23 | 3.53E-22 | -2.18E+00 | 1.72E-18 | 4.85E-18 | -1.86E+00 | 1.55E-13 | 5.70E-13 | -1.64E+00 | 1.06E-09 | 2.90E-09 | -1.39E+00 |
| HNRNPA1 | 2.40E-31 | 1.25E-30 | -1.71E+00 | 1.93E-23 | 8.33E-23 | -1.43E+00 | 4.34E-11 | 1.84E-10 | -1.70E+00 | 9.06E-08 | 3.46E-07 | -1.51E+00 | 9.48E-14 | 3.38E-13 | -1.64E+00 | 1.08E-05 | 3.06E-05 | -1.36E+00 | 1.12E-13 | [illegible] | -1.73E+00 | 2.05E-07 | 5.62E-07 | -1.53E+00 |

| | | | | | | | | | | | | | | | | | | | | | | | | |
|---|---|---|---|---|---|---|---|---|---|---|---|---|---|---|---|---|---|---|---|---|---|---|---|---|
| SDHC | 2.52E-24 | 1.31E-23 | -1.77E+00 | 2.00E-16 | 8.66E-16 | -1.32E+00 | 6.66E-16 | 2.83E-15 | -1.74E+00 | 1.54E-14 | 5.87E-14 | -1.45E+00 | 2.05E-10 | 7.30E-10 | -1.47E+00 | 4.63E-07 | 1.31E-06 | -1.23E+00 | 6.16E-15 | 2.26E-14 | -1.61E+00 | 1.10E-11 | 3.03E-11 | -1.40E+00 |
| AC114760.2 | 2.74E-24 | 1.43E-23 | 3.58E+00 | 6.73E-04 | 2.91E-03 | 2.26E+00 | 1.82E-13 | 7.74E-13 | 2.85E+00 | 1.46E-01 | 5.57E-01 | 1.47E+00 | 1.27E-14 | 4.52E-14 | 4.99E+00 | 3.34E-01 | 9.42E-01 | 1.47E+00 | 1.96E-03 | 7.18E-03 | 1.85E+00 | 1.05E-03 | 2.88E-03 | -1.68E+00 |
| WDR83OS | 2.91E-24 | 1.52E-23 | -1.41E+00 | 5.58E-20 | 2.41E-19 | -1.22E+00 | 4.67E-06 | 1.98E-05 | -1.30E+00 | 1.50E-05 | 5.73E-05 | -1.19E+00 | 1.76E-06 | 6.27E-06 | -1.27E+00 | 3.41E-04 | 9.63E-04 | -1.15E+00 | 5.25E-03 | 1.93E-02 | -1.11E+00 | 4.49E-02 | 1.23E-01 | -1.07E+00 |
| RPL5P17 | 3.00E-24 | 1.56E-23 | 2.37E+00 | 4.75E-08 | 2.05E-07 | 1.94E+00 | 1.63E-11 | 6.93E-11 | 2.36E+00 | 1.16E-04 | 4.41E-04 | 1.91E+00 | 2.48E-07 | 8.84E-07 | 2.26E+00 | 4.56E-03 | 1.29E-02 | 1.86E+00 | 2.69E-07 | 9.86E-07 | 2.07E+00 | 6.47E-03 | 1.78E-02 | 1.62E+00 |
| TAGLN2 | 3.30E-24 | 1.72E-23 | -1.67E+00 | 9.61E-14 | 4.15E-13 | -1.31E+00 | 1.46E-08 | 6.20E-08 | -1.60E+00 | 2.17E-04 | 8.29E-04 | -1.28E+00 | 8.23E-13 | 2.94E-12 | -1.67E+00 | 1.06E-07 | 2.98E-07 | -1.47E+00 | 1.08E-05 | 3.96E-05 | -1.37E+00 | 6.54E-04 | 1.80E-03 | -1.34E+00 |
| HSD17B4 | 3.50E-24 | 1.82E-23 | -1.70E+00 | 5.46E-10 | 2.36E-09 | -1.09E+00 | 1.73E-08 | 7.36E-08 | -1.45E+00 | 1.10E-04 | 4.19E-04 | -1.03E+00 | 2.26E-13 | 8.07E-13 | -1.68E+00 | 5.89E-04 | 1.66E-03 | -1.01E+00 | 5.38E-04 | 1.98E-03 | -1.08E+00 | 1.61E-01 | 4.42E-01 | 1.19E+00 |
| MDFI | 3.68E-24 | 1.92E-23 | -4.17E+00 | 6.34E-12 | 2.74E-11 | -1.71E+00 | 1.41E-28 | 5.98E-28 | -2.90E+00 | 1.29E-24 | 4.94E-24 | -2.09E+00 | 4.42E-04 | 1.58E-03 | -2.39E+00 | 1.06E-02 | 2.98E-02 | -1.44E+00 | 4.19E-04 | 1.54E-03 | -2.99E+00 | 5.70E-02 | 1.57E-01 | -1.32E+00 |
| LINC01829 | 3.96E-24 | 2.06E-23 | -3.19E+00 | 2.39E-17 | 1.03E-16 | -1.87E+00 | 2.10E-01 | 8.93E-01 | -1.00E+00 | 2.09E-01 | 7.97E-01 | -1.02E+00 | 1.38E-01 | 4.91E-01 | -1.33E+00 | 2.93E-01 | 8.25E-01 | 1.11E+00 | 2.30E-01 | 8.45E-01 | -1.11E+00 | 3.53E-01 | 9.70E-01 | -1.04E+00 |
| ISG15 | 3.96E-24 | 2.06E-23 | -2.18E+00 | 8.13E-28 | 3.51E-27 | -2.13E+00 | 1.45E-21 | 6.15E-21 | -2.48E+00 | 8.68E-25 | 3.31E-24 | -2.36E+00 | 4.03E-24 | 1.44E-23 | -4.19E+00 | 2.06E-20 | 5.81E-20 | -3.10E+00 | 1.22E-32 | 4.48E-32 | -3.91E+00 | 9.64E-28 | 2.65E-27 | -3.25E+00 |
| ZC3H13 | 4.01E-24 | 2.09E-23 | -1.69E+00 | 2.77E-06 | 1.20E-05 | -1.01E+00 | 1.59E-14 | 6.76E-14 | -1.77E+00 | 2.65E-06 | 1.01E-05 | -1.15E+00 | 6.23E-22 | 2.22E-21 | -1.96E+00 | 1.01E-08 | 2.84E-08 | -1.34E+00 | 1.18E-10 | 4.33E-10 | -1.60E+00 | 5.07E-03 | 1.39E-02 | -1.15E+00 |
| EFCAB14 | 4.03E-24 | 2.10E-23 | -1.91E+00 | 6.31E-11 | 2.73E-10 | -1.16E+00 | 2.34E-11 | 9.94E-11 | -1.80E+00 | 4.16E-05 | 1.59E-04 | -1.08E+00 | 6.09E-09 | 2.17E-08 | -1.69E+00 | 1.38E-06 | 3.88E-06 | -1.39E+00 | 6.21E-10 | 2.28E-09 | -1.45E+00 | 6.22E-05 | 1.71E-04 | -1.27E+00 |
| ADAR | 4.03E-24 | 2.10E-23 | -1.61E+00 | 4.30E-15 | 1.86E-14 | -1.28E+00 | 1.60E-09 | 6.80E-09 | -1.48E+00 | 1.46E-10 | 5.56E-10 | -1.39E+00 | 1.46E-14 | 5.19E-14 | -1.77E+00 | 1.02E-10 | 2.87E-10 | -1.55E+00 | 1.16E-13 | 4.27E-13 | -1.66E+00 | 2.65E-10 | 7.29E-10 | -1.62E+00 |
| POLR2J3 | 4.07E-24 | 2.12E-23 | -1.67E+00 | 5.91E-13 | 2.55E-12 | -1.19E+00 | 3.56E-09 | 1.51E-08 | -1.57E+00 | 5.39E-07 | 2.06E-06 | -1.27E+00 | 2.96E-09 | 1.06E-08 | -1.43E+00 | 3.91E-05 | 1.10E-04 | -1.12E+00 | 1.62E-12 | 5.93E-12 | -1.57E+00 | 1.10E-05 | 3.02E-05 | -1.21E+00 |
| ST13P19 | 4.55E-24 | 2.37E-23 | -1.92E+00 | 1.06E-20 | 4.57E-20 | -1.51E+00 | 1.63E-05 | 6.92E-05 | -1.39E+00 | 8.07E-06 | 3.08E-05 | -1.32E+00 | 4.27E-05 | 1.52E-04 | -1.19E+00 | 1.20E-03 | 3.39E-03 | 1.01E+00 | 2.28E-05 | 8.36E-05 | -1.25E+00 | 3.32E-04 | 9.12E-04 | -1.23E+00 |
| AC074033.1 | 4.59E-24 | 2.39E-23 | 1.89E+00 | 1.07E-03 | 4.64E-03 | 1.48E+00 | 2.57E-17 | 1.09E-16 | 1.87E+00 | 1.61E-05 | 6.15E-05 | 1.48E+00 | 1.60E-07 | 5.70E-07 | 1.78E+00 | 1.72E-02 | 4.85E-02 | 1.41E+00 | 2.88E-09 | 1.06E-08 | 1.71E+00 | 1.44E-04 | 3.95E-04 | 1.43E+00 |
| PGAM1 | 4.83E-24 | 2.51E-23 | -3.17E+00 | 1.48E-15 | 6.40E-15 | -1.68E+00 | 6.40E-21 | 2.72E-20 | -2.25E+00 | 4.61E-14 | 1.76E-13 | -1.50E+00 | 7.01E-12 | 2.50E-11 | -2.33E+00 | 6.71E-10 | 1.89E-09 | -1.87E+00 | 1.67E-28 | 6.13E-28 | -2.31E+00 | 4.34E-08 | 1.19E-07 | -1.23E+00 |
| NIP7 | 5.21E-24 | 2.72E-23 | -2.14E+00 | 8.93E-21 | 3.86E-20 | -1.75E+00 | 1.74E-10 | 7.41E-10 | -1.55E+00 | 1.42E-07 | 5.43E-07 | -1.17E+00 | 3.62E-09 | 1.29E-08 | -1.76E+00 | 1.32E-07 | 3.73E-07 | -1.55E+00 | 1.19E-09 | 4.36E-09 | -1.39E+00 | 4.94E-03 | 1.36E-02 | 1.03E+00 |
| HDAC1 | 5.35E-24 | 2.79E-23 | -1.60E+00 | 2.56E-13 | 1.11E-12 | -1.14E+00 | 1.15E-09 | 4.87E-09 | -1.47E+00 | 1.44E-06 | 5.49E-06 | -1.18E+00 | 1.34E-08 | 4.79E-08 | -1.28E+00 | 9.39E-04 | 2.65E-03 | 1.01E+00 | 1.91E-14 | 7.01E-14 | -1.59E+00 | 3.86E-03 | 1.06E-02 | -1.05E+00 |
| PAK2 | 5.40E-24 | 2.81E-23 | -1.56E+00 | 7.60E-13 | 3.29E-12 | -1.16E+00 | 8.38E-06 | 3.56E-05 | -1.34E+00 | 1.76E-02 | 6.74E-02 | -1.06E+00 | 2.04E-10 | 7.27E-10 | -1.32E+00 | 7.18E-08 | 2.02E-07 | -1.16E+00 | 4.43E-07 | 1.63E-06 | -1.29E+00 | 5.87E-04 | 1.61E-03 | -1.08E+00 |
| COX14 | 5.58E-24 | 2.91E-23 | -1.54E+00 | 4.12E-25 | 1.78E-24 | -1.42E+00 | 2.18E-03 | 9.26E-03 | -1.10E+00 | 1.90E-08 | 7.25E-08 | -1.24E+00 | 4.77E-08 | 1.70E-07 | -1.25E+00 | 1.09E-07 | 3.08E-07 | -1.20E+00 | 6.74E-03 | 2.47E-02 | -1.06E+00 | 3.14E-03 | 8.63E-03 | -1.11E+00 |
| QSER1 | 5.62E-24 | 2.93E-23 | -2.45E+00 | 1.36E-12 | 5.86E-12 | -1.39E+00 | 1.25E-10 | 5.30E-10 | -1.67E+00 | 3.67E-05 | 1.40E-04 | -1.12E+00 | 2.08E-10 | 7.42E-10 | -2.28E+00 | 1.38E-05 | 3.89E-05 | -1.42E+00 | 5.23E-14 | 1.92E-13 | -1.69E+00 | 7.19E-05 | 1.97E-04 | -1.05E+00 |
| RPL27 | 5.69E-24 | 2.97E-23 | 1.29E+00 | 2.00E-01 | 8.66E-01 | 1.02E+00 | 1.05E-11 | 4.46E-11 | 1.24E+00 | 2.59E-01 | 9.88E-01 | 1.02E+00 | 8.45E-10 | 3.01E-09 | 1.25E+00 | 3.95E-02 | 1.11E-01 | 1.07E+00 | 4.84E-05 | 1.78E-04 | 1.20E+00 | 3.40E-01 | 9.33E-01 | 1.02E+00 |
| GMPR | 6.37E-24 | 3.32E-23 | -3.18E+00 | 1.87E-14 | 8.07E-14 | -1.68E+00 | 7.97E-20 | 3.39E-19 | -2.58E+00 | 3.42E-11 | 1.31E-10 | -1.43E+00 | 2.48E-32 | 8.86E-32 | -2.33E+00 | 1.62E-12 | 4.57E-12 | -1.39E+00 | 3.43E-13 | 1.26E-12 | -1.30E+00 | 4.78E-06 | 1.31E-05 | -1.19E+00 |
| MIA3 | 6.49E-24 | 3.38E-23 | -1.76E+00 | 5.87E-12 | 2.54E-11 | -1.22E+00 | 6.80E-10 | 2.89E-09 | -1.45E+00 | 3.12E-07 | 1.19E-06 | -1.15E+00 | 1.28E-06 | 4.58E-06 | -1.30E+00 | 1.77E-03 | 4.98E-03 | -1.01E+00 | 1.88E-13 | 6.92E-13 | -1.57E+00 | 2.87E-06 | 7.88E-06 | -1.26E+00 |
| RPL10AP6 | 6.58E-24 | 3.43E-23 | -1.40E+00 | 8.55E-53 | 3.69E-52 | -1.66E+00 | 3.56E-11 | 1.51E-10 | -1.31E+00 | 6.69E-28 | 2.56E-27 | -1.58E+00 | 2.74E-10 | 9.78E-10 | -1.33E+00 | 2.73E-23 | 7.71E-23 | -1.57E+00 | 2.17E-21 | 7.95E-21 | -1.35E+00 | 3.98E-31 | 1.09E-30 | -1.61E+00 |
| PER1 | 6.85E-24 | 3.57E-23 | 3.31E+00 | 3.23E-05 | 1.40E-04 | 2.22E+00 | 1.07E-17 | 4.56E-17 | 3.24E+00 | 1.88E-07 | 7.18E-07 | 2.67E+00 | 3.43E-31 | 1.22E-30 | 6.19E+00 | 1.15E-23 | 3.23E-23 | 4.83E+00 | 1.43E-15 | 5.25E-15 | 2.95E+00 | 2.04E-09 | 5.59E-09 | 2.46E+00 |
| NDFIP1 | 7.01E-24 | 3.66E-23 | -1.61E+00 | 1.10E-18 | 4.77E-18 | -1.29E+00 | 2.24E-12 | 9.52E-12 | -1.59E+00 | 3.67E-12 | 1.40E-11 | -1.44E+00 | 3.43E-12 | 1.22E-11 | -1.46E+00 | 4.73E-10 | 1.33E-09 | -1.21E+00 | 2.12E-13 | 7.79E-13 | -1.75E+00 | 9.87E-12 | 2.71E-11 | -1.64E+00 |
| AC068522.1 | 9.18E-24 | 4.78E-23 | -1.64E+00 | 1.25E-43 | 5.41E-43 | -1.80E+00 | 1.43E-06 | 6.09E-06 | -1.49E+00 | 1.30E-15 | 4.95E-15 | -1.75E+00 | 1.29E-14 | 4.61E-14 | -1.63E+00 | 2.19E-20 | 6.16E-20 | -1.91E+00 | 7.10E-08 | 2.61E-07 | -1.45E+00 | 3.83E-09 | 1.05E-08 | -1.55E+00 |
| ANXA7 | 9.47E-24 | 4.93E-23 | -1.46E+00 | 1.39E-12 | 5.99E-12 | -1.09E+00 | 5.20E-06 | 2.21E-05 | -1.20E+00 | 2.07E-03 | 7.90E-03 | 1.06E+00 | 1.17E-07 | 4.17E-07 | -1.31E+00 | 8.29E-03 | 2.34E-02 | 1.03E+00 | 9.72E-04 | 3.57E-03 | -1.11E+00 | 7.56E-02 | 2.08E-01 | 1.03E+00 |
| ZMYM4 | 1.01E-23 | 5.24E-23 | -1.86E+00 | 4.37E-14 | 1.89E-13 | -1.26E+00 | 1.19E-15 | 5.06E-15 | -1.77E+00 | 1.05E-05 | 4.02E-05 | -1.08E+00 | 8.34E-05 | 2.97E-04 | -1.30E+00 | 1.14E-01 | 3.21E-01 | 1.19E+00 | 1.26E-10 | 4.62E-10 | -1.53E+00 | 5.98E-05 | 1.64E-04 | -1.16E+00 |
| SLC39A6 | 1.04E-23 | 5.40E-23 | -2.61E+00 | 2.75E-16 | 1.19E-15 | -1.60E+00 | 1.50E-19 | 6.38E-19 | -2.20E+00 | 6.44E-12 | 2.46E-11 | -1.42E+00 | 3.70E-07 | 1.32E-06 | -2.09E+00 | 1.97E-04 | 5.55E-04 | -1.34E+00 | 5.45E-10 | 2.00E-09 | -2.47E+00 | 4.15E-06 | 1.14E-05 | -1.67E+00 |
| MTIF3 | 1.06E-23 | 5.52E-23 | -2.02E+00 | 1.20E-13 | 5.18E-13 | -1.28E+00 | 1.35E-22 | 5.72E-22 | -2.10E+00 | 2.03E-17 | 7.74E-17 | -1.61E+00 | 8.23E-13 | 2.94E-12 | -1.91E+00 | 1.37E-09 | 3.87E-09 | -1.57E+00 | 4.03E-18 | 1.48E-17 | -1.86E+00 | 1.05E-09 | 2.89E-09 | -1.43E+00 |
| VDAC2 | 1.10E-23 | 5.75E-23 | -1.52E+00 | 9.78E-25 | 4.22E-24 | -1.42E+00 | 4.04E-06 | 1.72E-05 | -1.38E+00 | 2.58E-09 | 9.84E-09 | -1.39E+00 | 1.42E-09 | 5.05E-09 | -1.46E+00 | 2.26E-09 | 6.36E-09 | -1.37E+00 | 3.83E-08 | 1.40E-07 | -1.36E+00 | 5.64E-08 | 1.55E-07 | -1.36E+00 |
| RALA | 1.21E-23 | 6.31E-23 | -1.77E+00 | 1.12E-06 | 4.84E-06 | -1.03E+00 | 8.33E-08 | 3.54E-07 | -1.41E+00 | 6.76E-04 | 2.58E-03 | 1.01E+00 | 3.04E-16 | 1.08E-15 | -1.88E+00 | 4.45E-10 | 1.25E-09 | -1.34E+00 | 1.79E-05 | 6.58E-05 | -1.24E+00 | 2.67E-01 | 7.35E-01 | 1.22E+00 |
| TMSB10 | 1.21E-23 | 6.31E-23 | 1.46E+00 | 8.91E-02 | 3.85E-01 | 1.05E+00 | 3.98E-08 | 1.69E-07 | 1.26E+00 | 1.59E-01 | 6.07E-01 | -1.05E+00 | 3.51E-05 | 1.25E-04 | 1.24E+00 | 5.62E-02 | 1.59E-01 | -1.08E+00 | 8.56E-04 | 3.14E-03 | 1.26E+00 | 6.57E-02 | 1.81E-01 | -1.06E+00 |
| AC092017.1 | 1.21E-23 | 6.31E-23 | -1.82E+00 | 7.85E-28 | 3.39E-27 | -1.74E+00 | 2.55E-11 | 1.08E-10 | -1.43E+00 | 5.30E-19 | 2.03E-18 | -1.63E+00 | 3.06E-11 | 1.09E-10 | -1.74E+00 | 3.02E-08 | 8.51E-08 | -1.36E+00 | 2.40E-08 | 8.82E-08 | -1.31E+00 | 1.09E-06 | 3.00E-06 | -1.26E+00 |
| RPL12P35 | 1.27E-23 | 6.60E-23 | -1.44E+00 | 9.30E-55 | 4.02E-54 | -1.74E+00 | 2.36E-14 | 1.00E-13 | -1.40E+00 | 1.01E-28 | 3.84E-28 | -1.62E+00 | 1.28E-07 | 4.55E-07 | -1.34E+00 | 2.93E-20 | 8.25E-20 | -1.61E+00 | 3.18E-28 | 1.17E-27 | -1.45E+00 | 1.49E-34 | 4.08E-34 | -1.69E+00 |
| CST3 | 1.27E-23 | 6.60E-23 | -1.94E+00 | 5.02E-10 | 2.17E-09 | -1.31E+00 | 6.22E-02 | 2.64E-01 | -1.04E+00 | 1.95E-02 | 7.46E-02 | -1.05E+00 | 1.96E-02 | 7.00E-02 | 1.13E+00 | 1.99E-01 | 5.61E-01 | 1.25E+00 | 5.27E-02 | 1.93E-01 | -1.02E+00 | 1.81E-02 | 4.97E-02 | -1.20E+00 |
| ARPC1A | 1.27E-23 | 6.60E-23 | -1.86E+00 | 2.96E-17 | 1.28E-16 | -1.35E+00 | 1.94E-06 | 8.25E-06 | -1.33E+00 | 5.11E-04 | 1.95E-03 | -1.02E+00 | 2.28E-15 | 8.11E-15 | -1.81E+00 | 4.35E-10 | 1.23E-09 | -1.30E+00 | 3.06E-03 | 1.12E-02 | -1.09E+00 | 1.69E-01 | 4.65E-01 | 1.17E+00 |
| ERGIC2 | 1.40E-23 | 7.28E-23 | -1.84E+00 | 2.96E-20 | 1.28E-19 | -1.48E+00 | 2.23E-17 | 9.50E-17 | -1.84E+00 | 5.05E-14 | 1.93E-13 | -1.50E+00 | 5.62E-14 | 2.00E-13 | -1.89E+00 | 1.48E-10 | 4.17E-10 | -1.50E+00 | 1.19E-16 | 4.37E-16 | -1.61E+00 | 4.29E-09 | 1.18E-08 | -1.29E+00 |
| AC008813.1 | 1.41E-23 | 7.37E-23 | -1.46E+00 | 5.49E-36 | 2.37E-35 | -1.70E+00 | 1.03E-06 | 4.38E-06 | -1.31E+00 | 1.94E-20 | 7.42E-20 | -1.72E+00 | 4.99E-08 | 1.78E-07 | -1.23E+00 | 1.38E-13 | 3.89E-13 | -1.50E+00 | 1.26E-08 | 4.63E-08 | -1.34E+00 | 1.10E-11 | 3.03E-11 | -1.52E+00 |
| HNRNPH1 | 1.42E-23 | 7.42E-23 | -1.74E+00 | 1.67E-03 | 7.23E-03 | -1.08E+00 | 4.74E-11 | 2.01E-10 | -1.79E+00 | 1.54E-03 | 5.87E-03 | -1.17E+00 | 4.01E-18 | 1.43E-17 | -1.77E+00 | 2.81E-03 | 7.93E-03 | -1.11E+00 | 2.01E-20 | 7.37E-20 | -1.90E+00 | 9.75E-03 | 2.68E-02 | -1.10E+00 |
| PPAT | 1.49E-23 | 7.75E-23 | -2.42E+00 | 3.54E-11 | 1.53E-10 | -1.23E+00 | 2.35E-14 | 1.00E-13 | -1.79E+00 | 1.48E-06 | 5.65E-06 | -1.10E+00 | 9.87E-06 | 3.52E-05 | -1.82E+00 | 1.63E-02 | 4.61E-02 | -1.02E+00 | 1.35E-14 | 4.94E-14 | -1.97E+00 | 6.50E-03 | 1.79E-02 | -1.12E+00 |
| UROD | 1.50E-23 | 7.82E-23 | -1.53E+00 | 1.30E-11 | 5.63E-11 | -1.10E+00 | 3.76E-03 | 1.60E-02 | -1.11E+00 | 2.98E-02 | 1.14E-01 | 1.14E+00 | 5.29E-08 | 1.89E-07 | -1.32E+00 | 3.17E-01 | 8.93E-01 | 1.36E+00 | 4.17E-03 | 1.53E-02 | -1.26E+00 | 5.85E-02 | 1.61E-01 | -1.15E+00 |
| PRKDC | 1.76E-23 | 9.16E-23 | -1.60E+00 | 5.04E-07 | 2.18E-06 | 1.01E+00 | 1.03E-05 | 4.36E-05 | -1.23E+00 | 1.32E-01 | 5.04E-01 | 1.18E+00 | 1.17E-09 | 4.16E-09 | -1.31E+00 | 7.80E-03 | 2.20E-02 | 1.06E+00 | 7.60E-03 | 2.79E-02 | -1.12E+00 | 2.42E-01 | 6.65E-01 | 1.14E+00 |
| TAP2 | 1.79E-23 | 9.30E-23 | -2.84E+00 | 4.08E-14 | 1.76E-13 | -1.60E+00 | 3.82E-13 | 1.62E-12 | -1.72E+00 | 9.14E-12 | 3.49E-11 | -1.42E+00 | 1.92E-06 | 6.85E-06 | -1.83E+00 | 4.34E-04 | 1.22E-03 | -1.31E+00 | 4.25E-11 | 1.56E-10 | -1.86E+00 | 5.40E-04 | 1.48E-03 | -1.19E+00 |
| OIP5-AS1 | 1.85E-23 | 9.62E-23 | -1.61E+00 | 3.57E-07 | 1.54E-06 | 1.01E+00 | 7.40E-08 | 3.15E-07 | -1.37E+00 | 3.26E-02 | 1.25E-01 | 1.15E+00 | 1.22E-11 | 4.33E-11 | -1.48E+00 | 2.69E-03 | 7.58E-03 | 1.03E+00 | 9.36E-06 | 3.44E-05 | -1.35E+00 | 3.06E-01 | 8.40E-01 | 1.08E+00 |
| TBL1XR1 | 1.89E-23 | 9.86E-23 | -1.62E+00 | 3.69E-11 | 1.59E-10 | -1.14E+00 | 2.25E-06 | 9.54E-06 | -1.36E+00 | 3.15E-06 | 1.20E-05 | -1.20E+00 | 6.70E-05 | 2.39E-04 | -1.19E+00 | 2.43E-03 | 6.84E-03 | 1.02E+00 | 9.82E-10 | 3.60E-09 | -1.41E+00 | 2.15E-05 | 5.90E-05 | -1.21E+00 |
| AP006259.1 | 2.20E-23 | 1.15E-22 | -1.88E+00 | 1.52E-33 | 6.58E-33 | -2.46E+00 | 8.62E-12 | 3.66E-11 | -1.89E+00 | 2.05E-16 | 7.82E-16 | -2.25E+00 | 1.45E-06 | 5.16E-06 | -1.70E+00 | 3.27E-09 | 9.22E-09 | -2.20E+00 | 6.10E-13 | 2.24E-12 | -2.49E+00 | 2.20E-18 | 6.04E-18 | -4.06E+00 |
| RPL35 | 2.40E-23 | 1.25E-22 | 1.38E+00 | 6.40E-02 | 2.77E-01 | 1.07E+00 | 2.19E-22 | 9.32E-22 | 1.46E+00 | 2.04E-03 | 7.78E-03 | 1.14E+00 | 6.47E-12 | 2.31E-11 | 1.37E+00 | 4.55E-03 | 1.28E-02 | 1.13E+00 | 2.29E-13 | 8.41E-13 | 1.43E+00 | 5.30E-04 | 1.46E-03 | 1.20E+00 |
| TMBIM4 | 2.47E-23 | 1.29E-22 | -1.60E+00 | 4.62E-11 | 2.00E-10 | -1.14E+00 | 3.84E-11 | 1.63E-10 | -1.46E+00 | 5.31E-06 | 2.03E-05 | -1.08E+00 | 8.16E-15 | 2.91E-14 | -1.56E+00 | 1.96E-04 | 5.51E-04 | -1.06E+00 | 6.14E-11 | 2.25E-10 | -1.38E+00 | 4.72E-06 | 1.30E-05 | -1.16E+00 |
| HNRNPH2 | 2.62E-23 | 1.36E-22 | -1.77E+00 | 2.78E-11 | 1.20E-10 | -1.11E+00 | 9.49E-07 | 4.03E-06 | -1.38E+00 | 2.68E-04 | 1.02E-03 | -1.03E+00 | 1.15E-07 | 4.11E-07 | -1.30E+00 | 3.62E-02 | 1.02E-01 | 1.15E+00 | 3.68E-08 | 1.35E-07 | -1.28E+00 | 1.60E-01 | 4.38E-01 | 1.11E+00 |
| HNRNPD | 2.85E-23 | 1.49E-22 | -1.46E+00 | 5.43E-14 | 2.35E-13 | -1.16E+00 | 1.37E-03 | 5.83E-03 | -1.21E+00 | 4.81E-03 | 1.84E-02 | -1.14E+00 | 3.58E-10 | 1.28E-09 | -1.32E+00 | 6.69E-04 | 1.88E-03 | -1.06E+00 | 2.04E-05 | 7.49E-05 | -1.32E+00 | 1.90E-01 | 5.23E-01 | -1.03E+00 |
| EIF2AK1 | 2.91E-23 | 1.52E-22 | -1.63E+00 | 1.50E-12 | 6.50E-12 | -1.15E+00 | 1.13E-08 | 4.78E-08 | -1.33E+00 | 1.16E-06 | 4.42E-06 | -1.09E+00 | 9.88E-05 | 3.52E-04 | -1.07E+00 | 1.86E-01 | 5.24E-01 | 1.16E+00 | 2.25E-01 | 8.27E-01 | 1.19E+00 | 3.51E-01 | 9.63E-01 | 1.15E+00 |
| FKBP4 | 2.92E-23 | 1.52E-22 | -1.68E+00 | 1.51E-08 | 6.53E-08 | 1.02E+00 | 2.52E-06 | 1.07E-05 | -1.37E+00 | 1.47E-01 | 5.62E-01 | 1.16E+00 | 1.21E-20 | 4.31E-20 | -1.88E+00 | 3.74E-04 | 1.05E-03 | 1.12E+00 | 6.71E-11 | 2.46E-10 | -1.64E+00 | 3.12E-01 | 8.57E-01 | 1.19E+00 |
| BPTF | 2.97E-23 | 1.55E-22 | -1.54E+00 | 8.81E-07 | 3.81E-06 | -1.02E+00 | 1.45E-08 | 6.15E-08 | -1.42E+00 | 5.65E-04 | 2.16E-03 | -1.02E+00 | 2.61E-11 | 9.30E-11 | -1.45E+00 | 7.96E-07 | 2.24E-06 | -1.20E+00 | 2.16E-08 | 7.93E-08 | -1.37E+00 | 7.37E-04 | 2.03E-03 | -1.17E+00 |
| C5orf30 | 3.06E-23 | 1.59E-22 | 3.64E+00 | 2.23E-04 | 9.65E-04 | 2.35E+00 | 4.09E-06 | 1.74E-05 | 2.45E+00 | 9.68E-03 | 3.70E-02 | 2.03E+00 | 4.81E-02 | 1.72E-01 | 2.42E+00 | 2.28E-01 | 6.43E-01 | 1.36E+00 | 2.89E-03 | 1.06E-02 | 2.33E+00 | 8.02E-02 | 2.20E-01 | 1.70E+00 |
| RAB10 | 3.26E-23 | 1.70E-22 | -1.69E+00 | 1.69E-05 | 7.32E-05 | 1.15E+00 | 8.53E-12 | 3.62E-11 | -1.68E+00 | 9.44E-05 | 3.61E-04 | -1.08E+00 | 1.40E-24 | 4.98E-24 | -2.22E+00 | 6.19E-09 | 1.74E-08 | -1.20E+00 | 2.35E-18 | 8.61E-18 | -1.65E+00 | 1.76E-06 | 4.83E-06 | -1.24E+00 |
| DDX18 | 3.31E-23 | 1.72E-22 | -1.35E+00 | 6.07E-38 | 2.62E-37 | -1.61E+00 | 8.88E-04 | 3.77E-03 | -1.06E+00 | 4.48E-06 | 1.71E-05 | -1.11E+00 | 6.11E-09 | 2.18E-08 | -1.33E+00 | 3.96E-09 | 1.12E-08 | -1.38E+00 | 1.24E-06 | 4.53E-06 | -1.25E+00 | 5.79E-07 | 1.59E-06 | -1.29E+00 |
| GDI2 | 3.46E-23 | 1.80E-22 | -1.49E+00 | 3.10E-09 | 1.34E-08 | -1.15E+00 | 1.12E-07 | 4.74E-07 | -1.51E+00 | 2.14E-03 | 8.17E-03 | -1.19E+00 | 7.74E-13 | 2.76E-12 | -1.51E+00 | 1.68E-04 | 4.73E-04 | -1.16E+00 | 7.79E-11 | 2.86E-10 | -1.43E+00 | 1.57E-02 | 4.31E-02 | -1.12E+00 |
| SENP6 | 4.00E-23 | 2.09E-22 | -1.60E+00 | 6.39E-08 | 2.76E-07 | -1.04E+00 | 9.41E-11 | 4.00E-10 | -1.49E+00 | 1.93E-04 | 7.36E-04 | -1.05E+00 | 2.62E-05 | 9.35E-05 | -1.16E+00 | 1.64E-01 | 4.63E-01 | 1.13E+00 | 5.35E-05 | 1.96E-04 | -1.09E+00 | 1.06E-03 | 2.93E-03 | -1.03E+00 |
| PPP3CA | 4.16E-23 | 2.17E-22 | -1.59E+00 | 9.65E-06 | 4.17E-05 | 1.05E+00 | 8.30E-11 | 3.53E-10 | -1.47E+00 | 1.68E-05 | 6.43E-05 | -1.01E+00 | 8.52E-06 | 3.04E-05 | -1.25E+00 | 7.40E-02 | 2.09E-01 | 1.08E+00 | 5.47E-13 | 2.01E-12 | -1.64E+00 | 2.97E-04 | 8.16E-04 | -1.15E+00 |
| TRIM56 | 4.26E-23 | 2.22E-22 | -1.60E+00 | 9.52E-06 | 4.11E-05 | 1.06E+00 | 1.09E-14 | 4.62E-14 | -1.67E+00 | 1.57E-03 | 5.98E-03 | 1.01E+00 | 4.47E-07 | 1.59E-06 | -1.34E+00 | 9.88E-02 | 2.79E-01 | 1.16E+00 | 1.50E-10 | 5.49E-10 | -1.48E+00 | 7.99E-02 | 2.19E-01 | -1.01E+00 |
| AC244502.1 | 4.35E-23 | 2.27E-22 | 3.41E+00 | 4.45E-11 | 1.92E-10 | 2.72E+00 | 1.46E-01 | 6.20E-01 | 1.03E+00 | 1.38E-02 | 5.28E-02 | -1.05E+00 | 3.83E-17 | 1.37E-16 | -1.98E+00 | 3.52E-18 | 9.91E-18 | -2.12E+00 | 9.56E-11 | 3.51E-10 | -2.04E+00 | 7.09E-12 | 1.95E-11 | -2.32E+00 |
| TSEN34 | 4.36E-23 | 2.27E-22 | -1.59E+00 | 2.07E-10 | 8.94E-10 | -1.04E+00 | 1.23E-08 | 5.22E-08 | -1.30E+00 | 9.01E-05 | 3.44E-04 | -1.01E+00 | 2.49E-07 | 8.88E-07 | -1.61E+00 | 2.17E-02 | 6.11E-02 | 1.02E+00 | 1.72E-06 | 6.30E-06 | -1.27E+00 | 1.25E-02 | 3.44E-02 | -1.01E+00 |
| KEAP1 | 4.76E-23 | 2.48E-22 | -2.23E+00 | 2.35E-08 | 1.02E-07 | -1.06E+00 | 1.96E-09 | 8.34E-09 | -1.45E+00 | 4.44E-04 | 1.70E-03 | 1.06E+00 | 1.58E-15 | 5.62E-15 | -2.06E+00 | 1.45E-06 | 4.09E-06 | -1.11E+00 | 8.92E-25 | 3.27E-24 | -1.99E+00 | 8.39E-08 | 2.30E-07 | -1.27E+00 |
| FREM1 | 4.86E-23 | 2.53E-22 | -2.32E+00 | 1.81E-08 | 7.81E-08 | -1.20E+00 | 1.61E-07 | 6.84E-07 | -2.01E+00 | 4.88E-04 | 1.86E-03 | -1.18E+00 | 2.07E-13 | 7.38E-13 | -1.60E+00 | 4.27E-04 | 1.21E-03 | -1.11E+00 | 1.59E-11 | 5.84E-11 | -1.44E+00 | 8.01E-02 | 2.20E-01 | 1.04E+00 |
| AC021231.2 | 4.95E-23 | 2.58E-22 | -1.73E+00 | 3.96E-19 | 1.71E-18 | -1.45E+00 | 7.16E-12 | 3.04E-11 | -1.67E+00 | 2.88E-08 | 1.10E-07 | -1.27E+00 | 3.16E-24 | 1.13E-23 | -2.42E+00 | 2.08E-14 | 5.86E-14 | -1.60E+00 | 2.62E-22 | 9.63E-22 | -1.84E+00 | 2.30E-09 | 6.33E-09 | -1.41E+00 |
| CYTOR | 4.97E-23 | 2.59E-22 | -1.47E+00 | 1.47E-53 | 6.36E-53 | -2.02E+00 | 1.79E-15 | 7.61E-15 | -1.50E+00 | 3.51E-21 | 1.34E-20 | -1.66E+00 | 1.86E-04 | 6.62E-04 | -1.19E+00 | 1.24E-03 | 3.48E-03 | -1.22E+00 | 8.30E-02 | 3.05E-01 | -1.06E+00 | 1.73E-04 | 4.76E-04 | -1.56E+00 |
| RPL7 | 5.70E-23 | 2.97E-22 | 1.91E+00 | 9.47E-17 | 4.09E-16 | 1.91E+00 | 5.72E-18 | 2.43E-17 | 2.17E+00 | 1.76E-16 | 6.71E-16 | 2.19E+00 | 1.51E-08 | 5.37E-08 | 1.91E+00 | 5.46E-08 | 1.54E-07 | 1.88E+00 | 3.11E-08 | 1.14E-07 | 1.79E+00 | 1.13E-12 | 3.11E-12 | 2.02E+00 |
| RHOB | 5.73E-23 | 2.98E-22 | 3.23E+00 | 1.49E-16 | 6.42E-16 | 3.68E+00 | 1.11E-04 | 4.71E-04 | 2.20E+00 | 1.00E-05 | 3.82E-05 | 2.82E+00 | 1.06E-02 | 3.78E-02 | 2.06E+00 | 5.78E-02 | 1.63E-01 | 2.10E+00 | 1.01E-01 | 3.72E-01 | 2.28E+00 | 7.15E-02 | 1.96E-01 | 1.28E+00 |
| LAPTM4B | 5.77E-23 | 3.01E-22 | -1.75E+00 | 1.71E-01 | 7.37E-01 | 1.13E+00 | 8.24E-05 | 3.50E-04 | -1.31E+00 | 2.61E-01 | 9.98E-01 | 1.20E+00 | 3.22E-04 | 1.15E-03 | -1.07E+00 | 1.01E-03 | 2.84E-03 | 1.55E+00 | 9.71E-03 | 3.56E-02 | -1.15E+00 | 1.60E-01 | 4.38E-01 | 1.20E+00 |
| ANAPC5 | 6.21E-23 | 3.23E-22 | -1.50E+00 | 2.73E-12 | 1.18E-11 | -1.14E+00 | 8.05E-05 | 3.42E-04 | -1.19E+00 | 4.03E-03 | 1.54E-02 | 1.05E+00 | 2.25E-10 | 8.02E-10 | -1.38E+00 | 7.76E-04 | 2.19E-03 | -1.04E+00 | 5.90E-05 | 2.17E-04 | -1.13E+00 | 5.39E-02 | 1.48E-01 | 1.02E+00 |
| POLR2D | 6.35E-23 | 3.31E-22 | -2.22E+00 | 8.92E-15 | 3.86E-14 | -1.46E+00 | 1.14E-08 | 4.86E-08 | -1.74E+00 | 1.43E-04 | 5.46E-04 | -1.14E+00 | 8.82E-06 | 3.15E-05 | -1.48E+00 | 1.19E-03 | 3.36E-03 | -1.12E+00 | 1.64E-05 | 6.01E-05 | -1.37E+00 | 2.44E-02 | 6.71E-02 | 1.06E+00 |
| SMARCC1 | 6.35E-23 | 3.31E-22 | -1.39E+00 | 7.90E-11 | 3.41E-10 | -1.06E+00 | 1.02E-05 | 4.35E-05 | -1.26E+00 | 9.45E-04 | 3.61E-03 | -1.03E+00 | 3.73E-10 | 1.33E-09 | -1.36E+00 | 9.55E-07 | 2.69E-06 | -1.18E+00 | 5.06E-05 | 1.86E-04 | -1.27E+00 | 3.14E-02 | 8.62E-02 | -1.05E+00 |
| AC026369.3 | 6.66E-23 | 3.47E-22 | -1.58E+00 | 1.01E-14 | 4.38E-14 | -1.22E+00 | 4.48E-03 | 1.90E-02 | -1.11E+00 | 2.37E-06 | 9.06E-06 | -1.35E+00 | 1.45E-02 | 5.16E-02 | -1.54E+00 | 1.67E-02 | 4.71E-02 | -1.48E+00 | 1.50E-01 | 5.51E-01 | -1.35E+00 | 2.65E-01 | 7.28E-01 | -1.14E+00 |
| AL645608.4 | 6.83E-23 | 3.56E-22 | -1.60E+00 | 1.63E-44 | 7.04E-44 | -1.93E+00 | 2.54E-14 | 1.08E-13 | -1.69E+00 | 6.33E-28 | 2.42E-27 | -2.04E+00 | 6.10E-13 | 2.17E-12 | -1.54E+00 | 1.04E-22 | 2.93E-22 | -2.03E+00 | 1.06E-09 | 3.88E-09 | -1.52E+00 | 2.78E-23 | 7.62E-23 | -2.08E+00 |
| KDSR | 6.97E-23 | 3.63E-22 | -1.70E+00 | 5.36E-14 | 2.32E-13 | -1.24E+00 | 3.22E-10 | 1.37E-09 | -1.62E+00 | 2.48E-04 | 9.47E-04 | -1.03E+00 | 3.12E-09 | 1.11E-08 | -1.54E+00 | 5.82E-05 | 1.64E-04 | -1.06E+00 | 3.92E-14 | 1.44E-13 | -1.71E+00 | 2.81E-04 | 7.71E-04 | -1.12E+00 |
| NOL8 | 7.24E-23 | 3.77E-22 | -1.98E+00 | 9.15E-05 | 3.95E-04 | 1.07E+00 | 3.63E-09 | 1.54E-08 | -1.46E+00 | 1.89E-01 | 7.21E-01 | 1.35E+00 | 4.63E-14 | 1.65E-13 | -1.71E+00 | 1.73E-01 | 4.89E-01 | 1.24E+00 | 6.85E-18 | 2.51E-17 | -1.68E+00 | 2.38E-03 | 6.53E-03 | -1.07E+00 |
| H3F3B | 7.99E-23 | 4.16E-22 | 1.45E+00 | 2.25E-28 | 9.71E-28 | 1.56E+00 | 1.98E-12 | 8.43E-12 | 1.41E+00 | 4.39E-20 | 1.68E-19 | 1.59E+00 | 7.98E-06 | 2.84E-05 | 1.37E+00 | 6.90E-12 | 1.94E-11 | 1.65E+00 | 3.07E-05 | 1.13E-04 | 1.27E+00 | 1.02E-04 | 2.81E-04 | 1.21E+00 |
| MRPL9 | 7.99E-23 | 4.16E-22 | -1.93E+00 | 9.83E-14 | 4.25E-13 | -1.25E+00 | 2.36E-14 | 1.00E-13 | -1.70E+00 | 1.19E-12 | 4.56E-12 | -1.38E+00 | 6.29E-10 | 2.24E-09 | -1.57E+00 | 2.86E-05 | 8.06E-05 | -1.15E+00 | 3.94E-12 | 1.45E-11 | -1.62E+00 | 4.01E-04 | 1.10E-03 | -1.11E+00 |
| NLRC5 | 8.05E-23 | 4.19E-22 | -2.08E+00 | 1.18E-24 | 5.11E-24 | -1.96E+00 | 1.38E-17 | 5.87E-17 | -1.90E+00 | 9.79E-27 | 3.74E-26 | -2.37E+00 | 2.15E-12 | 7.65E-12 | -3.10E+00 | 2.84E-11 | 8.00E-11 | -2.42E+00 | 1.27E-28 | 4.66E-28 | -2.82E+00 | 5.03E-27 | 1.38E-26 | -2.97E+00 |
| MRPL4 | 8.11E-23 | 4.23E-22 | -1.56E+00 | 3.14E-12 | 1.36E-11 | -1.03E+00 | 5.78E-08 | 2.46E-07 | -1.25E+00 | 4.27E-06 | 1.63E-05 | -1.08E+00 | 2.88E-10 | 1.03E-09 | -1.52E+00 | 2.76E-03 | 7.79E-03 | 1.04E+00 | 3.72E-08 | 1.37E-07 | -1.26E+00 | 3.36E-01 | 9.22E-01 | 1.19E+00 |
| ARID3A | 9.02E-23 | 4.70E-22 | -2.28E+00 | 4.23E-17 | 1.83E-16 | -1.50E+00 | 1.28E-18 | 5.44E-18 | -2.55E+00 | 7.09E-09 | 2.71E-08 | -1.39E+00 | 2.59E-10 | 9.22E-10 | -2.21E+00 | 1.77E-08 | 4.99E-08 | -1.88E+00 | 5.64E-09 | 2.07E-08 | -1.83E+00 | 5.46E-08 | 1.50E-07 | -1.80E+00 |
| EIF4BP6 | 9.41E-23 | 4.90E-22 | -1.43E+00 | 6.98E-26 | 3.02E-25 | -1.40E+00 | 1.87E-08 | 7.93E-08 | -1.39E+00 | 4.46E-10 | 1.70E-09 | -1.33E+00 | 7.80E-08 | 2.78E-07 | -1.31E+00 | 3.07E-09 | 8.66E-09 | -1.35E+00 | 6.97E-06 | 2.56E-05 | -1.34E+00 | 3.48E-06 | 9.55E-06 | -1.28E+00 |
| PRNP | 9.80E-23 | 5.11E-22 | -1.78E+00 | 7.44E-17 | 3.21E-16 | -1.33E+00 | 7.70E-12 | 3.27E-11 | -1.56E+00 | 3.76E-11 | 1.44E-10 | -1.29E+00 | 2.31E-08 | 8.24E-08 | -2.03E+00 | 1.52E-06 | 4.28E-06 | -1.63E+00 | 1.72E-08 | 6.33E-08 | -1.68E+00 | 5.28E-04 | 1.45E-03 | -1.16E+00 |
| C20orf166-AS1 | 1.16E-22 | 6.04E-22 | -3.26E+00 | 7.09E-25 | 3.06E-24 | -3.97E+00 | 1.89E-08 | 8.05E-08 | -2.16E+00 | 2.97E-09 | 1.13E-08 | -2.44E+00 | 1.71E-26 | 6.10E-26 | -6.37E+00 | 4.86E-27 | 1.37E-26 | -5.59E+00 | 9.45E-07 | 3.47E-06 | -2.21E+00 | 9.01E-07 | 2.48E-06 | -2.57E+00 |
| PRKACB | 1.16E-22 | 6.06E-22 | -1.56E+00 | 2.22E-14 | 9.61E-14 | -1.25E+00 | 3.11E-07 | 1.32E-06 | -1.37E+00 | 1.18E-04 | 4.49E-04 | -1.09E+00 | 1.57E-02 | 5.59E-02 | 1.03E+00 | 4.66E-02 | 1.31E-01 | 1.08E+00 | 1.99E-01 | 7.32E-01 | 1.21E+00 | 1.34E-02 | 3.69E-02 | -1.02E+00 |
| RPS10P7 | 1.34E-22 | 7.00E-22 | -2.21E+00 | 2.21E-11 | 9.57E-11 | -1.24E+00 | 6.80E-28 | 2.89E-27 | -2.78E+00 | 4.89E-16 | 1.87E-15 | -1.56E+00 | 4.11E-10 | 1.47E-09 | -2.54E+00 | 1.08E-03 | 3.06E-03 | -1.10E+00 | 1.35E-15 | 4.95E-15 | -2.55E+00 | 8.29E-08 | 2.28E-07 | -1.52E+00 |
| TUBB | 1.37E-22 | 7.16E-22 | -1.43E+00 | 3.27E-06 | 1.41E-05 | -1.01E+00 | 4.31E-02 | 1.83E-01 | -1.04E+00 | 2.02E-01 | 7.70E-01 | 1.18E+00 | 6.52E-16 | 2.32E-15 | -1.80E+00 | 5.65E-06 | 1.59E-05 | -1.31E+00 | 6.38E-05 | 2.34E-04 | -1.17E+00 | 3.35E-01 | 9.21E-01 | 1.21E+00 |
| ACTN1 | 1.50E-22 | 7.84E-22 | -1.97E+00 | 1.45E-10 | 6.25E-10 | -1.18E+00 | 2.97E-11 | 1.26E-10 | -1.55E+00 | 1.28E-06 | 4.89E-06 | -1.17E+00 | 9.39E-23 | 3.35E-22 | -2.62E+00 | 1.10E-07 | 3.10E-07 | -1.36E+00 | 1.05E-07 | 3.86E-07 | -1.24E+00 | 1.32E-03 | 3.62E-03 | -1.13E+00 |
| PSME2P6 | 1.53E-22 | 7.99E-22 | -1.79E+00 | 1.76E-34 | 7.59E-34 | -2.26E+00 | 7.12E-12 | 3.03E-11 | -1.64E+00 | 2.64E-20 | 1.01E-19 | -1.83E+00 | 4.57E-12 | 1.63E-11 | -1.96E+00 | 7.05E-15 | 1.99E-14 | -2.29E+00 | 3.85E-27 | 1.41E-26 | -2.34E+00 | 3.34E-29 | 9.16E-29 | -2.51E+00 |
| ATP5A1 | 1.58E-22 | 8.25E-22 | -1.34E+00 | 1.52E-03 | 6.58E-03 | 1.02E+00 | 6.29E-03 | 2.67E-02 | -1.14E+00 | 1.42E-01 | 5.44E-01 | 1.02E+00 | 2.34E-07 | 8.33E-07 | -1.27E+00 | 8.20E-02 | 2.31E-01 | 1.03E+00 | 2.56E-04 | 9.39E-04 | -1.20E+00 | 2.19E-01 | 6.02E-01 | 1.00E+00 |
| SLTM | 1.66E-22 | 8.67E-22 | -1.57E+00 | 4.84E-18 | 2.09E-17 | -1.25E+00 | 6.39E-07 | 2.72E-06 | -1.37E+00 | 2.21E-04 | 8.42E-04 | -1.18E+00 | 5.99E-11 | 2.14E-10 | -1.42E+00 | 4.06E-07 | 1.14E-06 | -1.24E+00 | 2.02E-08 | 7.42E-08 | -1.35E+00 | 5.64E-06 | 1.55E-05 | -1.30E+00 |
| AL109618.1 | 1.88E-22 | 9.81E-22 | -1.90E+00 | 9.46E-20 | 4.09E-19 | -1.63E+00 | 5.00E-10 | 2.12E-09 | -1.50E+00 | 2.80E-13 | 1.07E-12 | -1.47E+00 | 1.57E-14 | 5.61E-14 | -1.98E+00 | 1.12E-12 | 3.16E-12 | -1.71E+00 | 9.53E-27 | 3.50E-26 | -2.07E+00 | 8.10E-17 | 2.23E-16 | -1.61E+00 |
| CTSO | 2.05E-22 | 1.07E-21 | -2.11E+00 | 3.60E-12 | 1.55E-11 | -1.22E+00 | 5.65E-08 | 2.40E-07 | -1.56E+00 | 2.67E-07 | 1.02E-06 | -1.34E+00 | 1.16E-02 | 4.13E-02 | -1.27E+00 | 7.56E-03 | 2.13E-02 | -1.33E+00 | 8.67E-06 | 3.18E-05 | -1.81E+00 | 1.70E-04 | 4.67E-04 | -1.53E+00 |
| EIF4H | 2.06E-22 | 1.08E-21 | -1.46E+00 | 1.78E-24 | 7.71E-24 | -1.37E+00 | 1.11E-06 | 4.73E-06 | -1.39E+00 | 2.06E-09 | 7.88E-09 | -1.33E+00 | 9.15E-11 | 3.26E-10 | -1.34E+00 | 8.93E-11 | 2.52E-10 | -1.35E+00 | 5.01E-06 | 1.84E-05 | -1.36E+00 | 3.03E-05 | 8.32E-05 | -1.27E+00 |
| DCP2 | 2.06E-22 | 1.08E-21 | -1.88E+00 | 8.85E-11 | 3.82E-10 | -1.18E+00 | 8.04E-07 | 3.42E-06 | -1.34E+00 | 4.97E-07 | 1.90E-06 | -1.20E+00 | 1.73E-11 | 6.15E-11 | -1.43E+00 | 2.60E-07 | 7.33E-07 | -1.15E+00 | 1.99E-03 | 7.31E-03 | -1.07E+00 | 2.44E-03 | 6.71E-03 | -1.01E+00 |
| ATP6V0B | 2.12E-22 | 1.10E-21 | -1.45E+00 | 2.15E-16 | 9.27E-16 | -1.22E+00 | 2.19E-03 | 9.29E-03 | -1.12E+00 | 2.87E-03 | 1.10E-02 | -1.07E+00 | 9.38E-07 | 3.35E-06 | -1.26E+00 | 1.30E-05 | 3.67E-05 | -1.14E+00 | 6.56E-04 | 2.41E-03 | -1.11E+00 | 1.35E-01 | 3.70E-01 | 1.11E+00 |
| CRBN | 2.13E-22 | 1.11E-21 | -1.49E+00 | 6.49E-06 | 2.80E-05 | 1.10E+00 | 3.48E-09 | 1.48E-08 | -1.46E+00 | 1.56E-03 | 5.96E-03 | 1.01E+00 | 1.27E-03 | 4.52E-03 | -1.09E+00 | 1.49E-01 | 4.20E-01 | 1.16E+00 | 3.54E-09 | 1.30E-08 | -1.30E+00 | 3.21E-05 | 8.83E-05 | -1.16E+00 |
| XRCC5 | 2.17E-22 | 1.13E-21 | -1.41E+00 | 1.17E-09 | 5.04E-09 | -1.08E+00 | 2.16E-04 | 9.17E-04 | -1.22E+00 | 2.14E-02 | 8.18E-02 | -1.02E+00 | 2.48E-13 | 8.85E-13 | -1.38E+00 | 2.82E-02 | 7.95E-02 | -1.02E+00 | 2.37E-04 | 8.68E-04 | -1.28E+00 | 2.09E-01 | 5.74E-01 | -1.04E+00 |
| TNFAIP3 | 2.29E-22 | 1.20E-21 | 4.56E+00 | 5.17E-02 | 2.23E-01 | 1.81E+00 | 5.94E-02 | 2.52E-01 | 2.28E+00 | 1.14E-08 | 4.36E-08 | -1.21E+00 | 5.69E-02 | 2.03E-01 | 3.15E+00 | 2.20E-06 | 6.21E-06 | -2.21E+00 | 4.88E-05 | 1.79E-04 | -1.28E+00 | 1.75E-08 | 4.82E-08 | -3.97E+00 |
| YWHAE | 2.29E-22 | 1.20E-21 | -1.59E+00 | 2.67E-08 | 1.16E-07 | -1.15E+00 | 8.63E-07 | 3.67E-06 | -1.44E+00 | 6.74E-03 | 2.57E-02 | -1.13E+00 | 2.46E-24 | 8.79E-24 | -1.90E+00 | 3.49E-06 | 9.84E-06 | -1.26E+00 | 4.39E-07 | 1.61E-06 | -1.44E+00 | 1.25E-01 | 3.42E-01 | -1.05E+00 |
| IL6R | 2.52E-22 | 1.32E-21 | -2.18E+00 | 4.36E-08 | 1.88E-07 | -1.14E+00 | 1.21E-21 | 5.14E-21 | -1.78E+00 | 1.03E-11 | 3.95E-11 | -1.21E+00 | 7.40E-03 | 2.64E-02 | -1.76E+00 | 4.14E-03 | 1.17E-02 | -2.29E+00 | 5.19E-02 | 1.91E-01 | -2.09E+00 | 1.81E-01 | 4.97E-01 | -1.24E+00 |
| RNPS1 | 2.58E-22 | 1.34E-21 | -1.40E+00 | 1.47E-09 | 6.34E-09 | -1.03E+00 | 2.16E-04 | 9.19E-04 | -1.22E+00 | 2.05E-02 | 7.84E-02 | 1.04E+00 | 5.42E-09 | 1.93E-08 | -1.36E+00 | 3.00E-03 | 8.46E-03 | -1.09E+00 | 8.30E-05 | 3.04E-04 | -1.25E+00 | 2.20E-01 | 6.05E-01 | 1.08E+00 |
| CCDC175 | 2.78E-22 | 1.45E-21 | -2.12E+00 | 1.19E-16 | 5.13E-16 | -1.62E+00 | 2.24E-08 | 9.51E-08 | -1.91E+00 | 7.10E-08 | 2.71E-07 | -1.68E+00 | 3.00E-03 | 1.07E-02 | -2.42E+00 | 1.32E-02 | 3.71E-02 | -1.93E+00 | 1.61E-01 | 5.90E-01 | 1.44E+00 | 3.53E-01 | 9.68E-01 | 1.15E+00 |
| SCML4 | 3.04E-22 | 1.58E-21 | -2.26E+00 | 1.78E-14 | 7.69E-14 | -1.34E+00 | 8.78E-15 | 3.73E-14 | -2.57E+00 | 5.05E-11 | 1.93E-10 | -1.76E+00 | 1.88E-02 | 6.70E-02 | -1.73E+00 | 5.38E-02 | 1.52E-01 | -1.36E+00 | 1.86E-01 | 6.81E-01 | -1.24E+00 | 2.56E-01 | 7.04E-01 | -1.15E+00 |
| ETNK1 | 3.08E-22 | 1.60E-21 | -1.53E+00 | 3.86E-05 | 1.67E-04 | 1.10E+00 | 6.17E-14 | 2.62E-13 | -1.68E+00 | 4.57E-05 | 1.74E-04 | -1.03E+00 | 1.71E-11 | 6.09E-11 | -1.51E+00 | 3.73E-02 | 1.05E-01 | 1.14E+00 | 3.05E-09 | 1.12E-08 | -1.43E+00 | 5.39E-03 | 1.48E-02 | -1.12E+00 |
| HDAC2 | 3.33E-22 | 1.73E-21 | -1.51E+00 | 7.55E-12 | 3.26E-11 | -1.11E+00 | 1.45E-06 | 6.15E-06 | -1.23E+00 | 6.26E-05 | 2.39E-04 | -1.03E+00 | 1.55E-13 | 5.52E-13 | -1.70E+00 | 1.05E-05 | 2.97E-05 | -1.25E+00 | 5.45E-10 | 2.00E-09 | -1.54E+00 | 3.36E-02 | 9.24E-02 | -1.11E+00 |
| KLF9 | 3.46E-22 | 1.81E-21 | 2.62E+00 | 2.18E-04 | 9.41E-04 | 2.00E+00 | 3.57E-09 | 1.52E-08 | 2.15E+00 | 3.14E-04 | 1.20E-03 | 1.90E+00 | 1.80E-10 | 6.40E-10 | 2.75E+00 | 6.43E-04 | 1.81E-03 | 1.92E+00 | 1.59E-05 | 5.82E-05 | 2.01E+00 | 2.84E-01 | 7.80E-01 | 1.41E+00 |
| BISPR | 3.87E-22 | 2.02E-21 | -1.75E+00 | 1.14E-16 | 4.91E-16 | -1.31E+00 | 3.93E-24 | 1.67E-23 | -2.69E+00 | 2.27E-17 | 8.69E-17 | -1.89E+00 | 9.38E-13 | 3.34E-12 | -2.09E+00 | 4.38E-11 | 1.24E-10 | -1.81E+00 | 1.43E-13 | 5.24E-13 | -2.81E+00 | 1.77E-10 | 4.87E-10 | -2.15E+00 |
| PDCD6 | 4.53E-22 | 2.36E-21 | -1.33E+00 | 5.94E-10 | 2.56E-09 | -1.04E+00 | 3.42E-04 | 1.45E-03 | -1.17E+00 | 7.49E-02 | 2.86E-01 | 1.04E+00 | 2.20E-10 | 7.86E-10 | -1.24E+00 | 5.12E-02 | 1.44E-01 | 1.13E+00 | 2.27E-03 | 8.34E-03 | -1.09E+00 | 3.10E-01 | 8.52E-01 | 1.10E+00 |
| FRG1CP | 4.53E-22 | 2.36E-21 | -1.93E+00 | 9.86E-19 | 4.26E-18 | -1.43E+00 | 2.87E-10 | 1.22E-09 | -1.69E+00 | 7.12E-10 | 2.72E-09 | -1.50E+00 | 2.42E-09 | 8.62E-09 | -1.82E+00 | 5.71E-07 | 1.61E-06 | -1.48E+00 | 7.20E-10 | 2.64E-09 | -1.46E+00 | 1.50E-06 | 4.12E-06 | -1.20E+00 |
| VEZF1 | 4.55E-22 | 2.37E-21 | -1.99E+00 | 1.59E-14 | 6.85E-14 | -1.36E+00 | 6.37E-15 | 2.71E-14 | -1.72E+00 | 4.75E-12 | 1.81E-11 | -1.34E+00 | 2.30E-13 | 8.21E-13 | -2.08E+00 | 4.61E-07 | 1.30E-06 | -1.35E+00 | 5.73E-17 | 2.10E-16 | -1.87E+00 | 1.07E-10 | 2.93E-10 | -1.51E+00 |
| AC025040.1 | 4.63E-22 | 2.41E-21 | -1.75E+00 | 1.22E-05 | 5.25E-05 | 1.17E+00 | 2.24E-21 | 9.50E-21 | -2.22E+00 | 4.92E-08 | 1.88E-07 | -1.14E+00 | 7.20E-02 | 2.57E-01 | 1.02E+00 | 8.12E-02 | 2.29E-01 | 1.08E+00 | 2.43E-01 | 8.94E-01 | 1.19E+00 | 3.51E-01 | 9.64E-01 | 1.22E+00 |
| HSP90AB2P | 5.59E-22 | 2.91E-21 | -1.52E+00 | 8.40E-35 | 3.63E-34 | -1.65E+00 | 3.60E-06 | 1.53E-05 | -1.45E+00 | 6.31E-10 | 2.41E-09 | -1.67E+00 | 3.39E-12 | 1.21E-11 | -1.60E+00 | 4.90E-10 | 1.38E-09 | -1.56E+00 | 6.30E-11 | 2.31E-10 | -1.59E+00 | 5.15E-07 | 1.41E-06 | -1.67E+00 |
| TOR1AIP2 | 5.89E-22 | 3.07E-21 | -1.55E+00 | 1.08E-21 | 4.68E-21 | -1.37E+00 | 8.00E-07 | 3.40E-06 | -1.45E+00 | 6.86E-07 | 2.62E-06 | -1.23E+00 | 2.37E-08 | 8.45E-08 | -1.43E+00 | 1.90E-06 | 5.37E-06 | -1.25E+00 | 1.12E-11 | 4.12E-11 | -1.59E+00 | 3.95E-05 | 1.08E-04 | -1.26E+00 |
| SBNO1 | 5.93E-22 | 3.09E-21 | -1.55E+00 | 6.77E-06 | 2.93E-05 | 1.09E+00 | 8.74E-13 | 3.71E-12 | -1.65E+00 | 5.77E-05 | 2.20E-04 | -1.09E+00 | 4.83E-08 | 1.72E-07 | -1.31E+00 | 5.41E-04 | 1.52E-03 | -1.01E+00 | 9.85E-07 | 3.62E-06 | -1.36E+00 | 1.11E-01 | 3.05E-01 | -1.01E+00 |
| HNRNPA0 | 6.05E-22 | 3.15E-21 | -1.48E+00 | 2.30E-12 | 9.92E-12 | -1.20E+00 | 4.29E-06 | 1.82E-05 | -1.37E+00 | 6.18E-04 | 2.36E-03 | -1.11E+00 | 2.01E-09 | 7.16E-09 | -1.48E+00 | 2.54E-04 | 7.15E-04 | -1.17E+00 | 1.85E-08 | 6.78E-08 | -1.37E+00 | 1.37E-02 | 3.75E-02 | -1.09E+00 |
| RGS18 | 6.12E-22 | 3.19E-21 | -3.36E+00 | 2.96E-17 | 1.28E-16 | -2.22E+00 | 1.01E-06 | 4.29E-06 | -1.88E+00 | 3.57E-04 | 1.36E-03 | -1.32E+00 | 4.51E-05 | 1.61E-04 | -1.59E+00 | 6.21E-03 | 1.75E-02 | -1.36E+00 | 1.07E-02 | 3.92E-02 | 1.83E+00 | 2.59E-01 | 7.12E-01 | 1.22E+00 |
| CD84 | 6.23E-22 | 3.25E-21 | -1.75E+00 | 8.77E-17 | 3.79E-16 | -1.38E+00 | 1.81E-12 | 7.70E-12 | -1.53E+00 | 6.60E-10 | 2.52E-09 | -1.19E+00 | 4.80E-06 | 1.71E-05 | -1.28E+00 | 1.66E-04 | 4.67E-04 | -1.23E+00 | 3.97E-06 | 1.46E-05 | -1.32E+00 | 2.86E-03 | 7.85E-03 | -1.22E+00 |
| SLC50A1 | 6.28E-22 | 3.27E-21 | -1.56E+00 | 4.00E-20 | 1.73E-19 | -1.41E+00 | 5.23E-16 | 2.22E-15 | -1.82E+00 | 3.13E-12 | 1.19E-11 | -1.39E+00 | 1.85E-11 | 6.60E-11 | -1.56E+00 | 6.03E-07 | 1.70E-06 | -1.18E+00 | 4.43E-04 | 1.63E-03 | -1.10E+00 | 1.20E-02 | 3.29E-02 | -1.02E+00 |
| RBPMS | 6.40E-22 | 3.34E-21 | -1.59E+00 | 6.20E-06 | 2.68E-05 | 1.06E+00 | 6.43E-08 | 2.73E-07 | -1.33E+00 | 2.04E-03 | 7.77E-03 | 1.05E+00 | 6.39E-02 | 2.28E-01 | -1.13E+00 | 1.86E-01 | 5.25E-01 | 1.16E+00 | 2.18E-01 | 7.99E-01 | -1.11E+00 | 2.56E-01 | 7.04E-01 | 1.61E+00 |
| HSD17B12 | 6.49E-22 | 3.38E-21 | -1.61E+00 | 2.38E-11 | 1.03E-10 | -1.11E+00 | 1.08E-07 | 4.59E-07 | -1.30E+00 | 3.13E-04 | 1.20E-03 | -1.04E+00 | 8.49E-11 | 3.03E-10 | -1.54E+00 | 1.42E-05 | 4.01E-05 | -1.17E+00 | 2.90E-08 | 1.06E-07 | -1.29E+00 | 1.52E-04 | 4.17E-04 | -1.11E+00 |
| MBOAT7 | 7.00E-22 | 3.65E-21 | -1.52E+00 | 1.84E-10 | 7.93E-10 | -1.09E+00 | 1.91E-06 | 8.11E-06 | -1.27E+00 | 1.71E-06 | 6.53E-06 | -1.16E+00 | 6.21E-12 | 2.21E-11 | -1.67E+00 | 1.77E-08 | 4.98E-08 | -1.38E+00 | 4.92E-05 | 1.81E-04 | -1.24E+00 | 1.24E-03 | 3.40E-03 | -1.09E+00 |
| UBXN4 | 8.09E-22 | 4.21E-21 | -1.50E+00 | 1.58E-10 | 6.84E-10 | -1.13E+00 | 4.36E-07 | 1.85E-06 | -1.41E+00 | 6.48E-05 | 2.48E-04 | -1.13E+00 | 1.82E-12 | 6.51E-12 | -1.56E+00 | 1.03E-06 | 2.90E-06 | -1.22E+00 | 2.30E-09 | 8.42E-09 | -1.47E+00 | 1.26E-02 | 3.45E-02 | -1.15E+00 |
| LAPTM4A | 8.12E-22 | 4.23E-21 | -1.42E+00 | 1.05E-11 | 4.52E-11 | -1.20E+00 | 6.91E-07 | 2.93E-06 | -1.31E+00 | 4.61E-08 | 1.76E-07 | -1.26E+00 | 1.86E-06 | 6.64E-06 | -1.43E+00 | 6.58E-08 | 1.86E-07 | -1.51E+00 | 1.92E-09 | 7.05E-09 | -1.37E+00 | 5.50E-06 | 1.51E-05 | -1.28E+00 |
| NXT1 | 8.29E-22 | 4.32E-21 | -1.65E+00 | 9.51E-38 | 4.11E-37 | -2.10E+00 | 1.13E-14 | 4.82E-14 | -1.61E+00 | 4.89E-18 | 1.87E-17 | -1.55E+00 | 8.45E-19 | 3.01E-18 | -2.07E+00 | 1.10E-21 | 3.09E-21 | -2.32E+00 | 7.00E-21 | 2.57E-20 | -2.00E+00 | 2.71E-11 | 7.43E-11 | -1.57E+00 |
| YWHAQ | 8.31E-22 | 4.33E-21 | -1.29E+00 | 8.49E-16 | 3.67E-15 | -1.13E+00 | 1.81E-03 | 7.71E-03 | -1.12E+00 | 2.94E-03 | 1.12E-02 | -1.06E+00 | 2.24E-06 | 7.99E-06 | -1.19E+00 | 1.90E-04 | 5.37E-04 | -1.11E+00 | 3.97E-03 | 1.46E-02 | -1.13E+00 | 2.98E-01 | 8.17E-01 | 1.06E+00 |
| STARD7 | 9.37E-22 | 4.89E-21 | -1.96E+00 | 4.69E-11 | 2.03E-10 | -1.31E+00 | 2.99E-09 | 1.27E-08 | -1.41E+00 | 4.09E-04 | 1.56E-03 | 1.03E+00 | 5.33E-12 | 1.90E-11 | -1.65E+00 | 8.48E-05 | 2.39E-04 | -1.08E+00 | 1.12E-05 | 4.13E-05 | -1.25E+00 | 2.19E-01 | 6.02E-01 | 1.14E+00 |
| BNIP3L | 9.63E-22 | 5.02E-21 | -1.48E+00 | 2.92E-13 | 1.26E-12 | -1.15E+00 | 2.61E-08 | 1.11E-07 | -1.47E+00 | 7.61E-09 | 2.91E-08 | -1.28E+00 | 1.27E-13 | 4.52E-13 | -1.64E+00 | 2.17E-05 | 6.12E-05 | -1.24E+00 | 3.77E-06 | 1.38E-05 | -1.34E+00 | 3.01E-04 | 8.27E-04 | -1.27E+00 |
| LRRC59 | 9.89E-22 | 5.15E-21 | -1.98E+00 | 1.39E-19 | 5.99E-19 | -1.71E+00 | 4.15E-07 | 1.76E-06 | -1.20E+00 | 2.43E-06 | 9.27E-06 | -1.03E+00 | 4.40E-09 | 1.57E-08 | -1.60E+00 | 1.49E-06 | 4.21E-06 | -1.36E+00 | 8.56E-07 | 3.14E-06 | -1.32E+00 | 2.75E-01 | 7.55E-01 | 1.11E+00 |
| ALKBH5 | 1.00E-21 | 5.23E-21 | -1.95E+00 | 1.29E-12 | 5.56E-12 | -1.25E+00 | 1.06E-11 | 4.51E-11 | -1.54E+00 | 2.26E-09 | 8.64E-09 | -1.19E+00 | 1.42E-04 | 5.07E-04 | -1.15E+00 | 7.06E-03 | 1.99E-02 | 1.12E+00 | 3.77E-10 | 1.38E-09 | -1.52E+00 | 3.26E-04 | 8.94E-04 | -1.10E+00 |
| SS18L2 | 1.04E-21 | 5.40E-21 | -1.58E+00 | 5.61E-15 | 2.42E-14 | -1.24E+00 | 1.66E-07 | 7.03E-07 | -1.38E+00 | 3.53E-05 | 1.35E-04 | -1.10E+00 | 2.23E-09 | 7.96E-09 | -1.13E+00 | 9.92E-06 | 2.80E-05 | 1.02E+00 | 3.01E-11 | 1.10E-10 | -1.45E+00 | 9.20E-05 | 2.53E-04 | -1.15E+00 |
| BEX3 | 1.09E-21 | 5.68E-21 | -1.42E+00 | 3.60E-07 | 1.55E-06 | -1.07E+00 | 4.84E-05 | 2.06E-04 | -1.29E+00 | 4.49E-04 | 1.71E-03 | -1.09E+00 | 7.50E-12 | 2.67E-11 | -1.54E+00 | 6.57E-05 | 1.85E-04 | -1.13E+00 | 1.16E-04 | 4.25E-04 | -1.14E+00 | 3.48E-02 | 9.57E-02 | -1.01E+00 |
| AL590867.2 | 1.13E-21 | 5.90E-21 | 1.97E+00 | 8.22E-22 | 3.55E-21 | 2.10E+00 | 3.57E-14 | 1.52E-13 | 1.87E+00 | 1.95E-16 | 7.43E-16 | 2.10E+00 | 5.14E-11 | 1.83E-10 | 1.93E+00 | 8.97E-20 | 2.53E-19 | 2.28E+00 | 2.37E-06 | 8.71E-06 | 1.65E+00 | 2.16E-15 | 5.93E-15 | 2.02E+00 |
| PHF20L1 | 1.24E-21 | 6.44E-21 | -1.64E+00 | 1.35E-15 | 5.84E-15 | -1.29E+00 | 1.13E-10 | 4.81E-10 | -1.57E+00 | 7.19E-08 | 2.74E-07 | -1.30E+00 | 7.48E-07 | 2.67E-06 | -1.30E+00 | 1.15E-06 | 3.25E-06 | -1.19E+00 | 4.99E-08 | 1.83E-07 | -1.38E+00 | 3.23E-05 | 8.88E-05 | -1.21E+00 |
| CAPN1 | 1.29E-21 | 6.70E-21 | -1.42E+00 | 3.69E-06 | 1.59E-05 | 1.13E+00 | 3.90E-05 | 1.66E-04 | -1.25E+00 | 5.43E-03 | 2.08E-02 | 1.04E+00 | 7.93E-06 | 2.83E-05 | -1.26E+00 | 5.05E-02 | 1.42E-01 | 1.03E+00 | 2.32E-02 | 8.52E-02 | -1.03E+00 | 1.13E-01 | 3.10E-01 | 1.04E+00 |
| SNX5 | 1.33E-21 | 6.95E-21 | -1.53E+00 | 7.32E-15 | 3.16E-14 | -1.20E+00 | 5.35E-10 | 2.27E-09 | -1.53E+00 | 6.90E-08 | 2.64E-07 | -1.23E+00 | 5.92E-06 | 2.11E-05 | -1.44E+00 | 6.98E-02 | 1.97E-01 | -1.02E+00 | 3.10E-06 | 1.14E-05 | -1.29E+00 | 1.62E-02 | 4.44E-02 | 1.01E+00 |
| RPL13A | 1.35E-21 | 7.03E-21 | 1.34E+00 | 1.91E-21 | 8.25E-21 | 1.35E+00 | 1.08E-11 | 4.59E-11 | 1.29E+00 | 3.99E-14 | 1.52E-13 | 1.33E+00 | 1.28E-11 | 4.57E-11 | 1.35E+00 | 2.93E-17 | 8.26E-17 | 1.43E+00 | 9.19E-06 | 3.37E-05 | 1.26E+00 | 3.57E-15 | 9.80E-15 | 1.35E+00 |
| RRP1B | 1.48E-21 | 7.73E-21 | -1.84E+00 | 6.68E-11 | 2.89E-10 | -1.18E+00 | 3.71E-04 | 1.58E-03 | -1.07E+00 | 1.34E-02 | 5.12E-02 | 1.25E+00 | 4.76E-05 | 1.70E-04 | -1.50E+00 | 1.01E-02 | 2.84E-02 | -1.05E+00 | 9.36E-08 | 3.44E-07 | -1.51E+00 | 5.34E-04 | 1.47E-03 | -1.19E+00 |
| ADAM8 | 1.53E-21 | 7.96E-21 | -1.63E+00 | 6.04E-14 | 2.61E-13 | -1.22E+00 | 2.87E-18 | 1.22E-17 | -2.11E+00 | 2.87E-10 | 1.10E-09 | -1.38E+00 | 2.79E-04 | 9.96E-04 | -1.54E+00 | 2.76E-05 | 7.79E-05 | -1.56E+00 | 1.03E-01 | 3.77E-01 | -1.02E+00 | 4.04E-02 | 1.11E-01 | -1.36E+00 |
| CTNNB1 | 1.54E-21 | 8.04E-21 | -1.72E+00 | 1.14E-11 | 4.91E-11 | -1.27E+00 | 1.06E-08 | 4.52E-08 | -1.71E+00 | 1.36E-07 | 5.19E-07 | -1.38E+00 | 1.67E-12 | 5.95E-12 | -1.63E+00 | 1.57E-05 | 4.41E-05 | -1.16E+00 | 3.33E-18 | 1.22E-17 | -2.02E+00 | 5.40E-06 | 1.48E-05 | -1.36E+00 |
| KIF5B | 1.57E-21 | 8.20E-21 | -1.43E+00 | 1.93E-22 | 8.35E-22 | -1.27E+00 | 1.88E-08 | 7.98E-08 | -1.43E+00 | 2.99E-09 | 1.14E-08 | -1.29E+00 | 4.39E-11 | 1.56E-10 | -1.38E+00 | 1.12E-09 | 3.16E-09 | -1.29E+00 | 9.50E-06 | 3.49E-05 | -1.20E+00 | 9.28E-03 | 2.55E-02 | -1.12E+00 |
| RASGEF1B | 1.58E-21 | 8.23E-21 | 3.41E+00 | 1.02E-04 | 4.40E-04 | -1.81E+00 | 5.43E-29 | 2.31E-28 | 5.07E+00 | 4.93E-02 | 1.88E-01 | -1.38E+00 | 1.27E-26 | 4.52E-26 | 8.94E+00 | 9.50E-02 | 2.68E-01 | -2.27E+00 | 1.81E-16 | 6.63E-16 | 7.60E+00 | 1.82E-01 | 5.00E-01 | -1.37E+00 |
| ABCF1 | 1.61E-21 | 8.38E-21 | -1.73E+00 | 1.91E-16 | 8.25E-16 | -1.27E+00 | 6.34E-10 | 2.69E-09 | -1.58E+00 | 4.55E-07 | 1.74E-06 | -1.22E+00 | 4.26E-10 | 1.52E-09 | -1.64E+00 | 2.03E-06 | 5.74E-06 | -1.27E+00 | 3.17E-09 | 1.16E-08 | -1.39E+00 | 7.47E-03 | 2.05E-02 | -1.04E+00 |
| SEPT11 | 1.63E-21 | 8.49E-21 | -1.63E+00 | 1.48E-06 | 6.41E-06 | -1.07E+00 | 3.65E-07 | 1.55E-06 | -1.43E+00 | 8.98E-02 | 3.43E-01 | 1.02E+00 | 5.01E-07 | 1.79E-06 | -1.34E+00 | 1.54E-02 | 4.34E-02 | -1.06E+00 | 8.66E-02 | 3.18E-01 | 1.01E+00 | 2.21E-01 | 6.06E-01 | 1.06E+00 |
| DKC1 | 1.77E-21 | 9.22E-21 | -1.64E+00 | 6.39E-04 | 2.76E-03 | 1.14E+00 | 2.33E-06 | 9.90E-06 | -1.28E+00 | 2.33E-01 | 8.90E-01 | 1.25E+00 | 1.82E-13 | 6.50E-13 | -1.58E+00 | 2.71E-01 | 7.63E-01 | 1.26E+00 | 1.88E-06 | 6.89E-06 | -1.32E+00 | 7.22E-02 | 1.98E-01 | 1.27E+00 |
| MAGEF1 | 2.00E-21 | 1.04E-20 | -2.07E+00 | 1.44E-07 | 6.24E-07 | -1.07E+00 | 4.18E-15 | 1.77E-14 | -1.85E+00 | 3.93E-09 | 1.50E-08 | -1.18E+00 | 2.31E-10 | 8.23E-10 | -1.62E+00 | 5.27E-02 | 1.49E-01 | 1.18E+00 | 4.29E-07 | 1.57E-06 | -1.34E+00 | 3.32E-03 | 9.11E-03 | -1.05E+00 |
| DDX24 | 2.01E-21 | 1.05E-20 | -1.51E+00 | 1.68E-19 | 7.27E-19 | -1.34E+00 | 3.01E-10 | 1.28E-09 | -1.52E+00 | 1.92E-12 | 7.32E-12 | -1.37E+00 | 5.40E-11 | 1.92E-10 | -1.47E+00 | 3.36E-11 | 9.48E-11 | -1.43E+00 | 8.06E-12 | 2.96E-11 | -1.61E+00 | 1.05E-04 | 2.89E-04 | -1.32E+00 |
| NDUFC1 | 2.02E-21 | 1.06E-20 | -1.53E+00 | 2.77E-23 | 1.20E-22 | -1.43E+00 | 7.80E-09 | 3.31E-08 | -1.43E+00 | 1.99E-15 | 7.61E-15 | -1.55E+00 | 5.01E-15 | 1.79E-14 | -1.71E+00 | 8.60E-11 | 2.42E-10 | -1.46E+00 | 9.27E-12 | 3.40E-11 | -1.52E+00 | 1.67E-08 | 4.59E-08 | -1.36E+00 |
| LYSMD2 | 2.06E-21 | 1.07E-20 | -1.50E+00 | 1.16E-09 | 5.02E-09 | -1.06E+00 | 5.67E-17 | 2.41E-16 | -1.72E+00 | 9.17E-06 | 3.50E-05 | -1.12E+00 | 3.15E-05 | 1.12E-04 | -1.31E+00 | 6.13E-03 | 1.73E-02 | -1.04E+00 | 1.42E-08 | 5.21E-08 | -1.27E+00 | 1.10E-04 | 3.03E-04 | -1.14E+00 |
| USP1 | 2.13E-21 | 1.11E-20 | -1.74E+00 | 1.19E-14 | 5.16E-14 | -1.26E+00 | 4.56E-03 | 1.94E-02 | -1.02E+00 | 1.96E-02 | 7.48E-02 | 1.09E+00 | 6.16E-12 | 2.20E-11 | -1.82E+00 | 8.00E-07 | 2.26E-06 | -1.30E+00 | 2.13E-09 | 7.83E-09 | -1.30E+00 | 2.34E-03 | 6.42E-03 | 1.05E+00 |
| NOC3L | 2.14E-21 | 1.11E-20 | -2.00E+00 | 6.82E-11 | 2.94E-10 | -1.28E+00 | 6.59E-12 | 2.80E-11 | -1.57E+00 | 6.39E-07 | 2.44E-06 | -1.09E+00 | 7.97E-14 | 2.84E-13 | -2.21E+00 | 7.81E-05 | 2.20E-04 | -1.18E+00 | 6.31E-16 | 2.32E-15 | -1.60E+00 | 7.25E-04 | 1.99E-03 | 1.04E+00 |
| SUDS3 | 2.18E-21 | 1.14E-20 | -1.89E+00 | 4.67E-16 | 2.02E-15 | -1.47E+00 | 2.24E-08 | 9.52E-08 | -1.43E+00 | 1.72E-08 | 6.59E-08 | -1.22E+00 | 1.68E-04 | 6.01E-04 | -1.29E+00 | 4.69E-05 | 1.32E-04 | -1.27E+00 | 3.86E-09 | 1.42E-08 | -1.31E+00 | 2.71E-12 | 7.44E-12 | -1.48E+00 |
| RBM8A | 2.33E-21 | 1.21E-20 | -1.36E+00 | 2.43E-20 | 1.05E-19 | -1.20E+00 | 2.42E-03 | 1.03E-02 | -1.09E+00 | 1.52E-03 | 5.81E-03 | -1.04E+00 | 1.76E-06 | 6.27E-06 | -1.17E+00 | 7.67E-06 | 2.16E-05 | -1.13E+00 | 2.87E-05 | 1.05E-04 | -1.18E+00 | 1.91E-02 | 5.25E-02 | -1.05E+00 |
| UBA7 | 2.33E-21 | 1.22E-20 | -1.74E+00 | 9.08E-16 | 3.92E-15 | -1.34E+00 | 1.57E-07 | 6.65E-07 | -1.53E+00 | 4.66E-07 | 1.78E-06 | -1.25E+00 | 9.43E-09 | 3.36E-08 | -1.34E+00 | 4.91E-09 | 1.38E-08 | -1.38E+00 | 2.46E-12 | 9.01E-12 | -1.49E+00 | 1.04E-08 | 2.86E-08 | -1.45E+00 |
| ABCE1 | 2.43E-21 | 1.27E-20 | -1.67E+00 | 6.39E-11 | 2.76E-10 | -1.12E+00 | 6.38E-07 | 2.71E-06 | -1.42E+00 | 3.35E-03 | 1.28E-02 | -1.00E+00 | 7.65E-08 | 2.73E-07 | -1.49E+00 | 1.17E-02 | 3.30E-02 | 1.10E+00 | 2.06E-12 | 7.57E-12 | -1.62E+00 | 2.08E-01 | 5.72E-01 | 1.08E+00 |
| APOL6 | 2.47E-21 | 1.29E-20 | -1.91E+00 | 5.29E-26 | 2.29E-25 | -1.90E+00 | 2.86E-20 | 1.21E-19 | -1.96E+00 | 1.16E-21 | 4.44E-21 | -1.84E+00 | 1.01E-12 | 3.59E-12 | -2.90E+00 | 1.78E-11 | 5.02E-11 | -2.33E+00 | 1.19E-18 | 4.35E-18 | -2.40E+00 | 7.37E-14 | 2.03E-13 | -2.00E+00 |
| CALM2 | 2.50E-21 | 1.30E-20 | -1.35E+00 | 2.17E-03 | 9.36E-03 | 1.39E+00 | 5.64E-04 | 2.40E-03 | -1.18E+00 | 2.16E-02 | 8.25E-02 | 1.25E+00 | 1.20E-12 | 4.28E-12 | -1.46E+00 | 3.61E-02 | 1.02E-01 | 1.24E+00 | 2.47E-05 | 9.05E-05 | -1.32E+00 | 1.38E-01 | 3.79E-01 | 1.12E+00 |
| MARCKS | 2.54E-21 | 1.32E-20 | -2.37E+00 | 2.53E-19 | 1.10E-18 | -2.07E+00 | 6.94E-26 | 2.95E-25 | -2.47E+00 | 9.58E-23 | 3.66E-22 | -2.29E+00 | 5.36E-25 | 1.91E-24 | -8.60E+00 | 2.62E-24 | 7.40E-24 | -5.79E+00 | 4.23E-27 | 1.55E-26 | -7.02E+00 | 8.40E-21 | 2.31E-20 | -3.63E+00 |
| PREX2 | 2.62E-21 | 1.36E-20 | -2.12E+00 | 7.59E-15 | 3.28E-14 | -1.46E+00 | 3.29E-02 | 1.40E-01 | -1.24E+00 | 2.67E-02 | 1.02E-01 | -1.10E+00 | 2.50E-01 | 8.90E-01 | 1.23E+00 | 3.55E-01 | 1.00E+00 | -1.00E+00 | 2.29E-01 | 8.40E-01 | -1.13E+00 | 3.08E-01 | 8.46E-01 | -1.06E+00 |
| HLA-DMA | 3.24E-21 | 1.69E-20 | -1.67E+00 | 1.78E-09 | 7.70E-09 | -1.23E+00 | 5.39E-10 | 2.29E-09 | -1.63E+00 | 1.37E-04 | 5.23E-04 | -1.24E+00 | 2.19E-18 | 7.80E-18 | -1.65E+00 | 1.37E-06 | 3.87E-06 | -1.16E+00 | 2.44E-27 | 8.95E-27 | -1.83E+00 | 5.82E-11 | 1.60E-10 | -1.42E+00 |
| APBB1IP | 3.36E-21 | 1.75E-20 | -1.58E+00 | 1.05E-11 | 4.56E-11 | -1.16E+00 | 1.39E-08 | 5.90E-08 | -1.44E+00 | 7.32E-09 | 2.80E-08 | -1.29E+00 | 3.68E-06 | 1.31E-05 | -1.19E+00 | 3.62E-05 | 1.02E-04 | -1.14E+00 | 4.36E-12 | 1.60E-11 | -1.76E+00 | 2.26E-07 | 6.21E-07 | -1.42E+00 |
| ZRANB2 | 4.06E-21 | 2.11E-20 | -1.39E+00 | 3.49E-03 | 1.51E-02 | 1.13E+00 | 1.52E-09 | 6.46E-09 | -1.49E+00 | 8.95E-04 | 3.42E-03 | -1.10E+00 | 5.25E-09 | 1.87E-08 | -1.29E+00 | 6.86E-03 | 1.93E-02 | -1.00E+00 | 1.87E-06 | 6.85E-06 | -1.27E+00 | 5.77E-02 | 1.59E-01 | -1.07E+00 |
| RPS2P46 | 4.71E-21 | 2.46E-20 | 1.35E+00 | 8.77E-72 | 3.79E-71 | 1.82E+00 | 4.82E-16 | 2.05E-15 | 1.42E+00 | 3.26E-46 | 1.25E-45 | 1.85E+00 | 7.49E-15 | 2.67E-14 | 1.45E+00 | 2.04E-46 | 5.75E-46 | 1.95E+00 | 1.02E-09 | 3.74E-09 | 1.41E+00 | 2.08E-42 | 5.72E-42 | 1.83E+00 |
| JUN | 4.89E-21 | 2.55E-20 | 2.12E+00 | 1.09E-01 | 4.72E-01 | 1.09E+00 | 8.55E-17 | 3.63E-16 | 2.09E+00 | 2.34E-01 | 8.94E-01 | 1.06E+00 | 6.05E-18 | 2.16E-17 | 2.16E+00 | 1.55E-02 | 4.37E-02 | 1.21E+00 | 1.56E-05 | 5.74E-05 | 1.64E+00 | 8.17E-04 | 2.25E-03 | -1.14E+00 |
| RPL17P36 | 4.95E-21 | 2.58E-20 | 1.35E+00 | 1.11E-49 | 4.78E-49 | 1.59E+00 | 2.12E-18 | 9.02E-18 | 1.40E+00 | 2.34E-37 | 8.94E-37 | 1.63E+00 | 1.66E-14 | 5.91E-14 | 1.41E+00 | 1.02E-30 | 2.87E-30 | 1.63E+00 | 1.22E-09 | 4.48E-09 | 1.35E+00 | 7.05E-35 | 1.94E-34 | 1.62E+00 |
| PSMB8-AS1 | 5.00E-21 | 2.61E-20 | -1.90E+00 | 2.59E-11 | 1.12E-10 | -1.22E+00 | 5.91E-11 | 2.51E-10 | -1.71E+00 | 3.52E-05 | 1.35E-04 | -1.15E+00 | 7.31E-17 | 2.60E-16 | -2.48E+00 | 6.47E-09 | 1.82E-08 | -1.34E+00 | 3.83E-23 | 1.41E-22 | -2.19E+00 | 1.58E-17 | 4.34E-17 | -1.77E+00 |
| ZNF146 | 5.55E-21 | 2.89E-20 | -1.85E+00 | 5.60E-13 | 2.42E-12 | -1.26E+00 | 1.07E-17 | 4.53E-17 | -2.00E+00 | 4.88E-09 | 1.87E-08 | -1.26E+00 | 4.33E-12 | 1.55E-11 | -2.16E+00 | 1.94E-02 | 5.47E-02 | 1.07E+00 | 1.44E-16 | 5.30E-16 | -1.79E+00 | 1.65E-04 | 4.53E-04 | -1.10E+00 |
| MRPL18 | 5.81E-21 | 3.03E-20 | -1.40E+00 | 3.46E-05 | 1.50E-04 | 1.15E+00 | 1.44E-03 | 6.14E-03 | -1.13E+00 | 1.91E-01 | 7.30E-01 | 1.24E+00 | 5.78E-16 | 2.06E-15 | -1.64E+00 | 2.78E-02 | 7.84E-02 | 1.17E+00 | 5.17E-04 | 1.90E-03 | -1.04E+00 | 3.21E-01 | 8.82E-01 | 1.21E+00 |
| C3orf80 | 6.09E-21 | 3.17E-20 | -1.73E+00 | 3.44E-05 | 1.49E-04 | 1.07E+00 | 1.12E-12 | 4.75E-12 | -1.92E+00 | 3.19E-03 | 1.22E-02 | 1.05E+00 | 6.32E-04 | 2.25E-03 | -1.35E+00 | 2.01E-02 | 5.66E-02 | -1.05E+00 | 5.68E-03 | 2.08E-02 | -1.22E+00 | 3.24E-01 | 8.90E-01 | 1.42E+00 |
| GOT2 | 6.13E-21 | 3.19E-20 | -2.26E+00 | 1.42E-10 | 6.14E-10 | -1.23E+00 | 1.92E-07 | 8.18E-07 | -1.35E+00 | 8.45E-07 | 3.23E-06 | -1.16E+00 | 1.89E-09 | 6.74E-09 | -2.27E+00 | 3.32E-04 | 9.37E-04 | -1.20E+00 | 2.51E-13 | 9.21E-13 | -1.71E+00 | 2.40E-04 | 6.58E-04 | -1.19E+00 |
| WDR1 | 6.31E-21 | 3.29E-20 | -1.73E+00 | 8.88E-08 | 3.84E-07 | -1.03E+00 | 3.44E-09 | 1.46E-08 | -1.50E+00 | 6.65E-06 | 2.54E-05 | -1.16E+00 | 2.00E-12 | 7.12E-12 | -1.44E+00 | 1.11E-02 | 3.12E-02 | 1.13E+00 | 1.29E-04 | 4.72E-04 | -1.19E+00 | 9.59E-02 | 2.63E-01 | 1.00E+00 |
| CYYR1 | 6.40E-21 | 3.34E-20 | -2.52E+00 | 9.66E-11 | 4.18E-10 | -1.36E+00 | 5.63E-05 | 2.39E-04 | -1.77E+00 | 2.25E-03 | 8.60E-03 | -1.15E+00 | 2.10E-01 | 7.49E-01 | 1.05E+00 | 2.91E-01 | 8.21E-01 | 1.09E+00 | 5.68E-02 | 2.09E-01 | -1.64E+00 | 2.31E-01 | 6.33E-01 | -1.02E+00 |
| KRCC1 | 6.60E-21 | 3.44E-20 | -1.72E+00 | 7.34E-13 | 3.17E-12 | -1.23E+00 | 1.75E-21 | 7.42E-21 | -2.45E+00 | 5.52E-11 | 2.11E-10 | -1.48E+00 | 9.54E-14 | 3.40E-13 | -2.07E+00 | 1.10E-07 | 3.11E-07 | -1.40E+00 | 1.44E-13 | 5.29E-13 | -1.73E+00 | 1.64E-07 | 4.51E-07 | -1.36E+00 |
| LAIR1 | 6.72E-21 | 3.50E-20 | -1.58E+00 | 1.59E-08 | 6.88E-08 | -1.05E+00 | 2.37E-08 | 1.00E-07 | -1.33E+00 | 9.52E-05 | 3.64E-04 | -1.00E+00 | 3.41E-06 | 1.21E-05 | -1.21E+00 | 1.89E-05 | 5.33E-05 | -1.22E+00 | 3.29E-03 | 1.21E-02 | -1.46E+00 | 8.48E-03 | 2.33E-02 | -1.41E+00 |
| TRA2A | 6.93E-21 | 3.61E-20 | -1.54E+00 | 6.14E-20 | 2.65E-19 | -1.37E+00 | 7.94E-08 | 3.38E-07 | -1.45E+00 | 2.73E-07 | 1.04E-06 | -1.31E+00 | 4.55E-13 | 1.62E-12 | -1.55E+00 | 1.68E-05 | 4.75E-05 | -1.23E+00 | 8.81E-09 | 3.23E-08 | -1.57E+00 | 1.47E-02 | 4.04E-02 | -1.18E+00 |
| SRP9 | 7.24E-21 | 3.77E-20 | -1.41E+00 | 9.83E-04 | 4.25E-03 | 1.09E+00 | 3.43E-04 | 1.46E-03 | -1.20E+00 | 4.27E-02 | 1.63E-01 | 1.02E+00 | 1.76E-17 | 6.27E-17 | -1.57E+00 | 7.99E-03 | 2.25E-02 | -1.04E+00 | 2.92E-04 | 1.07E-03 | -1.18E+00 | 3.25E-01 | 8.92E-01 | 1.11E+00 |
| RNF13 | 7.24E-21 | 3.77E-20 | -1.77E+00 | 2.30E-04 | 9.94E-04 | 1.18E+00 | 9.07E-10 | 3.85E-09 | -1.47E+00 | 8.45E-03 | 3.23E-02 | 1.10E+00 | 3.90E-12 | 1.39E-11 | -1.73E+00 | 5.92E-04 | 1.67E-03 | -1.00E+00 | 1.27E-14 | 4.65E-14 | -1.70E+00 | 7.23E-05 | 1.99E-04 | -1.23E+00 |
| CAV2 | 7.42E-21 | 3.87E-20 | -2.59E+00 | 2.54E-12 | 1.10E-11 | -1.45E+00 | 2.28E-11 | 9.67E-11 | -2.09E+00 | 4.94E-05 | 1.89E-04 | -1.14E+00 | 1.71E-13 | 6.11E-13 | -3.05E+00 | 4.26E-12 | 1.20E-11 | -2.67E+00 | 2.81E-02 | 1.03E-01 | 1.59E+00 | 6.63E-03 | 1.82E-02 | -1.01E+00 |
| AKAP9 | 7.64E-21 | 3.98E-20 | -1.53E+00 | 2.61E-16 | 1.13E-15 | -1.24E+00 | 3.72E-08 | 1.58E-07 | -1.51E+00 | 7.95E-06 | 3.04E-05 | -1.20E+00 | 1.28E-10 | 4.57E-10 | -1.56E+00 | 1.83E-07 | 5.17E-07 | -1.33E+00 | 2.43E-11 | 8.90E-11 | -1.53E+00 | 2.06E-07 | 5.67E-07 | -1.44E+00 |
| PPIL4 | 7.65E-21 | 3.99E-20 | -1.80E+00 | 9.29E-13 | 4.01E-12 | -1.22E+00 | 1.23E-11 | 5.23E-11 | -1.62E+00 | 7.01E-08 | 2.68E-07 | -1.19E+00 | 4.64E-13 | 1.65E-12 | -1.80E+00 | 1.38E-06 | 3.89E-06 | -1.25E+00 | 1.03E-10 | 3.79E-10 | -1.50E+00 | 7.37E-05 | 2.02E-04 | -1.16E+00 |
| AC020911.2 | 7.72E-21 | 4.02E-20 | 7.25E+00 | 2.73E-04 | 1.18E-03 | 3.56E+00 | 1.51E-08 | 6.41E-08 | 4.91E+00 | 3.01E-02 | 1.15E-01 | 2.40E+00 | 2.64E-03 | 9.43E-03 | 3.64E+00 | 3.33E-02 | 9.39E-02 | 3.04E+00 | 5.95E-02 | 2.18E-01 | 2.48E+00 | 2.49E-01 | 6.84E-01 | 1.61E+00 |
| SNTB1 | 7.82E-21 | 4.07E-20 | -2.00E+00 | 2.28E-15 | 9.86E-15 | -1.49E+00 | 4.05E-06 | 1.72E-05 | -1.40E+00 | 1.95E-03 | 7.46E-03 | -1.00E+00 | 9.80E-05 | 3.49E-04 | -1.25E+00 | 1.81E-05 | 5.10E-05 | -1.30E+00 | 1.22E-01 | 4.48E-01 | 1.07E+00 | 2.36E-01 | 6.48E-01 | 1.36E+00 |
| DDX6 | 8.45E-21 | 4.41E-20 | -1.56E+00 | 1.21E-14 | 5.21E-14 | -1.29E+00 | 1.73E-04 | 7.37E-04 | -1.35E+00 | 4.45E-04 | 1.70E-03 | -1.14E+00 | 4.54E-06 | 1.62E-05 | -1.29E+00 | 2.29E-08 | 6.45E-08 | -1.30E+00 | 5.38E-06 | 1.97E-05 | -1.34E+00 | 8.03E-04 | 2.21E-03 | -1.17E+00 |
| NDN | 8.51E-21 | 4.43E-20 | -1.80E+00 | 3.19E-05 | 1.38E-04 | 1.12E+00 | 5.52E-10 | 2.35E-09 | -1.49E+00 | 5.58E-03 | 2.13E-02 | 1.16E+00 | 1.49E-10 | 5.31E-10 | -1.69E+00 | 1.50E-02 | 4.22E-02 | 1.07E+00 | 1.52E-07 | 5.56E-07 | -1.28E+00 | 5.09E-02 | 1.40E-01 | 1.07E+00 |
| SMARCA5 | 8.72E-21 | 4.54E-20 | -1.46E+00 | 3.87E-17 | 1.67E-16 | -1.25E+00 | 6.22E-09 | 2.64E-08 | -1.40E+00 | 7.24E-08 | 2.77E-07 | -1.22E+00 | 5.65E-13 | 2.01E-12 | -1.59E+00 | 8.09E-07 | 2.28E-06 | -1.24E+00 | 3.36E-14 | 1.23E-13 | -1.58E+00 | 1.91E-05 | 5.26E-05 | -1.23E+00 |
| AL445433.1 | 8.78E-21 | 4.58E-20 | 2.45E+00 | 1.31E-02 | 5.68E-02 | 1.80E+00 | 5.82E-14 | 2.47E-13 | 2.59E+00 | 3.38E-04 | 1.29E-03 | 1.97E+00 | 6.78E-10 | 2.42E-09 | 2.40E+00 | 9.21E-04 | 2.60E-03 | 1.93E+00 | 6.24E-11 | 2.29E-10 | 2.25E+00 | 2.86E-04 | 7.86E-04 | 1.82E+00 |
| LIPA | 8.91E-21 | 4.64E-20 | -1.78E+00 | 4.74E-11 | 2.05E-10 | -1.20E+00 | 3.01E-07 | 1.28E-06 | -1.44E+00 | 5.10E-04 | 1.95E-03 | -1.01E+00 | 5.74E-14 | 2.05E-13 | -1.91E+00 | 2.68E-06 | 7.56E-06 | -1.18E+00 | 2.23E-04 | 8.20E-04 | -1.16E+00 | 1.64E-02 | 4.50E-02 | 1.05E+00 |
| CBX6 | 9.34E-21 | 4.87E-20 | -1.95E+00 | 4.06E-12 | 1.75E-11 | -1.31E+00 | 2.73E-16 | 1.16E-15 | -2.11E+00 | 6.82E-06 | 2.61E-05 | -1.15E+00 | 7.79E-11 | 2.78E-10 | -2.10E+00 | 3.44E-03 | 9.70E-03 | -1.09E+00 | 4.26E-09 | 1.56E-08 | -1.41E+00 | 2.32E-01 | 6.39E-01 | 1.10E+00 |
| GRINA | 9.96E-21 | 5.19E-20 | -1.82E+00 | 1.19E-14 | 5.12E-14 | -1.31E+00 | 6.72E-11 | 2.86E-10 | -1.67E+00 | 1.77E-07 | 6.77E-07 | -1.28E+00 | 8.65E-08 | 3.08E-07 | -1.76E+00 | 5.98E-07 | 1.69E-06 | -1.54E+00 | 2.45E-02 | 8.98E-02 | 1.07E+00 | 2.52E-02 | 6.93E-02 | -1.06E+00 |
| TBCA | 1.05E-20 | 5.47E-20 | -1.42E+00 | 2.86E-24 | 1.24E-23 | -1.36E+00 | 2.09E-07 | 8.86E-07 | -1.43E+00 | 8.23E-10 | 3.14E-09 | -1.44E+00 | 1.62E-16 | 5.78E-16 | -1.58E+00 | 2.06E-12 | 5.80E-12 | -1.35E+00 | 1.09E-06 | 4.01E-06 | -1.44E+00 | 3.90E-04 | 1.07E-03 | -1.39E+00 |
| NHP2P1 | 1.05E-20 | 5.47E-20 | -2.81E+00 | 8.98E-14 | 3.88E-13 | -1.65E+00 | 7.77E-19 | 3.30E-18 | -2.37E+00 | 1.21E-09 | 4.61E-09 | -1.33E+00 | 3.12E-09 | 1.11E-08 | -2.27E+00 | 1.61E-05 | 4.54E-05 | -1.30E+00 | 2.32E-25 | 8.51E-25 | -2.48E+00 | 1.87E-06 | 5.13E-06 | -1.17E+00 |
| CPT1A | 1.08E-20 | 5.61E-20 | -1.69E+00 | 4.29E-17 | 1.85E-16 | -1.34E+00 | 4.80E-13 | 2.04E-12 | -1.54E+00 | 8.78E-11 | 3.35E-10 | -1.26E+00 | 9.63E-12 | 3.43E-11 | -1.87E+00 | 2.09E-07 | 5.90E-07 | -1.30E+00 | 8.52E-14 | 3.13E-13 | -1.62E+00 | 4.56E-06 | 1.25E-05 | -1.17E+00 |
| CCT5 | 1.11E-20 | 5.77E-20 | -1.74E+00 | 7.27E-12 | 3.14E-11 | -1.23E+00 | 1.28E-12 | 5.42E-12 | -1.40E+00 | 2.11E-07 | 8.06E-07 | -1.09E+00 | 1.91E-15 | 6.81E-15 | -1.88E+00 | 3.76E-03 | 1.06E-02 | 1.03E+00 | 1.90E-11 | 6.97E-11 | -1.45E+00 | 1.98E-02 | 5.44E-02 | -1.06E+00 |
| HNRNPA3 | 1.29E-20 | 6.75E-20 | -1.42E+00 | 7.21E-09 | 3.11E-08 | -1.02E+00 | 2.60E-05 | 1.10E-04 | -1.34E+00 | 1.76E-01 | 6.71E-01 | 1.08E+00 | 6.16E-07 | 2.20E-06 | -1.15E+00 | 2.41E-02 | 6.79E-02 | 1.06E+00 | 4.69E-07 | 1.72E-06 | -1.32E+00 | 1.86E-01 | 5.10E-01 | 1.05E+00 |
| FAM96A | 1.40E-20 | 7.31E-20 | -1.63E+00 | 1.17E-18 | 5.06E-18 | -1.35E+00 | 8.40E-12 | 3.57E-11 | -1.49E+00 | 1.09E-12 | 4.15E-12 | -1.34E+00 | 2.33E-09 | 8.29E-09 | -1.65E+00 | 2.45E-06 | 6.91E-06 | -1.34E+00 | 1.80E-06 | 6.61E-06 | -1.22E+00 | 1.19E-06 | 3.26E-06 | -1.24E+00 |
| TPP1 | 1.45E-20 | 7.57E-20 | -1.72E+00 | 2.25E-10 | 9.71E-10 | -1.17E+00 | 2.94E-09 | 1.25E-08 | -1.43E+00 | 3.20E-04 | 1.22E-03 | -1.01E+00 | 1.46E-07 | 5.21E-07 | -1.44E+00 | 2.44E-04 | 6.88E-04 | -1.10E+00 | 9.22E-05 | 3.38E-04 | -1.06E+00 | 3.16E-03 | 8.68E-03 | 1.02E+00 |
| SPI1 | 1.55E-20 | 8.10E-20 | -1.60E+00 | 7.51E-12 | 3.25E-11 | -1.18E+00 | 1.74E-10 | 7.38E-10 | -1.56E+00 | 1.64E-05 | 6.26E-05 | -1.24E+00 | 1.01E-19 | 3.59E-19 | -1.93E+00 | 4.37E-13 | 1.23E-12 | -1.50E+00 | 2.96E-07 | 1.09E-06 | -1.36E+00 | 3.95E-05 | 1.08E-04 | -1.16E+00 |
| PYGO1 | 1.59E-20 | 8.28E-20 | -3.91E+00 | 4.71E-18 | 2.03E-17 | -3.02E+00 | 2.62E-21 | 1.11E-20 | -3.50E+00 | 2.47E-15 | 9.44E-15 | -2.04E+00 | 9.06E-02 | 3.23E-01 | -1.43E+00 | 1.45E-01 | 4.09E-01 | -1.36E+00 | 3.55E-07 | 1.30E-06 | -2.59E+00 | 8.77E-05 | 2.41E-04 | -1.80E+00 |
| ROCK1 | 1.65E-20 | 8.59E-20 | -1.45E+00 | 2.18E-11 | 9.41E-11 | -1.10E+00 | 1.79E-04 | 7.62E-04 | -1.27E+00 | 7.41E-06 | 2.83E-05 | -1.18E+00 | 2.28E-12 | 8.15E-12 | -1.55E+00 | 1.70E-06 | 4.78E-06 | -1.27E+00 | 2.80E-07 | 1.03E-06 | -1.34E+00 | 6.14E-05 | 1.69E-04 | -1.21E+00 |
| SLC9A3R1 | 1.70E-20 | 8.88E-20 | -1.68E+00 | 6.68E-08 | 2.88E-07 | -1.02E+00 | 1.73E-08 | 7.36E-08 | -1.37E+00 | 8.45E-06 | 3.23E-05 | -1.03E+00 | 2.66E-16 | 9.47E-16 | -2.07E+00 | 2.37E-06 | 6.68E-06 | -1.17E+00 | 5.08E-04 | 1.86E-03 | -1.17E+00 | 1.20E-02 | 3.30E-02 | -1.11E+00 |
| TMEM173 | 1.73E-20 | 9.02E-20 | -1.47E+00 | 1.59E-08 | 6.88E-08 | -1.01E+00 | 3.29E-12 | 1.40E-11 | -1.65E+00 | 1.64E-04 | 6.25E-04 | -1.12E+00 | 3.33E-09 | 1.19E-08 | -1.48E+00 | 5.65E-04 | 1.59E-03 | -1.14E+00 | 1.91E-02 | 7.01E-02 | -1.03E+00 | 2.15E-02 | 5.90E-02 | -1.04E+00 |
| HSPE1P4 | 1.84E-20 | 9.56E-20 | -1.51E+00 | 2.22E-19 | 9.58E-19 | -1.29E+00 | 4.11E-03 | 1.75E-02 | -1.16E+00 | 8.97E-05 | 3.43E-04 | -1.15E+00 | 7.91E-16 | 2.82E-15 | -1.71E+00 | 1.53E-07 | 4.30E-07 | -1.22E+00 | 2.40E-05 | 8.82E-05 | -1.17E+00 | 2.59E-01 | 7.10E-01 | -1.03E+00 |
| HLA-DQB1 | 1.86E-20 | 9.68E-20 | -1.70E+00 | 3.16E-20 | 1.36E-19 | -1.53E+00 | 7.43E-11 | 3.16E-10 | -1.69E+00 | 1.41E-10 | 5.39E-10 | -1.53E+00 | 5.27E-13 | 1.88E-12 | -1.72E+00 | 3.43E-13 | 9.68E-13 | -1.58E+00 | 1.37E-21 | 5.03E-21 | -2.48E+00 | 2.31E-13 | 6.35E-13 | -1.83E+00 |
| TNRC18 | 1.90E-20 | 9.92E-20 | -1.64E+00 | 6.12E-15 | 2.65E-14 | -1.24E+00 | 2.63E-07 | 1.12E-06 | -1.53E+00 | 1.43E-06 | 5.45E-06 | -1.18E+00 | 1.38E-08 | 4.93E-08 | -1.65E+00 | 6.50E-06 | 1.83E-05 | -1.25E+00 | 4.32E-08 | 1.59E-07 | -1.65E+00 | 2.96E-04 | 8.14E-04 | -1.29E+00 |
| PPP1CA | 1.91E-20 | 9.97E-20 | -1.36E+00 | 2.04E-07 | 8.83E-07 | 1.06E+00 | 3.05E-04 | 1.29E-03 | -1.16E+00 | 2.78E-02 | 1.06E-01 | 1.04E+00 | 6.41E-11 | 2.29E-10 | -1.46E+00 | 4.90E-04 | 1.38E-03 | -1.06E+00 | 7.44E-05 | 2.73E-04 | -1.24E+00 | 1.95E-01 | 5.35E-01 | 1.09E+00 |
| UQCRHL | 1.96E-20 | 1.02E-19 | -1.53E+00 | 4.93E-37 | 2.13E-36 | -1.60E+00 | 2.75E-06 | 1.17E-05 | -1.43E+00 | 9.18E-09 | 3.51E-08 | -1.53E+00 | 6.61E-16 | 2.36E-15 | -1.73E+00 | 1.13E-14 | 3.18E-14 | -1.68E+00 | 2.55E-09 | 9.36E-09 | -1.63E+00 | 2.42E-08 | 6.64E-08 | -1.64E+00 |
| ARPC1B | 2.04E-20 | 1.06E-19 | -1.68E+00 | 1.17E-18 | 5.04E-18 | -1.49E+00 | 1.48E-12 | 6.27E-12 | -1.52E+00 | 2.30E-21 | 8.80E-21 | -1.78E+00 | 6.96E-10 | 2.48E-09 | -1.54E+00 | 1.15E-14 | 3.25E-14 | -1.94E+00 | 4.72E-04 | 1.73E-03 | -1.06E+00 | 2.88E-09 | 7.90E-09 | -1.53E+00 |
| RPL10P16 | 2.44E-20 | 1.27E-19 | -1.30E+00 | 3.41E-28 | 1.47E-27 | -1.37E+00 | 1.31E-15 | 5.57E-15 | -1.31E+00 | 8.24E-18 | 3.15E-17 | -1.34E+00 | 2.32E-14 | 8.27E-14 | -1.35E+00 | 1.95E-24 | 5.51E-24 | -1.48E+00 | 4.41E-31 | 1.62E-30 | -1.37E+00 | 3.40E-29 | 9.35E-29 | -1.46E+00 |
| MTCO2P12 | 2.47E-20 | 1.29E-19 | -1.47E+00 | 1.44E-45 | 6.23E-45 | -2.31E+00 | 3.21E-08 | 1.36E-07 | -1.33E+00 | 3.14E-33 | 1.20E-32 | -2.25E+00 | 2.13E-14 | 7.61E-14 | -1.59E+00 | 4.86E-27 | 1.37E-26 | -2.20E+00 | 1.02E-09 | 3.75E-09 | -1.51E+00 | 1.64E-20 | 4.50E-20 | -2.13E+00 |
| CDC123 | 2.51E-20 | 1.31E-19 | -1.65E+00 | 2.84E-03 | 1.23E-02 | 1.22E+00 | 9.95E-11 | 4.23E-10 | -1.43E+00 | 8.30E-06 | 3.17E-05 | -1.06E+00 | 3.96E-19 | 1.41E-18 | -1.98E+00 | 1.64E-04 | 4.61E-04 | 1.05E+00 | 3.45E-06 | 1.27E-05 | -1.23E+00 | 2.22E-01 | 6.09E-01 | 1.12E+00 |
| PARP14 | 2.67E-20 | 1.39E-19 | -1.69E+00 | 1.61E-11 | 6.95E-11 | -1.28E+00 | 1.46E-17 | 6.19E-17 | -2.05E+00 | 1.84E-12 | 7.01E-12 | -1.60E+00 | 1.92E-22 | 6.85E-22 | -2.49E+00 | 3.62E-18 | 1.02E-17 | -2.03E+00 | 3.35E-20 | 1.23E-19 | -2.18E+00 | 3.48E-14 | 9.55E-14 | -1.94E+00 |
| PTBP3 | 2.68E-20 | 1.40E-19 | -1.51E+00 | 3.61E-12 | 1.56E-11 | -1.13E+00 | 5.75E-11 | 2.45E-10 | -1.59E+00 | 1.96E-06 | 7.48E-06 | -1.22E+00 | 5.22E-12 | 1.86E-11 | -1.41E+00 | 8.30E-09 | 2.34E-08 | -1.26E+00 | 1.10E-05 | 4.05E-05 | -1.32E+00 | 1.76E-03 | 4.83E-03 | -1.17E+00 |
| SPIN1 | 2.80E-20 | 1.46E-19 | -1.85E+00 | 8.84E-16 | 3.82E-15 | -1.42E+00 | 1.12E-10 | 4.77E-10 | -1.65E+00 | 3.91E-07 | 1.49E-06 | -1.18E+00 | 7.44E-10 | 2.65E-09 | -1.70E+00 | 9.43E-06 | 2.66E-05 | -1.26E+00 | 6.25E-06 | 2.30E-05 | -1.26E+00 | 2.56E-03 | 7.04E-03 | -1.10E+00 |
| TRIM58 | 2.82E-20 | 1.47E-19 | -1.97E+00 | 7.11E-13 | 3.07E-12 | -1.32E+00 | 3.92E-21 | 1.66E-20 | -2.52E+00 | 7.43E-12 | 2.84E-11 | -1.53E+00 | 4.30E-11 | 1.53E-10 | -1.51E+00 | 5.93E-05 | 1.67E-04 | -1.08E+00 | 6.42E-11 | 2.35E-10 | -1.17E+00 | 1.42E-07 | 3.90E-07 | -1.07E+00 |
| HACD3 | 2.88E-20 | 1.50E-19 | -1.94E+00 | 1.06E-09 | 4.59E-09 | -1.16E+00 | 1.28E-06 | 5.45E-06 | -1.29E+00 | 4.29E-03 | 1.64E-02 | 1.11E+00 | 3.09E-16 | 1.10E-15 | -2.27E+00 | 1.26E-05 | 3.55E-05 | -1.10E+00 | 2.05E-12 | 7.52E-12 | -1.45E+00 | 1.16E-01 | 3.20E-01 | 1.14E+00 |
| PHF5A | 2.92E-20 | 1.52E-19 | -1.46E+00 | 2.13E-17 | 9.21E-17 | -1.27E+00 | 4.23E-07 | 1.80E-06 | -1.28E+00 | 1.25E-07 | 4.77E-07 | -1.17E+00 | 3.75E-15 | 1.34E-14 | -1.71E+00 | 1.14E-08 | 3.21E-08 | -1.24E+00 | 1.60E-09 | 5.87E-09 | -1.39E+00 | 4.36E-04 | 1.20E-03 | -1.10E+00 |
| RPL29 | 2.98E-20 | 1.56E-19 | 1.22E+00 | 5.52E-09 | 2.38E-08 | 1.13E+00 | 9.77E-14 | 4.15E-13 | 1.23E+00 | 2.68E-07 | 1.02E-06 | 1.15E+00 | 1.99E-12 | 7.11E-12 | 1.25E+00 | 1.51E-08 | 4.25E-08 | 1.19E+00 | 9.04E-12 | 3.32E-11 | 1.26E+00 | 1.01E-06 | 2.77E-06 | 1.17E+00 |
| MKRN1 | 3.06E-20 | 1.59E-19 | -1.60E+00 | 2.17E-11 | 9.38E-11 | -1.11E+00 | 1.07E-09 | 4.53E-09 | -1.50E+00 | 1.29E-07 | 4.91E-07 | -1.18E+00 | 1.85E-11 | 6.60E-11 | -1.42E+00 | 1.91E-05 | 5.40E-05 | -1.05E+00 | 2.66E-08 | 9.75E-08 | -1.28E+00 | 1.08E-04 | 2.96E-04 | -1.12E+00 |
| ZDHHC4 | 3.22E-20 | 1.68E-19 | -1.95E+00 | 1.33E-13 | 5.76E-13 | -1.33E+00 | 8.72E-21 | 3.70E-20 | -2.35E+00 | 1.31E-12 | 5.01E-12 | -1.50E+00 | 1.14E-10 | 4.05E-10 | -1.72E+00 | 3.76E-05 | 1.06E-04 | -1.10E+00 | 1.26E-07 | 4.64E-07 | -1.37E+00 | 2.60E-04 | 7.14E-04 | -1.15E+00 |
| MRPL33 | 3.33E-20 | 1.73E-19 | -1.37E+00 | 2.02E-34 | 8.75E-34 | -1.64E+00 | 5.91E-05 | 2.51E-04 | -1.11E+00 | 4.50E-15 | 1.72E-14 | -1.52E+00 | 9.24E-09 | 3.30E-08 | -1.43E+00 | 1.52E-12 | 4.28E-12 | -1.60E+00 | 2.90E-06 | 1.06E-05 | -1.26E+00 | 2.40E-12 | 6.60E-12 | -1.58E+00 |
| CLIC1P1 | 3.34E-20 | 1.74E-19 | -2.69E+00 | 6.40E-11 | 2.77E-10 | -1.34E+00 | 2.39E-29 | 1.02E-28 | -3.34E+00 | 1.72E-16 | 6.55E-16 | -1.64E+00 | 4.63E-27 | 1.65E-26 | -4.11E+00 | 6.65E-17 | 1.87E-16 | -1.99E+00 | 1.60E-27 | 5.86E-27 | -3.20E+00 | 6.61E-09 | 1.82E-08 | -1.39E+00 |
| AP004242.1 | 3.50E-20 | 1.82E-19 | -2.57E+00 | 5.57E-18 | 2.41E-17 | -2.00E+00 | 4.45E-16 | 1.89E-15 | -2.47E+00 | 3.09E-12 | 1.18E-11 | -1.74E+00 | 7.64E-11 | 2.73E-10 | -2.22E+00 | 9.66E-08 | 2.72E-07 | -1.52E+00 | 8.17E-26 | 3.00E-25 | -2.34E+00 | 1.60E-19 | 4.41E-19 | -1.96E+00 |
| RTCB | 3.51E-20 | 1.83E-19 | -1.40E+00 | 2.94E-06 | 1.27E-05 | 1.11E+00 | 1.05E-04 | 4.45E-04 | -1.15E+00 | 1.65E-04 | 6.32E-04 | -1.01E+00 | 4.08E-03 | 1.46E-02 | -1.06E+00 | 2.00E-01 | 5.63E-01 | 1.22E+00 | 5.44E-04 | 2.00E-03 | -1.07E+00 | 2.48E-01 | 6.80E-01 | 1.16E+00 |
| DDX60 | 3.84E-20 | 2.00E-19 | -2.78E+00 | 3.88E-11 | 1.68E-10 | -1.47E+00 | 5.91E-30 | 2.51E-29 | -3.14E+00 | 8.84E-19 | 3.38E-18 | -1.81E+00 | 5.72E-10 | 2.04E-09 | -2.33E+00 | 3.25E-07 | 9.16E-07 | -1.67E+00 | 1.10E-25 | 4.04E-25 | -2.86E+00 | 3.17E-19 | 8.71E-19 | -2.27E+00 |
| OSBPL8 | 3.85E-20 | 2.01E-19 | -1.40E+00 | 2.89E-08 | 1.25E-07 | 1.01E+00 | 2.25E-06 | 9.55E-06 | -1.26E+00 | 3.34E-03 | 1.28E-02 | 1.04E+00 | 1.76E-09 | 6.28E-09 | -1.35E+00 | 4.55E-03 | 1.28E-02 | 1.01E+00 | 1.10E-05 | 4.04E-05 | -1.19E+00 | 1.39E-01 | 3.81E-01 | 1.07E+00 |
| STAT5A | 3.89E-20 | 2.03E-19 | -1.46E+00 | 7.42E-06 | 3.21E-05 | 1.07E+00 | 1.67E-10 | 7.11E-10 | -1.47E+00 | 3.09E-05 | 1.18E-04 | -1.04E+00 | 9.62E-09 | 3.43E-08 | -1.40E+00 | 2.46E-02 | 6.92E-02 | 1.06E+00 | 5.88E-06 | 2.16E-05 | -1.34E+00 | 4.37E-02 | 1.20E-01 | -1.07E+00 |
| KDM5A | 3.98E-20 | 2.07E-19 | -1.56E+00 | 3.76E-12 | 1.62E-11 | -1.14E+00 | 2.61E-08 | 1.11E-07 | -1.38E+00 | 1.84E-07 | 7.01E-07 | -1.18E+00 | 1.21E-06 | 4.32E-06 | -1.20E+00 | 1.65E-06 | 4.65E-06 | -1.18E+00 | 4.90E-07 | 1.80E-06 | -1.30E+00 | 2.01E-05 | 5.52E-05 | -1.23E+00 |
| MFSD2B | 4.04E-20 | 2.11E-19 | -1.48E+00 | 5.31E-31 | 2.29E-30 | -1.55E+00 | 4.29E-04 | 1.82E-03 | -1.27E+00 | 2.57E-09 | 9.82E-09 | -1.47E+00 | 4.92E-18 | 1.75E-17 | -1.70E+00 | 1.31E-17 | 3.70E-17 | -1.62E+00 | 1.65E-06 | 6.04E-06 | -1.44E+00 | 5.20E-04 | 1.43E-03 | -1.34E+00 |
| NOP58 | 4.12E-20 | 2.15E-19 | -1.37E+00 | 2.31E-21 | 9.99E-21 | -1.28E+00 | 7.85E-04 | 3.34E-03 | -1.16E+00 | 1.12E-05 | 4.28E-05 | -1.14E+00 | 3.13E-15 | 1.11E-14 | -1.54E+00 | 6.43E-05 | 1.81E-04 | -1.07E+00 | 1.68E-07 | 6.17E-07 | -1.38E+00 | 1.07E-01 | 2.94E-01 | -1.03E+00 |

| | | | | | | | | | | | | | | | | | | | | | | | | |
|---|---|---|---|---|---|---|---|---|---|---|---|---|---|---|---|---|---|---|---|---|---|---|---|---|
| ATP5F1 | 4.23E-20 | 2.21E-19 | -1.34E+00 | 5.74E-18 | 2.48E-17 | -1.16E+00 | 2.54E-05 | 1.08E-04 | -1.28E+00 | 7.10E-05 | 2.71E-04 | -1.22E+00 | 8.12E-09 | 2.90E-08 | -1.39E+00 | 2.89E-04 | 8.15E-04 | -1.06E+00 | 2.52E-06 | 9.25E-06 | -1.34E+00 | 1.37E-02 | 3.77E-02 | -1.12E+00 |
| EIF4G3 | 4.79E-20 | 2.50E-19 | -1.70E+00 | 7.85E-19 | 3.39E-18 | -1.48E+00 | 3.20E-10 | 1.36E-09 | -1.55E+00 | 1.85E-09 | 7.06E-09 | -1.26E+00 | 4.15E-08 | 1.48E-07 | -1.45E+00 | 3.15E-06 | 8.89E-06 | -1.23E+00 | 2.24E-01 | 8.20E-01 | 1.09E+00 | 5.39E-02 | 1.48E-01 | 1.08E+00 |
| TMEM136 | 4.91E-20 | 2.56E-19 | -1.85E+00 | 6.15E-13 | 2.66E-12 | -1.23E+00 | 6.51E-06 | 2.77E-05 | -1.78E+00 | 1.31E-03 | 5.00E-03 | -1.18E+00 | 2.63E-04 | 9.38E-04 | -1.93E+00 | 2.76E-03 | 7.79E-03 | -1.55E+00 | 1.17E-02 | 4.28E-02 | -1.51E+00 | 4.21E-02 | 1.16E-01 | -1.29E+00 |
| NOG | 5.12E-20 | 2.67E-19 | -2.60E+00 | 4.47E-08 | 1.93E-07 | -1.32E+00 | 8.73E-29 | 3.71E-28 | -4.05E+00 | 2.21E-20 | 8.42E-20 | -2.49E+00 | 2.41E-01 | 8.59E-01 | 1.66E+00 | 3.24E-01 | 9.15E-01 | 1.23E+00 | 2.16E-01 | 7.94E-01 | 1.46E+00 | 3.47E-01 | 9.54E-01 | -1.01E+00 |
| NCL | 5.17E-20 | 2.70E-19 | -1.46E+00 | 8.26E-12 | 3.57E-11 | -1.28E+00 | 1.81E-04 | 7.67E-04 | -1.28E+00 | 2.08E-02 | 7.95E-02 | -1.11E+00 | 3.46E-06 | 1.23E-05 | -1.31E+00 | 6.47E-03 | 1.82E-02 | -1.13E+00 | 7.88E-05 | 2.89E-04 | -1.14E+00 | 2.07E-01 | 5.69E-01 | 1.02E+00 |
| ANP32AP1 | 5.31E-20 | 2.77E-19 | -1.93E+00 | 1.62E-18 | 6.98E-18 | -1.62E+00 | 4.76E-09 | 2.02E-08 | -1.50E+00 | 4.94E-14 | 1.89E-13 | -1.54E+00 | 5.51E-19 | 1.97E-18 | -2.23E+00 | 2.69E-14 | 7.59E-14 | -1.64E+00 | 9.31E-28 | 3.42E-27 | -2.06E+00 | 5.21E-19 | 1.43E-18 | -1.66E+00 |
| AC098591.2 | 5.49E-20 | 2.86E-19 | -1.90E+00 | 5.27E-11 | 2.28E-10 | -1.22E+00 | 3.14E-12 | 1.33E-11 | -1.94E+00 | 3.05E-04 | 1.16E-03 | -1.10E+00 | 5.29E-10 | 1.89E-09 | -1.78E+00 | 4.48E-06 | 1.26E-05 | -1.22E+00 | 2.42E-11 | 8.87E-11 | -2.09E+00 | 4.31E-06 | 1.18E-05 | -1.35E+00 |
| CLDN15 | 5.62E-20 | 2.93E-19 | -1.49E+00 | 5.50E-06 | 2.38E-05 | 1.11E+00 | 1.35E-12 | 5.73E-12 | -1.79E+00 | 2.74E-05 | 1.05E-04 | -1.07E+00 | 4.75E-02 | 1.69E-01 | 1.04E+00 | 1.88E-01 | 5.30E-01 | 1.29E+00 | 2.60E-01 | 9.55E-01 | 1.40E+00 | 3.36E-01 | 9.22E-01 | 1.22E+00 |
| TYMP | 5.74E-20 | 2.99E-19 | -1.85E+00 | 1.05E-30 | 4.52E-30 | -2.29E+00 | 1.06E-09 | 4.49E-09 | -1.66E+00 | 3.85E-15 | 1.47E-14 | -1.91E+00 | 8.25E-12 | 2.94E-11 | -2.58E+00 | 1.24E-10 | 3.48E-10 | -2.15E+00 | 1.69E-04 | 6.21E-04 | -1.52E+00 | 1.79E-05 | 4.93E-05 | -1.82E+00 |
| AL512599.1 | 6.02E-20 | 3.14E-19 | -2.18E+00 | 7.89E-19 | 3.41E-18 | -1.82E+00 | 9.62E-12 | 4.09E-11 | -1.64E+00 | 2.68E-14 | 1.02E-13 | -1.59E+00 | 1.40E-15 | 4.98E-15 | -1.96E+00 | 1.91E-16 | 5.38E-16 | -1.87E+00 | 7.99E-10 | 2.93E-09 | -1.31E+00 | 1.04E-10 | 2.87E-10 | -1.59E+00 |
| RPS7P10 | 6.02E-20 | 3.14E-19 | -1.29E+00 | 7.05E-70 | 3.05E-69 | -1.64E+00 | 1.95E-09 | 8.28E-09 | -1.23E+00 | 1.08E-43 | 4.13E-43 | -1.61E+00 | 2.81E-15 | 1.00E-14 | -1.36E+00 | 4.62E-41 | 1.30E-40 | -1.66E+00 | 5.95E-23 | 2.18E-22 | -1.32E+00 | 8.06E-44 | 2.21E-43 | -1.64E+00 |
| ARL6IP1P2 | 6.25E-20 | 3.25E-19 | -1.63E+00 | 8.25E-17 | 3.56E-16 | -1.30E+00 | 8.40E-10 | 3.57E-09 | -1.52E+00 | 2.10E-08 | 8.01E-08 | -1.27E+00 | 3.17E-12 | 1.13E-11 | -1.78E+00 | 2.65E-10 | 7.46E-10 | -1.59E+00 | 6.39E-15 | 2.34E-14 | -1.65E+00 | 1.66E-12 | 4.56E-12 | -1.57E+00 |
| TMEM5 | 6.31E-20 | 3.29E-19 | -2.03E+00 | 3.98E-11 | 1.72E-10 | -1.27E+00 | 3.71E-08 | 1.58E-07 | -1.34E+00 | 7.05E-07 | 2.69E-06 | -1.12E+00 | 8.89E-05 | 3.17E-04 | -1.38E+00 | 3.55E-01 | 1.00E+00 | 1.37E+00 | 1.38E-08 | 5.07E-08 | -1.31E+00 | 9.71E-05 | 2.67E-04 | -1.12E+00 |
| AC068446.1 | 6.50E-20 | 3.39E-19 | -1.80E+00 | 4.47E-19 | 1.93E-18 | -1.55E+00 | 1.38E-24 | 5.84E-24 | -2.29E+00 | 2.02E-24 | 7.73E-24 | -1.98E+00 | 2.17E-09 | 7.72E-09 | -1.61E+00 | 1.39E-10 | 3.92E-10 | -1.75E+00 | 3.33E-20 | 1.22E-19 | -1.87E+00 | 7.29E-23 | 2.00E-22 | -1.94E+00 |
| EPHA1-AS1 | 6.55E-20 | 3.41E-19 | -1.84E+00 | 1.33E-15 | 5.73E-15 | -1.38E+00 | 1.47E-07 | 6.23E-07 | -1.65E+00 | 9.70E-07 | 3.71E-06 | -1.35E+00 | 2.33E-01 | 8.31E-01 | 1.10E+00 | 2.40E-01 | 6.76E-01 | -1.02E+00 | 9.43E-02 | 3.46E-01 | -1.37E+00 | 1.27E-01 | 3.50E-01 | -1.45E+00 |
| CHRM3 | 6.64E-20 | 3.46E-19 | -1.69E+00 | 6.16E-10 | 2.66E-09 | -1.13E+00 | 1.76E-11 | 7.50E-11 | -1.95E+00 | 1.80E-05 | 6.88E-05 | -1.14E+00 | 1.90E-01 | 6.78E-01 | 1.35E+00 | 3.08E-01 | 8.69E-01 | 1.53E+00 | 4.50E-02 | 1.65E-01 | -1.17E+00 | 2.08E-01 | 5.73E-01 | 1.16E+00 |
| SPTLC2 | 6.75E-20 | 3.52E-19 | -1.62E+00 | 2.38E-09 | 1.03E-08 | -1.07E+00 | 1.47E-06 | 6.23E-06 | -1.26E+00 | 8.32E-05 | 3.18E-04 | -1.03E+00 | 2.62E-07 | 9.35E-07 | -1.35E+00 | 5.23E-04 | 1.48E-03 | -1.07E+00 | 3.07E-06 | 1.13E-05 | -1.20E+00 | 1.10E-05 | 3.01E-05 | -1.18E+00 |
| TTC3 | 7.04E-20 | 3.67E-19 | -1.45E+00 | 2.08E-09 | 9.00E-09 | -1.10E+00 | 1.50E-06 | 6.38E-06 | -1.35E+00 | 7.34E-03 | 2.80E-02 | -1.01E+00 | 6.00E-10 | 2.14E-09 | -1.26E+00 | 7.80E-05 | 2.20E-04 | -1.00E+00 | 2.40E-07 | 8.82E-07 | -1.23E+00 | 5.85E-03 | 1.61E-02 | -1.04E+00 |
| LAMP1 | 7.22E-20 | 3.77E-19 | -1.46E+00 | 9.38E-11 | 4.05E-10 | -1.09E+00 | 3.82E-07 | 1.62E-06 | -1.31E+00 | 3.74E-06 | 1.43E-05 | -1.12E+00 | 1.05E-09 | 3.76E-09 | -1.29E+00 | 4.25E-06 | 1.20E-05 | -1.13E+00 | 3.46E-05 | 1.27E-04 | -1.19E+00 | 4.63E-02 | 1.27E-01 | 1.03E+00 |
| AC133435.1 | 7.47E-20 | 3.89E-19 | 2.06E+00 | 2.81E-12 | 1.22E-11 | 1.90E+00 | 1.84E-14 | 7.84E-14 | 2.14E+00 | 5.25E-09 | 2.01E-08 | 2.01E+00 | 1.02E-09 | 3.63E-09 | 1.98E+00 | 6.76E-08 | 1.91E-07 | 1.93E+00 | 9.95E-09 | 3.65E-08 | 1.86E+00 | 7.93E-09 | 2.18E-08 | 1.82E+00 |
| MPIG6B | 7.58E-20 | 3.95E-19 | -1.56E+00 | 8.86E-08 | 3.83E-07 | 1.01E+00 | 4.01E-08 | 1.70E-07 | -1.47E+00 | 5.93E-03 | 2.26E-02 | 1.02E+00 | 1.71E-06 | 6.09E-06 | -1.28E+00 | 8.78E-02 | 2.47E-01 | 1.07E+00 | 1.80E-04 | 6.61E-04 | -1.19E+00 | 1.83E-03 | 5.03E-03 | -1.26E+00 |
| CPNE8 | 7.98E-20 | 4.16E-19 | -1.73E+00 | 1.57E-10 | 6.78E-10 | -1.06E+00 | 5.15E-12 | 2.19E-11 | -1.58E+00 | 1.87E-09 | 7.14E-09 | -1.24E+00 | 7.89E-04 | 2.81E-03 | -1.33E+00 | 2.57E-02 | 7.24E-02 | -1.07E+00 | 4.16E-18 | 1.53E-17 | -2.24E+00 | 8.40E-10 | 2.31E-09 | -1.56E+00 |
| TMX1 | 8.01E-20 | 4.17E-19 | -1.54E+00 | 5.53E-11 | 2.39E-10 | -1.13E+00 | 1.18E-06 | 5.01E-06 | -1.18E+00 | 2.41E-04 | 9.20E-04 | 1.04E+00 | 2.49E-07 | 8.89E-07 | -1.45E+00 | 4.27E-02 | 1.20E-01 | 1.04E+00 | 1.28E-08 | 4.70E-08 | -1.41E+00 | 2.63E-02 | 7.23E-02 | -1.03E+00 |
| MYADM | 8.02E-20 | 4.18E-19 | 2.15E+00 | 9.02E-20 | 3.90E-19 | -1.29E+00 | 9.35E-11 | 3.97E-10 | 2.03E+00 | 5.16E-06 | 1.97E-05 | -1.09E+00 | 3.73E-17 | 1.33E-16 | 3.59E+00 | 1.31E-06 | 3.71E-06 | -1.08E+00 | 4.89E-27 | 1.79E-26 | 4.09E+00 | 5.88E-02 | 1.61E-01 | 1.10E+00 |
| ZFAND6 | 8.67E-20 | 4.52E-19 | -1.49E+00 | 1.14E-08 | 4.93E-08 | 1.03E+00 | 5.46E-11 | 2.32E-10 | -1.47E+00 | 4.42E-05 | 1.69E-04 | -1.02E+00 | 4.36E-14 | 1.55E-13 | -1.57E+00 | 1.17E-04 | 3.30E-04 | -1.05E+00 | 9.80E-11 | 3.60E-10 | -1.50E+00 | 4.78E-06 | 1.31E-05 | -1.27E+00 |
| RPS2 | 8.71E-20 | 4.54E-19 | 1.30E+00 | 3.07E-18 | 1.33E-17 | 1.29E+00 | 2.53E-16 | 1.07E-15 | 1.33E+00 | 3.90E-14 | 1.49E-13 | 1.31E+00 | 1.27E-12 | 4.51E-12 | 1.34E+00 | 9.08E-16 | 2.56E-15 | 1.38E+00 | 7.13E-10 | 2.62E-09 | 1.34E+00 | 9.94E-15 | 2.73E-14 | 1.34E+00 |
| ADSS | 8.91E-20 | 4.64E-19 | -1.67E+00 | 1.04E-13 | 4.51E-13 | -1.28E+00 | 3.83E-10 | 1.63E-09 | -1.44E+00 | 1.43E-07 | 5.48E-07 | -1.13E+00 | 1.68E-08 | 6.00E-08 | -1.42E+00 | 6.60E-04 | 1.86E-03 | 1.02E+00 | 2.24E-16 | 8.21E-16 | -1.62E+00 | 7.68E-07 | 2.11E-06 | -1.18E+00 |
| IPO9 | 9.06E-20 | 4.72E-19 | -2.00E+00 | 4.71E-08 | 2.03E-07 | -1.05E+00 | 4.56E-13 | 1.94E-12 | -1.79E+00 | 3.96E-03 | 1.51E-02 | 1.15E+00 | 6.22E-05 | 2.22E-04 | -1.34E+00 | 5.65E-02 | 1.59E-01 | 1.14E+00 | 9.24E-08 | 3.39E-07 | -1.56E+00 | 2.81E-01 | 7.72E-01 | 1.10E+00 |
| UBE2V1P2 | 9.10E-20 | 4.74E-19 | -1.88E+00 | 8.25E-26 | 3.56E-25 | -2.00E+00 | 5.97E-17 | 2.53E-16 | -1.80E+00 | 5.91E-21 | 2.26E-20 | -1.88E+00 | 3.14E-09 | 1.12E-08 | -1.62E+00 | 7.25E-11 | 2.05E-10 | -1.77E+00 | 1.63E-11 | 5.98E-11 | -1.58E+00 | 3.70E-12 | 1.02E-11 | -1.59E+00 |
| TMX2 | 9.24E-20 | 4.82E-19 | -1.84E+00 | 1.98E-12 | 8.57E-12 | -1.29E+00 | 1.17E-07 | 4.98E-07 | -1.39E+00 | 5.78E-05 | 2.21E-04 | -1.04E+00 | 7.06E-07 | 2.52E-06 | -1.24E+00 | 2.30E-03 | 6.49E-03 | 1.02E+00 | 3.23E-09 | 1.18E-08 | -1.39E+00 | 9.81E-04 | 2.69E-03 | -1.09E+00 |
| LETM1 | 9.26E-20 | 4.83E-19 | -2.15E+00 | 1.07E-10 | 4.60E-10 | -1.22E+00 | 9.37E-06 | 3.98E-05 | -1.43E+00 | 3.46E-02 | 1.32E-01 | 1.09E+00 | 4.30E-06 | 1.53E-05 | -1.55E+00 | 1.82E-02 | 5.13E-02 | 1.08E+00 | 1.07E-12 | 3.91E-12 | -1.74E+00 | 1.55E-01 | 4.26E-01 | 1.13E+00 |
| GATAD1 | 9.31E-20 | 4.85E-19 | -1.58E+00 | 3.06E-08 | 1.32E-07 | -1.06E+00 | 4.25E-09 | 1.81E-08 | -1.50E+00 | 5.66E-06 | 2.16E-05 | -1.19E+00 | 2.07E-04 | 7.39E-04 | -1.08E+00 | 1.58E-01 | 4.44E-01 | 1.23E+00 | 2.46E-10 | 9.01E-10 | -1.56E+00 | 1.32E-03 | 3.62E-03 | -1.08E+00 |
| TOMM22 | 9.39E-20 | 4.89E-19 | -1.35E+00 | 1.17E-12 | 5.07E-12 | -1.08E+00 | 2.72E-04 | 1.16E-03 | -1.16E+00 | 3.48E-03 | 1.33E-02 | -1.06E+00 | 8.31E-06 | 2.96E-05 | -1.18E+00 | 5.16E-04 | 1.46E-03 | -1.06E+00 | 2.01E-06 | 7.37E-06 | -1.23E+00 | 6.08E-02 | 1.67E-01 | -1.01E+00 |
| KDM5B | 9.48E-20 | 4.94E-19 | -1.60E+00 | 9.10E-14 | 3.93E-13 | -1.20E+00 | 1.98E-18 | 8.43E-18 | -1.94E+00 | 2.85E-09 | 1.09E-08 | -1.34E+00 | 1.02E-12 | 3.64E-12 | -1.68E+00 | 2.02E-14 | 5.70E-14 | -1.66E+00 | 1.35E-18 | 4.95E-18 | -1.83E+00 | 3.09E-13 | 8.47E-13 | -1.59E+00 |
| RBM6 | 1.03E-19 | 5.38E-19 | -1.67E+00 | 2.25E-07 | 9.73E-07 | -1.01E+00 | 2.83E-12 | 1.20E-11 | -1.62E+00 | 1.07E-06 | 4.08E-06 | -1.10E+00 | 8.48E-15 | 3.02E-14 | -1.58E+00 | 2.99E-07 | 8.44E-07 | -1.14E+00 | 1.79E-10 | 6.58E-10 | -1.55E+00 | 1.80E-03 | 4.95E-03 | -1.09E+00 |
| C14orf119 | 1.04E-19 | 5.40E-19 | -2.71E+00 | 2.87E-04 | 1.24E-03 | 1.05E+00 | 4.67E-31 | 1.98E-30 | -2.73E+00 | 3.86E-07 | 1.47E-06 | -1.04E+00 | 2.05E-14 | 7.30E-14 | -2.84E+00 | 3.20E-02 | 9.03E-02 | 1.27E+00 | 2.90E-23 | 1.07E-22 | -1.91E+00 | 1.05E-03 | 2.89E-03 | 1.01E+00 |
| TMEM159 | 1.05E-19 | 5.48E-19 | -1.54E+00 | 5.24E-08 | 2.26E-07 | 1.01E+00 | 1.51E-12 | 6.42E-12 | -1.75E+00 | 1.56E-06 | 5.95E-06 | -1.20E+00 | 1.00E-11 | 3.57E-11 | -1.58E+00 | 6.30E-09 | 1.77E-08 | -1.37E+00 | 3.46E-07 | 1.27E-06 | -1.44E+00 | 4.74E-03 | 1.30E-02 | -1.03E+00 |
| AC007238.1 | 1.08E-19 | 5.61E-19 | -1.62E+00 | 1.97E-18 | 8.50E-18 | -1.43E+00 | 1.92E-09 | 8.15E-09 | -1.41E+00 | 8.17E-20 | 3.12E-19 | -1.72E+00 | 2.25E-12 | 8.03E-12 | -1.67E+00 | 1.17E-11 | 3.29E-11 | -1.52E+00 | 1.10E-14 | 4.03E-14 | -1.58E+00 | 2.78E-16 | 7.65E-16 | -1.71E+00 |
| PDCD4 | 1.08E-19 | 5.61E-19 | -1.52E+00 | 8.59E-13 | 3.71E-12 | -1.13E+00 | 2.71E-10 | 1.15E-09 | -1.56E+00 | 4.64E-09 | 1.77E-08 | -1.30E+00 | 5.44E-09 | 1.94E-08 | -1.51E+00 | 1.06E-02 | 2.99E-02 | -1.05E+00 | 5.31E-05 | 1.95E-04 | -1.65E+00 | 6.06E-03 | 1.67E-02 | -1.37E+00 |
| ZNF32 | 1.08E-19 | 5.62E-19 | -1.67E+00 | 2.39E-14 | 1.03E-13 | -1.27E+00 | 1.80E-18 | 7.65E-18 | -1.87E+00 | 1.60E-09 | 6.13E-09 | -1.15E+00 | 1.08E-11 | 3.87E-11 | -1.62E+00 | 9.06E-07 | 2.55E-06 | -1.18E+00 | 2.68E-14 | 9.85E-14 | -1.88E+00 | 5.00E-08 | 1.37E-07 | -1.37E+00 |
| GTF2I | 1.09E-19 | 5.66E-19 | -1.69E+00 | 3.60E-14 | 1.55E-13 | -1.26E+00 | 5.52E-13 | 2.35E-12 | -1.60E+00 | 1.69E-08 | 6.44E-08 | -1.15E+00 | 6.04E-15 | 2.15E-14 | -1.77E+00 | 1.25E-11 | 3.54E-11 | -1.42E+00 | 4.90E-08 | 1.80E-07 | -1.39E+00 | 2.21E-08 | 6.08E-08 | -1.35E+00 |
| FLOT1 | 1.10E-19 | 5.74E-19 | -2.14E+00 | 2.58E-17 | 1.11E-16 | -1.76E+00 | 1.23E-09 | 5.24E-09 | -1.90E+00 | 1.73E-11 | 6.62E-11 | -2.00E+00 | 1.59E-10 | 5.66E-10 | -1.57E+00 | 5.57E-07 | 1.57E-06 | -1.32E+00 | 9.83E-07 | 3.61E-06 | -1.17E+00 | 6.18E-04 | 1.70E-03 | -1.04E+00 |
| BTN3A3 | 1.10E-19 | 5.75E-19 | -2.44E+00 | 3.04E-11 | 1.31E-10 | -1.42E+00 | 8.35E-15 | 3.55E-14 | -2.52E+00 | 1.21E-10 | 4.63E-10 | -1.78E+00 | 3.57E-14 | 1.27E-13 | -2.66E+00 | 3.92E-10 | 1.10E-09 | -1.75E+00 | 4.59E-32 | 1.69E-31 | -3.56E+00 | 6.72E-23 | 1.85E-22 | -2.41E+00 |
| ARF3 | 1.14E-19 | 5.95E-19 | -1.72E+00 | 1.67E-11 | 7.19E-11 | -1.15E+00 | 6.98E-08 | 2.96E-07 | -1.34E+00 | 9.24E-07 | 3.53E-06 | -1.14E+00 | 1.74E-08 | 6.19E-08 | -1.43E+00 | 3.28E-05 | 9.24E-05 | -1.12E+00 | 3.23E-05 | 1.19E-04 | -1.13E+00 | 3.04E-04 | 8.34E-04 | -1.12E+00 |
| CCNI | 1.20E-19 | 6.26E-19 | -1.30E+00 | 2.30E-02 | 9.94E-02 | -1.01E+00 | 3.98E-13 | 1.69E-12 | -1.35E+00 | 1.02E-02 | 3.89E-02 | -1.08E+00 | 8.05E-08 | 2.87E-07 | -1.25E+00 | 5.76E-02 | 1.62E-01 | -1.06E+00 | 4.64E-11 | 1.70E-10 | -1.28E+00 | 6.99E-03 | 1.92E-02 | -1.11E+00 |
| TRAF3IP3 | 1.20E-19 | 6.27E-19 | -2.18E+00 | 2.30E-03 | 9.93E-03 | 1.19E+00 | 2.26E-22 | 9.61E-22 | -2.47E+00 | 3.73E-09 | 1.42E-08 | -1.28E+00 | 2.93E-08 | 1.04E-07 | -2.01E+00 | 2.07E-02 | 5.82E-02 | 1.04E+00 | 6.14E-03 | 2.25E-02 | -1.47E+00 | 2.81E-01 | 7.72E-01 | 1.11E+00 |
| AC023512.1 | 1.29E-19 | 6.74E-19 | -2.70E+00 | 4.92E-18 | 2.12E-17 | -2.16E+00 | 3.28E-14 | 1.40E-13 | -2.09E+00 | 1.60E-12 | 6.12E-12 | -1.68E+00 | 8.93E-09 | 3.18E-08 | -1.76E+00 | 3.74E-09 | 1.05E-08 | -1.75E+00 | 1.09E-09 | 3.99E-09 | -1.59E+00 | 4.95E-04 | 1.36E-03 | -1.17E+00 |
| MESD | 1.29E-19 | 6.75E-19 | -1.45E+00 | 5.76E-06 | 2.49E-05 | 1.09E+00 | 1.05E-08 | 4.47E-08 | -1.33E+00 | 2.07E-04 | 7.89E-04 | 1.02E+00 | 8.00E-15 | 2.85E-14 | -1.89E+00 | 4.19E-05 | 1.18E-04 | -1.18E+00 | 1.87E-11 | 6.85E-11 | -1.48E+00 | 2.59E-02 | 7.12E-02 | -1.02E+00 |
| SNRPD3 | 1.39E-19 | 7.26E-19 | -1.27E+00 | 3.08E-14 | 1.33E-13 | -1.07E+00 | 9.41E-03 | 4.00E-02 | -1.07E+00 | 2.49E-02 | 9.50E-02 | 1.01E+00 | 5.47E-10 | 1.95E-09 | -1.31E+00 | 9.07E-05 | 2.56E-04 | -1.04E+00 | 3.41E-03 | 1.25E-02 | -1.14E+00 | 1.12E-01 | 3.09E-01 | -1.02E+00 |
| FTH1P10 | 1.47E-19 | 7.67E-19 | -1.45E+00 | 2.47E-15 | 1.07E-14 | -1.18E+00 | 2.06E-04 | 8.76E-04 | -1.18E+00 | 7.86E-02 | 3.00E-01 | 1.25E+00 | 8.33E-09 | 2.97E-08 | -1.41E+00 | 1.17E-03 | 3.29E-03 | -1.11E+00 | 1.47E-05 | 5.39E-05 | -1.35E+00 | 1.63E-01 | 4.47E-01 | 1.03E+00 |
| TMEM216 | 1.51E-19 | 7.85E-19 | -2.85E+00 | 5.82E-09 | 2.51E-08 | -1.27E+00 | 1.11E-12 | 4.72E-12 | -1.93E+00 | 8.19E-06 | 3.13E-05 | -1.10E+00 | 7.96E-10 | 2.84E-09 | -1.92E+00 | 3.64E-04 | 1.03E-03 | -1.18E+00 | 2.45E-12 | 8.99E-12 | -1.78E+00 | 1.82E-05 | 4.99E-05 | -1.18E+00 |
| PBXIP1 | 1.52E-19 | 7.90E-19 | -1.61E+00 | 1.54E-04 | 6.67E-04 | 1.10E+00 | 5.41E-09 | 2.30E-08 | -1.60E+00 | 1.54E-04 | 5.90E-04 | -1.10E+00 | 1.11E-12 | 3.94E-12 | -1.67E+00 | 1.57E-09 | 4.42E-09 | -1.37E+00 | 4.08E-04 | 1.50E-03 | -1.04E+00 | 9.35E-07 | 2.57E-06 | -1.47E+00 |
| PBX1 | 1.53E-19 | 7.95E-19 | -1.73E+00 | 5.63E-09 | 2.43E-08 | -1.12E+00 | 1.85E-09 | 7.87E-09 | -1.73E+00 | 5.51E-04 | 2.10E-03 | -1.04E+00 | 2.79E-12 | 9.95E-12 | -1.59E+00 | 2.45E-04 | 6.91E-04 | -1.19E+00 | 6.99E-02 | 2.56E-01 | 1.14E+00 | 2.68E-01 | 7.37E-01 | 1.32E+00 |
| AKR1C1 | 1.54E-19 | 8.02E-19 | 8.92E+00 | 1.36E-13 | 5.87E-13 | 7.38E+00 | 2.93E-03 | 1.25E-02 | 3.45E+00 | 1.99E-02 | 7.58E-02 | 2.96E+00 | 4.72E-05 | 1.68E-04 | 7.80E+00 | 2.38E-07 | 6.71E-07 | 1.06E+01 | 1.07E-05 | 3.93E-05 | 2.84E+00 | 8.45E-04 | 2.32E-03 | 2.56E+00 |
| SEPT7P8 | 1.59E-19 | 8.30E-19 | -2.08E+00 | 7.74E-20 | 3.35E-19 | -1.84E+00 | 4.97E-11 | 2.11E-10 | -1.64E+00 | 1.35E-11 | 5.15E-11 | -1.53E+00 | 1.10E-19 | 3.94E-19 | -2.19E+00 | 5.94E-17 | 1.67E-16 | -1.92E+00 | 9.04E-19 | 3.32E-18 | -1.73E+00 | 9.26E-18 | 2.54E-17 | -1.68E+00 |
| PLXDC2 | 1.65E-19 | 8.59E-19 | -1.61E+00 | 2.32E-07 | 1.00E-06 | -1.00E+00 | 1.24E-05 | 5.28E-05 | -1.30E+00 | 3.68E-03 | 1.41E-02 | 1.04E+00 | 7.52E-02 | 2.68E-01 | 1.15E+00 | 3.22E-01 | 9.07E-01 | 1.25E+00 | 2.09E-01 | 7.67E-01 | 1.01E+00 | 6.00E-02 | 1.65E-01 | 1.11E+00 |
| METTL2A | 1.67E-19 | 8.71E-19 | -2.00E+00 | 8.83E-16 | 3.81E-15 | -1.56E+00 | 4.27E-10 | 1.81E-09 | -1.59E+00 | 3.06E-06 | 1.17E-05 | -1.17E+00 | 7.97E-09 | 2.84E-08 | -1.71E+00 | 1.33E-04 | 3.74E-04 | -1.18E+00 | 1.50E-08 | 5.52E-08 | -1.42E+00 | 1.32E-03 | 3.62E-03 | -1.06E+00 |
| SNRPD2P1 | 1.86E-19 | 9.67E-19 | -1.94E+00 | 2.57E-25 | 1.11E-24 | -2.35E+00 | 1.36E-12 | 5.78E-12 | -1.65E+00 | 3.08E-21 | 1.18E-20 | -2.10E+00 | 3.02E-09 | 1.08E-08 | -1.73E+00 | 2.99E-13 | 8.42E-13 | -2.21E+00 | 1.92E-15 | 7.06E-15 | -1.85E+00 | 6.61E-16 | 1.82E-15 | -1.84E+00 |
| TSPYL1 | 1.87E-19 | 9.74E-19 | -1.57E+00 | 2.03E-15 | 8.76E-15 | -1.29E+00 | 1.29E-13 | 5.46E-13 | -1.87E+00 | 1.17E-10 | 4.48E-10 | -1.50E+00 | 9.43E-16 | 3.36E-15 | -1.69E+00 | 1.80E-07 | 5.08E-07 | -1.25E+00 | 4.71E-14 | 1.73E-13 | -1.70E+00 | 4.27E-05 | 1.17E-04 | -1.33E+00 |
| AC007387.1 | 1.91E-19 | 9.94E-19 | -1.58E+00 | 1.42E-30 | 6.15E-30 | -1.65E+00 | 4.32E-10 | 1.84E-09 | -1.37E+00 | 3.10E-15 | 1.18E-14 | -1.45E+00 | 1.95E-09 | 6.95E-09 | -1.33E+00 | 9.68E-10 | 2.73E-09 | -1.37E+00 | 4.70E-12 | 1.72E-11 | -1.58E+00 | 2.84E-10 | 7.80E-10 | -1.51E+00 |
| MRM2 | 1.92E-19 | 1.00E-18 | -2.52E+00 | 8.96E-08 | 3.87E-07 | -1.17E+00 | 3.06E-12 | 1.30E-11 | -1.87E+00 | 1.46E-03 | 5.58E-03 | 1.03E+00 | 2.59E-15 | 9.22E-15 | -2.75E+00 | 7.17E-06 | 2.02E-05 | -1.27E+00 | 4.65E-18 | 1.71E-17 | -1.69E+00 | 2.46E-04 | 6.75E-04 | -1.09E+00 |
| MAF1 | 1.98E-19 | 1.03E-18 | -1.37E+00 | 8.39E-07 | 3.63E-06 | 1.13E+00 | 2.82E-05 | 1.20E-04 | -1.23E+00 | 1.49E-04 | 5.71E-04 | -1.04E+00 | 3.54E-13 | 1.26E-12 | -1.38E+00 | 3.88E-04 | 1.09E-03 | -1.00E+00 | 2.05E-04 | 7.52E-04 | -1.16E+00 | 2.04E-02 | 5.61E-02 | -1.05E+00 |
| NNT-AS1 | 2.15E-19 | 1.12E-18 | -2.06E+00 | 3.66E-12 | 1.58E-11 | -1.33E+00 | 9.83E-12 | 4.18E-11 | -1.95E+00 | 6.34E-04 | 2.42E-03 | -1.10E+00 | 8.79E-17 | 3.13E-16 | -3.04E+00 | 1.92E-09 | 5.40E-09 | -1.72E+00 | 4.97E-20 | 1.82E-19 | -2.00E+00 | 3.08E-08 | 8.46E-08 | -1.33E+00 |
| AC145285.7 | 2.15E-19 | 1.12E-18 | -1.98E+00 | 1.64E-24 | 7.07E-24 | -2.34E+00 | 4.02E-12 | 1.71E-11 | -1.99E+00 | 8.36E-15 | 3.19E-14 | -2.20E+00 | 2.02E-12 | 7.19E-12 | -2.29E+00 | 1.01E-10 | 2.85E-10 | -1.77E+00 | 4.83E-19 | 1.77E-18 | -2.41E+00 | 1.07E-14 | 2.95E-14 | -1.85E+00 |
| ATP8B4 | 2.15E-19 | 1.12E-18 | -1.45E+00 | 1.29E-06 | 5.57E-06 | -1.01E+00 | 3.37E-18 | 1.43E-17 | -1.93E+00 | 1.30E-04 | 4.97E-04 | -1.13E+00 | 4.36E-06 | 1.56E-05 | -1.28E+00 | 3.42E-07 | 9.64E-07 | -1.38E+00 | 9.89E-02 | 3.63E-01 | -1.10E+00 | 2.60E-01 | 7.13E-01 | 1.11E+00 |
| RPS3AP6 | 2.31E-19 | 1.20E-18 | 1.26E+00 | 4.43E-03 | 1.91E-02 | 1.07E+00 | 2.65E-12 | 1.13E-11 | 1.25E+00 | 2.38E-03 | 9.10E-03 | 1.10E+00 | 5.68E-14 | 2.03E-13 | 1.31E+00 | 7.08E-03 | 2.00E-02 | 1.10E+00 | 1.55E-07 | 5.71E-07 | 1.24E+00 | 1.66E-01 | 4.57E-01 | 1.07E+00 |
| ITPRIPL2 | 2.32E-19 | 1.21E-18 | -2.03E+00 | 3.96E-10 | 1.71E-09 | -1.20E+00 | 1.10E-13 | 4.65E-13 | -2.08E+00 | 2.51E-05 | 9.57E-05 | -1.09E+00 | 5.86E-07 | 2.09E-06 | -2.68E+00 | 4.59E-05 | 1.29E-04 | -1.83E+00 | 4.47E-06 | 1.64E-05 | -2.75E+00 | 1.30E-04 | 3.57E-04 | -2.15E+00 |
| MIR181A1HG | 2.34E-19 | 1.22E-18 | 2.28E+00 | 2.55E-08 | 1.10E-07 | 1.92E+00 | 1.24E-02 | 5.26E-02 | 1.33E+00 | 2.86E-04 | 1.09E-03 | 1.69E+00 | 1.12E-06 | 3.99E-06 | 2.13E+00 | 1.27E-03 | 3.57E-03 | 1.78E+00 | 1.56E-01 | 5.73E-01 | 1.27E+00 | 2.49E-01 | 6.85E-01 | 1.28E+00 |
| RPL26P19 | 2.35E-19 | 1.23E-18 | -1.29E+00 | 1.24E-05 | 5.35E-05 | -1.10E+00 | 8.03E-07 | 3.41E-06 | -1.19E+00 | 3.13E-02 | 1.20E-01 | -1.05E+00 | 1.23E-05 | 4.37E-05 | -1.19E+00 | 3.31E-02 | 9.34E-02 | -1.07E+00 | 4.15E-14 | 1.52E-13 | -1.22E+00 | 2.67E-02 | 7.33E-02 | -1.06E+00 |
| RPS3AP47 | 2.35E-19 | 1.23E-18 | -1.44E+00 | 5.09E-05 | 2.20E-04 | -1.02E+00 | 9.22E-07 | 3.92E-06 | -1.38E+00 | 2.43E-01 | 9.30E-01 | 1.09E+00 | 1.11E-08 | 3.94E-08 | -1.35E+00 | 5.78E-03 | 1.63E-02 | -1.04E+00 | 5.14E-11 | 1.89E-10 | -1.39E+00 | 3.82E-02 | 1.05E-01 | 1.03E+00 |
| PBRM1 | 2.46E-19 | 1.28E-18 | -1.83E+00 | 2.31E-11 | 9.99E-11 | -1.20E+00 | 8.37E-15 | 3.56E-14 | -1.76E+00 | 1.22E-09 | 4.67E-09 | -1.28E+00 | 6.95E-20 | 2.48E-19 | -2.06E+00 | 4.94E-12 | 1.39E-11 | -1.54E+00 | 3.49E-12 | 1.28E-11 | -1.61E+00 | 1.89E-04 | 5.19E-04 | -1.24E+00 |
| C21orf33 | 2.50E-19 | 1.30E-18 | -1.40E+00 | 1.27E-13 | 5.49E-13 | -1.12E+00 | 5.65E-04 | 2.40E-03 | -1.13E+00 | 4.52E-03 | 1.73E-02 | 1.00E+00 | 6.41E-11 | 2.29E-10 | -1.45E+00 | 4.42E-04 | 1.25E-03 | -1.03E+00 | 3.47E-05 | 1.27E-04 | -1.20E+00 | 1.50E-02 | 4.11E-02 | -1.05E+00 |
| MFSD1 | 2.69E-19 | 1.40E-18 | -1.81E+00 | 5.26E-09 | 2.27E-08 | -1.13E+00 | 3.58E-11 | 1.52E-10 | -1.48E+00 | 2.58E-09 | 9.84E-09 | -1.24E+00 | 5.92E-11 | 2.11E-10 | -1.64E+00 | 2.15E-06 | 6.07E-06 | -1.21E+00 | 5.52E-12 | 2.03E-11 | -1.42E+00 | 1.28E-08 | 3.51E-08 | -1.37E+00 |
| GSTM3 | 2.70E-19 | 1.41E-18 | 3.72E+00 | 1.41E-06 | 6.10E-06 | 2.63E+00 | 2.36E-09 | 1.00E-08 | 2.74E+00 | 5.24E-03 | 2.00E-02 | 2.02E+00 | 2.81E-62 | 1.00E-61 | 3.02E+01 | 7.12E-44 | 2.01E-43 | 1.47E+01 | 9.05E-70 | 3.32E-69 | 1.33E+01 | 1.38E-47 | 3.78E-47 | 7.95E+00 |
| MCF2L | 2.73E-19 | 1.42E-18 | -2.95E+00 | 1.17E-15 | 5.04E-15 | -2.09E+00 | 1.08E-04 | 4.58E-04 | -1.79E+00 | 5.38E-04 | 2.05E-03 | -1.44E+00 | 1.70E-03 | 6.06E-03 | -2.76E+00 | 1.19E-02 | 3.36E-02 | -1.83E+00 | 3.75E-02 | 1.37E-01 | -1.79E+00 | 5.44E-02 | 1.50E-01 | -2.07E+00 |
| HLA-DQA1 | 2.74E-19 | 1.43E-18 | -1.91E+00 | 7.52E-07 | 3.25E-06 | -1.38E+00 | 6.72E-12 | 2.85E-11 | -1.88E+00 | 3.37E-06 | 1.29E-05 | -1.38E+00 | 4.19E-20 | 1.50E-19 | -1.99E+00 | 6.19E-14 | 1.75E-13 | -1.50E+00 | 7.63E-44 | 2.80E-43 | -3.17E+00 | 7.34E-19 | 2.02E-18 | -1.83E+00 |
| RPLP0P2 | 2.79E-19 | 1.46E-18 | -1.49E+00 | 2.30E-38 | 9.95E-38 | -1.68E+00 | 3.80E-07 | 1.62E-06 | -1.44E+00 | 1.05E-12 | 4.03E-12 | -1.61E+00 | 2.30E-09 | 8.21E-09 | -1.41E+00 | 3.23E-15 | 9.10E-15 | -1.70E+00 | 4.83E-11 | 1.77E-10 | -1.56E+00 | 2.52E-10 | 6.93E-10 | -1.62E+00 |
| MEAF6 | 2.91E-19 | 1.52E-18 | -1.37E+00 | 6.77E-09 | 2.92E-08 | 1.02E+00 | 2.77E-07 | 1.18E-06 | -1.27E+00 | 4.68E-04 | 1.79E-03 | 1.02E+00 | 5.08E-08 | 1.81E-07 | -1.32E+00 | 5.27E-03 | 1.48E-02 | 1.02E+00 | 1.51E-08 | 5.53E-08 | -1.31E+00 | 1.31E-02 | 3.60E-02 | -1.02E+00 |
| TIPARP | 2.97E-19 | 1.55E-18 | 2.96E+00 | 5.34E-09 | 2.31E-08 | -1.68E+00 | 2.31E-13 | 9.82E-13 | 2.86E+00 | 7.16E-07 | 2.73E-06 | -1.42E+00 | 2.17E-20 | 7.74E-20 | 3.61E+00 | 4.13E-05 | 1.17E-04 | -1.42E+00 | 4.90E-25 | 1.80E-24 | 4.43E+00 | 1.50E-03 | 4.13E-03 | -1.30E+00 |
| TRIP11 | 3.01E-19 | 1.57E-18 | -1.77E+00 | 5.65E-10 | 2.44E-09 | -1.18E+00 | 6.71E-13 | 2.85E-12 | -1.83E+00 | 1.11E-05 | 4.23E-05 | -1.17E+00 | 4.04E-17 | 1.44E-16 | -2.09E+00 | 2.02E-07 | 5.69E-07 | -1.26E+00 | 6.43E-10 | 2.36E-09 | -1.38E+00 | 2.64E-08 | 7.26E-08 | -1.42E+00 |
| RPSAP58 | 3.06E-19 | 1.59E-18 | 1.39E+00 | 3.24E-04 | 1.40E-03 | 1.15E+00 | 2.71E-16 | 1.15E-15 | 1.44E+00 | 5.87E-03 | 2.24E-02 | 1.13E+00 | 8.43E-13 | 3.01E-12 | 1.44E+00 | 3.61E-05 | 1.02E-04 | 1.22E+00 | 7.90E-10 | 2.90E-09 | 1.42E+00 | 2.36E-04 | 6.48E-04 | 1.21E+00 |
| DSC2 | 3.12E-19 | 1.63E-18 | -1.50E+00 | 1.28E-07 | 5.53E-07 | 1.04E+00 | 1.12E-12 | 4.75E-12 | -2.06E+00 | 8.82E-06 | 3.37E-05 | -1.31E+00 | 4.21E-02 | 1.50E-01 | 1.08E+00 | 7.45E-02 | 2.10E-01 | 1.17E+00 | 2.56E-01 | 9.38E-01 | 1.25E+00 | 3.56E-01 | 9.77E-01 | 1.19E+00 |
| NR1D2 | 3.15E-19 | 1.64E-18 | -1.99E+00 | 1.88E-19 | 8.12E-19 | -1.76E+00 | 1.41E-14 | 6.01E-14 | -1.93E+00 | 5.68E-15 | 2.17E-14 | -1.74E+00 | 6.20E-06 | 2.21E-05 | -1.71E+00 | 4.81E-04 | 1.36E-03 | -1.28E+00 | 1.77E-09 | 6.49E-09 | -1.52E+00 | 1.34E-10 | 3.67E-10 | -1.67E+00 |
| APOL2 | 3.17E-19 | 1.65E-18 | -2.58E+00 | 2.02E-05 | 8.73E-05 | -1.11E+00 | 3.95E-25 | 1.68E-24 | -3.33E+00 | 1.23E-12 | 4.70E-12 | -1.56E+00 | 5.17E-13 | 1.84E-12 | -2.65E+00 | 1.04E-07 | 2.95E-07 | -1.51E+00 | 2.71E-13 | 9.94E-13 | -2.66E+00 | 9.45E-06 | 2.60E-05 | -1.47E+00 |
| GULP1 | 3.19E-19 | 1.66E-18 | -2.74E+00 | 6.75E-17 | 2.92E-16 | -2.21E+00 | 4.45E-09 | 1.89E-08 | -3.00E+00 | 6.91E-05 | 2.64E-04 | -1.47E+00 | 5.34E-20 | 1.90E-19 | -3.11E+00 | 2.22E-13 | 6.26E-13 | -1.76E+00 | 1.89E-12 | 6.93E-12 | -2.11E+00 | 1.35E-03 | 3.70E-03 | -1.10E+00 |
| YTHDF2 | 3.31E-19 | 1.72E-18 | -1.39E+00 | 9.20E-22 | 3.98E-21 | -1.30E+00 | 5.73E-10 | 2.44E-09 | -1.45E+00 | 2.95E-05 | 1.13E-04 | -1.12E+00 | 1.61E-14 | 5.74E-14 | -1.65E+00 | 2.83E-08 | 7.99E-08 | -1.27E+00 | 5.11E-10 | 1.88E-09 | -1.48E+00 | 1.35E-04 | 3.71E-04 | -1.17E+00 |
| BAZ2A | 3.56E-19 | 1.85E-18 | -1.50E+00 | 3.72E-11 | 1.61E-10 | -1.07E+00 | 4.41E-12 | 1.87E-11 | -1.64E+00 | 3.72E-05 | 1.42E-04 | -1.11E+00 | 3.59E-05 | 1.28E-04 | -1.19E+00 | 2.44E-03 | 6.87E-03 | -1.00E+00 | 4.58E-05 | 1.68E-04 | -1.28E+00 | 5.93E-04 | 1.63E-03 | -1.18E+00 |
| Y_RNA | 3.67E-19 | 1.91E-18 | -1.75E+00 | 1.67E-14 | 7.22E-14 | -1.33E+00 | 2.09E-08 | 8.87E-08 | -1.41E+00 | 1.74E-05 | 6.65E-05 | -1.05E+00 | 2.25E-10 | 8.02E-10 | -1.50E+00 | 5.24E-07 | 1.48E-06 | -1.18E+00 | 8.52E-10 | 3.13E-09 | -1.51E+00 | 4.02E-08 | 1.10E-07 | -1.38E+00 |
| SDAD1 | 3.69E-19 | 1.92E-18 | -1.88E+00 | 2.91E-06 | 1.26E-05 | 1.02E+00 | 1.09E-14 | 4.62E-14 | -1.74E+00 | 1.51E-04 | 5.75E-04 | 1.08E+00 | 9.43E-09 | 3.36E-08 | -1.72E+00 | 2.93E-02 | 8.25E-02 | 1.17E+00 | 1.03E-10 | 3.76E-10 | -1.50E+00 | 1.05E-02 | 2.88E-02 | 1.05E+00 |
| SMIM14 | 3.79E-19 | 1.98E-18 | -3.57E+00 | 1.78E-09 | 7.67E-09 | -1.65E+00 | 2.45E-04 | 1.04E-03 | -1.63E+00 | 7.66E-03 | 2.93E-02 | -1.07E+00 | 4.56E-03 | 1.63E-02 | -1.59E+00 | 1.18E-02 | 3.32E-02 | -1.45E+00 | 1.44E-01 | 5.30E-01 | -1.13E+00 | 3.23E-01 | 8.87E-01 | 1.14E+00 |
| GIGYF2 | 3.84E-19 | 2.00E-18 | -1.83E+00 | 8.06E-13 | 3.48E-12 | -1.34E+00 | 5.00E-10 | 2.12E-09 | -1.66E+00 | 2.17E-06 | 8.28E-06 | -1.21E+00 | 1.01E-08 | 3.60E-08 | -1.52E+00 | 3.52E-04 | 9.92E-04 | -1.11E+00 | 1.51E-20 | 5.56E-20 | -1.98E+00 | 1.05E-09 | 2.89E-09 | -1.32E+00 |
| PDIA3P1 | 3.95E-19 | 2.06E-18 | -1.66E+00 | 5.17E-19 | 2.24E-18 | -1.50E+00 | 4.63E-10 | 1.97E-09 | -1.50E+00 | 4.27E-10 | 1.63E-09 | -1.29E+00 | 6.39E-14 | 2.28E-13 | -1.67E+00 | 2.49E-14 | 7.01E-14 | -1.61E+00 | 2.63E-08 | 9.67E-08 | -1.56E+00 | 3.32E-04 | 9.11E-04 | -1.38E+00 |
| CNBP | 3.95E-19 | 2.06E-18 | -1.28E+00 | 3.27E-02 | 1.41E-01 | 1.06E+00 | 7.69E-03 | 3.27E-02 | -1.13E+00 | 1.78E-01 | 6.78E-01 | 1.12E+00 | 5.29E-07 | 1.89E-06 | -1.17E+00 | 2.96E-01 | 8.35E-01 | 1.18E+00 | 7.07E-04 | 2.59E-03 | -1.16E+00 | 2.65E-01 | 7.29E-01 | 1.12E+00 |
| RARRES3 | 3.97E-19 | 2.07E-18 | -1.44E+00 | 3.34E-17 | 1.44E-16 | -1.28E+00 | 6.77E-07 | 2.87E-06 | -1.38E+00 | 4.37E-09 | 1.67E-08 | -1.39E+00 | 2.02E-07 | 7.20E-07 | -1.33E+00 | 3.08E-08 | 8.69E-08 | -1.29E+00 | 5.58E-09 | 2.05E-08 | -1.69E+00 | 5.41E-13 | 1.49E-12 | -1.95E+00 |
| ATP6V1A | 3.99E-19 | 2.08E-18 | -1.83E+00 | 3.03E-12 | 1.31E-11 | -1.27E+00 | 2.72E-12 | 1.16E-11 | -1.77E+00 | 9.29E-09 | 3.55E-08 | -1.40E+00 | 9.95E-13 | 3.55E-12 | -1.86E+00 | 5.31E-06 | 1.50E-05 | -1.14E+00 | 1.78E-03 | 6.52E-03 | -1.08E+00 | 1.00E-02 | 2.76E-02 | -1.10E+00 |
| BATF | 4.00E-19 | 2.08E-18 | -2.47E+00 | 1.17E-08 | 5.05E-08 | -1.17E+00 | 1.59E-16 | 6.76E-16 | -2.19E+00 | 7.38E-08 | 2.82E-07 | -1.26E+00 | 3.20E-07 | 1.14E-06 | -2.27E+00 | 1.41E-02 | 3.97E-02 | -1.10E+00 | 9.46E-09 | 3.47E-08 | -1.64E+00 | 1.91E-02 | 5.24E-02 | -1.00E+00 |
| BOD1 | 4.00E-19 | 2.08E-18 | -1.87E+00 | 1.49E-05 | 6.43E-05 | 1.05E+00 | 6.02E-11 | 2.56E-10 | -1.56E+00 | 1.87E-03 | 7.15E-03 | 1.06E+00 | 2.81E-09 | 1.00E-08 | -2.29E+00 | 4.03E-03 | 1.14E-02 | -1.11E+00 | 2.14E-13 | 7.85E-13 | -1.89E+00 | 7.85E-04 | 2.16E-03 | -1.04E+00 |
| SDCCAG8 | 4.01E-19 | 2.09E-18 | -1.52E+00 | 1.96E-10 | 8.47E-10 | -1.06E+00 | 1.66E-12 | 7.06E-12 | -1.69E+00 | 2.56E-08 | 9.79E-08 | -1.26E+00 | 4.82E-05 | 1.72E-04 | -1.13E+00 | 1.21E-03 | 3.42E-03 | 1.02E+00 | 1.67E-06 | 6.14E-06 | -1.39E+00 | 1.19E-06 | 3.28E-06 | -1.40E+00 |
| ZNRF1 | 4.32E-19 | 2.25E-18 | -1.38E+00 | 4.21E-07 | 1.82E-06 | 1.08E+00 | 6.03E-07 | 2.56E-06 | -1.24E+00 | 1.79E-03 | 6.84E-03 | 1.04E+00 | 1.68E-06 | 5.98E-06 | -1.19E+00 | 1.45E-03 | 4.10E-03 | -1.01E+00 | 2.09E-02 | 7.66E-02 | 1.09E+00 | 1.63E-01 | 4.47E-01 | 1.14E+00 |
| COPS3 | 4.34E-19 | 2.26E-18 | -1.37E+00 | 1.18E-19 | 5.12E-19 | -1.23E+00 | 1.19E-04 | 5.07E-04 | -1.15E+00 | 1.74E-06 | 6.65E-06 | -1.13E+00 | 4.96E-10 | 1.77E-09 | -1.31E+00 | 2.83E-03 | 7.99E-03 | 1.04E+00 | 7.06E-03 | 2.59E-02 | -1.06E+00 | 3.50E-01 | 9.62E-01 | 1.13E+00 |
| EBNA1BP2 | 4.72E-19 | 2.46E-18 | -1.41E+00 | 7.66E-12 | 3.31E-11 | -1.07E+00 | 3.70E-02 | 1.57E-01 | 1.07E+00 | 1.83E-01 | 6.98E-01 | 1.30E+00 | 2.28E-07 | 8.13E-07 | -1.34E+00 | 2.33E-02 | 6.57E-02 | 1.07E+00 | 5.83E-03 | 2.14E-02 | 1.10E+00 | 4.69E-02 | 1.29E-01 | 1.46E+00 |
| BCKDK | 4.74E-19 | 2.47E-18 | -1.94E+00 | 5.18E-18 | 2.24E-17 | -1.60E+00 | 2.89E-06 | 1.23E-05 | -1.25E+00 | 9.40E-06 | 3.59E-05 | -1.05E+00 | 9.55E-07 | 3.40E-06 | -1.59E+00 | 5.18E-05 | 1.46E-04 | -1.27E+00 | 1.39E-05 | 5.12E-05 | -1.16E+00 | 8.85E-03 | 2.43E-02 | 1.05E+00 |
| PTPN6 | 4.91E-19 | 2.56E-18 | -1.37E+00 | 2.35E-02 | 1.01E-01 | 1.19E+00 | 2.13E-05 | 9.04E-05 | -1.23E+00 | 3.61E-02 | 1.38E-01 | 1.09E+00 | 9.50E-04 | 3.39E-03 | -1.13E+00 | 3.49E-01 | 9.84E-01 | 1.19E+00 | 9.21E-03 | 3.38E-02 | -1.03E+00 | 3.03E-01 | 8.33E-01 | 1.14E+00 |
| GLUD1 | 4.97E-19 | 2.59E-18 | -1.38E+00 | 7.91E-16 | 3.42E-15 | -1.16E+00 | 1.43E-07 | 6.06E-07 | -1.41E+00 | 2.87E-03 | 1.10E-02 | -1.02E+00 | 2.09E-12 | 7.45E-12 | -1.79E+00 | 2.10E-05 | 5.93E-05 | -1.18E+00 | 8.92E-11 | 3.27E-10 | -1.44E+00 | 3.01E-05 | 8.28E-05 | -1.15E+00 |
| AC092641.1 | 5.10E-19 | 2.66E-18 | -3.04E+00 | 2.90E-12 | 1.25E-11 | -1.65E+00 | 1.52E-18 | 6.47E-18 | -3.00E+00 | 6.39E-11 | 2.44E-10 | -1.61E+00 | 1.02E-07 | 3.62E-07 | -2.57E+00 | 1.82E-04 | 5.13E-04 | -1.31E+00 | 2.25E-17 | 8.27E-17 | -2.78E+00 | 3.50E-10 | 9.60E-10 | -1.76E+00 |
| PPP2R1A | 5.21E-19 | 2.71E-18 | -1.58E+00 | 8.90E-11 | 3.85E-10 | -1.15E+00 | 3.61E-07 | 1.53E-06 | -1.32E+00 | 1.33E-07 | 5.09E-07 | -1.13E+00 | 2.12E-11 | 7.57E-11 | -1.56E+00 | 5.14E-05 | 1.45E-04 | -1.15E+00 | 1.56E-08 | 5.71E-08 | -1.40E+00 | 5.46E-03 | 1.50E-02 | -1.05E+00 |
| AFMID | 5.33E-19 | 2.78E-18 | -2.18E+00 | 4.14E-12 | 1.79E-11 | -1.37E+00 | 2.17E-16 | 9.24E-16 | -2.33E+00 | 3.88E-10 | 1.48E-09 | -1.51E+00 | 8.57E-11 | 3.05E-10 | -1.94E+00 | 3.13E-05 | 8.83E-05 | -1.19E+00 | 1.19E-17 | 4.38E-17 | -2.09E+00 | 1.15E-04 | 3.16E-04 | -1.17E+00 |
| CXXC5 | 5.50E-19 | 2.87E-18 | -1.43E+00 | 6.33E-03 | 2.73E-02 | 1.17E+00 | 1.39E-12 | 5.89E-12 | -1.67E+00 | 3.91E-04 | 1.49E-03 | -1.00E+00 | 1.25E-25 | 4.44E-25 | -2.23E+00 | 8.27E-05 | 2.33E-04 | -1.15E+00 | 1.60E-11 | 5.86E-11 | -1.63E+00 | 7.01E-03 | 1.92E-02 | -1.10E+00 |
| FBXO6 | 5.87E-19 | 3.06E-18 | -2.55E+00 | 6.89E-17 | 2.98E-16 | -1.94E+00 | 1.54E-13 | 6.56E-13 | -2.08E+00 | 1.15E-13 | 4.39E-13 | -1.91E+00 | 4.35E-04 | 1.55E-03 | -2.02E+00 | 1.17E-04 | 3.31E-04 | -2.85E+00 | 1.87E-06 | 6.87E-06 | -2.41E+00 | 1.89E-04 | 5.19E-04 | -1.76E+00 |
| RSAD2 | 5.99E-19 | 3.12E-18 | -3.39E+00 | 1.10E-17 | 4.75E-17 | -3.00E+00 | 1.01E-13 | 4.30E-13 | -2.97E+00 | 5.40E-13 | 2.06E-12 | -2.56E+00 | 7.79E-21 | 2.78E-20 | -3.18E+00 | 7.88E-23 | 2.22E-22 | -3.46E+00 | 7.02E-37 | 2.58E-36 | -4.31E+00 | 3.36E-37 | 9.23E-37 | -4.00E+00 |
| MRPL42 | 6.05E-19 | 3.15E-18 | -1.38E+00 | 5.93E-14 | 2.56E-13 | -1.08E+00 | 1.13E-06 | 4.80E-06 | -1.27E+00 | 1.20E-06 | 4.57E-06 | -1.13E+00 | 1.36E-09 | 4.85E-09 | -1.28E+00 | 5.30E-06 | 1.49E-05 | -1.10E+00 | 6.94E-04 | 2.55E-03 | -1.09E+00 | 3.37E-04 | 9.26E-04 | -1.08E+00 |
| ATF1 | 6.06E-19 | 3.16E-18 | -2.16E+00 | 5.37E-15 | 2.32E-14 | -1.62E+00 | 1.54E-13 | 6.52E-13 | -1.89E+00 | 1.76E-10 | 6.72E-10 | -1.44E+00 | 3.57E-09 | 1.27E-08 | -2.13E+00 | 1.41E-06 | 3.98E-06 | -1.45E+00 | 2.04E-29 | 7.47E-29 | -2.35E+00 | 4.60E-15 | 1.26E-14 | -1.57E+00 |
| PSMD12 | 6.10E-19 | 3.18E-18 | -1.52E+00 | 3.44E-16 | 1.49E-15 | -1.30E+00 | 1.45E-05 | 6.16E-05 | -1.27E+00 | 2.44E-06 | 9.31E-06 | -1.19E+00 | 1.08E-06 | 3.84E-06 | -1.20E+00 | 3.47E-08 | 9.77E-08 | -1.23E+00 | 6.82E-06 | 2.50E-05 | -1.28E+00 | 1.37E-03 | 3.76E-03 | -1.12E+00 |
| GANAB | 6.26E-19 | 3.26E-18 | -1.52E+00 | 2.04E-07 | 8.82E-07 | 1.03E+00 | 3.95E-06 | 1.68E-05 | -1.20E+00 | 1.33E-03 | 5.06E-03 | 1.12E+00 | 2.04E-11 | 7.27E-11 | -1.51E+00 | 2.47E-05 | 6.96E-05 | -1.13E+00 | 1.75E-06 | 6.42E-06 | -1.25E+00 | 4.30E-02 | 1.18E-01 | 1.08E+00 |
| SIL1 | 6.40E-19 | 3.33E-18 | -1.51E+00 | 2.90E-22 | 1.25E-21 | -1.42E+00 | 4.19E-04 | 1.78E-03 | -1.15E+00 | 4.29E-06 | 1.64E-05 | -1.07E+00 | 6.21E-05 | 2.21E-04 | 1.02E+00 | 8.37E-03 | 2.36E-02 | 1.15E+00 | 3.71E-05 | 1.36E-04 | -1.11E+00 | 2.35E-03 | 6.44E-03 | -1.06E+00 |
| TIMMDC1 | 6.43E-19 | 3.35E-18 | -1.46E+00 | 2.98E-09 | 1.29E-08 | -1.00E+00 | 3.84E-06 | 1.63E-05 | -1.27E+00 | 4.15E-04 | 1.59E-03 | -1.02E+00 | 2.19E-09 | 7.80E-09 | -1.37E+00 | 1.27E-03 | 3.59E-03 | 1.05E+00 | 1.14E-02 | 4.20E-02 | 1.01E+00 | 1.76E-01 | 4.84E-01 | 1.10E+00 |
| AP002381.1 | 6.47E-19 | 3.37E-18 | -1.32E+00 | 1.50E-20 | 6.49E-20 | -1.33E+00 | 2.14E-14 | 9.09E-14 | -1.34E+00 | 2.01E-14 | 7.67E-14 | -1.34E+00 | 4.26E-10 | 1.52E-09 | -1.30E+00 | 5.13E-08 | 1.45E-07 | -1.26E+00 | 6.60E-22 | 2.42E-21 | -1.36E+00 | 3.37E-19 | 9.26E-19 | -1.39E+00 |
| NDUFC2 | 6.84E-19 | 3.57E-18 | -1.34E+00 | 1.29E-37 | 5.59E-37 | -1.56E+00 | 7.60E-03 | 3.23E-02 | 1.03E+00 | 1.05E-10 | 4.01E-10 | -1.35E+00 | 1.03E-07 | 3.68E-07 | -1.34E+00 | 1.12E-10 | 3.16E-10 | -1.51E+00 | 9.17E-02 | 3.37E-01 | 1.07E+00 | 1.55E-03 | 4.25E-03 | -1.20E+00 |
| PSMB5 | 6.86E-19 | 3.58E-18 | -1.45E+00 | 2.46E-06 | 1.06E-05 | 1.04E+00 | 1.31E-08 | 5.58E-08 | -1.43E+00 | 8.88E-04 | 3.39E-03 | -1.08E+00 | 1.83E-11 | 6.52E-11 | -1.48E+00 | 2.82E-02 | 7.95E-02 | 1.08E+00 | 7.34E-07 | 2.69E-06 | -1.34E+00 | 3.64E-01 | 1.00E+00 | 1.09E+00 |
| MGA | 6.89E-19 | 3.59E-18 | -1.87E+00 | 1.57E-06 | 6.77E-06 | -1.01E+00 | 1.38E-17 | 5.88E-17 | -2.00E+00 | 5.31E-05 | 2.03E-04 | -1.07E+00 | 5.10E-16 | 1.82E-15 | -2.30E+00 | 5.07E-07 | 1.43E-06 | -1.21E+00 | 1.61E-15 | 5.90E-15 | -1.95E+00 | 7.25E-04 | 1.99E-03 | -1.10E+00 |
| CKAP2 | 7.34E-19 | 3.83E-18 | -1.43E+00 | 2.29E-21 | 9.91E-21 | -1.39E+00 | 3.37E-09 | 1.43E-08 | -1.37E+00 | 2.72E-09 | 1.04E-08 | -1.22E+00 | 1.84E-07 | 6.57E-07 | -1.43E+00 | 5.65E-06 | 1.59E-05 | -1.29E+00 | 8.94E-10 | 3.28E-09 | -1.47E+00 | 5.49E-07 | 1.51E-06 | -1.27E+00 |
| CTSZ | 7.48E-19 | 3.90E-18 | -1.88E+00 | 2.32E-08 | 1.00E-07 | -1.13E+00 | 7.53E-09 | 3.20E-08 | -1.42E+00 | 1.35E-06 | 5.17E-06 | -1.15E+00 | 1.88E-08 | 6.69E-08 | -1.63E+00 | 1.47E-05 | 4.14E-05 | -1.28E+00 | 1.96E-02 | 7.18E-02 | 1.48E+00 | 1.49E-01 | 4.09E-01 | 1.16E+00 |
| DHX33 | 7.71E-19 | 4.02E-18 | -2.35E+00 | 5.18E-13 | 2.24E-12 | -1.58E+00 | 6.67E-16 | 2.83E-15 | -1.95E+00 | 4.17E-08 | 1.59E-07 | -1.29E+00 | 9.94E-10 | 3.55E-09 | -2.24E+00 | 1.75E-06 | 4.92E-06 | -1.67E+00 | 2.48E-15 | 9.09E-15 | -1.72E+00 | 9.88E-04 | 2.71E-03 | -1.10E+00 |
| SPIRE1 | 7.87E-19 | 4.10E-18 | -3.11E+00 | 3.23E-12 | 1.39E-11 | -1.83E+00 | 2.35E-07 | 9.97E-07 | -2.14E+00 | 5.87E-06 | 2.24E-05 | -1.57E+00 | 9.36E-05 | 3.34E-04 | -2.06E+00 | 2.00E-04 | 5.63E-04 | -1.84E+00 | 4.52E-06 | 1.66E-05 | -2.57E+00 | 1.89E-02 | 5.18E-02 | -1.12E+00 |
| HERC5 | 8.13E-19 | 4.24E-18 | -2.21E+00 | 5.44E-14 | 2.35E-13 | -1.54E+00 | 2.45E-33 | 1.04E-32 | -2.35E+00 | 5.03E-35 | 1.92E-34 | -2.12E+00 | 5.09E-08 | 1.81E-07 | -3.27E+00 | 3.57E-09 | 1.01E-08 | -4.12E+00 | 8.17E-23 | 3.00E-22 | -5.26E+00 | 8.42E-19 | 2.31E-18 | -3.70E+00 |
| BNIP2 | 8.13E-19 | 4.24E-18 | -1.35E+00 | 1.86E-23 | 8.04E-23 | -1.31E+00 | 1.41E-03 | 6.01E-03 | -1.18E+00 | 1.86E-09 | 7.11E-09 | -1.34E+00 | 1.88E-03 | 6.72E-03 | -1.01E+00 | 8.43E-07 | 2.38E-06 | -1.20E+00 | 1.19E-02 | 4.37E-02 | 1.02E+00 | 2.67E-04 | 7.33E-04 | -1.14E+00 |
| TVP23A | 8.44E-19 | 4.40E-18 | -2.08E+00 | 1.27E-09 | 5.49E-09 | -1.20E+00 | 3.07E-13 | 1.31E-12 | -2.19E+00 | 4.55E-08 | 1.74E-07 | -1.34E+00 | 1.11E-04 | 3.95E-04 | -1.82E+00 | 5.05E-03 | 1.42E-02 | -1.25E+00 | 1.18E-02 | 4.33E-02 | -1.25E+00 | 5.80E-03 | 1.59E-02 | -1.40E+00 |
| CXorf38 | 8.54E-19 | 4.45E-18 | -1.84E+00 | 4.02E-14 | 1.74E-13 | -1.40E+00 | 2.71E-08 | 1.15E-07 | -1.60E+00 | 9.02E-06 | 3.45E-05 | -1.20E+00 | 1.04E-04 | 3.72E-04 | -1.19E+00 | 1.46E-05 | 4.10E-05 | -1.37E+00 | 2.81E-09 | 1.03E-08 | -1.32E+00 | 4.98E-06 | 1.37E-05 | -1.22E+00 |
| PPP4C | 9.02E-19 | 4.70E-18 | -1.54E+00 | 2.57E-08 | 1.11E-07 | -1.09E+00 | 5.44E-11 | 2.31E-10 | -1.41E+00 | 3.25E-06 | 1.24E-05 | -1.10E+00 | 1.69E-20 | 6.02E-20 | -1.85E+00 | 3.31E-05 | 9.32E-05 | -1.08E+00 | 4.40E-05 | 1.61E-04 | -1.18E+00 | 2.58E-01 | 7.09E-01 | 1.10E+00 |
| COPB1 | 9.11E-19 | 4.75E-18 | -1.54E+00 | 1.53E-09 | 6.59E-09 | -1.06E+00 | 3.51E-13 | 1.49E-12 | -1.53E+00 | 1.34E-08 | 5.10E-08 | -1.17E+00 | 3.25E-13 | 1.16E-12 | -1.62E+00 | 2.74E-05 | 7.73E-05 | -1.09E+00 | 1.97E-08 | 7.23E-08 | -1.36E+00 | 1.96E-03 | 5.38E-03 | -1.11E+00 |
| PLEKHB2 | 1.06E-18 | 5.50E-18 | -1.84E+00 | 2.97E-17 | 1.28E-16 | -1.57E+00 | 1.25E-09 | 5.30E-09 | -1.53E+00 | 1.48E-10 | 5.66E-10 | -1.41E+00 | 2.80E-16 | 9.97E-16 | -1.87E+00 | 4.34E-15 | 1.22E-14 | -1.62E+00 | 6.16E-15 | 2.26E-14 | -1.70E+00 | 1.77E-05 | 4.86E-05 | -1.24E+00 |
| RPS3AP4 | 1.06E-18 | 5.54E-18 | -1.47E+00 | 1.46E-09 | 6.31E-09 | -1.14E+00 | 3.00E-06 | 1.27E-05 | -1.41E+00 | 2.05E-03 | 7.83E-03 | -1.14E+00 | 6.62E-14 | 2.36E-13 | -1.55E+00 | 2.84E-06 | 8.02E-06 | -1.18E+00 | 2.15E-14 | 7.87E-14 | -1.58E+00 | 8.15E-03 | 2.24E-02 | -1.12E+00 |
| DOCK5 | 1.07E-18 | 5.60E-18 | -1.76E+00 | 1.35E-11 | 5.84E-11 | -1.27E+00 | 4.51E-08 | 1.92E-07 | -1.44E+00 | 3.50E-05 | 1.34E-04 | -1.07E+00 | 4.37E-07 | 1.56E-06 | -1.48E+00 | 2.37E-05 | 6.69E-05 | -1.28E+00 | 8.19E-11 | 3.01E-10 | -1.73E+00 | 2.53E-04 | 6.96E-04 | -1.27E+00 |
| ORAI2 | 1.07E-18 | 5.60E-18 | -1.54E+00 | 9.68E-07 | 4.18E-06 | 1.01E+00 | 4.83E-12 | 2.05E-11 | -1.72E+00 | 3.81E-03 | 1.45E-02 | 1.01E+00 | 1.03E-03 | 3.69E-03 | 1.02E+00 | 1.25E-01 | 3.53E-01 | 1.22E+00 | 4.55E-09 | 1.67E-08 | -1.29E+00 | 5.49E-02 | 1.51E-01 | 1.10E+00 |
| BMPR1B | 1.10E-18 | 5.73E-18 | -3.31E+00 | 1.28E-12 | 5.52E-12 | -2.08E+00 | 2.04E-04 | 8.67E-04 | -2.35E+00 | 1.07E-02 | 4.11E-02 | -1.39E+00 | 1.79E-01 | 6.38E-01 | -1.45E+00 | 2.92E-01 | 8.24E-01 | -1.13E+00 | 1.17E-01 | 4.31E-01 | -1.41E+00 | 1.78E-01 | 4.88E-01 | -1.43E+00 |
| TRIM24 | 1.10E-18 | 5.75E-18 | -1.52E+00 | 2.15E-09 | 9.30E-09 | -1.06E+00 | 1.43E-11 | 6.07E-11 | -1.65E+00 | 1.90E-06 | 7.26E-06 | -1.12E+00 | 3.55E-13 | 1.27E-12 | -1.44E+00 | 3.97E-05 | 1.12E-04 | -1.08E+00 | 5.08E-09 | 1.86E-08 | -1.35E+00 | 2.92E-02 | 8.01E-02 | 1.03E+00 |
| RAC2 | 1.11E-18 | 5.78E-18 | -1.43E+00 | 1.76E-03 | 7.62E-03 | -1.02E+00 | 8.32E-10 | 3.54E-09 | -1.51E+00 | 3.38E-02 | 1.29E-01 | -1.06E+00 | 1.09E-12 | 3.88E-12 | -1.60E+00 | 7.91E-04 | 2.23E-03 | -1.20E+00 | 1.60E-07 | 5.86E-07 | -1.30E+00 | 4.12E-02 | 1.13E-01 | -1.05E+00 |
| WAC-AS1 | 1.14E-18 | 5.94E-18 | -1.65E+00 | 4.36E-16 | 1.88E-15 | -1.33E+00 | 1.18E-06 | 5.02E-06 | -1.31E+00 | 9.72E-06 | 3.71E-05 | -1.15E+00 | 3.58E-10 | 1.28E-09 | -1.60E+00 | 7.04E-06 | 1.98E-05 | -1.26E+00 | 2.16E-07 | 7.93E-07 | -1.29E+00 | 3.66E-05 | 1.00E-04 | -1.22E+00 |
| MFSD1P1 | 1.14E-18 | 5.95E-18 | -3.42E+00 | 3.08E-17 | 1.33E-16 | -2.98E+00 | 1.14E-10 | 4.85E-10 | -2.12E+00 | 7.05E-16 | 2.69E-15 | -3.09E+00 | 6.80E-05 | 2.43E-04 | -1.71E+00 | 1.89E-08 | 5.31E-08 | -3.18E+00 | 3.85E-05 | 1.41E-04 | -1.71E+00 | 2.60E-04 | 7.15E-04 | -1.57E+00 |
| NOLC1 | 1.24E-18 | 6.44E-18 | -1.93E+00 | 5.51E-17 | 2.38E-16 | -1.60E+00 | 7.08E-06 | 3.01E-05 | -1.38E+00 | 1.36E-03 | 5.18E-03 | -1.06E+00 | 6.81E-09 | 2.43E-08 | -1.54E+00 | 1.63E-03 | 4.58E-03 | -1.02E+00 | 1.32E-06 | 4.84E-06 | -1.16E+00 | 3.41E-01 | 9.37E-01 | 1.30E+00 |
| ZNF397 | 1.27E-18 | 6.63E-18 | -1.81E+00 | 5.14E-02 | 2.22E-01 | 1.41E+00 | 5.32E-10 | 2.26E-09 | -1.93E+00 | 1.05E-01 | 4.01E-01 | 1.48E+00 | 9.23E-08 | 3.29E-07 | -1.55E+00 | 2.94E-01 | 8.28E-01 | 1.49E+00 | 2.39E-06 | 8.78E-06 | -1.55E+00 | 3.06E-01 | 8.40E-01 | 1.15E+00 |
| RGS2 | 1.27E-18 | 6.63E-18 | 4.16E+00 | 1.96E-01 | 8.45E-01 | 1.54E+00 | 9.59E-14 | 4.07E-13 | 3.14E+00 | 5.80E-02 | 2.21E-01 | 1.06E+00 | 1.04E-03 | 3.70E-03 | 3.86E+00 | 3.32E-03 | 9.37E-03 | -1.32E+00 | 2.52E-01 | 9.24E-01 | 1.65E+00 | 5.33E-03 | 1.46E-02 | -1.90E+00 |
| SSB | 1.31E-18 | 6.84E-18 | -1.34E+00 | 1.38E-05 | 5.96E-05 | 1.02E+00 | 1.97E-06 | 8.36E-06 | -1.37E+00 | 4.49E-03 | 1.72E-02 | -1.05E+00 | 3.15E-09 | 1.12E-08 | -1.39E+00 | 1.26E-01 | 3.56E-01 | 1.10E+00 | 1.31E-05 | 4.82E-05 | -1.26E+00 | 2.21E-01 | 6.06E-01 | 1.06E+00 |
| TAX1BP1 | 1.35E-18 | 7.06E-18 | -1.35E+00 | 8.76E-13 | 3.78E-12 | -1.11E+00 | 1.98E-08 | 8.42E-08 | -1.47E+00 | 2.53E-06 | 9.65E-06 | -1.20E+00 | 1.05E-06 | 3.74E-06 | -1.28E+00 | 5.52E-08 | 1.56E-07 | -1.29E+00 | 4.43E-05 | 1.63E-04 | -1.14E+00 | 8.79E-07 | 2.42E-06 | -1.26E+00 |
| CCDC126 | 1.36E-18 | 7.07E-18 | -2.26E+00 | 1.50E-17 | 6.49E-17 | -1.88E+00 | 3.21E-08 | 1.36E-07 | -2.09E+00 | 5.21E-07 | 1.99E-06 | -1.62E+00 | 1.95E-03 | 6.95E-03 | -1.15E+00 | 5.51E-04 | 1.55E-03 | -1.26E+00 | 2.28E-10 | 8.36E-10 | -1.92E+00 | 1.48E-08 | 4.08E-08 | -1.82E+00 |
| PRPF40A | 1.36E-18 | 7.07E-18 | -1.39E+00 | 5.90E-11 | 2.55E-10 | -1.11E+00 | 1.29E-05 | 5.47E-05 | -1.35E+00 | 6.34E-04 | 2.42E-03 | -1.16E+00 | 1.27E-13 | 4.52E-13 | -1.51E+00 | 2.23E-06 | 6.30E-06 | -1.25E+00 | 2.53E-04 | 9.30E-04 | -1.16E+00 | 1.39E-01 | 3.82E-01 | 1.01E+00 |
| BDP1 | 1.37E-18 | 7.15E-18 | -1.51E+00 | 1.38E-17 | 5.94E-17 | -1.32E+00 | 1.89E-05 | 8.03E-05 | -1.29E+00 | 2.59E-06 | 9.89E-06 | -1.22E+00 | 5.08E-06 | 1.81E-05 | -1.19E+00 | 1.77E-06 | 4.98E-06 | -1.17E+00 | 2.58E-05 | 9.47E-05 | -1.19E+00 | 1.88E-03 | 5.15E-03 | -1.12E+00 |
| FKBP1A | 1.43E-18 | 7.47E-18 | -1.47E+00 | 2.54E-16 | 1.10E-15 | -1.28E+00 | 4.22E-03 | 1.79E-02 | -1.12E+00 | 1.61E-03 | 6.14E-03 | -1.07E+00 | 1.11E-12 | 3.97E-12 | -1.47E+00 | 3.54E-07 | 9.97E-07 | -1.26E+00 | 8.90E-07 | 3.27E-06 | -1.28E+00 | 1.30E-02 | 3.56E-02 | -1.13E+00 |
| ST13P5 | 1.51E-18 | 7.89E-18 | -1.72E+00 | 1.02E-20 | 4.42E-20 | -1.68E+00 | 3.62E-06 | 1.54E-05 | -1.33E+00 | 1.92E-15 | 7.33E-15 | -1.71E+00 | 3.69E-06 | 1.32E-05 | -1.33E+00 | 4.77E-07 | 1.35E-06 | -1.43E+00 | 4.25E-06 | 1.56E-05 | -1.25E+00 | 4.02E-09 | 1.10E-08 | -1.44E+00 |
| SSTR2 | 1.57E-18 | 8.19E-18 | -1.45E+00 | 1.82E-33 | 7.87E-33 | -1.83E+00 | 4.24E-11 | 1.80E-10 | -1.40E+00 | 3.14E-27 | 1.20E-26 | -1.88E+00 | 1.05E-07 | 3.75E-07 | -1.26E+00 | 5.40E-17 | 1.52E-16 | -1.78E+00 | 3.50E-12 | 1.29E-11 | -1.61E+00 | 1.10E-21 | 3.01E-21 | -2.06E+00 |
| COX5AP2 | 1.59E-18 | 8.31E-18 | -1.75E+00 | 4.43E-22 | 1.91E-21 | -1.80E+00 | 1.03E-16 | 4.37E-16 | -1.71E+00 | 1.58E-25 | 6.03E-25 | -1.90E+00 | 2.76E-20 | 9.86E-20 | -2.22E+00 | 5.77E-21 | 1.63E-20 | -2.08E+00 | 3.02E-17 | 1.11E-16 | -1.75E+00 | 1.08E-18 | 2.97E-18 | -1.77E+00 |
| PTPN2P1 | 1.59E-18 | 8.31E-18 | -2.47E+00 | 5.36E-13 | 2.32E-12 | -1.64E+00 | 1.30E-11 | 5.52E-11 | -1.89E+00 | 9.98E-12 | 3.81E-11 | -1.73E+00 | 3.82E-03 | 1.36E-02 | -1.37E+00 | 6.75E-02 | 1.90E-01 | 1.00E+00 | 8.85E-06 | 3.25E-05 | -1.76E+00 | 7.27E-04 | 2.00E-03 | -1.32E+00 |
| FGFR1OP2 | 1.62E-18 | 8.42E-18 | -1.37E+00 | 3.86E-09 | 1.67E-08 | -1.06E+00 | 8.07E-06 | 3.43E-05 | -1.23E+00 | 1.32E-04 | 5.05E-04 | -1.03E+00 | 3.46E-05 | 1.23E-04 | -1.10E+00 | 1.14E-02 | 3.20E-02 | 1.10E+00 | 1.81E-01 | 6.66E-01 | 1.13E+00 | 1.99E-01 | 5.47E-01 | 1.03E+00 |
| PA2G4 | 1.63E-18 | 8.50E-18 | -1.32E+00 | 2.09E-15 | 9.04E-15 | -1.15E+00 | 7.89E-04 | 3.35E-03 | -1.17E+00 | 1.27E-02 | 4.84E-02 | -1.01E+00 | 2.65E-14 | 9.45E-14 | -1.61E+00 | 7.85E-06 | 2.21E-05 | -1.14E+00 | 2.10E-04 | 7.71E-04 | -1.20E+00 | 3.11E-01 | 8.55E-01 | 1.16E+00 |
| NIPBL | 1.66E-18 | 8.65E-18 | -1.39E+00 | 8.77E-09 | 3.79E-08 | -1.03E+00 | 1.88E-08 | 7.99E-08 | -1.49E+00 | 2.44E-07 | 9.32E-07 | -1.25E+00 | 2.84E-03 | 1.01E-02 | -1.11E+00 | 1.77E-02 | 5.00E-02 | 1.01E+00 | 8.10E-07 | 2.97E-06 | -1.24E+00 | 9.22E-04 | 2.53E-03 | -1.14E+00 |
| VPS13C | 1.68E-18 | 8.77E-18 | -1.43E+00 | 1.22E-10 | 5.29E-10 | -1.07E+00 | 4.83E-10 | 2.05E-09 | -1.50E+00 | 1.01E-04 | 3.87E-04 | -1.09E+00 | 2.78E-07 | 9.93E-07 | -1.32E+00 | 2.02E-07 | 5.69E-07 | -1.26E+00 | 1.59E-07 | 5.83E-07 | -1.41E+00 | 3.55E-06 | 9.74E-06 | -1.24E+00 |
| RAB5C | 1.70E-18 | 8.86E-18 | -1.51E+00 | 1.59E-10 | 6.89E-10 | -1.07E+00 | 6.59E-11 | 2.80E-10 | -1.50E+00 | 6.25E-05 | 2.39E-04 | -1.05E+00 | 8.91E-10 | 3.18E-09 | -1.38E+00 | 9.52E-06 | 2.68E-05 | -1.14E+00 | 1.09E-06 | 4.00E-06 | -1.25E+00 | 1.29E-02 | 3.53E-02 | 1.06E+00 |
| RPS19 | 1.77E-18 | 9.23E-18 | 1.26E+00 | 2.25E-01 | 9.74E-01 | 1.00E+00 | 1.71E-08 | 7.27E-08 | 1.19E+00 | 8.10E-02 | 3.09E-01 | -1.03E+00 | 4.83E-08 | 1.72E-07 | 1.22E+00 | 2.52E-01 | 7.11E-01 | 1.02E+00 | 1.44E-04 | 5.28E-04 | 1.21E+00 | 2.90E-01 | 7.97E-01 | 1.01E+00 |
| FOXP1 | 1.77E-18 | 9.23E-18 | -1.48E+00 | 3.24E-04 | 1.40E-03 | 1.05E+00 | 1.30E-08 | 5.54E-08 | -1.58E+00 | 7.98E-05 | 3.05E-04 | -1.12E+00 | 1.73E-11 | 6.17E-11 | -1.66E+00 | 1.07E-05 | 3.02E-05 | -1.37E+00 | 1.27E-13 | 4.67E-13 | -1.92E+00 | 1.18E-04 | 3.24E-04 | -1.39E+00 |
| KLHL23 | 1.77E-18 | 9.24E-18 | -1.59E+00 | 4.72E-12 | 2.04E-11 | -1.18E+00 | 6.36E-10 | 2.70E-09 | -1.54E+00 | 2.93E-06 | 1.12E-05 | -1.13E+00 | 3.72E-04 | 1.33E-03 | -1.33E+00 | 1.63E-02 | 4.61E-02 | 1.04E+00 | 2.47E-12 | 9.05E-12 | -1.72E+00 | 8.87E-04 | 2.44E-03 | -1.10E+00 |
| MTCYBP35 | 1.87E-18 | 9.73E-18 | -1.88E+00 | 9.27E-27 | 4.01E-26 | -2.31E+00 | 2.31E-20 | 9.83E-20 | -2.04E+00 | 2.69E-21 | 1.03E-20 | -1.91E+00 | 3.33E-11 | 1.19E-10 | -1.89E+00 | 1.52E-13 | 4.29E-13 | -2.10E+00 | 1.33E-21 | 4.88E-21 | -2.15E+00 | 1.24E-19 | 3.40E-19 | -1.96E+00 |
| ANGPT1 | 1.87E-18 | 9.73E-18 | -1.59E+00 | 1.05E-05 | 4.54E-05 | -1.01E+00 | 9.12E-04 | 3.88E-03 | -1.27E+00 | 1.61E-01 | 6.17E-01 | 1.17E+00 | 1.63E-06 | 5.83E-06 | -1.42E+00 | 6.93E-04 | 1.95E-03 | -1.16E+00 | 1.61E-01 | 5.90E-01 | 1.06E+00 | 2.28E-01 | 6.26E-01 | 1.17E+00 |
| AL121723.1 | 1.99E-18 | 1.04E-17 | -3.03E+00 | 1.62E-15 | 7.00E-15 | -2.00E+00 | 2.73E-07 | 1.16E-06 | -1.88E+00 | 1.80E-05 | 6.88E-05 | -1.36E+00 | 4.09E-08 | 1.46E-07 | -2.59E+00 | 7.40E-06 | 2.09E-05 | -1.70E+00 | 4.10E-06 | 1.51E-05 | -1.66E+00 | 2.22E-02 | 6.11E-02 | 1.02E+00 |
| WBP1L | 2.12E-18 | 1.11E-17 | -3.05E+00 | 5.49E-06 | 2.37E-05 | -1.03E+00 | 7.45E-11 | 3.17E-10 | -2.32E+00 | 3.52E-06 | 1.34E-05 | -1.38E+00 | 1.59E-10 | 5.68E-10 | -2.07E+00 | 1.18E-06 | 3.34E-06 | -1.30E+00 | 1.10E-09 | 4.04E-09 | -1.45E+00 | 3.58E-05 | 9.84E-05 | -1.12E+00 |
| SSPN | 2.18E-18 | 1.14E-17 | -3.16E+00 | 3.64E-06 | 1.57E-05 | -1.18E+00 | 6.12E-18 | 2.60E-17 | -3.34E+00 | 9.77E-11 | 3.73E-10 | -1.86E+00 | 7.18E-03 | 2.56E-02 | -2.63E+00 | 6.10E-02 | 1.72E-01 | -1.43E+00 | 4.80E-03 | 1.76E-02 | -2.09E+00 | 2.71E-01 | 7.45E-01 | 1.11E+00 |
| RPL21P2 | 2.22E-18 | 1.16E-17 | -1.46E+00 | 2.11E-31 | 9.13E-31 | -1.87E+00 | 4.53E-09 | 1.93E-08 | -1.36E+00 | 1.15E-20 | 4.39E-20 | -1.80E+00 | 1.75E-07 | 6.26E-07 | -1.21E+00 | 1.46E-17 | 4.13E-17 | -1.85E+00 | 4.77E-07 | 1.75E-06 | -1.25E+00 | 4.75E-17 | 1.31E-16 | -1.74E+00 |
| IWS1 | 2.26E-18 | 1.18E-17 | -1.72E+00 | 3.78E-14 | 1.63E-13 | -1.33E+00 | 1.39E-09 | 5.92E-09 | -1.52E+00 | 3.63E-05 | 1.39E-04 | -1.10E+00 | 1.35E-11 | 4.82E-11 | -1.94E+00 | 8.82E-06 | 2.49E-05 | -1.24E+00 | 2.92E-12 | 1.07E-11 | -1.65E+00 | 2.70E-03 | 7.42E-03 | -1.10E+00 |
| AC011979.1 | 2.31E-18 | 1.21E-17 | -1.47E+00 | 1.78E-47 | 7.70E-47 | -1.80E+00 | 8.07E-05 | 3.43E-04 | -1.37E+00 | 7.41E-16 | 2.83E-15 | -1.75E+00 | 9.91E-07 | 3.53E-06 | -1.37E+00 | 3.68E-14 | 1.04E-13 | -1.72E+00 | 1.70E-09 | 6.23E-09 | -1.45E+00 | 1.39E-14 | 3.81E-14 | -1.75E+00 |
| PSMC1P9 | 2.32E-18 | 1.21E-17 | -1.70E+00 | 3.89E-13 | 1.68E-12 | -1.26E+00 | 2.17E-12 | 9.20E-12 | -1.79E+00 | 3.46E-11 | 1.32E-10 | -1.48E+00 | 8.85E-18 | 3.15E-17 | -2.08E+00 | 2.33E-12 | 6.57E-12 | -1.57E+00 | 9.77E-14 | 3.59E-13 | -1.55E+00 | 3.46E-07 | 9.50E-07 | -1.20E+00 |
| REC8 | 2.41E-18 | 1.26E-17 | -1.68E+00 | 2.61E-16 | 1.13E-15 | -1.46E+00 | 9.76E-12 | 4.15E-11 | -1.63E+00 | 8.22E-13 | 3.14E-12 | -1.45E+00 | 2.98E-13 | 1.06E-12 | -1.77E+00 | 7.43E-12 | 2.09E-11 | -1.52E+00 | 8.39E-13 | 3.08E-12 | -1.74E+00 | 3.58E-06 | 9.83E-06 | -1.30E+00 |
| TNRC6C | 2.43E-18 | 1.26E-17 | -2.09E+00 | 4.67E-12 | 2.02E-11 | -1.31E+00 | 6.61E-07 | 2.81E-06 | -1.53E+00 | 1.09E-03 | 4.16E-03 | -1.04E+00 | 2.70E-07 | 9.61E-07 | -1.40E+00 | 1.03E-06 | 2.91E-06 | -1.23E+00 | 1.94E-10 | 7.13E-10 | -1.89E+00 | 4.28E-05 | 1.18E-04 | -1.25E+00 |
| PAFAH1B2 | 2.43E-18 | 1.27E-17 | -1.38E+00 | 4.70E-12 | 2.03E-11 | -1.12E+00 | 1.39E-06 | 5.89E-06 | -1.49E+00 | 2.99E-04 | 1.14E-03 | -1.17E+00 | 1.32E-10 | 4.72E-10 | -1.38E+00 | 3.04E-10 | 8.58E-10 | -1.39E+00 | 3.71E-10 | 1.36E-09 | -1.42E+00 | 9.69E-06 | 2.66E-05 | -1.26E+00 |
| ELN | 2.46E-18 | 1.28E-17 | -2.35E+00 | 7.16E-07 | 3.10E-06 | -1.16E+00 | 1.22E-01 | 5.17E-01 | -1.11E+00 | 1.92E-01 | 7.33E-01 | 1.16E+00 | 2.43E-01 | 8.67E-01 | 1.18E+00 | 3.09E-01 | 8.72E-01 | 1.53E+00 | 2.62E-01 | 9.63E-01 | 1.17E+00 | 2.09E-01 | 5.74E-01 | 1.53E+00 |
| AC073861.1 | 2.52E-18 | 1.31E-17 | 1.29E+00 | 1.39E-12 | 6.02E-12 | -1.23E+00 | 4.81E-15 | 2.04E-14 | 1.33E+00 | 6.38E-06 | 2.44E-05 | -1.17E+00 | 1.03E-09 | 3.69E-09 | 1.30E+00 | 4.30E-04 | 1.21E-03 | -1.15E+00 | 8.76E-06 | 3.21E-05 | 1.28E+00 | 8.19E-08 | 2.25E-07 | -1.19E+00 |
| CPNE1 | 2.55E-18 | 1.33E-17 | -1.42E+00 | 1.81E-07 | 7.81E-07 | -1.02E+00 | 6.96E-06 | 2.96E-05 | -1.38E+00 | 7.52E-03 | 2.87E-02 | -1.05E+00 | 1.16E-06 | 4.13E-06 | -1.31E+00 | 1.64E-05 | 4.63E-05 | -1.09E+00 | 1.75E-05 | 6.41E-05 | -1.20E+00 | 1.32E-02 | 3.64E-02 | 1.01E+00 |
| PPIAL4C | 2.57E-18 | 1.34E-17 | -2.92E+00 | 2.11E-12 | 9.13E-12 | -1.73E+00 | 4.50E-23 | 1.91E-22 | -2.81E+00 | 6.33E-17 | 2.42E-16 | -1.90E+00 | 2.57E-07 | 9.16E-07 | -2.11E+00 | 1.34E-04 | 3.79E-04 | -1.31E+00 | 1.94E-34 | 7.12E-34 | -3.06E+00 | 7.85E-17 | 2.16E-16 | -1.72E+00 |
| ISCU | 2.60E-18 | 1.35E-17 | -1.41E+00 | 4.40E-12 | 1.90E-11 | -1.08E+00 | 1.81E-04 | 7.69E-04 | -1.13E+00 | 5.55E-04 | 2.12E-03 | -1.00E+00 | 5.46E-10 | 1.95E-09 | -1.44E+00 | 3.49E-04 | 9.83E-04 | 1.02E+00 | 9.54E-07 | 3.50E-06 | -1.08E+00 | 7.80E-04 | 2.14E-03 | -1.03E+00 |
| ZNF518A | 2.70E-18 | 1.41E-17 | -1.66E+00 | 4.03E-15 | 1.74E-14 | -1.36E+00 | 1.27E-09 | 5.38E-09 | -1.47E+00 | 1.83E-06 | 6.97E-06 | -1.19E+00 | 1.20E-07 | 4.29E-07 | -1.40E+00 | 6.22E-06 | 1.75E-05 | -1.22E+00 | 1.06E-06 | 3.91E-06 | -1.21E+00 | 6.09E-04 | 1.67E-03 | -1.13E+00 |
| SMCO4 | 2.73E-18 | 1.42E-17 | -4.07E+00 | 1.06E-08 | 4.57E-08 | -1.51E+00 | 6.67E-27 | 2.83E-26 | -2.86E+00 | 9.29E-19 | 3.55E-18 | -1.89E+00 | 1.68E-01 | 5.98E-01 | -1.38E+00 | 2.97E-01 | 8.38E-01 | 1.06E+00 | 4.05E-06 | 1.49E-05 | -2.42E+00 | 4.18E-03 | 1.15E-02 | -1.39E+00 |
| SHISA5 | 2.76E-18 | 1.44E-17 | -1.55E+00 | 1.46E-12 | 6.30E-12 | -1.16E+00 | 6.71E-11 | 2.85E-10 | -1.64E+00 | 5.25E-08 | 2.01E-07 | -1.30E+00 | 1.98E-12 | 7.08E-12 | -1.67E+00 | 3.72E-08 | 1.05E-07 | -1.30E+00 | 3.22E-16 | 1.18E-15 | -1.66E+00 | 1.73E-10 | 4.75E-10 | -1.42E+00 |
| MACC1 | 2.79E-18 | 1.45E-17 | -1.97E+00 | 3.73E-13 | 1.61E-12 | -1.54E+00 | 1.03E-08 | 4.36E-08 | -1.71E+00 | 2.17E-06 | 8.28E-06 | -1.36E+00 | 1.20E-03 | 4.26E-03 | -1.57E+00 | 8.13E-04 | 2.29E-03 | -1.59E+00 | 1.09E-01 | 3.99E-01 | -1.05E+00 | 1.54E-01 | 4.24E-01 | -1.13E+00 |
| IBTK | 2.83E-18 | 1.47E-17 | -1.82E+00 | 2.20E-11 | 9.51E-11 | -1.17E+00 | 6.31E-12 | 2.68E-11 | -1.63E+00 | 2.69E-08 | 1.03E-07 | -1.20E+00 | 3.74E-03 | 1.33E-02 | -1.08E+00 | 8.31E-02 | 2.34E-01 | 1.13E+00 | 3.81E-07 | 1.40E-06 | -1.25E+00 | 4.27E-05 | 1.17E-04 | -1.11E+00 |
| MEST | 2.84E-18 | 1.48E-17 | -1.57E+00 | 5.71E-11 | 2.47E-10 | -1.18E+00 | 1.16E-09 | 4.91E-09 | -1.51E+00 | 2.02E-04 | 7.72E-04 | -1.02E+00 | 2.81E-04 | 1.00E-03 | -1.10E+00 | 3.19E-01 | 9.00E-01 | 1.31E+00 | 1.56E-03 | 5.72E-03 | -1.11E+00 | 3.32E-01 | 9.12E-01 | 1.22E+00 |
| RFLNB | 2.85E-18 | 1.49E-17 | -1.54E+00 | 1.34E-09 | 5.80E-09 | -1.18E+00 | 1.49E-03 | 6.32E-03 | -1.25E+00 | 1.41E-02 | 5.38E-02 | -1.06E+00 | 8.78E-05 | 3.13E-04 | -1.34E+00 | 7.45E-10 | 2.10E-09 | -1.75E+00 | 2.47E-04 | 9.06E-04 | 1.54E+00 | 1.05E-02 | 2.90E-02 | 1.43E+00 |
| CSK | 2.95E-18 | 1.54E-17 | -1.54E+00 | 3.39E-08 | 1.46E-07 | -1.02E+00 | 1.14E-14 | 4.85E-14 | -1.64E+00 | 1.20E-13 | 4.57E-13 | -1.43E+00 | 2.08E-10 | 7.41E-10 | -1.53E+00 | 1.15E-06 | 3.24E-06 | -1.28E+00 | 6.71E-05 | 2.46E-04 | -1.11E+00 | 2.89E-03 | 7.93E-03 | -1.02E+00 |
| RPL7P44 | 2.95E-18 | 1.54E-17 | -1.37E+00 | 7.27E-07 | 3.14E-06 | 1.06E+00 | 2.44E-07 | 1.04E-06 | -1.38E+00 | 4.86E-02 | 1.86E-01 | 1.07E+00 | 1.98E-09 | 7.07E-09 | -1.35E+00 | 8.08E-03 | 2.28E-02 | 1.07E+00 | 6.89E-11 | 2.53E-10 | -1.51E+00 | 9.94E-02 | 2.73E-01 | 1.04E+00 |
| SV2A | 2.99E-18 | 1.56E-17 | -2.01E+00 | 2.75E-09 | 1.19E-08 | -1.16E+00 | 3.62E-10 | 1.54E-09 | -1.96E+00 | 3.99E-04 | 1.52E-03 | -1.07E+00 | 4.70E-10 | 1.68E-09 | -1.94E+00 | 4.21E-09 | 1.19E-08 | -1.65E+00 | 2.42E-02 | 8.89E-02 | -1.22E+00 | 6.84E-02 | 1.88E-01 | -1.14E+00 |
| GNA15 | 3.04E-18 | 1.58E-17 | -1.43E+00 | 1.11E-10 | 4.82E-10 | -1.17E+00 | 3.98E-13 | 1.69E-12 | -1.62E+00 | 1.36E-05 | 5.19E-05 | -1.17E+00 | 1.33E-10 | 4.76E-10 | -1.40E+00 | 1.03E-07 | 2.91E-07 | -1.34E+00 | 4.61E-07 | 1.69E-06 | -1.31E+00 | 8.84E-02 | 2.43E-01 | 1.14E+00 |
| JPT1 | 3.12E-18 | 1.62E-17 | -1.59E+00 | 1.55E-15 | 6.70E-15 | -1.33E+00 | 1.39E-04 | 5.91E-04 | -1.15E+00 | 7.64E-05 | 2.92E-04 | 1.01E+00 | 1.31E-20 | 4.66E-20 | -1.83E+00 | 8.36E-09 | 2.36E-08 | -1.25E+00 | 4.43E-04 | 1.63E-03 | -1.11E+00 | 1.01E-02 | 2.78E-02 | -1.04E+00 |
| SLBP | 3.28E-18 | 1.71E-17 | -1.70E+00 | 1.87E-10 | 8.06E-10 | -1.14E+00 | 2.13E-06 | 9.04E-06 | -1.37E+00 | 5.31E-02 | 2.03E-01 | 1.09E+00 | 6.03E-15 | 2.15E-14 | -1.92E+00 | 1.12E-04 | 3.15E-04 | -1.10E+00 | 9.67E-06 | 3.55E-05 | -1.27E+00 | 3.26E-01 | 8.95E-01 | 1.15E+00 |
| GEMIN7 | 3.31E-18 | 1.73E-17 | -2.52E+00 | 9.24E-10 | 3.99E-09 | -1.38E+00 | 1.08E-16 | 4.60E-16 | -2.12E+00 | 6.14E-08 | 2.35E-07 | -1.18E+00 | 4.76E-05 | 1.70E-04 | -1.50E+00 | 1.36E-01 | 3.85E-01 | 1.26E+00 | 3.56E-10 | 1.31E-09 | -1.61E+00 | 1.24E-01 | 3.39E-01 | 1.12E+00 |
| ANKRD46 | 3.31E-18 | 1.73E-17 | -2.05E+00 | 1.22E-09 | 5.27E-09 | -1.17E+00 | 2.91E-10 | 1.24E-09 | -2.02E+00 | 3.17E-07 | 1.21E-06 | -1.39E+00 | 2.05E-05 | 7.32E-05 | -2.03E+00 | 1.69E-02 | 4.76E-02 | -1.03E+00 | 8.20E-07 | 3.01E-06 | -2.01E+00 | 4.41E-02 | 1.21E-01 | -1.09E+00 |
| IPO5 | 3.35E-18 | 1.75E-17 | -1.54E+00 | 3.00E-08 | 1.30E-07 | 1.03E+00 | 2.72E-05 | 1.16E-04 | -1.11E+00 | 3.69E-03 | 1.41E-02 | 1.10E+00 | 3.74E-07 | 1.33E-06 | -1.29E+00 | 1.44E-03 | 4.05E-03 | -1.01E+00 | 1.73E-07 | 6.35E-07 | -1.30E+00 | 1.96E-01 | 5.40E-01 | 1.10E+00 |
| COA4 | 3.42E-18 | 1.78E-17 | -1.41E+00 | 1.61E-08 | 6.96E-08 | -1.04E+00 | 1.02E-07 | 4.32E-07 | -1.34E+00 | 5.53E-03 | 2.11E-02 | 1.05E+00 | 6.95E-12 | 2.48E-11 | -1.50E+00 | 1.14E-03 | 3.21E-03 | 1.01E+00 | 3.34E-06 | 1.22E-05 | -1.28E+00 | 3.04E-01 | 8.35E-01 | 1.09E+00 |
| AK2 | 3.47E-18 | 1.81E-17 | -1.59E+00 | 3.71E-08 | 1.60E-07 | -1.03E+00 | 2.81E-06 | 1.20E-05 | -1.27E+00 | 2.14E-05 | 8.19E-05 | -1.10E+00 | 3.74E-12 | 1.33E-11 | -1.69E+00 | 9.09E-04 | 2.56E-03 | -1.03E+00 | 4.33E-14 | 1.59E-13 | -1.62E+00 | 1.54E-02 | 4.22E-02 | -1.05E+00 |
| JAK1 | 3.55E-18 | 1.85E-17 | -1.40E+00 | 7.88E-08 | 3.41E-07 | 1.01E+00 | 7.48E-08 | 3.18E-07 | -1.31E+00 | 6.08E-05 | 2.32E-04 | 1.03E+00 | 3.01E-05 | 1.07E-04 | -1.11E+00 | 1.55E-03 | 4.38E-03 | 1.03E+00 | 6.61E-05 | 2.43E-04 | -1.08E+00 | 2.09E-03 | 5.74E-03 | -1.04E+00 |
| PTEN | 3.60E-18 | 1.88E-17 | -1.46E+00 | 4.03E-12 | 1.74E-11 | -1.18E+00 | 7.64E-08 | 3.25E-07 | -1.45E+00 | 5.28E-06 | 2.02E-05 | -1.17E+00 | 1.21E-06 | 4.30E-06 | -1.22E+00 | 2.11E-04 | 5.95E-04 | -1.02E+00 | 8.47E-11 | 3.11E-10 | -1.46E+00 | 3.27E-06 | 8.98E-06 | -1.23E+00 |
| GSE1 | 3.68E-18 | 1.92E-17 | -1.49E+00 | 1.36E-08 | 5.90E-08 | -1.01E+00 | 1.83E-13 | 7.79E-13 | -1.61E+00 | 7.74E-05 | 2.96E-04 | 1.00E+00 | 3.58E-11 | 1.28E-10 | -1.43E+00 | 2.05E-05 | 5.77E-05 | -1.10E+00 | 3.35E-17 | 1.23E-16 | -1.87E+00 | 2.11E-06 | 5.81E-06 | -1.37E+00 |
| EIF3F | 3.95E-18 | 2.06E-17 | -1.42E+00 | 2.69E-16 | 1.16E-15 | -1.30E+00 | 2.12E-05 | 8.99E-05 | -1.33E+00 | 6.01E-05 | 2.30E-04 | -1.25E+00 | 1.36E-06 | 4.86E-06 | -1.33E+00 | 3.44E-04 | 9.70E-04 | -1.25E+00 | 6.79E-07 | 2.49E-06 | -1.30E+00 | 3.73E-06 | 1.02E-05 | -1.27E+00 |
| SLC3A2 | 3.96E-18 | 2.06E-17 | -1.42E+00 | 1.12E-22 | 4.85E-22 | -1.46E+00 | 4.30E-02 | 1.83E-01 | -1.03E+00 | 5.97E-05 | 2.28E-04 | -1.11E+00 | 3.02E-09 | 1.08E-08 | -1.31E+00 | 1.88E-07 | 5.29E-07 | -1.22E+00 | 6.62E-07 | 2.43E-06 | -1.21E+00 | 3.81E-03 | 1.05E-02 | -1.05E+00 |
| CENPCP1 | 3.99E-18 | 2.08E-17 | -1.90E+00 | 9.31E-20 | 4.02E-19 | -1.80E+00 | 5.84E-06 | 2.48E-05 | -1.41E+00 | 2.86E-08 | 1.09E-07 | -1.44E+00 | 7.11E-03 | 2.54E-02 | -1.03E+00 | 1.00E-03 | 2.82E-03 | -1.24E+00 | 4.35E-08 | 1.60E-07 | -1.41E+00 | 5.21E-10 | 1.43E-09 | -1.60E+00 |
| EIF1P5 | 4.08E-18 | 2.13E-17 | -1.46E+00 | 1.28E-38 | 5.51E-38 | -2.34E+00 | 1.90E-10 | 8.09E-10 | -1.49E+00 | 1.43E-19 | 5.47E-19 | -1.89E+00 | 2.73E-04 | 9.73E-04 | -1.02E+00 | 2.77E-09 | 7.81E-09 | -1.50E+00 | 2.36E-04 | 8.67E-04 | -1.06E+00 | 4.00E-13 | 1.10E-12 | -1.68E+00 |
| HSPD1 | 4.10E-18 | 2.14E-17 | -1.40E+00 | 2.38E-08 | 1.03E-07 | -1.14E+00 | 2.58E-04 | 1.10E-03 | -1.32E+00 | 9.29E-03 | 3.55E-02 | -1.17E+00 | 2.81E-15 | 1.00E-14 | -1.55E+00 | 8.19E-04 | 2.31E-03 | -1.13E+00 | 4.40E-07 | 1.62E-06 | -1.28E+00 | 1.68E-01 | 4.61E-01 | 1.00E+00 |
| ROCK2 | 4.19E-18 | 2.18E-17 | -1.95E+00 | 2.10E-06 | 9.08E-06 | -1.03E+00 | 2.29E-14 | 9.75E-14 | -1.80E+00 | 2.51E-05 | 9.60E-05 | -1.06E+00 | 3.71E-13 | 1.32E-12 | -1.79E+00 | 4.57E-08 | 1.29E-07 | -1.29E+00 | 4.03E-12 | 1.48E-11 | -1.20E+00 | 9.71E-03 | 2.67E-02 | -1.03E+00 |
| PRX | 4.21E-18 | 2.19E-17 | 8.32E+00 | 1.11E-07 | 4.82E-07 | 5.61E+00 | 6.36E-05 | 2.70E-04 | 3.39E+00 | 6.60E-02 | 2.52E-01 | 2.14E+00 | 1.60E-01 | 5.69E-01 | 2.28E+00 | 3.50E-01 | 9.87E-01 | 1.13E+00 | 2.46E-01 | 9.02E-01 | 1.06E+00 | 2.07E-01 | 5.69E-01 | -1.24E+00 |
| ARL1 | 4.28E-18 | 2.23E-17 | -1.68E+00 | 5.16E-11 | 2.23E-10 | -1.15E+00 | 5.95E-12 | 2.53E-11 | -1.58E+00 | 9.49E-07 | 3.62E-06 | -1.08E+00 | 7.76E-08 | 2.77E-07 | -1.51E+00 | 1.69E-03 | 4.78E-03 | -1.08E+00 | 4.01E-10 | 1.47E-09 | -1.44E+00 | 3.35E-04 | 9.20E-04 | -1.11E+00 |
| NIPSNAP2 | 4.33E-18 | 2.26E-17 | -1.40E+00 | 8.23E-08 | 3.56E-07 | 1.02E+00 | 6.12E-07 | 2.60E-06 | -1.30E+00 | 2.18E-05 | 8.34E-05 | -1.10E+00 | 1.65E-13 | 5.87E-13 | -1.51E+00 | 3.66E-06 | 1.03E-05 | -1.16E+00 | 8.06E-05 | 2.96E-04 | -1.14E+00 | 5.66E-04 | 1.56E-03 | -1.04E+00 |
| GRB2 | 4.54E-18 | 2.36E-17 | -1.53E+00 | 2.77E-18 | 1.20E-17 | -1.33E+00 | 1.30E-05 | 5.54E-05 | -1.27E+00 | 3.99E-09 | 1.53E-08 | -1.30E+00 | 3.10E-14 | 1.10E-13 | -1.60E+00 | 3.80E-11 | 1.07E-10 | -1.40E+00 | 2.11E-03 | 7.74E-03 | -1.10E+00 | 5.46E-04 | 1.50E-03 | -1.18E+00 |
| MYCBP2 | 4.60E-18 | 2.39E-17 | -1.43E+00 | 1.63E-04 | 7.04E-04 | 1.07E+00 | 8.44E-11 | 3.59E-10 | -1.57E+00 | 1.13E-04 | 4.30E-04 | -1.07E+00 | 7.51E-08 | 2.68E-07 | -1.38E+00 | 6.87E-04 | 1.94E-03 | -1.10E+00 | 1.13E-04 | 4.15E-04 | -1.23E+00 | 6.46E-02 | 1.77E-01 | 1.01E+00 |
| B4GALT1 | 4.77E-18 | 2.49E-17 | -2.05E+00 | 7.95E-12 | 3.44E-11 | -1.37E+00 | 3.50E-09 | 1.49E-08 | -1.63E+00 | 1.95E-08 | 7.46E-08 | -1.32E+00 | 8.86E-05 | 3.16E-04 | -1.34E+00 | 1.27E-04 | 3.58E-04 | -1.24E+00 | 4.85E-04 | 1.78E-03 | -1.17E+00 | 6.64E-02 | 1.82E-01 | 1.04E+00 |
| ITSN2 | 4.88E-18 | 2.54E-17 | -1.53E+00 | 7.55E-13 | 3.26E-12 | -1.16E+00 | 2.73E-08 | 1.16E-07 | -1.40E+00 | 3.97E-06 | 1.51E-05 | -1.07E+00 | 1.86E-07 | 6.65E-07 | -1.32E+00 | 6.50E-07 | 1.83E-06 | -1.24E+00 | 1.68E-11 | 6.17E-11 | -1.54E+00 | 8.16E-07 | 2.24E-06 | -1.26E+00 |
| ZNF467 | 4.90E-18 | 2.55E-17 | -1.84E+00 | 2.09E-15 | 9.04E-15 | -1.43E+00 | 4.22E-07 | 1.79E-06 | -1.84E+00 | 7.74E-07 | 2.96E-06 | -1.53E+00 | 1.07E-01 | 3.82E-01 | -1.42E+00 | 1.13E-01 | 3.19E-01 | -1.71E+00 | 2.68E-01 | 9.84E-01 | 1.05E+00 | 3.51E-01 | 9.65E-01 | 1.03E+00 |
| OAS3 | 4.99E-18 | 2.60E-17 | -2.58E+00 | 7.58E-18 | 3.27E-17 | -2.26E+00 | 1.14E-15 | 4.84E-15 | -2.40E+00 | 1.09E-12 | 4.16E-12 | -1.80E+00 | 2.61E-10 | 9.30E-10 | -7.30E+00 | 2.24E-10 | 6.32E-10 | -6.25E+00 | 8.02E-26 | 2.94E-25 | -4.92E+00 | 7.05E-21 | 1.94E-20 | -3.53E+00 |
| PLIN3 | 5.04E-18 | 2.63E-17 | -1.74E+00 | 4.66E-10 | 2.01E-09 | -1.21E+00 | 3.07E-15 | 1.31E-14 | -1.85E+00 | 8.86E-13 | 3.38E-12 | -1.38E+00 | 2.80E-12 | 1.00E-11 | -1.64E+00 | 1.06E-13 | 2.99E-13 | -1.63E+00 | 8.05E-15 | 2.96E-14 | -1.71E+00 | 1.45E-11 | 3.97E-11 | -1.52E+00 |
| TALDO1 | 5.04E-18 | 2.63E-17 | -1.26E+00 | 6.30E-03 | 2.72E-02 | -1.04E+00 | 7.11E-03 | 3.02E-02 | -1.09E+00 | 9.53E-02 | 3.64E-01 | 1.06E+00 | 3.60E-05 | 1.28E-04 | -1.20E+00 | 2.92E-01 | 8.22E-01 | -1.03E+00 | 2.38E-01 | 8.74E-01 | 1.05E+00 | 9.68E-03 | 2.66E-02 | 1.15E+00 |
| DDX21 | 5.07E-18 | 2.64E-17 | -1.40E+00 | 1.10E-34 | 4.74E-34 | -1.65E+00 | 1.24E-04 | 5.28E-04 | -1.17E+00 | 2.34E-06 | 8.92E-06 | -1.26E+00 | 1.89E-11 | 6.73E-11 | -1.48E+00 | 4.47E-12 | 1.26E-11 | -1.49E+00 | 1.37E-08 | 5.03E-08 | -1.44E+00 | 1.62E-03 | 4.45E-03 | -1.22E+00 |
| EIF1 | 5.11E-18 | 2.66E-17 | 1.21E+00 | 9.29E-02 | 4.01E-01 | 1.04E+00 | 2.44E-06 | 1.04E-05 | 1.12E+00 | 2.08E-01 | 7.93E-01 | 1.02E+00 | 1.17E-14 | 4.15E-14 | 1.27E+00 | 3.09E-04 | 8.70E-04 | 1.12E+00 | 1.73E-13 | 6.34E-13 | 1.24E+00 | 1.10E-02 | 3.01E-02 | 1.08E+00 |
| RSF1 | 5.14E-18 | 2.68E-17 | -1.46E+00 | 1.24E-14 | 5.35E-14 | -1.26E+00 | 4.33E-06 | 1.84E-05 | -1.24E+00 | 3.04E-06 | 1.16E-05 | -1.14E+00 | 3.01E-07 | 1.07E-06 | -1.30E+00 | 1.05E-04 | 2.97E-04 | -1.13E+00 | 6.51E-07 | 2.39E-06 | -1.32E+00 | 2.37E-04 | 6.51E-04 | -1.26E+00 |
| PHACTR1 | 5.53E-18 | 2.88E-17 | -1.52E+00 | 4.30E-20 | 1.86E-19 | -1.48E+00 | 8.63E-06 | 3.67E-05 | -1.29E+00 | 2.89E-07 | 1.10E-06 | -1.26E+00 | 9.80E-10 | 3.49E-09 | -1.63E+00 | 1.06E-11 | 2.98E-11 | -1.67E+00 | 4.53E-05 | 1.66E-04 | -1.26E+00 | 1.35E-04 | 3.72E-04 | -1.27E+00 |
| LGALS3BP | 5.63E-18 | 2.94E-17 | -2.39E+00 | 1.03E-12 | 4.44E-12 | -1.63E+00 | 6.22E-22 | 2.64E-21 | -2.92E+00 | 5.23E-17 | 2.00E-16 | -1.98E+00 | 1.03E-07 | 3.65E-07 | -4.59E+00 | 2.57E-06 | 7.23E-06 | -2.75E+00 | 3.25E-17 | 1.19E-16 | -3.62E+00 | 3.46E-12 | 9.50E-12 | -2.32E+00 |
| RPL3P2 | 5.64E-18 | 2.94E-17 | 2.58E+00 | 8.13E-04 | 3.51E-03 | 2.06E+00 | 7.23E-18 | 3.07E-17 | 2.51E+00 | 2.76E-05 | 1.05E-04 | 1.94E+00 | 6.91E-06 | 2.47E-05 | 2.27E+00 | 9.81E-02 | 2.77E-01 | 1.66E+00 | 3.28E-08 | 1.20E-07 | 2.37E+00 | 5.66E-04 | 1.56E-03 | 2.03E+00 |
| PRRC1 | 5.70E-18 | 2.97E-17 | -1.90E+00 | 8.28E-08 | 3.58E-07 | -1.08E+00 | 2.32E-07 | 9.87E-07 | -1.50E+00 | 3.57E-04 | 1.36E-03 | -1.02E+00 | 7.88E-15 | 2.81E-14 | -1.88E+00 | 5.45E-11 | 1.54E-10 | -1.48E+00 | 3.67E-16 | 1.35E-15 | -1.68E+00 | 1.75E-06 | 4.81E-06 | -1.17E+00 |
| GLG1 | 5.99E-18 | 3.12E-17 | -1.97E+00 | 3.65E-06 | 1.58E-05 | -1.04E+00 | 6.23E-14 | 2.65E-13 | -1.87E+00 | 4.43E-08 | 1.69E-07 | -1.29E+00 | 1.05E-04 | 3.75E-04 | -1.35E+00 | 1.98E-01 | 5.57E-01 | 1.36E+00 | 1.64E-05 | 6.00E-05 | -1.27E+00 | 3.30E-03 | 9.07E-03 | -1.12E+00 |
| RPS19P1 | 6.14E-18 | 3.20E-17 | 2.11E+00 | 1.31E-03 | 5.68E-03 | 1.69E+00 | 3.10E-10 | 1.32E-09 | 2.06E+00 | 9.07E-04 | 3.46E-03 | 1.64E+00 | 3.55E-04 | 1.27E-03 | 1.96E+00 | 7.64E-03 | 2.15E-02 | 1.70E+00 | 2.58E-09 | 9.46E-09 | 2.11E+00 | 7.69E-05 | 2.11E-04 | 1.77E+00 |
| C12orf75 | 6.24E-18 | 3.25E-17 | -1.86E+00 | 9.75E-21 | 4.21E-20 | -1.87E+00 | 7.05E-09 | 3.00E-08 | -1.35E+00 | 9.31E-14 | 3.56E-13 | -1.48E+00 | 3.05E-22 | 1.09E-21 | -5.25E+00 | 4.87E-27 | 1.37E-26 | -6.73E+00 | 1.14E-06 | 4.20E-06 | -1.21E+00 | 7.83E-06 | 2.15E-05 | -1.77E+00 |
| ZNF426 | 6.31E-18 | 3.29E-17 | -1.74E+00 | 1.42E-20 | 6.12E-20 | -1.68E+00 | 2.51E-05 | 1.07E-04 | -1.38E+00 | 6.07E-07 | 2.32E-06 | -1.35E+00 | 6.31E-07 | 2.25E-06 | -1.39E+00 | 9.09E-06 | 2.56E-05 | -1.23E+00 | 5.90E-03 | 2.17E-02 | -1.14E+00 | 9.69E-04 | 2.66E-03 | -1.20E+00 |
| KIZ | 6.31E-18 | 3.29E-17 | -1.68E+00 | 8.47E-17 | 3.66E-16 | -1.44E+00 | 7.91E-06 | 3.36E-05 | -1.39E+00 | 3.88E-05 | 1.48E-04 | -1.13E+00 | 2.18E-04 | 7.78E-04 | -1.20E+00 | 1.49E-02 | 4.19E-02 | 1.13E+00 | 7.40E-02 | 2.72E-01 | 1.04E+00 | 1.41E-02 | 3.86E-02 | -1.11E+00 |
| CSDE1 | 6.31E-18 | 3.29E-17 | -1.33E+00 | 2.38E-10 | 1.03E-09 | -1.15E+00 | 3.21E-04 | 1.36E-03 | -1.22E+00 | 1.28E-02 | 4.88E-02 | -1.06E+00 | 3.66E-06 | 1.31E-05 | -1.22E+00 | 4.06E-03 | 1.14E-02 | -1.12E+00 | 3.72E-05 | 1.37E-04 | -1.21E+00 | 1.04E-01 | 2.85E-01 | -1.07E+00 |
| PLD4 | 6.48E-18 | 3.38E-17 | -2.03E+00 | 1.62E-15 | 6.99E-15 | -1.56E+00 | 8.51E-09 | 3.61E-08 | -1.49E+00 | 4.00E-13 | 1.53E-12 | -1.60E+00 | 1.36E-01 | 4.87E-01 | -1.20E+00 | 2.67E-01 | 7.52E-01 | 1.09E+00 | 2.02E-01 | 7.40E-01 | -1.15E+00 | 3.03E-01 | 8.32E-01 | -1.11E+00 |
| TAOK1 | 6.58E-18 | 3.43E-17 | -1.62E+00 | 1.27E-08 | 5.50E-08 | -1.03E+00 | 6.96E-09 | 2.96E-08 | -1.56E+00 | 3.56E-06 | 1.36E-05 | -1.17E+00 | 1.06E-07 | 3.79E-07 | -1.43E+00 | 2.62E-04 | 7.37E-04 | -1.08E+00 | 2.58E-07 | 9.47E-07 | -1.09E+00 | 1.14E-04 | 3.12E-04 | -1.07E+00 |
| SUZ12P1 | 6.61E-18 | 3.44E-17 | -1.45E+00 | 5.48E-13 | 2.37E-12 | -1.12E+00 | 1.08E-11 | 4.58E-11 | -1.56E+00 | 6.81E-06 | 2.60E-05 | -1.10E+00 | 2.47E-09 | 8.79E-09 | -1.49E+00 | 1.89E-05 | 5.32E-05 | -1.19E+00 | 2.06E-09 | 7.56E-09 | -1.37E+00 | 7.64E-06 | 2.10E-05 | -1.25E+00 |
| RPS26P39 | 6.71E-18 | 3.50E-17 | -1.57E+00 | 1.73E-85 | 7.50E-85 | -2.82E+00 | 2.54E-08 | 1.08E-07 | -1.63E+00 | 4.86E-37 | 1.86E-36 | -2.71E+00 | 7.01E-13 | 2.50E-12 | -1.63E+00 | 8.58E-48 | 2.42E-47 | -2.84E+00 | 2.55E-09 | 9.37E-09 | -1.54E+00 | 5.49E-31 | 1.51E-30 | -2.65E+00 |

| | | | | | | | | | | | | | | | | | | | | | | | | |
|---|---|---|---|---|---|---|---|---|---|---|---|---|---|---|---|---|---|---|---|---|---|---|---|---|
| COPB2 | 6.76E-18 | 3.52E-17 | -1.66E+00 | 1.28E-05 | 5.52E-05 | 1.07E+00 | 1.07E-10 | 4.57E-10 | -1.42E+00 | 6.58E-05 | 2.51E-04 | -1.03E+00 | 3.44E-13 | 1.23E-12 | -1.92E+00 | 2.21E-03 | 6.24E-03 | -1.03E+00 | 5.23E-13 | 1.92E-12 | -1.65E+00 | 8.41E-04 | 2.31E-03 | -1.13E+00 |
| ATP2B1 | 6.78E-18 | 3.53E-17 | -1.67E+00 | 4.89E-09 | 2.11E-08 | -1.17E+00 | 1.57E-05 | 6.65E-05 | -1.42E+00 | 5.15E-04 | 1.97E-03 | -1.10E+00 | 1.29E-03 | 4.59E-03 | 1.04E+00 | 2.73E-03 | 7.70E-03 | 1.05E+00 | 1.15E-09 | 4.21E-09 | -1.67E+00 | 8.67E-02 | 2.38E-01 | 1.05E+00 |
| ZNF33B | 6.88E-18 | 3.58E-17 | -1.68E+00 | 2.99E-05 | 1.29E-04 | 1.14E+00 | 6.88E-09 | 2.92E-08 | -1.67E+00 | 3.43E-03 | 1.31E-02 | -1.01E+00 | 1.80E-08 | 6.43E-08 | -1.39E+00 | 1.11E-03 | 3.14E-03 | 1.02E+00 | 3.29E-05 | 1.21E-04 | -1.14E+00 | 1.76E-02 | 4.84E-02 | 1.03E+00 |
| GNAI2 | 6.99E-18 | 3.64E-17 | -1.34E+00 | 2.25E-05 | 9.73E-05 | 1.02E+00 | 1.38E-06 | 5.86E-06 | -1.40E+00 | 9.61E-03 | 3.67E-02 | -1.09E+00 | 4.62E-13 | 1.65E-12 | -1.56E+00 | 4.45E-06 | 1.25E-05 | -1.26E+00 | 9.31E-08 | 3.42E-07 | -1.38E+00 | 5.55E-03 | 1.53E-02 | -1.17E+00 |
| ANP32E | 7.08E-18 | 3.69E-17 | -1.54E+00 | 7.04E-09 | 3.04E-08 | -1.12E+00 | 1.16E-04 | 4.92E-04 | -1.21E+00 | 7.10E-03 | 2.71E-02 | 1.04E+00 | 1.04E-10 | 3.71E-10 | -1.37E+00 | 1.02E-04 | 2.88E-04 | -1.00E+00 | 5.25E-04 | 1.93E-03 | -1.18E+00 | 3.23E-01 | 8.87E-01 | 1.12E+00 |
| AKR1A1 | 7.19E-18 | 3.75E-17 | -1.35E+00 | 3.77E-08 | 1.63E-07 | -1.00E+00 | 3.30E-02 | 1.40E-01 | -1.03E+00 | 1.16E-01 | 4.44E-01 | 1.13E+00 | 9.80E-08 | 3.49E-07 | -1.18E+00 | 1.37E-03 | 3.87E-03 | 1.09E+00 | 1.84E-05 | 6.76E-05 | -1.15E+00 | 5.96E-02 | 1.64E-01 | 1.04E+00 |

| | | | | | | | | | | | | | | | | | | | | | | | | |
|---|---|---|---|---|---|---|---|---|---|---|---|---|---|---|---|---|---|---|---|---|---|---|---|---|
| PIGC | 2.60E-16 | 1.35E-15 | -1.51E+00 | 2.39E-06 | 1.03E-05 | 1.06E+00 | 9.06E-08 | 3.85E-07 | -1.38E+00 | 7.67E-03 | 2.93E-02 | 1.12E+00 | 6.47E-06 | 2.31E-05 | -1.29E+00 | 3.79E-02 | 1.07E-01 | 1.12E+00 | 6.56E-13 | 2.41E-12 | -1.63E+00 | 3.65E-03 | 1.00E-02 | -1.09E+00 |
| NAA20 | 2.61E-16 | 1.36E-15 | -1.35E+00 | 2.69E-05 | 1.16E-04 | 1.16E+00 | 3.58E-05 | 1.52E-04 | -1.17E+00 | 1.48E-03 | 5.64E-03 | 1.05E+00 | 5.38E-11 | 1.92E-10 | -1.55E+00 | 1.83E-02 | 5.15E-02 | 1.10E+00 | 1.44E-05 | 5.29E-05 | -1.09E+00 | 2.66E-01 | 7.29E-01 | 1.22E+00 |
| DNAJA2 | 2.63E-16 | 1.37E-15 | -1.46E+00 | 1.95E-12 | 8.44E-12 | -1.16E+00 | 4.95E-12 | 2.10E-11 | -1.49E+00 | 1.19E-07 | 4.53E-07 | -1.12E+00 | 2.28E-06 | 8.11E-06 | -1.15E+00 | 1.11E-02 | 3.14E-02 | 1.19E+00 | 6.65E-09 | 2.44E-08 | -1.33E+00 | 1.10E-02 | 3.03E-02 | -1.08E+00 |
| SNHG6 | 2.76E-16 | 1.44E-15 | 1.33E+00 | 2.80E-03 | 1.21E-02 | -1.07E+00 | 2.83E-09 | 1.20E-08 | 1.30E+00 | 5.97E-02 | 2.28E-01 | -1.04E+00 | 1.13E-10 | 4.04E-10 | 1.42E+00 | 4.11E-02 | 1.16E-01 | -1.05E+00 | 1.48E-11 | 5.44E-11 | 1.54E+00 | 1.06E-03 | 2.91E-03 | -1.07E+00 |
| AL133260.1 | 2.85E-16 | 1.49E-15 | 1.64E+00 | 3.97E-03 | 1.71E-02 | 1.05E+00 | 9.89E-13 | 4.20E-12 | 1.71E+00 | 1.75E-01 | 6.69E-01 | 1.13E+00 | 2.31E-07 | 8.22E-07 | 1.57E+00 | 1.16E-01 | 3.28E-01 | 1.06E+00 | 4.57E-06 | 1.68E-05 | 1.54E+00 | 1.21E-01 | 3.32E-01 | 1.02E+00 |
| AC090589.1 | 2.90E-16 | 1.51E-15 | -1.32E+00 | 4.58E-16 | 1.98E-15 | -1.21E+00 | 2.44E-08 | 1.03E-07 | -1.45E+00 | 1.52E-06 | 5.81E-06 | -1.24E+00 | 1.66E-10 | 5.92E-10 | -1.40E+00 | 2.80E-05 | 7.88E-05 | -1.14E+00 | 3.89E-10 | 1.43E-09 | -1.56E+00 | 2.26E-04 | 6.21E-04 | -1.22E+00 |
| AHCYL1 | 2.95E-16 | 1.54E-15 | -1.37E+00 | 1.70E-10 | 7.35E-10 | -1.07E+00 | 8.77E-06 | 3.73E-05 | -1.20E+00 | 5.00E-03 | 1.91E-02 | 1.07E+00 | 3.25E-05 | 1.16E-04 | -1.03E+00 | 1.06E-01 | 3.00E-01 | 1.28E+00 | 6.10E-04 | 2.24E-03 | -1.10E+00 | 1.55E-01 | 4.27E-01 | 1.11E+00 |
| KLHDC7B | 2.98E-16 | 1.55E-15 | -2.82E+00 | 6.66E-13 | 2.88E-12 | -2.03E+00 | 2.36E-63 | 1.00E-62 | -4.49E+00 | 3.44E-63 | 1.31E-62 | -3.87E+00 | 4.47E-03 | 1.60E-02 | -2.97E+00 | 2.08E-02 | 5.86E-02 | -1.87E+00 | 8.95E-02 | 3.29E-01 | -1.47E+00 | 1.12E-01 | 3.09E-01 | -1.60E+00 |
| COL4A3BP | 3.01E-16 | 1.57E-15 | -1.63E+00 | 1.16E-08 | 4.99E-08 | -1.12E+00 | 4.04E-09 | 1.72E-08 | -1.42E+00 | 5.10E-07 | 1.95E-06 | -1.15E+00 | 6.18E-04 | 2.20E-03 | -1.08E+00 | 3.26E-04 | 9.20E-04 | -1.09E+00 | 3.10E-06 | 1.14E-05 | -1.37E+00 | 3.84E-06 | 1.06E-05 | -1.38E+00 |
| CAPNS1 | 3.13E-16 | 1.63E-15 | -1.28E+00 | 3.82E-10 | 1.65E-09 | -1.05E+00 | 7.77E-08 | 3.30E-07 | -1.38E+00 | 1.76E-06 | 6.74E-06 | -1.16E+00 | 3.12E-11 | 1.11E-10 | -1.44E+00 | 2.54E-06 | 7.17E-06 | -1.20E+00 | 4.92E-04 | 1.81E-03 | -1.16E+00 | 2.64E-02 | 7.25E-02 | -1.08E+00 |
| SPECC1 | 3.27E-16 | 1.71E-15 | -1.79E+00 | 7.80E-07 | 3.37E-06 | -1.04E+00 | 1.93E-09 | 8.20E-09 | -1.70E+00 | 2.58E-07 | 9.87E-07 | -1.34E+00 | 4.31E-06 | 1.54E-05 | -1.34E+00 | 1.16E-03 | 3.28E-03 | -1.02E+00 | 3.79E-05 | 1.39E-04 | -1.12E+00 | 3.22E-03 | 8.84E-03 | -1.02E+00 |
| CRK | 3.36E-16 | 1.75E-15 | -2.18E+00 | 4.65E-07 | 2.01E-06 | -1.10E+00 | 9.50E-11 | 4.04E-10 | -1.86E+00 | 5.03E-07 | 1.92E-06 | -1.21E+00 | 2.42E-07 | 8.64E-07 | -1.72E+00 | 1.01E-06 | 2.85E-06 | -1.56E+00 | 2.24E-13 | 8.23E-13 | -2.01E+00 | 8.85E-07 | 2.43E-06 | -1.48E+00 |
| BRD7 | 3.42E-16 | 1.78E-15 | -1.44E+00 | 5.25E-12 | 2.27E-11 | -1.12E+00 | 3.23E-05 | 1.37E-04 | -1.19E+00 | 1.28E-07 | 4.88E-07 | -1.15E+00 | 2.08E-05 | 7.42E-05 | -1.11E+00 | 3.76E-05 | 1.06E-04 | -1.05E+00 | 9.74E-05 | 3.57E-04 | -1.14E+00 | 1.18E-06 | 3.25E-06 | -1.25E+00 |
| FAM181B | 3.44E-16 | 1.79E-15 | -3.17E+00 | 2.59E-10 | 1.12E-09 | -1.66E+00 | 6.78E-02 | 2.88E-01 | -1.38E+00 | 2.40E-01 | 9.16E-01 | 1.19E+00 | 1.77E-01 | 6.32E-01 | -1.45E+00 | 1.89E-01 | 5.33E-01 | -1.51E+00 | 8.54E-02 | 3.13E-01 | -1.24E+00 | 3.46E-01 | 9.51E-01 | 1.13E+00 |
| COPS7A | 3.46E-16 | 1.81E-15 | -1.83E+00 | 7.59E-11 | 3.28E-10 | -1.21E+00 | 9.60E-10 | 4.08E-09 | -1.67E+00 | 2.43E-06 | 9.29E-06 | -1.17E+00 | 3.91E-04 | 1.40E-03 | -1.36E+00 | 1.05E-03 | 2.97E-03 | -1.24E+00 | 1.30E-07 | 4.77E-07 | -1.47E+00 | 8.06E-04 | 2.21E-03 | -1.16E+00 |
| COX7A2P2 | 3.46E-16 | 1.81E-15 | -1.30E+00 | 5.12E-42 | 2.21E-41 | -1.80E+00 | 2.29E-04 | 9.74E-04 | -1.21E+00 | 2.76E-24 | 1.06E-23 | -1.97E+00 | 1.61E-07 | 5.74E-07 | -1.19E+00 | 1.43E-18 | 4.02E-18 | -1.74E+00 | 9.43E-04 | 3.46E-03 | -1.20E+00 | 7.62E-16 | 2.09E-15 | -1.84E+00 |
| TMEM87A | 3.53E-16 | 1.84E-15 | -1.51E+00 | 7.43E-14 | 3.21E-13 | -1.26E+00 | 8.80E-13 | 3.74E-12 | -1.57E+00 | 2.21E-12 | 8.44E-12 | -1.37E+00 | 3.89E-13 | 1.39E-12 | -1.56E+00 | 1.56E-09 | 4.40E-09 | -1.29E+00 | 2.37E-14 | 8.68E-14 | -1.62E+00 | 3.11E-08 | 8.53E-08 | -1.23E+00 |
| HSBP1L1 | 3.53E-16 | 1.84E-15 | -1.30E+00 | 1.36E-18 | 5.89E-18 | -1.28E+00 | 1.40E-04 | 5.95E-04 | -1.17E+00 | 8.69E-06 | 3.32E-05 | -1.16E+00 | 1.97E-08 | 7.04E-08 | -2.20E+00 | 3.74E-11 | 1.05E-10 | -2.56E+00 | 1.39E-02 | 5.09E-02 | -1.37E+00 | 2.85E-02 | 7.83E-02 | -1.42E+00 |
| HNRNPA1P48 | 3.54E-16 | 1.85E-15 | -1.44E+00 | 1.21E-16 | 5.25E-16 | -1.29E+00 | 2.95E-11 | 1.25E-10 | -1.56E+00 | 6.01E-12 | 2.29E-11 | -1.39E+00 | 1.92E-08 | 6.83E-08 | -1.46E+00 | 9.38E-04 | 2.64E-03 | -1.03E+00 | 5.90E-11 | 2.17E-10 | -1.59E+00 | 7.57E-06 | 2.08E-05 | -1.33E+00 |
| FAM46A | 3.55E-16 | 1.85E-15 | -1.69E+00 | 2.02E-12 | 8.73E-12 | -1.36E+00 | 1.91E-06 | 8.11E-06 | -1.18E+00 | 1.03E-04 | 3.94E-04 | 1.03E+00 | 1.66E-03 | 5.92E-03 | -1.15E+00 | 1.63E-06 | 4.59E-06 | -1.25E+00 | 7.29E-06 | 2.68E-05 | -1.28E+00 | 4.83E-04 | 1.33E-03 | -1.09E+00 |
| TMEM33 | 3.57E-16 | 1.86E-15 | -1.65E+00 | 4.70E-09 | 2.03E-08 | -1.17E+00 | 1.47E-06 | 6.23E-06 | -1.34E+00 | 1.61E-04 | 6.16E-04 | 1.07E+00 | 7.79E-08 | 2.78E-07 | -1.47E+00 | 8.84E-06 | 2.49E-05 | -1.27E+00 | 1.30E-15 | 4.77E-15 | -1.71E+00 | 1.23E-05 | 3.39E-05 | -1.15E+00 |
| DIAPH1 | 3.58E-16 | 1.87E-15 | -1.37E+00 | 1.29E-14 | 5.60E-14 | -1.21E+00 | 7.53E-04 | 3.20E-03 | -1.14E+00 | 4.12E-03 | 1.57E-02 | 1.07E+00 | 7.15E-02 | 2.55E-01 | 1.17E+00 | 2.70E-01 | 7.62E-01 | 1.52E+00 | 1.27E-02 | 4.67E-02 | 1.10E+00 | 2.11E-01 | 5.80E-01 | 1.00E+00 |
| DNAJC2 | 3.58E-16 | 1.87E-15 | -1.30E+00 | 5.54E-15 | 2.39E-14 | -1.15E+00 | 4.20E-04 | 1.78E-03 | -1.14E+00 | 1.16E-04 | 4.41E-04 | -1.02E+00 | 1.47E-02 | 5.23E-02 | 1.09E+00 | 3.32E-03 | 9.37E-03 | 1.04E+00 | 2.00E-01 | 7.33E-01 | 1.13E+00 | 3.59E-01 | 9.87E-01 | 1.14E+00 |
| LARP4 | 3.59E-16 | 1.87E-15 | -1.79E+00 | 9.42E-10 | 4.07E-09 | -1.18E+00 | 2.49E-12 | 1.06E-11 | -1.78E+00 | 1.16E-07 | 4.43E-07 | -1.28E+00 | 2.17E-05 | 7.74E-05 | -1.30E+00 | 7.21E-06 | 2.03E-05 | -1.33E+00 | 4.86E-07 | 1.78E-06 | -1.26E+00 | 7.85E-04 | 2.16E-03 | -1.02E+00 |
| PPP4R3B | 3.61E-16 | 1.88E-15 | -1.46E+00 | 7.48E-14 | 3.23E-13 | -1.25E+00 | 7.81E-10 | 3.32E-09 | -1.38E+00 | 1.10E-08 | 4.20E-08 | -1.21E+00 | 5.29E-10 | 1.89E-09 | -1.49E+00 | 5.03E-07 | 1.42E-06 | -1.25E+00 | 2.34E-06 | 8.58E-06 | -1.20E+00 | 5.86E-05 | 1.61E-04 | -1.10E+00 |
| ETV6 | 3.61E-16 | 1.88E-15 | -1.45E+00 | 2.66E-06 | 1.15E-05 | -1.05E+00 | 1.63E-08 | 6.90E-08 | -1.53E+00 | 1.12E-02 | 4.27E-02 | -1.08E+00 | 1.79E-12 | 6.37E-12 | -1.52E+00 | 1.86E-03 | 5.26E-03 | -1.07E+00 | 6.06E-10 | 2.23E-09 | -1.39E+00 | 1.05E-05 | 2.88E-05 | -1.23E+00 |
| CAPZA1 | 3.61E-16 | 1.88E-15 | -1.32E+00 | 5.31E-07 | 2.30E-06 | -1.00E+00 | 3.42E-04 | 1.45E-03 | -1.21E+00 | 8.56E-04 | 3.27E-03 | -1.07E+00 | 1.78E-09 | 6.35E-09 | -1.37E+00 | 2.85E-03 | 8.03E-03 | -1.02E+00 | 4.79E-06 | 1.76E-05 | -1.37E+00 | 3.83E-02 | 1.05E-01 | -1.13E+00 |
| BAX | 3.65E-16 | 1.90E-15 | -1.29E+00 | 1.26E-12 | 5.45E-12 | -1.11E+00 | 4.51E-03 | 1.92E-02 | -1.27E+00 | 4.84E-04 | 1.85E-03 | -1.15E+00 | 3.38E-04 | 1.20E-03 | -1.02E+00 | 9.78E-03 | 2.76E-02 | 1.10E+00 | 3.45E-04 | 1.26E-03 | -1.16E+00 | 6.67E-03 | 1.83E-02 | -1.09E+00 |
| RAPGEF2 | 3.72E-16 | 1.94E-15 | -1.80E+00 | 2.79E-09 | 1.21E-08 | -1.22E+00 | 4.08E-06 | 1.73E-05 | -1.54E+00 | 1.42E-04 | 5.44E-04 | -1.11E+00 | 4.47E-10 | 1.59E-09 | -1.31E+00 | 5.36E-07 | 1.51E-06 | -1.13E+00 | 3.80E-08 | 1.40E-07 | -1.38E+00 | 1.20E-03 | 3.29E-03 | -1.08E+00 |
| KLF13 | 3.85E-16 | 2.00E-15 | 2.32E+00 | 4.18E-02 | 1.81E-01 | 1.65E+00 | 7.37E-08 | 3.13E-07 | 1.97E+00 | 1.24E-03 | 4.75E-03 | 1.77E+00 | 6.22E-09 | 2.22E-08 | 2.63E+00 | 1.98E-01 | 5.59E-01 | 1.58E+00 | 2.27E-03 | 8.33E-03 | 1.82E+00 | 2.88E-01 | 7.91E-01 | 1.39E+00 |
| PPA2 | 3.85E-16 | 2.01E-15 | -1.63E+00 | 1.46E-11 | 6.32E-11 | -1.27E+00 | 2.77E-13 | 1.18E-12 | -1.60E+00 | 1.80E-08 | 6.89E-08 | -1.23E+00 | 2.30E-16 | 8.19E-16 | -2.08E+00 | 5.37E-08 | 1.51E-07 | -1.28E+00 | 3.07E-08 | 1.13E-07 | -1.45E+00 | 3.62E-03 | 9.95E-03 | -1.11E+00 |
| HSPA4L | 4.13E-16 | 2.15E-15 | -2.48E+00 | 5.78E-17 | 2.50E-16 | -2.29E+00 | 2.49E-09 | 1.06E-08 | -1.73E+00 | 1.60E-07 | 6.10E-07 | -1.40E+00 | 1.45E-01 | 5.16E-01 | -1.10E+00 | 1.87E-01 | 5.27E-01 | -1.05E+00 | 1.07E-02 | 3.92E-02 | -1.18E+00 | 1.48E-01 | 4.07E-01 | 1.08E+00 |
| AC009948.1 | 4.31E-16 | 2.25E-15 | -1.66E+00 | 5.95E-08 | 2.57E-07 | -1.09E+00 | 1.21E-07 | 5.15E-07 | -1.38E+00 | 6.19E-05 | 2.37E-04 | -1.06E+00 | 2.84E-09 | 1.01E-08 | -1.70E+00 | 1.16E-05 | 3.27E-05 | -1.26E+00 | 4.88E-06 | 1.79E-05 | -1.11E+00 | 5.61E-05 | 1.54E-04 | -1.12E+00 |
| BIRC6 | 4.32E-16 | 2.25E-15 | -1.40E+00 | 7.21E-06 | 3.12E-05 | 1.08E+00 | 2.22E-05 | 9.42E-05 | -1.20E+00 | 1.53E-04 | 5.84E-04 | -1.01E+00 | 3.95E-03 | 1.41E-02 | -1.03E+00 | 8.43E-03 | 2.38E-02 | 1.02E+00 | 5.64E-05 | 2.07E-04 | -1.13E+00 | 2.48E-02 | 6.81E-02 | 1.04E+00 |
| COX6A1 | 4.32E-16 | 2.25E-15 | -1.27E+00 | 6.34E-54 | 2.74E-53 | -2.02E+00 | 1.95E-02 | 8.27E-02 | -1.08E+00 | 5.17E-21 | 1.98E-20 | -1.81E+00 | 3.95E-10 | 1.41E-09 | -1.33E+00 | 4.60E-21 | 1.30E-20 | -1.70E+00 | 2.01E-03 | 7.37E-03 | -1.10E+00 | 1.05E-09 | 2.89E-09 | -1.58E+00 |
| KCNQ1OT1 | 4.37E-16 | 2.28E-15 | -1.84E+00 | 7.14E-08 | 3.08E-07 | -1.22E+00 | 4.82E-09 | 2.05E-08 | -1.94E+00 | 1.83E-05 | 6.97E-05 | -1.38E+00 | 1.98E-06 | 7.04E-06 | -1.42E+00 | 9.49E-04 | 2.68E-03 | -1.10E+00 | 4.31E-02 | 1.58E-01 | -1.11E+00 | 2.29E-01 | 6.30E-01 | 1.17E+00 |
| MT-ND4 | 4.39E-16 | 2.29E-15 | -1.22E+00 | 3.36E-10 | 1.45E-09 | -1.16E+00 | 2.12E-04 | 9.00E-04 | -1.11E+00 | 4.06E-02 | 1.55E-01 | -1.03E+00 | 6.77E-05 | 2.41E-04 | -1.13E+00 | 2.98E-01 | 8.39E-01 | 1.00E+00 | 3.77E-02 | 1.38E-01 | -1.02E+00 | 4.65E-03 | 1.28E-02 | 1.10E+00 |
| ZC3H6 | 4.58E-16 | 2.39E-15 | -1.99E+00 | 3.33E-15 | 1.44E-14 | -1.71E+00 | 3.18E-10 | 1.35E-09 | -1.89E+00 | 6.54E-13 | 2.50E-12 | -2.07E+00 | 1.93E-09 | 6.89E-09 | -2.18E+00 | 7.22E-12 | 2.03E-11 | -2.72E+00 | 5.90E-04 | 2.17E-03 | -1.18E+00 | 7.32E-06 | 2.01E-05 | -1.82E+00 |
| TSG101 | 4.62E-16 | 2.41E-15 | -1.52E+00 | 9.07E-11 | 3.92E-10 | -1.09E+00 | 2.69E-18 | 1.14E-17 | -1.84E+00 | 7.90E-14 | 3.02E-13 | -1.31E+00 | 3.76E-13 | 1.34E-12 | -1.80E+00 | 3.18E-05 | 8.97E-05 | -1.12E+00 | 2.15E-11 | 7.90E-11 | -1.36E+00 | 4.48E-06 | 1.23E-05 | -1.22E+00 |
| TNRC6B | 4.62E-16 | 2.41E-15 | -1.51E+00 | 4.54E-05 | 1.96E-04 | -1.04E+00 | 4.47E-08 | 1.90E-07 | -1.47E+00 | 2.06E-03 | 7.87E-03 | -1.05E+00 | 1.33E-11 | 4.74E-11 | -1.62E+00 | 2.35E-05 | 6.62E-05 | -1.22E+00 | 2.42E-15 | 8.90E-15 | -1.68E+00 | 2.10E-05 | 5.77E-05 | -1.18E+00 |
| BTN3A1 | 4.63E-16 | 2.41E-15 | -2.16E+00 | 3.05E-09 | 1.32E-08 | -1.29E+00 | 3.97E-13 | 1.69E-12 | -2.24E+00 | 1.17E-09 | 4.48E-09 | -1.56E+00 | 3.89E-11 | 1.39E-10 | -3.22E+00 | 2.18E-06 | 6.14E-06 | -1.66E+00 | 1.79E-16 | 6.58E-16 | -2.99E+00 | 9.51E-11 | 2.61E-10 | -2.15E+00 |
| S1PR2 | 4.67E-16 | 2.44E-15 | -2.06E+00 | 2.68E-18 | 1.16E-17 | -2.29E+00 | 1.57E-07 | 6.68E-07 | -1.84E+00 | 1.65E-09 | 6.30E-09 | -2.23E+00 | 6.33E-02 | 2.26E-01 | -1.18E+00 | 3.84E-02 | 1.08E-01 | -1.56E+00 | 4.80E-02 | 1.76E-01 | -1.26E+00 | 7.48E-02 | 2.05E-01 | -1.46E+00 |
| CEBPG | 4.69E-16 | 2.44E-15 | -1.58E+00 | 1.62E-26 | 7.02E-26 | -1.95E+00 | 3.81E-09 | 1.62E-08 | -1.38E+00 | 5.01E-17 | 1.91E-16 | -1.61E+00 | 1.29E-08 | 4.60E-08 | -1.35E+00 | 1.54E-14 | 4.35E-14 | -2.37E+00 | 2.72E-06 | 9.99E-06 | -1.29E+00 | 3.57E-05 | 9.80E-05 | -1.21E+00 |
| GCA | 4.71E-16 | 2.46E-15 | -1.72E+00 | 4.04E-03 | 1.75E-02 | 1.16E+00 | 3.16E-21 | 1.34E-20 | -2.07E+00 | 2.95E-07 | 1.13E-06 | -1.11E+00 | 2.28E-11 | 8.13E-11 | -1.89E+00 | 1.55E-04 | 4.36E-04 | -1.16E+00 | 4.84E-10 | 1.78E-09 | -1.54E+00 | 9.78E-05 | 2.69E-04 | -1.15E+00 |
| ZNF37A | 4.73E-16 | 2.47E-15 | -1.52E+00 | 1.20E-03 | 5.18E-03 | 1.16E+00 | 2.12E-09 | 9.01E-09 | -1.59E+00 | 1.01E-02 | 3.86E-02 | 1.01E+00 | 3.30E-06 | 1.18E-05 | -1.27E+00 | 3.51E-02 | 9.91E-02 | 1.16E+00 | 2.49E-05 | 9.14E-05 | -1.35E+00 | 1.51E-01 | 4.15E-01 | 1.01E+00 |
| EGLN1 | 4.73E-16 | 2.47E-15 | -2.07E+00 | 8.44E-15 | 3.64E-14 | -1.73E+00 | 3.95E-17 | 1.68E-16 | -1.78E+00 | 3.11E-23 | 1.19E-22 | -2.04E+00 | 9.36E-05 | 3.34E-04 | -1.81E+00 | 5.45E-06 | 1.54E-05 | -2.68E+00 | 1.55E-03 | 5.68E-03 | -1.84E+00 | 7.30E-04 | 2.00E-03 | -2.03E+00 |
| GTF2A1 | 4.77E-16 | 2.49E-15 | -2.58E+00 | 3.23E-12 | 1.40E-11 | -1.65E+00 | 1.12E-09 | 4.77E-09 | -1.96E+00 | 1.79E-05 | 6.84E-05 | -1.17E+00 | 7.63E-05 | 2.72E-04 | -1.54E+00 | 2.81E-02 | 7.93E-02 | 1.06E+00 | 1.79E-17 | 6.55E-17 | -1.67E+00 | 4.07E-06 | 1.12E-05 | -1.25E+00 |
| KIDINS220 | 4.91E-16 | 2.56E-15 | -1.60E+00 | 5.04E-06 | 2.18E-05 | 1.01E+00 | 6.23E-05 | 2.65E-04 | -1.26E+00 | 2.38E-03 | 9.10E-03 | 1.03E+00 | 5.22E-08 | 1.86E-07 | -1.53E+00 | 1.38E-05 | 3.89E-05 | -1.26E+00 | 1.14E-07 | 4.19E-07 | -1.31E+00 | 3.88E-05 | 1.07E-04 | -1.20E+00 |
| NR1H3 | 4.93E-16 | 2.57E-15 | -1.71E+00 | 1.99E-10 | 8.60E-10 | -1.18E+00 | 2.67E-09 | 1.14E-08 | -1.95E+00 | 1.82E-04 | 6.94E-04 | -1.07E+00 | 1.20E-02 | 4.29E-02 | -1.01E+00 | 4.27E-02 | 1.20E-01 | 1.12E+00 | 1.80E-02 | 6.62E-02 | -1.17E+00 | 2.44E-02 | 6.71E-02 | -1.10E+00 |
| ZBTB24 | 4.94E-16 | 2.57E-15 | -1.71E+00 | 1.30E-12 | 5.63E-12 | -1.33E+00 | 6.37E-11 | 2.71E-10 | -1.55E+00 | 1.06E-05 | 4.05E-05 | -1.08E+00 | 1.15E-06 | 4.09E-06 | -1.35E+00 | 4.65E-06 | 1.31E-05 | -1.33E+00 | 3.15E-05 | 1.16E-04 | -1.16E+00 | 8.47E-04 | 2.33E-03 | -1.02E+00 |
| NOB1 | 5.09E-16 | 2.65E-15 | -1.43E+00 | 6.70E-16 | 2.90E-15 | -1.32E+00 | 4.13E-11 | 1.75E-10 | -1.65E+00 | 5.78E-10 | 2.21E-09 | -1.43E+00 | 4.66E-15 | 1.66E-14 | -1.66E+00 | 4.38E-06 | 1.24E-05 | -1.14E+00 | 1.21E-11 | 4.43E-11 | -1.64E+00 | 2.10E-07 | 5.76E-07 | -1.47E+00 |
| HERC4 | 5.31E-16 | 2.77E-15 | -1.67E+00 | 7.36E-02 | 3.18E-01 | 1.35E+00 | 2.65E-14 | 1.12E-13 | -1.97E+00 | 1.39E-03 | 5.31E-03 | 1.04E+00 | 4.17E-09 | 1.49E-08 | -1.64E+00 | 4.84E-02 | 1.36E-01 | 1.10E+00 | 1.69E-11 | 6.21E-11 | -1.50E+00 | 2.87E-02 | 7.89E-02 | 1.07E+00 |
| USO1 | 5.50E-16 | 2.87E-15 | -1.52E+00 | 6.56E-08 | 2.84E-07 | -1.04E+00 | 1.57E-13 | 6.67E-13 | -1.65E+00 | 2.81E-09 | 1.07E-08 | -1.22E+00 | 1.78E-19 | 6.34E-19 | -1.98E+00 | 1.45E-09 | 4.09E-09 | -1.33E+00 | 8.25E-15 | 3.03E-14 | -1.81E+00 | 2.26E-05 | 6.21E-05 | -1.29E+00 |
| MYC | 5.56E-16 | 2.90E-15 | -2.12E+00 | 3.13E-07 | 1.35E-06 | -1.18E+00 | 7.78E-18 | 3.31E-17 | -2.43E+00 | 1.36E-01 | 5.20E-01 | 1.31E+00 | 8.10E-08 | 2.89E-07 | -1.81E+00 | 1.42E-02 | 4.00E-02 | 1.76E+00 | 3.80E-19 | 1.39E-18 | -2.98E+00 | 1.46E-01 | 4.01E-01 | -1.01E+00 |
| MYL6P3 | 5.57E-16 | 2.90E-15 | -1.57E+00 | 2.49E-15 | 1.07E-14 | -1.31E+00 | 8.70E-08 | 3.70E-07 | -1.44E+00 | 9.31E-09 | 3.56E-08 | -1.33E+00 | 3.09E-22 | 1.10E-21 | -2.06E+00 | 5.32E-25 | 1.50E-24 | -2.14E+00 | 5.58E-14 | 2.05E-13 | -1.72E+00 | 8.98E-12 | 2.47E-11 | -1.79E+00 |
| PLEK | 5.59E-16 | 2.91E-15 | -1.37E+00 | 2.69E-17 | 1.16E-16 | -1.34E+00 | 2.94E-08 | 1.25E-07 | -1.40E+00 | 9.44E-08 | 3.61E-07 | -1.20E+00 | 4.48E-02 | 1.60E-01 | -1.05E+00 | 2.38E-01 | 6.71E-01 | 1.10E+00 | 2.74E-16 | 1.01E-15 | 1.34E+00 | 4.19E-12 | 1.15E-11 | 1.29E+00 |
| INPP5D | 5.61E-16 | 2.92E-15 | -1.25E+00 | 2.40E-04 | 1.04E-03 | 1.21E+00 | 2.26E-05 | 9.60E-05 | -1.26E+00 | 7.05E-04 | 2.69E-03 | -1.01E+00 | 3.30E-03 | 1.18E-02 | -1.12E+00 | 1.55E-02 | 4.38E-02 | -1.05E+00 | 2.54E-10 | 9.33E-10 | -1.59E+00 | 1.49E-05 | 4.09E-05 | -1.20E+00 |
| GBP7 | 5.69E-16 | 2.96E-15 | -2.49E+00 | 1.18E-15 | 5.11E-15 | -2.23E+00 | 9.74E-09 | 4.14E-08 | -2.50E+00 | 3.05E-08 | 1.16E-07 | -2.23E+00 | 3.08E-02 | 1.10E-01 | -1.26E+00 | 1.75E-02 | 4.93E-02 | -1.65E+00 | 5.76E-07 | 2.12E-06 | -2.56E+00 | 7.55E-07 | 2.07E-06 | -2.80E+00 |
| ATAD2 | 5.86E-16 | 3.05E-15 | -1.58E+00 | 6.90E-27 | 2.98E-26 | -2.17E+00 | 8.46E-06 | 3.59E-05 | -1.16E+00 | 6.19E-14 | 2.36E-13 | -1.48E+00 | 3.27E-06 | 1.17E-05 | -1.39E+00 | 7.83E-11 | 2.21E-10 | -2.14E+00 | 2.04E-07 | 7.49E-07 | -1.28E+00 | 1.17E-09 | 3.22E-09 | -1.39E+00 |
| TM9SF3 | 5.95E-16 | 3.10E-15 | -1.38E+00 | 8.28E-18 | 3.58E-17 | -1.27E+00 | 4.98E-07 | 2.12E-06 | -1.26E+00 | 1.84E-07 | 7.04E-07 | -1.11E+00 | 1.29E-05 | 4.59E-05 | -1.14E+00 | 3.52E-04 | 9.93E-04 | -1.01E+00 | 2.51E-06 | 9.20E-06 | -1.21E+00 | 2.03E-06 | 5.58E-06 | -1.21E+00 |
| ZNF354B | 5.97E-16 | 3.11E-15 | -2.51E+00 | 8.62E-10 | 3.73E-09 | -1.53E+00 | 3.00E-07 | 1.27E-06 | -1.64E+00 | 9.76E-05 | 3.73E-04 | -1.14E+00 | 6.62E-03 | 2.36E-02 | -1.32E+00 | 7.36E-03 | 2.07E-02 | -1.24E+00 | 1.21E-04 | 4.46E-04 | -1.24E+00 | 1.16E-03 | 3.19E-03 | -1.14E+00 |
| CWC25 | 5.97E-16 | 3.11E-15 | -1.47E+00 | 6.87E-28 | 2.97E-27 | -1.83E+00 | 2.74E-07 | 1.16E-06 | -1.31E+00 | 3.97E-16 | 1.52E-15 | -1.49E+00 | 7.02E-07 | 2.50E-06 | -1.10E+00 | 6.62E-10 | 1.87E-09 | -1.39E+00 | 4.00E-07 | 1.47E-06 | -1.28E+00 | 4.77E-06 | 1.31E-05 | -1.30E+00 |
| BCL2A1 | 6.09E-16 | 3.17E-15 | -1.60E+00 | 2.73E-12 | 1.18E-11 | -1.21E+00 | 1.42E-02 | 6.02E-02 | -1.15E+00 | 2.38E-03 | 9.09E-03 | -1.15E+00 | 6.72E-06 | 2.40E-05 | -1.38E+00 | 8.12E-08 | 2.29E-07 | -1.69E+00 | 2.37E-04 | 8.71E-04 | -1.95E+00 | 9.02E-04 | 2.48E-03 | -1.91E+00 |
| SURP | 6.09E-16 | 3.17E-15 | -1.35E+00 | 1.10E-31 | 4.74E-31 | -1.74E+00 | 1.12E-04 | 4.76E-04 | -1.15E+00 | 1.38E-21 | 5.28E-21 | -1.70E+00 | 3.69E-11 | 1.32E-10 | -1.48E+00 | 3.39E-16 | 9.56E-16 | -1.75E+00 | 3.21E-03 | 1.18E-02 | -1.08E+00 | 2.41E-08 | 6.62E-08 | -1.36E+00 |
| EMC8 | 6.18E-16 | 3.22E-15 | -1.84E+00 | 8.24E-10 | 3.56E-09 | -1.22E+00 | 1.36E-04 | 5.78E-04 | -1.15E+00 | 4.89E-04 | 1.87E-03 | 1.03E+00 | 6.45E-09 | 2.30E-08 | -1.73E+00 | 3.51E-05 | 9.89E-05 | -1.16E+00 | 5.66E-04 | 2.08E-03 | -1.10E+00 | 2.16E-01 | 5.94E-01 | 1.18E+00 |
| TRIM44 | 6.25E-16 | 3.26E-15 | -1.41E+00 | 4.50E-09 | 1.95E-08 | -1.05E+00 | 2.82E-09 | 1.20E-08 | -1.45E+00 | 3.05E-05 | 1.16E-04 | -1.07E+00 | 1.80E-06 | 6.40E-06 | -1.21E+00 | 5.50E-04 | 1.55E-03 | -1.00E+00 | 1.71E-08 | 6.29E-08 | -1.33E+00 | 1.91E-03 | 5.24E-03 | -1.13E+00 |
| PSMA3-AS1 | 6.33E-16 | 3.30E-15 | -1.42E+00 | 1.80E-08 | 7.77E-08 | -1.04E+00 | 1.21E-10 | 5.16E-10 | -1.60E+00 | 4.43E-06 | 1.69E-05 | -1.24E+00 | 3.46E-06 | 1.23E-05 | -1.20E+00 | 1.81E-03 | 5.09E-03 | -1.00E+00 | 1.97E-06 | 7.25E-06 | -1.34E+00 | 1.15E-04 | 3.15E-04 | -1.24E+00 |
| PDCD10 | 6.45E-16 | 3.36E-15 | -1.28E+00 | 4.08E-07 | 1.77E-06 | 1.10E+00 | 4.09E-06 | 1.74E-05 | -1.22E+00 | 1.95E-02 | 7.45E-02 | 1.16E+00 | 2.36E-05 | 8.42E-05 | -1.08E+00 | 2.53E-01 | 7.13E-01 | 1.30E+00 | 5.67E-05 | 2.08E-04 | -1.13E+00 | 4.16E-02 | 1.14E-01 | -1.00E+00 |
| FTX | 6.53E-16 | 3.40E-15 | -1.33E+00 | 6.56E-02 | 2.84E-01 | 1.59E+00 | 1.06E-11 | 4.52E-11 | -1.63E+00 | 1.29E-01 | 4.93E-01 | 1.19E+00 | 1.22E-03 | 4.35E-03 | -1.05E+00 | 2.07E-01 | 5.84E-01 | 1.42E+00 | 3.22E-06 | 1.18E-05 | -1.18E+00 | 1.76E-01 | 4.84E-01 | 1.16E+00 |
| FAM83A | 6.54E-16 | 3.41E-15 | -1.83E+00 | 6.85E-05 | 2.96E-04 | -1.01E+00 | 1.86E-08 | 7.91E-08 | -2.33E+00 | 2.75E-02 | 1.05E-01 | -1.05E+00 | 2.53E-09 | 9.04E-09 | -1.50E+00 | 6.28E-05 | 1.77E-04 | -1.08E+00 | 4.94E-06 | 1.81E-05 | -1.93E+00 | 7.65E-02 | 2.10E-01 | -1.10E+00 |
| UBQLN2 | 6.54E-16 | 3.41E-15 | -1.66E+00 | 5.31E-12 | 2.29E-11 | -1.24E+00 | 1.17E-05 | 4.99E-05 | -1.34E+00 | 7.33E-05 | 2.80E-04 | -1.05E+00 | 5.89E-06 | 2.10E-05 | -1.50E+00 | 2.37E-03 | 6.68E-03 | -1.10E+00 | 1.09E-04 | 4.01E-04 | -1.32E+00 | 4.43E-02 | 1.22E-01 | 1.04E+00 |
| NR4A1 | 6.64E-16 | 3.46E-15 | 4.87E+00 | 1.54E-01 | 6.65E-01 | 1.12E+00 | 1.56E-07 | 6.62E-07 | 5.15E+00 | 1.67E-01 | 6.36E-01 | 1.95E+00 | 4.72E-09 | 1.68E-08 | 5.91E+00 | 1.17E-01 | 3.31E-01 | 2.26E+00 | 1.34E-05 | 4.91E-05 | 3.09E+00 | 2.19E-01 | 6.01E-01 | -1.03E+00 |
| AC245014.3 | 6.65E-16 | 3.46E-15 | 4.78E+00 | 1.61E-01 | 6.96E-01 | 1.41E+00 | 7.52E-07 | 3.19E-06 | 2.75E+00 | 8.18E-02 | 3.13E-01 | 1.10E+00 | 8.43E-02 | 3.01E-01 | 2.66E+00 | 9.48E-02 | 2.67E-01 | 2.35E+00 | 1.61E-01 | 5.90E-01 | 1.38E+00 | 1.30E-01 | 3.57E-01 | 1.05E+00 |
| CCDC115 | 6.65E-16 | 3.46E-15 | -1.23E+00 | 4.12E-08 | 1.78E-07 | 1.09E+00 | 3.13E-11 | 1.33E-10 | -1.40E+00 | 9.52E-06 | 3.64E-05 | -1.03E+00 | 4.53E-05 | 1.61E-04 | -1.17E+00 | 1.15E-01 | 3.23E-01 | 1.23E+00 | 2.49E-01 | 9.12E-01 | 1.18E+00 | 2.34E-01 | 6.43E-01 | 1.17E+00 |
| CHURC1 | 6.74E-16 | 3.51E-15 | -1.35E+00 | 4.72E-08 | 2.04E-07 | -1.04E+00 | 7.74E-08 | 3.29E-07 | -1.34E+00 | 5.39E-04 | 2.06E-03 | 1.06E+00 | 3.68E-06 | 1.31E-05 | -1.29E+00 | 4.93E-04 | 1.39E-03 | -1.05E+00 | 8.36E-02 | 3.07E-01 | 1.06E+00 | 1.90E-02 | 5.23E-02 | -1.00E+00 |
| RAD23A | 6.86E-16 | 3.58E-15 | -1.37E+00 | 4.13E-09 | 1.79E-08 | -1.00E+00 | 2.38E-10 | 1.01E-09 | -1.45E+00 | 1.23E-07 | 4.71E-07 | -1.16E+00 | 5.14E-08 | 1.83E-07 | -1.48E+00 | 2.61E-05 | 7.37E-05 | -1.24E+00 | 6.06E-09 | 2.23E-08 | -1.37E+00 | 1.07E-03 | 2.95E-03 | -1.05E+00 |
| BTG2 | 6.87E-16 | 3.58E-15 | 3.27E+00 | 4.77E-05 | 2.06E-04 | 1.24E+00 | 9.61E-07 | 4.08E-06 | 2.86E+00 | 5.57E-03 | 2.13E-02 | 1.14E+00 | 8.32E-15 | 2.97E-14 | 4.16E+00 | 1.04E-02 | 2.92E-02 | 1.31E+00 | 5.41E-04 | 1.99E-03 | 2.91E+00 | 1.64E-07 | 4.52E-07 | -1.64E+00 |
| FAM184A | 7.94E-16 | 4.14E-15 | -1.89E+00 | 1.14E-10 | 4.94E-10 | -1.23E+00 | 6.04E-07 | 2.57E-06 | -1.28E+00 | 5.94E-07 | 2.27E-06 | -1.13E+00 | 2.14E-01 | 7.65E-01 | 1.26E+00 | 2.12E-01 | 5.97E-01 | 1.26E+00 | 4.08E-03 | 1.50E-02 | -1.68E+00 | 8.56E-02 | 2.35E-01 | -1.14E+00 |
| PPIAP6 | 8.00E-16 | 4.17E-15 | -1.29E+00 | 1.60E-13 | 6.91E-13 | -1.12E+00 | 9.71E-05 | 4.13E-04 | -1.27E+00 | 2.04E-04 | 7.81E-04 | -1.15E+00 | 1.88E-09 | 6.70E-09 | -1.35E+00 | 1.19E-05 | 3.37E-05 | -1.08E+00 | 1.56E-08 | 5.74E-08 | -1.43E+00 | 1.69E-02 | 4.63E-02 | -1.12E+00 |
| HTATIP2 | 8.31E-16 | 4.33E-15 | -1.32E+00 | 1.03E-06 | 4.45E-06 | 1.07E+00 | 1.76E-05 | 7.48E-05 | -1.24E+00 | 7.10E-05 | 2.71E-04 | -1.00E+00 | 1.56E-03 | 5.58E-03 | -1.02E+00 | 6.03E-02 | 1.70E-01 | 1.22E+00 | 3.16E-02 | 1.16E-01 | 1.01E+00 | 6.05E-02 | 1.66E-01 | 1.02E+00 |
| LCN6 | 8.39E-16 | 4.37E-15 | 1.21E+01 | 6.74E-05 | 2.91E-04 | 6.06E+00 | 6.96E-05 | 2.96E-04 | 5.25E+00 | 4.25E-03 | 1.62E-02 | 4.22E+00 | 8.20E-08 | 2.93E-07 | 1.10E+01 | 2.60E-02 | 7.32E-02 | 5.04E+00 | 1.12E-01 | 4.10E-01 | 1.96E+00 | 3.07E-01 | 8.43E-01 | 1.27E+00 |
| CDC42 | 8.53E-16 | 4.44E-15 | -1.33E+00 | 1.19E-11 | 5.16E-11 | -1.16E+00 | 2.91E-09 | 1.24E-08 | -1.41E+00 | 7.88E-06 | 3.01E-05 | -1.22E+00 | 7.01E-08 | 2.50E-07 | -1.35E+00 | 6.82E-05 | 1.92E-04 | -1.23E+00 | 3.50E-08 | 1.28E-07 | -1.32E+00 | 7.36E-04 | 2.02E-03 | -1.19E+00 |
| STRAP | 8.59E-16 | 4.48E-15 | -1.23E+00 | 1.75E-16 | 7.54E-16 | -1.11E+00 | 9.48E-05 | 4.03E-04 | -1.14E+00 | 6.57E-04 | 2.51E-03 | -1.05E+00 | 2.21E-06 | 7.86E-06 | -1.22E+00 | 1.00E-05 | 2.82E-05 | -1.12E+00 | 3.61E-03 | 1.33E-02 | -1.09E+00 | 1.27E-01 | 3.48E-01 | 1.06E+00 |
| AC007318.1 | 8.63E-16 | 4.50E-15 | -1.82E+00 | 3.09E-08 | 1.34E-07 | -1.12E+00 | 1.67E-17 | 7.08E-17 | -1.80E+00 | 1.92E-10 | 7.33E-10 | -1.22E+00 | 4.25E-08 | 1.52E-07 | -1.62E+00 | 1.45E-03 | 4.09E-03 | -1.07E+00 | 2.24E-13 | 8.23E-13 | -1.80E+00 | 3.38E-04 | 9.30E-04 | -1.10E+00 |
| TRIM27 | 8.69E-16 | 4.53E-15 | -1.77E+00 | 1.72E-06 | 7.44E-06 | -1.03E+00 | 3.03E-05 | 1.29E-04 | -1.31E+00 | 6.79E-02 | 2.59E-01 | 1.17E+00 | 1.60E-14 | 5.70E-14 | -2.16E+00 | 2.50E-06 | 7.06E-06 | -1.25E+00 | 2.58E-19 | 9.47E-19 | -2.02E+00 | 8.54E-04 | 2.35E-03 | -1.04E+00 |
| NDUFV3 | 8.91E-16 | 4.64E-15 | -1.42E+00 | 4.69E-11 | 2.03E-10 | -1.13E+00 | 4.99E-09 | 2.12E-08 | -1.42E+00 | 3.04E-08 | 1.16E-07 | -1.22E+00 | 7.09E-14 | 2.53E-13 | -1.69E+00 | 4.90E-07 | 1.38E-06 | -1.24E+00 | 4.08E-10 | 1.50E-09 | -1.44E+00 | 9.49E-07 | 2.61E-06 | -1.25E+00 |
| GAB3 | 8.92E-16 | 4.65E-15 | -1.62E+00 | 1.33E-04 | 5.73E-04 | 1.07E+00 | 5.11E-16 | 2.17E-15 | -1.83E+00 | 2.43E-05 | 9.28E-05 | -1.03E+00 | 8.99E-07 | 3.20E-06 | -1.29E+00 | 1.10E-02 | 3.10E-02 | 1.05E+00 | 3.00E-10 | 1.10E-09 | -1.52E+00 | 4.58E-04 | 1.26E-03 | -1.07E+00 |
| HLA-DRB6 | 8.92E-16 | 4.65E-15 | -1.88E+00 | 1.36E-07 | 5.89E-07 | -1.41E+00 | 2.24E-10 | 9.53E-10 | -1.90E+00 | 1.29E-04 | 4.92E-04 | -1.38E+00 | 1.51E-39 | 5.39E-39 | -2.10E+00 | 3.38E-21 | 9.54E-21 | -1.70E+00 | 6.67E-65 | 2.45E-64 | -2.85E+00 | 4.81E-21 | 1.32E-20 | -1.79E+00 |
| RPIA | 8.96E-16 | 4.67E-15 | -1.52E+00 | 1.08E-09 | 4.68E-09 | -1.06E+00 | 5.09E-07 | 2.16E-06 | -1.24E+00 | 1.94E-06 | 7.42E-06 | -1.02E+00 | 1.44E-07 | 5.13E-07 | -1.51E+00 | 4.49E-03 | 1.26E-02 | -1.04E+00 | 4.54E-05 | 1.67E-04 | -1.20E+00 | 2.64E-02 | 7.25E-02 | 1.03E+00 |
| SEC62 | 9.02E-16 | 4.70E-15 | -1.26E+00 | 1.06E-20 | 4.59E-20 | -1.32E+00 | 1.33E-06 | 5.65E-06 | -1.25E+00 | 1.29E-08 | 4.92E-08 | -1.31E+00 | 1.49E-12 | 5.32E-12 | -1.49E+00 | 1.81E-13 | 5.09E-13 | -1.58E+00 | 3.08E-06 | 1.13E-05 | -1.28E+00 | 1.29E-07 | 3.54E-07 | -1.40E+00 |
| FEZ2 | 9.02E-16 | 4.70E-15 | -1.56E+00 | 1.29E-06 | 5.57E-06 | -1.00E+00 | 3.27E-08 | 1.39E-07 | -1.30E+00 | 3.17E-06 | 1.21E-05 | -1.06E+00 | 7.70E-11 | 2.75E-10 | -1.52E+00 | 9.30E-06 | 2.62E-05 | -1.17E+00 | 2.59E-05 | 9.50E-05 | -1.15E+00 | 4.91E-03 | 1.35E-02 | -1.03E+00 |
| XAF1 | 9.10E-16 | 4.74E-15 | -2.56E+00 | 4.73E-14 | 2.05E-13 | -2.10E+00 | 1.65E-35 | 7.00E-35 | -4.20E+00 | 5.40E-36 | 2.06E-35 | -3.98E+00 | 3.59E-19 | 1.28E-18 | -3.07E+00 | 6.98E-22 | 1.97E-21 | -3.18E+00 | 6.51E-29 | 2.39E-28 | -4.29E+00 | 2.41E-29 | 6.62E-29 | -4.84E+00 |
| CREB1 | 9.26E-16 | 4.83E-15 | -1.53E+00 | 3.00E-07 | 1.30E-06 | 1.01E+00 | 3.20E-11 | 1.36E-10 | -1.60E+00 | 2.31E-06 | 8.82E-06 | -1.14E+00 | 2.15E-11 | 7.68E-11 | -1.55E+00 | 2.87E-08 | 8.09E-08 | -1.33E+00 | 1.48E-11 | 5.43E-11 | -1.53E+00 | 3.83E-04 | 1.05E-03 | -1.08E+00 |
| TMEM109 | 9.44E-16 | 4.92E-15 | -1.87E+00 | 7.85E-05 | 3.39E-04 | -1.02E+00 | 7.73E-04 | 3.29E-03 | -1.17E+00 | 1.91E-02 | 7.31E-02 | 1.15E+00 | 9.20E-07 | 3.28E-06 | -1.47E+00 | 2.97E-02 | 8.37E-02 | 1.06E+00 | 5.18E-06 | 1.90E-05 | -1.22E+00 | 3.23E-01 | 8.87E-01 | 1.18E+00 |
| DNPEP | 9.55E-16 | 4.97E-15 | -1.45E+00 | 2.19E-12 | 9.48E-12 | -1.19E+00 | 6.36E-07 | 2.70E-06 | -1.18E+00 | 6.06E-07 | 2.31E-06 | -1.03E+00 | 8.04E-09 | 2.87E-08 | -1.35E+00 | 2.65E-05 | 7.48E-05 | -1.05E+00 | 3.10E-07 | 1.14E-06 | -1.21E+00 | 6.42E-03 | 1.76E-02 | -1.01E+00 |
| TRMT112 | 9.61E-16 | 5.01E-15 | -1.27E+00 | 1.29E-10 | 5.56E-10 | -1.08E+00 | 1.70E-02 | 7.22E-02 | -1.13E+00 | 7.36E-02 | 2.81E-01 | 1.02E+00 | 2.19E-07 | 7.80E-07 | -1.29E+00 | 2.52E-02 | 7.11E-02 | 1.00E+00 | 3.20E-05 | 1.17E-04 | -1.20E+00 | 1.68E-01 | 4.62E-01 | 1.03E+00 |
| WIPI2 | 9.78E-16 | 5.10E-15 | -1.36E+00 | 1.78E-08 | 7.68E-08 | 1.06E+00 | 1.60E-06 | 6.78E-06 | -1.25E+00 | 1.74E-04 | 6.63E-04 | -1.00E+00 | 3.01E-04 | 1.07E-03 | -1.05E+00 | 1.01E-02 | 2.84E-02 | 1.02E+00 | 9.59E-05 | 3.52E-04 | -1.20E+00 | 2.72E-02 | 7.48E-02 | -1.00E+00 |
| UFL1 | 9.88E-16 | 5.15E-15 | -1.58E+00 | 1.80E-13 | 7.79E-13 | -1.30E+00 | 1.36E-11 | 5.77E-11 | -1.59E+00 | 6.93E-12 | 2.65E-11 | -1.41E+00 | 3.44E-13 | 1.22E-12 | -1.88E+00 | 2.21E-06 | 6.23E-06 | -1.20E+00 | 1.90E-10 | 6.99E-10 | -1.63E+00 | 8.36E-06 | 2.30E-05 | -1.17E+00 |
| STX7 | 1.03E-15 | 5.35E-15 | -1.58E+00 | 2.41E-02 | 1.04E-01 | 1.36E+00 | 1.10E-11 | 4.66E-11 | -1.74E+00 | 8.05E-04 | 3.07E-03 | 1.04E+00 | 3.85E-12 | 1.37E-11 | -1.78E+00 | 1.49E-04 | 4.21E-04 | -1.02E+00 | 3.38E-11 | 1.24E-10 | -1.49E+00 | 3.32E-03 | 9.12E-03 | -1.00E+00 |
| TMEM192 | 1.04E-15 | 5.40E-15 | -1.70E+00 | 6.80E-10 | 2.94E-09 | -1.10E+00 | 2.17E-10 | 9.20E-10 | -1.60E+00 | 5.42E-06 | 2.07E-05 | -1.10E+00 | 1.31E-08 | 4.67E-08 | -1.73E+00 | 2.42E-04 | 6.82E-04 | -1.10E+00 | 3.92E-07 | 1.44E-06 | -1.31E+00 | 2.05E-03 | 5.64E-03 | -1.02E+00 |
| ATP5B | 1.04E-15 | 5.43E-15 | -1.30E+00 | 3.04E-04 | 1.31E-03 | -1.04E+00 | 9.12E-03 | 3.88E-02 | -1.13E+00 | 8.73E-02 | 3.34E-01 | -1.04E+00 | 3.81E-12 | 1.36E-11 | -1.40E+00 | 3.11E-02 | 8.76E-02 | -1.08E+00 | 8.17E-05 | 3.00E-04 | -1.23E+00 | 2.56E-01 | 7.03E-01 | -1.00E+00 |
| AAMP | 1.06E-15 | 5.51E-15 | -1.59E+00 | 2.19E-06 | 9.46E-06 | 1.03E+00 | 1.35E-06 | 5.73E-06 | -1.22E+00 | 2.58E-04 | 9.87E-04 | 1.02E+00 | 9.54E-09 | 3.40E-08 | -1.37E+00 | 2.47E-02 | 6.96E-02 | 1.21E+00 | 7.56E-05 | 2.77E-04 | -1.08E+00 | 2.70E-01 | 7.42E-01 | 1.26E+00 |
| TIAL1 | 1.09E-15 | 5.66E-15 | -1.39E+00 | 1.08E-07 | 4.65E-07 | 1.02E+00 | 7.41E-10 | 3.15E-09 | -1.49E+00 | 1.02E-04 | 3.91E-04 | -1.11E+00 | 1.28E-08 | 4.55E-08 | -1.40E+00 | 2.73E-03 | 7.70E-03 | -1.02E+00 | 5.18E-09 | 1.90E-08 | -1.40E+00 | 1.76E-03 | 4.83E-03 | -1.09E+00 |
| HP1BP3 | 1.10E-15 | 5.74E-15 | -1.34E+00 | 1.06E-05 | 4.56E-05 | 1.03E+00 | 1.50E-05 | 6.39E-05 | -1.23E+00 | 7.02E-03 | 2.68E-02 | 1.07E+00 | 8.75E-07 | 3.12E-06 | -1.20E+00 | 6.04E-03 | 1.70E-02 | 1.05E+00 | 3.65E-04 | 1.34E-03 | -1.19E+00 | 5.07E-03 | 1.39E-02 | -1.09E+00 |
| CCT8P1 | 1.11E-15 | 5.76E-15 | -1.73E+00 | 1.78E-19 | 7.70E-19 | -1.81E+00 | 1.18E-07 | 5.02E-07 | -1.49E+00 | 9.81E-12 | 3.75E-11 | -1.63E+00 | 1.44E-06 | 5.14E-06 | -1.42E+00 | 6.64E-08 | 1.87E-07 | -1.58E+00 | 6.51E-08 | 2.39E-07 | -1.37E+00 | 4.23E-17 | 1.16E-16 | -2.08E+00 |
| ZBTB11 | 1.12E-15 | 5.85E-15 | -1.93E+00 | 1.59E-21 | 6.86E-21 | -2.16E+00 | 6.39E-11 | 2.72E-10 | -1.49E+00 | 1.97E-19 | 7.53E-19 | -1.89E+00 | 3.38E-06 | 1.21E-05 | -1.38E+00 | 3.96E-09 | 1.12E-08 | -1.84E+00 | 4.18E-22 | 1.53E-21 | -2.22E+00 | 2.68E-22 | 7.36E-22 | -2.17E+00 |
| FAM89A | 1.14E-15 | 5.94E-15 | -2.56E+00 | 1.50E-13 | 6.46E-13 | -2.07E+00 | 1.05E-11 | 4.46E-11 | -2.00E+00 | 2.85E-11 | 1.09E-10 | -1.73E+00 | 1.32E-03 | 4.71E-03 | -2.84E+00 | 2.69E-03 | 7.58E-03 | -2.39E+00 | 7.10E-03 | 2.60E-02 | -1.59E+00 | 2.30E-02 | 6.32E-02 | -1.47E+00 |
| ZHX1 | 1.16E-15 | 6.06E-15 | -1.73E+00 | 9.72E-05 | 4.20E-04 | 1.12E+00 | 9.24E-18 | 3.93E-17 | -1.91E+00 | 1.90E-09 | 7.26E-09 | -1.18E+00 | 2.96E-11 | 1.05E-10 | -1.82E+00 | 4.47E-05 | 1.26E-04 | -1.14E+00 | 7.86E-11 | 2.89E-10 | -1.48E+00 | 2.36E-02 | 6.49E-02 | 1.01E+00 |
| TMEM57 | 1.16E-15 | 6.06E-15 | -1.88E+00 | 5.87E-12 | 2.54E-11 | -1.31E+00 | 2.08E-09 | 8.83E-09 | -1.59E+00 | 3.76E-07 | 1.44E-06 | -1.27E+00 | 8.79E-17 | 3.13E-16 | -1.94E+00 | 1.49E-09 | 4.19E-09 | -1.34E+00 | 6.63E-25 | 2.43E-24 | -2.06E+00 | 9.75E-14 | 2.68E-13 | -1.48E+00 |
| PLPBP | 1.16E-15 | 6.06E-15 | -1.53E+00 | 1.43E-03 | 6.18E-03 | 1.20E+00 | 1.49E-05 | 6.32E-05 | -1.27E+00 | 1.03E-01 | 3.95E-01 | 1.26E+00 | 2.36E-10 | 8.40E-10 | -1.60E+00 | 4.42E-03 | 1.25E-02 | 1.11E+00 | 8.46E-08 | 3.11E-07 | -1.41E+00 | 1.67E-01 | 4.57E-01 | 1.13E+00 |
| HLA-DOA | 1.20E-15 | 6.24E-15 | -1.48E+00 | 8.15E-09 | 3.52E-08 | -1.02E+00 | 1.16E-08 | 4.91E-08 | -1.39E+00 | 8.43E-05 | 3.22E-04 | 1.02E+00 | 1.45E-02 | 5.15E-02 | -1.62E+00 | 3.06E-02 | 8.62E-02 | -1.35E+00 | 4.97E-03 | 1.82E-02 | -2.14E+00 | 3.30E-02 | 9.05E-02 | -1.52E+00 |
| HIST1H4C | 1.22E-15 | 6.36E-15 | -1.31E+00 | 3.24E-61 | 1.40E-60 | -2.22E+00 | 2.98E-02 | 1.27E-01 | 1.41E+00 | 1.92E-14 | 7.34E-14 | -1.14E+00 | 1.90E-06 | 6.79E-06 | -1.28E+00 | 1.69E-24 | 4.77E-24 | -2.17E+00 | 4.95E-03 | 1.82E-02 | 1.09E+00 | 3.14E-11 | 8.62E-11 | -1.44E+00 |
| POGLUT1 | 1.24E-15 | 6.47E-15 | -2.07E+00 | 2.13E-05 | 9.21E-05 | -1.07E+00 | 3.22E-13 | 1.37E-12 | -1.83E+00 | 1.83E-03 | 7.01E-03 | 1.12E+00 | 1.58E-04 | 5.63E-04 | -1.53E+00 | 3.05E-01 | 8.60E-01 | 1.33E+00 | 7.11E-18 | 2.61E-17 | -2.28E+00 | 8.82E-06 | 2.42E-05 | -1.35E+00 |
| HAPLN3 | 1.25E-15 | 6.49E-15 | -2.29E+00 | 4.51E-14 | 1.95E-13 | -1.71E+00 | 1.76E-11 | 7.47E-11 | -2.73E+00 | 7.48E-08 | 2.86E-07 | -1.85E+00 | 2.72E-01 | 9.69E-01 | 1.16E+00 | 2.58E-01 | 7.26E-01 | -1.21E+00 | 1.19E-01 | 4.37E-01 | -1.06E+00 | 3.18E-01 | 8.72E-01 | 1.07E+00 |
| KLF15 | 1.25E-15 | 6.52E-15 | -2.51E+00 | 1.98E-14 | 8.57E-14 | -2.08E+00 | 3.62E-23 | 1.54E-22 | -3.42E+00 | 1.97E-20 | 7.52E-20 | -2.60E+00 | 6.72E-05 | 2.40E-04 | -1.60E+00 | 4.28E-06 | 1.21E-05 | -1.86E+00 | 1.79E-18 | 6.58E-18 | -2.44E+00 | 6.20E-16 | 1.70E-15 | -2.15E+00 |
| CD164 | 1.26E-15 | 6.57E-15 | -1.46E+00 | 3.76E-04 | 1.63E-03 | 1.41E+00 | 1.03E-03 | 4.37E-03 | -1.21E+00 | 8.83E-02 | 3.37E-01 | 1.30E+00 | 7.23E-12 | 2.58E-11 | -1.44E+00 | 5.56E-02 | 1.57E-01 | 1.09E+00 | 2.68E-07 | 9.82E-07 | -1.25E+00 | 2.94E-01 | 8.07E-01 | 1.14E+00 |
| TOM1 | 1.28E-15 | 6.68E-15 | -1.93E+00 | 1.74E-16 | 7.52E-16 | -1.86E+00 | 2.07E-12 | 8.82E-12 | -1.81E+00 | 3.57E-14 | 1.37E-13 | -1.85E+00 | 7.24E-09 | 2.58E-08 | -1.52E+00 | 4.50E-11 | 1.27E-10 | -1.70E+00 | 2.24E-04 | 8.23E-04 | -1.42E+00 | 1.97E-05 | 5.40E-05 | -1.69E+00 |
| RASSF3 | 1.29E-15 | 6.74E-15 | -3.09E+00 | 6.75E-09 | 2.92E-08 | -1.57E+00 | 5.57E-16 | 2.37E-15 | -2.50E+00 | 8.38E-12 | 3.20E-11 | -1.78E+00 | 3.42E-06 | 1.22E-05 | -2.65E+00 | 4.65E-04 | 1.31E-03 | -1.70E+00 | 2.75E-08 | 1.01E-07 | -1.73E+00 | 3.63E-06 | 9.98E-06 | -1.53E+00 |
| HSPA4 | 1.30E-15 | 6.79E-15 | -1.63E+00 | 3.07E-10 | 1.32E-09 | -1.17E+00 | 3.24E-10 | 1.38E-09 | -1.49E+00 | 1.57E-05 | 6.01E-05 | -1.02E+00 | 2.50E-09 | 8.93E-09 | -1.49E+00 | 1.54E-05 | 4.34E-05 | -1.10E+00 | 5.05E-07 | 1.85E-06 | -1.32E+00 | 6.57E-03 | 1.80E-02 | 1.03E+00 |
| VDAC3 | 1.31E-15 | 6.82E-15 | -1.22E+00 | 6.98E-06 | 3.02E-05 | 1.07E+00 | 3.00E-04 | 1.27E-03 | -1.08E+00 | 5.50E-03 | 2.10E-02 | 1.04E+00 | 2.24E-10 | 8.00E-10 | -1.43E+00 | 5.04E-02 | 1.42E-01 | 1.05E+00 | 7.29E-04 | 2.68E-03 | -1.10E+00 | 2.76E-01 | 7.58E-01 | 1.11E+00 |
| RPL7L1 | 1.31E-15 | 6.83E-15 | -1.29E+00 | 8.12E-08 | 3.51E-07 | 1.08E+00 | 8.63E-05 | 3.67E-04 | -1.22E+00 | 2.57E-02 | 9.80E-02 | 1.09E+00 | 1.51E-05 | 5.40E-05 | -1.10E+00 | 8.88E-02 | 2.50E-01 | 1.18E+00 | 6.55E-03 | 2.40E-02 | -1.08E+00 | 2.72E-01 | 7.46E-01 | 1.14E+00 |
| EEF1B2P1 | 1.32E-15 | 6.90E-15 | -1.66E+00 | 2.89E-29 | 1.25E-28 | -2.72E+00 | 4.72E-08 | 2.01E-07 | -1.41E+00 | 4.13E-23 | 1.58E-22 | -2.51E+00 | 5.59E-06 | 1.99E-05 | -1.22E+00 | 1.44E-12 | 4.07E-12 | -2.06E+00 | 3.32E-13 | 1.22E-12 | -1.66E+00 | 2.14E-23 | 5.87E-23 | -2.75E+00 |
| UBE3A | 1.33E-15 | 6.91E-15 | -1.43E+00 | 1.39E-13 | 6.00E-13 | -1.22E+00 | 2.68E-07 | 1.14E-06 | -1.31E+00 | 3.31E-07 | 1.26E-06 | -1.17E+00 | 4.68E-08 | 1.67E-07 | -1.25E+00 | 8.94E-05 | 2.52E-04 | -1.06E+00 | 2.79E-10 | 1.03E-09 | -1.37E+00 | 5.16E-05 | 1.42E-04 | -1.14E+00 |
| PSMD11 | 1.33E-15 | 6.93E-15 | -1.65E+00 | 8.57E-10 | 3.70E-09 | -1.18E+00 | 1.44E-07 | 6.14E-07 | -1.30E+00 | 6.90E-08 | 2.64E-07 | -1.12E+00 | 6.41E-10 | 2.29E-09 | -1.47E+00 | 4.12E-05 | 1.16E-04 | -1.13E+00 | 3.27E-10 | 1.20E-09 | -1.50E+00 | 1.00E-03 | 2.75E-03 | -1.12E+00 |
| RBPJ | 1.34E-15 | 6.98E-15 | -1.45E+00 | 1.46E-12 | 6.32E-12 | -1.18E+00 | 3.95E-10 | 1.68E-09 | -1.42E+00 | 3.31E-08 | 1.26E-07 | -1.15E+00 | 2.63E-06 | 9.39E-06 | -1.25E+00 | 5.99E-05 | 1.69E-04 | -1.06E+00 | 4.95E-12 | 1.82E-11 | -1.58E+00 | 2.33E-05 | 6.41E-05 | -1.24E+00 |
| NINJ1 | 1.34E-15 | 6.98E-15 | -1.84E+00 | 1.24E-15 | 5.35E-15 | -1.67E+00 | 7.95E-25 | 3.38E-24 | -2.35E+00 | 1.56E-23 | 5.97E-23 | -1.92E+00 | 3.56E-04 | 1.27E-03 | -2.39E+00 | 8.47E-05 | 2.39E-04 | -3.69E+00 | 4.18E-03 | 1.53E-02 | -1.60E+00 | 3.37E-03 | 9.25E-03 | -1.80E+00 |
| CSNK1A1L | 1.34E-15 | 7.00E-15 | -1.79E+00 | 2.17E-20 | 9.38E-20 | -2.21E+00 | 8.39E-07 | 3.56E-06 | -1.41E+00 | 7.29E-11 | 2.78E-10 | -1.66E+00 | 3.93E-05 | 1.40E-04 | -1.31E+00 | 2.51E-07 | 7.08E-07 | -1.64E+00 | 8.80E-15 | 3.23E-14 | -1.70E+00 | 2.95E-15 | 8.09E-15 | -1.82E+00 |
| RNF181 | 1.36E-15 | 7.10E-15 | -1.28E+00 | 5.26E-21 | 2.27E-20 | -1.30E+00 | 1.02E-03 | 4.33E-03 | -1.18E+00 | 1.02E-07 | 3.89E-07 | -1.22E+00 | 1.39E-09 | 4.96E-09 | -1.36E+00 | 9.88E-08 | 2.78E-07 | -1.27E+00 | 5.29E-03 | 1.94E-02 | -1.07E+00 | 1.96E-05 | 5.38E-05 | -1.26E+00 |
| RPS15AP12 | 1.41E-15 | 7.33E-15 | -1.53E+00 | 3.77E-48 | 1.63E-47 | -1.94E+00 | 3.67E-07 | 1.56E-06 | -1.47E+00 | 3.34E-23 | 1.27E-22 | -1.94E+00 | 3.97E-10 | 1.41E-09 | -1.50E+00 | 4.52E-21 | 1.27E-20 | -1.88E+00 | 1.55E-11 | 5.68E-11 | -1.51E+00 | 4.83E-25 | 1.33E-24 | -2.08E+00 |
| SLC16A9 | 1.42E-15 | 7.41E-15 | -2.98E+00 | 3.91E-10 | 1.69E-09 | -1.74E+00 | 3.37E-04 | 1.43E-03 | -1.44E+00 | 1.22E-04 | 4.64E-04 | -1.61E+00 | 1.90E-01 | 6.79E-01 | -1.17E+00 | 2.96E-01 | 8.35E-01 | 1.02E+00 | 2.24E-03 | 8.23E-03 | -1.51E+00 | 1.84E-02 | 5.05E-02 | -1.32E+00 |
| COQ2 | 1.43E-15 | 7.45E-15 | -2.05E+00 | 3.41E-13 | 1.47E-12 | -1.60E+00 | 4.01E-06 | 1.70E-05 | -1.54E+00 | 1.10E-06 | 4.20E-06 | -1.34E+00 | 2.84E-03 | 1.01E-02 | -1.57E+00 | 2.19E-02 | 6.18E-02 | -1.17E+00 | 3.63E-07 | 1.33E-06 | -1.41E+00 | 5.63E-06 | 1.55E-05 | -1.34E+00 |
| TMEM50B | 1.44E-15 | 7.48E-15 | -2.05E+00 | 5.26E-08 | 2.27E-07 | -1.10E+00 | 6.60E-10 | 2.80E-09 | -1.75E+00 | 9.52E-09 | 3.64E-08 | -1.43E+00 | 7.81E-06 | 2.78E-05 | -1.50E+00 | 5.68E-03 | 1.60E-02 | -1.04E+00 | 9.82E-06 | 3.61E-05 | -1.27E+00 | 3.84E-03 | 1.06E-02 | -1.10E+00 |
| LSM14A | 1.45E-15 | 7.57E-15 | -1.40E+00 | 9.79E-09 | 4.23E-08 | -1.07E+00 | 2.57E-05 | 1.09E-04 | -1.21E+00 | 3.24E-05 | 1.24E-04 | -1.04E+00 | 1.38E-09 | 4.91E-09 | -1.39E+00 | 5.22E-07 | 1.47E-06 | -1.23E+00 | 8.59E-03 | 3.15E-02 | -1.09E+00 | 2.96E-01 | 8.12E-01 | 1.05E+00 |
| GTF2H1 | 1.46E-15 | 7.59E-15 | -1.81E+00 | 1.22E-08 | 5.29E-08 | -1.15E+00 | 3.16E-06 | 1.34E-05 | -1.39E+00 | 1.05E-03 | 4.00E-03 | 1.01E+00 | 5.70E-09 | 2.03E-08 | -1.63E+00 | 2.59E-03 | 7.30E-03 | -1.01E+00 | 3.09E-06 | 1.13E-05 | -1.13E+00 | 1.85E-01 | 5.09E-01 | 1.20E+00 |
| CHMP3 | 1.46E-15 | 7.62E-15 | -1.73E+00 | 4.78E-12 | 2.07E-11 | -1.36E+00 | 3.12E-10 | 1.33E-09 | -1.50E+00 | 2.71E-12 | 1.04E-11 | -1.47E+00 | 6.15E-16 | 2.19E-15 | -1.88E+00 | 8.04E-11 | 2.27E-10 | -1.36E+00 | 3.65E-05 | 1.34E-04 | -1.17E+00 | 1.18E-03 | 3.23E-03 | -1.08E+00 |
| YARS | 1.47E-15 | 7.65E-15 | -1.54E+00 | 1.35E-11 | 5.83E-11 | -1.21E+00 | 1.36E-10 | 5.77E-10 | -1.47E+00 | 1.04E-09 | 3.96E-09 | -1.17E+00 | 1.08E-13 | 3.86E-13 | -1.49E+00 | 5.20E-05 | 1.47E-04 | -1.07E+00 | 2.02E-05 | 7.40E-05 | -1.25E+00 | 2.32E-01 | 6.36E-01 | 1.06E+00 |
| TUBA4A | 1.48E-15 | 7.69E-15 | -2.18E+00 | 3.11E-14 | 1.35E-13 | -1.94E+00 | 1.48E-02 | 6.28E-02 | -1.31E+00 | 1.75E-03 | 6.68E-03 | -1.47E+00 | 1.60E-01 | 5.71E-01 | 1.35E+00 | 2.90E-01 | 8.17E-01 | 1.37E+00 | 6.20E-04 | 2.28E-03 | 1.70E+00 | 8.30E-06 | 2.28E-05 | 1.04E+00 |
| GOPC | 1.49E-15 | 7.74E-15 | -1.57E+00 | 1.73E-09 | 7.49E-09 | -1.12E+00 | 3.34E-12 | 1.42E-11 | -1.68E+00 | 2.58E-06 | 9.86E-06 | -1.13E+00 | 3.92E-14 | 1.40E-13 | -1.75E+00 | 7.02E-10 | 1.98E-09 | -1.37E+00 | 1.36E-18 | 4.98E-18 | -1.79E+00 | 2.61E-07 | 7.16E-07 | -1.29E+00 |
| RFTN1 | 1.49E-15 | 7.78E-15 | -1.86E+00 | 3.05E-04 | 1.32E-03 | 1.17E+00 | 3.64E-04 | 1.55E-03 | -1.20E+00 | 2.46E-01 | 9.41E-01 | 1.50E+00 | 1.54E-03 | 5.48E-03 | -1.07E+00 | 3.12E-01 | 8.80E-01 | 1.48E+00 | 1.09E-01 | 4.01E-01 | 1.13E+00 | 2.92E-01 | 8.02E-01 | 1.32E+00 |
| MBP | 1.51E-15 | 7.86E-15 | -1.39E+00 | 1.98E-07 | 8.57E-07 | 1.01E+00 | 4.15E-14 | 1.76E-13 | -1.77E+00 | 3.61E-05 | 1.38E-04 | -1.13E+00 | 2.09E-14 | 7.45E-14 | -1.61E+00 | 1.07E-05 | 3.01E-05 | -1.16E+00 | 2.03E-13 | 7.44E-13 | -1.44E+00 | 3.86E-05 | 1.06E-04 | -1.20E+00 |
| ENTPD1 | 1.53E-15 | 7.96E-15 | -2.34E+00 | 8.66E-08 | 3.74E-07 | -1.26E+00 | 1.59E-08 | 6.76E-08 | -2.04E+00 | 3.14E-03 | 1.20E-02 | -1.05E+00 | 3.37E-05 | 1.20E-04 | -1.95E+00 | 1.70E-03 | 4.78E-03 | -1.34E+00 | 5.74E-02 | 2.11E-01 | -1.04E+00 | 2.18E-01 | 6.00E-01 | 1.44E+00 |
| LYRM1 | 1.55E-15 | 8.09E-15 | -1.46E+00 | 1.47E-17 | 6.35E-17 | -1.38E+00 | 2.40E-07 | 1.02E-06 | -1.33E+00 | 8.10E-08 | 3.09E-07 | -1.19E+00 | 1.97E-05 | 7.03E-05 | -1.29E+00 | 3.21E-06 | 9.06E-06 | -1.35E+00 | 2.58E-01 | 9.47E-01 | 1.26E+00 | 1.52E-01 | 4.16E-01 | 1.51E+00 |
| HSPA8 | 1.56E-15 | 8.14E-15 | -1.43E+00 | 1.54E-02 | 6.67E-02 | 1.04E+00 | 1.02E-02 | 4.32E-02 | -1.19E+00 | 2.62E-02 | 1.00E-01 | 1.22E+00 | 3.57E-10 | 1.27E-09 | -1.68E+00 | 9.00E-02 | 2.54E-01 | -1.07E+00 | 4.76E-06 | 1.75E-05 | -1.34E+00 | 6.08E-03 | 1.67E-02 | 1.26E+00 |
| HYLS1 | 1.59E-15 | 8.29E-15 | -3.71E+00 | 5.87E-11 | 2.53E-10 | -2.11E+00 | 4.25E-16 | 1.81E-15 | -2.31E+00 | 2.55E-13 | 9.76E-13 | -1.88E+00 | 1.76E-01 | 6.27E-01 | -1.11E+00 | 2.23E-01 | 6.29E-01 | -1.01E+00 | 6.65E-02 | 2.44E-01 | -1.22E+00 | 1.80E-01 | 4.95E-01 | 1.07E+00 |
| YIPF4 | 1.59E-15 | 8.31E-15 | -1.46E+00 | 3.19E-03 | 1.38E-02 | 1.15E+00 | 3.07E-06 | 1.31E-05 | -1.37E+00 | 2.87E-02 | 1.09E-01 | 1.04E+00 | 7.74E-12 | 2.76E-11 | -1.55E+00 | 4.79E-05 | 1.35E-04 | -1.15E+00 | 1.83E-07 | 6.72E-07 | -1.21E+00 | 4.50E-03 | 1.24E-02 | -1.02E+00 |
| SNRPE | 1.62E-15 | 8.43E-15 | -1.30E+00 | 4.33E-22 | 1.87E-21 | -1.39E+00 | 2.91E-04 | 1.24E-03 | -1.18E+00 | 6.28E-12 | 2.40E-11 | -1.38E+00 | 4.44E-10 | 1.58E-09 | -1.34E+00 | 1.41E-10 | 3.96E-10 | -1.29E+00 | 1.16E-05 | 4.27E-05 | -1.27E+00 | 1.18E-03 | 3.25E-03 | -1.25E+00 |
| HNRNPA1P4 | 1.64E-15 | 8.54E-15 | -1.61E+00 | 2.44E-06 | 1.05E-05 | 1.00E+00 | 9.50E-14 | 4.04E-13 | -1.67E+00 | 6.91E-08 | 2.64E-07 | -1.15E+00 | 1.10E-13 | 3.92E-13 | -1.74E+00 | 7.30E-04 | 2.06E-03 | -1.02E+00 | 5.10E-16 | 1.87E-15 | -1.91E+00 | 1.51E-03 | 4.15E-03 | -1.13E+00 |
| DNAJC8 | 1.67E-15 | 8.68E-15 | -1.29E+00 | 1.02E-12 | 4.42E-12 | -1.11E+00 | 4.81E-03 | 2.04E-02 | -1.05E+00 | 3.57E-04 | 1.36E-03 | -1.03E+00 | 2.21E-07 | 7.88E-07 | -1.15E+00 | 2.65E-05 | 7.48E-05 | -1.06E+00 | 2.60E-02 | 9.53E-02 | -1.05E+00 | 6.13E-02 | 1.68E-01 | -1.04E+00 |
| KRT18P62 | 1.67E-15 | 8.73E-15 | -2.02E+00 | 1.63E-12 | 7.06E-12 | -1.46E+00 | 2.21E-04 | 9.38E-04 | -1.41E+00 | 1.85E-04 | 7.06E-04 | -1.26E+00 | 3.54E-04 | 1.26E-03 | -1.67E+00 | 2.56E-03 | 7.22E-03 | -1.28E+00 | 2.45E-05 | 8.97E-05 | -1.56E+00 | 8.28E-05 | 2.27E-04 | -1.54E+00 |
| GLRX5 | 1.71E-15 | 8.89E-15 | -1.25E+00 | 7.37E-10 | 3.19E-09 | -1.01E+00 | 4.54E-02 | 1.93E-01 | 1.05E+00 | 5.59E-02 | 2.14E-01 | 1.18E+00 | 4.92E-05 | 1.75E-04 | -1.15E+00 | 2.84E-02 | 8.01E-02 | 1.05E+00 | 1.56E-01 | 5.73E-01 | 1.14E+00 | 1.20E-01 | 3.28E-01 | 1.28E+00 |
| ISCA1 | 1.71E-15 | 8.89E-15 | -1.60E+00 | 9.48E-20 | 4.10E-19 | -1.60E+00 | 1.39E-11 | 5.89E-11 | -1.64E+00 | 1.93E-10 | 7.36E-10 | -1.40E+00 | 1.10E-04 | 3.92E-04 | -1.07E+00 | 3.17E-06 | 8.95E-06 | -1.21E+00 | 6.30E-04 | 2.31E-03 | -1.13E+00 | 1.09E-05 | 2.99E-05 | -1.36E+00 |
| TCERG1 | 1.72E-15 | 8.99E-15 | -1.41E+00 | 2.45E-12 | 1.06E-11 | -1.23E+00 | 1.40E-08 | 5.93E-08 | -1.40E+00 | 2.47E-07 | 9.43E-07 | -1.19E+00 | 4.03E-11 | 1.44E-10 | -1.46E+00 | 9.34E-10 | 2.63E-09 | -1.26E+00 | 2.17E-08 | 7.95E-08 | -1.40E+00 | 5.72E-04 | 1.57E-03 | -1.14E+00 |
| SRRM2 | 1.72E-15 | 8.99E-15 | -1.33E+00 | 2.46E-09 | 1.06E-08 | -1.10E+00 | 1.34E-03 | 5.70E-03 | -1.17E+00 | 8.93E-02 | 3.41E-01 | 1.02E+00 | 1.44E-06 | 5.15E-06 | -1.21E+00 | 1.78E-02 | 5.02E-02 | -1.02E+00 | 2.56E-02 | 9.38E-02 | -1.01E+00 | 2.67E-01 | 7.33E-01 | 1.13E+00 |
| SSU72 | 1.74E-15 | 9.07E-15 | -1.25E+00 | 4.68E-12 | 2.02E-11 | -1.06E+00 | 6.71E-04 | 2.85E-03 | -1.14E+00 | 4.01E-02 | 1.53E-01 | 1.07E+00 | 1.80E-05 | 6.43E-05 | -1.02E+00 | 1.19E-01 | 3.37E-01 | 1.14E+00 | 4.51E-02 | 1.65E-01 | -1.02E+00 | 2.92E-01 | 8.02E-01 | 1.14E+00 |
| MLLT10 | 1.75E-15 | 9.11E-15 | -1.39E+00 | 5.08E-13 | 2.20E-12 | -1.14E+00 | 2.09E-08 | 8.87E-08 | -1.34E+00 | 2.57E-04 | 9.82E-04 | 1.01E+00 | 9.58E-13 | 3.42E-12 | -1.61E+00 | 5.06E-05 | 1.43E-04 | -1.04E+00 | 8.55E-12 | 3.14E-11 | -1.51E+00 | 4.91E-04 | 1.35E-03 | -1.10E+00 |
| NFIC | 1.80E-15 | 9.37E-15 | -1.61E+00 | 1.53E-08 | 6.62E-08 | -1.09E+00 | 7.34E-09 | 3.12E-08 | -1.42E+00 | 3.24E-04 | 1.24E-03 | -1.05E+00 | 1.43E-04 | 5.10E-04 | -1.35E+00 | 1.11E-02 | 3.12E-02 | -1.03E+00 | 5.98E-08 | 2.20E-07 | -1.45E+00 | 5.99E-03 | 1.65E-02 | -1.01E+00 |
| SMC4 | 1.82E-15 | 9.47E-15 | -1.47E+00 | 3.83E-03 | 1.66E-02 | 1.07E+00 | 1.88E-05 | 8.00E-05 | -1.12E+00 | 8.64E-04 | 3.30E-03 | 1.04E+00 | 2.92E-04 | 1.04E-03 | -1.04E+00 | 1.57E-02 | 4.43E-02 | 1.07E+00 | 1.37E-01 | 5.02E-01 | 1.10E+00 | 1.65E-01 | 4.54E-01 | 1.30E+00 |
| NPM1 | 1.82E-15 | 9.48E-15 | -1.34E+00 | 2.13E-04 | 9.18E-04 | -1.10E+00 | 7.54E-11 | 3.20E-10 | -1.38E+00 | 1.03E-04 | 3.95E-04 | -1.19E+00 | 1.68E-07 | 5.98E-07 | -1.33E+00 | 3.24E-01 | 9.12E-01 | 1.01E+00 | 1.05E-14 | 3.86E-14 | -1.33E+00 | 1.81E-02 | 4.97E-02 | -1.08E+00 |
| PRKCQ-AS1 | 1.86E-15 | 9.71E-15 | -1.52E+00 | 9.03E-04 | 3.90E-03 | 1.10E+00 | 5.98E-12 | 2.54E-11 | -1.63E+00 | 7.94E-04 | 3.03E-03 | 1.05E+00 | 1.81E-11 | 6.44E-11 | -1.62E+00 | 1.73E-01 | 4.88E-01 | 1.20E+00 | 1.52E-08 | 5.59E-08 | -1.37E+00 | 5.63E-02 | 1.55E-01 | 1.03E+00 |
| UBE2R2 | 1.87E-15 | 9.72E-15 | -1.42E+00 | 8.85E-11 | 3.82E-10 | -1.15E+00 | 9.85E-09 | 4.19E-08 | -1.41E+00 | 2.01E-05 | 7.67E-05 | -1.15E+00 | 1.01E-14 | 3.62E-14 | -1.60E+00 | 2.11E-05 | 5.96E-05 | -1.15E+00 | 7.21E-16 | 2.65E-15 | -1.74E+00 | 1.14E-05 | 3.12E-05 | -1.25E+00 |
| NDUFA3 | 1.88E-15 | 9.79E-15 | -1.50E+00 | 1.15E-25 | 4.96E-25 | -1.89E+00 | 2.45E-04 | 1.04E-03 | -1.11E+00 | 5.47E-18 | 2.09E-17 | -1.72E+00 | 1.01E-08 | 3.60E-08 | -1.39E+00 | 7.19E-12 | 2.03E-11 | -1.68E+00 | 1.32E-05 | 4.85E-05 | -1.16E+00 | 3.89E-15 | 1.07E-14 | -1.73E+00 |
| TTC17 | 1.88E-15 | 9.81E-15 | -1.51E+00 | 2.99E-07 | 1.29E-06 | -1.05E+00 | 3.30E-08 | 1.40E-07 | -1.44E+00 | 9.43E-04 | 3.60E-03 | 1.00E+00 | 1.14E-09 | 4.06E-09 | -1.48E+00 | 3.06E-03 | 8.64E-03 | 1.03E+00 | 3.14E-09 | 1.15E-08 | -1.31E+00 | 1.35E-03 | 3.70E-03 | -1.07E+00 |
| WAPL | 1.89E-15 | 9.86E-15 | -1.42E+00 | 4.02E-08 | 1.74E-07 | -1.01E+00 | 5.52E-10 | 2.34E-09 | -1.50E+00 | 7.91E-06 | 3.02E-05 | -1.09E+00 | 8.09E-09 | 2.89E-08 | -1.37E+00 | 1.46E-05 | 4.13E-05 | -1.11E+00 | 1.13E-07 | 4.13E-07 | -1.40E+00 | 3.22E-03 | 8.85E-03 | -1.10E+00 |
| SFXN3 | 1.90E-15 | 9.90E-15 | -1.78E+00 | 4.05E-05 | 1.75E-04 | 1.05E+00 | 9.50E-04 | 4.04E-03 | -1.27E+00 | 1.42E-02 | 5.41E-02 | 1.10E+00 | 1.48E-01 | 5.29E-01 | -1.15E+00 | 7.24E-02 | 2.04E-01 | -1.31E+00 | 1.57E-01 | 5.77E-01 | 1.65E+00 | 3.61E-01 | 9.91E-01 | 1.29E+00 |
| MPPED2 | 1.97E-15 | 1.03E-14 | -1.49E+00 | 8.40E-06 | 3.63E-05 | 1.09E+00 | 1.41E-04 | 6.00E-04 | -1.31E+00 | 1.58E-02 | 6.04E-02 | 1.13E+00 | 2.63E-01 | 9.37E-01 | 1.35E+00 | 9.39E-02 | 2.65E-01 | -1.07E+00 | 1.13E-01 | 4.15E-01 | 1.01E+00 | 1.80E-01 | 4.93E-01 | -1.04E+00 |
| PPP4R3A | 2.01E-15 | 1.05E-14 | -1.35E+00 | 1.68E-13 | 7.26E-13 | -1.21E+00 | 7.04E-08 | 2.99E-07 | -1.41E+00 | 2.95E-07 | 1.13E-06 | -1.21E+00 | 2.98E-09 | 1.06E-08 | -1.26E+00 | 7.54E-10 | 2.13E-09 | -1.25E+00 | 7.70E-05 | 2.82E-04 | -1.07E+00 | 4.59E-04 | 1.26E-03 | -1.04E+00 |
| ZNF638 | 2.10E-15 | 1.09E-14 | -1.44E+00 | 2.58E-04 | 1.12E-03 | 1.09E+00 | 1.41E-11 | 5.98E-11 | -1.66E+00 | 3.52E-04 | 1.34E-03 | -1.03E+00 | 3.77E-05 | 1.35E-04 | -1.19E+00 | 1.64E-01 | 4.61E-01 | 1.25E+00 | 1.15E-07 | 4.21E-07 | -1.45E+00 | 4.82E-03 | 1.32E-02 | -1.10E+00 |
| H1F0 | 2.13E-15 | 1.11E-14 | -1.59E+00 | 3.53E-05 | 1.53E-04 | -1.03E+00 | 8.73E-11 | 3.71E-10 | -1.94E+00 | 4.33E-04 | 1.65E-03 | -1.17E+00 | 1.73E-05 | 6.16E-05 | -1.26E+00 | 2.05E-01 | 5.77E-01 | 1.29E+00 | 2.56E-05 | 9.40E-05 | -1.28E+00 | 6.49E-05 | 1.78E-04 | -1.26E+00 |
| FOXK1 | 2.14E-15 | 1.11E-14 | -1.80E+00 | 1.27E-09 | 5.49E-09 | -1.22E+00 | 5.50E-13 | 2.34E-12 | -1.82E+00 | 1.27E-07 | 4.87E-07 | -1.18E+00 | 3.35E-10 | 1.19E-09 | -2.00E+00 | 3.19E-05 | 8.99E-05 | -1.18E+00 | 2.35E-11 | 8.63E-11 | -1.77E+00 | 6.99E-04 | 1.92E-03 | -1.07E+00 |
| SPATS2 | 2.15E-15 | 1.12E-14 | -1.50E+00 | 2.49E-08 | 1.08E-07 | -1.03E+00 | 8.06E-06 | 3.43E-05 | -1.38E+00 | 1.65E-03 | 6.29E-03 | -1.05E+00 | 3.80E-07 | 1.36E-06 | -1.28E+00 | 3.74E-02 | 1.05E-01 | 1.13E+00 | 1.01E-05 | 3.72E-05 | -1.15E+00 | 1.91E-03 | 5.24E-03 | -1.02E+00 |
| RPL35AP32 | 2.21E-15 | 1.15E-14 | -1.66E+00 | 9.30E-21 | 4.02E-20 | -1.94E+00 | 2.70E-07 | 1.15E-06 | -1.48E+00 | 3.10E-09 | 1.18E-08 | -1.52E+00 | 8.21E-04 | 2.93E-03 | -1.22E+00 | 2.00E-08 | 5.62E-08 | -2.25E+00 | 3.92E-07 | 1.44E-06 | -1.72E+00 | 5.42E-09 | 1.49E-08 | -1.94E+00 |
| WRB | 2.24E-15 | 1.17E-14 | -1.63E+00 | 1.69E-11 | 7.29E-11 | -1.25E+00 | 1.02E-03 | 4.32E-03 | -1.09E+00 | 1.85E-04 | 7.06E-04 | -1.00E+00 | 7.49E-09 | 2.67E-08 | -1.61E+00 | 2.11E-03 | 5.96E-03 | 1.02E+00 | 1.78E-03 | 6.55E-03 | 1.01E+00 | 1.66E-03 | 4.56E-03 | -1.17E+00 |
| CDIP1 | 2.25E-15 | 1.17E-14 | -1.70E+00 | 2.74E-08 | 1.18E-07 | -1.15E+00 | 1.04E-07 | 4.41E-07 | -1.42E+00 | 4.21E-05 | 1.61E-04 | -1.05E+00 | 7.42E-09 | 2.65E-08 | -1.48E+00 | 2.09E-06 | 5.88E-06 | -1.24E+00 | 2.67E-04 | 9.81E-04 | -1.04E+00 | 2.35E-04 | 6.45E-04 | -1.15E+00 |
| ACTA2 | 2.27E-15 | 1.18E-14 | -2.89E+00 | 8.05E-10 | 3.48E-09 | -1.63E+00 | 4.16E-15 | 1.77E-14 | -3.04E+00 | 6.80E-12 | 2.60E-11 | -2.14E+00 | 9.05E-02 | 3.23E-01 | -1.46E+00 | 3.41E-01 | 9.61E-01 | 1.34E+00 | 9.05E-05 | 3.32E-04 | -1.74E+00 | 8.89E-03 | 2.44E-02 | -1.36E+00 |
| DCAF16 | 2.29E-15 | 1.19E-14 | -1.57E+00 | 1.30E-12 | 5.62E-12 | -1.23E+00 | 1.83E-04 | 7.77E-04 | -1.14E+00 | 6.54E-04 | 2.50E-03 | 1.04E+00 | 1.29E-05 | 4.61E-05 | -1.28E+00 | 6.78E-04 | 1.91E-03 | -1.08E+00 | 6.28E-06 | 2.30E-05 | -1.34E+00 | 1.63E-02 | 4.48E-02 | 1.02E+00 |
| RPL21P28 | 2.31E-15 | 1.21E-14 | -1.39E+00 | 3.28E-47 | 1.42E-46 | -1.93E+00 | 6.54E-04 | 2.78E-03 | -1.31E+00 | 1.67E-15 | 6.39E-15 | -1.86E+00 | 3.35E-05 | 1.19E-04 | -1.24E+00 | 8.81E-18 | 2.48E-17 | -1.84E+00 | 2.97E-07 | 1.09E-06 | -1.36E+00 | 9.91E-20 | 2.72E-19 | -1.99E+00 |
| NDUFB9P2 | 2.35E-15 | 1.22E-14 | -1.73E+00 | 9.32E-13 | 4.03E-12 | -1.31E+00 | 3.49E-17 | 1.48E-16 | -1.93E+00 | 3.50E-17 | 1.34E-16 | -1.70E+00 | 2.41E-16 | 8.60E-16 | -2.27E+00 | 1.33E-08 | 3.75E-08 | -1.38E+00 | 7.90E-15 | 2.90E-14 | -1.68E+00 | 6.31E-09 | 1.73E-08 | -1.39E+00 |
| NUPL2 | 2.41E-15 | 1.25E-14 | -1.66E+00 | 2.94E-03 | 1.27E-02 | 1.23E+00 | 7.32E-09 | 3.11E-08 | -1.52E+00 | 1.86E-03 | 7.11E-03 | 1.03E+00 | 1.26E-12 | 4.49E-12 | -1.74E+00 | 3.66E-03 | 1.03E-02 | 1.14E+00 | 7.90E-14 | 2.90E-13 | -1.53E+00 | 2.05E-03 | 5.63E-03 | 1.02E+00 |
| RNF213 | 2.48E-15 | 1.29E-14 | -1.54E+00 | 1.85E-12 | 8.01E-12 | -1.28E+00 | 7.30E-17 | 3.10E-16 | -1.84E+00 | 5.10E-18 | 1.95E-17 | -1.69E+00 | 5.22E-10 | 1.86E-09 | -1.63E+00 | 4.23E-14 | 1.19E-13 | -1.88E+00 | 9.29E-10 | 3.41E-09 | -1.86E+00 | 5.19E-11 | 1.42E-10 | -2.02E+00 |
| PITPNB | 2.48E-15 | 1.29E-14 | -1.43E+00 | 3.70E-06 | 1.60E-05 | 1.08E+00 | 1.18E-13 | 5.00E-13 | -1.49E+00 | 1.31E-06 | 5.00E-06 | 1.02E+00 | 3.64E-14 | 1.30E-13 | -1.96E+00 | 1.05E-04 | 2.97E-04 | -1.09E+00 | 2.60E-11 | 9.54E-11 | -1.50E+00 | 1.35E-04 | 3.70E-04 | -1.05E+00 |
| HUWE1 | 2.52E-15 | 1.31E-14 | -1.43E+00 | 7.64E-07 | 3.30E-06 | -1.00E+00 | 1.89E-06 | 8.03E-06 | -1.26E+00 | 1.28E-02 | 4.88E-02 | 1.08E+00 | 2.68E-06 | 9.55E-06 | -1.20E+00 | 8.36E-02 | 2.36E-01 | 1.27E+00 | 3.34E-03 | 1.23E-02 | -1.12E+00 | 7.22E-02 | 1.98E-01 | 1.04E+00 |
| BUB3 | 2.53E-15 | 1.32E-14 | -1.40E+00 | 1.66E-03 | 7.18E-03 | 1.23E+00 | 2.38E-08 | 1.01E-07 | -1.25E+00 | 1.73E-03 | 6.60E-03 | 1.10E+00 | 3.74E-12 | 1.33E-11 | -1.62E+00 | 8.09E-02 | 2.28E-01 | 1.17E+00 | 2.89E-09 | 1.06E-08 | -1.40E+00 | 1.54E-01 | 4.24E-01 | 1.06E+00 |
| MYB | 2.54E-15 | 1.32E-14 | -1.32E+00 | 1.55E-03 | 6.68E-03 | 1.05E+00 | 4.65E-11 | 1.98E-10 | -1.68E+00 | 1.34E-03 | 5.10E-03 | -1.19E+00 | 6.50E-07 | 2.32E-06 | -1.49E+00 | 3.15E-01 | 8.89E-01 | 1.03E+00 | 3.65E-08 | 1.34E-07 | -1.46E+00 | 2.50E-01 | 6.87E-01 | 1.09E+00 |
| DDX52 | 2.55E-15 | 1.33E-14 | -1.54E+00 | 4.53E-14 | 1.96E-13 | -1.30E+00 | 2.72E-09 | 1.16E-08 | -1.47E+00 | 8.08E-07 | 3.09E-06 | -1.14E+00 | 6.01E-10 | 2.14E-09 | -1.36E+00 | 9.84E-09 | 2.77E-08 | -1.20E+00 | 3.80E-06 | 1.39E-05 | -1.21E+00 | 1.85E-04 | 5.09E-04 | -1.08E+00 |
| RHEB | 2.56E-15 | 1.33E-14 | -1.23E+00 | 1.13E-07 | 4.88E-07 | 1.10E+00 | 4.30E-04 | 1.83E-03 | -1.08E+00 | 2.51E-02 | 9.60E-02 | 1.16E+00 | 8.19E-08 | 2.92E-07 | -1.20E+00 | 9.30E-04 | 2.62E-03 | 1.04E+00 | 1.98E-03 | 7.26E-03 | -1.10E+00 | 1.10E-01 | 3.01E-01 | 1.00E+00 |
| PTGES3 | 2.56E-15 | 1.33E-14 | -1.26E+00 | 2.20E-01 | 9.51E-01 | 1.17E+00 | 5.28E-03 | 2.24E-02 | -1.13E+00 | 1.36E-01 | 5.18E-01 | 1.13E+00 | 2.49E-08 | 8.89E-08 | -1.27E+00 | 2.88E-01 | 8.12E-01 | 1.14E+00 | 1.72E-02 | 6.33E-02 | -1.08E+00 | 1.54E-02 | 4.22E-02 | 1.27E+00 |
| ANKRD10 | 2.56E-15 | 1.34E-14 | -1.43E+00 | 1.59E-06 | 6.85E-06 | -1.03E+00 | 4.04E-09 | 1.72E-08 | -1.43E+00 | 1.37E-04 | 5.23E-04 | -1.05E+00 | 1.05E-08 | 3.73E-08 | -1.35E+00 | 5.06E-04 | 1.43E-03 | -1.02E+00 | 8.73E-07 | 3.20E-06 | -1.31E+00 | 6.28E-02 | 1.73E-01 | -1.01E+00 |
| C1orf43 | 2.58E-15 | 1.34E-14 | -1.24E+00 | 4.36E-12 | 1.88E-11 | -1.08E+00 | 3.62E-08 | 1.54E-07 | -1.40E+00 | 1.67E-06 | 6.36E-06 | -1.19E+00 | 2.50E-10 | 8.91E-10 | -1.55E+00 | 3.14E-06 | 8.85E-06 | -1.22E+00 | 3.61E-06 | 1.33E-05 | -1.30E+00 | 1.35E-03 | 3.71E-03 | -1.10E+00 |
| RB1 | 2.63E-15 | 1.37E-14 | -1.52E+00 | 1.35E-08 | 5.85E-08 | -1.09E+00 | 8.32E-08 | 3.53E-07 | -1.37E+00 | 5.60E-06 | 2.14E-05 | -1.09E+00 | 7.87E-08 | 2.81E-07 | -1.25E+00 | 9.30E-05 | 2.62E-04 | 1.02E+00 | 2.46E-03 | 9.05E-03 | 1.03E+00 | 1.17E-02 | 3.22E-02 | 1.05E+00 |
| RPL32 | 2.67E-15 | 1.39E-14 | -1.18E+00 | 3.67E-34 | 1.59E-33 | -1.28E+00 | 4.32E-09 | 1.83E-08 | -1.16E+00 | 1.22E-16 | 4.65E-16 | -1.23E+00 | 4.28E-07 | 1.53E-06 | -1.16E+00 | 1.58E-11 | 4.45E-11 | -1.22E+00 | 1.49E-20 | 5.45E-20 | -1.21E+00 | 2.20E-22 | 6.04E-22 | -1.29E+00 |
| ABHD6 | 2.68E-15 | 1.40E-14 | -3.58E+00 | 4.26E-09 | 1.84E-08 | -1.73E+00 | 8.63E-03 | 3.67E-02 | -1.30E+00 | 9.22E-02 | 3.52E-01 | 1.12E+00 | 1.27E-04 | 4.52E-04 | -1.67E+00 | 3.94E-04 | 1.11E-03 | -1.62E+00 | 2.05E-03 | 7.54E-03 | -1.60E+00 | 6.78E-02 | 1.86E-01 | -1.16E+00 |
| KRIT1 | 2.69E-15 | 1.40E-14 | -1.54E+00 | 2.40E-09 | 1.04E-08 | -1.10E+00 | 5.74E-17 | 2.44E-16 | -1.99E+00 | 8.32E-13 | 3.18E-12 | -1.43E+00 | 7.13E-08 | 2.54E-07 | -1.55E+00 | 9.20E-05 | 2.59E-04 | -1.20E+00 | 1.75E-10 | 6.42E-10 | -1.51E+00 | 1.12E-07 | 3.07E-07 | -1.38E+00 |
| SLC35B2 | 2.73E-15 | 1.42E-14 | -2.40E+00 | 1.99E-09 | 8.60E-09 | -1.44E+00 | 7.00E-11 | 2.97E-10 | -1.64E+00 | 4.78E-09 | 1.83E-08 | -1.29E+00 | 8.61E-14 | 3.07E-13 | -2.37E+00 | 5.99E-09 | 1.69E-08 | -1.50E+00 | 9.39E-12 | 3.45E-11 | -1.66E+00 | 3.38E-04 | 9.28E-04 | -1.11E+00 |
| HPCAL1 | 2.80E-15 | 1.46E-14 | -1.36E+00 | 1.45E-10 | 6.26E-10 | -1.08E+00 | 8.33E-09 | 3.54E-08 | -1.33E+00 | 8.46E-07 | 3.23E-06 | -1.11E+00 | 1.61E-09 | 5.73E-09 | -1.53E+00 | 3.70E-05 | 1.04E-04 | -1.17E+00 | 2.57E-03 | 9.44E-03 | 1.13E+00 | 6.73E-03 | 1.85E-02 | 1.03E+00 |
| CU639417.1 | 2.95E-15 | 1.54E-14 | 5.30E+00 | 1.62E-01 | 6.98E-01 | 1.59E+00 | 5.35E-13 | 2.27E-12 | 6.10E+00 | 4.28E-02 | 1.63E-01 | 2.43E+00 | 7.18E-06 | 2.56E-05 | 3.75E+00 | 1.77E-01 | 4.99E-01 | 1.74E+00 | 1.54E-03 | 5.65E-03 | 2.45E+00 | 2.18E-01 | 5.98E-01 | 1.03E+00 |
| CTNNBL1 | 3.00E-15 | 1.56E-14 | -1.32E+00 | 4.11E-11 | 1.78E-10 | -1.08E+00 | 1.54E-05 | 6.56E-05 | -1.10E+00 | 3.65E-03 | 1.40E-02 | 1.10E+00 | 1.38E-01 | 4.91E-01 | 1.05E+00 | 3.30E-01 | 9.31E-01 | 1.10E+00 | 6.73E-02 | 2.47E-01 | -1.01E+00 | 2.94E-01 | 8.08E-01 | 1.08E+00 |
| GOLIM4 | 3.02E-15 | 1.57E-14 | -1.48E+00 | 7.87E-06 | 3.40E-05 | 1.08E+00 | 4.30E-12 | 1.83E-11 | -1.80E+00 | 1.66E-03 | 6.33E-03 | -1.02E+00 | 4.01E-09 | 1.43E-08 | -1.46E+00 | 1.29E-03 | 3.64E-03 | -1.04E+00 | 6.86E-10 | 2.52E-09 | -1.55E+00 | 9.63E-03 | 2.64E-02 | -1.11E+00 |
| AL078581.3 | 3.07E-15 | 1.60E-14 | -2.57E+00 | 1.10E-11 | 4.74E-11 | -1.71E+00 | 1.16E-04 | 4.91E-04 | -1.64E+00 | 8.83E-04 | 3.37E-03 | -1.28E+00 | 5.87E-02 | 2.09E-01 | -1.13E+00 | 1.20E-01 | 3.39E-01 | -1.02E+00 | 1.21E-04 | 4.45E-04 | -1.65E+00 | 1.12E-03 | 3.07E-03 | -1.42E+00 |
| GID8 | 3.17E-15 | 1.65E-14 | -1.42E+00 | 1.20E-08 | 5.17E-08 | -1.03E+00 | 2.88E-07 | 1.22E-06 | -1.30E+00 | 1.41E-06 | 5.38E-06 | -1.14E+00 | 2.03E-06 | 7.24E-06 | -1.25E+00 | 9.02E-04 | 2.54E-03 | 1.02E+00 | 4.14E-12 | 1.52E-11 | -1.56E+00 | 8.53E-04 | 2.34E-03 | -1.07E+00 |
| SLCO5A1 | 3.17E-15 | 1.65E-14 | -2.45E+00 | 2.20E-12 | 9.50E-12 | -1.58E+00 | 1.27E-04 | 5.40E-04 | -1.63E+00 | 7.27E-05 | 2.78E-04 | -1.61E+00 | 1.31E-02 | 4.65E-02 | -2.37E+00 | 2.45E-02 | 6.90E-02 | -2.04E+00 | 1.68E-01 | 6.18E-01 | 1.34E+00 | 2.51E-01 | 6.91E-01 | 1.34E+00 |
| USP10 | 3.18E-15 | 1.66E-14 | -1.71E+00 | 2.73E-08 | 1.18E-07 | -1.13E+00 | 2.33E-07 | 9.89E-07 | -1.32E+00 | 1.37E-05 | 5.24E-05 | -1.07E+00 | 2.26E-06 | 8.05E-06 | -1.36E+00 | 3.31E-04 | 9.34E-04 | -1.04E+00 | 2.40E-08 | 8.82E-08 | -1.37E+00 | 2.79E-02 | 7.66E-02 | 1.06E+00 |
| POLDIP2 | 3.20E-15 | 1.67E-14 | -1.43E+00 | 4.62E-12 | 2.00E-11 | -1.16E+00 | 2.40E-02 | 1.02E-01 | -1.03E+00 | 3.95E-04 | 1.51E-03 | -1.03E+00 | 1.44E-11 | 5.14E-11 | -1.70E+00 | 1.17E-02 | 3.30E-02 | 1.10E+00 | 4.75E-02 | 1.74E-01 | 1.05E+00 | 3.39E-01 | 9.32E-01 | 1.28E+00 |
| C6orf89 | 3.20E-15 | 1.67E-14 | -1.81E+00 | 2.94E-09 | 1.27E-08 | -1.23E+00 | 1.14E-05 | 4.84E-05 | -1.32E+00 | 1.83E-04 | 7.01E-04 | -1.09E+00 | 6.70E-04 | 2.39E-03 | -1.31E+00 | 2.13E-02 | 6.01E-02 | 1.02E+00 | 6.63E-11 | 2.43E-10 | -1.46E+00 | 1.71E-08 | 4.70E-08 | -1.31E+00 |
| CSNK2B | 3.23E-15 | 1.68E-14 | -1.33E+00 | 6.17E-17 | 2.66E-16 | -1.23E+00 | 9.29E-06 | 3.95E-05 | -1.35E+00 | 2.14E-08 | 8.16E-08 | -1.36E+00 | 1.21E-09 | 4.32E-09 | -1.33E+00 | 1.66E-07 | 4.69E-07 | -1.19E+00 | 1.99E-08 | 7.30E-08 | -1.45E+00 | 2.07E-04 | 5.68E-04 | -1.28E+00 |
| GPATCH4 | 3.24E-15 | 1.69E-14 | -1.91E+00 | 1.40E-06 | 6.05E-06 | -1.19E+00 | 1.17E-06 | 4.96E-06 | -1.45E+00 | 1.76E-02 | 6.72E-02 | -1.03E+00 | 2.47E-10 | 8.82E-10 | -1.88E+00 | 1.45E-02 | 4.10E-02 | 1.02E+00 | 2.96E-04 | 1.09E-03 | -1.13E+00 | 1.49E-01 | 4.09E-01 | 1.32E+00 |
| PPP4R2 | 3.24E-15 | 1.69E-14 | -1.50E+00 | 3.17E-10 | 1.37E-09 | -1.16E+00 | 7.35E-08 | 3.12E-07 | -1.40E+00 | 2.31E-07 | 8.84E-07 | -1.15E+00 | 2.90E-08 | 1.03E-07 | -1.54E+00 | 3.01E-08 | 8.48E-08 | -1.41E+00 | 1.92E-13 | 7.05E-13 | -1.53E+00 | 9.49E-08 | 2.61E-07 | -1.33E+00 |
| KDELR1 | 3.30E-15 | 1.72E-14 | -1.38E+00 | 2.38E-11 | 1.03E-10 | -1.18E+00 | 2.77E-06 | 1.18E-05 | -1.42E+00 | 2.17E-05 | 8.29E-05 | -1.28E+00 | 5.41E-11 | 1.93E-10 | -1.40E+00 | 1.92E-09 | 5.40E-09 | -1.35E+00 | 3.22E-05 | 1.18E-04 | -1.21E+00 | 9.65E-06 | 2.65E-05 | -1.21E+00 |
| ICE2 | 3.35E-15 | 1.75E-14 | -1.51E+00 | 5.68E-05 | 2.45E-04 | 1.08E+00 | 1.24E-07 | 5.26E-07 | -1.33E+00 | 1.08E-01 | 4.13E-01 | 1.32E+00 | 1.07E-13 | 3.83E-13 | -1.99E+00 | 1.22E-05 | 3.44E-05 | -1.22E+00 | 1.18E-06 | 4.33E-06 | -1.34E+00 | 5.53E-02 | 1.52E-01 | 1.05E+00 |
| PSMF1 | 3.36E-15 | 1.75E-14 | -1.33E+00 | 5.35E-13 | 2.31E-12 | -1.11E+00 | 1.44E-05 | 6.12E-05 | -1.27E+00 | 2.38E-04 | 9.09E-04 | -1.04E+00 | 2.75E-07 | 9.80E-07 | -1.17E+00 | 8.37E-05 | 2.36E-04 | 1.02E+00 | 1.03E-05 | 3.76E-05 | -1.22E+00 | 5.86E-03 | 1.61E-02 | -1.07E+00 |
| EREG | 3.43E-15 | 1.79E-14 | 2.59E+00 | 2.67E-09 | 1.16E-08 | 2.37E+00 | 4.70E-15 | 2.00E-14 | 2.47E+00 | 2.42E-12 | 9.24E-12 | 2.37E+00 | 2.39E-16 | 8.51E-16 | 3.39E+00 | 1.62E-12 | 4.57E-12 | 3.08E+00 | 1.07E-14 | 3.92E-14 | 3.00E+00 | 2.35E-09 | 6.47E-09 | 2.59E+00 |
| PGK1 | 3.47E-15 | 1.81E-14 | -1.34E+00 | 4.43E-11 | 1.92E-10 | -1.10E+00 | 1.21E-02 | 5.14E-02 | -1.15E+00 | 1.69E-01 | 6.46E-01 | 1.02E+00 | 2.94E-07 | 1.05E-06 | -1.32E+00 | 1.16E-03 | 3.26E-03 | -1.11E+00 | 3.06E-05 | 1.12E-04 | -1.18E+00 | 3.47E-01 | 9.52E-01 | 1.09E+00 |
| C4orf3 | 3.47E-15 | 1.81E-14 | -1.26E+00 | 1.38E-06 | 5.96E-06 | 1.02E+00 | 1.94E-04 | 8.24E-04 | -1.18E+00 | 1.36E-03 | 5.20E-03 | -1.03E+00 | 2.96E-07 | 1.05E-06 | -1.19E+00 | 1.08E-04 | 3.06E-04 | -1.05E+00 | 5.94E-03 | 2.18E-02 | -1.02E+00 | 2.89E-02 | 7.93E-02 | 1.01E+00 |
| FTH1P3 | 3.48E-15 | 1.81E-14 | -1.79E+00 | 4.91E-13 | 2.12E-12 | -1.52E+00 | 3.06E-07 | 1.30E-06 | -1.51E+00 | 1.29E-06 | 4.93E-06 | -1.32E+00 | 1.25E-11 | 4.44E-11 | -1.56E+00 | 4.86E-11 | 1.37E-10 | -1.42E+00 | 1.54E-23 | 5.64E-23 | -1.84E+00 | 4.31E-10 | 1.18E-09 | -1.31E+00 |
| UVRAG | 3.48E-15 | 1.81E-14 | -1.67E+00 | 1.62E-10 | 6.99E-10 | -1.24E+00 | 1.21E-17 | 5.15E-17 | -1.77E+00 | 4.45E-20 | 1.70E-19 | -1.77E+00 | 2.91E-08 | 1.04E-07 | -1.58E+00 | 5.46E-07 | 1.54E-06 | -1.47E+00 | 1.91E-09 | 7.00E-09 | -1.46E+00 | 1.28E-06 | 3.52E-06 | -1.21E+00 |
| RPS2P55 | 3.54E-15 | 1.85E-14 | 2.78E+00 | 1.36E-05 | 5.86E-05 | 2.43E+00 | 1.95E-12 | 8.27E-12 | 2.48E+00 | 1.06E-05 | 4.05E-05 | 2.22E+00 | 4.96E-08 | 1.77E-07 | 2.62E+00 | 6.70E-05 | 1.89E-04 | 2.31E+00 | 2.18E-10 | 8.00E-10 | 2.56E+00 | 4.33E-05 | 1.19E-04 | 2.11E+00 |
| AC133065.6 | 3.56E-15 | 1.85E-14 | -1.80E+00 | 3.62E-07 | 1.57E-06 | -1.08E+00 | 8.53E-10 | 3.62E-09 | -1.97E+00 | 3.21E-04 | 1.23E-03 | -1.08E+00 | 4.58E-05 | 1.63E-04 | -2.70E+00 | 7.31E-04 | 2.06E-03 | -1.75E+00 | 1.93E-02 | 7.07E-02 | -1.84E+00 | 1.28E-01 | 3.51E-01 | -1.28E+00 |
| HADHB | 3.57E-15 | 1.86E-14 | -1.27E+00 | 3.24E-06 | 1.40E-05 | 1.09E+00 | 8.90E-04 | 3.78E-03 | -1.18E+00 | 6.26E-04 | 2.39E-03 | -1.04E+00 | 6.86E-12 | 2.45E-11 | -1.38E+00 | 2.49E-04 | 7.01E-04 | -1.07E+00 | 1.59E-02 | 5.84E-02 | -1.01E+00 | 1.42E-01 | 3.91E-01 | 1.04E+00 |
| ODF3B | 3.59E-15 | 1.87E-14 | -1.42E+00 | 4.10E-32 | 1.77E-31 | -2.29E+00 | 1.44E-20 | 6.13E-20 | -1.87E+00 | 1.79E-40 | 6.85E-40 | -2.93E+00 | 2.80E-06 | 9.97E-06 | -2.65E+00 | 1.03E-05 | 2.90E-05 | -2.14E+00 | 1.08E-03 | 3.97E-03 | -2.49E+00 | 1.66E-03 | 4.55E-03 | -2.61E+00 |
| SESN3 | 3.59E-15 | 1.87E-14 | -1.46E+00 | 1.31E-13 | 5.65E-13 | -1.29E+00 | 3.89E-08 | 1.65E-07 | -1.70E+00 | 2.73E-10 | 1.04E-09 | -1.63E+00 | 2.27E-03 | 8.09E-03 | -1.02E+00 | 3.82E-07 | 1.08E-06 | -1.29E+00 | 3.89E-03 | 1.43E-02 | -1.11E+00 | 2.25E-08 | 6.18E-08 | -1.52E+00 |
| MIR4435-2HG | 3.64E-15 | 1.90E-14 | -1.31E+00 | 3.17E-17 | 1.37E-16 | -1.29E+00 | 6.57E-14 | 2.79E-13 | -1.66E+00 | 4.28E-10 | 1.63E-09 | -1.27E+00 | 1.81E-05 | 6.47E-05 | -1.27E+00 | 1.58E-02 | 4.44E-02 | -1.04E+00 | 6.99E-03 | 2.57E-02 | -1.12E+00 | 2.91E-03 | 8.00E-03 | -1.19E+00 |
| C14orf37 | 3.72E-15 | 1.94E-14 | -2.41E+00 | 2.93E-12 | 1.26E-11 | -1.81E+00 | 3.25E-05 | 1.38E-04 | -1.70E+00 | 3.89E-04 | 1.49E-03 | -1.25E+00 | 6.37E-05 | 2.27E-04 | -2.26E+00 | 7.65E-05 | 2.16E-04 | -2.14E+00 | 2.20E-01 | 8.07E-01 | -1.06E+00 | 3.42E-01 | 9.39E-01 | 1.21E+00 |
| AC104837.1 | 3.76E-15 | 1.96E-14 | -1.58E+00 | 1.27E-14 | 5.50E-14 | -1.35E+00 | 1.82E-11 | 7.73E-11 | -1.60E+00 | 3.30E-15 | 1.26E-14 | -1.65E+00 | 2.89E-10 | 1.03E-09 | -1.69E+00 | 3.62E-12 | 1.02E-11 | -1.79E+00 | 1.24E-15 | 4.54E-15 | -1.63E+00 | 1.09E-21 | 3.01E-21 | -1.84E+00 |
| PARVB | 3.78E-15 | 1.97E-14 | -1.37E+00 | 3.30E-07 | 1.43E-06 | 1.02E+00 | 2.27E-02 | 9.63E-02 | -1.02E+00 | 3.35E-02 | 1.28E-01 | 1.12E+00 | 2.21E-06 | 7.87E-06 | -1.21E+00 | 1.68E-02 | 4.73E-02 | 1.04E+00 | 1.40E-03 | 5.14E-03 | -1.20E+00 | 7.75E-02 | 2.13E-01 | -1.06E+00 |
| PRKAR1A | 3.81E-15 | 1.98E-14 | -1.38E+00 | 3.00E-11 | 1.30E-10 | -1.12E+00 | 3.88E-06 | 1.65E-05 | -1.27E+00 | 7.03E-06 | 2.68E-05 | -1.08E+00 | 2.39E-07 | 8.54E-07 | -1.18E+00 | 1.81E-04 | 5.09E-04 | -1.00E+00 | 3.77E-02 | 1.38E-01 | 1.05E+00 | 3.05E-03 | 8.38E-03 | -1.08E+00 |
| CACYBP | 3.82E-15 | 1.99E-14 | -1.29E+00 | 3.70E-06 | 1.60E-05 | 1.08E+00 | 5.11E-02 | 2.17E-01 | -1.02E+00 | 3.19E-02 | 1.22E-01 | 1.12E+00 | 1.77E-10 | 6.29E-10 | -1.34E+00 | 4.85E-05 | 1.37E-04 | -1.04E+00 | 1.08E-03 | 3.96E-03 | -1.10E+00 | 3.36E-01 | 9.22E-01 | 1.22E+00 |
| ATP6V1D | 3.84E-15 | 2.00E-14 | -1.50E+00 | 6.67E-13 | 2.88E-12 | -1.28E+00 | 7.69E-06 | 3.27E-05 | -1.30E+00 | 6.25E-06 | 2.39E-05 | -1.11E+00 | 3.03E-11 | 1.08E-10 | -1.77E+00 | 9.83E-09 | 2.77E-08 | -1.46E+00 | 3.16E-03 | 1.16E-02 | 1.04E+00 | 6.08E-02 | 1.67E-01 | 1.09E+00 |
| SUMO1 | 3.93E-15 | 2.05E-14 | -1.15E+00 | 7.07E-08 | 3.06E-07 | 1.03E+00 | 1.36E-03 | 5.76E-03 | -1.13E+00 | 8.11E-04 | 3.10E-03 | -1.03E+00 | 4.00E-09 | 1.43E-08 | -1.32E+00 | 1.05E-03 | 2.95E-03 | -1.02E+00 | 1.26E-03 | 4.64E-03 | -1.09E+00 | 2.56E-01 | 7.02E-01 | 1.07E+00 |
| LMNB1 | 4.00E-15 | 2.08E-14 | -2.48E+00 | 3.41E-10 | 1.47E-09 | -1.60E+00 | 5.81E-04 | 2.47E-03 | -1.06E+00 | 1.14E-02 | 4.34E-02 | 1.19E+00 | 3.47E-16 | 1.24E-15 | -3.67E+00 | 1.62E-08 | 4.56E-08 | -1.70E+00 | 2.19E-14 | 8.03E-14 | -1.56E+00 | 1.48E-01 | 4.07E-01 | 1.11E+00 |
| LRRC47 | 4.13E-15 | 2.15E-14 | -1.58E+00 | 3.34E-09 | 1.44E-08 | -1.09E+00 | 2.09E-12 | 8.90E-12 | -1.62E+00 | 5.98E-05 | 2.28E-04 | -1.06E+00 | 7.58E-06 | 2.70E-05 | -1.33E+00 | 3.98E-02 | 1.12E-01 | 1.15E+00 | 1.61E-10 | 5.91E-10 | -1.45E+00 | 1.58E-04 | 4.33E-04 | -1.04E+00 |
| LEPR | 4.20E-15 | 2.19E-14 | -1.95E+00 | 1.73E-05 | 7.46E-05 | -1.01E+00 | 5.91E-09 | 2.51E-08 | -1.84E+00 | 3.38E-04 | 1.29E-03 | -1.10E+00 | 1.59E-07 | 5.66E-07 | -1.64E+00 | 6.67E-02 | 1.88E-01 | 1.57E+00 | 1.27E-08 | 4.67E-08 | -1.40E+00 | 2.69E-01 | 7.38E-01 | 1.07E+00 |
| DHX9 | 4.24E-15 | 2.21E-14 | -1.35E+00 | 3.20E-07 | 1.38E-06 | -1.01E+00 | 9.28E-04 | 3.94E-03 | -1.11E+00 | 3.30E-03 | 1.26E-02 | 1.03E+00 | 7.97E-16 | 2.84E-15 | -1.41E+00 | 1.78E-08 | 5.02E-08 | -1.12E+00 | 3.20E-03 | 1.17E-02 | -1.06E+00 | 1.82E-01 | 5.01E-01 | 1.10E+00 |
| KAT6B | 4.31E-15 | 2.24E-14 | -1.30E+00 | 4.24E-13 | 1.83E-12 | -1.12E+00 | 3.10E-11 | 1.32E-10 | -1.44E+00 | 8.09E-08 | 3.09E-07 | -1.21E+00 | 1.04E-03 | 3.70E-03 | -1.03E+00 | 2.77E-03 | 7.80E-03 | -1.02E+00 | 1.54E-06 | 5.67E-06 | -1.35E+00 | 6.99E-07 | 1.92E-06 | -1.41E+00 |

| | | | | | | | | | | | | | | | | | | | | | | | | |
|---|---|---|---|---|---|---|---|---|---|---|---|---|---|---|---|---|---|---|---|---|---|---|---|---|
| ZFP90 | 4.40E-15 | 2.29E-14 | -1.69E+00 | 4.84E-09 | 2.09E-08 | -1.14E+00 | 9.47E-12 | 4.02E-11 | -1.67E+00 | 3.41E-05 | 1.30E-04 | -1.06E+00 | 2.96E-06 | 1.06E-05 | -1.59E+00 | 1.54E-04 | 4.34E-04 | -1.28E+00 | 8.04E-07 | 2.95E-06 | -1.42E+00 | 1.16E-03 | 3.20E-03 | -1.14E+00 |
| ST8SIA6 | 4.41E-15 | 2.30E-14 | -1.56E+00 | 6.68E-08 | 2.89E-07 | -1.08E+00 | 1.34E-04 | 5.69E-04 | -1.23E+00 | 9.08E-04 | 3.47E-03 | -1.01E+00 | 1.61E-12 | 5.72E-12 | -1.66E+00 | 2.76E-05 | 7.79E-05 | -1.28E+00 | 2.10E-04 | 7.69E-04 | -1.17E+00 | 8.27E-03 | 2.27E-02 | -1.09E+00 |
| MRPS27 | 4.51E-15 | 2.35E-14 | -1.36E+00 | 6.05E-09 | 2.61E-08 | -1.07E+00 | 5.28E-06 | 2.25E-05 | -1.30E+00 | 2.53E-05 | 9.65E-05 | -1.10E+00 | 4.64E-05 | 1.65E-04 | -1.24E+00 | 1.31E-01 | 3.69E-01 | 1.17E+00 | 2.59E-06 | 9.52E-06 | -1.21E+00 | 6.53E-02 | 1.79E-01 | 1.12E+00 |
| ZNF281 | 4.65E-15 | 2.42E-14 | -1.44E+00 | 7.05E-14 | 3.04E-13 | -1.24E+00 | 3.89E-11 | 1.65E-10 | -1.55E+00 | 2.26E-12 | 8.63E-12 | -1.46E+00 | 5.24E-07 | 1.87E-06 | -1.30E+00 | 2.52E-06 | 7.11E-06 | -1.30E+00 | 3.94E-06 | 1.44E-05 | -1.21E+00 | 6.48E-05 | 1.78E-04 | -1.11E+00 |
| WSB1 | 4.68E-15 | 2.44E-14 | -1.28E+00 | 9.90E-20 | 4.28E-19 | -1.32E+00 | 2.95E-07 | 1.25E-06 | -1.44E+00 | 2.36E-11 | 9.03E-11 | -1.53E+00 | 3.64E-10 | 1.30E-09 | -1.44E+00 | 1.38E-09 | 3.89E-09 | -1.45E+00 | 2.69E-13 | 9.89E-13 | -1.50E+00 | 1.41E-09 | 3.88E-09 | -1.42E+00 |
| SUCLG2 | 4.73E-15 | 2.47E-14 | -4.36E+00 | 9.68E-14 | 4.18E-13 | -3.20E+00 | 5.81E-05 | 2.47E-04 | -1.71E+00 | 2.32E-06 | 8.85E-06 | -1.97E+00 | 6.77E-10 | 2.41E-09 | -1.46E+00 | 1.45E-09 | 4.09E-09 | -1.33E+00 | 6.95E-07 | 2.55E-06 | -1.29E+00 | 1.76E-02 | 4.85E-02 | -1.05E+00 |
| SERBP1 | 4.74E-15 | 2.47E-14 | -1.31E+00 | 3.93E-10 | 1.70E-09 | -1.15E+00 | 2.56E-03 | 1.09E-02 | -1.16E+00 | 1.04E-02 | 3.97E-02 | -1.11E+00 | 7.60E-07 | 2.71E-06 | -1.19E+00 | 1.84E-02 | 5.17E-02 | 1.02E+00 | 2.76E-04 | 1.01E-03 | -1.22E+00 | 8.58E-02 | 2.36E-01 | -1.09E+00 |
| KXD1 | 4.79E-15 | 2.50E-14 | -1.42E+00 | 5.07E-10 | 2.19E-09 | -1.09E+00 | 2.72E-09 | 1.15E-08 | -1.35E+00 | 8.23E-04 | 3.14E-03 | 1.07E+00 | 2.64E-09 | 9.41E-09 | -1.43E+00 | 3.01E-04 | 8.49E-04 | -1.02E+00 | 3.28E-06 | 1.21E-05 | -1.25E+00 | 1.18E-01 | 3.25E-01 | 1.05E+00 |
| OCIAD1 | 4.85E-15 | 2.53E-14 | -1.34E+00 | 1.66E-05 | 7.16E-05 | 1.01E+00 | 5.12E-05 | 2.17E-04 | -1.23E+00 | 1.93E-03 | 7.37E-03 | -1.07E+00 | 2.18E-07 | 7.78E-07 | -1.31E+00 | 2.91E-04 | 8.20E-04 | -1.10E+00 | 1.94E-04 | 7.12E-04 | -1.22E+00 | 4.45E-02 | 1.22E-01 | -1.07E+00 |
| TCEA1P2 | 4.87E-15 | 2.54E-14 | -1.23E+00 | 2.36E-25 | 1.02E-24 | -1.33E+00 | 1.30E-04 | 5.52E-04 | -1.17E+00 | 7.44E-11 | 2.84E-10 | -1.34E+00 | 6.92E-03 | 2.47E-02 | 1.13E+00 | 8.64E-06 | 2.44E-05 | -1.08E+00 | 1.48E-01 | 5.43E-01 | 1.11E+00 | 9.80E-05 | 2.69E-04 | -1.26E+00 |
| GTF2H3 | 4.87E-15 | 2.54E-14 | -1.73E+00 | 5.68E-10 | 2.45E-09 | -1.23E+00 | 7.03E-08 | 2.99E-07 | -1.45E+00 | 1.66E-06 | 6.33E-06 | -1.21E+00 | 2.94E-04 | 1.05E-03 | -1.35E+00 | 2.39E-01 | 6.73E-01 | 1.31E+00 | 1.20E-05 | 4.41E-05 | -1.29E+00 | 8.66E-02 | 2.38E-01 | 1.10E+00 |
| NIPSNAP3A | 4.97E-15 | 2.59E-14 | -1.47E+00 | 2.17E-04 | 9.38E-04 | 1.12E+00 | 4.90E-09 | 2.08E-08 | -1.62E+00 | 9.43E-04 | 3.60E-03 | -1.05E+00 | 1.82E-12 | 6.50E-12 | -1.67E+00 | 1.60E-02 | 4.51E-02 | -1.07E+00 | 2.33E-08 | 8.55E-08 | -1.31E+00 | 5.77E-02 | 1.59E-01 | -1.01E+00 |
| LTC4S | 5.18E-15 | 2.70E-14 | -1.74E+00 | 1.07E-16 | 4.61E-16 | -1.69E+00 | 3.05E-04 | 1.29E-03 | -1.33E+00 | 3.75E-06 | 1.43E-05 | -1.36E+00 | 1.06E-07 | 3.77E-07 | -1.41E+00 | 2.08E-17 | 5.87E-17 | -2.03E+00 | 8.36E-02 | 3.07E-01 | 1.22E+00 | 3.23E-04 | 8.88E-04 | -1.34E+00 |
| YIPF3 | 5.34E-15 | 2.79E-14 | -1.38E+00 | 4.69E-06 | 2.03E-05 | 1.05E+00 | 1.80E-06 | 7.64E-06 | -1.35E+00 | 1.04E-05 | 3.95E-05 | -1.10E+00 | 5.35E-11 | 1.91E-10 | -1.38E+00 | 9.17E-06 | 2.58E-05 | -1.10E+00 | 2.00E-05 | 7.36E-05 | -1.16E+00 | 6.33E-04 | 1.74E-03 | -1.09E+00 |
| CCT3 | 5.35E-15 | 2.79E-14 | -1.22E+00 | 1.50E-01 | 6.49E-01 | 1.22E+00 | 9.79E-02 | 4.16E-01 | -1.02E+00 | 2.00E-01 | 7.63E-01 | 1.20E+00 | 1.50E-07 | 5.36E-07 | -1.18E+00 | 3.49E-01 | 9.84E-01 | 1.24E+00 | 9.74E-02 | 3.57E-01 | 1.06E+00 | 3.64E-02 | 1.00E-01 | 1.33E+00 |
| RUFY1 | 5.39E-15 | 2.81E-14 | -1.77E+00 | 1.14E-10 | 4.93E-10 | -1.28E+00 | 1.40E-08 | 5.94E-08 | -1.61E+00 | 2.93E-04 | 1.12E-03 | -1.05E+00 | 7.31E-06 | 2.60E-05 | -1.40E+00 | 3.23E-03 | 9.10E-03 | -1.05E+00 | 5.19E-09 | 1.91E-08 | 1.14E+00 | 4.05E-05 | 1.11E-04 | -1.11E+00 |
| PBDC1 | 5.44E-15 | 2.84E-14 | -1.76E+00 | 9.80E-07 | 4.23E-06 | -1.10E+00 | 9.44E-09 | 4.01E-08 | -1.41E+00 | 3.61E-06 | 1.38E-05 | -1.17E+00 | 1.99E-04 | 7.10E-04 | -1.25E+00 | 6.81E-03 | 1.92E-02 | 1.03E+00 | 1.90E-10 | 6.96E-10 | -1.38E+00 | 6.18E-03 | 1.70E-02 | 1.02E+00 |
| BIVM | 5.63E-15 | 2.93E-14 | -1.75E+00 | 1.47E-04 | 6.34E-04 | 1.11E+00 | 1.54E-09 | 6.54E-09 | -1.58E+00 | 1.51E-03 | 5.77E-03 | 1.02E+00 | 7.11E-06 | 2.54E-05 | -1.53E+00 | 2.48E-02 | 6.99E-02 | 1.05E+00 | 9.50E-06 | 3.49E-05 | -1.36E+00 | 7.91E-02 | 2.17E-01 | 1.03E+00 |
| CRTAP | 5.71E-15 | 2.98E-14 | -1.67E+00 | 1.63E-10 | 7.06E-10 | -1.23E+00 | 4.74E-14 | 2.01E-13 | -1.69E+00 | 4.85E-10 | 1.85E-09 | -1.29E+00 | 6.67E-09 | 2.38E-08 | -1.43E+00 | 1.42E-06 | 4.02E-06 | -1.24E+00 | 7.03E-08 | 2.58E-07 | -1.37E+00 | 2.82E-05 | 7.75E-05 | -1.16E+00 |
| PAK1IP1 | 5.76E-15 | 3.00E-14 | -1.91E+00 | 6.34E-10 | 2.74E-09 | -1.29E+00 | 8.44E-10 | 3.58E-09 | -1.48E+00 | 3.36E-05 | 1.28E-04 | -1.06E+00 | 6.87E-07 | 2.45E-06 | -1.50E+00 | 6.00E-04 | 1.69E-03 | -1.15E+00 | 3.68E-12 | 1.35E-11 | -1.46E+00 | 4.40E-02 | 1.21E-01 | 1.13E+00 |
| NBN | 5.80E-15 | 3.02E-14 | -1.46E+00 | 6.00E-13 | 2.59E-12 | -1.18E+00 | 4.12E-09 | 1.75E-08 | -1.41E+00 | 4.58E-08 | 1.75E-07 | -1.24E+00 | 3.09E-08 | 1.10E-07 | -1.63E+00 | 6.32E-06 | 1.78E-05 | -1.30E+00 | 3.86E-12 | 1.42E-11 | -1.61E+00 | 1.26E-06 | 3.47E-06 | -1.31E+00 |
| PCID2 | 5.81E-15 | 3.03E-14 | -1.53E+00 | 5.46E-06 | 2.36E-05 | -1.00E+00 | 6.22E-08 | 2.64E-07 | -1.40E+00 | 1.68E-03 | 6.41E-03 | 1.01E+00 | 6.04E-10 | 2.15E-09 | -1.49E+00 | 9.86E-04 | 2.78E-03 | 1.03E+00 | 3.06E-07 | 1.12E-06 | -1.31E+00 | 1.24E-01 | 3.40E-01 | 1.05E+00 |
| TRIM14 | 5.81E-15 | 3.03E-14 | -1.82E+00 | 2.62E-08 | 1.13E-07 | -1.20E+00 | 8.42E-17 | 3.58E-16 | -1.81E+00 | 2.06E-04 | 7.86E-04 | 1.01E+00 | 7.68E-06 | 2.74E-05 | -2.68E+00 | 5.16E-03 | 1.46E-02 | -1.34E+00 | 2.56E-13 | 9.41E-13 | -2.23E+00 | 1.06E-08 | 2.91E-08 | -1.59E+00 |
| PCMTD1P7 | 5.86E-15 | 3.06E-14 | -3.03E+00 | 7.95E-11 | 3.43E-10 | -1.79E+00 | 3.94E-05 | 1.68E-04 | -1.97E+00 | 3.12E-04 | 1.19E-03 | -1.46E+00 | 6.82E-06 | 2.43E-05 | -1.99E+00 | 9.70E-05 | 2.73E-04 | -1.57E+00 | 2.14E-05 | 7.87E-05 | -1.37E+00 | 1.47E-06 | 4.05E-06 | -1.99E+00 |
| RIF1 | 5.86E-15 | 3.06E-14 | -1.44E+00 | 1.26E-07 | 5.46E-07 | -1.02E+00 | 1.95E-09 | 8.29E-09 | -1.55E+00 | 8.12E-05 | 3.10E-04 | -1.13E+00 | 5.98E-17 | 2.13E-16 | -1.77E+00 | 3.25E-05 | 9.16E-05 | -1.10E+00 | 1.60E-14 | 5.86E-14 | -1.92E+00 | 4.39E-03 | 1.20E-02 | -1.20E+00 |
| SYK | 5.99E-15 | 3.12E-14 | -1.59E+00 | 2.07E-05 | 8.93E-05 | -1.01E+00 | 1.79E-08 | 7.59E-08 | -1.49E+00 | 9.01E-04 | 3.44E-03 | 1.01E+00 | 5.06E-08 | 1.80E-07 | -1.62E+00 | 1.25E-04 | 3.53E-04 | -1.19E+00 | 4.35E-03 | 1.60E-02 | -1.20E+00 | 5.02E-02 | 1.38E-01 | -1.08E+00 |
| ACSL3 | 6.02E-15 | 3.14E-14 | -1.77E+00 | 1.36E-13 | 5.90E-13 | -1.47E+00 | 2.83E-09 | 1.20E-08 | -1.68E+00 | 1.64E-05 | 6.27E-05 | -1.16E+00 | 5.09E-04 | 1.82E-03 | -1.06E+00 | 3.21E-05 | 9.06E-05 | -1.13E+00 | 4.05E-09 | 1.49E-08 | -1.38E+00 | 3.10E-04 | 8.53E-04 | -1.03E+00 |
| EP400NL | 6.21E-15 | 3.23E-14 | -1.65E+00 | 2.52E-11 | 1.09E-10 | -1.26E+00 | 1.87E-09 | 7.96E-09 | -1.54E+00 | 2.66E-13 | 1.02E-12 | -1.48E+00 | 1.24E-02 | 4.42E-02 | -1.05E+00 | 6.81E-02 | 1.92E-01 | 1.07E+00 | 1.65E-03 | 6.04E-03 | -1.08E+00 | 2.60E-03 | 7.15E-03 | -1.08E+00 |
| YPEL3 | 6.32E-15 | 3.29E-14 | -1.34E+00 | 4.67E-04 | 2.02E-03 | 1.14E+00 | 1.54E-11 | 6.56E-11 | -1.62E+00 | 2.14E-05 | 8.17E-05 | -1.12E+00 | 1.46E-06 | 5.22E-06 | -1.26E+00 | 6.21E-03 | 1.75E-02 | -1.00E+00 | 2.28E-10 | 8.37E-10 | -1.60E+00 | 1.29E-10 | 3.54E-10 | -1.56E+00 |
| SUPT16HP1 | 6.46E-15 | 3.37E-14 | -1.97E+00 | 2.49E-15 | 1.08E-14 | -1.80E+00 | 1.04E-10 | 4.41E-10 | -1.64E+00 | 1.24E-19 | 4.73E-19 | -1.72E+00 | 3.39E-05 | 1.21E-04 | -1.39E+00 | 3.51E-04 | 9.91E-04 | -1.31E+00 | 1.69E-07 | 6.19E-07 | -1.36E+00 | 2.01E-07 | 5.51E-07 | -1.35E+00 |
| PPDPF | 6.47E-15 | 3.37E-14 | -1.46E+00 | 3.72E-14 | 1.61E-13 | -1.31E+00 | 2.32E-03 | 9.86E-03 | -1.20E+00 | 4.09E-04 | 1.56E-03 | -1.13E+00 | 3.78E-10 | 1.35E-09 | -1.40E+00 | 1.34E-06 | 3.79E-06 | -1.30E+00 | 1.62E-05 | 5.93E-05 | -1.26E+00 | 7.42E-04 | 2.04E-03 | -1.28E+00 |
| GPATCH11 | 6.71E-15 | 3.50E-14 | -1.89E+00 | 1.52E-11 | 6.55E-11 | -1.54E+00 | 3.44E-08 | 1.46E-07 | -1.58E+00 | 3.69E-05 | 1.41E-04 | -1.18E+00 | 1.67E-05 | 5.95E-05 | -1.15E+00 | 3.75E-06 | 1.06E-05 | -1.25E+00 | 2.34E-03 | 8.60E-03 | -1.06E+00 | 3.94E-02 | 1.08E-01 | -1.01E+00 |
| PSIP1 | 7.04E-15 | 3.67E-14 | -1.28E+00 | 7.05E-04 | 3.05E-03 | 1.13E+00 | 1.30E-02 | 5.54E-02 | -1.03E+00 | 3.40E-02 | 1.30E-01 | 1.18E+00 | 6.29E-05 | 2.24E-04 | -1.03E+00 | 1.48E-02 | 4.16E-02 | 1.13E+00 | 1.54E-03 | 5.64E-03 | -1.00E+00 | 1.40E-01 | 3.84E-01 | 1.17E+00 |
| ZNF148 | 7.15E-15 | 3.72E-14 | -1.49E+00 | 7.07E-10 | 3.05E-09 | -1.16E+00 | 2.01E-08 | 8.53E-08 | -1.47E+00 | 1.83E-03 | 6.98E-03 | 1.01E+00 | 1.03E-06 | 3.66E-06 | -1.32E+00 | 7.42E-03 | 2.09E-02 | 1.02E+00 | 4.24E-15 | 1.56E-14 | -1.68E+00 | 1.25E-04 | 3.42E-04 | -1.16E+00 |
| RNF5 | 7.21E-15 | 3.76E-14 | -1.58E+00 | 2.40E-06 | 1.04E-05 | -1.00E+00 | 3.24E-08 | 1.38E-07 | -1.52E+00 | 1.19E-04 | 4.53E-04 | -1.07E+00 | 2.05E-14 | 7.31E-14 | -1.79E+00 | 5.65E-04 | 1.59E-03 | -1.04E+00 | 4.97E-11 | 1.82E-10 | -1.45E+00 | 4.72E-04 | 1.30E-03 | -1.09E+00 |
| AC026271.1 | 7.25E-15 | 3.78E-14 | -1.68E+00 | 1.70E-14 | 7.35E-14 | -1.47E+00 | 7.98E-06 | 3.39E-05 | -1.34E+00 | 1.86E-07 | 7.09E-07 | -1.29E+00 | 1.12E-08 | 4.01E-08 | -1.51E+00 | 2.17E-10 | 6.12E-10 | -1.60E+00 | 3.81E-16 | 1.40E-15 | -1.82E+00 | 2.84E-11 | 7.80E-11 | -1.54E+00 |
| H2AFV | 7.33E-15 | 3.82E-14 | -1.24E+00 | 1.40E-05 | 6.05E-05 | 1.00E+00 | 2.12E-03 | 9.00E-03 | -1.14E+00 | 1.63E-01 | 6.23E-01 | 1.09E+00 | 2.34E-02 | 8.33E-02 | -1.06E+00 | 1.40E-01 | 3.94E-01 | 1.05E+00 | 2.46E-02 | 9.04E-02 | -1.14E+00 | 2.11E-01 | 5.80E-01 | -1.01E+00 |
| MRPS7 | 7.33E-15 | 3.82E-14 | -1.38E+00 | 1.46E-08 | 6.31E-08 | -1.05E+00 | 2.52E-03 | 1.07E-02 | -1.05E+00 | 4.05E-04 | 1.55E-03 | 1.04E+00 | 5.96E-07 | 2.13E-06 | -1.21E+00 | 2.48E-03 | 6.99E-03 | 1.03E+00 | 5.24E-04 | 1.92E-03 | -1.04E+00 | 1.12E-01 | 3.07E-01 | 1.08E+00 |
| NDUFB6 | 7.73E-15 | 4.03E-14 | -1.38E+00 | 1.12E-12 | 4.82E-12 | -1.22E+00 | 5.84E-02 | 2.48E-01 | 1.03E+00 | 1.73E-03 | 6.62E-03 | 1.02E+00 | 2.79E-09 | 9.94E-09 | -1.41E+00 | 1.35E-05 | 3.80E-05 | -1.07E+00 | 2.31E-04 | 8.50E-04 | -1.10E+00 | 3.76E-04 | 1.03E-03 | -1.13E+00 |
| HIP1 | 7.73E-15 | 4.03E-14 | -1.64E+00 | 2.54E-10 | 1.10E-09 | -1.25E+00 | 6.40E-08 | 2.72E-07 | -1.39E+00 | 3.99E-04 | 1.52E-03 | 1.05E+00 | 1.66E-07 | 5.91E-07 | -1.72E+00 | 1.55E-05 | 4.36E-05 | -1.34E+00 | 2.06E-04 | 7.57E-04 | -1.45E+00 | 2.75E-03 | 7.56E-03 | -1.26E+00 |
| SDAD1P2 | 7.87E-15 | 4.10E-14 | -2.82E+00 | 7.57E-08 | 3.27E-07 | -1.42E+00 | 3.12E-13 | 1.33E-12 | -2.20E+00 | 3.02E-10 | 1.15E-09 | -1.56E+00 | 8.86E-11 | 3.16E-10 | -3.70E+00 | 1.34E-05 | 3.77E-05 | -1.50E+00 | 1.16E-15 | 4.24E-15 | -2.18E+00 | 1.35E-08 | 3.70E-08 | -1.60E+00 |
| PGAM1P8 | 7.87E-15 | 4.10E-14 | -2.13E+00 | 2.87E-10 | 1.24E-09 | -1.47E+00 | 1.27E-06 | 5.42E-06 | -1.28E+00 | 3.03E-09 | 1.16E-08 | -1.31E+00 | 1.57E-14 | 5.61E-14 | -2.07E+00 | 2.55E-12 | 7.18E-12 | -1.80E+00 | 1.69E-06 | 6.20E-06 | -1.18E+00 | 1.10E-03 | 3.02E-03 | -1.03E+00 |
| TIMM8B | 7.95E-15 | 4.14E-14 | -1.40E+00 | 5.75E-23 | 2.48E-22 | -1.56E+00 | 4.28E-04 | 1.82E-03 | -1.17E+00 | 3.18E-10 | 1.21E-09 | -1.35E+00 | 1.17E-13 | 4.17E-13 | -1.53E+00 | 3.62E-12 | 1.02E-11 | -1.48E+00 | 2.50E-04 | 9.16E-04 | -1.13E+00 | 1.07E-05 | 2.94E-05 | -1.33E+00 |
| COG4 | 8.21E-15 | 4.28E-14 | -1.61E+00 | 3.42E-09 | 1.48E-08 | -1.14E+00 | 1.57E-05 | 6.65E-05 | -1.25E+00 | 4.10E-06 | 1.57E-05 | -1.14E+00 | 4.55E-06 | 1.62E-05 | -1.25E+00 | 1.88E-04 | 5.30E-04 | -1.06E+00 | 5.51E-06 | 2.02E-05 | -1.16E+00 | 5.09E-06 | 1.40E-05 | -1.19E+00 |
| MYO5C | 8.22E-15 | 4.28E-14 | -1.84E+00 | 3.30E-05 | 1.42E-04 | 1.02E+00 | 3.38E-06 | 1.44E-05 | -1.51E+00 | 4.69E-06 | 1.79E-05 | -1.37E+00 | 2.39E-01 | 8.51E-01 | 1.46E+00 | 3.08E-01 | 8.67E-01 | 1.42E+00 | 2.16E-01 | 7.91E-01 | 1.34E+00 | 2.66E-01 | 7.30E-01 | 1.49E+00 |
| LUC7L2 | 8.35E-15 | 4.35E-14 | -1.34E+00 | 1.14E-09 | 4.93E-09 | -1.04E+00 | 3.58E-04 | 1.52E-03 | -1.12E+00 | 2.09E-03 | 7.97E-03 | 1.03E+00 | 4.41E-08 | 1.57E-07 | -1.47E+00 | 5.34E-03 | 1.51E-02 | 1.03E+00 | 7.47E-07 | 2.74E-06 | -1.25E+00 | 1.89E-02 | 5.18E-02 | 1.05E+00 |
| RPS3AP44 | 8.39E-15 | 4.37E-14 | -1.26E+00 | 4.82E-21 | 2.08E-20 | -1.31E+00 | 2.20E-07 | 9.33E-07 | -1.20E+00 | 1.73E-07 | 6.59E-07 | -1.21E+00 | 1.78E-05 | 6.35E-05 | -1.18E+00 | 4.89E-06 | 1.38E-05 | -1.20E+00 | 1.60E-18 | 5.87E-18 | -1.25E+00 | 1.25E-13 | 3.43E-13 | -1.28E+00 |
| PABPC1 | 8.43E-15 | 4.39E-14 | -1.21E+00 | 1.16E-01 | 5.00E-01 | -1.00E+00 | 6.62E-11 | 2.81E-10 | -1.23E+00 | 1.51E-01 | 5.76E-01 | -1.01E+00 | 1.40E-08 | 4.99E-08 | -1.21E+00 | 5.59E-02 | 1.58E-01 | -1.03E+00 | 6.24E-22 | 2.29E-21 | -1.33E+00 | 4.76E-02 | 1.31E-01 | -1.03E+00 |
| HIST1H2AC | 8.51E-15 | 4.43E-14 | -1.96E+00 | 1.24E-11 | 5.37E-11 | -1.45E+00 | 1.04E-02 | 4.41E-02 | -1.37E+00 | 2.80E-03 | 1.07E-02 | -1.28E+00 | 1.60E-06 | 5.69E-06 | -1.54E+00 | 1.23E-03 | 3.47E-03 | -1.03E+00 | 1.97E-04 | 7.22E-04 | 1.05E+00 | 1.39E-05 | 3.82E-05 | -1.30E+00 |
| SYPL1 | 8.79E-15 | 4.58E-14 | -1.34E+00 | 6.14E-02 | 2.65E-01 | 1.14E+00 | 7.08E-07 | 3.01E-06 | -1.36E+00 | 3.26E-02 | 1.24E-01 | 1.05E+00 | 4.15E-09 | 1.48E-08 | -1.20E+00 | 2.00E-01 | 5.65E-01 | 1.17E+00 | 4.52E-04 | 1.66E-03 | -1.17E+00 | 7.61E-02 | 2.09E-01 | 1.01E+00 |
| PRKAG2-AS1 | 8.88E-15 | 4.63E-14 | -2.23E+00 | 3.12E-06 | 1.35E-05 | -1.06E+00 | 4.36E-07 | 1.85E-06 | -1.41E+00 | 1.69E-03 | 6.46E-03 | 1.09E+00 | 2.21E-04 | 7.90E-04 | -2.21E+00 | 2.26E-02 | 6.38E-02 | -1.18E+00 | 8.59E-03 | 3.15E-02 | -1.25E+00 | 4.12E-02 | 1.13E-01 | 1.60E+00 |
| UBAC2 | 9.15E-15 | 4.77E-14 | -1.36E+00 | 1.33E-08 | 5.76E-08 | -1.01E+00 | 1.44E-05 | 6.11E-05 | -1.27E+00 | 5.66E-04 | 2.16E-03 | -1.06E+00 | 9.28E-10 | 3.31E-09 | -1.36E+00 | 4.95E-03 | 1.40E-02 | 1.11E+00 | 1.38E-07 | 5.06E-07 | -1.24E+00 | 3.12E-02 | 8.58E-02 | 1.04E+00 |
| LSM10 | 9.16E-15 | 4.77E-14 | -1.51E+00 | 1.21E-08 | 5.25E-08 | -1.05E+00 | 2.50E-06 | 1.06E-05 | -1.26E+00 | 2.06E-04 | 7.87E-04 | -1.01E+00 | 3.97E-11 | 1.41E-10 | -1.59E+00 | 7.03E-05 | 1.98E-04 | -1.08E+00 | 1.93E-05 | 7.08E-05 | -1.10E+00 | 6.09E-03 | 1.67E-02 | 1.03E+00 |
| DCLRE1C | 9.22E-15 | 4.81E-14 | -1.79E+00 | 6.42E-09 | 2.77E-08 | -1.17E+00 | 2.22E-11 | 9.45E-11 | -1.61E+00 | 2.98E-04 | 1.14E-03 | 1.05E+00 | 1.26E-09 | 4.50E-09 | -1.64E+00 | 9.27E-06 | 2.61E-05 | -1.21E+00 | 9.29E-08 | 3.41E-07 | -1.41E+00 | 9.35E-06 | 2.57E-05 | -1.22E+00 |
| CTSD | 9.31E-15 | 4.85E-14 | -1.37E+00 | 4.80E-09 | 2.07E-08 | -1.12E+00 | 8.40E-06 | 3.57E-05 | -1.23E+00 | 1.93E-04 | 7.39E-04 | -1.06E+00 | 9.13E-09 | 3.25E-08 | -1.43E+00 | 5.88E-09 | 1.66E-08 | -1.38E+00 | 3.02E-04 | 1.11E-03 | -1.06E+00 | 5.38E-02 | 1.48E-01 | 1.06E+00 |
| SELENOM | 9.37E-15 | 4.88E-14 | -2.04E+00 | 2.76E-11 | 1.19E-10 | -1.58E+00 | 1.26E-01 | 5.35E-01 | 1.44E+00 | 2.13E-01 | 8.14E-01 | 1.35E+00 | 1.97E-01 | 7.04E-01 | -1.04E+00 | 2.93E-01 | 8.27E-01 | -1.00E+00 | 1.82E-01 | 6.69E-01 | -1.14E+00 | 2.89E-01 | 7.94E-01 | -1.01E+00 |
| C1QTNF4 | 9.38E-15 | 4.89E-14 | -1.39E+00 | 1.58E-20 | 6.85E-20 | -1.57E+00 | 4.84E-11 | 2.06E-10 | -1.69E+00 | 2.25E-26 | 8.61E-26 | -2.26E+00 | 2.52E-01 | 8.99E-01 | 1.44E+00 | 3.39E-01 | 9.56E-01 | 1.19E+00 | 2.48E-01 | 9.11E-01 | 1.29E+00 | 3.35E-01 | 9.20E-01 | -1.08E+00 |
| ETF1 | 9.41E-15 | 4.91E-14 | -1.30E+00 | 5.20E-28 | 2.25E-27 | -1.70E+00 | 2.84E-03 | 1.21E-02 | -1.00E+00 | 1.68E-09 | 6.42E-09 | -1.21E+00 | 1.42E-09 | 5.06E-09 | -1.39E+00 | 1.23E-13 | 3.47E-13 | -1.56E+00 | 2.34E-04 | 8.59E-04 | -1.14E+00 | 3.02E-03 | 8.29E-03 | -1.09E+00 |
| ZBTB38 | 9.51E-15 | 4.95E-14 | -1.92E+00 | 7.52E-06 | 3.25E-05 | -1.03E+00 | 2.55E-11 | 1.08E-10 | -1.79E+00 | 2.96E-06 | 1.13E-05 | -1.21E+00 | 2.54E-04 | 9.06E-04 | -1.33E+00 | 5.61E-02 | 1.58E-01 | 1.01E+00 | 2.04E-04 | 7.49E-04 | -1.34E+00 | 3.10E-02 | 8.52E-02 | -1.03E+00 |
| PSMG3 | 9.64E-15 | 5.02E-14 | -1.40E+00 | 3.38E-12 | 1.46E-11 | -1.14E+00 | 2.93E-05 | 1.24E-04 | -1.23E+00 | 3.66E-03 | 1.40E-02 | 1.01E+00 | 1.54E-05 | 5.49E-05 | -1.09E+00 | 4.20E-03 | 1.18E-02 | 1.10E+00 | 2.60E-05 | 9.55E-05 | -1.20E+00 | 6.68E-02 | 1.84E-01 | 1.00E+00 |
| UFSP2 | 1.03E-14 | 5.35E-14 | -1.94E+00 | 1.96E-13 | 8.48E-13 | -1.64E+00 | 2.42E-10 | 1.03E-09 | -1.86E+00 | 1.56E-09 | 5.96E-09 | -1.55E+00 | 1.58E-05 | 5.64E-05 | -1.36E+00 | 3.96E-04 | 1.12E-03 | -1.15E+00 | 2.33E-05 | 8.55E-05 | -1.36E+00 | 3.06E-04 | 8.40E-04 | -1.16E+00 |
| MTDHP3 | 1.03E-14 | 5.37E-14 | -1.72E+00 | 1.62E-11 | 7.00E-11 | -1.40E+00 | 2.58E-05 | 1.09E-04 | -1.55E+00 | 1.63E-03 | 6.24E-03 | -1.15E+00 | 1.65E-01 | 5.87E-01 | 1.66E+00 | 4.13E-02 | 1.16E-01 | 2.01E+00 | 6.73E-02 | 2.47E-01 | 1.28E+00 | 4.41E-04 | 1.21E-03 | -1.11E+00 |
| DLG4 | 1.05E-14 | 5.49E-14 | -2.01E+00 | 4.27E-14 | 1.84E-13 | -1.71E+00 | 1.42E-06 | 6.05E-06 | -1.81E+00 | 2.97E-04 | 1.13E-03 | -1.23E+00 | 8.30E-06 | 2.96E-05 | -1.77E+00 | 9.78E-04 | 2.76E-03 | -1.26E+00 | 3.02E-02 | 1.11E-01 | -1.19E+00 | 1.14E-01 | 3.14E-01 | -1.04E+00 |
| TMF1 | 1.08E-14 | 5.64E-14 | -1.31E+00 | 9.53E-12 | 4.12E-11 | -1.14E+00 | 2.41E-05 | 1.02E-04 | -1.24E+00 | 7.39E-05 | 2.82E-04 | -1.12E+00 | 6.38E-02 | 2.28E-01 | 1.24E+00 | 1.68E-03 | 4.74E-03 | -1.04E+00 | 7.80E-03 | 2.86E-02 | 1.05E+00 | 4.55E-05 | 1.25E-04 | -1.11E+00 |
| ASPH | 1.10E-14 | 5.71E-14 | -1.43E+00 | 6.14E-11 | 2.65E-10 | -1.20E+00 | 8.63E-09 | 3.67E-08 | -1.43E+00 | 2.31E-04 | 8.81E-04 | -1.04E+00 | 5.36E-09 | 1.91E-08 | -1.84E+00 | 1.13E-06 | 3.19E-06 | -1.35E+00 | 1.02E-05 | 3.76E-05 | -1.24E+00 | 4.70E-05 | 1.29E-04 | -1.22E+00 |
| PPM1G | 1.10E-14 | 5.71E-14 | -1.33E+00 | 2.28E-16 | 9.86E-16 | -1.18E+00 | 1.08E-03 | 4.59E-03 | -1.15E+00 | 7.60E-05 | 2.90E-04 | -1.12E+00 | 2.61E-10 | 9.31E-10 | -1.43E+00 | 3.78E-05 | 1.07E-04 | -1.17E+00 | 6.39E-04 | 2.35E-03 | -1.14E+00 | 2.91E-01 | 8.00E-01 | 1.08E+00 |
| MTR | 1.11E-14 | 5.77E-14 | -1.67E+00 | 5.05E-06 | 2.18E-05 | -1.05E+00 | 1.28E-09 | 5.45E-09 | -1.45E+00 | 4.01E-04 | 1.53E-03 | 1.03E+00 | 1.29E-07 | 4.58E-07 | -1.48E+00 | 6.00E-03 | 1.69E-02 | -1.00E+00 | 8.80E-08 | 3.23E-07 | -1.43E+00 | 1.86E-03 | 5.11E-03 | -1.13E+00 |
| PSME1 | 1.12E-14 | 5.84E-14 | -1.34E+00 | 8.26E-10 | 3.57E-09 | -1.19E+00 | 7.90E-03 | 3.36E-02 | -1.16E+00 | 5.17E-03 | 1.98E-02 | -1.14E+00 | 1.10E-16 | 3.93E-16 | -1.54E+00 | 2.20E-05 | 6.19E-05 | -1.28E+00 | 1.17E-15 | 4.28E-15 | -1.65E+00 | 1.10E-06 | 3.01E-06 | -1.41E+00 |
| SERF1B | 1.13E-14 | 5.88E-14 | -1.76E+00 | 2.16E-11 | 9.33E-11 | -1.33E+00 | 5.18E-12 | 2.20E-11 | -1.57E+00 | 7.08E-11 | 2.70E-10 | -1.38E+00 | 2.88E-02 | 1.03E-01 | -1.12E+00 | 1.14E-01 | 3.20E-01 | 1.08E+00 | 1.04E-06 | 3.80E-06 | -1.46E+00 | 1.25E-05 | 3.43E-05 | -1.35E+00 |
| ZNF781 | 1.16E-14 | 6.05E-14 | -2.09E+00 | 1.27E-12 | 5.47E-12 | -1.58E+00 | 1.15E-07 | 4.91E-07 | -2.12E+00 | 4.38E-05 | 1.67E-04 | -1.37E+00 | 7.52E-02 | 2.68E-01 | 1.05E+00 | 1.80E-01 | 5.09E-01 | 1.15E+00 | 1.34E-02 | 4.92E-02 | -1.48E+00 | 4.70E-02 | 1.29E-01 | -1.37E+00 |
| HPS1 | 1.19E-14 | 6.19E-14 | -1.34E+00 | 2.07E-09 | 8.95E-09 | -1.02E+00 | 9.59E-05 | 4.07E-04 | -1.23E+00 | 1.10E-03 | 4.20E-03 | -1.00E+00 | 1.20E-04 | 4.26E-04 | -1.17E+00 | 6.52E-03 | 1.84E-02 | 1.04E+00 | 5.52E-08 | 2.03E-07 | -1.26E+00 | 2.09E-03 | 5.73E-03 | -1.08E+00 |
| DDX17 | 1.21E-14 | 6.31E-14 | -1.32E+00 | 1.98E-02 | 8.58E-02 | 1.30E+00 | 4.01E-13 | 1.71E-12 | -1.53E+00 | 1.11E-01 | 4.23E-01 | 1.14E+00 | 2.22E-05 | 7.93E-05 | -1.21E+00 | 3.03E-01 | 8.55E-01 | 1.09E+00 | 2.27E-06 | 8.35E-06 | -1.26E+00 | 2.09E-01 | 5.74E-01 | 1.09E+00 |
| RGCC | 1.21E-14 | 6.33E-14 | 4.95E+00 | 1.28E-02 | 5.54E-02 | -1.78E+00 | 9.83E-14 | 4.18E-13 | 2.50E+00 | 3.13E-03 | 1.20E-02 | -1.53E+00 | 9.89E-10 | 3.53E-09 | 8.63E+00 | 2.06E-01 | 5.80E-01 | -1.30E+00 | 5.19E-16 | 1.90E-15 | 1.21E+01 | 3.33E-01 | 9.15E-01 | 2.43E+00 |
| NUCKS1 | 1.23E-14 | 6.41E-14 | -1.32E+00 | 3.31E-06 | 1.43E-05 | -1.03E+00 | 5.60E-02 | 2.38E-01 | -1.03E+00 | 1.74E-01 | 6.64E-01 | 1.09E+00 | 8.22E-10 | 2.93E-09 | -1.29E+00 | 9.24E-03 | 2.60E-02 | 1.02E+00 | 1.09E-02 | 3.99E-02 | -1.20E+00 | 8.72E-02 | 2.40E-01 | 1.13E+00 |
| ECHDC1 | 1.27E-14 | 6.63E-14 | -1.47E+00 | 4.12E-08 | 1.78E-07 | -1.07E+00 | 8.50E-05 | 3.61E-04 | -1.17E+00 | 1.66E-04 | 6.33E-04 | -1.04E+00 | 4.62E-08 | 1.65E-07 | -1.40E+00 | 1.38E-03 | 3.89E-03 | 1.01E+00 | 7.58E-09 | 2.78E-08 | -1.28E+00 | 3.23E-02 | 8.87E-02 | 1.10E+00 |
| CRLF3 | 1.27E-14 | 6.63E-14 | -1.54E+00 | 2.81E-06 | 1.21E-05 | 1.03E+00 | 1.52E-08 | 6.44E-08 | -1.54E+00 | 6.85E-05 | 2.62E-04 | -1.05E+00 | 6.16E-08 | 2.20E-07 | -1.40E+00 | 1.28E-04 | 3.61E-04 | -1.07E+00 | 4.49E-05 | 1.65E-04 | -1.07E+00 | 4.45E-03 | 1.22E-02 | 1.01E+00 |
| CTSA | 1.28E-14 | 6.69E-14 | -1.82E+00 | 3.07E-11 | 1.32E-10 | -1.38E+00 | 9.21E-12 | 3.91E-11 | -1.53E+00 | 4.52E-06 | 1.73E-05 | -1.07E+00 | 2.33E-13 | 8.31E-13 | -1.74E+00 | 3.40E-07 | 9.58E-07 | -1.22E+00 | 2.45E-05 | 8.98E-05 | -1.08E+00 | 8.42E-03 | 2.31E-02 | -1.09E+00 |
| LNPEP | 1.30E-14 | 6.76E-14 | -1.54E+00 | 2.43E-03 | 1.05E-02 | 1.16E+00 | 3.62E-10 | 1.54E-09 | -1.49E+00 | 2.14E-04 | 8.17E-04 | 1.02E+00 | 1.47E-08 | 5.24E-08 | -1.43E+00 | 6.75E-03 | 1.90E-02 | 1.02E+00 | 2.45E-06 | 8.99E-06 | -1.30E+00 | 1.02E-01 | 2.79E-01 | 1.08E+00 |
| AL592114.1 | 1.31E-14 | 6.80E-14 | -1.28E+00 | 2.13E-40 | 9.19E-40 | -1.91E+00 | 1.88E-06 | 7.99E-06 | -1.22E+00 | 1.47E-23 | 5.60E-23 | -1.81E+00 | 3.58E-07 | 1.28E-06 | -1.14E+00 | 3.02E-17 | 8.52E-17 | -1.82E+00 | 1.61E-06 | 5.91E-06 | -1.29E+00 | 2.11E-15 | 5.78E-15 | -1.89E+00 |
| MRPS2 | 1.32E-14 | 6.87E-14 | -1.41E+00 | 8.55E-12 | 3.69E-11 | -1.15E+00 | 3.86E-08 | 1.64E-07 | -1.28E+00 | 4.91E-07 | 1.88E-06 | -1.12E+00 | 1.56E-12 | 5.54E-12 | -1.78E+00 | 3.92E-05 | 1.10E-04 | -1.17E+00 | 1.13E-08 | 4.16E-08 | -1.40E+00 | 6.43E-02 | 1.77E-01 | 1.03E+00 |
| ASB8 | 1.33E-14 | 6.93E-14 | -1.45E+00 | 1.85E-07 | 7.99E-07 | 1.01E+00 | 3.95E-10 | 1.68E-09 | -1.49E+00 | 3.63E-05 | 1.39E-04 | -1.05E+00 | 1.06E-07 | 3.79E-07 | -1.37E+00 | 1.75E-04 | 4.93E-04 | -1.12E+00 | 2.97E-08 | 1.09E-07 | -1.28E+00 | 2.88E-03 | 7.92E-03 | -1.01E+00 |
| IMMT | 1.33E-14 | 6.94E-14 | -1.44E+00 | 1.03E-05 | 4.43E-05 | 1.01E+00 | 2.04E-07 | 8.69E-07 | -1.28E+00 | 3.07E-04 | 1.17E-03 | -1.04E+00 | 4.74E-10 | 1.69E-09 | -1.43E+00 | 2.36E-03 | 6.66E-03 | 1.04E+00 | 1.53E-06 | 5.62E-06 | -1.24E+00 | 7.91E-03 | 2.17E-02 | 1.02E+00 |
| TXN2 | 1.37E-14 | 7.12E-14 | -1.29E+00 | 5.73E-09 | 2.48E-08 | -1.06E+00 | 3.43E-03 | 1.46E-02 | -1.09E+00 | 2.70E-03 | 1.03E-02 | 1.00E+00 | 2.88E-06 | 1.03E-05 | -1.19E+00 | 5.25E-03 | 1.48E-02 | 1.03E+00 | 7.37E-03 | 2.71E-02 | -1.06E+00 | 1.51E-01 | 4.14E-01 | 1.03E+00 |
| CTSW | 1.37E-14 | 7.14E-14 | -1.29E+00 | 2.17E-10 | 9.39E-10 | -1.08E+00 | 6.92E-05 | 2.94E-04 | -1.20E+00 | 1.29E-02 | 4.94E-02 | -1.05E+00 | 3.99E-06 | 1.42E-05 | -1.21E+00 | 1.94E-02 | 5.47E-02 | 1.01E+00 | 6.95E-02 | 2.55E-01 | 1.06E+00 | 2.45E-01 | 6.73E-01 | 1.08E+00 |
| RPL26P30 | 1.37E-14 | 7.16E-14 | -1.89E+00 | 1.64E-15 | 7.08E-15 | -1.79E+00 | 1.51E-14 | 6.40E-14 | -1.83E+00 | 5.77E-18 | 2.20E-17 | -1.79E+00 | 1.04E-06 | 3.71E-06 | -1.74E+00 | 2.11E-04 | 5.95E-04 | -1.29E+00 | 4.42E-11 | 1.62E-10 | -1.70E+00 | 1.06E-05 | 2.92E-05 | -1.33E+00 |
| CTR9 | 1.40E-14 | 7.31E-14 | -1.89E+00 | 4.67E-06 | 2.02E-05 | -1.13E+00 | 5.24E-15 | 2.23E-14 | -2.05E+00 | 7.50E-06 | 2.86E-05 | -1.14E+00 | 4.31E-13 | 1.54E-12 | -1.71E+00 | 2.91E-06 | 8.20E-06 | -1.17E+00 | 2.53E-09 | 9.29E-09 | -1.18E+00 | 3.96E-02 | 1.09E-01 | 1.06E+00 |
| FLI1 | 1.41E-14 | 7.36E-14 | -1.31E+00 | 7.04E-02 | 3.04E-01 | 1.35E+00 | 9.44E-09 | 4.01E-08 | -1.46E+00 | 6.69E-02 | 2.55E-01 | 1.16E+00 | 4.92E-06 | 1.76E-05 | -1.42E+00 | 5.91E-02 | 1.67E-01 | 1.02E+00 | 3.69E-06 | 1.35E-05 | -1.31E+00 | 1.24E-01 | 3.41E-01 | 1.13E+00 |
| BCL7B | 1.43E-14 | 7.43E-14 | -1.53E+00 | 1.80E-14 | 7.78E-14 | -1.40E+00 | 1.46E-09 | 6.18E-09 | -1.48E+00 | 2.63E-08 | 1.01E-07 | -1.28E+00 | 2.21E-08 | 7.88E-08 | -1.44E+00 | 2.62E-06 | 7.39E-06 | -1.18E+00 | 5.22E-10 | 1.92E-09 | -1.34E+00 | 4.53E-07 | 1.24E-06 | -1.27E+00 |
| PRCP | 1.45E-14 | 7.54E-14 | -1.41E+00 | 4.16E-10 | 1.80E-09 | -1.16E+00 | 1.25E-07 | 5.31E-07 | -1.28E+00 | 6.85E-05 | 2.62E-04 | 1.02E+00 | 1.44E-04 | 5.12E-04 | -1.28E+00 | 3.07E-03 | 8.67E-03 | -1.07E+00 | 3.29E-06 | 1.21E-05 | -1.16E+00 | 2.50E-03 | 6.86E-03 | -1.02E+00 |
| PSMD2 | 1.45E-14 | 7.54E-14 | -1.40E+00 | 2.12E-06 | 9.17E-06 | 1.05E+00 | 5.14E-04 | 2.19E-03 | -1.03E+00 | 2.22E-03 | 8.49E-03 | 1.19E+00 | 4.23E-02 | 1.51E-01 | 1.14E+00 | 3.24E-01 | 9.13E-01 | 1.36E+00 | 3.15E-02 | 1.16E-01 | 1.13E+00 | 1.73E-01 | 4.74E-01 | 1.33E+00 |
| SORD2P | 1.50E-14 | 7.80E-14 | -2.04E+00 | 1.51E-12 | 6.53E-12 | -1.63E+00 | 1.14E-08 | 4.86E-08 | -1.40E+00 | 5.78E-11 | 2.21E-10 | -1.41E+00 | 1.33E-08 | 4.74E-08 | -1.76E+00 | 4.45E-03 | 1.25E-02 | 1.00E+00 | 3.13E-10 | 1.15E-09 | -1.46E+00 | 1.16E-08 | 3.19E-08 | -1.38E+00 |
| RTL8B | 1.50E-14 | 7.83E-14 | -1.97E+00 | 7.25E-09 | 3.13E-08 | -1.29E+00 | 1.38E-04 | 5.86E-04 | -1.53E+00 | 1.15E-01 | 4.40E-01 | 1.25E+00 | 5.57E-03 | 1.99E-02 | -1.91E+00 | 2.68E-02 | 7.57E-02 | -1.34E+00 | 1.23E-02 | 4.52E-02 | -1.24E+00 | 1.52E-01 | 4.17E-01 | 1.15E+00 |
| LARS | 1.51E-14 | 7.88E-14 | -1.21E+00 | 3.04E-08 | 1.32E-07 | 1.03E+00 | 6.19E-04 | 2.63E-03 | -1.25E+00 | 1.01E-01 | 3.84E-01 | 1.05E+00 | 1.69E-10 | 6.02E-10 | -1.30E+00 | 2.33E-05 | 6.57E-05 | -1.02E+00 | 1.56E-05 | 5.73E-05 | -1.15E+00 | 1.09E-02 | 2.99E-02 | 1.02E+00 |
| SNX17 | 1.52E-14 | 7.90E-14 | -1.31E+00 | 6.93E-09 | 2.99E-08 | -1.03E+00 | 7.62E-07 | 3.24E-06 | -1.27E+00 | 3.20E-04 | 1.22E-03 | -1.04E+00 | 4.41E-11 | 1.57E-10 | -1.38E+00 | 6.88E-08 | 1.94E-07 | -1.11E+00 | 3.73E-03 | 1.37E-02 | -1.02E+00 | 2.69E-02 | 7.40E-02 | 1.02E+00 |
| ELOVL1 | 1.52E-14 | 7.92E-14 | -1.83E+00 | 1.27E-06 | 5.50E-06 | -1.11E+00 | 4.09E-11 | 1.74E-10 | -1.54E+00 | 5.52E-05 | 2.11E-04 | -1.02E+00 | 1.77E-15 | 6.30E-15 | -1.94E+00 | 9.80E-06 | 2.76E-05 | -1.05E+00 | 7.33E-16 | 2.69E-15 | -1.64E+00 | 6.65E-05 | 1.83E-04 | -1.10E+00 |
| TNFSF12 | 1.55E-14 | 8.06E-14 | -1.64E+00 | 4.61E-11 | 1.99E-10 | -1.27E+00 | 3.16E-06 | 1.34E-05 | -1.61E+00 | 1.54E-03 | 5.88E-03 | 1.03E+00 | 4.80E-11 | 1.71E-10 | -1.70E+00 | 3.91E-10 | 1.10E-09 | -1.55E+00 | 2.15E-03 | 7.89E-03 | -1.39E+00 | 2.78E-02 | 7.64E-02 | -1.12E+00 |
| ZC3H4 | 1.58E-14 | 8.23E-14 | -1.97E+00 | 3.81E-09 | 1.64E-08 | -1.21E+00 | 1.86E-12 | 7.88E-12 | -2.04E+00 | 1.30E-05 | 4.97E-05 | -1.19E+00 | 1.20E-10 | 4.29E-10 | -2.36E+00 | 1.60E-05 | 4.52E-05 | -1.38E+00 | 1.41E-18 | 5.19E-18 | -2.52E+00 | 1.57E-07 | 4.31E-07 | -1.33E+00 |
| DDX58 | 1.58E-14 | 8.24E-14 | -1.83E+00 | 4.16E-09 | 1.80E-08 | -1.21E+00 | 3.45E-12 | 1.47E-11 | -2.11E+00 | 3.44E-09 | 1.31E-08 | -1.52E+00 | 1.36E-13 | 4.84E-13 | -2.18E+00 | 3.75E-08 | 1.06E-07 | -1.36E+00 | 1.79E-09 | 6.58E-09 | -2.28E+00 | 8.87E-06 | 2.44E-05 | -1.67E+00 |
| RNMT | 1.58E-14 | 8.24E-14 | -1.29E+00 | 1.38E-27 | 5.95E-27 | -1.57E+00 | 3.74E-05 | 1.59E-04 | -1.35E+00 | 3.75E-11 | 1.43E-10 | -1.48E+00 | 1.84E-04 | 6.55E-04 | -1.02E+00 | 1.98E-10 | 5.59E-10 | -1.40E+00 | 3.16E-04 | 1.16E-03 | -1.21E+00 | 6.27E-05 | 1.72E-04 | -1.27E+00 |
| C5orf24 | 1.58E-14 | 8.24E-14 | -1.58E+00 | 4.54E-05 | 1.96E-04 | 1.05E+00 | 3.33E-11 | 1.41E-10 | -1.56E+00 | 3.94E-06 | 1.50E-05 | -1.09E+00 | 4.15E-05 | 1.48E-04 | -1.14E+00 | 1.93E-02 | 5.44E-02 | 1.05E+00 | 5.23E-09 | 1.92E-08 | -1.32E+00 | 1.20E-01 | 3.28E-01 | 1.10E+00 |
| KCNN4 | 1.58E-14 | 8.24E-14 | -1.82E+00 | 2.34E-05 | 1.01E-04 | -1.02E+00 | 8.42E-21 | 3.58E-20 | -2.03E+00 | 5.33E-12 | 2.04E-11 | -1.35E+00 | 5.76E-16 | 2.05E-15 | -1.73E+00 | 3.72E-06 | 1.05E-05 | -1.15E+00 | 1.09E-23 | 3.99E-23 | -2.16E+00 | 8.46E-08 | 2.32E-07 | -1.38E+00 |
| REV3L | 1.58E-14 | 8.24E-14 | -1.68E+00 | 3.50E-04 | 1.51E-03 | 1.09E+00 | 3.22E-13 | 1.37E-12 | -1.89E+00 | 1.01E-08 | 3.84E-08 | -1.34E+00 | 1.25E-11 | 4.46E-11 | -1.83E+00 | 3.13E-04 | 8.81E-04 | -1.06E+00 | 1.34E-11 | 4.90E-11 | -1.69E+00 | 6.58E-04 | 1.81E-03 | -1.20E+00 |
| ATXN2 | 1.67E-14 | 8.72E-14 | -1.49E+00 | 3.82E-04 | 1.65E-03 | 1.07E+00 | 9.91E-12 | 4.21E-11 | -1.60E+00 | 3.14E-04 | 1.20E-03 | 1.01E+00 | 3.50E-12 | 1.25E-11 | -1.82E+00 | 2.63E-04 | 7.42E-04 | -1.08E+00 | 6.24E-17 | 2.29E-16 | -1.93E+00 | 1.06E-08 | 2.91E-08 | -1.37E+00 |
| PPTC7 | 1.68E-14 | 8.76E-14 | -1.62E+00 | 1.06E-19 | 4.59E-19 | -1.78E+00 | 1.39E-06 | 5.91E-06 | -1.34E+00 | 1.37E-08 | 5.23E-08 | -1.29E+00 | 8.70E-07 | 3.10E-06 | -1.41E+00 | 3.91E-10 | 1.10E-09 | -1.95E+00 | 1.31E-07 | 4.80E-07 | -1.53E+00 | 4.76E-08 | 1.31E-07 | -1.51E+00 |
| CHTF8 | 1.70E-14 | 8.84E-14 | -1.80E+00 | 1.70E-06 | 7.33E-06 | -1.01E+00 | 1.90E-06 | 8.08E-06 | -1.39E+00 | 1.86E-04 | 7.12E-04 | -1.04E+00 | 2.62E-12 | 9.33E-12 | -1.93E+00 | 1.09E-05 | 3.06E-05 | -1.20E+00 | 3.80E-07 | 1.40E-06 | -1.20E+00 | 1.85E-02 | 5.07E-02 | 1.13E+00 |
| GHITM | 1.74E-14 | 9.07E-14 | -1.21E+00 | 1.45E-07 | 6.25E-07 | 1.07E+00 | 2.81E-04 | 1.19E-03 | -1.12E+00 | 1.52E-02 | 5.79E-02 | 1.04E+00 | 2.49E-07 | 8.88E-07 | -1.28E+00 | 3.82E-03 | 1.08E-02 | 1.01E+00 | 1.98E-04 | 7.28E-04 | -1.24E+00 | 1.82E-01 | 4.99E-01 | 1.04E+00 |
| ZNF385D | 1.75E-14 | 9.12E-14 | 2.07E+00 | 1.06E-07 | 4.58E-07 | 1.95E+00 | 4.14E-02 | 1.76E-01 | 1.35E+00 | 8.00E-03 | 3.06E-02 | 1.56E+00 | 9.19E-04 | 3.28E-03 | 1.81E+00 | 5.84E-02 | 1.65E-01 | 1.58E+00 | 6.49E-03 | 2.38E-02 | -1.13E+00 | 1.19E-02 | 3.27E-02 | -1.10E+00 |
| EGR2 | 1.82E-14 | 9.51E-14 | 1.23E+01 | 1.60E-01 | 6.90E-01 | 1.71E+00 | 6.52E-12 | 2.77E-11 | 8.96E+00 | 2.34E-01 | 8.93E-01 | 1.26E+00 | 2.20E-19 | 7.84E-19 | 2.28E+01 | 2.28E-01 | 6.44E-01 | 2.09E+00 | 1.55E-14 | 5.70E-14 | 1.26E+01 | 2.10E-01 | 5.78E-01 | 1.63E+00 |
| ZNF195 | 1.85E-14 | 9.62E-14 | -1.67E+00 | 5.21E-13 | 2.25E-12 | -1.41E+00 | 4.56E-16 | 1.94E-15 | -1.74E+00 | 1.43E-14 | 5.44E-14 | -1.48E+00 | 1.94E-07 | 6.92E-07 | -1.73E+00 | 6.33E-06 | 1.79E-05 | -1.47E+00 | 1.80E-12 | 6.61E-12 | -1.80E+00 | 1.61E-10 | 4.42E-10 | -1.69E+00 |
| TJP2 | 1.85E-14 | 9.66E-14 | -1.78E+00 | 1.18E-08 | 5.12E-08 | -1.13E+00 | 1.17E-08 | 4.98E-08 | -2.00E+00 | 8.19E-05 | 3.13E-04 | -1.30E+00 | 4.84E-08 | 1.73E-07 | -1.55E+00 | 6.30E-06 | 1.78E-05 | -1.23E+00 | 3.23E-02 | 1.19E-01 | 1.07E+00 | 3.17E-01 | 8.71E-01 | 1.34E+00 |
| PRKCI | 1.88E-14 | 9.82E-14 | -1.72E+00 | 4.44E-07 | 1.92E-06 | -1.07E+00 | 9.38E-09 | 3.99E-08 | -1.40E+00 | 2.10E-05 | 8.01E-05 | -1.03E+00 | 2.55E-08 | 9.10E-08 | -1.69E+00 | 3.32E-04 | 9.37E-04 | -1.12E+00 | 1.81E-07 | 6.65E-07 | -1.51E+00 | 2.51E-03 | 6.88E-03 | -1.15E+00 |
| LMNA | 1.89E-14 | 9.87E-14 | 2.58E+00 | 1.36E-01 | 5.86E-01 | 1.41E+00 | 3.30E-12 | 1.40E-11 | 2.79E+00 | 2.12E-01 | 8.11E-01 | 1.38E+00 | 1.09E-06 | 3.90E-06 | 2.17E+00 | 3.47E-01 | 9.77E-01 | 1.15E+00 | 1.90E-02 | 6.99E-02 | 1.65E+00 | 2.97E-03 | 8.16E-03 | -1.03E+00 |
| RAB11B | 1.97E-14 | 1.03E-13 | -1.39E+00 | 1.13E-11 | 4.89E-11 | -1.13E+00 | 3.02E-07 | 1.28E-06 | -1.34E+00 | 1.50E-05 | 5.72E-05 | -1.05E+00 | 9.28E-09 | 3.31E-08 | -1.39E+00 | 2.00E-05 | 5.63E-05 | -1.16E+00 | 1.10E-11 | 4.03E-11 | -1.55E+00 | 1.51E-05 | 4.16E-05 | -1.25E+00 |
| CALU | 2.00E-14 | 1.04E-13 | -1.66E+00 | 2.36E-09 | 1.02E-08 | -1.13E+00 | 3.81E-11 | 1.62E-10 | -1.53E+00 | 1.00E-05 | 3.82E-05 | -1.03E+00 | 4.03E-17 | 1.44E-16 | -1.85E+00 | 7.77E-10 | 2.19E-09 | -1.34E+00 | 1.14E-06 | 4.17E-06 | -1.37E+00 | 1.96E-03 | 5.38E-03 | -1.09E+00 |
| SND1 | 2.02E-14 | 1.05E-13 | -1.28E+00 | 1.07E-06 | 4.61E-06 | 1.00E+00 | 9.03E-06 | 3.84E-05 | -1.19E+00 | 5.50E-03 | 2.10E-02 | 1.04E+00 | 2.50E-09 | 8.93E-09 | -1.42E+00 | 7.87E-03 | 2.22E-02 | -1.03E+00 | 1.67E-05 | 6.12E-05 | -1.19E+00 | 8.27E-02 | 2.27E-01 | 1.09E+00 |
| WASHC5 | 2.04E-14 | 1.06E-13 | -1.86E+00 | 3.26E-10 | 1.41E-09 | -1.27E+00 | 4.24E-10 | 1.80E-09 | -1.64E+00 | 9.33E-07 | 3.56E-06 | -1.23E+00 | 1.63E-07 | 5.82E-07 | -1.56E+00 | 7.81E-03 | 2.20E-02 | 1.10E+00 | 1.61E-05 | 5.91E-05 | -1.26E+00 | 5.70E-03 | 1.57E-02 | -1.06E+00 |
| AFTPH | 2.04E-14 | 1.06E-13 | -1.39E+00 | 2.19E-07 | 9.45E-07 | -1.01E+00 | 2.25E-08 | 9.55E-08 | -1.42E+00 | 2.12E-05 | 8.11E-05 | -1.07E+00 | 2.75E-04 | 9.80E-04 | -1.19E+00 | 2.02E-02 | 5.69E-02 | 1.00E+00 | 1.22E-08 | 4.48E-08 | -1.16E+00 | 3.62E-04 | 9.95E-04 | -1.01E+00 |
| COX4I1P1 | 2.08E-14 | 1.09E-13 | -2.24E+00 | 1.31E-13 | 5.65E-13 | -2.01E+00 | 1.14E-10 | 4.86E-10 | -2.04E+00 | 2.82E-09 | 1.08E-08 | -1.66E+00 | 6.80E-10 | 2.42E-09 | -2.67E+00 | 1.21E-07 | 3.40E-07 | -2.03E+00 | 5.17E-18 | 1.90E-17 | -3.10E+00 | 1.43E-07 | 3.93E-07 | -1.45E+00 |
| PSENEN | 2.24E-14 | 1.17E-13 | -1.47E+00 | 2.76E-18 | 1.19E-17 | -1.55E+00 | 2.72E-09 | 1.16E-08 | -1.34E+00 | 1.46E-14 | 5.57E-14 | -1.50E+00 | 2.03E-09 | 7.25E-09 | -1.50E+00 | 2.88E-12 | 8.11E-12 | -1.61E+00 | 5.58E-04 | 2.05E-03 | -1.08E+00 | 1.13E-03 | 3.12E-03 | -1.06E+00 |
| LAMTOR1 | 2.25E-14 | 1.17E-13 | -1.26E+00 | 1.28E-04 | 5.51E-04 | 1.00E+00 | 4.18E-06 | 1.78E-05 | -1.33E+00 | 1.29E-02 | 4.92E-02 | -1.03E+00 | 3.54E-05 | 1.26E-04 | -1.26E+00 | 4.24E-02 | 1.20E-01 | -1.03E+00 | 1.14E-02 | 4.19E-02 | -1.03E+00 | 1.56E-01 | 4.28E-01 | -1.05E+00 |
| SLC25A36 | 2.26E-14 | 1.18E-13 | -1.27E+00 | 3.55E-23 | 1.53E-22 | -1.34E+00 | 1.35E-06 | 5.74E-06 | -1.31E+00 | 1.92E-09 | 7.35E-09 | -1.25E+00 | 2.50E-07 | 8.90E-07 | -1.20E+00 | 2.15E-10 | 6.06E-10 | -1.33E+00 | 2.11E-08 | 7.75E-08 | -1.39E+00 | 1.06E-05 | 2.91E-05 | -1.27E+00 |
| TERF2IP | 2.32E-14 | 1.21E-13 | -1.25E+00 | 9.29E-09 | 4.01E-08 | -1.01E+00 | 6.90E-07 | 2.93E-06 | -1.38E+00 | 6.98E-06 | 2.67E-05 | -1.15E+00 | 6.79E-11 | 2.42E-10 | -1.38E+00 | 1.23E-06 | 3.46E-06 | -1.14E+00 | 8.94E-06 | 3.28E-05 | -1.27E+00 | 1.34E-03 | 3.69E-03 | -1.19E+00 |
| G3BP1 | 2.36E-14 | 1.23E-13 | -1.27E+00 | 7.04E-07 | 3.04E-06 | 1.03E+00 | 1.33E-05 | 5.66E-05 | -1.22E+00 | 9.67E-02 | 3.69E-01 | 1.17E+00 | 3.43E-07 | 1.22E-06 | -1.14E+00 | 2.88E-02 | 8.12E-02 | 1.20E+00 | 1.01E-04 | 3.72E-04 | -1.23E+00 | 2.75E-01 | 7.57E-01 | 1.12E+00 |
| TRIM25 | 2.40E-14 | 1.25E-13 | -1.89E+00 | 3.55E-08 | 1.53E-07 | -1.25E+00 | 1.74E-13 | 7.41E-13 | -2.05E+00 | 7.90E-07 | 3.02E-06 | -1.26E+00 | 1.95E-05 | 6.95E-05 | -1.59E+00 | 3.60E-04 | 1.02E-03 | -1.41E+00 | 9.57E-13 | 3.51E-12 | -2.04E+00 | 1.46E-06 | 4.02E-06 | -1.36E+00 |
| BICD2 | 2.40E-14 | 1.25E-13 | -1.56E+00 | 4.26E-07 | 1.84E-06 | -1.07E+00 | 5.25E-05 | 2.23E-04 | -1.36E+00 | 2.68E-03 | 1.02E-02 | -1.03E+00 | 6.26E-07 | 2.23E-06 | -1.49E+00 | 1.54E-04 | 4.35E-04 | -1.25E+00 | 1.83E-09 | 6.72E-09 | -1.44E+00 | 1.06E-03 | 2.91E-03 | -1.12E+00 |
| LGALS9 | 2.42E-14 | 1.26E-13 | -1.35E+00 | 5.81E-10 | 2.51E-09 | -1.10E+00 | 2.90E-10 | 1.23E-09 | -1.65E+00 | 3.26E-09 | 1.25E-08 | -1.38E+00 | 3.92E-14 | 1.40E-13 | -1.81E+00 | 6.73E-13 | 1.90E-12 | -1.68E+00 | 8.64E-21 | 3.17E-20 | -1.94E+00 | 9.24E-15 | 2.54E-14 | -1.71E+00 |
| CPSF6 | 2.46E-14 | 1.28E-13 | -1.25E+00 | 8.56E-11 | 3.70E-10 | -1.04E+00 | 6.38E-04 | 2.71E-03 | -1.19E+00 | 3.86E-04 | 1.47E-03 | -1.05E+00 | 5.40E-08 | 1.93E-07 | -1.19E+00 | 1.92E-06 | 5.41E-06 | -1.10E+00 | 3.19E-04 | 1.17E-03 | -1.11E+00 | 2.16E-01 | 5.93E-01 | 1.13E+00 |
| TPI1 | 2.46E-14 | 1.28E-13 | -1.23E+00 | 5.51E-05 | 2.38E-04 | -1.02E+00 | 9.91E-02 | 4.21E-01 | -1.05E+00 | 1.61E-01 | 6.15E-01 | 1.03E+00 | 4.72E-05 | 1.68E-04 | -1.21E+00 | 2.70E-01 | 7.62E-01 | 1.07E+00 | 1.12E-01 | 4.10E-01 | -1.05E+00 | 3.83E-02 | 1.05E-01 | 1.18E+00 |
| PBX2 | 2.51E-14 | 1.31E-13 | -1.77E+00 | 8.03E-16 | 3.47E-15 | -1.80E+00 | 4.74E-09 | 2.02E-08 | -1.72E+00 | 9.85E-10 | 3.76E-09 | -1.61E+00 | 4.22E-08 | 1.51E-07 | -1.87E+00 | 7.34E-06 | 2.07E-05 | -1.34E+00 | 4.64E-12 | 1.70E-11 | -1.69E+00 | 2.25E-07 | 6.19E-07 | -1.24E+00 |
| CWC22 | 2.52E-14 | 1.31E-13 | -1.80E+00 | 2.39E-10 | 1.03E-09 | -1.30E+00 | 3.71E-17 | 1.58E-16 | -1.86E+00 | 3.01E-13 | 1.15E-12 | -1.44E+00 | 7.62E-07 | 2.72E-06 | -1.71E+00 | 1.57E-04 | 4.42E-04 | -1.22E+00 | 2.88E-08 | 1.06E-07 | -1.59E+00 | 1.96E-04 | 5.40E-04 | -1.19E+00 |
| KDM1A | 2.55E-14 | 1.33E-13 | -1.37E+00 | 1.68E-15 | 7.25E-15 | -1.23E+00 | 1.81E-06 | 7.69E-06 | -1.31E+00 | 4.40E-05 | 1.68E-04 | -1.12E+00 | 1.31E-08 | 4.67E-08 | -1.37E+00 | 3.89E-06 | 1.10E-05 | -1.14E+00 | 6.92E-06 | 2.54E-05 | -1.32E+00 | 1.09E-02 | 2.99E-02 | -1.13E+00 |
| BBX | 2.57E-14 | 1.34E-13 | -1.34E+00 | 2.57E-09 | 1.11E-08 | -1.07E+00 | 3.34E-08 | 1.42E-07 | -1.32E+00 | 3.00E-07 | 1.15E-06 | -1.11E+00 | 3.37E-07 | 1.20E-06 | -1.37E+00 | 7.60E-06 | 2.14E-05 | -1.28E+00 | 2.51E-08 | 9.21E-08 | -1.32E+00 | 4.76E-03 | 1.31E-02 | -1.08E+00 |
| GXYLT1 | 2.57E-14 | 1.34E-13 | -1.93E+00 | 3.65E-08 | 1.58E-07 | -1.20E+00 | 3.18E-11 | 1.35E-10 | -1.86E+00 | 6.25E-09 | 2.39E-08 | -1.39E+00 | 3.48E-08 | 1.24E-07 | -2.12E+00 | 7.67E-05 | 2.16E-04 | -1.41E+00 | 1.10E-10 | 4.04E-10 | -1.58E+00 | 3.09E-09 | 8.50E-09 | -1.42E+00 |
| PRPF38A | 2.59E-14 | 1.35E-13 | -1.33E+00 | 2.46E-12 | 1.06E-11 | -1.16E+00 | 4.72E-05 | 2.00E-04 | -1.14E+00 | 7.85E-06 | 3.00E-05 | -1.09E+00 | 3.62E-06 | 1.29E-05 | -1.07E+00 | 2.22E-04 | 6.26E-04 | 1.03E+00 | 1.95E-05 | 7.17E-05 | -1.13E+00 | 7.28E-05 | 2.00E-04 | -1.09E+00 |
| ATXN10 | 2.59E-14 | 1.35E-13 | -1.23E+00 | 6.92E-08 | 2.99E-07 | 1.08E+00 | 9.73E-05 | 4.13E-04 | -1.15E+00 | 1.63E-03 | 6.24E-03 | 1.01E+00 | 9.91E-07 | 3.53E-06 | -1.36E+00 | 4.07E-02 | 1.15E-01 | -1.07E+00 | 5.60E-04 | 2.05E-03 | -1.08E+00 | 2.20E-01 | 6.04E-01 | 1.14E+00 |
| C15orf61 | 2.72E-14 | 1.42E-13 | -1.54E+00 | 8.07E-13 | 3.49E-12 | -1.34E+00 | 2.57E-10 | 1.09E-09 | -1.48E+00 | 2.39E-13 | 9.15E-13 | -1.43E+00 | 7.28E-13 | 2.60E-12 | -1.82E+00 | 2.75E-13 | 7.75E-13 | -1.77E+00 | 1.76E-12 | 6.45E-12 | -1.57E+00 | 1.96E-08 | 5.38E-08 | -1.32E+00 |
| APPBP2 | 2.74E-14 | 1.43E-13 | -1.70E+00 | 7.15E-08 | 3.09E-07 | -1.09E+00 | 1.59E-12 | 6.74E-12 | -1.99E+00 | 1.85E-07 | 7.08E-07 | -1.32E+00 | 1.49E-07 | 5.32E-07 | -1.55E+00 | 7.85E-04 | 2.21E-03 | -1.14E+00 | 5.16E-14 | 1.89E-13 | -1.76E+00 | 1.66E-04 | 4.56E-04 | -1.15E+00 |
| PARP12 | 2.76E-14 | 1.44E-13 | -1.89E+00 | 2.72E-09 | 1.17E-08 | -1.30E+00 | 9.69E-13 | 4.12E-12 | -1.90E+00 | 1.14E-06 | 4.36E-06 | -1.18E+00 | 2.14E-10 | 7.64E-10 | -1.75E+00 | 4.98E-07 | 1.40E-06 | -1.29E+00 | 1.21E-25 | 4.43E-25 | -2.51E+00 | 4.72E-21 | 1.30E-20 | -2.10E+00 |
| DDX50 | 2.76E-14 | 1.44E-13 | -1.37E+00 | 4.86E-11 | 2.10E-10 | -1.11E+00 | 2.47E-07 | 1.05E-06 | -1.37E+00 | 4.82E-05 | 1.84E-04 | -1.14E+00 | 9.67E-07 | 3.45E-06 | -1.22E+00 | 4.99E-04 | 1.41E-03 | -1.00E+00 | 6.96E-04 | 2.55E-03 | -1.18E+00 | 1.54E-02 | 4.22E-02 | -1.04E+00 |
| MOSPD2 | 2.80E-14 | 1.46E-13 | -3.36E+00 | 1.97E-07 | 8.51E-07 | -1.58E+00 | 4.40E-04 | 1.87E-03 | -1.37E+00 | 2.02E-03 | 7.73E-03 | -1.11E+00 | 7.08E-03 | 2.52E-02 | -1.15E+00 | 5.38E-03 | 1.52E-02 | -1.23E+00 | 1.26E-06 | 4.62E-06 | -1.89E+00 | 2.35E-02 | 6.44E-02 | -1.06E+00 |
| MCOLN1 | 2.81E-14 | 1.47E-13 | -2.05E+00 | 1.37E-05 | 5.94E-05 | -1.04E+00 | 5.48E-17 | 2.33E-16 | -2.31E+00 | 7.08E-09 | 2.70E-08 | -1.33E+00 | 7.86E-06 | 2.80E-05 | -1.75E+00 | 3.01E-03 | 8.49E-03 | -1.19E+00 | 4.43E-03 | 1.63E-02 | -1.30E+00 | 1.78E-01 | 4.88E-01 | 1.17E+00 |
| MTDH | 2.81E-14 | 1.47E-13 | -1.27E+00 | 4.53E-03 | 1.96E-02 | 1.05E+00 | 8.81E-03 | 3.74E-02 | -1.09E+00 | 2.43E-01 | 9.27E-01 | 1.10E+00 | 5.62E-08 | 2.00E-07 | -1.33E+00 | 1.77E-02 | 4.98E-02 | -1.03E+00 | 3.05E-03 | 1.12E-02 | -1.18E+00 | 2.32E-01 | 6.38E-01 | -1.03E+00 |
| ARSD | 2.84E-14 | 1.48E-13 | -1.56E+00 | 6.05E-06 | 2.61E-05 | 1.03E+00 | 1.68E-11 | 7.12E-11 | -1.66E+00 | 1.74E-05 | 6.66E-05 | -1.03E+00 | 1.32E-15 | 4.69E-15 | -1.84E+00 | 2.65E-11 | 7.47E-11 | -1.52E+00 | 9.60E-04 | 3.52E-03 | -1.19E+00 | 4.06E-02 | 1.12E-01 | 1.06E+00 |
| TRAPPC3 | 2.85E-14 | 1.48E-13 | -1.36E+00 | 1.01E-06 | 4.35E-06 | 1.08E+00 | 3.51E-08 | 1.49E-07 | -1.35E+00 | 6.85E-05 | 2.62E-04 | -1.02E+00 | 8.42E-11 | 3.00E-10 | -1.45E+00 | 6.31E-04 | 1.78E-03 | 1.02E+00 | 1.81E-07 | 6.65E-07 | -1.28E+00 | 2.84E-02 | 7.81E-02 | 1.01E+00 |
| SLF2 | 2.86E-14 | 1.49E-13 | -2.10E+00 | 1.14E-03 | 4.92E-03 | 1.14E+00 | 5.91E-08 | 2.51E-07 | -1.66E+00 | 4.48E-04 | 1.71E-03 | -1.00E+00 | 4.67E-06 | 1.67E-05 | -1.92E+00 | 6.04E-03 | 1.70E-02 | -1.23E+00 | 9.84E-08 | 3.61E-07 | -1.40E+00 | 5.92E-03 | 1.63E-02 | 1.09E+00 |
| UBE2D3P1 | 2.88E-14 | 1.50E-13 | -1.56E+00 | 3.40E-18 | 1.47E-17 | -1.63E+00 | 1.66E-07 | 7.07E-07 | -1.35E+00 | 4.44E-13 | 1.70E-12 | -1.58E+00 | 1.05E-06 | 3.74E-06 | -1.29E+00 | 2.23E-10 | 6.28E-10 | -1.64E+00 | 2.80E-12 | 1.03E-11 | -1.49E+00 | 5.45E-18 | 1.50E-17 | -1.86E+00 |
| KIT | 2.89E-14 | 1.50E-13 | -1.67E+00 | 5.27E-07 | 2.28E-06 | -1.04E+00 | 6.21E-05 | 2.64E-04 | -1.22E+00 | 2.30E-02 | 8.77E-02 | 1.22E+00 | 3.32E-02 | 1.18E-01 | 1.79E+00 | 3.37E-01 | 9.49E-01 | 1.38E+00 | 2.15E-02 | 7.90E-02 | 1.63E+00 | 2.65E-02 | 7.29E-02 | 1.65E+00 |
| PRKX | 2.94E-14 | 1.53E-13 | -1.97E+00 | 1.64E-07 | 7.09E-07 | -1.14E+00 | 2.37E-08 | 1.00E-07 | -1.58E+00 | 1.16E-03 | 4.43E-03 | 1.05E+00 | 7.13E-05 | 2.54E-04 | -1.32E+00 | 6.49E-03 | 1.83E-02 | 1.05E+00 | 4.33E-11 | 1.59E-10 | -1.75E+00 | 2.46E-04 | 6.76E-04 | -1.16E+00 |
| SF3B1 | 3.03E-14 | 1.58E-13 | -1.34E+00 | 1.22E-22 | 5.29E-22 | -1.35E+00 | 2.88E-04 | 1.22E-03 | -1.22E+00 | 6.42E-06 | 2.45E-05 | -1.21E+00 | 1.21E-04 | 4.32E-04 | -1.16E+00 | 2.53E-05 | 7.13E-05 | -1.19E+00 | 7.67E-05 | 2.82E-04 | -1.22E+00 | 8.25E-03 | 2.27E-02 | -1.17E+00 |
| RPAP2 | 3.08E-14 | 1.61E-13 | -1.45E+00 | 2.88E-04 | 1.25E-03 | 1.14E+00 | 3.91E-09 | 1.66E-08 | -1.68E+00 | 6.53E-04 | 2.49E-03 | -1.05E+00 | 6.37E-05 | 2.27E-04 | -1.24E+00 | 2.03E-01 | 5.72E-01 | 1.22E+00 | 7.89E-06 | 2.89E-05 | -1.24E+00 | 9.70E-03 | 2.67E-02 | 1.02E+00 |
| CHRNA7 | 3.14E-14 | 1.64E-13 | -1.95E+00 | 7.08E-07 | 3.06E-06 | -1.15E+00 | 7.48E-05 | 3.18E-04 | -1.27E+00 | 2.71E-03 | 1.03E-02 | 1.10E+00 | 7.69E-02 | 2.74E-01 | -1.21E+00 | 9.02E-02 | 2.54E-01 | -1.28E+00 | 1.57E-02 | 5.76E-02 | -1.61E+00 | 6.80E-02 | 1.87E-01 | -1.35E+00 |
| ZCCHC17 | 3.22E-14 | 1.68E-13 | -1.57E+00 | 3.46E-06 | 1.49E-05 | 1.03E+00 | 1.38E-06 | 5.86E-06 | -1.23E+00 | 2.38E-03 | 9.10E-03 | 1.07E+00 | 3.35E-07 | 1.20E-06 | -1.43E+00 | 1.89E-03 | 5.32E-03 | 1.01E+00 | 4.78E-06 | 1.76E-05 | -1.14E+00 | 3.48E-03 | 9.55E-03 | -1.01E+00 |
| UBN2 | 3.22E-14 | 1.68E-13 | -1.55E+00 | 6.53E-06 | 2.82E-05 | 1.08E+00 | 1.25E-08 | 5.30E-08 | -1.56E+00 | 5.97E-04 | 2.28E-03 | -1.03E+00 | 2.17E-06 | 7.75E-06 | -1.38E+00 | 3.57E-04 | 1.01E-03 | -1.12E+00 | 1.20E-09 | 4.40E-09 | -1.45E+00 | 3.74E-05 | 1.03E-04 | -1.15E+00 |
| PCED1A | 3.26E-14 | 1.70E-13 | -1.72E+00 | 1.65E-12 | 7.14E-12 | -1.35E+00 | 3.62E-04 | 1.54E-03 | -1.25E+00 | 1.25E-02 | 4.79E-02 | 1.04E+00 | 2.82E-05 | 1.00E-04 | -1.26E+00 | 2.75E-04 | 7.76E-04 | -1.13E+00 | 4.90E-02 | 1.80E-01 | 1.04E+00 | 2.17E-01 | 5.95E-01 | 1.38E+00 |
| AC120049.1 | 3.29E-14 | 1.72E-13 | 3.37E+00 | 1.79E-02 | 7.74E-02 | 2.04E+00 | 1.12E-03 | 4.77E-03 | 2.41E+00 | 7.62E-02 | 2.91E-01 | 1.83E+00 | 6.50E-02 | 2.32E-01 | 3.36E+00 | 2.92E-01 | 8.22E-01 | 1.61E+00 | 1.32E-01 | 4.83E-01 | 1.89E+00 | 3.62E-01 | 9.95E-01 | 1.08E+00 |
| AAED1 | 3.36E-14 | 1.75E-13 | -1.34E+00 | 1.10E-09 | 4.74E-09 | -1.09E+00 | 7.00E-08 | 2.97E-07 | -1.52E+00 | 9.35E-07 | 3.57E-06 | -1.28E+00 | 1.29E-03 | 4.61E-03 | -1.00E+00 | 8.20E-05 | 2.31E-04 | -1.05E+00 | 8.87E-05 | 3.26E-04 | -1.17E+00 | 1.56E-05 | 4.29E-05 | -1.29E+00 |
| RTN3 | 3.51E-14 | 1.83E-13 | -1.29E+00 | 1.37E-13 | 5.91E-13 | -1.16E+00 | 5.75E-04 | 2.44E-03 | -1.15E+00 | 1.20E-04 | 4.60E-04 | -1.06E+00 | 2.54E-08 | 9.05E-08 | -1.30E+00 | 5.93E-03 | 1.67E-02 | -1.06E+00 | 5.32E-04 | 1.95E-03 | -1.18E+00 | 1.68E-02 | 4.60E-02 | -1.12E+00 |
| PNPT1 | 3.56E-14 | 1.86E-13 | -1.67E+00 | 1.60E-09 | 6.89E-09 | -1.18E+00 | 1.34E-07 | 5.69E-07 | -1.45E+00 | 1.24E-04 | 4.74E-04 | -1.12E+00 | 4.52E-07 | 1.61E-06 | -1.78E+00 | 1.42E-02 | 4.01E-02 | -1.07E+00 | 2.48E-13 | 9.11E-13 | -1.78E+00 | 3.20E-04 | 8.80E-04 | -1.25E+00 |
| WDR48 | 3.61E-14 | 1.88E-13 | -1.44E+00 | 2.31E-12 | 9.99E-12 | -1.22E+00 | 4.56E-12 | 1.94E-11 | -1.74E+00 | 1.56E-09 | 5.96E-09 | -1.31E+00 | 3.75E-08 | 1.34E-07 | -1.49E+00 | 5.85E-05 | 1.65E-04 | -1.16E+00 | 3.86E-10 | 1.42E-09 | -1.52E+00 | 2.95E-09 | 8.09E-09 | -1.49E+00 |
| EIF5A | 3.71E-14 | 1.93E-13 | -1.27E+00 | 9.28E-15 | 4.01E-14 | -1.11E+00 | 4.01E-02 | 1.70E-01 | -1.08E+00 | 1.61E-01 | 6.17E-01 | 1.08E+00 | 4.18E-06 | 1.49E-05 | -1.30E+00 | 2.51E-01 | 7.06E-01 | 1.14E+00 | 1.51E-02 | 5.54E-02 | -1.14E+00 | 1.83E-01 | 5.04E-01 | 1.16E+00 |
| PRR13 | 3.73E-14 | 1.95E-13 | -1.25E+00 | 3.54E-11 | 1.53E-10 | -1.08E+00 | 1.41E-04 | 5.98E-04 | -1.23E+00 | 9.42E-04 | 3.60E-03 | -1.10E+00 | 2.81E-07 | 1.00E-06 | -1.28E+00 | 1.31E-05 | 3.69E-05 | -1.12E+00 | 1.31E-03 | 4.80E-03 | -1.11E+00 | 6.53E-02 | 1.79E-01 | 1.03E+00 |
| LIG4 | 3.77E-14 | 1.96E-13 | -2.72E+00 | 3.57E-05 | 1.54E-04 | -1.06E+00 | 5.89E-19 | 2.50E-18 | -2.88E+00 | 5.15E-13 | 1.97E-12 | -1.81E+00 | 3.03E-07 | 1.08E-06 | -2.56E+00 | 1.23E-02 | 3.46E-02 | -1.00E+00 | 1.12E-09 | 4.11E-09 | -2.16E+00 | 4.77E-05 | 1.31E-04 | -1.38E+00 |
| HNRNPR | 3.79E-14 | 1.98E-13 | -1.28E+00 | 1.60E-02 | 6.93E-02 | 1.12E+00 | 7.81E-03 | 3.32E-02 | -1.06E+00 | 2.51E-01 | 9.58E-01 | 1.17E+00 | 8.68E-07 | 3.10E-06 | -1.22E+00 | 3.33E-01 | 9.39E-01 | 1.20E+00 | 2.67E-02 | 9.80E-02 | -1.07E+00 | 1.44E-01 | 3.95E-01 | 1.19E+00 |
| APIP | 3.80E-14 | 1.98E-13 | -1.40E+00 | 3.89E-11 | 1.68E-10 | -1.10E+00 | 9.45E-06 | 4.01E-05 | -1.25E+00 | 6.45E-07 | 2.46E-06 | -1.16E+00 | 2.02E-07 | 7.21E-07 | -1.60E+00 | 1.14E-02 | 3.21E-02 | -1.07E+00 | 4.65E-09 | 1.71E-08 | -1.42E+00 | 1.96E-03 | 5.38E-03 | -1.08E+00 |
| DAD1 | 3.84E-14 | 2.00E-13 | -1.23E+00 | 1.83E-12 | 7.89E-12 | -1.02E+00 | 1.93E-02 | 8.22E-02 | -1.09E+00 | 9.87E-02 | 3.77E-01 | 1.08E+00 | 1.22E-02 | 4.34E-02 | 1.02E+00 | 3.44E-01 | 9.69E-01 | 1.15E+00 | 9.62E-03 | 3.53E-02 | 1.40E+00 | 1.33E-02 | 3.66E-02 | 1.31E+00 |
| TFB2M | 3.87E-14 | 2.02E-13 | -1.63E+00 | 8.56E-14 | 3.70E-13 | -1.42E+00 | 4.48E-02 | 1.91E-01 | -1.01E+00 | 8.65E-02 | 3.30E-01 | 1.18E+00 | 3.24E-03 | 1.15E-02 | 1.06E+00 | 9.59E-03 | 2.70E-02 | 1.12E+00 | 3.80E-03 | 1.39E-02 | 1.07E+00 | 9.20E-02 | 2.53E-01 | 1.19E+00 |
| GRWD1 | 3.88E-14 | 2.02E-13 | -2.18E+00 | 8.65E-06 | 3.74E-05 | -1.09E+00 | 6.56E-14 | 2.79E-13 | -1.82E+00 | 1.64E-05 | 6.28E-05 | -1.09E+00 | 6.33E-11 | 2.26E-10 | -2.37E+00 | 5.29E-03 | 1.49E-02 | 1.01E+00 | 3.74E-22 | 1.37E-21 | -1.84E+00 | 1.67E-05 | 4.59E-05 | -1.14E+00 |
| BCCIP | 3.89E-14 | 2.03E-13 | -1.57E+00 | 2.04E-05 | 8.80E-05 | -1.01E+00 | 4.88E-05 | 2.07E-04 | -1.19E+00 | 4.99E-03 | 1.91E-02 | 1.09E+00 | 2.50E-02 | 8.90E-02 | 1.11E+00 | 1.84E-01 | 5.17E-01 | 1.30E+00 | 1.68E-06 | 6.16E-06 | -1.22E+00 | 4.23E-02 | 1.16E-01 | 1.11E+00 |
| ZNRD1 | 4.00E-14 | 2.09E-13 | -1.29E+00 | 3.69E-11 | 1.60E-10 | -1.11E+00 | 3.81E-06 | 1.62E-05 | -1.35E+00 | 2.13E-04 | 8.14E-04 | -1.08E+00 | 5.47E-10 | 1.95E-09 | -1.41E+00 | 7.69E-05 | 2.17E-04 | -1.10E+00 | 2.31E-09 | 8.49E-09 | -1.40E+00 | 6.75E-03 | 1.86E-02 | -1.06E+00 |
| CPXM1 | 4.05E-14 | 2.11E-13 | -1.41E+00 | 1.60E-14 | 6.90E-14 | -1.22E+00 | 5.22E-05 | 2.22E-04 | -1.23E+00 | 4.71E-06 | 1.80E-05 | -1.13E+00 | 2.43E-11 | 8.67E-11 | -1.60E+00 | 3.51E-09 | 9.90E-09 | -1.34E+00 | 3.46E-07 | 1.27E-06 | -1.41E+00 | 6.74E-04 | 1.85E-03 | -1.12E+00 |
| TRMT1L | 4.12E-14 | 2.15E-13 | -2.19E+00 | 2.49E-09 | 1.08E-08 | -1.38E+00 | 2.31E-09 | 9.83E-09 | -1.75E+00 | 1.66E-05 | 6.33E-05 | -1.18E+00 | 3.96E-09 | 1.41E-08 | -2.30E+00 | 1.24E-05 | 3.49E-05 | -1.50E+00 | 2.17E-20 | 7.97E-20 | -1.96E+00 | 9.94E-11 | 2.73E-10 | -1.49E+00 |
| GART | 4.15E-14 | 2.16E-13 | -1.53E+00 | 7.47E-07 | 3.23E-06 | 1.04E+00 | 6.30E-06 | 2.68E-05 | -1.30E+00 | 3.74E-02 | 1.43E-01 | 1.11E+00 | 1.17E-13 | 4.17E-13 | -2.16E+00 | 1.56E-03 | 4.39E-03 | -1.10E+00 | 1.08E-14 | 3.96E-14 | -1.55E+00 | 7.50E-02 | 2.06E-01 | 1.05E+00 |
| ACOT11 | 4.19E-14 | 2.19E-13 | -1.68E+00 | 3.54E-08 | 1.53E-07 | -1.17E+00 | 6.41E-03 | 2.72E-02 | -1.18E+00 | 1.22E-01 | 4.65E-01 | 1.21E+00 | 5.63E-07 | 2.01E-06 | -2.55E+00 | 9.71E-06 | 2.74E-05 | -2.00E+00 | 7.91E-02 | 2.90E-01 | -1.30E+00 | 3.58E-01 | 9.83E-01 | 1.17E+00 |
| EDEM3 | 4.29E-14 | 2.24E-13 | -1.99E+00 | 7.16E-09 | 3.10E-08 | -1.30E+00 | 1.74E-14 | 7.37E-14 | -1.90E+00 | 2.32E-05 | 8.86E-05 | -1.11E+00 | 7.61E-11 | 2.71E-10 | -2.34E+00 | 9.22E-06 | 2.60E-05 | -1.25E+00 | 7.24E-09 | 2.66E-08 | -1.69E+00 | 3.16E-03 | 8.69E-03 | -1.14E+00 |
| ARF1 | 4.30E-14 | 2.24E-13 | -1.21E+00 | 1.00E-07 | 4.34E-07 | 1.00E+00 | 3.69E-03 | 1.57E-02 | -1.14E+00 | 1.47E-01 | 5.63E-01 | 1.07E+00 | 1.12E-09 | 3.99E-09 | -1.30E+00 | 2.08E-02 | 5.88E-02 | -1.05E+00 | 1.40E-02 | 5.16E-02 | -1.14E+00 | 3.29E-01 | 9.04E-01 | 1.03E+00 |
| LTBR | 4.37E-14 | 2.28E-13 | -1.90E+00 | 2.22E-06 | 9.58E-06 | -1.14E+00 | 6.53E-05 | 2.78E-04 | -1.23E+00 | 9.05E-04 | 3.46E-03 | -1.00E+00 | 1.44E-04 | 5.13E-04 | -1.27E+00 | 2.46E-02 | 6.94E-02 | 1.02E+00 | 6.08E-04 | 2.23E-03 | -1.17E+00 | 7.37E-02 | 2.02E-01 | 1.05E+00 |
| PTGR1 | 4.45E-14 | 2.32E-13 | -2.42E+00 | 9.94E-07 | 4.30E-06 | -1.25E+00 | 1.87E-07 | 7.94E-07 | -1.63E+00 | 4.43E-06 | 1.69E-05 | -1.21E+00 | 3.09E-02 | 1.10E-01 | -1.41E+00 | 2.02E-01 | 5.68E-01 | 1.07E+00 | 7.41E-03 | 2.72E-02 | 1.06E+00 | 4.50E-02 | 1.24E-01 | 1.06E+00 |
| DHX58 | 4.54E-14 | 2.37E-13 | -2.02E+00 | 2.75E-13 | 1.19E-12 | -1.75E+00 | 1.77E-16 | 7.53E-16 | -2.31E+00 | 2.14E-13 | 8.16E-13 | -1.71E+00 | 4.31E-08 | 1.54E-07 | -2.00E+00 | 1.26E-08 | 3.56E-08 | -2.00E+00 | 2.26E-09 | 8.30E-09 | -1.88E+00 | 3.05E-06 | 8.39E-06 | -1.52E+00 |
| CP | 4.60E-14 | 2.40E-13 | -2.76E+00 | 5.11E-07 | 2.21E-06 | -1.36E+00 | 5.59E-20 | 2.38E-19 | -2.49E+00 | 2.03E-05 | 7.74E-05 | -1.25E+00 | 2.71E-01 | 9.66E-01 | 1.36E+00 | 2.39E-01 | 6.73E-01 | -1.34E+00 | 2.71E-01 | 9.94E-01 | 1.05E+00 | 3.53E-01 | 9.68E-01 | 1.08E+00 |
| CARD8-AS1 | 4.64E-14 | 2.42E-13 | -2.28E+00 | 2.50E-10 | 1.08E-09 | -1.54E+00 | 8.11E-07 | 3.45E-06 | -1.66E+00 | 1.01E-04 | 3.86E-04 | -1.15E+00 | 2.49E-04 | 8.87E-04 | -1.34E+00 | 1.56E-04 | 4.41E-04 | -1.43E+00 | 2.58E-07 | 9.48E-07 | -1.45E+00 | 3.99E-07 | 1.09E-06 | -1.62E+00 |
| CDCA7L | 4.65E-14 | 2.42E-13 | -1.68E+00 | 6.40E-12 | 2.76E-11 | -1.36E+00 | 2.43E-08 | 1.03E-07 | -1.34E+00 | 3.35E-10 | 1.28E-09 | -1.28E+00 | 2.97E-03 | 1.06E-02 | -1.61E+00 | 1.10E-03 | 3.09E-03 | -1.83E+00 | 1.21E-05 | 4.45E-05 | -1.69E+00 | 5.46E-02 | 1.50E-01 | -1.02E+00 |
| IER3IP1 | 4.70E-14 | 2.45E-13 | -1.59E+00 | 1.06E-06 | 4.56E-06 | -1.07E+00 | 3.37E-06 | 1.43E-05 | -1.23E+00 | 9.20E-06 | 3.52E-05 | -1.03E+00 | 1.34E-13 | 4.79E-13 | -1.82E+00 | 1.14E-05 | 3.20E-05 | -1.07E+00 | 1.04E-11 | 3.83E-11 | -1.42E+00 | 1.52E-04 | 4.18E-04 | -1.01E+00 |
| HERC2P2 | 4.71E-14 | 2.45E-13 | -1.74E+00 | 5.52E-07 | 2.38E-06 | -1.04E+00 | 1.36E-06 | 5.77E-06 | -1.48E+00 | 1.80E-05 | 6.87E-05 | -1.16E+00 | 2.28E-06 | 8.13E-06 | -1.48E+00 | 2.51E-01 | 7.09E-01 | 1.59E+00 | 6.09E-02 | 2.23E-01 | 1.06E+00 | 1.91E-03 | 5.25E-03 | -1.13E+00 |
| TMEM9B | 4.71E-14 | 2.46E-13 | -2.17E+00 | 2.57E-10 | 1.11E-09 | -1.49E+00 | 3.42E-10 | 1.45E-09 | -1.98E+00 | 2.02E-07 | 7.70E-07 | -1.48E+00 | 1.50E-06 | 5.33E-06 | -1.29E+00 | 1.36E-03 | 3.82E-03 | -1.02E+00 | 1.14E-07 | 4.18E-07 | -1.24E+00 | 2.26E-04 | 6.21E-04 | -1.09E+00 |
| NAALADL1 | 4.81E-14 | 2.51E-13 | -1.48E+00 | 5.57E-11 | 2.41E-10 | -1.23E+00 | 5.93E-11 | 2.52E-10 | -1.54E+00 | 5.00E-12 | 1.91E-11 | -1.36E+00 | 2.31E-06 | 8.22E-06 | -1.53E+00 | 7.83E-05 | 2.21E-04 | -1.19E+00 | 1.18E-01 | 4.33E-01 | 1.58E+00 | 1.37E-01 | 3.78E-01 | -1.07E+00 |
| DLD | 4.85E-14 | 2.53E-13 | -1.42E+00 | 3.93E-10 | 1.70E-09 | -1.11E+00 | 8.91E-09 | 3.79E-08 | -1.37E+00 | 5.20E-06 | 1.99E-05 | -1.13E+00 | 2.56E-11 | 9.12E-11 | -1.49E+00 | 1.58E-08 | 4.45E-08 | -1.28E+00 | 1.02E-10 | 3.75E-10 | -1.53E+00 | 2.51E-04 | 6.89E-04 | -1.14E+00 |
| HMGB1P26 | 4.88E-14 | 2.54E-13 | -1.75E+00 | 4.25E-21 | 1.84E-20 | -2.21E+00 | 1.10E-12 | 4.67E-12 | -1.62E+00 | 9.16E-26 | 3.50E-25 | -2.19E+00 | 4.42E-13 | 1.58E-12 | -2.09E+00 | 9.41E-14 | 2.65E-13 | -2.07E+00 | 4.05E-21 | 1.49E-20 | -1.87E+00 | 2.07E-22 | 5.69E-22 | -1.91E+00 |
| PMAIP1 | 4.88E-14 | 2.55E-13 | -1.22E+00 | 1.33E-50 | 5.74E-50 | -2.12E+00 | 5.37E-04 | 2.28E-03 | -1.17E+00 | 8.46E-18 | 3.23E-17 | -1.69E+00 | 1.06E-05 | 3.79E-05 | 1.07E+00 | 1.51E-16 | 4.26E-16 | -2.09E+00 | 2.91E-12 | 1.07E-11 | -1.58E+00 | 1.11E-19 | 3.06E-19 | -2.12E+00 |
| STAU1 | 4.93E-14 | 2.57E-13 | -1.43E+00 | 4.04E-08 | 1.75E-07 | -1.06E+00 | 1.08E-05 | 4.59E-05 | -1.20E+00 | 2.12E-04 | 8.10E-04 | -1.04E+00 | 7.73E-11 | 2.76E-10 | -1.48E+00 | 1.54E-05 | 4.34E-05 | -1.15E+00 | 2.73E-04 | 1.00E-03 | -1.14E+00 | 2.19E-04 | 6.02E-04 | -1.13E+00 |
| SPINT2 | 5.05E-14 | 2.63E-13 | -1.23E+00 | 1.36E-04 | 5.86E-04 | 1.00E+00 | 3.90E-04 | 1.66E-03 | -1.13E+00 | 1.81E-02 | 6.90E-02 | 1.01E+00 | 1.90E-04 | 6.77E-04 | -1.08E+00 | 1.42E-01 | 4.02E-01 | 1.11E+00 | 2.15E-01 | 7.90E-01 | 1.16E+00 | 3.17E-01 | 8.70E-01 | 1.12E+00 |
| ZNF605 | 5.07E-14 | 2.64E-13 | -2.19E+00 | 2.41E-07 | 1.04E-06 | -1.34E+00 | 2.37E-12 | 1.01E-11 | -2.19E+00 | 1.49E-07 | 5.69E-07 | -1.44E+00 | 3.30E-06 | 1.18E-05 | -1.54E+00 | 5.01E-03 | 1.41E-02 | 1.01E+00 | 7.66E-10 | 2.81E-09 | -1.86E+00 | 3.13E-06 | 8.60E-06 | -1.37E+00 |
| SMG1 | 5.07E-14 | 2.64E-13 | -1.35E+00 | 1.16E-08 | 4.99E-08 | -1.02E+00 | 2.50E-07 | 1.06E-06 | -1.39E+00 | 6.88E-05 | 2.63E-04 | -1.06E+00 | 4.81E-07 | 1.71E-06 | -1.24E+00 | 2.57E-07 | 7.25E-07 | -1.22E+00 | 8.50E-09 | 3.12E-08 | -1.43E+00 | 2.90E-05 | 7.97E-05 | -1.19E+00 |
| NPC2 | 5.09E-14 | 2.65E-13 | -1.36E+00 | 2.66E-12 | 1.15E-11 | -1.18E+00 | 3.77E-05 | 1.60E-04 | -1.23E+00 | 2.53E-06 | 9.68E-06 | -1.19E+00 | 7.85E-09 | 2.80E-08 | -1.30E+00 | 2.93E-09 | 8.27E-09 | -1.25E+00 | 6.54E-09 | 2.40E-08 | -1.37E+00 | 5.48E-07 | 1.51E-06 | -1.33E+00 |
| ZEB2 | 5.12E-14 | 2.67E-13 | -1.26E+00 | 5.32E-04 | 2.30E-03 | 1.06E+00 | 2.72E-09 | 1.16E-08 | -1.63E+00 | 1.63E-02 | 6.23E-02 | -1.12E+00 | 1.26E-02 | 4.49E-02 | -1.08E+00 | 2.93E-01 | 8.26E-01 | 1.06E+00 | 1.14E-02 | 4.19E-02 | -1.21E+00 | 1.21E-01 | 3.33E-01 | -1.04E+00 |
| HMGB1 | 5.38E-14 | 2.80E-13 | -1.35E+00 | 3.82E-08 | 1.65E-07 | -1.19E+00 | 3.12E-03 | 1.33E-02 | -1.11E+00 | 8.79E-05 | 3.36E-04 | -1.21E+00 | 7.10E-19 | 2.53E-18 | -1.64E+00 | 1.44E-09 | 4.06E-09 | -1.34E+00 | 1.42E-15 | 5.23E-15 | -1.44E+00 | 8.44E-06 | 2.32E-05 | -1.27E+00 |
| ZCCHC9 | 5.49E-14 | 2.86E-13 | -1.81E+00 | 4.35E-08 | 1.88E-07 | -1.21E+00 | 1.12E-12 | 4.76E-12 | -1.72E+00 | 8.46E-09 | 3.23E-08 | -1.24E+00 | 1.21E-10 | 4.32E-10 | -1.91E+00 | 8.47E-07 | 2.39E-06 | -1.46E+00 | 1.44E-05 | 5.28E-05 | -1.24E+00 | 3.78E-04 | 1.04E-03 | -1.15E+00 |
| TMX3 | 5.55E-14 | 2.89E-13 | -1.66E+00 | 6.16E-08 | 2.66E-07 | -1.12E+00 | 9.90E-12 | 4.21E-11 | -1.67E+00 | 2.59E-05 | 9.88E-05 | -1.03E+00 | 7.40E-13 | 2.64E-12 | -2.00E+00 | 7.89E-08 | 2.22E-07 | -1.45E+00 | 2.24E-11 | 8.23E-11 | -1.61E+00 | 3.55E-08 | 9.76E-08 | -1.36E+00 |
| HELZ | 5.57E-14 | 2.90E-13 | -1.33E+00 | 2.93E-12 | 1.27E-11 | -1.08E+00 | 1.47E-03 | 6.26E-03 | -1.14E+00 | 4.13E-04 | 1.58E-03 | 1.01E+00 | 2.80E-05 | 9.99E-05 | -1.03E+00 | 2.06E-05 | 5.80E-05 | -1.11E+00 | 9.12E-05 | 3.35E-04 | -1.02E+00 | 4.92E-04 | 1.35E-03 | -1.01E+00 |
| DEK | 5.66E-14 | 2.95E-13 | -1.30E+00 | 1.08E-06 | 4.67E-06 | -1.05E+00 | 2.23E-02 | 9.49E-02 | -1.07E+00 | 8.45E-02 | 3.23E-01 | 1.04E+00 | 1.21E-10 | 4.32E-10 | -1.34E+00 | 2.62E-05 | 7.37E-05 | -1.11E+00 | 2.66E-03 | 9.78E-03 | -1.22E+00 | 3.43E-01 | 9.41E-01 | 1.00E+00 |
| CEBPZ | 5.76E-14 | 3.00E-13 | -1.46E+00 | 7.12E-12 | 3.07E-11 | -1.23E+00 | 4.47E-04 | 1.90E-03 | -1.13E+00 | 3.43E-05 | 1.31E-04 | -1.08E+00 | 1.55E-06 | 5.52E-06 | -1.30E+00 | 5.17E-07 | 1.46E-06 | -1.39E+00 | 4.64E-10 | 1.70E-09 | -1.37E+00 | 6.14E-08 | 1.69E-07 | -1.32E+00 |
| ATP2B4 | 5.89E-14 | 3.07E-13 | -1.45E+00 | 5.86E-07 | 2.53E-06 | -1.07E+00 | 1.13E-06 | 4.78E-06 | -1.48E+00 | 4.39E-04 | 1.68E-03 | -1.05E+00 | 7.60E-06 | 2.71E-05 | -1.24E+00 | 1.46E-05 | 4.13E-05 | -1.18E+00 | 2.20E-07 | 8.07E-07 | -1.55E+00 | 3.75E-05 | 1.03E-04 | -1.28E+00 |
| COMMD7 | 5.97E-14 | 3.11E-13 | -1.22E+00 | 1.59E-06 | 6.88E-06 | 1.06E+00 | 1.43E-02 | 6.05E-02 | -1.06E+00 | 4.78E-02 | 1.83E-01 | 1.10E+00 | 1.24E-03 | 4.42E-03 | -1.12E+00 | 3.52E-02 | 9.93E-02 | 1.04E+00 | 8.31E-02 | 3.05E-01 | 1.02E+00 | 2.79E-01 | 7.67E-01 | 1.09E+00 |
| ILF2 | 6.04E-14 | 3.15E-13 | -1.31E+00 | 2.06E-11 | 8.91E-11 | -1.16E+00 | 1.94E-03 | 8.23E-03 | -1.17E+00 | 7.26E-03 | 2.77E-02 | -1.09E+00 | 4.37E-12 | 1.56E-11 | -1.47E+00 | 1.34E-04 | 3.78E-04 | -1.17E+00 | 2.64E-05 | 9.69E-05 | -1.24E+00 | 1.62E-01 | 4.44E-01 | -1.02E+00 |
| PDHB | 6.08E-14 | 3.17E-13 | -1.61E+00 | 2.41E-07 | 1.04E-06 | -1.06E+00 | 2.08E-12 | 8.84E-12 | -1.61E+00 | 6.00E-06 | 2.29E-05 | -1.16E+00 | 7.36E-16 | 2.62E-15 | -1.82E+00 | 9.09E-05 | 2.56E-04 | -1.02E+00 | 1.26E-14 | 4.63E-14 | -1.59E+00 | 4.03E-03 | 1.11E-02 | -1.02E+00 |
| EIF2AK4 | 6.16E-14 | 3.21E-13 | -1.46E+00 | 6.57E-08 | 2.84E-07 | -1.06E+00 | 1.72E-07 | 7.33E-07 | -1.34E+00 | 8.59E-06 | 3.28E-05 | -1.02E+00 | 8.87E-06 | 3.16E-05 | -1.16E+00 | 1.12E-04 | 3.16E-04 | -1.07E+00 | 1.44E-05 | 5.28E-05 | -1.20E+00 | 4.73E-02 | 1.30E-01 | 1.06E+00 |
| AIG1 | 6.19E-14 | 3.23E-13 | -1.46E+00 | 5.38E-13 | 2.32E-12 | -1.36E+00 | 1.83E-03 | 7.78E-03 | -1.12E+00 | 5.54E-06 | 2.12E-05 | -1.19E+00 | 1.55E-06 | 5.52E-06 | -1.17E+00 | 9.73E-03 | 2.74E-02 | 1.14E+00 | 1.30E-02 | 4.79E-02 | 1.07E+00 | 2.05E-04 | 5.63E-04 | -1.09E+00 |
| OTUD4 | 6.24E-14 | 3.25E-13 | -1.57E+00 | 1.80E-09 | 7.77E-09 | -1.08E+00 | 1.45E-15 | 6.18E-15 | -1.77E+00 | 5.79E-13 | 2.21E-12 | -1.41E+00 | 1.62E-04 | 5.79E-04 | -1.37E+00 | 5.40E-04 | 1.52E-03 | -1.22E+00 | 5.30E-14 | 1.95E-13 | -1.67E+00 | 2.49E-05 | 6.83E-05 | -1.18E+00 |
| SEL1L | 6.32E-14 | 3.30E-13 | -1.72E+00 | 1.45E-07 | 6.28E-07 | -1.11E+00 | 7.51E-07 | 3.19E-06 | -1.34E+00 | 5.23E-02 | 2.00E-01 | 1.24E+00 | 9.33E-06 | 3.33E-05 | -1.36E+00 | 4.25E-05 | 1.20E-04 | -1.17E+00 | 1.36E-06 | 4.99E-06 | -1.35E+00 | 1.04E-02 | 2.86E-02 | -1.02E+00 |
| AZIN1 | 6.46E-14 | 3.36E-13 | -1.43E+00 | 1.55E-13 | 6.70E-13 | -1.23E+00 | 1.20E-06 | 5.09E-06 | -1.35E+00 | 1.28E-04 | 4.90E-04 | -1.06E+00 | 1.29E-05 | 4.60E-05 | -1.11E+00 | 3.89E-06 | 1.10E-05 | -1.19E+00 | 1.29E-08 | 4.74E-08 | -1.28E+00 | 7.95E-05 | 2.18E-04 | -1.14E+00 |
| HCG18 | 6.46E-14 | 3.36E-13 | -1.59E+00 | 1.64E-04 | 7.08E-04 | 1.15E+00 | 4.51E-04 | 1.92E-03 | -1.17E+00 | 2.13E-01 | 8.13E-01 | 1.42E+00 | 1.09E-03 | 3.87E-03 | -1.18E+00 | 3.55E-01 | 1.00E+00 | 1.47E+00 | 8.21E-02 | 3.01E-01 | -1.04E+00 | 8.87E-02 | 2.44E-01 | 1.30E+00 |
| GNB4 | 6.63E-14 | 3.45E-13 | -1.44E+00 | 1.26E-12 | 5.43E-12 | -1.25E+00 | 9.00E-12 | 3.82E-11 | -1.63E+00 | 3.41E-08 | 1.30E-07 | -1.26E+00 | 1.50E-08 | 5.35E-08 | -1.53E+00 | 1.24E-05 | 3.48E-05 | -1.18E+00 | 6.46E-13 | 2.37E-12 | -1.67E+00 | 1.76E-06 | 4.82E-06 | -1.31E+00 |
| WFDC1 | 6.63E-14 | 3.45E-13 | -2.41E+00 | 4.03E-13 | 1.74E-12 | -2.11E+00 | 5.20E-10 | 2.21E-09 | -1.88E+00 | 1.17E-11 | 4.46E-11 | -1.97E+00 | 3.63E-03 | 1.29E-02 | -1.22E+00 | 3.27E-03 | 9.22E-03 | -1.35E+00 | 2.44E-02 | 8.96E-02 | 1.24E+00 | 2.48E-02 | 6.82E-02 | -1.18E+00 |
| TECR | 6.69E-14 | 3.49E-13 | -1.39E+00 | 3.90E-12 | 1.69E-11 | -1.20E+00 | 9.55E-07 | 4.06E-06 | -1.31E+00 | 4.04E-09 | 1.54E-08 | -1.20E+00 | 1.05E-12 | 3.74E-12 | -1.56E+00 | 1.39E-07 | 3.93E-07 | -1.20E+00 | 2.65E-06 | 9.74E-06 | -1.24E+00 | 1.19E-05 | 3.28E-05 | -1.15E+00 |
| ZFC3H1 | 6.75E-14 | 3.52E-13 | -1.39E+00 | 4.21E-05 | 1.82E-04 | 1.09E+00 | 8.24E-11 | 3.50E-10 | -1.55E+00 | 4.04E-06 | 1.54E-05 | -1.08E+00 | 1.43E-09 | 5.11E-09 | -1.48E+00 | 9.66E-08 | 2.72E-07 | -1.26E+00 | 5.58E-08 | 2.05E-07 | -1.37E+00 | 5.59E-04 | 1.54E-03 | -1.07E+00 |
| RFK | 6.82E-14 | 3.56E-13 | -1.55E+00 | 2.97E-15 | 1.28E-14 | -1.40E+00 | 9.84E-08 | 4.18E-07 | -1.24E+00 | 1.19E-06 | 4.56E-06 | -1.11E+00 | 1.22E-10 | 4.34E-10 | -1.57E+00 | 3.12E-05 | 8.79E-05 | -1.04E+00 | 1.35E-20 | 4.95E-20 | -2.10E+00 | 5.70E-13 | 1.57E-12 | -1.70E+00 |
| RPL12P4 | 6.87E-14 | 3.58E-13 | 1.30E+00 | 2.81E-13 | 1.21E-12 | 1.31E+00 | 1.20E-12 | 5.11E-12 | 1.36E+00 | 1.52E-13 | 5.79E-13 | 1.39E+00 | 5.12E-10 | 1.83E-09 | 1.40E+00 | 1.80E-12 | 5.07E-12 | 1.43E+00 | 1.01E-05 | 3.71E-05 | 1.36E+00 | 2.43E-13 | 6.69E-13 | 1.42E+00 |
| STAG3L3 | 6.91E-14 | 3.60E-13 | -1.33E+00 | 1.25E-07 | 5.39E-07 | 1.04E+00 | 1.39E-09 | 5.90E-09 | -1.47E+00 | 1.20E-04 | 4.58E-04 | -1.04E+00 | 9.62E-06 | 3.43E-05 | -1.29E+00 | 3.39E-05 | 9.55E-05 | -1.18E+00 | 5.76E-04 | 2.11E-03 | -1.13E+00 | 1.40E-01 | 3.85E-01 | 1.03E+00 |
| AC146507.2 | 6.96E-14 | 3.63E-13 | -2.59E+00 | 4.73E-04 | 2.04E-03 | 1.04E+00 | 7.59E-07 | 3.22E-06 | -1.76E+00 | 2.37E-02 | 9.03E-02 | 1.14E+00 | 6.64E-07 | 2.37E-06 | -2.29E+00 | 1.45E-02 | 4.08E-02 | -1.04E+00 | 2.04E-03 | 7.48E-03 | -1.28E+00 | 9.22E-02 | 2.53E-01 | 1.03E+00 |
| PRDX2 | 6.99E-14 | 3.64E-13 | -1.31E+00 | 2.99E-08 | 1.29E-07 | -1.05E+00 | 9.72E-03 | 4.13E-02 | -1.09E+00 | 2.40E-02 | 9.18E-02 | -1.03E+00 | 7.51E-16 | 2.68E-15 | -1.56E+00 | 1.08E-04 | 3.04E-04 | -1.10E+00 | 2.65E-05 | 9.71E-05 | -1.19E+00 | 1.21E-01 | 3.33E-01 | 1.06E+00 |
| AL513343.1 | 6.99E-14 | 3.64E-13 | -1.51E+00 | 8.79E-25 | 3.80E-24 | -2.10E+00 | 9.49E-05 | 4.03E-04 | -1.22E+00 | 2.47E-13 | 9.43E-13 | -1.81E+00 | 1.50E-06 | 5.35E-06 | -1.34E+00 | 1.95E-13 | 5.50E-13 | -2.15E+00 | 1.18E-05 | 4.34E-05 | -1.29E+00 | 1.70E-10 | 4.66E-10 | -1.78E+00 |
| LEO1 | 7.06E-14 | 3.68E-13 | -1.65E+00 | 7.96E-07 | 3.44E-06 | -1.04E+00 | 2.73E-15 | 1.16E-14 | -1.91E+00 | 8.27E-07 | 3.16E-06 | -1.17E+00 | 2.54E-12 | 9.06E-12 | -1.88E+00 | 5.73E-06 | 1.61E-05 | -1.17E+00 | 5.63E-18 | 2.07E-17 | -1.77E+00 | 1.07E-05 | 2.95E-05 | -1.20E+00 |
| TFG | 7.12E-14 | 3.71E-13 | -1.29E+00 | 5.88E-08 | 2.54E-07 | -1.02E+00 | 2.87E-08 | 1.22E-07 | -1.38E+00 | 2.01E-06 | 7.69E-06 | -1.10E+00 | 2.78E-11 | 9.92E-11 | -1.48E+00 | 2.20E-08 | 6.20E-08 | -1.26E+00 | 1.89E-13 | 6.94E-13 | -1.52E+00 | 6.21E-06 | 1.71E-05 | -1.27E+00 |
| CLCN3 | 7.15E-14 | 3.73E-13 | -2.53E+00 | 1.24E-06 | 5.34E-06 | -1.26E+00 | 1.86E-06 | 7.91E-06 | -2.04E+00 | 1.09E-02 | 4.17E-02 | -1.02E+00 | 2.09E-02 | 7.46E-02 | -1.50E+00 | 1.28E-01 | 3.60E-01 | -1.11E+00 | 3.80E-08 | 1.40E-07 | 1.11E+00 | 3.10E-05 | 8.51E-05 | 1.07E+00 |
| RABGAP1 | 7.16E-14 | 3.73E-13 | -1.39E+00 | 1.81E-02 | 7.80E-02 | 1.27E+00 | 4.77E-09 | 2.03E-08 | -1.48E+00 | 1.07E-03 | 4.10E-03 | 1.05E+00 | 9.36E-05 | 3.34E-04 | -1.23E+00 | 4.85E-02 | 1.37E-01 | 1.13E+00 | 1.53E-06 | 5.63E-06 | -1.21E+00 | 1.30E-04 | 3.58E-04 | -1.06E+00 |
| CTDSPL | 7.18E-14 | 3.74E-13 | -1.81E+00 | 3.06E-04 | 1.32E-03 | 1.05E+00 | 2.08E-07 | 8.82E-07 | -1.72E+00 | 9.83E-03 | 3.75E-02 | 1.04E+00 | 2.41E-02 | 8.59E-02 | -1.18E+00 | 3.28E-01 | 9.25E-01 | 1.41E+00 | 7.98E-06 | 2.93E-05 | -1.12E+00 | 2.81E-03 | 7.72E-03 | -1.07E+00 |
| MAN1A2 | 7.18E-14 | 3.74E-13 | -1.31E+00 | 7.23E-09 | 3.12E-08 | 1.02E+00 | 1.64E-10 | 6.95E-10 | -1.49E+00 | 3.34E-06 | 1.28E-05 | -1.12E+00 | 2.96E-06 | 1.05E-05 | -1.22E+00 | 2.55E-04 | 7.19E-04 | -1.06E+00 | 2.17E-06 | 7.97E-06 | -1.28E+00 | 1.78E-04 | 4.90E-04 | -1.09E+00 |
| RPL12P8 | 7.26E-14 | 3.79E-13 | -1.40E+00 | 5.26E-51 | 2.27E-50 | -2.07E+00 | 2.75E-06 | 1.17E-05 | -1.32E+00 | 2.00E-22 | 7.65E-22 | -1.88E+00 | 1.08E-05 | 3.86E-05 | -1.28E+00 | 5.12E-21 | 1.44E-20 | -1.96E+00 | 4.01E-07 | 1.47E-06 | -1.37E+00 | 2.05E-14 | 5.63E-14 | -1.90E+00 |
| COQ9 | 7.38E-14 | 3.85E-13 | -2.10E+00 | 1.06E-05 | 4.60E-05 | -1.08E+00 | 2.52E-14 | 1.07E-13 | -2.01E+00 | 2.40E-05 | 9.15E-05 | -1.09E+00 | 7.36E-11 | 2.62E-10 | -1.79E+00 | 2.99E-04 | 8.44E-04 | -1.02E+00 | 1.02E-16 | 3.73E-16 | -1.76E+00 | 4.22E-04 | 1.16E-03 | -1.03E+00 |
| SCARF1 | 7.39E-14 | 3.85E-13 | -1.63E+00 | 7.21E-09 | 3.12E-08 | -1.18E+00 | 2.86E-11 | 1.22E-10 | -2.09E+00 | 2.45E-04 | 9.37E-04 | -1.11E+00 | 2.00E-07 | 7.13E-07 | -2.12E+00 | 8.27E-06 | 2.33E-05 | -1.71E+00 | 3.89E-05 | 1.43E-04 | -1.95E+00 | 3.26E-04 | 8.96E-04 | -1.75E+00 |
| RMND5A | 7.46E-14 | 3.89E-13 | -1.32E+00 | 5.92E-18 | 2.56E-17 | -1.29E+00 | 4.12E-05 | 1.75E-04 | -1.21E+00 | 7.41E-08 | 2.83E-07 | -1.21E+00 | 2.53E-08 | 9.01E-08 | -1.25E+00 | 1.76E-09 | 4.96E-09 | -1.34E+00 | 2.73E-03 | 1.00E-02 | 1.01E+00 | 1.08E-05 | 2.97E-05 | -1.19E+00 |
| TOP2B | 7.47E-14 | 3.90E-13 | -1.34E+00 | 4.70E-08 | 2.03E-07 | -1.03E+00 | 3.31E-06 | 1.41E-05 | -1.34E+00 | 4.83E-04 | 1.84E-03 | -1.08E+00 | 4.17E-07 | 1.49E-06 | -1.31E+00 | 1.09E-03 | 3.06E-03 | -1.10E+00 | 1.95E-03 | 7.15E-03 | -1.03E+00 | 1.58E-03 | 4.35E-03 | -1.09E+00 |
| AC026150.1 | 7.54E-14 | 3.93E-13 | -2.51E+00 | 1.11E-11 | 4.78E-11 | -1.92E+00 | 1.37E-07 | 5.84E-07 | -2.24E+00 | 1.64E-05 | 6.27E-05 | -1.53E+00 | 6.49E-05 | 2.31E-04 | -2.07E+00 | 4.98E-05 | 1.40E-04 | -2.15E+00 | 2.56E-06 | 9.41E-06 | -1.68E+00 | 4.78E-05 | 1.31E-04 | -1.51E+00 |

| | | | | | | | | | | | | | | | | | | | | | | | | |
|---|---|---|---|---|---|---|---|---|---|---|---|---|---|---|---|---|---|---|---|---|---|---|---|---|
| ARFGAP2 | 7.64E-14 | 3.98E-13 | -1.34E+00 | 2.98E-12 | 1.29E-11 | -1.10E+00 | 6.11E-07 | 2.60E-06 | -1.31E+00 | 3.52E-05 | 1.35E-04 | -1.11E+00 | 4.99E-09 | 1.78E-08 | -1.34E+00 | 1.33E-04 | 3.75E-04 | -1.12E+00 | 8.77E-08 | 3.22E-07 | -1.22E+00 | 6.47E-03 | 1.78E-02 | 1.01E+00 |
| EPHX1 | 7.64E-14 | 3.98E-13 | -3.39E+00 | 3.18E-08 | 1.37E-07 | -1.63E+00 | 2.89E-03 | 1.23E-02 | -1.46E+00 | 1.43E-03 | 5.45E-03 | -1.35E+00 | 3.31E-02 | 1.18E-01 | -1.32E+00 | 3.34E-02 | 9.43E-02 | -1.44E+00 | 2.87E-06 | 1.05E-05 | -1.54E+00 | 3.69E-05 | 1.01E-04 | -1.51E+00 |
| SLC25A39 | 7.69E-14 | 4.01E-13 | -1.35E+00 | 6.74E-10 | 2.91E-09 | -1.03E+00 | 1.92E-08 | 8.15E-08 | -1.50E+00 | 1.57E-05 | 6.01E-05 | -1.12E+00 | 6.28E-11 | 2.24E-10 | -1.50E+00 | 2.14E-04 | 6.04E-04 | 1.05E+00 | 5.70E-08 | 2.09E-07 | -1.27E+00 | 9.31E-02 | 2.56E-01 | 1.13E+00 |
| AC025458.1 | 7.73E-14 | 4.03E-13 | -1.32E+00 | 1.15E-29 | 4.99E-29 | -1.74E+00 | 1.07E-07 | 4.55E-07 | -1.27E+00 | 5.96E-22 | 2.28E-21 | -1.92E+00 | 6.57E-05 | 2.34E-04 | -1.12E+00 | 9.34E-13 | 2.63E-12 | -1.65E+00 | 1.91E-08 | 6.99E-08 | -1.28E+00 | 6.24E-16 | 1.71E-15 | -1.79E+00 |
| ELK3 | 7.73E-14 | 4.03E-13 | -1.53E+00 | 3.40E-09 | 1.47E-08 | -1.13E+00 | 9.29E-08 | 3.95E-07 | -1.52E+00 | 5.55E-05 | 2.12E-04 | -1.17E+00 | 8.91E-03 | 3.18E-02 | -1.16E+00 | 6.48E-02 | 1.83E-01 | 1.11E+00 | 7.49E-14 | 2.75E-13 | -1.94E+00 | 2.58E-11 | 7.07E-11 | -1.81E+00 |
| SCO2 | 7.83E-14 | 4.08E-13 | -1.96E+00 | 1.02E-11 | 4.43E-11 | -1.58E+00 | 1.20E-13 | 5.10E-13 | -1.82E+00 | 2.36E-07 | 9.01E-07 | -1.25E+00 | 7.53E-12 | 2.68E-11 | -1.98E+00 | 1.17E-05 | 3.30E-05 | -1.31E+00 | 6.85E-19 | 2.51E-18 | -2.27E+00 | 5.02E-06 | 1.38E-05 | -1.32E+00 |
| DCAF12 | 8.06E-14 | 4.20E-13 | -2.17E+00 | 2.02E-05 | 8.73E-05 | -1.04E+00 | 3.97E-14 | 1.69E-13 | -2.06E+00 | 8.91E-07 | 3.40E-06 | -1.23E+00 | 5.28E-08 | 1.88E-07 | -2.44E+00 | 2.73E-03 | 7.70E-03 | -1.15E+00 | 6.28E-14 | 2.31E-13 | -1.93E+00 | 3.69E-05 | 1.01E-04 | -1.20E+00 |
| SFT2D2 | 8.10E-14 | 4.22E-13 | -1.35E+00 | 7.40E-12 | 3.20E-11 | -1.11E+00 | 6.47E-05 | 2.75E-04 | -1.35E+00 | 4.35E-05 | 1.66E-04 | -1.14E+00 | 1.18E-05 | 4.22E-05 | -1.25E+00 | 1.01E-03 | 2.85E-03 | -1.06E+00 | 5.67E-04 | 2.08E-03 | -1.25E+00 | 5.85E-02 | 1.61E-01 | -1.03E+00 |
| TTC14 | 8.14E-14 | 4.24E-13 | -1.30E+00 | 6.21E-02 | 2.68E-01 | 1.58E+00 | 3.01E-09 | 1.28E-08 | -1.42E+00 | 2.57E-01 | 9.82E-01 | 1.38E+00 | 9.66E-06 | 3.45E-05 | -1.10E+00 | 2.93E-01 | 8.26E-01 | 1.36E+00 | 6.93E-03 | 2.54E-02 | -1.01E+00 | 1.33E-01 | 3.66E-01 | 1.28E+00 |
| AHSA1 | 8.16E-14 | 4.25E-13 | -1.38E+00 | 6.25E-03 | 2.70E-02 | 1.28E+00 | 1.66E-02 | 7.06E-02 | 1.00E+00 | 9.93E-02 | 3.79E-01 | 1.43E+00 | 4.64E-12 | 1.66E-11 | -1.66E+00 | 5.00E-02 | 1.41E-01 | 1.12E+00 | 1.85E-03 | 6.77E-03 | -1.14E+00 | 1.98E-02 | 5.44E-02 | 1.42E+00 |
| NEIL2 | 8.28E-14 | 4.31E-13 | -1.86E+00 | 5.40E-08 | 2.33E-07 | -1.18E+00 | 1.99E-08 | 8.45E-08 | -1.62E+00 | 5.59E-06 | 2.14E-05 | -1.13E+00 | 1.52E-02 | 5.42E-02 | -1.04E+00 | 2.86E-01 | 8.05E-01 | 1.43E+00 | 8.91E-03 | 3.27E-02 | -1.14E+00 | 6.44E-02 | 1.77E-01 | 1.02E+00 |
| YTHDC1 | 8.33E-14 | 4.34E-13 | -1.30E+00 | 6.51E-23 | 2.81E-22 | -1.40E+00 | 3.71E-10 | 1.58E-09 | -1.43E+00 | 1.17E-15 | 4.48E-15 | -1.50E+00 | 1.02E-07 | 3.63E-07 | -1.23E+00 | 4.97E-09 | 1.40E-08 | -1.32E+00 | 7.40E-09 | 2.72E-08 | -1.38E+00 | 3.35E-13 | 9.21E-13 | -1.63E+00 |
| PURA | 8.33E-14 | 4.34E-13 | -1.57E+00 | 2.81E-07 | 1.22E-06 | -1.08E+00 | 4.55E-10 | 1.94E-09 | -1.75E+00 | 9.11E-04 | 3.48E-03 | -1.02E+00 | 2.62E-08 | 9.34E-08 | -1.73E+00 | 6.29E-05 | 1.77E-04 | -1.20E+00 | 1.95E-09 | 7.14E-09 | -1.39E+00 | 7.94E-04 | 2.18E-03 | -1.14E+00 |
| PSMA3 | 8.33E-14 | 4.34E-13 | -1.25E+00 | 6.62E-11 | 2.86E-10 | -1.01E+00 | 8.22E-05 | 3.49E-04 | -1.22E+00 | 9.13E-03 | 3.49E-02 | -1.01E+00 | 3.72E-10 | 1.33E-09 | -1.31E+00 | 7.80E-06 | 2.20E-05 | -1.07E+00 | 3.46E-06 | 1.27E-05 | -1.36E+00 | 5.66E-02 | 1.56E-01 | -1.03E+00 |
| SPG21 | 8.39E-14 | 4.37E-13 | -1.46E+00 | 1.27E-08 | 5.47E-08 | -1.06E+00 | 4.12E-11 | 1.75E-10 | -1.50E+00 | 1.88E-04 | 7.17E-04 | 1.01E+00 | 1.52E-07 | 5.42E-07 | -1.36E+00 | 4.55E-03 | 1.28E-02 | 1.07E+00 | 3.46E-07 | 1.27E-06 | -1.24E+00 | 2.87E-02 | 7.89E-02 | 1.15E+00 |
| SMARCD3 | 8.47E-14 | 4.41E-13 | -1.85E+00 | 3.30E-15 | 1.42E-14 | -1.81E+00 | 2.71E-09 | 1.15E-08 | -1.41E+00 | 7.65E-12 | 2.92E-11 | -1.42E+00 | 1.92E-01 | 6.86E-01 | 1.03E+00 | 2.28E-01 | 6.43E-01 | -1.02E+00 | 2.36E-01 | 8.65E-01 | -1.08E+00 | 3.19E-01 | 8.75E-01 | -1.06E+00 |
| HIC2 | 8.53E-14 | 4.45E-13 | -1.93E+00 | 2.20E-18 | 9.50E-18 | -2.19E+00 | 8.97E-12 | 3.81E-11 | -1.93E+00 | 1.45E-14 | 5.54E-14 | -2.07E+00 | 1.74E-04 | 6.21E-04 | -1.71E+00 | 3.85E-05 | 1.09E-04 | -1.90E+00 | 2.42E-11 | 8.87E-11 | -1.93E+00 | 1.34E-11 | 3.68E-11 | -2.03E+00 |
| UBE2W | 8.72E-14 | 4.55E-13 | -1.83E+00 | 4.58E-08 | 1.98E-07 | -1.22E+00 | 1.80E-10 | 7.63E-10 | -1.75E+00 | 1.29E-06 | 4.93E-06 | -1.17E+00 | 1.26E-07 | 4.48E-07 | -1.69E+00 | 2.39E-05 | 6.74E-05 | -1.34E+00 | 6.83E-11 | 2.51E-10 | -1.67E+00 | 5.58E-04 | 1.53E-03 | -1.06E+00 |
| ZC3H15 | 8.81E-14 | 4.59E-13 | -1.20E+00 | 1.06E-13 | 4.56E-13 | -1.11E+00 | 3.01E-02 | 1.28E-01 | -1.05E+00 | 2.13E-02 | 8.13E-02 | -1.01E+00 | 8.86E-05 | 3.16E-04 | -1.15E+00 | 5.40E-04 | 1.52E-03 | -1.12E+00 | 5.60E-02 | 2.06E-01 | -1.06E+00 | 2.80E-01 | 7.69E-01 | 1.02E+00 |
| GYG1 | 8.89E-14 | 4.63E-13 | -1.43E+00 | 8.28E-10 | 3.58E-09 | -1.09E+00 | 3.59E-11 | 1.53E-10 | -1.51E+00 | 1.20E-07 | 4.58E-07 | -1.12E+00 | 1.63E-08 | 5.82E-08 | -1.93E+00 | 5.61E-03 | 1.58E-02 | -1.08E+00 | 2.34E-05 | 8.58E-05 | -1.27E+00 | 2.96E-02 | 8.14E-02 | 1.04E+00 |
| VTA1 | 8.93E-14 | 4.65E-13 | -2.18E+00 | 6.80E-06 | 2.94E-05 | -1.16E+00 | 1.34E-10 | 5.68E-10 | -1.55E+00 | 8.34E-08 | 3.19E-07 | -1.20E+00 | 8.87E-12 | 3.16E-11 | -2.04E+00 | 1.72E-05 | 4.86E-05 | -1.26E+00 | 6.46E-23 | 2.37E-22 | -2.04E+00 | 4.67E-11 | 1.28E-10 | -1.41E+00 |
| PDS5B | 8.93E-14 | 4.65E-13 | -1.42E+00 | 4.53E-07 | 1.96E-06 | -1.00E+00 | 1.79E-07 | 7.63E-07 | -1.39E+00 | 1.22E-04 | 4.67E-04 | -1.09E+00 | 7.37E-06 | 2.63E-05 | -1.34E+00 | 4.70E-03 | 1.33E-02 | 1.00E+00 | 1.16E-07 | 4.27E-07 | -1.35E+00 | 7.31E-03 | 2.01E-02 | -1.04E+00 |
| SKIV2L2 | 8.94E-14 | 4.66E-13 | -1.53E+00 | 1.06E-08 | 4.57E-08 | -1.10E+00 | 5.73E-06 | 2.44E-05 | -1.26E+00 | 7.56E-06 | 2.89E-05 | -1.13E+00 | 7.52E-06 | 2.68E-05 | -1.20E+00 | 7.23E-04 | 2.04E-03 | 1.07E+00 | 8.83E-08 | 3.24E-07 | -1.39E+00 | 8.22E-03 | 2.26E-02 | -1.06E+00 |
| CLTA | 8.99E-14 | 4.68E-13 | -1.27E+00 | 3.29E-05 | 1.42E-04 | 1.05E+00 | 4.17E-04 | 1.77E-03 | -1.18E+00 | 5.28E-02 | 2.02E-01 | 1.02E+00 | 1.98E-09 | 7.07E-09 | -1.39E+00 | 1.89E-02 | 5.33E-02 | -1.02E+00 | 6.17E-03 | 2.27E-02 | -1.12E+00 | 1.67E-01 | 4.57E-01 | 1.12E+00 |
| WTAP | 9.00E-14 | 4.69E-13 | -1.22E+00 | 7.15E-08 | 3.09E-07 | 1.00E+00 | 1.37E-04 | 5.82E-04 | -1.29E+00 | 2.04E-03 | 7.78E-03 | -1.09E+00 | 1.17E-05 | 4.17E-05 | -1.07E+00 | 3.65E-03 | 1.03E-02 | 1.05E+00 | 1.54E-04 | 5.66E-04 | -1.21E+00 | 1.86E-02 | 5.11E-02 | -1.08E+00 |
| RPS20 | 9.03E-14 | 4.71E-13 | 1.29E+00 | 6.25E-15 | 2.70E-14 | 1.33E+00 | 2.13E-11 | 9.05E-11 | 1.31E+00 | 1.61E-17 | 6.16E-17 | 1.43E+00 | 2.39E-04 | 8.52E-04 | 1.23E+00 | 1.86E-10 | 5.24E-10 | 1.38E+00 | 1.49E-01 | 5.46E-01 | 1.13E+00 | 2.23E-09 | 6.13E-09 | 1.35E+00 |
| SLC35E2B | 9.05E-14 | 4.72E-13 | -1.55E+00 | 1.04E-05 | 4.48E-05 | 1.04E+00 | 2.15E-07 | 9.13E-07 | -1.46E+00 | 6.73E-04 | 2.57E-03 | -1.02E+00 | 6.80E-15 | 2.42E-14 | -2.05E+00 | 4.38E-05 | 1.23E-04 | -1.24E+00 | 1.39E-06 | 5.10E-06 | -1.30E+00 | 1.19E-04 | 3.26E-04 | -1.13E+00 |
| RRBP1 | 9.23E-14 | 4.81E-13 | -1.30E+00 | 2.04E-22 | 8.82E-22 | -1.40E+00 | 1.35E-04 | 5.73E-04 | -1.23E+00 | 5.77E-10 | 2.20E-09 | -1.47E+00 | 2.47E-04 | 8.79E-04 | -1.05E+00 | 1.94E-07 | 5.46E-07 | -1.22E+00 | 2.64E-05 | 9.69E-05 | -1.05E+00 | 1.45E-04 | 3.97E-04 | -1.12E+00 |
| ADO | 9.26E-14 | 4.83E-13 | -1.83E+00 | 6.76E-14 | 2.92E-13 | -1.59E+00 | 1.17E-07 | 4.96E-07 | -1.57E+00 | 5.74E-05 | 2.19E-04 | -1.13E+00 | 1.21E-06 | 4.30E-06 | -1.59E+00 | 4.05E-06 | 1.14E-05 | -1.51E+00 | 1.28E-07 | 4.71E-07 | -1.34E+00 | 6.09E-04 | 1.67E-03 | -1.01E+00 |
| TMED9 | 9.30E-14 | 4.84E-13 | -1.26E+00 | 5.15E-10 | 2.22E-09 | -1.01E+00 | 6.16E-03 | 2.62E-02 | -1.09E+00 | 1.40E-02 | 5.33E-02 | 1.02E+00 | 1.09E-10 | 3.89E-10 | -1.33E+00 | 7.64E-04 | 2.15E-03 | 1.03E+00 | 3.61E-05 | 1.32E-04 | -1.18E+00 | 2.09E-01 | 5.74E-01 | 1.08E+00 |
| CNIH4 | 9.30E-14 | 4.84E-13 | -1.95E+00 | 5.59E-10 | 2.42E-09 | -1.34E+00 | 2.32E-08 | 9.88E-08 | -1.43E+00 | 2.60E-09 | 9.93E-09 | -1.40E+00 | 4.08E-03 | 1.45E-02 | -1.05E+00 | 5.74E-02 | 1.62E-01 | 1.12E+00 | 2.69E-12 | 9.87E-12 | -1.64E+00 | 4.36E-05 | 1.20E-04 | -1.15E+00 |
| TGFB1 | 9.34E-14 | 4.87E-13 | -1.28E+00 | 2.44E-11 | 1.05E-10 | -1.13E+00 | 6.58E-03 | 2.80E-02 | -1.03E+00 | 2.38E-02 | 9.10E-02 | 1.14E+00 | 1.17E-08 | 4.16E-08 | -1.29E+00 | 1.53E-05 | 4.32E-05 | -1.22E+00 | 1.04E-01 | 3.84E-01 | 1.10E+00 | 6.94E-02 | 1.91E-01 | -1.05E+00 |
| GDF7 | 9.36E-14 | 4.88E-13 | -2.38E+00 | 1.34E-09 | 5.77E-09 | -1.65E+00 | 1.10E-02 | 4.68E-02 | -1.43E+00 | 7.58E-02 | 2.89E-01 | 1.04E+00 | 8.56E-02 | 3.05E-01 | -1.86E+00 | 1.45E-01 | 4.08E-01 | -1.57E+00 | 2.77E-02 | 1.02E-01 | -1.82E+00 | 3.06E-01 | 8.41E-01 | 1.07E+00 |
| RPL23AP75 | 9.48E-14 | 4.94E-13 | -1.26E+00 | 3.01E-27 | 1.30E-26 | -1.45E+00 | 1.51E-05 | 6.42E-05 | -1.31E+00 | 5.45E-10 | 2.08E-09 | -1.42E+00 | 1.73E-10 | 6.17E-10 | -1.36E+00 | 3.42E-10 | 9.63E-10 | -1.37E+00 | 1.05E-06 | 3.85E-06 | -1.35E+00 | 7.43E-06 | 2.04E-05 | -1.38E+00 |
| ATRAID | 9.54E-14 | 4.97E-13 | -1.20E+00 | 1.45E-08 | 6.26E-08 | 1.05E+00 | 8.26E-02 | 3.51E-01 | 1.00E+00 | 1.03E-01 | 3.93E-01 | 1.08E+00 | 7.39E-04 | 2.63E-03 | -1.00E+00 | 1.61E-01 | 4.53E-01 | 1.19E+00 | 4.14E-02 | 1.52E-01 | 1.04E+00 | 1.36E-01 | 3.72E-01 | 1.04E+00 |
| AC006504.5 | 9.55E-14 | 4.98E-13 | -2.74E+00 | 1.12E-06 | 4.86E-06 | -1.28E+00 | 1.53E-08 | 6.49E-08 | -2.25E+00 | 2.02E-03 | 7.72E-03 | -1.11E+00 | 8.12E-08 | 2.90E-07 | -2.81E+00 | 2.17E-03 | 6.11E-03 | -1.13E+00 | 4.00E-06 | 1.47E-05 | -2.12E+00 | 1.08E-01 | 2.97E-01 | 1.04E+00 |
| EMC2 | 9.58E-14 | 5.00E-13 | -1.63E+00 | 3.70E-07 | 1.60E-06 | -1.06E+00 | 3.80E-11 | 1.61E-10 | -1.78E+00 | 1.55E-04 | 5.91E-04 | -1.06E+00 | 5.30E-07 | 1.89E-06 | -1.63E+00 | 3.96E-04 | 1.12E-03 | -1.16E+00 | 2.68E-12 | 9.84E-12 | -1.52E+00 | 1.09E-04 | 2.98E-04 | -1.05E+00 |
| OGFR | 9.73E-14 | 5.07E-13 | -1.77E+00 | 1.11E-07 | 4.82E-07 | -1.13E+00 | 3.82E-09 | 1.62E-08 | -1.40E+00 | 6.45E-07 | 2.46E-06 | -1.08E+00 | 3.00E-03 | 1.07E-02 | -1.22E+00 | 2.76E-01 | 7.77E-01 | 1.24E+00 | 4.43E-11 | 1.63E-10 | -1.87E+00 | 9.01E-07 | 2.48E-06 | -1.40E+00 |
| BST1 | 9.85E-14 | 5.14E-13 | -1.84E+00 | 3.49E-13 | 1.51E-12 | -1.54E+00 | 1.76E-05 | 7.48E-05 | -1.24E+00 | 5.15E-04 | 1.97E-03 | 1.02E+00 | 2.37E-11 | 8.46E-11 | -2.10E+00 | 8.31E-09 | 2.34E-08 | -1.51E+00 | 1.11E-02 | 4.08E-02 | -1.08E+00 | 7.81E-03 | 2.15E-02 | -1.16E+00 |
| PPP2CA | 9.90E-14 | 5.16E-13 | -1.35E+00 | 4.71E-19 | 2.03E-18 | -1.49E+00 | 2.11E-05 | 8.98E-05 | -1.18E+00 | 1.58E-07 | 6.04E-07 | -1.20E+00 | 1.98E-08 | 7.07E-08 | -1.33E+00 | 7.36E-07 | 2.08E-06 | -1.30E+00 | 3.47E-13 | 1.27E-12 | -1.58E+00 | 7.62E-05 | 2.09E-04 | -1.21E+00 |
| ADPGK | 9.92E-14 | 5.17E-13 | -1.37E+00 | 1.98E-13 | 8.53E-13 | -1.19E+00 | 5.85E-09 | 2.49E-08 | -1.36E+00 | 1.04E-08 | 3.99E-08 | -1.17E+00 | 7.24E-06 | 2.58E-05 | -1.23E+00 | 3.74E-04 | 1.05E-03 | -1.09E+00 | 4.60E-13 | 1.69E-12 | -1.65E+00 | 1.08E-03 | 2.97E-03 | -1.02E+00 |
| U2SURP | 1.01E-13 | 5.25E-13 | -1.19E+00 | 9.68E-08 | 4.18E-07 | 1.03E+00 | 3.48E-04 | 1.48E-03 | -1.16E+00 | 3.25E-03 | 1.24E-02 | -1.00E+00 | 3.09E-05 | 1.10E-04 | -1.13E+00 | 1.23E-01 | 3.47E-01 | 1.14E+00 | 8.47E-05 | 3.11E-04 | -1.20E+00 | 9.63E-02 | 2.65E-01 | 1.04E+00 |
| HOOK3 | 1.01E-13 | 5.25E-13 | -1.49E+00 | 9.27E-06 | 4.01E-05 | 1.03E+00 | 3.48E-03 | 1.48E-02 | -1.15E+00 | 9.78E-02 | 3.73E-01 | 1.27E+00 | 6.39E-09 | 2.28E-08 | -1.40E+00 | 1.61E-05 | 4.54E-05 | -1.10E+00 | 4.84E-05 | 1.78E-04 | -1.19E+00 | 5.37E-05 | 1.48E-04 | -1.17E+00 |
| VAMP5 | 1.02E-13 | 5.29E-13 | -1.32E+00 | 7.26E-13 | 3.14E-12 | -1.22E+00 | 7.78E-04 | 3.30E-03 | -1.17E+00 | 6.70E-05 | 2.56E-04 | -1.13E+00 | 8.85E-03 | 3.15E-02 | -1.07E+00 | 7.48E-03 | 2.11E-02 | -1.14E+00 | 7.58E-09 | 2.78E-08 | -2.00E+00 | 1.26E-07 | 3.45E-07 | -1.88E+00 |
| NFATC1 | 1.02E-13 | 5.29E-13 | -1.72E+00 | 3.68E-09 | 1.59E-08 | -1.21E+00 | 1.26E-03 | 5.36E-03 | -1.18E+00 | 3.97E-03 | 1.52E-02 | 1.03E+00 | 3.55E-08 | 1.27E-07 | -1.68E+00 | 1.39E-05 | 3.91E-05 | -1.24E+00 | 1.41E-03 | 5.19E-03 | -1.06E+00 | 1.39E-01 | 3.81E-01 | 1.11E+00 |
| CD59 | 1.02E-13 | 5.30E-13 | -1.33E+00 | 1.59E-09 | 6.86E-09 | -1.04E+00 | 2.45E-02 | 1.04E-01 | -1.03E+00 | 1.34E-03 | 5.13E-03 | -1.01E+00 | 5.60E-03 | 2.00E-02 | 1.05E+00 | 1.00E-02 | 2.82E-02 | -1.07E+00 | 2.28E-01 | 8.36E-01 | 1.20E+00 | 2.13E-01 | 5.85E-01 | 1.12E+00 |
| CHMP5 | 1.02E-13 | 5.32E-13 | -1.27E+00 | 3.39E-14 | 1.46E-13 | -1.21E+00 | 1.14E-09 | 4.86E-09 | -1.43E+00 | 2.17E-09 | 8.29E-09 | -1.27E+00 | 2.29E-13 | 8.16E-13 | -1.55E+00 | 1.86E-10 | 5.25E-10 | -1.38E+00 | 4.55E-09 | 1.67E-08 | -1.28E+00 | 1.02E-08 | 2.80E-08 | -1.35E+00 |
| FNDC3A | 1.03E-13 | 5.34E-13 | -1.46E+00 | 1.72E-07 | 7.42E-07 | -1.02E+00 | 4.59E-06 | 1.95E-05 | -1.25E+00 | 1.28E-05 | 4.88E-05 | -1.10E+00 | 4.27E-04 | 1.52E-03 | -1.22E+00 | 5.00E-04 | 1.41E-03 | -1.25E+00 | 6.35E-08 | 2.33E-07 | -1.53E+00 | 2.21E-03 | 6.07E-03 | -1.15E+00 |
| RTF1 | 1.03E-13 | 5.35E-13 | -1.31E+00 | 6.70E-12 | 2.90E-11 | -1.15E+00 | 3.54E-04 | 1.50E-03 | -1.17E+00 | 2.13E-06 | 8.13E-06 | -1.21E+00 | 6.98E-07 | 2.49E-06 | -1.23E+00 | 1.67E-05 | 4.71E-05 | -1.10E+00 | 2.20E-03 | 8.07E-03 | -1.10E+00 | 1.24E-03 | 3.39E-03 | -1.14E+00 |
| ZYX | 1.03E-13 | 5.37E-13 | -1.27E+00 | 8.54E-11 | 3.69E-10 | -1.08E+00 | 3.56E-08 | 1.51E-07 | -1.42E+00 | 1.42E-13 | 5.41E-13 | -1.47E+00 | 3.48E-11 | 1.24E-10 | -1.65E+00 | 6.56E-12 | 1.85E-11 | -1.58E+00 | 2.52E-04 | 9.23E-04 | -1.44E+00 | 3.95E-03 | 1.09E-02 | -1.53E+00 |
| ALDH6A1 | 1.03E-13 | 5.37E-13 | -1.58E+00 | 7.66E-10 | 3.31E-09 | -1.18E+00 | 1.52E-10 | 6.47E-10 | -1.80E+00 | 1.15E-07 | 4.38E-07 | -1.29E+00 | 7.23E-15 | 2.58E-14 | -2.23E+00 | 1.16E-08 | 3.28E-08 | -1.33E+00 | 3.30E-13 | 1.21E-12 | -1.87E+00 | 1.91E-08 | 5.25E-08 | -1.37E+00 |
| TPRKBP1 | 1.04E-13 | 5.42E-13 | -1.82E+00 | 1.41E-10 | 6.08E-10 | -1.38E+00 | 1.66E-08 | 7.05E-08 | -1.66E+00 | 2.45E-08 | 9.37E-08 | -1.42E+00 | 4.63E-12 | 1.65E-11 | -2.40E+00 | 5.76E-08 | 1.62E-07 | -1.45E+00 | 7.27E-12 | 2.67E-11 | -1.77E+00 | 1.92E-08 | 5.27E-08 | -1.37E+00 |
| OASL | 1.04E-13 | 5.42E-13 | -4.20E+00 | 1.44E-13 | 6.23E-13 | -4.26E+00 | 5.52E-42 | 2.35E-41 | -7.03E+00 | 3.44E-43 | 1.31E-42 | -8.07E+00 | 8.24E-29 | 2.94E-28 | -7.06E+00 | 1.85E-29 | 5.21E-29 | -5.99E+00 | 5.30E-24 | 1.95E-23 | -5.01E+00 | 4.63E-20 | 1.27E-19 | -3.70E+00 |
| BICD1 | 1.05E-13 | 5.48E-13 | -1.62E+00 | 6.00E-07 | 2.59E-06 | -1.05E+00 | 1.90E-10 | 8.07E-10 | -1.65E+00 | 3.46E-04 | 1.32E-03 | 1.02E+00 | 1.09E-05 | 3.87E-05 | -1.76E+00 | 3.26E-03 | 9.18E-03 | -1.15E+00 | 8.25E-09 | 3.03E-08 | -1.79E+00 | 1.13E-03 | 3.10E-03 | -1.13E+00 |
| H3F3A | 1.06E-13 | 5.51E-13 | -1.26E+00 | 3.85E-14 | 1.66E-13 | -1.17E+00 | 4.38E-09 | 1.86E-08 | -1.42E+00 | 1.88E-07 | 7.18E-07 | -1.22E+00 | 2.43E-12 | 8.66E-12 | -1.47E+00 | 1.22E-10 | 3.44E-10 | -1.39E+00 | 5.60E-09 | 2.06E-08 | -1.43E+00 | 1.43E-06 | 3.94E-06 | -1.33E+00 |
| MPG | 1.06E-13 | 5.53E-13 | -2.86E+00 | 1.92E-09 | 8.29E-09 | -1.75E+00 | 3.86E-07 | 1.64E-06 | -1.79E+00 | 2.38E-06 | 9.08E-06 | -1.51E+00 | 3.58E-04 | 1.28E-03 | -2.63E+00 | 6.82E-03 | 1.92E-02 | -1.51E+00 | 3.27E-03 | 1.20E-02 | -1.50E+00 | 1.48E-01 | 4.07E-01 | 1.04E+00 |
| ACAA2 | 1.14E-13 | 5.96E-13 | -1.40E+00 | 7.32E-11 | 3.16E-10 | -1.18E+00 | 2.44E-09 | 1.03E-08 | -1.44E+00 | 2.96E-10 | 1.13E-09 | -1.34E+00 | 1.92E-06 | 6.83E-06 | -1.25E+00 | 1.25E-05 | 3.54E-05 | -1.18E+00 | 7.17E-09 | 2.63E-08 | -1.39E+00 | 5.33E-05 | 1.46E-04 | -1.14E+00 |
| USP39 | 1.15E-13 | 5.98E-13 | -1.71E+00 | 1.12E-08 | 4.85E-08 | -1.20E+00 | 1.98E-03 | 8.40E-03 | -1.09E+00 | 1.23E-04 | 4.72E-04 | -1.03E+00 | 2.01E-05 | 7.17E-05 | -1.46E+00 | 4.02E-03 | 1.13E-02 | -1.10E+00 | 1.44E-04 | 5.27E-04 | -1.02E+00 | 7.82E-03 | 2.15E-02 | 1.11E+00 |
| ANXA4 | 1.16E-13 | 6.03E-13 | -1.19E+00 | 2.06E-03 | 8.92E-03 | 1.24E+00 | 4.63E-05 | 1.97E-04 | -1.13E+00 | 8.82E-02 | 3.37E-01 | 1.24E+00 | 6.41E-14 | 2.29E-13 | -1.56E+00 | 1.31E-04 | 3.70E-04 | -1.01E+00 | 1.00E-02 | 3.67E-02 | -1.11E+00 | 3.60E-01 | 9.88E-01 | 1.20E+00 |
| CEP290 | 1.16E-13 | 6.06E-13 | -1.75E+00 | 2.62E-07 | 1.13E-06 | -1.22E+00 | 3.28E-07 | 1.39E-06 | -1.41E+00 | 1.64E-05 | 6.27E-05 | -1.11E+00 | 2.04E-07 | 7.26E-07 | -1.72E+00 | 1.09E-03 | 3.09E-03 | -1.15E+00 | 2.88E-05 | 1.06E-04 | -1.20E+00 | 2.00E-04 | 5.49E-04 | -1.19E+00 |
| NIPAL3 | 1.17E-13 | 6.11E-13 | -1.80E+00 | 7.71E-04 | 3.33E-03 | 1.17E+00 | 7.90E-10 | 3.36E-09 | -2.00E+00 | 3.75E-05 | 1.43E-04 | -1.27E+00 | 5.22E-07 | 1.86E-06 | -1.52E+00 | 1.38E-02 | 3.89E-02 | 1.13E+00 | 2.37E-09 | 8.68E-09 | -1.65E+00 | 1.36E-04 | 3.73E-04 | -1.21E+00 |
| CMPK2 | 1.18E-13 | 6.13E-13 | -2.55E+00 | 1.49E-09 | 6.44E-09 | -1.73E+00 | 2.60E-19 | 1.11E-18 | -3.10E+00 | 2.43E-11 | 9.30E-11 | -1.72E+00 | 2.26E-31 | 8.08E-31 | -2.89E+00 | 8.24E-25 | 2.32E-24 | -2.16E+00 | 1.71E-39 | 6.26E-39 | -2.74E+00 | 1.08E-18 | 2.97E-18 | -1.73E+00 |
| PRKCSH | 1.18E-13 | 6.15E-13 | -1.26E+00 | 5.43E-13 | 2.35E-12 | -1.13E+00 | 6.11E-05 | 2.60E-04 | -1.15E+00 | 9.02E-06 | 3.45E-05 | -1.07E+00 | 1.36E-03 | 4.86E-03 | -1.11E+00 | 2.07E-03 | 5.82E-03 | -1.05E+00 | 1.57E-05 | 5.77E-05 | -1.21E+00 | 3.09E-03 | 8.50E-03 | -1.05E+00 |
| RNF139 | 1.18E-13 | 6.16E-13 | -1.37E+00 | 1.73E-14 | 7.49E-14 | -1.31E+00 | 1.42E-05 | 6.04E-05 | -1.27E+00 | 5.84E-08 | 2.23E-07 | -1.29E+00 | 9.50E-06 | 3.39E-05 | -1.23E+00 | 1.75E-05 | 4.92E-05 | -1.17E+00 | 6.20E-06 | 2.27E-05 | -1.14E+00 | 1.02E-06 | 2.80E-06 | -1.27E+00 |
| FGFR1OP | 1.20E-13 | 6.24E-13 | -2.46E+00 | 2.94E-05 | 1.27E-04 | -1.18E+00 | 4.17E-10 | 1.77E-09 | -1.88E+00 | 2.14E-05 | 8.19E-05 | -1.14E+00 | 7.78E-03 | 2.78E-02 | -1.27E+00 | 1.52E-01 | 4.29E-01 | 1.07E+00 | 9.34E-07 | 3.43E-06 | -1.72E+00 | 2.44E-02 | 6.69E-02 | -1.02E+00 |
| ADPRHL2 | 1.20E-13 | 6.27E-13 | -1.85E+00 | 2.19E-09 | 9.46E-09 | -1.33E+00 | 1.73E-10 | 7.33E-10 | -1.59E+00 | 4.20E-08 | 1.60E-07 | -1.27E+00 | 1.65E-09 | 5.88E-09 | -2.13E+00 | 1.75E-04 | 4.94E-04 | -1.16E+00 | 1.27E-11 | 4.66E-11 | -1.61E+00 | 1.40E-03 | 3.85E-03 | -1.01E+00 |
| EXOC5 | 1.21E-13 | 6.28E-13 | -1.70E+00 | 6.50E-09 | 2.81E-08 | -1.31E+00 | 6.31E-05 | 2.68E-04 | -1.18E+00 | 1.16E-03 | 4.44E-03 | 1.04E+00 | 1.30E-07 | 4.63E-07 | -1.45E+00 | 5.56E-05 | 1.57E-04 | -1.22E+00 | 2.79E-08 | 1.02E-07 | -1.24E+00 | 1.88E-03 | 5.16E-03 | 1.01E+00 |
| DYM | 1.23E-13 | 6.40E-13 | -1.38E+00 | 1.50E-06 | 6.47E-06 | 1.08E+00 | 4.34E-05 | 1.84E-04 | -1.27E+00 | 2.44E-04 | 9.31E-04 | -1.07E+00 | 6.02E-06 | 2.15E-05 | -1.19E+00 | 7.50E-04 | 2.11E-03 | -1.04E+00 | 2.93E-04 | 1.07E-03 | -1.17E+00 | 9.99E-03 | 2.75E-02 | -1.09E+00 |
| NPM1P5 | 1.26E-13 | 6.59E-13 | -1.49E+00 | 3.49E-17 | 1.51E-16 | -1.63E+00 | 6.73E-13 | 2.86E-12 | -1.64E+00 | 3.90E-14 | 1.49E-13 | -1.52E+00 | 1.19E-08 | 4.26E-08 | -1.44E+00 | 8.05E-08 | 2.27E-07 | -1.31E+00 | 5.51E-09 | 2.02E-08 | -1.47E+00 | 1.62E-07 | 4.44E-07 | -1.43E+00 |
| TFAM | 1.29E-13 | 6.71E-13 | -1.36E+00 | 1.83E-22 | 7.92E-22 | -1.45E+00 | 9.41E-07 | 4.00E-06 | -1.19E+00 | 5.29E-08 | 2.02E-07 | -1.18E+00 | 2.19E-07 | 7.82E-07 | -1.20E+00 | 2.49E-08 | 7.02E-08 | -1.21E+00 | 3.27E-07 | 1.20E-06 | -1.30E+00 | 4.77E-04 | 1.31E-03 | -1.06E+00 |
| G3BP2 | 1.29E-13 | 6.74E-13 | -1.28E+00 | 5.66E-26 | 2.45E-25 | -1.48E+00 | 8.52E-05 | 3.62E-04 | -1.16E+00 | 1.26E-08 | 4.81E-08 | -1.29E+00 | 5.92E-06 | 2.11E-05 | -1.23E+00 | 1.45E-09 | 4.09E-09 | -1.37E+00 | 1.40E-03 | 5.16E-03 | -1.07E+00 | 4.16E-04 | 1.14E-03 | -1.12E+00 |
| S100PBP | 1.32E-13 | 6.86E-13 | -1.58E+00 | 1.08E-07 | 4.66E-07 | 1.00E+00 | 6.89E-06 | 2.93E-05 | -1.51E+00 | 3.57E-02 | 1.37E-01 | 1.10E+00 | 1.95E-14 | 6.94E-14 | -2.07E+00 | 6.76E-06 | 1.91E-05 | -1.21E+00 | 2.76E-08 | 1.01E-07 | -1.58E+00 | 6.12E-05 | 1.68E-04 | -1.19E+00 |
| ITGB1 | 1.32E-13 | 6.89E-13 | -1.50E+00 | 1.15E-17 | 4.99E-17 | -1.62E+00 | 1.27E-05 | 5.39E-05 | -1.17E+00 | 9.66E-15 | 3.69E-14 | -1.57E+00 | 1.23E-05 | 4.38E-05 | -1.16E+00 | 2.91E-08 | 8.19E-08 | -1.40E+00 | 3.66E-05 | 1.34E-04 | -1.13E+00 | 4.56E-14 | 1.25E-13 | -1.65E+00 |
| IVD | 1.32E-13 | 6.90E-13 | -1.52E+00 | 2.12E-04 | 9.18E-04 | 1.11E+00 | 3.64E-09 | 1.55E-08 | -1.41E+00 | 1.47E-03 | 5.61E-03 | 1.06E+00 | 3.72E-10 | 1.33E-09 | -1.40E+00 | 4.54E-06 | 1.28E-05 | -1.13E+00 | 1.06E-07 | 3.89E-07 | -1.31E+00 | 6.20E-03 | 1.70E-02 | 1.02E+00 |
| JADE1 | 1.36E-13 | 7.08E-13 | -1.62E+00 | 2.07E-14 | 8.95E-14 | -1.45E+00 | 2.50E-09 | 1.06E-08 | -1.46E+00 | 4.70E-11 | 1.80E-10 | -1.40E+00 | 3.30E-04 | 1.18E-03 | -1.22E+00 | 1.03E-04 | 2.91E-04 | -1.27E+00 | 5.68E-16 | 2.08E-15 | -1.96E+00 | 8.17E-09 | 2.24E-08 | -1.49E+00 |
| TMEM45A | 1.36E-13 | 7.11E-13 | -2.29E+00 | 1.35E-08 | 5.85E-08 | -1.35E+00 | 9.41E-06 | 4.00E-05 | -2.41E+00 | 2.32E-03 | 8.87E-03 | -1.29E+00 | 4.89E-04 | 1.74E-03 | -1.19E+00 | 1.86E-02 | 5.25E-02 | 1.06E+00 | 3.54E-08 | 1.30E-07 | -1.49E+00 | 1.51E-04 | 4.16E-04 | -1.23E+00 |
| PPP1R15A | 1.37E-13 | 7.14E-13 | 1.92E+00 | 9.78E-22 | 4.22E-21 | -1.27E+00 | 5.68E-10 | 2.41E-09 | 1.86E+00 | 3.05E-05 | 1.17E-04 | -1.13E+00 | 7.90E-07 | 2.82E-06 | 1.89E+00 | 6.92E-06 | 1.95E-05 | -1.17E+00 | 3.13E-04 | 1.15E-03 | 1.58E+00 | 3.74E-06 | 1.03E-05 | -1.29E+00 |
| NCBP3 | 1.37E-13 | 7.14E-13 | -1.47E+00 | 2.89E-08 | 1.25E-07 | -1.08E+00 | 2.55E-11 | 1.08E-10 | -1.58E+00 | 1.58E-10 | 6.05E-10 | -1.29E+00 | 1.39E-08 | 4.94E-08 | -1.30E+00 | 4.70E-07 | 1.32E-06 | -1.17E+00 | 1.09E-06 | 3.99E-06 | -1.38E+00 | 7.49E-06 | 2.06E-05 | -1.31E+00 |
| HLA-DPB1 | 1.39E-13 | 7.24E-13 | -1.86E+00 | 8.25E-06 | 3.56E-05 | -1.42E+00 | 2.44E-08 | 1.04E-07 | -1.79E+00 | 2.34E-03 | 8.95E-03 | -1.33E+00 | 3.96E-36 | 1.41E-35 | -2.18E+00 | 2.83E-26 | 7.99E-26 | -1.79E+00 | 1.78E-89 | 6.52E-89 | -3.07E+00 | 1.18E-30 | 3.23E-30 | -1.90E+00 |
| NIFK | 1.40E-13 | 7.28E-13 | -1.22E+00 | 1.37E-06 | 5.94E-06 | 1.07E+00 | 1.13E-08 | 4.78E-08 | -1.39E+00 | 5.10E-04 | 1.95E-03 | -1.00E+00 | 4.46E-14 | 1.59E-13 | -1.61E+00 | 3.40E-05 | 9.59E-05 | -1.12E+00 | 5.69E-12 | 2.09E-11 | -1.47E+00 | 4.45E-03 | 1.22E-02 | -1.07E+00 |
| NRDC | 1.40E-13 | 7.30E-13 | -1.36E+00 | 4.11E-08 | 1.77E-07 | -1.02E+00 | 6.01E-07 | 2.55E-06 | -1.35E+00 | 3.17E-03 | 1.21E-02 | 1.01E+00 | 1.62E-03 | 5.77E-03 | -1.01E+00 | 2.42E-02 | 6.83E-02 | 1.10E+00 | 2.91E-03 | 1.07E-02 | -1.10E+00 | 5.47E-03 | 1.50E-02 | -1.14E+00 |
| SLC30A5 | 1.40E-13 | 7.30E-13 | -1.56E+00 | 5.95E-10 | 2.57E-09 | -1.19E+00 | 3.79E-04 | 1.61E-03 | -1.09E+00 | 2.13E-04 | 8.12E-04 | 1.03E+00 | 2.96E-04 | 1.06E-03 | 1.00E+00 | 1.83E-02 | 5.16E-02 | 1.14E+00 | 2.02E-01 | 7.40E-01 | 1.14E+00 | 2.59E-01 | 7.11E-01 | 1.21E+00 |
| APOOL | 1.41E-13 | 7.33E-13 | -1.50E+00 | 1.33E-09 | 5.75E-09 | -1.17E+00 | 1.24E-02 | 5.28E-02 | -1.10E+00 | 1.02E-04 | 3.90E-04 | -1.10E+00 | 1.66E-04 | 5.93E-04 | -1.17E+00 | 3.74E-02 | 1.05E-01 | 1.16E+00 | 1.03E-02 | 3.77E-02 | 1.06E+00 | 1.39E-03 | 3.81E-03 | 1.01E+00 |
| POU2F2 | 1.42E-13 | 7.41E-13 | -1.56E+00 | 5.07E-07 | 2.19E-06 | -1.08E+00 | 3.22E-24 | 1.37E-23 | -2.18E+00 | 2.90E-17 | 1.11E-16 | -1.52E+00 | 6.64E-03 | 2.37E-02 | -1.52E+00 | 2.15E-02 | 6.05E-02 | -1.20E+00 | 1.88E-02 | 6.90E-02 | -1.92E+00 | 5.22E-02 | 1.43E-01 | -1.80E+00 |
| COMMD9 | 1.43E-13 | 7.45E-13 | -1.32E+00 | 8.31E-09 | 3.59E-08 | -1.02E+00 | 7.34E-07 | 3.12E-06 | -1.18E+00 | 5.80E-06 | 2.21E-05 | -1.06E+00 | 1.16E-11 | 4.12E-11 | -1.40E+00 | 5.50E-09 | 1.55E-08 | -1.23E+00 | 1.84E-05 | 6.74E-05 | -1.12E+00 | 3.60E-03 | 9.89E-03 | -1.01E+00 |
| GUCY1A3 | 1.45E-13 | 7.58E-13 | -1.39E+00 | 3.37E-05 | 1.46E-04 | 1.02E+00 | 4.07E-11 | 1.73E-10 | -1.43E+00 | 2.03E-04 | 7.77E-04 | 1.04E+00 | 4.14E-03 | 1.48E-02 | -1.02E+00 | 8.75E-02 | 2.47E-01 | 1.28E+00 | 7.53E-03 | 2.76E-02 | -1.07E+00 | 2.29E-02 | 6.29E-02 | -1.08E+00 |
| ARMCX3 | 1.46E-13 | 7.63E-13 | -1.30E+00 | 6.38E-09 | 2.76E-08 | -1.07E+00 | 1.21E-04 | 5.16E-04 | -1.23E+00 | 6.14E-06 | 2.34E-05 | -1.16E+00 | 3.11E-08 | 1.11E-07 | -1.32E+00 | 1.43E-04 | 4.04E-04 | -1.19E+00 | 2.60E-05 | 9.55E-05 | -1.04E+00 | 2.54E-02 | 6.99E-02 | 1.03E+00 |
| DCAF11 | 1.50E-13 | 7.83E-13 | -1.49E+00 | 2.59E-11 | 1.12E-10 | -1.21E+00 | 1.85E-07 | 7.85E-07 | -1.26E+00 | 2.25E-08 | 8.59E-08 | -1.18E+00 | 5.68E-08 | 2.03E-07 | -1.41E+00 | 1.20E-07 | 3.38E-07 | -1.29E+00 | 2.96E-04 | 1.09E-03 | -1.10E+00 | 2.17E-02 | 5.96E-02 | -1.01E+00 |
| PGS1 | 1.50E-13 | 7.83E-13 | -2.10E+00 | 4.53E-04 | 1.96E-03 | 1.04E+00 | 8.40E-08 | 3.57E-07 | -1.60E+00 | 3.09E-04 | 1.18E-03 | -1.08E+00 | 1.92E-10 | 6.84E-10 | -1.84E+00 | 1.04E-05 | 2.93E-05 | -1.26E+00 | 6.80E-07 | 2.50E-06 | -1.28E+00 | 5.27E-04 | 1.45E-03 | -1.06E+00 |
| SUMO2 | 1.51E-13 | 7.87E-13 | -1.25E+00 | 6.64E-08 | 2.87E-07 | -1.12E+00 | 6.17E-08 | 2.62E-07 | -1.33E+00 | 2.70E-05 | 1.03E-04 | -1.22E+00 | 3.15E-13 | 1.12E-12 | -1.43E+00 | 2.21E-07 | 6.24E-07 | -1.28E+00 | 4.75E-07 | 1.74E-06 | -1.28E+00 | 1.66E-03 | 4.56E-03 | -1.17E+00 |
| MTFR1L | 1.51E-13 | 7.88E-13 | -1.65E+00 | 5.15E-08 | 2.23E-07 | -1.14E+00 | 1.14E-03 | 4.85E-03 | -1.17E+00 | 1.70E-03 | 6.50E-03 | -1.01E+00 | 8.97E-06 | 3.20E-05 | -1.24E+00 | 1.84E-01 | 5.20E-01 | 1.17E+00 | 3.21E-08 | 1.18E-07 | -1.14E+00 | 3.29E-04 | 9.05E-04 | 1.03E+00 |
| FIP1L1 | 1.53E-13 | 7.98E-13 | -1.43E+00 | 1.10E-06 | 4.76E-06 | 1.04E+00 | 5.06E-07 | 2.15E-06 | -1.32E+00 | 2.26E-04 | 8.64E-04 | -1.00E+00 | 3.15E-07 | 1.12E-06 | -1.41E+00 | 4.72E-04 | 1.33E-03 | -1.08E+00 | 9.36E-14 | 3.44E-13 | -1.54E+00 | 7.23E-08 | 1.99E-07 | -1.27E+00 |
| TBC1D20 | 1.59E-13 | 8.28E-13 | -1.75E+00 | 1.67E-07 | 7.23E-07 | -1.15E+00 | 1.55E-05 | 6.59E-05 | -1.51E+00 | 1.01E-02 | 3.86E-02 | 1.04E+00 | 6.88E-03 | 2.45E-02 | -1.09E+00 | 1.22E-01 | 3.45E-01 | 1.17E+00 | 3.99E-05 | 1.47E-04 | -1.20E+00 | 5.73E-03 | 1.57E-02 | -1.01E+00 |
| XRCC6 | 1.64E-13 | 8.56E-13 | -1.14E+00 | 1.24E-04 | 5.35E-04 | 1.18E+00 | 5.81E-02 | 2.47E-01 | -1.00E+00 | 1.08E-01 | 4.12E-01 | 1.12E+00 | 1.91E-07 | 6.81E-07 | -1.18E+00 | 3.30E-02 | 9.29E-02 | 1.09E+00 | 7.67E-02 | 2.81E-01 | -1.04E+00 | 1.92E-01 | 5.26E-01 | 1.16E+00 |
| DNAJC4 | 1.64E-13 | 8.56E-13 | -1.33E+00 | 1.41E-10 | 6.09E-10 | -1.13E+00 | 3.74E-05 | 1.59E-04 | -1.23E+00 | 2.77E-06 | 1.06E-05 | -1.11E+00 | 1.33E-06 | 4.73E-06 | -1.20E+00 | 4.10E-08 | 1.15E-07 | -1.21E+00 | 6.90E-05 | 2.53E-04 | -1.07E+00 | 1.60E-04 | 4.40E-04 | -1.11E+00 |
| MBTD1 | 1.65E-13 | 8.58E-13 | -1.55E+00 | 1.97E-05 | 8.49E-05 | 1.02E+00 | 1.36E-09 | 5.78E-09 | -1.51E+00 | 4.69E-06 | 1.79E-05 | -1.10E+00 | 1.07E-05 | 3.83E-05 | -1.37E+00 | 7.71E-03 | 2.17E-02 | -1.05E+00 | 6.79E-06 | 2.49E-05 | -1.19E+00 | 7.57E-03 | 2.08E-02 | -1.02E+00 |
| GLOD4 | 1.68E-13 | 8.77E-13 | -1.42E+00 | 5.72E-04 | 2.47E-03 | 1.15E+00 | 5.92E-03 | 2.52E-02 | -1.05E+00 | 1.39E-01 | 5.31E-01 | 1.24E+00 | 7.08E-11 | 2.52E-10 | -1.63E+00 | 1.42E-02 | 4.00E-02 | 1.13E+00 | 1.08E-04 | 3.98E-04 | -1.07E+00 | 2.96E-01 | 8.14E-01 | 1.24E+00 |
| KIAA2026 | 1.69E-13 | 8.79E-13 | -1.55E+00 | 1.76E-06 | 7.61E-06 | 1.01E+00 | 3.68E-06 | 1.57E-05 | -1.31E+00 | 8.02E-06 | 3.06E-05 | -1.17E+00 | 3.76E-06 | 1.34E-05 | -1.29E+00 | 9.93E-07 | 2.80E-06 | -1.25E+00 | 4.54E-06 | 1.67E-05 | -1.35E+00 | 1.68E-05 | 4.62E-05 | -1.36E+00 |
| CNDP2 | 1.69E-13 | 8.79E-13 | -1.46E+00 | 7.87E-08 | 3.40E-07 | -1.05E+00 | 3.05E-06 | 1.29E-05 | -1.22E+00 | 2.80E-05 | 1.07E-04 | -1.00E+00 | 1.80E-07 | 6.41E-07 | -1.53E+00 | 8.91E-03 | 2.51E-02 | 1.01E+00 | 7.85E-07 | 2.88E-06 | -1.29E+00 | 6.38E-02 | 1.75E-01 | 1.09E+00 |
| TM2D1 | 1.69E-13 | 8.80E-13 | -1.30E+00 | 1.07E-05 | 4.61E-05 | 1.11E+00 | 4.02E-08 | 1.71E-07 | -1.44E+00 | 7.29E-03 | 2.79E-02 | 1.05E+00 | 2.57E-04 | 9.15E-04 | -1.12E+00 | 1.75E-02 | 4.93E-02 | 1.11E+00 | 2.82E-04 | 1.03E-03 | -1.12E+00 | 4.13E-02 | 1.13E-01 | 1.04E+00 |
| B3GNT2 | 1.70E-13 | 8.86E-13 | -1.56E+00 | 1.20E-10 | 5.18E-10 | -1.23E+00 | 1.19E-13 | 5.08E-13 | -1.91E+00 | 2.88E-12 | 1.10E-11 | -1.51E+00 | 6.79E-06 | 2.42E-05 | -1.17E+00 | 7.70E-05 | 2.17E-04 | -1.12E+00 | 6.12E-10 | 2.25E-09 | -1.27E+00 | 3.67E-14 | 1.01E-13 | -1.50E+00 |
| MZT1 | 1.70E-13 | 8.86E-13 | -1.53E+00 | 3.46E-13 | 1.49E-12 | -1.31E+00 | 2.17E-05 | 9.23E-05 | -1.17E+00 | 5.42E-06 | 2.07E-05 | -1.10E+00 | 4.57E-07 | 1.63E-06 | -1.33E+00 | 2.58E-05 | 7.26E-05 | -1.18E+00 | 6.55E-08 | 2.40E-07 | -1.29E+00 | 6.60E-06 | 1.81E-05 | -1.19E+00 |
| NIN | 1.71E-13 | 8.89E-13 | -1.51E+00 | 4.99E-10 | 2.16E-09 | -1.21E+00 | 7.99E-12 | 3.39E-11 | -1.52E+00 | 3.67E-07 | 1.40E-06 | -1.13E+00 | 1.63E-09 | 5.81E-09 | -1.59E+00 | 1.40E-07 | 3.96E-07 | -1.48E+00 | 1.35E-05 | 4.95E-05 | -1.20E+00 | 2.54E-03 | 6.98E-03 | -1.08E+00 |
| ZNF106 | 1.71E-13 | 8.93E-13 | -1.56E+00 | 2.73E-08 | 1.18E-07 | -1.15E+00 | 7.87E-05 | 3.34E-04 | -1.24E+00 | 9.73E-05 | 3.72E-04 | -1.10E+00 | 1.98E-07 | 7.06E-07 | -1.46E+00 | 1.77E-05 | 4.99E-05 | -1.19E+00 | 3.17E-07 | 1.16E-06 | -1.25E+00 | 8.88E-05 | 2.44E-04 | -1.21E+00 |
| ACAP2 | 1.71E-13 | 8.93E-13 | -1.35E+00 | 1.47E-07 | 6.33E-07 | -1.00E+00 | 1.20E-06 | 5.09E-06 | -1.31E+00 | 2.01E-05 | 7.66E-05 | -1.08E+00 | 2.68E-08 | 9.54E-08 | -1.33E+00 | 1.39E-07 | 3.93E-07 | -1.14E+00 | 7.64E-09 | 2.80E-08 | -1.43E+00 | 9.99E-05 | 2.74E-04 | -1.18E+00 |
| ATP2C1 | 1.73E-13 | 9.04E-13 | -1.31E+00 | 6.02E-25 | 2.60E-24 | -1.54E+00 | 1.05E-07 | 4.45E-07 | -1.41E+00 | 3.72E-10 | 1.42E-09 | -1.45E+00 | 1.72E-08 | 6.15E-08 | -1.45E+00 | 1.97E-10 | 5.56E-10 | -1.54E+00 | 1.78E-05 | 6.53E-05 | -1.23E+00 | 1.72E-05 | 4.74E-05 | -1.24E+00 |
| LSM3P3 | 1.75E-13 | 9.11E-13 | -2.30E+00 | 3.43E-11 | 1.48E-10 | -1.74E+00 | 9.62E-11 | 4.09E-10 | -1.74E+00 | 9.66E-12 | 3.69E-11 | -1.63E+00 | 2.96E-08 | 1.05E-07 | -1.75E+00 | 1.11E-06 | 3.12E-06 | -1.47E+00 | 7.26E-21 | 2.66E-20 | -2.25E+00 | 1.05E-09 | 2.88E-09 | -1.41E+00 |
| AC005480.1 | 1.75E-13 | 9.11E-13 | -2.50E+00 | 5.24E-13 | 2.26E-12 | -2.14E+00 | 3.48E-11 | 1.48E-10 | -2.30E+00 | 1.18E-11 | 4.52E-11 | -2.19E+00 | 5.08E-07 | 1.81E-06 | -2.41E+00 | 7.16E-07 | 2.02E-06 | -2.02E+00 | 2.34E-05 | 8.58E-05 | -1.56E+00 | 5.85E-08 | 1.61E-07 | -2.12E+00 |
| PAXIP1-AS2 | 1.75E-13 | 9.13E-13 | -1.89E+00 | 2.46E-07 | 1.06E-06 | -1.20E+00 | 6.44E-11 | 2.74E-10 | -2.13E+00 | 6.49E-08 | 2.48E-07 | -1.52E+00 | 1.28E-08 | 4.55E-08 | -1.94E+00 | 4.30E-07 | 1.21E-06 | -1.56E+00 | 1.58E-07 | 5.80E-07 | -1.67E+00 | 1.25E-06 | 3.42E-06 | -1.58E+00 |
| TLK1 | 1.78E-13 | 9.29E-13 | -1.31E+00 | 8.02E-07 | 3.47E-06 | 1.04E+00 | 3.72E-05 | 1.58E-04 | -1.35E+00 | 4.19E-03 | 1.60E-02 | -1.04E+00 | 1.92E-05 | 6.84E-05 | -1.20E+00 | 7.54E-04 | 2.13E-03 | -1.00E+00 | 2.55E-04 | 9.34E-04 | -1.06E+00 | 2.38E-02 | 6.52E-02 | 1.06E+00 |
| MAP1LC3B2 | 1.79E-13 | 9.31E-13 | -1.43E+00 | 3.76E-19 | 1.63E-18 | -1.65E+00 | 1.24E-09 | 5.26E-09 | -1.55E+00 | 1.42E-14 | 5.44E-14 | -1.70E+00 | 1.61E-09 | 5.73E-09 | -1.60E+00 | 4.56E-14 | 1.28E-13 | -1.86E+00 | 7.72E-06 | 2.83E-05 | -1.17E+00 | 1.19E-11 | 3.26E-11 | -1.74E+00 |
| AC021723.1 | 1.79E-13 | 9.31E-13 | -1.33E+00 | 6.32E-32 | 2.73E-31 | -2.07E+00 | 5.89E-07 | 2.50E-06 | -1.23E+00 | 5.90E-26 | 2.25E-25 | -2.10E+00 | 1.20E-05 | 4.29E-05 | -1.18E+00 | 2.60E-11 | 7.33E-11 | -1.75E+00 | 1.24E-07 | 4.55E-07 | -1.30E+00 | 6.45E-12 | 1.77E-11 | -1.68E+00 |
| NSMCE1 | 1.80E-13 | 9.41E-13 | -1.25E+00 | 3.50E-12 | 1.51E-11 | -1.11E+00 | 6.26E-05 | 2.66E-04 | -1.13E+00 | 1.52E-04 | 5.79E-04 | 1.01E+00 | 3.24E-06 | 1.16E-05 | -1.12E+00 | 2.11E-04 | 5.95E-04 | -1.02E+00 | 4.32E-02 | 1.59E-01 | 1.04E+00 | 7.73E-02 | 2.12E-01 | 1.05E+00 |
| TMEM59 | 1.81E-13 | 9.44E-13 | -1.27E+00 | 7.44E-05 | 3.21E-04 | 1.02E+00 | 2.41E-04 | 1.02E-03 | -1.18E+00 | 3.06E-02 | 1.17E-01 | 1.00E+00 | 1.73E-03 | 6.16E-03 | -1.11E+00 | 1.32E-01 | 3.71E-01 | 1.06E+00 | 1.07E-01 | 3.91E-01 | 1.03E+00 | 2.41E-01 | 6.62E-01 | 1.08E+00 |
| C19orf66 | 1.84E-13 | 9.59E-13 | -1.48E+00 | 2.96E-08 | 1.28E-07 | -1.12E+00 | 1.44E-15 | 6.11E-15 | -1.88E+00 | 1.15E-09 | 4.41E-09 | -1.37E+00 | 1.87E-12 | 6.67E-12 | -1.81E+00 | 9.40E-10 | 2.65E-09 | -1.61E+00 | 1.98E-08 | 7.25E-08 | -1.43E+00 | 1.86E-06 | 5.10E-06 | -1.31E+00 |
| BCAT2 | 1.86E-13 | 9.67E-13 | -1.97E+00 | 2.46E-05 | 1.06E-04 | 1.01E+00 | 3.69E-10 | 1.57E-09 | -1.71E+00 | 1.75E-05 | 6.70E-05 | -1.09E+00 | 1.34E-04 | 4.76E-04 | -1.97E+00 | 7.94E-02 | 2.24E-01 | 1.07E+00 | 1.98E-09 | 7.27E-09 | -1.54E+00 | 4.36E-04 | 1.20E-03 | -1.09E+00 |
| SLC25A6P2 | 1.86E-13 | 9.68E-13 | -2.03E+00 | 2.44E-11 | 1.05E-10 | -1.55E+00 | 2.80E-07 | 1.19E-06 | -1.71E+00 | 7.57E-05 | 2.89E-04 | -1.22E+00 | 2.58E-08 | 9.20E-08 | -1.88E+00 | 3.07E-07 | 8.65E-07 | -1.60E+00 | 3.32E-22 | 1.22E-21 | -2.69E+00 | 1.63E-15 | 4.48E-15 | -1.95E+00 |
| GTF2A2 | 1.86E-13 | 9.69E-13 | -1.30E+00 | 1.26E-14 | 5.43E-14 | -1.13E+00 | 5.21E-03 | 2.21E-02 | -1.10E+00 | 5.05E-02 | 1.93E-01 | 1.07E+00 | 1.71E-06 | 6.08E-06 | -1.15E+00 | 5.86E-04 | 1.65E-03 | -1.05E+00 | 1.77E-01 | 6.48E-01 | 1.06E+00 | 2.05E-01 | 5.64E-01 | -1.02E+00 |
| STAMBP | 1.89E-13 | 9.85E-13 | -1.38E+00 | 2.49E-11 | 1.07E-10 | -1.14E+00 | 3.05E-06 | 1.30E-05 | -1.32E+00 | 1.19E-05 | 4.53E-05 | -1.15E+00 | 1.87E-04 | 6.66E-04 | -1.13E+00 | 2.38E-04 | 6.70E-04 | -1.15E+00 | 4.65E-04 | 1.71E-03 | -1.18E+00 | 4.70E-04 | 1.29E-03 | -1.16E+00 |
| AGL | 1.93E-13 | 1.00E-12 | -1.90E+00 | 2.59E-08 | 1.12E-07 | -1.25E+00 | 9.70E-07 | 4.12E-06 | -1.44E+00 | 3.48E-05 | 1.33E-04 | -1.15E+00 | 4.30E-02 | 1.53E-01 | 1.01E+00 | 7.87E-02 | 2.22E-01 | 1.13E+00 | 1.25E-08 | 4.59E-08 | -1.48E+00 | 2.51E-07 | 6.89E-07 | -1.37E+00 |
| TWSG1 | 1.93E-13 | 1.00E-12 | -1.97E+00 | 2.77E-10 | 1.20E-09 | -1.55E+00 | 3.24E-04 | 1.38E-03 | -1.23E+00 | 2.90E-04 | 1.11E-03 | -1.18E+00 | 1.16E-03 | 4.13E-03 | -1.15E+00 | 4.82E-03 | 1.36E-02 | 1.02E+00 | 5.35E-08 | 1.96E-07 | -1.29E+00 | 8.44E-04 | 2.32E-03 | -1.12E+00 |
| PTGS1 | 1.93E-13 | 1.01E-12 | -1.69E+00 | 1.06E-05 | 4.56E-05 | -1.04E+00 | 3.66E-03 | 1.55E-02 | -1.24E+00 | 2.48E-01 | 9.48E-01 | 1.55E+00 | 7.67E-04 | 2.74E-03 | 1.02E+00 | 9.09E-02 | 2.56E-01 | 1.16E+00 | 2.70E-05 | 9.90E-05 | -1.04E+00 | 3.93E-03 | 1.08E-02 | 1.03E+00 |
| NKX3-1 | 1.95E-13 | 1.01E-12 | -2.15E+00 | 3.41E-14 | 1.47E-13 | -2.21E+00 | 9.43E-07 | 4.01E-06 | -2.18E+00 | 3.81E-04 | 1.46E-03 | -1.33E+00 | 1.17E-06 | 4.17E-06 | -2.26E+00 | 4.05E-05 | 1.14E-04 | -1.73E+00 | 3.21E-03 | 1.18E-02 | -1.55E+00 | 1.28E-03 | 3.53E-03 | -1.69E+00 |
| ASH1L | 1.98E-13 | 1.03E-12 | -1.34E+00 | 1.27E-11 | 5.48E-11 | -1.12E+00 | 4.25E-06 | 1.81E-05 | -1.20E+00 | 4.53E-08 | 1.73E-07 | -1.16E+00 | 1.31E-04 | 4.68E-04 | -1.08E+00 | 9.16E-06 | 2.58E-05 | -1.14E+00 | 1.42E-07 | 5.20E-07 | -1.18E+00 | 1.85E-11 | 5.07E-11 | -1.41E+00 |
| THAP11 | 1.98E-13 | 1.03E-12 | -1.72E+00 | 3.43E-04 | 1.48E-03 | 1.05E+00 | 5.67E-16 | 2.41E-15 | -1.75E+00 | 7.03E-07 | 2.69E-06 | -1.08E+00 | 2.96E-05 | 1.05E-04 | -1.60E+00 | 2.59E-01 | 7.30E-01 | 1.29E+00 | 1.85E-07 | 6.80E-07 | -1.31E+00 | 2.79E-02 | 7.66E-02 | 1.03E+00 |
| MRPS34 | 1.99E-13 | 1.04E-12 | -1.23E+00 | 2.01E-08 | 8.68E-08 | -1.01E+00 | 1.89E-02 | 8.05E-02 | -1.01E+00 | 4.32E-02 | 1.65E-01 | 1.17E+00 | 1.76E-11 | 6.27E-11 | -1.39E+00 | 6.30E-03 | 1.78E-02 | 1.10E+00 | 6.11E-03 | 2.24E-02 | 1.00E+00 | 2.31E-01 | 6.33E-01 | 1.25E+00 |
| RCC2 | 2.02E-13 | 1.05E-12 | -1.45E+00 | 3.87E-10 | 1.67E-09 | -1.12E+00 | 4.57E-11 | 1.94E-10 | -1.48E+00 | 1.56E-07 | 5.96E-07 | -1.16E+00 | 6.02E-07 | 2.15E-06 | -1.44E+00 | 4.87E-05 | 1.37E-04 | -1.18E+00 | 2.81E-09 | 1.03E-08 | -1.41E+00 | 5.97E-04 | 1.64E-03 | -1.11E+00 |
| NIFKP1 | 2.05E-13 | 1.07E-12 | -2.04E+00 | 1.15E-14 | 4.95E-14 | -2.04E+00 | 6.09E-15 | 2.59E-14 | -1.99E+00 | 1.54E-17 | 5.86E-17 | -1.97E+00 | 1.24E-08 | 4.42E-08 | -2.03E+00 | 1.68E-08 | 4.74E-08 | -1.91E+00 | 2.46E-10 | 9.02E-10 | -1.78E+00 | 5.26E-11 | 1.44E-10 | -1.75E+00 |
| RNF20 | 2.06E-13 | 1.07E-12 | -1.65E+00 | 1.87E-05 | 8.06E-05 | 1.02E+00 | 2.84E-07 | 1.21E-06 | -1.37E+00 | 1.90E-04 | 7.25E-04 | -1.05E+00 | 7.25E-08 | 2.59E-07 | -1.44E+00 | 3.95E-03 | 1.11E-02 | -1.02E+00 | 2.36E-07 | 8.67E-07 | -1.33E+00 | 9.07E-04 | 2.49E-03 | -1.04E+00 |
| AL590135.1 | 2.07E-13 | 1.08E-12 | -1.34E+00 | 3.39E-61 | 1.46E-60 | -2.19E+00 | 1.62E-04 | 6.87E-04 | -1.29E+00 | 1.85E-21 | 7.05E-21 | -1.95E+00 | 5.96E-08 | 2.13E-07 | -1.29E+00 | 8.69E-40 | 2.45E-39 | -2.43E+00 | 3.72E-06 | 1.37E-05 | -1.35E+00 | 4.90E-25 | 1.35E-24 | -2.25E+00 |
| SVIL | 2.07E-13 | 1.08E-12 | -1.75E+00 | 2.83E-06 | 1.22E-05 | -1.07E+00 | 2.38E-06 | 1.01E-05 | -1.53E+00 | 3.60E-03 | 1.38E-02 | 1.06E+00 | 6.76E-04 | 2.41E-03 | -1.80E+00 | 3.31E-03 | 9.32E-03 | -1.61E+00 | 7.56E-03 | 2.77E-02 | -1.69E+00 | 3.73E-02 | 1.03E-01 | -1.34E+00 |
| ZNF608 | 2.10E-13 | 1.09E-12 | -2.52E+00 | 7.07E-14 | 3.05E-13 | -2.39E+00 | 6.51E-05 | 2.76E-04 | -1.76E+00 | 1.10E-06 | 4.19E-06 | -1.98E+00 | 3.47E-04 | 1.24E-03 | -2.51E+00 | 1.16E-04 | 3.28E-04 | -3.00E+00 | 1.13E-01 | 4.16E-01 | -1.45E+00 | 1.73E-01 | 4.74E-01 | -1.47E+00 |
| TMEM230 | 2.10E-13 | 1.09E-12 | -1.25E+00 | 1.73E-11 | 7.49E-11 | -1.04E+00 | 2.13E-02 | 9.05E-02 | -1.04E+00 | 1.46E-01 | 5.60E-01 | 1.17E+00 | 2.24E-07 | 8.00E-07 | -1.19E+00 | 1.69E-04 | 4.76E-04 | -1.02E+00 | 9.45E-05 | 3.47E-04 | -1.16E+00 | 7.48E-02 | 2.05E-01 | 1.02E+00 |
| WDR26 | 2.10E-13 | 1.10E-12 | -1.49E+00 | 1.59E-16 | 6.88E-16 | -1.51E+00 | 2.86E-09 | 1.22E-08 | -1.60E+00 | 4.00E-14 | 1.53E-13 | -1.67E+00 | 2.44E-06 | 8.69E-06 | -1.19E+00 | 3.94E-10 | 1.11E-09 | -1.45E+00 | 1.79E-07 | 6.56E-07 | -1.25E+00 | 1.13E-12 | 3.09E-12 | -1.57E+00 |
| STRADB | 2.13E-13 | 1.11E-12 | -1.49E+00 | 7.29E-09 | 3.15E-08 | -1.11E+00 | 5.84E-06 | 2.48E-05 | -1.37E+00 | 1.11E-03 | 4.26E-03 | -1.02E+00 | 3.69E-06 | 1.31E-05 | -1.29E+00 | 6.06E-02 | 1.71E-01 | 1.17E+00 | 1.71E-11 | 6.26E-11 | -1.43E+00 | 1.83E-02 | 5.04E-02 | -1.03E+00 |
| ITGA4 | 2.13E-13 | 1.11E-12 | -1.25E+00 | 3.31E-02 | 1.43E-01 | 1.28E+00 | 1.02E-04 | 4.34E-04 | -1.25E+00 | 1.63E-01 | 6.23E-01 | 1.31E+00 | 7.64E-08 | 2.73E-07 | -1.14E+00 | 3.47E-01 | 9.78E-01 | 1.32E+00 | 2.31E-03 | 8.48E-03 | -1.04E+00 | 8.74E-02 | 2.40E-01 | 1.38E+00 |
| MAGED4B | 2.13E-13 | 1.11E-12 | -1.66E+00 | 4.46E-04 | 1.93E-03 | 1.17E+00 | 1.32E-03 | 5.60E-03 | -1.53E+00 | 1.06E-01 | 4.03E-01 | 1.18E+00 | 7.05E-03 | 2.51E-02 | -1.43E+00 | 2.11E-01 | 5.95E-01 | 1.54E+00 | 5.52E-03 | 2.03E-02 | -1.26E+00 | 1.96E-01 | 5.39E-01 | 1.19E+00 |
| EPS15 | 2.19E-13 | 1.14E-12 | -1.30E+00 | 9.73E-08 | 4.21E-07 | 1.02E+00 | 7.44E-07 | 3.16E-06 | -1.31E+00 | 8.91E-04 | 3.40E-03 | 1.01E+00 | 6.15E-05 | 2.19E-04 | -1.20E+00 | 9.46E-02 | 2.67E-01 | 1.10E+00 | 9.12E-06 | 3.35E-05 | -1.27E+00 | 7.56E-03 | 2.08E-02 | -1.12E+00 |
| PAPOLG | 2.22E-13 | 1.15E-12 | -1.92E+00 | 2.30E-08 | 9.93E-08 | -1.17E+00 | 1.24E-07 | 5.28E-07 | -1.73E+00 | 6.39E-04 | 2.44E-03 | -1.06E+00 | 2.43E-08 | 8.68E-08 | -2.53E+00 | 4.26E-05 | 1.20E-04 | -1.56E+00 | 2.90E-04 | 1.07E-03 | -1.57E+00 | 2.76E-01 | 7.58E-01 | 1.19E+00 |
| TGOLN2 | 2.23E-13 | 1.16E-12 | -1.36E+00 | 9.81E-13 | 4.24E-12 | -1.20E+00 | 2.26E-04 | 9.60E-04 | -1.22E+00 | 2.26E-02 | 8.65E-02 | 1.05E+00 | 1.09E-06 | 3.90E-06 | -1.13E+00 | 2.19E-05 | 6.19E-05 | -1.05E+00 | 3.77E-03 | 1.38E-02 | -1.06E+00 | 2.40E-02 | 6.60E-02 | -1.08E+00 |
| GBP2 | 2.29E-13 | 1.19E-12 | -1.34E+00 | 1.43E-09 | 6.16E-09 | -1.08E+00 | 1.17E-08 | 4.97E-08 | -1.51E+00 | 1.21E-10 | 4.63E-10 | -1.37E+00 | 5.79E-03 | 2.06E-02 | -1.18E+00 | 2.72E-04 | 7.65E-04 | -1.62E+00 | 1.65E-01 | 6.07E-01 | -1.03E+00 | 2.07E-01 | 5.68E-01 | -1.04E+00 |
| TOR3A | 2.34E-13 | 1.22E-12 | -1.72E+00 | 7.46E-08 | 3.22E-07 | -1.20E+00 | 3.21E-05 | 1.36E-04 | -1.11E+00 | 1.56E-03 | 5.95E-03 | 1.15E+00 | 7.55E-08 | 2.69E-07 | -1.81E+00 | 5.40E-04 | 1.52E-03 | -1.21E+00 | 1.74E-05 | 6.37E-05 | -1.21E+00 | 2.97E-02 | 8.16E-02 | 1.11E+00 |
| RPN1 | 2.34E-13 | 1.22E-12 | -1.33E+00 | 4.84E-10 | 2.09E-09 | -1.06E+00 | 2.13E-04 | 9.04E-04 | -1.10E+00 | 4.87E-02 | 1.86E-01 | 1.16E+00 | 6.63E-09 | 2.37E-08 | -1.42E+00 | 9.29E-05 | 2.62E-04 | -1.14E+00 | 2.67E-05 | 9.79E-05 | -1.25E+00 | 8.88E-02 | 2.44E-01 | 1.04E+00 |
| EIF3B | 2.39E-13 | 1.24E-12 | -1.20E+00 | 1.23E-08 | 5.32E-08 | 1.08E+00 | 1.61E-03 | 6.82E-03 | -1.07E+00 | 5.85E-02 | 2.23E-01 | 1.17E+00 | 1.18E-04 | 4.22E-04 | 1.02E+00 | 1.66E-01 | 4.69E-01 | 1.42E+00 | 1.21E-03 | 4.44E-03 | -1.07E+00 | 2.04E-01 | 5.59E-01 | 1.26E+00 |
| C11orf74 | 2.39E-13 | 1.24E-12 | -1.24E+00 | 1.68E-12 | 7.26E-12 | -1.08E+00 | 7.59E-06 | 3.23E-05 | -1.20E+00 | 1.43E-06 | 5.47E-06 | -1.11E+00 | 2.18E-03 | 7.76E-03 | -1.08E+00 | 9.42E-03 | 2.66E-02 | -1.03E+00 | 1.49E-03 | 5.49E-03 | -1.12E+00 | 8.19E-05 | 2.25E-04 | -1.19E+00 |
| NUB1 | 2.40E-13 | 1.25E-12 | -1.29E+00 | 7.53E-14 | 3.26E-13 | -1.21E+00 | 2.54E-07 | 1.08E-06 | -1.38E+00 | 2.13E-07 | 8.12E-07 | -1.20E+00 | 3.18E-07 | 1.13E-06 | -1.24E+00 | 2.15E-06 | 6.05E-06 | -1.11E+00 | 1.52E-08 | 5.59E-08 | -1.44E+00 | 3.44E-07 | 9.44E-07 | -1.35E+00 |
| DFFA | 2.42E-13 | 1.26E-12 | -1.55E+00 | 1.43E-05 | 6.19E-05 | -1.00E+00 | 4.47E-08 | 1.90E-07 | -1.38E+00 | 3.97E-04 | 1.52E-03 | 1.03E+00 | 1.15E-12 | 4.10E-12 | -1.77E+00 | 1.01E-04 | 2.86E-04 | -1.02E+00 | 2.51E-08 | 9.22E-08 | -1.38E+00 | 9.63E-03 | 2.65E-02 | 1.06E+00 |
| FAM96B | 2.45E-13 | 1.28E-12 | -1.27E+00 | 5.93E-09 | 2.56E-08 | -1.03E+00 | 8.50E-04 | 3.61E-03 | -1.09E+00 | 1.34E-04 | 5.11E-04 | -1.01E+00 | 2.39E-12 | 8.53E-12 | -1.29E+00 | 7.80E-08 | 2.20E-07 | -1.09E+00 | 3.43E-02 | 1.26E-01 | 1.08E+00 | 2.23E-02 | 6.13E-02 | 1.09E+00 |
| ESF1 | 2.48E-13 | 1.29E-12 | -1.36E+00 | 1.35E-08 | 5.85E-08 | -1.03E+00 | 1.07E-07 | 4.53E-07 | -1.26E+00 | 6.84E-08 | 2.61E-07 | -1.15E+00 | 8.28E-09 | 2.95E-08 | -1.54E+00 | 1.04E-05 | 2.95E-05 | -1.19E+00 | 2.61E-09 | 9.56E-09 | -1.35E+00 | 6.61E-05 | 1.82E-04 | -1.22E+00 |
| SRGAP3 | 2.49E-13 | 1.30E-12 | -2.10E+00 | 1.59E-09 | 6.88E-09 | -1.54E+00 | 7.99E-13 | 3.40E-12 | -2.00E+00 | 1.02E-13 | 3.89E-13 | -1.83E+00 | 7.26E-12 | 2.59E-11 | -2.55E+00 | 4.15E-12 | 1.17E-11 | -2.31E+00 | 1.20E-07 | 4.41E-07 | -3.37E+00 | 2.45E-05 | 6.72E-05 | -2.01E+00 |
| TAF3 | 2.53E-13 | 1.32E-12 | -1.81E+00 | 7.58E-08 | 3.28E-07 | -1.24E+00 | 1.05E-05 | 4.45E-05 | -1.42E+00 | 3.76E-05 | 1.43E-04 | -1.18E+00 | 1.73E-04 | 6.17E-04 | -1.38E+00 | 8.33E-04 | 2.35E-03 | -1.22E+00 | 1.12E-06 | 4.12E-06 | -1.34E+00 | 2.13E-03 | 5.86E-03 | -1.06E+00 |
| PRKAR2A | 2.57E-13 | 1.34E-12 | -1.67E+00 | 2.80E-05 | 1.21E-04 | 1.01E+00 | 1.66E-06 | 7.04E-06 | -1.35E+00 | 9.94E-04 | 3.80E-03 | 1.01E+00 | 1.24E-04 | 4.41E-04 | -1.18E+00 | 7.21E-03 | 2.03E-02 | 1.04E+00 | 2.07E-09 | 7.61E-09 | -1.49E+00 | 4.22E-03 | 1.16E-02 | 1.02E+00 |
| AC138409.2 | 2.59E-13 | 1.35E-12 | -1.47E+00 | 1.12E-06 | 4.86E-06 | -1.02E+00 | 1.85E-06 | 7.87E-06 | -1.36E+00 | 2.16E-04 | 8.27E-04 | -1.05E+00 | 9.52E-09 | 3.39E-08 | -1.58E+00 | 1.26E-06 | 3.55E-06 | -1.31E+00 | 9.84E-11 | 3.61E-10 | -1.71E+00 | 9.27E-07 | 2.55E-06 | -1.35E+00 |
| AC104978.1 | 2.62E-13 | 1.36E-12 | -1.38E+00 | 5.23E-57 | 2.26E-56 | -1.90E+00 | 8.38E-06 | 3.56E-05 | -1.29E+00 | 1.62E-22 | 6.17E-22 | -1.77E+00 | 3.88E-08 | 1.38E-07 | -1.38E+00 | 1.50E-22 | 4.22E-22 | -1.75E+00 | 3.68E-10 | 1.35E-09 | -1.39E+00 | 6.38E-26 | 1.75E-25 | -1.86E+00 |
| RAN | 2.71E-13 | 1.41E-12 | -1.28E+00 | 1.27E-02 | 5.49E-02 | 1.00E+00 | 8.89E-03 | 3.78E-02 | -1.14E+00 | 1.46E-01 | 5.58E-01 | 1.01E+00 | 1.16E-18 | 4.12E-18 | -1.55E+00 | 9.46E-03 | 2.67E-02 | -1.04E+00 | 2.41E-05 | 8.86E-05 | -1.21E+00 | 2.00E-01 | 5.48E-01 | 1.10E+00 |
| KDELR2 | 2.76E-13 | 1.44E-12 | -1.37E+00 | 4.85E-06 | 2.10E-05 | 1.03E+00 | 9.74E-04 | 4.14E-03 | -1.09E+00 | 2.24E-04 | 8.54E-04 | -1.02E+00 | 4.83E-11 | 1.72E-10 | -1.37E+00 | 1.57E-05 | 4.44E-05 | -1.10E+00 | 3.01E-03 | 1.10E-02 | -1.17E+00 | 1.25E-01 | 3.45E-01 | -1.03E+00 |
| MAF | 2.81E-13 | 1.46E-12 | -1.65E+00 | 1.26E-06 | 5.43E-06 | -1.03E+00 | 1.15E-03 | 4.87E-03 | -1.47E+00 | 5.25E-03 | 2.01E-02 | -1.08E+00 | 3.76E-03 | 1.34E-02 | -1.05E+00 | 4.50E-02 | 1.27E-01 | 1.08E+00 | 2.27E-06 | 8.35E-06 | -1.34E+00 | 5.94E-11 | 1.63E-10 | -1.78E+00 |
| WDR43 | 2.90E-13 | 1.51E-12 | -1.42E+00 | 2.35E-14 | 1.02E-13 | -1.33E+00 | 5.83E-04 | 2.48E-03 | -1.19E+00 | 2.46E-04 | 9.38E-04 | -1.09E+00 | 2.35E-07 | 8.37E-07 | -1.39E+00 | 2.14E-06 | 6.04E-06 | -1.19E+00 | 4.20E-07 | 1.54E-06 | -1.27E+00 | 2.59E-03 | 7.12E-03 | 1.02E+00 |
| RPS3 | 2.97E-13 | 1.55E-12 | -1.16E+00 | 7.07E-12 | 3.05E-11 | -1.15E+00 | 1.52E-06 | 6.45E-06 | -1.12E+00 | 1.42E-05 | 5.41E-05 | -1.11E+00 | 9.97E-03 | 3.56E-02 | -1.06E+00 | 9.27E-02 | 2.61E-01 | -1.04E+00 | 2.80E-05 | 1.03E-04 | -1.05E+00 | 6.05E-02 | 1.66E-01 | -1.04E+00 |
| LPP | 2.98E-13 | 1.55E-12 | -1.56E+00 | 2.99E-05 | 1.29E-04 | 1.05E+00 | 1.31E-06 | 5.55E-06 | -1.44E+00 | 1.87E-04 | 7.15E-04 | -1.11E+00 | 7.79E-02 | 2.78E-01 | 1.11E+00 | 2.18E-01 | 6.16E-01 | 1.14E+00 | 3.25E-07 | 1.19E-06 | -1.54E+00 | 1.64E-04 | 4.49E-04 | -1.26E+00 |
| EMC7 | 2.99E-13 | 1.56E-12 | -1.36E+00 | 1.99E-11 | 8.61E-11 | -1.15E+00 | 3.33E-03 | 1.42E-02 | -1.08E+00 | 4.50E-03 | 1.72E-02 | 1.04E+00 | 1.70E-06 | 6.06E-06 | -1.20E+00 | 2.80E-03 | 7.89E-03 | 1.03E+00 | 8.68E-04 | 3.19E-03 | -1.05E+00 | 2.96E-03 | 8.12E-03 | 1.01E+00 |
| LRRC61 | 3.01E-13 | 1.57E-12 | -2.04E+00 | 1.07E-06 | 4.64E-06 | -1.14E+00 | 1.63E-09 | 6.91E-09 | -1.80E+00 | 8.85E-05 | 3.38E-04 | -1.05E+00 | 4.77E-06 | 1.70E-05 | -1.81E+00 | 1.70E-01 | 4.80E-01 | 1.29E+00 | 3.04E-08 | 1.12E-07 | -1.65E+00 | 3.56E-02 | 9.77E-02 | 1.00E+00 |
| PIK3CG | 3.03E-13 | 1.58E-12 | -1.75E+00 | 9.52E-09 | 4.11E-08 | -1.25E+00 | 4.42E-09 | 1.88E-08 | -1.59E+00 | 1.15E-05 | 4.41E-05 | -1.16E+00 | 3.49E-04 | 1.24E-03 | -1.77E+00 | 8.04E-04 | 2.27E-03 | -1.55E+00 | 1.26E-05 | 4.64E-05 | -2.06E+00 | 5.13E-05 | 1.41E-04 | -2.07E+00 |
| NDUFA5 | 3.05E-13 | 1.59E-12 | -1.23E+00 | 1.84E-07 | 7.95E-07 | 1.03E+00 | 1.80E-02 | 7.63E-02 | -1.08E+00 | 1.06E-02 | 4.06E-02 | 1.00E+00 | 2.22E-04 | 7.93E-04 | -1.04E+00 | 1.76E-01 | 4.95E-01 | 1.20E+00 | 1.39E-01 | 5.09E-01 | 1.18E+00 | 2.99E-01 | 8.21E-01 | 1.24E+00 |
| UBC | 3.05E-13 | 1.59E-12 | 1.33E+00 | 6.98E-02 | 3.02E-01 | 1.02E+00 | 7.91E-11 | 3.36E-10 | 1.37E+00 | 2.74E-02 | 1.05E-01 | 1.15E+00 | 1.03E-05 | 3.69E-05 | 1.33E+00 | 1.51E-01 | 4.27E-01 | 1.00E+00 | 7.82E-06 | 2.87E-05 | 1.30E+00 | 2.93E-01 | 8.06E-01 | 1.03E+00 |
| LMAN2L | 3.11E-13 | 1.62E-12 | -2.43E+00 | 3.29E-08 | 1.42E-07 | -1.40E+00 | 2.47E-06 | 1.05E-05 | -1.60E+00 | 1.63E-03 | 6.24E-03 | -1.00E+00 | 4.62E-04 | 1.65E-03 | -1.63E+00 | 6.93E-02 | 1.95E-01 | 1.05E+00 | 4.58E-07 | 1.68E-06 | -1.55E+00 | 3.47E-03 | 9.54E-03 | -1.05E+00 |
| IFIT1 | 3.12E-13 | 1.63E-12 | -4.17E+00 | 7.14E-08 | 3.08E-07 | -1.84E+00 | 8.64E-21 | 3.67E-20 | -5.25E+00 | 7.70E-19 | 2.94E-18 | -3.94E+00 | 4.09E-18 | 1.46E-17 | -5.12E+00 | 4.98E-14 | 1.40E-13 | -2.98E+00 | 5.92E-35 | 2.17E-34 | -4.94E+00 | 1.53E-26 | 4.21E-26 | -3.23E+00 |
| YARS2 | 3.13E-13 | 1.63E-12 | -1.62E+00 | 3.96E-11 | 1.71E-10 | -1.33E+00 | 6.89E-10 | 2.93E-09 | -1.50E+00 | 3.43E-07 | 1.31E-06 | -1.15E+00 | 1.84E-07 | 6.57E-07 | -1.46E+00 | 2.12E-04 | 5.98E-04 | -1.08E+00 | 1.32E-07 | 4.84E-07 | -1.31E+00 | 9.69E-05 | 2.66E-04 | -1.14E+00 |
| BX890604.1 | 3.21E-13 | 1.67E-12 | -1.81E+00 | 6.56E-13 | 2.83E-12 | -1.63E+00 | 1.21E-04 | 5.14E-04 | -1.29E+00 | 2.10E-04 | 8.02E-04 | -1.13E+00 | 5.50E-03 | 1.96E-02 | -1.12E+00 | 1.43E-01 | 4.02E-01 | 1.28E+00 | 2.10E-03 | 7.71E-03 | -1.18E+00 | 6.64E-03 | 1.82E-02 | -1.21E+00 |
| TRBC1 | 3.26E-13 | 1.70E-12 | -2.16E+00 | 1.65E-10 | 7.14E-10 | -1.64E+00 | 6.36E-08 | 2.70E-07 | -1.93E+00 | 7.55E-08 | 2.88E-07 | -2.13E+00 | 1.71E-02 | 6.08E-02 | -1.41E+00 | 4.18E-02 | 1.18E-01 | -1.21E+00 | 2.18E-01 | 8.00E-01 | -1.00E+00 | 3.20E-01 | 8.79E-01 | 1.19E+00 |
| SRPK1 | 3.26E-13 | 1.70E-12 | -1.39E+00 | 1.89E-09 | 8.15E-09 | -1.10E+00 | 2.13E-06 | 9.07E-06 | -1.17E+00 | 5.02E-05 | 1.92E-04 | 1.01E+00 | 1.03E-06 | 3.69E-06 | -1.34E+00 | 3.23E-03 | 9.11E-03 | -1.06E+00 | 4.02E-06 | 1.47E-05 | -1.26E+00 | 1.96E-03 | 5.38E-03 | -1.06E+00 |
| MKRN2 | 3.27E-13 | 1.70E-12 | -1.69E+00 | 7.08E-09 | 3.06E-08 | -1.12E+00 | 1.68E-10 | 7.12E-10 | -1.53E+00 | 5.79E-06 | 2.21E-05 | -1.06E+00 | 8.84E-13 | 3.15E-12 | -1.58E+00 | 1.41E-07 | 3.98E-07 | -1.14E+00 | 2.04E-08 | 7.50E-08 | -1.41E+00 | 4.00E-02 | 1.10E-01 | -1.02E+00 |
| ZNF587 | 3.27E-13 | 1.70E-12 | -1.72E+00 | 8.85E-12 | 3.82E-11 | -1.48E+00 | 1.41E-07 | 5.99E-07 | -1.45E+00 | 1.06E-06 | 4.06E-06 | -1.30E+00 | 3.24E-06 | 1.15E-05 | -1.39E+00 | 8.96E-06 | 2.53E-05 | -1.34E+00 | 6.31E-11 | 2.31E-10 | -1.69E+00 | 3.62E-07 | 9.93E-07 | -1.30E+00 |
| ZNF74 | 3.27E-13 | 1.70E-12 | -2.50E+00 | 1.28E-09 | 5.55E-09 | -1.69E+00 | 1.53E-10 | 6.49E-10 | -1.81E+00 | 2.55E-07 | 9.76E-07 | -1.25E+00 | 1.83E-08 | 6.52E-08 | -2.30E+00 | 1.14E-04 | 3.21E-04 | -1.29E+00 | 9.80E-11 | 3.60E-10 | -2.20E+00 | 5.95E-04 | 1.63E-03 | -1.22E+00 |
| GNG5 | 3.27E-13 | 1.71E-12 | -1.17E+00 | 4.61E-14 | 1.99E-13 | -1.09E+00 | 5.20E-03 | 2.21E-02 | -1.14E+00 | 3.94E-04 | 1.51E-03 | -1.18E+00 | 5.08E-08 | 1.81E-07 | -1.37E+00 | 1.07E-03 | 3.01E-03 | -1.17E+00 | 2.74E-02 | 1.00E-01 | -1.13E+00 | 4.82E-02 | 1.32E-01 | -1.10E+00 |
| TUBAP2 | 3.29E-13 | 1.71E-12 | -1.32E+00 | 1.34E-11 | 5.80E-11 | -1.18E+00 | 5.44E-03 | 2.31E-02 | -1.18E+00 | 6.46E-02 | 2.47E-01 | -1.01E+00 | 3.48E-13 | 1.24E-12 | -1.62E+00 | 7.47E-08 | 2.11E-07 | -1.50E+00 | 1.85E-03 | 6.81E-03 | -1.21E+00 | 2.73E-01 | 7.49E-01 | 1.04E+00 |
| TMEM263 | 3.32E-13 | 1.73E-12 | -1.49E+00 | 2.62E-05 | 1.13E-04 | 1.03E+00 | 1.23E-06 | 5.21E-06 | -1.43E+00 | 9.32E-06 | 3.56E-05 | -1.16E+00 | 8.23E-13 | 2.94E-12 | -1.75E+00 | 1.53E-07 | 4.33E-07 | -1.20E+00 | 2.10E-06 | 7.70E-06 | -1.33E+00 | 2.41E-04 | 6.62E-04 | -1.14E+00 |
| RTL8A | 3.32E-13 | 1.73E-12 | -1.61E+00 | 8.50E-04 | 3.67E-03 | 1.11E+00 | 1.56E-09 | 6.63E-09 | -1.54E+00 | 1.03E-04 | 3.95E-04 | -1.00E+00 | 2.77E-08 | 9.89E-08 | -1.74E+00 | 1.29E-03 | 3.65E-03 | 1.01E+00 | 6.02E-07 | 2.21E-06 | -1.32E+00 | 6.36E-02 | 1.75E-01 | 1.09E+00 |
| STT3A | 3.33E-13 | 1.73E-12 | -1.83E+00 | 7.29E-06 | 3.15E-05 | -1.11E+00 | 7.73E-04 | 3.29E-03 | -1.25E+00 | 3.82E-02 | 1.46E-01 | 1.10E+00 | 1.48E-11 | 5.28E-11 | -2.07E+00 | 4.65E-05 | 1.31E-04 | -1.28E+00 | 1.41E-04 | 5.19E-04 | -1.30E+00 | 1.94E-01 | 5.32E-01 | 1.21E+00 |
| USP8 | 3.34E-13 | 1.74E-12 | -1.37E+00 | 5.57E-08 | 2.41E-07 | -1.04E+00 | 5.29E-07 | 2.25E-06 | -1.45E+00 | 4.08E-06 | 1.56E-05 | -1.21E+00 | 6.00E-05 | 2.14E-04 | -1.21E+00 | 2.78E-05 | 7.83E-05 | -1.24E+00 | 1.16E-07 | 4.27E-07 | -1.39E+00 | 1.36E-04 | 3.75E-04 | -1.24E+00 |
| GDE1 | 3.35E-13 | 1.75E-12 | -1.68E+00 | 5.31E-10 | 2.30E-09 | -1.26E+00 | 2.23E-11 | 9.49E-11 | -1.63E+00 | 2.45E-08 | 9.37E-08 | -1.31E+00 | 2.15E-11 | 7.68E-11 | -1.49E+00 | 1.25E-11 | 3.53E-11 | -1.49E+00 | 1.02E-12 | 3.76E-12 | -1.57E+00 | 5.20E-06 | 1.43E-05 | -1.13E+00 |
| AL031864.2 | 3.38E-13 | 1.76E-12 | -1.25E+00 | 8.46E-54 | 3.65E-53 | -2.29E+00 | 5.70E-07 | 2.42E-06 | -1.30E+00 | 6.61E-37 | 2.53E-36 | -2.39E+00 | 5.11E-11 | 1.82E-10 | -1.32E+00 | 9.36E-31 | 2.64E-30 | -2.24E+00 | 4.99E-05 | 1.83E-04 | -1.21E+00 | 3.03E-22 | 8.31E-22 | -2.13E+00 |
| PCMTD2 | 3.49E-13 | 1.82E-12 | -1.35E+00 | 4.01E-03 | 1.73E-02 | 1.29E+00 | 3.10E-09 | 1.32E-08 | -1.52E+00 | 3.70E-04 | 1.41E-03 | 1.01E+00 | 4.15E-03 | 1.48E-02 | -1.04E+00 | 2.74E-01 | 7.73E-01 | 1.38E+00 | 6.78E-06 | 2.49E-05 | -1.30E+00 | 1.96E-03 | 5.39E-03 | -1.08E+00 |
| TMEM106B | 3.49E-13 | 1.82E-12 | -1.33E+00 | 2.27E-05 | 9.80E-05 | 1.16E+00 | 1.45E-06 | 6.17E-06 | -1.30E+00 | 8.96E-03 | 3.42E-02 | 1.14E+00 | 8.07E-07 | 2.88E-06 | -1.28E+00 | 6.01E-03 | 1.70E-02 | 1.06E+00 | 3.40E-09 | 1.25E-08 | -1.30E+00 | 4.46E-04 | 1.22E-03 | -1.02E+00 |
| LIN7A | 3.55E-13 | 1.85E-12 | -2.34E+00 | 1.81E-12 | 7.80E-12 | -2.10E+00 | 5.05E-11 | 2.14E-10 | -1.71E+00 | 3.61E-13 | 1.38E-12 | -1.78E+00 | 5.88E-14 | 2.10E-13 | -2.02E+00 | 2.48E-22 | 6.99E-22 | -3.74E+00 | 8.08E-03 | 2.97E-02 | -2.03E+00 | 9.28E-02 | 2.55E-01 | -1.21E+00 |
| TRUB2 | 3.57E-13 | 1.86E-12 | -1.51E+00 | 1.88E-08 | 8.10E-08 | -1.13E+00 | 1.54E-08 | 6.56E-08 | -1.30E+00 | 7.78E-04 | 2.97E-03 | 1.07E+00 | 2.34E-06 | 8.35E-06 | -1.58E+00 | 1.86E-02 | 5.24E-02 | 1.03E+00 | 3.15E-10 | 1.16E-09 | -1.47E+00 | 2.83E-03 | 7.78E-03 | -1.02E+00 |
| GPX7 | 3.70E-13 | 1.93E-12 | -1.38E+00 | 5.41E-11 | 2.34E-10 | -1.16E+00 | 5.45E-08 | 2.32E-07 | -1.35E+00 | 1.01E-07 | 3.85E-07 | -1.19E+00 | 5.15E-11 | 1.84E-10 | -1.66E+00 | 2.81E-04 | 7.91E-04 | -1.11E+00 | 4.53E-06 | 1.66E-05 | -1.25E+00 | 1.44E-01 | 3.95E-01 | 1.12E+00 |
| BOLA2B | 3.74E-13 | 1.95E-12 | -1.28E+00 | 2.92E-24 | 1.26E-23 | -1.69E+00 | 5.62E-05 | 2.39E-04 | -1.12E+00 | 3.44E-14 | 1.31E-13 | -1.38E+00 | 7.41E-04 | 2.64E-03 | -1.08E+00 | 2.61E-07 | 7.35E-07 | -1.40E+00 | 5.66E-03 | 2.08E-02 | 1.01E+00 | 2.01E-05 | 5.53E-05 | -1.25E+00 |
| MARK3 | 3.88E-13 | 2.02E-12 | -1.40E+00 | 2.17E-04 | 9.37E-04 | 1.12E+00 | 2.27E-06 | 9.65E-06 | -1.32E+00 | 3.46E-04 | 1.32E-03 | 1.02E+00 | 8.25E-08 | 2.94E-07 | -1.40E+00 | 2.99E-03 | 8.43E-03 | -1.05E+00 | 3.16E-08 | 1.16E-07 | -1.41E+00 | 1.27E-05 | 3.48E-05 | -1.17E+00 |
| PHF11 | 3.89E-13 | 2.03E-12 | -1.45E+00 | 5.59E-08 | 2.42E-07 | -1.07E+00 | 2.41E-10 | 1.02E-09 | -1.44E+00 | 2.64E-07 | 1.01E-06 | -1.12E+00 | 8.38E-08 | 2.99E-07 | -1.44E+00 | 7.78E-06 | 2.19E-05 | -1.19E+00 | 7.21E-06 | 2.65E-05 | -1.30E+00 | 3.34E-05 | 9.17E-05 | -1.15E+00 |
| ZNF331 | 3.97E-13 | 2.07E-12 | -1.87E+00 | 4.90E-12 | 2.12E-11 | -1.57E+00 | 7.15E-08 | 3.04E-07 | -1.61E+00 | 5.86E-07 | 2.24E-06 | -1.37E+00 | 2.09E-07 | 7.47E-07 | -2.05E+00 | 6.75E-09 | 1.90E-08 | -2.61E+00 | 3.27E-10 | 1.20E-09 | -1.99E+00 | 6.60E-09 | 1.81E-08 | -1.94E+00 |
| SNAP23 | 3.97E-13 | 2.07E-12 | -1.56E+00 | 2.51E-08 | 1.08E-07 | -1.14E+00 | 1.20E-07 | 5.09E-07 | -1.46E+00 | 2.57E-10 | 9.82E-10 | -1.53E+00 | 6.80E-06 | 2.42E-05 | -1.29E+00 | 4.67E-04 | 1.32E-03 | -1.10E+00 | 4.32E-05 | 1.58E-04 | -1.04E+00 | 4.16E-06 | 1.14E-05 | -1.12E+00 |
| DNM3 | 3.98E-13 | 2.07E-12 | -1.85E+00 | 1.66E-08 | 7.19E-08 | -1.31E+00 | 5.76E-03 | 2.45E-02 | -1.41E+00 | 7.77E-02 | 2.97E-01 | 1.10E+00 | 8.42E-02 | 3.00E-01 | 1.00E+00 | 1.41E-01 | 3.98E-01 | 1.13E+00 | 1.82E-01 | 6.70E-01 | 1.81E+00 | 1.75E-01 | 4.81E-01 | 1.30E+00 |
| CHD9 | 3.98E-13 | 2.07E-12 | -1.24E+00 | 1.44E-07 | 6.21E-07 | -1.04E+00 | 7.99E-04 | 3.39E-03 | -1.26E+00 | 1.09E-03 | 4.17E-03 | -1.15E+00 | 1.58E-06 | 5.64E-06 | -1.27E+00 | 1.24E-05 | 3.48E-05 | -1.21E+00 | 6.95E-06 | 2.55E-05 | -1.26E+00 | 2.17E-05 | 5.97E-05 | -1.34E+00 |
| SMIM29 | 4.03E-13 | 2.10E-12 | -1.83E+00 | 1.30E-07 | 5.61E-07 | -1.18E+00 | 5.89E-04 | 2.50E-03 | -1.36E+00 | 1.59E-04 | 6.05E-04 | -1.27E+00 | 3.41E-07 | 1.22E-06 | -1.97E+00 | 6.28E-05 | 1.77E-04 | -1.50E+00 | 4.95E-06 | 1.82E-05 | -1.42E+00 | 9.59E-05 | 2.63E-04 | -1.30E+00 |
| ARPP19 | 4.06E-13 | 2.12E-12 | -1.26E+00 | 1.66E-10 | 7.18E-10 | -1.03E+00 | 3.04E-08 | 1.29E-07 | -1.31E+00 | 1.05E-05 | 4.00E-05 | -1.06E+00 | 1.70E-04 | 6.05E-04 | -1.08E+00 | 1.58E-05 | 4.46E-05 | -1.12E+00 | 1.95E-06 | 7.16E-06 | -1.23E+00 | 1.52E-02 | 4.17E-02 | -1.01E+00 |
| RAB14 | 4.10E-13 | 2.14E-12 | -1.26E+00 | 6.11E-09 | 2.64E-08 | -1.06E+00 | 2.43E-09 | 1.03E-08 | -1.42E+00 | 3.57E-08 | 1.36E-07 | -1.14E+00 | 8.14E-11 | 2.90E-10 | -1.50E+00 | 2.53E-04 | 7.12E-04 | -1.09E+00 | 1.19E-07 | 4.38E-07 | -1.38E+00 | 4.31E-04 | 1.18E-03 | -1.15E+00 |
| NUDT21 | 4.17E-13 | 2.17E-12 | -1.42E+00 | 1.99E-11 | 8.59E-11 | -1.19E+00 | 8.81E-04 | 3.75E-03 | -1.13E+00 | 2.54E-05 | 9.70E-05 | -1.08E+00 | 5.12E-05 | 1.83E-04 | -1.08E+00 | 9.44E-04 | 2.66E-03 | -1.07E+00 | 4.17E-02 | 1.53E-01 | 1.00E+00 | 5.89E-02 | 1.62E-01 | 1.12E+00 |
| TGIF2 | 4.21E-13 | 2.19E-12 | -1.48E+00 | 1.38E-11 | 5.97E-11 | -1.28E+00 | 9.01E-13 | 3.83E-12 | -1.62E+00 | 1.32E-12 | 5.03E-12 | -1.41E+00 | 3.40E-14 | 1.21E-13 | -1.68E+00 | 1.06E-15 | 2.98E-15 | -1.79E+00 | 2.17E-15 | 7.98E-15 | -2.38E+00 | 1.61E-06 | 4.41E-06 | -1.38E+00 |
| ERLIN2 | 4.22E-13 | 2.20E-12 | -1.51E+00 | 1.31E-12 | 5.65E-12 | -1.31E+00 | 2.57E-05 | 1.09E-04 | -1.28E+00 | 5.78E-05 | 2.21E-04 | -1.10E+00 | 1.45E-06 | 5.16E-06 | -1.48E+00 | 2.04E-05 | 5.76E-05 | -1.34E+00 | 6.57E-04 | 2.41E-03 | -1.18E+00 | 7.69E-03 | 2.11E-02 | -1.10E+00 |
| RPP78P | 4.33E-13 | 2.26E-12 | -2.17E+00 | 1.46E-05 | 6.32E-05 | -1.10E+00 | 3.22E-03 | 1.37E-02 | -1.08E+00 | 2.55E-01 | 9.74E-01 | 1.51E+00 | 9.33E-04 | 3.33E-03 | -1.29E+00 | 6.88E-02 | 1.94E-01 | 1.12E+00 | 2.13E-04 | 7.80E-04 | -1.41E+00 | 2.47E-01 | 6.80E-01 | 1.26E+00 |
| AL358176.3 | 4.34E-13 | 2.26E-12 | -1.42E+00 | 5.14E-60 | 2.22E-59 | -2.30E+00 | 2.08E-03 | 8.84E-03 | -1.28E+00 | 8.66E-22 | 3.31E-21 | -1.90E+00 | 2.13E-09 | 7.59E-09 | -1.39E+00 | 3.88E-27 | 1.09E-26 | -2.17E+00 | 1.07E-10 | 3.94E-10 | -1.58E+00 | 5.19E-27 | 1.42E-26 | -2.24E+00 |
| C8orf33 | 4.42E-13 | 2.31E-12 | -1.58E+00 | 1.96E-14 | 8.48E-14 | -1.48E+00 | 3.80E-02 | 1.61E-01 | -1.08E+00 | 6.45E-03 | 2.46E-02 | -1.02E+00 | 2.81E-05 | 1.00E-04 | -1.30E+00 | 2.12E-05 | 5.97E-05 | -1.25E+00 | 1.89E-04 | 6.92E-04 | -1.15E+00 | 1.04E-03 | 2.85E-03 | -1.06E+00 |
| MED13 | 4.51E-13 | 2.35E-12 | -1.46E+00 | 4.61E-07 | 1.99E-06 | 1.01E+00 | 1.40E-03 | 5.93E-03 | -1.22E+00 | 1.34E-04 | 5.12E-04 | -1.16E+00 | 3.07E-05 | 1.09E-04 | -1.18E+00 | 1.54E-05 | 4.34E-05 | -1.23E+00 | 5.01E-09 | 1.84E-08 | -1.44E+00 | 7.34E-06 | 2.02E-05 | -1.37E+00 |
| ZDHHC3 | 4.52E-13 | 2.35E-12 | -1.67E+00 | 2.49E-08 | 1.08E-07 | -1.16E+00 | 8.79E-06 | 3.73E-05 | -1.26E+00 | 4.59E-05 | 1.75E-04 | -1.08E+00 | 6.40E-03 | 2.28E-02 | 1.05E+00 | 7.21E-02 | 2.03E-01 | 1.30E+00 | 1.34E-06 | 4.90E-06 | -1.28E+00 | 3.42E-04 | 9.38E-04 | -1.07E+00 |
| SYNRG | 4.54E-13 | 2.37E-12 | -1.61E+00 | 2.78E-09 | 1.20E-08 | -1.20E+00 | 4.72E-12 | 2.01E-11 | -1.82E+00 | 1.32E-08 | 5.05E-08 | -1.39E+00 | 6.79E-06 | 2.42E-05 | -1.42E+00 | 2.43E-03 | 6.85E-03 | -1.10E+00 | 5.70E-15 | 2.09E-14 | -1.85E+00 | 4.94E-09 | 1.36E-08 | -1.46E+00 |
| DDHD1 | 4.56E-13 | 2.38E-12 | -1.34E+00 | 1.30E-13 | 5.63E-13 | -1.22E+00 | 1.37E-04 | 5.80E-04 | -1.17E+00 | 3.26E-06 | 1.24E-05 | -1.13E+00 | 1.92E-02 | 6.84E-02 | 1.13E+00 | 5.30E-04 | 1.49E-03 | -1.06E+00 | 1.68E-06 | 6.17E-06 | -1.33E+00 | 7.41E-07 | 2.04E-06 | -1.35E+00 |
| RBM3 | 4.58E-13 | 2.39E-12 | 1.41E+00 | 1.25E-02 | 5.39E-02 | 1.24E+00 | 8.71E-08 | 3.70E-07 | 1.35E+00 | 1.35E-04 | 5.14E-04 | 1.27E+00 | 5.20E-03 | 1.85E-02 | 1.25E+00 | 3.44E-01 | 9.70E-01 | 1.07E+00 | 3.54E-04 | 1.30E-03 | 1.22E+00 | 1.04E-04 | 2.87E-04 | 1.22E+00 |
| SLC1A6 | 4.59E-13 | 2.39E-12 | -1.90E+00 | 4.85E-07 | 2.10E-06 | -1.23E+00 | 7.71E-02 | 3.28E-01 | -1.14E+00 | 1.65E-01 | 6.30E-01 | 1.14E+00 | 2.02E-01 | 7.19E-01 | 1.01E+00 | 2.83E-01 | 7.98E-01 | 1.33E+00 | 2.62E-01 | 9.62E-01 | -1.02E+00 | 3.19E-01 | 8.77E-01 | 1.30E+00 |
| CMTM6 | 4.60E-13 | 2.40E-12 | -1.36E+00 | 1.37E-13 | 5.91E-13 | -1.25E+00 | 1.19E-05 | 5.08E-05 | -1.20E+00 | 7.34E-07 | 2.80E-06 | -1.14E+00 | 1.17E-06 | 4.17E-06 | -1.22E+00 | 4.59E-05 | 1.30E-04 | -1.16E+00 | 6.25E-04 | 2.29E-03 | -1.13E+00 | 1.53E-04 | 4.19E-04 | -1.11E+00 |
| TEX30 | 4.70E-13 | 2.45E-12 | -1.60E+00 | 2.52E-07 | 1.09E-06 | -1.08E+00 | 1.66E-08 | 7.05E-08 | -1.39E+00 | 8.56E-05 | 3.27E-04 | -1.02E+00 | 2.03E-04 | 7.24E-04 | -1.13E+00 | 1.61E-04 | 4.55E-04 | -1.22E+00 | 2.15E-08 | 7.88E-08 | -1.40E+00 | 5.47E-03 | 1.50E-02 | -1.08E+00 |
| COL2A1 | 4.70E-13 | 2.45E-12 | -2.36E+00 | 3.36E-13 | 1.45E-12 | -2.48E+00 | 1.31E-09 | 5.57E-09 | -2.42E+00 | 1.16E-10 | 4.42E-10 | -2.56E+00 | 2.64E-01 | 9.40E-01 | 1.10E+00 | 2.79E-01 | 7.88E-01 | -1.09E+00 | 2.17E-01 | 7.98E-01 | -1.09E+00 | 2.77E-01 | 7.62E-01 | -1.03E+00 |
| AC062028.2 | 4.79E-13 | 2.50E-12 | 2.55E+00 | 7.47E-04 | 3.23E-03 | 2.16E+00 | 2.98E-12 | 1.26E-11 | 2.33E+00 | 9.67E-08 | 3.69E-07 | 2.23E+00 | 2.73E-05 | 9.75E-05 | 2.30E+00 | 7.24E-04 | 2.04E-03 | 2.08E+00 | 4.89E-06 | 1.80E-05 | 2.06E+00 | 9.76E-05 | 2.68E-04 | 1.92E+00 |
| LRCH3 | 4.79E-13 | 2.50E-12 | -1.58E+00 | 4.53E-04 | 1.96E-03 | 1.16E+00 | 5.01E-09 | 2.13E-08 | -1.48E+00 | 6.45E-04 | 2.46E-03 | 1.08E+00 | 7.32E-05 | 2.61E-04 | -1.24E+00 | 6.81E-02 | 1.92E-01 | 1.26E+00 | 2.03E-05 | 7.46E-05 | -1.37E+00 | 4.72E-03 | 1.30E-02 | -1.07E+00 |
| ENTPD4 | 4.81E-13 | 2.51E-12 | -1.88E+00 | 2.65E-12 | 1.15E-11 | -1.67E+00 | 3.53E-11 | 1.50E-10 | -2.11E+00 | 2.21E-08 | 8.42E-08 | -1.62E+00 | 1.67E-06 | 5.94E-06 | -1.31E+00 | 5.08E-05 | 1.43E-04 | -1.17E+00 | 1.99E-07 | 7.31E-07 | -1.28E+00 | 1.71E-08 | 4.71E-08 | -1.39E+00 |
| ERCC6L2 | 4.88E-13 | 2.54E-12 | -1.77E+00 | 7.58E-09 | 3.28E-08 | -1.23E+00 | 1.41E-09 | 5.97E-09 | -1.61E+00 | 8.00E-08 | 3.06E-07 | -1.26E+00 | 1.73E-05 | 6.16E-05 | -1.45E+00 | 4.33E-04 | 1.22E-03 | -1.30E+00 | 4.28E-06 | 1.57E-05 | -1.28E+00 | 2.73E-04 | 7.51E-04 | -1.17E+00 |
| VBP1 | 5.00E-13 | 2.61E-12 | -1.23E+00 | 3.18E-10 | 1.38E-09 | -1.06E+00 | 3.35E-02 | 1.42E-01 | 1.09E+00 | 7.09E-03 | 2.71E-02 | 1.12E+00 | 2.55E-06 | 9.08E-06 | -1.35E+00 | 1.76E-03 | 4.96E-03 | -1.03E+00 | 1.68E-03 | 6.15E-03 | -1.02E+00 | 3.64E-01 | 1.00E+00 | 1.16E+00 |
| STXBP2 | 5.04E-13 | 2.63E-12 | -1.53E+00 | 5.19E-08 | 2.24E-07 | -1.12E+00 | 3.91E-09 | 1.66E-08 | -1.47E+00 | 8.35E-09 | 3.19E-08 | -1.20E+00 | 6.49E-12 | 2.32E-11 | -1.45E+00 | 4.55E-09 | 1.28E-08 | -1.23E+00 | 8.77E-07 | 3.22E-06 | -1.17E+00 | 2.34E-05 | 6.44E-05 | -1.16E+00 |
| HES1 | 5.05E-13 | 2.63E-12 | 8.51E+00 | 1.87E-01 | 8.09E-01 | 1.00E+00 | 4.41E-02 | 1.87E-01 | 2.35E+00 | 2.25E-01 | 8.61E-01 | -1.19E+00 | 2.73E-01 | 9.72E-01 | 1.28E+00 | 3.10E-01 | 8.75E-01 | 1.04E+00 | 2.06E-13 | 7.54E-13 | -1.75E+00 | 9.15E-16 | 2.51E-15 | -2.54E+00 |
| PSMC2 | 5.05E-13 | 2.63E-12 | -1.41E+00 | 6.12E-08 | 2.64E-07 | -1.03E+00 | 3.39E-05 | 1.44E-04 | -1.10E+00 | 3.08E-04 | 1.17E-03 | 1.05E+00 | 2.50E-07 | 8.90E-07 | -1.19E+00 | 8.41E-05 | 2.37E-04 | -1.03E+00 | 3.26E-04 | 1.20E-03 | -1.05E+00 | 1.29E-01 | 3.53E-01 | 1.13E+00 |
| ZDHHC2 | 5.09E-13 | 2.65E-12 | -1.36E+00 | 5.73E-04 | 2.47E-03 | 1.16E+00 | 1.83E-10 | 7.78E-10 | -1.62E+00 | 3.13E-05 | 1.20E-04 | -1.04E+00 | 4.03E-05 | 1.44E-04 | -1.14E+00 | 1.13E-01 | 3.18E-01 | 1.62E+00 | 8.45E-03 | 3.10E-02 | -1.16E+00 | 1.82E-01 | 5.00E-01 | 1.00E+00 |
| RBMX | 5.11E-13 | 2.66E-12 | -1.26E+00 | 4.94E-09 | 2.13E-08 | -1.11E+00 | 6.14E-04 | 2.61E-03 | -1.23E+00 | 2.56E-05 | 9.77E-05 | -1.24E+00 | 3.08E-05 | 1.10E-04 | -1.26E+00 | 6.49E-03 | 1.83E-02 | -1.09E+00 | 4.58E-04 | 1.68E-03 | -1.25E+00 | 2.56E-02 | 7.04E-02 | -1.13E+00 |
| MFAP1 | 5.13E-13 | 2.67E-12 | -1.88E+00 | 1.47E-03 | 6.35E-03 | 1.13E+00 | 9.91E-07 | 4.21E-06 | -1.53E+00 | 2.01E-02 | 7.69E-02 | 1.08E+00 | 1.28E-08 | 4.57E-08 | -1.64E+00 | 1.52E-02 | 4.28E-02 | 1.06E+00 | 3.87E-21 | 1.42E-20 | -1.83E+00 | 1.71E-04 | 4.70E-04 | -1.10E+00 |
| PSMD1 | 5.13E-13 | 2.67E-12 | -1.50E+00 | 1.03E-06 | 4.45E-06 | -1.03E+00 | 2.39E-04 | 1.01E-03 | -1.18E+00 | 6.07E-02 | 2.32E-01 | 1.12E+00 | 5.35E-09 | 1.91E-08 | -1.54E+00 | 3.59E-04 | 1.01E-03 | -1.08E+00 | 2.81E-05 | 1.03E-04 | -1.17E+00 | 3.32E-01 | 9.13E-01 | 1.18E+00 |
| CTNNA1 | 5.14E-13 | 2.68E-12 | -1.38E+00 | 1.85E-06 | 8.00E-06 | 1.05E+00 | 3.38E-06 | 1.44E-05 | -1.20E+00 | 1.32E-02 | 5.05E-02 | 1.13E+00 | 1.06E-07 | 3.79E-07 | -1.27E+00 | 6.69E-02 | 1.89E-01 | 1.16E+00 | 1.20E-03 | 4.41E-03 | -1.12E+00 | 2.11E-01 | 5.80E-01 | 1.09E+00 |
| POLR2L | 5.14E-13 | 2.68E-12 | -1.21E+00 | 3.60E-33 | 1.56E-32 | -1.57E+00 | 1.50E-02 | 6.38E-02 | -1.04E+00 | 2.35E-13 | 8.98E-13 | -1.51E+00 | 2.61E-06 | 9.29E-06 | -1.27E+00 | 8.42E-14 | 2.37E-13 | -1.59E+00 | 1.99E-03 | 7.30E-03 | -1.19E+00 | 3.39E-06 | 9.31E-06 | -1.54E+00 |
| SLC25A11 | 5.27E-13 | 2.75E-12 | -1.31E+00 | 6.65E-11 | 2.87E-10 | -1.09E+00 | 6.39E-05 | 2.71E-04 | -1.19E+00 | 3.97E-04 | 1.52E-03 | 1.00E+00 | 4.91E-09 | 1.75E-08 | -1.26E+00 | 1.16E-05 | 3.27E-05 | -1.08E+00 | 2.76E-03 | 1.01E-02 | -1.01E+00 | 1.87E-01 | 5.15E-01 | 1.23E+00 |
| NUDT22 | 5.34E-13 | 2.79E-12 | -3.12E+00 | 1.24E-07 | 5.37E-07 | -1.62E+00 | 1.08E-14 | 4.57E-14 | -2.81E+00 | 4.61E-11 | 1.76E-10 | -1.84E+00 | 8.08E-04 | 2.88E-03 | -1.55E+00 | 7.32E-03 | 2.06E-02 | -1.24E+00 | 1.56E-12 | 5.74E-12 | -1.95E+00 | 2.82E-05 | 7.74E-05 | -1.25E+00 |
| MORC1 | 5.37E-13 | 2.80E-12 | 8.27E+00 | 8.88E-03 | 3.84E-02 | 3.47E+00 | 1.32E-07 | 5.60E-07 | 4.86E+00 | 1.02E-01 | 3.88E-01 | 1.87E+00 | 1.91E-02 | 6.83E-02 | 4.22E+00 | 3.17E-01 | 8.93E-01 | 1.23E+00 | 2.55E-01 | 9.34E-01 | 1.01E+00 | 3.42E-01 | 9.40E-01 | -1.03E+00 |
| TRIAP1 | 5.41E-13 | 2.82E-12 | -1.47E+00 | 5.92E-11 | 2.56E-10 | -1.25E+00 | 1.26E-07 | 5.36E-07 | -1.25E+00 | 6.25E-06 | 2.39E-05 | -1.05E+00 | 1.11E-03 | 3.97E-03 | -1.04E+00 | 4.92E-03 | 1.39E-02 | 1.03E+00 | 9.47E-06 | 3.48E-05 | -1.20E+00 | 1.64E-02 | 4.51E-02 | 1.04E+00 |
| CREBBP | 5.41E-13 | 2.82E-12 | -1.60E+00 | 8.75E-06 | 3.78E-05 | -1.03E+00 | 1.29E-04 | 5.50E-04 | -1.34E+00 | 4.22E-03 | 1.61E-02 | -1.03E+00 | 8.83E-09 | 3.15E-08 | -1.54E+00 | 2.04E-05 | 5.75E-05 | -1.19E+00 | 4.76E-11 | 1.75E-10 | -1.70E+00 | 2.56E-06 | 7.04E-06 | -1.33E+00 |
| R3HDM2 | 5.41E-13 | 2.82E-12 | -1.35E+00 | 2.22E-14 | 9.57E-14 | -1.29E+00 | 5.08E-14 | 2.16E-13 | -1.73E+00 | 2.43E-17 | 9.29E-17 | -1.61E+00 | 7.15E-07 | 2.55E-06 | -1.26E+00 | 1.91E-08 | 5.37E-08 | -1.26E+00 | 5.93E-06 | 2.18E-05 | -1.28E+00 | 6.05E-08 | 1.66E-07 | -1.42E+00 |

| | | | | | | | | | | | | | | | | | | | | | | | | |
|---|---|---|---|---|---|---|---|---|---|---|---|---|---|---|---|---|---|---|---|---|---|---|---|---|
| FAM98B | 5.50E-13 | 2.86E-12 | -1.37E+00 | 2.00E-09 | 8.64E-09 | -1.11E+00 | 3.33E-07 | 1.41E-06 | -1.36E+00 | 6.39E-04 | 2.44E-03 | 1.01E+00 | 1.29E-04 | 4.59E-04 | -1.18E+00 | 3.09E-02 | 8.71E-02 | 1.05E+00 | 1.08E-06 | 3.97E-06 | -1.33E+00 | 1.79E-01 | 4.92E-01 | 1.14E+00 |
| DERL1 | 5.50E-13 | 2.86E-12 | -1.31E+00 | 8.88E-11 | 3.84E-10 | -1.12E+00 | 9.25E-05 | 3.93E-04 | -1.14E+00 | 5.28E-07 | 2.02E-06 | -1.12E+00 | 1.20E-06 | 4.30E-06 | -1.21E+00 | 9.60E-07 | 2.71E-06 | -1.18E+00 | 4.94E-09 | 1.81E-08 | -1.36E+00 | 4.76E-05 | 1.31E-04 | -1.09E+00 |
| ATG3 | 5.52E-13 | 2.88E-12 | -1.41E+00 | 5.36E-10 | 2.32E-09 | -1.07E+00 | 3.49E-07 | 1.48E-06 | -1.26E+00 | 2.77E-05 | 1.06E-04 | -1.04E+00 | 1.57E-06 | 5.58E-06 | -1.31E+00 | 1.17E-01 | 3.30E-01 | 1.27E+00 | 7.34E-09 | 2.69E-08 | -1.25E+00 | 6.91E-04 | 1.90E-03 | -1.01E+00 |
| PFDN2 | 5.53E-13 | 2.88E-12 | -1.30E+00 | 2.78E-11 | 1.20E-10 | -1.12E+00 | 3.38E-03 | 1.44E-02 | -1.09E+00 | 5.10E-04 | 1.95E-03 | -1.02E+00 | 1.86E-10 | 6.62E-10 | -1.44E+00 | 1.19E-03 | 3.35E-03 | -1.09E+00 | 3.03E-03 | 1.11E-02 | -1.07E+00 | 1.14E-01 | 3.12E-01 | 1.07E+00 |
| TIPRL | 5.59E-13 | 2.92E-12 | -1.30E+00 | 5.35E-10 | 2.31E-09 | -1.07E+00 | 5.75E-05 | 2.44E-04 | -1.10E+00 | 3.92E-07 | 1.50E-06 | -1.07E+00 | 4.52E-09 | 1.61E-08 | -1.24E+00 | 1.11E-05 | 3.12E-05 | -1.10E+00 | 3.69E-04 | 1.35E-03 | -1.11E+00 | 1.18E-03 | 3.24E-03 | -1.02E+00 |
| TM2D3 | 5.60E-13 | 2.92E-12 | -1.47E+00 | 3.41E-10 | 1.47E-09 | -1.19E+00 | 2.24E-07 | 9.54E-07 | -1.46E+00 | 1.93E-06 | 7.36E-06 | -1.21E+00 | 1.23E-05 | 4.39E-05 | -1.28E+00 | 5.91E-06 | 1.67E-05 | -1.39E+00 | 1.06E-10 | 3.89E-10 | -1.55E+00 | 9.28E-07 | 2.55E-06 | -1.26E+00 |
| UBTF | 5.60E-13 | 2.92E-12 | -1.44E+00 | 9.20E-03 | 3.97E-02 | 1.19E+00 | 2.12E-05 | 9.00E-05 | -1.23E+00 | 9.06E-02 | 3.46E-01 | 1.25E+00 | 1.65E-09 | 5.88E-09 | -1.43E+00 | 3.44E-03 | 9.69E-03 | 1.05E+00 | 3.33E-04 | 1.22E-03 | -1.14E+00 | 2.16E-01 | 5.92E-01 | 1.19E+00 |
| NOA1 | 5.75E-13 | 3.00E-12 | -1.27E+00 | 4.49E-09 | 1.94E-08 | -1.09E+00 | 2.32E-08 | 9.85E-08 | -1.42E+00 | 8.82E-08 | 3.37E-07 | -1.25E+00 | 8.21E-07 | 2.93E-06 | -1.25E+00 | 7.62E-05 | 2.15E-04 | -1.11E+00 | 1.22E-04 | 4.47E-04 | -1.20E+00 | 1.96E-03 | 5.39E-03 | -1.14E+00 |
| FXYD5 | 5.88E-13 | 3.06E-12 | -1.39E+00 | 5.09E-06 | 2.20E-05 | -1.18E+00 | 1.31E-05 | 5.57E-05 | -1.29E+00 | 2.53E-02 | 9.66E-02 | -1.05E+00 | 3.02E-05 | 1.08E-04 | -1.20E+00 | 1.07E-03 | 3.03E-03 | -1.11E+00 | 2.10E-04 | 7.72E-04 | -1.14E+00 | 8.21E-03 | 2.25E-02 | -1.06E+00 |
| COPS7B | 5.92E-13 | 3.08E-12 | -1.80E+00 | 6.79E-05 | 2.93E-04 | 1.03E+00 | 5.70E-12 | 2.42E-11 | -1.62E+00 | 1.23E-07 | 4.71E-07 | -1.19E+00 | 7.89E-06 | 2.81E-05 | -1.50E+00 | 3.62E-03 | 1.02E-02 | -1.03E+00 | 6.77E-10 | 2.49E-09 | -1.49E+00 | 8.31E-04 | 2.28E-03 | -1.07E+00 |
| MRPL16 | 6.01E-13 | 3.13E-12 | -1.39E+00 | 3.92E-03 | 1.69E-02 | 1.04E+00 | 8.24E-03 | 3.50E-02 | -1.04E+00 | 6.34E-02 | 2.42E-01 | 1.23E+00 | 3.29E-02 | 1.17E-01 | 1.43E+00 | 5.80E-06 | 1.63E-05 | 1.99E+00 | 7.41E-02 | 2.72E-01 | 1.49E+00 | 8.94E-05 | 2.46E-04 | 2.04E+00 |
| NDUFS3 | 6.12E-13 | 3.19E-12 | -1.19E+00 | 5.99E-08 | 2.59E-07 | 1.07E+00 | 1.32E-03 | 5.62E-03 | -1.08E+00 | 9.12E-03 | 3.48E-02 | 1.08E+00 | 1.01E-09 | 3.60E-09 | -1.34E+00 | 5.62E-02 | 1.59E-01 | 1.13E+00 | 1.85E-04 | 6.78E-04 | -1.19E+00 | 2.19E-01 | 6.03E-01 | 1.08E+00 |
| NME2P1 | 6.16E-13 | 3.21E-12 | -1.32E+00 | 2.56E-13 | 1.11E-12 | -1.29E+00 | 2.54E-03 | 1.08E-02 | -1.14E+00 | 5.07E-03 | 1.94E-02 | -1.12E+00 | 6.80E-09 | 2.42E-08 | -1.34E+00 | 1.41E-06 | 3.98E-06 | -1.26E+00 | 4.81E-11 | 1.77E-10 | -1.22E+00 | 1.45E-03 | 3.99E-03 | -1.13E+00 |
| HADHA | 6.20E-13 | 3.23E-12 | -1.25E+00 | 5.24E-07 | 2.27E-06 | -1.01E+00 | 6.43E-04 | 2.73E-03 | -1.14E+00 | 1.16E-02 | 4.45E-02 | 1.02E+00 | 1.91E-04 | 6.81E-04 | -1.08E+00 | 1.95E-02 | 5.49E-02 | 1.05E+00 | 3.27E-02 | 1.20E-01 | -1.05E+00 | 3.52E-01 | 9.67E-01 | 1.06E+00 |
| CHD8 | 6.33E-13 | 3.30E-12 | -1.42E+00 | 9.64E-09 | 4.16E-08 | -1.03E+00 | 1.54E-09 | 6.53E-09 | -1.42E+00 | 9.77E-07 | 3.73E-06 | -1.09E+00 | 2.70E-08 | 9.61E-08 | -1.36E+00 | 4.90E-06 | 1.38E-05 | -1.18E+00 | 3.21E-11 | 1.18E-10 | -1.45E+00 | 1.26E-09 | 3.45E-09 | -1.37E+00 |
| CCNT1 | 6.46E-13 | 3.37E-12 | -1.42E+00 | 1.07E-08 | 4.62E-08 | -1.04E+00 | 1.62E-07 | 6.89E-07 | -1.38E+00 | 9.66E-05 | 3.69E-04 | -1.08E+00 | 1.57E-04 | 5.60E-04 | -1.20E+00 | 2.86E-04 | 8.07E-04 | -1.18E+00 | 1.32E-08 | 4.84E-08 | -1.49E+00 | 6.22E-03 | 1.71E-02 | -1.07E+00 |
| BSPRY | 6.47E-13 | 3.37E-12 | -1.51E+00 | 3.22E-10 | 1.39E-09 | -1.22E+00 | 1.61E-10 | 6.85E-10 | -1.78E+00 | 3.21E-06 | 1.23E-05 | -1.21E+00 | 1.70E-01 | 6.06E-01 | 1.26E+00 | 7.36E-02 | 2.07E-01 | 1.72E+00 | 5.99E-02 | 2.20E-01 | 1.57E+00 | 2.65E-01 | 7.29E-01 | 1.47E+00 |
| BLOC1S2 | 6.54E-13 | 3.41E-12 | -1.40E+00 | 5.60E-16 | 2.42E-15 | -1.37E+00 | 1.53E-03 | 6.51E-03 | -1.10E+00 | 6.72E-07 | 2.57E-06 | -1.16E+00 | 5.47E-12 | 1.95E-11 | -1.55E+00 | 2.06E-08 | 5.81E-08 | -1.27E+00 | 6.33E-05 | 2.32E-04 | -1.12E+00 | 1.22E-04 | 3.34E-04 | -1.09E+00 |
| AC044849.2 | 6.59E-13 | 3.44E-12 | 2.75E+00 | 2.72E-07 | 1.18E-06 | -1.08E+00 | 7.95E-08 | 3.38E-07 | 2.04E+00 | 6.89E-06 | 2.63E-05 | -1.37E+00 | 7.49E-06 | 2.67E-05 | 2.58E+00 | 2.37E-06 | 6.68E-06 | -1.16E+00 | 3.24E-03 | 1.19E-02 | 1.95E+00 | 1.85E-05 | 5.07E-05 | -1.45E+00 |
| YWHABP2 | 6.61E-13 | 3.45E-12 | -2.27E+00 | 2.74E-09 | 1.18E-08 | -1.56E+00 | 1.01E-19 | 4.28E-19 | -2.87E+00 | 1.40E-11 | 5.33E-11 | -1.60E+00 | 1.67E-09 | 5.95E-09 | -3.13E+00 | 5.68E-05 | 1.60E-04 | -1.34E+00 | 3.45E-17 | 1.27E-16 | -2.44E+00 | 8.60E-08 | 2.36E-07 | -1.34E+00 |
| PPP2R5E | 6.66E-13 | 3.47E-12 | -1.38E+00 | 3.62E-08 | 1.57E-07 | -1.06E+00 | 8.20E-04 | 3.49E-03 | -1.11E+00 | 1.01E-03 | 3.86E-03 | 1.02E+00 | 3.08E-07 | 1.10E-06 | -1.24E+00 | 2.17E-04 | 6.11E-04 | 1.00E+00 | 1.49E-07 | 5.47E-07 | -1.29E+00 | 6.75E-04 | 1.85E-03 | -1.11E+00 |
| SYNE1 | 6.67E-13 | 3.48E-12 | -1.58E+00 | 3.97E-04 | 1.72E-03 | 1.08E+00 | 6.78E-08 | 2.88E-07 | -1.58E+00 | 1.48E-04 | 5.64E-04 | -1.13E+00 | 1.25E-05 | 4.47E-05 | -1.42E+00 | 5.26E-04 | 1.48E-03 | -1.11E+00 | 8.00E-04 | 2.94E-03 | -1.37E+00 | 1.22E-01 | 3.36E-01 | 1.03E+00 |
| PDIA3 | 6.69E-13 | 3.49E-12 | -1.29E+00 | 1.32E-06 | 5.70E-06 | 1.01E+00 | 3.28E-04 | 1.39E-03 | -1.15E+00 | 2.06E-03 | 7.88E-03 | 1.01E+00 | 6.71E-11 | 2.39E-10 | -1.49E+00 | 1.34E-03 | 3.79E-03 | -1.14E+00 | 5.66E-02 | 2.08E-01 | -1.11E+00 | 2.05E-01 | 5.64E-01 | 1.10E+00 |
| PPM1K | 6.69E-13 | 3.49E-12 | -1.67E+00 | 3.82E-04 | 1.65E-03 | -1.00E+00 | 2.90E-10 | 1.23E-09 | -1.56E+00 | 4.60E-04 | 1.76E-03 | -1.03E+00 | 8.76E-15 | 3.12E-14 | -1.91E+00 | 1.07E-07 | 3.02E-07 | -1.22E+00 | 1.53E-18 | 5.63E-18 | -2.12E+00 | 1.07E-08 | 2.95E-08 | -1.47E+00 |
| ACOT1 | 6.69E-13 | 3.49E-12 | -1.35E+00 | 7.06E-09 | 3.05E-08 | -1.04E+00 | 1.78E-04 | 7.58E-04 | -1.22E+00 | 3.70E-03 | 1.41E-02 | 1.02E+00 | 1.37E-08 | 4.90E-08 | -1.46E+00 | 3.66E-02 | 1.03E-01 | 1.16E+00 | 2.14E-04 | 7.84E-04 | -1.10E+00 | 4.63E-03 | 1.27E-02 | 1.02E+00 |
| TCOF1 | 6.77E-13 | 3.53E-12 | -1.33E+00 | 1.15E-11 | 4.96E-11 | -1.16E+00 | 8.68E-08 | 3.69E-07 | -1.37E+00 | 9.38E-04 | 3.58E-03 | 1.04E+00 | 8.47E-04 | 3.02E-03 | -1.21E+00 | 1.21E-01 | 3.42E-01 | 1.24E+00 | 1.96E-05 | 7.18E-05 | -1.21E+00 | 7.59E-02 | 2.08E-01 | 1.13E+00 |
| FAIM | 6.80E-13 | 3.54E-12 | -1.55E+00 | 5.87E-05 | 2.54E-04 | 1.08E+00 | 9.24E-09 | 3.92E-08 | -1.51E+00 | 3.86E-07 | 1.47E-06 | -1.18E+00 | 1.84E-09 | 6.55E-09 | -1.66E+00 | 1.76E-04 | 4.95E-04 | -1.08E+00 | 1.37E-05 | 5.01E-05 | -1.23E+00 | 3.81E-03 | 1.05E-02 | -1.02E+00 |
| DGKZ | 6.84E-13 | 3.56E-12 | -1.33E+00 | 1.37E-10 | 5.92E-10 | -1.14E+00 | 1.39E-09 | 5.92E-09 | -1.56E+00 | 9.97E-08 | 3.81E-07 | -1.28E+00 | 2.86E-05 | 1.02E-04 | -1.19E+00 | 1.63E-05 | 4.60E-05 | -1.19E+00 | 1.46E-05 | 5.37E-05 | -1.23E+00 | 1.15E-03 | 3.16E-03 | -1.15E+00 |
| RBMXL1 | 7.00E-13 | 3.65E-12 | -1.32E+00 | 9.51E-21 | 4.11E-20 | -1.49E+00 | 3.74E-09 | 1.59E-08 | -1.46E+00 | 1.32E-15 | 5.05E-15 | -1.53E+00 | 3.46E-12 | 1.23E-11 | -1.58E+00 | 9.41E-14 | 2.65E-13 | -1.60E+00 | 4.95E-13 | 1.82E-12 | -1.54E+00 | 6.02E-12 | 1.65E-11 | -1.54E+00 |
| ST3GAL2 | 7.01E-13 | 3.65E-12 | -1.65E+00 | 2.93E-06 | 1.27E-05 | -1.02E+00 | 3.10E-05 | 1.32E-04 | -1.48E+00 | 1.36E-01 | 5.19E-01 | 1.24E+00 | 1.82E-06 | 6.51E-06 | -1.70E+00 | 4.14E-02 | 1.17E-01 | 1.13E+00 | 2.66E-16 | 9.76E-16 | -2.10E+00 | 1.45E-09 | 3.98E-09 | -1.66E+00 |
| PDIA6 | 7.01E-13 | 3.66E-12 | -1.52E+00 | 3.39E-04 | 1.46E-03 | 1.11E+00 | 5.43E-03 | 2.31E-02 | -1.06E+00 | 2.06E-02 | 7.86E-02 | 1.13E+00 | 4.53E-09 | 1.62E-08 | -1.51E+00 | 1.68E-05 | 4.73E-05 | -1.27E+00 | 3.46E-03 | 1.27E-02 | -1.23E+00 | 2.39E-01 | 6.57E-01 | 1.04E+00 |
| ZNF136 | 7.08E-13 | 3.69E-12 | -1.52E+00 | 6.57E-12 | 2.84E-11 | -1.30E+00 | 3.86E-04 | 1.64E-03 | -1.14E+00 | 1.13E-04 | 4.33E-04 | -1.13E+00 | 1.38E-02 | 4.92E-02 | -1.04E+00 | 1.84E-03 | 5.17E-03 | -1.28E+00 | 6.25E-04 | 2.29E-03 | -1.10E+00 | 1.68E-05 | 4.62E-05 | -1.27E+00 |
| C9orf139 | 7.08E-13 | 3.69E-12 | -1.41E+00 | 8.19E-19 | 3.54E-18 | 2.71E+00 | 1.30E-10 | 5.52E-10 | -1.74E+00 | 1.95E-05 | 7.46E-05 | 2.13E+00 | 8.31E-03 | 2.96E-02 | -1.41E+00 | 9.13E-02 | 2.57E-01 | 1.09E+00 | 2.33E-01 | 8.57E-01 | 1.43E+00 | 1.84E-01 | 5.05E-01 | 1.98E+00 |
| RPS15AP38 | 7.10E-13 | 3.70E-12 | 1.45E+00 | 9.12E-02 | 3.94E-01 | 1.07E+00 | 2.64E-06 | 1.12E-05 | 1.37E+00 | 1.16E-01 | 4.42E-01 | 1.01E+00 | 6.00E-06 | 2.14E-05 | 1.41E+00 | 3.51E-01 | 9.88E-01 | 1.07E+00 | 1.07E-04 | 3.93E-04 | 1.37E+00 | 2.93E-01 | 8.05E-01 | 1.02E+00 |
| THAP5 | 7.10E-13 | 3.70E-12 | -1.88E+00 | 2.29E-06 | 9.89E-06 | -1.12E+00 | 1.12E-10 | 4.75E-10 | -1.96E+00 | 1.49E-05 | 5.69E-05 | -1.22E+00 | 4.41E-07 | 1.57E-06 | -1.73E+00 | 2.41E-03 | 6.79E-03 | -1.08E+00 | 2.59E-15 | 9.52E-15 | -1.91E+00 | 7.57E-06 | 2.08E-05 | -1.30E+00 |
| ETFRF1 | 7.16E-13 | 3.73E-12 | -1.29E+00 | 3.06E-15 | 1.32E-14 | -1.22E+00 | 2.46E-04 | 1.04E-03 | -1.17E+00 | 1.24E-07 | 4.75E-07 | -1.26E+00 | 1.31E-07 | 4.66E-07 | -1.36E+00 | 1.60E-07 | 4.52E-07 | -1.31E+00 | 1.38E-03 | 5.05E-03 | 1.02E+00 | 2.94E-04 | 8.07E-04 | -1.04E+00 |
| HSH2D | 7.29E-13 | 3.80E-12 | -1.29E+00 | 2.40E-09 | 1.04E-08 | -1.08E+00 | 6.18E-09 | 2.63E-08 | -1.63E+00 | 7.24E-04 | 2.77E-03 | -1.11E+00 | 2.15E-11 | 7.67E-11 | -1.47E+00 | 2.83E-08 | 7.98E-08 | -1.23E+00 | 5.37E-05 | 1.97E-04 | -1.26E+00 | 3.09E-04 | 8.49E-04 | -1.14E+00 |
| EIF4A1P10 | 7.31E-13 | 3.81E-12 | -1.37E+00 | 1.08E-08 | 4.67E-08 | -1.24E+00 | 1.26E-06 | 5.37E-06 | -1.39E+00 | 5.05E-05 | 1.93E-04 | -1.30E+00 | 1.03E-21 | 3.68E-21 | -1.61E+00 | 1.01E-08 | 2.86E-08 | -1.34E+00 | 9.06E-09 | 3.33E-08 | -1.33E+00 | 5.23E-03 | 1.44E-02 | -1.18E+00 |
| EXOSC2 | 7.36E-13 | 3.84E-12 | -1.85E+00 | 1.78E-07 | 7.69E-07 | -1.20E+00 | 3.03E-05 | 1.29E-04 | -1.20E+00 | 9.76E-03 | 3.73E-02 | 1.21E+00 | 1.42E-02 | 5.07E-02 | -1.07E+00 | 2.16E-01 | 6.08E-01 | 1.58E+00 | 1.50E-15 | 5.52E-15 | -1.54E+00 | 3.18E-05 | 8.74E-05 | 1.00E+00 |
| BAZ2B | 7.39E-13 | 3.85E-12 | -1.25E+00 | 3.27E-12 | 1.41E-11 | -1.13E+00 | 1.08E-09 | 4.60E-09 | -1.59E+00 | 7.08E-09 | 2.70E-08 | -1.39E+00 | 6.21E-05 | 2.21E-04 | -1.20E+00 | 4.41E-06 | 1.24E-05 | -1.24E+00 | 5.48E-11 | 2.01E-10 | -1.66E+00 | 5.53E-10 | 1.52E-09 | -1.63E+00 |
| FBXO7 | 7.45E-13 | 3.88E-12 | -1.15E+00 | 2.31E-06 | 1.00E-05 | 1.13E+00 | 3.39E-08 | 1.44E-07 | -1.20E+00 | 8.74E-06 | 3.34E-05 | 1.00E+00 | 1.65E-05 | 5.88E-05 | -1.24E+00 | 5.15E-02 | 1.45E-01 | 1.05E+00 | 4.71E-02 | 1.73E-01 | -1.04E+00 | 1.57E-01 | 4.31E-01 | 1.03E+00 |
| SNX20 | 7.47E-13 | 3.89E-12 | -1.72E+00 | 2.03E-05 | 8.78E-05 | 1.05E+00 | 3.17E-11 | 1.35E-10 | -1.87E+00 | 2.82E-08 | 1.08E-07 | -1.41E+00 | 3.73E-10 | 1.33E-09 | -2.38E+00 | 3.66E-06 | 1.03E-05 | -1.49E+00 | 6.41E-23 | 2.35E-22 | -2.90E+00 | 2.28E-09 | 6.25E-09 | -1.54E+00 |
| PGLYRP2 | 7.48E-13 | 3.90E-12 | -1.77E+00 | 9.57E-11 | 4.13E-10 | -1.35E+00 | 4.88E-02 | 2.07E-01 | -1.24E+00 | 1.12E-01 | 4.29E-01 | 1.02E+00 | 2.23E-01 | 7.96E-01 | -1.24E+00 | 3.42E-01 | 9.65E-01 | 1.07E+00 | 1.51E-01 | 5.56E-01 | -1.24E+00 | 2.00E-01 | 5.50E-01 | -1.25E+00 |
| LDHA | 7.66E-13 | 3.99E-12 | -1.33E+00 | 4.54E-05 | 1.96E-04 | -1.05E+00 | 1.65E-02 | 7.00E-02 | -1.12E+00 | 9.95E-02 | 3.80E-01 | -1.01E+00 | 2.53E-16 | 9.01E-16 | -1.48E+00 | 2.33E-02 | 6.56E-02 | 1.04E+00 | 5.89E-03 | 2.16E-02 | -1.06E+00 | 1.27E-01 | 3.49E-01 | 1.17E+00 |
| BAG2 | 7.71E-13 | 4.02E-12 | -1.67E+00 | 1.77E-08 | 7.64E-08 | -1.19E+00 | 3.38E-11 | 1.43E-10 | -1.72E+00 | 3.52E-09 | 1.34E-08 | -1.45E+00 | 2.78E-10 | 9.90E-10 | -1.88E+00 | 6.43E-07 | 1.81E-06 | -1.37E+00 | 3.37E-17 | 1.24E-16 | -1.85E+00 | 2.81E-12 | 7.72E-12 | -1.58E+00 |
| RHOT1 | 7.88E-13 | 4.11E-12 | -1.76E+00 | 3.13E-05 | 1.35E-04 | 1.05E+00 | 2.72E-11 | 1.16E-10 | -1.68E+00 | 1.42E-07 | 5.44E-07 | -1.20E+00 | 1.05E-06 | 3.76E-06 | -1.58E+00 | 4.33E-02 | 1.22E-01 | 1.16E+00 | 3.92E-15 | 1.44E-14 | -1.86E+00 | 6.91E-07 | 1.90E-06 | -1.31E+00 |
| ZBTB44 | 7.90E-13 | 4.12E-12 | -1.38E+00 | 2.84E-02 | 1.23E-01 | 1.30E+00 | 5.01E-09 | 2.13E-08 | -1.44E+00 | 4.85E-03 | 1.85E-02 | 1.06E+00 | 9.63E-12 | 3.43E-11 | -1.70E+00 | 3.67E-05 | 1.03E-04 | -1.16E+00 | 1.90E-07 | 6.98E-07 | -1.39E+00 | 1.99E-02 | 5.48E-02 | 1.02E+00 |
| STX3 | 7.95E-13 | 4.14E-12 | -1.89E+00 | 2.65E-09 | 1.14E-08 | -1.38E+00 | 5.77E-03 | 2.45E-02 | -1.21E+00 | 1.02E-02 | 3.90E-02 | -1.11E+00 | 9.74E-02 | 3.47E-01 | 2.03E+00 | 5.21E-02 | 1.47E-01 | 2.31E+00 | 1.54E-02 | 5.66E-02 | 2.91E+00 | 9.14E-03 | 2.51E-02 | 3.38E+00 |
| PSME2P2 | 7.95E-13 | 4.14E-12 | -1.30E+00 | 3.07E-16 | 1.33E-15 | -1.32E+00 | 5.42E-04 | 2.30E-03 | -1.24E+00 | 4.02E-07 | 1.54E-06 | -1.32E+00 | 2.14E-14 | 7.61E-14 | -1.70E+00 | 4.99E-17 | 1.41E-16 | -1.77E+00 | 7.01E-16 | 2.57E-15 | -1.87E+00 | 2.22E-11 | 6.09E-11 | -1.68E+00 |
| BCL11A | 7.96E-13 | 4.15E-12 | -1.35E+00 | 2.15E-01 | 9.28E-01 | 1.20E+00 | 2.53E-09 | 1.07E-08 | -1.72E+00 | 1.11E-01 | 4.23E-01 | -1.02E+00 | 1.22E-05 | 4.36E-05 | -1.14E+00 | 7.02E-02 | 1.98E-01 | 1.30E+00 | 1.60E-10 | 5.86E-10 | -1.60E+00 | 3.02E-02 | 8.29E-02 | -1.14E+00 |
| SMARCC2 | 8.11E-13 | 4.23E-12 | -1.35E+00 | 2.48E-10 | 1.07E-09 | -1.13E+00 | 5.26E-12 | 2.24E-11 | -1.49E+00 | 3.64E-10 | 1.39E-09 | -1.25E+00 | 4.09E-06 | 1.46E-05 | -1.24E+00 | 6.57E-07 | 1.85E-06 | -1.22E+00 | 6.71E-05 | 2.46E-04 | -1.18E+00 | 3.04E-04 | 8.34E-04 | -1.15E+00 |
| MPP6 | 8.19E-13 | 4.27E-12 | -1.54E+00 | 5.94E-07 | 2.57E-06 | -1.08E+00 | 1.40E-06 | 5.95E-06 | -1.39E+00 | 2.87E-04 | 1.10E-03 | -1.05E+00 | 6.87E-02 | 2.45E-01 | -1.04E+00 | 1.53E-01 | 4.32E-01 | 1.65E+00 | 5.06E-06 | 1.86E-05 | -1.33E+00 | 9.01E-02 | 2.47E-01 | 1.10E+00 |
| AP001992.1 | 8.26E-13 | 4.30E-12 | -1.51E+00 | 2.54E-17 | 1.10E-16 | -1.63E+00 | 1.55E-17 | 6.57E-17 | -2.09E+00 | 2.52E-19 | 9.64E-19 | -2.01E+00 | 2.94E-11 | 1.05E-10 | -1.86E+00 | 2.22E-12 | 6.26E-12 | -1.90E+00 | 5.31E-11 | 1.95E-10 | -1.70E+00 | 6.86E-10 | 1.88E-09 | -1.65E+00 |
| URM1 | 8.30E-13 | 4.32E-12 | -1.28E+00 | 3.52E-10 | 1.52E-09 | -1.02E+00 | 1.04E-05 | 4.42E-05 | -1.08E+00 | 7.20E-04 | 2.75E-03 | 1.14E+00 | 3.91E-07 | 1.39E-06 | -1.25E+00 | 7.04E-03 | 1.98E-02 | 1.07E+00 | 1.54E-05 | 5.65E-05 | -1.17E+00 | 9.11E-03 | 2.50E-02 | 1.01E+00 |
| TAP1 | 8.30E-13 | 4.32E-12 | -1.83E+00 | 3.01E-14 | 1.30E-13 | -1.90E+00 | 1.47E-15 | 6.24E-15 | -2.23E+00 | 1.86E-19 | 7.10E-19 | -2.59E+00 | 3.73E-08 | 1.33E-07 | -2.45E+00 | 6.01E-06 | 1.70E-05 | -1.70E+00 | 2.99E-14 | 1.10E-13 | -2.31E+00 | 8.70E-14 | 2.39E-13 | -2.16E+00 |
| ESCO1 | 8.36E-13 | 4.36E-12 | -1.56E+00 | 6.43E-07 | 2.78E-06 | -1.05E+00 | 3.22E-10 | 1.37E-09 | -1.56E+00 | 1.41E-07 | 5.38E-07 | -1.20E+00 | 2.83E-10 | 1.01E-09 | -1.51E+00 | 2.74E-09 | 7.71E-09 | -1.48E+00 | 6.12E-10 | 2.25E-09 | -1.45E+00 | 4.11E-07 | 1.13E-06 | -1.36E+00 |
| AC011495.1 | 8.58E-13 | 4.47E-12 | -1.67E+00 | 5.08E-13 | 2.20E-12 | -1.56E+00 | 1.00E-05 | 4.25E-05 | -1.42E+00 | 3.48E-07 | 1.33E-06 | -1.40E+00 | 6.61E-08 | 2.36E-07 | -1.87E+00 | 1.50E-10 | 4.22E-10 | -2.11E+00 | 1.70E-08 | 6.23E-08 | -1.58E+00 | 2.71E-11 | 7.43E-11 | -1.95E+00 |
| ATP13A3 | 8.60E-13 | 4.48E-12 | -1.64E+00 | 5.98E-11 | 2.58E-10 | -1.29E+00 | 1.83E-06 | 7.78E-06 | -1.30E+00 | 8.05E-07 | 3.07E-06 | -1.18E+00 | 1.33E-03 | 4.76E-03 | -1.17E+00 | 1.30E-04 | 3.68E-04 | -1.19E+00 | 1.75E-10 | 6.41E-10 | -1.53E+00 | 5.67E-04 | 1.56E-03 | -1.04E+00 |
| APLP2 | 8.63E-13 | 4.50E-12 | -1.36E+00 | 3.62E-06 | 1.56E-05 | -1.15E+00 | 2.56E-04 | 1.09E-03 | -1.31E+00 | 1.49E-02 | 5.70E-02 | -1.14E+00 | 4.56E-03 | 1.62E-02 | -1.13E+00 | 4.16E-02 | 1.17E-01 | 1.00E+00 | 1.64E-02 | 6.03E-02 | 1.03E+00 | 3.58E-01 | 9.83E-01 | 1.13E+00 |
| FTH1P16 | 8.75E-13 | 4.56E-12 | -1.73E+00 | 4.04E-13 | 1.75E-12 | -1.59E+00 | 5.36E-06 | 2.28E-05 | -1.39E+00 | 1.59E-06 | 6.08E-06 | -1.30E+00 | 8.98E-22 | 3.20E-21 | -1.79E+00 | 1.21E-24 | 3.40E-24 | -1.98E+00 | 4.41E-14 | 1.62E-13 | -1.53E+00 | 1.27E-14 | 3.48E-14 | -1.52E+00 |
| AC022182.2 | 8.76E-13 | 4.57E-12 | -2.59E+00 | 2.08E-12 | 8.97E-12 | -2.47E+00 | 3.41E-09 | 1.45E-08 | -2.14E+00 | 2.82E-09 | 1.08E-08 | -2.14E+00 | 9.54E-04 | 3.40E-03 | -2.60E+00 | 3.89E-03 | 1.10E-02 | -2.06E+00 | 2.91E-06 | 1.07E-05 | -2.13E+00 | 1.86E-04 | 5.10E-04 | -1.76E+00 |
| TAL1 | 8.78E-13 | 4.58E-12 | -1.82E+00 | 1.62E-07 | 6.98E-07 | -1.21E+00 | 3.96E-14 | 1.68E-13 | -2.07E+00 | 1.23E-07 | 4.70E-07 | -1.32E+00 | 1.53E-24 | 5.44E-24 | -2.19E+00 | 1.58E-03 | 4.45E-03 | 1.03E+00 | 2.74E-16 | 1.01E-15 | -1.43E+00 | 1.45E-06 | 3.99E-06 | -1.10E+00 |
| GTPBP10 | 8.80E-13 | 4.59E-12 | -1.97E+00 | 1.85E-07 | 7.99E-07 | -1.25E+00 | 9.01E-13 | 3.83E-12 | -1.98E+00 | 5.00E-06 | 1.91E-05 | -1.12E+00 | 2.06E-03 | 7.33E-03 | -1.21E+00 | 2.53E-02 | 7.13E-02 | 1.05E+00 | 4.29E-08 | 1.57E-07 | -1.52E+00 | 2.51E-10 | 6.89E-10 | -1.80E+00 |
| EWSR1 | 8.86E-13 | 4.62E-12 | -1.20E+00 | 5.97E-09 | 2.58E-08 | 1.00E+00 | 1.20E-02 | 5.10E-02 | -1.08E+00 | 2.45E-03 | 9.36E-03 | -1.00E+00 | 2.72E-06 | 9.69E-06 | -1.15E+00 | 4.55E-05 | 1.28E-04 | -1.06E+00 | 1.04E-03 | 3.82E-03 | -1.14E+00 | 1.39E-01 | 3.82E-01 | 1.02E+00 |
| UBN1 | 8.86E-13 | 4.62E-12 | -1.52E+00 | 2.71E-11 | 1.17E-10 | -1.26E+00 | 4.11E-08 | 1.75E-07 | -1.36E+00 | 7.92E-05 | 3.03E-04 | -1.01E+00 | 1.30E-07 | 4.63E-07 | -1.30E+00 | 1.66E-09 | 4.68E-09 | -1.35E+00 | 2.96E-07 | 1.09E-06 | -1.20E+00 | 1.27E-03 | 3.48E-03 | -1.03E+00 |
| DGCR6 | 8.88E-13 | 4.63E-12 | -1.81E+00 | 3.68E-16 | 1.59E-15 | -2.04E+00 | 1.73E-02 | 7.34E-02 | -1.02E+00 | 3.74E-05 | 1.43E-04 | -1.22E+00 | 8.33E-03 | 2.97E-02 | -1.53E+00 | 6.14E-03 | 1.73E-02 | -1.70E+00 | 2.02E-03 | 7.40E-03 | -1.07E+00 | 1.04E-06 | 2.86E-06 | -1.53E+00 |
| ATM | 8.91E-13 | 4.64E-12 | -1.30E+00 | 3.21E-03 | 1.39E-02 | 1.25E+00 | 3.26E-07 | 1.39E-06 | -1.46E+00 | 5.28E-04 | 2.02E-03 | 1.01E+00 | 5.09E-04 | 1.81E-03 | -1.24E+00 | 2.22E-01 | 6.26E-01 | 1.18E+00 | 7.97E-08 | 2.92E-07 | -1.49E+00 | 1.23E-02 | 3.38E-02 | -1.05E+00 |
| OGFOD1 | 8.95E-13 | 4.67E-12 | -1.78E+00 | 1.96E-08 | 8.46E-08 | -1.27E+00 | 1.69E-05 | 7.18E-05 | -1.24E+00 | 1.52E-02 | 5.79E-02 | 1.18E+00 | 1.81E-03 | 6.46E-03 | -1.05E+00 | 9.72E-02 | 2.74E-01 | 1.04E+00 | 5.40E-03 | 1.98E-02 | 1.11E+00 | 3.43E-01 | 9.42E-01 | 1.26E+00 |
| LPIN1 | 8.96E-13 | 4.67E-12 | -1.38E+00 | 8.35E-06 | 3.61E-05 | 1.03E+00 | 1.04E-05 | 4.41E-05 | -1.27E+00 | 1.04E-03 | 3.97E-03 | 1.05E+00 | 2.10E-10 | 7.49E-10 | -1.73E+00 | 3.06E-06 | 8.62E-06 | -1.30E+00 | 5.69E-11 | 2.09E-10 | -1.85E+00 | 6.35E-05 | 1.74E-04 | -1.31E+00 |
| NCSTN | 8.96E-13 | 4.67E-12 | -1.61E+00 | 1.12E-05 | 4.82E-05 | -1.03E+00 | 1.55E-07 | 6.58E-07 | -1.47E+00 | 1.26E-03 | 4.83E-03 | -1.02E+00 | 1.55E-02 | 5.51E-02 | -1.02E+00 | 1.82E-01 | 5.12E-01 | 1.25E+00 | 7.18E-06 | 2.64E-05 | -1.27E+00 | 3.85E-02 | 1.06E-01 | 1.06E+00 |
| PLD6 | 9.05E-13 | 4.72E-12 | -2.95E+00 | 2.68E-11 | 1.16E-10 | -2.39E+00 | 3.53E-09 | 1.50E-08 | -2.02E+00 | 2.37E-03 | 9.06E-03 | -1.01E+00 | 4.46E-04 | 1.59E-03 | -2.31E+00 | 1.18E-02 | 3.32E-02 | -1.43E+00 | 5.09E-09 | 1.87E-08 | -1.93E+00 | 1.09E-04 | 2.99E-04 | -1.23E+00 |
| GLTP | 9.10E-13 | 4.74E-12 | -1.64E+00 | 8.23E-08 | 3.55E-07 | -1.14E+00 | 7.62E-05 | 3.24E-04 | -1.23E+00 | 1.33E-04 | 5.08E-04 | -1.07E+00 | 4.36E-05 | 1.55E-04 | -1.20E+00 | 4.58E-04 | 1.29E-03 | -1.07E+00 | 4.37E-07 | 1.60E-06 | -1.27E+00 | 1.84E-03 | 5.05E-03 | -1.10E+00 |
| ING3 | 9.12E-13 | 4.75E-12 | -1.34E+00 | 2.36E-17 | 1.02E-16 | -1.38E+00 | 6.89E-11 | 2.93E-10 | -1.50E+00 | 1.08E-15 | 4.11E-15 | -1.58E+00 | 1.09E-03 | 3.89E-03 | -1.05E+00 | 1.02E-06 | 2.88E-06 | -1.28E+00 | 2.64E-10 | 9.70E-10 | -1.36E+00 | 6.19E-10 | 1.70E-09 | -1.40E+00 |
| RB1CC1 | 9.14E-13 | 4.76E-12 | -1.30E+00 | 1.64E-24 | 7.07E-24 | -1.50E+00 | 1.08E-05 | 4.61E-05 | -1.24E+00 | 2.47E-16 | 9.45E-16 | -1.68E+00 | 2.24E-06 | 7.99E-06 | -1.25E+00 | 5.34E-14 | 1.51E-13 | -1.59E+00 | 4.66E-08 | 1.71E-07 | -1.29E+00 | 3.24E-16 | 8.90E-16 | -1.73E+00 |
| VCP | 9.26E-13 | 4.82E-12 | -1.14E+00 | 1.12E-09 | 4.82E-09 | 1.01E+00 | 1.07E-02 | 4.55E-02 | -1.00E+00 | 5.01E-02 | 1.91E-01 | 1.15E+00 | 3.09E-07 | 1.10E-06 | -1.37E+00 | 5.71E-02 | 1.61E-01 | -1.00E+00 | 3.69E-04 | 1.36E-03 | -1.20E+00 | 1.38E-01 | 3.78E-01 | 1.04E+00 |
| PPP2R5C | 9.49E-13 | 4.94E-12 | -1.37E+00 | 2.86E-14 | 1.24E-13 | -1.31E+00 | 4.30E-04 | 1.83E-03 | -1.20E+00 | 1.29E-05 | 4.95E-05 | -1.19E+00 | 2.96E-06 | 1.05E-05 | -1.22E+00 | 1.54E-02 | 4.35E-02 | 1.13E+00 | 8.49E-07 | 3.12E-06 | -1.24E+00 | 1.69E-04 | 4.65E-04 | -1.18E+00 |
| CCDC66 | 9.69E-13 | 5.05E-12 | -1.67E+00 | 1.57E-04 | 6.78E-04 | 1.04E+00 | 1.01E-08 | 4.31E-08 | -1.55E+00 | 2.02E-04 | 7.72E-04 | -1.05E+00 | 2.98E-06 | 1.06E-05 | -1.58E+00 | 1.47E-03 | 4.13E-03 | -1.19E+00 | 2.86E-06 | 1.05E-05 | -1.40E+00 | 4.10E-02 | 1.13E-01 | 1.11E+00 |
| HDAC10 | 9.80E-13 | 5.11E-12 | -1.95E+00 | 2.47E-10 | 1.07E-09 | -1.47E+00 | 5.85E-11 | 2.48E-10 | -1.96E+00 | 3.15E-05 | 1.20E-04 | -1.12E+00 | 1.51E-06 | 5.39E-06 | -1.72E+00 | 2.81E-04 | 7.93E-04 | -1.23E+00 | 1.44E-09 | 5.27E-09 | -1.56E+00 | 4.92E-06 | 1.35E-05 | -1.20E+00 |
| AC012354.2 | 9.83E-13 | 5.12E-12 | -1.24E+00 | 7.17E-40 | 3.10E-39 | -1.70E+00 | 5.11E-04 | 2.17E-03 | -1.13E+00 | 1.27E-12 | 4.84E-12 | -1.46E+00 | 9.34E-09 | 3.33E-08 | -1.24E+00 | 3.73E-17 | 1.05E-16 | -1.57E+00 | 4.79E-04 | 1.76E-03 | -1.14E+00 | 1.47E-10 | 4.04E-10 | -1.56E+00 |
| PTMA | 9.84E-13 | 5.13E-12 | 1.27E+00 | 1.85E-07 | 7.98E-07 | 1.21E+00 | 7.54E-05 | 3.21E-04 | 1.17E+00 | 1.86E-04 | 7.12E-04 | 1.16E+00 | 4.26E-02 | 1.52E-01 | -1.06E+00 | 3.49E-01 | 9.85E-01 | 1.01E+00 | 1.50E-01 | 5.52E-01 | 1.01E+00 | 2.43E-04 | 6.68E-04 | 1.17E+00 |
| RNF44 | 9.85E-13 | 5.14E-12 | -1.69E+00 | 6.58E-10 | 2.84E-09 | -1.34E+00 | 9.19E-11 | 3.91E-10 | -1.70E+00 | 2.73E-09 | 1.04E-08 | -1.43E+00 | 2.42E-15 | 8.63E-15 | -1.75E+00 | 5.80E-15 | 1.64E-14 | -1.59E+00 | 8.29E-12 | 3.04E-11 | -1.55E+00 | 6.29E-11 | 1.73E-10 | -1.55E+00 |
| RMND1 | 9.88E-13 | 5.15E-12 | -1.61E+00 | 4.63E-09 | 2.00E-08 | -1.20E+00 | 1.65E-08 | 7.02E-08 | -1.55E+00 | 2.40E-08 | 9.17E-08 | -1.33E+00 | 3.33E-05 | 1.19E-04 | -1.46E+00 | 5.47E-03 | 1.54E-02 | -1.12E+00 | 4.36E-05 | 1.60E-04 | -1.31E+00 | 9.17E-04 | 2.52E-03 | -1.15E+00 |
| MFSD8 | 9.95E-13 | 5.18E-12 | -1.83E+00 | 6.22E-04 | 2.69E-03 | 1.19E+00 | 1.47E-06 | 6.23E-06 | -1.71E+00 | 6.01E-02 | 2.30E-01 | 1.19E+00 | 4.42E-06 | 1.58E-05 | -1.63E+00 | 1.85E-01 | 5.20E-01 | 1.32E+00 | 2.34E-05 | 8.57E-05 | -1.40E+00 | 1.86E-02 | 5.12E-02 | -1.07E+00 |
| TSPAN33 | 1.02E-12 | 5.32E-12 | -1.62E+00 | 7.85E-10 | 3.39E-09 | -1.24E+00 | 2.30E-04 | 9.77E-04 | -1.24E+00 | 1.07E-06 | 4.09E-06 | -1.33E+00 | 7.08E-03 | 2.52E-02 | -1.25E+00 | 3.85E-02 | 1.09E-01 | 1.01E+00 | 1.38E-01 | 5.07E-01 | 1.68E+00 | 3.21E-01 | 8.81E-01 | 1.13E+00 |
| PPIF | 1.04E-12 | 5.41E-12 | -2.03E+00 | 7.03E-11 | 3.04E-10 | -1.71E+00 | 9.09E-04 | 3.86E-03 | -1.18E+00 | 2.48E-06 | 9.47E-06 | -1.28E+00 | 1.66E-03 | 5.91E-03 | -1.12E+00 | 3.24E-02 | 9.14E-02 | 1.03E+00 | 3.06E-05 | 1.12E-04 | -1.00E+00 | 1.75E-02 | 4.80E-02 | -1.05E+00 |
| OBSL1 | 1.05E-12 | 5.48E-12 | -1.75E+00 | 1.39E-09 | 5.99E-09 | -1.36E+00 | 5.82E-05 | 2.47E-04 | -1.33E+00 | 8.37E-05 | 3.20E-04 | -1.13E+00 | 8.93E-05 | 3.18E-04 | -1.68E+00 | 3.39E-03 | 9.54E-03 | -1.26E+00 | 2.86E-04 | 1.05E-03 | -1.25E+00 | 1.27E-03 | 3.50E-03 | -1.11E+00 |
| XPC | 1.05E-12 | 5.48E-12 | -1.77E+00 | 2.09E-07 | 9.05E-07 | -1.18E+00 | 5.32E-10 | 2.26E-09 | -1.84E+00 | 3.72E-10 | 1.42E-09 | -1.62E+00 | 1.56E-04 | 5.56E-04 | -1.27E+00 | 1.44E-03 | 4.07E-03 | -1.10E+00 | 1.11E-04 | 4.08E-04 | -1.23E+00 | 5.52E-04 | 1.52E-03 | -1.14E+00 |
| ABHD17AP1 | 1.05E-12 | 5.48E-12 | -1.43E+00 | 2.10E-09 | 9.08E-09 | -1.13E+00 | 4.42E-05 | 1.88E-04 | -1.29E+00 | 2.98E-04 | 1.14E-03 | -1.11E+00 | 2.50E-06 | 8.91E-06 | -1.15E+00 | 1.05E-04 | 2.97E-04 | -1.07E+00 | 3.72E-02 | 1.37E-01 | 1.14E+00 | 3.38E-02 | 9.29E-02 | 1.11E+00 |
| CEP57 | 1.05E-12 | 5.49E-12 | -1.23E+00 | 7.18E-09 | 3.10E-08 | -1.03E+00 | 1.78E-03 | 7.55E-03 | -1.16E+00 | 8.78E-06 | 3.35E-05 | -1.16E+00 | 3.87E-08 | 1.38E-07 | -1.15E+00 | 1.26E-05 | 3.56E-05 | -1.03E+00 | 8.71E-07 | 3.20E-06 | -1.27E+00 | 6.51E-04 | 1.79E-03 | -1.17E+00 |
| AL158801.4 | 1.06E-12 | 5.54E-12 | -1.57E+00 | 1.23E-13 | 5.33E-13 | -1.38E+00 | 3.85E-08 | 1.64E-07 | -1.30E+00 | 5.96E-08 | 2.28E-07 | -1.18E+00 | 5.84E-12 | 2.08E-11 | -1.74E+00 | 2.74E-09 | 7.71E-09 | -1.37E+00 | 1.78E-15 | 6.52E-15 | -1.63E+00 | 3.19E-04 | 8.77E-04 | -1.02E+00 |
| SNRPEP7 | 1.06E-12 | 5.54E-12 | -2.44E+00 | 3.52E-10 | 1.52E-09 | -1.85E+00 | 3.38E-06 | 1.44E-05 | -1.56E+00 | 1.04E-07 | 3.96E-07 | -1.59E+00 | 2.24E-06 | 8.00E-06 | -2.42E+00 | 1.72E-05 | 4.85E-05 | -1.86E+00 | 2.64E-10 | 9.69E-10 | -1.81E+00 | 2.85E-06 | 7.83E-06 | -1.49E+00 |
| APMAP | 1.07E-12 | 5.58E-12 | -1.46E+00 | 1.06E-07 | 4.58E-07 | -1.09E+00 | 1.26E-09 | 5.37E-09 | -1.48E+00 | 4.58E-05 | 1.75E-04 | -1.03E+00 | 2.01E-06 | 7.15E-06 | -1.41E+00 | 8.42E-04 | 2.37E-03 | -1.15E+00 | 1.00E-05 | 3.67E-05 | -1.17E+00 | 1.77E-01 | 4.87E-01 | 1.17E+00 |
| USP11 | 1.08E-12 | 5.62E-12 | -1.35E+00 | 6.21E-06 | 2.68E-05 | 1.07E+00 | 1.02E-05 | 4.35E-05 | -1.25E+00 | 1.49E-04 | 5.68E-04 | 1.01E+00 | 9.80E-07 | 3.49E-06 | -1.34E+00 | 1.66E-04 | 4.67E-04 | -1.09E+00 | 1.67E-06 | 6.13E-06 | -1.29E+00 | 1.89E-03 | 5.20E-03 | -1.01E+00 |
| REPIN1 | 1.10E-12 | 5.74E-12 | -1.43E+00 | 1.31E-06 | 5.64E-06 | 1.01E+00 | 9.18E-04 | 3.90E-03 | -1.18E+00 | 1.93E-04 | 7.37E-04 | -1.07E+00 | 3.44E-10 | 1.23E-09 | -1.36E+00 | 1.11E-03 | 3.13E-03 | 1.08E+00 | 8.20E-07 | 3.01E-06 | -1.31E+00 | 2.09E-02 | 5.74E-02 | 1.02E+00 |
| PLA2G12A | 1.11E-12 | 5.78E-12 | -1.54E+00 | 5.05E-11 | 2.18E-10 | -1.26E+00 | 8.83E-08 | 3.75E-07 | -1.29E+00 | 5.38E-07 | 2.06E-06 | -1.13E+00 | 1.40E-06 | 4.99E-06 | -1.32E+00 | 5.30E-03 | 1.50E-02 | 1.07E+00 | 3.46E-07 | 1.27E-06 | -1.19E+00 | 6.94E-02 | 1.91E-01 | 1.12E+00 |
| GPBP1 | 1.12E-12 | 5.81E-12 | -1.26E+00 | 7.94E-10 | 3.43E-09 | -1.06E+00 | 1.85E-07 | 7.87E-07 | -1.34E+00 | 2.80E-04 | 1.07E-03 | -1.03E+00 | 7.25E-08 | 2.59E-07 | -1.20E+00 | 1.03E-06 | 2.90E-06 | -1.14E+00 | 1.74E-05 | 6.39E-05 | -1.24E+00 | 8.26E-04 | 2.27E-03 | -1.16E+00 |
| ARHGAP22 | 1.12E-12 | 5.84E-12 | -1.22E+00 | 1.61E-02 | 6.98E-02 | 1.42E+00 | 1.61E-06 | 6.85E-06 | -1.38E+00 | 5.95E-02 | 2.27E-01 | 1.19E+00 | 5.03E-09 | 1.79E-08 | -1.41E+00 | 2.47E-01 | 6.96E-01 | 1.35E+00 | 9.93E-08 | 3.65E-07 | -1.44E+00 | 2.08E-02 | 5.70E-02 | -1.00E+00 |
| P2RX7 | 1.13E-12 | 5.90E-12 | -1.91E+00 | 8.02E-11 | 3.46E-10 | -1.56E+00 | 1.58E-08 | 6.73E-08 | -1.99E+00 | 1.24E-03 | 4.72E-03 | -1.09E+00 | 3.67E-04 | 1.31E-03 | -1.70E+00 | 8.07E-02 | 2.27E-01 | 1.09E+00 | 2.93E-04 | 1.08E-03 | -1.49E+00 | 3.51E-03 | 9.63E-03 | -1.21E+00 |
| SIAH1 | 1.13E-12 | 5.91E-12 | -1.57E+00 | 2.94E-12 | 1.27E-11 | -1.42E+00 | 1.78E-07 | 7.57E-07 | -1.59E+00 | 2.73E-05 | 1.04E-04 | -1.16E+00 | 2.65E-09 | 9.46E-09 | -1.46E+00 | 1.17E-09 | 3.30E-09 | -1.33E+00 | 2.45E-11 | 8.97E-11 | -1.56E+00 | 2.29E-08 | 6.28E-08 | -1.29E+00 |
| SYNGR2 | 1.14E-12 | 5.92E-12 | -1.54E+00 | 1.45E-08 | 6.26E-08 | -1.20E+00 | 3.70E-10 | 1.57E-09 | -1.61E+00 | 7.64E-04 | 2.92E-03 | -1.04E+00 | 3.00E-15 | 1.07E-14 | -1.85E+00 | 1.45E-06 | 4.09E-06 | -1.27E+00 | 8.33E-08 | 3.06E-07 | -1.45E+00 | 6.27E-03 | 1.72E-02 | -1.08E+00 |
| TTC9C | 1.14E-12 | 5.94E-12 | -1.68E+00 | 9.44E-04 | 4.08E-03 | 1.18E+00 | 6.11E-09 | 2.60E-08 | -1.64E+00 | 1.20E-03 | 4.59E-03 | 1.01E+00 | 4.41E-11 | 1.57E-10 | -2.60E+00 | 3.28E-05 | 9.25E-05 | -1.31E+00 | 1.80E-07 | 6.59E-07 | -1.51E+00 | 5.17E-03 | 1.42E-02 | -1.07E+00 |
| ALG3 | 1.14E-12 | 5.95E-12 | -1.52E+00 | 4.60E-06 | 1.99E-05 | -1.07E+00 | 4.76E-05 | 2.02E-04 | -1.19E+00 | 4.11E-03 | 1.57E-02 | 1.09E+00 | 1.33E-07 | 4.76E-07 | -1.55E+00 | 6.19E-02 | 1.74E-01 | 1.13E+00 | 2.00E-06 | 7.33E-06 | -1.23E+00 | 7.30E-03 | 2.00E-02 | 1.10E+00 |
| SNRNP200 | 1.16E-12 | 6.03E-12 | -1.31E+00 | 2.13E-09 | 9.21E-09 | -1.11E+00 | 3.25E-05 | 1.38E-04 | -1.15E+00 | 1.03E-03 | 3.95E-03 | 1.05E+00 | 1.57E-03 | 5.58E-03 | 1.02E+00 | 1.61E-02 | 4.53E-02 | 1.14E+00 | 3.47E-03 | 1.27E-02 | -1.05E+00 | 3.12E-01 | 8.57E-01 | 1.13E+00 |
| TXLNG | 1.16E-12 | 6.05E-12 | -1.58E+00 | 8.63E-06 | 3.73E-05 | -1.00E+00 | 6.22E-08 | 2.64E-07 | -1.39E+00 | 1.01E-03 | 3.87E-03 | 1.09E+00 | 2.32E-05 | 8.26E-05 | -1.58E+00 | 1.43E-01 | 4.03E-01 | 1.14E+00 | 8.25E-12 | 3.03E-11 | -1.55E+00 | 3.93E-03 | 1.08E-02 | 1.05E+00 |
| OAF | 1.16E-12 | 6.06E-12 | -2.05E+00 | 4.63E-09 | 2.00E-08 | -1.37E+00 | 6.58E-16 | 2.79E-15 | -1.95E+00 | 1.97E-08 | 7.54E-08 | -1.26E+00 | 5.89E-09 | 2.10E-08 | -3.69E+00 | 4.71E-05 | 1.33E-04 | -1.70E+00 | 2.29E-07 | 8.41E-07 | -1.89E+00 | 5.01E-02 | 1.38E-01 | 1.04E+00 |
| TBCB | 1.16E-12 | 6.06E-12 | -1.20E+00 | 5.32E-09 | 2.30E-08 | 1.00E+00 | 1.67E-03 | 7.12E-03 | -1.07E+00 | 2.83E-03 | 1.08E-02 | 1.14E+00 | 7.97E-08 | 2.84E-07 | -1.28E+00 | 3.97E-04 | 1.12E-03 | 1.02E+00 | 2.59E-05 | 9.51E-05 | -1.17E+00 | 3.09E-02 | 8.50E-02 | 1.03E+00 |
| UQCR11 | 1.17E-12 | 6.12E-12 | -1.17E+00 | 6.55E-36 | 2.83E-35 | -1.66E+00 | 6.34E-02 | 2.70E-01 | 1.06E+00 | 2.13E-16 | 8.13E-16 | -1.59E+00 | 1.37E-06 | 4.90E-06 | -1.16E+00 | 1.14E-12 | 3.21E-12 | -1.50E+00 | 3.83E-02 | 1.41E-01 | 1.02E+00 | 8.97E-11 | 2.46E-10 | -1.54E+00 |
| RANBP2 | 1.17E-12 | 6.12E-12 | -1.47E+00 | 2.17E-10 | 9.37E-10 | -1.26E+00 | 1.67E-08 | 7.09E-08 | -1.53E+00 | 3.25E-06 | 1.24E-05 | -1.21E+00 | 1.24E-06 | 4.42E-06 | -1.29E+00 | 1.70E-06 | 4.78E-06 | -1.30E+00 | 2.75E-08 | 1.01E-07 | -1.31E+00 | 1.76E-05 | 4.84E-05 | -1.14E+00 |
| PPFIBP1 | 1.19E-12 | 6.22E-12 | -1.35E+00 | 1.71E-04 | 7.38E-04 | 1.14E+00 | 5.39E-08 | 2.29E-07 | -1.42E+00 | 2.20E-04 | 8.41E-04 | -1.04E+00 | 6.76E-04 | 2.41E-03 | -1.37E+00 | 5.51E-02 | 1.55E-01 | -1.00E+00 | 1.25E-07 | 4.59E-07 | -1.47E+00 | 6.50E-06 | 1.79E-05 | -1.37E+00 |
| POMGNT1 | 1.25E-12 | 6.53E-12 | -1.67E+00 | 1.18E-06 | 5.09E-06 | -1.06E+00 | 4.31E-04 | 1.83E-03 | -1.32E+00 | 1.52E-02 | 5.80E-02 | 1.02E+00 | 2.95E-06 | 1.05E-05 | -1.42E+00 | 2.17E-03 | 6.12E-03 | -1.04E+00 | 7.81E-05 | 2.87E-04 | -1.29E+00 | 1.76E-01 | 4.83E-01 | 1.15E+00 |
| UPF3A | 1.25E-12 | 6.53E-12 | -1.29E+00 | 5.15E-10 | 2.22E-09 | -1.01E+00 | 2.05E-06 | 8.69E-06 | -1.29E+00 | 1.57E-08 | 6.01E-08 | -1.20E+00 | 1.42E-13 | 5.08E-13 | -1.48E+00 | 3.58E-08 | 1.01E-07 | -1.26E+00 | 1.25E-06 | 4.57E-06 | -1.28E+00 | 1.53E-05 | 4.21E-05 | -1.26E+00 |
| NDUFS1 | 1.26E-12 | 6.59E-12 | -1.45E+00 | 2.87E-06 | 1.24E-05 | -1.02E+00 | 5.34E-09 | 2.27E-08 | -1.39E+00 | 3.81E-07 | 1.45E-06 | -1.15E+00 | 6.76E-10 | 2.41E-09 | -1.37E+00 | 8.09E-07 | 2.28E-06 | -1.07E+00 | 2.32E-05 | 8.53E-05 | -1.27E+00 | 4.95E-02 | 1.36E-01 | 1.02E+00 |
| VPS28 | 1.27E-12 | 6.64E-12 | -1.17E+00 | 2.61E-10 | 1.13E-09 | -1.04E+00 | 2.12E-03 | 8.99E-03 | -1.02E+00 | 1.44E-03 | 5.50E-03 | -1.01E+00 | 1.06E-04 | 3.78E-04 | -1.18E+00 | 1.06E-01 | 2.99E-01 | 1.05E+00 | 6.06E-02 | 2.23E-01 | 1.00E+00 | 4.06E-02 | 1.11E-01 | -1.07E+00 |
| SRRM1 | 1.30E-12 | 6.78E-12 | -1.26E+00 | 2.01E-24 | 8.69E-24 | -1.37E+00 | 1.32E-02 | 5.61E-02 | -1.14E+00 | 2.21E-05 | 8.45E-05 | -1.25E+00 | 8.21E-04 | 2.93E-03 | -1.18E+00 | 2.17E-08 | 6.11E-08 | -1.40E+00 | 1.38E-02 | 5.08E-02 | -1.11E+00 | 2.26E-02 | 6.19E-02 | -1.14E+00 |
| AC007537.1 | 1.30E-12 | 6.79E-12 | -1.29E+00 | 4.72E-25 | 2.04E-24 | -1.70E+00 | 4.81E-09 | 2.04E-08 | -1.43E+00 | 1.47E-23 | 5.60E-23 | -1.88E+00 | 3.16E-09 | 1.13E-08 | -1.53E+00 | 7.46E-19 | 2.10E-18 | -2.13E+00 | 2.52E-14 | 9.26E-14 | -1.59E+00 | 4.04E-22 | 1.11E-21 | -1.99E+00 |
| FUI | 1.32E-12 | 6.86E-12 | -1.44E+00 | 2.66E-08 | 1.15E-07 | -1.07E+00 | 4.25E-06 | 1.80E-05 | -1.23E+00 | 1.64E-04 | 6.25E-04 | -1.02E+00 | 3.86E-07 | 1.38E-06 | -1.33E+00 | 1.39E-04 | 3.92E-04 | -1.07E+00 | 2.23E-04 | 8.17E-04 | -1.15E+00 | 8.21E-03 | 2.25E-02 | -1.01E+00 |
| BLMH | 1.32E-12 | 6.87E-12 | -1.33E+00 | 1.10E-06 | 4.73E-06 | 1.00E+00 | 5.49E-08 | 2.33E-07 | -1.37E+00 | 1.80E-07 | 6.88E-07 | -1.15E+00 | 4.14E-08 | 1.47E-07 | -1.55E+00 | 1.06E-03 | 2.98E-03 | -1.08E+00 | 2.49E-08 | 9.12E-08 | -1.28E+00 | 2.96E-03 | 8.13E-03 | 1.03E+00 |
| ERLIN1 | 1.34E-12 | 6.97E-12 | -1.45E+00 | 9.00E-12 | 3.89E-11 | -1.25E+00 | 1.96E-09 | 8.33E-09 | -1.51E+00 | 4.07E-06 | 1.55E-05 | -1.10E+00 | 3.59E-02 | 1.28E-01 | 1.07E+00 | 1.06E-01 | 3.00E-01 | 1.19E+00 | 1.74E-06 | 6.38E-06 | -1.34E+00 | 6.98E-04 | 1.92E-03 | -1.12E+00 |
| COG2 | 1.34E-12 | 7.01E-12 | -1.90E+00 | 5.88E-07 | 2.54E-06 | -1.19E+00 | 1.97E-05 | 8.38E-05 | -1.23E+00 | 7.76E-05 | 2.96E-04 | -1.01E+00 | 1.21E-09 | 4.33E-09 | -2.31E+00 | 1.04E-03 | 2.94E-03 | -1.06E+00 | 7.32E-07 | 2.69E-06 | -1.46E+00 | 5.30E-04 | 1.46E-03 | -1.22E+00 |
| TXNL4A | 1.35E-12 | 7.05E-12 | -1.31E+00 | 4.59E-15 | 1.98E-14 | -1.24E+00 | 5.33E-03 | 2.26E-02 | -1.05E+00 | 3.70E-03 | 1.41E-02 | 1.01E+00 | 2.89E-09 | 1.03E-08 | -1.33E+00 | 9.06E-08 | 2.55E-07 | -1.26E+00 | 4.98E-06 | 1.83E-05 | -1.13E+00 | 5.02E-02 | 1.38E-01 | 1.03E+00 |
| PSMB2 | 1.37E-12 | 7.13E-12 | -1.27E+00 | 3.03E-09 | 1.31E-08 | -1.03E+00 | 4.76E-03 | 2.02E-02 | -1.08E+00 | 8.13E-04 | 3.10E-03 | -1.08E+00 | 3.12E-09 | 1.11E-08 | -1.35E+00 | 3.41E-04 | 9.61E-04 | -1.07E+00 | 2.36E-03 | 8.68E-03 | -1.18E+00 | 2.25E-01 | 6.18E-01 | 1.01E+00 |
| EPC2 | 1.42E-12 | 7.40E-12 | -1.56E+00 | 4.51E-11 | 1.95E-10 | -1.33E+00 | 3.64E-06 | 1.55E-05 | -1.32E+00 | 2.07E-06 | 7.90E-06 | -1.21E+00 | 1.72E-04 | 6.12E-04 | -1.14E+00 | 4.72E-05 | 1.33E-04 | -1.21E+00 | 1.03E-06 | 3.77E-06 | -1.31E+00 | 1.75E-06 | 4.80E-06 | -1.38E+00 |
| C17orf62 | 1.45E-12 | 7.57E-12 | -1.54E+00 | 1.00E-04 | 4.32E-04 | 1.02E+00 | 2.15E-10 | 9.15E-10 | -1.41E+00 | 2.64E-05 | 1.01E-04 | -1.01E+00 | 8.86E-14 | 3.16E-13 | -1.60E+00 | 3.03E-04 | 8.53E-04 | -1.01E+00 | 9.57E-09 | 3.51E-08 | -1.31E+00 | 2.46E-02 | 6.75E-02 | 1.09E+00 |
| UBL5P2 | 1.45E-12 | 7.57E-12 | -1.30E+00 | 3.26E-33 | 1.41E-32 | -1.94E+00 | 1.95E-03 | 8.30E-03 | -1.02E+00 | 2.80E-17 | 1.07E-16 | -1.62E+00 | 5.73E-07 | 2.04E-06 | -1.35E+00 | 5.59E-18 | 1.57E-17 | -1.98E+00 | 9.00E-06 | 3.30E-05 | -1.19E+00 | 9.87E-12 | 2.71E-11 | -1.54E+00 |
| CBFA2T3 | 1.46E-12 | 7.60E-12 | -1.41E+00 | 1.08E-05 | 4.68E-05 | 1.05E+00 | 1.83E-06 | 7.79E-06 | -1.49E+00 | 9.74E-02 | 3.72E-01 | 1.16E+00 | 5.72E-15 | 2.04E-14 | -1.96E+00 | 1.97E-04 | 5.54E-04 | -1.17E+00 | 5.80E-14 | 2.13E-13 | -1.51E+00 | 5.25E-06 | 1.44E-05 | -1.19E+00 |
| PPP1R16A | 1.46E-12 | 7.63E-12 | -1.74E+00 | 1.09E-07 | 4.69E-07 | -1.19E+00 | 6.04E-06 | 2.56E-05 | -1.58E+00 | 7.62E-04 | 2.91E-03 | -1.12E+00 | 6.24E-03 | 2.22E-02 | -1.32E+00 | 1.46E-01 | 4.13E-01 | 1.24E+00 | 4.98E-02 | 1.83E-01 | -1.03E+00 | 2.73E-01 | 7.50E-01 | 1.21E+00 |
| VKORC1 | 1.47E-12 | 7.64E-12 | -1.33E+00 | 1.84E-14 | 7.94E-14 | -1.29E+00 | 4.16E-05 | 1.77E-04 | -1.13E+00 | 8.28E-08 | 3.16E-07 | -1.11E+00 | 2.83E-09 | 1.01E-08 | -1.43E+00 | 6.26E-09 | 1.77E-08 | -1.35E+00 | 1.24E-04 | 4.54E-04 | -1.14E+00 | 5.21E-03 | 1.43E-02 | -1.06E+00 |
| DNAJC7 | 1.47E-12 | 7.65E-12 | -1.40E+00 | 9.45E-16 | 4.08E-15 | -1.32E+00 | 1.42E-08 | 6.04E-08 | -1.49E+00 | 5.13E-10 | 1.96E-09 | -1.42E+00 | 1.74E-08 | 6.19E-08 | -1.26E+00 | 1.03E-08 | 2.91E-08 | -1.25E+00 | 2.10E-07 | 7.69E-07 | -1.38E+00 | 9.66E-06 | 2.65E-05 | -1.25E+00 |
| CCZ1 | 1.47E-12 | 7.67E-12 | -1.50E+00 | 7.23E-10 | 3.12E-09 | -1.22E+00 | 7.72E-11 | 3.28E-10 | -1.57E+00 | 7.76E-11 | 2.96E-10 | -1.41E+00 | 2.21E-06 | 7.90E-06 | -1.43E+00 | 4.68E-08 | 1.32E-07 | -1.62E+00 | 1.43E-11 | 5.25E-11 | -1.59E+00 | 3.00E-08 | 8.25E-08 | -1.30E+00 |
| UHMK1 | 1.47E-12 | 7.67E-12 | -1.49E+00 | 2.96E-06 | 1.28E-05 | -1.03E+00 | 1.52E-12 | 6.44E-12 | -1.71E+00 | 8.44E-09 | 3.22E-08 | -1.28E+00 | 7.97E-14 | 2.84E-13 | -1.65E+00 | 1.94E-11 | 5.47E-11 | -1.36E+00 | 2.72E-07 | 9.98E-07 | -1.33E+00 | 1.32E-02 | 3.62E-02 | -1.04E+00 |
| POLE3 | 1.47E-12 | 7.68E-12 | -1.44E+00 | 1.95E-09 | 8.42E-09 | -1.14E+00 | 1.88E-09 | 7.97E-09 | -1.46E+00 | 2.80E-06 | 1.07E-05 | -1.20E+00 | 1.02E-10 | 3.62E-10 | -1.42E+00 | 2.84E-06 | 8.00E-06 | -1.11E+00 | 8.01E-06 | 2.94E-05 | -1.22E+00 | 1.11E-01 | 3.05E-01 | 1.06E+00 |
| WASHC2C | 1.47E-12 | 7.68E-12 | -1.67E+00 | 1.38E-06 | 5.97E-06 | -1.10E+00 | 1.64E-06 | 6.98E-06 | -1.27E+00 | 1.68E-08 | 6.43E-08 | -1.20E+00 | 1.18E-04 | 4.21E-04 | -1.21E+00 | 5.89E-05 | 1.66E-04 | -1.18E+00 | 4.14E-09 | 1.52E-08 | -1.33E+00 | 1.29E-07 | 3.53E-07 | -1.22E+00 |
| HADH | 1.48E-12 | 7.72E-12 | -1.39E+00 | 7.53E-11 | 3.25E-10 | -1.20E+00 | 6.95E-06 | 2.95E-05 | -1.19E+00 | 3.76E-07 | 1.44E-06 | -1.18E+00 | 1.32E-07 | 4.70E-07 | -1.36E+00 | 5.74E-03 | 1.62E-02 | 1.06E+00 | 5.55E-05 | 2.04E-04 | -1.12E+00 | 1.63E-01 | 4.49E-01 | 1.12E+00 |
| ACTL6A | 1.50E-12 | 7.84E-12 | -1.81E+00 | 3.37E-05 | 1.46E-04 | 1.02E+00 | 3.27E-05 | 1.39E-04 | -1.30E+00 | 6.42E-02 | 2.45E-01 | 1.17E+00 | 2.33E-07 | 8.30E-07 | -2.21E+00 | 4.23E-02 | 1.19E-01 | -1.01E+00 | 6.05E-15 | 2.22E-14 | -1.53E+00 | 4.02E-02 | 1.10E-01 | 1.07E+00 |
| RTL8C | 1.52E-12 | 7.92E-12 | -1.72E+00 | 1.20E-06 | 5.18E-06 | -1.08E+00 | 1.52E-05 | 6.44E-05 | -1.47E+00 | 4.51E-03 | 1.72E-02 | -1.01E+00 | 2.33E-09 | 8.30E-09 | -1.59E+00 | 7.52E-06 | 2.12E-05 | -1.20E+00 | 8.46E-07 | 3.11E-06 | -1.34E+00 | 8.47E-04 | 2.33E-03 | -1.11E+00 |
| BAG3 | 1.54E-12 | 8.04E-12 | -1.72E+00 | 4.04E-21 | 1.75E-20 | -2.93E+00 | 8.01E-13 | 3.40E-12 | -2.02E+00 | 6.20E-21 | 2.37E-20 | -3.08E+00 | 2.64E-02 | 9.40E-02 | -1.26E+00 | 4.82E-03 | 1.36E-02 | -1.79E+00 | 2.61E-01 | 9.59E-01 | 1.18E+00 | 7.93E-02 | 2.18E-01 | -1.38E+00 |
| STRN3 | 1.55E-12 | 8.10E-12 | -1.46E+00 | 2.37E-12 | 1.03E-11 | -1.25E+00 | 1.37E-07 | 5.84E-07 | -1.56E+00 | 2.08E-05 | 7.95E-05 | -1.21E+00 | 1.09E-06 | 3.89E-06 | -1.52E+00 | 3.42E-05 | 9.65E-05 | -1.21E+00 | 7.14E-11 | 2.62E-10 | -1.53E+00 | 2.23E-07 | 6.12E-07 | -1.34E+00 |
| TUG1 | 1.56E-12 | 8.13E-12 | -1.45E+00 | 1.07E-06 | 4.61E-06 | -1.03E+00 | 7.24E-06 | 3.07E-05 | -1.34E+00 | 4.76E-03 | 1.82E-02 | -1.01E+00 | 6.50E-11 | 2.32E-10 | -1.43E+00 | 5.06E-06 | 1.43E-05 | -1.10E+00 | 9.50E-11 | 3.49E-10 | -1.37E+00 | 4.21E-05 | 1.16E-04 | -1.07E+00 |
| MAPK1IP1L | 1.57E-12 | 8.20E-12 | -1.19E+00 | 1.79E-17 | 7.72E-17 | -1.19E+00 | 1.90E-04 | 8.05E-04 | -1.18E+00 | 5.98E-07 | 2.28E-06 | -1.18E+00 | 7.85E-09 | 2.80E-08 | -1.30E+00 | 9.57E-08 | 2.70E-07 | -1.27E+00 | 6.87E-07 | 2.52E-06 | -1.31E+00 | 2.64E-04 | 7.25E-04 | -1.18E+00 |
| SLC25A38 | 1.59E-12 | 8.27E-12 | -1.47E+00 | 8.11E-07 | 3.50E-06 | -1.03E+00 | 3.73E-04 | 1.59E-03 | -1.22E+00 | 9.50E-03 | 3.63E-02 | 1.10E+00 | 4.20E-08 | 1.50E-07 | -1.56E+00 | 5.88E-03 | 1.66E-02 | -1.02E+00 | 3.98E-07 | 1.46E-06 | -1.29E+00 | 4.58E-04 | 1.26E-03 | -1.11E+00 |
| AC108134.2 | 1.60E-12 | 8.36E-12 | -2.09E+00 | 3.61E-18 | 1.56E-17 | -2.94E+00 | 1.53E-03 | 6.48E-03 | -1.80E+00 | 9.94E-04 | 3.80E-03 | -1.70E+00 | 2.47E-01 | 8.80E-01 | 1.52E+00 | 3.49E-01 | 9.84E-01 | 1.30E+00 | 6.66E-02 | 2.44E-01 | -2.89E+00 | 2.49E-01 | 6.84E-01 | -1.07E+00 |
| TADA3 | 1.62E-12 | 8.46E-12 | -1.30E+00 | 7.85E-10 | 3.39E-09 | -1.09E+00 | 1.58E-08 | 6.70E-08 | -1.36E+00 | 1.18E-07 | 4.52E-07 | -1.15E+00 | 6.81E-11 | 2.43E-10 | -1.34E+00 | 7.85E-04 | 2.21E-03 | 1.00E+00 | 4.56E-03 | 1.67E-02 | -1.06E+00 | 6.44E-03 | 1.77E-02 | -1.02E+00 |
| SMC3 | 1.63E-12 | 8.48E-12 | -1.17E+00 | 1.79E-04 | 7.72E-04 | 1.16E+00 | 8.59E-02 | 3.65E-01 | 1.03E+00 | 2.30E-01 | 8.79E-01 | 1.24E+00 | 5.84E-09 | 2.08E-08 | -1.33E+00 | 2.61E-03 | 7.37E-03 | -1.01E+00 | 1.25E-04 | 4.59E-04 | -1.12E+00 | 1.64E-01 | 4.51E-01 | 1.07E+00 |
| MRPS16 | 1.63E-12 | 8.50E-12 | -1.38E+00 | 1.50E-10 | 6.47E-10 | -1.15E+00 | 6.20E-03 | 2.63E-02 | 1.02E+00 | 7.31E-04 | 2.79E-03 | 1.01E+00 | 2.47E-06 | 8.82E-06 | -1.15E+00 | 5.12E-05 | 1.44E-04 | -1.03E+00 | 8.30E-04 | 3.05E-03 | -1.11E+00 | 1.38E-02 | 3.79E-02 | -1.00E+00 |
| GNAQ | 1.63E-12 | 8.52E-12 | -1.29E+00 | 2.82E-12 | 1.22E-11 | -1.18E+00 | 1.39E-05 | 5.91E-05 | -1.27E+00 | 1.90E-09 | 7.27E-09 | -1.30E+00 | 6.23E-06 | 2.22E-05 | -1.39E+00 | 6.32E-05 | 1.78E-04 | -1.32E+00 | 3.93E-06 | 1.44E-05 | -1.36E+00 | 2.41E-03 | 6.61E-03 | -1.18E+00 |
| WAS | 1.64E-12 | 8.56E-12 | -1.26E+00 | 5.47E-09 | 2.36E-08 | -1.04E+00 | 2.10E-07 | 8.94E-07 | -1.34E+00 | 2.14E-06 | 8.19E-06 | -1.12E+00 | 3.18E-02 | 1.13E-01 | -1.01E+00 | 5.77E-02 | 1.63E-01 | -1.00E+00 | 4.37E-02 | 1.60E-01 | -1.03E+00 | 4.43E-02 | 1.22E-01 | -1.01E+00 |
| DAAM1 | 1.67E-12 | 8.71E-12 | -1.82E+00 | 7.48E-04 | 3.23E-03 | 1.01E+00 | 1.59E-07 | 6.76E-07 | -1.56E+00 | 4.19E-05 | 1.60E-04 | -1.19E+00 | 8.22E-02 | 2.93E-01 | 1.05E+00 | 1.48E-01 | 4.16E-01 | 1.65E+00 | 8.77E-04 | 3.22E-03 | -1.02E+00 | 2.05E-01 | 5.62E-01 | -1.12E+00 |
| GPX1 | 1.69E-12 | 8.82E-12 | -1.30E+00 | 6.06E-11 | 2.62E-10 | -1.23E+00 | 1.96E-07 | 8.33E-07 | -1.42E+00 | 3.59E-05 | 1.37E-04 | -1.30E+00 | 1.77E-06 | 6.31E-06 | -1.51E+00 | 3.17E-05 | 8.93E-05 | -1.51E+00 | 6.23E-04 | 2.29E-03 | -1.26E+00 | 9.50E-04 | 2.61E-03 | -1.37E+00 |
| SINHCAF | 1.74E-12 | 9.08E-12 | -1.29E+00 | 2.49E-17 | 1.08E-16 | -1.26E+00 | 8.69E-05 | 3.69E-04 | -1.31E+00 | 1.71E-06 | 6.53E-06 | -1.31E+00 | 5.53E-13 | 1.97E-12 | -1.58E+00 | 7.85E-16 | 2.21E-15 | -1.57E+00 | 1.32E-13 | 4.85E-13 | -1.80E+00 | 9.48E-10 | 2.60E-09 | -1.63E+00 |
| LRPAP1 | 1.74E-12 | 9.09E-12 | -1.20E+00 | 7.33E-11 | 3.17E-10 | -1.07E+00 | 6.77E-02 | 2.88E-01 | -1.01E+00 | 3.64E-02 | 1.39E-01 | 1.02E+00 | 3.49E-06 | 1.25E-05 | -1.10E+00 | 1.07E-05 | 3.01E-05 | -1.03E+00 | 8.48E-02 | 3.11E-01 | 1.16E+00 | 1.48E-01 | 4.05E-01 | 1.20E+00 |
| MTRNR2L12 | 1.75E-12 | 9.13E-12 | 1.33E+00 | 2.15E-21 | 9.30E-21 | -1.32E+00 | 1.33E-03 | 5.65E-03 | 1.19E+00 | 3.31E-14 | 1.27E-13 | -1.38E+00 | 5.74E-06 | 2.05E-05 | 1.33E+00 | 3.14E-06 | 8.85E-06 | -1.19E+00 | 7.64E-06 | 2.80E-05 | 1.29E+00 | 1.49E-08 | 4.10E-08 | -1.31E+00 |
| MOB1A | 1.75E-12 | 9.13E-12 | -1.21E+00 | 4.59E-15 | 1.98E-14 | -1.17E+00 | 1.23E-04 | 5.24E-04 | -1.23E+00 | 2.23E-05 | 8.53E-05 | -1.11E+00 | 3.76E-05 | 1.34E-04 | -1.15E+00 | 1.31E-04 | 3.68E-04 | -1.14E+00 | 2.63E-05 | 9.65E-05 | -1.29E+00 | 1.97E-02 | 5.42E-02 | -1.07E+00 |
| RAB2A | 1.79E-12 | 9.34E-12 | -1.21E+00 | 5.66E-09 | 2.45E-08 | -1.01E+00 | 2.27E-04 | 9.64E-04 | -1.30E+00 | 9.25E-06 | 3.53E-05 | -1.19E+00 | 7.71E-07 | 2.75E-06 | -1.18E+00 | 2.95E-05 | 8.33E-05 | -1.06E+00 | 5.25E-09 | 1.93E-08 | -1.41E+00 | 1.48E-05 | 4.05E-05 | -1.22E+00 |
| PTRHD1 | 1.79E-12 | 9.35E-12 | -1.23E+00 | 7.50E-25 | 3.24E-24 | -1.42E+00 | 3.42E-04 | 1.45E-03 | -1.13E+00 | 7.27E-07 | 2.78E-06 | -1.17E+00 | 5.52E-04 | 1.97E-03 | -1.05E+00 | 4.55E-05 | 1.28E-04 | -1.19E+00 | 1.32E-02 | 4.83E-02 | 1.02E+00 | 1.08E-03 | 2.97E-03 | -1.11E+00 |
| KHDRBS1 | 1.81E-12 | 9.43E-12 | -1.30E+00 | 1.73E-06 | 7.46E-06 | -1.09E+00 | 1.99E-05 | 8.46E-05 | -1.38E+00 | 4.70E-04 | 1.79E-03 | -1.21E+00 | 1.20E-09 | 4.29E-09 | -1.49E+00 | 7.79E-07 | 2.20E-06 | -1.31E+00 | 6.77E-08 | 2.49E-07 | -1.38E+00 | 7.08E-03 | 1.95E-02 | -1.12E+00 |
| IRAK4 | 1.81E-12 | 9.43E-12 | -1.59E+00 | 1.49E-07 | 6.44E-07 | -1.15E+00 | 5.46E-06 | 2.32E-05 | -1.46E+00 | 2.55E-03 | 9.75E-03 | -1.00E+00 | 1.64E-07 | 5.84E-07 | -1.44E+00 | 7.32E-05 | 2.06E-04 | -1.07E+00 | 1.92E-06 | 7.05E-06 | -1.30E+00 | 1.61E-03 | 4.43E-03 | -1.14E+00 |
| RTCA | 1.83E-12 | 9.52E-12 | -1.47E+00 | 1.49E-13 | 6.42E-13 | -1.33E+00 | 5.73E-07 | 2.43E-06 | -1.34E+00 | 1.54E-07 | 5.87E-07 | -1.25E+00 | 5.94E-09 | 2.12E-08 | -1.32E+00 | 7.27E-06 | 2.05E-05 | -1.07E+00 | 2.86E-06 | 1.05E-05 | -1.20E+00 | 6.77E-11 | 1.86E-10 | -1.46E+00 |
| SLC35E3 | 1.83E-12 | 9.52E-12 | -1.56E+00 | 9.49E-04 | 4.10E-03 | 1.10E+00 | 3.59E-05 | 1.52E-04 | -1.34E+00 | 2.63E-03 | 1.01E-02 | 1.06E+00 | 2.39E-08 | 8.51E-08 | -1.45E+00 | 1.31E-02 | 3.69E-02 | 1.19E+00 | 5.07E-04 | 1.86E-03 | -1.26E+00 | 2.65E-01 | 7.28E-01 | 1.16E+00 |
| NDUFS2 | 1.83E-12 | 9.53E-12 | -1.29E+00 | 2.49E-14 | 1.08E-13 | -1.24E+00 | 2.60E-08 | 1.10E-07 | -1.40E+00 | 7.08E-09 | 2.70E-08 | -1.22E+00 | 7.26E-15 | 2.59E-14 | -1.54E+00 | 9.88E-11 | 2.78E-10 | -1.23E+00 | 3.91E-08 | 1.43E-07 | -1.33E+00 | 3.31E-04 | 9.09E-04 | -1.16E+00 |
| AL033519.2 | 1.84E-12 | 9.56E-12 | -1.16E+00 | 5.64E-02 | 2.43E-01 | 1.27E+00 | 5.03E-04 | 2.14E-03 | -1.17E+00 | 2.02E-01 | 7.73E-01 | 1.29E+00 | 1.08E-10 | 3.86E-10 | -1.26E+00 | 9.64E-02 | 2.72E-01 | 1.21E+00 | 3.68E-05 | 1.35E-04 | -1.16E+00 | 1.22E-01 | 3.36E-01 | 1.21E+00 |
| HEXDC | 1.84E-12 | 9.57E-12 | -1.45E+00 | 4.98E-07 | 2.15E-06 | -1.02E+00 | 9.16E-10 | 3.89E-09 | -1.38E+00 | 9.76E-06 | 3.73E-05 | -1.04E+00 | 2.58E-06 | 9.19E-06 | -1.23E+00 | 4.12E-04 | 1.16E-03 | -1.05E+00 | 7.92E-06 | 2.91E-05 | -1.30E+00 | 2.07E-04 | 5.68E-04 | -1.13E+00 |
| CD2AP | 1.88E-12 | 9.79E-12 | -1.36E+00 | 1.80E-10 | 7.78E-10 | -1.11E+00 | 7.94E-08 | 3.38E-07 | -1.48E+00 | 4.18E-09 | 1.60E-08 | -1.36E+00 | 5.61E-09 | 2.00E-08 | -1.52E+00 | 3.24E-05 | 9.13E-05 | -1.15E+00 | 3.71E-15 | 1.36E-14 | -1.65E+00 | 3.23E-09 | 8.88E-09 | -1.36E+00 |
| VHL | 1.90E-12 | 9.89E-12 | -1.58E+00 | 2.12E-04 | 9.16E-04 | 1.04E+00 | 1.59E-09 | 6.76E-09 | -1.57E+00 | 1.37E-05 | 5.22E-05 | -1.10E+00 | 4.55E-05 | 1.62E-04 | -1.19E+00 | 9.52E-03 | 2.68E-02 | 1.11E+00 | 2.29E-07 | 8.39E-07 | -1.34E+00 | 5.48E-04 | 1.51E-03 | -1.09E+00 |
| USP2 | 1.91E-12 | 9.97E-12 | -3.75E+00 | 7.34E-09 | 3.17E-08 | -2.17E+00 | 2.11E-02 | 8.97E-02 | -1.24E+00 | 4.14E-02 | 1.58E-01 | -1.18E+00 | 2.48E-03 | 8.83E-03 | -1.91E+00 | 1.26E-03 | 3.55E-03 | -2.28E+00 | 2.29E-02 | 8.39E-02 | -1.77E+00 | 2.29E-01 | 6.30E-01 | -1.23E+00 |
| ZNF280B | 1.93E-12 | 1.01E-11 | -2.12E+00 | 2.08E-10 | 9.00E-10 | -1.78E+00 | 7.12E-03 | 3.02E-02 | -1.44E+00 | 2.30E-02 | 8.78E-02 | -1.11E+00 | 5.86E-02 | 2.09E-01 | -1.14E+00 | 8.21E-02 | 2.31E-01 | -1.12E+00 | 1.77E-06 | 6.51E-06 | -2.04E+00 | 1.80E-04 | 4.95E-04 | -1.63E+00 |
| CFAP97 | 1.95E-12 | 1.01E-11 | -1.32E+00 | 7.63E-08 | 3.30E-07 | -1.06E+00 | 1.18E-04 | 5.00E-04 | -1.19E+00 | 1.59E-02 | 6.09E-02 | 1.09E+00 | 1.37E-08 | 4.88E-08 | -1.31E+00 | 1.02E-04 | 2.87E-04 | -1.03E+00 | 3.23E-03 | 1.18E-02 | -1.11E+00 | 2.16E-01 | 5.94E-01 | 1.05E+00 |
| DMAC1 | 1.96E-12 | 1.02E-11 | -1.33E+00 | 2.48E-10 | 1.07E-09 | -1.12E+00 | 3.91E-02 | 1.66E-01 | 1.02E+00 | 3.06E-02 | 1.17E-01 | 1.14E+00 | 4.29E-05 | 1.53E-04 | -1.16E+00 | 5.66E-02 | 1.60E-01 | 1.14E+00 | 3.88E-04 | 1.43E-03 | -1.08E+00 | 1.57E-01 | 4.31E-01 | 1.14E+00 |
| CCDC43 | 1.98E-12 | 1.03E-11 | -1.48E+00 | 1.63E-08 | 7.02E-08 | -1.11E+00 | 5.43E-08 | 2.31E-07 | -1.34E+00 | 1.64E-06 | 6.27E-06 | -1.14E+00 | 2.23E-08 | 7.95E-08 | -1.58E+00 | 1.79E-04 | 5.03E-04 | -1.15E+00 | 2.62E-08 | 9.61E-08 | -1.51E+00 | 1.22E-02 | 3.35E-02 | -1.03E+00 |
| PLA2R1 | 1.99E-12 | 1.04E-11 | -2.36E+00 | 5.73E-06 | 2.48E-05 | -1.20E+00 | 1.36E-04 | 5.79E-04 | -2.14E+00 | 2.34E-03 | 8.95E-03 | -1.52E+00 | 1.88E-01 | 6.71E-01 | -1.22E+00 | 3.42E-01 | 9.64E-01 | 1.13E+00 | 2.26E-01 | 8.30E-01 | 1.16E+00 | 3.05E-01 | 8.39E-01 | 1.17E+00 |
| EIF5B | 1.99E-12 | 1.04E-11 | -1.36E+00 | 5.32E-08 | 2.30E-07 | -1.10E+00 | 3.35E-04 | 1.42E-03 | -1.20E+00 | 6.52E-03 | 2.49E-02 | -1.07E+00 | 8.35E-06 | 2.98E-05 | -1.30E+00 | 1.54E-03 | 4.33E-03 | -1.11E+00 | 1.86E-02 | 6.82E-02 | -1.06E+00 | 3.23E-01 | 8.88E-01 | 1.01E+00 |
| AC093014.1 | 1.99E-12 | 1.04E-11 | -1.28E+00 | 7.80E-23 | 3.37E-22 | -1.49E+00 | 1.03E-06 | 4.37E-06 | -1.35E+00 | 3.87E-12 | 1.48E-11 | -1.46E+00 | 3.78E-09 | 1.35E-08 | -1.35E+00 | 7.41E-13 | 2.09E-12 | -1.49E+00 | 8.69E-10 | 3.19E-09 | -1.40E+00 | 1.70E-11 | 4.68E-11 | -1.49E+00 |
| ATP6V1B2 | 2.00E-12 | 1.04E-11 | -1.67E+00 | 3.56E-05 | 1.54E-04 | 1.05E+00 | 3.84E-12 | 1.63E-11 | -1.82E+00 | 2.40E-07 | 9.17E-07 | -1.24E+00 | 2.94E-09 | 1.05E-08 | -1.89E+00 | 3.69E-04 | 1.04E-03 | -1.11E+00 | 3.07E-18 | 1.13E-17 | -1.93E+00 | 7.78E-08 | 2.14E-07 | -1.26E+00 |
| ATF3 | 2.04E-12 | 1.06E-11 | -1.14E+00 | 1.22E-76 | 5.27E-76 | -9.43E+00 | 4.79E-08 | 2.04E-07 | -1.29E+00 | 4.54E-60 | 1.73E-59 | -6.55E+00 | 1.04E-01 | 3.71E-01 | 1.74E+00 | 6.12E-06 | 1.72E-05 | -5.24E+00 | 6.30E-02 | 2.31E-01 | 1.90E+00 | 4.20E-13 | 1.15E-12 | -3.87E+00 |
| CYB5R3 | 2.05E-12 | 1.07E-11 | -1.38E+00 | 9.87E-07 | 4.27E-06 | -1.00E+00 | 4.33E-03 | 1.84E-02 | -1.03E+00 | 8.49E-03 | 3.24E-02 | 1.08E+00 | 1.75E-04 | 6.23E-04 | -1.12E+00 | 1.81E-04 | 5.09E-04 | -1.12E+00 | 5.57E-04 | 2.04E-03 | -1.08E+00 | 2.51E-03 | 6.88E-03 | -1.07E+00 |
| POGZ | 2.06E-12 | 1.07E-11 | -1.23E+00 | 7.06E-13 | 3.05E-12 | -1.14E+00 | 1.79E-07 | 7.61E-07 | -1.26E+00 | 1.56E-07 | 5.95E-07 | -1.22E+00 | 3.31E-06 | 1.18E-05 | -1.21E+00 | 1.31E-04 | 3.70E-04 | -1.12E+00 | 4.76E-09 | 1.75E-08 | -1.43E+00 | 2.01E-04 | 5.53E-04 | -1.18E+00 |
| NDUFA10 | 2.06E-12 | 1.07E-11 | -1.17E+00 | 3.22E-12 | 1.39E-11 | -1.04E+00 | 6.11E-04 | 2.59E-03 | -1.07E+00 | 9.66E-05 | 3.69E-04 | -1.03E+00 | 8.84E-02 | 3.15E-01 | 1.15E+00 | 3.44E-01 | 9.70E-01 | 1.28E+00 | 1.31E-02 | 4.81E-02 | 1.01E+00 | 1.44E-01 | 3.95E-01 | 1.07E+00 |
| VPS4A | 2.11E-12 | 1.10E-11 | -1.54E+00 | 2.70E-08 | 1.17E-07 | -1.13E+00 | 5.68E-06 | 2.41E-05 | -1.20E+00 | 4.89E-05 | 1.87E-04 | 1.03E+00 | 2.38E-09 | 8.49E-09 | -1.42E+00 | 4.36E-04 | 1.23E-03 | 1.02E+00 | 1.33E-05 | 4.88E-05 | -1.11E+00 | 3.04E-03 | 8.36E-03 | 1.00E+00 |
| PDXDC1 | 2.12E-12 | 1.10E-11 | -1.63E+00 | 9.65E-08 | 4.17E-07 | -1.10E+00 | 1.60E-05 | 6.79E-05 | -1.42E+00 | 3.23E-04 | 1.24E-03 | -1.14E+00 | 3.36E-07 | 1.20E-06 | -1.38E+00 | 1.89E-04 | 5.33E-04 | -1.09E+00 | 2.10E-20 | 7.71E-20 | -1.75E+00 | 2.92E-08 | 8.03E-08 | -1.16E+00 |
| BTK | 2.12E-12 | 1.11E-11 | -1.28E+00 | 3.12E-06 | 1.35E-05 | 1.05E+00 | 9.70E-09 | 4.12E-08 | -1.42E+00 | 5.93E-06 | 2.26E-05 | -1.05E+00 | 4.90E-08 | 1.75E-07 | -1.36E+00 | 1.46E-02 | 4.11E-02 | 1.08E+00 | 1.60E-05 | 5.86E-05 | -1.21E+00 | 5.16E-03 | 1.42E-02 | -1.05E+00 |
| MRPS36P1 | 2.15E-12 | 1.12E-11 | -1.66E+00 | 2.41E-15 | 1.04E-14 | -1.80E+00 | 3.26E-17 | 1.38E-16 | -1.96E+00 | 1.32E-21 | 5.06E-21 | -2.02E+00 | 1.30E-16 | 4.63E-16 | -2.19E+00 | 3.62E-19 | 1.02E-18 | -2.21E+00 | 2.33E-17 | 8.56E-17 | -1.82E+00 | 5.13E-21 | 1.41E-20 | -2.11E+00 |
| SPINK2 | 2.15E-12 | 1.12E-11 | 1.64E+00 | 1.65E-10 | 7.12E-10 | 1.63E+00 | 2.93E-06 | 1.24E-05 | 1.49E+00 | 2.60E-04 | 9.93E-04 | 1.37E+00 | 2.09E-07 | 7.45E-07 | 3.09E+00 | 6.21E-02 | 1.75E-01 | 1.98E+00 | 3.63E-03 | 1.33E-02 | 3.56E+00 | 9.44E-03 | 2.59E-02 | 3.41E+00 |
| TTC19 | 2.16E-12 | 1.12E-11 | -1.24E+00 | 8.96E-06 | 3.87E-05 | 1.07E+00 | 1.32E-02 | 5.60E-02 | -1.10E+00 | 1.27E-02 | 4.87E-02 | 1.06E+00 | 5.77E-05 | 2.06E-04 | -1.02E+00 | 1.09E-02 | 3.08E-02 | 1.12E+00 | 1.27E-03 | 4.66E-03 | -1.10E+00 | 1.71E-01 | 4.69E-01 | 1.06E+00 |
| SERPING1 | 2.18E-12 | 1.13E-11 | -1.38E+00 | 1.64E-05 | 7.08E-05 | 1.01E+00 | 1.51E-08 | 6.40E-08 | -1.58E+00 | 8.36E-06 | 3.19E-05 | -1.21E+00 | 3.25E-07 | 1.16E-06 | -1.48E+00 | 1.04E-05 | 2.94E-05 | -1.25E+00 | 5.39E-03 | 1.98E-02 | -1.06E+00 | 7.36E-02 | 2.02E-01 | 1.05E+00 |
| HDDC2 | 2.21E-12 | 1.15E-11 | -1.27E+00 | 1.97E-09 | 8.51E-09 | -1.04E+00 | 9.49E-04 | 4.03E-03 | 1.01E+00 | 1.26E-04 | 4.79E-04 | 1.01E+00 | 1.26E-03 | 4.50E-03 | -1.06E+00 | 1.18E-01 | 3.33E-01 | 1.19E+00 | 1.39E-01 | 5.09E-01 | 1.15E+00 | 2.90E-01 | 7.97E-01 | 1.22E+00 |
| FAM198B | 2.25E-12 | 1.17E-11 | -3.30E+00 | 1.57E-05 | 6.80E-05 | -1.42E+00 | 9.55E-08 | 4.06E-07 | -2.53E+00 | 4.76E-05 | 1.82E-04 | -1.67E+00 | 2.80E-01 | 1.00E+00 | 1.06E+00 | 3.28E-01 | 9.24E-01 | 1.29E+00 | 9.90E-02 | 3.63E-01 | -1.28E+00 | 2.19E-01 | 6.00E-01 | -1.10E+00 |
| FAM111A | 2.26E-12 | 1.18E-11 | -1.36E+00 | 2.88E-09 | 1.25E-08 | -1.05E+00 | 5.80E-07 | 2.46E-06 | -1.32E+00 | 4.02E-07 | 1.54E-06 | -1.24E+00 | 7.04E-09 | 2.51E-08 | -1.41E+00 | 3.37E-08 | 9.51E-08 | -1.34E+00 | 4.30E-05 | 1.58E-04 | -1.25E+00 | 6.90E-05 | 1.90E-04 | -1.14E+00 |
| HMGA2 | 2.27E-12 | 1.19E-11 | -1.50E+00 | 5.27E-03 | 2.28E-02 | 1.22E+00 | 1.56E-12 | 6.62E-12 | -2.02E+00 | 1.76E-04 | 6.71E-04 | 1.04E+00 | 5.04E-04 | 1.80E-03 | -1.67E+00 | 1.75E-02 | 4.93E-02 | -1.06E+00 | 1.41E-09 | 5.17E-09 | -2.91E+00 | 3.13E-04 | 8.60E-04 | -1.45E+00 |
| HNRNPL | 2.28E-12 | 1.19E-11 | -1.29E+00 | 7.06E-25 | 3.05E-24 | -1.62E+00 | 1.09E-05 | 4.62E-05 | -1.26E+00 | 5.41E-10 | 2.07E-09 | -1.36E+00 | 8.41E-08 | 3.00E-07 | -1.19E+00 | 3.09E-10 | 8.71E-10 | -1.34E+00 | 6.50E-14 | 2.39E-13 | -1.69E+00 | 7.01E-13 | 1.92E-12 | -1.66E+00 |
| MAN2B1 | 2.28E-12 | 1.19E-11 | -1.84E+00 | 2.01E-10 | 8.70E-10 | -1.55E+00 | 1.41E-15 | 6.00E-15 | -1.93E+00 | 3.58E-16 | 1.37E-15 | -1.68E+00 | 6.66E-08 | 2.37E-07 | -2.23E+00 | 2.21E-06 | 6.22E-06 | -1.68E+00 | 8.31E-08 | 3.05E-07 | -1.61E+00 | 1.30E-05 | 3.57E-05 | -1.41E+00 |
| GAS6-AS1 | 2.28E-12 | 1.19E-11 | -1.77E+00 | 6.91E-10 | 2.98E-09 | -1.33E+00 | 1.90E-06 | 8.05E-06 | -1.45E+00 | 3.83E-06 | 1.46E-05 | -1.27E+00 | 4.11E-04 | 1.46E-03 | -1.12E+00 | 5.58E-04 | 1.57E-03 | -1.10E+00 | 5.19E-02 | 1.91E-01 | -1.04E+00 | 3.20E-01 | 8.79E-01 | 1.33E+00 |
| MCTS2P | 2.29E-12 | 1.19E-11 | -1.41E+00 | 1.20E-19 | 5.16E-19 | -1.69E+00 | 2.91E-05 | 1.24E-04 | -1.15E+00 | 5.21E-14 | 1.99E-13 | -1.54E+00 | 3.76E-08 | 1.34E-07 | -1.46E+00 | 6.93E-11 | 1.95E-10 | -1.74E+00 | 2.06E-07 | 7.57E-07 | -1.30E+00 | 3.24E-11 | 8.90E-11 | -1.51E+00 |
| EZH2 | 2.30E-12 | 1.20E-11 | -1.70E+00 | 5.49E-13 | 2.37E-12 | -1.61E+00 | 3.53E-08 | 1.50E-07 | -1.37E+00 | 2.24E-08 | 8.56E-08 | -1.24E+00 | 2.31E-16 | 8.22E-16 | -2.23E+00 | 1.30E-12 | 3.65E-12 | -1.78E+00 | 7.71E-14 | 2.83E-13 | -1.60E+00 | 1.50E-08 | 4.13E-08 | -1.41E+00 |
| CSNK2A3 | 2.31E-12 | 1.20E-11 | -1.52E+00 | 5.50E-18 | 2.38E-17 | -1.82E+00 | 2.50E-03 | 1.06E-02 | -1.16E+00 | 1.84E-11 | 7.02E-11 | -1.66E+00 | 4.18E-04 | 1.49E-03 | -1.20E+00 | 8.19E-06 | 2.31E-05 | -1.48E+00 | 1.43E-12 | 5.25E-12 | -1.48E+00 | 1.73E-17 | 4.76E-17 | -1.68E+00 |
| TOMM20 | 2.31E-12 | 1.20E-11 | -1.28E+00 | 1.21E-17 | 5.22E-17 | -1.27E+00 | 7.32E-05 | 3.11E-04 | -1.33E+00 | 2.77E-05 | 1.06E-04 | -1.27E+00 | 7.27E-07 | 2.59E-06 | -1.27E+00 | 4.87E-05 | 1.37E-04 | -1.14E+00 | 9.12E-08 | 3.35E-07 | -1.40E+00 | 4.24E-05 | 1.17E-04 | -1.35E+00 |
| SAR1A | 2.31E-12 | 1.21E-11 | -1.35E+00 | 2.85E-09 | 1.23E-08 | -1.03E+00 | 1.10E-03 | 4.66E-03 | -1.13E+00 | 1.79E-03 | 6.82E-03 | 1.05E+00 | 1.65E-06 | 5.89E-06 | -1.24E+00 | 5.13E-02 | 1.45E-01 | 1.15E+00 | 5.57E-02 | 2.04E-01 | -1.10E+00 | 2.15E-01 | 5.90E-01 | 1.00E+00 |
| NUDT5 | 2.33E-12 | 1.21E-11 | -1.24E+00 | 2.25E-03 | 9.73E-03 | 1.26E+00 | 1.19E-04 | 5.04E-04 | -1.15E+00 | 3.80E-02 | 1.45E-01 | 1.11E+00 | 5.94E-08 | 2.12E-07 | -1.34E+00 | 3.45E-02 | 9.72E-02 | 1.16E+00 | 8.15E-08 | 2.99E-07 | -1.37E+00 | 1.34E-01 | 3.69E-01 | 1.10E+00 |
| SRSF1 | 2.33E-12 | 1.21E-11 | -1.27E+00 | 6.06E-08 | 2.62E-07 | 1.05E+00 | 4.66E-05 | 1.98E-04 | -1.18E+00 | 9.67E-02 | 3.69E-01 | 1.20E+00 | 5.01E-10 | 1.79E-09 | -1.42E+00 | 1.35E-03 | 3.80E-03 | 1.02E+00 | 1.73E-05 | 6.37E-05 | -1.23E+00 | 3.46E-01 | 9.52E-01 | 1.18E+00 |
| SKAP1 | 2.36E-12 | 1.23E-11 | -1.38E+00 | 7.86E-06 | 3.40E-05 | 1.02E+00 | 1.58E-01 | 6.69E-01 | 1.14E+00 | 1.23E-01 | 4.71E-01 | 1.29E+00 | 1.68E-01 | 5.98E-01 | 1.52E+00 | 2.70E-01 | 7.62E-01 | 1.41E+00 | 9.14E-03 | 3.35E-02 | 1.86E+00 | 1.24E-01 | 3.41E-01 | 1.59E+00 |
| GPBP1L1 | 2.37E-12 | 1.23E-11 | -1.29E+00 | 1.54E-14 | 6.64E-14 | -1.24E+00 | 1.27E-05 | 5.40E-05 | -1.19E+00 | 3.86E-05 | 1.47E-04 | -1.07E+00 | 2.44E-04 | 8.71E-04 | -1.07E+00 | 8.98E-05 | 2.53E-04 | -1.12E+00 | 1.76E-09 | 6.47E-09 | -1.42E+00 | 6.13E-05 | 1.68E-04 | -1.16E+00 |
| PDS5A | 2.46E-12 | 1.28E-11 | -1.28E+00 | 7.80E-05 | 3.37E-04 | 1.15E+00 | 1.34E-05 | 5.70E-05 | -1.19E+00 | 3.88E-02 | 1.48E-01 | 1.17E+00 | 1.70E-06 | 6.07E-06 | -1.30E+00 | 4.27E-03 | 1.20E-02 | -1.00E+00 | 2.13E-07 | 7.83E-07 | -1.16E+00 | 5.65E-03 | 1.55E-02 | 1.07E+00 |
| SLK | 2.48E-12 | 1.29E-11 | -1.36E+00 | 7.90E-08 | 3.42E-07 | -1.04E+00 | 1.50E-03 | 6.36E-03 | -1.03E+00 | 1.08E-03 | 4.13E-03 | 1.07E+00 | 2.39E-03 | 8.54E-03 | 1.03E+00 | 1.69E-02 | 4.75E-02 | 1.08E+00 | 1.20E-04 | 4.41E-04 | -1.08E+00 | 1.51E-03 | 4.14E-03 | 1.03E+00 |
| AHCY | 2.48E-12 | 1.29E-11 | -1.28E+00 | 1.50E-07 | 6.48E-07 | -1.02E+00 | 1.11E-04 | 4.72E-04 | -1.17E+00 | 1.03E-03 | 3.95E-03 | -1.03E+00 | 8.73E-07 | 3.11E-06 | -1.22E+00 | 5.22E-04 | 1.47E-03 | 1.02E+00 | 2.86E-04 | 1.05E-03 | -1.10E+00 | 3.63E-01 | 9.97E-01 | 1.18E+00 |
| EXT2 | 2.48E-12 | 1.29E-11 | -1.88E+00 | 4.96E-07 | 2.15E-06 | -1.20E+00 | 6.03E-04 | 2.56E-03 | -1.14E+00 | 1.18E-02 | 4.50E-02 | 1.11E+00 | 1.33E-04 | 4.76E-04 | -1.26E+00 | 7.23E-03 | 2.04E-02 | 1.03E+00 | 8.05E-03 | 2.95E-02 | -1.05E+00 | 4.03E-02 | 1.11E-01 | 1.01E+00 |
| GPALPP1 | 2.51E-12 | 1.31E-11 | -1.48E+00 | 3.89E-03 | 1.68E-02 | 1.20E+00 | 1.73E-06 | 7.36E-06 | -1.42E+00 | 1.19E-02 | 4.55E-02 | 1.09E+00 | 2.22E-06 | 7.91E-06 | -1.28E+00 | 9.00E-03 | 2.54E-02 | 1.14E+00 | 4.64E-04 | 1.70E-03 | -1.12E+00 | 1.14E-02 | 3.14E-02 | -1.03E+00 |
| UBE2G2 | 2.52E-12 | 1.31E-11 | -1.32E+00 | 5.78E-05 | 2.50E-04 | 1.03E+00 | 2.43E-02 | 1.03E-01 | -1.02E+00 | 2.23E-01 | 8.51E-01 | 1.30E+00 | 9.68E-03 | 3.45E-02 | -1.02E+00 | 2.92E-01 | 8.22E-01 | 1.29E+00 | 1.10E-02 | 4.04E-02 | -1.00E+00 | 3.44E-01 | 9.44E-01 | 1.27E+00 |
| IRF1 | 2.56E-12 | 1.34E-11 | -1.39E+00 | 2.39E-91 | 1.03E-90 | -3.68E+00 | 5.57E-03 | 2.37E-02 | -1.21E+00 | 1.31E-38 | 4.99E-38 | -3.20E+00 | 3.07E-25 | 1.09E-24 | -2.31E+00 | 5.45E-56 | 1.54E-55 | -5.15E+00 | 4.17E-36 | 1.53E-35 | -3.08E+00 | 6.64E-74 | 1.82E-73 | -6.27E+00 |
| OCLN | 2.57E-12 | 1.34E-11 | -1.78E+00 | 4.44E-07 | 1.92E-06 | -1.15E+00 | 5.64E-11 | 2.40E-10 | -1.87E+00 | 3.00E-08 | 1.15E-07 | -1.41E+00 | 1.93E-05 | 6.87E-05 | -1.59E+00 | 3.76E-05 | 1.06E-04 | -1.57E+00 | 3.97E-04 | 1.46E-03 | -1.68E+00 | 2.22E-02 | 6.10E-02 | -1.24E+00 |
| RPL36AP41 | 2.61E-12 | 1.36E-11 | -1.19E+00 | 1.15E-42 | 4.98E-42 | -1.97E+00 | 1.07E-09 | 4.53E-09 | -1.30E+00 | 1.70E-25 | 6.51E-25 | -1.86E+00 | 4.87E-10 | 1.74E-09 | -1.35E+00 | 1.07E-20 | 3.00E-20 | -1.89E+00 | 8.53E-08 | 3.13E-07 | -1.28E+00 | 1.76E-22 | 4.82E-22 | -2.02E+00 |
| TM2D2 | 2.62E-12 | 1.37E-11 | -1.71E+00 | 6.63E-06 | 2.87E-05 | -1.04E+00 | 1.01E-11 | 4.30E-11 | -1.68E+00 | 2.59E-07 | 9.89E-07 | -1.22E+00 | 5.55E-06 | 1.98E-05 | -1.38E+00 | 1.96E-03 | 5.52E-03 | -1.03E+00 | 4.62E-06 | 1.70E-05 | -1.24E+00 | 9.35E-03 | 2.57E-02 | 1.07E+00 |
| SLC9A7 | 2.66E-12 | 1.38E-11 | -1.39E+00 | 6.18E-09 | 2.67E-08 | -1.08E+00 | 2.61E-04 | 1.11E-03 | -1.11E+00 | 2.54E-05 | 9.70E-05 | -1.05E+00 | 2.24E-03 | 7.99E-03 | -1.11E+00 | 2.37E-03 | 6.68E-03 | -1.05E+00 | 1.93E-05 | 7.10E-05 | -1.19E+00 | 1.18E-03 | 3.24E-03 | -1.07E+00 |
| APOL3 | 2.67E-12 | 1.39E-11 | -1.64E+00 | 4.44E-05 | 1.92E-04 | -1.01E+00 | 5.94E-14 | 2.52E-13 | -1.97E+00 | 1.59E-07 | 6.07E-07 | -1.23E+00 | 4.87E-06 | 1.74E-05 | -2.43E+00 | 2.11E-03 | 5.96E-03 | -1.34E+00 | 1.49E-28 | 5.48E-28 | -2.82E+00 | 6.40E-19 | 1.76E-18 | -2.04E+00 |
| GINM1 | 2.68E-12 | 1.40E-11 | -1.50E+00 | 2.81E-12 | 1.22E-11 | -1.33E+00 | 1.02E-10 | 4.32E-10 | -1.49E+00 | 1.90E-11 | 7.26E-11 | -1.32E+00 | 7.55E-06 | 2.69E-05 | -1.32E+00 | 1.20E-07 | 3.37E-07 | -1.42E+00 | 2.06E-10 | 7.57E-10 | -1.42E+00 | 1.51E-07 | 4.14E-07 | -1.32E+00 |
| ZBTB20 | 2.69E-12 | 1.40E-11 | -1.51E+00 | 1.90E-04 | 8.20E-04 | -1.08E+00 | 1.19E-10 | 5.06E-10 | -1.77E+00 | 5.67E-05 | 2.16E-04 | -1.14E+00 | 3.89E-09 | 1.39E-08 | -1.49E+00 | 8.33E-11 | 2.35E-10 | -1.43E+00 | 4.65E-14 | 1.71E-13 | -1.90E+00 | 8.05E-04 | 2.21E-03 | -1.24E+00 |
| ILF3 | 2.72E-12 | 1.42E-11 | -1.19E+00 | 3.01E-10 | 1.30E-09 | -1.02E+00 | 1.71E-02 | 7.26E-02 | -1.07E+00 | 1.29E-02 | 4.94E-02 | 1.03E+00 | 3.45E-09 | 1.23E-08 | -1.29E+00 | 1.95E-04 | 5.51E-04 | -1.08E+00 | 1.38E-05 | 5.08E-05 | -1.25E+00 | 1.47E-01 | 4.03E-01 | -1.00E+00 |
| MAP3K20 | 2.73E-12 | 1.42E-11 | -1.55E+00 | 9.02E-07 | 3.90E-06 | -1.10E+00 | 2.20E-07 | 9.34E-07 | -1.36E+00 | 1.99E-03 | 7.60E-03 | 1.10E+00 | 8.59E-08 | 3.06E-07 | -1.76E+00 | 2.25E-04 | 6.34E-04 | -1.23E+00 | 4.64E-06 | 1.70E-05 | -1.25E+00 | 1.94E-01 | 5.33E-01 | 1.19E+00 |
| ZNF639 | 2.74E-12 | 1.43E-11 | -1.40E+00 | 2.74E-11 | 1.18E-10 | -1.23E+00 | 5.41E-03 | 2.30E-02 | -1.13E+00 | 5.52E-05 | 2.11E-04 | -1.16E+00 | 1.42E-03 | 5.06E-03 | -1.03E+00 | 6.02E-04 | 1.70E-03 | -1.02E+00 | 4.43E-06 | 1.62E-05 | -1.10E+00 | 1.51E-02 | 4.14E-02 | 1.04E+00 |
| LSM6 | 2.74E-12 | 1.43E-11 | -1.27E+00 | 3.63E-09 | 1.57E-08 | -1.06E+00 | 2.97E-04 | 1.26E-03 | -1.10E+00 | 7.59E-03 | 2.90E-02 | 1.06E+00 | 4.78E-11 | 1.70E-10 | -1.55E+00 | 5.46E-04 | 1.54E-03 | -1.01E+00 | 9.06E-06 | 3.32E-05 | -1.23E+00 | 8.58E-02 | 2.36E-01 | 1.06E+00 |
| UBIAD1 | 2.75E-12 | 1.43E-11 | -2.02E+00 | 2.22E-08 | 9.60E-08 | -1.37E+00 | 1.14E-09 | 4.84E-09 | -1.73E+00 | 5.65E-06 | 2.16E-05 | -1.20E+00 | 1.32E-10 | 4.71E-10 | -2.28E+00 | 2.65E-07 | 7.48E-07 | -1.58E+00 | 6.89E-07 | 2.53E-06 | -1.60E+00 | 8.43E-03 | 2.31E-02 | -1.01E+00 |
| KIF21A | 2.78E-12 | 1.45E-11 | -2.06E+00 | 2.75E-06 | 1.19E-05 | -1.17E+00 | 6.94E-22 | 2.95E-21 | -2.43E+00 | 8.35E-10 | 3.19E-09 | -1.37E+00 | 6.29E-10 | 2.24E-09 | -4.02E+00 | 2.05E-05 | 5.79E-05 | -1.85E+00 | 6.31E-10 | 2.32E-09 | -2.08E+00 | 8.60E-02 | 2.36E-01 | 1.11E+00 |
| TIMP3 | 2.79E-12 | 1.45E-11 | -2.02E+00 | 5.91E-07 | 2.55E-06 | -1.34E+00 | 1.86E-03 | 7.89E-03 | -1.50E+00 | 6.28E-02 | 2.40E-01 | 1.08E+00 | 3.81E-11 | 1.36E-10 | -1.03E+00 | 1.31E-05 | 3.70E-05 | -1.07E+00 | 2.60E-01 | 9.53E-01 | 2.34E+00 | 4.58E-02 | 1.26E-01 | 2.09E+00 |
| UQCRQ | 2.80E-12 | 1.46E-11 | -1.19E+00 | 1.12E-36 | 4.85E-36 | -1.54E+00 | 2.35E-01 | 1.00E+00 | 1.12E+00 | 3.44E-05 | 1.32E-04 | -1.18E+00 | 1.35E-06 | 4.81E-06 | -1.23E+00 | 6.03E-12 | 1.70E-11 | -1.47E+00 | 5.76E-02 | 2.11E-01 | -1.04E+00 | 9.77E-03 | 2.68E-02 | -1.23E+00 |
| CSPP1 | 2.83E-12 | 1.47E-11 | -1.68E+00 | 2.09E-09 | 9.03E-09 | -1.27E+00 | 6.23E-08 | 2.65E-07 | -1.44E+00 | 5.21E-06 | 1.99E-05 | -1.17E+00 | 5.59E-07 | 1.99E-06 | -1.53E+00 | 2.15E-05 | 6.07E-05 | -1.35E+00 | 8.80E-09 | 3.23E-08 | -1.61E+00 | 2.98E-04 | 8.18E-04 | -1.11E+00 |
| UBA1 | 2.84E-12 | 1.48E-11 | -1.43E+00 | 2.89E-09 | 1.25E-08 | -1.11E+00 | 6.80E-08 | 2.89E-07 | -1.29E+00 | 1.20E-07 | 4.57E-07 | -1.16E+00 | 6.73E-08 | 2.40E-07 | -1.48E+00 | 2.47E-04 | 6.96E-04 | -1.13E+00 | 4.87E-09 | 1.79E-08 | -1.30E+00 | 6.98E-05 | 1.92E-04 | -1.12E+00 |
| ZNF252P | 2.84E-12 | 1.48E-11 | -1.99E+00 | 3.58E-09 | 1.55E-08 | -1.51E+00 | 3.08E-04 | 1.31E-03 | -1.80E+00 | 1.12E-01 | 4.29E-01 | 1.16E+00 | 6.74E-07 | 2.40E-06 | -2.34E+00 | 5.55E-04 | 1.56E-03 | -1.44E+00 | 2.26E-08 | 8.28E-08 | -1.85E+00 | 6.61E-03 | 1.82E-02 | -1.09E+00 |
| EEF1A1P3 | 2.84E-12 | 1.48E-11 | -1.62E+00 | 2.42E-11 | 1.05E-10 | -1.34E+00 | 2.09E-07 | 8.86E-07 | -1.46E+00 | 4.18E-09 | 1.60E-08 | -1.40E+00 | 6.19E-07 | 2.21E-06 | -1.68E+00 | 1.48E-06 | 4.17E-06 | -1.41E+00 | 1.22E-04 | 4.48E-04 | -1.41E+00 | 2.07E-04 | 5.68E-04 | -1.35E+00 |
| SAFB | 2.86E-12 | 1.49E-11 | -1.36E+00 | 6.82E-06 | 2.95E-05 | 1.00E+00 | 2.26E-04 | 9.61E-04 | -1.13E+00 | 3.16E-04 | 1.21E-03 | 1.01E+00 | 1.37E-06 | 4.88E-06 | -1.25E+00 | 4.31E-03 | 1.21E-02 | 1.04E+00 | 6.10E-10 | 2.24E-09 | -1.37E+00 | 1.61E-05 | 4.41E-05 | -1.16E+00 |
| FXN | 2.87E-12 | 1.50E-11 | -1.54E+00 | 1.81E-06 | 7.80E-06 | -1.05E+00 | 1.03E-09 | 4.37E-09 | -1.53E+00 | 2.10E-06 | 8.01E-06 | -1.17E+00 | 6.19E-04 | 2.21E-03 | -1.21E+00 | 1.19E-01 | 3.34E-01 | 1.44E+00 | 2.92E-10 | 1.07E-09 | -1.39E+00 | 8.75E-03 | 2.40E-02 | -1.11E+00 |
| ADIPOR1 | 2.88E-12 | 1.50E-11 | -1.45E+00 | 9.14E-10 | 3.95E-09 | -1.18E+00 | 1.23E-05 | 5.24E-05 | -1.26E+00 | 5.22E-04 | 1.99E-03 | 1.00E+00 | 6.83E-05 | 2.44E-04 | -1.11E+00 | 1.78E-04 | 5.02E-04 | -1.09E+00 | 5.68E-11 | 2.08E-10 | -1.27E+00 | 2.86E-04 | 7.86E-04 | -1.04E+00 |
| EIF4E | 2.88E-12 | 1.50E-11 | -1.72E+00 | 5.56E-08 | 2.40E-07 | -1.24E+00 | 2.37E-10 | 1.01E-09 | -1.57E+00 | 9.50E-05 | 3.63E-04 | -1.01E+00 | 1.25E-04 | 4.46E-04 | -1.41E+00 | 1.83E-02 | 5.17E-02 | -1.05E+00 | 1.39E-08 | 5.11E-08 | -1.26E+00 | 1.32E-03 | 3.63E-03 | 1.01E+00 |
| TMEM65 | 2.92E-12 | 1.52E-11 | -1.42E+00 | 7.28E-08 | 3.15E-07 | -1.06E+00 | 8.25E-03 | 3.51E-02 | -1.20E+00 | 1.23E-02 | 4.71E-02 | -1.02E+00 | 1.65E-01 | 5.88E-01 | 1.49E+00 | 2.18E-03 | 6.14E-03 | 1.97E+00 | 1.37E-01 | 5.01E-01 | 1.24E+00 | 1.80E-01 | 4.94E-01 | 1.28E+00 |
| MSRB2 | 2.97E-12 | 1.55E-11 | -1.34E+00 | 6.50E-12 | 2.81E-11 | -1.21E+00 | 9.58E-05 | 4.07E-04 | -1.18E+00 | 1.21E-06 | 4.61E-06 | -1.19E+00 | 3.75E-10 | 1.34E-09 | -1.46E+00 | 3.67E-09 | 1.04E-08 | -1.32E+00 | 1.54E-02 | 5.64E-02 | 1.04E+00 | 6.24E-04 | 1.71E-03 | -1.07E+00 |
| SCP2 | 3.04E-12 | 1.58E-11 | -1.19E+00 | 7.03E-06 | 3.04E-05 | 1.10E+00 | 1.43E-04 | 6.09E-04 | -1.15E+00 | 2.65E-03 | 1.01E-02 | -1.01E+00 | 2.35E-08 | 8.39E-08 | -1.24E+00 | 2.57E-03 | 7.26E-03 | 1.01E+00 | 9.63E-03 | 3.54E-02 | 1.00E+00 | 6.06E-02 | 1.67E-01 | 1.01E+00 |

| | | | | | | | | | | | | | | | | | | | | | | | |
|---|---|---|---|---|---|---|---|---|---|---|---|---|---|---|---|---|---|---|---|---|---|---|---|
| AC078817.1 | 3.04E-12 | 1.58E-11 | -1.32E+00 | 4.32E-38 | 1.87E-37 | -1.86E+00 | 5.52E-06 | 2.34E-05 | -1.22E+00 | 2.83E-25 | 1.08E-24 | -1.88E+00 | 2.74E-08 | 9.78E-08 | -1.25E+00 | 2.72E-20 | 7.66E-20 | -1.79E+00 | 2.56E-06 | 9.41E-06 | -1.16E+00 | 5.90E-16 | [illegible] |
| HOMER2 | 3.06E-12 | 1.60E-11 | -2.03E+00 | 1.49E-07 | 6.45E-07 | -1.31E+00 | 1.39E-08 | 5.90E-08 | -1.58E+00 | 5.82E-07 | 2.22E-06 | -1.22E+00 | 1.33E-14 | 4.74E-14 | -2.69E+00 | 2.24E-07 | 6.33E-07 | -1.35E+00 | 6.27E-11 | 2.30E-10 | -1.46E+00 | 3.13E-05 | [illegible] |
| IDS | 3.06E-12 | 1.60E-11 | 2.05E+00 | 1.35E-05 | 5.81E-05 | 1.00E+00 | 2.13E-09 | 9.05E-09 | 2.07E+00 | 1.89E-02 | 7.23E-02 | 1.03E+00 | 7.45E-17 | 2.66E-16 | 3.45E+00 | 3.15E-02 | 8.88E-02 | 1.27E+00 | 2.41E-12 | 8.85E-12 | 2.86E+00 | 3.62E-01 | [illegible] |
| DHRS7 | 3.09E-12 | 1.61E-11 | -1.23E+00 | 1.46E-09 | 6.31E-09 | -1.05E+00 | 1.91E-07 | 8.13E-07 | -1.34E+00 | 5.39E-07 | 2.06E-06 | -1.19E+00 | 5.98E-07 | 2.13E-06 | -1.21E+00 | 2.30E-05 | 6.48E-05 | -1.04E+00 | 4.51E-04 | 1.66E-03 | -1.15E+00 | 3.14E-05 | [illegible] |
| HIST2H2AA4 | 3.12E-12 | 1.63E-11 | 2.03E+00 | 3.22E-03 | 1.39E-02 | 1.65E+00 | 5.48E-04 | 2.33E-03 | 1.64E+00 | 1.36E-02 | 5.21E-02 | 2.06E+00 | 5.12E-09 | 1.82E-08 | 2.68E+00 | 8.07E-04 | 2.27E-03 | 2.24E+00 | 5.77E-02 | 2.12E-01 | 1.83E+00 | 2.22E-03 | [illegible] |
| GTF3C4 | 3.14E-12 | 1.64E-11 | -1.92E+00 | 2.87E-10 | 1.24E-09 | -1.51E+00 | 1.70E-07 | 7.20E-07 | -1.48E+00 | 4.51E-08 | 1.72E-07 | -1.41E+00 | 1.43E-05 | 5.11E-05 | -1.56E+00 | 1.69E-04 | 4.76E-04 | -1.33E+00 | 9.59E-06 | 3.52E-05 | -1.35E+00 | 2.08E-03 | [illegible] |
| PARP10 | 3.14E-12 | 1.64E-11 | -1.54E+00 | 1.60E-10 | 6.90E-10 | -1.35E+00 | 1.00E-08 | 4.27E-08 | -1.50E+00 | 1.02E-09 | 3.88E-09 | -1.46E+00 | 3.49E-07 | 1.25E-06 | -1.57E+00 | 2.93E-09 | 8.27E-09 | -1.97E+00 | 1.76E-09 | 6.47E-09 | -1.87E+00 | 2.01E-08 | [illegible] |
| DALRD3 | 3.15E-12 | 1.64E-11 | -1.66E+00 | 3.60E-10 | 1.56E-09 | -1.34E+00 | 5.65E-09 | 2.40E-08 | -1.55E+00 | 1.20E-08 | 4.58E-08 | -1.38E+00 | 2.14E-10 | 7.64E-10 | -2.09E+00 | 1.50E-05 | 4.23E-05 | -1.31E+00 | 6.10E-04 | 2.24E-03 | -1.24E+00 | 9.97E-03 | [illegible] |
| HIF1AN | 3.23E-12 | 1.68E-11 | -1.67E+00 | 1.54E-07 | 6.66E-07 | -1.10E+00 | 1.27E-06 | 5.39E-06 | -1.46E+00 | 4.89E-04 | 1.87E-03 | -1.01E+00 | 1.34E-07 | 4.77E-07 | -1.88E+00 | 1.04E-04 | 2.92E-04 | -1.27E+00 | 2.19E-05 | 8.04E-05 | -1.25E+00 | 2.84E-04 | [illegible] |
| CDIPT | 3.26E-12 | 1.70E-11 | -1.35E+00 | 8.15E-07 | 3.52E-06 | 1.03E+00 | 4.84E-08 | 2.06E-07 | -1.40E+00 | 3.57E-04 | 1.36E-03 | 1.03E+00 | 1.21E-12 | 4.32E-12 | -1.55E+00 | 3.04E-06 | 8.56E-06 | -1.14E+00 | 1.91E-06 | 6.99E-06 | -1.19E+00 | 8.71E-04 | [illegible] |
| PFKFB2 | 3.30E-12 | 1.72E-11 | 3.52E+00 | 5.69E-10 | 2.46E-09 | 3.91E+00 | 3.47E-07 | 1.47E-06 | 2.66E+00 | 1.70E-05 | 6.51E-05 | 2.85E+00 | 4.64E-02 | 1.65E-01 | 2.20E+00 | 3.27E-01 | 9.22E-01 | 1.39E+00 | 1.54E-01 | 5.67E-01 | 1.63E+00 | 1.78E-01 | [illegible] |
| ACADM | 3.30E-12 | 1.72E-11 | -1.32E+00 | 3.65E-07 | 1.58E-06 | 1.00E+00 | 3.80E-05 | 1.62E-04 | -1.28E+00 | 6.67E-05 | 2.55E-04 | -1.15E+00 | 1.80E-10 | 6.43E-10 | -1.55E+00 | 3.58E-05 | 1.01E-04 | -1.15E+00 | 7.06E-06 | 2.59E-05 | -1.28E+00 | 7.76E-04 | [illegible] |
| CFL2 | 3.30E-12 | 1.72E-11 | -1.64E+00 | 6.45E-07 | 2.79E-06 | -1.10E+00 | 7.05E-04 | 3.00E-03 | -1.28E+00 | 7.55E-03 | 2.88E-02 | -1.00E+00 | 1.06E-02 | 3.77E-02 | -1.13E+00 | 2.56E-01 | 7.20E-01 | 1.43E+00 | 1.53E-02 | 5.61E-02 | -1.20E+00 | 7.87E-02 | [illegible] |
| TTC1 | 3.33E-12 | 1.74E-11 | -1.25E+00 | 3.87E-08 | 1.67E-07 | -1.01E+00 | 1.20E-06 | 5.11E-06 | -1.24E+00 | 6.02E-10 | 2.30E-09 | -1.22E+00 | 1.37E-07 | 4.89E-07 | -1.33E+00 | 1.74E-03 | 4.90E-03 | -1.02E+00 | 3.00E-04 | 1.10E-03 | -1.19E+00 | 1.92E-02 | [illegible] |
| LYRM7 | 3.35E-12 | 1.75E-11 | -1.57E+00 | 2.65E-05 | 1.15E-04 | 1.07E+00 | 4.81E-10 | 2.04E-09 | -1.66E+00 | 6.01E-04 | 2.29E-03 | -1.03E+00 | 3.47E-14 | 1.24E-13 | -2.01E+00 | 8.07E-06 | 2.27E-05 | -1.16E+00 | 1.28E-06 | 4.70E-06 | -1.30E+00 | 9.77E-02 | [illegible] |
| MRPL32 | 3.39E-12 | 1.76E-11 | -1.31E+00 | 7.28E-11 | 3.14E-10 | -1.13E+00 | 2.26E-08 | 9.59E-08 | -1.32E+00 | 5.74E-05 | 2.19E-04 | -1.11E+00 | 4.64E-13 | 1.65E-12 | -1.59E+00 | 4.35E-05 | 1.23E-04 | -1.10E+00 | 1.41E-12 | 5.16E-12 | -1.59E+00 | 1.27E-04 | [illegible] |
| ECH1 | 3.43E-12 | 1.79E-11 | -1.25E+00 | 4.21E-07 | 1.82E-06 | 1.00E+00 | 2.46E-04 | 1.05E-03 | -1.23E+00 | 2.74E-04 | 1.05E-03 | -1.13E+00 | 3.43E-11 | 1.22E-10 | -1.41E+00 | 2.01E-02 | 5.66E-02 | 1.03E+00 | 2.15E-02 | 7.88E-02 | -1.12E+00 | 3.24E-01 | [illegible] |
| RCE1 | 3.46E-12 | 1.80E-11 | -1.92E+00 | 2.40E-08 | 1.04E-07 | -1.30E+00 | 2.42E-10 | 1.03E-09 | -1.83E+00 | 1.96E-06 | 7.49E-06 | -1.18E+00 | 6.73E-08 | 2.40E-07 | -1.84E+00 | 3.40E-06 | 9.59E-06 | -1.51E+00 | 6.44E-17 | 2.36E-16 | -2.26E+00 | 5.11E-08 | [illegible] |
| FKBP8 | 3.47E-12 | 1.81E-11 | -1.22E+00 | 1.80E-05 | 7.79E-05 | 1.06E+00 | 1.46E-02 | 6.20E-02 | -1.11E+00 | 1.76E-02 | 6.71E-02 | 1.03E+00 | 1.27E-07 | 4.53E-07 | -1.16E+00 | 9.42E-03 | 2.66E-02 | 1.08E+00 | 1.94E-02 | 7.10E-02 | -1.07E+00 | 2.38E-01 | [illegible] |
| NR0B1 | 3.48E-12 | 1.81E-11 | 9.20E+00 | 2.21E-16 | 9.54E-16 | 1.28E+01 | 1.25E-03 | 5.31E-03 | 4.09E+00 | 3.83E-06 | 1.46E-05 | 6.88E+00 | 1.23E-01 | 4.39E-01 | 2.65E+00 | 1.35E-01 | 3.82E-01 | 2.68E+00 | 1.01E-01 | 3.70E-01 | 1.86E+00 | 1.60E-01 | [illegible] |
| ROMO1 | 3.55E-12 | 1.85E-11 | -1.20E+00 | 1.36E-33 | 5.86E-33 | -1.78E+00 | 3.34E-04 | 1.42E-03 | -1.09E+00 | 2.84E-20 | 1.08E-19 | -1.81E+00 | 1.70E-13 | 6.05E-13 | -1.42E+00 | 1.07E-24 | 3.03E-24 | -2.02E+00 | 7.39E-04 | 2.71E-03 | -1.08E+00 | 1.82E-08 | [illegible] |
| LSM2 | 3.56E-12 | 1.86E-11 | -1.28E+00 | 2.64E-05 | 1.14E-04 | -1.05E+00 | 1.04E-04 | 4.42E-04 | -1.32E+00 | 3.29E-03 | 1.26E-02 | -1.16E+00 | 2.07E-09 | 7.40E-09 | -1.38E+00 | 1.39E-02 | 3.93E-02 | -1.02E+00 | 3.48E-05 | 1.28E-04 | -1.23E+00 | 5.46E-02 | [illegible] |
| FRAT2 | 3.56E-12 | 1.86E-11 | -1.68E+00 | 4.46E-13 | 1.93E-12 | -1.60E+00 | 9.51E-10 | 4.04E-09 | -1.44E+00 | 8.69E-10 | 3.32E-09 | -1.42E+00 | 1.40E-09 | 4.98E-09 | -1.61E+00 | 2.28E-11 | 6.42E-11 | -1.55E+00 | 9.15E-19 | 3.36E-18 | -1.91E+00 | 1.77E-10 | [illegible] |
| HSPB1P1 | 3.65E-12 | 1.90E-11 | -1.37E+00 | 6.66E-05 | 2.88E-04 | 1.12E+00 | 4.96E-05 | 2.11E-04 | -1.17E+00 | 2.86E-05 | 1.09E-04 | -1.01E+00 | 8.49E-05 | 3.03E-04 | -1.30E+00 | 1.85E-02 | 5.23E-02 | 1.04E+00 | 1.51E-03 | 5.53E-03 | 1.06E+00 | 2.00E-03 | [illegible] |
| STX16 | 3.66E-12 | 1.91E-11 | -1.35E+00 | 4.47E-09 | 1.93E-08 | -1.05E+00 | 4.70E-06 | 2.00E-05 | -1.39E+00 | 9.66E-05 | 3.69E-04 | -1.11E+00 | 2.86E-07 | 1.02E-06 | -1.27E+00 | 7.94E-04 | 2.24E-03 | 1.01E+00 | 4.88E-05 | 1.79E-04 | -1.25E+00 | 1.90E-03 | [illegible] |
| NUDC | 3.75E-12 | 1.95E-11 | -1.14E+00 | 3.77E-11 | 1.63E-10 | -1.05E+00 | 2.32E-02 | 9.87E-02 | -1.01E+00 | 1.74E-02 | 6.65E-02 | 1.06E+00 | 4.26E-08 | 1.52E-07 | -1.28E+00 | 1.93E-02 | 5.44E-02 | 1.05E+00 | 2.80E-02 | 1.03E-01 | 1.02E+00 | 2.76E-01 | [illegible] |
| ATP6AP1 | 3.75E-12 | 1.96E-11 | -1.30E+00 | 5.54E-08 | 2.39E-07 | 1.01E+00 | 2.47E-07 | 1.05E-06 | -1.18E+00 | 7.10E-07 | 2.71E-06 | -1.03E+00 | 1.31E-06 | 4.69E-06 | -1.22E+00 | 6.78E-07 | 1.91E-06 | -1.18E+00 | 1.14E-05 | 4.18E-05 | -1.26E+00 | 3.74E-04 | [illegible] |
| ELP2 | 3.78E-12 | 1.97E-11 | -1.31E+00 | 1.35E-04 | 5.85E-04 | 1.09E+00 | 1.16E-06 | 4.94E-06 | -1.26E+00 | 1.86E-04 | 7.10E-04 | -1.03E+00 | 1.75E-06 | 6.24E-06 | -1.20E+00 | 1.30E-04 | 3.67E-04 | 1.00E+00 | 1.38E-11 | 5.08E-11 | -1.47E+00 | 3.33E-05 | [illegible] |
| ASNSD1 | 3.79E-12 | 1.98E-11 | -1.38E+00 | 8.75E-14 | 3.78E-13 | -1.31E+00 | 1.42E-04 | 6.04E-04 | -1.16E+00 | 1.73E-05 | 6.60E-05 | -1.02E+00 | 3.72E-04 | 1.33E-03 | -1.08E+00 | 3.04E-04 | 8.58E-04 | -1.08E+00 | 3.93E-09 | 1.44E-08 | -1.34E+00 | 1.85E-05 | [illegible] |
| TMEM248 | 3.80E-12 | 1.98E-11 | -1.43E+00 | 4.29E-05 | 1.85E-04 | 1.05E+00 | 2.49E-05 | 1.06E-04 | -1.21E+00 | 1.27E-03 | 4.84E-03 | 1.03E+00 | 7.99E-06 | 2.85E-05 | -1.19E+00 | 2.97E-05 | 8.38E-05 | -1.13E+00 | 1.34E-06 | 4.91E-06 | -1.11E+00 | 9.75E-03 | [illegible] |
| WASF4P | 3.86E-12 | 2.01E-11 | 3.31E+00 | 1.11E-04 | 4.79E-04 | 2.78E+00 | 1.55E-11 | 6.57E-11 | 3.05E+00 | 4.14E-03 | 1.58E-02 | 2.15E+00 | 1.62E-04 | 5.77E-04 | 2.62E+00 | 5.19E-02 | 1.46E-01 | 2.01E+00 | 1.47E-05 | 5.38E-05 | 2.43E+00 | 3.22E-02 | [illegible] |
| NAV1 | 3.98E-12 | 2.07E-11 | -1.74E+00 | 2.27E-06 | 9.81E-06 | -1.03E+00 | 5.44E-07 | 2.31E-06 | -2.08E+00 | 2.66E-03 | 1.01E-02 | -1.17E+00 | 6.62E-05 | 2.36E-04 | -2.24E+00 | 8.43E-03 | 2.38E-02 | -1.23E+00 | 8.49E-02 | 3.12E-01 | -1.42E+00 | 3.33E-01 | [illegible] |
| AASDHPPT | 3.98E-12 | 2.07E-11 | -1.48E+00 | 3.69E-09 | 1.59E-08 | -1.20E+00 | 5.19E-11 | 2.21E-10 | -1.48E+00 | 4.33E-06 | 1.65E-05 | -1.02E+00 | 1.59E-14 | 5.67E-14 | -2.20E+00 | 6.96E-06 | 1.96E-05 | -1.23E+00 | 7.77E-21 | 2.85E-20 | -1.72E+00 | 3.68E-09 | [illegible] |
| PAWR | 3.98E-12 | 2.08E-11 | -1.26E+00 | 1.19E-10 | 5.14E-10 | -1.08E+00 | 1.09E-06 | 4.65E-06 | -1.31E+00 | 1.06E-05 | 4.06E-05 | -1.09E+00 | 5.48E-04 | 1.95E-03 | -1.22E+00 | 8.97E-03 | 2.53E-02 | 1.03E+00 | 1.28E-02 | 4.69E-02 | -1.26E+00 | 2.65E-01 | [illegible] |
| ELK4 | 4.04E-12 | 2.11E-11 | -1.53E+00 | 3.38E-08 | 1.46E-07 | -1.15E+00 | 2.28E-06 | 9.68E-06 | -1.35E+00 | 8.88E-03 | 3.39E-02 | 1.14E+00 | 6.08E-04 | 2.17E-03 | -1.16E+00 | 1.93E-03 | 5.44E-03 | -1.15E+00 | 1.33E-02 | 4.87E-02 | -1.04E+00 | 3.85E-04 | [illegible] |
| AC087633.2 | 4.11E-12 | 2.14E-11 | -1.23E+00 | 9.79E-09 | 4.23E-08 | -1.10E+00 | 1.57E-04 | 6.67E-04 | -1.18E+00 | 8.95E-03 | 3.42E-02 | -1.01E+00 | 4.17E-03 | 1.49E-02 | -1.11E+00 | 4.59E-03 | 1.29E-02 | -1.10E+00 | 3.26E-06 | 1.20E-05 | -1.14E+00 | 6.65E-03 | [illegible] |
| ITGA6 | 4.13E-12 | 2.15E-11 | -1.92E+00 | 6.50E-07 | 2.81E-06 | -1.16E+00 | 1.83E-03 | 7.76E-03 | -1.18E+00 | 3.94E-03 | 1.51E-02 | -1.00E+00 | 2.55E-01 | 9.08E-01 | 1.38E+00 | 2.91E-01 | 8.21E-01 | -1.08E+00 | 2.67E-01 | 9.80E-01 | 1.11E+00 | 7.48E-02 | [illegible] |
| EI24 | 4.13E-12 | 2.15E-11 | -1.30E+00 | 7.66E-07 | 3.31E-06 | 1.03E+00 | 2.25E-05 | 9.57E-05 | -1.23E+00 | 2.92E-02 | 1.12E-01 | 1.11E+00 | 3.62E-11 | 1.29E-10 | -1.56E+00 | 3.28E-06 | 9.24E-06 | -1.11E+00 | 1.28E-08 | 4.70E-08 | -1.39E+00 | 1.93E-02 | [illegible] |
| DYNC1H1 | 4.16E-12 | 2.17E-11 | -1.28E+00 | 1.82E-06 | 7.87E-06 | 1.08E+00 | 5.99E-05 | 2.54E-04 | -1.15E+00 | 2.97E-03 | 1.14E-02 | 1.11E+00 | 1.75E-02 | 6.25E-02 | 1.17E+00 | 1.78E-02 | 5.03E-02 | 1.08E+00 | 6.95E-04 | 2.55E-03 | -1.09E+00 | 5.55E-04 | [illegible] |

[illegible]

| | | | | | | | | | | | | | | | | | | | | | | | | |
|---|---|---|---|---|---|---|---|---|---|---|---|---|---|---|---|---|---|---|---|---|---|---|---|---|
| JAKMIP2 | 1.58E-11 | 8.22E-11 | -1.56E+00 | 6.04E-09 | 2.61E-08 | -1.17E+00 | 7.02E-18 | 2.98E-17 | -2.27E+00 | 1.54E-09 | 5.89E-09 | -1.31E+00 | 6.94E-13 | 2.48E-12 | -2.06E+00 | 1.27E-10 | 3.57E-10 | -1.66E+00 | 7.88E-04 | 2.89E-03 | -1.31E+00 | 1.30E-02 | 3.58E-02 | -1.14E+00 |
| PIM2 | 1.59E-11 | 8.28E-11 | -1.54E+00 | 3.03E-13 | 1.31E-12 | -1.55E+00 | 1.53E-11 | 6.49E-11 | -1.96E+00 | 9.10E-12 | 3.48E-11 | -1.90E+00 | 1.52E-03 | 5.42E-03 | -1.11E+00 | 9.11E-08 | 2.57E-07 | -1.73E+00 | 6.12E-03 | 2.25E-02 | -1.23E+00 | 3.16E-12 | 8.69E-12 | -2.36E+00 |
| MINPP1 | 1.60E-11 | 8.35E-11 | -2.96E+00 | 4.11E-06 | 1.78E-05 | -1.43E+00 | 2.45E-13 | 1.04E-12 | -3.11E+00 | 5.54E-04 | 2.11E-03 | -1.11E+00 | 3.14E-20 | 1.12E-19 | -1.50E+00 | 1.24E-01 | 3.50E-01 | 1.25E+00 | 4.31E-15 | 1.58E-14 | -1.62E+00 | 1.68E-01 | 4.62E-01 | 1.18E+00 |
| AP2M1 | 1.62E-11 | 8.43E-11 | -1.11E+00 | 5.47E-02 | 2.36E-01 | 1.24E+00 | 9.94E-03 | 4.22E-02 | -1.08E+00 | 2.13E-01 | 8.13E-01 | 1.11E+00 | 1.60E-03 | 5.72E-03 | 1.00E+00 | 3.33E-01 | 9.37E-01 | 1.26E+00 | 1.35E-01 | 4.95E-01 | 1.01E+00 | 2.56E-01 | 7.04E-01 | 1.13E+00 |
| PSMG2 | 1.62E-11 | 8.45E-11 | -1.14E+00 | 2.37E-05 | 1.02E-04 | 1.10E+00 | 1.09E-05 | 4.64E-05 | -1.34E+00 | 2.04E-02 | 7.80E-02 | -1.02E+00 | 7.10E-10 | 2.53E-09 | -1.35E+00 | 3.06E-02 | 8.63E-02 | 1.12E+00 | 6.30E-07 | 2.31E-06 | -1.25E+00 | 6.54E-03 | 1.80E-02 | -1.01E+00 |
| FBP1 | 1.63E-11 | 8.48E-11 | -3.07E+00 | 8.47E-11 | 3.66E-10 | -2.34E+00 | 9.13E-13 | 3.88E-12 | -2.36E+00 | 2.27E-15 | 8.67E-15 | -2.44E+00 | 8.64E-09 | 3.08E-08 | -1.88E+00 | 7.95E-10 | 2.24E-09 | -2.05E+00 | 2.56E-04 | 9.41E-04 | -1.85E+00 | 4.59E-03 | 1.26E-02 | -1.44E+00 |
| SPAG9 | 1.63E-11 | 8.49E-11 | -1.26E+00 | 2.13E-19 | 9.22E-19 | -1.35E+00 | 1.33E-03 | 5.64E-03 | -1.06E+00 | 1.15E-09 | 4.41E-09 | -1.34E+00 | 5.01E-05 | 1.79E-04 | -1.12E+00 | 2.66E-11 | 7.49E-11 | -1.59E+00 | 2.14E-09 | 7.84E-09 | -1.32E+00 | 5.41E-12 | 1.49E-11 | -1.70E+00 |
| CCDC88A | 1.63E-11 | 8.49E-11 | -1.34E+00 | 2.47E-13 | 1.07E-12 | -1.30E+00 | 4.96E-05 | 2.11E-04 | -1.22E+00 | 3.59E-08 | 1.37E-07 | -1.28E+00 | 5.19E-07 | 1.85E-06 | -1.24E+00 | 1.40E-07 | 3.94E-07 | -1.27E+00 | 5.17E-07 | 1.90E-06 | -1.27E+00 | 6.91E-08 | 1.90E-07 | -1.31E+00 |
| HSPA9P1 | 1.64E-11 | 8.52E-11 | -1.93E+00 | 1.55E-13 | 6.71E-13 | -2.12E+00 | 1.99E-12 | 8.48E-12 | -1.91E+00 | 2.96E-12 | 1.13E-11 | -1.70E+00 | 1.99E-07 | 7.10E-07 | -1.94E+00 | 1.06E-08 | 3.00E-08 | -2.17E+00 | 7.25E-14 | 2.66E-13 | -1.81E+00 | 2.76E-11 | 7.58E-11 | -1.59E+00 |
| CEP350 | 1.65E-11 | 8.59E-11 | -1.33E+00 | 2.07E-10 | 8.95E-10 | -1.14E+00 | 9.12E-10 | 3.87E-09 | -1.46E+00 | 3.94E-08 | 1.50E-07 | -1.25E+00 | 5.34E-06 | 1.90E-05 | -1.28E+00 | 1.00E-05 | 2.83E-05 | -1.26E+00 | 1.34E-07 | 4.91E-07 | -1.28E+00 | 1.29E-03 | 3.54E-03 | -1.15E+00 |
| TRIM11 | 1.65E-11 | 8.60E-11 | -1.84E+00 | 2.39E-06 | 1.03E-05 | -1.11E+00 | 1.93E-04 | 8.18E-04 | -1.32E+00 | 9.66E-04 | 3.69E-03 | -1.05E+00 | 5.91E-12 | 2.11E-11 | -2.20E+00 | 7.71E-08 | 2.17E-07 | -1.59E+00 | 1.16E-11 | 4.27E-11 | -1.86E+00 | 1.46E-03 | 4.01E-03 | -1.07E+00 |
| NEK1 | 1.67E-11 | 8.72E-11 | -1.73E+00 | 8.59E-08 | 3.71E-07 | -1.16E+00 | 4.78E-09 | 2.03E-08 | -1.85E+00 | 1.10E-04 | 4.21E-04 | -1.18E+00 | 8.61E-04 | 3.07E-03 | -1.53E+00 | 4.93E-02 | 1.39E-01 | -1.02E+00 | 7.52E-09 | 2.76E-08 | -1.63E+00 | 4.87E-07 | 1.34E-06 | -1.40E+00 |
| EPHX2 | 1.67E-11 | 8.73E-11 | -1.64E+00 | 6.41E-09 | 2.77E-08 | -1.28E+00 | 1.12E-02 | 4.75E-02 | -1.16E+00 | 3.91E-02 | 1.49E-01 | 1.03E+00 | 4.38E-03 | 1.56E-02 | -1.72E+00 | 1.06E-01 | 2.98E-01 | -1.01E+00 | 7.71E-07 | 2.83E-06 | -1.42E+00 | 2.85E-08 | 7.82E-08 | -1.64E+00 |
| GIGYF1 | 1.68E-11 | 8.74E-11 | -1.33E+00 | 1.28E-09 | 5.55E-09 | -1.03E+00 | 2.64E-04 | 1.12E-03 | -1.27E+00 | 8.74E-03 | 3.34E-02 | 1.11E+00 | 1.76E-04 | 6.28E-04 | -1.17E+00 | 1.65E-03 | 4.65E-03 | 1.01E+00 | 5.35E-10 | 1.97E-09 | -1.51E+00 | 9.30E-04 | 2.56E-03 | -1.05E+00 |
| NOL11 | 1.75E-11 | 9.12E-11 | -1.27E+00 | 5.86E-06 | 2.53E-05 | 1.09E+00 | 1.24E-08 | 5.29E-08 | -1.38E+00 | 2.74E-03 | 1.04E-02 | 1.07E+00 | 4.54E-02 | 1.62E-01 | 1.19E+00 | 2.25E-01 | 6.35E-01 | 1.36E+00 | 4.59E-07 | 1.69E-06 | -1.25E+00 | 1.67E-01 | 4.58E-01 | 1.12E+00 |
| SKAP2 | 1.76E-11 | 9.18E-11 | -1.30E+00 | 1.00E-02 | 4.33E-02 | 1.21E+00 | 7.01E-07 | 2.98E-06 | -1.42E+00 | 1.86E-03 | 7.09E-03 | -1.02E+00 | 1.73E-11 | 6.17E-11 | -1.57E+00 | 2.28E-05 | 6.44E-05 | -1.09E+00 | 8.72E-09 | 3.20E-08 | -1.42E+00 | 1.09E-03 | 2.99E-03 | -1.07E+00 |
| ODC1 | 1.80E-11 | 9.38E-11 | -1.39E+00 | 1.96E-18 | 8.47E-18 | -1.54E+00 | 1.71E-04 | 7.26E-04 | -1.02E+00 | 6.64E-08 | 2.54E-07 | -1.13E+00 | 1.27E-07 | 4.54E-07 | -1.29E+00 | 6.50E-06 | 1.83E-05 | -1.17E+00 | 7.66E-02 | 2.81E-01 | 1.11E+00 | 1.37E-01 | 3.76E-01 | 1.31E+00 |
| AC016700.1 | 1.81E-11 | 9.44E-11 | -2.29E+00 | 1.92E-09 | 8.31E-09 | -1.73E+00 | 1.05E-28 | 4.48E-28 | -3.40E+00 | 3.65E-18 | 1.39E-17 | -1.90E+00 | 3.55E-08 | 1.27E-07 | -2.25E+00 | 2.63E-07 | 7.40E-07 | -1.93E+00 | 7.92E-12 | 2.91E-11 | -2.03E+00 | 5.40E-07 | 1.48E-06 | -1.44E+00 |
| SMIM10L1 | 1.81E-11 | 9.44E-11 | -1.27E+00 | 5.22E-08 | 2.26E-07 | -1.05E+00 | 5.51E-07 | 2.34E-06 | -1.22E+00 | 1.29E-03 | 4.93E-03 | 1.06E+00 | 1.59E-06 | 5.68E-06 | -1.15E+00 | 5.66E-04 | 1.60E-03 | 1.00E+00 | 2.53E-02 | 9.27E-02 | -1.06E+00 | 1.83E-01 | 5.04E-01 | 1.04E+00 |
| SORT1 | 1.84E-11 | 9.61E-11 | -1.97E+00 | 6.92E-08 | 2.99E-07 | -1.24E+00 | 2.38E-03 | 1.01E-02 | -1.41E+00 | 1.09E-02 | 4.16E-02 | -1.18E+00 | 4.31E-04 | 1.54E-03 | -2.08E+00 | 9.27E-03 | 2.61E-02 | -1.28E+00 | 1.36E-01 | 5.00E-01 | 1.12E+00 | 1.92E-01 | 5.27E-01 | 1.08E+00 |
| LRRN1 | 1.84E-11 | 9.61E-11 | -2.69E+00 | 1.71E-02 | 7.39E-02 | 1.16E+00 | 1.76E-02 | 7.48E-02 | -1.46E+00 | 1.73E-01 | 6.61E-01 | 1.50E+00 | 8.52E-02 | 3.04E-01 | -1.06E+00 | 1.16E-01 | 3.28E-01 | 2.23E+00 | 2.52E-01 | 9.25E-01 | 1.05E+00 | 1.67E-01 | 4.58E-01 | 1.74E+00 |
| UBE2D3 | 1.85E-11 | 9.63E-11 | -1.15E+00 | 1.07E-10 | 4.62E-10 | -1.08E+00 | 4.35E-03 | 1.85E-02 | -1.13E+00 | 5.00E-03 | 1.91E-02 | -1.05E+00 | 2.73E-07 | 9.72E-07 | -1.32E+00 | 2.85E-05 | 8.03E-05 | -1.17E+00 | 5.65E-03 | 2.07E-02 | -1.10E+00 | 2.57E-02 | 7.06E-02 | -1.08E+00 |
| ARID1B | 1.87E-11 | 9.74E-11 | -1.31E+00 | 6.25E-06 | 2.70E-05 | -1.06E+00 | 2.93E-05 | 1.25E-04 | -1.41E+00 | 2.62E-02 | 9.99E-02 | -1.10E+00 | 1.19E-06 | 4.25E-06 | -1.22E+00 | 2.82E-04 | 7.96E-04 | -1.12E+00 | 3.28E-14 | 1.20E-13 | -1.59E+00 | 2.24E-05 | 6.15E-05 | -1.25E+00 |
| MRPL3 | 1.88E-11 | 9.77E-11 | -1.23E+00 | 1.98E-07 | 8.57E-07 | 1.01E+00 | 2.15E-02 | 9.13E-02 | -1.06E+00 | 3.56E-02 | 1.36E-01 | 1.10E+00 | 1.66E-08 | 5.92E-08 | -1.30E+00 | 1.45E-02 | 4.08E-02 | 1.14E+00 | 4.31E-06 | 1.58E-05 | -1.16E+00 | 3.30E-01 | 9.07E-01 | 1.20E+00 |
| SCAMP1 | 1.89E-11 | 9.86E-11 | -1.48E+00 | 3.86E-07 | 1.67E-06 | -1.07E+00 | 1.49E-18 | 6.34E-18 | -1.83E+00 | 1.10E-13 | 4.21E-13 | -1.36E+00 | 7.61E-11 | 2.71E-10 | -1.59E+00 | 9.22E-06 | 2.60E-05 | -1.10E+00 | 4.95E-13 | 1.82E-12 | -1.79E+00 | 3.63E-07 | 9.98E-07 | -1.37E+00 |
| ATIC | 1.90E-11 | 9.89E-11 | -1.21E+00 | 1.77E-07 | 7.64E-07 | 1.02E+00 | 9.62E-05 | 4.09E-04 | -1.14E+00 | 1.27E-03 | 4.84E-03 | 1.03E+00 | 8.82E-13 | 3.14E-12 | -1.43E+00 | 1.08E-03 | 3.03E-03 | 1.08E+00 | 7.51E-06 | 2.75E-05 | -1.27E+00 | 2.45E-02 | 6.72E-02 | 1.01E+00 |
| RNF170 | 1.90E-11 | 9.90E-11 | -1.94E+00 | 1.24E-02 | 5.35E-02 | 1.30E+00 | 8.13E-11 | 3.46E-10 | -2.18E+00 | 5.02E-02 | 1.92E-01 | 1.16E+00 | 1.97E-04 | 7.04E-04 | -1.51E+00 | 2.81E-01 | 7.91E-01 | 1.31E+00 | 2.98E-10 | 1.10E-09 | -1.73E+00 | 5.86E-03 | 1.61E-02 | 1.02E+00 |
| ARAP2 | 1.96E-11 | 1.02E-10 | -1.47E+00 | 4.82E-09 | 2.08E-08 | -1.12E+00 | 2.00E-08 | 8.48E-08 | -1.62E+00 | 1.68E-06 | 6.40E-06 | -1.25E+00 | 2.00E-01 | 7.14E-01 | 1.03E+00 | 6.85E-02 | 1.93E-01 | -1.45E+00 | 7.34E-02 | 2.70E-01 | -1.49E+00 | 1.22E-01 | 3.34E-01 | -1.30E+00 |
| RPL10P12 | 1.96E-11 | 1.02E-10 | -1.37E+00 | 1.94E-35 | 8.36E-35 | -1.70E+00 | 4.18E-05 | 1.78E-04 | -1.32E+00 | 6.35E-16 | 2.43E-15 | -1.71E+00 | 3.41E-06 | 1.22E-05 | -1.40E+00 | 4.02E-17 | 1.13E-16 | -1.83E+00 | 2.31E-08 | 8.46E-08 | -1.39E+00 | 2.12E-19 | 5.81E-19 | -1.82E+00 |
| TACC3 | 1.97E-11 | 1.03E-10 | -1.23E+00 | 3.70E-04 | 1.60E-03 | 1.21E+00 | 1.89E-03 | 8.01E-03 | -1.09E+00 | 2.05E-01 | 7.84E-01 | 1.27E+00 | 3.91E-06 | 1.40E-05 | -1.27E+00 | 2.22E-05 | 6.25E-05 | -1.24E+00 | 3.86E-03 | 1.42E-02 | -1.08E+00 | 2.97E-01 | 8.17E-01 | 1.15E+00 |
| [illegible] | [illegible] | [illegible] | [illegible] | [illegible] | [illegible] | [illegible] | [illegible] | [illegible] | [illegible] | [illegible] | [illegible] | [illegible] | [illegible] | [illegible] | [illegible] | [illegible] | [illegible] | [illegible] | [illegible] | [illegible] | [illegible] | [illegible] | [illegible] | [illegible] |

| | | | | | | | | | | | | | | | | | | | | | | | | |
|---|---|---|---|---|---|---|---|---|---|---|---|---|---|---|---|---|---|---|---|---|---|---|---|---|
| RPL30P14 | 5.55E-11 | 2.89E-10 | -1.18E+00 | 4.75E-48 | 2.05E-47 | -1.94E+00 | 4.19E-03 | 1.78E-02 | -1.09E+00 | 1.11E-24 | 4.26E-24 | -1.98E+00 | 8.66E-03 | 3.09E-02 | -1.02E+00 | 7.24E-16 | 2.04E-15 | -1.80E+00 | 1.63E-06 | 5.98E-06 | -1.24E+00 | 3.29E-19 | 9.03E-19 | -1.94E+00 |
| MALL | 5.59E-11 | 2.92E-10 | -2.42E+00 | 5.74E-07 | 2.48E-06 | -1.45E+00 | 7.08E-06 | 3.01E-05 | -1.93E+00 | 9.52E-03 | 3.64E-02 | -1.10E+00 | 5.81E-02 | 2.07E-01 | -1.70E+00 | 1.90E-01 | 5.35E-01 | -1.12E+00 | 4.85E-08 | 1.78E-07 | -1.96E+00 | 3.84E-04 | 1.06E-03 | -1.13E+00 |
| SELENOK | 5.61E-11 | 2.92E-10 | -1.14E+00 | 3.80E-16 | 1.64E-15 | -1.28E+00 | 2.61E-03 | 1.11E-02 | -1.12E+00 | 1.19E-06 | 4.53E-06 | -1.24E+00 | 3.69E-07 | 1.31E-06 | -1.17E+00 | 9.91E-16 | 2.79E-15 | -1.66E+00 | 3.86E-03 | 1.42E-02 | -1.12E+00 | 1.33E-05 | 3.65E-05 | -1.40E+00 |
| IL16 | 5.62E-11 | 2.93E-10 | -1.47E+00 | 2.14E-04 | 9.23E-04 | 1.09E+00 | 2.62E-11 | 1.11E-10 | -1.75E+00 | 5.81E-06 | 2.22E-05 | -1.09E+00 | 8.77E-05 | 3.13E-04 | -1.90E+00 | 5.39E-03 | 1.52E-02 | -1.31E+00 | 7.92E-04 | 2.91E-03 | -1.39E+00 | 3.58E-03 | 9.83E-03 | -1.30E+00 |
| KIAA0141 | 5.69E-11 | 2.96E-10 | -1.34E+00 | 2.07E-08 | 8.93E-08 | -1.06E+00 | 2.00E-04 | 8.48E-04 | -1.16E+00 | 2.08E-03 | 7.96E-03 | 1.02E+00 | 1.14E-09 | 4.06E-09 | -1.44E+00 | 4.15E-07 | 1.17E-06 | -1.20E+00 | 1.54E-04 | 5.66E-04 | -1.19E+00 | 2.11E-02 | 5.78E-02 | 1.06E+00 |
| PLS3 | 5.70E-11 | 2.97E-10 | -2.06E+00 | 1.35E-05 | 5.82E-05 | -1.19E+00 | 2.19E-03 | 9.30E-03 | -1.19E+00 | 6.04E-02 | 2.31E-01 | 1.16E+00 | 1.13E-02 | 4.04E-02 | -1.10E+00 | 8.25E-02 | 2.32E-01 | 1.10E+00 | 1.21E-01 | 4.45E-01 | 1.63E+00 | 1.65E-01 | 4.52E-01 | 1.61E+00 |
| GUCY1B3 | 5.77E-11 | 3.01E-10 | -1.42E+00 | 5.36E-03 | 2.32E-02 | 1.22E+00 | 1.28E-03 | 5.43E-03 | -1.30E+00 | 2.05E-01 | 7.83E-01 | 1.31E+00 | 4.36E-02 | 1.55E-01 | 1.01E+00 | 3.02E-01 | 8.51E-01 | 1.37E+00 | 2.65E-01 | 9.74E-01 | 1.01E+00 | 2.76E-01 | 7.59E-01 | -1.07E+00 |
| MARCH5 | 5.79E-11 | 3.02E-10 | -1.36E+00 | 1.51E-08 | 6.54E-08 | -1.08E+00 | 3.93E-05 | 1.67E-04 | -1.32E+00 | 5.74E-05 | 2.19E-04 | -1.17E+00 | 7.93E-07 | 2.83E-06 | -1.30E+00 | 1.41E-04 | 3.98E-04 | -1.11E+00 | 1.13E-06 | 4.15E-06 | -1.30E+00 | 1.13E-05 | 3.11E-05 | -1.24E+00 |
| ID2 | 5.83E-11 | 3.04E-10 | 3.01E+00 | 1.80E-12 | 7.80E-12 | -1.58E+00 | 5.48E-03 | 2.33E-02 | 1.20E+00 | 9.87E-19 | 3.77E-18 | -3.53E+00 | 2.03E-01 | 7.25E-01 | 1.13E+00 | 1.46E-03 | 4.12E-03 | -3.42E+00 | 1.79E-03 | 6.57E-03 | 3.47E+00 | 2.17E-02 | 5.97E-02 | 1.40E+00 |
| TMEM51 | 5.91E-11 | 3.08E-10 | -2.62E+00 | 9.49E-05 | 4.10E-04 | -1.11E+00 | 3.03E-09 | 1.29E-08 | -2.32E+00 | 1.01E-04 | 3.84E-04 | -1.30E+00 | 8.46E-03 | 3.02E-02 | -2.40E+00 | 3.92E-02 | 1.11E-01 | -1.69E+00 | 9.10E-02 | 3.34E-01 | -1.38E+00 | 2.83E-01 | 7.77E-01 | -1.01E+00 |
| NDUFA11 | 5.92E-11 | 3.08E-10 | -1.17E+00 | 3.84E-33 | 1.66E-32 | -1.57E+00 | 2.37E-03 | 1.00E-02 | -1.14E+00 | 1.22E-16 | 4.65E-16 | -1.62E+00 | 1.66E-08 | 5.91E-08 | -1.42E+00 | 2.71E-16 | 7.65E-16 | -1.76E+00 | 4.59E-04 | 1.69E-03 | -1.13E+00 | 1.13E-10 | 3.09E-10 | -1.52E+00 |
| INTS10 | 5.94E-11 | 3.10E-10 | -1.23E+00 | 1.13E-08 | 4.89E-08 | 1.00E+00 | 1.07E-08 | 4.54E-08 | -1.35E+00 | 3.07E-09 | 1.17E-08 | -1.19E+00 | 3.94E-02 | 1.40E-01 | 1.18E+00 | 1.50E-01 | 4.23E-01 | 1.39E+00 | 5.09E-05 | 1.87E-04 | -1.08E+00 | 1.25E-04 | 3.42E-04 | -1.10E+00 |
| ZNF562 | 5.96E-11 | 3.10E-10 | -1.49E+00 | 1.32E-08 | 5.72E-08 | -1.21E+00 | 1.82E-02 | 7.73E-02 | 1.04E+00 | 4.23E-03 | 1.61E-02 | 1.14E+00 | 1.30E-05 | 4.63E-05 | -1.11E+00 | 1.85E-04 | 5.23E-04 | -1.01E+00 | 1.19E-08 | 4.38E-08 | -1.54E+00 | 1.64E-04 | 4.52E-04 | -1.22E+00 |
| PIAS1 | 5.98E-11 | 3.11E-10 | -1.41E+00 | 1.14E-05 | 4.95E-05 | 1.00E+00 | 1.16E-04 | 4.92E-04 | -1.14E+00 | 1.13E-02 | 4.32E-02 | 1.13E+00 | 7.11E-05 | 2.53E-04 | -1.12E+00 | 1.38E-04 | 3.89E-04 | -1.06E+00 | 1.67E-06 | 6.13E-06 | -1.30E+00 | 1.84E-02 | 5.06E-02 | 1.03E+00 |
| ARF4 | 6.10E-11 | 3.18E-10 | -1.08E+00 | 6.25E-15 | 2.70E-14 | -1.11E+00 | 2.67E-03 | 1.13E-02 | -1.01E+00 | 9.52E-04 | 3.64E-03 | 1.05E+00 | 5.44E-07 | 1.94E-06 | -1.23E+00 | 2.57E-02 | 7.24E-02 | 1.09E+00 | 4.02E-05 | 1.48E-04 | -1.10E+00 | 1.86E-01 | 5.12E-01 | 1.04E+00 |
| USP28 | 6.10E-11 | 3.18E-10 | -2.33E+00 | 5.70E-06 | 2.46E-05 | -1.24E+00 | 3.24E-06 | 1.38E-05 | -1.54E+00 | 9.82E-04 | 3.75E-03 | -1.08E+00 | 4.44E-02 | 1.58E-01 | -1.23E+00 | 9.38E-02 | 2.64E-01 | -1.20E+00 | 6.56E-04 | 2.41E-03 | -1.64E+00 | 8.31E-02 | 2.28E-01 | -1.10E+00 |
| MED31 | 6.13E-11 | 3.19E-10 | -2.49E+00 | 6.64E-09 | 2.87E-08 | -1.85E+00 | 4.35E-06 | 1.85E-05 | -1.77E+00 | 3.50E-04 | 1.34E-03 | -1.31E+00 | 4.41E-02 | 1.57E-01 | -1.04E+00 | 9.60E-02 | 2.71E-01 | 1.13E+00 | 7.20E-10 | 2.64E-09 | -1.50E+00 | 2.21E-06 | 6.08E-06 | -1.30E+00 |
| TUBB2B | 6.14E-11 | 3.20E-10 | -1.43E+00 | 2.21E-09 | 9.56E-09 | -1.21E+00 | 8.80E-05 | 3.74E-04 | -1.08E+00 | 5.37E-03 | 2.05E-02 | 1.32E+00 | 1.67E-10 | 5.97E-10 | -1.76E+00 | 5.88E-09 | 1.66E-08 | -1.53E+00 | 1.79E-05 | 6.56E-05 | -1.26E+00 | 1.29E-03 | 3.53E-03 | -1.05E+00 |
| IPO7 | 6.14E-11 | 3.20E-10 | -1.38E+00 | 9.60E-05 | 4.15E-04 | 1.11E+00 | 1.00E-09 | 4.26E-09 | -1.45E+00 | 1.91E-05 | 7.30E-05 | -1.05E+00 | 3.40E-12 | 1.21E-11 | -1.72E+00 | 5.84E-07 | 1.65E-06 | -1.23E+00 | 1.31E-12 | 4.80E-12 | -1.62E+00 | 7.11E-04 | 1.95E-03 | -1.18E+00 |
| AKR1B1 | 6.15E-11 | 3.20E-10 | -1.26E+00 | 1.12E-06 | 4.86E-06 | 1.00E+00 | 3.69E-03 | 1.57E-02 | -1.08E+00 | 2.19E-04 | 8.36E-04 | -1.04E+00 | 1.49E-08 | 5.33E-08 | -1.49E+00 | 7.45E-05 | 2.10E-04 | -1.14E+00 | 9.52E-04 | 3.49E-03 | -1.09E+00 | 1.17E-01 | 3.22E-01 | 1.10E+00 |
| HACD4 | 6.16E-11 | 3.21E-10 | -1.20E+00 | 1.55E-08 | 6.70E-08 | 1.01E+00 | 8.78E-06 | 3.73E-05 | -1.24E+00 | 1.07E-05 | 4.08E-05 | -1.09E+00 | 2.20E-05 | 7.83E-05 | -1.19E+00 | 3.61E-03 | 1.02E-02 | 1.04E+00 | 1.22E-07 | 4.48E-07 | -1.29E+00 | 3.79E-05 | 1.04E-04 | -1.17E+00 |
| SLC35B4 | 6.17E-11 | 3.21E-10 | -1.88E+00 | 4.41E-07 | 1.91E-06 | -1.27E+00 | 1.02E-09 | 4.33E-09 | -1.67E+00 | 1.29E-04 | 4.92E-04 | -1.06E+00 | 4.77E-05 | 1.70E-04 | -1.51E+00 | 9.38E-03 | 2.64E-02 | 1.00E+00 | 9.75E-10 | 3.58E-09 | -1.68E+00 | 1.84E-02 | 5.05E-02 | -1.01E+00 |
| ATAD1 | 6.31E-11 | 3.29E-10 | -1.54E+00 | 9.56E-07 | 4.13E-06 | -1.08E+00 | 1.36E-07 | 5.78E-07 | -1.39E+00 | 6.29E-06 | 2.40E-05 | -1.09E+00 | 1.68E-06 | 5.98E-06 | -1.53E+00 | 1.77E-04 | 4.98E-04 | -1.11E+00 | 3.98E-17 | 1.46E-16 | -1.93E+00 | 2.90E-08 | 7.98E-08 | -1.33E+00 |
| C19orf54 | 6.36E-11 | 3.32E-10 | -2.07E+00 | 1.04E-08 | 4.50E-08 | -1.56E+00 | 3.14E-06 | 1.33E-05 | -1.53E+00 | 5.59E-08 | 2.13E-07 | -1.59E+00 | 3.27E-06 | 1.17E-05 | -2.00E+00 | 5.49E-05 | 1.55E-04 | -1.74E+00 | 1.07E-05 | 3.93E-05 | -1.91E+00 | 3.60E-05 | 9.87E-05 | -1.87E+00 |
| NME3 | 6.37E-11 | 3.32E-10 | -1.17E+00 | 3.14E-11 | 1.36E-10 | -1.07E+00 | 6.07E-04 | 2.58E-03 | -1.16E+00 | 3.59E-03 | 1.37E-02 | 1.04E+00 | 5.56E-03 | 1.98E-02 | -1.05E+00 | 1.32E-01 | 3.72E-01 | 1.16E+00 | 4.98E-03 | 1.83E-02 | -1.02E+00 | 7.48E-02 | 2.05E-01 | 1.15E+00 |
| MIEN1 | 6.37E-11 | 3.32E-10 | -1.23E+00 | 2.74E-05 | 1.18E-04 | 1.11E+00 | 1.59E-08 | 6.77E-08 | -1.36E+00 | 4.33E-06 | 1.65E-05 | -1.07E+00 | 1.23E-06 | 4.39E-06 | -1.14E+00 | 2.92E-03 | 8.23E-03 | 1.04E+00 | 1.94E-03 | 7.13E-03 | 1.03E+00 | 1.49E-01 | 4.10E-01 | 1.17E+00 |
| CUEDC2 | 6.37E-11 | 3.32E-10 | -1.19E+00 | 4.33E-07 | 1.87E-06 | 1.03E+00 | 1.97E-02 | 8.39E-02 | -1.04E+00 | 2.86E-02 | 1.09E-01 | 1.08E+00 | 1.92E-06 | 6.85E-06 | -1.21E+00 | 3.04E-04 | 8.58E-04 | -1.02E+00 | 9.79E-02 | 3.59E-01 | 1.07E+00 | 2.71E-01 | 7.44E-01 | 1.23E+00 |
| MDM2 | 6.38E-11 | 3.32E-10 | -1.61E+00 | 1.12E-09 | 4.82E-09 | -1.35E+00 | 4.64E-06 | 1.97E-05 | -1.50E+00 | 7.85E-09 | 3.00E-08 | -1.53E+00 | 2.57E-04 | 9.15E-04 | -1.07E+00 | 3.19E-04 | 9.00E-04 | -1.05E+00 | 2.03E-08 | 7.47E-08 | -1.41E+00 | 1.35E-08 | 3.72E-08 | -1.45E+00 |
| FBXO21 | 6.44E-11 | 3.36E-10 | -1.68E+00 | 3.63E-09 | 1.57E-08 | -1.27E+00 | 1.26E-03 | 5.37E-03 | -1.26E+00 | 1.54E-03 | 5.88E-03 | -1.13E+00 | 7.03E-06 | 2.51E-05 | -2.22E+00 | 6.53E-04 | 1.84E-03 | -1.37E+00 | 2.37E-05 | 8.71E-05 | -1.17E+00 | 4.17E-03 | 1.15E-02 | 1.06E+00 |
| MARC1 | 6.46E-11 | 3.37E-10 | -2.26E+00 | 3.40E-09 | 1.47E-08 | -1.79E+00 | 7.01E-05 | 2.98E-04 | -1.09E+00 | 3.89E-05 | 1.49E-04 | 1.01E+00 | 4.18E-03 | 1.49E-02 | -2.30E+00 | 2.43E-02 | 6.84E-02 | -1.52E+00 | 5.87E-03 | 2.16E-02 | -1.81E+00 | 3.56E-02 | 9.79E-02 | -1.35E+00 |
| DDA1 | 6.46E-11 | 3.37E-10 | -1.34E+00 | 9.26E-14 | 4.00E-13 | -1.39E+00 | 1.51E-05 | 6.42E-05 | -1.26E+00 | 1.74E-08 | 6.64E-08 | -1.30E+00 | 1.74E-06 | 6.19E-06 | -1.28E+00 | 9.80E-05 | 2.76E-04 | -1.16E+00 | 1.63E-06 | 5.98E-06 | -1.15E+00 | 1.17E-04 | 3.22E-04 | -1.08E+00 |
| USP48 | 6.53E-11 | 3.40E-10 | -1.34E+00 | 2.77E-05 | 1.20E-04 | 1.03E+00 | 4.66E-10 | 1.98E-09 | -1.64E+00 | 2.09E-05 | 7.98E-05 | -1.14E+00 | 5.70E-15 | 2.03E-14 | -1.75E+00 | 2.96E-08 | 8.35E-08 | -1.36E+00 | 6.82E-12 | 2.50E-11 | -1.61E+00 | 1.71E-04 | 4.70E-04 | -1.21E+00 |
| MFN1 | 6.56E-11 | 3.42E-10 | -1.62E+00 | 1.55E-06 | 6.70E-06 | -1.13E+00 | 1.19E-08 | 5.06E-08 | -1.68E+00 | 1.37E-06 | 5.24E-06 | -1.25E+00 | 7.24E-07 | 2.58E-06 | -1.48E+00 | 5.36E-06 | 1.51E-05 | -1.32E+00 | 3.27E-11 | 1.20E-10 | -1.84E+00 | 2.61E-05 | 7.16E-05 | -1.21E+00 |
| SCAMP1-AS1 | 6.58E-11 | 3.43E-10 | -1.29E+00 | 2.65E-06 | 1.15E-05 | 1.06E+00 | 3.73E-05 | 1.59E-04 | -1.24E+00 | 1.82E-04 | 6.94E-04 | -1.07E+00 | 1.40E-07 | 5.01E-07 | -1.33E+00 | 6.28E-04 | 1.77E-03 | 1.03E+00 | 1.95E-01 | 7.16E-01 | 1.13E+00 | 9.56E-02 | 2.63E-01 | 1.36E+00 |
| AC114491.1 | 6.62E-11 | 3.45E-10 | -1.27E+00 | 3.47E-22 | 1.50E-21 | -1.33E+00 | 4.07E-02 | 1.73E-01 | -1.12E+00 | 7.11E-06 | 2.72E-05 | -1.23E+00 | 9.88E-05 | 3.52E-04 | -1.25E+00 | 1.65E-06 | 4.64E-06 | -1.29E+00 | 2.94E-02 | 1.08E-01 | -1.11E+00 | 1.68E-02 | 4.62E-02 | -1.21E+00 |
| COLGALT1 | 6.64E-11 | 3.46E-10 | -2.10E+00 | 6.46E-06 | 2.79E-05 | -1.23E+00 | 3.20E-09 | 1.36E-08 | -1.56E+00 | 4.07E-03 | 1.56E-02 | -1.01E+00 | 1.74E-03 | 6.20E-03 | -1.48E+00 | 2.64E-01 | 7.44E-01 | 1.33E+00 | 2.97E-07 | 1.09E-06 | -1.46E+00 | 3.43E-01 | 9.41E-01 | 1.22E+00 |
| [illegible] | | | | | | | | | | | | | | | | | | | | | | | | |

| | | | | | | | | | | | | | | | | | | | | | | | | |
|---|---|---|---|---|---|---|---|---|---|---|---|---|---|---|---|---|---|---|---|---|---|---|---|---|
| DUSP23 | 1.94E-10 | 1.01E-09 | -1.18E+00 | 3.43E-07 | 1.48E-06 | 1.03E+00 | 5.41E-03 | 2.30E-02 | -1.04E+00 | 1.30E-02 | 4.96E-02 | 1.21E+00 | 2.41E-07 | 8.58E-07 | -1.19E+00 | 1.16E-03 | 3.27E-03 | 1.13E+00 | 3.89E-03 | 1.43E-02 | -1.02E+00 | 8.68E-02 | 2.38E-01 | 1.04E+00 |
| NSMCE3 | 2.01E-10 | 1.05E-09 | -1.23E+00 | 4.47E-10 | 1.93E-09 | -1.13E+00 | 6.00E-10 | 2.55E-09 | -1.53E+00 | 2.22E-06 | 8.49E-06 | -1.13E+00 | 7.26E-05 | 2.59E-04 | -1.13E+00 | 5.06E-04 | 1.43E-03 | -1.09E+00 | 7.76E-08 | 2.85E-07 | 1.00E+00 | 5.45E-03 | 1.50E-02 | 1.01E+00 |
| NUP214 | 2.01E-10 | 1.05E-09 | -1.28E+00 | 9.00E-08 | 3.89E-07 | -1.06E+00 | 2.17E-06 | 9.23E-06 | -1.21E+00 | 3.43E-05 | 1.31E-04 | 1.05E+00 | 2.12E-01 | 7.57E-01 | 1.20E+00 | 3.06E-01 | 8.64E-01 | 1.29E+00 | 4.88E-06 | 1.79E-05 | -1.25E+00 | 3.94E-04 | 1.08E-03 | -1.12E+00 |
| SEC31A | 2.01E-10 | 1.05E-09 | -1.23E+00 | 7.73E-05 | 3.34E-04 | 1.04E+00 | 2.85E-05 | 1.21E-04 | -1.23E+00 | 8.24E-05 | 3.15E-04 | -1.05E+00 | 7.19E-07 | 2.56E-06 | -1.32E+00 | 3.29E-03 | 9.29E-03 | -1.03E+00 | 2.57E-07 | 9.42E-07 | -1.33E+00 | 5.71E-02 | 1.57E-01 | -1.07E+00 |
| BET1 | 2.04E-10 | 1.06E-09 | -1.74E+00 | 7.50E-07 | 3.24E-06 | -1.19E+00 | 6.25E-10 | 2.66E-09 | -1.71E+00 | 1.24E-05 | 4.72E-05 | -1.12E+00 | 8.52E-12 | 3.04E-11 | -2.43E+00 | 1.49E-06 | 4.19E-06 | -1.42E+00 | 2.46E-12 | 9.01E-12 | -1.32E+00 | 6.27E-07 | 1.72E-06 | -1.29E+00 |
| EXOC1 | 2.04E-10 | 1.06E-09 | -1.34E+00 | 8.93E-07 | 3.86E-06 | -1.02E+00 | 5.72E-06 | 2.43E-05 | -1.26E+00 | 4.19E-03 | 1.60E-02 | 1.07E+00 | 9.55E-07 | 3.40E-06 | -1.48E+00 | 3.62E-03 | 1.02E-02 | -1.04E+00 | 1.11E-07 | 4.09E-07 | -1.25E+00 | 3.02E-04 | 8.30E-04 | -1.12E+00 |
| PEF1 | 2.05E-10 | 1.07E-09 | -1.28E+00 | 3.96E-05 | 1.71E-04 | 1.07E+00 | 1.91E-07 | 8.12E-07 | -1.30E+00 | 7.57E-05 | 2.89E-04 | 1.00E+00 | 5.68E-08 | 2.03E-07 | -1.24E+00 | 2.34E-03 | 6.61E-03 | 1.10E+00 | 1.22E-06 | 4.48E-06 | -1.21E+00 | 1.03E-01 | 2.83E-01 | 1.12E+00 |
| ITPR1 | 2.06E-10 | 1.07E-09 | -1.28E+00 | 5.12E-08 | 2.21E-07 | -1.00E+00 | 1.09E-06 | 4.62E-06 | -1.42E+00 | 4.80E-06 | 1.83E-05 | -1.15E+00 | 5.67E-07 | 2.02E-06 | -1.50E+00 | 3.18E-06 | 8.96E-06 | -1.40E+00 | 6.49E-12 | 2.38E-11 | -1.65E+00 | 3.35E-13 | 9.21E-13 | -1.77E+00 |
| EME2 | 2.07E-10 | 1.08E-09 | -2.08E+00 | 1.04E-06 | 4.50E-06 | -1.36E+00 | 1.46E-04 | 6.19E-04 | -1.47E+00 | 1.71E-04 | 6.53E-04 | -1.24E+00 | 5.92E-09 | 2.11E-08 | -2.07E+00 | 4.39E-08 | 1.24E-07 | -1.77E+00 | 5.30E-03 | 1.94E-02 | -1.25E+00 | 3.74E-03 | 1.03E-02 | -1.32E+00 |
| BRD9 | 2.09E-10 | 1.09E-09 | -1.88E+00 | 7.30E-08 | 3.15E-07 | -1.40E+00 | 1.26E-08 | 5.34E-08 | -1.46E+00 | 1.77E-11 | 6.75E-11 | -1.47E+00 | 2.19E-04 | 7.83E-04 | -1.34E+00 | 1.59E-03 | 4.49E-03 | -1.16E+00 | 1.17E-03 | 4.29E-03 | -1.12E+00 | 4.40E-03 | 1.21E-02 | -1.10E+00 |
| CDC37 | 2.09E-10 | 1.09E-09 | -1.12E+00 | 1.01E-10 | 4.36E-10 | -1.01E+00 | 8.21E-03 | 3.49E-02 | -1.03E+00 | 2.90E-04 | 1.11E-03 | 1.02E+00 | 8.07E-05 | 2.88E-04 | -1.14E+00 | 6.08E-04 | 1.71E-03 | -1.04E+00 | 7.44E-02 | 2.73E-01 | 1.04E+00 | 1.82E-01 | 5.00E-01 | 1.07E+00 |
| N4BP2 | 2.12E-10 | 1.11E-09 | -1.26E+00 | 1.15E-10 | 4.98E-10 | -1.11E+00 | 9.96E-06 | 4.23E-05 | -1.30E+00 | 7.91E-08 | 3.02E-07 | -1.22E+00 | 5.20E-10 | 1.86E-09 | -1.59E+00 | 2.53E-09 | 7.13E-09 | -1.45E+00 | 1.46E-05 | 5.35E-05 | -1.45E+00 | 2.06E-05 | 5.66E-05 | -1.33E+00 |
| ZMYM3 | 2.13E-10 | 1.11E-09 | -1.71E+00 | 4.84E-05 | 2.09E-04 | -1.06E+00 | 1.03E-15 | 4.36E-15 | -1.88E+00 | 2.36E-11 | 9.03E-11 | -1.42E+00 | 4.85E-04 | 1.73E-03 | -1.42E+00 | 1.32E-02 | 3.73E-02 | -1.08E+00 | 1.07E-04 | 3.94E-04 | -1.34E+00 | 5.75E-03 | 1.58E-02 | -1.13E+00 |
| DLAT | 2.13E-10 | 1.11E-09 | -2.38E+00 | 8.46E-06 | 3.66E-05 | -1.32E+00 | 4.38E-11 | 1.86E-10 | -1.91E+00 | 3.82E-07 | 1.46E-06 | -1.28E+00 | 2.71E-04 | 9.65E-04 | -1.47E+00 | 1.01E-01 | 2.86E-01 | 1.25E+00 | 5.32E-08 | 1.95E-07 | -1.40E+00 | 1.73E-02 | 4.75E-02 | 1.00E+00 |
| NENF | 2.15E-10 | 1.12E-09 | -1.18E+00 | 5.47E-15 | 2.36E-14 | -1.13E+00 | 6.47E-02 | 2.75E-01 | 1.09E+00 | 1.91E-02 | 7.29E-02 | 1.10E+00 | 5.75E-02 | 2.05E-01 | 1.08E+00 | 1.57E-01 | 4.44E-01 | 1.11E+00 | 1.16E-01 | 4.25E-01 | 1.19E+00 | 6.25E-02 | 1.72E-01 | 1.21E+00 |
| PPIE | 2.16E-10 | 1.12E-09 | -1.13E+00 | 3.88E-07 | 1.68E-06 | 1.07E+00 | 4.88E-04 | 2.08E-03 | -1.11E+00 | 1.77E-03 | 6.78E-03 | 1.04E+00 | 1.25E-06 | 4.44E-06 | -1.19E+00 | 1.83E-02 | 5.17E-02 | 1.13E+00 | 9.36E-04 | 3.44E-03 | -1.09E+00 | 3.86E-02 | 1.06E-01 | 1.04E+00 |
| SUZ12 | 2.17E-10 | 1.13E-09 | -1.33E+00 | 1.11E-06 | 4.80E-06 | -1.01E+00 | 3.26E-02 | 1.39E-01 | 1.08E+00 | 1.87E-02 | 7.14E-02 | 1.14E+00 | 8.47E-04 | 3.02E-03 | -1.07E+00 | 9.60E-02 | 2.71E-01 | 1.11E+00 | 1.41E-04 | 5.18E-04 | -1.18E+00 | 5.30E-02 | 1.46E-01 | 1.00E+00 |
| DSG2 | 2.17E-10 | 1.13E-09 | -1.29E+00 | 3.57E-04 | 1.54E-03 | 1.17E+00 | 2.69E-07 | 1.14E-06 | -1.40E+00 | 2.01E-03 | 7.69E-03 | 1.05E+00 | 1.20E-01 | 4.29E-01 | 1.00E+00 | 2.87E-01 | 8.09E-01 | 1.29E+00 | 2.35E-01 | 8.63E-01 | 1.32E+00 | 4.01E-02 | 1.10E-01 | 1.87E+00 |
| SLC25A3 | 2.19E-10 | 1.14E-09 | -1.17E+00 | 2.09E-04 | 9.03E-04 | -1.03E+00 | 3.66E-02 | 1.55E-01 | -1.06E+00 | 5.71E-02 | 2.18E-01 | -1.06E+00 | 5.83E-02 | 2.08E-01 | 1.03E+00 | 2.70E-01 | 7.62E-01 | 1.09E+00 | 2.69E-01 | 9.87E-01 | 1.06E+00 | 3.47E-01 | 9.54E-01 | 1.08E+00 |
| ATP2A3 | 2.19E-10 | 1.14E-09 | -1.35E+00 | 5.01E-09 | 2.17E-08 | -1.13E+00 | 2.10E-03 | 8.93E-03 | -1.06E+00 | 1.83E-02 | 6.99E-02 | 1.15E+00 | 6.24E-08 | 2.23E-07 | -1.44E+00 | 2.73E-08 | 7.69E-08 | -1.41E+00 | 1.90E-01 | 6.97E-01 | 1.11E+00 | 2.53E-01 | 6.94E-01 | 1.04E+00 |
| GOLGA8R | 2.19E-10 | 1.14E-09 | -1.70E+00 | 1.49E-11 | 6.42E-11 | -1.64E+00 | 3.19E-06 | 1.36E-05 | -1.80E+00 | 3.13E-06 | 1.19E-05 | -1.60E+00 | 2.39E-07 | 8.54E-07 | -2.30E+00 | 2.13E-06 | 6.02E-06 | -1.86E+00 | 1.21E-02 | 4.44E-02 | -1.20E+00 | 1.57E-04 | 4.30E-04 | -1.76E+00 |
| TRIR | 2.22E-10 | 1.16E-09 | -1.18E+00 | 4.92E-05 | 2.13E-04 | -1.01E+00 | 1.91E-02 | 8.12E-02 | -1.08E+00 | 1.21E-01 | 4.62E-01 | 1.02E+00 | 7.39E-06 | 2.64E-05 | -1.22E+00 | 9.98E-03 | 2.81E-02 | -1.05E+00 | 3.01E-04 | 1.10E-03 | -1.16E+00 | 9.43E-02 | 2.59E-01 | -1.07E+00 |
| FPGS | 2.22E-10 | 1.16E-09 | -1.40E+00 | 8.16E-09 | 3.52E-08 | -1.18E+00 | 3.66E-08 | 1.55E-07 | -1.30E+00 | 4.05E-09 | 1.55E-08 | -1.20E+00 | 4.54E-04 | 1.62E-03 | -1.09E+00 | 1.73E-02 | 4.88E-02 | 1.13E+00 | 1.05E-02 | 3.87E-02 | -1.03E+00 | 2.03E-01 | 5.57E-01 | 1.11E+00 |
| AGGF1 | 2.24E-10 | 1.17E-09 | -1.44E+00 | 1.13E-06 | 4.86E-06 | -1.08E+00 | 1.05E-12 | 4.46E-12 | -1.77E+00 | 9.49E-08 | 3.63E-07 | -1.21E+00 | 1.37E-05 | 4.88E-05 | -1.40E+00 | 6.21E-03 | 1.75E-02 | -1.01E+00 | 5.95E-08 | 2.19E-07 | -1.43E+00 | 4.94E-02 | 1.36E-01 | 1.06E+00 |
| NLGN1 | 2.24E-10 | 1.17E-09 | -2.53E+00 | 1.33E-09 | 5.73E-09 | -2.03E+00 | 6.69E-02 | 2.84E-01 | -1.34E+00 | 1.30E-01 | 4.98E-01 | -1.03E+00 | 2.27E-01 | 8.11E-01 | 1.38E+00 | 2.47E-02 | 6.96E-02 | 2.59E+00 | 2.68E-01 | 9.84E-01 | 1.16E+00 | 8.17E-02 | 2.25E-01 | 1.72E+00 |
| DAB2 | 2.24E-10 | 1.17E-09 | -1.66E+00 | 2.18E-05 | 9.42E-05 | -1.13E+00 | 7.72E-06 | 3.28E-05 | -1.61E+00 | 1.82E-03 | 6.96E-03 | -1.04E+00 | 1.62E-02 | 5.78E-02 | -1.00E+00 | 1.49E-01 | 4.21E-01 | 1.21E+00 | 4.75E-02 | 1.74E-01 | 1.62E+00 | 3.26E-01 | 8.96E-01 | 1.37E+00 |
| AGTRAP | 2.25E-10 | 1.17E-09 | -1.44E+00 | 7.03E-09 | 3.04E-08 | -1.18E+00 | 4.38E-09 | 1.86E-08 | -1.28E+00 | 1.35E-13 | 5.16E-13 | -1.42E+00 | 6.76E-02 | 2.41E-01 | -1.56E+00 | 1.49E-01 | 4.21E-01 | -1.23E+00 | 9.96E-02 | 3.66E-01 | -1.08E+00 | 3.06E-01 | 8.42E-01 | 1.15E+00 |
| PRPF8 | 2.26E-10 | 1.18E-09 | -1.21E+00 | 1.10E-02 | 4.75E-02 | 1.25E+00 | 8.82E-03 | 3.75E-02 | 1.02E+00 | 1.85E-01 | 7.08E-01 | 1.28E+00 | 9.40E-03 | 3.35E-02 | 1.06E+00 | 6.25E-02 | 1.76E-01 | 1.22E+00 | 2.01E-01 | 7.38E-01 | 1.16E+00 | 3.07E-01 | 8.45E-01 | 1.32E+00 |
| GABPB1-AS1 | 2.26E-10 | 1.18E-09 | -1.35E+00 | 8.98E-02 | 3.88E-01 | 1.26E+00 | 2.51E-09 | 1.07E-08 | -1.53E+00 | 9.48E-02 | 3.62E-01 | 1.05E+00 | 9.75E-03 | 3.48E-02 | 1.08E+00 | 3.92E-02 | 1.10E-01 | 1.51E+00 | 1.17E-04 | 4.29E-04 | -1.23E+00 | 1.39E-01 | 3.81E-01 | -1.04E+00 |
| ANKRD26 | 2.27E-10 | 1.19E-09 | -1.66E+00 | 3.92E-08 | 1.70E-07 | -1.30E+00 | 2.32E-15 | 9.85E-15 | -2.07E+00 | 1.41E-13 | 5.40E-13 | -1.69E+00 | 9.94E-04 | 3.55E-03 | -1.85E+00 | 1.72E-02 | 4.85E-02 | -1.30E+00 | 1.31E-11 | 4.81E-11 | -1.80E+00 | 9.68E-13 | 2.66E-12 | -2.01E+00 |
| TTLL7 | 2.28E-10 | 1.19E-09 | -1.60E+00 | 2.44E-04 | 1.06E-03 | 1.05E+00 | 4.97E-12 | 2.11E-11 | -1.95E+00 | 4.50E-09 | 1.72E-08 | -1.39E+00 | 1.88E-14 | 6.72E-14 | -2.61E+00 | 2.26E-15 | 6.36E-15 | -2.60E+00 | 2.06E-02 | 7.57E-02 | -1.04E+00 | 6.54E-02 | 1.80E-01 | -1.11E+00 |
| GATM | 2.29E-10 | 1.20E-09 | -2.27E+00 | 6.24E-04 | 2.69E-03 | -1.02E+00 | 2.99E-11 | 1.27E-10 | -1.79E+00 | 4.50E-04 | 1.72E-03 | 1.00E+00 | 2.50E-02 | 8.91E-02 | -1.65E+00 | 1.52E-01 | 4.28E-01 | 1.10E+00 | 7.93E-04 | 2.91E-03 | -1.71E+00 | 2.72E-01 | 7.46E-01 | 1.16E+00 |
| USP33 | 2.31E-10 | 1.21E-09 | -1.43E+00 | 1.21E-05 | 5.24E-05 | -1.03E+00 | 1.45E-10 | 6.15E-10 | -1.57E+00 | 1.31E-05 | 5.00E-05 | -1.07E+00 | 2.62E-09 | 9.34E-09 | -1.52E+00 | 2.06E-04 | 5.81E-04 | -1.12E+00 | 2.01E-07 | 7.38E-07 | -1.20E+00 | 3.84E-04 | 1.06E-03 | -1.07E+00 |
| MCCC2 | 2.34E-10 | 1.22E-09 | -1.60E+00 | 3.37E-05 | 1.46E-04 | 1.01E+00 | 6.03E-04 | 2.56E-03 | -1.20E+00 | 1.05E-02 | 4.01E-02 | 1.07E+00 | 1.47E-06 | 5.25E-06 | -1.53E+00 | 6.59E-04 | 1.86E-03 | -1.10E+00 | 3.87E-07 | 1.42E-06 | -1.35E+00 | 1.48E-03 | 4.07E-03 | -1.10E+00 |
| TPRG1L | 2.34E-10 | 1.22E-09 | -1.81E+00 | 9.59E-05 | 4.14E-04 | -1.02E+00 | 1.20E-08 | 5.09E-08 | -1.96E+00 | 1.38E-04 | 5.26E-04 | -1.19E+00 | 1.38E-11 | 4.91E-11 | -2.16E+00 | 4.65E-06 | 1.31E-05 | -1.25E+00 | 4.37E-08 | 1.60E-07 | -1.36E+00 | 9.31E-07 | 2.56E-06 | -1.33E+00 |
| RPS15 | 2.35E-10 | 1.22E-09 | -1.16E+00 | 1.28E-57 | 5.51E-57 | -1.48E+00 | 1.69E-03 | 7.19E-03 | -1.09E+00 | 6.54E-31 | 2.50E-30 | -1.43E+00 | 1.49E-07 | 5.32E-07 | -1.19E+00 | 1.78E-30 | 5.02E-30 | -1.49E+00 | 1.59E-08 | 5.84E-08 | -1.11E+00 | 3.09E-28 | 8.50E-28 | -1.39E+00 |
| MOB4 | 2.36E-10 | 1.23E-09 | -1.41E+00 | 5.61E-13 | 2.42E-12 | -1.38E+00 | 4.57E-05 | 1.94E-04 | -1.25E+00 | 6.53E-05 | 2.50E-04 | -1.07E+00 | 3.55E-10 | 1.27E-09 | -1.67E+00 | 8.34E-14 | 2.35E-13 | -1.91E+00 | 8.09E-08 | 2.97E-07 | -1.34E+00 | 8.04E-07 | 2.21E-06 | -1.26E+00 |
| MSRB3 | 2.36E-10 | 1.23E-09 | -1.23E+00 | 1.01E-03 | 4.35E-03 | 1.15E+00 | 5.17E-06 | 2.20E-05 | -1.38E+00 | 2.52E-03 | 9.63E-03 | 1.01E+00 | 5.44E-04 | 1.94E-03 | -1.16E+00 | 1.24E-02 | 3.50E-02 | 1.09E+00 | 2.50E-01 | 9.17E-01 | 1.20E+00 | 1.48E-01 | 4.07E-01 | 1.39E+00 |
| RBM25 | 2.36E-10 | 1.23E-09 | -1.26E+00 | 7.03E-10 | 3.04E-09 | -1.18E+00 | 5.22E-05 | 2.22E-04 | -1.28E+00 | 7.48E-05 | 2.86E-04 | -1.19E+00 | 4.79E-06 | 1.71E-05 | -1.28E+00 | 8.51E-04 | 2.40E-03 | -1.17E+00 | 2.81E-06 | 1.03E-05 | -1.32E+00 | 1.58E-02 | 4.35E-02 | -1.14E+00 |
| MAP2K5 | 2.37E-10 | 1.23E-09 | -1.54E+00 | 2.14E-07 | 9.23E-07 | -1.19E+00 | 6.71E-13 | 2.85E-12 | -1.67E+00 | 1.67E-10 | 6.38E-10 | -1.42E+00 | 2.88E-09 | 1.03E-08 | -1.83E+00 | 4.85E-06 | 1.37E-05 | -1.26E+00 | 1.84E-07 | 6.76E-07 | -1.41E+00 | 3.14E-06 | 8.61E-06 | -1.30E+00 |
| RPL13P5 | 2.39E-10 | 1.24E-09 | -1.81E+00 | 3.01E-09 | 1.30E-08 | -1.55E+00 | 1.34E-04 | 5.71E-04 | -1.44E+00 | 3.69E-05 | 1.41E-04 | -1.43E+00 | 2.47E-02 | 8.80E-02 | -1.04E+00 | 1.61E-02 | 4.54E-02 | -1.08E+00 | 7.91E-10 | 2.90E-09 | -1.67E+00 | 3.03E-09 | 8.34E-09 | -1.67E+00 |
| DIS3L | 2.40E-10 | 1.25E-09 | -1.71E+00 | 1.81E-04 | 7.84E-04 | 1.01E+00 | 1.39E-06 | 5.91E-06 | -1.38E+00 | 2.13E-03 | 8.15E-03 | 1.04E+00 | 6.98E-07 | 2.49E-06 | -1.69E+00 | 8.25E-03 | 2.32E-02 | -1.00E+00 | 3.55E-14 | 1.30E-13 | -1.74E+00 | 1.20E-06 | 3.31E-06 | -1.27E+00 |
| WASF1 | 2.42E-10 | 1.26E-09 | -1.42E+00 | 5.46E-06 | 2.36E-05 | -1.01E+00 | 3.81E-11 | 1.62E-10 | -1.62E+00 | 6.20E-10 | 2.37E-09 | -1.36E+00 | 3.26E-02 | 1.16E-01 | -1.14E+00 | 3.54E-01 | 9.97E-01 | 1.32E+00 | 1.14E-01 | 4.18E-01 | 1.09E+00 | 2.59E-01 | 7.12E-01 | 1.08E+00 |
| AC012065.5 | 2.42E-10 | 1.26E-09 | -1.70E+00 | 1.55E-08 | 6.70E-08 | -1.39E+00 | 2.51E-07 | 1.07E-06 | -1.59E+00 | 5.20E-05 | 1.99E-04 | -1.21E+00 | 6.11E-08 | 2.18E-07 | -2.21E+00 | 6.27E-04 | 1.77E-03 | -1.25E+00 | 5.55E-14 | 2.04E-13 | -1.75E+00 | 2.64E-06 | 7.25E-06 | -1.23E+00 |
| ANKRD17 | 2.43E-10 | 1.27E-09 | -1.31E+00 | 9.49E-07 | 4.10E-06 | -1.03E+00 | 1.54E-09 | 6.55E-09 | -1.40E+00 | 4.78E-06 | 1.82E-05 | -1.06E+00 | 4.93E-11 | 1.76E-10 | -1.54E+00 | 1.65E-03 | 4.66E-03 | 1.01E+00 | 1.46E-05 | 5.36E-05 | -1.25E+00 | 1.01E-01 | 2.77E-01 | 1.06E+00 |
| UBE2V2 | 2.43E-10 | 1.27E-09 | -1.20E+00 | 1.06E-03 | 4.60E-03 | 1.20E+00 | 1.13E-04 | 4.79E-04 | -1.10E+00 | 1.50E-02 | 5.73E-02 | 1.14E+00 | 3.84E-07 | 1.37E-06 | -1.28E+00 | 1.99E-01 | 5.61E-01 | 1.33E+00 | 5.37E-04 | 1.97E-03 | -1.13E+00 | 1.97E-01 | 5.41E-01 | 1.13E+00 |
| ZZZ3 | 2.44E-10 | 1.27E-09 | -1.55E+00 | 1.81E-06 | 7.84E-06 | -1.09E+00 | 3.97E-11 | 1.69E-10 | -1.50E+00 | 2.33E-07 | 8.89E-07 | -1.13E+00 | 4.63E-04 | 1.65E-03 | -1.30E+00 | 5.06E-04 | 1.43E-03 | -1.22E+00 | 5.37E-04 | 1.97E-03 | -1.10E+00 | 2.73E-03 | 7.49E-03 | -1.10E+00 |
| HAX1 | 2.46E-10 | 1.28E-09 | -1.11E+00 | 1.25E-06 | 5.41E-06 | 1.12E+00 | 9.81E-05 | 4.17E-04 | -1.18E+00 | 4.55E-04 | 1.74E-03 | -1.04E+00 | 1.51E-06 | 5.37E-06 | -1.18E+00 | 1.53E-05 | 4.32E-05 | -1.10E+00 | 4.67E-03 | 1.72E-02 | 1.01E+00 | 2.17E-02 | 5.96E-02 | 1.09E+00 |
| DNAH14 | 2.47E-10 | 1.29E-09 | -1.50E+00 | 3.04E-08 | 1.31E-07 | -1.18E+00 | 2.90E-04 | 1.23E-03 | -1.28E+00 | 5.11E-04 | 1.95E-03 | -1.11E+00 | 4.58E-06 | 1.63E-05 | -1.46E+00 | 1.54E-04 | 4.35E-04 | -1.15E+00 | 3.28E-06 | 1.21E-05 | -1.32E+00 | 1.35E-02 | 3.70E-02 | 1.04E+00 |
| GSDMD | 2.48E-10 | 1.29E-09 | -1.22E+00 | 3.59E-07 | 1.55E-06 | -1.02E+00 | 8.96E-04 | 3.81E-03 | -1.10E+00 | 2.90E-05 | 1.11E-04 | -1.03E+00 | 3.97E-11 | 1.42E-10 | -1.49E+00 | 8.25E-07 | 2.33E-06 | -1.12E+00 | 9.32E-04 | 3.42E-03 | -1.04E+00 | 1.31E-03 | 3.60E-03 | -1.02E+00 |
| ZNF280D | 2.48E-10 | 1.29E-09 | -1.26E+00 | 4.86E-06 | 2.10E-05 | 1.06E+00 | 4.35E-09 | 1.85E-08 | -1.43E+00 | 2.47E-05 | 9.42E-05 | -1.09E+00 | 8.29E-08 | 2.96E-07 | -1.34E+00 | 3.99E-05 | 1.13E-04 | -1.15E+00 | 1.81E-05 | 6.65E-05 | -1.22E+00 | 1.13E-03 | 3.10E-03 | -1.08E+00 |
| RPS14P8 | 2.49E-10 | 1.30E-09 | -1.21E+00 | 3.74E-09 | 1.62E-08 | -1.09E+00 | 1.71E-07 | 7.25E-07 | -1.30E+00 | 1.28E-09 | 4.87E-09 | -1.22E+00 | 2.54E-08 | 9.05E-08 | -1.46E+00 | 6.92E-04 | 1.95E-03 | -1.02E+00 | 5.76E-06 | 2.11E-05 | -1.30E+00 | 9.14E-06 | 2.51E-05 | -1.26E+00 |
| ERP44 | 2.50E-10 | 1.30E-09 | -1.21E+00 | 4.52E-08 | 1.95E-07 | -1.00E+00 | 9.99E-04 | 4.25E-03 | -1.08E+00 | 9.43E-04 | 3.60E-03 | 1.04E+00 | 4.30E-07 | 1.53E-06 | -1.25E+00 | 4.40E-04 | 1.24E-03 | -1.02E+00 | 2.23E-02 | 8.19E-02 | 1.06E+00 | 3.47E-01 | 9.52E-01 | 1.19E+00 |
| PIK3CB | 2.50E-10 | 1.30E-09 | -1.64E+00 | 2.49E-05 | 1.08E-04 | -1.15E+00 | 1.97E-06 | 8.36E-06 | -1.57E+00 | 2.09E-03 | 8.00E-03 | -1.04E+00 | 1.42E-01 | 5.08E-01 | 1.39E+00 | 1.64E-01 | 4.63E-01 | 1.27E+00 | 1.40E-01 | 5.16E-01 | 1.33E+00 | 7.57E-03 | 2.08E-02 | 1.04E+00 |
| CHP1 | 2.50E-10 | 1.30E-09 | -1.49E+00 | 1.86E-05 | 8.05E-05 | 1.02E+00 | 3.12E-06 | 1.33E-05 | -1.40E+00 | 1.96E-03 | 7.50E-03 | 1.00E+00 | 3.05E-07 | 1.09E-06 | -1.36E+00 | 3.92E-02 | 1.10E-01 | 1.05E+00 | 3.33E-07 | 1.22E-06 | -1.30E+00 | 2.96E-03 | 8.14E-03 | -1.09E+00 |
| ANXA11 | 2.52E-10 | 1.31E-09 | -1.22E+00 | 5.29E-09 | 2.29E-08 | -1.02E+00 | 7.22E-07 | 3.07E-06 | -1.27E+00 | 1.02E-04 | 3.88E-04 | -1.08E+00 | 1.26E-09 | 4.50E-09 | -1.27E+00 | 4.34E-06 | 1.22E-05 | -1.09E+00 | 8.80E-06 | 3.23E-05 | -1.20E+00 | 7.40E-03 | 2.03E-02 | -1.07E+00 |
| RPP14 | 2.54E-10 | 1.32E-09 | -1.72E+00 | 1.58E-08 | 6.82E-08 | -1.40E+00 | 2.22E-10 | 9.43E-10 | -1.65E+00 | 1.65E-07 | 6.30E-07 | -1.20E+00 | 9.27E-06 | 3.30E-05 | -1.67E+00 | 6.59E-05 | 1.86E-04 | -1.41E+00 | 7.10E-08 | 2.61E-07 | -1.63E+00 | 2.70E-05 | 7.43E-05 | -1.28E+00 |
| LINC02399 | 2.55E-10 | 1.33E-09 | 3.11E+00 | 1.77E-01 | 7.65E-01 | 1.52E+00 | 5.20E-05 | 2.21E-04 | 2.64E+00 | 8.31E-02 | 3.17E-01 | 1.62E+00 | 2.10E-01 | 7.50E-01 | 1.67E+00 | 2.34E-01 | 6.61E-01 | -1.20E+00 | 2.45E-01 | 9.00E-01 | 1.25E+00 | 3.10E-01 | 8.52E-01 | 1.36E+00 |
| PAPD7 | 2.55E-10 | 1.33E-09 | -2.30E+00 | 4.28E-09 | 1.85E-08 | -1.90E+00 | 1.68E-04 | 7.16E-04 | -1.38E+00 | 2.41E-04 | 9.21E-04 | -1.28E+00 | 2.55E-03 | 9.11E-03 | -1.42E+00 | 1.11E-03 | 3.13E-03 | -1.73E+00 | 4.23E-07 | 1.55E-06 | -1.72E+00 | 2.19E-05 | 6.03E-05 | -1.55E+00 |
| ZNF420 | 2.57E-10 | 1.34E-09 | -2.05E+00 | 2.75E-06 | 1.19E-05 | -1.27E+00 | 8.26E-06 | 3.51E-05 | -1.52E+00 | 3.67E-03 | 1.40E-02 | 1.00E+00 | 1.28E-04 | 4.58E-04 | -2.07E+00 | 1.57E-02 | 4.44E-02 | -1.21E+00 | 1.10E-09 | 4.03E-09 | -2.03E+00 | 3.95E-06 | 1.09E-05 | -1.38E+00 |
| PARP3 | 2.58E-10 | 1.34E-09 | -2.10E+00 | 2.59E-09 | 1.12E-08 | -1.77E+00 | 1.72E-03 | 7.32E-03 | -1.38E+00 | 1.29E-03 | 4.94E-03 | -1.27E+00 | 1.27E-02 | 4.52E-02 | -1.41E+00 | 6.71E-03 | 1.89E-02 | -1.47E+00 | 2.29E-05 | 8.42E-05 | -2.07E+00 | 2.04E-03 | 5.61E-03 | -1.43E+00 |
| IDH3B | 2.59E-10 | 1.35E-09 | -1.28E+00 | 2.02E-06 | 8.74E-06 | -1.01E+00 | 2.95E-05 | 1.25E-04 | -1.06E+00 | 5.07E-06 | 1.94E-05 | -1.01E+00 | 1.66E-05 | 5.93E-05 | -1.23E+00 | 4.22E-03 | 1.19E-02 | 1.03E+00 | 1.10E-02 | 4.04E-02 | 1.04E+00 | 4.22E-02 | 1.16E-01 | 1.12E+00 |
| CTBP1 | 2.62E-10 | 1.36E-09 | -1.51E+00 | 1.49E-06 | 6.43E-06 | -1.08E+00 | 8.27E-08 | 3.52E-07 | -1.51E+00 | 3.38E-04 | 1.29E-03 | -1.06E+00 | 2.17E-09 | 7.72E-09 | -1.70E+00 | 6.52E-06 | 1.84E-05 | -1.26E+00 | 1.02E-09 | 3.74E-09 | -1.62E+00 | 1.20E-03 | 3.29E-03 | -1.16E+00 |
| TLNRD1 | 2.63E-10 | 1.37E-09 | -1.76E+00 | 8.15E-09 | 3.52E-08 | -1.43E+00 | 8.16E-04 | 3.47E-03 | -1.18E+00 | 4.92E-03 | 1.88E-02 | -1.02E+00 | 8.42E-11 | 3.00E-10 | -2.45E+00 | 7.87E-07 | 2.22E-06 | -1.60E+00 | 1.14E-10 | 4.19E-10 | -1.86E+00 | 1.25E-05 | 3.44E-05 | -1.47E+00 |
| LYAR | 2.65E-10 | 1.38E-09 | -1.65E+00 | 6.94E-05 | 3.00E-04 | -1.04E+00 | 6.66E-08 | 2.83E-07 | -1.33E+00 | 7.42E-06 | 2.83E-05 | -1.08E+00 | 2.31E-13 | 8.25E-13 | -2.23E+00 | 3.94E-03 | 1.11E-02 | 1.03E+00 | 3.93E-10 | 1.44E-09 | -1.38E+00 | 1.48E-02 | 4.08E-02 | 1.03E+00 |
| RPARP-AS1 | 2.66E-10 | 1.38E-09 | -1.34E+00 | 2.96E-06 | 1.28E-05 | 1.06E+00 | 3.85E-06 | 1.63E-05 | -1.25E+00 | 5.71E-06 | 2.18E-05 | -1.07E+00 | 8.57E-04 | 3.05E-03 | -1.15E+00 | 1.53E-02 | 4.32E-02 | 1.07E+00 | 3.32E-03 | 1.22E-02 | -1.15E+00 | 6.15E-02 | 1.69E-01 | 1.05E+00 |
| STN1 | 2.66E-10 | 1.39E-09 | -1.28E+00 | 1.53E-06 | 6.60E-06 | -1.02E+00 | 2.49E-03 | 1.06E-02 | -1.13E+00 | 5.90E-05 | 2.25E-04 | -1.13E+00 | 5.88E-03 | 2.10E-02 | -1.00E+00 | 1.09E-02 | 3.07E-02 | 1.08E+00 | 7.10E-02 | 2.61E-01 | 1.22E+00 | 8.65E-02 | 2.38E-01 | 1.20E+00 |
| DYNC1I2P1 | 2.66E-10 | 1.39E-09 | -1.64E+00 | 2.27E-07 | 9.81E-07 | -1.22E+00 | 3.40E-08 | 1.44E-07 | -1.59E+00 | 2.30E-06 | 8.79E-06 | -1.21E+00 | 4.16E-07 | 1.48E-06 | -1.75E+00 | 1.10E-05 | 3.11E-05 | -1.39E+00 | 7.53E-08 | 2.76E-07 | -1.38E+00 | 4.82E-08 | 1.32E-07 | -1.40E+00 |
| TUFM | 2.66E-10 | 1.39E-09 | -1.17E+00 | 1.02E-08 | 4.42E-08 | -1.05E+00 | 4.04E-02 | 1.71E-01 | -1.03E+00 | 1.49E-02 | 5.71E-02 | -1.03E+00 | 1.68E-04 | 6.00E-04 | -1.11E+00 | 4.18E-02 | 1.18E-01 | 1.14E+00 | 6.94E-02 | 2.55E-01 | 1.03E+00 | 1.37E-01 | 3.75E-01 | 1.21E+00 |
| CTCF | 2.67E-10 | 1.39E-09 | -1.35E+00 | 4.83E-06 | 2.09E-05 | -1.01E+00 | 5.00E-12 | 2.12E-11 | -1.51E+00 | 2.13E-06 | 8.13E-06 | -1.03E+00 | 3.30E-11 | 1.18E-10 | -1.45E+00 | 7.24E-05 | 2.04E-04 | -1.03E+00 | 9.38E-12 | 3.44E-11 | -1.52E+00 | 4.09E-04 | 1.12E-03 | -1.13E+00 |
| PPM1H | 2.67E-10 | 1.39E-09 | -1.77E+00 | 1.72E-06 | 7.44E-06 | -1.15E+00 | 3.45E-06 | 1.46E-05 | -1.49E+00 | 3.01E-05 | 1.15E-04 | -1.27E+00 | 7.69E-19 | 2.74E-18 | -1.71E+00 | 6.96E-20 | 1.96E-19 | -1.67E+00 | 3.72E-08 | 1.37E-07 | -1.59E+00 | 6.57E-06 | 1.80E-05 | -1.43E+00 |
| MBNL1-AS1 | 2.68E-10 | 1.40E-09 | -1.74E+00 | 1.04E-03 | 4.48E-03 | 2.05E+00 | 8.81E-05 | 3.74E-04 | -1.58E+00 | 2.98E-02 | 1.14E-01 | 1.82E+00 | 4.12E-02 | 1.47E-01 | -1.60E+00 | 6.32E-02 | 1.78E-01 | 1.80E+00 | 1.52E-02 | 5.59E-02 | -1.29E+00 | 2.08E-02 | 5.72E-02 | 1.83E+00 |
| RNF219 | 2.68E-10 | 1.40E-09 | -1.76E+00 | 4.36E-11 | 1.88E-10 | -1.78E+00 | 7.05E-07 | 2.99E-06 | -1.55E+00 | 1.21E-05 | 4.64E-05 | -1.21E+00 | 1.75E-05 | 6.25E-05 | -1.74E+00 | 1.27E-03 | 3.58E-03 | -1.24E+00 | 6.64E-09 | 2.44E-08 | -1.66E+00 | 6.85E-04 | 1.88E-03 | -1.06E+00 |
| AF127577.4 | 2.70E-10 | 1.41E-09 | -2.07E+00 | 1.19E-03 | 5.14E-03 | 1.04E+00 | 4.19E-07 | 1.78E-06 | -2.21E+00 | 4.52E-02 | 1.73E-01 | 1.16E+00 | 1.83E-01 | 6.54E-01 | 1.08E+00 | 2.69E-01 | 7.58E-01 | 1.13E+00 | 2.38E-01 | 8.75E-01 | 1.24E+00 | 2.59E-01 | 7.12E-01 | 1.27E+00 |
| RPLP0 | 2.71E-10 | 1.41E-09 | -1.19E+00 | 2.33E-08 | 1.01E-07 | -1.16E+00 | 3.97E-08 | 1.69E-07 | -1.21E+00 | 3.01E-04 | 1.15E-03 | -1.12E+00 | 1.20E-08 | 4.29E-08 | -1.23E+00 | 1.60E-04 | 4.51E-04 | -1.16E+00 | 1.78E-20 | 6.55E-20 | -1.28E+00 | 1.51E-05 | 4.16E-05 | -1.14E+00 |
| GOLGA7 | 2.71E-10 | 1.41E-09 | -1.25E+00 | 8.72E-06 | 3.77E-05 | 1.03E+00 | 7.53E-05 | 3.20E-04 | -1.25E+00 | 2.34E-04 | 8.95E-04 | -1.08E+00 | 2.85E-08 | 1.01E-07 | -1.39E+00 | 1.60E-03 | 4.52E-03 | -1.01E+00 | 7.57E-09 | 2.78E-08 | -1.36E+00 | 6.99E-04 | 1.92E-03 | -1.07E+00 |
| FES | 2.72E-10 | 1.42E-09 | -1.21E+00 | 1.10E-09 | 4.75E-09 | -1.04E+00 | 6.15E-09 | 2.61E-08 | -1.36E+00 | 8.23E-11 | 3.15E-10 | -1.21E+00 | 2.84E-02 | 1.01E-01 | 1.15E+00 | 3.48E-01 | 9.80E-01 | 1.39E+00 | 1.60E-01 | 5.89E-01 | 1.13E+00 | 2.37E-01 | 6.50E-01 | 1.22E+00 |
| CRIP2 | 2.73E-10 | 1.42E-09 | 4.00E+00 | 1.35E-03 | 5.82E-03 | 2.65E+00 | 1.66E-04 | 7.04E-04 | 3.04E+00 | 1.90E-02 | 7.26E-02 | 2.61E+00 | 2.00E-01 | 7.12E-01 | 1.65E+00 | 3.29E-01 | 9.29E-01 | 1.18E+00 | 1.82E-01 | 6.67E-01 | 2.05E+00 | 3.06E-01 | 8.40E-01 | -1.16E+00 |
| PTPRE | 2.73E-10 | 1.42E-09 | -1.49E+00 | 4.23E-24 | 1.83E-23 | -2.54E+00 | 1.49E-03 | 6.31E-03 | -1.11E+00 | 6.04E-19 | 2.31E-18 | -2.27E+00 | 3.86E-08 | 1.37E-07 | -1.67E+00 | 1.05E-06 | 2.95E-06 | -1.57E+00 | 3.85E-03 | 1.41E-02 | -1.48E+00 | 8.67E-03 | 2.38E-02 | -1.49E+00 |
| HOXA10 | 2.73E-10 | 1.42E-09 | -1.56E+00 | 1.79E-02 | 7.72E-02 | 1.20E+00 | 1.72E-05 | 7.32E-05 | -1.37E+00 | 2.65E-02 | 1.01E-01 | 1.17E+00 | 6.17E-02 | 2.20E-01 | -1.02E+00 | 2.03E-01 | 5.73E-01 | 1.49E+00 | 1.60E-04 | 5.88E-04 | -1.13E+00 | 3.53E-02 | 9.70E-02 | 1.07E+00 |
| PLRG1 | 2.75E-10 | 1.43E-09 | -1.32E+00 | 1.81E-04 | 7.81E-04 | 1.10E+00 | 8.69E-04 | 3.69E-03 | -1.23E+00 | 2.33E-02 | 8.91E-02 | 1.06E+00 | 8.93E-07 | 3.19E-06 | -1.20E+00 | 1.33E-01 | 3.76E-01 | 1.30E+00 | 2.25E-04 | 8.25E-04 | -1.14E+00 | 3.39E-01 | 9.32E-01 | 1.20E+00 |
| CD302 | 2.82E-10 | 1.47E-09 | -1.43E+00 | 6.05E-10 | 2.62E-09 | -1.23E+00 | 4.34E-06 | 1.84E-05 | -1.31E+00 | 3.67E-09 | 1.40E-08 | -1.36E+00 | 6.75E-03 | 2.41E-02 | -1.09E+00 | 2.20E-04 | 6.22E-04 | -1.29E+00 | 3.35E-04 | 1.23E-03 | -1.32E+00 | 6.01E-02 | 1.65E-01 | 1.04E+00 |
| LDAH | 2.83E-10 | 1.48E-09 | -1.73E+00 | 9.25E-04 | 4.00E-03 | 1.03E+00 | 3.09E-06 | 1.31E-05 | -1.59E+00 | 4.15E-02 | 1.59E-01 | 1.10E+00 | 5.17E-06 | 1.84E-05 | -1.68E+00 | 7.43E-03 | 2.10E-02 | -1.00E+00 | 7.79E-07 | 2.86E-06 | -1.30E+00 | 1.26E-03 | 3.45E-03 | -1.02E+00 |
| MYCBP | 2.84E-10 | 1.48E-09 | -2.09E+00 | 3.86E-06 | 1.67E-05 | -1.28E+00 | 1.74E-06 | 7.41E-06 | -1.34E+00 | 2.58E-05 | 9.87E-05 | -1.10E+00 | 3.12E-06 | 1.11E-05 | -1.91E+00 | 2.21E-03 | 6.24E-03 | -1.18E+00 | 2.44E-13 | 8.95E-13 | -1.79E+00 | 9.72E-06 | 2.67E-05 | -1.20E+00 |
| CNOT6L | 2.86E-10 | 1.49E-09 | -1.24E+00 | 5.04E-07 | 2.18E-06 | 1.05E+00 | 6.68E-07 | 2.84E-06 | -1.37E+00 | 8.82E-05 | 3.37E-04 | -1.09E+00 | 2.69E-08 | 9.61E-08 | -1.39E+00 | 1.16E-04 | 3.28E-04 | -1.14E+00 | 2.32E-09 | 8.52E-09 | -1.52E+00 | 5.01E-05 | 1.38E-04 | -1.23E+00 |
| SBDS | 2.86E-10 | 1.49E-09 | -1.14E+00 | 9.17E-48 | 3.96E-47 | -2.21E+00 | 2.99E-06 | 1.27E-05 | -1.31E+00 | 2.89E-27 | 1.10E-26 | -2.35E+00 | 3.47E-02 | 1.24E-01 | 1.10E+00 | 2.32E-17 | 6.53E-17 | -2.00E+00 | 5.97E-08 | 2.19E-07 | -1.15E+00 | 4.59E-23 | 1.26E-22 | -1.91E+00 |
| DNAJC10 | 2.87E-10 | 1.50E-09 | -1.43E+00 | 2.03E-03 | 8.77E-03 | 1.17E+00 | 3.14E-04 | 1.34E-03 | -1.29E+00 | 3.24E-02 | 1.24E-01 | 1.11E+00 | 2.34E-05 | 8.35E-05 | -1.30E+00 | 1.27E-01 | 3.59E-01 | 1.20E+00 | 1.34E-04 | 4.90E-04 | -1.12E+00 | 4.12E-02 | 1.13E-01 | 1.07E+00 |
| METTL23 | 2.87E-10 | 1.50E-09 | -1.24E+00 | 8.90E-06 | 3.85E-05 | 1.04E+00 | 8.89E-06 | 3.78E-05 | -1.25E+00 | 1.47E-03 | 5.62E-03 | 1.04E+00 | 1.75E-09 | 6.26E-09 | -1.30E+00 | 1.97E-03 | 5.56E-03 | 1.12E+00 | 4.82E-06 | 1.77E-05 | -1.13E+00 | 7.82E-03 | 2.15E-02 | 1.03E+00 |
| NEK7 | 2.87E-10 | 1.50E-09 | -1.48E+00 | 4.70E-07 | 2.03E-06 | -1.11E+00 | 1.09E-06 | 4.62E-06 | -1.38E+00 | 5.81E-06 | 2.22E-05 | -1.18E+00 | 2.14E-04 | 7.61E-04 | -1.49E+00 | 2.12E-03 | 5.97E-03 | -1.27E+00 | 7.22E-06 | 2.65E-05 | -1.22E+00 | 1.87E-04 | 5.14E-04 | -1.10E+00 |
| MON2 | 2.90E-10 | 1.51E-09 | -1.52E+00 | 5.53E-05 | 2.39E-04 | 1.01E+00 | 6.45E-11 | 2.74E-10 | -1.62E+00 | 1.43E-06 | 5.45E-06 | -1.10E+00 | 8.87E-05 | 3.16E-04 | -1.37E+00 | 2.86E-02 | 8.07E-02 | 1.02E+00 | 1.79E-08 | 6.56E-08 | -1.49E+00 | 8.79E-06 | 2.42E-05 | -1.28E+00 |
| RNF138 | 2.91E-10 | 1.52E-09 | -1.30E+00 | 3.19E-13 | 1.38E-12 | -1.32E+00 | 3.42E-04 | 1.45E-03 | -1.09E+00 | 4.56E-07 | 1.74E-06 | -1.12E+00 | 1.06E-05 | 3.77E-05 | -1.14E+00 | 1.20E-05 | 3.38E-05 | -1.17E+00 | 1.57E-05 | 5.77E-05 | -1.20E+00 | 1.03E-02 | 2.82E-02 | -1.02E+00 |
| STK4 | 2.93E-10 | 1.53E-09 | -1.16E+00 | 1.16E-07 | 5.01E-07 | 1.01E+00 | 1.52E-04 | 6.46E-04 | -1.10E+00 | 2.39E-02 | 9.11E-02 | 1.10E+00 | 2.64E-01 | 9.42E-01 | 1.36E+00 | 9.18E-02 | 2.59E-01 | 1.11E+00 | 4.07E-02 | 1.49E-01 | 1.12E+00 | 5.88E-04 | 1.61E-03 | -1.14E+00 |
| STOM | 2.95E-10 | 1.54E-09 | -1.28E+00 | 8.09E-05 | 3.49E-04 | 1.13E+00 | 1.22E-04 | 5.20E-04 | -1.21E+00 | 3.35E-02 | 1.28E-01 | 1.15E+00 | 6.38E-02 | 2.28E-01 | -1.08E+00 | 1.86E-01 | 5.24E-01 | 1.31E+00 | 4.25E-03 | 1.56E-02 | -1.07E+00 | 7.89E-03 | 2.17E-02 | -1.26E+00 |
| GRASP | 2.95E-10 | 1.54E-09 | 4.50E+00 | 2.07E-01 | 8.95E-01 | 1.34E+00 | 2.26E-04 | 9.59E-04 | 3.13E+00 | 1.95E-01 | 7.43E-01 | 1.37E+00 | 6.53E-03 | 2.33E-02 | 3.60E+00 | 3.15E-01 | 8.89E-01 | 1.22E+00 | 7.67E-02 | 2.82E-01 | 1.83E+00 | 2.99E-01 | 8.22E-01 | 1.01E+00 |
| WIPF1 | 2.96E-10 | 1.54E-09 | -1.23E+00 | 3.29E-06 | 1.42E-05 | 1.06E+00 | 6.80E-03 | 2.89E-02 | -1.15E+00 | 6.81E-02 | 2.60E-01 | 1.06E+00 | 2.13E-06 | 7.60E-06 | -1.15E+00 | 4.20E-07 | 1.18E-06 | -1.19E+00 | 3.19E-02 | 1.17E-01 | 1.03E+00 | 1.20E-02 | 3.28E-02 | -1.15E+00 |
| CYP4V2 | 2.98E-10 | 1.55E-09 | -1.67E+00 | 1.64E-07 | 7.09E-07 | -1.17E+00 | 3.64E-09 | 1.55E-08 | -1.82E+00 | 8.31E-06 | 3.17E-05 | -1.20E+00 | 2.67E-05 | 9.52E-05 | -1.67E+00 | 2.43E-03 | 6.85E-03 | -1.16E+00 | 3.71E-05 | 1.36E-04 | -1.45E+00 | 1.55E-03 | 4.26E-03 | -1.26E+00 |
| OSBPL9 | 3.03E-10 | 1.58E-09 | -1.38E+00 | 5.83E-08 | 2.52E-07 | -1.10E+00 | 4.70E-03 | 1.99E-02 | -1.02E+00 | 1.09E-04 | 4.17E-04 | -1.02E+00 | 1.12E-06 | 4.00E-06 | -1.18E+00 | 1.85E-05 | 5.21E-05 | -1.07E+00 | 3.70E-09 | 1.36E-08 | -1.31E+00 | 5.09E-07 | 1.40E-06 | -1.26E+00 |
| MAGEH1 | 3.03E-10 | 1.58E-09 | -1.75E+00 | 6.64E-04 | 2.87E-03 | 1.05E+00 | 7.23E-06 | 3.07E-05 | -1.58E+00 | 8.62E-03 | 3.29E-02 | 1.02E+00 | 1.07E-08 | 3.81E-08 | -1.83E+00 | 4.21E-03 | 1.19E-02 | 1.00E+00 | 1.35E-05 | 4.96E-05 | -1.28E+00 | 2.44E-01 | 6.71E-01 | 1.25E+00 |
| GNL3 | 3.05E-10 | 1.59E-09 | -1.31E+00 | 3.31E-02 | 1.43E-01 | 1.31E+00 | 2.92E-07 | 1.24E-06 | -1.36E+00 | 4.63E-03 | 1.77E-02 | 1.02E+00 | 2.17E-06 | 7.75E-06 | -1.27E+00 | 9.37E-02 | 2.64E-01 | 1.16E+00 | 2.12E-06 | 7.76E-06 | -1.30E+00 | 3.19E-01 | 8.77E-01 | 1.16E+00 |
| SPPL2B | 3.07E-10 | 1.60E-09 | -1.19E+00 | 1.76E-07 | 7.62E-07 | 1.00E+00 | 1.80E-04 | 7.67E-04 | -1.10E+00 | 9.77E-03 | 3.73E-02 | 1.11E+00 | 2.50E-03 | 8.93E-03 | -1.00E+00 | 1.28E-02 | 3.60E-02 | 1.09E+00 | 2.19E-02 | 8.04E-02 | 1.07E+00 | 2.68E-02 | 7.35E-02 | 1.07E+00 |
| PARD6B | 3.07E-10 | 1.60E-09 | -1.74E+00 | 2.10E-13 | 9.08E-13 | -1.89E+00 | 3.33E-10 | 1.42E-09 | -2.09E+00 | 2.21E-07 | 8.45E-07 | -1.47E+00 | 6.57E-06 | 2.34E-05 | -2.19E+00 | 8.41E-07 | 2.37E-06 | -2.57E+00 | 3.16E-10 | 1.16E-09 | -2.73E+00 | 2.15E-08 | 5.91E-08 | -2.29E+00 |
| FBXW9 | 3.09E-10 | 1.61E-09 | -1.32E+00 | 2.08E-14 | 9.00E-14 | -1.44E+00 | 4.10E-04 | 1.74E-03 | -1.18E+00 | 2.72E-09 | 1.04E-08 | -1.50E+00 | 6.37E-05 | 2.27E-04 | -1.38E+00 | 3.22E-05 | 9.08E-05 | -1.41E+00 | 7.25E-02 | 2.66E-01 | 1.09E+00 | 2.83E-04 | 7.76E-04 | -1.32E+00 |
| ATP6V0D1 | 3.10E-10 | 1.62E-09 | -1.56E+00 | 2.78E-08 | 1.20E-07 | -1.22E+00 | 1.28E-07 | 5.44E-07 | -1.40E+00 | 2.52E-07 | 9.64E-07 | -1.17E+00 | 5.64E-14 | 2.01E-13 | -1.62E+00 | 2.03E-10 | 5.71E-10 | -1.28E+00 | 3.14E-06 | 1.15E-05 | -1.11E+00 | 3.19E-05 | 8.77E-05 | -1.11E+00 |
| PRELID3B | 3.12E-10 | 1.63E-09 | -1.27E+00 | 6.58E-12 | 2.84E-11 | -1.14E+00 | 3.70E-09 | 1.57E-08 | -1.32E+00 | 1.54E-08 | 5.88E-08 | -1.25E+00 | 4.43E-05 | 1.58E-04 | -1.06E+00 | 2.09E-05 | 5.89E-05 | -1.03E+00 | 2.84E-04 | 1.04E-03 | -1.10E+00 | 2.44E-03 | 6.71E-03 | -1.10E+00 |
| IFI27L2 | 3.13E-10 | 1.63E-09 | -1.22E+00 | 1.13E-24 | 4.87E-24 | -1.65E+00 | 2.22E-02 | 9.42E-02 | 1.05E+00 | 4.66E-15 | 1.78E-14 | -1.56E+00 | 1.93E-03 | 6.87E-03 | -1.08E+00 | 1.68E-08 | 4.74E-08 | -1.73E+00 | 5.73E-03 | 2.10E-02 | 1.42E+00 | 3.71E-03 | 1.02E-02 | -1.24E+00 |
| RRP15 | 3.14E-10 | 1.63E-09 | -1.52E+00 | 1.43E-07 | 6.16E-07 | -1.15E+00 | 4.90E-05 | 2.08E-04 | -1.28E+00 | 4.51E-06 | 1.72E-05 | -1.19E+00 | 2.49E-05 | 8.89E-05 | -1.33E+00 | 1.24E-04 | 3.50E-04 | -1.15E+00 | 1.86E-08 | 6.83E-08 | -1.41E+00 | 1.16E-07 | 3.18E-07 | -1.23E+00 |
| GPATCH8 | 3.16E-10 | 1.65E-09 | -1.26E+00 | 1.04E-07 | 4.51E-07 | -1.07E+00 | 4.79E-05 | 2.04E-04 | -1.20E+00 | 5.08E-04 | 1.94E-03 | -1.02E+00 | 4.16E-07 | 1.48E-06 | -1.23E+00 | 1.40E-04 | 3.95E-04 | -1.09E+00 | 1.94E-06 | 7.11E-06 | -1.15E+00 | 7.38E-05 | 2.03E-04 | -1.13E+00 |
| WDR82 | 3.17E-10 | 1.65E-09 | -1.31E+00 | 3.69E-12 | 1.59E-11 | -1.23E+00 | 3.61E-06 | 1.53E-05 | -1.20E+00 | 4.05E-06 | 1.55E-05 | -1.13E+00 | 8.44E-05 | 3.01E-04 | -1.05E+00 | 3.15E-04 | 8.88E-04 | 1.04E+00 | 1.21E-07 | 4.44E-07 | -1.27E+00 | 4.72E-04 | 1.30E-03 | -1.13E+00 |
| MIAT | 3.20E-10 | 1.67E-09 | -1.68E+00 | 1.34E-06 | 5.79E-06 | -1.22E+00 | 5.45E-04 | 2.32E-03 | -2.00E+00 | 3.10E-02 | 1.18E-01 | -1.18E+00 | 3.18E-02 | 1.13E-01 | -1.79E+00 | 1.68E-01 | 4.73E-01 | -1.01E+00 | 1.25E-01 | 4.57E-01 | -1.28E+00 | 2.55E-01 | 7.00E-01 | -1.07E+00 |
| CHCHD3 | 3.20E-10 | 1.67E-09 | -1.11E+00 | 1.47E-11 | 6.35E-11 | -1.02E+00 | 5.80E-02 | 2.46E-01 | 1.11E+00 | 1.61E-02 | 6.16E-02 | 1.11E+00 | 5.57E-06 | 1.99E-05 | -1.10E+00 | 6.71E-04 | 1.89E-03 | 1.03E+00 | 6.54E-03 | 2.40E-02 | 1.08E+00 | 1.76E-01 | 4.84E-01 | 1.18E+00 |
| BCL7A | 3.22E-10 | 1.68E-09 | -1.37E+00 | 2.30E-15 | 9.96E-15 | -1.40E+00 | 1.40E-04 | 5.94E-04 | -1.17E+00 | 2.68E-05 | 1.02E-04 | -1.10E+00 | 3.00E-07 | 1.07E-06 | -1.53E+00 | 6.08E-05 | 1.71E-04 | -1.21E+00 | 1.18E-08 | 4.34E-08 | -1.51E+00 | 1.21E-04 | 3.32E-04 | -1.17E+00 |
| MTHFD1 | 3.26E-10 | 1.70E-09 | -1.43E+00 | 3.91E-07 | 1.69E-06 | -1.11E+00 | 7.60E-06 | 3.23E-05 | -1.20E+00 | 5.81E-04 | 2.22E-03 | 1.06E+00 | 7.65E-06 | 2.73E-05 | -1.26E+00 | 1.96E-01 | 5.51E-01 | 1.44E+00 | 3.19E-05 | 1.17E-04 | -1.12E+00 | 3.31E-02 | 9.10E-02 | 1.45E+00 |
| OSBPL1A | 3.30E-10 | 1.72E-09 | -1.39E+00 | 5.10E-07 | 2.20E-06 | -1.08E+00 | 7.01E-06 | 2.98E-05 | -1.31E+00 | 1.13E-03 | 4.30E-03 | 1.01E+00 | 1.24E-09 | 4.41E-09 | -1.76E+00 | 1.08E-06 | 3.03E-06 | -1.27E+00 | 1.13E-07 | 4.13E-07 | -1.52E+00 | 9.73E-03 | 2.67E-02 | 1.03E+00 |
| ETFA | 3.30E-10 | 1.72E-09 | -1.16E+00 | 7.37E-09 | 3.18E-08 | 1.01E+00 | 4.05E-03 | 1.72E-02 | -1.01E+00 | 2.18E-03 | 8.31E-03 | 1.07E+00 | 3.12E-05 | 1.11E-04 | -1.20E+00 | 1.23E-03 | 3.47E-03 | -1.05E+00 | 1.09E-01 | 3.99E-01 | 1.20E+00 | 2.01E-01 | 5.51E-01 | 1.14E+00 |
| C2orf88 | 3.31E-10 | 1.72E-09 | -1.70E+00 | 2.70E-03 | 1.17E-02 | 1.11E+00 | 1.39E-04 | 5.92E-04 | -1.45E+00 | 1.31E-01 | 5.02E-01 | 1.41E+00 | 1.85E-01 | 6.60E-01 | 1.28E+00 | 8.45E-04 | 2.38E-03 | 1.73E+00 | 1.01E-01 | 3.69E-01 | 1.46E+00 | 1.67E-02 | 4.60E-02 | 1.26E+00 |
| MRPL10 | 3.31E-10 | 1.73E-09 | -1.32E+00 | 2.87E-07 | 1.24E-06 | -1.06E+00 | 6.05E-06 | 2.57E-05 | -1.27E+00 | 1.10E-04 | 4.21E-04 | -1.05E+00 | 1.23E-12 | 4.38E-12 | -1.57E+00 | 5.35E-05 | 1.51E-04 | -1.07E+00 | 6.79E-06 | 2.49E-05 | -1.23E+00 | 1.76E-05 | 4.84E-05 | -1.15E+00 |
| TMEM223 | 3.35E-10 | 1.75E-09 | -1.36E+00 | 1.91E-03 | 8.27E-03 | 1.18E+00 | 1.36E-07 | 5.79E-07 | -1.37E+00 | 3.46E-02 | 1.32E-01 | 1.18E+00 | 4.11E-08 | 1.47E-07 | -1.71E+00 | 2.14E-02 | 6.03E-02 | 1.04E+00 | 7.05E-08 | 2.59E-07 | -1.43E+00 | 1.53E-01 | 4.20E-01 | 1.11E+00 |
| DYRK4 | 3.41E-10 | 1.78E-09 | -1.43E+00 | 4.36E-06 | 1.89E-05 | -1.05E+00 | 4.10E-07 | 1.74E-06 | -1.38E+00 | 4.24E-07 | 1.62E-06 | -1.22E+00 | 1.11E-08 | 3.96E-08 | -1.60E+00 | 4.96E-03 | 1.40E-02 | 1.06E+00 | 6.58E-07 | 2.42E-06 | -1.29E+00 | 2.84E-02 | 7.79E-02 | 1.03E+00 |
| NOL4L | 3.43E-10 | 1.79E-09 | -1.17E+00 | 3.75E-07 | 1.62E-06 | 1.03E+00 | 1.68E-05 | 7.13E-05 | -1.26E+00 | 1.67E-06 | 6.39E-06 | -1.15E+00 | 3.01E-11 | 1.07E-10 | -1.40E+00 | 2.22E-07 | 6.26E-07 | -1.23E+00 | 1.68E-16 | 6.17E-16 | -1.48E+00 | 2.49E-12 | 6.84E-12 | -1.46E+00 |
| E2F4 | 3.45E-10 | 1.80E-09 | -1.60E+00 | 2.06E-05 | 8.92E-05 | -1.02E+00 | 7.18E-13 | 3.05E-12 | -1.67E+00 | 8.46E-06 | 3.23E-05 | -1.05E+00 | 9.96E-09 | 3.55E-08 | -1.84E+00 | 4.27E-03 | 1.20E-02 | 1.02E+00 | 6.96E-08 | 2.55E-07 | -1.43E+00 | 1.15E-02 | 3.17E-02 | -1.06E+00 |
| TRAPPC11 | 3.45E-10 | 1.80E-09 | -1.40E+00 | 2.37E-04 | 1.02E-03 | 1.10E+00 | 3.60E-04 | 1.53E-03 | -1.22E+00 | 9.50E-02 | 3.63E-01 | 1.27E+00 | 6.68E-03 | 2.38E-02 | 1.05E+00 | 1.07E-01 | 3.01E-01 | 1.35E+00 | 4.92E-05 | 1.81E-04 | -1.20E+00 | 4.34E-02 | 1.19E-01 | 1.10E+00 |
| SCN9A | 3.45E-10 | 1.80E-09 | -1.48E+00 | 8.95E-03 | 3.87E-02 | 1.14E+00 | 1.30E-16 | 5.51E-16 | -2.20E+00 | 1.94E-07 | 7.41E-07 | -1.29E+00 | 1.27E-01 | 4.54E-01 | 1.18E+00 | 3.14E-01 | 8.84E-01 | 1.44E+00 | 2.79E-04 | 1.03E-03 | -1.33E+00 | 3.59E-04 | 9.86E-04 | -1.41E+00 |
| ISG20L2 | 3.51E-10 | 1.83E-09 | -1.49E+00 | 2.26E-08 | 9.77E-08 | -1.26E+00 | 4.49E-06 | 1.91E-05 | -1.27E+00 | 6.07E-06 | 2.32E-05 | -1.14E+00 | 3.84E-08 | 1.37E-07 | -1.45E+00 | 2.25E-05 | 6.34E-05 | -1.17E+00 | 2.23E-10 | 8.20E-10 | -1.68E+00 | 5.03E-03 | 1.38E-02 | -1.10E+00 |
| CEP104 | 3.52E-10 | 1.83E-09 | -1.71E+00 | 6.87E-04 | 2.97E-03 | 1.02E+00 | 4.31E-07 | 1.83E-06 | -1.44E+00 | 2.28E-04 | 8.70E-04 | -1.07E+00 | 1.50E-06 | 5.34E-06 | -1.94E+00 | 6.52E-04 | 1.84E-03 | -1.18E+00 | 1.80E-09 | 6.59E-09 | -1.43E+00 | 3.91E-07 | 1.07E-06 | -1.38E+00 |
| KRR1 | 3.55E-10 | 1.85E-09 | -1.23E+00 | 2.57E-16 | 1.11E-15 | -1.29E+00 | 2.49E-03 | 1.06E-02 | -1.17E+00 | 3.14E-06 | 1.20E-05 | -1.18E+00 | 1.18E-04 | 4.22E-04 | -1.05E+00 | 4.46E-06 | 1.26E-05 | -1.15E+00 | 4.33E-06 | 1.59E-05 | -1.11E+00 | 6.45E-03 | 1.77E-02 | 1.05E+00 |
| GTF2H5 | 3.56E-10 | 1.86E-09 | -1.31E+00 | 1.32E-08 | 5.70E-08 | -1.07E+00 | 4.08E-04 | 1.73E-03 | -1.10E+00 | 4.80E-06 | 1.83E-05 | -1.09E+00 | 3.15E-09 | 1.12E-08 | -1.51E+00 | 1.48E-04 | 4.17E-04 | -1.08E+00 | 1.22E-06 | 4.47E-06 | -1.26E+00 | 4.24E-05 | 1.17E-04 | -1.18E+00 |
| CGGBP1 | 3.59E-10 | 1.87E-09 | -1.13E+00 | 4.43E-06 | 1.91E-05 | 1.12E+00 | 2.08E-08 | 8.82E-08 | -1.28E+00 | 2.44E-03 | 9.34E-03 | 1.04E+00 | 2.53E-10 | 9.02E-10 | -1.33E+00 | 5.29E-05 | 1.49E-04 | -1.06E+00 | 1.85E-05 | 6.77E-05 | -1.26E+00 | 4.95E-03 | 1.36E-02 | -1.11E+00 |
| LETMD1 | 3.65E-10 | 1.90E-09 | -1.26E+00 | 1.40E-08 | 6.07E-08 | -1.10E+00 | 1.11E-05 | 4.72E-05 | -1.35E+00 | 1.18E-05 | 4.50E-05 | -1.15E+00 | 2.55E-03 | 9.09E-03 | 1.03E+00 | 1.60E-02 | 4.50E-02 | 1.13E+00 | 1.04E-04 | 3.81E-04 | -1.09E+00 | 6.49E-05 | 1.78E-04 | -1.20E+00 |
| POLR2I | 3.66E-10 | 1.91E-09 | -1.22E+00 | 1.38E-19 | 5.98E-19 | -1.39E+00 | 1.73E-02 | 7.34E-02 | 1.09E+00 | 2.60E-08 | 9.94E-08 | -1.12E+00 | 1.17E-08 | 4.18E-08 | -1.25E+00 | 3.77E-11 | 1.06E-10 | -1.36E+00 | 1.57E-03 | 5.78E-03 | -1.01E+00 | 2.34E-06 | 6.42E-06 | -1.24E+00 |
| NSUN4 | 3.66E-10 | 1.91E-09 | -1.34E+00 | 3.10E-11 | 1.34E-10 | -1.24E+00 | 9.30E-05 | 3.95E-04 | -1.24E+00 | 3.39E-05 | 1.29E-04 | -1.11E+00 | 2.82E-05 | 1.01E-04 | -1.31E+00 | 1.34E-05 | 3.78E-05 | -1.36E+00 | 2.01E-06 | 7.39E-06 | -1.39E+00 | 4.71E-03 | 1.29E-02 | -1.13E+00 |
| ANKIB1 | 3.66E-10 | 1.91E-09 | -1.54E+00 | 1.22E-04 | 5.27E-04 | 1.01E+00 | 8.35E-15 | 3.55E-14 | -1.94E+00 | 2.07E-11 | 7.92E-11 | -1.43E+00 | 2.90E-07 | 1.03E-06 | -1.54E+00 | 2.33E-06 | 6.56E-06 | -1.35E+00 | 1.24E-10 | 4.57E-10 | -1.72E+00 | 5.47E-05 | 1.50E-04 | -1.22E+00 |
| PNMA6A | 3.67E-10 | 1.91E-09 | -1.89E+00 | 4.22E-11 | 1.83E-10 | -1.85E+00 | 2.73E-03 | 1.16E-02 | -1.12E+00 | 1.61E-04 | 6.15E-04 | -1.11E+00 | 1.98E-01 | 7.06E-01 | 1.12E+00 | 1.46E-01 | 4.11E-01 | 1.04E+00 | 3.97E-02 | 1.46E-01 | -1.22E+00 | 3.35E-01 | 9.21E-01 | 1.29E+00 |
| PRPF4 | 3.67E-10 | 1.91E-09 | -1.54E+00 | 4.54E-06 | 1.96E-05 | -1.08E+00 | 1.81E-11 | 7.70E-11 | -1.58E+00 | 4.18E-06 | 1.60E-05 | -1.14E+00 | 1.97E-05 | 7.04E-05 | -1.49E+00 | 4.54E-02 | 1.28E-01 | 1.10E+00 | 1.69E-13 | 6.21E-13 | -1.61E+00 | 3.23E-02 | 8.88E-02 | 1.12E+00 |
| DR1 | 3.67E-10 | 1.91E-09 | -1.20E+00 | 7.94E-08 | 3.43E-07 | -1.03E+00 | 2.02E-04 | 8.56E-04 | -1.10E+00 | 1.05E-03 | 4.01E-03 | 1.07E+00 | 4.94E-05 | 1.76E-04 | -1.04E+00 | 4.34E-04 | 1.22E-03 | 1.04E+00 | 2.29E-03 | 8.41E-03 | -1.06E+00 | 4.90E-03 | 1.35E-02 | -1.05E+00 |
| FAM241A | 3.69E-10 | 1.92E-09 | -1.44E+00 | 5.26E-09 | 2.27E-08 | -1.26E+00 | 1.17E-06 | 4.99E-06 | -1.38E+00 | 2.03E-04 | 7.77E-04 | -1.06E+00 | 1.35E-05 | 4.80E-05 | -1.29E+00 | 6.39E-04 | 1.80E-03 | -1.01E+00 | 3.38E-08 | 1.24E-07 | -1.41E+00 | 5.21E-04 | 1.43E-03 | -1.06E+00 |
| ZNF546 | 3.72E-10 | 1.94E-09 | -1.98E+00 | 2.64E-02 | 1.14E-01 | 1.19E+00 | 1.05E-05 | 4.46E-05 | -2.12E+00 | 2.74E-02 | 1.05E-01 | 1.04E+00 | 3.83E-03 | 1.37E-02 | -1.36E+00 | 2.41E-01 | 6.81E-01 | 1.29E+00 | 4.18E-02 | 1.53E-01 | -1.12E+00 | 3.56E-01 | 9.79E-01 | 1.19E+00 |
| C5orf42 | 3.74E-10 | 1.95E-09 | -1.46E+00 | 1.03E-03 | 4.43E-03 | 1.08E+00 | 2.51E-04 | 1.06E-03 | -1.34E+00 | 1.16E-01 | 4.44E-01 | 1.15E+00 | 3.06E-03 | 1.09E-02 | -1.21E+00 | 2.04E-01 | 5.75E-01 | 1.24E+00 | 3.16E-04 | 1.16E-03 | -1.21E+00 | 1.00E-01 | 2.75E-01 | 1.18E+00 |
| CISD2 | 3.76E-10 | 1.96E-09 | -1.43E+00 | 6.00E-06 | 2.59E-05 | -1.02E+00 | 6.07E-08 | 2.58E-07 | -1.35E+00 | 1.84E-08 | 7.03E-08 | -1.24E+00 | 2.48E-10 | 8.84E-10 | -1.62E+00 | 6.39E-07 | 1.80E-06 | -1.24E+00 | 1.20E-06 | 4.39E-06 | -1.20E+00 | 3.56E-05 | 9.79E-05 | -1.11E+00 |
| RILP | 3.78E-10 | 1.97E-09 | -1.53E+00 | 4.21E-05 | 1.82E-04 | 1.02E+00 | 2.02E-03 | 8.56E-03 | -1.22E+00 | 8.97E-03 | 3.42E-02 | 1.02E+00 | 2.19E-08 | 7.82E-08 | -1.52E+00 | 2.87E-03 | 8.08E-03 | -1.04E+00 | 1.58E-14 | 5.81E-14 | -1.31E+00 | 5.47E-10 | 1.50E-09 | -1.35E+00 |
| NHP2 | 3.78E-10 | 1.97E-09 | -1.78E+00 | 3.66E-07 | 1.58E-06 | -1.32E+00 | 1.36E-13 | 5.77E-13 | -2.14E+00 | 7.29E-10 | 2.78E-09 | -1.45E+00 | 2.45E-07 | 8.75E-07 | -1.91E+00 | 7.29E-04 | 2.06E-03 | -1.16E+00 | 8.75E-11 | 3.21E-10 | -1.70E+00 | 9.51E-05 | 2.61E-04 | -1.22E+00 |
| SLC35C2 | 3.80E-10 | 1.98E-09 | -1.38E+00 | 3.14E-05 | 1.36E-04 | 1.05E+00 | 3.62E-07 | 1.54E-06 | -1.31E+00 | 8.86E-05 | 3.38E-04 | 1.03E+00 | 3.74E-12 | 1.33E-11 | -1.67E+00 | 6.91E-06 | 1.95E-05 | -1.08E+00 | 4.40E-05 | 1.62E-04 | -1.14E+00 | 4.54E-02 | 1.25E-01 | 1.10E+00 |
| ZFAND5 | 3.82E-10 | 1.99E-09 | -1.13E+00 | 5.60E-29 | 2.42E-28 | -1.59E+00 | 2.13E-07 | 9.07E-07 | -1.34E+00 | 8.89E-12 | 3.40E-11 | -1.46E+00 | 4.48E-08 | 1.60E-07 | -1.23E+00 | 1.19E-11 | 3.36E-11 | -1.43E+00 | 7.01E-09 | 2.57E-08 | -1.32E+00 | 2.18E-07 | 5.99E-07 | -1.27E+00 |
| SNORD3B-1 | 3.83E-10 | 1.99E-09 | -1.22E+00 | 4.37E-43 | 1.89E-42 | -3.46E+00 | 2.30E-03 | 9.76E-03 | -1.09E+00 | 1.08E-19 | 4.14E-19 | -3.12E+00 | 7.21E-09 | 2.57E-08 | -1.97E+00 | 1.69E-09 | 4.78E-09 | -2.22E+00 | 4.08E-12 | 1.50E-11 | -1.97E+00 | 1.28E-11 | 3.52E-11 | -2.08E+00 |
| MGAT1 | 3.85E-10 | 2.01E-09 | -1.30E+00 | 7.25E-04 | 3.13E-03 | 1.17E+00 | 8.80E-08 | 3.74E-07 | -1.35E+00 | 1.12E-02 | 4.29E-02 | 1.14E+00 | 1.77E-07 | 6.32E-07 | -1.53E+00 | 1.51E-02 | 4.25E-02 | 1.09E+00 | 2.85E-04 | 1.04E-03 | -1.19E+00 | 2.89E-01 | 7.95E-01 | 1.16E+00 |
| NFYC | 3.87E-10 | 2.02E-09 | -1.45E+00 | 1.70E-07 | 7.35E-07 | -1.10E+00 | 1.13E-06 | 4.79E-06 | -1.64E+00 | 5.81E-04 | 2.22E-03 | -1.21E+00 | 6.51E-06 | 2.32E-05 | -1.65E+00 | 6.25E-03 | 1.76E-02 | -1.11E+00 | 4.02E-18 | 1.48E-17 | -1.88E+00 | 8.17E-11 | 2.24E-10 | -1.47E+00 |
| DYNLRB1 | 3.88E-10 | 2.02E-09 | -1.10E+00 | 6.37E-06 | 2.75E-05 | 1.09E+00 | 1.57E-02 | 6.68E-02 | 1.01E+00 | 8.54E-03 | 3.26E-02 | 1.09E+00 | 6.99E-06 | 2.49E-05 | -1.11E+00 | 5.38E-02 | 1.52E-01 | 1.06E+00 | 1.90E-01 | 6.96E-01 | 1.14E+00 | 2.01E-01 | 5.52E-01 | 1.09E+00 |
| SLX4IP | 3.91E-10 | 2.04E-09 | -1.82E+00 | 2.30E-06 | 9.95E-06 | -1.22E+00 | 9.30E-12 | 3.95E-11 | -1.84E+00 | 2.45E-09 | 9.35E-09 | -1.45E+00 | 2.04E-05 | 7.27E-05 | -1.32E+00 | 5.91E-03 | 1.67E-02 | -1.01E+00 | 1.14E-07 | 4.17E-07 | -1.31E+00 | 3.36E-05 | 9.22E-05 | -1.19E+00 |
| LGALS8 | 3.93E-10 | 2.05E-09 | -1.47E+00 | 2.01E-08 | 8.69E-08 | -1.22E+00 | 1.68E-06 | 7.15E-06 | -1.43E+00 | 3.07E-05 | 1.17E-04 | -1.18E+00 | 8.06E-10 | 2.87E-09 | -1.76E+00 | 2.38E-06 | 6.72E-06 | -1.27E+00 | 9.91E-17 | 3.64E-16 | -1.60E+00 | 1.56E-13 | 4.27E-13 | -1.58E+00 |
| PYCARD | 3.93E-10 | 2.05E-09 | -1.20E+00 | 6.34E-11 | 2.74E-10 | -1.09E+00 | 2.39E-02 | 1.02E-01 | -1.07E+00 | 1.01E-02 | 3.86E-02 | -1.08E+00 | 1.81E-10 | 6.45E-10 | -1.44E+00 | 5.46E-07 | 1.54E-06 | -1.31E+00 | 1.64E-05 | 6.01E-05 | -1.21E+00 | 4.43E-04 | 1.22E-03 | -1.12E+00 |
| GCDH | 3.94E-10 | 2.06E-09 | -1.50E+00 | 5.12E-06 | 2.21E-05 | -1.07E+00 | 1.63E-03 | 6.93E-03 | -1.10E+00 | 1.28E-03 | 4.90E-03 | 1.04E+00 | 6.98E-04 | 2.49E-03 | -1.23E+00 | 1.56E-02 | 4.40E-02 | -1.01E+00 | 3.75E-04 | 1.38E-03 | -1.06E+00 | 1.22E-02 | 3.34E-02 | 1.08E+00 |
| SMIM20 | 3.94E-10 | 2.06E-09 | -1.20E+00 | 1.45E-10 | 6.25E-10 | -1.09E+00 | 7.72E-04 | 3.28E-03 | -1.10E+00 | 9.42E-06 | 3.60E-05 | -1.10E+00 | 2.35E-08 | 8.38E-08 | -1.32E+00 | 2.52E-04 | 7.11E-04 | 1.02E+00 | 1.19E-02 | 4.36E-02 | 1.02E+00 | 9.64E-03 | 2.65E-02 | 1.01E+00 |
| SCCPDH | 3.95E-10 | 2.06E-09 | -1.14E+00 | 3.79E-14 | 1.64E-13 | -1.12E+00 | 8.66E-02 | 3.68E-01 | 1.03E+00 | 1.34E-02 | 5.11E-02 | 1.03E+00 | 1.99E-05 | 7.10E-05 | -1.12E+00 | 3.30E-02 | 9.29E-02 | 1.05E+00 | 2.00E-01 | 7.34E-01 | 1.19E+00 | 2.43E-01 | 6.66E-01 | 1.33E+00 |
| CDC42EP3 | 3.96E-10 | 2.07E-09 | -1.41E+00 | 4.75E-10 | 2.05E-09 | -1.26E+00 | 5.61E-03 | 2.39E-02 | -1.16E+00 | 2.00E-02 | 7.65E-02 | 1.04E+00 | 2.00E-07 | 7.12E-07 | -1.27E+00 | 8.18E-03 | 2.31E-02 | 1.06E+00 | 5.73E-05 | 2.10E-04 | 1.02E+00 | 7.11E-04 | 1.95E-03 | -1.02E+00 |
| CSPG4P12 | 3.97E-10 | 2.07E-09 | -2.48E+00 | 1.71E-08 | 7.37E-08 | -1.92E+00 | 9.87E-07 | 4.19E-06 | -2.21E+00 | 5.25E-06 | 2.00E-05 | -1.74E+00 | 1.06E-05 | 3.77E-05 | -2.04E+00 | 1.15E-06 | 3.23E-06 | -2.46E+00 | 6.11E-11 | 2.24E-10 | -2.20E+00 | 1.97E-06 | 5.41E-06 | -1.35E+00 |
| EMC4 | 3.99E-10 | 2.08E-09 | -1.16E+00 | 1.67E-05 | 7.21E-05 | 1.15E+00 | 2.98E-02 | 1.27E-01 | 1.07E+00 | 2.05E-01 | 7.82E-01 | 1.30E+00 | 1.00E-12 | 3.57E-12 | -1.53E+00 | 4.78E-05 | 1.35E-04 | -1.11E+00 | 7.72E-04 | 2.83E-03 | -1.12E+00 | 2.23E-01 | 6.11E-01 | 1.14E+00 |
| RPL23AP58 | 3.99E-10 | 2.08E-09 | -1.22E+00 | 1.64E-20 | 7.10E-20 | -1.49E+00 | 7.88E-06 | 3.35E-05 | -1.25E+00 | 1.06E-16 | 4.05E-16 | -1.46E+00 | 6.00E-07 | 2.14E-06 | -1.20E+00 | 3.93E-12 | 1.11E-11 | -1.39E+00 | 1.26E-07 | 4.63E-07 | -1.32E+00 | 1.61E-10 | 4.43E-10 | -1.52E+00 |
| C1QBP | 4.05E-10 | 2.11E-09 | -1.15E+00 | 1.13E-07 | 4.87E-07 | -1.01E+00 | 1.89E-01 | 8.03E-01 | 1.03E+00 | 2.04E-01 | 7.78E-01 | 1.06E+00 | 5.89E-05 | 2.10E-04 | -1.12E+00 | 2.62E-01 | 7.39E-01 | 1.13E+00 | 3.24E-02 | 1.19E-01 | 1.03E+00 | 1.59E-01 | 4.36E-01 | 1.20E+00 |
| HIKESHI | 4.09E-10 | 2.13E-09 | -1.12E+00 | 2.62E-09 | 1.13E-08 | 1.02E+00 | 3.73E-03 | 1.58E-02 | -1.02E+00 | 9.88E-03 | 3.77E-02 | 1.10E+00 | 2.15E-07 | 7.66E-07 | -1.31E+00 | 1.53E-03 | 4.32E-03 | -1.06E+00 | 5.66E-03 | 2.08E-02 | -1.07E+00 | 2.92E-02 | 8.02E-02 | -1.00E+00 |
| RCHY1 | 4.14E-10 | 2.16E-09 | -1.44E+00 | 3.94E-05 | 1.70E-04 | 1.04E+00 | 8.05E-05 | 3.42E-04 | -1.21E+00 | 2.48E-02 | 9.47E-02 | 1.16E+00 | 3.39E-05 | 1.21E-04 | -1.39E+00 | 4.32E-02 | 1.22E-01 | 1.11E+00 | 2.76E-04 | 1.01E-03 | 1.00E+00 | 2.60E-03 | 7.14E-03 | -1.11E+00 |
| MRPS26 | 4.17E-10 | 2.17E-09 | -1.33E+00 | 5.44E-05 | 2.35E-04 | 1.03E+00 | 2.50E-05 | 1.06E-04 | -1.08E+00 | 1.41E-03 | 5.40E-03 | 1.10E+00 | 1.14E-17 | 4.06E-17 | -1.71E+00 | 9.35E-04 | 2.64E-03 | 1.07E+00 | 3.85E-03 | 1.41E-02 | -1.09E+00 | 1.11E-01 | 3.05E-01 | 1.31E+00 |
| NOL9 | 4.20E-10 | 2.19E-09 | -1.68E+00 | 2.87E-05 | 1.24E-04 | -1.06E+00 | 1.09E-04 | 4.63E-04 | -1.31E+00 | 5.84E-02 | 2.23E-01 | 1.13E+00 | 4.17E-04 | 1.49E-03 | -1.32E+00 | 7.74E-03 | 2.18E-02 | -1.08E+00 | 3.38E-06 | 1.24E-05 | -1.27E+00 | 8.34E-05 | 2.29E-04 | -1.13E+00 |
| HDLBP | 4.25E-10 | 2.21E-09 | -1.28E+00 | 3.47E-06 | 1.50E-05 | 1.01E+00 | 1.24E-04 | 5.25E-04 | -1.13E+00 | 2.68E-02 | 1.02E-01 | 1.17E+00 | 1.47E-07 | 5.24E-07 | -1.36E+00 | 7.99E-06 | 2.25E-05 | -1.16E+00 | 1.23E-03 | 4.50E-03 | -1.12E+00 | 1.77E-01 | 4.87E-01 | 1.11E+00 |
| NUDT19 | 4.26E-10 | 2.22E-09 | -1.73E+00 | 1.84E-08 | 7.94E-08 | -1.37E+00 | 1.29E-08 | 5.49E-08 | -1.91E+00 | 5.35E-05 | 2.04E-04 | -1.17E+00 | 3.99E-10 | 1.42E-09 | -1.72E+00 | 5.80E-06 | 1.63E-05 | -1.18E+00 | 5.75E-22 | 2.11E-21 | -1.87E+00 | 2.26E-09 | 6.20E-09 | -1.32E+00 |
| EIF3M | 4.28E-10 | 2.23E-09 | -1.20E+00 | 8.74E-03 | 3.77E-02 | -1.00E+00 | 1.28E-03 | 5.43E-03 | -1.19E+00 | 1.58E-01 | 6.02E-01 | -1.00E+00 | 8.75E-05 | 3.12E-04 | -1.20E+00 | 2.04E-01 | 5.75E-01 | 1.02E+00 | 7.16E-08 | 2.63E-07 | -1.23E+00 | 2.39E-01 | 6.57E-01 | 1.04E+00 |
| HOXA9 | 4.28E-10 | 2.23E-09 | -1.29E+00 | 1.83E-01 | 7.92E-01 | 1.44E+00 | 6.43E-05 | 2.73E-04 | -1.25E+00 | 1.00E-01 | 3.84E-01 | 1.39E+00 | 3.42E-06 | 1.22E-05 | -1.51E+00 | 3.66E-03 | 1.03E-02 | -1.07E+00 | 5.96E-07 | 2.19E-06 | -1.57E+00 | 2.31E-02 | 6.33E-02 | 1.00E+00 |
| SCARB2 | 4.28E-10 | 2.23E-09 | -1.31E+00 | 4.33E-05 | 1.87E-04 | 1.06E+00 | 2.41E-07 | 1.02E-06 | -1.44E+00 | 4.62E-07 | 1.76E-06 | -1.26E+00 | 2.39E-06 | 8.51E-06 | -1.69E+00 | 9.37E-06 | 2.64E-05 | -1.48E+00 | 4.02E-06 | 1.48E-05 | -1.49E+00 | 4.47E-07 | 1.23E-06 | -1.69E+00 |
| SUCLA2 | 4.31E-10 | 2.24E-09 | -1.53E+00 | 3.72E-03 | 1.61E-02 | 1.18E+00 | 1.38E-07 | 5.84E-07 | -1.44E+00 | 1.91E-05 | 7.30E-05 | -1.12E+00 | 2.06E-06 | 7.34E-06 | -1.35E+00 | 3.55E-02 | 1.00E-01 | 1.16E+00 | 1.72E-07 | 6.31E-07 | -1.36E+00 | 7.42E-05 | 2.04E-04 | -1.21E+00 |
| ACSL5 | 4.31E-10 | 2.25E-09 | -1.70E+00 | 1.72E-05 | 7.41E-05 | -1.10E+00 | 3.51E-09 | 1.49E-08 | -1.58E+00 | 4.07E-05 | 1.55E-04 | -1.13E+00 | 2.68E-05 | 9.55E-05 | -1.45E+00 | 7.38E-02 | 2.08E-01 | 1.22E+00 | 5.38E-09 | 1.97E-08 | -1.43E+00 | 3.03E-04 | 8.33E-04 | -1.16E+00 |
| GCLM | 4.34E-10 | 2.26E-09 | -1.56E+00 | 1.53E-09 | 6.62E-09 | -1.40E+00 | 6.52E-12 | 2.77E-11 | -1.73E+00 | 6.83E-10 | 2.61E-09 | -1.32E+00 | 1.08E-08 | 3.86E-08 | -1.89E+00 | 8.26E-07 | 2.33E-06 | -1.46E+00 | 2.28E-14 | 8.35E-14 | -1.63E+00 | 8.16E-07 | 2.24E-06 | -1.21E+00 |
| PIP5K1B | 4.34E-10 | 2.26E-09 | -1.86E+00 | 1.12E-09 | 4.85E-09 | -1.74E+00 | 1.06E-05 | 4.49E-05 | -1.71E+00 | 6.74E-05 | 2.57E-04 | -1.34E+00 | 1.95E-07 | 6.95E-07 | -1.39E+00 | 5.14E-06 | 1.45E-05 | -1.25E+00 | 5.18E-19 | 1.90E-18 | -1.51E+00 | 2.49E-15 | 6.83E-15 | -1.51E+00 |
| GTF2IRD2 | 4.40E-10 | 2.29E-09 | -1.61E+00 | 8.94E-04 | 3.86E-03 | 1.09E+00 | 2.34E-03 | 9.95E-03 | -1.20E+00 | 3.42E-02 | 1.31E-01 | 1.09E+00 | 8.21E-04 | 2.93E-03 | -1.10E+00 | 1.80E-01 | 5.08E-01 | 1.34E+00 | 3.61E-03 | 1.32E-02 | -1.07E+00 | 2.50E-03 | 6.88E-03 | 1.00E+00 |
| PCYOX1 | 4.40E-10 | 2.30E-09 | -1.76E+00 | 2.63E-04 | 1.14E-03 | -1.01E+00 | 3.42E-05 | 1.45E-04 | -1.45E+00 | 7.20E-02 | 2.75E-01 | 1.20E+00 | 1.40E-07 | 4.97E-07 | -1.89E+00 | 3.37E-05 | 9.49E-05 | -1.37E+00 | 8.91E-05 | 3.27E-04 | -1.33E+00 | 2.57E-02 | 7.06E-02 | -1.03E+00 |
| NUMB | 4.43E-10 | 2.31E-09 | -1.42E+00 | 2.14E-04 | 9.25E-04 | 1.09E+00 | 1.60E-05 | 6.79E-05 | -1.29E+00 | 4.17E-03 | 1.59E-02 | 1.08E+00 | 1.93E-08 | 6.89E-08 | -1.45E+00 | 7.19E-05 | 2.03E-04 | -1.12E+00 | 9.14E-06 | 3.36E-05 | -1.23E+00 | 1.05E-04 | 2.87E-04 | -1.16E+00 |
| ZNF302 | 4.49E-10 | 2.34E-09 | -1.20E+00 | 3.59E-02 | 1.55E-01 | 1.34E+00 | 1.89E-08 | 8.01E-08 | -1.33E+00 | 6.68E-03 | 2.55E-02 | 1.14E+00 | 1.28E-03 | 4.56E-03 | -1.00E+00 | 3.31E-01 | 9.33E-01 | 1.37E+00 | 1.12E-05 | 4.10E-05 | -1.22E+00 | 1.61E-02 | 4.43E-02 | 1.02E+00 |
| PAIP1 | 4.50E-10 | 2.34E-09 | -1.31E+00 | 2.10E-08 | 9.09E-08 | -1.07E+00 | 5.26E-05 | 2.23E-04 | -1.30E+00 | 1.43E-04 | 5.45E-04 | -1.09E+00 | 6.60E-07 | 2.35E-06 | -1.44E+00 | 6.17E-06 | 1.74E-05 | -1.36E+00 | 9.13E-09 | 3.35E-08 | -1.51E+00 | 1.98E-06 | 5.45E-06 | -1.29E+00 |
| NRROS | 4.50E-10 | 2.35E-09 | -1.31E+00 | 1.46E-04 | 6.31E-04 | 1.09E+00 | 1.75E-08 | 7.43E-08 | -1.47E+00 | 4.37E-02 | 1.67E-01 | 1.19E+00 | 3.27E-02 | 1.16E-01 | 1.25E+00 | 4.41E-02 | 1.24E-01 | 1.39E+00 | 3.95E-06 | 1.45E-05 | -1.20E+00 | 6.33E-03 | 1.74E-02 | 1.03E+00 |
| UBR2 | 4.51E-10 | 2.35E-09 | -1.33E+00 | 1.74E-04 | 7.52E-04 | 1.05E+00 | 5.94E-06 | 2.52E-05 | -1.27E+00 | 3.82E-04 | 1.46E-03 | 1.01E+00 | 1.50E-03 | 5.34E-03 | -1.06E+00 | 1.31E-03 | 3.70E-03 | 1.01E+00 | 5.14E-05 | 1.89E-04 | -1.18E+00 | 5.59E-05 | 1.54E-04 | -1.18E+00 |
| DDX42 | 4.53E-10 | 2.36E-09 | -1.28E+00 | 4.48E-07 | 1.94E-06 | -1.02E+00 | 9.94E-07 | 4.22E-06 | -1.22E+00 | 1.00E-04 | 3.82E-04 | 1.04E+00 | 4.13E-06 | 1.47E-05 | -1.15E+00 | 2.46E-04 | 6.94E-04 | -1.08E+00 | 2.26E-06 | 8.29E-06 | -1.27E+00 | 7.79E-03 | 2.14E-02 | -1.03E+00 |
| MTMR12 | 4.54E-10 | 2.36E-09 | -1.96E+00 | 5.47E-07 | 2.36E-06 | -1.28E+00 | 2.33E-03 | 9.91E-03 | -1.28E+00 | 3.03E-03 | 1.16E-02 | -1.08E+00 | 3.32E-07 | 1.18E-06 | -2.13E+00 | 5.37E-05 | 1.51E-04 | -1.44E+00 | 1.45E-07 | 5.33E-07 | -1.39E+00 | 6.54E-08 | 1.80E-07 | -1.62E+00 |
| TRNT1 | 4.55E-10 | 2.37E-09 | -1.34E+00 | 4.20E-05 | 1.82E-04 | 1.02E+00 | 7.65E-11 | 3.25E-10 | -1.55E+00 | 3.22E-04 | 1.23E-03 | 1.02E+00 | 5.11E-07 | 1.82E-06 | -1.43E+00 | 5.71E-04 | 1.61E-03 | -1.11E+00 | 1.24E-07 | 4.56E-07 | -1.38E+00 | 1.88E-01 | 5.17E-01 | 1.12E+00 |
| OSER1-AS1 | 4.55E-10 | 2.37E-09 | -1.62E+00 | 3.93E-06 | 1.70E-05 | -1.15E+00 | 1.83E-02 | 7.79E-02 | -1.31E+00 | 1.36E-01 | 5.19E-01 | 1.12E+00 | 2.04E-05 | 7.27E-05 | -1.57E+00 | 1.64E-02 | 4.63E-02 | 1.03E+00 | 4.33E-02 | 1.59E-01 | -1.07E+00 | 2.47E-01 | 6.79E-01 | 1.15E+00 |
| COL24A1 | 4.58E-10 | 2.39E-09 | -1.47E+00 | 1.12E-09 | 4.82E-09 | -1.26E+00 | 1.58E-08 | 6.70E-08 | -1.61E+00 | 4.77E-07 | 1.82E-06 | -1.38E+00 | 4.33E-02 | 1.54E-01 | -1.17E+00 | 5.90E-03 | 1.66E-02 | -1.84E+00 | 8.20E-02 | 3.01E-01 | 2.11E+00 | 2.56E-01 | 7.04E-01 | 1.08E+00 |
| TNKS2 | 4.58E-10 | 2.39E-09 | -1.51E+00 | 1.21E-07 | 5.21E-07 | -1.16E+00 | 9.04E-09 | 3.84E-08 | -1.46E+00 | 1.48E-05 | 5.67E-05 | -1.10E+00 | 1.93E-05 | 6.87E-05 | -1.31E+00 | 3.47E-04 | 9.78E-04 | -1.10E+00 | 5.78E-16 | 2.12E-15 | -1.85E+00 | 1.34E-10 | 3.67E-10 | -1.56E+00 |
| TNFAIP8L1 | 4.58E-10 | 2.39E-09 | -1.65E+00 | 1.02E-07 | 4.40E-07 | -1.27E+00 | 2.20E-07 | 9.35E-07 | -1.85E+00 | 1.74E-05 | 6.66E-05 | -1.32E+00 | 2.64E-07 | 9.41E-07 | -1.73E+00 | 1.19E-04 | 3.35E-04 | -1.27E+00 | 3.06E-13 | 1.12E-12 | -1.84E+00 | 1.83E-09 | 5.03E-09 | -1.75E+00 |
| DUSP15 | 4.58E-10 | 2.39E-09 | -1.65E+00 | 4.17E-06 | 1.80E-05 | -1.10E+00 | 1.22E-07 | 5.17E-07 | -1.50E+00 | 7.34E-07 | 2.80E-06 | -1.23E+00 | 1.30E-02 | 4.65E-02 | -1.21E+00 | 1.47E-02 | 4.15E-02 | -1.27E+00 | 1.71E-01 | 6.27E-01 | -1.24E+00 | 3.59E-01 | 9.85E-01 | 1.22E+00 |
| MRPL44 | 4.60E-10 | 2.40E-09 | -1.28E+00 | 2.48E-09 | 1.07E-08 | -1.10E+00 | 4.98E-07 | 2.11E-06 | -1.32E+00 | 2.40E-05 | 9.16E-05 | -1.06E+00 | 3.46E-12 | 1.23E-11 | -1.79E+00 | 8.52E-08 | 2.40E-07 | -1.24E+00 | 2.51E-13 | 9.21E-13 | -1.48E+00 | 3.80E-05 | 1.05E-04 | -1.11E+00 |
| FKBP3 | 4.65E-10 | 2.42E-09 | -1.37E+00 | 9.99E-10 | 4.32E-09 | -1.24E+00 | 1.98E-02 | 8.40E-02 | -1.06E+00 | 3.48E-04 | 1.33E-03 | -1.05E+00 | 8.15E-07 | 2.91E-06 | -1.22E+00 | 7.02E-07 | 1.98E-06 | -1.19E+00 | 1.42E-04 | 5.22E-04 | -1.18E+00 | 1.26E-03 | 3.45E-03 | -1.13E+00 |
| RDH11 | 4.66E-10 | 2.43E-09 | -1.18E+00 | 1.54E-05 | 6.64E-05 | 1.13E+00 | 1.44E-03 | 6.13E-03 | -1.09E+00 | 4.28E-02 | 1.64E-01 | 1.19E+00 | 6.15E-12 | 2.19E-11 | -1.42E+00 | 4.03E-04 | 1.14E-03 | -1.02E+00 | 9.47E-06 | 3.48E-05 | -1.08E+00 | 9.49E-02 | 2.61E-01 | 1.16E+00 |
| KLRG1 | 4.66E-10 | 2.43E-09 | -1.52E+00 | 3.34E-07 | 1.44E-06 | -1.17E+00 | 1.58E-07 | 6.72E-07 | -1.50E+00 | 2.60E-07 | 9.92E-07 | -1.31E+00 | 1.52E-02 | 5.43E-02 | -1.01E+00 | 2.62E-07 | 7.38E-07 | -1.53E+00 | 2.89E-04 | 1.06E-03 | -1.34E+00 | 1.10E-03 | 3.01E-03 | -1.32E+00 |
| ATF6 | 4.70E-10 | 2.45E-09 | -1.34E+00 | 3.85E-04 | 1.67E-03 | 1.07E+00 | 6.42E-03 | 2.73E-02 | -1.03E+00 | 1.51E-01 | 5.76E-01 | 1.34E+00 | 3.42E-09 | 1.22E-08 | -1.43E+00 | 5.43E-06 | 1.53E-05 | -1.11E+00 | 4.04E-06 | 1.48E-05 | -1.27E+00 | 1.06E-02 | 2.90E-02 | -1.02E+00 |
| CMTM8 | 4.70E-10 | 2.45E-09 | -1.76E+00 | 6.48E-05 | 2.80E-04 | -1.11E+00 | 2.30E-03 | 9.77E-03 | -1.46E+00 | 1.11E-02 | 4.26E-02 | -1.06E+00 | 1.68E-01 | 6.01E-01 | 1.23E+00 | 2.41E-01 | 6.79E-01 | 1.64E+00 | 9.27E-02 | 3.40E-01 | -1.12E+00 | 7.34E-02 | 2.02E-01 | 1.52E+00 |
| INTS8 | 4.71E-10 | 2.45E-09 | -1.50E+00 | 7.43E-07 | 3.21E-06 | -1.15E+00 | 6.96E-03 | 2.96E-02 | -1.09E+00 | 1.01E-02 | 3.87E-02 | 1.05E+00 | 7.76E-06 | 2.77E-05 | -1.36E+00 | 4.08E-04 | 1.15E-03 | -1.08E+00 | 7.38E-07 | 2.71E-06 | -1.50E+00 | 2.49E-02 | 6.84E-02 | 1.04E+00 |
| PHLDA1 | 4.79E-10 | 2.50E-09 | 5.35E+00 | 6.49E-02 | 2.80E-01 | 2.08E+00 | 1.37E-05 | 5.84E-05 | 3.41E+00 | 2.47E-01 | 9.43E-01 | 1.11E+00 | 1.43E-01 | 5.09E-01 | 1.90E+00 | 2.95E-01 | 8.31E-01 | 1.34E+00 | 1.07E-01 | 3.92E-01 | 1.71E+00 | 3.59E-01 | 9.87E-01 | 1.09E+00 |
| MPLKIP | 4.81E-10 | 2.51E-09 | -1.15E+00 | 1.15E-02 | 4.96E-02 | 1.21E+00 | 2.27E-03 | 9.65E-03 | -1.09E+00 | 1.76E-02 | 6.71E-02 | 1.06E+00 | 8.66E-06 | 3.09E-05 | -1.28E+00 | 2.34E-01 | 6.60E-01 | 1.07E+00 | 2.30E-02 | 8.44E-02 | -1.03E+00 | 9.90E-02 | 2.72E-01 | 1.23E+00 |
| RP9P | 4.84E-10 | 2.52E-09 | -1.71E+00 | 8.14E-08 | 3.52E-07 | -1.35E+00 | 1.30E-04 | 5.51E-04 | -1.15E+00 | 3.07E-05 | 1.17E-04 | -1.09E+00 | 6.75E-06 | 2.41E-05 | -1.41E+00 | 3.14E-04 | 8.86E-04 | -1.16E+00 | 1.69E-06 | 6.20E-06 | -1.15E+00 | 3.12E-06 | 8.58E-06 | -1.12E+00 |
| AL158801.1 | 4.91E-10 | 2.56E-09 | -2.68E+00 | 1.07E-07 | 4.64E-07 | -1.81E+00 | 1.12E-04 | 4.75E-04 | -1.55E+00 | 1.91E-05 | 7.31E-05 | -1.60E+00 | 8.59E-03 | 3.06E-02 | -1.10E+00 | 1.18E-03 | 3.34E-03 | -1.38E+00 | 2.87E-06 | 1.05E-05 | -1.84E+00 | 1.95E-05 | 5.35E-05 | -1.58E+00 |
| DCTD | 4.91E-10 | 2.56E-09 | -1.22E+00 | 2.54E-12 | 1.10E-11 | -1.17E+00 | 8.10E-05 | 3.44E-04 | -1.23E+00 | 2.37E-04 | 9.06E-04 | -1.06E+00 | 2.72E-05 | 9.68E-05 | -1.11E+00 | 2.18E-03 | 6.15E-03 | 1.03E+00 | 1.18E-03 | 4.32E-03 | -1.04E+00 | 4.45E-02 | 1.22E-01 | 1.12E+00 |
| FARSA | 4.92E-10 | 2.57E-09 | -1.47E+00 | 4.20E-03 | 1.81E-02 | 1.06E+00 | 4.77E-09 | 2.03E-08 | -1.40E+00 | 9.52E-04 | 3.64E-03 | -1.03E+00 | 9.68E-15 | 3.45E-14 | -1.62E+00 | 4.55E-04 | 1.28E-03 | 1.06E+00 | 1.09E-08 | 4.00E-08 | -1.34E+00 | 3.34E-01 | 9.18E-01 | 1.16E+00 |
| TRIB3 | 4.93E-10 | 2.57E-09 | -1.84E+00 | 4.27E-04 | 1.85E-03 | 1.00E+00 | 2.12E-04 | 9.00E-04 | -1.26E+00 | 9.04E-02 | 3.45E-01 | 1.25E+00 | 1.99E-02 | 7.10E-02 | -1.45E+00 | 2.81E-01 | 7.92E-01 | 1.27E+00 | 2.30E-01 | 8.43E-01 | 1.81E+00 | 6.30E-02 | 1.73E-01 | 2.00E+00 |
| ITFG2 | 4.93E-10 | 2.57E-09 | -1.72E+00 | 2.83E-06 | 1.22E-05 | -1.19E+00 | 2.43E-07 | 1.03E-06 | -1.41E+00 | 1.02E-06 | 3.91E-06 | -1.20E+00 | 2.84E-05 | 1.01E-04 | -1.24E+00 | 4.91E-04 | 1.38E-03 | -1.06E+00 | 1.42E-07 | 5.22E-07 | -1.57E+00 | 8.68E-03 | 2.39E-02 | -1.05E+00 |
| EP300 | 4.98E-10 | 2.59E-09 | -1.17E+00 | 2.14E-12 | 9.23E-12 | -1.13E+00 | 2.06E-07 | 8.74E-07 | -1.38E+00 | 1.15E-15 | 4.40E-15 | -1.60E+00 | 6.56E-05 | 2.34E-04 | -1.16E+00 | 5.41E-13 | 1.52E-12 | -1.85E+00 | 2.19E-07 | 8.03E-07 | -1.25E+00 | 7.21E-08 | 1.98E-07 | -1.34E+00 |
| NCKAP1L | 5.00E-10 | 2.61E-09 | -1.20E+00 | 1.53E-04 | 6.63E-04 | 1.10E+00 | 1.01E-05 | 4.30E-05 | -1.37E+00 | 4.01E-05 | 1.53E-04 | -1.10E+00 | 3.01E-04 | 1.07E-03 | -1.03E+00 | 1.13E-03 | 3.18E-03 | 1.06E+00 | 4.76E-03 | 1.75E-02 | -1.17E+00 | 7.20E-02 | 1.98E-01 | -1.02E+00 |
| SUGT1 | 5.01E-10 | 2.61E-09 | -1.19E+00 | 1.65E-03 | 7.15E-03 | 1.17E+00 | 5.22E-04 | 2.22E-03 | -1.10E+00 | 6.36E-03 | 2.43E-02 | 1.10E+00 | 1.97E-12 | 7.02E-12 | -1.29E+00 | 3.44E-04 | 9.70E-04 | -1.01E+00 | 4.63E-06 | 1.70E-05 | -1.24E+00 | 2.74E-02 | 7.52E-02 | -1.01E+00 |
| BTF3L4 | 5.02E-10 | 2.62E-09 | -1.30E+00 | 2.78E-08 | 1.20E-07 | -1.10E+00 | 3.41E-06 | 1.45E-05 | -1.20E+00 | 2.15E-05 | 8.20E-05 | -1.02E+00 | 2.49E-08 | 8.89E-08 | -1.29E+00 | 2.15E-06 | 6.07E-06 | -1.16E+00 | 7.37E-06 | 2.71E-05 | -1.21E+00 | 1.89E-02 | 5.19E-02 | -1.02E+00 |
| KMT5A | 5.06E-10 | 2.63E-09 | -1.70E+00 | 1.29E-07 | 5.57E-07 | -1.28E+00 | 2.83E-07 | 1.20E-06 | -1.52E+00 | 1.80E-09 | 6.88E-09 | -1.43E+00 | 5.96E-08 | 2.13E-07 | -1.98E+00 | 8.98E-05 | 2.53E-04 | -1.33E+00 | 4.10E-15 | 1.51E-14 | -2.05E+00 | 1.36E-07 | 3.74E-07 | -1.51E+00 |
| PORCN | 5.07E-10 | 2.64E-09 | -2.04E+00 | 5.18E-05 | 2.24E-04 | -1.11E+00 | 6.42E-03 | 2.73E-02 | -1.39E+00 | 3.11E-02 | 1.19E-01 | -1.01E+00 | 1.28E-02 | 4.55E-02 | -1.35E+00 | 2.74E-02 | 7.72E-02 | -1.30E+00 | 1.52E-05 | 5.59E-05 | -2.08E+00 | 1.29E-03 | 3.55E-03 | -1.60E+00 |
| TMEM258 | 5.10E-10 | 2.66E-09 | -1.28E+00 | 8.44E-35 | 3.65E-34 | -1.58E+00 | 2.39E-02 | 1.02E-01 | -1.14E+00 | 2.63E-09 | 1.01E-08 | -1.52E+00 | 9.16E-09 | 3.27E-08 | -1.33E+00 | 1.65E-12 | 4.66E-12 | -1.61E+00 | 2.16E-04 | 7.92E-04 | -1.23E+00 | 1.80E-09 | 4.95E-09 | -1.58E+00 |
| TAF1 | 5.10E-10 | 2.66E-09 | -1.57E+00 | 9.16E-04 | 3.96E-03 | 1.06E+00 | 1.16E-09 | 4.91E-09 | -1.64E+00 | 2.28E-05 | 8.70E-05 | -1.05E+00 | 2.48E-03 | 8.84E-03 | -1.14E+00 | 7.17E-02 | 2.02E-01 | 1.12E+00 | 1.19E-03 | 4.37E-03 | -1.13E+00 | 1.12E-01 | 3.07E-01 | 1.10E+00 |
| CTDSP2 | 5.19E-10 | 2.70E-09 | -1.26E+00 | 2.19E-03 | 9.48E-03 | 1.19E+00 | 2.39E-07 | 1.02E-06 | -1.45E+00 | 5.27E-04 | 2.01E-03 | -1.02E+00 | 2.54E-06 | 9.07E-06 | -1.10E+00 | 7.31E-03 | 2.06E-02 | 1.15E+00 | 6.15E-09 | 2.26E-08 | -1.39E+00 | 1.97E-06 | 5.41E-06 | -1.22E+00 |
| NAA38 | 5.20E-10 | 2.71E-09 | -1.18E+00 | 1.57E-38 | 6.77E-38 | -1.66E+00 | 9.94E-02 | 4.22E-01 | 1.01E+00 | 1.68E-08 | 6.41E-08 | -1.43E+00 | 5.80E-05 | 2.07E-04 | -1.25E+00 | 2.36E-06 | 6.65E-06 | -1.39E+00 | 1.50E-01 | 5.49E-01 | 1.03E+00 | 8.06E-04 | 2.21E-03 | -1.37E+00 |
| RPL34P33 | 5.21E-10 | 2.72E-09 | -1.15E+00 | 3.19E-40 | 1.38E-39 | -1.87E+00 | 9.10E-04 | 3.87E-03 | -1.05E+00 | 1.38E-19 | 5.28E-19 | -1.78E+00 | 2.46E-06 | 8.78E-06 | -1.15E+00 | 1.24E-23 | 3.48E-23 | -2.06E+00 | 4.78E-05 | 1.75E-04 | -1.19E+00 | 6.95E-18 | 1.91E-17 | -2.05E+00 |
| SDF2 | 5.28E-10 | 2.75E-09 | -1.26E+00 | 7.58E-06 | 3.27E-05 | 1.04E+00 | 1.11E-03 | 4.71E-03 | -1.03E+00 | 3.41E-02 | 1.30E-01 | 1.21E+00 | 1.66E-05 | 5.93E-05 | -1.25E+00 | 6.11E-02 | 1.72E-01 | 1.15E+00 | 1.42E-04 | 5.22E-04 | -1.08E+00 | 1.69E-02 | 4.64E-02 | 1.05E+00 |
| IFI16 | 5.29E-10 | 2.76E-09 | -1.25E+00 | 1.08E-03 | 4.66E-03 | 1.07E+00 | 3.38E-06 | 1.44E-05 | -1.44E+00 | 3.75E-05 | 1.43E-04 | -1.16E+00 | 2.16E-03 | 7.71E-03 | -1.13E+00 | 3.13E-01 | 8.82E-01 | 1.29E+00 | 1.02E-16 | 3.73E-16 | -1.78E+00 | 6.44E-11 | 1.77E-10 | -1.52E+00 |
| KCTD20 | 5.33E-10 | 2.78E-09 | -1.49E+00 | 2.10E-04 | 9.09E-04 | 1.06E+00 | 7.87E-05 | 3.34E-04 | -1.24E+00 | 3.37E-03 | 1.29E-02 | 1.06E+00 | 2.57E-06 | 9.17E-06 | -1.24E+00 | 4.32E-04 | 1.22E-03 | -1.05E+00 | 7.06E-06 | 2.59E-05 | -1.11E+00 | 2.86E-04 | 7.85E-04 | -1.01E+00 |
| DSEL | 5.38E-10 | 2.80E-09 | -2.34E+00 | 2.35E-09 | 1.01E-08 | -2.28E+00 | 2.30E-02 | 9.77E-02 | -1.34E+00 | 5.21E-02 | 1.99E-01 | -1.18E+00 | 1.71E-02 | 6.11E-02 | -1.45E+00 | 1.78E-02 | 5.03E-02 | -1.71E+00 | 2.60E-04 | 9.54E-04 | -1.68E+00 | 2.95E-04 | 8.09E-04 | -1.79E+00 |
| ERAP1 | 5.40E-10 | 2.81E-09 | -1.40E+00 | 3.27E-05 | 1.41E-04 | 1.07E+00 | 5.59E-06 | 2.38E-05 | -1.46E+00 | 7.73E-03 | 2.95E-02 | 1.02E+00 | 1.32E-06 | 4.72E-06 | -1.52E+00 | 1.20E-04 | 3.38E-04 | -1.24E+00 | 6.89E-06 | 2.53E-05 | -1.38E+00 | 4.72E-03 | 1.30E-02 | -1.05E+00 |
| POGK | 5.43E-10 | 2.83E-09 | -1.55E+00 | 5.55E-06 | 2.40E-05 | -1.08E+00 | 1.48E-11 | 6.28E-11 | -1.74E+00 | 5.31E-05 | 2.03E-04 | 1.01E+00 | 4.18E-04 | 1.49E-03 | -1.50E+00 | 1.26E-01 | 3.54E-01 | 1.11E+00 | 4.55E-11 | 1.67E-10 | -1.72E+00 | 1.33E-03 | 3.65E-03 | -1.10E+00 |
| FH | 5.46E-10 | 2.85E-09 | -1.36E+00 | 9.79E-07 | 4.23E-06 | -1.02E+00 | 6.96E-04 | 2.96E-03 | -1.10E+00 | 1.29E-04 | 4.94E-04 | -1.02E+00 | 3.46E-12 | 1.23E-11 | -1.67E+00 | 2.07E-05 | 5.83E-05 | -1.07E+00 | 4.48E-05 | 1.64E-04 | -1.15E+00 | 1.60E-01 | 4.39E-01 | 1.21E+00 |
| EIF6 | 5.47E-10 | 2.85E-09 | -1.21E+00 | 2.07E-07 | 8.94E-07 | -1.05E+00 | 2.98E-03 | 1.27E-02 | -1.03E+00 | 4.32E-04 | 1.65E-03 | 1.00E+00 | 2.90E-05 | 1.03E-04 | -1.13E+00 | 8.36E-04 | 2.36E-03 | -1.03E+00 | 4.19E-03 | 1.54E-02 | -1.06E+00 | 5.43E-02 | 1.49E-01 | 1.02E+00 |
| SDHAF1 | 5.49E-10 | 2.86E-09 | -1.68E+00 | 3.07E-04 | 1.33E-03 | 1.00E+00 | 1.08E-11 | 4.60E-11 | -1.79E+00 | 3.91E-03 | 1.49E-02 | 1.14E+00 | 2.27E-09 | 8.09E-09 | -2.01E+00 | 5.23E-04 | 1.47E-03 | -1.16E+00 | 2.91E-20 | 1.07E-19 | -1.94E+00 | 1.11E-07 | 3.04E-07 | -1.21E+00 |
| PPP1R11 | 5.56E-10 | 2.90E-09 | -1.21E+00 | 1.19E-08 | 5.15E-08 | -1.03E+00 | 2.91E-06 | 1.24E-05 | -1.33E+00 | 3.49E-05 | 1.33E-04 | -1.06E+00 | 1.56E-04 | 5.55E-04 | -1.06E+00 | 1.44E-03 | 4.05E-03 | 1.05E+00 | 2.99E-02 | 1.10E-01 | 1.01E+00 | 3.25E-01 | 8.92E-01 | 1.21E+00 |
| HNRNPC | 5.56E-10 | 2.90E-09 | -1.27E+00 | 7.17E-14 | 3.10E-13 | -1.30E+00 | 1.35E-04 | 5.74E-04 | -1.26E+00 | 4.69E-05 | 1.79E-04 | -1.27E+00 | 6.28E-07 | 2.24E-06 | -1.37E+00 | 9.95E-04 | 2.80E-03 | -1.20E+00 | 1.87E-07 | 6.86E-07 | -1.37E+00 | 6.58E-04 | 1.81E-03 | -1.24E+00 |
| GLB1 | 5.57E-10 | 2.90E-09 | -1.56E+00 | 7.51E-04 | 3.24E-03 | 1.10E+00 | 5.91E-13 | 2.51E-12 | -1.59E+00 | 1.88E-10 | 7.20E-10 | -1.26E+00 | 9.41E-08 | 3.35E-07 | -1.49E+00 | 2.85E-03 | 8.02E-03 | 1.03E+00 | 1.36E-08 | 5.00E-08 | -1.43E+00 | 1.01E-04 | 2.78E-04 | -1.09E+00 |
| EPDR1 | 5.57E-10 | 2.90E-09 | -2.12E+00 | 3.40E-04 | 1.47E-03 | -1.03E+00 | 5.25E-12 | 2.23E-11 | -1.91E+00 | 8.66E-06 | 3.31E-05 | -1.08E+00 | 3.38E-10 | 1.21E-09 | -2.22E+00 | 1.45E-02 | 4.10E-02 | -1.02E+00 | 1.61E-15 | 5.91E-15 | -1.49E+00 | 3.75E-09 | 1.03E-08 | -1.23E+00 |
| PSMB6 | 5.59E-10 | 2.91E-09 | -1.19E+00 | 2.89E-04 | 1.25E-03 | 1.11E+00 | 2.45E-02 | 1.04E-01 | 1.08E+00 | 4.31E-02 | 1.65E-01 | 1.10E+00 | 5.44E-09 | 1.94E-08 | -1.21E+00 | 4.37E-05 | 1.23E-04 | -1.02E+00 | 7.90E-02 | 2.90E-01 | -1.05E+00 | 3.18E-01 | 8.73E-01 | 1.10E+00 |
| RHOG | 5.62E-10 | 2.93E-09 | -1.14E+00 | 4.50E-06 | 1.94E-05 | 1.05E+00 | 2.30E-09 | 9.76E-09 | -1.49E+00 | 3.52E-06 | 1.34E-05 | -1.24E+00 | 1.52E-10 | 5.41E-10 | -1.48E+00 | 5.65E-05 | 1.59E-04 | -1.23E+00 | 4.32E-11 | 1.58E-10 | -1.60E+00 | 3.35E-05 | 9.21E-05 | -1.30E+00 |
| COPG2 | 5.63E-10 | 2.93E-09 | -1.68E+00 | 1.37E-06 | 5.94E-06 | -1.19E+00 | 2.33E-14 | 9.92E-14 | -1.91E+00 | 1.00E-13 | 3.83E-13 | -1.65E+00 | 2.02E-09 | 7.22E-09 | -1.95E+00 | 2.29E-06 | 6.47E-06 | -1.32E+00 | 2.12E-11 | 7.76E-11 | -1.68E+00 | 6.83E-07 | 1.88E-06 | -1.31E+00 |
| FSTL1 | 5.65E-10 | 2.94E-09 | -1.49E+00 | 5.56E-04 | 2.40E-03 | 1.15E+00 | 3.23E-03 | 1.37E-02 | -1.09E+00 | 3.92E-03 | 1.50E-02 | 1.09E+00 | 2.42E-03 | 8.64E-03 | -1.82E+00 | 1.15E-02 | 3.24E-02 | -1.25E+00 | 6.11E-02 | 2.24E-01 | -1.19E+00 | 1.78E-01 | 4.90E-01 | 1.10E+00 |
| TIMM10B | 5.66E-10 | 2.95E-09 | -1.29E+00 | 7.75E-08 | 3.35E-07 | -1.05E+00 | 3.60E-06 | 1.53E-05 | -1.45E+00 | 3.34E-03 | 1.27E-02 | -1.01E+00 | 2.88E-06 | 1.03E-05 | -1.24E+00 | 2.19E-02 | 6.16E-02 | 1.13E+00 | 4.23E-05 | 1.55E-04 | -1.18E+00 | 3.28E-01 | 9.00E-01 | 1.29E+00 |
| PRKAB2 | 5.73E-10 | 2.99E-09 | -1.68E+00 | 4.39E-06 | 1.90E-05 | -1.19E+00 | 1.25E-04 | 5.32E-04 | -1.24E+00 | 3.61E-03 | 1.38E-02 | 1.06E+00 | 1.61E-01 | 5.73E-01 | 1.18E+00 | 5.01E-03 | 1.41E-02 | -1.18E+00 | 2.66E-05 | 9.78E-05 | -1.36E+00 | 6.19E-04 | 1.70E-03 | -1.31E+00 |
| KIN | 5.76E-10 | 3.00E-09 | -1.39E+00 | 4.13E-03 | 1.78E-02 | 1.18E+00 | 4.44E-06 | 1.89E-05 | -1.37E+00 | 1.13E-02 | 4.31E-02 | 1.09E+00 | 3.31E-05 | 1.18E-04 | -1.16E+00 | 1.33E-02 | 3.76E-02 | 1.11E+00 | 4.06E-03 | 1.49E-02 | -1.07E+00 | 2.74E-02 | 7.54E-02 | 1.07E+00 |
| UBBP4 | 5.79E-10 | 3.02E-09 | -1.18E+00 | 3.51E-09 | 1.52E-08 | 1.00E+00 | 7.80E-02 | 3.31E-01 | -1.02E+00 | 8.69E-02 | 3.32E-01 | 1.16E+00 | 6.39E-04 | 2.28E-03 | -1.02E+00 | 2.97E-03 | 8.37E-03 | 1.10E+00 | 1.61E-01 | 5.91E-01 | 1.16E+00 | 3.32E-01 | 9.11E-01 | 1.23E+00 |

| | | | | | | | | | | | | | | | | | | | | | | | | |
|---|---|---|---|---|---|---|---|---|---|---|---|---|---|---|---|---|---|---|---|---|---|---|---|---|
| AC008124.1 | 5.80E-10 | 3.02E-09 | -1.14E+00 | 7.45E-07 | 3.22E-06 | 1.11E+00 | 3.31E-02 | 1.41E-01 | 1.05E+00 | 6.19E-02 | 2.37E-01 | 1.23E+00 | 4.18E-04 | 1.49E-03 | -1.09E+00 | 4.86E-02 | 1.37E-01 | 1.13E+00 | 1.79E-03 | 6.58E-03 | -1.06E+00 | 1.20E-03 | 3.29E-03 | -1.07E+00 |
| RMDN3 | 5.84E-10 | 3.05E-09 | -1.62E+00 | 1.89E-07 | 8.16E-07 | -1.27E+00 | 3.27E-04 | 1.39E-03 | -1.32E+00 | 1.83E-02 | 6.99E-02 | 1.09E+00 | 2.87E-03 | 1.02E-02 | -1.17E+00 | 1.19E-02 | 3.35E-02 | -1.01E+00 | 4.09E-15 | 1.50E-14 | -1.68E+00 | 3.34E-08 | 9.17E-08 | -1.33E+00 |
| NAPEPLD | 5.86E-10 | 3.06E-09 | -1.91E+00 | 2.17E-01 | 9.37E-01 | 1.49E+00 | 1.05E-04 | 4.45E-04 | -1.52E+00 | 1.97E-02 | 7.52E-02 | 1.87E+00 | 1.08E-02 | 3.86E-02 | -1.24E+00 | 1.18E-02 | 3.34E-02 | 1.97E+00 | 3.51E-11 | 1.29E-10 | -1.72E+00 | 9.70E-03 | 2.67E-02 | 1.01E+00 |
| MMP28 | 5.92E-10 | 3.09E-09 | -1.49E+00 | 3.28E-05 | 1.42E-04 | 1.00E+00 | 2.20E-02 | 9.33E-02 | -1.35E+00 | 4.33E-02 | 1.66E-01 | -1.09E+00 | 1.80E-02 | 6.43E-02 | -2.63E+00 | 5.42E-02 | 1.53E-01 | -1.58E+00 | 9.12E-02 | 3.35E-01 | -1.65E+00 | 1.93E-01 | 5.29E-01 | -1.39E+00 |
| MTND2P28 | 5.96E-10 | 3.10E-09 | -1.21E+00 | 2.42E-13 | 1.05E-12 | -1.19E+00 | 3.95E-05 | 1.68E-04 | -1.26E+00 | 1.90E-06 | 7.24E-06 | -1.23E+00 | 1.66E-04 | 5.91E-04 | -1.09E+00 | 4.72E-03 | 1.33E-02 | -1.06E+00 | 1.44E-01 | 5.30E-01 | 1.07E+00 | 1.85E-01 | 5.09E-01 | 1.07E+00 |
| FABP5P2 | 5.96E-10 | 3.11E-09 | -1.26E+00 | 3.57E-07 | 1.54E-06 | -1.04E+00 | 3.22E-03 | 1.37E-02 | -1.10E+00 | 4.06E-02 | 1.55E-01 | 1.10E+00 | 2.44E-08 | 8.71E-08 | -1.38E+00 | 5.46E-03 | 1.54E-02 | -1.00E+00 | 1.48E-02 | 5.45E-02 | 1.01E+00 | 3.96E-02 | 1.09E-01 | 1.38E+00 |
| RPS6KA5 | 5.98E-10 | 3.12E-09 | -1.53E+00 | 1.09E-02 | 4.72E-02 | 1.25E+00 | 3.45E-05 | 1.46E-04 | -1.33E+00 | 2.60E-01 | 9.94E-01 | 1.45E+00 | 1.70E-07 | 6.06E-07 | -1.66E+00 | 2.23E-07 | 6.28E-07 | -1.37E+00 | 1.58E-06 | 5.79E-06 | -1.55E+00 | 2.55E-02 | 7.01E-02 | -1.01E+00 |
| WDSUB1 | 5.99E-10 | 3.12E-09 | -1.40E+00 | 1.23E-05 | 5.33E-05 | 1.05E+00 | 1.69E-06 | 7.19E-06 | -1.40E+00 | 5.47E-06 | 2.09E-05 | -1.18E+00 | 3.98E-05 | 1.42E-04 | -1.42E+00 | 1.30E-02 | 3.68E-02 | 1.05E+00 | 5.44E-08 | 2.00E-07 | -1.35E+00 | 5.57E-04 | 1.53E-03 | -1.02E+00 |
| ZFP36L2 | 5.99E-10 | 3.12E-09 | 1.49E+00 | 5.41E-51 | 2.34E-50 | 2.61E+00 | 1.30E-03 | 5.54E-03 | 1.32E+00 | 6.68E-28 | 2.55E-27 | 2.49E+00 | 1.60E-33 | 5.71E-33 | 2.75E+00 | 1.85E-63 | 5.23E-63 | 3.71E+00 | 3.20E-12 | 1.17E-11 | 1.83E+00 | 9.37E-36 | 2.57E-35 | 2.73E+00 |
| UTP3 | 6.03E-10 | 3.14E-09 | -1.41E+00 | 3.22E-04 | 1.39E-03 | 1.07E+00 | 1.61E-07 | 6.82E-07 | -1.36E+00 | 1.38E-02 | 5.26E-02 | 1.17E+00 | 2.02E-03 | 7.21E-03 | -1.13E+00 | 3.20E-01 | 9.03E-01 | 1.44E+00 | 3.06E-09 | 1.12E-08 | -1.44E+00 | 1.39E-01 | 3.82E-01 | 1.11E+00 |
| KANSL1 | 6.06E-10 | 3.16E-09 | -1.21E+00 | 9.91E-04 | 4.28E-03 | 1.14E+00 | 3.44E-05 | 1.46E-04 | -1.31E+00 | 2.83E-04 | 1.08E-03 | -1.10E+00 | 1.72E-06 | 6.13E-06 | -1.14E+00 | 6.49E-05 | 1.83E-04 | -1.05E+00 | 1.88E-06 | 6.89E-06 | -1.29E+00 | 4.97E-04 | 1.37E-03 | -1.14E+00 |
| C11orf49 | 6.08E-10 | 3.17E-09 | -1.55E+00 | 2.63E-03 | 1.14E-02 | 1.12E+00 | 8.39E-06 | 3.56E-05 | -1.32E+00 | 2.62E-04 | 9.99E-04 | -1.04E+00 | 6.66E-05 | 2.37E-04 | -1.14E+00 | 2.26E-02 | 6.38E-02 | 1.16E+00 | 3.02E-04 | 1.11E-03 | -1.12E+00 | 5.94E-02 | 1.63E-01 | 1.07E+00 |
| MCM3AP-AS1 | 6.12E-10 | 3.19E-09 | -1.77E+00 | 1.18E-03 | 5.12E-03 | 1.08E+00 | 4.57E-07 | 1.94E-06 | -1.94E+00 | 5.10E-02 | 1.95E-01 | 1.08E+00 | 1.13E-03 | 4.04E-03 | -1.59E+00 | 2.54E-02 | 7.16E-02 | -1.13E+00 | 8.02E-03 | 2.95E-02 | -1.23E+00 | 9.12E-02 | 2.50E-01 | 1.05E+00 |
| CCDC85B | 6.13E-10 | 3.19E-09 | -1.13E+00 | 9.12E-13 | 3.94E-12 | -1.07E+00 | 1.21E-01 | 5.15E-01 | 1.01E+00 | 9.66E-02 | 3.69E-01 | 1.09E+00 | 6.74E-04 | 2.40E-03 | -1.04E+00 | 1.04E-01 | 2.93E-01 | 1.12E+00 | 6.40E-02 | 2.35E-01 | 1.01E+00 | 1.43E-01 | 3.93E-01 | 1.18E+00 |
| NUP62 | 6.14E-10 | 3.20E-09 | -1.69E+00 | 3.16E-05 | 1.37E-04 | -1.11E+00 | 1.60E-14 | 6.79E-14 | -1.69E+00 | 5.31E-07 | 2.03E-06 | -1.08E+00 | 2.37E-11 | 8.46E-11 | -1.92E+00 | 2.02E-04 | 5.68E-04 | -1.02E+00 | 1.11E-21 | 4.09E-21 | -1.85E+00 | 5.83E-05 | 1.60E-04 | -1.09E+00 |
| MCFD2 | 6.18E-10 | 3.22E-09 | -1.29E+00 | 1.69E-07 | 7.30E-07 | -1.06E+00 | 1.03E-03 | 4.40E-03 | -1.12E+00 | 9.56E-06 | 3.65E-05 | -1.08E+00 | 1.09E-04 | 3.89E-04 | -1.19E+00 | 5.73E-04 | 1.62E-03 | -1.10E+00 | 1.12E-03 | 4.13E-03 | -1.06E+00 | 4.55E-03 | 1.25E-02 | -1.04E+00 |
| PROM1 | 6.20E-10 | 3.23E-09 | -1.21E+00 | 3.46E-03 | 1.49E-02 | 1.29E+00 | 2.07E-05 | 8.78E-05 | -1.29E+00 | 5.13E-02 | 1.96E-01 | 1.16E+00 | 2.00E-01 | 7.12E-01 | 1.71E+00 | 3.24E-01 | 9.12E-01 | 1.46E+00 | 2.71E-01 | 9.93E-01 | 1.08E+00 | 3.22E-01 | 8.85E-01 | 1.32E+00 |
| NAT6 | 6.24E-10 | 3.25E-09 | -1.58E+00 | 2.05E-07 | 8.86E-07 | -1.19E+00 | 2.42E-07 | 1.03E-06 | -1.55E+00 | 1.93E-04 | 7.37E-04 | -1.04E+00 | 1.15E-09 | 4.11E-09 | -1.24E+00 | 1.06E-03 | 3.00E-03 | 1.09E+00 | 9.05E-09 | 3.32E-08 | -1.37E+00 | 3.95E-08 | 1.09E-07 | -1.36E+00 |
| UBE2B | 6.28E-10 | 3.27E-09 | -1.21E+00 | 7.99E-07 | 3.45E-06 | -1.02E+00 | 3.54E-06 | 1.50E-05 | -1.24E+00 | 7.94E-06 | 3.03E-05 | -1.10E+00 | 7.48E-07 | 2.67E-06 | -1.18E+00 | 5.85E-04 | 1.65E-03 | -1.02E+00 | 2.88E-08 | 1.06E-07 | -1.23E+00 | 5.82E-06 | 1.60E-05 | -1.22E+00 |
| NIPSNAP1 | 6.31E-10 | 3.29E-09 | -1.29E+00 | 1.70E-11 | 7.35E-11 | -1.21E+00 | 1.54E-08 | 6.55E-08 | -1.37E+00 | 4.07E-12 | 1.55E-11 | -1.34E+00 | 4.17E-06 | 1.49E-05 | -1.21E+00 | 7.18E-06 | 2.02E-05 | -1.20E+00 | 4.52E-07 | 1.66E-06 | -1.35E+00 | 4.78E-04 | 1.31E-03 | -1.17E+00 |
| AC020898.1 | 6.31E-10 | 3.29E-09 | 1.63E+00 | 6.13E-33 | 2.65E-32 | 2.29E+00 | 2.19E-06 | 9.31E-06 | 1.54E+00 | 1.20E-23 | 4.57E-23 | 2.36E+00 | 1.11E-06 | 3.96E-06 | 1.61E+00 | 8.15E-21 | 2.30E-20 | 2.21E+00 | 1.26E-05 | 4.61E-05 | 1.58E+00 | 2.54E-20 | 6.96E-20 | 2.16E+00 |
| TERF1P5 | 6.32E-10 | 3.29E-09 | -1.81E+00 | 1.01E-07 | 4.37E-07 | -1.38E+00 | 1.00E-07 | 4.26E-07 | -1.60E+00 | 9.43E-06 | 3.60E-05 | -1.22E+00 | 3.28E-08 | 1.17E-07 | -1.92E+00 | 1.50E-06 | 4.22E-06 | -1.63E+00 | 4.77E-17 | 1.75E-16 | -2.02E+00 | 2.56E-11 | 7.02E-11 | -1.56E+00 |
| DCAF10 | 6.34E-10 | 3.30E-09 | -1.53E+00 | 2.46E-03 | 1.06E-02 | 1.20E+00 | 1.16E-11 | 4.94E-11 | -1.82E+00 | 3.58E-05 | 1.37E-04 | -1.03E+00 | 4.55E-08 | 1.62E-07 | -1.60E+00 | 1.49E-04 | 4.20E-04 | -1.05E+00 | 3.36E-14 | 1.23E-13 | -1.80E+00 | 3.62E-04 | 9.95E-04 | -1.15E+00 |
| SPTBN1 | 6.34E-10 | 3.31E-09 | -1.26E+00 | 1.71E-05 | 7.40E-05 | 1.03E+00 | 1.13E-02 | 4.82E-02 | 1.05E+00 | 3.06E-02 | 1.17E-01 | 1.21E+00 | 3.24E-03 | 1.16E-02 | -1.05E+00 | 6.10E-03 | 1.72E-02 | -1.03E+00 | 2.16E-04 | 7.91E-04 | -1.46E+00 | 8.94E-03 | 2.46E-02 | -1.14E+00 |
| SPG11 | 6.37E-10 | 3.32E-09 | -1.32E+00 | 5.52E-05 | 2.38E-04 | 1.12E+00 | 2.56E-13 | 1.09E-12 | -1.65E+00 | 2.46E-06 | 9.38E-06 | -1.07E+00 | 7.93E-09 | 2.83E-08 | -1.61E+00 | 8.20E-03 | 2.31E-02 | 1.06E+00 | 1.21E-08 | 4.45E-08 | -1.38E+00 | 1.22E-03 | 3.35E-03 | -1.16E+00 |
| TIMM23 | 6.46E-10 | 3.37E-09 | -1.56E+00 | 6.02E-08 | 2.60E-07 | -1.29E+00 | 1.13E-04 | 4.82E-04 | -1.16E+00 | 6.77E-07 | 2.59E-06 | -1.19E+00 | 5.55E-04 | 1.98E-03 | -1.48E+00 | 1.66E-03 | 4.67E-03 | -1.26E+00 | 2.60E-05 | 9.53E-05 | -1.14E+00 | 3.30E-03 | 9.07E-03 | -1.04E+00 |
| NPR3 | 6.48E-10 | 3.38E-09 | -1.35E+00 | 9.17E-02 | 3.96E-01 | 1.12E+00 | 1.86E-04 | 7.92E-04 | -1.32E+00 | 1.16E-01 | 4.43E-01 | 1.07E+00 | 7.54E-05 | 2.69E-04 | -1.37E+00 | 6.83E-02 | 1.93E-01 | -1.05E+00 | 2.65E-01 | 9.74E-01 | 1.01E+00 | 1.43E-05 | 3.94E-05 | 1.62E+00 |
| SPON2 | 6.50E-10 | 3.39E-09 | 3.00E+00 | 3.74E-02 | 1.62E-01 | 1.89E+00 | 2.08E-05 | 8.85E-05 | 2.01E+00 | 2.42E-01 | 9.25E-01 | 1.48E+00 | 1.18E-02 | 4.22E-02 | 2.66E+00 | 3.30E-01 | 9.30E-01 | 1.21E+00 | 1.42E-01 | 5.22E-01 | 1.89E+00 | 3.48E-01 | 9.56E-01 | 1.13E+00 |
| GSTK1 | 6.51E-10 | 3.39E-09 | -1.24E+00 | 1.96E-08 | 8.45E-08 | -1.07E+00 | 8.63E-05 | 3.67E-04 | -1.23E+00 | 1.57E-02 | 5.98E-02 | -1.01E+00 | 3.09E-06 | 1.10E-05 | -1.10E+00 | 1.24E-04 | 3.50E-04 | -1.05E+00 | 6.29E-03 | 2.31E-02 | 1.01E+00 | 1.16E-01 | 3.20E-01 | 1.08E+00 |
| TRNAU1AP | 6.61E-10 | 3.45E-09 | -1.30E+00 | 3.31E-05 | 1.43E-04 | 1.06E+00 | 3.77E-05 | 1.60E-04 | -1.23E+00 | 6.63E-04 | 2.53E-03 | -1.05E+00 | 7.52E-06 | 2.68E-05 | -1.33E+00 | 3.74E-04 | 1.05E-03 | -1.12E+00 | 9.85E-03 | 3.61E-02 | -1.09E+00 | 4.62E-02 | 1.27E-01 | 1.04E+00 |
| HMG20B | 6.74E-10 | 3.51E-09 | -1.23E+00 | 7.37E-08 | 3.18E-07 | -1.01E+00 | 2.61E-05 | 1.11E-04 | -1.18E+00 | 1.38E-04 | 5.29E-04 | 1.01E+00 | 7.17E-11 | 2.56E-10 | -1.50E+00 | 5.99E-07 | 1.69E-06 | -1.25E+00 | 2.95E-04 | 1.08E-03 | -1.15E+00 | 3.69E-02 | 1.01E-01 | 1.02E+00 |
| CDK11B | 6.81E-10 | 3.55E-09 | -1.40E+00 | 1.62E-09 | 6.98E-09 | -1.26E+00 | 1.28E-04 | 5.44E-04 | -1.14E+00 | 9.98E-07 | 3.81E-06 | -1.17E+00 | 2.68E-08 | 9.54E-08 | -1.40E+00 | 1.40E-07 | 3.96E-07 | -1.31E+00 | 3.41E-13 | 1.25E-12 | -1.60E+00 | 6.45E-12 | 1.77E-11 | -1.56E+00 |
| LTN1 | 6.81E-10 | 3.55E-09 | -1.41E+00 | 7.84E-06 | 3.39E-05 | -1.00E+00 | 1.44E-09 | 6.14E-09 | -1.63E+00 | 2.89E-05 | 1.10E-04 | -1.11E+00 | 4.53E-09 | 1.62E-08 | -1.45E+00 | 5.07E-06 | 1.43E-05 | -1.23E+00 | 3.14E-05 | 1.15E-04 | -1.13E+00 | 3.85E-03 | 1.06E-02 | -1.02E+00 |
| ENAH | 6.81E-10 | 3.55E-09 | -1.45E+00 | 2.68E-03 | 1.16E-02 | 1.21E+00 | 4.55E-16 | 1.93E-15 | -2.04E+00 | 5.94E-06 | 2.27E-05 | -1.08E+00 | 2.21E-04 | 7.87E-04 | -2.15E+00 | 3.88E-02 | 1.09E-01 | -1.07E+00 | 1.20E-05 | 4.39E-05 | -1.37E+00 | 2.11E-01 | 5.80E-01 | 1.25E+00 |
| PMVK | 6.91E-10 | 3.60E-09 | -1.15E+00 | 1.25E-07 | 5.39E-07 | 1.02E+00 | 1.22E-04 | 5.19E-04 | -1.18E+00 | 8.40E-05 | 3.21E-04 | -1.03E+00 | 3.83E-04 | 1.37E-03 | 1.05E+00 | 9.67E-04 | 2.72E-03 | 1.09E+00 | 2.09E-01 | 7.68E-01 | 1.20E+00 | 7.86E-02 | 2.16E-01 | 1.12E+00 |
| NFIB | 6.94E-10 | 3.62E-09 | -3.09E+00 | 1.75E-09 | 7.56E-09 | -3.02E+00 | 7.73E-02 | 3.28E-01 | -1.34E+00 | 7.31E-02 | 2.79E-01 | -1.38E+00 | 6.62E-02 | 2.36E-01 | -1.44E+00 | 8.59E-02 | 2.42E-01 | -1.44E+00 | 7.20E-10 | 2.64E-09 | -1.83E+00 | 3.01E-12 | 8.28E-12 | -2.81E+00 |

| | | | | | | | | | | | | | | | | | | | | | |
|---|---|---|---|---|---|---|---|---|---|---|---|---|---|---|---|---|---|---|---|---|---|
| C12orf76 | 1.75E-09 | 9.14E-09 | -1.26E+00 | 4.15E-06 | 1.79E-05 | 1.08E+00 | 1.96E-07 | 8.33E-07 | -1.42E+00 | 2.36E-05 | 9.03E-05 | -1.10E+00 | 1.31E-05 | 4.68E-05 | -1.08E+00 | 1.97E-03 | 5.54E-03 | 1.09E+00 | 4.42E-03 | 1.62E-02 | 1.07E+00 | 1.00E-01 | 2.75E-01 | 1.08E+00 |
| AL020997.1 | 1.76E-09 | 9.17E-09 | -1.84E+00 | 2.60E-12 | 1.12E-11 | -2.31E+00 | 1.85E-09 | 7.85E-09 | -1.55E+00 | 2.10E-17 | 8.02E-17 | -1.87E+00 | 9.94E-09 | 3.54E-08 | -1.93E+00 | 9.44E-10 | 2.66E-09 | -1.98E+00 | 1.14E-20 | 4.19E-20 | -1.93E+00 | 4.82E-19 | 1.32E-18 | -1.94E+00 |
| WIPF2 | 1.76E-09 | 9.20E-09 | -1.55E+00 | 4.34E-06 | 1.88E-05 | -1.07E+00 | 8.88E-07 | 3.77E-06 | -1.50E+00 | 2.32E-05 | 8.87E-05 | -1.16E+00 | 2.60E-07 | 9.27E-07 | -1.49E+00 | 3.00E-07 | 8.46E-07 | -1.50E+00 | 5.58E-05 | 2.05E-04 | -1.18E+00 | 3.10E-04 | 8.50E-04 | -1.19E+00 |
| FAM234B | 1.77E-09 | 9.23E-09 | -1.84E+00 | 2.46E-06 | 1.06E-05 | -1.19E+00 | 4.67E-10 | 1.99E-09 | -2.12E+00 | 1.25E-04 | 4.78E-04 | -1.18E+00 | 1.33E-01 | 4.73E-01 | -1.12E+00 | 3.43E-01 | 9.68E-01 | 1.15E+00 | 3.44E-07 | 1.26E-06 | -1.70E+00 | 3.09E-04 | 8.49E-04 | -1.29E+00 |
| DNAJC19P9 | 1.77E-09 | 9.23E-09 | -1.21E+00 | 1.34E-12 | 5.79E-12 | -1.14E+00 | 7.73E-04 | 3.29E-03 | -1.19E+00 | 2.09E-04 | 7.98E-04 | -1.09E+00 | 1.06E-07 | 3.80E-07 | -1.40E+00 | 2.34E-06 | 6.61E-06 | -1.30E+00 | 2.09E-07 | 7.65E-07 | -1.30E+00 | 2.49E-06 | 6.83E-06 | -1.22E+00 |
| ATP6V1E1 | 1.78E-09 | 9.30E-09 | -1.45E+00 | 4.28E-05 | 1.85E-04 | 1.06E+00 | 1.45E-04 | 6.16E-04 | -1.21E+00 | 6.16E-05 | 2.35E-04 | -1.11E+00 | 1.41E-07 | 5.01E-07 | -1.38E+00 | 6.61E-04 | 1.86E-03 | -1.01E+00 | 7.43E-06 | 2.73E-05 | -1.25E+00 | 3.66E-03 | 1.01E-02 | -1.02E+00 |
| SPCS2P4 | 1.79E-09 | 9.34E-09 | -1.21E+00 | 1.39E-13 | 6.00E-13 | -1.22E+00 | 6.59E-04 | 2.80E-03 | -1.08E+00 | 4.35E-07 | 1.66E-06 | -1.16E+00 | 7.04E-08 | 2.51E-07 | -1.33E+00 | 1.99E-08 | 5.61E-08 | -1.28E+00 | 1.10E-02 | 4.04E-02 | -1.03E+00 | 1.28E-04 | 3.51E-04 | -1.27E+00 |
| ATP5S | 1.80E-09 | 9.36E-09 | -1.24E+00 | 1.21E-05 | 5.22E-05 | 1.05E+00 | 1.85E-05 | 7.84E-05 | -1.32E+00 | 2.81E-03 | 1.07E-02 | 1.03E+00 | 8.06E-07 | 2.87E-06 | -1.10E+00 | 3.96E-04 | 1.12E-03 | 1.07E+00 | 9.72E-04 | 3.57E-03 | -1.05E+00 | 3.25E-02 | 8.92E-02 | 1.10E+00 |
| TULP4 | 1.80E-09 | 9.37E-09 | -1.71E+00 | 6.96E-05 | 3.01E-04 | -1.03E+00 | 3.06E-02 | 1.30E-01 | -1.17E+00 | 7.88E-02 | 3.01E-01 | 1.18E+00 | 7.58E-06 | 2.70E-05 | -1.53E+00 | 1.27E-04 | 3.59E-04 | -1.15E+00 | 4.35E-04 | 1.60E-03 | -1.51E+00 | 6.51E-02 | 1.79E-01 | -1.15E+00 |
| AC098934.3 | 1.80E-09 | 9.38E-09 | -1.20E+00 | 1.16E-50 | 5.01E-50 | -1.85E+00 | 8.25E-05 | 3.51E-04 | -1.33E+00 | 2.52E-25 | 9.64E-25 | -2.03E+00 | 1.79E-06 | 6.38E-06 | -1.37E+00 | 3.48E-17 | 9.81E-17 | -1.89E+00 | 1.35E-06 | 4.94E-06 | -1.33E+00 | 1.21E-17 | 3.32E-17 | -1.98E+00 |
| SP100 | 1.80E-09 | 9.38E-09 | -1.09E+00 | 9.73E-06 | 4.20E-05 | 1.11E+00 | 2.82E-05 | 1.20E-04 | -1.28E+00 | 2.50E-06 | 9.56E-06 | -1.15E+00 | 9.04E-09 | 3.22E-08 | -1.46E+00 | 1.21E-05 | 3.42E-05 | -1.29E+00 | 4.49E-08 | 1.65E-07 | -1.46E+00 | 1.53E-10 | 4.22E-10 | -1.50E+00 |
| MNT | 1.80E-09 | 9.40E-09 | -1.39E+00 | 1.06E-10 | 4.57E-10 | -1.40E+00 | 3.78E-15 | 1.61E-14 | -1.88E+00 | 3.63E-13 | 1.39E-12 | -1.51E+00 | 6.14E-06 | 2.19E-05 | -1.71E+00 | 8.09E-07 | 2.28E-06 | -1.94E+00 | 4.55E-08 | 1.67E-07 | -1.71E+00 | 3.65E-02 | 1.00E-01 | -1.06E+00 |
| MSI2 | 1.81E-09 | 9.45E-09 | -1.42E+00 | 5.67E-03 | 2.45E-02 | -1.05E+00 | 1.80E-06 | 7.64E-06 | -1.48E+00 | 4.64E-03 | 1.77E-02 | -1.14E+00 | 2.16E-15 | 7.71E-15 | -1.72E+00 | 7.18E-14 | 2.03E-13 | -1.56E+00 | 1.08E-16 | 3.96E-16 | -1.62E+00 | 3.23E-07 | 8.87E-07 | -1.31E+00 |
| HSPB11 | 1.82E-09 | 9.50E-09 | -1.24E+00 | 4.48E-11 | 1.94E-10 | -1.17E+00 | 1.72E-01 | 7.33E-01 | 1.19E+00 | 6.79E-03 | 2.59E-02 | 1.09E+00 | 1.04E-05 | 3.70E-05 | -1.17E+00 | 2.48E-05 | 7.00E-05 | -1.11E+00 | 2.23E-01 | 8.18E-01 | 1.12E+00 | 1.39E-01 | 3.81E-01 | 1.08E+00 |
| FTH1P15 | 1.82E-09 | 9.51E-09 | -1.23E+00 | 1.88E-02 | 8.12E-02 | 1.38E+00 | 1.06E-03 | 4.51E-03 | -1.14E+00 | 2.03E-01 | 7.74E-01 | 1.33E+00 | 7.20E-07 | 2.57E-06 | -1.29E+00 | 1.96E-01 | 5.52E-01 | 1.24E+00 | 6.52E-07 | 2.39E-06 | -1.30E+00 | 2.89E-01 | 7.93E-01 | 1.30E+00 |
| TMEM50A | 1.84E-09 | 9.60E-09 | -1.13E+00 | 1.20E-08 | 5.18E-08 | 1.02E+00 | 8.89E-03 | 3.78E-02 | 1.02E+00 | 6.64E-04 | 2.53E-03 | 1.05E+00 | 4.58E-05 | 1.63E-04 | -1.10E+00 | 2.22E-01 | 6.26E-01 | 1.16E+00 | 1.59E-01 | 5.85E-01 | 1.15E+00 | 1.29E-01 | 3.55E-01 | 1.24E+00 |
| EXOSC7 | 1.84E-09 | 9.61E-09 | -1.24E+00 | 4.10E-04 | 1.77E-03 | 1.17E+00 | 2.30E-03 | 9.78E-03 | -1.02E+00 | 8.88E-02 | 3.39E-01 | 1.29E+00 | 1.34E-03 | 4.77E-03 | -1.02E+00 | 3.30E-01 | 9.29E-01 | 1.53E+00 | 9.20E-05 | 3.38E-04 | -1.08E+00 | 3.49E-01 | 9.58E-01 | 1.29E+00 |
| C4orf46 | 1.86E-09 | 9.70E-09 | -1.95E+00 | 1.08E-07 | 4.67E-07 | -1.51E+00 | 2.49E-06 | 1.06E-05 | -1.45E+00 | 5.30E-06 | 2.03E-05 | -1.20E+00 | 5.09E-03 | 1.81E-02 | -1.70E+00 | 5.92E-03 | 1.67E-02 | -1.68E+00 | 9.87E-11 | 3.62E-10 | -1.71E+00 | 3.24E-07 | 8.89E-07 | -1.36E+00 |
| CEP63 | 1.88E-09 | 9.82E-09 | -1.30E+00 | 1.17E-06 | 5.04E-06 | -1.02E+00 | 1.57E-06 | 6.65E-06 | -1.30E+00 | 4.30E-06 | 1.64E-05 | -1.15E+00 | 1.04E-10 | 3.71E-10 | -1.51E+00 | 2.06E-09 | 5.80E-09 | -1.27E+00 | 1.16E-04 | 4.26E-04 | -1.07E+00 | 4.91E-04 | 1.35E-03 | -1.06E+00 |
| CLK3 | 1.90E-09 | 9.92E-09 | -1.26E+00 | 4.93E-10 | 2.13E-09 | -1.09E+00 | 1.85E-06 | 7.84E-06 | -1.29E+00 | 2.24E-06 | 8.54E-06 | -1.13E+00 | 4.55E-06 | 1.62E-05 | -1.44E+00 | 2.54E-04 | 7.15E-04 | -1.19E+00 | 4.47E-05 | 1.64E-04 | -1.09E+00 | 1.43E-05 | 3.94E-05 | -1.28E+00 |
| ZNF738 | 1.90E-09 | 9.92E-09 | -1.85E+00 | 6.17E-03 | 2.67E-02 | 1.10E+00 | 1.79E-06 | 7.61E-06 | -1.72E+00 | 7.05E-02 | 2.69E-01 | 1.05E+00 | 1.80E-03 | 6.43E-03 | -1.07E+00 | 3.31E-01 | 9.32E-01 | 1.42E+00 | 1.62E-02 | 5.93E-02 | -1.12E+00 | 2.24E-01 | 6.16E-01 | 1.23E+00 |
| CXorf21 | 1.93E-09 | 1.01E-08 | -1.55E+00 | 6.99E-05 | 3.02E-04 | 1.02E+00 | 1.17E-15 | 4.97E-15 | -2.07E+00 | 8.73E-08 | 3.34E-07 | -1.24E+00 | 1.11E-04 | 3.96E-04 | -1.28E+00 | 4.55E-03 | 1.28E-02 | -1.20E+00 | 6.91E-02 | 2.54E-01 | -1.07E+00 | 2.11E-01 | 5.80E-01 | 1.05E+00 |
| RGS14 | 1.96E-09 | 1.02E-08 | -1.33E+00 | 1.43E-05 | 6.17E-05 | 1.06E+00 | 2.53E-08 | 1.07E-07 | -1.42E+00 | 2.46E-05 | 9.39E-05 | -1.01E+00 | 1.27E-14 | 4.52E-14 | -2.35E+00 | 2.72E-07 | 7.68E-07 | -1.32E+00 | 2.67E-04 | 9.79E-04 | -1.46E+00 | 2.52E-01 | 6.93E-01 | 1.13E+00 |
| CHSY1 | 1.96E-09 | 1.02E-08 | -1.71E+00 | 1.81E-04 | 7.81E-04 | -1.07E+00 | 2.39E-06 | 1.02E-05 | -1.54E+00 | 4.77E-06 | 1.82E-05 | -1.34E+00 | 6.80E-03 | 2.42E-02 | -1.08E+00 | 1.45E-02 | 4.10E-02 | -1.07E+00 | 5.49E-06 | 2.01E-05 | -1.46E+00 | 2.76E-05 | 7.58E-05 | -1.39E+00 |
| SUN2 | 1.96E-09 | 1.02E-08 | -1.81E+00 | 1.66E-03 | 7.16E-03 | 1.08E+00 | 4.95E-06 | 2.10E-05 | -1.51E+00 | 3.25E-04 | 1.24E-03 | -1.11E+00 | 2.53E-04 | 9.03E-04 | -1.88E+00 | 1.87E-02 | 5.28E-02 | -1.12E+00 | 1.87E-02 | 6.87E-02 | -1.16E+00 | 8.53E-02 | 2.34E-01 | -1.08E+00 |
| AIMP1 | 1.96E-09 | 1.02E-08 | -1.15E+00 | 2.38E-14 | 1.03E-13 | -1.16E+00 | 7.28E-04 | 3.09E-03 | -1.08E+00 | 2.16E-04 | 8.26E-04 | -1.05E+00 | 1.75E-05 | 6.25E-05 | -1.12E+00 | 3.53E-05 | 9.95E-05 | -1.12E+00 | 8.25E-03 | 3.03E-02 | 1.00E+00 | 1.89E-02 | 5.19E-02 | -1.01E+00 |
| RBFOX2 | 1.99E-09 | 1.04E-08 | -1.91E+00 | 1.14E-05 | 4.91E-05 | -1.23E+00 | 1.58E-04 | 6.73E-04 | -1.48E+00 | 5.47E-03 | 2.09E-02 | -1.06E+00 | 2.36E-03 | 8.40E-03 | -1.60E+00 | 8.82E-03 | 2.49E-02 | -1.37E+00 | 1.36E-05 | 4.98E-05 | -1.70E+00 | 1.22E-01 | 3.35E-01 | 1.09E+00 |
| FBXL5 | 1.99E-09 | 1.04E-08 | -1.30E+00 | 9.66E-04 | 4.17E-03 | 1.20E+00 | 3.39E-10 | 1.44E-09 | -1.56E+00 | 4.84E-07 | 1.85E-06 | -1.21E+00 | 7.31E-06 | 2.61E-05 | -1.15E+00 | 2.47E-05 | 6.96E-05 | -1.12E+00 | 5.03E-05 | 1.85E-04 | -1.25E+00 | 3.62E-02 | 9.95E-02 | 1.00E+00 |
| NDUFAF3 | 1.99E-09 | 1.04E-08 | -1.15E+00 | 7.36E-09 | 3.18E-08 | 1.01E+00 | 7.77E-02 | 3.30E-01 | 1.02E+00 | 2.42E-01 | 9.24E-01 | 1.25E+00 | 1.47E-09 | 5.24E-09 | -1.24E+00 | 3.72E-03 | 1.05E-02 | 1.07E+00 | 6.03E-02 | 2.21E-01 | 1.05E+00 | 2.86E-01 | 7.87E-01 | 1.04E+00 |
| AC138393.1 | 2.01E-09 | 1.05E-08 | -1.74E+00 | 6.18E-07 | 2.67E-06 | -1.26E+00 | 3.28E-04 | 1.39E-03 | -1.34E+00 | 3.75E-04 | 1.43E-03 | -1.21E+00 | 1.39E-05 | 4.94E-05 | -1.44E+00 | 1.50E-03 | 4.23E-03 | -1.17E+00 | 4.14E-11 | 1.52E-10 | -1.92E+00 | 1.27E-07 | 3.48E-07 | -1.48E+00 |
| IKZF5 | 2.01E-09 | 1.05E-08 | -1.72E+00 | 7.83E-06 | 3.38E-05 | -1.10E+00 | 6.69E-04 | 2.84E-03 | -1.27E+00 | 9.40E-04 | 3.59E-03 | -1.09E+00 | 4.30E-02 | 1.53E-01 | 1.05E+00 | 7.40E-02 | 2.09E-01 | 1.12E+00 | 6.24E-05 | 2.29E-04 | -1.22E+00 | 1.06E-04 | 2.92E-04 | -1.27E+00 |
| EGLN3 | 2.01E-09 | 1.05E-08 | -1.62E+00 | 1.00E-03 | 4.33E-03 | 1.01E+00 | 1.37E-01 | 5.82E-01 | -1.06E+00 | 9.73E-02 | 3.72E-01 | 1.77E+00 | 2.72E-01 | 9.69E-01 | 1.04E+00 | 1.79E-01 | 5.05E-01 | 1.08E+00 | 1.53E-02 | 5.61E-02 | 1.97E+00 | 3.36E-02 | 9.24E-02 | 1.58E+00 |
| SCAMP3 | 2.02E-09 | 1.05E-08 | -1.73E+00 | 1.45E-03 | 6.26E-03 | 1.09E+00 | 7.60E-08 | 3.23E-07 | -1.45E+00 | 2.28E-05 | 8.69E-05 | -1.04E+00 | 3.00E-13 | 1.07E-12 | -2.62E+00 | 1.08E-06 | 3.04E-06 | -1.41E+00 | 1.29E-06 | 4.74E-06 | -1.38E+00 | 5.85E-04 | 1.61E-03 | -1.15E+00 |
| SMG1P3 | 2.06E-09 | 1.07E-08 | -1.34E+00 | 3.45E-11 | 1.49E-10 | -1.33E+00 | 4.29E-09 | 1.82E-08 | -1.48E+00 | 1.51E-10 | 5.78E-10 | -1.41E+00 | 2.90E-03 | 1.03E-02 | 1.01E+00 | 1.12E-03 | 3.14E-03 | -1.01E+00 | 1.45E-03 | 5.32E-03 | 1.04E+00 | 3.58E-07 | 9.84E-07 | -1.25E+00 |
| FAM204BP | 2.06E-09 | 1.08E-08 | -1.56E+00 | 2.28E-09 | 9.86E-09 | -1.46E+00 | 8.09E-04 | 3.44E-03 | -1.17E+00 | 5.67E-04 | 2.17E-03 | -1.05E+00 | 1.43E-05 | 5.09E-05 | -1.31E+00 | 4.05E-06 | 1.14E-05 | -1.48E+00 | 2.67E-06 | 9.82E-06 | -1.36E+00 | 4.27E-05 | 1.17E-04 | -1.25E+00 |
| RGPD8 | 2.06E-09 | 1.08E-08 | -1.88E+00 | 5.04E-06 | 2.18E-05 | -1.23E+00 | 4.04E-02 | 1.72E-01 | -1.02E+00 | 4.51E-03 | 1.72E-02 | -1.12E+00 | 1.70E-01 | 6.07E-01 | 1.31E+00 | 5.77E-02 | 1.63E-01 | -1.03E+00 | 6.07E-02 | 2.23E-01 | -1.04E+00 | 2.00E-01 | 5.50E-01 | 1.16E+00 |
| LIN7C | 2.06E-09 | 1.08E-08 | -1.43E+00 | 6.06E-11 | 2.62E-10 | -1.32E+00 | 1.57E-08 | 6.67E-08 | -1.56E+00 | 2.42E-06 | 9.25E-06 | -1.17E+00 | 1.30E-07 | 4.63E-07 | -1.47E+00 | 1.50E-08 | 4.23E-08 | -1.44E+00 | 2.29E-09 | 8.42E-09 | -1.41E+00 | 1.12E-06 | 3.06E-06 | -1.21E+00 |
| SMIM10 | 2.08E-09 | 1.08E-08 | -1.49E+00 | 1.81E-05 | 7.83E-05 | -1.02E+00 | 5.08E-02 | 2.16E-01 | 1.03E+00 | 2.47E-01 | 9.45E-01 | 1.37E+00 | 1.18E-05 | 4.20E-05 | -1.53E+00 | 3.43E-01 | 9.66E-01 | 1.40E+00 | 2.20E-04 | 8.09E-04 | -1.33E+00 | 5.74E-02 | 1.58E-01 | -1.05E+00 |
| PDRG1 | 2.08E-09 | 1.09E-08 | -1.19E+00 | 7.23E-15 | 3.13E-14 | -1.38E+00 | 3.42E-04 | 1.45E-03 | -1.12E+00 | 1.56E-09 | 5.96E-09 | -1.33E+00 | 3.78E-08 | 1.35E-07 | -1.33E+00 | 8.56E-07 | 2.41E-06 | -1.23E+00 | 5.40E-04 | 1.98E-03 | -1.05E+00 | 2.39E-05 | 6.56E-05 | -1.01E+00 |
| WDR61 | 2.09E-09 | 1.09E-08 | -1.21E+00 | 1.22E-04 | 5.27E-04 | 1.16E+00 | 5.62E-05 | 2.39E-04 | -1.11E+00 | 2.55E-03 | 9.73E-03 | 1.11E+00 | 1.77E-08 | 6.32E-08 | -1.37E+00 | 2.82E-03 | 7.95E-03 | 1.03E+00 | 2.85E-05 | 1.05E-04 | -1.14E+00 | 4.06E-02 | 1.12E-01 | 1.11E+00 |
| EXOC3 | 2.09E-09 | 1.09E-08 | -1.46E+00 | 3.72E-05 | 1.61E-04 | -1.03E+00 | 8.26E-05 | 3.51E-04 | -1.34E+00 | 2.05E-02 | 7.85E-02 | 1.09E+00 | 6.92E-05 | 2.47E-04 | -1.34E+00 | 3.08E-03 | 8.69E-03 | -1.11E+00 | 3.86E-04 | 1.42E-03 | -1.05E+00 | 1.73E-02 | 4.75E-02 | 1.05E+00 |
| MUT | 2.10E-09 | 1.09E-08 | -1.76E+00 | 1.98E-05 | 8.56E-05 | -1.08E+00 | 3.25E-08 | 1.38E-07 | -1.70E+00 | 1.73E-03 | 6.62E-03 | -1.03E+00 | 7.18E-07 | 2.56E-06 | -1.80E+00 | 6.13E-03 | 1.73E-02 | -1.05E+00 | 2.49E-22 | 9.13E-22 | -2.01E+00 | 2.45E-09 | 6.73E-09 | -1.31E+00 |
| MARK2 | 2.10E-09 | 1.09E-08 | -1.55E+00 | 1.55E-04 | 6.71E-04 | 1.02E+00 | 5.20E-10 | 2.21E-09 | -1.61E+00 | 5.84E-08 | 2.23E-07 | -1.22E+00 | 1.49E-09 | 5.31E-09 | -1.53E+00 | 8.22E-08 | 2.32E-07 | -1.34E+00 | 9.71E-08 | 3.56E-07 | -1.43E+00 | 4.95E-05 | 1.36E-04 | -1.23E+00 |
| NAP1L4P1 | 2.12E-09 | 1.10E-08 | -1.68E+00 | 1.44E-14 | 6.22E-14 | -2.53E+00 | 9.41E-07 | 4.00E-06 | -1.46E+00 | 1.07E-09 | 4.09E-09 | -1.70E+00 | 1.86E-05 | 6.64E-05 | -1.30E+00 | 5.58E-09 | 1.57E-08 | -2.14E+00 | 5.43E-07 | 1.99E-06 | -1.30E+00 | 1.12E-10 | 3.08E-10 | -1.68E+00 |
| RALGDS | 2.12E-09 | 1.10E-08 | -1.52E+00 | 9.72E-08 | 4.20E-07 | -1.25E+00 | 4.81E-09 | 2.04E-08 | -1.65E+00 | 1.76E-07 | 6.74E-07 | -1.30E+00 | 2.62E-04 | 9.33E-04 | -1.29E+00 | 5.40E-04 | 1.52E-03 | -1.33E+00 | 2.04E-01 | 7.49E-01 | 1.41E+00 | 1.67E-01 | 4.59E-01 | 1.54E+00 |
| TMEM165 | 2.12E-09 | 1.10E-08 | -1.10E+00 | 1.52E-02 | 6.57E-02 | 1.28E+00 | 2.86E-06 | 1.21E-05 | -1.23E+00 | 1.09E-01 | 4.15E-01 | 1.20E+00 | 1.76E-02 | 6.26E-02 | 1.18E+00 | 3.03E-01 | 8.54E-01 | 1.47E+00 | 9.06E-04 | 3.33E-03 | -1.18E+00 | 1.49E-01 | 4.10E-01 | 1.09E+00 |
| MPHOSPH8 | 2.12E-09 | 1.10E-08 | -1.13E+00 | 7.89E-02 | 3.41E-01 | 1.36E+00 | 4.71E-06 | 2.00E-05 | -1.28E+00 | 6.39E-02 | 2.44E-01 | 1.17E+00 | 6.35E-07 | 2.27E-06 | -1.25E+00 | 9.11E-02 | 2.57E-01 | 1.15E+00 | 3.40E-03 | 1.25E-02 | -1.06E+00 | 2.24E-01 | 6.15E-01 | 1.15E+00 |
| ZNF814 | 2.14E-09 | 1.11E-08 | -1.22E+00 | 4.44E-07 | 1.92E-06 | 1.03E+00 | 6.06E-06 | 2.57E-05 | -1.32E+00 | 3.17E-06 | 1.21E-05 | -1.16E+00 | 3.23E-03 | 1.15E-02 | -1.02E+00 | 1.67E-02 | 4.70E-02 | 1.11E+00 | 3.18E-04 | 1.17E-03 | -1.21E+00 | 5.88E-03 | 1.61E-02 | -1.03E+00 |
| INSIG2 | 2.14E-09 | 1.11E-08 | -1.49E+00 | 7.91E-06 | 3.42E-05 | -1.01E+00 | 9.06E-06 | 3.85E-05 | -1.53E+00 | 1.91E-03 | 7.30E-03 | -1.01E+00 | 1.30E-05 | 4.64E-05 | -1.19E+00 | 6.59E-04 | 1.86E-03 | 1.05E+00 | 5.93E-09 | 2.18E-08 | -1.45E+00 | 8.88E-04 | 2.44E-03 | -1.14E+00 |
| HCFC1R1 | 2.14E-09 | 1.11E-08 | -1.32E+00 | 1.13E-08 | 4.88E-08 | -1.17E+00 | 1.11E-05 | 4.71E-05 | -1.30E+00 | 7.93E-05 | 3.03E-04 | -1.10E+00 | 3.98E-03 | 1.42E-02 | -1.01E+00 | 6.68E-03 | 1.88E-02 | 1.02E+00 | 3.12E-06 | 1.14E-05 | -1.33E+00 | 2.75E-06 | 7.57E-06 | -1.34E+00 |
| A2M | 2.18E-09 | 1.13E-08 | -1.51E+00 | 2.07E-06 | 8.94E-06 | -1.11E+00 | 4.65E-06 | 1.98E-05 | -1.61E+00 | 1.08E-04 | 4.14E-04 | -1.24E+00 | 1.42E-03 | 5.05E-03 | -1.52E+00 | 1.15E-02 | 3.23E-02 | -1.28E+00 | 3.51E-02 | 1.29E-01 | -1.51E+00 | 3.10E-01 | 8.52E-01 | 1.10E+00 |
| COQ10B | 2.18E-09 | 1.14E-08 | -1.18E+00 | 5.75E-23 | 2.48E-22 | -1.69E+00 | 5.77E-05 | 2.45E-04 | -1.23E+00 | 7.96E-15 | 3.04E-14 | -1.82E+00 | 2.22E-05 | 7.92E-05 | -1.08E+00 | 1.93E-20 | 5.44E-20 | -2.16E+00 | 1.90E-06 | 6.97E-06 | -1.23E+00 | 6.50E-14 | 1.79E-13 | -1.76E+00 |
| JMJD1C | 2.21E-09 | 1.15E-08 | -1.14E+00 | 9.89E-16 | 4.27E-15 | -1.17E+00 | 1.28E-02 | 5.43E-02 | -1.02E+00 | 3.06E-09 | 1.17E-08 | -1.32E+00 | 1.49E-04 | 5.31E-04 | -1.05E+00 | 4.16E-06 | 1.17E-05 | -1.21E+00 | 4.11E-07 | 1.51E-06 | -1.29E+00 | 3.39E-09 | 9.31E-09 | -1.46E+00 |
| ITPKB | 2.22E-09 | 1.16E-08 | -1.62E+00 | 5.65E-06 | 2.44E-05 | -1.18E+00 | 1.28E-06 | 5.44E-06 | -1.81E+00 | 1.23E-03 | 4.69E-03 | -1.18E+00 | 4.49E-03 | 1.60E-02 | -1.68E+00 | 1.92E-01 | 5.41E-01 | 1.03E+00 | 1.37E-02 | 5.04E-02 | -1.02E+00 | 5.81E-02 | 1.60E-01 | 1.01E+00 |
| AK3 | 2.23E-09 | 1.16E-08 | -1.22E+00 | 5.19E-10 | 2.24E-09 | -1.10E+00 | 5.61E-06 | 2.38E-05 | -1.23E+00 | 1.21E-05 | 4.63E-05 | -1.05E+00 | 2.22E-03 | 7.92E-03 | -1.04E+00 | 1.78E-02 | 5.02E-02 | 1.03E+00 | 2.73E-05 | 1.00E-04 | -1.11E+00 | 2.02E-03 | 5.54E-03 | -1.05E+00 |
| SRP54 | 2.24E-09 | 1.17E-08 | -1.32E+00 | 3.34E-05 | 1.45E-04 | 1.03E+00 | 6.00E-10 | 2.55E-09 | -1.52E+00 | 1.97E-05 | 7.53E-05 | -1.10E+00 | 1.64E-08 | 5.84E-08 | -1.34E+00 | 7.27E-03 | 2.05E-02 | 1.10E+00 | 5.94E-07 | 2.18E-06 | -1.25E+00 | 6.07E-03 | 1.67E-02 | -1.05E+00 |
| AP2B1 | 2.25E-09 | 1.17E-08 | -1.39E+00 | 3.08E-04 | 1.33E-03 | 1.09E+00 | 2.22E-05 | 9.43E-05 | -1.25E+00 | 1.19E-03 | 4.53E-03 | 1.01E+00 | 1.67E-04 | 5.96E-04 | -1.17E+00 | 1.37E-04 | 3.87E-04 | -1.11E+00 | 6.80E-08 | 2.50E-07 | -1.32E+00 | 1.25E-05 | 3.42E-05 | -1.18E+00 |
| NDUFAB1 | 2.26E-09 | 1.18E-08 | -1.17E+00 | 4.59E-05 | 1.98E-04 | 1.11E+00 | 2.10E-02 | 8.94E-02 | -1.02E+00 | 1.09E-02 | 4.16E-02 | 1.03E+00 | 2.28E-08 | 8.13E-08 | -1.24E+00 | 7.18E-03 | 2.02E-02 | 1.06E+00 | 1.46E-02 | 5.37E-02 | -1.05E+00 | 3.24E-01 | 8.89E-01 | 1.13E+00 |
| DERL2 | 2.27E-09 | 1.18E-08 | -1.30E+00 | 5.81E-06 | 2.51E-05 | 1.00E+00 | 1.96E-03 | 8.31E-03 | -1.06E+00 | 7.25E-05 | 2.77E-04 | -1.03E+00 | 4.41E-08 | 1.57E-07 | -1.26E+00 | 6.42E-05 | 1.81E-04 | -1.06E+00 | 5.94E-03 | 2.18E-02 | -1.07E+00 | 4.65E-02 | 1.28E-01 | -1.03E+00 |
| UAP1 | 2.29E-09 | 1.19E-08 | -1.49E+00 | 9.79E-06 | 4.23E-05 | -1.07E+00 | 1.20E-06 | 5.11E-06 | -1.27E+00 | 1.84E-06 | 7.01E-06 | -1.08E+00 | 7.45E-07 | 2.66E-06 | -1.86E+00 | 1.50E-02 | 4.23E-02 | -1.04E+00 | 7.67E-07 | 2.81E-06 | -1.35E+00 | 7.12E-03 | 1.96E-02 | -1.02E+00 |
| UBE2G1 | 2.29E-09 | 1.19E-08 | -1.34E+00 | 5.52E-09 | 2.38E-08 | -1.23E+00 | 1.27E-06 | 5.41E-06 | -1.52E+00 | 3.62E-06 | 1.38E-05 | -1.24E+00 | 4.55E-04 | 1.62E-03 | -1.27E+00 | 2.00E-03 | 5.62E-03 | -1.15E+00 | 1.45E-07 | 5.31E-07 | -1.44E+00 | 9.62E-04 | 2.64E-03 | -1.10E+00 |
| CLINT1 | 2.32E-09 | 1.21E-08 | -1.20E+00 | 4.85E-12 | 2.10E-11 | -1.19E+00 | 3.35E-05 | 1.43E-04 | -1.30E+00 | 6.73E-08 | 2.57E-07 | -1.28E+00 | 1.97E-11 | 7.04E-11 | -1.47E+00 | 3.28E-13 | 9.26E-13 | -1.51E+00 | 3.13E-07 | 1.15E-06 | -1.35E+00 | 1.85E-05 | 5.07E-05 | -1.25E+00 |
| DNAJB11 | 2.33E-09 | 1.21E-08 | -1.28E+00 | 1.46E-13 | 6.29E-13 | -1.36E+00 | 2.10E-02 | 8.92E-02 | 1.06E+00 | 1.94E-04 | 7.41E-04 | -1.03E+00 | 7.42E-09 | 2.64E-08 | -1.44E+00 | 5.86E-09 | 1.65E-08 | -1.42E+00 | 1.96E-04 | 7.18E-04 | -1.26E+00 | 2.54E-04 | 6.97E-04 | -1.31E+00 |
| SLC39A4 | 2.33E-09 | 1.21E-08 | -1.61E+00 | 6.27E-06 | 2.71E-05 | -1.09E+00 | 1.62E-07 | 6.90E-07 | -1.42E+00 | 3.23E-04 | 1.24E-03 | -1.03E+00 | 3.80E-10 | 1.35E-09 | -1.39E+00 | 5.50E-06 | 1.55E-05 | -1.03E+00 | 1.52E-04 | 5.58E-04 | -1.12E+00 | 8.28E-02 | 2.28E-01 | 1.06E+00 |
| P3H1 | 2.37E-09 | 1.23E-08 | -1.59E+00 | 1.29E-05 | 5.60E-05 | -1.09E+00 | 4.65E-06 | 1.98E-05 | -1.23E+00 | 4.10E-06 | 1.57E-05 | -1.11E+00 | 3.49E-09 | 1.25E-08 | -1.60E+00 | 3.83E-06 | 1.08E-05 | -1.25E+00 | 1.39E-08 | 5.11E-08 | -1.51E+00 | 1.44E-03 | 3.96E-03 | -1.16E+00 |
| SCAF4 | 2.41E-09 | 1.26E-08 | -1.28E+00 | 8.27E-09 | 3.57E-08 | -1.20E+00 | 6.91E-07 | 2.94E-06 | -1.26E+00 | 4.50E-05 | 1.72E-04 | -1.06E+00 | 6.00E-03 | 2.14E-02 | 1.02E+00 | 7.14E-03 | 2.01E-02 | -1.01E+00 | 4.64E-05 | 1.70E-04 | -1.13E+00 | 2.96E-04 | 8.13E-04 | -1.06E+00 |
| XRCC1 | 2.45E-09 | 1.28E-08 | -1.52E+00 | 1.01E-07 | 4.37E-07 | -1.26E+00 | 1.20E-18 | 5.08E-18 | -2.38E+00 | 7.18E-18 | 2.74E-17 | -1.95E+00 | 4.71E-06 | 1.68E-05 | -1.47E+00 | 3.40E-04 | 9.57E-04 | -1.16E+00 | 1.40E-11 | 5.13E-11 | -1.88E+00 | 1.50E-06 | 4.12E-06 | -1.49E+00 |
| LINC00969 | 2.46E-09 | 1.28E-08 | -1.29E+00 | 6.75E-04 | 2.92E-03 | 1.14E+00 | 2.15E-07 | 9.15E-07 | -1.30E+00 | 1.11E-04 | 4.24E-04 | -1.03E+00 | 7.39E-08 | 2.63E-07 | -1.28E+00 | 2.50E-04 | 7.04E-04 | -1.03E+00 | 2.56E-06 | 9.39E-06 | -1.23E+00 | 1.67E-04 | 4.58E-04 | -1.17E+00 |
| MTAP | 2.47E-09 | 1.29E-08 | -1.49E+00 | 1.71E-04 | 7.38E-04 | 1.05E+00 | 1.43E-06 | 6.09E-06 | -1.34E+00 | 3.89E-03 | 1.48E-02 | 1.08E+00 | 3.97E-07 | 1.41E-06 | -1.43E+00 | 4.03E-03 | 1.14E-02 | -1.04E+00 | 1.47E-12 | 5.39E-12 | -1.55E+00 | 1.56E-02 | 4.30E-02 | 1.06E+00 |
| M6PR | 2.48E-09 | 1.29E-08 | -1.49E+00 | 1.78E-07 | 7.70E-07 | -1.16E+00 | 1.56E-12 | 6.62E-12 | -1.56E+00 | 5.16E-08 | 1.97E-07 | -1.21E+00 | 1.19E-07 | 4.26E-07 | -1.38E+00 | 7.77E-04 | 2.19E-03 | -1.07E+00 | 9.46E-11 | 3.47E-10 | -1.61E+00 | 2.29E-04 | 6.28E-04 | -1.18E+00 |
| B3GNT5 | 2.48E-09 | 1.29E-08 | -1.64E+00 | 5.97E-09 | 2.58E-08 | -1.39E+00 | 6.05E-06 | 2.57E-05 | -1.44E+00 | 1.92E-05 | 7.34E-05 | -1.23E+00 | 1.66E-03 | 5.91E-03 | -1.48E+00 | 2.11E-03 | 5.95E-03 | -1.50E+00 | 8.06E-07 | 2.96E-06 | -2.53E+00 | 5.39E-04 | 1.48E-03 | -1.52E+00 |
| AC008771.1 | 2.50E-09 | 1.30E-08 | -2.14E+00 | 6.39E-05 | 2.76E-04 | -1.12E+00 | 9.53E-02 | 4.05E-01 | -1.04E+00 | 1.89E-01 | 7.21E-01 | 1.20E+00 | 5.27E-04 | 1.88E-03 | -2.56E+00 | 8.45E-03 | 2.38E-02 | -1.52E+00 | 1.02E-03 | 3.76E-03 | -1.50E+00 | 2.90E-02 | 7.97E-02 | -1.10E+00 |
| DDB1 | 2.51E-09 | 1.31E-08 | -1.23E+00 | 2.92E-06 | 1.26E-05 | 1.00E+00 | 1.27E-04 | 5.42E-04 | -1.17E+00 | 5.64E-03 | 2.15E-02 | 1.01E+00 | 1.05E-05 | 3.73E-05 | -1.19E+00 | 1.48E-02 | 4.16E-02 | 1.08E+00 | 1.08E-04 | 3.98E-04 | -1.18E+00 | 2.57E-02 | 7.06E-02 | -1.03E+00 |
| ATP1B1 | 2.52E-09 | 1.31E-08 | -1.22E+00 | 5.06E-06 | 2.19E-05 | 1.07E+00 | 3.68E-05 | 1.56E-04 | -1.22E+00 | 3.72E-03 | 1.42E-02 | 1.04E+00 | 2.05E-01 | 7.30E-01 | 1.51E+00 | 2.12E-01 | 5.98E-01 | 1.40E+00 | 2.53E-01 | 9.28E-01 | 1.26E+00 | 9.69E-02 | 2.66E-01 | 1.21E+00 |
| NUP160 | 2.52E-09 | 1.31E-08 | -1.45E+00 | 1.50E-04 | 6.47E-04 | 1.05E+00 | 2.38E-02 | 1.01E-01 | 1.02E+00 | 1.78E-01 | 6.80E-01 | 1.32E+00 | 2.70E-03 | 9.62E-03 | -1.19E+00 | 1.50E-01 | 4.23E-01 | 1.20E+00 | 6.26E-06 | 2.30E-05 | -1.21E+00 | 8.70E-02 | 2.39E-01 | 1.16E+00 |
| SMAD2 | 2.52E-09 | 1.32E-08 | -1.30E+00 | 4.74E-03 | 2.05E-02 | 1.19E+00 | 6.95E-09 | 2.95E-08 | -1.47E+00 | 8.66E-03 | 3.31E-02 | 1.10E+00 | 4.20E-05 | 1.50E-04 | -1.27E+00 | 5.40E-02 | 1.52E-01 | 1.06E+00 | 1.36E-06 | 5.00E-06 | -1.36E+00 | 2.69E-03 | 7.40E-03 | -1.12E+00 |
| LCOR | 2.54E-09 | 1.32E-08 | -1.28E+00 | 3.31E-06 | 1.43E-05 | 1.01E+00 | 7.78E-04 | 3.31E-03 | -1.16E+00 | 1.22E-03 | 4.65E-03 | -1.01E+00 | 3.72E-04 | 1.33E-03 | -1.18E+00 | 1.78E-04 | 5.02E-04 | -1.19E+00 | 9.29E-10 | 3.41E-09 | -1.27E+00 | 1.32E-06 | 3.63E-06 | -1.15E+00 |
| MGST1 | 2.55E-09 | 1.33E-08 | -1.25E+00 | 1.32E-02 | 5.71E-02 | 1.07E+00 | 9.48E-05 | 4.03E-04 | -1.34E+00 | 2.34E-01 | 8.95E-01 | 1.08E+00 | 3.07E-05 | 1.09E-04 | -1.30E+00 | 3.37E-02 | 9.51E-02 | 1.03E+00 | 1.43E-03 | 5.25E-03 | -1.04E+00 | 7.12E-03 | 1.95E-02 | 1.46E+00 |
| POT1 | 2.56E-09 | 1.34E-08 | -1.64E+00 | 1.44E-04 | 6.24E-04 | 1.07E+00 | 1.12E-02 | 4.75E-02 | -1.14E+00 | 2.46E-02 | 9.39E-02 | 1.03E+00 | 2.52E-03 | 9.00E-03 | -1.14E+00 | 2.72E-02 | 7.68E-02 | 1.12E+00 | 3.03E-08 | 1.11E-07 | -1.32E+00 | 6.57E-06 | 1.81E-05 | -1.20E+00 |
| DYNC2LI1 | 2.61E-09 | 1.36E-08 | -1.38E+00 | 5.06E-08 | 2.19E-07 | -1.18E+00 | 1.11E-04 | 4.73E-04 | -1.27E+00 | 4.50E-04 | 1.72E-03 | -1.08E+00 | 4.31E-04 | 1.54E-03 | -1.19E+00 | 3.56E-03 | 1.00E-02 | 1.01E+00 | 5.97E-04 | 2.19E-03 | -1.14E+00 | 4.19E-03 | 1.15E-02 | 1.00E+00 |
| COG7 | 2.62E-09 | 1.36E-08 | -1.96E+00 | 2.57E-05 | 1.11E-04 | -1.17E+00 | 1.53E-06 | 6.52E-06 | -1.63E+00 | 4.25E-03 | 1.62E-02 | 1.05E+00 | 2.45E-03 | 8.74E-03 | -1.43E+00 | 2.06E-02 | 5.80E-02 | -1.10E+00 | 1.82E-05 | 6.69E-05 | -1.41E+00 | 1.52E-01 | 4.17E-01 | 1.15E+00 |
| ZNF85 | 2.64E-09 | 1.38E-08 | -1.40E+00 | 2.64E-09 | 1.14E-08 | -1.31E+00 | 3.06E-14 | 1.30E-13 | -1.72E+00 | 1.00E-13 | 3.83E-13 | -1.56E+00 | 1.67E-07 | 5.97E-07 | -1.64E+00 | 1.65E-06 | 4.65E-06 | -1.40E+00 | 7.45E-05 | 2.73E-04 | -1.06E+00 | 5.44E-06 | 1.50E-05 | -1.24E+00 |
| GOLGA8A | 2.65E-09 | 1.38E-08 | -1.23E+00 | 8.26E-03 | 3.57E-02 | 1.13E+00 | 3.42E-04 | 1.45E-03 | -1.26E+00 | 2.26E-01 | 8.62E-01 | 1.12E+00 | 2.55E-03 | 9.09E-03 | -1.01E+00 | 1.64E-01 | 4.62E-01 | 1.20E+00 | 7.31E-02 | 2.68E-01 | 1.17E+00 | 2.03E-01 | 5.57E-01 | 1.25E+00 |
| HPF1 | 2.66E-09 | 1.38E-08 | -1.34E+00 | 3.13E-04 | 1.35E-03 | 1.12E+00 | 8.81E-08 | 3.74E-07 | -1.30E+00 | 3.04E-07 | 1.16E-06 | -1.08E+00 | 2.67E-06 | 9.52E-06 | -1.52E+00 | 3.33E-02 | 9.38E-02 | 1.04E+00 | 1.80E-07 | 6.61E-07 | -1.35E+00 | 2.27E-03 | 6.22E-03 | -1.12E+00 |
| METTL2B | 2.66E-09 | 1.39E-08 | -1.76E+00 | 1.13E-05 | 4.88E-05 | -1.11E+00 | 7.19E-06 | 3.06E-05 | -1.37E+00 | 9.08E-04 | 3.47E-03 | 1.02E+00 | 1.37E-04 | 4.90E-04 | -1.41E+00 | 3.79E-03 | 1.07E-02 | -1.10E+00 | 4.52E-06 | 1.66E-05 | -1.27E+00 | 4.39E-03 | 1.21E-02 | -1.02E+00 |
| TMEM126A | 2.66E-09 | 1.39E-08 | -1.17E+00 | 1.30E-08 | 5.64E-08 | -1.08E+00 | 2.52E-04 | 1.07E-03 | -1.19E+00 | 1.41E-04 | 5.40E-04 | -1.07E+00 | 2.97E-07 | 1.06E-06 | -1.16E+00 | 5.04E-06 | 1.42E-05 | -1.08E+00 | 1.32E-03 | 4.85E-03 | 1.00E+00 | 4.28E-03 | 1.18E-02 | 1.01E+00 |
| ABRAXAS1 | 2.69E-09 | 1.40E-08 | -1.31E+00 | 1.26E-09 | 5.43E-09 | -1.20E+00 | 2.90E-07 | 1.23E-06 | -1.44E+00 | 1.32E-06 | 5.05E-06 | -1.19E+00 | 8.99E-07 | 3.20E-06 | -1.21E+00 | 4.34E-07 | 1.22E-06 | -1.23E+00 | 4.99E-04 | 1.83E-03 | -1.11E+00 | 4.18E-05 | 1.15E-04 | -1.25E+00 |
| MRNIP | 2.69E-09 | 1.40E-08 | -1.60E+00 | 8.95E-07 | 3.87E-06 | -1.28E+00 | 9.68E-06 | 4.11E-05 | -1.78E+00 | 1.21E-03 | 4.62E-03 | -1.22E+00 | 3.53E-03 | 1.26E-02 | -1.43E+00 | 2.40E-02 | 6.77E-02 | -1.10E+00 | 2.95E-07 | 1.08E-06 | -2.00E+00 | 6.34E-05 | 1.74E-04 | -1.59E+00 |
| GPAA1 | 2.71E-09 | 1.41E-08 | -1.42E+00 | 2.05E-07 | 8.87E-07 | -1.16E+00 | 6.95E-02 | 2.95E-01 | 1.07E+00 | 1.88E-03 | 7.19E-03 | 1.03E+00 | 3.20E-02 | 1.14E-01 | 1.14E+00 | 4.51E-02 | 1.27E-01 | 1.23E+00 | 1.71E-02 | 6.26E-02 | 1.06E+00 | 1.29E-01 | 3.54E-01 | 1.15E+00 |
| CAPZB | 2.72E-09 | 1.42E-08 | -1.09E+00 | 2.60E-04 | 1.12E-03 | 1.14E+00 | 9.81E-03 | 4.17E-02 | -1.08E+00 | 1.07E-01 | 4.09E-01 | 1.12E+00 | 6.50E-12 | 2.32E-11 | -1.34E+00 | 6.96E-06 | 1.96E-05 | -1.11E+00 | 1.37E-03 | 5.04E-03 | -1.08E+00 | 6.31E-02 | 1.73E-01 | 1.02E+00 |
| RPS3AP37 | 2.73E-09 | 1.42E-08 | -1.71E+00 | 7.09E-12 | 3.06E-11 | -2.08E+00 | 3.44E-04 | 1.46E-03 | -1.47E+00 | 1.30E-05 | 4.97E-05 | -1.83E+00 | 4.81E-04 | 1.72E-03 | -1.66E+00 | 2.37E-05 | 6.69E-05 | -2.11E+00 | 3.00E-07 | 1.10E-06 | -1.96E+00 | 4.12E-06 | 1.13E-05 | -1.79E+00 |
| RSAD1 | 2.73E-09 | 1.42E-08 | -1.61E+00 | 1.68E-05 | 7.24E-05 | -1.10E+00 | 1.47E-02 | 6.24E-02 | -1.24E+00 | 1.33E-01 | 5.09E-01 | 1.13E+00 | 4.29E-04 | 1.53E-03 | -1.29E+00 | 2.46E-02 | 6.92E-02 | 1.00E+00 | 8.66E-05 | 3.18E-04 | -1.31E+00 | 6.97E-02 | 1.92E-01 | 1.06E+00 |
| AGPAT5 | 2.73E-09 | 1.43E-08 | -1.50E+00 | 1.08E-04 | 4.66E-04 | 1.02E+00 | 3.65E-10 | 1.55E-09 | -1.62E+00 | 6.07E-04 | 2.32E-03 | -1.01E+00 | 2.97E-23 | 1.06E-22 | -2.38E+00 | 1.93E-12 | 5.45E-12 | -1.49E+00 | 4.94E-14 | 1.81E-13 | -1.63E+00 | 6.17E-05 | 1.69E-04 | -1.11E+00 |
| RPL23AP25 | 2.74E-09 | 1.43E-08 | -2.01E+00 | 3.77E-07 | 1.63E-06 | -1.49E+00 | 1.66E-12 | 7.06E-12 | -2.28E+00 | 2.88E-09 | 1.10E-08 | -1.72E+00 | 1.66E-05 | 5.91E-05 | -2.25E+00 | 2.57E-04 | 7.24E-04 | -1.76E+00 | 2.75E-03 | 1.01E-02 | -1.26E+00 | 3.54E-02 | 9.71E-02 | -1.05E+00 |
| COPS4 | 2.77E-09 | 1.44E-08 | -1.24E+00 | 3.31E-04 | 1.43E-03 | 1.19E+00 | 8.73E-07 | 3.71E-06 | -1.28E+00 | 3.04E-05 | 1.16E-04 | -1.08E+00 | 1.72E-07 | 6.13E-07 | -1.25E+00 | 4.79E-04 | 1.35E-03 | 1.06E+00 | 2.55E-06 | 9.35E-06 | -1.15E+00 | 1.86E-03 | 5.11E-03 | 1.02E+00 |
| EMC1 | 2.80E-09 | 1.46E-08 | -1.48E+00 | 4.82E-07 | 2.08E-06 | -1.11E+00 | 5.50E-09 | 2.34E-08 | -1.50E+00 | 8.00E-08 | 3.06E-07 | -1.28E+00 | 3.44E-03 | 1.23E-02 | -1.09E+00 | 1.39E-02 | 3.93E-02 | 1.08E+00 | 2.81E-03 | 1.03E-02 | -1.06E+00 | 1.34E-01 | 3.67E-01 | 1.19E+00 |
| STXBP3 | 2.80E-09 | 1.46E-08 | -1.25E+00 | 2.38E-03 | 1.03E-02 | 1.12E+00 | 3.26E-08 | 1.39E-07 | -1.35E+00 | 9.99E-06 | 3.82E-05 | -1.06E+00 | 1.08E-10 | 3.85E-10 | -1.56E+00 | 7.94E-07 | 2.24E-06 | -1.25E+00 | 1.50E-05 | 5.50E-05 | -1.12E+00 | 6.96E-04 | 1.91E-03 | -1.09E+00 |
| GGPS1 | 2.80E-09 | 1.46E-08 | -1.36E+00 | 8.89E-03 | 3.84E-02 | 1.23E+00 | 1.02E-07 | 4.35E-07 | -1.29E+00 | 9.86E-04 | 3.77E-03 | 1.04E+00 | 1.96E-03 | 7.00E-03 | -1.10E+00 | 2.29E-01 | 6.45E-01 | 1.32E+00 | 1.90E-03 | 6.96E-03 | -1.21E+00 | 1.04E-01 | 2.85E-01 | 1.11E+00 |
| RPL23A | 2.80E-09 | 1.46E-08 | -1.13E+00 | 1.77E-11 | 7.64E-11 | -1.07E+00 | 6.33E-04 | 2.69E-03 | -1.16E+00 | 1.85E-05 | 7.07E-05 | -1.13E+00 | 1.55E-06 | 5.52E-06 | -1.20E+00 | 9.74E-06 | 2.75E-05 | -1.19E+00 | 4.17E-07 | 1.53E-06 | -1.24E+00 | 2.94E-04 | 8.08E-04 | -1.12E+00 |
| DRAM1 | 2.81E-09 | 1.47E-08 | -1.41E+00 | 2.18E-05 | 9.40E-05 | 1.02E+00 | 2.49E-09 | 1.06E-08 | -1.56E+00 | 4.38E-09 | 1.67E-08 | -1.30E+00 | 1.00E-02 | 3.58E-02 | -1.37E+00 | 4.13E-02 | 1.16E-01 | -1.13E+00 | 3.15E-07 | 1.16E-06 | -1.75E+00 | 9.98E-05 | 2.74E-04 | -1.42E+00 |
| LAGE3P1 | 2.82E-09 | 1.47E-08 | -2.13E+00 | 2.70E-08 | 1.17E-07 | -1.77E+00 | 4.03E-11 | 1.71E-10 | -2.13E+00 | 4.97E-11 | 1.90E-10 | -1.96E+00 | 5.69E-07 | 2.03E-06 | -2.38E+00 | 2.89E-04 | 8.15E-04 | -1.44E+00 | 1.65E-10 | 6.05E-10 | -1.90E+00 | 3.10E-08 | 8.53E-08 | -1.69E+00 |
| BCL3 | 2.82E-09 | 1.47E-08 | -1.54E+00 | 2.71E-20 | 1.17E-19 | -3.62E+00 | 3.25E-07 | 1.38E-06 | -1.81E+00 | 4.18E-16 | 1.60E-15 | -4.23E+00 | 9.21E-07 | 3.28E-06 | -2.38E+00 | 7.89E-08 | 2.22E-07 | -3.13E+00 | 5.13E-02 | 1.88E-01 | -1.25E+00 | 2.68E-02 | 7.35E-02 | -1.99E+00 |
| PRPSAP1 | 2.87E-09 | 1.49E-08 | -1.30E+00 | 3.01E-04 | 1.30E-03 | 1.10E+00 | 8.90E-04 | 3.78E-03 | -1.10E+00 | 2.05E-03 | 7.81E-03 | 1.05E+00 | 2.30E-09 | 8.19E-09 | -1.48E+00 | 6.86E-06 | 1.93E-05 | -1.13E+00 | 3.39E-03 | 1.25E-02 | -1.10E+00 | 3.04E-01 | 8.35E-01 | 1.18E+00 |
| AC092329.4 | 2.88E-09 | 1.50E-08 | -1.54E+00 | 2.81E-09 | 1.21E-08 | -1.44E+00 | 2.34E-03 | 9.93E-03 | -1.27E+00 | 1.38E-02 | 5.29E-02 | 1.03E+00 | 4.61E-02 | 1.64E-01 | -1.03E+00 | 2.41E-02 | 6.78E-02 | -1.04E+00 | 1.68E-01 | 6.17E-01 | 1.22E+00 | 9.43E-03 | 2.59E-02 | -1.20E+00 |
| ALDH1B1 | 2.88E-09 | 1.50E-08 | -2.16E+00 | 4.56E-05 | 1.97E-04 | -1.19E+00 | 1.07E-03 | 4.55E-03 | -1.24E+00 | 5.52E-03 | 2.11E-02 | -1.05E+00 | 6.33E-06 | 2.26E-05 | -2.51E+00 | 1.98E-02 | 5.59E-02 | -1.18E+00 | 1.88E-11 | 6.89E-11 | -1.77E+00 | 3.44E-03 | 9.45E-03 | -1.01E+00 |
| CCDC42 | 2.88E-09 | 1.50E-08 | -1.28E+00 | 8.47E-05 | 3.66E-04 | 1.03E+00 | 2.41E-04 | 1.02E-03 | -1.26E+00 | 1.56E-02 | 5.95E-02 | 1.09E+00 | 5.43E-02 | 1.94E-01 | 1.15E+00 | 1.94E-01 | 5.48E-01 | 1.59E+00 | 3.00E-02 | 1.10E-01 | 1.53E+00 | 3.14E-02 | 8.64E-02 | 1.50E+00 |
| PTMS | 2.89E-09 | 1.50E-08 | -1.23E+00 | 1.01E-02 | 4.38E-02 | 1.04E+00 | 5.09E-09 | 2.16E-08 | -1.32E+00 | 3.07E-02 | 1.17E-01 | 1.04E+00 | 1.93E-06 | 6.89E-06 | -1.15E+00 | 6.84E-05 | 1.93E-04 | -1.03E+00 | 9.53E-04 | 3.50E-03 | -1.12E+00 | 4.31E-02 | 1.19E-01 | -1.01E+00 |
| RAMP1 | 2.89E-09 | 1.51E-08 | 1.83E+00 | 2.43E-05 | 1.05E-04 | 1.77E+00 | 1.14E-07 | 4.83E-07 | 1.74E+00 | 2.51E-05 | 9.59E-05 | 1.69E+00 | 1.46E-09 | 5.21E-09 | 2.81E+00 | 6.67E-04 | 1.88E-03 | 2.28E+00 | 8.94E-33 | 3.28E-32 | 6.51E+00 | 3.70E-20 | 1.02E-19 | 4.93E+00 |
| TWF2 | 2.89E-09 | 1.51E-08 | -1.17E+00 | 5.79E-07 | 2.50E-06 | 1.02E+00 | 5.23E-03 | 2.22E-02 | -1.03E+00 | 2.76E-03 | 1.06E-02 | 1.05E+00 | 4.83E-08 | 1.72E-07 | -1.27E+00 | 1.03E-07 | 2.92E-07 | -1.23E+00 | 2.17E-01 | 7.96E-01 | 1.28E+00 | 3.04E-01 | 8.34E-01 | 1.22E+00 |
| FBXW2 | 2.89E-09 | 1.51E-08 | -1.57E+00 | 3.95E-06 | 1.71E-05 | -1.14E+00 | 5.24E-13 | 2.23E-12 | -1.60E+00 | 1.81E-05 | 6.91E-05 | 1.03E+00 | 2.46E-08 | 8.79E-08 | -1.55E+00 | 6.05E-05 | 1.70E-04 | -1.13E+00 | 1.30E-07 | 4.78E-07 | -1.32E+00 | 2.73E-03 | 7.51E-03 | -1.03E+00 |
| NDUFA1 | 2.94E-09 | 1.53E-08 | -1.11E+00 | 4.69E-35 | 2.03E-34 | -1.51E+00 | 1.42E-02 | 6.04E-02 | -1.04E+00 | 3.24E-15 | 1.24E-14 | -1.46E+00 | 7.22E-05 | 2.57E-04 | -1.13E+00 | 1.19E-13 | 3.37E-13 | -1.50E+00 | 6.05E-03 | 2.22E-02 | -1.12E+00 | 4.14E-11 | 1.14E-10 | -1.57E+00 |
| AC242426.3 | 2.95E-09 | 1.54E-08 | -2.27E+00 | 4.15E-05 | 1.79E-04 | -1.25E+00 | 4.80E-04 | 2.04E-03 | -1.76E+00 | 9.60E-03 | 3.67E-02 | -1.14E+00 | 1.10E-01 | 3.92E-01 | -1.02E+00 | 9.31E-02 | 2.62E-01 | -1.08E+00 | 2.69E-01 | 9.86E-01 | 1.36E+00 | 1.20E-01 | 3.29E-01 | -1.12E+00 |
| TMEM140 | 2.97E-09 | 1.55E-08 | -1.60E+00 | 7.30E-06 | 3.16E-05 | -1.15E+00 | 9.26E-06 | 3.94E-05 | -1.76E+00 | 4.66E-06 | 1.78E-05 | -1.73E+00 | 2.26E-02 | 8.05E-02 | -1.40E+00 | 3.41E-02 | 9.62E-02 | -1.32E+00 | 8.79E-02 | 3.23E-01 | 2.00E+00 | 2.48E-02 | 6.82E-02 | 1.39E+00 |
| KLHL42 | 2.99E-09 | 1.56E-08 | -1.37E+00 | 3.39E-06 | 1.46E-05 | -1.08E+00 | 1.38E-11 | 5.88E-11 | -1.52E+00 | 3.46E-09 | 1.32E-08 | -1.21E+00 | 1.17E-07 | 4.18E-07 | -1.51E+00 | 5.66E-07 | 1.60E-06 | -1.43E+00 | 2.50E-08 | 9.16E-08 | -1.32E+00 | 1.46E-05 | 4.01E-05 | -1.12E+00 |
| MUM1 | 2.99E-09 | 1.56E-08 | -1.41E+00 | 1.13E-02 | 4.89E-02 | 1.34E+00 | 1.08E-05 | 4.59E-05 | -1.40E+00 | 5.08E-02 | 1.94E-01 | 1.22E+00 | 1.56E-05 | 5.57E-05 | -1.27E+00 | 1.41E-02 | 3.98E-02 | 1.16E+00 | 6.47E-08 | 2.38E-07 | -1.52E+00 | 4.17E-02 | 1.15E-01 | 1.10E+00 |
| LRRK1 | 3.00E-09 | 1.56E-08 | -1.46E+00 | 6.81E-05 | 2.94E-04 | 1.04E+00 | 6.27E-04 | 2.66E-03 | -1.29E+00 | 8.61E-02 | 3.29E-01 | 1.17E+00 | 3.49E-03 | 1.24E-02 | -1.41E+00 | 3.44E-03 | 9.71E-03 | -1.25E+00 | 2.49E-03 | 9.14E-03 | -1.50E+00 | 1.02E-01 | 2.81E-01 | -1.06E+00 |
| RAC1 | 3.02E-09 | 1.57E-08 | -1.10E+00 | 1.77E-02 | 7.64E-02 | 1.30E+00 | 6.84E-02 | 2.91E-01 | 1.06E+00 | 9.14E-02 | 3.49E-01 | 1.23E+00 | 1.32E-06 | 4.71E-06 | -1.27E+00 | 1.80E-03 | 5.08E-03 | 1.01E+00 | 4.53E-02 | 1.66E-01 | -1.05E+00 | 5.46E-02 | 1.50E-01 | 1.04E+00 |
| LAGE3 | 3.04E-09 | 1.58E-08 | -1.11E+00 | 1.07E-07 | 4.63E-07 | 1.06E+00 | 1.88E-01 | 7.99E-01 | 1.15E+00 | 1.14E-01 | 4.35E-01 | 1.21E+00 | 3.14E-06 | 1.12E-05 | -1.18E+00 | 7.99E-03 | 2.25E-02 | 1.08E+00 | 1.62E-01 | 5.95E-01 | 1.16E+00 | 9.46E-02 | 2.60E-01 | 1.35E+00 |
| FAM241B | 3.04E-09 | 1.59E-08 | -1.94E+00 | 1.82E-09 | 7.87E-09 | -1.76E+00 | 1.06E-02 | 4.51E-02 | -1.27E+00 | 5.95E-04 | 2.27E-03 | -1.60E+00 | 4.21E-02 | 1.50E-01 | -1.31E+00 | 3.27E-02 | 9.22E-02 | -1.47E+00 | 2.57E-02 | 9.44E-02 | -1.32E+00 | 8.34E-02 | 2.29E-01 | -1.05E+00 |
| C1orf35 | 3.04E-09 | 1.59E-08 | -1.23E+00 | 6.47E-04 | 2.80E-03 | 1.18E+00 | 9.98E-06 | 4.24E-05 | -1.21E+00 | 1.17E-04 | 4.46E-04 | -1.04E+00 | 1.54E-05 | 5.49E-05 | -1.45E+00 | 1.54E-02 | 4.34E-02 | 1.00E+00 | 1.00E-10 | 3.68E-10 | -1.56E+00 | 5.23E-03 | 1.44E-02 | -1.08E+00 |
| AOAH | 3.06E-09 | 1.60E-08 | -1.24E+00 | 3.22E-04 | 1.39E-03 | 1.09E+00 | 1.02E-05 | 4.33E-05 | -1.41E+00 | 9.69E-03 | 3.70E-02 | 1.04E+00 | 4.96E-03 | 1.77E-02 | -1.11E+00 | 7.58E-02 | 2.14E-01 | 1.12E+00 | 8.73E-02 | 3.20E-01 | 1.48E+00 | 1.80E-01 | 4.94E-01 | 1.24E+00 |
| CNPPD1 | 3.07E-09 | 1.60E-08 | -1.42E+00 | 2.57E-05 | 1.11E-04 | 1.03E+00 | 8.83E-10 | 3.75E-09 | -1.50E+00 | 1.46E-06 | 5.58E-06 | -1.11E+00 | 2.62E-04 | 9.35E-04 | -1.29E+00 | 4.71E-02 | 1.33E-01 | 1.03E+00 | 1.83E-05 | 6.72E-05 | -1.22E+00 | 6.31E-03 | 1.73E-02 | -1.09E+00 |
| FAH | 3.07E-09 | 1.60E-08 | -1.31E+00 | 1.36E-11 | 5.87E-11 | -1.22E+00 | 8.64E-06 | 3.67E-05 | -1.29E+00 | 1.50E-04 | 5.74E-04 | -1.07E+00 | 4.61E-05 | 1.64E-04 | -1.33E+00 | 3.46E-03 | 9.76E-03 | -1.04E+00 | 1.11E-03 | 4.06E-03 | -1.07E+00 | 1.37E-01 | 3.76E-01 | 1.19E+00 |
| CNOT6 | 3.07E-09 | 1.60E-08 | -1.47E+00 | 7.62E-06 | 3.29E-05 | -1.03E+00 | 2.77E-05 | 1.18E-04 | -1.26E+00 | 4.37E-04 | 1.67E-03 | -1.07E+00 | 5.23E-05 | 1.87E-04 | -1.40E+00 | 3.11E-03 | 8.76E-03 | -1.03E+00 | 2.02E-08 | 7.42E-08 | -1.44E+00 | 1.07E-03 | 2.93E-03 | -1.05E+00 |
| EFTUD2 | 3.07E-09 | 1.60E-08 | -1.34E+00 | 2.44E-05 | 1.05E-04 | -1.03E+00 | 9.05E-09 | 3.85E-08 | -1.29E+00 | 4.31E-06 | 1.65E-05 | -1.02E+00 | 1.05E-05 | 3.74E-05 | -1.20E+00 | 2.17E-02 | 6.11E-02 | 1.13E+00 | 5.36E-06 | 1.97E-05 | -1.25E+00 | 5.09E-02 | 1.40E-01 | 1.05E+00 |
| NANS | 3.07E-09 | 1.60E-08 | -1.15E+00 | 1.24E-19 | 5.37E-19 | -1.30E+00 | 8.48E-03 | 3.60E-02 | -1.00E+00 | 6.57E-05 | 2.51E-04 | -1.04E+00 | 1.33E-06 | 4.73E-06 | -1.22E+00 | 1.95E-08 | 5.51E-08 | -1.40E+00 | 8.23E-04 | 3.02E-03 | -1.07E+00 | 2.27E-02 | 6.24E-02 | -1.04E+00 |
| NASP | 3.07E-09 | 1.60E-08 | -1.18E+00 | 9.20E-08 | 3.97E-07 | -1.06E+00 | 1.68E-01 | 7.14E-01 | 1.00E+00 | 1.09E-01 | 4.17E-01 | -1.01E+00 | 4.38E-07 | 1.56E-06 | -1.18E+00 | 2.43E-05 | 6.84E-05 | -1.10E+00 | 5.91E-02 | 2.17E-01 | -1.05E+00 | 3.36E-01 | 9.22E-01 | 1.05E+00 |
| MRPS9 | 3.07E-09 | 1.60E-08 | -1.31E+00 | 2.54E-05 | 1.10E-04 | 1.03E+00 | 6.23E-10 | 2.65E-09 | -1.40E+00 | 1.13E-08 | 4.33E-08 | -1.22E+00 | 2.21E-10 | 7.88E-10 | -1.77E+00 | 1.07E-03 | 3.02E-03 | -1.03E+00 | 2.63E-09 | 9.64E-09 | -1.34E+00 | 1.71E-05 | 4.68E-05 | -1.14E+00 |
| AC244034.1 | 3.08E-09 | 1.60E-08 | -1.65E+00 | 6.66E-09 | 2.88E-08 | -1.47E+00 | 2.06E-05 | 8.77E-05 | -1.50E+00 | 2.89E-05 | 1.10E-04 | -1.26E+00 | 4.46E-03 | 1.59E-02 | -1.27E+00 | 1.44E-02 | 4.05E-02 | -1.13E+00 | 2.50E-05 | 9.18E-05 | -1.46E+00 | 1.82E-04 | 5.00E-04 | -1.29E+00 |
| FAM118A | 3.09E-09 | 1.61E-08 | -1.15E+00 | 2.60E-03 | 1.12E-02 | 1.21E+00 | 6.78E-04 | 2.88E-03 | -1.14E+00 | 9.77E-03 | 3.73E-02 | 1.09E+00 | 4.89E-06 | 1.74E-05 | -1.15E+00 | 2.65E-01 | 7.46E-01 | 1.46E+00 | 4.08E-05 | 1.50E-04 | -1.22E+00 | 4.24E-02 | 1.17E-01 | -1.00E+00 |
| MED21 | 3.09E-09 | 1.61E-08 | -1.27E+00 | 5.29E-10 | 2.29E-09 | -1.18E+00 | 8.46E-13 | 3.59E-12 | -1.63E+00 | 2.41E-08 | 9.19E-08 | -1.23E+00 | 2.23E-06 | 7.97E-06 | -1.29E+00 | 1.94E-03 | 5.48E-03 | -1.02E+00 | 4.46E-13 | 1.64E-12 | -1.54E+00 | 8.35E-05 | 2.29E-04 | -1.15E+00 |
| HADHAP2 | 3.09E-09 | 1.61E-08 | -1.73E+00 | 3.82E-12 | 1.65E-11 | -2.18E+00 | 1.08E-07 | 4.58E-07 | -1.75E+00 | 3.53E-07 | 1.35E-06 | -1.47E+00 | 6.09E-04 | 2.17E-03 | -1.29E+00 | 1.58E-04 | 4.46E-04 | -1.43E+00 | 9.45E-10 | 3.47E-09 | -1.91E+00 | 8.85E-08 | 2.43E-07 | -1.76E+00 |
| MRPL46 | 3.10E-09 | 1.61E-08 | -1.56E+00 | 1.76E-06 | 7.61E-06 | -1.16E+00 | 3.36E-07 | 1.43E-06 | -1.35E+00 | 6.81E-08 | 2.60E-07 | -1.21E+00 | 1.04E-06 | 3.69E-06 | -1.54E+00 | 3.85E-04 | 1.09E-03 | -1.17E+00 | 1.72E-12 | 6.32E-12 | -1.48E+00 | 1.58E-07 | 4.35E-07 | -1.24E+00 |
| SMC5 | 3.10E-09 | 1.62E-08 | -1.22E+00 | 2.06E-07 | 8.88E-07 | -1.01E+00 | 8.20E-03 | 3.49E-02 | -1.00E+00 | 1.11E-02 | 4.23E-02 | 1.10E+00 | 9.94E-04 | 3.55E-03 | 1.00E+00 | 2.47E-03 | 6.97E-03 | 1.00E+00 | 1.85E-02 | 6.80E-02 | 1.02E+00 | 3.25E-02 | 8.93E-02 | 1.03E+00 |
| MRPL20 | 3.12E-09 | 1.63E-08 | -1.13E+00 | 8.84E-07 | 3.82E-06 | 1.11E+00 | 4.27E-03 | 1.81E-02 | -1.07E+00 | 9.41E-03 | 3.59E-02 | 1.03E+00 | 9.13E-09 | 3.25E-08 | -1.21E+00 | 1.45E-03 | 4.10E-03 | 1.06E+00 | 4.59E-03 | 1.68E-02 | -1.04E+00 | 2.83E-01 | 7.78E-01 | 1.23E+00 |
| MBD5 | 3.15E-09 | 1.64E-08 | -1.24E+00 | 2.30E-05 | 9.92E-05 | 1.08E+00 | 6.79E-06 | 2.88E-05 | -1.30E+00 | 7.63E-04 | 2.92E-03 | -1.00E+00 | 9.98E-03 | 3.56E-02 | 1.02E+00 | 3.35E-01 | 9.46E-01 | 1.39E+00 | 1.06E-05 | 3.89E-05 | -1.26E+00 | 1.13E-02 | 3.09E-02 | -1.05E+00 |
| AC107075.1 | 3.15E-09 | 1.64E-08 | -1.23E+00 | 4.91E-02 | 2.12E-01 | -1.03E+00 | 2.46E-05 | 1.04E-04 | -1.19E+00 | 1.65E-01 | 6.30E-01 | 1.00E+00 | 2.37E-03 | 8.45E-03 | -1.13E+00 | 1.28E-01 | 3.60E-01 | -1.04E+00 | 6.52E-12 | 2.39E-11 | -1.22E+00 | 2.28E-03 | 6.26E-03 | -1.08E+00 |
| UBXN2A | 3.15E-09 | 1.64E-08 | -1.23E+00 | 1.71E-14 | 7.38E-14 | -1.44E+00 | 1.29E-02 | 5.50E-02 | -1.05E+00 | 2.67E-03 | 1.02E-02 | 1.02E+00 | 4.01E-04 | 1.43E-03 | -1.07E+00 | 1.62E-05 | 4.58E-05 | -1.20E+00 | 1.90E-04 | 6.99E-04 | -1.06E+00 | 5.92E-06 | 1.63E-05 | -1.18E+00 |
| POLR2E | 3.16E-09 | 1.65E-08 | -1.18E+00 | 1.12E-06 | 4.86E-06 | 1.05E+00 | 2.22E-03 | 9.41E-03 | -1.10E+00 | 5.69E-03 | 2.17E-02 | 1.05E+00 | 8.82E-06 | 3.15E-05 | -1.22E+00 | 9.87E-02 | 2.78E-01 | 1.12E+00 | 1.06E-02 | 3.89E-02 | -1.04E+00 | 1.39E-01 | 3.81E-01 | 1.21E+00 |
| FAR2 | 3.16E-09 | 1.65E-08 | -1.46E+00 | 3.42E-06 | 1.48E-05 | -1.01E+00 | 3.73E-05 | 1.58E-04 | -1.37E+00 | 4.56E-03 | 1.74E-02 | -1.02E+00 | 4.16E-09 | 1.48E-08 | -1.97E+00 | 4.58E-07 | 1.29E-06 | -1.59E+00 | 8.97E-07 | 3.29E-06 | -1.43E+00 | 1.16E-02 | 3.18E-02 | -1.03E+00 |
| TSFM | 3.16E-09 | 1.65E-08 | -1.56E+00 | 5.02E-03 | 2.17E-02 | 1.15E+00 | 5.96E-05 | 2.53E-04 | -1.08E+00 | 1.08E-02 | 4.12E-02 | 1.19E+00 | 1.15E-04 | 4.10E-04 | -1.47E+00 | 3.29E-01 | 9.27E-01 | 1.45E+00 | 9.63E-09 | 3.53E-08 | -1.27E+00 | 2.57E-01 | 7.05E-01 | 1.26E+00 |
| ACTR10 | 3.21E-09 | 1.67E-08 | -1.29E+00 | 1.92E-05 | 8.31E-05 | 1.03E+00 | 1.17E-05 | 4.95E-05 | -1.24E+00 | 6.81E-04 | 2.60E-03 | -1.02E+00 | 1.85E-11 | 6.58E-11 | -1.45E+00 | 2.12E-05 | 5.99E-05 | -1.04E+00 | 2.04E-09 | 7.50E-09 | -1.41E+00 | 3.78E-04 | 1.04E-03 | -1.20E+00 |
| MIEF1 | 3.23E-09 | 1.68E-08 | -1.61E+00 | 5.34E-05 | 2.31E-04 | -1.05E+00 | 4.23E-07 | 1.80E-06 | -1.58E+00 | 2.55E-02 | 9.75E-02 | 1.07E+00 | 1.16E-08 | 4.13E-08 | -2.06E+00 | 1.47E-03 | 4.15E-03 | -1.18E+00 | 4.71E-10 | 1.73E-09 | -1.62E+00 | 2.14E-02 | 5.88E-02 | 1.08E+00 |
| SETDB2 | 3.23E-09 | 1.68E-08 | -1.50E+00 | 2.40E-03 | 1.04E-02 | 1.14E+00 | 7.98E-06 | 3.39E-05 | -1.25E+00 | 3.66E-04 | 1.40E-03 | 1.04E+00 | 1.55E-03 | 5.51E-03 | -1.21E+00 | 2.31E-02 | 6.52E-02 | 1.02E+00 | 2.24E-08 | 8.21E-08 | -1.56E+00 | 4.05E-03 | 1.11E-02 | -1.10E+00 |
| ASB9 | 3.24E-09 | 1.69E-08 | -1.70E+00 | 1.11E-07 | 4.78E-07 | -1.35E+00 | 1.47E-01 | 6.23E-01 | 1.18E+00 | 1.38E-01 | 5.27E-01 | 1.09E+00 | 3.90E-03 | 1.39E-02 | -1.21E+00 | 5.82E-03 | 1.64E-02 | -1.06E+00 | 5.08E-04 | 1.86E-03 | -1.33E+00 | 4.18E-02 | 1.15E-01 | 1.03E+00 |
| EPN1 | 3.25E-09 | 1.69E-08 | -1.28E+00 | 4.57E-07 | 1.97E-06 | -1.03E+00 | 1.53E-03 | 6.51E-03 | -1.12E+00 | 1.04E-03 | 3.98E-03 | -1.03E+00 | 7.45E-04 | 2.66E-03 | -1.13E+00 | 1.26E-02 | 3.55E-02 | 1.04E+00 | 7.19E-05 | 2.64E-04 | -1.17E+00 | 6.57E-03 | 1.80E-02 | 1.00E+00 |
| PCNX4 | 3.25E-09 | 1.69E-08 | -1.41E+00 | 4.80E-02 | 2.07E-01 | 1.32E+00 | 5.52E-10 | 2.35E-09 | -1.55E+00 | 1.23E-02 | 4.71E-02 | 1.20E+00 | 7.17E-05 | 2.56E-04 | -1.17E+00 | 1.86E-01 | 5.24E-01 | 1.36E+00 | 3.49E-04 | 1.28E-03 | -1.14E+00 | 4.97E-02 | 1.37E-01 | 1.44E+00 |
| VTI1B | 3.28E-09 | 1.71E-08 | -1.75E+00 | 2.41E-08 | 1.04E-07 | -1.46E+00 | 7.99E-12 | 3.40E-11 | -1.79E+00 | 3.89E-14 | 1.49E-13 | -1.76E+00 | 8.87E-12 | 3.16E-11 | -2.20E+00 | 5.46E-12 | 1.54E-11 | -1.93E+00 | 2.92E-09 | 1.07E-08 | -1.43E+00 | 4.43E-08 | 1.22E-07 | -1.37E+00 |
| PPIA | 3.30E-09 | 1.72E-08 | 1.54E+00 | 2.79E-10 | 1.21E-09 | 1.61E+00 | 9.62E-05 | 4.09E-04 | 1.46E+00 | 1.40E-05 | 5.34E-05 | 1.55E+00 | 2.55E-01 | 9.09E-01 | 1.13E+00 | 1.12E-03 | 3.17E-03 | 1.40E+00 | 5.52E-03 | 2.02E-02 | 1.27E+00 | 6.15E-06 | 1.69E-05 | 1.53E+00 |
| LIN54 | 3.30E-09 | 1.72E-08 | -1.85E+00 | 7.09E-07 | 3.06E-06 | -1.30E+00 | 8.41E-02 | 3.57E-01 | 1.11E+00 | 5.70E-02 | 2.18E-01 | 1.18E+00 | 2.97E-03 | 1.06E-02 | -1.27E+00 | 2.19E-02 | 6.16E-02 | -1.16E+00 | 6.63E-05 | 2.43E-04 | -1.19E+00 | 8.34E-05 | 2.29E-04 | -1.32E+00 |
| RBM48 | 3.33E-09 | 1.74E-08 | -1.28E+00 | 5.95E-11 | 2.57E-10 | -1.26E+00 | 1.20E-03 | 5.10E-03 | -1.10E+00 | 1.84E-03 | 7.01E-03 | -1.01E+00 | 9.92E-02 | 3.54E-01 | 1.36E+00 | 7.62E-03 | 2.15E-02 | -1.01E+00 | 2.38E-02 | 8.75E-02 | 1.01E+00 | 2.66E-03 | 7.30E-03 | -1.06E+00 |
| APBA3 | 3.34E-09 | 1.74E-08 | -2.01E+00 | 1.21E-05 | 5.24E-05 | -1.26E+00 | 1.56E-04 | 6.61E-04 | -1.56E+00 | 1.31E-05 | 5.00E-05 | -1.62E+00 | 6.18E-02 | 2.20E-01 | 1.02E+00 | 7.66E-02 | 2.16E-01 | 1.00E+00 | 7.33E-05 | 2.69E-04 | -1.36E+00 | 3.57E-02 | 9.79E-02 | 1.06E+00 |
| RRM1 | 3.34E-09 | 1.74E-08 | -1.78E+00 | 4.94E-06 | 2.14E-05 | -1.22E+00 | 9.06E-03 | 3.85E-02 | -1.02E+00 | 2.00E-04 | 7.63E-04 | 1.08E+00 | 4.38E-08 | 1.56E-07 | -1.94E+00 | 1.47E-04 | 4.14E-04 | -1.19E+00 | 2.44E-05 | 8.96E-05 | -1.25E+00 | 8.86E-02 | 2.43E-01 | 1.07E+00 |
| SEH1L | 3.35E-09 | 1.75E-08 | -1.43E+00 | 9.89E-08 | 4.27E-07 | -1.19E+00 | 3.25E-08 | 1.38E-07 | -1.45E+00 | 4.21E-05 | 1.61E-04 | -1.08E+00 | 1.32E-02 | 4.69E-02 | 1.01E+00 | 2.10E-01 | 5.91E-01 | 1.27E+00 | 9.49E-08 | 3.48E-07 | -1.35E+00 | 1.97E-02 | 5.40E-02 | 1.06E+00 |
| CCAR1 | 3.38E-09 | 1.76E-08 | -1.17E+00 | 9.79E-06 | 4.23E-05 | 1.05E+00 | 1.95E-06 | 8.26E-06 | -1.38E+00 | 4.10E-05 | 1.57E-04 | -1.16E+00 | 2.54E-05 | 9.06E-05 | -1.34E+00 | 1.03E-01 | 2.91E-01 | 1.05E+00 | 2.50E-09 | 9.18E-09 | -1.47E+00 | 3.79E-03 | 1.04E-02 | -1.10E+00 |
| NAAA | 3.38E-09 | 1.76E-08 | -1.64E+00 | 1.03E-02 | 4.44E-02 | 1.21E+00 | 6.11E-06 | 2.60E-05 | -1.38E+00 | 3.87E-03 | 1.48E-02 | 1.00E+00 | 6.64E-03 | 2.37E-02 | -1.30E+00 | 9.51E-02 | 2.68E-01 | 1.05E+00 | 1.15E-04 | 4.21E-04 | -1.43E+00 | 9.25E-02 | 2.54E-01 | 1.15E+00 |
| PIKFYVE | 3.39E-09 | 1.76E-08 | -1.41E+00 | 7.09E-04 | 3.06E-03 | 1.12E+00 | 7.98E-05 | 3.39E-04 | -1.22E+00 | 3.65E-03 | 1.39E-02 | 1.09E+00 | 2.18E-04 | 7.76E-04 | -1.11E+00 | 7.46E-03 | 2.10E-02 | 1.00E+00 | 1.00E-10 | 3.67E-10 | -1.43E+00 | 3.68E-05 | 1.01E-04 | -1.04E+00 |
| PTPN1 | 3.39E-09 | 1.77E-08 | -1.28E+00 | 4.72E-12 | 2.04E-11 | -1.24E+00 | 6.67E-04 | 2.83E-03 | -1.07E+00 | 5.82E-05 | 2.22E-04 | -1.09E+00 | 3.21E-04 | 1.15E-03 | -1.07E+00 | 1.69E-03 | 4.76E-03 | 1.06E+00 | 2.51E-06 | 9.21E-06 | -1.23E+00 | 2.29E-06 | 6.28E-06 | -1.26E+00 |
| RPL12P32 | 3.39E-09 | 1.77E-08 | -1.26E+00 | 1.17E-67 | 5.07E-67 | -2.19E+00 | 4.74E-03 | 2.02E-02 | -1.18E+00 | 4.99E-23 | 1.90E-22 | -1.93E+00 | 1.47E-03 | 5.23E-03 | -1.15E+00 | 3.62E-19 | 1.02E-18 | -1.91E+00 | 1.10E-03 | 4.04E-03 | -1.17E+00 | 5.91E-17 | 1.62E-16 | -1.91E+00 |
| MANBA | 3.39E-09 | 1.77E-08 | -1.28E+00 | 1.16E-05 | 5.01E-05 | -1.00E+00 | 1.10E-07 | 4.68E-07 | -1.44E+00 | 1.80E-06 | 6.87E-06 | -1.19E+00 | 3.11E-06 | 1.11E-05 | -1.23E+00 | 4.43E-06 | 1.25E-05 | -1.15E+00 | 1.01E-04 | 3.71E-04 | -1.26E+00 | 1.06E-04 | 2.92E-04 | -1.24E+00 |
| SLAIN2 | 3.44E-09 | 1.79E-08 | -1.40E+00 | 6.29E-07 | 2.72E-06 | -1.12E+00 | 3.06E-05 | 1.30E-04 | -1.27E+00 | 4.61E-07 | 1.76E-06 | -1.25E+00 | 8.06E-06 | 2.87E-05 | -1.18E+00 | 7.15E-06 | 2.02E-05 | -1.21E+00 | 1.21E-02 | 4.43E-02 | 1.05E+00 | 4.69E-04 | 1.29E-03 | -1.15E+00 |
| TMEM185A | 3.44E-09 | 1.79E-08 | -1.71E+00 | 7.89E-04 | 3.41E-03 | -1.02E+00 | 3.41E-04 | 1.45E-03 | -1.33E+00 | 3.98E-03 | 1.52E-02 | -1.04E+00 | 1.48E-01 | 5.27E-01 | 1.18E+00 | 2.68E-01 | 7.55E-01 | 1.31E+00 | 5.92E-04 | 2.17E-03 | -1.39E+00 | 6.36E-03 | 1.75E-02 | -1.21E+00 |
| DNPH1 | 3.47E-09 | 1.81E-08 | -1.11E+00 | 1.11E-07 | 4.80E-07 | 1.04E+00 | 1.06E-01 | 4.49E-01 | 1.10E+00 | 2.26E-02 | 8.62E-02 | 1.11E+00 | 4.29E-07 | 1.53E-06 | -1.18E+00 | 2.65E-01 | 7.46E-01 | 1.38E+00 | 9.52E-03 | 3.49E-02 | -1.03E+00 | 2.31E-01 | 6.35E-01 | 1.19E+00 |
| CHMP4B | 3.47E-09 | 1.81E-08 | -1.34E+00 | 3.16E-09 | 1.37E-08 | -1.15E+00 | 1.12E-05 | 4.77E-05 | -1.25E+00 | 8.09E-09 | 3.09E-08 | -1.25E+00 | 1.21E-05 | 4.31E-05 | -1.21E+00 | 7.14E-05 | 2.01E-04 | -1.07E+00 | 1.18E-06 | 4.34E-06 | -1.14E+00 | 1.92E-07 | 5.27E-07 | -1.24E+00 |
| BAG5 | 3.48E-09 | 1.81E-08 | -1.24E+00 | 6.73E-15 | 2.91E-14 | -1.46E+00 | 2.24E-06 | 9.50E-06 | -1.28E+00 | 1.84E-07 | 7.01E-07 | -1.17E+00 | 1.62E-06 | 5.78E-06 | -1.20E+00 | 2.43E-10 | 6.86E-10 | -1.42E+00 | 3.09E-04 | 1.13E-03 | -1.06E+00 | 2.78E-06 | 7.63E-06 | -1.18E+00 |
| ATR | 3.49E-09 | 1.82E-08 | -1.92E+00 | 5.70E-05 | 2.46E-04 | -1.18E+00 | 8.54E-05 | 3.63E-04 | -1.33E+00 | 1.43E-01 | 5.47E-01 | 1.30E+00 | 1.93E-02 | 6.87E-02 | -1.07E+00 | 1.24E-01 | 3.51E-01 | 1.16E+00 | 1.12E-05 | 4.11E-05 | -1.20E+00 | 2.41E-02 | 6.63E-02 | 1.03E+00 |
| EIF4E2 | 3.49E-09 | 1.82E-08 | -1.28E+00 | 4.56E-08 | 1.97E-07 | -1.14E+00 | 8.48E-04 | 3.60E-03 | -1.14E+00 | 9.66E-05 | 3.69E-04 | -1.06E+00 | 1.48E-09 | 5.29E-09 | -1.43E+00 | 1.03E-05 | 2.89E-05 | -1.13E+00 | 3.36E-06 | 1.23E-05 | -1.16E+00 | 3.78E-04 | 1.04E-03 | -1.08E+00 |
| CEACAM4 | 3.50E-09 | 1.82E-08 | 2.94E+00 | 1.22E-03 | 5.28E-03 | 2.63E+00 | 1.07E-09 | 4.53E-09 | 2.74E+00 | 2.82E-02 | 1.08E-01 | 1.89E+00 | 2.24E-01 | 7.98E-01 | 1.70E+00 | 3.37E-01 | 9.49E-01 | 1.20E+00 | 2.67E-01 | 9.78E-01 | 1.05E+00 | 3.42E-01 | 9.41E-01 | 1.10E+00 |
| EML2 | 3.51E-09 | 1.83E-08 | -2.01E+00 | 4.95E-05 | 2.14E-04 | -1.16E+00 | 5.02E-06 | 2.13E-05 | -1.76E+00 | 2.61E-05 | 9.97E-05 | -1.34E+00 | 3.85E-04 | 1.37E-03 | -1.63E+00 | 5.38E-02 | 1.52E-01 | 1.11E+00 | 7.85E-09 | 2.88E-08 | -1.79E+00 | 3.84E-06 | 1.06E-05 | -1.45E+00 |
| SELENOS | 3.52E-09 | 1.84E-08 | -1.31E+00 | 2.94E-08 | 1.27E-07 | -1.17E+00 | 3.26E-03 | 1.39E-02 | -1.07E+00 | 7.88E-06 | 3.01E-05 | -1.13E+00 | 3.71E-13 | 1.32E-12 | -1.58E+00 | 7.90E-10 | 2.23E-09 | -1.41E+00 | 2.10E-08 | 7.70E-08 | -1.38E+00 | 2.35E-07 | 6.45E-07 | -1.43E+00 |
| PAN3 | 3.55E-09 | 1.85E-08 | -1.25E+00 | 1.68E-06 | 7.24E-06 | 1.02E+00 | 3.77E-05 | 1.60E-04 | -1.30E+00 | 8.91E-04 | 3.40E-03 | -1.03E+00 | 5.00E-05 | 1.78E-04 | -1.09E+00 | 5.46E-02 | 1.54E-01 | 1.14E+00 | 2.09E-05 | 7.68E-05 | -1.22E+00 | 1.84E-03 | 5.05E-03 | -1.06E+00 |
| MARCH6 | 3.56E-09 | 1.86E-08 | -1.20E+00 | 4.19E-08 | 1.81E-07 | -1.07E+00 | 5.75E-04 | 2.45E-03 | -1.21E+00 | 1.22E-03 | 4.67E-03 | -1.09E+00 | 2.05E-04 | 7.32E-04 | -1.13E+00 | 1.53E-04 | 4.31E-04 | -1.11E+00 | 1.22E-04 | 4.49E-04 | -1.26E+00 | 3.82E-04 | 1.05E-03 | -1.19E+00 |
| RNF34 | 3.58E-09 | 1.87E-08 | -1.65E+00 | 8.23E-04 | 3.56E-03 | 1.06E+00 | 6.35E-09 | 2.70E-08 | -1.93E+00 | 2.69E-04 | 1.03E-03 | -1.11E+00 | 5.31E-10 | 1.89E-09 | -2.35E+00 | 1.46E-04 | 4.11E-04 | -1.17E+00 | 5.94E-29 | 2.18E-28 | -2.58E+00 | 1.13E-10 | 3.10E-10 | -1.40E+00 |
| ARMC10 | 3.59E-09 | 1.87E-08 | -1.40E+00 | 5.82E-07 | 2.51E-06 | -1.06E+00 | 4.65E-06 | 1.98E-05 | -1.30E+00 | 1.69E-03 | 6.44E-03 | 1.01E+00 | 1.03E-06 | 3.67E-06 | -1.26E+00 | 1.61E-03 | 4.55E-03 | 1.09E+00 | 1.04E-05 | 3.83E-05 | -1.10E+00 | 3.22E-03 | 8.84E-03 | 1.01E+00 |
| ARHGAP17 | 3.60E-09 | 1.87E-08 | -1.80E+00 | 7.68E-05 | 3.32E-04 | -1.15E+00 | 9.23E-08 | 3.92E-07 | -1.45E+00 | 5.89E-06 | 2.25E-05 | -1.17E+00 | 1.62E-02 | 5.77E-02 | -1.06E+00 | 5.76E-02 | 1.63E-01 | 1.03E+00 | 1.06E-03 | 3.89E-03 | -1.12E+00 | 4.78E-02 | 1.31E-01 | 1.04E+00 |
| CD27-AS1 | 3.60E-09 | 1.87E-08 | -1.50E+00 | 2.35E-06 | 1.02E-05 | -1.11E+00 | 4.89E-07 | 2.08E-06 | -1.59E+00 | 9.34E-06 | 3.57E-05 | -1.32E+00 | 1.86E-04 | 6.62E-04 | -1.46E+00 | 6.19E-02 | 1.75E-01 | 1.08E+00 | 2.21E-08 | 8.11E-08 | -1.32E+00 | 1.21E-06 | 3.32E-06 | -1.30E+00 |
| FTH1P20 | 3.60E-09 | 1.88E-08 | -1.29E+00 | 1.30E-21 | 5.62E-21 | -1.42E+00 | 1.32E-05 | 5.60E-05 | -1.23E+00 | 8.65E-07 | 3.30E-06 | -1.24E+00 | 6.24E-06 | 2.23E-05 | -1.29E+00 | 7.40E-11 | 2.09E-10 | -1.51E+00 | 3.24E-07 | 1.19E-06 | -1.29E+00 | 2.91E-08 | 7.98E-08 | -1.35E+00 |
| PGM1 | 3.61E-09 | 1.88E-08 | -1.36E+00 | 1.11E-05 | 4.79E-05 | -1.02E+00 | 1.34E-07 | 5.71E-07 | -1.31E+00 | 1.11E-03 | 4.24E-03 | 1.07E+00 | 1.87E-02 | 6.66E-02 | 1.01E+00 | 1.16E-01 | 3.28E-01 | 1.14E+00 | 8.49E-02 | 3.12E-01 | 1.19E+00 | 2.41E-01 | 6.62E-01 | 1.27E+00 |
| SMC1A | 3.64E-09 | 1.90E-08 | -1.45E+00 | 1.97E-05 | 8.50E-05 | -1.06E+00 | 9.26E-03 | 3.93E-02 | -1.07E+00 | 1.78E-02 | 6.78E-02 | 1.10E+00 | 1.28E-02 | 4.55E-02 | 1.10E+00 | 3.39E-01 | 9.56E-01 | 1.43E+00 | 1.48E-01 | 5.45E-01 | 1.15E+00 | 2.64E-01 | 7.25E-01 | 1.25E+00 |
| DUBR | 3.64E-09 | 1.90E-08 | -1.99E+00 | 3.93E-07 | 1.70E-06 | -1.52E+00 | 3.91E-07 | 1.66E-06 | -2.11E+00 | 6.22E-04 | 2.38E-03 | -1.25E+00 | 9.97E-04 | 3.55E-03 | -1.40E+00 | 2.41E-02 | 6.79E-02 | -1.04E+00 | 1.33E-02 | 4.87E-02 | 1.05E+00 | 1.55E-02 | 4.25E-02 | -1.02E+00 |
| AL360270.2 | 3.65E-09 | 1.90E-08 | 4.24E+00 | 7.41E-07 | 3.20E-06 | 4.11E+00 | 8.30E-05 | 3.52E-04 | 3.86E+00 | 2.26E-03 | 8.64E-03 | 3.26E+00 | 7.26E-03 | 2.59E-02 | 2.78E+00 | 2.02E-02 | 5.69E-02 | 2.33E+00 | 3.11E-02 | 1.14E-01 | 2.23E+00 | 3.54E-01 | 9.74E-01 | 1.36E+00 |
| RBFA | 3.66E-09 | 1.91E-08 | -1.26E+00 | 9.72E-08 | 4.20E-07 | -1.05E+00 | 9.67E-03 | 4.11E-02 | -1.00E+00 | 1.17E-03 | 4.49E-03 | 1.05E+00 | 1.20E-06 | 4.26E-06 | -1.34E+00 | 1.88E-03 | 5.29E-03 | -1.01E+00 | 6.43E-05 | 2.36E-04 | -1.15E+00 | 3.17E-02 | 8.72E-02 | 1.05E+00 |
| ERP29 | 3.66E-09 | 1.91E-08 | -1.20E+00 | 1.79E-06 | 7.73E-06 | -1.08E+00 | 7.17E-04 | 3.05E-03 | -1.20E+00 | 1.97E-03 | 7.53E-03 | -1.10E+00 | 4.54E-04 | 1.62E-03 | -1.09E+00 | 2.53E-02 | 7.13E-02 | 1.01E+00 | 3.19E-03 | 1.17E-02 | -1.04E+00 | 8.90E-03 | 2.45E-02 | -1.09E+00 |
| LRRC2 | 3.67E-09 | 1.91E-08 | -2.08E+00 | 4.43E-04 | 1.91E-03 | -1.09E+00 | 3.67E-08 | 1.56E-07 | -2.29E+00 | 3.55E-04 | 1.36E-03 | -1.33E+00 | 9.70E-05 | 3.46E-04 | -1.89E+00 | 2.04E-03 | 5.76E-03 | -1.30E+00 | 2.99E-11 | 1.10E-10 | -3.00E+00 | 3.25E-06 | 8.93E-06 | -1.75E+00 |
| HLA-DPB2 | 3.67E-09 | 1.91E-08 | -1.78E+00 | 1.79E-04 | 7.74E-04 | -1.03E+00 | 9.47E-05 | 4.02E-04 | -1.75E+00 | 1.16E-01 | 4.44E-01 | 1.12E+00 | 1.34E-03 | 4.77E-03 | -2.02E+00 | 1.11E-02 | 3.13E-02 | -1.34E+00 | 1.28E-01 | 4.69E-01 | -1.41E+00 | 3.10E-01 | 8.51E-01 | 1.25E+00 |
| TEX264 | 3.67E-09 | 1.91E-08 | -1.15E+00 | 2.15E-04 | 9.29E-04 | 1.16E+00 | 1.03E-03 | 4.36E-03 | -1.10E+00 | 7.66E-03 | 2.93E-02 | 1.09E+00 | 2.56E-07 | 9.11E-07 | -1.17E+00 | 2.31E-03 | 6.51E-03 | 1.10E+00 | 1.77E-02 | 6.49E-02 | 1.06E+00 | 9.12E-02 | 2.50E-01 | 1.05E+00 |
| CCNL1 | 3.68E-09 | 1.92E-08 | 1.78E+00 | 4.57E-08 | 1.97E-07 | -1.06E+00 | 7.05E-04 | 3.00E-03 | 1.45E+00 | 4.27E-05 | 1.63E-04 | -1.17E+00 | 7.73E-02 | 2.76E-01 | 1.39E+00 | 1.80E-05 | 5.06E-05 | -1.10E+00 | 1.02E-01 | 3.74E-01 | 1.16E+00 | 1.17E-06 | 3.21E-06 | -1.26E+00 |
| SERINC1 | 3.79E-09 | 1.98E-08 | -1.17E+00 | 4.07E-09 | 1.76E-08 | -1.04E+00 | 2.90E-04 | 1.23E-03 | -1.19E+00 | 4.61E-04 | 1.76E-03 | -1.00E+00 | 4.64E-07 | 1.65E-06 | -1.28E+00 | 6.69E-06 | 1.89E-05 | -1.12E+00 | 1.71E-05 | 6.28E-05 | -1.22E+00 | 1.88E-04 | 5.17E-04 | -1.12E+00 |
| DGUOK | 3.81E-09 | 1.99E-08 | -1.12E+00 | 4.73E-09 | 2.04E-08 | 1.01E+00 | 8.54E-04 | 3.63E-03 | -1.09E+00 | 4.42E-03 | 1.69E-02 | 1.08E+00 | 1.46E-05 | 5.21E-05 | -1.15E+00 | 1.34E-03 | 3.77E-03 | 1.01E+00 | 2.93E-02 | 1.07E-01 | 1.08E+00 | 2.04E-01 | 5.61E-01 | 1.14E+00 |
| TOR1A | 3.82E-09 | 1.99E-08 | -1.42E+00 | 1.08E-03 | 4.66E-03 | 1.11E+00 | 1.07E-05 | 4.57E-05 | -1.17E+00 | 1.20E-05 | 4.56E-05 | -1.02E+00 | 2.46E-08 | 8.77E-08 | -1.44E+00 | 1.41E-05 | 3.98E-05 | -1.15E+00 | 3.43E-10 | 1.26E-09 | -1.57E+00 | 2.70E-03 | 7.41E-03 | -1.09E+00 |
| AL365258.2 | 3.82E-09 | 1.99E-08 | -1.33E+00 | 8.07E-55 | 3.49E-54 | -2.04E+00 | 6.11E-04 | 2.60E-03 | -1.25E+00 | 4.02E-17 | 1.54E-16 | -1.82E+00 | 1.47E-05 | 5.23E-05 | -1.29E+00 | 1.99E-19 | 5.61E-19 | -1.83E+00 | 1.58E-06 | 5.79E-06 | -1.34E+00 | 5.81E-16 | 1.60E-15 | -1.87E+00 |
| KIAA1683 | 3.84E-09 | 2.00E-08 | 4.08E+00 | 1.44E-01 | 6.24E-01 | 1.31E+00 | 7.22E-06 | 3.07E-05 | 3.02E+00 | 1.84E-01 | 7.04E-01 | 1.14E+00 | 2.44E-01 | 8.70E-01 | 1.71E+00 | 6.97E-02 | 1.96E-01 | -1.10E+00 | 5.43E-02 | 1.99E-01 | 2.05E+00 | 3.39E-01 | 9.31E-01 | 1.17E+00 |
| EIF3C | 3.84E-09 | 2.00E-08 | -1.09E+00 | 1.61E-06 | 6.96E-06 | 1.06E+00 | 2.11E-01 | 8.96E-01 | 1.06E+00 | 2.62E-01 | 1.00E+00 | 1.19E+00 | 6.52E-04 | 2.33E-03 | -1.06E+00 | 9.55E-02 | 2.69E-01 | 1.10E+00 | 4.87E-03 | 1.79E-02 | -1.03E+00 | 1.52E-01 | 4.19E-01 | 1.21E+00 |
| TCHP | 3.86E-09 | 2.01E-08 | -1.59E+00 | 2.60E-05 | 1.13E-04 | -1.11E+00 | 7.24E-03 | 3.08E-02 | -1.14E+00 | 3.32E-02 | 1.27E-01 | 1.08E+00 | 1.14E-04 | 4.07E-04 | -1.39E+00 | 1.52E-02 | 4.27E-02 | -1.07E+00 | 1.11E-04 | 4.08E-04 | -1.15E+00 | 1.87E-02 | 5.13E-02 | 1.04E+00 |
| COA6 | 3.88E-09 | 2.02E-08 | -1.25E+00 | 2.68E-06 | 1.16E-05 | 1.01E+00 | 2.97E-10 | 1.26E-09 | -1.37E+00 | 1.73E-07 | 6.62E-07 | -1.06E+00 | 1.17E-09 | 4.16E-09 | -1.57E+00 | 1.00E-03 | 2.82E-03 | 1.06E+00 | 1.14E-08 | 4.17E-08 | -1.31E+00 | 1.66E-03 | 4.57E-03 | -1.03E+00 |
| CCDC25 | 3.88E-09 | 2.02E-08 | -1.24E+00 | 5.84E-08 | 2.52E-07 | -1.05E+00 | 8.71E-03 | 3.70E-02 | 1.01E+00 | 3.34E-05 | 1.28E-04 | 1.03E+00 | 1.87E-05 | 6.68E-05 | -1.12E+00 | 4.97E-03 | 1.40E-02 | 1.03E+00 | 4.69E-03 | 1.72E-02 | -1.10E+00 | 2.17E-03 | 5.97E-03 | -1.13E+00 |
| BRWD1 | 3.89E-09 | 2.03E-08 | -1.16E+00 | 3.37E-05 | 1.46E-04 | 1.13E+00 | 1.04E-04 | 4.40E-04 | -1.11E+00 | 8.07E-03 | 3.08E-02 | 1.12E+00 | 1.88E-02 | 6.69E-02 | 1.14E+00 | 1.98E-01 | 5.59E-01 | 1.35E+00 | 1.36E-02 | 4.99E-02 | 1.09E+00 | 2.55E-03 | 6.99E-03 | -1.04E+00 |
| ING5 | 3.89E-09 | 2.03E-08 | -1.20E+00 | 1.56E-08 | 6.75E-08 | -1.07E+00 | 2.95E-06 | 1.25E-05 | -1.22E+00 | 1.84E-06 | 7.02E-06 | -1.06E+00 | 4.29E-04 | 1.53E-03 | -1.14E+00 | 9.09E-03 | 2.56E-02 | 1.06E+00 | 2.25E-02 | 8.27E-02 | 1.16E+00 | 5.20E-02 | 1.43E-01 | 1.12E+00 |
| ZFYVE27 | 3.90E-09 | 2.03E-08 | -1.58E+00 | 2.55E-08 | 1.10E-07 | -1.34E+00 | 3.05E-06 | 1.29E-05 | -1.46E+00 | 1.64E-05 | 6.27E-05 | -1.17E+00 | 3.24E-04 | 1.15E-03 | -1.26E+00 | 4.42E-03 | 1.25E-02 | -1.06E+00 | 1.08E-11 | 3.96E-11 | -1.67E+00 | 1.89E-08 | 5.20E-08 | -1.51E+00 |
| OR2X1P | 3.91E-09 | 2.04E-08 | -1.96E+00 | 1.03E-05 | 4.46E-05 | -1.27E+00 | 6.20E-04 | 2.63E-03 | -1.83E+00 | 1.10E-02 | 4.19E-02 | -1.22E+00 | 4.25E-04 | 1.52E-03 | -1.97E+00 | 4.26E-02 | 1.20E-01 | -1.09E+00 | 2.28E-02 | 8.37E-02 | -1.32E+00 | 9.64E-02 | 2.65E-01 | -1.18E+00 |
| CYHR1 | 3.92E-09 | 2.04E-08 | -1.32E+00 | 1.80E-06 | 7.79E-06 | -1.01E+00 | 6.37E-07 | 2.71E-06 | -1.39E+00 | 2.63E-03 | 1.01E-02 | 1.10E+00 | 5.69E-10 | 2.03E-09 | -1.54E+00 | 6.01E-06 | 1.70E-05 | -1.10E+00 | 1.89E-03 | 6.92E-03 | -1.04E+00 | 9.09E-02 | 2.50E-01 | 1.10E+00 |
| ACO1 | 3.93E-09 | 2.05E-08 | -1.59E+00 | 1.87E-04 | 8.10E-04 | -1.03E+00 | 2.47E-07 | 1.05E-06 | -1.59E+00 | 4.71E-04 | 1.80E-03 | -1.10E+00 | 3.78E-05 | 1.35E-04 | -1.32E+00 | 6.31E-02 | 1.78E-01 | 1.17E+00 | 3.55E-03 | 1.30E-02 | -1.07E+00 | 2.35E-01 | 6.47E-01 | 1.14E+00 |
| YME1L1 | 3.94E-09 | 2.05E-08 | -1.10E+00 | 1.59E-22 | 6.86E-22 | -1.37E+00 | 2.88E-04 | 1.22E-03 | -1.20E+00 | 1.36E-07 | 5.18E-07 | -1.29E+00 | 7.45E-10 | 2.66E-09 | -1.42E+00 | 8.90E-12 | 2.51E-11 | -1.47E+00 | 1.18E-09 | 4.34E-09 | -1.50E+00 | 2.19E-09 | 6.01E-09 | -1.53E+00 |
| RTP4 | 3.94E-09 | 2.05E-08 | -1.67E+00 | 1.47E-06 | 6.37E-06 | -1.24E+00 | 1.82E-15 | 7.72E-15 | -2.32E+00 | 1.63E-12 | 6.22E-12 | -1.74E+00 | 3.24E-13 | 1.16E-12 | -3.03E+00 | 1.91E-08 | 5.37E-08 | -1.74E+00 | 2.74E-24 | 1.01E-23 | -3.14E+00 | 6.58E-14 | 1.81E-13 | -1.94E+00 |
| KLHL36 | 3.96E-09 | 2.06E-08 | -1.41E+00 | 1.19E-05 | 5.15E-05 | -1.08E+00 | 3.23E-06 | 1.37E-05 | -1.43E+00 | 3.05E-06 | 1.16E-05 | -1.33E+00 | 4.53E-04 | 1.62E-03 | -1.15E+00 | 3.76E-03 | 1.06E-02 | -1.13E+00 | 1.01E-04 | 3.72E-04 | -1.18E+00 | 4.95E-05 | 1.36E-04 | -1.22E+00 |
| ELAVL1 | 3.98E-09 | 2.07E-08 | -1.28E+00 | 4.68E-05 | 2.02E-04 | 1.07E+00 | 2.78E-07 | 1.18E-06 | -1.18E+00 | 2.54E-07 | 9.70E-07 | -1.09E+00 | 6.72E-06 | 2.40E-05 | -1.22E+00 | 9.55E-02 | 2.69E-01 | 1.12E+00 | 2.18E-04 | 8.00E-04 | -1.10E+00 | 4.79E-02 | 1.31E-01 | 1.06E+00 |
| SF3B4 | 4.01E-09 | 2.09E-08 | -1.30E+00 | 1.09E-06 | 4.73E-06 | 1.01E+00 | 2.29E-06 | 9.75E-06 | -1.35E+00 | 2.34E-05 | 8.92E-05 | -1.13E+00 | 1.02E-07 | 3.64E-07 | -1.38E+00 | 1.30E-03 | 3.67E-03 | 1.02E+00 | 5.47E-06 | 2.01E-05 | -1.27E+00 | 1.52E-02 | 4.17E-02 | -1.01E+00 |
| PRPF39 | 4.02E-09 | 2.10E-08 | -1.45E+00 | 2.50E-05 | 1.08E-04 | -1.02E+00 | 1.75E-02 | 7.44E-02 | -1.08E+00 | 3.02E-02 | 1.15E-01 | 1.13E+00 | 1.59E-02 | 5.67E-02 | -1.01E+00 | 8.64E-02 | 2.43E-01 | 1.13E+00 | 1.60E-05 | 5.87E-05 | -1.12E+00 | 4.21E-03 | 1.16E-02 | 1.02E+00 |
| HCK | 4.09E-09 | 2.13E-08 | -2.14E+00 | 3.65E-08 | 1.58E-07 | -1.68E+00 | 4.26E-27 | 1.81E-26 | -2.58E+00 | 2.25E-19 | 8.60E-19 | -1.73E+00 | 2.05E-05 | 7.30E-05 | -2.16E+00 | 2.85E-05 | 8.04E-05 | -2.12E+00 | 3.50E-02 | 1.28E-01 | -2.13E+00 | 7.76E-02 | 2.13E-01 | -1.92E+00 |
| CSNK1A1P1 | 4.11E-09 | 2.14E-08 | -2.49E+00 | 3.60E-06 | 1.56E-05 | -1.54E+00 | 1.32E-07 | 5.59E-07 | -1.73E+00 | 1.60E-06 | 6.13E-06 | -1.38E+00 | 7.20E-07 | 2.57E-06 | -1.92E+00 | 6.79E-05 | 1.92E-04 | -1.41E+00 | 5.52E-08 | 2.03E-07 | -1.93E+00 | 2.47E-06 | 6.77E-06 | -1.53E+00 |
| RFT1 | 4.14E-09 | 2.16E-08 | -1.65E+00 | 7.02E-05 | 3.03E-04 | -1.02E+00 | 6.29E-09 | 2.67E-08 | -1.56E+00 | 2.20E-06 | 8.41E-06 | -1.16E+00 | 9.30E-06 | 3.31E-05 | -1.47E+00 | 4.25E-02 | 1.20E-01 | 1.15E+00 | 4.24E-11 | 1.56E-10 | -1.49E+00 | 5.58E-03 | 1.53E-02 | -1.01E+00 |
| DERA | 4.16E-09 | 2.17E-08 | -1.25E+00 | 3.36E-06 | 1.45E-05 | 1.03E+00 | 1.35E-02 | 5.72E-02 | 1.04E+00 | 2.04E-04 | 7.81E-04 | 1.03E+00 | 5.77E-06 | 2.06E-05 | -1.45E+00 | 4.06E-05 | 1.14E-04 | -1.32E+00 | 2.13E-08 | 7.81E-08 | -1.27E+00 | 5.32E-07 | 1.46E-06 | -1.26E+00 |
| AC026124.1 | 4.16E-09 | 2.17E-08 | -1.26E+00 | 2.83E-02 | 1.22E-01 | 1.30E+00 | 2.13E-04 | 9.04E-04 | -1.26E+00 | 1.64E-01 | 6.27E-01 | 1.26E+00 | 3.32E-02 | 1.18E-01 | -1.26E+00 | 3.53E-01 | 9.95E-01 | 1.28E+00 | 2.70E-01 | 9.93E-01 | 1.28E+00 | 3.37E-01 | 9.26E-01 | 1.19E+00 |
| HOOK1 | 4.18E-09 | 2.18E-08 | -1.66E+00 | 2.58E-09 | 1.12E-08 | -1.59E+00 | 1.93E-05 | 8.22E-05 | -1.48E+00 | 1.69E-03 | 6.45E-03 | -1.08E+00 | 7.94E-04 | 2.83E-03 | -1.58E+00 | 6.09E-03 | 1.72E-02 | -1.20E+00 | 1.13E-01 | 4.14E-01 | 1.09E+00 | 3.44E-01 | 9.44E-01 | 1.26E+00 |
| MTIF2 | 4.21E-09 | 2.19E-08 | -1.43E+00 | 2.45E-08 | 1.06E-07 | -1.19E+00 | 1.40E-07 | 5.94E-07 | -1.41E+00 | 7.83E-08 | 2.99E-07 | -1.24E+00 | 1.87E-04 | 6.69E-04 | -1.14E+00 | 7.60E-03 | 2.14E-02 | 1.04E+00 | 4.79E-06 | 1.76E-05 | -1.24E+00 | 2.47E-03 | 6.79E-03 | -1.06E+00 |
| PRKRA | 4.21E-09 | 2.19E-08 | -1.17E+00 | 5.16E-09 | 2.23E-08 | -1.06E+00 | 1.38E-05 | 5.84E-05 | -1.18E+00 | 9.74E-05 | 3.72E-04 | -1.01E+00 | 7.80E-06 | 2.78E-05 | -1.27E+00 | 2.44E-04 | 6.88E-04 | -1.13E+00 | 1.75E-03 | 6.41E-03 | -1.17E+00 | 2.46E-01 | 6.77E-01 | 1.10E+00 |
| ALPK1 | 4.24E-09 | 2.21E-08 | -1.40E+00 | 3.83E-08 | 1.66E-07 | -1.17E+00 | 2.49E-09 | 1.06E-08 | -1.67E+00 | 2.92E-06 | 1.12E-05 | -1.22E+00 | 6.10E-05 | 2.17E-04 | -1.14E+00 | 1.61E-03 | 4.53E-03 | 1.03E+00 | 7.88E-05 | 2.89E-04 | -1.29E+00 | 3.32E-06 | 9.13E-06 | -1.35E+00 |
| TOMM70 | 4.27E-09 | 2.23E-08 | -1.40E+00 | 1.48E-04 | 6.40E-04 | 1.05E+00 | 2.20E-05 | 9.37E-05 | -1.23E+00 | 1.91E-04 | 7.31E-04 | -1.02E+00 | 7.01E-06 | 2.50E-05 | -1.46E+00 | 4.06E-02 | 1.14E-01 | 1.11E+00 | 3.36E-12 | 1.23E-11 | -1.56E+00 | 5.51E-07 | 1.51E-06 | -1.26E+00 |
| GGCT | 4.31E-09 | 2.24E-08 | -1.10E+00 | 4.39E-10 | 1.90E-09 | -1.04E+00 | 2.19E-02 | 9.32E-02 | -1.03E+00 | 1.60E-03 | 6.11E-03 | -1.02E+00 | 6.51E-07 | 2.32E-06 | -1.23E+00 | 4.40E-03 | 1.24E-02 | 1.06E+00 | 1.32E-02 | 4.84E-02 | -1.07E+00 | 3.04E-01 | 8.35E-01 | 1.12E+00 |
| MRPL12 | 4.31E-09 | 2.25E-08 | -1.15E+00 | 4.12E-08 | 1.78E-07 | -1.02E+00 | 7.24E-03 | 3.08E-02 | 1.02E+00 | 1.82E-03 | 6.96E-03 | 1.02E+00 | 6.74E-09 | 2.40E-08 | -1.41E+00 | 1.74E-03 | 4.90E-03 | 1.03E+00 | 1.59E-04 | 5.84E-04 | -1.05E+00 | 2.78E-01 | 7.63E-01 | 1.30E+00 |
| PTCD3 | 4.35E-09 | 2.27E-08 | -1.28E+00 | 1.46E-05 | 6.31E-05 | -1.02E+00 | 4.18E-05 | 1.78E-04 | -1.29E+00 | 4.99E-04 | 1.90E-03 | -1.03E+00 | 8.77E-03 | 3.13E-02 | 1.01E+00 | 8.54E-02 | 2.41E-01 | 1.13E+00 | 3.81E-07 | 1.40E-06 | -1.24E+00 | 3.32E-03 | 9.12E-03 | 1.03E+00 |
| TRMT5 | 4.36E-09 | 2.27E-08 | -1.67E+00 | 1.43E-05 | 6.16E-05 | -1.05E+00 | 5.81E-11 | 2.47E-10 | -1.63E+00 | 1.80E-09 | 6.87E-09 | -1.31E+00 | 3.20E-06 | 1.14E-05 | -1.75E+00 | 2.98E-03 | 8.41E-03 | -1.05E+00 | 1.00E-14 | 3.67E-14 | -1.73E+00 | 5.52E-08 | 1.52E-07 | -1.36E+00 |
| RPS15AP11 | 4.37E-09 | 2.28E-08 | -1.39E+00 | 4.47E-51 | 1.93E-50 | -2.05E+00 | 9.62E-05 | 4.09E-04 | -1.37E+00 | 8.88E-27 | 3.39E-26 | -2.20E+00 | 1.74E-07 | 6.20E-07 | -1.45E+00 | 1.28E-25 | 3.60E-25 | -2.04E+00 | 9.64E-07 | 3.54E-06 | -1.35E+00 | 1.57E-19 | 4.32E-19 | -1.95E+00 |
| PPA1 | 4.39E-09 | 2.29E-08 | -1.22E+00 | 3.68E-06 | 1.59E-05 | -1.10E+00 | 4.40E-02 | 1.87E-01 | -1.10E+00 | 7.65E-02 | 2.92E-01 | -1.06E+00 | 1.26E-09 | 4.50E-09 | -1.27E+00 | 7.06E-02 | 1.99E-01 | 1.07E+00 | 3.17E-03 | 1.16E-02 | 1.05E+00 | 3.33E-02 | 9.13E-02 | 1.32E+00 |
| ZKSCAN8 | 4.39E-09 | 2.29E-08 | -1.41E+00 | 3.24E-05 | 1.40E-04 | 1.01E+00 | 7.19E-06 | 3.05E-05 | -1.47E+00 | 2.68E-03 | 1.02E-02 | -1.02E+00 | 1.04E-02 | 3.69E-02 | -1.02E+00 | 3.12E-02 | 8.79E-02 | 1.02E+00 | 3.42E-06 | 1.25E-05 | -1.27E+00 | 4.01E-06 | 1.10E-05 | -1.30E+00 |
| SLC5A3 | 4.40E-09 | 2.29E-08 | -1.49E+00 | 2.04E-05 | 8.80E-05 | -1.12E+00 | 2.29E-07 | 9.73E-07 | -1.45E+00 | 6.38E-02 | 2.44E-01 | 1.23E+00 | 5.05E-03 | 1.80E-02 | -1.04E+00 | 4.88E-02 | 1.38E-01 | 1.05E+00 | 1.54E-02 | 5.65E-02 | 1.02E+00 | 7.44E-02 | 2.04E-01 | 1.08E+00 |
| SELENOW | 4.42E-09 | 2.30E-08 | -1.08E+00 | 5.90E-07 | 2.55E-06 | 1.10E+00 | 3.82E-02 | 1.62E-01 | 1.03E+00 | 1.36E-02 | 5.18E-02 | 1.10E+00 | 4.06E-03 | 1.45E-02 | 1.06E+00 | 2.17E-01 | 6.12E-01 | 1.25E+00 | 1.14E-01 | 4.20E-01 | 1.10E+00 | 3.39E-01 | 9.31E-01 | 1.16E+00 |
| LSM12 | 4.42E-09 | 2.30E-08 | -1.25E+00 | 8.23E-10 | 3.55E-09 | -1.21E+00 | 5.99E-05 | 2.54E-04 | -1.18E+00 | 1.93E-08 | 7.35E-08 | -1.26E+00 | 8.70E-04 | 3.10E-03 | -1.00E+00 | 4.70E-04 | 1.32E-03 | -1.04E+00 | 5.23E-06 | 1.92E-05 | -1.19E+00 | 1.86E-06 | 5.10E-06 | -1.25E+00 |
| YTHDF1 | 4.42E-09 | 2.31E-08 | -1.70E+00 | 5.59E-07 | 2.41E-06 | -1.31E+00 | 3.15E-06 | 1.34E-05 | -1.47E+00 | 2.16E-03 | 8.24E-03 | 1.01E+00 | 3.54E-05 | 1.26E-04 | -1.27E+00 | 7.60E-03 | 2.14E-02 | 1.09E+00 | 4.60E-10 | 1.69E-09 | -1.34E+00 | 1.02E-04 | 2.80E-04 | -1.08E+00 |
| MLLT6 | 4.42E-09 | 2.31E-08 | -1.33E+00 | 2.70E-08 | 1.16E-07 | -1.15E+00 | 9.92E-05 | 4.21E-04 | -1.20E+00 | 1.28E-05 | 4.87E-05 | -1.10E+00 | 9.24E-05 | 3.29E-04 | -1.22E+00 | 1.26E-02 | 3.55E-02 | 1.05E+00 | 1.85E-05 | 6.80E-05 | -1.30E+00 | 9.16E-03 | 2.52E-02 | -1.02E+00 |
| NQO2 | 4.46E-09 | 2.32E-08 | -1.53E+00 | 6.12E-08 | 2.64E-07 | -1.25E+00 | 1.32E-02 | 5.63E-02 | -1.09E+00 | 3.13E-02 | 1.20E-01 | 1.11E+00 | 8.58E-06 | 3.06E-05 | -1.21E+00 | 1.96E-04 | 5.52E-04 | -1.08E+00 | 1.23E-06 | 4.50E-06 | -1.13E+00 | 6.89E-03 | 1.89E-02 | 1.02E+00 |
| PLEKHA5 | 4.48E-09 | 2.33E-08 | -1.30E+00 | 3.33E-06 | 1.44E-05 | 1.01E+00 | 5.20E-06 | 2.21E-05 | -1.29E+00 | 4.76E-04 | 1.82E-03 | -1.02E+00 | 1.98E-05 | 7.07E-05 | -1.43E+00 | 1.08E-02 | 3.03E-02 | -1.05E+00 | 6.91E-06 | 2.54E-05 | -1.20E+00 | 1.08E-03 | 2.97E-03 | -1.13E+00 |

| | | | | | | | | | | | | | | | | | | | | | | | | |
|---|---|---|---|---|---|---|---|---|---|---|---|---|---|---|---|---|---|---|---|---|---|---|---|---|
| ADAMTSL4-AS1 | 4.49E-09 | 2.34E-08 | 3.33E+00 | 9.12E-03 | 3.94E-02 | 2.34E+00 | 4.46E-04 | 1.90E-03 | 2.24E+00 | 1.29E-02 | 4.92E-02 | 2.18E+00 | 3.38E-02 | 1.20E-01 | 2.65E+00 | 1.79E-01 | 5.04E-01 | 1.87E+00 | 5.61E-02 | 2.06E-01 | 1.93E+00 | 1.66E-01 | 4.56E-01 | 1.76E+00 |
| PANX1 | 4.52E-09 | 2.35E-08 | -2.72E+00 | 1.89E-09 | 8.16E-09 | -2.78E+00 | 3.40E-06 | 1.44E-05 | -1.79E+00 | 8.66E-07 | 3.31E-06 | -1.63E+00 | 7.70E-03 | 2.75E-02 | -1.80E+00 | 1.84E-02 | 5.19E-02 | -1.60E+00 | 1.02E-05 | 3.76E-05 | 1.02E+00 | 3.06E-08 | 8.40E-08 | -1.77E+00 |
| FBXW8 | 4.55E-09 | 2.37E-08 | -1.90E+00 | 5.67E-06 | 2.45E-05 | -1.26E+00 | 5.81E-05 | 2.47E-04 | -1.53E+00 | 9.72E-03 | 3.71E-02 | -1.03E+00 | 1.30E-03 | 4.65E-03 | -1.38E+00 | 5.90E-02 | 1.66E-01 | 1.08E+00 | 4.29E-07 | 1.57E-06 | -1.48E+00 | 5.14E-03 | 1.41E-02 | -1.03E+00 |
| NBAS | 4.57E-09 | 2.38E-08 | -1.44E+00 | 6.77E-06 | 2.92E-05 | -1.03E+00 | 1.11E-05 | 4.72E-05 | -1.24E+00 | 1.37E-03 | 5.25E-03 | 1.02E+00 | 7.25E-04 | 2.59E-03 | -1.05E+00 | 3.85E-02 | 1.09E-01 | 1.11E+00 | 2.91E-04 | 1.07E-03 | -1.09E+00 | 4.83E-02 | 1.33E-01 | 1.13E+00 |
| B4GAT1 | 4.59E-09 | 2.39E-08 | -2.27E+00 | 3.84E-05 | 1.66E-04 | -1.25E+00 | 8.18E-08 | 3.48E-07 | -1.49E+00 | 3.35E-10 | 1.28E-09 | -1.55E+00 | 2.26E-02 | 8.07E-02 | -1.52E+00 | 1.93E-01 | 5.44E-01 | 1.14E+00 | 4.74E-07 | 1.74E-06 | -2.11E+00 | 1.19E-03 | 3.27E-03 | -1.31E+00 |
| NOP10 | 4.59E-09 | 2.39E-08 | -1.11E+00 | 9.49E-15 | 4.10E-14 | -1.20E+00 | 5.36E-02 | 2.28E-01 | -1.04E+00 | 2.72E-04 | 1.04E-03 | -1.09E+00 | 3.20E-06 | 1.14E-05 | -1.19E+00 | 5.00E-05 | 1.41E-04 | -1.08E+00 | 3.44E-04 | 1.26E-03 | -1.23E+00 | 3.43E-02 | 9.41E-02 | -1.04E+00 |
| PIGS | 4.59E-09 | 2.39E-08 | -1.51E+00 | 1.90E-08 | 8.21E-08 | -1.31E+00 | 1.92E-03 | 8.17E-03 | -1.18E+00 | 2.94E-02 | 1.12E-01 | 1.13E+00 | 1.67E-05 | 5.94E-05 | -1.33E+00 | 2.12E-03 | 5.98E-03 | -1.04E+00 | 3.24E-09 | 1.19E-08 | -1.33E+00 | 6.34E-05 | 1.74E-04 | -1.07E+00 |
| RPL23AP82 | 4.60E-09 | 2.40E-08 | -2.27E+00 | 1.87E-06 | 8.08E-06 | -1.50E+00 | 6.80E-06 | 2.89E-05 | -1.65E+00 | 1.43E-06 | 5.46E-06 | -1.57E+00 | 4.30E-03 | 1.53E-02 | -1.35E+00 | 3.85E-03 | 1.09E-02 | -1.54E+00 | 1.75E-02 | 6.44E-02 | -1.24E+00 | 1.59E-01 | 4.37E-01 | 1.06E+00 |
| ZNF12 | 4.61E-09 | 2.40E-08 | -1.22E+00 | 4.76E-10 | 2.05E-09 | -1.18E+00 | 1.25E-08 | 5.31E-08 | -1.63E+00 | 1.91E-06 | 7.31E-06 | -1.30E+00 | 1.17E-09 | 4.17E-09 | -1.59E+00 | 1.96E-11 | 5.51E-11 | -1.70E+00 | 1.06E-09 | 3.88E-09 | -1.68E+00 | 8.81E-07 | 2.42E-06 | -1.37E+00 |
| HDGF | 4.61E-09 | 2.40E-08 | -1.22E+00 | 5.83E-08 | 2.52E-07 | -1.08E+00 | 1.41E-03 | 6.01E-03 | -1.06E+00 | 2.63E-03 | 1.01E-02 | 1.08E+00 | 3.02E-06 | 1.08E-05 | -1.20E+00 | 1.36E-03 | 3.85E-03 | -1.01E+00 | 4.04E-03 | 1.48E-02 | -1.09E+00 | 2.85E-01 | 7.84E-01 | 1.09E+00 |
| CFAP36 | 4.63E-09 | 2.41E-08 | -1.16E+00 | 5.67E-11 | 2.45E-10 | -1.12E+00 | 8.81E-04 | 3.74E-03 | -1.14E+00 | 4.22E-04 | 1.61E-03 | -1.05E+00 | 6.42E-04 | 2.29E-03 | -1.17E+00 | 2.16E-03 | 6.09E-03 | -1.06E+00 | 2.57E-03 | 9.41E-03 | 1.04E+00 | 1.17E-03 | 3.20E-03 | 1.01E+00 |
| RASSF2 | 4.64E-09 | 2.42E-08 | -1.64E+00 | 1.62E-07 | 7.01E-07 | -1.32E+00 | 3.64E-03 | 1.55E-02 | -1.14E+00 | 8.64E-04 | 3.30E-03 | -1.10E+00 | 1.69E-03 | 6.03E-03 | -1.32E+00 | 1.82E-03 | 5.14E-03 | -1.41E+00 | 2.11E-06 | 7.73E-06 | -2.10E+00 | 1.95E-05 | 5.35E-05 | -1.85E+00 |
| UCP2 | 4.70E-09 | 2.45E-08 | -1.36E+00 | 1.44E-02 | 6.20E-02 | 1.15E+00 | 1.97E-06 | 8.38E-06 | -1.37E+00 | 6.73E-04 | 2.57E-03 | -1.03E+00 | 1.44E-04 | 5.12E-04 | -1.23E+00 | 3.30E-01 | 9.30E-01 | 1.26E+00 | 6.39E-02 | 2.35E-01 | 1.04E+00 | 3.21E-01 | 8.81E-01 | 1.26E+00 |
| CPNE2 | 4.72E-09 | 2.46E-08 | -1.75E+00 | 1.23E-06 | 5.33E-06 | -1.33E+00 | 8.00E-04 | 3.40E-03 | -1.21E+00 | 8.82E-03 | 3.37E-02 | 1.06E+00 | 6.49E-02 | 2.32E-01 | -1.38E+00 | 1.41E-01 | 3.98E-01 | -1.11E+00 | 1.48E-02 | 5.44E-02 | -1.28E+00 | 7.73E-02 | 2.12E-01 | 1.01E+00 |
| PPM1F | 4.73E-09 | 2.47E-08 | -1.12E+00 | 2.11E-05 | 9.13E-05 | 1.15E+00 | 3.37E-02 | 1.43E-01 | 1.14E+00 | 1.23E-01 | 4.70E-01 | 1.39E+00 | 1.79E-02 | 6.38E-02 | -1.24E+00 | 1.54E-01 | 4.33E-01 | 1.12E+00 | 5.24E-04 | 1.92E-03 | -1.65E+00 | 1.63E-02 | 4.47E-02 | -1.33E+00 |
| APBB1 | 4.75E-09 | 2.47E-08 | -1.63E+00 | 7.01E-07 | 3.03E-06 | -1.19E+00 | 1.79E-01 | 7.62E-01 | 1.27E+00 | 2.54E-01 | 9.72E-01 | 1.29E+00 | 1.95E-01 | 6.97E-01 | 1.15E+00 | 2.12E-01 | 5.97E-01 | 1.13E+00 | 2.04E-01 | 7.47E-01 | 1.15E+00 | 2.45E-01 | 6.73E-01 | 1.10E+00 |
| CDK13 | 4.75E-09 | 2.47E-08 | -1.31E+00 | 6.95E-05 | 3.00E-04 | 1.05E+00 | 2.42E-08 | 1.03E-07 | -1.56E+00 | 3.08E-04 | 1.18E-03 | -1.00E+00 | 4.80E-10 | 1.71E-09 | -1.52E+00 | 3.93E-07 | 1.11E-06 | -1.18E+00 | 5.95E-11 | 2.18E-10 | -1.53E+00 | 3.17E-07 | 8.71E-07 | -1.43E+00 |
| VASP | 4.76E-09 | 2.48E-08 | -1.15E+00 | 7.30E-08 | 3.15E-07 | 1.03E+00 | 1.03E-06 | 4.38E-06 | -1.22E+00 | 2.59E-06 | 9.87E-06 | -1.12E+00 | 2.38E-05 | 8.48E-05 | -1.17E+00 | 2.33E-03 | 6.58E-03 | -1.06E+00 | 5.94E-04 | 2.18E-03 | -1.17E+00 | 4.95E-02 | 1.36E-01 | -1.08E+00 |
| LRRC41 | 4.79E-09 | 2.49E-08 | -1.20E+00 | 2.85E-04 | 1.23E-03 | 1.11E+00 | 1.42E-08 | 6.04E-08 | -1.29E+00 | 1.46E-07 | 5.57E-07 | -1.12E+00 | 4.10E-07 | 1.46E-06 | -1.32E+00 | 1.01E-02 | 2.86E-02 | 1.09E+00 | 3.76E-07 | 1.38E-06 | -1.32E+00 | 3.04E-04 | 8.35E-04 | -1.07E+00 |
| AL512844.2 | 4.80E-09 | 2.50E-08 | -2.00E+00 | 1.57E-09 | 6.77E-09 | -2.05E+00 | 1.83E-08 | 7.78E-08 | -1.79E+00 | 2.25E-10 | 8.60E-10 | -1.84E+00 | 1.24E-04 | 4.43E-04 | -1.75E+00 | 1.68E-03 | 4.73E-03 | -1.37E+00 | 1.91E-10 | 7.00E-10 | -1.70E+00 | 6.89E-12 | 1.89E-11 | -1.92E+00 |
| ATG9A | 4.82E-09 | 2.51E-08 | -1.83E+00 | 8.95E-08 | 3.87E-07 | -1.46E+00 | 9.00E-03 | 3.82E-02 | -1.20E+00 | 5.85E-02 | 2.23E-01 | 1.08E+00 | 3.61E-03 | 1.29E-02 | -1.27E+00 | 1.92E-02 | 5.42E-02 | -1.05E+00 | 6.66E-06 | 2.45E-05 | -1.13E+00 | 1.94E-05 | 5.34E-05 | -1.20E+00 |
| AP006222.1 | 4.83E-09 | 2.51E-08 | -2.13E+00 | 6.29E-06 | 2.72E-05 | -1.32E+00 | 1.80E-10 | 7.63E-10 | -2.19E+00 | 9.20E-11 | 3.51E-10 | -2.10E+00 | 1.75E-05 | 6.22E-05 | -2.61E+00 | 1.02E-03 | 2.86E-03 | -1.52E+00 | 4.92E-08 | 1.81E-07 | -2.37E+00 | 7.08E-05 | 1.94E-04 | -1.50E+00 |
| MARS | 4.83E-09 | 2.52E-08 | -1.50E+00 | 2.38E-04 | 1.03E-03 | 1.01E+00 | 8.79E-02 | 3.73E-01 | -1.01E+00 | 2.07E-01 | 7.91E-01 | 1.26E+00 | 1.13E-04 | 4.03E-04 | -1.43E+00 | 4.07E-03 | 1.15E-02 | -1.17E+00 | 6.22E-06 | 2.28E-05 | -1.30E+00 | 1.14E-01 | 3.12E-01 | 1.15E+00 |
| TMEM19 | 4.91E-09 | 2.56E-08 | -1.57E+00 | 4.46E-06 | 1.93E-05 | -1.08E+00 | 1.23E-05 | 5.24E-05 | -1.57E+00 | 4.28E-03 | 1.63E-02 | -1.00E+00 | 2.55E-03 | 9.08E-03 | -1.40E+00 | 1.80E-01 | 5.06E-01 | 1.30E+00 | 3.48E-04 | 1.28E-03 | -1.39E+00 | 5.44E-02 | 1.49E-01 | -1.01E+00 |
| NREP | 4.91E-09 | 2.56E-08 | -1.27E+00 | 1.05E-07 | 4.55E-07 | -1.04E+00 | 3.30E-11 | 1.40E-10 | -1.65E+00 | 2.56E-14 | 9.79E-14 | -1.56E+00 | 8.90E-11 | 3.17E-10 | -1.60E+00 | 6.87E-12 | 1.94E-11 | -1.67E+00 | 4.63E-13 | 1.70E-12 | -1.81E+00 | 8.02E-10 | 2.20E-09 | -1.55E+00 |
| TCEAL9 | 4.91E-09 | 2.56E-08 | -1.27E+00 | 7.80E-07 | 3.37E-06 | -1.07E+00 | 2.23E-02 | 9.46E-02 | 1.06E+00 | 6.58E-03 | 2.51E-02 | 1.14E+00 | 2.35E-05 | 8.39E-05 | -1.23E+00 | 2.00E-03 | 5.63E-03 | 1.02E+00 | 3.54E-08 | 1.30E-07 | -1.22E+00 | 3.26E-04 | 8.94E-04 | -1.07E+00 |
| ALDH18A1 | 4.96E-09 | 2.59E-08 | -1.51E+00 | 2.23E-05 | 9.62E-05 | -1.01E+00 | 3.95E-06 | 1.68E-05 | -1.26E+00 | 7.80E-04 | 2.98E-03 | 1.02E+00 | 3.02E-06 | 1.08E-05 | -1.46E+00 | 5.69E-03 | 1.60E-02 | 1.05E+00 | 4.98E-05 | 1.83E-04 | -1.21E+00 | 9.22E-02 | 2.53E-01 | 1.43E+00 |
| ATF6B | 4.97E-09 | 2.59E-08 | -1.19E+00 | 5.20E-03 | 2.24E-02 | 1.22E+00 | 3.30E-03 | 1.40E-02 | -1.07E+00 | 9.52E-03 | 3.64E-02 | 1.10E+00 | 1.48E-03 | 5.27E-03 | 1.01E+00 | 3.75E-03 | 1.06E-02 | 1.04E+00 | 1.58E-02 | 5.80E-02 | -1.06E+00 | 8.44E-02 | 2.32E-01 | 1.05E+00 |
| DDHD2 | 4.97E-09 | 2.59E-08 | -1.49E+00 | 3.63E-06 | 1.57E-05 | -1.09E+00 | 4.75E-06 | 2.02E-05 | -1.35E+00 | 9.64E-05 | 3.68E-04 | -1.07E+00 | 5.75E-06 | 2.05E-05 | -1.53E+00 | 4.47E-03 | 1.26E-02 | -1.01E+00 | 9.66E-03 | 3.55E-02 | 1.04E+00 | 3.53E-01 | 9.71E-01 | 1.33E+00 |
| CCDC28A | 4.98E-09 | 2.60E-08 | -1.28E+00 | 1.04E-05 | 4.49E-05 | 1.01E+00 | 2.05E-07 | 8.71E-07 | -1.36E+00 | 3.08E-07 | 1.17E-06 | -1.19E+00 | 1.09E-11 | 3.89E-11 | -1.64E+00 | 2.07E-05 | 5.85E-05 | -1.06E+00 | 3.76E-07 | 1.38E-06 | -1.22E+00 | 3.77E-07 | 1.04E-06 | -1.19E+00 |
| DEF6 | 4.99E-09 | 2.60E-08 | -1.25E+00 | 9.76E-07 | 4.22E-06 | -1.02E+00 | 3.70E-08 | 1.57E-07 | -1.43E+00 | 1.42E-08 | 5.41E-08 | -1.31E+00 | 9.88E-13 | 3.52E-12 | -1.78E+00 | 4.40E-08 | 1.24E-07 | -1.33E+00 | 2.60E-09 | 9.54E-09 | -1.36E+00 | 2.82E-08 | 7.75E-08 | -1.35E+00 |
| MOCS2 | 4.99E-09 | 2.60E-08 | -1.61E+00 | 2.73E-05 | 1.18E-04 | -1.06E+00 | 8.19E-11 | 3.48E-10 | -1.72E+00 | 4.80E-09 | 1.83E-08 | -1.42E+00 | 1.30E-06 | 4.63E-06 | -1.53E+00 | 1.30E-04 | 3.68E-04 | -1.18E+00 | 9.32E-06 | 3.42E-05 | -1.12E+00 | 3.86E-04 | 1.06E-03 | -1.07E+00 |
| MAP1LC3B | 5.00E-09 | 2.61E-08 | -1.17E+00 | 2.60E-14 | 1.12E-13 | -1.24E+00 | 1.63E-01 | 6.93E-01 | 1.11E+00 | 1.18E-03 | 4.52E-03 | -1.06E+00 | 1.29E-05 | 4.62E-05 | -1.07E+00 | 1.64E-07 | 4.61E-07 | -1.20E+00 | 3.85E-04 | 1.41E-03 | 1.01E+00 | 2.22E-04 | 6.09E-04 | -1.16E+00 |
| TSR2 | 5.01E-09 | 2.61E-08 | -1.19E+00 | 6.70E-06 | 2.89E-05 | 1.07E+00 | 1.04E-01 | 4.42E-01 | 1.13E+00 | 4.43E-02 | 1.69E-01 | 1.19E+00 | 3.90E-04 | 1.39E-03 | 1.03E+00 | 3.76E-03 | 1.06E-02 | 1.09E+00 | 2.66E-02 | 9.78E-02 | 1.05E+00 | 3.31E-01 | 9.08E-01 | 1.20E+00 |
| CLDN5 | 5.03E-09 | 2.62E-08 | -2.40E+00 | 2.66E-02 | 1.15E-01 | 1.17E+00 | 2.67E-04 | 1.13E-03 | -1.52E+00 | 3.85E-03 | 1.47E-02 | -1.26E+00 | 5.65E-02 | 2.02E-01 | -2.21E+00 | 1.11E-01 | 3.13E-01 | -1.74E+00 | 1.69E-01 | 6.22E-01 | 2.45E+00 | 1.43E-02 | 3.94E-02 | -1.86E+00 |
| CEP85L | 5.04E-09 | 2.63E-08 | -1.37E+00 | 2.80E-11 | 1.21E-10 | -1.40E+00 | 5.04E-04 | 2.14E-03 | -1.18E+00 | 7.47E-07 | 2.85E-06 | -1.33E+00 | 2.95E-04 | 1.05E-03 | -1.21E+00 | 5.92E-05 | 1.67E-04 | -1.38E+00 | 1.39E-11 | 5.12E-11 | -1.82E+00 | 4.16E-12 | 1.14E-11 | -1.82E+00 |
| AP1G1 | 5.04E-09 | 2.63E-08 | -1.31E+00 | 1.54E-08 | 6.65E-08 | -1.23E+00 | 9.17E-05 | 3.89E-04 | -1.27E+00 | 3.86E-07 | 1.47E-06 | -1.28E+00 | 7.66E-03 | 2.73E-02 | -1.02E+00 | 1.12E-04 | 3.15E-04 | -1.23E+00 | 6.92E-07 | 2.54E-06 | -1.34E+00 | 1.20E-06 | 3.29E-06 | -1.34E+00 |
| MTHFD2 | 5.04E-09 | 2.63E-08 | -1.60E+00 | 5.62E-08 | 2.43E-07 | -1.32E+00 | 1.08E-07 | 4.57E-07 | -1.31E+00 | 3.72E-06 | 1.42E-05 | -1.06E+00 | 2.18E-16 | 7.78E-16 | -1.83E+00 | 9.66E-08 | 2.72E-07 | -1.27E+00 | 5.67E-17 | 2.08E-16 | -1.79E+00 | 6.70E-08 | 1.84E-07 | -1.36E+00 |
| SMN2 | 5.08E-09 | 2.65E-08 | -1.22E+00 | 2.57E-09 | 1.11E-08 | -1.14E+00 | 1.19E-08 | 5.04E-08 | -1.26E+00 | 9.52E-09 | 3.64E-08 | -1.15E+00 | 8.76E-07 | 3.13E-06 | -1.28E+00 | 5.56E-05 | 1.57E-04 | -1.03E+00 | 1.34E-10 | 4.91E-10 | -1.44E+00 | 7.79E-05 | 2.14E-04 | -1.15E+00 |
| THAP3 | 5.09E-09 | 2.65E-08 | -1.83E+00 | 1.54E-04 | 6.67E-04 | -1.02E+00 | 3.96E-06 | 1.68E-05 | -1.52E+00 | 9.35E-04 | 3.57E-03 | -1.08E+00 | 3.22E-06 | 1.15E-05 | -1.91E+00 | 6.23E-03 | 1.75E-02 | -1.13E+00 | 3.82E-08 | 1.40E-07 | -1.62E+00 | 1.54E-02 | 4.23E-02 | -1.05E+00 |
| TADA2A | 5.11E-09 | 2.66E-08 | -2.03E+00 | 8.56E-06 | 3.70E-05 | -1.31E+00 | 1.58E-08 | 6.72E-08 | -1.57E+00 | 2.83E-03 | 1.08E-02 | 1.01E+00 | 2.52E-04 | 8.97E-04 | -1.43E+00 | 1.71E-02 | 4.82E-02 | -1.04E+00 | 9.82E-14 | 3.60E-13 | -1.69E+00 | 3.20E-04 | 8.78E-04 | -1.06E+00 |
| MED24 | 5.13E-09 | 2.67E-08 | -2.00E+00 | 7.89E-04 | 3.41E-03 | -1.03E+00 | 3.28E-08 | 1.40E-07 | -1.75E+00 | 3.82E-04 | 1.46E-03 | -1.03E+00 | 1.12E-10 | 4.01E-10 | -2.01E+00 | 3.93E-06 | 1.11E-05 | -1.23E+00 | 2.69E-12 | 9.87E-12 | -1.65E+00 | 1.05E-05 | 2.90E-05 | -1.15E+00 |
| KDM1B | 5.13E-09 | 2.67E-08 | -1.37E+00 | 4.36E-09 | 1.88E-08 | -1.20E+00 | 1.77E-05 | 7.51E-05 | -1.25E+00 | 2.29E-05 | 8.75E-05 | -1.10E+00 | 2.80E-04 | 1.00E-03 | -1.23E+00 | 4.77E-04 | 1.35E-03 | -1.31E+00 | 5.71E-04 | 2.10E-03 | -1.36E+00 | 3.51E-03 | 9.64E-03 | -1.17E+00 |
| TXNP5 | 5.13E-09 | 2.68E-08 | -1.21E+00 | 5.28E-14 | 2.28E-13 | -1.28E+00 | 6.64E-03 | 2.82E-02 | -1.11E+00 | 2.49E-04 | 9.52E-04 | -1.09E+00 | 6.45E-09 | 2.30E-08 | -1.30E+00 | 1.44E-06 | 4.07E-06 | -1.14E+00 | 1.76E-06 | 6.46E-06 | -1.20E+00 | 4.34E-04 | 1.19E-03 | -1.12E+00 |
| HDAC3 | 5.16E-09 | 2.69E-08 | -1.30E+00 | 7.82E-07 | 3.38E-06 | -1.02E+00 | 7.19E-05 | 3.06E-04 | -1.16E+00 | 1.86E-05 | 7.11E-05 | -1.06E+00 | 1.49E-06 | 5.30E-06 | -1.09E+00 | 1.43E-05 | 4.03E-05 | -1.04E+00 | 1.88E-02 | 6.90E-02 | 1.09E+00 | 4.97E-02 | 1.37E-01 | 1.16E+00 |
| MRPL51 | 5.16E-09 | 2.69E-08 | -1.06E+00 | 1.79E-08 | 7.73E-08 | 1.03E+00 | 8.92E-02 | 3.79E-01 | 1.04E+00 | 9.68E-02 | 3.70E-01 | 1.09E+00 | 1.31E-06 | 4.67E-06 | -1.18E+00 | 2.38E-02 | 6.72E-02 | 1.05E+00 | 1.89E-01 | 6.94E-01 | 1.04E+00 | 2.86E-01 | 7.87E-01 | 1.16E+00 |
| CARMIL1 | 5.16E-09 | 2.69E-08 | -1.64E+00 | 1.42E-05 | 6.13E-05 | -1.03E+00 | 1.36E-06 | 5.78E-06 | -2.09E+00 | 9.07E-04 | 3.46E-03 | -1.16E+00 | 5.98E-13 | 2.13E-12 | -1.88E+00 | 5.64E-06 | 1.59E-05 | -1.19E+00 | 7.52E-04 | 2.76E-03 | -1.22E+00 | 1.12E-01 | 3.09E-01 | 1.06E+00 |
| C1R | 5.20E-09 | 2.71E-08 | -1.88E+00 | 1.88E-08 | 8.11E-08 | -1.72E+00 | 1.20E-03 | 5.11E-03 | -1.43E+00 | 1.85E-03 | 7.07E-03 | -1.30E+00 | 1.27E-04 | 4.54E-04 | -2.10E+00 | 3.33E-04 | 9.38E-04 | -1.86E+00 | 3.66E-02 | 1.34E-01 | -1.26E+00 | 1.75E-01 | 4.79E-01 | 1.04E+00 |
| OXA1L | 5.20E-09 | 2.71E-08 | -1.15E+00 | 1.56E-03 | 6.75E-03 | 1.20E+00 | 2.43E-05 | 1.03E-04 | -1.18E+00 | 2.68E-04 | 1.02E-03 | -1.02E+00 | 4.86E-08 | 1.73E-07 | -1.20E+00 | 7.34E-04 | 2.07E-03 | 1.06E+00 | 4.74E-04 | 1.74E-03 | -1.05E+00 | 1.06E-02 | 2.90E-02 | 1.06E+00 |
| RAB3GAP1 | 5.21E-09 | 2.71E-08 | -1.39E+00 | 3.51E-07 | 1.52E-06 | -1.09E+00 | 1.21E-07 | 5.14E-07 | -1.44E+00 | 3.45E-07 | 1.32E-06 | -1.24E+00 | 7.18E-06 | 2.56E-05 | -1.30E+00 | 3.58E-03 | 1.01E-02 | -1.00E+00 | 7.14E-04 | 2.62E-03 | -1.12E+00 | 9.43E-04 | 2.59E-03 | -1.09E+00 |
| SEC23IP | 5.21E-09 | 2.71E-08 | -1.68E+00 | 2.47E-04 | 1.07E-03 | -1.03E+00 | 9.20E-08 | 3.91E-07 | -1.59E+00 | 1.28E-03 | 4.89E-03 | -1.04E+00 | 1.21E-05 | 4.32E-05 | -1.47E+00 | 4.17E-03 | 1.17E-02 | -1.11E+00 | 1.76E-07 | 6.47E-07 | -1.34E+00 | 1.31E-02 | 3.61E-02 | 1.10E+00 |
| COMMD8 | 5.22E-09 | 2.72E-08 | -1.42E+00 | 7.44E-05 | 3.21E-04 | 1.00E+00 | 2.67E-06 | 1.14E-05 | -1.14E+00 | 9.49E-08 | 3.63E-07 | -1.16E+00 | 1.74E-07 | 6.20E-07 | -1.44E+00 | 1.12E-05 | 3.15E-05 | -1.22E+00 | 1.93E-06 | 7.09E-06 | -1.18E+00 | 3.47E-06 | 9.53E-06 | -1.13E+00 |
| RPS24 | 5.22E-09 | 2.72E-08 | -1.16E+00 | 9.06E-48 | 3.92E-47 | -1.45E+00 | 1.67E-05 | 7.10E-05 | -1.14E+00 | 4.50E-24 | 1.72E-23 | -1.38E+00 | 2.75E-05 | 9.79E-05 | -1.15E+00 | 2.24E-27 | 6.32E-27 | -1.46E+00 | 7.52E-21 | 2.76E-20 | -1.25E+00 | 1.03E-38 | 2.82E-38 | -1.50E+00 |
| AATF | 5.22E-09 | 2.72E-08 | -1.09E+00 | 5.24E-13 | 2.26E-12 | -1.13E+00 | 3.81E-03 | 1.62E-02 | 1.03E+00 | 1.30E-03 | 4.97E-03 | 1.12E+00 | 8.89E-06 | 3.17E-05 | -1.20E+00 | 1.62E-02 | 4.57E-02 | 1.04E+00 | 3.10E-03 | 1.14E-02 | -1.02E+00 | 2.09E-02 | 5.74E-02 | 1.05E+00 |
| CREB3L4 | 5.23E-09 | 2.73E-08 | -1.28E+00 | 1.80E-05 | 7.79E-05 | 1.08E+00 | 7.77E-08 | 3.30E-07 | -1.36E+00 | 2.33E-06 | 8.91E-06 | -1.14E+00 | 2.08E-08 | 7.40E-08 | -1.45E+00 | 4.39E-05 | 1.24E-04 | -1.13E+00 | 4.94E-08 | 1.81E-07 | -1.36E+00 | 2.50E-06 | 6.87E-06 | -1.24E+00 |
| PPP3CB | 5.28E-09 | 2.75E-08 | -1.66E+00 | 4.05E-05 | 1.75E-04 | -1.11E+00 | 2.18E-12 | 9.27E-12 | -1.97E+00 | 5.94E-10 | 2.27E-09 | -1.41E+00 | 3.50E-02 | 1.25E-01 | 1.05E+00 | 6.40E-02 | 1.80E-01 | 1.11E+00 | 1.20E-07 | 4.39E-07 | -1.33E+00 | 3.10E-04 | 8.53E-04 | -1.06E+00 |
| HOXB3 | 5.33E-09 | 2.78E-08 | -1.33E+00 | 9.12E-03 | 3.94E-02 | 1.25E+00 | 3.54E-05 | 1.50E-04 | -1.27E+00 | 6.89E-02 | 2.63E-01 | 1.31E+00 | 7.65E-02 | 2.73E-01 | -1.04E+00 | 1.82E-01 | 5.14E-01 | 1.54E+00 | 1.23E-03 | 4.50E-03 | -1.32E+00 | 2.38E-01 | 6.55E-01 | 1.20E+00 |
| TNPO1 | 5.36E-09 | 2.79E-08 | -1.18E+00 | 4.59E-09 | 1.98E-08 | -1.07E+00 | 4.21E-06 | 1.79E-05 | -1.16E+00 | 1.64E-05 | 6.28E-05 | -1.04E+00 | 9.64E-09 | 3.44E-08 | -1.42E+00 | 8.86E-08 | 2.50E-07 | -1.25E+00 | 3.77E-08 | 1.39E-07 | -1.45E+00 | 1.44E-03 | 3.95E-03 | -1.11E+00 |
| HIGD2A | 5.42E-09 | 2.82E-08 | -1.23E+00 | 2.79E-11 | 1.20E-10 | -1.18E+00 | 1.64E-03 | 6.96E-03 | -1.14E+00 | 1.00E-03 | 3.83E-03 | -1.09E+00 | 6.52E-07 | 2.33E-06 | -1.29E+00 | 8.95E-04 | 2.52E-03 | -1.09E+00 | 8.94E-06 | 3.28E-05 | -1.19E+00 | 5.48E-03 | 1.51E-02 | -1.09E+00 |
| COMMD5 | 5.43E-09 | 2.83E-08 | -1.63E+00 | 2.24E-06 | 9.69E-06 | -1.21E+00 | 2.01E-04 | 8.55E-04 | -1.34E+00 | 7.00E-02 | 2.67E-01 | 1.19E+00 | 1.67E-05 | 5.94E-05 | -1.52E+00 | 5.52E-03 | 1.56E-02 | -1.03E+00 | 1.06E-11 | 3.88E-11 | -1.75E+00 | 1.20E-02 | 3.31E-02 | -1.04E+00 |
| LIAS | 5.48E-09 | 2.86E-08 | -1.26E+00 | 2.31E-07 | 9.99E-07 | -1.07E+00 | 2.04E-03 | 8.66E-03 | -1.07E+00 | 6.24E-05 | 2.38E-04 | -1.07E+00 | 1.67E-05 | 5.95E-05 | -1.20E+00 | 1.93E-03 | 5.44E-03 | 1.01E+00 | 2.42E-03 | 8.87E-03 | -1.02E+00 | 1.96E-02 | 5.39E-02 | 1.01E+00 |
| CCND3 | 5.48E-09 | 2.86E-08 | 1.85E+00 | 2.02E-13 | 8.73E-13 | 2.19E+00 | 2.99E-02 | 1.27E-01 | 1.47E+00 | 1.42E-03 | 5.41E-03 | 1.85E+00 | 9.90E-14 | 3.53E-13 | 2.74E+00 | 5.14E-10 | 1.45E-09 | 2.36E+00 | 6.15E-10 | 2.26E-09 | 1.55E+00 | 3.80E-04 | 1.04E-03 | 1.17E+00 |
| HMGN2 | 5.51E-09 | 2.87E-08 | -1.22E+00 | 6.76E-08 | 2.92E-07 | -1.05E+00 | 1.79E-03 | 7.61E-03 | -1.04E+00 | 2.93E-06 | 1.12E-05 | -1.08E+00 | 4.99E-04 | 1.78E-03 | 1.02E+00 | 1.05E-03 | 2.95E-03 | 1.04E+00 | 5.61E-04 | 2.06E-03 | -1.08E+00 | 1.17E-03 | 3.22E-03 | -1.14E+00 |
| MPV17L | 5.51E-09 | 2.87E-08 | -1.70E+00 | 7.16E-05 | 3.09E-04 | -1.02E+00 | 2.62E-03 | 1.11E-02 | -1.18E+00 | 1.69E-03 | 6.45E-03 | -1.04E+00 | 2.00E-03 | 7.13E-03 | -1.26E+00 | 2.41E-02 | 6.79E-02 | 1.02E+00 | 3.86E-10 | 1.42E-09 | -1.52E+00 | 1.47E-04 | 4.04E-04 | -1.19E+00 |
| ZMPSTE24 | 5.55E-09 | 2.89E-08 | -2.03E+00 | 1.53E-04 | 6.62E-04 | -1.09E+00 | 4.60E-04 | 1.96E-03 | -1.30E+00 | 2.20E-03 | 8.40E-03 | 1.03E+00 | 1.87E-05 | 6.67E-05 | -1.48E+00 | 5.15E-04 | 1.45E-03 | -1.16E+00 | 3.65E-05 | 1.34E-04 | -1.22E+00 | 6.19E-02 | 1.70E-01 | 1.06E+00 |
| PPP2R3C | 5.56E-09 | 2.90E-08 | -1.17E+00 | 2.62E-08 | 1.13E-07 | 1.02E+00 | 2.07E-01 | 8.79E-01 | 1.09E+00 | 1.62E-03 | 6.20E-03 | 1.01E+00 | 4.00E-02 | 1.42E-01 | 1.10E+00 | 5.06E-02 | 1.43E-01 | 1.14E+00 | 1.55E-03 | 5.70E-03 | -1.04E+00 | 1.05E-03 | 2.88E-03 | -1.07E+00 |
| TANK | 5.58E-09 | 2.91E-08 | -1.18E+00 | 3.40E-10 | 1.47E-09 | -1.14E+00 | 3.49E-09 | 1.48E-08 | -1.37E+00 | 2.33E-07 | 8.90E-07 | -1.13E+00 | 2.69E-03 | 9.59E-03 | -1.09E+00 | 3.59E-04 | 1.01E-03 | -1.17E+00 | 2.62E-07 | 9.61E-07 | -1.35E+00 | 2.35E-04 | 6.45E-04 | -1.02E+00 |
| EMSY | 5.63E-09 | 2.93E-08 | -1.67E+00 | 2.87E-05 | 1.24E-04 | -1.14E+00 | 2.06E-08 | 8.76E-08 | -1.66E+00 | 3.11E-06 | 1.19E-05 | -1.31E+00 | 6.65E-02 | 2.37E-01 | 1.02E+00 | 2.34E-01 | 6.61E-01 | 1.32E+00 | 4.67E-06 | 1.71E-05 | -1.31E+00 | 6.37E-04 | 1.75E-03 | -1.16E+00 |
| AP1M1 | 5.67E-09 | 2.96E-08 | -1.30E+00 | 2.84E-04 | 1.23E-03 | 1.10E+00 | 4.21E-09 | 1.79E-08 | -1.40E+00 | 7.43E-09 | 2.84E-08 | -1.19E+00 | 4.83E-06 | 1.72E-05 | -1.35E+00 | 4.27E-04 | 1.21E-03 | -1.13E+00 | 8.84E-05 | 3.24E-04 | -1.21E+00 | 6.11E-03 | 1.68E-02 | -1.03E+00 |
| KIFAP3 | 5.72E-09 | 2.98E-08 | -1.38E+00 | 2.27E-08 | 9.82E-08 | -1.25E+00 | 1.31E-04 | 5.56E-04 | -1.31E+00 | 2.58E-07 | 9.85E-07 | -1.41E+00 | 1.14E-10 | 4.08E-10 | -1.84E+00 | 1.83E-06 | 5.17E-06 | -1.32E+00 | 2.81E-14 | 1.03E-13 | -1.55E+00 | 5.98E-21 | 1.64E-20 | -2.00E+00 |
| ASAH1 | 5.73E-09 | 2.99E-08 | -1.18E+00 | 8.77E-06 | 3.79E-05 | 1.04E+00 | 5.38E-04 | 2.28E-03 | -1.13E+00 | 4.37E-05 | 1.67E-04 | -1.06E+00 | 4.96E-07 | 1.77E-06 | -1.25E+00 | 7.52E-07 | 2.12E-06 | -1.24E+00 | 9.21E-03 | 3.38E-02 | -1.02E+00 | 1.28E-02 | 3.51E-02 | -1.03E+00 |
| PP7080 | 5.76E-09 | 3.00E-08 | -2.34E+00 | 1.25E-04 | 5.42E-04 | -1.22E+00 | 3.91E-05 | 1.66E-04 | -1.91E+00 | 5.43E-03 | 2.08E-02 | -1.16E+00 | 8.62E-08 | 3.07E-07 | -2.00E+00 | 2.96E-05 | 8.35E-05 | -1.50E+00 | 3.73E-07 | 1.37E-06 | -1.66E+00 | 9.03E-02 | 2.48E-01 | 1.08E+00 |
| UBE2Z | 5.76E-09 | 3.00E-08 | -1.25E+00 | 2.47E-06 | 1.07E-05 | 1.02E+00 | 8.47E-06 | 3.60E-05 | -1.28E+00 | 4.22E-04 | 1.61E-03 | -1.02E+00 | 8.72E-03 | 3.11E-02 | 1.06E+00 | 1.03E-02 | 2.89E-02 | 1.07E+00 | 5.96E-04 | 2.19E-03 | -1.10E+00 | 1.38E-01 | 3.79E-01 | 1.16E+00 |
| MRPL50 | 5.80E-09 | 3.02E-08 | -1.26E+00 | 2.26E-09 | 9.77E-09 | -1.22E+00 | 1.22E-06 | 5.19E-06 | -1.29E+00 | 2.89E-04 | 1.10E-03 | -1.03E+00 | 2.60E-02 | 9.27E-02 | -1.04E+00 | 3.21E-01 | 9.05E-01 | 1.40E+00 | 4.87E-06 | 1.79E-05 | -1.28E+00 | 2.95E-01 | 8.09E-01 | 1.20E+00 |
| STK17A | 5.84E-09 | 3.04E-08 | -1.11E+00 | 8.93E-20 | 3.86E-19 | -1.55E+00 | 2.82E-07 | 1.20E-06 | -1.34E+00 | 2.63E-16 | 1.00E-15 | -1.81E+00 | 3.44E-09 | 1.23E-08 | -1.29E+00 | 1.16E-15 | 3.26E-15 | -1.69E+00 | 1.85E-11 | 6.80E-11 | -1.46E+00 | 3.19E-15 | 8.75E-15 | -1.64E+00 |
| SMNDC1 | 5.85E-09 | 3.05E-08 | -1.18E+00 | 1.39E-09 | 6.01E-09 | -1.12E+00 | 7.58E-06 | 3.22E-05 | -1.23E+00 | 9.70E-06 | 3.70E-05 | -1.08E+00 | 1.10E-04 | 3.92E-04 | -1.07E+00 | 1.91E-06 | 5.39E-06 | -1.20E+00 | 3.44E-10 | 1.26E-09 | -1.34E+00 | 1.52E-06 | 4.17E-06 | -1.21E+00 |
| FZD7 | 5.90E-09 | 3.07E-08 | -2.25E+00 | 4.83E-06 | 2.09E-05 | -1.41E+00 | 5.45E-03 | 2.31E-02 | -1.34E+00 | 6.16E-02 | 2.35E-01 | 1.09E+00 | 1.66E-02 | 5.93E-02 | -1.38E+00 | 3.63E-02 | 1.02E-01 | -1.22E+00 | 2.17E-05 | 7.98E-05 | -2.25E+00 | 8.76E-03 | 2.41E-02 | -1.40E+00 |
| RPL41P5 | 5.93E-09 | 3.09E-08 | -1.41E+00 | 2.00E-13 | 8.63E-13 | -1.84E+00 | 8.32E-10 | 3.54E-09 | -1.63E+00 | 2.90E-15 | 1.11E-14 | -1.92E+00 | 3.73E-04 | 1.33E-03 | -1.47E+00 | 1.10E-04 | 3.11E-04 | -1.54E+00 | 4.43E-10 | 1.63E-09 | -1.81E+00 | 1.34E-10 | 3.68E-10 | -2.09E+00 |
| ACSF3 | 5.93E-09 | 3.09E-08 | -1.25E+00 | 2.25E-05 | 9.71E-05 | 1.03E+00 | 6.39E-07 | 2.72E-06 | -1.34E+00 | 4.50E-04 | 1.72E-03 | 1.01E+00 | 1.61E-03 | 5.76E-03 | -1.09E+00 | 4.50E-02 | 1.27E-01 | 1.23E+00 | 7.39E-07 | 2.71E-06 | -1.41E+00 | 1.61E-02 | 4.43E-02 | 1.00E+00 |
| LARP6 | 5.95E-09 | 3.10E-08 | -2.04E+00 | 1.41E-07 | 6.09E-07 | -1.58E+00 | 4.66E-02 | 1.98E-01 | -1.04E+00 | 7.43E-04 | 2.84E-03 | -1.24E+00 | 3.41E-03 | 1.22E-02 | -1.83E+00 | 1.04E-02 | 2.94E-02 | -1.57E+00 | 1.88E-04 | 6.89E-04 | -1.66E+00 | 1.38E-03 | 3.80E-03 | -1.49E+00 |
| GON7 | 5.96E-09 | 3.10E-08 | -1.48E+00 | 9.91E-08 | 4.28E-07 | -1.24E+00 | 2.49E-03 | 1.06E-02 | -1.02E+00 | 5.39E-03 | 2.06E-02 | 1.09E+00 | 1.53E-09 | 5.44E-09 | -1.99E+00 | 1.75E-06 | 4.93E-06 | -1.48E+00 | 2.99E-06 | 1.10E-05 | -1.22E+00 | 8.42E-03 | 2.31E-02 | 1.01E+00 |
| RPA1 | 5.99E-09 | 3.12E-08 | -1.44E+00 | 1.69E-03 | 7.31E-03 | 1.15E+00 | 2.52E-04 | 1.07E-03 | -1.16E+00 | 2.59E-02 | 9.89E-02 | 1.15E+00 | 5.19E-07 | 1.85E-06 | -1.44E+00 | 4.17E-03 | 1.17E-02 | -1.01E+00 | 9.16E-05 | 3.36E-04 | -1.12E+00 | 6.06E-02 | 1.66E-01 | 1.47E+00 |
| SUPT16H | 6.00E-09 | 3.13E-08 | -1.25E+00 | 5.85E-05 | 2.53E-04 | 1.03E+00 | 1.66E-05 | 7.05E-05 | -1.16E+00 | 2.27E-03 | 8.69E-03 | 1.12E+00 | 1.23E-05 | 4.38E-05 | -1.26E+00 | 1.92E-01 | 5.42E-01 | 1.17E+00 | 1.53E-02 | 5.63E-02 | 1.01E+00 | 1.74E-01 | 4.78E-01 | 1.30E+00 |
| AF131215.6 | 6.05E-09 | 3.15E-08 | -2.10E+00 | 1.30E-03 | 5.64E-03 | -1.09E+00 | 5.04E-04 | 2.14E-03 | -1.52E+00 | 1.53E-01 | 5.84E-01 | 1.25E+00 | 1.93E-02 | 6.89E-02 | -1.05E+00 | 1.37E-01 | 3.85E-01 | 1.28E+00 | 1.54E-04 | 5.65E-04 | -1.54E+00 | 8.58E-02 | 2.36E-01 | -1.01E+00 |
| AP001157.1 | 6.05E-09 | 3.16E-08 | -1.57E+00 | 1.14E-01 | 4.92E-01 | 1.39E+00 | 1.90E-06 | 8.09E-06 | -1.51E+00 | 5.81E-04 | 2.22E-03 | -1.11E+00 | 4.32E-04 | 1.54E-03 | -1.29E+00 | 2.99E-02 | 8.43E-02 | 1.05E+00 | 1.50E-03 | 5.49E-03 | -1.41E+00 | 2.22E-01 | 6.10E-01 | 1.23E+00 |
| HK1 | 6.10E-09 | 3.18E-08 | -1.40E+00 | 2.28E-06 | 9.84E-06 | -1.05E+00 | 1.58E-08 | 6.72E-08 | -1.45E+00 | 2.27E-06 | 8.67E-06 | -1.16E+00 | 2.99E-05 | 1.07E-04 | -1.21E+00 | 5.64E-02 | 1.59E-01 | 1.24E+00 | 2.08E-10 | 7.64E-10 | -1.28E+00 | 4.92E-03 | 1.35E-02 | -1.03E+00 |
| BOLA3 | 6.10E-09 | 3.18E-08 | -1.47E+00 | 4.55E-08 | 1.97E-07 | -1.22E+00 | 6.48E-04 | 2.75E-03 | -1.01E+00 | 8.00E-10 | 3.06E-09 | -1.32E+00 | 1.37E-03 | 4.90E-03 | -1.29E+00 | 5.10E-04 | 1.44E-03 | -1.38E+00 | 1.91E-07 | 7.02E-07 | -1.27E+00 | 2.50E-08 | 6.87E-08 | -1.33E+00 |
| AGPAT1 | 6.12E-09 | 3.19E-08 | -1.90E+00 | 1.59E-07 | 6.86E-07 | -1.47E+00 | 8.44E-08 | 3.59E-07 | -2.13E+00 | 3.04E-04 | 1.16E-03 | -1.29E+00 | 2.77E-02 | 9.86E-02 | -1.13E+00 | 2.25E-01 | 6.35E-01 | 1.20E+00 | 4.95E-11 | 1.82E-10 | -1.99E+00 | 2.44E-05 | 6.69E-05 | -1.26E+00 |
| AC093909.1 | 6.13E-09 | 3.20E-08 | -1.64E+00 | 5.64E-11 | 2.44E-10 | -1.73E+00 | 2.55E-04 | 1.08E-03 | -1.34E+00 | 6.44E-07 | 2.46E-06 | -1.66E+00 | 9.07E-08 | 3.24E-07 | -1.66E+00 | 3.28E-06 | 9.24E-06 | -1.36E+00 | 2.50E-10 | 9.17E-10 | -1.47E+00 | 2.31E-13 | 6.35E-13 | -1.80E+00 |
| EIF2B1 | 6.14E-09 | 3.20E-08 | -1.20E+00 | 1.21E-05 | 5.23E-05 | 1.09E+00 | 9.11E-06 | 3.87E-05 | -1.15E+00 | 1.06E-02 | 4.07E-02 | 1.20E+00 | 3.73E-03 | 1.33E-02 | 1.06E+00 | 2.08E-01 | 5.87E-01 | 1.33E+00 | 4.11E-03 | 1.51E-02 | -1.10E+00 | 2.87E-01 | 7.88E-01 | 1.17E+00 |
| MAP7D3 | 6.17E-09 | 3.22E-08 | -1.16E+00 | 4.82E-10 | 2.08E-09 | -1.12E+00 | 2.82E-05 | 1.20E-04 | -1.16E+00 | 1.57E-06 | 5.98E-06 | -1.14E+00 | 7.25E-08 | 2.59E-07 | -1.30E+00 | 6.13E-06 | 1.73E-05 | -1.20E+00 | 5.58E-05 | 2.05E-04 | -1.18E+00 | 1.05E-02 | 2.89E-02 | -1.03E+00 |
| TMEM182 | 6.17E-09 | 3.22E-08 | -1.95E+00 | 1.19E-07 | 5.15E-07 | -1.57E+00 | 2.36E-02 | 1.00E-01 | -1.24E+00 | 1.70E-02 | 6.51E-02 | -1.08E+00 | 5.02E-05 | 1.79E-04 | -1.70E+00 | 4.98E-03 | 1.40E-02 | -1.16E+00 | 2.43E-01 | 8.90E-01 | 1.23E+00 | 1.53E-01 | 4.20E-01 | 1.05E+00 |
| ORC6 | 6.17E-09 | 3.22E-08 | -2.67E+00 | 7.77E-08 | 3.36E-07 | -2.10E+00 | 6.98E-03 | 2.97E-02 | -1.00E+00 | 3.06E-04 | 1.17E-03 | 1.08E+00 | 1.57E-10 | 5.61E-10 | -3.40E+00 | 1.13E-11 | 3.18E-11 | -3.70E+00 | 8.80E-09 | 3.23E-08 | -1.33E+00 | 1.03E-01 | 2.83E-01 | 1.15E+00 |
| GAN | 6.17E-09 | 3.22E-08 | -1.59E+00 | 1.50E-04 | 6.47E-04 | 1.05E+00 | 5.65E-06 | 2.40E-05 | -1.74E+00 | 6.65E-02 | 2.54E-01 | 1.10E+00 | 1.18E-08 | 4.21E-08 | -2.02E+00 | 7.95E-05 | 2.24E-04 | -1.21E+00 | 3.09E-07 | 1.13E-06 | -1.84E+00 | 3.26E-03 | 8.95E-03 | -1.13E+00 |
| EBAG9 | 6.18E-09 | 3.22E-08 | -1.24E+00 | 3.69E-06 | 1.59E-05 | 1.08E+00 | 1.55E-04 | 6.57E-04 | -1.21E+00 | 7.59E-03 | 2.90E-02 | 1.02E+00 | 4.19E-05 | 1.49E-04 | -1.05E+00 | 1.65E-02 | 4.66E-02 | 1.15E+00 | 8.44E-04 | 3.10E-03 | -1.01E+00 | 1.12E-02 | 3.08E-02 | 1.09E+00 |
| MRVI1 | 6.23E-09 | 3.25E-08 | -3.18E+00 | 1.78E-07 | 7.68E-07 | -2.50E+00 | 3.13E-02 | 1.33E-01 | -1.36E+00 | 8.54E-02 | 3.26E-01 | -1.11E+00 | 2.41E-01 | 8.60E-01 | 1.19E+00 | 1.53E-01 | 4.32E-01 | 1.59E+00 | 6.67E-02 | 2.45E-01 | 1.19E+00 | 7.31E-02 | 2.01E-01 | -1.82E+00 |
| PAPD4 | 6.23E-09 | 3.25E-08 | -1.27E+00 | 5.11E-07 | 2.21E-06 | -1.03E+00 | 4.04E-12 | 1.72E-11 | -1.61E+00 | 1.14E-07 | 4.34E-07 | -1.21E+00 | 9.28E-05 | 3.31E-04 | -1.14E+00 | 2.86E-06 | 8.07E-06 | -1.26E+00 | 6.90E-08 | 2.53E-07 | -1.29E+00 | 8.97E-07 | 2.46E-06 | -1.19E+00 |
| ENO1 | 6.24E-09 | 3.25E-08 | 1.30E+00 | 8.16E-19 | 3.53E-18 | 1.49E+00 | 6.63E-07 | 2.82E-06 | 1.31E+00 | 5.48E-09 | 2.09E-08 | 1.39E+00 | 1.82E-01 | 6.48E-01 | 1.11E+00 | 2.78E-03 | 7.84E-03 | 1.23E+00 | 2.70E-01 | 9.90E-01 | 1.08E+00 | 2.33E-05 | 6.41E-05 | 1.29E+00 |
| NPM1P27 | 6.24E-09 | 3.25E-08 | -1.14E+00 | 2.09E-08 | 9.03E-08 | -1.05E+00 | 2.61E-02 | 1.11E-01 | -1.08E+00 | 3.79E-02 | 1.45E-01 | -1.03E+00 | 5.93E-05 | 2.12E-04 | -1.15E+00 | 1.35E-01 | 3.81E-01 | 1.03E+00 | 6.21E-04 | 2.28E-03 | -1.11E+00 | 7.35E-02 | 2.02E-01 | -1.05E+00 |
| POMC | 6.28E-09 | 3.27E-08 | -3.09E+00 | 8.55E-10 | 3.69E-09 | -3.33E+00 | 8.64E-03 | 3.67E-02 | -1.19E+00 | 8.21E-08 | 3.14E-07 | -2.19E+00 | 3.27E-03 | 1.17E-02 | -1.66E+00 | 6.61E-04 | 1.86E-03 | -2.06E+00 | 1.50E-01 | 5.49E-01 | -1.26E+00 | 2.90E-01 | 7.97E-01 | -1.09E+00 |
| MCCC1 | 6.33E-09 | 3.30E-08 | -1.33E+00 | 6.30E-07 | 2.72E-06 | -1.08E+00 | 1.71E-06 | 7.27E-06 | -1.43E+00 | 2.29E-05 | 8.74E-05 | -1.22E+00 | 3.22E-02 | 1.15E-01 | 1.11E+00 | 2.58E-01 | 7.29E-01 | 1.55E+00 | 1.30E-05 | 4.78E-05 | -1.13E+00 | 7.29E-04 | 2.00E-03 | -1.06E+00 |
| HMGN5 | 6.34E-09 | 3.30E-08 | -1.42E+00 | 1.11E-07 | 4.78E-07 | -1.15E+00 | 6.28E-03 | 2.67E-02 | -1.07E+00 | 1.20E-06 | 4.58E-06 | -1.15E+00 | 1.49E-03 | 5.30E-03 | -1.12E+00 | 5.54E-02 | 1.56E-01 | 1.22E+00 | 2.16E-01 | 7.92E-01 | 1.16E+00 | 7.75E-02 | 2.13E-01 | 1.07E+00 |
| RUVBL1 | 6.34E-09 | 3.31E-08 | -1.27E+00 | 9.63E-06 | 4.16E-05 | 1.04E+00 | 7.65E-03 | 3.25E-02 | -1.01E+00 | 5.17E-02 | 1.97E-01 | 1.18E+00 | 1.37E-12 | 4.89E-12 | -1.48E+00 | 2.22E-02 | 6.25E-02 | 1.17E+00 | 2.10E-04 | 7.71E-04 | -1.05E+00 | 2.76E-01 | 7.57E-01 | 1.24E+00 |
| SULT1C4 | 6.35E-09 | 3.31E-08 | -1.20E+00 | 5.31E-06 | 2.30E-05 | 1.00E+00 | 1.06E-02 | 4.51E-02 | -1.21E+00 | 3.02E-02 | 1.15E-01 | 1.02E+00 | 1.19E-01 | 4.24E-01 | 1.15E+00 | 1.64E-01 | 4.61E-01 | 1.24E+00 | 2.10E-01 | 7.71E-01 | 1.35E+00 | 3.28E-01 | 9.01E-01 | 1.35E+00 |
| SNX1 | 6.38E-09 | 3.32E-08 | -1.48E+00 | 2.52E-03 | 1.09E-02 | 1.08E+00 | 3.01E-12 | 1.28E-11 | -1.55E+00 | 1.44E-07 | 5.51E-07 | -1.15E+00 | 5.13E-07 | 1.83E-06 | -1.50E+00 | 1.45E-02 | 4.08E-02 | 1.05E+00 | 9.13E-11 | 3.35E-10 | -1.55E+00 | 3.40E-04 | 9.34E-04 | -1.20E+00 |
| METTL21A | 6.38E-09 | 3.33E-08 | -1.28E+00 | 6.13E-07 | 2.65E-06 | -1.04E+00 | 7.54E-04 | 3.20E-03 | -1.16E+00 | 4.80E-02 | 1.83E-01 | 1.18E+00 | 7.62E-07 | 2.72E-06 | -1.36E+00 | 7.96E-04 | 2.24E-03 | -1.05E+00 | 2.53E-09 | 9.29E-09 | -1.37E+00 | 3.49E-02 | 9.59E-02 | 1.10E+00 |
| VPS13B | 6.40E-09 | 3.33E-08 | -1.59E+00 | 8.41E-06 | 3.63E-05 | -1.15E+00 | 4.90E-05 | 2.08E-04 | -1.32E+00 | 1.64E-03 | 6.26E-03 | -1.03E+00 | 2.96E-05 | 1.05E-04 | -1.20E+00 | 4.64E-04 | 1.31E-03 | -1.07E+00 | 2.62E-02 | 9.60E-02 | 1.07E+00 | 3.12E-02 | 8.56E-02 | 1.19E+00 |
| NUP37 | 6.40E-09 | 3.33E-08 | -1.39E+00 | 9.57E-05 | 4.13E-04 | 1.02E+00 | 1.07E-09 | 4.53E-09 | -1.48E+00 | 5.92E-05 | 2.26E-04 | -1.09E+00 | 1.87E-08 | 6.67E-08 | -1.57E+00 | 6.42E-04 | 1.81E-03 | -1.04E+00 | 3.11E-10 | 1.14E-09 | -1.40E+00 | 4.39E-04 | 1.21E-03 | -1.09E+00 |
| MIGA2 | 6.46E-09 | 3.37E-08 | -1.65E+00 | 2.17E-06 | 9.37E-06 | -1.25E+00 | 4.24E-03 | 1.80E-02 | -1.20E+00 | 6.82E-02 | 2.60E-01 | 1.17E+00 | 2.71E-06 | 9.66E-06 | -1.59E+00 | 3.25E-03 | 9.17E-03 | -1.01E+00 | 2.88E-07 | 1.06E-06 | -1.10E+00 | 1.36E-04 | 3.75E-04 | -1.10E+00 |
| PHF23 | 6.49E-09 | 3.38E-08 | -2.06E+00 | 2.77E-04 | 1.20E-03 | -1.21E+00 | 9.92E-09 | 4.21E-08 | -1.91E+00 | 1.03E-02 | 3.93E-02 | 1.16E+00 | 2.32E-04 | 8.26E-04 | -1.63E+00 | 1.21E-01 | 3.42E-01 | -1.02E+00 | 3.35E-15 | 1.23E-14 | -1.70E+00 | 1.91E-05 | 5.25E-05 | -1.05E+00 |
| KCNE3 | 6.50E-09 | 3.39E-08 | -2.24E+00 | 1.64E-05 | 7.07E-05 | -1.39E+00 | 9.17E-29 | 3.90E-28 | -2.59E+00 | 6.39E-24 | 2.44E-23 | -1.93E+00 | 4.40E-07 | 1.57E-06 | -3.65E+00 | 1.05E-06 | 2.95E-06 | -3.19E+00 | 2.92E-03 | 1.07E-02 | -1.08E+00 | 8.76E-03 | 2.41E-02 | -1.45E+00 |
| RPS7P1 | 6.51E-09 | 3.39E-08 | -1.12E+00 | 4.10E-02 | 1.77E-01 | 1.04E+00 | 2.51E-05 | 1.07E-04 | -1.11E+00 | 5.79E-03 | 2.21E-02 | 1.07E+00 | 1.01E-08 | 3.60E-08 | -1.18E+00 | 3.07E-01 | 8.67E-01 | 1.01E+00 | 1.89E-12 | 6.93E-12 | -1.15E+00 | 6.08E-02 | 1.67E-01 | 1.04E+00 |
| PPP1R2 | 6.54E-09 | 3.41E-08 | -1.19E+00 | 8.35E-13 | 3.61E-12 | -1.16E+00 | 2.51E-04 | 1.07E-03 | -1.08E+00 | 6.62E-07 | 2.53E-06 | -1.13E+00 | 3.35E-05 | 1.19E-04 | -1.10E+00 | 2.51E-09 | 7.07E-09 | -1.31E+00 | 6.54E-05 | 2.40E-04 | -1.07E+00 | 2.62E-07 | 7.19E-07 | -1.27E+00 |
| AAMDC | 6.55E-09 | 3.41E-08 | -1.30E+00 | 9.62E-10 | 4.16E-09 | -1.19E+00 | 1.11E-04 | 4.70E-04 | -1.19E+00 | 5.39E-09 | 2.06E-08 | -1.36E+00 | 1.69E-04 | 6.02E-04 | 1.01E+00 | 9.49E-05 | 2.68E-04 | -1.05E+00 | 2.67E-06 | 9.82E-06 | -1.14E+00 | 5.70E-08 | 1.57E-07 | -1.27E+00 |
| RPS2P7 | 6.55E-09 | 3.41E-08 | 2.63E+00 | 2.21E-01 | 9.55E-01 | 1.52E+00 | 1.44E-09 | 6.14E-09 | 2.50E+00 | 1.99E-01 | 7.59E-01 | 1.37E+00 | 9.07E-05 | 3.24E-04 | 2.51E+00 | 1.67E-01 | 4.71E-01 | 1.64E+00 | 9.01E-09 | 3.31E-08 | 2.50E+00 | 9.45E-03 | 2.60E-02 | 1.84E+00 |
| MAEA | 6.59E-09 | 3.43E-08 | -1.53E+00 | 1.63E-02 | 7.03E-02 | 1.27E+00 | 5.49E-08 | 2.33E-07 | -1.40E+00 | 3.94E-04 | 1.51E-03 | 1.05E+00 | 3.92E-08 | 1.40E-07 | -1.57E+00 | 3.92E-04 | 1.10E-03 | -1.10E+00 | 5.29E-15 | 1.94E-14 | -1.71E+00 | 6.30E-03 | 1.73E-02 | -1.02E+00 |
| RNPEP | 6.59E-09 | 3.43E-08 | -1.53E+00 | 1.45E-07 | 6.28E-07 | -1.31E+00 | 9.31E-10 | 3.95E-09 | -1.43E+00 | 2.27E-10 | 8.66E-10 | -1.35E+00 | 4.58E-17 | 1.63E-16 | -2.08E+00 | 6.99E-13 | 1.97E-12 | -1.66E+00 | 1.02E-08 | 3.73E-08 | -1.36E+00 | 1.62E-04 | 4.44E-04 | -1.08E+00 |
| EEF1A1P14 | 6.62E-09 | 3.45E-08 | -1.44E+00 | 5.37E-14 | 2.32E-13 | -1.69E+00 | 3.29E-07 | 1.40E-06 | -1.38E+00 | 3.35E-08 | 1.28E-07 | -1.33E+00 | 1.27E-02 | 4.51E-02 | 1.00E+00 | 1.21E-03 | 3.41E-03 | -1.22E+00 | 2.03E-10 | 7.43E-10 | -1.75E+00 | 2.36E-06 | 6.48E-06 | -1.34E+00 |
| SRP72 | 6.69E-09 | 3.49E-08 | -1.12E+00 | 5.52E-05 | 2.38E-04 | 1.06E+00 | 1.50E-02 | 6.37E-02 | -1.09E+00 | 6.02E-02 | 2.30E-01 | -1.01E+00 | 4.78E-06 | 1.70E-05 | -1.17E+00 | 2.44E-03 | 6.88E-03 | -1.00E+00 | 2.36E-02 | 8.68E-02 | -1.05E+00 | 2.00E-01 | 5.49E-01 | 1.01E+00 |
| BTN2A2 | 6.69E-09 | 3.49E-08 | -1.63E+00 | 1.35E-07 | 5.81E-07 | -1.31E+00 | 9.59E-08 | 4.08E-07 | -1.79E+00 | 8.98E-07 | 3.43E-06 | -1.46E+00 | 5.16E-04 | 1.84E-03 | -1.38E+00 | 1.69E-03 | 4.78E-03 | -1.35E+00 | 7.44E-04 | 2.73E-03 | -1.37E+00 | 3.17E-03 | 8.70E-03 | -1.29E+00 |
| HNRNPAB | 6.76E-09 | 3.52E-08 | -1.14E+00 | 2.70E-07 | 1.17E-06 | -1.03E+00 | 8.89E-02 | 3.78E-01 | 1.06E+00 | 3.21E-02 | 1.23E-01 | 1.04E+00 | 1.23E-03 | 4.40E-03 | -1.17E+00 | 2.48E-03 | 6.99E-03 | -1.10E+00 | 6.33E-02 | 2.32E-01 | 1.01E+00 | 2.44E-02 | 6.69E-01 | 1.04E+00 |
| FBXO22 | 6.77E-09 | 3.53E-08 | -1.15E+00 | 3.43E-04 | 1.48E-03 | 1.16E+00 | 6.31E-10 | 2.68E-09 | -1.53E+00 | 2.95E-06 | 1.13E-05 | -1.12E+00 | 1.93E-04 | 6.87E-04 | -1.17E+00 | 9.56E-03 | 2.69E-02 | -1.04E+00 | 5.14E-07 | 1.88E-06 | -1.36E+00 | 2.66E-02 | 7.32E-02 | -1.05E+00 |
| PEX19 | 6.78E-09 | 3.53E-08 | -1.52E+00 | 4.64E-08 | 2.01E-07 | -1.36E+00 | 1.73E-06 | 7.34E-06 | -1.38E+00 | 1.24E-07 | 4.72E-07 | -1.31E+00 | 6.78E-06 | 2.42E-05 | -1.31E+00 | 8.43E-04 | 2.38E-03 | -1.07E+00 | 1.94E-10 | 7.13E-10 | -1.38E+00 | 2.81E-07 | 7.72E-07 | -1.21E+00 |
| SH3BGRL | 6.84E-09 | 3.57E-08 | -1.17E+00 | 1.63E-05 | 7.03E-05 | -1.03E+00 | 2.20E-02 | 9.34E-02 | -1.09E+00 | 2.12E-01 | 8.08E-01 | 1.07E+00 | 1.64E-01 | 5.86E-01 | 1.07E+00 | 6.55E-02 | 1.85E-01 | 1.23E+00 | 4.49E-02 | 1.65E-01 | -1.00E+00 | 3.15E-01 | 8.66E-01 | 1.04E+00 |
| NFAT5 | 6.84E-09 | 3.57E-08 | -1.21E+00 | 2.11E-05 | 9.11E-05 | 1.11E+00 | 3.22E-02 | 1.37E-01 | -1.03E+00 | 1.30E-03 | 4.97E-03 | -1.05E+00 | 4.47E-04 | 1.59E-03 | -1.04E+00 | 1.50E-03 | 4.23E-03 | 1.03E+00 | 6.56E-04 | 2.41E-03 | -1.05E+00 | 1.87E-06 | 5.15E-06 | -1.35E+00 |
| ZNF598 | 6.85E-09 | 3.57E-08 | -1.83E+00 | 1.31E-04 | 5.65E-04 | -1.05E+00 | 2.96E-06 | 1.26E-05 | -1.39E+00 | 7.75E-06 | 2.96E-05 | -1.16E+00 | 3.44E-02 | 1.23E-01 | -1.18E+00 | 3.21E-01 | 9.06E-01 | 1.45E+00 | 1.12E-03 | 4.11E-03 | -1.22E+00 | 1.96E-01 | 5.39E-01 | 1.14E+00 |
| POLR2K | 6.86E-09 | 3.57E-08 | -1.13E+00 | 2.91E-09 | 1.26E-08 | -1.02E+00 | 1.43E-03 | 6.08E-03 | -1.10E+00 | 1.14E-02 | 4.37E-02 | 1.07E+00 | 9.42E-07 | 3.36E-06 | -1.15E+00 | 1.70E-02 | 4.80E-02 | 1.14E+00 | 8.98E-05 | 3.29E-04 | -1.12E+00 | 1.40E-02 | 3.85E-02 | 1.01E+00 |
| RPL34P31 | 6.88E-09 | 3.58E-08 | -1.18E+00 | 2.89E-44 | 1.25E-43 | -1.81E+00 | 7.56E-04 | 3.21E-03 | -1.10E+00 | 3.59E-21 | 1.37E-20 | -1.68E+00 | 3.05E-04 | 1.09E-03 | -1.10E+00 | 1.07E-20 | 3.01E-20 | -1.81E+00 | 1.79E-05 | 6.57E-05 | -1.19E+00 | 2.87E-18 | 7.87E-18 | -1.98E+00 |
| COX15 | 6.98E-09 | 3.64E-08 | -1.44E+00 | 3.05E-05 | 1.32E-04 | 1.00E+00 | 1.12E-05 | 4.78E-05 | -1.29E+00 | 2.17E-05 | 8.29E-05 | -1.15E+00 | 4.43E-08 | 1.58E-07 | -1.35E+00 | 1.74E-05 | 4.91E-05 | -1.11E+00 | 1.09E-08 | 4.00E-08 | -1.22E+00 | 4.80E-06 | 1.32E-05 | -1.19E+00 |
| RPL28 | 6.99E-09 | 3.64E-08 | -1.13E+00 | 9.06E-34 | 3.92E-33 | -1.30E+00 | 2.73E-05 | 1.16E-04 | -1.11E+00 | 5.20E-20 | 1.99E-19 | -1.28E+00 | 6.01E-08 | 2.14E-07 | -1.17E+00 | 3.62E-17 | 1.02E-16 | -1.29E+00 | 1.80E-10 | 6.61E-10 | -1.13E+00 | 4.60E-17 | 1.26E-16 | -1.25E+00 |
| CREBL2 | 7.03E-09 | 3.66E-08 | -1.66E+00 | 1.09E-07 | 4.71E-07 | -1.39E+00 | 6.04E-04 | 2.57E-03 | -1.56E+00 | 2.58E-03 | 9.84E-03 | -1.24E+00 | 1.67E-06 | 5.95E-06 | -2.41E+00 | 1.45E-04 | 4.09E-04 | -1.49E+00 | 2.69E-11 | 9.86E-11 | -1.96E+00 | 6.02E-08 | 1.65E-07 | -1.65E+00 |
| DNTTIP2 | 7.04E-09 | 3.67E-08 | -1.00E+00 | 2.13E-22 | 9.19E-22 | -1.26E+00 | 4.20E-02 | 1.79E-01 | -1.00E+00 | 4.30E-06 | 1.64E-05 | -1.12E+00 | 1.34E-04 | 4.79E-04 | 1.11E+00 | 4.87E-07 | 1.37E-06 | -1.10E+00 | 6.40E-02 | 2.35E-01 | 1.08E+00 | 2.91E-02 | 7.99E-02 | 1.04E+00 |
| C1orf21 | 7.06E-09 | 3.68E-08 | -1.19E+00 | 9.80E-05 | 4.23E-04 | 1.08E+00 | 1.15E-05 | 4.87E-05 | -1.10E+00 | 3.38E-04 | 1.29E-03 | 1.02E+00 | 1.73E-04 | 6.15E-04 | -1.14E+00 | 6.00E-02 | 1.69E-01 | 1.10E+00 | 1.27E-05 | 4.65E-05 | -1.26E+00 | 9.34E-03 | 2.57E-02 | -1.03E+00 |
| LLPH | 7.07E-09 | 3.68E-08 | -1.12E+00 | 7.07E-12 | 3.06E-11 | -1.13E+00 | 9.06E-07 | 3.85E-06 | -1.15E+00 | 3.86E-07 | 1.47E-06 | -1.03E+00 | 2.64E-07 | 9.41E-07 | -1.21E+00 | 6.01E-06 | 1.70E-05 | -1.13E+00 | 5.76E-05 | 2.11E-04 | -1.08E+00 | 1.07E-05 | 2.93E-05 | -1.20E+00 |
| NOP56 | 7.09E-09 | 3.70E-08 | -1.15E+00 | 1.45E-01 | 6.27E-01 | 1.32E+00 | 6.04E-02 | 2.57E-01 | 1.01E+00 | 1.93E-01 | 7.37E-01 | 1.23E+00 | 1.27E-05 | 4.52E-05 | -1.16E+00 | 4.89E-02 | 1.38E-01 | 1.52E+00 | 5.97E-04 | 2.19E-03 | -1.20E+00 | 1.47E-02 | 4.03E-02 | 1.36E+00 |
| SETD2 | 7.10E-09 | 3.70E-08 | -1.20E+00 | 3.53E-07 | 1.53E-06 | -1.03E+00 | 1.37E-07 | 5.84E-07 | -1.44E+00 | 2.57E-05 | 9.82E-05 | -1.11E+00 | 7.19E-04 | 2.56E-03 | -1.12E+00 | 8.93E-04 | 2.52E-03 | -1.13E+00 | 2.61E-07 | 9.57E-07 | -1.26E+00 | 1.14E-04 | 3.14E-04 | -1.07E+00 |
| PIK3AP1 | 7.12E-09 | 3.71E-08 | -1.99E+00 | 2.02E-06 | 8.73E-06 | -1.39E+00 | 2.02E-13 | 8.59E-13 | -2.28E+00 | 1.44E-09 | 5.48E-09 | -1.60E+00 | 5.26E-11 | 1.87E-10 | -1.84E+00 | 4.18E-08 | 1.18E-07 | -1.41E+00 | 1.51E-03 | 5.54E-03 | -1.08E+00 | 9.88E-03 | 2.71E-02 | -1.06E+00 |
| CMAS | 7.12E-09 | 3.71E-08 | -1.28E+00 | 3.82E-03 | 1.65E-02 | 1.22E+00 | 2.31E-05 | 9.79E-05 | -1.16E+00 | 1.66E-03 | 6.34E-03 | 1.12E+00 | 6.07E-08 | 2.16E-07 | -1.25E+00 | 1.74E-03 | 4.91E-03 | 1.09E+00 | 2.09E-08 | 7.67E-08 | -1.25E+00 | 6.89E-03 | 1.89E-02 | -1.18E+00 |
| RPSAP54 | 7.24E-09 | 3.77E-08 | -1.81E+00 | 2.46E-04 | 1.06E-03 | -1.04E+00 | 3.48E-17 | 1.48E-16 | -2.33E+00 | 6.54E-08 | 2.50E-07 | -1.23E+00 | 1.55E-05 | 5.53E-05 | -1.49E+00 | 3.45E-02 | 9.72E-02 | 1.15E+00 | 2.77E-10 | 1.02E-09 | -1.93E+00 | 2.39E-03 | 6.57E-03 | -1.06E+00 |
| TIMM44 | 7.29E-09 | 3.80E-08 | -1.40E+00 | 3.53E-12 | 1.53E-11 | -1.46E+00 | 1.17E-06 | 4.95E-06 | -1.43E+00 | 2.08E-06 | 7.95E-06 | -1.20E+00 | 4.05E-05 | 1.44E-04 | -1.42E+00 | 3.82E-03 | 1.08E-02 | -1.02E+00 | 8.74E-06 | 3.21E-05 | -1.26E+00 | 1.27E-02 | 3.48E-02 | -1.05E+00 |
| EDEM1 | 7.29E-09 | 3.80E-08 | -1.70E+00 | 7.69E-07 | 3.32E-06 | -1.32E+00 | 5.86E-05 | 2.49E-04 | -1.38E+00 | 2.59E-07 | 9.89E-07 | -1.43E+00 | 6.55E-07 | 2.34E-06 | -1.72E+00 | 1.64E-07 | 4.61E-07 | -1.70E+00 | 4.34E-02 | 1.59E-01 | -1.10E+00 | 5.80E-02 | 1.59E-01 | -1.16E+00 |
| PKP4 | 7.38E-09 | 3.85E-08 | -1.83E+00 | 1.53E-03 | 6.62E-03 | 1.03E+00 | 3.76E-07 | 1.60E-06 | -1.52E+00 | 3.02E-02 | 1.15E-01 | 1.15E+00 | 3.28E-02 | 1.17E-01 | -1.29E+00 | 2.58E-01 | 7.27E-01 | 1.32E+00 | 1.24E-04 | 4.53E-04 | -1.30E+00 | 3.00E-01 | 8.23E-01 | 1.31E+00 |
| MRE11 | 7.39E-09 | 3.85E-08 | -1.32E+00 | 6.38E-05 | 2.76E-04 | 1.03E+00 | 1.28E-05 | 5.44E-05 | -1.24E+00 | 5.65E-03 | 2.16E-02 | -1.02E+00 | 4.55E-05 | 1.62E-04 | -1.16E+00 | 5.30E-02 | 1.50E-01 | 1.18E+00 | 3.75E-06 | 1.37E-05 | -1.23E+00 | 1.19E-01 | 3.27E-01 | 1.04E+00 |
| HNRNPDL | 7.40E-09 | 3.86E-08 | -1.22E+00 | 3.39E-21 | 1.46E-20 | -1.35E+00 | 1.31E-02 | 5.58E-02 | -1.17E+00 | 1.41E-05 | 5.39E-05 | -1.33E+00 | 6.92E-08 | 2.47E-07 | -1.23E+00 | 9.43E-11 | 2.66E-10 | -1.38E+00 | 7.35E-05 | 2.70E-04 | -1.30E+00 | 3.86E-05 | 1.06E-04 | -1.35E+00 |
| DENR | 7.42E-09 | 3.86E-08 | -1.05E+00 | 9.09E-10 | 3.93E-09 | 1.00E+00 | 3.52E-04 | 1.49E-03 | -1.09E+00 | 4.49E-07 | 1.72E-06 | -1.11E+00 | 5.94E-08 | 2.12E-07 | -1.28E+00 | 2.93E-06 | 8.27E-06 | -1.18E+00 | 1.79E-04 | 6.56E-04 | -1.07E+00 | 2.07E-04 | 5.69E-04 | -1.14E+00 |
| CEP41 | 7.43E-09 | 3.87E-08 | -1.69E+00 | 8.06E-05 | 3.48E-04 | -1.11E+00 | 2.26E-07 | 9.62E-07 | -1.31E+00 | 2.87E-05 | 1.10E-04 | -1.04E+00 | 2.08E-04 | 7.40E-04 | -1.42E+00 | 2.98E-03 | 8.40E-03 | -1.18E+00 | 3.39E-08 | 1.24E-07 | -1.44E+00 | 5.26E-05 | 1.45E-04 | -1.11E+00 |
| MRPL55 | 7.43E-09 | 3.87E-08 | -1.15E+00 | 2.00E-10 | 8.65E-10 | -1.12E+00 | 1.78E-01 | 7.56E-01 | 1.16E+00 | 9.73E-02 | 3.72E-01 | 1.20E+00 | 3.88E-04 | 1.38E-03 | -1.05E+00 | 4.62E-02 | 1.30E-01 | 1.15E+00 | 1.09E-01 | 3.99E-01 | 1.20E+00 | 5.65E-02 | 1.55E-01 | 1.17E+00 |
| F11R | 7.43E-09 | 3.87E-08 | -1.43E+00 | 4.30E-06 | 1.86E-05 | -1.09E+00 | 6.77E-13 | 2.88E-12 | -1.91E+00 | 5.13E-13 | 1.96E-12 | -1.69E+00 | 3.78E-05 | 1.35E-04 | -1.40E+00 | 2.49E-05 | 7.01E-05 | -1.38E+00 | 1.53E-02 | 5.61E-02 | 1.18E+00 | 1.96E-02 | 5.39E-02 | -1.00E+00 |
| ACSL1 | 7.49E-09 | 3.90E-08 | -1.83E+00 | 1.35E-05 | 5.85E-05 | -1.22E+00 | 1.66E-08 | 7.05E-08 | -1.71E+00 | 3.52E-05 | 1.34E-04 | -1.19E+00 | 1.40E-05 | 4.98E-05 | -1.40E+00 | 1.17E-01 | 3.29E-01 | 1.35E+00 | 1.63E-07 | 5.99E-07 | -1.37E+00 | 1.70E-03 | 4.67E-03 | -1.08E+00 |
| IFNGR2 | 7.57E-09 | 3.94E-08 | -1.38E+00 | 3.26E-05 | 1.41E-04 | 1.02E+00 | 1.25E-09 | 5.30E-09 | -1.45E+00 | 1.36E-08 | 5.18E-08 | -1.22E+00 | 6.37E-08 | 2.27E-07 | -1.56E+00 | 2.44E-04 | 6.87E-04 | -1.16E+00 | 1.00E-10 | 3.68E-10 | -1.53E+00 | 3.39E-05 | 9.32E-05 | -1.15E+00 |
| OGT | 7.58E-09 | 3.95E-08 | -1.16E+00 | 1.04E-01 | 4.47E-01 | 1.49E+00 | 3.22E-07 | 1.37E-06 | -1.34E+00 | 2.22E-01 | 8.48E-01 | 1.33E+00 | 8.90E-03 | 3.17E-02 | 1.04E+00 | 1.01E-01 | 2.85E-01 | 1.55E+00 | 1.30E-02 | 4.79E-02 | -1.05E+00 | 2.89E-01 | 7.94E-01 | 1.26E+00 |
| TRIM52 | 7.58E-09 | 3.95E-08 | -1.41E+00 | 1.15E-03 | 4.95E-03 | 1.16E+00 | 1.98E-04 | 8.40E-04 | -1.25E+00 | 3.01E-03 | 1.15E-02 | 1.02E+00 | 5.43E-03 | 1.94E-02 | -1.14E+00 | 1.79E-01 | 5.05E-01 | 1.23E+00 | 2.04E-03 | 7.48E-03 | -1.11E+00 | 3.23E-02 | 8.86E-02 | -1.01E+00 |
| RBM27 | 7.63E-09 | 3.98E-08 | -1.54E+00 | 6.27E-06 | 2.71E-05 | -1.16E+00 | 1.47E-05 | 6.26E-05 | -1.27E+00 | 9.81E-05 | 3.75E-04 | -1.10E+00 | 5.23E-11 | 1.87E-10 | -1.83E+00 | 5.49E-08 | 1.55E-07 | -1.39E+00 | 7.70E-08 | 2.83E-07 | -1.54E+00 | 2.29E-02 | 6.29E-02 | 1.03E+00 |
| MAP1A | 7.64E-09 | 3.98E-08 | -1.57E+00 | 1.74E-04 | 7.51E-04 | -1.06E+00 | 6.57E-11 | 2.79E-10 | -1.68E+00 | 4.71E-09 | 1.80E-08 | -1.35E+00 | 5.08E-04 | 1.81E-03 | -1.01E+00 | 1.31E-03 | 3.68E-03 | 1.02E+00 | 2.69E-01 | 9.88E-01 | 1.43E+00 | 7.92E-02 | 2.18E-01 | 1.09E+00 |
| MRPL14 | 7.68E-09 | 4.00E-08 | -1.13E+00 | 6.53E-07 | 2.82E-06 | 1.10E+00 | 1.22E-03 | 5.19E-03 | -1.13E+00 | 9.56E-03 | 3.65E-02 | 1.07E+00 | 3.51E-10 | 1.25E-09 | -1.45E+00 | 8.12E-03 | 2.29E-02 | 1.10E+00 | 1.00E-05 | 3.68E-05 | -1.18E+00 | 3.23E-01 | 8.86E-01 | 1.14E+00 |
| LXN | 7.69E-09 | 4.01E-08 | -1.34E+00 | 3.91E-04 | 1.69E-03 | 1.11E+00 | 6.01E-05 | 2.55E-04 | -1.32E+00 | 7.02E-04 | 2.68E-03 | -1.05E+00 | 2.26E-12 | 8.06E-12 | -1.49E+00 | 3.71E-02 | 1.05E-01 | 1.28E+00 | 1.04E-05 | 3.81E-05 | 1.07E+00 | 1.07E-01 | 2.95E-01 | 1.42E+00 |
| RSPRY1 | 7.73E-09 | 4.03E-08 | -1.36E+00 | 1.46E-11 | 6.32E-11 | -1.47E+00 | 7.03E-09 | 2.99E-08 | -1.50E+00 | 1.14E-07 | 4.36E-07 | -1.27E+00 | 1.13E-06 | 4.04E-06 | -1.62E+00 | 2.23E-04 | 6.28E-04 | -1.21E+00 | 2.32E-09 | 8.53E-09 | -1.66E+00 | 7.86E-08 | 2.16E-07 | -1.51E+00 |
| AJ009632.2 | 7.82E-09 | 4.07E-08 | -1.26E+00 | 1.12E-01 | 4.82E-01 | 1.41E+00 | 1.89E-01 | 8.04E-01 | 1.29E+00 | 3.75E-02 | 1.43E-01 | 2.06E+00 | 1.89E-01 | 6.73E-01 | -1.13E+00 | 2.96E-01 | 8.34E-01 | 1.08E+00 | 2.67E-01 | 9.80E-01 | 1.08E+00 | 2.42E-01 | 6.65E-01 | 1.69E+00 |
| SNX27 | 7.83E-09 | 4.08E-08 | -1.25E+00 | 3.94E-08 | 1.70E-07 | -1.08E+00 | 6.09E-08 | 2.59E-07 | -1.38E+00 | 1.45E-07 | 5.53E-07 | -1.20E+00 | 7.12E-05 | 2.54E-04 | -1.27E+00 | 2.65E-03 | 7.46E-03 | -1.02E+00 | 1.36E-08 | 4.99E-08 | -1.46E+00 | 1.76E-04 | 4.83E-04 | -1.15E+00 |
| PTPN18 | 7.84E-09 | 4.08E-08 | -1.22E+00 | 6.88E-05 | 2.97E-04 | 1.11E+00 | 9.59E-08 | 4.07E-07 | -1.33E+00 | 5.88E-06 | 2.25E-05 | -1.07E+00 | 1.41E-03 | 5.02E-03 | -1.11E+00 | 5.81E-02 | 1.64E-01 | 1.12E+00 | 3.75E-03 | 1.37E-02 | 1.14E+00 | 8.18E-02 | 2.25E-01 | 1.17E+00 |
| WDFY1 | 7.87E-09 | 4.10E-08 | -1.33E+00 | 8.89E-07 | 3.84E-06 | -1.16E+00 | 1.47E-05 | 6.25E-05 | -1.49E+00 | 2.20E-05 | 8.40E-05 | -1.22E+00 | 1.59E-06 | 5.68E-06 | -1.43E+00 | 3.42E-05 | 9.65E-05 | -1.23E+00 | 3.02E-07 | 1.11E-06 | -1.43E+00 | 5.10E-05 | 1.40E-04 | -1.18E+00 |
| HTT | 7.90E-09 | 4.12E-08 | -1.21E+00 | 3.43E-08 | 1.48E-07 | -1.13E+00 | 7.14E-02 | 3.04E-01 | 1.04E+00 | 2.12E-03 | 8.09E-03 | -1.02E+00 | 1.29E-03 | 4.60E-03 | 1.08E+00 | 7.09E-03 | 2.00E-02 | 1.14E+00 | 2.62E-01 | 9.63E-01 | 1.37E+00 | 1.09E-01 | 2.98E-01 | 1.10E+00 |
| ZADH2 | 7.92E-09 | 4.13E-08 | -1.40E+00 | 1.07E-05 | 4.63E-05 | 1.01E+00 | 4.71E-09 | 2.00E-08 | -1.47E+00 | 1.16E-08 | 4.43E-08 | -1.24E+00 | 9.43E-09 | 3.36E-08 | -1.35E+00 | 4.78E-06 | 1.35E-05 | -1.14E+00 | 1.13E-06 | 4.13E-06 | -1.28E+00 | 3.64E-03 | 1.00E-02 | -1.05E+00 |
| SEC63P1 | 7.92E-09 | 4.13E-08 | -1.56E+00 | 8.96E-10 | 3.87E-09 | -1.57E+00 | 4.74E-09 | 2.02E-08 | -1.67E+00 | 7.28E-08 | 2.78E-07 | -1.32E+00 | 2.68E-07 | 9.57E-07 | -1.55E+00 | 4.60E-07 | 1.30E-06 | -1.46E+00 | 1.83E-11 | 6.73E-11 | -1.52E+00 | 2.83E-09 | 7.78E-09 | -1.39E+00 |
| PPME1 | 7.93E-09 | 4.13E-08 | -1.54E+00 | 6.72E-07 | 2.90E-06 | -1.22E+00 | 1.38E-05 | 5.88E-05 | -1.39E+00 | 3.08E-04 | 1.18E-03 | -1.11E+00 | 3.19E-05 | 1.14E-04 | -1.34E+00 | 4.44E-04 | 1.25E-03 | -1.17E+00 | 7.45E-07 | 2.73E-06 | -1.26E+00 | 1.27E-04 | 3.49E-04 | -1.16E+00 |
| POLR2G | 7.95E-09 | 4.14E-08 | -1.07E+00 | 1.16E-07 | 5.00E-07 | 1.08E+00 | 9.18E-05 | 3.90E-04 | -1.12E+00 | 5.10E-05 | 1.95E-04 | -1.02E+00 | 6.75E-06 | 2.41E-05 | -1.16E+00 | 1.09E-02 | 3.07E-02 | 1.00E+00 | 2.07E-01 | 7.61E-01 | 1.24E+00 | 3.55E-01 | 9.74E-01 | 1.26E+00 |
| ALS2 | 7.95E-09 | 4.14E-08 | -1.46E+00 | 1.68E-02 | 7.25E-02 | 1.28E+00 | 2.33E-04 | 9.92E-04 | -1.24E+00 | 1.99E-01 | 7.62E-01 | 1.55E+00 | 5.31E-07 | 1.89E-06 | -1.45E+00 | 4.10E-04 | 1.16E-03 | 1.02E+00 | 1.02E-03 | 3.76E-03 | -1.31E+00 | 5.83E-02 | 1.60E-01 | 1.05E+00 |
| WNK1 | 7.97E-09 | 4.16E-08 | -1.13E+00 | 1.79E-04 | 7.75E-04 | 1.15E+00 | 7.00E-08 | 2.97E-07 | -1.33E+00 | 3.39E-03 | 1.30E-02 | 1.10E+00 | 5.53E-07 | 1.97E-06 | -1.20E+00 | 1.42E-03 | 3.99E-03 | -1.01E+00 | 1.41E-09 | 5.17E-09 | -1.29E+00 | 1.90E-03 | 5.21E-03 | -1.02E+00 |
| CYTL1 | 8.00E-09 | 4.17E-08 | -1.33E+00 | 1.91E-02 | 8.24E-02 | -1.02E+00 | 3.72E-05 | 1.58E-04 | -1.33E+00 | 1.34E-02 | 5.12E-02 | -1.03E+00 | 1.04E-01 | 3.71E-01 | -1.01E+00 | 6.93E-02 | 1.95E-01 | 1.24E+00 | 1.62E-04 | 5.95E-04 | -1.20E+00 | 2.13E-01 | 5.85E-01 | 1.15E+00 |
| FAM192A | 8.03E-09 | 4.18E-08 | -1.39E+00 | 7.09E-05 | 3.06E-04 | 1.02E+00 | 1.23E-06 | 5.23E-06 | -1.21E+00 | 4.08E-05 | 1.56E-04 | 1.00E+00 | 7.68E-08 | 2.74E-07 | -1.19E+00 | 7.08E-05 | 2.00E-04 | 1.02E+00 | 4.31E-04 | 1.58E-03 | -1.07E+00 | 8.15E-04 | 2.24E-03 | -1.07E+00 |
| COPE | 8.08E-09 | 4.21E-08 | -1.08E+00 | 1.02E-09 | 4.43E-09 | 1.01E+00 | 1.31E-01 | 5.56E-01 | 1.13E+00 | 3.51E-02 | 1.34E-01 | 1.07E+00 | 1.12E-06 | 4.00E-06 | -1.16E+00 | 5.58E-04 | 1.57E-03 | 1.02E+00 | 2.47E-01 | 9.05E-01 | 1.14E+00 | 2.10E-01 | 5.77E-01 | 1.18E+00 |
| ZMYND8 | 8.10E-09 | 4.22E-08 | -1.19E+00 | 3.14E-04 | 1.36E-03 | 1.14E+00 | 2.42E-10 | 1.03E-09 | -1.47E+00 | 1.75E-04 | 6.70E-04 | -1.04E+00 | 1.89E-06 | 6.73E-06 | -1.27E+00 | 5.83E-02 | 1.64E-01 | 1.08E+00 | 1.83E-07 | 6.72E-07 | -1.41E+00 | 1.84E-02 | 5.05E-02 | -1.11E+00 |
| VPS29 | 8.10E-09 | 4.22E-08 | -1.12E+00 | 7.52E-07 | 3.25E-06 | 1.10E+00 | 6.45E-02 | 2.74E-01 | 1.08E+00 | 3.28E-02 | 1.25E-01 | 1.16E+00 | 1.40E-05 | 4.98E-05 | -1.22E+00 | 1.06E-02 | 3.00E-02 | -1.02E+00 | 1.18E-01 | 4.31E-01 | -1.01E+00 | 3.35E-01 | 9.20E-01 | 1.10E+00 |
| MCM3AP | 8.10E-09 | 4.22E-08 | -1.62E+00 | 1.55E-04 | 6.71E-04 | -1.02E+00 | 1.70E-06 | 7.22E-06 | -1.50E+00 | 8.92E-03 | 3.41E-02 | 1.11E+00 | 3.82E-03 | 1.36E-02 | -1.18E+00 | 8.18E-02 | 2.31E-01 | 1.10E+00 | 2.03E-02 | 7.43E-02 | -1.11E+00 | 5.70E-02 | 1.57E-01 | 1.08E+00 |
| MAGOH | 8.11E-09 | 4.23E-08 | -1.02E+00 | 1.80E-09 | 7.78E-09 | 1.02E+00 | 2.05E-02 | 8.70E-02 | 1.01E+00 | 1.16E-03 | 4.44E-03 | 1.02E+00 | 8.79E-06 | 3.14E-05 | -1.11E+00 | 7.78E-05 | 2.19E-04 | -1.04E+00 | 1.52E-04 | 5.57E-04 | -1.13E+00 | 5.04E-02 | 1.38E-01 | 1.04E+00 |
| ZNF521 | 8.11E-09 | 4.23E-08 | -1.36E+00 | 9.06E-02 | 3.91E-01 | 1.32E+00 | 4.43E-04 | 1.88E-03 | -1.21E+00 | 2.22E-01 | 8.48E-01 | 1.40E+00 | 1.77E-15 | 6.31E-15 | -2.00E+00 | 5.43E-06 | 1.53E-05 | -1.18E+00 | 2.07E-12 | 7.59E-12 | -1.95E+00 | 3.19E-04 | 8.77E-04 | -1.25E+00 |
| AC010997.6 | 8.15E-09 | 4.25E-08 | -1.58E+00 | 1.15E-05 | 4.95E-05 | -1.10E+00 | 3.78E-15 | 1.61E-14 | -1.89E+00 | 7.18E-07 | 2.74E-06 | -1.14E+00 | 4.64E-04 | 1.66E-03 | -1.28E+00 | 1.05E-02 | 2.97E-02 | -1.05E+00 | 4.82E-09 | 1.77E-08 | -2.47E+00 | 6.48E-04 | 1.78E-03 | -1.37E+00 |
| LPCAT2 | 8.16E-09 | 4.25E-08 | -1.23E+00 | 1.77E-09 | 7.66E-09 | -1.11E+00 | 2.37E-04 | 1.01E-03 | -1.08E+00 | 8.43E-05 | 3.22E-04 | -1.03E+00 | 6.50E-05 | 2.32E-04 | -1.13E+00 | 1.94E-10 | 5.47E-10 | -1.55E+00 | 1.80E-02 | 6.62E-02 | -1.20E+00 | 1.26E-01 | 3.45E-01 | 1.01E+00 |
| FAM102A | 8.17E-09 | 4.26E-08 | -1.85E+00 | 4.77E-08 | 2.06E-07 | -1.62E+00 | 1.09E-03 | 4.62E-03 | -1.67E+00 | 1.85E-02 | 7.05E-02 | -1.13E+00 | 1.73E-03 | 6.16E-03 | -1.89E+00 | 4.05E-03 | 1.14E-02 | -1.62E+00 | 1.15E-01 | 4.24E-01 | -1.14E+00 | 1.72E-01 | 4.72E-01 | -1.13E+00 |
| KYAT3 | 8.19E-09 | 4.27E-08 | -1.22E+00 | 3.65E-09 | 1.58E-08 | -1.11E+00 | 3.27E-06 | 1.39E-05 | -1.34E+00 | 1.81E-06 | 6.91E-06 | -1.21E+00 | 2.93E-06 | 1.05E-05 | -1.26E+00 | 2.52E-02 | 7.10E-02 | 1.07E+00 | 3.72E-08 | 1.37E-07 | -1.31E+00 | 9.50E-05 | 2.61E-04 | -1.12E+00 |
| MZB1 | 8.19E-09 | 4.27E-08 | 2.12E+00 | 4.32E-06 | 1.87E-05 | 2.08E+00 | 5.16E-02 | 2.19E-01 | 1.53E+00 | 1.87E-01 | 7.15E-01 | 1.37E+00 | 9.18E-02 | 3.27E-01 | -1.10E+00 | 1.72E-01 | 4.84E-01 | 1.01E+00 | 2.33E-01 | 8.57E-01 | 1.44E+00 | 3.10E-01 | 8.52E-01 | 1.39E+00 |
| SLC25A5P2 | 8.20E-09 | 4.27E-08 | -3.13E+00 | 9.23E-06 | 3.99E-05 | -1.63E+00 | 7.78E-15 | 3.30E-14 | -2.97E+00 | 6.05E-10 | 2.31E-09 | -1.78E+00 | 3.43E-03 | 1.22E-02 | -2.27E+00 | 6.12E-02 | 1.72E-01 | -1.24E+00 | 3.72E-13 | 1.36E-12 | -2.78E+00 | 1.17E-06 | 3.21E-06 | -1.55E+00 |
| CPVL | 8.20E-09 | 4.27E-08 | -1.19E+00 | 2.55E-08 | 1.10E-07 | -1.11E+00 | 1.01E-04 | 4.28E-04 | -1.28E+00 | 2.40E-02 | 9.18E-02 | -1.01E+00 | 4.07E-05 | 1.45E-04 | -1.06E+00 | 8.94E-02 | 2.52E-01 | 1.23E+00 | 5.48E-03 | 2.01E-02 | -1.03E+00 | 3.03E-01 | 8.32E-01 | 1.25E+00 |
| TARS | 8.21E-09 | 4.28E-08 | -1.23E+00 | 6.99E-06 | 3.02E-05 | 1.03E+00 | 2.86E-03 | 1.22E-02 | -1.07E+00 | 5.21E-04 | 1.99E-03 | 1.03E+00 | 5.65E-06 | 2.01E-05 | -1.25E+00 | 1.16E-03 | 3.26E-03 | 1.03E+00 | 1.91E-05 | 7.02E-05 | -1.15E+00 | 6.79E-02 | 1.86E-01 | 1.14E+00 |
| PTMAP8 | 8.22E-09 | 4.28E-08 | 2.29E+00 | 3.57E-04 | 1.54E-03 | 2.17E+00 | 3.58E-09 | 1.52E-08 | 2.03E+00 | 3.64E-05 | 1.39E-04 | 2.04E+00 | 1.12E-02 | 4.00E-02 | 1.98E+00 | 5.87E-03 | 1.66E-02 | 2.01E+00 | 9.42E-04 | 3.46E-03 | 1.70E+00 | 1.22E-03 | 3.36E-03 | 1.81E+00 |
| TSN | 8.28E-09 | 4.32E-08 | -1.16E+00 | 1.09E-03 | 4.72E-03 | 1.16E+00 | 1.86E-04 | 7.92E-04 | -1.19E+00 | 9.11E-03 | 3.48E-02 | 1.06E+00 | 4.16E-05 | 1.48E-04 | -1.14E+00 | 3.46E-02 | 9.76E-02 | 1.13E+00 | 5.30E-06 | 1.94E-05 | -1.16E+00 | 3.56E-01 | 9.77E-01 | 1.22E+00 |
| MRPL17 | 8.32E-09 | 4.34E-08 | -1.76E+00 | 1.23E-04 | 5.32E-04 | -1.09E+00 | 1.04E-07 | 4.44E-07 | -1.65E+00 | 2.90E-02 | 1.11E-01 | 1.06E+00 | 3.43E-04 | 1.22E-03 | -1.49E+00 | 1.36E-01 | 3.83E-01 | 1.12E+00 | 4.41E-13 | 1.62E-12 | -1.66E+00 | 1.93E-02 | 5.31E-02 | -1.02E+00 |
| VAMP3 | 8.35E-09 | 4.35E-08 | -1.28E+00 | 3.91E-11 | 1.69E-10 | -1.31E+00 | 1.06E-03 | 4.50E-03 | -1.21E+00 | 3.52E-04 | 1.34E-03 | -1.04E+00 | 5.51E-06 | 1.97E-05 | -1.22E+00 | 1.90E-06 | 5.36E-06 | -1.31E+00 | 4.67E-06 | 1.71E-05 | -1.27E+00 | 2.73E-04 | 7.49E-04 | -1.26E+00 |
| NF1 | 8.41E-09 | 4.38E-08 | -1.28E+00 | 2.20E-02 | 9.51E-02 | 1.29E+00 | 3.38E-02 | 1.44E-01 | -1.04E+00 | 2.29E-02 | 8.76E-02 | 1.11E+00 | 6.41E-05 | 2.28E-04 | -1.28E+00 | 8.47E-03 | 2.39E-02 | 1.07E+00 | 5.51E-07 | 2.02E-06 | -1.47E+00 | 5.44E-05 | 1.49E-04 | -1.29E+00 |
| AL513477.1 | 8.41E-09 | 4.38E-08 | -2.09E+00 | 1.75E-09 | 7.56E-09 | -2.22E+00 | 1.64E-02 | 6.96E-02 | -1.22E+00 | 8.45E-04 | 3.23E-03 | -1.55E+00 | 9.20E-13 | 3.28E-12 | -2.91E+00 | 1.34E-12 | 3.79E-12 | -2.74E+00 | 1.71E-07 | 6.28E-07 | -1.50E+00 | 5.52E-13 | 1.52E-12 | -2.08E+00 |
| PLOD2 | 8.42E-09 | 4.39E-08 | -1.44E+00 | 1.16E-04 | 5.02E-04 | 1.02E+00 | 2.71E-04 | 1.15E-03 | -1.20E+00 | 6.43E-05 | 2.45E-04 | -1.12E+00 | 3.93E-03 | 1.40E-02 | -1.16E+00 | 1.51E-01 | 4.26E-01 | 1.17E+00 | 1.75E-02 | 6.44E-02 | 1.01E+00 | 7.70E-02 | 2.11E-01 | 1.08E+00 |
| SLC16A3 | 8.50E-09 | 4.43E-08 | -1.74E+00 | 1.47E-07 | 6.34E-07 | -1.40E+00 | 1.86E-07 | 7.92E-07 | -1.59E+00 | 2.70E-05 | 1.03E-04 | -1.17E+00 | 7.90E-05 | 2.81E-04 | -1.46E+00 | 1.68E-05 | 4.74E-05 | -1.72E+00 | 7.16E-03 | 2.63E-02 | -1.03E+00 | 8.17E-03 | 2.24E-02 | -1.55E+00 |
| TMEM123 | 8.60E-09 | 4.48E-08 | -1.11E+00 | 4.88E-05 | 2.11E-04 | 1.07E+00 | 3.56E-03 | 1.51E-02 | -1.13E+00 | 2.45E-02 | 9.35E-02 | 1.02E+00 | 1.04E-06 | 3.69E-06 | -1.05E+00 | 2.20E-04 | 6.22E-04 | 1.03E+00 | 4.56E-03 | 1.67E-02 | -1.02E+00 | 1.73E-01 | 4.74E-01 | 1.12E+00 |
| AC009053.1 | 8.60E-09 | 4.48E-08 | -2.00E+00 | 7.93E-07 | 3.43E-06 | -1.50E+00 | 2.70E-02 | 1.15E-01 | -1.07E+00 | 6.10E-03 | 2.33E-02 | -1.20E+00 | 1.43E-02 | 5.10E-02 | -1.38E+00 | 5.22E-02 | 1.47E-01 | -1.10E+00 | 3.66E-03 | 1.34E-02 | -1.33E+00 | 5.64E-02 | 1.55E-01 | -1.13E+00 |
| SLFN12 | 8.62E-09 | 4.49E-08 | -1.88E+00 | 9.61E-09 | 4.15E-08 | -1.90E+00 | 7.59E-07 | 3.22E-06 | -2.34E+00 | 8.67E-07 | 3.31E-06 | -1.94E+00 | 1.65E-12 | 5.87E-12 | -2.51E+00 | 2.51E-08 | 7.07E-08 | -1.63E+00 | 3.24E-09 | 1.19E-08 | -2.06E+00 | 3.02E-06 | 8.28E-06 | -1.61E+00 |
| LZTFL1 | 8.63E-09 | 4.50E-08 | -1.47E+00 | 4.50E-07 | 1.95E-06 | -1.19E+00 | 6.01E-11 | 2.55E-10 | -1.57E+00 | 4.52E-09 | 1.73E-08 | -1.32E+00 | 1.76E-05 | 6.27E-05 | -1.33E+00 | 6.80E-04 | 1.92E-03 | -1.16E+00 | 2.26E-04 | 8.30E-04 | -1.08E+00 | 9.66E-03 | 2.65E-02 | 1.02E+00 |
| IRAK1 | 8.63E-09 | 4.50E-08 | -1.25E+00 | 1.39E-04 | 6.00E-04 | 1.12E+00 | 8.81E-03 | 3.74E-02 | 1.01E+00 | 3.42E-03 | 1.31E-02 | 1.10E+00 | 4.25E-06 | 1.51E-05 | -1.20E+00 | 1.19E-03 | 3.35E-03 | 1.02E+00 | 5.42E-02 | 1.99E-01 | 1.05E+00 | 3.47E-01 | 9.53E-01 | 1.27E+00 |
| AUP1 | 8.64E-09 | 4.50E-08 | -1.27E+00 | 8.14E-04 | 3.52E-03 | 1.05E+00 | 2.59E-04 | 1.10E-03 | -1.17E+00 | 8.88E-04 | 3.39E-03 | -1.06E+00 | 1.36E-06 | 4.85E-06 | -1.13E+00 | 4.01E-03 | 1.13E-02 | 1.10E+00 | 1.38E-04 | 5.07E-04 | -1.07E+00 | 1.72E-01 | 4.72E-01 | 1.17E+00 |
| NKRF | 8.65E-09 | 4.51E-08 | -1.83E+00 | 6.53E-08 | 2.82E-07 | -1.52E+00 | 1.63E-08 | 6.94E-08 | -1.81E+00 | 2.13E-07 | 8.12E-07 | -1.45E+00 | 4.97E-03 | 1.77E-02 | -1.41E+00 | 6.03E-02 | 1.70E-01 | -1.01E+00 | 5.45E-09 | 2.00E-08 | -1.98E+00 | 1.89E-03 | 5.18E-03 | -1.17E+00 |
| RIOK1 | 8.70E-09 | 4.53E-08 | -1.51E+00 | 6.03E-04 | 2.60E-03 | 1.06E+00 | 3.77E-07 | 1.60E-06 | -1.43E+00 | 6.72E-04 | 2.57E-03 | 1.04E+00 | 1.25E-06 | 4.46E-06 | -1.61E+00 | 5.20E-02 | 1.47E-01 | 1.09E+00 | 1.44E-14 | 5.30E-14 | -1.61E+00 | 9.11E-03 | 2.50E-02 | 1.01E+00 |
| ATP5I2 | 8.70E-09 | 4.54E-08 | -1.09E+00 | 5.70E-30 | 2.46E-29 | -1.38E+00 | 1.32E-01 | 5.63E-01 | 1.06E+00 | 3.10E-10 | 1.18E-09 | -1.31E+00 | 5.90E-07 | 2.11E-06 | -1.19E+00 | 2.74E-12 | 7.71E-12 | -1.36E+00 | 2.13E-02 | 7.84E-02 | -1.11E+00 | 4.46E-04 | 1.23E-03 | -1.35E+00 |
| RNASEL | 8.73E-09 | 4.55E-08 | -2.19E+00 | 3.64E-05 | 1.57E-04 | -1.30E+00 | 1.84E-04 | 7.80E-04 | -1.52E+00 | 7.50E-04 | 2.86E-03 | -1.23E+00 | 4.41E-12 | 1.57E-11 | -2.94E+00 | 3.14E-08 | 8.84E-08 | -1.77E+00 | 3.08E-07 | 1.13E-06 | -1.82E+00 | 2.76E-04 | 7.57E-04 | -1.30E+00 |
| LIMD1 | 8.81E-09 | 4.59E-08 | -1.93E+00 | 7.28E-02 | 3.15E-01 | 1.19E+00 | 8.28E-11 | 3.52E-10 | -1.94E+00 | 3.72E-03 | 1.42E-02 | 1.12E+00 | 6.96E-07 | 2.48E-06 | -1.89E+00 | 4.86E-03 | 1.37E-02 | -1.07E+00 | 2.05E-04 | 7.51E-04 | -1.18E+00 | 3.45E-02 | 9.48E-02 | -1.03E+00 |
| SLC26A6 | 8.83E-09 | 4.60E-08 | -2.75E+00 | 2.02E-06 | 8.71E-06 | -1.73E+00 | 3.97E-02 | 1.69E-01 | -1.11E+00 | 2.57E-01 | 9.80E-01 | 1.31E+00 | 1.20E-03 | 4.26E-03 | -1.84E+00 | 3.02E-02 | 8.51E-02 | -1.09E+00 | 7.55E-03 | 2.77E-02 | -1.84E+00 | 4.79E-02 | 1.31E-01 | -1.47E+00 |
| QDPR | 8.84E-09 | 4.61E-08 | -1.34E+00 | 8.46E-06 | 3.66E-05 | -1.03E+00 | 3.22E-04 | 1.37E-03 | -1.12E+00 | 1.75E-05 | 6.68E-05 | -1.00E+00 | 1.15E-11 | 4.10E-11 | -1.93E+00 | 6.11E-05 | 1.72E-04 | -1.19E+00 | 1.56E-04 | 5.71E-04 | -1.06E+00 | 1.86E-01 | 5.11E-01 | 1.18E+00 |
| ARAF | 8.92E-09 | 4.65E-08 | -1.46E+00 | 2.60E-09 | 1.12E-08 | -1.47E+00 | 2.06E-03 | 8.76E-03 | -1.23E+00 | 2.23E-04 | 8.52E-04 | -1.13E+00 | 1.56E-03 | 5.55E-03 | -1.10E+00 | 1.60E-02 | 4.50E-02 | 1.05E+00 | 4.35E-05 | 1.60E-04 | -1.32E+00 | 1.07E-02 | 2.95E-02 | -1.05E+00 |
| DCXR | 8.98E-09 | 4.68E-08 | -1.22E+00 | 5.16E-07 | 2.23E-06 | 1.01E+00 | 8.39E-04 | 3.56E-03 | -1.02E+00 | 5.77E-02 | 2.20E-01 | 1.22E+00 | 6.46E-04 | 2.30E-03 | -1.12E+00 | 1.20E-02 | 3.40E-02 | 1.03E+00 | 4.71E-02 | 1.73E-01 | 1.08E+00 | 2.63E-02 | 7.22E-02 | 1.39E+00 |
| PCBP2 | 9.06E-09 | 4.72E-08 | -1.16E+00 | 6.14E-02 | 2.65E-01 | 1.05E+00 | 1.34E-05 | 5.68E-05 | -1.26E+00 | 4.82E-02 | 1.84E-01 | -1.07E+00 | 9.70E-07 | 3.46E-06 | -1.19E+00 | 5.48E-02 | 1.54E-01 | -1.02E+00 | 1.44E-05 | 5.29E-05 | -1.19E+00 | 6.88E-02 | 1.89E-01 | -1.05E+00 |
| LPGAT1 | 9.07E-09 | 4.73E-08 | -1.28E+00 | 2.64E-10 | 1.14E-09 | -1.19E+00 | 5.01E-06 | 2.13E-05 | -1.37E+00 | 8.00E-07 | 3.06E-06 | -1.28E+00 | 6.02E-05 | 2.15E-04 | -1.01E+00 | 8.16E-07 | 2.30E-06 | -1.16E+00 | 1.78E-03 | 6.54E-03 | -1.02E+00 | 1.93E-04 | 5.31E-04 | -1.14E+00 |
| NOM1 | 9.08E-09 | 4.73E-08 | -1.90E+00 | 1.03E-06 | 4.46E-06 | -1.42E+00 | 1.08E-03 | 4.61E-03 | -1.19E+00 | 1.90E-02 | 7.25E-02 | 1.09E+00 | 3.76E-03 | 1.34E-02 | -1.20E+00 | 1.64E-02 | 4.61E-02 | -1.03E+00 | 2.31E-09 | 8.48E-09 | -1.48E+00 | 4.03E-03 | 1.11E-02 | -1.04E+00 |
| SLC38A7 | 9.20E-09 | 4.80E-08 | -1.61E+00 | 1.29E-09 | 5.58E-09 | -1.51E+00 | 2.01E-03 | 8.56E-03 | -1.30E+00 | 4.25E-03 | 1.62E-02 | -1.03E+00 | 1.87E-03 | 6.65E-03 | -1.43E+00 | 3.18E-02 | 8.96E-02 | -1.04E+00 | 4.67E-06 | 1.71E-05 | -1.63E+00 | 2.64E-03 | 7.26E-03 | -1.15E+00 |
| TRPC2 | 9.21E-09 | 4.80E-08 | -1.52E+00 | 1.17E-05 | 5.06E-05 | -1.08E+00 | 1.11E-01 | 4.72E-01 | 1.12E+00 | 5.09E-02 | 1.94E-01 | 1.15E+00 | 8.13E-02 | 2.90E-01 | -1.18E+00 | 1.63E-01 | 4.59E-01 | 1.09E+00 | 1.68E-03 | 6.15E-03 | 2.65E+00 | 1.60E-01 | 4.40E-01 | 1.74E+00 |
| VRK3 | 9.24E-09 | 4.82E-08 | -1.36E+00 | 1.05E-07 | 4.53E-07 | -1.21E+00 | 1.74E-03 | 7.38E-03 | -1.22E+00 | 1.76E-03 | 6.74E-03 | -1.14E+00 | 2.41E-04 | 8.59E-04 | -1.16E+00 | 6.19E-03 | 1.75E-02 | 1.05E+00 | 2.59E-04 | 9.52E-04 | -1.18E+00 | 3.05E-03 | 8.39E-03 | -1.03E+00 |
| MARCH2 | 9.26E-09 | 4.83E-08 | -1.34E+00 | 1.30E-04 | 5.62E-04 | 1.09E+00 | 4.62E-02 | 1.96E-01 | 1.01E+00 | 6.96E-02 | 2.66E-01 | 1.13E+00 | 1.28E-04 | 4.56E-04 | -1.12E+00 | 1.76E-04 | 4.96E-04 | -1.13E+00 | 1.82E-04 | 6.69E-04 | 1.10E+00 | 1.82E-04 | 5.00E-04 | -1.20E+00 |
| CCDC146 | 9.32E-09 | 4.86E-08 | -2.14E+00 | 1.36E-08 | 5.86E-08 | -2.00E+00 | 1.20E-03 | 5.09E-03 | -1.20E+00 | 1.71E-04 | 6.52E-04 | -1.19E+00 | 4.53E-02 | 1.62E-01 | -1.13E+00 | 1.28E-01 | 3.62E-01 | 1.12E+00 | 1.65E-03 | 6.06E-03 | -1.35E+00 | 4.51E-04 | 1.24E-03 | -1.80E+00 |
| INTS7 | 9.38E-09 | 4.89E-08 | -1.98E+00 | 4.56E-04 | 1.97E-03 | -1.04E+00 | 9.76E-12 | 4.15E-11 | -1.82E+00 | 4.82E-05 | 1.84E-04 | -1.02E+00 | 2.19E-10 | 7.82E-10 | -2.53E+00 | 6.92E-03 | 1.95E-02 | 1.01E+00 | 6.67E-13 | 2.45E-12 | -1.52E+00 | 1.13E-03 | 3.09E-03 | -1.01E+00 |
| EXD2 | 9.44E-09 | 4.92E-08 | -1.79E+00 | 4.79E-04 | 2.07E-03 | -1.05E+00 | 4.72E-06 | 2.00E-05 | -1.49E+00 | 8.85E-06 | 3.38E-05 | -1.29E+00 | 3.26E-04 | 1.16E-03 | -1.45E+00 | 9.16E-03 | 2.58E-02 | -1.11E+00 | 2.81E-06 | 1.03E-05 | -1.41E+00 | 9.08E-04 | 2.49E-03 | -1.18E+00 |
| TUSC2 | 9.45E-09 | 4.93E-08 | -1.71E+00 | 8.91E-06 | 3.85E-05 | -1.20E+00 | 2.37E-09 | 1.01E-08 | -1.77E+00 | 2.77E-06 | 1.06E-05 | -1.24E+00 | 4.80E-06 | 1.71E-05 | -1.42E+00 | 3.13E-05 | 8.82E-05 | -1.31E+00 | 1.06E-07 | 3.89E-07 | -1.35E+00 | 4.53E-03 | 1.24E-02 | -1.06E+00 |
| CMC1 | 9.45E-09 | 4.93E-08 | -1.31E+00 | 1.35E-05 | 5.83E-05 | 1.02E+00 | 4.59E-03 | 1.95E-02 | -1.08E+00 | 4.06E-05 | 1.55E-04 | -1.10E+00 | 7.40E-05 | 2.64E-04 | -1.23E+00 | 4.14E-02 | 1.17E-01 | 1.15E+00 | 5.79E-05 | 2.13E-04 | -1.14E+00 | 8.76E-02 | 2.41E-01 | 1.05E+00 |
| PABPN1 | 9.54E-09 | 4.97E-08 | -1.11E+00 | 4.46E-02 | 1.93E-01 | 1.21E+00 | 5.91E-05 | 2.51E-04 | -1.14E+00 | 1.49E-02 | 5.70E-02 | 1.10E+00 | 2.64E-06 | 9.41E-06 | -1.15E+00 | 2.49E-02 | 7.03E-02 | 1.11E+00 | 3.41E-02 | 1.25E-01 | -1.06E+00 | 1.64E-01 | 4.50E-01 | 1.13E+00 |
| TCEA2 | 9.55E-09 | 4.98E-08 | -1.19E+00 | 5.98E-04 | 2.58E-03 | 1.13E+00 | 1.48E-04 | 6.27E-04 | -1.14E+00 | 4.34E-04 | 1.66E-03 | 1.07E+00 | 1.19E-04 | 4.26E-04 | -1.15E+00 | 2.28E-03 | 6.44E-03 | 1.02E+00 | 2.36E-01 | 8.65E-01 | 1.31E+00 | 3.58E-01 | 9.85E-01 | 1.05E+00 |
| TNPO3 | 9.56E-09 | 4.98E-08 | -1.28E+00 | 4.89E-05 | 2.11E-04 | 1.08E+00 | 4.20E-05 | 1.79E-04 | -1.17E+00 | 5.32E-04 | 2.03E-03 | 1.02E+00 | 7.06E-07 | 2.52E-06 | -1.31E+00 | 5.21E-05 | 1.47E-04 | -1.13E+00 | 2.61E-10 | 9.57E-10 | -1.42E+00 | 6.64E-04 | 1.82E-03 | 1.00E+00 |
| ARL4A | 9.56E-09 | 4.98E-08 | 2.14E+00 | 5.81E-02 | 2.51E-01 | 1.45E+00 | 7.09E-03 | 3.01E-02 | 1.69E+00 | 4.81E-03 | 1.84E-02 | 1.18E+00 | 6.56E-13 | 2.34E-12 | 3.76E+00 | 1.38E-05 | 3.88E-05 | 2.83E+00 | 1.86E-12 | 6.83E-12 | 2.72E+00 | 4.25E-07 | 1.17E-06 | 2.47E+00 |
| DMTF1 | 9.57E-09 | 4.99E-08 | -1.21E+00 | 8.58E-04 | 3.71E-03 | 1.18E+00 | 2.46E-07 | 1.04E-06 | -1.28E+00 | 4.46E-05 | 1.70E-04 | -1.03E+00 | 1.99E-04 | 7.10E-04 | -1.20E+00 | 3.78E-02 | 1.07E-01 | 1.04E+00 | 2.23E-03 | 8.19E-03 | -1.12E+00 | 1.88E-03 | 5.15E-03 | -1.13E+00 |
| AL136370.1 | 9.72E-09 | 5.07E-08 | -1.90E+00 | 3.12E-05 | 1.35E-04 | -1.20E+00 | 9.69E-03 | 4.12E-02 | -1.42E+00 | 4.11E-02 | 1.57E-01 | -1.03E+00 | 2.12E-03 | 7.56E-03 | -1.54E+00 | 9.24E-02 | 2.60E-01 | 1.17E+00 | 1.18E-08 | 4.32E-08 | -1.61E+00 | 6.93E-04 | 1.90E-03 | -1.13E+00 |
| NUP88 | 9.77E-09 | 5.09E-08 | -1.36E+00 | 3.85E-04 | 1.66E-03 | 1.11E+00 | 2.25E-03 | 9.57E-03 | -1.06E+00 | 1.40E-03 | 5.35E-03 | 1.08E+00 | 1.10E-03 | 3.93E-03 | -1.11E+00 | 9.83E-03 | 2.77E-02 | 1.03E+00 | 2.72E-03 | 9.98E-03 | -1.02E+00 | 1.50E-02 | 4.12E-02 | 1.03E+00 |
| ZNF383 | 9.79E-09 | 5.10E-08 | -1.45E+00 | 3.15E-07 | 1.36E-06 | -1.18E+00 | 1.02E-03 | 4.34E-03 | -1.22E+00 | 7.65E-04 | 2.92E-03 | -1.08E+00 | 4.89E-03 | 1.74E-02 | -1.25E+00 | 3.73E-02 | 1.05E-01 | -1.01E+00 | 1.93E-02 | 7.10E-02 | -1.20E+00 | 1.40E-02 | 3.85E-02 | -1.18E+00 |
| PDIK1L | 9.81E-09 | 5.11E-08 | -1.34E+00 | 9.80E-09 | 4.24E-08 | -1.23E+00 | 4.95E-02 | 2.10E-01 | 1.06E+00 | 3.36E-03 | 1.29E-02 | -1.04E+00 | 2.61E-02 | 9.32E-02 | 1.08E+00 | 2.29E-02 | 6.47E-02 | 1.18E+00 | 3.00E-06 | 1.10E-05 | -1.39E+00 | 1.32E-02 | 3.63E-02 | -1.03E+00 |
| POLR2H | 9.85E-09 | 5.13E-08 | -1.21E+00 | 8.97E-05 | 3.88E-04 | 1.08E+00 | 5.20E-04 | 2.21E-03 | -1.12E+00 | 3.57E-04 | 1.36E-03 | -1.01E+00 | 5.28E-09 | 1.88E-08 | -1.28E+00 | 8.54E-05 | 2.41E-04 | -1.00E+00 | 1.93E-07 | 7.07E-07 | -1.30E+00 | 2.48E-02 | 6.81E-02 | 1.02E+00 |
| MRPL15 | 9.91E-09 | 5.16E-08 | -1.13E+00 | 2.46E-03 | 1.07E-02 | 1.20E+00 | 9.85E-04 | 4.19E-03 | -1.09E+00 | 1.77E-02 | 6.78E-02 | 1.14E+00 | 1.88E-04 | 6.70E-04 | -1.16E+00 | 1.76E-01 | 4.95E-01 | 1.51E+00 | 2.37E-09 | 8.69E-09 | -1.27E+00 | 1.37E-01 | 3.77E-01 | 1.11E+00 |
| TMEM38B | 9.97E-09 | 5.19E-08 | -1.23E+00 | 1.80E-06 | 7.78E-06 | 1.02E+00 | 1.27E-06 | 5.41E-06 | -1.40E+00 | 4.98E-08 | 1.90E-07 | -1.34E+00 | 5.13E-05 | 1.83E-04 | -1.75E+00 | 6.79E-05 | 1.92E-04 | -1.62E+00 | 4.17E-06 | 1.53E-05 | -1.28E+00 | 1.12E-03 | 3.07E-03 | -1.00E+00 |
| TMEM219 | 1.00E-08 | 5.21E-08 | -1.09E+00 | 1.49E-08 | 6.46E-08 | -1.01E+00 | 7.72E-04 | 3.28E-03 | -1.11E+00 | 1.06E-03 | 4.05E-03 | 1.02E+00 | 2.92E-06 | 1.04E-05 | -1.14E+00 | 9.05E-06 | 2.55E-05 | -1.11E+00 | 5.57E-02 | 2.05E-01 | 1.06E+00 | 2.01E-02 | 5.52E-02 | -1.06E+00 |

| | | | | | | | | | | | | | | | | | | | | | | | | |
|---|---|---|---|---|---|---|---|---|---|---|---|---|---|---|---|---|---|---|---|---|---|---|---|---|
| MYD88 | 1.01E-08 | 5.29E-08 | -1.55E+00 | 1.83E-06 | 7.89E-06 | -1.16E+00 | 3.74E-03 | 1.59E-02 | -1.22E+00 | 8.55E-03 | 3.27E-02 | -1.01E+00 | 1.31E-07 | 4.66E-07 | -1.53E+00 | 2.28E-05 | 6.42E-05 | -1.20E+00 | 4.24E-08 | 1.56E-07 | -1.54E+00 | 9.36E-05 | 2.57E-04 | -1.22E+00 |
| FUCA1 | 1.02E-08 | 5.31E-08 | -2.24E+00 | 3.10E-04 | 1.34E-03 | -1.07E+00 | 8.91E-07 | 3.79E-06 | -1.90E+00 | 2.57E-06 | 9.82E-06 | -1.56E+00 | 3.48E-08 | 1.24E-07 | -2.45E+00 | 9.85E-04 | 2.78E-03 | -1.17E+00 | 8.96E-15 | 3.29E-14 | -1.70E+00 | 1.69E-07 | 4.65E-07 | -1.28E+00 |
| ERLEC1 | 1.02E-08 | 5.34E-08 | -1.41E+00 | 9.33E-05 | 4.03E-04 | -1.02E+00 | 7.20E-06 | 3.06E-05 | -1.26E+00 | 2.49E-04 | 9.53E-04 | -1.07E+00 | 2.63E-10 | 9.38E-10 | -1.75E+00 | 1.81E-07 | 5.10E-07 | -1.39E+00 | 2.57E-05 | 9.44E-05 | -1.27E+00 | 1.17E-03 | 3.20E-03 | -1.16E+00 |
| CRELD1 | 1.02E-08 | 5.34E-08 | -1.84E+00 | 3.69E-03 | 1.59E-02 | 1.04E+00 | 7.61E-05 | 3.23E-04 | -1.56E+00 | 1.83E-03 | 6.99E-03 | -1.12E+00 | 6.65E-02 | 2.37E-01 | 1.00E+00 | 1.84E-01 | 5.19E-01 | 1.24E+00 | 1.80E-02 | 6.60E-02 | -1.02E+00 | 8.85E-02 | 2.43E-01 | 1.04E+00 |
| ETHE1 | 1.03E-08 | 5.35E-08 | -1.17E+00 | 2.89E-08 | 1.25E-07 | -1.03E+00 | 4.91E-03 | 2.08E-02 | -1.09E+00 | 6.14E-05 | 2.34E-04 | -1.05E+00 | 3.86E-05 | 1.38E-04 | -1.15E+00 | 1.11E-04 | 3.12E-04 | -1.08E+00 | 1.03E-03 | 3.80E-03 | -1.05E+00 | 5.44E-04 | 1.49E-03 | -1.04E+00 |
| DCTN5 | 1.03E-08 | 5.36E-08 | -1.38E+00 | 6.48E-03 | 2.80E-02 | 1.20E+00 | 2.38E-07 | 1.01E-06 | -1.42E+00 | 5.08E-04 | 1.94E-03 | 1.00E+00 | 8.29E-06 | 2.96E-05 | -1.21E+00 | 2.41E-03 | 6.79E-03 | 1.04E+00 | 3.82E-17 | 1.40E-16 | -1.72E+00 | 1.11E-03 | 3.04E-03 | -1.05E+00 |
| UBE2J2 | 1.04E-08 | 5.40E-08 | -1.31E+00 | 9.07E-04 | 3.92E-03 | 1.12E+00 | 1.82E-04 | 7.72E-04 | -1.19E+00 | 1.79E-03 | 6.85E-03 | 1.04E+00 | 1.02E-07 | 3.63E-07 | -1.26E+00 | 2.25E-03 | 6.36E-03 | 1.13E+00 | 5.66E-08 | 2.08E-07 | -1.25E+00 | 3.64E-04 | 1.00E-03 | -1.06E+00 |
| RAVER2 | 1.04E-08 | 5.42E-08 | -1.42E+00 | 3.38E-06 | 1.46E-05 | -1.10E+00 | 4.53E-09 | 1.93E-08 | -1.45E+00 | 9.84E-07 | 3.76E-06 | -1.15E+00 | 3.91E-04 | 1.39E-03 | -1.49E+00 | 1.06E-02 | 3.00E-02 | -1.08E+00 | 5.99E-04 | 2.20E-03 | -1.30E+00 | 3.66E-02 | 1.00E-01 | -1.01E+00 |
| MRPS18B | 1.04E-08 | 5.43E-08 | -1.24E+00 | 5.89E-03 | 2.55E-02 | 1.15E+00 | 3.52E-07 | 1.50E-06 | -1.40E+00 | 4.03E-03 | 1.54E-02 | -1.03E+00 | 5.51E-12 | 1.97E-11 | -1.39E+00 | 8.49E-02 | 2.39E-01 | 1.26E+00 | 2.36E-06 | 8.68E-06 | -1.22E+00 | 3.73E-02 | 1.02E-01 | 1.05E+00 |
| DUSP14 | 1.05E-08 | 5.46E-08 | -1.58E+00 | 1.07E-05 | 4.62E-05 | -1.10E+00 | 3.16E-05 | 1.34E-04 | -1.34E+00 | 2.60E-04 | 9.94E-04 | -1.07E+00 | 2.12E-04 | 7.57E-04 | -1.41E+00 | 1.97E-03 | 5.55E-03 | -1.26E+00 | 1.78E-03 | 6.52E-03 | -1.05E+00 | 2.02E-02 | 5.55E-02 | -1.05E+00 |
| NDUFV1 | 1.05E-08 | 5.50E-08 | -1.09E+00 | 3.33E-11 | 1.44E-10 | -1.04E+00 | 1.13E-01 | 4.82E-01 | 1.11E+00 | 1.08E-02 | 4.13E-02 | 1.07E+00 | 3.93E-05 | 1.40E-04 | -1.07E+00 | 7.51E-05 | 2.12E-04 | -1.06E+00 | 1.96E-02 | 7.19E-02 | 1.10E+00 | 2.14E-01 | 5.89E-01 | 1.23E+00 |
| AC005785.2 | 1.06E-08 | 5.51E-08 | -1.89E+00 | 2.16E-12 | 9.32E-12 | -2.78E+00 | 4.99E-11 | 2.12E-10 | -2.24E+00 | 2.00E-11 | 7.62E-11 | -2.09E+00 | 8.16E-04 | 2.91E-03 | -1.30E+00 | 1.43E-05 | 4.03E-05 | -2.04E+00 | 3.54E-06 | 1.30E-05 | -1.54E+00 | 4.86E-04 | 1.34E-03 | -1.33E+00 |
| FUT8 | 1.06E-08 | 5.51E-08 | -1.74E+00 | 3.62E-04 | 1.56E-03 | -1.02E+00 | 2.29E-06 | 9.74E-06 | -1.43E+00 | 8.27E-04 | 3.16E-03 | -1.00E+00 | 4.83E-08 | 1.72E-07 | -1.62E+00 | 1.77E-03 | 4.98E-03 | -1.02E+00 | 3.56E-11 | 1.31E-10 | -1.41E+00 | 6.56E-05 | 1.80E-04 | -1.20E+00 |
| RPSAP53 | 1.06E-08 | 5.52E-08 | -1.83E+00 | 6.21E-04 | 2.68E-03 | 1.00E+00 | 1.08E-14 | 4.57E-14 | -2.07E+00 | 2.05E-07 | 7.82E-07 | -1.19E+00 | 6.06E-05 | 2.16E-04 | -1.53E+00 | 1.84E-02 | 5.19E-02 | 1.01E+00 | 2.94E-06 | 1.08E-05 | -1.55E+00 | 2.83E-01 | 7.76E-01 | 1.24E+00 |
| WDR86 | 1.06E-08 | 5.54E-08 | -1.78E+00 | 2.09E-02 | 9.03E-02 | 1.22E+00 | 2.27E-06 | 9.65E-06 | -1.76E+00 | 1.52E-05 | 5.80E-05 | -1.50E+00 | 2.45E-01 | 8.72E-01 | 1.37E+00 | 9.81E-02 | 2.76E-01 | 2.12E+00 | 2.28E-01 | 8.37E-01 | 1.37E+00 | 2.50E-01 | 6.86E-01 | 1.56E+00 |
| CD74 | 1.07E-08 | 5.57E-08 | -1.60E+00 | 8.27E-03 | 3.57E-02 | -1.19E+00 | 8.13E-07 | 3.45E-06 | -1.64E+00 | 1.52E-02 | 5.80E-02 | -1.22E+00 | 3.30E-24 | 1.18E-23 | -2.00E+00 | 9.24E-18 | 2.61E-17 | -1.59E+00 | 4.17E-111 | 1.53E-110 | -3.03E+00 | 1.51E-26 | 4.16E-26 | -1.77E+00 |
| WDR3 | 1.07E-08 | 5.58E-08 | -1.70E+00 | 2.82E-03 | 1.22E-02 | 1.13E+00 | 1.50E-06 | 6.36E-06 | -1.42E+00 | 7.33E-02 | 2.80E-01 | 1.20E+00 | 1.87E-05 | 6.67E-05 | -1.61E+00 | 8.87E-02 | 2.50E-01 | 1.20E+00 | 1.90E-14 | 6.98E-14 | -1.71E+00 | 7.32E-04 | 2.01E-03 | -1.03E+00 |
| ANKRD13A | 1.08E-08 | 5.62E-08 | -1.20E+00 | 3.25E-06 | 1.40E-05 | 1.01E+00 | 2.46E-05 | 1.05E-04 | -1.21E+00 | 2.99E-05 | 1.14E-04 | -1.11E+00 | 1.11E-03 | 3.97E-03 | -1.05E+00 | 7.21E-03 | 2.03E-02 | 1.05E+00 | 2.65E-05 | 9.71E-05 | -1.07E+00 | 1.58E-06 | 4.33E-06 | -1.24E+00 |
| LANCL1 | 1.08E-08 | 5.63E-08 | -1.45E+00 | 6.98E-05 | 3.02E-04 | -1.04E+00 | 8.47E-06 | 3.60E-05 | -1.26E+00 | 7.21E-05 | 2.76E-04 | -1.07E+00 | 1.26E-04 | 4.48E-04 | -1.19E+00 | 1.24E-02 | 3.49E-02 | 1.09E+00 | 8.80E-10 | 3.23E-09 | -1.50E+00 | 6.84E-05 | 1.88E-04 | -1.17E+00 |
| MAPRE3 | 1.08E-08 | 5.63E-08 | -2.80E+00 | 3.53E-06 | 1.52E-05 | -1.88E+00 | 1.02E-02 | 4.33E-02 | -1.56E+00 | 8.30E-03 | 3.17E-02 | -1.68E+00 | 1.77E-08 | 6.29E-08 | -1.91E+00 | 1.20E-07 | 3.38E-07 | -1.58E+00 | 2.51E-01 | 9.22E-01 | 1.30E+00 | 1.98E-01 | 5.43E-01 | 1.00E+00 |
| TAS2R14 | 1.09E-08 | 5.71E-08 | -2.05E+00 | 1.94E-04 | 8.40E-04 | -1.12E+00 | 4.57E-06 | 1.94E-05 | -2.00E+00 | 3.40E-04 | 1.30E-03 | -1.32E+00 | 4.53E-02 | 1.62E-01 | -1.09E+00 | 1.87E-01 | 5.28E-01 | 1.16E+00 | 8.68E-08 | 3.18E-07 | -2.01E+00 | 4.44E-04 | 1.22E-03 | -1.32E+00 |
| ZNF260 | 1.10E-08 | 5.71E-08 | -1.72E+00 | 3.10E-05 | 1.34E-04 | -1.14E+00 | 4.20E-05 | 1.78E-04 | -1.40E+00 | 5.08E-04 | 1.94E-03 | -1.10E+00 | 5.69E-04 | 2.03E-03 | -1.41E+00 | 6.12E-04 | 1.73E-03 | -1.36E+00 | 1.96E-05 | 7.21E-05 | -1.28E+00 | 1.27E-04 | 3.49E-04 | -1.30E+00 |
| PTPRA | 1.10E-08 | 5.71E-08 | -1.34E+00 | 6.18E-06 | 2.67E-05 | 1.01E+00 | 3.57E-12 | 1.52E-11 | -1.59E+00 | 4.80E-09 | 1.83E-08 | -1.18E+00 | 1.36E-04 | 4.85E-04 | -1.25E+00 | 1.81E-02 | 5.10E-02 | 1.03E+00 | 1.44E-03 | 5.29E-03 | -1.11E+00 | 4.58E-03 | 1.26E-02 | -1.03E+00 |
| VAV1 | 1.10E-08 | 5.76E-08 | -1.29E+00 | 1.61E-04 | 6.96E-04 | 1.13E+00 | 6.55E-06 | 2.78E-05 | -1.35E+00 | 3.82E-05 | 1.46E-04 | -1.17E+00 | 1.86E-07 | 6.63E-07 | -1.74E+00 | 4.35E-04 | 1.23E-03 | -1.17E+00 | 6.10E-04 | 2.24E-03 | -1.18E+00 | 1.43E-03 | 3.93E-03 | -1.04E+00 |
| DDX23 | 1.11E-08 | 5.79E-08 | -1.44E+00 | 9.47E-04 | 4.09E-03 | 1.11E+00 | 3.79E-03 | 1.61E-02 | -1.14E+00 | 1.14E-01 | 4.35E-01 | 1.27E+00 | 2.62E-07 | 9.34E-07 | -1.37E+00 | 5.64E-04 | 1.59E-03 | 1.01E+00 | 2.76E-04 | 1.01E-03 | -1.07E+00 | 2.06E-01 | 5.67E-01 | 1.36E+00 |
| CCDC90B | 1.12E-08 | 5.82E-08 | -1.11E+00 | 1.38E-05 | 5.97E-05 | 1.07E+00 | 9.37E-05 | 3.98E-04 | -1.18E+00 | 8.83E-06 | 3.37E-05 | -1.13E+00 | 1.79E-08 | 6.39E-08 | -1.28E+00 | 4.66E-03 | 1.31E-02 | 1.04E+00 | 1.97E-03 | 7.21E-03 | -1.07E+00 | 1.71E-03 | 4.68E-03 | -1.12E+00 |
| PDGFD | 1.13E-08 | 5.88E-08 | -1.42E+00 | 2.10E-06 | 9.08E-06 | -1.12E+00 | 3.35E-04 | 1.43E-03 | -1.24E+00 | 5.85E-04 | 2.23E-03 | -1.07E+00 | 2.76E-01 | 9.82E-01 | 1.49E+00 | 2.85E-01 | 8.04E-01 | 1.28E+00 | 1.60E-01 | 5.88E-01 | 1.46E+00 | 3.09E-01 | 8.49E-01 | 1.11E+00 |
| SRFBP1 | 1.13E-08 | 5.89E-08 | -1.22E+00 | 9.60E-07 | 4.15E-06 | -1.05E+00 | 2.45E-06 | 1.04E-05 | -1.20E+00 | 1.69E-04 | 6.45E-04 | 1.04E+00 | 1.70E-01 | 6.06E-01 | 1.28E+00 | 3.55E-01 | 1.00E+00 | 1.31E+00 | 3.88E-02 | 1.43E-01 | 1.10E+00 | 1.15E-01 | 3.16E-01 | 1.14E+00 |
| TTN | 1.13E-08 | 5.89E-08 | -1.23E+00 | 1.27E-02 | 5.50E-02 | 1.26E+00 | 7.58E-07 | 3.22E-06 | -1.41E+00 | 1.92E-03 | 7.34E-03 | 1.04E+00 | 1.58E-02 | 5.64E-02 | -1.11E+00 | 1.36E-01 | 3.83E-01 | 1.15E+00 | 1.25E-01 | 4.59E-01 | 1.18E+00 | 2.76E-01 | 7.59E-01 | 1.24E+00 |
| OTULIN | 1.15E-08 | 5.98E-08 | -1.64E+00 | 4.22E-06 | 1.82E-05 | -1.23E+00 | 1.71E-05 | 7.28E-05 | -1.32E+00 | 1.28E-04 | 4.88E-04 | -1.10E+00 | 4.59E-02 | 1.64E-01 | 1.05E+00 | 7.76E-02 | 2.19E-01 | 1.13E+00 | 1.23E-08 | 4.53E-08 | -1.31E+00 | 1.37E-08 | 3.77E-08 | -1.34E+00 |
| DMAP1 | 1.15E-08 | 6.01E-08 | -1.19E+00 | 8.25E-05 | 3.57E-04 | 1.16E+00 | 4.30E-03 | 1.83E-02 | -1.03E+00 | 2.61E-06 | 9.97E-06 | -1.08E+00 | 1.91E-03 | 6.82E-03 | 1.02E+00 | 2.72E-02 | 7.65E-02 | 1.18E+00 | 5.03E-02 | 1.85E-01 | 1.10E+00 | 1.60E-02 | 4.39E-02 | 1.02E+00 |
| IRF5 | 1.16E-08 | 6.03E-08 | -1.54E+00 | 1.92E-05 | 8.28E-05 | -1.07E+00 | 3.68E-10 | 1.57E-09 | -1.64E+00 | 6.86E-06 | 2.62E-05 | -1.11E+00 | 1.22E-05 | 4.35E-05 | -1.36E+00 | 1.89E-04 | 5.34E-04 | -1.20E+00 | 1.47E-04 | 5.38E-04 | -1.47E+00 | 2.48E-03 | 6.82E-03 | -1.18E+00 |
| SRP72P1 | 1.16E-08 | 6.04E-08 | -1.74E+00 | 6.82E-10 | 2.95E-09 | -1.80E+00 | 7.44E-15 | 3.16E-14 | -2.09E+00 | 5.54E-14 | 2.11E-13 | -1.75E+00 | 6.83E-03 | 2.43E-02 | -1.22E+00 | 1.69E-02 | 4.76E-02 | -1.08E+00 | 9.74E-14 | 3.57E-13 | -1.79E+00 | 1.47E-11 | 4.05E-11 | -1.56E+00 |
| ACOT9 | 1.16E-08 | 6.07E-08 | -1.58E+00 | 2.43E-05 | 1.05E-04 | -1.06E+00 | 3.65E-04 | 1.55E-03 | -1.20E+00 | 2.07E-03 | 7.90E-03 | 1.03E+00 | 5.26E-10 | 1.87E-09 | -1.56E+00 | 5.67E-06 | 1.60E-05 | -1.15E+00 | 5.29E-06 | 1.94E-05 | -1.18E+00 | 3.99E-04 | 1.10E-03 | -1.06E+00 |
| AC241952.1 | 1.17E-08 | 6.07E-08 | -1.97E+00 | 2.06E-07 | 8.88E-07 | -1.62E+00 | 1.89E-02 | 8.03E-02 | -1.28E+00 | 1.95E-02 | 7.43E-02 | -1.17E+00 | 5.22E-02 | 1.86E-01 | -1.13E+00 | 2.89E-02 | 8.15E-02 | -1.57E+00 | 1.27E-01 | 4.68E-01 | 2.16E+00 | 1.08E-01 | 2.97E-01 | 1.08E+00 |
| TPD52L2 | 1.17E-08 | 6.08E-08 | -1.34E+00 | 4.91E-04 | 2.12E-03 | 1.14E+00 | 3.27E-04 | 1.39E-03 | -1.16E+00 | 3.91E-02 | 1.49E-01 | 1.16E+00 | 3.48E-07 | 1.24E-06 | -1.40E+00 | 2.98E-02 | 8.40E-02 | 1.13E+00 | 2.22E-05 | 8.16E-05 | -1.14E+00 | 3.04E-03 | 8.36E-03 | 1.03E+00 |
| SNAPC5 | 1.17E-08 | 6.09E-08 | -1.44E+00 | 1.53E-06 | 6.61E-06 | -1.14E+00 | 1.68E-05 | 7.14E-05 | -1.25E+00 | 2.67E-03 | 1.02E-02 | 1.09E+00 | 1.31E-03 | 4.67E-03 | -1.20E+00 | 1.40E-01 | 3.96E-01 | 1.20E+00 | 1.13E-04 | 4.14E-04 | -1.15E+00 | 1.08E-01 | 2.96E-01 | 1.17E+00 |
| PRPS1 | 1.18E-08 | 6.14E-08 | -1.37E+00 | 3.21E-07 | 1.39E-06 | -1.10E+00 | 2.93E-05 | 1.25E-04 | -1.23E+00 | 3.46E-05 | 1.32E-04 | -1.05E+00 | 2.54E-06 | 9.07E-06 | -1.18E+00 | 1.98E-03 | 5.59E-03 | 1.10E+00 | 1.07E-05 | 3.92E-05 | -1.17E+00 | 8.63E-02 | 2.37E-01 | 1.06E+00 |
| TSEN15 | 1.18E-08 | 6.15E-08 | -1.53E+00 | 3.32E-04 | 1.43E-03 | 1.03E+00 | 2.24E-08 | 9.51E-08 | -1.42E+00 | 1.47E-05 | 5.62E-05 | -1.07E+00 | 6.47E-07 | 2.31E-06 | -1.51E+00 | 8.67E-03 | 2.44E-02 | 1.06E+00 | 3.36E-09 | 1.23E-08 | -1.26E+00 | 9.64E-03 | 2.65E-02 | 1.16E+00 |
| PSMA6 | 1.19E-08 | 6.18E-08 | -1.81E+00 | 3.22E-02 | 1.39E-01 | 1.69E+00 | 3.30E-09 | 1.40E-08 | -1.79E+00 | 6.96E-02 | 2.66E-01 | 1.23E+00 | 3.35E-03 | 1.20E-02 | -1.25E+00 | 6.74E-02 | 1.90E-01 | 1.11E+00 | 4.70E-06 | 1.72E-05 | -1.49E+00 | 1.85E-03 | 5.08E-03 | -1.10E+00 |
| GBGT1 | 1.19E-08 | 6.19E-08 | -1.75E+00 | 4.95E-08 | 2.14E-07 | -1.50E+00 | 3.93E-05 | 1.67E-04 | -1.53E+00 | 7.44E-05 | 2.84E-04 | -1.36E+00 | 5.21E-08 | 1.86E-07 | -1.60E+00 | 8.55E-03 | 2.41E-02 | 1.22E+00 | 1.13E-06 | 4.15E-06 | -1.22E+00 | 2.54E-03 | 6.99E-03 | -1.04E+00 |
| TNIP1 | 1.19E-08 | 6.20E-08 | -1.19E+00 | 4.06E-06 | 1.76E-05 | 1.00E+00 | 6.87E-08 | 2.92E-07 | -1.33E+00 | 1.85E-06 | 7.07E-06 | -1.12E+00 | 5.15E-07 | 1.84E-06 | -1.24E+00 | 2.91E-03 | 8.22E-03 | 1.06E+00 | 1.39E-07 | 5.12E-07 | -1.28E+00 | 3.86E-04 | 1.06E-03 | -1.06E+00 |
| EIF1AXP1 | 1.20E-08 | 6.25E-08 | -1.31E+00 | 6.32E-07 | 2.73E-06 | -1.10E+00 | 5.18E-04 | 2.20E-03 | -1.11E+00 | 1.95E-03 | 7.46E-03 | 1.11E+00 | 8.90E-04 | 3.17E-03 | -1.26E+00 | 3.95E-02 | 1.11E-01 | 1.10E+00 | 2.29E-11 | 8.40E-11 | -1.44E+00 | 4.06E-05 | 1.11E-04 | -1.09E+00 |
| LRRC8A | 1.20E-08 | 6.25E-08 | -1.56E+00 | 6.00E-12 | 2.59E-11 | -1.83E+00 | 9.83E-09 | 4.18E-08 | -1.44E+00 | 2.76E-17 | 1.06E-16 | -1.90E+00 | 6.42E-04 | 2.29E-03 | -1.23E+00 | 3.49E-04 | 9.83E-04 | -1.38E+00 | 1.52E-06 | 5.58E-06 | -1.53E+00 | 4.95E-06 | 1.36E-05 | -1.42E+00 |
| LPCAT4 | 1.20E-08 | 6.26E-08 | -1.80E+00 | 7.00E-05 | 3.03E-04 | -1.12E+00 | 1.13E-04 | 4.78E-04 | -1.56E+00 | 1.33E-02 | 5.09E-02 | 1.02E+00 | 1.98E-04 | 7.06E-04 | -1.99E+00 | 2.05E-03 | 5.78E-03 | -1.45E+00 | 9.11E-04 | 3.34E-03 | -1.47E+00 | 4.71E-02 | 1.29E-01 | -1.04E+00 |
| GGA1 | 1.20E-08 | 6.26E-08 | -1.28E+00 | 1.31E-05 | 5.65E-05 | 1.04E+00 | 5.06E-05 | 2.15E-04 | -1.17E+00 | 1.84E-05 | 7.04E-05 | -1.12E+00 | 3.68E-04 | 1.31E-03 | -1.08E+00 | 4.92E-03 | 1.39E-02 | 1.06E+00 | 1.71E-02 | 6.27E-02 | 1.03E+00 | 2.36E-01 | 6.48E-01 | 1.29E+00 |
| IRF2BP2 | 1.21E-08 | 6.31E-08 | -1.03E+00 | 1.17E-01 | 5.03E-01 | 1.39E+00 | 2.42E-02 | 1.03E-01 | -1.02E+00 | 3.73E-02 | 1.43E-01 | 1.42E+00 | 1.84E-01 | 6.56E-01 | 1.27E+00 | 8.81E-02 | 2.48E-01 | 1.48E+00 | 1.61E-01 | 5.91E-01 | 1.12E+00 | 2.28E-01 | 6.26E-01 | 1.25E+00 |
| IKBIP | 1.21E-08 | 6.31E-08 | -1.25E+00 | 2.81E-05 | 1.21E-04 | 1.02E+00 | 8.43E-06 | 3.58E-05 | -1.25E+00 | 2.37E-03 | 9.07E-03 | 1.03E+00 | 6.39E-05 | 2.28E-04 | -1.25E+00 | 3.49E-02 | 9.85E-02 | 1.11E+00 | 4.47E-06 | 1.64E-05 | -1.19E+00 | 2.74E-03 | 7.54E-03 | -1.02E+00 |
| THEM6 | 1.21E-08 | 6.33E-08 | -1.45E+00 | 2.26E-05 | 9.78E-05 | -1.00E+00 | 1.49E-02 | 6.33E-02 | -1.05E+00 | 1.38E-01 | 5.27E-01 | 1.31E+00 | 4.17E-04 | 1.49E-03 | -1.30E+00 | 1.32E-01 | 3.72E-01 | 1.34E+00 | 1.24E-05 | 4.53E-05 | -1.23E+00 | 1.23E-01 | 3.38E-01 | 1.19E+00 |
| SLC20A1 | 1.22E-08 | 6.35E-08 | -1.46E+00 | 3.32E-10 | 1.43E-09 | -1.49E+00 | 1.10E-05 | 4.66E-05 | -1.35E+00 | 2.90E-05 | 1.11E-04 | -1.15E+00 | 1.65E-02 | 5.88E-02 | -1.04E+00 | 1.68E-02 | 4.74E-02 | -1.13E+00 | 6.26E-07 | 2.30E-06 | -1.26E+00 | 3.45E-03 | 9.47E-03 | 1.04E+00 |
| VPS53 | 1.22E-08 | 6.36E-08 | -1.48E+00 | 6.14E-04 | 2.65E-03 | 1.01E+00 | 6.60E-05 | 2.80E-04 | -1.39E+00 | 6.68E-03 | 2.55E-02 | 1.03E+00 | 1.22E-04 | 4.34E-04 | -1.22E+00 | 1.33E-02 | 3.75E-02 | 1.08E+00 | 2.64E-05 | 9.68E-05 | -1.18E+00 | 8.74E-04 | 2.40E-03 | -1.09E+00 |
| RAB1A | 1.22E-08 | 6.36E-08 | -1.14E+00 | 3.91E-10 | 1.69E-09 | -1.12E+00 | 2.20E-07 | 9.34E-07 | -1.25E+00 | 1.20E-12 | 4.60E-12 | -1.34E+00 | 2.81E-11 | 1.00E-10 | -1.44E+00 | 6.45E-14 | 1.82E-13 | -1.57E+00 | 7.24E-09 | 2.66E-08 | -1.29E+00 | 3.08E-07 | 8.47E-07 | -1.37E+00 |
| UBE2D1 | 1.22E-08 | 6.36E-08 | -1.37E+00 | 1.71E-06 | 7.37E-06 | -1.06E+00 | 2.35E-09 | 1.00E-08 | -1.65E+00 | 5.50E-05 | 2.10E-04 | -1.10E+00 | 2.26E-08 | 8.07E-08 | -1.88E+00 | 1.25E-04 | 3.53E-04 | -1.24E+00 | 3.63E-13 | 1.33E-12 | -1.93E+00 | 1.12E-02 | 3.08E-02 | -1.05E+00 |
| DRAP1 | 1.22E-08 | 6.36E-08 | -1.14E+00 | 1.19E-12 | 5.16E-12 | -1.09E+00 | 9.06E-05 | 3.85E-04 | -1.25E+00 | 1.43E-04 | 5.47E-04 | -1.19E+00 | 2.45E-02 | 8.73E-02 | -1.10E+00 | 2.23E-02 | 6.28E-02 | -1.11E+00 | 3.03E-02 | 1.11E-01 | -1.10E+00 | 4.09E-02 | 1.12E-01 | -1.14E+00 |
| NSD3 | 1.22E-08 | 6.37E-08 | -1.15E+00 | 4.92E-07 | 2.13E-06 | -1.01E+00 | 4.00E-05 | 1.70E-04 | -1.23E+00 | 4.86E-04 | 1.86E-03 | -1.09E+00 | 6.58E-03 | 2.35E-02 | 1.08E+00 | 8.77E-04 | 2.47E-03 | 1.06E+00 | 2.49E-03 | 9.12E-03 | -1.11E+00 | 4.97E-02 | 1.36E-01 | -1.09E+00 |
| CALM2P2 | 1.23E-08 | 6.40E-08 | -1.68E+00 | 2.68E-10 | 1.16E-09 | -1.82E+00 | 1.41E-02 | 5.97E-02 | -1.27E+00 | 2.27E-02 | 8.67E-02 | -1.07E+00 | 4.30E-02 | 1.53E-01 | -1.08E+00 | 2.81E-02 | 7.92E-02 | -1.12E+00 | 1.25E-03 | 4.57E-03 | -1.10E+00 | 8.07E-07 | 2.22E-06 | -1.37E+00 |
| CXorf40A | 1.23E-08 | 6.40E-08 | -1.57E+00 | 3.99E-05 | 1.72E-04 | 1.01E+00 | 3.16E-07 | 1.34E-06 | -1.39E+00 | 6.78E-06 | 2.59E-05 | -1.13E+00 | 7.74E-07 | 2.76E-06 | -1.67E+00 | 3.46E-03 | 9.77E-03 | -1.01E+00 | 1.29E-08 | 4.74E-08 | -1.38E+00 | 1.29E-04 | 3.54E-04 | -1.09E+00 |
| C21orf59 | 1.23E-08 | 6.41E-08 | -1.28E+00 | 5.45E-08 | 2.36E-07 | -1.06E+00 | 1.95E-06 | 8.29E-06 | -1.19E+00 | 7.31E-10 | 2.79E-09 | -1.20E+00 | 9.67E-06 | 3.45E-05 | -1.36E+00 | 1.00E-04 | 2.82E-04 | -1.21E+00 | 9.72E-04 | 3.57E-03 | -1.08E+00 | 3.37E-02 | 9.25E-02 | 1.06E+00 |
| ACBD6 | 1.23E-08 | 6.42E-08 | -1.13E+00 | 1.19E-09 | 5.16E-09 | -1.02E+00 | 1.08E-02 | 4.59E-02 | -1.00E+00 | 1.40E-03 | 5.33E-03 | 1.07E+00 | 4.33E-03 | 1.54E-02 | 1.08E+00 | 1.03E-01 | 2.91E-01 | 1.21E+00 | 1.25E-03 | 4.58E-03 | -1.05E+00 | 1.09E-01 | 2.99E-01 | 1.10E+00 |
| PREB | 1.23E-08 | 6.43E-08 | -1.89E+00 | 3.20E-05 | 1.38E-04 | -1.21E+00 | 1.65E-06 | 7.02E-06 | -1.37E+00 | 9.82E-06 | 3.75E-05 | -1.15E+00 | 1.92E-06 | 6.83E-06 | -1.62E+00 | 5.22E-03 | 1.47E-02 | -1.03E+00 | 1.30E-09 | 4.78E-09 | -1.43E+00 | 1.43E-03 | 3.93E-03 | -1.04E+00 |
| RPP38 | 1.24E-08 | 6.44E-08 | -1.43E+00 | 2.19E-09 | 9.47E-09 | -1.38E+00 | 1.91E-06 | 8.12E-06 | -1.37E+00 | 3.31E-07 | 1.26E-06 | -1.31E+00 | 1.99E-01 | 7.10E-01 | 1.21E+00 | 2.86E-01 | 8.05E-01 | 1.22E+00 | 2.05E-12 | 7.52E-12 | -1.72E+00 | 7.31E-12 | 2.01E-11 | -1.63E+00 |
| RBM28 | 1.24E-08 | 6.47E-08 | -1.20E+00 | 7.74E-06 | 3.34E-05 | -1.01E+00 | 1.73E-06 | 7.33E-06 | -1.26E+00 | 1.58E-03 | 6.04E-03 | 1.09E+00 | 2.37E-02 | 8.46E-02 | 1.14E+00 | 2.86E-01 | 8.07E-01 | 1.35E+00 | 3.22E-07 | 1.18E-06 | -1.30E+00 | 1.50E-02 | 4.12E-02 | 1.06E+00 |
| MMACHC | 1.25E-08 | 6.50E-08 | -2.30E+00 | 5.43E-05 | 2.34E-04 | -1.25E+00 | 1.74E-04 | 7.40E-04 | -1.71E+00 | 1.72E-02 | 6.59E-02 | -1.11E+00 | 5.76E-03 | 2.05E-02 | -1.69E+00 | 1.33E-01 | 3.75E-01 | 1.19E+00 | 2.76E-02 | 1.01E-01 | -1.32E+00 | 3.09E-01 | 8.50E-01 | 1.06E+00 |
| RNF150 | 1.25E-08 | 6.52E-08 | -1.66E+00 | 2.99E-08 | 1.29E-07 | -1.41E+00 | 2.75E-03 | 1.17E-02 | -1.30E+00 | 4.44E-03 | 1.70E-02 | -1.11E+00 | 8.50E-02 | 3.03E-01 | -1.16E+00 | 1.28E-01 | 3.61E-01 | -1.19E+00 | 5.92E-03 | 2.17E-02 | -1.77E+00 | 1.01E-01 | 2.77E-01 | -1.08E+00 |
| KLHL12 | 1.26E-08 | 6.58E-08 | -1.74E+00 | 5.19E-07 | 2.24E-06 | -1.28E+00 | 4.17E-14 | 1.77E-13 | -2.08E+00 | 2.39E-11 | 9.15E-11 | -1.62E+00 | 2.78E-09 | 9.91E-09 | -2.46E+00 | 1.70E-04 | 4.79E-04 | -1.27E+00 | 1.92E-09 | 7.04E-09 | -1.49E+00 | 5.37E-03 | 1.47E-02 | -1.02E+00 |
| C14orf2 | 1.26E-08 | 6.58E-08 | -1.15E+00 | 7.15E-21 | 3.09E-20 | -1.29E+00 | 4.84E-02 | 2.06E-01 | -1.06E+00 | 7.18E-07 | 2.74E-06 | -1.37E+00 | 2.12E-02 | 7.55E-02 | -1.05E+00 | 1.55E-04 | 4.38E-04 | -1.18E+00 | 6.96E-02 | 2.56E-01 | -1.05E+00 | 4.17E-04 | 1.15E-03 | -1.30E+00 |
| FAM110A | 1.28E-08 | 6.70E-08 | -1.86E+00 | 4.68E-04 | 2.02E-03 | 1.00E+00 | 5.08E-07 | 2.16E-06 | -1.98E+00 | 6.46E-03 | 2.47E-02 | -1.01E+00 | 3.03E-17 | 1.08E-16 | -1.86E+00 | 7.64E-10 | 2.15E-09 | -1.35E+00 | 2.28E-02 | 8.37E-02 | 1.15E+00 | 1.40E-01 | 3.85E-01 | 1.22E+00 |
| TXN | 1.29E-08 | 6.72E-08 | 1.55E+00 | 6.36E-07 | 2.75E-06 | 1.56E+00 | 2.41E-12 | 1.02E-11 | 1.73E+00 | 6.54E-13 | 2.50E-12 | 1.80E+00 | 6.39E-04 | 2.28E-03 | 1.49E+00 | 1.24E-05 | 3.48E-05 | 1.59E+00 | 6.82E-06 | 2.50E-05 | 1.52E+00 | 1.68E-07 | 4.60E-07 | 1.63E+00 |
| CKAP4 | 1.29E-08 | 6.73E-08 | -3.15E+00 | 1.16E-03 | 5.02E-03 | -1.23E+00 | 2.00E-05 | 8.48E-05 | -1.60E+00 | 6.51E-04 | 2.49E-03 | -1.15E+00 | 1.11E-02 | 3.97E-02 | -1.39E+00 | 1.46E-01 | 4.12E-01 | 1.14E+00 | 9.99E-06 | 3.67E-05 | -1.49E+00 | 1.50E-02 | 4.12E-02 | -1.03E+00 |
| MYL6B | 1.29E-08 | 6.74E-08 | -1.06E+00 | 2.58E-09 | 1.12E-08 | 1.01E+00 | 3.71E-02 | 1.58E-01 | -1.01E+00 | 1.70E-04 | 6.50E-04 | -1.06E+00 | 5.67E-06 | 2.02E-05 | -1.22E+00 | 3.59E-03 | 1.01E-02 | -1.04E+00 | 1.23E-01 | 4.53E-01 | 1.14E+00 | 1.20E-01 | 3.31E-01 | 1.07E+00 |
| BHLHE41 | 1.30E-08 | 6.76E-08 | -1.87E+00 | 1.99E-06 | 8.61E-06 | -1.43E+00 | 4.70E-02 | 2.00E-01 | -1.06E+00 | 1.88E-02 | 7.19E-02 | 1.95E+00 | 1.30E-01 | 4.64E-01 | 2.36E+00 | 3.03E-01 | 8.54E-01 | 1.48E+00 | 1.75E-01 | 6.42E-01 | 1.41E+00 | 3.21E-01 | 8.81E-01 | 1.19E+00 |
| TSEN2 | 1.31E-08 | 6.85E-08 | -1.72E+00 | 4.18E-03 | 1.81E-02 | 1.11E+00 | 2.79E-05 | 1.18E-04 | -1.38E+00 | 2.55E-01 | 9.75E-01 | 1.38E+00 | 1.70E-04 | 6.07E-04 | -1.58E+00 | 8.56E-02 | 2.41E-01 | 1.22E+00 | 1.95E-07 | 7.18E-07 | -1.59E+00 | 3.26E-01 | 8.94E-01 | 1.29E+00 |
| HNRNPUL2 | 1.33E-08 | 6.91E-08 | -1.33E+00 | 4.79E-07 | 2.07E-06 | -1.06E+00 | 1.44E-04 | 6.11E-04 | -1.19E+00 | 4.93E-02 | 1.88E-01 | 1.20E+00 | 2.56E-08 | 9.12E-08 | -1.53E+00 | 7.77E-05 | 2.19E-04 | -1.17E+00 | 2.42E-08 | 8.88E-08 | -1.32E+00 | 1.09E-03 | 2.99E-03 | 1.02E+00 |
| MSH2 | 1.33E-08 | 6.91E-08 | -1.39E+00 | 1.25E-03 | 5.42E-03 | 1.09E+00 | 6.86E-07 | 2.91E-06 | -1.29E+00 | 6.33E-05 | 2.42E-04 | 1.00E+00 | 1.16E-03 | 4.14E-03 | -1.13E+00 | 1.81E-01 | 5.11E-01 | 1.27E+00 | 1.64E-09 | 6.03E-09 | -1.39E+00 | 2.43E-02 | 6.67E-02 | 1.02E+00 |
| ABHD10 | 1.33E-08 | 6.91E-08 | -1.70E+00 | 1.01E-04 | 4.36E-04 | -1.10E+00 | 9.41E-13 | 4.00E-12 | -1.67E+00 | 1.75E-08 | 6.69E-08 | -1.23E+00 | 7.01E-06 | 2.50E-05 | -1.59E+00 | 5.30E-04 | 1.49E-03 | -1.16E+00 | 4.15E-09 | 1.52E-08 | -1.43E+00 | 1.79E-06 | 4.91E-06 | -1.26E+00 |
| PQBP1 | 1.33E-08 | 6.93E-08 | -1.14E+00 | 4.47E-07 | 1.93E-06 | -1.00E+00 | 3.75E-04 | 1.59E-03 | -1.19E+00 | 8.62E-03 | 3.29E-02 | 1.03E+00 | 4.66E-04 | 1.66E-03 | -1.04E+00 | 1.39E-02 | 3.92E-02 | 1.10E+00 | 2.74E-03 | 1.01E-02 | -1.03E+00 | 7.42E-02 | 2.04E-01 | 1.07E+00 |
| USP42 | 1.33E-08 | 6.96E-08 | -1.62E+00 | 6.42E-10 | 2.78E-09 | -1.63E+00 | 2.28E-16 | 9.69E-16 | -1.94E+00 | 1.98E-21 | 7.55E-21 | -2.11E+00 | 2.88E-05 | 1.03E-04 | -1.36E+00 | 4.49E-08 | 1.27E-07 | -1.97E+00 | 2.82E-08 | 1.04E-07 | -1.57E+00 | 7.92E-12 | 2.17E-11 | -1.89E+00 |
| CPSF2 | 1.34E-08 | 6.98E-08 | -1.54E+00 | 2.81E-03 | 1.21E-02 | 1.10E+00 | 1.82E-06 | 7.75E-06 | -1.31E+00 | 3.82E-03 | 1.46E-02 | 1.07E+00 | 3.60E-07 | 1.28E-06 | -1.54E+00 | 2.36E-03 | 6.65E-03 | 1.07E+00 | 6.26E-07 | 2.30E-06 | -1.26E+00 | 1.37E-01 | 3.76E-01 | 1.17E+00 |
| GPR12 | 1.34E-08 | 7.00E-08 | -2.19E+00 | 3.21E-08 | 1.39E-07 | -1.88E+00 | 2.38E-25 | 1.01E-24 | -2.62E+00 | 8.52E-16 | 3.25E-15 | -1.53E+00 | 1.32E-01 | 4.69E-01 | -1.54E+00 | 2.74E-01 | 7.72E-01 | -1.04E+00 | 2.60E-01 | 9.53E-01 | 1.19E+00 | 3.64E-01 | 1.00E+00 | 1.07E+00 |
| XXYLT1 | 1.35E-08 | 7.03E-08 | -1.52E+00 | 2.66E-09 | 1.15E-08 | -1.48E+00 | 2.31E-06 | 9.80E-06 | -1.37E+00 | 1.29E-06 | 4.93E-06 | -1.18E+00 | 1.98E-04 | 7.06E-04 | -1.53E+00 | 9.08E-03 | 2.56E-02 | -1.12E+00 | 1.10E-06 | 4.03E-06 | -1.39E+00 | 5.22E-02 | 1.44E-01 | 1.09E+00 |
| FKTN | 1.36E-08 | 7.08E-08 | -1.94E+00 | 1.42E-02 | 6.16E-02 | 1.21E+00 | 7.81E-06 | 3.32E-05 | -1.65E+00 | 1.08E-01 | 4.12E-01 | 1.17E+00 | 4.27E-03 | 1.52E-02 | -1.43E+00 | 8.02E-02 | 2.26E-01 | 1.01E+00 | 7.51E-03 | 2.76E-02 | -1.35E+00 | 2.88E-01 | 7.91E-01 | 1.10E+00 |
| RANGRF | 1.36E-08 | 7.08E-08 | -1.62E+00 | 2.31E-06 | 1.00E-05 | -1.18E+00 | 3.87E-10 | 1.64E-09 | -1.68E+00 | 5.76E-09 | 2.20E-08 | -1.34E+00 | 1.01E-04 | 3.61E-04 | -1.48E+00 | 3.77E-02 | 1.06E-01 | 1.03E+00 | 1.03E-15 | 3.76E-15 | -1.63E+00 | 2.43E-08 | 6.68E-08 | -1.23E+00 |
| PQLC2 | 1.36E-08 | 7.09E-08 | -2.17E+00 | 2.58E-05 | 1.11E-04 | -1.31E+00 | 8.07E-05 | 3.43E-04 | -1.59E+00 | 2.64E-03 | 1.01E-02 | -1.15E+00 | 1.75E-01 | 6.25E-01 | -1.16E+00 | 2.20E-01 | 6.20E-01 | -1.18E+00 | 1.74E-02 | 6.37E-02 | -1.19E+00 | 3.25E-02 | 8.93E-02 | -1.23E+00 |
| MAGED1 | 1.36E-08 | 7.09E-08 | -1.29E+00 | 8.91E-06 | 3.85E-05 | -1.01E+00 | 5.31E-05 | 2.25E-04 | -1.18E+00 | 2.33E-05 | 8.88E-05 | -1.09E+00 | 2.80E-07 | 1.00E-06 | -1.35E+00 | 2.22E-04 | 6.25E-04 | 1.00E+00 | 7.95E-02 | 2.92E-01 | 1.03E+00 | 2.48E-01 | 6.82E-01 | 1.23E+00 |
| FBXW11 | 1.36E-08 | 7.09E-08 | -1.50E+00 | 1.54E-08 | 6.65E-08 | -1.33E+00 | 2.65E-03 | 1.12E-02 | -1.25E+00 | 1.51E-03 | 5.78E-03 | -1.11E+00 | 2.97E-05 | 1.06E-04 | -1.29E+00 | 5.99E-06 | 1.69E-05 | -1.42E+00 | 4.16E-07 | 1.53E-06 | -1.46E+00 | 8.33E-04 | 2.29E-03 | -1.16E+00 |
| ADGRG5 | 1.36E-08 | 7.10E-08 | -1.52E+00 | 3.74E-06 | 1.62E-05 | -1.18E+00 | 6.17E-14 | 2.62E-13 | -1.72E+00 | 9.45E-10 | 3.61E-09 | -1.24E+00 | 3.33E-06 | 1.19E-05 | -4.23E+00 | 1.23E-05 | 3.46E-05 | -3.49E+00 | 1.77E-03 | 6.51E-03 | -2.71E+00 | 2.24E-03 | 6.16E-03 | -2.86E+00 |
| STK3 | 1.36E-08 | 7.10E-08 | -1.41E+00 | 2.45E-07 | 1.06E-06 | -1.19E+00 | 4.04E-05 | 1.71E-04 | -1.32E+00 | 1.50E-03 | 5.72E-03 | -1.02E+00 | 2.01E-05 | 7.17E-05 | -1.33E+00 | 2.40E-03 | 6.76E-03 | -1.11E+00 | 1.91E-06 | 7.01E-06 | -1.31E+00 | 5.17E-07 | 1.42E-06 | -1.43E+00 |
| DDT | 1.37E-08 | 7.14E-08 | -1.21E+00 | 9.23E-14 | 3.99E-13 | -1.21E+00 | 2.13E-02 | 9.04E-02 | -1.12E+00 | 4.88E-04 | 1.86E-03 | -1.16E+00 | 2.21E-08 | 7.88E-08 | -1.36E+00 | 4.60E-05 | 1.30E-04 | -1.17E+00 | 1.74E-05 | 6.37E-05 | -1.28E+00 | 4.85E-04 | 1.33E-03 | -1.17E+00 |
| AC009318.1 | 1.38E-08 | 7.17E-08 | -2.27E+00 | 1.07E-06 | 4.63E-06 | -1.62E+00 | 1.30E-05 | 5.53E-05 | -1.94E+00 | 7.07E-04 | 2.70E-03 | -1.40E+00 | 8.84E-02 | 3.15E-01 | 1.00E+00 | 2.61E-01 | 7.36E-01 | 1.25E+00 | 3.54E-04 | 1.30E-03 | -1.88E+00 | 5.72E-02 | 1.57E-01 | -1.16E+00 |
| ACSS2 | 1.38E-08 | 7.17E-08 | -1.54E+00 | 7.95E-06 | 3.44E-05 | -1.10E+00 | 7.60E-05 | 3.23E-04 | -1.32E+00 | 5.18E-06 | 1.98E-05 | -1.25E+00 | 6.71E-02 | 2.39E-01 | 1.05E+00 | 1.38E-01 | 3.89E-01 | 1.20E+00 | 1.35E-07 | 4.97E-07 | -1.38E+00 | 1.85E-04 | 5.09E-04 | -1.14E+00 |
| PIH1D1 | 1.38E-08 | 7.18E-08 | -1.18E+00 | 4.05E-06 | 1.75E-05 | 1.08E+00 | 9.58E-02 | 4.07E-01 | 1.19E+00 | 2.45E-02 | 9.38E-02 | 1.23E+00 | 5.12E-05 | 1.83E-04 | -1.15E+00 | 1.15E-02 | 3.24E-02 | 1.08E+00 | 9.78E-02 | 3.59E-01 | 1.12E+00 | 3.27E-01 | 8.98E-01 | 1.19E+00 |
| PIA2 | 1.38E-08 | 7.18E-08 | -1.21E+00 | 1.45E-06 | 6.28E-06 | 1.03E+00 | 9.26E-07 | 3.93E-06 | -1.25E+00 | 2.07E-06 | 7.91E-06 | -1.09E+00 | 7.64E-06 | 2.73E-05 | -1.18E+00 | 6.96E-06 | 1.96E-05 | -1.18E+00 | 1.18E-04 | 4.35E-04 | -1.17E+00 | 6.74E-03 | 1.85E-02 | -1.11E+00 |
| ASL | 1.38E-08 | 7.21E-08 | -1.93E+00 | 2.44E-08 | 1.05E-07 | -1.65E+00 | 9.41E-02 | 4.00E-01 | 1.08E+00 | 3.87E-03 | 1.48E-02 | -1.19E+00 | 2.91E-08 | 1.04E-07 | -1.56E+00 | 3.74E-09 | 1.06E-08 | -1.72E+00 | 2.74E-02 | 1.01E-01 | 1.10E+00 | 3.52E-06 | 9.66E-06 | -1.22E+00 |
| DNAJC15 | 1.39E-08 | 7.22E-08 | -1.07E+00 | 3.75E-11 | 1.62E-10 | -1.01E+00 | 4.18E-02 | 1.78E-01 | 1.07E+00 | 1.59E-04 | 6.08E-04 | -1.03E+00 | 2.66E-07 | 9.50E-07 | -1.22E+00 | 4.79E-05 | 1.35E-04 | -1.09E+00 | 9.53E-02 | 3.50E-01 | 1.08E+00 | 7.61E-02 | 2.09E-01 | 1.03E+00 |
| RBBP5 | 1.39E-08 | 7.24E-08 | -1.76E+00 | 4.83E-05 | 2.09E-04 | -1.14E+00 | 2.74E-06 | 1.17E-05 | -1.69E+00 | 4.14E-03 | 1.58E-02 | -1.08E+00 | 1.48E-05 | 5.29E-05 | -1.94E+00 | 9.38E-03 | 2.65E-02 | -1.15E+00 | 8.45E-07 | 3.10E-06 | -1.34E+00 | 1.40E-02 | 3.85E-02 | 1.00E+00 |
| AL136126.1 | 1.39E-08 | 7.25E-08 | -1.19E+00 | 9.51E-15 | 4.11E-14 | -1.23E+00 | 3.66E-05 | 1.55E-04 | -1.15E+00 | 7.52E-08 | 2.87E-07 | -1.19E+00 | 1.97E-06 | 7.02E-06 | -1.15E+00 | 3.80E-05 | 1.07E-04 | -1.05E+00 | 1.41E-10 | 5.19E-10 | -1.40E+00 | 1.06E-06 | 2.91E-06 | -1.28E+00 |
| ZNF664 | 1.39E-08 | 7.26E-08 | -1.63E+00 | 4.78E-05 | 2.07E-04 | -1.08E+00 | 3.26E-05 | 1.38E-04 | -1.39E+00 | 1.21E-04 | 4.62E-04 | -1.13E+00 | 1.42E-03 | 5.06E-03 | -1.16E+00 | 2.68E-03 | 7.56E-03 | -1.08E+00 | 4.48E-04 | 1.65E-03 | -1.04E+00 | 7.76E-02 | 2.13E-01 | 1.16E+00 |
| ACE | 1.39E-08 | 7.26E-08 | -1.56E+00 | 1.71E-07 | 7.40E-07 | -1.28E+00 | 2.08E-01 | 8.84E-01 | 1.02E+00 | 1.81E-01 | 6.93E-01 | 1.02E+00 | 1.27E-01 | 4.52E-01 | -1.47E+00 | 2.25E-01 | 6.34E-01 | -1.24E+00 | 1.63E-01 | 5.97E-01 | -1.34E+00 | 2.18E-01 | 5.99E-01 | -1.30E+00 |
| GCFC2 | 1.39E-08 | 7.27E-08 | -1.21E+00 | 6.38E-06 | 2.76E-05 | 1.05E+00 | 6.08E-02 | 2.58E-01 | 1.01E+00 | 2.51E-02 | 9.60E-02 | 1.12E+00 | 3.01E-04 | 1.07E-03 | -1.17E+00 | 1.80E-02 | 5.08E-02 | 1.09E+00 | 1.54E-02 | 5.65E-02 | 1.02E+00 | 9.53E-02 | 2.62E-01 | 1.14E+00 |
| NELFE | 1.40E-08 | 7.29E-08 | -1.19E+00 | 3.71E-09 | 1.60E-08 | -1.05E+00 | 8.34E-04 | 3.54E-03 | -1.00E+00 | 2.16E-04 | 8.27E-04 | 1.08E+00 | 8.47E-07 | 3.02E-06 | -1.14E+00 | 2.62E-04 | 7.40E-04 | 1.04E+00 | 1.06E-02 | 3.88E-02 | 1.02E+00 | 3.61E-01 | 9.92E-01 | 1.26E+00 |
| CTSF | 1.42E-08 | 7.40E-08 | -1.12E+00 | 1.20E-04 | 5.18E-04 | 1.15E+00 | 5.80E-04 | 2.46E-03 | -1.13E+00 | 3.56E-02 | 1.36E-01 | 1.17E+00 | 1.30E-01 | 4.64E-01 | 1.30E+00 | 1.48E-01 | 4.18E-01 | 1.27E+00 | 1.14E-03 | 4.20E-03 | -1.07E+00 | 2.12E-02 | 5.83E-02 | 1.07E+00 |
| KIF3A | 1.42E-08 | 7.40E-08 | -1.86E+00 | 3.73E-05 | 1.61E-04 | -1.20E+00 | 3.64E-05 | 1.55E-04 | -1.44E+00 | 2.70E-05 | 1.03E-04 | -1.25E+00 | 1.63E-12 | 5.80E-12 | -2.68E+00 | 6.06E-10 | 1.71E-09 | -1.85E+00 | 1.17E-08 | 4.28E-08 | -2.12E+00 | 6.91E-07 | 1.90E-06 | -1.76E+00 |
| MATR3 | 1.42E-08 | 7.42E-08 | -1.12E+00 | 5.29E-05 | 2.29E-04 | 1.03E+00 | 5.57E-03 | 2.37E-02 | -1.14E+00 | 5.39E-02 | 2.06E-01 | -1.00E+00 | 9.87E-02 | 3.52E-01 | 1.08E+00 | 1.91E-02 | 5.39E-02 | 1.03E+00 | 3.52E-02 | 1.29E-01 | -1.03E+00 | 3.27E-01 | 8.99E-01 | 1.04E+00 |
| ZNF618 | 1.43E-08 | 7.43E-08 | -1.44E+00 | 6.34E-06 | 2.74E-05 | -1.06E+00 | 9.88E-07 | 4.20E-06 | -1.49E+00 | 6.33E-05 | 2.42E-04 | -1.17E+00 | 1.55E-09 | 5.54E-09 | -1.63E+00 | 1.00E-07 | 2.82E-07 | -1.38E+00 | 1.28E-11 | 4.70E-11 | -1.76E+00 | 3.62E-09 | 9.94E-09 | -1.53E+00 |
| RSL1D1 | 1.44E-08 | 7.51E-08 | -1.24E+00 | 5.86E-03 | 2.53E-02 | 1.00E+00 | 3.66E-04 | 1.56E-03 | -1.21E+00 | 1.20E-01 | 4.59E-01 | 1.03E+00 | 1.07E-09 | 3.81E-09 | -1.28E+00 | 1.66E-02 | 4.67E-02 | -1.04E+00 | 1.24E-08 | 4.55E-08 | -1.25E+00 | 3.53E-02 | 9.69E-02 | -1.01E+00 |
| RNF41 | 1.45E-08 | 7.54E-08 | -1.46E+00 | 3.02E-04 | 1.31E-03 | -1.01E+00 | 2.51E-07 | 1.07E-06 | -1.39E+00 | 8.02E-05 | 3.06E-04 | -1.00E+00 | 1.69E-08 | 6.01E-08 | -1.35E+00 | 2.84E-06 | 8.00E-06 | -1.16E+00 | 1.71E-04 | 6.27E-04 | -1.24E+00 | 1.45E-01 | 3.99E-01 | 1.11E+00 |
| MAP3K5 | 1.45E-08 | 7.55E-08 | -1.36E+00 | 6.36E-07 | 2.75E-06 | -1.06E+00 | 4.74E-06 | 2.02E-05 | -1.53E+00 | 2.69E-03 | 1.03E-02 | -1.07E+00 | 4.99E-05 | 1.78E-04 | -1.17E+00 | 3.01E-04 | 8.48E-04 | -1.12E+00 | 1.38E-12 | 5.07E-12 | -1.44E+00 | 1.11E-03 | 3.05E-03 | -1.01E+00 |
| MACF1 | 1.45E-08 | 7.57E-08 | -1.14E+00 | 3.42E-02 | 1.48E-01 | 1.21E+00 | 1.03E-03 | 4.36E-03 | -1.17E+00 | 7.53E-02 | 2.88E-01 | 1.10E+00 | 1.90E-04 | 6.79E-04 | -1.15E+00 | 3.09E-02 | 8.71E-02 | 1.04E+00 | 1.72E-04 | 6.30E-04 | -1.05E+00 | 1.52E-01 | 4.16E-01 | 1.09E+00 |
| MXI1 | 1.47E-08 | 7.64E-08 | -1.33E+00 | 2.21E-06 | 9.54E-06 | -1.09E+00 | 2.76E-05 | 1.17E-04 | -1.33E+00 | 2.10E-04 | 8.01E-04 | -1.10E+00 | 2.48E-03 | 8.83E-03 | -1.04E+00 | 1.63E-02 | 4.59E-02 | 1.13E+00 | 6.50E-05 | 2.39E-04 | -1.27E+00 | 4.04E-03 | 1.11E-02 | -1.07E+00 |
| CUL4B | 1.47E-08 | 7.68E-08 | -1.43E+00 | 5.03E-04 | 2.17E-03 | 1.07E+00 | 1.18E-08 | 5.01E-08 | -1.70E+00 | 1.81E-06 | 6.90E-06 | -1.30E+00 | 5.71E-04 | 2.04E-03 | -1.29E+00 | 1.80E-02 | 5.08E-02 | -1.01E+00 | 2.48E-08 | 9.11E-08 | -1.45E+00 | 8.16E-07 | 2.24E-06 | -1.41E+00 |
| MRPL35 | 1.49E-08 | 7.74E-08 | -1.62E+00 | 1.84E-05 | 7.93E-05 | -1.09E+00 | 1.48E-05 | 6.28E-05 | -1.29E+00 | 7.62E-05 | 2.91E-04 | -1.04E+00 | 1.52E-05 | 5.42E-05 | -1.43E+00 | 6.57E-03 | 1.85E-02 | -1.00E+00 | 1.10E-05 | 4.05E-05 | -1.20E+00 | 1.95E-01 | 5.37E-01 | 1.14E+00 |
| PARN | 1.49E-08 | 7.75E-08 | -1.40E+00 | 2.67E-06 | 1.15E-05 | -1.16E+00 | 5.02E-06 | 2.13E-05 | -1.36E+00 | 5.08E-04 | 1.94E-03 | -1.06E+00 | 6.11E-06 | 2.18E-05 | -1.38E+00 | 5.37E-06 | 1.51E-05 | -1.36E+00 | 7.91E-08 | 2.90E-07 | -1.38E+00 | 4.86E-05 | 1.34E-04 | -1.20E+00 |
| [illegible] | [illegible] | [illegible] | [illegible] | [illegible] | [illegible] | [illegible] | [illegible] | [illegible] | [illegible] | [illegible] | [illegible] | [illegible] | [illegible] | [illegible] | [illegible] | [illegible] | [illegible] | [illegible] | [illegible] | [illegible] | [illegible] | [illegible] | [illegible] | [illegible] |
| USP18 | 1.54E-08 | 8.05E-08 | -2.93E+00 | 4.41E-06 | 1.91E-05 | -1.70E+00 | 2.83E-17 | 1.20E-16 | -3.60E+00 | 6.26E-16 | 2.39E-15 | -2.89E+00 | 1.23E-15 | 4.38E-15 | -5.12E+00 | 9.24E-18 | 2.61E-17 | -6.39E+00 | 3.49E-35 | 1.28E-34 | -2.97E+00 | 7.95E-32 | 2.18E-31 | -2.57E+00 |
| AC008894.3 | 1.56E-08 | 8.13E-08 | 4.31E+00 | 1.26E-01 | 5.44E-01 | 1.09E+00 | 2.89E-07 | 1.23E-06 | 3.43E+00 | 2.27E-01 | 8.68E-01 | 1.38E+00 | 2.22E-03 | 7.91E-03 | 4.05E+00 | 2.25E-01 | 6.35E-01 | 1.94E+00 | 1.78E-03 | 6.54E-03 | 2.71E+00 | 1.81E-01 | 4.97E-01 | -1.15E+00 |
| HIST1H4K | 1.57E-08 | 8.17E-08 | -1.44E+00 | 2.59E-17 | 1.12E-16 | -2.71E+00 | 1.78E-01 | 7.57E-01 | 1.16E+00 | 6.74E-06 | 2.57E-05 | -1.62E+00 | 8.24E-03 | 2.94E-02 | -1.00E+00 | 4.64E-06 | 1.31E-05 | -1.80E+00 | 2.61E-08 | 9.58E-08 | -1.36E+00 | 4.30E-16 | 1.18E-15 | -2.45E+00 |
| DHPS | 1.59E-08 | 8.31E-08 | -1.14E+00 | 1.22E-08 | 5.28E-08 | -1.04E+00 | 1.67E-06 | 7.08E-06 | -1.32E+00 | 3.03E-05 | 1.16E-04 | -1.12E+00 | 1.19E-02 | 4.23E-02 | -1.08E+00 | 1.94E-01 | 5.47E-01 | 1.10E+00 | 1.42E-03 | 5.22E-03 | -1.02E+00 | 1.67E-02 | 4.57E-02 | 1.04E+00 |
| RNF149 | 1.60E-08 | 8.31E-08 | -1.24E+00 | 1.10E-06 | 4.76E-06 | -1.00E+00 | 5.10E-09 | 2.17E-08 | -1.51E+00 | 9.99E-07 | 3.82E-06 | -1.19E+00 | 1.28E-04 | 4.56E-04 | -1.26E+00 | 6.66E-05 | 1.88E-04 | -1.38E+00 | 1.79E-11 | 6.56E-11 | -1.77E+00 | 1.91E-06 | 5.25E-06 | -1.36E+00 |
| ICK | 1.60E-08 | 8.36E-08 | -1.76E+00 | 2.70E-09 | 1.17E-08 | -1.77E+00 | 3.99E-05 | 1.70E-04 | -1.45E+00 | 1.21E-04 | 4.62E-04 | -1.24E+00 | 1.09E-06 | 3.88E-06 | -1.74E+00 | 8.74E-05 | 2.47E-04 | -1.39E+00 | 1.18E-04 | 4.33E-04 | -1.74E+00 | 2.67E-02 | 7.35E-02 | -1.11E+00 |
| ARHGEF2 | 1.61E-08 | 8.37E-08 | -1.19E+00 | 1.59E-04 | 6.86E-04 | 1.07E+00 | 1.85E-07 | 7.87E-07 | -1.40E+00 | 6.16E-04 | 2.35E-03 | 1.00E+00 | 7.09E-11 | 2.53E-10 | -1.48E+00 | 2.45E-05 | 6.91E-05 | -1.15E+00 | 2.13E-09 | 7.83E-09 | -1.33E+00 | 8.98E-04 | 2.47E-03 | -1.07E+00 |
| CDA5 | 1.61E-08 | 8.37E-08 | -1.23E+00 | 1.66E-06 | 7.19E-06 | 1.01E+00 | 1.43E-02 | 6.08E-02 | -1.05E+00 | 3.20E-04 | 1.22E-03 | -1.05E+00 | 6.83E-04 | 2.44E-03 | -1.09E+00 | 4.21E-03 | 1.19E-02 | 1.03E+00 | 1.27E-03 | 4.65E-03 | -1.11E+00 | 3.39E-02 | 9.32E-02 | 1.02E+00 |
| SYTL4 | 1.62E-08 | 8.42E-08 | -1.71E+00 | 2.53E-04 | 1.09E-03 | -1.11E+00 | 2.21E-02 | 9.38E-02 | -1.43E+00 | 2.52E-01 | 9.63E-01 | 1.30E+00 | 2.66E-03 | 9.49E-03 | -1.73E+00 | 4.10E-02 | 1.15E-01 | -1.31E+00 | 1.88E-05 | 6.91E-05 | -1.30E+00 | 3.29E-03 | 9.03E-03 | -1.07E+00 |
| ATP5A1P3 | 1.63E-08 | 8.47E-08 | -1.94E+00 | 1.13E-07 | 4.87E-07 | -1.65E+00 | 3.35E-11 | 1.42E-10 | -1.79E+00 | 1.25E-14 | 4.78E-14 | -2.02E+00 | 3.28E-03 | 1.17E-02 | -1.26E+00 | 3.80E-03 | 1.07E-02 | -1.26E+00 | 6.95E-10 | 2.55E-09 | -1.82E+00 | 2.71E-09 | 7.44E-09 | -1.77E+00 |
| USP37 | 1.63E-08 | 8.47E-08 | -2.07E+00 | 4.56E-05 | 1.97E-04 | -1.26E+00 | 7.69E-03 | 3.27E-02 | -1.14E+00 | 8.38E-03 | 3.20E-02 | -1.04E+00 | 1.38E-06 | 4.91E-06 | -2.20E+00 | 3.19E-07 | 8.98E-07 | -2.34E+00 | 1.23E-05 | 4.50E-05 | -1.30E+00 | 2.71E-03 | 7.44E-03 | -1.02E+00 |
| LRRCC1 | 1.63E-08 | 8.50E-08 | -1.45E+00 | 2.20E-06 | 9.50E-06 | -1.17E+00 | 5.89E-07 | 2.50E-06 | -1.35E+00 | 1.71E-05 | 6.54E-05 | -1.16E+00 | 1.53E-03 | 5.47E-03 | -1.48E+00 | 9.52E-02 | 2.68E-01 | 1.13E+00 | 8.49E-11 | 3.12E-10 | -1.43E+00 | 2.91E-03 | 7.99E-03 | -1.05E+00 |
| C18orf21 | 1.63E-08 | 8.51E-08 | -1.31E+00 | 3.96E-11 | 1.71E-10 | -1.30E+00 | 1.66E-02 | 7.04E-02 | -1.03E+00 | 7.22E-06 | 2.76E-05 | -1.28E+00 | 4.02E-04 | 1.43E-03 | -1.15E+00 | 1.08E-04 | 3.04E-04 | -1.25E+00 | 1.19E-04 | 4.37E-04 | -1.17E+00 | 1.55E-04 | 4.26E-04 | -1.15E+00 |
| CXXC1 | 1.63E-08 | 8.52E-08 | -1.33E+00 | 8.58E-05 | 3.71E-04 | 1.05E+00 | 2.00E-04 | 8.52E-04 | -1.14E+00 | 1.21E-04 | 4.64E-04 | -1.08E+00 | 4.46E-03 | 1.59E-02 | -1.15E+00 | 3.91E-02 | 1.10E-01 | 1.03E+00 | 5.06E-05 | 1.86E-04 | -1.10E+00 | 2.70E-02 | 7.41E-02 | 1.10E+00 |
| AGAP4 | 1.66E-08 | 8.63E-08 | -1.64E+00 | 1.28E-04 | 5.53E-04 | -1.06E+00 | 3.81E-12 | 1.62E-11 | -1.74E+00 | 1.73E-09 | 6.60E-09 | -1.30E+00 | 1.14E-04 | 4.05E-04 | -1.36E+00 | 1.05E-02 | 2.97E-02 | -1.08E+00 | 1.26E-10 | 4.61E-10 | -1.97E+00 | 4.96E-09 | 1.36E-08 | -1.85E+00 |
| ATAT1 | 1.66E-08 | 8.67E-08 | -1.47E+00 | 7.61E-06 | 3.29E-05 | -1.03E+00 | 3.80E-04 | 1.61E-03 | -1.47E+00 | 5.92E-02 | 2.26E-01 | 1.08E+00 | 9.08E-04 | 3.24E-03 | -1.23E+00 | 4.49E-02 | 1.26E-01 | 1.15E+00 | 2.32E-04 | 8.52E-04 | -1.35E+00 | 3.93E-02 | 1.08E-01 | -1.03E+00 |
| RFWD2 | 1.66E-08 | 8.67E-08 | -1.27E+00 | 7.88E-09 | 3.40E-08 | -1.22E+00 | 5.20E-04 | 2.21E-03 | -1.18E+00 | 1.37E-04 | 5.24E-04 | -1.06E+00 | 1.42E-05 | 5.06E-05 | -1.15E+00 | 2.78E-06 | 7.83E-06 | -1.17E+00 | 1.02E-04 | 3.73E-04 | -1.15E+00 | 6.65E-04 | 1.83E-03 | -1.13E+00 |
| CYFIP1 | 1.67E-08 | 8.71E-08 | -1.38E+00 | 2.24E-03 | 9.67E-03 | 1.10E+00 | 2.10E-03 | 8.90E-03 | -1.10E+00 | 3.87E-02 | 1.48E-01 | 1.18E+00 | 1.04E-03 | 3.70E-03 | -1.25E+00 | 7.51E-02 | 2.12E-01 | 1.12E+00 | 2.22E-05 | 8.14E-05 | -1.19E+00 | 2.11E-03 | 5.78E-03 | -1.14E+00 |
| DGCR6L | 1.69E-08 | 8.80E-08 | -1.13E+00 | 6.36E-11 | 2.75E-10 | -1.11E+00 | 2.13E-01 | 9.05E-01 | 1.22E+00 | 6.71E-02 | 2.56E-01 | 1.20E+00 | 2.97E-04 | 1.06E-03 | -1.04E+00 | 5.20E-05 | 1.47E-04 | -1.09E+00 | 1.23E-01 | 4.53E-01 | 1.28E+00 | 2.50E-01 | 6.86E-01 | 1.36E+00 |
| ASPHD2 | 1.70E-08 | 8.87E-08 | -1.94E+00 | 1.02E-07 | 4.41E-07 | -1.52E+00 | 1.43E-07 | 6.07E-07 | -1.96E+00 | 4.63E-06 | 1.77E-05 | -1.60E+00 | 2.86E-03 | 1.02E-02 | -2.19E+00 | 1.65E-02 | 4.65E-02 | -1.60E+00 | 8.30E-02 | 3.05E-01 | -1.12E+00 | 2.95E-02 | 8.09E-02 | -1.47E+00 |
| MLXIP | 1.71E-08 | 8.91E-08 | -1.14E+00 | 3.35E-03 | 1.45E-02 | 1.21E+00 | 7.36E-06 | 3.13E-05 | -1.26E+00 | 8.90E-04 | 3.40E-03 | 1.02E+00 | 6.88E-07 | 2.45E-06 | -1.20E+00 | 8.62E-04 | 2.43E-03 | -1.08E+00 | 4.59E-05 | 1.68E-04 | -1.11E+00 | 3.40E-03 | 9.33E-03 | 1.06E+00 |
| NME6 | 1.72E-08 | 8.98E-08 | -1.54E+00 | 3.27E-04 | 1.41E-03 | -1.01E+00 | 2.54E-12 | 1.08E-11 | -1.75E+00 | 3.27E-09 | 1.25E-08 | -1.30E+00 | 1.56E-08 | 5.56E-08 | -2.14E+00 | 2.52E-04 | 7.10E-04 | -1.25E+00 | 1.44E-09 | 5.27E-09 | -1.42E+00 | 1.15E-05 | 3.16E-05 | -1.22E+00 |
| CDC42EP4 | 1.72E-08 | 8.98E-08 | -2.36E+00 | 3.13E-09 | 1.35E-08 | -2.59E+00 | 2.07E-03 | 8.79E-03 | -1.69E+00 | 1.08E-03 | 4.12E-03 | -1.75E+00 | 2.47E-07 | 8.80E-07 | -2.60E+00 | 1.41E-06 | 3.97E-06 | -2.22E+00 | 6.82E-26 | 2.50E-25 | -2.02E+00 | 2.16E-23 | 5.94E-23 | -1.79E+00 |
| NUP93 | 1.74E-08 | 9.09E-08 | -1.39E+00 | 4.31E-03 | 1.86E-02 | 1.23E+00 | 8.69E-03 | 3.69E-02 | -1.04E+00 | 6.19E-02 | 2.37E-01 | 1.26E+00 | 4.87E-07 | 1.73E-06 | -1.53E+00 | 3.79E-03 | 1.07E-02 | 1.03E+00 | 3.69E-05 | 1.36E-04 | -1.19E+00 | 3.12E-01 | 8.58E-01 | 1.29E+00 |
| DHX30 | 1.75E-08 | 9.13E-08 | -1.42E+00 | 3.99E-06 | 1.73E-05 | -1.07E+00 | 2.44E-04 | 1.04E-03 | -1.15E+00 | 1.98E-03 | 7.58E-03 | 1.09E+00 | 2.60E-06 | 9.27E-06 | -1.37E+00 | 9.30E-05 | 2.62E-04 | -1.22E+00 | 1.63E-03 | 5.99E-03 | -1.11E+00 | 7.71E-02 | 2.12E-01 | 1.07E+00 |
| RAB5A | 1.75E-08 | 9.13E-08 | -1.23E+00 | 3.33E-08 | 1.44E-07 | -1.16E+00 | 2.64E-03 | 1.12E-02 | -1.06E+00 | 4.45E-04 | 1.70E-03 | 1.01E+00 | 8.42E-04 | 3.00E-03 | 1.01E+00 | 4.17E-04 | 1.18E-03 | -1.02E+00 | 4.50E-05 | 1.65E-04 | -1.20E+00 | 8.96E-04 | 2.46E-03 | -1.08E+00 |
| CHST13 | 1.75E-08 | 9.13E-08 | -1.17E+00 | 2.61E-03 | 1.13E-02 | 1.24E+00 | 3.67E-05 | 1.56E-04 | -1.20E+00 | 5.77E-04 | 2.20E-03 | 1.03E+00 | 1.28E-06 | 4.58E-06 | -1.29E+00 | 5.18E-02 | 1.46E-01 | 1.14E+00 | 3.03E-02 | 1.11E-01 | 1.07E+00 | 3.13E-01 | 8.60E-01 | 1.35E+00 |
| ADNP | 1.77E-08 | 9.21E-08 | -1.30E+00 | 2.38E-07 | 1.03E-06 | -1.02E+00 | 1.13E-05 | 4.80E-05 | -1.45E+00 | 7.94E-06 | 3.03E-05 | -1.21E+00 | 4.40E-07 | 1.57E-06 | -1.30E+00 | 9.14E-04 | 2.58E-03 | 1.02E+00 | 1.16E-10 | 4.27E-10 | -1.67E+00 | 2.49E-04 | 6.85E-04 | -1.15E+00 |
| CAP1 | 1.77E-08 | 9.21E-08 | -1.64E+00 | 5.76E-08 | 2.49E-07 | -1.46E+00 | 1.98E-04 | 8.43E-04 | -1.39E+00 | 5.93E-05 | 2.27E-04 | -1.37E+00 | 2.14E-03 | 7.64E-03 | -1.14E+00 | 2.52E-05 | 7.10E-05 | -1.51E+00 | 4.04E-02 | 1.48E-01 | 1.05E+00 | 3.01E-03 | 8.27E-03 | -1.24E+00 |
| OGFOD3 | 1.77E-08 | 9.25E-08 | -1.41E+00 | 4.78E-08 | 2.06E-07 | -1.22E+00 | 5.86E-02 | 2.49E-01 | 1.07E+00 | 4.04E-03 | 1.54E-02 | 1.09E+00 | 3.51E-03 | 1.25E-02 | -1.06E+00 | 2.19E-02 | 6.17E-02 | 1.00E+00 | 3.47E-04 | 1.28E-03 | -1.08E+00 | 5.63E-02 | 1.55E-01 | 1.11E+00 |
| ZFX | 1.78E-08 | 9.26E-08 | -1.25E+00 | 1.85E-07 | 7.99E-07 | -1.09E+00 | 4.28E-05 | 1.82E-04 | -1.26E+00 | 4.44E-05 | 1.70E-04 | -1.13E+00 | 7.37E-05 | 2.63E-04 | -1.24E+00 | 1.31E-06 | 3.69E-06 | -1.39E+00 | 1.94E-08 | 7.13E-08 | -1.53E+00 | 2.92E-04 | 8.01E-04 | -1.19E+00 |
| IMPACT | 1.80E-08 | 9.36E-08 | -1.29E+00 | 6.00E-06 | 2.59E-05 | 1.01E+00 | 2.40E-09 | 1.02E-08 | -1.54E+00 | 5.58E-09 | 2.13E-08 | -1.40E+00 | 2.25E-12 | 8.03E-12 | -1.64E+00 | 1.61E-07 | 4.53E-07 | -1.24E+00 | 2.06E-03 | 7.55E-03 | -1.12E+00 | 7.35E-03 | 2.02E-02 | -1.08E+00 |
| ALPP | 1.82E-08 | 9.50E-08 | -1.88E+00 | 5.23E-05 | 2.26E-04 | -1.18E+00 | 1.91E-01 | 8.12E-01 | 1.01E+00 | 2.42E-01 | 9.25E-01 | 1.21E+00 | 1.19E-02 | 4.25E-02 | -1.57E+00 | 9.27E-03 | 2.61E-02 | -1.82E+00 | 3.23E-03 | 1.19E-02 | -2.06E+00 | 5.04E-02 | 1.38E-01 | -1.53E+00 |
| ZNF609 | 1.82E-08 | 9.50E-08 | -1.11E+00 | 4.51E-04 | 1.95E-03 | 1.17E+00 | 2.27E-04 | 9.64E-04 | -1.11E+00 | 4.72E-03 | 1.80E-02 | 1.13E+00 | 2.08E-07 | 7.42E-07 | -1.33E+00 | 1.02E-06 | 2.88E-06 | -1.24E+00 | 1.30E-11 | 4.77E-11 | -1.84E+00 | 2.21E-08 | 6.07E-08 | -1.50E+00 |
| RNF141 | 1.82E-08 | 9.50E-08 | -1.34E+00 | 2.60E-05 | 1.12E-04 | -1.00E+00 | 3.73E-05 | 1.58E-04 | -1.28E+00 | 1.10E-06 | 4.18E-06 | -1.23E+00 | 5.30E-10 | 1.89E-09 | -1.60E+00 | 2.12E-06 | 5.99E-06 | -1.21E+00 | 3.95E-06 | 1.45E-05 | -1.20E+00 | 6.46E-03 | 1.77E-02 | 1.02E+00 |
| NRBP1 | 1.83E-08 | 9.56E-08 | -1.46E+00 | 1.62E-07 | 6.98E-07 | -1.20E+00 | 3.21E-09 | 1.36E-08 | -1.62E+00 | 4.04E-08 | 1.54E-07 | -1.35E+00 | 1.37E-12 | 4.90E-12 | -1.53E+00 | 1.95E-08 | 5.49E-08 | -1.19E+00 | 6.09E-11 | 2.24E-10 | -1.44E+00 | 7.57E-05 | 2.08E-04 | -1.13E+00 |
| CCDC184 | 1.83E-08 | 9.56E-08 | 4.20E+00 | 1.82E-01 | 7.86E-01 | -1.13E+00 | 1.61E-04 | 6.86E-04 | 2.99E+00 | 2.19E-01 | 8.35E-01 | -1.05E+00 | 4.92E-03 | 1.76E-02 | 4.07E+00 | 3.37E-01 | 9.49E-01 | 1.01E+00 | 2.84E-04 | 1.04E-03 | 2.94E+00 | 3.39E-01 | 9.31E-01 | 1.09E+00 |
| RNF166 | 1.84E-08 | 9.57E-08 | -1.40E+00 | 1.22E-06 | 5.25E-06 | -1.13E+00 | 1.62E-04 | 6.90E-04 | -1.25E+00 | 5.85E-04 | 2.23E-03 | -1.03E+00 | 3.56E-12 | 1.27E-11 | -1.68E+00 | 9.28E-10 | 2.62E-09 | -1.47E+00 | 3.58E-06 | 1.31E-05 | -1.22E+00 | 4.55E-03 | 1.25E-02 | -1.02E+00 |
| QSOX1 | 1.84E-08 | 9.62E-08 | -1.43E+00 | 4.99E-08 | 2.16E-07 | -1.24E+00 | 8.58E-06 | 3.64E-05 | -1.20E+00 | 2.13E-07 | 8.14E-07 | -1.11E+00 | 3.02E-11 | 1.08E-10 | -1.62E+00 | 4.70E-08 | 1.32E-07 | -1.33E+00 | 3.14E-07 | 1.15E-06 | -1.51E+00 | 5.61E-06 | 1.54E-05 | -1.39E+00 |
| ARID2 | 1.85E-08 | 9.64E-08 | -1.20E+00 | 1.90E-04 | 8.22E-04 | 1.15E+00 | 3.15E-04 | 1.34E-03 | -1.25E+00 | 7.86E-04 | 3.00E-03 | -1.02E+00 | 9.22E-03 | 3.29E-02 | 1.05E+00 | 1.23E-02 | 3.46E-02 | 1.07E+00 | 1.63E-07 | 5.98E-07 | -1.43E+00 | 5.27E-04 | 1.45E-03 | -1.17E+00 |
| ARV1 | 1.86E-08 | 9.68E-08 | -1.67E+00 | 8.34E-05 | 3.60E-04 | -1.10E+00 | 2.59E-06 | 1.10E-05 | -1.40E+00 | 6.69E-04 | 2.56E-03 | -1.02E+00 | 2.37E-07 | 8.44E-07 | -1.54E+00 | 1.12E-01 | 3.16E-01 | 1.28E+00 | 4.39E-07 | 1.61E-06 | -1.29E+00 | 2.15E-02 | 5.90E-02 | -1.06E+00 |
| GAA | 1.86E-08 | 9.69E-08 | -1.39E+00 | 9.79E-10 | 4.23E-09 | -1.42E+00 | 4.09E-04 | 1.74E-03 | -1.23E+00 | 1.31E-05 | 5.00E-05 | -1.18E+00 | 5.36E-04 | 1.91E-03 | -1.39E+00 | 2.09E-03 | 5.91E-03 | -1.18E+00 | 2.38E-01 | 8.74E-01 | 1.26E+00 | 2.50E-01 | 6.88E-01 | 1.44E+00 |
| FAM136A | 1.87E-08 | 9.73E-08 | -1.14E+00 | 9.40E-09 | 4.06E-08 | -1.04E+00 | 4.29E-02 | 1.82E-01 | 1.01E+00 | 6.31E-02 | 2.41E-01 | 1.12E+00 | 1.42E-09 | 5.07E-09 | -1.35E+00 | 3.44E-03 | 9.70E-03 | 1.03E+00 | 1.35E-03 | 4.94E-03 | 1.02E+00 | 3.58E-01 | 9.83E-01 | 1.29E+00 |
| [illegible] | [illegible] | [illegible] | [illegible] | [illegible] | [illegible] | [illegible] | [illegible] | [illegible] | [illegible] | [illegible] | [illegible] | [illegible] | [illegible] | [illegible] | [illegible] | [illegible] | [illegible] | [illegible] | [illegible] | [illegible] | [illegible] | [illegible] | [illegible] | [illegible] |
| LAMTOR2 | 1.88E-08 | 9.80E-08 | -1.07E+00 | 5.43E-07 | 2.35E-06 | 1.04E+00 | 8.86E-02 | 3.76E-01 | 1.07E+00 | 6.35E-04 | 2.42E-03 | 1.02E+00 | 1.38E-05 | 4.92E-05 | -1.19E+00 | 8.57E-04 | 2.42E-03 | -1.03E+00 | 9.61E-03 | 3.53E-02 | -1.02E+00 | 3.05E-01 | 8.37E-01 | 1.08E+00 |
| RXRB | 1.89E-08 | 9.83E-08 | -1.51E+00 | 3.21E-05 | 1.39E-04 | -1.03E+00 | 1.83E-06 | 7.75E-06 | -1.29E+00 | 6.83E-03 | 2.61E-02 | 1.15E+00 | 1.86E-06 | 6.65E-06 | -1.65E+00 | 6.24E-04 | 1.76E-03 | -1.17E+00 | 7.21E-07 | 2.65E-06 | -1.42E+00 | 1.12E-02 | 3.08E-02 | -1.02E+00 |
| SHQ1 | 1.89E-08 | 9.84E-08 | -2.05E+00 | 3.69E-03 | 1.59E-02 | -1.02E+00 | 9.92E-08 | 4.21E-07 | -1.73E+00 | 1.01E-02 | 3.86E-02 | 1.04E+00 | 1.05E-06 | 3.74E-06 | -2.48E+00 | 1.19E-02 | 3.35E-02 | -1.02E+00 | 8.76E-10 | 3.22E-09 | -1.73E+00 | 2.24E-01 | 6.16E-01 | 1.16E+00 |
| ACTR3 | 1.89E-08 | 9.85E-08 | -1.20E+00 | 6.85E-05 | 2.96E-04 | 1.08E+00 | 1.81E-03 | 7.69E-03 | -1.13E+00 | 1.91E-03 | 7.29E-03 | -1.07E+00 | 2.03E-10 | 7.23E-10 | -1.59E+00 | 1.62E-05 | 4.58E-05 | -1.23E+00 | 4.16E-07 | 1.53E-06 | -1.41E+00 | 4.64E-02 | 1.28E-01 | -1.08E+00 |
| UGGT1 | 1.92E-08 | 9.99E-08 | -1.70E+00 | 6.93E-05 | 3.00E-04 | -1.10E+00 | 2.66E-05 | 1.13E-04 | -1.17E+00 | 5.98E-05 | 2.29E-04 | -1.01E+00 | 6.28E-07 | 2.24E-06 | -1.40E+00 | 4.29E-03 | 1.21E-02 | 1.09E+00 | 1.02E-07 | 3.76E-07 | -1.26E+00 | 2.72E-04 | 7.48E-04 | -1.13E+00 |
| CA13 | 1.92E-08 | 1.00E-07 | -1.76E+00 | 3.83E-03 | 1.65E-02 | 1.09E+00 | 1.32E-01 | 5.61E-01 | -1.05E+00 | 1.98E-01 | 7.55E-01 | 1.34E+00 | 1.13E-01 | 4.04E-01 | -1.30E+00 | 2.27E-01 | 6.40E-01 | -1.05E+00 | 2.09E-02 | 7.69E-02 | -1.50E+00 | 6.73E-02 | 1.85E-01 | -1.23E+00 |
| TSSC4 | 1.93E-08 | 1.00E-07 | -1.19E+00 | 6.15E-09 | 2.66E-08 | -1.10E+00 | 3.83E-04 | 1.63E-03 | -1.05E+00 | 4.62E-05 | 1.77E-04 | -1.02E+00 | 6.05E-06 | 2.16E-05 | -1.12E+00 | 4.78E-06 | 1.35E-05 | -1.13E+00 | 1.89E-03 | 6.94E-03 | 1.07E+00 | 4.72E-03 | 1.30E-02 | 1.08E+00 |
| TMED3 | 1.93E-08 | 1.00E-07 | -1.12E+00 | 5.68E-09 | 2.45E-08 | 1.01E+00 | 1.55E-03 | 6.57E-03 | -1.11E+00 | 2.54E-02 | 9.71E-02 | 1.14E+00 | 2.97E-04 | 1.06E-03 | -1.07E+00 | 1.20E-03 | 3.37E-03 | 1.06E+00 | 1.22E-01 | 4.46E-01 | 1.08E+00 | 2.72E-01 | 7.46E-01 | 1.20E+00 |
| ZNF667 | 1.93E-08 | 1.01E-07 | -1.25E+00 | 9.81E-10 | 4.24E-09 | -1.21E+00 | 3.21E-04 | 1.36E-03 | -1.22E+00 | 2.20E-04 | 8.39E-04 | -1.14E+00 | 9.96E-02 | 3.55E-01 | 1.27E+00 | 2.40E-02 | 6.77E-02 | 1.07E+00 | 2.24E-03 | 8.23E-03 | -1.15E+00 | 2.04E-02 | 5.61E-02 | -1.07E+00 |
| PCTP | 1.95E-08 | 1.01E-07 | -2.30E+00 | 5.99E-05 | 2.59E-04 | -1.23E+00 | 1.84E-06 | 7.84E-06 | -1.63E+00 | 1.69E-03 | 6.45E-03 | -1.08E+00 | 2.59E-13 | 9.22E-13 | -1.77E+00 | 5.35E-11 | 1.51E-10 | -1.45E+00 | 1.37E-32 | 5.03E-32 | -2.04E+00 | 5.60E-15 | 1.54E-14 | -1.38E+00 |
| TMEM80 | 1.95E-08 | 1.02E-07 | -1.46E+00 | 1.50E-03 | 6.47E-03 | 1.14E+00 | 1.04E-06 | 4.41E-06 | -1.57E+00 | 7.25E-03 | 2.77E-02 | 1.08E+00 | 5.09E-04 | 1.82E-03 | -1.22E+00 | 2.03E-01 | 5.72E-01 | 1.37E+00 | 3.95E-04 | 1.45E-03 | -1.21E+00 | 2.43E-03 | 6.68E-03 | -1.10E+00 |
| LCP2 | 1.96E-08 | 1.02E-07 | -1.18E+00 | 5.08E-05 | 2.20E-04 | 1.04E+00 | 2.12E-07 | 9.02E-07 | -1.45E+00 | 3.69E-06 | 1.41E-05 | -1.23E+00 | 9.14E-12 | 3.26E-11 | -1.49E+00 | 2.12E-08 | 5.98E-08 | -1.22E+00 | 3.43E-04 | 1.26E-03 | -1.19E+00 | 2.16E-03 | 5.93E-03 | -1.14E+00 |
| ZNF644 | 1.96E-08 | 1.02E-07 | -1.13E+00 | 2.02E-16 | 8.71E-16 | -1.27E+00 | 2.55E-05 | 1.08E-04 | -1.19E+00 | 5.39E-07 | 2.06E-06 | -1.23E+00 | 2.58E-08 | 9.20E-08 | -1.21E+00 | 1.78E-11 | 5.01E-11 | -1.31E+00 | 6.95E-05 | 2.55E-04 | -1.20E+00 | 3.26E-05 | 8.95E-05 | -1.19E+00 |
| NAGPA | 1.96E-08 | 1.02E-07 | -1.38E+00 | 1.76E-06 | 7.61E-06 | -1.10E+00 | 1.28E-04 | 5.44E-04 | -1.34E+00 | 2.88E-07 | 1.10E-06 | -1.32E+00 | 6.35E-07 | 2.27E-06 | -1.67E+00 | 1.04E-03 | 2.93E-03 | -1.14E+00 | 7.12E-07 | 2.61E-06 | -1.34E+00 | 8.95E-03 | 2.46E-02 | 1.02E+00 |
| LRBA | 1.96E-08 | 1.02E-07 | -1.17E+00 | 5.31E-02 | 2.29E-01 | 1.45E+00 | 8.72E-04 | 3.70E-03 | -1.03E+00 | 1.79E-01 | 6.84E-01 | 1.42E+00 | 3.39E-04 | 1.21E-03 | -1.12E+00 | 9.45E-02 | 2.66E-01 | 1.19E+00 | 2.51E-01 | 9.22E-01 | 1.32E+00 | 3.58E-01 | 9.83E-01 | 1.18E+00 |
| GLIPR2 | 1.96E-08 | 1.02E-07 | -1.25E+00 | 2.20E-04 | 9.51E-04 | 1.08E+00 | 4.93E-13 | 2.09E-12 | -1.66E+00 | 3.80E-10 | 1.45E-09 | -1.38E+00 | 1.49E-06 | 5.33E-06 | -1.74E+00 | 1.59E-04 | 4.49E-04 | -1.28E+00 | 4.44E-06 | 1.63E-05 | -1.25E+00 | 4.94E-04 | 1.36E-03 | -1.12E+00 |
| SLC22A17 | 1.96E-08 | 1.02E-07 | -1.67E+00 | 1.23E-05 | 5.30E-05 | -1.19E+00 | 2.96E-03 | 1.26E-02 | -1.45E+00 | 3.40E-02 | 1.30E-01 | 1.02E+00 | 4.88E-02 | 1.74E-01 | 1.09E+00 | 3.39E-01 | 9.56E-01 | 1.36E+00 | 1.16E-01 | 4.26E-01 | 1.46E+00 | 8.34E-02 | 2.29E-01 | 1.07E+00 |
| RITA1 | 1.98E-08 | 1.03E-07 | -1.75E+00 | 2.94E-04 | 1.27E-03 | 1.01E+00 | 1.63E-06 | 6.92E-06 | -1.47E+00 | 6.51E-04 | 2.49E-03 | -1.03E+00 | 1.36E-04 | 4.86E-04 | -1.33E+00 | 2.13E-02 | 6.01E-02 | 1.07E+00 | 5.29E-05 | 1.94E-04 | -1.20E+00 | 3.10E-01 | 8.51E-01 | 1.32E+00 |
| GALNT1 | 1.99E-08 | 1.04E-07 | -1.16E+00 | 2.69E-06 | 1.16E-05 | 1.08E+00 | 6.44E-05 | 2.74E-04 | -1.21E+00 | 6.84E-05 | 2.61E-04 | -1.08E+00 | 9.04E-09 | 3.22E-08 | -1.32E+00 | 3.33E-09 | 9.38E-09 | -1.30E+00 | 2.54E-05 | 9.33E-05 | -1.22E+00 | 1.87E-03 | 5.13E-03 | -1.09E+00 |
| ZC3H14 | 1.99E-08 | 1.04E-07 | -1.60E+00 | 2.82E-06 | 1.22E-05 | -1.25E+00 | 3.45E-04 | 1.47E-03 | -1.09E+00 | 3.43E-05 | 1.31E-04 | -1.04E+00 | 2.61E-05 | 9.30E-05 | -1.61E+00 | 1.01E-02 | 2.84E-02 | -1.01E+00 | 1.47E-06 | 5.39E-06 | -1.36E+00 | 5.97E-06 | 1.64E-05 | -1.27E+00 |
| CYP20A1 | 1.99E-08 | 1.04E-07 | -1.48E+00 | 2.40E-05 | 1.04E-04 | -1.06E+00 | 1.28E-05 | 5.45E-05 | -1.35E+00 | 6.04E-05 | 2.31E-04 | -1.12E+00 | 1.65E-06 | 5.89E-06 | -1.58E+00 | 2.53E-04 | 7.13E-04 | -1.26E+00 | 3.62E-06 | 1.33E-05 | -1.21E+00 | 8.49E-04 | 2.33E-03 | -1.15E+00 |
| AL512844.1 | 2.00E-08 | 1.04E-07 | -1.39E+00 | 3.33E-12 | 1.44E-11 | -1.49E+00 | 1.90E-11 | 8.07E-11 | -1.55E+00 | 2.43E-16 | 9.29E-16 | -1.67E+00 | 2.14E-04 | 7.62E-04 | -1.14E+00 | 1.92E-06 | 5.41E-06 | -1.44E+00 | 2.63E-07 | 9.65E-07 | -1.09E+00 | 1.88E-11 | 5.16E-11 | -1.52E+00 |
| NONO | 2.01E-08 | 1.05E-07 | -1.13E+00 | 2.52E-08 | 1.09E-07 | -1.02E+00 | 1.18E-04 | 5.00E-04 | -1.28E+00 | 8.27E-04 | 3.16E-03 | -1.15E+00 | 1.62E-08 | 5.78E-08 | -1.26E+00 | 1.23E-03 | 3.46E-03 | -1.05E+00 | 3.67E-04 | 1.35E-03 | -1.26E+00 | 6.75E-03 | 1.85E-02 | -1.16E+00 |
| WDFY3 | 2.01E-08 | 1.05E-07 | -2.62E+00 | 1.47E-07 | 6.36E-07 | -2.19E+00 | 4.20E-05 | 1.78E-04 | -1.82E+00 | 2.42E-04 | 9.26E-04 | -1.45E+00 | 1.80E-03 | 6.43E-03 | -2.39E+00 | 2.41E-03 | 6.80E-03 | -2.26E+00 | 6.03E-13 | 2.21E-12 | -2.79E+00 | 2.05E-13 | 5.63E-13 | -3.13E+00 |
| EMILIN2 | 2.02E-08 | 1.05E-07 | -1.48E+00 | 4.26E-05 | 1.84E-04 | -1.10E+00 | 3.06E-06 | 1.30E-05 | -1.28E+00 | 5.79E-04 | 2.21E-03 | 1.03E+00 | 1.93E-04 | 6.87E-04 | -1.37E+00 | 6.56E-04 | 1.85E-03 | -1.30E+00 | 5.78E-08 | 2.12E-07 | -1.44E+00 | 1.66E-05 | 4.56E-05 | -1.25E+00 |
| YIF1B | 2.02E-08 | 1.05E-07 | -1.61E+00 | 7.75E-09 | 3.35E-08 | -1.54E+00 | 3.93E-06 | 1.67E-05 | -1.40E+00 | 1.31E-06 | 5.01E-06 | -1.26E+00 | 2.70E-11 | 9.63E-11 | -1.52E+00 | 1.25E-06 | 3.52E-06 | -1.13E+00 | 1.31E-06 | 4.81E-06 | -1.14E+00 | 2.39E-04 | 6.55E-04 | -1.04E+00 |
| NTPCR | 2.03E-08 | 1.06E-07 | -1.19E+00 | 4.45E-09 | 1.92E-08 | -1.03E+00 | 4.24E-03 | 1.80E-02 | -1.04E+00 | 4.85E-04 | 1.85E-03 | 1.01E+00 | 3.56E-10 | 1.27E-09 | -1.63E+00 | 1.10E-04 | 3.09E-04 | -1.31E+00 | 5.26E-04 | 1.93E-03 | -1.11E+00 | 4.91E-02 | 1.35E-01 | 1.04E+00 |
| CAV1 | 2.03E-08 | 1.06E-07 | -2.02E+00 | 1.76E-03 | 7.60E-03 | -1.22E+00 | 4.36E-06 | 1.85E-05 | -1.67E+00 | 1.88E-01 | 7.20E-01 | 1.32E+00 | 1.28E-01 | 4.55E-01 | -1.42E+00 | 8.10E-02 | 2.28E-01 | -2.95E+00 | 2.14E-01 | 7.86E-01 | -1.03E+00 | 2.44E-01 | 6.69E-01 | 1.12E+00 |
| ERO1A | 2.04E-08 | 1.06E-07 | -1.32E+00 | 3.22E-04 | 1.39E-03 | 1.09E+00 | 2.17E-03 | 9.21E-03 | -1.04E+00 | 4.46E-03 | 1.70E-02 | 1.11E+00 | 9.54E-04 | 3.40E-03 | -1.37E+00 | 1.30E-03 | 3.67E-03 | -1.26E+00 | 4.23E-08 | 1.55E-07 | -1.64E+00 | 4.97E-04 | 1.37E-03 | -1.10E+00 |
| MOAP1 | 2.04E-08 | 1.06E-07 | -1.14E+00 | 9.90E-18 | 4.28E-17 | -1.46E+00 | 4.92E-06 | 2.09E-05 | -1.28E+00 | 1.74E-08 | 6.66E-08 | -1.26E+00 | 2.73E-03 | 9.73E-03 | -1.09E+00 | 2.01E-02 | 5.67E-02 | 1.07E+00 | 2.22E-07 | 8.13E-07 | -1.34E+00 | 3.71E-04 | 1.02E-03 | -1.18E+00 |
| AC254562.1 | 2.04E-08 | 1.06E-07 | -2.26E+00 | 1.65E-05 | 7.14E-05 | -1.38E+00 | 4.03E-03 | 1.71E-02 | -1.50E+00 | 8.13E-02 | 3.11E-01 | 1.05E+00 | 3.18E-03 | 1.14E-02 | -1.95E+00 | 1.41E-01 | 3.97E-01 | -1.00E+00 | 5.79E-02 | 2.12E-01 | -1.14E+00 | 3.06E-01 | 8.40E-01 | 1.18E+00 |
| WDR77 | 2.04E-08 | 1.06E-07 | -1.61E+00 | 5.03E-03 | 2.18E-02 | 1.18E+00 | 7.67E-07 | 3.26E-06 | -1.51E+00 | 1.11E-03 | 4.24E-03 | 1.01E+00 | 2.06E-05 | 7.36E-05 | -1.74E+00 | 2.47E-02 | 6.96E-02 | -1.03E+00 | 6.30E-11 | 2.31E-10 | -1.64E+00 | 4.41E-02 | 1.21E-01 | 1.05E+00 |
| ACOT8 | 2.05E-08 | 1.07E-07 | -1.82E+00 | 2.78E-06 | 1.20E-05 | -1.31E+00 | 5.40E-07 | 2.30E-06 | -1.55E+00 | 4.88E-05 | 1.86E-04 | -1.18E+00 | 3.11E-06 | 1.11E-05 | -1.46E+00 | 9.79E-05 | 2.76E-04 | -1.17E+00 | 7.15E-04 | 2.62E-03 | -1.13E+00 | 2.08E-02 | 5.71E-02 | 1.03E+00 |
| CDC27 | 2.06E-08 | 1.07E-07 | -1.41E+00 | 1.45E-07 | 6.27E-07 | -1.17E+00 | 2.98E-06 | 1.27E-05 | -1.33E+00 | 1.15E-06 | 4.38E-06 | -1.21E+00 | 3.27E-04 | 1.17E-03 | -1.15E+00 | 3.33E-03 | 9.37E-03 | -1.03E+00 | 8.45E-22 | 3.10E-21 | -1.59E+00 | 7.58E-10 | 2.08E-09 | -1.38E+00 |
| LRMP | 2.06E-08 | 1.08E-07 | -1.41E+00 | 2.02E-05 | 8.74E-05 | -1.03E+00 | 1.64E-09 | 6.97E-09 | -1.61E+00 | 2.28E-09 | 8.69E-09 | -1.45E+00 | 3.91E-10 | 1.39E-09 | -1.66E+00 | 2.98E-06 | 8.40E-06 | -1.27E+00 | 1.20E-15 | 4.40E-15 | -1.80E+00 | 2.37E-11 | 6.52E-11 | -1.49E+00 |
| DTWD1 | 2.07E-08 | 1.08E-07 | -1.34E+00 | 2.43E-02 | 1.05E-01 | 1.27E+00 | 2.38E-03 | 1.01E-02 | -1.04E+00 | 2.36E-01 | 9.01E-01 | 1.44E+00 | 2.38E-06 | 8.47E-06 | -1.17E+00 | 2.12E-01 | 5.97E-01 | 1.41E+00 | 3.11E-03 | 1.14E-02 | -1.05E+00 | 1.43E-02 | 3.93E-02 | 1.03E+00 |
| PSMA4 | 2.08E-08 | 1.09E-07 | -1.13E+00 | 1.16E-07 | 4.99E-07 | 1.00E+00 | 5.68E-02 | 2.42E-01 | 1.00E+00 | 9.54E-02 | 3.64E-01 | 1.05E+00 | 2.41E-09 | 8.59E-09 | -1.30E+00 | 5.49E-03 | 1.55E-02 | -1.03E+00 | 2.98E-03 | 1.09E-02 | -1.12E+00 | 3.24E-01 | 8.90E-01 | 1.08E+00 |
| SIGIRR | 2.09E-08 | 1.09E-07 | -1.19E+00 | 6.91E-05 | 2.99E-04 | 1.06E+00 | 2.00E-08 | 8.48E-08 | -1.37E+00 | 4.94E-06 | 1.89E-05 | -1.07E+00 | 3.31E-06 | 1.18E-05 | -1.26E+00 | 8.93E-04 | 2.52E-03 | 1.00E+00 | 3.90E-02 | 1.43E-01 | -1.03E+00 | 7.21E-03 | 1.98E-02 | -1.11E+00 |
| OSBPL7 | 2.09E-08 | 1.09E-07 | -1.90E+00 | 3.89E-04 | 1.68E-03 | -1.02E+00 | 2.22E-04 | 9.42E-04 | -1.50E+00 | 1.68E-02 | 6.43E-02 | -1.03E+00 | 1.08E-02 | 3.83E-02 | -1.08E+00 | 1.73E-02 | 4.88E-02 | -1.21E+00 | 6.00E-04 | 2.20E-03 | -1.37E+00 | 3.76E-02 | 1.03E-01 | -1.03E+00 |
| RNASEH2C | 2.09E-08 | 1.09E-07 | -1.09E+00 | 1.57E-08 | 6.78E-08 | -1.00E+00 | 1.26E-01 | 5.33E-01 | 1.17E+00 | 1.01E-01 | 3.88E-01 | 1.04E+00 | 9.22E-03 | 3.29E-02 | 1.09E+00 | 4.64E-02 | 1.31E-01 | 1.16E+00 | 5.09E-02 | 1.87E-01 | 1.31E+00 | 7.36E-03 | 2.02E-02 | 1.41E+00 |
| MDM4 | 2.09E-08 | 1.09E-07 | -1.14E+00 | 2.88E-06 | 1.24E-05 | -1.01E+00 | 3.69E-04 | 1.57E-03 | -1.25E+00 | 1.73E-03 | 6.59E-03 | -1.05E+00 | 2.85E-04 | 1.02E-03 | -1.19E+00 | 1.71E-03 | 4.82E-03 | -1.12E+00 | 3.04E-07 | 1.12E-06 | -1.34E+00 | 8.22E-04 | 2.26E-03 | -1.16E+00 |
| ARL2 | 2.10E-08 | 1.09E-07 | -1.08E+00 | 1.29E-05 | 5.55E-05 | 1.10E+00 | 2.31E-01 | 9.79E-01 | 1.12E+00 | 1.87E-01 | 7.14E-01 | 1.15E+00 | 2.54E-05 | 9.06E-05 | -1.08E+00 | 1.13E-01 | 3.20E-01 | 1.16E+00 | 2.58E-01 | 9.46E-01 | 1.21E+00 | 7.39E-02 | 2.03E-01 | 1.36E+00 |
| MCAT | 2.13E-08 | 1.11E-07 | -1.79E+00 | 2.70E-04 | 1.17E-03 | -1.09E+00 | 7.61E-05 | 3.23E-04 | -1.15E+00 | 4.01E-04 | 1.53E-03 | 1.01E+00 | 6.98E-07 | 2.49E-06 | -1.64E+00 | 2.14E-04 | 6.05E-04 | -1.19E+00 | 4.99E-07 | 1.83E-06 | -1.37E+00 | 3.77E-02 | 1.04E-01 | 1.05E+00 |
| C20orf27 | 2.13E-08 | 1.11E-07 | -1.16E+00 | 6.51E-08 | 2.81E-07 | -1.01E+00 | 2.34E-03 | 9.93E-03 | -1.04E+00 | 5.71E-04 | 2.18E-03 | 1.05E+00 | 7.53E-05 | 2.69E-04 | -1.16E+00 | 5.65E-04 | 1.59E-03 | -1.07E+00 | 3.21E-04 | 1.18E-03 | -1.09E+00 | 1.62E-01 | 4.44E-01 | 1.12E+00 |
| ANKRD36C | 2.14E-08 | 1.12E-07 | -1.16E+00 | 5.97E-06 | 2.58E-05 | 1.06E+00 | 4.65E-05 | 1.98E-04 | -1.18E+00 | 7.77E-05 | 2.97E-04 | -1.01E+00 | 5.65E-04 | 2.01E-03 | -1.05E+00 | 6.13E-04 | 1.73E-03 | -1.04E+00 | 1.47E-04 | 5.40E-04 | -1.19E+00 | 6.13E-04 | 1.68E-03 | -1.14E+00 |
| BLVRB | 2.15E-08 | 1.12E-07 | -1.23E+00 | 1.44E-05 | 6.20E-05 | 1.05E+00 | 1.04E-01 | 4.44E-01 | 1.07E+00 | 3.94E-02 | 1.51E-01 | 1.15E+00 | 5.22E-03 | 1.86E-02 | 1.22E+00 | 1.69E-01 | 4.76E-01 | 1.96E+00 | 1.93E-01 | 7.09E-01 | 1.16E+00 | 3.40E-01 | 9.35E-01 | 1.11E+00 |
| NDUFAF7 | 2.15E-08 | 1.12E-07 | -1.37E+00 | 1.77E-06 | 7.67E-06 | -1.10E+00 | 1.07E-06 | 4.55E-06 | -1.50E+00 | 5.54E-05 | 2.11E-04 | -1.14E+00 | 4.20E-06 | 1.50E-05 | -1.39E+00 | 1.50E-03 | 4.23E-03 | -1.05E+00 | 8.02E-03 | 2.95E-02 | -1.07E+00 | 4.71E-02 | 1.29E-01 | 1.01E+00 |
| DNMT1 | 2.15E-08 | 1.12E-07 | -1.21E+00 | 1.83E-04 | 7.91E-04 | 1.11E+00 | 1.15E-01 | 4.89E-01 | 1.08E+00 | 1.76E-01 | 6.71E-01 | 1.31E+00 | 1.84E-06 | 6.57E-06 | -1.27E+00 | 1.36E-03 | 3.85E-03 | 1.04E+00 | 5.19E-03 | 1.91E-02 | -1.02E+00 | 7.94E-02 | 2.18E-01 | 1.33E+00 |
| SUMF1 | 2.18E-08 | 1.13E-07 | -1.70E+00 | 2.19E-08 | 9.48E-08 | -1.54E+00 | 8.97E-04 | 3.81E-03 | -1.24E+00 | 2.63E-03 | 1.00E-02 | -1.03E+00 | 1.06E-02 | 3.77E-02 | -1.14E+00 | 6.43E-02 | 1.81E-01 | 1.00E+00 | 3.63E-03 | 1.33E-02 | -1.16E+00 | 2.88E-01 | 7.90E-01 | 1.20E+00 |
| MCTP2 | 2.18E-08 | 1.14E-07 | -1.19E+00 | 1.01E-08 | 4.35E-08 | -1.14E+00 | 2.64E-02 | 1.12E-01 | 1.05E+00 | 5.15E-06 | 1.97E-05 | -1.11E+00 | 1.71E-01 | 6.11E-01 | 1.42E+00 | 6.14E-03 | 1.73E-02 | 1.04E+00 | 9.36E-03 | 3.43E-02 | -1.01E+00 | 1.66E-02 | 4.56E-02 | -1.04E+00 |
| POLR3E | 2.18E-08 | 1.14E-07 | -1.48E+00 | 3.09E-03 | 1.33E-02 | 1.12E+00 | 1.19E-05 | 5.08E-05 | -1.51E+00 | 2.98E-02 | 1.14E-01 | 1.03E+00 | 1.33E-05 | 4.72E-05 | -1.17E+00 | 2.53E-03 | 7.13E-03 | -1.07E+00 | 7.70E-13 | 2.83E-12 | -1.61E+00 | 7.38E-04 | 2.03E-03 | -1.07E+00 |
| SNHG14 | 2.20E-08 | 1.15E-07 | -1.42E+00 | 3.05E-06 | 1.32E-05 | -1.03E+00 | 2.46E-06 | 1.04E-05 | -1.54E+00 | 8.55E-06 | 3.26E-05 | -1.34E+00 | 9.65E-07 | 3.44E-06 | -1.34E+00 | 1.22E-04 | 3.45E-04 | -1.20E+00 | 1.28E-04 | 4.68E-04 | -1.22E+00 | 5.56E-05 | 1.53E-04 | -1.21E+00 |
| KIF5A | 2.21E-08 | 1.15E-07 | -2.24E+00 | 8.79E-03 | 3.80E-02 | 1.03E+00 | 3.25E-04 | 1.38E-03 | -2.12E+00 | 3.21E-02 | 1.23E-01 | -1.04E+00 | 6.05E-04 | 2.16E-03 | -1.73E+00 | 3.42E-02 | 9.64E-02 | -1.12E+00 | 1.99E-02 | 7.31E-02 | -1.52E+00 | 3.55E-01 | 9.74E-01 | 1.16E+00 |
| TKFC | 2.25E-08 | 1.17E-07 | -1.43E+00 | 1.79E-08 | 7.73E-08 | -1.30E+00 | 5.54E-02 | 2.35E-01 | -1.01E+00 | 1.56E-02 | 5.96E-02 | 1.00E+00 | 4.36E-04 | 1.55E-03 | -1.19E+00 | 1.01E-02 | 2.85E-02 | 1.02E+00 | 4.03E-03 | 1.48E-02 | -1.06E+00 | 6.70E-04 | 1.84E-03 | -1.04E+00 |
| UBE3B | 2.26E-08 | 1.18E-07 | -1.79E+00 | 4.10E-06 | 1.77E-05 | -1.27E+00 | 1.09E-04 | 4.63E-04 | -1.48E+00 | 2.18E-02 | 8.32E-02 | 1.05E+00 | 5.17E-03 | 1.84E-02 | -1.09E+00 | 8.43E-02 | 2.38E-01 | 1.22E+00 | 1.04E-06 | 3.80E-06 | -1.43E+00 | 7.92E-03 | 2.18E-02 | -1.02E+00 |
| ATP5HP4 | 2.28E-08 | 1.19E-07 | -1.15E+00 | 2.57E-16 | 1.11E-15 | -1.27E+00 | 4.21E-04 | 1.79E-03 | -1.12E+00 | 1.51E-10 | 5.78E-10 | -1.34E+00 | 3.02E-06 | 1.08E-05 | -1.15E+00 | 5.25E-10 | 1.48E-09 | -1.37E+00 | 5.76E-05 | 2.11E-04 | -1.16E+00 | 9.89E-06 | 2.72E-05 | -1.24E+00 |
| RAP1B | 2.29E-08 | 1.19E-07 | -1.03E+00 | 2.62E-13 | 1.13E-12 | -1.09E+00 | 1.35E-03 | 5.75E-03 | 1.03E+00 | 3.43E-05 | 1.31E-04 | -1.07E+00 | 1.62E-02 | 5.76E-02 | 1.01E+00 | 1.81E-02 | 5.11E-02 | 1.00E+00 | 4.09E-02 | 1.50E-01 | 1.02E+00 | 1.09E-04 | 2.99E-04 | -1.40E+00 |
| POLR1H | 2.29E-08 | 1.19E-07 | -1.85E+00 | 6.30E-06 | 2.72E-05 | -1.23E+00 | 1.86E-02 | 7.89E-02 | -1.13E+00 | 1.78E-01 | 6.79E-01 | 1.45E+00 | 5.44E-06 | 1.94E-05 | -1.97E+00 | 1.56E-02 | 4.39E-02 | -1.02E+00 | 3.30E-06 | 1.21E-05 | -1.22E+00 | 1.87E-01 | 5.14E-01 | 1.30E+00 |
| AC016734.1 | 2.30E-08 | 1.20E-07 | -1.29E+00 | 2.35E-04 | 1.01E-03 | 1.12E+00 | 6.36E-04 | 2.70E-03 | -1.13E+00 | 1.82E-03 | 6.96E-03 | 1.06E+00 | 4.42E-11 | 1.58E-10 | -1.71E+00 | 1.03E-03 | 2.91E-03 | 1.09E+00 | 7.94E-04 | 2.91E-03 | -1.11E+00 | 3.33E-01 | 9.14E-01 | 1.22E+00 |
| C21orf58 | 2.30E-08 | 1.20E-07 | -1.50E+00 | 5.35E-05 | 2.31E-04 | -1.03E+00 | 7.72E-02 | 3.28E-01 | 1.18E+00 | 1.35E-01 | 5.17E-01 | 1.35E+00 | 1.77E-04 | 6.31E-04 | -1.41E+00 | 7.30E-03 | 2.06E-02 | -1.10E+00 | 3.43E-02 | 1.26E-01 | 1.30E+00 | 5.98E-02 | 1.64E-01 | 1.12E+00 |
| RHPN1 | 2.31E-08 | 1.20E-07 | -1.23E+00 | 6.64E-08 | 2.87E-07 | -1.09E+00 | 1.90E-04 | 8.07E-04 | -1.05E+00 | 2.83E-05 | 1.08E-04 | -1.05E+00 | 2.47E-01 | 8.79E-01 | 1.02E+00 | 3.11E-01 | 8.77E-01 | 1.08E+00 | 2.36E-01 | 8.65E-01 | -1.09E+00 | 2.51E-01 | 6.88E-01 | -1.28E+00 |
| DLGAP4 | 2.32E-08 | 1.21E-07 | -1.29E+00 | 2.31E-04 | 1.00E-03 | 1.10E+00 | 5.54E-08 | 2.35E-07 | -1.58E+00 | 2.94E-08 | 1.12E-07 | -1.36E+00 | 3.63E-05 | 1.29E-04 | -1.03E+00 | 1.74E-05 | 4.89E-05 | -1.15E+00 | 1.30E-03 | 4.79E-03 | -1.02E+00 | 1.91E-02 | 5.26E-02 | 1.12E+00 |
| BCAS4 | 2.34E-08 | 1.22E-07 | -1.12E+00 | 3.59E-08 | 1.55E-07 | 1.04E+00 | 4.37E-06 | 1.86E-05 | -1.16E+00 | 1.84E-06 | 7.01E-06 | -1.05E+00 | 2.43E-05 | 8.67E-05 | -1.15E+00 | 7.77E-05 | 2.19E-04 | -1.04E+00 | 9.94E-02 | 3.65E-01 | -1.72E+00 | 2.28E-01 | 6.27E-01 | -1.49E+00 |
| ERAL1 | 2.35E-08 | 1.22E-07 | -1.54E+00 | 3.79E-05 | 1.64E-04 | -1.08E+00 | 1.74E-10 | 7.38E-10 | -1.47E+00 | 1.41E-07 | 5.38E-07 | -1.12E+00 | 4.73E-09 | 1.69E-08 | -1.73E+00 | 1.92E-02 | 5.42E-02 | 1.14E+00 | 3.04E-21 | 1.11E-20 | -1.83E+00 | 2.87E-05 | 7.89E-05 | -1.10E+00 |
| BX322639.1 | 2.35E-08 | 1.22E-07 | -1.70E+00 | 2.97E-07 | 1.28E-06 | -1.35E+00 | 9.20E-04 | 3.91E-03 | -1.39E+00 | 2.29E-04 | 8.74E-04 | -1.32E+00 | 5.79E-03 | 2.06E-02 | -1.31E+00 | 3.55E-02 | 1.00E-01 | -1.01E+00 | 4.69E-09 | 1.72E-08 | -1.76E+00 | 6.69E-05 | 1.84E-04 | -1.24E+00 |
| G2E3 | 2.35E-08 | 1.23E-07 | -1.31E+00 | 4.37E-07 | 1.89E-06 | -1.16E+00 | 4.75E-05 | 2.02E-04 | -1.39E+00 | 6.03E-04 | 2.30E-03 | -1.04E+00 | 3.38E-04 | 1.20E-03 | -1.10E+00 | 2.41E-03 | 6.79E-03 | -1.15E+00 | 5.57E-07 | 2.04E-06 | -1.41E+00 | 1.46E-05 | 4.00E-05 | -1.32E+00 |
| C1GALT1C1 | 2.37E-08 | 1.24E-07 | -2.28E+00 | 1.17E-02 | 5.07E-02 | 1.15E+00 | 3.17E-11 | 1.34E-10 | -1.90E+00 | 5.49E-06 | 2.10E-05 | -1.23E+00 | 6.20E-13 | 2.21E-12 | -2.68E+00 | 3.72E-07 | 1.05E-06 | -1.44E+00 | 7.22E-09 | 2.65E-08 | -1.66E+00 | 8.56E-02 | 2.35E-01 | 1.09E+00 |
| SIVA1 | 2.38E-08 | 1.24E-07 | -1.04E+00 | 5.94E-13 | 2.57E-12 | -1.07E+00 | 1.25E-02 | 5.31E-02 | 1.31E+00 | 1.32E-01 | 5.03E-01 | 1.15E+00 | 1.54E-05 | 5.49E-05 | -1.05E+00 | 2.64E-04 | 7.43E-04 | 1.02E+00 | 1.92E-01 | 7.05E-01 | 1.18E+00 | 2.55E-01 | 7.01E-01 | 1.26E+00 |
| TMEM69 | 2.40E-08 | 1.25E-07 | -1.41E+00 | 2.09E-04 | 9.01E-04 | 1.03E+00 | 3.34E-11 | 1.42E-10 | -1.68E+00 | 1.12E-08 | 4.27E-08 | -1.29E+00 | 3.07E-09 | 1.10E-08 | -1.83E+00 | 9.43E-06 | 2.66E-05 | -1.30E+00 | 5.68E-16 | 2.08E-15 | -1.99E+00 | 3.19E-07 | 8.77E-07 | -1.40E+00 |
| FKBP1C | 2.42E-08 | 1.26E-07 | -1.22E+00 | 6.97E-05 | 3.01E-04 | 1.87E+00 | 2.26E-01 | 9.62E-01 | 1.24E+00 | 2.38E-05 | 9.11E-05 | 2.14E+00 | 9.60E-08 | 3.42E-07 | -1.26E+00 | 1.91E-01 | 5.38E-01 | 1.71E+00 | 1.20E-01 | 4.41E-01 | 1.14E+00 | 4.64E-06 | 1.28E-05 | 1.96E+00 |

| | | | | | | | | | | | | | | | | | | | | | | | | |
|---|---|---|---|---|---|---|---|---|---|---|---|---|---|---|---|---|---|---|---|---|---|---|---|---|
| CLPX | 2.42E-08 | 1.26E-07 | -1.21E+00 | 1.83E-03 | 7.91E-03 | 1.22E+00 | 2.04E-02 | 8.67E-02 | -1.02E+00 | 1.45E-01 | 5.53E-01 | 1.25E+00 | 1.13E-06 | 4.01E-06 | -1.37E+00 | 4.87E-03 | 1.37E-02 | 1.01E+00 | 6.71E-02 | 2.46E-01 | 1.12E+00 | 3.48E-01 | 9.56E-01 | 1.26E+00 |
| MSH6 | 2.42E-08 | 1.26E-07 | -1.37E+00 | 7.43E-05 | 3.21E-04 | 1.00E+00 | 2.87E-04 | 1.22E-03 | -1.10E+00 | 4.16E-02 | 1.59E-01 | 1.21E+00 | 4.13E-05 | 1.47E-04 | -1.32E+00 | 4.04E-03 | 1.14E-02 | -1.08E+00 | 1.19E-07 | 4.36E-07 | -1.28E+00 | 3.02E-02 | 8.30E-02 | 1.09E+00 |
| YIPF5 | 2.42E-08 | 1.26E-07 | -1.23E+00 | 3.64E-06 | 1.57E-05 | 1.02E+00 | 1.46E-07 | 6.20E-07 | -1.34E+00 | 2.74E-07 | 1.05E-06 | -1.18E+00 | 2.21E-06 | 7.89E-06 | -1.29E+00 | 4.28E-03 | 1.21E-02 | 1.03E+00 | 5.07E-08 | 1.86E-07 | -1.45E+00 | 4.97E-04 | 1.36E-03 | -1.13E+00 |
| PPIAP22 | 2.44E-08 | 1.27E-07 | -1.22E+00 | 6.48E-03 | 2.80E-02 | 1.27E+00 | 1.75E-07 | 7.43E-07 | -1.30E+00 | 1.58E-02 | 6.02E-02 | 1.17E+00 | 1.94E-09 | 6.91E-09 | -1.43E+00 | 4.94E-02 | 1.39E-01 | 1.19E+00 | 1.94E-11 | 7.12E-11 | -1.48E+00 | 2.49E-01 | 6.84E-01 | 1.22E+00 |
| MYH10 | 2.45E-08 | 1.28E-07 | -1.36E+00 | 2.17E-04 | 9.36E-04 | 1.08E+00 | 6.36E-02 | 2.70E-01 | -1.04E+00 | 2.54E-01 | 9.71E-01 | 1.37E+00 | 6.33E-09 | 2.26E-08 | -1.48E+00 | 2.20E-03 | 6.21E-03 | -1.02E+00 | 1.63E-08 | 5.99E-08 | -1.34E+00 | 1.86E-03 | 5.10E-03 | -1.05E+00 |
| RMI1 | 2.45E-08 | 1.28E-07 | -2.47E+00 | 5.00E-04 | 2.16E-03 | -1.14E+00 | 5.44E-08 | 2.31E-07 | -1.67E+00 | 1.14E-03 | 4.34E-03 | 1.03E+00 | 9.19E-07 | 3.28E-06 | -2.38E+00 | 5.34E-04 | 1.51E-03 | -1.38E+00 | 2.89E-10 | 1.06E-09 | -1.82E+00 | 9.94E-05 | 2.73E-04 | -1.14E+00 |
| UTP14A | 2.46E-08 | 1.28E-07 | -1.38E+00 | 4.67E-06 | 2.02E-05 | -1.10E+00 | 1.02E-05 | 4.34E-05 | -1.22E+00 | 1.55E-03 | 5.90E-03 | 1.12E+00 | 8.49E-07 | 3.03E-06 | -1.38E+00 | 1.93E-03 | 5.45E-03 | -1.01E+00 | 5.95E-05 | 2.18E-04 | -1.15E+00 | 2.05E-02 | 5.62E-02 | 1.10E+00 |
| AC116917.1 | 2.46E-08 | 1.28E-07 | -1.40E+00 | 2.79E-03 | 1.20E-02 | 1.18E+00 | 5.12E-03 | 2.18E-02 | -1.20E+00 | 1.09E-01 | 4.15E-01 | 1.25E+00 | 2.41E-01 | 8.61E-01 | -1.04E+00 | 3.37E-01 | 9.50E-01 | 1.18E+00 | 9.72E-02 | 3.57E-01 | 1.07E+00 | 1.67E-01 | 4.59E-01 | -1.18E+00 |
| UBAP2 | 2.49E-08 | 1.30E-07 | -1.24E+00 | 8.11E-04 | 3.51E-03 | 1.11E+00 | 8.02E-04 | 3.41E-03 | -1.14E+00 | 4.90E-03 | 1.87E-02 | 1.03E+00 | 8.08E-05 | 2.88E-04 | -1.20E+00 | 1.53E-02 | 4.31E-02 | 1.02E+00 | 1.22E-02 | 4.49E-02 | -1.05E+00 | 6.07E-02 | 1.67E-01 | 1.10E+00 |
| DDX1 | 2.50E-08 | 1.30E-07 | -1.20E+00 | 1.11E-03 | 4.80E-03 | 1.20E+00 | 2.96E-03 | 1.26E-02 | 1.00E+00 | 4.76E-02 | 1.82E-01 | 1.24E+00 | 3.09E-04 | 1.10E-03 | -1.13E+00 | 8.07E-02 | 2.27E-01 | 1.18E+00 | 2.21E-02 | 8.10E-02 | 1.05E+00 | 8.21E-02 | 2.25E-01 | 1.36E+00 |
| TMEM30A | 2.51E-08 | 1.31E-07 | -1.52E+00 | 2.77E-06 | 1.20E-05 | -1.23E+00 | 2.59E-04 | 1.10E-03 | -1.25E+00 | 3.52E-03 | 1.34E-02 | -1.03E+00 | 3.86E-03 | 1.38E-02 | -1.19E+00 | 2.18E-03 | 6.14E-03 | -1.16E+00 | 5.44E-05 | 2.00E-04 | -1.26E+00 | 4.08E-04 | 1.12E-03 | -1.16E+00 |
| C14orf39 | 2.52E-08 | 1.31E-07 | -2.03E+00 | 1.15E-08 | 4.99E-08 | -2.17E+00 | 7.58E-02 | 3.22E-01 | -1.17E+00 | 1.34E-02 | 5.12E-02 | -1.59E+00 | 1.02E-01 | 3.63E-01 | -1.27E+00 | 2.92E-01 | 8.24E-01 | -1.02E+00 | 9.10E-04 | 3.34E-03 | -1.41E+00 | 1.59E-05 | 4.36E-05 | -2.09E+00 |
| FAU | 2.52E-08 | 1.32E-07 | -1.09E+00 | 1.49E-57 | 6.44E-57 | -1.28E+00 | 1.60E-04 | 6.81E-04 | -1.07E+00 | 1.47E-35 | 5.62E-35 | -1.26E+00 | 1.34E-03 | 4.79E-03 | -1.06E+00 | 1.33E-35 | 3.76E-35 | -1.30E+00 | 2.67E-04 | 9.78E-04 | -1.05E+00 | 3.61E-32 | 9.92E-32 | -1.24E+00 |
| DESI2 | 2.54E-08 | 1.32E-07 | -1.27E+00 | 1.83E-10 | 7.92E-10 | -1.29E+00 | 2.52E-04 | 1.07E-03 | -1.15E+00 | 5.00E-06 | 1.91E-05 | -1.12E+00 | 1.03E-02 | 3.67E-02 | 1.06E+00 | 1.22E-03 | 3.43E-03 | -1.09E+00 | 1.53E-03 | 5.61E-03 | -1.13E+00 | 2.13E-02 | 5.85E-02 | -1.01E+00 |
| CIAO1 | 2.55E-08 | 1.33E-07 | -1.28E+00 | 9.69E-06 | 4.19E-05 | -1.04E+00 | 1.92E-07 | 8.17E-07 | -1.39E+00 | 2.44E-05 | 9.31E-05 | -1.08E+00 | 2.41E-08 | 8.58E-08 | -1.40E+00 | 4.04E-05 | 1.14E-04 | -1.17E+00 | 4.93E-08 | 1.81E-07 | -1.27E+00 | 1.29E-02 | 3.53E-02 | 1.03E+00 |
| HHLA3 | 2.55E-08 | 1.33E-07 | -1.57E+00 | 3.99E-09 | 1.72E-08 | -1.52E+00 | 9.29E-02 | 3.95E-01 | 1.00E+00 | 2.86E-02 | 1.09E-01 | -1.11E+00 | 1.07E-04 | 3.83E-04 | -1.70E+00 | 1.65E-04 | 4.66E-04 | -1.51E+00 | 9.17E-07 | 3.37E-06 | -1.37E+00 | 2.75E-06 | 7.55E-06 | -1.38E+00 |
| DAZAP2 | 2.56E-08 | 1.34E-07 | -1.19E+00 | 2.39E-05 | 1.03E-04 | 1.03E+00 | 1.66E-04 | 7.03E-04 | -1.18E+00 | 4.06E-02 | 1.55E-01 | 1.08E+00 | 7.11E-05 | 2.53E-04 | -1.12E+00 | 9.42E-03 | 2.66E-02 | 1.03E+00 | 1.72E-04 | 6.32E-04 | -1.13E+00 | 1.93E-02 | 5.30E-02 | -1.01E+00 |
| FAM133A | 2.57E-08 | 1.34E-07 | -2.12E+00 | 9.79E-07 | 4.23E-06 | -1.69E+00 | 1.14E-01 | 4.83E-01 | -1.30E+00 | 6.52E-02 | 2.49E-01 | -1.26E+00 | 2.30E-01 | 8.19E-01 | 1.36E+00 | 2.94E-01 | 8.30E-01 | 1.92E+00 | 2.69E-01 | 9.88E-01 | -1.01E+00 | 3.62E-01 | 9.93E-01 | -1.01E+00 |
| GALK1 | 2.57E-08 | 1.34E-07 | -1.27E+00 | 5.59E-06 | 2.42E-05 | 1.00E+00 | 2.65E-03 | 1.12E-02 | 1.45E+00 | 3.89E-02 | 1.49E-01 | 1.60E+00 | 9.94E-04 | 3.55E-03 | -1.27E+00 | 5.67E-03 | 1.60E-02 | -1.10E+00 | 2.63E-01 | 9.64E-01 | 1.28E+00 | 3.12E-01 | 8.57E-01 | 1.51E+00 |
| ERG | 2.59E-08 | 1.35E-07 | -1.20E+00 | 1.63E-03 | 7.04E-03 | 1.14E+00 | 7.02E-10 | 2.98E-09 | -1.45E+00 | 4.45E-05 | 1.70E-04 | -1.03E+00 | 1.90E-04 | 6.77E-04 | -1.05E+00 | 3.13E-02 | 8.83E-02 | 1.16E+00 | 2.99E-04 | 1.10E-03 | -1.20E+00 | 4.50E-03 | 1.24E-02 | -1.13E+00 |
| THOC5 | 2.61E-08 | 1.36E-07 | -1.46E+00 | 5.45E-05 | 2.36E-04 | -1.05E+00 | 5.30E-06 | 2.25E-05 | -1.31E+00 | 4.94E-07 | 1.89E-06 | -1.17E+00 | 2.99E-06 | 1.07E-05 | -1.35E+00 | 8.81E-05 | 2.48E-04 | -1.11E+00 | 6.78E-05 | 2.49E-04 | -1.20E+00 | 2.26E-02 | 6.21E-02 | 1.04E+00 |
| MINDY2 | 2.62E-08 | 1.37E-07 | -1.32E+00 | 3.05E-05 | 1.32E-04 | 1.05E+00 | 1.77E-06 | 7.53E-06 | -1.51E+00 | 2.88E-04 | 1.10E-03 | -1.19E+00 | 1.96E-03 | 6.98E-03 | -1.10E+00 | 3.97E-02 | 1.12E-01 | 1.14E+00 | 1.47E-06 | 5.40E-06 | -1.12E+00 | 1.16E-04 | 3.19E-04 | -1.23E+00 |
| ABHD3 | 2.67E-08 | 1.39E-07 | -1.54E+00 | 1.21E-06 | 5.23E-06 | -1.20E+00 | 5.76E-02 | 2.45E-01 | -1.01E+00 | 8.12E-02 | 3.10E-01 | 1.12E+00 | 7.88E-04 | 2.81E-03 | -1.33E+00 | 1.02E-03 | 2.87E-03 | -1.28E+00 | 2.99E-05 | 1.10E-04 | -1.46E+00 | 3.88E-03 | 1.07E-02 | -1.13E+00 |
| SMCHD1 | 2.67E-08 | 1.39E-07 | -1.10E+00 | 3.63E-11 | 1.57E-10 | -1.14E+00 | 5.22E-06 | 2.22E-05 | -1.20E+00 | 1.25E-07 | 4.77E-07 | -1.13E+00 | 1.45E-06 | 5.16E-06 | -1.15E+00 | 6.14E-12 | 1.73E-11 | -1.42E+00 | 1.67E-06 | 6.14E-06 | -1.23E+00 | 9.54E-07 | 2.62E-06 | -1.28E+00 |
| GLRX3 | 2.69E-08 | 1.40E-07 | -1.26E+00 | 1.55E-07 | 6.70E-07 | -1.09E+00 | 1.14E-05 | 4.85E-05 | -1.17E+00 | 6.27E-05 | 2.40E-04 | -1.00E+00 | 3.43E-06 | 1.22E-05 | -1.25E+00 | 1.17E-02 | 3.31E-02 | 1.15E+00 | 5.08E-06 | 1.86E-05 | -1.25E+00 | 5.33E-04 | 1.46E-03 | -1.10E+00 |
| ZBTB1 | 2.70E-08 | 1.40E-07 | -1.18E+00 | 6.61E-18 | 2.86E-17 | -1.47E+00 | 1.17E-07 | 4.98E-07 | -1.43E+00 | 5.17E-11 | 1.98E-10 | -1.53E+00 | 6.78E-04 | 2.42E-03 | -1.09E+00 | 3.68E-08 | 1.04E-07 | -1.51E+00 | 2.50E-10 | 9.19E-10 | -1.57E+00 | 5.29E-10 | 1.45E-09 | -1.60E+00 |
| FAM92A | 2.70E-08 | 1.41E-07 | -1.32E+00 | 3.17E-04 | 1.37E-03 | 1.10E+00 | 3.51E-09 | 1.49E-08 | -1.53E+00 | 2.28E-07 | 8.70E-07 | -1.25E+00 | 4.14E-07 | 1.48E-06 | -1.59E+00 | 1.81E-05 | 5.11E-05 | -1.34E+00 | 5.18E-05 | 1.90E-04 | -1.21E+00 | 5.25E-02 | 1.44E-01 | 1.08E+00 |
| PRMT3 | 2.70E-08 | 1.41E-07 | -1.76E+00 | 8.41E-04 | 3.63E-03 | 1.01E+00 | 4.43E-05 | 1.88E-04 | -1.37E+00 | 5.16E-05 | 1.97E-04 | -1.18E+00 | 7.89E-07 | 2.81E-06 | -1.86E+00 | 2.57E-03 | 7.24E-03 | -1.18E+00 | 4.64E-10 | 1.70E-09 | -1.52E+00 | 6.90E-05 | 1.90E-04 | -1.10E+00 |
| RNF214 | 2.71E-08 | 1.41E-07 | -1.23E+00 | 3.80E-09 | 1.64E-08 | -1.21E+00 | 3.43E-08 | 1.46E-07 | -1.69E+00 | 1.18E-06 | 4.51E-06 | -1.37E+00 | 1.31E-02 | 4.68E-02 | -1.04E+00 | 2.17E-02 | 6.12E-02 | 1.01E+00 | 3.17E-05 | 1.16E-04 | -1.29E+00 | 2.44E-06 | 6.70E-06 | -1.41E+00 |
| TMEM267 | 2.71E-08 | 1.41E-07 | -1.61E+00 | 6.13E-05 | 2.65E-04 | -1.08E+00 | 1.42E-04 | 6.03E-04 | -1.38E+00 | 1.58E-04 | 6.04E-04 | -1.15E+00 | 2.35E-02 | 8.38E-02 | -1.06E+00 | 9.08E-02 | 2.56E-01 | 1.14E+00 | 3.77E-08 | 1.38E-07 | -1.94E+00 | 1.32E-03 | 3.62E-03 | -1.19E+00 |
| BEX4 | 2.71E-08 | 1.41E-07 | -1.16E+00 | 5.81E-04 | 2.51E-03 | 1.11E+00 | 2.41E-05 | 1.02E-04 | -1.26E+00 | 5.65E-03 | 2.16E-02 | 1.04E+00 | 1.43E-09 | 5.09E-09 | -1.43E+00 | 2.37E-02 | 6.68E-02 | -1.02E+00 | 5.42E-03 | 1.99E-02 | -1.13E+00 | 2.65E-01 | 7.29E-01 | 1.06E+00 |
| RNASEH2B | 2.71E-08 | 1.41E-07 | -1.16E+00 | 8.35E-04 | 3.61E-03 | 1.13E+00 | 1.74E-03 | 7.40E-03 | -1.22E+00 | 2.35E-02 | 8.97E-02 | 1.01E+00 | 4.49E-10 | 1.60E-09 | -1.32E+00 | 3.66E-03 | 1.03E-02 | 1.03E+00 | 3.04E-08 | 1.12E-07 | -1.21E+00 | 9.52E-02 | 2.61E-01 | 1.06E+00 |
| PABPC4 | 2.71E-08 | 1.41E-07 | -1.20E+00 | 4.10E-12 | 1.77E-11 | -1.20E+00 | 8.18E-04 | 3.48E-03 | -1.07E+00 | 3.48E-07 | 1.33E-06 | -1.10E+00 | 8.33E-05 | 2.97E-04 | -1.06E+00 | 7.81E-05 | 2.20E-04 | -1.09E+00 | 5.14E-03 | 1.89E-02 | -1.09E+00 | 2.03E-01 | 5.57E-01 | 1.10E+00 |
| TMEM41B | 2.72E-08 | 1.42E-07 | -1.27E+00 | 1.04E-08 | 4.50E-08 | -1.16E+00 | 1.20E-06 | 5.08E-06 | -1.50E+00 | 5.41E-07 | 2.07E-06 | -1.31E+00 | 1.58E-03 | 5.62E-03 | -1.20E+00 | 6.40E-03 | 1.80E-02 | -1.05E+00 | 3.13E-04 | 1.15E-03 | -1.10E+00 | 1.78E-03 | 4.90E-03 | -1.06E+00 |
| C1orf50 | 2.73E-08 | 1.42E-07 | -2.03E+00 | 1.14E-06 | 4.92E-06 | -1.58E+00 | 1.66E-03 | 7.04E-03 | -1.41E+00 | 3.34E-02 | 1.28E-01 | 1.03E+00 | 1.24E-03 | 4.42E-03 | -1.66E+00 | 2.74E-02 | 7.74E-02 | -1.16E+00 | 7.47E-08 | 2.74E-07 | -1.64E+00 | 6.30E-06 | 1.73E-05 | -1.41E+00 |
| ZNF792 | 2.74E-08 | 1.43E-07 | -1.83E+00 | 4.22E-05 | 1.82E-04 | -1.14E+00 | 4.17E-08 | 1.77E-07 | -1.78E+00 | 1.57E-04 | 6.01E-04 | -1.06E+00 | 7.18E-06 | 2.56E-05 | -1.51E+00 | 2.20E-05 | 6.19E-05 | -1.38E+00 | 5.47E-02 | 2.01E-01 | 1.16E+00 | 1.43E-01 | 3.93E-01 | 1.18E+00 |
| GATB | 2.75E-08 | 1.43E-07 | -1.47E+00 | 2.21E-07 | 9.54E-07 | -1.24E+00 | 9.08E-07 | 3.86E-06 | -1.25E+00 | 2.10E-11 | 8.02E-11 | -1.48E+00 | 9.81E-05 | 3.50E-04 | -1.38E+00 | 1.59E-02 | 4.49E-02 | 1.01E+00 | 2.40E-09 | 8.81E-09 | -1.50E+00 | 1.98E-06 | 5.44E-06 | -1.33E+00 |
| SRGAP2C | 2.81E-08 | 1.46E-07 | -1.66E+00 | 1.89E-06 | 8.18E-06 | -1.26E+00 | 3.28E-06 | 1.39E-05 | -1.64E+00 | 2.13E-04 | 8.14E-04 | -1.21E+00 | 3.23E-07 | 1.15E-06 | -1.68E+00 | 3.49E-08 | 9.85E-08 | -2.15E+00 | 7.73E-06 | 2.84E-05 | -1.45E+00 | 4.63E-06 | 1.27E-05 | -1.63E+00 |
| AC073063.1 | 2.83E-08 | 1.48E-07 | -1.42E+00 | 1.54E-06 | 6.66E-06 | -1.07E+00 | 1.85E-08 | 7.85E-08 | -1.45E+00 | 1.90E-06 | 7.26E-06 | -1.10E+00 | 9.05E-09 | 3.23E-08 | -1.54E+00 | 1.06E-06 | 2.99E-06 | -1.29E+00 | 6.28E-11 | 2.31E-10 | -1.50E+00 | 4.18E-04 | 1.15E-03 | -1.08E+00 |
| PPIL1 | 2.83E-08 | 1.48E-07 | -1.90E+00 | 1.04E-04 | 4.51E-04 | -1.17E+00 | 1.17E-10 | 4.99E-10 | -1.53E+00 | 6.49E-05 | 2.48E-04 | 1.03E+00 | 1.84E-12 | 6.57E-12 | -2.52E+00 | 1.15E-04 | 3.25E-04 | -1.09E+00 | 9.32E-17 | 3.42E-16 | -1.51E+00 | 3.40E-04 | 9.34E-04 | 1.04E+00 |
| AC090204.1 | 2.84E-08 | 1.48E-07 | -1.35E+00 | 2.47E-08 | 1.07E-07 | -1.21E+00 | 5.55E-05 | 2.36E-04 | -1.30E+00 | 3.01E-05 | 1.15E-04 | -1.19E+00 | 4.68E-05 | 1.67E-04 | -1.42E+00 | 9.32E-06 | 2.63E-05 | -1.44E+00 | 3.39E-02 | 1.25E-01 | -1.03E+00 | 3.72E-02 | 1.02E-01 | -1.01E+00 |
| TBC1D7 | 2.86E-08 | 1.49E-07 | -1.43E+00 | 3.70E-07 | 1.60E-06 | -1.21E+00 | 3.47E-03 | 1.47E-02 | -1.09E+00 | 1.56E-04 | 5.97E-04 | -1.14E+00 | 2.22E-05 | 7.93E-05 | -1.28E+00 | 8.09E-04 | 2.28E-03 | -1.10E+00 | 9.89E-06 | 3.63E-05 | -1.14E+00 | 1.62E-04 | 4.44E-04 | -1.10E+00 |
| TRAF3IP2 | 2.86E-08 | 1.49E-07 | -1.34E+00 | 1.54E-04 | 6.66E-04 | 1.04E+00 | 9.64E-03 | 4.09E-02 | -1.05E+00 | 1.35E-03 | 5.15E-03 | -1.02E+00 | 8.56E-05 | 3.05E-04 | -1.76E+00 | 6.67E-03 | 1.88E-02 | -1.22E+00 | 4.50E-06 | 1.65E-05 | -1.50E+00 | 8.76E-04 | 2.41E-03 | -1.19E+00 |
| CDC73 | 2.90E-08 | 1.51E-07 | -1.21E+00 | 1.64E-07 | 7.07E-07 | -1.01E+00 | 4.25E-07 | 1.81E-06 | -1.32E+00 | 1.09E-06 | 4.18E-06 | -1.14E+00 | 2.13E-03 | 7.60E-03 | -1.07E+00 | 2.15E-04 | 6.07E-04 | -1.11E+00 | 9.32E-10 | 3.42E-09 | -1.41E+00 | 4.86E-08 | 1.34E-07 | -1.37E+00 |
| GFM1 | 2.91E-08 | 1.52E-07 | -1.46E+00 | 5.07E-05 | 2.19E-04 | -1.06E+00 | 1.73E-05 | 7.33E-05 | -1.30E+00 | 5.78E-03 | 2.21E-02 | 1.10E+00 | 3.36E-05 | 1.20E-04 | -1.31E+00 | 1.61E-02 | 4.54E-02 | 1.07E+00 | 2.44E-05 | 8.96E-05 | -1.30E+00 | 1.39E-02 | 3.81E-02 | -1.05E+00 |
| MRPL48 | 2.92E-08 | 1.52E-07 | -1.34E+00 | 9.35E-06 | 4.04E-05 | -1.01E+00 | 4.70E-03 | 1.99E-02 | -1.08E+00 | 1.64E-02 | 6.28E-02 | 1.10E+00 | 5.62E-06 | 2.00E-05 | -1.24E+00 | 7.55E-03 | 2.13E-02 | 1.14E+00 | 3.46E-03 | 1.27E-02 | -1.05E+00 | 1.98E-01 | 5.43E-01 | 1.14E+00 |
| NYNRIN | 2.92E-08 | 1.52E-07 | -1.61E+00 | 8.76E-03 | 3.79E-02 | 1.09E+00 | 8.40E-07 | 3.57E-06 | -1.78E+00 | 3.91E-04 | 1.49E-03 | -1.17E+00 | 6.79E-02 | 2.42E-01 | -1.15E+00 | 1.92E-01 | 5.40E-01 | 1.08E+00 | 8.34E-02 | 3.06E-01 | -1.14E+00 | 1.57E-01 | 4.30E-01 | -1.00E+00 |
| HPS3 | 2.96E-08 | 1.54E-07 | -1.88E+00 | 6.70E-06 | 2.89E-05 | -1.28E+00 | 2.44E-02 | 1.04E-01 | -1.10E+00 | 1.58E-01 | 6.04E-01 | 1.19E+00 | 1.33E-03 | 4.74E-03 | -1.71E+00 | 1.90E-03 | 5.36E-03 | -1.63E+00 | 1.56E-03 | 5.71E-03 | -1.47E+00 | 3.01E-02 | 8.28E-02 | -1.18E+00 |
| ZNF395 | 2.97E-08 | 1.55E-07 | -1.27E+00 | 8.58E-04 | 3.71E-03 | 1.15E+00 | 1.06E-06 | 4.51E-06 | -1.34E+00 | 1.19E-03 | 4.55E-03 | 1.05E+00 | 4.39E-03 | 1.57E-02 | -1.13E+00 | 8.59E-02 | 2.42E-01 | 1.10E+00 | 5.31E-07 | 1.95E-06 | -1.46E+00 | 1.35E-03 | 3.71E-03 | -1.17E+00 |
| GPATCH2L | 2.97E-08 | 1.55E-07 | -1.13E+00 | 1.20E-08 | 5.18E-08 | -1.03E+00 | 4.48E-05 | 1.90E-04 | -1.27E+00 | 1.38E-06 | 5.26E-06 | -1.24E+00 | 6.30E-04 | 2.25E-03 | -1.08E+00 | 7.76E-04 | 2.19E-03 | -1.12E+00 | 3.04E-05 | 1.11E-04 | -1.19E+00 | 8.57E-05 | 2.35E-04 | -1.20E+00 |
| ACTR6 | 2.97E-08 | 1.55E-07 | -1.30E+00 | 1.91E-04 | 8.27E-04 | 1.09E+00 | 4.38E-06 | 1.86E-05 | -1.32E+00 | 8.57E-05 | 3.27E-04 | -1.10E+00 | 7.45E-04 | 2.66E-03 | -1.04E+00 | 3.39E-02 | 9.57E-02 | 1.15E+00 | 1.11E-04 | 4.07E-04 | -1.16E+00 | 4.90E-03 | 1.35E-02 | -1.03E+00 |
| AL354702.1 | 2.97E-08 | 1.55E-07 | -2.06E+00 | 7.40E-10 | 3.20E-09 | -2.38E+00 | 1.19E-02 | 5.07E-02 | -1.18E+00 | 1.48E-03 | 5.67E-03 | -1.20E+00 | 1.18E-07 | 4.22E-07 | -2.78E+00 | 6.48E-07 | 1.83E-06 | -2.80E+00 | 6.12E-18 | 2.24E-17 | -2.36E+00 | 5.22E-14 | 1.43E-13 | -1.91E+00 |
| RENBP | 2.98E-08 | 1.55E-07 | -1.75E+00 | 2.03E-09 | 8.77E-09 | -1.84E+00 | 1.83E-03 | 7.77E-03 | -1.27E+00 | 7.17E-05 | 2.74E-04 | -1.51E+00 | 3.42E-06 | 1.22E-05 | -1.84E+00 | 6.87E-05 | 1.94E-04 | -1.53E+00 | 1.41E-07 | 5.16E-07 | -1.36E+00 | 4.56E-12 | 1.25E-11 | -2.04E+00 |
| PDHX | 3.00E-08 | 1.56E-07 | -1.52E+00 | 3.46E-06 | 1.49E-05 | -1.17E+00 | 6.93E-05 | 2.94E-04 | -1.24E+00 | 1.33E-03 | 5.09E-03 | 1.04E+00 | 9.34E-06 | 3.33E-05 | -1.23E+00 | 9.09E-02 | 2.56E-01 | 1.17E+00 | 7.83E-11 | 2.87E-10 | -1.52E+00 | 3.11E-08 | 8.53E-08 | -1.35E+00 |
| HACD1 | 3.01E-08 | 1.57E-07 | -1.23E+00 | 1.47E-05 | 6.34E-05 | -1.02E+00 | 7.57E-04 | 3.22E-03 | -1.11E+00 | 2.67E-02 | 1.02E-01 | 1.18E+00 | 5.45E-08 | 1.94E-07 | -1.25E+00 | 5.36E-02 | 1.51E-01 | 1.18E+00 | 5.47E-02 | 2.01E-01 | 1.04E+00 | 1.14E-01 | 3.12E-01 | 1.13E+00 |
| UGGT2 | 3.02E-08 | 1.58E-07 | -1.67E+00 | 2.18E-06 | 9.42E-06 | -1.24E+00 | 2.84E-06 | 1.21E-05 | -1.53E+00 | 3.48E-06 | 1.33E-05 | -1.33E+00 | 6.53E-04 | 2.33E-03 | -1.47E+00 | 5.40E-02 | 1.52E-01 | -1.01E+00 | 7.73E-04 | 2.84E-03 | -1.36E+00 | 1.36E-02 | 3.74E-02 | -1.14E+00 |
| RHNO1 | 3.02E-08 | 1.58E-07 | -1.39E+00 | 4.44E-04 | 1.92E-03 | 1.08E+00 | 9.31E-04 | 3.96E-03 | -1.29E+00 | 1.82E-02 | 6.94E-02 | 1.08E+00 | 1.59E-07 | 5.67E-07 | -1.66E+00 | 1.56E-05 | 4.41E-05 | -1.39E+00 | 1.95E-07 | 7.16E-07 | -1.33E+00 | 8.81E-04 | 2.42E-03 | -1.03E+00 |
| CTSB | 3.02E-08 | 1.58E-07 | -1.59E+00 | 3.45E-04 | 1.49E-03 | -1.03E+00 | 3.35E-06 | 1.42E-05 | -1.41E+00 | 6.67E-06 | 2.55E-05 | -1.22E+00 | 1.17E-10 | 4.18E-10 | -1.52E+00 | 1.69E-03 | 4.78E-03 | 1.04E+00 | 4.10E-10 | 1.50E-09 | -1.44E+00 | 2.11E-03 | 5.80E-03 | -1.00E+00 |
| CERS6 | 3.04E-08 | 1.59E-07 | -1.26E+00 | 1.37E-10 | 5.93E-10 | -1.32E+00 | 1.77E-03 | 7.53E-03 | -1.10E+00 | 2.85E-06 | 1.09E-05 | -1.24E+00 | 6.08E-06 | 2.17E-05 | -1.22E+00 | 6.47E-07 | 1.82E-06 | -1.35E+00 | 8.38E-08 | 3.08E-07 | -1.44E+00 | 5.11E-07 | 1.40E-06 | -1.39E+00 |
| TTC8 | 3.11E-08 | 1.62E-07 | -2.13E+00 | 3.11E-06 | 1.34E-05 | -1.43E+00 | 1.59E-11 | 6.76E-11 | -2.38E+00 | 2.01E-08 | 7.68E-08 | -1.88E+00 | 2.06E-05 | 7.35E-05 | -2.31E+00 | 3.86E-03 | 1.09E-02 | -1.28E+00 | 7.66E-08 | 2.81E-07 | -1.55E+00 | 5.07E-05 | 1.39E-04 | -1.20E+00 |
| KIAA0586 | 3.13E-08 | 1.63E-07 | -1.46E+00 | 1.46E-06 | 6.30E-06 | -1.14E+00 | 8.65E-05 | 3.68E-04 | -1.26E+00 | 3.40E-04 | 1.30E-03 | 1.00E+00 | 7.20E-04 | 2.57E-03 | -1.11E+00 | 1.84E-03 | 5.18E-03 | -1.09E+00 | 3.57E-07 | 1.31E-06 | -1.22E+00 | 8.01E-04 | 2.20E-03 | -1.05E+00 |
| DACH1 | 3.14E-08 | 1.64E-07 | -1.32E+00 | 5.70E-04 | 2.46E-03 | 1.06E+00 | 4.61E-05 | 1.96E-04 | -1.38E+00 | 9.52E-05 | 3.64E-04 | -1.17E+00 | 1.78E-04 | 6.34E-04 | -1.34E+00 | 1.26E-03 | 3.56E-03 | -1.18E+00 | 5.50E-05 | 2.02E-04 | -1.52E+00 | 1.20E-02 | 3.30E-02 | -1.22E+00 |
| BLOC1S4 | 3.18E-08 | 1.66E-07 | -1.20E+00 | 1.20E-06 | 5.18E-06 | -1.05E+00 | 1.17E-09 | 4.96E-09 | -1.50E+00 | 2.68E-07 | 1.02E-06 | -1.21E+00 | 2.06E-06 | 7.36E-06 | -1.22E+00 | 2.90E-05 | 8.17E-05 | -1.10E+00 | 5.94E-10 | 2.18E-09 | -1.45E+00 | 2.51E-05 | 6.90E-05 | -1.17E+00 |
| INIP | 3.19E-08 | 1.66E-07 | -1.83E+00 | 2.31E-05 | 9.98E-05 | -1.28E+00 | 1.72E-09 | 7.29E-09 | -1.80E+00 | 1.60E-04 | 6.10E-04 | -1.08E+00 | 2.34E-07 | 8.35E-07 | -1.56E+00 | 4.90E-05 | 1.38E-04 | -1.17E+00 | 5.02E-21 | 1.84E-20 | -2.30E+00 | 4.47E-07 | 1.23E-06 | -1.29E+00 |
| MAPK3 | 3.23E-08 | 1.68E-07 | -1.36E+00 | 1.46E-04 | 6.29E-04 | 1.03E+00 | 2.63E-02 | 1.12E-01 | 1.04E+00 | 8.76E-03 | 3.35E-02 | 1.13E+00 | 3.24E-03 | 1.16E-02 | 1.01E+00 | 3.98E-03 | 1.12E-02 | -1.01E+00 | 1.13E-02 | 4.16E-02 | 1.02E+00 | 3.08E-02 | 8.47E-02 | -1.01E+00 |
| AC092745.3 | 3.23E-08 | 1.69E-07 | -2.18E+00 | 7.37E-07 | 3.18E-06 | -1.61E+00 | 1.04E-17 | 4.41E-17 | -2.80E+00 | 6.50E-15 | 2.48E-14 | -2.08E+00 | 1.03E-13 | 3.68E-13 | -2.82E+00 | 2.66E-11 | 7.49E-11 | -2.00E+00 | 1.11E-15 | 4.07E-15 | -2.15E+00 | 1.00E-08 | 2.76E-08 | -1.44E+00 |
| HSPA9 | 3.24E-08 | 1.69E-07 | -1.08E+00 | 2.26E-04 | 9.77E-04 | 1.11E+00 | 3.36E-02 | 1.43E-01 | -1.07E+00 | 4.36E-02 | 1.67E-01 | 1.07E+00 | 1.19E-08 | 4.26E-08 | -1.35E+00 | 1.04E-03 | 2.93E-03 | 1.03E+00 | 2.45E-05 | 8.99E-05 | -1.20E+00 | 3.41E-01 | 9.36E-01 | 1.23E+00 |
| LIMCH1 | 3.24E-08 | 1.69E-07 | -1.53E+00 | 1.03E-02 | 4.43E-02 | 1.22E+00 | 1.81E-01 | 7.69E-01 | 1.29E+00 | 6.33E-02 | 2.42E-01 | 1.72E+00 | 2.24E-01 | 7.99E-01 | 1.48E+00 | 3.37E-01 | 9.51E-01 | 1.12E+00 | 2.35E-01 | 8.61E-01 | -1.11E+00 | 3.47E-01 | 9.53E-01 | 1.01E+00 |
| FAM57A | 3.24E-08 | 1.69E-07 | -2.15E+00 | 6.93E-07 | 2.99E-06 | -1.73E+00 | 3.67E-05 | 1.56E-04 | -1.65E+00 | 1.03E-05 | 3.92E-05 | -1.47E+00 | 5.31E-03 | 1.89E-02 | -1.86E+00 | 1.48E-02 | 4.18E-02 | -1.50E+00 | 1.66E-05 | 6.09E-05 | -1.65E+00 | 2.62E-02 | 7.21E-02 | -1.00E+00 |
| SP1 | 3.26E-08 | 1.70E-07 | -1.39E+00 | 3.56E-04 | 1.54E-03 | 1.04E+00 | 1.34E-10 | 5.68E-10 | -1.57E+00 | 4.29E-08 | 1.64E-07 | -1.28E+00 | 1.64E-04 | 5.83E-04 | -1.29E+00 | 4.73E-02 | 1.33E-01 | 1.03E+00 | 1.57E-06 | 5.78E-06 | -1.42E+00 | 8.35E-04 | 2.29E-03 | -1.21E+00 |
| TXNDC17 | 3.29E-08 | 1.71E-07 | -1.18E+00 | 2.00E-06 | 8.64E-06 | 1.03E+00 | 6.66E-03 | 2.83E-02 | 1.04E+00 | 5.89E-03 | 2.25E-02 | 1.12E+00 | 5.00E-10 | 1.78E-09 | -1.46E+00 | 2.00E-05 | 5.63E-05 | -1.09E+00 | 8.67E-04 | 3.18E-03 | -1.10E+00 | 2.55E-02 | 7.01E-02 | 1.02E+00 |
| TMEM131 | 3.30E-08 | 1.72E-07 | -1.23E+00 | 1.13E-04 | 4.89E-04 | 1.09E+00 | 8.78E-05 | 3.73E-04 | -1.25E+00 | 3.10E-04 | 1.19E-03 | -1.05E+00 | 7.67E-06 | 2.74E-05 | -1.28E+00 | 1.02E-03 | 2.89E-03 | -1.04E+00 | 9.98E-06 | 3.66E-05 | -1.31E+00 | 3.28E-04 | 9.00E-04 | -1.11E+00 |
| YEATS4 | 3.31E-08 | 1.73E-07 | -1.36E+00 | 1.58E-06 | 6.81E-06 | -1.11E+00 | 5.67E-06 | 2.41E-05 | -1.06E+00 | 6.22E-08 | 2.37E-07 | -1.10E+00 | 1.33E-10 | 4.76E-10 | -1.78E+00 | 3.54E-05 | 9.98E-05 | -1.21E+00 | 3.16E-06 | 1.16E-05 | -1.11E+00 | 9.74E-03 | 2.67E-02 | -1.01E+00 |
| SUCLG1 | 3.31E-08 | 1.73E-07 | -1.17E+00 | 3.08E-06 | 1.33E-05 | 1.04E+00 | 4.47E-04 | 1.90E-03 | -1.11E+00 | 4.24E-05 | 1.62E-04 | -1.05E+00 | 2.10E-07 | 7.50E-07 | -1.19E+00 | 4.69E-03 | 1.32E-02 | 1.10E+00 | 1.82E-02 | 6.70E-02 | -1.02E+00 | 2.26E-01 | 6.21E-01 | 1.10E+00 |
| DLL3 | 3.34E-08 | 1.74E-07 | -2.06E+00 | 1.03E-03 | 4.43E-03 | -1.07E+00 | 1.59E-03 | 6.74E-03 | -1.43E+00 | 9.32E-02 | 3.56E-01 | 1.24E+00 | 1.86E-01 | 6.65E-01 | -1.04E+00 | 2.86E-01 | 8.05E-01 | 1.05E+00 | 2.68E-01 | 9.85E-01 | 1.01E+00 | 1.36E-01 | 3.75E-01 | 1.44E+00 |
| BUD23 | 3.35E-08 | 1.75E-07 | -1.26E+00 | 6.98E-07 | 3.02E-06 | -1.04E+00 | 2.68E-06 | 1.14E-05 | -1.22E+00 | 2.66E-06 | 1.02E-05 | -1.08E+00 | 9.20E-03 | 3.28E-02 | 1.02E+00 | 8.11E-02 | 2.29E-01 | 1.22E+00 | 6.66E-03 | 2.44E-02 | 1.02E+00 | 2.74E-01 | 7.53E-01 | 1.11E+00 |
| YPEL1 | 3.36E-08 | 1.75E-07 | -1.75E+00 | 1.25E-04 | 5.38E-04 | -1.08E+00 | 1.01E-06 | 4.27E-06 | -1.55E+00 | 9.22E-06 | 3.52E-05 | -1.26E+00 | 1.41E-06 | 5.02E-06 | -1.94E+00 | 6.35E-06 | 1.79E-05 | -1.60E+00 | 3.77E-05 | 1.38E-04 | -1.72E+00 | 2.44E-03 | 6.71E-03 | -1.36E+00 |
| LIMA1 | 3.36E-08 | 1.75E-07 | -1.35E+00 | 7.62E-07 | 3.29E-06 | -1.20E+00 | 4.78E-07 | 2.03E-06 | -1.38E+00 | 3.11E-06 | 1.19E-05 | -1.19E+00 | 1.48E-08 | 5.26E-08 | -1.85E+00 | 4.21E-04 | 1.19E-03 | -1.23E+00 | 9.07E-14 | 3.33E-13 | -1.65E+00 | 5.79E-17 | 1.59E-16 | -1.69E+00 |
| NDUFAF1 | 3.36E-08 | 1.75E-07 | -1.68E+00 | 4.89E-04 | 2.11E-03 | -1.00E+00 | 2.48E-03 | 1.05E-02 | -1.20E+00 | 2.01E-03 | 7.66E-03 | -1.04E+00 | 1.08E-05 | 3.87E-05 | -1.56E+00 | 8.33E-03 | 2.35E-02 | -1.11E+00 | 6.98E-05 | 2.56E-04 | -1.10E+00 | 3.95E-03 | 1.09E-02 | -1.00E+00 |
| CNOT6LP1 | 3.37E-08 | 1.75E-07 | -1.30E+00 | 8.25E-09 | 3.57E-08 | -1.30E+00 | 1.06E-03 | 4.52E-03 | -1.25E+00 | 1.58E-04 | 6.03E-04 | -1.28E+00 | 3.34E-02 | 1.19E-01 | -1.11E+00 | 3.02E-03 | 8.52E-03 | -1.46E+00 | 2.27E-04 | 8.33E-04 | -1.23E+00 | 2.52E-05 | 6.93E-05 | -1.44E+00 |
| EXTL2 | 3.38E-08 | 1.76E-07 | -1.47E+00 | 3.18E-03 | 1.37E-02 | 1.22E+00 | 2.25E-05 | 9.57E-05 | -1.25E+00 | 5.33E-04 | 2.04E-03 | 1.02E+00 | 1.40E-11 | 4.99E-11 | -1.87E+00 | 1.05E-03 | 2.97E-03 | -1.01E+00 | 3.40E-17 | 1.25E-16 | -1.86E+00 | 5.27E-05 | 1.45E-04 | -1.18E+00 |
| GNB5 | 3.38E-08 | 1.76E-07 | -1.35E+00 | 1.02E-05 | 4.40E-05 | 1.06E+00 | 2.88E-07 | 1.22E-06 | -1.36E+00 | 1.14E-10 | 4.35E-10 | -1.39E+00 | 6.16E-05 | 2.20E-04 | -1.46E+00 | 4.28E-06 | 1.21E-05 | -1.94E+00 | 2.19E-01 | 8.05E-01 | 1.26E+00 | 2.05E-01 | 5.64E-01 | -1.19E+00 |
| PIK3C3 | 3.42E-08 | 1.78E-07 | -1.29E+00 | 2.63E-02 | 1.13E-01 | 1.31E+00 | 7.22E-06 | 3.07E-05 | -1.28E+00 | 1.85E-02 | 7.06E-02 | 1.16E+00 | 2.11E-05 | 7.53E-05 | -1.16E+00 | 1.22E-03 | 3.44E-03 | 1.02E+00 | 8.36E-08 | 3.07E-07 | -1.37E+00 | 8.45E-04 | 2.32E-03 | -1.03E+00 |
| LINC00662 | 3.42E-08 | 1.78E-07 | -1.29E+00 | 4.52E-04 | 1.95E-03 | 1.11E+00 | 5.77E-04 | 2.45E-03 | -1.16E+00 | 1.28E-02 | 4.87E-02 | 1.13E+00 | 4.51E-04 | 1.61E-03 | -1.30E+00 | 1.47E-02 | 4.15E-02 | -1.07E+00 | 5.83E-04 | 2.14E-03 | -1.44E+00 | 6.43E-02 | 1.77E-01 | 1.02E+00 |
| HLA-DPA1 | 3.43E-08 | 1.79E-07 | -1.52E+00 | 2.02E-02 | 8.75E-02 | -1.15E+00 | 7.43E-08 | 3.16E-07 | -1.62E+00 | 2.34E-02 | 8.92E-02 | -1.16E+00 | 6.20E-21 | 2.21E-20 | -1.92E+00 | 8.15E-21 | 2.30E-20 | -1.63E+00 | 4.98E-63 | 1.83E-62 | -2.33E+00 | 8.13E-19 | 2.23E-18 | -1.56E+00 |
| CNKSR1 | 3.43E-08 | 1.79E-07 | -2.33E+00 | 3.48E-04 | 1.50E-03 | -1.20E+00 | 2.51E-05 | 1.07E-04 | -2.02E+00 | 6.80E-04 | 2.60E-03 | -1.45E+00 | 1.87E-03 | 6.68E-03 | -2.25E+00 | 1.37E-02 | 3.85E-02 | -1.55E+00 | 8.11E-02 | 2.98E-01 | -1.16E+00 | 1.60E-01 | 4.40E-01 | -1.06E+00 |
| SKP1 | 3.44E-08 | 1.79E-07 | -1.16E+00 | 1.93E-11 | 8.36E-11 | -1.20E+00 | 1.22E-01 | 5.19E-01 | -1.02E+00 | 1.01E-03 | 3.86E-03 | -1.12E+00 | 6.32E-04 | 2.25E-03 | -1.17E+00 | 1.74E-04 | 4.91E-04 | -1.14E+00 | 2.29E-01 | 8.39E-01 | -1.01E+00 | 4.44E-02 | 1.22E-01 | -1.12E+00 |
| GOSR1 | 3.45E-08 | 1.80E-07 | -1.16E+00 | 8.00E-09 | 3.46E-08 | -1.09E+00 | 1.24E-06 | 5.26E-06 | -1.36E+00 | 1.76E-04 | 6.72E-04 | -1.10E+00 | 1.38E-02 | 4.91E-02 | 1.07E+00 | 1.25E-02 | 3.53E-02 | 1.10E+00 | 3.81E-07 | 1.40E-06 | -1.30E+00 | 1.39E-05 | 3.82E-05 | -1.21E+00 |
| ANKRD50 | 3.45E-08 | 1.80E-07 | -1.75E+00 | 7.88E-08 | 3.41E-07 | -1.55E+00 | 2.52E-04 | 1.07E-03 | -1.30E+00 | 3.73E-04 | 1.43E-03 | -1.21E+00 | 1.61E-01 | 5.73E-01 | 1.10E+00 | 4.78E-02 | 1.35E-01 | -1.10E+00 | 1.32E-02 | 4.83E-02 | -1.07E+00 | 1.90E-04 | 5.22E-04 | -1.34E+00 |
| Z83843.1 | 3.46E-08 | 1.80E-07 | -1.33E+00 | 7.10E-02 | 3.07E-01 | 1.75E+00 | 1.30E-06 | 5.53E-06 | -1.62E+00 | 2.13E-02 | 8.13E-02 | 1.21E+00 | 9.39E-03 | 3.35E-02 | -1.04E+00 | 3.31E-01 | 9.32E-01 | 1.38E+00 | 4.38E-04 | 1.61E-03 | -1.45E+00 | 5.33E-02 | 1.46E-01 | 1.00E+00 |
| SUPT6H | 3.46E-08 | 1.80E-07 | -1.24E+00 | 3.06E-07 | 1.32E-06 | -1.09E+00 | 2.30E-05 | 9.76E-05 | -1.27E+00 | 2.53E-04 | 9.66E-04 | -1.05E+00 | 8.87E-06 | 3.16E-05 | -1.20E+00 | 5.20E-05 | 1.47E-04 | -1.11E+00 | 7.75E-03 | 2.84E-02 | 1.01E+00 | 2.81E-02 | 7.71E-02 | 1.07E+00 |
| ZBTB17 | 3.49E-08 | 1.82E-07 | -1.48E+00 | 1.36E-06 | 5.89E-06 | -1.17E+00 | 1.57E-06 | 6.66E-06 | -1.53E+00 | 7.60E-05 | 2.90E-04 | -1.17E+00 | 5.17E-11 | 1.84E-10 | -1.76E+00 | 5.14E-10 | 1.45E-09 | -1.58E+00 | 3.09E-04 | 1.14E-03 | -1.27E+00 | 1.20E-02 | 3.30E-02 | -1.08E+00 |
| CD37 | 3.50E-08 | 1.82E-07 | -1.30E+00 | 2.51E-03 | 1.08E-02 | -1.09E+00 | 8.89E-12 | 3.78E-11 | -1.50E+00 | 4.91E-06 | 1.88E-05 | -1.23E+00 | 1.20E-04 | 4.27E-04 | -1.24E+00 | 2.59E-04 | 7.29E-04 | -1.18E+00 | 3.80E-08 | 1.40E-07 | -1.21E+00 | 7.30E-05 | 2.01E-04 | -1.14E+00 |
| UNC93B1 | 3.50E-08 | 1.83E-07 | -1.91E+00 | 3.24E-08 | 1.40E-07 | -1.79E+00 | 1.36E-06 | 5.78E-06 | -1.59E+00 | 8.73E-06 | 3.34E-05 | -1.41E+00 | 5.29E-08 | 1.89E-07 | -2.04E+00 | 1.72E-05 | 4.86E-05 | -1.47E+00 | 7.00E-08 | 2.57E-07 | -1.68E+00 | 9.39E-07 | 2.58E-06 | -1.51E+00 |
| CNOT4 | 3.51E-08 | 1.83E-07 | -1.16E+00 | 4.13E-06 | 1.78E-05 | 1.05E+00 | 1.08E-05 | 4.61E-05 | -1.21E+00 | 1.76E-08 | 6.74E-08 | -1.16E+00 | 2.05E-02 | 7.30E-02 | 1.04E+00 | 4.92E-04 | 1.39E-03 | -1.08E+00 | 1.37E-04 | 5.02E-04 | -1.13E+00 | 7.65E-07 | 2.10E-06 | -1.30E+00 |
| SLC15A2 | 3.53E-08 | 1.84E-07 | -1.79E+00 | 1.81E-02 | 7.83E-02 | 1.25E+00 | 5.21E-08 | 2.21E-07 | -1.85E+00 | 1.92E-01 | 7.33E-01 | 1.33E+00 | 7.38E-02 | 2.63E-01 | -1.26E+00 | 1.94E-01 | 5.48E-01 | 1.43E+00 | 5.25E-02 | 1.93E-01 | -1.02E+00 | 3.58E-01 | 9.82E-01 | 1.30E+00 |
| MUL1 | 3.53E-08 | 1.84E-07 | -1.67E+00 | 3.84E-07 | 1.66E-06 | -1.44E+00 | 1.17E-10 | 4.98E-10 | -1.63E+00 | 5.80E-10 | 2.22E-09 | -1.41E+00 | 1.07E-08 | 3.81E-08 | -1.91E+00 | 1.17E-05 | 3.28E-05 | -1.26E+00 | 1.14E-07 | 4.20E-07 | -1.26E+00 | 3.42E-05 | 9.39E-05 | -1.09E+00 |
| NEPRO | 3.54E-08 | 1.85E-07 | -1.37E+00 | 4.26E-07 | 1.84E-06 | -1.08E+00 | 5.35E-11 | 2.27E-10 | -1.47E+00 | 1.31E-08 | 5.02E-08 | -1.22E+00 | 1.26E-09 | 4.50E-09 | -1.42E+00 | 2.30E-08 | 6.48E-08 | -1.26E+00 | 5.62E-08 | 2.06E-07 | -1.30E+00 | 1.05E-02 | 2.89E-02 | 1.02E+00 |
| ZNF655 | 3.55E-08 | 1.85E-07 | -1.22E+00 | 5.69E-05 | 2.46E-04 | 1.08E+00 | 2.10E-06 | 8.93E-06 | -1.26E+00 | 1.50E-05 | 5.73E-05 | -1.06E+00 | 3.63E-03 | 1.29E-02 | -1.03E+00 | 8.32E-03 | 2.35E-02 | 1.01E+00 | 1.30E-03 | 4.77E-03 | -1.06E+00 | 9.58E-02 | 2.63E-01 | 1.04E+00 |
| ATF7IP | 3.57E-08 | 1.86E-07 | -1.21E+00 | 1.06E-03 | 4.58E-03 | 1.09E+00 | 6.67E-09 | 2.84E-08 | -1.60E+00 | 4.01E-04 | 1.53E-03 | -1.08E+00 | 1.84E-06 | 6.57E-06 | -1.26E+00 | 4.09E-03 | 1.15E-02 | 1.04E+00 | 1.97E-06 | 7.22E-06 | -1.22E+00 | 1.90E-05 | 5.21E-05 | -1.21E+00 |
| SAR1B | 3.58E-08 | 1.86E-07 | -1.20E+00 | 3.82E-09 | 1.65E-08 | -1.15E+00 | 4.92E-06 | 2.09E-05 | -1.30E+00 | 7.31E-06 | 2.79E-05 | -1.13E+00 | 7.72E-06 | 2.75E-05 | -1.29E+00 | 6.57E-04 | 1.85E-03 | -1.10E+00 | 1.77E-09 | 6.50E-09 | -1.36E+00 | 3.85E-04 | 1.06E-03 | -1.06E+00 |
| COQ8B | 3.59E-08 | 1.87E-07 | -1.36E+00 | 6.35E-08 | 2.75E-07 | -1.19E+00 | 1.09E-10 | 4.64E-10 | -1.49E+00 | 2.39E-10 | 9.13E-10 | -1.36E+00 | 6.56E-05 | 2.34E-04 | -1.17E+00 | 9.34E-04 | 2.63E-03 | -1.00E+00 | 1.15E-07 | 4.23E-07 | -1.42E+00 | 1.19E-06 | 3.28E-06 | -1.32E+00 |
| AC091544.3 | 3.61E-08 | 1.88E-07 | -1.90E+00 | 1.44E-07 | 6.22E-07 | -1.59E+00 | 1.57E-11 | 6.68E-11 | -2.06E+00 | 5.67E-10 | 2.17E-09 | -1.60E+00 | 4.78E-10 | 1.70E-09 | -3.49E+00 | 1.20E-07 | 3.38E-07 | -2.11E+00 | 1.07E-22 | 3.94E-22 | -2.36E+00 | 3.34E-16 | 9.16E-16 | -1.83E+00 |
| KMT2D | 3.61E-08 | 1.88E-07 | -1.31E+00 | 4.29E-11 | 1.85E-10 | -1.40E+00 | 1.12E-06 | 4.75E-06 | -1.29E+00 | 2.98E-10 | 1.14E-09 | -1.48E+00 | 1.73E-07 | 6.16E-07 | -1.50E+00 | 4.52E-10 | 1.28E-09 | -1.71E+00 | 4.75E-05 | 1.74E-04 | -1.42E+00 | 1.05E-04 | 2.89E-04 | -1.31E+00 |
| CALHM2 | 3.65E-08 | 1.90E-07 | -2.16E+00 | 6.18E-05 | 2.67E-04 | -1.38E+00 | 3.36E-04 | 1.43E-03 | -1.51E+00 | 2.35E-04 | 8.97E-04 | -1.41E+00 | 3.31E-02 | 1.18E-01 | -1.58E+00 | 1.11E-01 | 3.13E-01 | -1.12E+00 | 6.89E-02 | 2.53E-01 | -1.38E+00 | 1.25E-01 | 3.45E-01 | -1.24E+00 |
| TBC1D9B | 3.66E-08 | 1.91E-07 | -1.39E+00 | 2.72E-04 | 1.17E-03 | 1.04E+00 | 4.66E-03 | 1.98E-02 | 1.00E+00 | 8.81E-02 | 3.36E-01 | 1.37E+00 | 1.23E-02 | 4.40E-02 | -1.02E+00 | 6.08E-02 | 1.72E-01 | 1.08E+00 | 7.61E-02 | 2.79E-01 | 1.13E+00 | 3.31E-01 | 9.10E-01 | 1.28E+00 |
| CEP95 | 3.68E-08 | 1.92E-07 | -1.27E+00 | 7.72E-08 | 3.34E-07 | -1.12E+00 | 1.40E-04 | 5.93E-04 | -1.20E+00 | 1.41E-05 | 5.38E-05 | -1.12E+00 | 6.69E-06 | 2.38E-05 | -1.30E+00 | 1.46E-03 | 4.11E-03 | -1.11E+00 | 5.06E-04 | 1.86E-03 | -1.29E+00 | 1.46E-02 | 4.01E-02 | -1.20E+00 |
| TTC37 | 3.68E-08 | 1.92E-07 | -1.19E+00 | 9.50E-06 | 4.11E-05 | 1.05E+00 | 1.40E-07 | 5.97E-07 | -1.22E+00 | 1.58E-04 | 6.02E-04 | 1.02E+00 | 1.95E-08 | 6.95E-08 | -1.17E+00 | 5.36E-06 | 1.51E-05 | -1.04E+00 | 2.58E-05 | 9.47E-05 | -1.25E+00 | 2.44E-04 | 6.70E-04 | -1.12E+00 |
| PGAP2 | 3.70E-08 | 1.93E-07 | -1.39E+00 | 4.19E-05 | 1.81E-04 | -1.00E+00 | 5.87E-06 | 2.50E-05 | -1.28E+00 | 4.21E-06 | 1.61E-05 | -1.13E+00 | 1.05E-02 | 3.74E-02 | 1.09E+00 | 1.05E-01 | 2.95E-01 | 1.23E+00 | 9.53E-05 | 3.50E-04 | -1.20E+00 | 4.26E-03 | 1.17E-02 | 1.03E+00 |
| ENTPD6 | 3.75E-08 | 1.95E-07 | -2.05E+00 | 1.47E-04 | 6.35E-04 | -1.39E+00 | 2.66E-10 | 1.13E-09 | -1.98E+00 | 6.17E-09 | 2.36E-08 | -1.55E+00 | 1.98E-02 | 7.07E-02 | -1.36E+00 | 2.97E-01 | 8.36E-01 | 1.32E+00 | 1.02E-05 | 3.74E-05 | -1.63E+00 | 7.97E-02 | 2.19E-01 | 1.08E+00 |
| PHF21A | 3.75E-08 | 1.96E-07 | -1.34E+00 | 2.79E-02 | 1.20E-01 | 1.28E+00 | 8.02E-09 | 3.41E-08 | -1.42E+00 | 2.59E-04 | 9.90E-04 | -1.02E+00 | 4.24E-04 | 1.51E-03 | -1.21E+00 | 5.74E-02 | 1.62E-01 | 1.08E+00 | 6.47E-04 | 2.38E-03 | -1.15E+00 | 1.42E-02 | 3.91E-02 | 1.04E+00 |
| AC068775.1 | 3.77E-08 | 1.96E-07 | -1.22E+00 | 6.15E-11 | 2.66E-10 | -1.26E+00 | 4.48E-07 | 1.91E-06 | -1.39E+00 | 2.48E-09 | 9.49E-09 | -1.39E+00 | 1.80E-03 | 6.41E-03 | -1.02E+00 | 1.40E-04 | 3.95E-04 | -1.17E+00 | 3.01E-05 | 1.10E-04 | -1.21E+00 | 1.11E-05 | 3.06E-05 | -1.22E+00 |
| C11orf68 | 3.78E-08 | 1.97E-07 | -1.56E+00 | 5.31E-03 | 2.30E-02 | 1.18E+00 | 2.36E-08 | 1.00E-07 | -1.65E+00 | 2.17E-05 | 8.28E-05 | -1.14E+00 | 4.48E-10 | 1.60E-09 | -1.82E+00 | 3.50E-03 | 9.88E-03 | 1.05E+00 | 2.79E-15 | 1.02E-14 | -1.59E+00 | 3.56E-10 | 9.78E-10 | -1.32E+00 |
| SPRYD4 | 3.81E-08 | 1.99E-07 | -1.46E+00 | 1.42E-05 | 6.13E-05 | -1.03E+00 | 3.42E-10 | 1.45E-09 | -1.90E+00 | 1.98E-06 | 7.55E-06 | -1.25E+00 | 1.41E-05 | 5.04E-05 | -1.40E+00 | 1.57E-02 | 4.42E-02 | 1.07E+00 | 4.43E-06 | 1.62E-05 | -1.32E+00 | 8.46E-02 | 2.32E-01 | 1.03E+00 |
| SNRPB2 | 3.81E-08 | 1.99E-07 | -1.11E+00 | 1.18E-17 | 5.12E-17 | -1.18E+00 | 3.03E-03 | 1.29E-02 | -1.08E+00 | 4.76E-05 | 1.82E-04 | -1.16E+00 | 1.39E-04 | 4.96E-04 | -1.02E+00 | 5.66E-04 | 1.60E-03 | 1.00E+00 | 2.27E-03 | 8.34E-03 | -1.09E+00 | 1.85E-02 | 5.07E-02 | -1.04E+00 |
| PUM3 | 3.82E-08 | 1.99E-07 | -1.26E+00 | 2.39E-07 | 1.03E-06 | -1.07E+00 | 1.60E-06 | 6.79E-06 | -1.27E+00 | 1.30E-04 | 4.95E-04 | 1.02E+00 | 2.47E-04 | 8.82E-04 | -1.18E+00 | 1.30E-01 | 3.67E-01 | 1.23E+00 | 2.32E-10 | 8.50E-10 | -1.40E+00 | 4.53E-03 | 1.24E-02 | -1.03E+00 |
| CITED2 | 3.84E-08 | 2.00E-07 | 2.41E+00 | 3.14E-07 | 1.36E-06 | 2.69E+00 | 8.90E-04 | 3.78E-03 | 1.72E+00 | 1.94E-04 | 7.41E-04 | 2.02E+00 | 2.29E-02 | 8.18E-02 | 1.63E+00 | 1.50E-06 | 4.23E-06 | 2.27E+00 | 3.14E-06 | 1.15E-05 | 2.09E+00 | 5.07E-07 | 1.39E-06 | 2.29E+00 |
| TEKT4 | 3.84E-08 | 2.00E-07 | -2.18E+00 | 9.72E-08 | 4.20E-07 | -1.67E+00 | 7.69E-05 | 3.27E-04 | -1.57E+00 | 8.77E-06 | 3.35E-05 | -1.55E+00 | 2.17E-03 | 7.74E-03 | -1.39E+00 | 5.58E-02 | 1.57E-01 | 1.00E+00 | 5.63E-04 | 2.07E-03 | -1.18E+00 | 3.50E-06 | 9.63E-06 | -1.44E+00 |
| FRY | 3.86E-08 | 2.01E-07 | -1.31E+00 | 4.11E-08 | 1.78E-07 | -1.20E+00 | 3.40E-07 | 1.44E-06 | -1.65E+00 | 3.71E-04 | 1.42E-03 | -1.14E+00 | 5.02E-06 | 1.79E-05 | -1.30E+00 | 1.11E-06 | 3.14E-06 | -1.43E+00 | 8.51E-03 | 3.12E-02 | -1.24E+00 | 3.64E-02 | 1.00E-01 | -1.14E+00 |
| DHX36 | 3.90E-08 | 2.03E-07 | -1.13E+00 | 5.84E-11 | 2.52E-10 | -1.11E+00 | 1.47E-03 | 6.24E-03 | -1.10E+00 | 1.45E-03 | 5.56E-03 | -1.00E+00 | 1.17E-02 | 4.19E-02 | 1.02E+00 | 8.47E-03 | 2.39E-02 | 1.09E+00 | 3.02E-04 | 1.11E-03 | -1.16E+00 | 1.24E-02 | 3.40E-02 | -1.05E+00 |
| TSPAN2 | 3.90E-08 | 2.03E-07 | -1.37E+00 | 1.02E-02 | 4.39E-02 | 1.09E+00 | 4.58E-04 | 1.94E-03 | -1.30E+00 | 5.91E-02 | 2.26E-01 | 1.06E+00 | 3.76E-03 | 1.34E-02 | -1.35E+00 | 1.18E-01 | 3.34E-01 | -1.01E+00 | 8.01E-02 | 2.94E-01 | 1.06E+00 | 3.50E-01 | 9.62E-01 | 1.31E+00 |
| MLC1 | 3.93E-08 | 2.05E-07 | -1.24E+00 | 1.19E-04 | 5.14E-04 | 1.08E+00 | 4.95E-06 | 2.10E-05 | -1.19E+00 | 1.60E-04 | 6.11E-04 | 1.02E+00 | 4.00E-03 | 1.43E-02 | -1.02E+00 | 1.82E-01 | 5.14E-01 | 1.35E+00 | 2.72E-01 | 1.00E+00 | 1.28E+00 | 3.03E-01 | 8.32E-01 | 1.25E+00 |
| SNX6 | 3.94E-08 | 2.05E-07 | -1.11E+00 | 7.82E-09 | 3.38E-08 | -1.03E+00 | 3.46E-06 | 1.47E-05 | -1.32E+00 | 2.14E-05 | 8.18E-05 | -1.13E+00 | 3.30E-06 | 1.18E-05 | -1.11E+00 | 3.09E-04 | 8.71E-04 | -1.04E+00 | 4.15E-04 | 1.52E-03 | -1.18E+00 | 1.47E-04 | 4.04E-04 | -1.16E+00 |
| TRPM7 | 4.02E-08 | 2.10E-07 | -1.29E+00 | 3.93E-02 | 1.70E-01 | 1.35E+00 | 1.80E-06 | 7.64E-06 | -1.34E+00 | 9.54E-03 | 3.64E-02 | 1.08E+00 | 1.14E-02 | 4.08E-02 | -1.07E+00 | 1.26E-01 | 3.54E-01 | 1.13E+00 | 7.70E-06 | 2.83E-05 | -1.35E+00 | 7.43E-02 | 2.04E-01 | 1.06E+00 |
| KLHL9 | 4.03E-08 | 2.10E-07 | -1.27E+00 | 5.22E-05 | 2.26E-04 | 1.04E+00 | 1.09E-05 | 4.64E-05 | -1.43E+00 | 4.27E-03 | 1.63E-02 | 1.02E+00 | 1.07E-05 | 3.82E-05 | -1.39E+00 | 2.95E-04 | 8.31E-04 | -1.18E+00 | 5.44E-07 | 1.99E-06 | -1.51E+00 | 7.12E-05 | 1.96E-04 | -1.32E+00 |
| DOCK11P1 | 4.09E-08 | 2.13E-07 | -2.80E+00 | 3.05E-05 | 1.32E-04 | -1.57E+00 | 5.87E-05 | 2.50E-04 | -1.96E+00 | 8.22E-04 | 3.14E-03 | -1.47E+00 | 5.23E-05 | 1.87E-04 | -1.66E+00 | 5.35E-04 | 1.51E-03 | -1.50E+00 | 1.61E-04 | 5.91E-04 | -1.96E+00 | 1.58E-02 | 4.34E-02 | -1.31E+00 |
| CEP135 | 4.11E-08 | 2.14E-07 | -1.24E+00 | 7.30E-06 | 3.15E-05 | -1.00E+00 | 3.65E-06 | 1.55E-05 | -1.31E+00 | 2.43E-04 | 9.27E-04 | -1.00E+00 | 1.09E-03 | 3.89E-03 | -1.17E+00 | 2.54E-03 | 7.17E-03 | -1.08E+00 | 1.75E-02 | 6.41E-02 | -1.09E+00 | 3.57E-01 | 9.81E-01 | 1.21E+00 |
| KANTR | 4.11E-08 | 2.14E-07 | -1.61E+00 | 2.16E-01 | 9.32E-01 | 1.45E+00 | 4.65E-04 | 1.97E-03 | -1.39E+00 | 2.53E-01 | 9.66E-01 | 1.31E+00 | 3.50E-02 | 1.25E-01 | 1.03E+00 | 3.00E-01 | 8.46E-01 | 1.65E+00 | 2.87E-06 | 1.05E-05 | -1.78E+00 | 2.14E-02 | 5.87E-02 | -1.10E+00 |
| AC069236.1 | 4.11E-08 | 2.14E-07 | 3.90E+00 | 6.05E-04 | 2.61E-03 | 3.11E+00 | 1.17E-03 | 4.98E-03 | 1.96E+00 | 2.01E-01 | 7.68E-01 | 1.44E+00 | 2.17E-01 | 7.74E-01 | 1.16E+00 | 2.32E-01 | 6.54E-01 | 1.67E+00 | 2.39E-02 | 8.75E-02 | -1.07E+00 | 1.72E-01 | 4.72E-01 | 1.13E+00 |
| TMEM141 | 4.12E-08 | 2.15E-07 | -1.39E+00 | 4.08E-09 | 1.76E-08 | -1.30E+00 | 8.32E-03 | 3.54E-02 | 1.00E+00 | 8.60E-05 | 3.29E-04 | -1.02E+00 | 1.59E-10 | 5.66E-10 | -1.40E+00 | 1.50E-05 | 4.24E-05 | -1.03E+00 | 1.08E-01 | 3.98E-01 | 1.07E+00 | 3.02E-02 | 8.29E-02 | -1.00E+00 |
| HNRNPA2B1 | 4.14E-08 | 2.16E-07 | -1.12E+00 | 3.92E-02 | 1.69E-01 | 1.11E+00 | 2.23E-01 | 9.46E-01 | 1.02E+00 | 2.33E-03 | 8.90E-03 | 1.15E+00 | 3.38E-04 | 1.20E-03 | -1.13E+00 | 8.73E-02 | 2.46E-01 | 1.07E+00 | 1.96E-03 | 7.18E-03 | -1.10E+00 | 1.36E-02 | 3.74E-02 | 1.10E+00 |
| LAS1L | 4.14E-08 | 2.16E-07 | -1.32E+00 | 4.19E-05 | 1.81E-04 | 1.02E+00 | 7.13E-02 | 3.03E-01 | 1.03E+00 | 1.69E-02 | 6.44E-02 | 1.07E+00 | 1.68E-05 | 6.00E-05 | -1.38E+00 | 1.69E-03 | 4.75E-03 | -1.12E+00 | 1.11E-05 | 4.08E-05 | -1.28E+00 | 5.38E-02 | 1.48E-01 | 1.03E+00 |
| ANKFY1 | 4.20E-08 | 2.19E-07 | -1.86E+00 | 7.26E-05 | 3.14E-04 | -1.08E+00 | 6.52E-06 | 2.77E-05 | -1.61E+00 | 2.29E-02 | 8.73E-02 | 1.05E+00 | 2.93E-03 | 1.04E-02 | -1.28E+00 | 2.48E-02 | 6.99E-02 | 1.01E+00 | 1.71E-06 | 6.29E-06 | -1.60E+00 | 3.51E-03 | 9.64E-03 | -1.12E+00 |
| TMC8 | 4.24E-08 | 2.21E-07 | -1.19E+00 | 1.08E-04 | 4.68E-04 | 1.10E+00 | 1.06E-05 | 4.51E-05 | -1.32E+00 | 7.19E-04 | 2.75E-03 | 1.04E+00 | 1.76E-07 | 6.28E-07 | -1.42E+00 | 9.35E-07 | 2.64E-06 | -1.21E+00 | 6.03E-10 | 2.21E-09 | -1.51E+00 | 4.10E-05 | 1.13E-04 | -1.18E+00 |
| VAMP2 | 4.26E-08 | 2.22E-07 | -1.12E+00 | 1.14E-03 | 4.94E-03 | 1.11E+00 | 1.46E-08 | 6.19E-08 | -1.28E+00 | 3.45E-05 | 1.32E-04 | -1.05E+00 | 8.00E-04 | 2.85E-03 | -1.04E+00 | 5.18E-03 | 1.46E-02 | 1.01E+00 | 1.16E-04 | 4.27E-04 | -1.14E+00 | 8.61E-04 | 2.36E-03 | -1.14E+00 |
| SLC38A5 | 4.26E-08 | 2.22E-07 | -1.54E+00 | 5.67E-06 | 2.45E-05 | -1.18E+00 | 2.96E-04 | 1.26E-03 | -1.19E+00 | 1.48E-02 | 5.65E-02 | 1.13E+00 | 2.76E-04 | 9.84E-04 | -1.78E+00 | 1.05E-01 | 2.95E-01 | 1.19E+00 | 5.56E-11 | 2.04E-10 | -1.72E+00 | 2.31E-02 | 6.36E-02 | 1.05E+00 |
| GNPDA2 | 4.26E-08 | 2.22E-07 | -1.28E+00 | 4.52E-11 | 1.95E-10 | -1.40E+00 | 2.38E-04 | 1.01E-03 | -1.31E+00 | 2.21E-03 | 8.43E-03 | -1.05E+00 | 1.29E-04 | 4.61E-04 | -1.22E+00 | 3.56E-03 | 1.00E-02 | -1.01E+00 | 5.43E-02 | 1.99E-01 | 1.19E+00 | 9.71E-02 | 2.67E-01 | 1.19E+00 |
| PNO1 | 4.29E-08 | 2.24E-07 | -1.35E+00 | 9.09E-08 | 3.93E-07 | -1.19E+00 | 3.11E-02 | 1.32E-01 | -1.04E+00 | 8.26E-04 | 3.16E-03 | -1.10E+00 | 1.55E-03 | 5.54E-03 | -1.17E+00 | 2.25E-02 | 6.34E-02 | 1.03E+00 | 1.72E-02 | 6.31E-02 | -1.06E+00 | 2.88E-01 | 7.91E-01 | 1.13E+00 |
| NCOA3 | 4.30E-08 | 2.24E-07 | -1.30E+00 | 1.06E-02 | 4.58E-02 | 1.26E+00 | 4.19E-06 | 1.78E-05 | -1.30E+00 | 4.68E-04 | 1.79E-03 | 1.06E+00 | 2.94E-07 | 1.05E-06 | -1.42E+00 | 8.41E-05 | 2.37E-04 | -1.14E+00 | 1.79E-07 | 6.57E-07 | -1.62E+00 | 1.24E-04 | 3.41E-04 | -1.20E+00 |
| RPS17 | 4.32E-08 | 2.25E-07 | -1.16E+00 | 9.62E-28 | 4.16E-27 | -1.40E+00 | 4.48E-04 | 1.91E-03 | -1.13E+00 | 3.37E-13 | 1.29E-12 | -1.34E+00 | 1.18E-06 | 4.22E-06 | -1.19E+00 | 4.99E-08 | 1.41E-07 | -1.27E+00 | 3.40E-09 | 1.25E-08 | -1.18E+00 | 3.86E-07 | 1.06E-06 | -1.22E+00 |
| HNRNPKP2 | 4.34E-08 | 2.26E-07 | -1.47E+00 | 8.14E-12 | 3.52E-11 | -1.77E+00 | 1.39E-03 | 5.89E-03 | -1.12E+00 | 2.33E-06 | 8.90E-06 | -1.27E+00 | 8.08E-05 | 2.88E-04 | -1.16E+00 | 6.31E-08 | 1.78E-07 | -1.54E+00 | 1.00E-06 | 3.67E-06 | -1.30E+00 | 2.06E-13 | 5.65E-13 | -1.70E+00 |
| CASP8AP2 | 4.34E-08 | 2.26E-07 | -1.36E+00 | 3.36E-04 | 1.45E-03 | 1.02E+00 | 3.68E-05 | 1.56E-04 | -1.40E+00 | 1.36E-02 | 5.18E-02 | -1.01E+00 | 9.83E-03 | 3.50E-02 | -1.16E+00 | 1.30E-01 | 3.66E-01 | 1.19E+00 | 2.47E-09 | 9.05E-09 | -1.51E+00 | 2.21E-01 | 6.07E-01 | 1.03E+00 |
| ZNF836 | 4.34E-08 | 2.26E-07 | -1.85E+00 | 2.94E-06 | 1.27E-05 | -1.40E+00 | 5.58E-03 | 2.37E-02 | -1.27E+00 | 1.10E-03 | 4.21E-03 | -1.43E+00 | 2.72E-02 | 9.69E-02 | -1.10E+00 | 4.35E-02 | 1.23E-01 | -1.07E+00 | 6.91E-03 | 2.54E-02 | -1.38E+00 | 1.17E-03 | 3.20E-03 | -1.80E+00 |
| SART3 | 4.34E-08 | 2.26E-07 | -1.55E+00 | 1.03E-05 | 4.46E-05 | -1.09E+00 | 9.62E-10 | 4.09E-09 | -1.74E+00 | 2.14E-03 | 8.18E-03 | 1.02E+00 | 1.21E-03 | 4.32E-03 | -1.21E+00 | 8.20E-02 | 2.31E-01 | 1.14E+00 | 1.71E-17 | 6.27E-17 | -1.88E+00 | 3.41E-05 | 9.38E-05 | -1.14E+00 |
| ESYT1 | 4.34E-08 | 2.26E-07 | -1.20E+00 | 4.85E-08 | 2.10E-07 | -1.03E+00 | 3.33E-05 | 1.41E-04 | -1.18E+00 | 9.98E-04 | 3.81E-03 | 1.08E+00 | 1.10E-05 | 3.92E-05 | -1.26E+00 | 1.46E-04 | 4.13E-04 | -1.09E+00 | 4.90E-03 | 1.80E-02 | -1.13E+00 | 4.19E-02 | 1.15E-01 | -1.03E+00 |
| KHSRP | 4.38E-08 | 2.28E-07 | -1.20E+00 | 9.92E-07 | 4.29E-06 | -1.06E+00 | 1.31E-02 | 5.56E-02 | -1.07E+00 | 1.25E-02 | 4.76E-02 | 1.11E+00 | 4.56E-05 | 1.63E-04 | -1.12E+00 | 1.92E-03 | 5.42E-03 | 1.06E+00 | 2.45E-13 | 8.97E-13 | -1.57E+00 | 1.30E-03 | 3.57E-03 | -1.01E+00 |
| PARP11 | 4.38E-08 | 2.28E-07 | -1.44E+00 | 1.02E-05 | 4.41E-05 | -1.08E+00 | 8.16E-15 | 3.47E-14 | -2.21E+00 | 3.34E-09 | 1.28E-08 | -1.43E+00 | 6.46E-06 | 2.30E-05 | -1.56E+00 | 4.55E-03 | 1.28E-02 | -1.11E+00 | 6.06E-05 | 2.22E-04 | -1.15E+00 | 7.88E-05 | 2.17E-04 | -1.15E+00 |
| AL157938.3 | 4.41E-08 | 2.30E-07 | -1.73E+00 | 3.64E-04 | 1.57E-03 | -1.02E+00 | 9.02E-08 | 3.83E-07 | -1.63E+00 | 1.11E-07 | 4.23E-07 | -1.48E+00 | 8.92E-05 | 3.18E-04 | -1.93E+00 | 1.45E-02 | 4.09E-02 | -1.18E+00 | 8.49E-11 | 3.12E-10 | -2.41E+00 | 4.24E-06 | 1.16E-05 | -1.59E+00 |
| RPS15AP30 | 4.46E-08 | 2.32E-07 | -1.71E+00 | 8.54E-09 | 3.69E-08 | -1.60E+00 | 6.94E-05 | 2.95E-04 | -1.38E+00 | 2.92E-07 | 1.12E-06 | -1.48E+00 | 1.28E-01 | 4.58E-01 | -1.95E+00 | 2.56E-01 | 7.20E-01 | -1.34E+00 | 4.93E-02 | 1.81E-01 | -1.69E+00 | 4.62E-02 | 1.27E-01 | -1.98E+00 |
| NR5A1 | 4.47E-08 | 2.33E-07 | 4.51E+00 | 7.42E-05 | 3.21E-04 | 4.01E+00 | 2.31E-02 | 9.81E-02 | 2.04E+00 | 7.74E-04 | 2.96E-03 | 3.18E+00 | 5.67E-02 | 2.02E-01 | 2.65E+00 | 5.36E-02 | 1.51E-01 | 2.97E+00 | 1.29E-01 | 4.75E-01 | 1.81E+00 | 1.21E-01 | 3.32E-01 | 1.83E+00 |
| XPOT | 4.49E-08 | 2.34E-07 | -1.38E+00 | 5.51E-04 | 2.38E-03 | 1.06E+00 | 3.93E-09 | 1.67E-08 | -1.46E+00 | 7.23E-05 | 2.76E-04 | -1.04E+00 | 8.70E-04 | 3.10E-03 | -1.10E+00 | 2.84E-03 | 8.02E-03 | -1.01E+00 | 1.73E-08 | 6.36E-08 | -1.38E+00 | 3.44E-02 | 9.46E-02 | 1.10E+00 |
| TMPO | 4.51E-08 | 2.35E-07 | -1.16E+00 | 1.75E-06 | 7.55E-06 | 1.02E+00 | 9.53E-03 | 4.05E-02 | 1.00E+00 | 3.44E-03 | 1.31E-02 | 1.10E+00 | 1.18E-10 | 4.20E-10 | -1.47E+00 | 6.29E-09 | 1.77E-08 | -1.40E+00 | 6.36E-06 | 2.33E-05 | -1.23E+00 | 5.28E-03 | 1.45E-02 | -1.09E+00 |
| CD81 | 4.51E-08 | 2.35E-07 | 1.84E+00 | 8.92E-05 | 3.85E-04 | 1.89E+00 | 1.35E-09 | 5.72E-09 | 1.82E+00 | 8.75E-05 | 3.34E-04 | 1.68E+00 | 2.02E-02 | 7.20E-02 | 1.90E+00 | 2.99E-01 | 8.44E-01 | 1.57E+00 | 3.69E-06 | 1.35E-05 | 2.12E+00 | 3.17E-05 | 8.71E-05 | 1.93E+00 |
| NSD1 | 4.51E-08 | 2.35E-07 | -1.16E+00 | 7.83E-06 | 3.38E-05 | 1.04E+00 | 5.66E-07 | 2.40E-06 | -1.34E+00 | 2.29E-05 | 8.73E-05 | -1.09E+00 | 1.13E-02 | 4.03E-02 | -1.03E+00 | 5.89E-04 | 1.66E-03 | -1.11E+00 | 9.87E-05 | 3.62E-04 | -1.10E+00 | 7.76E-04 | 2.13E-03 | -1.06E+00 |
| PRKAA2 | 4.55E-08 | 2.37E-07 | -1.72E+00 | 5.89E-06 | 2.55E-05 | -1.35E+00 | 7.79E-05 | 3.31E-04 | -2.07E+00 | 2.21E-02 | 8.45E-02 | 1.03E+00 | 3.97E-02 | 1.42E-01 | -1.56E+00 | 1.13E-01 | 3.19E-01 | -1.10E+00 | 1.79E-01 | 6.56E-01 | -1.03E+00 | 2.84E-01 | 7.80E-01 | -1.10E+00 |
| JAM2 | 4.56E-08 | 2.37E-07 | -1.47E+00 | 5.34E-04 | 2.31E-03 | 1.05E+00 | 2.47E-06 | 1.05E-05 | -1.54E+00 | 2.51E-05 | 9.60E-05 | -1.35E+00 | 6.37E-02 | 2.27E-01 | -1.46E+00 | 1.85E-01 | 5.20E-01 | -1.01E+00 | 2.43E-01 | 8.91E-01 | 1.03E+00 | 2.67E-01 | 7.34E-01 | -1.05E+00 |
| INPP5B | 4.61E-08 | 2.40E-07 | -1.41E+00 | 2.57E-05 | 1.11E-04 | -1.03E+00 | 2.38E-08 | 1.01E-07 | -1.58E+00 | 2.60E-07 | 9.92E-07 | -1.34E+00 | 2.89E-04 | 1.03E-03 | -1.19E+00 | 3.68E-03 | 1.04E-02 | -1.02E+00 | 3.64E-04 | 1.34E-03 | -1.16E+00 | 4.42E-05 | 1.21E-04 | -1.34E+00 |
| CHCHD1 | 4.61E-08 | 2.40E-07 | -1.09E+00 | 6.16E-05 | 2.66E-04 | 1.14E+00 | 9.43E-02 | 4.01E-01 | 1.05E+00 | 6.52E-02 | 2.49E-01 | 1.15E+00 | 2.36E-04 | 8.43E-04 | -1.14E+00 | 1.55E-02 | 4.37E-02 | 1.03E+00 | 1.90E-01 | 6.96E-01 | 1.16E+00 | 2.84E-01 | 7.80E-01 | 1.21E+00 |
| SH3KBP1 | 4.64E-08 | 2.42E-07 | -1.25E+00 | 6.38E-07 | 2.75E-06 | -1.03E+00 | 2.31E-04 | 9.82E-04 | -1.33E+00 | 5.12E-05 | 1.96E-04 | -1.16E+00 | 1.53E-07 | 5.46E-07 | -1.38E+00 | 7.55E-04 | 2.13E-03 | -1.11E+00 | 4.64E-10 | 1.70E-09 | -1.56E+00 | 1.42E-07 | 3.91E-07 | -1.42E+00 |
| AKT3 | 4.65E-08 | 2.42E-07 | -1.16E+00 | 1.61E-06 | 6.97E-06 | 1.07E+00 | 4.60E-02 | 1.95E-01 | 1.00E+00 | 8.61E-02 | 3.29E-01 | 1.21E+00 | 3.48E-05 | 1.24E-04 | -1.21E+00 | 1.14E-04 | 3.22E-04 | -1.11E+00 | 6.77E-03 | 2.48E-02 | -1.27E+00 | 3.05E-02 | 8.37E-02 | -1.13E+00 |
| DDX51 | 4.70E-08 | 2.45E-07 | -1.72E+00 | 2.67E-06 | 1.15E-05 | -1.22E+00 | 2.96E-03 | 1.26E-02 | -1.11E+00 | 5.86E-03 | 2.24E-02 | 1.15E+00 | 2.01E-04 | 7.17E-04 | -1.31E+00 | 1.25E-02 | 3.53E-02 | 1.07E+00 | 4.86E-04 | 1.78E-03 | -1.17E+00 | 8.84E-02 | 2.43E-01 | 1.10E+00 |
| SRSF10 | 4.71E-08 | 2.46E-07 | -1.11E+00 | 2.99E-16 | 1.29E-15 | -1.15E+00 | 1.40E-01 | 5.96E-01 | 1.03E+00 | 1.77E-03 | 6.77E-03 | -1.04E+00 | 7.93E-05 | 2.83E-04 | -1.08E+00 | 4.28E-06 | 1.21E-05 | -1.14E+00 | 6.87E-03 | 2.52E-02 | -1.09E+00 | 1.17E-01 | 3.22E-01 | -1.01E+00 |
| MLH3 | 4.72E-08 | 2.46E-07 | -1.42E+00 | 3.54E-03 | 1.53E-02 | 1.16E+00 | 7.59E-11 | 3.22E-10 | -1.89E+00 | 1.67E-04 | 6.39E-04 | -1.07E+00 | 2.50E-05 | 8.90E-05 | -1.35E+00 | 6.33E-03 | 1.78E-02 | 1.01E+00 | 1.15E-06 | 4.23E-06 | -1.26E+00 | 1.17E-01 | 3.20E-01 | 1.03E+00 |
| ZP3 | 4.72E-08 | 2.46E-07 | -1.38E+00 | 4.21E-08 | 1.82E-07 | -1.20E+00 | 1.74E-10 | 7.38E-10 | -1.49E+00 | 2.10E-06 | 8.04E-06 | -1.10E+00 | 1.82E-03 | 6.49E-03 | -1.11E+00 | 3.27E-04 | 9.22E-04 | -1.23E+00 | 4.38E-06 | 1.61E-05 | -1.39E+00 | 6.07E-05 | 1.67E-04 | -1.33E+00 |
| ARHGAP18 | 4.75E-08 | 2.47E-07 | -1.82E+00 | 6.68E-08 | 2.89E-07 | -1.53E+00 | 9.40E-10 | 3.99E-09 | -2.03E+00 | 2.17E-11 | 8.30E-11 | -2.25E+00 | 7.29E-06 | 2.60E-05 | -1.33E+00 | 6.81E-06 | 1.92E-05 | -1.35E+00 | 1.59E-03 | 5.83E-03 | 1.04E+00 | 3.10E-10 | 8.51E-10 | -1.38E+00 |
| PAK4 | 4.75E-08 | 2.48E-07 | -1.86E+00 | 5.34E-06 | 2.31E-05 | -1.43E+00 | 3.30E-04 | 1.40E-03 | -1.50E+00 | 2.06E-04 | 7.89E-04 | -1.34E+00 | 8.58E-04 | 3.06E-03 | -1.46E+00 | 8.33E-03 | 2.35E-02 | -1.15E+00 | 3.37E-07 | 1.24E-06 | -1.52E+00 | 6.13E-04 | 1.68E-03 | -1.24E+00 |
| AC127164.1 | 4.77E-08 | 2.49E-07 | -3.37E+00 | 1.20E-02 | 5.17E-02 | -1.08E+00 | 2.16E-03 | 9.19E-03 | -1.55E+00 | 3.65E-02 | 1.40E-01 | -1.01E+00 | 2.64E-04 | 9.43E-04 | -2.36E+00 | 7.73E-03 | 2.18E-02 | -1.34E+00 | 3.02E-03 | 1.11E-02 | -1.39E+00 | 1.87E-01 | 5.13E-01 | 1.03E+00 |
| XIAP | 4.77E-08 | 2.49E-07 | -1.28E+00 | 5.33E-08 | 2.30E-07 | -1.12E+00 | 3.92E-08 | 1.67E-07 | -1.50E+00 | 1.90E-06 | 7.26E-06 | -1.17E+00 | 2.76E-03 | 9.83E-03 | -1.29E+00 | 6.93E-03 | 1.95E-02 | -1.23E+00 | 1.76E-10 | 6.47E-10 | -1.59E+00 | 1.14E-08 | 3.12E-08 | -1.53E+00 |
| SLC35A1 | 4.80E-08 | 2.50E-07 | -1.58E+00 | 5.92E-03 | 2.56E-02 | 1.21E+00 | 1.93E-12 | 8.21E-12 | -1.84E+00 | 7.64E-05 | 2.92E-04 | -1.01E+00 | 2.89E-05 | 1.03E-04 | -1.58E+00 | 1.25E-01 | 3.54E-01 | 1.23E+00 | 2.12E-05 | 7.79E-05 | -1.53E+00 | 4.96E-02 | 1.36E-01 | -1.05E+00 |
| KIAA0368 | 4.80E-08 | 2.50E-07 | -1.33E+00 | 3.94E-07 | 1.70E-06 | -1.14E+00 | 2.05E-02 | 8.70E-02 | -1.00E+00 | 1.42E-01 | 5.43E-01 | 1.31E+00 | 6.06E-03 | 2.16E-02 | 1.01E+00 | 1.09E-01 | 3.07E-01 | 1.24E+00 | 4.43E-04 | 1.63E-03 | -1.12E+00 | 7.81E-03 | 2.14E-02 | 1.01E+00 |
| SCFD1 | 4.81E-08 | 2.51E-07 | -1.09E+00 | 3.86E-06 | 1.67E-05 | 1.09E+00 | 1.77E-04 | 7.53E-04 | -1.13E+00 | 3.60E-03 | 1.38E-02 | 1.08E+00 | 2.05E-04 | 7.32E-04 | -1.20E+00 | 7.70E-02 | 2.17E-01 | 1.09E+00 | 3.13E-03 | 1.15E-02 | -1.07E+00 | 5.33E-03 | 1.46E-02 | -1.11E+00 |
| INTS11 | 4.85E-08 | 2.53E-07 | -1.09E+00 | 2.66E-08 | 1.15E-07 | 1.04E+00 | 1.92E-05 | 8.17E-05 | -1.18E+00 | 1.72E-04 | 6.59E-04 | 1.02E+00 | 4.06E-06 | 1.45E-05 | -1.01E+00 | 5.85E-04 | 1.65E-03 | 1.12E+00 | 1.36E-06 | 4.99E-06 | -1.14E+00 | 1.16E-01 | 3.20E-01 | 1.16E+00 |
| TBK1 | 4.87E-08 | 2.54E-07 | -1.68E+00 | 7.49E-05 | 3.24E-04 | -1.11E+00 | 3.30E-07 | 1.40E-06 | -1.83E+00 | 7.10E-05 | 2.71E-04 | -1.31E+00 | 8.99E-07 | 3.21E-06 | -1.41E+00 | 6.09E-05 | 1.72E-04 | -1.21E+00 | 3.01E-14 | 1.10E-13 | -1.70E+00 | 4.43E-06 | 1.22E-05 | -1.25E+00 |
| EIF1 | 4.87E-08 | 2.54E-07 | -1.85E+00 | 1.32E-04 | 5.71E-04 | -1.08E+00 | 1.26E-04 | 5.33E-04 | -1.28E+00 | 8.03E-03 | 3.07E-02 | 1.07E+00 | 1.42E-04 | 5.06E-04 | -1.32E+00 | 5.23E-04 | 1.48E-03 | -1.19E+00 | 9.19E-07 | 3.37E-06 | -1.26E+00 | 2.66E-02 | 7.30E-02 | 1.04E+00 |
| P3H4 | 4.87E-08 | 2.54E-07 | -1.74E+00 | 1.81E-06 | 7.82E-06 | -1.33E+00 | 1.85E-05 | 7.86E-05 | -1.36E+00 | 1.23E-04 | 4.71E-04 | -1.10E+00 | 2.05E-05 | 7.30E-05 | -1.70E+00 | 2.28E-03 | 6.42E-03 | -1.17E+00 | 1.76E-09 | 6.47E-09 | -1.72E+00 | 1.88E-03 | 5.18E-03 | -1.12E+00 |
| HSP90AA1 | 4.89E-08 | 2.55E-07 | -1.15E+00 | 3.66E-02 | 1.58E-01 | 1.12E+00 | 1.07E-01 | 4.54E-01 | 1.04E+00 | 1.25E-08 | 4.77E-08 | 1.31E+00 | 6.97E-03 | 2.49E-02 | -1.11E+00 | 1.43E-02 | 4.04E-02 | 1.14E+00 | 1.30E-01 | 4.76E-01 | -1.00E+00 | 7.99E-15 | 2.19E-14 | 1.46E+00 |
| NAMPTP1 | 4.89E-08 | 2.55E-07 | -1.25E+00 | 1.55E-15 | 6.71E-15 | -1.63E+00 | 3.12E-03 | 1.33E-02 | -1.13E+00 | 6.26E-05 | 2.39E-04 | -1.17E+00 | 6.53E-06 | 2.33E-05 | -1.30E+00 | 2.17E-02 | 6.12E-02 | -1.03E+00 | 1.65E-11 | 6.05E-11 | -1.75E+00 | 7.49E-05 | 2.06E-04 | -1.25E+00 |
| PPARA | 4.89E-08 | 2.55E-07 | -1.78E+00 | 1.94E-04 | 8.37E-04 | -1.05E+00 | 2.16E-03 | 9.18E-03 | -1.34E+00 | 4.63E-02 | 1.77E-01 | 1.16E+00 | 3.09E-03 | 1.10E-02 | -1.25E+00 | 2.41E-02 | 6.79E-02 | 1.09E+00 | 1.05E-13 | 3.84E-13 | -2.15E+00 | 2.63E-08 | 7.21E-08 | -1.39E+00 |
| RNF10 | 4.90E-08 | 2.55E-07 | -1.16E+00 | 4.96E-13 | 2.15E-12 | -1.21E+00 | 1.19E-04 | 5.04E-04 | -1.13E+00 | 7.55E-04 | 2.88E-03 | -1.00E+00 | 8.30E-02 | 2.96E-01 | 1.19E+00 | 2.46E-02 | 6.93E-02 | 1.10E+00 | 2.64E-03 | 9.70E-03 | -1.11E+00 | 3.11E-04 | 8.54E-04 | -1.18E+00 |
| TRIM37 | 4.90E-08 | 2.55E-07 | -1.67E+00 | 5.24E-07 | 2.27E-06 | -1.30E+00 | 7.76E-06 | 3.30E-05 | -1.44E+00 | 3.42E-03 | 1.31E-02 | 1.02E+00 | 5.94E-12 | 2.12E-11 | -2.43E+00 | 2.48E-08 | 7.00E-08 | -1.49E+00 | 3.53E-14 | 1.29E-13 | -1.76E+00 | 6.60E-07 | 1.81E-06 | -1.25E+00 |
| DSTYK | 4.93E-08 | 2.57E-07 | -1.26E+00 | 9.91E-08 | 4.28E-07 | -1.14E+00 | 6.53E-07 | 2.77E-06 | -1.38E+00 | 7.06E-08 | 2.70E-07 | -1.27E+00 | 7.37E-03 | 2.63E-02 | 1.02E+00 | 2.59E-05 | 7.31E-05 | -1.26E+00 | 3.42E-10 | 1.25E-09 | -1.57E+00 | 1.42E-07 | 3.91E-07 | -1.52E+00 |
| CUX1 | 4.93E-08 | 2.57E-07 | -1.14E+00 | 1.87E-05 | 8.08E-05 | 1.04E+00 | 4.43E-06 | 1.88E-05 | -1.29E+00 | 5.11E-04 | 1.95E-03 | -1.05E+00 | 3.56E-03 | 1.27E-02 | 1.05E+00 | 2.04E-04 | 5.76E-04 | -1.10E+00 | 2.65E-04 | 9.72E-04 | -1.20E+00 | 3.64E-05 | 1.00E-04 | -1.26E+00 |
| BRD7P5 | 4.93E-08 | 2.57E-07 | -1.63E+00 | 2.25E-06 | 9.73E-06 | -1.30E+00 | 1.27E-04 | 5.39E-04 | -1.43E+00 | 2.59E-04 | 9.89E-04 | -1.20E+00 | 6.23E-03 | 2.22E-02 | -1.48E+00 | 1.85E-02 | 5.21E-02 | -1.33E+00 | 4.98E-04 | 1.83E-03 | -1.18E+00 | 1.87E-03 | 5.13E-03 | -1.10E+00 |
| ZNF321P | 4.95E-08 | 2.58E-07 | -1.61E+00 | 1.27E-03 | 5.49E-03 | 1.08E+00 | 4.36E-04 | 1.85E-03 | -1.39E+00 | 7.32E-02 | 2.80E-01 | 1.27E+00 | 7.11E-02 | 2.54E-01 | -1.11E+00 | 1.18E-01 | 3.31E-01 | -1.03E+00 | 2.21E-03 | 8.10E-03 | -1.47E+00 | 4.44E-02 | 1.22E-01 | -1.18E+00 |
| ZNF672 | 4.95E-08 | 2.58E-07 | -1.53E+00 | 6.28E-06 | 2.72E-05 | -1.26E+00 | 2.16E-11 | 9.16E-11 | -1.69E+00 | 6.86E-08 | 2.62E-07 | -1.16E+00 | 1.41E-04 | 5.04E-04 | -1.47E+00 | 1.90E-03 | 5.34E-03 | -1.19E+00 | 2.37E-08 | 8.70E-08 | -1.55E+00 | 8.05E-03 | 2.21E-02 | -1.03E+00 |
| CHML | 4.97E-08 | 2.59E-07 | -1.41E+00 | 5.62E-05 | 2.43E-04 | -1.02E+00 | 8.05E-02 | 3.42E-01 | -1.02E+00 | 4.65E-02 | 1.78E-01 | 1.51E+00 | 4.31E-04 | 1.54E-03 | -1.27E+00 | 3.01E-02 | 8.49E-02 | 1.03E+00 | 1.67E-03 | 6.12E-03 | -1.16E+00 | 1.21E-02 | 3.33E-02 | -1.11E+00 |
| GOLGA3 | 4.99E-08 | 2.60E-07 | -1.41E+00 | 2.30E-05 | 9.94E-05 | -1.03E+00 | 2.27E-06 | 9.63E-06 | -1.32E+00 | 2.94E-03 | 1.12E-02 | 1.10E+00 | 2.91E-05 | 1.04E-04 | -1.31E+00 | 4.18E-02 | 1.18E-01 | 1.09E+00 | 1.55E-05 | 5.69E-05 | -1.31E+00 | 2.97E-02 | 8.17E-02 | -1.02E+00 |
| RPL11 | 5.00E-08 | 2.61E-07 | 1.11E+00 | 1.17E-02 | 5.06E-02 | 1.05E+00 | 1.93E-10 | 8.21E-10 | 1.17E+00 | 1.18E-04 | 4.49E-04 | 1.10E+00 | 1.19E-07 | 4.24E-07 | 1.16E+00 | 4.31E-02 | 1.21E-01 | 1.05E+00 | 2.00E-05 | 7.34E-05 | 1.15E+00 | 1.26E-03 | 3.47E-03 | 1.07E+00 |
| CHD6 | 5.00E-08 | 2.61E-07 | -1.16E+00 | 4.08E-07 | 1.77E-06 | 1.02E+00 | 8.25E-06 | 3.51E-05 | -1.24E+00 | 5.58E-07 | 2.13E-06 | -1.11E+00 | 7.79E-07 | 2.78E-06 | -1.13E+00 | 7.67E-08 | 2.16E-07 | -1.09E+00 | 9.67E-08 | 3.55E-07 | -1.36E+00 | 9.28E-09 | 2.55E-08 | -1.39E+00 |
| ZNF589 | 5.02E-08 | 2.61E-07 | -1.85E+00 | 9.64E-06 | 4.17E-05 | -1.31E+00 | 7.66E-02 | 3.25E-01 | 1.06E+00 | 1.83E-01 | 7.00E-01 | 1.57E+00 | 1.26E-02 | 4.50E-02 | -1.08E+00 | 7.79E-03 | 2.20E-02 | -1.19E+00 | 4.85E-03 | 1.78E-02 | -1.43E+00 | 1.55E-01 | 4.26E-01 | 1.01E+00 |
| [illegible] | [illegible] | [illegible] | [illegible] | [illegible] | [illegible] | [illegible] | [illegible] | [illegible] | [illegible] | [illegible] | [illegible] | [illegible] | [illegible] | [illegible] | [illegible] | [illegible] | [illegible] | [illegible] | [illegible] | [illegible] | [illegible] | [illegible] | [illegible] | [illegible] |

| | | | | | | | | | | | | | | | | | | | | | | | | |
|---|---|---|---|---|---|---|---|---|---|---|---|---|---|---|---|---|---|---|---|---|---|---|---|---|
| SLC46A3 | 5.72E-08 | 2.98E-07 | -1.68E+00 | 1.54E-04 | 6.68E-04 | -1.14E+00 | 1.57E-07 | 6.65E-07 | -1.88E+00 | 2.50E-04 | 9.56E-04 | -1.19E+00 | 1.15E-02 | 4.11E-02 | -1.58E+00 | 5.98E-02 | 1.68E-01 | -1.20E+00 | 3.60E-02 | 1.32E-01 | -1.11E+00 | 2.59E-01 | 7.12E-01 | 1.13E+00 |
| POLR3F | 5.75E-08 | 3.00E-07 | -1.71E+00 | 9.15E-05 | 3.95E-04 | -1.12E+00 | 1.26E-05 | 5.34E-05 | -1.34E+00 | 1.28E-04 | 4.88E-04 | -1.05E+00 | 9.53E-04 | 3.40E-03 | -1.30E+00 | 4.92E-03 | 1.39E-02 | -1.09E+00 | 1.58E-07 | 5.81E-07 | -1.57E+00 | 3.77E-05 | 1.03E-04 | -1.31E+00 |
| UBE2E3 | 5.77E-08 | 3.01E-07 | -1.09E+00 | 2.59E-07 | 1.12E-06 | 1.04E+00 | 2.97E-05 | 1.26E-04 | -1.18E+00 | 5.22E-04 | 1.99E-03 | -1.03E+00 | 3.37E-06 | 1.20E-05 | -1.08E+00 | 2.64E-04 | 7.45E-04 | 1.01E+00 | 3.30E-04 | 1.21E-03 | -1.09E+00 | 2.26E-03 | 6.21E-03 | -1.09E+00 |
| HSP90B3P | 5.78E-08 | 3.01E-07 | -2.06E+00 | 4.36E-06 | 1.88E-05 | -1.48E+00 | 2.83E-08 | 1.20E-07 | -1.74E+00 | 5.14E-09 | 1.96E-08 | -1.67E+00 | 6.21E-05 | 2.21E-04 | -1.81E+00 | 3.68E-05 | 1.04E-04 | -2.02E+00 | 1.53E-07 | 5.61E-07 | -1.65E+00 | 1.37E-05 | 3.76E-05 | -1.38E+00 |
| JMY | 5.78E-08 | 3.01E-07 | -1.20E+00 | 7.37E-10 | 3.19E-09 | -1.21E+00 | 2.67E-04 | 1.13E-03 | -1.33E+00 | 4.80E-04 | 1.83E-03 | -1.12E+00 | 1.30E-03 | 4.62E-03 | -1.04E+00 | 3.05E-05 | 8.59E-05 | -1.14E+00 | 1.69E-04 | 6.19E-04 | -1.25E+00 | 1.01E-03 | 2.76E-03 | -1.18E+00 |
| SPRT | 5.80E-08 | 3.02E-07 | -1.27E+00 | 7.99E-06 | 3.45E-05 | -1.00E+00 | 1.81E-02 | 7.67E-02 | -1.13E+00 | 1.11E-02 | 4.23E-02 | 1.05E+00 | 2.66E-07 | 9.48E-07 | -1.36E+00 | 6.27E-04 | 1.77E-03 | -1.03E+00 | 4.08E-02 | 1.50E-01 | 1.06E+00 | 3.34E-01 | 9.17E-01 | 1.23E+00 |
| HERC6 | 5.83E-08 | 3.04E-07 | -1.43E+00 | 9.15E-08 | 3.95E-07 | -1.31E+00 | 3.46E-12 | 1.47E-11 | -2.19E+00 | 1.12E-09 | 4.30E-09 | -1.62E+00 | 3.10E-07 | 1.10E-06 | -2.55E+00 | 4.79E-06 | 1.35E-05 | -1.86E+00 | 2.47E-17 | 9.06E-17 | -2.81E+00 | 6.59E-16 | 1.81E-15 | -2.78E+00 |
| RNVU1-19 | 5.86E-08 | 3.05E-07 | 3.30E+00 | 1.57E-01 | 6.79E-01 | 1.21E+00 | 5.16E-03 | 2.19E-02 | 1.96E+00 | 5.14E-02 | 1.97E-01 | -1.01E+00 | 8.82E-02 | 3.15E-01 | -1.19E+00 | 2.88E-01 | 8.11E-01 | 1.23E+00 | 1.43E-02 | 5.26E-02 | -1.45E+00 | 8.41E-02 | 2.31E-01 | -1.11E+00 |
| TLR2 | 5.86E-08 | 3.05E-07 | -2.67E+00 | 2.59E-04 | 1.12E-03 | -1.25E+00 | 1.82E-02 | 7.71E-02 | -1.19E+00 | 2.34E-04 | 8.95E-04 | -1.45E+00 | 1.05E-02 | 3.73E-02 | -1.82E+00 | 8.85E-03 | 2.49E-02 | -1.88E+00 | 1.70E-05 | 6.24E-05 | -2.95E+00 | 1.08E-03 | 2.95E-03 | -1.93E+00 |
| RBCK1 | 5.91E-08 | 3.08E-07 | -1.08E+00 | 1.24E-07 | 5.38E-07 | 1.04E+00 | 3.54E-05 | 1.50E-04 | -1.20E+00 | 1.21E-07 | 4.63E-07 | -1.25E+00 | 1.31E-08 | 4.67E-08 | -1.23E+00 | 1.14E-06 | 3.20E-06 | -1.06E+00 | 1.59E-04 | 5.85E-04 | -1.06E+00 | 1.40E-03 | 3.85E-03 | -1.11E+00 |
| ABL2 | 5.94E-08 | 3.10E-07 | -1.48E+00 | 9.22E-03 | 3.99E-02 | 1.16E+00 | 1.05E-07 | 4.46E-07 | -1.56E+00 | 8.62E-05 | 3.29E-04 | -1.08E+00 | 6.38E-05 | 2.28E-04 | -1.48E+00 | 5.70E-03 | 1.61E-02 | -1.07E+00 | 1.39E-07 | 5.12E-07 | -1.66E+00 | 1.11E-03 | 3.06E-03 | -1.20E+00 |
| AC078899.1 | 5.96E-08 | 3.10E-07 | -1.68E+00 | 1.46E-09 | 6.32E-09 | -1.80E+00 | 6.39E-07 | 2.72E-06 | -1.57E+00 | 1.66E-08 | 6.33E-08 | -1.69E+00 | 3.38E-03 | 1.20E-02 | -1.23E+00 | 6.04E-05 | 1.70E-04 | -1.66E+00 | 5.69E-03 | 2.09E-02 | -1.10E+00 | 1.41E-02 | 3.87E-02 | -1.04E+00 |
| NAA50 | 5.96E-08 | 3.11E-07 | -1.15E+00 | 5.86E-09 | 2.53E-08 | -1.07E+00 | 1.11E-03 | 4.70E-03 | -1.03E+00 | 4.71E-04 | 1.80E-03 | 1.03E+00 | 2.61E-07 | 9.30E-07 | -1.34E+00 | 1.85E-04 | 5.23E-04 | -1.07E+00 | 9.51E-06 | 3.49E-05 | -1.22E+00 | 1.32E-01 | 3.63E-01 | 1.06E+00 |
| NLRC3 | 5.97E-08 | 3.11E-07 | -1.50E+00 | 2.36E-04 | 1.02E-03 | 1.01E+00 | 5.10E-03 | 2.17E-02 | -1.19E+00 | 1.26E-02 | 4.80E-02 | -1.04E+00 | 1.56E-05 | 5.56E-05 | -1.44E+00 | 1.57E-04 | 4.41E-04 | -1.21E+00 | 2.18E-02 | 8.00E-02 | -1.11E+00 | 1.10E-01 | 3.02E-01 | 1.04E+00 |
| SQOR | 5.98E-08 | 3.12E-07 | -1.22E+00 | 1.05E-07 | 4.55E-07 | -1.07E+00 | 1.69E-04 | 7.20E-04 | -1.17E+00 | 1.05E-08 | 3.99E-08 | -1.22E+00 | 2.23E-03 | 7.95E-03 | -1.61E+00 | 4.86E-03 | 1.37E-02 | -1.59E+00 | 1.14E-01 | 4.19E-01 | -1.12E+00 | 6.59E-02 | 1.81E-01 | -1.32E+00 |
| NUCB2 | 6.00E-08 | 3.13E-07 | -1.22E+00 | 3.57E-03 | 1.54E-02 | -1.04E+00 | 1.86E-01 | 7.90E-01 | -1.03E+00 | 1.36E-01 | 5.19E-01 | -1.06E+00 | 3.66E-06 | 1.30E-05 | -1.11E+00 | 5.47E-03 | 1.54E-02 | -1.03E+00 | 2.71E-01 | 9.96E-01 | 1.10E+00 | 3.51E-01 | 9.64E-01 | 1.09E+00 |
| SPATA2D | 6.01E-08 | 3.13E-07 | -1.45E+00 | 9.33E-07 | 4.03E-06 | -1.13E+00 | 7.75E-05 | 3.29E-04 | -1.33E+00 | 6.62E-07 | 2.53E-06 | -1.50E+00 | 8.03E-02 | 2.86E-01 | -1.05E+00 | 2.41E-02 | 6.79E-02 | -1.47E+00 | 1.45E-01 | 5.33E-01 | 1.05E+00 | 2.81E-01 | 7.71E-01 | 1.18E+00 |
| AKIRIN1 | 6.03E-08 | 3.14E-07 | -1.17E+00 | 8.47E-11 | 3.66E-10 | -1.15E+00 | 1.72E-11 | 7.29E-11 | -1.59E+00 | 1.42E-14 | 5.41E-14 | -1.48E+00 | 6.99E-11 | 2.49E-10 | -1.29E+00 | 3.49E-13 | 9.83E-13 | -1.31E+00 | 1.05E-06 | 3.86E-06 | -1.20E+00 | 1.65E-05 | 4.53E-05 | -1.16E+00 |
| SEC61A1 | 6.04E-08 | 3.15E-07 | -1.42E+00 | 5.00E-05 | 2.16E-04 | -1.07E+00 | 1.76E-03 | 7.49E-03 | -1.02E+00 | 4.05E-03 | 1.55E-02 | 1.11E+00 | 9.42E-06 | 3.36E-05 | -1.33E+00 | 1.31E-04 | 3.69E-04 | -1.19E+00 | 8.87E-05 | 3.26E-04 | -1.06E+00 | 2.99E-01 | 8.22E-01 | 1.23E+00 |
| SORD | 6.04E-08 | 3.15E-07 | -1.58E+00 | 1.02E-04 | 4.41E-04 | -1.08E+00 | 3.44E-02 | 1.46E-01 | -1.03E+00 | 1.73E-01 | 6.59E-01 | 1.28E+00 | 3.18E-02 | 1.13E-01 | -1.02E+00 | 2.14E-01 | 6.03E-01 | 1.64E+00 | 3.36E-05 | 1.23E-04 | -1.25E+00 | 3.37E-01 | 9.26E-01 | 1.24E+00 |
| FUNDC1 | 6.05E-08 | 3.15E-07 | -1.32E+00 | 6.58E-07 | 2.84E-06 | -1.14E+00 | 1.11E-05 | 4.72E-05 | -1.25E+00 | 4.47E-05 | 1.71E-04 | -1.06E+00 | 1.65E-07 | 5.87E-07 | -1.59E+00 | 1.22E-03 | 3.45E-03 | -1.14E+00 | 4.36E-03 | 1.60E-02 | -1.04E+00 | 6.30E-02 | 1.73E-01 | 1.10E+00 |
| GPR157 | 6.07E-08 | 3.16E-07 | -2.25E+00 | 1.02E-03 | 4.42E-03 | -1.10E+00 | 5.60E-04 | 2.38E-03 | -1.53E+00 | 1.73E-02 | 6.63E-02 | 1.04E+00 | 1.51E-02 | 5.38E-02 | -1.42E+00 | 4.28E-02 | 1.21E-01 | -1.27E+00 | 9.15E-02 | 3.36E-01 | -1.42E+00 | 1.37E-01 | 3.77E-01 | -1.37E+00 |
| DBP | 6.07E-08 | 3.16E-07 | -1.59E+00 | 7.52E-04 | 3.25E-03 | -1.01E+00 | 7.93E-10 | 3.37E-09 | -1.68E+00 | 1.15E-05 | 4.40E-05 | -1.12E+00 | 3.43E-11 | 1.22E-10 | -2.51E+00 | 6.79E-06 | 1.91E-05 | -1.34E+00 | 2.98E-19 | 1.09E-18 | -3.10E+00 | 7.21E-09 | 1.98E-08 | -1.64E+00 |
| PRDX6 | 6.11E-08 | 3.19E-07 | -1.17E+00 | 5.74E-06 | 2.48E-05 | -1.06E+00 | 1.22E-01 | 5.18E-01 | -1.01E+00 | 2.18E-01 | 8.33E-01 | 1.05E+00 | 6.13E-03 | 2.19E-02 | -1.08E+00 | 6.75E-02 | 1.90E-01 | -1.03E+00 | 1.52E-01 | 5.56E-01 | 1.03E+00 | 3.63E-01 | 9.98E-01 | 1.04E+00 |
| NHLRC3 | 6.14E-08 | 3.20E-07 | -1.03E+00 | 9.14E-02 | 3.95E-01 | 1.40E+00 | 1.63E-04 | 6.94E-04 | -1.14E+00 | 2.41E-01 | 9.20E-01 | 1.35E+00 | 1.09E-05 | 3.89E-05 | -1.05E+00 | 7.25E-02 | 2.04E-01 | 1.32E+00 | 1.11E-01 | 4.08E-01 | 1.09E+00 | 7.69E-02 | 2.11E-01 | 1.36E+00 |
| HEMK1 | 6.17E-08 | 3.22E-07 | -1.45E+00 | 8.73E-05 | 3.77E-04 | -1.01E+00 | 4.75E-06 | 2.02E-05 | -1.45E+00 | 1.20E-05 | 4.59E-05 | -1.28E+00 | 1.55E-04 | 5.52E-04 | -1.37E+00 | 1.34E-03 | 3.77E-03 | -1.15E+00 | 1.92E-04 | 7.04E-04 | -1.22E+00 | 1.24E-02 | 3.40E-02 | -1.02E+00 |
| TCEANC2 | 6.19E-08 | 3.22E-07 | -1.51E+00 | 1.11E-02 | 4.78E-02 | 1.25E+00 | 1.22E-06 | 5.20E-06 | -1.88E+00 | 9.77E-02 | 3.73E-01 | 1.12E+00 | 2.27E-08 | 8.08E-08 | -2.21E+00 | 2.89E-04 | 8.15E-04 | -1.20E+00 | 6.44E-05 | 2.36E-04 | -1.39E+00 | 5.25E-03 | 1.44E-02 | -1.17E+00 |
| KANSL1L | 6.20E-08 | 3.23E-07 | -1.38E+00 | 2.03E-03 | 8.77E-03 | 1.12E+00 | 1.70E-06 | 7.21E-06 | -1.49E+00 | 2.06E-04 | 7.86E-04 | -1.10E+00 | 3.31E-02 | 1.18E-01 | 1.05E+00 | 5.98E-02 | 1.68E-01 | 1.14E+00 | 1.98E-05 | 7.25E-05 | -1.43E+00 | 4.04E-03 | 1.11E-02 | -1.13E+00 |
| SERF2 | 6.21E-08 | 3.24E-07 | -1.13E+00 | 8.39E-17 | 3.63E-16 | -1.23E+00 | 1.55E-01 | 6.59E-01 | 1.02E+00 | 1.33E-07 | 5.07E-07 | -1.21E+00 | 2.03E-03 | 7.24E-03 | -1.10E+00 | 3.44E-05 | 9.69E-05 | -1.18E+00 | 4.15E-04 | 1.52E-03 | -1.12E+00 | 6.68E-09 | 1.84E-08 | -1.28E+00 |
| RPS27 | 6.27E-08 | 3.27E-07 | -1.14E+00 | 4.17E-12 | 1.80E-11 | -1.13E+00 | 4.30E-02 | 1.83E-01 | 1.02E+00 | 5.69E-06 | 2.17E-05 | -1.13E+00 | 7.61E-03 | 2.71E-02 | 1.04E+00 | 3.61E-04 | 1.02E-03 | -1.04E+00 | 3.64E-02 | 1.34E-01 | 1.01E+00 | 3.26E-07 | 8.95E-07 | -1.26E+00 |
| NSMCE4A | 6.28E-08 | 3.27E-07 | -1.21E+00 | 8.33E-05 | 3.60E-04 | 1.13E+00 | 2.04E-02 | 8.66E-02 | -1.01E+00 | 3.59E-02 | 1.37E-01 | 1.21E+00 | 8.45E-10 | 3.01E-09 | -1.49E+00 | 9.23E-05 | 2.60E-04 | -1.01E+00 | 5.68E-05 | 2.08E-04 | -1.19E+00 | 2.45E-01 | 6.74E-01 | 1.24E+00 |
| REXO5 | 6.28E-08 | 3.27E-07 | -1.99E+00 | 5.66E-07 | 2.44E-06 | -1.58E+00 | 2.83E-06 | 1.20E-05 | -1.39E+00 | 1.17E-06 | 4.46E-06 | -1.33E+00 | 6.26E-15 | 2.23E-14 | -2.14E+00 | 1.23E-19 | 3.46E-19 | -2.39E+00 | 2.79E-02 | 1.02E-01 | -1.01E+00 | 2.88E-03 | 7.92E-03 | -1.27E+00 |
| RFX7 | 6.29E-08 | 3.28E-07 | -1.49E+00 | 1.96E-05 | 8.49E-05 | -1.07E+00 | 3.03E-08 | 1.29E-07 | -1.57E+00 | 2.67E-04 | 1.02E-03 | -1.03E+00 | 7.40E-08 | 2.64E-07 | -1.77E+00 | 1.54E-03 | 4.35E-03 | -1.08E+00 | 2.02E-16 | 7.41E-16 | -1.94E+00 | 6.94E-08 | 1.91E-07 | -1.34E+00 |
| KRT8 | 6.32E-08 | 3.30E-07 | -1.12E+00 | 1.39E-02 | 6.02E-02 | 1.28E+00 | 9.28E-02 | 3.94E-01 | 1.06E+00 | 2.19E-01 | 8.38E-01 | 1.45E+00 | 3.91E-04 | 1.40E-03 | -1.25E+00 | 4.38E-04 | 1.23E-03 | -1.28E+00 | 1.96E-01 | 7.21E-01 | 1.17E+00 | 3.20E-01 | 8.79E-01 | 1.27E+00 |
| DCUN1D5 | 6.33E-08 | 3.30E-07 | -1.26E+00 | 1.28E-08 | 5.55E-08 | -1.22E+00 | 5.67E-09 | 2.41E-08 | -1.29E+00 | 1.60E-11 | 6.13E-11 | -1.29E+00 | 8.54E-07 | 3.05E-06 | -1.31E+00 | 2.72E-06 | 7.68E-06 | -1.22E+00 | 2.16E-07 | 7.93E-07 | -1.23E+00 | 4.49E-03 | 1.23E-02 | -1.04E+00 |
| TGFBR1 | 6.35E-08 | 3.31E-07 | -1.53E+00 | 3.16E-07 | 1.37E-06 | -1.31E+00 | 8.53E-09 | 3.63E-08 | -1.60E+00 | 1.24E-09 | 4.73E-09 | -1.47E+00 | 1.46E-02 | 5.21E-02 | 1.03E+00 | 2.49E-04 | 7.02E-04 | -1.51E+00 | 2.11E-01 | 7.73E-01 | 1.11E+00 | 3.26E-01 | 8.96E-01 | 1.21E+00 |
| DYNLT1 | 6.36E-08 | 3.32E-07 | -1.07E+00 | 1.32E-08 | 5.72E-08 | -1.03E+00 | 4.95E-07 | 2.10E-06 | -1.32E+00 | 1.02E-05 | 3.88E-05 | -1.12E+00 | 1.58E-03 | 5.65E-03 | 1.17E+00 | 3.50E-07 | 9.86E-07 | -1.21E+00 | 4.21E-11 | 1.55E-10 | -1.59E+00 | 5.63E-06 | 1.55E-05 | -1.27E+00 |
| RPS6KA6 | 6.36E-08 | 3.32E-07 | -1.87E+00 | 6.46E-04 | 2.79E-03 | -1.04E+00 | 4.61E-02 | 1.96E-01 | -1.18E+00 | 8.40E-02 | 3.21E-01 | 1.01E+00 | 2.29E-01 | 8.16E-01 | -1.16E+00 | 3.27E-01 | 9.23E-01 | 1.01E+00 | 2.05E-01 | 7.52E-01 | 1.04E+00 | 2.00E-01 | 5.49E-01 | -1.16E+00 |
| H3F3AP4 | 6.42E-08 | 3.34E-07 | -1.23E+00 | 2.57E-07 | 1.11E-06 | -1.17E+00 | 6.33E-05 | 2.69E-04 | -1.29E+00 | 4.35E-03 | 1.66E-02 | -1.17E+00 | 1.39E-10 | 4.95E-10 | -1.48E+00 | 1.73E-07 | 4.89E-07 | -1.34E+00 | 1.11E-09 | 4.07E-09 | -1.45E+00 | 5.54E-07 | 1.52E-06 | -1.34E+00 |
| LRRC57 | 6.52E-08 | 3.40E-07 | -1.26E+00 | 9.93E-04 | 4.29E-03 | 1.14E+00 | 7.05E-05 | 2.99E-04 | -1.34E+00 | 7.33E-03 | 2.80E-02 | 1.06E+00 | 1.57E-04 | 5.58E-04 | -1.14E+00 | 2.39E-02 | 6.74E-02 | 1.12E+00 | 2.29E-05 | 8.42E-05 | -1.36E+00 | 2.67E-02 | 7.32E-02 | 1.00E+00 |
| ZNRF3 | 6.53E-08 | 3.40E-07 | -2.55E+00 | 6.41E-05 | 2.77E-04 | -1.43E+00 | 1.23E-07 | 5.21E-07 | -2.36E+00 | 7.34E-07 | 2.80E-06 | -1.93E+00 | 8.54E-02 | 3.04E-01 | -1.51E+00 | 3.03E-01 | 8.53E-01 | 1.00E+00 | 6.91E-02 | 2.54E-01 | 1.17E+00 | 2.33E-01 | 6.41E-01 | 1.14E+00 |
| RACK1 | 6.55E-08 | 3.41E-07 | 1.12E+00 | 2.05E-10 | 8.84E-10 | 1.15E+00 | 1.20E-07 | 5.08E-07 | 1.16E+00 | 3.80E-12 | 1.45E-11 | 1.22E+00 | 4.83E-06 | 1.72E-05 | 1.15E+00 | 2.20E-08 | 6.20E-08 | 1.19E+00 | 2.69E-05 | 9.86E-05 | 1.15E+00 | 2.41E-09 | 6.61E-09 | 1.19E+00 |
| CWF19L2 | 6.59E-08 | 3.43E-07 | -1.36E+00 | 7.09E-05 | 3.06E-04 | -1.00E+00 | 2.46E-06 | 1.04E-05 | -1.41E+00 | 1.91E-04 | 7.31E-04 | -1.11E+00 | 8.97E-04 | 3.20E-03 | -1.41E+00 | 8.26E-03 | 2.33E-02 | -1.21E+00 | 2.66E-03 | 9.77E-03 | -1.08E+00 | 1.68E-03 | 4.60E-03 | -1.20E+00 |
| B3GALNT2 | 6.61E-08 | 3.44E-07 | -1.71E+00 | 1.26E-07 | 5.44E-07 | -1.51E+00 | 5.00E-08 | 2.13E-07 | -1.74E+00 | 4.78E-05 | 1.83E-04 | -1.22E+00 | 7.34E-11 | 2.62E-10 | -1.99E+00 | 1.98E-08 | 5.58E-08 | -1.62E+00 | 2.48E-06 | 9.09E-06 | -1.30E+00 | 1.89E-06 | 5.18E-06 | -1.31E+00 |
| TESC | 6.61E-08 | 3.45E-07 | -1.53E+00 | 3.50E-10 | 1.51E-09 | -1.77E+00 | 1.40E-03 | 5.94E-03 | -1.03E+00 | 1.30E-07 | 4.96E-07 | -1.28E+00 | 1.91E-37 | 6.82E-37 | -2.93E+00 | 3.04E-22 | 8.58E-22 | -1.99E+00 | 2.05E-30 | 7.54E-30 | -1.81E+00 | 4.81E-14 | 1.32E-13 | -1.52E+00 |
| RIPK1 | 6.61E-08 | 3.45E-07 | -1.53E+00 | 3.84E-05 | 1.66E-04 | -1.05E+00 | 3.35E-08 | 1.42E-07 | -1.70E+00 | 1.30E-06 | 4.95E-06 | -1.29E+00 | 1.62E-04 | 5.78E-04 | -1.64E+00 | 5.51E-02 | 1.55E-01 | -1.01E+00 | 2.02E-05 | 7.40E-05 | -1.38E+00 | 3.03E-02 | 8.31E-02 | -1.00E+00 |
| TMEM14C | 6.64E-08 | 3.46E-07 | -1.09E+00 | 3.21E-04 | 1.39E-03 | 1.09E+00 | 9.38E-02 | 3.99E-01 | 1.02E+00 | 2.01E-01 | 7.69E-01 | 1.14E+00 | 4.96E-05 | 1.77E-04 | 1.01E+00 | 2.65E-01 | 7.46E-01 | 1.51E+00 | 2.52E-01 | 9.25E-01 | 1.04E+00 | 2.07E-01 | 5.69E-01 | -1.01E+00 |
| CDK2AP2 | 6.66E-08 | 3.47E-07 | -1.47E+00 | 2.19E-03 | 9.46E-03 | 1.08E+00 | 1.32E-07 | 5.61E-07 | -1.35E+00 | 1.20E-05 | 4.57E-05 | -1.09E+00 | 2.12E-17 | 7.56E-17 | -2.41E+00 | 2.05E-11 | 5.79E-11 | -1.73E+00 | 7.44E-07 | 2.73E-06 | -1.54E+00 | 5.24E-03 | 1.44E-02 | -1.20E+00 |
| SLC4A1AP | 6.66E-08 | 3.47E-07 | -1.43E+00 | 1.91E-05 | 8.24E-05 | -1.11E+00 | 3.19E-05 | 1.35E-04 | -1.35E+00 | 2.12E-04 | 8.10E-04 | -1.10E+00 | 3.30E-07 | 1.18E-06 | -1.56E+00 | 2.05E-06 | 5.77E-06 | -1.39E+00 | 7.28E-09 | 2.67E-08 | -1.35E+00 | 1.06E-05 | 2.92E-05 | -1.16E+00 |
| CC2D1B | 6.67E-08 | 3.47E-07 | -1.60E+00 | 2.66E-05 | 1.15E-04 | -1.16E+00 | 7.42E-02 | 3.15E-01 | -1.11E+00 | 5.76E-02 | 2.20E-01 | -1.00E+00 | 4.18E-03 | 1.49E-02 | -1.27E+00 | 8.34E-03 | 2.35E-02 | -1.22E+00 | 1.04E-01 | 3.83E-01 | 1.02E+00 | 1.62E-01 | 4.44E-01 | 1.05E+00 |
| RAD51C | 6.75E-08 | 3.52E-07 | -1.32E+00 | 4.48E-05 | 1.94E-04 | 1.03E+00 | 7.38E-04 | 3.13E-03 | -1.05E+00 | 3.73E-04 | 1.43E-03 | 1.06E+00 | 2.28E-05 | 8.15E-05 | -1.30E+00 | 1.41E-03 | 3.98E-03 | -1.06E+00 | 1.47E-04 | 5.39E-04 | -1.08E+00 | 2.57E-01 | 7.05E-01 | 1.23E+00 |
| FAM110B | 6.75E-08 | 3.52E-07 | -1.91E+00 | 7.74E-06 | 3.34E-05 | -1.31E+00 | 3.00E-02 | 1.27E-01 | -1.27E+00 | 7.22E-02 | 2.76E-01 | -1.09E+00 | 3.44E-02 | 1.23E-01 | -1.68E+00 | 1.16E-01 | 3.26E-01 | -1.11E+00 | 2.57E-01 | 9.45E-01 | 1.13E+00 | 2.31E-01 | 6.34E-01 | 1.59E+00 |
| COX8A | 6.81E-08 | 3.55E-07 | -1.14E+00 | 3.13E-19 | 1.35E-18 | -1.25E+00 | 2.20E-01 | 9.34E-01 | 1.14E+00 | 3.59E-04 | 1.37E-03 | -1.12E+00 | 6.03E-04 | 2.15E-03 | -1.10E+00 | 2.14E-05 | 6.04E-05 | -1.17E+00 | 1.96E-01 | 7.20E-01 | 1.03E+00 | 6.34E-02 | 1.74E-01 | -1.13E+00 |
| KIAA1147 | 6.82E-08 | 3.55E-07 | -1.23E+00 | 3.01E-05 | 1.30E-04 | 1.03E+00 | 1.55E-05 | 6.57E-05 | -1.19E+00 | 8.10E-05 | 3.09E-04 | 1.01E+00 | 2.06E-04 | 7.33E-04 | -1.12E+00 | 1.00E-03 | 2.83E-03 | -1.07E+00 | 3.17E-02 | 1.16E-01 | 1.03E+00 | 1.02E-01 | 2.81E-01 | 1.15E+00 |
| EGFEM1P | 6.82E-08 | 3.55E-07 | -1.26E+00 | 2.12E-01 | 9.17E-01 | 1.50E+00 | 1.15E-07 | 4.90E-07 | -1.49E+00 | 4.32E-04 | 1.65E-03 | 1.05E+00 | 4.46E-02 | 1.59E-01 | 2.74E+00 | 1.39E-02 | 3.92E-02 | 3.37E+00 | 2.56E-01 | 9.40E-01 | 1.08E+00 | 2.35E-01 | 6.46E-01 | 1.33E+00 |
| IMPDH1 | 6.86E-08 | 3.58E-07 | -1.64E+00 | 1.38E-04 | 5.94E-04 | -1.03E+00 | 1.03E-09 | 4.36E-09 | -1.45E+00 | 1.55E-04 | 5.93E-04 | 1.04E+00 | 3.94E-04 | 1.40E-03 | -1.24E+00 | 8.54E-03 | 2.41E-02 | -1.08E+00 | 6.91E-02 | 2.54E-01 | 1.10E+00 | 5.99E-02 | 1.65E-01 | 1.55E+00 |
| FAM229B | 6.87E-08 | 3.58E-07 | -1.43E+00 | 2.55E-08 | 1.10E-07 | -1.30E+00 | 6.43E-04 | 2.73E-03 | -1.20E+00 | 2.07E-05 | 7.89E-05 | -1.27E+00 | 1.71E-02 | 6.09E-02 | 1.07E+00 | 1.93E-01 | 5.44E-01 | 1.38E+00 | 2.25E-01 | 8.24E-01 | 1.24E+00 | 1.49E-01 | 4.09E-01 | 1.38E+00 |
| C11orf54 | 6.90E-08 | 3.60E-07 | -1.30E+00 | 3.74E-03 | 1.62E-02 | 1.23E+00 | 1.37E-07 | 5.84E-07 | -1.46E+00 | 2.02E-04 | 7.72E-04 | -1.02E+00 | 3.30E-05 | 1.18E-04 | -1.18E+00 | 9.98E-03 | 2.81E-02 | 1.14E+00 | 2.67E-04 | 9.79E-04 | -1.06E+00 | 4.89E-02 | 1.34E-01 | 1.07E+00 |
| PDE4D | 6.90E-08 | 3.60E-07 | -1.29E+00 | 4.03E-07 | 1.74E-06 | -1.07E+00 | 2.54E-04 | 1.08E-03 | -1.23E+00 | 1.56E-06 | 5.95E-06 | -1.31E+00 | 3.91E-03 | 1.40E-02 | -1.22E+00 | 2.00E-03 | 5.62E-03 | -1.32E+00 | 7.28E-02 | 2.67E-01 | 1.03E+00 | 1.43E-02 | 3.93E-02 | -1.17E+00 |
| NFKB2 | 6.91E-08 | 3.60E-07 | -1.28E+00 | 2.07E-16 | 8.95E-16 | -2.09E+00 | 2.21E-09 | 9.38E-09 | -1.56E+00 | 8.45E-13 | 3.23E-12 | -2.03E+00 | 2.28E-04 | 8.11E-04 | -1.69E+00 | 1.72E-04 | 4.86E-04 | -1.74E+00 | 5.98E-11 | 2.19E-10 | -2.08E+00 | 7.08E-14 | 1.94E-13 | -3.01E+00 |
| GTF3C5 | 6.91E-08 | 3.60E-07 | -1.43E+00 | 2.57E-06 | 1.11E-05 | -1.14E+00 | 4.10E-10 | 1.74E-09 | -1.53E+00 | 4.51E-06 | 1.72E-05 | -1.10E+00 | 3.02E-06 | 1.08E-05 | -1.53E+00 | 7.01E-03 | 1.98E-02 | -1.09E+00 | 2.29E-09 | 8.39E-09 | -1.29E+00 | 2.96E-05 | 8.12E-05 | -1.06E+00 |
| MAP4K2 | 6.95E-08 | 3.62E-07 | -1.18E+00 | 3.63E-02 | 1.57E-01 | 1.35E+00 | 9.40E-04 | 3.99E-03 | -1.13E+00 | 5.94E-02 | 2.27E-01 | 1.24E+00 | 2.79E-05 | 9.96E-05 | -1.59E+00 | 3.51E-04 | 9.88E-04 | -1.22E+00 | 2.71E-01 | 9.93E-01 | 1.28E+00 | 2.45E-01 | 6.73E-01 | 1.45E+00 |
| AL359976.1 | 6.99E-08 | 3.64E-07 | -1.76E+00 | 1.54E-10 | 6.64E-10 | -2.23E+00 | 2.56E-07 | 1.09E-06 | -1.73E+00 | 3.76E-11 | 1.44E-10 | -2.46E+00 | 1.51E-05 | 5.39E-05 | -2.13E+00 | 3.46E-05 | 9.75E-05 | -1.93E+00 | 2.30E-09 | 8.42E-09 | -1.69E+00 | 2.17E-12 | 5.97E-12 | -2.27E+00 |
| PCLO | 6.99E-08 | 3.64E-07 | -1.85E+00 | 4.68E-04 | 2.02E-03 | -1.16E+00 | 5.70E-02 | 2.42E-01 | -1.27E+00 | 6.83E-02 | 2.61E-01 | -1.20E+00 | 2.69E-01 | 9.59E-01 | 1.20E+00 | 3.40E-01 | 9.57E-01 | 1.49E+00 | 2.23E-01 | 8.19E-01 | 1.03E+00 | 3.58E-01 | 9.82E-01 | 1.08E+00 |
| CBX5 | 7.00E-08 | 3.65E-07 | -1.07E+00 | 1.06E-04 | 4.58E-04 | 1.16E+00 | 5.78E-03 | 2.46E-02 | -1.05E+00 | 7.09E-03 | 2.71E-02 | 1.10E+00 | 5.51E-09 | 1.97E-08 | -1.33E+00 | 2.78E-04 | 7.83E-04 | -1.03E+00 | 2.67E-01 | 9.80E-01 | 1.08E+00 | 9.57E-02 | 2.63E-01 | 1.23E+00 |
| SHTN1 | 7.04E-08 | 3.67E-07 | -2.40E+00 | 1.06E-04 | 4.58E-04 | -1.40E+00 | 5.92E-02 | 2.51E-01 | -1.08E+00 | 2.07E-01 | 7.90E-01 | 1.22E+00 | 5.70E-02 | 2.03E-01 | -1.11E+00 | 5.48E-02 | 1.55E-01 | -1.13E+00 | 1.98E-04 | 7.25E-04 | -1.35E+00 | 3.42E-02 | 9.40E-02 | -1.08E+00 |
| GUCD1 | 7.04E-08 | 3.67E-07 | -1.49E+00 | 4.06E-06 | 1.75E-05 | -1.10E+00 | 2.63E-03 | 1.12E-02 | -1.16E+00 | 5.56E-03 | 2.12E-02 | 1.05E+00 | 2.03E-05 | 7.24E-05 | -1.32E+00 | 3.32E-04 | 9.37E-04 | -1.07E+00 | 1.12E-04 | 4.10E-04 | -1.18E+00 | 5.22E-02 | 1.43E-01 | 1.09E+00 |
| B3GNT8 | 7.10E-08 | 3.70E-07 | -1.64E+00 | 1.23E-05 | 5.31E-05 | -1.20E+00 | 2.48E-11 | 1.05E-10 | -2.25E+00 | 4.12E-05 | 1.57E-04 | -1.25E+00 | 1.47E-06 | 5.23E-06 | -5.04E+00 | 1.46E-04 | 4.11E-04 | -2.26E+00 | 2.59E-01 | 9.52E-01 | 1.05E+00 | 2.61E-01 | 7.18E-01 | 1.42E+00 |
| AC012150.2 | 7.12E-08 | 3.71E-07 | -2.71E+00 | 1.16E-06 | 5.02E-06 | -2.06E+00 | 6.76E-04 | 2.87E-03 | -1.56E+00 | 9.16E-05 | 3.50E-04 | -1.68E+00 | 9.41E-05 | 3.36E-04 | -1.88E+00 | 5.36E-04 | 1.51E-03 | -1.48E+00 | 1.17E-01 | 4.30E-01 | -1.57E+00 | 1.93E-01 | 5.30E-01 | -1.45E+00 |
| ORC4 | 7.14E-08 | 3.72E-07 | -1.15E+00 | 2.12E-03 | 9.16E-03 | 1.27E+00 | 6.67E-06 | 2.83E-05 | -1.27E+00 | 3.70E-03 | 1.41E-02 | 1.10E+00 | 2.82E-04 | 1.00E-03 | 1.02E+00 | 2.51E-01 | 7.08E-01 | 1.41E+00 | 2.50E-07 | 9.18E-07 | -1.25E+00 | 1.57E-03 | 4.31E-03 | -1.05E+00 |
| DHRS4 | 7.17E-08 | 3.74E-07 | -1.16E+00 | 2.10E-10 | 9.07E-10 | -1.16E+00 | 1.66E-04 | 7.05E-04 | -1.17E+00 | 4.34E-07 | 1.66E-06 | -1.22E+00 | 5.65E-07 | 2.01E-06 | -1.37E+00 | 4.92E-07 | 1.39E-06 | -1.28E+00 | 7.68E-02 | 2.82E-01 | 1.15E+00 | 6.86E-02 | 1.88E-01 | 1.09E+00 |
| WASF3 | 7.17E-08 | 3.74E-07 | -2.25E+00 | 2.62E-05 | 1.13E-04 | -1.46E+00 | 4.12E-02 | 1.75E-01 | -1.09E+00 | 1.61E-02 | 6.17E-02 | -1.18E+00 | 2.11E-02 | 7.51E-02 | -1.37E+00 | 2.77E-02 | 7.82E-02 | -1.47E+00 | 1.33E-04 | 4.87E-04 | -1.20E+00 | 1.88E-02 | 5.17E-02 | -1.01E+00 |
| CGRRF1 | 7.25E-08 | 3.78E-07 | -1.36E+00 | 3.81E-06 | 1.65E-05 | -1.10E+00 | 8.87E-04 | 3.77E-03 | -1.39E+00 | 8.08E-04 | 3.09E-03 | -1.16E+00 | 8.96E-05 | 3.20E-04 | -1.30E+00 | 6.60E-05 | 1.86E-04 | -1.22E+00 | 7.77E-03 | 2.85E-02 | 1.20E+00 | 8.49E-05 | 2.33E-04 | -1.15E+00 |
| TRAPPC12 | 7.30E-08 | 3.80E-07 | -1.17E+00 | 7.01E-05 | 3.03E-04 | 1.10E+00 | 1.69E-06 | 7.18E-06 | -1.23E+00 | 1.52E-03 | 5.79E-03 | 1.10E+00 | 1.19E-04 | 4.26E-04 | -1.30E+00 | 4.46E-02 | 1.26E-01 | 1.06E+00 | 4.56E-03 | 1.67E-02 | -1.14E+00 | 7.95E-02 | 2.18E-01 | 1.04E+00 |
| METAP1D | 7.31E-08 | 3.81E-07 | -1.07E+00 | 4.54E-05 | 1.96E-04 | 1.14E+00 | 1.29E-11 | 5.48E-11 | -1.66E+00 | 1.62E-04 | 6.18E-04 | 1.01E+00 | 1.81E-03 | 6.44E-03 | -1.15E+00 | 3.38E-01 | 9.51E-01 | 1.40E+00 | 2.14E-05 | 7.87E-05 | -1.22E+00 | 6.18E-02 | 1.70E-01 | 1.09E+00 |
| F2RL2 | 7.31E-08 | 3.81E-07 | -1.26E+00 | 2.79E-04 | 1.21E-03 | 1.10E+00 | 2.93E-05 | 1.24E-04 | -1.30E+00 | 7.50E-03 | 2.86E-02 | 1.08E+00 | 3.53E-08 | 1.26E-07 | -1.60E+00 | 1.05E-05 | 2.97E-05 | -1.22E+00 | 1.54E-05 | 5.66E-05 | -1.68E+00 | 3.07E-02 | 8.44E-02 | -1.02E+00 |
| MAP2K7 | 7.31E-08 | 3.81E-07 | -1.33E+00 | 4.37E-05 | 1.89E-04 | -1.03E+00 | 1.16E-04 | 4.92E-04 | -1.31E+00 | 3.17E-03 | 1.21E-02 | -1.04E+00 | 1.56E-04 | 5.55E-04 | -1.18E+00 | 5.84E-04 | 1.65E-03 | -1.05E+00 | 1.88E-06 | 6.89E-06 | -1.29E+00 | 5.92E-04 | 1.63E-03 | -1.05E+00 |
| SPRYD7 | 7.31E-08 | 3.81E-07 | -1.47E+00 | 9.56E-07 | 4.13E-06 | -1.24E+00 | 7.17E-07 | 3.05E-06 | -1.62E+00 | 2.72E-05 | 1.04E-04 | -1.22E+00 | 3.22E-03 | 1.15E-02 | -1.31E+00 | 8.90E-03 | 2.51E-02 | -1.13E+00 | 1.64E-10 | 6.00E-10 | -1.51E+00 | 5.03E-09 | 1.38E-08 | -1.33E+00 |
| ANKRD10-IT1 | 7.31E-08 | 3.81E-07 | -1.79E+00 | 2.57E-03 | 1.11E-02 | 1.02E+00 | 1.36E-11 | 5.77E-11 | -2.13E+00 | 1.55E-04 | 5.92E-04 | -1.06E+00 | 2.48E-07 | 8.83E-07 | -1.96E+00 | 1.20E-03 | 3.37E-03 | -1.15E+00 | 2.54E-05 | 9.34E-05 | -1.48E+00 | 7.97E-02 | 2.19E-01 | 1.03E+00 |
| WDR91 | 7.33E-08 | 3.82E-07 | -1.50E+00 | 9.71E-04 | 4.20E-03 | 1.13E+00 | 1.43E-06 | 6.06E-06 | -1.77E+00 | 1.28E-03 | 4.88E-03 | -1.12E+00 | 9.18E-04 | 3.27E-03 | -1.54E+00 | 5.64E-02 | 1.59E-01 | 1.01E+00 | 3.40E-05 | 1.25E-04 | -1.40E+00 | 5.93E-03 | 1.63E-02 | -1.08E+00 |
| RPL26P37 | 7.34E-08 | 3.82E-07 | 2.52E+00 | 3.78E-02 | 1.63E-01 | 1.27E+00 | 5.03E-06 | 2.14E-05 | 2.19E+00 | 7.62E-02 | 2.91E-01 | 1.16E+00 | 7.21E-02 | 2.57E-01 | 1.66E+00 | 2.39E-02 | 6.73E-02 | 1.01E+00 | 1.48E-06 | 5.42E-06 | 2.37E+00 | 1.88E-01 | 5.17E-01 | 1.42E+00 |
| DENND4C | 7.37E-08 | 3.84E-07 | -1.35E+00 | 2.19E-05 | 9.45E-05 | -1.05E+00 | 9.74E-05 | 4.14E-04 | -1.24E+00 | 3.66E-03 | 1.40E-02 | 1.05E+00 | 2.43E-04 | 8.65E-04 | -1.22E+00 | 2.30E-03 | 6.49E-03 | -1.09E+00 | 6.50E-05 | 2.39E-04 | -1.23E+00 | 5.27E-04 | 1.45E-03 | -1.13E+00 |
| NPTN | 7.38E-08 | 3.85E-07 | -1.85E+00 | 9.50E-07 | 4.11E-06 | -1.43E+00 | 4.47E-07 | 1.90E-06 | -1.39E+00 | 1.41E-06 | 5.38E-06 | -1.20E+00 | 1.40E-02 | 4.98E-02 | -1.05E+00 | 5.41E-02 | 1.52E-01 | 1.11E+00 | 3.81E-03 | 1.40E-02 | -1.07E+00 | 4.07E-02 | 1.12E-01 | 1.08E+00 |
| SHE | 7.38E-08 | 3.85E-07 | -2.05E+00 | 7.22E-04 | 3.12E-03 | -1.17E+00 | 2.94E-06 | 1.25E-05 | -1.85E+00 | 9.30E-05 | 3.55E-04 | -1.58E+00 | 3.20E-02 | 1.14E-01 | -1.73E+00 | 4.66E-02 | 1.31E-01 | -1.62E+00 | 1.42E-01 | 5.23E-01 | -1.24E+00 | 2.91E-01 | 7.99E-01 | -1.03E+00 |
| THUMPD1 | 7.39E-08 | 3.85E-07 | -1.02E+00 | 1.82E-10 | 7.84E-10 | -1.04E+00 | 2.59E-04 | 1.10E-03 | -1.06E+00 | 2.80E-04 | 1.07E-03 | 1.02E+00 | 1.20E-01 | 4.27E-01 | 1.28E+00 | 2.21E-01 | 6.22E-01 | 1.50E+00 | 1.09E-01 | 4.00E-01 | 1.18E+00 | 2.24E-01 | 6.15E-01 | 1.06E+00 |
| PPID | 7.40E-08 | 3.86E-07 | -1.28E+00 | 3.25E-06 | 1.40E-05 | -1.04E+00 | 2.23E-02 | 9.47E-02 | -1.02E+00 | 7.87E-04 | 3.01E-03 | -1.02E+00 | 5.91E-04 | 2.11E-03 | -1.21E+00 | 2.99E-03 | 8.43E-03 | -1.09E+00 | 1.59E-03 | 5.84E-03 | -1.09E+00 | 7.20E-03 | 1.98E-02 | -1.10E+00 |
| CLIP1 | 7.40E-08 | 3.86E-07 | -1.25E+00 | 2.94E-05 | 1.27E-04 | 1.01E+00 | 2.44E-05 | 1.04E-04 | -1.28E+00 | 2.80E-05 | 1.07E-04 | -1.12E+00 | 2.27E-05 | 8.09E-05 | -1.34E+00 | 8.38E-05 | 2.36E-04 | -1.23E+00 | 2.21E-08 | 8.12E-08 | -1.50E+00 | 1.01E-04 | 2.77E-04 | -1.22E+00 |
| DDX10 | 7.44E-08 | 3.88E-07 | -1.40E+00 | 3.52E-02 | 1.52E-01 | 1.28E+00 | 8.10E-06 | 3.44E-05 | -1.28E+00 | 3.45E-02 | 1.32E-01 | 1.16E+00 | 1.65E-05 | 5.87E-05 | -1.41E+00 | 2.38E-01 | 6.70E-01 | 1.35E+00 | 1.17E-05 | 4.30E-05 | -1.21E+00 | 1.13E-01 | 3.12E-01 | 1.44E+00 |
| ZC2HC1A | 7.50E-08 | 3.91E-07 | -1.26E+00 | 2.11E-05 | 9.10E-05 | 1.00E+00 | 6.20E-08 | 2.63E-07 | -1.72E+00 | 2.03E-05 | 7.74E-05 | -1.25E+00 | 8.38E-05 | 2.99E-04 | -1.73E+00 | 2.26E-04 | 6.37E-04 | -1.64E+00 | 1.52E-03 | 5.59E-03 | -1.52E+00 | 2.50E-03 | 6.88E-03 | -1.43E+00 |
| SIN3A | 7.52E-08 | 3.92E-07 | -1.24E+00 | 3.03E-09 | 1.31E-08 | -1.24E+00 | 3.09E-07 | 1.31E-06 | -1.40E+00 | 1.31E-06 | 5.00E-06 | -1.17E+00 | 5.28E-08 | 1.88E-07 | -1.34E+00 | 1.83E-08 | 5.16E-08 | -1.34E+00 | 7.06E-09 | 2.59E-08 | -1.39E+00 | 1.32E-05 | 3.63E-05 | -1.22E+00 |
| SPAST | 7.53E-08 | 3.92E-07 | -1.48E+00 | 1.19E-04 | 5.12E-04 | 1.03E+00 | 3.11E-10 | 1.32E-09 | -1.88E+00 | 1.24E-05 | 4.73E-05 | -1.16E+00 | 3.16E-06 | 1.13E-05 | -1.46E+00 | 9.17E-03 | 2.59E-02 | 1.13E+00 | 6.28E-09 | 2.31E-08 | -1.64E+00 | 5.59E-03 | 1.54E-02 | -1.04E+00 |
| OSTF1 | 7.55E-08 | 3.94E-07 | -1.12E+00 | 3.66E-06 | 1.58E-05 | 1.06E+00 | 7.73E-04 | 3.28E-03 | -1.11E+00 | 2.04E-05 | 7.81E-05 | -1.08E+00 | 9.92E-07 | 3.54E-06 | -1.27E+00 | 5.14E-05 | 1.45E-04 | -1.07E+00 | 2.08E-04 | 7.63E-04 | -1.11E+00 | 5.78E-06 | 1.59E-05 | -1.23E+00 |
| SNRPA1 | 7.55E-08 | 3.94E-07 | -1.14E+00 | 2.31E-09 | 9.98E-09 | -1.05E+00 | 7.25E-03 | 3.08E-02 | 1.02E+00 | 1.01E-04 | 3.86E-04 | -1.04E+00 | 2.10E-05 | 7.47E-05 | -1.15E+00 | 5.16E-04 | 1.45E-03 | -1.07E+00 | 1.70E-04 | 6.24E-04 | -1.09E+00 | 8.33E-02 | 2.29E-01 | 1.05E+00 |
| AP005431.1 | 7.55E-08 | 3.94E-07 | -1.35E+00 | 1.20E-10 | 5.19E-10 | -1.47E+00 | 3.57E-04 | 1.52E-03 | -1.18E+00 | 7.04E-07 | 2.69E-06 | -1.27E+00 | 1.77E-07 | 6.31E-07 | -1.44E+00 | 2.17E-10 | 6.12E-10 | -1.80E+00 | 2.24E-09 | 8.21E-09 | -1.91E+00 | 4.95E-09 | 1.36E-08 | -1.81E+00 |
| GLT8D1 | 7.55E-08 | 3.94E-07 | -1.41E+00 | 3.83E-02 | 1.66E-01 | 1.27E+00 | 2.64E-07 | 1.12E-06 | -1.32E+00 | 5.36E-04 | 2.05E-03 | 1.03E+00 | 4.82E-06 | 1.72E-05 | -1.30E+00 | 1.99E-02 | 5.61E-02 | 1.19E+00 | 1.78E-04 | 6.54E-04 | -1.11E+00 | 1.44E-01 | 3.95E-01 | 1.20E+00 |
| AC096664.2 | 7.56E-08 | 3.94E-07 | 2.42E+00 | 7.36E-02 | 3.18E-01 | 1.81E+00 | 3.46E-10 | 1.47E-09 | 2.57E+00 | 1.09E-04 | 4.17E-04 | 2.24E+00 | 1.85E-04 | 6.59E-04 | 2.47E+00 | 1.90E-03 | 5.35E-03 | 2.32E+00 | 1.15E-06 | 4.23E-06 | 2.25E+00 | 1.15E-03 | 3.17E-03 | 2.15E+00 |
| FCF1 | 7.58E-08 | 3.95E-07 | -1.70E+00 | 1.34E-04 | 5.80E-04 | -1.10E+00 | 8.18E-07 | 3.48E-06 | -1.58E+00 | 4.43E-05 | 1.69E-04 | -1.26E+00 | 4.73E-06 | 1.69E-05 | -1.52E+00 | 1.01E-04 | 2.84E-04 | -1.31E+00 | 2.16E-04 | 7.94E-04 | -1.45E+00 | 9.52E-03 | 2.61E-02 | -1.19E+00 |
| FBXO3 | 7.60E-08 | 3.96E-07 | -1.17E+00 | 1.12E-04 | 4.82E-04 | 1.10E+00 | 2.36E-06 | 1.00E-05 | -1.27E+00 | 9.62E-05 | 3.68E-04 | -1.09E+00 | 1.74E-04 | 6.21E-04 | -1.25E+00 | 4.44E-04 | 1.25E-03 | -1.23E+00 | 2.51E-04 | 9.20E-04 | -1.10E+00 | 1.29E-02 | 3.53E-02 | 1.00E+00 |
| RGS12 | 7.63E-08 | 3.98E-07 | -1.58E+00 | 8.40E-05 | 3.63E-04 | 1.04E+00 | 5.28E-03 | 2.24E-02 | -1.11E+00 | 1.95E-02 | 7.45E-02 | 1.13E+00 | 3.28E-05 | 1.17E-04 | -1.64E+00 | 4.67E-05 | 1.32E-04 | -1.58E+00 | 8.40E-03 | 3.08E-02 | -1.22E+00 | 1.02E-01 | 2.81E-01 | -1.01E+00 |
| CYC1 | 7.66E-08 | 3.99E-07 | -1.08E+00 | 1.97E-07 | 8.50E-07 | 1.06E+00 | 1.03E-01 | 4.36E-01 | 1.10E+00 | 1.46E-02 | 5.59E-02 | 1.08E+00 | 3.58E-06 | 1.28E-05 | -1.12E+00 | 1.02E-02 | 2.88E-02 | 1.12E+00 | 8.91E-02 | 3.27E-01 | -1.00E+00 | 2.44E-01 | 6.69E-01 | 1.20E+00 |
| UQCRFS1 | 7.67E-08 | 4.00E-07 | -1.11E+00 | 1.72E-15 | 7.44E-15 | -1.15E+00 | 6.14E-02 | 2.61E-01 | 1.02E+00 | 2.94E-04 | 1.12E-03 | -1.08E+00 | 9.43E-04 | 3.36E-03 | -1.07E+00 | 1.26E-03 | 3.54E-03 | -1.11E+00 | 9.90E-03 | 3.63E-02 | -1.06E+00 | 9.61E-02 | 2.64E-01 | 1.00E+00 |
| DCTN2 | 7.67E-08 | 4.00E-07 | -1.07E+00 | 2.68E-08 | 1.16E-07 | 1.04E+00 | 8.17E-03 | 3.47E-02 | -1.08E+00 | 1.08E-03 | 4.14E-03 | 1.01E+00 | 1.12E-06 | 3.98E-06 | -1.09E+00 | 4.43E-03 | 1.25E-02 | 1.10E+00 | 2.95E-03 | 1.08E-02 | 1.02E+00 | 5.35E-03 | 1.47E-02 | -1.01E+00 |
| SEC63 | 7.69E-08 | 4.01E-07 | -1.29E+00 | 1.93E-07 | 8.34E-07 | -1.09E+00 | 8.24E-06 | 3.50E-05 | -1.24E+00 | 2.84E-06 | 1.09E-05 | -1.16E+00 | 1.10E-07 | 3.91E-07 | -1.38E+00 | 8.20E-05 | 2.31E-04 | -1.06E+00 | 2.01E-02 | 7.38E-02 | -1.01E+00 | 3.35E-01 | 9.20E-01 | 1.19E+00 |
| PTGES3P1 | 7.72E-08 | 4.03E-07 | -1.15E+00 | 3.12E-14 | 1.35E-13 | -1.31E+00 | 5.31E-04 | 2.26E-03 | -1.07E+00 | 1.97E-08 | 7.53E-08 | -1.17E+00 | 6.53E-09 | 2.33E-08 | -1.38E+00 | 3.07E-09 | 8.65E-09 | -1.38E+00 | 6.77E-03 | 2.48E-02 | -1.04E+00 | 9.39E-05 | 2.58E-04 | -1.25E+00 |
| QARS | 7.73E-08 | 4.03E-07 | -1.04E+00 | 1.14E-07 | 4.92E-07 | 1.07E+00 | 3.96E-03 | 1.68E-02 | -1.04E+00 | 2.03E-03 | 7.74E-03 | 1.06E+00 | 2.85E-04 | 1.02E-03 | -1.01E+00 | 6.37E-03 | 1.80E-02 | 1.10E+00 | 1.28E-02 | 4.69E-02 | -1.03E+00 | 3.06E-01 | 8.40E-01 | 1.12E+00 |
| MGRN1 | 7.80E-08 | 4.07E-07 | -1.17E+00 | 9.15E-09 | 3.95E-08 | -1.15E+00 | 2.69E-06 | 1.14E-05 | -1.16E+00 | 1.04E-06 | 3.98E-06 | -1.05E+00 | 6.29E-04 | 2.24E-03 | -1.03E+00 | 2.59E-05 | 7.30E-05 | -1.21E+00 | 1.05E-08 | 3.86E-08 | -1.56E+00 | 1.59E-09 | 4.36E-09 | -1.72E+00 |
| WHAMMP2 | 7.81E-08 | 4.07E-07 | -1.40E+00 | 1.07E-04 | 4.64E-04 | 1.00E+00 | 8.66E-07 | 3.68E-06 | -1.45E+00 | 4.02E-05 | 1.54E-04 | -1.09E+00 | 2.09E-03 | 7.46E-03 | -1.18E+00 | 9.27E-03 | 2.61E-02 | 1.01E+00 | 2.97E-03 | 1.09E-02 | 1.10E+00 | 2.20E-03 | 6.05E-03 | -1.08E+00 |
| RUNX1 | 7.81E-08 | 4.07E-07 | -1.11E+00 | 2.09E-07 | 9.02E-07 | -1.00E+00 | 7.17E-03 | 3.05E-02 | -1.14E+00 | 1.35E-02 | 5.16E-02 | 1.01E+00 | 1.81E-08 | 6.46E-08 | -1.36E+00 | 2.17E-08 | 6.12E-08 | -1.35E+00 | 1.95E-07 | 7.16E-07 | -1.29E+00 | 1.55E-06 | 4.25E-06 | -1.24E+00 |
| PTBP1 | 7.87E-08 | 4.10E-07 | -1.09E+00 | 2.60E-08 | 1.12E-07 | 1.02E+00 | 1.43E-03 | 6.09E-03 | -1.11E+00 | 2.47E-04 | 9.42E-04 | -1.05E+00 | 1.33E-06 | 4.73E-06 | -1.22E+00 | 2.49E-03 | 7.02E-03 | 1.01E+00 | 1.26E-05 | 4.62E-05 | -1.18E+00 | 2.22E-02 | 6.10E-02 | 1.01E+00 |
| CASP10 | 7.87E-08 | 4.10E-07 | -1.16E+00 | 1.51E-10 | 6.52E-10 | -1.22E+00 | 1.26E-02 | 5.34E-02 | -1.03E+00 | 6.32E-05 | 2.41E-04 | -1.15E+00 | 2.93E-02 | 1.05E-01 | 1.06E+00 | 6.21E-03 | 1.75E-02 | -1.09E+00 | 8.93E-03 | 3.28E-02 | -1.04E+00 | 3.59E-05 | 9.87E-05 | -1.23E+00 |
| DPAGT1 | 7.94E-08 | 4.14E-07 | -1.67E+00 | 2.22E-04 | 9.61E-04 | -1.06E+00 | 1.85E-02 | 7.88E-02 | -1.12E+00 | 1.38E-01 | 5.26E-01 | 1.24E+00 | 6.27E-08 | 2.23E-07 | -2.13E+00 | 1.88E-02 | 5.31E-02 | 1.03E+00 | 5.70E-05 | 2.09E-04 | -1.21E+00 | 1.32E-01 | 3.62E-01 | 1.12E+00 |
| AC084824.5 | 7.94E-08 | 4.14E-07 | -1.50E+00 | 3.35E-07 | 1.45E-06 | -1.39E+00 | 4.43E-13 | 1.88E-12 | -1.70E+00 | 2.92E-13 | 1.12E-12 | -1.50E+00 | 2.73E-03 | 9.72E-03 | -1.48E+00 | 8.31E-04 | 2.34E-03 | -1.85E+00 | 1.15E-08 | 4.23E-08 | -1.82E+00 | 9.09E-06 | 2.50E-05 | -1.49E+00 |
| AL139100.1 | 7.97E-08 | 4.15E-07 | -1.09E+00 | 6.73E-02 | 2.91E-01 | 1.73E+00 | 4.34E-03 | 1.84E-02 | 1.02E+00 | 2.82E-05 | 1.08E-04 | 2.16E+00 | 6.98E-03 | 2.49E-02 | 1.09E+00 | 2.45E-03 | 6.92E-03 | 2.16E+00 | 2.41E-03 | 8.83E-03 | -1.09E+00 | 8.39E-06 | 2.30E-05 | 1.97E+00 |
| PPP2R1B | 7.98E-08 | 4.16E-07 | -1.88E+00 | 1.99E-07 | 8.61E-07 | -1.61E+00 | 3.84E-12 | 1.63E-11 | -2.08E+00 | 2.35E-08 | 8.99E-08 | -1.44E+00 | 1.11E-02 | 3.96E-02 | -1.16E+00 | 2.42E-03 | 6.83E-03 | -1.33E+00 | 5.97E-10 | 2.19E-09 | -1.47E+00 | 4.67E-03 | 1.28E-02 | 1.04E+00 |
| MARCH8 | 7.98E-08 | 4.16E-07 | -1.46E+00 | 2.69E-04 | 1.16E-03 | 1.01E+00 | 8.04E-05 | 3.42E-04 | -1.58E+00 | 3.86E-03 | 1.47E-02 | -1.07E+00 | 9.95E-04 | 3.55E-03 | -1.15E+00 | 2.70E-01 | 7.61E-01 | 1.32E+00 | 4.77E-04 | 1.75E-03 | -1.18E+00 | 1.65E-02 | 4.52E-02 | 1.01E+00 |
| ACADL | 8.02E-08 | 4.18E-07 | 4.04E+00 | 1.08E-02 | 4.65E-02 | 2.47E+00 | 1.64E-03 | 6.96E-03 | 2.38E+00 | 5.47E-02 | 2.09E-01 | 1.80E+00 | 2.29E-01 | 8.17E-01 | 1.24E+00 | 3.33E-01 | 9.39E-01 | 1.09E+00 | 1.32E-01 | 4.84E-01 | 1.41E+00 | 2.90E-01 | 7.96E-01 | 1.26E+00 |
| BZW2 | 8.06E-08 | 4.20E-07 | -1.04E+00 | 1.43E-05 | 6.16E-05 | 1.07E+00 | 2.53E-02 | 1.07E-01 | -1.08E+00 | 5.85E-02 | 2.23E-01 | 1.10E+00 | 2.00E-04 | 7.13E-04 | -1.08E+00 | 1.72E-03 | 4.85E-03 | 1.02E+00 | 3.46E-02 | 1.27E-01 | 1.01E+00 | 2.87E-01 | 7.87E-01 | 1.12E+00 |
| AL357054.4 | 8.06E-08 | 4.20E-07 | -1.82E+00 | 6.90E-06 | 2.98E-05 | -1.49E+00 | 9.55E-07 | 4.06E-06 | -1.96E+00 | 8.44E-05 | 3.22E-04 | -1.53E+00 | 5.84E-02 | 2.08E-01 | -1.47E+00 | 8.09E-02 | 2.28E-01 | -1.39E+00 | 1.82E-02 | 6.68E-02 | -1.48E+00 | 7.05E-02 | 1.94E-01 | -1.41E+00 |
| DIDO1 | 8.07E-08 | 4.21E-07 | -1.19E+00 | 3.62E-02 | 1.57E-01 | 1.43E+00 | 6.67E-08 | 2.83E-07 | -1.52E+00 | 2.21E-03 | 8.46E-03 | 1.00E+00 | 6.54E-05 | 2.33E-04 | -1.16E+00 | 1.31E-01 | 3.70E-01 | 1.27E+00 | 2.93E-04 | 1.08E-03 | -1.19E+00 | 5.08E-02 | 1.40E-01 | 1.09E+00 |
| AC103769.1 | 8.10E-08 | 4.22E-07 | -1.71E+00 | 9.01E-02 | 3.89E-01 | 1.40E+00 | 8.86E-05 | 3.77E-04 | -1.60E+00 | 2.29E-02 | 8.74E-02 | 1.04E+00 | 2.17E-01 | 7.75E-01 | 1.28E+00 | 2.00E-01 | 5.65E-01 | 1.71E+00 | 2.16E-04 | 7.91E-04 | -1.89E+00 | 9.04E-02 | 2.48E-01 | -1.09E+00 |
| SLC39A7 | 8.11E-08 | 4.23E-07 | -1.46E+00 | 7.46E-06 | 3.22E-05 | -1.15E+00 | 2.24E-03 | 9.52E-03 | -1.11E+00 | 6.11E-03 | 2.33E-02 | 1.02E+00 | 3.22E-02 | 1.15E-01 | 1.08E+00 | 3.71E-02 | 1.05E-01 | 1.12E+00 | 1.46E-04 | 5.35E-04 | -1.25E+00 | 6.17E-05 | 1.69E-04 | -1.22E+00 |
| QRICH1 | 8.14E-08 | 4.24E-07 | -1.33E+00 | 2.08E-05 | 8.98E-05 | -1.06E+00 | 2.53E-03 | 1.07E-02 | -1.06E+00 | 1.71E-03 | 6.53E-03 | 1.09E+00 | 1.61E-03 | 5.76E-03 | -1.31E+00 | 8.30E-02 | 2.34E-01 | 1.04E+00 | 1.92E-13 | 7.05E-13 | -1.78E+00 | 1.34E-07 | 3.68E-07 | -1.31E+00 |
| JTB | 8.15E-08 | 4.25E-07 | -1.07E+00 | 4.23E-06 | 1.83E-05 | 1.06E+00 | 1.10E-01 | 4.67E-01 | 1.05E+00 | 1.90E-01 | 7.25E-01 | 1.14E+00 | 8.71E-03 | 3.10E-02 | 1.03E+00 | 1.20E-01 | 3.37E-01 | 1.12E+00 | 2.36E-01 | 8.64E-01 | 1.06E+00 | 5.42E-02 | 1.49E-01 | 1.22E+00 |
| NDUFA13 | 8.19E-08 | 4.27E-07 | -1.75E+00 | 1.50E-07 | 6.48E-07 | -1.48E+00 | 4.38E-03 | 1.86E-02 | -1.13E+00 | 9.72E-06 | 3.71E-05 | -1.36E+00 | 1.02E-04 | 3.64E-04 | -1.15E+00 | 4.31E-09 | 1.21E-08 | -1.83E+00 | 6.30E-09 | 2.31E-08 | -1.40E+00 | 1.08E-10 | 2.97E-10 | -1.59E+00 |
| NAPRT | 8.22E-08 | 4.29E-07 | -1.20E+00 | 4.97E-06 | 2.15E-05 | 1.01E+00 | 3.20E-04 | 1.36E-03 | -1.07E+00 | 7.57E-04 | 2.89E-03 | 1.04E+00 | 5.36E-06 | 1.91E-05 | -1.16E+00 | 1.27E-03 | 3.57E-03 | 1.03E+00 | 2.38E-03 | 8.73E-03 | -1.08E+00 | 1.64E-03 | 4.50E-03 | -1.04E+00 |
| PPP2R5B | 8.23E-08 | 4.29E-07 | -2.11E+00 | 2.32E-09 | 1.00E-08 | -2.47E+00 | 5.43E-10 | 2.31E-09 | -2.17E+00 | 8.51E-11 | 3.25E-10 | -2.34E+00 | 2.96E-03 | 1.06E-02 | -1.44E+00 | 3.24E-02 | 9.12E-02 | -1.07E+00 | 1.26E-02 | 4.63E-02 | -1.09E+00 | 2.19E-03 | 6.02E-03 | -1.31E+00 |
| ATP5I | 8.23E-08 | 4.29E-07 | -1.11E+00 | 1.52E-27 | 6.55E-27 | -1.91E+00 | 2.51E-02 | 1.07E-01 | 1.03E+00 | 1.22E-21 | 4.65E-21 | -1.98E+00 | 6.61E-03 | 2.36E-02 | 1.07E+00 | 2.25E-13 | 6.34E-13 | -1.72E+00 | 3.28E-06 | 1.20E-05 | -1.10E+00 | 5.29E-22 | 1.45E-21 | -1.99E+00 |
| IL10RA | 8.27E-08 | 4.31E-07 | -1.76E+00 | 2.81E-08 | 1.21E-07 | -1.79E+00 | 2.07E-07 | 8.81E-07 | -2.00E+00 | 2.96E-10 | 1.13E-09 | -2.84E+00 | 1.72E-01 | 6.12E-01 | -1.09E+00 | 7.52E-02 | 2.12E-01 | -1.80E+00 | 2.37E-02 | 8.70E-02 | -1.79E+00 | 1.34E-02 | 3.67E-02 | -2.53E+00 |
| CRNKL1 | 8.27E-08 | 4.31E-07 | -1.55E+00 | 1.03E-05 | 4.44E-05 | -1.19E+00 | 9.91E-07 | 4.21E-06 | -1.43E+00 | 2.66E-06 | 1.01E-05 | -1.23E+00 | 1.92E-05 | 6.85E-05 | -1.43E+00 | 6.18E-04 | 1.74E-03 | -1.28E+00 | 2.35E-15 | 8.61E-15 | -1.88E+00 | 5.54E-07 | 1.52E-06 | -1.24E+00 |
| INTS3 | 8.32E-08 | 4.33E-07 | -1.46E+00 | 1.48E-03 | 6.40E-03 | 1.06E+00 | 2.90E-06 | 1.23E-05 | -1.52E+00 | 1.48E-03 | 5.67E-03 | 1.02E+00 | 3.45E-05 | 1.23E-04 | -1.41E+00 | 5.40E-03 | 1.52E-02 | -1.05E+00 | 1.94E-03 | 7.14E-03 | -1.43E+00 | 2.61E-02 | 7.17E-02 | -1.20E+00 |
| SDF4 | 8.41E-08 | 4.38E-07 | -1.21E+00 | 7.51E-05 | 3.24E-04 | 1.05E+00 | 5.99E-07 | 2.55E-06 | -1.23E+00 | 1.67E-06 | 6.38E-06 | -1.07E+00 | 1.55E-09 | 5.54E-09 | -1.42E+00 | 1.38E-03 | 3.89E-03 | 1.04E+00 | 6.94E-03 | 2.55E-02 | -1.00E+00 | 5.45E-02 | 1.50E-01 | 1.09E+00 |
| ZNF799 | 8.41E-08 | 4.38E-07 | -2.60E+00 | 3.49E-06 | 1.51E-05 | -1.82E+00 | 6.67E-07 | 2.83E-06 | -2.35E+00 | 1.30E-03 | 4.96E-03 | -1.30E+00 | 1.28E-06 | 4.58E-06 | -2.06E+00 | 3.32E-07 | 9.35E-07 | -2.24E+00 | 5.01E-08 | 1.84E-07 | -2.30E+00 | 1.12E-03 | 3.08E-03 | -1.40E+00 |
| USP36 | 8.41E-08 | 4.38E-07 | -1.48E+00 | 2.50E-06 | 1.08E-05 | -1.16E+00 | 3.77E-03 | 1.60E-02 | -1.08E+00 | 3.20E-05 | 1.22E-04 | -1.14E+00 | 2.34E-03 | 8.34E-03 | -1.14E+00 | 2.66E-02 | 7.50E-02 | 1.07E+00 | 8.08E-09 | 2.97E-08 | -1.61E+00 | 9.58E-02 | 2.63E-01 | 1.17E+00 |
| PPP1CC | 8.43E-08 | 4.39E-07 | -1.17E+00 | 2.81E-05 | 1.22E-04 | 1.03E+00 | 1.13E-05 | 4.80E-05 | -1.28E+00 | 3.82E-05 | 1.46E-04 | -1.14E+00 | 1.04E-11 | 3.72E-11 | -1.40E+00 | 2.99E-04 | 8.44E-04 | -1.06E+00 | 1.31E-02 | 4.82E-02 | -1.16E+00 | 2.69E-01 | 7.38E-01 | 1.04E+00 |
| RPL18P13 | 8.44E-08 | 4.40E-07 | -1.16E+00 | 2.35E-67 | 1.02E-66 | -2.09E+00 | 2.37E-02 | 1.01E-01 | -1.16E+00 | 1.24E-21 | 4.75E-21 | -1.93E+00 | 6.67E-04 | 2.38E-03 | -1.11E+00 | 3.66E-24 | 1.03E-23 | -1.81E+00 | 1.52E-04 | 5.58E-04 | -1.19E+00 | 4.05E-18 | 1.11E-17 | -1.84E+00 |
| CENPF | 8.45E-08 | 4.41E-07 | -2.65E+00 | 3.92E-06 | 1.70E-05 | -1.69E+00 | 2.29E-04 | 9.73E-04 | -1.13E+00 | 6.12E-04 | 2.34E-03 | 1.22E+00 | 5.94E-04 | 2.12E-03 | -1.49E+00 | 1.20E-01 | 3.39E-01 | 1.31E+00 | 4.25E-07 | 1.56E-06 | -1.15E+00 | 6.35E-04 | 1.74E-03 | 1.08E+00 |
| VPS39 | 8.46E-08 | 4.41E-07 | -1.43E+00 | 8.23E-05 | 3.56E-04 | 1.02E+00 | 7.60E-02 | 3.23E-01 | 1.09E+00 | 6.28E-02 | 2.40E-01 | 1.23E+00 | 9.88E-05 | 3.52E-04 | -1.18E+00 | 4.65E-04 | 1.31E-03 | -1.14E+00 | 9.40E-02 | 3.45E-01 | 1.17E+00 | 3.12E-03 | 8.56E-03 | -1.11E+00 |
| ERG28 | 8.52E-08 | 4.44E-07 | -1.15E+00 | 1.39E-04 | 6.01E-04 | 1.18E+00 | 5.26E-03 | 2.23E-02 | -1.09E+00 | 1.26E-01 | 4.81E-01 | 1.22E+00 | 8.26E-06 | 2.95E-05 | -1.26E+00 | 1.35E-01 | 3.79E-01 | 1.24E+00 | 1.91E-02 | 7.02E-02 | 1.04E+00 | 1.42E-01 | 3.90E-01 | 1.25E+00 |
| UBE2J1 | 8.53E-08 | 4.44E-07 | -1.18E+00 | 3.66E-05 | 1.58E-04 | 1.08E+00 | 3.72E-03 | 1.58E-02 | -1.09E+00 | 6.26E-05 | 2.39E-04 | -1.06E+00 | 3.00E-09 | 1.07E-08 | -1.42E+00 | 3.26E-10 | 9.18E-10 | -1.44E+00 | 9.92E-04 | 3.64E-03 | -1.30E+00 | 6.36E-03 | 1.75E-02 | -1.22E+00 |
| EBLN3P | 8.54E-08 | 4.45E-07 | -1.26E+00 | 5.30E-05 | 2.29E-04 | 1.00E+00 | 7.04E-10 | 2.99E-09 | -1.40E+00 | 9.59E-05 | 3.66E-04 | 1.01E+00 | 1.40E-05 | 4.98E-05 | -1.21E+00 | 4.89E-03 | 1.38E-02 | 1.10E+00 | 2.64E-07 | 9.68E-07 | -1.30E+00 | 6.33E-04 | 1.74E-03 | -1.12E+00 |
| BNIP3 | 8.59E-08 | 4.47E-07 | -1.21E+00 | 4.18E-05 | 1.81E-04 | 1.05E+00 | 1.95E-07 | 8.31E-07 | -1.40E+00 | 1.71E-04 | 6.53E-04 | -1.07E+00 | 4.19E-05 | 1.49E-04 | -1.28E+00 | 2.82E-03 | 7.95E-03 | 1.00E+00 | 3.18E-05 | 1.17E-04 | -1.13E+00 | 3.52E-03 | 9.68E-03 | -1.03E+00 |
| MBD6 | 8.59E-08 | 4.48E-07 | -1.27E+00 | 9.75E-09 | 4.21E-08 | -1.23E+00 | 2.78E-02 | 1.18E-01 | -1.03E+00 | 1.19E-04 | 4.53E-04 | -1.21E+00 | 4.69E-02 | 1.67E-01 | 1.01E+00 | 1.23E-02 | 3.48E-02 | -1.10E+00 | 4.89E-05 | 1.79E-04 | -1.29E+00 | 4.92E-06 | 1.35E-05 | -1.35E+00 |
| KMT5B | 8.62E-08 | 4.49E-07 | -1.20E+00 | 1.15E-06 | 4.96E-06 | 1.00E+00 | 3.26E-09 | 1.39E-08 | -1.45E+00 | 3.23E-08 | 1.23E-07 | -1.22E+00 | 6.19E-07 | 2.21E-06 | -1.29E+00 | 7.71E-07 | 2.17E-06 | -1.27E+00 | 1.73E-09 | 6.34E-09 | -1.40E+00 | 6.35E-09 | 1.75E-08 | -1.48E+00 |
| UQCRC1 | 8.63E-08 | 4.50E-07 | -1.19E+00 | 1.44E-08 | 6.23E-08 | -1.08E+00 | 1.75E-02 | 7.44E-02 | -1.02E+00 | 2.92E-05 | 1.11E-04 | -1.11E+00 | 4.47E-06 | 1.59E-05 | -1.13E+00 | 4.93E-05 | 1.39E-04 | -1.04E+00 | 1.77E-03 | 6.48E-03 | -1.05E+00 | 1.39E-01 | 3.83E-01 | 1.11E+00 |
| MRPL13 | 8.67E-08 | 4.52E-07 | -1.19E+00 | 3.90E-03 | 1.69E-02 | 1.19E+00 | 3.98E-05 | 1.69E-04 | -1.18E+00 | 1.03E-06 | 3.94E-06 | -1.14E+00 | 1.74E-07 | 6.20E-07 | -1.35E+00 | 6.86E-02 | 1.93E-01 | 1.18E+00 | 2.73E-08 | 1.00E-07 | -1.34E+00 | 3.79E-02 | 1.04E-01 | -1.01E+00 |
| TRAPPC4 | 8.69E-08 | 4.53E-07 | -1.30E+00 | 2.25E-05 | 9.70E-05 | 1.02E+00 | 8.19E-02 | 3.48E-01 | 1.10E+00 | 2.05E-01 | 7.82E-01 | 1.47E+00 | 1.30E-05 | 4.64E-05 | -1.21E+00 | 2.92E-02 | 8.22E-02 | 1.11E+00 | 2.86E-04 | 1.05E-03 | -1.04E+00 | 2.78E-02 | 7.63E-02 | 1.13E+00 |
| NPEPPS | 8.69E-08 | 4.53E-07 | -1.32E+00 | 1.66E-02 | 7.17E-02 | 1.23E+00 | 4.94E-08 | 2.10E-07 | -1.58E+00 | 9.98E-04 | 3.81E-03 | -1.07E+00 | 6.76E-08 | 2.41E-07 | -1.59E+00 | 1.31E-04 | 3.69E-04 | -1.14E+00 | 2.10E-07 | 7.73E-07 | -1.40E+00 | 1.36E-02 | 3.74E-02 | -1.08E+00 |
| HMGB2 | 8.77E-08 | 4.57E-07 | 1.66E+00 | 1.41E-01 | 6.11E-01 | 1.41E+00 | 1.25E-09 | 5.31E-09 | 2.05E+00 | 5.93E-03 | 2.27E-02 | 1.62E+00 | 7.67E-02 | 2.73E-01 | 1.12E+00 | 7.55E-02 | 2.13E-01 | 1.13E+00 | 1.14E-01 | 4.19E-01 | 1.08E+00 | 3.21E-01 | 8.83E-01 | 1.10E+00 |
| ATP5J2P4 | 8.81E-08 | 4.59E-07 | -2.19E+00 | 8.31E-07 | 3.59E-06 | -1.71E+00 | 9.47E-03 | 4.02E-02 | -1.13E+00 | 2.68E-06 | 1.02E-05 | -1.65E+00 | 1.53E-03 | 5.44E-03 | -1.52E+00 | 1.72E-03 | 4.84E-03 | -1.36E+00 | 1.60E-08 | 5.87E-08 | -1.48E+00 | 4.21E-12 | 1.16E-11 | -1.84E+00 |
| C16orf52 | 8.88E-08 | 4.63E-07 | -1.38E+00 | 5.53E-04 | 2.39E-03 | 1.10E+00 | 1.02E-08 | 4.33E-08 | -1.76E+00 | 5.10E-07 | 1.95E-06 | -1.43E+00 | 1.42E-07 | 5.05E-07 | -1.48E+00 | 1.74E-05 | 4.89E-05 | -1.13E+00 | 1.06E-05 | 3.87E-05 | -1.40E+00 | 5.07E-04 | 1.39E-03 | -1.22E+00 |
| DYNC1LI1 | 8.88E-08 | 4.63E-07 | -1.21E+00 | 7.93E-07 | 3.43E-06 | -1.03E+00 | 8.26E-06 | 3.51E-05 | -1.23E+00 | 1.19E-08 | 4.54E-08 | -1.24E+00 | 5.35E-07 | 1.91E-06 | -1.38E+00 | 2.65E-04 | 7.47E-04 | -1.18E+00 | 4.67E-08 | 1.72E-07 | -1.35E+00 | 5.70E-07 | 1.57E-06 | -1.34E+00 |
| RBM18 | 8.91E-08 | 4.65E-07 | -1.41E+00 | 1.61E-08 | 6.96E-08 | -1.36E+00 | 4.34E-08 | 1.84E-07 | -1.54E+00 | 4.85E-07 | 1.85E-06 | -1.29E+00 | 7.04E-05 | 2.51E-04 | -1.30E+00 | 1.41E-04 | 3.97E-04 | -1.12E+00 | 1.62E-09 | 5.94E-09 | -1.49E+00 | 1.48E-02 | 4.08E-02 | 1.05E+00 |
| OS9 | 8.95E-08 | 4.67E-07 | -1.39E+00 | 2.82E-05 | 1.22E-04 | -1.05E+00 | 1.58E-04 | 6.73E-04 | -1.27E+00 | 1.96E-05 | 7.50E-05 | -1.17E+00 | 1.89E-04 | 6.75E-04 | -1.15E+00 | 4.18E-04 | 1.18E-03 | -1.08E+00 | 1.78E-03 | 6.53E-03 | -1.16E+00 | 1.99E-03 | 5.48E-03 | -1.16E+00 |
| ECHS1 | 8.96E-08 | 4.67E-07 | -1.08E+00 | 2.22E-05 | 9.57E-05 | 1.06E+00 | 1.75E-01 | 7.43E-01 | 1.11E+00 | 1.40E-02 | 5.35E-02 | 1.09E+00 | 2.30E-11 | 8.21E-11 | -1.20E+00 | 3.63E-03 | 1.02E-02 | 1.09E+00 | 8.26E-02 | 3.03E-01 | 1.04E+00 | 3.03E-01 | 8.32E-01 | 1.17E+00 |
| MAD2L2 | 8.99E-08 | 4.69E-07 | -1.20E+00 | 7.79E-09 | 3.37E-08 | -1.12E+00 | 8.32E-02 | 3.54E-01 | 1.12E+00 | 1.70E-04 | 6.48E-04 | -1.01E+00 | 1.08E-02 | 3.83E-02 | -1.08E+00 | 7.99E-02 | 2.25E-01 | 1.12E+00 | 2.52E-02 | 9.25E-02 | 1.16E+00 | 6.10E-02 | 1.68E-01 | 1.19E+00 |
| VPS26A | 9.01E-08 | 4.70E-07 | -1.14E+00 | 1.27E-05 | 5.51E-05 | 1.08E+00 | 4.80E-07 | 2.04E-06 | -1.22E+00 | 8.67E-06 | 3.31E-05 | -1.09E+00 | 1.20E-10 | 4.29E-10 | -1.38E+00 | 2.16E-10 | 6.08E-10 | -1.29E+00 | 3.58E-06 | 1.31E-05 | -1.27E+00 | 5.31E-03 | 1.46E-02 | -1.05E+00 |
| CNOT2 | 9.01E-08 | 4.70E-07 | -1.06E+00 | 1.01E-09 | 4.37E-09 | -1.05E+00 | 2.32E-02 | 9.87E-02 | -1.01E+00 | 4.68E-04 | 1.79E-03 | -1.01E+00 | 4.00E-05 | 1.43E-04 | -1.02E+00 | 1.62E-05 | 4.56E-05 | -1.04E+00 | 2.16E-04 | 7.92E-04 | -1.22E+00 | 8.33E-05 | 2.29E-04 | -1.20E+00 |
| NUFIP1 | 9.07E-08 | 4.73E-07 | -1.55E+00 | 4.22E-07 | 1.82E-06 | -1.25E+00 | 1.29E-07 | 5.48E-07 | -1.39E+00 | 1.01E-11 | 3.88E-11 | -1.49E+00 | 2.02E-05 | 7.19E-05 | -1.53E+00 | 6.06E-04 | 1.71E-03 | -1.33E+00 | 1.08E-12 | 3.97E-12 | -1.64E+00 | 2.73E-08 | 7.51E-08 | -1.33E+00 |
| NADK | 9.08E-08 | 4.73E-07 | -1.42E+00 | 2.97E-03 | 1.28E-02 | 1.12E+00 | 7.46E-07 | 3.17E-06 | -1.57E+00 | 2.31E-03 | 8.83E-03 | -1.06E+00 | 7.27E-04 | 2.59E-03 | -1.36E+00 | 5.63E-02 | 1.59E-01 | 1.04E+00 | 6.33E-06 | 2.32E-05 | -1.21E+00 | 6.61E-02 | 1.81E-01 | 1.10E+00 |
| OAZ2 | 9.12E-08 | 4.75E-07 | -1.12E+00 | 2.55E-03 | 1.10E-02 | 1.25E+00 | 1.22E-03 | 5.19E-03 | -1.10E+00 | 7.60E-02 | 2.90E-01 | 1.20E+00 | 7.95E-03 | 2.83E-02 | 1.20E+00 | 1.26E-01 | 3.56E-01 | 1.37E+00 | 4.52E-02 | 1.66E-01 | 1.04E+00 | 8.56E-02 | 2.35E-01 | 1.14E+00 |
| TBPL1 | 9.14E-08 | 4.76E-07 | -1.38E+00 | 6.25E-06 | 2.70E-05 | -1.12E+00 | 6.66E-08 | 2.83E-07 | -1.48E+00 | 6.67E-06 | 2.55E-05 | -1.12E+00 | 3.10E-12 | 1.11E-11 | -1.53E+00 | 9.19E-06 | 2.59E-05 | -1.07E+00 | 4.26E-14 | 1.56E-13 | -1.45E+00 | 1.08E-06 | 2.98E-06 | -1.28E+00 |
| DDX27 | 9.17E-08 | 4.78E-07 | -1.24E+00 | 7.09E-08 | 3.07E-07 | -1.07E+00 | 2.64E-06 | 1.12E-05 | -1.22E+00 | 1.64E-05 | 6.26E-05 | -1.04E+00 | 2.50E-07 | 8.90E-07 | -1.26E+00 | 1.24E-03 | 3.49E-03 | 1.06E+00 | 4.40E-06 | 1.62E-05 | -1.22E+00 | 9.55E-07 | 2.62E-06 | -1.15E+00 |
| PGAP3 | 9.18E-08 | 4.79E-07 | -1.93E+00 | 1.66E-06 | 7.19E-06 | -1.46E+00 | 1.21E-01 | 5.15E-01 | 1.11E+00 | 2.16E-01 | 8.25E-01 | 1.40E+00 | 1.63E-01 | 5.82E-01 | 1.14E+00 | 1.37E-01 | 3.85E-01 | 1.02E+00 | 1.93E-01 | 7.09E-01 | 1.19E+00 | 2.33E-01 | 6.39E-01 | 1.22E+00 |
| BABAM1 | 9.24E-08 | 4.81E-07 | -1.16E+00 | 6.66E-07 | 2.88E-06 | 1.01E+00 | 3.30E-05 | 1.40E-04 | -1.21E+00 | 7.33E-06 | 2.80E-05 | -1.12E+00 | 2.38E-10 | 8.48E-10 | -1.45E+00 | 5.99E-06 | 1.69E-05 | -1.07E+00 | 6.28E-06 | 2.31E-05 | -1.32E+00 | 1.23E-02 | 3.39E-02 | -1.08E+00 |
| FAM221A | 9.27E-08 | 4.83E-07 | -1.54E+00 | 9.22E-05 | 3.98E-04 | -1.06E+00 | 1.38E-05 | 5.86E-05 | -1.54E+00 | 2.34E-04 | 8.95E-04 | -1.22E+00 | 8.27E-04 | 2.95E-03 | -1.42E+00 | 8.84E-03 | 2.49E-02 | -1.10E+00 | 6.87E-02 | 2.52E-01 | -1.09E+00 | 1.08E-01 | 2.97E-01 | -1.02E+00 |
| ARL6IP5 | 9.27E-08 | 4.83E-07 | -1.08E+00 | 5.25E-02 | 2.27E-01 | 1.29E+00 | 1.41E-04 | 6.00E-04 | -1.16E+00 | 2.28E-01 | 8.69E-01 | 1.14E+00 | 2.54E-01 | 9.04E-01 | 1.30E+00 | 3.00E-02 | 8.47E-02 | 1.36E+00 | 5.45E-03 | 2.00E-02 | 1.41E+00 | 9.89E-03 | 2.72E-02 | 1.30E+00 |
| EHD2 | 9.30E-08 | 4.85E-07 | -1.41E+00 | 2.16E-01 | 9.32E-01 | 1.41E+00 | 1.04E-02 | 4.42E-02 | -1.08E+00 | 1.99E-01 | 7.61E-01 | 1.48E+00 | 5.17E-07 | 1.84E-06 | -1.89E+00 | 4.42E-03 | 1.25E-02 | -1.07E+00 | 8.10E-07 | 2.97E-06 | -1.49E+00 | 9.89E-06 | 2.72E-05 | -1.36E+00 |
| CORO1B | 9.32E-08 | 4.86E-07 | -1.20E+00 | 1.33E-05 | 5.73E-05 | 1.07E+00 | 1.41E-07 | 6.00E-07 | -1.43E+00 | 5.54E-06 | 2.12E-05 | -1.08E+00 | 1.69E-04 | 6.03E-04 | -1.15E+00 | 1.15E-02 | 3.25E-02 | 1.06E+00 | 2.83E-02 | 1.04E-01 | 1.09E+00 | 5.79E-02 | 1.59E-01 | 1.05E+00 |
| BEND4 | 9.32E-08 | 4.86E-07 | -1.41E+00 | 2.44E-05 | 1.06E-04 | -1.02E+00 | 8.74E-13 | 3.71E-12 | -1.98E+00 | 4.31E-07 | 1.65E-06 | -1.25E+00 | 6.60E-07 | 2.35E-06 | -1.69E+00 | 8.31E-05 | 2.34E-04 | -1.27E+00 | 1.56E-08 | 5.71E-08 | -1.82E+00 | 1.35E-03 | 3.72E-03 | -1.23E+00 |
| LRSAM1 | 9.32E-08 | 4.86E-07 | -1.75E+00 | 9.50E-04 | 4.11E-03 | -1.05E+00 | 1.80E-03 | 7.66E-03 | -1.26E+00 | 3.43E-03 | 1.31E-02 | -1.01E+00 | 1.65E-06 | 5.87E-06 | -1.63E+00 | 3.29E-03 | 9.29E-03 | -1.04E+00 | 2.94E-02 | 1.08E-01 | -1.20E+00 | 1.30E-01 | 3.57E-01 | 1.04E+00 |
| PPM1D | 9.32E-08 | 4.86E-07 | -1.54E+00 | 1.12E-09 | 4.86E-09 | -1.64E+00 | 1.37E-06 | 5.83E-06 | -1.44E+00 | 3.80E-11 | 1.45E-10 | -1.88E+00 | 1.06E-02 | 3.78E-02 | -1.11E+00 | 4.71E-05 | 1.33E-04 | -1.62E+00 | 1.43E-03 | 5.25E-03 | -1.08E+00 | 5.83E-05 | 1.60E-04 | -1.34E+00 |
| RPL12P33 | 9.35E-08 | 4.87E-07 | -1.23E+00 | 2.56E-19 | 1.10E-18 | -1.90E+00 | 4.02E-04 | 1.71E-03 | -1.16E+00 | 3.36E-13 | 1.28E-12 | -1.85E+00 | 9.23E-04 | 3.29E-03 | -1.09E+00 | 3.15E-08 | 8.89E-08 | -1.75E+00 | 3.13E-05 | 1.15E-04 | -1.09E+00 | 7.16E-13 | 1.97E-12 | -1.72E+00 |
| PCNA | 9.39E-08 | 4.89E-07 | -1.63E+00 | 8.48E-03 | 3.66E-02 | 1.08E+00 | 1.00E-02 | 4.25E-02 | 1.28E+00 | 9.74E-03 | 3.72E-02 | 1.55E+00 | 1.74E-05 | 6.19E-05 | -1.64E+00 | 3.01E-01 | 8.47E-01 | 1.34E+00 | 2.29E-06 | 8.42E-06 | -1.07E+00 | 1.64E-03 | 4.50E-03 | 1.55E+00 |
| SF3B3 | 9.41E-08 | 4.91E-07 | -1.27E+00 | 5.56E-02 | 2.40E-01 | 1.37E+00 | 1.05E-07 | 4.46E-07 | -1.26E+00 | 1.14E-02 | 4.34E-02 | 1.21E+00 | 2.48E-05 | 8.86E-05 | -1.24E+00 | 1.16E-01 | 3.28E-01 | 1.45E+00 | 6.04E-08 | 2.22E-07 | -1.47E+00 | 3.18E-01 | 8.75E-01 | 1.06E+00 |
| ANKRD20A9P | 9.43E-08 | 4.91E-07 | -1.57E+00 | 1.68E-07 | 7.28E-07 | -1.45E+00 | 1.96E-04 | 8.34E-04 | -1.49E+00 | 8.52E-04 | 3.25E-03 | -1.28E+00 | 2.00E-01 | 7.14E-01 | 1.50E+00 | 2.23E-01 | 6.28E-01 | -1.04E+00 | 2.37E-01 | 8.69E-01 | 1.16E+00 | 6.79E-02 | 1.87E-01 | -1.30E+00 |
| ZBTB8OS | 9.44E-08 | 4.92E-07 | -1.46E+00 | 8.08E-05 | 3.49E-04 | -1.03E+00 | 7.67E-04 | 3.26E-03 | -1.12E+00 | 3.54E-05 | 1.35E-04 | -1.12E+00 | 2.70E-03 | 9.63E-03 | -1.03E+00 | 2.14E-02 | 6.03E-02 | 1.11E+00 | 1.27E-05 | 4.67E-05 | -1.15E+00 | 1.62E-08 | 4.46E-08 | -1.30E+00 |
| SRSF2 | 9.44E-08 | 4.92E-07 | -1.08E+00 | 6.42E-08 | 2.77E-07 | 1.04E+00 | 1.43E-01 | 6.09E-01 | 1.01E+00 | 9.49E-02 | 3.62E-01 | 1.20E+00 | 1.86E-06 | 6.65E-06 | -1.23E+00 | 3.22E-03 | 9.08E-03 | -1.02E+00 | 1.02E-01 | 3.75E-01 | 1.03E+00 | 1.63E-02 | 4.47E-02 | 1.33E+00 |
| CBR4 | 9.50E-08 | 4.95E-07 | -1.41E+00 | 6.44E-02 | 2.78E-01 | 1.35E+00 | 1.37E-07 | 5.81E-07 | -1.62E+00 | 4.02E-02 | 1.54E-01 | 1.21E+00 | 3.10E-05 | 1.10E-04 | -1.49E+00 | 1.65E-01 | 4.66E-01 | 1.27E+00 | 1.20E-07 | 4.39E-07 | -1.34E+00 | 1.77E-01 | 4.86E-01 | 1.20E+00 |
| HEATR6 | 9.50E-08 | 4.95E-07 | -1.91E+00 | 2.30E-02 | 9.95E-02 | 1.19E+00 | 3.40E-04 | 1.45E-03 | -1.46E+00 | 5.75E-02 | 2.20E-01 | 1.15E+00 | 1.27E-06 | 4.53E-06 | -1.41E+00 | 1.08E-04 | 3.06E-04 | -1.19E+00 | 1.25E-09 | 4.58E-09 | -1.64E+00 | 5.16E-03 | 1.42E-02 | -1.04E+00 |
| YIF1A | 9.52E-08 | 4.96E-07 | -1.23E+00 | 9.22E-08 | 3.98E-07 | -1.10E+00 | 3.80E-06 | 1.61E-05 | -1.20E+00 | 1.38E-03 | 5.29E-03 | 1.09E+00 | 5.50E-06 | 1.96E-05 | -1.13E+00 | 6.15E-02 | 1.73E-01 | 1.25E+00 | 1.05E-03 | 3.85E-03 | -1.03E+00 | 5.29E-02 | 1.45E-01 | 1.15E+00 |
| TPM1 | 9.55E-08 | 4.98E-07 | -1.21E+00 | 1.26E-01 | 5.46E-01 | 1.42E+00 | 1.04E-01 | 4.40E-01 | 1.10E+00 | 1.95E-01 | 7.44E-01 | 1.61E+00 | 2.84E-02 | 1.01E-01 | 1.16E+00 | 2.16E-01 | 6.08E-01 | 1.49E+00 | 5.25E-03 | 1.93E-02 | -1.12E+00 | 5.80E-02 | 1.59E-01 | -1.13E+00 |
| HMGN1 | 9.66E-08 | 5.03E-07 | -1.15E+00 | 6.61E-12 | 2.85E-11 | -1.13E+00 | 1.49E-01 | 6.33E-01 | 1.01E+00 | 2.07E-02 | 7.90E-02 | -1.07E+00 | 2.35E-05 | 8.37E-05 | -1.18E+00 | 3.49E-04 | 9.83E-04 | -1.15E+00 | 1.11E-01 | 4.08E-01 | 1.12E+00 | 2.92E-01 | 8.01E-01 | 1.02E+00 |
| MICAL1 | 9.76E-08 | 5.09E-07 | -1.12E+00 | 9.73E-08 | 4.21E-07 | -1.03E+00 | 1.62E-02 | 6.88E-02 | 1.01E+00 | 2.48E-02 | 9.47E-02 | 1.20E+00 | 5.47E-02 | 1.95E-01 | 1.18E+00 | 2.64E-03 | 7.44E-03 | -1.09E+00 | 1.69E-01 | 6.19E-01 | 1.35E+00 | 1.84E-01 | 5.06E-01 | 1.40E+00 |
| WBP11 | 9.78E-08 | 5.10E-07 | -1.07E+00 | 3.01E-09 | 1.30E-08 | -1.04E+00 | 5.12E-03 | 2.18E-02 | -1.07E+00 | 4.91E-04 | 1.87E-03 | 1.00E+00 | 3.94E-02 | 1.40E-01 | 1.14E+00 | 4.25E-02 | 1.20E-01 | 1.11E+00 | 5.45E-03 | 2.00E-02 | -1.01E+00 | 8.30E-02 | 2.28E-01 | 1.03E+00 |
| HPS4 | 9.79E-08 | 5.10E-07 | -1.17E+00 | 1.96E-05 | 8.48E-05 | 1.06E+00 | 1.72E-02 | 7.31E-02 | 1.04E+00 | 1.38E-02 | 5.28E-02 | 1.18E+00 | 1.96E-02 | 6.97E-02 | 1.16E+00 | 1.89E-01 | 5.31E-01 | 1.42E+00 | 7.08E-03 | 2.60E-02 | 1.06E+00 | 1.98E-01 | 5.44E-01 | 1.33E+00 |
| NKAIN2 | 9.81E-08 | 5.11E-07 | -1.31E+00 | 4.90E-03 | 2.12E-02 | 1.16E+00 | 1.40E-01 | 5.94E-01 | 1.14E+00 | 2.58E-01 | 9.84E-01 | 1.50E+00 | 4.39E-02 | 1.56E-01 | 1.10E+00 | 1.07E-01 | 3.02E-01 | 1.28E+00 | 6.48E-03 | 2.38E-02 | -1.34E+00 | 3.04E-02 | 8.35E-02 | -1.17E+00 |
| F2R | 9.85E-08 | 5.13E-07 | -1.20E+00 | 1.23E-03 | 5.34E-03 | 1.11E+00 | 2.55E-07 | 1.08E-06 | -1.42E+00 | 2.41E-03 | 9.20E-03 | -1.02E+00 | 6.32E-04 | 2.25E-03 | 1.03E+00 | 2.56E-06 | 7.22E-06 | -1.10E+00 | 1.21E-01 | 4.45E-01 | 1.08E+00 | 6.89E-02 | 1.89E-01 | 1.00E+00 |
| CALCOCO1 | 9.96E-08 | 5.19E-07 | -1.45E+00 | 5.87E-03 | 2.54E-02 | 1.13E+00 | 1.44E-08 | 6.11E-08 | -1.64E+00 | 3.53E-05 | 1.35E-04 | -1.20E+00 | 6.74E-05 | 2.40E-04 | -1.18E+00 | 7.71E-02 | 2.17E-01 | 1.22E+00 | 7.15E-05 | 2.63E-04 | -1.18E+00 | 8.83E-03 | 2.42E-02 | -1.03E+00 |
| TFAP4 | 9.99E-08 | 5.21E-07 | -1.60E+00 | 1.23E-02 | 5.31E-02 | 1.14E+00 | 3.59E-15 | 1.52E-14 | -1.77E+00 | 5.02E-06 | 1.92E-05 | -1.03E+00 | 2.64E-06 | 9.42E-06 | -2.29E+00 | 1.11E-01 | 3.12E-01 | 1.15E+00 | 2.90E-16 | 1.07E-15 | -2.17E+00 | 4.69E-04 | 1.29E-03 | -1.08E+00 |
| ZNF677 | 1.01E-07 | 5.25E-07 | -1.29E+00 | 1.23E-03 | 5.32E-03 | 1.11E+00 | 6.71E-06 | 2.85E-05 | -1.46E+00 | 3.03E-05 | 1.16E-04 | -1.15E+00 | 1.03E-03 | 3.66E-03 | -1.09E+00 | 5.63E-02 | 1.59E-01 | 1.16E+00 | 1.37E-02 | 5.03E-02 | -1.14E+00 | 2.27E-01 | 6.24E-01 | 1.15E+00 |
| U47924.2 | 1.01E-07 | 5.26E-07 | 3.37E+00 | 3.52E-04 | 1.52E-03 | -2.03E+00 | 2.24E-08 | 9.52E-08 | 2.86E+00 | 6.61E-07 | 2.53E-06 | -2.67E+00 | 1.91E-01 | 6.82E-01 | 1.33E+00 | 8.57E-02 | 2.42E-01 | -1.29E+00 | 3.93E-02 | 1.44E-01 | -1.06E+00 | 7.12E-03 | 1.95E-02 | -1.55E+00 |
| AP3D1 | 1.02E-07 | 5.31E-07 | -1.28E+00 | 8.28E-03 | 3.58E-02 | 1.29E+00 | 2.09E-04 | 8.88E-04 | -1.12E+00 | 8.89E-04 | 3.39E-03 | 1.08E+00 | 5.97E-07 | 2.13E-06 | -1.34E+00 | 2.57E-03 | 7.25E-03 | 1.12E+00 | 6.42E-04 | 2.36E-03 | -1.19E+00 | 3.22E-01 | 8.84E-01 | 1.19E+00 |
| PTPN2 | 1.02E-07 | 5.31E-07 | -1.12E+00 | 6.86E-07 | 2.97E-06 | 1.09E+00 | 2.16E-04 | 9.20E-04 | -1.11E+00 | 2.40E-05 | 9.18E-05 | -1.04E+00 | 1.26E-05 | 4.48E-05 | -1.18E+00 | 1.45E-03 | 4.08E-03 | 1.00E+00 | 3.13E-06 | 1.15E-05 | -1.22E+00 | 1.12E-04 | 3.08E-04 | -1.18E+00 |
| PREP | 1.02E-07 | 5.31E-07 | -1.64E+00 | 2.60E-04 | 1.13E-03 | -1.02E+00 | 2.41E-08 | 1.02E-07 | -1.55E+00 | 2.56E-04 | 9.78E-04 | -1.03E+00 | 3.51E-08 | 1.25E-07 | -2.18E+00 | 5.86E-03 | 1.65E-02 | -1.01E+00 | 3.16E-09 | 1.16E-08 | -1.64E+00 | 1.73E-02 | 4.76E-02 | 1.03E+00 |
| NPLOC4 | 1.02E-07 | 5.31E-07 | -1.35E+00 | 3.10E-06 | 1.34E-05 | -1.11E+00 | 7.70E-04 | 3.27E-03 | -1.14E+00 | 1.58E-03 | 6.03E-03 | -1.01E+00 | 1.56E-03 | 5.58E-03 | -1.10E+00 | 2.00E-03 | 5.64E-03 | -1.10E+00 | 9.42E-05 | 3.46E-04 | -1.13E+00 | 1.04E-02 | 2.85E-02 | 1.01E+00 |
| CEP19 | 1.02E-07 | 5.31E-07 | -1.81E+00 | 6.26E-03 | 2.71E-02 | 1.07E+00 | 2.86E-05 | 1.22E-04 | -1.53E+00 | 1.47E-03 | 5.60E-03 | -1.01E+00 | 8.01E-04 | 2.86E-03 | -1.42E+00 | 5.86E-02 | 1.65E-01 | 1.07E+00 | 1.42E-02 | 5.23E-02 | -1.12E+00 | 2.00E-01 | 5.50E-01 | 1.15E+00 |
| VPS41 | 1.02E-07 | 5.33E-07 | -1.28E+00 | 1.00E-03 | 4.33E-03 | 1.03E+00 | 5.05E-07 | 2.15E-06 | -1.27E+00 | 3.00E-06 | 1.15E-05 | -1.10E+00 | 3.55E-11 | 1.27E-10 | -1.76E+00 | 6.10E-08 | 1.72E-07 | -1.41E+00 | 1.99E-07 | 7.31E-07 | -1.38E+00 | 1.78E-08 | 4.88E-08 | -1.46E+00 |
| BCAP29 | 1.03E-07 | 5.35E-07 | -1.18E+00 | 2.20E-05 | 9.52E-05 | 1.07E+00 | 2.29E-04 | 9.72E-04 | -1.23E+00 | 2.26E-03 | 8.63E-03 | -1.05E+00 | 3.76E-07 | 1.34E-06 | -1.30E+00 | 3.49E-04 | 9.83E-04 | -1.06E+00 | 2.12E-09 | 7.77E-09 | -1.25E+00 | 1.42E-05 | 3.91E-05 | -1.14E+00 |
| SAP30L | 1.03E-07 | 5.37E-07 | -1.85E+00 | 4.40E-03 | 1.90E-02 | 1.06E+00 | 8.89E-04 | 3.78E-03 | -1.21E+00 | 1.43E-01 | 5.45E-01 | 1.27E+00 | 4.59E-09 | 1.64E-08 | -2.50E+00 | 3.95E-06 | 1.11E-05 | -1.63E+00 | 1.08E-04 | 3.97E-04 | -1.18E+00 | 2.69E-01 | 7.38E-01 | 1.22E+00 |
| ATL3 | 1.03E-07 | 5.38E-07 | -1.35E+00 | 1.04E-02 | 4.51E-02 | 1.19E+00 | 1.95E-06 | 8.26E-06 | -1.29E+00 | 1.53E-02 | 5.83E-02 | 1.17E+00 | 6.55E-05 | 2.34E-04 | -1.16E+00 | 1.45E-02 | 4.09E-02 | 1.18E+00 | 1.35E-05 | 4.95E-05 | -1.29E+00 | 3.57E-01 | 9.79E-01 | 1.14E+00 |
| RAB29 | 1.03E-07 | 5.39E-07 | -1.54E+00 | 1.76E-03 | 7.62E-03 | 1.09E+00 | 3.73E-03 | 1.59E-02 | -1.37E+00 | 9.73E-02 | 3.71E-01 | 1.19E+00 | 1.11E-01 | 3.97E-01 | -1.11E+00 | 1.47E-01 | 4.14E-01 | 1.62E+00 | 3.86E-04 | 1.42E-03 | -1.88E+00 | 3.39E-02 | 9.31E-02 | -1.24E+00 |
| FZD3 | 1.04E-07 | 5.42E-07 | -1.71E+00 | 5.88E-08 | 2.54E-07 | -1.55E+00 | 1.15E-07 | 4.88E-07 | -1.97E+00 | 9.12E-06 | 3.48E-05 | -1.46E+00 | 2.26E-10 | 8.06E-10 | -1.76E+00 | 7.55E-06 | 2.13E-05 | -1.12E+00 | 1.98E-17 | 7.25E-17 | -1.93E+00 | 1.03E-11 | 2.83E-11 | -1.49E+00 |
| BMPR2 | 1.04E-07 | 5.42E-07 | -1.53E+00 | 4.68E-04 | 2.02E-03 | -1.01E+00 | 5.37E-05 | 2.28E-04 | -1.71E+00 | 7.06E-03 | 2.70E-02 | -1.02E+00 | 1.26E-04 | 4.48E-04 | -2.00E+00 | 7.06E-04 | 1.99E-03 | -1.61E+00 | 8.11E-05 | 2.98E-04 | -1.47E+00 | 1.15E-04 | 3.15E-04 | -1.59E+00 |
| NUDT10 | 1.04E-07 | 5.44E-07 | -1.75E+00 | 3.66E-03 | 1.58E-02 | 1.09E+00 | 1.18E-06 | 5.02E-06 | -1.49E+00 | 2.56E-03 | 9.78E-03 | 1.03E+00 | 5.73E-03 | 2.04E-02 | -1.28E+00 | 8.72E-02 | 2.46E-01 | 1.10E+00 | 1.40E-01 | 5.14E-01 | -1.21E+00 | 3.62E-01 | 9.95E-01 | 1.17E+00 |
| AC009244.1 | 1.05E-07 | 5.45E-07 | -1.31E+00 | 4.20E-04 | 1.81E-03 | 1.12E+00 | 2.25E-05 | 9.54E-05 | -1.41E+00 | 2.30E-02 | 8.80E-02 | 1.09E+00 | 8.56E-06 | 3.05E-05 | -1.33E+00 | 1.54E-03 | 4.33E-03 | 1.00E+00 | 9.00E-06 | 3.30E-05 | -1.29E+00 | 2.62E-01 | 7.20E-01 | 1.24E+00 |
| ICA1L | 1.05E-07 | 5.46E-07 | -1.61E+00 | 5.09E-06 | 2.20E-05 | -1.17E+00 | 7.61E-04 | 3.23E-03 | -1.31E+00 | 2.35E-04 | 8.98E-04 | -1.09E+00 | 3.77E-03 | 1.35E-02 | -1.24E+00 | 7.23E-03 | 2.04E-02 | -1.11E+00 | 2.42E-05 | 8.89E-05 | -1.31E+00 | 1.52E-04 | 4.18E-04 | -1.13E+00 |
| KIAA1109 | 1.05E-07 | 5.49E-07 | -1.20E+00 | 2.31E-04 | 9.98E-04 | 1.09E+00 | 8.59E-05 | 3.65E-04 | -1.25E+00 | 5.59E-05 | 2.14E-04 | -1.12E+00 | 2.18E-03 | 7.77E-03 | -1.04E+00 | 5.64E-04 | 1.59E-03 | -1.21E+00 | 8.91E-06 | 3.27E-05 | -1.27E+00 | 3.32E-04 | 9.11E-04 | -1.21E+00 |
| ZNHIT6 | 1.05E-07 | 5.49E-07 | -1.30E+00 | 1.26E-04 | 5.44E-04 | 1.04E+00 | 7.03E-14 | 2.99E-13 | -1.66E+00 | 6.47E-08 | 2.47E-07 | -1.23E+00 | 3.48E-10 | 1.24E-09 | -1.36E+00 | 7.44E-09 | 2.10E-08 | -1.33E+00 | 6.48E-05 | 2.38E-04 | -1.28E+00 | 2.79E-02 | 7.66E-02 | -1.05E+00 |
| CTDSP1 | 1.07E-07 | 5.56E-07 | -1.17E+00 | 7.66E-07 | 3.31E-06 | -1.04E+00 | 9.81E-06 | 4.17E-05 | -1.20E+00 | 6.76E-05 | 2.58E-04 | -1.04E+00 | 2.31E-07 | 8.24E-07 | -1.23E+00 | 2.68E-04 | 7.55E-04 | 1.00E+00 | 7.87E-08 | 2.89E-07 | -1.30E+00 | 1.27E-03 | 3.48E-03 | -1.10E+00 |
| SNAP47 | 1.07E-07 | 5.56E-07 | -1.55E+00 | 7.80E-03 | 3.37E-02 | 1.15E+00 | 1.67E-03 | 7.12E-03 | -1.30E+00 | 3.43E-02 | 1.31E-01 | 1.06E+00 | 1.39E-01 | 4.94E-01 | 1.39E+00 | 1.48E-01 | 4.17E-01 | 1.80E+00 | 2.90E-02 | 1.07E-01 | 1.69E+00 | 4.22E-02 | 1.16E-01 | 1.68E+00 |
| UTP23 | 1.07E-07 | 5.56E-07 | -1.23E+00 | 7.27E-15 | 3.14E-14 | -1.44E+00 | 5.17E-10 | 2.20E-09 | -1.45E+00 | 5.12E-15 | 1.96E-14 | -1.57E+00 | 1.81E-02 | 6.46E-02 | 1.03E+00 | 1.99E-04 | 5.60E-04 | -1.26E+00 | 1.24E-08 | 4.56E-08 | -1.42E+00 | 1.02E-06 | 2.80E-06 | -1.28E+00 |
| TMED10 | 1.07E-07 | 5.57E-07 | -1.02E+00 | 2.81E-07 | 1.22E-06 | 1.07E+00 | 1.44E-02 | 6.14E-02 | -1.04E+00 | 6.49E-03 | 2.48E-02 | 1.03E+00 | 2.71E-12 | 9.65E-12 | -1.40E+00 | 4.69E-07 | 1.32E-06 | -1.17E+00 | 7.93E-04 | 2.91E-03 | -1.07E+00 | 1.67E-01 | 4.60E-01 | 1.15E+00 |
| USP47 | 1.08E-07 | 5.62E-07 | -1.15E+00 | 4.45E-05 | 1.92E-04 | 1.08E+00 | 5.69E-06 | 2.42E-05 | -1.29E+00 | 2.44E-06 | 9.31E-06 | -1.13E+00 | 1.55E-08 | 5.54E-08 | -1.43E+00 | 5.46E-05 | 1.54E-04 | -1.11E+00 | 6.66E-08 | 2.45E-07 | -1.26E+00 | 6.07E-04 | 1.67E-03 | -1.10E+00 |
| ATF2 | 1.08E-07 | 5.63E-07 | -1.22E+00 | 3.98E-05 | 1.72E-04 | 1.13E+00 | 2.18E-06 | 9.27E-06 | -1.43E+00 | 1.33E-05 | 5.09E-05 | -1.14E+00 | 7.38E-07 | 2.63E-06 | -1.38E+00 | 3.16E-04 | 8.92E-04 | -1.11E+00 | 1.86E-06 | 6.82E-06 | -1.25E+00 | 4.80E-03 | 1.32E-02 | -1.06E+00 |
| CTBS | 1.08E-07 | 5.64E-07 | -1.20E+00 | 2.62E-05 | 1.13E-04 | 1.02E+00 | 1.89E-04 | 8.03E-04 | -1.20E+00 | 3.01E-03 | 1.15E-02 | 1.01E+00 | 5.54E-03 | 1.98E-02 | 1.08E+00 | 2.29E-02 | 6.45E-02 | 1.13E+00 | 3.22E-02 | 1.18E-01 | 1.08E+00 | 2.75E-02 | 7.55E-02 | -1.02E+00 |
| RPL18P11 | 1.09E-07 | 5.66E-07 | -1.14E+00 | 5.63E-39 | 2.43E-38 | -2.12E+00 | 7.17E-03 | 3.05E-02 | -1.10E+00 | 2.01E-21 | 7.67E-21 | -2.05E+00 | 2.38E-06 | 8.47E-06 | -1.15E+00 | 7.64E-22 | 2.15E-21 | -2.10E+00 | 1.52E-03 | 5.57E-03 | -1.09E+00 | 2.50E-16 | 6.86E-16 | -1.86E+00 |
| ZNF765 | 1.09E-07 | 5.66E-07 | -1.64E+00 | 1.24E-05 | 5.34E-05 | -1.24E+00 | 1.48E-07 | 6.27E-07 | -1.82E+00 | 1.19E-05 | 4.54E-05 | -1.38E+00 | 3.07E-03 | 1.09E-02 | -1.29E+00 | 7.70E-02 | 2.17E-01 | 1.11E+00 | 2.85E-08 | 1.05E-07 | -1.44E+00 | 8.94E-04 | 2.46E-03 | -1.04E+00 |
| ACADS | 1.09E-07 | 5.68E-07 | -1.29E+00 | 1.28E-06 | 5.53E-06 | -1.07E+00 | 5.90E-03 | 2.51E-02 | -1.05E+00 | 4.07E-05 | 1.56E-04 | -1.12E+00 | 1.18E-03 | 4.21E-03 | -1.18E+00 | 3.63E-02 | 1.02E-01 | 1.12E+00 | 1.23E-02 | 4.53E-02 | -1.03E+00 | 2.41E-03 | 6.62E-03 | -1.09E+00 |
| LINC02001 | 1.09E-07 | 5.69E-07 | -1.10E+00 | 3.49E-07 | 1.51E-06 | 1.02E+00 | 1.48E-01 | 6.31E-01 | 1.30E+00 | 1.81E-01 | 6.90E-01 | 1.36E+00 | 1.97E-02 | 7.01E-02 | 1.27E+00 | 1.30E-01 | 3.67E-01 | 1.37E+00 | 1.99E-01 | 7.32E-01 | 1.25E+00 | 1.34E-01 | 3.67E-01 | 1.09E+00 |
| THRA | 1.09E-07 | 5.69E-07 | -1.36E+00 | 9.81E-05 | 4.24E-04 | 1.04E+00 | 3.55E-03 | 1.51E-02 | -1.16E+00 | 2.75E-03 | 1.05E-02 | -1.06E+00 | 1.61E-04 | 5.75E-04 | -1.34E+00 | 1.24E-03 | 3.50E-03 | -1.17E+00 | 3.45E-08 | 1.27E-07 | -1.32E+00 | 3.71E-07 | 1.02E-06 | -1.35E+00 |
| RPF2P1 | 1.10E-07 | 5.71E-07 | -1.50E+00 | 3.10E-06 | 1.34E-05 | -1.28E+00 | 1.28E-08 | 5.42E-08 | -1.62E+00 | 7.52E-08 | 2.87E-07 | -1.32E+00 | 4.16E-10 | 1.48E-09 | -2.77E+00 | 2.12E-05 | 5.97E-05 | -1.43E+00 | 3.46E-14 | 1.27E-13 | -1.85E+00 | 1.11E-09 | 3.06E-09 | -1.46E+00 |
| USF1 | 1.10E-07 | 5.71E-07 | -1.75E+00 | 2.07E-07 | 8.94E-07 | -1.58E+00 | 3.27E-06 | 1.39E-05 | -1.68E+00 | 2.61E-04 | 9.95E-04 | -1.20E+00 | 2.14E-09 | 7.62E-09 | -1.81E+00 | 1.20E-06 | 3.39E-06 | -1.36E+00 | 2.41E-04 | 8.86E-04 | -1.36E+00 | 5.65E-02 | 1.55E-01 | 1.02E+00 |
| NFATC2 | 1.10E-07 | 5.74E-07 | -1.25E+00 | 2.80E-02 | 1.21E-01 | 1.37E+00 | 1.98E-04 | 8.41E-04 | -1.38E+00 | 3.84E-02 | 1.47E-01 | 1.17E+00 | 4.04E-06 | 1.44E-05 | -1.43E+00 | 1.85E-03 | 5.23E-03 | 1.01E+00 | 2.62E-07 | 9.61E-07 | -1.75E+00 | 1.20E-03 | 3.29E-03 | -1.20E+00 |
| TIMP1 | 1.10E-07 | 5.75E-07 | -1.16E+00 | 3.39E-08 | 1.47E-07 | -1.03E+00 | 9.29E-02 | 3.95E-01 | 1.10E+00 | 1.10E-01 | 4.19E-01 | 1.15E+00 | 4.69E-03 | 1.67E-02 | -1.10E+00 | 1.77E-03 | 4.99E-03 | -1.16E+00 | 6.78E-02 | 2.49E-01 | 1.12E+00 | 3.14E-02 | 8.63E-02 | 1.06E+00 |
| ROCK1P1 | 1.10E-07 | 5.75E-07 | -1.98E+00 | 8.82E-06 | 3.81E-05 | -1.46E+00 | 2.28E-08 | 9.68E-08 | -1.91E+00 | 8.19E-10 | 3.13E-09 | -1.91E+00 | 1.96E-06 | 7.00E-06 | -2.53E+00 | 5.10E-04 | 1.44E-03 | -1.45E+00 | 2.56E-11 | 9.40E-11 | -2.31E+00 | 1.72E-08 | 4.73E-08 | -1.93E+00 |
| POLB | 1.10E-07 | 5.75E-07 | -1.43E+00 | 3.82E-09 | 1.65E-08 | -1.40E+00 | 4.25E-05 | 1.81E-04 | -1.30E+00 | 7.38E-08 | 2.82E-07 | -1.51E+00 | 7.46E-05 | 2.66E-04 | -1.27E+00 | 1.03E-06 | 2.91E-06 | -1.69E+00 | 8.43E-03 | 3.09E-02 | -1.19E+00 | 7.10E-04 | 1.95E-03 | -1.43E+00 |
| BCL9 | 1.10E-07 | 5.75E-07 | -1.84E+00 | 6.41E-05 | 2.77E-04 | -1.25E+00 | 3.26E-04 | 1.39E-03 | -1.59E+00 | 4.95E-02 | 1.89E-01 | 1.10E+00 | 1.99E-02 | 7.11E-02 | -1.04E+00 | 3.09E-02 | 8.72E-02 | -1.02E+00 | 5.85E-06 | 2.15E-05 | -2.41E+00 | 1.43E-02 | 3.93E-02 | -1.26E+00 |
| COPS2 | 1.10E-07 | 5.76E-07 | -1.18E+00 | 1.36E-10 | 5.87E-10 | -1.16E+00 | 1.93E-04 | 8.20E-04 | -1.16E+00 | 1.31E-06 | 5.00E-06 | -1.18E+00 | 3.39E-04 | 1.21E-03 | -1.12E+00 | 1.31E-05 | 3.68E-05 | -1.19E+00 | 9.17E-05 | 3.37E-04 | -1.10E+00 | 3.95E-05 | 1.08E-04 | -1.13E+00 |
| UACA | 1.11E-07 | 5.77E-07 | -1.75E+00 | 7.61E-06 | 3.29E-05 | -1.40E+00 | 1.52E-09 | 6.44E-09 | -1.81E+00 | 3.40E-09 | 1.30E-08 | -1.51E+00 | 4.14E-05 | 1.48E-04 | -1.92E+00 | 3.57E-02 | 1.01E-01 | 1.10E+00 | 2.67E-18 | 9.78E-18 | -2.18E+00 | 3.35E-09 | 9.19E-09 | -1.58E+00 |
| TAF5 | 1.12E-07 | 5.82E-07 | -2.30E+00 | 2.26E-06 | 9.75E-06 | -1.64E+00 | 5.29E-06 | 2.25E-05 | -1.86E+00 | 2.20E-03 | 8.41E-03 | -1.02E+00 | 3.45E-05 | 1.23E-04 | -3.14E+00 | 5.78E-03 | 1.63E-02 | -1.39E+00 | 8.52E-08 | 3.13E-07 | -1.46E+00 | 3.02E-03 | 8.31E-03 | -1.13E+00 |
| HCLS1 | 1.12E-07 | 5.85E-07 | -1.12E+00 | 8.27E-08 | 3.58E-07 | 1.02E+00 | 1.75E-04 | 7.44E-04 | -1.18E+00 | 4.16E-02 | 1.59E-01 | 1.16E+00 | 2.23E-08 | 7.96E-08 | -1.23E+00 | 7.09E-05 | 2.00E-04 | 1.02E+00 | 3.48E-03 | 1.28E-02 | -1.02E+00 | 2.51E-01 | 6.89E-01 | 1.18E+00 |
| MAGOHB | 1.13E-07 | 5.87E-07 | -1.24E+00 | 1.08E-03 | 4.65E-03 | 1.18E+00 | 4.74E-03 | 2.01E-02 | -1.01E+00 | 9.57E-04 | 3.66E-03 | 1.09E+00 | 9.95E-09 | 3.55E-08 | -1.95E+00 | 5.09E-03 | 1.43E-02 | -1.05E+00 | 5.74E-07 | 2.11E-06 | -1.26E+00 | 1.47E-01 | 4.04E-01 | 1.14E+00 |

| | | | | | | | | | | | | | | | | | | | | | | | | |
|---|---|---|---|---|---|---|---|---|---|---|---|---|---|---|---|---|---|---|---|---|---|---|---|---|
| SLC16A7 | 1.13E-07 | 5.87E-07 | -1.52E+00 | 4.70E-05 | 2.03E-04 | -1.14E+00 | 2.70E-06 | 1.15E-05 | -1.48E+00 | 1.83E-02 | 6.97E-02 | 1.13E+00 | 1.58E-03 | 5.62E-03 | -1.40E+00 | 1.87E-02 | 5.26E-02 | -1.05E+00 | 7.28E-04 | 2.67E-03 | -1.42E+00 | 2.55E-02 | 7.01E-02 | -1.15E+00 |
| DDX49 | 1.13E-07 | 5.90E-07 | -1.16E+00 | 1.33E-05 | 5.74E-05 | 1.04E+00 | 9.26E-05 | 3.93E-04 | -1.07E+00 | 8.44E-06 | 3.22E-05 | -1.04E+00 | 1.29E-04 | 4.61E-04 | -1.12E+00 | 4.50E-02 | 1.27E-01 | 1.10E+00 | 1.06E-01 | 3.87E-01 | 1.12E+00 | 1.59E-01 | 4.37E-01 | 1.36E+00 |
| FAM222B | 1.14E-07 | 5.94E-07 | -1.45E+00 | 6.31E-07 | 2.72E-06 | -1.18E+00 | 1.44E-04 | 6.13E-04 | -1.26E+00 | 2.23E-04 | 8.52E-04 | -1.15E+00 | 3.84E-04 | 1.37E-03 | -1.50E+00 | 3.80E-02 | 1.07E-01 | 1.05E+00 | 9.81E-09 | 3.60E-08 | -2.04E+00 | 3.04E-04 | 8.34E-04 | -1.35E+00 |
| AP000766.1 | 1.15E-07 | 5.97E-07 | -1.78E+00 | 1.29E-03 | 5.58E-03 | 1.01E+00 | 7.10E-02 | 3.02E-01 | -1.20E+00 | 1.33E-01 | 5.07E-01 | 1.15E+00 | 2.06E-01 | 7.36E-01 | 1.26E+00 | 1.52E-01 | 4.30E-01 | 1.09E+00 | 2.93E-02 | 1.07E-01 | -1.06E+00 | 7.29E-02 | 2.00E-01 | -1.03E+00 |
| DAP | 1.15E-07 | 5.98E-07 | -1.28E+00 | 9.99E-07 | 4.32E-06 | -1.07E+00 | 4.36E-03 | 1.85E-02 | -1.15E+00 | 4.65E-04 | 1.78E-03 | -1.05E+00 | 7.61E-06 | 2.71E-05 | -1.25E+00 | 3.16E-02 | 8.90E-02 | 1.19E+00 | 1.39E-02 | 5.09E-02 | 1.06E+00 | 2.91E-02 | 7.98E-02 | -1.04E+00 |
| FGF7 | 1.16E-07 | 6.02E-07 | -1.94E+00 | 1.54E-03 | 6.67E-03 | 1.02E+00 | 5.43E-04 | 2.31E-03 | -1.49E+00 | 9.52E-02 | 3.63E-01 | 1.21E+00 | 7.46E-03 | 2.66E-02 | -1.27E+00 | 1.52E-02 | 4.29E-02 | -1.11E+00 | 1.20E-03 | 4.42E-03 | -1.50E+00 | 1.22E-02 | 3.35E-02 | -1.26E+00 |
| UTP18 | 1.16E-07 | 6.02E-07 | -1.27E+00 | 4.63E-09 | 2.00E-08 | -1.18E+00 | 2.71E-06 | 1.15E-05 | -1.20E+00 | 1.15E-08 | 4.38E-08 | -1.09E+00 | 8.15E-07 | 2.91E-06 | -1.54E+00 | 5.04E-04 | 1.42E-03 | -1.11E+00 | 1.03E-08 | 3.79E-08 | -1.35E+00 | 1.35E-03 | 3.70E-03 | -1.06E+00 |
| EHMT1 | 1.16E-07 | 6.03E-07 | -1.17E+00 | 9.61E-06 | 4.15E-05 | 1.09E+00 | 1.30E-06 | 5.54E-06 | -1.46E+00 | 4.06E-03 | 1.55E-02 | -1.06E+00 | 1.06E-05 | 3.79E-05 | -1.20E+00 | 4.28E-03 | 1.21E-02 | 1.00E+00 | 2.77E-08 | 1.02E-07 | -1.41E+00 | 4.14E-02 | 1.14E-01 | 1.06E+00 |
| ARHGEF6 | 1.16E-07 | 6.03E-07 | -1.14E+00 | 1.61E-08 | 6.96E-08 | -1.01E+00 | 4.24E-05 | 1.80E-04 | -1.26E+00 | 9.77E-04 | 3.73E-03 | -1.02E+00 | 5.19E-06 | 1.85E-05 | -1.33E+00 | 2.65E-06 | 7.46E-06 | -1.30E+00 | 1.18E-02 | 4.33E-02 | -1.09E+00 | 2.49E-01 | 6.84E-01 | 1.14E+00 |
| ARMT1 | 1.17E-07 | 6.08E-07 | -2.17E+00 | 1.33E-05 | 5.73E-05 | -1.51E+00 | 3.36E-02 | 1.43E-01 | -1.10E+00 | 3.17E-02 | 1.21E-01 | 1.05E+00 | 6.23E-05 | 2.22E-04 | -1.88E+00 | 7.47E-03 | 2.11E-02 | -1.21E+00 | 2.56E-09 | 9.41E-09 | -1.47E+00 | 1.77E-05 | 4.86E-05 | -1.17E+00 |
| SP4 | 1.17E-07 | 6.11E-07 | -1.56E+00 | 4.61E-05 | 1.99E-04 | -1.10E+00 | 6.16E-03 | 2.62E-02 | -1.19E+00 | 2.74E-02 | 1.05E-01 | 1.08E+00 | 8.84E-06 | 3.15E-05 | -1.72E+00 | 4.56E-06 | 1.29E-05 | -1.77E+00 | 5.72E-06 | 2.10E-05 | -1.68E+00 | 3.67E-03 | 1.01E-02 | -1.14E+00 |
| PCCB | 1.17E-07 | 6.11E-07 | -1.51E+00 | 5.59E-09 | 2.42E-08 | -1.48E+00 | 9.39E-04 | 3.99E-03 | -1.19E+00 | 2.08E-08 | 7.93E-08 | -1.40E+00 | 2.83E-03 | 1.01E-02 | -1.14E+00 | 1.61E-01 | 4.53E-01 | 1.21E+00 | 3.55E-06 | 1.30E-05 | -1.23E+00 | 1.82E-03 | 4.99E-03 | -1.02E+00 |
| HSPA5 | 1.17E-07 | 6.11E-07 | -1.19E+00 | 1.37E-15 | 5.91E-15 | -1.29E+00 | 2.68E-02 | 1.14E-01 | 1.07E+00 | 1.27E-02 | 4.85E-02 | 1.05E+00 | 1.41E-09 | 5.03E-09 | -1.65E+00 | 5.84E-12 | 1.65E-11 | -1.78E+00 | 2.10E-05 | 7.71E-05 | -1.40E+00 | 7.47E-02 | 2.05E-01 | -1.18E+00 |
| RASAL2 | 1.17E-07 | 6.12E-07 | -2.58E+00 | 6.65E-05 | 2.87E-04 | -1.44E+00 | 1.92E-10 | 8.17E-10 | -2.19E+00 | 1.78E-09 | 6.81E-09 | -1.65E+00 | 3.23E-04 | 1.15E-03 | -2.09E+00 | 3.85E-04 | 1.09E-03 | -2.36E+00 | 3.59E-04 | 1.32E-03 | -1.36E+00 | 8.61E-03 | 2.36E-02 | -1.20E+00 |
| MYEF2 | 1.17E-07 | 6.12E-07 | -1.53E+00 | 9.96E-10 | 4.30E-09 | -1.42E+00 | 6.19E-03 | 2.63E-02 | -1.14E+00 | 4.33E-04 | 1.65E-03 | -1.10E+00 | 3.90E-03 | 1.39E-02 | -1.06E+00 | 5.15E-05 | 1.45E-04 | -1.45E+00 | 3.92E-02 | 1.44E-01 | -1.10E+00 | 8.93E-03 | 2.45E-02 | -1.18E+00 |
| TUSC1 | 1.18E-07 | 6.13E-07 | -1.29E+00 | 8.78E-06 | 3.79E-05 | -1.09E+00 | 1.76E-03 | 7.49E-03 | -1.10E+00 | 1.14E-02 | 4.36E-02 | 1.13E+00 | 3.26E-04 | 1.16E-03 | -1.31E+00 | 9.00E-03 | 2.54E-02 | -1.15E+00 | 5.44E-06 | 1.99E-05 | -1.18E+00 | 6.53E-02 | 1.79E-01 | 1.15E+00 |
| SETD1B | 1.18E-07 | 6.13E-07 | -1.62E+00 | 1.86E-04 | 8.06E-04 | -1.06E+00 | 4.86E-03 | 2.06E-02 | -1.21E+00 | 6.97E-02 | 2.66E-01 | 1.10E+00 | 4.11E-07 | 1.47E-06 | -1.63E+00 | 9.94E-04 | 2.80E-03 | -1.14E+00 | 1.27E-05 | 4.66E-05 | -1.51E+00 | 1.04E-01 | 2.87E-01 | 1.07E+00 |
| AC092017.2 | 1.18E-07 | 6.16E-07 | -1.17E+00 | 1.09E-62 | 4.70E-62 | -2.14E+00 | 2.51E-04 | 1.07E-03 | -1.24E+00 | 1.76E-42 | 6.72E-42 | -2.49E+00 | 1.92E-04 | 6.84E-04 | -1.16E+00 | 4.60E-24 | 1.30E-23 | -2.08E+00 | 5.93E-05 | 2.18E-04 | -1.32E+00 | 1.49E-23 | 4.09E-23 | -2.18E+00 |
| FYB1 | 1.18E-07 | 6.16E-07 | -2.92E+00 | 6.73E-08 | 2.91E-07 | -2.60E+00 | 5.92E-20 | 2.51E-19 | -3.14E+00 | 1.21E-22 | 4.62E-22 | -3.03E+00 | 1.20E-28 | 4.26E-28 | -2.11E+00 | 4.22E-30 | 1.19E-29 | -1.93E+00 | 3.55E-02 | 1.30E-01 | -1.11E+00 | 3.90E-07 | 1.07E-06 | -1.55E+00 |
| IPO8 | 1.18E-07 | 6.16E-07 | -1.63E+00 | 3.73E-04 | 1.61E-03 | -1.04E+00 | 2.00E-03 | 8.50E-03 | -1.13E+00 | 5.33E-03 | 2.04E-02 | 1.02E+00 | 6.36E-07 | 2.27E-06 | -1.62E+00 | 5.67E-05 | 1.60E-04 | -1.26E+00 | 3.41E-04 | 1.25E-03 | -1.22E+00 | 9.31E-02 | 2.56E-01 | 1.03E+00 |
| MCTP1 | 1.18E-07 | 6.16E-07 | -1.39E+00 | 1.66E-08 | 7.15E-08 | -1.27E+00 | 8.98E-04 | 3.82E-03 | -1.15E+00 | 4.89E-04 | 1.87E-03 | 1.00E+00 | 2.66E-01 | 9.48E-01 | 1.37E+00 | 2.68E-01 | 7.56E-01 | 1.32E+00 | 7.96E-02 | 2.92E-01 | 1.37E+00 | 1.63E-02 | 4.47E-02 | -1.16E+00 |
| MKNK2 | 1.18E-07 | 6.16E-07 | -1.31E+00 | 1.85E-06 | 8.00E-06 | -1.04E+00 | 6.84E-08 | 2.91E-07 | -1.44E+00 | 3.45E-06 | 1.32E-05 | -1.11E+00 | 7.99E-04 | 2.85E-03 | -1.20E+00 | 4.21E-03 | 1.19E-02 | -1.11E+00 | 1.44E-03 | 5.29E-03 | -1.13E+00 | 3.87E-05 | 1.06E-04 | -1.36E+00 |
| VSIR | 1.18E-07 | 6.17E-07 | 1.75E+00 | 1.54E-05 | 6.65E-05 | 1.85E+00 | 5.93E-03 | 2.52E-02 | 1.49E+00 | 4.28E-02 | 1.64E-01 | 1.52E+00 | 8.90E-04 | 3.17E-03 | 1.12E+00 | 9.45E-06 | 2.67E-05 | -1.00E+00 | 2.09E-01 | 7.66E-01 | 1.48E+00 | 1.97E-01 | 5.42E-01 | 1.08E+00 |
| RBL2 | 1.19E-07 | 6.18E-07 | -1.84E+00 | 2.03E-05 | 8.76E-05 | -1.27E+00 | 9.47E-04 | 4.02E-03 | -1.60E+00 | 2.31E-02 | 8.83E-02 | -1.11E+00 | 4.23E-11 | 1.51E-10 | -1.97E+00 | 4.30E-11 | 1.21E-10 | -1.77E+00 | 2.46E-05 | 9.03E-05 | -1.31E+00 | 2.64E-03 | 7.24E-03 | -1.12E+00 |
| MANBAL | 1.19E-07 | 6.19E-07 | -1.55E+00 | 8.54E-04 | 3.69E-03 | -1.01E+00 | 5.50E-06 | 2.34E-05 | -1.70E+00 | 5.70E-02 | 2.18E-01 | 1.07E+00 | 2.34E-03 | 8.33E-03 | -1.38E+00 | 4.81E-02 | 1.36E-01 | 1.09E+00 | 1.49E-04 | 5.46E-04 | -1.44E+00 | 7.73E-03 | 2.12E-02 | -1.18E+00 |
| LINC00694 | 1.19E-07 | 6.23E-07 | -2.41E+00 | 5.21E-04 | 2.25E-03 | -1.31E+00 | 9.85E-03 | 4.19E-02 | -1.51E+00 | 3.65E-02 | 1.39E-01 | -1.16E+00 | 1.20E-11 | 4.29E-11 | -3.17E+00 | 1.84E-08 | 5.18E-08 | -2.06E+00 | 2.84E-03 | 1.04E-02 | -1.69E+00 | 1.60E-03 | 4.40E-03 | -2.17E+00 |
| THEM4 | 1.20E-07 | 6.25E-07 | -1.20E+00 | 5.95E-07 | 2.57E-06 | -1.04E+00 | 3.93E-06 | 1.67E-05 | -1.38E+00 | 5.37E-07 | 2.05E-06 | -1.28E+00 | 5.27E-04 | 1.88E-03 | -1.09E+00 | 1.05E-04 | 2.97E-04 | -1.14E+00 | 1.67E-03 | 6.12E-03 | -1.09E+00 | 1.21E-02 | 3.34E-02 | 1.03E+00 |
| SOS1 | 1.20E-07 | 6.25E-07 | -1.40E+00 | 3.68E-05 | 1.59E-04 | -1.03E+00 | 8.98E-10 | 3.82E-09 | -1.54E+00 | 4.00E-06 | 1.53E-05 | -1.11E+00 | 1.20E-08 | 4.27E-08 | -1.76E+00 | 5.78E-03 | 1.63E-02 | 1.06E+00 | 2.57E-11 | 9.45E-11 | -1.69E+00 | 5.83E-06 | 1.60E-05 | -1.21E+00 |
| PDCL | 1.21E-07 | 6.29E-07 | -1.19E+00 | 6.59E-05 | 2.85E-04 | 1.09E+00 | 5.82E-06 | 2.47E-05 | -1.25E+00 | 1.84E-03 | 7.03E-03 | 1.03E+00 | 7.70E-05 | 2.74E-04 | -1.23E+00 | 8.93E-04 | 2.52E-03 | -1.06E+00 | 9.96E-04 | 3.66E-03 | -1.07E+00 | 8.54E-03 | 2.34E-02 | -1.01E+00 |
| B4GALT6 | 1.21E-07 | 6.32E-07 | -1.43E+00 | 6.00E-03 | 2.59E-02 | 1.19E+00 | 4.94E-08 | 2.10E-07 | -1.49E+00 | 1.19E-05 | 4.53E-05 | -1.10E+00 | 1.63E-02 | 5.82E-02 | -1.28E+00 | 3.56E-02 | 1.00E-01 | -1.21E+00 | 6.78E-04 | 2.49E-03 | -1.64E+00 | 1.52E-01 | 4.17E-01 | -1.00E+00 |
| HSPE1P2 | 1.22E-07 | 6.34E-07 | -1.59E+00 | 4.08E-03 | 1.76E-02 | 1.15E+00 | 4.26E-02 | 1.81E-01 | 1.03E+00 | 7.81E-02 | 2.98E-01 | 1.68E+00 | 2.27E-05 | 8.10E-05 | -1.66E+00 | 3.47E-01 | 9.78E-01 | 1.47E+00 | 4.24E-05 | 1.56E-04 | -1.10E+00 | 3.22E-04 | 8.85E-04 | 1.92E+00 |
| PAPOLA | 1.22E-07 | 6.34E-07 | -1.11E+00 | 4.82E-05 | 2.08E-04 | 1.05E+00 | 4.68E-03 | 1.99E-02 | -1.07E+00 | 5.97E-03 | 2.28E-02 | -1.00E+00 | 2.29E-04 | 8.15E-04 | -1.00E+00 | 1.35E-03 | 3.80E-03 | 1.02E+00 | 1.32E-02 | 4.86E-02 | -1.07E+00 | 6.40E-02 | 1.76E-01 | -1.06E+00 |
| EIF2D | 1.22E-07 | 6.38E-07 | -1.09E+00 | 3.21E-07 | 1.39E-06 | 1.03E+00 | 5.43E-05 | 2.31E-04 | -1.14E+00 | 1.91E-05 | 7.29E-05 | 1.01E+00 | 1.09E-05 | 3.87E-05 | -1.12E+00 | 1.45E-03 | 4.08E-03 | 1.07E+00 | 6.23E-03 | 2.28E-02 | 1.03E+00 | 3.05E-02 | 8.37E-02 | 1.08E+00 |
| SLC29A3 | 1.23E-07 | 6.41E-07 | -2.25E+00 | 4.74E-05 | 2.05E-04 | -1.39E+00 | 1.05E-13 | 4.48E-13 | -2.30E+00 | 6.13E-09 | 2.34E-08 | -1.45E+00 | 6.29E-06 | 2.24E-05 | -1.77E+00 | 6.61E-05 | 1.86E-04 | -1.41E+00 | 2.24E-03 | 8.21E-03 | -1.56E+00 | 4.34E-02 | 1.19E-01 | -1.19E+00 |
| CMC2 | 1.24E-07 | 6.47E-07 | -1.11E+00 | 1.06E-11 | 4.57E-11 | -1.15E+00 | 1.03E-02 | 4.37E-02 | -1.01E+00 | 1.83E-04 | 6.98E-04 | -1.03E+00 | 3.30E-03 | 1.18E-02 | 1.08E+00 | 2.77E-03 | 7.81E-03 | 1.04E+00 | 1.82E-03 | 6.67E-03 | -1.10E+00 | 1.87E-02 | 5.12E-02 | 1.03E+00 |
| PGRMC2 | 1.24E-07 | 6.47E-07 | -1.10E+00 | 4.05E-06 | 1.75E-05 | -1.03E+00 | 2.30E-09 | 9.75E-09 | -1.39E+00 | 2.61E-05 | 9.98E-05 | -1.03E+00 | 2.65E-06 | 9.47E-06 | -1.23E+00 | 4.07E-04 | 1.15E-03 | -1.02E+00 | 9.37E-09 | 3.44E-08 | -1.13E+00 | 2.40E-04 | 6.59E-04 | -1.06E+00 |
| CIB1 | 1.25E-07 | 6.50E-07 | -1.09E+00 | 1.31E-06 | 5.67E-06 | 1.07E+00 | 5.79E-03 | 2.46E-02 | -1.02E+00 | 4.86E-03 | 1.86E-02 | 1.09E+00 | 1.10E-04 | 3.91E-04 | 1.05E+00 | 1.47E-02 | 4.15E-02 | 1.23E+00 | 1.44E-01 | 5.27E-01 | 1.16E+00 | 3.48E-01 | 9.55E-01 | 1.22E+00 |
| ZDHHC21 | 1.25E-07 | 6.51E-07 | -1.28E+00 | 2.66E-03 | 1.15E-02 | 1.20E+00 | 1.10E-09 | 4.66E-09 | -1.54E+00 | 6.00E-05 | 2.29E-04 | -1.03E+00 | 1.22E-03 | 4.37E-03 | -1.32E+00 | 1.52E-02 | 4.29E-02 | -1.01E+00 | 3.60E-07 | 1.32E-06 | -1.75E+00 | 2.75E-04 | 7.54E-04 | -1.41E+00 |
| CLGN | 1.25E-07 | 6.53E-07 | -2.08E+00 | 2.35E-04 | 1.01E-03 | -1.17E+00 | 1.97E-02 | 8.38E-02 | -1.17E+00 | 5.80E-03 | 2.22E-02 | -1.05E+00 | 4.87E-03 | 1.74E-02 | -1.92E+00 | 4.08E-02 | 1.15E-01 | -1.25E+00 | 2.25E-06 | 8.27E-06 | -1.57E+00 | 1.22E-02 | 3.35E-02 | -1.08E+00 |
| MGME1 | 1.25E-07 | 6.53E-07 | -1.34E+00 | 2.09E-05 | 9.01E-05 | -1.07E+00 | 1.06E-06 | 4.52E-06 | -1.47E+00 | 2.00E-03 | 7.62E-03 | 1.05E+00 | 4.65E-05 | 1.66E-04 | -1.04E+00 | 1.17E-03 | 3.30E-03 | 1.01E+00 | 6.30E-06 | 2.31E-05 | -1.21E+00 | 1.93E-04 | 5.30E-04 | -1.12E+00 |
| VEZT | 1.25E-07 | 6.53E-07 | -1.23E+00 | 3.36E-07 | 1.45E-06 | -1.08E+00 | 1.56E-03 | 6.64E-03 | -1.11E+00 | 6.30E-04 | 2.41E-03 | -1.02E+00 | 3.31E-03 | 1.18E-02 | -1.03E+00 | 1.33E-04 | 3.74E-04 | -1.14E+00 | 1.27E-03 | 4.67E-03 | -1.13E+00 | 6.14E-05 | 1.69E-04 | -1.16E+00 |
| TFDP1 | 1.25E-07 | 6.54E-07 | -1.32E+00 | 5.45E-08 | 2.36E-07 | -1.19E+00 | 2.76E-02 | 1.17E-01 | 1.07E+00 | 2.94E-04 | 1.12E-03 | 1.07E+00 | 8.27E-05 | 2.95E-04 | -1.42E+00 | 3.05E-03 | 8.61E-03 | -1.10E+00 | 4.11E-05 | 1.51E-04 | -1.16E+00 | 8.50E-02 | 2.33E-01 | 1.10E+00 |
| KCTD6 | 1.26E-07 | 6.55E-07 | -1.81E+00 | 5.17E-06 | 2.23E-05 | -1.29E+00 | 2.10E-10 | 8.91E-10 | -2.26E+00 | 8.83E-09 | 3.37E-08 | -1.75E+00 | 2.24E-06 | 7.99E-06 | -1.92E+00 | 9.46E-06 | 2.67E-05 | -1.65E+00 | 6.27E-07 | 2.30E-06 | -1.69E+00 | 3.16E-04 | 8.68E-04 | -1.26E+00 |
| MYL12BP1 | 1.26E-07 | 6.55E-07 | -1.82E+00 | 6.99E-09 | 3.02E-08 | -2.00E+00 | 7.55E-07 | 3.21E-06 | -1.67E+00 | 1.32E-07 | 5.04E-07 | -1.72E+00 | 6.45E-04 | 2.30E-03 | -1.84E+00 | 1.07E-03 | 3.01E-03 | -1.57E+00 | 2.69E-05 | 9.87E-05 | -1.52E+00 | 8.12E-07 | 2.23E-06 | -1.79E+00 |
| LMO7 | 1.26E-07 | 6.55E-07 | -1.92E+00 | 1.49E-02 | 6.43E-02 | 1.05E+00 | 1.05E-04 | 4.44E-04 | -1.63E+00 | 3.45E-02 | 1.32E-01 | 1.03E+00 | 1.75E-03 | 6.25E-03 | -1.68E+00 | 6.20E-02 | 1.75E-01 | -1.03E+00 | 1.64E-05 | 6.01E-05 | -1.97E+00 | 1.45E-02 | 3.98E-02 | -1.13E+00 |
| CDK9 | 1.26E-07 | 6.55E-07 | -1.27E+00 | 2.38E-04 | 1.03E-03 | 1.10E+00 | 1.90E-05 | 8.05E-05 | -1.25E+00 | 3.22E-06 | 1.23E-05 | -1.15E+00 | 1.29E-02 | 4.60E-02 | 1.12E+00 | 2.36E-03 | 6.66E-03 | 1.02E+00 | 1.41E-03 | 5.17E-03 | -1.07E+00 | 9.63E-03 | 2.65E-02 | 1.01E+00 |
| APOBEC3G | 1.26E-07 | 6.57E-07 | -1.13E+00 | 1.18E-04 | 5.10E-04 | 1.12E+00 | 1.19E-07 | 5.04E-07 | -1.37E+00 | 6.63E-05 | 2.53E-04 | -1.06E+00 | 1.47E-10 | 5.24E-10 | -1.78E+00 | 9.88E-07 | 2.78E-06 | -1.36E+00 | 1.61E-09 | 5.91E-09 | -1.55E+00 | 3.99E-06 | 1.10E-05 | -1.32E+00 |
| ULK4P2 | 1.26E-07 | 6.58E-07 | -1.44E+00 | 1.17E-05 | 5.07E-05 | -1.08E+00 | 1.16E-05 | 4.92E-05 | -1.48E+00 | 5.25E-04 | 2.01E-03 | -1.10E+00 | 6.77E-04 | 2.41E-03 | -1.37E+00 | 1.19E-02 | 3.37E-02 | -1.10E+00 | 3.88E-03 | 1.43E-02 | -1.16E+00 | 8.95E-04 | 2.46E-03 | -1.27E+00 |
| ZSCAN29 | 1.26E-07 | 6.59E-07 | -1.61E+00 | 3.30E-04 | 1.43E-03 | -1.05E+00 | 1.79E-05 | 7.61E-05 | -1.51E+00 | 5.20E-03 | 1.99E-02 | 1.07E+00 | 1.01E-03 | 3.59E-03 | -1.54E+00 | 7.58E-02 | 2.14E-01 | 1.16E+00 | 1.38E-02 | 5.07E-02 | -1.12E+00 | 2.84E-01 | 7.80E-01 | 1.16E+00 |
| AC134050.1 | 1.27E-07 | 6.59E-07 | -2.01E+00 | 2.48E-05 | 1.07E-04 | -1.40E+00 | 2.55E-05 | 1.08E-04 | -1.52E+00 | 8.66E-05 | 3.31E-04 | -1.35E+00 | 1.50E-02 | 5.35E-02 | -1.43E+00 | 1.35E-01 | 3.82E-01 | 1.05E+00 | 1.55E-06 | 5.69E-06 | -1.53E+00 | 2.80E-04 | 7.68E-04 | -1.24E+00 |
| ZNF493 | 1.27E-07 | 6.60E-07 | -1.22E+00 | 6.85E-02 | 2.96E-01 | 1.33E+00 | 1.96E-07 | 8.35E-07 | -1.31E+00 | 1.06E-02 | 4.04E-02 | 1.17E+00 | 3.48E-04 | 1.24E-03 | -1.11E+00 | 2.59E-01 | 7.29E-01 | 1.37E+00 | 1.64E-02 | 6.00E-02 | -1.01E+00 | 3.35E-01 | 9.19E-01 | 1.29E+00 |
| C20orf194 | 1.27E-07 | 6.64E-07 | -1.33E+00 | 3.85E-04 | 1.66E-03 | 1.08E+00 | 5.93E-07 | 2.52E-06 | -1.50E+00 | 2.85E-06 | 1.09E-05 | -1.22E+00 | 5.81E-04 | 2.07E-03 | -1.11E+00 | 4.01E-04 | 1.13E-03 | -1.14E+00 | 1.53E-08 | 5.62E-08 | -1.86E+00 | 6.99E-04 | 1.92E-03 | -1.20E+00 |
| RFX2 | 1.28E-07 | 6.65E-07 | -1.48E+00 | 7.12E-07 | 3.08E-06 | -1.19E+00 | 1.69E-06 | 7.16E-06 | -1.29E+00 | 2.40E-05 | 9.18E-05 | -1.06E+00 | 6.07E-02 | 2.16E-01 | 1.06E+00 | 1.61E-02 | 4.54E-02 | -1.01E+00 | 4.89E-04 | 1.79E-03 | -1.41E+00 | 1.20E-02 | 3.29E-02 | -1.15E+00 |
| PMS2P5 | 1.28E-07 | 6.65E-07 | -2.02E+00 | 4.80E-07 | 2.07E-06 | -1.80E+00 | 1.91E-04 | 8.10E-04 | -1.77E+00 | 5.32E-03 | 2.03E-02 | -1.15E+00 | 2.24E-01 | 8.00E-01 | 1.29E+00 | 1.61E-01 | 4.54E-01 | 1.11E+00 | 2.58E-05 | 9.49E-05 | -1.51E+00 | 2.16E-03 | 5.93E-03 | -1.26E+00 |
| METTL6 | 1.28E-07 | 6.66E-07 | -1.63E+00 | 2.36E-04 | 1.02E-03 | -1.07E+00 | 1.04E-06 | 4.42E-06 | -1.54E+00 | 2.63E-04 | 1.00E-03 | -1.15E+00 | 1.31E-03 | 4.65E-03 | -1.26E+00 | 1.20E-01 | 3.39E-01 | 1.21E+00 | 2.07E-02 | 7.58E-02 | 1.03E+00 | 8.98E-02 | 2.47E-01 | 1.19E+00 |
| EIF3I | 1.28E-07 | 6.66E-07 | -1.14E+00 | 3.36E-11 | 1.45E-10 | -1.13E+00 | 9.81E-04 | 4.17E-03 | -1.19E+00 | 3.05E-05 | 1.17E-04 | -1.12E+00 | 5.75E-06 | 2.05E-05 | -1.19E+00 | 8.70E-03 | 2.45E-02 | -1.04E+00 | 1.37E-02 | 5.03E-02 | -1.05E+00 | 3.35E-01 | 9.20E-01 | 1.08E+00 |
| EXOSC5 | 1.28E-07 | 6.67E-07 | -1.26E+00 | 1.84E-05 | 7.97E-05 | -1.01E+00 | 1.28E-02 | 5.42E-02 | -1.01E+00 | 2.66E-02 | 1.01E-01 | 1.18E+00 | 7.69E-07 | 2.74E-06 | -1.43E+00 | 6.65E-02 | 1.87E-01 | 1.16E+00 | 4.12E-09 | 1.51E-08 | -1.38E+00 | 9.44E-02 | 2.59E-01 | 1.09E+00 |
| PSRC1 | 1.28E-07 | 6.67E-07 | -2.10E+00 | 7.33E-08 | 3.17E-07 | -2.08E+00 | 3.19E-05 | 1.36E-04 | -1.62E+00 | 9.59E-04 | 3.66E-03 | -1.17E+00 | 1.26E-01 | 4.50E-01 | -1.29E+00 | 2.29E-01 | 6.46E-01 | -1.07E+00 | 1.15E-01 | 4.23E-01 | -1.44E+00 | 3.63E-01 | 9.97E-01 | 1.10E+00 |
| ZNF542P | 1.28E-07 | 6.68E-07 | -1.96E+00 | 1.56E-05 | 6.73E-05 | -1.47E+00 | 1.04E-10 | 4.42E-10 | -2.29E+00 | 1.06E-06 | 4.06E-06 | -1.40E+00 | 3.81E-06 | 1.36E-05 | -1.93E+00 | 5.18E-05 | 1.46E-04 | -1.69E+00 | 9.48E-03 | 3.48E-02 | 1.11E+00 | 2.69E-01 | 7.39E-01 | 1.51E+00 |
| IP6K2 | 1.29E-07 | 6.72E-07 | -1.12E+00 | 2.43E-15 | 1.05E-14 | -1.22E+00 | 5.03E-07 | 2.14E-06 | -1.30E+00 | 1.85E-10 | 7.08E-10 | -1.32E+00 | 1.90E-04 | 6.78E-04 | -1.18E+00 | 1.68E-08 | 4.74E-08 | -1.42E+00 | 2.12E-09 | 7.78E-09 | -1.54E+00 | 1.69E-10 | 4.64E-10 | -1.61E+00 |
| DNAJC3-AS1 | 1.30E-07 | 6.78E-07 | -1.73E+00 | 7.60E-04 | 3.28E-03 | -1.10E+00 | 1.42E-05 | 6.04E-05 | -1.80E+00 | 1.74E-03 | 6.63E-03 | -1.21E+00 | 6.86E-04 | 2.45E-03 | -1.57E+00 | 1.70E-02 | 4.81E-02 | -1.13E+00 | 2.82E-03 | 1.03E-02 | -1.19E+00 | 2.78E-01 | 7.65E-01 | 1.20E+00 |
| ATP10D | 1.30E-07 | 6.78E-07 | -1.89E+00 | 6.52E-04 | 2.82E-03 | -1.09E+00 | 1.06E-02 | 4.52E-02 | -1.31E+00 | 1.46E-01 | 5.57E-01 | 1.17E+00 | 1.83E-02 | 6.54E-02 | 1.02E+00 | 4.84E-02 | 1.37E-01 | 1.09E+00 | 1.28E-02 | 4.69E-02 | -1.43E+00 | 1.11E-01 | 3.05E-01 | -1.11E+00 |
| CHST10 | 1.30E-07 | 6.79E-07 | -1.95E+00 | 1.93E-03 | 8.33E-03 | -1.02E+00 | 4.95E-05 | 2.10E-04 | -1.55E+00 | 1.01E-03 | 3.86E-03 | -1.20E+00 | 1.45E-04 | 5.17E-04 | -1.73E+00 | 2.05E-03 | 5.77E-03 | -1.27E+00 | 2.28E-07 | 8.35E-07 | -1.56E+00 | 5.53E-04 | 1.52E-03 | -1.15E+00 |
| TIMM17A | 1.30E-07 | 6.79E-07 | -1.30E+00 | 2.50E-08 | 1.08E-07 | -1.17E+00 | 1.83E-02 | 7.77E-02 | 1.02E+00 | 2.44E-03 | 9.34E-03 | 1.06E+00 | 6.23E-09 | 2.22E-08 | -1.53E+00 | 2.56E-06 | 7.22E-06 | -1.20E+00 | 1.84E-05 | 6.76E-05 | -1.25E+00 | 4.24E-03 | 1.17E-02 | -1.03E+00 |
| SVOPL | 1.31E-07 | 6.83E-07 | -1.69E+00 | 6.56E-07 | 2.84E-06 | -1.48E+00 | 1.67E-06 | 7.11E-06 | -1.60E+00 | 5.06E-06 | 1.93E-05 | -1.30E+00 | 1.37E-10 | 4.87E-10 | -1.32E+00 | 1.36E-05 | 3.83E-05 | -1.09E+00 | 5.80E-03 | 2.13E-02 | 1.01E+00 | 1.24E-02 | 3.40E-02 | -1.01E+00 |
| EFNB1 | 1.34E-07 | 6.96E-07 | -1.83E+00 | 2.63E-05 | 1.14E-04 | -1.34E+00 | 9.48E-07 | 4.03E-06 | -1.85E+00 | 1.14E-04 | 4.34E-04 | -1.41E+00 | 1.44E-01 | 5.15E-01 | -1.40E+00 | 1.63E-01 | 4.60E-01 | -1.58E+00 | 1.19E-01 | 4.38E-01 | 1.93E+00 | 2.41E-01 | 6.61E-01 | 1.62E+00 |
| GDI2P2 | 1.34E-07 | 6.98E-07 | -1.78E+00 | 6.65E-06 | 2.87E-05 | -1.32E+00 | 3.01E-16 | 1.28E-15 | -2.25E+00 | 2.67E-17 | 1.02E-16 | -2.19E+00 | 7.71E-08 | 2.75E-07 | -2.49E+00 | 1.10E-05 | 3.09E-05 | -1.59E+00 | 1.79E-08 | 6.55E-08 | -1.81E+00 | 1.28E-04 | 3.51E-04 | -1.25E+00 |
| OSTM1 | 1.34E-07 | 7.00E-07 | -2.36E+00 | 4.06E-05 | 1.75E-04 | -1.42E+00 | 7.71E-04 | 3.28E-03 | -1.62E+00 | 3.00E-03 | 1.15E-02 | -1.30E+00 | 6.65E-04 | 2.37E-03 | -1.67E+00 | 5.04E-02 | 1.42E-01 | 1.03E+00 | 2.32E-08 | 8.51E-08 | -1.51E+00 | 4.22E-05 | 1.16E-04 | -1.30E+00 |
| TMCO3 | 1.34E-07 | 7.01E-07 | -1.29E+00 | 2.10E-03 | 9.06E-03 | 1.15E+00 | 2.06E-02 | 8.76E-02 | -1.06E+00 | 3.63E-03 | 1.39E-02 | -1.00E+00 | 1.30E-03 | 4.62E-03 | -1.09E+00 | 5.72E-05 | 1.61E-04 | -1.24E+00 | 1.82E-03 | 6.69E-03 | -1.04E+00 | 2.60E-03 | 7.13E-03 | -1.07E+00 |
| APH1A | 1.35E-07 | 7.02E-07 | -1.09E+00 | 1.46E-06 | 6.31E-06 | 1.12E+00 | 1.90E-05 | 8.08E-05 | -1.13E+00 | 4.73E-04 | 1.81E-03 | 1.05E+00 | 4.03E-07 | 1.44E-06 | -1.21E+00 | 3.98E-04 | 1.12E-03 | 1.01E+00 | 1.63E-02 | 5.99E-02 | 1.02E+00 | 1.01E-01 | 2.78E-01 | 1.11E+00 |
| TMEM205 | 1.35E-07 | 7.02E-07 | -1.11E+00 | 1.20E-08 | 5.18E-08 | -1.03E+00 | 2.12E-01 | 9.01E-01 | 1.03E+00 | 2.06E-02 | 7.86E-02 | 1.01E+00 | 5.90E-04 | 2.10E-03 | -1.09E+00 | 2.25E-02 | 6.34E-02 | 1.14E+00 | 2.60E-01 | 9.53E-01 | 1.22E+00 | 3.11E-01 | 8.54E-01 | 1.33E+00 |
| SURF6 | 1.35E-07 | 7.03E-07 | -1.58E+00 | 5.18E-04 | 2.24E-03 | -1.02E+00 | 1.51E-03 | 6.41E-03 | -1.12E+00 | 4.20E-02 | 1.60E-01 | 1.24E+00 | 1.55E-04 | 5.51E-04 | -1.16E+00 | 1.72E-02 | 4.85E-02 | 1.11E+00 | 1.83E-10 | 6.71E-10 | -1.50E+00 | 7.41E-03 | 2.04E-02 | 1.03E+00 |
| GADD45GIP1 | 1.35E-07 | 7.05E-07 | 1.00E+00 | 2.03E-04 | 8.79E-04 | 1.12E+00 | 1.65E-01 | 6.99E-01 | 1.09E+00 | 1.84E-01 | 7.03E-01 | 1.14E+00 | 1.26E-04 | 4.49E-04 | -1.10E+00 | 9.92E-02 | 2.80E-01 | 1.11E+00 | 9.71E-03 | 3.57E-02 | -1.02E+00 | 3.30E-01 | 9.08E-01 | 1.17E+00 |
| PAAF1 | 1.36E-07 | 7.06E-07 | -1.40E+00 | 2.65E-02 | 1.15E-01 | 1.24E+00 | 1.10E-04 | 4.66E-04 | -1.20E+00 | 7.97E-02 | 3.04E-01 | 1.34E+00 | 1.94E-05 | 6.92E-05 | -1.28E+00 | 1.38E-01 | 3.89E-01 | 1.35E+00 | 1.32E-03 | 4.86E-03 | -1.09E+00 | 3.18E-01 | 8.74E-01 | 1.21E+00 |
| AKAP1 | 1.36E-07 | 7.09E-07 | -1.93E+00 | 2.49E-02 | 1.07E-01 | 1.14E+00 | 5.57E-07 | 2.37E-06 | -1.56E+00 | 1.27E-01 | 4.83E-01 | 1.27E+00 | 1.17E-02 | 4.16E-02 | -1.54E+00 | 2.94E-01 | 8.29E-01 | 1.53E+00 | 1.74E-07 | 6.40E-07 | -1.46E+00 | 2.47E-01 | 6.79E-01 | 1.13E+00 |
| C22orf15 | 1.36E-07 | 7.10E-07 | -2.05E+00 | 1.39E-04 | 6.02E-04 | -1.29E+00 | 4.95E-03 | 2.10E-02 | -1.38E+00 | 1.95E-02 | 7.45E-02 | -1.10E+00 | 2.72E-01 | 9.69E-01 | 1.24E+00 | 3.10E-01 | 8.74E-01 | 1.34E+00 | 2.57E-01 | 9.42E-01 | 1.28E+00 | 1.60E-01 | 4.39E-01 | 1.44E+00 |
| PDE6B | 1.36E-07 | 7.10E-07 | -1.40E+00 | 4.97E-04 | 2.15E-03 | 1.03E+00 | 3.76E-08 | 1.60E-07 | -1.72E+00 | 1.27E-05 | 4.86E-05 | -1.21E+00 | 1.06E-05 | 3.78E-05 | -1.47E+00 | 3.92E-03 | 1.11E-02 | 1.00E+00 | 3.66E-03 | 1.34E-02 | -1.05E+00 | 1.63E-02 | 4.47E-02 | 1.03E+00 |
| PTPN22 | 1.36E-07 | 7.10E-07 | -1.33E+00 | 1.29E-09 | 5.58E-09 | -1.43E+00 | 5.52E-05 | 2.34E-04 | -1.46E+00 | 5.48E-04 | 2.09E-03 | -1.21E+00 | 3.24E-04 | 1.15E-03 | -1.11E+00 | 2.13E-04 | 6.00E-04 | -1.19E+00 | 1.61E-02 | 5.90E-02 | 1.04E+00 | 4.10E-02 | 1.13E-01 | 1.02E+00 |
| DPM3 | 1.37E-07 | 7.13E-07 | -1.29E+00 | 8.73E-14 | 3.77E-13 | -1.44E+00 | 7.46E-05 | 3.17E-04 | -1.14E+00 | 2.43E-17 | 9.29E-17 | -1.59E+00 | 3.78E-04 | 1.35E-03 | -1.11E+00 | 4.33E-06 | 1.22E-05 | -1.27E+00 | 7.76E-05 | 2.85E-04 | -1.12E+00 | 6.24E-10 | 1.71E-09 | -1.44E+00 |
| HDAC5 | 1.38E-07 | 7.20E-07 | -1.18E+00 | 3.82E-09 | 1.65E-08 | -1.18E+00 | 6.02E-05 | 2.56E-04 | -1.28E+00 | 4.98E-07 | 1.90E-06 | -1.35E+00 | 3.62E-02 | 1.29E-01 | 1.07E+00 | 1.86E-02 | 5.24E-02 | 1.05E+00 | 1.37E-01 | 5.01E-01 | 1.36E+00 | 9.55E-04 | 2.62E-03 | -1.10E+00 |
| NT5M | 1.38E-07 | 7.20E-07 | -1.06E+00 | 2.32E-10 | 1.00E-09 | -1.04E+00 | 5.77E-03 | 2.45E-02 | -1.03E+00 | 2.17E-02 | 8.30E-02 | 1.10E+00 | 5.43E-03 | 1.93E-02 | 1.07E+00 | 3.70E-02 | 1.04E-01 | 1.15E+00 | 2.55E-01 | 9.35E-01 | 1.33E+00 | 7.94E-02 | 2.18E-01 | 1.24E+00 |
| SECISBP2 | 1.38E-07 | 7.21E-07 | -1.13E+00 | 3.98E-06 | 1.72E-05 | -1.00E+00 | 3.59E-05 | 1.52E-04 | -1.13E+00 | 5.71E-04 | 2.18E-03 | 1.08E+00 | 3.88E-06 | 1.38E-05 | -1.16E+00 | 2.85E-04 | 8.04E-04 | -1.04E+00 | 1.16E-05 | 4.24E-05 | -1.24E+00 | 1.91E-02 | 5.26E-02 | -1.02E+00 |
| B4GALT7 | 1.38E-07 | 7.21E-07 | -1.55E+00 | 1.24E-04 | 5.38E-04 | -1.13E+00 | 2.06E-03 | 8.76E-03 | -1.11E+00 | 1.28E-02 | 4.87E-02 | 1.13E+00 | 4.58E-03 | 1.63E-02 | 1.03E+00 | 2.62E-02 | 7.38E-02 | 1.15E+00 | 3.10E-02 | 1.14E-01 | 1.10E+00 | 2.96E-02 | 8.13E-02 | 1.11E+00 |
| ZNF330 | 1.39E-07 | 7.24E-07 | -1.25E+00 | 3.26E-07 | 1.41E-06 | -1.12E+00 | 1.35E-04 | 5.73E-04 | -1.26E+00 | 8.49E-04 | 3.24E-03 | -1.05E+00 | 1.05E-05 | 3.75E-05 | -1.30E+00 | 1.50E-03 | 4.23E-03 | -1.06E+00 | 3.93E-09 | 1.44E-08 | -1.40E+00 | 3.07E-06 | 8.44E-06 | -1.28E+00 |
| LINC00891 | 1.39E-07 | 7.24E-07 | 2.97E+00 | 2.25E-15 | 9.72E-15 | 4.99E+00 | 8.15E-02 | 3.46E-01 | 1.51E+00 | 5.06E-03 | 1.93E-02 | 2.96E+00 | 2.21E-02 | 7.87E-02 | 4.51E+00 | 1.56E-05 | 4.39E-05 | 9.81E+00 | 1.27E-01 | 4.68E-01 | 1.30E+00 | 2.12E-01 | 5.82E-01 | 1.36E+00 |
| FAM220A | 1.40E-07 | 7.27E-07 | -1.75E+00 | 5.88E-07 | 2.54E-06 | -1.51E+00 | 6.43E-08 | 2.73E-07 | -2.12E+00 | 1.29E-04 | 4.93E-04 | -1.28E+00 | 5.38E-02 | 1.92E-01 | -1.27E+00 | 2.55E-01 | 7.20E-01 | 1.10E+00 | 1.42E-03 | 5.20E-03 | -1.44E+00 | 1.24E-03 | 3.39E-03 | -1.52E+00 |
| USP32 | 1.40E-07 | 7.28E-07 | -1.63E+00 | 2.17E-03 | 9.40E-03 | 1.02E+00 | 5.32E-03 | 2.26E-02 | -1.09E+00 | 1.19E-03 | 4.54E-03 | -1.02E+00 | 2.07E-09 | 7.40E-09 | -2.02E+00 | 7.84E-05 | 2.21E-04 | -1.20E+00 | 1.25E-05 | 4.59E-05 | -1.34E+00 | 1.42E-02 | 3.89E-02 | -1.04E+00 |
| MAP2K3 | 1.40E-07 | 7.28E-07 | -1.21E+00 | 4.22E-04 | 1.82E-03 | 1.15E+00 | 6.76E-08 | 2.87E-07 | -1.24E+00 | 2.87E-05 | 1.10E-04 | 1.04E+00 | 1.66E-08 | 5.91E-08 | -1.37E+00 | 1.07E-04 | 3.00E-04 | -1.07E+00 | 8.77E-05 | 3.22E-04 | -1.13E+00 | 1.49E-02 | 4.08E-02 | 1.05E+00 |
| PXK | 1.40E-07 | 7.32E-07 | -1.34E+00 | 1.08E-05 | 4.68E-05 | -1.04E+00 | 3.51E-08 | 1.49E-07 | -1.38E+00 | 4.77E-04 | 1.82E-03 | 1.08E+00 | 4.72E-04 | 1.68E-03 | -1.32E+00 | 1.52E-02 | 4.27E-02 | -1.05E+00 | 4.83E-07 | 1.77E-06 | -1.30E+00 | 3.00E-02 | 8.24E-02 | 1.03E+00 |
| AC110994.2 | 1.41E-07 | 7.33E-07 | -1.69E+00 | 1.79E-07 | 7.75E-07 | -1.57E+00 | 2.38E-03 | 1.01E-02 | -1.37E+00 | 8.53E-04 | 3.26E-03 | -1.40E+00 | 9.72E-03 | 3.47E-02 | -1.19E+00 | 6.68E-03 | 1.88E-02 | -1.27E+00 | 2.10E-02 | 7.72E-02 | -1.34E+00 | 9.48E-02 | 2.60E-01 | -1.13E+00 |
| EEF1A1P22 | 1.41E-07 | 7.34E-07 | -1.32E+00 | 2.06E-12 | 8.92E-12 | -1.55E+00 | 2.38E-04 | 1.01E-03 | -1.14E+00 | 1.40E-06 | 5.35E-06 | -1.19E+00 | 7.07E-03 | 2.52E-02 | 1.09E+00 | 1.56E-04 | 4.41E-04 | -1.18E+00 | 1.72E-05 | 6.32E-05 | -1.41E+00 | 1.42E-05 | 3.89E-05 | -1.40E+00 |
| FKBP9 | 1.42E-07 | 7.42E-07 | -4.16E+00 | 8.72E-06 | 3.77E-05 | -2.50E+00 | 1.71E-01 | 7.27E-01 | -1.01E+00 | 1.01E-01 | 3.86E-01 | -1.14E+00 | 4.71E-03 | 1.68E-02 | -1.30E+00 | 3.27E-02 | 9.22E-02 | -1.08E+00 | 1.42E-06 | 5.21E-06 | -1.50E+00 | 1.22E-07 | 3.36E-07 | -1.60E+00 |
| NME7 | 1.43E-07 | 7.43E-07 | -1.32E+00 | 7.84E-06 | 3.39E-05 | -1.05E+00 | 6.81E-06 | 2.89E-05 | -1.30E+00 | 1.32E-04 | 5.06E-04 | -1.03E+00 | 1.11E-05 | 3.97E-05 | -1.43E+00 | 2.15E-02 | 6.06E-02 | 1.03E+00 | 2.55E-04 | 9.38E-04 | -1.10E+00 | 2.40E-02 | 6.58E-02 | 1.10E+00 |
| MRPL36 | 1.44E-07 | 7.50E-07 | -1.17E+00 | 8.46E-06 | 3.66E-05 | 1.07E+00 | 2.43E-05 | 1.03E-04 | -1.20E+00 | 9.00E-04 | 3.44E-03 | -1.01E+00 | 1.35E-06 | 4.81E-06 | -1.29E+00 | 9.63E-04 | 2.72E-03 | -1.02E+00 | 6.27E-02 | 2.30E-01 | 1.10E+00 | 2.97E-01 | 8.14E-01 | 1.18E+00 |
| CARF | 1.45E-07 | 7.53E-07 | -1.43E+00 | 2.03E-01 | 8.77E-01 | 1.53E+00 | 3.05E-07 | 1.30E-06 | -2.07E+00 | 3.76E-02 | 1.43E-01 | 1.16E+00 | 8.42E-02 | 3.00E-01 | -1.06E+00 | 3.14E-01 | 8.86E-01 | 1.34E+00 | 3.42E-04 | 1.25E-03 | -1.25E+00 | 1.41E-01 | 3.86E-01 | 1.12E+00 |
| CAPN7 | 1.45E-07 | 7.57E-07 | -1.27E+00 | 5.52E-13 | 2.39E-12 | -1.58E+00 | 4.69E-04 | 1.99E-03 | -1.27E+00 | 4.59E-04 | 1.75E-03 | -1.11E+00 | 7.65E-06 | 2.73E-05 | -1.53E+00 | 7.23E-08 | 2.04E-07 | -1.67E+00 | 1.78E-10 | 6.53E-10 | -1.59E+00 | 2.79E-10 | 7.65E-10 | -1.85E+00 |
| AC006453.2 | 1.46E-07 | 7.63E-07 | -1.26E+00 | 9.00E-10 | 3.89E-09 | -1.32E+00 | 3.29E-06 | 1.40E-05 | -1.31E+00 | 4.61E-09 | 1.76E-08 | -1.36E+00 | 6.43E-05 | 2.29E-04 | -1.34E+00 | 8.57E-05 | 2.42E-04 | -1.24E+00 | 5.63E-03 | 2.07E-02 | -1.01E+00 | 6.84E-05 | 1.88E-04 | -1.18E+00 |
| SUCO | 1.47E-07 | 7.66E-07 | -1.36E+00 | 1.65E-11 | 7.13E-11 | -1.41E+00 | 2.12E-02 | 9.02E-02 | 1.12E+00 | 1.02E-05 | 3.89E-05 | -1.07E+00 | 1.85E-04 | 6.59E-04 | -1.18E+00 | 1.07E-05 | 3.01E-05 | -1.29E+00 | 1.08E-10 | 3.98E-10 | -1.66E+00 | 1.26E-09 | 3.47E-09 | -1.63E+00 |
| SEPT1 | 1.48E-07 | 7.71E-07 | -1.33E+00 | 3.25E-07 | 1.40E-06 | -1.14E+00 | 1.63E-08 | 6.92E-08 | -1.56E+00 | 9.53E-10 | 3.64E-09 | -1.52E+00 | 1.54E-03 | 5.48E-03 | -1.38E+00 | 1.06E-03 | 2.97E-03 | -1.43E+00 | 6.87E-02 | 2.52E-01 | -1.09E+00 | 5.18E-02 | 1.42E-01 | -1.15E+00 |
| ATMIN | 1.48E-07 | 7.72E-07 | -1.63E+00 | 1.19E-04 | 5.14E-04 | -1.05E+00 | 1.12E-04 | 4.76E-04 | -1.33E+00 | 6.01E-04 | 2.30E-03 | -1.03E+00 | 1.91E-05 | 6.80E-05 | -1.35E+00 | 1.25E-05 | 3.51E-05 | -1.37E+00 | 1.54E-04 | 5.67E-04 | -1.16E+00 | 2.27E-04 | 6.23E-04 | -1.22E+00 |
| ZNF594 | 1.49E-07 | 7.75E-07 | -1.86E+00 | 4.91E-04 | 2.12E-03 | -1.12E+00 | 1.95E-06 | 8.26E-06 | -2.31E+00 | 9.73E-03 | 3.72E-02 | -1.09E+00 | 5.00E-03 | 1.78E-02 | -1.67E+00 | 1.51E-01 | 4.26E-01 | 1.09E+00 | 1.44E-07 | 5.27E-07 | -2.44E+00 | 7.60E-03 | 2.09E-02 | -1.17E+00 |
| MIF4GD | 1.49E-07 | 7.77E-07 | -1.25E+00 | 6.64E-07 | 2.87E-06 | -1.05E+00 | 5.71E-04 | 2.43E-03 | -1.04E+00 | 3.21E-05 | 1.23E-04 | 1.05E+00 | 1.58E-04 | 5.63E-04 | -1.25E+00 | 3.09E-02 | 8.71E-02 | 1.09E+00 | 2.54E-03 | 9.31E-03 | -1.07E+00 | 5.10E-02 | 1.40E-01 | 1.04E+00 |
| AIFM2 | 1.49E-07 | 7.78E-07 | -2.13E+00 | 5.87E-06 | 2.53E-05 | -1.62E+00 | 2.41E-02 | 1.02E-01 | -1.33E+00 | 3.35E-02 | 1.28E-01 | -1.20E+00 | 2.09E-01 | 7.46E-01 | 1.11E+00 | 1.77E-01 | 5.00E-01 | -1.10E+00 | 1.43E-01 | 5.23E-01 | -1.03E+00 | 5.66E-02 | 1.55E-01 | -1.22E+00 |
| WDR36 | 1.50E-07 | 7.79E-07 | -1.42E+00 | 1.86E-06 | 8.02E-06 | -1.20E+00 | 3.51E-08 | 1.49E-07 | -1.50E+00 | 3.21E-06 | 1.23E-05 | -1.16E+00 | 9.60E-05 | 3.42E-04 | -1.30E+00 | 2.16E-03 | 6.10E-03 | -1.08E+00 | 3.71E-10 | 1.36E-09 | -1.62E+00 | 1.23E-02 | 3.37E-02 | -1.06E+00 |
| LINC00854 | 1.50E-07 | 7.79E-07 | -2.03E+00 | 5.30E-03 | 2.29E-02 | 1.04E+00 | 1.55E-06 | 6.60E-06 | -1.87E+00 | 2.33E-04 | 8.89E-04 | -1.27E+00 | 1.57E-04 | 5.59E-04 | -1.81E+00 | 3.65E-05 | 1.03E-04 | -2.04E+00 | 2.57E-01 | 9.43E-01 | 1.08E+00 | 2.14E-01 | 5.87E-01 | -1.02E+00 |
| KIAA0930 | 1.50E-07 | 7.82E-07 | -1.62E+00 | 3.70E-04 | 1.60E-03 | -1.06E+00 | 6.13E-04 | 2.61E-03 | -1.12E+00 | 4.36E-04 | 1.67E-03 | 1.05E+00 | 5.45E-04 | 1.94E-03 | -1.47E+00 | 7.10E-03 | 2.00E-02 | -1.13E+00 | 1.13E-02 | 4.13E-02 | -1.15E+00 | 3.22E-01 | 8.84E-01 | 1.32E+00 |
| APC | 1.50E-07 | 7.83E-07 | -1.37E+00 | 7.15E-05 | 3.09E-04 | -1.02E+00 | 3.77E-08 | 1.60E-07 | -1.47E+00 | 1.95E-06 | 7.47E-06 | -1.13E+00 | 1.17E-05 | 4.18E-05 | -1.28E+00 | 3.54E-05 | 9.97E-05 | -1.15E+00 | 3.24E-06 | 1.19E-05 | -1.18E+00 | 8.22E-06 | 2.26E-05 | -1.22E+00 |
| UQCC2 | 1.52E-07 | 7.92E-07 | -1.21E+00 | 5.18E-13 | 2.24E-12 | -1.28E+00 | 4.80E-02 | 2.04E-01 | 1.08E+00 | 6.67E-06 | 2.55E-05 | -1.13E+00 | 8.97E-09 | 3.20E-08 | -1.39E+00 | 2.13E-08 | 6.00E-08 | -1.29E+00 | 1.53E-04 | 5.63E-04 | -1.19E+00 | 6.84E-02 | 1.88E-01 | -1.06E+00 |
| AL356275.1 | 1.52E-07 | 7.93E-07 | -1.57E+00 | 1.21E-02 | 5.24E-02 | 1.18E+00 | 1.89E-02 | 8.01E-02 | -1.22E+00 | 2.43E-01 | 9.29E-01 | 1.42E+00 | 1.08E-01 | 3.84E-01 | -1.07E+00 | 2.81E-01 | 7.93E-01 | 1.63E+00 | 1.42E-01 | 5.21E-01 | 1.09E+00 | 3.35E-01 | 9.21E-01 | 1.30E+00 |
| RFX5 | 1.52E-07 | 7.94E-07 | -1.48E+00 | 1.73E-08 | 7.46E-08 | -1.46E+00 | 2.73E-05 | 1.16E-04 | -1.42E+00 | 2.72E-05 | 1.04E-04 | -1.29E+00 | 2.69E-04 | 9.59E-04 | -1.29E+00 | 6.96E-03 | 1.96E-02 | -1.02E+00 | 2.75E-07 | 1.01E-06 | -1.52E+00 | 9.50E-08 | 2.61E-07 | -1.55E+00 |
| RASGEF1A | 1.52E-07 | 7.94E-07 | -1.56E+00 | 2.65E-06 | 1.14E-05 | -1.25E+00 | 2.61E-04 | 1.11E-03 | -1.39E+00 | 3.01E-03 | 1.15E-02 | -1.10E+00 | 2.24E-01 | 7.98E-01 | 1.40E+00 | 3.34E-01 | 9.42E-01 | 1.13E+00 | 9.20E-02 | 3.38E-01 | -1.26E+00 | 1.73E-01 | 4.74E-01 | -1.01E+00 |
| GTF2E2 | 1.53E-07 | 7.96E-07 | -1.61E+00 | 2.82E-04 | 1.22E-03 | -1.01E+00 | 2.41E-16 | 1.03E-15 | -2.15E+00 | 2.60E-13 | 9.93E-13 | -1.63E+00 | 1.35E-05 | 4.81E-05 | -1.64E+00 | 6.27E-05 | 1.77E-04 | -1.43E+00 | 3.22E-16 | 1.18E-15 | -1.74E+00 | 6.01E-06 | 1.65E-05 | -1.12E+00 |
| AC007993.3 | 1.55E-07 | 8.06E-07 | -1.33E+00 | 8.80E-07 | 3.80E-06 | -1.15E+00 | 3.85E-03 | 1.63E-02 | -1.04E+00 | 2.86E-05 | 1.09E-04 | -1.04E+00 | 1.59E-04 | 5.66E-04 | -1.70E+00 | 7.89E-03 | 2.22E-02 | -1.26E+00 | 2.07E-02 | 7.58E-02 | -1.00E+00 | 1.30E-02 | 3.57E-02 | -1.06E+00 |
| ATXN2L | 1.55E-07 | 8.09E-07 | -1.17E+00 | 5.64E-12 | 2.44E-11 | -1.29E+00 | 5.82E-11 | 2.47E-10 | -1.43E+00 | 8.88E-13 | 3.39E-12 | -1.37E+00 | 2.50E-06 | 8.92E-06 | -1.33E+00 | 2.46E-06 | 6.92E-06 | -1.42E+00 | 1.56E-11 | 5.74E-11 | -1.59E+00 | 4.35E-11 | 1.19E-10 | -1.55E+00 |
| CHM | 1.56E-07 | 8.14E-07 | -1.41E+00 | 2.45E-05 | 1.06E-04 | -1.02E+00 | 1.30E-10 | 5.54E-10 | -1.74E+00 | 4.32E-07 | 1.65E-06 | -1.27E+00 | 4.22E-04 | 1.50E-03 | -1.52E+00 | 4.29E-02 | 1.21E-01 | 1.03E+00 | 2.00E-12 | 7.33E-12 | -1.98E+00 | 1.90E-08 | 5.21E-08 | -1.61E+00 |
| ZNF791 | 1.56E-07 | 8.16E-07 | -1.30E+00 | 5.64E-11 | 2.44E-10 | -1.43E+00 | 4.66E-05 | 1.98E-04 | -1.16E+00 | 8.22E-08 | 3.14E-07 | -1.26E+00 | 2.30E-02 | 8.20E-02 | 1.08E+00 | 1.30E-04 | 3.67E-04 | -1.20E+00 | 3.91E-05 | 1.44E-04 | -1.17E+00 | 2.96E-05 | 8.12E-05 | -1.25E+00 |
| POLR2A | 1.57E-07 | 8.17E-07 | -1.18E+00 | 4.07E-08 | 1.76E-07 | -1.10E+00 | 1.16E-04 | 4.92E-04 | -1.20E+00 | 5.00E-05 | 1.91E-04 | -1.11E+00 | 8.21E-04 | 2.93E-03 | -1.04E+00 | 1.52E-04 | 4.27E-04 | -1.10E+00 | 2.17E-04 | 7.96E-04 | -1.06E+00 | 3.87E-04 | 1.06E-03 | -1.07E+00 |
| USP51 | 1.58E-07 | 8.22E-07 | -1.53E+00 | 3.34E-05 | 1.44E-04 | -1.21E+00 | 1.11E-05 | 4.72E-05 | -1.48E+00 | 3.00E-03 | 1.15E-02 | -1.01E+00 | 2.04E-03 | 7.26E-03 | -1.33E+00 | 3.37E-03 | 9.51E-03 | -1.23E+00 | 1.41E-01 | 5.16E-01 | -1.01E+00 | 2.43E-01 | 6.66E-01 | 1.06E+00 |
| TERF2 | 1.59E-07 | 8.29E-07 | -1.28E+00 | 1.66E-06 | 7.17E-06 | -1.06E+00 | 1.50E-05 | 6.39E-05 | -1.20E+00 | 2.26E-03 | 8.63E-03 | 1.11E+00 | 1.13E-04 | 4.02E-04 | -1.15E+00 | 1.81E-02 | 5.11E-02 | 1.14E+00 | 1.00E-06 | 3.69E-06 | -1.39E+00 | 5.07E-06 | 1.39E-05 | -1.27E+00 |
| PLEKHF1 | 1.59E-07 | 8.30E-07 | -2.41E+00 | 1.65E-04 | 7.11E-04 | -1.31E+00 | 8.98E-04 | 3.81E-03 | -1.37E+00 | 6.08E-02 | 2.32E-01 | -1.02E+00 | 2.38E-02 | 8.47E-02 | -1.28E+00 | 1.23E-02 | 3.47E-02 | -1.61E+00 | 2.06E-02 | 7.55E-02 | -1.51E+00 | 5.01E-02 | 1.38E-01 | -1.44E+00 |
| AK6 | 1.60E-07 | 8.35E-07 | -1.14E+00 | 5.11E-05 | 2.21E-04 | 1.09E+00 | 1.82E-03 | 7.74E-03 | -1.09E+00 | 2.28E-02 | 8.70E-02 | 1.11E+00 | 8.95E-11 | 3.19E-10 | -1.46E+00 | 2.04E-03 | 5.74E-03 | 1.06E+00 | 2.73E-04 | 1.00E-03 | -1.09E+00 | 1.42E-01 | 3.91E-01 | 1.15E+00 |
| FLAD1 | 1.61E-07 | 8.37E-07 | -1.42E+00 | 4.63E-05 | 2.00E-04 | -1.07E+00 | 1.84E-04 | 7.83E-04 | -1.32E+00 | 8.10E-03 | 3.10E-02 | 1.07E+00 | 1.34E-03 | 4.76E-03 | -1.20E+00 | 1.28E-02 | 3.61E-02 | -1.00E+00 | 7.24E-04 | 2.66E-03 | -1.08E+00 | 8.09E-02 | 2.22E-01 | 1.10E+00 |
| CLIP3 | 1.61E-07 | 8.39E-07 | -1.44E+00 | 4.37E-06 | 1.89E-05 | -1.32E+00 | 1.66E-01 | 7.06E-01 | 1.09E+00 | 2.01E-01 | 7.68E-01 | 1.19E+00 | 1.94E-08 | 6.93E-08 | -2.16E+00 | 2.12E-06 | 5.99E-06 | -1.66E+00 | 4.53E-02 | 1.66E-01 | -1.48E+00 | 6.47E-02 | 1.78E-01 | -1.55E+00 |
| OMA1 | 1.61E-07 | 8.41E-07 | -1.33E+00 | 1.34E-02 | 5.81E-02 | 1.21E+00 | 7.23E-18 | 3.07E-17 | -1.90E+00 | 3.14E-13 | 1.20E-12 | -1.33E+00 | 3.11E-11 | 1.11E-10 | -1.92E+00 | 1.44E-04 | 4.06E-04 | -1.17E+00 | 9.96E-14 | 3.65E-13 | -1.91E+00 | 1.57E-04 | 4.32E-04 | -1.14E+00 |
| HINT3 | 1.62E-07 | 8.44E-07 | -1.24E+00 | 1.42E-06 | 6.15E-06 | -1.10E+00 | 2.83E-05 | 1.20E-04 | -1.32E+00 | 3.65E-05 | 1.40E-04 | -1.18E+00 | 3.48E-04 | 1.24E-03 | -1.10E+00 | 9.50E-04 | 2.68E-03 | -1.09E+00 | 5.20E-08 | 1.91E-07 | -1.62E+00 | 8.87E-06 | 2.44E-05 | -1.37E+00 |
| CCP110 | 1.63E-07 | 8.48E-07 | -1.36E+00 | 3.50E-04 | 1.51E-03 | 1.03E+00 | 1.76E-07 | 7.49E-07 | -1.50E+00 | 6.42E-05 | 2.45E-04 | -1.16E+00 | 4.59E-09 | 1.63E-08 | -1.86E+00 | 5.71E-05 | 1.61E-04 | -1.26E+00 | 9.85E-04 | 3.62E-03 | -1.06E+00 | 4.09E-02 | 1.12E-01 | 1.13E+00 |
| GTF2F1 | 1.63E-07 | 8.49E-07 | -1.12E+00 | 2.57E-07 | 1.11E-06 | -1.03E+00 | 5.00E-05 | 2.12E-04 | -1.18E+00 | 3.09E-08 | 1.18E-07 | -1.22E+00 | 6.09E-05 | 2.17E-04 | -1.06E+00 | 4.09E-03 | 1.15E-02 | 1.04E+00 | 4.32E-03 | 1.59E-02 | -1.03E+00 | 1.45E-03 | 3.98E-03 | -1.09E+00 |
| TMEM115 | 1.63E-07 | 8.49E-07 | -1.58E+00 | 2.47E-03 | 1.07E-02 | 1.06E+00 | 3.68E-09 | 1.56E-08 | -1.67E+00 | 2.41E-04 | 9.21E-04 | -1.06E+00 | 2.23E-07 | 7.96E-07 | -1.97E+00 | 9.92E-03 | 2.80E-02 | -1.07E+00 | 8.50E-10 | 3.12E-09 | -1.48E+00 | 1.81E-05 | 4.98E-05 | -1.14E+00 |
| ABHD12B | 1.64E-07 | 8.56E-07 | -2.07E+00 | 8.51E-05 | 3.68E-04 | -1.29E+00 | 9.18E-23 | 3.90E-22 | -2.48E+00 | 3.77E-21 | 1.44E-20 | -2.12E+00 | 6.02E-03 | 2.15E-02 | -2.08E+00 | 3.52E-02 | 9.91E-02 | -1.36E+00 | 1.71E-01 | 6.26E-01 | -1.28E+00 | 2.60E-01 | 7.16E-01 | -1.17E+00 |
| TMEM250 | 1.65E-07 | 8.59E-07 | -1.99E+00 | 1.48E-04 | 6.41E-04 | -1.21E+00 | 8.09E-09 | 3.44E-08 | -1.71E+00 | 4.54E-06 | 1.73E-05 | -1.18E+00 | 5.69E-10 | 2.03E-09 | -2.02E+00 | 2.21E-05 | 6.24E-05 | -1.20E+00 | 2.62E-09 | 9.63E-09 | -1.32E+00 | 2.24E-03 | 6.14E-03 | -1.01E+00 |
| SH3GLB1 | 1.65E-07 | 8.60E-07 | -1.09E+00 | 2.20E-08 | 9.52E-08 | -1.07E+00 | 8.70E-08 | 3.69E-07 | -1.38E+00 | 5.58E-07 | 2.13E-06 | -1.21E+00 | 6.34E-07 | 2.26E-06 | -1.26E+00 | 5.27E-06 | 1.48E-05 | -1.16E+00 | 1.30E-06 | 4.76E-06 | -1.23E+00 | 1.76E-06 | 4.82E-06 | -1.22E+00 |
| Z82217.1 | 1.65E-07 | 8.60E-07 | -1.67E+00 | 1.05E-07 | 4.55E-07 | -1.62E+00 | 3.18E-05 | 1.35E-04 | -1.86E+00 | 1.36E-03 | 5.18E-03 | -1.26E+00 | 4.91E-02 | 1.75E-01 | -2.10E+00 | 1.90E-01 | 5.35E-01 | -1.48E+00 | 1.09E-02 | 4.00E-02 | -1.54E+00 | 4.99E-02 | 1.37E-01 | -1.14E+00 |
| LINC00847 | 1.65E-07 | 8.62E-07 | -1.57E+00 | 3.00E-03 | 1.30E-02 | 1.05E+00 | 6.73E-09 | 2.86E-08 | -2.08E+00 | 3.47E-05 | 1.33E-04 | -1.31E+00 | 3.18E-07 | 1.13E-06 | -1.97E+00 | 3.19E-02 | 8.98E-02 | 1.07E+00 | 7.67E-10 | 2.82E-09 | -1.89E+00 | 6.07E-04 | 1.67E-03 | -1.14E+00 |
| AC090617.3 | 1.66E-07 | 8.65E-07 | -1.62E+00 | 1.33E-08 | 5.74E-08 | -1.66E+00 | 1.21E-02 | 5.14E-02 | -1.12E+00 | 5.19E-03 | 1.98E-02 | -1.02E+00 | 4.34E-05 | 1.55E-04 | -1.49E+00 | 2.65E-05 | 7.47E-05 | -1.45E+00 | 4.98E-20 | 1.83E-19 | -1.83E+00 | 7.47E-13 | 2.05E-12 | -1.47E+00 |
| DNAJC30 | 1.66E-07 | 8.65E-07 | -1.52E+00 | 4.42E-08 | 1.91E-07 | -1.47E+00 | 7.03E-05 | 2.99E-04 | -1.36E+00 | 2.37E-02 | 9.03E-02 | 1.13E+00 | 1.84E-03 | 6.54E-03 | -1.30E+00 | 5.60E-02 | 1.58E-01 | 1.05E+00 | 2.53E-06 | 9.28E-06 | -1.24E+00 | 4.64E-02 | 1.28E-01 | 1.14E+00 |
| TRMT61A | 1.66E-07 | 8.67E-07 | -1.74E+00 | 3.48E-03 | 1.50E-02 | 1.03E+00 | 3.03E-04 | 1.29E-03 | -1.14E+00 | 7.58E-03 | 2.89E-02 | 1.18E+00 | 1.19E-04 | 4.24E-04 | -1.17E+00 | 1.03E-01 | 2.91E-01 | 1.30E+00 | 3.85E-04 | 1.41E-03 | -1.16E+00 | 3.00E-01 | 8.25E-01 | 1.28E+00 |
| KLF11 | 1.67E-07 | 8.69E-07 | -2.26E+00 | 8.90E-09 | 3.84E-08 | -2.71E+00 | 9.07E-02 | 3.85E-01 | -1.10E+00 | 2.21E-02 | 8.43E-02 | -1.27E+00 | 1.66E-03 | 5.92E-03 | -1.36E+00 | 1.21E-04 | 3.42E-04 | -1.80E+00 | 1.26E-02 | 4.63E-02 | -1.05E+00 | 6.09E-04 | 1.67E-03 | -1.26E+00 |
| CRCP | 1.67E-07 | 8.70E-07 | -1.31E+00 | 2.54E-07 | 1.10E-06 | -1.10E+00 | 3.29E-07 | 1.40E-06 | -1.35E+00 | 9.49E-07 | 3.62E-06 | -1.16E+00 | 7.39E-03 | 2.63E-02 | -1.15E+00 | 8.48E-05 | 2.39E-04 | -1.33E+00 | 5.53E-04 | 2.03E-03 | -1.08E+00 | 1.54E-03 | 4.24E-03 | -1.03E+00 |
| EIF3CL | 1.69E-07 | 8.79E-07 | -1.15E+00 | 7.53E-16 | 3.25E-15 | -1.40E+00 | 1.72E-02 | 7.31E-02 | -1.02E+00 | 1.73E-06 | 6.63E-06 | -1.12E+00 | 5.23E-04 | 1.86E-03 | -1.03E+00 | 3.32E-06 | 9.35E-06 | -1.18E+00 | 7.80E-02 | 2.86E-01 | 1.07E+00 | 6.01E-05 | 1.65E-04 | -1.19E+00 |
| AC019205.2 | 1.69E-07 | 8.79E-07 | -1.70E+00 | 3.43E-05 | 1.48E-04 | -1.20E+00 | 1.56E-03 | 6.64E-03 | -1.32E+00 | 7.96E-02 | 3.04E-01 | 1.22E+00 | 8.16E-02 | 2.91E-01 | 1.07E+00 | 3.47E-01 | 9.77E-01 | 1.44E+00 | 7.64E-03 | 2.80E-02 | -1.13E+00 | 7.25E-02 | 1.99E-01 | 1.09E+00 |
| MRGBP | 1.69E-07 | 8.81E-07 | -1.74E+00 | 3.95E-04 | 1.71E-03 | -1.09E+00 | 1.55E-06 | 6.58E-06 | -1.51E+00 | 8.75E-05 | 3.34E-04 | -1.16E+00 | 6.08E-06 | 2.17E-05 | -1.71E+00 | 1.87E-03 | 5.27E-03 | -1.14E+00 | 8.91E-09 | 3.27E-08 | -1.38E+00 | 5.71E-06 | 1.57E-05 | -1.26E+00 |
| DENND2D | 1.69E-07 | 8.82E-07 | -1.36E+00 | 9.24E-06 | 3.99E-05 | -1.11E+00 | 4.17E-06 | 1.77E-05 | -1.66E+00 | 6.25E-03 | 2.39E-02 | -1.08E+00 | 9.42E-06 | 3.36E-05 | -1.67E+00 | 7.79E-04 | 2.19E-03 | -1.22E+00 | 8.05E-02 | 2.95E-01 | 1.22E+00 | 2.38E-01 | 6.55E-01 | 1.30E+00 |
| WARS | 1.69E-07 | 8.83E-07 | -2.13E+00 | 1.55E-05 | 6.69E-05 | -1.48E+00 | 2.15E-08 | 9.16E-08 | -1.59E+00 | 2.43E-08 | 9.28E-08 | -1.39E+00 | 6.21E-05 | 2.21E-04 | -1.67E+00 | 8.98E-04 | 2.53E-03 | -1.33E+00 | 3.73E-06 | 1.37E-05 | -1.42E+00 | 1.21E-04 | 3.33E-04 | -1.29E+00 |
| USB1 | 1.69E-07 | 8.83E-07 | -1.55E+00 | 2.18E-05 | 9.42E-05 | -1.16E+00 | 6.66E-03 | 2.83E-02 | -1.06E+00 | 2.97E-02 | 1.14E-01 | 1.18E+00 | 1.57E-03 | 5.60E-03 | -1.05E+00 | 2.01E-02 | 5.68E-02 | 1.13E+00 | 9.12E-06 | 3.35E-05 | -1.16E+00 | 1.49E-02 | 4.08E-02 | 1.12E+00 |
| SNHG17 | 1.72E-07 | 8.98E-07 | -1.28E+00 | 3.14E-06 | 1.35E-05 | -1.04E+00 | 1.80E-05 | 7.66E-05 | -1.19E+00 | 6.00E-05 | 2.29E-04 | -1.06E+00 | 3.23E-03 | 1.15E-02 | -1.01E+00 | 4.33E-02 | 1.22E-01 | 1.16E+00 | 2.29E-01 | 8.42E-01 | 1.31E+00 | 2.10E-01 | 5.78E-01 | 1.20E+00 |
| TTC5 | 1.74E-07 | 9.06E-07 | -1.28E+00 | 2.74E-03 | 1.18E-02 | 1.15E+00 | 1.25E-07 | 5.32E-07 | -1.46E+00 | 7.43E-03 | 2.84E-02 | 1.12E+00 | 4.06E-04 | 1.45E-03 | -1.09E+00 | 4.38E-02 | 1.23E-01 | 1.17E+00 | 1.02E-05 | 3.75E-05 | -1.20E+00 | 7.12E-04 | 1.95E-03 | -1.08E+00 |
| RUBCN | 1.76E-07 | 9.19E-07 | -1.39E+00 | 7.17E-05 | 3.10E-04 | -1.01E+00 | 7.98E-02 | 3.39E-01 | 1.06E+00 | 1.16E-01 | 4.42E-01 | 1.23E+00 | 1.48E-02 | 5.27E-02 | -1.25E+00 | 3.77E-02 | 1.06E-01 | -1.08E+00 | 2.10E-02 | 7.71E-02 | 1.04E+00 | 1.23E-02 | 3.37E-02 | -1.05E+00 |
| BEX1 | 1.77E-07 | 9.20E-07 | -1.12E+00 | 7.52E-07 | 3.25E-06 | -1.01E+00 | 1.07E-01 | 4.55E-01 | 1.33E+00 | 9.06E-02 | 3.46E-01 | 1.39E+00 | 2.85E-07 | 1.02E-06 | -1.04E+00 | 6.84E-04 | 1.93E-03 | 1.18E+00 | 1.39E-02 | 5.12E-02 | 1.08E+00 | 1.31E-01 | 3.59E-01 | 1.54E+00 |
| L3MBTL3 | 1.78E-07 | 9.27E-07 | -1.48E+00 | 1.13E-05 | 4.89E-05 | -1.14E+00 | 1.72E-02 | 7.30E-02 | -1.00E+00 | 4.22E-02 | 1.61E-01 | 1.23E+00 | 6.96E-04 | 2.48E-03 | -1.54E+00 | 1.13E-02 | 3.17E-02 | -1.20E+00 | 1.61E-04 | 5.90E-04 | -1.24E+00 | 5.77E-03 | 1.58E-02 | -1.16E+00 |
| ALDOA | 1.79E-07 | 9.34E-07 | -1.18E+00 | 1.52E-06 | 6.57E-06 | -1.08E+00 | 1.95E-01 | 8.29E-01 | 1.04E+00 | 1.28E-01 | 4.89E-01 | 1.08E+00 | 1.23E-01 | 4.38E-01 | -1.01E+00 | 2.02E-01 | 5.69E-01 | 1.01E+00 | 1.02E-02 | 3.75E-02 | 1.17E+00 | 1.05E-04 | 2.89E-04 | 1.25E+00 |
| ARL4D | 1.82E-07 | 9.48E-07 | -2.00E+00 | 1.33E-06 | 5.73E-06 | -1.59E+00 | 2.91E-03 | 1.24E-02 | -1.23E+00 | 1.73E-03 | 6.61E-03 | -1.29E+00 | 5.12E-05 | 1.83E-04 | -2.95E+00 | 1.69E-03 | 4.76E-03 | -1.66E+00 | 1.38E-02 | 5.06E-02 | -1.45E+00 | 2.76E-04 | 7.57E-04 | -2.43E+00 |
| MIS18A | 1.82E-07 | 9.48E-07 | -1.48E+00 | 2.81E-03 | 1.21E-02 | 1.10E+00 | 6.32E-04 | 2.68E-03 | -1.11E+00 | 1.26E-02 | 4.82E-02 | 1.15E+00 | 1.20E-06 | 4.30E-06 | -1.71E+00 | 2.76E-04 | 7.79E-04 | -1.22E+00 | 1.51E-07 | 5.53E-07 | -1.35E+00 | 2.14E-01 | 5.89E-01 | 1.14E+00 |
| MED1 | 1.83E-07 | 9.51E-07 | -1.36E+00 | 7.49E-06 | 3.24E-05 | -1.09E+00 | 3.08E-10 | 1.31E-09 | -1.53E+00 | 2.92E-07 | 1.12E-06 | -1.12E+00 | 9.41E-04 | 3.35E-03 | -1.08E+00 | 5.94E-03 | 1.68E-02 | 1.01E+00 | 8.03E-11 | 2.95E-10 | -1.52E+00 | 3.23E-04 | 8.88E-04 | -1.16E+00 |
| RNFT1 | 1.83E-07 | 9.52E-07 | -1.54E+00 | 3.64E-04 | 1.57E-03 | -1.08E+00 | 1.37E-06 | 5.84E-06 | -1.60E+00 | 4.15E-04 | 1.58E-03 | -1.13E+00 | 5.38E-10 | 1.92E-09 | -1.66E+00 | 9.67E-05 | 2.72E-04 | -1.09E+00 | 7.55E-02 | 2.77E-01 | 1.09E+00 | 2.99E-01 | 8.21E-01 | 1.30E+00 |
| DDX60L | 1.83E-07 | 9.56E-07 | -1.40E+00 | 2.57E-05 | 1.11E-04 | -1.05E+00 | 1.67E-06 | 7.11E-06 | -1.45E+00 | 2.84E-06 | 1.08E-05 | -1.27E+00 | 1.57E-08 | 5.60E-08 | -1.92E+00 | 1.37E-04 | 3.87E-04 | -1.29E+00 | 4.22E-10 | 1.55E-09 | -1.97E+00 | 5.07E-07 | 1.39E-06 | -1.60E+00 |
| RBM7 | 1.83E-07 | 9.56E-07 | -1.07E+00 | 1.96E-10 | 8.47E-10 | -1.07E+00 | 1.03E-05 | 4.37E-05 | -1.30E+00 | 4.92E-10 | 1.88E-09 | -1.42E+00 | 1.53E-05 | 5.44E-05 | -1.18E+00 | 1.68E-07 | 4.75E-07 | -1.30E+00 | 1.05E-05 | 3.86E-05 | -1.13E+00 | 1.15E-05 | 3.16E-05 | -1.18E+00 |
| TUBE1 | 1.84E-07 | 9.61E-07 | -1.89E+00 | 7.55E-04 | 3.26E-03 | -1.08E+00 | 7.00E-05 | 2.97E-04 | -1.56E+00 | 9.28E-03 | 3.54E-02 | 1.02E+00 | 1.12E-02 | 3.99E-02 | -1.16E+00 | 8.51E-02 | 2.40E-01 | 1.11E+00 | 2.40E-01 | 8.80E-01 | 1.22E+00 | 3.45E-01 | 9.48E-01 | 1.16E+00 |
| SEC24A | 1.85E-07 | 9.64E-07 | -1.91E+00 | 3.06E-06 | 1.32E-05 | -1.35E+00 | 4.76E-05 | 2.02E-04 | -1.44E+00 | 1.58E-04 | 6.02E-04 | -1.14E+00 | 1.76E-05 | 6.29E-05 | -1.41E+00 | 1.16E-04 | 3.28E-04 | -1.32E+00 | 9.68E-07 | 3.55E-06 | -1.43E+00 | 1.04E-04 | 2.85E-04 | -1.18E+00 |
| GUSBP2 | 1.86E-07 | 9.67E-07 | -1.55E+00 | 4.54E-04 | 1.96E-03 | -1.05E+00 | 2.66E-05 | 1.13E-04 | -1.33E+00 | 3.65E-04 | 1.39E-03 | -1.04E+00 | 1.39E-06 | 4.95E-06 | -1.64E+00 | 9.16E-03 | 2.58E-02 | 1.05E+00 | 2.18E-05 | 8.01E-05 | -1.43E+00 | 6.28E-06 | 1.73E-05 | -1.60E+00 |
| PRR4 | 1.86E-07 | 9.67E-07 | -1.10E+00 | 1.56E-01 | 6.73E-01 | 1.55E+00 | 1.25E-05 | 5.32E-05 | -1.21E+00 | 6.16E-03 | 2.35E-02 | 1.14E+00 | 5.32E-03 | 1.90E-02 | -1.01E+00 | 1.56E-01 | 4.39E-01 | 1.24E+00 | 1.65E-05 | 6.04E-05 | -1.27E+00 | 1.90E-01 | 5.22E-01 | 1.15E+00 |
| LIN52 | 1.87E-07 | 9.75E-07 | -2.03E+00 | 1.63E-05 | 7.03E-05 | -1.33E+00 | 1.01E-06 | 4.29E-06 | -1.93E+00 | 1.30E-04 | 4.96E-04 | -1.38E+00 | 4.11E-03 | 1.47E-02 | -1.51E+00 | 3.06E-02 | 8.64E-02 | -1.17E+00 | 8.87E-09 | 3.26E-08 | -2.16E+00 | 3.32E-03 | 9.13E-03 | -1.19E+00 |
| DTD1 | 1.88E-07 | 9.80E-07 | -1.07E+00 | 3.49E-10 | 1.51E-09 | -1.06E+00 | 8.98E-02 | 3.82E-01 | 1.07E+00 | 4.08E-03 | 1.56E-02 | 1.06E+00 | 4.58E-02 | 1.63E-01 | 1.04E+00 | 4.08E-02 | 1.15E-01 | 1.06E+00 | 5.09E-02 | 1.87E-01 | 1.35E+00 | 3.33E-01 | 9.14E-01 | 1.24E+00 |
| AKR1C4 | 1.88E-07 | 9.81E-07 | 4.41E+00 | 1.95E-02 | 8.42E-02 | 2.45E+00 | 1.79E-02 | 7.62E-02 | 2.23E+00 | 1.15E-01 | 4.41E-01 | 1.75E+00 | 2.02E-03 | 7.22E-03 | 4.47E+00 | 6.72E-02 | 1.90E-01 | 2.75E+00 | 2.38E-03 | 8.74E-03 | 2.49E+00 | 7.80E-02 | 2.14E-01 | 1.96E+00 |
| SPDYE11 | 1.89E-07 | 9.85E-07 | -2.39E+00 | 2.96E-04 | 1.28E-03 | -1.31E+00 | 1.73E-02 | 7.33E-02 | -1.26E+00 | 9.49E-02 | 3.63E-01 | 1.05E+00 | 8.56E-02 | 3.05E-01 | -1.13E+00 | 3.19E-01 | 9.01E-01 | 1.30E+00 | 4.18E-03 | 1.53E-02 | -1.32E+00 | 6.65E-03 | 1.83E-02 | -1.33E+00 |
| UBAC1 | 1.90E-07 | 9.88E-07 | -1.12E+00 | 9.44E-06 | 4.08E-05 | 1.07E+00 | 5.65E-03 | 2.40E-02 | -1.02E+00 | 6.89E-05 | 2.63E-04 | 1.00E+00 | 1.61E-08 | 5.74E-08 | -1.27E+00 | 4.96E-05 | 1.40E-04 | 1.07E+00 | 1.33E-01 | 4.87E-01 | 1.17E+00 | 2.59E-02 | 7.13E-02 | 1.02E+00 |
| C12orf29 | 1.90E-07 | 9.88E-07 | -1.34E+00 | 2.45E-05 | 1.06E-04 | 1.02E+00 | 1.79E-04 | 7.62E-04 | -1.40E+00 | 8.74E-05 | 3.34E-04 | -1.20E+00 | 6.39E-09 | 2.28E-08 | -2.17E+00 | 7.78E-05 | 2.19E-04 | -1.29E+00 | 2.35E-11 | 8.63E-11 | -1.62E+00 | 1.41E-03 | 3.88E-03 | -1.10E+00 |
| SPTLC3 | 1.90E-07 | 9.90E-07 | -1.58E+00 | 8.71E-03 | 3.76E-02 | 1.12E+00 | 1.06E-04 | 4.50E-04 | -1.46E+00 | 5.44E-03 | 2.08E-02 | -1.01E+00 | 4.28E-04 | 1.53E-03 | -1.38E+00 | 5.69E-03 | 1.60E-02 | -1.21E+00 | 3.58E-02 | 1.32E-01 | -1.16E+00 | 8.20E-02 | 2.25E-01 | -1.07E+00 |
| SMPDL3B | 1.90E-07 | 9.90E-07 | -1.43E+00 | 8.19E-07 | 3.54E-06 | -1.17E+00 | 2.01E-04 | 8.56E-04 | -1.37E+00 | 2.79E-03 | 1.06E-02 | -1.04E+00 | 5.52E-02 | 1.97E-01 | -1.10E+00 | 1.59E-01 | 4.49E-01 | 1.09E+00 | 5.64E-02 | 2.07E-01 | -1.37E+00 | 3.34E-01 | 9.17E-01 | 1.11E+00 |
| HIBCH | 1.90E-07 | 9.91E-07 | -1.25E+00 | 5.39E-02 | 2.33E-01 | 1.32E+00 | 4.50E-07 | 1.91E-06 | -1.35E+00 | 5.54E-03 | 2.11E-02 | 1.11E+00 | 1.46E-06 | 5.19E-06 | -1.37E+00 | 8.06E-02 | 2.27E-01 | 1.19E+00 | 6.93E-06 | 2.54E-05 | -1.21E+00 | 6.50E-02 | 1.78E-01 | 1.07E+00 |
| IPMK | 1.90E-07 | 9.92E-07 | -1.80E+00 | 7.62E-06 | 3.29E-05 | -1.37E+00 | 2.07E-05 | 8.78E-05 | -1.52E+00 | 1.52E-06 | 5.81E-06 | -1.54E+00 | 3.08E-03 | 1.10E-02 | -1.37E+00 | 1.28E-02 | 3.61E-02 | -1.16E+00 | 3.49E-08 | 1.28E-07 | -1.89E+00 | 2.94E-05 | 8.08E-05 | -1.46E+00 |
| INO80C | 1.90E-07 | 9.92E-07 | -1.40E+00 | 2.12E-06 | 9.17E-06 | -1.16E+00 | 2.59E-05 | 1.10E-04 | -1.30E+00 | 1.64E-05 | 6.26E-05 | -1.09E+00 | 1.62E-14 | 5.76E-14 | -1.83E+00 | 2.21E-06 | 6.23E-06 | -1.25E+00 | 1.39E-21 | 5.11E-21 | -1.71E+00 | 9.60E-11 | 2.64E-10 | -1.30E+00 |
| ECHDC2 | 1.91E-07 | 9.93E-07 | -1.44E+00 | 6.38E-07 | 2.75E-06 | -1.31E+00 | 2.33E-06 | 9.90E-06 | -1.41E+00 | 4.02E-06 | 1.54E-05 | -1.30E+00 | 3.76E-09 | 1.34E-08 | -2.94E+00 | 5.50E-09 | 1.55E-08 | -2.62E+00 | 2.90E-05 | 1.06E-04 | -2.35E+00 | 1.83E-04 | 5.03E-04 | -2.06E+00 |
| FBA1 | 1.92E-07 | 1.00E-06 | -2.01E+00 | 2.85E-04 | 1.23E-03 | -1.14E+00 | 1.91E-11 | 8.10E-11 | -1.93E+00 | 3.68E-06 | 1.40E-05 | -1.16E+00 | 1.31E-03 | 4.67E-03 | -1.28E+00 | 3.10E-02 | 8.73E-02 | 1.08E+00 | 4.57E-06 | 1.68E-05 | -1.43E+00 | 6.57E-03 | 1.80E-02 | -1.06E+00 |
| ZNF718 | 1.94E-07 | 1.01E-06 | 3.04E+00 | 2.63E-07 | 1.14E-06 | 3.45E+00 | 2.16E-01 | 9.19E-01 | 1.30E+00 | 1.20E-01 | 4.57E-01 | 1.62E+00 | 5.05E-02 | 1.80E-01 | 2.28E+00 | 6.88E-02 | 1.94E-01 | 2.21E+00 | 2.27E-02 | 8.32E-02 | 1.83E+00 | 1.87E-02 | 5.12E-02 | 1.99E+00 |
| C16orf87 | 1.94E-07 | 1.01E-06 | -1.04E+00 | 1.08E-21 | 4.68E-21 | -1.42E+00 | 3.98E-04 | 1.69E-03 | -1.15E+00 | 4.76E-13 | 1.82E-12 | -1.44E+00 | 6.43E-06 | 2.29E-05 | -1.21E+00 | 2.10E-11 | 5.91E-11 | -1.54E+00 | 2.09E-06 | 7.66E-06 | -1.27E+00 | 4.39E-11 | 1.21E-10 | -1.59E+00 |
| TPD52 | 1.94E-07 | 1.01E-06 | -1.22E+00 | 1.09E-04 | 4.71E-04 | 1.08E+00 | 1.86E-01 | 7.90E-01 | 1.26E+00 | 2.43E-01 | 9.28E-01 | 1.56E+00 | 1.90E-02 | 6.77E-02 | -1.02E+00 | 1.13E-01 | 3.18E-01 | 1.13E+00 | 1.41E-03 | 5.17E-03 | -1.16E+00 | 2.39E-01 | 6.56E-01 | 1.17E+00 |
| C1orf54 | 1.95E-07 | 1.01E-06 | -1.23E+00 | 1.27E-07 | 5.47E-07 | -1.10E+00 | 1.18E-04 | 5.00E-04 | -1.19E+00 | 5.59E-06 | 2.14E-05 | -1.13E+00 | 2.28E-03 | 8.12E-03 | -1.22E+00 | 1.32E-03 | 3.73E-03 | -1.26E+00 | 1.84E-02 | 6.76E-02 | -1.17E+00 | 6.89E-02 | 1.89E-01 | -1.10E+00 |
| CCPG1 | 1.95E-07 | 1.02E-06 | -1.52E+00 | 1.23E-05 | 5.32E-05 | -1.21E+00 | 6.84E-10 | 2.91E-09 | -1.91E+00 | 7.84E-09 | 2.99E-08 | -1.52E+00 | 4.25E-03 | 1.51E-02 | -1.29E+00 | 6.65E-02 | 1.87E-01 | 1.12E+00 | 4.73E-03 | 1.74E-02 | -1.06E+00 | 1.16E-02 | 3.18E-02 | -1.09E+00 |
| IFT27 | 1.95E-07 | 1.02E-06 | -1.45E+00 | 1.45E-07 | 6.27E-07 | -1.28E+00 | 1.90E-07 | 8.06E-07 | -1.48E+00 | 4.85E-06 | 1.85E-05 | -1.12E+00 | 4.26E-07 | 1.52E-06 | -1.50E+00 | 1.26E-04 | 3.56E-04 | -1.20E+00 | 1.34E-06 | 4.90E-06 | -1.29E+00 | 6.42E-04 | 1.76E-03 | -1.13E+00 |
| DCAF8 | 1.95E-07 | 1.02E-06 | -1.36E+00 | 6.43E-04 | 2.78E-03 | 1.04E+00 | 3.70E-06 | 1.57E-05 | -1.46E+00 | 4.84E-05 | 1.85E-04 | -1.12E+00 | 2.51E-06 | 8.96E-06 | -1.62E+00 | 2.30E-02 | 6.49E-02 | 1.09E+00 | 7.68E-04 | 2.82E-03 | -1.31E+00 | 2.69E-02 | 7.39E-02 | -1.04E+00 |
| NUP43 | 1.96E-07 | 1.02E-06 | -1.71E+00 | 1.31E-01 | 5.65E-01 | 1.78E+00 | 5.17E-12 | 2.20E-11 | -1.94E+00 | 9.88E-03 | 3.77E-02 | 1.16E+00 | 4.90E-05 | 1.75E-04 | -1.66E+00 | 8.95E-02 | 2.52E-01 | 1.71E+00 | 1.21E-11 | 4.42E-11 | -1.77E+00 | 3.67E-02 | 1.01E-01 | 1.10E+00 |
| CDC42BPA | 1.96E-07 | 1.02E-06 | -1.29E+00 | 6.38E-03 | 2.76E-02 | 1.23E+00 | 8.11E-05 | 3.45E-04 | -1.29E+00 | 1.23E-02 | 4.68E-02 | 1.06E+00 | 2.93E-04 | 1.05E-03 | -1.09E+00 | 1.04E-01 | 2.92E-01 | 1.28E+00 | 4.89E-03 | 1.80E-02 | 1.05E+00 | 1.40E-01 | 3.85E-01 | 1.12E+00 |
| SLC30A6 | 1.97E-07 | 1.03E-06 | -1.74E+00 | 4.65E-05 | 2.01E-04 | -1.17E+00 | 5.36E-08 | 2.28E-07 | -1.82E+00 | 1.27E-03 | 4.85E-03 | -1.01E+00 | 1.07E-05 | 3.83E-05 | -1.96E+00 | 2.73E-02 | 7.70E-02 | 1.01E+00 | 1.44E-08 | 5.29E-08 | -1.79E+00 | 8.16E-02 | 2.24E-01 | 1.14E+00 |
| ZNF839 | 1.97E-07 | 1.03E-06 | -1.98E+00 | 1.05E-04 | 4.55E-04 | -1.29E+00 | 2.19E-02 | 9.30E-02 | -1.27E+00 | 8.91E-02 | 3.40E-01 | 1.11E+00 | 1.51E-02 | 5.38E-02 | -1.38E+00 | 2.09E-02 | 5.88E-02 | -1.40E+00 | 9.17E-02 | 3.37E-01 | -1.13E+00 | 5.10E-02 | 1.40E-01 | -1.23E+00 |
| ZMIZ1 | 1.98E-07 | 1.03E-06 | -1.23E+00 | 3.65E-06 | 1.58E-05 | -1.04E+00 | 2.29E-07 | 9.72E-07 | -1.36E+00 | 1.05E-07 | 4.00E-07 | -1.21E+00 | 1.75E-09 | 6.24E-09 | -1.41E+00 | 1.45E-10 | 4.08E-10 | -1.38E+00 | 4.72E-07 | 1.73E-06 | -1.37E+00 | 3.74E-05 | 1.03E-04 | -1.27E+00 |
| HELB | 1.98E-07 | 1.03E-06 | -1.25E+00 | 3.05E-07 | 1.32E-06 | -1.09E+00 | 2.26E-07 | 9.60E-07 | -1.59E+00 | 2.19E-05 | 8.36E-05 | -1.18E+00 | 1.14E-05 | 4.05E-05 | -1.73E+00 | 2.03E-03 | 5.73E-03 | -1.14E+00 | 2.37E-06 | 8.70E-06 | -1.74E+00 | 5.59E-03 | 1.53E-02 | -1.29E+00 |
| CAMKK2 | 1.98E-07 | 1.03E-06 | -1.40E+00 | 3.86E-04 | 1.67E-03 | -1.01E+00 | 2.15E-06 | 9.16E-06 | -1.26E+00 | 5.35E-02 | 2.04E-01 | 1.25E+00 | 2.91E-06 | 1.04E-05 | -1.42E+00 | 2.94E-03 | 8.29E-03 | -1.00E+00 | 2.43E-04 | 8.93E-04 | -1.07E+00 | 1.97E-01 | 5.41E-01 | 1.23E+00 |
| SNAI3 | 1.99E-07 | 1.04E-06 | -2.05E+00 | 1.10E-04 | 4.75E-04 | -1.31E+00 | 1.09E-21 | 4.64E-21 | -2.52E+00 | 2.42E-14 | 9.26E-14 | -1.53E+00 | 3.62E-05 | 1.29E-04 | -2.41E+00 | 2.80E-04 | 7.89E-04 | -1.65E+00 | 4.56E-02 | 1.67E-01 | -1.95E+00 | 2.57E-01 | 7.05E-01 | -1.12E+00 |
| KDM2A | 1.99E-07 | 1.04E-06 | -1.09E+00 | 1.83E-13 | 7.89E-13 | -1.27E+00 | 2.58E-03 | 1.10E-02 | -1.02E+00 | 2.54E-11 | 9.72E-11 | -1.38E+00 | 7.11E-05 | 2.53E-04 | -1.08E+00 | 1.40E-08 | 3.94E-08 | -1.37E+00 | 6.73E-04 | 2.47E-03 | -1.11E+00 | 2.65E-07 | 7.27E-07 | -1.40E+00 |
| TMX4 | 2.00E-07 | 1.04E-06 | -1.06E+00 | 8.58E-10 | 3.71E-09 | -1.08E+00 | 1.61E-03 | 6.85E-03 | -1.14E+00 | 1.01E-05 | 3.87E-05 | -1.19E+00 | 1.06E-02 | 3.77E-02 | 1.11E+00 | 4.13E-04 | 1.16E-03 | -1.02E+00 | 1.02E-02 | 3.75E-02 | 1.14E+00 | 1.66E-04 | 4.55E-04 | -1.06E+00 |
| TMEM9 | 2.00E-07 | 1.04E-06 | -1.23E+00 | 9.40E-08 | 4.06E-07 | -1.09E+00 | 3.70E-04 | 1.57E-03 | -1.13E+00 | 3.21E-03 | 1.23E-02 | 1.08E+00 | 1.41E-06 | 5.01E-06 | -1.18E+00 | 3.41E-07 | 9.62E-07 | -1.18E+00 | 5.61E-05 | 2.06E-04 | -1.19E+00 | 1.71E-03 | 4.69E-03 | -1.10E+00 |
| TARS2 | 2.00E-07 | 1.04E-06 | -1.45E+00 | 2.76E-04 | 1.19E-03 | 1.05E+00 | 1.18E-04 | 5.00E-04 | -1.37E+00 | 5.09E-05 | 1.94E-04 | -1.26E+00 | 3.78E-05 | 1.35E-04 | -1.47E+00 | 3.41E-03 | 9.61E-03 | -1.11E+00 | 2.44E-10 | 8.95E-10 | -1.56E+00 | 4.01E-05 | 1.10E-04 | -1.15E+00 |
| GOLGA5 | 2.00E-07 | 1.04E-06 | -1.66E+00 | 5.89E-05 | 2.55E-04 | -1.16E+00 | 6.10E-05 | 2.59E-04 | -1.50E+00 | 9.86E-02 | 3.76E-01 | 1.20E+00 | 1.47E-06 | 5.24E-06 | -1.47E+00 | 1.62E-03 | 4.57E-03 | -1.03E+00 | 8.23E-06 | 3.02E-05 | -1.25E+00 | 6.42E-06 | 1.76E-05 | -1.27E+00 |
| EXOSC1 | 2.01E-07 | 1.05E-06 | -1.13E+00 | 2.20E-04 | 9.51E-04 | 1.16E+00 | 6.64E-02 | 2.82E-01 | 1.05E+00 | 4.46E-03 | 1.71E-02 | 1.06E+00 | 4.55E-06 | 1.62E-05 | -1.23E+00 | 1.69E-03 | 4.77E-03 | -1.01E+00 | 6.53E-03 | 2.40E-02 | 1.00E+00 | 1.49E-02 | 4.10E-02 | 1.01E+00 |
| TAF4B | 2.02E-07 | 1.05E-06 | -1.41E+00 | 9.12E-07 | 3.94E-06 | -1.19E+00 | 2.17E-02 | 9.24E-02 | -1.18E+00 | 4.46E-02 | 1.70E-01 | -1.01E+00 | 1.21E-02 | 4.30E-02 | -1.18E+00 | 6.03E-02 | 1.70E-01 | -1.02E+00 | 7.79E-14 | 2.86E-13 | -1.88E+00 | 2.61E-07 | 7.16E-07 | -1.29E+00 |
| RCOR3 | 2.02E-07 | 1.05E-06 | -1.22E+00 | 1.58E-04 | 6.81E-04 | 1.06E+00 | 6.21E-05 | 2.64E-04 | -1.26E+00 | 1.98E-03 | 7.58E-03 | -1.03E+00 | 1.83E-03 | 6.53E-03 | 1.01E+00 | 5.23E-02 | 1.48E-01 | 1.14E+00 | 2.39E-06 | 8.78E-06 | -1.11E+00 | 1.85E-04 | 5.07E-04 | -1.06E+00 |
| SVIP | 2.04E-07 | 1.07E-06 | -1.10E+00 | 8.40E-07 | 3.63E-06 | 1.05E+00 | 7.06E-02 | 3.00E-01 | 1.08E+00 | 7.12E-03 | 2.72E-02 | 1.09E+00 | 8.32E-03 | 2.97E-02 | 1.08E+00 | 6.20E-04 | 1.75E-03 | -1.05E+00 | 1.93E-01 | 7.10E-01 | 1.22E+00 | 3.06E-01 | 8.40E-01 | 1.17E+00 |
| XPO7 | 2.06E-07 | 1.07E-06 | -1.47E+00 | 7.25E-05 | 3.13E-04 | -1.13E+00 | 2.53E-04 | 1.08E-03 | -1.25E+00 | 1.02E-02 | 3.88E-02 | 1.10E+00 | 4.96E-07 | 1.77E-06 | -1.50E+00 | 1.08E-04 | 3.06E-04 | -1.20E+00 | 3.46E-07 | 1.27E-06 | -1.41E+00 | 1.88E-05 | 5.17E-05 | -1.17E+00 |
| FAHD2A | 2.07E-07 | 1.08E-06 | -1.32E+00 | 3.65E-08 | 1.58E-07 | -1.22E+00 | 8.72E-05 | 3.71E-04 | -1.23E+00 | 3.14E-09 | 1.20E-08 | -1.34E+00 | 1.77E-05 | 6.31E-05 | -1.42E+00 | 8.82E-04 | 2.49E-03 | -1.12E+00 | 2.41E-06 | 8.84E-06 | -1.39E+00 | 6.10E-04 | 1.68E-03 | -1.11E+00 |
| AC087257.1 | 2.07E-07 | 1.08E-06 | -1.71E+00 | 2.21E-03 | 9.53E-03 | 1.06E+00 | 7.98E-02 | 3.39E-01 | -1.05E+00 | 2.26E-01 | 8.63E-01 | 1.50E+00 | 1.59E-01 | 5.66E-01 | 1.05E+00 | 2.36E-01 | 6.65E-01 | 1.19E+00 | 2.40E-06 | 8.81E-06 | -2.70E+00 | 3.35E-04 | 9.20E-04 | -1.84E+00 |
| AC233699.1 | 2.09E-07 | 1.09E-06 | -2.12E+00 | 9.34E-04 | 4.03E-03 | -1.12E+00 | 2.35E-04 | 1.00E-03 | -1.79E+00 | 3.76E-02 | 1.44E-01 | -1.03E+00 | 8.95E-05 | 3.19E-04 | -1.69E+00 | 1.07E-05 | 3.01E-05 | -2.04E+00 | 2.32E-05 | 8.52E-05 | -1.78E+00 | 9.11E-03 | 2.50E-02 | -1.10E+00 |
| CEP152 | 2.09E-07 | 1.09E-06 | -1.51E+00 | 8.31E-04 | 3.59E-03 | 1.09E+00 | 6.77E-04 | 2.88E-03 | -1.19E+00 | 6.43E-03 | 2.45E-02 | 1.03E+00 | 7.05E-06 | 2.51E-05 | -1.79E+00 | 4.04E-03 | 1.14E-02 | -1.05E+00 | 4.17E-10 | 1.53E-09 | -1.52E+00 | 2.23E-03 | 6.14E-03 | -1.08E+00 |
| CACTIN | 2.11E-07 | 1.10E-06 | -1.78E+00 | 3.11E-04 | 1.34E-03 | -1.14E+00 | 2.35E-01 | 9.98E-01 | 1.17E+00 | 1.78E-01 | 6.79E-01 | 1.42E+00 | 1.78E-01 | 6.36E-01 | 1.35E+00 | 3.42E-01 | 9.64E-01 | 1.56E+00 | 7.64E-03 | 2.80E-02 | -1.11E+00 | 2.52E-01 | 6.91E-01 | 1.19E+00 |
| CAMLG | 2.12E-07 | 1.11E-06 | -1.09E+00 | 1.28E-05 | 5.52E-05 | 1.04E+00 | 5.48E-03 | 2.33E-02 | -1.14E+00 | 4.20E-03 | 1.60E-02 | -1.06E+00 | 4.41E-13 | 1.57E-12 | -1.78E+00 | 1.52E-08 | 4.29E-08 | -1.56E+00 | 5.23E-05 | 1.92E-04 | -1.21E+00 | 2.65E-03 | 7.28E-03 | -1.13E+00 |
| AL033519.1 | 2.13E-07 | 1.11E-06 | 4.60E+00 | 3.57E-02 | 1.54E-01 | 2.35E+00 | 1.26E-04 | 5.36E-04 | 2.53E+00 | 1.97E-01 | 7.51E-01 | 1.43E+00 | 7.70E-03 | 2.75E-02 | 3.84E+00 | 1.43E-01 | 4.04E-01 | 2.18E+00 | 1.42E-01 | 5.20E-01 | 1.49E+00 | 3.39E-01 | 9.31E-01 | 1.19E+00 |
| ACCS | 2.13E-07 | 1.11E-06 | -1.43E+00 | 1.03E-06 | 4.47E-06 | -1.21E+00 | 2.13E-03 | 9.04E-03 | -1.30E+00 | 4.09E-02 | 1.56E-01 | 1.07E+00 | 3.33E-03 | 1.19E-02 | -1.20E+00 | 2.95E-02 | 8.32E-02 | 1.07E+00 | 2.36E-02 | 8.65E-02 | 1.12E+00 | 9.11E-02 | 2.50E-01 | 1.15E+00 |
| VWA5A | 2.14E-07 | 1.11E-06 | -1.55E+00 | 8.13E-03 | 3.51E-02 | 1.20E+00 | 8.21E-03 | 3.49E-02 | -1.34E+00 | 3.01E-02 | 1.15E-01 | -1.05E+00 | 5.07E-06 | 1.81E-05 | -3.48E+00 | 1.48E-06 | 4.16E-06 | -3.66E+00 | 1.50E-01 | 5.52E-01 | 1.74E+00 | 2.06E-02 | 5.65E-02 | -1.18E+00 |
| SYDE2 | 2.14E-07 | 1.11E-06 | -1.68E+00 | 2.32E-05 | 1.00E-04 | -1.20E+00 | 9.09E-06 | 3.86E-05 | -1.77E+00 | 8.84E-04 | 3.38E-03 | -1.18E+00 | 5.58E-05 | 1.99E-04 | -1.83E+00 | 7.81E-03 | 2.20E-02 | -1.23E+00 | 4.62E-05 | 1.70E-04 | -1.38E+00 | 7.86E-03 | 2.16E-02 | -1.18E+00 |
| RHOBTB1 | 2.14E-07 | 1.11E-06 | -1.82E+00 | 4.83E-06 | 2.09E-05 | -1.44E+00 | 1.42E-06 | 6.03E-06 | -1.48E+00 | 2.93E-06 | 1.12E-05 | -1.35E+00 | 2.17E-06 | 7.75E-06 | -2.24E+00 | 7.23E-05 | 2.04E-04 | -1.70E+00 | 6.01E-04 | 2.21E-03 | -1.10E+00 | 5.62E-03 | 1.54E-02 | -1.11E+00 |
| NME1 | 2.15E-07 | 1.12E-06 | -1.09E+00 | 1.60E-05 | 6.93E-05 | 1.07E+00 | 2.28E-01 | 9.69E-01 | 1.16E+00 | 1.90E-01 | 7.26E-01 | 1.24E+00 | 1.85E-03 | 6.61E-03 | -1.15E+00 | 3.02E-01 | 8.50E-01 | 1.26E+00 | 5.10E-02 | 1.87E-01 | 1.17E+00 | 5.20E-03 | 1.43E-02 | 1.55E+00 |
| DPY19L2 | 2.15E-07 | 1.12E-06 | -1.73E+00 | 1.86E-02 | 8.03E-02 | 1.20E+00 | 8.08E-05 | 3.43E-04 | -1.33E+00 | 5.87E-03 | 2.24E-02 | 1.07E+00 | 1.16E-03 | 4.15E-03 | -1.19E+00 | 8.57E-02 | 2.42E-01 | 1.12E+00 | 4.18E-06 | 1.53E-05 | -1.38E+00 | 3.52E-02 | 9.66E-02 | 1.07E+00 |
| COQ4 | 2.15E-07 | 1.12E-06 | -1.15E+00 | 1.83E-08 | 7.89E-08 | -1.05E+00 | 4.92E-05 | 2.09E-04 | -1.10E+00 | 1.49E-06 | 5.70E-06 | -1.06E+00 | 1.10E-07 | 3.91E-07 | -1.34E+00 | 9.99E-06 | 2.82E-05 | -1.09E+00 | 2.37E-03 | 8.69E-03 | -1.03E+00 | 1.01E-03 | 2.76E-03 | -1.07E+00 |
| DVL2 | 2.16E-07 | 1.12E-06 | -1.30E+00 | 1.97E-04 | 8.50E-04 | 1.12E+00 | 1.11E-02 | 4.72E-02 | -1.13E+00 | 3.93E-02 | 1.50E-01 | 1.08E+00 | 2.15E-02 | 7.67E-02 | -1.02E+00 | 1.30E-01 | 3.67E-01 | 1.20E+00 | 1.38E-02 | 5.05E-02 | -1.04E+00 | 2.18E-01 | 5.99E-01 | 1.13E+00 |
| WDR89 | 2.16E-07 | 1.12E-06 | -1.59E+00 | 8.74E-08 | 3.77E-07 | -1.49E+00 | 2.08E-03 | 8.85E-03 | -1.22E+00 | 9.17E-03 | 3.50E-02 | -1.00E+00 | 1.54E-01 | 5.50E-01 | 1.23E+00 | 9.01E-02 | 2.54E-01 | 1.12E+00 | 8.68E-09 | 3.18E-08 | -1.61E+00 | 2.25E-05 | 6.18E-05 | -1.25E+00 |
| LRRC27 | 2.16E-07 | 1.13E-06 | -1.54E+00 | 1.04E-03 | 4.51E-03 | 1.03E+00 | 4.75E-05 | 2.02E-04 | -1.56E+00 | 1.09E-04 | 4.14E-04 | -1.22E+00 | 2.38E-10 | 8.48E-10 | -2.15E+00 | 2.84E-08 | 7.99E-08 | -1.62E+00 | 3.50E-08 | 1.28E-07 | -1.70E+00 | 1.92E-08 | 5.26E-08 | -1.85E+00 |
| CARD6 | 2.16E-07 | 1.13E-06 | -1.87E+00 | 4.82E-05 | 2.08E-04 | -1.36E+00 | 3.04E-08 | 1.29E-07 | -1.91E+00 | 2.56E-05 | 9.77E-05 | -1.20E+00 | 2.05E-10 | 7.30E-10 | -2.64E+00 | 2.12E-06 | 5.99E-06 | -1.57E+00 | 9.16E-09 | 3.36E-08 | -1.96E+00 | 4.96E-06 | 1.36E-05 | -1.52E+00 |
| PSMD4 | 2.16E-07 | 1.13E-06 | -1.10E+00 | 3.67E-08 | 1.59E-07 | 1.01E+00 | 8.39E-04 | 3.57E-03 | -1.12E+00 | 9.02E-06 | 3.45E-05 | -1.13E+00 | 4.64E-05 | 1.66E-04 | -1.16E+00 | 2.89E-04 | 8.15E-04 | -1.05E+00 | 1.09E-04 | 4.01E-04 | -1.14E+00 | 1.06E-03 | 2.92E-03 | -1.11E+00 |
| MTHFR | 2.17E-07 | 1.13E-06 | -1.53E+00 | 2.22E-03 | 9.59E-03 | 1.01E+00 | 2.19E-06 | 9.32E-06 | -1.54E+00 | 7.80E-03 | 2.98E-02 | 1.11E+00 | 8.43E-05 | 3.01E-04 | -1.22E+00 | 5.64E-02 | 1.59E-01 | 1.17E+00 | 5.77E-10 | 2.12E-09 | -1.52E+00 | 5.28E-03 | 1.45E-02 | -1.09E+00 |
| MPHOSPH9 | 2.17E-07 | 1.13E-06 | -1.28E+00 | 3.12E-03 | 1.35E-02 | 1.09E+00 | 1.33E-02 | 5.65E-02 | -1.08E+00 | 8.43E-02 | 3.22E-01 | 1.22E+00 | 1.18E-04 | 4.20E-04 | -1.10E+00 | 2.80E-02 | 7.90E-02 | 1.13E+00 | 8.38E-07 | 3.07E-06 | -1.33E+00 | 1.17E-02 | 3.20E-02 | -1.01E+00 |
| GPSM3 | 2.18E-07 | 1.14E-06 | -1.17E+00 | 9.57E-07 | 4.14E-06 | 1.00E+00 | 2.71E-06 | 1.15E-05 | -1.35E+00 | 4.98E-06 | 1.90E-05 | -1.25E+00 | 1.34E-04 | 4.76E-04 | -1.25E+00 | 4.19E-04 | 1.18E-03 | -1.13E+00 | 4.76E-02 | 1.75E-01 | 1.03E+00 | 5.48E-03 | 1.51E-02 | -1.07E+00 |
| ZMAT1 | 2.19E-07 | 1.14E-06 | -1.29E+00 | 1.17E-03 | 5.06E-03 | 1.12E+00 | 1.75E-06 | 7.45E-06 | -1.45E+00 | 1.09E-05 | 4.16E-05 | -1.24E+00 | 3.28E-03 | 1.17E-02 | -1.08E+00 | 6.93E-02 | 1.95E-01 | 1.09E+00 | 3.31E-02 | 1.21E-01 | -1.03E+00 | 2.83E-01 | 7.77E-01 | 1.27E+00 |
| HEATR1 | 2.19E-07 | 1.14E-06 | -1.42E+00 | 3.70E-08 | 1.60E-07 | -1.43E+00 | 1.12E-05 | 4.76E-05 | -1.43E+00 | 5.28E-05 | 2.02E-04 | -1.22E+00 | 4.59E-04 | 1.64E-03 | -1.38E+00 | 3.06E-03 | 8.61E-03 | -1.29E+00 | 1.50E-04 | 5.51E-04 | -1.13E+00 | 4.59E-03 | 1.26E-02 | -1.00E+00 |
| CHMP2B | 2.20E-07 | 1.14E-06 | -1.16E+00 | 2.34E-09 | 1.01E-08 | -1.08E+00 | 5.65E-04 | 2.40E-03 | -1.06E+00 | 5.13E-05 | 1.96E-04 | -1.01E+00 | 4.67E-06 | 1.67E-05 | -1.13E+00 | 5.65E-07 | 1.59E-06 | -1.22E+00 | 2.22E-03 | 8.16E-03 | -1.02E+00 | 4.99E-03 | 1.37E-02 | 1.03E+00 |
| POM121C | 2.22E-07 | 1.16E-06 | -1.41E+00 | 6.10E-04 | 2.63E-03 | 1.02E+00 | 2.46E-02 | 1.04E-01 | 1.06E+00 | 8.73E-02 | 3.33E-01 | 1.32E+00 | 1.62E-03 | 5.77E-03 | -1.23E+00 | 1.07E-01 | 3.02E-01 | 1.11E+00 | 8.63E-03 | 3.17E-02 | -1.09E+00 | 3.29E-01 | 9.04E-01 | 1.20E+00 |
| IL10RB | 2.25E-07 | 1.17E-06 | -1.76E+00 | 8.59E-06 | 3.71E-05 | -1.43E+00 | 4.92E-04 | 2.09E-03 | -1.36E+00 | 9.89E-04 | 3.78E-03 | -1.16E+00 | 5.46E-14 | 1.95E-13 | -2.03E+00 | 1.09E-09 | 3.07E-09 | -1.35E+00 | 8.84E-13 | 3.24E-12 | -1.78E+00 | 2.90E-07 | 7.97E-07 | -1.37E+00 |
| SLC25A17 | 2.25E-07 | 1.17E-06 | -2.07E+00 | 3.92E-05 | 1.69E-04 | -1.46E+00 | 7.99E-03 | 3.40E-02 | -1.12E+00 | 1.99E-01 | 7.62E-01 | 1.35E+00 | 1.32E-03 | 4.71E-03 | -1.75E+00 | 4.49E-02 | 1.27E-01 | -1.19E+00 | 1.71E-08 | 6.29E-08 | -1.42E+00 | 4.41E-02 | 1.21E-01 | 1.10E+00 |
| RRN3P1 | 2.25E-07 | 1.17E-06 | -1.28E+00 | 7.34E-06 | 3.17E-05 | -1.07E+00 | 9.60E-07 | 4.08E-06 | -1.34E+00 | 2.36E-07 | 9.00E-07 | -1.24E+00 | 1.72E-06 | 6.15E-06 | -1.28E+00 | 7.08E-06 | 1.99E-05 | -1.16E+00 | 1.26E-06 | 4.61E-06 | -1.42E+00 | 1.62E-02 | 4.46E-02 | -1.02E+00 |
| SNAPC2 | 2.27E-07 | 1.18E-06 | -1.39E+00 | 4.48E-07 | 1.94E-06 | -1.21E+00 | 3.86E-03 | 1.64E-02 | -1.13E+00 | 1.92E-03 | 7.33E-03 | 1.01E+00 | 2.69E-03 | 9.60E-03 | -1.09E+00 | 2.11E-02 | 5.96E-02 | 1.06E+00 | 8.05E-03 | 2.96E-02 | -1.05E+00 | 1.60E-01 | 4.40E-01 | 1.08E+00 |
| CBWD1 | 2.27E-07 | 1.19E-06 | -1.10E+00 | 4.18E-05 | 1.80E-04 | 1.13E+00 | 7.00E-04 | 2.97E-03 | -1.18E+00 | 2.05E-02 | 7.83E-02 | 1.11E+00 | 5.55E-03 | 1.98E-02 | 1.02E+00 | 1.65E-01 | 4.65E-01 | 1.29E+00 | 4.33E-04 | 1.59E-03 | -1.10E+00 | 2.27E-01 | 6.23E-01 | 1.14E+00 |
| ANKH | 2.28E-07 | 1.19E-06 | -1.60E+00 | 3.38E-04 | 1.46E-03 | -1.09E+00 | 1.67E-02 | 7.12E-02 | -1.01E+00 | 1.54E-01 | 5.87E-01 | 1.41E+00 | 4.41E-02 | 1.57E-01 | -1.25E+00 | 2.53E-01 | 7.13E-01 | 1.21E+00 | 3.23E-03 | 1.18E-02 | -1.20E+00 | 9.17E-02 | 2.52E-01 | 1.01E+00 |
| IST1 | 2.28E-07 | 1.19E-06 | -1.18E+00 | 5.16E-04 | 2.23E-03 | 1.15E+00 | 7.20E-08 | 3.06E-07 | -1.39E+00 | 3.00E-05 | 1.15E-04 | -1.05E+00 | 1.79E-05 | 6.38E-05 | -1.24E+00 | 1.60E-02 | 4.50E-02 | 1.11E+00 | 7.87E-07 | 2.89E-06 | -1.27E+00 | 1.55E-02 | 4.26E-02 | 1.02E+00 |
| PIN4P1 | 2.29E-07 | 1.19E-06 | -1.24E+00 | 2.54E-10 | 1.10E-09 | -1.41E+00 | 7.10E-05 | 3.02E-04 | -1.27E+00 | 6.65E-08 | 2.54E-07 | -1.39E+00 | 3.22E-04 | 1.15E-03 | -1.19E+00 | 3.76E-08 | 1.06E-07 | -1.79E+00 | 8.31E-05 | 3.05E-04 | -1.09E+00 | 5.63E-09 | 1.55E-08 | -1.38E+00 |

| | | | | | | | | | | | | | | | | | | | | | | | | |
|---|---|---|---|---|---|---|---|---|---|---|---|---|---|---|---|---|---|---|---|---|---|---|---|---|
| AC022400.7 | 2.29E-07 | 1.19E-06 | -1.07E+00 | 1.23E-13 | 5.30E-13 | -1.25E+00 | 1.04E-03 | 4.43E-03 | -1.15E+00 | 9.90E-09 | 3.78E-08 | -1.36E+00 | 3.32E-03 | 1.18E-02 | -1.00E+00 | 2.10E-05 | 5.91E-05 | -1.30E+00 | 1.11E-01 | 4.09E-01 | 1.18E+00 | 1.03E-02 | 2.83E-02 | -1.03E+00 |
| SACS | 2.30E-07 | 1.20E-06 | -1.39E+00 | 2.74E-05 | 1.18E-04 | -1.04E+00 | 2.01E-04 | 8.54E-04 | -1.28E+00 | 5.41E-03 | 2.07E-02 | 1.07E+00 | 1.79E-03 | 6.40E-03 | -1.46E+00 | 1.52E-01 | 4.27E-01 | 1.14E+00 | 1.97E-09 | 7.24E-09 | -1.67E+00 | 1.52E-05 | 4.17E-05 | -1.29E+00 |
| NFKBIE | 2.30E-07 | 1.20E-06 | -1.14E+00 | 6.40E-20 | 2.77E-19 | -2.15E+00 | 1.21E-10 | 5.16E-10 | -1.54E+00 | 5.56E-15 | 2.13E-14 | -1.81E+00 | 7.50E-07 | 2.67E-06 | -2.36E+00 | 1.94E-07 | 5.48E-07 | -2.35E+00 | 6.94E-11 | 2.55E-10 | -2.24E+00 | 1.37E-07 | 3.76E-07 | -1.87E+00 |
| STXBP5 | 2.30E-07 | 1.20E-06 | -1.31E+00 | 7.32E-04 | 3.16E-03 | 1.11E+00 | 6.43E-07 | 2.73E-06 | -1.48E+00 | 8.50E-04 | 3.25E-03 | -1.01E+00 | 3.58E-16 | 1.28E-15 | -1.80E+00 | 7.05E-10 | 1.99E-09 | -1.36E+00 | 1.69E-17 | 6.20E-17 | -1.74E+00 | 3.22E-08 | 8.84E-08 | -1.37E+00 |
| SNRNP27 | 2.30E-07 | 1.20E-06 | -1.39E+00 | 5.49E-08 | 2.37E-07 | -1.31E+00 | 7.12E-03 | 3.03E-02 | -1.13E+00 | 1.93E-03 | 7.38E-03 | -1.09E+00 | 2.50E-07 | 8.92E-07 | -1.38E+00 | 3.60E-05 | 1.02E-04 | -1.18E+00 | 5.09E-10 | 1.87E-09 | -1.38E+00 | 4.56E-07 | 1.25E-06 | -1.24E+00 |
| AC068888.2 | 2.31E-07 | 1.21E-06 | -2.44E+00 | 4.32E-04 | 1.87E-03 | -1.27E+00 | 1.73E-08 | 7.36E-08 | -2.38E+00 | 4.22E-04 | 1.61E-03 | -1.24E+00 | 5.37E-02 | 1.91E-01 | -1.17E+00 | 1.78E-02 | 5.03E-02 | -1.37E+00 | 3.48E-04 | 1.28E-03 | -1.74E+00 | 7.09E-02 | 1.95E-01 | -1.02E+00 |
| SRP68 | 2.33E-07 | 1.21E-06 | -1.23E+00 | 3.27E-03 | 1.41E-02 | 1.15E+00 | 1.96E-03 | 8.32E-03 | -1.15E+00 | 2.69E-02 | 1.03E-01 | 1.11E+00 | 5.99E-09 | 2.14E-08 | -1.52E+00 | 1.43E-03 | 4.02E-03 | 1.01E+00 | 2.42E-07 | 8.89E-07 | -1.28E+00 | 6.59E-02 | 1.81E-01 | 1.15E+00 |
| DHX29 | 2.34E-07 | 1.22E-06 | -1.11E+00 | 2.68E-07 | 1.16E-06 | -1.02E+00 | 4.17E-03 | 1.77E-02 | -1.05E+00 | 3.60E-03 | 1.37E-02 | 1.08E+00 | 9.70E-05 | 3.46E-04 | -1.06E+00 | 5.44E-05 | 1.53E-04 | -1.04E+00 | 1.03E-02 | 3.79E-02 | 1.01E+00 | 3.30E-02 | 9.06E-02 | 1.05E+00 |
| VIPAS39 | 2.34E-07 | 1.22E-06 | -1.46E+00 | 2.48E-03 | 1.07E-02 | 1.10E+00 | 1.31E-07 | 5.56E-07 | -1.51E+00 | 3.41E-04 | 1.30E-03 | -1.03E+00 | 3.55E-08 | 1.27E-07 | -1.70E+00 | 2.71E-05 | 7.64E-05 | -1.20E+00 | 1.88E-04 | 6.88E-04 | -1.07E+00 | 2.32E-03 | 6.36E-03 | -1.00E+00 |
| THOC3 | 2.34E-07 | 1.22E-06 | -1.32E+00 | 2.27E-03 | 9.79E-03 | 1.16E+00 | 1.97E-07 | 8.37E-07 | -1.37E+00 | 3.45E-04 | 1.32E-03 | -1.03E+00 | 1.02E-04 | 3.63E-04 | -1.27E+00 | 1.58E-02 | 4.46E-02 | 1.11E+00 | 1.03E-10 | 3.76E-10 | -1.41E+00 | 2.50E-03 | 6.88E-03 | 1.00E+00 |
| SHOC2 | 2.34E-07 | 1.22E-06 | -1.12E+00 | 3.97E-15 | 1.71E-14 | -1.34E+00 | 1.34E-06 | 5.71E-06 | -1.25E+00 | 6.50E-15 | 2.48E-14 | -1.53E+00 | 2.08E-04 | 7.43E-04 | -1.22E+00 | 1.04E-07 | 2.95E-07 | -1.48E+00 | 4.11E-07 | 1.51E-06 | -1.28E+00 | 3.01E-08 | 8.26E-08 | -1.49E+00 |
| ARMC1 | 2.34E-07 | 1.22E-06 | -1.28E+00 | 7.35E-05 | 3.18E-04 | 1.04E+00 | 3.03E-06 | 1.29E-05 | -1.35E+00 | 1.19E-05 | 4.53E-05 | -1.17E+00 | 4.76E-18 | 1.70E-17 | -2.33E+00 | 4.24E-10 | 1.20E-09 | -1.37E+00 | 1.08E-20 | 3.96E-20 | -1.89E+00 | 2.73E-08 | 7.50E-08 | -1.29E+00 |
| CCT5P2 | 2.36E-07 | 1.23E-06 | -2.28E+00 | 2.59E-06 | 1.12E-05 | -1.82E+00 | 6.10E-05 | 2.59E-04 | -1.68E+00 | 4.92E-07 | 1.88E-06 | -1.95E+00 | 8.52E-08 | 3.04E-07 | -2.73E+00 | 7.71E-08 | 2.17E-07 | -2.72E+00 | 6.66E-09 | 2.44E-08 | -1.73E+00 | 6.80E-09 | 1.87E-08 | -1.70E+00 |
| TIMM50 | 2.36E-07 | 1.23E-06 | -1.33E+00 | 9.31E-06 | 4.02E-05 | -1.10E+00 | 5.38E-03 | 2.28E-02 | -1.09E+00 | 1.54E-03 | 5.88E-03 | -1.00E+00 | 1.67E-08 | 5.94E-08 | -1.50E+00 | 2.82E-03 | 7.94E-03 | 1.08E+00 | 3.26E-13 | 1.20E-12 | -1.45E+00 | 2.73E-04 | 7.51E-04 | -1.02E+00 |
| TMEM234 | 2.37E-07 | 1.23E-06 | -1.69E+00 | 2.81E-06 | 1.21E-05 | -1.39E+00 | 1.91E-03 | 8.10E-03 | -1.24E+00 | 1.87E-03 | 7.16E-03 | -1.09E+00 | 1.61E-03 | 5.74E-03 | -1.46E+00 | 6.64E-02 | 1.87E-01 | -1.01E+00 | 1.05E-01 | 3.86E-01 | -1.06E+00 | 3.61E-01 | 9.92E-01 | 1.12E+00 |
| FAM171B | 2.37E-07 | 1.24E-06 | -1.41E+00 | 2.53E-04 | 1.09E-03 | -1.02E+00 | 3.96E-07 | 1.68E-06 | -1.76E+00 | 1.42E-04 | 5.43E-04 | -1.22E+00 | 2.29E-01 | 8.15E-01 | 1.03E+00 | 2.84E-01 | 8.01E-01 | 1.33E+00 | 2.26E-01 | 8.31E-01 | 1.27E+00 | 3.47E-01 | 9.54E-01 | 1.30E+00 |
| AL445363.3 | 2.38E-07 | 1.24E-06 | -1.58E+00 | 3.41E-04 | 1.47E-03 | -1.04E+00 | 5.85E-06 | 2.49E-05 | -1.60E+00 | 2.48E-05 | 9.47E-05 | -1.35E+00 | 3.64E-04 | 1.30E-03 | -1.87E+00 | 6.51E-03 | 1.83E-02 | -1.36E+00 | 1.88E-02 | 6.91E-02 | -1.02E+00 | 2.21E-02 | 6.07E-02 | -1.13E+00 |
| GMEB1 | 2.40E-07 | 1.25E-06 | -1.50E+00 | 5.81E-05 | 2.51E-04 | -1.13E+00 | 3.65E-06 | 1.55E-05 | -1.49E+00 | 2.53E-03 | 9.65E-03 | -1.03E+00 | 1.07E-06 | 3.82E-06 | -1.54E+00 | 7.08E-04 | 2.00E-03 | -1.10E+00 | 1.05E-11 | 3.84E-11 | -1.72E+00 | 8.08E-07 | 2.22E-06 | -1.34E+00 |
| AMFR | 2.41E-07 | 1.26E-06 | -1.43E+00 | 4.20E-08 | 1.81E-07 | -1.31E+00 | 1.77E-06 | 7.52E-06 | -1.42E+00 | 4.46E-05 | 1.70E-04 | -1.08E+00 | 8.28E-02 | 2.95E-01 | 1.21E+00 | 1.11E-01 | 3.14E-01 | 1.16E+00 | 1.91E-01 | 7.00E-01 | 1.29E+00 | 2.04E-01 | 5.60E-01 | 1.26E+00 |
| ZNF232 | 2.42E-07 | 1.26E-06 | -1.70E+00 | 9.98E-05 | 4.31E-04 | -1.14E+00 | 3.72E-06 | 1.58E-05 | -1.54E+00 | 5.93E-04 | 2.27E-03 | -1.10E+00 | 5.09E-04 | 1.81E-03 | -1.93E+00 | 1.07E-02 | 3.01E-02 | -1.29E+00 | 1.14E-02 | 4.19E-02 | -1.14E+00 | 2.89E-02 | 7.93E-02 | -1.15E+00 |
| TWISTNB | 2.42E-07 | 1.26E-06 | -1.08E+00 | 3.69E-11 | 1.60E-10 | -1.08E+00 | 7.30E-04 | 3.10E-03 | -1.09E+00 | 3.96E-05 | 1.51E-04 | -1.05E+00 | 1.56E-03 | 5.55E-03 | 1.20E+00 | 1.31E-05 | 3.69E-05 | -1.00E+00 | 3.77E-02 | 1.38E-01 | 1.11E+00 | 3.05E-02 | 8.37E-02 | 1.09E+00 |
| FAF2 | 2.42E-07 | 1.26E-06 | -1.41E+00 | 5.37E-07 | 2.32E-06 | -1.23E+00 | 3.80E-06 | 1.61E-05 | -1.29E+00 | 4.50E-07 | 1.72E-06 | -1.18E+00 | 1.39E-04 | 4.97E-04 | -1.19E+00 | 2.81E-04 | 7.93E-04 | -1.14E+00 | 1.92E-06 | 7.03E-06 | -1.28E+00 | 1.25E-03 | 3.43E-03 | -1.10E+00 |
| CRLS1 | 2.43E-07 | 1.27E-06 | -1.39E+00 | 5.05E-04 | 2.18E-03 | 1.03E+00 | 2.64E-11 | 1.12E-10 | -1.51E+00 | 8.98E-06 | 3.43E-05 | -1.07E+00 | 3.67E-17 | 1.31E-16 | -2.15E+00 | 1.56E-07 | 4.39E-07 | -1.28E+00 | 2.58E-12 | 9.48E-12 | -1.42E+00 | 8.00E-03 | 2.20E-02 | 1.06E+00 |
| PARP4 | 2.43E-07 | 1.27E-06 | -1.25E+00 | 9.04E-02 | 3.91E-01 | 1.28E+00 | 5.14E-04 | 2.19E-03 | -1.09E+00 | 1.18E-01 | 4.49E-01 | 1.36E+00 | 1.41E-06 | 5.04E-06 | -1.27E+00 | 1.57E-02 | 4.42E-02 | 1.09E+00 | 2.39E-05 | 8.78E-05 | -1.21E+00 | 5.53E-03 | 1.52E-02 | -1.07E+00 |
| RORA | 2.46E-07 | 1.28E-06 | -1.79E+00 | 5.01E-04 | 2.17E-03 | -1.10E+00 | 1.87E-01 | 7.96E-01 | -1.02E+00 | 2.08E-01 | 7.94E-01 | 1.22E+00 | 3.01E-04 | 1.07E-03 | -1.77E+00 | 6.09E-03 | 1.72E-02 | -1.38E+00 | 1.03E-02 | 3.79E-02 | -1.86E+00 | 2.57E-01 | 7.05E-01 | 1.08E+00 |
| CHCHD7 | 2.47E-07 | 1.29E-06 | -1.03E+00 | 1.02E-05 | 4.42E-05 | 1.13E+00 | 6.52E-04 | 2.77E-03 | -1.11E+00 | 2.55E-03 | 9.74E-03 | -1.00E+00 | 1.22E-03 | 4.36E-03 | -1.03E+00 | 1.26E-02 | 3.56E-02 | 1.05E+00 | 2.51E-02 | 9.19E-02 | 1.45E+00 | 2.05E-01 | 5.64E-01 | 1.30E+00 |
| RBM12B | 2.49E-07 | 1.30E-06 | -1.41E+00 | 1.51E-02 | 6.53E-02 | 1.13E+00 | 2.11E-11 | 8.96E-11 | -1.69E+00 | 1.09E-06 | 4.16E-06 | -1.08E+00 | 1.20E-05 | 4.28E-05 | -1.23E+00 | 1.00E-03 | 2.83E-03 | -1.03E+00 | 6.45E-14 | 2.37E-13 | -1.79E+00 | 1.16E-06 | 3.19E-06 | -1.27E+00 |
| C1QBPP2 | 2.51E-07 | 1.31E-06 | -2.00E+00 | 1.47E-06 | 6.34E-06 | -1.63E+00 | 4.74E-09 | 2.02E-08 | -1.62E+00 | 1.01E-10 | 3.85E-10 | -1.59E+00 | 7.97E-05 | 2.84E-04 | -1.82E+00 | 5.58E-04 | 1.57E-03 | -1.44E+00 | 6.27E-13 | 2.30E-12 | -1.76E+00 | 1.58E-08 | 4.34E-08 | -1.39E+00 |
| GSTO1 | 2.52E-07 | 1.31E-06 | -1.03E+00 | 5.26E-07 | 2.27E-06 | 1.07E+00 | 1.21E-01 | 5.13E-01 | 1.05E+00 | 3.36E-02 | 1.28E-01 | 1.05E+00 | 4.72E-06 | 1.68E-05 | -1.04E+00 | 1.54E-02 | 4.35E-02 | 1.12E+00 | 1.65E-01 | 6.05E-01 | 1.29E+00 | 2.51E-01 | 6.88E-01 | 1.14E+00 |
| LINC00526 | 2.53E-07 | 1.32E-06 | -1.48E+00 | 2.81E-05 | 1.21E-04 | -1.14E+00 | 5.00E-03 | 2.12E-02 | -1.31E+00 | 6.51E-02 | 2.49E-01 | 1.07E+00 | 5.10E-02 | 1.82E-01 | -1.07E+00 | 1.67E-01 | 4.70E-01 | 1.63E+00 | 4.32E-02 | 1.58E-01 | -1.01E+00 | 3.23E-01 | 8.86E-01 | 1.21E+00 |
| B3GALNT1 | 2.54E-07 | 1.32E-06 | -1.51E+00 | 1.26E-02 | 5.46E-02 | 1.24E+00 | 2.68E-06 | 1.14E-05 | -1.64E+00 | 7.88E-02 | 3.01E-01 | 1.23E+00 | 2.12E-07 | 7.57E-07 | -2.07E+00 | 5.69E-03 | 1.60E-02 | -1.16E+00 | 8.83E-04 | 3.24E-03 | -1.18E+00 | 3.56E-01 | 9.78E-01 | 1.25E+00 |
| ERCC1 | 2.55E-07 | 1.33E-06 | -1.12E+00 | 9.30E-06 | 4.02E-05 | 1.10E+00 | 3.08E-02 | 1.31E-01 | -1.02E+00 | 4.20E-03 | 1.60E-02 | 1.05E+00 | 1.25E-04 | 4.44E-04 | -1.06E+00 | 7.66E-02 | 2.16E-01 | 1.20E+00 | 1.52E-02 | 5.59E-02 | 1.00E+00 | 2.47E-01 | 6.79E-01 | 1.10E+00 |
| ATOX1 | 2.55E-07 | 1.33E-06 | -1.07E+00 | 2.31E-21 | 1.00E-20 | -1.34E+00 | 1.96E-01 | 8.32E-01 | 1.03E+00 | 1.66E-07 | 6.35E-07 | -1.29E+00 | 1.14E-05 | 4.07E-05 | -1.11E+00 | 4.07E-15 | 1.15E-14 | -1.43E+00 | 3.72E-02 | 1.37E-01 | 1.01E+00 | 9.38E-05 | 2.58E-04 | -1.19E+00 |
| TRIM38 | 2.56E-07 | 1.33E-06 | -1.22E+00 | 6.08E-04 | 2.63E-03 | 1.08E+00 | 1.75E-04 | 7.45E-04 | -1.26E+00 | 5.86E-05 | 2.24E-04 | -1.12E+00 | 1.13E-03 | 4.05E-03 | -1.04E+00 | 4.95E-04 | 1.39E-03 | -1.09E+00 | 3.97E-03 | 1.46E-02 | -1.01E+00 | 2.08E-04 | 5.71E-04 | -1.15E+00 |
| GLE1 | 2.56E-07 | 1.33E-06 | -1.58E+00 | 3.58E-04 | 1.55E-03 | -1.07E+00 | 5.28E-12 | 2.24E-11 | -1.78E+00 | 4.27E-08 | 1.63E-07 | -1.26E+00 | 2.97E-05 | 1.06E-04 | -1.65E+00 | 6.62E-02 | 1.87E-01 | 1.10E+00 | 9.29E-08 | 3.41E-07 | -1.21E+00 | 4.71E-03 | 1.29E-02 | 1.04E+00 |
| AC004801.1 | 2.56E-07 | 1.33E-06 | -2.11E+00 | 5.74E-06 | 2.48E-05 | -1.58E+00 | 2.72E-07 | 1.16E-06 | -2.13E+00 | 6.21E-04 | 2.37E-03 | -1.24E+00 | 2.23E-04 | 7.95E-04 | -1.94E+00 | 1.61E-03 | 4.53E-03 | -1.60E+00 | 5.52E-04 | 2.02E-03 | -1.53E+00 | 1.02E-02 | 2.79E-02 | -1.38E+00 |
| GPCPD1 | 2.56E-07 | 1.33E-06 | -1.14E+00 | 3.18E-07 | 1.37E-06 | -1.05E+00 | 1.10E-05 | 4.68E-05 | -1.33E+00 | 5.41E-06 | 2.07E-05 | -1.24E+00 | 6.97E-03 | 2.48E-02 | -1.05E+00 | 3.36E-04 | 9.47E-04 | -1.27E+00 | 1.41E-05 | 5.18E-05 | -1.21E+00 | 2.35E-07 | 6.45E-07 | -1.42E+00 |
| SYTL1 | 2.58E-07 | 1.34E-06 | -1.14E+00 | 1.10E-07 | 4.77E-07 | -1.03E+00 | 3.00E-06 | 1.28E-05 | -1.29E+00 | 1.63E-05 | 6.21E-05 | -1.06E+00 | 1.23E-06 | 4.39E-06 | -1.39E+00 | 1.21E-04 | 3.41E-04 | -1.07E+00 | 9.61E-02 | 3.53E-01 | 1.44E+00 | 3.02E-01 | 8.29E-01 | 1.20E+00 |
| RAB1B | 2.58E-07 | 1.34E-06 | -2.00E+00 | 1.06E-05 | 4.56E-05 | -1.45E+00 | 2.35E-06 | 1.00E-05 | -1.91E+00 | 8.12E-05 | 3.10E-04 | -1.36E+00 | 8.28E-07 | 2.95E-06 | -1.80E+00 | 9.77E-08 | 2.75E-07 | -1.99E+00 | 6.36E-22 | 2.33E-21 | -2.84E+00 | 1.18E-11 | 3.24E-11 | -1.67E+00 |
| ATF4 | 2.59E-07 | 1.35E-06 | 1.97E+00 | 3.99E-02 | 1.72E-01 | 1.71E+00 | 3.82E-03 | 1.62E-02 | 1.57E+00 | 1.26E-01 | 4.81E-01 | 1.52E+00 | 2.58E-02 | 9.18E-02 | 1.75E+00 | 2.96E-01 | 8.35E-01 | 1.55E+00 | 5.43E-04 | 1.99E-03 | 1.85E+00 | 4.53E-02 | 1.24E-01 | 1.60E+00 |
| THAP12 | 2.59E-07 | 1.35E-06 | -1.32E+00 | 8.22E-07 | 3.55E-06 | -1.19E+00 | 8.25E-03 | 3.50E-02 | -1.08E+00 | 1.50E-03 | 5.74E-03 | -1.02E+00 | 7.32E-06 | 2.61E-05 | -1.27E+00 | 1.79E-05 | 5.04E-05 | -1.19E+00 | 6.49E-07 | 2.38E-06 | -1.34E+00 | 2.45E-07 | 6.73E-07 | -1.38E+00 |
| B3GAT3 | 2.60E-07 | 1.36E-06 | -1.20E+00 | 5.91E-07 | 2.56E-06 | -1.08E+00 | 3.93E-03 | 1.67E-02 | 1.04E+00 | 1.95E-03 | 7.43E-03 | 1.14E+00 | 4.29E-07 | 1.53E-06 | -1.20E+00 | 3.92E-04 | 1.10E-03 | 1.07E+00 | 3.44E-03 | 1.26E-02 | -1.06E+00 | 3.34E-01 | 9.19E-01 | 1.29E+00 |
| CDKN2C | 2.60E-07 | 1.36E-06 | -3.65E+00 | 2.91E-06 | 1.26E-05 | -2.64E+00 | 5.81E-12 | 2.47E-11 | -3.32E+00 | 2.39E-11 | 9.15E-11 | -2.98E+00 | 1.17E-27 | 4.17E-27 | -8.17E+00 | 7.89E-27 | 2.22E-26 | -5.81E+00 | 2.43E-56 | 8.93E-56 | -2.83E+00 | 1.80E-53 | 4.93E-53 | -3.06E+00 |
| N4BP2L1 | 2.62E-07 | 1.36E-06 | -1.16E+00 | 1.10E-03 | 4.77E-03 | 1.14E+00 | 3.50E-06 | 1.49E-05 | -1.26E+00 | 1.07E-03 | 4.08E-03 | 1.04E+00 | 3.02E-06 | 1.08E-05 | -1.33E+00 | 2.90E-02 | 8.18E-02 | 1.09E+00 | 5.59E-07 | 2.05E-06 | -1.38E+00 | 4.28E-02 | 1.18E-01 | -1.00E+00 |
| TERF1 | 2.62E-07 | 1.36E-06 | -1.18E+00 | 3.52E-03 | 1.52E-02 | 1.16E+00 | 3.57E-03 | 1.52E-02 | -1.13E+00 | 2.73E-02 | 1.04E-01 | 1.12E+00 | 3.00E-03 | 1.07E-02 | -1.11E+00 | 2.24E-01 | 6.31E-01 | 1.27E+00 | 5.95E-05 | 2.18E-04 | -1.20E+00 | 3.54E-03 | 9.72E-03 | -1.05E+00 |
| TUFT1 | 2.63E-07 | 1.37E-06 | -1.77E+00 | 2.68E-12 | 1.16E-11 | -2.91E+00 | 1.15E-09 | 4.87E-09 | -2.21E+00 | 4.11E-14 | 1.57E-13 | -3.54E+00 | 1.18E-01 | 4.20E-01 | 1.03E+00 | 6.22E-02 | 1.75E-01 | -1.52E+00 | 2.03E-01 | 7.45E-01 | 1.23E+00 | 2.90E-01 | 7.96E-01 | 1.30E+00 |
| TMEM268 | 2.66E-07 | 1.39E-06 | -1.44E+00 | 2.22E-04 | 9.58E-04 | 1.00E+00 | 1.44E-02 | 6.11E-02 | -1.05E+00 | 3.62E-02 | 1.38E-01 | 1.14E+00 | 2.31E-01 | 8.24E-01 | 1.23E+00 | 3.23E-01 | 9.12E-01 | 1.31E+00 | 7.40E-03 | 2.72E-02 | -1.78E+00 | 1.15E-01 | 3.17E-01 | -1.14E+00 |
| THYN1 | 2.67E-07 | 1.39E-06 | -1.08E+00 | 2.22E-05 | 9.58E-05 | 1.09E+00 | 6.14E-04 | 2.61E-03 | -1.07E+00 | 5.38E-03 | 2.06E-02 | 1.13E+00 | 4.53E-09 | 1.61E-08 | -1.32E+00 | 5.10E-05 | 1.44E-04 | -1.06E+00 | 8.60E-06 | 3.16E-05 | -1.13E+00 | 2.20E-03 | 6.05E-03 | 1.01E+00 |
| RPP30 | 2.68E-07 | 1.40E-06 | -1.23E+00 | 2.32E-03 | 1.00E-02 | 1.13E+00 | 7.75E-05 | 3.29E-04 | -1.15E+00 | 4.18E-04 | 1.60E-03 | 1.06E+00 | 4.88E-06 | 1.74E-05 | -1.27E+00 | 9.24E-03 | 2.60E-02 | 1.01E+00 | 2.83E-05 | 1.04E-04 | -1.14E+00 | 1.17E-01 | 3.22E-01 | 1.14E+00 |
| ARRDC2 | 2.68E-07 | 1.40E-06 | -1.33E+00 | 2.15E-02 | 9.30E-02 | 1.24E+00 | 6.19E-07 | 2.63E-06 | -1.40E+00 | 3.26E-04 | 1.24E-03 | 1.10E+00 | 3.40E-07 | 1.21E-06 | -1.94E+00 | 1.05E-04 | 2.97E-04 | -1.34E+00 | 1.22E-08 | 4.47E-08 | -1.89E+00 | 1.42E-03 | 3.89E-03 | -1.25E+00 |
| ZNF333 | 2.68E-07 | 1.40E-06 | -1.36E+00 | 5.25E-03 | 2.27E-02 | 1.15E+00 | 7.69E-06 | 3.27E-05 | -1.36E+00 | 6.84E-04 | 2.61E-03 | 1.04E+00 | 9.04E-03 | 3.22E-02 | 1.01E+00 | 6.13E-02 | 1.73E-01 | 1.12E+00 | 1.05E-04 | 3.84E-04 | -1.18E+00 | 6.26E-03 | 1.72E-02 | -1.04E+00 |
| LINC00339 | 2.69E-07 | 1.40E-06 | -1.36E+00 | 2.38E-03 | 1.03E-02 | 1.12E+00 | 4.77E-12 | 2.03E-11 | -1.79E+00 | 4.84E-08 | 1.85E-07 | -1.33E+00 | 4.44E-07 | 1.58E-06 | -1.78E+00 | 8.42E-04 | 2.37E-03 | -1.20E+00 | 4.46E-09 | 1.64E-08 | -1.46E+00 | 1.79E-06 | 4.92E-06 | -1.23E+00 |
| LIX1L | 2.69E-07 | 1.40E-06 | -1.31E+00 | 8.74E-03 | 3.78E-02 | 1.21E+00 | 8.25E-05 | 3.51E-04 | -1.20E+00 | 2.28E-03 | 8.71E-03 | 1.09E+00 | 4.26E-05 | 1.52E-04 | -1.21E+00 | 7.93E-03 | 2.24E-02 | 1.10E+00 | 4.96E-07 | 1.82E-06 | -1.39E+00 | 6.85E-03 | 1.88E-02 | -1.04E+00 |
| RPP25L | 2.75E-07 | 1.43E-06 | -1.20E+00 | 4.90E-03 | 2.12E-02 | 1.22E+00 | 7.21E-13 | 3.06E-12 | -1.51E+00 | 1.06E-07 | 4.06E-07 | -1.11E+00 | 1.75E-09 | 6.23E-09 | -1.69E+00 | 2.15E-04 | 6.05E-04 | -1.07E+00 | 6.85E-14 | 2.51E-13 | -1.69E+00 | 1.20E-03 | 3.29E-03 | -1.06E+00 |
| LDOC1 | 2.75E-07 | 1.43E-06 | -1.30E+00 | 1.15E-06 | 4.96E-06 | -1.14E+00 | 4.35E-04 | 1.85E-03 | -1.35E+00 | 1.26E-02 | 4.81E-02 | 1.08E+00 | 6.64E-03 | 2.37E-02 | -1.44E+00 | 8.83E-02 | 2.49E-01 | 1.05E+00 | 7.04E-02 | 2.59E-01 | -1.08E+00 | 1.61E-01 | 4.42E-01 | 1.02E+00 |
| ERCC4 | 2.75E-07 | 1.43E-06 | -1.95E+00 | 4.40E-03 | 1.90E-02 | 1.04E+00 | 5.50E-06 | 2.34E-05 | -1.77E+00 | 1.31E-03 | 4.99E-03 | -1.13E+00 | 2.66E-04 | 9.50E-04 | -1.98E+00 | 2.83E-02 | 7.98E-02 | -1.03E+00 | 9.17E-17 | 3.36E-16 | -2.67E+00 | 4.89E-07 | 1.34E-06 | -1.45E+00 |
| FOPNL | 2.78E-07 | 1.45E-06 | -1.20E+00 | 1.46E-06 | 6.33E-06 | -1.06E+00 | 6.23E-08 | 2.65E-07 | -1.32E+00 | 9.09E-07 | 3.47E-06 | -1.13E+00 | 1.31E-07 | 4.67E-07 | -1.31E+00 | 2.70E-06 | 7.61E-06 | -1.26E+00 | 6.29E-06 | 2.31E-05 | -1.20E+00 | 3.83E-04 | 1.05E-03 | -1.09E+00 |
| SH2B2 | 2.78E-07 | 1.45E-06 | -2.39E+00 | 3.92E-05 | 1.69E-04 | -1.57E+00 | 8.49E-06 | 3.61E-05 | -1.45E+00 | 4.10E-07 | 1.57E-06 | -1.54E+00 | 8.56E-02 | 3.05E-01 | -1.31E+00 | 6.42E-02 | 1.81E-01 | -1.72E+00 | 1.84E-01 | 6.75E-01 | -1.33E+00 | 3.23E-01 | 8.86E-01 | -1.08E+00 |
| CAMK1D | 2.78E-07 | 1.45E-06 | -1.22E+00 | 5.11E-04 | 2.21E-03 | 1.13E+00 | 4.09E-02 | 1.74E-01 | -1.08E+00 | 1.17E-01 | 4.46E-01 | 1.19E+00 | 3.51E-04 | 1.25E-03 | -1.17E+00 | 1.07E-02 | 3.03E-02 | 1.02E+00 | 1.77E-06 | 6.49E-06 | -1.45E+00 | 1.03E-02 | 2.84E-02 | -1.08E+00 |
| FHIT | 2.79E-07 | 1.45E-06 | -1.18E+00 | 8.21E-07 | 3.55E-06 | -1.04E+00 | 6.58E-02 | 2.80E-01 | -1.01E+00 | 7.87E-02 | 3.01E-01 | 1.13E+00 | 8.67E-03 | 3.09E-02 | -1.03E+00 | 2.58E-01 | 7.27E-01 | 1.30E+00 | 2.15E-01 | 7.90E-01 | 1.19E+00 | 6.76E-03 | 1.86E-02 | 1.58E+00 |
| YTHDF3 | 2.79E-07 | 1.45E-06 | -1.12E+00 | 1.30E-15 | 5.61E-15 | -1.40E+00 | 5.22E-05 | 2.22E-04 | -1.23E+00 | 8.57E-11 | 3.27E-10 | -1.49E+00 | 1.69E-02 | 6.03E-02 | 1.04E+00 | 5.78E-07 | 1.63E-06 | -1.44E+00 | 8.41E-07 | 3.09E-06 | -1.24E+00 | 1.01E-09 | 2.76E-09 | -1.45E+00 |
| MAPKAP1 | 2.79E-07 | 1.45E-06 | -1.27E+00 | 4.26E-04 | 1.84E-03 | 1.08E+00 | 4.05E-05 | 1.72E-04 | -1.13E+00 | 2.07E-03 | 7.92E-03 | 1.10E+00 | 9.21E-04 | 3.28E-03 | -1.05E+00 | 3.61E-02 | 1.02E-01 | 1.13E+00 | 1.94E-03 | 7.10E-03 | 1.03E+00 | 3.15E-01 | 8.66E-01 | 1.26E+00 |
| TBP | 2.79E-07 | 1.46E-06 | -1.83E+00 | 2.56E-03 | 1.11E-02 | 1.02E+00 | 2.92E-07 | 1.24E-06 | -1.49E+00 | 2.74E-04 | 1.05E-03 | -1.06E+00 | 3.67E-06 | 1.31E-05 | -1.43E+00 | 1.57E-04 | 4.42E-04 | -1.24E+00 | 7.33E-09 | 2.69E-08 | -1.56E+00 | 2.65E-05 | 7.27E-05 | -1.22E+00 |
| KIAA1549 | 2.79E-07 | 1.46E-06 | -1.64E+00 | 8.64E-05 | 3.73E-04 | -1.12E+00 | 4.37E-05 | 1.85E-04 | -1.55E+00 | 7.44E-02 | 2.84E-01 | 1.20E+00 | 9.99E-03 | 3.56E-02 | -1.27E+00 | 1.37E-01 | 3.86E-01 | 1.06E+00 | 3.96E-05 | 1.45E-04 | -1.61E+00 | 4.97E-03 | 1.36E-02 | -1.16E+00 |
| LOXL1-AS1 | 2.80E-07 | 1.46E-06 | -1.36E+00 | 6.27E-05 | 2.71E-04 | -1.06E+00 | 1.98E-06 | 8.41E-06 | -1.65E+00 | 1.11E-04 | 4.24E-04 | -1.25E+00 | 2.31E-01 | 8.25E-01 | 1.01E+00 | 2.62E-01 | 7.38E-01 | -1.02E+00 | 2.31E-01 | 8.49E-01 | -1.12E+00 | 3.52E-01 | 9.68E-01 | 1.09E+00 |
| PUS3 | 2.80E-07 | 1.46E-06 | -1.41E+00 | 1.11E-04 | 4.80E-04 | -1.06E+00 | 5.40E-05 | 2.29E-04 | -1.42E+00 | 1.99E-03 | 7.58E-03 | -1.05E+00 | 5.21E-07 | 1.86E-06 | -1.74E+00 | 1.19E-04 | 3.35E-04 | -1.25E+00 | 3.39E-06 | 1.25E-05 | -1.56E+00 | 3.62E-04 | 9.94E-04 | -1.41E+00 |
| RPL26P27 | 2.81E-07 | 1.46E-06 | -1.59E+00 | 1.13E-09 | 4.87E-09 | -2.02E+00 | 4.64E-03 | 1.97E-02 | -1.29E+00 | 1.44E-04 | 5.48E-04 | -1.53E+00 | 1.04E-01 | 3.72E-01 | -1.10E+00 | 2.70E-01 | 7.61E-01 | 1.19E+00 | 4.18E-02 | 1.54E-01 | -1.04E+00 | 2.55E-02 | 7.00E-02 | -1.21E+00 |
| MORF4L1P1 | 2.81E-07 | 1.46E-06 | -1.07E+00 | 1.52E-25 | 6.55E-25 | -1.64E+00 | 2.29E-05 | 9.73E-05 | -1.16E+00 | 6.39E-18 | 2.44E-17 | -1.61E+00 | 2.48E-06 | 8.85E-06 | -1.22E+00 | 4.41E-14 | 1.24E-13 | -1.69E+00 | 1.09E-04 | 4.01E-04 | -1.15E+00 | 3.92E-14 | 1.08E-13 | -1.59E+00 |
| RPL8 | 2.81E-07 | 1.47E-06 | 1.13E+00 | 3.80E-18 | 1.64E-17 | 1.23E+00 | 5.61E-11 | 2.38E-10 | 1.21E+00 | 3.54E-16 | 1.35E-15 | 1.26E+00 | 8.06E-07 | 2.87E-06 | 1.18E+00 | 3.73E-21 | 1.05E-20 | 1.34E+00 | 6.15E-15 | 2.26E-14 | 1.29E+00 | 7.27E-29 | 2.00E-28 | 1.39E+00 |
| CHMP6 | 2.81E-07 | 1.47E-06 | -1.42E+00 | 3.19E-05 | 1.38E-04 | -1.06E+00 | 7.85E-12 | 3.34E-11 | -2.08E+00 | 1.74E-06 | 6.65E-06 | -1.27E+00 | 5.74E-06 | 2.05E-05 | -1.80E+00 | 4.33E-03 | 1.22E-02 | -1.07E+00 | 4.46E-07 | 1.64E-06 | -1.47E+00 | 7.18E-06 | 1.97E-05 | -1.31E+00 |
| MBTPS1 | 2.82E-07 | 1.47E-06 | -1.21E+00 | 3.30E-06 | 1.43E-05 | 1.02E+00 | 5.34E-09 | 2.27E-08 | -1.46E+00 | 6.75E-06 | 2.58E-05 | -1.04E+00 | 3.21E-03 | 1.14E-02 | -1.02E+00 | 1.91E-02 | 5.38E-02 | 1.16E+00 | 2.69E-02 | 9.87E-02 | 1.04E+00 | 2.30E-01 | 6.32E-01 | 1.15E+00 |
| TMEM243 | 2.82E-07 | 1.47E-06 | -1.08E+00 | 4.02E-05 | 1.74E-04 | 1.09E+00 | 1.17E-03 | 4.98E-03 | -1.14E+00 | 3.49E-04 | 1.33E-03 | -1.07E+00 | 2.03E-04 | 7.25E-04 | -1.11E+00 | 4.06E-05 | 1.14E-04 | -1.19E+00 | 1.43E-01 | 5.26E-01 | 1.07E+00 | 4.62E-02 | 1.27E-01 | -1.04E+00 |
| POLM | 2.82E-07 | 1.47E-06 | -1.19E+00 | 2.10E-04 | 9.08E-04 | 1.14E+00 | 9.27E-03 | 3.94E-02 | -1.11E+00 | 1.01E-01 | 3.86E-01 | 1.29E+00 | 2.34E-01 | 8.34E-01 | 1.22E+00 | 1.21E-01 | 3.42E-01 | 1.58E+00 | 2.04E-05 | 7.50E-05 | -1.53E+00 | 1.30E-02 | 3.57E-02 | -1.10E+00 |
| UBE2Q2P1 | 2.83E-07 | 1.47E-06 | -1.21E+00 | 1.77E-05 | 7.65E-05 | 1.00E+00 | 2.77E-03 | 1.18E-02 | -1.08E+00 | 3.04E-03 | 1.16E-02 | 1.06E+00 | 5.81E-02 | 2.07E-01 | 1.25E+00 | 9.57E-03 | 2.70E-02 | 1.08E+00 | 1.00E-01 | 3.69E-01 | 1.17E+00 | 5.42E-02 | 1.49E-01 | -1.04E+00 |
| AC244669.1 | 2.84E-07 | 1.48E-06 | -1.37E+00 | 6.88E-08 | 2.97E-07 | -1.38E+00 | 3.29E-08 | 1.40E-07 | -1.81E+00 | 1.40E-11 | 5.33E-11 | -2.12E+00 | 2.13E-03 | 7.59E-03 | -1.11E+00 | 1.83E-03 | 5.15E-03 | -1.23E+00 | 2.58E-03 | 9.46E-03 | -1.26E+00 | 1.47E-03 | 4.04E-03 | -1.41E+00 |
| AC139426.3 | 2.84E-07 | 1.48E-06 | -2.02E+00 | 1.31E-02 | 5.68E-02 | 1.04E+00 | 8.19E-08 | 3.48E-07 | -2.25E+00 | 2.90E-04 | 1.11E-03 | -1.25E+00 | 2.98E-02 | 1.06E-01 | -2.25E+00 | 4.74E-02 | 1.33E-01 | -2.11E+00 | 1.11E-01 | 4.08E-01 | -1.28E+00 | 2.36E-01 | 6.50E-01 | -1.11E+00 |
| CREB3L2 | 2.85E-07 | 1.49E-06 | -1.15E+00 | 7.24E-05 | 3.13E-04 | 1.08E+00 | 2.11E-02 | 8.96E-02 | -1.02E+00 | 1.60E-02 | 6.12E-02 | 1.08E+00 | 6.88E-13 | 2.45E-12 | -1.78E+00 | 2.49E-07 | 7.01E-07 | -1.26E+00 | 7.34E-12 | 2.69E-11 | -1.65E+00 | 1.32E-03 | 3.62E-03 | -1.01E+00 |
| HIST1H4I | 2.85E-07 | 1.49E-06 | -1.84E+00 | 9.45E-05 | 4.08E-04 | -1.25E+00 | 3.36E-06 | 1.43E-05 | -1.97E+00 | 3.21E-04 | 1.23E-03 | -1.35E+00 | 2.47E-02 | 8.82E-02 | -1.69E+00 | 2.14E-01 | 6.04E-01 | -1.01E+00 | 3.72E-08 | 1.36E-07 | -1.96E+00 | 1.88E-06 | 5.16E-06 | -1.58E+00 |
| SEC13 | 2.87E-07 | 1.49E-06 | -1.19E+00 | 1.01E-06 | 4.36E-06 | -1.03E+00 | 3.25E-02 | 1.38E-01 | 1.05E+00 | 4.26E-02 | 1.63E-01 | 1.11E+00 | 4.38E-08 | 1.56E-07 | -1.27E+00 | 3.04E-05 | 8.58E-05 | -1.05E+00 | 2.65E-05 | 9.72E-05 | -1.16E+00 | 9.45E-03 | 2.60E-02 | 1.02E+00 |
| SPIN2B | 2.87E-07 | 1.50E-06 | -1.32E+00 | 1.70E-06 | 7.33E-06 | -1.12E+00 | 1.23E-07 | 5.24E-07 | -1.54E+00 | 2.06E-05 | 7.88E-05 | -1.15E+00 | 1.27E-05 | 4.52E-05 | -1.44E+00 | 1.44E-04 | 4.06E-04 | -1.17E+00 | 1.66E-04 | 6.08E-04 | -1.29E+00 | 4.79E-03 | 1.32E-02 | -1.08E+00 |
| ZBTB25 | 2.88E-07 | 1.50E-06 | -1.37E+00 | 3.03E-06 | 1.31E-05 | -1.14E+00 | 1.54E-02 | 6.53E-02 | -1.11E+00 | 6.88E-03 | 2.63E-02 | 1.03E+00 | 2.89E-04 | 1.03E-03 | -1.22E+00 | 6.02E-02 | 1.70E-01 | 1.13E+00 | 2.37E-06 | 8.68E-06 | -1.58E+00 | 2.64E-02 | 7.24E-02 | -1.03E+00 |
| LINC01220 | 2.89E-07 | 1.51E-06 | 4.14E+00 | 1.46E-01 | 6.29E-01 | 1.53E+00 | 4.66E-05 | 1.98E-04 | 2.60E+00 | 2.11E-01 | 8.06E-01 | 1.02E+00 | 6.61E-02 | 2.36E-01 | 2.20E+00 | 3.31E-01 | 9.34E-01 | 1.23E+00 | 4.51E-02 | 1.65E-01 | 1.84E+00 | 2.82E-01 | 7.75E-01 | 1.28E+00 |
| MGLL | 2.89E-07 | 1.51E-06 | -1.13E+00 | 2.65E-05 | 1.14E-04 | 1.07E+00 | 1.08E-02 | 4.57E-02 | -1.04E+00 | 7.12E-05 | 2.72E-04 | -1.02E+00 | 1.08E-03 | 3.87E-03 | -1.08E+00 | 1.43E-02 | 4.03E-02 | 1.10E+00 | 1.18E-01 | 4.32E-01 | 1.09E+00 | 1.20E-02 | 3.30E-02 | -1.06E+00 |
| POLR3GL | 2.90E-07 | 1.51E-06 | -1.08E+00 | 2.71E-04 | 1.17E-03 | 1.14E+00 | 1.69E-04 | 7.17E-04 | -1.22E+00 | 1.36E-04 | 5.20E-04 | -1.07E+00 | 1.68E-03 | 5.99E-03 | -1.01E+00 | 8.85E-03 | 2.49E-02 | 1.05E+00 | 1.69E-02 | 6.22E-02 | 1.06E+00 | 1.15E-02 | 3.16E-02 | 1.09E+00 |
| LACTB | 2.92E-07 | 1.52E-06 | -1.27E+00 | 5.89E-06 | 2.54E-05 | -1.04E+00 | 4.04E-06 | 1.72E-05 | -1.35E+00 | 5.64E-06 | 2.16E-05 | -1.19E+00 | 5.06E-09 | 1.80E-08 | -1.70E+00 | 8.73E-05 | 2.46E-04 | -1.19E+00 | 9.23E-04 | 3.39E-03 | -1.24E+00 | 1.82E-04 | 5.00E-04 | -1.39E+00 |
| URGCP | 2.92E-07 | 1.52E-06 | -1.58E+00 | 1.38E-05 | 5.94E-05 | -1.27E+00 | 1.15E-06 | 4.87E-06 | -1.70E+00 | 2.77E-03 | 1.06E-02 | -1.00E+00 | 2.63E-03 | 9.36E-03 | -1.37E+00 | 1.27E-01 | 3.59E-01 | 1.26E+00 | 3.26E-09 | 1.20E-08 | -1.54E+00 | 1.81E-04 | 4.96E-04 | -1.15E+00 |
| ZNF566 | 2.96E-07 | 1.54E-06 | -1.70E+00 | 2.08E-03 | 8.99E-03 | 1.06E+00 | 1.51E-05 | 6.42E-05 | -1.78E+00 | 3.07E-02 | 1.17E-01 | 1.02E+00 | 4.91E-03 | 1.75E-02 | -1.17E+00 | 1.23E-02 | 3.46E-02 | -1.09E+00 | 2.86E-07 | 1.05E-06 | -1.66E+00 | 2.01E-04 | 5.52E-04 | -1.35E+00 |
| DOK3 | 2.96E-07 | 1.54E-06 | -1.51E+00 | 1.65E-05 | 7.12E-05 | -1.11E+00 | 2.23E-10 | 9.46E-10 | -1.96E+00 | 4.50E-07 | 1.72E-06 | -1.41E+00 | 7.63E-04 | 2.72E-03 | -1.58E+00 | 2.16E-02 | 6.10E-02 | -1.14E+00 | 9.80E-03 | 3.60E-02 | -1.12E+00 | 5.02E-03 | 1.38E-02 | -1.37E+00 |
| FRG1HP | 2.96E-07 | 1.54E-06 | -1.12E+00 | 3.58E-06 | 1.55E-05 | 1.03E+00 | 1.55E-05 | 6.59E-05 | -1.21E+00 | 8.48E-04 | 3.24E-03 | 1.02E+00 | 3.33E-06 | 1.19E-05 | -1.13E+00 | 6.79E-07 | 1.91E-06 | -1.14E+00 | 4.51E-02 | 1.66E-01 | 1.12E+00 | 8.24E-03 | 2.26E-02 | 1.00E+00 |
| ACTR2 | 2.97E-07 | 1.55E-06 | -1.13E+00 | 1.85E-07 | 8.01E-07 | -1.04E+00 | 1.24E-03 | 5.27E-03 | -1.11E+00 | 4.52E-03 | 1.73E-02 | -1.02E+00 | 7.13E-07 | 2.54E-06 | -1.24E+00 | 7.43E-06 | 2.10E-05 | -1.27E+00 | 2.98E-03 | 1.09E-02 | -1.17E+00 | 2.85E-02 | 7.83E-02 | -1.09E+00 |
| PLCB2 | 2.98E-07 | 1.55E-06 | -1.32E+00 | 5.13E-06 | 2.22E-05 | -1.11E+00 | 3.84E-06 | 1.63E-05 | -1.47E+00 | 7.87E-04 | 3.01E-03 | -1.08E+00 | 4.01E-04 | 1.43E-03 | -1.16E+00 | 9.34E-04 | 2.63E-03 | -1.13E+00 | 1.93E-04 | 7.07E-04 | -1.37E+00 | 1.87E-02 | 5.13E-02 | -1.09E+00 |
| ABCC5 | 2.99E-07 | 1.56E-06 | -1.61E+00 | 1.22E-03 | 5.29E-03 | -1.00E+00 | 1.38E-04 | 5.85E-04 | -1.55E+00 | 7.08E-02 | 2.71E-01 | 1.11E+00 | 1.72E-10 | 6.14E-10 | -1.55E+00 | 9.72E-04 | 2.74E-03 | -1.02E+00 | 8.08E-18 | 2.96E-17 | -1.68E+00 | 5.56E-08 | 1.53E-07 | -1.23E+00 |
| ABHD14A | 2.99E-07 | 1.56E-06 | -1.20E+00 | 1.10E-03 | 4.75E-03 | 1.19E+00 | 4.38E-02 | 1.86E-01 | 1.07E+00 | 5.03E-02 | 1.92E-01 | 1.27E+00 | 2.29E-02 | 8.16E-02 | 1.01E+00 | 1.80E-01 | 5.07E-01 | 1.20E+00 | 4.76E-03 | 1.75E-02 | 1.03E+00 | 1.09E-01 | 2.98E-01 | 1.12E+00 |
| ZNRD1ASP | 3.00E-07 | 1.56E-06 | -2.16E+00 | 1.95E-01 | 8.44E-01 | 1.40E+00 | 7.03E-06 | 2.99E-05 | -2.42E+00 | 9.97E-02 | 3.81E-01 | 1.13E+00 | 1.93E-05 | 6.87E-05 | -2.03E+00 | 9.87E-02 | 2.78E-01 | 1.08E+00 | 1.82E-04 | 6.69E-04 | -1.88E+00 | 2.76E-01 | 7.58E-01 | 1.24E+00 |
| OSBPL6 | 3.00E-07 | 1.56E-06 | -2.88E+00 | 1.08E-05 | 4.69E-05 | -1.92E+00 | 1.25E-02 | 5.31E-02 | -1.60E+00 | 6.00E-02 | 2.29E-01 | -1.13E+00 | 3.34E-03 | 1.19E-02 | -1.78E+00 | 1.08E-01 | 3.04E-01 | 1.05E+00 | 2.98E-56 | 1.09E-55 | -3.76E+00 | 1.00E-21 | 2.75E-21 | -1.66E+00 |
| ARMC6 | 3.03E-07 | 1.58E-06 | -1.53E+00 | 1.34E-04 | 5.80E-04 | -1.02E+00 | 5.82E-06 | 2.47E-05 | -1.31E+00 | 1.15E-03 | 4.40E-03 | 1.05E+00 | 1.79E-04 | 6.40E-04 | -1.33E+00 | 1.61E-01 | 4.55E-01 | 1.30E+00 | 5.71E-11 | 2.10E-10 | -1.41E+00 | 9.71E-04 | 2.67E-03 | 1.03E+00 |
| SMIM8 | 3.04E-07 | 1.58E-06 | -1.25E+00 | 1.91E-03 | 8.26E-03 | 1.21E+00 | 1.01E-07 | 4.29E-07 | -1.49E+00 | 1.09E-03 | 4.15E-03 | 1.00E+00 | 4.88E-06 | 1.74E-05 | -1.31E+00 | 7.07E-02 | 1.99E-01 | 1.15E+00 | 3.90E-06 | 1.43E-05 | -1.36E+00 | 2.91E-02 | 7.99E-02 | -1.03E+00 |
| SLC25A24 | 3.05E-07 | 1.59E-06 | -1.51E+00 | 2.43E-04 | 1.05E-03 | -1.07E+00 | 4.07E-05 | 1.73E-04 | -1.31E+00 | 1.24E-05 | 4.72E-05 | -1.22E+00 | 2.15E-04 | 7.66E-04 | -1.34E+00 | 9.30E-04 | 2.62E-03 | -1.23E+00 | 1.98E-09 | 7.26E-09 | -1.66E+00 | 6.80E-07 | 1.87E-06 | -1.40E+00 |
| MAST1 | 3.06E-07 | 1.60E-06 | -1.84E+00 | 2.35E-05 | 1.02E-04 | -1.32E+00 | 4.15E-07 | 1.76E-06 | -1.91E+00 | 2.23E-05 | 8.52E-05 | -1.34E+00 | 6.04E-06 | 2.15E-05 | -1.80E+00 | 1.60E-04 | 4.52E-04 | -1.48E+00 | 4.97E-03 | 1.82E-02 | -1.35E+00 | 3.10E-02 | 8.53E-02 | -1.13E+00 |
| EIF4G2 | 3.09E-07 | 1.61E-06 | -1.03E+00 | 7.31E-11 | 3.16E-10 | -1.00E+00 | 1.05E-02 | 4.45E-02 | -1.07E+00 | 3.46E-03 | 1.32E-02 | -1.05E+00 | 2.05E-01 | 7.31E-01 | 1.24E+00 | 1.43E-01 | 4.02E-01 | 1.21E+00 | 1.66E-01 | 6.10E-01 | 1.19E+00 | 1.20E-01 | 3.29E-01 | 1.17E+00 |
| SYT11 | 3.10E-07 | 1.62E-06 | -1.28E+00 | 6.22E-05 | 2.69E-04 | -1.04E+00 | 1.11E-03 | 4.72E-03 | -1.21E+00 | 4.59E-03 | 1.75E-02 | -1.03E+00 | 1.41E-03 | 5.02E-03 | -1.15E+00 | 3.09E-02 | 8.71E-02 | 1.12E+00 | 7.40E-02 | 2.72E-01 | 1.14E+00 | 4.16E-02 | 1.14E-01 | 1.01E+00 |
| DPP7 | 3.10E-07 | 1.62E-06 | -1.03E+00 | 1.07E-08 | 4.64E-08 | -1.02E+00 | 1.41E-01 | 5.99E-01 | 1.18E+00 | 9.16E-02 | 3.50E-01 | 1.16E+00 | 7.41E-06 | 2.64E-05 | -1.09E+00 | 4.39E-05 | 1.24E-04 | -1.00E+00 | 1.11E-01 | 4.07E-01 | 1.31E+00 | 4.44E-02 | 1.22E-01 | 1.44E+00 |
| HDGFL3 | 3.10E-07 | 1.62E-06 | -1.10E+00 | 4.06E-08 | 1.75E-07 | -1.05E+00 | 4.57E-02 | 1.94E-01 | 1.01E+00 | 7.39E-03 | 2.82E-02 | 1.08E+00 | 3.49E-05 | 1.24E-04 | -1.17E+00 | 1.02E-06 | 2.88E-06 | -1.29E+00 | 4.86E-06 | 1.78E-05 | -1.20E+00 | 1.57E-04 | 4.32E-04 | -1.12E+00 |
| PLA2G6 | 3.10E-07 | 1.62E-06 | -1.79E+00 | 2.35E-04 | 1.01E-03 | -1.12E+00 | 1.10E-04 | 4.66E-04 | -1.66E+00 | 8.36E-04 | 3.19E-03 | -1.30E+00 | 2.43E-01 | 8.66E-01 | 1.42E+00 | 2.47E-01 | 6.95E-01 | 1.70E+00 | 2.74E-06 | 1.00E-05 | -1.50E+00 | 3.93E-08 | 1.08E-07 | -1.67E+00 |
| BEND7 | 3.11E-07 | 1.62E-06 | -1.23E+00 | 1.50E-05 | 6.50E-05 | 1.02E+00 | 3.37E-03 | 1.43E-02 | -1.10E+00 | 4.85E-03 | 1.85E-02 | 1.01E+00 | 4.61E-02 | 1.64E-01 | -1.41E+00 | 3.00E-01 | 8.47E-01 | 1.25E+00 | 2.17E-01 | 7.97E-01 | 1.03E+00 | 3.55E-01 | 9.76E-01 | 1.27E+00 |
| TUBGCP5 | 3.12E-07 | 1.62E-06 | -1.75E+00 | 1.61E-04 | 6.96E-04 | -1.14E+00 | 2.63E-03 | 1.12E-02 | -1.14E+00 | 2.14E-04 | 8.16E-04 | -1.10E+00 | 4.25E-02 | 1.51E-01 | 1.11E+00 | 6.85E-03 | 1.93E-02 | -1.04E+00 | 7.15E-03 | 2.63E-02 | -1.11E+00 | 9.05E-02 | 2.49E-01 | 1.13E+00 |
| AC026362.2 | 3.12E-07 | 1.62E-06 | -2.52E+00 | 1.25E-03 | 5.40E-03 | -1.27E+00 | 1.09E-03 | 4.63E-03 | -2.19E+00 | 1.40E-01 | 5.37E-01 | 1.14E+00 | 1.60E-06 | 5.70E-06 | -2.22E+00 | 2.95E-01 | 8.30E-01 | 1.32E+00 | 1.15E-08 | 4.23E-08 | -2.21E+00 | 7.63E-02 | 2.10E-01 | 1.00E+00 |
| SECISBP2L | 3.12E-07 | 1.63E-06 | -1.35E+00 | 1.41E-05 | 6.11E-05 | -1.08E+00 | 4.40E-03 | 1.87E-02 | -1.15E+00 | 2.78E-04 | 1.06E-03 | -1.22E+00 | 2.41E-01 | 8.61E-01 | 1.32E+00 | 4.28E-02 | 1.21E-01 | 1.08E+00 | 6.05E-07 | 2.22E-06 | -1.23E+00 | 1.73E-05 | 4.75E-05 | -1.26E+00 |
| SULT1A1 | 3.12E-07 | 1.63E-06 | -1.20E+00 | 7.55E-04 | 3.26E-03 | 1.22E+00 | 2.00E-04 | 8.49E-04 | -1.21E+00 | 1.55E-03 | 5.94E-03 | -1.00E+00 | 1.32E-05 | 4.71E-05 | -1.18E+00 | 5.90E-02 | 1.66E-01 | 1.20E+00 | 3.58E-05 | 1.31E-04 | -1.13E+00 | 2.07E-03 | 5.70E-03 | -1.11E+00 |
| IMPA1 | 3.13E-07 | 1.63E-06 | -1.26E+00 | 7.06E-04 | 3.05E-03 | 1.17E+00 | 7.62E-11 | 3.24E-10 | -1.65E+00 | 3.40E-07 | 1.30E-06 | -1.21E+00 | 5.78E-06 | 2.06E-05 | -1.26E+00 | 1.84E-05 | 5.20E-05 | -1.19E+00 | 1.67E-08 | 6.15E-08 | -1.55E+00 | 1.38E-05 | 3.78E-05 | -1.33E+00 |
| FECH | 3.13E-07 | 1.63E-06 | -1.59E+00 | 9.56E-06 | 4.13E-05 | -1.25E+00 | 3.60E-05 | 1.53E-04 | -1.32E+00 | 3.26E-04 | 1.25E-03 | -1.06E+00 | 2.81E-12 | 1.00E-11 | -1.69E+00 | 1.95E-04 | 5.51E-04 | -1.06E+00 | 3.36E-07 | 1.23E-06 | -1.26E+00 | 3.09E-04 | 8.49E-04 | -1.10E+00 |
| DEPTOR | 3.13E-07 | 1.63E-06 | -1.23E+00 | 1.18E-06 | 5.12E-06 | -1.09E+00 | 3.08E-07 | 1.31E-06 | -1.38E+00 | 5.48E-07 | 2.09E-06 | -1.22E+00 | 1.95E-30 | 6.96E-30 | -2.53E+00 | 1.43E-17 | 4.03E-17 | -1.65E+00 | 2.61E-15 | 9.57E-15 | -2.18E+00 | 7.26E-06 | 1.99E-05 | -1.42E+00 |
| TOB1 | 3.16E-07 | 1.65E-06 | -1.09E+00 | 8.84E-26 | 3.82E-25 | -2.07E+00 | 7.19E-09 | 3.05E-08 | -1.45E+00 | 3.90E-23 | 1.49E-22 | -2.49E+00 | 2.08E-06 | 7.41E-06 | -1.83E+00 | 1.97E-10 | 5.56E-10 | -3.36E+00 | 4.23E-11 | 1.55E-10 | -1.61E+00 | 3.75E-16 | 1.03E-15 | -2.72E+00 |
| EHD4 | 3.16E-07 | 1.65E-06 | -1.77E+00 | 2.96E-09 | 1.28E-08 | -2.25E+00 | 6.66E-05 | 2.83E-04 | -1.59E+00 | 6.12E-04 | 2.34E-03 | -1.28E+00 | 3.63E-09 | 1.29E-08 | -1.95E+00 | 1.40E-09 | 3.95E-09 | -2.14E+00 | 1.50E-03 | 5.49E-03 | -1.09E+00 | 1.05E-02 | 2.88E-02 | -1.05E+00 |
| NOTCH1 | 3.20E-07 | 1.67E-06 | -1.50E+00 | 2.59E-03 | 1.12E-02 | 1.08E+00 | 1.55E-09 | 6.60E-09 | -1.64E+00 | 3.27E-04 | 1.25E-03 | 1.05E+00 | 2.54E-02 | 9.05E-02 | -1.51E+00 | 1.02E-01 | 2.87E-01 | -1.00E+00 | 3.06E-02 | 1.12E-01 | -1.39E+00 | 2.51E-01 | 6.89E-01 | 1.09E+00 |
| FAM8A1 | 3.21E-07 | 1.67E-06 | -1.22E+00 | 3.45E-04 | 1.49E-03 | 1.09E+00 | 6.87E-06 | 2.92E-05 | -1.51E+00 | 6.58E-04 | 2.51E-03 | -1.09E+00 | 1.70E-02 | 6.06E-02 | 1.06E+00 | 3.42E-01 | 9.65E-01 | 1.30E+00 | 7.03E-05 | 2.58E-04 | -1.14E+00 | 1.46E-03 | 4.01E-03 | -1.08E+00 |
| OARD1 | 3.24E-07 | 1.69E-06 | -1.03E+00 | 3.12E-09 | 1.35E-08 | 1.03E+00 | 8.07E-07 | 3.43E-06 | -1.26E+00 | 1.37E-05 | 5.22E-05 | -1.08E+00 | 1.98E-06 | 7.07E-06 | -1.13E+00 | 1.80E-03 | 5.06E-03 | 1.05E+00 | 7.28E-04 | 2.67E-03 | -1.06E+00 | 4.83E-03 | 1.33E-02 | -1.06E+00 |
| BCAS3 | 3.25E-07 | 1.69E-06 | -1.46E+00 | 7.76E-04 | 3.35E-03 | 1.11E+00 | 7.40E-04 | 3.14E-03 | -1.32E+00 | 1.85E-02 | 7.06E-02 | -1.00E+00 | 2.08E-05 | 7.40E-05 | -1.47E+00 | 7.59E-03 | 2.14E-02 | 1.01E+00 | 3.71E-04 | 1.36E-03 | -1.44E+00 | 9.61E-03 | 2.64E-02 | -1.17E+00 |
| AP3B1 | 3.30E-07 | 1.72E-06 | -1.28E+00 | 9.93E-06 | 4.29E-05 | -1.03E+00 | 7.38E-09 | 3.14E-08 | -1.38E+00 | 4.89E-08 | 1.87E-07 | -1.18E+00 | 1.88E-05 | 6.70E-05 | -1.15E+00 | 2.42E-04 | 6.83E-04 | -1.07E+00 | 6.29E-07 | 2.31E-06 | -1.30E+00 | 8.38E-04 | 2.30E-03 | -1.07E+00 |
| PLEKHA1 | 3.30E-07 | 1.72E-06 | -1.55E+00 | 1.52E-02 | 6.55E-02 | 1.18E+00 | 5.71E-03 | 2.42E-02 | -1.25E+00 | 1.15E-01 | 4.38E-01 | 1.15E+00 | 2.69E-02 | 9.61E-02 | -1.16E+00 | 3.41E-01 | 9.61E-01 | 1.56E+00 | 3.47E-04 | 1.27E-03 | -1.56E+00 | 6.04E-02 | 1.66E-01 | -1.07E+00 |
| TRAF7 | 3.30E-07 | 1.72E-06 | -1.54E+00 | 3.21E-06 | 1.39E-05 | -1.28E+00 | 9.97E-04 | 4.24E-03 | -1.18E+00 | 1.13E-06 | 4.31E-06 | -1.27E+00 | 6.64E-08 | 2.37E-07 | -1.53E+00 | 2.96E-05 | 8.35E-05 | -1.17E+00 | 2.20E-08 | 8.06E-08 | -1.29E+00 | 2.35E-04 | 6.46E-04 | -1.05E+00 |
| ZCCHC10 | 3.30E-07 | 1.72E-06 | -1.22E+00 | 2.63E-07 | 1.14E-06 | -1.07E+00 | 1.52E-11 | 6.45E-11 | -1.82E+00 | 8.35E-10 | 3.19E-09 | -1.45E+00 | 4.49E-04 | 1.60E-03 | -1.31E+00 | 8.92E-03 | 2.51E-02 | -1.01E+00 | 1.91E-13 | 7.03E-13 | -1.75E+00 | 1.08E-07 | 2.98E-07 | -1.36E+00 |
| TOR2A | 3.31E-07 | 1.73E-06 | -1.47E+00 | 1.16E-05 | 5.01E-05 | -1.15E+00 | 8.13E-07 | 3.46E-06 | -1.30E+00 | 1.79E-07 | 6.82E-07 | -1.17E+00 | 1.23E-08 | 4.40E-08 | -1.84E+00 | 4.57E-04 | 1.29E-03 | -1.15E+00 | 2.52E-10 | 9.24E-10 | -1.38E+00 | 4.31E-08 | 1.18E-07 | -1.21E+00 |
| NAGK | 3.32E-07 | 1.73E-06 | -1.29E+00 | 7.20E-04 | 3.11E-03 | 1.09E+00 | 1.61E-09 | 6.86E-09 | -1.79E+00 | 4.80E-05 | 1.83E-04 | -1.19E+00 | 2.40E-07 | 8.55E-07 | -1.75E+00 | 4.61E-04 | 1.30E-03 | -1.22E+00 | 1.19E-06 | 4.35E-06 | -1.51E+00 | 9.34E-03 | 2.56E-02 | -1.05E+00 |
| MRPS22 | 3.32E-07 | 1.73E-06 | -1.14E+00 | 2.67E-06 | 1.16E-05 | 1.01E+00 | 8.65E-06 | 3.67E-05 | -1.18E+00 | 6.16E-06 | 2.35E-05 | -1.06E+00 | 4.79E-05 | 1.71E-04 | -1.25E+00 | 1.64E-02 | 4.62E-02 | 1.08E+00 | 4.43E-16 | 1.63E-15 | -1.53E+00 | 5.80E-06 | 1.59E-05 | -1.09E+00 |
| CCDC122 | 3.34E-07 | 1.74E-06 | -1.56E+00 | 5.98E-06 | 2.58E-05 | -1.26E+00 | 3.05E-03 | 1.30E-02 | -1.24E+00 | 1.27E-02 | 4.87E-02 | -1.02E+00 | 4.20E-04 | 1.50E-03 | -1.34E+00 | 5.06E-03 | 1.43E-02 | -1.13E+00 | 4.26E-05 | 1.56E-04 | -1.33E+00 | 5.43E-04 | 1.49E-03 | -1.22E+00 |
| ADA | 3.36E-07 | 1.75E-06 | -1.13E+00 | 3.07E-10 | 1.33E-09 | -1.13E+00 | 1.95E-02 | 8.30E-02 | -1.03E+00 | 1.46E-05 | 5.59E-05 | -1.11E+00 | 1.76E-06 | 6.28E-06 | -1.28E+00 | 3.96E-06 | 1.12E-05 | -1.25E+00 | 5.16E-07 | 1.90E-06 | -1.45E+00 | 1.51E-04 | 4.16E-04 | -1.26E+00 |
| SNHG7 | 3.38E-07 | 1.76E-06 | 1.49E+00 | 1.28E-03 | 5.52E-03 | 1.39E+00 | 2.05E-02 | 8.72E-02 | -1.01E+00 | 5.27E-02 | 2.01E-01 | 1.05E+00 | 5.13E-04 | 1.83E-03 | 1.42E+00 | 3.53E-02 | 9.94E-02 | 1.29E+00 | 8.62E-05 | 3.16E-04 | 1.53E+00 | 2.65E-01 | 7.29E-01 | 1.15E+00 |
| PARD3B | 3.38E-07 | 1.76E-06 | -1.69E+00 | 2.42E-05 | 1.04E-04 | -1.17E+00 | 2.03E-02 | 8.64E-02 | -1.15E+00 | 1.53E-01 | 5.83E-01 | 1.29E+00 | 6.32E-02 | 2.25E-01 | -1.19E+00 | 1.72E-01 | 4.85E-01 | 1.15E+00 | 2.71E-02 | 9.96E-02 | -1.14E+00 | 3.59E-01 | 9.87E-01 | 1.29E+00 |
| TRIQK | 3.40E-07 | 1.77E-06 | -1.23E+00 | 5.99E-05 | 2.59E-04 | 1.03E+00 | 1.86E-04 | 7.92E-04 | -1.31E+00 | 1.09E-03 | 4.18E-03 | -1.01E+00 | 6.46E-06 | 2.30E-05 | -1.59E+00 | 5.31E-04 | 1.50E-03 | -1.24E+00 | 4.09E-04 | 1.50E-03 | -1.29E+00 | 1.01E-02 | 2.78E-02 | -1.14E+00 |
| GPR35 | 3.42E-07 | 1.78E-06 | -2.05E+00 | 2.09E-04 | 9.02E-04 | -1.23E+00 | 4.81E-04 | 2.04E-03 | -1.29E+00 | 1.48E-04 | 5.65E-04 | -1.30E+00 | 2.36E-01 | 8.43E-01 | 1.22E+00 | 3.23E-01 | 9.11E-01 | 1.10E+00 | 2.40E-01 | 8.81E-01 | 1.02E+00 | 2.79E-01 | 7.68E-01 | -1.24E+00 |
| TUBGCP2 | 3.42E-07 | 1.78E-06 | -1.12E+00 | 1.61E-04 | 6.97E-04 | 1.13E+00 | 7.19E-07 | 3.06E-06 | -1.27E+00 | 3.48E-09 | 1.33E-08 | -1.20E+00 | 1.06E-04 | 3.77E-04 | -1.13E+00 | 4.60E-05 | 1.30E-04 | -1.03E+00 | 2.41E-04 | 8.84E-04 | -1.02E+00 | 5.55E-04 | 1.52E-03 | 1.01E+00 |
| NIT2 | 3.43E-07 | 1.79E-06 | -1.09E+00 | 1.29E-07 | 5.58E-07 | 1.00E+00 | 3.85E-04 | 1.64E-03 | -1.11E+00 | 6.32E-03 | 2.41E-02 | 1.05E+00 | 1.33E-04 | 4.75E-04 | -1.08E+00 | 1.04E-04 | 2.94E-04 | -1.06E+00 | 9.36E-02 | 3.44E-01 | 1.13E+00 | 3.24E-01 | 8.89E-01 | 1.23E+00 |
| DGAT1 | 3.45E-07 | 1.80E-06 | -1.33E+00 | 3.73E-06 | 1.61E-05 | -1.12E+00 | 2.95E-04 | 1.25E-03 | -1.24E+00 | 1.23E-03 | 4.69E-03 | 1.00E+00 | 6.04E-04 | 2.15E-03 | -1.49E+00 | 2.71E-03 | 7.63E-03 | -1.25E+00 | 2.34E-02 | 8.60E-02 | -1.01E+00 | 5.40E-02 | 1.48E-01 | 1.02E+00 |
| RANBP6 | 3.45E-07 | 1.80E-06 | -1.24E+00 | 2.18E-09 | 9.43E-09 | -1.35E+00 | 5.82E-04 | 2.47E-03 | -1.30E+00 | 8.54E-05 | 3.26E-04 | -1.25E+00 | 1.11E-02 | 3.95E-02 | -1.03E+00 | 6.74E-04 | 1.90E-03 | -1.24E+00 | 3.55E-07 | 1.30E-06 | -1.42E+00 | 1.46E-06 | 4.00E-06 | -1.30E+00 |
| RPS3P6 | 3.46E-07 | 1.80E-06 | 2.82E+00 | 1.38E-02 | 5.97E-02 | 2.32E+00 | 3.73E-04 | 1.59E-03 | 2.14E+00 | 3.47E-02 | 1.33E-01 | 1.86E+00 | 4.52E-02 | 1.61E-01 | 2.36E+00 | 6.92E-02 | 1.95E-01 | 2.26E+00 | 2.05E-02 | 7.51E-02 | 1.92E+00 | 4.37E-02 | 1.20E-01 | 1.89E+00 |
| RPP8 | 3.48E-07 | 1.82E-06 | -1.26E+00 | 5.48E-06 | 2.37E-05 | -1.04E+00 | 8.93E-05 | 3.79E-04 | -1.33E+00 | 2.67E-04 | 1.02E-03 | -1.11E+00 | 4.54E-02 | 1.62E-01 | 1.07E+00 | 1.68E-01 | 4.73E-01 | 1.17E+00 | 8.12E-07 | 2.98E-06 | -1.23E+00 | 4.31E-04 | 1.18E-03 | -1.20E+00 |
| TNIP2 | 3.50E-07 | 1.83E-06 | -1.43E+00 | 1.21E-05 | 5.25E-05 | -1.09E+00 | 4.29E-03 | 1.82E-02 | -1.08E+00 | 2.45E-03 | 9.37E-03 | -1.00E+00 | 2.27E-05 | 8.09E-05 | -1.36E+00 | 2.61E-02 | 7.37E-02 | 1.12E+00 | 1.40E-02 | 5.13E-02 | -1.05E+00 | 9.42E-02 | 2.59E-01 | 1.09E+00 |
| BPHL | 3.51E-07 | 1.83E-06 | -1.82E+00 | 1.77E-05 | 7.67E-05 | -1.32E+00 | 1.42E-04 | 6.02E-04 | -1.50E+00 | 1.44E-03 | 5.49E-03 | -1.18E+00 | 5.14E-02 | 1.83E-01 | -1.05E+00 | 1.63E-01 | 4.61E-01 | 1.11E+00 | 1.92E-03 | 7.03E-03 | -1.26E+00 | 6.27E-02 | 1.72E-01 | -1.05E+00 |
| RNF216P1 | 3.53E-07 | 1.84E-06 | -1.54E+00 | 3.20E-05 | 1.38E-04 | -1.18E+00 | 9.92E-10 | 4.21E-09 | -1.76E+00 | 4.42E-06 | 1.69E-05 | -1.17E+00 | 4.05E-07 | 1.44E-06 | -1.92E+00 | 8.44E-06 | 2.38E-05 | -1.54E+00 | 5.95E-13 | 2.18E-12 | -1.89E+00 | 3.25E-06 | 8.92E-06 | -1.29E+00 |
| GAR1 | 3.54E-07 | 1.84E-06 | -1.21E+00 | 9.55E-04 | 4.13E-03 | 1.13E+00 | 2.41E-04 | 1.03E-03 | -1.16E+00 | 1.05E-02 | 4.01E-02 | 1.11E+00 | 8.68E-10 | 3.09E-09 | -1.47E+00 | 4.94E-03 | 1.39E-02 | 1.14E+00 | 6.45E-07 | 2.37E-06 | -1.27E+00 | 2.09E-01 | 5.74E-01 | 1.13E+00 |
| NOP14 | 3.54E-07 | 1.84E-06 | -1.13E+00 | 2.43E-06 | 1.05E-05 | 1.02E+00 | 7.19E-03 | 3.05E-02 | -1.07E+00 | 1.19E-02 | 4.54E-02 | 1.12E+00 | 8.67E-06 | 3.09E-05 | -1.19E+00 | 4.80E-04 | 1.35E-03 | -1.04E+00 | 1.34E-04 | 4.92E-04 | -1.11E+00 | 5.32E-02 | 1.46E-01 | 1.10E+00 |
| NFATC2IP | 3.54E-07 | 1.85E-06 | -1.25E+00 | 1.18E-04 | 5.09E-04 | 1.05E+00 | 1.50E-04 | 6.36E-04 | -1.28E+00 | 4.75E-02 | 1.81E-01 | 1.07E+00 | 4.91E-08 | 1.75E-07 | -1.37E+00 | 1.26E-03 | 3.55E-03 | -1.05E+00 | 1.35E-08 | 4.97E-08 | -1.42E+00 | 9.73E-04 | 2.67E-03 | -1.11E+00 |
| RPL36AP45 | 3.55E-07 | 1.85E-06 | 2.35E+00 | 6.93E-02 | 3.00E-01 | 1.80E+00 | 3.36E-04 | 1.43E-03 | 1.80E+00 | 1.81E-01 | 6.90E-01 | 1.54E+00 | 5.98E-02 | 2.13E-01 | 1.69E+00 | 2.71E-01 | 7.63E-01 | 1.39E+00 | 9.61E-05 | 3.53E-04 | 2.04E+00 | 1.59E-03 | 4.37E-03 | 1.88E+00 |
| KIAA1143 | 3.55E-07 | 1.85E-06 | -1.27E+00 | 2.97E-09 | 1.28E-08 | -1.39E+00 | 7.82E-03 | 3.32E-02 | -1.04E+00 | 8.40E-03 | 3.21E-02 | 1.10E+00 | 2.64E-02 | 9.43E-02 | -1.03E+00 | 3.87E-03 | 1.09E-02 | -1.21E+00 | 6.59E-05 | 2.42E-04 | -1.18E+00 | 4.63E-03 | 1.27E-02 | -1.00E+00 |
| SMARCAD1 | 3.55E-07 | 1.85E-06 | -1.41E+00 | 4.82E-04 | 2.08E-03 | -1.04E+00 | 2.13E-07 | 9.07E-07 | -1.37E+00 | 3.72E-03 | 1.42E-02 | 1.17E+00 | 7.39E-08 | 2.63E-07 | -1.46E+00 | 4.83E-05 | 1.36E-04 | -1.23E+00 | 3.51E-08 | 1.29E-07 | -1.29E+00 | 2.70E-03 | 7.42E-03 | -1.03E+00 |
| MTRR | 3.56E-07 | 1.85E-06 | -1.72E+00 | 5.62E-04 | 2.43E-03 | -1.11E+00 | 9.14E-04 | 3.89E-03 | -1.14E+00 | 2.48E-05 | 9.47E-05 | -1.32E+00 | 1.28E-01 | 4.56E-01 | 1.11E+00 | 2.65E-01 | 7.46E-01 | 1.47E+00 | 2.16E-04 | 7.91E-04 | -1.32E+00 | 6.75E-03 | 1.85E-02 | -1.13E+00 |
| DZIP3 | 3.56E-07 | 1.86E-06 | -1.57E+00 | 6.28E-06 | 2.71E-05 | -1.28E+00 | 8.39E-05 | 3.56E-04 | -1.69E+00 | 6.03E-04 | 2.30E-03 | -1.33E+00 | 1.66E-04 | 5.92E-04 | -1.50E+00 | 2.16E-03 | 6.08E-03 | -1.21E+00 | 5.45E-03 | 2.00E-02 | -1.31E+00 | 2.32E-02 | 6.38E-02 | -1.20E+00 |
| SLC12A6 | 3.57E-07 | 1.86E-06 | -1.28E+00 | 1.73E-02 | 7.46E-02 | 1.22E+00 | 1.05E-02 | 4.44E-02 | -1.05E+00 | 4.61E-02 | 1.76E-01 | 1.22E+00 | 1.86E-02 | 6.62E-02 | 1.06E+00 | 2.57E-01 | 7.24E-01 | 1.27E+00 | 7.73E-04 | 2.84E-03 | -1.04E+00 | 1.55E-02 | 4.25E-02 | 1.12E+00 |
| CHRFAM7A | 3.58E-07 | 1.86E-06 | -1.42E+00 | 1.39E-04 | 6.00E-04 | 1.01E+00 | 1.83E-03 | 7.77E-03 | -1.17E+00 | 9.21E-03 | 3.52E-02 | 1.10E+00 | 5.27E-04 | 1.88E-03 | -1.75E+00 | 1.40E-03 | 3.96E-03 | -1.56E+00 | 5.05E-03 | 1.85E-02 | -1.46E+00 | 4.63E-03 | 1.27E-02 | -1.47E+00 |
| AL132989.1 | 3.58E-07 | 1.86E-06 | -1.43E+00 | 1.88E-07 | 8.14E-07 | -1.30E+00 | 1.57E-03 | 6.67E-03 | -1.19E+00 | 2.28E-03 | 8.72E-03 | -1.02E+00 | 2.98E-02 | 1.06E-01 | -1.07E+00 | 4.58E-02 | 1.29E-01 | -1.05E+00 | 3.82E-04 | 1.40E-03 | -1.39E+00 | 1.66E-03 | 4.56E-03 | -1.29E+00 |
| ZCCHC6 | 3.61E-07 | 1.88E-06 | -1.20E+00 | 4.31E-07 | 1.86E-06 | -1.12E+00 | 1.29E-08 | 5.47E-08 | -1.71E+00 | 4.28E-06 | 1.63E-05 | -1.23E+00 | 1.79E-03 | 6.40E-03 | -1.12E+00 | 2.90E-03 | 8.17E-03 | -1.12E+00 | 1.24E-07 | 4.53E-07 | -1.18E+00 | 4.83E-06 | 1.33E-05 | -1.32E+00 |
| INPP4A | 3.62E-07 | 1.89E-06 | -1.85E+00 | 3.29E-03 | 1.42E-02 | -1.03E+00 | 3.79E-03 | 1.61E-02 | -1.29E+00 | 1.91E-02 | 7.29E-02 | 1.04E+00 | 1.46E-06 | 5.21E-06 | -2.05E+00 | 3.55E-05 | 1.00E-04 | -1.67E+00 | 8.54E-06 | 3.13E-05 | -1.59E+00 | 1.39E-03 | 3.81E-03 | -1.19E+00 |
| GABARAPL1 | 3.62E-07 | 1.89E-06 | -1.45E+00 | 1.87E-10 | 8.08E-10 | -1.78E+00 | 2.08E-04 | 8.85E-04 | -1.57E+00 | 8.98E-06 | 3.43E-05 | -1.78E+00 | 6.25E-07 | 2.23E-06 | -1.33E+00 | 6.94E-11 | 1.96E-10 | -2.24E+00 | 5.44E-12 | 2.00E-11 | -2.01E+00 | 5.67E-12 | 1.56E-11 | -2.18E+00 |
| TMEM101 | 3.63E-07 | 1.89E-06 | -1.12E+00 | 6.98E-05 | 3.02E-04 | 1.10E+00 | 1.09E-03 | 4.64E-03 | -1.08E+00 | 1.03E-04 | 3.95E-04 | -1.04E+00 | 1.55E-06 | 5.52E-06 | -1.25E+00 | 3.29E-02 | 9.28E-02 | 1.18E+00 | 4.78E-06 | 1.76E-05 | -1.31E+00 | 2.18E-02 | 6.00E-02 | -1.02E+00 |
| WDR12 | 3.63E-07 | 1.89E-06 | -1.53E+00 | 1.94E-04 | 8.38E-04 | -1.05E+00 | 7.27E-07 | 3.09E-06 | -1.33E+00 | 6.50E-03 | 2.48E-02 | 1.13E+00 | 1.92E-05 | 6.84E-05 | -1.57E+00 | 2.51E-01 | 7.09E-01 | 1.34E+00 | 3.93E-22 | 1.44E-21 | -1.88E+00 | 6.52E-03 | 1.79E-02 | -1.00E+00 |
| CASP9 | 3.64E-07 | 1.90E-06 | -1.82E+00 | 6.06E-04 | 2.62E-03 | -1.12E+00 | 7.88E-06 | 3.35E-05 | -1.59E+00 | 7.95E-07 | 3.04E-06 | -1.48E+00 | 3.09E-10 | 1.10E-09 | -2.34E+00 | 1.64E-07 | 4.61E-07 | -1.64E+00 | 1.25E-04 | 4.57E-04 | -1.42E+00 | 5.50E-04 | 1.51E-03 | -1.28E+00 |
| BLOC1S5 | 3.64E-07 | 1.90E-06 | -1.53E+00 | 1.67E-02 | 7.20E-02 | 1.17E+00 | 3.24E-04 | 1.38E-03 | -1.25E+00 | 4.41E-02 | 1.69E-01 | 1.17E+00 | 2.53E-04 | 9.03E-04 | -1.47E+00 | 3.47E-01 | 9.78E-01 | 1.46E+00 | 1.36E-05 | 4.98E-05 | -1.20E+00 | 6.71E-03 | 1.84E-02 | 1.00E+00 |
| NKIRAS1 | 3.64E-07 | 1.90E-06 | -1.51E+00 | 9.08E-04 | 3.92E-03 | 1.03E+00 | 2.28E-09 | 9.70E-09 | -1.74E+00 | 2.00E-06 | 7.64E-06 | -1.20E+00 | 4.15E-02 | 1.48E-01 | -1.03E+00 | 1.58E-01 | 4.45E-01 | 1.16E+00 | 7.72E-03 | 2.83E-02 | -1.11E+00 | 3.15E-02 | 8.65E-02 | -1.10E+00 |
| TTTY15 | 3.65E-07 | 1.90E-06 | -1.19E+00 | 8.36E-05 | 3.61E-04 | 1.03E+00 | 1.21E-04 | 5.13E-04 | -1.22E+00 | 1.14E-03 | 4.35E-03 | -1.03E+00 | 3.86E-04 | 1.37E-03 | -1.19E+00 | 2.28E-03 | 6.44E-03 | -1.12E+00 | 1.32E-04 | 4.84E-04 | -1.20E+00 | 1.79E-03 | 4.92E-03 | -1.08E+00 |
| TMEM127 | 3.66E-07 | 1.91E-06 | -1.58E+00 | 3.68E-04 | 1.59E-03 | -1.02E+00 | 1.31E-04 | 5.58E-04 | -1.50E+00 | 2.61E-03 | 9.99E-03 | -1.09E+00 | 1.31E-02 | 4.67E-02 | -1.10E+00 | 4.51E-03 | 1.27E-02 | -1.26E+00 | 4.59E-04 | 1.69E-03 | -1.26E+00 | 3.86E-02 | 1.06E-01 | 1.02E+00 |
| AC103987.1 | 3.66E-07 | 1.91E-06 | -2.04E+00 | 1.16E-03 | 5.02E-03 | -1.10E+00 | 1.72E-05 | 7.31E-05 | -1.88E+00 | 1.01E-03 | 3.85E-03 | -1.28E+00 | 8.74E-02 | 3.12E-01 | -1.01E+00 | 9.17E-02 | 2.59E-01 | -1.08E+00 | 1.44E-02 | 5.28E-02 | -1.53E+00 | 2.41E-01 | 6.61E-01 | 1.05E+00 |
| ZFP64 | 3.68E-07 | 1.92E-06 | -1.91E+00 | 7.15E-05 | 3.09E-04 | -1.37E+00 | 8.99E-08 | 3.82E-07 | -2.07E+00 | 1.84E-04 | 7.04E-04 | -1.28E+00 | 2.31E-07 | 8.22E-07 | -2.38E+00 | 6.80E-05 | 1.92E-04 | -1.52E+00 | 6.29E-07 | 2.31E-06 | -1.61E+00 | 5.22E-05 | 1.43E-04 | -1.21E+00 |
| MIER3 | 3.68E-07 | 1.92E-06 | -1.63E+00 | 1.48E-07 | 6.41E-07 | -1.47E+00 | 4.57E-06 | 1.94E-05 | -1.64E+00 | 1.95E-04 | 7.45E-04 | -1.17E+00 | 5.94E-02 | 2.12E-01 | -1.09E+00 | 1.60E-02 | 4.50E-02 | -1.32E+00 | 8.92E-07 | 3.27E-06 | -1.37E+00 | 1.33E-06 | 3.65E-06 | -1.56E+00 |
| ADH5 | 3.71E-07 | 1.94E-06 | 1.00E+00 | 5.16E-02 | 2.23E-01 | 1.39E+00 | 3.97E-03 | 1.69E-02 | -1.01E+00 | 6.82E-02 | 2.60E-01 | 1.24E+00 | 7.41E-05 | 2.64E-04 | -1.09E+00 | 2.80E-01 | 7.89E-01 | 1.30E+00 | 1.43E-03 | 5.24E-03 | -1.06E+00 | 9.68E-02 | 2.66E-01 | 1.08E+00 |
| SPEF2 | 3.71E-07 | 1.94E-06 | -1.31E+00 | 3.73E-05 | 1.61E-04 | 1.03E+00 | 2.32E-05 | 9.86E-05 | -1.51E+00 | 1.94E-02 | 7.42E-02 | 1.06E+00 | 3.30E-02 | 1.18E-01 | -1.20E+00 | 2.14E-01 | 6.03E-01 | 1.11E+00 | 1.72E-02 | 6.31E-02 | -1.07E+00 | 8.59E-03 | 2.36E-02 | -1.13E+00 |
| C8orf58 | 3.76E-07 | 1.96E-06 | -1.69E+00 | 4.80E-04 | 2.07E-03 | -1.03E+00 | 3.36E-04 | 1.43E-03 | -1.27E+00 | 2.46E-05 | 9.41E-05 | -1.31E+00 | 7.12E-02 | 2.54E-01 | 1.11E+00 | 3.46E-02 | 9.75E-02 | 1.00E+00 | 8.38E-03 | 3.07E-02 | -1.06E+00 | 6.16E-03 | 1.69E-02 | -1.17E+00 |
| RTN2 | 3.78E-07 | 1.97E-06 | -1.44E+00 | 2.31E-03 | 9.98E-03 | 1.10E+00 | 6.58E-04 | 2.80E-03 | -1.64E+00 | 2.51E-02 | 9.58E-02 | -1.05E+00 | 1.20E-03 | 4.29E-03 | -1.14E+00 | 3.06E-02 | 8.62E-02 | 1.12E+00 | 2.61E-01 | 9.59E-01 | 1.38E+00 | 1.91E-01 | 5.25E-01 | 1.24E+00 |
| WDR17 | 3.81E-07 | 1.98E-06 | -1.69E+00 | 5.59E-06 | 2.42E-05 | -1.32E+00 | 2.76E-02 | 1.17E-01 | -1.08E+00 | 9.30E-02 | 3.55E-01 | 1.18E+00 | 3.54E-02 | 1.26E-01 | -1.56E+00 | 2.28E-01 | 6.43E-01 | 1.11E+00 | 6.94E-03 | 2.55E-02 | -1.25E+00 | 3.24E-02 | 8.90E-02 | -1.18E+00 |
| PLTP | 3.81E-07 | 1.98E-06 | -1.53E+00 | 5.59E-06 | 2.41E-05 | -1.29E+00 | 8.41E-06 | 3.57E-05 | -1.29E+00 | 2.16E-04 | 8.24E-04 | 1.01E+00 | 8.95E-05 | 3.19E-04 | -1.30E+00 | 2.08E-03 | 5.86E-03 | -1.16E+00 | 1.24E-04 | 4.53E-04 | -1.27E+00 | 3.13E-02 | 8.60E-02 | -1.05E+00 |
| AC060780.1 | 3.81E-07 | 1.99E-06 | -1.37E+00 | 1.56E-01 | 6.76E-01 | 1.40E+00 | 1.05E-04 | 4.46E-04 | -1.38E+00 | 6.02E-02 | 2.30E-01 | 1.15E+00 | 6.30E-02 | 2.25E-01 | 1.24E+00 | 3.36E-01 | 9.48E-01 | 1.60E+00 | 1.21E-04 | 4.44E-04 | -1.12E+00 | 6.69E-03 | 1.84E-02 | -1.05E+00 |
| MAT2A | 3.82E-07 | 1.99E-06 | -1.35E+00 | 1.58E-01 | 6.81E-01 | 1.54E+00 | 2.58E-08 | 1.10E-07 | -1.59E+00 | 6.90E-02 | 2.64E-01 | 1.72E+00 | 1.25E-05 | 4.47E-05 | -1.29E+00 | 1.25E-01 | 3.51E-01 | 1.27E+00 | 4.74E-04 | 1.74E-03 | -1.09E+00 | 1.33E-01 | 3.67E-01 | 1.56E+00 |
| AC099560.2 | 3.82E-07 | 1.99E-06 | 1.29E+00 | 1.26E-35 | 5.43E-35 | -1.44E+00 | 5.94E-06 | 2.52E-05 | 1.29E+00 | 3.38E-13 | 1.29E-12 | -1.43E+00 | 1.30E-05 | 4.63E-05 | 1.37E+00 | 6.91E-09 | 1.95E-08 | -1.30E+00 | 2.06E-06 | 7.58E-06 | 1.40E+00 | 2.14E-12 | 5.88E-12 | -1.41E+00 |
| GPR89A | 3.86E-07 | 2.01E-06 | -1.15E+00 | 1.39E-05 | 6.01E-05 | 1.05E+00 | 1.93E-05 | 8.18E-05 | -1.35E+00 | 1.95E-06 | 7.44E-06 | -1.24E+00 | 3.49E-05 | 1.25E-04 | -1.34E+00 | 1.88E-04 | 5.30E-04 | -1.21E+00 | 4.46E-06 | 1.64E-05 | -1.27E+00 | 9.36E-04 | 2.57E-03 | -1.08E+00 |
| ST6GAL1 | 3.86E-07 | 2.01E-06 | -1.32E+00 | 6.53E-03 | 2.82E-02 | 1.23E+00 | 1.13E-06 | 4.80E-06 | -1.67E+00 | 1.34E-03 | 5.11E-03 | -1.08E+00 | 3.87E-05 | 1.38E-04 | -1.65E+00 | 3.20E-01 | 9.03E-01 | 1.30E+00 | 5.30E-11 | 1.95E-10 | -1.67E+00 | 8.64E-03 | 2.37E-02 | 1.03E+00 |
| IL17RC | 3.86E-07 | 2.01E-06 | -1.37E+00 | 8.87E-06 | 3.83E-05 | -1.14E+00 | 3.99E-02 | 1.70E-01 | -1.02E+00 | 4.69E-03 | 1.79E-02 | -1.05E+00 | 2.65E-03 | 9.46E-03 | -1.68E+00 | 6.99E-03 | 1.97E-02 | -1.43E+00 | 2.30E-03 | 8.45E-03 | -1.11E+00 | 5.28E-03 | 1.45E-02 | -1.10E+00 |
| AC244154.1 | 3.87E-07 | 2.02E-06 | -1.75E+00 | 2.62E-05 | 1.13E-04 | -1.24E+00 | 1.59E-01 | 6.77E-01 | 1.12E+00 | 2.31E-01 | 8.81E-01 | 1.36E+00 | 3.19E-03 | 1.14E-02 | -1.51E+00 | 3.96E-03 | 1.11E-02 | -1.44E+00 | 6.73E-02 | 2.47E-01 | -1.03E+00 | 6.30E-02 | 1.73E-01 | -1.03E+00 |
| CASP4 | 3.90E-07 | 2.03E-06 | -1.13E+00 | 5.81E-06 | 2.51E-05 | -1.04E+00 | 4.14E-03 | 1.76E-02 | -1.08E+00 | 6.06E-03 | 2.31E-02 | 1.04E+00 | 1.65E-04 | 5.89E-04 | 1.00E+00 | 3.52E-03 | 9.91E-03 | 1.04E+00 | 2.79E-06 | 1.02E-05 | -1.25E+00 | 1.27E-04 | 3.50E-04 | -1.16E+00 |
| NECAB3 | 3.91E-07 | 2.04E-06 | -1.63E+00 | 1.07E-05 | 4.62E-05 | -1.26E+00 | 6.96E-09 | 2.96E-08 | -1.76E+00 | 1.57E-06 | 5.99E-06 | -1.23E+00 | 3.00E-02 | 1.07E-01 | -1.42E+00 | 7.90E-02 | 2.23E-01 | -1.17E+00 | 2.55E-06 | 9.37E-06 | -1.39E+00 | 8.78E-03 | 2.41E-02 | -1.21E+00 |
| IMP4 | 3.91E-07 | 2.04E-06 | -1.13E+00 | 9.26E-05 | 4.00E-04 | 1.08E+00 | 6.38E-05 | 2.71E-04 | -1.20E+00 | 4.72E-04 | 1.80E-03 | -1.03E+00 | 1.85E-04 | 6.59E-04 | -1.08E+00 | 1.97E-01 | 5.57E-01 | 1.28E+00 | 9.09E-04 | 3.34E-03 | -1.06E+00 | 2.48E-01 | 6.81E-01 | 1.18E+00 |
| KIF1B | 3.91E-07 | 2.04E-06 | -1.41E+00 | 1.36E-03 | 5.89E-03 | 1.04E+00 | 5.50E-05 | 2.34E-04 | -1.32E+00 | 3.19E-04 | 1.22E-03 | -1.05E+00 | 3.04E-03 | 1.08E-02 | -1.19E+00 | 1.76E-02 | 4.97E-02 | -1.03E+00 | 3.37E-05 | 1.24E-04 | -1.38E+00 | 1.82E-03 | 5.01E-03 | -1.16E+00 |
| TCAF1 | 3.94E-07 | 2.05E-06 | -1.28E+00 | 1.03E-07 | 4.45E-07 | -1.12E+00 | 8.22E-07 | 3.49E-06 | -1.36E+00 | 2.41E-06 | 9.22E-06 | -1.21E+00 | 3.03E-08 | 1.08E-07 | -1.48E+00 | 2.44E-04 | 6.89E-04 | -1.11E+00 | 1.88E-10 | 6.90E-10 | -1.45E+00 | 6.73E-07 | 1.85E-06 | -1.35E+00 |
| ZNF7 | 3.94E-07 | 2.05E-06 | -1.45E+00 | 6.92E-04 | 2.99E-03 | 1.06E+00 | 7.05E-03 | 3.00E-02 | -1.15E+00 | 2.77E-02 | 1.06E-01 | 1.12E+00 | 1.18E-07 | 4.21E-07 | -1.65E+00 | 3.75E-04 | 1.06E-03 | -1.10E+00 | 1.11E-03 | 4.07E-03 | -1.17E+00 | 4.01E-02 | 1.10E-01 | 1.05E+00 |
| SEL1L3 | 3.94E-07 | 2.05E-06 | -1.51E+00 | 2.46E-05 | 1.06E-04 | -1.08E+00 | 1.39E-06 | 5.89E-06 | -1.53E+00 | 4.16E-06 | 1.59E-05 | -1.28E+00 | 2.57E-01 | 9.15E-01 | 1.42E+00 | 2.69E-01 | 7.58E-01 | -1.05E+00 | 2.37E-01 | 8.71E-01 | 1.29E+00 | 1.19E-01 | 3.26E-01 | -1.12E+00 |
| ZNF667-AS1 | 3.96E-07 | 2.06E-06 | -1.17E+00 | 2.27E-04 | 9.80E-04 | 1.14E+00 | 4.82E-02 | 2.05E-01 | 1.09E+00 | 2.22E-01 | 8.48E-01 | 1.48E+00 | 7.49E-04 | 2.67E-03 | -1.11E+00 | 5.18E-02 | 1.46E-01 | 1.11E+00 | 8.72E-06 | 3.20E-05 | -1.25E+00 | 1.55E-02 | 4.25E-02 | 1.00E+00 |
| BIRC2 | 3.98E-07 | 2.08E-06 | -1.09E+00 | 7.25E-09 | 3.13E-08 | -1.08E+00 | 2.69E-06 | 1.14E-05 | -1.26E+00 | 8.93E-07 | 3.41E-06 | -1.12E+00 | 1.89E-04 | 6.74E-04 | 1.03E+00 | 4.56E-05 | 1.29E-04 | -1.08E+00 | 9.46E-07 | 3.47E-06 | -1.34E+00 | 6.80E-04 | 1.87E-03 | -1.19E+00 |
| AC093155.2 | 3.99E-07 | 2.08E-06 | -2.29E+00 | 8.44E-07 | 3.65E-06 | -1.99E+00 | 1.52E-03 | 6.47E-03 | -1.29E+00 | 6.12E-04 | 2.34E-03 | -1.26E+00 | 4.05E-04 | 1.44E-03 | -1.59E+00 | 2.25E-05 | 6.35E-05 | -2.13E+00 | 1.70E-07 | 6.23E-07 | -1.55E+00 | 8.42E-08 | 2.31E-07 | -1.72E+00 |
| SOCS4 | 4.00E-07 | 2.08E-06 | -1.35E+00 | 4.35E-09 | 1.88E-08 | -1.36E+00 | 2.56E-05 | 1.09E-04 | -1.21E+00 | 1.56E-08 | 5.98E-08 | -1.43E+00 | 3.32E-02 | 1.18E-01 | -1.01E+00 | 2.53E-02 | 7.14E-02 | -1.04E+00 | 5.73E-04 | 2.10E-03 | -1.11E+00 | 1.00E-04 | 2.75E-04 | -1.25E+00 |
| ZNF620 | 4.01E-07 | 2.09E-06 | -1.60E+00 | 7.43E-07 | 3.21E-06 | -1.38E+00 | 7.40E-08 | 3.15E-07 | -1.76E+00 | 1.18E-06 | 4.52E-06 | -1.42E+00 | 7.52E-05 | 2.68E-04 | -1.62E+00 | 4.42E-04 | 1.25E-03 | -1.36E+00 | 2.86E-09 | 1.05E-08 | -1.82E+00 | 2.35E-07 | 6.45E-07 | -1.67E+00 |
| HDGFL2 | 4.04E-07 | 2.10E-06 | -1.11E+00 | 7.07E-05 | 3.06E-04 | 1.09E+00 | 5.27E-03 | 2.24E-02 | -1.03E+00 | 1.40E-03 | 5.34E-03 | 1.06E+00 | 9.24E-05 | 3.29E-04 | -1.01E+00 | 7.17E-04 | 2.02E-03 | 1.09E+00 | 1.00E-01 | 3.67E-01 | 1.14E+00 | 6.36E-02 | 1.75E-01 | 1.15E+00 |
| GUSB | 4.04E-07 | 2.11E-06 | -1.16E+00 | 8.23E-08 | 3.56E-07 | -1.07E+00 | 6.68E-02 | 2.84E-01 | 1.15E+00 | 6.49E-02 | 2.48E-01 | 1.24E+00 | 2.60E-05 | 9.27E-05 | -1.10E+00 | 5.49E-03 | 1.55E-02 | 1.09E+00 | 1.42E-01 | 5.22E-01 | 1.12E+00 | 2.29E-01 | 6.29E-01 | 1.13E+00 |
| CELF1 | 4.05E-07 | 2.11E-06 | -1.14E+00 | 4.73E-05 | 2.05E-04 | 1.11E+00 | 8.48E-09 | 3.60E-08 | -1.38E+00 | 1.35E-06 | 5.17E-06 | -1.10E+00 | 7.63E-05 | 2.72E-04 | -1.11E+00 | 1.93E-04 | 5.44E-04 | -1.09E+00 | 3.24E-11 | 1.19E-10 | -1.62E+00 | 6.39E-04 | 1.76E-03 | -1.18E+00 |
| DUSP10 | 4.07E-07 | 2.12E-06 | 1.93E+00 | 4.78E-04 | 2.07E-03 | 1.05E+00 | 1.33E-04 | 5.66E-04 | 1.78E+00 | 5.37E-02 | 2.05E-01 | 1.27E+00 | 1.53E-01 | 5.45E-01 | 1.79E+00 | 1.50E-03 | 4.23E-03 | 1.01E+00 | 2.10E-01 | 7.69E-01 | 1.41E+00 | 2.79E-02 | 7.66E-02 | -1.13E+00 |
| FMC1 | 4.08E-07 | 2.12E-06 | -1.24E+00 | 8.15E-13 | 3.52E-12 | -1.65E+00 | 4.24E-03 | 1.80E-02 | -1.13E+00 | 1.56E-12 | 5.95E-12 | -1.72E+00 | 1.93E-06 | 6.88E-06 | -1.38E+00 | 1.38E-08 | 3.89E-08 | -1.72E+00 | 6.98E-04 | 2.56E-03 | -1.06E+00 | 1.20E-05 | 3.30E-05 | -1.21E+00 |
| BBS7 | 4.08E-07 | 2.13E-06 | -1.80E+00 | 7.14E-05 | 3.09E-04 | -1.28E+00 | 1.82E-03 | 7.74E-03 | -1.23E+00 | 1.72E-04 | 6.58E-04 | -1.22E+00 | 4.73E-02 | 1.69E-01 | -1.33E+00 | 1.09E-01 | 3.08E-01 | -1.07E+00 | 4.24E-11 | 1.56E-10 | -1.64E+00 | 6.59E-08 | 1.81E-07 | -1.36E+00 |
| EFR3A | 4.11E-07 | 2.14E-06 | -1.49E+00 | 2.12E-05 | 9.14E-05 | -1.13E+00 | 7.81E-12 | 3.32E-11 | -1.81E+00 | 1.93E-10 | 7.38E-10 | -1.44E+00 | 3.80E-07 | 1.35E-06 | -1.82E+00 | 1.17E-08 | 3.30E-08 | -2.05E+00 | 2.11E-08 | 7.75E-08 | -1.53E+00 | 2.14E-04 | 5.89E-04 | -1.11E+00 |
| RPL10P6 | 4.11E-07 | 2.14E-06 | -1.13E+00 | 1.17E-04 | 5.07E-04 | 1.10E+00 | 1.93E-04 | 8.21E-04 | -1.11E+00 | 1.14E-04 | 4.37E-04 | 1.13E+00 | 1.15E-05 | 4.09E-05 | -1.16E+00 | 1.25E-01 | 3.51E-01 | 1.04E+00 | 1.57E-12 | 5.75E-12 | -1.17E+00 | 3.06E-01 | 8.40E-01 | 1.03E+00 |
| UQCRC2 | 4.12E-07 | 2.15E-06 | -1.09E+00 | 3.45E-05 | 1.49E-04 | 1.07E+00 | 3.05E-03 | 1.30E-02 | -1.09E+00 | 6.38E-02 | 2.44E-01 | 1.06E+00 | 5.98E-05 | 2.13E-04 | -1.17E+00 | 7.01E-03 | 1.98E-02 | 1.01E+00 | 7.91E-03 | 2.90E-02 | -1.06E+00 | 1.39E-01 | 3.82E-01 | 1.07E+00 |
| STK26 | 4.12E-07 | 2.15E-06 | -1.14E+00 | 4.48E-07 | 1.93E-06 | -1.04E+00 | 3.31E-06 | 1.41E-05 | -1.19E+00 | 4.30E-06 | 1.64E-05 | -1.11E+00 | 7.70E-05 | 2.74E-04 | -1.12E+00 | 3.07E-04 | 8.64E-04 | -1.04E+00 | 3.08E-03 | 1.13E-02 | -1.06E+00 | 7.45E-03 | 2.05E-02 | 1.01E+00 |
| ICMT | 4.13E-07 | 2.15E-06 | -2.13E+00 | 6.60E-04 | 2.85E-03 | -1.18E+00 | 4.30E-04 | 1.83E-03 | -1.34E+00 | 3.07E-02 | 1.17E-01 | 1.18E+00 | 4.61E-04 | 1.64E-03 | -2.22E+00 | 3.71E-02 | 1.05E-01 | -1.23E+00 | 8.41E-05 | 3.09E-04 | -1.19E+00 | 1.00E-01 | 2.75E-01 | 1.18E+00 |
| SLC27A1 | 4.20E-07 | 2.19E-06 | -1.43E+00 | 3.61E-04 | 1.56E-03 | -1.03E+00 | 1.52E-03 | 6.45E-03 | -1.10E+00 | 1.53E-02 | 5.83E-02 | 1.21E+00 | 2.55E-04 | 9.11E-04 | -1.76E+00 | 2.44E-02 | 6.88E-02 | -1.08E+00 | 4.67E-02 | 1.71E-01 | -1.12E+00 | 1.09E-01 | 2.99E-01 | -1.03E+00 |
| IMPA2 | 4.20E-07 | 2.19E-06 | -1.10E+00 | 2.12E-06 | 9.16E-06 | 1.08E+00 | 2.93E-03 | 1.24E-02 | -1.01E+00 | 4.65E-02 | 1.78E-01 | 1.20E+00 | 7.23E-04 | 2.58E-03 | -1.05E+00 | 1.75E-01 | 4.94E-01 | 1.27E+00 | 2.20E-01 | 8.07E-01 | 1.25E+00 | 4.44E-02 | 1.22E-01 | 1.48E+00 |
| SMARCD2 | 4.23E-07 | 2.21E-06 | -1.29E+00 | 4.46E-04 | 1.93E-03 | 1.10E+00 | 1.82E-05 | 7.75E-05 | -1.21E+00 | 2.46E-03 | 9.40E-03 | 1.06E+00 | 5.56E-05 | 1.98E-04 | -1.22E+00 | 1.39E-01 | 3.91E-01 | 1.31E+00 | 3.58E-10 | 1.31E-09 | -1.34E+00 | 2.51E-04 | 6.90E-04 | -1.04E+00 |
| GCLC | 4.24E-07 | 2.21E-06 | -1.49E+00 | 1.61E-04 | 6.94E-04 | -1.02E+00 | 3.42E-10 | 1.45E-09 | -1.68E+00 | 5.06E-10 | 1.93E-09 | -1.47E+00 | 7.13E-08 | 2.54E-07 | -1.61E+00 | 6.59E-02 | 1.86E-01 | 1.22E+00 | 1.19E-25 | 4.38E-25 | -2.04E+00 | 8.44E-09 | 2.32E-08 | -1.33E+00 |
| FCRLB | 4.29E-07 | 2.24E-06 | -2.14E+00 | 3.66E-05 | 1.58E-04 | -1.49E+00 | 1.23E-02 | 5.23E-02 | -1.64E+00 | 3.68E-02 | 1.41E-01 | -1.26E+00 | 2.96E-02 | 1.05E-01 | -1.41E+00 | 3.40E-01 | 9.58E-01 | 1.31E+00 | 8.24E-03 | 3.02E-02 | -1.19E+00 | 9.67E-02 | 2.66E-01 | 1.10E+00 |
| UTY | 4.30E-07 | 2.24E-06 | -1.15E+00 | 8.85E-06 | 3.82E-05 | 1.02E+00 | 2.61E-09 | 1.11E-08 | -1.44E+00 | 1.12E-06 | 4.30E-06 | -1.11E+00 | 9.13E-09 | 3.25E-08 | -1.24E+00 | 4.88E-08 | 1.38E-07 | -1.14E+00 | 1.82E-09 | 6.67E-09 | -1.33E+00 | 2.34E-06 | 6.44E-06 | -1.20E+00 |
| COX17 | 4.31E-07 | 2.25E-06 | -1.35E+00 | 5.36E-09 | 2.31E-08 | -1.43E+00 | 1.04E-04 | 4.41E-04 | -1.17E+00 | 2.96E-11 | 1.13E-10 | -1.54E+00 | 4.52E-04 | 1.61E-03 | -1.15E+00 | 4.59E-04 | 1.29E-03 | -1.16E+00 | 2.05E-04 | 7.52E-04 | -1.14E+00 | 1.14E-05 | 3.14E-05 | -1.25E+00 |
| SLC25A46 | 4.33E-07 | 2.26E-06 | -1.26E+00 | 9.60E-05 | 4.15E-04 | 1.06E+00 | 1.75E-06 | 7.44E-06 | -1.32E+00 | 9.11E-04 | 3.48E-03 | 1.02E+00 | 8.58E-04 | 3.06E-03 | -1.26E+00 | 3.39E-02 | 9.55E-02 | 1.02E+00 | 8.08E-09 | 2.97E-08 | -1.32E+00 | 1.85E-03 | 5.08E-03 | 1.05E+00 |
| TCAIM | 4.33E-07 | 2.26E-06 | -1.37E+00 | 3.32E-02 | 1.44E-01 | 1.23E+00 | 4.38E-12 | 1.86E-11 | -1.64E+00 | 2.08E-09 | 7.93E-09 | -1.27E+00 | 2.21E-06 | 7.89E-06 | -1.37E+00 | 1.06E-03 | 3.00E-03 | -1.09E+00 | 3.67E-07 | 1.35E-06 | -1.40E+00 | 7.41E-04 | 2.03E-03 | -1.10E+00 |
| FIBP | 4.34E-07 | 2.26E-06 | -1.05E+00 | 2.12E-09 | 9.16E-09 | -1.03E+00 | 6.52E-03 | 2.77E-02 | -1.02E+00 | 7.46E-05 | 2.85E-04 | -1.04E+00 | 1.38E-04 | 4.91E-04 | -1.02E+00 | 2.29E-04 | 6.47E-04 | -1.01E+00 | 1.27E-01 | 4.67E-01 | 1.18E+00 | 5.04E-02 | 1.38E-01 | 1.05E+00 |
| IER5 | 4.36E-07 | 2.27E-06 | 2.24E+00 | 6.93E-10 | 2.99E-09 | -1.73E+00 | 1.40E-01 | 5.96E-01 | 1.47E+00 | 1.15E-07 | 4.38E-07 | -1.52E+00 | 5.26E-07 | 1.88E-06 | 2.88E+00 | 8.65E-02 | 2.44E-01 | 1.41E+00 | 6.91E-02 | 2.54E-01 | 1.56E+00 | 6.75E-09 | 1.85E-08 | -1.36E+00 |
| NDUFAF4 | 4.37E-07 | 2.28E-06 | -1.29E+00 | 1.32E-05 | 5.71E-05 | -1.04E+00 | 2.68E-06 | 1.14E-05 | -1.37E+00 | 1.42E-03 | 5.43E-03 | 1.00E+00 | 1.22E-03 | 4.35E-03 | -1.19E+00 | 1.59E-01 | 4.49E-01 | 1.17E+00 | 1.40E-22 | 5.15E-22 | -1.87E+00 | 3.43E-05 | 9.42E-05 | -1.09E+00 |
| SMAD4 | 4.38E-07 | 2.28E-06 | -1.16E+00 | 4.92E-08 | 2.13E-07 | -1.11E+00 | 5.63E-05 | 2.39E-04 | -1.28E+00 | 1.06E-02 | 4.04E-02 | 1.03E+00 | 1.21E-05 | 4.32E-05 | -1.22E+00 | 1.20E-03 | 3.40E-03 | -1.02E+00 | 4.14E-03 | 1.52E-02 | 1.06E+00 | 4.90E-03 | 1.35E-02 | 1.02E+00 |
| PTCD2 | 4.38E-07 | 2.28E-06 | -1.73E+00 | 1.36E-02 | 5.86E-02 | 1.16E+00 | 5.98E-07 | 2.54E-06 | -1.76E+00 | 2.63E-02 | 1.00E-01 | 1.15E+00 | 2.56E-03 | 9.14E-03 | -1.54E+00 | 3.36E-01 | 9.48E-01 | 1.34E+00 | 1.67E-07 | 6.15E-07 | -1.64E+00 | 6.08E-02 | 1.67E-01 | 1.15E+00 |
| SF3B5 | 4.39E-07 | 2.29E-06 | -1.07E+00 | 1.37E-08 | 5.90E-08 | -1.02E+00 | 7.55E-02 | 3.21E-01 | 1.22E+00 | 2.42E-01 | 9.25E-01 | 1.16E+00 | 5.14E-03 | 1.83E-02 | -1.05E+00 | 2.88E-01 | 8.12E-01 | 1.12E+00 | 1.66E-01 | 6.09E-01 | 1.07E+00 | 2.18E-01 | 6.00E-01 | 1.13E+00 |
| PTER | 4.41E-07 | 2.30E-06 | -1.38E+00 | 7.48E-02 | 3.23E-01 | 1.30E+00 | 1.14E-04 | 4.83E-04 | -1.39E+00 | 1.95E-02 | 7.46E-02 | 1.07E+00 | 2.54E-07 | 9.05E-07 | -1.78E+00 | 1.30E-03 | 3.67E-03 | -1.01E+00 | 1.53E-04 | 5.62E-04 | -1.29E+00 | 9.37E-02 | 2.57E-01 | 1.13E+00 |
| NAT10 | 4.44E-07 | 2.32E-06 | -1.30E+00 | 2.39E-05 | 1.03E-04 | -1.05E+00 | 1.53E-07 | 6.52E-07 | -1.39E+00 | 1.06E-03 | 4.04E-03 | 1.04E+00 | 2.50E-08 | 8.91E-08 | -1.59E+00 | 1.93E-03 | 5.43E-03 | 1.05E+00 | 1.43E-17 | 5.26E-17 | -1.81E+00 | 3.21E-06 | 8.81E-06 | -1.18E+00 |
| UTP6 | 4.45E-07 | 2.32E-06 | -1.13E+00 | 5.65E-04 | 2.44E-03 | 1.18E+00 | 1.15E-07 | 4.91E-07 | -1.37E+00 | 8.96E-04 | 3.42E-03 | 1.01E+00 | 3.94E-07 | 1.40E-06 | -1.48E+00 | 6.22E-04 | 1.75E-03 | -1.05E+00 | 2.54E-10 | 9.30E-10 | -1.34E+00 | 1.52E-02 | 4.17E-02 | 1.11E+00 |
| WSB2 | 4.47E-07 | 2.33E-06 | -1.90E+00 | 5.35E-05 | 2.31E-04 | -1.26E+00 | 1.02E-02 | 4.34E-02 | -1.15E+00 | 5.75E-03 | 2.20E-02 | -1.06E+00 | 2.46E-06 | 8.77E-06 | -1.72E+00 | 5.80E-07 | 1.63E-06 | -1.79E+00 | 3.00E-10 | 1.10E-09 | -1.83E+00 | 8.15E-07 | 2.24E-06 | -1.54E+00 |
| PDCD5P1 | 4.47E-07 | 2.33E-06 | -1.08E+00 | 8.53E-20 | 3.68E-19 | -1.47E+00 | 1.05E-02 | 4.47E-02 | -1.09E+00 | 1.22E-09 | 4.66E-09 | -1.45E+00 | 1.90E-08 | 6.76E-08 | -1.50E+00 | 2.44E-10 | 6.89E-10 | -1.57E+00 | 6.84E-05 | 2.51E-04 | -1.17E+00 | 1.23E-09 | 3.37E-09 | -1.37E+00 |
| TRIM47 | 4.50E-07 | 2.34E-06 | -1.64E+00 | 3.57E-02 | 1.54E-01 | 1.21E+00 | 6.11E-03 | 2.60E-02 | -1.15E+00 | 4.49E-02 | 1.72E-01 | 1.15E+00 | 1.70E-01 | 6.06E-01 | -1.23E+00 | 2.69E-01 | 7.57E-01 | -1.02E+00 | 2.53E-01 | 9.30E-01 | 1.07E+00 | 3.64E-01 | 1.00E+00 | 1.12E+00 |
| DNAJC12 | 4.50E-07 | 2.35E-06 | -1.48E+00 | 3.62E-03 | 1.56E-02 | 1.08E+00 | 5.91E-09 | 2.51E-08 | -1.53E+00 | 3.38E-07 | 1.29E-06 | -1.27E+00 | 5.73E-15 | 2.04E-14 | -2.39E+00 | 4.41E-05 | 1.24E-04 | -1.16E+00 | 2.59E-15 | 9.51E-15 | -1.64E+00 | 1.47E-05 | 4.03E-05 | -1.14E+00 |
| EIF1P3 | 4.51E-07 | 2.35E-06 | -1.75E+00 | 1.99E-07 | 8.60E-07 | -1.71E+00 | 2.48E-09 | 1.05E-08 | -1.91E+00 | 3.49E-11 | 1.33E-10 | -2.08E+00 | 2.53E-04 | 9.02E-04 | -1.39E+00 | 1.76E-05 | 4.96E-05 | -1.61E+00 | 4.26E-06 | 1.56E-05 | -1.32E+00 | 4.20E-09 | 1.15E-08 | -1.77E+00 |
| LINS1 | 4.53E-07 | 2.36E-06 | -2.07E+00 | 1.32E-03 | 5.72E-03 | -1.10E+00 | 2.47E-08 | 1.05E-07 | -2.16E+00 | 3.13E-03 | 1.20E-02 | -1.06E+00 | 1.75E-03 | 6.23E-03 | -1.48E+00 | 2.08E-01 | 5.87E-01 | 1.34E+00 | 1.03E-06 | 3.79E-06 | -1.45E+00 | 1.81E-03 | 4.98E-03 | -1.12E+00 |
| IKZF1 | 4.53E-07 | 2.36E-06 | 1.80E+00 | 1.80E-04 | 7.76E-04 | 1.79E+00 | 7.43E-03 | 3.16E-02 | 1.44E+00 | 5.66E-02 | 2.16E-01 | 1.46E+00 | 6.37E-04 | 2.27E-03 | 1.10E+00 | 1.61E-02 | 4.53E-02 | 1.13E+00 | 8.02E-06 | 2.94E-05 | -1.19E+00 | 8.21E-03 | 2.25E-02 | 1.01E+00 |
| TTC38 | 4.56E-07 | 2.38E-06 | -1.60E+00 | 1.44E-06 | 6.21E-06 | -1.36E+00 | 1.22E-02 | 5.17E-02 | -1.10E+00 | 3.16E-02 | 1.21E-01 | 1.07E+00 | 6.55E-02 | 2.34E-01 | 1.10E+00 | 3.50E-02 | 9.87E-02 | 1.08E+00 | 3.47E-05 | 1.27E-04 | -1.18E+00 | 4.58E-02 | 1.26E-01 | 1.09E+00 |
| AK4 | 4.57E-07 | 2.38E-06 | -2.27E+00 | 8.96E-07 | 3.87E-06 | -2.15E+00 | 3.71E-29 | 1.58E-28 | -2.78E+00 | 1.66E-20 | 6.36E-20 | -1.80E+00 | 3.35E-03 | 1.19E-02 | -1.62E+00 | 4.96E-03 | 1.40E-02 | -1.49E+00 | 1.04E-04 | 3.82E-04 | -1.63E+00 | 4.23E-04 | 1.16E-03 | -1.55E+00 |
| DSCR3 | 4.64E-07 | 2.42E-06 | -1.22E+00 | 5.25E-04 | 2.27E-03 | 1.12E+00 | 6.11E-09 | 2.60E-08 | -1.39E+00 | 9.94E-12 | 3.80E-11 | -1.34E+00 | 3.21E-04 | 1.15E-03 | -1.27E+00 | 1.30E-03 | 3.67E-03 | -1.14E+00 | 2.24E-04 | 8.23E-04 | -1.11E+00 | 3.18E-03 | 8.74E-03 | -1.01E+00 |
| ALKBH8 | 4.65E-07 | 2.43E-06 | -2.02E+00 | 1.10E-04 | 4.75E-04 | -1.32E+00 | 2.54E-07 | 1.08E-06 | -1.77E+00 | 2.29E-05 | 8.76E-05 | -1.28E+00 | 3.40E-04 | 1.21E-03 | -2.26E+00 | 1.16E-02 | 3.28E-02 | -1.36E+00 | 2.85E-04 | 1.04E-03 | -1.49E+00 | 1.06E-03 | 2.90E-03 | -1.42E+00 |
| RPS26P13 | 4.65E-07 | 2.43E-06 | -1.41E+00 | 3.58E-12 | 1.55E-11 | -2.08E+00 | 5.14E-03 | 2.19E-02 | -1.19E+00 | 1.55E-07 | 5.92E-07 | -1.99E+00 | 8.43E-02 | 3.00E-01 | 1.03E+00 | 1.86E-03 | 5.26E-03 | -1.77E+00 | 9.89E-02 | 3.63E-01 | 1.06E+00 | 6.04E-05 | 1.66E-04 | -1.74E+00 |
| AC011405.1 | 4.68E-07 | 2.44E-06 | -2.16E+00 | 3.04E-04 | 1.31E-03 | -1.27E+00 | 8.49E-02 | 3.61E-01 | -1.07E+00 | 2.15E-01 | 8.20E-01 | 1.27E+00 | 1.17E-01 | 4.17E-01 | 1.04E+00 | 3.55E-01 | 1.00E+00 | 1.26E+00 | 1.66E-06 | 6.09E-06 | -1.83E+00 | 1.01E-04 | 2.78E-04 | -1.49E+00 |
| ULK3 | 4.70E-07 | 2.45E-06 | -1.38E+00 | 1.25E-05 | 5.40E-05 | -1.09E+00 | 5.08E-03 | 2.16E-02 | -1.18E+00 | 4.60E-03 | 1.76E-02 | -1.01E+00 | 5.04E-05 | 1.80E-04 | -1.33E+00 | 1.41E-02 | 3.98E-02 | 1.06E+00 | 1.73E-06 | 6.34E-06 | -1.13E+00 | 3.29E-03 | 9.03E-03 | -1.05E+00 |

| | | | | | | | | | | | | | | | | | | | | | | | | |
|---|---|---|---|---|---|---|---|---|---|---|---|---|---|---|---|---|---|---|---|---|---|---|---|---|
| TBXAS1 | 4.70E-07 | 2.45E-06 | -1.10E+00 | 1.38E-01 | 5.98E-01 | 1.44E+00 | 2.81E-03 | 1.19E-02 | -1.10E+00 | 9.18E-02 | 3.51E-01 | 1.25E+00 | 3.22E-03 | 1.15E-02 | 1.01E+00 | 1.78E-01 | 5.03E-01 | 1.45E+00 | 6.22E-02 | 2.28E-01 | -1.04E+00 | 1.19E-01 | 3.28E-01 | 1.14E+00 |
| DUS4L | 4.71E-07 | 2.45E-06 | -1.65E+00 | 2.73E-04 | 1.18E-03 | -1.13E+00 | 1.46E-02 | 6.20E-02 | -1.04E+00 | 9.75E-04 | 3.73E-03 | -1.11E+00 | 7.11E-04 | 2.53E-03 | -1.39E+00 | 2.57E-02 | 7.24E-02 | 1.03E+00 | 4.31E-02 | 1.58E-01 | -1.02E+00 | 1.91E-01 | 5.25E-01 | 1.14E+00 |
| ZNF559 | 4.72E-07 | 2.46E-06 | -1.55E+00 | 6.69E-04 | 2.89E-03 | -1.04E+00 | 5.36E-08 | 2.28E-07 | -1.76E+00 | 1.05E-04 | 4.01E-04 | -1.17E+00 | 1.06E-05 | 3.77E-05 | -1.73E+00 | 2.27E-03 | 6.41E-03 | -1.21E+00 | 2.17E-06 | 7.97E-06 | -1.58E+00 | 7.86E-03 | 2.16E-02 | -1.05E+00 |
| AL031729.1 | 4.73E-07 | 2.47E-06 | -1.07E+00 | 1.44E-46 | 6.23E-46 | -1.74E+00 | 3.04E-02 | 1.29E-01 | -1.00E+00 | 5.56E-17 | 2.13E-16 | -1.64E+00 | 3.78E-08 | 1.35E-07 | -1.26E+00 | 1.10E-23 | 3.10E-23 | -1.90E+00 | 7.70E-03 | 2.83E-02 | -1.10E+00 | 1.30E-11 | 3.57E-11 | -1.70E+00 |
| DPF2 | 4.77E-07 | 2.48E-06 | -1.26E+00 | 1.49E-02 | 6.44E-02 | 1.23E+00 | 4.98E-04 | 2.12E-03 | -1.14E+00 | 1.03E-02 | 3.95E-02 | 1.13E+00 | 2.28E-02 | 8.12E-02 | 1.02E+00 | 1.27E-01 | 3.57E-01 | 1.55E+00 | 5.45E-02 | 2.00E-01 | 1.12E+00 | 4.90E-02 | 1.35E-01 | 1.11E+00 |
| UNC50 | 4.79E-07 | 2.49E-06 | -1.30E+00 | 1.94E-03 | 8.39E-03 | 1.17E+00 | 1.56E-06 | 6.61E-06 | -1.47E+00 | 7.64E-04 | 2.92E-03 | -1.06E+00 | 4.04E-06 | 1.44E-05 | -1.42E+00 | 9.09E-03 | 2.56E-02 | 1.03E+00 | 2.33E-09 | 8.56E-09 | -1.47E+00 | 1.12E-02 | 3.07E-02 | 1.03E+00 |
| NSUN2 | 4.81E-07 | 2.51E-06 | -1.51E+00 | 2.40E-04 | 1.04E-03 | -1.04E+00 | 1.86E-03 | 7.90E-03 | -1.16E+00 | 5.53E-02 | 2.11E-01 | 1.15E+00 | 4.59E-04 | 1.64E-03 | -1.37E+00 | 5.80E-02 | 1.63E-01 | 1.21E+00 | 6.71E-07 | 2.46E-06 | -1.27E+00 | 1.67E-02 | 4.60E-02 | 1.00E+00 |
| TPM3 | 4.83E-07 | 2.52E-06 | -1.16E+00 | 9.69E-13 | 4.19E-12 | -1.20E+00 | 1.19E-01 | 5.04E-01 | -1.00E+00 | 5.89E-02 | 2.25E-01 | -1.01E+00 | 4.25E-04 | 1.52E-03 | -1.09E+00 | 1.87E-04 | 5.26E-04 | -1.18E+00 | 1.39E-01 | 5.09E-01 | -1.09E+00 | 2.67E-01 | 7.33E-01 | -1.08E+00 |
| USP25 | 4.84E-07 | 2.52E-06 | -1.52E+00 | 5.66E-05 | 2.45E-04 | -1.14E+00 | 8.81E-04 | 3.75E-03 | -1.14E+00 | 3.53E-03 | 1.35E-02 | 1.10E+00 | 4.06E-06 | 1.45E-05 | -1.73E+00 | 5.76E-04 | 1.62E-03 | -1.32E+00 | 3.79E-07 | 1.39E-06 | -1.35E+00 | 3.75E-04 | 1.03E-03 | -1.09E+00 |
| FPGT | 4.84E-07 | 2.52E-06 | -1.90E+00 | 5.98E-03 | 2.58E-02 | 1.06E+00 | 1.82E-06 | 7.75E-06 | -2.38E+00 | 1.94E-02 | 7.42E-02 | -1.03E+00 | 2.39E-03 | 8.54E-03 | -1.83E+00 | 6.19E-02 | 1.74E-01 | -1.17E+00 | 5.80E-05 | 2.13E-04 | -1.66E+00 | 6.51E-02 | 1.79E-01 | 1.06E+00 |
| NIPSNAP3B | 4.84E-07 | 2.52E-06 | -1.67E+00 | 1.47E-04 | 6.36E-04 | -1.15E+00 | 7.81E-03 | 3.32E-02 | -1.18E+00 | 3.34E-02 | 1.28E-01 | 1.06E+00 | 6.75E-02 | 2.41E-01 | 1.04E+00 | 1.15E-01 | 3.25E-01 | 1.65E+00 | 6.58E-02 | 2.42E-01 | 1.00E+00 | 1.07E-01 | 2.94E-01 | 1.04E+00 |
| SEC23A | 4.86E-07 | 2.53E-06 | -1.24E+00 | 1.84E-06 | 7.93E-06 | -1.08E+00 | 1.13E-09 | 4.82E-09 | -1.64E+00 | 1.82E-04 | 6.93E-04 | -1.05E+00 | 5.56E-06 | 1.98E-05 | -1.24E+00 | 5.73E-06 | 1.62E-05 | -1.22E+00 | 1.04E-08 | 3.83E-08 | -1.40E+00 | 1.50E-05 | 4.11E-05 | -1.25E+00 |
| DGKD | 4.86E-07 | 2.53E-06 | -1.51E+00 | 2.76E-02 | 1.19E-01 | 1.26E+00 | 2.93E-02 | 1.24E-01 | 1.02E+00 | 9.75E-03 | 3.73E-02 | -1.02E+00 | 4.92E-03 | 1.75E-02 | -1.09E+00 | 7.34E-03 | 2.07E-02 | -1.01E+00 | 7.24E-02 | 2.66E-01 | 1.81E+00 | 1.33E-01 | 3.66E-01 | 1.37E+00 |
| SEC11A | 4.87E-07 | 2.54E-06 | -1.07E+00 | 2.38E-05 | 1.03E-04 | 1.06E+00 | 6.01E-02 | 2.55E-01 | -1.03E+00 | 1.29E-02 | 4.94E-02 | 1.00E+00 | 1.26E-02 | 4.51E-02 | -1.09E+00 | 6.33E-02 | 1.79E-01 | -1.06E+00 | 1.49E-02 | 5.48E-02 | -1.03E+00 | 1.73E-01 | 4.74E-01 | -1.01E+00 |
| PAXBP1-AS1 | 4.90E-07 | 2.56E-06 | -1.74E+00 | 9.52E-06 | 4.11E-05 | -1.31E+00 | 4.57E-02 | 1.94E-01 | -1.03E+00 | 4.59E-02 | 1.75E-01 | 1.07E+00 | 3.74E-02 | 1.33E-01 | -1.19E+00 | 7.25E-02 | 2.04E-01 | -1.07E+00 | 1.60E-01 | 5.86E-01 | 1.62E+00 | 2.66E-01 | 7.30E-01 | 1.07E+00 |
| MAP4 | 4.93E-07 | 2.57E-06 | -1.20E+00 | 1.04E-02 | 4.51E-02 | 1.22E+00 | 2.04E-02 | 8.66E-02 | 1.01E+00 | 1.62E-01 | 6.19E-01 | 1.32E+00 | 1.50E-03 | 5.35E-03 | -1.06E+00 | 2.44E-02 | 6.88E-02 | 1.06E+00 | 8.31E-05 | 3.05E-04 | -1.18E+00 | 6.26E-03 | 1.72E-02 | -1.01E+00 |
| SAMM50 | 4.94E-07 | 2.57E-06 | -1.14E+00 | 2.32E-05 | 1.00E-04 | 1.07E+00 | 1.07E-05 | 4.56E-05 | -1.16E+00 | 4.12E-06 | 1.58E-05 | -1.08E+00 | 1.52E-06 | 5.42E-06 | -1.39E+00 | 1.76E-02 | 4.96E-02 | 1.02E+00 | 4.44E-04 | 1.63E-03 | -1.06E+00 | 1.34E-03 | 3.69E-03 | -1.05E+00 |
| PTS | 4.94E-07 | 2.57E-06 | -1.26E+00 | 7.97E-15 | 3.44E-14 | -2.01E+00 | 6.53E-06 | 2.77E-05 | -1.38E+00 | 1.72E-16 | 6.58E-16 | -2.54E+00 | 1.40E-07 | 4.99E-07 | -1.18E+00 | 2.79E-16 | 7.87E-16 | -2.26E+00 | 1.56E-08 | 5.71E-08 | -1.50E+00 | 7.82E-12 | 2.15E-11 | -1.71E+00 |
| SLC25A33 | 4.94E-07 | 2.58E-06 | -1.27E+00 | 1.44E-11 | 6.22E-11 | -1.60E+00 | 2.49E-06 | 1.06E-05 | -1.24E+00 | 1.80E-09 | 6.87E-09 | -1.32E+00 | 1.85E-02 | 6.58E-02 | 1.11E+00 | 5.17E-04 | 1.46E-03 | -1.12E+00 | 7.83E-04 | 2.87E-03 | -1.11E+00 | 2.16E-03 | 5.93E-03 | -1.12E+00 |
| CYP51A1P2 | 4.97E-07 | 2.59E-06 | -1.60E+00 | 1.06E-06 | 4.59E-06 | -1.41E+00 | 1.34E-04 | 5.70E-04 | -1.54E+00 | 9.52E-03 | 3.64E-02 | -1.07E+00 | 1.38E-01 | 4.93E-01 | 1.20E+00 | 2.59E-01 | 7.30E-01 | 1.25E+00 | 2.69E-03 | 9.87E-03 | -1.25E+00 | 3.09E-02 | 8.49E-02 | -1.13E+00 |
| CCDC169 | 4.98E-07 | 2.60E-06 | -1.97E+00 | 2.67E-04 | 1.16E-03 | -1.22E+00 | 2.02E-07 | 8.59E-07 | -2.65E+00 | 2.33E-03 | 8.88E-03 | -1.22E+00 | 1.03E-06 | 3.67E-06 | -2.58E+00 | 1.52E-03 | 4.29E-03 | -1.26E+00 | 1.38E-06 | 5.08E-06 | -1.50E+00 | 3.94E-02 | 1.08E-01 | 1.08E+00 |
| C19orf12 | 4.99E-07 | 2.60E-06 | -1.51E+00 | 5.54E-05 | 2.39E-04 | -1.14E+00 | 5.41E-02 | 2.30E-01 | -1.06E+00 | 1.14E-01 | 4.34E-01 | 1.50E+00 | 1.76E-05 | 6.27E-05 | -1.55E+00 | 3.65E-03 | 1.03E-02 | -1.10E+00 | 1.86E-14 | 6.82E-14 | -1.85E+00 | 8.75E-09 | 2.40E-08 | -1.34E+00 |
| APPL2 | 5.01E-07 | 2.61E-06 | -1.43E+00 | 1.76E-06 | 7.61E-06 | -1.26E+00 | 7.53E-04 | 3.20E-03 | -1.27E+00 | 1.70E-04 | 6.50E-04 | -1.22E+00 | 6.54E-05 | 2.33E-04 | -2.53E+00 | 6.64E-04 | 1.87E-03 | -1.88E+00 | 9.74E-03 | 3.58E-02 | -1.02E+00 | 3.59E-03 | 9.87E-03 | -1.46E+00 |
| PHAX | 5.01E-07 | 2.61E-06 | -1.17E+00 | 1.33E-08 | 5.76E-08 | -1.13E+00 | 8.42E-04 | 3.58E-03 | -1.15E+00 | 4.42E-03 | 1.69E-02 | 1.04E+00 | 3.98E-05 | 1.42E-04 | -1.27E+00 | 1.69E-02 | 4.77E-02 | 1.03E+00 | 9.89E-04 | 3.63E-03 | -1.11E+00 | 1.52E-02 | 4.19E-02 | -1.01E+00 |
| SF3A1 | 5.02E-07 | 2.62E-06 | -1.15E+00 | 3.31E-05 | 1.43E-04 | 1.10E+00 | 1.68E-06 | 7.13E-06 | -1.29E+00 | 2.63E-04 | 1.01E-03 | -1.08E+00 | 2.52E-03 | 8.99E-03 | -1.01E+00 | 9.27E-04 | 2.61E-03 | 1.03E+00 | 1.52E-05 | 5.57E-05 | -1.20E+00 | 1.93E-02 | 5.30E-02 | 1.04E+00 |
| PRRSL | 5.04E-07 | 2.63E-06 | -1.98E+00 | 8.40E-06 | 3.63E-05 | -1.42E+00 | 3.16E-09 | 1.34E-08 | -2.24E+00 | 4.55E-07 | 1.74E-06 | -1.60E+00 | 4.47E-02 | 1.59E-01 | -1.51E+00 | 1.58E-01 | 4.46E-01 | -1.20E+00 | 2.79E-02 | 1.03E-01 | -1.36E+00 | 2.10E-01 | 5.76E-01 | 1.12E+00 |
| AASS | 5.05E-07 | 2.63E-06 | -1.33E+00 | 2.07E-03 | 8.96E-03 | 1.16E+00 | 1.40E-04 | 5.93E-04 | -1.61E+00 | 3.75E-03 | 1.43E-02 | -1.06E+00 | 2.64E-02 | 9.42E-02 | -1.08E+00 | 6.92E-02 | 1.95E-01 | 1.01E+00 | 1.62E-04 | 5.94E-04 | -1.57E+00 | 1.55E-02 | 4.27E-02 | -1.17E+00 |
| ZNF507 | 5.06E-07 | 2.64E-06 | -1.40E+00 | 4.04E-06 | 1.75E-05 | -1.14E+00 | 2.11E-02 | 8.98E-02 | -1.02E+00 | 1.02E-02 | 3.90E-02 | 1.06E+00 | 1.68E-01 | 6.00E-01 | 1.24E+00 | 2.82E-01 | 7.96E-01 | 1.59E+00 | 1.57E-02 | 5.75E-02 | 1.08E+00 | 4.30E-02 | 1.18E-01 | 1.15E+00 |
| EID2 | 5.07E-07 | 2.64E-06 | -1.44E+00 | 2.20E-05 | 9.50E-05 | -1.09E+00 | 9.36E-10 | 3.98E-09 | -1.54E+00 | 1.96E-09 | 7.48E-09 | -1.32E+00 | 1.06E-04 | 3.77E-04 | -1.49E+00 | 2.24E-02 | 6.31E-02 | 1.08E+00 | 8.37E-07 | 3.07E-06 | -1.28E+00 | 7.18E-04 | 1.97E-03 | -1.03E+00 |
| NR8F2P3 | 5.08E-07 | 2.65E-06 | -1.33E+00 | 9.98E-14 | 4.31E-13 | -2.31E+00 | 3.27E-05 | 1.39E-04 | -1.37E+00 | 4.13E-10 | 1.58E-09 | -1.75E+00 | 2.09E-02 | 7.46E-02 | 1.07E+00 | 6.10E-05 | 1.72E-04 | -1.56E+00 | 9.34E-05 | 3.43E-04 | -1.11E+00 | 1.75E-09 | 4.82E-09 | -1.74E+00 |
| CLPTM1 | 5.08E-07 | 2.65E-06 | -1.49E+00 | 1.62E-02 | 7.00E-02 | 1.24E+00 | 4.38E-04 | 1.86E-03 | -1.20E+00 | 6.86E-02 | 2.62E-01 | 1.20E+00 | 1.82E-05 | 6.48E-05 | -1.33E+00 | 2.53E-03 | 7.13E-03 | -1.01E+00 | 1.36E-04 | 4.99E-04 | -1.18E+00 | 1.59E-02 | 4.36E-02 | 1.00E+00 |
| CHUK | 5.08E-07 | 2.65E-06 | -1.63E+00 | 2.91E-07 | 1.26E-06 | -1.53E+00 | 1.24E-07 | 5.27E-07 | -1.66E+00 | 5.89E-05 | 2.25E-04 | -1.22E+00 | 1.79E-03 | 6.37E-03 | -1.44E+00 | 1.68E-03 | 4.74E-03 | -1.46E+00 | 2.68E-07 | 9.85E-07 | -1.59E+00 | 1.10E-03 | 3.01E-03 | -1.21E+00 |
| MAN1B1 | 5.10E-07 | 2.66E-06 | -1.35E+00 | 3.51E-06 | 1.52E-05 | -1.08E+00 | 2.12E-03 | 9.02E-03 | -1.14E+00 | 1.98E-02 | 7.55E-02 | 1.09E+00 | 3.41E-06 | 1.21E-05 | -1.32E+00 | 3.12E-05 | 8.80E-05 | -1.15E+00 | 9.31E-05 | 3.42E-04 | -1.26E+00 | 7.78E-03 | 2.14E-02 | -1.00E+00 |
| ZBTB20-AS3 | 5.10E-07 | 2.66E-06 | -2.16E+00 | 3.37E-04 | 1.45E-03 | -1.23E+00 | 3.91E-04 | 1.66E-03 | -2.31E+00 | 1.30E-01 | 4.98E-01 | 1.11E+00 | 5.60E-02 | 2.00E-01 | -1.26E+00 | 2.10E-01 | 5.91E-01 | 1.04E+00 | 1.73E-01 | 6.35E-01 | -1.09E+00 | 2.11E-01 | 5.79E-01 | 1.28E+00 |
| ZNF273 | 5.10E-07 | 2.66E-06 | -1.30E+00 | 6.32E-09 | 2.73E-08 | -1.33E+00 | 3.03E-05 | 1.29E-04 | -1.32E+00 | 2.23E-06 | 8.53E-06 | -1.29E+00 | 4.76E-03 | 1.70E-02 | -1.07E+00 | 4.43E-02 | 1.25E-01 | 1.16E+00 | 3.67E-07 | 1.35E-06 | -1.23E+00 | 1.50E-05 | 4.11E-05 | -1.23E+00 |
| RIPOR3 | 5.10E-07 | 2.66E-06 | -1.90E+00 | 2.13E-04 | 9.22E-04 | -1.17E+00 | 7.18E-02 | 3.05E-01 | -1.32E+00 | 1.94E-01 | 7.42E-01 | 1.19E+00 | 2.05E-10 | 7.30E-10 | -1.35E+00 | 1.86E-02 | 5.24E-02 | 1.15E+00 | 3.62E-19 | 1.33E-18 | -1.31E+00 | 5.53E-07 | 1.52E-06 | -1.15E+00 |
| TCEA1 | 5.11E-07 | 2.66E-06 | -1.13E+00 | 6.20E-07 | 2.68E-06 | 1.00E+00 | 1.87E-03 | 7.96E-03 | -1.08E+00 | 8.82E-06 | 3.37E-05 | -1.04E+00 | 8.74E-06 | 3.11E-05 | -1.22E+00 | 2.72E-04 | 7.66E-04 | -1.05E+00 | 8.51E-05 | 3.12E-04 | -1.17E+00 | 8.63E-05 | 2.37E-04 | -1.16E+00 |
| LRP10 | 5.17E-07 | 2.69E-06 | -1.50E+00 | 4.56E-05 | 1.97E-04 | -1.14E+00 | 5.17E-06 | 2.20E-05 | -1.67E+00 | 8.04E-06 | 3.07E-05 | -1.44E+00 | 4.94E-03 | 1.76E-02 | 1.05E+00 | 4.59E-04 | 1.29E-03 | -1.09E+00 | 7.42E-03 | 2.72E-02 | 1.11E+00 | 4.06E-03 | 1.12E-02 | 1.02E+00 |
| DPH5 | 5.19E-07 | 2.70E-06 | -1.17E+00 | 2.85E-08 | 1.23E-07 | -1.10E+00 | 1.87E-04 | 7.96E-04 | -1.15E+00 | 4.38E-04 | 1.67E-03 | -1.01E+00 | 1.30E-02 | 4.65E-02 | 1.02E+00 | 2.78E-02 | 7.85E-02 | 1.17E+00 | 1.27E-02 | 4.67E-02 | -1.05E+00 | 1.37E-01 | 3.78E-01 | 1.13E+00 |
| CSNK2A1 | 5.20E-07 | 2.71E-06 | -1.24E+00 | 3.38E-03 | 1.46E-02 | 1.12E+00 | 2.19E-03 | 9.32E-03 | -1.11E+00 | 2.37E-02 | 9.04E-02 | 1.17E+00 | 3.45E-05 | 1.23E-04 | -1.16E+00 | 1.25E-01 | 3.51E-01 | 1.25E+00 | 8.10E-08 | 2.97E-07 | -1.30E+00 | 7.64E-03 | 2.10E-02 | 1.01E+00 |
| AC005336.3 | 5.23E-07 | 2.73E-06 | -2.23E+00 | 1.26E-02 | 5.46E-02 | 1.03E+00 | 8.73E-03 | 3.71E-02 | -1.36E+00 | 8.01E-02 | 3.06E-01 | 1.63E+00 | 1.24E-01 | 4.40E-01 | 1.03E+00 | 2.06E-01 | 5.80E-01 | 1.77E+00 | 2.42E-01 | 8.87E-01 | -1.01E+00 | 1.71E-01 | 4.69E-01 | 1.54E+00 |
| ELOBP2 | 5.24E-07 | 2.73E-06 | -1.38E+00 | 4.49E-15 | 1.94E-14 | -2.61E+00 | 2.14E-02 | 9.08E-02 | -1.05E+00 | 3.40E-07 | 1.30E-06 | -1.63E+00 | 2.61E-07 | 9.30E-07 | -1.54E+00 | 3.15E-08 | 8.87E-08 | -1.61E+00 | 1.66E-06 | 6.09E-06 | -1.29E+00 | 2.29E-14 | 6.30E-14 | -1.95E+00 |
| RAB3GAP2 | 5.24E-07 | 2.73E-06 | -1.19E+00 | 1.03E-03 | 4.47E-03 | 1.18E+00 | 3.92E-04 | 1.67E-03 | -1.22E+00 | 5.08E-04 | 1.94E-03 | 1.01E+00 | 2.60E-04 | 9.25E-04 | -1.21E+00 | 4.58E-03 | 1.29E-02 | 1.02E+00 | 2.90E-03 | 1.06E-02 | -1.10E+00 | 1.47E-01 | 4.05E-01 | 1.09E+00 |
| LRRC8C-DT | 5.25E-07 | 2.74E-06 | -1.42E+00 | 6.82E-05 | 2.95E-04 | -1.11E+00 | 2.24E-01 | 9.51E-01 | 1.17E+00 | 1.62E-02 | 6.20E-02 | -1.04E+00 | 8.65E-02 | 3.09E-01 | 1.20E+00 | 1.64E-02 | 4.63E-02 | -1.12E+00 | 4.23E-02 | 1.55E-01 | -1.15E+00 | 1.28E-02 | 3.51E-02 | -1.52E+00 |
| OAZ1 | 5.25E-07 | 2.74E-06 | -1.11E+00 | 8.73E-02 | 3.77E-01 | 1.10E+00 | 1.76E-01 | 7.49E-01 | 1.06E+00 | 1.07E-02 | 4.08E-02 | 1.15E+00 | 3.24E-04 | 1.15E-03 | -1.09E+00 | 2.96E-01 | 8.36E-01 | 1.05E+00 | 1.20E-04 | 4.40E-04 | 1.35E+00 | 7.64E-03 | 2.10E-02 | 1.16E+00 |
| P2RX4 | 5.25E-07 | 2.74E-06 | -1.17E+00 | 6.72E-07 | 2.91E-06 | -1.04E+00 | 3.23E-05 | 1.37E-04 | -1.27E+00 | 2.87E-05 | 1.10E-04 | -1.12E+00 | 4.58E-03 | 1.63E-02 | -1.02E+00 | 3.04E-03 | 8.56E-03 | -1.08E+00 | 6.03E-04 | 2.21E-03 | -1.29E+00 | 3.27E-04 | 8.99E-04 | -1.41E+00 |
| NFIA | 5.29E-07 | 2.76E-06 | -1.12E+00 | 2.13E-01 | 9.21E-01 | 1.41E+00 | 1.84E-02 | 7.80E-02 | -1.05E+00 | 1.56E-01 | 5.96E-01 | 1.53E+00 | 1.35E-03 | 4.82E-03 | -1.21E+00 | 2.57E-01 | 7.26E-01 | 1.33E+00 | 1.67E-10 | 6.14E-10 | -1.75E+00 | 4.24E-04 | 1.16E-03 | -1.30E+00 |
| PSMD14 | 5.29E-07 | 2.76E-06 | -1.21E+00 | 1.74E-04 | 7.52E-04 | 1.09E+00 | 7.20E-06 | 3.06E-05 | -1.12E+00 | 3.08E-04 | 1.18E-03 | 1.08E+00 | 8.53E-05 | 3.04E-04 | -1.19E+00 | 1.31E-01 | 3.70E-01 | 1.15E+00 | 1.47E-02 | 5.39E-02 | 1.05E+00 | 1.56E-01 | 4.28E-01 | 1.33E+00 |
| METTL3 | 5.30E-07 | 2.76E-06 | -1.38E+00 | 3.59E-06 | 1.55E-05 | -1.19E+00 | 5.94E-07 | 2.52E-06 | -1.58E+00 | 1.93E-07 | 7.37E-07 | -1.39E+00 | 8.38E-06 | 2.99E-05 | -1.42E+00 | 6.02E-04 | 1.70E-03 | -1.15E+00 | 3.24E-09 | 1.19E-08 | -1.64E+00 | 9.18E-06 | 2.52E-05 | -1.35E+00 |
| ARHGAP23 | 5.30E-07 | 2.76E-06 | -2.05E+00 | 7.03E-05 | 3.04E-04 | -1.37E+00 | 6.73E-04 | 2.86E-03 | -1.44E+00 | 5.57E-04 | 2.13E-03 | -1.33E+00 | 3.65E-02 | 1.30E-01 | -1.44E+00 | 3.43E-02 | 9.68E-02 | -1.64E+00 | 1.39E-01 | 5.09E-01 | -1.18E+00 | 2.27E-01 | 6.23E-01 | -1.08E+00 |
| GKAP1 | 5.31E-07 | 2.77E-06 | -1.10E+00 | 1.27E-06 | 5.48E-06 | 1.02E+00 | 2.36E-06 | 1.00E-05 | -1.21E+00 | 3.02E-07 | 1.15E-06 | -1.10E+00 | 1.78E-05 | 6.34E-05 | -1.19E+00 | 4.58E-04 | 1.29E-03 | -1.07E+00 | 2.01E-03 | 7.38E-03 | -1.04E+00 | 3.57E-04 | 9.80E-04 | -1.20E+00 |
| SAMD10 | 5.33E-07 | 2.78E-06 | -1.47E+00 | 4.99E-04 | 2.15E-03 | 1.02E+00 | 1.25E-08 | 5.31E-08 | -1.88E+00 | 6.80E-05 | 2.60E-04 | -1.19E+00 | 9.62E-03 | 3.43E-02 | -1.74E+00 | 1.03E-01 | 2.89E-01 | 1.00E+00 | 2.94E-03 | 1.08E-02 | -1.68E+00 | 8.42E-02 | 2.31E-01 | -1.16E+00 |
| SUPV3L1 | 5.33E-07 | 2.78E-06 | -1.35E+00 | 5.41E-09 | 2.34E-08 | -1.35E+00 | 1.82E-05 | 7.75E-05 | -1.31E+00 | 1.70E-08 | 6.49E-08 | -1.36E+00 | 6.89E-06 | 2.46E-05 | -1.45E+00 | 9.67E-06 | 2.73E-05 | -1.37E+00 | 5.96E-07 | 2.19E-06 | -1.25E+00 | 2.87E-03 | 7.88E-03 | -1.03E+00 |
| NBR2 | 5.33E-07 | 2.78E-06 | -1.93E+00 | 2.63E-05 | 1.14E-04 | -1.44E+00 | 1.40E-04 | 5.93E-04 | -1.71E+00 | 2.30E-02 | 8.80E-02 | -1.01E+00 | 1.60E-01 | 5.69E-01 | 1.29E+00 | 3.16E-01 | 8.91E-01 | 1.59E+00 | 1.50E-02 | 5.51E-02 | -1.15E+00 | 2.15E-01 | 5.90E-01 | 1.19E+00 |
| SMAP2 | 5.38E-07 | 2.80E-06 | 2.07E+00 | 4.44E-04 | 1.92E-03 | 2.04E+00 | 1.91E-08 | 8.13E-08 | 2.12E+00 | 2.83E-04 | 1.08E-03 | 1.88E+00 | 3.30E-03 | 1.18E-02 | 2.02E+00 | 2.78E-01 | 7.84E-01 | 1.71E+00 | 1.22E-03 | 4.49E-03 | 1.63E+00 | 1.34E-01 | 3.69E-01 | 1.14E+00 |
| RPE | 5.40E-07 | 2.82E-06 | -1.52E+00 | 3.79E-02 | 1.64E-01 | 1.32E+00 | 1.42E-06 | 6.02E-06 | -1.44E+00 | 2.14E-04 | 8.19E-04 | -1.06E+00 | 3.49E-07 | 1.25E-06 | -1.91E+00 | 4.78E-04 | 1.35E-03 | -1.15E+00 | 3.85E-05 | 1.41E-04 | -1.26E+00 | 2.49E-01 | 6.85E-01 | 1.21E+00 |
| ATXN7L1 | 5.40E-07 | 2.82E-06 | -1.47E+00 | 9.49E-06 | 4.10E-05 | -1.20E+00 | 3.66E-05 | 1.56E-04 | -1.51E+00 | 7.95E-03 | 3.04E-02 | 1.06E+00 | 1.20E-05 | 4.28E-05 | -1.60E+00 | 1.30E-03 | 3.67E-03 | -1.20E+00 | 8.76E-06 | 3.22E-05 | -1.39E+00 | 1.54E-04 | 4.22E-04 | -1.29E+00 |
| IFRD2 | 5.40E-07 | 2.82E-06 | -2.06E+00 | 8.78E-05 | 3.80E-04 | -1.36E+00 | 6.01E-05 | 2.55E-04 | -1.67E+00 | 1.23E-02 | 4.71E-02 | -1.03E+00 | 1.54E-04 | 5.50E-04 | -2.06E+00 | 1.45E-03 | 4.08E-03 | -1.47E+00 | 1.30E-10 | 4.76E-10 | -1.58E+00 | 2.92E-04 | 8.03E-04 | -1.11E+00 |
| ODF2 | 5.40E-07 | 2.82E-06 | -1.31E+00 | 2.35E-03 | 1.02E-02 | 1.13E+00 | 1.46E-04 | 6.20E-04 | -1.20E+00 | 6.86E-05 | 2.62E-04 | -1.06E+00 | 3.92E-02 | 1.40E-01 | 1.05E+00 | 2.81E-01 | 7.93E-01 | 1.30E+00 | 3.42E-05 | 1.25E-04 | -1.22E+00 | 1.51E-01 | 4.14E-01 | 1.09E+00 |
| MT1F | 5.40E-07 | 2.82E-06 | -1.18E+00 | 6.79E-14 | 2.93E-13 | -1.63E+00 | 1.29E-08 | 5.47E-08 | -1.52E+00 | 9.55E-16 | 3.65E-15 | -2.15E+00 | 1.46E-07 | 5.22E-07 | -1.33E+00 | 2.19E-11 | 6.16E-11 | -1.56E+00 | 1.89E-01 | 6.95E-01 | 1.20E+00 | 1.44E-03 | 3.97E-03 | -1.35E+00 |
| FASTKD2 | 5.45E-07 | 2.84E-06 | -1.21E+00 | 1.33E-05 | 5.74E-05 | 1.04E+00 | 9.67E-08 | 4.11E-07 | -1.33E+00 | 5.38E-07 | 2.06E-06 | -1.11E+00 | 5.94E-03 | 2.12E-02 | 1.04E+00 | 4.05E-04 | 1.14E-03 | -1.11E+00 | 4.74E-09 | 1.74E-08 | -1.42E+00 | 1.06E-04 | 2.91E-04 | -1.11E+00 |
| CYP51A1 | 5.45E-07 | 2.84E-06 | -1.97E+00 | 1.40E-04 | 6.06E-04 | -1.19E+00 | 1.04E-01 | 4.44E-01 | -1.03E+00 | 7.66E-02 | 2.93E-01 | 1.69E+00 | 1.94E-01 | 6.92E-01 | 1.06E+00 | 2.51E-01 | 7.08E-01 | 1.45E+00 | 3.98E-02 | 1.46E-01 | -1.14E+00 | 2.70E-01 | 7.42E-01 | 1.12E+00 |
| MYSM1 | 5.49E-07 | 2.86E-06 | -1.21E+00 | 8.84E-07 | 3.82E-06 | -1.06E+00 | 4.34E-05 | 1.85E-04 | -1.15E+00 | 3.45E-05 | 1.32E-04 | -1.05E+00 | 1.49E-03 | 5.30E-03 | -1.00E+00 | 4.06E-03 | 1.14E-02 | 1.05E+00 | 1.50E-03 | 5.49E-03 | -1.04E+00 | 1.42E-02 | 3.90E-02 | -1.01E+00 |
| AC016065.1 | 5.52E-07 | 2.88E-06 | -1.44E+00 | 6.03E-04 | 2.60E-03 | 1.00E+00 | 3.39E-07 | 1.44E-06 | -1.42E+00 | 1.47E-05 | 5.60E-05 | -1.12E+00 | 5.41E-05 | 1.93E-04 | -1.38E+00 | 3.10E-02 | 8.75E-02 | 1.12E+00 | 1.18E-05 | 4.34E-05 | -1.41E+00 | 2.94E-03 | 8.08E-03 | -1.09E+00 |
| GAPVD1 | 5.52E-07 | 2.88E-06 | -1.16E+00 | 3.63E-06 | 1.57E-05 | -1.01E+00 | 1.91E-02 | 8.11E-02 | 1.01E+00 | 7.18E-04 | 2.74E-03 | 1.02E+00 | 9.23E-08 | 3.29E-07 | -1.24E+00 | 1.16E-07 | 3.26E-07 | -1.21E+00 | 2.75E-03 | 1.01E-02 | -1.12E+00 | 2.96E-03 | 8.13E-03 | -1.13E+00 |
| TWNK | 5.54E-07 | 2.89E-06 | -2.32E+00 | 1.12E-03 | 4.83E-03 | -1.15E+00 | 1.93E-04 | 8.19E-04 | -1.61E+00 | 1.66E-01 | 6.35E-01 | 1.29E+00 | 1.54E-01 | 5.51E-01 | 1.11E+00 | 1.50E-01 | 4.24E-01 | 1.92E+00 | 8.57E-10 | 3.15E-09 | -1.88E+00 | 1.48E-02 | 4.05E-02 | 1.03E+00 |
| CYP7B1 | 5.56E-07 | 2.90E-06 | -2.25E+00 | 1.09E-05 | 4.72E-05 | -1.70E+00 | 5.65E-05 | 2.40E-04 | -1.66E+00 | 1.00E-04 | 3.82E-04 | -1.40E+00 | 1.60E-10 | 5.72E-10 | -1.88E+00 | 5.03E-08 | 1.42E-07 | -1.42E+00 | 6.88E-08 | 2.52E-07 | -1.38E+00 | 9.40E-10 | 2.58E-09 | -1.52E+00 |
| DICER1 | 5.56E-07 | 2.90E-06 | -1.20E+00 | 5.89E-06 | 2.55E-05 | -1.00E+00 | 5.41E-09 | 2.30E-08 | -1.45E+00 | 1.24E-03 | 4.75E-03 | 1.04E+00 | 8.87E-08 | 3.16E-07 | -1.43E+00 | 9.66E-06 | 2.72E-05 | -1.19E+00 | 6.87E-12 | 2.52E-11 | -1.51E+00 | 1.44E-04 | 3.95E-04 | -1.10E+00 |
| CARD16 | 5.59E-07 | 2.91E-06 | -1.06E+00 | 5.51E-10 | 2.38E-09 | -1.07E+00 | 6.85E-03 | 2.91E-02 | -1.05E+00 | 8.57E-09 | 3.28E-08 | -1.32E+00 | 3.26E-04 | 1.16E-03 | -1.24E+00 | 6.60E-04 | 1.86E-03 | -1.18E+00 | 1.50E-10 | 5.51E-10 | -1.36E+00 | 1.59E-09 | 4.37E-09 | -1.43E+00 |
| HMGCL | 5.64E-07 | 2.94E-06 | -1.40E+00 | 2.91E-02 | 1.26E-01 | 1.28E+00 | 2.96E-06 | 1.26E-05 | -1.43E+00 | 2.30E-04 | 8.77E-04 | -1.05E+00 | 4.39E-06 | 1.56E-05 | -1.26E+00 | 1.54E-01 | 4.34E-01 | 1.42E+00 | 1.77E-03 | 6.50E-03 | 1.02E+00 | 1.26E-03 | 3.47E-03 | -1.04E+00 |
| B3GNTL1 | 5.64E-07 | 2.94E-06 | -1.60E+00 | 1.45E-04 | 6.26E-04 | -1.13E+00 | 5.76E-05 | 2.45E-04 | -1.30E+00 | 1.72E-03 | 6.56E-03 | -1.01E+00 | 3.43E-06 | 1.22E-05 | -1.88E+00 | 2.52E-03 | 7.09E-03 | -1.10E+00 | 1.39E-04 | 5.09E-04 | -1.62E+00 | 2.22E-02 | 6.09E-02 | -1.17E+00 |
| SPATA13 | 5.65E-07 | 2.94E-06 | -1.36E+00 | 9.41E-09 | 4.07E-08 | -1.41E+00 | 1.01E-04 | 4.28E-04 | -1.30E+00 | 2.67E-04 | 1.02E-03 | -1.05E+00 | 1.33E-01 | 4.76E-01 | 1.01E+00 | 7.84E-02 | 2.21E-01 | -1.12E+00 | 7.62E-05 | 2.80E-04 | -1.48E+00 | 1.45E-02 | 3.99E-02 | -1.17E+00 |
| ZNF483 | 5.66E-07 | 2.95E-06 | -1.85E+00 | 5.36E-06 | 2.32E-05 | -1.46E+00 | 1.40E-01 | 5.94E-01 | 1.03E+00 | 7.32E-02 | 2.80E-01 | -1.05E+00 | 2.14E-02 | 7.62E-02 | -1.49E+00 | 1.26E-02 | 3.55E-02 | -1.76E+00 | 6.05E-02 | 2.22E-01 | -1.19E+00 | 2.12E-02 | 5.82E-02 | -1.38E+00 |
| AL355987.4 | 5.66E-07 | 2.95E-06 | -2.21E+00 | 3.67E-02 | 1.59E-01 | 1.16E+00 | 8.22E-03 | 3.49E-02 | -1.24E+00 | 2.48E-01 | 9.47E-01 | 1.42E+00 | 2.11E-02 | 7.51E-02 | -1.04E+00 | 1.61E-01 | 4.53E-01 | 1.32E+00 | 4.88E-02 | 1.79E-01 | -1.05E+00 | 2.38E-02 | 6.55E-02 | 1.69E+00 |
| PFDN6 | 5.67E-07 | 2.96E-06 | -1.11E+00 | 2.28E-05 | 9.86E-05 | 1.11E+00 | 1.10E-02 | 4.65E-02 | -1.01E+00 | 2.59E-03 | 9.90E-03 | 1.07E+00 | 1.16E-07 | 4.14E-07 | -1.31E+00 | 4.97E-03 | 1.40E-02 | 1.08E+00 | 2.41E-04 | 8.85E-04 | -1.06E+00 | 7.64E-02 | 2.10E-01 | 1.08E+00 |
| RPA3 | 5.68E-07 | 2.96E-06 | -1.12E+00 | 1.72E-06 | 7.41E-06 | -1.03E+00 | 1.15E-01 | 4.90E-01 | 1.10E+00 | 2.47E-03 | 9.43E-03 | 1.03E+00 | 1.69E-07 | 6.04E-07 | -1.50E+00 | 4.00E-05 | 1.13E-04 | -1.20E+00 | 4.25E-02 | 1.56E-01 | 1.05E+00 | 3.24E-01 | 8.89E-01 | 1.17E+00 |
| AC008105.3 | 5.68E-07 | 2.96E-06 | -1.13E+00 | 1.66E-01 | 7.16E-01 | 1.42E+00 | 2.87E-03 | 1.22E-02 | -1.12E+00 | 3.63E-02 | 1.39E-01 | 1.17E+00 | 3.52E-06 | 1.26E-05 | -1.30E+00 | 3.02E-02 | 8.52E-02 | 1.16E+00 | 2.34E-01 | 8.59E-01 | 1.17E+00 | 1.81E-01 | 4.97E-01 | 1.38E+00 |
| PNISR | 5.69E-07 | 2.96E-06 | -1.14E+00 | 8.54E-11 | 3.69E-10 | 1.37E+00 | 2.91E-06 | 1.24E-05 | -1.18E+00 | 6.95E-04 | 2.66E-03 | 1.24E+00 | 1.78E-03 | 6.34E-03 | -1.07E+00 | 2.25E-03 | 6.35E-03 | 1.25E+00 | 7.11E-05 | 2.61E-04 | -1.06E+00 | 2.56E-03 | 7.03E-03 | 1.23E+00 |
| ITM2A | 5.73E-07 | 2.99E-06 | -1.09E+00 | 3.03E-02 | 1.31E-01 | 1.27E+00 | 1.03E-02 | 4.37E-02 | -1.13E+00 | 6.44E-02 | 2.46E-01 | 1.03E+00 | 1.01E-05 | 3.59E-05 | -1.08E+00 | 2.78E-02 | 7.84E-02 | 1.13E+00 | 2.51E-04 | 9.21E-04 | -1.10E+00 | 1.47E-02 | 4.03E-02 | -1.00E+00 |
| JCHAIN | 5.73E-07 | 2.99E-06 | -1.41E+01 | 1.87E-07 | 8.10E-07 | -1.73E+01 | 1.50E-01 | 6.39E-01 | -1.36E+00 | 3.83E-02 | 1.46E-01 | -2.13E+00 | 2.80E-01 | 1.00E+00 | 1.00E+00 | 3.55E-01 | 1.00E+00 | 1.00E+00 | 2.72E-01 | 1.00E+00 | 1.00E+00 | 3.64E-01 | 1.00E+00 | 1.00E+00 |
| MCEE | 5.75E-07 | 2.99E-06 | -1.30E+00 | 2.68E-05 | 1.16E-04 | -1.01E+00 | 1.83E-03 | 7.76E-03 | -1.18E+00 | 1.57E-02 | 5.99E-02 | 1.07E+00 | 1.14E-04 | 4.05E-04 | -1.19E+00 | 2.68E-03 | 7.56E-03 | 1.04E+00 | 5.33E-04 | 1.95E-03 | -1.07E+00 | 4.38E-04 | 1.20E-03 | -1.13E+00 |
| DPY19L4 | 5.75E-07 | 3.00E-06 | -1.34E+00 | 1.68E-02 | 7.26E-02 | 1.30E+00 | 1.93E-05 | 8.20E-05 | -1.38E+00 | 8.72E-03 | 3.33E-02 | 1.02E+00 | 4.15E-06 | 1.48E-05 | -1.62E+00 | 3.43E-04 | 9.67E-04 | -1.25E+00 | 8.40E-05 | 3.08E-04 | -1.25E+00 | 3.98E-02 | 1.09E-01 | 1.08E+00 |
| SLC22A16 | 5.79E-07 | 3.02E-06 | -1.29E+00 | 3.62E-04 | 1.57E-03 | 1.07E+00 | 2.95E-03 | 1.25E-02 | -1.12E+00 | 1.97E-04 | 7.51E-04 | -1.05E+00 | 3.38E-03 | 1.20E-02 | -1.47E+00 | 2.76E-03 | 7.79E-03 | -1.51E+00 | 1.95E-03 | 7.17E-03 | -1.74E+00 | 1.22E-01 | 3.34E-01 | -1.03E+00 |
| MTA3 | 5.80E-07 | 3.02E-06 | -1.79E+00 | 7.65E-05 | 3.31E-04 | -1.19E+00 | 1.27E-03 | 5.40E-03 | -1.13E+00 | 1.81E-05 | 6.90E-05 | -1.23E+00 | 8.28E-04 | 2.95E-03 | -2.01E+00 | 7.99E-03 | 2.25E-02 | -1.51E+00 | 9.28E-09 | 3.40E-08 | -1.92E+00 | 4.73E-05 | 1.30E-04 | -1.36E+00 |
| ABCF2 | 5.80E-07 | 3.02E-06 | -1.39E+00 | 5.03E-05 | 2.17E-04 | -1.05E+00 | 1.25E-03 | 5.31E-03 | -1.16E+00 | 1.80E-01 | 6.86E-01 | 1.45E+00 | 6.37E-04 | 2.27E-03 | -1.19E+00 | 2.18E-01 | 6.16E-01 | 1.39E+00 | 9.43E-05 | 3.46E-04 | -1.07E+00 | 2.11E-01 | 5.80E-01 | 1.35E+00 |
| C1orf228 | 5.81E-07 | 3.03E-06 | 1.36E+00 | 3.82E-06 | 1.65E-05 | 1.41E+00 | 5.26E-02 | 2.24E-01 | 1.13E+00 | 7.29E-03 | 2.78E-02 | 1.28E+00 | 9.39E-02 | 3.35E-01 | 1.00E+00 | 8.66E-04 | 2.44E-03 | -1.12E+00 | 1.17E-02 | 4.28E-02 | 1.15E+00 | 4.66E-02 | 1.28E-01 | 1.12E+00 |
| B4GALT5 | 5.81E-07 | 3.03E-06 | -1.83E+00 | 2.81E-06 | 1.21E-05 | -1.56E+00 | 8.62E-04 | 3.66E-03 | -1.37E+00 | 2.98E-03 | 1.14E-02 | -1.18E+00 | 1.04E-02 | 3.70E-02 | -1.22E+00 | 7.77E-03 | 2.19E-02 | -1.19E+00 | 5.16E-04 | 1.89E-03 | -1.30E+00 | 6.29E-06 | 1.73E-05 | -1.59E+00 |
| IPO5P1 | 5.81E-07 | 3.03E-06 | -1.57E+00 | 7.19E-07 | 3.11E-06 | -1.51E+00 | 5.72E-05 | 2.43E-04 | -1.61E+00 | 1.69E-04 | 6.44E-04 | -1.39E+00 | 3.01E-05 | 1.07E-04 | -1.52E+00 | 3.16E-05 | 8.92E-05 | -1.54E+00 | 4.70E-04 | 1.73E-03 | -1.53E+00 | 1.00E-03 | 2.75E-03 | -1.66E+00 |
| GPR137 | 5.82E-07 | 3.03E-06 | -1.46E+00 | 1.08E-07 | 4.68E-07 | -1.46E+00 | 4.44E-06 | 1.89E-05 | -1.39E+00 | 4.99E-09 | 1.91E-08 | -1.59E+00 | 2.17E-05 | 7.74E-05 | -1.55E+00 | 1.21E-06 | 3.40E-06 | -1.77E+00 | 2.44E-02 | 8.95E-02 | 1.01E+00 | 7.07E-02 | 1.94E-01 | 1.02E+00 |
| CDC34 | 5.83E-07 | 3.04E-06 | -1.28E+00 | 1.96E-06 | 8.45E-06 | -1.12E+00 | 8.43E-05 | 3.58E-04 | -1.18E+00 | 7.70E-06 | 2.94E-05 | -1.11E+00 | 4.99E-07 | 1.78E-06 | -1.44E+00 | 9.21E-05 | 2.60E-04 | -1.14E+00 | 1.11E-05 | 4.06E-05 | -1.15E+00 | 4.68E-03 | 1.29E-02 | 1.02E+00 |
| AL133243.3 | 5.83E-07 | 3.04E-06 | -1.41E+00 | 1.19E-01 | 5.16E-01 | 1.38E+00 | 4.25E-04 | 1.81E-03 | -1.30E+00 | 1.25E-01 | 4.78E-01 | 1.29E+00 | 1.54E-03 | 5.48E-03 | -1.18E+00 | 2.45E-01 | 6.92E-01 | 1.37E+00 | 4.09E-04 | 1.50E-03 | -1.23E+00 | 1.61E-01 | 4.41E-01 | 1.14E+00 |
| SMIM30 | 5.86E-07 | 3.05E-06 | -1.04E+00 | 8.01E-06 | 3.46E-05 | 1.13E+00 | 5.61E-05 | 2.38E-04 | -1.14E+00 | 8.54E-05 | 3.26E-04 | 1.04E+00 | 3.52E-11 | 1.26E-10 | -1.59E+00 | 3.50E-03 | 9.87E-03 | 1.04E+00 | 3.76E-03 | 1.38E-02 | 1.00E+00 | 4.43E-02 | 1.22E-01 | 1.09E+00 |
| TYK2 | 5.89E-07 | 3.07E-06 | -1.40E+00 | 4.44E-03 | 1.92E-02 | 1.17E+00 | 1.46E-02 | 6.21E-02 | -1.13E+00 | 1.77E-02 | 6.75E-02 | 1.08E+00 | 4.85E-04 | 1.73E-03 | -1.27E+00 | 2.32E-01 | 6.54E-01 | 1.39E+00 | 2.14E-04 | 7.84E-04 | -1.25E+00 | 1.05E-01 | 2.88E-01 | 1.06E+00 |
| CU459211.1 | 5.90E-07 | 3.07E-06 | -1.32E+00 | 9.69E-04 | 4.19E-03 | 1.09E+00 | 1.32E-04 | 5.60E-04 | -1.21E+00 | 1.58E-02 | 6.03E-02 | 1.12E+00 | 4.19E-04 | 1.49E-03 | -1.22E+00 | 6.73E-03 | 1.90E-02 | -1.02E+00 | 1.52E-04 | 5.58E-04 | -1.13E+00 | 6.60E-02 | 1.81E-01 | 1.17E+00 |
| FGF14-AS2 | 5.91E-07 | 3.08E-06 | -1.90E+00 | 3.62E-05 | 1.56E-04 | -1.36E+00 | 1.16E-03 | 4.91E-03 | -1.26E+00 | 6.30E-04 | 2.41E-03 | -1.15E+00 | 9.53E-03 | 3.40E-02 | -1.22E+00 | 2.52E-02 | 7.10E-02 | -1.14E+00 | 2.56E-01 | 9.41E-01 | 1.28E+00 | 3.07E-01 | 8.43E-01 | 1.34E+00 |
| KIAA1191 | 5.92E-07 | 3.09E-06 | -1.35E+00 | 7.08E-03 | 3.06E-02 | 1.16E+00 | 1.16E-05 | 4.92E-05 | -1.45E+00 | 4.45E-04 | 1.70E-03 | -1.09E+00 | 1.98E-04 | 7.08E-04 | -1.37E+00 | 7.79E-03 | 2.20E-02 | -1.08E+00 | 2.10E-01 | 7.72E-01 | 1.29E+00 | 3.34E-01 | 9.17E-01 | 1.32E+00 |
| ARL3 | 5.95E-07 | 3.10E-06 | -1.21E+00 | 1.73E-04 | 7.47E-04 | 1.13E+00 | 5.83E-02 | 2.48E-01 | 1.08E+00 | 9.75E-03 | 3.72E-02 | 1.10E+00 | 3.40E-03 | 1.21E-02 | -1.10E+00 | 7.71E-03 | 2.17E-02 | -1.02E+00 | 9.15E-02 | 3.36E-01 | 1.14E+00 | 2.12E-01 | 5.83E-01 | 1.21E+00 |
| MRI1 | 5.95E-07 | 3.10E-06 | -1.85E+00 | 7.04E-05 | 3.04E-04 | -1.35E+00 | 3.54E-05 | 1.50E-04 | -1.53E+00 | 4.78E-02 | 1.82E-01 | 1.13E+00 | 1.08E-03 | 3.85E-03 | -1.90E+00 | 2.78E-02 | 7.84E-02 | -1.16E+00 | 4.46E-09 | 1.64E-08 | -1.68E+00 | 1.82E-06 | 4.99E-06 | -1.42E+00 |
| AP002387.1 | 5.96E-07 | 3.11E-06 | -1.21E+00 | 7.53E-05 | 3.25E-04 | 1.04E+00 | 1.48E-02 | 6.28E-02 | -1.07E+00 | 3.10E-02 | 1.18E-01 | 1.10E+00 | 2.27E-01 | 8.08E-01 | 1.56E+00 | 4.66E-02 | 1.32E-01 | 2.18E+00 | 1.74E-01 | 6.38E-01 | 1.19E+00 | 2.70E-01 | 7.40E-01 | 1.27E+00 |
| DGKE | 5.96E-07 | 3.11E-06 | -1.18E+00 | 1.16E-04 | 5.01E-04 | 1.13E+00 | 2.46E-05 | 1.05E-04 | -1.31E+00 | 1.14E-03 | 4.35E-03 | -1.06E+00 | 1.99E-01 | 7.11E-01 | 1.30E+00 | 1.48E-01 | 4.16E-01 | 1.07E+00 | 4.51E-03 | 1.66E-02 | 1.84E+00 | 3.62E-01 | 9.95E-01 | 1.13E+00 |
| IRF2BPL | 6.00E-07 | 3.13E-06 | -1.36E+00 | 5.96E-11 | 2.57E-10 | -1.88E+00 | 3.59E-07 | 1.53E-06 | -1.65E+00 | 1.38E-10 | 5.26E-10 | -1.97E+00 | 5.06E-07 | 1.81E-06 | -1.71E+00 | 1.50E-08 | 4.23E-08 | -1.85E+00 | 2.80E-10 | 1.03E-09 | -1.81E+00 | 1.65E-10 | 4.53E-10 | -1.88E+00 |
| ABHD14B | 6.00E-07 | 3.13E-06 | -1.04E+00 | 8.78E-05 | 3.80E-04 | 1.14E+00 | 2.78E-02 | 1.18E-01 | -1.02E+00 | 5.91E-02 | 2.26E-01 | 1.18E+00 | 6.58E-02 | 2.35E-01 | 1.20E+00 | 2.33E-01 | 6.56E-01 | 1.37E+00 | 2.21E-01 | 8.12E-01 | 1.16E+00 | 1.90E-01 | 5.22E-01 | 1.26E+00 |
| C12orf49 | 6.02E-07 | 3.14E-06 | -2.02E+00 | 7.56E-03 | 3.27E-02 | 1.02E+00 | 8.13E-03 | 3.45E-02 | -1.28E+00 | 1.46E-01 | 5.58E-01 | 1.16E+00 | 2.76E-04 | 9.83E-04 | -1.42E+00 | 1.91E-02 | 5.38E-02 | -1.00E+00 | 1.03E-06 | 3.79E-06 | -1.44E+00 | 5.46E-05 | 1.50E-04 | -1.33E+00 |
| ORMDL2 | 6.04E-07 | 3.15E-06 | -1.44E+00 | 3.26E-04 | 1.41E-03 | 1.04E+00 | 1.22E-05 | 5.17E-05 | -1.34E+00 | 1.35E-03 | 5.15E-03 | 1.01E+00 | 4.73E-08 | 1.69E-07 | -1.68E+00 | 1.07E-02 | 3.01E-02 | 1.06E+00 | 2.38E-04 | 8.72E-04 | -1.18E+00 | 5.13E-02 | 1.41E-01 | 1.08E+00 |
| AF279873.2 | 6.05E-07 | 3.15E-06 | 2.09E+00 | 2.61E-03 | 1.13E-02 | 1.99E+00 | 1.03E-07 | 4.36E-07 | 2.17E+00 | 1.32E-04 | 5.06E-04 | 2.14E+00 | 1.09E-01 | 3.90E-01 | 1.66E+00 | 1.55E-01 | 4.38E-01 | 1.66E+00 | 2.68E-04 | 9.84E-04 | 1.80E+00 | 2.58E-03 | 7.08E-03 | 1.84E+00 |
| PSMC3 | 6.05E-07 | 3.16E-06 | -1.08E+00 | 1.10E-07 | 4.76E-07 | -1.01E+00 | 2.29E-02 | 9.75E-02 | 1.06E+00 | 1.28E-03 | 4.90E-03 | 1.12E+00 | 3.10E-04 | 1.10E-03 | -1.08E+00 | 1.14E-02 | 3.23E-02 | 1.07E+00 | 7.83E-02 | 2.88E-01 | 1.08E+00 | 2.02E-01 | 5.55E-01 | 1.14E+00 |
| MCM3 | 6.07E-07 | 3.16E-06 | -1.15E+00 | 1.57E-01 | 6.77E-01 | 1.49E+00 | 1.13E-01 | 4.81E-01 | 1.18E+00 | 2.86E-02 | 1.09E-01 | 1.42E+00 | 4.04E-05 | 1.44E-04 | -1.35E+00 | 9.22E-02 | 2.60E-01 | 1.28E+00 | 1.55E-04 | 5.68E-04 | -1.14E+00 | 3.22E-02 | 8.84E-02 | 1.34E+00 |
| SNX14 | 6.07E-07 | 3.16E-06 | -1.26E+00 | 7.18E-03 | 3.10E-02 | 1.18E+00 | 1.21E-07 | 5.15E-07 | -1.29E+00 | 6.53E-04 | 2.49E-03 | 1.06E+00 | 1.43E-05 | 5.11E-05 | -1.12E+00 | 8.33E-05 | 2.35E-04 | -1.00E+00 | 8.63E-09 | 3.17E-08 | -1.43E+00 | 2.58E-06 | 7.10E-06 | -1.29E+00 |
| ZNF780A | 6.08E-07 | 3.17E-06 | -2.24E+00 | 1.91E-05 | 8.25E-05 | -1.67E+00 | 2.43E-06 | 1.03E-05 | -2.21E+00 | 4.57E-06 | 1.74E-05 | -1.95E+00 | 2.26E-01 | 8.08E-01 | 1.14E+00 | 3.19E-01 | 8.99E-01 | 1.17E+00 | 1.12E-03 | 4.13E-03 | -1.30E+00 | 2.02E-03 | 5.55E-03 | -1.38E+00 |
| METAP2 | 6.09E-07 | 3.18E-06 | -1.03E+00 | 1.76E-08 | 7.61E-08 | 1.00E+00 | 9.83E-04 | 4.18E-03 | -1.16E+00 | 3.81E-07 | 1.45E-06 | -1.23E+00 | 2.28E-06 | 8.13E-06 | -1.23E+00 | 1.95E-04 | 5.51E-04 | -1.14E+00 | 4.24E-05 | 1.55E-04 | -1.18E+00 | 9.52E-05 | 2.62E-04 | -1.19E+00 |
| DARS | 6.14E-07 | 3.20E-06 | -1.11E+00 | 5.35E-02 | 2.31E-01 | 1.21E+00 | 7.99E-04 | 3.39E-03 | -1.16E+00 | 1.22E-01 | 4.67E-01 | 1.14E+00 | 1.49E-07 | 5.31E-07 | -1.09E+00 | 1.22E-01 | 3.44E-01 | 1.30E+00 | 3.28E-03 | 1.20E-02 | -1.13E+00 | 3.61E-01 | 9.91E-01 | 1.09E+00 |
| ANKRD65 | 6.18E-07 | 3.22E-06 | -1.42E+00 | 1.51E-03 | 6.54E-03 | 1.03E+00 | 4.86E-03 | 2.06E-02 | -1.06E+00 | 4.16E-03 | 1.59E-02 | 1.07E+00 | 2.72E-02 | 9.69E-02 | -1.24E+00 | 2.10E-02 | 5.92E-02 | -1.37E+00 | 2.46E-01 | 9.01E-01 | 1.02E+00 | 3.21E-01 | 8.82E-01 | 1.31E+00 |
| RBX1 | 6.20E-07 | 3.23E-06 | -1.08E+00 | 8.17E-05 | 3.53E-04 | 1.10E+00 | 7.61E-02 | 3.23E-01 | 1.04E+00 | 2.03E-02 | 7.75E-02 | 1.10E+00 | 2.12E-09 | 7.56E-09 | -1.22E+00 | 3.31E-03 | 9.32E-03 | 1.09E+00 | 2.13E-01 | 7.83E-01 | 1.10E+00 | 3.43E-01 | 9.43E-01 | 1.06E+00 |
| DDIT4 | 6.20E-07 | 3.23E-06 | 2.97E+00 | 4.54E-45 | 1.96E-44 | 8.33E+00 | 1.31E-02 | 5.57E-02 | 1.65E+00 | 5.31E-18 | 2.03E-17 | 4.70E+00 | 2.13E-01 | 7.61E-01 | 1.62E+00 | 3.19E-03 | 8.99E-03 | 2.54E+00 | 1.99E-02 | 7.31E-02 | 2.36E+00 | 1.93E-03 | 5.29E-03 | 3.39E+00 |
| ANKRD36BP2 | 6.20E-07 | 3.23E-06 | -2.02E+00 | 3.40E-03 | 1.47E-02 | 1.03E+00 | 9.62E-06 | 4.09E-05 | -1.60E+00 | 8.42E-05 | 3.22E-04 | -1.37E+00 | 7.60E-03 | 2.71E-02 | -1.41E+00 | 5.28E-02 | 1.49E-01 | -1.03E+00 | 1.97E-06 | 7.25E-06 | -1.58E+00 | 5.53E-06 | 1.52E-05 | -1.64E+00 |
| MPP5 | 6.21E-07 | 3.24E-06 | -1.19E+00 | 6.58E-11 | 2.84E-10 | -1.38E+00 | 1.05E-06 | 4.46E-06 | -1.43E+00 | 2.07E-08 | 7.90E-08 | -1.47E+00 | 1.39E-02 | 4.96E-02 | -1.03E+00 | 3.45E-02 | 9.72E-02 | 1.07E+00 | 8.12E-04 | 2.98E-03 | -1.22E+00 | 3.51E-04 | 9.65E-04 | -1.39E+00 |
| ITPR2 | 6.23E-07 | 3.25E-06 | -1.13E+00 | 3.75E-04 | 1.62E-03 | 1.13E+00 | 5.34E-09 | 2.27E-08 | -1.66E+00 | 3.46E-04 | 1.32E-03 | -1.19E+00 | 7.12E-03 | 2.54E-02 | -1.05E+00 | 7.34E-02 | 2.07E-01 | 1.17E+00 | 1.56E-04 | 5.74E-04 | -1.24E+00 | 3.38E-03 | 9.27E-03 | -1.05E+00 |
| AC211433.1 | 6.25E-07 | 3.26E-06 | -1.93E+00 | 6.02E-03 | 2.60E-02 | 1.04E+00 | 1.85E-01 | 7.87E-01 | 1.07E+00 | 1.98E-01 | 7.55E-01 | 1.10E+00 | 1.20E-01 | 4.29E-01 | 1.12E+00 | 2.10E-01 | 5.93E-01 | 1.24E+00 | 1.95E-02 | 7.17E-02 | -1.19E+00 | 1.80E-02 | 4.96E-02 | -1.28E+00 |
| CPTP | 6.27E-07 | 3.27E-06 | -1.48E+00 | 1.20E-04 | 5.17E-04 | -1.12E+00 | 7.94E-04 | 3.37E-03 | -1.50E+00 | 1.82E-02 | 6.95E-02 | 1.02E+00 | 4.02E-04 | 1.43E-03 | -1.65E+00 | 2.18E-01 | 6.15E-01 | 1.40E+00 | 9.43E-05 | 3.46E-04 | -1.16E+00 | 1.39E-02 | 3.83E-02 | -1.01E+00 |
| KLHL28 | 6.30E-07 | 3.29E-06 | -1.36E+00 | 4.05E-07 | 1.75E-06 | -1.28E+00 | 6.25E-02 | 2.66E-01 | 1.01E+00 | 1.43E-02 | 5.46E-02 | 1.04E+00 | 1.21E-10 | 4.32E-10 | -1.76E+00 | 3.61E-08 | 1.02E-07 | -1.47E+00 | 1.85E-03 | 6.79E-03 | -1.13E+00 | 3.50E-03 | 9.61E-03 | -1.08E+00 |
| TMEM220 | 6.32E-07 | 3.30E-06 | -1.02E+00 | 1.83E-03 | 7.89E-03 | 1.31E+00 | 6.48E-05 | 2.75E-04 | -1.15E+00 | 3.41E-03 | 1.30E-02 | 1.11E+00 | 1.38E-02 | 4.93E-02 | 1.05E+00 | 3.06E-01 | 8.63E-01 | 1.42E+00 | 6.90E-02 | 2.53E-01 | 1.09E+00 | 2.98E-01 | 8.19E-01 | 1.28E+00 |
| GCNT2 | 6.34E-07 | 3.30E-06 | -1.15E+00 | 3.54E-03 | 1.53E-02 | 1.19E+00 | 3.85E-10 | 1.64E-09 | -1.83E+00 | 5.38E-07 | 2.06E-06 | -1.27E+00 | 6.02E-04 | 2.15E-03 | -1.20E+00 | 4.86E-04 | 1.37E-03 | -1.22E+00 | 2.34E-05 | 8.60E-05 | -1.43E+00 | 1.22E-04 | 3.35E-04 | -1.30E+00 |
| AC105233.5 | 6.34E-07 | 3.30E-06 | -2.32E+00 | 1.05E-02 | 4.55E-02 | -1.02E+00 | 4.31E-04 | 1.83E-03 | -1.38E+00 | 3.52E-03 | 1.35E-02 | -1.05E+00 | 1.35E-01 | 4.82E-01 | -1.03E+00 | 2.34E-01 | 6.59E-01 | 1.62E+00 | 1.38E-05 | 5.07E-05 | -1.89E+00 | 3.49E-03 | 9.58E-03 | -1.30E+00 |
| LAT2 | 6.40E-07 | 3.34E-06 | -1.17E+00 | 2.01E-07 | 8.67E-07 | -1.06E+00 | 1.92E-05 | 8.15E-05 | -1.29E+00 | 9.26E-06 | 3.54E-05 | -1.14E+00 | 6.94E-08 | 2.47E-07 | -1.26E+00 | 4.21E-07 | 1.19E-06 | -1.22E+00 | 5.85E-02 | 2.15E-01 | -1.08E+00 | 9.22E-02 | 2.53E-01 | -1.06E+00 |
| RPS24P8 | 6.41E-07 | 3.34E-06 | 1.68E+00 | 2.91E-03 | 1.26E-02 | 1.23E+00 | 1.56E-05 | 6.61E-05 | 1.61E+00 | 2.14E-01 | 8.17E-01 | 1.18E+00 | 7.19E-03 | 2.56E-02 | 1.63E+00 | 3.54E-02 | 9.97E-02 | 1.19E+00 | 7.46E-04 | 2.74E-03 | 1.55E+00 | 1.52E-01 | 4.16E-01 | 1.12E+00 |
| SDCBP | 6.44E-07 | 3.35E-06 | -1.05E+00 | 5.15E-04 | 2.23E-03 | 1.19E+00 | 1.02E-04 | 4.34E-04 | -1.27E+00 | 3.73E-03 | 1.42E-02 | -1.02E+00 | 8.84E-03 | 3.15E-02 | 1.14E+00 | 1.08E-02 | 3.06E-02 | 1.05E+00 | 7.23E-03 | 2.65E-02 | 1.01E+00 | 1.58E-04 | 4.33E-04 | -1.13E+00 |
| EVA1B | 6.53E-07 | 3.40E-06 | -1.15E+00 | 3.59E-12 | 1.55E-11 | -1.22E+00 | 1.16E-05 | 4.94E-05 | -1.27E+00 | 2.88E-05 | 1.10E-04 | -1.17E+00 | 5.15E-08 | 1.84E-07 | -1.26E+00 | 1.72E-09 | 4.85E-09 | -1.34E+00 | 2.67E-01 | 9.80E-01 | 1.47E+00 | 5.17E-02 | 1.42E-01 | 1.13E+00 |
| CALCRL | 6.55E-07 | 3.41E-06 | -1.17E+00 | 4.10E-04 | 1.77E-03 | 1.10E+00 | 5.37E-12 | 2.28E-11 | -1.95E+00 | 2.75E-10 | 1.05E-09 | -1.59E+00 | 1.66E-03 | 5.92E-03 | -1.23E+00 | 8.33E-05 | 2.35E-04 | -1.49E+00 | 8.62E-02 | 3.16E-01 | -1.12E+00 | 5.71E-02 | 1.57E-01 | -1.22E+00 |
| AL158206.1 | 6.64E-07 | 3.46E-06 | -2.14E+00 | 2.28E-04 | 9.83E-04 | -1.43E+00 | 8.83E-03 | 3.75E-02 | -1.53E+00 | 7.57E-03 | 2.89E-02 | -1.45E+00 | 6.88E-06 | 2.45E-05 | -2.56E+00 | 5.73E-05 | 1.62E-04 | -2.23E+00 | 1.09E-05 | 4.01E-05 | -1.98E+00 | 1.24E-04 | 3.41E-04 | -1.91E+00 |
| MSL3 | 6.65E-07 | 3.46E-06 | -1.39E+00 | 2.94E-04 | 1.27E-03 | -1.02E+00 | 3.25E-04 | 1.38E-03 | -1.18E+00 | 6.81E-03 | 2.60E-02 | 1.03E+00 | 2.82E-02 | 1.00E-01 | 1.11E+00 | 2.69E-01 | 7.59E-01 | 1.40E+00 | 4.62E-05 | 1.70E-04 | -1.16E+00 | 4.75E-02 | 1.30E-01 | 1.10E+00 |
| ZDHHC12 | 6.66E-07 | 3.47E-06 | -1.37E+00 | 6.17E-05 | 2.67E-04 | -1.02E+00 | 4.36E-05 | 1.85E-04 | -1.15E+00 | 2.39E-06 | 9.14E-06 | -1.10E+00 | 1.28E-05 | 4.55E-05 | -1.14E+00 | 2.72E-04 | 7.66E-04 | -1.02E+00 | 1.86E-03 | 6.81E-03 | 1.09E+00 | 8.73E-03 | 2.40E-02 | 1.02E+00 |
| RPUSD3 | 6.67E-07 | 3.48E-06 | -1.24E+00 | 4.32E-04 | 1.87E-03 | 1.09E+00 | 8.64E-06 | 3.67E-05 | -1.21E+00 | 4.29E-06 | 1.64E-05 | -1.12E+00 | 4.25E-08 | 1.51E-07 | -1.41E+00 | 1.82E-02 | 5.14E-02 | 1.22E+00 | 8.34E-04 | 3.06E-03 | -1.04E+00 | 1.16E-01 | 3.20E-01 | 1.07E+00 |
| RPAP1 | 6.68E-07 | 3.48E-06 | -1.61E+00 | 4.05E-04 | 1.75E-03 | -1.12E+00 | 1.50E-01 | 6.35E-01 | 1.05E+00 | 2.51E-01 | 9.60E-01 | 1.25E+00 | 3.35E-04 | 1.19E-03 | -1.69E+00 | 3.51E-02 | 9.91E-02 | 1.05E+00 | 1.05E-03 | 3.87E-03 | -1.16E+00 | 9.26E-03 | 2.54E-02 | -1.11E+00 |
| WBP2 | 6.69E-07 | 3.49E-06 | -1.18E+00 | 6.79E-04 | 2.94E-03 | 1.11E+00 | 1.54E-07 | 6.55E-07 | -1.47E+00 | 9.97E-04 | 3.81E-03 | -1.00E+00 | 1.95E-09 | 6.97E-09 | -1.45E+00 | 1.25E-06 | 3.53E-06 | -1.18E+00 | 1.43E-05 | 5.25E-05 | -1.10E+00 | 2.07E-04 | 5.68E-04 | -1.10E+00 |
| THAP4 | 6.75E-07 | 3.52E-06 | -1.32E+00 | 6.33E-05 | 2.74E-04 | -1.04E+00 | 3.89E-03 | 1.65E-02 | -1.11E+00 | 1.50E-02 | 5.73E-02 | 1.10E+00 | 9.70E-03 | 3.46E-02 | -1.21E+00 | 6.25E-02 | 1.76E-01 | 1.04E+00 | 4.93E-04 | 1.81E-03 | 1.00E+00 | 7.62E-03 | 2.09E-02 | 1.07E+00 |
| TPT1 | 6.76E-07 | 3.52E-06 | 1.08E+00 | 3.84E-22 | 1.66E-21 | 1.17E+00 | 4.58E-02 | 1.94E-01 | 1.04E+00 | 7.72E-18 | 2.95E-17 | 1.20E+00 | 4.76E-05 | 1.70E-04 | 1.11E+00 | 1.19E-02 | 3.35E-02 | 1.06E+00 | 1.14E-02 | 4.19E-02 | 1.07E+00 | 1.53E-01 | 4.22E-01 | 1.02E+00 |
| NDUFA6-AS1 | 6.77E-07 | 3.53E-06 | -1.70E+00 | 1.87E-05 | 8.08E-05 | -1.37E+00 | 2.05E-01 | 8.70E-01 | 1.22E+00 | 2.02E-01 | 7.73E-01 | 1.36E+00 | 1.40E-03 | 4.98E-03 | -1.45E+00 | 5.21E-03 | 1.47E-02 | -1.30E+00 | 3.33E-03 | 1.22E-02 | -1.27E+00 | 3.06E-02 | 8.41E-02 | -1.11E+00 |
| AC090602.1 | 6.77E-07 | 3.53E-06 | -1.02E+00 | 1.66E-22 | 7.17E-22 | -1.31E+00 | 9.42E-04 | 4.00E-03 | -1.06E+00 | 6.93E-11 | 2.65E-10 | -1.34E+00 | 3.35E-02 | 1.19E-01 | 1.07E+00 | 7.66E-09 | 2.16E-08 | -1.32E+00 | 6.02E-03 | 2.21E-02 | -1.03E+00 | 2.67E-08 | 7.32E-08 | -1.42E+00 |
| CCDC137 | 6.82E-07 | 3.55E-06 | -1.45E+00 | 6.70E-04 | 2.90E-03 | 1.01E+00 | 1.53E-05 | 6.49E-05 | -1.33E+00 | 1.96E-03 | 7.47E-03 | 1.02E+00 | 2.33E-05 | 8.30E-05 | -1.25E+00 | 8.51E-03 | 2.40E-02 | 1.03E+00 | 4.26E-08 | 1.56E-07 | -1.31E+00 | 6.18E-04 | 1.70E-03 | -1.05E+00 |
| PRDX5 | 6.85E-07 | 3.57E-06 | -1.06E+00 | 2.47E-05 | 1.07E-04 | 1.10E+00 | 4.89E-02 | 2.08E-01 | 1.06E+00 | 7.27E-02 | 2.78E-01 | 1.15E+00 | 4.74E-06 | 1.69E-05 | -1.10E+00 | 7.28E-03 | 2.05E-02 | 1.07E+00 | 2.50E-01 | 9.19E-01 | 1.15E+00 | 1.77E-01 | 4.86E-01 | 1.26E+00 |
| CD151 | 6.86E-07 | 3.57E-06 | -1.22E+00 | 1.27E-04 | 5.50E-04 | 1.04E+00 | 7.27E-04 | 3.09E-03 | -1.07E+00 | 8.41E-02 | 3.21E-01 | 1.28E+00 | 3.02E-03 | 1.08E-02 | -1.21E+00 | 9.75E-03 | 2.75E-02 | -1.03E+00 | 1.92E-01 | 7.06E-01 | 1.43E+00 | 1.54E-01 | 4.24E-01 | 1.30E+00 |
| PDZD11 | 6.89E-07 | 3.59E-06 | -1.53E+00 | 3.54E-04 | 1.53E-03 | -1.04E+00 | 1.17E-10 | 4.98E-10 | -1.71E+00 | 1.79E-07 | 6.84E-07 | -1.24E+00 | 5.85E-03 | 2.09E-02 | -1.49E+00 | 1.47E-01 | 4.14E-01 | 1.09E+00 | 6.94E-09 | 2.55E-08 | -1.59E+00 | 1.86E-04 | 5.11E-04 | -1.16E+00 |
| THOC1 | 6.90E-07 | 3.60E-06 | -1.23E+00 | 3.04E-05 | 1.32E-04 | -1.02E+00 | 5.54E-08 | 2.35E-07 | -1.45E+00 | 1.32E-06 | 5.03E-06 | -1.19E+00 | 8.31E-04 | 2.96E-03 | -1.16E+00 | 8.06E-02 | 2.27E-01 | 1.05E+00 | 4.48E-05 | 1.64E-04 | -1.21E+00 | 2.06E-01 | 5.66E-01 | 1.11E+00 |
| AL158801.5 | 6.91E-07 | 3.60E-06 | -1.06E+00 | 8.38E-12 | 3.62E-11 | -1.14E+00 | 1.64E-04 | 6.97E-04 | -1.12E+00 | 4.98E-11 | 1.90E-10 | -1.27E+00 | 2.56E-11 | 9.13E-11 | -1.58E+00 | 1.20E-07 | 3.38E-07 | -1.33E+00 | 2.85E-06 | 1.05E-05 | -1.26E+00 | 5.59E-05 | 1.54E-04 | -1.20E+00 |
| RIN2 | 6.92E-07 | 3.61E-06 | -1.61E+00 | 1.03E-04 | 4.43E-04 | -1.18E+00 | 1.34E-02 | 5.70E-02 | -1.50E+00 | 7.77E-02 | 2.97E-01 | -1.07E+00 | 6.26E-05 | 2.23E-04 | -2.04E+00 | 1.09E-05 | 3.08E-05 | -2.21E+00 | 1.91E-02 | 7.01E-02 | -1.54E+00 | 1.33E-01 | 3.66E-01 | -1.01E+00 |
| CCT6P3 | 6.93E-07 | 3.61E-06 | -1.51E+00 | 8.75E-06 | 3.78E-05 | -1.25E+00 | 3.10E-07 | 1.32E-06 | -1.57E+00 | 8.59E-04 | 3.28E-03 | -1.03E+00 | 3.44E-03 | 1.23E-02 | -1.07E+00 | 3.11E-02 | 8.78E-02 | 1.17E+00 | 1.73E-01 | 6.36E-01 | 1.36E+00 | 1.60E-01 | 4.40E-01 | 1.11E+00 |
| FLT3 | 7.00E-07 | 3.65E-06 | 1.93E+00 | 1.25E-14 | 5.38E-14 | 2.54E+00 | 1.19E-05 | 5.06E-05 | 1.60E+00 | 1.84E-07 | 7.04E-07 | 2.02E+00 | 3.33E-05 | 1.19E-04 | 4.35E+00 | 4.84E-02 | 1.37E-01 | 2.70E+00 | 8.47E-03 | 3.11E-02 | 5.04E+00 | 2.73E-03 | 7.50E-03 | 6.91E+00 |
| UTP20 | 7.05E-07 | 3.67E-06 | -1.54E+00 | 3.15E-03 | 1.36E-02 | 1.10E+00 | 1.44E-04 | 6.11E-04 | -1.22E+00 | 5.63E-02 | 2.15E-01 | 1.27E+00 | 3.29E-04 | 1.17E-03 | -1.73E+00 | 1.72E-02 | 4.83E-02 | -1.15E+00 | 7.90E-05 | 2.90E-04 | -1.36E+00 | 3.26E-01 | 8.95E-01 | 1.23E+00 |
| PTDSS1 | 7.05E-07 | 3.67E-06 | -1.14E+00 | 1.19E-04 | 5.14E-04 | 1.07E+00 | 8.69E-04 | 3.69E-03 | -1.11E+00 | 1.02E-01 | 3.88E-01 | 1.18E+00 | 1.46E-05 | 5.22E-05 | -1.09E+00 | 5.01E-02 | 1.41E-01 | 1.21E+00 | 5.54E-02 | 2.03E-01 | -1.00E+00 | 1.80E-01 | 4.94E-01 | 1.08E+00 |
| SURF4 | 7.05E-07 | 3.67E-06 | -1.10E+00 | 4.70E-10 | 2.03E-09 | -1.13E+00 | 1.22E-02 | 5.20E-02 | -1.01E+00 | 2.22E-03 | 8.48E-03 | 1.03E+00 | 2.11E-07 | 7.54E-07 | -1.30E+00 | 8.52E-08 | 2.40E-07 | -1.35E+00 | 4.77E-03 | 1.75E-02 | -1.13E+00 | 6.93E-02 | 1.90E-01 | -1.05E+00 |
| AKR7A2 | 7.05E-07 | 3.68E-06 | -1.13E+00 | 2.43E-06 | 1.05E-05 | 1.03E+00 | 2.01E-01 | 8.52E-01 | 1.21E+00 | 2.11E-02 | 8.04E-02 | 1.21E+00 | 4.87E-03 | 1.74E-02 | -1.01E+00 | 4.62E-03 | 1.30E-02 | 1.03E+00 | 1.96E-01 | 7.18E-01 | 1.25E+00 | 1.96E-01 | 5.37E-01 | 1.24E+00 |
| DIP2A | 7.06E-07 | 3.68E-06 | -1.22E+00 | 5.33E-03 | 2.30E-02 | 1.21E+00 | 6.18E-13 | 2.63E-12 | -2.03E+00 | 1.64E-06 | 6.27E-06 | -1.18E+00 | 1.86E-07 | 6.65E-07 | -1.78E+00 | 6.87E-04 | 1.94E-03 | -1.18E+00 | 1.46E-04 | 5.37E-04 | -1.21E+00 | 2.64E-03 | 7.24E-03 | -1.11E+00 |
| TRAF3IP2-AS1 | 7.09E-07 | 3.69E-06 | -1.26E+00 | 1.28E-04 | 5.51E-04 | -1.00E+00 | 6.81E-05 | 2.89E-04 | -1.16E+00 | 1.13E-04 | 4.30E-04 | 1.04E+00 | 4.30E-02 | 1.53E-01 | 1.11E+00 | 3.18E-01 | 8.96E-01 | 1.40E+00 | 6.02E-02 | 2.21E-01 | 1.02E+00 | 2.45E-01 | 6.74E-01 | 1.14E+00 |
| CDC16 | 7.10E-07 | 3.70E-06 | -1.15E+00 | 3.00E-04 | 1.30E-03 | 1.11E+00 | 9.86E-06 | 4.19E-05 | -1.24E+00 | 1.97E-04 | 7.53E-04 | -1.05E+00 | 8.71E-08 | 3.10E-07 | -1.36E+00 | 1.95E-06 | 5.50E-06 | -1.20E+00 | 4.60E-05 | 1.69E-04 | -1.18E+00 | 1.91E-02 | 5.23E-02 | 1.00E+00 |
| AP000787.1 | 7.11E-07 | 3.71E-06 | -1.59E+00 | 4.75E-04 | 2.05E-03 | -1.09E+00 | 9.95E-06 | 4.23E-05 | -1.88E+00 | 2.60E-04 | 9.95E-04 | -1.39E+00 | 1.49E-03 | 5.32E-03 | -1.39E+00 | 4.25E-03 | 1.20E-02 | -1.23E+00 | 2.17E-02 | 7.97E-02 | -1.25E+00 | 1.57E-02 | 4.30E-02 | -1.41E+00 |
| SMARCD1 | 7.11E-07 | 3.71E-06 | -1.18E+00 | 3.67E-07 | 1.59E-06 | -1.10E+00 | 3.02E-06 | 1.28E-05 | -1.24E+00 | 5.31E-08 | 2.03E-07 | -1.28E+00 | 1.39E-05 | 4.94E-05 | -1.18E+00 | 3.00E-06 | 8.47E-06 | -1.24E+00 | 1.78E-06 | 6.55E-06 | -1.17E+00 | 2.83E-08 | 7.78E-08 | -1.29E+00 |
| PCBP1-AS1 | 7.12E-07 | 3.71E-06 | -1.20E+00 | 1.16E-02 | 5.01E-02 | 1.20E+00 | 7.84E-07 | 3.33E-06 | -1.40E+00 | 1.20E-03 | 4.57E-03 | 1.03E+00 | 8.29E-06 | 2.96E-05 | -1.25E+00 | 1.07E-03 | 3.02E-03 | 1.00E+00 | 8.50E-02 | 3.12E-01 | 1.05E+00 | 1.97E-01 | 5.41E-01 | 1.17E+00 |
| PPP1R3D | 7.18E-07 | 3.74E-06 | -1.66E+00 | 8.30E-06 | 3.59E-05 | -1.45E+00 | 3.07E-03 | 1.30E-02 | -1.35E+00 | 7.63E-03 | 2.92E-02 | -1.08E+00 | 2.49E-04 | 8.86E-04 | -1.38E+00 | 7.56E-05 | 2.13E-04 | -1.48E+00 | 2.24E-02 | 8.22E-02 | -1.12E+00 | 8.47E-02 | 2.33E-01 | 1.02E+00 |
| LRMDA | 7.19E-07 | 3.75E-06 | -1.10E+00 | 3.45E-10 | 1.49E-09 | -1.16E+00 | 6.50E-03 | 2.76E-02 | -1.00E+00 | 2.71E-04 | 1.04E-03 | -1.03E+00 | 1.11E-05 | 3.97E-05 | -1.28E+00 | 8.97E-06 | 2.53E-05 | -1.22E+00 | 7.59E-02 | 2.79E-01 | 1.06E+00 | 2.24E-01 | 6.15E-01 | 1.17E+00 |
| ZSWIM6 | 7.19E-07 | 3.75E-06 | -1.56E+00 | 1.29E-02 | 5.57E-02 | 1.20E+00 | 1.85E-06 | 7.88E-06 | -1.69E+00 | 5.48E-04 | 2.09E-03 | -1.13E+00 | 2.31E-05 | 8.25E-05 | -1.65E+00 | 3.27E-03 | 9.21E-03 | -1.15E+00 | 4.84E-07 | 1.78E-06 | -1.90E+00 | 1.77E-03 | 4.85E-03 | -1.23E+00 |
| DEGS2 | 7.21E-07 | 3.76E-06 | -2.84E+00 | 4.94E-05 | 2.13E-04 | -1.73E+00 | 1.14E-02 | 4.84E-02 | -1.88E+00 | 6.46E-02 | 2.47E-01 | -1.11E+00 | 3.29E-04 | 1.17E-03 | -1.78E+00 | 4.68E-02 | 1.32E-01 | 1.53E+00 | 4.66E-15 | 1.71E-14 | -1.50E+00 | 3.51E-07 | 9.65E-07 | -1.21E+00 |
| MYO18A | 7.22E-07 | 3.76E-06 | -1.20E+00 | 1.80E-04 | 7.78E-04 | 1.07E+00 | 1.97E-09 | 8.38E-09 | -1.59E+00 | 3.13E-04 | 1.20E-03 | -1.05E+00 | 7.55E-03 | 2.69E-02 | -1.29E+00 | 4.09E-02 | 1.15E-01 | -1.01E+00 | 1.92E-01 | 7.03E-01 | 1.32E+00 | 2.83E-01 | 7.77E-01 | 1.32E+00 |
| C1orf109 | 7.23E-07 | 3.77E-06 | -1.91E+00 | 7.09E-05 | 3.07E-04 | -1.39E+00 | 1.83E-05 | 7.77E-05 | -1.68E+00 | 5.56E-06 | 2.12E-05 | -1.57E+00 | 1.24E-01 | 4.43E-01 | 1.29E+00 | 3.51E-02 | 9.88E-02 | 1.04E+00 | 8.95E-13 | 3.29E-12 | -1.55E+00 | 4.19E-05 | 1.15E-04 | -1.12E+00 |
| EEF1B2P3 | 7.25E-07 | 3.78E-06 | 1.30E+00 | 4.93E-27 | 2.13E-26 | 1.68E+00 | 2.03E-13 | 8.65E-13 | 1.50E+00 | 1.93E-33 | 7.39E-33 | 1.93E+00 | 5.91E-08 | 2.11E-07 | 1.45E+00 | 1.28E-28 | 3.62E-28 | 1.96E+00 | 9.79E-10 | 3.59E-09 | 1.49E+00 | 2.02E-39 | 5.55E-39 | 2.02E+00 |
| PUM1 | 7.25E-07 | 3.78E-06 | -1.10E+00 | 3.97E-03 | 1.72E-02 | 1.23E+00 | 9.95E-10 | 4.23E-09 | -1.45E+00 | 1.68E-03 | 6.41E-03 | 1.03E+00 | 1.78E-08 | 6.34E-08 | -1.36E+00 | 2.95E-03 | 8.31E-03 | -1.02E+00 | 9.42E-08 | 3.46E-07 | -1.36E+00 | 5.62E-02 | 1.54E-01 | 1.03E+00 |
| G6PD | 7.26E-07 | 3.78E-06 | -1.13E+00 | 2.29E-05 | 9.88E-05 | 1.09E+00 | 3.62E-04 | 1.54E-03 | -1.16E+00 | 8.18E-03 | 3.12E-02 | 1.15E+00 | 1.78E-06 | 6.35E-06 | -1.22E+00 | 7.61E-07 | 2.14E-06 | -1.20E+00 | 5.45E-05 | 2.00E-04 | -1.17E+00 | 1.29E-02 | 3.53E-02 | 1.02E+00 |
| LONRF2 | 7.27E-07 | 3.79E-06 | -2.00E+00 | 1.37E-07 | 5.91E-07 | -2.16E+00 | 8.04E-06 | 3.42E-05 | -1.70E+00 | 5.29E-06 | 2.02E-05 | -1.82E+00 | 1.19E-02 | 4.23E-02 | -1.32E+00 | 3.80E-03 | 1.07E-02 | -1.62E+00 | 2.71E-02 | 9.95E-02 | -1.18E+00 | 3.32E-03 | 9.11E-03 | -1.76E+00 |
| AC013356.1 | 7.30E-07 | 3.80E-06 | -1.60E+00 | 3.34E-08 | 1.45E-07 | -1.70E+00 | 3.45E-02 | 1.46E-01 | -1.02E+00 | 3.08E-04 | 1.17E-03 | -1.33E+00 | 2.29E-01 | 8.16E-01 | 1.30E+00 | 3.39E-01 | 9.54E-01 | 1.50E+00 | 3.26E-04 | 1.20E-03 | -1.61E+00 | 2.31E-05 | 6.33E-05 | -1.73E+00 |
| LINC01132 | 7.30E-07 | 3.80E-06 | 3.52E+00 | 1.92E-01 | 8.28E-01 | 1.37E+00 | 2.25E-02 | 9.55E-02 | 1.85E+00 | 2.10E-01 | 8.01E-01 | 1.09E+00 | 6.73E-02 | 2.40E-01 | 1.99E+00 | 3.45E-01 | 9.74E-01 | 1.20E+00 | 1.40E-01 | 5.15E-01 | 1.61E+00 | 3.06E-01 | 8.40E-01 | 1.22E+00 |
| RFXANK | 7.32E-07 | 3.81E-06 | -1.11E+00 | 4.02E-06 | 1.74E-05 | 1.01E+00 | 1.74E-04 | 7.37E-04 | -1.07E+00 | 7.15E-04 | 2.73E-03 | 1.09E+00 | 2.32E-06 | 8.28E-06 | -1.18E+00 | 2.66E-04 | 7.50E-04 | 1.05E+00 | 1.90E-04 | 6.97E-04 | -1.08E+00 | 6.64E-02 | 1.82E-01 | 1.03E+00 |
| KCTD5 | 7.34E-07 | 3.82E-06 | -1.56E+00 | 1.14E-03 | 4.94E-03 | -1.05E+00 | 2.10E-05 | 8.93E-05 | -1.44E+00 | 1.57E-04 | 5.98E-04 | -1.11E+00 | 6.78E-04 | 2.42E-03 | -1.29E+00 | 2.66E-02 | 7.51E-02 | 1.02E+00 | 3.92E-12 | 1.44E-11 | -1.67E+00 | 2.87E-05 | 7.87E-05 | -1.13E+00 |
| SLC16A1-AS1 | 7.34E-07 | 3.83E-06 | -1.54E+00 | 5.77E-02 | 2.49E-01 | 1.27E+00 | 1.42E-02 | 6.04E-02 | -1.21E+00 | 9.70E-02 | 3.71E-01 | 1.40E+00 | 4.12E-03 | 1.47E-02 | -1.26E+00 | 1.19E-01 | 3.35E-01 | 1.27E+00 | 2.23E-07 | 8.19E-07 | -1.71E+00 | 2.83E-02 | 7.79E-02 | -1.10E+00 |
| BRD4 | 7.38E-07 | 3.85E-06 | -1.07E+00 | 7.66E-06 | 3.31E-05 | 1.07E+00 | 4.87E-02 | 2.07E-01 | 1.05E+00 | 1.55E-02 | 5.91E-02 | 1.13E+00 | 2.02E-02 | 7.19E-02 | 1.08E+00 | 1.84E-01 | 5.20E-01 | 1.18E+00 | 8.88E-03 | 3.26E-02 | -1.03E+00 | 1.24E-01 | 3.40E-01 | 1.07E+00 |
| FAM177A1 | 7.39E-07 | 3.85E-06 | -1.18E+00 | 2.62E-11 | 1.13E-10 | -1.27E+00 | 4.69E-03 | 1.99E-02 | -1.05E+00 | 9.44E-05 | 3.60E-04 | -1.08E+00 | 7.32E-05 | 2.61E-04 | -1.08E+00 | 3.97E-05 | 1.12E-04 | -1.15E+00 | 2.62E-08 | 9.62E-08 | -1.27E+00 | 2.78E-08 | 7.63E-08 | -1.39E+00 |
| PLCG2 | 7.39E-07 | 3.85E-06 | -1.33E+00 | 1.27E-04 | 5.51E-04 | -1.02E+00 | 6.42E-04 | 2.73E-03 | -1.15E+00 | 1.12E-02 | 4.29E-02 | 1.12E+00 | 7.70E-05 | 2.74E-04 | -1.63E+00 | 4.97E-03 | 1.40E-02 | -1.11E+00 | 4.65E-03 | 1.70E-02 | -1.24E+00 | 2.22E-02 | 6.09E-02 | -1.12E+00 |
| CHPT1 | 7.39E-07 | 3.85E-06 | -1.07E+00 | 1.87E-06 | 8.10E-06 | 1.07E+00 | 1.18E-02 | 5.03E-02 | 1.01E+00 | 2.02E-03 | 7.70E-03 | 1.08E+00 | 3.26E-07 | 1.16E-06 | -1.28E+00 | 4.54E-06 | 1.28E-05 | -1.19E+00 | 3.37E-02 | 1.24E-01 | 1.05E+00 | 2.65E-01 | 7.27E-01 | 1.14E+00 |
| ZNF324B | 7.39E-07 | 3.85E-06 | -2.13E+00 | 1.20E-04 | 5.18E-04 | -1.29E+00 | 3.98E-07 | 1.69E-06 | -1.87E+00 | 2.73E-05 | 1.04E-04 | -1.34E+00 | 9.41E-02 | 3.36E-01 | -1.04E+00 | 3.10E-01 | 8.75E-01 | 1.58E+00 | 5.53E-02 | 2.03E-01 | -1.40E+00 | 3.37E-01 | 9.25E-01 | 1.15E+00 |
| AC105285.1 | 7.41E-07 | 3.86E-06 | -1.59E+00 | 1.50E-05 | 6.48E-05 | -1.21E+00 | 6.92E-02 | 2.94E-01 | -1.04E+00 | 1.45E-01 | 5.52E-01 | 1.15E+00 | 2.67E-01 | 9.52E-01 | 1.31E+00 | 5.23E-02 | 1.47E-01 | 2.26E+00 | 2.58E-01 | 9.46E-01 | 1.21E+00 | 2.29E-01 | 6.28E-01 | 1.41E+00 |
| EIF2A | 7.42E-07 | 3.87E-06 | -1.06E+00 | 6.51E-06 | 2.81E-05 | 1.03E+00 | 4.61E-04 | 1.96E-03 | -1.21E+00 | 1.20E-03 | 4.57E-03 | -1.10E+00 | 1.67E-02 | 5.95E-02 | 1.01E+00 | 3.16E-01 | 8.91E-01 | 1.14E+00 | 6.66E-04 | 2.44E-03 | -1.11E+00 | 9.60E-02 | 2.64E-01 | -1.02E+00 |
| DOCK7 | 7.47E-07 | 3.89E-06 | -1.28E+00 | 7.95E-05 | 3.43E-04 | 1.02E+00 | 2.87E-05 | 1.22E-04 | -1.31E+00 | 5.54E-04 | 2.12E-03 | -1.04E+00 | 2.49E-02 | 8.87E-02 | -1.14E+00 | 8.55E-02 | 2.41E-01 | 1.10E+00 | 1.37E-16 | 5.03E-16 | -1.82E+00 | 3.34E-10 | 9.17E-10 | -1.49E+00 |
| ANO10 | 7.49E-07 | 3.90E-06 | -1.32E+00 | 7.20E-06 | 3.11E-05 | -1.14E+00 | 5.83E-03 | 2.48E-02 | -1.16E+00 | 4.17E-02 | 1.59E-01 | 1.10E+00 | 6.77E-03 | 2.42E-02 | -1.23E+00 | 1.07E-01 | 3.02E-01 | 1.13E+00 | 1.00E-03 | 3.68E-03 | -1.11E+00 | 2.59E-04 | 7.13E-04 | -1.15E+00 |
| GTF3C2 | 7.51E-07 | 3.91E-06 | -1.40E+00 | 3.23E-05 | 1.39E-04 | -1.09E+00 | 2.20E-05 | 9.36E-05 | -1.30E+00 | 6.16E-03 | 2.35E-02 | 1.10E+00 | 5.72E-05 | 2.04E-04 | -1.32E+00 | 5.86E-04 | 1.65E-03 | -1.09E+00 | 1.53E-04 | 5.60E-04 | -1.24E+00 | 4.80E-02 | 1.32E-01 | 1.09E+00 |
| FHL1 | 7.52E-07 | 3.92E-06 | -1.14E+00 | 5.59E-03 | 2.42E-02 | 1.13E+00 | 3.05E-03 | 1.30E-02 | -1.06E+00 | 5.92E-03 | 2.26E-02 | 1.08E+00 | 1.44E-01 | 5.13E-01 | -1.04E+00 | 3.08E-01 | 8.67E-01 | 1.15E+00 | 1.69E-01 | 6.19E-01 | 1.44E+00 | 3.08E-01 | 8.46E-01 | 1.13E+00 |
| USP15 | 7.54E-07 | 3.93E-06 | -1.14E+00 | 3.01E-06 | 1.30E-05 | -1.00E+00 | 1.49E-04 | 6.35E-04 | -1.20E+00 | 1.28E-03 | 4.90E-03 | -1.01E+00 | 1.97E-05 | 7.02E-05 | -1.24E+00 | 2.04E-04 | 5.75E-04 | -1.16E+00 | 1.14E-03 | 4.19E-03 | -1.16E+00 | 1.22E-03 | 3.36E-03 | -1.29E+00 |
| ADD1 | 7.55E-07 | 3.93E-06 | -1.10E+00 | 5.05E-02 | 2.18E-01 | 1.32E+00 | 1.87E-03 | 7.93E-03 | -1.13E+00 | 9.51E-02 | 3.63E-01 | 1.20E+00 | 4.36E-05 | 1.55E-04 | -1.15E+00 | 1.87E-02 | 5.27E-02 | 1.08E+00 | 2.06E-03 | 7.55E-03 | -1.05E+00 | 1.36E-02 | 3.73E-02 | 1.04E+00 |
| DOCK1 | 7.60E-07 | 3.96E-06 | -1.33E+00 | 3.66E-02 | 1.58E-01 | 1.30E+00 | 8.48E-07 | 3.60E-06 | -1.66E+00 | 2.12E-02 | 8.10E-02 | 1.12E+00 | 1.96E-01 | 6.99E-01 | 1.11E+00 | 2.96E-01 | 8.34E-01 | 1.53E+00 | 2.41E-02 | 8.85E-02 | -1.30E+00 | 3.31E-01 | 9.10E-01 | 1.27E+00 |
| EFCAB13 | 7.65E-07 | 3.99E-06 | -1.23E+00 | 7.23E-03 | 3.12E-02 | 1.32E+00 | 1.17E-02 | 4.96E-02 | -1.13E+00 | 4.32E-02 | 1.65E-01 | 1.04E+00 | 6.57E-02 | 2.34E-01 | 1.15E+00 | 2.18E-01 | 6.15E-01 | 1.37E+00 | 2.31E-02 | 8.49E-02 | -1.01E+00 | 9.39E-02 | 2.58E-01 | 1.17E+00 |
| NIFK-AS1 | 7.67E-07 | 4.00E-06 | -1.25E+00 | 3.16E-02 | 1.37E-01 | 1.27E+00 | 7.87E-06 | 3.35E-05 | -1.28E+00 | 4.52E-03 | 1.73E-02 | 1.11E+00 | 3.33E-04 | 1.19E-03 | -1.20E+00 | 1.26E-01 | 3.55E-01 | 1.15E+00 | 6.04E-04 | 2.22E-03 | -1.10E+00 | 3.15E-01 | 8.65E-01 | 1.26E+00 |
| DPM2 | 7.67E-07 | 4.00E-06 | -1.17E+00 | 1.93E-03 | 8.34E-03 | 1.19E+00 | 1.14E-05 | 4.85E-05 | -1.16E+00 | 1.05E-04 | 4.01E-04 | -1.00E+00 | 1.39E-05 | 4.97E-05 | -1.22E+00 | 7.38E-03 | 2.08E-02 | 1.09E+00 | 5.72E-05 | 2.10E-04 | -1.08E+00 | 1.74E-02 | 4.79E-02 | 1.09E+00 |
| SNX29 | 7.67E-07 | 4.00E-06 | -1.36E+00 | 2.81E-06 | 1.22E-05 | -1.14E+00 | 9.44E-06 | 4.01E-05 | -1.38E+00 | 2.51E-04 | 9.60E-04 | -1.09E+00 | 1.58E-04 | 5.65E-04 | -1.55E+00 | 2.17E-03 | 6.12E-03 | -1.23E+00 | 4.26E-02 | 1.56E-01 | 1.05E+00 | 1.14E-01 | 3.12E-01 | 1.12E+00 |
| WDR59 | 7.69E-07 | 4.01E-06 | -1.22E+00 | 2.35E-02 | 1.02E-01 | 1.27E+00 | 3.95E-04 | 1.68E-03 | -1.45E+00 | 6.69E-02 | 2.56E-01 | 1.12E+00 | 4.31E-08 | 1.54E-07 | -1.25E+00 | 5.99E-03 | 1.69E-02 | 1.21E+00 | 8.46E-03 | 3.10E-02 | -1.07E+00 | 3.35E-01 | 9.20E-01 | 1.21E+00 |
| ZNF428 | 7.70E-07 | 4.01E-06 | -1.07E+00 | 3.13E-07 | 1.35E-06 | 1.05E+00 | 1.49E-01 | 6.32E-01 | 1.00E+00 | 2.01E-02 | 7.66E-02 | 1.03E+00 | 3.70E-06 | 1.32E-05 | -1.20E+00 | 4.89E-04 | 1.38E-03 | -1.08E+00 | 5.96E-02 | 2.19E-01 | 1.08E+00 | 2.73E-01 | 7.50E-01 | 1.12E+00 |
| PDCL3 | 7.73E-07 | 4.03E-06 | -1.28E+00 | 5.54E-05 | 2.39E-04 | -1.02E+00 | 1.49E-05 | 6.32E-05 | -1.22E+00 | 6.12E-04 | 2.34E-03 | 1.06E+00 | 7.60E-06 | 2.71E-05 | -1.15E+00 | 1.19E-03 | 3.36E-03 | 1.07E+00 | 8.50E-07 | 3.12E-06 | -1.28E+00 | 1.93E-02 | 5.29E-02 | 1.00E+00 |
| AC127024.8 | 7.81E-07 | 4.07E-06 | -1.72E+00 | 9.34E-06 | 4.04E-05 | -1.39E+00 | 5.23E-05 | 2.22E-04 | -1.46E+00 | 6.92E-03 | 2.64E-02 | 1.02E+00 | 4.26E-07 | 1.52E-06 | -1.98E+00 | 1.13E-04 | 3.20E-04 | -1.35E+00 | 1.63E-07 | 5.98E-07 | -1.55E+00 | 3.49E-03 | 9.60E-03 | -1.11E+00 |
| C2 | 7.82E-07 | 4.07E-06 | -2.22E+00 | 5.12E-05 | 2.21E-04 | -1.54E+00 | 5.51E-08 | 2.34E-07 | -2.47E+00 | 2.64E-05 | 1.01E-04 | -1.70E+00 | 2.36E-03 | 8.40E-03 | -2.27E+00 | 5.16E-03 | 1.45E-02 | -1.98E+00 | 2.47E-02 | 9.05E-02 | -1.54E+00 | 2.36E-02 | 6.48E-02 | -1.81E+00 |
| DYNC1LI2 | 7.83E-07 | 4.08E-06 | -1.32E+00 | 4.78E-06 | 2.07E-05 | -1.10E+00 | 8.80E-05 | 3.74E-04 | -1.21E+00 | 2.27E-04 | 8.68E-04 | 1.00E+00 | 4.08E-04 | 1.46E-03 | -1.17E+00 | 1.30E-06 | 3.66E-06 | -1.43E+00 | 2.58E-08 | 9.46E-08 | -1.17E+00 | 3.07E-06 | 8.44E-06 | -1.11E+00 |
| FEM1B | 7.86E-07 | 4.09E-06 | -1.22E+00 | 6.73E-11 | 2.91E-10 | -1.37E+00 | 9.26E-07 | 3.93E-06 | -1.30E+00 | 5.40E-09 | 2.06E-08 | -1.32E+00 | 9.57E-06 | 3.41E-05 | -1.33E+00 | 1.74E-08 | 4.91E-08 | -1.47E+00 | 4.03E-03 | 1.48E-02 | -1.15E+00 | 3.01E-02 | 8.27E-02 | -1.01E+00 |
| ALOX12-AS1 | 7.88E-07 | 4.11E-06 | -1.44E+00 | 6.01E-04 | 2.60E-03 | -1.03E+00 | 1.09E-05 | 4.62E-05 | -1.51E+00 | 7.62E-04 | 2.91E-03 | -1.14E+00 | 8.13E-04 | 2.90E-03 | -1.28E+00 | 2.82E-02 | 7.95E-02 | 1.06E+00 | 1.77E-04 | 6.48E-04 | -1.29E+00 | 6.58E-03 | 1.81E-02 | -1.01E+00 |
| NFIL3 | 7.95E-07 | 4.14E-06 | 2.60E+00 | 1.87E-01 | 8.07E-01 | 1.39E+00 | 1.36E-01 | 5.78E-01 | 1.35E+00 | 2.58E-03 | 9.85E-03 | -1.11E+00 | 2.94E-02 | 1.05E-01 | 2.13E+00 | 2.01E-02 | 5.68E-02 | 1.13E+00 | 3.82E-02 | 1.40E-01 | 1.70E+00 | 6.57E-03 | 1.80E-02 | -1.49E+00 |
| KAT8 | 7.99E-07 | 4.17E-06 | -1.13E+00 | 1.23E-03 | 5.31E-03 | 1.15E+00 | 1.00E-04 | 4.27E-04 | -1.17E+00 | 3.58E-03 | 1.37E-02 | 1.06E+00 | 2.93E-07 | 1.04E-06 | -1.36E+00 | 3.95E-03 | 1.11E-02 | 1.01E+00 | 2.02E-06 | 7.41E-06 | -1.27E+00 | 1.37E-03 | 3.76E-03 | -1.06E+00 |
| AMPD2 | 7.99E-07 | 4.17E-06 | -1.22E+00 | 6.49E-08 | 2.80E-07 | -1.15E+00 | 7.01E-05 | 2.98E-04 | -1.20E+00 | 6.15E-06 | 2.35E-05 | -1.14E+00 | 5.00E-03 | 1.78E-02 | -1.01E+00 | 3.17E-01 | 8.95E-01 | 1.53E+00 | 1.28E-02 | 4.68E-02 | -1.01E+00 | 3.05E-03 | 8.37E-03 | -1.08E+00 |
| IFIT5 | 8.00E-07 | 4.17E-06 | -1.34E+00 | 6.58E-05 | 2.84E-04 | -1.07E+00 | 6.39E-08 | 2.71E-07 | -1.68E+00 | 2.70E-05 | 1.03E-04 | -1.25E+00 | 5.53E-08 | 1.97E-07 | -1.73E+00 | 5.29E-08 | 1.49E-07 | -1.63E+00 | 5.06E-10 | 1.86E-09 | -1.67E+00 | 2.36E-08 | 6.48E-08 | -1.51E+00 |
| HGSNAT | 8.05E-07 | 4.19E-06 | -1.24E+00 | 2.94E-02 | 1.27E-01 | 1.23E+00 | 1.92E-05 | 8.17E-05 | -1.14E+00 | 4.16E-03 | 1.59E-02 | 1.15E+00 | 2.20E-02 | 7.84E-02 | 1.10E+00 | 2.38E-01 | 6.70E-01 | 1.60E+00 | 7.01E-03 | 2.57E-02 | 1.05E+00 | 1.48E-01 | 4.05E-01 | 1.18E+00 |
| PTX3 | 8.06E-07 | 4.20E-06 | 2.37E+00 | 1.44E-06 | 6.21E-06 | -1.17E+00 | 2.73E-04 | 1.16E-03 | 1.88E+00 | 4.50E-04 | 1.72E-03 | 1.06E+00 | 9.01E-03 | 3.21E-02 | -1.41E+00 | 1.17E-03 | 3.31E-03 | -3.74E+00 | 1.76E-01 | 6.45E-01 | 1.08E+00 | 1.35E-01 | 3.72E-01 | -1.92E+00 |
| CLEC2B | 8.14E-07 | 4.24E-06 | 1.71E+00 | 2.47E-02 | 1.07E-01 | 1.14E+00 | 1.35E-02 | 5.75E-02 | 1.51E+00 | 1.91E-03 | 7.30E-03 | -1.04E+00 | 2.38E-01 | 8.49E-01 | 1.27E+00 | 1.48E-04 | 4.18E-04 | -1.17E+00 | 8.75E-05 | 3.21E-04 | -1.03E+00 | 6.61E-14 | 1.81E-13 | -1.56E+00 |
| COPS9 | 8.14E-07 | 4.24E-06 | -1.03E+00 | 1.65E-24 | 7.13E-24 | -1.32E+00 | 2.05E-01 | 8.71E-01 | 1.09E+00 | 8.87E-08 | 3.39E-07 | -1.29E+00 | 7.26E-03 | 2.59E-02 | -1.02E+00 | 3.68E-07 | 1.04E-06 | -1.28E+00 | 5.27E-02 | 1.94E-01 | 1.03E+00 | 2.23E-06 | 6.13E-06 | -1.38E+00 |
| BAG1 | 8.14E-07 | 4.24E-06 | -1.09E+00 | 1.31E-04 | 5.65E-04 | 1.10E+00 | 2.79E-02 | 1.18E-01 | -1.02E+00 | 1.80E-02 | 6.87E-02 | 1.07E+00 | 1.92E-02 | 6.84E-02 | 1.11E+00 | 9.17E-02 | 2.59E-01 | 1.24E+00 | 4.28E-02 | 1.57E-01 | 1.40E+00 | 3.69E-02 | 1.01E-01 | 1.43E+00 |
| AC018645.2 | 8.15E-07 | 4.25E-06 | -2.16E+00 | 3.55E-06 | 1.53E-05 | -1.83E+00 | 2.92E-05 | 1.24E-04 | -1.87E+00 | 7.35E-04 | 2.81E-03 | -1.30E+00 | 6.29E-04 | 2.24E-03 | -1.85E+00 | 2.03E-03 | 5.73E-03 | -1.62E+00 | 5.90E-04 | 2.16E-03 | -1.33E+00 | 1.40E-02 | 3.84E-02 | -1.13E+00 |
| AP5M1 | 8.18E-07 | 4.26E-06 | -1.30E+00 | 1.92E-03 | 8.29E-03 | 1.10E+00 | 4.73E-05 | 2.01E-04 | -1.23E+00 | 4.54E-02 | 1.74E-01 | 1.19E+00 | 2.09E-05 | 7.44E-05 | -1.11E+00 | 4.34E-02 | 1.22E-01 | 1.30E+00 | 5.61E-07 | 2.06E-06 | -1.43E+00 | 2.73E-04 | 7.49E-04 | -1.20E+00 |
| SLFN5 | 8.19E-07 | 4.27E-06 | -3.98E+00 | 3.33E-06 | 1.44E-05 | -3.21E+00 | 9.30E-04 | 3.95E-03 | -2.65E+00 | 6.35E-03 | 2.43E-02 | -1.93E+00 | 4.73E-09 | 1.69E-08 | -2.76E+00 | 1.95E-09 | 5.49E-09 | -2.60E+00 | 6.12E-70 | 2.25E-69 | -7.30E+00 | 1.77E-56 | 4.85E-56 | -4.42E+00 |
| HIST1H1D | 8.19E-07 | 4.27E-06 | 1.62E+00 | 1.45E-02 | 6.28E-02 | 1.51E+00 | 1.27E-04 | 5.39E-04 | 1.49E+00 | 1.17E-02 | 4.46E-02 | 1.40E+00 | 2.60E-01 | 9.28E-01 | 1.19E+00 | 1.78E-02 | 5.02E-02 | 1.01E+00 | 4.50E-02 | 1.65E-01 | 1.21E+00 | 5.09E-02 | 1.40E-01 | 1.14E+00 |
| RPL7P15 | 8.20E-07 | 4.27E-06 | -1.05E+00 | 1.06E-01 | 4.56E-01 | 1.47E+00 | 8.04E-07 | 3.42E-06 | -1.28E+00 | 1.63E-01 | 6.21E-01 | 1.31E+00 | 8.11E-03 | 2.89E-02 | 1.04E+00 | 8.92E-02 | 2.51E-01 | 1.54E+00 | 1.89E-04 | 6.94E-04 | -1.21E+00 | 4.59E-02 | 1.26E-01 | 1.41E+00 |
| AC120114.4 | 8.20E-07 | 4.27E-06 | -1.64E+00 | 1.48E-05 | 6.38E-05 | -1.23E+00 | 9.84E-07 | 4.18E-06 | -1.53E+00 | 1.22E-09 | 4.65E-09 | -1.92E+00 | 7.01E-07 | 2.50E-06 | -2.20E+00 | 4.21E-05 | 1.19E-04 | -1.50E+00 | 1.18E-02 | 4.33E-02 | -1.66E+00 | 6.88E-02 | 1.89E-01 | -1.25E+00 |
| EID2B | 8.20E-07 | 4.27E-06 | -1.44E+00 | 2.13E-05 | 9.20E-05 | -1.21E+00 | 6.23E-02 | 2.65E-01 | -1.08E+00 | 9.59E-02 | 3.66E-01 | 1.12E+00 | 2.80E-01 | 9.97E-01 | 1.15E+00 | 1.24E-01 | 3.49E-01 | -1.02E+00 | 1.06E-02 | 3.90E-02 | -1.17E+00 | 3.97E-02 | 1.09E-01 | -1.06E+00 |
| CASD1 | 8.29E-07 | 4.32E-06 | -1.22E+00 | 1.80E-05 | 7.76E-05 | -1.01E+00 | 1.10E-06 | 4.68E-06 | -1.39E+00 | 1.37E-03 | 5.22E-03 | 1.01E+00 | 5.61E-09 | 2.00E-08 | -1.81E+00 | 5.08E-05 | 1.43E-04 | -1.27E+00 | 8.49E-06 | 3.12E-05 | -1.31E+00 | 2.05E-03 | 5.64E-03 | -1.05E+00 |
| SNX19 | 8.30E-07 | 4.33E-06 | -1.44E+00 | 8.00E-05 | 3.45E-04 | -1.05E+00 | 2.15E-04 | 9.15E-04 | -1.31E+00 | 1.01E-03 | 3.85E-03 | -1.03E+00 | 4.55E-03 | 1.62E-02 | -1.05E+00 | 4.97E-03 | 1.40E-02 | -1.12E+00 | 7.85E-09 | 2.88E-08 | -1.64E+00 | 2.08E-03 | 5.72E-03 | -1.09E+00 |
| PPIP5K2 | 8.31E-07 | 4.33E-06 | -1.08E+00 | 9.40E-06 | 4.06E-05 | 1.04E+00 | 7.03E-05 | 2.99E-04 | -1.16E+00 | 2.98E-05 | 1.14E-04 | -1.08E+00 | 4.26E-03 | 1.52E-02 | -1.01E+00 | 3.91E-02 | 1.10E-01 | 1.05E+00 | 4.75E-03 | 1.74E-02 | -1.04E+00 | 7.22E-02 | 1.98E-01 | 1.02E+00 |
| DDAH1 | 8.34E-07 | 4.35E-06 | -1.68E+00 | 5.66E-04 | 2.45E-03 | -1.03E+00 | 4.87E-08 | 2.07E-07 | -1.46E+00 | 6.69E-06 | 2.56E-05 | -1.14E+00 | 4.47E-02 | 1.59E-01 | -1.32E+00 | 2.34E-01 | 6.60E-01 | 1.13E+00 | 1.23E-03 | 4.53E-03 | -1.38E+00 | 2.62E-01 | 7.20E-01 | 1.23E+00 |
| UBE2Q1 | 8.34E-07 | 4.35E-06 | -1.39E+00 | 2.12E-05 | 9.17E-05 | -1.06E+00 | 2.91E-04 | 1.24E-03 | -1.26E+00 | 1.48E-02 | 5.64E-02 | 1.10E+00 | 5.16E-04 | 1.84E-03 | -1.14E+00 | 7.72E-04 | 2.18E-03 | -1.15E+00 | 3.78E-03 | 1.39E-02 | -1.05E+00 | 5.80E-02 | 1.59E-01 | 1.05E+00 |
| SAMD4B | 8.35E-07 | 4.35E-06 | -1.22E+00 | 6.92E-08 | 2.99E-07 | -1.13E+00 | 1.00E-02 | 4.26E-02 | -1.06E+00 | 1.74E-04 | 6.64E-04 | -1.10E+00 | 6.56E-05 | 2.34E-04 | -1.06E+00 | 2.79E-05 | 7.88E-05 | -1.12E+00 | 1.44E-02 | 5.30E-02 | -1.10E+00 | 8.03E-03 | 2.21E-02 | -1.07E+00 |
| PPT2 | 8.37E-07 | 4.36E-06 | -1.95E+00 | 2.23E-04 | 9.65E-04 | -1.25E+00 | 6.43E-08 | 2.73E-07 | -1.80E+00 | 4.91E-07 | 1.88E-06 | -1.54E+00 | 2.09E-03 | 7.46E-03 | -1.58E+00 | 1.68E-02 | 4.74E-02 | -1.18E+00 | 1.95E-07 | 7.18E-07 | -1.31E+00 | 2.56E-07 | 7.02E-07 | -1.32E+00 |
| UBR5 | 8.39E-07 | 4.37E-06 | -1.11E+00 | 1.81E-04 | 7.80E-04 | 1.08E+00 | 1.26E-03 | 5.37E-03 | -1.14E+00 | 1.91E-03 | 7.30E-03 | 1.07E+00 | 5.33E-05 | 1.90E-04 | -1.25E+00 | 3.87E-03 | 1.09E-02 | -1.05E+00 | 1.97E-05 | 7.22E-05 | -1.27E+00 | 3.12E-03 | 8.58E-03 | -1.08E+00 |
| RAP1A | 8.43E-07 | 4.39E-06 | -1.12E+00 | 1.49E-04 | 6.44E-04 | 1.11E+00 | 8.71E-07 | 3.70E-06 | -1.31E+00 | 4.24E-06 | 1.62E-05 | -1.11E+00 | 9.46E-08 | 3.37E-07 | -1.24E+00 | 3.36E-04 | 9.47E-04 | 1.04E+00 | 1.61E-05 | 5.89E-05 | -1.10E+00 | 1.45E-02 | 3.98E-02 | -1.00E+00 |
| MAGT1 | 8.52E-07 | 4.44E-06 | -1.20E+00 | 1.38E-03 | 5.94E-03 | 1.21E+00 | 2.49E-03 | 1.06E-02 | -1.09E+00 | 1.31E-02 | 5.01E-02 | 1.11E+00 | 4.27E-03 | 1.52E-02 | 1.03E+00 | 5.20E-03 | 1.47E-02 | 1.03E+00 | 2.49E-06 | 9.16E-06 | -1.39E+00 | 4.87E-02 | 1.34E-01 | -1.06E+00 |
| TMEM168 | 8.55E-07 | 4.45E-06 | -1.26E+00 | 2.28E-04 | 9.85E-04 | 1.04E+00 | 3.60E-06 | 1.53E-05 | -1.47E+00 | 3.40E-04 | 1.30E-03 | -1.09E+00 | 1.82E-03 | 6.50E-03 | -1.18E+00 | 5.89E-02 | 1.66E-01 | 1.18E+00 | 4.21E-03 | 1.54E-02 | -1.16E+00 | 1.77E-01 | 4.86E-01 | 1.21E+00 |
| CD47 | 8.56E-07 | 4.46E-06 | -1.10E+00 | 1.46E-02 | 6.30E-02 | 1.24E+00 | 5.09E-02 | 2.16E-01 | -1.03E+00 | 1.12E-01 | 4.27E-01 | 1.17E+00 | 3.95E-07 | 1.41E-06 | -1.30E+00 | 1.11E-01 | 3.12E-01 | 1.24E+00 | 2.30E-09 | 8.42E-09 | -1.25E+00 | 6.42E-04 | 1.76E-03 | -1.04E+00 |
| FAM20B | 8.61E-07 | 4.49E-06 | -1.45E+00 | 9.27E-04 | 4.01E-03 | 1.07E+00 | 1.34E-05 | 5.68E-05 | -1.39E+00 | 2.84E-02 | 1.09E-01 | 1.11E+00 | 5.27E-04 | 1.88E-03 | -1.28E+00 | 1.45E-02 | 4.08E-02 | 1.01E+00 | 2.74E-10 | 1.01E-09 | -1.45E+00 | 1.39E-02 | 3.83E-02 | 1.06E+00 |
| GTPBP1 | 8.61E-07 | 4.49E-06 | -1.44E+00 | 1.21E-05 | 5.22E-05 | -1.20E+00 | 2.29E-02 | 9.75E-02 | -1.04E+00 | 1.82E-02 | 6.95E-02 | 1.03E+00 | 1.97E-03 | 7.01E-03 | -1.12E+00 | 7.72E-04 | 2.18E-03 | -1.20E+00 | 7.24E-03 | 2.66E-02 | -1.01E+00 | 4.94E-03 | 1.36E-02 | -1.05E+00 |
| SDHB | 8.64E-07 | 4.50E-06 | -1.17E+00 | 4.75E-04 | 2.05E-03 | 1.09E+00 | 5.93E-04 | 2.52E-03 | -1.04E+00 | 3.92E-05 | 1.50E-04 | -1.06E+00 | 1.41E-03 | 5.02E-03 | -1.05E+00 | 2.93E-02 | 8.25E-02 | 1.11E+00 | 5.79E-11 | 2.13E-10 | -1.40E+00 | 2.16E-03 | 5.93E-03 | -1.01E+00 |
| ANKRD28 | 8.68E-07 | 4.52E-06 | 1.30E+00 | 4.11E-07 | 1.78E-06 | 1.43E+00 | 1.68E-04 | 7.14E-04 | 1.25E+00 | 8.22E-08 | 3.14E-07 | 1.51E+00 | 5.83E-26 | 2.08E-25 | 2.39E+00 | 8.37E-21 | 2.36E-20 | 2.16E+00 | 4.68E-55 | 1.72E-54 | 3.15E+00 | 3.99E-34 | 1.10E-33 | 2.54E+00 |
| ZKSCAN3 | 8.70E-07 | 4.53E-06 | -1.72E+00 | 6.11E-04 | 2.64E-03 | -1.05E+00 | 1.37E-08 | 5.81E-08 | -2.40E+00 | 5.20E-05 | 1.99E-04 | -1.47E+00 | 9.08E-05 | 3.24E-04 | -1.56E+00 | 2.08E-01 | 5.85E-01 | 1.24E+00 | 2.26E-15 | 8.31E-15 | -2.73E+00 | 5.70E-07 | 1.56E-06 | -1.61E+00 |
| PRC1 | 8.70E-07 | 4.54E-06 | -1.49E+00 | 3.16E-07 | 1.36E-06 | -1.44E+00 | 1.56E-01 | 6.61E-01 | 1.23E+00 | 9.80E-03 | 3.74E-02 | 1.16E+00 | 3.17E-03 | 1.13E-02 | -1.07E+00 | 6.01E-04 | 1.69E-03 | -1.19E+00 | 4.26E-05 | 1.56E-04 | -1.32E+00 | 7.67E-03 | 2.11E-02 | -1.15E+00 |

| | | | | | | | | | | | | | | | | | | | | | | | | |
|---|---|---|---|---|---|---|---|---|---|---|---|---|---|---|---|---|---|---|---|---|---|---|---|---|
| RUNX3 | 8.70E-07 | 4.54E-06 | -1.13E+00 | 1.59E-05 | 6.88E-05 | 1.02E+00 | 7.57E-02 | 3.22E-01 | 1.05E+00 | 1.26E-03 | 4.80E-03 | -1.00E+00 | 4.71E-04 | 1.68E-03 | -1.73E+00 | 1.90E-03 | 5.34E-03 | -1.58E+00 | 5.58E-02 | 2.05E-01 | -1.32E+00 | 3.37E-01 | 9.27E-01 | 1.23E+00 |
| ARHGEF35 | 8.76E-07 | 4.57E-06 | -1.68E+00 | 2.52E-06 | 1.09E-05 | -1.51E+00 | 1.72E-06 | 7.32E-06 | -1.71E+00 | 9.79E-08 | 3.74E-07 | -1.79E+00 | 1.02E-01 | 3.65E-01 | 1.75E+00 | 2.10E-01 | 5.91E-01 | -1.10E+00 | 1.67E-01 | 6.13E-01 | -1.06E+00 | 1.89E-01 | 5.19E-01 | -1.34E+00 |
| STX5 | 8.80E-07 | 4.59E-06 | -1.16E+00 | 2.28E-05 | 9.85E-05 | 1.07E+00 | 2.11E-05 | 8.98E-05 | -1.24E+00 | 5.74E-08 | 2.19E-07 | -1.23E+00 | 2.31E-06 | 8.22E-06 | -1.15E+00 | 5.09E-05 | 1.43E-04 | -1.03E+00 | 1.76E-03 | 6.46E-03 | -1.01E+00 | 1.02E-03 | 2.79E-03 | -1.02E+00 |
| ENG | 8.83E-07 | 4.60E-06 | -1.23E+00 | 3.54E-03 | 1.53E-02 | 1.20E+00 | 8.52E-09 | 3.62E-08 | -1.48E+00 | 3.80E-06 | 1.45E-05 | -1.04E+00 | 1.51E-03 | 5.38E-03 | -1.50E+00 | 9.06E-02 | 2.55E-01 | 1.10E+00 | 3.00E-16 | 1.10E-15 | -1.79E+00 | 3.83E-07 | 1.05E-06 | -1.20E+00 |
| PNRC2 | 8.84E-07 | 4.61E-06 | -1.10E+00 | 3.99E-15 | 1.72E-14 | -1.25E+00 | 8.33E-09 | 3.54E-08 | -1.41E+00 | 4.03E-13 | 1.54E-12 | -1.43E+00 | 2.74E-07 | 9.77E-07 | -1.25E+00 | 4.21E-07 | 1.19E-06 | -1.24E+00 | 1.61E-04 | 5.90E-04 | -1.24E+00 | 1.15E-05 | 3.16E-05 | -1.28E+00 |
| ADGRE5 | 8.87E-07 | 4.62E-06 | 2.24E+00 | 1.49E-03 | 6.42E-03 | 1.15E+00 | 9.95E-02 | 4.23E-01 | 1.48E+00 | 2.28E-04 | 8.71E-04 | 1.04E+00 | 2.02E-01 | 7.19E-01 | 1.47E+00 | 1.50E-02 | 4.23E-02 | -1.12E+00 | 8.89E-02 | 3.26E-01 | 1.89E+00 | 2.47E-01 | 6.78E-01 | 1.16E+00 |
| AL353622.1 | 8.93E-07 | 4.65E-06 | -1.49E+00 | 4.98E-03 | 2.15E-02 | 1.13E+00 | 1.46E-06 | 6.19E-06 | -2.01E+00 | 3.22E-05 | 1.23E-04 | -1.50E+00 | 8.88E-02 | 3.17E-01 | 1.10E+00 | 3.10E-01 | 8.74E-01 | 1.48E+00 | 4.46E-02 | 1.64E-01 | -1.21E+00 | 2.34E-01 | 6.44E-01 | 1.02E+00 |
| SSSCA1 | 8.94E-07 | 4.66E-06 | -1.24E+00 | 3.58E-04 | 1.55E-03 | 1.03E+00 | 1.20E-01 | 5.11E-01 | 1.12E+00 | 1.49E-02 | 5.70E-02 | 1.14E+00 | 9.54E-03 | 3.40E-02 | 1.12E+00 | 7.94E-02 | 2.24E-01 | 1.24E+00 | 1.96E-05 | 7.21E-05 | -1.05E+00 | 4.17E-02 | 1.15E-01 | 1.07E+00 |
| MBOAT2 | 8.94E-07 | 4.66E-06 | -1.26E+00 | 2.03E-02 | 8.77E-02 | 1.21E+00 | 1.34E-05 | 5.70E-05 | -1.41E+00 | 1.89E-02 | 7.21E-02 | 1.11E+00 | 3.24E-06 | 1.16E-05 | -1.11E+00 | 2.32E-01 | 6.54E-01 | 1.48E+00 | 2.45E-02 | 9.00E-02 | 1.16E+00 | 2.88E-01 | 7.92E-01 | 1.37E+00 |
| STK10 | 8.96E-07 | 4.67E-06 | -1.21E+00 | 4.99E-03 | 2.15E-02 | 1.19E+00 | 3.22E-04 | 1.37E-03 | -1.22E+00 | 7.69E-02 | 2.94E-01 | 1.13E+00 | 1.09E-03 | 3.88E-03 | -1.22E+00 | 5.64E-02 | 1.59E-01 | 1.08E+00 | 1.78E-04 | 6.53E-04 | -1.19E+00 | 7.28E-03 | 2.00E-02 | -1.03E+00 |
| NBPF12 | 8.96E-07 | 4.67E-06 | -1.55E+00 | 1.34E-02 | 5.79E-02 | 1.20E+00 | 3.40E-06 | 1.44E-05 | -1.63E+00 | 3.59E-04 | 1.37E-03 | -1.09E+00 | 4.34E-03 | 1.55E-02 | -1.23E+00 | 2.62E-02 | 7.38E-02 | -1.05E+00 | 1.06E-06 | 3.88E-06 | -1.26E+00 | 6.98E-05 | 1.92E-04 | -1.21E+00 |
| PTPN12 | 8.97E-07 | 4.68E-06 | -1.33E+00 | 6.08E-07 | 2.63E-06 | -1.17E+00 | 1.29E-04 | 5.48E-04 | -1.21E+00 | 9.35E-05 | 3.57E-04 | -1.08E+00 | 1.98E-04 | 7.06E-04 | -1.17E+00 | 2.38E-03 | 6.70E-03 | -1.03E+00 | 6.30E-04 | 2.31E-03 | -1.12E+00 | 2.85E-03 | 7.84E-03 | -1.29E+00 |
| Z98749.2 | 9.01E-07 | 4.70E-06 | -1.47E+00 | 6.13E-02 | 2.65E-01 | 1.26E+00 | 4.44E-08 | 1.89E-07 | -1.85E+00 | 4.22E-04 | 1.61E-03 | -1.07E+00 | 3.04E-03 | 1.08E-02 | -1.23E+00 | 1.22E-01 | 3.43E-01 | 1.11E+00 | 2.41E-02 | 8.84E-02 | -1.29E+00 | 3.06E-01 | 8.40E-01 | 1.16E+00 |
| CHMP2A | 9.02E-07 | 4.70E-06 | -1.03E+00 | 5.46E-11 | 2.36E-10 | -1.03E+00 | 2.11E-01 | 8.95E-01 | 1.14E+00 | 9.09E-03 | 3.47E-02 | 1.06E+00 | 2.10E-08 | 7.48E-08 | -1.10E+00 | 1.23E-05 | 3.47E-05 | -1.03E+00 | 1.93E-01 | 7.09E-01 | 1.26E+00 | 3.17E-01 | 8.70E-01 | 1.20E+00 |
| COPS6 | 9.05E-07 | 4.71E-06 | -1.01E+00 | 1.30E-03 | 5.63E-03 | 1.23E+00 | 2.31E-02 | 9.81E-02 | 1.03E+00 | 1.24E-02 | 4.74E-02 | 1.13E+00 | 8.62E-03 | 3.07E-02 | 1.13E+00 | 5.22E-02 | 1.47E-01 | 1.27E+00 | 1.32E-01 | 4.85E-01 | 1.12E+00 | 8.81E-02 | 2.42E-01 | 1.33E+00 |
| ZNF511 | 9.07E-07 | 4.73E-06 | -1.08E+00 | 6.58E-07 | 2.84E-06 | 1.01E+00 | 1.05E-02 | 4.47E-02 | 1.01E+00 | 1.40E-03 | 5.35E-03 | 1.08E+00 | 8.72E-05 | 3.11E-04 | -1.10E+00 | 2.76E-02 | 7.78E-02 | 1.15E+00 | 2.65E-02 | 9.72E-02 | 1.06E+00 | 2.41E-01 | 6.62E-01 | 1.15E+00 |
| NNT | 9.07E-07 | 4.73E-06 | -1.30E+00 | 6.02E-04 | 2.60E-03 | 1.04E+00 | 1.13E-09 | 4.80E-09 | -1.50E+00 | 2.07E-07 | 7.92E-07 | -1.16E+00 | 8.79E-06 | 3.13E-05 | -1.24E+00 | 1.53E-03 | 4.33E-03 | 1.03E+00 | 1.23E-06 | 4.51E-06 | -1.31E+00 | 5.61E-05 | 1.54E-04 | -1.18E+00 |
| SNUPN | 9.09E-07 | 4.73E-06 | -1.57E+00 | 3.05E-07 | 1.32E-06 | -1.50E+00 | 1.30E-08 | 5.50E-08 | -1.52E+00 | 2.83E-10 | 1.08E-09 | -1.50E+00 | 1.15E-06 | 4.08E-06 | -1.61E+00 | 3.61E-05 | 1.02E-04 | -1.27E+00 | 3.21E-04 | 1.18E-03 | -1.14E+00 | 2.93E-06 | 8.04E-06 | -1.33E+00 |
| ACTR1B | 9.19E-07 | 4.79E-06 | -1.19E+00 | 5.14E-06 | 2.22E-05 | -1.04E+00 | 8.63E-05 | 3.67E-04 | -1.23E+00 | 7.86E-06 | 3.00E-05 | -1.17E+00 | 2.39E-04 | 8.52E-04 | -1.34E+00 | 7.64E-04 | 2.15E-03 | -1.21E+00 | 5.04E-07 | 1.85E-06 | -1.42E+00 | 1.85E-05 | 5.07E-05 | -1.25E+00 |
| C1orf56 | 9.20E-07 | 4.79E-06 | -1.62E+00 | 9.81E-05 | 4.24E-04 | -1.18E+00 | 5.38E-04 | 2.28E-03 | -1.45E+00 | 3.19E-03 | 1.22E-02 | -1.12E+00 | 2.32E-02 | 8.28E-02 | -1.24E+00 | 2.13E-01 | 6.01E-01 | 1.13E+00 | 1.20E-03 | 4.39E-03 | -1.38E+00 | 2.26E-02 | 6.21E-02 | -1.09E+00 |
| SMG6 | 9.22E-07 | 4.81E-06 | -1.34E+00 | 1.23E-07 | 5.32E-07 | -1.29E+00 | 9.95E-06 | 4.23E-05 | -1.32E+00 | 5.84E-08 | 2.23E-07 | -1.35E+00 | 2.66E-03 | 9.48E-03 | 1.04E+00 | 5.48E-04 | 1.54E-03 | -1.09E+00 | 1.58E-02 | 5.79E-02 | 1.11E+00 | 7.92E-04 | 2.18E-03 | -1.04E+00 |
| SMS | 9.27E-07 | 4.83E-06 | -1.08E+00 | 1.53E-09 | 6.60E-09 | -1.04E+00 | 3.85E-02 | 1.63E-01 | -1.05E+00 | 5.92E-02 | 2.26E-01 | 1.04E+00 | 1.17E-07 | 4.18E-07 | -1.15E+00 | 8.79E-05 | 2.48E-04 | 1.03E+00 | 4.99E-05 | 1.83E-04 | -1.19E+00 | 1.56E-02 | 4.27E-02 | -1.07E+00 |
| SORBS3 | 9.27E-07 | 4.83E-06 | -1.13E+00 | 4.78E-05 | 2.06E-04 | 1.03E+00 | 1.66E-04 | 7.03E-04 | -1.22E+00 | 1.40E-03 | 5.34E-03 | 1.02E+00 | 1.58E-01 | 5.64E-01 | -1.04E+00 | 1.86E-01 | 5.24E-01 | -1.02E+00 | 2.35E-01 | 8.61E-01 | 1.31E+00 | 3.10E-01 | 8.52E-01 | 1.29E+00 |
| EIF1B-AS1 | 9.28E-07 | 4.83E-06 | -1.52E+00 | 8.06E-06 | 3.48E-05 | -1.26E+00 | 1.47E-03 | 6.23E-03 | -1.45E+00 | 1.69E-03 | 6.45E-03 | -1.22E+00 | 4.37E-05 | 1.56E-04 | -1.87E+00 | 2.11E-04 | 5.95E-04 | -1.59E+00 | 2.93E-02 | 1.08E-01 | -1.15E+00 | 2.55E-02 | 7.01E-02 | -1.13E+00 |
| CYP4F22 | 9.31E-07 | 4.85E-06 | -1.87E+00 | 4.71E-04 | 2.04E-03 | -1.17E+00 | 1.13E-02 | 4.80E-02 | -1.34E+00 | 2.59E-01 | 9.90E-01 | 1.28E+00 | 1.47E-01 | 5.26E-01 | -1.47E+00 | 2.98E-01 | 8.41E-01 | -1.02E+00 | 2.02E-01 | 7.43E-01 | -1.13E+00 | 3.46E-01 | 9.49E-01 | 1.05E+00 |
| PLA2G15 | 9.34E-07 | 4.87E-06 | -1.69E+00 | 1.81E-04 | 7.83E-04 | -1.21E+00 | 6.07E-05 | 2.58E-04 | -1.64E+00 | 7.00E-03 | 2.67E-02 | -1.01E+00 | 2.80E-06 | 9.98E-06 | -1.91E+00 | 1.50E-06 | 4.22E-06 | -1.94E+00 | 1.57E-05 | 5.77E-05 | -1.40E+00 | 1.69E-03 | 4.65E-03 | -1.13E+00 |
| POLH | 9.35E-07 | 4.87E-06 | -1.70E+00 | 4.64E-06 | 2.00E-05 | -1.44E+00 | 2.07E-03 | 8.79E-03 | -1.44E+00 | 1.27E-02 | 4.85E-02 | -1.07E+00 | 4.52E-02 | 1.61E-01 | -1.16E+00 | 3.55E-01 | 1.00E+00 | 1.34E+00 | 1.99E-05 | 7.30E-05 | -1.46E+00 | 1.42E-02 | 3.90E-02 | -1.12E+00 |
| INTU | 9.37E-07 | 4.88E-06 | -1.50E+00 | 5.60E-07 | 2.42E-06 | -1.27E+00 | 1.15E-06 | 4.87E-06 | -1.56E+00 | 4.95E-05 | 1.89E-04 | -1.07E+00 | 6.70E-05 | 2.39E-04 | -1.53E+00 | 8.00E-03 | 2.25E-02 | -1.03E+00 | 8.76E-10 | 3.22E-09 | -1.98E+00 | 9.45E-04 | 2.60E-03 | -1.25E+00 |
| GTPBP3 | 9.41E-07 | 4.91E-06 | -1.41E+00 | 2.62E-04 | 1.13E-03 | 1.00E+00 | 5.29E-04 | 2.25E-03 | -1.21E+00 | 5.58E-05 | 2.13E-04 | -1.18E+00 | 1.11E-03 | 3.97E-03 | -1.24E+00 | 9.35E-03 | 2.64E-02 | -1.03E+00 | 2.17E-10 | 7.97E-10 | -1.64E+00 | 4.68E-04 | 1.29E-03 | -1.11E+00 |
| SNAP29 | 9.41E-07 | 4.91E-06 | -1.30E+00 | 1.61E-04 | 6.95E-04 | -1.04E+00 | 1.52E-04 | 6.45E-04 | -1.25E+00 | 1.05E-04 | 4.01E-04 | -1.16E+00 | 2.23E-05 | 7.94E-05 | -1.27E+00 | 3.00E-03 | 8.45E-03 | 1.01E+00 | 1.92E-06 | 7.05E-06 | -1.18E+00 | 6.29E-04 | 1.73E-03 | -1.08E+00 |
| OR2A20P | 9.48E-07 | 4.94E-06 | -1.60E+00 | 2.20E-05 | 9.51E-05 | -1.28E+00 | 7.09E-02 | 3.01E-01 | -1.05E+00 | 4.74E-02 | 1.81E-01 | -1.05E+00 | 1.82E-01 | 6.49E-01 | -1.14E+00 | 2.08E-01 | 5.87E-01 | -1.17E+00 | 2.36E-01 | 8.67E-01 | 1.20E+00 | 3.23E-01 | 8.87E-01 | 1.02E+00 |
| TNFRSF10A | 9.55E-07 | 4.98E-06 | -1.70E+00 | 1.03E-05 | 4.46E-05 | -1.43E+00 | 6.49E-06 | 2.76E-05 | -1.69E+00 | 3.75E-06 | 1.43E-05 | -1.59E+00 | 2.18E-03 | 7.79E-03 | -1.60E+00 | 3.07E-04 | 8.64E-04 | -2.21E+00 | 2.76E-04 | 1.01E-03 | -2.13E+00 | 4.32E-04 | 1.19E-03 | -1.98E+00 |
| TIMM21 | 9.62E-07 | 5.01E-06 | -1.86E+00 | 1.55E-03 | 6.69E-03 | -1.01E+00 | 1.41E-05 | 5.98E-05 | -1.36E+00 | 1.32E-04 | 5.04E-04 | -1.12E+00 | 1.28E-02 | 4.55E-02 | -1.49E+00 | 1.59E-01 | 4.48E-01 | 1.03E+00 | 7.30E-07 | 2.68E-06 | -1.41E+00 | 1.11E-03 | 3.04E-03 | -1.09E+00 |
| FAM98C | 9.62E-07 | 5.01E-06 | -1.39E+00 | 4.15E-04 | 1.79E-03 | 1.06E+00 | 7.32E-05 | 3.11E-04 | -1.61E+00 | 1.25E-04 | 4.76E-04 | -1.40E+00 | 1.30E-03 | 4.65E-03 | -1.27E+00 | 1.28E-01 | 3.62E-01 | 1.28E+00 | 2.58E-02 | 9.48E-02 | -1.08E+00 | 4.70E-02 | 1.29E-01 | -1.05E+00 |
| SALL4 | 9.67E-07 | 5.04E-06 | -2.01E+00 | 1.24E-04 | 5.36E-04 | -1.42E+00 | 4.18E-05 | 1.78E-04 | -1.58E+00 | 5.31E-04 | 2.03E-03 | -1.30E+00 | 1.10E-01 | 3.94E-01 | -1.10E+00 | 3.40E-01 | 9.59E-01 | 1.41E+00 | 9.30E-02 | 3.41E-01 | -1.36E+00 | 8.80E-02 | 2.42E-01 | -1.56E+00 |
| ZMYM6 | 9.70E-07 | 5.06E-06 | -1.38E+00 | 1.16E-04 | 5.00E-04 | -1.02E+00 | 8.45E-05 | 3.59E-04 | -1.32E+00 | 1.03E-03 | 3.94E-03 | -1.08E+00 | 1.11E-01 | 3.97E-01 | 1.14E+00 | 1.91E-01 | 5.38E-01 | 1.30E+00 | 5.20E-03 | 1.91E-02 | -1.48E+00 | 7.32E-03 | 2.01E-02 | -1.47E+00 |
| RGPD6 | 9.71E-07 | 5.06E-06 | -1.37E+00 | 3.38E-03 | 1.46E-02 | 1.09E+00 | 3.34E-03 | 1.42E-02 | -1.14E+00 | 2.49E-02 | 9.52E-02 | 1.16E+00 | 1.31E-01 | 4.67E-01 | 1.25E+00 | 2.21E-01 | 6.22E-01 | 1.40E+00 | 4.84E-04 | 1.77E-03 | -1.37E+00 | 1.10E-02 | 3.02E-02 | -1.19E+00 |
| EDEM2 | 9.73E-07 | 5.07E-06 | -1.23E+00 | 6.31E-05 | 2.73E-04 | 1.04E+00 | 1.11E-03 | 4.72E-03 | -1.12E+00 | 7.39E-05 | 2.82E-04 | -1.04E+00 | 1.19E-09 | 4.26E-09 | -1.43E+00 | 2.71E-07 | 7.64E-07 | -1.24E+00 | 2.34E-03 | 8.60E-03 | -1.04E+00 | 3.85E-02 | 1.06E-01 | 1.11E+00 |
| TIMM22 | 9.73E-07 | 5.07E-06 | -1.43E+00 | 2.45E-04 | 1.06E-03 | -1.03E+00 | 2.79E-03 | 1.18E-02 | -1.16E+00 | 1.82E-02 | 6.95E-02 | 1.14E+00 | 1.92E-10 | 6.85E-10 | -1.86E+00 | 8.92E-04 | 2.51E-03 | -1.05E+00 | 9.09E-09 | 3.33E-08 | -1.23E+00 | 3.57E-03 | 9.81E-03 | 1.00E+00 |
| NR4A2 | 9.76E-07 | 5.09E-06 | 3.32E+00 | 3.64E-02 | 1.57E-01 | -1.19E+00 | 1.73E-02 | 7.36E-02 | 2.27E+00 | 2.24E-02 | 8.54E-02 | -1.63E+00 | 5.60E-11 | 2.00E-10 | 5.36E+00 | 9.12E-02 | 2.57E-01 | 1.07E+00 | 7.17E-11 | 2.63E-10 | 2.91E+00 | 1.72E-04 | 4.72E-04 | -1.57E+00 |
| SEMA4C | 9.79E-07 | 5.10E-06 | -1.51E+00 | 2.47E-05 | 1.07E-04 | -1.23E+00 | 4.67E-07 | 1.98E-06 | -1.93E+00 | 5.00E-06 | 1.91E-05 | -1.60E+00 | 1.12E-02 | 4.00E-02 | -1.19E+00 | 3.17E-02 | 8.94E-02 | -1.09E+00 | 5.57E-06 | 2.05E-05 | -1.64E+00 | 5.65E-02 | 1.55E-01 | -1.07E+00 |
| COX7CP1 | 9.80E-07 | 5.11E-06 | -1.14E+00 | 8.44E-36 | 3.65E-35 | -1.65E+00 | 1.45E-04 | 6.16E-04 | -1.17E+00 | 2.86E-19 | 1.09E-18 | -1.70E+00 | 1.09E-04 | 3.90E-04 | -1.22E+00 | 1.43E-13 | 4.04E-13 | -1.59E+00 | 7.84E-03 | 2.88E-02 | -1.08E+00 | 1.28E-12 | 3.51E-12 | -1.67E+00 |
| NSUN7 | 9.81E-07 | 5.11E-06 | -1.48E+00 | 1.31E-07 | 5.65E-07 | -1.45E+00 | 5.37E-04 | 2.28E-03 | -1.39E+00 | 2.38E-04 | 9.10E-04 | -1.40E+00 | 2.08E-01 | 7.41E-01 | 1.29E+00 | 2.23E-01 | 6.29E-01 | 1.09E+00 | 2.71E-01 | 9.93E-01 | 1.11E+00 | 2.00E-01 | 5.48E-01 | 1.42E+00 |
| SDHAF3 | 9.89E-07 | 5.16E-06 | -1.18E+00 | 1.82E-04 | 7.85E-04 | 1.14E+00 | 1.35E-03 | 5.72E-03 | -1.04E+00 | 2.02E-02 | 7.70E-02 | 1.16E+00 | 5.02E-03 | 1.79E-02 | 1.01E+00 | 7.18E-02 | 2.03E-01 | 1.26E+00 | 3.77E-02 | 1.38E-01 | 1.20E+00 | 1.80E-01 | 4.94E-01 | 1.41E+00 |
| METTL9 | 9.96E-07 | 5.19E-06 | -1.02E+00 | 1.20E-05 | 5.18E-05 | 1.09E+00 | 3.23E-02 | 1.37E-01 | -1.04E+00 | 4.11E-02 | 1.57E-01 | 1.05E+00 | 9.62E-08 | 3.43E-07 | -1.13E+00 | 3.68E-04 | 1.04E-03 | 1.02E+00 | 2.04E-03 | 7.51E-03 | -1.17E+00 | 4.91E-02 | 1.35E-01 | -1.05E+00 |
| PON2 | 9.96E-07 | 5.19E-06 | -1.15E+00 | 7.35E-04 | 3.18E-03 | 1.13E+00 | 3.46E-07 | 1.47E-06 | -1.27E+00 | 3.27E-04 | 1.25E-03 | 1.08E+00 | 7.71E-04 | 2.75E-03 | 1.07E+00 | 2.73E-03 | 7.70E-03 | 1.11E+00 | 6.88E-02 | 2.52E-01 | 1.64E+00 | 1.46E-01 | 4.01E-01 | 1.37E+00 |
| MROH6 | 9.97E-07 | 5.20E-06 | -1.31E+00 | 7.11E-06 | 3.07E-05 | -1.13E+00 | 1.06E-05 | 4.51E-05 | -1.53E+00 | 2.83E-04 | 1.08E-03 | -1.26E+00 | 9.67E-02 | 3.45E-01 | 1.00E+00 | 9.79E-02 | 2.76E-01 | -1.04E+00 | 1.10E-02 | 4.04E-02 | -1.60E+00 | 1.53E-02 | 4.20E-02 | -1.61E+00 |
| RPSAP8 | 9.98E-07 | 5.20E-06 | 2.08E+00 | 1.01E-01 | 4.35E-01 | 1.64E+00 | 1.33E-08 | 5.66E-08 | 1.99E+00 | 1.06E-01 | 4.06E-01 | 1.54E+00 | 3.43E-02 | 1.22E-01 | 1.78E+00 | 2.91E-01 | 8.20E-01 | 1.54E+00 | 2.49E-05 | 9.14E-05 | 2.00E+00 | 3.24E-02 | 8.89E-02 | 1.61E+00 |
| BSG | 9.98E-07 | 5.20E-06 | -1.09E+00 | 1.06E-03 | 4.59E-03 | 1.15E+00 | 8.13E-04 | 3.46E-03 | -1.19E+00 | 1.49E-02 | 5.67E-02 | 1.02E+00 | 6.81E-08 | 2.43E-07 | -1.25E+00 | 1.26E-03 | 3.56E-03 | -1.02E+00 | 4.35E-04 | 1.60E-03 | -1.23E+00 | 2.09E-01 | 5.74E-01 | 1.06E+00 |
| ZNF451 | 1.00E-06 | 5.22E-06 | -1.03E+00 | 3.88E-18 | 1.68E-17 | -1.30E+00 | 3.41E-04 | 1.45E-03 | -1.13E+00 | 4.53E-09 | 1.73E-08 | -1.26E+00 | 7.06E-02 | 2.52E-01 | 1.09E+00 | 1.21E-04 | 3.41E-04 | -1.14E+00 | 1.96E-05 | 7.21E-05 | -1.22E+00 | 1.19E-10 | 3.26E-10 | -1.53E+00 |
| PPP2R3A | 1.00E-06 | 5.23E-06 | -1.65E+00 | 5.23E-06 | 2.26E-05 | -1.34E+00 | 8.92E-04 | 3.79E-03 | -1.39E+00 | 5.46E-03 | 2.08E-02 | -1.12E+00 | 3.37E-02 | 1.20E-01 | -1.21E+00 | 1.11E-01 | 3.13E-01 | 1.00E+00 | 2.83E-02 | 1.04E-01 | -1.40E+00 | 7.77E-02 | 2.14E-01 | -1.22E+00 |
| GGA2 | 1.00E-06 | 5.23E-06 | -2.00E+00 | 7.88E-05 | 3.40E-04 | -1.42E+00 | 3.66E-07 | 1.56E-06 | -1.37E+00 | 1.23E-05 | 4.71E-05 | -1.08E+00 | 5.85E-03 | 2.09E-02 | -1.21E+00 | 6.18E-02 | 1.74E-01 | 1.12E+00 | 3.37E-05 | 1.24E-04 | -1.17E+00 | 5.60E-02 | 1.54E-01 | 1.12E+00 |
| ZNF8 | 1.00E-06 | 5.23E-06 | -1.46E+00 | 3.24E-06 | 1.40E-05 | -1.34E+00 | 1.23E-02 | 5.21E-02 | -1.15E+00 | 1.20E-03 | 4.60E-03 | -1.14E+00 | 3.89E-04 | 1.39E-03 | -1.26E+00 | 3.91E-03 | 1.10E-02 | -1.06E+00 | 5.42E-07 | 1.99E-06 | -1.52E+00 | 9.87E-06 | 2.71E-05 | -1.41E+00 |
| RELA | 1.00E-06 | 5.23E-06 | -1.13E+00 | 4.88E-06 | 2.11E-05 | -1.00E+00 | 1.38E-04 | 5.84E-04 | -1.22E+00 | 3.20E-04 | 1.22E-03 | -1.01E+00 | 4.03E-02 | 1.44E-01 | 1.13E+00 | 2.27E-01 | 6.39E-01 | 1.25E+00 | 1.88E-07 | 6.91E-07 | -1.26E+00 | 1.90E-05 | 5.22E-05 | -1.10E+00 |
| AC012618.1 | 1.01E-06 | 5.27E-06 | -1.29E+00 | 1.23E-81 | 5.30E-81 | -2.47E+00 | 6.88E-03 | 2.92E-02 | -1.22E+00 | 7.53E-27 | 2.88E-26 | -2.25E+00 | 1.55E-03 | 5.52E-03 | -1.26E+00 | 5.80E-27 | 1.64E-26 | -2.21E+00 | 5.35E-06 | 1.96E-05 | -1.33E+00 | 2.29E-26 | 6.30E-26 | -2.29E+00 |
| TMEM42 | 1.01E-06 | 5.28E-06 | -1.34E+00 | 3.11E-05 | 1.35E-04 | -1.08E+00 | 1.70E-06 | 7.24E-06 | -1.37E+00 | 3.58E-06 | 1.37E-05 | -1.16E+00 | 4.22E-05 | 1.50E-04 | -1.29E+00 | 3.67E-02 | 1.04E-01 | 1.14E+00 | 1.33E-01 | 4.87E-01 | 1.15E+00 | 4.81E-02 | 1.32E-01 | 1.11E+00 |
| C20orf96 | 1.01E-06 | 5.29E-06 | -1.36E+00 | 8.74E-04 | 3.78E-03 | 1.02E+00 | 2.48E-06 | 1.05E-05 | -1.41E+00 | 2.61E-05 | 9.96E-05 | -1.15E+00 | 4.77E-07 | 1.70E-06 | -1.54E+00 | 4.67E-03 | 1.32E-02 | -1.02E+00 | 6.78E-04 | 2.49E-03 | -1.26E+00 | 2.94E-03 | 8.07E-03 | -1.27E+00 |
| MKNK1 | 1.02E-06 | 5.31E-06 | -1.17E+00 | 1.10E-05 | 4.76E-05 | -1.04E+00 | 5.51E-03 | 2.34E-02 | -1.06E+00 | 6.05E-04 | 2.31E-03 | -1.01E+00 | 5.63E-03 | 2.01E-02 | -1.04E+00 | 4.22E-03 | 1.19E-02 | -1.04E+00 | 3.23E-02 | 1.19E-01 | 1.15E+00 | 4.74E-02 | 1.30E-01 | 1.06E+00 |
| RHBDD2 | 1.02E-06 | 5.31E-06 | -1.62E+00 | 1.15E-05 | 4.98E-05 | -1.28E+00 | 8.29E-12 | 3.52E-11 | -1.85E+00 | 6.93E-11 | 2.65E-10 | -1.60E+00 | 1.12E-08 | 4.00E-08 | -1.36E+00 | 7.42E-10 | 2.09E-09 | -1.59E+00 | 5.70E-04 | 2.09E-03 | -1.21E+00 | 2.50E-02 | 6.87E-02 | 1.02E+00 |
| MPHOSPH10 | 1.02E-06 | 5.32E-06 | -1.17E+00 | 2.00E-01 | 8.63E-01 | 1.46E+00 | 3.70E-07 | 1.57E-06 | -1.26E+00 | 2.60E-01 | 9.92E-01 | 1.30E+00 | 7.85E-05 | 2.80E-04 | -1.04E+00 | 2.24E-01 | 6.31E-01 | 1.32E+00 | 3.10E-08 | 1.14E-07 | -1.35E+00 | 2.98E-01 | 8.18E-01 | 1.19E+00 |
| CLHC1 | 1.02E-06 | 5.33E-06 | -1.92E+00 | 5.57E-03 | 2.41E-02 | -1.04E+00 | 3.01E-05 | 1.28E-04 | -1.86E+00 | 5.35E-03 | 2.04E-02 | -1.15E+00 | 1.49E-03 | 5.32E-03 | -1.48E+00 | 1.69E-01 | 4.77E-01 | 1.26E+00 | 3.79E-04 | 1.39E-03 | -1.38E+00 | 1.30E-01 | 3.57E-01 | 1.14E+00 |
| POLDIP3 | 1.03E-06 | 5.35E-06 | -1.26E+00 | 6.34E-07 | 2.74E-06 | -1.14E+00 | 3.18E-05 | 1.35E-04 | -1.19E+00 | 1.54E-04 | 5.87E-04 | 1.00E+00 | 9.47E-06 | 3.38E-05 | -1.24E+00 | 1.27E-05 | 3.57E-05 | -1.18E+00 | 3.04E-10 | 1.11E-09 | -1.44E+00 | 1.04E-03 | 2.85E-03 | -1.04E+00 |
| SPOCK2 | 1.03E-06 | 5.35E-06 | 2.24E+00 | 3.47E-06 | 1.50E-05 | 2.43E+00 | 1.45E-01 | 6.14E-01 | 1.26E+00 | 1.09E-01 | 4.17E-01 | 1.16E+00 | 1.90E-01 | 6.76E-01 | 1.62E+00 | 1.89E-01 | 5.33E-01 | 1.76E+00 | 1.93E-01 | 7.09E-01 | 1.60E+00 | 2.47E-01 | 6.78E-01 | 1.60E+00 |
| NR2F6 | 1.03E-06 | 5.36E-06 | -1.67E+00 | 8.31E-04 | 3.59E-03 | -1.06E+00 | 4.08E-06 | 1.74E-05 | -1.52E+00 | 4.92E-04 | 1.88E-03 | -1.08E+00 | 1.21E-04 | 4.30E-04 | -1.55E+00 | 1.05E-01 | 2.96E-01 | 1.21E+00 | 4.97E-07 | 1.83E-06 | -1.12E+00 | 5.96E-03 | 1.64E-02 | 1.05E+00 |
| EEF1A1P33 | 1.04E-06 | 5.42E-06 | -1.36E+00 | 1.24E-07 | 5.35E-07 | -1.35E+00 | 1.82E-04 | 7.72E-04 | -1.31E+00 | 1.46E-06 | 5.59E-06 | -1.42E+00 | 2.39E-02 | 8.52E-02 | -1.00E+00 | 7.61E-03 | 2.14E-02 | -1.03E+00 | 2.16E-01 | 7.92E-01 | 1.17E+00 | 2.32E-03 | 6.36E-03 | -1.31E+00 |
| TBC1D15 | 1.04E-06 | 5.42E-06 | -1.15E+00 | 8.91E-08 | 3.85E-07 | -1.07E+00 | 2.95E-04 | 1.25E-03 | -1.04E+00 | 3.68E-05 | 1.41E-04 | 1.02E+00 | 6.41E-02 | 2.28E-01 | 1.26E+00 | 1.03E-01 | 2.90E-01 | 1.24E+00 | 1.26E-01 | 4.63E-01 | 1.41E+00 | 6.27E-02 | 1.72E-01 | 1.09E+00 |
| TMEM251 | 1.04E-06 | 5.43E-06 | -1.46E+00 | 3.69E-03 | 1.59E-02 | 1.09E+00 | 2.82E-05 | 1.20E-04 | -1.46E+00 | 3.16E-02 | 1.21E-01 | 1.11E+00 | 2.62E-12 | 9.33E-12 | -2.07E+00 | 7.14E-05 | 2.01E-04 | -1.20E+00 | 9.45E-06 | 3.47E-05 | -1.15E+00 | 1.64E-02 | 4.52E-02 | 1.10E+00 |
| MTCH2 | 1.05E-06 | 5.47E-06 | -1.21E+00 | 6.25E-03 | 2.70E-02 | 1.16E+00 | 2.53E-07 | 1.07E-06 | -1.30E+00 | 2.55E-04 | 9.75E-04 | 1.01E+00 | 2.76E-08 | 9.85E-08 | -1.36E+00 | 1.05E-04 | 2.95E-04 | -1.03E+00 | 6.63E-05 | 2.43E-04 | -1.11E+00 | 2.97E-01 | 8.15E-01 | 1.31E+00 |
| MICU1 | 1.06E-06 | 5.50E-06 | -1.31E+00 | 3.58E-06 | 1.55E-05 | -1.13E+00 | 5.51E-05 | 2.34E-04 | -1.32E+00 | 3.52E-06 | 1.35E-05 | -1.33E+00 | 6.50E-06 | 2.32E-05 | -1.28E+00 | 4.10E-06 | 1.16E-05 | -1.29E+00 | 2.15E-07 | 7.89E-07 | -1.16E+00 | 1.02E-11 | 2.80E-11 | -1.45E+00 |
| TMEM161B | 1.06E-06 | 5.54E-06 | -1.51E+00 | 1.04E-02 | 4.50E-02 | 1.18E+00 | 3.57E-08 | 1.52E-07 | -1.65E+00 | 3.76E-05 | 1.44E-04 | -1.11E+00 | 8.27E-04 | 2.95E-03 | -1.23E+00 | 1.34E-03 | 3.78E-03 | -1.17E+00 | 4.80E-04 | 1.76E-03 | -1.12E+00 | 4.07E-02 | 1.12E-01 | 1.04E+00 |
| TPP2 | 1.07E-06 | 5.56E-06 | -1.17E+00 | 6.27E-06 | 2.71E-05 | -1.03E+00 | 3.46E-05 | 1.47E-04 | -1.23E+00 | 6.31E-05 | 2.41E-04 | -1.04E+00 | 1.70E-05 | 6.08E-05 | -1.08E+00 | 2.74E-05 | 7.72E-05 | -1.09E+00 | 6.91E-02 | 2.54E-01 | 1.09E+00 | 8.78E-03 | 2.41E-02 | -1.07E+00 |
| HLA-E | 1.07E-06 | 5.56E-06 | -1.24E+00 | 2.13E-02 | 9.19E-02 | -1.07E+00 | 1.66E-07 | 7.07E-07 | -1.37E+00 | 4.92E-04 | 1.88E-03 | -1.19E+00 | 1.42E-08 | 5.06E-08 | -1.38E+00 | 7.02E-10 | 1.98E-09 | -1.38E+00 | 1.06E-27 | 3.89E-27 | -1.61E+00 | 5.39E-29 | 1.48E-28 | -1.68E+00 |
| PRIM2 | 1.07E-06 | 5.57E-06 | -1.44E+00 | 8.30E-05 | 3.59E-04 | -1.07E+00 | 1.08E-03 | 4.59E-03 | -1.13E+00 | 2.52E-03 | 9.63E-03 | 1.03E+00 | 2.58E-03 | 9.19E-03 | -1.26E+00 | 5.30E-03 | 1.50E-02 | -1.15E+00 | 1.12E-01 | 4.12E-01 | 1.09E+00 | 3.00E-01 | 8.23E-01 | 1.22E+00 |
| NBPF14 | 1.07E-06 | 5.57E-06 | -1.32E+00 | 1.55E-03 | 6.68E-03 | 1.11E+00 | 1.68E-11 | 7.15E-11 | -1.75E+00 | 6.02E-06 | 2.30E-05 | -1.14E+00 | 4.84E-06 | 1.72E-05 | -1.21E+00 | 4.92E-04 | 1.39E-03 | -1.02E+00 | 7.30E-07 | 2.68E-06 | -1.28E+00 | 1.86E-03 | 5.12E-03 | -1.11E+00 |
| IL1B | 1.07E-06 | 5.57E-06 | -1.04E+00 | 1.38E-01 | 5.95E-01 | 1.50E+00 | 6.94E-10 | 2.95E-09 | -1.74E+00 | 3.37E-02 | 1.29E-01 | 1.11E+00 | 4.55E-03 | 1.62E-02 | 1.15E+00 | 1.70E-03 | 4.79E-03 | 1.97E+00 | 2.19E-02 | 8.03E-02 | 1.04E+00 | 3.51E-01 | 9.63E-01 | 1.18E+00 |
| BTF3P8 | 1.07E-06 | 5.58E-06 | 2.14E+00 | 8.52E-09 | 3.68E-08 | 2.73E+00 | 9.60E-06 | 4.08E-05 | 2.10E+00 | 2.69E-06 | 1.03E-05 | 2.50E+00 | 3.76E-05 | 1.34E-04 | 2.73E+00 | 3.81E-07 | 1.07E-06 | 3.17E+00 | 3.91E-06 | 1.44E-05 | 2.34E+00 | 7.22E-10 | 1.98E-09 | 2.88E+00 |
| NUDT11 | 1.07E-06 | 5.58E-06 | -1.36E+00 | 4.27E-03 | 1.85E-02 | 1.13E+00 | 1.87E-07 | 7.94E-07 | -1.44E+00 | 2.94E-04 | 1.12E-03 | -1.07E+00 | 7.13E-02 | 2.54E-01 | 1.17E+00 | 1.94E-01 | 5.47E-01 | 1.83E+00 | 2.43E-01 | 8.92E-01 | 1.38E+00 | 3.36E-02 | 9.24E-02 | 2.09E+00 |
| HLA-L | 1.07E-06 | 5.59E-06 | -1.68E+00 | 6.64E-04 | 2.87E-03 | -1.07E+00 | 2.09E-02 | 8.89E-02 | -1.26E+00 | 4.13E-02 | 1.58E-01 | -1.08E+00 | 3.57E-02 | 1.27E-01 | -1.52E+00 | 6.09E-02 | 1.72E-01 | -1.45E+00 | 7.70E-02 | 2.82E-01 | -1.27E+00 | 6.79E-02 | 1.86E-01 | -1.45E+00 |
| LYSMD3 | 1.08E-06 | 5.61E-06 | -1.27E+00 | 9.19E-13 | 3.97E-12 | -1.69E+00 | 6.31E-07 | 2.68E-06 | -1.38E+00 | 1.39E-11 | 5.33E-11 | -1.72E+00 | 1.94E-03 | 6.93E-03 | 1.01E+00 | 2.26E-05 | 6.37E-05 | -1.21E+00 | 3.58E-06 | 1.31E-05 | -1.29E+00 | 3.16E-07 | 8.69E-07 | -1.53E+00 |
| ARHGEF9 | 1.08E-06 | 5.62E-06 | -1.28E+00 | 5.61E-08 | 2.42E-07 | -1.33E+00 | 6.27E-07 | 2.67E-06 | -1.50E+00 | 3.21E-05 | 1.23E-04 | -1.18E+00 | 7.80E-05 | 2.78E-04 | -1.31E+00 | 1.93E-02 | 5.44E-02 | 1.01E+00 | 2.62E-04 | 9.60E-04 | -1.26E+00 | 7.48E-03 | 2.05E-02 | -1.11E+00 |
| CCZ1B | 1.08E-06 | 5.64E-06 | -1.24E+00 | 4.33E-05 | 1.87E-04 | 1.03E+00 | 3.39E-04 | 1.44E-03 | -1.30E+00 | 6.30E-03 | 2.41E-02 | 1.02E+00 | 1.59E-04 | 5.67E-04 | -1.43E+00 | 3.76E-03 | 1.06E-02 | -1.07E+00 | 3.24E-02 | 1.19E-01 | -1.05E+00 | 1.14E-01 | 3.12E-01 | 1.05E+00 |
| DPP3 | 1.09E-06 | 5.68E-06 | -1.36E+00 | 7.47E-07 | 3.23E-06 | -1.26E+00 | 6.27E-05 | 2.66E-04 | -1.27E+00 | 9.30E-06 | 3.55E-05 | -1.18E+00 | 7.18E-05 | 2.56E-04 | -1.28E+00 | 1.97E-04 | 5.55E-04 | -1.21E+00 | 1.67E-02 | 6.14E-02 | 1.12E+00 | 2.15E-02 | 5.91E-02 | 1.11E+00 |
| RNF217 | 1.09E-06 | 5.69E-06 | -1.24E+00 | 1.13E-02 | 4.88E-02 | 1.21E+00 | 4.32E-03 | 1.83E-02 | -1.29E+00 | 1.98E-01 | 7.54E-01 | 1.28E+00 | 6.43E-03 | 2.29E-02 | 1.09E+00 | 6.06E-02 | 1.71E-01 | 1.21E+00 | 7.54E-02 | 2.77E-01 | 1.11E+00 | 2.11E-02 | 5.78E-02 | 1.05E+00 |
| ABO | 1.10E-06 | 5.71E-06 | -1.12E+00 | 4.93E-02 | 2.13E-01 | 1.28E+00 | 9.21E-03 | 3.91E-02 | -1.06E+00 | 1.32E-01 | 5.05E-01 | 1.26E+00 | 1.56E-01 | 5.58E-01 | 1.43E+00 | 2.18E-02 | 6.16E-02 | 1.84E+00 | 1.29E-02 | 4.73E-02 | 1.02E+00 | 2.87E-01 | 7.90E-01 | 1.29E+00 |
| SAC3D1 | 1.10E-06 | 5.72E-06 | -1.41E+00 | 1.29E-04 | 5.56E-04 | -1.03E+00 | 2.95E-02 | 1.25E-01 | -1.12E+00 | 6.89E-02 | 2.63E-01 | 1.08E+00 | 1.41E-08 | 5.02E-08 | -2.21E+00 | 1.77E-03 | 4.98E-03 | -1.08E+00 | 3.09E-05 | 1.13E-04 | -1.16E+00 | 2.56E-02 | 7.02E-02 | 1.09E+00 |
| ITGA2 | 1.10E-06 | 5.75E-06 | -1.57E+00 | 5.97E-04 | 2.58E-03 | -1.03E+00 | 5.51E-06 | 2.34E-05 | -2.18E+00 | 1.03E-04 | 3.94E-04 | -1.49E+00 | 1.99E-01 | 7.11E-01 | 2.05E+00 | 1.17E-01 | 3.30E-01 | 3.04E+00 | 6.52E-03 | 2.39E-02 | 1.71E+00 | 2.47E-01 | 6.78E-01 | -1.02E+00 |
| TRIM33 | 1.10E-06 | 5.75E-06 | -1.14E+00 | 2.03E-03 | 8.77E-03 | 1.15E+00 | 2.23E-02 | 9.49E-02 | -1.02E+00 | 1.76E-02 | 6.71E-02 | 1.14E+00 | 4.17E-05 | 1.49E-04 | -1.13E+00 | 2.93E-03 | 8.26E-03 | -1.03E+00 | 3.66E-10 | 1.34E-09 | -1.38E+00 | 1.02E-05 | 2.81E-05 | -1.15E+00 |
| GORASP2 | 1.10E-06 | 5.75E-06 | -1.19E+00 | 3.48E-04 | 1.50E-03 | 1.06E+00 | 7.11E-04 | 3.02E-03 | -1.18E+00 | 3.38E-03 | 1.29E-02 | 1.04E+00 | 1.03E-07 | 3.68E-07 | -1.33E+00 | 6.65E-03 | 1.87E-02 | 1.07E+00 | 8.22E-07 | 3.02E-06 | -1.19E+00 | 6.86E-04 | 1.88E-03 | -1.07E+00 |
| CBX3P9 | 1.11E-06 | 5.79E-06 | -1.53E+00 | 1.70E-08 | 7.36E-08 | -1.71E+00 | 2.35E-04 | 9.98E-04 | -1.29E+00 | 5.74E-05 | 2.19E-04 | -1.22E+00 | 1.72E-10 | 6.14E-10 | -1.84E+00 | 7.72E-11 | 2.18E-10 | -1.79E+00 | 5.66E-09 | 2.08E-08 | -1.54E+00 | 1.84E-07 | 5.06E-07 | -1.46E+00 |
| AC024619.4 | 1.11E-06 | 5.80E-06 | 1.94E+00 | 4.48E-02 | 1.94E-01 | 1.71E+00 | 7.77E-08 | 3.30E-07 | 2.23E+00 | 3.98E-03 | 1.52E-02 | 1.86E+00 | 9.81E-04 | 3.50E-03 | 1.91E+00 | 4.82E-02 | 1.36E-01 | 1.67E+00 | 1.52E-11 | 5.58E-11 | 2.07E+00 | 2.85E-09 | 7.82E-09 | 1.99E+00 |
| ZNF48 | 1.11E-06 | 5.81E-06 | -1.45E+00 | 8.46E-04 | 3.66E-03 | 1.02E+00 | 6.70E-06 | 2.84E-05 | -1.45E+00 | 1.34E-03 | 5.14E-03 | 1.02E+00 | 5.99E-04 | 2.14E-03 | -1.42E+00 | 5.83E-02 | 1.64E-01 | 1.09E+00 | 3.01E-04 | 1.10E-03 | -1.33E+00 | 1.35E-02 | 3.71E-02 | -1.08E+00 |
| ZNF184 | 1.12E-06 | 5.81E-06 | -1.34E+00 | 1.17E-11 | 5.07E-11 | -1.79E+00 | 6.31E-06 | 2.68E-05 | -1.37E+00 | 7.33E-13 | 2.80E-12 | -1.86E+00 | 2.68E-07 | 9.56E-07 | -1.63E+00 | 5.31E-14 | 1.50E-13 | -2.72E+00 | 4.80E-06 | 1.76E-05 | -1.68E+00 | 1.09E-05 | 3.00E-05 | -1.61E+00 |
| SNRNP48 | 1.12E-06 | 5.81E-06 | -1.17E+00 | 7.42E-05 | 3.21E-04 | 1.05E+00 | 4.91E-05 | 2.09E-04 | -1.26E+00 | 3.82E-06 | 1.46E-05 | -1.13E+00 | 2.11E-06 | 7.54E-06 | -1.46E+00 | 2.19E-03 | 6.19E-03 | -1.15E+00 | 7.43E-09 | 2.73E-08 | -1.36E+00 | 2.34E-04 | 6.44E-04 | -1.10E+00 |
| ZC3HC1 | 1.12E-06 | 5.83E-06 | -1.69E+00 | 3.54E-03 | 1.53E-02 | 1.08E+00 | 7.38E-10 | 3.14E-09 | -1.82E+00 | 1.02E-06 | 3.91E-06 | -1.27E+00 | 3.23E-05 | 1.15E-04 | -1.88E+00 | 5.97E-02 | 1.68E-01 | -1.04E+00 | 4.90E-07 | 1.80E-06 | -1.46E+00 | 5.32E-03 | 1.46E-02 | -1.07E+00 |
| DDX41 | 1.12E-06 | 5.83E-06 | -1.37E+00 | 5.28E-02 | 2.28E-01 | 1.31E+00 | 6.11E-07 | 2.60E-06 | -1.40E+00 | 6.83E-05 | 2.61E-04 | -1.07E+00 | 6.82E-03 | 2.43E-02 | -1.15E+00 | 2.62E-01 | 7.39E-01 | 1.23E+00 | 3.02E-03 | 1.11E-02 | 1.02E+00 | 2.60E-01 | 7.14E-01 | 1.24E+00 |
| OGDH | 1.12E-06 | 5.83E-06 | -1.42E+00 | 6.64E-04 | 2.87E-03 | -1.00E+00 | 3.92E-05 | 1.67E-04 | -1.39E+00 | 6.26E-03 | 2.39E-02 | -1.01E+00 | 2.68E-03 | 9.54E-03 | -1.13E+00 | 1.22E-01 | 3.43E-01 | 1.20E+00 | 5.46E-04 | 2.01E-03 | -1.19E+00 | 2.18E-01 | 5.99E-01 | 1.21E+00 |
| RPL5P4 | 1.13E-06 | 5.87E-06 | -1.12E+00 | 1.55E-04 | 6.71E-04 | -1.10E+00 | 1.50E-02 | 6.36E-02 | -1.06E+00 | 8.71E-02 | 3.33E-01 | -1.03E+00 | 1.51E-03 | 5.37E-03 | -1.11E+00 | 1.26E-02 | 3.55E-02 | -1.08E+00 | 1.94E-07 | 7.13E-07 | -1.13E+00 | 6.14E-03 | 1.69E-02 | -1.09E+00 |
| PPP6C | 1.13E-06 | 5.88E-06 | -1.23E+00 | 2.92E-06 | 1.26E-05 | -1.07E+00 | 1.32E-07 | 5.60E-07 | -1.45E+00 | 3.40E-06 | 1.30E-05 | -1.21E+00 | 7.53E-10 | 2.68E-09 | -1.52E+00 | 1.03E-05 | 2.89E-05 | -1.13E+00 | 2.37E-09 | 8.68E-09 | -1.23E+00 | 7.91E-05 | 2.17E-04 | -1.08E+00 |
| ATXN1 | 1.13E-06 | 5.89E-06 | -1.27E+00 | 2.07E-06 | 8.92E-06 | -1.13E+00 | 3.69E-06 | 1.57E-05 | -1.28E+00 | 5.76E-07 | 2.20E-06 | -1.24E+00 | 2.58E-02 | 9.22E-02 | -1.01E+00 | 3.36E-01 | 9.46E-01 | 1.28E+00 | 2.21E-02 | 8.10E-02 | -1.01E+00 | 1.34E-01 | 3.69E-01 | 1.01E+00 |
| AL031587.6 | 1.13E-06 | 5.89E-06 | -1.46E+00 | 4.65E-05 | 2.01E-04 | -1.19E+00 | 4.52E-08 | 1.92E-07 | -1.71E+00 | 4.24E-07 | 1.62E-06 | -1.38E+00 | 5.69E-04 | 2.03E-03 | -1.21E+00 | 7.94E-04 | 2.24E-03 | -1.21E+00 | 7.92E-04 | 2.91E-03 | -1.38E+00 | 8.07E-03 | 2.22E-02 | -1.25E+00 |
| AC009779.2 | 1.13E-06 | 5.91E-06 | -1.86E+00 | 2.63E-05 | 1.13E-04 | -1.50E+00 | 1.26E-03 | 5.34E-03 | -1.63E+00 | 4.06E-03 | 1.55E-02 | -1.28E+00 | 5.43E-08 | 1.94E-07 | -2.58E+00 | 6.35E-06 | 1.79E-05 | -1.72E+00 | 4.97E-06 | 1.82E-05 | -1.39E+00 | 6.60E-04 | 1.81E-03 | -1.27E+00 |
| CLN5 | 1.13E-06 | 5.91E-06 | -1.16E+00 | 1.56E-03 | 6.75E-03 | 1.16E+00 | 7.31E-04 | 3.11E-03 | -1.11E+00 | 2.31E-03 | 8.84E-03 | 1.05E+00 | 3.82E-03 | 1.36E-02 | 1.01E+00 | 5.17E-03 | 1.46E-02 | 1.09E+00 | 6.68E-03 | 2.45E-02 | -1.11E+00 | 2.72E-01 | 7.48E-01 | 1.16E+00 |
| ATP8B3 | 1.13E-06 | 5.91E-06 | -1.77E+00 | 6.89E-07 | 2.98E-06 | -1.71E+00 | 1.30E-09 | 5.53E-09 | -2.05E+00 | 5.60E-11 | 2.14E-10 | -2.31E+00 | 3.03E-06 | 1.08E-05 | -2.22E+00 | 7.20E-04 | 2.03E-03 | -1.49E+00 | 1.95E-03 | 7.17E-03 | -1.72E+00 | 5.88E-03 | 1.62E-02 | -1.55E+00 |
| DNAJC21 | 1.15E-06 | 5.98E-06 | -1.12E+00 | 8.29E-06 | 3.58E-05 | 1.06E+00 | 1.93E-03 | 8.20E-03 | -1.02E+00 | 1.05E-04 | 4.01E-04 | -1.00E+00 | 4.43E-02 | 1.58E-01 | 1.11E+00 | 5.40E-02 | 1.52E-01 | 1.17E+00 | 1.99E-02 | 7.31E-02 | 1.08E+00 | 1.55E-01 | 4.25E-01 | 1.14E+00 |
| UNG | 1.15E-06 | 5.99E-06 | -1.55E+00 | 5.25E-04 | 2.27E-03 | -1.07E+00 | 1.58E-02 | 6.72E-02 | -1.06E+00 | 5.45E-03 | 2.08E-02 | 1.11E+00 | 4.57E-11 | 1.63E-10 | -2.78E+00 | 1.00E-05 | 2.82E-05 | -1.49E+00 | 4.29E-07 | 1.57E-06 | -1.29E+00 | 1.49E-01 | 4.09E-01 | 1.43E+00 |
| CD2BP2 | 1.15E-06 | 6.00E-06 | -1.27E+00 | 4.52E-04 | 1.95E-03 | 1.06E+00 | 2.03E-07 | 8.61E-07 | -1.32E+00 | 3.84E-08 | 1.47E-07 | -1.19E+00 | 1.46E-07 | 5.19E-07 | -1.53E+00 | 4.09E-03 | 1.15E-02 | -1.04E+00 | 8.34E-07 | 3.06E-06 | -1.20E+00 | 5.33E-04 | 1.46E-03 | -1.06E+00 |
| DDTL | 1.15E-06 | 6.00E-06 | -1.48E+00 | 1.42E-06 | 6.12E-06 | -1.36E+00 | 4.51E-06 | 1.92E-05 | -1.40E+00 | 2.03E-07 | 7.77E-07 | -1.45E+00 | 5.08E-06 | 1.81E-05 | -1.91E+00 | 7.72E-05 | 2.18E-04 | -1.49E+00 | 3.28E-06 | 1.20E-05 | -1.43E+00 | 8.11E-07 | 2.23E-06 | -1.47E+00 |
| LITAF | 1.15E-06 | 6.02E-06 | -1.12E+00 | 1.36E-04 | 5.86E-04 | 1.12E+00 | 1.67E-08 | 7.10E-08 | -1.21E+00 | 6.40E-04 | 2.45E-03 | 1.14E+00 | 2.77E-03 | 9.87E-03 | -1.18E+00 | 1.21E-03 | 3.40E-03 | -1.16E+00 | 7.40E-02 | 2.71E-01 | -1.35E+00 | 2.11E-01 | 5.79E-01 | -1.10E+00 |
| AC008915.2 | 1.16E-06 | 6.03E-06 | -1.77E+00 | 4.73E-05 | 2.04E-04 | -1.30E+00 | 1.85E-02 | 7.84E-02 | -1.22E+00 | 1.27E-02 | 4.87E-02 | -1.13E+00 | 3.55E-04 | 1.26E-03 | -1.34E+00 | 5.68E-04 | 1.60E-03 | -1.34E+00 | 2.24E-02 | 8.23E-02 | -1.11E+00 | 8.54E-03 | 2.35E-02 | -1.26E+00 |
| STK33 | 1.16E-06 | 6.03E-06 | -1.29E+00 | 1.44E-06 | 6.24E-06 | -1.15E+00 | 8.66E-06 | 3.68E-05 | -1.33E+00 | 1.56E-08 | 5.97E-08 | -1.43E+00 | 3.55E-04 | 1.27E-03 | -1.76E+00 | 2.54E-04 | 7.16E-04 | -1.76E+00 | 4.80E-03 | 1.76E-02 | -1.94E+00 | 1.44E-01 | 3.96E-01 | -1.03E+00 |
| DESI1 | 1.16E-06 | 6.03E-06 | -1.25E+00 | 3.45E-07 | 1.49E-06 | -1.17E+00 | 2.87E-05 | 1.22E-04 | -1.26E+00 | 4.17E-04 | 1.59E-03 | -1.03E+00 | 2.07E-03 | 7.37E-03 | -1.48E+00 | 2.18E-02 | 6.14E-02 | -1.15E+00 | 4.14E-02 | 1.52E-01 | -1.08E+00 | 1.56E-01 | 4.30E-01 | 1.04E+00 |
| PLSCR4 | 1.16E-06 | 6.04E-06 | -1.44E+00 | 2.06E-04 | 8.89E-04 | -1.10E+00 | 1.00E-03 | 4.26E-03 | -1.36E+00 | 1.71E-02 | 6.51E-02 | 1.07E+00 | 5.10E-03 | 1.82E-02 | -1.42E+00 | 9.31E-02 | 2.62E-01 | 1.11E+00 | 3.32E-04 | 1.22E-03 | -1.31E+00 | 1.05E-03 | 2.89E-03 | -1.23E+00 |
| PLD3 | 1.17E-06 | 6.11E-06 | -1.11E+00 | 7.67E-04 | 3.31E-03 | 1.21E+00 | 2.69E-02 | 1.15E-01 | 1.04E+00 | 1.59E-01 | 6.07E-01 | 1.37E+00 | 7.33E-04 | 2.61E-03 | -1.09E+00 | 1.56E-02 | 4.40E-02 | 1.06E+00 | 2.96E-03 | 1.09E-02 | -1.02E+00 | 7.86E-03 | 2.16E-02 | 1.02E+00 |
| RN7SL587P | 1.18E-06 | 6.13E-06 | -1.87E+00 | 1.10E-05 | 4.74E-05 | -1.52E+00 | 3.28E-04 | 1.39E-03 | -1.50E+00 | 2.95E-05 | 1.13E-04 | -1.62E+00 | 2.64E-01 | 9.41E-01 | 1.19E+00 | 2.06E-01 | 5.82E-01 | -1.19E+00 | 1.25E-01 | 4.57E-01 | -1.03E+00 | 2.25E-02 | 6.19E-02 | -1.37E+00 |
| ZNF320 | 1.19E-06 | 6.18E-06 | -1.80E+00 | 6.90E-04 | 2.98E-03 | -1.09E+00 | 3.73E-07 | 1.59E-06 | -1.55E+00 | 1.33E-05 | 5.08E-05 | -1.21E+00 | 2.35E-03 | 8.39E-03 | -1.32E+00 | 2.29E-01 | 6.46E-01 | 1.30E+00 | 9.40E-07 | 3.45E-06 | -1.61E+00 | 3.43E-04 | 9.42E-04 | -1.35E+00 |
| PGBD2 | 1.19E-06 | 6.20E-06 | -1.98E+00 | 5.29E-06 | 2.29E-05 | -1.70E+00 | 5.13E-03 | 2.18E-02 | -1.76E+00 | 2.95E-02 | 1.13E-01 | -1.30E+00 | 1.44E-10 | 5.13E-10 | -2.47E+00 | 1.01E-07 | 2.84E-07 | -1.78E+00 | 9.72E-07 | 3.57E-06 | -1.77E+00 | 5.07E-04 | 1.39E-03 | -1.31E+00 |
| ELOVL5 | 1.20E-06 | 6.25E-06 | -1.11E+00 | 1.52E-05 | 6.55E-05 | 1.05E+00 | 3.92E-04 | 1.66E-03 | -1.06E+00 | 4.24E-04 | 1.62E-03 | 1.03E+00 | 9.14E-03 | 3.26E-02 | 1.05E+00 | 7.16E-02 | 2.02E-01 | 1.16E+00 | 3.04E-07 | 1.12E-06 | -1.26E+00 | 6.04E-05 | 1.66E-04 | -1.17E+00 |
| CAST | 1.20E-06 | 6.26E-06 | -1.13E+00 | 5.68E-03 | 2.45E-02 | 1.14E+00 | 5.02E-07 | 2.13E-06 | -1.25E+00 | 3.76E-03 | 1.44E-02 | 1.08E+00 | 1.50E-07 | 5.35E-07 | -1.24E+00 | 1.28E-03 | 3.60E-03 | -1.00E+00 | 8.35E-04 | 3.07E-03 | -1.15E+00 | 3.66E-02 | 1.01E-01 | -1.03E+00 |
| ACBD3 | 1.20E-06 | 6.27E-06 | -1.27E+00 | 5.68E-09 | 2.45E-08 | -1.30E+00 | 1.74E-02 | 7.41E-02 | 1.07E+00 | 1.70E-03 | 6.51E-03 | 1.11E+00 | 1.10E-04 | 3.93E-04 | -1.12E+00 | 1.44E-04 | 4.06E-04 | -1.08E+00 | 5.58E-06 | 2.05E-05 | -1.24E+00 | 4.12E-05 | 1.13E-04 | -1.19E+00 |
| PMS1 | 1.21E-06 | 6.28E-06 | -1.39E+00 | 8.97E-04 | 3.88E-03 | 1.04E+00 | 1.64E-09 | 6.96E-09 | -1.45E+00 | 2.94E-07 | 1.12E-06 | -1.14E+00 | 2.41E-05 | 8.58E-05 | -1.76E+00 | 9.62E-03 | 2.71E-02 | -1.09E+00 | 5.40E-07 | 1.98E-06 | -1.47E+00 | 5.44E-03 | 1.50E-02 | -1.10E+00 |
| GRAMD1B | 1.21E-06 | 6.29E-06 | -1.58E+00 | 6.13E-05 | 2.65E-04 | -1.17E+00 | 2.69E-04 | 1.14E-03 | -1.25E+00 | 1.41E-04 | 5.38E-04 | -1.18E+00 | 1.07E-01 | 3.81E-01 | -1.52E+00 | 8.47E-02 | 2.39E-01 | -1.74E+00 | 6.74E-02 | 2.47E-01 | -1.44E+00 | 1.34E-01 | 3.67E-01 | -1.36E+00 |
| POLR2C | 1.21E-06 | 6.29E-06 | -1.13E+00 | 9.03E-11 | 3.90E-10 | -1.14E+00 | 1.42E-06 | 6.03E-06 | -1.13E+00 | 8.10E-07 | 3.09E-06 | -1.07E+00 | 6.65E-05 | 2.37E-04 | -1.17E+00 | 3.37E-03 | 9.51E-03 | -1.02E+00 | 1.40E-03 | 5.15E-03 | -1.07E+00 | 3.94E-02 | 1.08E-01 | 1.04E+00 |
| ALDH3B1 | 1.21E-06 | 6.30E-06 | -1.19E+00 | 1.46E-08 | 6.30E-08 | -1.20E+00 | 1.20E-04 | 5.12E-04 | -1.19E+00 | 6.80E-05 | 2.60E-04 | -1.11E+00 | 2.36E-10 | 8.43E-10 | -1.55E+00 | 2.64E-07 | 7.45E-07 | -1.28E+00 | 3.18E-03 | 1.17E-02 | -1.06E+00 | 2.41E-02 | 6.62E-02 | 1.01E+00 |
| UBE2Q2P2 | 1.21E-06 | 6.31E-06 | -1.72E+00 | 4.23E-06 | 1.83E-05 | -1.38E+00 | 1.20E-03 | 5.12E-03 | -1.52E+00 | 5.38E-04 | 2.05E-03 | -1.50E+00 | 6.21E-05 | 2.21E-04 | -1.78E+00 | 8.75E-03 | 2.47E-02 | -1.10E+00 | 1.55E-07 | 5.70E-07 | -1.39E+00 | 2.65E-07 | 7.27E-07 | -1.36E+00 |
| BMS1P1 | 1.21E-06 | 6.33E-06 | -1.35E+00 | 4.51E-06 | 1.95E-05 | -1.24E+00 | 2.23E-06 | 9.47E-06 | -1.57E+00 | 1.48E-06 | 5.67E-06 | -1.37E+00 | 1.37E-09 | 4.88E-09 | -1.67E+00 | 7.43E-05 | 2.09E-04 | -1.12E+00 | 1.35E-03 | 4.97E-03 | -1.20E+00 | 1.50E-02 | 4.12E-02 | -1.05E+00 |
| PPIH | 1.22E-06 | 6.36E-06 | -1.17E+00 | 3.49E-04 | 1.51E-03 | 1.12E+00 | 3.90E-02 | 1.66E-01 | -1.02E+00 | 2.41E-05 | 9.22E-05 | -1.05E+00 | 5.42E-06 | 1.93E-05 | -1.30E+00 | 2.54E-03 | 7.16E-03 | -1.03E+00 | 3.27E-04 | 1.20E-03 | -1.16E+00 | 1.00E-01 | 2.75E-01 | 1.11E+00 |
| DMXL2 | 1.23E-06 | 6.40E-06 | -1.72E+00 | 9.70E-04 | 4.19E-03 | -1.08E+00 | 4.40E-05 | 1.87E-04 | -1.55E+00 | 1.91E-06 | 7.28E-06 | -1.72E+00 | 1.72E-01 | 6.15E-01 | -1.37E+00 | 1.95E-01 | 5.49E-01 | -1.61E+00 | 9.71E-02 | 3.56E-01 | -1.53E+00 | 2.37E-01 | 6.52E-01 | -1.07E+00 |
| PRKAG2 | 1.24E-06 | 6.48E-06 | -1.34E+00 | 5.98E-05 | 2.59E-04 | -1.05E+00 | 2.38E-13 | 1.01E-12 | -1.56E+00 | 7.88E-09 | 3.01E-08 | -1.14E+00 | 3.00E-07 | 1.07E-06 | -1.68E+00 | 1.98E-04 | 5.59E-04 | -1.25E+00 | 4.60E-06 | 1.69E-05 | -1.48E+00 | 1.49E-03 | 4.08E-03 | -1.16E+00 |
| GAPDHP40 | 1.26E-06 | 6.55E-06 | -1.13E+00 | 4.27E-08 | 1.84E-07 | -1.17E+00 | 3.21E-02 | 1.36E-01 | -1.11E+00 | 7.16E-02 | 2.74E-01 | -1.07E+00 | 8.32E-07 | 2.97E-06 | -1.29E+00 | 5.65E-05 | 1.59E-04 | -1.23E+00 | 2.29E-04 | 8.41E-04 | -1.14E+00 | 1.23E-02 | 3.38E-02 | -1.08E+00 |
| ELOC | 1.26E-06 | 6.56E-06 | -1.21E+00 | 6.03E-06 | 2.61E-05 | -1.07E+00 | 5.64E-04 | 2.39E-03 | -1.07E+00 | 6.25E-05 | 2.39E-04 | -1.03E+00 | 7.45E-04 | 2.66E-03 | -1.10E+00 | 5.48E-02 | 1.55E-01 | 1.10E+00 | 8.47E-03 | 3.11E-02 | -1.02E+00 | 2.60E-02 | 7.15E-02 | 1.05E+00 |
| AC004865.2 | 1.26E-06 | 6.58E-06 | -1.13E+00 | 6.31E-08 | 2.72E-07 | -1.06E+00 | 5.90E-03 | 2.51E-02 | -1.16E+00 | 4.66E-04 | 1.78E-03 | -1.17E+00 | 2.04E-02 | 7.28E-02 | 1.14E+00 | 3.10E-02 | 8.74E-02 | 1.13E+00 | 1.34E-01 | 4.91E-01 | 1.64E+00 | 3.07E-01 | 8.44E-01 | 1.29E+00 |
| TSC22D4 | 1.26E-06 | 6.58E-06 | -1.11E+00 | 4.72E-02 | 2.04E-01 | 1.35E+00 | 5.92E-07 | 2.52E-06 | -1.38E+00 | 3.15E-03 | 1.20E-02 | 1.05E+00 | 4.68E-03 | 1.67E-02 | 1.13E+00 | 1.50E-01 | 4.24E-01 | 1.36E+00 | 1.54E-01 | 5.66E-01 | 1.32E+00 | 1.78E-01 | 4.88E-01 | 1.36E+00 |
| BIN2 | 1.27E-06 | 6.59E-06 | -1.15E+00 | 1.25E-03 | 5.41E-03 | 1.11E+00 | 1.73E-02 | 7.34E-02 | -1.02E+00 | 1.81E-01 | 6.90E-01 | 1.24E+00 | 1.66E-02 | 5.90E-02 | 1.07E+00 | 1.43E-02 | 4.03E-02 | 1.13E+00 | 4.93E-04 | 1.81E-03 | 1.71E+00 | 1.70E-02 | 4.68E-02 | 1.44E+00 |
| TMEM231 | 1.27E-06 | 6.61E-06 | -2.49E+00 | 3.89E-04 | 1.68E-03 | -1.49E+00 | 1.54E-04 | 6.54E-04 | -1.84E+00 | 2.78E-04 | 1.06E-03 | -1.65E+00 | 3.77E-02 | 1.34E-01 | -1.07E+00 | 7.53E-02 | 2.12E-01 | -1.07E+00 | 1.30E-02 | 4.77E-02 | 1.02E+00 | 5.36E-02 | 1.47E-01 | 1.02E+00 |
| ZNF576 | 1.27E-06 | 6.62E-06 | -1.37E+00 | 7.58E-05 | 3.28E-04 | -1.02E+00 | 2.77E-04 | 1.18E-03 | -1.38E+00 | 4.66E-03 | 1.78E-02 | -1.08E+00 | 2.53E-04 | 9.01E-04 | -1.44E+00 | 3.45E-02 | 9.71E-02 | 1.03E+00 | 2.37E-05 | 8.68E-05 | -1.28E+00 | 7.16E-03 | 1.97E-02 | 1.03E+00 |
| SLC25A20 | 1.27E-06 | 6.63E-06 | -1.62E+00 | 1.17E-04 | 5.04E-04 | -1.17E+00 | 3.14E-05 | 1.34E-04 | -1.57E+00 | 6.31E-05 | 2.41E-04 | -1.29E+00 | 5.57E-09 | 1.99E-08 | -1.66E+00 | 7.57E-06 | 2.14E-05 | -1.21E+00 | 5.89E-12 | 2.16E-11 | -1.63E+00 | 1.72E-04 | 4.72E-04 | -1.08E+00 |
| NLN | 1.27E-06 | 6.63E-06 | -1.71E+00 | 6.13E-03 | 2.65E-02 | 1.09E+00 | 8.26E-15 | 3.51E-14 | -2.00E+00 | 8.15E-07 | 3.11E-06 | -1.11E+00 | 4.57E-08 | 1.63E-07 | -2.61E+00 | 2.67E-02 | 7.54E-02 | 1.01E+00 | 1.07E-07 | 3.93E-07 | -1.51E+00 | 6.20E-02 | 1.70E-01 | 1.13E+00 |
| TRAPPC8 | 1.27E-06 | 6.64E-06 | -1.68E+00 | 3.73E-03 | 1.61E-02 | 1.03E+00 | 1.68E-05 | 7.15E-05 | -1.33E+00 | 1.77E-03 | 6.76E-03 | 1.09E+00 | 1.17E-02 | 4.17E-02 | -1.16E+00 | 6.39E-02 | 1.80E-01 | 1.03E+00 | 1.52E-05 | 5.59E-05 | -1.42E+00 | 3.17E-03 | 8.70E-03 | -1.11E+00 |
| TBC1D10B | 1.28E-06 | 6.66E-06 | -1.58E+00 | 1.38E-03 | 5.96E-03 | -1.02E+00 | 2.84E-11 | 1.21E-10 | -1.68E+00 | 5.94E-10 | 2.27E-09 | -1.38E+00 | 2.57E-05 | 9.15E-05 | -1.62E+00 | 9.14E-03 | 2.58E-02 | -1.11E+00 | 3.32E-07 | 1.22E-06 | -1.22E+00 | 4.68E-05 | 1.28E-04 | -1.23E+00 |
| PSMA2 | 1.28E-06 | 6.67E-06 | -1.26E+00 | 2.48E-04 | 1.07E-03 | 1.02E+00 | 1.39E-06 | 5.91E-06 | -1.25E+00 | 7.52E-09 | 2.87E-08 | -1.21E+00 | 2.95E-08 | 1.05E-07 | -1.32E+00 | 1.50E-06 | 4.23E-06 | -1.17E+00 | 3.92E-04 | 1.44E-03 | -1.05E+00 | 2.99E-04 | 8.23E-04 | -1.07E+00 |
| CASP3 | 1.28E-06 | 6.69E-06 | -1.28E+00 | 6.72E-02 | 2.90E-01 | 1.68E+00 | 1.06E-08 | 4.49E-08 | -1.58E+00 | 1.47E-01 | 5.60E-01 | 1.45E+00 | 2.04E-02 | 7.29E-02 | 1.25E+00 | 1.96E-04 | 5.54E-04 | 2.07E+00 | 1.19E-02 | 4.37E-02 | 1.01E+00 | 9.66E-02 | 2.65E-01 | 1.43E+00 |
| SH3PXD2A | 1.28E-06 | 6.69E-06 | -1.41E+00 | 5.43E-03 | 2.35E-02 | 1.07E+00 | 1.77E-09 | 7.53E-09 | -1.64E+00 | 5.27E-07 | 2.01E-06 | -1.19E+00 | 6.82E-04 | 2.43E-03 | -1.22E+00 | 4.21E-02 | 1.19E-01 | 1.02E+00 | 2.35E-11 | 8.62E-11 | -1.62E+00 | 3.55E-05 | 9.76E-05 | -1.14E+00 |
| KLHL8 | 1.28E-06 | 6.69E-06 | -1.31E+00 | 6.69E-06 | 2.89E-05 | -1.15E+00 | 1.53E-03 | 6.51E-03 | -1.16E+00 | 2.28E-07 | 8.72E-07 | -1.41E+00 | 1.31E-04 | 4.65E-04 | -1.28E+00 | 3.52E-04 | 9.91E-04 | -1.21E+00 | 1.52E-07 | 5.58E-07 | -1.41E+00 | 1.88E-06 | 5.16E-06 | -1.34E+00 |
| ESR1 | 1.28E-06 | 6.70E-06 | -1.78E+00 | 6.18E-07 | 2.67E-06 | -1.70E+00 | 3.31E-02 | 1.41E-01 | -1.13E+00 | 1.96E-03 | 7.51E-03 | -1.43E+00 | 2.00E-01 | 7.13E-01 | 1.31E+00 | 3.32E-01 | 9.37E-01 | 1.05E+00 | 6.89E-02 | 2.53E-01 | -1.41E+00 | 1.13E-01 | 3.12E-01 | -1.37E+00 |
| RUVBL2 | 1.28E-06 | 6.70E-06 | -1.07E+00 | 1.35E-04 | 5.83E-04 | 1.11E+00 | 2.60E-02 | 1.10E-01 | 1.06E+00 | 5.25E-03 | 2.00E-02 | 1.10E+00 | 4.51E-07 | 1.61E-06 | -1.21E+00 | 5.60E-03 | 1.58E-02 | 1.15E+00 | 2.43E-04 | 8.93E-04 | -1.04E+00 | 2.19E-01 | 6.03E-01 | 1.22E+00 |
| CRYGD | 1.29E-06 | 6.70E-06 | -1.07E+00 | 1.62E-18 | 7.00E-18 | -1.39E+00 | 3.95E-02 | 1.68E-01 | 1.02E+00 | 6.14E-05 | 2.35E-04 | -1.11E+00 | 1.95E-06 | 6.96E-06 | -1.36E+00 | 4.11E-10 | 1.16E-09 | -1.68E+00 | 6.11E-02 | 2.24E-01 | -1.07E+00 | 2.14E-02 | 5.88E-02 | -1.22E+00 |
| TGS1 | 1.29E-06 | 6.71E-06 | -1.09E+00 | 1.61E-14 | 6.98E-14 | -1.42E+00 | 3.34E-05 | 1.42E-04 | -1.14E+00 | 3.55E-08 | 1.36E-07 | -1.28E+00 | 6.22E-03 | 2.22E-02 | 1.11E+00 | 3.91E-04 | 1.10E-03 | -1.08E+00 | 3.82E-06 | 1.40E-05 | -1.28E+00 | 6.45E-03 | 1.77E-02 | -1.04E+00 |
| HMGB1P11 | 1.29E-06 | 6.72E-06 | -1.37E+00 | 1.43E-11 | 6.19E-11 | -1.87E+00 | 1.72E-09 | 7.29E-09 | -1.36E+00 | 1.24E-19 | 4.73E-19 | -1.90E+00 | 3.50E-15 | 1.25E-14 | -2.11E+00 | 2.25E-17 | 6.33E-17 | -2.22E+00 | 3.26E-11 | 1.20E-10 | -1.43E+00 | 2.05E-21 | 5.62E-21 | -2.01E+00 |
| SLC35E1 | 1.30E-06 | 6.77E-06 | -1.32E+00 | 9.01E-07 | 3.90E-06 | -1.18E+00 | 3.47E-05 | 1.48E-04 | -1.28E+00 | 1.48E-05 | 5.67E-05 | -1.19E+00 | 1.73E-06 | 6.19E-06 | -1.88E+00 | 4.18E-04 | 1.18E-03 | -1.36E+00 | 2.27E-06 | 8.35E-06 | -1.37E+00 | 2.55E-03 | 7.00E-03 | -1.11E+00 |
| MAP1S | 1.30E-06 | 6.77E-06 | -1.61E+00 | 2.77E-02 | 1.20E-01 | 1.23E+00 | 1.88E-06 | 7.97E-06 | -1.63E+00 | 3.10E-02 | 1.18E-01 | 1.12E+00 | 2.23E-06 | 7.94E-06 | -1.80E+00 | 4.51E-02 | 1.27E-01 | 1.19E+00 | 2.66E-08 | 9.77E-08 | -1.82E+00 | 7.05E-04 | 1.94E-03 | -1.17E+00 |
| RTN4R | 1.31E-06 | 6.83E-06 | -2.27E+00 | 1.15E-04 | 4.95E-04 | -1.57E+00 | 5.76E-07 | 2.45E-06 | -1.57E+00 | 4.05E-03 | 1.55E-02 | 1.07E+00 | 9.36E-02 | 3.34E-01 | -1.49E+00 | 9.84E-02 | 2.77E-01 | -1.64E+00 | 1.31E-01 | 4.80E-01 | -1.32E+00 | 2.27E-01 | 6.24E-01 | -1.22E+00 |
| BRF1 | 1.32E-06 | 6.85E-06 | -1.30E+00 | 8.37E-05 | 3.62E-04 | -1.02E+00 | 3.24E-04 | 1.38E-03 | -1.26E+00 | 3.93E-04 | 1.50E-03 | -1.10E+00 | 1.14E-03 | 4.06E-03 | -1.25E+00 | 4.04E-03 | 1.14E-02 | -1.07E+00 | 8.27E-07 | 3.03E-06 | -1.32E+00 | 7.65E-06 | 2.10E-05 | -1.36E+00 |
| NUP98 | 1.32E-06 | 6.86E-06 | -1.10E+00 | 5.62E-07 | 2.43E-06 | -1.03E+00 | 5.88E-03 | 2.50E-02 | -1.05E+00 | 7.51E-03 | 2.87E-02 | 1.03E+00 | 1.98E-01 | 7.07E-01 | 1.31E+00 | 3.51E-01 | 9.88E-01 | 1.43E+00 | 6.94E-08 | 2.55E-07 | -1.27E+00 | 5.12E-04 | 1.41E-03 | -1.12E+00 |
| AC079601.2 | 1.32E-06 | 6.86E-06 | -1.29E+00 | 1.25E-11 | 5.39E-11 | -1.78E+00 | 3.17E-05 | 1.35E-04 | -1.31E+00 | 2.09E-11 | 7.98E-11 | -2.01E+00 | 1.04E-04 | 3.72E-04 | -1.37E+00 | 9.47E-07 | 2.67E-06 | -1.85E+00 | 2.72E-03 | 1.00E-02 | -1.35E+00 | 2.95E-05 | 8.11E-05 | -1.69E+00 |
| AC005154.1 | 1.32E-06 | 6.88E-06 | -1.31E+00 | 1.11E-02 | 4.80E-02 | 1.32E+00 | 8.53E-11 | 3.62E-10 | -2.06E+00 | 2.35E-03 | 8.97E-03 | -1.00E+00 | 1.34E-07 | 4.77E-07 | -1.85E+00 | 1.40E-03 | 3.95E-03 | -1.14E+00 | 7.18E-04 | 2.64E-03 | -1.20E+00 | 7.80E-02 | 2.14E-01 | 1.14E+00 |
| ANKRD35 | 1.32E-06 | 6.88E-06 | -1.76E+00 | 2.27E-08 | 9.80E-08 | -2.16E+00 | 3.56E-02 | 1.51E-01 | -1.13E+00 | 1.90E-03 | 7.27E-03 | -1.32E+00 | 1.31E-03 | 4.67E-03 | -1.29E+00 | 2.76E-03 | 7.79E-03 | -1.22E+00 | 6.44E-02 | 2.36E-01 | 1.64E+00 | 1.93E-01 | 5.30E-01 | 1.23E+00 |
| MLST8 | 1.32E-06 | 6.89E-06 | -1.29E+00 | 1.46E-07 | 6.32E-07 | -1.23E+00 | 1.91E-06 | 8.13E-06 | -1.25E+00 | 3.68E-06 | 1.41E-05 | -1.09E+00 | 6.48E-08 | 2.31E-07 | -1.38E+00 | 6.06E-07 | 1.71E-06 | -1.29E+00 | 1.07E-04 | 3.94E-04 | -1.12E+00 | 4.45E-03 | 1.22E-02 | 1.00E+00 |
| CD44 | 1.32E-06 | 6.90E-06 | 1.51E+00 | 5.27E-02 | 2.28E-01 | 1.23E+00 | 3.42E-03 | 1.45E-02 | 1.36E+00 | 1.26E-01 | 4.80E-01 | 1.24E+00 | 3.08E-14 | 1.10E-13 | 2.42E+00 | 2.90E-01 | 8.16E-01 | 1.42E+00 | 8.37E-02 | 3.07E-01 | 1.42E+00 | 2.26E-02 | 6.22E-02 | 1.07E+00 |
| RPRD1A | 1.33E-06 | 6.91E-06 | -1.18E+00 | 3.63E-03 | 1.57E-02 | 1.20E+00 | 7.19E-03 | 3.06E-02 | -1.05E+00 | 8.33E-02 | 3.18E-01 | 1.29E+00 | 1.45E-02 | 5.17E-02 | 1.00E+00 | 3.34E-02 | 9.43E-02 | 1.11E+00 | 9.91E-05 | 3.64E-04 | -1.22E+00 | 1.51E-02 | 4.16E-02 | 1.02E+00 |
| TMS84XP4 | 1.33E-06 | 6.92E-06 | -1.09E+00 | 1.43E-03 | 6.20E-03 | 1.19E+00 | 4.23E-07 | 1.80E-06 | -1.47E+00 | 6.47E-02 | 2.47E-01 | 1.06E+00 | 6.74E-16 | 2.40E-15 | -1.99E+00 | 4.10E-10 | 1.16E-09 | -1.54E+00 | 7.18E-04 | 2.64E-03 | -1.94E+00 | 3.11E-01 | 8.55E-01 | -1.39E+00 |
| RING1 | 1.34E-06 | 6.98E-06 | -1.20E+00 | 3.44E-04 | 1.49E-03 | 1.12E+00 | 4.96E-06 | 2.11E-05 | -1.23E+00 | 1.59E-03 | 6.06E-03 | 1.10E+00 | 2.35E-08 | 8.39E-08 | -1.52E+00 | 6.34E-05 | 1.79E-04 | -1.07E+00 | 2.10E-05 | 7.72E-05 | -1.27E+00 | 4.21E-02 | 1.16E-01 | 1.04E+00 |
| AL513365.1 | 1.34E-06 | 6.99E-06 | -1.08E+00 | 8.95E-25 | 3.87E-24 | -1.71E+00 | 8.43E-02 | 3.58E-01 | 1.10E+00 | 9.89E-11 | 3.78E-10 | -1.45E+00 | 3.04E-06 | 1.08E-05 | -1.11E+00 | 7.09E-15 | 2.00E-14 | -1.77E+00 | 1.06E-06 | 3.89E-06 | -1.16E+00 | 9.01E-18 | 2.48E-17 | -1.77E+00 |
| ZCCHC2 | 1.35E-06 | 7.02E-06 | -1.44E+00 | 8.98E-09 | 3.88E-08 | -1.48E+00 | 2.05E-07 | 8.72E-07 | -1.74E+00 | 7.59E-08 | 2.90E-07 | -1.61E+00 | 2.73E-03 | 9.72E-03 | -1.05E+00 | 6.23E-06 | 1.76E-05 | -1.52E+00 | 1.16E-07 | 4.27E-07 | -1.46E+00 | 1.14E-07 | 3.12E-07 | -1.57E+00 |
| TRPC4AP | 1.35E-06 | 7.03E-06 | -1.27E+00 | 1.10E-04 | 4.75E-04 | 1.02E+00 | 2.77E-05 | 1.18E-04 | -1.22E+00 | 1.17E-06 | 4.48E-06 | -1.13E+00 | 2.38E-08 | 8.50E-08 | -1.41E+00 | 3.02E-07 | 8.51E-07 | -1.25E+00 | 1.77E-08 | 6.50E-08 | -1.35E+00 | 9.44E-05 | 2.59E-04 | -1.11E+00 |
| ARFIP1 | 1.35E-06 | 7.04E-06 | -1.54E+00 | 8.16E-05 | 3.53E-04 | -1.11E+00 | 5.85E-04 | 2.49E-03 | -1.36E+00 | 3.34E-02 | 1.28E-01 | 1.07E+00 | 3.53E-05 | 1.26E-04 | -1.37E+00 | 3.19E-04 | 9.00E-04 | -1.14E+00 | 7.66E-12 | 2.81E-11 | -1.43E+00 | 4.29E-05 | 1.18E-04 | -1.14E+00 |
| PRKAB1 | 1.35E-06 | 7.05E-06 | -1.41E+00 | 9.81E-05 | 4.24E-04 | -1.06E+00 | 8.11E-05 | 3.45E-04 | -1.32E+00 | 1.86E-03 | 7.11E-03 | 1.03E+00 | 1.53E-07 | 5.46E-07 | -1.60E+00 | 1.53E-04 | 4.31E-04 | -1.13E+00 | 1.14E-08 | 4.19E-08 | -1.39E+00 | 1.13E-06 | 3.11E-06 | -1.35E+00 |
| TMEM138 | 1.35E-06 | 7.06E-06 | -1.59E+00 | 2.03E-03 | 8.77E-03 | -1.04E+00 | 3.03E-02 | 1.29E-01 | -1.03E+00 | 1.96E-01 | 7.50E-01 | 1.33E+00 | 1.92E-01 | 6.84E-01 | 1.39E+00 | 3.00E-01 | 8.46E-01 | 1.60E+00 | 1.07E-04 | 3.94E-04 | -1.17E+00 | 1.12E-01 | 3.08E-01 | 1.10E+00 |
| MCF2L2 | 1.36E-06 | 7.06E-06 | -1.60E+00 | 3.96E-05 | 1.71E-04 | -1.30E+00 | 3.66E-02 | 1.56E-01 | -1.03E+00 | 1.94E-02 | 7.43E-02 | -1.06E+00 | 2.56E-01 | 9.14E-01 | 1.75E+00 | 1.53E-01 | 4.32E-01 | -1.18E+00 | 8.76E-04 | 3.22E-03 | -1.78E+00 | 3.54E-03 | 9.72E-03 | -1.59E+00 |
| OPA1 | 1.36E-06 | 7.08E-06 | -1.20E+00 | 6.00E-10 | 2.59E-09 | -1.26E+00 | 8.19E-04 | 3.48E-03 | -1.22E+00 | 1.47E-05 | 5.62E-05 | -1.21E+00 | 6.49E-02 | 2.31E-01 | 1.24E+00 | 1.58E-03 | 4.46E-03 | 1.04E+00 | 7.63E-04 | 2.80E-03 | -1.03E+00 | 9.88E-03 | 2.71E-02 | -1.02E+00 |
| SENP2 | 1.36E-06 | 7.08E-06 | -1.34E+00 | 1.98E-07 | 8.56E-07 | -1.30E+00 | 9.15E-07 | 3.89E-06 | -1.31E+00 | 1.96E-11 | 7.50E-11 | -1.40E+00 | 2.35E-02 | 8.37E-02 | -1.06E+00 | 2.88E-02 | 8.13E-02 | 1.01E+00 | 5.11E-04 | 1.88E-03 | -1.27E+00 | 7.55E-03 | 2.07E-02 | -1.18E+00 |
| FTLP3 | 1.36E-06 | 7.09E-06 | 1.92E+00 | 2.58E-28 | 1.11E-27 | 3.29E+00 | 2.15E-05 | 9.12E-05 | 1.77E+00 | 1.20E-14 | 4.57E-14 | 2.91E+00 | 1.30E-03 | 4.62E-03 | 1.98E+00 | 9.69E-09 | 2.73E-08 | 2.83E+00 | 4.25E-05 | 1.56E-04 | 1.83E+00 | 2.92E-21 | 8.02E-21 | 3.03E+00 |
| AC073857.1 | 1.36E-06 | 7.09E-06 | -2.32E+00 | 2.93E-04 | 1.26E-03 | -1.44E+00 | 4.08E-06 | 1.73E-05 | -2.01E+00 | 5.21E-05 | 1.99E-04 | -1.63E+00 | 3.38E-03 | 1.20E-02 | -1.53E+00 | 3.27E-02 | 9.21E-02 | -1.07E+00 | 1.96E-06 | 7.19E-06 | -1.61E+00 | 3.82E-05 | 1.05E-04 | -1.33E+00 |
| KLHL3 | 1.36E-06 | 7.11E-06 | -1.40E+00 | 1.18E-04 | 5.08E-04 | -1.07E+00 | 1.79E-01 | 7.61E-01 | 1.20E+00 | 1.10E-01 | 4.19E-01 | 1.23E+00 | 2.57E-01 | 9.16E-01 | 1.21E+00 | 3.45E-01 | 9.72E-01 | 1.31E+00 | 6.89E-03 | 2.53E-02 | -1.50E+00 | 1.05E-02 | 2.87E-02 | -1.75E+00 |
| LCORL | 1.37E-06 | 7.12E-06 | -1.33E+00 | 8.02E-05 | 3.46E-04 | -1.09E+00 | 4.05E-04 | 1.72E-03 | -1.22E+00 | 2.37E-02 | 9.06E-02 | 1.16E+00 | 1.10E-02 | 3.94E-02 | 1.01E+00 | 1.15E-01 | 3.23E-01 | 1.20E+00 | 8.01E-05 | 2.94E-04 | -1.18E+00 | 1.17E-02 | 3.22E-02 | 1.02E+00 |
| MON1B | 1.37E-06 | 7.14E-06 | -1.50E+00 | 3.77E-03 | 1.63E-02 | 1.05E+00 | 6.05E-08 | 2.57E-07 | -1.70E+00 | 1.94E-03 | 7.39E-03 | -1.00E+00 | 5.55E-07 | 1.98E-06 | -1.57E+00 | 1.45E-03 | 4.08E-03 | -1.02E+00 | 1.92E-04 | 7.04E-04 | -1.22E+00 | 2.32E-01 | 6.37E-01 | 1.25E+00 |
| TAF7 | 1.37E-06 | 7.14E-06 | -1.12E+00 | 2.19E-04 | 9.46E-04 | 1.05E+00 | 1.43E-04 | 6.09E-04 | -1.20E+00 | 3.09E-02 | 1.18E-01 | 1.01E+00 | 4.30E-04 | 1.53E-03 | -1.13E+00 | 1.67E-03 | 4.71E-03 | -1.00E+00 | 9.71E-03 | 3.57E-02 | -1.11E+00 | 2.08E-01 | 5.72E-01 | -1.01E+00 |
| AL136531.3 | 1.37E-06 | 7.16E-06 | 2.89E+00 | 1.45E-01 | 6.26E-01 | -1.07E+00 | 1.82E-05 | 7.75E-05 | 3.73E+00 | 1.38E-01 | 5.25E-01 | 1.64E+00 | 9.70E-04 | 3.46E-03 | 5.53E+00 | 1.75E-01 | 4.94E-01 | 1.92E+00 | 3.58E-02 | 1.31E-01 | 2.09E+00 | 2.82E-01 | 7.75E-01 | 1.35E+00 |
| ANXA2R | 1.38E-06 | 7.17E-06 | 2.19E+00 | 5.25E-02 | 2.27E-01 | 1.30E+00 | 2.95E-03 | 1.25E-02 | 1.67E+00 | 7.72E-03 | 2.95E-02 | -1.03E+00 | 1.85E-01 | 6.60E-01 | 1.38E+00 | 4.80E-02 | 1.35E-01 | 1.05E+00 | 5.46E-03 | 2.00E-02 | 1.90E+00 | 2.76E-01 | 7.58E-01 | 1.14E+00 |
| USP16 | 1.39E-06 | 7.22E-06 | -1.10E+00 | 3.08E-14 | 1.33E-13 | -1.29E+00 | 1.98E-04 | 8.42E-04 | -1.06E+00 | 1.50E-07 | 5.73E-07 | -1.14E+00 | 1.84E-05 | 6.56E-05 | -1.05E+00 | 3.16E-10 | 8.91E-10 | -1.32E+00 | 4.14E-03 | 1.52E-02 | 1.01E+00 | 6.68E-08 | 1.83E-07 | -1.29E+00 |
| ACAT1 | 1.39E-06 | 7.26E-06 | -1.27E+00 | 2.94E-02 | 1.27E-01 | 1.27E+00 | 2.42E-03 | 1.03E-02 | -1.05E+00 | 6.90E-04 | 2.64E-03 | -1.03E+00 | 2.63E-05 | 9.39E-05 | -1.25E+00 | 6.65E-02 | 1.87E-01 | 1.24E+00 | 1.07E-04 | 3.94E-04 | -1.18E+00 | 2.41E-02 | 6.62E-02 | 1.04E+00 |
| CCDC18 | 1.39E-06 | 7.27E-06 | -1.37E+00 | 5.23E-03 | 2.26E-02 | 1.06E+00 | 1.85E-09 | 7.85E-09 | -1.52E+00 | 1.96E-07 | 7.49E-07 | -1.16E+00 | 3.44E-10 | 1.23E-09 | -1.77E+00 | 5.16E-04 | 1.45E-03 | -1.06E+00 | 1.17E-03 | 4.30E-03 | -1.19E+00 | 3.69E-02 | 1.01E-01 | -1.08E+00 |
| ZNF121 | 1.40E-06 | 7.28E-06 | -1.08E+00 | 1.05E-08 | 4.53E-08 | -1.05E+00 | 1.32E-06 | 5.61E-06 | -1.26E+00 | 3.85E-09 | 1.47E-08 | -1.19E+00 | 9.38E-02 | 3.35E-01 | 1.13E+00 | 1.11E-01 | 3.13E-01 | 1.18E+00 | 4.76E-09 | 1.75E-08 | -1.42E+00 | 2.00E-05 | 5.50E-05 | -1.24E+00 |
| CLIP2 | 1.40E-06 | 7.28E-06 | -1.53E+00 | 9.84E-04 | 4.25E-03 | -1.01E+00 | 7.09E-08 | 3.01E-07 | -1.60E+00 | 2.25E-06 | 8.60E-06 | -1.27E+00 | 6.48E-10 | 2.31E-09 | -1.67E+00 | 1.40E-06 | 3.94E-06 | -1.30E+00 | 1.25E-01 | 4.57E-01 | 1.20E+00 | 9.90E-02 | 2.72E-01 | 1.02E+00 |
| NECAP1 | 1.40E-06 | 7.28E-06 | -1.10E+00 | 3.17E-12 | 1.37E-11 | -1.31E+00 | 1.06E-01 | 4.52E-01 | 1.17E+00 | 1.11E-04 | 4.24E-04 | -1.07E+00 | 1.26E-01 | 4.50E-01 | 1.33E+00 | 2.57E-04 | 7.24E-04 | -1.08E+00 | 2.36E-07 | 8.67E-07 | -1.29E+00 | 3.79E-10 | 1.04E-09 | -1.44E+00 |
| RAB16 | 1.40E-06 | 7.28E-06 | -1.16E+00 | 2.07E-04 | 8.96E-04 | 1.07E+00 | 1.64E-01 | 6.98E-01 | 1.04E+00 | 1.25E-01 | 4.76E-01 | 1.21E+00 | 3.18E-08 | 1.13E-07 | -1.49E+00 | 1.85E-03 | 5.22E-03 | -1.01E+00 | 3.98E-05 | 1.46E-04 | -1.22E+00 | 2.46E-01 | 6.76E-01 | 1.25E+00 |
| FRYL | 1.40E-06 | 7.29E-06 | -1.14E+00 | 2.93E-03 | 1.27E-02 | 1.16E+00 | 2.64E-06 | 1.12E-05 | -1.35E+00 | 5.80E-04 | 2.22E-03 | -1.01E+00 | 2.18E-05 | 7.79E-05 | -1.40E+00 | 9.54E-03 | 2.69E-02 | -1.02E+00 | 5.02E-05 | 1.84E-04 | -1.31E+00 | 4.32E-03 | 1.19E-02 | -1.09E+00 |
| SSFA2 | 1.40E-06 | 7.31E-06 | -1.20E+00 | 4.99E-06 | 2.16E-05 | -1.03E+00 | 1.50E-07 | 6.38E-07 | -1.27E+00 | 2.22E-08 | 8.49E-08 | -1.26E+00 | 8.01E-03 | 2.85E-02 | 1.04E+00 | 2.49E-03 | 7.02E-03 | -1.02E+00 | 2.59E-04 | 9.49E-04 | -1.07E+00 | 1.25E-05 | 3.42E-05 | -1.19E+00 |
| ZNF326 | 1.40E-06 | 7.31E-06 | -1.08E+00 | 2.49E-06 | 1.08E-05 | 1.01E+00 | 4.51E-02 | 1.92E-01 | 1.01E+00 | 7.33E-03 | 2.80E-02 | 1.05E+00 | 5.37E-03 | 1.92E-02 | 1.07E+00 | 2.19E-02 | 6.16E-02 | 1.12E+00 | 4.48E-02 | 1.64E-01 | 1.08E+00 | 3.42E-02 | 9.39E-02 | -1.01E+00 |
| SLC9A3R2 | 1.40E-06 | 7.32E-06 | -1.66E+00 | 3.10E-04 | 1.34E-03 | -1.13E+00 | 2.28E-01 | 9.70E-01 | 1.21E+00 | 2.33E-01 | 8.90E-01 | 1.36E+00 | 1.12E-01 | 4.01E-01 | 2.28E+00 | 8.00E-02 | 2.26E-01 | 2.50E+00 | 9.69E-03 | 3.56E-02 | 3.27E+00 | 1.88E-02 | 5.15E-02 | 2.18E+00 |
| PDAP1 | 1.41E-06 | 7.32E-06 | -1.05E+00 | 1.44E-05 | 6.21E-05 | 1.10E+00 | 1.74E-01 | 7.40E-01 | 1.08E+00 | 6.54E-02 | 2.50E-01 | 1.13E+00 | 1.88E-04 | 6.70E-04 | -1.08E+00 | 2.19E-02 | 6.18E-02 | 1.15E+00 | 3.96E-02 | 1.45E-01 | 1.02E+00 | 3.34E-01 | 9.17E-01 | 1.13E+00 |
| ZNF267 | 1.41E-06 | 7.33E-06 | -1.10E+00 | 3.31E-14 | 1.43E-13 | -1.47E+00 | 1.91E-02 | 8.11E-02 | 1.04E+00 | 1.65E-05 | 6.31E-05 | -1.21E+00 | 1.00E-01 | 3.58E-01 | 1.35E+00 | 3.91E-05 | 1.10E-04 | -1.24E+00 | 1.57E-04 | 5.75E-04 | 1.16E+00 | 5.84E-06 | 1.60E-05 | -1.14E+00 |
| AKIP1 | 1.41E-06 | 7.35E-06 | -1.45E+00 | 2.64E-06 | 1.14E-05 | -1.26E+00 | 1.41E-04 | 5.97E-04 | -1.27E+00 | 4.52E-04 | 1.73E-03 | -1.08E+00 | 8.50E-04 | 3.03E-03 | -1.24E+00 | 1.83E-02 | 5.17E-02 | 1.03E+00 | 2.14E-06 | 7.85E-06 | -1.24E+00 | 5.12E-05 | 1.41E-04 | -1.09E+00 |
| KNOP1 | 1.41E-06 | 7.36E-06 | -1.73E+00 | 2.49E-03 | 1.08E-02 | 1.00E+00 | 2.33E-08 | 9.89E-08 | -1.63E+00 | 1.42E-04 | 5.44E-04 | -1.12E+00 | 5.71E-05 | 2.04E-04 | -1.67E+00 | 8.05E-02 | 2.27E-01 | 1.08E+00 | 1.71E-07 | 6.27E-07 | -1.47E+00 | 8.93E-03 | 2.45E-02 | -1.06E+00 |
| SLX1B-SULT1A4 | 1.41E-06 | 7.37E-06 | -1.80E+00 | 8.38E-05 | 3.62E-04 | -1.22E+00 | 1.04E-03 | 4.42E-03 | -1.33E+00 | 9.19E-03 | 3.51E-02 | -1.03E+00 | 4.47E-02 | 1.59E-01 | -1.13E+00 | 1.65E-01 | 4.64E-01 | 1.13E+00 | 1.05E-23 | 3.85E-23 | -2.51E+00 | 9.05E-14 | 2.49E-13 | -1.74E+00 |
| MAP3K4 | 1.43E-06 | 7.45E-06 | -1.33E+00 | 1.85E-04 | 8.00E-04 | -1.00E+00 | 2.43E-03 | 1.03E-02 | -1.20E+00 | 4.29E-02 | 1.64E-01 | 1.10E+00 | 4.01E-08 | 1.43E-07 | -1.81E+00 | 1.84E-05 | 5.17E-05 | -1.27E+00 | 5.43E-09 | 1.99E-08 | -1.50E+00 | 4.24E-06 | 1.16E-05 | -1.29E+00 |
| ADAMTSL4 | 1.44E-06 | 7.48E-06 | -1.71E+00 | 1.95E-07 | 8.41E-07 | -1.72E+00 | 1.44E-03 | 6.11E-03 | -1.32E+00 | 8.87E-04 | 3.39E-03 | -1.21E+00 | 1.92E-01 | 6.83E-01 | 1.04E+00 | 1.87E-01 | 5.28E-01 | -1.25E+00 | 2.26E-01 | 8.30E-01 | 1.28E+00 | 3.20E-01 | 8.80E-01 | 1.00E+00 |
| AC020951.1 | 1.44E-06 | 7.51E-06 | -1.58E+00 | 3.56E-02 | 1.54E-01 | 1.25E+00 | 1.17E-05 | 4.96E-05 | -1.51E+00 | 2.53E-02 | 9.66E-02 | 1.11E+00 | 6.06E-03 | 2.16E-02 | -1.41E+00 | 2.54E-01 | 7.16E-01 | 1.23E+00 | 1.06E-02 | 3.90E-02 | -1.34E+00 | 2.15E-01 | 5.91E-01 | 1.06E+00 |
| PDPR | 1.44E-06 | 7.51E-06 | -1.48E+00 | 1.21E-02 | 5.24E-02 | 1.14E+00 | 5.21E-07 | 2.22E-06 | -1.32E+00 | 5.12E-03 | 1.96E-02 | 1.17E+00 | 4.06E-07 | 1.45E-06 | -1.59E+00 | 7.11E-04 | 2.00E-03 | -1.12E+00 | 1.38E-08 | 5.06E-08 | -1.52E+00 | 2.95E-02 | 8.11E-02 | -1.01E+00 |
| BEND6 | 1.45E-06 | 7.55E-06 | -1.61E+00 | 2.21E-06 | 9.55E-06 | -1.45E+00 | 4.11E-05 | 1.75E-04 | -1.50E+00 | 1.15E-04 | 4.41E-04 | -1.20E+00 | 2.61E-06 | 9.29E-06 | -2.15E+00 | 3.63E-04 | 1.02E-03 | -1.26E+00 | 2.09E-01 | 7.65E-01 | 1.20E+00 | 2.76E-01 | 7.57E-01 | 1.11E+00 |
| GTPBP8 | 1.45E-06 | 7.56E-06 | -1.18E+00 | 3.46E-07 | 1.49E-06 | -1.12E+00 | 8.96E-07 | 3.81E-06 | -1.21E+00 | 4.36E-08 | 1.67E-07 | -1.16E+00 | 3.94E-06 | 1.41E-05 | -1.23E+00 | 1.25E-03 | 3.52E-03 | -1.04E+00 | 1.77E-06 | 6.49E-06 | -1.18E+00 | 4.18E-03 | 1.15E-02 | -1.06E+00 |
| CHD1L | 1.45E-06 | 7.56E-06 | -1.16E+00 | 2.39E-05 | 1.03E-04 | 1.01E+00 | 2.06E-04 | 8.76E-04 | -1.18E+00 | 8.87E-04 | 3.39E-03 | 1.00E+00 | 5.43E-07 | 1.94E-06 | -1.21E+00 | 6.34E-04 | 1.79E-03 | 1.02E+00 | 5.41E-03 | 1.98E-02 | 1.00E+00 | 2.87E-01 | 7.88E-01 | 1.18E+00 |
| CKAP5 | 1.45E-06 | 7.57E-06 | -1.70E+00 | 2.42E-02 | 1.05E-01 | 1.12E+00 | 1.94E-04 | 8.22E-04 | -1.22E+00 | 2.09E-01 | 8.00E-01 | 1.34E+00 | 2.17E-10 | 7.75E-10 | -1.80E+00 | 1.03E-03 | 2.91E-03 | 1.00E+00 | 7.85E-11 | 2.88E-10 | -1.37E+00 | 1.06E-02 | 2.92E-02 | 1.04E+00 |
| MTA2 | 1.46E-06 | 7.59E-06 | -1.38E+00 | 1.44E-05 | 6.20E-05 | -1.16E+00 | 2.99E-06 | 1.27E-05 | -1.40E+00 | 1.98E-06 | 7.55E-06 | -1.19E+00 | 1.32E-05 | 4.69E-05 | -1.33E+00 | 6.06E-04 | 1.71E-03 | -1.14E+00 | 1.01E-04 | 3.71E-04 | -1.18E+00 | 4.79E-03 | 1.32E-02 | -1.10E+00 |
| CERS4 | 1.46E-06 | 7.59E-06 | -1.27E+00 | 1.84E-03 | 7.97E-03 | 1.16E+00 | 2.12E-02 | 9.00E-02 | -1.06E+00 | 7.75E-03 | 2.96E-02 | 1.01E+00 | 1.05E-03 | 3.74E-03 | -1.23E+00 | 2.53E-03 | 7.13E-03 | -1.12E+00 | 2.21E-01 | 8.12E-01 | 1.27E+00 | 4.37E-02 | 1.20E-01 | -1.09E+00 |
| UBE2E1 | 1.46E-06 | 7.61E-06 | -1.06E+00 | 3.11E-03 | 1.35E-02 | 1.20E+00 | 3.55E-04 | 1.51E-03 | -1.15E+00 | 2.46E-04 | 9.38E-04 | -1.01E+00 | 3.02E-07 | 1.08E-06 | -1.25E+00 | 1.10E-02 | 3.10E-02 | 1.04E+00 | 3.97E-04 | 1.46E-03 | -1.16E+00 | 2.76E-04 | 7.57E-04 | -1.08E+00 |
| SCAPER | 1.47E-06 | 7.65E-06 | -1.09E+00 | 2.35E-06 | 1.02E-05 | 1.01E+00 | 8.26E-02 | 3.51E-01 | 1.05E+00 | 1.50E-01 | 5.72E-01 | 1.23E+00 | 1.71E-03 | 6.10E-03 | -1.27E+00 | 1.54E-02 | 4.34E-02 | -1.13E+00 | 1.20E-01 | 4.42E-01 | 1.16E+00 | 5.79E-02 | 1.59E-01 | -1.06E+00 |
| ARHGAP11B | 1.47E-06 | 7.67E-06 | -1.73E+00 | 2.84E-05 | 1.23E-04 | -1.31E+00 | 1.11E-02 | 4.74E-02 | -1.03E+00 | 7.60E-03 | 2.90E-02 | 1.09E+00 | 8.84E-05 | 3.15E-04 | -1.98E+00 | 2.54E-03 | 7.15E-03 | -1.35E+00 | 2.06E-08 | 7.56E-08 | -1.69E+00 | 9.47E-06 | 2.60E-05 | -1.40E+00 |
| CLUAP1 | 1.47E-06 | 7.67E-06 | -1.43E+00 | 9.46E-03 | 4.09E-02 | 1.17E+00 | 1.11E-08 | 4.71E-08 | -1.65E+00 | 9.81E-03 | 3.75E-02 | 1.13E+00 | 5.33E-05 | 1.90E-04 | -1.76E+00 | 8.23E-02 | 2.32E-01 | 1.09E+00 | 3.21E-06 | 1.18E-05 | -1.58E+00 | 1.86E-01 | 5.10E-01 | 1.13E+00 |
| PLEKHA2 | 1.48E-06 | 7.69E-06 | -1.28E+00 | 3.79E-05 | 1.64E-04 | -1.03E+00 | 1.21E-03 | 5.15E-03 | -1.21E+00 | 2.77E-04 | 1.06E-03 | -1.14E+00 | 2.78E-05 | 9.92E-05 | -1.24E+00 | 4.98E-06 | 1.40E-05 | -1.22E+00 | 5.29E-04 | 1.94E-03 | -1.03E+00 | 9.68E-05 | 2.66E-04 | -1.13E+00 |
| SLC25A32 | 1.48E-06 | 7.70E-06 | -1.26E+00 | 6.90E-09 | 2.98E-08 | -1.29E+00 | 5.20E-06 | 2.21E-05 | -1.40E+00 | 4.95E-08 | 1.89E-07 | -1.30E+00 | 9.96E-02 | 3.55E-01 | 1.30E+00 | 1.44E-02 | 4.06E-02 | 1.00E+00 | 2.62E-06 | 9.61E-06 | -1.44E+00 | 1.14E-04 | 3.13E-04 | -1.23E+00 |
| NLK | 1.49E-06 | 7.75E-06 | -1.53E+00 | 5.32E-05 | 2.30E-04 | -1.13E+00 | 3.36E-07 | 1.43E-06 | -1.72E+00 | 8.82E-06 | 3.37E-05 | -1.30E+00 | 6.60E-24 | 2.35E-23 | -1.81E+00 | 2.75E-13 | 7.75E-13 | -1.33E+00 | 1.10E-17 | 4.04E-17 | -1.40E+00 | 3.30E-10 | 9.08E-10 | -1.19E+00 |
| MSN | 1.49E-06 | 7.76E-06 | -1.03E+00 | 8.47E-02 | 3.66E-01 | 1.25E+00 | 9.24E-02 | 3.93E-01 | 1.04E+00 | 8.26E-02 | 3.16E-01 | 1.30E+00 | 1.99E-02 | 7.08E-02 | 1.14E+00 | 1.47E-01 | 4.14E-01 | 1.17E+00 | 1.77E-01 | 6.51E-01 | 1.19E+00 | 2.91E-01 | 7.99E-01 | 1.21E+00 |
| NBL1 | 1.49E-06 | 7.77E-06 | -1.10E+00 | 4.43E-04 | 1.91E-03 | 1.09E+00 | 7.56E-02 | 3.21E-01 | 1.14E+00 | 1.18E-01 | 4.52E-01 | 1.28E+00 | 1.20E-01 | 4.28E-01 | 2.76E+00 | 2.87E-01 | 8.10E-01 | 1.81E+00 | 9.29E-03 | 3.41E-02 | 4.05E+00 | 4.56E-02 | 1.25E-01 | 3.60E+00 |
| RPL5P34 | 1.50E-06 | 7.83E-06 | 2.28E+00 | 1.42E-02 | 6.16E-02 | 1.99E+00 | 1.43E-08 | 6.06E-08 | 2.40E+00 | 8.23E-04 | 3.14E-03 | 2.13E+00 | 2.64E-04 | 9.41E-04 | 2.60E+00 | 1.47E-02 | 4.13E-02 | 2.24E+00 | 6.44E-07 | 2.36E-06 | 2.33E+00 | 2.94E-03 | 8.08E-03 | 2.05E+00 |
| TRMT11 | 1.50E-06 | 7.84E-06 | -1.26E+00 | 1.66E-02 | 7.15E-02 | 1.27E+00 | 4.26E-05 | 1.81E-04 | -1.39E+00 | 8.73E-02 | 3.34E-01 | 1.19E+00 | 6.21E-07 | 2.21E-06 | -1.38E+00 | 3.47E-05 | 9.79E-05 | -1.20E+00 | 1.37E-08 | 5.02E-08 | -1.49E+00 | 1.70E-02 | 4.66E-02 | 1.00E+00 |
| ZNF263 | 1.51E-06 | 7.85E-06 | -1.38E+00 | 4.76E-05 | 2.06E-04 | -1.07E+00 | 3.34E-06 | 1.42E-05 | -1.55E+00 | 7.23E-05 | 2.76E-04 | -1.13E+00 | 1.77E-05 | 6.33E-05 | -1.37E+00 | 6.66E-05 | 1.88E-04 | -1.27E+00 | 1.75E-11 | 6.41E-11 | -1.76E+00 | 2.18E-06 | 6.00E-06 | -1.34E+00 |
| RNF2 | 1.51E-06 | 7.87E-06 | -1.17E+00 | 1.75E-05 | 7.56E-05 | -1.03E+00 | 5.61E-08 | 2.38E-07 | -1.36E+00 | 9.48E-05 | 3.62E-04 | -1.02E+00 | 9.54E-05 | 3.40E-04 | -1.23E+00 | 8.40E-03 | 2.37E-02 | 1.06E+00 | 1.05E-07 | 3.84E-07 | -1.29E+00 | 3.40E-03 | 9.34E-03 | 1.00E+00 |
| AC092324.1 | 1.51E-06 | 7.88E-06 | -1.26E+00 | 9.18E-49 | 3.97E-48 | -1.93E+00 | 2.98E-03 | 1.27E-02 | -1.25E+00 | 4.94E-21 | 1.89E-20 | -1.98E+00 | 4.04E-04 | 1.44E-03 | -1.21E+00 | 3.30E-27 | 9.30E-27 | -2.04E+00 | 1.01E-04 | 3.71E-04 | -1.22E+00 | 8.00E-18 | 2.20E-17 | -1.86E+00 |
| MED10 | 1.53E-06 | 7.98E-06 | -1.01E+00 | 8.79E-06 | 3.80E-05 | 1.11E+00 | 9.11E-02 | 3.87E-01 | 1.16E+00 | 2.80E-03 | 1.07E-02 | 1.12E+00 | 5.82E-04 | 2.07E-03 | -1.05E+00 | 7.21E-04 | 2.03E-03 | 1.00E+00 | 2.33E-01 | 8.57E-01 | 1.23E+00 | 8.80E-02 | 2.42E-01 | 1.07E+00 |
| NOL8P1 | 1.53E-06 | 7.98E-06 | -2.31E+00 | 9.23E-06 | 3.99E-05 | -1.83E+00 | 1.41E-09 | 6.00E-09 | -2.49E+00 | 5.61E-06 | 2.14E-05 | -1.51E+00 | 6.86E-12 | 2.45E-11 | -3.26E+00 | 3.10E-07 | 8.74E-07 | -1.64E+00 | 3.75E-26 | 1.38E-25 | -2.64E+00 | 2.55E-17 | 7.00E-17 | -1.83E+00 |
| RPL36 | 1.53E-06 | 7.98E-06 | -1.10E+00 | 1.14E-118 | 4.91E-118 | -1.64E+00 | 5.09E-02 | 2.16E-01 | -1.03E+00 | 3.49E-68 | 1.33E-67 | -1.58E+00 | 5.53E-05 | 1.97E-04 | -1.12E+00 | 2.45E-57 | 6.91E-57 | -1.59E+00 | 8.86E-05 | 3.25E-04 | -1.06E+00 | 1.31E-59 | 3.59E-59 | -1.54E+00 |
| AC016747.1 | 1.54E-06 | 8.01E-06 | -1.66E+00 | 3.15E-04 | 1.36E-03 | -1.14E+00 | 2.64E-03 | 1.12E-02 | -1.34E+00 | 2.05E-03 | 7.81E-03 | -1.22E+00 | 1.92E-01 | 6.84E-01 | 1.28E+00 | 1.41E-01 | 3.97E-01 | 1.15E+00 | 1.78E-01 | 6.54E-01 | 1.10E+00 | 2.78E-01 | 7.65E-01 | 1.19E+00 |
| DHRS4-AS1 | 1.54E-06 | 8.01E-06 | -1.45E+00 | 9.23E-03 | 3.99E-02 | 1.18E+00 | 8.26E-13 | 3.51E-12 | -1.94E+00 | 2.66E-08 | 1.02E-07 | -1.28E+00 | 2.86E-03 | 1.02E-02 | -1.25E+00 | 2.76E-01 | 7.79E-01 | 1.33E+00 | 9.98E-07 | 3.66E-06 | -1.36E+00 | 6.71E-04 | 1.84E-03 | -1.05E+00 |
| POLG2 | 1.54E-06 | 8.02E-06 | -1.30E+00 | 1.41E-04 | 6.09E-04 | 1.01E+00 | 3.37E-07 | 1.43E-06 | -1.40E+00 | 4.58E-04 | 1.75E-03 | 1.02E+00 | 3.62E-02 | 1.29E-01 | -1.02E+00 | 2.25E-01 | 6.36E-01 | 1.16E+00 | 6.25E-05 | 2.29E-04 | -1.26E+00 | 3.90E-02 | 1.07E-01 | 1.03E+00 |
| CYB5R4 | 1.55E-06 | 8.07E-06 | -1.13E+00 | 1.14E-05 | 4.94E-05 | -1.01E+00 | 1.89E-05 | 8.03E-05 | -1.23E+00 | 1.66E-05 | 6.33E-05 | -1.09E+00 | 6.61E-05 | 2.36E-04 | -1.03E+00 | 5.53E-05 | 1.56E-04 | -1.05E+00 | 2.78E-05 | 1.02E-04 | -1.18E+00 | 6.60E-07 | 1.81E-06 | -1.28E+00 |
| DCUN1D4 | 1.55E-06 | 8.07E-06 | -1.31E+00 | 5.93E-03 | 2.56E-02 | 1.20E+00 | 4.08E-08 | 1.73E-07 | -1.63E+00 | 1.68E-04 | 6.42E-04 | -1.06E+00 | 2.25E-03 | 8.03E-03 | -1.32E+00 | 1.17E-01 | 3.29E-01 | 1.10E+00 | 1.83E-03 | 6.73E-03 | -1.12E+00 | 3.06E-01 | 8.41E-01 | 1.51E+00 |
| HMBOX1 | 1.55E-06 | 8.09E-06 | -1.15E+00 | 7.03E-03 | 3.04E-02 | 1.22E+00 | 2.00E-06 | 8.48E-06 | -1.33E+00 | 5.65E-03 | 2.16E-02 | 1.06E+00 | 1.49E-06 | 5.30E-06 | -1.18E+00 | 1.23E-03 | 3.48E-03 | 1.09E+00 | 1.25E-15 | 4.57E-15 | -1.78E+00 | 1.30E-07 | 3.57E-07 | -1.32E+00 |
| ZNF626 | 1.55E-06 | 8.10E-06 | -1.71E+00 | 1.03E-04 | 4.43E-04 | -1.24E+00 | 2.05E-04 | 8.73E-04 | -1.45E+00 | 1.47E-04 | 5.62E-04 | -1.23E+00 | 8.23E-06 | 2.93E-05 | -1.98E+00 | 1.72E-03 | 4.85E-03 | -1.25E+00 | 4.80E-04 | 1.76E-03 | -1.38E+00 | 2.80E-03 | 7.69E-03 | -1.36E+00 |
| EFHC2 | 1.56E-06 | 8.12E-06 | -1.67E+00 | 9.87E-06 | 4.27E-05 | -1.37E+00 | 1.31E-07 | 5.58E-07 | -1.80E+00 | 1.32E-06 | 5.05E-06 | -1.48E+00 | 2.14E-13 | 7.64E-13 | -1.95E+00 | 2.13E-14 | 6.00E-14 | -2.05E+00 | 4.00E-05 | 1.47E-04 | -1.19E+00 | 1.36E-04 | 3.73E-04 | -1.25E+00 |
| CD33 | 1.56E-06 | 8.12E-06 | -1.73E+00 | 1.78E-08 | 7.71E-08 | -1.82E+00 | 1.11E-01 | 4.72E-01 | 1.22E+00 | 8.44E-05 | 3.23E-04 | -1.03E+00 | 3.24E-17 | 1.16E-16 | -2.35E+00 | 1.39E-19 | 3.92E-19 | -2.38E+00 | 2.28E-14 | 8.38E-14 | -2.54E+00 | 1.08E-14 | 2.96E-14 | -2.52E+00 |
| SFXN5 | 1.57E-06 | 8.17E-06 | -1.26E+00 | 1.44E-02 | 6.22E-02 | 1.30E+00 | 7.40E-04 | 3.15E-03 | -1.19E+00 | 1.90E-02 | 7.24E-02 | 1.13E+00 | 5.27E-02 | 1.88E-01 | -1.05E+00 | 2.49E-01 | 7.01E-01 | 1.23E+00 | 6.46E-02 | 2.37E-01 | 1.03E+00 | 2.34E-01 | 6.44E-01 | 1.16E+00 |
| ZMAT3 | 1.58E-06 | 8.21E-06 | -2.27E+00 | 3.01E-05 | 1.30E-04 | -1.74E+00 | 2.47E-07 | 1.05E-06 | -2.00E+00 | 5.61E-06 | 2.14E-05 | -1.49E+00 | 3.86E-05 | 1.38E-04 | -1.62E+00 | 3.49E-02 | 9.83E-02 | 1.04E+00 | 1.20E-10 | 4.40E-10 | -1.56E+00 | 3.08E-09 | 8.46E-09 | -1.53E+00 |

| | | | | | | | | | | | | | | | | | | | | | | | | |
|---|---|---|---|---|---|---|---|---|---|---|---|---|---|---|---|---|---|---|---|---|---|---|---|---|
| RPL7AP53 | 1.58E-06 | 8.23E-06 | -1.34E+00 | 4.60E-12 | 1.99E-11 | -2.40E+00 | 1.62E-01 | 6.90E-01 | 1.34E+00 | 1.18E-02 | 4.51E-02 | -1.12E+00 | 6.76E-02 | 2.41E-01 | 1.00E+00 | 3.16E-03 | 8.92E-03 | -1.65E+00 | 6.96E-04 | 2.55E-03 | -1.30E+00 | 3.79E-04 | 1.04E-03 | -1.40E+00 |
| NELFCD | 1.58E-06 | 8.24E-06 | -1.09E+00 | 6.06E-03 | 2.62E-02 | 1.21E+00 | 3.19E-03 | 1.35E-02 | -1.10E+00 | 6.72E-03 | 2.57E-02 | 1.07E+00 | 5.12E-05 | 1.82E-04 | -1.13E+00 | 1.92E-02 | 5.42E-02 | 1.18E+00 | 4.60E-02 | 1.69E-01 | -1.03E+00 | 6.13E-02 | 1.68E-01 | -1.01E+00 |
| ATL2 | 1.58E-06 | 8.24E-06 | -1.17E+00 | 1.90E-08 | 8.19E-08 | -1.26E+00 | 1.36E-04 | 5.76E-04 | -1.17E+00 | 1.24E-06 | 4.74E-06 | -1.18E+00 | 5.37E-02 | 1.92E-01 | 1.13E+00 | 8.49E-04 | 2.39E-03 | -1.19E+00 | 6.18E-06 | 2.27E-05 | -1.28E+00 | 3.58E-06 | 9.83E-06 | -1.37E+00 |
| GRPEL1 | 1.58E-06 | 8.26E-06 | -1.45E+00 | 2.71E-04 | 1.17E-03 | -1.06E+00 | 4.68E-10 | 1.99E-09 | -1.46E+00 | 4.66E-08 | 1.78E-07 | -1.20E+00 | 3.21E-09 | 1.14E-08 | -1.74E+00 | 8.92E-04 | 2.51E-03 | -1.04E+00 | 2.24E-07 | 8.24E-07 | -1.29E+00 | 1.41E-02 | 3.87E-02 | 1.02E+00 |
| SLC31A1 | 1.59E-06 | 8.27E-06 | -1.75E+00 | 2.08E-07 | 8.98E-07 | -1.97E+00 | 2.46E-04 | 1.05E-03 | -1.20E+00 | 1.27E-05 | 4.85E-05 | -1.20E+00 | 5.07E-03 | 1.81E-02 | -1.35E+00 | 1.37E-03 | 3.87E-03 | -1.50E+00 | 4.00E-09 | 1.47E-08 | -1.56E+00 | 1.98E-06 | 5.43E-06 | -1.33E+00 |
| RIT1 | 1.59E-06 | 8.29E-06 | -1.17E+00 | 1.02E-11 | 4.39E-11 | -1.36E+00 | 4.04E-03 | 1.72E-02 | -1.04E+00 | 2.68E-07 | 1.02E-06 | -1.25E+00 | 5.32E-05 | 1.90E-04 | -1.22E+00 | 5.37E-07 | 1.51E-06 | -1.45E+00 | 7.05E-05 | 2.59E-04 | -1.14E+00 | 1.83E-05 | 5.04E-05 | -1.21E+00 |
| CEPT1 | 1.60E-06 | 8.32E-06 | -1.33E+00 | 1.54E-03 | 6.67E-03 | 1.09E+00 | 3.37E-03 | 1.43E-02 | -1.13E+00 | 2.19E-02 | 8.38E-02 | 1.10E+00 | 1.59E-03 | 5.66E-03 | -1.15E+00 | 2.03E-03 | 5.71E-03 | -1.23E+00 | 7.83E-02 | 2.87E-01 | -1.06E+00 | 2.18E-01 | 5.99E-01 | 1.04E+00 |
| COPS5 | 1.60E-06 | 8.33E-06 | -1.05E+00 | 3.75E-03 | 1.62E-02 | 1.20E+00 | 3.76E-04 | 1.60E-03 | -1.23E+00 | 6.85E-03 | 2.62E-02 | 1.00E+00 | 8.07E-10 | 2.88E-09 | -1.36E+00 | 2.44E-04 | 6.87E-04 | -1.09E+00 | 6.69E-03 | 2.46E-02 | -1.05E+00 | 1.19E-01 | 3.27E-01 | 1.08E+00 |
| AP003352.1 | 1.60E-06 | 8.33E-06 | -1.60E+00 | 1.18E-05 | 5.08E-05 | -1.23E+00 | 1.85E-03 | 7.84E-03 | -1.41E+00 | 5.69E-02 | 2.17E-01 | 1.10E+00 | 1.38E-04 | 4.93E-04 | -1.67E+00 | 2.21E-02 | 6.24E-02 | -1.10E+00 | 2.41E-02 | 8.85E-02 | -1.13E+00 | 3.41E-01 | 9.36E-01 | 1.34E+00 |
| L3MBTL4-AS1 | 1.60E-06 | 8.36E-06 | -1.54E+00 | 5.57E-04 | 2.40E-03 | -1.06E+00 | 1.69E-03 | 7.17E-03 | -1.21E+00 | 8.46E-03 | 3.23E-02 | -1.03E+00 | 2.54E-01 | 9.06E-01 | 1.22E+00 | 2.59E-01 | 7.31E-01 | 1.14E+00 | 4.41E-02 | 1.62E-01 | -1.19E+00 | 3.37E-01 | 9.25E-01 | 1.38E+00 |
| BTBD7 | 1.61E-06 | 8.39E-06 | -1.19E+00 | 3.95E-05 | 1.71E-04 | 1.03E+00 | 6.19E-09 | 2.63E-08 | -1.50E+00 | 5.49E-07 | 2.10E-06 | -1.21E+00 | 1.60E-04 | 5.71E-04 | -1.19E+00 | 5.11E-06 | 1.44E-05 | -1.36E+00 | 9.64E-07 | 3.54E-06 | -1.46E+00 | 6.86E-04 | 1.88E-03 | -1.22E+00 |
| LYPLA2 | 1.62E-06 | 8.43E-06 | -1.33E+00 | 1.03E-03 | 4.45E-03 | 1.12E+00 | 3.47E-06 | 1.47E-05 | -1.34E+00 | 1.37E-03 | 5.22E-03 | 1.03E+00 | 2.08E-12 | 7.43E-12 | -1.90E+00 | 1.12E-07 | 3.16E-07 | -1.29E+00 | 6.89E-08 | 2.53E-07 | -1.22E+00 | 7.66E-04 | 2.10E-03 | -1.03E+00 |
| MRPL49 | 1.62E-06 | 8.45E-06 | -1.22E+00 | 5.31E-06 | 2.30E-05 | -1.08E+00 | 1.26E-03 | 5.35E-03 | -1.09E+00 | 4.57E-03 | 1.74E-02 | 1.09E+00 | 2.28E-06 | 8.13E-06 | -1.39E+00 | 2.21E-03 | 6.24E-03 | -1.02E+00 | 1.61E-03 | 5.92E-03 | -1.05E+00 | 2.68E-01 | 7.36E-01 | 1.14E+00 |
| STAG3L2 | 1.62E-06 | 8.45E-06 | -1.10E+00 | 4.07E-03 | 1.76E-02 | 1.26E+00 | 1.88E-02 | 8.00E-02 | 1.06E+00 | 1.93E-01 | 7.36E-01 | 1.35E+00 | 1.92E-01 | 6.85E-01 | 1.39E+00 | 3.52E-01 | 9.92E-01 | 1.49E+00 | 1.22E-01 | 4.48E-01 | 1.52E+00 | 1.24E-02 | 3.41E-02 | 1.74E+00 |
| IKZF3 | 1.62E-06 | 8.46E-06 | 7.18E+00 | 2.10E-05 | 9.09E-05 | 7.77E+00 | 1.35E-02 | 5.73E-02 | 2.45E+00 | 8.14E-04 | 3.11E-03 | 3.72E+00 | 2.02E-04 | 7.21E-04 | 4.80E+00 | 2.48E-07 | 6.99E-07 | 7.23E+00 | 1.34E-05 | 4.90E-05 | 3.19E+00 | 1.63E-12 | 4.48E-12 | 4.81E+00 |
| OXR1 | 1.63E-06 | 8.47E-06 | -1.15E+00 | 9.78E-04 | 4.22E-03 | 1.13E+00 | 2.84E-07 | 1.21E-06 | -1.38E+00 | 4.70E-04 | 1.79E-03 | -1.07E+00 | 2.03E-09 | 7.25E-09 | -1.67E+00 | 1.21E-04 | 3.42E-04 | -1.15E+00 | 1.10E-08 | 4.04E-08 | -1.45E+00 | 3.62E-06 | 9.94E-06 | -1.24E+00 |
| BRAP | 1.63E-06 | 8.50E-06 | -1.40E+00 | 4.04E-05 | 1.74E-04 | -1.16E+00 | 1.40E-04 | 5.96E-04 | -1.24E+00 | 7.63E-04 | 2.92E-03 | -1.05E+00 | 2.24E-02 | 8.00E-02 | -1.02E+00 | 8.53E-02 | 2.41E-01 | 1.17E+00 | 4.52E-08 | 1.66E-07 | -1.42E+00 | 3.76E-04 | 1.03E-03 | -1.08E+00 |
| MIXL1 | 1.63E-06 | 8.51E-06 | -1.74E+00 | 4.44E-07 | 1.92E-06 | -1.73E+00 | 5.24E-03 | 2.23E-02 | -1.57E+00 | 5.87E-02 | 2.24E-01 | -1.03E+00 | 4.65E-03 | 1.66E-02 | -2.88E+00 | 1.99E-02 | 5.61E-02 | -2.11E+00 | 1.35E-01 | 4.95E-01 | 1.44E+00 | 2.60E-01 | 7.13E-01 | 1.21E+00 |
| NPM3 | 1.63E-06 | 8.51E-06 | -1.29E+00 | 1.24E-03 | 5.35E-03 | 1.07E+00 | 3.51E-02 | 1.49E-01 | 1.06E+00 | 1.20E-02 | 4.59E-02 | 1.13E+00 | 1.55E-03 | 5.54E-03 | -1.08E+00 | 5.98E-02 | 1.68E-01 | 1.23E+00 | 2.00E-05 | 7.33E-05 | -1.11E+00 | 1.26E-01 | 3.45E-01 | 1.18E+00 |
| KIAA0040 | 1.64E-06 | 8.53E-06 | -1.53E+00 | 1.04E-05 | 4.48E-05 | -1.28E+00 | 1.32E-08 | 5.62E-08 | -1.69E+00 | 4.34E-07 | 1.66E-06 | -1.42E+00 | 6.49E-11 | 2.31E-10 | -2.07E+00 | 7.27E-06 | 2.05E-05 | -1.30E+00 | 2.82E-22 | 1.03E-21 | -2.29E+00 | 1.39E-09 | 3.81E-09 | -1.35E+00 |
| TESMIN | 1.64E-06 | 8.53E-06 | -1.11E+00 | 8.09E-06 | 3.50E-05 | 1.08E+00 | 3.42E-10 | 1.45E-09 | -1.46E+00 | 5.81E-13 | 2.22E-12 | -1.37E+00 | 4.38E-06 | 1.56E-05 | -1.27E+00 | 2.37E-05 | 6.69E-05 | -1.19E+00 | 1.82E-04 | 6.69E-04 | -1.22E+00 | 2.34E-02 | 6.44E-02 | -1.09E+00 |
| APTX | 1.64E-06 | 8.56E-06 | -1.27E+00 | 3.90E-04 | 1.68E-03 | 1.09E+00 | 5.30E-04 | 2.25E-03 | -1.14E+00 | 1.10E-04 | 4.22E-04 | -1.03E+00 | 6.38E-05 | 2.28E-04 | -1.12E+00 | 1.59E-03 | 4.48E-03 | -1.02E+00 | 3.18E-03 | 1.17E-02 | 1.03E+00 | 1.21E-02 | 3.31E-02 | 1.05E+00 |
| KRT18P10 | 1.65E-06 | 8.60E-06 | 3.66E+00 | 2.40E-03 | 1.04E-02 | 2.94E+00 | 2.98E-03 | 1.27E-02 | 2.22E+00 | 4.25E-02 | 1.62E-01 | 2.00E+00 | 1.27E-01 | 4.54E-01 | 1.96E+00 | 3.02E-01 | 8.52E-01 | 1.36E+00 | 2.50E-01 | 9.18E-01 | 1.21E+00 | 3.22E-01 | 8.84E-01 | 1.21E+00 |
| METAP1 | 1.67E-06 | 8.68E-06 | -1.44E+00 | 4.81E-03 | 2.08E-02 | 1.07E+00 | 6.70E-09 | 2.85E-08 | -1.47E+00 | 1.25E-04 | 4.76E-04 | 1.01E+00 | 2.52E-07 | 8.99E-07 | -1.72E+00 | 1.29E-03 | 3.64E-03 | -1.07E+00 | 1.15E-18 | 4.21E-18 | -1.86E+00 | 1.06E-05 | 2.91E-05 | -1.09E+00 |
| SLC29A1 | 1.67E-06 | 8.69E-06 | -1.88E+00 | 2.69E-03 | 1.16E-02 | -1.01E+00 | 1.22E-07 | 5.20E-07 | -1.61E+00 | 8.55E-03 | 3.27E-02 | 1.03E+00 | 1.18E-06 | 4.22E-06 | -2.40E+00 | 3.68E-05 | 1.04E-04 | -1.70E+00 | 1.62E-03 | 5.96E-03 | -1.14E+00 | 2.50E-01 | 6.88E-01 | 1.37E+00 |
| ISCA1P1 | 1.67E-06 | 8.69E-06 | -1.68E+00 | 1.13E-05 | 4.89E-05 | -1.43E+00 | 7.79E-05 | 3.31E-04 | -1.47E+00 | 1.88E-06 | 7.19E-06 | -1.69E+00 | 1.45E-04 | 5.17E-04 | -1.49E+00 | 2.39E-04 | 6.74E-04 | -1.37E+00 | 6.15E-09 | 2.26E-08 | -1.45E+00 | 1.54E-06 | 4.22E-06 | -1.49E+00 |
| ACSL4 | 1.67E-06 | 8.73E-06 | -1.43E+00 | 3.12E-04 | 1.35E-03 | 1.02E+00 | 4.38E-09 | 1.86E-08 | -1.62E+00 | 2.23E-05 | 8.50E-05 | -1.12E+00 | 8.86E-37 | 3.16E-36 | -3.12E+00 | 3.17E-18 | 8.95E-18 | -1.91E+00 | 6.09E-25 | 2.23E-24 | -2.19E+00 | 4.07E-11 | 1.12E-10 | -1.47E+00 |
| MEGF9 | 1.68E-06 | 8.73E-06 | -1.46E+00 | 1.51E-04 | 6.51E-04 | -1.10E+00 | 8.02E-04 | 3.41E-03 | -1.33E+00 | 4.41E-06 | 1.69E-05 | -1.52E+00 | 6.15E-04 | 2.19E-03 | -1.49E+00 | 6.34E-04 | 1.79E-03 | -1.41E+00 | 4.83E-03 | 1.77E-02 | -1.33E+00 | 6.11E-02 | 1.68E-01 | -1.15E+00 |
| GNPNAT1 | 1.68E-06 | 8.78E-06 | -1.47E+00 | 1.79E-06 | 7.72E-06 | -1.39E+00 | 1.30E-07 | 5.54E-07 | -1.47E+00 | 6.09E-08 | 2.33E-07 | -1.31E+00 | 1.11E-02 | 3.96E-02 | -1.35E+00 | 5.55E-02 | 1.56E-01 | -1.04E+00 | 8.38E-05 | 3.08E-04 | -1.22E+00 | 2.96E-02 | 8.13E-02 | 1.06E+00 |
| GALNT2 | 1.68E-06 | 8.78E-06 | 2.11E+00 | 6.99E-04 | 3.02E-03 | 2.10E+00 | 2.15E-01 | 9.15E-01 | 1.35E+00 | 2.49E-01 | 9.51E-01 | 1.49E+00 | 7.41E-02 | 2.64E-01 | 1.72E+00 | 6.71E-03 | 1.89E-02 | 1.94E+00 | 1.56E-01 | 5.71E-01 | 1.19E+00 | 1.91E-02 | 5.24E-02 | 1.08E+00 |
| CEP78 | 1.70E-06 | 8.86E-06 | -1.41E+00 | 1.63E-04 | 7.04E-04 | -1.08E+00 | 2.46E-05 | 1.05E-04 | -1.32E+00 | 7.21E-04 | 2.75E-03 | 1.01E+00 | 1.45E-05 | 5.15E-05 | -1.84E+00 | 1.95E-03 | 5.49E-03 | -1.27E+00 | 6.49E-07 | 2.38E-06 | -1.45E+00 | 8.08E-03 | 2.22E-02 | -1.10E+00 |
| ABHD18 | 1.70E-06 | 8.88E-06 | -1.41E+00 | 2.60E-04 | 1.12E-03 | -1.00E+00 | 8.07E-04 | 3.43E-03 | -1.22E+00 | 1.49E-01 | 5.68E-01 | 1.32E+00 | 6.53E-06 | 2.33E-05 | -1.35E+00 | 2.05E-04 | 5.77E-04 | -1.21E+00 | 3.52E-02 | 1.29E-01 | 1.07E+00 | 3.53E-01 | 9.69E-01 | 1.42E+00 |
| NDUFAF6 | 1.71E-06 | 8.94E-06 | -1.16E+00 | 6.46E-02 | 2.79E-01 | 1.42E+00 | 1.91E-03 | 8.10E-03 | -1.11E+00 | 2.46E-01 | 9.41E-01 | 1.31E+00 | 9.51E-04 | 3.39E-03 | -1.07E+00 | 1.57E-01 | 4.42E-01 | 1.33E+00 | 6.12E-04 | 2.25E-03 | -1.10E+00 | 1.88E-01 | 5.16E-01 | 1.18E+00 |
| THUMPD2 | 1.72E-06 | 8.96E-06 | -1.22E+00 | 3.01E-05 | 1.30E-04 | 1.05E+00 | 1.07E-03 | 4.56E-03 | -1.26E+00 | 1.42E-05 | 5.44E-05 | -1.24E+00 | 3.22E-03 | 1.15E-02 | -1.02E+00 | 3.76E-03 | 1.06E-02 | -1.01E+00 | 3.30E-03 | 1.21E-02 | 1.11E+00 | 3.82E-03 | 1.05E-02 | -1.15E+00 |
| ERP27 | 1.72E-06 | 8.96E-06 | -1.82E+00 | 3.26E-05 | 1.41E-04 | -1.41E+00 | 1.26E-03 | 5.37E-03 | -1.59E+00 | 2.61E-02 | 9.97E-02 | -1.06E+00 | 2.20E-01 | 7.83E-01 | 1.09E+00 | 2.66E-01 | 7.51E-01 | 1.55E+00 | 5.14E-03 | 1.89E-02 | -2.02E+00 | 1.07E-01 | 2.93E-01 | -1.27E+00 |
| CLN3 | 1.73E-06 | 9.00E-06 | -1.75E+00 | 2.63E-06 | 1.14E-05 | -1.53E+00 | 6.44E-09 | 2.74E-08 | -2.12E+00 | 5.52E-08 | 2.11E-07 | -1.72E+00 | 3.87E-05 | 1.38E-04 | -1.71E+00 | 6.51E-04 | 1.84E-03 | -1.34E+00 | 6.60E-07 | 2.42E-06 | -1.85E+00 | 5.03E-04 | 1.38E-03 | -1.33E+00 |
| NOP2 | 1.73E-06 | 9.01E-06 | -1.77E+00 | 1.69E-03 | 7.28E-03 | -1.01E+00 | 1.37E-11 | 5.82E-11 | -1.89E+00 | 3.11E-08 | 1.19E-07 | -1.35E+00 | 2.69E-03 | 9.60E-03 | -1.38E+00 | 4.39E-02 | 1.24E-01 | -1.08E+00 | 5.71E-07 | 2.10E-06 | -1.38E+00 | 5.88E-02 | 1.61E-01 | 1.09E+00 |
| RPS27L | 1.73E-06 | 9.02E-06 | -1.04E+00 | 1.85E-08 | 8.00E-08 | 1.04E+00 | 5.45E-03 | 2.32E-02 | -1.20E+00 | 1.21E-05 | 4.61E-05 | -1.23E+00 | 1.13E-04 | 4.02E-04 | -1.07E+00 | 8.69E-03 | 2.45E-02 | 1.01E+00 | 3.29E-02 | 1.21E-01 | -1.09E+00 | 8.13E-04 | 2.23E-03 | -1.35E+00 |
| CEP89 | 1.73E-06 | 9.02E-06 | -1.67E+00 | 7.78E-04 | 3.36E-03 | -1.06E+00 | 1.89E-05 | 8.03E-05 | -1.42E+00 | 5.33E-05 | 2.04E-04 | -1.21E+00 | 1.77E-03 | 6.30E-03 | -1.27E+00 | 4.70E-03 | 1.33E-02 | -1.20E+00 | 5.63E-05 | 2.07E-04 | -1.28E+00 | 4.61E-05 | 1.27E-04 | -1.44E+00 |
| N4BP2L2 | 1.74E-06 | 9.06E-06 | -1.13E+00 | 1.33E-01 | 5.73E-01 | 1.26E+00 | 1.43E-05 | 6.08E-05 | -1.26E+00 | 2.58E-01 | 9.85E-01 | 1.14E+00 | 7.32E-05 | 2.61E-04 | -1.10E+00 | 2.53E-01 | 7.15E-01 | 1.16E+00 | 7.45E-04 | 2.73E-03 | -1.10E+00 | 3.01E-01 | 8.26E-01 | 1.10E+00 |
| PTPRC | 1.74E-06 | 9.08E-06 | -1.22E+00 | 7.10E-03 | 3.07E-02 | 1.05E+00 | 4.21E-03 | 1.79E-02 | -1.16E+00 | 7.17E-02 | 2.74E-01 | 1.00E+00 | 3.23E-04 | 1.15E-03 | -1.04E+00 | 4.85E-06 | 1.37E-05 | -1.14E+00 | 8.91E-05 | 3.27E-04 | -1.12E+00 | 2.84E-16 | 7.80E-16 | -1.56E+00 |
| EML4 | 1.74E-06 | 9.08E-06 | -1.34E+00 | 6.96E-05 | 3.01E-04 | -1.02E+00 | 1.03E-04 | 4.37E-04 | -1.27E+00 | 3.22E-05 | 1.23E-04 | -1.10E+00 | 9.59E-05 | 3.42E-04 | -1.18E+00 | 7.06E-06 | 1.99E-05 | -1.37E+00 | 2.04E-09 | 7.49E-09 | -1.45E+00 | 9.98E-05 | 2.74E-04 | -1.14E+00 |
| PCF11 | 1.74E-06 | 9.08E-06 | -1.07E+00 | 1.74E-04 | 7.52E-04 | 1.10E+00 | 4.31E-04 | 1.83E-03 | -1.12E+00 | 1.11E-04 | 4.26E-04 | -1.09E+00 | 1.37E-02 | 4.89E-02 | -1.01E+00 | 3.33E-02 | 9.39E-02 | 1.04E+00 | 1.46E-05 | 5.34E-05 | -1.15E+00 | 1.96E-02 | 5.38E-02 | -1.01E+00 |
| TMEM170B | 1.75E-06 | 9.10E-06 | -1.70E+00 | 1.98E-06 | 8.55E-06 | -1.65E+00 | 1.14E-01 | 4.83E-01 | 1.08E+00 | 5.39E-03 | 2.06E-02 | -1.05E+00 | 5.98E-03 | 2.13E-02 | -1.52E+00 | 1.77E-03 | 4.99E-03 | -1.84E+00 | 1.10E-03 | 4.04E-03 | -1.54E+00 | 5.33E-03 | 1.46E-02 | -1.80E+00 |
| IKZF4 | 1.75E-06 | 9.14E-06 | -1.85E+00 | 1.41E-03 | 6.08E-03 | -1.18E+00 | 5.89E-06 | 2.50E-05 | -1.74E+00 | 6.34E-05 | 2.42E-04 | -1.39E+00 | 3.29E-05 | 1.17E-04 | -2.52E+00 | 4.08E-03 | 1.15E-02 | -1.39E+00 | 6.25E-08 | 2.30E-07 | -1.82E+00 | 1.56E-05 | 4.29E-05 | -1.44E+00 |
| TMEM97 | 1.76E-06 | 9.16E-06 | -1.26E+00 | 3.50E-03 | 1.51E-02 | 1.13E+00 | 1.52E-03 | 6.46E-03 | -1.01E+00 | 1.36E-02 | 5.18E-02 | 1.28E+00 | 1.54E-07 | 5.51E-07 | -1.43E+00 | 7.69E-05 | 2.17E-04 | -1.06E+00 | 1.58E-07 | 5.79E-07 | -1.40E+00 | 2.90E-01 | 7.97E-01 | 1.31E+00 |
| AF131215.4 | 1.76E-06 | 9.17E-06 | -1.54E+00 | 7.76E-05 | 3.35E-04 | -1.14E+00 | 8.55E-07 | 3.63E-06 | -1.99E+00 | 1.20E-04 | 4.57E-04 | -1.38E+00 | 3.69E-03 | 1.31E-02 | -1.56E+00 | 2.38E-01 | 6.71E-01 | 1.20E+00 | 5.29E-05 | 1.94E-04 | -1.83E+00 | 2.38E-02 | 6.54E-02 | -1.19E+00 |
| PQLC3 | 1.76E-06 | 9.17E-06 | -1.31E+00 | 7.71E-04 | 3.33E-03 | 1.02E+00 | 3.68E-07 | 1.57E-06 | -1.40E+00 | 1.36E-05 | 5.19E-05 | -1.13E+00 | 1.41E-05 | 5.02E-05 | -1.17E+00 | 5.95E-05 | 1.68E-04 | -1.08E+00 | 1.07E-01 | 3.91E-01 | 1.09E+00 | 4.83E-02 | 1.33E-01 | -1.00E+00 |
| CCS | 1.76E-06 | 9.19E-06 | -1.10E+00 | 1.69E-02 | 7.32E-02 | 1.27E+00 | 1.72E-03 | 7.29E-03 | -1.11E+00 | 5.91E-03 | 2.26E-02 | 1.14E+00 | 6.09E-04 | 2.17E-03 | 1.04E+00 | 6.79E-02 | 1.91E-01 | 1.24E+00 | 2.17E-01 | 7.97E-01 | 1.36E+00 | 3.21E-01 | 8.82E-01 | 1.31E+00 |
| AP2S1 | 1.77E-06 | 9.21E-06 | 1.01E+00 | 2.38E-06 | 1.03E-05 | 1.13E+00 | 1.65E-01 | 7.02E-01 | 1.12E+00 | 1.67E-02 | 6.37E-02 | 1.09E+00 | 1.71E-06 | 6.11E-06 | -1.12E+00 | 6.20E-04 | 1.75E-03 | 1.02E+00 | 7.51E-02 | 2.76E-01 | 1.06E+00 | 9.22E-02 | 2.53E-01 | 1.04E+00 |
| RPUSD1 | 1.77E-06 | 9.21E-06 | -1.45E+00 | 7.58E-04 | 3.28E-03 | -1.03E+00 | 1.96E-02 | 8.33E-02 | -1.05E+00 | 9.68E-02 | 3.70E-01 | 1.16E+00 | 1.45E-03 | 5.17E-03 | -1.15E+00 | 6.45E-02 | 1.82E-01 | 1.16E+00 | 9.79E-06 | 3.59E-05 | -1.20E+00 | 2.63E-02 | 7.21E-02 | 1.10E+00 |
| JMJD8 | 1.77E-06 | 9.22E-06 | -2.06E+00 | 1.02E-05 | 4.42E-05 | -1.71E+00 | 4.40E-02 | 1.87E-01 | -1.11E+00 | 1.73E-02 | 6.62E-02 | -1.10E+00 | 9.54E-03 | 3.40E-02 | -1.69E+00 | 4.46E-02 | 1.26E-01 | -1.26E+00 | 8.24E-03 | 3.02E-02 | -1.14E+00 | 2.89E-02 | 7.93E-02 | -1.05E+00 |
| PLXNC1 | 1.78E-06 | 9.26E-06 | -1.41E+00 | 4.40E-04 | 1.90E-03 | -1.02E+00 | 9.78E-07 | 4.16E-06 | -1.48E+00 | 6.53E-05 | 2.49E-04 | -1.11E+00 | 5.92E-16 | 2.11E-15 | -2.07E+00 | 6.43E-05 | 1.81E-04 | -1.19E+00 | 3.37E-32 | 1.24E-31 | -2.89E+00 | 3.44E-05 | 9.44E-05 | -1.28E+00 |
| METTL1 | 1.79E-06 | 9.31E-06 | -1.79E+00 | 2.84E-05 | 1.23E-04 | -1.38E+00 | 1.52E-03 | 6.44E-03 | -1.23E+00 | 2.22E-02 | 8.47E-02 | 1.06E+00 | 1.70E-03 | 6.07E-03 | -1.67E+00 | 1.45E-01 | 4.09E-01 | 1.05E+00 | 1.68E-06 | 6.17E-06 | -1.31E+00 | 8.13E-02 | 2.23E-01 | 1.13E+00 |
| EXOC6 | 1.79E-06 | 9.31E-06 | -1.15E+00 | 5.56E-04 | 2.40E-03 | 1.14E+00 | 3.84E-03 | 1.63E-02 | -1.01E+00 | 4.47E-02 | 1.71E-01 | 1.26E+00 | 6.62E-04 | 2.36E-03 | -1.20E+00 | 1.08E-02 | 3.04E-02 | 1.03E+00 | 7.91E-06 | 2.90E-05 | -1.16E+00 | 9.88E-02 | 2.71E-01 | 1.11E+00 |
| RPS2P4 | 1.79E-06 | 9.33E-06 | 1.34E+00 | 2.01E-01 | 8.71E-01 | 1.13E+00 | 1.04E-05 | 4.42E-05 | 1.38E+00 | 8.90E-02 | 3.40E-01 | 1.14E+00 | 7.23E-04 | 2.58E-03 | 1.43E+00 | 5.14E-02 | 1.45E-01 | 1.26E+00 | 5.94E-04 | 2.18E-03 | 1.42E+00 | 1.63E-01 | 4.48E-01 | 1.17E+00 |
| AC023794.3 | 1.79E-06 | 9.33E-06 | -1.43E+00 | 6.19E-05 | 2.67E-04 | -1.20E+00 | 5.37E-05 | 2.28E-04 | -1.57E+00 | 5.89E-05 | 2.25E-04 | -1.41E+00 | 2.02E-01 | 7.19E-01 | -1.22E+00 | 3.41E-01 | 9.60E-01 | 1.14E+00 | 8.91E-02 | 3.27E-01 | -1.30E+00 | 1.59E-01 | 4.38E-01 | -1.26E+00 |
| TIGD7 | 1.79E-06 | 9.33E-06 | -2.75E+00 | 8.13E-04 | 3.51E-03 | -1.55E+00 | 1.28E-02 | 5.43E-02 | -1.32E+00 | 4.29E-02 | 1.64E-01 | -1.12E+00 | 2.57E-03 | 9.16E-03 | -1.75E+00 | 9.83E-03 | 2.77E-02 | -1.52E+00 | 5.70E-02 | 2.09E-01 | -1.12E+00 | 2.18E-01 | 5.98E-01 | 1.04E+00 |
| SSBP3 | 1.79E-06 | 9.34E-06 | -1.14E+00 | 6.16E-03 | 2.66E-02 | 1.14E+00 | 2.05E-06 | 8.71E-06 | -1.29E+00 | 2.64E-05 | 1.01E-04 | -1.08E+00 | 2.16E-06 | 7.70E-06 | -1.27E+00 | 2.65E-04 | 7.47E-04 | -1.07E+00 | 6.97E-06 | 2.56E-05 | -1.32E+00 | 8.07E-05 | 2.22E-04 | -1.25E+00 |
| LINC01232 | 1.80E-06 | 9.39E-06 | -1.75E+00 | 1.75E-05 | 7.55E-05 | -1.45E+00 | 7.89E-06 | 3.35E-05 | -1.75E+00 | 1.16E-05 | 4.45E-05 | -1.76E+00 | 4.05E-02 | 1.44E-01 | -1.29E+00 | 4.11E-02 | 1.16E-01 | -1.34E+00 | 1.82E-06 | 6.67E-06 | -1.92E+00 | 1.66E-06 | 4.56E-06 | -2.02E+00 |
| APOOP5 | 1.81E-06 | 9.42E-06 | -2.40E+00 | 1.58E-04 | 6.84E-04 | -1.56E+00 | 8.28E-06 | 3.52E-05 | -1.73E+00 | 3.14E-06 | 1.20E-05 | -1.77E+00 | 5.38E-04 | 1.92E-03 | -1.99E+00 | 1.99E-02 | 5.61E-02 | -1.19E+00 | 7.52E-04 | 2.76E-03 | -1.46E+00 | 5.48E-03 | 1.51E-02 | -1.28E+00 |
| NSRP1 | 1.81E-06 | 9.44E-06 | -1.08E+00 | 4.67E-05 | 2.02E-04 | 1.06E+00 | 2.43E-03 | 1.03E-02 | -1.14E+00 | 1.22E-02 | 4.65E-02 | 1.04E+00 | 2.84E-04 | 1.01E-03 | -1.07E+00 | 2.02E-02 | 5.68E-02 | 1.04E+00 | 1.53E-04 | 5.60E-04 | -1.06E+00 | 4.85E-04 | 1.33E-03 | -1.09E+00 |
| PES1 | 1.81E-06 | 9.44E-06 | -1.34E+00 | 5.54E-04 | 2.39E-03 | 1.02E+00 | 3.12E-05 | 1.33E-04 | -1.18E+00 | 2.24E-04 | 8.57E-04 | 1.05E+00 | 1.39E-03 | 4.95E-03 | -1.32E+00 | 4.23E-02 | 1.19E-01 | -1.00E+00 | 1.14E-06 | 4.17E-06 | -1.35E+00 | 4.66E-04 | 1.28E-03 | -1.07E+00 |
| KRT18P39 | 1.82E-06 | 9.47E-06 | 3.42E+00 | 1.61E-04 | 6.98E-04 | 3.33E+00 | 3.46E-03 | 1.47E-02 | 1.99E+00 | 1.59E-01 | 6.09E-01 | 1.55E+00 | 1.58E-01 | 5.62E-01 | 1.93E+00 | 3.44E-01 | 9.71E-01 | 1.27E+00 | 1.17E-01 | 4.30E-01 | 2.00E+00 | 1.25E-01 | 3.45E-01 | 1.87E+00 |
| LINC00116 | 1.83E-06 | 9.52E-06 | -1.34E+00 | 7.70E-06 | 3.33E-05 | -1.22E+00 | 5.18E-12 | 2.20E-11 | -1.79E+00 | 2.28E-09 | 8.72E-09 | -1.33E+00 | 4.67E-07 | 1.67E-06 | -1.70E+00 | 5.61E-04 | 1.58E-03 | -1.22E+00 | 8.51E-17 | 3.12E-16 | -1.32E+00 | 2.14E-13 | 5.88E-13 | -1.56E+00 |
| WFDC2 | 1.83E-06 | 9.53E-06 | -1.66E+00 | 3.98E-04 | 1.72E-03 | -1.17E+00 | 7.90E-03 | 3.36E-02 | -1.38E+00 | 2.26E-02 | 8.63E-02 | -1.15E+00 | 8.31E-02 | 2.96E-01 | -1.62E+00 | 1.46E-01 | 4.11E-01 | -1.31E+00 | 1.86E-01 | 6.82E-01 | -1.21E+00 | 3.38E-01 | 9.28E-01 | 1.12E+00 |
| COX7A2L | 1.83E-06 | 9.54E-06 | -1.07E+00 | 1.44E-06 | 6.24E-06 | -1.00E+00 | 5.78E-02 | 2.45E-01 | -1.04E+00 | 4.96E-02 | 1.89E-01 | 1.00E+00 | 3.96E-09 | 1.41E-08 | -1.20E+00 | 1.54E-04 | 4.35E-04 | -1.08E+00 | 1.12E-01 | 4.11E-01 | 1.04E+00 | 2.91E-01 | 7.99E-01 | 1.10E+00 |
| GCNT1 | 1.83E-06 | 9.56E-06 | -1.28E+00 | 6.89E-05 | 2.98E-04 | -1.02E+00 | 1.33E-08 | 5.67E-08 | -1.64E+00 | 4.11E-07 | 1.57E-06 | -1.28E+00 | 1.34E-06 | 4.77E-06 | -2.63E+00 | 7.94E-03 | 2.24E-02 | -1.16E+00 | 1.50E-12 | 5.50E-12 | -1.72E+00 | 7.75E-06 | 2.13E-05 | -1.26E+00 |
| NAPB | 1.84E-06 | 9.57E-06 | -1.47E+00 | 7.02E-02 | 3.03E-01 | 1.37E+00 | 2.90E-07 | 1.23E-06 | -2.01E+00 | 1.70E-02 | 6.50E-02 | -1.02E+00 | 4.90E-06 | 1.75E-05 | -1.54E+00 | 9.25E-04 | 2.61E-03 | -1.16E+00 | 1.74E-02 | 6.37E-02 | -1.06E+00 | 9.95E-02 | 2.73E-01 | 1.01E+00 |
| TPRA1 | 1.84E-06 | 9.61E-06 | -1.55E+00 | 3.47E-04 | 1.50E-03 | -1.08E+00 | 1.73E-03 | 7.33E-03 | -1.24E+00 | 1.52E-01 | 5.79E-01 | 1.29E+00 | 9.59E-08 | 3.42E-07 | -2.24E+00 | 1.08E-04 | 3.04E-04 | -1.33E+00 | 9.32E-09 | 3.42E-08 | -1.54E+00 | 1.76E-04 | 4.82E-04 | -1.25E+00 |
| PUM2 | 1.85E-06 | 9.63E-06 | -1.23E+00 | 1.30E-03 | 5.62E-03 | 1.11E+00 | 8.93E-10 | 3.80E-09 | -1.57E+00 | 4.61E-04 | 1.76E-03 | -1.01E+00 | 1.08E-03 | 3.85E-03 | -1.03E+00 | 1.88E-02 | 5.30E-02 | 1.08E+00 | 7.29E-07 | 2.68E-06 | -1.18E+00 | 1.24E-05 | 3.42E-05 | -1.12E+00 |
| NADK2 | 1.86E-06 | 9.71E-06 | -1.52E+00 | 1.44E-05 | 6.24E-05 | -1.32E+00 | 2.87E-07 | 1.22E-06 | -1.50E+00 | 5.19E-05 | 1.98E-04 | -1.07E+00 | 4.66E-05 | 1.66E-04 | -1.30E+00 | 1.59E-01 | 4.48E-01 | 1.27E+00 | 4.67E-05 | 1.72E-04 | -1.25E+00 | 1.75E-02 | 4.80E-02 | 1.02E+00 |
| RAB6A | 1.86E-06 | 9.71E-06 | -1.15E+00 | 5.04E-06 | 2.18E-05 | -1.03E+00 | 5.90E-09 | 2.50E-08 | -1.43E+00 | 4.01E-09 | 1.53E-08 | -1.35E+00 | 2.30E-05 | 8.20E-05 | -1.16E+00 | 4.57E-04 | 1.29E-03 | -1.16E+00 | 9.79E-06 | 3.59E-05 | -1.27E+00 | 2.49E-05 | 6.83E-05 | -1.23E+00 |
| OR7C1 | 1.87E-06 | 9.75E-06 | -1.55E+00 | 8.91E-06 | 3.85E-05 | -1.32E+00 | 2.75E-10 | 1.17E-09 | -2.20E+00 | 9.11E-09 | 3.48E-08 | -1.68E+00 | 3.17E-04 | 1.13E-03 | -1.83E+00 | 1.82E-02 | 5.14E-02 | -1.27E+00 | 1.97E-15 | 7.25E-15 | -2.14E+00 | 1.15E-06 | 3.16E-06 | -1.17E+00 |
| GSTO2 | 1.87E-06 | 9.76E-06 | -2.07E+00 | 1.28E-04 | 5.53E-04 | -1.42E+00 | 1.29E-01 | 5.49E-01 | 1.08E+00 | 3.35E-02 | 1.28E-01 | -1.05E+00 | 2.55E-01 | 9.10E-01 | 1.23E+00 | 3.51E-01 | 9.89E-01 | 1.29E+00 | 2.18E-01 | 8.01E-01 | 1.10E+00 | 3.06E-01 | 8.40E-01 | 1.21E+00 |
| COMMD2 | 1.88E-06 | 9.78E-06 | -1.10E+00 | 1.55E-03 | 6.70E-03 | 1.17E+00 | 1.46E-02 | 6.21E-02 | 1.02E+00 | 2.68E-02 | 1.02E-01 | 1.16E+00 | 7.99E-04 | 2.85E-03 | -1.03E+00 | 2.68E-02 | 7.56E-02 | 1.11E+00 | 1.42E-05 | 5.22E-05 | -1.12E+00 | 2.21E-04 | 6.08E-04 | -1.05E+00 |
| BPNT1 | 1.88E-06 | 9.80E-06 | -1.50E+00 | 1.27E-04 | 5.50E-04 | -1.07E+00 | 2.05E-04 | 8.71E-04 | -1.36E+00 | 3.42E-03 | 1.31E-02 | -1.05E+00 | 1.44E-08 | 5.14E-08 | -2.02E+00 | 3.51E-03 | 9.89E-03 | 1.02E+00 | 5.97E-13 | 2.19E-12 | -1.99E+00 | 3.89E-06 | 1.07E-05 | -1.31E+00 |
| HENMT1 | 1.88E-06 | 9.81E-06 | -1.07E+00 | 3.56E-05 | 1.54E-04 | 1.09E+00 | 8.91E-07 | 3.79E-06 | -1.24E+00 | 4.48E-06 | 1.71E-05 | -1.01E+00 | 1.54E-03 | 5.51E-03 | -1.12E+00 | 2.10E-02 | 5.91E-02 | 1.07E+00 | 8.10E-02 | 2.97E-01 | 1.44E+00 | 6.74E-03 | 1.85E-02 | 1.67E+00 |
| MCM9 | 1.89E-06 | 9.83E-06 | -1.38E+00 | 1.93E-06 | 8.35E-06 | -1.27E+00 | 2.02E-04 | 8.56E-04 | -1.34E+00 | 1.48E-04 | 5.67E-04 | -1.21E+00 | 2.63E-03 | 9.38E-03 | -1.16E+00 | 1.62E-03 | 4.58E-03 | -1.19E+00 | 2.99E-07 | 1.10E-06 | -1.36E+00 | 4.12E-06 | 1.13E-05 | -1.39E+00 |
| SCLY | 1.89E-06 | 9.87E-06 | -2.17E+00 | 1.03E-03 | 4.44E-03 | -1.20E+00 | 6.36E-03 | 2.70E-02 | -1.24E+00 | 2.90E-02 | 1.11E-01 | 1.06E+00 | 8.13E-02 | 2.90E-01 | -1.18E+00 | 3.08E-01 | 8.68E-01 | 1.39E+00 | 7.20E-03 | 2.64E-02 | -1.26E+00 | 2.83E-01 | 7.77E-01 | 1.26E+00 |
| CHD3 | 1.89E-06 | 9.87E-06 | -1.24E+00 | 8.08E-05 | 3.49E-04 | -1.04E+00 | 2.65E-05 | 1.13E-04 | -1.22E+00 | 1.57E-05 | 6.01E-05 | -1.14E+00 | 3.87E-04 | 1.38E-03 | -1.04E+00 | 4.62E-05 | 1.30E-04 | -1.10E+00 | 4.21E-06 | 1.55E-05 | -1.11E+00 | 2.38E-05 | 6.53E-05 | -1.14E+00 |
| AC093433.1 | 1.90E-06 | 9.90E-06 | -1.60E+00 | 8.08E-07 | 3.49E-06 | -1.48E+00 | 1.28E-05 | 5.43E-05 | -1.41E+00 | 5.68E-05 | 2.17E-04 | -1.22E+00 | 4.68E-04 | 1.67E-03 | -1.39E+00 | 1.43E-02 | 4.03E-02 | -1.06E+00 | 4.60E-07 | 1.69E-06 | -1.48E+00 | 6.10E-04 | 1.68E-03 | -1.22E+00 |
| TSPYL4 | 1.91E-06 | 9.93E-06 | -1.07E+00 | 5.29E-08 | 2.28E-07 | -1.14E+00 | 8.98E-03 | 3.81E-02 | -1.00E+00 | 6.12E-05 | 2.34E-04 | -1.12E+00 | 4.53E-02 | 1.61E-01 | 1.12E+00 | 6.32E-04 | 1.78E-03 | -1.10E+00 | 2.22E-03 | 8.14E-03 | -1.09E+00 | 3.90E-05 | 1.07E-04 | -1.33E+00 |
| MFHAS1 | 1.91E-06 | 9.94E-06 | -1.99E+00 | 1.68E-03 | 7.25E-03 | -1.16E+00 | 1.06E-10 | 4.49E-10 | -1.99E+00 | 1.51E-06 | 5.76E-06 | -1.25E+00 | 1.00E-02 | 3.58E-02 | -1.37E+00 | 6.56E-02 | 1.85E-01 | -1.09E+00 | 8.86E-11 | 3.25E-10 | -1.77E+00 | 1.41E-06 | 3.87E-06 | -1.35E+00 |
| GATA2 | 1.91E-06 | 9.95E-06 | -1.23E+00 | 1.58E-02 | 6.84E-02 | 1.09E+00 | 6.92E-08 | 2.94E-07 | -1.44E+00 | 2.08E-01 | 7.94E-01 | 1.16E+00 | 2.34E-10 | 8.35E-10 | -1.61E+00 | 2.97E-04 | 8.37E-04 | -1.21E+00 | 2.30E-10 | 8.44E-10 | -1.45E+00 | 8.58E-04 | 2.36E-03 | -1.10E+00 |
| SSNA1 | 1.91E-06 | 9.98E-06 | -1.07E+00 | 3.60E-05 | 1.56E-04 | 1.10E+00 | 4.92E-03 | 2.09E-02 | -1.02E+00 | 1.94E-02 | 7.43E-02 | 1.12E+00 | 4.88E-06 | 1.74E-05 | -1.13E+00 | 1.00E-02 | 2.83E-02 | 1.06E+00 | 4.59E-02 | 1.68E-01 | 1.10E+00 | 5.67E-02 | 1.56E-01 | 1.10E+00 |
| WASHC3 | 1.92E-06 | 1.00E-05 | -1.09E+00 | 3.70E-04 | 1.60E-03 | 1.12E+00 | 1.47E-03 | 6.23E-03 | -1.08E+00 | 3.22E-03 | 1.23E-02 | 1.04E+00 | 6.64E-05 | 2.37E-04 | -1.17E+00 | 1.46E-03 | 4.11E-03 | 1.02E+00 | 1.74E-02 | 6.39E-02 | 1.18E+00 | 1.16E-02 | 3.19E-02 | 1.09E+00 |
| LPAR2 | 1.92E-06 | 1.00E-05 | -1.29E+00 | 4.55E-06 | 1.97E-05 | -1.07E+00 | 2.43E-08 | 1.03E-07 | -1.77E+00 | 6.25E-06 | 2.39E-05 | -1.28E+00 | 4.98E-02 | 1.78E-01 | 1.01E+00 | 4.08E-02 | 1.15E-01 | -1.09E+00 | 1.48E-01 | 5.42E-01 | -1.13E+00 | 2.16E-01 | 5.92E-01 | -1.06E+00 |
| AC091563.1 | 1.93E-06 | 1.00E-05 | -2.08E+00 | 2.19E-04 | 9.48E-04 | -1.28E+00 | 3.01E-03 | 1.28E-02 | -1.56E+00 | 6.46E-02 | 2.47E-01 | 1.15E+00 | 8.98E-33 | 3.20E-32 | -4.18E+00 | 3.86E-21 | 1.09E-20 | -2.10E+00 | 1.31E-07 | 4.81E-07 | -3.12E+00 | 2.48E-04 | 6.82E-04 | -1.75E+00 |
| SRD5A3 | 1.93E-06 | 1.01E-05 | -1.28E+00 | 7.61E-03 | 3.29E-02 | 1.21E+00 | 2.21E-05 | 9.37E-05 | -1.21E+00 | 5.97E-03 | 2.28E-02 | 1.08E+00 | 1.64E-03 | 5.85E-03 | -1.09E+00 | 1.48E-01 | 4.17E-01 | 1.24E+00 | 2.52E-03 | 9.26E-03 | -1.17E+00 | 2.57E-01 | 7.05E-01 | 1.15E+00 |
| AC006001.3 | 1.93E-06 | 1.01E-05 | -1.19E+00 | 4.87E-07 | 2.11E-06 | -1.08E+00 | 1.76E-06 | 7.49E-06 | -1.43E+00 | 2.89E-05 | 1.10E-04 | -1.10E+00 | 2.57E-04 | 9.15E-04 | -1.17E+00 | 2.09E-02 | 5.89E-02 | 1.10E+00 | 1.45E-05 | 5.30E-05 | -1.23E+00 | 5.88E-03 | 1.62E-02 | -1.06E+00 |
| BAK1 | 1.96E-06 | 1.02E-05 | -1.36E+00 | 1.40E-05 | 6.03E-05 | -1.05E+00 | 7.94E-17 | 3.37E-16 | -2.22E+00 | 3.60E-13 | 1.38E-12 | -1.59E+00 | 2.19E-10 | 7.82E-10 | -1.98E+00 | 1.52E-07 | 4.30E-07 | -1.53E+00 | 4.74E-16 | 1.74E-15 | -1.59E+00 | 3.96E-09 | 1.09E-08 | -1.27E+00 |
| NR1H2 | 1.96E-06 | 1.02E-05 | -1.10E+00 | 1.07E-03 | 4.61E-03 | 1.13E+00 | 1.65E-03 | 6.99E-03 | -1.10E+00 | 1.28E-02 | 4.90E-02 | 1.15E+00 | 8.43E-03 | 3.01E-02 | 1.01E+00 | 1.19E-01 | 3.37E-01 | 1.22E+00 | 2.24E-05 | 8.23E-05 | -1.07E+00 | 6.97E-04 | 1.92E-03 | -1.09E+00 |
| UQCC3 | 1.97E-06 | 1.02E-05 | -1.25E+00 | 1.14E-04 | 4.92E-04 | 1.02E+00 | 9.89E-04 | 4.20E-03 | -1.11E+00 | 1.49E-04 | 5.68E-04 | -1.03E+00 | 3.11E-06 | 1.11E-05 | -1.46E+00 | 1.85E-03 | 5.21E-03 | -1.08E+00 | 7.77E-06 | 2.85E-05 | -1.20E+00 | 2.73E-03 | 7.50E-03 | -1.04E+00 |
| LRRFIP2 | 1.97E-06 | 1.03E-05 | -1.09E+00 | 8.53E-07 | 3.68E-06 | -1.01E+00 | 3.35E-04 | 1.43E-03 | -1.16E+00 | 1.55E-03 | 5.92E-03 | -1.05E+00 | 8.82E-02 | 3.14E-01 | 1.08E+00 | 8.30E-02 | 2.34E-01 | 1.08E+00 | 2.40E-01 | 8.82E-01 | 1.33E+00 | 6.55E-02 | 1.80E-01 | 1.02E+00 |
| AL139274.2 | 1.98E-06 | 1.03E-05 | -1.16E+00 | 7.73E-09 | 3.34E-08 | -1.24E+00 | 5.58E-02 | 2.37E-01 | -1.15E+00 | 9.32E-03 | 3.56E-02 | -1.20E+00 | 6.84E-03 | 2.44E-02 | -1.22E+00 | 3.35E-02 | 9.43E-02 | -1.02E+00 | 1.23E-01 | 4.51E-01 | -1.02E+00 | 3.91E-02 | 1.07E-01 | -1.13E+00 |
| ZNF378P | 1.98E-06 | 1.03E-05 | -1.30E+00 | 6.12E-02 | 2.64E-01 | 1.22E+00 | 1.10E-05 | 4.68E-05 | -1.36E+00 | 3.29E-02 | 1.26E-01 | 1.16E+00 | 1.34E-05 | 4.76E-05 | -1.42E+00 | 1.79E-04 | 5.04E-04 | -1.20E+00 | 6.47E-08 | 2.38E-07 | -1.28E+00 | 1.82E-06 | 5.01E-06 | -1.21E+00 |
| AC239868.2 | 1.99E-06 | 1.04E-05 | -1.49E+00 | 2.19E-05 | 9.46E-05 | 2.43E+00 | 6.40E-07 | 2.72E-06 | -2.17E+00 | 1.94E-01 | 7.43E-01 | 1.36E+00 | 3.65E-03 | 1.30E-02 | -1.44E+00 | 1.33E-02 | 3.76E-02 | 2.18E+00 | 7.64E-02 | 2.80E-01 | -1.12E+00 | 1.60E-02 | 4.41E-02 | 2.15E+00 |
| MAPRE1 | 1.99E-06 | 1.04E-05 | -1.01E+00 | 1.02E-11 | 4.43E-11 | -1.09E+00 | 5.40E-02 | 2.30E-01 | 1.07E+00 | 7.77E-03 | 2.97E-02 | 1.10E+00 | 7.68E-02 | 2.74E-01 | 1.64E+00 | 4.84E-02 | 1.37E-01 | 1.19E+00 | 3.17E-02 | 1.17E-01 | 1.45E+00 | 2.46E-01 | 6.77E-01 | 1.32E+00 |
| DNAJA3 | 2.00E-06 | 1.04E-05 | -1.40E+00 | 5.34E-02 | 2.31E-01 | 1.28E+00 | 2.26E-05 | 9.61E-05 | -1.37E+00 | 9.68E-03 | 3.70E-02 | 1.09E+00 | 6.87E-04 | 2.45E-03 | -1.43E+00 | 3.17E-01 | 8.93E-01 | 1.44E+00 | 2.57E-16 | 9.42E-16 | -1.83E+00 | 2.51E-04 | 6.90E-04 | -1.07E+00 |
| TAZ | 2.00E-06 | 1.04E-05 | -1.23E+00 | 7.55E-07 | 3.26E-06 | -1.12E+00 | 1.38E-04 | 5.88E-04 | -1.15E+00 | 3.40E-05 | 1.30E-04 | -1.09E+00 | 1.94E-02 | 6.90E-02 | -1.05E+00 | 7.55E-02 | 2.13E-01 | 1.12E+00 | 3.15E-02 | 1.16E-01 | 1.03E+00 | 5.35E-02 | 1.47E-01 | 1.04E+00 |
| ARHGEF3 | 2.01E-06 | 1.05E-05 | -1.26E+00 | 4.44E-03 | 1.92E-02 | 1.18E+00 | 2.76E-07 | 1.17E-06 | -1.60E+00 | 1.65E-06 | 6.31E-06 | -1.28E+00 | 1.12E-02 | 3.99E-02 | -1.25E+00 | 2.02E-01 | 5.68E-01 | 1.18E+00 | 5.56E-07 | 2.04E-06 | -1.55E+00 | 2.19E-04 | 6.03E-04 | -1.29E+00 |
| GNA11 | 2.01E-06 | 1.05E-05 | -1.81E+00 | 1.58E-04 | 6.81E-04 | -1.34E+00 | 3.38E-03 | 1.43E-02 | -1.37E+00 | 1.68E-02 | 6.41E-02 | -1.05E+00 | 2.10E-01 | 7.51E-01 | 1.01E+00 | 3.46E-01 | 9.75E-01 | 1.32E+00 | 6.21E-02 | 2.28E-01 | -1.14E+00 | 1.74E-01 | 4.77E-01 | 1.04E+00 |
| IFT74 | 2.01E-06 | 1.05E-05 | -1.07E+00 | 5.70E-08 | 2.46E-07 | -1.06E+00 | 4.28E-03 | 1.82E-02 | -1.12E+00 | 3.03E-03 | 1.16E-02 | 1.05E+00 | 5.25E-06 | 1.87E-05 | -1.40E+00 | 3.53E-02 | 9.95E-02 | 1.08E+00 | 6.27E-04 | 2.30E-03 | -1.10E+00 | 2.36E-03 | 6.49E-03 | -1.07E+00 |
| AP3M1 | 2.03E-06 | 1.06E-05 | -1.25E+00 | 2.17E-03 | 9.37E-03 | 1.12E+00 | 1.52E-03 | 6.47E-03 | -1.16E+00 | 1.25E-01 | 4.77E-01 | 1.31E+00 | 5.52E-05 | 1.97E-04 | -1.27E+00 | 9.50E-04 | 2.68E-03 | -1.04E+00 | 1.04E-04 | 3.82E-04 | -1.10E+00 | 9.71E-03 | 2.67E-02 | 1.10E+00 |
| TYSND1 | 2.04E-06 | 1.06E-05 | -1.41E+00 | 1.89E-03 | 8.17E-03 | 1.05E+00 | 5.47E-07 | 2.33E-06 | -1.51E+00 | 3.67E-04 | 1.40E-03 | -1.02E+00 | 5.20E-04 | 1.85E-03 | -1.61E+00 | 2.50E-02 | 7.05E-02 | 1.01E+00 | 5.43E-06 | 1.99E-05 | -1.70E+00 | 3.35E-03 | 9.21E-03 | -1.20E+00 |
| CNTROB | 2.05E-06 | 1.07E-05 | -1.37E+00 | 1.81E-04 | 7.84E-04 | 1.02E+00 | 1.79E-02 | 7.62E-02 | -1.05E+00 | 1.93E-02 | 7.38E-02 | 1.05E+00 | 1.56E-02 | 5.58E-02 | -1.20E+00 | 2.73E-01 | 7.70E-01 | 1.34E+00 | 1.42E-02 | 5.22E-02 | -1.10E+00 | 2.85E-01 | 7.82E-01 | 1.22E+00 |
| SH2B3 | 2.05E-06 | 1.07E-05 | -1.57E+00 | 2.29E-05 | 9.88E-05 | -1.29E+00 | 3.32E-06 | 1.41E-05 | -1.67E+00 | 3.52E-04 | 1.34E-03 | -1.10E+00 | 2.55E-08 | 9.08E-08 | -1.85E+00 | 5.62E-07 | 1.58E-06 | -1.56E+00 | 1.15E-07 | 4.22E-07 | -1.26E+00 | 8.49E-05 | 2.33E-04 | -1.14E+00 |
| AC110373.1 | 2.06E-06 | 1.07E-05 | -2.19E+00 | 3.79E-03 | 1.64E-02 | -1.14E+00 | 7.60E-06 | 3.23E-05 | -1.67E+00 | 1.70E-03 | 6.50E-03 | -1.08E+00 | 1.22E-02 | 4.34E-02 | -1.48E+00 | 2.01E-01 | 5.67E-01 | 1.18E+00 | 2.89E-07 | 1.06E-06 | -1.47E+00 | 3.31E-04 | 9.09E-04 | -1.10E+00 |
| RHBDD1 | 2.06E-06 | 1.07E-05 | -1.83E+00 | 1.75E-03 | 7.55E-03 | -1.07E+00 | 3.16E-06 | 1.34E-05 | -1.67E+00 | 6.25E-05 | 2.39E-04 | -1.19E+00 | 7.23E-03 | 2.58E-02 | -1.51E+00 | 1.24E-01 | 3.51E-01 | 1.08E+00 | 3.27E-04 | 1.20E-03 | -1.36E+00 | 4.54E-02 | 1.25E-01 | -1.09E+00 |
| DAXX | 2.06E-06 | 1.07E-05 | -1.35E+00 | 2.81E-02 | 1.22E-01 | 1.28E+00 | 3.64E-09 | 1.55E-08 | -1.53E+00 | 6.38E-07 | 2.44E-06 | -1.21E+00 | 1.26E-06 | 4.50E-06 | -1.70E+00 | 1.38E-02 | 3.88E-02 | -1.08E+00 | 3.61E-06 | 1.32E-05 | -1.32E+00 | 9.84E-02 | 2.70E-01 | 1.07E+00 |
| PPP1R16B | 2.06E-06 | 1.07E-05 | -1.56E+00 | 5.69E-05 | 2.46E-04 | -1.17E+00 | 2.20E-01 | 9.36E-01 | 1.07E+00 | 1.54E-01 | 5.89E-01 | 1.40E+00 | 2.76E-01 | 9.85E-01 | 1.26E+00 | 1.99E-01 | 5.61E-01 | -1.18E+00 | 1.80E-01 | 6.60E-01 | 1.32E+00 | 3.36E-01 | 9.23E-01 | -1.02E+00 |
| HNRNPA3P1 | 2.06E-06 | 1.07E-05 | -1.36E+00 | 8.47E-11 | 3.66E-10 | -1.84E+00 | 2.43E-09 | 1.03E-08 | -1.66E+00 | 8.01E-09 | 3.06E-08 | -1.48E+00 | 7.50E-06 | 2.67E-05 | -1.61E+00 | 6.23E-07 | 1.76E-06 | -1.76E+00 | 6.16E-11 | 2.26E-10 | -1.64E+00 | 1.70E-10 | 4.67E-10 | -1.85E+00 |
| ST6GALNAC6 | 2.07E-06 | 1.08E-05 | -1.32E+00 | 3.82E-04 | 1.65E-03 | 1.05E+00 | 6.89E-04 | 2.93E-03 | -1.29E+00 | 2.30E-02 | 8.79E-02 | 1.07E+00 | 2.65E-07 | 9.44E-07 | -1.31E+00 | 1.41E-04 | 3.97E-04 | -1.12E+00 | 1.11E-03 | 4.09E-03 | 1.01E+00 | 6.90E-03 | 1.89E-02 | 1.10E+00 |
| TEKT4P2 | 2.08E-06 | 1.08E-05 | -1.85E+00 | 6.45E-03 | 2.79E-02 | -1.01E+00 | 3.64E-06 | 1.55E-05 | -1.47E+00 | 1.02E-04 | 3.88E-04 | -1.12E+00 | 7.48E-07 | 2.67E-06 | -1.79E+00 | 8.80E-02 | 2.48E-01 | 1.19E+00 | 1.08E-11 | 3.97E-11 | -1.51E+00 | 1.36E-03 | 3.72E-03 | 1.06E+00 |
| SLC27A5 | 2.08E-06 | 1.08E-05 | -1.18E+00 | 3.04E-06 | 1.31E-05 | -1.08E+00 | 1.18E-03 | 5.01E-03 | -1.08E+00 | 6.16E-05 | 2.35E-04 | -1.06E+00 | 8.47E-06 | 3.02E-05 | -1.45E+00 | 5.79E-04 | 1.63E-03 | -1.15E+00 | 3.02E-03 | 1.11E-02 | -1.09E+00 | 9.94E-02 | 2.73E-01 | 1.05E+00 |
| SVBP | 2.09E-06 | 1.09E-05 | -1.15E+00 | 5.89E-06 | 2.54E-05 | 1.03E+00 | 1.13E-05 | 4.79E-05 | -1.25E+00 | 7.55E-07 | 2.88E-06 | -1.19E+00 | 5.47E-14 | 1.95E-13 | -1.63E+00 | 1.31E-07 | 3.70E-07 | -1.16E+00 | 9.47E-07 | 3.48E-06 | -1.18E+00 | 8.24E-05 | 2.26E-04 | -1.13E+00 |
| CBFA2T2 | 2.10E-06 | 1.09E-05 | -1.24E+00 | 6.30E-05 | 2.72E-04 | -1.02E+00 | 1.29E-03 | 5.50E-03 | -1.36E+00 | 3.30E-02 | 1.26E-01 | 1.05E+00 | 1.43E-02 | 5.11E-02 | -1.21E+00 | 2.16E-01 | 6.09E-01 | 1.20E+00 | 1.97E-06 | 7.25E-06 | -1.66E+00 | 4.04E-04 | 1.11E-03 | -1.36E+00 |
| ZNF271P | 2.11E-06 | 1.10E-05 | -1.20E+00 | 5.24E-07 | 2.27E-06 | -1.12E+00 | 5.65E-06 | 2.40E-05 | -1.39E+00 | 1.21E-04 | 4.62E-04 | -1.13E+00 | 2.33E-03 | 8.29E-03 | 1.00E+00 | 1.26E-03 | 3.56E-03 | -1.03E+00 | 1.44E-02 | 5.27E-02 | 1.05E+00 | 1.03E-02 | 2.82E-02 | 1.03E+00 |
| MAD2L1 | 2.11E-06 | 1.10E-05 | -1.28E+00 | 1.10E-05 | 4.75E-05 | -1.08E+00 | 1.96E-01 | 8.31E-01 | 1.19E+00 | 1.94E-02 | 7.40E-02 | 1.12E+00 | 1.13E-06 | 4.04E-06 | -1.56E+00 | 1.04E-05 | 2.92E-05 | -1.34E+00 | 7.78E-02 | 2.85E-01 | 1.07E+00 | 6.66E-02 | 1.83E-01 | 1.35E+00 |
| GOLGA8N | 2.12E-06 | 1.11E-05 | -1.28E+00 | 4.24E-06 | 1.83E-05 | -1.14E+00 | 3.26E-06 | 1.38E-05 | -1.32E+00 | 7.75E-07 | 2.96E-06 | -1.21E+00 | 3.68E-04 | 1.31E-03 | -1.37E+00 | 3.44E-03 | 9.69E-03 | -1.14E+00 | 4.36E-06 | 1.60E-05 | -1.41E+00 | 1.29E-06 | 3.54E-06 | -1.60E+00 |
| METTL26 | 2.13E-06 | 1.11E-05 | -1.01E+00 | 2.56E-07 | 1.11E-06 | 1.08E+00 | 1.19E-01 | 5.04E-01 | 1.18E+00 | 2.04E-01 | 7.81E-01 | 1.20E+00 | 1.39E-04 | 4.96E-04 | -1.06E+00 | 3.44E-04 | 9.69E-04 | 1.02E+00 | 2.04E-02 | 7.50E-02 | -1.00E+00 | 3.49E-01 | 9.59E-01 | 1.15E+00 |
| PIGN | 2.14E-06 | 1.11E-05 | -1.20E+00 | 6.04E-03 | 2.61E-02 | 1.29E+00 | 5.99E-05 | 2.54E-04 | -1.35E+00 | 6.57E-02 | 2.51E-01 | 1.19E+00 | 4.13E-04 | 1.47E-03 | -1.14E+00 | 5.82E-03 | 1.64E-02 | 1.04E+00 | 3.45E-04 | 1.26E-03 | -1.08E+00 | 2.67E-03 | 7.34E-03 | -1.08E+00 |
| DBF4P1 | 2.14E-06 | 1.12E-05 | -1.66E+00 | 4.93E-07 | 2.13E-06 | -1.77E+00 | 1.92E-04 | 8.14E-04 | -1.34E+00 | 3.81E-05 | 1.46E-04 | -1.29E+00 | 1.73E-03 | 6.18E-03 | -1.17E+00 | 3.06E-05 | 8.63E-05 | -1.67E+00 | 1.63E-17 | 6.00E-17 | -1.73E+00 | 6.47E-12 | 1.78E-11 | -1.36E+00 |
| CBR1 | 2.15E-06 | 1.12E-05 | -1.10E+00 | 1.26E-01 | 5.42E-01 | 1.47E+00 | 2.61E-05 | 1.11E-04 | -1.21E+00 | 4.13E-03 | 1.58E-02 | 1.10E+00 | 1.01E-13 | 3.61E-13 | -1.77E+00 | 4.12E-03 | 1.16E-02 | 1.08E+00 | 1.05E-03 | 3.86E-03 | -1.05E+00 | 3.53E-01 | 9.69E-01 | 1.32E+00 |
| WDR11 | 2.15E-06 | 1.12E-05 | -1.21E+00 | 6.47E-02 | 2.80E-01 | 1.33E+00 | 3.92E-06 | 1.67E-05 | -1.24E+00 | 3.87E-03 | 1.48E-02 | 1.12E+00 | 8.41E-06 | 3.00E-05 | -1.32E+00 | 4.90E-04 | 1.38E-03 | -1.07E+00 | 8.28E-05 | 3.04E-04 | -1.12E+00 | 1.59E-02 | 4.36E-02 | 1.02E+00 |
| GNB2 | 2.15E-06 | 1.12E-05 | 1.02E+00 | 2.65E-04 | 1.15E-03 | 1.23E+00 | 5.99E-02 | 2.55E-01 | 1.10E+00 | 9.41E-02 | 3.59E-01 | 1.26E+00 | 4.99E-05 | 1.78E-04 | -1.03E+00 | 7.29E-02 | 2.06E-01 | 1.21E+00 | 1.05E-01 | 3.85E-01 | 1.13E+00 | 1.67E-01 | 4.58E-01 | 1.29E+00 |
| FARP2 | 2.15E-06 | 1.12E-05 | -1.58E+00 | 2.37E-05 | 1.02E-04 | -1.31E+00 | 1.92E-03 | 8.16E-03 | -1.26E+00 | 3.10E-03 | 1.18E-02 | -1.11E+00 | 9.90E-02 | 3.53E-01 | 1.02E+00 | 3.17E-01 | 8.95E-01 | 1.33E+00 | 1.77E-05 | 6.51E-05 | -1.50E+00 | 4.11E-05 | 1.13E-04 | -1.57E+00 |
| POLD3 | 2.15E-06 | 1.12E-05 | -1.51E+00 | 1.90E-03 | 8.21E-03 | 1.03E+00 | 4.86E-03 | 2.06E-02 | -1.07E+00 | 4.08E-02 | 1.56E-01 | 1.16E+00 | 5.64E-03 | 2.01E-02 | -1.28E+00 | 7.78E-02 | 2.19E-01 | 1.02E+00 | 1.01E-05 | 3.70E-05 | -1.33E+00 | 2.01E-01 | 5.53E-01 | 1.15E+00 |
| THG1L | 2.15E-06 | 1.12E-05 | -1.29E+00 | 1.43E-05 | 6.17E-05 | -1.07E+00 | 7.20E-07 | 3.06E-06 | -1.27E+00 | 3.97E-07 | 1.52E-06 | -1.19E+00 | 4.98E-03 | 1.78E-02 | -1.09E+00 | 1.01E-01 | 2.84E-01 | 1.16E+00 | 1.55E-06 | 5.69E-06 | -1.33E+00 | 2.24E-02 | 6.15E-02 | -1.01E+00 |
| HNRNPA1L2 | 2.16E-06 | 1.13E-05 | -1.11E+00 | 2.34E-11 | 1.01E-10 | -1.25E+00 | 3.94E-07 | 1.68E-06 | -1.23E+00 | 1.63E-12 | 6.23E-12 | -1.38E+00 | 4.72E-05 | 1.68E-04 | -1.10E+00 | 3.03E-07 | 8.55E-07 | -1.26E+00 | 2.26E-07 | 8.30E-07 | -1.21E+00 | 8.69E-08 | 2.39E-07 | -1.30E+00 |
| MYDGF | 2.16E-06 | 1.13E-05 | -1.02E+00 | 8.13E-04 | 3.51E-03 | 1.14E+00 | 1.93E-01 | 8.21E-01 | 1.24E+00 | 2.52E-01 | 9.63E-01 | 1.24E+00 | 1.14E-07 | 4.05E-07 | -1.27E+00 | 4.84E-04 | 1.36E-03 | -1.11E+00 | 2.42E-01 | 8.88E-01 | 1.13E+00 | 2.08E-02 | 5.70E-02 | 1.27E+00 |
| C4orf48 | 2.17E-06 | 1.13E-05 | -1.08E+00 | 1.34E-15 | 5.77E-15 | -1.34E+00 | 4.92E-02 | 2.09E-01 | -1.00E+00 | 2.98E-07 | 1.14E-06 | -1.26E+00 | 5.33E-06 | 1.90E-05 | -1.27E+00 | 6.29E-13 | 1.77E-12 | -1.71E+00 | 2.12E-02 | 7.79E-02 | 1.07E+00 | 3.40E-03 | 9.35E-03 | -1.16E+00 |
| AC009996.1 | 2.17E-06 | 1.13E-05 | -1.84E+00 | 3.68E-06 | 1.59E-05 | -1.63E+00 | 2.49E-05 | 1.06E-04 | -1.54E+00 | 9.73E-08 | 3.72E-07 | -2.04E+00 | 7.50E-04 | 2.68E-03 | -1.99E+00 | 6.36E-02 | 1.79E-01 | -1.21E+00 | 1.63E-03 | 5.97E-03 | -1.71E+00 | 1.51E-02 | 4.14E-02 | -1.44E+00 |
| CNTRL | 2.17E-06 | 1.13E-05 | -1.15E+00 | 4.67E-05 | 2.02E-04 | 1.02E+00 | 3.75E-06 | 1.59E-05 | -1.27E+00 | 1.69E-06 | 6.45E-06 | -1.08E+00 | 7.01E-09 | 2.50E-08 | -1.45E+00 | 9.66E-09 | 2.72E-08 | -1.27E+00 | 5.64E-08 | 2.07E-07 | -1.34E+00 | 2.79E-06 | 7.67E-06 | -1.28E+00 |
| TEX10 | 2.17E-06 | 1.13E-05 | -1.17E+00 | 9.81E-04 | 4.24E-03 | 1.15E+00 | 8.73E-10 | 3.71E-09 | -1.46E+00 | 9.06E-04 | 3.46E-03 | 1.05E+00 | 1.62E-07 | 5.78E-07 | -1.46E+00 | 5.21E-04 | 1.47E-03 | -1.11E+00 | 1.22E-04 | 4.46E-04 | -1.20E+00 | 5.35E-02 | 1.47E-01 | 1.02E+00 |
| FAM215B | 2.19E-06 | 1.14E-05 | -1.23E+00 | 1.61E-04 | 6.95E-04 | 1.02E+00 | 5.12E-07 | 2.17E-06 | -1.37E+00 | 4.18E-05 | 1.60E-04 | -1.10E+00 | 2.75E-10 | 9.82E-10 | -1.63E+00 | 3.04E-10 | 8.58E-10 | -1.56E+00 | 4.20E-05 | 1.54E-04 | -1.51E+00 | 2.61E-02 | 7.16E-02 | -1.04E+00 |
| UBE3C | 2.19E-06 | 1.14E-05 | -1.31E+00 | 5.18E-04 | 2.24E-03 | 1.04E+00 | 5.93E-08 | 2.52E-07 | -1.45E+00 | 2.01E-04 | 7.67E-04 | -1.03E+00 | 1.36E-05 | 4.86E-05 | -1.34E+00 | 4.94E-03 | 1.39E-02 | -1.04E+00 | 2.10E-08 | 7.72E-08 | -1.43E+00 | 1.66E-02 | 4.56E-02 | 1.02E+00 |
| TOMM40 | 2.20E-06 | 1.15E-05 | -1.08E+00 | 5.85E-06 | 2.53E-05 | 1.08E+00 | 1.47E-01 | 6.25E-01 | 1.16E+00 | 1.49E-01 | 5.67E-01 | 1.30E+00 | 1.05E-03 | 3.74E-03 | -1.06E+00 | 2.57E-01 | 7.26E-01 | 1.29E+00 | 1.10E-02 | 4.02E-02 | 1.02E+00 | 4.11E-02 | 1.13E-01 | 1.39E+00 |
| MRPS10 | 2.20E-06 | 1.15E-05 | -1.09E+00 | 5.71E-05 | 2.47E-04 | 1.13E+00 | 1.64E-03 | 6.97E-03 | -1.08E+00 | 2.96E-02 | 1.13E-01 | 1.18E+00 | 7.45E-07 | 2.66E-06 | -1.22E+00 | 1.06E-03 | 2.97E-03 | 1.03E+00 | 1.11E-02 | 4.09E-02 | -1.05E+00 | 2.71E-01 | 7.45E-01 | 1.28E+00 |
| AC006970.1 | 2.21E-06 | 1.15E-05 | -1.45E+00 | 1.07E-02 | 4.64E-02 | 1.13E+00 | 8.91E-09 | 3.79E-08 | -1.85E+00 | 8.93E-04 | 3.41E-03 | -1.06E+00 | 2.37E-05 | 8.46E-05 | -2.49E+00 | 2.16E-03 | 6.10E-03 | -1.44E+00 | 4.49E-05 | 1.65E-04 | -1.51E+00 | 3.31E-03 | 9.08E-03 | -1.22E+00 |
| PCLAF | 2.22E-06 | 1.15E-05 | -1.65E+00 | 1.77E-11 | 7.66E-11 | -2.13E+00 | 5.77E-02 | 2.45E-01 | 1.03E+00 | 5.28E-08 | 2.02E-07 | -1.18E+00 | 1.87E-10 | 6.66E-10 | -2.18E+00 | 3.68E-08 | 1.04E-07 | -1.75E+00 | 1.63E-05 | 5.98E-05 | -1.15E+00 | 5.38E-03 | 1.48E-02 | 1.01E+00 |
| TRIM66 | 2.22E-06 | 1.15E-05 | -1.71E+00 | 1.33E-05 | 5.73E-05 | -1.42E+00 | 8.41E-04 | 3.58E-03 | -1.45E+00 | 2.17E-03 | 8.28E-03 | -1.22E+00 | 3.32E-02 | 1.18E-01 | -1.17E+00 | 3.48E-01 | 9.80E-01 | 1.31E+00 | 1.32E-02 | 4.84E-02 | -1.15E+00 | 6.28E-02 | 1.72E-01 | -1.03E+00 |
| CTDP1 | 2.22E-06 | 1.16E-05 | -1.35E+00 | 1.55E-06 | 6.68E-06 | -1.27E+00 | 1.08E-08 | 4.59E-08 | -1.53E+00 | 5.27E-11 | 2.01E-10 | -1.47E+00 | 7.26E-05 | 2.59E-04 | -1.49E+00 | 4.97E-04 | 1.40E-03 | -1.33E+00 | 5.52E-04 | 2.03E-03 | -1.31E+00 | 1.43E-01 | 3.92E-01 | 1.13E+00 |
| RPS6P25 | 2.22E-06 | 1.16E-05 | -1.17E+00 | 6.47E-54 | 2.80E-53 | -1.68E+00 | 4.62E-04 | 1.96E-03 | -1.15E+00 | 1.89E-27 | 7.20E-27 | -1.63E+00 | 3.97E-05 | 1.42E-04 | -1.21E+00 | 9.87E-25 | 2.78E-24 | -1.65E+00 | 1.11E-12 | 4.09E-12 | -1.24E+00 | 1.87E-31 | 5.14E-31 | -1.68E+00 |
| UBE2M | 2.22E-06 | 1.16E-05 | 1.90E+00 | 1.81E-01 | 7.84E-01 | 1.45E+00 | 2.94E-04 | 1.25E-03 | 1.66E+00 | 1.22E-01 | 4.65E-01 | 1.25E+00 | 7.37E-03 | 2.63E-02 | 1.76E+00 | 3.27E-01 | 9.21E-01 | 1.41E+00 | 1.78E-09 | 6.52E-09 | 2.08E+00 | 6.77E-03 | 1.86E-02 | 1.53E+00 |
| TMEM237 | 2.23E-06 | 1.16E-05 | -1.80E+00 | 3.75E-06 | 1.62E-05 | -1.47E+00 | 1.68E-01 | 7.14E-01 | 1.10E+00 | 2.08E-01 | 7.96E-01 | 1.28E+00 | 8.69E-04 | 3.10E-03 | -1.73E+00 | 1.15E-02 | 3.23E-02 | -1.21E+00 | 2.86E-06 | 1.05E-05 | -1.28E+00 | 2.16E-04 | 5.92E-04 | -1.12E+00 |
| SDAD1P1 | 2.23E-06 | 1.16E-05 | -1.28E+00 | 1.79E-05 | 7.73E-05 | -1.08E+00 | 1.17E-03 | 4.99E-03 | -1.38E+00 | 1.09E-02 | 4.16E-02 | -1.04E+00 | 1.20E-03 | 4.30E-03 | -1.26E+00 | 4.01E-03 | 1.13E-02 | -1.14E+00 | 1.57E-01 | 5.78E-01 | 1.09E+00 | 1.39E-01 | 3.82E-01 | 1.05E+00 |
| IGBP1 | 2.23E-06 | 1.16E-05 | -1.09E+00 | 1.54E-04 | 6.66E-04 | 1.08E+00 | 3.32E-03 | 1.41E-02 | -1.09E+00 | 1.41E-03 | 5.38E-03 | 1.00E+00 | 3.35E-03 | 1.20E-02 | -1.03E+00 | 2.00E-01 | 5.64E-01 | 1.14E+00 | 1.72E-02 | 6.30E-02 | 1.03E+00 | 6.44E-02 | 1.77E-01 | 1.05E+00 |
| PFDN1 | 2.23E-06 | 1.16E-05 | -1.04E+00 | 8.03E-09 | 3.47E-08 | -1.06E+00 | 2.28E-02 | 9.71E-02 | -1.00E+00 | 1.20E-03 | 4.57E-03 | -1.02E+00 | 3.92E-04 | 1.40E-03 | -1.06E+00 | 1.34E-02 | 3.78E-02 | 1.07E+00 | 1.94E-05 | 7.13E-05 | -1.07E+00 | 1.47E-04 | 4.03E-04 | -1.06E+00 |
| FYN | 2.24E-06 | 1.17E-05 | -1.25E+00 | 1.30E-02 | 5.62E-02 | 1.26E+00 | 6.71E-04 | 2.85E-03 | -1.18E+00 | 5.91E-03 | 2.26E-02 | 1.09E+00 | 3.89E-07 | 1.39E-06 | -1.87E+00 | 1.04E-02 | 2.92E-02 | -1.05E+00 | 6.74E-05 | 2.47E-04 | -1.04E+00 | 6.75E-03 | 1.85E-02 | 1.04E+00 |
| ZNF160 | 2.24E-06 | 1.17E-05 | -1.39E+00 | 1.08E-05 | 4.65E-05 | -1.16E+00 | 3.65E-03 | 1.55E-02 | -1.11E+00 | 3.82E-03 | 1.46E-02 | -1.01E+00 | 4.57E-02 | 1.63E-01 | 1.04E+00 | 6.74E-02 | 1.90E-01 | 1.19E+00 | 3.96E-02 | 1.45E-01 | -1.03E+00 | 5.98E-03 | 1.64E-02 | -1.11E+00 |
| AGGF1P1 | 2.25E-06 | 1.17E-05 | -1.69E+00 | 1.26E-01 | 5.43E-01 | 1.41E+00 | 1.26E-04 | 5.34E-04 | -1.60E+00 | 2.59E-01 | 9.90E-01 | 1.43E+00 | 1.36E-02 | 4.85E-02 | -1.32E+00 | 3.51E-01 | 9.91E-01 | 1.36E+00 | 1.70E-01 | 6.25E-01 | 1.23E+00 | 1.33E-01 | 3.64E-01 | 1.62E+00 |
| KLHL20 | 2.25E-06 | 1.18E-05 | -1.33E+00 | 1.44E-04 | 6.21E-04 | -1.01E+00 | 5.61E-06 | 2.38E-05 | -1.40E+00 | 5.60E-05 | 2.14E-04 | -1.15E+00 | 2.19E-02 | 7.80E-02 | 1.10E+00 | 1.09E-02 | 3.09E-02 | -1.01E+00 | 4.22E-06 | 1.55E-05 | -1.28E+00 | 1.54E-06 | 4.22E-06 | -1.37E+00 |
| STRN4 | 2.26E-06 | 1.18E-05 | -1.43E+00 | 4.69E-07 | 2.02E-06 | -1.25E+00 | 5.08E-04 | 2.16E-03 | -1.20E+00 | 4.77E-03 | 1.82E-02 | 1.02E+00 | 4.91E-05 | 1.75E-04 | -1.34E+00 | 9.43E-04 | 2.66E-03 | -1.08E+00 | 6.10E-10 | 2.24E-09 | -1.50E+00 | 1.95E-04 | 5.36E-04 | -1.15E+00 |
| AC023051.1 | 2.26E-06 | 1.18E-05 | -1.46E+00 | 1.76E-03 | 7.62E-03 | 1.08E+00 | 5.09E-05 | 2.16E-04 | -1.55E+00 | 6.31E-04 | 2.41E-03 | -1.24E+00 | 1.59E-01 | 5.69E-01 | 1.15E+00 | 2.69E-01 | 7.57E-01 | 1.59E+00 | 2.00E-05 | 7.34E-05 | -1.77E+00 | 1.57E-02 | 4.31E-02 | -1.12E+00 |
| RABEP1 | 2.26E-06 | 1.18E-05 | -1.23E+00 | 4.38E-05 | 1.89E-04 | -1.02E+00 | 7.19E-06 | 3.05E-05 | -1.34E+00 | 1.28E-04 | 4.87E-04 | -1.07E+00 | 1.26E-03 | 4.51E-03 | -1.15E+00 | 5.70E-02 | 1.61E-01 | 1.11E+00 | 2.60E-05 | 9.55E-05 | -1.19E+00 | 2.73E-01 | 7.49E-01 | 1.18E+00 |
| SLC2A10 | 2.26E-06 | 1.18E-05 | -2.11E+00 | 1.66E-04 | 7.16E-04 | -1.49E+00 | 1.22E-02 | 5.17E-02 | -1.36E+00 | 8.16E-03 | 3.12E-02 | -1.27E+00 | 3.55E-08 | 1.27E-07 | -1.98E+00 | 1.16E-03 | 3.27E-03 | -1.06E+00 | 3.77E-03 | 1.38E-02 | -1.14E+00 | 1.21E-01 | 3.33E-01 | 1.14E+00 |
| AC233279.1 | 2.27E-06 | 1.18E-05 | -1.22E+00 | 3.67E-07 | 1.59E-06 | -1.17E+00 | 1.44E-03 | 6.14E-03 | -1.09E+00 | 2.68E-04 | 1.02E-03 | -1.06E+00 | 2.26E-08 | 8.06E-08 | -1.58E+00 | 7.33E-06 | 2.07E-05 | -1.32E+00 | 5.64E-10 | 2.07E-09 | -1.38E+00 | 5.33E-07 | 1.47E-06 | -1.21E+00 |
| FMO1 | 2.28E-06 | 1.19E-05 | -1.51E+00 | 3.30E-09 | 1.42E-08 | -1.75E+00 | 1.28E-03 | 5.45E-03 | -1.02E+00 | 1.61E-12 | 6.15E-12 | -1.47E+00 | 7.80E-09 | 2.78E-08 | -1.67E+00 | 2.27E-09 | 6.41E-09 | -1.75E+00 | 1.71E-11 | 6.29E-11 | -1.55E+00 | 3.41E-15 | 9.35E-15 | -1.68E+00 |
| SMAD5-AS1 | 2.28E-06 | 1.19E-05 | -1.86E+00 | 1.21E-05 | 5.25E-05 | -1.58E+00 | 5.48E-06 | 2.33E-05 | -1.78E+00 | 1.21E-06 | 4.60E-06 | -1.83E+00 | 4.11E-02 | 1.47E-01 | -1.10E+00 | 8.63E-03 | 2.43E-02 | -1.52E+00 | 4.58E-02 | 1.68E-01 | -1.07E+00 | 1.07E-02 | 2.95E-02 | -1.23E+00 |
| SGPP1 | 2.28E-06 | 1.19E-05 | -2.43E+00 | 1.33E-04 | 5.75E-04 | -1.58E+00 | 1.16E-03 | 4.91E-03 | -1.55E+00 | 4.57E-03 | 1.75E-02 | -1.23E+00 | 2.06E-02 | 7.36E-02 | -1.11E+00 | 1.67E-01 | 4.72E-01 | 1.73E+00 | 6.82E-19 | 2.50E-18 | -1.83E+00 | 3.25E-08 | 8.92E-08 | -1.37E+00 |
| HS6ST1 | 2.30E-06 | 1.20E-05 | -1.84E+00 | 2.59E-03 | 1.12E-02 | -1.08E+00 | 4.94E-05 | 2.10E-04 | -1.54E+00 | 2.59E-04 | 9.88E-04 | -1.29E+00 | 4.72E-08 | 1.68E-07 | -1.94E+00 | 4.29E-05 | 1.21E-04 | -1.35E+00 | 2.46E-11 | 9.04E-11 | -2.19E+00 | 1.03E-07 | 2.83E-07 | -1.75E+00 |
| EXOSC8 | 2.30E-06 | 1.20E-05 | -1.10E+00 | 1.34E-02 | 5.81E-02 | 1.29E+00 | 4.56E-02 | 1.94E-01 | 1.15E+00 | 7.40E-03 | 2.83E-02 | 1.19E+00 | 1.25E-03 | 4.45E-03 | -1.01E+00 | 1.44E-01 | 4.07E-01 | 1.28E+00 | 1.26E-03 | 4.64E-03 | -1.02E+00 | 2.91E-01 | 7.99E-01 | 1.26E+00 |
| NDUFB2 | 2.31E-06 | 1.20E-05 | -1.06E+00 | 1.60E-14 | 6.90E-14 | -1.17E+00 | 1.46E-01 | 6.21E-01 | 1.09E+00 | 1.86E-03 | 7.11E-03 | -1.04E+00 | 1.07E-03 | 3.83E-03 | -1.10E+00 | 5.72E-05 | 1.61E-04 | -1.15E+00 | 1.28E-01 | 4.71E-01 | 1.07E+00 | 4.88E-02 | 1.34E-01 | -1.07E+00 |
| CHCHD2 | 2.34E-06 | 1.22E-05 | -1.13E+00 | 2.22E-11 | 9.60E-11 | -1.18E+00 | 1.05E-01 | 4.46E-01 | -1.03E+00 | 3.58E-03 | 1.37E-02 | -1.12E+00 | 6.61E-06 | 2.36E-05 | -1.21E+00 | 7.89E-07 | 2.23E-06 | -1.23E+00 | 1.27E-03 | 4.66E-03 | -1.13E+00 | 3.31E-02 | 9.08E-02 | -1.11E+00 |
| WDCP | 2.35E-06 | 1.22E-05 | -2.20E+00 | 3.15E-04 | 1.36E-03 | -1.33E+00 | 4.81E-13 | 2.04E-12 | -2.82E+00 | 4.30E-06 | 1.64E-05 | -1.32E+00 | 2.00E-03 | 7.14E-03 | -1.59E+00 | 5.10E-02 | 1.44E-01 | -1.06E+00 | 6.87E-06 | 2.52E-05 | -1.67E+00 | 8.31E-02 | 2.28E-01 | 1.00E+00 |
| KPNA1 | 2.35E-06 | 1.22E-05 | -1.38E+00 | 1.28E-04 | 5.54E-04 | -1.06E+00 | 1.58E-04 | 6.73E-04 | -1.36E+00 | 2.09E-03 | 7.98E-03 | -1.09E+00 | 2.24E-03 | 7.98E-03 | -1.10E+00 | 3.09E-02 | 8.72E-02 | 1.09E+00 | 1.41E-05 | 5.17E-05 | -1.27E+00 | 1.09E-03 | 2.99E-03 | -1.07E+00 |
| C12orf4 | 2.36E-06 | 1.23E-05 | -1.58E+00 | 3.59E-05 | 1.55E-04 | -1.33E+00 | 3.25E-02 | 1.38E-01 | -1.05E+00 | 1.34E-01 | 5.12E-01 | 1.19E+00 | 3.08E-03 | 1.10E-02 | -1.44E+00 | 1.06E-02 | 2.99E-02 | -1.16E+00 | 1.59E-03 | 5.85E-03 | -1.09E+00 | 2.29E-02 | 6.28E-02 | -1.03E+00 |
| EMD | 2.36E-06 | 1.23E-05 | -1.18E+00 | 6.91E-10 | 2.99E-09 | -1.25E+00 | 1.29E-03 | 5.49E-03 | -1.13E+00 | 2.83E-04 | 1.08E-03 | -1.09E+00 | 2.24E-07 | 7.98E-07 | -1.30E+00 | 6.08E-06 | 1.71E-05 | -1.21E+00 | 1.07E-04 | 3.94E-04 | -1.14E+00 | 1.87E-02 | 5.13E-02 | -1.06E+00 |
| SNAPC1 | 2.37E-06 | 1.24E-05 | -1.58E+00 | 1.48E-05 | 6.40E-05 | -1.37E+00 | 7.20E-04 | 3.06E-03 | -1.30E+00 | 9.60E-05 | 3.67E-04 | -1.26E+00 | 6.65E-03 | 2.37E-02 | -1.21E+00 | 5.17E-03 | 1.46E-02 | -1.14E+00 | 4.29E-08 | 1.57E-07 | -1.57E+00 | 1.07E-06 | 2.93E-06 | -1.31E+00 |
| AC007041.1 | 2.37E-06 | 1.24E-05 | -1.87E+00 | 8.13E-04 | 3.51E-03 | -1.16E+00 | 1.87E-08 | 7.96E-08 | -1.63E+00 | 6.91E-08 | 2.64E-07 | -1.35E+00 | 4.28E-08 | 1.53E-07 | -1.97E+00 | 3.82E-04 | 1.08E-03 | -1.08E+00 | 1.14E-26 | 4.18E-26 | -2.49E+00 | 5.46E-14 | 1.50E-13 | -1.56E+00 |
| TMOD3 | 2.38E-06 | 1.24E-05 | -1.08E+00 | 1.03E-05 | 4.44E-05 | 1.06E+00 | 3.40E-03 | 1.44E-02 | -1.05E+00 | 1.88E-03 | 7.19E-03 | 1.03E+00 | 1.23E-04 | 4.37E-04 | -1.16E+00 | 9.90E-04 | 2.79E-03 | -1.09E+00 | 1.83E-05 | 6.72E-05 | -1.13E+00 | 1.15E-04 | 3.15E-04 | -1.13E+00 |
| RAD50 | 2.38E-06 | 1.24E-05 | -1.62E+00 | 2.51E-05 | 1.08E-04 | -1.33E+00 | 1.84E-02 | 7.83E-02 | -1.11E+00 | 1.31E-02 | 5.02E-02 | -1.03E+00 | 8.10E-04 | 2.89E-03 | -1.47E+00 | 1.80E-02 | 5.07E-02 | -1.10E+00 | 9.28E-03 | 3.41E-02 | -1.03E+00 | 1.18E-02 | 3.24E-02 | -1.04E+00 |
| CCDC71 | 2.38E-06 | 1.24E-05 | -1.93E+00 | 1.19E-03 | 5.15E-03 | -1.10E+00 | 2.44E-07 | 1.04E-06 | -1.85E+00 | 3.05E-05 | 1.17E-04 | -1.38E+00 | 5.52E-02 | 1.97E-01 | -1.19E+00 | 3.42E-01 | 9.64E-01 | 1.45E+00 | 1.77E-19 | 6.49E-19 | -2.35E+00 | 1.27E-07 | 3.48E-07 | -1.31E+00 |
| RPL7AP66 | 2.38E-06 | 1.24E-05 | 1.34E+00 | 8.92E-24 | 3.85E-23 | 1.73E+00 | 7.53E-10 | 3.20E-09 | 1.49E+00 | 1.27E-18 | 4.85E-18 | 1.79E+00 | 1.58E-06 | 5.63E-06 | 1.51E+00 | 3.45E-15 | 9.71E-15 | 1.81E+00 | 3.10E-08 | 1.14E-07 | 1.56E+00 | 1.85E-26 | 5.07E-26 | 1.97E+00 |
| KBTBD11 | 2.39E-06 | 1.24E-05 | -1.56E+00 | 2.31E-05 | 9.98E-05 | -1.27E+00 | 1.45E-09 | 6.18E-09 | -1.39E+00 | 8.04E-07 | 3.07E-06 | 1.01E+00 | 2.43E-01 | 8.68E-01 | 1.00E+00 | 3.06E-01 | 8.63E-01 | 1.37E+00 | 1.07E-01 | 3.94E-01 | 1.77E+00 | 1.72E-01 | 4.72E-01 | 1.71E+00 |
| HERPUD1 | 2.39E-06 | 1.25E-05 | -1.35E+00 | 1.80E-12 | 7.76E-12 | -1.51E+00 | 5.60E-02 | 2.38E-01 | 1.09E+00 | 4.64E-06 | 1.77E-05 | -1.11E+00 | 3.43E-07 | 1.22E-06 | -1.43E+00 | 1.82E-15 | 5.14E-15 | -1.75E+00 | 4.04E-03 | 1.48E-02 | -1.20E+00 | 7.80E-08 | 2.14E-07 | -1.71E+00 |
| PATL1 | 2.40E-06 | 1.25E-05 | -1.51E+00 | 4.50E-04 | 1.95E-03 | -1.00E+00 | 4.74E-03 | 2.02E-02 | -1.22E+00 | 4.78E-03 | 1.82E-02 | -1.05E+00 | 5.32E-03 | 1.90E-02 | -1.39E+00 | 4.75E-03 | 1.34E-02 | -1.40E+00 | 1.96E-04 | 7.19E-04 | -1.20E+00 | 1.39E-02 | 3.81E-02 | -1.02E+00 |
| AP001972.5 | 2.42E-06 | 1.26E-05 | -1.31E+00 | 3.64E-05 | 1.57E-04 | -1.13E+00 | 5.97E-03 | 2.54E-02 | -1.14E+00 | 3.90E-03 | 1.49E-02 | -1.03E+00 | 6.80E-05 | 2.43E-04 | -1.55E+00 | 3.71E-03 | 1.05E-02 | -1.09E+00 | 2.41E-03 | 8.86E-03 | -1.18E+00 | 4.55E-02 | 1.25E-01 | 1.00E+00 |
| PRMT2 | 2.42E-06 | 1.26E-05 | -1.05E+00 | 1.55E-01 | 6.72E-01 | 1.54E+00 | 6.89E-04 | 2.93E-03 | -1.19E+00 | 6.13E-02 | 2.34E-01 | 1.17E+00 | 1.92E-04 | 6.83E-04 | -1.14E+00 | 3.45E-02 | 9.74E-02 | 1.04E+00 | 7.84E-04 | 2.88E-03 | -1.05E+00 | 2.59E-04 | 7.13E-04 | -1.09E+00 |
| EYA3 | 2.42E-06 | 1.26E-05 | -1.41E+00 | 1.49E-03 | 6.42E-03 | 1.07E+00 | 2.56E-06 | 1.09E-05 | -1.54E+00 | 1.04E-03 | 3.96E-03 | -1.00E+00 | 7.52E-06 | 2.68E-05 | -1.94E+00 | 7.27E-03 | 2.05E-02 | -1.15E+00 | 5.30E-11 | 1.95E-10 | -1.83E+00 | 8.18E-03 | 2.25E-02 | -1.05E+00 |
| FAM124B | 2.43E-06 | 1.27E-05 | 2.23E+00 | 7.75E-03 | 3.35E-02 | 2.13E+00 | 3.70E-02 | 1.57E-01 | 1.53E+00 | 7.72E-02 | 2.95E-01 | 1.64E+00 | 1.85E-06 | 6.61E-06 | 2.96E+00 | 1.31E-05 | 3.69E-05 | 3.04E+00 | 2.21E-04 | 8.10E-04 | 1.97E+00 | 2.36E-04 | 6.48E-04 | 2.09E+00 |
| YIPF6 | 2.44E-06 | 1.27E-05 | -1.26E+00 | 5.44E-03 | 2.35E-02 | 1.20E+00 | 3.02E-06 | 1.28E-05 | -1.64E+00 | 1.75E-03 | 6.69E-03 | -1.17E+00 | 2.07E-05 | 7.38E-05 | -1.26E+00 | 2.46E-03 | 6.93E-03 | 1.01E+00 | 2.21E-07 | 8.10E-07 | -1.31E+00 | 5.70E-05 | 1.57E-04 | -1.14E+00 |
| SCMH1 | 2.45E-06 | 1.28E-05 | -1.28E+00 | 3.81E-03 | 1.65E-02 | 1.18E+00 | 4.06E-02 | 1.73E-01 | -1.03E+00 | 9.78E-02 | 3.74E-01 | 1.20E+00 | 4.78E-02 | 1.71E-01 | 1.04E+00 | 8.65E-03 | 2.44E-02 | -1.26E+00 | 1.05E-01 | 3.84E-01 | 1.02E+00 | 1.14E-01 | 3.12E-01 | 1.60E+00 |
| ZNF580 | 2.46E-06 | 1.28E-05 | -1.14E+00 | 4.39E-04 | 1.90E-03 | 1.12E+00 | 2.17E-06 | 9.23E-06 | -1.31E+00 | 3.33E-03 | 1.27E-02 | 1.06E+00 | 4.86E-02 | 1.73E-01 | 1.07E+00 | 3.02E-01 | 8.51E-01 | 1.35E+00 | 2.91E-04 | 1.07E-03 | -1.08E+00 | 1.38E-02 | 3.79E-02 | 1.04E+00 |
| HACL1 | 2.47E-06 | 1.29E-05 | -1.35E+00 | 3.39E-05 | 1.46E-04 | -1.07E+00 | 2.11E-02 | 8.98E-02 | -1.12E+00 | 5.00E-03 | 1.91E-02 | -1.10E+00 | 3.81E-06 | 1.36E-05 | -1.45E+00 | 3.02E-02 | 8.53E-02 | 1.12E+00 | 2.40E-02 | 8.80E-02 | 1.01E+00 | 3.29E-01 | 9.03E-01 | 1.20E+00 |
| KCTD15 | 2.47E-06 | 1.29E-05 | -1.57E+00 | 8.70E-05 | 3.76E-04 | -1.12E+00 | 5.47E-05 | 2.32E-04 | -1.49E+00 | 7.91E-03 | 3.02E-02 | -1.04E+00 | 1.36E-03 | 4.87E-03 | -1.36E+00 | 2.02E-02 | 5.68E-02 | -1.02E+00 | 9.74E-03 | 3.57E-02 | -1.22E+00 | 1.96E-01 | 5.39E-01 | 1.08E+00 |
| ZMYND11 | 2.47E-06 | 1.29E-05 | -1.36E+00 | 1.53E-04 | 6.62E-04 | -1.04E+00 | 1.94E-06 | 8.23E-06 | -1.47E+00 | 7.09E-04 | 2.71E-03 | -1.04E+00 | 1.86E-06 | 6.65E-06 | -1.53E+00 | 1.71E-03 | 4.83E-03 | -1.18E+00 | 5.73E-11 | 2.10E-10 | -1.47E+00 | 1.77E-05 | 4.86E-05 | -1.08E+00 |
| ALDH5A1 | 2.49E-06 | 1.30E-05 | -1.35E+00 | 3.14E-03 | 1.36E-02 | 1.12E+00 | 3.52E-06 | 1.50E-05 | -1.30E+00 | 9.86E-04 | 3.77E-03 | 1.02E+00 | 2.79E-02 | 9.93E-02 | -1.02E+00 | 2.07E-01 | 5.83E-01 | 1.34E+00 | 9.34E-04 | 3.43E-03 | -1.03E+00 | 1.23E-02 | 3.37E-02 | -1.00E+00 |
| ZNF548 | 2.50E-06 | 1.30E-05 | -1.15E+00 | 3.93E-13 | 1.70E-12 | -1.66E+00 | 5.57E-03 | 2.37E-02 | -1.10E+00 | 5.23E-06 | 2.00E-05 | -1.38E+00 | 3.71E-03 | 1.32E-02 | 1.00E+00 | 3.54E-04 | 9.98E-04 | -1.30E+00 | 6.31E-03 | 2.32E-02 | -1.16E+00 | 2.03E-04 | 5.57E-04 | -1.35E+00 |
| FLVCR1 | 2.51E-06 | 1.31E-05 | -1.28E+00 | 5.79E-04 | 2.50E-03 | 1.01E+00 | 1.27E-04 | 5.40E-04 | -1.29E+00 | 5.23E-03 | 2.00E-02 | 1.06E+00 | 4.09E-03 | 1.46E-02 | -1.25E+00 | 5.04E-02 | 1.42E-01 | 1.09E+00 | 7.84E-03 | 2.88E-02 | -1.13E+00 | 1.00E-01 | 2.75E-01 | 1.08E+00 |
| HSPG2 | 2.51E-06 | 1.31E-05 | -1.25E+00 | 8.43E-03 | 3.64E-02 | 1.20E+00 | 1.24E-07 | 5.28E-07 | -1.81E+00 | 6.35E-05 | 2.42E-04 | -1.21E+00 | 4.16E-07 | 1.48E-06 | -1.48E+00 | 1.10E-03 | 3.10E-03 | -1.05E+00 | 2.89E-10 | 1.06E-09 | -2.93E+00 | 6.81E-07 | 1.87E-06 | -2.08E+00 |
| TMCC1-AS1 | 2.51E-06 | 1.31E-05 | -1.40E+00 | 5.48E-03 | 2.37E-02 | 1.16E+00 | 6.26E-09 | 2.66E-08 | -1.67E+00 | 5.53E-05 | 2.11E-04 | -1.02E+00 | 6.27E-05 | 2.24E-04 | -1.37E+00 | 2.88E-02 | 8.13E-02 | 1.08E+00 | 6.47E-09 | 2.37E-08 | -1.65E+00 | 3.26E-02 | 8.94E-02 | 1.08E+00 |
| LRRC37A4P | 2.52E-06 | 1.31E-05 | -1.27E+00 | 3.67E-02 | 1.59E-01 | 1.32E+00 | 3.18E-10 | 1.35E-09 | -1.75E+00 | 2.30E-07 | 8.79E-07 | -1.34E+00 | 5.86E-08 | 2.09E-07 | -1.97E+00 | 1.05E-05 | 2.96E-05 | -1.36E+00 | 1.25E-08 | 4.58E-08 | -2.84E+00 | 1.15E-05 | 3.16E-05 | -1.97E+00 |
| AL160400.1 | 2.52E-06 | 1.31E-05 | 2.38E+00 | 5.10E-05 | 2.20E-04 | 2.45E+00 | 1.38E-01 | 5.87E-01 | 1.54E+00 | 2.62E-01 | 1.00E+00 | 1.56E+00 | 1.32E-02 | 4.70E-02 | 1.02E+00 | 8.34E-02 | 2.35E-01 | 1.12E+00 | 2.46E-01 | 9.02E-01 | 1.26E+00 | 4.52E-03 | 1.24E-02 | -1.00E+00 |
| DLEU1 | 2.53E-06 | 1.32E-05 | -1.33E+00 | 4.69E-06 | 2.03E-05 | -1.23E+00 | 2.63E-06 | 1.12E-05 | -1.45E+00 | 1.97E-07 | 7.54E-07 | -1.54E+00 | 9.41E-06 | 3.36E-05 | -1.68E+00 | 5.61E-05 | 1.58E-04 | -1.47E+00 | 7.61E-05 | 2.79E-04 | -1.38E+00 | 4.01E-03 | 1.10E-02 | -1.12E+00 |
| SPINT1 | 2.54E-06 | 1.32E-05 | -1.20E+00 | 2.74E-04 | 1.18E-03 | 1.10E+00 | 1.29E-08 | 5.47E-08 | -1.53E+00 | 4.55E-08 | 1.74E-07 | -1.35E+00 | 4.89E-06 | 1.74E-05 | -1.41E+00 | 9.52E-06 | 2.68E-05 | -1.28E+00 | 5.44E-06 | 2.00E-05 | -1.69E+00 | 6.83E-05 | 1.88E-04 | -1.51E+00 |
| CD46P1 | 2.54E-06 | 1.32E-05 | -1.62E+00 | 4.08E-08 | 1.76E-07 | -1.92E+00 | 1.38E-03 | 5.84E-03 | -1.46E+00 | 2.02E-03 | 7.71E-03 | -1.33E+00 | 3.96E-03 | 1.41E-02 | -1.37E+00 | 1.19E-02 | 3.37E-02 | -1.24E+00 | 8.29E-03 | 3.04E-02 | -1.07E+00 | 3.35E-03 | 9.20E-03 | -1.22E+00 |
| DDX28 | 2.55E-06 | 1.33E-05 | -1.30E+00 | 2.82E-03 | 1.22E-02 | 1.16E+00 | 1.00E-04 | 4.25E-04 | -1.21E+00 | 1.58E-02 | 6.03E-02 | 1.12E+00 | 1.30E-04 | 4.63E-04 | -1.28E+00 | 2.66E-02 | 7.51E-02 | 1.11E+00 | 6.26E-06 | 2.30E-05 | -1.27E+00 | 2.50E-03 | 6.88E-03 | -1.14E+00 |
| LINC02160 | 2.56E-06 | 1.34E-05 | -1.45E+00 | 1.66E-02 | 7.17E-02 | 1.19E+00 | 6.48E-03 | 2.75E-02 | -1.25E+00 | 7.36E-02 | 2.81E-01 | 1.10E+00 | 2.27E-01 | 8.08E-01 | 1.65E+00 | 3.12E-01 | 8.80E-01 | 1.46E+00 | 1.73E-01 | 6.36E-01 | 1.51E+00 | 2.44E-01 | 6.71E-01 | 1.48E+00 |
| PRMT7 | 2.59E-06 | 1.35E-05 | -1.42E+00 | 6.28E-04 | 2.71E-03 | 1.02E+00 | 1.93E-06 | 8.19E-06 | -1.44E+00 | 1.84E-03 | 7.02E-03 | 1.01E+00 | 2.09E-03 | 7.45E-03 | -1.06E+00 | 2.58E-02 | 7.28E-02 | 1.17E+00 | 9.70E-04 | 3.56E-03 | -1.09E+00 | 2.35E-01 | 6.47E-01 | 1.24E+00 |
| ALAS1 | 2.59E-06 | 1.35E-05 | -1.71E+00 | 5.36E-04 | 2.32E-03 | -1.22E+00 | 1.50E-11 | 6.36E-11 | -1.82E+00 | 4.74E-11 | 1.81E-10 | -1.50E+00 | 1.24E-14 | 4.41E-14 | -2.33E+00 | 1.20E-12 | 3.39E-12 | -1.96E+00 | 8.39E-06 | 3.08E-05 | -1.48E+00 | 2.08E-04 | 5.71E-04 | -1.29E+00 |
| JAK2 | 2.62E-06 | 1.36E-05 | -1.45E+00 | 3.84E-04 | 1.66E-03 | -1.05E+00 | 9.80E-08 | 4.17E-07 | -1.68E+00 | 1.11E-04 | 4.23E-04 | -1.16E+00 | 2.99E-02 | 1.07E-01 | 1.08E+00 | 1.14E-02 | 3.22E-02 | -1.05E+00 | 1.66E-03 | 6.08E-03 | 1.06E+00 | 1.30E-03 | 3.57E-03 | -1.05E+00 |
| PADI4 | 2.62E-06 | 1.36E-05 | -1.13E+00 | 5.88E-05 | 2.54E-04 | 1.04E+00 | 2.22E-02 | 9.41E-02 | -1.07E+00 | 5.18E-04 | 1.98E-03 | -1.15E+00 | 5.71E-02 | 2.04E-01 | -1.14E+00 | 2.56E-01 | 7.20E-01 | 1.12E+00 | 2.38E-03 | 8.75E-03 | 2.46E+00 | 3.30E-02 | 9.06E-02 | 1.93E+00 |
| BRAF | 2.62E-06 | 1.36E-05 | -1.01E+00 | 1.30E-07 | 5.62E-07 | 1.03E+00 | 6.27E-04 | 2.66E-03 | -1.09E+00 | 2.27E-07 | 8.68E-07 | -1.23E+00 | 7.72E-02 | 2.75E-01 | 1.19E+00 | 7.98E-03 | 2.25E-02 | 1.06E+00 | 6.10E-02 | 2.24E-01 | 1.03E+00 | 9.31E-03 | 2.56E-02 | -1.03E+00 |
| DCAF13 | 2.62E-06 | 1.37E-05 | -1.10E+00 | 1.88E-03 | 8.13E-03 | 1.16E+00 | 1.44E-02 | 6.12E-02 | 1.04E+00 | 1.60E-01 | 6.13E-01 | 1.38E+00 | 3.22E-02 | 1.15E-01 | 1.09E+00 | 8.17E-02 | 2.30E-01 | 1.52E+00 | 7.07E-04 | 2.59E-03 | -1.03E+00 | 2.50E-01 | 6.88E-01 | 1.28E+00 |
| OTUD6B-AS1 | 2.63E-06 | 1.37E-05 | -1.13E+00 | 2.78E-13 | 1.20E-12 | -1.32E+00 | 1.64E-03 | 6.97E-03 | 1.00E+00 | 3.94E-05 | 1.51E-04 | -1.07E+00 | 1.14E-03 | 4.08E-03 | 1.01E+00 | 2.26E-05 | 6.37E-05 | -1.16E+00 | 1.28E-06 | 4.70E-06 | -1.20E+00 | 2.15E-08 | 5.90E-08 | -1.40E+00 |
| LRCH1 | 2.65E-06 | 1.38E-05 | -1.26E+00 | 2.30E-06 | 9.92E-06 | -1.12E+00 | 1.54E-04 | 6.54E-04 | -1.23E+00 | 3.03E-05 | 1.16E-04 | -1.21E+00 | 1.62E-04 | 5.78E-04 | -1.46E+00 | 9.48E-03 | 2.67E-02 | -1.02E+00 | 1.09E-07 | 4.00E-07 | -1.72E+00 | 4.99E-06 | 1.37E-05 | -1.46E+00 |
| BCL7C | 2.65E-06 | 1.38E-05 | -1.10E+00 | 8.30E-06 | 3.59E-05 | 1.06E+00 | 1.92E-02 | 8.15E-02 | 1.14E+00 | 4.55E-05 | 1.74E-04 | 1.04E+00 | 4.74E-02 | 1.69E-01 | 1.15E+00 | 1.67E-01 | 4.72E-01 | 1.21E+00 | 1.48E-01 | 5.42E-01 | 1.16E+00 | 3.07E-01 | 8.45E-01 | 1.20E+00 |
| DZIP1 | 2.66E-06 | 1.39E-05 | -2.37E+00 | 9.67E-03 | 4.18E-02 | -1.06E+00 | 6.92E-08 | 2.94E-07 | -2.34E+00 | 1.82E-03 | 6.94E-03 | -1.19E+00 | 7.00E-03 | 2.49E-02 | -1.73E+00 | 9.80E-03 | 2.76E-02 | -1.69E+00 | 2.67E-02 | 9.81E-02 | -1.27E+00 | 1.02E-01 | 2.79E-01 | -1.05E+00 |
| Z95152.1 | 2.67E-06 | 1.39E-05 | 2.25E+00 | 2.11E-01 | 9.13E-01 | 1.37E+00 | 1.27E-04 | 5.37E-04 | 2.21E+00 | 4.81E-02 | 1.84E-01 | 1.84E+00 | 1.70E-07 | 6.07E-07 | 4.02E+00 | 9.17E-02 | 2.59E-01 | 2.11E+00 | 1.77E-09 | 6.49E-09 | 3.25E+00 | 5.96E-02 | 1.64E-01 | 1.84E+00 |
| HIGD1A | 2.67E-06 | 1.39E-05 | -1.23E+00 | 3.80E-06 | 1.64E-05 | -1.13E+00 | 3.74E-04 | 1.59E-03 | -1.18E+00 | 7.96E-04 | 3.04E-03 | 1.02E+00 | 4.37E-08 | 1.56E-07 | -1.42E+00 | 1.67E-03 | 4.69E-03 | 1.01E+00 | 1.07E-12 | 3.91E-12 | -1.42E+00 | 1.13E-08 | 3.11E-08 | -1.35E+00 |
| TOX4 | 2.67E-06 | 1.39E-05 | -1.01E+00 | 5.27E-16 | 2.28E-15 | -1.20E+00 | 3.50E-04 | 1.49E-03 | -1.11E+00 | 3.15E-10 | 1.20E-09 | -1.27E+00 | 7.60E-04 | 2.71E-03 | -1.03E+00 | 3.30E-06 | 9.30E-06 | -1.20E+00 | 6.49E-02 | 2.38E-01 | 1.10E+00 | 1.53E-03 | 4.20E-03 | -1.12E+00 |
| CICP27 | 2.68E-06 | 1.39E-05 | -1.63E+00 | 1.23E-04 | 5.31E-04 | -1.21E+00 | 2.66E-06 | 1.13E-05 | -1.62E+00 | 2.60E-07 | 9.92E-07 | -1.56E+00 | 7.20E-02 | 2.57E-01 | 1.04E+00 | 1.39E-01 | 3.91E-01 | 1.20E+00 | 9.40E-03 | 3.45E-02 | -1.32E+00 | 1.38E-02 | 3.80E-02 | -1.35E+00 |
| PNMA1 | 2.68E-06 | 1.40E-05 | -1.24E+00 | 3.86E-04 | 1.67E-03 | 1.06E+00 | 6.54E-02 | 2.78E-01 | 1.08E+00 | 1.23E-01 | 4.68E-01 | 1.29E+00 | 1.09E-04 | 3.90E-04 | -1.29E+00 | 1.20E-03 | 3.39E-03 | -1.16E+00 | 7.14E-04 | 2.62E-03 | 1.11E+00 | 7.79E-02 | 2.14E-01 | 1.20E+00 |
| NGLY1 | 2.68E-06 | 1.40E-05 | -1.14E+00 | 5.68E-05 | 2.46E-04 | 1.06E+00 | 1.20E-05 | 5.09E-05 | -1.25E+00 | 3.31E-08 | 1.26E-07 | -1.24E+00 | 8.88E-04 | 3.16E-03 | 1.00E+00 | 7.61E-03 | 2.14E-02 | 1.05E+00 | 9.24E-04 | 3.39E-03 | -1.10E+00 | 2.02E-02 | 5.54E-02 | -1.02E+00 |
| AC108693.1 | 2.69E-06 | 1.40E-05 | -1.68E+00 | 1.53E-04 | 6.63E-04 | -1.24E+00 | 6.89E-03 | 2.93E-02 | -1.18E+00 | 1.60E-01 | 6.09E-01 | 1.32E+00 | 6.49E-02 | 2.31E-01 | 1.05E+00 | 1.20E-01 | 3.37E-01 | 1.08E+00 | 1.11E-02 | 4.06E-02 | -1.21E+00 | 2.05E-01 | 5.64E-01 | 1.11E+00 |
| GSEC | 2.69E-06 | 1.40E-05 | -1.87E+00 | 3.92E-06 | 1.69E-05 | -1.73E+00 | 3.18E-13 | 1.35E-12 | -2.35E+00 | 8.37E-07 | 3.20E-06 | -1.27E+00 | 5.93E-10 | 2.11E-09 | -3.13E+00 | 5.02E-08 | 1.42E-07 | -2.06E+00 | 2.99E-08 | 1.10E-07 | -2.28E+00 | 1.04E-03 | 2.85E-03 | -1.31E+00 |
| TRIM68 | 2.70E-06 | 1.41E-05 | -1.72E+00 | 4.95E-03 | 2.14E-02 | 1.04E+00 | 5.14E-04 | 2.18E-03 | -1.42E+00 | 4.70E-03 | 1.80E-02 | -1.15E+00 | 1.06E-06 | 3.79E-06 | -2.45E+00 | 7.84E-04 | 2.21E-03 | -1.39E+00 | 1.53E-04 | 5.63E-04 | -1.39E+00 | 5.81E-03 | 1.59E-02 | -1.14E+00 |
| TPT1-AS1 | 2.71E-06 | 1.41E-05 | -1.28E+00 | 8.75E-02 | 3.78E-01 | 1.52E+00 | 1.80E-07 | 7.63E-07 | -1.48E+00 | 1.19E-01 | 4.53E-01 | 1.32E+00 | 3.79E-05 | 1.35E-04 | -1.44E+00 | 2.57E-01 | 7.24E-01 | 1.28E+00 | 2.71E-04 | 9.95E-04 | -1.37E+00 | 3.23E-01 | 8.88E-01 | 1.31E+00 |
| VSTM4 | 2.72E-06 | 1.42E-05 | 3.13E+00 | 4.73E-03 | 2.04E-02 | 2.53E+00 | 3.05E-07 | 1.30E-06 | 4.08E+00 | 1.24E-04 | 4.75E-04 | 3.92E+00 | 1.53E-02 | 5.44E-02 | 3.68E+00 | 1.64E-01 | 4.61E-01 | 2.28E+00 | 2.40E-03 | 8.81E-03 | 2.80E+00 | 5.65E-02 | 1.55E-01 | 2.22E+00 |
| AKT2 | 2.72E-06 | 1.42E-05 | -1.16E+00 | 5.21E-06 | 2.25E-05 | -1.06E+00 | 8.10E-05 | 3.44E-04 | -1.16E+00 | 7.77E-06 | 2.97E-05 | -1.13E+00 | 1.54E-04 | 5.51E-04 | -1.11E+00 | 1.53E-04 | 4.32E-04 | -1.03E+00 | 8.63E-06 | 3.17E-05 | -1.28E+00 | 9.31E-04 | 2.56E-03 | -1.14E+00 |
| MTX2 | 2.74E-06 | 1.43E-05 | -1.16E+00 | 1.51E-05 | 6.51E-05 | 1.02E+00 | 1.26E-04 | 5.33E-04 | -1.20E+00 | 3.81E-05 | 1.46E-04 | -1.06E+00 | 2.54E-07 | 9.07E-07 | -1.57E+00 | 1.48E-03 | 4.18E-03 | -1.06E+00 | 8.25E-15 | 3.03E-14 | -1.59E+00 | 2.60E-06 | 7.13E-06 | -1.16E+00 |
| MPHOSPH6 | 2.75E-06 | 1.43E-05 | -1.23E+00 | 6.64E-06 | 2.87E-05 | -1.07E+00 | 6.74E-03 | 2.87E-02 | -1.05E+00 | 1.13E-03 | 4.30E-03 | 1.04E+00 | 9.06E-06 | 3.23E-05 | -1.51E+00 | 2.13E-02 | 6.00E-02 | 1.02E+00 | 2.33E-05 | 8.54E-05 | -1.08E+00 | 1.14E-01 | 3.14E-01 | 1.14E+00 |
| RASGRP4 | 2.75E-06 | 1.43E-05 | -1.44E+00 | 5.18E-03 | 2.24E-02 | 1.07E+00 | 1.55E-03 | 6.57E-03 | -1.27E+00 | 1.69E-02 | 6.44E-02 | 1.07E+00 | 3.43E-10 | 1.22E-09 | -2.08E+00 | 2.38E-08 | 6.72E-08 | -1.65E+00 | 2.99E-02 | 1.10E-01 | -1.25E+00 | 8.97E-02 | 2.46E-01 | -1.11E+00 |
| SLCO3A1 | 2.75E-06 | 1.43E-05 | -1.40E+00 | 5.17E-06 | 2.23E-05 | -1.28E+00 | 3.27E-08 | 1.39E-07 | -1.41E+00 | 1.85E-14 | 7.07E-14 | -1.67E+00 | 2.93E-07 | 1.05E-06 | -1.50E+00 | 1.20E-10 | 3.38E-10 | -2.05E+00 | 8.21E-02 | 3.01E-01 | -1.04E+00 | 9.62E-02 | 2.64E-01 | 1.01E+00 |
| HPRT1 | 2.76E-06 | 1.44E-05 | -1.01E+00 | 9.47E-05 | 4.09E-04 | 1.12E+00 | 3.40E-02 | 1.44E-01 | 1.06E+00 | 3.89E-02 | 1.49E-01 | 1.16E+00 | 4.26E-07 | 1.52E-06 | -1.28E+00 | 1.37E-01 | 3.86E-01 | 1.22E+00 | 2.94E-02 | 1.08E-01 | 1.02E+00 | 5.46E-02 | 1.50E-01 | 1.34E+00 |
| DTNBP1 | 2.76E-06 | 1.44E-05 | -1.12E+00 | 2.99E-03 | 1.29E-02 | 1.16E+00 | 3.89E-05 | 1.65E-04 | -1.25E+00 | 4.55E-05 | 1.74E-04 | -1.06E+00 | 1.57E-04 | 5.58E-04 | -1.20E+00 | 1.16E-03 | 3.26E-03 | -1.05E+00 | 1.50E-02 | 5.51E-02 | -1.01E+00 | 7.27E-02 | 2.00E-01 | 1.12E+00 |
| GMPR2 | 2.76E-06 | 1.44E-05 | -1.10E+00 | 2.79E-04 | 1.20E-03 | 1.17E+00 | 2.96E-11 | 1.26E-10 | -1.59E+00 | 2.76E-06 | 1.05E-05 | -1.15E+00 | 1.10E-05 | 3.93E-05 | -1.22E+00 | 2.29E-03 | 6.46E-03 | -1.02E+00 | 8.54E-03 | 3.13E-02 | 1.03E+00 | 2.36E-01 | 6.47E-01 | 1.15E+00 |
| PNN | 2.76E-06 | 1.44E-05 | -1.09E+00 | 1.87E-01 | 8.07E-01 | 1.15E+00 | 3.94E-05 | 1.67E-04 | -1.29E+00 | 1.98E-01 | 7.56E-01 | -1.01E+00 | 4.39E-02 | 1.56E-01 | -1.03E+00 | 3.02E-02 | 8.52E-02 | 1.24E+00 | 1.24E-03 | 4.54E-03 | -1.12E+00 | 6.66E-02 | 1.83E-01 | 1.14E+00 |
| HMOX2 | 2.76E-06 | 1.44E-05 | -1.22E+00 | 1.32E-02 | 5.72E-02 | 1.26E+00 | 1.07E-07 | 4.56E-07 | -1.26E+00 | 1.23E-04 | 4.69E-04 | 1.02E+00 | 1.52E-01 | 5.43E-01 | 1.20E+00 | 2.42E-01 | 6.83E-01 | 1.34E+00 | 3.56E-05 | 1.31E-04 | -1.38E+00 | 4.51E-02 | 1.24E-01 | 1.06E+00 |

| | | | | | | | | | | | | | | | | | | | | | | | | |
|---|---|---|---|---|---|---|---|---|---|---|---|---|---|---|---|---|---|---|---|---|---|---|---|---|
| STAC3 | 2.76E-06 | 1.44E-05 | -2.73E+00 | 2.74E-05 | 1.18E-04 | -1.97E+00 | 1.63E-07 | 6.91E-07 | -1.67E+00 | 3.39E-10 | 1.30E-09 | -1.96E+00 | 4.04E-06 | 1.44E-05 | -1.86E+00 | 3.93E-07 | 1.11E-06 | -2.06E+00 | 9.09E-03 | 3.34E-02 | -1.25E+00 | 6.49E-03 | 1.78E-02 | -1.34E+00 |
| LAMTOR3 | 2.76E-06 | 1.44E-05 | -1.14E+00 | 1.42E-04 | 6.15E-04 | 1.11E+00 | 9.77E-05 | 4.15E-04 | -1.25E+00 | 7.91E-04 | 3.02E-03 | 1.00E+00 | 2.55E-03 | 9.09E-03 | -1.02E+00 | 3.86E-02 | 1.09E-01 | 1.15E+00 | 5.89E-08 | 2.16E-07 | -1.31E+00 | 2.30E-03 | 6.32E-03 | -1.04E+00 |
| GPR137C | 2.77E-06 | 1.44E-05 | -1.70E+00 | 6.84E-07 | 2.96E-06 | -1.80E+00 | 1.45E-03 | 6.16E-03 | -1.51E+00 | 8.33E-04 | 3.18E-03 | -1.54E+00 | 2.88E-08 | 1.03E-07 | -3.23E+00 | 1.03E-06 | 2.91E-06 | -1.70E+00 | 8.09E-04 | 2.97E-03 | -1.94E+00 | 6.57E-03 | 1.80E-02 | -1.63E+00 |
| VIRMA | 2.77E-06 | 1.44E-05 | -1.23E+00 | 1.30E-03 | 5.60E-03 | 1.11E+00 | 3.52E-05 | 1.50E-04 | -1.35E+00 | 1.30E-06 | 4.96E-06 | -1.20E+00 | 1.74E-04 | 6.21E-04 | -1.22E+00 | 5.53E-04 | 1.56E-03 | -1.17E+00 | 6.93E-05 | 2.54E-04 | -1.18E+00 | 1.55E-03 | 4.26E-03 | -1.05E+00 |
| ITPRIP | 2.77E-06 | 1.44E-05 | -1.45E+00 | 1.30E-09 | 5.62E-09 | -2.17E+00 | 1.52E-06 | 6.48E-06 | -1.66E+00 | 3.85E-08 | 1.47E-07 | -1.84E+00 | 2.86E-02 | 1.02E-01 | -1.04E+00 | 8.49E-04 | 2.39E-03 | -1.81E+00 | 2.83E-02 | 1.04E-01 | 1.03E+00 | 2.12E-03 | 5.81E-03 | -1.35E+00 |
| CASP2 | 2.77E-06 | 1.44E-05 | -1.07E+00 | 5.30E-02 | 2.29E-01 | 1.33E+00 | 7.63E-02 | 3.24E-01 | 1.02E+00 | 2.49E-01 | 9.51E-01 | 1.32E+00 | 4.71E-05 | 1.68E-04 | -1.02E+00 | 5.82E-05 | 1.64E-04 | 1.01E+00 | 2.95E-06 | 1.08E-05 | -1.26E+00 | 1.98E-01 | 5.43E-01 | 1.20E+00 |
| DUSP12 | 2.77E-06 | 1.44E-05 | -1.25E+00 | 4.99E-02 | 2.16E-01 | 1.32E+00 | 7.09E-03 | 3.01E-02 | 1.00E+00 | 7.85E-02 | 3.00E-01 | 1.26E+00 | 4.64E-03 | 1.65E-02 | 1.04E+00 | 2.50E-01 | 7.05E-01 | 1.43E+00 | 1.98E-01 | 7.28E-01 | 1.27E+00 | 2.68E-01 | 7.36E-01 | 1.37E+00 |
| RBMS2 | 2.78E-06 | 1.45E-05 | -1.48E+00 | 1.92E-05 | 8.30E-05 | -1.25E+00 | 1.05E-05 | 4.45E-05 | -1.33E+00 | 2.28E-05 | 8.69E-05 | -1.11E+00 | 1.75E-01 | 6.24E-01 | 2.02E+00 | 2.82E-01 | 7.96E-01 | 1.56E+00 | 2.71E-01 | 9.93E-01 | 1.17E+00 | 3.58E-01 | 9.85E-01 | 1.12E+00 |
| GNPTG | 2.79E-06 | 1.46E-05 | -1.22E+00 | 8.96E-05 | 3.87E-04 | -1.01E+00 | 6.98E-03 | 2.96E-02 | -1.18E+00 | 1.79E-03 | 6.85E-03 | -1.10E+00 | 3.33E-06 | 1.19E-05 | -1.37E+00 | 5.12E-07 | 1.44E-06 | -1.49E+00 | 1.54E-05 | 5.64E-05 | -1.24E+00 | 1.09E-05 | 3.00E-05 | -1.32E+00 |
| AC007406.3 | 2.81E-06 | 1.46E-05 | -1.66E+00 | 1.83E-03 | 7.89E-03 | -1.09E+00 | 7.29E-04 | 3.10E-03 | -1.64E+00 | 1.04E-01 | 3.95E-01 | 1.19E+00 | 1.84E-02 | 6.54E-02 | -1.64E+00 | 5.38E-02 | 1.52E-01 | -1.20E+00 | 1.87E-02 | 6.88E-02 | -1.55E+00 | 8.12E-02 | 2.23E-01 | -1.15E+00 |
| AC007938.3 | 2.82E-06 | 1.47E-05 | -2.18E+00 | 1.88E-02 | 8.13E-02 | 1.02E+00 | 5.25E-03 | 2.23E-02 | -1.55E+00 | 3.89E-02 | 1.49E-01 | -1.09E+00 | 2.47E-01 | 8.80E-01 | 1.31E+00 | 2.65E-01 | 7.46E-01 | 1.16E+00 | 1.74E-02 | 6.37E-02 | -1.42E+00 | 2.13E-01 | 5.86E-01 | 1.03E+00 |
| CCDC82 | 2.84E-06 | 1.48E-05 | -1.22E+00 | 4.18E-05 | 1.81E-04 | 1.00E+00 | 1.01E-04 | 4.28E-04 | -1.27E+00 | 2.08E-04 | 7.96E-04 | -1.12E+00 | 2.68E-06 | 9.55E-06 | -1.34E+00 | 6.72E-04 | 1.90E-03 | -1.05E+00 | 1.32E-11 | 4.85E-11 | -1.55E+00 | 1.93E-05 | 5.31E-05 | -1.18E+00 |
| RPS20P14 | 2.84E-06 | 1.48E-05 | -1.04E+00 | 3.12E-09 | 1.35E-08 | -1.02E+00 | 5.21E-02 | 2.21E-01 | -1.01E+00 | 8.73E-04 | 3.33E-03 | -1.01E+00 | 2.64E-03 | 9.42E-03 | -1.01E+00 | 1.20E-03 | 3.37E-03 | 1.02E+00 | 5.09E-06 | 1.87E-05 | -1.14E+00 | 1.12E-03 | 3.07E-03 | -1.02E+00 |
| DOCK8 | 2.85E-06 | 1.48E-05 | -1.11E+00 | 7.02E-03 | 3.03E-02 | 1.24E+00 | 1.60E-06 | 6.79E-06 | -1.38E+00 | 1.72E-04 | 6.57E-04 | -1.10E+00 | 2.89E-07 | 1.03E-06 | -1.24E+00 | 1.04E-04 | 2.93E-04 | -1.08E+00 | 1.34E-03 | 4.91E-03 | -1.06E+00 | 2.66E-02 | 7.32E-02 | 1.02E+00 |
| RUNDC1 | 2.85E-06 | 1.49E-05 | -1.56E+00 | 3.15E-05 | 1.36E-04 | -1.25E+00 | 4.54E-05 | 1.93E-04 | -1.36E+00 | 6.41E-05 | 2.45E-04 | -1.22E+00 | 1.09E-02 | 3.88E-02 | -1.24E+00 | 3.13E-03 | 8.81E-03 | -1.54E+00 | 4.19E-04 | 1.54E-03 | -1.26E+00 | 1.70E-05 | 4.67E-05 | -1.61E+00 |
| SYNGAP1 | 2.85E-06 | 1.49E-05 | -1.27E+00 | 2.30E-04 | 9.94E-04 | 1.04E+00 | 7.37E-13 | 3.13E-12 | -1.75E+00 | 5.51E-07 | 2.11E-06 | -1.14E+00 | 5.70E-03 | 2.03E-02 | -1.16E+00 | 7.27E-02 | 2.05E-01 | 1.10E+00 | 1.81E-07 | 6.63E-07 | -1.46E+00 | 4.70E-04 | 1.29E-03 | -1.13E+00 |
| CCDC34 | 2.88E-06 | 1.50E-05 | -1.16E+00 | 1.50E-05 | 6.48E-05 | -1.01E+00 | 4.21E-02 | 1.79E-01 | 1.03E+00 | 1.57E-03 | 6.00E-03 | 1.02E+00 | 8.32E-04 | 2.97E-03 | -1.22E+00 | 2.90E-02 | 8.19E-02 | 1.06E+00 | 1.51E-04 | 5.54E-04 | -1.10E+00 | 8.86E-02 | 2.43E-01 | 1.02E+00 |
| ZNF486 | 2.88E-06 | 1.50E-05 | -1.49E+00 | 9.21E-05 | 3.98E-04 | -1.14E+00 | 6.13E-05 | 2.61E-04 | -1.51E+00 | 1.05E-04 | 4.00E-04 | -1.30E+00 | 5.52E-02 | 1.97E-01 | 1.03E+00 | 1.81E-02 | 5.11E-02 | -1.16E+00 | 2.10E-03 | 7.70E-03 | -1.31E+00 | 6.57E-03 | 1.80E-02 | -1.27E+00 |
| PIP5K1C | 2.89E-06 | 1.50E-05 | -1.56E+00 | 4.15E-03 | 1.79E-02 | 1.07E+00 | 3.47E-02 | 1.47E-01 | -1.03E+00 | 2.51E-01 | 9.61E-01 | 1.39E+00 | 1.73E-01 | 6.16E-01 | 1.30E+00 | 1.59E-01 | 4.47E-01 | 1.13E+00 | 2.61E-01 | 9.57E-01 | 1.49E+00 | 3.39E-01 | 9.31E-01 | 1.51E+00 |
| DYNLT3 | 2.89E-06 | 1.51E-05 | -1.15E+00 | 1.27E-07 | 5.48E-07 | -1.10E+00 | 3.75E-04 | 1.59E-03 | -1.10E+00 | 3.99E-04 | 1.52E-03 | -1.03E+00 | 2.05E-04 | 7.31E-04 | -1.03E+00 | 5.86E-05 | 1.65E-04 | -1.13E+00 | 5.23E-06 | 1.92E-05 | -1.22E+00 | 2.79E-05 | 7.66E-05 | -1.27E+00 |
| TAF1B | 2.90E-06 | 1.51E-05 | -1.44E+00 | 4.68E-06 | 2.02E-05 | -1.27E+00 | 1.06E-04 | 4.52E-04 | -1.22E+00 | 9.46E-05 | 3.61E-04 | -1.19E+00 | 7.69E-06 | 2.74E-05 | -1.13E+00 | 1.88E-04 | 5.30E-04 | -1.10E+00 | 1.48E-09 | 5.44E-09 | -1.47E+00 | 1.14E-04 | 3.12E-04 | -1.12E+00 |
| RBM10 | 2.90E-06 | 1.51E-05 | -1.32E+00 | 5.95E-04 | 2.57E-03 | 1.03E+00 | 5.80E-03 | 2.46E-02 | -1.05E+00 | 1.90E-03 | 7.28E-03 | 1.05E+00 | 3.88E-04 | 1.38E-03 | -1.13E+00 | 1.43E-02 | 4.02E-02 | 1.08E+00 | 1.34E-06 | 4.92E-06 | -1.21E+00 | 2.41E-03 | 6.62E-03 | -1.03E+00 |
| CETN3 | 2.91E-06 | 1.52E-05 | -1.44E+00 | 1.33E-05 | 5.73E-05 | -1.18E+00 | 1.46E-17 | 6.19E-17 | -2.14E+00 | 3.81E-10 | 1.45E-09 | -1.29E+00 | 1.20E-04 | 4.29E-04 | -2.00E+00 | 5.06E-02 | 1.43E-01 | -1.04E+00 | 1.38E-15 | 5.06E-15 | -2.01E+00 | 6.36E-06 | 1.75E-05 | -1.22E+00 |
| LINC01184 | 2.91E-06 | 1.52E-05 | -1.12E+00 | 1.53E-03 | 6.60E-03 | 1.22E+00 | 2.04E-05 | 8.65E-05 | -1.38E+00 | 1.21E-03 | 4.61E-03 | -1.10E+00 | 1.21E-03 | 4.33E-03 | -1.21E+00 | 6.28E-02 | 1.77E-01 | 1.07E+00 | 2.31E-05 | 8.49E-05 | -1.24E+00 | 2.91E-02 | 8.00E-02 | 1.09E+00 |
| CD55 | 2.91E-06 | 1.52E-05 | -1.04E+00 | 2.52E-10 | 1.09E-09 | -1.10E+00 | 1.60E-04 | 6.79E-04 | -1.11E+00 | 1.84E-07 | 7.01E-07 | -1.11E+00 | 7.55E-07 | 2.69E-06 | -1.25E+00 | 1.66E-05 | 4.69E-05 | -1.20E+00 | 2.59E-06 | 9.51E-06 | -1.10E+00 | 5.59E-07 | 1.54E-06 | -1.27E+00 |
| SELP | 2.91E-06 | 1.52E-05 | -2.91E+00 | 2.62E-05 | 1.13E-04 | -2.04E+00 | 5.72E-02 | 2.43E-01 | -1.45E+00 | 8.49E-02 | 3.24E-01 | -1.24E+00 | 3.80E-02 | 1.35E-01 | -2.67E+00 | 2.19E-02 | 6.19E-02 | -2.94E+00 | 1.26E-03 | 4.64E-03 | -1.16E+00 | 2.97E-04 | 8.16E-04 | -1.95E+00 |
| EMP1 | 2.92E-06 | 1.52E-05 | 1.98E+00 | 1.38E-01 | 5.97E-01 | 1.52E+00 | 2.20E-01 | 9.34E-01 | 1.14E+00 | 2.68E-04 | 1.02E-03 | -1.05E+00 | 5.13E-03 | 1.83E-02 | 3.24E+00 | 1.84E-01 | 5.17E-01 | -1.24E+00 | 6.14E-02 | 2.25E-01 | 3.08E+00 | 3.31E-01 | 9.10E-01 | 1.43E+00 |
| AC068587.6 | 2.93E-06 | 1.53E-05 | -1.46E+00 | 1.34E-04 | 5.78E-04 | -1.14E+00 | 4.98E-06 | 2.12E-05 | -1.37E+00 | 1.98E-04 | 7.55E-04 | -1.12E+00 | 9.27E-05 | 3.31E-04 | -1.64E+00 | 5.42E-04 | 1.53E-03 | -1.33E+00 | 5.36E-06 | 1.97E-05 | -2.17E+00 | 3.60E-03 | 9.90E-03 | -1.36E+00 |
| AL121603.2 | 2.95E-06 | 1.54E-05 | -1.33E+00 | 4.19E-09 | 1.81E-08 | -1.52E+00 | 5.79E-03 | 2.46E-02 | -1.17E+00 | 4.15E-03 | 1.59E-02 | -1.13E+00 | 5.75E-04 | 2.05E-03 | -1.42E+00 | 4.91E-04 | 1.38E-03 | -1.44E+00 | 3.73E-04 | 1.37E-03 | -1.48E+00 | 3.38E-04 | 9.29E-04 | -1.62E+00 |
| PCDHGA6 | 2.96E-06 | 1.54E-05 | -1.36E+00 | 3.05E-03 | 1.32E-02 | 1.13E+00 | 1.81E-04 | 7.68E-04 | -1.40E+00 | 8.25E-03 | 3.15E-02 | -1.03E+00 | 2.22E-03 | 7.90E-03 | -1.45E+00 | 3.03E-02 | 8.55E-02 | -1.13E+00 | 2.11E-01 | 7.74E-01 | -1.03E+00 | 3.33E-01 | 9.14E-01 | 1.24E+00 |
| CCNJ | 2.97E-06 | 1.55E-05 | -1.69E+00 | 9.53E-06 | 4.12E-05 | -1.45E+00 | 1.99E-05 | 8.47E-05 | -1.67E+00 | 3.02E-05 | 1.15E-04 | -1.38E+00 | 9.83E-03 | 3.50E-02 | -1.05E+00 | 1.91E-03 | 5.39E-03 | -1.20E+00 | 5.47E-07 | 2.01E-06 | -1.45E+00 | 1.39E-03 | 3.83E-03 | -1.11E+00 |
| TTC4 | 2.98E-06 | 1.55E-05 | -1.85E+00 | 1.17E-04 | 5.05E-04 | -1.37E+00 | 7.07E-04 | 3.00E-03 | -1.44E+00 | 1.79E-05 | 6.84E-05 | -1.78E+00 | 3.91E-04 | 1.39E-03 | -1.75E+00 | 1.74E-03 | 4.91E-03 | -1.44E+00 | 2.57E-02 | 9.42E-02 | -1.14E+00 | 2.96E-02 | 8.12E-02 | -1.09E+00 |
| AVPI1 | 3.00E-06 | 1.56E-05 | -1.79E+00 | 3.69E-05 | 1.59E-04 | -1.41E+00 | 7.57E-03 | 3.22E-02 | -1.35E+00 | 1.39E-02 | 5.31E-02 | -1.17E+00 | 3.94E-03 | 1.41E-02 | -1.31E+00 | 1.11E-03 | 3.14E-03 | -1.52E+00 | 2.25E-05 | 8.27E-05 | -1.41E+00 | 3.83E-04 | 1.05E-03 | -1.25E+00 |
| HSD3B7 | 3.01E-06 | 1.57E-05 | -1.62E+00 | 2.17E-03 | 9.37E-03 | -1.02E+00 | 3.35E-05 | 1.42E-04 | -1.78E+00 | 1.71E-02 | 6.52E-02 | -1.02E+00 | 2.58E-08 | 9.21E-08 | -2.62E+00 | 4.60E-04 | 1.30E-03 | -1.23E+00 | 1.75E-05 | 6.44E-05 | -1.55E+00 | 2.20E-03 | 6.04E-03 | -1.35E+00 |
| SFR1 | 3.01E-06 | 1.57E-05 | -1.23E+00 | 5.90E-06 | 2.55E-05 | -1.09E+00 | 2.20E-04 | 9.36E-04 | -1.23E+00 | 6.66E-04 | 2.55E-03 | -1.07E+00 | 1.01E-03 | 3.59E-03 | -1.24E+00 | 2.33E-01 | 6.56E-01 | 1.25E+00 | 2.42E-05 | 8.90E-05 | -1.21E+00 | 6.82E-03 | 1.87E-02 | -1.11E+00 |
| DPP10 | 3.01E-06 | 1.57E-05 | -1.80E+00 | 1.37E-04 | 5.90E-04 | -1.31E+00 | 7.60E-04 | 3.23E-03 | -1.80E+00 | 1.71E-01 | 6.52E-01 | 1.34E+00 | 3.11E-09 | 1.11E-08 | -2.16E+00 | 1.22E-04 | 3.43E-04 | -1.20E+00 | 5.11E-06 | 1.88E-05 | -1.47E+00 | 2.12E-04 | 5.82E-04 | -1.23E+00 |
| RPL15P3 | 3.02E-06 | 1.57E-05 | 1.78E+00 | 3.98E-05 | 1.72E-04 | 1.92E+00 | 1.60E-05 | 6.80E-05 | 1.65E+00 | 7.11E-06 | 2.72E-05 | 1.88E+00 | 1.56E-04 | 5.57E-04 | 1.90E+00 | 3.89E-08 | 1.10E-07 | 2.32E+00 | 8.70E-06 | 3.19E-05 | 1.85E+00 | 8.97E-10 | 2.46E-09 | 2.12E+00 |
| SIMC1 | 3.02E-06 | 1.58E-05 | -1.58E+00 | 3.74E-03 | 1.61E-02 | 1.04E+00 | 1.91E-03 | 8.11E-03 | -1.18E+00 | 7.46E-04 | 2.85E-03 | -1.17E+00 | 2.50E-03 | 8.93E-03 | -1.56E+00 | 1.46E-02 | 4.11E-02 | -1.16E+00 | 1.85E-01 | 6.80E-01 | 1.06E+00 | 2.45E-01 | 6.73E-01 | 1.11E+00 |
| UBE2O | 3.03E-06 | 1.58E-05 | -1.50E+00 | 2.89E-03 | 1.25E-02 | 1.01E+00 | 1.14E-05 | 4.85E-05 | -1.44E+00 | 2.32E-02 | 8.87E-02 | 1.14E+00 | 3.70E-07 | 1.32E-06 | -1.63E+00 | 1.31E-02 | 3.69E-02 | 1.10E+00 | 3.25E-11 | 1.19E-10 | -1.75E+00 | 8.03E-06 | 2.21E-05 | -1.27E+00 |
| DAPP1 | 3.04E-06 | 1.58E-05 | -1.28E+00 | 2.37E-03 | 1.02E-02 | 1.11E+00 | 1.37E-08 | 5.84E-08 | -1.93E+00 | 5.52E-06 | 2.11E-05 | -1.39E+00 | 6.06E-03 | 2.16E-02 | -1.56E+00 | 4.99E-03 | 1.41E-02 | -1.62E+00 | 6.53E-02 | 2.40E-01 | 1.55E+00 | 9.16E-03 | 2.52E-02 | -1.04E+00 |
| SLC12A5 | 3.04E-06 | 1.59E-05 | -1.39E+00 | 5.33E-09 | 2.30E-08 | -1.77E+00 | 6.54E-03 | 2.78E-02 | -1.34E+00 | 3.54E-07 | 1.35E-06 | -2.28E+00 | 1.91E-01 | 6.82E-01 | 1.04E+00 | 1.68E-01 | 4.75E-01 | -1.04E+00 | 2.02E-01 | 7.40E-01 | -1.04E+00 | 1.67E-01 | 4.58E-01 | -1.16E+00 |
| CCDC91 | 3.05E-06 | 1.59E-05 | -1.15E+00 | 2.00E-05 | 8.64E-05 | 1.02E+00 | 7.07E-04 | 3.01E-03 | -1.10E+00 | 1.64E-02 | 6.27E-02 | 1.17E+00 | 1.13E-03 | 4.02E-03 | -1.11E+00 | 3.89E-04 | 1.10E-03 | -1.14E+00 | 1.67E-01 | 6.13E-01 | 1.08E+00 | 2.82E-01 | 7.75E-01 | 1.19E+00 |
| RPSAP52 | 3.06E-06 | 1.59E-05 | -1.92E+00 | 2.09E-04 | 9.02E-04 | 2.36E+00 | 5.58E-02 | 2.37E-01 | -1.28E+00 | 1.89E-02 | 7.21E-02 | 1.89E+00 | 9.07E-02 | 3.23E-01 | -1.25E+00 | 2.65E-01 | 7.46E-01 | 1.35E+00 | 4.93E-02 | 1.81E-01 | -1.37E+00 | 2.36E-01 | 6.48E-01 | 1.48E+00 |
| GALNT11 | 3.06E-06 | 1.60E-05 | -1.23E+00 | 9.56E-07 | 4.13E-06 | -1.20E+00 | 6.94E-05 | 2.95E-04 | -1.29E+00 | 1.12E-05 | 4.28E-05 | -1.27E+00 | 8.41E-07 | 3.00E-06 | -1.54E+00 | 1.76E-06 | 4.97E-06 | -1.43E+00 | 3.08E-03 | 1.13E-02 | 1.04E+00 | 4.37E-02 | 1.20E-01 | 1.08E+00 |
| FUCA2 | 3.07E-06 | 1.60E-05 | -1.49E+00 | 9.70E-06 | 4.19E-05 | -1.26E+00 | 2.80E-04 | 1.19E-03 | -1.23E+00 | 3.66E-04 | 1.40E-03 | -1.03E+00 | 4.40E-07 | 1.57E-06 | -1.58E+00 | 1.16E-05 | 3.27E-05 | -1.31E+00 | 1.22E-06 | 4.47E-06 | -1.23E+00 | 8.02E-06 | 2.20E-05 | -1.10E+00 |
| UCK2 | 3.07E-06 | 1.60E-05 | -1.41E+00 | 4.09E-05 | 1.77E-04 | -1.14E+00 | 7.38E-05 | 3.14E-04 | -1.11E+00 | 3.83E-04 | 1.46E-03 | 1.02E+00 | 8.36E-03 | 2.98E-02 | -1.17E+00 | 8.25E-02 | 2.33E-01 | 1.04E+00 | 8.73E-06 | 3.21E-05 | -1.28E+00 | 4.02E-02 | 1.11E-01 | 1.09E+00 |
| MRPS30 | 3.07E-06 | 1.60E-05 | -1.02E+00 | 4.92E-07 | 2.13E-06 | 1.02E+00 | 8.32E-04 | 3.54E-03 | -1.12E+00 | 9.51E-03 | 3.63E-02 | 1.10E+00 | 1.62E-07 | 5.79E-07 | -1.27E+00 | 1.30E-03 | 3.66E-03 | 1.03E+00 | 3.53E-05 | 1.30E-04 | -1.08E+00 | 1.47E-02 | 4.04E-02 | 1.12E+00 |
| TAF12 | 3.08E-06 | 1.60E-05 | -1.11E+00 | 3.20E-04 | 1.38E-03 | 1.14E+00 | 4.20E-03 | 1.79E-02 | -1.07E+00 | 1.66E-03 | 6.33E-03 | 1.02E+00 | 4.40E-06 | 1.57E-05 | -1.30E+00 | 1.98E-02 | 5.59E-02 | 1.10E+00 | 2.67E-03 | 9.79E-03 | -1.01E+00 | 4.81E-02 | 1.32E-01 | 1.12E+00 |
| PTRH2 | 3.08E-06 | 1.61E-05 | -1.55E+00 | 3.28E-04 | 1.42E-03 | -1.13E+00 | 1.16E-09 | 4.93E-09 | -1.69E+00 | 2.93E-06 | 1.12E-05 | -1.26E+00 | 6.71E-03 | 2.39E-02 | -1.13E+00 | 2.18E-02 | 6.15E-02 | -1.03E+00 | 2.07E-09 | 7.59E-09 | -1.64E+00 | 1.18E-04 | 3.24E-04 | -1.19E+00 |
| RAB40B | 3.08E-06 | 1.61E-05 | -1.49E+00 | 5.09E-03 | 2.20E-02 | 1.08E+00 | 1.23E-08 | 5.21E-08 | -1.95E+00 | 5.66E-05 | 2.16E-04 | -1.19E+00 | 4.95E-04 | 1.77E-03 | -1.61E+00 | 1.73E-03 | 4.87E-03 | -1.54E+00 | 3.84E-04 | 1.41E-03 | -1.38E+00 | 3.99E-02 | 1.10E-01 | 1.01E+00 |
| ATG2B | 3.10E-06 | 1.61E-05 | -1.45E+00 | 8.37E-05 | 3.62E-04 | -1.12E+00 | 1.16E-04 | 4.93E-04 | -1.42E+00 | 3.38E-03 | 1.29E-02 | -1.02E+00 | 6.25E-02 | 2.23E-01 | -1.02E+00 | 1.50E-01 | 4.23E-01 | 1.18E+00 | 2.47E-07 | 9.06E-07 | -1.72E+00 | 8.58E-04 | 2.36E-03 | -1.17E+00 |
| ARPC5 | 3.10E-06 | 1.62E-05 | -1.02E+00 | 3.06E-02 | 1.32E-01 | 1.42E+00 | 2.26E-01 | 9.61E-01 | 1.13E+00 | 1.06E-02 | 4.05E-02 | 1.42E+00 | 7.54E-06 | 2.69E-05 | -1.12E+00 | 2.05E-01 | 5.78E-01 | 1.18E+00 | 2.21E-01 | 8.12E-01 | 1.02E+00 | 1.18E-01 | 3.25E-01 | 1.18E+00 |
| SDE2 | 3.11E-06 | 1.62E-05 | -1.04E+00 | 1.08E-24 | 4.66E-24 | -1.97E+00 | 2.58E-05 | 1.10E-04 | -1.13E+00 | 3.99E-16 | 1.53E-15 | -1.75E+00 | 2.88E-02 | 1.03E-01 | 1.23E+00 | 2.23E-06 | 6.28E-06 | -1.30E+00 | 1.58E-04 | 5.79E-04 | -1.06E+00 | 1.81E-13 | 4.99E-13 | -1.71E+00 |
| PDP1 | 3.11E-06 | 1.62E-05 | -1.36E+00 | 2.07E-07 | 8.95E-07 | -1.44E+00 | 8.76E-05 | 3.72E-04 | -1.50E+00 | 9.26E-06 | 3.54E-05 | -1.45E+00 | 2.37E-11 | 8.46E-11 | -2.29E+00 | 7.16E-13 | 2.02E-12 | -2.66E+00 | 2.01E-06 | 7.38E-06 | -1.93E+00 | 3.04E-06 | 8.35E-06 | -1.89E+00 |
| GPS1 | 3.11E-06 | 1.62E-05 | -1.18E+00 | 1.38E-06 | 5.97E-06 | -1.05E+00 | 2.63E-03 | 1.12E-02 | -1.00E+00 | 8.14E-03 | 3.11E-02 | 1.18E+00 | 3.98E-06 | 1.42E-05 | -1.33E+00 | 5.39E-03 | 1.52E-02 | 1.03E+00 | 3.78E-02 | 1.39E-01 | 1.11E+00 | 2.19E-01 | 6.01E-01 | 1.40E+00 |
| DUSP19 | 3.11E-06 | 1.62E-05 | -1.82E+00 | 1.63E-04 | 7.06E-04 | -1.30E+00 | 8.90E-04 | 3.78E-03 | -1.58E+00 | 7.02E-03 | 2.68E-02 | -1.24E+00 | 5.51E-04 | 1.96E-03 | -1.90E+00 | 6.41E-02 | 1.81E-01 | -1.03E+00 | 4.29E-11 | 1.57E-10 | -2.24E+00 | 1.91E-08 | 5.25E-08 | -1.75E+00 |
| BTBD10 | 3.11E-06 | 1.62E-05 | -1.68E+00 | 2.69E-04 | 1.16E-03 | -1.19E+00 | 3.14E-09 | 1.34E-08 | -1.77E+00 | 9.36E-09 | 3.58E-08 | -1.51E+00 | 3.72E-07 | 1.33E-06 | -2.13E+00 | 1.81E-05 | 5.10E-05 | -1.59E+00 | 6.71E-09 | 2.46E-08 | -1.43E+00 | 3.11E-07 | 8.55E-07 | -1.52E+00 |
| MAPK11 | 3.12E-06 | 1.63E-05 | -1.48E+00 | 7.24E-06 | 3.13E-05 | -1.28E+00 | 1.18E-02 | 5.03E-02 | -1.26E+00 | 1.98E-02 | 7.55E-02 | -1.11E+00 | 1.51E-01 | 5.39E-01 | 1.02E+00 | 1.33E-01 | 3.75E-01 | -1.15E+00 | 1.56E-01 | 5.71E-01 | 1.61E+00 | 3.58E-01 | 9.83E-01 | 1.24E+00 |
| PHF1 | 3.13E-06 | 1.63E-05 | -1.21E+00 | 9.59E-05 | 4.14E-04 | -1.02E+00 | 4.57E-06 | 1.94E-05 | -1.46E+00 | 4.96E-05 | 1.89E-04 | -1.17E+00 | 1.08E-04 | 3.85E-04 | -1.34E+00 | 1.51E-04 | 4.26E-04 | -1.34E+00 | 3.71E-04 | 1.36E-03 | -1.20E+00 | 5.57E-03 | 1.53E-02 | -1.09E+00 |
| HMG20A | 3.13E-06 | 1.63E-05 | -1.31E+00 | 1.61E-06 | 6.94E-06 | -1.20E+00 | 4.67E-04 | 1.99E-03 | -1.37E+00 | 1.93E-02 | 7.37E-02 | 1.10E+00 | 7.03E-02 | 2.51E-01 | 1.12E+00 | 2.42E-01 | 6.81E-01 | 1.43E+00 | 6.83E-03 | 2.51E-02 | -1.09E+00 | 1.06E-02 | 2.91E-02 | -1.12E+00 |
| ZBTB16 | 3.13E-06 | 1.63E-05 | 1.88E+00 | 1.92E-01 | 8.30E-01 | 1.40E+00 | 8.90E-02 | 3.78E-01 | 1.30E+00 | 3.61E-02 | 1.38E-01 | 1.15E+00 | 7.01E-06 | 2.50E-05 | 1.99E+00 | 2.70E-01 | 7.62E-01 | 1.31E+00 | 3.11E-06 | 1.14E-05 | 2.04E+00 | 1.59E-03 | 4.38E-03 | 1.64E+00 |
| TMED7 | 3.14E-06 | 1.64E-05 | -1.15E+00 | 4.19E-06 | 1.81E-05 | -1.02E+00 | 2.60E-03 | 1.11E-02 | -1.02E+00 | 1.53E-03 | 5.84E-03 | 1.03E+00 | 2.27E-03 | 8.08E-03 | -1.07E+00 | 8.56E-04 | 2.41E-03 | -1.04E+00 | 7.48E-06 | 2.74E-05 | -1.31E+00 | 6.07E-03 | 1.67E-02 | -1.14E+00 |
| NSD2 | 3.15E-06 | 1.64E-05 | -1.31E+00 | 6.87E-05 | 2.97E-04 | -1.09E+00 | 1.59E-02 | 6.76E-02 | -1.02E+00 | 1.19E-02 | 4.54E-02 | 1.16E+00 | 7.56E-05 | 2.70E-04 | -1.15E+00 | 1.09E-02 | 3.06E-02 | 1.13E+00 | 5.17E-13 | 1.90E-12 | -1.50E+00 | 3.53E-08 | 9.69E-08 | -1.21E+00 |
| NUBPL | 3.15E-06 | 1.64E-05 | -1.44E+00 | 2.14E-03 | 9.23E-03 | 1.02E+00 | 5.22E-06 | 2.22E-05 | -1.51E+00 | 1.36E-04 | 5.19E-04 | -1.09E+00 | 3.01E-04 | 1.07E-03 | -1.31E+00 | 2.08E-02 | 5.87E-02 | 1.10E+00 | 8.65E-05 | 3.17E-04 | -1.31E+00 | 4.72E-02 | 1.30E-01 | 1.10E+00 |
| VPS36 | 3.17E-06 | 1.65E-05 | -1.11E+00 | 2.76E-04 | 1.19E-03 | 1.12E+00 | 7.61E-06 | 3.23E-05 | -1.22E+00 | 4.41E-02 | 1.69E-01 | 1.14E+00 | 1.27E-04 | 4.53E-04 | -1.14E+00 | 2.37E-03 | 6.69E-03 | -1.01E+00 | 8.41E-06 | 3.09E-05 | -1.26E+00 | 1.57E-03 | 4.32E-03 | -1.13E+00 |
| INSR | 3.19E-06 | 1.66E-05 | -1.36E+00 | 2.92E-03 | 1.26E-02 | 1.10E+00 | 2.27E-03 | 9.65E-03 | -1.18E+00 | 1.31E-02 | 5.02E-02 | 1.12E+00 | 3.59E-02 | 1.28E-01 | 1.56E+00 | 2.88E-02 | 8.12E-02 | 1.69E+00 | 1.86E-02 | 6.84E-02 | -1.07E+00 | 1.24E-01 | 3.41E-01 | 1.37E+00 |
| HIF1A | 3.19E-06 | 1.66E-05 | -1.01E+00 | 7.29E-06 | 3.15E-05 | 1.11E+00 | 1.88E-02 | 7.99E-02 | 1.06E+00 | 4.09E-03 | 1.56E-02 | 1.09E+00 | 1.76E-01 | 6.29E-01 | 1.49E+00 | 3.35E-01 | 9.45E-01 | 1.27E+00 | 4.21E-02 | 1.55E-01 | 1.10E+00 | 3.09E-04 | 8.49E-04 | -1.17E+00 |
| HSCB | 3.20E-06 | 1.67E-05 | -1.10E+00 | 2.64E-05 | 1.14E-04 | 1.06E+00 | 1.35E-03 | 5.74E-03 | -1.03E+00 | 9.17E-04 | 3.50E-03 | 1.09E+00 | 5.81E-06 | 2.07E-05 | -1.20E+00 | 8.02E-04 | 2.26E-03 | 1.05E+00 | 8.88E-02 | 3.26E-01 | 1.21E+00 | 4.06E-02 | 1.12E-01 | 1.08E+00 |
| XKR8 | 3.20E-06 | 1.67E-05 | -2.07E+00 | 7.78E-05 | 3.36E-04 | -1.48E+00 | 2.72E-08 | 1.15E-07 | -1.95E+00 | 7.87E-05 | 3.01E-04 | -1.28E+00 | 3.25E-03 | 1.16E-02 | -1.77E+00 | 4.66E-02 | 1.31E-01 | -1.16E+00 | 5.50E-08 | 2.02E-07 | -1.75E+00 | 1.60E-04 | 4.41E-04 | -1.19E+00 |
| RNF175 | 3.20E-06 | 1.67E-05 | -2.88E+00 | 3.15E-05 | 1.36E-04 | -2.17E+00 | 1.75E-10 | 7.42E-10 | -2.50E+00 | 7.47E-07 | 2.85E-06 | -1.55E+00 | 2.24E-01 | 7.98E-01 | -1.04E+00 | 2.51E-01 | 7.08E-01 | 1.82E+00 | 1.98E-03 | 7.26E-03 | 2.32E+00 | 1.22E-01 | 3.34E-01 | 1.55E+00 |
| MAD1L1 | 3.21E-06 | 1.67E-05 | -1.22E+00 | 9.35E-03 | 4.04E-02 | 1.25E+00 | 1.46E-02 | 6.19E-02 | -1.14E+00 | 3.04E-02 | 1.16E-01 | 1.08E+00 | 2.13E-06 | 7.60E-06 | -1.59E+00 | 3.81E-05 | 1.07E-04 | -1.32E+00 | 5.02E-04 | 1.84E-03 | -1.08E+00 | 1.80E-01 | 4.95E-01 | 1.18E+00 |
| LSM14B | 3.21E-06 | 1.67E-05 | -1.56E+00 | 3.49E-04 | 1.51E-03 | -1.09E+00 | 1.56E-02 | 6.63E-02 | -1.09E+00 | 3.00E-02 | 1.15E-01 | 1.08E+00 | 1.32E-04 | 4.72E-04 | -1.61E+00 | 6.58E-02 | 1.85E-01 | 1.00E+00 | 7.32E-04 | 2.69E-03 | -1.02E+00 | 1.61E-03 | 4.41E-03 | -1.03E+00 |
| SNX3 | 3.22E-06 | 1.68E-05 | -1.10E+00 | 9.49E-02 | 4.10E-01 | 1.11E+00 | 5.43E-04 | 2.31E-03 | -1.18E+00 | 1.67E-01 | 6.38E-01 | 1.05E+00 | 6.13E-06 | 2.18E-05 | -1.02E+00 | 2.56E-01 | 7.22E-01 | 1.20E+00 | 3.05E-02 | 1.12E-01 | -1.03E+00 | 2.59E-01 | 7.10E-01 | -1.01E+00 |
| EIF2S2P4 | 3.24E-06 | 1.69E-05 | -1.08E+00 | 1.18E-13 | 5.10E-13 | -1.47E+00 | 1.05E-02 | 4.46E-02 | -1.06E+00 | 2.04E-12 | 7.79E-12 | -1.59E+00 | 1.76E-03 | 6.27E-03 | -1.12E+00 | 1.18E-05 | 3.34E-05 | -1.47E+00 | 1.15E-05 | 4.21E-05 | -1.15E+00 | 6.22E-15 | 1.71E-14 | -1.84E+00 |
| UBE2V1P1 | 3.27E-06 | 1.70E-05 | -1.45E+00 | 1.81E-09 | 7.83E-09 | -2.06E+00 | 9.45E-14 | 4.02E-13 | -2.16E+00 | 1.87E-16 | 7.14E-16 | -2.33E+00 | 1.26E-04 | 4.51E-04 | -1.60E+00 | 2.62E-05 | 7.37E-05 | -2.10E+00 | 7.04E-08 | 2.58E-07 | -1.57E+00 | 3.29E-11 | 9.04E-11 | -2.48E+00 |
| TGFBRAP1 | 3.27E-06 | 1.71E-05 | -1.14E+00 | 2.60E-03 | 1.12E-02 | 1.15E+00 | 8.16E-06 | 3.47E-05 | -1.25E+00 | 1.53E-03 | 5.86E-03 | 1.07E+00 | 3.31E-02 | 1.18E-01 | 1.09E+00 | 3.05E-01 | 8.59E-01 | 1.38E+00 | 2.80E-02 | 1.03E-01 | -1.02E+00 | 2.12E-01 | 5.83E-01 | 1.16E+00 |
| IGSF6 | 3.27E-06 | 1.71E-05 | -2.31E+00 | 9.47E-05 | 4.09E-04 | -1.65E+00 | 2.38E-05 | 1.01E-04 | -2.08E+00 | 1.99E-06 | 7.58E-06 | -2.09E+00 | 8.21E-08 | 2.93E-07 | -2.36E+00 | 4.58E-08 | 1.29E-07 | -2.40E+00 | 5.06E-08 | 1.86E-07 | -2.28E+00 | 1.39E-05 | 3.83E-05 | -1.69E+00 |
| FTSJ3 | 3.27E-06 | 1.71E-05 | -1.26E+00 | 8.92E-04 | 3.85E-03 | 1.10E+00 | 3.70E-04 | 1.57E-03 | -1.19E+00 | 5.36E-04 | 2.05E-03 | 1.00E+00 | 2.33E-04 | 8.32E-04 | -1.25E+00 | 4.10E-02 | 1.16E-01 | 1.12E+00 | 8.47E-06 | 3.11E-05 | -1.23E+00 | 1.40E-01 | 3.86E-01 | 1.19E+00 |
| STK38 | 3.28E-06 | 1.71E-05 | -1.43E+00 | 6.12E-02 | 2.64E-01 | 1.30E+00 | 1.27E-09 | 5.42E-09 | -1.70E+00 | 7.38E-05 | 2.82E-04 | -1.07E+00 | 2.63E-05 | 9.39E-05 | -1.50E+00 | 4.52E-03 | 1.27E-02 | -1.13E+00 | 1.17E-07 | 4.30E-07 | -1.55E+00 | 4.49E-04 | 1.23E-03 | -1.24E+00 |
| ANKZF1 | 3.28E-06 | 1.71E-05 | -1.26E+00 | 6.48E-03 | 2.80E-02 | 1.12E+00 | 4.82E-03 | 2.05E-02 | -1.11E+00 | 1.51E-02 | 5.76E-02 | 1.10E+00 | 1.04E-04 | 3.71E-04 | -1.25E+00 | 3.64E-02 | 1.03E-01 | 1.07E+00 | 2.34E-07 | 8.60E-07 | -1.35E+00 | 9.39E-05 | 2.58E-04 | -1.19E+00 |
| APOPT1 | 3.31E-06 | 1.72E-05 | -1.16E+00 | 3.43E-05 | 1.48E-04 | 1.02E+00 | 9.39E-02 | 3.99E-01 | 1.09E+00 | 6.16E-02 | 2.35E-01 | 1.23E+00 | 3.62E-03 | 1.29E-02 | 1.08E+00 | 3.23E-03 | 9.10E-03 | 1.06E+00 | 4.80E-03 | 1.76E-02 | 1.02E+00 | 1.12E-01 | 3.07E-01 | 1.12E+00 |
| SRBD1 | 3.31E-06 | 1.72E-05 | -1.24E+00 | 2.49E-05 | 1.08E-04 | -1.02E+00 | 4.55E-14 | 1.93E-13 | -1.77E+00 | 3.68E-11 | 1.41E-10 | -1.39E+00 | 1.49E-08 | 5.33E-08 | -1.48E+00 | 3.08E-07 | 8.67E-07 | -1.31E+00 | 1.24E-11 | 4.56E-11 | -1.52E+00 | 4.30E-11 | 1.18E-10 | -1.64E+00 |
| SLC7A6OS | 3.31E-06 | 1.72E-05 | -1.54E+00 | 2.57E-05 | 1.11E-04 | -1.29E+00 | 1.37E-04 | 5.84E-04 | -1.56E+00 | 6.60E-03 | 2.52E-02 | -1.13E+00 | 1.03E-01 | 3.66E-01 | 1.05E+00 | 1.25E-01 | 3.52E-01 | 1.02E+00 | 7.92E-06 | 2.91E-05 | -1.40E+00 | 4.58E-05 | 1.26E-04 | -1.35E+00 |
| AL035078.1 | 3.32E-06 | 1.73E-05 | -1.85E+00 | 1.56E-07 | 6.75E-07 | -2.35E+00 | 1.71E-01 | 7.27E-01 | 1.27E+00 | 4.12E-02 | 1.57E-01 | -1.19E+00 | 6.48E-04 | 2.31E-03 | -1.58E+00 | 2.95E-06 | 8.31E-06 | -2.89E+00 | 1.43E-06 | 5.24E-06 | -1.68E+00 | 5.90E-11 | 1.62E-10 | -2.61E+00 |
| FBXW4P1 | 3.34E-06 | 1.74E-05 | 2.77E+00 | 7.10E-04 | 3.07E-03 | 2.66E+00 | 1.24E-07 | 5.28E-07 | 2.84E+00 | 3.39E-04 | 1.29E-03 | 2.62E+00 | 2.78E-02 | 9.92E-02 | 2.43E+00 | 2.07E-01 | 5.82E-01 | 1.78E+00 | 9.78E-02 | 3.59E-01 | 1.64E+00 | 2.32E-01 | 6.37E-01 | -1.05E+00 |
| RPL5P8 | 3.34E-06 | 1.74E-05 | 2.18E+00 | 1.06E-03 | 4.60E-03 | 2.22E+00 | 3.13E-06 | 1.33E-05 | 2.06E+00 | 5.72E-04 | 2.18E-03 | 2.08E+00 | 1.03E-02 | 3.67E-02 | 1.84E+00 | 1.28E-03 | 3.60E-03 | 2.07E+00 | 2.87E-04 | 1.05E-03 | 1.94E+00 | 5.71E-04 | 1.57E-03 | 2.08E+00 |
| ATP5LP4 | 3.36E-06 | 1.75E-05 | 2.33E+00 | 1.13E-01 | 4.90E-01 | 1.68E+00 | 1.40E-06 | 5.96E-06 | 2.41E+00 | 1.28E-02 | 4.87E-02 | 1.98E+00 | 1.53E-01 | 5.47E-01 | 1.47E+00 | 2.91E-01 | 8.21E-01 | 1.27E+00 | 7.74E-05 | 2.84E-04 | 2.01E+00 | 8.30E-02 | 2.28E-01 | 1.53E+00 |
| AARSD1 | 3.36E-06 | 1.75E-05 | -1.66E+00 | 3.86E-03 | 1.67E-02 | 1.04E+00 | 3.58E-02 | 1.52E-01 | -1.11E+00 | 2.37E-01 | 9.06E-01 | 1.35E+00 | 1.10E-01 | 3.91E-01 | 1.19E+00 | 1.03E-01 | 2.90E-01 | 1.17E+00 | 1.56E-01 | 5.72E-01 | 1.04E+00 | 3.35E-01 | 9.21E-01 | 1.20E+00 |
| CASP8 | 3.37E-06 | 1.76E-05 | -1.04E+00 | 3.51E-03 | 1.52E-02 | 1.23E+00 | 5.51E-05 | 2.34E-04 | -1.13E+00 | 1.27E-03 | 4.87E-03 | 1.05E+00 | 1.78E-03 | 6.33E-03 | 1.00E+00 | 5.75E-04 | 1.62E-03 | -1.03E+00 | 2.64E-03 | 9.68E-03 | -1.09E+00 | 1.23E-01 | 3.37E-01 | 1.10E+00 |
| ATP5C1 | 3.38E-06 | 1.76E-05 | -1.08E+00 | 5.46E-03 | 2.36E-02 | 1.06E+00 | 3.82E-02 | 1.62E-01 | -1.07E+00 | 5.84E-02 | 2.23E-01 | -1.01E+00 | 2.84E-03 | 1.01E-02 | -1.09E+00 | 1.74E-01 | 4.89E-01 | 1.03E+00 | 2.05E-02 | 7.52E-02 | -1.05E+00 | 2.51E-01 | 6.88E-01 | 1.03E+00 |
| SMG7 | 3.38E-06 | 1.76E-05 | -1.22E+00 | 2.72E-04 | 1.18E-03 | 1.05E+00 | 7.54E-04 | 3.21E-03 | -1.17E+00 | 1.80E-04 | 6.88E-04 | -1.02E+00 | 3.30E-06 | 1.18E-05 | -1.34E+00 | 1.63E-04 | 4.59E-04 | -1.17E+00 | 2.35E-06 | 8.63E-06 | -1.27E+00 | 3.65E-05 | 1.00E-04 | -1.21E+00 |
| PCSK7 | 3.39E-06 | 1.77E-05 | -1.23E+00 | 2.34E-09 | 1.01E-08 | -1.32E+00 | 7.86E-05 | 3.34E-04 | -1.30E+00 | 5.34E-04 | 2.04E-03 | -1.08E+00 | 3.50E-03 | 1.25E-02 | -1.13E+00 | 3.05E-03 | 8.59E-03 | -1.13E+00 | 4.99E-05 | 1.83E-04 | -1.38E+00 | 1.92E-05 | 5.29E-05 | -1.38E+00 |
| NOS1AP | 3.40E-06 | 1.77E-05 | -2.59E+00 | 5.94E-06 | 2.57E-05 | -2.47E+00 | 2.28E-01 | 9.67E-01 | 1.01E+00 | 2.50E-01 | 9.56E-01 | 1.16E+00 | 2.56E-01 | 9.12E-01 | -1.21E+00 | 3.22E-01 | 9.09E-01 | -1.19E+00 | 1.87E-03 | 6.86E-03 | -1.02E+00 | 1.70E-04 | 4.67E-04 | -1.17E+00 |
| RAB11B-AS1 | 3.40E-06 | 1.77E-05 | -1.74E+00 | 3.11E-06 | 1.34E-05 | -1.69E+00 | 2.33E-02 | 9.89E-02 | -1.13E+00 | 1.17E-02 | 4.49E-02 | -1.05E+00 | 6.09E-04 | 2.17E-03 | -1.85E+00 | 3.94E-03 | 1.11E-02 | -1.44E+00 | 2.35E-03 | 8.62E-03 | -1.16E+00 | 2.47E-05 | 6.78E-05 | -1.46E+00 |
| BCLAF1P2 | 3.40E-06 | 1.77E-05 | -1.12E+00 | 2.18E-15 | 9.44E-15 | -1.36E+00 | 1.60E-02 | 6.80E-02 | -1.02E+00 | 8.04E-07 | 3.07E-06 | -1.21E+00 | 2.08E-03 | 7.42E-03 | -1.09E+00 | 5.93E-05 | 1.67E-04 | -1.19E+00 | 1.17E-05 | 4.30E-05 | -1.15E+00 | 6.66E-06 | 1.83E-05 | -1.16E+00 |
| LMBR1 | 3.45E-06 | 1.80E-05 | -1.50E+00 | 1.37E-06 | 5.94E-06 | -1.43E+00 | 3.76E-04 | 1.60E-03 | -1.20E+00 | 6.06E-04 | 2.31E-03 | -1.07E+00 | 6.07E-03 | 2.16E-02 | -1.10E+00 | 3.65E-02 | 1.03E-01 | 1.10E+00 | 4.08E-03 | 1.50E-02 | -1.11E+00 | 4.63E-02 | 1.27E-01 | 1.04E+00 |
| ZBTB49 | 3.46E-06 | 1.80E-05 | -1.97E+00 | 1.81E-05 | 7.82E-05 | -1.64E+00 | 9.41E-04 | 4.00E-03 | -1.47E+00 | 3.26E-03 | 1.24E-02 | -1.22E+00 | 1.60E-02 | 5.69E-02 | -1.60E+00 | 5.66E-02 | 1.60E-01 | -1.18E+00 | 1.40E-03 | 5.15E-03 | -1.37E+00 | 2.25E-02 | 6.18E-02 | -1.15E+00 |
| CRKL | 3.46E-06 | 1.80E-05 | -1.58E+00 | 1.78E-03 | 7.71E-03 | -1.05E+00 | 1.74E-08 | 7.38E-08 | -1.58E+00 | 6.63E-07 | 2.53E-06 | -1.24E+00 | 9.82E-04 | 3.50E-03 | -1.41E+00 | 1.19E-02 | 3.35E-02 | -1.14E+00 | 1.20E-09 | 4.40E-09 | -1.31E+00 | 1.21E-04 | 3.32E-04 | -1.11E+00 |
| RBMX2P5 | 3.46E-06 | 1.80E-05 | -1.29E+00 | 1.66E-07 | 7.15E-07 | -1.26E+00 | 8.22E-03 | 3.49E-02 | -1.10E+00 | 5.69E-04 | 2.17E-03 | -1.09E+00 | 2.69E-03 | 9.60E-03 | -1.20E+00 | 2.46E-03 | 6.92E-03 | -1.19E+00 | 4.89E-03 | 1.80E-02 | 1.05E+00 | 1.34E-05 | 3.68E-05 | -1.36E+00 |
| WASL | 3.47E-06 | 1.81E-05 | -1.17E+00 | 9.66E-10 | 4.17E-09 | -1.34E+00 | 8.13E-04 | 3.45E-03 | -1.12E+00 | 2.98E-07 | 1.14E-06 | -1.38E+00 | 1.41E-02 | 5.04E-02 | -1.04E+00 | 4.02E-03 | 1.13E-02 | -1.26E+00 | 7.86E-07 | 2.88E-06 | -1.39E+00 | 1.34E-11 | 3.68E-11 | -1.94E+00 |
| SERPINE2 | 3.47E-06 | 1.81E-05 | -1.05E+00 | 2.27E-03 | 9.80E-03 | 1.22E+00 | 8.25E-02 | 3.50E-01 | 1.13E+00 | 2.05E-01 | 7.84E-01 | 1.45E+00 | 1.01E-05 | 3.62E-05 | -1.08E+00 | 1.42E-01 | 4.00E-01 | 1.28E+00 | 2.70E-04 | 9.91E-04 | -1.13E+00 | 1.24E-01 | 3.40E-01 | 1.06E+00 |
| AC116533.1 | 3.48E-06 | 1.81E-05 | -1.11E+00 | 1.14E-121 | 4.93E-121 | -1.78E+00 | 1.78E-03 | 7.55E-03 | -1.09E+00 | 1.03E-67 | 3.95E-67 | -1.70E+00 | 7.01E-05 | 2.50E-04 | -1.14E+00 | 4.80E-55 | 1.35E-54 | -1.71E+00 | 2.03E-07 | 7.46E-07 | -1.14E+00 | 1.47E-79 | 4.04E-79 | -1.78E+00 |
| SPATA2 | 3.49E-06 | 1.82E-05 | -1.50E+00 | 7.22E-14 | 3.12E-13 | -3.28E+00 | 2.24E-04 | 9.50E-04 | -1.31E+00 | 4.36E-12 | 1.67E-11 | -2.39E+00 | 2.60E-05 | 9.28E-05 | -1.52E+00 | 1.87E-08 | 5.26E-08 | -3.01E+00 | 1.66E-05 | 6.10E-05 | -1.33E+00 | 5.28E-14 | 1.45E-13 | -2.85E+00 |
| FAM193A | 3.49E-06 | 1.82E-05 | -1.26E+00 | 9.23E-04 | 3.99E-03 | 1.10E+00 | 6.38E-06 | 2.71E-05 | -1.41E+00 | 3.80E-04 | 1.45E-03 | -1.04E+00 | 6.03E-07 | 2.15E-06 | -1.59E+00 | 9.82E-05 | 2.77E-04 | -1.29E+00 | 1.84E-07 | 6.74E-07 | -1.53E+00 | 6.13E-03 | 1.68E-02 | -1.09E+00 |
| UBALD1 | 3.49E-06 | 1.82E-05 | -1.42E+00 | 2.75E-04 | 1.19E-03 | -1.01E+00 | 5.98E-10 | 2.54E-09 | -1.76E+00 | 1.92E-06 | 7.32E-06 | -1.23E+00 | 1.90E-05 | 6.79E-05 | -1.73E+00 | 1.54E-03 | 4.35E-03 | -1.25E+00 | 4.89E-12 | 1.79E-11 | -1.73E+00 | 5.63E-06 | 1.55E-05 | -1.22E+00 |
| JARID2 | 3.50E-06 | 1.82E-05 | -1.15E+00 | 5.04E-09 | 2.18E-08 | -1.15E+00 | 1.04E-02 | 4.44E-02 | -1.06E+00 | 1.07E-05 | 4.10E-05 | -1.16E+00 | 1.43E-01 | 5.09E-01 | 1.35E+00 | 1.15E-04 | 3.25E-04 | -1.07E+00 | 3.80E-03 | 1.40E-02 | -1.02E+00 | 8.88E-04 | 2.44E-03 | -1.13E+00 |
| BRD2 | 3.50E-06 | 1.82E-05 | -1.06E+00 | 5.60E-08 | 2.42E-07 | 1.01E+00 | 8.64E-04 | 3.67E-03 | -1.19E+00 | 4.34E-04 | 1.66E-03 | -1.05E+00 | 5.63E-06 | 2.01E-05 | -1.13E+00 | 5.30E-07 | 1.49E-06 | -1.15E+00 | 4.29E-06 | 1.58E-05 | -1.17E+00 | 7.14E-03 | 1.96E-02 | -1.04E+00 |
| CEP164 | 3.51E-06 | 1.83E-05 | -1.48E+00 | 3.43E-04 | 1.48E-03 | -1.11E+00 | 1.21E-04 | 5.15E-04 | -1.26E+00 | 1.28E-03 | 4.87E-03 | -1.05E+00 | 6.47E-03 | 2.31E-02 | -1.23E+00 | 1.01E-01 | 2.84E-01 | 1.15E+00 | 1.38E-08 | 5.08E-08 | -1.73E+00 | 2.22E-04 | 6.10E-04 | -1.24E+00 |
| MRPL58 | 3.52E-06 | 1.83E-05 | -1.20E+00 | 4.31E-04 | 1.86E-03 | 1.07E+00 | 3.41E-06 | 1.45E-05 | -1.18E+00 | 4.31E-06 | 1.65E-05 | -1.05E+00 | 1.36E-04 | 4.87E-04 | -1.18E+00 | 1.84E-02 | 5.20E-02 | 1.14E+00 | 5.47E-02 | 2.01E-01 | 1.06E+00 | 3.15E-01 | 8.66E-01 | 1.31E+00 |
| CAB39 | 3.53E-06 | 1.84E-05 | -1.29E+00 | 6.09E-04 | 2.63E-03 | 1.06E+00 | 2.66E-04 | 1.13E-03 | -1.20E+00 | 5.89E-04 | 2.25E-03 | -1.06E+00 | 2.75E-04 | 9.82E-04 | -1.05E+00 | 1.32E-01 | 3.72E-01 | 1.22E+00 | 1.05E-04 | 3.85E-04 | -1.08E+00 | 1.31E-01 | 3.59E-01 | 1.14E+00 |
| ago-04 | 3.54E-06 | 1.84E-05 | -1.32E+00 | 1.70E-04 | 7.34E-04 | -1.02E+00 | 9.44E-13 | 4.01E-12 | -1.69E+00 | 2.61E-12 | 9.96E-12 | -1.39E+00 | 1.93E-02 | 6.89E-02 | -1.02E+00 | 9.40E-03 | 2.65E-02 | -1.12E+00 | 8.86E-06 | 3.25E-05 | -1.43E+00 | 3.88E-06 | 1.07E-05 | -1.53E+00 |
| MRPL45P2 | 3.54E-06 | 1.85E-05 | -1.42E+00 | 3.01E-02 | 1.30E-01 | 1.28E+00 | 2.70E-05 | 1.15E-04 | -1.47E+00 | 1.10E-03 | 4.18E-03 | -1.08E+00 | 2.00E-05 | 7.13E-05 | -1.68E+00 | 4.57E-03 | 1.29E-02 | -1.05E+00 | 7.18E-03 | 2.64E-02 | -1.15E+00 | 1.83E-01 | 5.03E-01 | 1.19E+00 |
| RIDA | 3.55E-06 | 1.85E-05 | -1.17E+00 | 9.74E-06 | 4.21E-05 | -1.08E+00 | 2.24E-05 | 9.50E-05 | -1.28E+00 | 9.19E-07 | 3.51E-06 | -1.23E+00 | 2.26E-02 | 8.07E-02 | 1.11E+00 | 5.61E-02 | 1.58E-01 | 1.18E+00 | 6.12E-05 | 2.25E-04 | -1.17E+00 | 6.62E-03 | 1.82E-02 | -1.02E+00 |
| THNSL1 | 3.55E-06 | 1.85E-05 | -1.98E+00 | 3.22E-03 | 1.39E-02 | -1.07E+00 | 5.52E-10 | 2.34E-09 | -2.61E+00 | 4.98E-04 | 1.90E-03 | -1.20E+00 | 4.97E-02 | 1.77E-01 | -1.58E+00 | 1.94E-01 | 5.47E-01 | 1.61E+00 | 2.25E-11 | 8.26E-11 | -1.99E+00 | 3.67E-03 | 1.01E-02 | -1.07E+00 |
| WDR70 | 3.55E-06 | 1.85E-05 | -1.28E+00 | 3.96E-04 | 1.71E-03 | 1.05E+00 | 1.77E-04 | 7.51E-04 | -1.20E+00 | 1.50E-05 | 5.73E-05 | -1.14E+00 | 3.48E-03 | 1.24E-02 | -1.13E+00 | 1.19E-02 | 3.36E-02 | -1.00E+00 | 3.29E-04 | 1.21E-03 | -1.08E+00 | 3.80E-04 | 1.04E-03 | -1.12E+00 |
| MFSD3 | 3.55E-06 | 1.85E-05 | -1.75E+00 | 1.19E-03 | 5.13E-03 | -1.10E+00 | 1.27E-06 | 5.38E-06 | -1.59E+00 | 3.54E-05 | 1.35E-04 | -1.20E+00 | 1.06E-05 | 3.78E-05 | -1.86E+00 | 2.23E-02 | 6.29E-02 | 1.01E+00 | 1.29E-05 | 4.74E-05 | -1.26E+00 | 1.98E-03 | 5.44E-03 | -1.05E+00 |
| CYB5R1 | 3.57E-06 | 1.86E-05 | -1.21E+00 | 8.99E-03 | 3.88E-02 | 1.25E+00 | 6.25E-04 | 2.66E-03 | -1.16E+00 | 2.34E-03 | 8.93E-03 | 1.05E+00 | 3.03E-03 | 1.08E-02 | -1.08E+00 | 7.84E-02 | 2.21E-01 | 1.15E+00 | 1.35E-01 | 4.96E-01 | 1.13E+00 | 7.07E-02 | 1.94E-01 | 1.05E+00 |
| PRR5 | 3.58E-06 | 1.87E-05 | -1.11E+00 | 9.45E-06 | 4.08E-05 | 1.08E+00 | 4.89E-03 | 2.08E-02 | -1.07E+00 | 1.00E-01 | 3.84E-01 | 1.26E+00 | 2.10E-09 | 7.47E-09 | -1.77E+00 | 1.99E-05 | 5.60E-05 | -1.19E+00 | 3.21E-03 | 1.18E-02 | -1.17E+00 | 3.11E-01 | 8.55E-01 | 1.28E+00 |
| HS3ST3B1 | 3.60E-06 | 1.87E-05 | -1.82E+00 | 4.10E-05 | 1.77E-04 | -1.48E+00 | 1.25E-07 | 5.31E-07 | -1.67E+00 | 1.51E-06 | 5.76E-06 | -1.51E+00 | 2.26E-01 | 8.06E-01 | -1.29E+00 | 3.43E-01 | 9.66E-01 | -1.09E+00 | 1.29E-01 | 4.74E-01 | -1.22E+00 | 1.35E-01 | 3.70E-01 | -1.48E+00 |
| HELZ2 | 3.61E-06 | 1.88E-05 | -1.48E+00 | 3.82E-06 | 1.65E-05 | -1.44E+00 | 1.31E-12 | 5.57E-12 | -2.45E+00 | 7.33E-17 | 2.80E-16 | -3.26E+00 | 5.55E-06 | 1.98E-05 | -2.16E+00 | 1.03E-07 | 2.90E-07 | -3.45E+00 | 7.57E-06 | 2.78E-05 | -1.79E+00 | 1.27E-07 | 3.48E-07 | -3.76E+00 |
| SEC23B | 3.62E-06 | 1.89E-05 | -1.38E+00 | 3.98E-03 | 1.72E-02 | 1.14E+00 | 3.84E-03 | 1.63E-02 | -1.09E+00 | 4.70E-02 | 1.79E-01 | 1.19E+00 | 3.04E-05 | 1.08E-04 | -1.54E+00 | 8.76E-02 | 2.47E-01 | 1.16E+00 | 1.67E-11 | 6.13E-11 | -1.62E+00 | 3.64E-04 | 1.00E-03 | -1.05E+00 |
| ST8SIA6-AS1 | 3.63E-06 | 1.89E-05 | -1.53E+00 | 3.25E-04 | 1.40E-03 | -1.06E+00 | 1.82E-02 | 7.75E-02 | -1.30E+00 | 1.74E-01 | 6.64E-01 | 1.18E+00 | 2.85E-06 | 1.02E-05 | -1.38E+00 | 9.44E-03 | 2.66E-02 | -1.01E+00 | 1.28E-05 | 4.69E-05 | -1.31E+00 | 2.45E-02 | 6.73E-02 | 1.04E+00 |
| ZNF568 | 3.63E-06 | 1.89E-05 | -1.63E+00 | 1.84E-02 | 7.96E-02 | 1.17E+00 | 1.02E-03 | 4.32E-03 | -1.36E+00 | 2.33E-02 | 8.88E-02 | 1.02E+00 | 5.57E-05 | 1.99E-04 | -1.50E+00 | 5.57E-03 | 1.57E-02 | -1.16E+00 | 6.34E-05 | 2.33E-04 | -1.47E+00 | 1.06E-04 | 2.91E-04 | -1.45E+00 |
| GBE1 | 3.65E-06 | 1.90E-05 | -1.42E+00 | 6.33E-04 | 2.73E-03 | -1.06E+00 | 2.09E-01 | 8.89E-01 | 1.17E+00 | 2.40E-01 | 9.19E-01 | 1.34E+00 | 1.52E-04 | 5.43E-04 | -1.47E+00 | 2.35E-02 | 6.64E-02 | 1.00E+00 | 7.50E-03 | 2.75E-02 | -1.13E+00 | 4.56E-03 | 1.25E-02 | -1.08E+00 |
| PPM1A | 3.67E-06 | 1.91E-05 | -1.21E+00 | 1.66E-05 | 7.18E-05 | -1.02E+00 | 2.53E-05 | 1.07E-04 | -1.22E+00 | 2.60E-05 | 9.92E-05 | -1.02E+00 | 5.81E-04 | 2.07E-03 | -1.08E+00 | 7.40E-02 | 2.09E-01 | 1.18E+00 | 3.11E-10 | 1.14E-09 | -1.45E+00 | 1.02E-05 | 2.80E-05 | -1.20E+00 |
| PDE4DIP | 3.67E-06 | 1.91E-05 | -1.35E+00 | 2.69E-02 | 1.16E-01 | 1.30E+00 | 7.10E-04 | 3.02E-03 | -1.46E+00 | 1.26E-01 | 4.82E-01 | 1.11E+00 | 2.96E-02 | 1.05E-01 | -1.10E+00 | 1.27E-02 | 3.57E-02 | -1.24E+00 | 2.71E-01 | 9.95E-01 | 1.67E+00 | 3.63E-01 | 9.98E-01 | 1.37E+00 |
| ZNF783 | 3.68E-06 | 1.92E-05 | -1.41E+00 | 1.02E-03 | 4.40E-03 | -1.02E+00 | 1.23E-05 | 5.24E-05 | -1.81E+00 | 4.26E-03 | 1.63E-02 | -1.05E+00 | 3.21E-05 | 1.14E-04 | -2.06E+00 | 1.29E-03 | 3.63E-03 | -1.30E+00 | 4.75E-02 | 1.74E-01 | -1.39E+00 | 1.54E-01 | 4.23E-01 | 1.35E+00 |
| AGGF1P10 | 3.68E-06 | 1.92E-05 | -1.57E+00 | 3.73E-03 | 1.61E-02 | 1.05E+00 | 4.27E-05 | 1.82E-04 | -1.86E+00 | 7.80E-03 | 2.98E-02 | -1.11E+00 | 6.98E-04 | 2.49E-03 | -1.53E+00 | 3.41E-02 | 9.61E-02 | -1.06E+00 | 8.49E-05 | 3.12E-04 | -1.54E+00 | 5.99E-03 | 1.64E-02 | -1.21E+00 |
| ANKRD27 | 3.70E-06 | 1.93E-05 | -1.52E+00 | 3.05E-05 | 1.32E-04 | -1.26E+00 | 1.18E-13 | 5.01E-13 | -2.08E+00 | 2.75E-07 | 1.05E-06 | -1.30E+00 | 1.79E-13 | 6.39E-13 | -1.78E+00 | 1.68E-04 | 4.73E-04 | -1.12E+00 | 1.01E-14 | 3.72E-14 | -1.55E+00 | 3.08E-05 | 8.47E-05 | -1.14E+00 |
| LPCAT1 | 3.71E-06 | 1.93E-05 | -1.47E+00 | 7.35E-04 | 3.17E-03 | -1.03E+00 | 3.32E-10 | 1.41E-09 | -1.80E+00 | 1.46E-05 | 5.58E-05 | -1.20E+00 | 8.61E-06 | 3.07E-05 | -2.91E+00 | 3.47E-04 | 9.77E-04 | -1.69E+00 | 5.55E-04 | 2.04E-03 | -1.52E+00 | 3.35E-02 | 9.21E-02 | -1.06E+00 |
| CTDSPL2 | 3.71E-06 | 1.93E-05 | -1.15E+00 | 6.07E-06 | 2.62E-05 | 1.03E+00 | 8.46E-04 | 3.60E-03 | -1.06E+00 | 9.23E-04 | 3.53E-03 | 1.05E+00 | 7.51E-03 | 2.68E-02 | 1.08E+00 | 2.27E-02 | 6.40E-02 | 1.18E+00 | 2.78E-05 | 1.02E-04 | -1.11E+00 | 1.89E-04 | 5.19E-04 | -1.10E+00 |
| ANKRD11 | 3.75E-06 | 1.96E-05 | -1.11E+00 | 8.93E-04 | 3.86E-03 | 1.04E+00 | 1.29E-04 | 5.48E-04 | -1.16E+00 | 3.78E-04 | 1.44E-03 | 1.01E+00 | 1.11E-02 | 3.97E-02 | -1.05E+00 | 3.43E-02 | 9.68E-02 | 1.03E+00 | 1.87E-05 | 6.87E-05 | -1.24E+00 | 3.24E-01 | 8.91E-01 | 1.09E+00 |
| MMS22L | 3.75E-06 | 1.96E-05 | -1.24E+00 | 1.57E-02 | 6.77E-02 | 1.20E+00 | 2.92E-03 | 1.24E-02 | -1.14E+00 | 4.94E-02 | 1.89E-01 | 1.18E+00 | 3.85E-04 | 1.37E-03 | -1.25E+00 | 3.30E-02 | 9.31E-02 | 1.05E+00 | 2.29E-09 | 8.42E-09 | -1.29E+00 | 1.62E-02 | 4.45E-02 | 1.08E+00 |
| TMEM39B | 3.79E-06 | 1.98E-05 | -1.28E+00 | 2.59E-03 | 1.12E-02 | 1.19E+00 | 5.19E-04 | 2.20E-03 | -1.13E+00 | 2.58E-04 | 9.85E-04 | -1.10E+00 | 4.53E-05 | 1.62E-04 | -1.30E+00 | 4.04E-04 | 1.14E-03 | -1.14E+00 | 4.34E-04 | 1.59E-03 | -1.11E+00 | 1.04E-02 | 2.86E-02 | 1.00E+00 |
| AC008560.1 | 3.80E-06 | 1.98E-05 | -1.52E+00 | 1.21E-06 | 5.25E-06 | -1.44E+00 | 3.73E-03 | 1.59E-02 | -1.28E+00 | 6.23E-05 | 2.38E-04 | -1.38E+00 | 1.43E-04 | 5.10E-04 | -1.41E+00 | 2.00E-03 | 5.65E-03 | -1.12E+00 | 1.52E-04 | 5.56E-04 | -1.10E+00 | 8.49E-05 | 2.33E-04 | -1.17E+00 |
| AC012618.3 | 3.82E-06 | 1.99E-05 | -1.88E+00 | 5.75E-04 | 2.49E-03 | -1.16E+00 | 1.11E-04 | 4.70E-04 | -1.86E+00 | 2.73E-02 | 1.04E-01 | -1.11E+00 | 3.69E-03 | 1.31E-02 | -2.06E+00 | 5.18E-02 | 1.46E-01 | -1.39E+00 | 2.41E-01 | 8.83E-01 | 1.14E+00 | 3.48E-01 | 9.55E-01 | 1.20E+00 |
| WHAMMP3 | 3.82E-06 | 1.99E-05 | -1.20E+00 | 6.95E-03 | 3.01E-02 | 1.19E+00 | 3.40E-08 | 1.45E-07 | -1.53E+00 | 1.87E-06 | 7.15E-06 | -1.18E+00 | 1.04E-02 | 3.69E-02 | -1.01E+00 | 1.02E-02 | 2.88E-02 | 1.05E+00 | 1.96E-02 | 7.20E-02 | 1.10E+00 | 1.13E-02 | 3.10E-02 | -1.07E+00 |
| AC015883.1 | 3.82E-06 | 1.99E-05 | -1.29E+00 | 3.34E-04 | 1.44E-03 | -1.01E+00 | 5.73E-07 | 2.43E-06 | -1.70E+00 | 1.71E-05 | 6.54E-05 | -1.30E+00 | 1.50E-10 | 5.33E-10 | -2.37E+00 | 2.65E-09 | 7.48E-09 | -1.96E+00 | 1.40E-05 | 5.13E-05 | -2.11E+00 | 1.09E-03 | 2.98E-03 | -1.55E+00 |
| NDUFA6 | 3.85E-06 | 2.00E-05 | -1.10E+00 | 6.31E-12 | 2.73E-11 | -1.23E+00 | 4.86E-02 | 2.07E-01 | 1.05E+00 | 6.07E-05 | 2.32E-04 | -1.07E+00 | 1.39E-07 | 4.96E-07 | -1.26E+00 | 1.19E-06 | 3.34E-06 | -1.18E+00 | 7.99E-02 | 2.93E-01 | 1.20E+00 | 2.36E-01 | 6.48E-01 | 1.14E+00 |
| RRAGA | 3.85E-06 | 2.01E-05 | -1.12E+00 | 2.87E-04 | 1.24E-03 | 1.11E+00 | 5.99E-08 | 2.54E-07 | -1.28E+00 | 8.11E-08 | 3.10E-07 | -1.07E+00 | 7.97E-08 | 2.84E-07 | -1.32E+00 | 1.05E-03 | 2.95E-03 | 1.04E+00 | 6.21E-11 | 2.28E-10 | -1.54E+00 | 1.56E-06 | 4.29E-06 | -1.26E+00 |
| NOSIP | 3.86E-06 | 2.01E-05 | 1.01E+00 | 2.05E-04 | 8.85E-04 | 1.19E+00 | 1.37E-03 | 5.82E-03 | -1.13E+00 | 3.96E-04 | 1.51E-03 | -1.04E+00 | 2.20E-06 | 7.85E-06 | -1.14E+00 | 1.30E-02 | 3.67E-02 | 1.06E+00 | 9.49E-03 | 3.48E-02 | 1.07E+00 | 2.48E-01 | 6.80E-01 | 1.16E+00 |
| A2MP1 | 3.88E-06 | 2.02E-05 | -2.10E+00 | 1.23E-03 | 5.34E-03 | -1.18E+00 | 7.27E-04 | 3.09E-03 | -1.42E+00 | 1.88E-03 | 7.20E-03 | -1.21E+00 | 2.61E-03 | 9.31E-03 | -1.77E+00 | 9.64E-03 | 2.72E-02 | -1.39E+00 | 1.66E-01 | 6.08E-01 | 1.42E+00 | 3.03E-01 | 8.33E-01 | 1.29E+00 |
| CHCHD2P6 | 3.89E-06 | 2.03E-05 | -1.47E+00 | 5.52E-06 | 2.39E-05 | -1.35E+00 | 5.20E-07 | 2.21E-06 | -1.44E+00 | 9.55E-10 | 3.65E-09 | -1.68E+00 | 5.19E-03 | 1.85E-02 | -1.39E+00 | 1.26E-03 | 3.56E-03 | -1.76E+00 | 2.57E-10 | 9.44E-10 | -1.80E+00 | 5.70E-07 | 1.57E-06 | -1.35E+00 |
| MTERF3 | 3.90E-06 | 2.03E-05 | -1.65E+00 | 6.77E-03 | 2.92E-02 | 1.04E+00 | 3.85E-08 | 1.64E-07 | -1.74E+00 | 4.45E-04 | 1.70E-03 | -1.06E+00 | 6.20E-04 | 2.21E-03 | -1.51E+00 | 3.84E-03 | 1.08E-02 | -1.27E+00 | 4.13E-09 | 1.52E-08 | -1.68E+00 | 6.28E-03 | 1.72E-02 | -1.04E+00 |
| FP236241.1 | 3.90E-06 | 2.03E-05 | -1.80E+00 | 1.53E-02 | 6.62E-02 | 1.16E+00 | 1.85E-09 | 7.85E-09 | -1.88E+00 | 1.00E-01 | 3.84E-01 | 1.20E+00 | 7.88E-08 | 2.81E-07 | -1.84E+00 | 5.52E-02 | 1.56E-01 | 1.15E+00 | 8.91E-13 | 3.27E-12 | -1.52E+00 | 1.34E-01 | 3.67E-01 | 1.11E+00 |
| PRR14L | 3.90E-06 | 2.03E-05 | -1.18E+00 | 5.14E-09 | 2.22E-08 | -1.34E+00 | 1.27E-07 | 5.39E-07 | -1.45E+00 | 8.21E-08 | 3.14E-07 | -1.28E+00 | 1.68E-05 | 5.98E-05 | -1.23E+00 | 4.60E-05 | 1.30E-04 | -1.25E+00 | 1.99E-06 | 7.31E-06 | -1.29E+00 | 1.22E-07 | 3.35E-07 | -1.49E+00 |
| IDH3G | 3.90E-06 | 2.04E-05 | -1.02E+00 | 1.19E-04 | 5.16E-04 | 1.17E+00 | 7.44E-04 | 3.16E-03 | -1.12E+00 | 2.95E-03 | 1.12E-02 | 1.00E+00 | 6.07E-09 | 2.16E-08 | -1.29E+00 | 3.93E-07 | 1.11E-06 | -1.18E+00 | 6.61E-04 | 2.43E-03 | -1.08E+00 | 6.12E-02 | 1.68E-01 | 1.10E+00 |
| SEZ6L2 | 3.93E-06 | 2.05E-05 | -1.53E+00 | 2.46E-04 | 1.06E-03 | -1.12E+00 | 3.65E-03 | 1.55E-02 | -1.83E+00 | 8.90E-02 | 3.40E-01 | 1.00E+00 | 7.50E-02 | 2.68E-01 | -1.30E+00 | 1.75E-01 | 4.94E-01 | -1.09E+00 | 2.66E-01 | 9.76E-01 | 1.13E+00 | 3.04E-01 | 8.34E-01 | 1.01E+00 |
| PNPLA2 | 3.93E-06 | 2.05E-05 | -1.20E+00 | 2.43E-06 | 1.05E-05 | -1.07E+00 | 6.11E-04 | 2.60E-03 | -1.17E+00 | 2.71E-04 | 1.04E-03 | -1.06E+00 | 7.58E-05 | 2.70E-04 | -1.25E+00 | 4.24E-03 | 1.19E-02 | -1.00E+00 | 1.66E-04 | 6.11E-04 | -1.06E+00 | 9.34E-04 | 2.57E-03 | -1.06E+00 |
| TMEM117 | 3.94E-06 | 2.05E-05 | -2.59E+00 | 3.91E-05 | 1.69E-04 | -2.02E+00 | 2.06E-01 | 8.75E-01 | -1.05E+00 | 1.41E-01 | 5.38E-01 | -1.21E+00 | 1.84E-01 | 6.57E-01 | 1.04E+00 | 3.55E-01 | 9.99E-01 | 1.32E+00 | 2.03E-06 | 7.47E-06 | -1.73E+00 | 3.93E-03 | 1.08E-02 | -1.13E+00 |
| STX10 | 3.94E-06 | 2.06E-05 | -1.08E+00 | 4.85E-06 | 2.10E-05 | 1.08E+00 | 3.37E-04 | 1.43E-03 | -1.10E+00 | 1.88E-04 | 7.18E-04 | 1.00E+00 | 1.14E-03 | 4.08E-03 | -1.03E+00 | 3.27E-02 | 9.22E-02 | 1.12E+00 | 1.84E-01 | 6.77E-01 | 1.11E+00 | 2.97E-01 | 8.15E-01 | 1.22E+00 |
| RWDD2B | 3.98E-06 | 2.07E-05 | -1.47E+00 | 1.26E-05 | 5.45E-05 | -1.28E+00 | 4.38E-06 | 1.86E-05 | -1.52E+00 | 7.05E-04 | 2.69E-03 | -1.12E+00 | 1.62E-03 | 5.76E-03 | -1.30E+00 | 1.03E-01 | 2.91E-01 | 1.16E+00 | 5.58E-04 | 2.05E-03 | -1.09E+00 | 1.66E-02 | 4.56E-02 | -1.04E+00 |
| SEPT2 | 3.98E-06 | 2.08E-05 | -1.08E+00 | 4.74E-09 | 2.05E-08 | -1.05E+00 | 2.33E-06 | 9.90E-06 | -1.20E+00 | 1.55E-06 | 5.94E-06 | -1.04E+00 | 5.48E-02 | 1.95E-01 | 1.14E+00 | 1.85E-01 | 5.21E-01 | 1.19E+00 | 8.40E-02 | 3.08E-01 | 1.03E+00 | 1.30E-01 | 3.58E-01 | 1.08E+00 |
| ZMAT5 | 3.98E-06 | 2.08E-05 | -1.90E+00 | 5.89E-04 | 2.54E-03 | -1.20E+00 | 3.37E-03 | 1.43E-02 | -1.67E+00 | 3.08E-02 | 1.18E-01 | -1.15E+00 | 3.09E-09 | 1.10E-08 | -3.47E+00 | 5.14E-07 | 1.45E-06 | -2.26E+00 | 3.57E-07 | 1.31E-06 | -1.61E+00 | 9.49E-06 | 2.61E-05 | -1.71E+00 |
| GID4 | 3.99E-06 | 2.08E-05 | -1.31E+00 | 3.59E-05 | 1.55E-04 | -1.13E+00 | 1.81E-05 | 7.68E-05 | -1.51E+00 | 4.68E-04 | 1.79E-03 | -1.16E+00 | 5.90E-05 | 2.10E-04 | -1.57E+00 | 6.38E-04 | 1.80E-03 | -1.34E+00 | 7.11E-05 | 2.61E-04 | -1.37E+00 | 3.60E-02 | 9.88E-02 | 1.01E+00 |
| KANK1 | 3.99E-06 | 2.08E-05 | -1.59E+00 | 2.03E-01 | 8.77E-01 | 1.30E+00 | 6.77E-04 | 2.87E-03 | -1.44E+00 | 5.03E-02 | 1.92E-01 | 1.15E+00 | 1.91E-03 | 6.82E-03 | -1.30E+00 | 3.18E-01 | 8.96E-01 | 1.47E+00 | 2.14E-05 | 7.87E-05 | -1.61E+00 | 1.03E-01 | 2.82E-01 | -1.01E+00 |
| TMEM98-AS1 | 4.00E-06 | 2.08E-05 | -1.77E+00 | 1.67E-02 | 7.20E-02 | 1.11E+00 | 1.53E-06 | 6.52E-06 | -2.09E+00 | 1.59E-03 | 6.08E-03 | -1.18E+00 | 4.27E-04 | 1.52E-03 | -2.39E+00 | 1.97E-02 | 5.54E-02 | -1.28E+00 | 3.45E-03 | 1.27E-02 | -1.30E+00 | 8.60E-03 | 2.36E-02 | -1.13E+00 |
| ADI1 | 4.02E-06 | 2.09E-05 | -1.10E+00 | 1.10E-04 | 4.76E-04 | 1.10E+00 | 5.25E-02 | 2.23E-01 | 1.08E+00 | 9.15E-03 | 3.50E-02 | 1.08E+00 | 2.47E-04 | 8.81E-04 | -1.09E+00 | 2.33E-02 | 6.56E-02 | 1.05E+00 | 3.11E-02 | 1.14E-01 | 1.10E+00 | 4.18E-02 | 1.15E-01 | 1.14E+00 |
| TRAPPC2 | 4.02E-06 | 2.09E-05 | -1.24E+00 | 2.37E-02 | 1.02E-01 | 1.23E+00 | 2.93E-05 | 1.25E-04 | -1.46E+00 | 1.54E-02 | 5.89E-02 | 1.03E+00 | 3.21E-05 | 1.14E-04 | -1.05E+00 | 6.19E-02 | 1.75E-01 | 1.28E+00 | 2.58E-01 | 9.48E-01 | 1.21E+00 | 3.10E-01 | 8.50E-01 | 1.23E+00 |
| RBBP6 | 4.02E-06 | 2.10E-05 | -1.05E+00 | 1.55E-20 | 6.72E-20 | -1.25E+00 | 2.49E-02 | 1.06E-01 | -1.01E+00 | 3.74E-08 | 1.43E-07 | -1.21E+00 | 1.70E-03 | 6.05E-03 | -1.03E+00 | 4.35E-11 | 1.23E-10 | -1.45E+00 | 1.87E-02 | 6.85E-02 | 1.01E+00 | 8.42E-05 | 2.31E-04 | -1.18E+00 |
| RALGPS2 | 4.03E-06 | 2.10E-05 | -1.12E+00 | 3.26E-07 | 1.41E-06 | -1.07E+00 | 3.12E-06 | 1.33E-05 | -1.33E+00 | 2.00E-07 | 7.63E-07 | -1.27E+00 | 7.00E-04 | 2.50E-03 | -1.16E+00 | 1.35E-03 | 3.82E-03 | -1.13E+00 | 4.28E-04 | 1.57E-03 | -1.22E+00 | 2.76E-04 | 7.59E-04 | -1.20E+00 |
| JRKL | 4.03E-06 | 2.10E-05 | -1.50E+00 | 4.53E-03 | 1.96E-02 | 1.01E+00 | 4.45E-02 | 1.89E-01 | -1.05E+00 | 1.95E-01 | 7.44E-01 | 1.30E+00 | 6.25E-04 | 2.23E-03 | -1.48E+00 | 7.41E-03 | 2.09E-02 | -1.23E+00 | 2.84E-04 | 1.04E-03 | -1.29E+00 | 3.21E-02 | 8.82E-02 | 1.07E+00 |
| TPMT | 4.03E-06 | 2.10E-05 | -1.18E+00 | 6.84E-06 | 2.95E-05 | -1.05E+00 | 9.17E-04 | 3.89E-03 | -1.20E+00 | 4.05E-04 | 1.55E-03 | -1.06E+00 | 2.09E-06 | 7.47E-06 | -1.41E+00 | 3.76E-04 | 1.06E-03 | -1.17E+00 | 3.85E-06 | 1.41E-05 | -1.19E+00 | 2.41E-03 | 6.62E-03 | 1.04E+00 |
| PIK3CD | 4.04E-06 | 2.11E-05 | -1.31E+00 | 1.28E-03 | 5.54E-03 | 1.03E+00 | 1.42E-08 | 6.04E-08 | -1.71E+00 | 2.07E-04 | 7.89E-04 | -1.13E+00 | 5.48E-04 | 1.96E-03 | -1.76E+00 | 1.98E-02 | 5.59E-02 | -1.16E+00 | 1.63E-03 | 5.99E-03 | -1.95E+00 | 5.91E-02 | 1.62E-01 | -1.14E+00 |
| POU2F1 | 4.06E-06 | 2.12E-05 | -1.07E+00 | 2.47E-04 | 1.07E-03 | 1.17E+00 | 6.15E-10 | 2.61E-09 | -1.58E+00 | 1.88E-07 | 7.18E-07 | -1.19E+00 | 8.13E-06 | 2.90E-05 | -1.25E+00 | 1.67E-03 | 4.71E-03 | -1.02E+00 | 7.49E-07 | 2.75E-06 | -1.48E+00 | 2.09E-02 | 5.73E-02 | -1.00E+00 |
| TOMM34 | 4.06E-06 | 2.12E-05 | -1.40E+00 | 2.36E-05 | 1.02E-04 | -1.16E+00 | 3.58E-06 | 1.52E-05 | -1.37E+00 | 9.06E-08 | 3.46E-07 | -1.34E+00 | 2.75E-05 | 9.82E-05 | -1.49E+00 | 6.00E-05 | 1.69E-04 | -1.39E+00 | 9.83E-08 | 3.61E-07 | -1.29E+00 | 7.60E-06 | 2.09E-05 | -1.23E+00 |
| THAP1 | 4.08E-06 | 2.13E-05 | -1.28E+00 | 1.78E-05 | 7.70E-05 | -1.14E+00 | 6.09E-06 | 2.59E-05 | -1.31E+00 | 9.79E-08 | 3.74E-07 | -1.31E+00 | 2.88E-09 | 1.03E-08 | -1.78E+00 | 2.33E-09 | 6.57E-09 | -1.83E+00 | 4.95E-07 | 1.82E-06 | -1.54E+00 | 1.16E-04 | 3.19E-04 | -1.35E+00 |
| FMR1 | 4.10E-06 | 2.14E-05 | -1.27E+00 | 6.68E-05 | 2.89E-04 | -1.03E+00 | 1.92E-05 | 8.18E-05 | -1.40E+00 | 2.85E-04 | 1.09E-03 | -1.09E+00 | 1.48E-03 | 5.26E-03 | -1.19E+00 | 3.98E-03 | 1.12E-02 | -1.09E+00 | 8.38E-05 | 3.07E-04 | -1.24E+00 | 2.11E-02 | 5.80E-02 | -1.07E+00 |
| CCM2 | 4.13E-06 | 2.15E-05 | -1.16E+00 | 1.09E-01 | 4.70E-01 | 1.34E+00 | 1.67E-06 | 7.08E-06 | -1.28E+00 | 1.88E-03 | 7.18E-03 | 1.10E+00 | 2.51E-05 | 8.96E-05 | -1.05E+00 | 1.95E-02 | 5.49E-02 | 1.12E+00 | 2.76E-08 | 1.01E-07 | -1.32E+00 | 6.95E-05 | 1.91E-04 | -1.12E+00 |
| ERCC5 | 4.14E-06 | 2.16E-05 | -1.53E+00 | 1.63E-05 | 7.05E-05 | -1.33E+00 | 5.14E-05 | 2.19E-04 | -1.61E+00 | 2.89E-03 | 1.10E-02 | -1.08E+00 | 1.23E-02 | 4.37E-02 | -1.40E+00 | 3.22E-02 | 9.06E-02 | -1.26E+00 | 1.21E-02 | 4.43E-02 | -1.19E+00 | 3.41E-03 | 9.37E-03 | -1.45E+00 |
| SLC33A1 | 4.16E-06 | 2.17E-05 | -1.54E+00 | 1.15E-01 | 4.99E-01 | 1.26E+00 | 2.52E-06 | 1.07E-05 | -1.55E+00 | 1.01E-01 | 3.87E-01 | 1.30E+00 | 4.84E-07 | 1.73E-06 | -1.79E+00 | 9.54E-03 | 2.69E-02 | -1.02E+00 | 4.76E-04 | 1.75E-03 | -1.14E+00 | 3.49E-01 | 9.58E-01 | 1.33E+00 |
| FITM2 | 4.16E-06 | 2.17E-05 | -2.22E+00 | 1.01E-04 | 4.34E-04 | -1.59E+00 | 5.76E-02 | 2.45E-01 | 1.64E+00 | 4.70E-02 | 1.79E-01 | 1.94E+00 | 2.05E-01 | 7.32E-01 | 1.08E+00 | 2.13E-01 | 6.02E-01 | 1.58E+00 | 2.45E-03 | 8.98E-03 | -1.53E+00 | 1.21E-01 | 3.33E-01 | -1.12E+00 |
| ABCC4 | 4.17E-06 | 2.17E-05 | -1.80E+00 | 1.96E-04 | 8.47E-04 | -1.28E+00 | 1.78E-04 | 7.58E-04 | -1.43E+00 | 1.43E-02 | 5.45E-02 | 1.02E+00 | 8.57E-22 | 3.05E-21 | -1.73E+00 | 4.58E-07 | 1.29E-06 | -1.23E+00 | 1.26E-07 | 4.64E-07 | -1.23E+00 | 3.08E-03 | 8.46E-03 | -1.14E+00 |
| NRAS | 4.18E-06 | 2.18E-05 | -1.19E+00 | 2.25E-06 | 9.71E-06 | -1.08E+00 | 1.94E-03 | 8.24E-03 | -1.13E+00 | 3.42E-04 | 1.31E-03 | -1.09E+00 | 1.48E-08 | 5.28E-08 | -1.47E+00 | 1.87E-09 | 5.28E-09 | -1.48E+00 | 2.14E-04 | 7.85E-04 | -1.08E+00 | 1.39E-03 | 3.83E-03 | -1.03E+00 |
| ARHGAP1 | 4.19E-06 | 2.18E-05 | -1.18E+00 | 3.90E-03 | 1.69E-02 | 1.18E+00 | 1.39E-06 | 5.92E-06 | -1.43E+00 | 2.09E-05 | 7.99E-05 | -1.13E+00 | 3.02E-06 | 1.08E-05 | -1.59E+00 | 2.83E-03 | 7.98E-03 | -1.18E+00 | 6.28E-06 | 2.31E-05 | -1.30E+00 | 1.00E-03 | 2.75E-03 | -1.18E+00 |
| ZNF200 | 4.20E-06 | 2.19E-05 | -1.63E+00 | 2.39E-05 | 1.03E-04 | -1.44E+00 | 2.15E-05 | 9.13E-05 | -1.86E+00 | 1.60E-03 | 6.11E-03 | -1.28E+00 | 2.13E-02 | 7.58E-02 | -1.84E+00 | 2.78E-01 | 7.84E-01 | 1.17E+00 | 1.69E-06 | 6.22E-06 | -1.70E+00 | 4.61E-03 | 1.27E-02 | -1.12E+00 |
| PPIAP3 | 4.20E-06 | 2.19E-05 | -1.13E+00 | 3.48E-05 | 1.51E-04 | 1.08E+00 | 1.01E-02 | 4.29E-02 | 1.03E+00 | 1.12E-02 | 4.29E-02 | 1.16E+00 | 1.57E-06 | 5.60E-06 | -1.32E+00 | 1.13E-03 | 3.19E-03 | -1.07E+00 | 1.76E-06 | 6.47E-06 | -1.21E+00 | 2.98E-02 | 8.17E-02 | 1.06E+00 |
| PIAS4 | 4.23E-06 | 2.20E-05 | -1.73E+00 | 3.48E-03 | 1.50E-02 | -1.03E+00 | 1.00E-06 | 4.26E-06 | -1.74E+00 | 3.07E-04 | 1.17E-03 | -1.10E+00 | 1.03E-04 | 3.67E-04 | -1.51E+00 | 6.46E-04 | 1.82E-03 | -1.30E+00 | 8.10E-02 | 2.97E-01 | -1.07E+00 | 1.09E-01 | 2.99E-01 | 1.48E+00 |
| USF3 | 4.24E-06 | 2.21E-05 | -1.13E+00 | 2.35E-09 | 1.01E-08 | -1.23E+00 | 1.53E-05 | 6.49E-05 | -1.33E+00 | 6.12E-08 | 2.34E-07 | -1.39E+00 | 5.87E-05 | 2.09E-04 | -1.16E+00 | 3.49E-09 | 9.84E-09 | -1.56E+00 | 1.66E-05 | 6.10E-05 | -1.32E+00 | 3.00E-07 | 8.25E-07 | -1.54E+00 |
| MEIOC | 4.25E-06 | 2.21E-05 | -2.04E+00 | 7.50E-05 | 3.24E-04 | -1.43E+00 | 4.31E-04 | 1.83E-03 | -1.43E+00 | 2.95E-06 | 1.13E-05 | -1.69E+00 | 9.82E-03 | 3.50E-02 | -1.46E+00 | 1.52E-02 | 4.28E-02 | -1.45E+00 | 1.21E-04 | 4.43E-04 | -1.46E+00 | 4.46E-03 | 1.22E-02 | -1.16E+00 |
| KANSL2 | 4.26E-06 | 2.22E-05 | -1.19E+00 | 1.41E-06 | 6.11E-06 | -1.14E+00 | 1.53E-03 | 6.49E-03 | -1.19E+00 | 8.65E-05 | 3.31E-04 | -1.19E+00 | 1.92E-04 | 6.84E-04 | -1.11E+00 | 3.42E-05 | 9.65E-05 | -1.29E+00 | 1.31E-05 | 4.81E-05 | -1.26E+00 | 7.85E-04 | 2.16E-03 | -1.13E+00 |
| FASTKD1 | 4.27E-06 | 2.22E-05 | -1.41E+00 | 4.97E-03 | 2.15E-02 | 1.15E+00 | 2.05E-06 | 8.69E-06 | -1.55E+00 | 3.51E-04 | 1.34E-03 | -1.09E+00 | 1.13E-05 | 4.04E-05 | -1.80E+00 | 2.76E-02 | 7.77E-02 | 1.01E+00 | 5.63E-06 | 2.07E-05 | -1.23E+00 | 5.26E-02 | 1.45E-01 | 1.11E+00 |
| LFNG | 4.27E-06 | 2.22E-05 | -1.73E+00 | 2.96E-04 | 1.28E-03 | -1.31E+00 | 9.62E-05 | 4.09E-04 | -1.68E+00 | 3.34E-03 | 1.28E-02 | -1.25E+00 | 9.82E-02 | 3.50E-01 | -1.13E+00 | 1.06E-01 | 2.99E-01 | -1.27E+00 | 1.37E-01 | 5.03E-01 | -1.31E+00 | 1.57E-01 | 4.30E-01 | -1.40E+00 |
| AEBP1 | 4.27E-06 | 2.23E-05 | -1.37E+00 | 5.61E-03 | 2.42E-02 | 1.18E+00 | 2.21E-04 | 9.39E-04 | -1.16E+00 | 7.52E-03 | 2.87E-02 | 1.10E+00 | 1.25E-01 | 4.44E-01 | 1.11E+00 | 2.36E-01 | 6.66E-01 | 1.20E+00 | 2.23E-02 | 8.17E-02 | -1.32E+00 | 3.28E-01 | 9.01E-01 | 1.17E+00 |
| STMP1 | 4.27E-06 | 2.23E-05 | -1.05E+00 | 6.87E-02 | 2.97E-01 | 1.16E+00 | 6.24E-03 | 2.65E-02 | -1.12E+00 | 2.49E-01 | 9.50E-01 | 1.09E+00 | 1.37E-02 | 4.87E-02 | -1.11E+00 | 2.01E-01 | 5.67E-01 | 1.01E+00 | 2.07E-01 | 7.59E-01 | 1.06E+00 | 3.43E-01 | 9.41E-01 | 1.04E+00 |
| SLC39A11 | 4.30E-06 | 2.24E-05 | -1.85E+00 | 9.13E-03 | 3.94E-02 | -1.04E+00 | 1.02E-06 | 4.34E-06 | -1.68E+00 | 1.80E-03 | 6.87E-03 | -1.04E+00 | 2.53E-05 | 9.03E-05 | -1.99E+00 | 2.01E-01 | 5.67E-01 | 1.20E+00 | 8.58E-14 | 3.15E-13 | -2.02E+00 | 3.92E-07 | 1.08E-06 | -1.45E+00 |
| POLD2 | 4.31E-06 | 2.25E-05 | -1.10E+00 | 4.80E-05 | 2.08E-04 | 1.06E+00 | 6.49E-02 | 2.76E-01 | 1.05E+00 | 1.18E-01 | 4.50E-01 | 1.26E+00 | 1.93E-05 | 6.89E-05 | -1.21E+00 | 8.35E-02 | 2.35E-01 | 1.20E+00 | 7.66E-02 | 2.81E-01 | 1.02E+00 | 2.72E-01 | 7.46E-01 | 1.21E+00 |
| ACTBP2 | 4.33E-06 | 2.26E-05 | -1.64E+00 | 5.75E-05 | 2.49E-04 | -1.26E+00 | 3.83E-02 | 1.63E-01 | -1.05E+00 | 5.60E-03 | 2.14E-02 | -1.10E+00 | 6.08E-02 | 2.17E-01 | -1.20E+00 | 1.82E-01 | 5.14E-01 | 1.10E+00 | 6.08E-15 | 2.23E-14 | -1.88E+00 | 2.07E-10 | 5.70E-10 | -1.77E+00 |
| ARHGEF17 | 4.38E-06 | 2.28E-05 | -1.25E+00 | 3.10E-02 | 1.34E-01 | 1.28E+00 | 3.71E-02 | 1.58E-01 | -1.09E+00 | 1.38E-01 | 5.26E-01 | 1.19E+00 | 2.54E-01 | 9.06E-01 | 1.16E+00 | 3.42E-01 | 9.64E-01 | 1.40E+00 | 2.40E-01 | 8.83E-01 | 1.24E+00 | 1.98E-01 | 5.43E-01 | 1.74E+00 |
| FAM66C | 4.40E-06 | 2.29E-05 | -1.49E+00 | 3.00E-03 | 1.30E-02 | 1.04E+00 | 1.01E-03 | 4.29E-03 | -1.70E+00 | 4.22E-02 | 1.61E-01 | -1.05E+00 | 1.02E-01 | 3.65E-01 | -1.01E+00 | 7.38E-02 | 2.08E-01 | -1.18E+00 | 5.52E-02 | 2.03E-01 | -1.21E+00 | 2.20E-01 | 6.05E-01 | -1.03E+00 |
| TARSL2 | 4.41E-06 | 2.30E-05 | -1.15E+00 | 9.55E-09 | 4.13E-08 | -1.23E+00 | 1.34E-04 | 5.69E-04 | -1.22E+00 | 7.06E-07 | 2.70E-06 | -1.31E+00 | 2.63E-01 | 9.37E-01 | 1.41E+00 | 3.61E-04 | 1.02E-03 | -1.12E+00 | 1.39E-01 | 5.12E-01 | 1.29E+00 | 3.08E-02 | 8.45E-02 | 1.05E+00 |
| FIZ1 | 4.44E-06 | 2.32E-05 | -1.43E+00 | 6.72E-07 | 2.91E-06 | -1.46E+00 | 6.21E-03 | 2.64E-02 | -1.18E+00 | 2.71E-03 | 1.03E-02 | -1.12E+00 | 1.07E-01 | 3.82E-01 | -1.01E+00 | 3.54E-02 | 9.97E-02 | -1.32E+00 | 2.11E-03 | 7.75E-03 | -1.23E+00 | 2.05E-02 | 5.62E-02 | -1.13E+00 |
| GTPBP4 | 4.44E-06 | 2.32E-05 | -1.23E+00 | 6.60E-04 | 2.85E-03 | 1.07E+00 | 7.37E-06 | 3.13E-05 | -1.26E+00 | 1.64E-05 | 6.28E-05 | -1.08E+00 | 1.14E-06 | 4.07E-06 | -1.27E+00 | 8.82E-04 | 2.49E-03 | -1.03E+00 | 3.76E-06 | 1.38E-05 | -1.21E+00 | 8.15E-03 | 2.24E-02 | 1.01E+00 |
| ARMCX2 | 4.45E-06 | 2.32E-05 | -1.50E+00 | 4.33E-02 | 1.87E-01 | 1.22E+00 | 1.14E-05 | 4.85E-05 | -1.58E+00 | 1.35E-03 | 5.15E-03 | -1.12E+00 | 3.15E-02 | 1.12E-01 | -1.70E+00 | 2.27E-01 | 6.39E-01 | 1.07E+00 | 1.39E-02 | 5.11E-02 | -1.44E+00 | 1.23E-01 | 3.38E-01 | -1.00E+00 |
| WDR54 | 4.47E-06 | 2.33E-05 | -1.13E+00 | 5.48E-05 | 2.37E-04 | 1.01E+00 | 1.21E-01 | 5.15E-01 | 1.12E+00 | 1.05E-01 | 4.01E-01 | 1.28E+00 | 3.64E-03 | 1.30E-02 | 1.01E+00 | 1.40E-02 | 3.94E-02 | 1.11E+00 | 6.99E-02 | 2.57E-01 | 1.25E+00 | 3.60E-01 | 9.89E-01 | 1.41E+00 |
| AP5S1 | 4.48E-06 | 2.33E-05 | -1.57E+00 | 2.17E-03 | 9.36E-03 | 1.01E+00 | 5.52E-06 | 2.35E-05 | -1.52E+00 | 9.25E-04 | 3.53E-03 | -1.12E+00 | 1.36E-03 | 4.86E-03 | -1.69E+00 | 3.00E-02 | 8.47E-02 | -1.17E+00 | 8.33E-08 | 3.06E-07 | -1.55E+00 | 3.84E-08 | 1.05E-07 | -1.70E+00 |
| HMCES | 4.49E-06 | 2.34E-05 | -1.16E+00 | 1.71E-04 | 7.38E-04 | 1.05E+00 | 1.61E-02 | 6.86E-02 | -1.08E+00 | 3.01E-03 | 1.15E-02 | 1.02E+00 | 9.18E-03 | 3.27E-02 | -1.10E+00 | 1.62E-01 | 4.57E-01 | 1.23E+00 | 2.52E-02 | 9.26E-02 | 1.03E+00 | 2.48E-01 | 6.81E-01 | 1.23E+00 |
| TMTC3 | 4.50E-06 | 2.34E-05 | -1.30E+00 | 1.88E-06 | 8.14E-06 | -1.25E+00 | 8.84E-06 | 3.76E-05 | -1.65E+00 | 2.15E-06 | 8.21E-06 | -1.61E+00 | 1.27E-02 | 4.52E-02 | -1.11E+00 | 2.74E-02 | 7.73E-02 | -1.03E+00 | 3.85E-04 | 1.41E-03 | -1.40E+00 | 9.25E-02 | 2.54E-01 | -1.01E+00 |
| SAT2 | 4.52E-06 | 2.36E-05 | -1.05E+00 | 3.96E-05 | 1.71E-04 | 1.07E+00 | 2.67E-04 | 1.13E-03 | -1.16E+00 | 7.34E-03 | 2.80E-02 | 1.10E+00 | 1.16E-02 | 4.12E-02 | 1.08E+00 | 3.11E-02 | 8.76E-02 | 1.12E+00 | 1.09E-01 | 4.01E-01 | 1.27E+00 | 4.62E-02 | 1.27E-01 | 1.13E+00 |
| WDR41 | 4.53E-06 | 2.36E-05 | -1.19E+00 | 1.50E-04 | 6.47E-04 | 1.03E+00 | 7.14E-03 | 3.03E-02 | -1.11E+00 | 7.35E-04 | 2.81E-03 | -1.00E+00 | 1.82E-03 | 6.50E-03 | -1.05E+00 | 5.83E-03 | 1.64E-02 | 1.02E+00 | 5.63E-07 | 2.07E-06 | -1.26E+00 | 3.62E-03 | 9.95E-03 | -1.08E+00 |
| TAF2 | 4.54E-06 | 2.37E-05 | -1.43E+00 | 2.25E-05 | 9.72E-05 | -1.26E+00 | 8.21E-08 | 3.49E-07 | -1.60E+00 | 6.80E-08 | 2.60E-07 | -1.44E+00 | 2.73E-04 | 9.73E-04 | -1.30E+00 | 4.72E-03 | 1.33E-02 | -1.05E+00 | 4.98E-05 | 1.83E-04 | -1.24E+00 | 3.84E-03 | 1.06E-02 | -1.05E+00 |
| DOCK11 | 4.55E-06 | 2.37E-05 | -1.26E+00 | 4.74E-02 | 2.05E-01 | 1.27E+00 | 4.69E-06 | 1.99E-05 | -1.38E+00 | 1.40E-03 | 5.36E-03 | 1.01E+00 | 3.13E-06 | 1.12E-05 | -1.22E+00 | 1.08E-04 | 3.04E-04 | -1.03E+00 | 2.08E-06 | 7.64E-06 | -1.21E+00 | 4.98E-04 | 1.37E-03 | -1.09E+00 |
| AL021368.2 | 4.56E-06 | 2.38E-05 | -1.59E+00 | 1.14E-02 | 4.94E-02 | 1.05E+00 | 6.86E-06 | 2.92E-05 | -1.60E+00 | 9.03E-04 | 3.45E-03 | -1.12E+00 | 7.37E-03 | 2.63E-02 | -1.02E+00 | 2.14E-02 | 6.04E-02 | 1.03E+00 | 3.89E-02 | 1.43E-01 | -1.08E+00 | 9.22E-02 | 2.53E-01 | 1.63E+00 |
| MARF1 | 4.57E-06 | 2.38E-05 | -1.27E+00 | 4.53E-04 | 1.96E-03 | 1.00E+00 | 1.17E-06 | 4.95E-06 | -1.40E+00 | 6.86E-07 | 2.62E-06 | -1.19E+00 | 1.28E-02 | 4.57E-02 | -1.10E+00 | 5.48E-02 | 1.55E-01 | 1.01E+00 | 8.78E-05 | 3.22E-04 | -1.36E+00 | 1.65E-04 | 4.52E-04 | -1.33E+00 |
| VPS54 | 4.63E-06 | 2.41E-05 | -1.59E+00 | 1.40E-02 | 6.07E-02 | 1.07E+00 | 1.55E-02 | 6.58E-02 | -1.11E+00 | 1.47E-01 | 5.62E-01 | 1.25E+00 | 1.18E-02 | 4.20E-02 | -1.47E+00 | 3.23E-01 | 9.11E-01 | 1.35E+00 | 1.16E-02 | 4.27E-02 | -1.02E+00 | 2.13E-01 | 5.85E-01 | 1.16E+00 |
| FNBP1 | 4.70E-06 | 2.45E-05 | -1.13E+00 | 4.39E-06 | 1.90E-05 | -1.06E+00 | 1.32E-02 | 5.62E-02 | -1.11E+00 | 1.71E-04 | 6.52E-04 | -1.14E+00 | 2.61E-10 | 9.31E-10 | -1.26E+00 | 4.72E-14 | 1.33E-13 | -1.48E+00 | 8.25E-15 | 3.03E-14 | -1.49E+00 | 3.10E-13 | 8.51E-13 | -1.55E+00 |
| TTC13 | 4.71E-06 | 2.45E-05 | -2.31E+00 | 8.21E-03 | 3.55E-02 | -1.07E+00 | 2.70E-03 | 1.15E-02 | -1.30E+00 | 2.43E-02 | 9.27E-02 | 1.09E+00 | 2.21E-01 | 7.89E-01 | 1.07E+00 | 1.35E-01 | 3.79E-01 | 2.01E+00 | 2.90E-02 | 1.06E-01 | -1.24E+00 | 2.69E-01 | 7.39E-01 | 1.12E+00 |
| C2orf49 | 4.83E-06 | 2.51E-05 | -1.34E+00 | 8.89E-03 | 3.84E-02 | 1.14E+00 | 5.40E-09 | 2.29E-08 | -1.73E+00 | 2.70E-07 | 1.03E-06 | -1.39E+00 | 3.53E-04 | 1.26E-03 | -1.22E+00 | 3.53E-04 | 9.96E-04 | -1.21E+00 | 8.25E-09 | 3.03E-08 | -1.51E+00 | 2.44E-06 | 6.70E-06 | -1.37E+00 |
| MRPS6 | 4.85E-06 | 2.53E-05 | -1.11E+00 | 1.04E-01 | 4.47E-01 | 1.36E+00 | 2.61E-02 | 1.11E-01 | -1.00E+00 | 1.53E-01 | 5.85E-01 | 1.24E+00 | 7.39E-06 | 2.64E-05 | -1.32E+00 | 2.29E-01 | 6.45E-01 | 1.29E+00 | 2.02E-01 | 7.42E-01 | 1.27E+00 | 8.80E-05 | 2.42E-04 | 1.64E+00 |
| WDR45 | 4.85E-06 | 2.53E-05 | -1.12E+00 | 1.55E-04 | 6.71E-04 | 1.08E+00 | 6.67E-03 | 2.84E-02 | -1.03E+00 | 5.75E-02 | 2.20E-01 | 1.28E+00 | 2.53E-04 | 9.01E-04 | -1.11E+00 | 4.53E-05 | 1.28E-04 | -1.15E+00 | 4.51E-03 | 1.65E-02 | 1.04E+00 | 7.05E-03 | 1.94E-02 | 1.07E+00 |
| AC008870.2 | 4.89E-06 | 2.55E-05 | -2.30E+00 | 4.01E-03 | 1.73E-02 | -1.17E+00 | 9.21E-02 | 3.91E-01 | -1.14E+00 | 8.79E-02 | 3.36E-01 | 1.69E+00 | 4.08E-02 | 1.46E-01 | -1.36E+00 | 2.90E-01 | 8.19E-01 | 1.44E+00 | 4.52E-05 | 1.66E-04 | -2.41E+00 | 7.32E-02 | 2.01E-01 | -1.05E+00 |
| DCTN3 | 4.89E-06 | 2.55E-05 | -1.03E+00 | 1.21E-06 | 5.23E-06 | 1.07E+00 | 3.52E-02 | 1.50E-01 | 1.04E+00 | 1.57E-03 | 6.01E-03 | 1.06E+00 | 4.20E-05 | 1.50E-04 | -1.05E+00 | 1.53E-03 | 4.32E-03 | 1.06E+00 | 2.56E-01 | 9.40E-01 | 1.20E+00 | 3.54E-01 | 9.72E-01 | 1.17E+00 |
| STK19 | 4.90E-06 | 2.55E-05 | -1.17E+00 | 1.73E-05 | 7.46E-05 | 1.02E+00 | 4.12E-05 | 1.75E-04 | -1.22E+00 | 2.31E-04 | 8.81E-04 | -1.07E+00 | 5.49E-03 | 1.96E-02 | 1.05E+00 | 3.00E-03 | 8.45E-03 | 1.02E+00 | 9.79E-02 | 3.59E-01 | 1.20E+00 | 5.20E-02 | 1.43E-01 | 1.11E+00 |
| TCEAL8 | 4.93E-06 | 2.57E-05 | -1.09E+00 | 1.16E-03 | 5.00E-03 | 1.16E+00 | 7.44E-05 | 3.16E-04 | -1.11E+00 | 1.19E-02 | 4.53E-02 | 1.14E+00 | 5.78E-08 | 2.06E-07 | -1.34E+00 | 5.99E-02 | 1.69E-01 | 1.20E+00 | 4.94E-03 | 1.81E-02 | -1.04E+00 | 1.36E-01 | 3.73E-01 | 1.14E+00 |
| CLASRP | 4.94E-06 | 2.57E-05 | -1.22E+00 | 3.27E-06 | 1.42E-05 | -1.13E+00 | 2.80E-07 | 1.19E-06 | -1.38E+00 | 4.93E-06 | 1.88E-05 | -1.07E+00 | 5.94E-05 | 2.12E-04 | -1.21E+00 | 6.70E-05 | 1.89E-04 | -1.10E+00 | 4.67E-04 | 1.71E-03 | -1.13E+00 | 6.80E-04 | 1.87E-03 | -1.15E+00 |
| BAG4 | 4.94E-06 | 2.57E-05 | -1.30E+00 | 2.39E-04 | 1.03E-03 | -1.02E+00 | 2.77E-04 | 1.18E-03 | -1.24E+00 | 5.17E-04 | 1.98E-03 | -1.05E+00 | 8.37E-05 | 2.98E-04 | -1.30E+00 | 3.96E-04 | 1.12E-03 | -1.19E+00 | 3.22E-07 | 1.18E-06 | -1.40E+00 | 3.32E-07 | 9.13E-07 | -1.33E+00 |
| ERC1 | 4.95E-06 | 2.58E-05 | -1.28E+00 | 5.20E-04 | 2.25E-03 | 1.05E+00 | 5.00E-04 | 2.12E-03 | -1.24E+00 | 6.03E-03 | 2.30E-02 | 1.02E+00 | 1.25E-06 | 4.45E-06 | -1.37E+00 | 3.24E-06 | 9.12E-06 | -1.39E+00 | 2.98E-05 | 1.09E-04 | -1.23E+00 | 7.05E-04 | 1.94E-03 | -1.16E+00 |
| AL391058.1 | 4.95E-06 | 2.58E-05 | -2.00E+00 | 1.27E-03 | 5.50E-03 | -1.20E+00 | 7.44E-14 | 3.16E-13 | -2.63E+00 | 1.48E-09 | 5.67E-09 | -1.80E+00 | 7.61E-06 | 2.71E-05 | -2.17E+00 | 1.03E-04 | 2.91E-04 | -1.78E+00 | 4.09E-09 | 1.50E-08 | -1.88E+00 | 2.60E-04 | 7.15E-04 | -1.22E+00 |
| ATP13A1 | 4.95E-06 | 2.58E-05 | -1.29E+00 | 1.99E-05 | 8.59E-05 | -1.08E+00 | 2.15E-05 | 9.12E-05 | -1.33E+00 | 2.17E-05 | 8.29E-05 | -1.19E+00 | 2.23E-05 | 7.96E-05 | -1.41E+00 | 3.45E-03 | 9.74E-03 | -1.03E+00 | 1.47E-06 | 5.39E-06 | -1.30E+00 | 2.95E-03 | 8.09E-03 | -1.04E+00 |
| AC078883.2 | 4.97E-06 | 2.59E-05 | -1.75E+00 | 1.25E-03 | 5.40E-03 | -1.12E+00 | 3.61E-03 | 1.53E-02 | -1.32E+00 | 1.06E-01 | 4.07E-01 | 1.30E+00 | 2.16E-01 | 7.70E-01 | 1.52E+00 | 3.29E-01 | 9.26E-01 | 1.37E+00 | 1.69E-01 | 6.21E-01 | 2.25E+00 | 1.17E-01 | 3.22E-01 | 1.06E+00 |
| AC243772.3 | 4.97E-06 | 2.59E-05 | -1.96E+00 | 1.24E-02 | 5.35E-02 | 1.02E+00 | 2.30E-05 | 9.76E-05 | -2.04E+00 | 3.25E-03 | 1.24E-02 | -1.31E+00 | 1.93E-01 | 6.88E-01 | 1.60E+00 | 3.23E-01 | 9.10E-01 | 1.35E+00 | 1.13E-01 | 4.15E-01 | 1.51E+00 | 2.89E-01 | 7.95E-01 | 1.17E+00 |
| AC136628.3 | 4.99E-06 | 2.60E-05 | -2.14E+00 | 1.46E-02 | 6.31E-02 | 1.02E+00 | 2.88E-03 | 1.23E-02 | -1.51E+00 | 7.55E-02 | 2.88E-01 | 1.07E+00 | 1.46E-01 | 5.19E-01 | 1.07E+00 | 3.27E-01 | 9.21E-01 | 1.51E+00 | 2.20E-02 | 8.08E-02 | -1.09E+00 | 2.61E-01 | 7.17E-01 | 1.17E+00 |
| CRYBG1 | 5.01E-06 | 2.61E-05 | -1.85E+00 | 2.08E-05 | 8.97E-05 | -1.56E+00 | 1.75E-06 | 7.42E-06 | -1.37E+00 | 1.51E-10 | 5.78E-10 | -1.50E+00 | 1.94E-01 | 6.92E-01 | -1.12E+00 | 2.38E-01 | 6.72E-01 | -1.06E+00 | 8.34E-02 | 3.06E-01 | -1.38E+00 | 9.93E-02 | 2.73E-01 | -1.51E+00 |
| UCHL1 | 5.02E-06 | 2.62E-05 | -1.26E+00 | 1.11E-02 | 4.80E-02 | 1.18E+00 | 2.06E-06 | 8.74E-06 | -1.45E+00 | 1.72E-01 | 6.56E-01 | 1.50E+00 | 1.24E-01 | 4.43E-01 | 1.70E+00 | 5.97E-02 | 1.68E-01 | 2.18E+00 | 2.46E-01 | 9.04E-01 | 1.21E+00 | 7.00E-02 | 1.92E-01 | 2.50E+00 |
| CNST | 5.03E-06 | 2.62E-05 | -1.01E+00 | 2.77E-06 | 1.20E-05 | 1.11E+00 | 1.83E-02 | 7.78E-02 | 1.07E+00 | 7.43E-02 | 2.84E-01 | 1.29E+00 | 1.44E-03 | 5.15E-03 | 1.66E+00 | 2.58E-01 | 7.28E-01 | 1.33E+00 | 1.33E-10 | 4.89E-10 | 2.13E+00 | 1.03E-03 | 2.83E-03 | 1.61E+00 |
| PSMD6 | 5.05E-06 | 2.63E-05 | -1.07E+00 | 3.94E-04 | 1.70E-03 | 1.14E+00 | 1.42E-04 | 6.02E-04 | -1.10E+00 | 1.06E-05 | 4.04E-05 | -1.07E+00 | 5.21E-05 | 1.86E-04 | -1.12E+00 | 2.38E-02 | 6.70E-02 | 1.07E+00 | 5.51E-02 | 2.02E-01 | -1.04E+00 | 2.75E-01 | 7.55E-01 | 1.06E+00 |
| SMIM27 | 5.08E-06 | 2.65E-05 | -1.07E+00 | 1.42E-21 | 6.15E-21 | -1.72E+00 | 1.03E-02 | 4.37E-02 | -1.00E+00 | 1.66E-10 | 6.33E-10 | -1.51E+00 | 2.14E-03 | 7.64E-03 | 1.01E+00 | 2.68E-12 | 7.56E-12 | -1.55E+00 | 1.44E-04 | 5.28E-04 | -1.05E+00 | 1.11E-10 | 3.06E-10 | -1.52E+00 |
| ZNF529 | 5.08E-06 | 2.65E-05 | -1.24E+00 | 2.21E-06 | 9.56E-06 | -1.13E+00 | 4.89E-06 | 2.08E-05 | -1.38E+00 | 1.01E-04 | 3.87E-04 | -1.06E+00 | 1.17E-03 | 4.17E-03 | -1.09E+00 | 1.13E-02 | 3.18E-02 | 1.04E+00 | 5.72E-02 | 2.10E-01 | -1.05E+00 | 1.37E-02 | 3.75E-02 | -1.12E+00 |
| CAPZA2 | 5.09E-06 | 2.65E-05 | -1.01E+00 | 9.62E-03 | 4.16E-02 | 1.20E+00 | 3.79E-03 | 1.61E-02 | -1.06E+00 | 4.46E-04 | 1.70E-03 | -1.03E+00 | 3.00E-06 | 1.07E-05 | -1.17E+00 | 1.70E-02 | 4.80E-02 | 1.12E+00 | 2.84E-03 | 1.04E-02 | -1.11E+00 | 1.10E-01 | 3.01E-01 | -1.07E+00 |
| TFR2 | 5.10E-06 | 2.66E-05 | -1.29E+00 | 5.43E-05 | 2.34E-04 | -1.03E+00 | 1.79E-04 | 7.61E-04 | -1.20E+00 | 5.97E-03 | 2.28E-02 | 1.07E+00 | 1.87E-05 | 6.67E-05 | -1.05E+00 | 1.08E-01 | 3.04E-01 | 1.41E+00 | 6.32E-03 | 2.32E-02 | -1.08E+00 | 1.40E-01 | 3.84E-01 | 1.07E+00 |
| SMC2 | 5.11E-06 | 2.66E-05 | -1.16E+00 | 2.24E-03 | 9.67E-03 | 1.14E+00 | 4.35E-03 | 1.85E-02 | 1.02E+00 | 4.47E-02 | 1.71E-01 | 1.27E+00 | 2.38E-05 | 8.48E-05 | -1.51E+00 | 1.41E-01 | 3.96E-01 | 1.17E+00 | 7.64E-05 | 2.80E-04 | -1.17E+00 | 3.54E-01 | 9.72E-01 | 1.16E+00 |
| AC004889.1 | 5.11E-06 | 2.67E-05 | -1.46E+00 | 2.95E-05 | 1.27E-04 | -1.22E+00 | 5.20E-03 | 2.21E-02 | -1.08E+00 | 5.02E-05 | 1.92E-04 | -1.48E+00 | 1.06E-01 | 3.76E-01 | -1.32E+00 | 1.25E-01 | 3.53E-01 | -1.38E+00 | 1.91E-01 | 7.01E-01 | -1.05E+00 | 3.08E-01 | 8.47E-01 | 1.06E+00 |

| | | | | | | | | | | | | | | | | | | | | | | | |
|---|---|---|---|---|---|---|---|---|---|---|---|---|---|---|---|---|---|---|---|---|---|---|---|
| CFAP57 | 5.12E-06 | 2.67E-05 | -1.99E+00 | 1.44E-02 | 6.23E-02 | -1.08E+00 | 5.38E-03 | 2.29E-02 | -1.64E+00 | 2.27E-01 | 8.66E-01 | 1.22E+00 | 1.13E-01 | 4.04E-01 | -1.17E+00 | 1.06E-01 | 2.99E-01 | 1.89E+00 | 2.10E-01 | 7.69E-01 | 1.04E+00 | 2.85E-01 | 7.82E-01 | 1.34E+00 |
| PTMAP5 | 5.13E-06 | 2.67E-05 | -1.15E+00 | 2.33E-25 | 1.01E-24 | -1.41E+00 | 3.08E-09 | 1.31E-08 | -1.27E+00 | 4.08E-21 | 1.56E-20 | -1.47E+00 | 4.97E-22 | 1.77E-21 | -1.53E+00 | 2.93E-32 | 8.27E-32 | -1.70E+00 | 3.37E-32 | 1.24E-31 | -1.54E+00 | 1.23E-26 | 3.38E-26 | -1.54E+00 |
| NDRG3 | 5.14E-06 | 2.68E-05 | -1.52E+00 | 1.55E-03 | 6.70E-03 | -1.03E+00 | 1.92E-04 | 8.14E-04 | -1.24E+00 | 6.93E-05 | 2.65E-04 | -1.21E+00 | 3.36E-04 | 1.20E-03 | -1.36E+00 | 1.34E-02 | 3.78E-02 | 1.03E+00 | 4.01E-07 | 1.47E-06 | -1.46E+00 | 1.77E-05 | 4.87E-05 | -1.24E+00 |
| ZNF141 | 5.15E-06 | 2.69E-05 | -1.21E+00 | 1.97E-02 | 8.49E-02 | 1.26E+00 | 1.01E-03 | 4.29E-03 | -1.26E+00 | 8.27E-02 | 3.16E-01 | 1.14E+00 | 2.78E-02 | 9.93E-02 | 1.04E+00 | 3.35E-01 | 9.45E-01 | 1.43E+00 | 1.82E-01 | 6.69E-01 | 1.23E+00 | 1.08E-01 | 2.97E-01 | 1.51E+00 |
| RN7SL263P | 5.15E-06 | 2.69E-05 | 3.48E+00 | 3.44E-03 | 1.49E-02 | 2.83E+00 | 2.03E-04 | 8.63E-04 | 2.62E+00 | 1.91E-02 | 7.29E-02 | 2.17E+00 | 1.22E-02 | 4.34E-02 | 3.38E+00 | 1.31E-01 | 3.70E-01 | 2.17E+00 | 1.24E-01 | 4.54E-01 | 1.67E+00 | 2.81E-01 | 7.72E-01 | -1.16E+00 |
| SOWAHD | 5.16E-06 | 2.69E-05 | -2.11E+00 | 4.75E-03 | 2.05E-02 | -1.10E+00 | 1.54E-11 | 6.56E-11 | -2.37E+00 | 6.28E-07 | 2.40E-06 | -1.45E+00 | 2.08E-02 | 7.43E-02 | -1.77E+00 | 1.73E-02 | 4.87E-02 | -2.03E+00 | 2.09E-01 | 7.67E-01 | -1.18E+00 | 3.29E-01 | 9.03E-01 | -1.04E+00 |
| ADAM28 | 5.16E-06 | 2.69E-05 | -1.06E+00 | 2.40E-04 | 1.04E-03 | 1.16E+00 | 8.59E-03 | 3.65E-02 | 1.01E+00 | 1.15E-03 | 4.40E-03 | -1.03E+00 | 2.27E-02 | 8.10E-02 | 1.02E+00 | 1.31E-03 | 3.70E-03 | -1.26E+00 | 1.38E-01 | 5.07E-01 | -1.18E+00 | 2.38E-01 | 6.55E-01 | -1.05E+00 |
| NOL12 | 5.17E-06 | 2.70E-05 | -1.47E+00 | 5.35E-06 | 2.31E-05 | -1.34E+00 | 1.13E-04 | 4.80E-04 | -1.34E+00 | 2.08E-05 | 7.95E-05 | -1.33E+00 | 1.74E-01 | 6.20E-01 | 1.17E+00 | 2.88E-01 | 8.12E-01 | 1.39E+00 | 2.00E-05 | 7.35E-05 | -1.26E+00 | 2.49E-03 | 6.83E-03 | -1.05E+00 |
| MTO1 | 5.20E-06 | 2.71E-05 | -1.22E+00 | 1.90E-04 | 8.23E-04 | 1.03E+00 | 3.32E-04 | 1.41E-03 | -1.28E+00 | 1.22E-04 | 4.67E-04 | -1.17E+00 | 2.04E-02 | 7.26E-02 | -1.31E+00 | 1.45E-01 | 4.09E-01 | 1.11E+00 | 1.15E-04 | 4.21E-04 | -1.23E+00 | 3.21E-04 | 8.81E-04 | -1.17E+00 |
| HTATSF1 | 5.21E-06 | 2.72E-05 | 1.00E+00 | 1.06E-04 | 4.59E-04 | 1.11E+00 | 2.09E-02 | 8.88E-02 | -1.02E+00 | 2.77E-03 | 1.06E-02 | 1.02E+00 | 5.05E-08 | 1.80E-07 | -1.30E+00 | 2.53E-04 | 7.13E-04 | -1.02E+00 | 8.58E-05 | 3.15E-04 | -1.17E+00 | 2.86E-02 | 7.87E-02 | -1.03E+00 |
| C9orf43 | 5.22E-06 | 2.72E-05 | -1.02E+00 | 2.09E-07 | 9.05E-07 | 1.07E+00 | 2.22E-06 | 9.42E-06 | -1.32E+00 | 3.48E-07 | 1.33E-06 | -1.21E+00 | 6.57E-02 | 2.34E-01 | 1.20E+00 | 9.20E-03 | 2.59E-02 | 1.01E+00 | 1.04E-01 | 3.82E-01 | 1.62E+00 | 1.50E-01 | 4.12E-01 | 1.62E+00 |
| CABIN1 | 5.22E-06 | 2.72E-05 | -1.23E+00 | 1.07E-03 | 4.61E-03 | 1.08E+00 | 1.45E-04 | 6.14E-04 | -1.14E+00 | 5.78E-04 | 2.21E-03 | 1.00E+00 | 4.88E-02 | 1.74E-01 | 1.05E+00 | 1.96E-01 | 5.52E-01 | 1.31E+00 | 2.42E-02 | 8.87E-02 | -1.06E+00 | 3.45E-01 | 9.48E-01 | 1.32E+00 |
| TP53RK | 5.23E-06 | 2.73E-05 | -1.52E+00 | 2.03E-03 | 8.77E-03 | 1.01E+00 | 1.39E-09 | 5.92E-09 | -1.81E+00 | 1.64E-04 | 6.27E-04 | -1.09E+00 | 1.16E-08 | 4.15E-08 | -1.53E+00 | 6.92E-04 | 1.95E-03 | -1.02E+00 | 2.13E-18 | 7.81E-18 | -1.77E+00 | 3.50E-10 | 9.60E-10 | -1.22E+00 |
| SEPHS2 | 5.24E-06 | 2.73E-05 | 1.01E+00 | 1.39E-08 | 5.99E-08 | -1.04E+00 | 7.29E-07 | 3.10E-06 | -1.25E+00 | 3.59E-05 | 1.37E-04 | -1.06E+00 | 2.03E-06 | 7.24E-06 | -1.21E+00 | 1.59E-03 | 4.49E-03 | -1.03E+00 | 8.10E-10 | 2.97E-09 | -1.39E+00 | 1.87E-02 | 5.13E-02 | -1.04E+00 |
| DHX57 | 5.25E-06 | 2.74E-05 | -1.33E+00 | 4.35E-04 | 1.88E-03 | 1.03E+00 | 1.38E-04 | 5.87E-04 | -1.36E+00 | 5.61E-04 | 2.14E-03 | -1.14E+00 | 1.23E-02 | 4.38E-02 | -1.07E+00 | 1.35E-01 | 3.81E-01 | 1.19E+00 | 5.48E-03 | 2.01E-02 | -1.03E+00 | 2.39E-02 | 6.55E-02 | 1.01E+00 |
| TMA16P2 | 5.26E-06 | 2.74E-05 | -1.61E+00 | 1.27E-05 | 5.48E-05 | -1.29E+00 | 1.82E-02 | 7.75E-02 | -1.11E+00 | 1.65E-03 | 6.29E-03 | -1.16E+00 | 2.07E-05 | 7.39E-05 | -1.77E+00 | 6.23E-05 | 1.76E-04 | -1.60E+00 | 2.91E-08 | 1.07E-07 | -1.56E+00 | 2.30E-06 | 6.32E-06 | -1.27E+00 |
| USP49 | 5.26E-06 | 2.74E-05 | -1.60E+00 | 2.33E-04 | 1.01E-03 | -1.27E+00 | 1.23E-05 | 5.24E-05 | -1.39E+00 | 7.39E-04 | 2.82E-03 | -1.03E+00 | 1.10E-01 | 3.93E-01 | -1.02E+00 | 3.23E-01 | 9.11E-01 | 1.30E+00 | 5.76E-06 | 2.11E-05 | -1.54E+00 | 3.21E-04 | 8.81E-04 | -1.20E+00 |
| SQLE | 5.27E-06 | 2.75E-05 | -1.14E+00 | 4.33E-02 | 1.87E-01 | 1.31E+00 | 2.41E-03 | 1.02E-02 | -1.09E+00 | 2.17E-01 | 8.31E-01 | 1.37E+00 | 3.31E-04 | 1.18E-03 | -1.15E+00 | 1.54E-01 | 4.34E-01 | 1.62E+00 | 1.82E-01 | 6.67E-01 | 1.08E+00 | 5.25E-04 | 1.44E-03 | 1.70E+00 |
| UBL4A | 5.28E-06 | 2.75E-05 | -1.53E+00 | 5.12E-02 | 2.21E-01 | 1.27E+00 | 7.34E-09 | 3.12E-08 | -1.62E+00 | 2.44E-03 | 9.33E-03 | 1.05E+00 | 2.15E-09 | 7.66E-09 | -1.86E+00 | 2.00E-03 | 5.63E-03 | -1.02E+00 | 2.04E-18 | 7.48E-18 | -1.49E+00 | 3.34E-04 | 9.19E-04 | -1.04E+00 |
| POFUT2 | 5.31E-06 | 2.77E-05 | -1.63E+00 | 2.45E-05 | 1.06E-04 | -1.36E+00 | 5.90E-07 | 2.51E-06 | -1.78E+00 | 1.07E-06 | 4.09E-06 | -1.53E+00 | 1.58E-04 | 5.63E-04 | -1.65E+00 | 1.45E-02 | 4.09E-02 | -1.12E+00 | 1.63E-01 | 5.97E-01 | 1.19E+00 | 5.78E-02 | 1.59E-01 | -1.00E+00 |
| C1orf27 | 5.37E-06 | 2.80E-05 | -1.20E+00 | 1.80E-08 | 7.77E-08 | -1.19E+00 | 2.49E-03 | 1.06E-02 | -1.13E+00 | 4.09E-06 | 1.56E-05 | -1.20E+00 | 4.70E-06 | 1.68E-05 | -1.34E+00 | 1.58E-07 | 4.45E-07 | -1.37E+00 | 5.87E-04 | 2.15E-03 | -1.14E+00 | 3.17E-05 | 8.70E-05 | -1.23E+00 |
| METTL7A | 5.37E-06 | 2.80E-05 | 1.87E+00 | 3.53E-07 | 1.52E-06 | 2.23E+00 | 2.98E-02 | 1.27E-01 | 1.27E+00 | 1.54E-02 | 5.89E-02 | 1.47E+00 | 2.16E-01 | 7.71E-01 | 1.34E+00 | 2.96E-01 | 8.34E-01 | 1.51E+00 | 7.12E-03 | 2.61E-02 | -1.02E+00 | 7.85E-02 | 2.16E-01 | 1.09E+00 |
| ZBTB21 | 5.38E-06 | 2.81E-05 | -1.05E+00 | 9.83E-18 | 4.25E-17 | -1.65E+00 | 7.88E-03 | 3.35E-02 | -1.02E+00 | 1.55E-12 | 5.93E-12 | -1.74E+00 | 7.28E-03 | 2.60E-02 | -1.04E+00 | 4.10E-07 | 1.16E-06 | -2.00E+00 | 5.54E-09 | 2.03E-08 | -1.83E+00 | 1.03E-12 | 2.82E-12 | -2.47E+00 |
| PZP | 5.39E-06 | 2.81E-05 | -1.58E+00 | 3.82E-03 | 1.65E-02 | 1.02E+00 | 9.57E-05 | 4.07E-04 | -1.98E+00 | 1.09E-03 | 4.16E-03 | -1.48E+00 | 1.07E-03 | 3.80E-03 | -1.37E+00 | 7.22E-03 | 2.04E-02 | -1.11E+00 | 1.18E-01 | 4.32E-01 | -1.11E+00 | 1.97E-01 | 5.40E-01 | -1.02E+00 |
| VDAC1P5 | 5.40E-06 | 2.82E-05 | -1.51E+00 | 8.10E-11 | 3.50E-10 | -2.73E+00 | 1.32E-05 | 5.60E-05 | -1.41E+00 | 1.21E-13 | 4.62E-13 | -2.45E+00 | 1.43E-06 | 5.10E-06 | -1.73E+00 | 3.51E-11 | 9.90E-11 | -3.32E+00 | 2.06E-06 | 7.55E-06 | -1.36E+00 | 1.89E-15 | 5.18E-15 | -2.46E+00 |
| TNFRSF21 | 5.42E-06 | 2.82E-05 | -1.48E+00 | 1.43E-06 | 6.17E-06 | -1.42E+00 | 1.61E-08 | 6.84E-08 | -1.96E+00 | 1.88E-08 | 7.19E-08 | -1.83E+00 | 2.01E-04 | 7.18E-04 | -2.15E+00 | 6.22E-04 | 1.75E-03 | -1.78E+00 | 3.33E-12 | 1.22E-11 | -2.64E+00 | 3.20E-08 | 8.78E-08 | -2.00E+00 |
| ZNF414 | 5.44E-06 | 2.83E-05 | -1.19E+00 | 1.26E-05 | 5.45E-05 | -1.00E+00 | 7.83E-06 | 3.33E-05 | -1.30E+00 | 3.08E-07 | 1.17E-06 | -1.25E+00 | 2.36E-04 | 8.40E-04 | -1.41E+00 | 4.28E-03 | 1.21E-02 | -1.05E+00 | 1.51E-07 | 5.53E-07 | -1.38E+00 | 5.40E-07 | 1.48E-06 | -1.46E+00 |
| AC009126.1 | 5.44E-06 | 2.84E-05 | -1.31E+00 | 7.53E-05 | 3.25E-04 | -1.07E+00 | 2.30E-02 | 9.76E-02 | -1.10E+00 | 1.53E-03 | 5.83E-03 | -1.32E+00 | 2.01E-01 | 7.18E-01 | 1.27E+00 | 3.22E-01 | 9.07E-01 | 1.47E+00 | 6.39E-02 | 2.35E-01 | 1.09E+00 | 1.40E-01 | 3.84E-01 | 1.06E+00 |
| TINF2 | 5.45E-06 | 2.84E-05 | -1.14E+00 | 9.33E-06 | 4.03E-05 | -1.00E+00 | 3.53E-04 | 1.50E-03 | -1.23E+00 | 5.47E-02 | 2.09E-01 | 1.10E+00 | 6.24E-05 | 2.22E-04 | -1.15E+00 | 3.75E-03 | 1.06E-02 | 1.06E+00 | 1.94E-03 | 7.14E-03 | 1.00E+00 | 6.73E-03 | 1.85E-02 | 1.09E+00 |
| DYNLL2 | 5.45E-06 | 2.84E-05 | -1.15E+00 | 3.34E-12 | 1.44E-11 | -1.51E+00 | 1.01E-02 | 4.29E-02 | -1.04E+00 | 1.77E-03 | 6.76E-03 | -1.02E+00 | 6.57E-04 | 2.34E-03 | -1.03E+00 | 7.22E-06 | 2.04E-05 | -1.28E+00 | 2.38E-07 | 8.75E-07 | -1.13E+00 | 1.92E-05 | 5.29E-05 | -1.09E+00 |
| MEX3A | 5.46E-06 | 2.84E-05 | -2.22E+00 | 5.00E-03 | 2.16E-02 | -1.10E+00 | 6.36E-07 | 2.70E-06 | -2.20E+00 | 1.81E-06 | 6.92E-06 | -1.81E+00 | 6.21E-09 | 2.21E-08 | -3.01E+00 | 2.90E-07 | 8.18E-07 | -2.15E+00 | 7.40E-08 | 2.72E-07 | -2.18E+00 | 2.75E-04 | 7.56E-04 | -1.46E+00 |
| HRH2 | 5.47E-06 | 2.85E-05 | -2.51E+00 | 7.01E-05 | 3.03E-04 | -1.86E+00 | 1.46E-08 | 6.19E-08 | -1.69E+00 | 1.52E-07 | 5.82E-07 | -1.46E+00 | 7.13E-02 | 2.54E-01 | -1.47E+00 | 1.28E-01 | 3.61E-01 | -1.40E+00 | 9.35E-04 | 3.43E-03 | -2.59E+00 | 5.60E-03 | 1.54E-02 | -2.22E+00 |
| AL355488.1 | 5.47E-06 | 2.85E-05 | -2.88E+00 | 1.41E-01 | 6.08E-01 | 1.58E+00 | 3.11E-06 | 1.32E-05 | -2.89E+00 | 6.97E-02 | 2.66E-01 | 1.61E+00 | 3.35E-05 | 1.19E-04 | -4.08E+00 | 2.30E-01 | 6.49E-01 | 1.11E+00 | 9.78E-06 | 3.59E-05 | -2.05E+00 | 2.46E-02 | 6.75E-02 | 1.71E+00 |
| CANT1 | 5.47E-06 | 2.85E-05 | -1.53E+00 | 3.32E-03 | 1.43E-02 | 1.02E+00 | 3.35E-07 | 1.42E-06 | -1.77E+00 | 1.64E-04 | 6.28E-04 | -1.14E+00 | 1.86E-05 | 6.64E-05 | -1.75E+00 | 2.32E-04 | 6.53E-04 | -1.47E+00 | 7.53E-18 | 2.76E-17 | -2.33E+00 | 1.53E-09 | 4.21E-09 | -1.49E+00 |
| TP53BP2 | 5.49E-06 | 2.86E-05 | -1.34E+00 | 1.99E-09 | 8.62E-09 | -1.54E+00 | 4.61E-04 | 1.96E-03 | -1.19E+00 | 6.91E-06 | 2.64E-05 | -1.25E+00 | 1.33E-06 | 4.73E-06 | -1.48E+00 | 4.40E-05 | 1.24E-04 | -1.23E+00 | 1.08E-08 | 3.98E-08 | -1.45E+00 | 4.90E-06 | 1.35E-05 | -1.36E+00 |
| FUZ | 5.50E-06 | 2.86E-05 | -1.30E+00 | 5.71E-06 | 2.47E-05 | -1.13E+00 | 1.49E-09 | 6.35E-09 | -1.54E+00 | 3.72E-09 | 1.42E-08 | -1.30E+00 | 3.39E-05 | 1.21E-04 | -1.66E+00 | 5.77E-03 | 1.63E-02 | -1.07E+00 | 6.38E-04 | 2.34E-03 | -1.21E+00 | 1.66E-03 | 4.57E-03 | -1.12E+00 |
| MTMR4 | 5.50E-06 | 2.87E-05 | -1.17E+00 | 1.13E-02 | 4.88E-02 | 1.21E+00 | 6.68E-02 | 2.84E-01 | 1.02E+00 | 2.51E-01 | 9.58E-01 | 1.37E+00 | 1.10E-02 | 3.91E-02 | 1.10E+00 | 9.89E-03 | 2.79E-02 | 1.10E+00 | 3.08E-07 | 1.13E-06 | -1.36E+00 | 5.20E-04 | 1.43E-03 | -1.14E+00 |
| ATP5D | 5.51E-06 | 2.87E-05 | -1.15E+00 | 3.52E-06 | 1.52E-05 | -1.02E+00 | 5.27E-03 | 2.24E-02 | -1.16E+00 | 5.39E-02 | 2.06E-01 | -1.05E+00 | 1.12E-06 | 3.99E-06 | -1.16E+00 | 1.14E-01 | 3.21E-01 | 1.05E+00 | 1.87E-02 | 6.87E-02 | -1.08E+00 | 3.51E-01 | 9.63E-01 | 1.06E+00 |
| EMID1 | 5.56E-06 | 2.90E-05 | -1.08E+00 | 1.17E-05 | 5.07E-05 | 1.11E+00 | 7.73E-02 | 3.28E-01 | 1.18E+00 | 1.36E-01 | 5.19E-01 | 1.32E+00 | 1.86E-03 | 6.63E-03 | 1.05E+00 | 2.76E-03 | 7.79E-03 | 1.03E+00 | 8.17E-03 | 3.00E-02 | 1.01E+00 | 3.19E-01 | 8.76E-01 | 1.28E+00 |
| NBPF20 | 5.62E-06 | 2.93E-05 | -1.76E+00 | 9.01E-04 | 3.89E-03 | -1.09E+00 | 2.60E-02 | 1.11E-01 | -1.26E+00 | 4.10E-02 | 1.56E-01 | -1.03E+00 | 3.08E-03 | 1.10E-02 | -1.42E+00 | 5.08E-02 | 1.43E-01 | -1.08E+00 | 8.60E-08 | 3.16E-07 | -2.19E+00 | 4.70E-04 | 1.29E-03 | -1.38E+00 |
| LRRC40 | 5.62E-06 | 2.93E-05 | -1.27E+00 | 2.75E-04 | 1.19E-03 | 1.02E+00 | 4.67E-04 | 1.98E-03 | -1.21E+00 | 2.87E-03 | 1.09E-02 | -1.00E+00 | 3.70E-03 | 1.32E-02 | -1.11E+00 | 9.97E-03 | 2.81E-02 | -1.08E+00 | 6.23E-04 | 2.29E-03 | -1.05E+00 | 1.70E-02 | 4.66E-02 | 1.09E+00 |
| HECA | 5.64E-06 | 2.94E-05 | 1.80E+00 | 1.52E-02 | 6.56E-02 | 1.08E+00 | 1.41E-06 | 5.98E-06 | 2.15E+00 | 2.31E-01 | 8.83E-01 | 1.45E+00 | 1.89E-01 | 6.74E-01 | 1.69E+00 | 7.76E-04 | 2.19E-03 | -1.15E+00 | 2.27E-01 | 8.32E-01 | 1.28E+00 | 4.28E-03 | 1.18E-02 | -1.19E+00 |
| ATP5J2P5 | 5.65E-06 | 2.94E-05 | -1.70E+00 | 6.27E-06 | 2.71E-05 | -1.55E+00 | 4.70E-02 | 2.00E-01 | -1.00E+00 | 1.95E-07 | 7.45E-07 | -1.69E+00 | 2.36E-08 | 8.41E-08 | -2.17E+00 | 2.15E-08 | 6.06E-08 | -2.05E+00 | 3.95E-12 | 1.45E-11 | -1.60E+00 | 1.37E-18 | 3.76E-18 | -2.12E+00 |
| XYLT2 | 5.68E-06 | 2.96E-05 | -1.62E+00 | 7.29E-04 | 3.15E-03 | -1.04E+00 | 2.58E-08 | 1.10E-07 | -1.84E+00 | 2.46E-04 | 9.40E-04 | -1.17E+00 | 2.40E-03 | 8.56E-03 | -1.72E+00 | 8.07E-02 | 2.28E-01 | -1.02E+00 | 1.34E-11 | 4.93E-11 | -1.81E+00 | 1.92E-07 | 5.28E-07 | -1.85E+00 |
| ZNF614 | 5.69E-06 | 2.97E-05 | -1.54E+00 | 9.30E-08 | 4.02E-07 | -1.72E+00 | 1.41E-01 | 6.01E-01 | 1.20E+00 | 2.05E-02 | 7.82E-02 | 1.02E+00 | 4.37E-02 | 1.56E-01 | 1.02E+00 | 1.09E-02 | 3.06E-02 | -1.21E+00 | 1.23E-01 | 4.51E-01 | 1.12E+00 | 2.74E-02 | 7.53E-02 | -1.07E+00 |
| CCDC28B | 5.70E-06 | 2.97E-05 | -1.18E+00 | 1.47E-04 | 6.36E-04 | 1.05E+00 | 2.18E-03 | 9.27E-03 | -1.10E+00 | 4.16E-04 | 1.59E-03 | -1.00E+00 | 5.93E-10 | 2.11E-09 | -1.47E+00 | 4.01E-05 | 1.13E-04 | -1.09E+00 | 1.68E-02 | 6.17E-02 | 1.12E+00 | 2.76E-02 | 7.58E-02 | 1.08E+00 |
| MBNL2 | 5.70E-06 | 2.97E-05 | -1.25E+00 | 1.26E-05 | 5.43E-05 | -1.13E+00 | 3.41E-06 | 1.45E-05 | -1.69E+00 | 4.71E-03 | 1.80E-02 | -1.13E+00 | 5.12E-09 | 1.83E-08 | -1.72E+00 | 5.97E-05 | 1.68E-04 | -1.17E+00 | 9.21E-14 | 3.38E-13 | -1.90E+00 | 5.22E-11 | 1.43E-10 | -1.73E+00 |
| ITGB2-AS1 | 5.71E-06 | 2.97E-05 | -1.49E+00 | 4.44E-06 | 1.92E-05 | -1.40E+00 | 1.07E-03 | 4.53E-03 | -1.32E+00 | 7.84E-04 | 2.99E-03 | -1.30E+00 | 1.98E-01 | 7.06E-01 | 1.83E+00 | 3.05E-01 | 8.60E-01 | 1.14E+00 | 2.29E-01 | 8.40E-01 | 1.15E+00 | 3.22E-01 | 8.84E-01 | -1.10E+00 |
| NBEAL1 | 5.71E-06 | 2.98E-05 | -1.21E+00 | 1.19E-07 | 5.15E-07 | -1.21E+00 | 4.06E-06 | 1.73E-05 | -1.55E+00 | 3.23E-06 | 1.23E-05 | -1.37E+00 | 3.20E-05 | 1.14E-04 | -1.45E+00 | 5.97E-04 | 1.68E-03 | -1.26E+00 | 8.28E-05 | 3.04E-04 | -1.32E+00 | 1.20E-05 | 3.30E-05 | -1.43E+00 |
| WDR60 | 5.71E-06 | 2.98E-05 | -1.16E+00 | 1.68E-03 | 7.26E-03 | 1.06E+00 | 3.15E-06 | 1.34E-05 | -1.29E+00 | 1.64E-05 | 6.25E-05 | -1.10E+00 | 1.31E-03 | 4.67E-03 | -1.14E+00 | 3.76E-04 | 1.06E-03 | -1.19E+00 | 1.50E-02 | 5.50E-02 | -1.06E+00 | 7.10E-03 | 1.95E-02 | -1.06E+00 |
| SMARCE1P5 | 5.72E-06 | 2.98E-05 | -1.23E+00 | 1.09E-07 | 4.71E-07 | -1.29E+00 | 3.83E-07 | 1.63E-06 | -1.44E+00 | 2.63E-14 | 1.00E-13 | -2.10E+00 | 5.53E-10 | 1.97E-09 | -1.75E+00 | 2.72E-11 | 7.67E-11 | -1.92E+00 | 5.51E-09 | 2.02E-08 | -1.43E+00 | 2.57E-13 | 7.06E-13 | -1.71E+00 |
| TGDS | 5.78E-06 | 3.01E-05 | -1.65E+00 | 1.51E-01 | 6.52E-01 | 1.37E+00 | 4.92E-07 | 2.09E-06 | -1.72E+00 | 2.77E-02 | 1.06E-01 | 1.16E+00 | 9.82E-04 | 3.50E-03 | -1.81E+00 | 4.93E-02 | 1.39E-01 | -1.04E+00 | 9.15E-08 | 3.36E-07 | -1.95E+00 | 4.37E-02 | 1.20E-01 | -1.04E+00 |
| AC090825.1 | 5.80E-06 | 3.02E-05 | -1.76E+00 | 5.23E-03 | 2.26E-02 | 1.06E+00 | 8.79E-02 | 3.73E-01 | -1.07E+00 | 1.94E-01 | 7.41E-01 | 1.19E+00 | 9.55E-03 | 3.41E-02 | -1.56E+00 | 7.62E-02 | 2.15E-01 | -1.18E+00 | 4.51E-02 | 1.66E-01 | -1.15E+00 | 3.49E-01 | 9.59E-01 | 1.32E+00 |
| CERS5 | 5.81E-06 | 3.03E-05 | -1.17E+00 | 5.99E-03 | 2.59E-02 | 1.21E+00 | 4.08E-04 | 1.73E-03 | -1.07E+00 | 4.00E-05 | 1.53E-04 | -1.01E+00 | 3.46E-04 | 1.23E-03 | -1.07E+00 | 3.04E-02 | 8.56E-02 | 1.25E+00 | 4.99E-02 | 1.83E-01 | -1.02E+00 | 9.86E-02 | 2.71E-01 | 1.04E+00 |
| RSL24D1 | 5.82E-06 | 3.03E-05 | -1.15E+00 | 2.78E-07 | 1.20E-06 | -1.10E+00 | 4.31E-03 | 1.83E-02 | -1.13E+00 | 2.53E-03 | 9.67E-03 | -1.12E+00 | 9.44E-05 | 3.37E-04 | -1.09E+00 | 1.18E-02 | 3.33E-02 | -1.01E+00 | 1.24E-04 | 4.54E-04 | -1.12E+00 | 4.43E-03 | 1.22E-02 | -1.11E+00 |
| SMARCAL1 | 5.84E-06 | 3.04E-05 | -1.30E+00 | 1.47E-04 | 6.36E-04 | -1.01E+00 | 6.58E-05 | 2.80E-04 | -1.39E+00 | 3.47E-03 | 1.32E-02 | 1.00E+00 | 1.93E-07 | 6.87E-07 | -1.40E+00 | 3.18E-05 | 8.97E-05 | -1.12E+00 | 2.62E-13 | 9.61E-13 | -1.60E+00 | 1.76E-06 | 4.83E-06 | -1.24E+00 |
| IGFBP2 | 5.84E-06 | 3.05E-05 | -1.41E+00 | 1.08E-07 | 4.66E-07 | -1.60E+00 | 2.50E-02 | 1.06E-01 | 1.31E+00 | 3.91E-04 | 1.49E-03 | 1.10E+00 | 3.06E-05 | 1.09E-04 | -1.85E+00 | 2.25E-05 | 6.33E-05 | -1.91E+00 | 6.42E-04 | 2.36E-03 | -1.24E+00 | 1.66E-01 | 4.55E-01 | 1.17E+00 |
| ZNF506 | 5.85E-06 | 3.05E-05 | -1.17E+00 | 3.13E-04 | 1.35E-03 | 1.06E+00 | 5.31E-06 | 2.26E-05 | -1.40E+00 | 9.19E-04 | 3.51E-03 | -1.04E+00 | 5.34E-07 | 1.91E-06 | -1.32E+00 | 1.60E-06 | 4.51E-06 | -1.22E+00 | 3.16E-03 | 1.16E-02 | -1.12E+00 | 3.85E-03 | 1.06E-02 | -1.10E+00 |
| C5orf51 | 5.88E-06 | 3.07E-05 | -1.29E+00 | 1.28E-05 | 5.51E-05 | -1.12E+00 | 6.30E-07 | 2.68E-06 | -1.69E+00 | 2.57E-04 | 9.83E-04 | -1.17E+00 | 1.64E-06 | 5.84E-06 | -1.65E+00 | 9.62E-05 | 2.71E-04 | -1.26E+00 | 1.19E-06 | 4.37E-06 | -1.42E+00 | 1.93E-03 | 5.31E-03 | -1.09E+00 |
| CHST11 | 5.93E-06 | 3.09E-05 | -1.06E+00 | 1.22E-11 | 5.27E-11 | -1.13E+00 | 3.98E-03 | 1.69E-02 | -1.01E+00 | 4.21E-05 | 1.61E-04 | -1.08E+00 | 1.60E-02 | 5.71E-02 | 1.17E+00 | 8.37E-05 | 2.36E-04 | -1.05E+00 | 2.56E-06 | 9.38E-06 | -1.15E+00 | 2.28E-03 | 6.26E-03 | -1.00E+00 |
| ARL5A | 5.93E-06 | 3.09E-05 | -1.14E+00 | 3.69E-07 | 1.59E-06 | -1.18E+00 | 4.57E-05 | 1.94E-04 | -1.23E+00 | 1.57E-03 | 5.99E-03 | 1.07E+00 | 1.57E-08 | 5.60E-08 | -1.71E+00 | 1.41E-06 | 3.99E-06 | -1.38E+00 | 1.93E-05 | 7.08E-05 | -1.21E+00 | 1.75E-02 | 4.81E-02 | 1.07E+00 |
| MIGA1 | 5.93E-06 | 3.09E-05 | -1.29E+00 | 1.26E-02 | 5.45E-02 | 1.20E+00 | 5.84E-03 | 2.48E-02 | -1.04E+00 | 1.69E-02 | 6.45E-02 | 1.15E+00 | 1.16E-01 | 4.15E-01 | 1.15E+00 | 1.97E-01 | 5.56E-01 | 1.17E+00 | 3.66E-02 | 1.34E-01 | 1.12E+00 | 1.31E-01 | 3.59E-01 | 1.41E+00 |
| CLK4 | 5.94E-06 | 3.09E-05 | -1.13E+00 | 7.09E-04 | 3.06E-03 | 1.07E+00 | 1.17E-03 | 4.99E-03 | -1.13E+00 | 1.09E-02 | 4.17E-02 | 1.05E+00 | 5.97E-05 | 2.13E-04 | -1.08E+00 | 9.73E-04 | 2.74E-03 | 1.03E+00 | 5.26E-03 | 1.93E-02 | -1.02E+00 | 6.58E-02 | 1.81E-01 | 1.05E+00 |
| ECE2 | 5.94E-06 | 3.09E-05 | -2.00E+00 | 2.26E-03 | 9.75E-03 | -1.14E+00 | 1.89E-04 | 8.04E-04 | -1.16E+00 | 1.80E-02 | 6.87E-02 | 1.19E+00 | 1.59E-03 | 5.68E-03 | -1.78E+00 | 2.98E-01 | 8.40E-01 | 1.49E+00 | 2.17E-03 | 7.96E-03 | -1.17E+00 | 1.78E-02 | 4.89E-02 | 1.83E+00 |
| CUL1 | 5.96E-06 | 3.11E-05 | -1.18E+00 | 1.13E-03 | 4.87E-03 | 1.08E+00 | 1.73E-07 | 7.37E-07 | -1.30E+00 | 8.87E-04 | 3.39E-03 | 1.07E+00 | 1.27E-06 | 4.54E-06 | -1.44E+00 | 1.49E-02 | 4.19E-02 | 1.10E+00 | 4.43E-11 | 1.63E-10 | -1.37E+00 | 1.62E-04 | 4.45E-04 | -1.05E+00 |
| UBE4A | 5.96E-06 | 3.11E-05 | -1.19E+00 | 1.47E-04 | 6.37E-04 | 1.06E+00 | 3.80E-05 | 1.62E-04 | -1.22E+00 | 3.54E-05 | 1.35E-04 | -1.09E+00 | 4.67E-03 | 1.66E-02 | 1.01E+00 | 7.06E-04 | 1.99E-03 | -1.04E+00 | 4.46E-03 | 1.64E-02 | -1.00E+00 | 8.94E-03 | 2.46E-02 | -1.00E+00 |
| DNAJC18 | 5.97E-06 | 3.11E-05 | -1.25E+00 | 4.13E-06 | 1.79E-05 | -1.09E+00 | 1.22E-05 | 5.20E-05 | -1.47E+00 | 1.70E-04 | 6.50E-04 | -1.14E+00 | 1.57E-04 | 5.58E-04 | -1.30E+00 | 9.75E-04 | 2.75E-03 | -1.12E+00 | 8.86E-03 | 3.25E-02 | -1.18E+00 | 2.44E-02 | 6.69E-02 | -1.16E+00 |
| FLCN | 5.97E-06 | 3.11E-05 | -1.59E+00 | 1.95E-04 | 8.43E-04 | -1.13E+00 | 7.18E-04 | 3.05E-03 | -1.43E+00 | 1.28E-03 | 4.89E-03 | -1.18E+00 | 2.08E-05 | 7.42E-05 | -1.39E+00 | 8.52E-04 | 2.40E-03 | -1.10E+00 | 6.84E-08 | 2.51E-07 | -1.73E+00 | 1.16E-04 | 3.19E-04 | -1.38E+00 |
| RC3H1 | 6.02E-06 | 3.14E-05 | -1.04E+00 | 5.84E-08 | 2.52E-07 | -1.06E+00 | 5.57E-02 | 2.37E-01 | 1.01E+00 | 2.80E-04 | 1.07E-03 | 1.00E+00 | 7.39E-05 | 2.64E-04 | -1.04E+00 | 1.10E-07 | 3.11E-07 | -1.27E+00 | 1.41E-02 | 5.19E-02 | 1.01E+00 | 1.03E-05 | 2.83E-05 | -1.22E+00 |
| PIGU | 6.02E-06 | 3.14E-05 | -1.60E+00 | 1.22E-02 | 5.27E-02 | 1.09E+00 | 1.95E-03 | 8.28E-03 | -1.24E+00 | 6.32E-03 | 2.41E-02 | 1.01E+00 | 1.46E-02 | 5.21E-02 | -1.55E+00 | 2.03E-01 | 5.71E-01 | 1.24E+00 | 1.63E-07 | 5.98E-07 | -1.81E+00 | 8.91E-02 | 2.45E-01 | 1.06E+00 |
| TMEM120B | 6.02E-06 | 3.14E-05 | -1.61E+00 | 4.45E-03 | 1.92E-02 | 1.00E+00 | 5.16E-03 | 2.19E-02 | -1.22E+00 | 3.83E-02 | 1.46E-01 | 1.10E+00 | 4.29E-02 | 1.53E-01 | -1.11E+00 | 2.92E-01 | 8.22E-01 | 1.56E+00 | 3.19E-02 | 1.17E-01 | 1.02E+00 | 2.38E-01 | 6.55E-01 | 1.19E+00 |
| TMEM144 | 6.07E-06 | 3.16E-05 | -1.64E+00 | 1.93E-03 | 8.32E-03 | -1.07E+00 | 6.95E-05 | 2.95E-04 | -1.41E+00 | 1.34E-03 | 5.13E-03 | -1.09E+00 | 5.71E-02 | 2.04E-01 | -1.41E+00 | 1.48E-01 | 4.18E-01 | -1.09E+00 | 1.10E-05 | 4.05E-05 | -1.81E+00 | 1.36E-02 | 3.73E-02 | -1.17E+00 |
| FXR1 | 6.10E-06 | 3.18E-05 | -1.05E+00 | 1.99E-09 | 8.60E-09 | -1.01E+00 | 1.26E-03 | 5.36E-03 | -1.21E+00 | 3.94E-05 | 1.51E-04 | -1.16E+00 | 3.00E-03 | 1.07E-02 | -1.00E+00 | 7.76E-06 | 2.19E-05 | -1.09E+00 | 1.13E-04 | 4.15E-04 | -1.14E+00 | 6.30E-06 | 1.73E-05 | -1.27E+00 |
| SHMT2 | 6.12E-06 | 3.19E-05 | -1.21E+00 | 3.71E-05 | 1.60E-04 | -1.02E+00 | 2.64E-03 | 1.12E-02 | -1.04E+00 | 1.88E-03 | 7.18E-03 | 1.05E+00 | 2.34E-06 | 8.33E-06 | -1.25E+00 | 1.24E-03 | 3.49E-03 | 1.00E+00 | 9.87E-03 | 3.62E-02 | 1.08E+00 | 2.09E-01 | 5.74E-01 | 1.22E+00 |
| MTMR2 | 6.12E-06 | 3.19E-05 | -1.26E+00 | 1.20E-04 | 5.20E-04 | 1.03E+00 | 3.39E-05 | 1.44E-04 | -1.24E+00 | 1.12E-05 | 4.28E-05 | -1.15E+00 | 9.37E-04 | 3.34E-03 | -1.15E+00 | 5.85E-02 | 1.65E-01 | 1.17E+00 | 2.42E-06 | 8.89E-06 | -1.20E+00 | 5.33E-04 | 1.46E-03 | -1.02E+00 |
| CA8 | 6.19E-06 | 3.22E-05 | -1.48E+00 | 4.73E-02 | 2.04E-01 | 1.26E+00 | 2.58E-04 | 1.10E-03 | -1.38E+00 | 6.33E-04 | 2.42E-03 | -1.11E+00 | 1.28E-01 | 4.55E-01 | -1.55E+00 | 2.50E-01 | 7.06E-01 | -1.05E+00 | 8.71E-03 | 3.20E-02 | -2.18E+00 | 3.84E-02 | 1.06E-01 | -1.60E+00 |
| SEC24C | 6.19E-06 | 3.23E-05 | -1.63E+00 | 1.01E-04 | 4.38E-04 | -1.27E+00 | 6.38E-07 | 2.71E-06 | -1.82E+00 | 2.42E-06 | 9.25E-06 | -1.56E+00 | 9.89E-05 | 3.53E-04 | -1.41E+00 | 7.30E-06 | 2.06E-05 | -1.65E+00 | 6.11E-05 | 2.24E-04 | -1.38E+00 | 1.75E-05 | 4.82E-05 | -1.52E+00 |
| NUMA1 | 6.20E-06 | 3.23E-05 | -1.33E+00 | 3.98E-04 | 1.72E-03 | 1.00E+00 | 3.00E-02 | 1.27E-01 | -1.09E+00 | 3.38E-02 | 1.29E-01 | 1.11E+00 | 3.63E-02 | 1.30E-01 | 1.09E+00 | 1.27E-01 | 3.58E-01 | 1.29E+00 | 4.54E-03 | 1.67E-02 | -1.13E+00 | 1.25E-02 | 3.43E-02 | -1.08E+00 |
| RNF135 | 6.20E-06 | 3.23E-05 | -1.25E+00 | 2.70E-03 | 1.17E-02 | 1.08E+00 | 6.25E-05 | 2.65E-04 | -1.30E+00 | 4.07E-03 | 1.56E-02 | -1.04E+00 | 8.02E-03 | 2.86E-02 | -1.18E+00 | 2.18E-01 | 6.15E-01 | 1.21E+00 | 2.81E-05 | 1.03E-04 | -1.38E+00 | 2.20E-02 | 6.03E-02 | 1.02E+00 |
| RAB3IP | 6.21E-06 | 3.24E-05 | -1.21E+00 | 7.34E-04 | 3.17E-03 | 1.11E+00 | 4.51E-07 | 1.91E-06 | -1.39E+00 | 3.81E-05 | 1.45E-04 | -1.10E+00 | 6.35E-03 | 2.27E-02 | -1.22E+00 | 6.50E-02 | 1.83E-01 | 1.06E+00 | 2.00E-03 | 7.35E-03 | -1.25E+00 | 2.00E-01 | 5.51E-01 | 1.14E+00 |
| CRYZ | 6.23E-06 | 3.24E-05 | -1.61E+00 | 1.05E-01 | 4.53E-01 | 1.64E+00 | 2.17E-06 | 9.22E-06 | -1.63E+00 | 3.57E-02 | 1.36E-01 | 1.24E+00 | 1.98E-08 | 7.04E-08 | -2.22E+00 | 6.62E-03 | 1.87E-02 | 1.00E+00 | 1.89E-02 | 6.95E-02 | -1.24E+00 | 3.11E-01 | 8.55E-01 | 1.20E+00 |
| LRRC28 | 6.23E-06 | 3.25E-05 | -1.14E+00 | 3.97E-03 | 1.71E-02 | 1.23E+00 | 3.08E-04 | 1.31E-03 | -1.27E+00 | 7.18E-04 | 2.74E-03 | -1.03E+00 | 1.18E-03 | 4.22E-03 | -1.16E+00 | 1.73E-01 | 4.88E-01 | 1.27E+00 | 1.51E-05 | 5.55E-05 | -1.33E+00 | 2.50E-05 | 6.87E-05 | -1.39E+00 |
| SPRYD3 | 6.24E-06 | 3.25E-05 | -1.89E+00 | 4.41E-04 | 1.91E-03 | -1.22E+00 | 7.65E-03 | 3.25E-02 | -1.37E+00 | 1.26E-01 | 4.82E-01 | 1.17E+00 | 4.21E-03 | 1.50E-02 | -1.49E+00 | 5.17E-02 | 1.46E-01 | 1.02E+00 | 3.23E-07 | 1.19E-06 | -1.79E+00 | 2.20E-02 | 6.04E-02 | -1.13E+00 |
| HOOK2 | 6.25E-06 | 3.26E-05 | -1.23E+00 | 5.99E-03 | 2.59E-02 | 1.22E+00 | 8.63E-06 | 3.67E-05 | -1.28E+00 | 3.13E-04 | 1.20E-03 | 1.02E+00 | 8.57E-08 | 3.05E-07 | -1.32E+00 | 4.80E-04 | 1.35E-03 | -1.00E+00 | 3.84E-04 | 1.41E-03 | -1.12E+00 | 1.87E-02 | 5.12E-02 | 1.01E+00 |
| UBE3D | 6.28E-06 | 3.27E-05 | -1.27E+00 | 3.71E-02 | 1.61E-01 | 1.27E+00 | 2.29E-06 | 9.72E-06 | -1.51E+00 | 5.53E-04 | 2.11E-03 | -1.10E+00 | 1.64E-03 | 5.85E-03 | -1.60E+00 | 1.40E-01 | 3.94E-01 | 1.05E+00 | 1.51E-05 | 5.56E-05 | -1.68E+00 | 5.68E-02 | 1.56E-01 | -1.08E+00 |
| ARL10 | 6.28E-06 | 3.27E-05 | -1.48E+00 | 2.45E-02 | 1.06E-01 | 1.19E+00 | 8.45E-08 | 3.59E-07 | -1.84E+00 | 3.69E-02 | 1.41E-01 | 1.11E+00 | 1.03E-04 | 3.67E-04 | -1.81E+00 | 8.25E-02 | 2.33E-01 | 1.12E+00 | 3.24E-02 | 1.19E-01 | -1.21E+00 | 2.84E-01 | 7.81E-01 | 1.42E+00 |
| NFKBIB | 6.28E-06 | 3.27E-05 | -1.22E+00 | 1.76E-08 | 7.58E-08 | -1.37E+00 | 9.34E-06 | 3.97E-05 | -1.25E+00 | 1.31E-08 | 5.01E-08 | -1.36E+00 | 1.80E-05 | 6.43E-05 | -1.53E+00 | 4.60E-04 | 1.30E-03 | -1.22E+00 | 6.16E-10 | 2.26E-09 | -1.42E+00 | 1.54E-06 | 4.24E-06 | -1.29E+00 |
| RAB22A | 6.29E-06 | 3.28E-05 | -1.30E+00 | 2.49E-05 | 1.08E-04 | -1.13E+00 | 4.69E-05 | 1.99E-04 | -1.44E+00 | 7.33E-04 | 2.80E-03 | -1.09E+00 | 1.37E-06 | 4.89E-06 | -1.34E+00 | 8.55E-08 | 2.41E-07 | -1.46E+00 | 1.11E-04 | 4.06E-04 | -1.25E+00 | 8.00E-04 | 2.20E-03 | -1.13E+00 |
| CCDC171 | 6.30E-06 | 3.28E-05 | -1.13E+00 | 1.33E-07 | 5.76E-07 | -1.10E+00 | 1.66E-11 | 7.06E-11 | -1.47E+00 | 1.33E-12 | 5.07E-12 | -1.39E+00 | 9.31E-04 | 3.32E-03 | -1.08E+00 | 5.05E-05 | 1.42E-04 | -1.18E+00 | 4.29E-03 | 1.58E-02 | -1.27E+00 | 2.61E-02 | 7.17E-02 | -1.13E+00 |
| GALNT5 | 6.31E-06 | 3.29E-05 | -1.84E+00 | 8.22E-03 | 3.55E-02 | 1.02E+00 | 6.17E-06 | 2.62E-05 | -1.79E+00 | 6.31E-04 | 2.41E-03 | -1.16E+00 | 2.38E-01 | 8.48E-01 | 1.37E+00 | 1.24E-01 | 3.49E-01 | 2.17E+00 | 1.79E-01 | 6.56E-01 | 1.40E+00 | 1.77E-01 | 4.87E-01 | 1.66E+00 |
| SNF8 | 6.32E-06 | 3.29E-05 | 1.01E+00 | 5.32E-11 | 2.30E-10 | -1.02E+00 | 2.11E-01 | 8.95E-01 | 1.16E+00 | 8.76E-03 | 3.35E-02 | 1.07E+00 | 2.55E-02 | 9.08E-02 | 1.11E+00 | 4.34E-02 | 1.22E-01 | 1.09E+00 | 2.33E-01 | 8.53E-01 | 1.21E+00 | 2.84E-01 | 7.81E-01 | 1.20E+00 |
| SEMA4B | 6.32E-06 | 3.30E-05 | -2.66E+00 | 1.09E-03 | 4.69E-03 | -1.56E+00 | 1.00E-02 | 4.26E-02 | -1.40E+00 | 6.82E-02 | 2.61E-01 | 1.03E+00 | 1.11E-01 | 3.97E-01 | -1.17E+00 | 2.86E-01 | 8.06E-01 | 1.23E+00 | 2.03E-01 | 7.44E-01 | 1.16E+00 | 2.39E-01 | 6.56E-01 | 1.39E+00 |
| ASGR1 | 6.33E-06 | 3.30E-05 | -1.75E+00 | 1.37E-06 | 5.94E-06 | -1.75E+00 | 7.03E-05 | 2.99E-04 | -1.53E+00 | 4.25E-05 | 1.62E-04 | -1.44E+00 | 6.24E-02 | 2.22E-01 | -1.19E+00 | 9.12E-02 | 2.57E-01 | -1.07E+00 | 9.22E-03 | 3.39E-02 | -1.23E+00 | 2.24E-01 | 6.16E-01 | 1.18E+00 |
| ZNF131 | 6.33E-06 | 3.30E-05 | -1.03E+00 | 1.31E-07 | 5.66E-07 | -1.03E+00 | 8.69E-04 | 3.69E-03 | -1.14E+00 | 9.77E-04 | 3.73E-03 | -1.03E+00 | 5.26E-04 | 1.88E-03 | -1.03E+00 | 1.11E-03 | 3.13E-03 | -1.00E+00 | 8.61E-04 | 3.16E-03 | -1.12E+00 | 1.14E-02 | 3.13E-02 | -1.01E+00 |
| ARRDC1 | 6.38E-06 | 3.32E-05 | -1.09E+00 | 1.20E-02 | 5.17E-02 | 1.25E+00 | 2.50E-03 | 1.06E-02 | -1.03E+00 | 8.70E-03 | 3.32E-02 | 1.14E+00 | 6.24E-04 | 2.23E-03 | -1.15E+00 | 1.25E-02 | 3.51E-02 | 1.02E+00 | 6.24E-03 | 2.29E-02 | 1.03E+00 | 5.58E-02 | 1.53E-01 | 1.21E+00 |
| RGS10 | 6.38E-06 | 3.32E-05 | -1.10E+00 | 4.33E-07 | 1.87E-06 | -1.09E+00 | 2.02E-04 | 8.58E-04 | -1.22E+00 | 4.59E-06 | 1.75E-05 | -1.25E+00 | 8.14E-06 | 2.90E-05 | -1.44E+00 | 1.79E-03 | 5.04E-03 | -1.19E+00 | 2.08E-02 | 7.64E-02 | 1.40E+00 | 5.88E-02 | 1.62E-01 | -1.02E+00 |
| YY1AP1 | 6.39E-06 | 3.33E-05 | -1.25E+00 | 5.05E-06 | 2.18E-05 | -1.14E+00 | 2.44E-06 | 1.03E-05 | -1.22E+00 | 5.25E-10 | 2.00E-09 | -1.35E+00 | 2.87E-06 | 1.02E-05 | -1.32E+00 | 8.95E-06 | 2.52E-05 | -1.27E+00 | 2.27E-03 | 8.32E-03 | -1.09E+00 | 1.68E-04 | 4.61E-04 | -1.15E+00 |
| C22orf39 | 6.40E-06 | 3.34E-05 | -1.14E+00 | 1.08E-04 | 4.66E-04 | 1.06E+00 | 5.07E-02 | 2.15E-01 | -1.00E+00 | 3.08E-02 | 1.18E-01 | 1.05E+00 | 3.17E-03 | 1.13E-02 | -1.01E+00 | 1.48E-02 | 4.16E-02 | 1.07E+00 | 2.36E-05 | 8.67E-05 | -1.11E+00 | 8.08E-04 | 2.22E-03 | -1.05E+00 |
| PLCE1 | 6.41E-06 | 3.34E-05 | -1.72E+00 | 5.64E-03 | 2.44E-02 | 1.00E+00 | 1.97E-02 | 8.38E-02 | -1.27E+00 | 2.39E-01 | 9.12E-01 | 1.34E+00 | 5.21E-02 | 1.86E-01 | -1.20E+00 | 1.71E-01 | 4.83E-01 | 1.10E+00 | 1.35E-02 | 4.96E-02 | -1.33E+00 | 2.11E-01 | 5.80E-01 | 1.08E+00 |
| SLC39A14 | 6.42E-06 | 3.35E-05 | -2.47E+00 | 6.69E-03 | 2.89E-02 | -1.11E+00 | 7.53E-06 | 3.20E-05 | -1.76E+00 | 1.89E-02 | 7.20E-02 | 1.01E+00 | 1.70E-02 | 6.07E-02 | -1.73E+00 | 2.58E-01 | 7.28E-01 | 1.19E+00 | 4.54E-07 | 1.67E-06 | -1.30E+00 | 1.33E-02 | 3.66E-02 | 1.05E+00 |
| ZNF264 | 6.42E-06 | 3.35E-05 | -1.15E+00 | 6.14E-06 | 2.65E-05 | -1.01E+00 | 1.37E-08 | 5.82E-08 | -1.61E+00 | 1.94E-07 | 7.43E-07 | -1.29E+00 | 1.38E-06 | 4.91E-06 | -1.87E+00 | 1.77E-04 | 4.98E-04 | -1.41E+00 | 1.48E-06 | 5.44E-06 | -1.43E+00 | 1.15E-05 | 3.15E-05 | -1.41E+00 |
| TAF5L | 6.42E-06 | 3.35E-05 | -1.45E+00 | 2.43E-07 | 1.05E-06 | -1.52E+00 | 6.42E-04 | 2.73E-03 | -1.33E+00 | 9.90E-06 | 3.78E-05 | -1.56E+00 | 2.30E-04 | 8.21E-04 | -1.53E+00 | 2.02E-04 | 5.68E-04 | -1.62E+00 | 3.99E-12 | 1.46E-11 | -1.87E+00 | 1.95E-09 | 5.36E-09 | -1.61E+00 |
| RGS19 | 6.42E-06 | 3.35E-05 | -1.14E+00 | 1.11E-02 | 4.78E-02 | 1.10E+00 | 9.57E-03 | 4.07E-02 | -1.03E+00 | 1.46E-01 | 5.58E-01 | 1.17E+00 | 6.68E-09 | 2.38E-08 | -1.53E+00 | 1.82E-08 | 5.13E-08 | -1.42E+00 | 8.81E-03 | 3.23E-02 | -1.02E+00 | 1.24E-01 | 3.42E-01 | 1.28E+00 |
| RBMX2P3 | 6.44E-06 | 3.35E-05 | -1.54E+00 | 4.66E-07 | 2.01E-06 | -1.64E+00 | 1.15E-04 | 4.88E-04 | -1.53E+00 | 4.67E-05 | 1.78E-04 | -1.50E+00 | 3.86E-06 | 1.38E-05 | -1.69E+00 | 2.16E-05 | 6.08E-05 | -1.51E+00 | 2.69E-04 | 9.86E-04 | -1.32E+00 | 2.54E-06 | 6.96E-06 | -1.60E+00 |
| TCIRG1 | 6.44E-06 | 3.36E-05 | -1.23E+00 | 1.23E-04 | 5.31E-04 | 1.01E+00 | 1.07E-03 | 4.55E-03 | -1.34E+00 | 2.71E-04 | 1.04E-03 | -1.23E+00 | 8.48E-04 | 3.02E-03 | -1.33E+00 | 3.11E-04 | 8.77E-04 | -1.43E+00 | 5.75E-06 | 2.11E-05 | -1.44E+00 | 1.27E-05 | 3.48E-05 | -1.54E+00 |
| RAB28 | 6.45E-06 | 3.36E-05 | -1.60E+00 | 1.93E-03 | 8.33E-03 | -1.03E+00 | 5.10E-05 | 2.17E-04 | -1.36E+00 | 6.23E-04 | 2.38E-03 | -1.03E+00 | 1.10E-04 | 3.93E-04 | -1.39E+00 | 2.71E-04 | 7.63E-04 | -1.21E+00 | 6.58E-02 | 2.42E-01 | 1.08E+00 | 2.60E-01 | 7.15E-01 | 1.14E+00 |
| KCTD3 | 6.45E-06 | 3.36E-05 | -1.27E+00 | 7.26E-04 | 3.14E-03 | 1.03E+00 | 4.55E-06 | 1.93E-05 | -1.37E+00 | 2.99E-04 | 1.14E-03 | -1.00E+00 | 4.39E-03 | 1.57E-02 | -1.26E+00 | 2.51E-01 | 7.09E-01 | 1.31E+00 | 3.13E-04 | 1.15E-03 | -1.30E+00 | 2.26E-01 | 6.22E-01 | 1.11E+00 |
| DNM2 | 6.47E-06 | 3.37E-05 | -1.17E+00 | 2.22E-05 | 9.58E-05 | -1.07E+00 | 5.79E-03 | 2.46E-02 | 1.03E+00 | 1.09E-03 | 4.15E-03 | 1.12E+00 | 2.24E-05 | 7.98E-05 | -1.30E+00 | 2.98E-02 | 8.40E-02 | 1.08E+00 | 1.04E-04 | 3.83E-04 | -1.09E+00 | 2.20E-02 | 6.04E-02 | 1.05E+00 |
| RGPD4 | 6.47E-06 | 3.37E-05 | -1.45E+00 | 6.14E-07 | 2.65E-06 | -1.48E+00 | 8.94E-04 | 3.80E-03 | -1.24E+00 | 9.09E-07 | 3.47E-06 | -1.55E+00 | 1.25E-03 | 4.47E-03 | -1.36E+00 | 1.54E-05 | 4.34E-05 | -2.10E+00 | 1.16E-02 | 4.26E-02 | -1.12E+00 | 5.40E-05 | 1.48E-04 | -1.50E+00 |
| COX6B1 | 6.47E-06 | 3.37E-05 | -1.17E+00 | 2.10E-25 | 9.07E-25 | -1.42E+00 | 2.75E-02 | 1.17E-01 | -1.10E+00 | 2.83E-06 | 1.08E-05 | -1.33E+00 | 1.06E-04 | 3.78E-04 | -1.17E+00 | 4.74E-10 | 1.34E-09 | -1.30E+00 | 3.36E-02 | 1.23E-01 | -1.08E+00 | 2.22E-04 | 6.11E-04 | -1.26E+00 |
| SLC7A1 | 6.48E-06 | 3.38E-05 | -1.52E+00 | 4.74E-03 | 2.05E-02 | 1.06E+00 | 2.04E-03 | 8.66E-03 | -1.12E+00 | 5.09E-02 | 1.95E-01 | 1.18E+00 | 2.97E-03 | 1.06E-02 | -1.50E+00 | 2.53E-01 | 7.13E-01 | 1.18E+00 | 3.80E-05 | 1.39E-04 | -1.35E+00 | 3.53E-01 | 9.70E-01 | 1.32E+00 |
| AC007383.2 | 6.51E-06 | 3.39E-05 | -1.74E+00 | 8.90E-03 | 3.85E-02 | 1.02E+00 | 1.14E-07 | 4.85E-07 | -2.14E+00 | 3.80E-06 | 1.45E-05 | -1.55E+00 | 1.03E-04 | 3.67E-04 | -2.08E+00 | 1.00E-01 | 2.82E-01 | 1.07E+00 | 1.49E-04 | 5.47E-04 | -1.51E+00 | 1.01E-01 | 2.77E-01 | 1.09E+00 |
| METTL16 | 6.52E-06 | 3.40E-05 | -1.22E+00 | 3.19E-06 | 1.38E-05 | -1.10E+00 | 8.44E-03 | 3.59E-02 | 1.01E+00 | 1.71E-03 | 6.53E-03 | 1.05E+00 | 1.38E-04 | 4.90E-04 | -1.23E+00 | 1.02E-02 | 2.88E-02 | 1.04E+00 | 9.08E-02 | 3.33E-01 | 1.10E+00 | 1.50E-01 | 4.13E-01 | 1.14E+00 |
| TTC28 | 6.55E-06 | 3.41E-05 | -1.54E+00 | 1.15E-02 | 4.96E-02 | 1.09E+00 | 2.12E-03 | 9.02E-03 | -1.43E+00 | 1.28E-01 | 4.90E-01 | 1.22E+00 | 1.20E-01 | 4.27E-01 | 1.06E+00 | 1.11E-01 | 3.12E-01 | 1.86E+00 | 2.48E-06 | 9.09E-06 | -1.63E+00 | 2.83E-03 | 7.77E-03 | -1.18E+00 |
| RABL2A | 6.55E-06 | 3.42E-05 | -1.39E+00 | 1.12E-02 | 4.83E-02 | 1.11E+00 | 1.00E-06 | 4.27E-06 | -1.49E+00 | 6.12E-04 | 2.34E-03 | -1.04E+00 | 1.65E-04 | 5.87E-04 | -1.32E+00 | 3.89E-03 | 1.10E-02 | -1.05E+00 | 3.55E-03 | 1.30E-02 | -1.15E+00 | 1.29E-01 | 3.54E-01 | 1.15E+00 |
| CPOX | 6.56E-06 | 3.42E-05 | -1.73E+00 | 3.50E-03 | 1.51E-02 | 1.04E+00 | 1.89E-10 | 8.03E-10 | -1.79E+00 | 2.54E-08 | 9.69E-08 | -1.28E+00 | 2.38E-03 | 8.48E-03 | -1.33E+00 | 3.32E-01 | 9.36E-01 | 1.46E+00 | 9.47E-37 | 3.48E-36 | -2.26E+00 | 2.16E-15 | 5.93E-15 | -1.40E+00 |
| EIF2S3 | 6.56E-06 | 3.42E-05 | -1.11E+00 | 2.07E-01 | 8.96E-01 | 1.16E+00 | 2.95E-03 | 1.25E-02 | -1.15E+00 | 1.93E-01 | 7.39E-01 | 1.07E+00 | 1.78E-02 | 6.36E-02 | -1.01E+00 | 1.91E-01 | 5.38E-01 | 1.09E+00 | 8.98E-05 | 3.30E-04 | -1.09E+00 | 3.06E-01 | 8.42E-01 | 1.12E+00 |
| MARC2 | 6.59E-06 | 3.43E-05 | -1.33E+00 | 1.81E-05 | 7.80E-05 | -1.17E+00 | 1.37E-06 | 5.84E-06 | -1.38E+00 | 1.81E-07 | 6.90E-07 | -1.36E+00 | 3.03E-03 | 1.08E-02 | -1.80E+00 | 1.93E-03 | 5.45E-03 | -1.97E+00 | 5.64E-02 | 2.07E-01 | -1.43E+00 | 2.95E-01 | 8.10E-01 | 1.37E+00 |
| AL109811.3 | 6.59E-06 | 3.43E-05 | -2.00E+00 | 8.03E-02 | 3.47E-01 | 1.74E+00 | 4.41E-05 | 1.88E-04 | -2.07E+00 | 1.81E-01 | 6.91E-01 | 1.18E+00 | 3.91E-03 | 1.39E-02 | -1.77E+00 | 3.40E-01 | 9.57E-01 | 1.10E+00 | 2.83E-03 | 1.04E-02 | -1.35E+00 | 3.08E-01 | 8.45E-01 | 1.18E+00 |
| ARSB | 6.61E-06 | 3.45E-05 | -1.43E+00 | 1.35E-03 | 5.81E-03 | 1.02E+00 | 1.69E-02 | 7.18E-02 | -1.17E+00 | 1.94E-01 | 7.42E-01 | 1.38E+00 | 1.39E-01 | 4.94E-01 | 1.00E+00 | 1.70E-01 | 4.80E-01 | 1.06E+00 | 7.00E-02 | 2.57E-01 | 1.01E+00 | 1.47E-01 | 4.05E-01 | 1.68E+00 |
| SERPINB1 | 6.61E-06 | 3.45E-05 | -1.23E+00 | 1.24E-02 | 5.34E-02 | -1.11E+00 | 3.71E-07 | 1.57E-06 | -1.35E+00 | 8.61E-04 | 3.29E-03 | -1.17E+00 | 8.17E-04 | 2.91E-03 | -1.17E+00 | 8.36E-03 | 2.36E-02 | -1.09E+00 | 5.63E-02 | 2.07E-01 | 1.02E+00 | 1.36E-02 | 3.75E-02 | -1.03E+00 |
| COIL | 6.62E-06 | 3.45E-05 | -1.14E+00 | 2.57E-08 | 1.11E-07 | -1.14E+00 | 1.03E-04 | 4.38E-04 | -1.12E+00 | 5.77E-08 | 2.20E-07 | -1.20E+00 | 8.51E-03 | 3.03E-02 | -1.00E+00 | 6.44E-04 | 1.82E-03 | -1.12E+00 | 3.41E-05 | 1.25E-04 | -1.08E+00 | 3.04E-08 | 8.36E-08 | -1.25E+00 |
| NUP153 | 6.62E-06 | 3.45E-05 | -1.28E+00 | 1.39E-05 | 5.99E-05 | -1.18E+00 | 8.35E-03 | 3.55E-02 | -1.07E+00 | 5.18E-06 | 1.98E-05 | -1.23E+00 | 1.98E-02 | 7.06E-02 | 1.10E+00 | 2.05E-02 | 5.79E-02 | 1.17E+00 | 1.22E-06 | 4.48E-06 | -1.31E+00 | 1.49E-03 | 4.08E-03 | -1.10E+00 |
| ORC2 | 6.62E-06 | 3.45E-05 | -1.29E+00 | 4.45E-03 | 1.92E-02 | 1.17E+00 | 1.74E-05 | 7.39E-05 | -1.37E+00 | 4.69E-05 | 1.79E-04 | -1.20E+00 | 1.70E-06 | 6.05E-06 | -1.55E+00 | 1.07E-03 | 3.01E-03 | -1.09E+00 | 6.74E-08 | 2.47E-07 | -1.54E+00 | 2.38E-02 | 6.55E-02 | 1.03E+00 |
| HGH1 | 6.64E-06 | 3.46E-05 | -1.74E+00 | 4.27E-03 | 1.85E-02 | -1.01E+00 | 3.31E-06 | 1.41E-05 | -1.59E+00 | 9.40E-03 | 3.59E-02 | 1.07E+00 | 6.71E-02 | 2.39E-01 | -1.06E+00 | 3.16E-01 | 8.92E-01 | 1.37E+00 | 3.96E-06 | 1.45E-05 | -1.37E+00 | 1.66E-01 | 4.57E-01 | 1.22E+00 |
| TCEAL4 | 6.66E-06 | 3.47E-05 | -1.09E+00 | 6.62E-06 | 2.86E-05 | 1.00E+00 | 9.02E-03 | 3.83E-02 | -1.06E+00 | 1.10E-03 | 4.20E-03 | -1.06E+00 | 2.16E-05 | 7.70E-05 | -1.18E+00 | 1.86E-04 | 5.25E-04 | -1.15E+00 | 1.41E-02 | 5.19E-02 | -1.04E+00 | 1.54E-02 | 4.24E-02 | -1.08E+00 |
| STARD10 | 6.67E-06 | 3.48E-05 | -1.16E+00 | 2.78E-06 | 1.20E-05 | -1.04E+00 | 1.91E-04 | 8.12E-04 | -1.32E+00 | 7.76E-06 | 2.96E-05 | -1.26E+00 | 1.27E-02 | 4.53E-02 | 1.02E+00 | 3.29E-02 | 9.28E-02 | 1.18E+00 | 1.83E-01 | 6.73E-01 | 1.12E+00 | 7.82E-03 | 2.15E-02 | -1.32E+00 |
| SMUG1 | 6.68E-06 | 3.48E-05 | -1.64E+00 | 8.97E-04 | 3.88E-03 | -1.07E+00 | 2.23E-01 | 9.46E-01 | 1.04E+00 | 8.78E-02 | 3.35E-01 | 1.46E+00 | 3.78E-03 | 1.35E-02 | -1.11E+00 | 6.99E-02 | 1.97E-01 | 1.16E+00 | 8.49E-04 | 3.11E-03 | -1.05E+00 | 3.04E-03 | 8.35E-03 | 1.01E+00 |
| DHX15 | 6.69E-06 | 3.49E-05 | -1.13E+00 | 3.90E-05 | 1.69E-04 | 1.01E+00 | 6.56E-02 | 2.79E-01 | -1.07E+00 | 3.32E-02 | 1.27E-01 | 1.05E+00 | 2.31E-07 | 8.22E-07 | -1.39E+00 | 4.71E-04 | 1.33E-03 | -1.12E+00 | 4.48E-04 | 1.64E-03 | -1.10E+00 | 1.16E-02 | 3.18E-02 | 1.03E+00 |
| PDSS2 | 6.71E-06 | 3.50E-05 | -1.50E+00 | 5.07E-04 | 2.19E-03 | -1.08E+00 | 3.04E-05 | 1.29E-04 | -1.35E+00 | 6.23E-05 | 2.38E-04 | -1.18E+00 | 2.86E-05 | 1.02E-04 | -1.60E+00 | 1.72E-03 | 4.86E-03 | -1.13E+00 | 3.72E-04 | 1.36E-03 | -1.19E+00 | 4.21E-03 | 1.16E-02 | -1.21E+00 |
| NFYB | 6.72E-06 | 3.50E-05 | -1.31E+00 | 4.49E-04 | 1.94E-03 | 1.03E+00 | 1.55E-04 | 6.60E-04 | -1.17E+00 | 1.55E-04 | 5.93E-04 | -1.09E+00 | 5.70E-06 | 2.03E-05 | -1.52E+00 | 4.90E-05 | 1.38E-04 | -1.41E+00 | 4.97E-05 | 1.82E-04 | -1.18E+00 | 1.22E-02 | 3.35E-02 | 1.01E+00 |
| FMNL1 | 6.76E-06 | 3.52E-05 | -1.07E+00 | 7.39E-04 | 3.19E-03 | 1.10E+00 | 1.51E-03 | 6.42E-03 | -1.10E+00 | 5.21E-03 | 1.99E-02 | 1.05E+00 | 1.70E-02 | 6.07E-02 | 1.18E+00 | 7.39E-02 | 2.08E-01 | 1.23E+00 | 1.02E-01 | 3.75E-01 | 1.11E+00 | 1.29E-02 | 3.54E-02 | -1.00E+00 |
| ECHDC3 | 6.76E-06 | 3.52E-05 | -1.56E+00 | 1.55E-03 | 6.72E-03 | -1.01E+00 | 1.10E-06 | 4.68E-06 | -1.49E+00 | 8.55E-07 | 3.27E-06 | -1.30E+00 | 4.02E-04 | 1.43E-03 | -2.01E+00 | 3.28E-02 | 9.25E-02 | -1.12E+00 | 1.72E-02 | 6.30E-02 | -1.04E+00 | 9.76E-02 | 2.68E-01 | 1.03E+00 |
| AC099343.3 | 6.76E-06 | 3.52E-05 | -1.21E+00 | 2.11E-05 | 9.11E-05 | -1.11E+00 | 3.76E-05 | 1.60E-04 | -1.33E+00 | 5.10E-04 | 1.95E-03 | -1.14E+00 | 1.14E-03 | 4.07E-03 | -1.32E+00 | 7.81E-02 | 2.20E-01 | 1.16E+00 | 7.60E-03 | 2.79E-02 | -1.08E+00 | 1.12E-02 | 3.09E-02 | -1.09E+00 |
| ZNF500 | 6.79E-06 | 3.54E-05 | -1.52E+00 | 4.31E-02 | 1.86E-01 | 1.23E+00 | 6.66E-07 | 2.83E-06 | -1.78E+00 | 6.84E-04 | 2.61E-03 | -1.16E+00 | 3.31E-04 | 1.18E-03 | -1.66E+00 | 1.92E-02 | 5.41E-02 | -1.11E+00 | 5.44E-08 | 2.00E-07 | -1.66E+00 | 1.83E-06 | 5.04E-06 | -1.42E+00 |
| NFIX | 6.80E-06 | 3.55E-05 | -1.15E+00 | 6.27E-03 | 2.71E-02 | 1.24E+00 | 4.51E-06 | 1.91E-05 | -1.41E+00 | 2.23E-03 | 8.54E-03 | -1.00E+00 | 3.27E-08 | 1.17E-07 | -1.67E+00 | 5.20E-04 | 1.47E-03 | -1.11E+00 | 3.62E-08 | 1.33E-07 | -1.40E+00 | 1.30E-04 | 3.58E-04 | -1.19E+00 |
| CCDC130 | 6.81E-06 | 3.55E-05 | -1.28E+00 | 3.09E-04 | 1.34E-03 | 1.02E+00 | 1.85E-09 | 7.85E-09 | -1.55E+00 | 8.27E-09 | 3.16E-08 | -1.38E+00 | 7.83E-03 | 2.79E-02 | -1.08E+00 | 7.16E-02 | 2.02E-01 | 1.11E+00 | 2.02E-04 | 7.41E-04 | -1.17E+00 | 3.58E-03 | 9.84E-03 | -1.08E+00 |
| EFNA4 | 6.86E-06 | 3.57E-05 | -1.76E+00 | 9.57E-04 | 4.14E-03 | -1.22E+00 | 7.11E-10 | 3.02E-09 | -2.13E+00 | 7.73E-06 | 2.95E-05 | -1.25E+00 | 3.14E-05 | 1.12E-04 | -2.02E+00 | 6.70E-04 | 1.89E-03 | -1.48E+00 | 2.39E-03 | 8.77E-03 | -1.72E+00 | 1.40E-01 | 3.84E-01 | -1.02E+00 |
| MORC3 | 6.87E-06 | 3.58E-05 | -1.08E+00 | 3.28E-07 | 1.42E-06 | -1.08E+00 | 7.79E-06 | 3.31E-05 | -1.24E+00 | 5.74E-05 | 2.19E-04 | -1.11E+00 | 3.85E-06 | 1.37E-05 | -1.19E+00 | 7.35E-08 | 2.07E-07 | -1.34E+00 | 1.73E-08 | 6.35E-08 | -1.19E+00 | 1.48E-10 | 4.07E-10 | -1.38E+00 |
| GALNS | 6.88E-06 | 3.59E-05 | -1.64E+00 | 1.54E-03 | 6.65E-03 | -1.11E+00 | 5.13E-06 | 2.18E-05 | -1.40E+00 | 8.35E-07 | 3.19E-06 | -1.37E+00 | 6.55E-03 | 2.33E-02 | -1.14E+00 | 1.08E-01 | 3.04E-01 | 1.25E+00 | 1.60E-03 | 5.88E-03 | -1.33E+00 | 6.47E-03 | 1.78E-02 | -1.16E+00 |
| TMEM41A | 6.89E-06 | 3.59E-05 | -1.29E+00 | 9.08E-06 | 3.93E-05 | -1.12E+00 | 2.46E-06 | 1.04E-05 | -1.38E+00 | 2.95E-05 | 1.13E-04 | -1.08E+00 | 2.37E-01 | 8.44E-01 | 1.24E+00 | 3.05E-01 | 8.61E-01 | 1.32E+00 | 4.30E-07 | 1.58E-06 | -1.21E+00 | 1.13E-04 | 3.11E-04 | -1.07E+00 |
| MGEA5 | 6.94E-06 | 3.62E-05 | -1.03E+00 | 3.79E-02 | 1.64E-01 | 1.30E+00 | 1.18E-03 | 5.01E-03 | -1.09E+00 | 4.83E-03 | 1.84E-02 | 1.08E+00 | 2.77E-03 | 9.88E-03 | 1.01E+00 | 9.60E-02 | 2.71E-01 | 1.13E+00 | 6.53E-04 | 2.40E-03 | -1.09E+00 | 9.16E-03 | 2.52E-02 | 1.02E+00 |
| FAXDC2 | 6.94E-06 | 3.62E-05 | -2.08E+00 | 8.58E-04 | 3.71E-03 | -1.30E+00 | 6.24E-02 | 2.65E-01 | -1.22E+00 | 1.57E-01 | 6.01E-01 | 1.19E+00 | 6.49E-02 | 2.31E-01 | -1.08E+00 | 3.09E-02 | 8.71E-02 | -1.81E+00 | 1.89E-03 | 6.94E-03 | 1.43E+00 | 7.81E-06 | 2.15E-05 | -1.08E+00 |
| ATL1 | 6.94E-06 | 3.62E-05 | -1.31E+00 | 2.56E-03 | 1.11E-02 | 1.13E+00 | 9.83E-04 | 4.18E-03 | -1.43E+00 | 4.62E-03 | 1.77E-02 | -1.18E+00 | 2.95E-06 | 1.05E-05 | -1.91E+00 | 1.32E-02 | 3.73E-02 | 1.00E+00 | 1.00E-03 | 3.68E-03 | -1.24E+00 | 1.41E-02 | 3.87E-02 | -1.14E+00 |
| LINC01251 | 6.96E-06 | 3.63E-05 | 3.18E+00 | 6.37E-02 | 2.75E-01 | 1.68E+00 | 6.45E-03 | 2.74E-02 | 1.92E+00 | 1.27E-01 | 4.84E-01 | 1.45E+00 | 3.78E-02 | 1.35E-01 | 2.48E+00 | 1.17E-01 | 3.30E-01 | 2.17E+00 | 1.50E-01 | 5.50E-01 | 1.34E+00 | 3.03E-01 | 8.33E-01 | 1.22E+00 |
| KIAA0753 | 6.96E-06 | 3.63E-05 | -1.66E+00 | 2.16E-03 | 9.35E-03 | -1.07E+00 | 4.13E-08 | 1.76E-07 | -1.84E+00 | 1.71E-04 | 6.54E-04 | -1.14E+00 | 5.22E-04 | 1.86E-03 | -1.43E+00 | 3.26E-03 | 9.19E-03 | -1.22E+00 | 2.46E-02 | 9.02E-02 | -1.16E+00 | 1.30E-01 | 3.58E-01 | -1.07E+00 |
| PCYOX1L | 6.98E-06 | 3.64E-05 | -1.62E+00 | 1.72E-03 | 7.42E-03 | -1.10E+00 | 4.76E-07 | 2.02E-06 | -1.42E+00 | 1.14E-04 | 4.37E-04 | -1.03E+00 | 2.79E-09 | 9.94E-09 | -2.09E+00 | 2.43E-06 | 6.86E-06 | -1.42E+00 | 9.91E-07 | 3.64E-06 | -1.36E+00 | 1.84E-03 | 5.05E-03 | -1.06E+00 |
| CAAP1 | 6.98E-06 | 3.64E-05 | -1.31E+00 | 5.85E-05 | 2.53E-04 | -1.04E+00 | 8.35E-08 | 3.55E-07 | -1.41E+00 | 1.26E-04 | 4.83E-04 | -1.00E+00 | 3.16E-03 | 1.13E-02 | -1.21E+00 | 3.15E-02 | 8.88E-02 | -1.05E+00 | 3.04E-07 | 1.12E-06 | -1.29E+00 | 2.85E-04 | 7.83E-04 | -1.02E+00 |
| C19orf24 | 6.98E-06 | 3.64E-05 | -1.19E+00 | 5.63E-05 | 2.43E-04 | -1.02E+00 | 5.83E-03 | 2.48E-02 | -1.01E+00 | 3.00E-03 | 1.15E-02 | 1.15E+00 | 1.36E-06 | 4.86E-06 | -1.41E+00 | 1.06E-03 | 2.99E-03 | -1.05E+00 | 9.81E-04 | 3.60E-03 | -1.13E+00 | 1.01E-01 | 2.78E-01 | 1.11E+00 |
| RNF6 | 7.04E-06 | 3.67E-05 | -1.19E+00 | 9.67E-09 | 4.18E-08 | -1.35E+00 | 1.51E-03 | 6.40E-03 | -1.12E+00 | 1.49E-06 | 5.70E-06 | -1.23E+00 | 4.63E-02 | 1.65E-01 | 1.17E+00 | 9.70E-08 | 2.74E-07 | -1.49E+00 | 7.76E-04 | 2.85E-03 | 1.05E+00 | 4.63E-03 | 1.27E-02 | 1.05E+00 |
| RPL21P10 | 7.05E-06 | 3.68E-05 | -1.13E+00 | 8.16E-13 | 3.53E-12 | -1.45E+00 | 2.30E-05 | 9.79E-05 | -1.14E+00 | 3.37E-15 | 1.29E-14 | -1.55E+00 | 3.88E-04 | 1.38E-03 | -1.23E+00 | 2.40E-06 | 6.77E-06 | -1.42E+00 | 2.88E-04 | 1.06E-03 | -1.10E+00 | 7.22E-08 | 1.98E-07 | -1.49E+00 |
| LYL1 | 7.06E-06 | 3.68E-05 | -1.10E+00 | 7.92E-03 | 3.42E-02 | 1.03E+00 | 3.43E-04 | 1.46E-03 | -1.20E+00 | 1.27E-01 | 4.87E-01 | -1.03E+00 | 5.42E-10 | 1.93E-09 | -1.45E+00 | 4.55E-03 | 1.28E-02 | -1.18E+00 | 3.48E-03 | 1.28E-02 | 1.06E+00 | 1.63E-02 | 4.47E-02 | -1.00E+00 |
| TBCK | 7.06E-06 | 3.68E-05 | -1.49E+00 | 6.28E-03 | 2.71E-02 | 1.08E+00 | 1.11E-10 | 4.73E-10 | -1.89E+00 | 3.99E-05 | 1.52E-04 | -1.10E+00 | 5.26E-05 | 1.88E-04 | -1.59E+00 | 1.26E-02 | 3.56E-02 | -1.14E+00 | 6.14E-11 | 2.25E-10 | -1.59E+00 | 1.87E-05 | 5.14E-05 | -1.13E+00 |
| ARL17A | 7.08E-06 | 3.69E-05 | -1.42E+00 | 1.69E-04 | 7.30E-04 | -1.02E+00 | 1.05E-04 | 4.46E-04 | -1.38E+00 | 1.61E-03 | 6.16E-03 | -1.07E+00 | 8.75E-05 | 3.12E-04 | -1.31E+00 | 7.79E-03 | 2.20E-02 | 1.05E+00 | 2.34E-05 | 8.57E-05 | -1.53E+00 | 1.14E-01 | 3.12E-01 | 1.13E+00 |
| FAHD1 | 7.08E-06 | 3.69E-05 | -1.29E+00 | 2.42E-02 | 1.05E-01 | 1.26E+00 | 2.61E-04 | 1.11E-03 | -1.24E+00 | 2.24E-01 | 8.57E-01 | 1.42E+00 | 1.27E-03 | 4.53E-03 | -1.21E+00 | 1.30E-01 | 3.66E-01 | 1.26E+00 | 8.16E-06 | 2.99E-05 | -1.27E+00 | 2.07E-01 | 5.69E-01 | 1.22E+00 |
| ELOVL6 | 7.09E-06 | 3.69E-05 | -1.48E+00 | 1.04E-04 | 4.50E-04 | -1.16E+00 | 3.16E-03 | 1.34E-02 | -1.27E+00 | 3.89E-03 | 1.49E-02 | 1.00E+00 | 4.29E-07 | 1.53E-06 | -1.63E+00 | 3.16E-04 | 8.92E-04 | -1.12E+00 | 4.71E-12 | 1.73E-11 | -1.57E+00 | 2.16E-03 | 5.93E-03 | -1.04E+00 |
| RPS6KB1 | 7.09E-06 | 3.69E-05 | -1.17E+00 | 2.69E-06 | 1.16E-05 | -1.08E+00 | 1.99E-06 | 8.45E-06 | -1.35E+00 | 8.10E-08 | 3.09E-07 | -1.26E+00 | 2.29E-02 | 8.18E-02 | 1.14E+00 | 3.54E-02 | 9.97E-02 | 1.16E+00 | 3.23E-06 | 1.18E-05 | -1.20E+00 | 2.49E-05 | 6.83E-05 | -1.19E+00 |
| ELAC2 | 7.10E-06 | 3.70E-05 | -1.32E+00 | 3.70E-03 | 1.60E-02 | 1.09E+00 | 1.02E-05 | 4.34E-05 | -1.22E+00 | 1.38E-03 | 5.29E-03 | 1.02E+00 | 1.56E-08 | 5.56E-08 | -1.93E+00 | 3.06E-02 | 8.63E-02 | 1.05E+00 | 1.26E-06 | 4.62E-06 | -1.19E+00 | 1.00E-01 | 2.76E-01 | 1.22E+00 |
| BRCA2 | 7.11E-06 | 3.71E-05 | -1.38E+00 | 1.31E-07 | 5.68E-07 | -1.56E+00 | 1.31E-01 | 5.55E-01 | 1.12E+00 | 2.67E-05 | 1.02E-04 | -1.16E+00 | 3.06E-02 | 1.09E-01 | -1.49E+00 | 1.92E-02 | 5.40E-02 | -1.80E+00 | 3.00E-03 | 1.10E-02 | 1.01E+00 | 1.61E-01 | 4.43E-01 | 1.18E+00 |
| MRPL19 | 7.11E-06 | 3.71E-05 | -1.15E+00 | 1.94E-04 | 8.40E-04 | 1.12E+00 | 2.54E-02 | 1.08E-01 | -1.05E+00 | 2.53E-01 | 9.65E-01 | 1.27E+00 | 2.00E-06 | 7.14E-06 | -1.18E+00 | 1.34E-05 | 3.79E-05 | -1.06E+00 | 3.21E-04 | 1.18E-03 | -1.08E+00 | 5.46E-02 | 1.50E-01 | 1.12E+00 |
| ZCCHC7 | 7.12E-06 | 3.71E-05 | -1.07E+00 | 2.32E-04 | 1.00E-03 | 1.16E+00 | 3.76E-07 | 1.60E-06 | -1.19E+00 | 1.98E-03 | 7.56E-03 | 1.12E+00 | 2.29E-05 | 8.15E-05 | -1.20E+00 | 9.89E-02 | 2.79E-01 | 1.24E+00 | 5.54E-06 | 2.03E-05 | -1.24E+00 | 1.71E-03 | 4.70E-03 | 1.01E+00 |
| KLHDC10 | 7.13E-06 | 3.72E-05 | -1.26E+00 | 5.58E-03 | 2.41E-02 | 1.11E+00 | 4.95E-03 | 2.10E-02 | -1.11E+00 | 2.47E-03 | 9.44E-03 | 1.04E+00 | 8.11E-05 | 2.89E-04 | -1.34E+00 | 1.24E-05 | 3.48E-05 | -1.52E+00 | 5.35E-03 | 1.97E-02 | -1.04E+00 | 2.37E-02 | 6.51E-02 | -1.03E+00 |
| RBMXP2 | 7.14E-06 | 3.72E-05 | -1.20E+00 | 1.36E-08 | 5.89E-08 | -1.34E+00 | 8.49E-07 | 3.61E-06 | -1.29E+00 | 2.50E-16 | 9.54E-16 | -1.79E+00 | 1.30E-02 | 4.65E-02 | -1.02E+00 | 4.32E-04 | 1.22E-03 | -1.26E+00 | 2.20E-07 | 8.08E-07 | -1.50E+00 | 6.53E-11 | 1.79E-10 | -1.68E+00 |
| SUSD1 | 7.16E-06 | 3.73E-05 | -1.64E+00 | 1.15E-04 | 4.95E-04 | -1.28E+00 | 9.20E-03 | 3.91E-02 | -1.12E+00 | 6.67E-03 | 2.55E-02 | -1.12E+00 | 3.51E-10 | 1.25E-09 | -1.87E+00 | 1.69E-04 | 4.75E-04 | -1.12E+00 | 1.90E-05 | 6.98E-05 | 1.12E+00 | 2.15E-02 | 5.91E-02 | 1.11E+00 |
| PRTFDC1 | 7.18E-06 | 3.74E-05 | -1.40E+00 | 1.41E-04 | 6.09E-04 | -1.08E+00 | 3.74E-02 | 1.59E-01 | -1.01E+00 | 3.41E-02 | 1.30E-01 | 1.10E+00 | 6.24E-04 | 2.23E-03 | -1.70E+00 | 2.59E-04 | 7.29E-04 | -1.95E+00 | 3.07E-03 | 1.13E-02 | -1.18E+00 | 1.49E-03 | 4.09E-03 | -1.24E+00 |
| LRTOMT | 7.19E-06 | 3.75E-05 | -1.76E+00 | 6.10E-02 | 2.64E-01 | 1.18E+00 | 1.63E-08 | 6.91E-08 | -2.31E+00 | 3.10E-03 | 1.18E-02 | 1.01E+00 | 8.56E-03 | 3.05E-02 | -1.32E+00 | 2.44E-01 | 6.87E-01 | 1.24E+00 | 3.29E-05 | 1.21E-04 | -1.62E+00 | 2.14E-01 | 5.89E-01 | 1.17E+00 |
| TMOD2 | 7.22E-06 | 3.76E-05 | -1.18E+00 | 2.45E-03 | 1.06E-02 | 1.17E+00 | 2.78E-03 | 1.18E-02 | -1.16E+00 | 2.16E-01 | 8.26E-01 | 1.43E+00 | 1.84E-02 | 6.57E-02 | -1.27E+00 | 7.13E-02 | 2.01E-01 | -1.13E+00 | 1.75E-01 | 6.41E-01 | -1.04E+00 | 2.19E-01 | 6.03E-01 | 1.69E+00 |
| WIZ | 7.22E-06 | 3.76E-05 | -1.45E+00 | 5.21E-02 | 2.25E-01 | 1.25E+00 | 5.18E-05 | 2.20E-04 | -1.43E+00 | 8.03E-03 | 3.07E-02 | -1.01E+00 | 9.75E-04 | 3.48E-03 | -1.30E+00 | 7.26E-03 | 2.05E-02 | -1.05E+00 | 1.07E-02 | 3.93E-02 | -1.13E+00 | 8.27E-02 | 2.27E-01 | 1.45E+00 |
| GPX4 | 7.23E-06 | 3.77E-05 | -1.06E+00 | 3.36E-04 | 1.45E-03 | 1.02E+00 | 1.97E-01 | 8.39E-01 | -1.01E+00 | 1.56E-01 | 5.97E-01 | 1.01E+00 | 2.81E-03 | 1.00E-02 | -1.14E+00 | 3.02E-01 | 8.50E-01 | 1.04E+00 | 2.02E-01 | 7.41E-01 | 1.04E+00 | 3.30E-01 | 9.06E-01 | 1.03E+00 |
| MAPK14 | 7.24E-06 | 3.78E-05 | -1.02E+00 | 2.64E-08 | 1.14E-07 | -1.02E+00 | 1.99E-04 | 8.45E-04 | -1.08E+00 | 2.01E-06 | 7.68E-06 | -1.10E+00 | 2.08E-01 | 7.43E-01 | 1.25E+00 | 2.32E-02 | 6.54E-02 | 1.04E+00 | 2.33E-01 | 8.56E-01 | 1.26E+00 | 6.13E-02 | 1.68E-01 | 1.02E+00 |
| PSMC4 | 7.26E-06 | 3.78E-05 | -1.06E+00 | 1.51E-04 | 6.53E-04 | 1.10E+00 | 1.25E-02 | 5.30E-02 | 1.04E+00 | 6.22E-02 | 2.38E-01 | 1.18E+00 | 1.70E-05 | 6.07E-05 | -1.19E+00 | 4.81E-03 | 1.36E-02 | -1.00E+00 | 3.08E-02 | 1.13E-01 | 1.02E+00 | 3.35E-01 | 9.21E-01 | 1.19E+00 |
| OSTC | 7.27E-06 | 3.79E-05 | -1.10E+00 | 2.97E-02 | 1.28E-01 | 1.13E+00 | 4.46E-02 | 1.89E-01 | -1.05E+00 | 2.12E-01 | 8.09E-01 | 1.11E+00 | 4.55E-08 | 1.62E-07 | -1.33E+00 | 3.55E-02 | 1.00E-01 | -1.02E+00 | 6.46E-04 | 2.37E-03 | -1.13E+00 | 3.49E-01 | 9.60E-01 | 1.08E+00 |
| TMEM242 | 7.27E-06 | 3.79E-05 | -1.08E+00 | 4.78E-04 | 2.07E-03 | 1.10E+00 | 1.30E-02 | 5.54E-02 | 1.02E+00 | 1.24E-02 | 4.74E-02 | 1.14E+00 | 1.31E-01 | 4.68E-01 | 1.22E+00 | 2.10E-01 | 5.93E-01 | 1.25E+00 | 2.33E-01 | 8.55E-01 | 1.22E+00 | 1.99E-01 | 5.47E-01 | 1.22E+00 |
| PIGL | 7.27E-06 | 3.79E-05 | -1.52E+00 | 1.22E-02 | 5.26E-02 | 1.19E+00 | 8.70E-08 | 3.69E-07 | -1.60E+00 | 1.34E-05 | 5.11E-05 | -1.16E+00 | 6.73E-04 | 2.40E-03 | -1.35E+00 | 2.14E-02 | 6.04E-02 | -1.00E+00 | 2.04E-04 | 7.48E-04 | -1.33E+00 | 8.36E-03 | 2.30E-02 | -1.05E+00 |
| ARFRP1 | 7.28E-06 | 3.79E-05 | -1.17E+00 | 4.54E-03 | 1.96E-02 | 1.17E+00 | 1.20E-04 | 5.09E-04 | -1.23E+00 | 3.50E-02 | 1.34E-01 | 1.15E+00 | 2.21E-06 | 7.87E-06 | -1.34E+00 | 1.10E-01 | 3.10E-01 | 1.23E+00 | 9.77E-04 | 3.59E-03 | 1.03E+00 | 2.02E-01 | 5.54E-01 | 1.25E+00 |
| GTF2H2C | 7.29E-06 | 3.80E-05 | -1.27E+00 | 2.37E-03 | 1.02E-02 | 1.09E+00 | 4.51E-06 | 1.91E-05 | -1.46E+00 | 1.07E-05 | 4.10E-05 | -1.30E+00 | 8.34E-04 | 2.97E-03 | -1.15E+00 | 2.02E-02 | 5.68E-02 | 1.07E+00 | 1.76E-07 | 6.45E-07 | -1.42E+00 | 1.64E-07 | 4.51E-07 | -1.50E+00 |
| ZNF43 | 7.29E-06 | 3.80E-05 | -1.06E+00 | 7.15E-05 | 3.09E-04 | 1.06E+00 | 2.01E-02 | 8.54E-02 | 1.03E+00 | 1.20E-01 | 4.58E-01 | 1.20E+00 | 1.74E-04 | 6.21E-04 | -1.17E+00 | 5.30E-03 | 1.50E-02 | -1.01E+00 | 8.26E-04 | 3.03E-03 | -1.09E+00 | 3.10E-03 | 8.50E-03 | -1.10E+00 |
| RAB34 | 7.31E-06 | 3.81E-05 | -1.02E+00 | 1.60E-05 | 6.90E-05 | 1.09E+00 | 2.36E-02 | 1.00E-01 | -1.07E+00 | 2.83E-02 | 1.08E-01 | 1.07E+00 | 9.89E-06 | 3.52E-05 | -1.13E+00 | 2.18E-06 | 6.14E-06 | -1.17E+00 | 1.30E-02 | 4.79E-02 | 1.08E+00 | 2.29E-01 | 6.30E-01 | 1.22E+00 |
| FBXL4 | 7.31E-06 | 3.81E-05 | -1.28E+00 | 3.15E-03 | 1.36E-02 | 1.09E+00 | 1.25E-05 | 5.32E-05 | -1.34E+00 | 1.11E-03 | 4.22E-03 | 1.03E+00 | 1.28E-04 | 4.56E-04 | -1.19E+00 | 3.86E-03 | 1.09E-02 | 1.02E+00 | 1.52E-03 | 5.59E-03 | -1.11E+00 | 1.67E-02 | 4.57E-02 | 1.02E+00 |
| CCDC191 | 7.34E-06 | 3.82E-05 | -1.70E+00 | 4.31E-05 | 1.86E-04 | -1.37E+00 | 4.83E-02 | 2.05E-01 | -1.11E+00 | 1.18E-02 | 4.50E-02 | -1.07E+00 | 2.23E-01 | 7.94E-01 | 1.33E+00 | 2.50E-01 | 7.06E-01 | 1.20E+00 | 2.37E-01 | 8.70E-01 | 1.08E+00 | 8.53E-02 | 2.34E-01 | -1.17E+00 |
| OSGEP | 7.36E-06 | 3.83E-05 | -1.06E+00 | 7.95E-06 | 3.43E-05 | 1.04E+00 | 9.83E-04 | 4.18E-03 | 1.00E+00 | 2.96E-04 | 1.13E-03 | 1.08E+00 | 1.40E-06 | 4.98E-06 | -1.26E+00 | 1.49E-04 | 4.20E-04 | -1.12E+00 | 4.08E-02 | 1.50E-01 | 1.05E+00 | 2.41E-02 | 6.63E-02 | -1.01E+00 |
| LINC00674 | 7.37E-06 | 3.84E-05 | -1.33E+00 | 1.47E-04 | 6.33E-04 | -1.14E+00 | 1.21E-10 | 5.15E-10 | -2.08E+00 | 2.48E-07 | 9.46E-07 | -1.38E+00 | 4.44E-06 | 1.58E-05 | -1.60E+00 | 6.72E-05 | 1.90E-04 | -1.36E+00 | 1.04E-02 | 3.81E-02 | -1.06E+00 | 2.04E-02 | 5.60E-02 | -1.00E+00 |
| AC016027.1 | 7.40E-06 | 3.85E-05 | -1.96E+00 | 1.86E-03 | 8.04E-03 | -1.19E+00 | 6.39E-07 | 2.71E-06 | -1.83E+00 | 2.67E-04 | 1.02E-03 | -1.22E+00 | 5.83E-04 | 2.08E-03 | -1.75E+00 | 2.90E-02 | 8.16E-02 | 1.02E+00 | 4.25E-03 | 1.56E-02 | -1.56E+00 | 8.24E-02 | 2.26E-01 | -1.10E+00 |
| UBA6-AS1 | 7.42E-06 | 3.87E-05 | -1.45E+00 | 6.03E-04 | 2.60E-03 | -1.04E+00 | 2.71E-02 | 1.15E-01 | -1.24E+00 | 1.06E-01 | 4.05E-01 | 1.08E+00 | 1.15E-01 | 4.09E-01 | 1.01E+00 | 3.07E-01 | 8.64E-01 | 1.20E+00 | 7.61E-07 | 2.79E-06 | -1.36E+00 | 4.81E-08 | 1.32E-07 | -1.63E+00 |
| AP2A1 | 7.44E-06 | 3.88E-05 | -1.31E+00 | 2.48E-05 | 1.07E-04 | -1.08E+00 | 4.37E-07 | 1.86E-06 | -1.32E+00 | 4.99E-10 | 1.91E-09 | -1.43E+00 | 1.48E-06 | 5.27E-06 | -1.28E+00 | 4.71E-07 | 1.33E-06 | -1.32E+00 | 2.65E-08 | 9.73E-08 | -1.41E+00 | 1.26E-04 | 3.47E-04 | -1.16E+00 |
| AC073130.3 | 7.44E-06 | 3.88E-05 | 2.70E+00 | 7.47E-02 | 3.23E-01 | 1.30E+00 | 5.40E-04 | 2.29E-03 | 2.28E+00 | 3.46E-02 | 1.32E-01 | 1.07E+00 | 2.35E-01 | 8.39E-01 | 1.46E+00 | 1.73E-02 | 4.88E-02 | -1.39E+00 | 6.80E-04 | 2.49E-03 | 2.40E+00 | 7.99E-02 | 2.20E-01 | -1.13E+00 |
| NEDD4 | 7.44E-06 | 3.88E-05 | -1.23E+00 | 1.05E-02 | 4.54E-02 | 1.22E+00 | 9.31E-05 | 3.96E-04 | -1.24E+00 | 1.25E-03 | 4.77E-03 | 1.02E+00 | 5.06E-02 | 1.80E-01 | 1.07E+00 | 4.37E-02 | 1.23E-01 | 1.01E+00 | 4.94E-03 | 1.81E-02 | -1.07E+00 | 1.79E-02 | 4.92E-02 | -1.01E+00 |
| WAC | 7.45E-06 | 3.88E-05 | -1.04E+00 | 6.15E-04 | 2.66E-03 | 1.10E+00 | 2.89E-03 | 1.23E-02 | -1.07E+00 | 4.79E-04 | 1.83E-03 | 1.03E+00 | 4.28E-08 | 1.53E-07 | -1.19E+00 | 1.57E-05 | 4.41E-05 | -1.03E+00 | 9.56E-08 | 3.51E-07 | -1.33E+00 | 5.92E-04 | 1.63E-03 | -1.17E+00 |
| HAAO | 7.45E-06 | 3.88E-05 | -1.67E+00 | 1.20E-04 | 5.18E-04 | -1.24E+00 | 1.97E-04 | 8.36E-04 | -1.30E+00 | 5.78E-04 | 2.21E-03 | -1.16E+00 | 2.28E-05 | 8.12E-05 | -1.59E+00 | 9.60E-04 | 2.71E-03 | -1.21E+00 | 3.75E-04 | 1.38E-03 | -1.58E+00 | 3.00E-03 | 8.24E-03 | -1.32E+00 |
| DAPK1-IT1 | 7.45E-06 | 3.89E-05 | -1.53E+00 | 1.35E-02 | 5.82E-02 | 1.13E+00 | 2.38E-02 | 1.01E-01 | -1.29E+00 | 1.48E-01 | 5.67E-01 | 1.17E+00 | 2.26E-02 | 8.05E-02 | -1.08E+00 | 3.48E-01 | 9.80E-01 | 1.45E+00 | 1.27E-02 | 4.64E-02 | -1.92E+00 | 2.20E-01 | 6.04E-01 | 1.44E+00 |
| NARS2 | 7.46E-06 | 3.89E-05 | -1.29E+00 | 7.83E-04 | 3.38E-03 | 1.03E+00 | 2.91E-05 | 1.24E-04 | -1.42E+00 | 8.27E-03 | 3.16E-02 | 1.04E+00 | 2.97E-04 | 1.06E-03 | -1.78E+00 | 7.12E-02 | 2.01E-01 | -1.01E+00 | 3.56E-10 | 1.31E-09 | -1.66E+00 | 8.43E-02 | 2.32E-01 | 1.14E+00 |
| MRPL41 | 7.46E-06 | 3.89E-05 | -1.11E+00 | 1.57E-06 | 6.79E-06 | -1.03E+00 | 8.47E-02 | 3.60E-01 | 1.03E+00 | 3.00E-03 | 1.15E-02 | 1.01E+00 | 9.73E-05 | 3.47E-04 | -1.13E+00 | 3.29E-04 | 9.28E-04 | -1.10E+00 | 7.84E-02 | 2.88E-01 | 1.05E+00 | 1.21E-01 | 3.34E-01 | 1.08E+00 |
| MIIP | 7.49E-06 | 3.90E-05 | -1.23E+00 | 8.49E-04 | 3.67E-03 | 1.07E+00 | 1.45E-03 | 6.16E-03 | -1.18E+00 | 5.72E-04 | 2.18E-03 | -1.01E+00 | 3.14E-02 | 1.12E-01 | 1.03E+00 | 6.19E-02 | 1.74E-01 | 1.18E+00 | 1.54E-05 | 5.67E-05 | -1.17E+00 | 5.26E-03 | 1.44E-02 | 1.00E+00 |
| TFB1M | 7.50E-06 | 3.91E-05 | -1.40E+00 | 3.94E-02 | 1.70E-01 | 1.24E+00 | 1.27E-08 | 5.39E-08 | -1.69E+00 | 1.54E-03 | 5.88E-03 | 1.02E+00 | 1.58E-05 | 5.62E-05 | -1.48E+00 | 2.03E-02 | 5.73E-02 | 1.01E+00 | 6.26E-11 | 2.30E-10 | -1.89E+00 | 1.37E-03 | 3.75E-03 | -1.17E+00 |
| AC068620.4 | 7.52E-06 | 3.92E-05 | -1.59E+00 | 4.18E-05 | 1.81E-04 | -1.36E+00 | 5.02E-03 | 2.13E-02 | -1.22E+00 | 1.42E-03 | 5.44E-03 | -1.30E+00 | 2.06E-01 | 7.36E-01 | 1.49E+00 | 1.09E-01 | 3.06E-01 | -1.03E+00 | 5.90E-02 | 2.17E-01 | -1.12E+00 | 9.28E-02 | 2.55E-01 | -1.14E+00 |
| TRMT2A | 7.56E-06 | 3.94E-05 | -1.26E+00 | 1.57E-05 | 6.79E-05 | -1.10E+00 | 8.60E-03 | 3.65E-02 | -1.03E+00 | 1.23E-02 | 4.70E-02 | 1.17E+00 | 1.59E-02 | 5.68E-02 | -1.18E+00 | 1.39E-01 | 3.90E-01 | 1.14E+00 | 5.11E-02 | 1.88E-01 | 1.14E+00 | 2.38E-01 | 6.55E-01 | 1.36E+00 |
| NUS1 | 7.58E-06 | 3.95E-05 | -1.43E+00 | 6.62E-09 | 2.86E-08 | -1.68E+00 | 3.98E-04 | 1.69E-03 | -1.21E+00 | 1.74E-05 | 6.63E-05 | -1.23E+00 | 1.27E-09 | 4.52E-09 | -2.02E+00 | 7.78E-09 | 2.19E-08 | -1.64E+00 | 1.53E-05 | 5.62E-05 | -1.38E+00 | 1.10E-03 | 3.03E-03 | -1.17E+00 |
| SLC27A3 | 7.58E-06 | 3.95E-05 | -1.51E+00 | 3.32E-04 | 1.44E-03 | -1.07E+00 | 1.12E-08 | 4.77E-08 | -1.56E+00 | 3.14E-06 | 1.20E-05 | -1.20E+00 | 2.95E-04 | 1.05E-03 | -1.53E+00 | 2.30E-02 | 6.48E-02 | -1.02E+00 | 1.06E-01 | 3.87E-01 | 1.15E+00 | 4.76E-02 | 1.31E-01 | -1.01E+00 |
| LYSMD1 | 7.60E-06 | 3.96E-05 | -2.07E+00 | 8.47E-04 | 3.66E-03 | -1.31E+00 | 1.02E-05 | 4.33E-05 | -2.30E+00 | 1.28E-03 | 4.87E-03 | -1.42E+00 | 6.70E-02 | 2.39E-01 | -1.46E+00 | 1.41E-01 | 3.98E-01 | -1.22E+00 | 8.51E-03 | 3.12E-02 | -1.31E+00 | 2.55E-02 | 7.00E-02 | -1.15E+00 |
| RAD1 | 7.62E-06 | 3.97E-05 | -1.46E+00 | 1.99E-04 | 8.58E-04 | -1.11E+00 | 1.90E-06 | 8.09E-06 | -1.39E+00 | 1.15E-03 | 4.40E-03 | 1.01E+00 | 2.83E-06 | 1.01E-05 | -1.48E+00 | 3.32E-04 | 9.37E-04 | -1.13E+00 | 5.07E-06 | 1.86E-05 | -1.25E+00 | 1.35E-01 | 3.70E-01 | 1.14E+00 |
| ZNF91 | 7.64E-06 | 3.98E-05 | -1.07E+00 | 1.73E-04 | 7.46E-04 | 1.11E+00 | 1.34E-06 | 5.69E-06 | -1.28E+00 | 6.56E-05 | 2.51E-04 | -1.05E+00 | 5.23E-04 | 1.86E-03 | -1.10E+00 | 2.28E-04 | 6.43E-04 | -1.15E+00 | 3.54E-02 | 1.30E-01 | 1.07E+00 | 1.85E-02 | 5.08E-02 | 1.07E+00 |
| PEX2 | 7.66E-06 | 3.99E-05 | -1.14E+00 | 2.69E-05 | 1.16E-04 | 1.07E+00 | 1.45E-03 | 6.16E-03 | -1.10E+00 | 5.67E-04 | 2.16E-03 | -1.02E+00 | 1.95E-03 | 6.95E-03 | -1.02E+00 | 9.52E-02 | 2.68E-01 | 1.21E+00 | 3.49E-06 | 1.28E-05 | -1.24E+00 | 5.18E-05 | 1.42E-04 | -1.17E+00 |
| MED15 | 7.68E-06 | 4.00E-05 | -1.28E+00 | 1.21E-03 | 5.22E-03 | 1.06E+00 | 4.36E-03 | 1.85E-02 | -1.05E+00 | 3.66E-03 | 1.40E-02 | 1.08E+00 | 1.66E-04 | 5.92E-04 | -1.31E+00 | 2.17E-01 | 6.12E-01 | 1.30E+00 | 6.89E-03 | 2.53E-02 | 1.01E+00 | 6.30E-02 | 1.73E-01 | 1.13E+00 |
| ROGDI | 7.68E-06 | 4.00E-05 | -1.42E+00 | 3.32E-03 | 1.43E-02 | 1.03E+00 | 1.94E-04 | 8.24E-04 | -1.34E+00 | 2.32E-05 | 8.87E-05 | -1.26E+00 | 6.01E-04 | 2.14E-03 | -1.57E+00 | 2.73E-02 | 7.69E-02 | 1.00E+00 | 2.49E-01 | 9.15E-01 | 1.41E+00 | 9.30E-02 | 2.55E-01 | 1.07E+00 |
| HINT2 | 7.69E-06 | 4.01E-05 | -1.01E+00 | 2.50E-08 | 1.08E-07 | -1.02E+00 | 4.90E-03 | 2.08E-02 | 1.05E+00 | 9.59E-10 | 3.66E-09 | -1.21E+00 | 5.00E-02 | 1.78E-01 | 1.13E+00 | 1.92E-03 | 5.42E-03 | -1.06E+00 | 4.31E-02 | 1.58E-01 | 1.14E+00 | 4.91E-04 | 1.35E-03 | -1.08E+00 |
| MMGT1 | 7.75E-06 | 4.04E-05 | -1.15E+00 | 2.17E-07 | 9.36E-07 | -1.20E+00 | 4.59E-06 | 1.95E-05 | -1.25E+00 | 1.58E-07 | 6.02E-07 | -1.22E+00 | 7.82E-02 | 2.79E-01 | 1.16E+00 | 6.44E-03 | 1.82E-02 | -1.06E+00 | 2.58E-03 | 9.45E-03 | -1.11E+00 | 6.44E-03 | 1.77E-02 | -1.03E+00 |
| ABHD13 | 7.77E-06 | 4.05E-05 | -1.33E+00 | 9.98E-06 | 4.31E-05 | -1.21E+00 | 1.61E-07 | 6.82E-07 | -1.60E+00 | 1.94E-06 | 7.41E-06 | -1.29E+00 | 3.29E-06 | 1.17E-05 | -1.78E+00 | 3.05E-05 | 8.59E-05 | -1.53E+00 | 1.19E-09 | 4.36E-09 | -1.60E+00 | 9.94E-09 | 2.73E-08 | -1.55E+00 |
| SMG9 | 7.77E-06 | 4.05E-05 | -1.42E+00 | 1.28E-03 | 5.52E-03 | -1.03E+00 | 1.22E-03 | 5.19E-03 | -1.20E+00 | 4.65E-03 | 1.77E-02 | 1.03E+00 | 8.86E-04 | 3.16E-03 | -1.49E+00 | 2.06E-01 | 5.82E-01 | 1.18E+00 | 4.69E-05 | 1.72E-04 | -1.22E+00 | 1.23E-01 | 3.38E-01 | 1.15E+00 |
| FAM224B | 7.78E-06 | 4.06E-05 | -1.17E+00 | 4.11E-04 | 1.78E-03 | 1.11E+00 | 1.11E-04 | 4.73E-04 | -1.49E+00 | 3.46E-05 | 1.32E-04 | -1.46E+00 | 1.53E-02 | 5.47E-02 | -1.53E+00 | 1.29E-02 | 3.64E-02 | -1.53E+00 | 1.25E-01 | 4.58E-01 | 1.02E+00 | 4.53E-02 | 1.24E-01 | -1.49E+00 |
| CYBRD1 | 7.79E-06 | 4.06E-05 | -1.13E+00 | 4.22E-03 | 1.82E-02 | 1.24E+00 | 6.21E-03 | 2.64E-02 | -1.10E+00 | 1.34E-01 | 5.10E-01 | 1.22E+00 | 2.35E-02 | 8.37E-02 | -1.05E+00 | 6.99E-02 | 1.97E-01 | 1.11E+00 | 3.27E-03 | 1.20E-02 | -1.17E+00 | 2.58E-02 | 7.10E-02 | -1.01E+00 |
| PSMD13 | 7.80E-06 | 4.06E-05 | 1.04E+00 | 6.38E-05 | 2.75E-04 | 1.18E+00 | 2.17E-01 | 9.23E-01 | 1.09E+00 | 2.62E-01 | 1.00E+00 | 1.25E+00 | 1.43E-04 | 5.08E-04 | -1.03E+00 | 7.13E-03 | 2.01E-02 | 1.10E+00 | 1.35E-01 | 4.97E-01 | 1.13E+00 | 2.67E-01 | 7.33E-01 | 1.24E+00 |
| GCH1 | 7.80E-06 | 4.07E-05 | -1.38E+00 | 1.08E-04 | 4.68E-04 | -1.15E+00 | 9.73E-16 | 4.14E-15 | -2.33E+00 | 1.74E-15 | 6.64E-15 | -1.79E+00 | 1.64E-02 | 5.86E-02 | -1.97E+00 | 9.41E-02 | 2.65E-01 | -1.23E+00 | 3.16E-03 | 1.16E-02 | -1.49E+00 | 2.86E-03 | 7.84E-03 | -1.69E+00 |
| SGF29 | 7.82E-06 | 4.08E-05 | -1.15E+00 | 1.82E-03 | 7.88E-03 | 1.11E+00 | 3.75E-06 | 1.59E-05 | -1.39E+00 | 1.42E-03 | 5.41E-03 | 1.01E+00 | 9.60E-08 | 3.42E-07 | -1.43E+00 | 2.78E-06 | 7.82E-06 | -1.31E+00 | 9.82E-08 | 3.60E-07 | -1.38E+00 | 2.92E-04 | 8.02E-04 | -1.18E+00 |
| STYX | 7.87E-06 | 4.10E-05 | -1.34E+00 | 1.19E-05 | 5.12E-05 | -1.17E+00 | 3.77E-06 | 1.60E-05 | -1.42E+00 | 2.88E-06 | 1.10E-05 | -1.25E+00 | 5.46E-04 | 1.95E-03 | -1.29E+00 | 1.85E-03 | 5.21E-03 | -1.22E+00 | 3.36E-06 | 1.23E-05 | -1.40E+00 | 1.23E-06 | 3.39E-06 | -1.40E+00 |
| REXO4 | 7.89E-06 | 4.11E-05 | -1.16E+00 | 6.47E-03 | 2.79E-02 | 1.19E+00 | 1.42E-04 | 6.02E-04 | -1.25E+00 | 3.26E-04 | 1.25E-03 | -1.04E+00 | 3.33E-02 | 1.19E-01 | 1.06E+00 | 7.81E-02 | 2.20E-01 | 1.16E+00 | 2.12E-03 | 7.77E-03 | -1.05E+00 | 4.20E-03 | 1.15E-02 | 1.00E+00 |
| AC087385.2 | 7.91E-06 | 4.12E-05 | -1.27E+00 | 1.83E-01 | 7.93E-01 | 1.50E+00 | 5.70E-04 | 2.42E-03 | -1.20E+00 | 1.38E-01 | 5.28E-01 | 1.53E+00 | 3.07E-04 | 1.09E-03 | -1.29E+00 | 2.63E-01 | 7.42E-01 | 1.46E+00 | 1.22E-03 | 4.48E-03 | -1.07E+00 | 2.34E-02 | 6.42E-02 | 1.76E+00 |
| ZNF512B | 7.96E-06 | 4.15E-05 | -1.42E+00 | 3.00E-04 | 1.30E-03 | -1.12E+00 | 1.55E-03 | 6.59E-03 | -1.28E+00 | 2.64E-03 | 1.01E-02 | -1.05E+00 | 2.62E-01 | 9.36E-01 | 1.41E+00 | 1.96E-01 | 5.54E-01 | 1.80E+00 | 1.61E-02 | 5.91E-02 | -1.10E+00 | 3.11E-02 | 8.54E-02 | 1.01E+00 |
| CENPN | 7.97E-06 | 4.15E-05 | -1.81E+00 | 2.03E-06 | 8.77E-06 | -1.96E+00 | 2.21E-02 | 9.38E-02 | -1.06E+00 | 6.43E-06 | 2.46E-05 | -1.35E+00 | 7.63E-05 | 2.72E-04 | -1.81E+00 | 3.96E-04 | 1.12E-03 | -1.54E+00 | 3.99E-04 | 1.46E-03 | -1.14E+00 | 4.48E-04 | 1.23E-03 | -1.10E+00 |
| RPL3P1 | 8.01E-06 | 4.18E-05 | -1.61E+00 | 1.13E-06 | 4.90E-06 | -1.85E+00 | 2.39E-04 | 1.02E-03 | -1.66E+00 | 4.21E-03 | 1.61E-02 | -1.21E+00 | 1.61E-02 | 5.72E-02 | -1.31E+00 | 9.90E-03 | 2.79E-02 | -1.38E+00 | 6.58E-04 | 2.42E-03 | -1.40E+00 | 1.60E-04 | 4.40E-04 | -1.74E+00 |
| EIF4A2 | 8.02E-06 | 4.18E-05 | -1.10E+00 | 1.98E-03 | 8.57E-03 | 1.05E+00 | 2.01E-03 | 8.56E-03 | -1.13E+00 | 2.05E-01 | 7.84E-01 | 1.06E+00 | 1.06E-06 | 3.76E-06 | -1.21E+00 | 1.68E-01 | 4.73E-01 | 1.08E+00 | 1.88E-03 | 6.91E-03 | -1.15E+00 | 1.01E-01 | 2.78E-01 | -1.05E+00 |
| PIK3C2B | 8.02E-06 | 4.18E-05 | -1.49E+00 | 2.91E-03 | 1.26E-02 | 1.01E+00 | 1.20E-02 | 5.11E-02 | -1.24E+00 | 6.27E-02 | 2.40E-01 | 1.07E+00 | 6.20E-03 | 2.21E-02 | -2.03E+00 | 1.18E-02 | 3.33E-02 | -1.53E+00 | 1.59E-01 | 5.82E-01 | -1.09E+00 | 1.35E-01 | 3.70E-01 | -1.29E+00 |
| C16orf45 | 8.03E-06 | 4.19E-05 | -1.30E+00 | 3.19E-04 | 1.38E-03 | 1.00E+00 | 8.43E-03 | 3.58E-02 | -1.22E+00 | 8.88E-04 | 3.39E-03 | -1.37E+00 | 1.12E-01 | 3.99E-01 | -1.03E+00 | 2.45E-01 | 6.90E-01 | 1.17E+00 | 1.69E-06 | 6.22E-06 | -1.53E+00 | 2.91E-05 | 8.00E-05 | -1.29E+00 |
| KAT7 | 8.03E-06 | 4.19E-05 | -1.12E+00 | 1.17E-07 | 5.04E-07 | -1.09E+00 | 2.58E-04 | 1.10E-03 | -1.22E+00 | 2.64E-06 | 1.01E-05 | -1.21E+00 | 1.95E-05 | 6.95E-05 | -1.25E+00 | 3.37E-07 | 9.49E-07 | -1.35E+00 | 1.39E-07 | 5.11E-07 | -1.37E+00 | 8.28E-09 | 2.27E-08 | -1.47E+00 |
| SNHG20 | 8.10E-06 | 4.22E-05 | -1.49E+00 | 3.75E-02 | 1.62E-01 | 1.21E+00 | 8.60E-03 | 3.65E-02 | -1.28E+00 | 2.19E-01 | 8.37E-01 | 1.28E+00 | 3.31E-03 | 1.18E-02 | -1.28E+00 | 3.05E-01 | 8.59E-01 | 1.50E+00 | 5.04E-02 | 1.85E-01 | 1.12E+00 | 6.96E-03 | 1.91E-02 | 1.91E+00 |
| TNRC6A | 8.10E-06 | 4.22E-05 | -1.15E+00 | 1.32E-07 | 5.72E-07 | -1.19E+00 | 3.65E-05 | 1.55E-04 | -1.28E+00 | 6.63E-05 | 2.53E-04 | -1.10E+00 | 5.43E-04 | 1.94E-03 | -1.04E+00 | 5.62E-04 | 1.58E-03 | -1.05E+00 | 1.27E-05 | 4.66E-05 | -1.13E+00 | 1.43E-08 | 3.92E-08 | -1.46E+00 |
| ALG10 | 8.11E-06 | 4.23E-05 | -1.43E+00 | 1.91E-06 | 8.25E-06 | -1.39E+00 | 9.16E-03 | 3.89E-02 | -1.20E+00 | 5.63E-04 | 2.15E-03 | -1.35E+00 | 2.04E-01 | 7.28E-01 | 1.19E+00 | 2.84E-01 | 8.01E-01 | 1.20E+00 | 3.26E-02 | 1.20E-01 | -1.03E+00 | 5.82E-02 | 1.60E-01 | 1.01E+00 |
| LENG8 | 8.14E-06 | 4.24E-05 | -1.17E+00 | 1.04E-02 | 4.48E-02 | 1.78E+00 | 7.58E-03 | 3.22E-02 | -1.02E+00 | 1.19E-01 | 4.56E-01 | 1.66E+00 | 7.17E-05 | 2.56E-04 | -1.14E+00 | 1.66E-01 | 4.68E-01 | 1.29E+00 | 8.97E-03 | 3.29E-02 | 1.02E+00 | 1.09E-01 | 2.98E-01 | 1.49E+00 |
| MEGF8 | 8.16E-06 | 4.25E-05 | -1.52E+00 | 4.93E-03 | 2.13E-02 | 1.10E+00 | 2.46E-02 | 1.05E-01 | -1.02E+00 | 6.85E-02 | 2.62E-01 | 1.16E+00 | 2.06E-03 | 7.33E-03 | -1.31E+00 | 5.97E-02 | 1.68E-01 | 1.01E+00 | 8.34E-05 | 3.06E-04 | -1.43E+00 | 2.47E-02 | 6.80E-02 | -1.11E+00 |
| LINC00982 | 8.18E-06 | 4.26E-05 | -1.39E+00 | 3.21E-05 | 1.39E-04 | -1.19E+00 | 1.89E-02 | 8.02E-02 | -1.27E+00 | 1.43E-01 | 5.46E-01 | 1.12E+00 | 3.29E-02 | 1.17E-01 | -1.08E+00 | 2.89E-02 | 8.14E-02 | -1.18E+00 | 1.96E-01 | 7.18E-01 | 1.39E+00 | 1.39E-01 | 3.82E-01 | 1.84E+00 |
| BUD31 | 8.18E-06 | 4.26E-05 | -1.34E+00 | 3.66E-07 | 1.58E-06 | -1.40E+00 | 5.71E-09 | 2.43E-08 | -1.54E+00 | 1.66E-12 | 6.33E-12 | -1.67E+00 | 9.95E-09 | 3.55E-08 | -1.51E+00 | 1.25E-08 | 3.52E-08 | -1.60E+00 | 2.72E-07 | 9.97E-07 | -1.32E+00 | 2.74E-09 | 7.52E-09 | -1.47E+00 |
| CCNG2 | 8.20E-06 | 4.27E-05 | -1.20E+00 | 2.95E-09 | 1.27E-08 | -1.36E+00 | 3.20E-10 | 1.36E-09 | -1.69E+00 | 1.39E-11 | 5.33E-11 | -1.66E+00 | 1.69E-07 | 6.03E-07 | -1.52E+00 | 1.31E-09 | 3.70E-09 | -1.82E+00 | 1.23E-04 | 4.52E-04 | -1.70E+00 | 2.17E-05 | 5.96E-05 | -1.87E+00 |
| PSME2 | 8.23E-06 | 4.29E-05 | -1.06E+00 | 2.43E-03 | 1.05E-02 | 1.23E+00 | 1.09E-01 | 4.61E-01 | 1.19E+00 | 3.45E-02 | 1.32E-01 | 1.45E+00 | 3.92E-03 | 1.40E-02 | 1.04E+00 | 3.01E-01 | 8.50E-01 | 1.56E+00 | 4.62E-07 | 1.70E-06 | -1.25E+00 | 2.11E-01 | 5.80E-01 | 1.29E+00 |

| | | | | | | | | | | | | | | | | | | | | | | | | |
|---|---|---|---|---|---|---|---|---|---|---|---|---|---|---|---|---|---|---|---|---|---|---|---|---|
| NDEL1 | 8.23E-06 | 4.29E-05 | -1.27E+00 | 2.02E-06 | 8.71E-06 | -1.26E+00 | 1.36E-05 | 5.79E-05 | -1.38E+00 | 1.44E-04 | 5.49E-04 | -1.10E+00 | 3.32E-03 | 1.18E-02 | -1.06E+00 | 8.06E-02 | 2.27E-01 | 1.09E+00 | 2.77E-08 | 1.02E-07 | -1.22E+00 | 1.49E-08 | 4.08E-08 | -1.32E+00 |
| CDCA4 | 8.23E-06 | 4.29E-05 | -1.80E+00 | 6.73E-04 | 2.91E-03 | -1.20E+00 | 1.50E-07 | 6.37E-07 | -1.46E+00 | 4.87E-04 | 1.86E-03 | -1.03E+00 | 8.17E-11 | 2.91E-10 | -3.13E+00 | 3.77E-08 | 1.06E-07 | -1.89E+00 | 2.54E-18 | 9.33E-18 | -1.79E+00 | 7.38E-06 | 2.03E-05 | -1.14E+00 |
| COCH | 8.24E-06 | 4.30E-05 | -1.31E+00 | 9.80E-04 | 4.23E-03 | 1.01E+00 | 2.18E-01 | 9.25E-01 | 1.11E+00 | 2.20E-01 | 8.40E-01 | 1.33E+00 | 2.50E-04 | 8.91E-04 | -1.37E+00 | 9.73E-03 | 2.74E-02 | 1.07E+00 | 1.03E-02 | 3.76E-02 | 1.29E+00 | 9.61E-06 | 2.64E-05 | 1.68E+00 |
| C1RL | 8.24E-06 | 4.30E-05 | -1.18E+00 | 2.28E-02 | 9.87E-02 | 1.22E+00 | 2.06E-07 | 8.74E-07 | -1.47E+00 | 2.99E-04 | 1.14E-03 | -1.03E+00 | 4.97E-05 | 1.77E-04 | -1.30E+00 | 1.25E-02 | 3.51E-02 | 1.14E+00 | 8.20E-02 | 3.01E-01 | 1.05E+00 | 1.60E-01 | 4.38E-01 | 1.00E+00 |
| COMMD1 | 8.27E-06 | 4.31E-05 | -1.12E+00 | 1.00E-04 | 4.32E-04 | 1.06E+00 | 2.12E-04 | 8.99E-04 | -1.06E+00 | 1.85E-05 | 7.07E-05 | 1.01E+00 | 5.37E-05 | 1.92E-04 | -1.16E+00 | 1.99E-02 | 5.62E-02 | 1.09E+00 | 1.77E-02 | 6.50E-02 | 1.14E+00 | 5.91E-02 | 1.62E-01 | 1.19E+00 |
| GPANK1 | 8.29E-06 | 4.32E-05 | -1.08E+00 | 9.62E-08 | 4.15E-07 | -1.06E+00 | 2.03E-04 | 8.62E-04 | -1.17E+00 | 1.91E-07 | 7.31E-07 | -1.21E+00 | 2.18E-03 | 7.77E-03 | -1.10E+00 | 2.18E-02 | 6.15E-02 | -1.02E+00 | 1.10E-07 | 4.03E-07 | -1.33E+00 | 1.95E-05 | 5.36E-05 | -1.19E+00 |
| CCND2 | 8.31E-06 | 4.33E-05 | -1.09E+00 | 1.45E-13 | 6.25E-13 | -1.18E+00 | 6.42E-03 | 2.73E-02 | -1.13E+00 | 3.50E-02 | 1.34E-01 | 1.00E+00 | 1.81E-05 | 6.44E-05 | -1.17E+00 | 4.83E-05 | 1.36E-04 | -1.08E+00 | 6.82E-06 | 2.50E-05 | -1.24E+00 | 2.97E-03 | 8.15E-03 | -1.10E+00 |
| ZNF92 | 8.36E-06 | 4.36E-05 | -1.04E+00 | 1.36E-13 | 5.87E-13 | -1.34E+00 | 6.57E-04 | 2.79E-03 | -1.16E+00 | 2.18E-08 | 8.34E-08 | -1.34E+00 | 5.84E-03 | 2.08E-02 | 1.03E+00 | 4.37E-06 | 1.23E-05 | -1.26E+00 | 8.21E-06 | 3.01E-05 | -1.16E+00 | 8.11E-11 | 2.23E-10 | -1.53E+00 |
| UBE2Q2 | 8.41E-06 | 4.38E-05 | -1.13E+00 | 5.13E-10 | 2.21E-09 | -1.32E+00 | 1.99E-04 | 8.44E-04 | -1.11E+00 | 3.36E-04 | 1.28E-03 | 1.06E+00 | 3.25E-04 | 1.16E-03 | -1.07E+00 | 4.22E-07 | 1.19E-06 | -1.32E+00 | 3.75E-05 | 1.38E-04 | -1.10E+00 | 3.12E-06 | 8.58E-06 | -1.25E+00 |
| AC007686.3 | 8.44E-06 | 4.40E-05 | -1.46E+00 | 1.57E-05 | 6.78E-05 | -1.26E+00 | 3.80E-03 | 1.61E-02 | -1.72E+00 | 3.85E-02 | 1.47E-01 | -1.20E+00 | 8.21E-04 | 2.93E-03 | -1.80E+00 | 4.02E-02 | 1.13E-01 | -1.14E+00 | 6.43E-02 | 2.36E-01 | -1.27E+00 | 2.58E-01 | 7.08E-01 | 1.09E+00 |
| IER3 | 8.48E-06 | 4.42E-05 | 2.90E+00 | 8.16E-04 | 3.52E-03 | -2.37E+00 | 1.43E-03 | 6.07E-03 | 2.92E+00 | 7.24E-02 | 2.77E-01 | -1.36E+00 | 1.75E-01 | 6.26E-01 | 1.96E+00 | 3.40E-01 | 9.59E-01 | 1.04E+00 | 6.95E-02 | 2.55E-01 | 1.49E+00 | 2.07E-01 | 5.68E-01 | -1.53E+00 |
| AL592148.3 | 8.48E-06 | 4.42E-05 | -1.89E+00 | 1.86E-04 | 8.03E-04 | -1.50E+00 | 1.29E-01 | 5.48E-01 | -1.05E+00 | 2.14E-01 | 8.17E-01 | 1.43E+00 | 9.89E-04 | 3.53E-03 | -1.71E+00 | 1.79E-03 | 5.03E-03 | -1.60E+00 | 3.27E-02 | 1.20E-01 | -1.11E+00 | 7.25E-03 | 1.99E-02 | -1.21E+00 |
| PPARGC1B | 8.51E-06 | 4.43E-05 | -1.72E+00 | 1.93E-03 | 8.35E-03 | -1.05E+00 | 1.33E-12 | 5.64E-12 | -1.82E+00 | 8.21E-06 | 3.14E-05 | -1.05E+00 | 8.32E-02 | 2.97E-01 | -1.16E+00 | 3.20E-01 | 9.03E-01 | 1.43E+00 | 1.79E-08 | 6.59E-08 | -2.55E+00 | 1.27E-03 | 3.49E-03 | -1.23E+00 |
| ERI2 | 8.52E-06 | 4.44E-05 | -1.87E+00 | 7.21E-04 | 3.12E-03 | -1.24E+00 | 2.53E-03 | 1.08E-02 | -1.24E+00 | 2.49E-02 | 9.52E-02 | 1.10E+00 | 6.96E-08 | 2.48E-07 | -1.94E+00 | 8.54E-06 | 2.41E-05 | -1.37E+00 | 3.16E-03 | 1.16E-02 | -1.12E+00 | 1.38E-01 | 3.78E-01 | 1.15E+00 |
| MESTIT1 | 8.54E-06 | 4.45E-05 | -1.77E+00 | 2.00E-02 | 8.63E-02 | 1.08E+00 | 1.74E-03 | 7.38E-03 | -1.55E+00 | 2.57E-01 | 9.83E-01 | 1.24E+00 | 1.30E-02 | 4.65E-02 | -1.34E+00 | 3.46E-01 | 9.74E-01 | 1.40E+00 | 2.49E-01 | 9.15E-01 | 1.27E+00 | 4.46E-02 | 1.23E-01 | 1.52E+00 |
| MT-ND6 | 8.56E-06 | 4.46E-05 | -1.13E+00 | 2.20E-12 | 9.49E-12 | -1.20E+00 | 7.47E-02 | 3.18E-01 | -1.03E+00 | 1.80E-04 | 6.86E-04 | -1.14E+00 | 3.62E-03 | 1.29E-02 | -1.07E+00 | 1.06E-03 | 2.98E-03 | -1.07E+00 | 2.37E-01 | 8.71E-01 | 1.05E+00 | 3.23E-01 | 8.88E-01 | 1.03E+00 |
| CEP44 | 8.58E-06 | 4.47E-05 | -1.16E+00 | 2.42E-03 | 1.04E-02 | 1.16E+00 | 1.11E-08 | 4.70E-08 | -1.61E+00 | 7.46E-05 | 2.85E-04 | -1.07E+00 | 8.05E-04 | 2.87E-03 | -1.20E+00 | 1.01E-02 | 2.84E-02 | 1.02E+00 | 1.52E-05 | 5.59E-05 | -1.25E+00 | 3.10E-04 | 8.51E-04 | -1.18E+00 |
| NFKBID | 8.58E-06 | 4.47E-05 | 2.24E+00 | 2.52E-05 | 1.09E-04 | -1.29E+00 | 5.87E-02 | 2.49E-01 | 1.56E+00 | 1.89E-06 | 7.22E-06 | -1.44E+00 | 2.76E-01 | 9.85E-01 | 1.57E+00 | 1.19E-01 | 3.37E-01 | -1.00E+00 | 1.35E-01 | 4.94E-01 | 1.03E+00 | 1.22E-02 | 3.35E-02 | -1.61E+00 |
| CDC23 | 8.65E-06 | 4.51E-05 | -1.56E+00 | 2.70E-05 | 1.17E-04 | -1.34E+00 | 6.71E-04 | 2.85E-03 | -1.34E+00 | 2.20E-03 | 8.42E-03 | -1.14E+00 | 1.34E-03 | 4.78E-03 | -1.32E+00 | 4.25E-04 | 1.20E-03 | -1.30E+00 | 1.23E-08 | 4.50E-08 | -1.60E+00 | 9.05E-04 | 2.49E-03 | -1.19E+00 |
| AC093462.1 | 8.65E-06 | 4.51E-05 | 2.43E+00 | 2.05E-01 | 8.84E-01 | 1.38E+00 | 6.21E-03 | 2.64E-02 | 1.94E+00 | 1.37E-01 | 5.22E-01 | 1.07E+00 | 1.41E-01 | 5.03E-01 | 1.74E+00 | 3.11E-01 | 8.77E-01 | 1.27E+00 | 4.38E-02 | 1.61E-01 | 2.19E+00 | 3.99E-02 | 1.09E-01 | 2.25E+00 |
| DYRK1B | 8.67E-06 | 4.52E-05 | -1.49E+00 | 4.74E-04 | 2.05E-03 | -1.12E+00 | 1.10E-06 | 4.67E-06 | -1.54E+00 | 1.66E-06 | 6.33E-06 | -1.38E+00 | 3.46E-06 | 1.24E-05 | -1.39E+00 | 8.41E-05 | 2.37E-04 | -1.22E+00 | 9.83E-02 | 3.61E-01 | -1.03E+00 | 1.34E-01 | 3.67E-01 | -1.12E+00 |
| AC026471.1 | 8.72E-06 | 4.55E-05 | -1.21E+00 | 7.87E-04 | 3.40E-03 | 1.12E+00 | 7.96E-05 | 3.38E-04 | -1.29E+00 | 6.12E-03 | 2.34E-02 | 1.07E+00 | 1.34E-02 | 4.77E-02 | -1.08E+00 | 1.64E-01 | 4.62E-01 | 1.11E+00 | 1.07E-02 | 3.94E-02 | -1.22E+00 | 1.14E-01 | 3.12E-01 | 1.01E+00 |
| ZNF804A | 8.73E-06 | 4.55E-05 | -1.40E+00 | 4.03E-03 | 1.74E-02 | 1.09E+00 | 6.16E-09 | 2.62E-08 | -1.84E+00 | 2.64E-06 | 1.01E-05 | -1.33E+00 | 2.77E-11 | 9.87E-11 | -1.79E+00 | 7.39E-08 | 2.08E-07 | -1.35E+00 | 1.04E-23 | 3.82E-23 | -2.92E+00 | 1.99E-07 | 5.46E-07 | -1.35E+00 |
| ALAD | 8.75E-06 | 4.56E-05 | -1.47E+00 | 3.92E-04 | 1.69E-03 | -1.08E+00 | 4.79E-06 | 2.03E-05 | -1.52E+00 | 1.98E-04 | 7.55E-04 | -1.14E+00 | 2.34E-01 | 8.33E-01 | 1.52E+00 | 2.29E-05 | 6.46E-05 | 2.56E+00 | 1.04E-01 | 3.81E-01 | 1.11E+00 | 1.85E-01 | 5.09E-01 | 1.37E+00 |
| WDR92 | 8.78E-06 | 4.58E-05 | -1.60E+00 | 1.14E-01 | 4.94E-01 | 1.39E+00 | 5.83E-07 | 2.48E-06 | -1.97E+00 | 8.88E-03 | 3.39E-02 | -1.03E+00 | 5.64E-07 | 2.01E-06 | -1.44E+00 | 2.98E-03 | 8.40E-03 | 1.00E+00 | 3.19E-08 | 1.17E-07 | -1.38E+00 | 2.72E-02 | 7.48E-02 | 1.12E+00 |
| ATXN8OS | 8.78E-06 | 4.58E-05 | -1.16E+00 | 1.87E-08 | 8.08E-08 | -1.17E+00 | 8.22E-06 | 3.49E-05 | -1.46E+00 | 6.56E-10 | 2.51E-09 | -1.59E+00 | 1.38E-06 | 4.92E-06 | -1.47E+00 | 1.32E-09 | 3.72E-09 | -1.75E+00 | 8.60E-08 | 3.15E-07 | -1.48E+00 | 5.63E-08 | 1.55E-07 | -1.52E+00 |
| ZNF236 | 8.80E-06 | 4.59E-05 | -1.38E+00 | 5.01E-04 | 2.17E-03 | -1.07E+00 | 9.66E-03 | 4.10E-02 | -1.14E+00 | 3.83E-02 | 1.46E-01 | 1.11E+00 | 2.89E-02 | 1.03E-01 | -1.11E+00 | 7.02E-02 | 1.98E-01 | 1.07E+00 | 1.14E-04 | 4.20E-04 | -1.26E+00 | 3.23E-03 | 8.88E-03 | -1.12E+00 |
| FBXO48 | 8.80E-06 | 4.59E-05 | -2.12E+00 | 1.16E-03 | 5.02E-03 | -1.25E+00 | 2.29E-01 | 9.74E-01 | 1.23E+00 | 9.15E-02 | 3.50E-01 | -1.11E+00 | 9.99E-02 | 3.56E-01 | -1.16E+00 | 6.59E-02 | 1.86E-01 | -1.26E+00 | 5.03E-02 | 1.85E-01 | 1.05E+00 | 9.94E-03 | 2.73E-02 | -1.33E+00 |
| SPDYE5 | 8.85E-06 | 4.61E-05 | -1.59E+00 | 3.48E-07 | 1.50E-06 | -1.90E+00 | 4.75E-06 | 2.02E-05 | -1.74E+00 | 2.91E-05 | 1.11E-04 | -1.49E+00 | 8.53E-04 | 3.04E-03 | -2.01E+00 | 4.91E-03 | 1.38E-02 | -1.51E+00 | 2.08E-05 | 7.64E-05 | -1.79E+00 | 8.31E-04 | 2.28E-03 | -1.44E+00 |
| TNIK | 8.88E-06 | 4.63E-05 | -1.36E+00 | 6.98E-06 | 3.02E-05 | -1.24E+00 | 2.92E-03 | 1.24E-02 | -1.29E+00 | 1.89E-03 | 7.23E-03 | -1.16E+00 | 1.21E-01 | 4.31E-01 | 1.30E+00 | 1.54E-02 | 4.33E-02 | 1.11E+00 | 8.32E-04 | 3.06E-03 | 1.05E+00 | 4.48E-07 | 1.23E-06 | -1.16E+00 |
| GTF3C6 | 8.88E-06 | 4.63E-05 | -1.03E+00 | 1.20E-08 | 5.19E-08 | 1.02E+00 | 6.42E-02 | 2.73E-01 | 1.04E+00 | 1.03E-02 | 3.92E-02 | 1.08E+00 | 2.46E-08 | 8.78E-08 | -1.32E+00 | 2.09E-02 | 5.88E-02 | 1.03E+00 | 1.68E-02 | 6.16E-02 | -1.01E+00 | 2.19E-01 | 6.02E-01 | 1.05E+00 |
| DYRK2 | 8.88E-06 | 4.63E-05 | -1.23E+00 | 9.14E-07 | 3.95E-06 | -1.19E+00 | 2.25E-06 | 9.55E-06 | -1.36E+00 | 1.97E-08 | 7.53E-08 | -1.41E+00 | 1.42E-01 | 5.06E-01 | 1.14E+00 | 2.33E-01 | 6.57E-01 | 1.16E+00 | 1.51E-05 | 5.56E-05 | -1.33E+00 | 3.70E-08 | 1.02E-07 | -1.55E+00 |
| AC005838.2 | 8.90E-06 | 4.64E-05 | -1.41E+00 | 1.84E-03 | 7.97E-03 | 1.03E+00 | 6.66E-03 | 2.83E-02 | -1.30E+00 | 1.70E-02 | 6.51E-02 | -1.09E+00 | 4.76E-03 | 1.70E-02 | -1.05E+00 | 1.40E-02 | 3.96E-02 | -1.05E+00 | 4.51E-06 | 1.65E-05 | -1.20E+00 | 2.88E-06 | 7.92E-06 | -1.23E+00 |
| RCOR2 | 8.92E-06 | 4.65E-05 | -1.85E+00 | 5.73E-05 | 2.47E-04 | -1.63E+00 | 1.59E-02 | 6.74E-02 | -1.38E+00 | 5.91E-02 | 2.26E-01 | -1.11E+00 | 1.37E-01 | 4.88E-01 | -1.18E+00 | 1.03E-01 | 2.90E-01 | -1.35E+00 | 9.32E-05 | 3.42E-04 | -1.73E+00 | 1.59E-06 | 4.38E-06 | -2.48E+00 |
| PRR3 | 8.92E-06 | 4.65E-05 | -1.42E+00 | 2.50E-03 | 1.08E-02 | -1.02E+00 | 5.65E-03 | 2.40E-02 | -1.10E+00 | 5.88E-02 | 2.25E-01 | 1.28E+00 | 1.23E-07 | 4.40E-07 | -1.56E+00 | 1.08E-04 | 3.06E-04 | -1.14E+00 | 2.61E-06 | 9.57E-06 | -1.58E+00 | 2.60E-02 | 7.14E-02 | -1.04E+00 |
| C1orf52 | 8.95E-06 | 4.66E-05 | -1.08E+00 | 6.36E-09 | 2.75E-08 | -1.11E+00 | 9.43E-05 | 4.00E-04 | -1.20E+00 | 4.63E-06 | 1.77E-05 | -1.19E+00 | 5.83E-04 | 2.08E-03 | -1.07E+00 | 3.25E-04 | 9.15E-04 | -1.12E+00 | 1.53E-07 | 5.62E-07 | -1.19E+00 | 1.15E-09 | 3.17E-09 | -1.31E+00 |
| BCLAF3 | 8.95E-06 | 4.66E-05 | -1.56E+00 | 1.03E-02 | 4.45E-02 | 1.07E+00 | 7.31E-05 | 3.11E-04 | -1.97E+00 | 8.54E-03 | 3.26E-02 | -1.15E+00 | 3.20E-05 | 1.14E-04 | -1.58E+00 | 2.74E-03 | 7.73E-03 | -1.26E+00 | 3.86E-03 | 1.42E-02 | -1.23E+00 | 3.25E-03 | 8.92E-03 | -1.22E+00 |
| MORF4L2 | 9.03E-06 | 4.71E-05 | -1.01E+00 | 4.77E-03 | 2.06E-02 | 1.25E+00 | 3.17E-03 | 1.35E-02 | 1.00E+00 | 8.91E-02 | 3.41E-01 | 1.29E+00 | 9.80E-05 | 3.49E-04 | -1.08E+00 | 2.39E-01 | 6.73E-01 | 1.27E+00 | 3.84E-05 | 1.41E-04 | -1.16E+00 | 2.61E-01 | 7.16E-01 | 1.17E+00 |
| HILPDA | 9.04E-06 | 4.71E-05 | -2.20E+00 | 5.18E-04 | 2.24E-03 | -1.41E+00 | 2.39E-10 | 1.02E-09 | -1.89E+00 | 3.59E-05 | 1.37E-04 | -1.10E+00 | 8.81E-09 | 3.14E-08 | -2.75E+00 | 3.89E-06 | 1.10E-05 | -1.69E+00 | 4.76E-07 | 1.75E-06 | -1.60E+00 | 4.88E-02 | 1.34E-01 | 1.03E+00 |
| RPGRIP1L | 9.05E-06 | 4.72E-05 | -1.60E+00 | 2.25E-06 | 9.74E-06 | -1.60E+00 | 7.12E-05 | 3.03E-04 | -1.49E+00 | 1.39E-03 | 5.32E-03 | -1.09E+00 | 8.81E-04 | 3.14E-03 | -1.12E+00 | 2.60E-04 | 7.33E-04 | -1.40E+00 | 2.86E-02 | 1.05E-01 | 1.08E+00 | 1.24E-01 | 3.42E-01 | 1.10E+00 |
| RPS4XP11 | 9.06E-06 | 4.72E-05 | -1.15E+00 | 7.74E-54 | 3.35E-53 | -1.67E+00 | 1.79E-02 | 7.60E-02 | -1.08E+00 | 7.15E-26 | 2.73E-25 | -1.54E+00 | 1.59E-01 | 5.66E-01 | -1.02E+00 | 3.23E-17 | 9.10E-17 | -1.47E+00 | 1.41E-06 | 5.17E-06 | -1.09E+00 | 1.96E-36 | 5.38E-36 | -1.62E+00 |
| TRMT44 | 9.07E-06 | 4.72E-05 | -1.43E+00 | 3.52E-04 | 1.52E-03 | -1.06E+00 | 2.13E-06 | 9.07E-06 | -1.53E+00 | 6.75E-05 | 2.58E-04 | -1.11E+00 | 1.15E-03 | 4.10E-03 | -1.47E+00 | 7.79E-02 | 2.20E-01 | 1.06E+00 | 4.52E-07 | 1.66E-06 | -1.55E+00 | 4.66E-05 | 1.28E-04 | -1.29E+00 |
| PHF7 | 9.08E-06 | 4.73E-05 | -1.40E+00 | 5.86E-03 | 2.53E-02 | 1.08E+00 | 7.50E-09 | 3.19E-08 | -1.62E+00 | 2.07E-06 | 7.91E-06 | -1.22E+00 | 2.32E-03 | 8.28E-03 | -1.21E+00 | 7.79E-02 | 2.20E-01 | 1.09E+00 | 8.08E-07 | 2.97E-06 | -1.46E+00 | 2.84E-02 | 7.81E-02 | -1.02E+00 |
| UTP14C | 9.08E-06 | 4.73E-05 | -1.76E+00 | 1.04E-03 | 4.51E-03 | -1.16E+00 | 1.25E-05 | 5.32E-05 | -1.55E+00 | 5.80E-03 | 2.22E-02 | 1.01E+00 | 2.24E-02 | 7.99E-02 | -1.04E+00 | 1.71E-01 | 4.81E-01 | 1.72E+00 | 4.93E-11 | 1.81E-10 | -1.76E+00 | 1.24E-05 | 3.40E-05 | -1.32E+00 |
| RPS3AP12 | 9.08E-06 | 4.73E-05 | -1.30E+00 | 2.54E-06 | 1.10E-05 | -1.25E+00 | 1.76E-01 | 7.47E-01 | 1.19E+00 | 9.40E-02 | 3.59E-01 | 1.12E+00 | 6.31E-02 | 2.25E-01 | 1.11E+00 | 4.04E-03 | 1.14E-02 | -1.38E+00 | 3.34E-02 | 1.23E-01 | -1.03E+00 | 1.14E-01 | 3.12E-01 | 1.08E+00 |
| BAG6 | 9.10E-06 | 4.75E-05 | -1.07E+00 | 6.59E-06 | 2.85E-05 | 1.04E+00 | 1.70E-04 | 7.23E-04 | -1.13E+00 | 2.51E-05 | 9.57E-05 | -1.04E+00 | 3.00E-03 | 1.07E-02 | 1.05E+00 | 2.35E-02 | 6.62E-02 | 1.21E+00 | 7.14E-03 | 2.62E-02 | 1.00E+00 | 3.70E-02 | 1.02E-01 | 1.12E+00 |
| FDPS | 9.14E-06 | 4.76E-05 | -1.10E+00 | 8.11E-04 | 3.50E-03 | 1.16E+00 | 5.51E-03 | 2.34E-02 | -1.13E+00 | 2.76E-02 | 1.05E-01 | 1.13E+00 | 4.26E-05 | 1.52E-04 | -1.12E+00 | 7.77E-03 | 2.19E-02 | 1.15E+00 | 1.56E-04 | 5.74E-04 | -1.21E+00 | 1.66E-01 | 4.55E-01 | 1.04E+00 |
| RER1 | 9.16E-06 | 4.77E-05 | -1.12E+00 | 9.54E-09 | 4.12E-08 | -1.09E+00 | 5.58E-04 | 2.37E-03 | -1.10E+00 | 8.66E-05 | 3.31E-04 | -1.05E+00 | 2.24E-06 | 8.00E-06 | -1.21E+00 | 2.04E-03 | 5.76E-03 | 1.01E+00 | 5.12E-02 | 1.88E-01 | 1.01E+00 | 8.93E-03 | 2.45E-02 | -1.03E+00 |
| IKBKE | 9.17E-06 | 4.78E-05 | -1.33E+00 | 1.06E-08 | 4.58E-08 | -1.64E+00 | 2.36E-06 | 1.00E-05 | -1.44E+00 | 6.67E-06 | 2.55E-05 | -1.17E+00 | 9.51E-03 | 3.39E-02 | -1.25E+00 | 2.94E-02 | 8.29E-02 | -1.15E+00 | 5.83E-02 | 2.14E-01 | -1.02E+00 | 5.07E-02 | 1.39E-01 | -1.10E+00 |
| NAA15 | 9.17E-06 | 4.78E-05 | -1.05E+00 | 2.23E-05 | 9.64E-05 | 1.08E+00 | 7.05E-07 | 3.00E-06 | -1.31E+00 | 5.31E-06 | 2.03E-05 | -1.12E+00 | 1.55E-04 | 5.52E-04 | -1.08E+00 | 1.06E-02 | 3.00E-02 | 1.12E+00 | 1.67E-08 | 6.15E-08 | -1.51E+00 | 6.50E-04 | 1.79E-03 | -1.13E+00 |
| SHPK | 9.17E-06 | 4.78E-05 | -1.93E+00 | 1.22E-04 | 5.29E-04 | -1.51E+00 | 8.75E-03 | 3.72E-02 | -1.34E+00 | 8.38E-03 | 3.20E-02 | -1.24E+00 | 2.76E-03 | 9.86E-03 | -1.66E+00 | 4.21E-02 | 1.19E-01 | -1.13E+00 | 1.60E-03 | 5.86E-03 | -1.40E+00 | 3.30E-01 | 9.08E-01 | 1.25E+00 |
| CDK7 | 9.20E-06 | 4.79E-05 | -1.53E+00 | 1.20E-02 | 5.17E-02 | 1.07E+00 | 3.97E-05 | 1.69E-04 | -1.42E+00 | 2.02E-05 | 7.73E-05 | -1.34E+00 | 9.28E-03 | 3.31E-02 | -1.15E+00 | 2.26E-02 | 6.36E-02 | -1.05E+00 | 3.45E-07 | 1.27E-06 | -1.35E+00 | 3.14E-06 | 8.64E-06 | -1.22E+00 |
| RN7SKP230 | 9.20E-06 | 4.80E-05 | 3.32E+00 | 1.72E-02 | 7.43E-02 | 2.51E+00 | 1.27E-01 | 5.38E-01 | 1.44E+00 | 2.03E-01 | 7.74E-01 | 1.24E+00 | 3.92E-02 | 1.40E-01 | 2.11E+00 | 9.03E-02 | 2.55E-01 | 2.08E+00 | 7.44E-02 | 2.73E-01 | 1.56E+00 | 2.08E-01 | 5.73E-01 | 1.46E+00 |
| EIF2B5 | 9.22E-06 | 4.80E-05 | -1.10E+00 | 4.78E-05 | 2.06E-04 | 1.06E+00 | 4.42E-02 | 1.88E-01 | 1.04E+00 | 3.06E-02 | 1.17E-01 | 1.20E+00 | 1.12E-09 | 4.00E-09 | -1.44E+00 | 8.95E-06 | 2.52E-05 | -1.10E+00 | 5.25E-02 | 1.93E-01 | -1.01E+00 | 2.81E-01 | 7.71E-01 | 1.13E+00 |
| BPGM | 9.24E-06 | 4.82E-05 | -1.40E+00 | 8.56E-03 | 3.70E-02 | 1.08E+00 | 3.19E-02 | 1.35E-01 | -1.06E+00 | 6.31E-02 | 2.41E-01 | 1.15E+00 | 1.23E-10 | 4.39E-10 | -1.93E+00 | 5.74E-05 | 1.62E-04 | -1.06E+00 | 3.06E-06 | 1.12E-05 | -1.25E+00 | 3.38E-04 | 9.29E-04 | -1.10E+00 |
| PIR | 9.24E-06 | 4.82E-05 | -1.53E+00 | 1.11E-04 | 4.80E-04 | -1.17E+00 | 9.74E-06 | 4.14E-05 | -1.86E+00 | 5.79E-04 | 2.21E-03 | -1.30E+00 | 1.43E-02 | 5.08E-02 | -1.17E+00 | 3.02E-01 | 8.50E-01 | 1.35E+00 | 1.84E-01 | 6.74E-01 | 1.40E+00 | 6.47E-02 | 1.78E-01 | 1.41E+00 |
| AC018607.1 | 9.28E-06 | 4.84E-05 | -1.98E+00 | 2.01E-04 | 8.69E-04 | -1.53E+00 | 2.81E-03 | 1.19E-02 | -1.49E+00 | 4.14E-03 | 1.58E-02 | -1.33E+00 | 7.94E-04 | 2.83E-03 | -1.75E+00 | 3.06E-02 | 8.63E-02 | -1.12E+00 | 2.15E-02 | 7.91E-02 | -1.38E+00 | 8.59E-02 | 2.36E-01 | -1.19E+00 |
| FUT11 | 9.28E-06 | 4.84E-05 | -1.64E+00 | 1.09E-03 | 4.69E-03 | -1.11E+00 | 5.76E-08 | 2.45E-07 | -1.87E+00 | 6.07E-06 | 2.32E-05 | -1.33E+00 | 4.33E-05 | 1.54E-04 | -1.85E+00 | 9.47E-04 | 2.67E-03 | -1.35E+00 | 9.46E-07 | 3.47E-06 | -1.97E+00 | 5.11E-03 | 1.40E-02 | -1.21E+00 |
| PAQR8 | 9.29E-06 | 4.84E-05 | -1.98E+00 | 1.15E-01 | 4.97E-01 | 1.24E+00 | 3.23E-10 | 1.37E-09 | -2.52E+00 | 1.42E-03 | 5.43E-03 | -1.08E+00 | 1.05E-02 | 3.73E-02 | -2.39E+00 | 2.98E-01 | 8.39E-01 | 1.68E+00 | 5.25E-14 | 1.93E-13 | -2.77E+00 | 1.38E-03 | 3.80E-03 | -1.11E+00 |
| PPIAP31 | 9.34E-06 | 4.87E-05 | -1.77E+00 | 1.52E-06 | 6.55E-06 | -1.95E+00 | 1.87E-07 | 7.93E-07 | -1.83E+00 | 4.77E-07 | 1.82E-06 | -1.58E+00 | 2.45E-03 | 8.73E-03 | -1.48E+00 | 2.87E-03 | 8.10E-03 | -1.52E+00 | 1.30E-10 | 4.77E-10 | -1.84E+00 | 1.05E-08 | 2.88E-08 | -1.62E+00 |
| CARD8 | 9.36E-06 | 4.88E-05 | -1.06E+00 | 5.17E-04 | 2.23E-03 | 1.19E+00 | 4.86E-05 | 2.06E-04 | -1.32E+00 | 8.69E-04 | 3.32E-03 | -1.05E+00 | 3.03E-12 | 1.08E-11 | -1.44E+00 | 2.35E-10 | 6.61E-10 | -1.27E+00 | 3.52E-09 | 1.29E-08 | -1.38E+00 | 2.25E-07 | 6.19E-07 | -1.28E+00 |
| CAPN10 | 9.36E-06 | 4.88E-05 | -1.40E+00 | 1.98E-03 | 8.56E-03 | 1.03E+00 | 3.19E-03 | 1.35E-02 | -1.18E+00 | 1.28E-02 | 4.90E-02 | 1.04E+00 | 7.48E-04 | 2.67E-03 | -1.49E+00 | 2.66E-02 | 7.48E-02 | -1.04E+00 | 1.92E-04 | 7.04E-04 | -1.29E+00 | 7.39E-02 | 2.03E-01 | 1.16E+00 |
| MAML2 | 9.40E-06 | 4.90E-05 | -1.32E+00 | 1.07E-04 | 4.63E-04 | -1.09E+00 | 3.68E-02 | 1.56E-01 | -1.11E+00 | 2.80E-04 | 1.07E-03 | -1.44E+00 | 1.50E-01 | 5.35E-01 | 1.08E+00 | 5.25E-02 | 1.48E-01 | -1.23E+00 | 1.05E-01 | 3.87E-01 | 1.07E+00 | 3.98E-02 | 1.09E-01 | -1.22E+00 |
| AC018797.2 | 9.43E-06 | 4.91E-05 | -2.19E+00 | 1.60E-03 | 6.90E-03 | -1.22E+00 | 7.20E-03 | 3.06E-02 | -1.62E+00 | 4.34E-02 | 1.66E-01 | -1.18E+00 | 2.71E-02 | 9.67E-02 | -1.40E+00 | 2.84E-01 | 8.02E-01 | 1.24E+00 | 1.17E-07 | 4.30E-07 | -1.97E+00 | 3.72E-02 | 1.02E-01 | -1.06E+00 |
| RNF185 | 9.43E-06 | 4.92E-05 | -1.79E+00 | 3.63E-04 | 1.57E-03 | -1.29E+00 | 3.30E-06 | 1.40E-05 | -1.89E+00 | 1.15E-02 | 4.40E-02 | -1.06E+00 | 4.63E-03 | 1.65E-02 | -1.44E+00 | 2.45E-02 | 6.91E-02 | -1.19E+00 | 6.19E-09 | 2.27E-08 | -2.41E+00 | 7.08E-04 | 1.95E-03 | -1.35E+00 |
| TSTA3 | 9.49E-06 | 4.95E-05 | -1.20E+00 | 8.39E-06 | 3.62E-05 | -1.06E+00 | 9.21E-03 | 3.91E-02 | -1.04E+00 | 1.48E-03 | 5.64E-03 | -1.01E+00 | 8.10E-05 | 2.89E-04 | -1.22E+00 | 1.11E-03 | 3.12E-03 | -1.07E+00 | 5.86E-02 | 2.15E-01 | 1.16E+00 | 1.48E-01 | 4.08E-01 | 1.10E+00 |
| ZKSCAN7 | 9.52E-06 | 4.96E-05 | -1.79E+00 | 3.88E-04 | 1.68E-03 | -1.29E+00 | 6.93E-09 | 2.94E-08 | -2.03E+00 | 8.86E-05 | 3.38E-04 | -1.19E+00 | 1.20E-03 | 4.26E-03 | -1.54E+00 | 1.78E-02 | 5.02E-02 | -1.15E+00 | 1.47E-04 | 5.38E-04 | -1.47E+00 | 5.25E-03 | 1.44E-02 | -1.27E+00 |
| SPCS1 | 9.53E-06 | 4.97E-05 | -1.10E+00 | 3.89E-09 | 1.68E-08 | -1.05E+00 | 1.02E-02 | 4.35E-02 | -1.06E+00 | 3.97E-02 | 1.52E-01 | -1.02E+00 | 2.59E-02 | 9.23E-02 | -1.01E+00 | 2.43E-02 | 6.84E-02 | 1.05E+00 | 1.90E-01 | 6.97E-01 | 1.11E+00 | 1.91E-01 | 5.24E-01 | 1.19E+00 |
| RAD51-AS1 | 9.55E-06 | 4.98E-05 | -1.31E+00 | 1.71E-01 | 7.40E-01 | 1.53E+00 | 4.43E-04 | 1.88E-03 | -1.54E+00 | 2.20E-01 | 8.41E-01 | 1.37E+00 | 1.92E-01 | 6.83E-01 | 1.33E+00 | 1.45E-02 | 4.09E-02 | 2.00E+00 | 1.62E-02 | 5.94E-02 | -1.11E+00 | 3.47E-01 | 9.53E-01 | 1.15E+00 |
| AC010468.1 | 9.57E-06 | 4.99E-05 | 1.69E+00 | 9.33E-25 | 4.03E-24 | 2.53E+00 | 5.61E-06 | 2.38E-05 | 1.69E+00 | 4.02E-19 | 1.54E-18 | 2.63E+00 | 2.06E-03 | 7.35E-03 | 1.64E+00 | 2.38E-16 | 6.72E-16 | 2.42E+00 | 3.06E-03 | 1.12E-02 | 1.62E+00 | 5.91E-16 | 1.62E-15 | 2.42E+00 |
| AC098935.1 | 9.59E-06 | 5.00E-05 | -1.14E+00 | 1.37E-12 | 5.93E-12 | -1.72E+00 | 6.54E-03 | 2.78E-02 | -1.04E+00 | 2.71E-06 | 1.04E-05 | -1.43E+00 | 1.14E-02 | 4.08E-02 | -1.07E+00 | 8.94E-04 | 2.52E-03 | -1.35E+00 | 2.79E-05 | 1.02E-04 | -1.40E+00 | 2.14E-08 | 5.87E-08 | -1.90E+00 |
| GFM2 | 9.60E-06 | 5.00E-05 | -1.82E+00 | 4.54E-04 | 1.96E-03 | -1.21E+00 | 5.65E-04 | 2.40E-03 | -1.43E+00 | 9.98E-03 | 3.81E-02 | -1.06E+00 | 8.06E-06 | 2.87E-05 | -1.59E+00 | 1.94E-04 | 5.47E-04 | -1.30E+00 | 1.73E-06 | 6.35E-06 | -1.32E+00 | 1.16E-03 | 3.19E-03 | -1.09E+00 |
| CENPC | 9.68E-06 | 5.05E-05 | -1.11E+00 | 1.17E-05 | 5.06E-05 | 1.04E+00 | 7.24E-03 | 3.08E-02 | -1.08E+00 | 4.40E-06 | 1.68E-05 | -1.22E+00 | 1.52E-05 | 5.42E-05 | -1.09E+00 | 2.03E-03 | 5.71E-03 | 1.07E+00 | 3.46E-03 | 1.27E-02 | -1.02E+00 | 1.91E-02 | 5.23E-02 | 1.00E+00 |
| FAM13A | 9.69E-06 | 5.05E-05 | -1.08E+00 | 1.15E-02 | 4.95E-02 | 1.32E+00 | 1.94E-04 | 8.23E-04 | -1.15E+00 | 1.03E-04 | 3.95E-04 | 1.01E+00 | 9.88E-04 | 3.52E-03 | -1.37E+00 | 9.00E-02 | 2.54E-01 | 1.09E+00 | 1.47E-03 | 5.38E-03 | -1.22E+00 | 4.23E-02 | 1.16E-01 | -1.01E+00 |
| DDRGK1 | 9.69E-06 | 5.05E-05 | -1.10E+00 | 3.11E-05 | 1.35E-04 | 1.06E+00 | 1.20E-03 | 5.11E-03 | -1.16E+00 | 1.05E-04 | 4.03E-04 | -1.07E+00 | 1.62E-06 | 5.78E-06 | -1.13E+00 | 3.06E-06 | 8.63E-06 | -1.04E+00 | 4.59E-02 | 1.68E-01 | 1.08E+00 | 1.79E-02 | 4.91E-02 | 1.07E+00 |
| FTLP2 | 9.70E-06 | 5.05E-05 | 2.61E+00 | 2.59E-08 | 1.12E-07 | 3.32E+00 | 2.55E-03 | 1.08E-02 | 1.89E+00 | 4.04E-11 | 1.54E-10 | 3.44E+00 | 2.17E-01 | 7.73E-01 | 1.60E+00 | 2.13E-03 | 6.00E-03 | 2.31E+00 | 7.38E-04 | 2.71E-03 | 2.19E+00 | 9.90E-10 | 2.72E-09 | 3.47E+00 |
| ITGB2 | 9.78E-06 | 5.10E-05 | -1.05E+00 | 2.74E-09 | 1.18E-08 | -1.04E+00 | 6.91E-02 | 2.94E-01 | 1.08E+00 | 2.26E-03 | 8.64E-03 | -1.02E+00 | 3.39E-05 | 1.21E-04 | -1.22E+00 | 3.11E-05 | 8.76E-05 | -1.24E+00 | 3.99E-03 | 1.46E-02 | -1.07E+00 | 4.88E-04 | 1.34E-03 | -1.19E+00 |
| DTX2P1-UPK3BP1- | 9.81E-06 | 5.11E-05 | -1.67E+00 | 4.74E-05 | 2.05E-04 | -1.44E+00 | 7.34E-09 | 3.12E-08 | -2.09E+00 | 5.32E-07 | 2.03E-06 | -1.57E+00 | 1.84E-02 | 6.58E-02 | -1.35E+00 | 5.58E-02 | 1.57E-01 | -1.10E+00 | 2.76E-02 | 1.01E-01 | -1.08E+00 | 5.78E-02 | 1.59E-01 | 1.02E+00 |
| ZNF671 | 9.83E-06 | 5.12E-05 | -1.52E+00 | 5.53E-03 | 2.39E-02 | 1.09E+00 | 6.36E-05 | 2.70E-04 | -1.53E+00 | 4.25E-03 | 1.62E-02 | -1.01E+00 | 1.31E-01 | 4.66E-01 | 1.06E+00 | 1.40E-01 | 3.96E-01 | 1.06E+00 | 6.10E-04 | 2.24E-03 | -1.35E+00 | 9.75E-03 | 2.68E-02 | -1.19E+00 |
| SOAT1 | 9.87E-06 | 5.14E-05 | -1.42E+00 | 1.59E-03 | 6.86E-03 | 1.01E+00 | 7.13E-04 | 3.03E-03 | -1.26E+00 | 8.43E-04 | 3.22E-03 | -1.13E+00 | 4.38E-05 | 1.56E-04 | -1.87E+00 | 8.48E-05 | 2.39E-04 | -1.67E+00 | 2.37E-08 | 8.68E-08 | -1.62E+00 | 1.11E-03 | 3.05E-03 | -1.11E+00 |
| LINC02256 | 9.88E-06 | 5.15E-05 | -1.43E+00 | 4.66E-06 | 2.02E-05 | -1.45E+00 | 4.17E-07 | 1.77E-06 | -2.13E+00 | 2.45E-06 | 9.38E-06 | -1.69E+00 | 2.98E-06 | 1.06E-05 | -2.06E+00 | 2.09E-04 | 5.88E-04 | -1.45E+00 | 2.64E-01 | 9.68E-01 | 1.22E+00 | 3.43E-01 | 9.42E-01 | 1.19E+00 |
| RAB9B | 9.94E-06 | 5.18E-05 | -1.85E+00 | 1.55E-01 | 6.71E-01 | 1.36E+00 | 2.94E-02 | 1.25E-01 | -1.27E+00 | 1.38E-01 | 5.25E-01 | 1.54E+00 | 3.57E-04 | 1.27E-03 | -2.05E+00 | 2.69E-01 | 7.59E-01 | 1.38E+00 | 8.04E-03 | 2.95E-02 | -1.32E+00 | 1.09E-01 | 2.99E-01 | 1.09E+00 |
| UQCC1 | 9.96E-06 | 5.19E-05 | -1.58E+00 | 3.11E-02 | 1.34E-01 | 1.16E+00 | 2.69E-04 | 1.14E-03 | -1.28E+00 | 4.18E-02 | 1.60E-01 | 1.16E+00 | 2.41E-03 | 8.58E-03 | -1.20E+00 | 1.47E-01 | 4.14E-01 | 1.20E+00 | 5.44E-04 | 2.00E-03 | -1.06E+00 | 3.49E-01 | 9.59E-01 | 1.31E+00 |
| FAM126A | 9.97E-06 | 5.20E-05 | -1.38E+00 | 1.02E-02 | 4.39E-02 | 1.09E+00 | 2.46E-08 | 1.04E-07 | -1.69E+00 | 8.30E-04 | 3.17E-03 | -1.01E+00 | 6.94E-05 | 2.47E-04 | -1.48E+00 | 4.58E-03 | 1.29E-02 | -1.14E+00 | 1.94E-02 | 7.11E-02 | 1.00E+00 | 2.85E-01 | 7.82E-01 | 1.24E+00 |
| UTRN | 1.00E-05 | 5.22E-05 | -1.10E+00 | 2.09E-05 | 9.04E-05 | 1.05E+00 | 1.02E-07 | 4.33E-07 | -1.38E+00 | 2.99E-08 | 1.14E-07 | -1.26E+00 | 4.42E-03 | 1.58E-02 | -1.06E+00 | 3.03E-06 | 8.55E-06 | -1.40E+00 | 1.36E-03 | 5.00E-03 | -1.25E+00 | 6.73E-04 | 1.85E-03 | -1.23E+00 |
| ELL2 | 1.01E-05 | 5.26E-05 | -1.32E+00 | 1.55E-10 | 6.68E-10 | -1.59E+00 | 2.32E-02 | 9.85E-02 | 1.00E+00 | 3.88E-05 | 1.48E-04 | -1.21E+00 | 5.02E-03 | 1.79E-02 | -1.07E+00 | 5.10E-06 | 1.44E-05 | -1.51E+00 | 3.25E-09 | 1.19E-08 | -1.93E+00 | 5.46E-09 | 1.50E-08 | -1.85E+00 |
| ANAPC11 | 1.01E-05 | 5.28E-05 | -1.03E+00 | 1.42E-10 | 6.12E-10 | -1.05E+00 | 1.98E-01 | 8.40E-01 | 1.19E+00 | 5.49E-02 | 2.10E-01 | 1.15E+00 | 1.21E-05 | 4.31E-05 | -1.13E+00 | 1.25E-05 | 3.54E-05 | -1.11E+00 | 2.58E-01 | 9.46E-01 | 1.05E+00 | 2.24E-01 | 6.15E-01 | 1.07E+00 |
| LINC01006 | 1.01E-05 | 5.28E-05 | -1.84E+00 | 4.46E-03 | 1.93E-02 | -1.10E+00 | 1.22E-07 | 5.20E-07 | -2.28E+00 | 1.62E-04 | 6.18E-04 | -1.32E+00 | 2.32E-07 | 8.26E-07 | -1.72E+00 | 2.31E-04 | 6.52E-04 | -1.17E+00 | 6.87E-04 | 2.52E-03 | -1.25E+00 | 2.31E-01 | 6.33E-01 | 1.13E+00 |
| CCDC9 | 1.01E-05 | 5.29E-05 | -1.51E+00 | 1.50E-07 | 6.48E-07 | -1.74E+00 | 7.04E-02 | 2.99E-01 | 1.06E+00 | 5.99E-02 | 2.29E-01 | 1.07E+00 | 1.84E-02 | 6.57E-02 | -1.12E+00 | 7.44E-03 | 2.10E-02 | -1.27E+00 | 7.76E-04 | 2.85E-03 | -1.13E+00 | 2.76E-05 | 7.59E-05 | -1.53E+00 |
| LINC00667 | 1.01E-05 | 5.29E-05 | -1.11E+00 | 3.50E-02 | 1.51E-01 | 1.35E+00 | 1.51E-04 | 6.42E-04 | -1.22E+00 | 1.87E-02 | 7.14E-02 | 1.19E+00 | 5.31E-05 | 1.89E-04 | -1.08E+00 | 2.17E-02 | 6.12E-02 | 1.17E+00 | 8.20E-05 | 3.01E-04 | -1.08E+00 | 1.38E-02 | 3.79E-02 | 1.08E+00 |
| AC010632.2 | 1.02E-05 | 5.29E-05 | -1.90E+00 | 4.79E-03 | 2.07E-02 | -1.11E+00 | 4.58E-06 | 1.95E-05 | -1.70E+00 | 6.10E-04 | 2.33E-03 | -1.19E+00 | 9.53E-03 | 3.40E-02 | -1.39E+00 | 1.12E-02 | 3.17E-02 | -1.37E+00 | 4.48E-03 | 1.64E-02 | -1.65E+00 | 1.49E-01 | 4.10E-01 | 1.01E+00 |
| C11orf71 | 1.02E-05 | 5.30E-05 | -1.17E+00 | 4.93E-03 | 2.13E-02 | 1.18E+00 | 1.50E-08 | 6.38E-08 | -1.66E+00 | 9.89E-07 | 3.78E-06 | -1.31E+00 | 3.08E-02 | 1.10E-01 | 1.01E+00 | 2.13E-01 | 6.01E-01 | 1.34E+00 | 6.67E-04 | 2.45E-03 | -1.27E+00 | 1.36E-02 | 3.73E-02 | -1.08E+00 |
| NFU1 | 1.02E-05 | 5.30E-05 | -1.06E+00 | 3.71E-06 | 1.60E-05 | 1.01E+00 | 1.20E-02 | 5.11E-02 | -1.00E+00 | 1.99E-02 | 7.58E-02 | 1.11E+00 | 6.14E-05 | 2.19E-04 | -1.11E+00 | 7.23E-03 | 2.04E-02 | 1.12E+00 | 1.92E-01 | 7.06E-01 | 1.15E+00 | 3.59E-03 | 9.87E-03 | -1.03E+00 |
| SLC25A1 | 1.02E-05 | 5.31E-05 | -1.12E+00 | 9.05E-03 | 3.91E-02 | 1.18E+00 | 1.72E-03 | 7.30E-03 | -1.02E+00 | 2.32E-03 | 8.86E-03 | 1.10E+00 | 1.51E-04 | 5.37E-04 | -1.24E+00 | 1.09E-02 | 3.08E-02 | 1.00E+00 | 9.91E-02 | 3.64E-01 | 1.39E+00 | 6.76E-03 | 1.86E-02 | 1.64E+00 |
| AL132656.3 | 1.02E-05 | 5.32E-05 | -1.99E+00 | 5.90E-05 | 2.55E-04 | -1.65E+00 | 2.16E-02 | 9.18E-02 | -1.39E+00 | 2.08E-01 | 7.96E-01 | 1.21E+00 | 4.72E-04 | 1.68E-03 | -1.46E+00 | 1.34E-03 | 3.77E-03 | -1.35E+00 | 1.93E-02 | 7.10E-02 | -1.24E+00 | 3.28E-01 | 9.02E-01 | 1.21E+00 |
| TRAF3IP1 | 1.02E-05 | 5.32E-05 | -1.46E+00 | 1.88E-05 | 8.12E-05 | -1.38E+00 | 3.82E-02 | 1.62E-01 | -1.11E+00 | 5.51E-02 | 2.10E-01 | 1.07E+00 | 2.35E-03 | 8.37E-03 | -1.31E+00 | 1.65E-02 | 4.66E-02 | -1.11E+00 | 1.74E-01 | 6.40E-01 | 1.12E+00 | 2.86E-01 | 7.86E-01 | 1.21E+00 |
| ZDHHC7 | 1.02E-05 | 5.32E-05 | -1.67E+00 | 5.56E-03 | 2.40E-02 | -1.00E+00 | 1.07E-07 | 4.57E-07 | -1.88E+00 | 8.76E-07 | 3.35E-06 | -1.62E+00 | 2.16E-06 | 7.72E-06 | -1.88E+00 | 4.51E-03 | 1.27E-02 | -1.06E+00 | 3.05E-12 | 1.12E-11 | -1.48E+00 | 3.59E-06 | 9.86E-06 | -1.15E+00 |
| AC006299.1 | 1.02E-05 | 5.33E-05 | -2.13E+00 | 1.84E-03 | 7.96E-03 | -1.28E+00 | 1.61E-01 | 6.85E-01 | -1.01E+00 | 2.52E-01 | 9.62E-01 | 1.26E+00 | 1.92E-01 | 6.83E-01 | 1.03E+00 | 1.92E-01 | 5.41E-01 | -1.15E+00 | 1.90E-01 | 6.98E-01 | 1.27E+00 | 3.36E-01 | 9.24E-01 | 1.11E+00 |
| OTUB1 | 1.02E-05 | 5.33E-05 | -1.12E+00 | 5.26E-03 | 2.27E-02 | 1.18E+00 | 8.10E-07 | 3.44E-06 | -1.28E+00 | 7.74E-06 | 2.95E-05 | -1.04E+00 | 7.64E-08 | 2.73E-07 | -1.38E+00 | 1.62E-06 | 4.56E-06 | -1.17E+00 | 3.64E-07 | 1.34E-06 | -1.34E+00 | 7.56E-04 | 2.08E-03 | -1.10E+00 |
| ZNF574 | 1.03E-05 | 5.34E-05 | -1.25E+00 | 3.13E-11 | 1.35E-10 | -1.94E+00 | 2.20E-08 | 9.33E-08 | -1.50E+00 | 8.32E-14 | 3.18E-13 | -1.88E+00 | 1.28E-04 | 4.58E-04 | -1.49E+00 | 1.86E-02 | 5.24E-02 | 1.03E+00 | 1.12E-10 | 4.10E-10 | -1.73E+00 | 1.09E-06 | 2.99E-06 | -1.36E+00 |
| SPDL1 | 1.03E-05 | 5.35E-05 | -1.70E+00 | 3.05E-04 | 1.32E-03 | -1.26E+00 | 1.64E-07 | 6.99E-07 | -1.74E+00 | 2.57E-04 | 9.80E-04 | -1.06E+00 | 1.72E-03 | 6.13E-03 | -2.10E+00 | 5.18E-02 | 1.46E-01 | -1.23E+00 | 5.00E-10 | 1.83E-09 | -1.90E+00 | 4.34E-04 | 1.19E-03 | -1.14E+00 |
| KIF9 | 1.03E-05 | 5.35E-05 | -1.08E+00 | 7.04E-10 | 3.04E-09 | -1.20E+00 | 8.15E-05 | 3.46E-04 | -1.14E+00 | 1.58E-08 | 6.02E-08 | -1.29E+00 | 9.36E-06 | 3.34E-05 | -1.21E+00 | 2.74E-07 | 7.71E-07 | -1.30E+00 | 4.78E-04 | 1.76E-03 | -1.10E+00 | 6.22E-07 | 1.71E-06 | -1.32E+00 |
| SLFN11 | 1.03E-05 | 5.35E-05 | -1.37E+00 | 3.98E-04 | 1.72E-03 | -1.07E+00 | 1.56E-03 | 6.62E-03 | -1.25E+00 | 7.82E-02 | 2.99E-01 | 1.19E+00 | 7.26E-05 | 2.59E-04 | -1.60E+00 | 4.60E-02 | 1.30E-01 | 1.19E+00 | 7.89E-20 | 2.90E-19 | -1.90E+00 | 1.18E-07 | 3.24E-07 | -1.15E+00 |
| ZNF480 | 1.03E-05 | 5.37E-05 | -1.19E+00 | 8.29E-08 | 3.58E-07 | -1.22E+00 | 2.04E-07 | 8.66E-07 | -1.47E+00 | 3.19E-08 | 1.22E-07 | -1.41E+00 | 2.69E-02 | 9.61E-02 | 1.00E+00 | 5.77E-03 | 1.63E-02 | -1.06E+00 | 6.62E-09 | 2.43E-08 | -1.47E+00 | 3.23E-09 | 8.88E-09 | -1.44E+00 |
| GDI1 | 1.03E-05 | 5.39E-05 | -1.53E+00 | 3.30E-02 | 1.43E-01 | 1.20E+00 | 4.42E-06 | 1.88E-05 | -1.66E+00 | 5.30E-03 | 2.02E-02 | -1.05E+00 | 1.15E-05 | 4.12E-05 | -1.64E+00 | 2.00E-03 | 5.63E-03 | -1.18E+00 | 7.67E-04 | 2.81E-03 | -1.11E+00 | 1.36E-01 | 3.74E-01 | 1.21E+00 |
| AVL9 | 1.04E-05 | 5.42E-05 | -1.13E+00 | 5.46E-05 | 2.36E-04 | -1.01E+00 | 1.39E-04 | 5.89E-04 | -1.33E+00 | 6.11E-02 | 2.33E-01 | 1.16E+00 | 2.00E-04 | 7.12E-04 | -1.36E+00 | 3.53E-02 | 9.94E-02 | 1.06E+00 | 8.84E-04 | 3.25E-03 | -1.18E+00 | 7.54E-03 | 2.07E-02 | -1.08E+00 |
| AC008895.1 | 1.04E-05 | 5.43E-05 | -2.04E+00 | 7.03E-04 | 3.04E-03 | -1.34E+00 | 3.18E-08 | 1.35E-07 | -2.94E+00 | 1.81E-04 | 6.92E-04 | -1.41E+00 | 7.94E-02 | 2.83E-01 | -1.17E+00 | 3.53E-01 | 9.95E-01 | 1.29E+00 | 5.53E-04 | 2.03E-03 | -1.80E+00 | 1.50E-01 | 4.11E-01 | -1.01E+00 |
| RTN4 | 1.04E-05 | 5.44E-05 | -1.11E+00 | 1.26E-03 | 5.45E-03 | 1.07E+00 | 1.92E-02 | 8.16E-02 | -1.04E+00 | 6.28E-02 | 2.40E-01 | 1.04E+00 | 7.56E-07 | 2.70E-06 | -1.27E+00 | 1.70E-03 | 4.80E-03 | -1.16E+00 | 2.95E-02 | 1.08E-01 | 1.08E+00 | 8.73E-02 | 2.40E-01 | -1.01E+00 |
| SEPHS1 | 1.05E-05 | 5.46E-05 | -1.45E+00 | 2.43E-04 | 1.05E-03 | -1.11E+00 | 5.97E-02 | 2.54E-01 | 1.04E+00 | 2.70E-03 | 1.03E-02 | -1.00E+00 | 3.14E-03 | 1.12E-02 | -1.18E+00 | 4.93E-02 | 1.39E-01 | 1.13E+00 | 1.14E-04 | 4.20E-04 | -1.10E+00 | 4.21E-03 | 1.16E-02 | -1.07E+00 |
| PRR13P5 | 1.05E-05 | 5.47E-05 | -1.63E+00 | 3.12E-06 | 1.35E-05 | -1.68E+00 | 1.03E-06 | 4.37E-06 | -1.70E+00 | 8.87E-09 | 3.39E-08 | -1.97E+00 | 1.06E-10 | 3.77E-10 | -1.94E+00 | 1.42E-12 | 4.01E-12 | -2.15E+00 | 1.03E-06 | 3.77E-06 | -1.58E+00 | 1.92E-08 | 5.28E-08 | -2.13E+00 |
| MPND | 1.06E-05 | 5.50E-05 | -1.20E+00 | 7.14E-09 | 3.09E-08 | -1.47E+00 | 3.61E-03 | 1.53E-02 | -1.06E+00 | 2.50E-05 | 9.57E-05 | -1.21E+00 | 1.69E-05 | 6.03E-05 | -1.35E+00 | 5.46E-06 | 1.54E-05 | -1.42E+00 | 1.13E-06 | 4.13E-06 | -1.29E+00 | 4.25E-04 | 1.17E-03 | -1.20E+00 |
| ACTR8 | 1.06E-05 | 5.53E-05 | -1.42E+00 | 3.70E-04 | 1.60E-03 | -1.09E+00 | 7.62E-07 | 3.24E-06 | -1.64E+00 | 1.04E-03 | 3.99E-03 | -1.06E+00 | 1.85E-02 | 6.61E-02 | -1.03E+00 | 9.65E-02 | 2.72E-01 | 1.11E+00 | 4.89E-12 | 1.79E-11 | -1.53E+00 | 1.16E-05 | 3.18E-05 | -1.13E+00 |
| SPPL3 | 1.06E-05 | 5.53E-05 | -1.19E+00 | 5.48E-03 | 2.37E-02 | 1.13E+00 | 3.63E-02 | 1.54E-01 | 1.05E+00 | 9.03E-03 | 3.45E-02 | 1.14E+00 | 1.23E-04 | 4.40E-04 | -1.28E+00 | 9.39E-04 | 2.65E-03 | -1.04E+00 | 2.22E-05 | 8.16E-05 | -1.26E+00 | 2.68E-02 | 7.35E-02 | 1.01E+00 |
| RIC8A | 1.06E-05 | 5.53E-05 | -1.26E+00 | 2.98E-04 | 1.29E-03 | -1.06E+00 | 2.14E-06 | 9.10E-06 | -1.31E+00 | 5.10E-05 | 1.95E-04 | -1.07E+00 | 5.13E-05 | 1.83E-04 | -1.22E+00 | 5.43E-02 | 1.53E-01 | 1.12E+00 | 7.38E-05 | 2.71E-04 | -1.06E+00 | 1.10E-02 | 3.03E-02 | 1.10E+00 |
| LRP11 | 1.06E-05 | 5.54E-05 | -2.21E+00 | 1.43E-04 | 6.18E-04 | -1.75E+00 | 2.74E-04 | 1.16E-03 | -1.52E+00 | 8.52E-06 | 3.25E-05 | -1.98E+00 | 1.72E-01 | 6.11E-01 | -1.18E+00 | 2.59E-01 | 7.29E-01 | -1.06E+00 | 4.07E-07 | 1.49E-06 | -1.77E+00 | 9.43E-03 | 2.59E-02 | -1.04E+00 |
| HS1BP3 | 1.06E-05 | 5.54E-05 | -1.33E+00 | 1.90E-03 | 8.19E-03 | 1.06E+00 | 5.50E-03 | 2.34E-02 | -1.20E+00 | 7.63E-02 | 2.91E-01 | 1.21E+00 | 3.12E-02 | 1.11E-01 | 1.04E+00 | 1.17E-01 | 3.31E-01 | 1.13E+00 | 2.44E-04 | 8.94E-04 | -1.26E+00 | 8.88E-05 | 2.44E-04 | -1.45E+00 |
| ZNF518B | 1.07E-05 | 5.55E-05 | -1.50E+00 | 2.58E-04 | 1.11E-03 | -1.09E+00 | 3.74E-03 | 1.59E-02 | -1.46E+00 | 7.96E-03 | 3.04E-02 | -1.21E+00 | 1.26E-03 | 4.51E-03 | -1.22E+00 | 2.20E-03 | 6.21E-03 | -1.27E+00 | 7.06E-03 | 2.59E-02 | -1.06E+00 | 2.85E-02 | 7.84E-02 | -1.03E+00 |
| MAP3K13 | 1.07E-05 | 5.55E-05 | 1.00E+00 | 8.08E-07 | 3.49E-06 | 1.06E+00 | 2.18E-04 | 9.28E-04 | -1.14E+00 | 2.21E-03 | 8.45E-03 | 1.07E+00 | 1.61E-02 | 5.74E-02 | 1.02E+00 | 2.49E-03 | 7.01E-03 | -1.09E+00 | 1.10E-03 | 4.02E-03 | -1.07E+00 | 9.07E-04 | 2.49E-03 | -1.05E+00 |
| TRIP6 | 1.07E-05 | 5.56E-05 | -1.12E+00 | 1.85E-04 | 7.98E-04 | 1.05E+00 | 5.02E-04 | 2.13E-03 | -1.05E+00 | 8.12E-05 | 3.10E-04 | 1.06E+00 | 9.21E-02 | 3.29E-01 | 2.04E+00 | 1.25E-02 | 3.52E-02 | 3.00E+00 | 8.31E-03 | 3.05E-02 | 1.77E+00 | 6.49E-02 | 1.78E-01 | 1.64E+00 |
| PDCD2L | 1.07E-05 | 5.56E-05 | -1.78E+00 | 4.66E-05 | 2.02E-04 | -1.47E+00 | 3.98E-07 | 1.69E-06 | -1.95E+00 | 3.67E-04 | 1.40E-03 | -1.22E+00 | 1.35E-04 | 4.81E-04 | -1.86E+00 | 1.21E-02 | 3.40E-02 | -1.20E+00 | 4.81E-12 | 1.76E-11 | -1.96E+00 | 3.21E-05 | 8.83E-05 | -1.25E+00 |
| RPS6KA4 | 1.07E-05 | 5.60E-05 | -1.34E+00 | 2.46E-02 | 1.06E-01 | 1.19E+00 | 7.36E-09 | 3.13E-08 | -1.66E+00 | 1.09E-05 | 4.16E-05 | -1.23E+00 | 1.86E-01 | 6.65E-01 | 1.09E+00 | 3.10E-01 | 8.75E-01 | 1.56E+00 | 9.90E-04 | 3.63E-03 | -1.29E+00 | 5.33E-02 | 1.46E-01 | 1.07E+00 |
| PSPC1P1 | 1.08E-05 | 5.62E-05 | -1.48E+00 | 1.23E-07 | 5.30E-07 | -1.69E+00 | 8.09E-06 | 3.44E-05 | -1.50E+00 | 1.97E-08 | 7.52E-08 | -1.76E+00 | 1.91E-05 | 6.80E-05 | -2.08E+00 | 4.16E-04 | 1.17E-03 | -1.67E+00 | 3.58E-11 | 1.31E-10 | -1.61E+00 | 7.78E-09 | 2.14E-08 | -1.49E+00 |
| PDCD11 | 1.09E-05 | 5.67E-05 | -1.37E+00 | 2.09E-03 | 9.04E-03 | 1.05E+00 | 2.54E-02 | 1.08E-01 | -1.05E+00 | 1.38E-01 | 5.28E-01 | 1.44E+00 | 2.11E-02 | 7.53E-02 | -1.27E+00 | 7.16E-02 | 2.02E-01 | 1.74E+00 | 1.90E-03 | 6.96E-03 | -1.04E+00 | 3.60E-01 | 9.89E-01 | 1.36E+00 |
| PCNT | 1.09E-05 | 5.68E-05 | -1.17E+00 | 1.04E-02 | 4.49E-02 | 1.21E+00 | 8.07E-03 | 3.43E-02 | -1.05E+00 | 1.66E-01 | 6.34E-01 | 1.27E+00 | 7.02E-04 | 2.50E-03 | -1.32E+00 | 6.78E-02 | 1.91E-01 | 1.14E+00 | 9.19E-03 | 3.37E-02 | 1.00E+00 | 3.91E-04 | 1.07E-03 | -1.19E+00 |
| TMEM189 | 1.09E-05 | 5.68E-05 | -1.44E+00 | 2.31E-03 | 9.97E-03 | 1.01E+00 | 5.55E-04 | 2.36E-03 | -1.23E+00 | 3.94E-02 | 1.50E-01 | 1.19E+00 | 6.14E-06 | 2.19E-05 | -1.67E+00 | 9.56E-05 | 2.69E-04 | -1.42E+00 | 2.83E-03 | 1.04E-02 | -1.02E+00 | 1.56E-01 | 4.28E-01 | 1.23E+00 |
| ZNF713 | 1.09E-05 | 5.70E-05 | -1.75E+00 | 7.11E-03 | 3.07E-02 | 1.06E+00 | 1.25E-03 | 5.30E-03 | -1.48E+00 | 1.57E-03 | 6.01E-03 | -1.37E+00 | 1.32E-06 | 4.71E-06 | -1.95E+00 | 8.41E-08 | 2.37E-07 | -2.66E+00 | 5.18E-07 | 1.90E-06 | -1.88E+00 | 5.70E-07 | 1.57E-06 | -1.93E+00 |
| TNFAIP2 | 1.10E-05 | 5.72E-05 | -1.09E+00 | 8.17E-03 | 3.53E-02 | 1.27E+00 | 1.07E-07 | 4.55E-07 | -1.43E+00 | 4.80E-05 | 1.83E-04 | -1.11E+00 | 3.66E-03 | 1.31E-02 | -1.45E+00 | 2.00E-03 | 5.63E-03 | -1.34E+00 | 7.85E-02 | 2.88E-01 | -1.87E+00 | 1.13E-01 | 3.12E-01 | -1.77E+00 |
| SCD1 | 1.10E-05 | 5.72E-05 | -1.17E+00 | 3.89E-02 | 1.68E-01 | 1.29E+00 | 1.26E-03 | 5.36E-03 | -1.23E+00 | 1.67E-01 | 6.40E-01 | 1.19E+00 | 2.54E-03 | 9.05E-03 | -1.03E+00 | 2.27E-01 | 6.39E-01 | 1.35E+00 | 1.39E-03 | 5.08E-03 | -1.11E+00 | 1.66E-01 | 4.55E-01 | 1.16E+00 |
| TMEM2 | 1.10E-05 | 5.73E-05 | -1.12E+00 | 6.33E-09 | 2.74E-08 | -1.30E+00 | 5.38E-03 | 2.28E-02 | -1.24E+00 | 1.77E-06 | 6.78E-06 | -1.93E+00 | 7.38E-03 | 2.63E-02 | 1.05E+00 | 9.55E-02 | 2.69E-01 | 1.20E+00 | 3.04E-06 | 1.11E-05 | -1.27E+00 | 5.76E-09 | 1.58E-08 | -1.56E+00 |
| CCNQ | 1.10E-05 | 5.73E-05 | -1.46E+00 | 1.41E-02 | 6.11E-02 | 1.10E+00 | 1.27E-06 | 5.38E-06 | -1.25E+00 | 8.29E-06 | 3.17E-05 | -1.06E+00 | 1.27E-04 | 4.52E-04 | -1.46E+00 | 1.40E-02 | 3.94E-02 | -1.01E+00 | 4.32E-05 | 1.59E-04 | -1.25E+00 | 1.80E-01 | 4.95E-01 | 1.21E+00 |
| ACVR1 | 1.10E-05 | 5.73E-05 | -1.70E+00 | 2.72E-04 | 1.18E-03 | -1.31E+00 | 8.34E-04 | 3.54E-03 | -1.49E+00 | 1.90E-03 | 7.26E-03 | -1.33E+00 | 3.21E-10 | 1.14E-09 | -2.11E+00 | 1.14E-04 | 3.21E-04 | -1.22E+00 | 6.89E-22 | 2.53E-21 | -2.18E+00 | 4.53E-11 | 1.24E-10 | -1.51E+00 |
| KIF22 | 1.10E-05 | 5.75E-05 | -1.37E+00 | 1.56E-05 | 6.73E-05 | -1.27E+00 | 1.96E-03 | 8.32E-03 | -1.09E+00 | 3.64E-03 | 1.39E-02 | 1.05E+00 | 6.15E-05 | 2.19E-04 | -1.37E+00 | 3.95E-05 | 1.11E-04 | -1.39E+00 | 6.19E-05 | 2.27E-04 | -1.12E+00 | 4.59E-04 | 1.26E-03 | -1.03E+00 |
| PXMP4 | 1.10E-05 | 5.75E-05 | -1.72E+00 | 1.76E-02 | 7.60E-02 | 1.14E+00 | 7.39E-06 | 3.14E-05 | -1.64E+00 | 3.25E-04 | 1.24E-03 | -1.13E+00 | 2.61E-05 | 9.30E-05 | -1.79E+00 | 8.04E-03 | 2.27E-02 | -1.08E+00 | 1.34E-05 | 4.92E-05 | -1.11E+00 | 3.43E-02 | 9.43E-02 | 1.16E+00 |
| MAN1B1-AS1 | 1.11E-05 | 5.78E-05 | -1.32E+00 | 4.29E-05 | 1.85E-04 | -1.16E+00 | 2.41E-05 | 1.02E-04 | -1.46E+00 | 4.41E-06 | 1.68E-05 | -1.40E+00 | 1.34E-04 | 4.78E-04 | -1.93E+00 | 1.21E-03 | 3.42E-03 | -1.45E+00 | 6.33E-07 | 2.32E-06 | -1.68E+00 | 1.15E-04 | 3.17E-04 | -1.42E+00 |
| GLCE | 1.11E-05 | 5.81E-05 | -1.41E+00 | 6.03E-04 | 2.61E-03 | 1.02E+00 | 1.37E-06 | 5.84E-06 | -1.56E+00 | 2.51E-03 | 9.60E-03 | -1.02E+00 | 1.08E-03 | 3.84E-03 | -1.44E+00 | 2.66E-02 | 7.48E-02 | -1.02E+00 | 1.23E-03 | 4.52E-03 | -1.22E+00 | 2.13E-02 | 5.84E-02 | -1.08E+00 |
| KIAA1456 | 1.12E-05 | 5.83E-05 | -2.06E+00 | 4.13E-05 | 1.78E-04 | -1.87E+00 | 2.12E-07 | 9.00E-07 | -2.05E+00 | 1.39E-07 | 5.33E-07 | -2.07E+00 | 7.12E-03 | 2.54E-02 | -1.73E+00 | 6.35E-03 | 1.79E-02 | -1.93E+00 | 2.75E-08 | 1.01E-07 | -1.97E+00 | 3.58E-05 | 9.84E-05 | -1.41E+00 |
| C6orf203 | 1.13E-05 | 5.86E-05 | -1.63E+00 | 9.76E-02 | 4.22E-01 | 1.32E+00 | 1.06E-09 | 4.49E-09 | -1.72E+00 | 1.54E-05 | 5.88E-05 | -1.14E+00 | 3.98E-05 | 1.42E-04 | -1.52E+00 | 4.81E-02 | 1.36E-01 | 1.21E+00 | 1.17E-14 | 4.29E-14 | -1.77E+00 | 6.06E-04 | 1.66E-03 | -1.07E+00 |
| MRPL40 | 1.13E-05 | 5.87E-05 | -1.05E+00 | 1.76E-04 | 7.59E-04 | 1.12E+00 | 1.73E-02 | 7.35E-02 | 1.08E+00 | 2.99E-04 | 1.14E-03 | 1.07E+00 | 6.92E-06 | 2.47E-05 | -1.15E+00 | 5.90E-03 | 1.66E-02 | 1.12E+00 | 3.82E-02 | 1.40E-01 | 1.18E+00 | 1.88E-01 | 5.16E-01 | 1.24E+00 |
| AC008079.2 | 1.13E-05 | 5.87E-05 | -2.09E+00 | 2.28E-04 | 9.83E-04 | -1.45E+00 | 3.17E-04 | 1.35E-03 | -1.73E+00 | 2.37E-02 | 9.07E-02 | -1.10E+00 | 1.24E-01 | 4.41E-01 | 1.02E+00 | 3.16E-01 | 8.90E-01 | 1.36E+00 | 4.33E-03 | 1.59E-02 | -1.21E+00 | 8.64E-03 | 2.37E-02 | -1.26E+00 |
| PHF3 | 1.13E-05 | 5.87E-05 | -1.04E+00 | 8.70E-07 | 3.76E-06 | 1.03E+00 | 1.20E-04 | 5.10E-04 | -1.23E+00 | 1.28E-05 | 4.89E-05 | -1.15E+00 | 9.81E-05 | 3.50E-04 | -1.05E+00 | 1.05E-04 | 2.96E-04 | -1.09E+00 | 3.04E-07 | 1.12E-06 | -1.26E+00 | 1.23E-06 | 3.38E-06 | -1.21E+00 |
| KBTBD7 | 1.13E-05 | 5.88E-05 | -1.47E+00 | 1.47E-03 | 6.36E-03 | -1.07E+00 | 1.01E-03 | 4.28E-03 | -1.39E+00 | 3.91E-02 | 1.49E-01 | 1.02E+00 | 1.25E-04 | 4.47E-04 | -1.57E+00 | 8.81E-04 | 2.48E-03 | -1.34E+00 | 7.13E-04 | 2.62E-03 | -1.32E+00 | 3.71E-03 | 1.02E-02 | -1.18E+00 |
| TSHZ1 | 1.13E-05 | 5.89E-05 | -1.52E+00 | 4.19E-03 | 1.81E-02 | 1.05E+00 | 2.57E-06 | 1.09E-05 | -1.69E+00 | 4.72E-02 | 1.80E-01 | 1.06E+00 | 2.69E-02 | 9.61E-02 | -1.32E+00 | 3.55E-01 | 1.00E+00 | 1.45E+00 | 9.49E-07 | 3.48E-06 | -1.96E+00 | 1.32E-01 | 3.62E-01 | 1.10E+00 |
| SLC43A3 | 1.13E-05 | 5.89E-05 | -1.86E+00 | 1.54E-03 | 6.67E-03 | -1.13E+00 | 3.16E-05 | 1.34E-04 | -1.33E+00 | 1.86E-03 | 7.11E-03 | -1.03E+00 | 9.58E-19 | 3.42E-18 | -2.53E+00 | 1.63E-10 | 4.59E-10 | -1.40E+00 | 3.32E-13 | 1.22E-12 | -1.45E+00 | 7.36E-02 | 2.02E-01 | 1.05E+00 |
| EDARADD | 1.13E-05 | 5.91E-05 | -1.96E+00 | 8.77E-05 | 3.79E-04 | -1.55E+00 | 9.96E-06 | 4.23E-05 | -2.26E+00 | 4.47E-04 | 1.71E-03 | -1.54E+00 | 6.05E-04 | 2.16E-03 | -1.11E+00 | 1.03E-02 | 2.90E-02 | 1.03E+00 | 4.69E-07 | 1.72E-06 | -1.84E+00 | 2.72E-05 | 7.48E-05 | -1.54E+00 |
| XPNPEP1 | 1.14E-05 | 5.92E-05 | -1.22E+00 | 6.56E-08 | 2.83E-07 | -1.22E+00 | 6.81E-05 | 2.90E-04 | -1.13E+00 | 4.13E-06 | 1.58E-05 | -1.06E+00 | 7.94E-03 | 2.83E-02 | -1.06E+00 | 8.97E-03 | 2.53E-02 | -1.02E+00 | 7.20E-04 | 2.64E-03 | 1.05E+00 | 2.55E-02 | 7.00E-02 | 1.08E+00 |
| AC011899.2 | 1.14E-05 | 5.94E-05 | -1.71E+00 | 3.55E-03 | 1.53E-02 | -1.07E+00 | 1.35E-03 | 5.74E-03 | -1.30E+00 | 8.58E-04 | 3.28E-03 | -1.30E+00 | 2.36E-06 | 8.42E-06 | -3.48E+00 | 3.47E-05 | 9.79E-05 | -2.41E+00 | 1.99E-01 | 7.29E-01 | -1.12E+00 | 3.19E-01 | 8.75E-01 | 1.03E+00 |
| AZI2 | 1.14E-05 | 5.96E-05 | -1.35E+00 | 4.95E-05 | 2.14E-04 | -1.14E+00 | 1.11E-03 | 4.72E-03 | -1.18E+00 | 5.83E-06 | 2.23E-05 | -1.26E+00 | 3.42E-03 | 1.22E-02 | -1.31E+00 | 4.03E-02 | 1.14E-01 | -1.02E+00 | 2.84E-04 | 1.04E-03 | -1.26E+00 | 1.32E-04 | 3.62E-04 | -1.39E+00 |
| AL365436.1 | 1.15E-05 | 5.98E-05 | -1.43E+00 | 9.34E-08 | 4.03E-07 | -1.72E+00 | 2.31E-01 | 9.83E-01 | 1.07E+00 | 7.67E-03 | 2.93E-02 | -1.44E+00 | 3.58E-02 | 1.28E-01 | -1.20E+00 | 3.59E-03 | 1.01E-02 | -1.76E+00 | 3.84E-02 | 1.41E-01 | -1.21E+00 | 1.60E-02 | 4.40E-02 | -1.39E+00 |
| SLC25A40 | 1.16E-05 | 6.05E-05 | -1.50E+00 | 2.93E-03 | 1.26E-02 | 1.01E+00 | 8.63E-05 | 3.67E-04 | -1.41E+00 | 1.43E-02 | 5.45E-02 | 1.08E+00 | 6.63E-03 | 2.36E-02 | -1.09E+00 | 2.48E-02 | 7.00E-02 | 1.06E+00 | 9.50E-05 | 3.49E-04 | -1.32E+00 | 6.92E-02 | 1.90E-01 | 1.06E+00 |
| PPWD1 | 1.16E-05 | 6.06E-05 | -1.25E+00 | 1.21E-03 | 5.23E-03 | 1.13E+00 | 3.60E-09 | 1.53E-08 | -1.53E+00 | 2.73E-05 | 1.04E-04 | -1.03E+00 | 5.22E-04 | 1.86E-03 | -1.27E+00 | 1.21E-02 | 3.42E-02 | -1.05E+00 | 6.27E-09 | 2.30E-08 | -1.49E+00 | 1.48E-09 | 4.05E-09 | -1.56E+00 |
| FAM204A | 1.18E-05 | 6.13E-05 | -1.05E+00 | 3.42E-05 | 1.48E-04 | 1.07E+00 | 7.29E-03 | 3.10E-02 | 1.00E+00 | 6.81E-04 | 2.60E-03 | 1.02E+00 | 6.59E-02 | 2.35E-01 | 1.16E+00 | 9.45E-04 | 2.66E-03 | 1.01E+00 | 1.13E-01 | 4.14E-01 | 1.18E+00 | 2.26E-02 | 6.22E-02 | 1.07E+00 |
| SDHA | 1.18E-05 | 6.13E-05 | -1.14E+00 | 4.65E-03 | 2.01E-02 | 1.19E+00 | 2.72E-03 | 1.16E-02 | -1.14E+00 | 2.76E-04 | 1.05E-03 | -1.06E+00 | 3.63E-05 | 1.29E-04 | -1.30E+00 | 4.68E-03 | 1.32E-02 | -1.03E+00 | 5.65E-03 | 2.07E-02 | -1.04E+00 | 3.53E-01 | 9.70E-01 | 1.14E+00 |
| KIAA1551 | 1.18E-05 | 6.15E-05 | -1.09E+00 | 4.73E-03 | 2.04E-02 | 1.22E+00 | 9.70E-03 | 4.12E-02 | -1.04E+00 | 8.18E-02 | 3.13E-01 | 1.23E+00 | 2.42E-07 | 8.65E-07 | -1.27E+00 | 4.01E-08 | 1.13E-07 | -1.25E+00 | 1.77E-08 | 6.51E-08 | -1.36E+00 | 1.06E-03 | 2.91E-03 | -1.21E+00 |
| PRMT6 | 1.18E-05 | 6.16E-05 | -1.63E+00 | 4.33E-03 | 1.87E-02 | -1.02E+00 | 5.33E-02 | 2.26E-01 | -1.03E+00 | 7.70E-02 | 2.94E-01 | 1.70E+00 | 2.11E-05 | 7.51E-05 | -1.85E+00 | 3.21E-02 | 9.05E-02 | -1.02E+00 | 4.19E-07 | 1.54E-06 | -1.57E+00 | 1.87E-01 | 5.14E-01 | 1.19E+00 |
| FBXL3 | 1.18E-05 | 6.17E-05 | -1.16E+00 | 2.90E-06 | 1.25E-05 | -1.13E+00 | 2.59E-04 | 1.10E-03 | -1.20E+00 | 6.31E-08 | 2.41E-07 | -1.35E+00 | 6.39E-04 | 2.28E-03 | -1.19E+00 | 2.25E-04 | 6.34E-04 | -1.31E+00 | 1.73E-08 | 6.34E-08 | -1.26E+00 | 1.64E-06 | 4.49E-06 | -1.25E+00 |
| MTERF1 | 1.19E-05 | 6.22E-05 | -1.47E+00 | 4.59E-04 | 1.98E-03 | -1.17E+00 | 4.40E-02 | 1.87E-01 | -1.08E+00 | 2.27E-01 | 8.66E-01 | 1.28E+00 | 1.11E-01 | 3.97E-01 | 1.22E+00 | 1.86E-01 | 5.25E-01 | 1.23E+00 | 8.39E-03 | 3.08E-02 | -1.12E+00 | 2.65E-01 | 7.29E-01 | 1.19E+00 |
| PRR15 | 1.20E-05 | 6.23E-05 | -1.36E+00 | 2.09E-01 | 9.03E-01 | 1.59E+00 | 2.91E-02 | 1.24E-01 | -1.20E+00 | 1.90E-01 | 7.27E-01 | 1.44E+00 | 4.33E-02 | 1.54E-01 | -1.45E+00 | 3.22E-01 | 9.07E-01 | 1.39E+00 | 7.38E-03 | 2.71E-02 | -1.18E+00 | 2.06E-01 | 5.66E-01 | 1.25E+00 |
| SCAI | 1.20E-05 | 6.27E-05 | -1.08E+00 | 1.57E-01 | 6.79E-01 | 1.43E+00 | 4.20E-05 | 1.79E-04 | -1.24E+00 | 4.38E-03 | 1.67E-02 | 1.07E+00 | 4.24E-02 | 1.51E-01 | 1.18E+00 | 1.50E-01 | 4.23E-01 | 1.69E+00 | 4.18E-04 | 1.54E-03 | -1.11E+00 | 2.56E-01 | 7.04E-01 | 1.25E+00 |
| LYRM2 | 1.21E-05 | 6.29E-05 | -1.09E+00 | 5.50E-06 | 2.38E-05 | 1.02E+00 | 4.44E-06 | 1.89E-05 | -1.20E+00 | 1.88E-04 | 7.17E-04 | -1.04E+00 | 8.48E-04 | 3.02E-03 | -1.06E+00 | 3.59E-03 | 1.01E-02 | -1.03E+00 | 4.11E-02 | 1.51E-01 | 1.02E+00 | 7.79E-02 | 2.14E-01 | 1.02E+00 |
| SUOX | 1.21E-05 | 6.29E-05 | -1.50E+00 | 1.34E-02 | 5.81E-02 | 1.13E+00 | 6.75E-04 | 2.87E-03 | -1.53E+00 | 9.60E-02 | 3.67E-01 | 1.15E+00 | 1.05E-03 | 3.75E-03 | -1.80E+00 | 7.19E-02 | 2.03E-01 | 1.04E+00 | 1.36E-08 | 4.99E-08 | -1.83E+00 | 9.73E-03 | 2.67E-02 | -1.07E+00 |
| ZNF692 | 1.21E-05 | 6.29E-05 | -1.24E+00 | 6.76E-04 | 2.92E-03 | 1.06E+00 | 1.74E-08 | 7.39E-08 | -1.63E+00 | 1.00E-04 | 3.82E-04 | -1.11E+00 | 1.29E-08 | 4.59E-08 | -1.72E+00 | 1.52E-05 | 4.28E-05 | -1.26E+00 | 9.09E-08 | 3.34E-07 | -1.46E+00 | 4.33E-05 | 1.19E-04 | -1.20E+00 |
| RAB42 | 1.21E-05 | 6.30E-05 | -1.85E+00 | 8.79E-04 | 3.80E-03 | -1.24E+00 | 1.54E-02 | 6.55E-02 | -1.28E+00 | 2.17E-01 | 8.30E-01 | 1.29E+00 | 2.11E-01 | 7.51E-01 | 1.02E+00 | 2.64E-01 | 7.44E-01 | 1.56E+00 | 1.15E-01 | 4.21E-01 | -1.20E+00 | 3.45E-01 | 9.48E-01 | 1.10E+00 |
| MTFR1 | 1.21E-05 | 6.33E-05 | -1.67E+00 | 7.77E-04 | 3.36E-03 | -1.10E+00 | 6.34E-07 | 2.69E-06 | -1.72E+00 | 1.29E-03 | 4.92E-03 | 1.01E+00 | 1.74E-06 | 6.20E-06 | -1.73E+00 | 2.89E-04 | 8.16E-04 | -1.24E+00 | 2.29E-07 | 8.39E-07 | -1.61E+00 | 4.97E-02 | 1.37E-01 | 1.02E+00 |
| DDX59 | 1.23E-05 | 6.40E-05 | -1.28E+00 | 9.66E-07 | 4.18E-06 | -1.27E+00 | 2.83E-02 | 1.20E-01 | -1.15E+00 | 3.24E-04 | 1.24E-03 | -1.34E+00 | 5.17E-12 | 1.84E-11 | -2.05E+00 | 1.79E-10 | 5.05E-10 | -1.75E+00 | 3.60E-04 | 1.32E-03 | -1.28E+00 | 7.54E-05 | 2.07E-04 | -1.51E+00 |
| GNL3L | 1.23E-05 | 6.41E-05 | -1.24E+00 | 6.07E-05 | 2.62E-04 | -1.09E+00 | 1.34E-03 | 5.68E-03 | -1.22E+00 | 1.69E-03 | 6.46E-03 | -1.03E+00 | 2.31E-03 | 8.25E-03 | -1.33E+00 | 7.99E-03 | 2.25E-02 | -1.13E+00 | 6.09E-03 | 2.23E-02 | -1.07E+00 | 8.77E-03 | 2.41E-02 | -1.02E+00 |
| HK2 | 1.23E-05 | 6.42E-05 | -1.58E+00 | 3.67E-05 | 1.59E-04 | -1.33E+00 | 1.26E-04 | 5.34E-04 | -1.30E+00 | 1.11E-02 | 4.23E-02 | 1.11E+00 | 6.88E-05 | 2.45E-04 | -1.96E+00 | 1.09E-03 | 3.08E-03 | -1.56E+00 | 4.17E-04 | 1.53E-03 | -1.59E+00 | 6.01E-03 | 1.65E-02 | -1.33E+00 |
| ZNF204P | 1.23E-05 | 6.42E-05 | -1.36E+00 | 7.44E-03 | 3.21E-02 | 1.12E+00 | 1.91E-03 | 8.10E-03 | -1.66E+00 | 1.02E-02 | 3.89E-02 | -1.24E+00 | 4.14E-03 | 1.48E-02 | -1.75E+00 | 2.22E-01 | 6.27E-01 | 1.27E+00 | 1.47E-02 | 5.39E-02 | -1.46E+00 | 3.35E-02 | 9.19E-02 | -1.26E+00 |
| VPS72 | 1.23E-05 | 6.42E-05 | -1.16E+00 | 1.81E-02 | 7.80E-02 | 1.27E+00 | 2.85E-09 | 1.21E-08 | -1.39E+00 | 2.67E-04 | 1.02E-03 | 1.03E+00 | 4.95E-11 | 1.77E-10 | -1.66E+00 | 8.25E-03 | 2.33E-02 | 1.11E+00 | 2.17E-12 | 7.95E-12 | -1.50E+00 | 2.64E-03 | 7.24E-03 | -1.04E+00 |
| R3HDM4 | 1.24E-05 | 6.44E-05 | -1.10E+00 | 4.30E-05 | 1.86E-04 | 1.06E+00 | 4.61E-08 | 1.96E-07 | -1.43E+00 | 9.58E-05 | 3.66E-04 | -1.04E+00 | 3.03E-04 | 1.08E-03 | -1.17E+00 | 1.14E-02 | 3.21E-02 | 1.07E+00 | 7.39E-03 | 2.71E-02 | 1.13E+00 | 2.98E-03 | 8.19E-03 | -1.06E+00 |
| SH3BP2 | 1.24E-05 | 6.47E-05 | -1.30E+00 | 1.46E-04 | 6.31E-04 | 1.02E+00 | 1.54E-05 | 6.54E-05 | -1.40E+00 | 7.84E-04 | 2.99E-03 | -1.09E+00 | 3.06E-03 | 1.09E-02 | -1.19E+00 | 3.00E-03 | 8.44E-03 | -1.17E+00 | 6.46E-07 | 2.37E-06 | -1.43E+00 | 9.30E-04 | 2.56E-03 | -1.16E+00 |
| ZNF33A | 1.25E-05 | 6.49E-05 | -1.16E+00 | 1.10E-03 | 4.77E-03 | 1.06E+00 | 1.04E-10 | 4.42E-10 | -1.64E+00 | 3.49E-10 | 1.33E-09 | -1.44E+00 | 7.35E-10 | 2.62E-09 | -1.52E+00 | 4.55E-06 | 1.28E-05 | -1.23E+00 | 1.59E-07 | 5.85E-07 | -1.44E+00 | 7.09E-05 | 1.95E-04 | -1.24E+00 |
| ZNF514 | 1.25E-05 | 6.50E-05 | -1.19E+00 | 2.65E-02 | 1.15E-01 | 1.27E+00 | 2.07E-07 | 8.80E-07 | -1.67E+00 | 5.34E-03 | 2.04E-02 | 1.08E+00 | 4.41E-03 | 1.57E-02 | -1.15E+00 | 1.13E-01 | 3.20E-01 | 1.18E+00 | 6.53E-02 | 2.40E-01 | -1.00E+00 | 1.69E-01 | 4.63E-01 | 1.46E+00 |
| MXD4 | 1.25E-05 | 6.50E-05 | -1.01E+00 | 2.12E-08 | 9.16E-08 | -1.01E+00 | 1.41E-08 | 6.01E-08 | -1.33E+00 | 2.90E-09 | 1.11E-08 | -1.27E+00 | 1.41E-06 | 5.01E-06 | -1.15E+00 | 3.24E-06 | 9.13E-06 | -1.17E+00 | 7.17E-09 | 2.63E-08 | -1.49E+00 | 5.93E-09 | 1.63E-08 | -1.57E+00 |
| NUFIP2 | 1.25E-05 | 6.50E-05 | 1.09E+00 | 2.42E-18 | 1.05E-17 | -1.19E+00 | 4.96E-03 | 2.11E-02 | -1.03E+00 | 7.05E-10 | 2.69E-09 | -1.29E+00 | 6.04E-03 | 2.15E-02 | 1.08E+00 | 3.56E-10 | 1.00E-09 | -1.35E+00 | 6.24E-03 | 2.29E-02 | -1.04E+00 | 9.12E-04 | 2.51E-03 | -1.12E+00 |
| NDUFA5P11 | 1.25E-05 | 6.51E-05 | -1.65E+00 | 9.44E-08 | 4.08E-07 | -2.19E+00 | 1.41E-02 | 6.00E-02 | -1.12E+00 | 1.43E-03 | 5.46E-03 | -1.21E+00 | 2.11E-03 | 7.52E-03 | -1.45E+00 | 1.47E-02 | 4.13E-02 | -1.07E+00 | 1.48E-01 | 5.44E-01 | 1.09E+00 | 4.04E-03 | 1.11E-02 | -1.27E+00 |
| CD300A | 1.25E-05 | 6.51E-05 | -1.10E+00 | 1.44E-01 | 6.24E-01 | 1.39E+00 | 6.01E-04 | 2.55E-03 | -1.09E+00 | 2.22E-01 | 8.46E-01 | 1.33E+00 | 2.83E-04 | 1.01E-03 | -1.15E+00 | 4.18E-03 | 1.18E-02 | -1.03E+00 | 1.14E-01 | 4.18E-01 | 1.05E+00 | 2.85E-01 | 7.84E-01 | 1.16E+00 |
| XPNPEP3 | 1.25E-05 | 6.51E-05 | -1.22E+00 | 1.11E-03 | 4.79E-03 | 1.05E+00 | 8.98E-03 | 3.82E-02 | -1.07E+00 | 3.79E-04 | 1.45E-03 | -1.01E+00 | 4.11E-07 | 1.46E-06 | -1.60E+00 | 2.82E-05 | 7.95E-05 | -1.34E+00 | 2.52E-01 | 9.27E-01 | 1.19E+00 | 3.55E-01 | 9.75E-01 | 1.30E+00 |
| LINC01278 | 1.25E-05 | 6.52E-05 | -1.12E+00 | 7.56E-03 | 3.26E-02 | 1.21E+00 | 2.12E-04 | 9.00E-04 | -1.34E+00 | 1.02E-01 | 3.89E-01 | 1.25E+00 | 2.75E-01 | 9.79E-01 | 1.39E+00 | 7.82E-02 | 2.21E-01 | 1.80E+00 | 2.04E-02 | 7.49E-02 | -1.05E+00 | 1.49E-01 | 4.09E-01 | 1.17E+00 |
| EHHADH | 1.26E-05 | 6.55E-05 | -1.74E+00 | 9.33E-02 | 4.03E-01 | 1.34E+00 | 4.36E-04 | 1.85E-03 | -1.47E+00 | 4.14E-03 | 1.58E-02 | -1.17E+00 | 5.69E-03 | 2.03E-02 | -1.46E+00 | 1.06E-01 | 2.99E-01 | 1.00E+00 | 1.17E-07 | 4.28E-07 | -1.66E+00 | 2.44E-06 | 6.70E-06 | -1.49E+00 |
| YPEL2 | 1.26E-05 | 6.59E-05 | -1.16E+00 | 6.40E-03 | 2.76E-02 | 1.20E+00 | 5.32E-10 | 2.26E-09 | -1.57E+00 | 9.07E-07 | 3.46E-06 | -1.17E+00 | 6.31E-06 | 2.25E-05 | -1.31E+00 | 9.94E-03 | 2.80E-02 | 1.04E+00 | 1.70E-04 | 6.24E-04 | -1.21E+00 | 1.82E-04 | 5.00E-04 | -1.16E+00 |
| SAMD13 | 1.27E-05 | 6.60E-05 | -1.37E+00 | 4.70E-05 | 2.03E-04 | -1.16E+00 | 4.09E-06 | 1.74E-05 | -1.48E+00 | 1.09E-05 | 4.16E-05 | -1.27E+00 | 6.31E-02 | 2.25E-01 | 1.13E+00 | 2.66E-02 | 7.50E-02 | 1.08E+00 | 1.21E-01 | 4.45E-01 | -1.00E+00 | 1.03E-01 | 2.82E-01 | -1.05E+00 |
| TTBK2 | 1.27E-05 | 6.60E-05 | -1.35E+00 | 2.62E-05 | 1.13E-04 | -1.19E+00 | 1.16E-09 | 4.94E-09 | -1.80E+00 | 1.58E-07 | 6.05E-07 | -1.45E+00 | 3.18E-02 | 1.13E-01 | 1.14E+00 | 1.13E-01 | 3.17E-01 | 1.23E+00 | 3.38E-03 | 1.24E-02 | -1.23E+00 | 8.01E-04 | 2.20E-03 | -1.43E+00 |
| PHIP | 1.27E-05 | 6.61E-05 | -1.08E+00 | 6.73E-04 | 2.91E-03 | 1.10E+00 | 8.21E-02 | 3.49E-01 | 1.05E+00 | 3.80E-02 | 1.45E-01 | 1.08E+00 | 7.40E-03 | 2.64E-02 | 1.06E+00 | 6.27E-05 | 1.77E-04 | -1.05E+00 | 1.34E-04 | 4.90E-04 | -1.17E+00 | 8.25E-03 | 2.27E-02 | -1.12E+00 |
| CSNK1G1 | 1.27E-05 | 6.62E-05 | -1.25E+00 | 3.28E-04 | 1.42E-03 | 1.02E+00 | 1.14E-05 | 4.85E-05 | -1.26E+00 | 1.71E-03 | 6.54E-03 | 1.07E+00 | 4.93E-05 | 1.76E-04 | -1.37E+00 | 3.31E-05 | 9.33E-05 | -1.26E+00 | 1.42E-07 | 5.20E-07 | -1.42E+00 | 1.17E-03 | 3.21E-03 | -1.13E+00 |
| ATP5EP2 | 1.27E-05 | 6.62E-05 | -1.11E+00 | 4.60E-52 | 1.99E-51 | -1.75E+00 | 2.98E-02 | 1.27E-01 | -1.06E+00 | 3.30E-21 | 1.26E-20 | -1.75E+00 | 4.20E-03 | 1.50E-02 | -1.15E+00 | 1.72E-20 | 4.84E-20 | -1.83E+00 | 2.05E-02 | 7.53E-02 | -1.14E+00 | 1.28E-12 | 3.51E-12 | -1.91E+00 |
| NBDY | 1.27E-05 | 6.63E-05 | -1.01E+00 | 2.85E-05 | 1.23E-04 | 1.09E+00 | 2.02E-01 | 8.59E-01 | 1.19E+00 | 2.52E-01 | 9.63E-01 | 1.28E+00 | 6.92E-03 | 2.47E-02 | 1.12E+00 | 3.27E-02 | 9.23E-02 | 1.10E+00 | 1.93E-01 | 7.09E-01 | 1.22E+00 | 1.89E-01 | 5.19E-01 | 1.22E+00 |
| FASTK | 1.27E-05 | 6.63E-05 | -1.17E+00 | 1.51E-03 | 6.54E-03 | 1.10E+00 | 2.82E-07 | 1.20E-06 | -1.42E+00 | 1.25E-04 | 4.78E-04 | -1.07E+00 | 1.68E-06 | 5.98E-06 | -1.48E+00 | 3.38E-05 | 9.52E-05 | -1.32E+00 | 3.38E-08 | 1.24E-07 | -1.33E+00 | 9.03E-08 | 2.48E-07 | -1.31E+00 |
| RIMKLB | 1.27E-05 | 6.63E-05 | -1.17E+00 | 2.74E-03 | 1.18E-02 | 1.16E+00 | 2.01E-04 | 8.53E-04 | -1.23E+00 | 3.35E-03 | 1.28E-02 | 1.04E+00 | 1.18E-02 | 4.22E-02 | 1.08E+00 | 3.42E-03 | 9.63E-03 | 1.01E+00 | 2.33E-03 | 8.54E-03 | 1.02E+00 | 4.18E-03 | 1.15E-02 | -1.03E+00 |
| MAPKAPK5-AS1 | 1.27E-05 | 6.64E-05 | -1.08E+00 | 3.93E-03 | 1.70E-02 | 1.19E+00 | 1.45E-04 | 6.17E-04 | -1.13E+00 | 2.26E-03 | 8.65E-03 | 1.06E+00 | 1.05E-03 | 3.74E-03 | -1.04E+00 | 9.18E-02 | 2.59E-01 | 1.21E+00 | 1.18E-01 | 4.35E-01 | 1.15E+00 | 1.89E-01 | 5.19E-01 | 1.22E+00 |
| TCTN3 | 1.28E-05 | 6.66E-05 | -1.12E+00 | 4.21E-03 | 1.82E-02 | 1.21E+00 | 5.44E-07 | 2.31E-06 | -1.36E+00 | 1.02E-03 | 3.91E-03 | 1.02E+00 | 7.79E-05 | 2.78E-04 | -1.33E+00 | 4.62E-02 | 1.30E-01 | 1.13E+00 | 5.77E-03 | 2.12E-02 | -1.07E+00 | 5.43E-02 | 1.49E-01 | 1.11E+00 |
| ZNF543 | 1.28E-05 | 6.70E-05 | -1.23E+00 | 1.35E-05 | 5.84E-05 | -1.17E+00 | 3.54E-03 | 1.51E-02 | -1.27E+00 | 3.09E-03 | 1.18E-02 | -1.19E+00 | 1.39E-02 | 4.96E-02 | 1.01E+00 | 1.34E-03 | 3.79E-03 | -1.31E+00 | 3.24E-03 | 1.19E-02 | -1.37E+00 | 8.54E-04 | 2.35E-03 | -1.60E+00 |
| DST | 1.29E-05 | 6.70E-05 | -1.02E+00 | 6.46E-04 | 2.79E-03 | 1.18E+00 | 1.33E-04 | 5.65E-04 | -1.18E+00 | 1.95E-02 | 7.44E-02 | 1.04E+00 | 1.45E-06 | 5.18E-06 | -1.39E+00 | 6.81E-06 | 1.92E-05 | -1.33E+00 | 4.52E-06 | 1.66E-05 | -1.51E+00 | 4.02E-03 | 1.10E-02 | -1.19E+00 |
| AARS | 1.29E-05 | 6.71E-05 | -1.70E+00 | 1.31E-01 | 5.66E-01 | 1.31E+00 | 2.64E-03 | 1.12E-02 | -1.42E+00 | 2.01E-01 | 7.68E-01 | 1.22E+00 | 2.01E-04 | 7.15E-04 | -1.57E+00 | 3.65E-02 | 1.03E-01 | -1.03E+00 | 6.93E-05 | 2.54E-04 | -1.19E+00 | 2.14E-01 | 5.89E-01 | 1.08E+00 |
| AC023157.2 | 1.29E-05 | 6.72E-05 | -1.96E+00 | 1.24E-02 | 5.38E-02 | 1.09E+00 | 1.22E-04 | 5.17E-04 | -1.52E+00 | 1.24E-03 | 4.75E-03 | -1.12E+00 | 3.71E-02 | 1.32E-01 | -1.37E+00 | 2.69E-01 | 7.57E-01 | 1.29E+00 | 1.86E-08 | 6.83E-08 | -2.28E+00 | 4.26E-05 | 1.17E-04 | -1.59E+00 |
| LONP2 | 1.29E-05 | 6.72E-05 | -1.07E+00 | 1.06E-02 | 4.57E-02 | 1.32E+00 | 3.09E-02 | 1.31E-01 | -1.03E+00 | 1.84E-01 | 7.04E-01 | 1.42E+00 | 1.06E-03 | 3.78E-03 | -1.06E+00 | 3.42E-01 | 9.64E-01 | 1.31E+00 | 8.34E-05 | 3.06E-04 | -1.32E+00 | 4.00E-02 | 1.10E-01 | -1.04E+00 |
| SIGLEC14 | 1.29E-05 | 6.72E-05 | -1.85E+00 | 9.62E-06 | 4.16E-05 | -1.73E+00 | 7.63E-02 | 3.24E-01 | -1.04E+00 | 2.12E-03 | 8.12E-03 | -1.47E+00 | 6.74E-04 | 2.40E-03 | -1.79E+00 | 2.84E-03 | 8.01E-03 | -1.40E+00 | 8.49E-02 | 3.12E-01 | -1.50E+00 | 1.58E-01 | 4.35E-01 | -1.34E+00 |
| GRHPR | 1.29E-05 | 6.74E-05 | -1.03E+00 | 4.44E-05 | 1.92E-04 | 1.11E+00 | 8.83E-02 | 3.75E-01 | 1.08E+00 | 1.11E-02 | 4.23E-02 | 1.07E+00 | 1.80E-03 | 6.40E-03 | -1.00E+00 | 9.59E-03 | 2.70E-02 | 1.05E+00 | 1.34E-01 | 4.92E-01 | 1.19E+00 | 2.95E-01 | 8.12E-01 | 1.21E+00 |
| LMAN2 | 1.29E-05 | 6.74E-05 | -1.01E+00 | 7.52E-06 | 3.25E-05 | 1.12E+00 | 2.36E-02 | 1.00E-01 | -1.01E+00 | 1.15E-02 | 4.40E-02 | 1.06E+00 | 1.14E-03 | 4.08E-03 | -1.06E+00 | 2.12E-02 | 5.97E-02 | 1.11E+00 | 3.88E-02 | 1.42E-01 | 1.01E+00 | 2.55E-01 | 7.00E-01 | 1.13E+00 |
| ABI3BP | 1.29E-05 | 6.74E-05 | -1.38E+00 | 3.73E-05 | 1.61E-04 | -1.36E+00 | 7.05E-02 | 2.99E-01 | 1.58E+00 | 1.81E-01 | 6.92E-01 | 1.30E+00 | 2.80E-01 | 9.99E-01 | 1.06E+00 | 2.90E-01 | 8.17E-01 | 1.30E+00 | 2.36E-01 | 8.67E-01 | 1.05E+00 | 3.09E-01 | 8.49E-01 | 1.04E+00 |
| CSTB | 1.30E-05 | 6.75E-05 | -1.01E+00 | 1.15E-05 | 4.95E-05 | 1.07E+00 | 2.23E-01 | 9.46E-01 | 1.18E+00 | 2.60E-01 | 9.92E-01 | 1.21E+00 | 1.36E-07 | 4.84E-07 | -1.16E+00 | 1.31E-03 | 3.71E-03 | -1.01E+00 | 8.25E-03 | 3.03E-02 | -1.01E+00 | 2.23E-02 | 6.14E-02 | 1.02E+00 |
| ADAL | 1.30E-05 | 6.77E-05 | -1.41E+00 | 1.66E-03 | 7.18E-03 | 1.02E+00 | 3.67E-03 | 1.56E-02 | -1.19E+00 | 6.88E-03 | 2.63E-02 | -1.04E+00 | 1.13E-04 | 4.02E-04 | -1.34E+00 | 6.75E-03 | 1.90E-02 | -1.06E+00 | 2.13E-02 | 7.82E-02 | 1.03E+00 | 3.15E-01 | 8.65E-01 | 1.27E+00 |
| PPP3CC | 1.30E-05 | 6.77E-05 | -1.30E+00 | 3.68E-06 | 1.59E-05 | -1.27E+00 | 8.47E-04 | 3.60E-03 | -1.34E+00 | 1.68E-04 | 6.43E-04 | -1.30E+00 | 2.65E-02 | 9.44E-02 | -1.06E+00 | 1.21E-02 | 3.40E-02 | -1.14E+00 | 1.54E-04 | 5.66E-04 | -1.35E+00 | 1.35E-05 | 3.72E-05 | -1.66E+00 |
| CRISPLD1 | 1.30E-05 | 6.79E-05 | -1.50E+00 | 3.27E-02 | 1.41E-01 | 1.19E+00 | 4.73E-02 | 2.01E-01 | -1.11E+00 | 1.54E-01 | 5.88E-01 | 1.20E+00 | 1.45E-05 | 5.16E-05 | -1.54E+00 | 5.81E-05 | 1.64E-04 | -1.44E+00 | 4.47E-03 | 1.64E-02 | -1.21E+00 | 1.30E-02 | 3.56E-02 | -1.11E+00 |
| MAGI2-AS3 | 1.30E-05 | 6.79E-05 | -1.10E+00 | 1.96E-02 | 8.47E-02 | 1.25E+00 | 1.08E-03 | 4.57E-03 | -1.22E+00 | 3.83E-02 | 1.46E-01 | 1.12E+00 | 4.33E-02 | 1.54E-01 | 1.14E+00 | 2.54E-01 | 7.15E-01 | 1.41E+00 | 1.10E-01 | 4.02E-01 | 1.10E+00 | 2.66E-01 | 7.31E-01 | 1.24E+00 |
| SESTD1 | 1.31E-05 | 6.80E-05 | -1.24E+00 | 2.26E-02 | 9.75E-02 | 1.24E+00 | 3.95E-05 | 1.68E-04 | -1.32E+00 | 3.01E-07 | 1.15E-06 | -1.27E+00 | 2.42E-01 | 8.64E-01 | 1.19E+00 | 1.10E-01 | 3.09E-01 | -1.02E+00 | 6.01E-03 | 2.21E-02 | 1.01E+00 | 4.11E-03 | 1.13E-02 | -1.01E+00 |
| AL357060.2 | 1.31E-05 | 6.82E-05 | 2.40E+00 | 4.10E-02 | 1.77E-01 | 2.00E+00 | 5.68E-03 | 2.41E-02 | 1.89E+00 | 2.56E-01 | 9.79E-01 | 1.35E+00 | 2.61E-01 | 9.31E-01 | 1.51E+00 | 1.08E-01 | 3.04E-01 | -1.02E+00 | 1.79E-01 | 6.56E-01 | 1.06E+00 | 2.03E-01 | 5.58E-01 | -1.03E+00 |
| ZNF581 | 1.31E-05 | 6.82E-05 | -1.09E+00 | 3.04E-04 | 1.31E-03 | 1.10E+00 | 3.60E-06 | 1.53E-05 | -1.29E+00 | 1.73E-05 | 6.61E-05 | -1.12E+00 | 1.33E-07 | 4.76E-07 | -1.32E+00 | 4.94E-06 | 1.39E-05 | -1.13E+00 | 2.95E-05 | 1.08E-04 | -1.19E+00 | 1.07E-03 | 2.93E-03 | -1.12E+00 |
| VPS45 | 1.31E-05 | 6.84E-05 | -1.49E+00 | 1.86E-03 | 8.03E-03 | -1.03E+00 | 2.43E-03 | 1.03E-02 | -1.21E+00 | 3.43E-02 | 1.31E-01 | 1.12E+00 | 5.86E-03 | 2.09E-02 | -1.21E+00 | 1.70E-01 | 4.79E-01 | 1.28E+00 | 2.05E-05 | 7.52E-05 | -1.25E+00 | 1.26E-02 | 3.47E-02 | -1.00E+00 |
| PTAR1 | 1.31E-05 | 6.85E-05 | -1.00E+00 | 6.82E-03 | 2.95E-02 | 1.29E+00 | 1.74E-03 | 7.39E-03 | -1.06E+00 | 6.26E-03 | 2.39E-02 | 1.06E+00 | 2.71E-01 | 9.67E-01 | 1.38E+00 | 2.53E-01 | 7.14E-01 | 1.36E+00 | 3.03E-02 | 1.11E-01 | -1.02E+00 | 2.08E-01 | 5.70E-01 | 1.10E+00 |
| GDPD1 | 1.32E-05 | 6.86E-05 | -1.47E+00 | 1.15E-03 | 4.99E-03 | -1.05E+00 | 1.07E-04 | 4.56E-04 | -1.96E+00 | 4.30E-03 | 1.64E-02 | -1.25E+00 | 4.57E-06 | 1.63E-05 | -1.39E+00 | 2.37E-01 | 6.68E-01 | 1.37E+00 | 1.32E-29 | 4.85E-29 | -2.28E+00 | 3.04E-10 | 8.35E-10 | -1.36E+00 |
| STARD4-AS1 | 1.32E-05 | 6.88E-05 | -1.17E+00 | 1.15E-01 | 4.95E-01 | 1.37E+00 | 1.57E-06 | 6.65E-06 | -1.33E+00 | 4.39E-04 | 1.68E-03 | 1.02E+00 | 1.54E-05 | 5.50E-05 | -1.17E+00 | 7.57E-03 | 2.13E-02 | 1.11E+00 | 5.07E-07 | 1.86E-06 | -1.93E+00 | 1.15E-02 | 3.15E-02 | -1.12E+00 |
| LRFN1 | 1.33E-05 | 6.91E-05 | -1.71E+00 | 1.20E-02 | 5.17E-02 | 1.09E+00 | 6.69E-04 | 2.84E-03 | -1.52E+00 | 3.06E-02 | 1.17E-01 | 1.00E+00 | 1.19E-02 | 4.23E-02 | -1.89E+00 | 4.19E-02 | 1.18E-01 | -1.52E+00 | 2.52E-02 | 9.24E-02 | -1.33E+00 | 7.85E-02 | 2.16E-01 | -1.10E+00 |
| DACT1 | 1.33E-05 | 6.94E-05 | -1.65E+00 | 7.86E-05 | 3.40E-04 | -1.37E+00 | 1.43E-01 | 6.10E-01 | 1.24E+00 | 1.52E-01 | 5.79E-01 | 1.46E+00 | 1.52E-01 | 5.43E-01 | -1.09E+00 | 1.29E-01 | 3.64E-01 | -1.47E+00 | 2.57E-01 | 9.43E-01 | 1.51E+00 | 1.92E-01 | 5.27E-01 | 1.06E+00 |
| AL590682.1 | 1.33E-05 | 6.95E-05 | -1.10E+00 | 1.30E-61 | 5.61E-61 | -2.06E+00 | 6.89E-02 | 2.93E-01 | -1.09E+00 | 2.47E-22 | 9.44E-22 | -2.00E+00 | 7.81E-05 | 2.79E-04 | -1.16E+00 | 1.41E-24 | 3.98E-24 | -1.92E+00 | 1.31E-04 | 4.80E-04 | -1.27E+00 | 1.95E-19 | 5.35E-19 | -2.08E+00 |
| FBXO11 | 1.34E-05 | 6.96E-05 | -1.14E+00 | 5.28E-10 | 2.28E-09 | -1.38E+00 | 3.99E-02 | 1.69E-01 | 1.00E+00 | 7.19E-03 | 2.75E-02 | 1.04E+00 | 3.48E-02 | 1.24E-01 | 1.07E+00 | 1.35E-02 | 3.82E-02 | 1.01E+00 | 1.02E-02 | 3.74E-02 | -1.02E+00 | 3.94E-03 | 1.08E-02 | -1.11E+00 |
| SLC52A3 | 1.34E-05 | 6.97E-05 | -2.02E+00 | 4.00E-03 | 1.73E-02 | -1.18E+00 | 1.81E-04 | 7.67E-04 | -1.77E+00 | 1.92E-03 | 7.35E-03 | -1.39E+00 | 4.17E-03 | 1.49E-02 | -2.00E+00 | 3.41E-02 | 9.61E-02 | -1.31E+00 | 7.82E-02 | 2.87E-01 | -1.12E+00 | 1.34E-01 | 3.67E-01 | 1.66E+00 |
| PSMD8P1 | 1.34E-05 | 6.98E-05 | -1.73E+00 | 5.09E-06 | 2.20E-05 | -1.68E+00 | 4.46E-04 | 1.89E-03 | -1.39E+00 | 1.01E-05 | 3.87E-05 | -1.55E+00 | 6.35E-02 | 2.26E-01 | -1.02E+00 | 6.98E-02 | 1.97E-01 | -1.08E+00 | 1.97E-03 | 7.22E-03 | -1.16E+00 | 4.39E-03 | 1.20E-02 | -1.20E+00 |
| PPP2CB | 1.34E-05 | 6.98E-05 | -1.22E+00 | 1.50E-03 | 6.47E-03 | 1.09E+00 | 6.74E-06 | 2.86E-05 | -1.50E+00 | 5.74E-05 | 2.19E-04 | -1.23E+00 | 6.58E-10 | 2.34E-09 | -1.87E+00 | 6.41E-05 | 1.81E-04 | -1.15E+00 | 2.60E-15 | 9.54E-15 | -1.89E+00 | 5.22E-09 | 1.43E-08 | -1.49E+00 |
| EBLN2 | 1.34E-05 | 6.99E-05 | -1.31E+00 | 1.92E-05 | 8.28E-05 | -1.23E+00 | 3.03E-02 | 1.29E-01 | 1.03E+00 | 4.46E-03 | 1.70E-02 | -1.01E+00 | 3.77E-03 | 1.35E-02 | -1.14E+00 | 1.71E-03 | 4.82E-03 | -1.21E+00 | 9.17E-03 | 3.37E-02 | -1.16E+00 | 1.17E-02 | 3.21E-02 | -1.20E+00 |
| TAPT1-AS1 | 1.34E-05 | 7.01E-05 | -1.57E+00 | 8.00E-04 | 3.46E-03 | -1.07E+00 | 4.96E-03 | 2.11E-02 | -1.26E+00 | 3.90E-02 | 1.49E-01 | 1.07E+00 | 3.06E-03 | 1.09E-02 | -2.18E+00 | 2.41E-01 | 6.81E-01 | 1.10E+00 | 1.35E-01 | 4.97E-01 | 1.06E+00 | 1.59E-01 | 4.38E-01 | 1.12E+00 |
| PEX11B | 1.35E-05 | 7.02E-05 | -1.95E+00 | 2.15E-01 | 9.30E-01 | 1.34E+00 | 1.54E-07 | 6.53E-07 | -1.93E+00 | 2.51E-02 | 9.60E-02 | 1.17E+00 | 5.49E-07 | 1.96E-06 | -2.24E+00 | 1.18E-02 | 3.32E-02 | -1.01E+00 | 1.32E-08 | 4.85E-08 | -1.75E+00 | 1.38E-02 | 3.79E-02 | -1.02E+00 |
| SNORA25 | 1.35E-05 | 7.04E-05 | -1.83E+00 | 3.12E-04 | 1.35E-03 | -1.40E+00 | 4.83E-10 | 2.05E-09 | -2.80E+00 | 1.55E-07 | 5.91E-07 | -1.84E+00 | 1.61E-05 | 5.73E-05 | -2.49E+00 | 7.02E-05 | 1.98E-04 | -2.00E+00 | 6.08E-04 | 2.23E-03 | -1.76E+00 | 2.04E-03 | 5.61E-03 | -1.64E+00 |
| SLC35C1 | 1.36E-05 | 7.07E-05 | -1.75E+00 | 1.98E-03 | 8.57E-03 | -1.12E+00 | 2.40E-09 | 1.02E-08 | -1.90E+00 | 7.55E-07 | 2.88E-06 | -1.32E+00 | 1.95E-01 | 6.95E-01 | 1.14E+00 | 2.76E-01 | 7.79E-01 | 1.21E+00 | 4.24E-04 | 1.56E-03 | -1.54E+00 | 2.18E-02 | 6.00E-02 | -1.15E+00 |
| GALC | 1.36E-05 | 7.07E-05 | -1.87E+00 | 2.08E-04 | 8.98E-04 | -1.39E+00 | 3.71E-04 | 1.58E-03 | -1.46E+00 | 3.75E-03 | 1.43E-02 | -1.06E+00 | 2.03E-07 | 7.23E-07 | -1.93E+00 | 8.55E-04 | 2.41E-03 | -1.28E+00 | 3.14E-08 | 1.15E-07 | -1.22E+00 | 5.28E-11 | 1.45E-10 | -1.32E+00 |
| AL359075.1 | 1.36E-05 | 7.08E-05 | -1.77E+00 | 1.57E-03 | 6.76E-03 | -1.12E+00 | 5.91E-03 | 2.51E-02 | -1.35E+00 | 1.67E-02 | 6.38E-02 | -1.07E+00 | 4.14E-03 | 1.48E-02 | -1.51E+00 | 3.13E-01 | 8.82E-01 | 1.33E+00 | 8.92E-02 | 3.27E-01 | -1.13E+00 | 3.63E-01 | 9.98E-01 | 1.21E+00 |
| APBB2 | 1.36E-05 | 7.09E-05 | -1.76E+00 | 1.24E-03 | 5.37E-03 | -1.19E+00 | 3.19E-04 | 1.35E-03 | -1.56E+00 | 1.10E-03 | 4.19E-03 | -1.43E+00 | 8.23E-02 | 2.94E-01 | -1.48E+00 | 1.08E-01 | 3.05E-01 | -1.55E+00 | 2.49E-01 | 9.15E-01 | 1.09E+00 | 2.04E-01 | 5.61E-01 | 1.39E+00 |
| CTNS | 1.36E-05 | 7.09E-05 | -2.05E+00 | 1.35E-02 | 5.82E-02 | -1.05E+00 | 6.26E-05 | 2.66E-04 | -1.68E+00 | 2.94E-02 | 1.12E-01 | 1.09E+00 | 1.66E-12 | 5.93E-12 | -2.02E+00 | 2.98E-06 | 8.40E-06 | -1.22E+00 | 2.33E-01 | 8.56E-01 | 1.31E+00 | 3.06E-01 | 8.39E-01 | 1.45E+00 |
| AC114495.2 | 1.36E-05 | 7.09E-05 | -1.80E+00 | 1.75E-04 | 7.55E-04 | -1.40E+00 | 4.12E-02 | 1.75E-01 | -1.28E+00 | 3.35E-02 | 1.28E-01 | -1.16E+00 | 1.28E-03 | 4.57E-03 | -1.50E+00 | 2.63E-02 | 7.42E-02 | -1.14E+00 | 3.93E-03 | 1.44E-02 | -1.44E+00 | 3.54E-01 | 9.72E-01 | 1.14E+00 |
| SNW1 | 1.36E-05 | 7.11E-05 | 1.05E+00 | 2.13E-03 | 9.22E-03 | 1.33E+00 | 1.50E-03 | 6.36E-03 | -1.05E+00 | 8.43E-04 | 3.22E-03 | 1.04E+00 | 1.43E-05 | 5.10E-05 | -1.05E+00 | 1.10E-01 | 3.11E-01 | 1.33E+00 | 7.90E-02 | 2.90E-01 | 1.03E+00 | 2.34E-01 | 6.44E-01 | 1.13E+00 |
| CACNA1A | 1.37E-05 | 7.13E-05 | -1.20E+00 | 1.55E-04 | 6.70E-04 | 1.01E+00 | 8.40E-06 | 3.57E-05 | -1.44E+00 | 1.71E-05 | 6.54E-05 | -1.19E+00 | 7.44E-07 | 2.65E-06 | -1.58E+00 | 4.06E-04 | 1.14E-03 | -1.16E+00 | 2.84E-04 | 1.04E-03 | -1.29E+00 | 3.10E-03 | 8.52E-03 | -1.18E+00 |
| COL6A2 | 1.37E-05 | 7.14E-05 | -1.09E+00 | 9.39E-03 | 4.06E-02 | 1.29E+00 | 8.03E-02 | 3.41E-01 | 1.06E+00 | 1.80E-01 | 6.88E-01 | 1.53E+00 | 3.80E-02 | 1.36E-01 | -1.13E+00 | 7.99E-03 | 2.25E-02 | -1.27E+00 | 1.71E-01 | 6.28E-01 | 1.52E+00 | 3.06E-01 | 8.41E-01 | 1.41E+00 |
| KIF13A | 1.37E-05 | 7.15E-05 | -1.09E+00 | 1.24E-05 | 5.34E-05 | 1.01E+00 | 3.22E-03 | 1.37E-02 | -1.22E+00 | 4.14E-03 | 1.58E-02 | -1.04E+00 | 2.77E-06 | 9.88E-06 | -1.37E+00 | 1.50E-04 | 4.22E-04 | -1.20E+00 | 1.96E-10 | 7.18E-10 | -1.55E+00 | 1.03E-07 | 2.82E-07 | -1.41E+00 |

| | | | | | | | | | | | | | | | | | | | | | | | | |
|---|---|---|---|---|---|---|---|---|---|---|---|---|---|---|---|---|---|---|---|---|---|---|---|---|
| AC118344.2 | 1.37E-05 | 7.16E-05 | -1.66E+00 | 3.61E-04 | 1.56E-03 | -1.22E+00 | 8.07E-02 | 3.43E-01 | -1.15E+00 | 1.19E-01 | 4.56E-01 | 1.17E+00 | 3.06E-03 | 1.09E-02 | -1.59E+00 | 7.45E-02 | 2.10E-01 | -1.18E+00 | 8.31E-04 | 3.05E-03 | -1.86E+00 | 9.07E-03 | 2.49E-02 | -1.33E+00 |
| CLN8 | 1.38E-05 | 7.18E-05 | -1.21E+00 | 2.55E-04 | 1.10E-03 | -1.02E+00 | 2.74E-02 | 1.17E-01 | 1.02E+00 | 1.94E-01 | 7.40E-01 | 1.34E+00 | 8.22E-05 | 2.93E-04 | -1.58E+00 | 1.30E-04 | 3.68E-04 | -1.52E+00 | 1.11E-07 | 4.06E-07 | -1.56E+00 | 1.59E-04 | 4.37E-04 | -1.31E+00 |
| IFFO1 | 1.38E-05 | 7.18E-05 | -1.30E+00 | 5.98E-04 | 2.58E-03 | 1.00E+00 | 1.27E-02 | 5.40E-02 | -1.16E+00 | 3.28E-02 | 1.25E-01 | 1.02E+00 | 2.09E-04 | 7.45E-04 | -1.34E+00 | 5.26E-03 | 1.48E-02 | 1.02E+00 | 5.18E-02 | 1.90E-01 | -1.01E+00 | 7.35E-02 | 2.02E-01 | -1.07E+00 |
| FCHSD2 | 1.38E-05 | 7.18E-05 | -1.22E+00 | 1.32E-03 | 5.71E-03 | 1.07E+00 | 1.22E-06 | 5.18E-06 | -1.43E+00 | 4.28E-06 | 1.64E-05 | -1.18E+00 | 4.31E-03 | 1.54E-02 | -1.12E+00 | 1.07E-02 | 3.03E-02 | -1.01E+00 | 3.07E-06 | 1.13E-05 | -1.38E+00 | 5.20E-04 | 1.43E-03 | -1.13E+00 |
| MED8 | 1.38E-05 | 7.21E-05 | -1.19E+00 | 2.93E-02 | 1.27E-01 | 1.30E+00 | 9.15E-05 | 3.89E-04 | -1.18E+00 | 2.72E-04 | 1.04E-03 | -1.03E+00 | 4.83E-07 | 1.72E-06 | -1.38E+00 | 1.58E-05 | 4.46E-05 | -1.20E+00 | 1.77E-09 | 6.48E-09 | -1.39E+00 | 1.94E-04 | 5.33E-04 | -1.18E+00 |
| IK | 1.39E-05 | 7.26E-05 | 1.02E+00 | 5.11E-03 | 2.21E-02 | 1.33E+00 | 2.28E-01 | 9.69E-01 | 1.23E+00 | 2.15E-01 | 8.21E-01 | 1.34E+00 | 1.22E-02 | 4.36E-02 | 1.03E+00 | 2.09E-01 | 5.88E-01 | 1.29E+00 | 1.57E-01 | 5.75E-01 | 1.18E+00 | 3.51E-01 | 9.65E-01 | 1.23E+00 |
| UBA6 | 1.39E-05 | 7.26E-05 | -1.18E+00 | 2.30E-04 | 9.93E-04 | 1.04E+00 | 2.62E-04 | 1.11E-03 | -1.16E+00 | 3.53E-06 | 1.35E-05 | -1.09E+00 | 2.97E-04 | 1.06E-03 | -1.02E+00 | 7.58E-03 | 2.14E-02 | 1.11E+00 | 5.51E-08 | 2.02E-07 | -1.26E+00 | 3.64E-05 | 9.99E-05 | -1.15E+00 |
| FAM120C | 1.40E-05 | 7.31E-05 | -1.28E+00 | 4.90E-04 | 2.12E-03 | -1.00E+00 | 1.15E-02 | 4.91E-02 | -1.17E+00 | 1.73E-01 | 6.61E-01 | 1.23E+00 | 2.07E-04 | 7.39E-04 | -1.63E+00 | 3.88E-02 | 1.09E-01 | -1.06E+00 | 9.35E-04 | 3.43E-03 | -1.15E+00 | 6.93E-02 | 1.90E-01 | 1.07E+00 |
| PABPC1P10 | 1.41E-05 | 7.33E-05 | -1.36E+00 | 1.10E-07 | 4.76E-07 | -1.69E+00 | 1.36E-12 | 5.78E-12 | -2.21E+00 | 2.47E-11 | 9.45E-11 | -1.83E+00 | 2.87E-05 | 1.02E-04 | -1.38E+00 | 4.72E-06 | 1.33E-05 | -1.67E+00 | 2.47E-03 | 9.06E-03 | -1.55E+00 | 1.14E-02 | 3.14E-02 | -1.41E+00 |
| SERAC1 | 1.41E-05 | 7.33E-05 | -1.61E+00 | 1.14E-03 | 4.94E-03 | -1.16E+00 | 2.30E-06 | 9.78E-06 | -1.76E+00 | 5.08E-05 | 1.94E-04 | -1.36E+00 | 8.25E-04 | 2.94E-03 | -1.21E+00 | 1.63E-02 | 4.58E-02 | 1.09E+00 | 3.12E-02 | 1.15E-01 | 1.02E+00 | 3.85E-02 | 1.06E-01 | 1.04E+00 |
| TDRD3 | 1.41E-05 | 7.33E-05 | -1.21E+00 | 9.08E-05 | 3.92E-04 | 1.03E+00 | 1.76E-02 | 7.49E-02 | 1.10E+00 | 1.06E-02 | 4.05E-02 | 1.17E+00 | 2.70E-03 | 9.62E-03 | -1.16E+00 | 4.28E-02 | 1.21E-01 | 1.12E+00 | 9.69E-07 | 3.56E-06 | -1.44E+00 | 1.40E-03 | 3.86E-03 | -1.20E+00 |
| KDELC2 | 1.41E-05 | 7.35E-05 | -1.61E+00 | 5.48E-06 | 2.37E-05 | -1.56E+00 | 1.87E-02 | 7.93E-02 | -1.02E+00 | 1.97E-05 | 7.54E-05 | -1.12E+00 | 1.20E-03 | 4.26E-03 | -1.80E+00 | 2.45E-02 | 6.91E-02 | -1.08E+00 | 6.01E-10 | 2.21E-09 | -1.68E+00 | 2.24E-03 | 6.15E-03 | -1.08E+00 |
| ZNF76 | 1.41E-05 | 7.37E-05 | -1.34E+00 | 1.39E-03 | 6.02E-03 | -1.01E+00 | 3.56E-03 | 1.51E-02 | -1.19E+00 | 8.82E-03 | 3.37E-02 | 1.02E+00 | 3.45E-05 | 1.23E-04 | -1.39E+00 | 4.19E-03 | 1.18E-02 | 1.07E+00 | 6.98E-04 | 2.56E-03 | -1.18E+00 | 5.65E-02 | 1.55E-01 | 1.07E+00 |
| CUTC | 1.41E-05 | 7.37E-05 | -1.06E+00 | 1.48E-03 | 6.41E-03 | 1.15E+00 | 1.05E-03 | 4.44E-03 | -1.06E+00 | 1.85E-03 | 7.08E-03 | 1.05E+00 | 4.98E-05 | 1.78E-04 | -1.06E+00 | 3.08E-03 | 8.69E-03 | 1.09E+00 | 5.43E-02 | 1.99E-01 | 1.04E+00 | 1.57E-01 | 4.30E-01 | 1.18E+00 |
| STAM | 1.42E-05 | 7.40E-05 | -1.12E+00 | 2.92E-08 | 1.26E-07 | -1.24E+00 | 5.55E-03 | 2.36E-02 | -1.04E+00 | 4.81E-04 | 1.84E-03 | -1.13E+00 | 4.45E-02 | 1.59E-01 | 1.17E+00 | 9.92E-06 | 2.80E-05 | -1.15E+00 | 2.21E-02 | 8.10E-02 | 1.20E+00 | 1.65E-05 | 4.54E-05 | -1.05E+00 |
| SCYL2 | 1.43E-05 | 7.44E-05 | -1.22E+00 | 4.49E-06 | 1.94E-05 | -1.11E+00 | 2.67E-05 | 1.13E-04 | -1.29E+00 | 3.78E-06 | 1.44E-05 | -1.23E+00 | 2.56E-07 | 9.13E-07 | -1.53E+00 | 2.24E-05 | 6.31E-05 | -1.24E+00 | 7.70E-08 | 2.83E-07 | -1.25E+00 | 8.71E-07 | 2.39E-06 | -1.34E+00 |
| LRIG2 | 1.43E-05 | 7.44E-05 | -1.28E+00 | 2.25E-02 | 9.72E-02 | 1.28E+00 | 3.28E-07 | 1.40E-06 | -1.78E+00 | 5.79E-04 | 2.21E-03 | -1.14E+00 | 1.73E-01 | 6.17E-01 | 1.35E+00 | 1.29E-01 | 3.63E-01 | 1.22E+00 | 2.66E-01 | 9.75E-01 | 1.34E+00 | 2.06E-01 | 5.65E-01 | 1.28E+00 |
| GLIDR | 1.43E-05 | 7.44E-05 | -1.65E+00 | 4.43E-02 | 1.91E-01 | 1.18E+00 | 2.67E-06 | 1.13E-05 | -1.88E+00 | 9.26E-04 | 3.54E-03 | -1.16E+00 | 8.23E-05 | 2.94E-04 | -1.30E+00 | 2.48E-01 | 7.00E-01 | 1.48E+00 | 2.33E-05 | 8.56E-05 | -1.50E+00 | 3.05E-01 | 8.39E-01 | 1.27E+00 |
| DCUN1D3 | 1.43E-05 | 7.47E-05 | -1.57E+00 | 7.84E-05 | 3.39E-04 | -1.34E+00 | 7.53E-02 | 3.20E-01 | -1.10E+00 | 4.58E-02 | 1.75E-01 | -1.08E+00 | 1.45E-01 | 5.16E-01 | 1.09E+00 | 5.64E-02 | 1.59E-01 | -1.34E+00 | 8.40E-02 | 3.08E-01 | -1.24E+00 | 2.62E-01 | 7.20E-01 | -1.02E+00 |
| GGCX | 1.43E-05 | 7.47E-05 | -1.30E+00 | 2.18E-05 | 9.41E-05 | -1.19E+00 | 4.71E-07 | 2.00E-06 | -1.41E+00 | 2.95E-07 | 1.13E-06 | -1.25E+00 | 2.57E-02 | 9.16E-02 | -1.05E+00 | 1.16E-02 | 3.28E-02 | -1.13E+00 | 8.40E-05 | 3.08E-04 | -1.29E+00 | 1.42E-05 | 3.90E-05 | -1.24E+00 |
| KLF3-AS1 | 1.44E-05 | 7.48E-05 | -1.24E+00 | 2.21E-01 | 9.54E-01 | 1.45E+00 | 3.65E-02 | 1.55E-01 | -1.15E+00 | 2.96E-02 | 1.13E-01 | 1.83E+00 | 2.28E-01 | 8.11E-01 | 1.37E+00 | 1.84E-01 | 5.19E-01 | 1.54E+00 | 9.04E-02 | 3.32E-01 | -1.01E+00 | 2.70E-01 | 7.41E-01 | 1.12E+00 |
| TMEM256 | 1.44E-05 | 7.49E-05 | -1.06E+00 | 5.73E-22 | 2.47E-21 | -1.40E+00 | 1.98E-04 | 8.43E-04 | -1.14E+00 | 1.76E-16 | 6.71E-16 | -1.59E+00 | 8.98E-05 | 3.20E-04 | -1.22E+00 | 1.57E-15 | 4.42E-15 | -1.74E+00 | 1.93E-02 | 7.07E-02 | -1.00E+00 | 1.10E-07 | 3.02E-07 | -1.40E+00 |
| ASCC1 | 1.44E-05 | 7.50E-05 | -1.19E+00 | 1.83E-03 | 7.89E-03 | 1.17E+00 | 2.64E-06 | 1.12E-05 | -1.30E+00 | 1.39E-03 | 5.29E-03 | 1.03E+00 | 8.34E-05 | 2.97E-04 | -1.22E+00 | 1.21E-02 | 3.40E-02 | 1.02E+00 | 5.80E-04 | 2.13E-03 | -1.12E+00 | 9.28E-03 | 2.55E-02 | -1.04E+00 |
| CARS2 | 1.44E-05 | 7.50E-05 | -1.06E+00 | 2.30E-05 | 9.94E-05 | 1.06E+00 | 8.41E-02 | 3.58E-01 | 1.08E+00 | 2.51E-01 | 9.58E-01 | 1.34E+00 | 3.50E-04 | 1.25E-03 | -1.01E+00 | 2.79E-02 | 7.86E-02 | 1.23E+00 | 4.26E-02 | 1.56E-01 | -1.07E+00 | 2.44E-01 | 6.70E-01 | 1.24E+00 |
| NVL | 1.44E-05 | 7.53E-05 | -1.55E+00 | 7.39E-04 | 3.19E-03 | -1.12E+00 | 2.89E-04 | 1.23E-03 | -1.41E+00 | 2.34E-03 | 8.95E-03 | -1.09E+00 | 8.54E-06 | 3.05E-05 | -1.57E+00 | 1.47E-03 | 4.15E-03 | -1.18E+00 | 3.95E-03 | 1.45E-02 | -1.07E+00 | 3.14E-02 | 8.62E-02 | -1.04E+00 |
| GTF2H2 | 1.45E-05 | 7.53E-05 | -1.11E+00 | 9.82E-02 | 4.24E-01 | 1.35E+00 | 9.94E-07 | 4.22E-06 | -1.24E+00 | 1.57E-02 | 6.00E-02 | 1.19E+00 | 1.32E-07 | 4.70E-07 | -1.34E+00 | 7.70E-02 | 2.17E-01 | 1.20E+00 | 2.73E-09 | 1.00E-08 | -1.30E+00 | 4.20E-02 | 1.15E-01 | 1.13E+00 |
| SZT2 | 1.45E-05 | 7.56E-05 | -1.20E+00 | 3.39E-03 | 1.47E-02 | 1.11E+00 | 1.46E-03 | 6.22E-03 | -1.22E+00 | 9.27E-02 | 3.54E-01 | 1.22E+00 | 2.06E-01 | 7.35E-01 | 1.30E+00 | 2.46E-01 | 6.93E-01 | 1.32E+00 | 2.14E-05 | 7.87E-05 | -1.35E+00 | 6.02E-03 | 1.65E-02 | -1.02E+00 |
| ALYREF | 1.45E-05 | 7.56E-05 | -1.01E+00 | 7.33E-07 | 3.17E-06 | -1.00E+00 | 2.31E-01 | 9.83E-01 | 1.20E+00 | 8.37E-02 | 3.20E-01 | 1.27E+00 | 1.53E-02 | 5.46E-02 | 1.03E+00 | 1.26E-01 | 3.54E-01 | 1.11E+00 | 6.89E-03 | 2.53E-02 | 1.09E+00 | 1.67E-01 | 4.60E-01 | 1.42E+00 |
| RPS2P35 | 1.46E-05 | 7.58E-05 | 1.85E+00 | 4.76E-08 | 2.06E-07 | 2.26E+00 | 6.26E-08 | 2.66E-07 | 1.92E+00 | 1.04E-07 | 3.97E-07 | 2.23E+00 | 4.68E-03 | 1.67E-02 | 1.87E+00 | 1.07E-05 | 3.01E-05 | 2.21E+00 | 1.12E-06 | 4.11E-06 | 1.95E+00 | 3.22E-11 | 8.86E-11 | 2.44E+00 |
| ALDH3A2 | 1.46E-05 | 7.59E-05 | -1.25E+00 | 8.62E-03 | 3.72E-02 | 1.18E+00 | 1.06E-02 | 4.50E-02 | -1.05E+00 | 3.49E-02 | 1.33E-01 | 1.18E+00 | 5.65E-05 | 2.01E-04 | -1.40E+00 | 2.49E-03 | 7.02E-03 | -1.11E+00 | 3.09E-02 | 1.13E-01 | 1.09E+00 | 1.23E-01 | 3.38E-01 | 1.52E+00 |
| MAPK9 | 1.46E-05 | 7.62E-05 | -1.25E+00 | 2.84E-04 | 1.23E-03 | -1.02E+00 | 2.79E-09 | 1.19E-08 | -1.45E+00 | 1.14E-06 | 4.36E-06 | -1.15E+00 | 3.28E-02 | 1.17E-01 | -1.11E+00 | 1.05E-01 | 2.95E-01 | 1.10E+00 | 7.19E-04 | 2.64E-03 | -1.13E+00 | 9.40E-03 | 2.58E-02 | -1.04E+00 |
| PDXDC2P-NPIPB14 | 1.46E-05 | 7.63E-05 | -1.60E+00 | 8.33E-05 | 3.60E-04 | -1.37E+00 | 1.34E-11 | 5.71E-11 | -2.11E+00 | 1.01E-12 | 3.87E-12 | -2.28E+00 | 1.45E-14 | 5.18E-14 | -2.96E+00 | 1.73E-10 | 4.89E-10 | -1.83E+00 | 1.12E-07 | 4.12E-07 | -2.12E+00 | 6.84E-04 | 1.88E-03 | -1.45E+00 |
| RABEPK | 1.47E-05 | 7.64E-05 | -1.41E+00 | 6.11E-03 | 2.64E-02 | 1.06E+00 | 1.02E-03 | 4.32E-03 | -1.20E+00 | 3.26E-02 | 1.24E-01 | 1.15E+00 | 1.04E-04 | 3.70E-04 | -1.63E+00 | 2.22E-01 | 6.27E-01 | 1.30E+00 | 2.87E-05 | 1.05E-04 | -1.23E+00 | 2.39E-01 | 6.57E-01 | 1.28E+00 |
| EGR4 | 1.47E-05 | 7.64E-05 | 5.17E+00 | 1.49E-01 | 6.45E-01 | -1.17E+00 | 1.89E-06 | 8.01E-06 | 5.82E+00 | 1.66E-01 | 6.35E-01 | -1.11E+00 | 9.01E-02 | 3.21E-01 | 2.23E+00 | 3.55E-01 | 1.00E+00 | 1.00E+00 | 2.41E-02 | 8.84E-02 | 2.62E+00 | 3.35E-01 | 9.19E-01 | -1.03E+00 |
| PSTPIP1 | 1.47E-05 | 7.64E-05 | -1.15E+00 | 1.78E-08 | 7.71E-08 | -1.24E+00 | 1.46E-06 | 6.20E-06 | -1.42E+00 | 6.19E-09 | 2.36E-08 | -1.58E+00 | 3.42E-03 | 1.22E-02 | -1.86E+00 | 7.63E-03 | 2.15E-02 | -1.62E+00 | 9.92E-02 | 3.64E-01 | -1.71E+00 | 2.35E-01 | 6.46E-01 | -1.26E+00 |
| RPS24P13 | 1.48E-05 | 7.70E-05 | -1.35E+00 | 7.21E-09 | 3.12E-08 | -1.81E+00 | 3.48E-04 | 1.48E-03 | -1.29E+00 | 8.57E-09 | 3.27E-08 | -2.03E+00 | 1.85E-07 | 6.59E-07 | -2.28E+00 | 1.35E-07 | 3.81E-07 | -2.22E+00 | 4.93E-03 | 1.81E-02 | -1.37E+00 | 2.59E-03 | 7.12E-03 | -1.46E+00 |
| AC011472.2 | 1.48E-05 | 7.74E-05 | 3.53E+00 | 1.33E-01 | 5.73E-01 | 1.87E+00 | 7.93E-03 | 3.37E-02 | 1.76E+00 | 2.62E-01 | 1.00E+00 | 1.53E+00 | 8.52E-02 | 3.04E-01 | 2.12E+00 | 1.86E-01 | 5.25E-01 | 1.98E+00 | 1.09E-01 | 3.99E-01 | 1.95E+00 | 3.62E-01 | 9.93E-01 | 1.07E+00 |
| FCER1G | 1.48E-05 | 7.74E-05 | -1.62E+00 | 1.07E-04 | 4.63E-04 | -1.11E+00 | 5.75E-02 | 2.44E-01 | -1.03E+00 | 3.13E-02 | 1.20E-01 | -1.12E+00 | 3.00E-11 | 1.07E-10 | -1.85E+00 | 5.26E-15 | 1.48E-14 | -2.20E+00 | 5.93E-02 | 2.18E-01 | -1.01E+00 | 2.01E-03 | 5.52E-03 | -1.65E+00 |
| NCAPD2 | 1.49E-05 | 7.74E-05 | -1.77E+00 | 1.69E-03 | 7.31E-03 | -1.24E+00 | 1.87E-02 | 7.94E-02 | -1.11E+00 | 8.02E-03 | 3.06E-02 | 1.01E+00 | 4.50E-03 | 1.60E-02 | -1.32E+00 | 1.74E-02 | 4.91E-02 | -1.16E+00 | 1.24E-10 | 4.56E-10 | -1.54E+00 | 2.87E-04 | 7.87E-04 | -1.15E+00 |
| SERTAD1 | 1.49E-05 | 7.75E-05 | 2.08E+00 | 3.85E-06 | 1.66E-05 | -1.54E+00 | 1.06E-05 | 4.49E-05 | 2.44E+00 | 1.31E-02 | 5.00E-02 | -1.21E+00 | 1.08E-01 | 3.86E-01 | 1.54E+00 | 3.28E-04 | 9.24E-04 | -1.98E+00 | 2.72E-01 | 9.99E-01 | 1.41E+00 | 2.92E-08 | 8.01E-08 | -2.23E+00 |
| AL512343.1 | 1.50E-05 | 7.80E-05 | -1.58E+00 | 6.57E-06 | 2.84E-05 | -1.61E+00 | 1.16E-04 | 4.93E-04 | -1.40E+00 | 2.24E-05 | 8.55E-05 | -1.34E+00 | 5.12E-05 | 1.83E-04 | -1.66E+00 | 2.80E-04 | 7.89E-04 | -1.51E+00 | 6.23E-14 | 2.29E-13 | -1.62E+00 | 5.18E-14 | 1.42E-13 | -1.63E+00 |
| AC099524.1 | 1.50E-05 | 7.83E-05 | -2.26E+00 | 8.81E-06 | 3.81E-05 | -2.15E+00 | 8.24E-02 | 3.50E-01 | -1.42E+00 | 1.07E-01 | 4.11E-01 | -1.31E+00 | 9.28E-08 | 3.31E-07 | -2.59E+00 | 1.31E-03 | 3.68E-03 | -1.27E+00 | 7.65E-04 | 2.81E-03 | -1.35E+00 | 1.43E-02 | 3.93E-02 | -1.13E+00 |
| TNPO2 | 1.50E-05 | 7.83E-05 | -1.27E+00 | 7.89E-04 | 3.41E-03 | 1.03E+00 | 1.20E-04 | 5.09E-04 | -1.20E+00 | 3.57E-04 | 1.36E-03 | -1.04E+00 | 9.43E-03 | 3.36E-02 | 1.05E+00 | 5.66E-02 | 1.59E-01 | 1.16E+00 | 1.04E-04 | 3.83E-04 | -1.27E+00 | 9.34E-03 | 2.56E-02 | -1.03E+00 |
| MRPL21 | 1.51E-05 | 7.88E-05 | -1.01E+00 | 9.91E-07 | 4.28E-06 | 1.04E+00 | 2.20E-01 | 9.36E-01 | 1.18E+00 | 4.90E-02 | 1.87E-01 | 1.18E+00 | 2.48E-04 | 8.85E-04 | 1.01E+00 | 6.53E-04 | 1.84E-03 | 1.05E+00 | 6.87E-02 | 2.52E-01 | 1.10E+00 | 1.77E-01 | 4.85E-01 | 1.11E+00 |
| ZNF766 | 1.51E-05 | 7.88E-05 | -1.08E+00 | 1.57E-12 | 6.78E-12 | -1.35E+00 | 5.44E-04 | 2.31E-03 | -1.21E+00 | 5.33E-07 | 2.03E-06 | -1.27E+00 | 8.12E-05 | 2.90E-04 | -1.19E+00 | 3.36E-04 | 9.48E-04 | -1.06E+00 | 5.56E-07 | 2.04E-06 | -1.30E+00 | 1.56E-07 | 4.27E-07 | -1.39E+00 |
| ASB13 | 1.51E-05 | 7.89E-05 | -1.60E+00 | 1.08E-02 | 4.67E-02 | 1.07E+00 | 2.29E-08 | 9.72E-08 | -1.98E+00 | 2.64E-02 | 1.01E-01 | 1.16E+00 | 1.39E-02 | 4.96E-02 | -1.54E+00 | 2.98E-01 | 8.40E-01 | 1.31E+00 | 1.22E-03 | 4.48E-03 | -1.32E+00 | 2.55E-02 | 7.01E-02 | -1.08E+00 |
| BLOC1S1 | 1.52E-05 | 7.90E-05 | -1.09E+00 | 1.92E-22 | 8.31E-22 | -1.65E+00 | 4.52E-02 | 1.92E-01 | 1.11E+00 | 1.51E-14 | 5.78E-14 | -1.37E+00 | 3.97E-03 | 1.42E-02 | 1.10E+00 | 8.78E-12 | 2.48E-11 | -1.41E+00 | 1.00E-01 | 3.68E-01 | 1.20E+00 | 1.34E-07 | 3.68E-07 | -1.30E+00 |
| APEX2 | 1.52E-05 | 7.92E-05 | -1.75E+00 | 2.13E-02 | 9.22E-02 | 1.11E+00 | 2.08E-04 | 8.83E-04 | -1.33E+00 | 2.87E-02 | 1.09E-01 | 1.15E+00 | 4.37E-05 | 1.56E-04 | -1.85E+00 | 2.52E-02 | 7.11E-02 | -1.02E+00 | 1.33E-08 | 4.90E-08 | -1.36E+00 | 2.02E-01 | 5.55E-01 | 1.17E+00 |
| HACE1 | 1.52E-05 | 7.92E-05 | -1.45E+00 | 6.12E-03 | 2.64E-02 | 1.09E+00 | 3.09E-03 | 1.31E-02 | -1.21E+00 | 6.91E-03 | 2.64E-02 | 1.02E+00 | 5.28E-05 | 1.88E-04 | -1.74E+00 | 6.90E-04 | 1.95E-03 | -1.44E+00 | 4.81E-04 | 1.77E-03 | -1.45E+00 | 1.89E-03 | 5.20E-03 | -1.25E+00 |
| LRRC14 | 1.53E-05 | 7.96E-05 | -2.21E+00 | 7.67E-05 | 3.31E-04 | -1.73E+00 | 7.68E-03 | 3.26E-02 | -1.43E+00 | 1.83E-01 | 7.00E-01 | 1.15E+00 | 1.67E-03 | 5.94E-03 | -1.75E+00 | 8.38E-02 | 2.36E-01 | 1.03E+00 | 1.08E-05 | 3.95E-05 | -1.78E+00 | 2.61E-02 | 7.18E-02 | -1.02E+00 |
| TRIM32 | 1.53E-05 | 7.97E-05 | -1.76E+00 | 3.26E-03 | 1.41E-02 | -1.01E+00 | 2.47E-05 | 1.05E-04 | -1.60E+00 | 1.86E-03 | 7.09E-03 | -1.07E+00 | 1.18E-03 | 4.22E-03 | -1.71E+00 | 1.22E-03 | 3.43E-03 | -1.75E+00 | 1.46E-03 | 5.36E-03 | -1.45E+00 | 1.76E-01 | 4.84E-01 | 1.16E+00 |
| FLYWCH2 | 1.53E-05 | 7.97E-05 | -1.11E+00 | 1.14E-03 | 4.92E-03 | 1.17E+00 | 7.14E-04 | 3.03E-03 | -1.10E+00 | 6.05E-02 | 2.31E-01 | 1.24E+00 | 5.08E-06 | 1.81E-05 | -1.20E+00 | 8.08E-03 | 2.28E-02 | 1.13E+00 | 2.29E-07 | 8.41E-07 | -1.24E+00 | 6.03E-03 | 1.66E-02 | 1.02E+00 |
| TMEM14B | 1.53E-05 | 7.98E-05 | 1.01E+00 | 1.32E-06 | 5.72E-06 | 1.07E+00 | 1.30E-01 | 5.52E-01 | 1.12E+00 | 1.21E-01 | 4.63E-01 | 1.10E+00 | 5.67E-05 | 2.02E-04 | -1.04E+00 | 1.55E-01 | 4.36E-01 | 1.27E+00 | 6.97E-02 | 2.56E-01 | 1.04E+00 | 2.52E-01 | 6.91E-01 | 1.02E+00 |
| NMRAL1 | 1.53E-05 | 7.98E-05 | -1.06E+00 | 1.88E-05 | 8.12E-05 | 1.04E+00 | 1.15E-01 | 4.91E-01 | 1.11E+00 | 4.22E-04 | 1.61E-03 | 1.00E+00 | 6.53E-05 | 2.33E-04 | -1.08E+00 | 3.22E-02 | 9.07E-02 | 1.11E+00 | 1.18E-02 | 4.32E-02 | 1.07E+00 | 3.26E-01 | 8.95E-01 | 1.35E+00 |
| CLIP4 | 1.53E-05 | 7.98E-05 | -1.18E+00 | 3.25E-04 | 1.41E-03 | 1.05E+00 | 2.91E-04 | 1.23E-03 | -1.26E+00 | 7.39E-02 | 2.82E-01 | 1.23E+00 | 1.33E-01 | 4.75E-01 | 1.18E+00 | 1.27E-02 | 3.58E-02 | -1.20E+00 | 1.21E-01 | 4.44E-01 | 1.09E+00 | 6.90E-02 | 1.90E-01 | 1.04E+00 |
| ZNF22 | 1.53E-05 | 7.98E-05 | -1.02E+00 | 1.56E-01 | 6.73E-01 | 1.41E+00 | 4.08E-03 | 1.73E-02 | -1.09E+00 | 7.41E-02 | 2.83E-01 | 1.21E+00 | 1.56E-03 | 5.56E-03 | -1.14E+00 | 7.48E-02 | 2.11E-01 | 1.18E+00 | 1.87E-02 | 6.88E-02 | -1.02E+00 | 3.29E-01 | 9.03E-01 | 1.19E+00 |
| TLE4 | 1.53E-05 | 7.98E-05 | 2.88E+00 | 7.71E-02 | 3.33E-01 | -1.19E+00 | 1.56E-10 | 6.65E-10 | 2.54E+00 | 2.72E-02 | 1.04E-01 | -1.14E+00 | 7.91E-05 | 2.82E-04 | 2.72E+00 | 1.15E-01 | 3.23E-01 | -1.34E+00 | 1.08E-01 | 3.98E-01 | 1.45E+00 | 4.71E-02 | 1.29E-01 | -2.39E+00 |
| CDKN3 | 1.53E-05 | 7.98E-05 | -2.36E+00 | 9.51E-05 | 4.11E-04 | -1.96E+00 | 2.00E-01 | 8.48E-01 | 1.29E+00 | 8.50E-02 | 3.25E-01 | 1.19E+00 | 2.28E-04 | 8.12E-04 | -2.74E+00 | 2.28E-03 | 6.43E-03 | -1.72E+00 | 2.39E-01 | 8.77E-01 | -1.12E+00 | 4.59E-03 | 1.26E-02 | 1.06E+00 |
| TRIM8 | 1.54E-05 | 8.00E-05 | -1.17E+00 | 1.87E-03 | 8.07E-03 | 1.12E+00 | 3.09E-07 | 1.31E-06 | -1.49E+00 | 2.51E-04 | 9.57E-04 | -1.02E+00 | 3.85E-02 | 1.37E-01 | 1.21E+00 | 2.01E-02 | 5.67E-02 | 1.18E+00 | 4.25E-03 | 1.56E-02 | 1.02E+00 | 1.21E-02 | 3.34E-02 | -1.09E+00 |
| POMT1 | 1.54E-05 | 8.03E-05 | -1.33E+00 | 2.28E-04 | 9.86E-04 | -1.02E+00 | 7.32E-04 | 3.11E-03 | -1.24E+00 | 1.90E-03 | 7.25E-03 | -1.11E+00 | 1.66E-03 | 5.91E-03 | -1.46E+00 | 4.00E-02 | 1.13E-01 | -1.06E+00 | 5.48E-03 | 2.01E-02 | -1.29E+00 | 1.57E-01 | 4.32E-01 | 1.09E+00 |
| PLA2G4A | 1.54E-05 | 8.04E-05 | -1.45E+00 | 6.10E-06 | 2.63E-05 | -1.33E+00 | 3.94E-14 | 1.68E-13 | -2.06E+00 | 2.13E-15 | 8.15E-15 | -1.86E+00 | 3.24E-06 | 1.16E-05 | -1.92E+00 | 6.63E-06 | 1.87E-05 | -1.85E+00 | 2.14E-04 | 7.84E-04 | -1.67E+00 | 7.73E-02 | 2.12E-01 | -1.03E+00 |
| H6PD | 1.54E-05 | 8.04E-05 | -1.35E+00 | 8.42E-06 | 3.64E-05 | -1.24E+00 | 2.66E-06 | 1.13E-05 | -1.80E+00 | 6.62E-05 | 2.53E-04 | -1.32E+00 | 1.80E-01 | 6.42E-01 | 1.10E+00 | 1.41E-01 | 3.97E-01 | 1.07E+00 | 3.33E-03 | 1.22E-02 | -1.43E+00 | 2.40E-01 | 6.60E-01 | 1.08E+00 |
| SOCS2 | 1.54E-05 | 8.05E-05 | 1.50E+00 | 5.75E-12 | 2.48E-11 | 1.95E+00 | 2.26E-01 | 9.62E-01 | 1.35E+00 | 4.60E-03 | 1.76E-02 | 1.69E+00 | 8.60E-04 | 3.07E-03 | 2.02E+00 | 7.43E-02 | 2.09E-01 | 1.38E+00 | 2.26E-02 | 8.28E-02 | 1.76E+00 | 2.09E-01 | 5.73E-01 | 1.37E+00 |
| PI4KB | 1.54E-05 | 8.05E-05 | -1.15E+00 | 9.43E-04 | 4.07E-03 | 1.14E+00 | 1.65E-04 | 7.01E-04 | -1.18E+00 | 4.15E-04 | 1.59E-03 | 1.02E+00 | 7.28E-03 | 2.59E-02 | 1.00E+00 | 1.87E-02 | 5.27E-02 | 1.04E+00 | 1.61E-04 | 5.91E-04 | -1.11E+00 | 8.74E-04 | 2.40E-03 | -1.07E+00 |
| ZNF233 | 1.55E-05 | 8.06E-05 | -1.82E+00 | 2.92E-02 | 1.26E-01 | 1.07E+00 | 8.23E-02 | 3.49E-01 | -1.14E+00 | 2.46E-01 | 9.38E-01 | 1.26E+00 | 1.15E-01 | 4.09E-01 | -1.37E+00 | 3.55E-01 | 1.00E+00 | 1.13E+00 | 2.16E-02 | 7.93E-02 | -1.34E+00 | 3.49E-01 | 9.58E-01 | 1.29E+00 |
| AC010435.1 | 1.55E-05 | 8.09E-05 | -1.42E+00 | 5.61E-08 | 2.43E-07 | -1.71E+00 | 4.13E-03 | 1.75E-02 | -1.15E+00 | 3.35E-05 | 1.28E-04 | -1.34E+00 | 1.98E-05 | 7.06E-05 | -1.42E+00 | 1.40E-07 | 3.94E-07 | -1.79E+00 | 1.80E-04 | 6.59E-04 | -1.23E+00 | 9.62E-04 | 2.64E-03 | -1.14E+00 |
| FGGY | 1.56E-05 | 8.12E-05 | -1.19E+00 | 8.23E-05 | 3.56E-04 | -1.04E+00 | 1.25E-01 | 5.32E-01 | 1.03E+00 | 2.13E-02 | 8.13E-02 | -1.03E+00 | 2.27E-04 | 8.09E-04 | -1.17E+00 | 5.49E-05 | 1.55E-04 | -1.25E+00 | 2.59E-01 | 9.50E-01 | 1.19E+00 | 3.34E-01 | 9.17E-01 | 1.24E+00 |
| ARPC3 | 1.56E-05 | 8.14E-05 | 1.34E+00 | 4.50E-04 | 1.94E-03 | 1.35E+00 | 3.22E-07 | 1.37E-06 | 1.41E+00 | 8.75E-05 | 3.34E-04 | 1.33E+00 | 5.39E-04 | 1.92E-03 | 1.32E+00 | 1.39E-02 | 3.93E-02 | 1.23E+00 | 1.78E-11 | 6.52E-11 | 1.57E+00 | 1.84E-06 | 5.05E-06 | 1.44E+00 |
| C1orf115 | 1.56E-05 | 8.15E-05 | -1.65E+00 | 5.28E-02 | 2.28E-01 | 1.09E+00 | 6.92E-02 | 2.94E-01 | -1.25E+00 | 1.52E-01 | 5.82E-01 | 1.00E+00 | 2.15E-01 | 7.67E-01 | -1.20E+00 | 2.90E-01 | 8.18E-01 | 1.31E+00 | 4.25E-02 | 1.56E-01 | -1.75E+00 | 4.25E-02 | 1.17E-01 | -2.04E+00 |
| MLLT3 | 1.57E-05 | 8.19E-05 | -1.22E+00 | 1.75E-01 | 7.55E-01 | 1.16E+00 | 8.10E-05 | 3.44E-04 | -1.01E+00 | 5.01E-03 | 1.91E-02 | 1.31E+00 | 2.60E-03 | 9.27E-03 | 1.47E+00 | 2.05E-04 | 5.78E-04 | 1.67E+00 | 3.69E-02 | 1.35E-01 | 1.06E+00 | 1.19E-02 | 3.28E-02 | -1.07E+00 |
| F13A1 | 1.57E-05 | 8.20E-05 | -3.78E+00 | 9.15E-06 | 3.95E-05 | -3.98E+00 | 9.19E-44 | 3.90E-43 | -4.39E+00 | 9.69E-18 | 3.70E-17 | -1.92E+00 | 2.00E-01 | 7.13E-01 | -1.12E+00 | 2.33E-01 | 6.56E-01 | -2.18E+00 | 5.09E-02 | 1.87E-01 | 1.12E+00 | 4.20E-03 | 1.15E-02 | -1.90E+00 |
| IL6ST | 1.57E-05 | 8.20E-05 | -1.23E+00 | 2.21E-10 | 9.56E-10 | -1.47E+00 | 1.33E-03 | 5.65E-03 | -1.16E+00 | 3.00E-07 | 1.15E-06 | -1.58E+00 | 1.63E-01 | 5.81E-01 | 1.66E+00 | 8.10E-02 | 2.28E-01 | -1.26E+00 | 5.82E-03 | 2.14E-02 | 1.82E+00 | 1.16E-04 | 3.17E-04 | -1.09E+00 |
| PNPLA8 | 1.57E-05 | 8.20E-05 | -1.13E+00 | 4.07E-11 | 1.76E-10 | -1.39E+00 | 1.14E-06 | 4.83E-06 | -1.27E+00 | 1.09E-13 | 4.17E-13 | -1.57E+00 | 2.03E-04 | 7.24E-04 | -1.06E+00 | 1.29E-07 | 3.64E-07 | -1.42E+00 | 4.46E-07 | 1.64E-06 | -1.29E+00 | 2.09E-10 | 5.75E-10 | -1.59E+00 |
| DPH7 | 1.57E-05 | 8.21E-05 | -1.25E+00 | 1.38E-03 | 5.96E-03 | 1.13E+00 | 8.11E-06 | 3.44E-05 | -1.26E+00 | 1.85E-04 | 7.05E-04 | -1.00E+00 | 7.48E-10 | 2.67E-09 | -1.61E+00 | 1.21E-04 | 3.41E-04 | -1.11E+00 | 6.65E-02 | 2.44E-01 | 1.11E+00 | 2.52E-01 | 6.91E-01 | 1.29E+00 |
| POLL | 1.58E-05 | 8.21E-05 | -1.13E+00 | 5.59E-04 | 2.41E-03 | 1.12E+00 | 1.83E-04 | 7.77E-04 | -1.22E+00 | 1.33E-02 | 5.07E-02 | 1.09E+00 | 7.70E-02 | 2.75E-01 | 1.07E+00 | 2.39E-01 | 6.74E-01 | 1.32E+00 | 3.27E-02 | 1.20E-01 | -1.04E+00 | 1.74E-02 | 4.79E-02 | -1.02E+00 |
| S100Z | 1.58E-05 | 8.21E-05 | -1.57E+00 | 1.50E-01 | 6.47E-01 | 1.26E+00 | 3.53E-07 | 1.50E-06 | -1.62E+00 | 1.80E-06 | 6.87E-06 | -1.38E+00 | 1.56E-01 | 5.55E-01 | -1.10E+00 | 2.44E-01 | 6.87E-01 | 1.67E+00 | 1.95E-01 | 7.17E-01 | -1.11E+00 | 3.20E-01 | 8.80E-01 | 1.09E+00 |
| PPP1R14B | 1.58E-05 | 8.23E-05 | 1.67E+00 | 1.78E-01 | 7.69E-01 | 1.30E+00 | 4.47E-08 | 1.90E-07 | 1.76E+00 | 3.75E-03 | 1.43E-02 | 1.38E+00 | 2.03E-01 | 7.25E-01 | 1.25E+00 | 2.78E-02 | 7.85E-02 | 1.14E+00 | 8.28E-02 | 3.04E-01 | 1.49E+00 | 1.10E-01 | 3.03E-01 | 1.34E+00 |
| C19orf53 | 1.58E-05 | 8.26E-05 | -1.03E+00 | 1.93E-09 | 8.35E-09 | -1.05E+00 | 3.75E-02 | 1.59E-01 | 1.01E+00 | 6.57E-04 | 2.51E-03 | -1.07E+00 | 6.10E-04 | 2.17E-03 | 1.01E+00 | 4.00E-03 | 1.13E-02 | 1.06E+00 | 9.53E-02 | 3.50E-01 | 1.06E+00 | 3.01E-01 | 8.28E-01 | 1.09E+00 |
| HPS5 | 1.58E-05 | 8.26E-05 | -1.43E+00 | 3.53E-03 | 1.52E-02 | 1.03E+00 | 3.94E-05 | 1.67E-04 | -1.40E+00 | 3.71E-03 | 1.42E-02 | -1.02E+00 | 4.25E-05 | 1.51E-04 | -1.44E+00 | 4.37E-04 | 1.23E-03 | -1.24E+00 | 1.24E-05 | 4.56E-05 | -1.41E+00 | 3.68E-05 | 1.01E-04 | -1.39E+00 |
| NDUFA2 | 1.58E-05 | 8.26E-05 | -1.07E+00 | 1.16E-15 | 5.02E-15 | -1.25E+00 | 5.87E-02 | 2.50E-01 | 1.10E+00 | 6.89E-06 | 2.63E-05 | -1.12E+00 | 6.92E-06 | 2.47E-05 | -1.09E+00 | 3.94E-05 | 1.11E-04 | -1.09E+00 | 1.40E-01 | 5.16E-01 | 1.12E+00 | 2.49E-03 | 6.85E-03 | -1.14E+00 |
| SLITRK4 | 1.59E-05 | 8.26E-05 | -1.45E+00 | 1.61E-01 | 6.97E-01 | 1.38E+00 | 1.11E-09 | 4.73E-09 | -1.75E+00 | 1.25E-01 | 4.78E-01 | 1.37E+00 | 2.39E-01 | 8.52E-01 | 1.36E+00 | 2.63E-01 | 7.41E-01 | 2.41E+00 | 2.28E-01 | 8.38E-01 | 1.28E+00 | 2.98E-01 | 8.18E-01 | 1.40E+00 |
| SIPA1L3 | 1.59E-05 | 8.27E-05 | -1.40E+00 | 2.82E-05 | 1.22E-04 | -1.25E+00 | 6.87E-03 | 2.92E-02 | -1.18E+00 | 3.19E-03 | 1.22E-02 | -1.08E+00 | 2.21E-01 | 7.88E-01 | 1.14E+00 | 6.89E-02 | 1.94E-01 | -1.22E+00 | 1.58E-02 | 5.80E-02 | -1.17E+00 | 5.38E-02 | 1.48E-01 | -1.08E+00 |
| AP000347.1 | 1.59E-05 | 8.27E-05 | -1.19E+00 | 4.09E-03 | 1.77E-02 | 1.14E+00 | 9.31E-03 | 3.96E-02 | -1.05E+00 | 3.07E-02 | 1.17E-01 | 1.16E+00 | 2.43E-02 | 8.65E-02 | -1.04E+00 | 8.45E-02 | 2.38E-01 | 1.14E+00 | 4.59E-03 | 1.69E-02 | -1.24E+00 | 1.79E-01 | 4.92E-01 | 1.10E+00 |
| MEF2C-AS1 | 1.59E-05 | 8.30E-05 | -1.32E+00 | 1.38E-01 | 5.95E-01 | 1.42E+00 | 2.94E-03 | 1.25E-02 | -1.19E+00 | 4.48E-02 | 1.71E-01 | 1.17E+00 | 6.99E-06 | 2.49E-05 | -1.46E+00 | 1.13E-02 | 3.19E-02 | 1.06E+00 | 3.05E-07 | 1.12E-06 | -1.49E+00 | 2.98E-04 | 8.18E-04 | -1.21E+00 |
| IL1R1 | 1.60E-05 | 8.32E-05 | -1.21E+00 | 8.15E-03 | 3.52E-02 | 1.12E+00 | 1.21E-01 | 5.14E-01 | -1.02E+00 | 2.56E-01 | 9.77E-01 | 1.13E+00 | 2.59E-01 | 9.23E-01 | 1.45E+00 | 3.21E-01 | 9.06E-01 | 1.53E+00 | 2.53E-01 | 9.28E-01 | -1.03E+00 | 3.54E-01 | 9.73E-01 | 1.35E+00 |
| UMPS | 1.60E-05 | 8.33E-05 | -1.27E+00 | 2.71E-03 | 1.17E-02 | 1.06E+00 | 7.94E-06 | 3.37E-05 | -1.35E+00 | 2.98E-02 | 1.14E-01 | 1.19E+00 | 2.40E-03 | 8.57E-03 | -1.23E+00 | 1.68E-01 | 4.73E-01 | 1.17E+00 | 8.27E-12 | 3.04E-11 | -1.44E+00 | 1.31E-01 | 3.60E-01 | 1.19E+00 |
| ZNF24 | 1.60E-05 | 8.35E-05 | -1.07E+00 | 2.42E-11 | 1.05E-10 | -1.15E+00 | 5.96E-06 | 2.53E-05 | -1.29E+00 | 2.34E-09 | 8.95E-09 | -1.33E+00 | 2.17E-04 | 7.75E-04 | -1.04E+00 | 2.42E-04 | 6.82E-04 | -1.01E+00 | 1.16E-02 | 4.27E-02 | -1.04E+00 | 9.20E-05 | 2.53E-04 | -1.16E+00 |
| ZSWIM7 | 1.60E-05 | 8.36E-05 | -1.06E+00 | 1.07E-02 | 4.60E-02 | 1.28E+00 | 2.80E-04 | 1.19E-03 | -1.21E+00 | 5.07E-03 | 1.94E-02 | 1.05E+00 | 3.98E-05 | 1.42E-04 | -1.20E+00 | 1.52E-02 | 4.27E-02 | 1.05E+00 | 1.43E-04 | 5.24E-04 | -1.09E+00 | 9.84E-03 | 2.70E-02 | 1.03E+00 |
| ZCCHC4 | 1.61E-05 | 8.37E-05 | -1.52E+00 | 3.20E-04 | 1.38E-03 | -1.12E+00 | 2.69E-03 | 1.14E-02 | -1.30E+00 | 3.99E-03 | 1.53E-02 | -1.10E+00 | 1.43E-01 | 5.10E-01 | -1.10E+00 | 1.18E-01 | 3.34E-01 | -1.29E+00 | 1.50E-05 | 5.51E-05 | -1.61E+00 | 1.79E-02 | 4.91E-02 | -1.06E+00 |
| PANK2 | 1.62E-05 | 8.45E-05 | -1.29E+00 | 6.68E-04 | 2.89E-03 | 1.02E+00 | 3.76E-07 | 1.60E-06 | -1.47E+00 | 1.90E-05 | 7.24E-05 | -1.17E+00 | 1.21E-03 | 4.32E-03 | -1.08E+00 | 2.74E-01 | 7.71E-01 | 1.34E+00 | 6.60E-10 | 2.42E-09 | -1.30E+00 | 3.14E-07 | 8.62E-07 | -1.21E+00 |
| CDH26 | 1.62E-05 | 8.46E-05 | -1.26E+00 | 4.92E-05 | 2.13E-04 | -1.10E+00 | 1.45E-04 | 6.18E-04 | -1.19E+00 | 2.37E-07 | 9.04E-07 | -1.28E+00 | 4.82E-04 | 1.72E-03 | -2.48E+00 | 1.45E-03 | 4.08E-03 | -1.87E+00 | 8.11E-04 | 2.98E-03 | -1.24E+00 | 3.46E-04 | 9.49E-04 | -1.35E+00 |
| WASHC2A | 1.63E-05 | 8.49E-05 | -1.15E+00 | 5.38E-03 | 2.32E-02 | 1.25E+00 | 2.85E-02 | 1.21E-01 | 1.02E+00 | 5.70E-02 | 2.18E-01 | 1.30E+00 | 3.06E-03 | 1.09E-02 | -1.16E+00 | 7.77E-02 | 2.19E-01 | 1.13E+00 | 9.41E-02 | 3.45E-01 | 1.27E+00 | 2.54E-01 | 6.98E-01 | 1.34E+00 |
| KIAA1841 | 1.63E-05 | 8.50E-05 | -1.89E+00 | 4.24E-03 | 1.83E-02 | -1.12E+00 | 4.46E-06 | 1.89E-05 | -1.77E+00 | 1.17E-02 | 4.47E-02 | 1.07E+00 | 7.46E-04 | 2.66E-03 | -1.64E+00 | 8.08E-03 | 2.28E-02 | -1.18E+00 | 8.21E-06 | 3.01E-05 | -1.54E+00 | 4.42E-03 | 1.21E-02 | -1.09E+00 |
| TRAK1 | 1.63E-05 | 8.50E-05 | -1.38E+00 | 8.24E-03 | 3.56E-02 | 1.10E+00 | 4.76E-05 | 2.02E-04 | -1.33E+00 | 2.46E-02 | 9.39E-02 | 1.14E+00 | 1.80E-04 | 6.40E-04 | -1.34E+00 | 1.00E-01 | 2.83E-01 | 1.21E+00 | 7.33E-06 | 2.69E-05 | -1.53E+00 | 1.22E-01 | 3.36E-01 | 1.12E+00 |
| AC092821.3 | 1.63E-05 | 8.51E-05 | -1.85E+00 | 2.65E-03 | 1.15E-02 | -1.16E+00 | 7.37E-03 | 3.13E-02 | -1.18E+00 | 8.58E-03 | 3.28E-02 | -1.11E+00 | 9.71E-02 | 3.46E-01 | -1.04E+00 | 1.52E-01 | 4.28E-01 | 1.06E+00 | 7.63E-02 | 2.80E-01 | -1.18E+00 | 1.85E-01 | 5.09E-01 | 1.02E+00 |
| SLC38A2 | 1.63E-05 | 8.52E-05 | -1.00E+00 | 2.81E-05 | 1.22E-04 | 1.11E+00 | 1.44E-01 | 6.13E-01 | 1.06E+00 | 1.70E-01 | 6.49E-01 | 1.36E+00 | 1.42E-03 | 5.06E-03 | -1.04E+00 | 2.20E-04 | 6.20E-04 | -1.03E+00 | 4.52E-06 | 1.66E-05 | -1.27E+00 | 9.05E-03 | 2.49E-02 | -1.01E+00 |
| AC011444.2 | 1.64E-05 | 8.55E-05 | 3.25E+00 | 7.10E-02 | 3.07E-01 | -1.14E+00 | 2.05E-04 | 8.71E-04 | 2.77E+00 | 2.55E-01 | 9.74E-01 | 1.28E+00 | 3.26E-02 | 1.16E-01 | 2.50E+00 | 8.38E-02 | 2.36E-01 | 2.20E+00 | 1.88E-02 | 6.89E-02 | 2.01E+00 | 3.41E-03 | 9.38E-03 | 2.82E+00 |
| GCC1 | 1.65E-05 | 8.58E-05 | -1.27E+00 | 1.23E-09 | 5.30E-09 | -1.69E+00 | 1.49E-02 | 6.32E-02 | -1.04E+00 | 7.45E-06 | 2.85E-05 | -1.36E+00 | 6.38E-05 | 2.28E-04 | -1.24E+00 | 8.06E-06 | 2.27E-05 | -1.48E+00 | 2.61E-03 | 9.56E-03 | -1.15E+00 | 1.46E-05 | 4.01E-05 | -1.64E+00 |
| Z97192.3 | 1.65E-05 | 8.58E-05 | -1.30E+00 | 1.15E-01 | 4.98E-01 | 1.34E+00 | 1.41E-06 | 5.99E-06 | -1.77E+00 | 9.93E-02 | 3.79E-01 | 1.20E+00 | 5.09E-03 | 1.81E-02 | -1.08E+00 | 2.93E-01 | 8.27E-01 | 1.48E+00 | 1.58E-03 | 5.80E-03 | -1.60E+00 | 1.73E-01 | 4.76E-01 | 1.10E+00 |
| C11orf21 | 1.65E-05 | 8.61E-05 | -1.22E+00 | 3.99E-06 | 1.72E-05 | -1.20E+00 | 7.58E-05 | 3.22E-04 | -1.35E+00 | 7.62E-04 | 2.91E-03 | -1.05E+00 | 3.19E-05 | 1.14E-04 | -1.42E+00 | 4.38E-05 | 1.24E-04 | -1.34E+00 | 4.75E-03 | 1.74E-02 | 1.10E+00 | 9.34E-04 | 2.56E-03 | -1.28E+00 |
| CCDC6 | 1.67E-05 | 8.68E-05 | -1.08E+00 | 3.13E-11 | 1.35E-10 | -1.23E+00 | 1.00E-02 | 4.25E-02 | 1.07E+00 | 8.26E-05 | 3.16E-04 | -1.04E+00 | 2.51E-01 | 8.93E-01 | 1.37E+00 | 1.64E-02 | 4.63E-02 | -1.06E+00 | 1.53E-01 | 5.62E-01 | 1.25E+00 | 1.09E-02 | 2.98E-02 | -1.04E+00 |
| ZNF600 | 1.67E-05 | 8.69E-05 | -1.40E+00 | 2.07E-05 | 8.94E-05 | -1.25E+00 | 1.33E-02 | 5.64E-02 | -1.14E+00 | 1.13E-02 | 4.32E-02 | 1.00E+00 | 1.48E-02 | 5.26E-02 | 1.02E+00 | 8.56E-03 | 2.41E-02 | -1.05E+00 | 1.93E-03 | 7.08E-03 | -1.36E+00 | 3.44E-03 | 9.45E-03 | -1.25E+00 |
| RARS | 1.67E-05 | 8.71E-05 | -1.08E+00 | 1.19E-03 | 5.14E-03 | 1.17E+00 | 1.29E-06 | 5.48E-06 | -1.28E+00 | 2.81E-03 | 1.07E-02 | 1.07E+00 | 1.59E-05 | 5.65E-05 | -1.30E+00 | 3.56E-02 | 1.00E-01 | 1.09E+00 | 8.20E-08 | 3.01E-07 | -1.28E+00 | 6.22E-02 | 1.71E-01 | 1.11E+00 |
| EIPR1 | 1.67E-05 | 8.71E-05 | -1.21E+00 | 3.67E-04 | 1.59E-03 | 1.04E+00 | 1.64E-01 | 6.99E-01 | 1.20E+00 | 2.25E-03 | 8.60E-03 | 1.11E+00 | 7.99E-06 | 2.85E-05 | -1.38E+00 | 1.56E-03 | 4.39E-03 | 1.01E+00 | 3.67E-03 | 1.35E-02 | -1.02E+00 | 9.12E-02 | 2.51E-01 | 1.12E+00 |
| ZNF586 | 1.68E-05 | 8.74E-05 | -1.39E+00 | 2.00E-05 | 8.66E-05 | -1.25E+00 | 1.80E-07 | 7.63E-07 | -1.44E+00 | 8.32E-09 | 3.18E-08 | -1.38E+00 | 2.36E-04 | 8.43E-04 | -1.23E+00 | 2.22E-03 | 6.26E-03 | -1.04E+00 | 1.32E-04 | 4.84E-04 | -1.32E+00 | 6.82E-03 | 1.87E-02 | -1.07E+00 |
| PMF1 | 1.68E-05 | 8.74E-05 | -1.02E+00 | 7.23E-09 | 3.12E-08 | -1.01E+00 | 2.02E-01 | 8.57E-01 | 1.14E+00 | 2.02E-02 | 7.71E-02 | 1.18E+00 | 3.46E-02 | 1.23E-01 | 1.14E+00 | 2.20E-02 | 6.21E-02 | 1.11E+00 | 1.73E-01 | 6.35E-01 | 1.22E+00 | 1.97E-01 | 5.40E-01 | 1.13E+00 |
| OST4 | 1.68E-05 | 8.74E-05 | -1.12E+00 | 2.51E-23 | 1.08E-22 | -1.38E+00 | 8.21E-02 | 3.49E-01 | -1.06E+00 | 5.95E-09 | 2.27E-08 | -1.34E+00 | 3.64E-03 | 1.30E-02 | -1.12E+00 | 3.27E-05 | 9.23E-05 | -1.32E+00 | 1.96E-01 | 7.19E-01 | 1.06E+00 | 7.44E-04 | 2.04E-03 | -1.28E+00 |
| BTBD19 | 1.68E-05 | 8.75E-05 | 2.28E+00 | 5.35E-03 | 2.31E-02 | -1.82E+00 | 1.60E-01 | 6.80E-01 | 1.37E+00 | 2.99E-03 | 1.14E-02 | -1.85E+00 | 5.23E-02 | 1.86E-01 | 2.09E+00 | 1.06E-01 | 3.00E-01 | -1.19E+00 | 5.24E-02 | 1.92E-01 | 2.22E+00 | 1.19E-01 | 3.27E-01 | 1.20E+00 |
| HLA-DRB1 | 1.69E-05 | 8.78E-05 | -1.25E+00 | 3.39E-02 | 1.46E-01 | 1.01E+00 | 2.83E-02 | 1.20E-01 | -1.15E+00 | 2.52E-01 | 9.62E-01 | 1.07E+00 | 3.89E-16 | 1.39E-15 | -1.39E+00 | 9.13E-09 | 2.57E-08 | -1.15E+00 | 1.45E-43 | 5.33E-43 | -1.99E+00 | 5.63E-12 | 1.55E-11 | -1.30E+00 |
| HNRNPH3 | 1.69E-05 | 8.79E-05 | 1.01E+00 | 1.99E-06 | 8.59E-06 | 1.06E+00 | 1.18E-01 | 5.00E-01 | 1.21E+00 | 1.66E-01 | 6.34E-01 | 1.21E+00 | 3.27E-03 | 1.17E-02 | 1.08E+00 | 3.96E-02 | 1.12E-01 | 1.16E+00 | 1.07E-01 | 3.92E-01 | 1.11E+00 | 5.84E-02 | 1.61E-01 | 1.18E+00 |
| AL450998.1 | 1.69E-05 | 8.80E-05 | -1.22E+00 | 1.44E-57 | 6.20E-57 | -1.89E+00 | 2.30E-03 | 9.76E-03 | -1.21E+00 | 1.44E-15 | 5.49E-15 | -1.77E+00 | 9.45E-04 | 3.37E-03 | -1.20E+00 | 4.53E-13 | 1.28E-12 | -1.62E+00 | 2.77E-07 | 1.02E-06 | -1.31E+00 | 1.75E-13 | 4.81E-13 | -1.77E+00 |
| AC008026.1 | 1.69E-05 | 8.83E-05 | 1.61E+00 | 2.96E-04 | 1.28E-03 | 1.70E+00 | 7.98E-04 | 3.39E-03 | 1.52E+00 | 4.86E-04 | 1.86E-03 | 1.67E+00 | 4.05E-02 | 1.44E-01 | 1.57E+00 | 1.19E-02 | 3.35E-02 | 1.67E+00 | 3.95E-02 | 1.45E-01 | 1.52E+00 | 2.50E-03 | 6.86E-03 | 1.73E+00 |
| PPP1R21 | 1.69E-05 | 8.83E-05 | -1.47E+00 | 6.19E-03 | 2.68E-02 | 1.07E+00 | 2.50E-06 | 1.06E-05 | -1.53E+00 | 4.48E-03 | 1.71E-02 | -1.03E+00 | 6.27E-05 | 2.23E-04 | -1.70E+00 | 1.42E-02 | 4.01E-02 | -1.19E+00 | 8.63E-03 | 3.17E-02 | -1.04E+00 | 5.60E-02 | 1.54E-01 | 1.09E+00 |
| NAA30 | 1.70E-05 | 8.87E-05 | -1.27E+00 | 5.77E-05 | 2.49E-04 | -1.06E+00 | 2.09E-07 | 8.88E-07 | -1.26E+00 | 1.28E-07 | 4.90E-07 | -1.16E+00 | 8.89E-02 | 3.17E-01 | 1.06E+00 | 4.08E-02 | 1.15E-01 | -1.09E+00 | 1.60E-05 | 5.87E-05 | -1.35E+00 | 2.80E-03 | 7.68E-03 | -1.09E+00 |
| LINC00869 | 1.70E-05 | 8.87E-05 | -1.15E+00 | 2.90E-03 | 1.25E-02 | 1.16E+00 | 9.88E-07 | 4.20E-06 | -1.27E+00 | 2.49E-05 | 9.51E-05 | -1.02E+00 | 4.02E-03 | 1.43E-02 | -1.21E+00 | 3.55E-02 | 9.99E-02 | 1.03E+00 | 3.65E-05 | 1.34E-04 | -1.22E+00 | 5.31E-04 | 1.46E-03 | -1.06E+00 |
| DPP8 | 1.70E-05 | 8.87E-05 | -1.29E+00 | 4.50E-03 | 1.95E-02 | 1.12E+00 | 9.00E-03 | 3.82E-02 | -1.11E+00 | 1.28E-02 | 4.90E-02 | 1.07E+00 | 2.88E-02 | 1.03E-01 | -1.06E+00 | 1.91E-01 | 5.39E-01 | 1.19E+00 | 2.90E-02 | 1.07E-01 | 1.05E+00 | 4.42E-03 | 1.21E-02 | -1.05E+00 |
| MAGI3 | 1.71E-05 | 8.89E-05 | -1.86E+00 | 2.50E-04 | 1.08E-03 | -1.46E+00 | 1.33E-08 | 5.63E-08 | -2.36E+00 | 7.49E-07 | 2.86E-06 | -1.77E+00 | 1.33E-01 | 4.73E-01 | -1.16E+00 | 9.14E-02 | 2.58E-01 | -1.55E+00 | 6.60E-03 | 2.42E-02 | -1.43E+00 | 1.78E-02 | 4.88E-02 | -1.27E+00 |
| CNOT1 | 1.71E-05 | 8.89E-05 | -1.21E+00 | 3.39E-07 | 1.47E-06 | -1.26E+00 | 2.81E-03 | 1.19E-02 | -1.07E+00 | 1.17E-06 | 4.47E-06 | -1.32E+00 | 6.23E-04 | 2.22E-03 | -1.14E+00 | 3.97E-05 | 1.12E-04 | -1.30E+00 | 4.14E-03 | 1.52E-02 | -1.06E+00 | 3.39E-04 | 9.32E-04 | -1.25E+00 |
| AC104389.3 | 1.71E-05 | 8.90E-05 | -1.49E+00 | 4.22E-07 | 1.83E-06 | -1.66E+00 | 4.90E-02 | 2.08E-01 | 1.03E+00 | 7.84E-06 | 2.99E-05 | -1.69E+00 | 1.50E-03 | 5.34E-03 | -1.45E+00 | 4.92E-04 | 1.39E-03 | -1.62E+00 | 1.82E-03 | 6.67E-03 | -1.16E+00 | 1.84E-07 | 5.07E-07 | -2.01E+00 |
| RAD18 | 1.71E-05 | 8.92E-05 | -1.35E+00 | 8.29E-03 | 3.58E-02 | 1.09E+00 | 1.29E-01 | 5.49E-01 | 1.24E+00 | 1.15E-01 | 4.41E-01 | 1.55E+00 | 1.40E-02 | 5.01E-02 | -1.13E+00 | 9.13E-02 | 2.57E-01 | 1.09E+00 | 2.90E-02 | 1.06E-01 | 1.08E+00 | 8.65E-02 | 2.38E-01 | 1.13E+00 |
| VSIG10 | 1.71E-05 | 8.93E-05 | -1.37E+00 | 6.83E-03 | 2.95E-02 | 1.05E+00 | 3.66E-06 | 1.56E-05 | -1.41E+00 | 4.26E-03 | 1.63E-02 | 1.07E+00 | 4.81E-05 | 1.72E-04 | -1.71E+00 | 1.30E-03 | 3.67E-03 | -1.36E+00 | 9.32E-07 | 3.42E-06 | -1.50E+00 | 1.09E-04 | 3.01E-04 | -1.28E+00 |
| RAD52 | 1.72E-05 | 8.97E-05 | -1.27E+00 | 6.97E-04 | 3.01E-03 | 1.03E+00 | 8.40E-07 | 3.57E-06 | -1.61E+00 | 7.53E-04 | 2.88E-03 | -1.04E+00 | 8.53E-04 | 3.04E-03 | -1.54E+00 | 3.07E-02 | 8.65E-02 | -1.04E+00 | 1.28E-07 | 4.69E-07 | -1.65E+00 | 1.86E-02 | 5.10E-02 | -1.02E+00 |
| RPL37A | 1.72E-05 | 8.97E-05 | 1.11E+00 | 2.93E-68 | 1.27E-67 | -1.55E+00 | 1.15E-04 | 4.89E-04 | 1.12E+00 | 1.09E-41 | 4.16E-41 | -1.52E+00 | 6.84E-03 | 2.44E-02 | 1.10E+00 | 1.09E-27 | 3.06E-27 | -1.46E+00 | 4.82E-02 | 1.77E-01 | 1.03E+00 | 7.73E-45 | 2.12E-44 | -1.52E+00 |
| AC245060.6 | 1.73E-05 | 8.99E-05 | -2.05E+00 | 7.57E-02 | 3.27E-01 | 1.31E+00 | 1.94E-05 | 8.26E-05 | -2.16E+00 | 2.45E-02 | 9.35E-02 | -1.06E+00 | 2.98E-03 | 1.06E-02 | -1.51E+00 | 1.70E-01 | 4.78E-01 | 1.14E+00 | 1.92E-02 | 7.06E-02 | -1.43E+00 | 3.42E-01 | 9.39E-01 | 1.21E+00 |
| SPATA2L | 1.73E-05 | 9.02E-05 | -1.42E+00 | 4.57E-03 | 1.97E-02 | 1.09E+00 | 2.89E-04 | 1.23E-03 | -1.29E+00 | 6.04E-05 | 2.31E-04 | -1.31E+00 | 1.02E-01 | 3.63E-01 | 1.83E+00 | 3.51E-01 | 9.89E-01 | 1.37E+00 | 1.35E-01 | 4.96E-01 | 1.01E+00 | 2.14E-01 | 5.86E-01 | 1.15E+00 |
| TAPBP | 1.74E-05 | 9.07E-05 | -1.11E+00 | 5.13E-05 | 2.22E-04 | 1.01E+00 | 1.21E-02 | 5.13E-02 | -1.07E+00 | 5.35E-03 | 2.04E-02 | -1.03E+00 | 6.78E-08 | 2.42E-07 | -1.19E+00 | 1.50E-08 | 4.23E-08 | -1.19E+00 | 1.54E-04 | 5.67E-04 | -1.34E+00 | 1.77E-03 | 4.86E-03 | -1.24E+00 |
| RNF8 | 1.74E-05 | 9.09E-05 | -1.02E+00 | 2.00E-02 | 8.63E-02 | 1.35E+00 | 1.49E-04 | 6.32E-04 | -1.21E+00 | 2.92E-02 | 1.12E-01 | 1.15E+00 | 1.30E-03 | 4.62E-03 | -1.20E+00 | 9.16E-02 | 2.58E-01 | 1.21E+00 | 2.86E-04 | 1.05E-03 | -1.07E+00 | 1.91E-01 | 5.25E-01 | 1.26E+00 |
| AC108449.2 | 1.74E-05 | 9.09E-05 | -1.34E+00 | 2.15E-01 | 9.31E-01 | 1.49E+00 | 3.15E-05 | 1.34E-04 | -1.52E+00 | 6.82E-02 | 2.60E-01 | 1.19E+00 | 1.15E-04 | 4.10E-04 | -1.19E+00 | 1.45E-01 | 4.09E-01 | 1.35E+00 | 6.01E-05 | 2.21E-04 | -1.30E+00 | 8.41E-03 | 2.31E-02 | -1.06E+00 |
| TSPAN17 | 1.75E-05 | 9.10E-05 | -1.16E+00 | 7.93E-05 | 3.43E-04 | -1.02E+00 | 5.20E-03 | 2.21E-02 | -1.07E+00 | 7.39E-05 | 2.82E-04 | -1.19E+00 | 8.01E-03 | 2.86E-02 | -1.15E+00 | 3.30E-02 | 9.29E-02 | 1.07E+00 | 6.38E-03 | 2.34E-02 | -1.15E+00 | 4.02E-02 | 1.10E-01 | 1.02E+00 |
| CAMTA2 | 1.75E-05 | 9.10E-05 | -1.47E+00 | 3.29E-04 | 1.42E-03 | -1.13E+00 | 9.60E-04 | 4.08E-03 | -1.61E+00 | 1.67E-02 | 6.37E-02 | -1.13E+00 | 1.99E-01 | 7.10E-01 | -1.03E+00 | 1.90E-01 | 5.35E-01 | -1.10E+00 | 2.40E-01 | 8.80E-01 | 1.21E+00 | 1.42E-01 | 3.89E-01 | 1.00E+00 |
| FAM117B | 1.75E-05 | 9.13E-05 | -1.17E+00 | 1.48E-05 | 6.38E-05 | -1.07E+00 | 7.52E-02 | 3.19E-01 | -1.05E+00 | 1.72E-04 | 6.57E-04 | -1.26E+00 | 2.71E-03 | 9.65E-03 | -1.25E+00 | 1.76E-02 | 4.96E-02 | -1.14E+00 | 1.15E-03 | 4.22E-03 | -1.16E+00 | 2.69E-03 | 7.38E-03 | -1.13E+00 |
| TMEM177 | 1.76E-05 | 9.15E-05 | -1.71E+00 | 1.33E-02 | 5.73E-02 | 1.13E+00 | 3.75E-08 | 1.59E-07 | -2.19E+00 | 2.98E-02 | 1.14E-01 | 1.15E+00 | 2.93E-03 | 1.04E-02 | -1.46E+00 | 1.64E-01 | 4.63E-01 | 1.21E+00 | 2.32E-08 | 8.51E-08 | -1.68E+00 | 1.19E-01 | 3.26E-01 | 1.05E+00 |
| FAM200B | 1.76E-05 | 9.15E-05 | -1.02E+00 | 1.39E-08 | 6.02E-08 | -1.07E+00 | 1.48E-01 | 6.28E-01 | 1.16E+00 | 1.15E-03 | 4.40E-03 | -1.02E+00 | 7.38E-02 | 2.63E-01 | 1.23E+00 | 6.31E-03 | 1.78E-02 | -1.02E+00 | 2.64E-02 | 9.70E-02 | 1.09E+00 | 3.60E-04 | 9.89E-04 | -1.19E+00 |
| DLEU2L | 1.76E-05 | 9.16E-05 | -1.84E+00 | 1.52E-02 | 6.57E-02 | 1.06E+00 | 1.32E-03 | 5.63E-03 | -1.40E+00 | 2.30E-03 | 8.78E-03 | -1.18E+00 | 1.56E-03 | 5.58E-03 | -2.00E+00 | 7.81E-02 | 2.20E-01 | 1.01E+00 | 1.61E-01 | 5.91E-01 | -1.06E+00 | 2.12E-01 | 5.81E-01 | 1.33E+00 |
| TIMP2 | 1.77E-05 | 9.21E-05 | -1.12E+00 | 4.37E-03 | 1.89E-02 | 1.21E+00 | 2.12E-04 | 9.00E-04 | -1.09E+00 | 4.15E-04 | 1.59E-03 | 1.04E+00 | 6.83E-02 | 2.44E-01 | 1.01E+00 | 4.32E-02 | 1.22E-01 | -1.13E+00 | 2.65E-01 | 9.72E-01 | 1.23E+00 | 3.39E-01 | 9.30E-01 | 1.38E+00 |
| GOLGB1 | 1.77E-05 | 9.25E-05 | -1.11E+00 | 1.13E-04 | 4.88E-04 | -1.03E+00 | 4.75E-06 | 2.02E-05 | -1.24E+00 | 4.21E-06 | 1.61E-05 | -1.24E+00 | 3.55E-03 | 1.26E-02 | -1.02E+00 | 2.44E-03 | 6.88E-03 | 1.05E+00 | 1.23E-01 | 4.50E-01 | 1.14E+00 | 6.19E-02 | 1.70E-01 | 1.08E+00 |
| PIK3R1 | 1.78E-05 | 9.25E-05 | 1.64E+00 | 1.13E-01 | 4.89E-01 | 1.46E+00 | 5.02E-03 | 2.13E-02 | 1.40E+00 | 2.51E-01 | 9.58E-01 | 1.28E+00 | 8.40E-03 | 2.99E-02 | 1.72E+00 | 1.78E-02 | 5.03E-02 | 1.05E+00 | 2.34E-02 | 8.60E-02 | 1.08E+00 | 2.21E-03 | 6.08E-03 | 1.01E+00 |
| CCNB1IP1 | 1.78E-05 | 9.26E-05 | -1.02E+00 | 3.42E-04 | 1.48E-03 | 1.10E+00 | 9.43E-03 | 4.01E-02 | -1.05E+00 | 2.70E-03 | 1.03E-02 | -1.00E+00 | 6.96E-02 | 2.48E-01 | 1.08E+00 | 1.37E-01 | 3.87E-01 | 1.19E+00 | 2.61E-01 | 9.59E-01 | 1.23E+00 | 2.64E-01 | 7.25E-01 | 1.15E+00 |
| HOXB5 | 1.78E-05 | 9.30E-05 | -1.06E+00 | 1.24E-02 | 5.35E-02 | 1.25E+00 | 7.25E-04 | 3.08E-03 | -1.39E+00 | 2.61E-01 | 9.98E-01 | 1.29E+00 | 3.03E-03 | 1.08E-02 | -1.43E+00 | 1.52E-01 | 4.29E-01 | 1.07E+00 | 1.65E-01 | 6.05E-01 | 1.06E+00 | 2.10E-01 | 5.76E-01 | 1.52E+00 |
| SCLT1 | 1.79E-05 | 9.32E-05 | -1.34E+00 | 5.79E-03 | 2.50E-02 | 1.03E+00 | 3.35E-02 | 1.42E-01 | 1.04E+00 | 6.00E-02 | 2.29E-01 | 1.18E+00 | 2.98E-07 | 1.06E-06 | -1.57E+00 | 2.40E-04 | 6.76E-04 | -1.22E+00 | 3.26E-04 | 1.20E-03 | -1.10E+00 | 1.53E-03 | 4.20E-03 | -1.09E+00 |
| ZNF431 | 1.80E-05 | 9.37E-05 | -1.07E+00 | 4.06E-06 | 1.75E-05 | 1.01E+00 | 2.59E-04 | 1.10E-03 | -1.24E+00 | 1.68E-04 | 6.40E-04 | -1.10E+00 | 4.51E-04 | 1.61E-03 | 1.04E+00 | 6.28E-04 | 1.77E-03 | 1.13E+00 | 9.14E-02 | 3.36E-01 | 1.37E+00 | 2.07E-01 | 5.67E-01 | 1.39E+00 |
| AL049873.1 | 1.80E-05 | 9.39E-05 | -1.19E+00 | 4.61E-55 | 1.99E-54 | -1.82E+00 | 1.98E-03 | 8.40E-03 | -1.15E+00 | 3.50E-27 | 1.34E-26 | -1.72E+00 | 1.04E-01 | 3.72E-01 | -1.05E+00 | 1.51E-16 | 4.26E-16 | -1.58E+00 | 1.79E-05 | 6.56E-05 | -1.16E+00 | 1.98E-25 | 5.44E-25 | -1.69E+00 |
| AC019069.1 | 1.80E-05 | 9.40E-05 | -1.95E+00 | 4.08E-04 | 1.76E-03 | -1.40E+00 | 8.47E-08 | 3.60E-07 | -1.87E+00 | 3.94E-05 | 1.51E-04 | -1.27E+00 | 1.28E-03 | 4.55E-03 | -2.00E+00 | 1.29E-01 | 3.64E-01 | 1.20E+00 | 6.74E-05 | 2.47E-04 | -1.62E+00 | 2.97E-01 | 8.15E-01 | 1.35E+00 |
| FBXL18 | 1.81E-05 | 9.44E-05 | -1.49E+00 | 4.76E-06 | 2.06E-05 | -1.61E+00 | 7.12E-05 | 3.02E-04 | -1.65E+00 | 2.33E-05 | 8.92E-05 | -1.69E+00 | 3.19E-04 | 1.14E-03 | -1.82E+00 | 1.16E-03 | 3.28E-03 | -1.65E+00 | 1.35E-03 | 4.94E-03 | -1.69E+00 | 1.52E-02 | 4.17E-02 | -1.36E+00 |
| SULT1A2 | 1.82E-05 | 9.50E-05 | -1.77E+00 | 4.56E-04 | 1.97E-03 | -1.23E+00 | 8.77E-06 | 3.73E-05 | -2.06E+00 | 4.03E-03 | 1.54E-02 | -1.12E+00 | 1.61E-18 | 5.74E-18 | -3.45E+00 | 1.03E-12 | 2.92E-12 | -1.93E+00 | 3.21E-29 | 1.18E-28 | -2.07E+00 | 1.79E-24 | 4.92E-24 | -1.91E+00 |
| EXPH5 | 1.83E-05 | 9.52E-05 | -1.44E+00 | 4.81E-04 | 2.08E-03 | -1.11E+00 | 6.72E-07 | 2.85E-06 | -1.85E+00 | 2.37E-03 | 9.06E-03 | -1.07E+00 | 3.05E-03 | 1.09E-02 | -2.01E+00 | 9.31E-02 | 2.62E-01 | 1.03E+00 | 5.59E-04 | 2.05E-03 | -1.53E+00 | 4.02E-03 | 1.10E-02 | -1.36E+00 |
| MSRA | 1.83E-05 | 9.55E-05 | -1.15E+00 | 9.02E-04 | 3.90E-03 | 1.08E+00 | 1.12E-01 | 4.74E-01 | 1.06E+00 | 2.64E-02 | 1.01E-01 | 1.03E+00 | 2.02E-01 | 7.21E-01 | 1.32E+00 | 3.43E-02 | 9.68E-02 | 1.12E+00 | 9.21E-03 | 3.38E-02 | 1.03E+00 | 4.96E-02 | 1.36E-01 | 1.12E+00 |
| HSD17B8 | 1.83E-05 | 9.55E-05 | -1.16E+00 | 1.23E-05 | 5.31E-05 | -1.06E+00 | 4.56E-04 | 1.94E-03 | -1.02E+00 | 8.10E-07 | 3.09E-06 | -1.09E+00 | 1.48E-03 | 5.26E-03 | -1.25E+00 | 4.45E-02 | 1.25E-01 | 1.07E+00 | 2.42E-02 | 8.89E-02 | -1.03E+00 | 1.15E-02 | 3.16E-02 | -1.06E+00 |
| ZNF30 | 1.83E-05 | 9.56E-05 | -1.80E+00 | 5.57E-04 | 2.41E-03 | -1.28E+00 | 5.67E-05 | 2.41E-04 | -1.80E+00 | 3.31E-03 | 1.26E-02 | -1.11E+00 | 1.70E-02 | 6.06E-02 | -1.44E+00 | 1.05E-01 | 2.95E-01 | -1.11E+00 | 4.75E-03 | 1.74E-02 | -1.25E+00 | 6.07E-02 | 1.67E-01 | -1.11E+00 |
| FAM76A | 1.83E-05 | 9.56E-05 | -1.35E+00 | 1.86E-02 | 8.03E-02 | 1.23E+00 | 1.22E-08 | 5.20E-08 | -1.90E+00 | 6.03E-06 | 2.30E-05 | -1.30E+00 | 1.09E-07 | 3.89E-07 | -1.96E+00 | 2.37E-03 | 6.68E-03 | -1.14E+00 | 4.29E-08 | 1.57E-07 | -1.87E+00 | 2.59E-04 | 7.13E-04 | -1.35E+00 |
| TMEM135 | 1.85E-05 | 9.63E-05 | -1.33E+00 | 4.01E-03 | 1.73E-02 | 1.07E+00 | 2.51E-05 | 1.07E-04 | -1.37E+00 | 1.51E-03 | 5.76E-03 | -1.06E+00 | 4.85E-04 | 1.73E-03 | -1.51E+00 | 1.59E-03 | 4.48E-03 | -1.23E+00 | 1.96E-06 | 7.20E-06 | -1.10E+00 | 4.97E-03 | 1.36E-02 | -1.17E+00 |
| AC010642.2 | 1.85E-05 | 9.66E-05 | -1.31E+00 | 8.93E-08 | 3.86E-07 | -1.51E+00 | 1.71E-04 | 7.26E-04 | -1.31E+00 | 7.09E-08 | 2.71E-07 | -1.77E+00 | 2.24E-02 | 7.99E-02 | 1.03E+00 | 3.02E-02 | 8.53E-02 | 1.08E+00 | 8.04E-03 | 2.95E-02 | -1.02E+00 | 9.04E-04 | 2.48E-03 | -1.14E+00 |
| NOTCH4 | 1.86E-05 | 9.70E-05 | 3.28E+00 | 1.15E-01 | 4.95E-01 | 1.11E+00 | 1.07E-02 | 4.56E-02 | 1.76E+00 | 2.05E-02 | 7.84E-02 | -1.26E+00 | 2.37E-01 | 8.46E-01 | 1.66E+00 | 1.89E-01 | 5.34E-01 | 1.10E+00 | 2.22E-01 | 8.15E-01 | 1.24E+00 | 1.20E-01 | 3.30E-01 | -1.12E+00 |
| CHID1 | 1.86E-05 | 9.71E-05 | -1.11E+00 | 9.57E-05 | 4.13E-04 | 1.06E+00 | 1.47E-02 | 6.22E-02 | 1.05E+00 | 7.62E-02 | 2.91E-01 | 1.24E+00 | 1.19E-02 | 4.23E-02 | -1.01E+00 | 1.12E-01 | 3.17E-01 | 1.15E+00 | 4.85E-02 | 1.78E-01 | 1.07E+00 | 2.12E-01 | 5.83E-01 | 1.34E+00 |
| MMRN1 | 1.87E-05 | 9.74E-05 | -1.03E+00 | 2.15E-01 | 9.31E-01 | 1.50E+00 | 7.17E-02 | 3.04E-01 | -1.01E+00 | 1.34E-01 | 5.12E-01 | 1.49E+00 | 1.54E-01 | 5.48E-01 | 1.17E+00 | 1.85E-01 | 5.21E-01 | 1.12E+00 | 1.08E-02 | 3.97E-02 | 1.16E+00 | 1.84E-01 | 5.04E-01 | -1.20E+00 |
| LCLAT1 | 1.88E-05 | 9.80E-05 | -1.42E+00 | 1.00E-02 | 4.33E-02 | 1.19E+00 | 2.82E-04 | 1.20E-03 | -1.24E+00 | 2.94E-03 | 1.12E-02 | -1.01E+00 | 1.92E-07 | 6.86E-07 | -1.94E+00 | 2.24E-04 | 6.31E-04 | -1.23E+00 | 1.41E-14 | 5.18E-14 | -1.70E+00 | 6.94E-07 | 1.91E-06 | -1.27E+00 |
| POMP | 1.88E-05 | 9.81E-05 | -1.01E+00 | 1.57E-06 | 6.76E-06 | 1.04E+00 | 1.88E-01 | 7.97E-01 | 1.19E+00 | 4.25E-02 | 1.62E-01 | 1.08E+00 | 2.57E-06 | 9.18E-06 | -1.16E+00 | 3.06E-06 | 8.62E-06 | -1.14E+00 | 9.64E-02 | 3.54E-01 | 1.07E+00 | 1.84E-01 | 5.06E-01 | 1.02E+00 |
| AC016876.1 | 1.89E-05 | 9.83E-05 | -1.48E+00 | 9.59E-05 | 4.14E-04 | -1.25E+00 | 1.59E-01 | 6.77E-01 | 1.09E+00 | 4.24E-02 | 1.62E-01 | -1.02E+00 | 2.02E-02 | 7.21E-02 | -1.17E+00 | 9.66E-02 | 2.72E-01 | 1.03E+00 | 3.37E-02 | 1.24E-01 | -1.05E+00 | 1.68E-01 | 4.62E-01 | 1.22E+00 |
| DIXDC1 | 1.89E-05 | 9.87E-05 | -2.11E+00 | 4.15E-04 | 1.79E-03 | -1.56E+00 | 1.96E-09 | 8.35E-09 | -2.18E+00 | 2.58E-06 | 9.86E-06 | -1.47E+00 | 1.83E-02 | 6.51E-02 | -1.74E+00 | 3.19E-02 | 8.99E-02 | -1.46E+00 | 2.21E-08 | 8.09E-08 | -2.33E+00 | 3.67E-03 | 1.01E-02 | -1.09E+00 |
| ago-01 | 1.89E-05 | 9.87E-05 | -1.11E+00 | 1.12E-03 | 4.83E-03 | 1.16E+00 | 1.72E-04 | 7.32E-04 | -1.16E+00 | 2.51E-02 | 9.60E-02 | 1.13E+00 | 2.30E-01 | 8.20E-01 | 1.54E+00 | 3.17E-01 | 8.94E-01 | 1.50E+00 | 7.31E-03 | 2.68E-02 | 1.19E+00 | 7.20E-04 | 1.98E-03 | 1.04E+00 |
| GEMIN5 | 1.89E-05 | 9.87E-05 | -1.68E+00 | 1.06E-01 | 4.59E-01 | 1.33E+00 | 2.94E-06 | 1.25E-05 | -1.52E+00 | 2.44E-03 | 9.32E-03 | 1.02E+00 | 7.98E-03 | 2.84E-02 | -1.56E+00 | 3.15E-01 | 8.89E-01 | 1.40E+00 | 1.61E-05 | 5.92E-05 | -1.35E+00 | 1.30E-01 | 3.56E-01 | 1.49E+00 |
| ZFP91 | 1.90E-05 | 9.91E-05 | -1.19E+00 | 5.17E-08 | 2.23E-07 | -1.30E+00 | 3.33E-07 | 1.41E-06 | -1.44E+00 | 1.40E-12 | 5.36E-12 | -1.69E+00 | 2.53E-03 | 9.02E-03 | -1.07E+00 | 1.41E-04 | 3.99E-04 | -1.34E+00 | 4.43E-12 | 1.63E-11 | -1.53E+00 | 6.97E-10 | 1.91E-09 | -1.42E+00 |
| AKIRIN2 | 1.90E-05 | 9.91E-05 | -1.04E+00 | 2.78E-06 | 1.20E-05 | 1.04E+00 | 6.19E-05 | 2.63E-04 | -1.20E+00 | 5.35E-07 | 2.04E-06 | -1.20E+00 | 6.56E-05 | 2.34E-04 | -1.17E+00 | 9.59E-06 | 2.70E-05 | -1.20E+00 | 3.57E-05 | 1.31E-04 | -1.14E+00 | 9.74E-07 | 2.67E-06 | -1.23E+00 |
| NAA16 | 1.90E-05 | 9.92E-05 | -1.22E+00 | 3.46E-05 | 1.49E-04 | -1.14E+00 | 2.25E-02 | 9.54E-02 | 1.01E+00 | 3.21E-05 | 1.23E-04 | -1.14E+00 | 1.44E-02 | 5.15E-02 | 1.04E+00 | 6.34E-03 | 1.79E-02 | -1.05E+00 | 2.06E-03 | 7.57E-03 | -1.11E+00 | 4.87E-04 | 1.34E-03 | -1.20E+00 |
| SLC35A4 | 1.90E-05 | 9.92E-05 | -1.20E+00 | 1.23E-03 | 5.32E-03 | 1.08E+00 | 9.03E-04 | 3.84E-03 | -1.07E+00 | 5.62E-03 | 2.15E-02 | 1.22E+00 | 1.47E-03 | 5.24E-03 | -1.32E+00 | 9.52E-02 | 2.68E-01 | 1.09E+00 | 2.52E-02 | 9.26E-02 | 1.05E+00 | 3.11E-01 | 8.54E-01 | 1.31E+00 |
| TMEM183A | 1.92E-05 | 9.98E-05 | -1.21E+00 | 2.46E-07 | 1.06E-06 | -1.26E+00 | 4.91E-05 | 2.08E-04 | -1.37E+00 | 8.01E-09 | 3.06E-08 | -1.52E+00 | 6.58E-03 | 2.35E-02 | 1.07E+00 | 1.04E-04 | 2.93E-04 | -1.13E+00 | 3.83E-03 | 1.41E-02 | -1.05E+00 | 1.79E-05 | 4.91E-05 | -1.22E+00 |
| PAXBP1 | 1.92E-05 | 9.99E-05 | -1.12E+00 | 3.56E-07 | 1.54E-06 | -1.12E+00 | 2.38E-04 | 1.01E-03 | -1.13E+00 | 8.66E-06 | 3.31E-05 | -1.09E+00 | 1.44E-04 | 5.13E-04 | -1.18E+00 | 2.66E-04 | 7.49E-04 | -1.22E+00 | 1.23E-06 | 4.52E-06 | -1.18E+00 | 1.51E-04 | 4.13E-04 | -1.16E+00 |
| SIK3 | 1.92E-05 | 1.00E-04 | -1.16E+00 | 9.02E-07 | 3.90E-06 | -1.08E+00 | 3.98E-04 | 1.69E-03 | -1.19E+00 | 3.94E-04 | 1.50E-03 | -1.10E+00 | 1.62E-01 | 5.79E-01 | 1.39E+00 | 4.20E-02 | 1.18E-01 | 1.03E+00 | 1.01E-01 | 3.70E-01 | 1.09E+00 | 4.77E-03 | 1.31E-02 | -1.12E+00 |
| SH3BP5L | 1.92E-05 | 1.00E-04 | -1.88E+00 | 8.02E-03 | 3.47E-02 | -1.03E+00 | 8.35E-06 | 3.55E-05 | -2.08E+00 | 3.58E-03 | 1.37E-02 | -1.26E+00 | 3.03E-03 | 1.08E-02 | -1.61E+00 | 8.20E-02 | 2.31E-01 | -1.00E+00 | 1.51E-03 | 5.53E-03 | -1.47E+00 | 2.55E-01 | 7.02E-01 | 1.15E+00 |
| DTD2 | 1.93E-05 | 1.00E-04 | -1.24E+00 | 2.27E-04 | 9.82E-04 | -1.06E+00 | 8.08E-06 | 3.43E-05 | -1.35E+00 | 7.76E-03 | 2.96E-02 | 1.06E+00 | 7.70E-05 | 2.74E-04 | -1.64E+00 | 9.62E-02 | 2.71E-01 | 1.20E+00 | 2.86E-05 | 1.05E-04 | -1.25E+00 | 4.83E-02 | 1.33E-01 | 1.07E+00 |
| ZNF385A | 1.93E-05 | 1.01E-04 | -1.88E+00 | 3.66E-04 | 1.58E-03 | -1.41E+00 | 2.25E-02 | 9.55E-02 | -1.27E+00 | 3.64E-02 | 1.39E-01 | -1.07E+00 | 1.92E-04 | 6.84E-04 | -1.68E+00 | 8.51E-03 | 2.40E-02 | -1.18E+00 | 2.88E-08 | 1.06E-07 | -1.13E+00 | 5.47E-16 | 1.50E-15 | -1.39E+00 |
| RPAP3 | 1.93E-05 | 1.01E-04 | -1.19E+00 | 4.40E-05 | 1.90E-04 | -1.05E+00 | 2.54E-05 | 1.08E-04 | -1.26E+00 | 4.45E-03 | 1.70E-02 | 1.09E+00 | 1.90E-07 | 6.79E-07 | -1.43E+00 | 1.88E-04 | 5.30E-04 | -1.07E+00 | 7.67E-06 | 2.81E-05 | -1.22E+00 | 1.01E-03 | 2.79E-03 | -1.10E+00 |
| RPL23P2 | 1.93E-05 | 1.01E-04 | -1.45E+00 | 2.26E-04 | 9.75E-04 | -1.10E+00 | 8.18E-07 | 3.48E-06 | -1.60E+00 | 1.47E-02 | 5.62E-02 | 1.08E+00 | 5.48E-05 | 1.96E-04 | -1.51E+00 | 1.55E-02 | 4.36E-02 | 1.05E+00 | 3.47E-05 | 1.27E-04 | -1.38E+00 | 2.33E-02 | 6.39E-02 | 1.00E+00 |
| FAM189B | 1.93E-05 | 1.01E-04 | -1.35E+00 | 1.79E-03 | 7.75E-03 | 1.03E+00 | 1.77E-04 | 7.54E-04 | -1.44E+00 | 1.92E-02 | 7.35E-02 | 1.01E+00 | 3.04E-03 | 1.08E-02 | -1.40E+00 | 4.83E-02 | 1.36E-01 | -1.03E+00 | 4.79E-04 | 1.76E-03 | 1.04E+00 | 1.51E-04 | 4.14E-04 | -1.16E+00 |
| SKP2 | 1.94E-05 | 1.01E-04 | -1.42E+00 | 2.04E-04 | 8.80E-04 | -1.16E+00 | 3.37E-04 | 1.43E-03 | -1.20E+00 | 9.38E-04 | 3.58E-03 | -1.01E+00 | 2.57E-02 | 9.16E-02 | -1.20E+00 | 2.90E-01 | 8.16E-01 | 1.40E+00 | 1.87E-02 | 6.88E-02 | -1.06E+00 | 2.11E-01 | 5.81E-01 | 1.14E+00 |
| PSMA6P1 | 1.95E-05 | 1.02E-04 | 1.76E+00 | 1.10E-01 | 4.73E-01 | 1.54E+00 | 2.16E-07 | 9.18E-07 | 2.02E+00 | 1.09E-01 | 4.15E-01 | 1.52E+00 | 7.07E-02 | 2.52E-01 | 1.18E+00 | 4.15E-02 | 1.17E-01 | 1.15E+00 | 3.11E-03 | 1.14E-02 | 1.51E+00 | 1.35E-01 | 3.70E-01 | 1.29E+00 |
| RIOK2 | 1.96E-05 | 1.02E-04 | -1.24E+00 | 2.50E-04 | 1.08E-03 | -1.01E+00 | 7.67E-04 | 3.26E-03 | -1.13E+00 | 6.72E-06 | 2.57E-05 | -1.11E+00 | 7.51E-05 | 2.68E-04 | -1.21E+00 | 1.12E-03 | 3.14E-03 | -1.04E+00 | 7.09E-14 | 2.60E-13 | -1.74E+00 | 2.87E-06 | 7.89E-06 | -1.25E+00 |
| PTGIR | 1.96E-05 | 1.02E-04 | -2.45E+00 | 5.86E-04 | 2.53E-03 | -1.61E+00 | 6.56E-06 | 2.79E-05 | -3.12E+00 | 3.88E-03 | 1.48E-02 | -1.47E+00 | 2.00E-02 | 7.14E-02 | -2.05E+00 | 3.06E-02 | 8.63E-02 | -1.95E+00 | 1.99E-01 | 7.30E-01 | 3.42E+00 | 1.09E-01 | 2.99E-01 | 1.43E+00 |
| DPYD | 1.96E-05 | 1.02E-04 | -1.21E+00 | 1.48E-02 | 6.38E-02 | 1.21E+00 | 2.57E-04 | 1.09E-03 | -1.26E+00 | 3.33E-02 | 1.27E-01 | 1.19E+00 | 1.72E-03 | 6.15E-03 | -2.57E+00 | 3.11E-02 | 8.77E-02 | -1.41E+00 | 3.74E-05 | 1.37E-04 | -1.95E+00 | 9.74E-04 | 2.67E-03 | -1.54E+00 |
| ARMCX1 | 1.97E-05 | 1.03E-04 | -1.06E+00 | 1.51E-06 | 6.53E-06 | 1.01E+00 | 5.87E-03 | 2.49E-02 | -1.03E+00 | 2.84E-05 | 1.08E-04 | -1.12E+00 | 1.70E-04 | 6.06E-04 | -1.12E+00 | 5.49E-04 | 1.55E-03 | -1.10E+00 | 5.61E-06 | 2.06E-05 | -1.29E+00 | 7.38E-04 | 2.03E-03 | -1.17E+00 |
| GNB1 | 1.98E-05 | 1.03E-04 | -1.00E+00 | 6.85E-03 | 2.96E-02 | 1.22E+00 | 6.81E-02 | 2.89E-01 | 1.02E+00 | 4.64E-02 | 1.77E-01 | 1.08E+00 | 1.53E-03 | 5.45E-03 | -1.00E+00 | 8.38E-03 | 2.36E-02 | 1.08E+00 | 1.90E-05 | 6.96E-05 | -1.21E+00 | 4.13E-03 | 1.13E-02 | -1.11E+00 |
| AL136298.3 | 1.99E-05 | 1.03E-04 | -1.30E+00 | 8.03E-11 | 3.47E-10 | -1.83E+00 | 5.56E-09 | 2.36E-08 | -1.35E+00 | 1.52E-16 | 5.79E-16 | -1.81E+00 | 3.09E-04 | 1.10E-03 | -1.25E+00 | 1.54E-05 | 4.35E-05 | -1.49E+00 | 2.30E-04 | 8.45E-04 | -1.13E+00 | 2.85E-09 | 7.82E-09 | -1.52E+00 |
| GTF3C3 | 2.00E-05 | 1.04E-04 | -1.14E+00 | 1.18E-02 | 5.08E-02 | 1.22E+00 | 1.97E-03 | 8.37E-03 | -1.08E+00 | 3.74E-02 | 1.43E-01 | 1.20E+00 | 8.59E-04 | 3.06E-03 | -1.11E+00 | 2.64E-02 | 7.44E-02 | 1.11E+00 | 8.63E-06 | 3.17E-05 | -1.23E+00 | 6.50E-04 | 1.78E-03 | -1.12E+00 |
| PSMA7 | 2.00E-05 | 1.04E-04 | 1.01E+00 | 3.54E-02 | 1.53E-01 | 1.15E+00 | 4.96E-02 | 2.11E-01 | 1.17E+00 | 8.48E-02 | 3.24E-01 | 1.19E+00 | 2.11E-03 | 7.53E-03 | -1.07E+00 | 2.83E-01 | 7.99E-01 | 1.11E+00 | 8.15E-02 | 2.99E-01 | 1.14E+00 | 4.92E-03 | 1.35E-02 | 1.29E+00 |
| ZFYVE16 | 2.01E-05 | 1.05E-04 | -1.29E+00 | 7.54E-03 | 3.26E-02 | 1.08E+00 | 9.33E-06 | 3.96E-05 | -1.39E+00 | 3.81E-02 | 1.46E-01 | 1.18E+00 | 4.25E-04 | 1.51E-03 | -1.15E+00 | 4.10E-03 | 1.16E-02 | 1.02E+00 | 2.38E-08 | 8.72E-08 | -1.79E+00 | 3.14E-06 | 8.61E-06 | -1.64E+00 |
| SMIM19 | 2.01E-05 | 1.05E-04 | 1.01E+00 | 4.76E-09 | 2.05E-08 | -1.05E+00 | 2.49E-02 | 1.06E-01 | 1.30E+00 | 9.08E-02 | 3.47E-01 | 1.19E+00 | 7.98E-04 | 2.85E-03 | -1.09E+00 | 2.04E-03 | 5.75E-03 | -1.05E+00 | 1.85E-01 | 6.81E-01 | 1.25E+00 | 8.54E-02 | 2.35E-01 | 1.10E+00 |
| CCDC93 | 2.01E-05 | 1.05E-04 | -1.26E+00 | 7.43E-05 | 3.21E-04 | -1.07E+00 | 3.42E-03 | 1.45E-02 | -1.11E+00 | 1.85E-05 | 7.08E-05 | -1.14E+00 | 8.20E-04 | 2.92E-03 | -1.22E+00 | 2.54E-05 | 7.16E-05 | -1.53E+00 | 6.48E-03 | 2.38E-02 | -1.04E+00 | 8.15E-03 | 2.24E-02 | -1.02E+00 |
| FAM122B | 2.01E-05 | 1.05E-04 | -1.37E+00 | 6.98E-03 | 3.02E-02 | 1.14E+00 | 5.08E-05 | 2.16E-04 | -1.39E+00 | 1.78E-03 | 6.80E-03 | -1.03E+00 | 1.65E-02 | 5.90E-02 | -1.01E+00 | 1.79E-01 | 5.04E-01 | 1.22E+00 | 3.41E-06 | 1.25E-05 | -1.39E+00 | 6.30E-04 | 1.73E-03 | -1.10E+00 |
| GRK2 | 2.02E-05 | 1.05E-04 | -1.03E+00 | 3.89E-03 | 1.68E-02 | 1.22E+00 | 6.37E-08 | 2.71E-07 | -1.30E+00 | 4.88E-07 | 1.86E-06 | -1.08E+00 | 5.35E-04 | 1.91E-03 | 1.03E+00 | 3.92E-05 | 1.10E-04 | -1.05E+00 | 1.61E-04 | 5.90E-04 | -1.04E+00 | 2.98E-06 | 8.18E-06 | -1.16E+00 |
| AKAP11 | 2.02E-05 | 1.05E-04 | -1.22E+00 | 5.90E-04 | 2.55E-03 | 1.01E+00 | 2.91E-03 | 1.23E-02 | -1.25E+00 | 7.02E-02 | 2.68E-01 | 1.18E+00 | 8.82E-03 | 3.14E-02 | -1.17E+00 | 1.46E-02 | 4.11E-02 | -1.11E+00 | 4.97E-04 | 1.82E-03 | -1.11E+00 | 5.32E-04 | 1.46E-03 | -1.10E+00 |
| ASRGL1 | 2.02E-05 | 1.05E-04 | -1.55E+00 | 8.56E-08 | 3.70E-07 | -2.01E+00 | 1.64E-03 | 6.97E-03 | 1.00E+00 | 1.00E-07 | 3.82E-07 | -1.16E+00 | 7.09E-14 | 2.53E-13 | -2.06E+00 | 6.66E-14 | 1.88E-13 | -1.84E+00 | 3.38E-03 | 1.24E-02 | -1.02E+00 | 4.71E-02 | 1.29E-01 | 1.10E+00 |
| OTUD6B | 2.02E-05 | 1.06E-04 | -1.24E+00 | 1.67E-04 | 7.24E-04 | -1.01E+00 | 3.65E-05 | 1.55E-04 | -1.35E+00 | 4.62E-03 | 1.76E-02 | -1.02E+00 | 1.07E-03 | 3.80E-03 | -1.20E+00 | 1.47E-02 | 4.13E-02 | 1.00E+00 | 1.55E-08 | 5.68E-08 | -1.49E+00 | 2.37E-04 | 6.50E-04 | -1.18E+00 |
| NT5C3A | 2.02E-05 | 1.06E-04 | 1.01E+00 | 2.77E-04 | 1.20E-03 | 1.19E+00 | 3.58E-02 | 1.52E-01 | 1.05E+00 | 2.83E-02 | 1.08E-01 | 1.14E+00 | 1.26E-02 | 4.50E-02 | 1.14E+00 | 1.79E-01 | 5.04E-01 | 1.31E+00 | 1.98E-01 | 7.26E-01 | 1.38E+00 | 1.60E-01 | 4.38E-01 | 1.21E+00 |
| TICAM1 | 2.03E-05 | 1.06E-04 | -1.91E+00 | 1.86E-05 | 8.03E-05 | -1.80E+00 | 3.01E-05 | 1.28E-04 | -1.75E+00 | 1.02E-04 | 3.88E-04 | -1.52E+00 | 1.04E-03 | 3.70E-03 | -2.29E+00 | 8.16E-02 | 2.30E-01 | -1.04E+00 | 2.82E-15 | 1.03E-14 | -2.64E+00 | 1.61E-11 | 4.43E-11 | -1.95E+00 |
| C5orf63 | 2.04E-05 | 1.06E-04 | -1.61E+00 | 1.70E-02 | 7.33E-02 | 1.06E+00 | 5.65E-04 | 2.40E-03 | -1.43E+00 | 2.05E-02 | 7.84E-02 | 1.18E+00 | 2.15E-04 | 7.68E-04 | -1.27E+00 | 7.98E-02 | 2.25E-01 | 1.21E+00 | 1.31E-07 | 4.80E-07 | -1.49E+00 | 6.46E-03 | 1.78E-02 | -1.01E+00 |
| DGCR8 | 2.04E-05 | 1.06E-04 | -1.34E+00 | 5.01E-03 | 2.16E-02 | 1.05E+00 | 2.35E-03 | 1.00E-02 | -1.21E+00 | 2.55E-01 | 9.75E-01 | 1.52E+00 | 5.77E-06 | 2.06E-05 | -1.71E+00 | 2.35E-04 | 6.64E-04 | -1.33E+00 | 4.63E-06 | 1.70E-05 | -1.45E+00 | 2.17E-01 | 5.95E-01 | 1.20E+00 |
| PPIG | 2.04E-05 | 1.07E-04 | -1.01E+00 | 4.52E-03 | 1.95E-02 | 1.15E+00 | 1.14E-02 | 4.86E-02 | -1.04E+00 | 1.73E-02 | 6.60E-02 | 1.02E+00 | 6.97E-07 | 2.48E-06 | -1.23E+00 | 2.00E-03 | 5.65E-03 | -1.00E+00 | 4.01E-03 | 1.47E-02 | -1.06E+00 | 1.08E-01 | 2.96E-01 | -1.00E+00 |
| AC104027.1 | 2.06E-05 | 1.07E-04 | -1.90E+00 | 4.06E-04 | 1.76E-03 | -1.49E+00 | 2.23E-08 | 9.49E-08 | -2.12E+00 | 3.27E-06 | 1.25E-05 | -1.48E+00 | 2.35E-07 | 8.37E-07 | -2.41E+00 | 1.62E-04 | 4.56E-04 | -1.40E+00 | 1.10E-14 | 4.03E-14 | -2.61E+00 | 3.70E-07 | 1.02E-06 | -1.51E+00 |
| SIRPA | 2.06E-05 | 1.07E-04 | -1.56E+00 | 8.00E-04 | 3.46E-03 | -1.13E+00 | 3.87E-05 | 1.64E-04 | -1.53E+00 | 4.45E-04 | 1.70E-03 | -1.23E+00 | 8.05E-02 | 2.87E-01 | -1.80E+00 | 1.18E-01 | 3.32E-01 | -1.63E+00 | 9.84E-02 | 3.61E-01 | -1.59E+00 | 2.36E-01 | 6.49E-01 | -1.27E+00 |
| DBF4B | 2.06E-05 | 1.07E-04 | -1.99E+00 | 6.27E-03 | 2.71E-02 | -1.13E+00 | 2.08E-02 | 8.85E-02 | -1.08E+00 | 4.84E-02 | 1.85E-01 | 1.11E+00 | 1.82E-01 | 6.47E-01 | 1.11E+00 | 2.31E-01 | 6.51E-01 | 1.74E+00 | 9.49E-03 | 3.48E-02 | -1.11E+00 | 2.70E-01 | 7.42E-01 | 1.27E+00 |
| NOMO1 | 2.07E-05 | 1.08E-04 | -1.40E+00 | 6.36E-07 | 2.75E-06 | -1.59E+00 | 5.78E-06 | 2.46E-05 | -1.45E+00 | 1.58E-04 | 6.02E-04 | -1.10E+00 | 5.07E-06 | 1.81E-05 | -1.65E+00 | 2.27E-05 | 6.40E-05 | -1.40E+00 | 3.86E-06 | 1.42E-05 | -1.48E+00 | 4.59E-04 | 1.26E-03 | -1.20E+00 |
| R3HDM1 | 2.07E-05 | 1.08E-04 | -1.08E+00 | 5.94E-04 | 2.57E-03 | 1.15E+00 | 4.47E-04 | 1.90E-03 | -1.11E+00 | 2.12E-03 | 8.10E-03 | 1.04E+00 | 7.67E-05 | 2.73E-04 | -1.21E+00 | 2.28E-02 | 6.44E-02 | 1.07E+00 | 1.05E-03 | 3.85E-03 | -1.01E+00 | 7.36E-02 | 2.02E-01 | 1.07E+00 |
| NPM1P29 | 2.07E-05 | 1.08E-04 | -1.29E+00 | 3.59E-04 | 1.55E-03 | -1.10E+00 | 3.84E-02 | 1.63E-01 | 1.01E+00 | 1.86E-02 | 7.10E-02 | 1.08E+00 | 1.84E-03 | 6.55E-03 | -1.26E+00 | 4.61E-03 | 1.30E-02 | -1.18E+00 | 2.54E-03 | 9.31E-03 | -1.26E+00 | 1.18E-01 | 3.25E-01 | 1.17E+00 |
| AL137802.1 | 2.08E-05 | 1.08E-04 | -1.58E+00 | 5.60E-05 | 2.42E-04 | -1.36E+00 | 6.35E-05 | 2.70E-04 | -1.39E+00 | 4.55E-07 | 1.74E-06 | -1.69E+00 | 2.27E-02 | 8.09E-02 | -1.31E+00 | 1.01E-02 | 2.84E-02 | -1.71E+00 | 3.63E-04 | 1.33E-03 | -1.45E+00 | 1.98E-07 | 5.43E-07 | -2.35E+00 |
| P4HB | 2.08E-05 | 1.08E-04 | -1.10E+00 | 2.37E-03 | 1.02E-02 | 1.06E+00 | 2.92E-02 | 1.24E-01 | 1.03E+00 | 1.36E-01 | 5.20E-01 | 1.12E+00 | 5.18E-11 | 1.85E-10 | -1.58E+00 | 4.03E-05 | 1.14E-04 | -1.25E+00 | 6.12E-02 | 2.25E-01 | -1.05E+00 | 1.95E-01 | 5.36E-01 | 1.11E+00 |
| DDX19B | 2.08E-05 | 1.09E-04 | -1.13E+00 | 2.25E-06 | 9.70E-06 | -1.10E+00 | 2.19E-03 | 9.33E-03 | -1.04E+00 | 1.28E-06 | 4.90E-06 | -1.17E+00 | 3.94E-03 | 1.40E-02 | 1.02E+00 | 1.54E-02 | 4.34E-02 | 1.07E+00 | 5.42E-06 | 1.99E-05 | -1.16E+00 | 1.12E-04 | 3.09E-04 | -1.15E+00 |
| LINC00987 | 2.08E-05 | 1.09E-04 | -1.51E+00 | 8.59E-03 | 3.71E-02 | 1.11E+00 | 2.51E-03 | 1.07E-02 | -1.39E+00 | 8.21E-02 | 3.14E-01 | 1.10E+00 | 1.30E-03 | 4.63E-03 | -1.47E+00 | 2.96E-02 | 8.34E-02 | -1.08E+00 | 1.62E-01 | 5.95E-01 | -1.09E+00 | 2.56E-01 | 7.04E-01 | 1.32E+00 |
| VAMP4 | 2.10E-05 | 1.09E-04 | -1.21E+00 | 1.70E-02 | 7.36E-02 | 1.24E+00 | 5.55E-07 | 2.36E-06 | -1.43E+00 | 1.01E-03 | 3.86E-03 | 1.01E+00 | 6.50E-06 | 2.32E-05 | -1.37E+00 | 9.21E-03 | 2.59E-02 | 1.06E+00 | 1.72E-03 | 6.31E-03 | 1.04E+00 | 1.87E-01 | 5.13E-01 | 1.28E+00 |
| KRTCAP2 | 2.10E-05 | 1.09E-04 | -1.75E+00 | 5.97E-03 | 2.58E-02 | -1.00E+00 | 1.92E-05 | 8.15E-05 | -1.57E+00 | 3.00E-03 | 1.15E-02 | -1.04E+00 | 2.00E-02 | 7.13E-02 | -1.53E+00 | 2.55E-01 | 7.20E-01 | 1.21E+00 | 5.57E-02 | 2.04E-01 | 1.00E+00 | 1.14E-02 | 3.14E-02 | -1.24E+00 |
| BRI3BP | 2.11E-05 | 1.10E-04 | -1.66E+00 | 3.27E-04 | 1.41E-03 | -1.09E+00 | 6.19E-09 | 2.63E-08 | -1.63E+00 | 9.52E-05 | 3.64E-04 | -1.15E+00 | 4.12E-03 | 1.47E-02 | -1.61E+00 | 3.40E-02 | 9.59E-02 | -1.13E+00 | 2.48E-12 | 9.12E-12 | -1.70E+00 | 4.33E-03 | 1.19E-02 | 1.02E+00 |
| JMJD4 | 2.11E-05 | 1.10E-04 | -1.38E+00 | 9.69E-04 | 4.19E-03 | 1.02E+00 | 3.65E-04 | 1.55E-03 | -1.19E+00 | 1.08E-02 | 4.13E-02 | 1.16E+00 | 4.04E-05 | 1.44E-04 | -1.34E+00 | 2.96E-03 | 8.36E-03 | -1.04E+00 | 2.27E-01 | 8.34E-01 | 1.21E+00 | 7.22E-02 | 1.98E-01 | 1.39E+00 |
| AAK1 | 2.12E-05 | 1.11E-04 | -1.09E+00 | 6.34E-07 | 2.74E-06 | -1.08E+00 | 4.02E-03 | 1.71E-02 | -1.13E+00 | 4.43E-05 | 1.69E-04 | -1.17E+00 | 9.66E-06 | 3.44E-05 | -1.19E+00 | 1.08E-06 | 3.03E-06 | -1.23E+00 | 9.57E-05 | 3.51E-04 | -1.30E+00 | 1.72E-04 | 4.72E-04 | -1.24E+00 |
| CLASP2 | 2.13E-05 | 1.11E-04 | -1.21E+00 | 1.65E-03 | 7.12E-03 | 1.06E+00 | 1.55E-04 | 6.59E-04 | -1.25E+00 | 1.36E-03 | 5.18E-03 | 1.02E+00 | 2.77E-06 | 9.88E-06 | -1.42E+00 | 1.62E-04 | 4.56E-04 | -1.25E+00 | 4.35E-05 | 1.60E-04 | -1.27E+00 | 9.88E-03 | 2.71E-02 | -1.01E+00 |
| STYXL1 | 2.14E-05 | 1.12E-04 | -1.18E+00 | 1.03E-04 | 4.46E-04 | 1.01E+00 | 4.81E-02 | 2.04E-01 | -1.03E+00 | 2.13E-02 | 8.15E-02 | 1.08E+00 | 1.10E-03 | 3.93E-03 | -1.12E+00 | 4.35E-02 | 1.23E-01 | 1.07E+00 | 1.34E-01 | 4.92E-01 | 1.20E+00 | 1.57E-01 | 4.30E-01 | 1.41E+00 |
| SAYSD1 | 2.15E-05 | 1.12E-04 | -1.40E+00 | 7.86E-03 | 3.40E-02 | 1.15E+00 | 7.22E-07 | 3.07E-06 | -1.54E+00 | 5.65E-04 | 2.16E-03 | -1.04E+00 | 8.72E-06 | 3.11E-05 | -1.92E+00 | 2.79E-03 | 7.86E-03 | -1.14E+00 | 5.15E-02 | 1.89E-01 | 1.11E+00 | 3.61E-01 | 9.92E-01 | 1.20E+00 |
| LTB | 2.16E-05 | 1.13E-04 | 1.92E+00 | 7.90E-11 | 3.41E-10 | 2.67E+00 | 1.04E-02 | 4.41E-02 | -1.06E+00 | 6.36E-04 | 2.43E-03 | 1.35E+00 | 2.16E-01 | 7.69E-01 | 1.34E+00 | 2.19E-01 | 6.17E-01 | 1.00E+00 | 2.61E-01 | 9.56E-01 | 1.46E+00 | 2.53E-01 | 6.96E-01 | 1.11E+00 |
| GPC3 | 2.16E-05 | 1.13E-04 | -2.27E+00 | 5.49E-05 | 2.37E-04 | -1.91E+00 | 5.06E-08 | 2.15E-07 | -1.48E+00 | 4.95E-07 | 1.89E-06 | -1.29E+00 | 3.31E-02 | 1.18E-01 | -1.49E+00 | 2.45E-02 | 6.90E-02 | -2.15E+00 | 2.60E-01 | 9.56E-01 | 1.12E+00 | 3.38E-01 | 9.29E-01 | 1.23E+00 |
| INSIG1 | 2.17E-05 | 1.13E-04 | -1.02E+00 | 9.97E-05 | 4.31E-04 | 1.14E+00 | 9.11E-03 | 3.87E-02 | -1.23E+00 | 1.11E-01 | 4.23E-01 | 1.14E+00 | 2.52E-01 | 8.99E-01 | 1.60E+00 | 8.22E-02 | 2.32E-01 | 1.98E+00 | 1.49E-01 | 5.45E-01 | 1.17E+00 | 1.82E-03 | 5.00E-03 | 1.67E+00 |
| FAM118B | 2.17E-05 | 1.13E-04 | -1.84E+00 | 1.95E-01 | 8.42E-01 | 1.48E+00 | 7.89E-06 | 3.35E-05 | -1.52E+00 | 8.47E-03 | 3.23E-02 | 1.07E+00 | 7.55E-06 | 2.69E-05 | -2.08E+00 | 3.53E-04 | 9.96E-04 | -1.44E+00 | 1.40E-02 | 5.13E-02 | -1.24E+00 | 1.21E-01 | 3.32E-01 | -1.00E+00 |
| WWP1 | 2.18E-05 | 1.13E-04 | -1.11E+00 | 9.39E-04 | 4.06E-03 | 1.16E+00 | 3.95E-10 | 1.68E-09 | -1.61E+00 | 1.28E-08 | 4.89E-08 | -1.32E+00 | 9.89E-03 | 3.53E-02 | -1.03E+00 | 6.06E-03 | 1.71E-02 | -1.13E+00 | 8.36E-02 | 3.07E-01 | 1.04E+00 | 3.21E-01 | 8.83E-01 | 1.16E+00 |
| ALG13 | 2.18E-05 | 1.13E-04 | -1.05E+00 | 8.43E-12 | 3.64E-11 | -1.21E+00 | 7.70E-05 | 3.27E-04 | -1.14E+00 | 1.40E-08 | 5.34E-08 | -1.26E+00 | 3.93E-03 | 1.40E-02 | 1.05E+00 | 2.49E-03 | 7.02E-03 | 1.01E+00 | 3.47E-04 | 1.27E-03 | -1.13E+00 | 3.62E-05 | 9.95E-05 | -1.25E+00 |
| TMEM167A | 2.18E-05 | 1.14E-04 | -1.00E+00 | 7.62E-06 | 3.29E-05 | 1.12E+00 | 3.62E-02 | 1.54E-01 | 1.04E+00 | 5.83E-02 | 2.23E-01 | 1.19E+00 | 1.53E-02 | 5.45E-02 | 1.12E+00 | 9.34E-02 | 2.63E-01 | 1.22E+00 | 3.62E-02 | 1.33E-01 | 1.03E+00 | 5.22E-02 | 1.44E-01 | 1.03E+00 |
| POLR2M | 2.18E-05 | 1.14E-04 | -1.23E+00 | 6.75E-03 | 2.92E-02 | 1.17E+00 | 8.72E-05 | 3.70E-04 | -1.31E+00 | 4.54E-03 | 1.73E-02 | 1.07E+00 | 5.54E-06 | 1.98E-05 | -1.32E+00 | 2.88E-03 | 8.12E-03 | 1.07E+00 | 1.48E-10 | 5.43E-10 | -1.57E+00 | 1.29E-06 | 3.54E-06 | -1.24E+00 |
| DHODH | 2.18E-05 | 1.14E-04 | -1.39E+00 | 1.76E-04 | 7.59E-04 | -1.16E+00 | 3.70E-03 | 1.57E-02 | -1.15E+00 | 3.33E-02 | 1.27E-01 | 1.10E+00 | 3.63E-03 | 1.29E-02 | -1.47E+00 | 2.92E-01 | 8.23E-01 | 1.35E+00 | 3.04E-07 | 1.12E-06 | -1.33E+00 | 1.49E-02 | 4.10E-02 | 1.09E+00 |
| CCDC92 | 2.18E-05 | 1.14E-04 | -1.27E+00 | 1.30E-03 | 5.60E-03 | 1.07E+00 | 3.22E-02 | 1.37E-01 | -1.02E+00 | 1.84E-01 | 7.04E-01 | 1.51E+00 | 2.77E-01 | 9.88E-01 | 1.48E+00 | 1.04E-01 | 2.93E-01 | 1.86E+00 | 1.07E-01 | 3.94E-01 | 1.13E+00 | 2.13E-01 | 5.85E-01 | 1.15E+00 |
| AC079089.1 | 2.18E-05 | 1.14E-04 | -1.67E+00 | 8.28E-04 | 3.58E-03 | -1.22E+00 | 1.57E-10 | 6.66E-10 | -2.11E+00 | 1.16E-08 | 4.41E-08 | -1.63E+00 | 1.48E-01 | 5.29E-01 | -1.44E+00 | 3.14E-01 | 8.85E-01 | 1.09E+00 | 2.49E-01 | 9.13E-01 | 1.07E+00 | 3.01E-01 | 8.28E-01 | 1.17E+00 |
| AC023161.2 | 2.19E-05 | 1.14E-04 | -1.83E+00 | 1.98E-04 | 8.56E-04 | -1.43E+00 | 2.72E-11 | 1.16E-10 | -2.83E+00 | 6.54E-07 | 2.50E-06 | -1.62E+00 | 3.36E-07 | 1.20E-06 | -2.70E+00 | 1.09E-05 | 3.07E-05 | -1.87E+00 | 2.86E-06 | 1.05E-05 | -2.06E+00 | 1.87E-04 | 5.14E-04 | -1.59E+00 |
| TP53BP1 | 2.20E-05 | 1.15E-04 | -1.22E+00 | 1.16E-08 | 5.03E-08 | -1.36E+00 | 1.06E-05 | 4.50E-05 | -1.36E+00 | 1.26E-05 | 4.80E-05 | -1.22E+00 | 9.48E-10 | 3.38E-09 | -1.64E+00 | 1.28E-07 | 3.60E-07 | -1.35E+00 | 1.82E-07 | 6.69E-07 | -1.43E+00 | 2.88E-07 | 7.91E-07 | -1.45E+00 |
| GDAP2 | 2.21E-05 | 1.15E-04 | -1.35E+00 | 4.24E-03 | 1.83E-02 | 1.10E+00 | 1.37E-07 | 5.82E-07 | -1.71E+00 | 6.40E-05 | 2.44E-04 | -1.08E+00 | 2.55E-08 | 9.08E-08 | -1.85E+00 | 9.68E-06 | 2.73E-05 | -1.37E+00 | 2.65E-05 | 9.72E-05 | -1.29E+00 | 3.06E-03 | 8.41E-03 | -1.07E+00 |
| CHRM3-AS2 | 2.21E-05 | 1.15E-04 | -1.24E+00 | 1.82E-03 | 7.87E-03 | 1.06E+00 | 1.15E-02 | 4.90E-02 | -1.17E+00 | 7.18E-03 | 2.74E-02 | -1.08E+00 | 3.87E-08 | 1.38E-07 | 3.52E+00 | 7.50E-04 | 2.11E-03 | 2.35E+00 | 7.85E-04 | 2.88E-03 | 1.78E+00 | 1.88E-01 | 5.18E-01 | 1.38E+00 |
| EEF2 | 2.22E-05 | 1.16E-04 | -1.10E+00 | 8.13E-04 | 3.51E-03 | 1.10E+00 | 1.66E-02 | 7.04E-02 | -1.05E+00 | 1.29E-04 | 4.93E-04 | 1.16E+00 | 2.43E-01 | 8.67E-01 | 1.00E+00 | 1.63E-03 | 4.60E-03 | 1.13E+00 | 3.30E-02 | 1.21E-01 | -1.02E+00 | 2.06E-03 | 5.66E-03 | 1.13E+00 |
| UBAP1 | 2.23E-05 | 1.16E-04 | -1.15E+00 | 8.07E-08 | 3.49E-07 | -1.36E+00 | 7.44E-04 | 3.16E-03 | -1.17E+00 | 1.09E-07 | 4.17E-07 | -1.47E+00 | 1.12E-02 | 3.98E-02 | -1.06E+00 | 3.68E-03 | 1.04E-02 | -1.15E+00 | 4.16E-10 | 1.53E-09 | -1.61E+00 | 4.30E-09 | 1.18E-08 | -1.47E+00 |

| | | | | | | | | | | | | | | | | | | | | | | | | |
|---|---|---|---|---|---|---|---|---|---|---|---|---|---|---|---|---|---|---|---|---|---|---|---|---|
| AC105749.1 | 2.23E-05 | 1.16E-04 | -1.56E+00 | 6.02E-04 | 2.60E-03 | -1.12E+00 | 1.36E-02 | 5.80E-02 | -1.45E+00 | 8.14E-02 | 3.11E-01 | 1.04E+00 | 1.28E-11 | 4.57E-11 | -3.07E+00 | 2.37E-10 | 6.67E-10 | -2.50E+00 | 1.13E-07 | 4.14E-07 | -2.91E+00 | 1.87E-04 | 5.14E-04 | -1.79E+00 |
| ASB16-AS1 | 2.23E-05 | 1.16E-04 | -1.49E+00 | 6.64E-03 | 2.87E-02 | 1.05E+00 | 4.31E-05 | 1.83E-04 | -1.40E+00 | 3.97E-03 | 1.52E-02 | -1.01E+00 | 6.66E-07 | 2.38E-06 | -2.05E+00 | 3.38E-04 | 9.53E-04 | -1.24E+00 | 2.00E-05 | 7.35E-05 | -1.35E+00 | 1.37E-03 | 3.76E-03 | -1.25E+00 |
| PTPN4 | 2.24E-05 | 1.17E-04 | -1.09E+00 | 3.68E-04 | 1.59E-03 | 1.07E+00 | 5.43E-08 | 2.31E-07 | -1.30E+00 | 1.79E-07 | 6.82E-07 | -1.19E+00 | 8.34E-03 | 2.97E-02 | -1.07E+00 | 1.47E-01 | 4.14E-01 | 1.28E+00 | 2.78E-05 | 1.02E-04 | -1.22E+00 | 1.29E-03 | 3.55E-03 | -1.12E+00 |
| ZRANB3 | 2.24E-05 | 1.17E-04 | -1.31E+00 | 8.29E-05 | 3.58E-04 | -1.14E+00 | 5.43E-06 | 2.31E-05 | -1.29E+00 | 1.35E-06 | 5.17E-06 | -1.25E+00 | 2.09E-02 | 7.47E-02 | 1.04E+00 | 1.08E-02 | 3.05E-02 | 1.00E+00 | 1.45E-06 | 5.30E-06 | -1.37E+00 | 1.82E-03 | 5.01E-03 | -1.11E+00 |
| IL17D | 2.24E-05 | 1.17E-04 | -1.85E+00 | 3.44E-03 | 1.49E-02 | -1.05E+00 | 2.10E-01 | 8.94E-01 | 1.21E+00 | 2.33E-01 | 8.91E-01 | 1.34E+00 | 7.57E-02 | 2.70E-01 | -1.67E+00 | 1.41E-01 | 3.99E-01 | -1.30E+00 | 2.39E-01 | 8.78E-01 | 1.12E+00 | 2.47E-01 | 6.80E-01 | 1.34E+00 |
| CCT8 | 2.25E-05 | 1.17E-04 | -1.03E+00 | 1.18E-02 | 5.10E-02 | 1.15E+00 | 1.26E-01 | 5.35E-01 | 1.02E+00 | 2.02E-01 | 7.70E-01 | 1.08E+00 | 5.23E-05 | 1.87E-04 | -1.07E+00 | 2.18E-01 | 6.15E-01 | 1.12E+00 | 8.32E-02 | 3.05E-01 | 1.01E+00 | 2.68E-01 | 7.35E-01 | 1.12E+00 |
| ANKRD36 | 2.26E-05 | 1.18E-04 | -1.18E+00 | 6.72E-02 | 2.90E-01 | 1.20E+00 | 3.84E-06 | 1.63E-05 | -1.24E+00 | 9.75E-03 | 3.73E-02 | 1.09E+00 | 2.37E-05 | 8.43E-05 | -1.18E+00 | 3.47E-04 | 9.78E-04 | -1.07E+00 | 1.59E-05 | 5.82E-05 | -1.21E+00 | 3.85E-02 | 1.06E-01 | -1.02E+00 |
| EIF3H | 2.26E-05 | 1.18E-04 | -1.12E+00 | 8.83E-05 | 3.81E-04 | -1.07E+00 | 1.86E-02 | 7.89E-02 | -1.09E+00 | 1.63E-02 | 6.24E-02 | -1.06E+00 | 1.29E-02 | 4.61E-02 | -1.07E+00 | 1.66E-01 | 4.69E-01 | 1.02E+00 | 8.00E-06 | 2.93E-05 | -1.14E+00 | 2.86E-02 | 7.87E-02 | -1.06E+00 |
| KATNA1 | 2.26E-05 | 1.18E-04 | -1.22E+00 | 3.30E-03 | 1.43E-02 | 1.18E+00 | 6.34E-04 | 2.69E-03 | -1.12E+00 | 1.72E-03 | 6.56E-03 | 1.02E+00 | 7.95E-05 | 2.83E-04 | -1.29E+00 | 7.09E-03 | 2.00E-02 | 1.01E+00 | 2.13E-05 | 7.80E-05 | -1.13E+00 | 2.11E-04 | 5.78E-04 | -1.09E+00 |
| TSR1 | 2.27E-05 | 1.18E-04 | -1.41E+00 | 8.81E-04 | 3.81E-03 | -1.00E+00 | 1.33E-06 | 5.67E-06 | -1.38E+00 | 2.69E-04 | 1.03E-03 | -1.01E+00 | 1.13E-05 | 4.04E-05 | -1.39E+00 | 3.58E-05 | 1.01E-04 | -1.28E+00 | 9.16E-03 | 3.36E-02 | -1.03E+00 | 3.82E-02 | 1.05E-01 | 1.15E+00 |
| ZNF780B | 2.27E-05 | 1.18E-04 | -1.38E+00 | 2.90E-02 | 1.25E-01 | 1.29E+00 | 5.64E-04 | 2.39E-03 | -1.39E+00 | 4.43E-02 | 1.69E-01 | 1.18E+00 | 1.00E-01 | 3.58E-01 | 1.15E+00 | 2.14E-01 | 6.02E-01 | 1.70E+00 | 4.36E-05 | 1.60E-04 | -1.59E+00 | 1.35E-03 | 3.70E-03 | -1.29E+00 |
| HIPK2 | 2.28E-05 | 1.19E-04 | -1.17E+00 | 4.14E-04 | 1.79E-03 | 1.10E+00 | 3.57E-02 | 1.52E-01 | -1.04E+00 | 1.47E-02 | 5.63E-02 | 1.03E+00 | 1.33E-01 | 4.75E-01 | 1.57E+00 | 3.40E-01 | 9.59E-01 | 1.43E+00 | 1.75E-01 | 6.41E-01 | 1.16E+00 | 3.08E-01 | 8.46E-01 | 1.19E+00 |
| REPS1 | 2.28E-05 | 1.19E-04 | -1.10E+00 | 5.98E-04 | 2.59E-03 | 1.12E+00 | 3.03E-03 | 1.29E-02 | -1.05E+00 | 5.04E-04 | 1.93E-03 | 1.02E+00 | 4.94E-02 | 1.76E-01 | 1.19E+00 | 2.75E-02 | 7.75E-02 | 1.19E+00 | 1.39E-01 | 5.11E-01 | 1.15E+00 | 2.66E-01 | 7.29E-01 | 1.28E+00 |
| RIOX2 | 2.28E-05 | 1.19E-04 | -1.60E+00 | 2.55E-02 | 1.10E-01 | 1.19E+00 | 1.92E-04 | 8.18E-04 | -1.28E+00 | 2.44E-03 | 9.31E-03 | 1.02E+00 | 5.83E-04 | 2.08E-03 | -1.77E+00 | 3.44E-02 | 9.71E-02 | -1.11E+00 | 4.27E-06 | 1.57E-05 | -1.39E+00 | 5.91E-03 | 1.62E-02 | -1.05E+00 |
| EXOC4 | 2.28E-05 | 1.19E-04 | -1.05E+00 | 8.82E-07 | 3.81E-06 | 1.01E+00 | 3.51E-04 | 1.49E-03 | -1.08E+00 | 8.59E-04 | 3.28E-03 | 1.09E+00 | 6.57E-05 | 2.34E-04 | -1.15E+00 | 5.17E-04 | 1.46E-03 | 1.01E+00 | 3.24E-03 | 1.19E-02 | -1.02E+00 | 1.26E-02 | 3.47E-02 | 1.06E+00 |
| ZNF771 | 2.29E-05 | 1.19E-04 | -1.35E+00 | 2.32E-03 | 1.00E-02 | 1.05E+00 | 1.52E-03 | 6.47E-03 | -1.24E+00 | 2.05E-03 | 7.82E-03 | -1.09E+00 | 2.35E-01 | 8.36E-01 | 1.12E+00 | 2.43E-01 | 6.84E-01 | 1.55E+00 | 1.28E-02 | 4.69E-02 | -1.19E+00 | 2.79E-02 | 7.67E-02 | -1.18E+00 |
| GAMT | 2.29E-05 | 1.19E-04 | -1.03E+00 | 2.30E-05 | 9.96E-05 | 1.08E+00 | 1.84E-01 | 7.80E-01 | 1.23E+00 | 7.45E-02 | 2.84E-01 | 1.28E+00 | 9.92E-04 | 3.54E-03 | -1.11E+00 | 3.56E-03 | 1.00E-02 | 1.00E+00 | 2.55E-01 | 9.36E-01 | 1.27E+00 | 1.49E-01 | 4.10E-01 | 1.39E+00 |
| AC008736.1 | 2.29E-05 | 1.19E-04 | -2.15E+00 | 5.12E-04 | 2.21E-03 | -1.46E+00 | 1.29E-02 | 5.48E-02 | -1.41E+00 | 6.30E-03 | 2.41E-02 | -1.45E+00 | 8.66E-02 | 3.09E-01 | 1.10E+00 | 2.19E-01 | 6.17E-01 | 1.27E+00 | 5.58E-04 | 2.05E-03 | -1.05E+00 | 6.86E-04 | 1.88E-03 | -1.16E+00 |
| TMUB2 | 2.29E-05 | 1.20E-04 | -1.10E+00 | 9.34E-06 | 4.03E-05 | -1.03E+00 | 4.83E-08 | 2.05E-07 | -1.38E+00 | 7.10E-08 | 2.71E-07 | -1.23E+00 | 1.79E-04 | 6.39E-04 | -1.08E+00 | 1.57E-05 | 4.43E-05 | -1.14E+00 | 2.35E-03 | 8.61E-03 | -1.11E+00 | 8.43E-02 | 2.31E-01 | 1.04E+00 |
| FAM104B | 2.30E-05 | 1.20E-04 | -1.26E+00 | 1.30E-02 | 5.63E-02 | 1.21E+00 | 1.14E-05 | 4.85E-05 | -1.23E+00 | 1.26E-03 | 4.83E-03 | 1.02E+00 | 1.85E-03 | 6.60E-03 | -1.09E+00 | 2.14E-01 | 6.02E-01 | 1.36E+00 | 1.14E-01 | 4.19E-01 | 1.13E+00 | 7.90E-02 | 2.17E-01 | 1.39E+00 |
| SIRT6 | 2.30E-05 | 1.20E-04 | -1.42E+00 | 7.86E-04 | 3.40E-03 | -1.08E+00 | 1.70E-03 | 7.23E-03 | -1.22E+00 | 1.32E-02 | 5.05E-02 | 1.10E+00 | 4.22E-04 | 1.51E-03 | -1.26E+00 | 1.49E-02 | 4.20E-02 | -1.01E+00 | 9.69E-04 | 3.56E-03 | -1.10E+00 | 6.33E-03 | 1.74E-02 | -1.01E+00 |
| ZNF714 | 2.30E-05 | 1.20E-04 | -1.21E+00 | 3.63E-04 | 1.57E-03 | 1.06E+00 | 1.87E-04 | 7.94E-04 | -1.28E+00 | 3.34E-04 | 1.28E-03 | -1.08E+00 | 3.32E-04 | 1.18E-03 | -1.27E+00 | 5.57E-03 | 1.57E-02 | -1.00E+00 | 1.23E-06 | 4.53E-06 | -1.19E+00 | 1.35E-04 | 3.71E-04 | -1.03E+00 |
| AKR1C6P | 2.31E-05 | 1.20E-04 | 4.26E+00 | 1.49E-07 | 6.44E-07 | 5.95E+00 | 4.15E-02 | 1.76E-01 | 1.76E+00 | 4.31E-02 | 1.65E-01 | 1.95E+00 | 1.29E-01 | 4.59E-01 | 1.79E+00 | 1.30E-01 | 3.66E-01 | 2.18E+00 | 2.63E-01 | 9.64E-01 | 1.22E+00 | 3.58E-01 | 9.83E-01 | 1.08E+00 |
| KDM4C | 2.31E-05 | 1.21E-04 | -1.16E+00 | 1.76E-04 | 7.58E-04 | 1.01E+00 | 1.12E-03 | 4.75E-03 | -1.16E+00 | 6.12E-03 | 2.34E-02 | 1.07E+00 | 1.59E-02 | 5.67E-02 | 1.03E+00 | 1.51E-02 | 4.26E-02 | 1.05E+00 | 2.23E-03 | 8.17E-03 | -1.10E+00 | 5.73E-04 | 1.57E-03 | -1.23E+00 |
| AMER1 | 2.31E-05 | 1.21E-04 | -1.74E+00 | 2.43E-05 | 1.05E-04 | -1.61E+00 | 6.77E-04 | 2.88E-03 | -1.44E+00 | 1.87E-02 | 7.15E-02 | -1.08E+00 | 4.85E-04 | 1.73E-03 | -2.63E+00 | 4.75E-03 | 1.34E-02 | -1.76E+00 | 3.75E-03 | 1.38E-02 | -1.38E+00 | 6.79E-02 | 1.87E-01 | -1.06E+00 |
| AC108718.1 | 2.32E-05 | 1.21E-04 | -1.77E+00 | 3.02E-04 | 1.31E-03 | -1.37E+00 | 1.57E-02 | 6.68E-02 | -1.48E+00 | 7.83E-03 | 2.99E-02 | -1.66E+00 | 1.98E-01 | 7.07E-01 | 1.01E+00 | 3.26E-01 | 9.20E-01 | 1.27E+00 | 2.21E-02 | 8.10E-02 | -1.69E+00 | 5.11E-02 | 1.40E-01 | -1.66E+00 |
| ZNF658B | 2.32E-05 | 1.21E-04 | -1.86E+00 | 1.52E-04 | 6.59E-04 | -1.54E+00 | 1.09E-03 | 4.63E-03 | -1.40E+00 | 5.77E-04 | 2.20E-03 | -1.49E+00 | 6.28E-02 | 2.24E-01 | -1.04E+00 | 4.75E-02 | 1.34E-01 | -1.28E+00 | 2.81E-03 | 1.03E-02 | -1.54E+00 | 2.36E-02 | 6.49E-02 | -1.25E+00 |
| CDC42BPB | 2.32E-05 | 1.21E-04 | -1.69E+00 | 1.60E-03 | 6.92E-03 | -1.09E+00 | 7.23E-03 | 3.07E-02 | -1.63E+00 | 1.29E-02 | 4.94E-02 | -1.40E+00 | 2.10E-03 | 7.49E-03 | -1.47E+00 | 1.46E-02 | 4.11E-02 | -1.28E+00 | 1.86E-02 | 6.82E-02 | 1.31E+00 | 1.36E-02 | 3.73E-02 | -1.04E+00 |
| HLA-F-AS1 | 2.33E-05 | 1.21E-04 | -1.65E+00 | 2.38E-02 | 1.03E-01 | 1.04E+00 | 6.69E-02 | 2.84E-01 | -1.07E+00 | 1.99E-01 | 7.60E-01 | 1.27E+00 | 8.15E-02 | 2.91E-01 | -1.18E+00 | 1.09E-01 | 3.08E-01 | -1.13E+00 | 1.17E-01 | 4.28E-01 | -1.16E+00 | 3.27E-01 | 8.99E-01 | 1.07E+00 |
| BEX5 | 2.33E-05 | 1.21E-04 | -1.17E+00 | 2.78E-06 | 1.20E-05 | -1.15E+00 | 1.72E-01 | 7.32E-01 | -1.09E+00 | 1.09E-02 | 4.16E-02 | -1.35E+00 | 1.93E-02 | 6.88E-02 | 1.99E+00 | 9.07E-02 | 2.56E-01 | 1.95E+00 | 3.75E-03 | 1.38E-02 | 2.45E+00 | 2.56E-02 | 7.04E-02 | 2.11E+00 |
| AXL | 2.33E-05 | 1.22E-04 | -1.52E+00 | 2.16E-01 | 9.33E-01 | 1.25E+00 | 9.68E-08 | 4.11E-07 | -2.44E+00 | 3.13E-03 | 1.20E-02 | -1.17E+00 | 2.50E-01 | 8.93E-01 | -1.06E+00 | 3.27E-01 | 9.22E-01 | 1.01E+00 | 1.18E-01 | 4.35E-01 | -1.61E+00 | 2.21E-01 | 6.08E-01 | -1.24E+00 |
| AL158071.2 | 2.33E-05 | 1.22E-04 | 3.08E+00 | 1.09E-01 | 4.69E-01 | 1.45E+00 | 7.81E-02 | 3.32E-01 | 1.48E+00 | 1.67E-01 | 6.37E-01 | 1.24E+00 | 4.13E-03 | 1.47E-02 | 3.57E+00 | 3.55E-01 | 1.00E+00 | -1.00E+00 | 1.33E-01 | 4.88E-01 | 1.49E+00 | 3.31E-01 | 9.09E-01 | -1.01E+00 |
| C6orf47 | 2.33E-05 | 1.22E-04 | -1.55E+00 | 1.05E-03 | 4.56E-03 | -1.17E+00 | 2.40E-04 | 1.02E-03 | -1.89E+00 | 4.21E-02 | 1.61E-01 | -1.02E+00 | 1.08E-02 | 3.87E-02 | -1.50E+00 | 1.67E-01 | 4.72E-01 | 1.09E+00 | 2.88E-03 | 1.06E-02 | -1.19E+00 | 9.56E-02 | 2.63E-01 | -1.06E+00 |
| NME8 | 2.34E-05 | 1.22E-04 | -1.79E+00 | 6.25E-03 | 2.70E-02 | 1.02E+00 | 1.09E-04 | 4.65E-04 | -1.93E+00 | 3.26E-04 | 1.25E-03 | -1.53E+00 | 3.74E-12 | 1.33E-11 | -1.55E+00 | 2.92E-09 | 8.24E-09 | -1.23E+00 | 2.27E-01 | 8.32E-01 | 1.20E+00 | 1.78E-01 | 4.88E-01 | -1.08E+00 |
| MIEF2 | 2.34E-05 | 1.22E-04 | -1.48E+00 | 7.18E-04 | 3.10E-03 | -1.08E+00 | 4.54E-05 | 1.93E-04 | -1.86E+00 | 1.76E-02 | 6.72E-02 | -1.03E+00 | 1.12E-01 | 3.98E-01 | 1.02E+00 | 1.50E-01 | 4.24E-01 | 1.01E+00 | 1.91E-09 | 7.02E-09 | -1.98E+00 | 1.85E-04 | 5.07E-04 | -1.28E+00 |
| STRN | 2.35E-05 | 1.22E-04 | -1.26E+00 | 9.45E-05 | 4.08E-04 | -1.08E+00 | 5.18E-04 | 2.20E-03 | -1.19E+00 | 2.51E-02 | 9.59E-02 | 1.14E+00 | 1.33E-01 | 4.75E-01 | 1.29E+00 | 1.09E-01 | 3.07E-01 | 1.19E+00 | 6.17E-02 | 2.27E-01 | 1.09E+00 | 1.54E-01 | 4.24E-01 | 1.05E+00 |
| AC138470.1 | 2.35E-05 | 1.22E-04 | -1.78E+00 | 4.07E-02 | 1.76E-01 | 1.06E+00 | 1.84E-01 | 7.82E-01 | 1.30E+00 | 8.07E-03 | 3.08E-02 | 2.24E+00 | 1.80E-01 | 6.42E-01 | 1.23E+00 | 1.26E-01 | 3.54E-01 | 1.97E+00 | 2.69E-02 | 9.88E-02 | -1.34E+00 | 1.17E-01 | 3.22E-01 | -1.06E+00 |
| ATG10 | 2.36E-05 | 1.23E-04 | -1.20E+00 | 1.85E-02 | 7.98E-02 | 1.24E+00 | 2.44E-07 | 1.04E-06 | -1.33E+00 | 1.91E-07 | 7.31E-07 | -1.19E+00 | 1.03E-08 | 3.66E-08 | -1.55E+00 | 1.12E-04 | 3.16E-04 | -1.05E+00 | 8.15E-11 | 2.99E-10 | -1.57E+00 | 9.30E-06 | 2.56E-05 | -1.17E+00 |
| SEM1P1 | 2.36E-05 | 1.23E-04 | -1.57E+00 | 1.95E-06 | 8.43E-06 | -1.77E+00 | 1.19E-04 | 5.07E-04 | -1.40E+00 | 1.57E-07 | 5.99E-07 | -1.77E+00 | 5.43E-02 | 1.94E-01 | -1.08E+00 | 1.89E-03 | 5.32E-03 | -1.82E+00 | 3.34E-02 | 1.23E-01 | -1.03E+00 | 2.85E-02 | 7.84E-02 | -1.10E+00 |
| MAFG | 2.36E-05 | 1.23E-04 | -1.04E+00 | 1.78E-11 | 7.69E-11 | -1.32E+00 | 1.85E-03 | 7.84E-03 | -1.08E+00 | 7.14E-08 | 2.73E-07 | -1.25E+00 | 2.40E-04 | 8.55E-04 | 1.03E+00 | 2.20E-08 | 6.20E-08 | -1.35E+00 | 5.24E-05 | 1.92E-04 | -1.05E+00 | 1.00E-06 | 2.75E-06 | -1.15E+00 |
| DANT2 | 2.37E-05 | 1.23E-04 | -1.54E+00 | 5.88E-02 | 2.54E-01 | 1.23E+00 | 8.85E-09 | 3.76E-08 | -1.69E+00 | 1.82E-07 | 6.97E-07 | -1.48E+00 | 9.51E-02 | 3.39E-01 | -1.26E+00 | 2.35E-01 | 6.63E-01 | 1.14E+00 | 1.73E-01 | 6.36E-01 | 1.04E+00 | 3.15E-01 | 8.65E-01 | 1.08E+00 |
| UIMC1 | 2.37E-05 | 1.23E-04 | -1.03E+00 | 9.04E-05 | 3.91E-04 | 1.10E+00 | 2.14E-05 | 9.08E-05 | -1.24E+00 | 2.64E-05 | 1.01E-04 | -1.10E+00 | 1.60E-03 | 5.71E-03 | -1.01E+00 | 4.33E-02 | 1.22E-01 | 1.17E+00 | 1.33E-05 | 4.88E-05 | -1.17E+00 | 1.44E-04 | 3.96E-04 | -1.09E+00 |
| FBL | 2.37E-05 | 1.24E-04 | -1.05E+00 | 3.17E-04 | 1.37E-03 | 1.08E+00 | 1.91E-01 | 8.13E-01 | 1.05E+00 | 1.84E-01 | 7.04E-01 | 1.10E+00 | 3.09E-06 | 1.10E-05 | -1.08E+00 | 2.51E-02 | 7.06E-02 | 1.18E+00 | 6.69E-04 | 2.46E-03 | -1.07E+00 | 3.28E-01 | 9.00E-01 | 1.13E+00 |
| FTH1P4 | 2.37E-05 | 1.24E-04 | -1.09E+00 | 7.93E-09 | 3.43E-08 | -1.07E+00 | 9.08E-03 | 3.86E-02 | 1.04E+00 | 3.98E-04 | 1.52E-03 | 1.06E+00 | 1.87E-02 | 6.67E-02 | -1.02E+00 | 1.77E-04 | 4.98E-04 | -1.13E+00 | 2.56E-02 | 9.40E-02 | -1.02E+00 | 9.85E-03 | 2.71E-02 | -1.10E+00 |
| RTL6 | 2.38E-05 | 1.24E-04 | -1.61E+00 | 9.71E-05 | 4.19E-04 | -1.28E+00 | 1.01E-03 | 4.30E-03 | -1.34E+00 | 1.11E-04 | 4.24E-04 | -1.34E+00 | 2.06E-02 | 7.36E-02 | -1.54E+00 | 2.96E-01 | 8.35E-01 | 1.22E+00 | 4.97E-03 | 1.82E-02 | -1.34E+00 | 1.28E-02 | 3.51E-02 | -1.31E+00 |
| DENND3 | 2.38E-05 | 1.24E-04 | -1.24E+00 | 3.86E-15 | 1.67E-14 | -2.37E+00 | 1.21E-04 | 5.13E-04 | -1.15E+00 | 9.79E-19 | 3.74E-18 | -1.93E+00 | 4.95E-03 | 1.77E-02 | 1.04E+00 | 2.33E-10 | 6.56E-10 | -2.27E+00 | 3.22E-05 | 1.18E-04 | -1.29E+00 | 1.31E-07 | 3.60E-07 | -2.08E+00 |
| SATB1-AS1 | 2.39E-05 | 1.25E-04 | -1.06E+00 | 2.73E-04 | 1.18E-03 | 1.10E+00 | 2.02E-04 | 8.59E-04 | -1.18E+00 | 1.20E-03 | 4.57E-03 | 1.02E+00 | 3.42E-02 | 1.22E-01 | 2.49E+00 | 3.05E-01 | 8.60E-01 | 1.76E+00 | 2.71E-02 | 9.93E-02 | -1.13E+00 | 8.65E-02 | 2.38E-01 | -1.05E+00 |
| ENTPD5 | 2.41E-05 | 1.26E-04 | -1.56E+00 | 9.90E-05 | 4.28E-04 | -1.28E+00 | 5.67E-05 | 2.41E-04 | -1.50E+00 | 7.18E-05 | 2.74E-04 | -1.37E+00 | 2.50E-04 | 8.90E-04 | -1.39E+00 | 2.62E-03 | 7.39E-03 | -1.11E+00 | 6.81E-04 | 2.50E-03 | -1.38E+00 | 1.97E-02 | 5.40E-02 | -1.22E+00 |
| EPM2AIP1 | 2.41E-05 | 1.26E-04 | -1.10E+00 | 1.42E-01 | 6.15E-01 | 1.38E+00 | 3.29E-06 | 1.40E-05 | -1.23E+00 | 1.27E-01 | 4.86E-01 | 1.34E+00 | 3.57E-03 | 1.27E-02 | 1.08E+00 | 3.47E-01 | 9.78E-01 | 1.42E+00 | 2.36E-03 | 8.65E-03 | -1.06E+00 | 3.26E-01 | 8.96E-01 | 1.31E+00 |
| RBM8B | 2.41E-05 | 1.26E-04 | 2.12E+00 | 2.27E-01 | 9.79E-01 | 1.46E+00 | 1.97E-07 | 8.37E-07 | 2.31E+00 | 1.60E-02 | 6.10E-02 | 1.87E+00 | 7.23E-04 | 2.58E-03 | 2.10E+00 | 4.90E-02 | 1.38E-01 | 1.74E+00 | 1.51E-03 | 5.53E-03 | 1.96E+00 | 7.71E-03 | 2.12E-02 | 1.84E+00 |
| PRRG4 | 2.41E-05 | 1.26E-04 | -1.12E+00 | 8.88E-11 | 3.84E-10 | -1.47E+00 | 1.06E-06 | 4.51E-06 | -1.22E+00 | 3.38E-10 | 1.29E-09 | -1.28E+00 | 6.37E-03 | 2.27E-02 | -1.52E+00 | 1.00E-04 | 2.82E-04 | -2.41E+00 | 1.17E-02 | 4.30E-02 | -1.43E+00 | 7.64E-02 | 2.10E-01 | -1.20E+00 |
| TMEM214 | 2.42E-05 | 1.26E-04 | -1.57E+00 | 2.58E-02 | 1.12E-01 | 1.16E+00 | 8.86E-05 | 3.76E-04 | -1.36E+00 | 1.86E-04 | 7.12E-04 | -1.23E+00 | 5.47E-06 | 1.95E-05 | -1.42E+00 | 4.07E-04 | 1.15E-03 | -1.16E+00 | 1.56E-01 | 5.73E-01 | 1.09E+00 | 1.22E-01 | 3.36E-01 | 1.22E+00 |
| AC006509.1 | 2.43E-05 | 1.26E-04 | -1.18E+00 | 2.04E-13 | 8.83E-13 | -1.96E+00 | 4.37E-06 | 1.86E-05 | -1.34E+00 | 1.06E-11 | 4.06E-11 | -1.93E+00 | 1.27E-02 | 4.52E-02 | 1.05E+00 | 3.29E-05 | 9.27E-05 | -1.46E+00 | 7.35E-09 | 2.70E-08 | -1.48E+00 | 7.51E-16 | 2.06E-15 | -2.05E+00 |
| MRPS17 | 2.43E-05 | 1.26E-04 | -1.57E+00 | 8.12E-04 | 3.51E-03 | -1.18E+00 | 6.52E-06 | 2.77E-05 | -1.47E+00 | 5.17E-04 | 1.97E-03 | 1.01E+00 | 1.21E-04 | 4.32E-04 | -1.51E+00 | 1.20E-02 | 3.38E-02 | -1.00E+00 | 8.38E-08 | 3.08E-07 | -1.50E+00 | 7.89E-04 | 2.17E-03 | -1.05E+00 |
| POLR3GP2 | 2.45E-05 | 1.28E-04 | -2.46E+00 | 2.35E-04 | 1.02E-03 | -1.83E+00 | 1.93E-01 | 8.19E-01 | 1.08E+00 | 2.05E-01 | 7.84E-01 | 1.16E+00 | 1.50E-02 | 5.33E-02 | -1.87E+00 | 2.99E-01 | 8.44E-01 | 1.23E+00 | 5.79E-03 | 2.13E-02 | -1.30E+00 | 2.61E-01 | 7.18E-01 | 1.12E+00 |
| POM121 | 2.45E-05 | 1.28E-04 | -1.29E+00 | 3.28E-06 | 1.42E-05 | -1.26E+00 | 7.96E-03 | 3.38E-02 | -1.19E+00 | 4.26E-03 | 1.63E-02 | -1.05E+00 | 2.58E-03 | 9.18E-03 | -1.17E+00 | 3.50E-03 | 9.87E-03 | -1.14E+00 | 2.71E-03 | 9.94E-03 | -1.15E+00 | 4.84E-02 | 1.33E-01 | 1.06E+00 |
| YPEL5 | 2.46E-05 | 1.28E-04 | 1.98E+00 | 1.56E-07 | 6.74E-07 | 1.04E+00 | 1.36E-01 | 5.80E-01 | 1.48E+00 | 3.80E-09 | 1.45E-08 | -1.19E+00 | 1.32E-01 | 4.72E-01 | 1.27E+00 | 8.54E-09 | 2.41E-08 | -1.40E+00 | 4.95E-02 | 1.82E-01 | 1.19E+00 | 1.72E-08 | 4.72E-08 | -1.31E+00 |
| RPL5P24 | 2.47E-05 | 1.28E-04 | 2.06E+00 | 5.72E-03 | 2.47E-02 | 2.00E+00 | 8.54E-07 | 3.63E-06 | 2.14E+00 | 2.01E-03 | 7.69E-03 | 2.04E+00 | 1.34E-07 | 4.79E-07 | 2.69E+00 | 7.25E-04 | 2.04E-03 | 2.28E+00 | 1.15E-08 | 4.22E-08 | 2.48E+00 | 3.23E-07 | 8.88E-07 | 2.35E+00 |
| FBXL15 | 2.47E-05 | 1.29E-04 | -1.14E+00 | 2.99E-03 | 1.29E-02 | 1.15E+00 | 1.22E-04 | 5.18E-04 | -1.21E+00 | 1.63E-02 | 6.23E-02 | 1.08E+00 | 2.13E-04 | 7.61E-04 | -1.09E+00 | 4.58E-02 | 1.29E-01 | 1.22E+00 | 3.41E-04 | 1.25E-03 | -1.09E+00 | 7.73E-03 | 2.12E-02 | 1.03E+00 |
| AHCTF1 | 2.47E-05 | 1.29E-04 | -1.23E+00 | 3.33E-07 | 1.44E-06 | -1.35E+00 | 2.57E-05 | 1.09E-04 | -1.27E+00 | 1.23E-10 | 4.71E-10 | -1.60E+00 | 9.85E-08 | 3.51E-07 | -1.88E+00 | 2.02E-08 | 5.71E-08 | -1.82E+00 | 3.11E-15 | 1.14E-14 | -1.93E+00 | 1.82E-11 | 5.01E-11 | -1.55E+00 |
| NEGR1 | 2.47E-05 | 1.29E-04 | -1.58E+00 | 2.54E-05 | 1.10E-04 | -1.50E+00 | 2.10E-04 | 8.91E-04 | -1.63E+00 | 7.67E-04 | 2.93E-03 | -1.33E+00 | 1.64E-01 | 5.85E-01 | 1.77E+00 | 2.72E-01 | 7.66E-01 | 1.50E+00 | 7.25E-02 | 2.66E-01 | -1.53E+00 | 1.38E-01 | 3.79E-01 | -1.42E+00 |
| COX11 | 2.48E-05 | 1.29E-04 | -1.03E+00 | 8.44E-05 | 3.65E-04 | 1.09E+00 | 1.17E-02 | 4.95E-02 | 1.03E+00 | 2.06E-03 | 7.87E-03 | 1.08E+00 | 1.04E-04 | 3.69E-04 | -1.15E+00 | 3.93E-04 | 1.11E-03 | -1.02E+00 | 1.96E-02 | 7.19E-02 | 1.02E+00 | 2.08E-02 | 5.71E-02 | 1.08E+00 |
| CCDC80 | 2.49E-05 | 1.30E-04 | -1.94E+00 | 2.47E-05 | 1.07E-04 | -2.73E+00 | 1.22E-01 | 5.17E-01 | 1.15E+00 | 3.33E-02 | 1.27E-01 | -1.51E+00 | 1.73E-01 | 6.19E-01 | -1.23E+00 | 2.65E-01 | 7.47E-01 | -1.04E+00 | 2.18E-01 | 7.98E-01 | -1.05E+00 | 2.18E-01 | 6.00E-01 | -1.17E+00 |
| NFS1 | 2.50E-05 | 1.30E-04 | -1.59E+00 | 4.58E-03 | 1.98E-02 | -1.02E+00 | 5.58E-04 | 2.37E-03 | -1.41E+00 | 2.32E-02 | 8.87E-02 | 1.05E+00 | 3.64E-03 | 1.30E-02 | -1.24E+00 | 1.30E-01 | 3.68E-01 | 1.23E+00 | 1.32E-06 | 4.86E-06 | -1.60E+00 | 1.07E-02 | 2.93E-02 | -1.03E+00 |
| GATC | 2.51E-05 | 1.31E-04 | -1.30E+00 | 2.74E-02 | 1.19E-01 | 1.29E+00 | 3.76E-05 | 1.60E-04 | -1.40E+00 | 9.50E-02 | 3.63E-01 | 1.16E+00 | 2.37E-03 | 8.45E-03 | -1.10E+00 | 6.91E-02 | 1.95E-01 | 1.18E+00 | 9.43E-07 | 3.46E-06 | -1.35E+00 | 3.62E-03 | 9.94E-03 | -1.04E+00 |
| ZBTB22 | 2.52E-05 | 1.31E-04 | -1.38E+00 | 3.03E-04 | 1.31E-03 | -1.11E+00 | 7.66E-07 | 3.25E-06 | -1.93E+00 | 3.24E-05 | 1.24E-04 | -1.31E+00 | 2.08E-04 | 7.41E-04 | -1.27E+00 | 8.36E-03 | 2.36E-02 | -1.01E+00 | 2.55E-06 | 9.37E-06 | -1.54E+00 | 1.48E-03 | 4.05E-03 | -1.15E+00 |
| AP000487.1 | 2.53E-05 | 1.32E-04 | -2.36E+00 | 1.64E-02 | 7.10E-02 | -1.13E+00 | 5.78E-02 | 2.46E-01 | -1.28E+00 | 6.25E-02 | 2.39E-01 | -1.22E+00 | 7.91E-02 | 2.82E-01 | -1.37E+00 | 3.41E-01 | 9.63E-01 | 1.25E+00 | 1.08E-01 | 3.98E-01 | 1.33E+00 | 1.89E-01 | 5.20E-01 | 1.51E+00 |
| PIGBOS1 | 2.53E-05 | 1.32E-04 | -1.11E+00 | 4.67E-04 | 2.02E-03 | 1.09E+00 | 2.47E-03 | 1.05E-02 | -1.04E+00 | 6.84E-03 | 2.61E-02 | 1.08E+00 | 7.11E-10 | 2.53E-09 | -1.42E+00 | 6.84E-06 | 1.93E-05 | -1.05E+00 | 5.24E-04 | 1.92E-03 | -1.10E+00 | 1.21E-01 | 3.33E-01 | 1.08E+00 |
| ZNHIT1 | 2.53E-05 | 1.32E-04 | 1.02E+00 | 4.00E-05 | 1.73E-04 | 1.13E+00 | 1.50E-01 | 6.38E-01 | 1.10E+00 | 2.19E-02 | 8.37E-02 | 1.12E+00 | 1.71E-05 | 6.09E-05 | -1.11E+00 | 2.75E-03 | 7.74E-03 | 1.01E+00 | 2.62E-02 | 9.63E-02 | 1.05E+00 | 8.35E-02 | 2.29E-01 | 1.08E+00 |
| ZNF329 | 2.53E-05 | 1.32E-04 | -1.35E+00 | 8.80E-07 | 3.80E-06 | -1.44E+00 | 1.77E-04 | 7.54E-04 | -1.33E+00 | 1.28E-05 | 4.90E-05 | -1.37E+00 | 3.90E-02 | 1.39E-01 | 1.09E+00 | 5.15E-03 | 1.45E-02 | -1.19E+00 | 1.37E-01 | 5.01E-01 | 1.16E+00 | 2.79E-02 | 7.68E-02 | -1.09E+00 |
| NTNG2 | 2.53E-05 | 1.32E-04 | -1.62E+00 | 4.88E-05 | 2.11E-04 | -1.48E+00 | 2.08E-18 | 8.85E-18 | -2.64E+00 | 6.72E-17 | 2.57E-16 | -2.16E+00 | 2.88E-02 | 1.03E-01 | -2.02E+00 | 5.20E-02 | 1.47E-01 | -1.74E+00 | 8.73E-02 | 3.20E-01 | -1.41E+00 | 1.07E-01 | 2.93E-01 | -1.42E+00 |
| MGAT4A | 2.54E-05 | 1.33E-04 | -1.12E+00 | 2.78E-11 | 1.20E-10 | -1.26E+00 | 1.23E-01 | 5.24E-01 | 1.08E+00 | 2.51E-03 | 9.58E-03 | -1.01E+00 | 1.39E-02 | 4.94E-02 | -1.11E+00 | 2.43E-04 | 6.86E-04 | -1.53E+00 | 5.48E-03 | 2.01E-02 | -1.31E+00 | 2.54E-03 | 6.97E-03 | -1.58E+00 |
| MECP2 | 2.55E-05 | 1.33E-04 | -1.21E+00 | 6.26E-05 | 2.71E-04 | -1.07E+00 | 3.80E-05 | 1.61E-04 | -1.22E+00 | 1.24E-03 | 4.73E-03 | 1.05E+00 | 6.86E-02 | 2.44E-01 | 1.11E+00 | 1.61E-02 | 4.54E-02 | -1.02E+00 | 1.05E-05 | 3.85E-05 | -1.15E+00 | 1.39E-05 | 3.83E-05 | -1.18E+00 |
| ST13P4 | 2.55E-05 | 1.33E-04 | -1.21E+00 | 6.95E-03 | 3.00E-02 | 1.19E+00 | 8.35E-05 | 3.55E-04 | -1.24E+00 | 3.95E-04 | 1.51E-03 | -1.02E+00 | 1.27E-02 | 4.53E-02 | 1.04E+00 | 5.29E-02 | 1.49E-01 | 1.20E+00 | 6.58E-04 | 2.42E-03 | -1.16E+00 | 2.17E-02 | 5.96E-02 | 1.05E+00 |
| AP002812.1 | 2.55E-05 | 1.33E-04 | -1.69E+00 | 1.60E-04 | 6.92E-04 | -1.41E+00 | 4.16E-03 | 1.77E-02 | -1.43E+00 | 4.22E-03 | 1.61E-02 | -1.32E+00 | 1.21E-02 | 4.33E-02 | -1.62E+00 | 7.43E-02 | 2.09E-01 | -1.12E+00 | 3.83E-05 | 1.40E-04 | -1.76E+00 | 9.21E-03 | 2.53E-02 | -1.21E+00 |
| SETSIP | 2.55E-05 | 1.33E-04 | -1.34E+00 | 4.72E-05 | 2.04E-04 | -1.15E+00 | 7.48E-06 | 3.18E-05 | -1.35E+00 | 8.04E-09 | 3.07E-08 | -1.54E+00 | 7.76E-04 | 2.77E-03 | -1.52E+00 | 2.23E-04 | 6.28E-04 | -1.58E+00 | 6.97E-11 | 2.56E-10 | -1.69E+00 | 1.05E-09 | 2.88E-09 | -1.60E+00 |
| ANKRD18B | 2.57E-05 | 1.34E-04 | -1.40E+00 | 6.48E-06 | 2.80E-05 | -1.28E+00 | 3.98E-05 | 1.69E-04 | -1.31E+00 | 4.51E-04 | 1.72E-03 | -1.11E+00 | 8.20E-03 | 2.92E-02 | -1.65E+00 | 1.13E-01 | 3.20E-01 | -1.16E+00 | 3.16E-07 | 1.16E-06 | -1.39E+00 | 6.37E-03 | 1.75E-02 | -1.06E+00 |
| SBNO2 | 2.58E-05 | 1.34E-04 | -1.45E+00 | 4.41E-04 | 1.91E-03 | -1.19E+00 | 1.35E-03 | 5.74E-03 | -1.32E+00 | 1.63E-02 | 6.21E-02 | 1.01E+00 | 1.06E-03 | 3.77E-03 | -1.41E+00 | 2.65E-02 | 7.48E-02 | 1.01E+00 | 6.11E-04 | 2.24E-03 | -1.72E+00 | 4.36E-02 | 1.20E-01 | -1.11E+00 |
| BCL2L13 | 2.58E-05 | 1.34E-04 | -1.13E+00 | 4.28E-05 | 1.85E-04 | 1.01E+00 | 6.84E-10 | 2.91E-09 | -1.49E+00 | 6.68E-09 | 2.55E-08 | -1.21E+00 | 1.09E-04 | 3.88E-04 | -1.23E+00 | 1.40E-03 | 3.94E-03 | -1.14E+00 | 4.44E-04 | 1.63E-03 | -1.05E+00 | 7.73E-04 | 2.12E-03 | -1.05E+00 |
| MIR3142HG | 2.58E-05 | 1.34E-04 | -1.67E+00 | 7.83E-02 | 3.38E-01 | 1.28E+00 | 2.21E-01 | 9.40E-01 | 1.11E+00 | 3.81E-02 | 1.46E-01 | 2.02E+00 | 5.78E-02 | 2.06E-01 | -1.36E+00 | 8.25E-02 | 2.33E-01 | -1.31E+00 | 2.66E-01 | 9.77E-01 | 1.07E+00 | 2.78E-01 | 7.64E-01 | -1.15E+00 |
| SFXN1 | 2.58E-05 | 1.35E-04 | -1.14E+00 | 2.12E-05 | 9.16E-05 | -1.05E+00 | 3.82E-03 | 1.62E-02 | -1.14E+00 | 1.76E-03 | 6.71E-03 | -1.09E+00 | 1.30E-03 | 4.64E-03 | -1.26E+00 | 2.32E-03 | 6.55E-03 | -1.28E+00 | 1.94E-10 | 7.13E-10 | -1.58E+00 | 1.29E-03 | 3.54E-03 | -1.05E+00 |
| TCEAL5 | 2.59E-05 | 1.35E-04 | -1.24E+00 | 2.03E-06 | 8.79E-06 | -1.32E+00 | 2.02E-01 | 8.56E-01 | 1.21E+00 | 2.51E-01 | 9.60E-01 | 1.28E+00 | 2.22E-01 | 7.90E-01 | 1.07E+00 | 1.78E-01 | 5.03E-01 | -1.03E+00 | 2.29E-01 | 8.42E-01 | 1.09E+00 | 2.55E-01 | 7.00E-01 | 1.43E+00 |
| NPEPL1 | 2.59E-05 | 1.35E-04 | -1.23E+00 | 1.88E-04 | 8.12E-04 | -1.06E+00 | 1.34E-01 | 5.70E-01 | 1.05E+00 | 2.22E-01 | 8.50E-01 | 1.38E+00 | 1.31E-01 | 4.66E-01 | 1.18E+00 | 1.28E-01 | 3.62E-01 | 1.17E+00 | 1.41E-04 | 5.16E-04 | -1.50E+00 | 9.99E-03 | 2.75E-02 | -1.11E+00 |
| NPM1P25 | 2.59E-05 | 1.35E-04 | -1.32E+00 | 2.43E-07 | 1.05E-06 | -1.55E+00 | 1.89E-05 | 8.03E-05 | -1.32E+00 | 7.97E-09 | 3.04E-08 | -1.52E+00 | 1.06E-02 | 3.79E-02 | -1.11E+00 | 7.73E-03 | 2.18E-02 | -1.12E+00 | 1.37E-03 | 5.01E-03 | -1.16E+00 | 1.22E-03 | 3.35E-03 | -1.22E+00 |
| PUS1 | 2.60E-05 | 1.35E-04 | -1.39E+00 | 5.81E-04 | 2.51E-03 | -1.02E+00 | 4.98E-05 | 2.12E-04 | -1.28E+00 | 1.27E-02 | 4.86E-02 | 1.15E+00 | 2.01E-03 | 7.17E-03 | -1.25E+00 | 1.67E-01 | 4.71E-01 | 1.31E+00 | 1.72E-03 | 6.30E-03 | -1.13E+00 | 2.97E-01 | 8.17E-01 | 1.35E+00 |
| PHB | 2.60E-05 | 1.36E-04 | -1.87E+00 | 6.89E-04 | 2.98E-03 | -1.44E+00 | 1.60E-08 | 6.80E-08 | -1.88E+00 | 1.19E-05 | 4.54E-05 | -1.31E+00 | 2.00E-02 | 7.14E-02 | -1.61E+00 | 1.31E-01 | 3.70E-01 | -1.02E+00 | 3.29E-07 | 1.21E-06 | -1.69E+00 | 2.47E-02 | 6.77E-02 | -1.05E+00 |
| DTL | 2.61E-05 | 1.36E-04 | -2.81E+00 | 1.02E-04 | 4.41E-04 | -2.17E+00 | 2.31E-01 | 9.82E-01 | 1.15E+00 | 2.33E-01 | 8.91E-01 | 1.34E+00 | 1.27E-08 | 4.53E-08 | -1.92E+00 | 1.02E-05 | 2.88E-05 | -1.51E+00 | 2.30E-06 | 8.45E-06 | -1.13E+00 | 2.81E-01 | 7.72E-01 | 1.25E+00 |
| SSH1 | 2.62E-05 | 1.37E-04 | -1.81E+00 | 1.78E-03 | 7.68E-03 | -1.26E+00 | 6.22E-09 | 2.64E-08 | -2.20E+00 | 2.56E-06 | 9.78E-06 | -1.46E+00 | 3.92E-05 | 1.40E-04 | -1.83E+00 | 1.63E-02 | 4.59E-02 | -1.09E+00 | 1.50E-06 | 5.49E-06 | -1.61E+00 | 2.29E-04 | 6.30E-04 | -1.31E+00 |
| NFKBIL1 | 2.64E-05 | 1.37E-04 | -1.09E+00 | 1.07E-03 | 4.62E-03 | 1.09E+00 | 7.20E-05 | 3.06E-04 | -1.23E+00 | 2.95E-05 | 1.13E-04 | -1.14E+00 | 5.60E-06 | 2.00E-05 | -1.22E+00 | 2.92E-04 | 8.22E-04 | -1.03E+00 | 6.77E-05 | 2.49E-04 | -1.11E+00 | 6.86E-04 | 1.88E-03 | -1.09E+00 |
| MIR155HG | 2.65E-05 | 1.38E-04 | -1.08E+00 | 1.83E-03 | 7.90E-03 | 1.21E+00 | 1.18E-03 | 5.02E-03 | -1.13E+00 | 2.00E-01 | 7.63E-01 | 1.50E+00 | 3.59E-02 | 1.28E-01 | 1.13E+00 | 4.44E-02 | 1.25E-01 | 1.09E+00 | 1.15E-01 | 4.21E-01 | 1.14E+00 | 2.06E-01 | 5.65E-01 | 1.14E+00 |
| C1GALT1 | 2.67E-05 | 1.39E-04 | -1.06E+00 | 1.27E-13 | 5.49E-13 | -1.40E+00 | 5.91E-03 | 2.51E-02 | -1.08E+00 | 2.38E-08 | 9.08E-08 | -1.37E+00 | 2.18E-08 | 7.76E-08 | -1.24E+00 | 1.96E-14 | 5.53E-14 | -1.75E+00 | 7.51E-06 | 2.75E-05 | -1.19E+00 | 4.12E-08 | 1.13E-07 | -1.33E+00 |
| C1QTNF6 | 2.68E-05 | 1.40E-04 | -1.57E+00 | 2.73E-03 | 1.18E-02 | -1.08E+00 | 1.73E-02 | 7.35E-02 | -1.35E+00 | 1.60E-01 | 6.10E-01 | 1.14E+00 | 3.48E-03 | 1.24E-02 | -1.74E+00 | 3.39E-02 | 9.56E-02 | -1.10E+00 | 7.74E-04 | 2.84E-03 | -1.60E+00 | 2.17E-03 | 5.95E-03 | -1.62E+00 |
| CEP126 | 2.69E-05 | 1.40E-04 | -1.40E+00 | 6.80E-03 | 2.94E-02 | 1.03E+00 | 7.09E-07 | 3.01E-06 | -1.58E+00 | 1.54E-05 | 5.89E-05 | -1.33E+00 | 2.40E-03 | 8.56E-03 | -1.79E+00 | 1.39E-02 | 3.91E-02 | -1.37E+00 | 3.89E-02 | 1.43E-01 | -1.19E+00 | 2.49E-01 | 6.83E-01 | 1.17E+00 |
| TRIM16 | 2.70E-05 | 1.41E-04 | -1.20E+00 | 2.68E-05 | 1.16E-04 | -1.08E+00 | 5.57E-08 | 2.37E-07 | -1.46E+00 | 6.81E-06 | 2.60E-05 | -1.09E+00 | 2.30E-03 | 8.19E-03 | -1.08E+00 | 5.42E-02 | 1.53E-01 | 1.27E+00 | 9.63E-05 | 3.53E-04 | -1.29E+00 | 8.60E-03 | 2.36E-02 | -1.06E+00 |
| KANK2 | 2.71E-05 | 1.41E-04 | -1.49E+00 | 2.65E-03 | 1.15E-02 | 1.01E+00 | 1.64E-04 | 6.95E-04 | -1.27E+00 | 5.06E-02 | 1.93E-01 | 1.17E+00 | 1.27E-02 | 4.52E-02 | -1.69E+00 | 2.61E-02 | 7.36E-02 | -1.45E+00 | 8.77E-06 | 3.22E-05 | -1.54E+00 | 4.14E-02 | 1.14E-01 | 1.07E+00 |
| C18orf54 | 2.71E-05 | 1.41E-04 | -1.74E+00 | 1.04E-01 | 4.50E-01 | 1.35E+00 | 7.09E-05 | 3.01E-04 | -1.61E+00 | 1.92E-02 | 7.34E-02 | 1.08E+00 | 3.15E-04 | 1.12E-03 | -2.65E+00 | 1.19E-02 | 3.37E-02 | -1.53E+00 | 1.03E-03 | 3.79E-03 | -1.44E+00 | 1.45E-01 | 3.98E-01 | 1.05E+00 |
| BORCS8 | 2.73E-05 | 1.42E-04 | -1.23E+00 | 3.10E-04 | 1.34E-03 | -1.00E+00 | 1.65E-04 | 6.99E-04 | -1.31E+00 | 1.08E-03 | 4.14E-03 | -1.07E+00 | 4.16E-07 | 1.48E-06 | -1.44E+00 | 3.55E-05 | 1.00E-04 | -1.17E+00 | 3.27E-02 | 1.20E-01 | 1.05E+00 | 5.39E-02 | 1.48E-01 | 1.03E+00 |
| TMEM126B | 2.74E-05 | 1.43E-04 | -1.14E+00 | 2.76E-02 | 1.19E-01 | 1.25E+00 | 1.27E-04 | 5.37E-04 | -1.18E+00 | 5.40E-04 | 2.06E-03 | 1.02E+00 | 3.34E-02 | 1.19E-01 | 1.10E+00 | 3.24E-01 | 9.14E-01 | 1.37E+00 | 2.15E-02 | 7.88E-02 | 1.09E+00 | 2.28E-01 | 6.25E-01 | 1.17E+00 |
| LRRC37A3 | 2.74E-05 | 1.43E-04 | -1.33E+00 | 5.51E-05 | 2.38E-04 | -1.22E+00 | 1.82E-02 | 7.72E-02 | -1.06E+00 | 3.11E-02 | 1.19E-01 | 1.11E+00 | 2.69E-03 | 9.60E-03 | -1.35E+00 | 1.23E-02 | 3.47E-02 | -1.08E+00 | 2.04E-02 | 7.48E-02 | -1.22E+00 | 1.62E-01 | 4.45E-01 | 1.07E+00 |
| ALDH7A1P1 | 2.74E-05 | 1.43E-04 | -1.83E+00 | 1.33E-03 | 5.75E-03 | -1.19E+00 | 1.94E-03 | 8.26E-03 | -1.30E+00 | 1.45E-03 | 5.53E-03 | -1.14E+00 | 6.32E-02 | 2.25E-01 | -1.14E+00 | 7.98E-02 | 2.25E-01 | -1.13E+00 | 5.09E-03 | 1.87E-02 | -1.23E+00 | 8.59E-02 | 2.36E-01 | 1.01E+00 |
| ERBIN | 2.74E-05 | 1.43E-04 | -1.10E+00 | 2.65E-03 | 1.15E-02 | 1.16E+00 | 1.64E-04 | 6.95E-04 | -1.19E+00 | 3.08E-04 | 1.18E-03 | -1.08E+00 | 1.22E-03 | 4.35E-03 | -1.07E+00 | 4.92E-02 | 1.39E-01 | 1.10E+00 | 1.22E-02 | 4.49E-02 | 1.06E+00 | 7.85E-02 | 2.16E-01 | 1.15E+00 |
| SERPINI1 | 2.75E-05 | 1.43E-04 | -1.69E+00 | 2.56E-04 | 1.11E-03 | -1.35E+00 | 9.54E-02 | 4.05E-01 | -1.14E+00 | 1.47E-01 | 5.62E-01 | 1.05E+00 | 1.87E-02 | 6.67E-02 | 1.06E+00 | 1.22E-01 | 3.45E-01 | 1.29E+00 | 6.42E-04 | 2.36E-03 | 1.08E+00 | 4.30E-03 | 1.18E-02 | 1.09E+00 |
| NAP1L3 | 2.75E-05 | 1.43E-04 | -1.25E+00 | 2.84E-03 | 1.23E-02 | 1.12E+00 | 6.77E-03 | 2.88E-02 | -1.03E+00 | 8.10E-02 | 3.10E-01 | 1.30E+00 | 1.06E-01 | 3.79E-01 | 1.14E+00 | 2.86E-01 | 8.05E-01 | 1.58E+00 | 9.17E-02 | 3.37E-01 | 1.51E+00 | 1.00E-01 | 2.75E-01 | 1.71E+00 |
| TMEM71 | 2.75E-05 | 1.43E-04 | -1.33E+00 | 5.94E-03 | 2.57E-02 | 1.05E+00 | 3.39E-02 | 1.44E-01 | -1.11E+00 | 1.37E-01 | 5.22E-01 | 1.15E+00 | 2.21E-01 | 7.86E-01 | 1.49E+00 | 3.04E-01 | 8.57E-01 | 1.11E+00 | 2.53E-01 | 9.30E-01 | -1.03E+00 | 2.85E-01 | 7.81E-01 | -1.15E+00 |
| ANKRD39 | 2.75E-05 | 1.43E-04 | -1.37E+00 | 1.04E-04 | 4.49E-04 | -1.17E+00 | 1.68E-03 | 7.15E-03 | -1.03E+00 | 1.67E-06 | 6.36E-06 | -1.19E+00 | 3.57E-08 | 1.27E-07 | -1.76E+00 | 8.55E-08 | 2.41E-07 | -1.57E+00 | 2.09E-07 | 7.68E-07 | -1.47E+00 | 1.35E-08 | 3.70E-08 | -1.51E+00 |
| DCBLD2 | 2.75E-05 | 1.44E-04 | -1.33E+00 | 2.29E-03 | 9.91E-03 | 1.02E+00 | 1.35E-01 | 5.76E-01 | 1.15E+00 | 1.84E-02 | 7.04E-02 | 1.65E+00 | 6.33E-04 | 2.26E-03 | -1.46E+00 | 8.24E-02 | 2.32E-01 | 1.15E+00 | 1.36E-03 | 5.00E-03 | -1.20E+00 | 5.58E-02 | 1.53E-01 | 1.10E+00 |
| MT1XP1 | 2.76E-05 | 1.44E-04 | -1.44E+00 | 5.23E-06 | 2.26E-05 | -1.68E+00 | 1.93E-05 | 8.18E-05 | -1.37E+00 | 1.25E-08 | 4.78E-08 | -1.65E+00 | 2.07E-03 | 7.39E-03 | 1.03E+00 | 5.98E-06 | 1.69E-05 | -1.18E+00 | 7.33E-06 | 2.69E-05 | -1.08E+00 | 3.22E-17 | 8.84E-17 | -2.15E+00 |
| ARL14EP | 2.76E-05 | 1.44E-04 | -1.01E+00 | 7.07E-07 | 3.06E-06 | 1.01E+00 | 1.09E-04 | 4.65E-04 | -1.11E+00 | 1.17E-02 | 4.48E-02 | 1.15E+00 | 5.42E-10 | 1.93E-09 | -1.28E+00 | 1.76E-08 | 4.97E-08 | -1.20E+00 | 8.63E-07 | 3.17E-06 | -1.15E+00 | 3.24E-04 | 8.90E-04 | -1.03E+00 |
| ZNF767P | 2.78E-05 | 1.45E-04 | -1.59E+00 | 1.03E-02 | 4.46E-02 | 1.04E+00 | 2.40E-03 | 1.02E-02 | -1.39E+00 | 8.93E-02 | 3.41E-01 | 1.61E+00 | 2.89E-04 | 1.03E-03 | -1.50E+00 | 3.23E-02 | 9.10E-02 | 1.07E+00 | 4.59E-02 | 1.69E-01 | -1.07E+00 | 3.25E-01 | 8.92E-01 | 1.33E+00 |
| MLYCD | 2.78E-05 | 1.45E-04 | -1.40E+00 | 1.23E-02 | 5.32E-02 | 1.18E+00 | 1.23E-05 | 5.22E-05 | -1.51E+00 | 8.40E-02 | 3.21E-01 | 1.30E+00 | 3.04E-02 | 1.08E-01 | -1.08E+00 | 1.26E-01 | 3.54E-01 | 1.07E+00 | 5.40E-07 | 1.98E-06 | -1.43E+00 | 8.75E-02 | 2.40E-01 | 1.08E+00 |
| LSM5 | 2.79E-05 | 1.45E-04 | 1.01E+00 | 6.06E-11 | 2.62E-10 | 1.03E+00 | 1.77E-01 | 7.54E-01 | 1.14E+00 | 2.19E-02 | 8.36E-02 | 1.11E+00 | 2.50E-03 | 8.91E-03 | -1.05E+00 | 6.73E-02 | 1.90E-01 | 1.05E+00 | 4.62E-02 | 1.70E-01 | 1.01E+00 | 1.67E-01 | 4.58E-01 | 1.05E+00 |
| HMGN2P46 | 2.79E-05 | 1.45E-04 | -1.32E+00 | 1.53E-08 | 6.60E-08 | -1.95E+00 | 1.29E-03 | 5.49E-03 | -1.10E+00 | 8.76E-13 | 3.34E-12 | -1.85E+00 | 1.63E-03 | 5.83E-03 | -1.33E+00 | 2.00E-05 | 5.63E-05 | -2.03E+00 | 1.11E-04 | 4.06E-04 | -1.13E+00 | 1.03E-06 | 2.84E-06 | -1.31E+00 |
| PDP2 | 2.80E-05 | 1.46E-04 | -1.73E+00 | 2.40E-04 | 1.04E-03 | -1.39E+00 | 2.08E-04 | 8.84E-04 | -1.67E+00 | 3.27E-03 | 1.25E-02 | -1.15E+00 | 2.26E-04 | 8.06E-04 | -2.56E+00 | 1.33E-02 | 3.74E-02 | -1.31E+00 | 2.47E-04 | 9.05E-04 | -1.51E+00 | 5.95E-04 | 1.64E-03 | -1.56E+00 |
| DHX8 | 2.80E-05 | 1.46E-04 | -1.29E+00 | 3.58E-04 | 1.55E-03 | -1.08E+00 | 8.75E-05 | 3.72E-04 | -1.18E+00 | 4.90E-06 | 1.87E-05 | -1.20E+00 | 2.80E-01 | 1.00E+00 | 1.48E+00 | 5.08E-02 | 1.43E-01 | 1.12E+00 | 3.88E-02 | 1.42E-01 | 1.07E+00 | 9.94E-04 | 2.73E-03 | -1.12E+00 |
| RIC8B | 2.81E-05 | 1.46E-04 | -1.80E+00 | 6.03E-02 | 2.60E-01 | 1.11E+00 | 2.74E-02 | 1.17E-01 | -1.17E+00 | 2.58E-01 | 9.85E-01 | 1.27E+00 | 3.42E-06 | 1.22E-05 | -1.99E+00 | 1.25E-02 | 3.51E-02 | 1.02E+00 | 4.67E-06 | 1.71E-05 | -1.73E+00 | 7.54E-03 | 2.07E-02 | -1.20E+00 |
| HCFC1 | 2.81E-05 | 1.47E-04 | -1.15E+00 | 5.26E-05 | 2.27E-04 | -1.08E+00 | 3.49E-02 | 1.48E-01 | -1.01E+00 | 1.52E-02 | 5.80E-02 | 1.07E+00 | 2.43E-04 | 8.67E-04 | -1.15E+00 | 7.41E-03 | 2.09E-02 | 1.11E+00 | 2.56E-06 | 9.41E-06 | -1.29E+00 | 1.81E-02 | 4.96E-02 | 1.06E+00 |
| RPL10P4 | 2.81E-05 | 1.47E-04 | -1.26E+00 | 9.65E-04 | 4.17E-03 | 1.02E+00 | 1.91E-02 | 8.12E-02 | -1.02E+00 | 2.31E-04 | 8.84E-04 | -1.16E+00 | 8.43E-07 | 3.00E-06 | -1.47E+00 | 1.54E-10 | 4.33E-10 | -1.80E+00 | 4.53E-05 | 1.66E-04 | -1.28E+00 | 1.37E-05 | 3.75E-05 | -1.39E+00 |
| AC093909.6 | 2.82E-05 | 1.47E-04 | -1.84E+00 | 1.60E-04 | 6.89E-04 | -1.44E+00 | 1.82E-03 | 7.74E-03 | -1.31E+00 | 1.20E-03 | 4.59E-03 | -1.25E+00 | 8.51E-03 | 3.04E-02 | -1.19E+00 | 1.98E-03 | 5.58E-03 | -1.47E+00 | 3.73E-05 | 1.37E-04 | -1.50E+00 | 3.21E-03 | 8.81E-03 | -1.28E+00 |
| SEPT5 | 2.83E-05 | 1.47E-04 | -2.00E+00 | 1.84E-03 | 7.97E-03 | -1.25E+00 | 5.91E-03 | 2.51E-02 | -1.65E+00 | 5.07E-02 | 1.94E-01 | -1.07E+00 | 2.34E-01 | 8.35E-01 | 1.25E+00 | 2.99E-01 | 8.42E-01 | 1.11E+00 | 2.34E-01 | 8.60E-01 | 1.29E+00 | 1.11E-01 | 3.05E-01 | 1.55E+00 |
| NKTR | 2.83E-05 | 1.47E-04 | -1.04E+00 | 8.69E-03 | 3.76E-02 | 1.11E+00 | 2.02E-03 | 8.60E-03 | -1.14E+00 | 4.11E-03 | 1.57E-02 | -1.06E+00 | 1.82E-01 | 6.48E-01 | 1.23E+00 | 3.53E-01 | 9.94E-01 | 1.17E+00 | 2.83E-03 | 1.04E-02 | -1.09E+00 | 1.50E-01 | 4.12E-01 | -1.03E+00 |
| GGTA1P | 2.83E-05 | 1.47E-04 | -1.03E+00 | 6.74E-08 | 2.91E-07 | -1.04E+00 | 7.32E-04 | 3.11E-03 | -1.13E+00 | 2.92E-04 | 1.12E-03 | -1.09E+00 | 1.92E-05 | 6.84E-05 | -1.66E+00 | 9.71E-07 | 2.74E-06 | -1.96E+00 | 6.65E-02 | 2.44E-01 | 1.06E+00 | 1.35E-01 | 3.71E-01 | -1.14E+00 |
| WDYHV1 | 2.84E-05 | 1.48E-04 | -1.17E+00 | 1.47E-03 | 6.33E-03 | 1.14E+00 | 6.57E-06 | 2.79E-05 | -1.28E+00 | 1.25E-04 | 4.76E-04 | -1.01E+00 | 1.39E-05 | 4.94E-05 | -1.37E+00 | 6.10E-03 | 1.72E-02 | 1.01E+00 | 3.43E-03 | 1.26E-02 | -1.14E+00 | 4.22E-03 | 1.16E-02 | -1.09E+00 |
| COL4A5 | 2.84E-05 | 1.48E-04 | -1.67E+00 | 7.42E-04 | 3.21E-03 | -1.21E+00 | 1.87E-04 | 7.93E-04 | -1.25E+00 | 4.38E-04 | 1.67E-03 | -1.06E+00 | 1.50E-03 | 5.35E-03 | -1.25E+00 | 8.75E-03 | 2.47E-02 | -1.11E+00 | 3.95E-04 | 1.45E-03 | -1.54E+00 | 1.36E-02 | 3.73E-02 | -1.13E+00 |
| CNOT11 | 2.84E-05 | 1.48E-04 | -1.19E+00 | 2.45E-04 | 1.06E-03 | 1.05E+00 | 4.51E-04 | 1.92E-03 | -1.14E+00 | 1.19E-03 | 4.56E-03 | 1.07E+00 | 6.63E-09 | 2.36E-08 | -1.58E+00 | 9.23E-04 | 2.60E-03 | -1.04E+00 | 4.62E-05 | 1.70E-04 | -1.23E+00 | 1.43E-01 | 3.93E-01 | 1.09E+00 |
| HTRA2 | 2.84E-05 | 1.48E-04 | -1.64E+00 | 7.69E-04 | 3.32E-03 | -1.22E+00 | 1.12E-03 | 4.75E-03 | -1.40E+00 | 2.77E-04 | 1.06E-03 | -1.27E+00 | 7.38E-08 | 2.63E-07 | -2.14E+00 | 1.45E-05 | 4.10E-05 | -1.50E+00 | 9.75E-04 | 3.58E-03 | -1.15E+00 | 2.15E-02 | 5.90E-02 | -1.01E+00 |
| TRIM41 | 2.85E-05 | 1.49E-04 | -1.18E+00 | 7.19E-06 | 3.11E-05 | -1.13E+00 | 8.06E-04 | 3.42E-03 | -1.17E+00 | 2.22E-07 | 8.48E-07 | -1.38E+00 | 8.11E-03 | 2.89E-02 | -1.09E+00 | 1.33E-02 | 3.76E-02 | -1.10E+00 | 1.28E-03 | 4.69E-03 | -1.27E+00 | 4.81E-03 | 1.32E-02 | -1.28E+00 |
| TP53TG1 | 2.85E-05 | 1.49E-04 | -1.06E+00 | 9.33E-10 | 4.03E-09 | -1.22E+00 | 1.08E-05 | 4.57E-05 | -1.26E+00 | 4.89E-13 | 1.87E-12 | -1.55E+00 | 9.89E-05 | 3.53E-04 | -1.04E+00 | 9.16E-08 | 2.58E-07 | -1.29E+00 | 1.64E-02 | 6.02E-02 | 1.07E+00 | 4.70E-06 | 1.29E-05 | -1.29E+00 |
| HOMEZ | 2.85E-05 | 1.49E-04 | -1.44E+00 | 2.82E-03 | 1.22E-02 | 1.03E+00 | 2.32E-08 | 9.85E-08 | -1.95E+00 | 3.20E-03 | 1.22E-02 | -1.04E+00 | 4.49E-04 | 1.60E-03 | -1.51E+00 | 1.36E-02 | 3.83E-02 | -1.07E+00 | 4.13E-07 | 1.52E-06 | -1.69E+00 | 2.39E-02 | 6.57E-02 | 1.06E+00 |
| CDK19 | 2.87E-05 | 1.49E-04 | -1.13E+00 | 1.46E-02 | 6.32E-02 | 1.21E+00 | 4.47E-06 | 1.90E-05 | -1.49E+00 | 1.39E-03 | 5.29E-03 | -1.08E+00 | 1.91E-08 | 6.83E-08 | -1.38E+00 | 4.68E-03 | 1.32E-02 | 1.06E+00 | 4.68E-08 | 1.72E-07 | -1.42E+00 | 1.06E-03 | 2.92E-03 | -1.01E+00 |
| AC011712.1 | 2.88E-05 | 1.50E-04 | -1.89E+00 | 1.32E-02 | 5.72E-02 | -1.01E+00 | 7.33E-13 | 3.12E-12 | -2.35E+00 | 3.27E-09 | 1.25E-08 | -1.58E+00 | 2.01E-06 | 7.16E-06 | -2.88E+00 | 5.14E-04 | 1.45E-03 | -1.53E+00 | 3.11E-10 | 1.14E-09 | -2.26E+00 | 1.29E-03 | 3.54E-03 | -1.22E+00 |
| TRAF2 | 2.89E-05 | 1.51E-04 | -1.38E+00 | 1.51E-03 | 6.54E-03 | -1.03E+00 | 1.99E-03 | 8.45E-03 | -1.11E+00 | 2.21E-03 | 8.44E-03 | 1.02E+00 | 2.18E-03 | 7.76E-03 | -1.35E+00 | 2.48E-02 | 6.99E-02 | -1.02E+00 | 3.58E-10 | 1.31E-09 | -1.49E+00 | 1.18E-03 | 3.24E-03 | 1.04E+00 |
| ZNF883 | 2.89E-05 | 1.51E-04 | -1.33E+00 | 2.05E-02 | 8.86E-02 | 1.20E+00 | 3.03E-06 | 1.29E-05 | -1.43E+00 | 4.24E-04 | 1.62E-03 | -1.06E+00 | 3.27E-04 | 1.17E-03 | -1.49E+00 | 6.72E-02 | 1.89E-01 | 1.12E+00 | 3.84E-02 | 1.41E-01 | -1.09E+00 | 4.25E-02 | 1.17E-01 | 1.54E+00 |
| NARF | 2.89E-05 | 1.51E-04 | -1.04E+00 | 1.06E-05 | 4.58E-05 | 1.06E+00 | 2.17E-04 | 9.20E-04 | -1.10E+00 | 1.51E-04 | 5.75E-04 | 1.01E+00 | 2.65E-08 | 9.43E-08 | -1.21E+00 | 8.25E-04 | 2.33E-03 | 1.14E+00 | 6.60E-03 | 2.42E-02 | -1.08E+00 | 9.65E-03 | 2.65E-02 | -1.06E+00 |
| ZDHHC9 | 2.89E-05 | 1.51E-04 | -1.51E+00 | 6.00E-03 | 2.59E-02 | 1.02E+00 | 3.67E-05 | 1.56E-04 | -1.33E+00 | 4.91E-03 | 1.88E-02 | 1.05E+00 | 2.14E-09 | 7.64E-09 | -1.95E+00 | 1.81E-05 | 5.11E-05 | -1.24E+00 | 2.12E-05 | 7.77E-05 | -1.47E+00 | 5.00E-02 | 1.37E-01 | 1.00E+00 |
| PPP1R9A | 2.90E-05 | 1.51E-04 | -1.32E+00 | 7.40E-03 | 3.20E-02 | 1.11E+00 | 6.08E-04 | 2.58E-03 | -1.36E+00 | 3.30E-04 | 1.26E-03 | -1.21E+00 | 6.74E-02 | 2.40E-01 | -1.11E+00 | 4.39E-02 | 1.24E-01 | -1.09E+00 | 1.20E-01 | 4.42E-01 | 1.10E+00 | 1.43E-01 | 3.93E-01 | 1.07E+00 |
| LINC02470 | 2.90E-05 | 1.51E-04 | -1.66E+00 | 7.02E-05 | 3.03E-04 | -1.46E+00 | 1.59E-02 | 6.76E-02 | -1.34E+00 | 2.47E-01 | 9.45E-01 | 1.30E+00 | 7.32E-08 | 2.61E-07 | -3.08E+00 | 2.56E-06 | 7.23E-06 | -2.18E+00 | 5.56E-02 | 2.04E-01 | 1.76E+00 | 1.68E-01 | 4.61E-01 | 1.48E+00 |
| USP34 | 2.90E-05 | 1.51E-04 | 1.04E+00 | 2.26E-06 | 9.76E-06 | 1.08E+00 | 2.58E-05 | 1.10E-04 | -1.23E+00 | 4.50E-05 | 1.72E-04 | -1.13E+00 | 3.46E-03 | 1.23E-02 | 1.09E+00 | 6.82E-04 | 1.92E-03 | -1.01E+00 | 3.29E-03 | 1.21E-02 | -1.03E+00 | 2.14E-05 | 5.87E-05 | -1.05E+00 |
| AL139174.1 | 2.90E-05 | 1.51E-04 | -1.54E+00 | 1.45E-08 | 6.26E-08 | -2.64E+00 | 8.48E-03 | 3.60E-02 | -1.16E+00 | 9.81E-05 | 3.75E-04 | -1.42E+00 | 1.11E-02 | 3.96E-02 | -1.30E+00 | 1.17E-03 | 3.30E-03 | -1.78E+00 | 1.86E-01 | 6.84E-01 | 1.15E+00 | 3.38E-04 | 9.29E-04 | -1.72E+00 |
| HERC2 | 2.90E-05 | 1.51E-04 | -1.09E+00 | 1.23E-03 | 5.31E-03 | 1.14E+00 | 4.76E-04 | 2.02E-03 | -1.08E+00 | 2.15E-03 | 8.21E-03 | 1.07E+00 | 1.32E-01 | 4.70E-01 | 1.26E+00 | 2.52E-01 | 7.09E-01 | 1.40E+00 | 1.53E-02 | 5.61E-02 | 1.07E+00 | 7.55E-02 | 2.07E-01 | 1.08E+00 |
| POU5F2 | 2.91E-05 | 1.52E-04 | -1.43E+00 | 3.59E-04 | 1.55E-03 | -1.16E+00 | 1.45E-05 | 6.17E-05 | -1.40E+00 | 2.30E-04 | 8.80E-04 | -1.14E+00 | 7.42E-03 | 2.64E-02 | -1.21E+00 | 1.08E-02 | 3.06E-02 | -1.19E+00 | 1.08E-01 | 3.95E-01 | -1.07E+00 | 1.06E-01 | 2.91E-01 | -1.01E+00 |
| CERCAM | 2.91E-05 | 1.52E-04 | -1.53E+00 | 6.12E-04 | 2.64E-03 | -1.08E+00 | 1.37E-11 | 5.83E-11 | -1.79E+00 | 2.84E-06 | 1.08E-05 | -1.16E+00 | 1.37E-01 | 4.90E-01 | 1.04E+00 | 1.98E-01 | 5.59E-01 | 1.18E+00 | 1.16E-01 | 4.25E-01 | 1.02E+00 | 3.94E-02 | 1.08E-01 | 1.82E+00 |
| FMNL2 | 2.92E-05 | 1.52E-04 | -1.40E+00 | 6.65E-03 | 2.87E-02 | 1.12E+00 | 8.12E-03 | 3.45E-02 | -1.30E+00 | 2.46E-01 | 9.39E-01 | 1.35E+00 | 7.61E-02 | 2.71E-01 | -1.13E+00 | 1.46E-01 | 4.12E-01 | -1.01E+00 | 1.65E-01 | 6.06E-01 | -1.12E+00 | 1.87E-01 | 5.14E-01 | 1.50E+00 |
| AC016747.3 | 2.92E-05 | 1.52E-04 | -1.91E+00 | 2.69E-03 | 1.16E-02 | -1.23E+00 | 1.55E-01 | 6.59E-01 | 1.08E+00 | 1.73E-01 | 6.62E-01 | 1.20E+00 | 6.39E-03 | 2.28E-02 | -1.57E+00 | 1.03E-02 | 2.89E-02 | -1.46E+00 | 2.08E-04 | 7.63E-04 | -1.44E+00 | 4.34E-02 | 1.19E-01 | -1.01E+00 |
| TIAM2 | 2.92E-05 | 1.52E-04 | -1.62E+00 | 9.91E-02 | 4.28E-01 | 1.31E+00 | 1.05E-03 | 4.47E-03 | -1.63E+00 | 1.50E-01 | 5.74E-01 | 1.29E+00 | 4.94E-03 | 1.76E-02 | -1.44E+00 | 2.60E-01 | 7.32E-01 | 1.30E+00 | 4.58E-02 | 1.68E-01 | -1.05E+00 | 3.05E-01 | 8.38E-01 | 1.26E+00 |
| SLC41A3 | 2.93E-05 | 1.53E-04 | -1.19E+00 | 6.72E-03 | 2.91E-02 | 1.16E+00 | 1.33E-05 | 5.66E-05 | -1.24E+00 | 1.08E-03 | 4.14E-03 | 1.02E+00 | 1.72E-03 | 6.14E-03 | -1.19E+00 | 1.03E-01 | 2.90E-01 | 1.17E+00 | 5.07E-03 | 1.86E-02 | -1.04E+00 | 1.52E-01 | 4.19E-01 | 1.15E+00 |
| UPF3B | 2.93E-05 | 1.53E-04 | -1.12E+00 | 2.03E-02 | 8.76E-02 | 1.13E+00 | 1.70E-05 | 7.20E-05 | -1.18E+00 | 5.97E-04 | 2.28E-03 | -1.01E+00 | 7.18E-09 | 2.56E-08 | -1.44E+00 | 7.79E-03 | 2.19E-02 | -1.04E+00 | 3.12E-02 | 1.14E-01 | -1.02E+00 | 2.78E-01 | 7.65E-01 | 1.05E+00 |
| NOL10 | 2.95E-05 | 1.54E-04 | -1.22E+00 | 1.63E-03 | 7.03E-03 | 1.09E+00 | 1.55E-06 | 6.60E-06 | -1.23E+00 | 6.17E-07 | 2.36E-06 | -1.10E+00 | 4.46E-03 | 1.59E-02 | 1.07E+00 | 2.60E-01 | 7.33E-01 | 1.46E+00 | 5.10E-06 | 1.87E-05 | -1.14E+00 | 5.01E-02 | 1.38E-01 | 1.10E+00 |
| MPRIP-AS1 | 2.95E-05 | 1.54E-04 | -2.14E+00 | 9.88E-03 | 4.27E-02 | -1.19E+00 | 7.73E-06 | 3.28E-05 | -2.29E+00 | 2.99E-04 | 1.14E-03 | -1.65E+00 | 6.41E-11 | 2.29E-10 | -3.23E+00 | 1.09E-08 | 3.06E-08 | -2.05E+00 | 1.54E-02 | 5.64E-02 | -1.29E+00 | 1.07E-01 | 2.95E-01 | -1.03E+00 |
| GLYR1 | 2.96E-05 | 1.54E-04 | -1.29E+00 | 2.84E-02 | 1.23E-01 | 1.18E+00 | 6.16E-06 | 2.62E-05 | -1.45E+00 | 5.57E-03 | 2.13E-02 | 1.05E+00 | 3.92E-09 | 1.40E-08 | -1.45E+00 | 2.07E-05 | 5.83E-05 | -1.05E+00 | 1.31E-07 | 4.81E-07 | -1.30E+00 | 2.18E-02 | 5.99E-02 | 1.05E+00 |
| CTNNBIP1 | 2.96E-05 | 1.55E-04 | -1.11E+00 | 6.15E-06 | 2.66E-05 | -1.04E+00 | 8.01E-06 | 3.40E-05 | -1.30E+00 | 1.21E-07 | 4.63E-07 | -1.25E+00 | 9.65E-03 | 3.44E-02 | -1.02E+00 | 3.68E-02 | 1.04E-01 | 1.09E+00 | 6.06E-07 | 2.22E-06 | -1.34E+00 | 4.11E-06 | 1.13E-05 | -1.25E+00 |
| CCT7 | 2.96E-05 | 1.55E-04 | -1.03E+00 | 1.55E-03 | 6.68E-03 | 1.13E+00 | 2.00E-01 | 8.52E-01 | 1.09E+00 | 1.32E-01 | 5.03E-01 | 1.22E+00 | 4.70E-06 | 1.68E-05 | -1.17E+00 | 2.37E-01 | 6.69E-01 | 1.14E+00 | 1.03E-01 | 3.80E-01 | 1.04E+00 | 1.92E-01 | 5.27E-01 | 1.24E+00 |
| TMEM245 | 2.96E-05 | 1.55E-04 | -1.16E+00 | 3.55E-03 | 1.53E-02 | 1.17E+00 | 9.12E-07 | 3.87E-06 | -1.50E+00 | 6.45E-05 | 2.46E-04 | -1.11E+00 | 1.73E-06 | 6.16E-06 | -1.83E+00 | 7.77E-02 | 2.19E-01 | 1.08E+00 | 8.76E-15 | 3.22E-14 | -1.82E+00 | 6.80E-06 | 1.87E-05 | -1.30E+00 |
| CIRBP | 2.97E-05 | 1.55E-04 | 1.15E+00 | 1.70E-03 | 7.33E-03 | 1.11E+00 | 1.30E-07 | 5.53E-07 | 1.28E+00 | 4.42E-04 | 1.69E-03 | 1.20E+00 | 2.66E-07 | 9.50E-07 | 1.41E+00 | 1.40E-01 | 3.94E-01 | 1.14E+00 | 2.86E-11 | 1.05E-10 | 1.43E+00 | 4.23E-04 | 1.16E-03 | 1.21E+00 |
| MRPS36 | 2.97E-05 | 1.55E-04 | 1.01E+00 | 1.36E-08 | 5.86E-08 | -1.04E+00 | 4.90E-03 | 2.08E-02 | -1.03E+00 | 6.44E-08 | 2.46E-07 | -1.23E+00 | 1.41E-06 | 5.03E-06 | -1.07E+00 | 6.46E-09 | 1.82E-08 | -1.31E+00 | 3.15E-05 | 1.16E-04 | -1.08E+00 | 1.89E-06 | 5.20E-06 | -1.20E+00 |
| PPM1L | 2.98E-05 | 1.55E-04 | -1.34E+00 | 9.43E-03 | 4.07E-02 | 1.14E+00 | 9.95E-06 | 4.23E-05 | -1.41E+00 | 1.00E-03 | 3.82E-03 | -1.01E+00 | 6.50E-03 | 2.32E-02 | -1.05E+00 | 2.44E-01 | 6.89E-01 | 1.36E+00 | 1.39E-01 | 5.10E-01 | 1.12E+00 | 1.60E-01 | 4.40E-01 | 1.47E+00 |
| AP001469.3 | 2.98E-05 | 1.55E-04 | -1.70E+00 | 1.95E-02 | 8.42E-02 | 1.03E+00 | 8.65E-03 | 3.68E-02 | -1.50E+00 | 9.21E-02 | 3.52E-01 | 1.03E+00 | 5.21E-03 | 1.86E-02 | -1.65E+00 | 2.36E-01 | 6.65E-01 | 1.25E+00 | 1.09E-04 | 4.01E-04 | -1.70E+00 | 3.19E-03 | 8.78E-03 | -1.26E+00 |
| MRPS5 | 2.98E-05 | 1.55E-04 | -1.06E+00 | 4.35E-03 | 1.88E-02 | 1.21E+00 | 8.98E-06 | 3.82E-05 | -1.24E+00 | 3.18E-07 | 1.22E-06 | -1.15E+00 | 2.80E-05 | 9.97E-05 | -1.20E+00 | 1.50E-02 | 4.24E-02 | 1.06E+00 | 3.39E-10 | 1.24E-09 | -1.38E+00 | 1.00E-03 | 2.75E-03 | -1.05E+00 |
| EIF2B4 | 2.99E-05 | 1.56E-04 | -1.31E+00 | 1.05E-02 | 4.54E-02 | 1.10E+00 | 5.12E-07 | 2.18E-06 | -1.39E+00 | 5.94E-06 | 2.27E-05 | -1.11E+00 | 1.29E-01 | 4.60E-01 | 1.20E+00 | 3.16E-01 | 8.90E-01 | 1.38E+00 | 2.01E-05 | 7.37E-05 | -1.25E+00 | 9.21E-02 | 2.53E-01 | 1.13E+00 |
| TAB2 | 2.99E-05 | 1.56E-04 | -1.00E+00 | 4.29E-03 | 1.86E-02 | 1.24E+00 | 4.49E-03 | 1.91E-02 | -1.09E+00 | 3.51E-02 | 1.34E-01 | 1.14E+00 | 4.49E-02 | 1.60E-01 | 1.29E+00 | 1.42E-01 | 4.00E-01 | 1.36E+00 | 2.21E-01 | 8.09E-01 | 1.23E+00 | 3.01E-01 | 8.26E-01 | 1.15E+00 |
| SAP30BP | 2.99E-05 | 1.56E-04 | -1.14E+00 | 2.12E-03 | 9.16E-03 | 1.13E+00 | 1.51E-06 | 6.41E-06 | -1.24E+00 | 3.76E-07 | 1.43E-06 | -1.15E+00 | 1.79E-02 | 6.40E-02 | 1.01E+00 | 1.95E-01 | 5.49E-01 | 1.25E+00 | 1.24E-08 | 4.56E-08 | -1.32E+00 | 1.60E-04 | 4.40E-04 | -1.18E+00 |
| RAB27A | 2.99E-05 | 1.56E-04 | -1.32E+00 | 3.65E-05 | 1.58E-04 | -1.13E+00 | 4.27E-05 | 1.81E-04 | -1.11E+00 | 3.54E-05 | 1.35E-04 | -1.07E+00 | 9.14E-05 | 3.26E-04 | -1.65E+00 | 6.31E-05 | 1.78E-04 | -1.72E+00 | 4.57E-02 | 1.68E-01 | 1.04E+00 | 3.56E-01 | 9.79E-01 | 1.34E+00 |
| CCAR2 | 2.99E-05 | 1.56E-04 | -1.21E+00 | 3.12E-04 | 1.35E-03 | -1.04E+00 | 2.30E-03 | 9.79E-03 | -1.21E+00 | 2.61E-03 | 9.97E-03 | -1.01E+00 | 3.76E-03 | 1.34E-02 | -1.20E+00 | 4.82E-02 | 1.36E-01 | 1.09E+00 | 3.56E-05 | 1.31E-04 | -1.30E+00 | 2.25E-02 | 6.18E-02 | -1.03E+00 |
| RPL32P3 | 2.99E-05 | 1.56E-04 | -1.18E+00 | 5.24E-03 | 2.26E-02 | 1.90E+00 | 1.27E-05 | 5.39E-05 | -1.47E+00 | 2.60E-01 | 9.92E-01 | 1.48E+00 | 1.35E-02 | 4.80E-02 | 1.07E+00 | 6.19E-02 | 1.74E-01 | 1.73E+00 | 2.42E-01 | 8.87E-01 | 1.26E+00 | 9.99E-03 | 2.75E-02 | 1.61E+00 |
| KLF7-IT1 | 3.00E-05 | 1.56E-04 | 2.83E+00 | 2.05E-03 | 8.86E-03 | 2.51E+00 | 4.69E-03 | 1.99E-02 | 2.03E+00 | 3.12E-02 | 1.19E-01 | 2.23E+00 | 1.43E-01 | 5.09E-01 | 2.02E+00 | 3.27E-01 | 9.22E-01 | 1.41E+00 | 1.79E-01 | 6.58E-01 | 1.25E+00 | 2.51E-01 | 6.89E-01 | -1.24E+00 |
| AC127502.2 | 3.01E-05 | 1.57E-04 | -1.79E+00 | 5.07E-03 | 2.19E-02 | -1.07E+00 | 1.61E-06 | 6.83E-06 | -1.93E+00 | 8.20E-05 | 3.13E-04 | -1.37E+00 | 3.01E-05 | 1.07E-04 | -2.45E+00 | 2.68E-03 | 7.55E-03 | -1.35E+00 | 3.59E-03 | 1.32E-02 | -1.55E+00 | 1.69E-01 | 4.63E-01 | -1.10E+00 |
| NISCH | 3.01E-05 | 1.57E-04 | -1.34E+00 | 3.34E-03 | 1.44E-02 | 1.03E+00 | 7.50E-03 | 3.19E-02 | -1.06E+00 | 1.45E-02 | 5.52E-02 | 1.13E+00 | 1.78E-03 | 6.36E-03 | -1.07E+00 | 1.42E-01 | 4.00E-01 | 1.39E+00 | 9.12E-06 | 3.35E-05 | -1.22E+00 | 4.91E-02 | 1.35E-01 | 1.13E+00 |
| VWDE | 3.01E-05 | 1.57E-04 | -1.32E+00 | 2.96E-02 | 1.28E-01 | 1.22E+00 | 8.62E-03 | 3.66E-02 | -1.18E+00 | 2.58E-01 | 9.84E-01 | 1.43E+00 | 1.43E-01 | 5.11E-01 | 1.79E+00 | 3.39E-03 | 9.56E-03 | 2.97E+00 | 2.30E-03 | 8.43E-03 | 1.74E+00 | 1.59E-02 | 4.36E-02 | 1.61E+00 |
| GLIPR1 | 3.02E-05 | 1.57E-04 | -1.07E+00 | 1.17E-02 | 5.07E-02 | 1.28E+00 | 1.35E-02 | 5.72E-02 | -1.05E+00 | 8.17E-04 | 3.12E-03 | 1.01E+00 | 2.48E-01 | 8.83E-01 | 1.17E+00 | 3.29E-01 | 9.26E-01 | 1.26E+00 | 1.43E-01 | 5.26E-01 | -1.12E+00 | 3.48E-01 | 9.56E-01 | 1.27E+00 |
| ARID4A | 3.03E-05 | 1.58E-04 | 1.01E+00 | 6.43E-14 | 2.78E-13 | -1.22E+00 | 7.58E-03 | 3.22E-02 | -1.17E+00 | 3.99E-14 | 1.52E-13 | -1.58E+00 | 7.82E-03 | 2.79E-02 | -1.08E+00 | 1.01E-07 | 2.86E-07 | -1.52E+00 | 5.12E-05 | 1.88E-04 | -1.14E+00 | 5.01E-10 | 1.38E-09 | -1.49E+00 |
| S1PR4 | 3.03E-05 | 1.58E-04 | 1.04E+00 | 1.15E-01 | 4.99E-01 | 1.36E+00 | 1.34E-08 | 5.69E-08 | -1.37E+00 | 4.78E-03 | 1.82E-02 | 1.10E+00 | 3.63E-06 | 1.29E-05 | -1.22E+00 | 1.32E-01 | 3.73E-01 | 1.23E+00 | 9.05E-08 | 3.32E-07 | -1.35E+00 | 2.72E-02 | 7.46E-02 | -1.04E+00 |
| C6orf106 | 3.05E-05 | 1.59E-04 | -1.20E+00 | 1.26E-04 | 5.44E-04 | -1.00E+00 | 4.21E-03 | 1.79E-02 | -1.13E+00 | 4.71E-03 | 1.80E-02 | 1.02E+00 | 6.69E-05 | 2.38E-04 | -1.19E+00 | 6.61E-03 | 1.86E-02 | -1.00E+00 | 3.68E-03 | 1.35E-02 | 1.04E+00 | 5.67E-03 | 1.56E-02 | 1.01E+00 |
| PRR14 | 3.07E-05 | 1.60E-04 | -1.07E+00 | 4.37E-07 | 1.89E-06 | -1.06E+00 | 1.98E-03 | 8.43E-03 | -1.13E+00 | 1.36E-04 | 5.19E-04 | -1.12E+00 | 1.67E-05 | 5.94E-05 | -1.43E+00 | 4.16E-04 | 1.17E-03 | -1.18E+00 | 1.55E-02 | 5.69E-02 | -1.01E+00 | 4.68E-03 | 1.28E-02 | -1.07E+00 |
| BID | 3.07E-05 | 1.60E-04 | -1.10E+00 | 8.42E-05 | 3.64E-04 | 1.06E+00 | 9.32E-02 | 3.96E-01 | 1.04E+00 | 1.76E-02 | 6.71E-02 | 1.12E+00 | 1.74E-05 | 6.19E-05 | -1.20E+00 | 1.40E-02 | 3.95E-02 | 1.07E+00 | 1.36E-03 | 5.01E-03 | -1.00E+00 | 1.66E-02 | 4.57E-02 | 1.05E+00 |
| SDR42E1 | 3.07E-05 | 1.60E-04 | -1.84E+00 | 2.08E-04 | 9.00E-04 | -1.43E+00 | 6.77E-04 | 2.88E-03 | -1.27E+00 | 3.97E-04 | 1.52E-03 | -1.31E+00 | 2.12E-01 | 7.57E-01 | -1.05E+00 | 3.49E-01 | 9.84E-01 | 1.24E+00 | 1.02E-01 | 3.73E-01 | -1.09E+00 | 1.18E-01 | 3.25E-01 | -1.10E+00 |
| PLBD2 | 3.08E-05 | 1.60E-04 | -1.36E+00 | 1.39E-04 | 5.98E-04 | -1.16E+00 | 1.35E-05 | 5.75E-05 | -1.50E+00 | 8.86E-04 | 3.39E-03 | -1.13E+00 | 2.81E-02 | 1.00E-01 | 1.01E+00 | 1.26E-01 | 3.56E-01 | 1.22E+00 | 9.19E-03 | 3.37E-02 | -1.11E+00 | 3.40E-02 | 9.34E-02 | -1.16E+00 |
| DZIP1L | 3.08E-05 | 1.60E-04 | -1.68E+00 | 9.90E-05 | 4.28E-04 | -1.36E+00 | 1.37E-01 | 5.84E-01 | 1.01E+00 | 5.69E-02 | 2.17E-01 | -1.10E+00 | 2.69E-01 | 9.58E-01 | 1.31E+00 | 3.39E-01 | 9.56E-01 | 1.11E+00 | 6.80E-03 | 2.50E-02 | -1.76E+00 | 2.05E-02 | 5.62E-02 | -1.69E+00 |
| KCNK5 | 3.08E-05 | 1.61E-04 | -1.45E+00 | 4.03E-06 | 1.74E-05 | -1.52E+00 | 6.25E-04 | 2.66E-03 | -1.57E+00 | 3.85E-03 | 1.47E-02 | -1.19E+00 | 2.96E-03 | 1.05E-02 | -1.20E+00 | 7.36E-02 | 2.07E-01 | 1.11E+00 | 2.26E-09 | 8.30E-09 | -1.54E+00 | 2.98E-04 | 8.18E-04 | -1.10E+00 |
| AL591866.1 | 3.09E-05 | 1.61E-04 | -1.33E+00 | 8.32E-10 | 3.60E-09 | -2.30E+00 | 3.83E-02 | 1.63E-01 | -1.11E+00 | 3.67E-03 | 1.40E-02 | -1.31E+00 | 2.61E-02 | 9.30E-02 | -1.30E+00 | 4.12E-03 | 1.16E-02 | -1.63E+00 | 1.23E-01 | 4.50E-01 | -1.01E+00 | 1.90E-02 | 5.22E-02 | -1.35E+00 |
| HMGN1P38 | 3.11E-05 | 1.62E-04 | -1.15E+00 | 1.91E-10 | 8.26E-10 | -1.69E+00 | 8.31E-02 | 3.53E-01 | 1.07E+00 | 2.57E-09 | 9.83E-09 | -2.07E+00 | 7.28E-03 | 2.59E-02 | -1.10E+00 | 1.58E-06 | 4.45E-06 | -1.88E+00 | 1.30E-01 | 4.76E-01 | 1.29E+00 | 1.66E-07 | 4.56E-07 | -1.60E+00 |
| NDST1 | 3.14E-05 | 1.63E-04 | -1.67E+00 | 7.25E-03 | 3.13E-02 | -1.02E+00 | 4.19E-06 | 1.78E-05 | -1.65E+00 | 8.48E-06 | 3.24E-05 | -1.41E+00 | 3.47E-02 | 1.24E-01 | -1.38E+00 | 4.81E-02 | 1.36E-01 | -1.29E+00 | 1.60E-01 | 5.86E-01 | 1.01E+00 | 2.81E-01 | 7.73E-01 | 1.41E+00 |
| AC100861.1 | 3.14E-05 | 1.63E-04 | -1.60E+00 | 2.37E-04 | 1.02E-03 | -1.24E+00 | 5.21E-07 | 2.21E-06 | -2.23E+00 | 1.44E-04 | 5.49E-04 | -1.44E+00 | 4.97E-03 | 1.77E-02 | -2.04E+00 | 9.18E-02 | 2.59E-01 | -1.15E+00 | 2.47E-05 | 9.05E-05 | -2.04E+00 | 1.15E-03 | 3.17E-03 | -1.52E+00 |
| AC087241.4 | 3.14E-05 | 1.64E-04 | -1.64E+00 | 1.52E-02 | 6.55E-02 | 1.06E+00 | 2.45E-02 | 1.04E-01 | -1.24E+00 | 1.77E-01 | 6.75E-01 | 1.27E+00 | 2.59E-01 | 9.25E-01 | 1.12E+00 | 4.46E-02 | 1.26E-01 | 1.83E+00 | 8.75E-03 | 3.21E-02 | -1.30E+00 | 1.94E-02 | 5.34E-02 | -1.27E+00 |
| C11orf58 | 3.15E-05 | 1.64E-04 | -1.00E+00 | 1.05E-04 | 4.54E-04 | 1.07E+00 | 7.90E-02 | 3.36E-01 | -1.03E+00 | 9.19E-02 | 3.51E-01 | 1.02E+00 | 3.36E-02 | 1.20E-01 | 1.02E+00 | 2.39E-01 | 6.72E-01 | 1.08E+00 | 7.53E-03 | 2.76E-02 | -1.07E+00 | 1.36E-01 | 3.75E-01 | -1.00E+00 |
| GPATCH1 | 3.15E-05 | 1.64E-04 | -1.52E+00 | 1.85E-02 | 7.99E-02 | 1.10E+00 | 6.30E-04 | 2.68E-03 | -1.39E+00 | 2.57E-02 | 9.81E-02 | 1.02E+00 | 2.42E-03 | 8.62E-03 | -1.29E+00 | 1.43E-01 | 4.03E-01 | 1.32E+00 | 1.16E-08 | 4.24E-08 | -1.71E+00 | 1.99E-02 | 5.48E-02 | -1.05E+00 |
| KIF3B | 3.15E-05 | 1.64E-04 | -1.88E+00 | 1.46E-02 | 6.30E-02 | -1.04E+00 | 2.86E-05 | 1.22E-04 | -1.59E+00 | 1.90E-03 | 7.24E-03 | -1.11E+00 | 3.30E-06 | 1.18E-05 | -2.08E+00 | 9.72E-04 | 2.74E-03 | -1.33E+00 | 8.76E-06 | 3.22E-05 | -1.74E+00 | 1.56E-03 | 4.29E-03 | -1.30E+00 |
| RNF113A | 3.15E-05 | 1.64E-04 | -1.11E+00 | 2.80E-04 | 1.21E-03 | 1.05E+00 | 1.61E-04 | 6.83E-04 | -1.26E+00 | 8.85E-04 | 3.38E-03 | -1.06E+00 | 1.09E-04 | 3.90E-04 | -1.25E+00 | 2.54E-03 | 7.15E-03 | -1.11E+00 | 9.62E-05 | 3.53E-04 | -1.19E+00 | 5.70E-03 | 1.56E-02 | -1.04E+00 |
| NAB1 | 3.15E-05 | 1.64E-04 | -1.20E+00 | 2.77E-04 | 1.20E-03 | 1.03E+00 | 1.73E-04 | 7.36E-04 | -1.26E+00 | 2.88E-04 | 1.10E-03 | -1.07E+00 | 7.51E-03 | 2.68E-02 | -1.08E+00 | 4.16E-02 | 1.17E-01 | 1.01E+00 | 5.15E-11 | 1.89E-10 | -1.75E+00 | 3.59E-07 | 9.85E-07 | -1.39E+00 |
| CPQ | 3.16E-05 | 1.65E-04 | -1.19E+00 | 1.88E-04 | 8.11E-04 | -1.02E+00 | 1.10E-01 | 4.67E-01 | 1.05E+00 | 5.07E-02 | 1.94E-01 | 1.02E+00 | 9.88E-05 | 3.52E-04 | -1.24E+00 | 1.78E-02 | 5.00E-02 | 1.14E+00 | 1.25E-03 | 4.59E-03 | -1.06E+00 | 4.44E-03 | 1.22E-02 | -1.05E+00 |
| DNAJC28 | 3.16E-05 | 1.65E-04 | -2.24E+00 | 1.41E-02 | 6.10E-02 | -1.18E+00 | 9.27E-02 | 3.94E-01 | -1.22E+00 | 2.33E-01 | 8.89E-01 | 1.11E+00 | 2.07E-03 | 7.39E-03 | -1.98E+00 | 5.69E-03 | 1.60E-02 | -1.71E+00 | 2.43E-01 | 8.94E-01 | 1.05E+00 | 3.63E-01 | 9.96E-01 | 1.15E+00 |
| BBS2 | 3.16E-05 | 1.65E-04 | -1.43E+00 | 1.14E-02 | 4.93E-02 | 1.09E+00 | 3.74E-05 | 1.59E-04 | -1.59E+00 | 2.79E-02 | 1.07E-01 | 1.08E+00 | 9.94E-03 | 3.54E-02 | -1.03E+00 | 3.18E-01 | 8.97E-01 | 1.48E+00 | 2.26E-02 | 8.28E-02 | -1.13E+00 | 1.69E-01 | 4.63E-01 | 1.18E+00 |
| IRAK3 | 3.17E-05 | 1.65E-04 | 1.74E+00 | 6.94E-03 | 3.00E-02 | 1.87E+00 | 2.86E-03 | 1.22E-02 | 1.54E+00 | 1.06E-02 | 4.05E-02 | 1.59E+00 | 3.78E-05 | 1.35E-04 | 2.11E+00 | 1.58E-01 | 4.45E-01 | 1.60E+00 | 2.67E-02 | 9.78E-02 | 1.54E+00 | 9.49E-02 | 2.61E-01 | 1.43E+00 |
| IAH1 | 3.18E-05 | 1.66E-04 | -1.03E+00 | 1.19E-05 | 5.12E-05 | 1.05E+00 | 2.74E-04 | 1.17E-03 | -1.01E+00 | 6.49E-06 | 2.48E-05 | -1.00E+00 | 1.76E-05 | 6.26E-05 | -1.08E+00 | 3.22E-03 | 9.08E-03 | 1.10E+00 | 1.78E-04 | 6.53E-04 | -1.14E+00 | 1.28E-01 | 3.52E-01 | 1.13E+00 |
| SPSB2 | 3.18E-05 | 1.66E-04 | -1.41E+00 | 1.04E-02 | 4.50E-02 | 1.10E+00 | 2.93E-08 | 1.25E-07 | -1.64E+00 | 7.79E-06 | 2.98E-05 | -1.26E+00 | 8.47E-05 | 3.02E-04 | -1.31E+00 | 8.93E-03 | 2.52E-02 | 1.09E+00 | 5.87E-03 | 2.16E-02 | -1.07E+00 | 2.81E-02 | 7.73E-02 | -1.01E+00 |
| RARA | 3.19E-05 | 1.66E-04 | -1.27E+00 | 2.29E-04 | 9.87E-04 | -1.06E+00 | 1.23E-05 | 5.22E-05 | -1.45E+00 | 4.95E-07 | 1.89E-06 | -1.47E+00 | 4.41E-03 | 1.57E-02 | -1.07E+00 | 4.90E-04 | 1.38E-03 | -1.22E+00 | 4.44E-04 | 1.63E-03 | -1.31E+00 | 2.12E-05 | 5.81E-05 | -1.50E+00 |
| CCDC138 | 3.19E-05 | 1.66E-04 | -1.33E+00 | 1.22E-04 | 5.26E-04 | -1.17E+00 | 3.55E-03 | 1.51E-02 | -1.12E+00 | 6.50E-02 | 2.48E-01 | 1.22E+00 | 1.84E-01 | 6.57E-01 | 1.23E+00 | 2.01E-01 | 5.66E-01 | 1.86E+00 | 3.01E-03 | 1.11E-02 | -1.15E+00 | 2.93E-01 | 8.06E-01 | 1.25E+00 |
| H2AFZP3 | 3.19E-05 | 1.66E-04 | -1.86E+00 | 2.71E-04 | 1.17E-03 | -1.48E+00 | 6.44E-11 | 2.74E-10 | -1.87E+00 | 4.40E-07 | 1.68E-06 | -1.20E+00 | 3.32E-04 | 1.18E-03 | -2.48E+00 | 4.99E-03 | 1.41E-02 | -1.63E+00 | 4.11E-19 | 1.51E-18 | -2.68E+00 | 8.38E-11 | 2.30E-10 | -1.80E+00 |
| AC092128.1 | 3.19E-05 | 1.66E-04 | -1.08E+00 | 8.79E-12 | 3.80E-11 | -1.53E+00 | 3.95E-02 | 1.68E-01 | 1.07E+00 | 9.43E-07 | 3.60E-06 | -1.31E+00 | 7.26E-02 | 2.59E-01 | 1.23E+00 | 1.65E-03 | 4.66E-03 | -1.04E+00 | 1.16E-02 | 4.27E-02 | 1.10E+00 | 2.26E-08 | 6.21E-08 | -1.46E+00 |
| KATNB1 | 3.20E-05 | 1.67E-04 | -1.28E+00 | 2.31E-02 | 1.00E-01 | 1.22E+00 | 3.00E-04 | 1.27E-03 | -1.18E+00 | 5.14E-03 | 1.96E-02 | 1.16E+00 | 2.74E-04 | 9.79E-04 | -1.48E+00 | 2.92E-02 | 8.22E-02 | 1.01E+00 | 1.92E-03 | 7.06E-03 | -1.03E+00 | 2.41E-01 | 6.62E-01 | 1.29E+00 |
| TRBC2 | 3.21E-05 | 1.67E-04 | -1.02E+00 | 2.72E-02 | 1.18E-01 | 1.25E+00 | 5.82E-05 | 2.47E-04 | -1.14E+00 | 1.47E-03 | 5.63E-03 | 1.04E+00 | 5.86E-02 | 2.09E-01 | 1.11E+00 | 8.47E-02 | 2.39E-01 | 1.14E+00 | 1.45E-01 | 5.31E-01 | 1.00E+00 | 2.29E-01 | 6.28E-01 | 1.30E+00 |
| MRPS15 | 3.22E-05 | 1.68E-04 | 1.04E+00 | 1.09E-07 | 4.70E-07 | 1.05E+00 | 1.44E-01 | 6.12E-01 | 1.17E+00 | 1.84E-01 | 7.04E-01 | 1.16E+00 | 2.39E-05 | 8.51E-05 | -1.13E+00 | 7.59E-03 | 2.14E-02 | -1.01E+00 | 4.62E-02 | 1.70E-01 | -1.03E+00 | 3.00E-01 | 8.23E-01 | 1.12E+00 |
| ATP6V0E2 | 3.22E-05 | 1.68E-04 | -1.40E+00 | 9.99E-05 | 4.32E-04 | -1.18E+00 | 2.75E-04 | 1.17E-03 | -1.48E+00 | 5.56E-03 | 2.12E-02 | -1.12E+00 | 5.61E-03 | 2.00E-02 | -1.99E+00 | 9.84E-02 | 2.77E-01 | -1.01E+00 | 2.09E-06 | 7.68E-06 | -1.70E+00 | 1.98E-02 | 5.44E-02 | -1.08E+00 |
| BYSL | 3.23E-05 | 1.68E-04 | -1.87E+00 | 1.68E-03 | 7.25E-03 | -1.29E+00 | 7.76E-04 | 3.30E-03 | -1.14E+00 | 9.88E-05 | 3.77E-04 | -1.07E+00 | 1.19E-04 | 4.23E-04 | -1.93E+00 | 1.32E-02 | 3.73E-02 | -1.15E+00 | 1.26E-18 | 4.63E-18 | -1.72E+00 | 8.61E-12 | 2.37E-11 | -1.34E+00 |
| ARHGAP12 | 3.25E-05 | 1.69E-04 | -1.50E+00 | 2.31E-05 | 9.99E-05 | -1.46E+00 | 2.18E-03 | 9.27E-03 | -1.87E+00 | 7.42E-03 | 2.84E-02 | -1.46E+00 | 2.33E-01 | 8.32E-01 | 1.52E+00 | 3.35E-01 | 9.43E-01 | 1.29E+00 | 3.16E-06 | 1.16E-05 | -1.66E+00 | 7.55E-06 | 2.07E-05 | -1.64E+00 |
| SPATC1L | 3.25E-05 | 1.69E-04 | -1.03E+00 | 2.58E-04 | 1.11E-03 | 1.11E+00 | 2.37E-04 | 1.01E-03 | -1.10E+00 | 3.90E-03 | 1.49E-02 | 1.11E+00 | 2.27E-05 | 8.10E-05 | -1.38E+00 | 5.21E-03 | 1.47E-02 | -1.09E+00 | 9.94E-03 | 3.65E-02 | 1.01E+00 | 1.46E-01 | 4.00E-01 | 1.17E+00 |
| USP20 | 3.25E-05 | 1.70E-04 | -1.35E+00 | 1.19E-03 | 5.13E-03 | -1.01E+00 | 5.04E-04 | 2.14E-03 | -1.32E+00 | 4.51E-03 | 1.72E-02 | -1.00E+00 | 2.54E-12 | 9.07E-12 | -2.11E+00 | 1.22E-04 | 3.43E-04 | -1.21E+00 | 3.12E-09 | 1.15E-08 | 1.06E+00 | 6.81E-08 | 1.87E-07 | -1.36E+00 |
| IMMP1L | 3.29E-05 | 1.71E-04 | -1.09E+00 | 2.34E-09 | 1.01E-08 | -1.26E+00 | 1.18E-01 | 5.00E-01 | 1.26E+00 | 8.73E-04 | 3.33E-03 | 1.04E+00 | 3.48E-03 | 1.24E-02 | 1.08E+00 | 5.15E-03 | 1.45E-02 | 1.10E+00 | 4.29E-05 | 1.57E-04 | -1.07E+00 | 7.29E-07 | 2.00E-06 | -1.22E+00 |
| NACC1 | 3.29E-05 | 1.71E-04 | -1.51E+00 | 2.84E-04 | 1.23E-03 | -1.21E+00 | 3.65E-02 | 1.55E-01 | -1.04E+00 | 1.12E-01 | 4.26E-01 | 1.31E+00 | 2.67E-01 | 9.53E-01 | 1.24E+00 | 1.99E-01 | 5.62E-01 | 1.44E+00 | 2.45E-02 | 9.00E-02 | 1.11E+00 | 1.67E-01 | 4.60E-01 | 1.25E+00 |
| NBPF8 | 3.29E-05 | 1.71E-04 | -1.46E+00 | 4.15E-04 | 1.79E-03 | -1.14E+00 | 3.37E-07 | 1.43E-06 | -1.66E+00 | 1.43E-05 | 5.46E-05 | -1.29E+00 | 9.05E-06 | 3.23E-05 | -1.76E+00 | 4.36E-04 | 1.23E-03 | -1.37E+00 | 7.34E-04 | 2.69E-03 | -1.38E+00 | 1.42E-02 | 3.91E-02 | -1.27E+00 |
| ECI1 | 3.29E-05 | 1.71E-04 | -1.02E+00 | 5.96E-05 | 2.57E-04 | 1.10E+00 | 4.07E-02 | 1.73E-01 | 1.34E+00 | 1.70E-01 | 6.49E-01 | 1.36E+00 | 8.85E-05 | 3.16E-04 | -1.08E+00 | 3.71E-03 | 1.05E-02 | 1.06E+00 | 5.24E-02 | 1.92E-01 | 1.11E+00 | 2.09E-01 | 5.74E-01 | 1.35E+00 |
| CCDC112 | 3.30E-05 | 1.72E-04 | -1.14E+00 | 4.64E-05 | 2.01E-04 | -1.01E+00 | 4.36E-05 | 1.85E-04 | -1.18E+00 | 7.38E-08 | 2.82E-07 | -1.16E+00 | 4.73E-09 | 1.69E-08 | -1.38E+00 | 4.12E-04 | 1.16E-03 | -1.03E+00 | 5.25E-07 | 1.93E-06 | -1.28E+00 | 1.68E-05 | 4.61E-05 | -1.19E+00 |
| FHL3 | 3.31E-05 | 1.73E-04 | -1.00E+00 | 1.10E-05 | 4.76E-05 | 1.07E+00 | 9.23E-07 | 3.92E-06 | -1.27E+00 | 4.07E-06 | 1.56E-05 | -1.10E+00 | 1.56E-03 | 5.57E-03 | 1.01E+00 | 1.11E-04 | 3.12E-04 | -1.02E+00 | 4.23E-02 | 1.55E-01 | 1.15E+00 | 1.17E-02 | 3.20E-02 | -1.03E+00 |
| CCDC14 | 3.32E-05 | 1.73E-04 | -1.14E+00 | 2.46E-03 | 1.06E-02 | 1.11E+00 | 7.18E-06 | 3.05E-05 | -1.22E+00 | 6.77E-04 | 2.59E-03 | 1.02E+00 | 3.70E-04 | 1.32E-03 | -1.10E+00 | 3.51E-03 | 9.89E-03 | -1.02E+00 | 2.96E-03 | 1.09E-02 | -1.13E+00 | 1.25E-01 | 3.42E-01 | 1.00E+00 |
| PET117 | 3.32E-05 | 1.73E-04 | -1.34E+00 | 8.86E-04 | 3.83E-03 | -1.06E+00 | 1.03E-01 | 4.36E-01 | -1.00E+00 | 1.70E-01 | 6.48E-01 | 1.22E+00 | 1.48E-01 | 5.26E-01 | 1.03E+00 | 1.45E-01 | 4.09E-01 | 1.04E+00 | 6.63E-03 | 2.43E-02 | -1.15E+00 | 1.91E-01 | 5.26E-01 | 1.12E+00 |
| ZNF425 | 3.33E-05 | 1.73E-04 | -1.93E+00 | 6.32E-03 | 2.73E-02 | -1.16E+00 | 1.95E-09 | 8.29E-09 | -2.62E+00 | 6.69E-06 | 2.55E-05 | -1.58E+00 | 4.01E-05 | 1.43E-04 | -3.40E+00 | 3.55E-02 | 1.00E-01 | -1.16E+00 | 7.83E-04 | 2.87E-03 | -1.87E+00 | 4.30E-02 | 1.18E-01 | -1.17E+00 |
| RNF125 | 3.34E-05 | 1.74E-04 | -1.04E+00 | 4.52E-06 | 1.95E-05 | 1.06E+00 | 2.64E-04 | 1.12E-03 | -1.11E+00 | 7.96E-07 | 3.04E-06 | -1.11E+00 | 9.18E-02 | 3.27E-01 | 1.08E+00 | 8.03E-03 | 2.26E-02 | -1.11E+00 | 1.75E-01 | 6.42E-01 | 1.06E+00 | 2.72E-01 | 7.46E-01 | 1.11E+00 |
| NAT8 | 3.34E-05 | 1.74E-04 | -2.28E+00 | 1.80E-03 | 7.76E-03 | -1.41E+00 | 9.87E-06 | 4.19E-05 | -2.05E+00 | 1.78E-04 | 6.80E-04 | -1.54E+00 | 7.59E-04 | 2.71E-03 | -1.88E+00 | 5.77E-03 | 1.63E-02 | -1.42E+00 | 2.74E-06 | 1.00E-05 | -1.39E+00 | 8.74E-06 | 2.40E-05 | -1.59E+00 |
| ANO5 | 3.34E-05 | 1.74E-04 | -1.94E+00 | 7.69E-02 | 3.32E-01 | 1.25E+00 | 1.63E-02 | 6.91E-02 | -1.20E+00 | 1.99E-01 | 7.58E-01 | 1.23E+00 | 1.91E-01 | 6.80E-01 | 1.08E+00 | 1.82E-01 | 5.13E-01 | 2.04E+00 | 7.40E-02 | 2.71E-01 | -1.11E+00 | 1.49E-01 | 4.09E-01 | 1.05E+00 |
| HHIP-AS1 | 3.34E-05 | 1.74E-04 | -1.50E+00 | 1.36E-06 | 5.88E-06 | -1.95E+00 | 1.86E-01 | 7.91E-01 | 1.27E+00 | 1.29E-01 | 4.95E-01 | 1.05E+00 | 2.20E-01 | 7.85E-01 | 1.33E+00 | 3.50E-01 | 9.88E-01 | 1.51E+00 | 2.07E-01 | 7.61E-01 | -1.03E+00 | 2.11E-01 | 5.79E-01 | -1.04E+00 |
| NUDT15 | 3.35E-05 | 1.74E-04 | -1.31E+00 | 2.10E-04 | 9.08E-04 | -1.06E+00 | 1.48E-07 | 6.28E-07 | -1.42E+00 | 8.74E-08 | 3.34E-07 | -1.27E+00 | 1.54E-03 | 5.50E-03 | -1.51E+00 | 7.78E-03 | 2.19E-02 | -1.25E+00 | 1.47E-02 | 5.40E-02 | -1.06E+00 | 4.33E-02 | 1.19E-01 | 1.06E+00 |
| ARSA | 3.36E-05 | 1.75E-04 | -1.69E+00 | 1.06E-04 | 4.59E-04 | -1.42E+00 | 2.36E-02 | 1.00E-01 | -1.10E+00 | 6.26E-02 | 2.39E-01 | 1.15E+00 | 8.85E-03 | 3.15E-02 | -1.24E+00 | 2.09E-01 | 5.88E-01 | 1.18E+00 | 6.99E-02 | 2.57E-01 | 1.02E+00 | 2.04E-01 | 5.61E-01 | 1.16E+00 |
| GBA2 | 3.36E-05 | 1.75E-04 | -1.46E+00 | 1.98E-02 | 8.57E-02 | 1.20E+00 | 2.93E-02 | 1.25E-01 | -1.17E+00 | 1.66E-01 | 6.35E-01 | 1.56E+00 | 1.47E-01 | 5.22E-01 | 1.23E+00 | 1.96E-01 | 5.53E-01 | 1.23E+00 | 6.24E-02 | 2.29E-01 | 1.00E+00 | 1.07E-01 | 2.94E-01 | 1.03E+00 |
| COX4I1 | 3.37E-05 | 1.76E-04 | 1.12E+00 | 2.45E-03 | 1.06E-02 | 1.11E+00 | 2.75E-04 | 1.17E-03 | 1.14E+00 | 1.13E-02 | 4.33E-02 | 1.10E+00 | 8.48E-02 | 3.02E-01 | 1.06E+00 | 5.85E-02 | 1.65E-01 | 1.08E+00 | 6.45E-02 | 2.37E-01 | 1.10E+00 | 5.51E-02 | 1.51E-01 | 1.09E+00 |
| REL | 3.38E-05 | 1.76E-04 | -1.05E+00 | 2.10E-16 | 9.08E-16 | -1.38E+00 | 1.55E-04 | 6.58E-04 | -1.26E+00 | 9.77E-11 | 3.73E-10 | -1.45E+00 | 1.56E-05 | 5.54E-05 | -1.23E+00 | 1.08E-09 | 3.04E-09 | -1.68E+00 | 4.09E-10 | 1.50E-09 | -1.58E+00 | 2.89E-13 | 7.93E-13 | -1.81E+00 |
| MANF | 3.38E-05 | 1.76E-04 | -1.23E+00 | 9.57E-02 | 4.14E-01 | 1.21E+00 | 2.05E-01 | 8.70E-01 | 1.16E+00 | 4.48E-02 | 1.71E-01 | 1.47E+00 | 4.24E-05 | 1.51E-04 | -1.48E+00 | 2.81E-02 | 7.92E-02 | -1.08E+00 | 1.51E-02 | 5.54E-02 | -1.09E+00 | 3.22E-01 | 8.83E-01 | 1.06E+00 |
| FAM149B1 | 3.39E-05 | 1.77E-04 | -1.44E+00 | 6.94E-05 | 3.00E-04 | -1.21E+00 | 3.49E-07 | 1.48E-06 | -1.41E+00 | 4.12E-05 | 1.57E-04 | -1.04E+00 | 1.75E-05 | 6.26E-05 | -1.30E+00 | 2.37E-03 | 6.67E-03 | 1.01E+00 | 1.02E-05 | 3.76E-05 | -1.22E+00 | 7.38E-02 | 2.03E-01 | 1.06E+00 |
| NUBP1 | 3.40E-05 | 1.77E-04 | -1.76E+00 | 1.26E-02 | 5.43E-02 | -1.04E+00 | 8.61E-08 | 3.66E-07 | -2.21E+00 | 3.58E-05 | 1.37E-04 | -1.46E+00 | 3.53E-02 | 1.26E-01 | -1.19E+00 | 7.74E-02 | 2.18E-01 | 1.04E+00 | 1.36E-02 | 5.00E-02 | -1.37E+00 | 8.95E-02 | 2.46E-01 | -1.07E+00 |
| SPATA6 | 3.40E-05 | 1.77E-04 | -1.11E+00 | 5.84E-04 | 2.53E-03 | 1.07E+00 | 5.80E-07 | 2.46E-06 | -1.37E+00 | 1.20E-05 | 4.58E-05 | -1.10E+00 | 1.16E-02 | 4.12E-02 | -1.03E+00 | 4.71E-03 | 1.33E-02 | -1.25E+00 | 1.38E-01 | 5.06E-01 | -1.04E+00 | 2.89E-01 | 7.93E-01 | 1.20E+00 |
| RHOC | 3.40E-05 | 1.77E-04 | -1.09E+00 | 9.30E-07 | 4.02E-06 | -1.02E+00 | 1.25E-01 | 5.32E-01 | 1.08E+00 | 1.39E-01 | 5.30E-01 | 1.21E+00 | 3.82E-04 | 1.36E-03 | 1.06E+00 | 7.34E-06 | 2.07E-05 | -1.11E+00 | 1.68E-03 | 6.15E-03 | 1.96E+00 | 2.93E-01 | 8.05E-01 | 1.32E+00 |
| TMEM102 | 3.41E-05 | 1.77E-04 | -1.48E+00 | 2.97E-03 | 1.28E-02 | 1.02E+00 | 1.35E-09 | 5.72E-09 | -2.20E+00 | 1.77E-04 | 6.77E-04 | -1.18E+00 | 1.53E-05 | 5.45E-05 | -2.05E+00 | 1.22E-02 | 3.45E-02 | -1.10E+00 | 2.62E-07 | 9.62E-07 | -1.86E+00 | 1.95E-03 | 5.36E-03 | -1.24E+00 |
| TMED8 | 3.41E-05 | 1.77E-04 | -1.07E+00 | 2.96E-02 | 1.28E-01 | 1.30E+00 | 2.38E-07 | 1.01E-06 | -1.48E+00 | 2.98E-02 | 1.14E-01 | 1.07E+00 | 6.73E-05 | 2.40E-04 | -1.24E+00 | 3.97E-02 | 1.12E-01 | 1.13E+00 | 2.22E-01 | 8.16E-01 | 1.21E+00 | 2.98E-01 | 8.18E-01 | 1.32E+00 |
| PGGT1B | 3.41E-05 | 1.77E-04 | -1.16E+00 | 5.68E-02 | 2.45E-01 | 1.36E+00 | 1.77E-08 | 7.52E-08 | -1.56E+00 | 1.12E-05 | 4.29E-05 | -1.15E+00 | 2.15E-06 | 7.66E-06 | -1.37E+00 | 2.61E-03 | 7.36E-03 | 1.02E+00 | 9.40E-11 | 3.45E-10 | -1.58E+00 | 1.40E-06 | 3.83E-06 | -1.24E+00 |
| GOLGA8B | 3.41E-05 | 1.78E-04 | -1.04E+00 | 6.49E-05 | 2.80E-04 | 1.12E+00 | 4.01E-02 | 1.71E-01 | -1.03E+00 | 2.01E-01 | 7.68E-01 | 1.24E+00 | 1.33E-02 | 4.74E-02 | 1.18E+00 | 1.47E-02 | 4.13E-02 | 1.22E+00 | 1.33E-01 | 4.89E-01 | 1.45E+00 | 2.82E-02 | 7.75E-02 | 1.61E+00 |

| | | | | | | | | | | | | | | | | | | | | | | | | |
|---|---|---|---|---|---|---|---|---|---|---|---|---|---|---|---|---|---|---|---|---|---|---|---|---|
| CCDC18-AS1 | 3.41E-05 | 1.78E-04 | -1.09E+00 | 1.16E-02 | 5.02E-02 | 1.19E+00 | 3.66E-09 | 1.55E-08 | -1.40E+00 | 5.15E-03 | 1.97E-02 | 1.11E+00 | 8.66E-02 | 3.09E-01 | 1.12E+00 | 3.35E-01 | 9.43E-01 | 1.18E+00 | 4.48E-02 | 1.64E-01 | 1.16E+00 | 6.80E-02 | 1.87E-01 | 1.21E+00 |
| EPHB6 | 3.42E-05 | 1.78E-04 | -1.16E+00 | 1.73E-04 | 7.46E-04 | 1.01E+00 | 1.14E-05 | 4.84E-05 | -1.27E+00 | 2.03E-08 | 7.77E-08 | -1.32E+00 | 1.96E-03 | 6.98E-03 | -1.46E+00 | 7.17E-04 | 2.02E-03 | -1.51E+00 | 1.61E-03 | 5.91E-03 | -1.46E+00 | 1.09E-02 | 2.98E-02 | -1.27E+00 |
| SEC22C | 3.42E-05 | 1.78E-04 | -1.06E+00 | 3.31E-05 | 1.43E-04 | 1.05E+00 | 1.95E-03 | 8.27E-03 | -1.09E+00 | 4.75E-04 | 1.81E-03 | -1.02E+00 | 1.22E-01 | 4.33E-01 | 1.31E+00 | 4.17E-03 | 1.17E-02 | 1.01E+00 | 5.23E-03 | 1.92E-02 | -1.01E+00 | 9.69E-06 | 2.66E-05 | -1.17E+00 |
| IQSEC3 | 3.43E-05 | 1.79E-04 | -1.32E+00 | 2.02E-05 | 8.73E-05 | -1.21E+00 | 4.83E-05 | 2.05E-04 | -1.44E+00 | 3.56E-04 | 1.36E-03 | -1.13E+00 | 2.01E-02 | 7.17E-02 | -1.21E+00 | 7.93E-03 | 2.23E-02 | -1.55E+00 | 3.47E-03 | 1.27E-02 | -1.23E+00 | 3.92E-02 | 1.08E-01 | -1.05E+00 |
| SH3GL3 | 3.45E-05 | 1.80E-04 | -2.06E+00 | 5.73E-04 | 2.48E-03 | -1.43E+00 | 3.52E-02 | 1.49E-01 | -1.39E+00 | 2.12E-01 | 8.12E-01 | 1.10E+00 | 2.25E-03 | 8.02E-03 | -1.73E+00 | 4.01E-02 | 1.13E-01 | -1.08E+00 | 5.19E-04 | 1.90E-03 | -1.37E+00 | 1.81E-01 | 4.98E-01 | 1.28E+00 |
| MED14 | 3.45E-05 | 1.80E-04 | -1.31E+00 | 2.01E-04 | 8.68E-04 | -1.12E+00 | 8.27E-08 | 3.52E-07 | -1.58E+00 | 8.29E-07 | 3.17E-06 | -1.34E+00 | 9.23E-05 | 3.29E-04 | -1.33E+00 | 4.71E-03 | 1.33E-02 | 1.00E+00 | 6.50E-05 | 2.39E-04 | -1.27E+00 | 1.70E-02 | 4.68E-02 | -1.12E+00 |
| PPFIA3 | 3.45E-05 | 1.80E-04 | -1.54E+00 | 2.57E-04 | 1.11E-03 | -1.33E+00 | 1.30E-02 | 5.52E-02 | -1.33E+00 | 9.59E-02 | 3.67E-01 | 1.07E+00 | 1.92E-01 | 6.83E-01 | 1.17E+00 | 1.65E-01 | 4.64E-01 | -1.11E+00 | 1.53E-01 | 5.61E-01 | 1.20E+00 | 2.99E-01 | 8.21E-01 | 1.46E+00 |
| DNAL1 | 3.45E-05 | 1.80E-04 | -1.64E+00 | 9.43E-04 | 4.07E-03 | -1.15E+00 | 5.63E-02 | 2.39E-01 | -1.01E+00 | 1.47E-01 | 5.61E-01 | 1.18E+00 | 2.41E-04 | 8.60E-04 | -1.51E+00 | 5.06E-03 | 1.43E-02 | -1.18E+00 | 1.07E-04 | 3.94E-04 | -1.61E+00 | 1.56E-02 | 4.30E-02 | -1.19E+00 |
| ANKRD54 | 3.46E-05 | 1.80E-04 | -1.01E+00 | 8.73E-12 | 3.77E-11 | -1.26E+00 | 3.78E-02 | 1.61E-01 | 1.08E+00 | 3.59E-05 | 1.37E-04 | -1.06E+00 | 2.11E-02 | 7.53E-02 | 1.08E+00 | 6.79E-07 | 1.91E-06 | -1.28E+00 | 2.10E-01 | 7.70E-01 | 1.33E+00 | 1.13E-01 | 3.10E-01 | 1.17E+00 |
| GMFB | 3.46E-05 | 1.80E-04 | -1.07E+00 | 2.90E-04 | 1.25E-03 | 1.08E+00 | 4.84E-02 | 2.06E-01 | 1.03E+00 | 5.54E-03 | 2.12E-02 | 1.04E+00 | 3.14E-02 | 1.12E-01 | 1.10E+00 | 9.05E-02 | 2.55E-01 | 1.23E+00 | 9.78E-04 | 3.59E-03 | -1.09E+00 | 5.58E-04 | 1.53E-03 | -1.14E+00 |
| RHEBL1 | 3.46E-05 | 1.80E-04 | -2.81E+00 | 1.80E-03 | 7.76E-03 | -1.55E+00 | 1.60E-03 | 6.78E-03 | -1.64E+00 | 6.38E-03 | 2.44E-02 | -1.29E+00 | 9.87E-06 | 3.52E-05 | -2.91E+00 | 5.67E-03 | 1.60E-02 | -1.39E+00 | 1.55E-02 | 5.71E-02 | -1.28E+00 | 1.35E-01 | 3.71E-01 | 1.44E+00 |
| TOR1B | 3.46E-05 | 1.80E-04 | -1.70E+00 | 2.67E-02 | 1.15E-01 | 1.14E+00 | 6.63E-05 | 2.82E-04 | -1.48E+00 | 9.96E-03 | 3.80E-02 | -1.02E+00 | 1.30E-07 | 4.63E-07 | -3.33E+00 | 7.82E-03 | 2.20E-02 | -1.20E+00 | 3.24E-08 | 1.19E-07 | -1.75E+00 | 5.73E-03 | 1.57E-02 | -1.09E+00 |
| VPS13A | 3.47E-05 | 1.81E-04 | -1.10E+00 | 1.37E-01 | 5.93E-01 | 1.35E+00 | 1.03E-03 | 4.37E-03 | -1.11E+00 | 1.19E-02 | 4.55E-02 | 1.13E+00 | 5.09E-07 | 1.81E-06 | -1.29E+00 | 1.22E-03 | 3.44E-03 | -1.05E+00 | 9.17E-05 | 3.36E-04 | -1.09E+00 | 3.09E-02 | 8.49E-02 | 1.10E+00 |
| NCKAP5L | 3.48E-05 | 1.82E-04 | -1.20E+00 | 4.12E-06 | 1.78E-05 | -1.19E+00 | 1.44E-04 | 6.10E-04 | -1.29E+00 | 2.85E-06 | 1.09E-05 | -1.39E+00 | 1.07E-01 | 3.82E-01 | 1.12E+00 | 1.70E-02 | 4.78E-02 | -1.10E+00 | 1.17E-03 | 4.29E-03 | -1.48E+00 | 1.31E-02 | 3.60E-02 | -1.20E+00 |
| AC009120.2 | 3.51E-05 | 1.83E-04 | -2.24E+00 | 1.13E-02 | 4.86E-02 | -1.13E+00 | 4.42E-03 | 1.88E-02 | -1.36E+00 | 6.26E-02 | 2.39E-01 | 1.17E+00 | 1.77E-06 | 6.31E-06 | -3.64E+00 | 4.28E-02 | 1.21E-01 | 1.06E+00 | 1.36E-04 | 4.98E-04 | -1.58E+00 | 5.19E-02 | 1.43E-01 | 1.01E+00 |
| ARAP1 | 3.51E-05 | 1.83E-04 | -2.28E+00 | 1.52E-02 | 6.57E-02 | -1.15E+00 | 9.84E-04 | 4.18E-03 | -1.73E+00 | 6.20E-02 | 2.37E-01 | 1.01E+00 | 1.51E-04 | 5.40E-04 | -1.79E+00 | 3.96E-03 | 1.11E-02 | -1.17E+00 | 1.64E-03 | 6.01E-03 | -1.38E+00 | 1.28E-02 | 3.51E-02 | -1.17E+00 |
| EEF1A1P16 | 3.52E-05 | 1.83E-04 | -1.01E+00 | 8.41E-16 | 3.63E-15 | -1.42E+00 | 4.28E-03 | 1.82E-02 | -1.08E+00 | 5.14E-09 | 1.96E-08 | -1.42E+00 | 8.64E-03 | 3.08E-02 | 1.03E+00 | 6.45E-08 | 1.82E-07 | -1.48E+00 | 7.71E-03 | 2.83E-02 | -1.06E+00 | 7.33E-08 | 2.01E-07 | -1.55E+00 |
| NAE1 | 3.53E-05 | 1.84E-04 | -1.01E+00 | 2.71E-02 | 1.17E-01 | 1.36E+00 | 4.05E-02 | 1.72E-01 | 1.04E+00 | 6.13E-02 | 2.34E-01 | 1.25E+00 | 1.11E-03 | 3.97E-03 | -1.07E+00 | 1.84E-01 | 5.19E-01 | 1.24E+00 | 3.99E-02 | 1.46E-01 | 1.07E+00 | 2.36E-01 | 6.47E-01 | 1.30E+00 |
| FRG1DP | 3.53E-05 | 1.84E-04 | -1.23E+00 | 1.12E-06 | 4.86E-06 | -1.26E+00 | 1.64E-02 | 6.95E-02 | -1.22E+00 | 3.72E-04 | 1.42E-03 | -1.43E+00 | 1.95E-03 | 6.94E-03 | -1.25E+00 | 2.71E-05 | 7.63E-05 | -1.65E+00 | 2.55E-02 | 9.35E-02 | -1.12E+00 | 2.84E-02 | 7.81E-02 | -1.06E+00 |
| NAA25 | 3.53E-05 | 1.84E-04 | -1.10E+00 | 1.76E-04 | 7.62E-04 | 1.08E+00 | 9.59E-07 | 4.07E-06 | -1.29E+00 | 6.59E-06 | 2.52E-05 | -1.13E+00 | 2.33E-04 | 8.32E-04 | -1.26E+00 | 3.53E-02 | 9.96E-02 | 1.10E+00 | 8.61E-07 | 3.16E-06 | -1.31E+00 | 9.98E-04 | 2.74E-03 | -1.09E+00 |
| DOCK2 | 3.53E-05 | 1.84E-04 | 1.01E+00 | 6.79E-03 | 2.94E-02 | 1.31E+00 | 2.05E-01 | 8.71E-01 | 1.05E+00 | 2.27E-01 | 8.65E-01 | 1.26E+00 | 2.67E-03 | 9.51E-03 | 1.00E+00 | 3.63E-02 | 1.02E-01 | 1.13E+00 | 1.43E-01 | 5.23E-01 | 1.19E+00 | 3.27E-01 | 8.99E-01 | 1.27E+00 |
| AC010469.1 | 3.55E-05 | 1.85E-04 | 1.75E+00 | 1.66E-01 | 7.16E-01 | 1.48E+00 | 5.39E-08 | 2.29E-07 | 2.22E+00 | 1.91E-03 | 7.31E-03 | 2.00E+00 | 1.76E-06 | 6.28E-06 | 2.64E+00 | 8.01E-04 | 2.26E-03 | 2.34E+00 | 1.25E-06 | 4.58E-06 | 1.94E+00 | 2.21E-02 | 6.06E-02 | 1.62E+00 |
| NPM2 | 3.55E-05 | 1.85E-04 | -1.05E+00 | 5.38E-07 | 2.32E-06 | -1.04E+00 | 2.82E-04 | 1.20E-03 | -1.12E+00 | 3.56E-05 | 1.36E-04 | -1.11E+00 | 8.98E-02 | 3.20E-01 | -1.29E+00 | 8.37E-02 | 2.36E-01 | -1.40E+00 | 7.23E-02 | 2.65E-01 | -1.35E+00 | 2.28E-01 | 6.26E-01 | 1.07E+00 |
| SS18L1 | 3.57E-05 | 1.86E-04 | -1.36E+00 | 1.16E-07 | 5.00E-07 | -1.55E+00 | 1.66E-03 | 7.04E-03 | -1.23E+00 | 6.38E-03 | 2.44E-02 | 1.00E+00 | 1.85E-03 | 6.61E-03 | -1.56E+00 | 4.39E-03 | 1.24E-02 | -1.45E+00 | 4.75E-06 | 1.74E-05 | -1.60E+00 | 3.59E-03 | 9.85E-03 | -1.28E+00 |
| PEPD | 3.59E-05 | 1.87E-04 | -1.05E+00 | 2.51E-04 | 1.09E-03 | 1.12E+00 | 7.84E-04 | 3.33E-03 | -1.13E+00 | 2.78E-02 | 1.06E-01 | 1.11E+00 | 6.64E-05 | 2.37E-04 | -1.18E+00 | 5.26E-04 | 1.48E-03 | -1.05E+00 | 6.65E-02 | 2.44E-01 | 1.15E+00 | 3.11E-01 | 8.54E-01 | 1.13E+00 |
| ZXDC | 3.59E-05 | 1.87E-04 | -1.31E+00 | 1.59E-03 | 6.86E-03 | 1.01E+00 | 5.73E-05 | 2.44E-04 | -1.30E+00 | 2.60E-03 | 9.95E-03 | -1.01E+00 | 6.66E-02 | 2.37E-01 | 1.02E+00 | 1.13E-01 | 3.19E-01 | 1.13E+00 | 1.86E-03 | 6.83E-03 | -1.23E+00 | 6.16E-02 | 1.69E-01 | -1.05E+00 |
| PAXIP1-AS1 | 3.59E-05 | 1.87E-04 | -1.00E+00 | 1.38E-01 | 5.95E-01 | 1.50E+00 | 6.46E-02 | 2.74E-01 | 1.12E+00 | 2.54E-02 | 9.69E-02 | 1.77E+00 | 3.20E-02 | 1.14E-01 | 1.75E+00 | 4.47E-04 | 1.26E-03 | 2.37E+00 | 9.67E-02 | 3.55E-01 | 1.41E+00 | 2.97E-04 | 8.15E-04 | 1.80E+00 |
| PEX1 | 3.60E-05 | 1.88E-04 | -1.22E+00 | 3.11E-03 | 1.35E-02 | 1.12E+00 | 6.71E-03 | 2.85E-02 | -1.11E+00 | 1.70E-02 | 6.49E-02 | 1.07E+00 | 8.26E-02 | 2.95E-01 | 1.20E+00 | 1.35E-01 | 3.81E-01 | 1.19E+00 | 5.32E-04 | 1.95E-03 | -1.12E+00 | 2.85E-04 | 7.82E-04 | -1.24E+00 |
| AL137100.1 | 3.60E-05 | 1.88E-04 | -1.46E+00 | 3.51E-04 | 1.52E-03 | -1.18E+00 | 4.91E-10 | 2.09E-09 | -1.87E+00 | 2.58E-10 | 9.87E-10 | -1.72E+00 | 3.90E-05 | 1.39E-04 | -2.12E+00 | 1.27E-03 | 3.57E-03 | -1.43E+00 | 9.86E-05 | 3.62E-04 | -1.41E+00 | 1.46E-03 | 4.01E-03 | -1.21E+00 |
| PELO | 3.61E-05 | 1.88E-04 | -1.45E+00 | 1.03E-03 | 4.46E-03 | -1.08E+00 | 5.59E-05 | 2.37E-04 | -1.36E+00 | 4.45E-04 | 1.70E-03 | -1.13E+00 | 1.39E-04 | 4.96E-04 | -1.31E+00 | 4.59E-05 | 1.29E-04 | -1.47E+00 | 2.28E-05 | 8.36E-05 | -1.27E+00 | 3.86E-05 | 1.06E-04 | -1.38E+00 |
| FEN1 | 3.61E-05 | 1.88E-04 | -2.23E+00 | 4.51E-03 | 1.95E-02 | -1.22E+00 | 2.15E-01 | 9.13E-01 | 1.28E+00 | 2.06E-01 | 7.88E-01 | 1.40E+00 | 2.56E-02 | 9.14E-02 | -1.34E+00 | 1.90E-01 | 5.36E-01 | 1.16E+00 | 7.60E-05 | 2.79E-04 | -1.19E+00 | 4.71E-04 | 1.29E-03 | 1.61E+00 |
| GAPDHP65 | 3.62E-05 | 1.89E-04 | -1.10E+00 | 2.93E-04 | 1.26E-03 | 1.04E+00 | 5.87E-03 | 2.49E-02 | -1.16E+00 | 1.58E-01 | 6.03E-01 | 1.10E+00 | 2.78E-08 | 9.90E-08 | -1.19E+00 | 1.91E-03 | 5.38E-03 | 1.10E+00 | 2.42E-04 | 8.89E-04 | -1.17E+00 | 2.30E-01 | 6.32E-01 | 1.13E+00 |
| LTV1 | 3.62E-05 | 1.89E-04 | -1.11E+00 | 6.80E-06 | 2.94E-05 | -1.09E+00 | 1.17E-04 | 4.97E-04 | -1.15E+00 | 5.25E-09 | 2.00E-08 | -1.28E+00 | 5.04E-03 | 1.80E-02 | 1.05E+00 | 1.34E-03 | 3.77E-03 | -1.07E+00 | 4.96E-08 | 1.82E-07 | -1.21E+00 | 1.66E-02 | 4.56E-02 | 1.06E+00 |
| CFAP20 | 3.62E-05 | 1.89E-04 | -1.05E+00 | 3.09E-15 | 1.34E-14 | -1.51E+00 | 1.64E-04 | 6.97E-04 | -1.20E+00 | 2.40E-11 | 9.17E-11 | -1.49E+00 | 1.11E-02 | 3.97E-02 | 1.11E+00 | 8.94E-09 | 2.52E-08 | -1.44E+00 | 3.53E-03 | 1.30E-02 | -1.00E+00 | 3.72E-04 | 1.02E-03 | -1.13E+00 |
| C7orf26 | 3.62E-05 | 1.89E-04 | -1.23E+00 | 1.34E-02 | 5.79E-02 | 1.21E+00 | 5.47E-05 | 2.32E-04 | -1.26E+00 | 2.12E-03 | 8.10E-03 | -1.00E+00 | 1.64E-07 | 5.85E-07 | -1.94E+00 | 1.08E-02 | 3.04E-02 | -1.00E+00 | 4.42E-05 | 1.62E-04 | -1.32E+00 | 9.82E-02 | 2.70E-01 | 1.17E+00 |
| PDE6G | 3.64E-05 | 1.89E-04 | -1.12E+00 | 4.44E-05 | 1.92E-04 | 1.03E+00 | 7.98E-04 | 3.39E-03 | -1.16E+00 | 9.73E-03 | 3.72E-02 | 1.06E+00 | 4.57E-06 | 1.63E-05 | -1.37E+00 | 3.92E-04 | 1.10E-03 | -1.15E+00 | 2.31E-01 | 8.46E-01 | 1.31E+00 | 2.59E-01 | 7.13E-01 | 1.37E+00 |
| MOGS | 3.64E-05 | 1.89E-04 | -1.16E+00 | 2.70E-02 | 1.16E-01 | 1.30E+00 | 3.32E-04 | 1.41E-03 | -1.19E+00 | 2.62E-02 | 1.00E-01 | 1.20E+00 | 4.36E-04 | 1.55E-03 | -1.33E+00 | 2.35E-01 | 6.63E-01 | 1.59E+00 | 4.82E-05 | 1.77E-04 | -1.17E+00 | 1.57E-01 | 4.30E-01 | 1.22E+00 |
| ARL17B | 3.64E-05 | 1.90E-04 | -1.77E+00 | 6.61E-04 | 2.85E-03 | -1.24E+00 | 1.31E-04 | 5.56E-04 | -1.71E+00 | 8.91E-04 | 3.40E-03 | -1.28E+00 | 8.88E-05 | 3.17E-04 | -2.22E+00 | 5.63E-02 | 1.59E-01 | -1.02E+00 | 2.08E-06 | 7.64E-06 | -1.73E+00 | 6.08E-02 | 1.67E-01 | 1.01E+00 |
| LMNB2 | 3.66E-05 | 1.91E-04 | -1.53E+00 | 5.32E-04 | 2.30E-03 | -1.15E+00 | 8.96E-02 | 3.81E-01 | 1.16E+00 | 2.24E-01 | 8.54E-01 | 1.59E+00 | 8.00E-04 | 2.85E-03 | -1.77E+00 | 1.26E-01 | 3.55E-01 | 1.17E+00 | 2.04E-02 | 7.49E-02 | 1.04E+00 | 1.31E-02 | 3.60E-02 | 1.76E+00 |
| COMMD10 | 3.66E-05 | 1.91E-04 | -1.16E+00 | 2.46E-03 | 1.06E-02 | 1.13E+00 | 1.16E-04 | 4.93E-04 | -1.10E+00 | 5.34E-04 | 2.04E-03 | 1.04E+00 | 1.14E-02 | 4.08E-02 | -1.03E+00 | 2.88E-01 | 8.11E-01 | 1.35E+00 | 1.08E-08 | 3.96E-08 | -1.43E+00 | 9.55E-03 | 2.62E-02 | 1.03E+00 |
| SPCS2 | 3.68E-05 | 1.92E-04 | -1.10E+00 | 1.53E-03 | 6.60E-03 | 1.16E+00 | 3.65E-03 | 1.55E-02 | -1.10E+00 | 8.17E-04 | 3.12E-03 | -1.01E+00 | 3.23E-07 | 1.15E-06 | -1.35E+00 | 8.35E-04 | 2.35E-03 | -1.06E+00 | 1.83E-06 | 6.73E-06 | -1.12E+00 | 2.24E-03 | 6.16E-03 | 1.02E+00 |
| LINC01140 | 3.68E-05 | 1.92E-04 | -2.12E+00 | 8.19E-05 | 3.54E-04 | -1.96E+00 | 1.07E-05 | 4.55E-05 | -1.95E+00 | 3.49E-04 | 1.33E-03 | -1.48E+00 | 2.74E-24 | 9.78E-24 | -3.72E+00 | 5.03E-24 | 1.42E-23 | -3.26E+00 | 4.54E-06 | 1.67E-05 | -3.08E+00 | 1.00E-03 | 2.75E-03 | -1.83E+00 |
| ZFP30 | 3.68E-05 | 1.92E-04 | -1.38E+00 | 4.36E-02 | 1.88E-01 | 1.18E+00 | 1.62E-02 | 6.87E-02 | -1.04E+00 | 2.57E-01 | 9.83E-01 | 1.42E+00 | 2.07E-01 | 7.37E-01 | 1.14E+00 | 2.24E-01 | 6.30E-01 | 1.55E+00 | 3.50E-02 | 1.29E-01 | -1.11E+00 | 1.06E-01 | 2.91E-01 | 1.10E+00 |
| SUPT7L | 3.69E-05 | 1.92E-04 | -1.07E+00 | 1.51E-03 | 6.51E-03 | 1.12E+00 | 3.35E-04 | 1.42E-03 | -1.20E+00 | 3.20E-04 | 1.22E-03 | -1.03E+00 | 1.46E-04 | 5.20E-04 | -1.23E+00 | 5.61E-03 | 1.58E-02 | -1.02E+00 | 2.92E-06 | 1.07E-05 | -1.26E+00 | 9.14E-05 | 2.51E-04 | -1.15E+00 |
| EVC2 | 3.69E-05 | 1.92E-04 | -1.37E+00 | 2.47E-03 | 1.07E-02 | 1.06E+00 | 1.07E-05 | 4.57E-05 | -1.60E+00 | 3.13E-04 | 1.19E-03 | -1.26E+00 | 2.02E-02 | 7.20E-02 | -1.72E+00 | 2.27E-02 | 6.40E-02 | -1.82E+00 | 1.58E-02 | 5.78E-02 | -1.88E+00 | 1.88E-01 | 5.18E-01 | -1.11E+00 |
| TCTN2 | 3.70E-05 | 1.93E-04 | -1.78E+00 | 2.42E-03 | 1.05E-02 | -1.14E+00 | 4.64E-06 | 1.97E-05 | -1.62E+00 | 3.96E-05 | 1.51E-04 | -1.34E+00 | 1.46E-04 | 5.19E-04 | -2.17E+00 | 5.21E-03 | 1.47E-02 | -1.54E+00 | 2.28E-03 | 8.39E-03 | -1.70E+00 | 1.67E-01 | 4.59E-01 | -1.00E+00 |
| POR | 3.71E-05 | 1.93E-04 | -1.49E+00 | 8.21E-02 | 3.55E-01 | 1.30E+00 | 1.54E-02 | 6.56E-02 | -1.13E+00 | 1.73E-02 | 6.60E-02 | 1.03E+00 | 5.78E-02 | 2.06E-01 | 1.01E+00 | 2.28E-01 | 6.42E-01 | 1.58E+00 | 2.84E-02 | 1.04E-01 | -1.11E+00 | 1.16E-01 | 3.20E-01 | 1.51E+00 |
| TMCO6 | 3.71E-05 | 1.93E-04 | -1.65E+00 | 3.30E-03 | 1.43E-02 | -1.12E+00 | 2.48E-04 | 1.05E-03 | -1.45E+00 | 9.91E-03 | 3.79E-02 | 1.03E+00 | 1.75E-04 | 6.22E-04 | -1.83E+00 | 8.14E-02 | 2.29E-01 | 1.09E+00 | 4.46E-05 | 1.64E-04 | -1.31E+00 | 6.96E-03 | 1.91E-02 | -1.01E+00 |
| GANC | 3.71E-05 | 1.93E-04 | -1.32E+00 | 9.33E-04 | 4.03E-03 | 1.00E+00 | 6.34E-08 | 2.69E-07 | -1.86E+00 | 2.88E-04 | 1.10E-03 | -1.13E+00 | 2.49E-02 | 8.87E-02 | -1.25E+00 | 2.25E-01 | 6.35E-01 | 1.11E+00 | 5.83E-06 | 2.14E-05 | -1.66E+00 | 1.50E-03 | 4.13E-03 | -1.31E+00 |
| HEXDC-IT1 | 3.72E-05 | 1.94E-04 | -1.45E+00 | 5.11E-04 | 2.21E-03 | -1.11E+00 | 2.95E-03 | 1.25E-02 | -1.33E+00 | 2.62E-02 | 1.00E-01 | 1.00E+00 | 2.45E-01 | 8.73E-01 | 1.11E+00 | 2.85E-01 | 8.04E-01 | 1.15E+00 | 2.64E-02 | 9.70E-02 | -1.38E+00 | 2.44E-01 | 6.69E-01 | 1.10E+00 |
| AC064850.1 | 3.72E-05 | 1.94E-04 | -1.84E+00 | 6.15E-05 | 2.66E-04 | -1.68E+00 | 3.14E-02 | 1.33E-01 | -1.24E+00 | 4.89E-02 | 1.87E-01 | -1.04E+00 | 1.66E-02 | 5.90E-02 | -1.38E+00 | 4.01E-02 | 1.13E-01 | -1.32E+00 | 8.15E-03 | 2.99E-02 | -1.43E+00 | 4.44E-02 | 1.22E-01 | -1.33E+00 |
| FIGNL1 | 3.72E-05 | 1.94E-04 | -1.84E+00 | 3.18E-04 | 1.37E-03 | -1.53E+00 | 3.25E-03 | 1.38E-02 | -1.32E+00 | 7.09E-03 | 2.71E-02 | -1.14E+00 | 4.70E-03 | 1.67E-02 | -2.11E+00 | 3.75E-02 | 1.06E-01 | -1.36E+00 | 1.16E-09 | 4.26E-09 | -1.56E+00 | 2.92E-08 | 8.01E-08 | -1.48E+00 |
| PDE5A | 3.72E-05 | 1.94E-04 | -1.51E+00 | 1.76E-03 | 7.62E-03 | -1.09E+00 | 1.50E-02 | 6.37E-02 | -1.19E+00 | 3.32E-02 | 1.27E-01 | -1.04E+00 | 6.73E-03 | 2.40E-02 | -1.35E+00 | 1.25E-02 | 3.51E-02 | -1.13E+00 | 7.96E-02 | 2.92E-01 | 1.30E+00 | 3.30E-01 | 9.06E-01 | -1.06E+00 |
| ENOSF1 | 3.73E-05 | 1.94E-04 | -1.04E+00 | 9.32E-06 | 4.03E-05 | 1.06E+00 | 4.57E-04 | 1.94E-03 | -1.09E+00 | 1.18E-04 | 4.49E-04 | 1.06E+00 | 1.64E-05 | 5.85E-05 | -1.30E+00 | 4.58E-02 | 1.29E-01 | 1.13E+00 | 8.68E-09 | 3.19E-08 | -1.33E+00 | 1.29E-03 | 3.53E-03 | -1.06E+00 |
| SNX10 | 3.73E-05 | 1.95E-04 | -1.18E+00 | 2.18E-02 | 9.41E-02 | 1.22E+00 | 8.32E-02 | 3.53E-01 | 1.05E+00 | 9.85E-03 | 3.76E-02 | 1.06E+00 | 2.90E-04 | 1.03E-03 | -1.26E+00 | 2.40E-04 | 6.77E-04 | -1.33E+00 | 1.39E-02 | 5.11E-02 | -1.16E+00 | 1.44E-01 | 3.94E-01 | 1.00E+00 |
| GPAM | 3.75E-05 | 1.95E-04 | -1.52E+00 | 1.55E-03 | 6.70E-03 | -1.05E+00 | 2.21E-02 | 9.38E-02 | -1.19E+00 | 2.35E-01 | 8.97E-01 | 1.37E+00 | 1.36E-04 | 4.85E-04 | -1.49E+00 | 2.65E-04 | 7.47E-04 | -1.48E+00 | 1.86E-06 | 6.81E-06 | -1.71E+00 | 8.87E-07 | 2.44E-06 | -1.74E+00 |
| PCOLCE2 | 3.76E-05 | 1.96E-04 | -1.68E+00 | 3.12E-02 | 1.35E-01 | 1.08E+00 | 3.15E-02 | 1.34E-01 | -1.11E+00 | 3.55E-02 | 1.36E-01 | 1.69E+00 | 2.10E-01 | 7.49E-01 | -1.11E+00 | 2.83E-01 | 7.97E-01 | 1.56E+00 | 7.49E-02 | 2.75E-01 | -1.19E+00 | 2.68E-01 | 7.36E-01 | 1.45E+00 |
| DCK | 3.77E-05 | 1.96E-04 | -1.11E+00 | 3.29E-03 | 1.42E-02 | 1.13E+00 | 2.25E-04 | 9.58E-04 | -1.16E+00 | 8.75E-05 | 3.34E-04 | -1.01E+00 | 4.62E-05 | 1.65E-04 | -1.24E+00 | 1.41E-02 | 3.97E-02 | 1.10E+00 | 5.85E-10 | 2.15E-09 | -1.33E+00 | 6.22E-04 | 1.71E-03 | -1.07E+00 |
| AL133338.1 | 3.77E-05 | 1.96E-04 | -1.57E+00 | 1.01E-01 | 4.37E-01 | 1.23E+00 | 1.37E-02 | 5.80E-02 | -1.32E+00 | 1.92E-01 | 7.34E-01 | 1.27E+00 | 2.36E-03 | 8.40E-03 | -1.57E+00 | 5.34E-02 | 1.50E-01 | 1.02E+00 | 1.63E-01 | 5.96E-01 | 1.18E+00 | 3.56E-02 | 9.78E-02 | 1.76E+00 |
| MTERF4 | 3.77E-05 | 1.96E-04 | -1.19E+00 | 1.88E-02 | 8.12E-02 | 1.20E+00 | 4.66E-02 | 1.98E-01 | 1.08E+00 | 2.54E-02 | 9.69E-02 | 1.23E+00 | 6.62E-02 | 2.36E-01 | 1.25E+00 | 2.38E-02 | 6.70E-02 | 1.13E+00 | 2.30E-01 | 8.43E-01 | 1.28E+00 | 2.28E-01 | 6.27E-01 | 1.22E+00 |
| SH3YL1 | 3.77E-05 | 1.96E-04 | -1.07E+00 | 7.15E-02 | 3.09E-01 | 1.37E+00 | 1.34E-03 | 5.70E-03 | -1.13E+00 | 2.06E-01 | 7.87E-01 | 1.29E+00 | 1.10E-02 | 3.91E-02 | -1.12E+00 | 3.30E-01 | 9.29E-01 | 1.45E+00 | 9.02E-02 | 3.31E-01 | 1.06E+00 | 3.52E-01 | 9.66E-01 | 1.25E+00 |
| TSC2 | 3.79E-05 | 1.98E-04 | -1.40E+00 | 1.28E-03 | 5.54E-03 | -1.05E+00 | 1.24E-02 | 5.29E-02 | -1.08E+00 | 9.21E-02 | 3.52E-01 | 1.24E+00 | 3.78E-05 | 1.35E-04 | -1.63E+00 | 1.86E-02 | 5.24E-02 | 1.05E+00 | 3.61E-03 | 1.32E-02 | -1.11E+00 | 1.41E-01 | 3.87E-01 | 1.11E+00 |
| GALT | 3.80E-05 | 1.98E-04 | -1.20E+00 | 1.14E-03 | 4.92E-03 | 1.12E+00 | 1.81E-05 | 7.68E-05 | -1.30E+00 | 1.53E-04 | 5.84E-04 | -1.05E+00 | 1.27E-03 | 4.51E-03 | -1.20E+00 | 3.04E-02 | 8.57E-02 | 1.06E+00 | 6.59E-02 | 2.42E-01 | 1.10E+00 | 9.00E-02 | 2.47E-01 | 1.11E+00 |
| AKNA | 3.81E-05 | 1.98E-04 | -1.02E+00 | 2.43E-02 | 1.05E-01 | 1.32E+00 | 2.62E-03 | 1.11E-02 | -1.06E+00 | 2.08E-01 | 7.95E-01 | 1.35E+00 | 1.41E-03 | 5.02E-03 | -1.28E+00 | 5.57E-03 | 1.57E-02 | -1.13E+00 | 1.29E-02 | 4.72E-02 | -1.47E+00 | 7.22E-02 | 1.98E-01 | -1.15E+00 |
| AGTPBP1 | 3.83E-05 | 1.99E-04 | -1.21E+00 | 2.04E-07 | 8.80E-07 | -1.35E+00 | 1.68E-06 | 7.15E-06 | -1.31E+00 | 1.14E-06 | 4.36E-06 | -1.19E+00 | 1.03E-01 | 3.69E-01 | -1.04E+00 | 1.01E-01 | 2.84E-01 | -1.07E+00 | 1.92E-04 | 7.06E-04 | -1.21E+00 | 3.77E-05 | 1.04E-04 | -1.38E+00 |
| ALKBH3 | 3.83E-05 | 1.99E-04 | -1.17E+00 | 2.14E-06 | 9.27E-06 | -1.22E+00 | 1.92E-02 | 8.16E-02 | -1.01E+00 | 5.31E-03 | 2.03E-02 | 1.07E+00 | 1.96E-06 | 7.00E-06 | -1.56E+00 | 4.58E-04 | 1.29E-03 | -1.21E+00 | 1.28E-02 | 4.69E-02 | 1.03E+00 | 1.41E-02 | 3.87E-02 | 1.04E+00 |
| CFP | 3.83E-05 | 2.00E-04 | -1.16E+00 | 2.09E-02 | 9.04E-02 | 1.26E+00 | 1.90E-04 | 8.09E-04 | -1.23E+00 | 1.17E-03 | 4.46E-03 | -1.06E+00 | 2.55E-04 | 9.10E-04 | -1.69E+00 | 3.62E-05 | 1.02E-04 | -2.47E+00 | 4.70E-12 | 1.72E-11 | -1.44E+00 | 1.96E-10 | 5.39E-10 | -1.67E+00 |
| FKBP10 | 3.83E-05 | 2.00E-04 | -1.66E+00 | 1.41E-04 | 6.07E-04 | -1.34E+00 | 1.58E-04 | 6.69E-04 | -1.52E+00 | 3.80E-04 | 1.45E-03 | -1.35E+00 | 1.20E-04 | 4.26E-04 | -2.08E+00 | 3.08E-04 | 8.68E-04 | -1.86E+00 | 7.24E-02 | 2.66E-01 | 1.11E+00 | 2.91E-01 | 8.00E-01 | 1.26E+00 |
| AC017007.3 | 3.83E-05 | 2.00E-04 | -1.33E+00 | 1.35E-01 | 5.82E-01 | 1.42E+00 | 1.26E-02 | 5.34E-02 | -1.13E+00 | 8.52E-02 | 3.25E-01 | 1.61E+00 | 5.14E-02 | 1.83E-01 | -1.06E+00 | 1.64E-01 | 4.61E-01 | 1.76E+00 | 4.95E-03 | 1.82E-02 | -1.26E+00 | 3.38E-01 | 9.29E-01 | 1.17E+00 |
| DYNC1I2 | 3.84E-05 | 2.00E-04 | -1.32E+00 | 1.88E-03 | 8.14E-03 | -1.02E+00 | 7.32E-04 | 3.11E-03 | -1.30E+00 | 4.39E-05 | 1.68E-04 | -1.29E+00 | 3.57E-02 | 1.27E-01 | -1.05E+00 | 1.61E-01 | 4.54E-01 | 1.28E+00 | 4.62E-02 | 1.69E-01 | -1.02E+00 | 5.12E-03 | 1.41E-02 | -1.16E+00 |
| AF127577.2 | 3.85E-05 | 2.01E-04 | -1.39E+00 | 2.64E-02 | 1.14E-01 | 1.62E+00 | 2.37E-05 | 1.01E-04 | -1.54E+00 | 5.08E-03 | 1.94E-02 | 1.02E+00 | 1.89E-01 | 6.74E-01 | 1.24E+00 | 3.49E-01 | 9.85E-01 | 1.51E+00 | 1.50E-02 | 5.51E-02 | -1.35E+00 | 8.24E-02 | 2.26E-01 | -1.06E+00 |
| TSPAN31 | 3.85E-05 | 2.01E-04 | -1.24E+00 | 2.19E-05 | 9.48E-05 | -1.18E+00 | 2.71E-03 | 1.15E-02 | -1.21E+00 | 4.49E-02 | 1.71E-01 | 1.18E+00 | 6.54E-02 | 2.33E-01 | 1.14E+00 | 1.36E-02 | 3.82E-02 | -1.04E+00 | 8.98E-02 | 3.30E-01 | 1.10E+00 | 2.98E-01 | 8.18E-01 | 1.42E+00 |
| ARL9 | 3.86E-05 | 2.01E-04 | -1.36E+00 | 3.47E-05 | 1.50E-04 | -1.28E+00 | 1.42E-05 | 6.04E-05 | -1.34E+00 | 1.39E-07 | 5.31E-07 | -1.55E+00 | 5.09E-02 | 1.81E-01 | -1.50E+00 | 7.87E-02 | 2.22E-01 | -1.27E+00 | 1.48E-02 | 5.42E-02 | -1.42E+00 | 8.11E-03 | 2.23E-02 | -1.65E+00 |
| RNF180 | 3.87E-05 | 2.02E-04 | -1.28E+00 | 1.28E-05 | 5.54E-05 | -1.35E+00 | 6.02E-02 | 2.56E-01 | -1.27E+00 | 1.49E-02 | 5.68E-02 | -1.67E+00 | 2.64E-01 | 9.43E-01 | 1.33E+00 | 1.91E-01 | 5.38E-01 | -1.01E+00 | 1.95E-01 | 7.16E-01 | -1.03E+00 | 3.25E-01 | 8.91E-01 | 1.11E+00 |
| GOT1 | 3.88E-05 | 2.02E-04 | -1.44E+00 | 1.23E-02 | 5.33E-02 | 1.09E+00 | 4.68E-03 | 1.99E-02 | -1.10E+00 | 1.85E-02 | 7.05E-02 | 1.11E+00 | 1.05E-06 | 3.75E-06 | -1.67E+00 | 1.23E-02 | 3.46E-02 | -1.04E+00 | 3.39E-08 | 1.24E-07 | -1.27E+00 | 2.31E-03 | 6.35E-03 | 1.02E+00 |
| GMDS-AS1 | 3.90E-05 | 2.03E-04 | -1.14E+00 | 2.57E-03 | 1.11E-02 | 1.12E+00 | 2.60E-06 | 1.10E-05 | -1.50E+00 | 2.35E-05 | 8.98E-05 | -1.25E+00 | 2.51E-03 | 8.94E-03 | -1.21E+00 | 9.92E-02 | 2.80E-01 | 1.13E+00 | 4.82E-02 | 1.77E-01 | 1.03E+00 | 1.11E-01 | 3.06E-01 | 1.11E+00 |
| SLC30A4 | 3.90E-05 | 2.03E-04 | -1.89E+00 | 1.61E-03 | 6.98E-03 | -1.25E+00 | 1.76E-01 | 7.48E-01 | 1.26E+00 | 8.58E-02 | 3.28E-01 | 1.65E+00 | 1.08E-01 | 3.84E-01 | -1.35E+00 | 2.99E-01 | 8.43E-01 | 1.15E+00 | 5.78E-02 | 2.12E-01 | -1.14E+00 | 8.89E-02 | 2.44E-01 | -1.16E+00 |
| CD40 | 3.90E-05 | 2.03E-04 | -1.80E+00 | 6.47E-03 | 2.80E-02 | -1.04E+00 | 3.76E-02 | 1.60E-01 | -1.63E+00 | 5.70E-02 | 2.18E-01 | -1.30E+00 | 2.74E-01 | 9.78E-01 | 1.55E+00 | 6.64E-03 | 1.87E-02 | 2.60E+00 | 1.34E-01 | 4.93E-01 | 1.27E+00 | 5.10E-03 | 1.40E-02 | 1.05E+00 |
| NIT1 | 3.92E-05 | 2.04E-04 | -1.17E+00 | 1.31E-02 | 5.66E-02 | 1.25E+00 | 9.44E-06 | 4.01E-05 | -1.28E+00 | 7.36E-03 | 2.81E-02 | 1.07E+00 | 9.06E-09 | 3.23E-08 | -1.55E+00 | 3.15E-03 | 8.88E-03 | 1.06E+00 | 1.18E-04 | 4.32E-04 | -1.14E+00 | 1.64E-01 | 4.52E-01 | 1.19E+00 |
| PSME4 | 3.93E-05 | 2.05E-04 | -1.09E+00 | 3.07E-10 | 1.33E-09 | -1.28E+00 | 4.84E-06 | 2.06E-05 | -1.16E+00 | 2.77E-11 | 1.06E-10 | -1.31E+00 | 2.11E-02 | 7.53E-02 | 1.05E+00 | 1.03E-05 | 2.90E-05 | -1.18E+00 | 6.26E-07 | 2.30E-06 | -1.34E+00 | 8.74E-06 | 2.40E-05 | -1.23E+00 |
| MAP4K5 | 3.93E-05 | 2.05E-04 | -1.07E+00 | 7.36E-07 | 3.18E-06 | -1.07E+00 | 1.86E-05 | 7.90E-05 | -1.18E+00 | 5.52E-08 | 2.11E-07 | -1.19E+00 | 6.56E-05 | 2.34E-04 | -1.03E+00 | 4.27E-08 | 1.20E-07 | -1.26E+00 | 8.08E-05 | 2.96E-04 | -1.06E+00 | 5.01E-05 | 1.38E-04 | -1.26E+00 |
| TCP1P3 | 3.95E-05 | 2.06E-04 | -1.39E+00 | 1.32E-06 | 5.71E-06 | -1.54E+00 | 1.19E-05 | 5.07E-05 | -1.47E+00 | 3.09E-05 | 1.18E-04 | -1.25E+00 | 1.29E-04 | 4.60E-04 | -1.54E+00 | 9.60E-05 | 2.71E-04 | -1.62E+00 | 5.92E-02 | 2.17E-01 | 1.00E+00 | 9.11E-02 | 2.50E-01 | 1.04E+00 |
| MKLN1 | 3.97E-05 | 2.07E-04 | -1.03E+00 | 7.64E-03 | 3.30E-02 | 1.26E+00 | 3.15E-04 | 1.34E-03 | -1.10E+00 | 5.75E-03 | 2.20E-02 | 1.14E+00 | 3.22E-02 | 1.15E-01 | 1.07E+00 | 2.62E-02 | 7.39E-02 | 1.07E+00 | 7.03E-04 | 2.58E-03 | -1.13E+00 | 6.90E-02 | 1.90E-01 | 1.00E+00 |
| TFRC | 3.98E-05 | 2.08E-04 | 1.03E+00 | 3.81E-12 | 1.65E-11 | -1.17E+00 | 3.18E-07 | 1.35E-06 | -1.36E+00 | 1.28E-06 | 4.91E-06 | -1.19E+00 | 1.35E-04 | 4.80E-04 | -1.13E+00 | 1.80E-01 | 5.06E-01 | 1.25E+00 | 6.68E-04 | 2.45E-03 | -1.28E+00 | 1.05E-01 | 2.88E-01 | 1.05E+00 |
| SUGP2 | 3.99E-05 | 2.08E-04 | -1.01E+00 | 2.06E-01 | 8.88E-01 | 1.46E+00 | 3.02E-03 | 1.28E-02 | -1.04E+00 | 7.58E-02 | 2.90E-01 | 1.23E+00 | 5.39E-02 | 1.92E-01 | 1.12E+00 | 3.00E-01 | 8.46E-01 | 1.33E+00 | 2.27E-02 | 8.35E-02 | 1.06E+00 | 2.16E-01 | 5.92E-01 | 1.36E+00 |
| ZSWIM5 | 3.99E-05 | 2.08E-04 | -2.07E+00 | 1.13E-02 | 4.87E-02 | -1.10E+00 | 5.41E-04 | 2.30E-03 | -1.60E+00 | 8.79E-04 | 3.36E-03 | -1.45E+00 | 2.15E-01 | 7.66E-01 | 1.02E+00 | 2.72E-01 | 7.67E-01 | 1.43E+00 | 9.82E-04 | 3.60E-03 | -1.66E+00 | 3.01E-02 | 8.28E-02 | -1.21E+00 |
| CCNH | 3.99E-05 | 2.08E-04 | 1.08E+00 | 4.42E-03 | 1.91E-02 | 1.26E+00 | 3.58E-05 | 1.52E-04 | -1.12E+00 | 1.36E-04 | 5.20E-04 | -1.01E+00 | 8.92E-02 | 3.18E-01 | 1.15E+00 | 5.79E-02 | 1.63E-01 | 1.09E+00 | 6.42E-03 | 2.36E-02 | -1.03E+00 | 4.93E-03 | 1.36E-02 | -1.02E+00 |
| BICRAL | 3.99E-05 | 2.08E-04 | -1.56E+00 | 5.69E-03 | 2.46E-02 | -1.03E+00 | 2.23E-03 | 9.47E-03 | -1.19E+00 | 1.40E-01 | 5.36E-01 | 1.30E+00 | 2.11E-02 | 7.54E-02 | -1.07E+00 | 2.28E-01 | 6.42E-01 | 1.38E+00 | 1.96E-04 | 7.20E-04 | -1.41E+00 | 1.59E-01 | 4.37E-01 | 1.09E+00 |
| PEX5 | 3.99E-05 | 2.08E-04 | -1.41E+00 | 1.86E-03 | 8.03E-03 | -1.00E+00 | 2.58E-04 | 1.10E-03 | -1.29E+00 | 1.14E-02 | 4.35E-02 | 1.14E+00 | 6.61E-03 | 2.36E-02 | -1.31E+00 | 9.38E-02 | 2.64E-01 | 1.08E+00 | 2.89E-05 | 1.06E-04 | -1.32E+00 | 4.57E-02 | 1.26E-01 | 1.09E+00 |
| EIF4BP7 | 4.00E-05 | 2.09E-04 | 1.02E+00 | 4.04E-03 | 1.74E-02 | 1.30E+00 | 5.96E-02 | 2.53E-01 | 1.10E+00 | 2.50E-01 | 9.55E-01 | 1.38E+00 | 1.17E-03 | 4.17E-03 | 1.01E+00 | 7.18E-02 | 2.02E-01 | 1.27E+00 | 2.79E-02 | 1.02E-01 | 1.09E+00 | 3.60E-01 | 9.90E-01 | 1.37E+00 |
| BHLHB9 | 4.02E-05 | 2.09E-04 | -1.62E+00 | 5.90E-02 | 2.55E-01 | 1.13E+00 | 1.89E-02 | 8.03E-02 | -1.22E+00 | 6.44E-03 | 2.46E-02 | -1.20E+00 | 2.00E-02 | 7.13E-02 | -1.61E+00 | 2.32E-01 | 6.54E-01 | 1.10E+00 | 9.80E-02 | 3.60E-01 | -1.06E+00 | 1.08E-01 | 2.97E-01 | -1.11E+00 |
| NR2C2AP | 4.02E-05 | 2.09E-04 | -1.57E+00 | 1.69E-03 | 7.30E-03 | -1.22E+00 | 1.37E-04 | 5.84E-04 | -1.27E+00 | 5.02E-03 | 1.92E-02 | 1.04E+00 | 7.13E-02 | 2.54E-01 | -1.14E+00 | 2.80E-01 | 7.90E-01 | 1.22E+00 | 1.57E-08 | 5.75E-08 | -1.41E+00 | 2.17E-03 | 5.97E-03 | -1.01E+00 |
| AC096775.1 | 4.03E-05 | 2.10E-04 | -1.53E+00 | 8.15E-06 | 3.52E-05 | -1.58E+00 | 1.61E-02 | 6.86E-02 | -1.46E+00 | 4.72E-02 | 1.80E-01 | -1.20E+00 | 2.80E-01 | 9.99E-01 | 1.36E+00 | 3.18E-01 | 8.96E-01 | 1.46E+00 | 2.37E-01 | 8.69E-01 | 1.25E+00 | 1.66E-01 | 4.57E-01 | 1.61E+00 |
| PIM3 | 4.03E-05 | 2.10E-04 | -1.01E+00 | 7.53E-08 | 3.25E-07 | -1.10E+00 | 7.91E-08 | 3.36E-07 | -1.39E+00 | 2.05E-06 | 7.82E-06 | -1.17E+00 | 1.17E-14 | 4.15E-14 | -2.76E+00 | 7.61E-13 | 2.15E-12 | -2.23E+00 | 3.26E-15 | 1.20E-14 | -1.94E+00 | 6.35E-11 | 1.74E-10 | -1.60E+00 |
| PHF20 | 4.03E-05 | 2.10E-04 | 1.00E+00 | 6.70E-06 | 2.90E-05 | 1.07E+00 | 2.22E-02 | 9.45E-02 | 1.01E+00 | 2.56E-03 | 9.79E-03 | 1.03E+00 | 6.28E-02 | 2.24E-01 | 1.06E+00 | 3.02E-02 | 8.51E-02 | 1.06E+00 | 4.06E-02 | 1.49E-01 | 1.00E+00 | 3.31E-01 | 9.09E-01 | 1.12E+00 |
| EEF1GP1 | 4.04E-05 | 2.11E-04 | -1.63E+00 | 1.97E-05 | 8.52E-05 | -1.76E+00 | 1.62E-01 | 6.89E-01 | 1.03E+00 | 2.53E-01 | 9.68E-01 | 1.31E+00 | 2.77E-02 | 9.88E-02 | -1.16E+00 | 4.78E-03 | 1.35E-02 | -1.53E+00 | 5.75E-03 | 2.11E-02 | -1.26E+00 | 2.24E-02 | 6.15E-02 | -1.25E+00 |
| SUMO3 | 4.04E-05 | 2.11E-04 | 1.08E+00 | 8.04E-05 | 3.47E-04 | 1.14E+00 | 6.49E-02 | 2.76E-01 | 1.07E+00 | 6.46E-02 | 2.47E-01 | 1.12E+00 | 2.06E-02 | 7.34E-02 | 1.13E+00 | 9.97E-02 | 2.81E-01 | 1.17E+00 | 3.44E-02 | 1.26E-01 | 1.37E+00 | 9.16E-02 | 2.52E-01 | 1.30E+00 |
| AC006141.1 | 4.05E-05 | 2.11E-04 | -1.31E+00 | 4.00E-02 | 1.73E-01 | 1.16E+00 | 9.81E-07 | 4.17E-06 | -1.61E+00 | 1.30E-03 | 4.96E-03 | -1.02E+00 | 6.16E-04 | 2.20E-03 | -1.23E+00 | 4.87E-02 | 1.37E-01 | 1.10E+00 | 4.98E-07 | 1.83E-06 | -2.08E+00 | 4.02E-02 | 1.11E-01 | -1.13E+00 |
| FGL2 | 4.05E-05 | 2.11E-04 | -2.14E+00 | 3.24E-03 | 1.40E-02 | -1.04E+00 | 2.19E-03 | 9.29E-03 | -1.65E+00 | 2.22E-02 | 8.49E-02 | -1.41E+00 | 1.73E-01 | 6.16E-01 | -1.13E+00 | 1.58E-01 | 4.45E-01 | -1.32E+00 | 2.24E-03 | 8.23E-03 | -2.50E+00 | 1.77E-02 | 4.87E-02 | -1.87E+00 |
| AC127502.1 | 4.06E-05 | 2.12E-04 | -1.06E+00 | 1.51E-06 | 6.51E-06 | -1.10E+00 | 7.89E-06 | 3.35E-05 | -1.25E+00 | 9.31E-07 | 3.56E-06 | -1.21E+00 | 7.82E-04 | 2.79E-03 | -1.32E+00 | 9.02E-05 | 2.54E-04 | -1.52E+00 | 5.58E-04 | 2.05E-03 | -1.05E+00 | 9.71E-05 | 2.67E-04 | -1.19E+00 |
| CNKSR3 | 4.06E-05 | 2.12E-04 | -1.54E+00 | 7.84E-03 | 3.39E-02 | 1.14E+00 | 2.16E-05 | 9.18E-05 | -1.77E+00 | 8.14E-04 | 3.11E-03 | -1.19E+00 | 9.68E-09 | 3.45E-08 | -1.40E+00 | 1.03E-06 | 2.91E-06 | -1.11E+00 | 4.58E-07 | 1.68E-06 | -2.04E+00 | 5.15E-03 | 1.41E-02 | -1.05E+00 |
| PRKACA | 4.07E-05 | 2.12E-04 | -1.42E+00 | 1.13E-03 | 4.90E-03 | -1.10E+00 | 4.16E-03 | 1.77E-02 | -1.17E+00 | 9.93E-03 | 3.79E-02 | -1.01E+00 | 1.06E-02 | 3.76E-02 | -1.22E+00 | 2.92E-02 | 8.22E-02 | -1.04E+00 | 2.61E-02 | 9.56E-02 | -1.00E+00 | 2.28E-01 | 6.27E-01 | 1.21E+00 |
| PRKAG1 | 4.07E-05 | 2.12E-04 | -1.10E+00 | 3.07E-03 | 1.33E-02 | 1.16E+00 | 2.59E-05 | 1.10E-04 | -1.26E+00 | 1.95E-04 | 7.43E-04 | -1.03E+00 | 1.39E-03 | 4.96E-03 | 1.03E+00 | 2.46E-01 | 6.94E-01 | 1.44E+00 | 1.32E-05 | 4.85E-05 | -1.11E+00 | 3.17E-02 | 8.70E-02 | 1.12E+00 |
| MFSD11 | 4.08E-05 | 2.13E-04 | -1.31E+00 | 7.61E-03 | 3.29E-02 | 1.07E+00 | 1.12E-05 | 4.75E-05 | -1.56E+00 | 6.69E-05 | 2.56E-04 | -1.32E+00 | 1.54E-07 | 5.50E-07 | -1.99E+00 | 2.20E-04 | 6.21E-04 | -1.28E+00 | 3.16E-10 | 1.16E-09 | -1.61E+00 | 8.14E-08 | 2.24E-07 | -1.41E+00 |
| SH3GL1 | 4.09E-05 | 2.13E-04 | -1.33E+00 | 3.63E-03 | 1.57E-02 | 1.09E+00 | 9.49E-04 | 4.03E-03 | -1.25E+00 | 1.36E-02 | 5.20E-02 | 1.00E+00 | 2.80E-04 | 1.00E-03 | -1.26E+00 | 2.16E-02 | 6.09E-02 | 1.08E+00 | 9.84E-09 | 3.61E-08 | -1.35E+00 | 1.51E-03 | 4.15E-03 | 1.01E+00 |
| KIAA1324L | 4.09E-05 | 2.13E-04 | -1.17E+00 | 1.91E-04 | 8.25E-04 | 1.05E+00 | 5.76E-03 | 2.45E-02 | -1.04E+00 | 9.40E-02 | 3.59E-01 | 1.30E+00 | 1.62E-03 | 5.77E-03 | -1.08E+00 | 7.34E-02 | 2.07E-01 | 1.21E+00 | 6.24E-07 | 2.29E-06 | -1.42E+00 | 1.65E-05 | 4.52E-05 | -1.36E+00 |
| ZNF135 | 4.10E-05 | 2.13E-04 | -1.56E+00 | 1.78E-04 | 7.71E-04 | -1.30E+00 | 1.08E-04 | 4.60E-04 | -1.34E+00 | 4.36E-04 | 1.67E-03 | -1.20E+00 | 5.19E-03 | 1.85E-02 | -1.26E+00 | 1.29E-02 | 3.63E-02 | -1.10E+00 | 4.17E-04 | 1.53E-03 | -1.51E+00 | 1.10E-03 | 3.02E-03 | -1.48E+00 |
| ADK | 4.10E-05 | 2.14E-04 | -1.01E+00 | 2.74E-04 | 1.19E-03 | 1.11E+00 | 4.74E-03 | 2.02E-02 | -1.03E+00 | 1.95E-04 | 7.45E-04 | 1.04E+00 | 5.91E-02 | 2.11E-01 | 1.09E+00 | 2.97E-01 | 8.36E-01 | 1.30E+00 | 1.16E-03 | 4.26E-03 | -1.08E+00 | 7.86E-02 | 2.16E-01 | 1.12E+00 |
| UPF2 | 4.10E-05 | 2.14E-04 | 1.01E+00 | 5.22E-09 | 2.26E-08 | -1.05E+00 | 5.31E-04 | 2.26E-03 | -1.07E+00 | 3.14E-07 | 1.20E-06 | -1.18E+00 | 1.51E-04 | 5.39E-04 | -1.09E+00 | 2.56E-07 | 7.22E-07 | -1.32E+00 | 2.02E-04 | 7.42E-04 | -1.19E+00 | 6.31E-08 | 1.73E-07 | -1.43E+00 |
| CCDC61 | 4.11E-05 | 2.14E-04 | -1.57E+00 | 3.59E-03 | 1.55E-02 | -1.04E+00 | 4.55E-06 | 1.93E-05 | -1.39E+00 | 1.10E-05 | 4.21E-05 | -1.17E+00 | 9.67E-05 | 3.45E-04 | -1.57E+00 | 6.45E-03 | 1.82E-02 | -1.08E+00 | 1.59E-02 | 5.85E-02 | -1.09E+00 | 5.37E-02 | 1.47E-01 | -1.07E+00 |
| PSMD8 | 4.11E-05 | 2.14E-04 | 1.01E+00 | 1.62E-05 | 7.00E-05 | 1.13E+00 | 2.13E-01 | 9.05E-01 | 1.13E+00 | 1.81E-01 | 6.93E-01 | 1.21E+00 | 2.74E-02 | 9.77E-02 | 1.09E+00 | 2.76E-01 | 7.78E-01 | 1.26E+00 | 1.21E-01 | 4.45E-01 | 1.20E+00 | 2.86E-02 | 7.87E-02 | 1.34E+00 |
| EFEMP2 | 4.11E-05 | 2.14E-04 | -1.29E+00 | 4.29E-04 | 1.85E-03 | -1.02E+00 | 3.09E-02 | 1.31E-01 | -1.05E+00 | 3.60E-03 | 1.38E-02 | -1.11E+00 | 7.82E-03 | 2.79E-02 | -1.15E+00 | 2.71E-03 | 7.64E-03 | -1.36E+00 | 1.49E-01 | 5.48E-01 | 1.18E+00 | 7.10E-02 | 1.95E-01 | 1.10E+00 |
| MAMDC2 | 4.13E-05 | 2.15E-04 | -1.34E+00 | 3.69E-02 | 1.59E-01 | 1.26E+00 | 8.62E-10 | 3.66E-09 | -1.51E+00 | 1.49E-03 | 5.68E-03 | 1.11E+00 | 6.06E-02 | 2.16E-01 | -1.22E+00 | 2.61E-02 | 7.36E-02 | -1.95E+00 | 2.56E-01 | 9.38E-01 | 1.27E+00 | 3.61E-01 | 9.92E-01 | 1.22E+00 |
| CLN6 | 4.15E-05 | 2.16E-04 | -1.63E+00 | 6.64E-03 | 2.87E-02 | -1.07E+00 | 1.70E-04 | 7.21E-04 | -1.39E+00 | 8.96E-03 | 3.42E-02 | 1.05E+00 | 1.61E-01 | 5.74E-01 | 1.21E+00 | 2.09E-01 | 5.89E-01 | 1.92E+00 | 1.19E-05 | 4.36E-05 | -1.40E+00 | 7.02E-02 | 1.93E-01 | 1.09E+00 |
| PRAM1 | 4.15E-05 | 2.16E-04 | 1.02E+00 | 3.08E-04 | 1.33E-03 | 1.15E+00 | 5.42E-10 | 2.30E-09 | -1.42E+00 | 4.13E-06 | 1.58E-05 | -1.09E+00 | 2.28E-01 | 8.14E-01 | 1.18E+00 | 1.87E-01 | 5.28E-01 | -1.43E+00 | 2.46E-01 | 9.04E-01 | 1.15E+00 | 3.42E-01 | 9.40E-01 | 1.09E+00 |
| AGAP2 | 4.16E-05 | 2.17E-04 | -1.22E+00 | 3.48E-04 | 1.50E-03 | 1.01E+00 | 3.93E-07 | 1.67E-06 | -1.60E+00 | 2.47E-07 | 9.43E-07 | -1.31E+00 | 5.00E-03 | 1.78E-02 | -1.26E+00 | 5.22E-03 | 1.47E-02 | -1.23E+00 | 1.06E-03 | 3.88E-03 | -1.35E+00 | 6.01E-03 | 1.65E-02 | -1.35E+00 |
| MAU2 | 4.18E-05 | 2.18E-04 | -1.13E+00 | 1.76E-02 | 7.60E-02 | 1.26E+00 | 8.30E-05 | 3.53E-04 | -1.27E+00 | 4.25E-03 | 1.62E-02 | 1.08E+00 | 5.16E-05 | 1.84E-04 | -1.24E+00 | 7.87E-03 | 2.22E-02 | -1.01E+00 | 4.11E-04 | 1.51E-03 | -1.21E+00 | 3.23E-02 | 8.87E-02 | 1.03E+00 |
| USP7 | 4.18E-05 | 2.18E-04 | -1.14E+00 | 7.20E-05 | 3.11E-04 | -1.01E+00 | 7.75E-05 | 3.29E-04 | -1.16E+00 | 8.07E-06 | 3.08E-05 | -1.10E+00 | 2.59E-03 | 9.23E-03 | 1.09E+00 | 6.85E-04 | 1.93E-03 | -1.02E+00 | 2.13E-06 | 7.81E-06 | -1.27E+00 | 3.69E-04 | 1.01E-03 | -1.07E+00 |
| EIF3FP3 | 4.18E-05 | 2.18E-04 | 1.51E+00 | 8.45E-05 | 3.65E-04 | 1.72E+00 | 2.18E-04 | 9.26E-04 | 1.55E+00 | 1.96E-05 | 7.49E-05 | 1.77E+00 | 2.48E-01 | 8.83E-01 | 1.41E+00 | 1.37E-03 | 3.87E-03 | 1.84E+00 | 8.25E-03 | 3.03E-02 | 1.51E+00 | 2.08E-04 | 5.72E-04 | 1.72E+00 |
| MBD1 | 4.19E-05 | 2.18E-04 | -1.26E+00 | 8.40E-03 | 3.63E-02 | 1.13E+00 | 4.42E-03 | 1.88E-02 | -1.05E+00 | 2.01E-02 | 7.69E-02 | 1.16E+00 | 1.64E-10 | 5.86E-10 | -1.94E+00 | 3.12E-05 | 8.79E-05 | -1.16E+00 | 1.23E-09 | 4.52E-09 | -1.38E+00 | 4.12E-05 | 1.13E-04 | -1.09E+00 |
| SSR2 | 4.22E-05 | 2.20E-04 | -1.06E+00 | 7.54E-02 | 3.26E-01 | 1.10E+00 | 4.58E-02 | 1.95E-01 | -1.07E+00 | 2.48E-01 | 9.48E-01 | 1.05E+00 | 1.35E-04 | 4.80E-04 | -1.25E+00 | 7.06E-02 | 1.99E-01 | -1.08E+00 | 1.05E-02 | 3.86E-02 | -1.02E+00 | 3.15E-01 | 8.67E-01 | 1.09E+00 |
| RMDN2 | 4.22E-05 | 2.20E-04 | -2.13E+00 | 6.87E-05 | 2.97E-04 | -2.03E+00 | 1.56E-02 | 6.63E-02 | -1.43E+00 | 4.58E-03 | 1.75E-02 | -1.62E+00 | 1.25E-07 | 4.46E-07 | -1.50E+00 | 7.99E-08 | 2.25E-07 | -1.48E+00 | 2.90E-04 | 1.07E-03 | -1.10E+00 | 1.25E-06 | 3.42E-06 | -1.32E+00 |
| MALAT1 | 4.22E-05 | 2.20E-04 | 1.13E+00 | 1.04E-08 | 4.51E-08 | 1.21E+00 | 2.99E-02 | 1.27E-01 | 1.11E+00 | 2.70E-05 | 1.03E-04 | 1.22E+00 | 1.12E-04 | 4.00E-04 | 1.24E+00 | 1.23E-02 | 3.45E-02 | 1.12E+00 | 4.01E-04 | 1.47E-03 | 1.35E+00 | 1.59E-04 | 4.37E-04 | 1.28E+00 |
| DDX20 | 4.23E-05 | 2.21E-04 | -1.19E+00 | 8.76E-05 | 3.79E-04 | -1.07E+00 | 3.45E-05 | 1.47E-04 | -1.38E+00 | 3.94E-04 | 1.50E-03 | -1.08E+00 | 2.85E-03 | 1.02E-02 | -1.01E+00 | 9.66E-04 | 2.72E-03 | -1.18E+00 | 7.27E-03 | 2.67E-02 | -1.12E+00 | 2.29E-01 | 6.28E-01 | 1.14E+00 |
| HOXA5 | 4.24E-05 | 2.21E-04 | -1.17E+00 | 1.74E-01 | 7.53E-01 | 1.43E+00 | 4.22E-04 | 1.79E-03 | -1.27E+00 | 2.42E-02 | 9.23E-02 | 1.06E+00 | 1.92E-01 | 6.85E-01 | 1.27E+00 | 2.38E-01 | 6.71E-01 | 1.52E+00 | 1.47E-01 | 5.39E-01 | -1.04E+00 | 3.62E-01 | 9.94E-01 | 1.16E+00 |
| CD58 | 4.25E-05 | 2.21E-04 | -1.25E+00 | 1.55E-02 | 6.71E-02 | 1.20E+00 | 4.43E-03 | 1.88E-02 | -1.15E+00 | 1.93E-04 | 7.37E-04 | -1.12E+00 | 2.51E-05 | 8.96E-05 | -1.71E+00 | 1.06E-01 | 2.99E-01 | 1.24E+00 | 3.19E-05 | 1.17E-04 | -1.19E+00 | 2.89E-01 | 7.93E-01 | 1.35E+00 |
| SERPINE1 | 4.25E-05 | 2.21E-04 | 4.08E+00 | 1.21E-01 | 5.23E-01 | 1.82E+00 | 1.84E-01 | 7.82E-01 | 1.28E+00 | 4.56E-02 | 1.74E-01 | -1.26E+00 | 2.26E-04 | 8.07E-04 | 5.38E+00 | 1.63E-01 | 4.60E-01 | -1.50E+00 | 4.74E-12 | 1.74E-11 | 1.12E+01 | 1.59E-01 | 4.37E-01 | 3.76E+00 |
| SPATA5L1 | 4.25E-05 | 2.22E-04 | -1.79E+00 | 2.84E-02 | 1.23E-01 | 1.16E+00 | 1.71E-02 | 7.27E-02 | -1.33E+00 | 2.55E-01 | 9.74E-01 | 1.22E+00 | 7.58E-02 | 2.70E-01 | -1.20E+00 | 2.60E-01 | 7.33E-01 | 1.28E+00 | 1.26E-10 | 4.63E-10 | -1.93E+00 | 3.32E-04 | 9.12E-04 | -1.12E+00 |
| SUPT5H | 4.26E-05 | 2.22E-04 | -1.10E+00 | 4.98E-06 | 2.15E-05 | -1.07E+00 | 3.65E-02 | 1.55E-01 | 1.08E+00 | 1.11E-01 | 4.25E-01 | 1.29E+00 | 1.18E-02 | 4.22E-02 | 1.08E+00 | 2.60E-03 | 7.33E-03 | 1.02E+00 | 1.08E-01 | 3.98E-01 | 1.13E+00 | 2.83E-01 | 7.76E-01 | 1.25E+00 |
| BSDC1 | 4.27E-05 | 2.23E-04 | 1.02E+00 | 7.71E-12 | 3.33E-11 | -1.17E+00 | 4.09E-03 | 1.74E-02 | -1.05E+00 | 2.00E-06 | 7.66E-06 | -1.20E+00 | 1.02E-01 | 3.65E-01 | 1.21E+00 | 8.64E-03 | 2.43E-02 | 1.05E+00 | 4.43E-12 | 1.63E-11 | -1.50E+00 | 7.08E-14 | 1.94E-13 | -1.66E+00 |
| ZEB1 | 4.28E-05 | 2.23E-04 | -1.05E+00 | 1.91E-06 | 8.24E-06 | -1.05E+00 | 2.26E-01 | 9.62E-01 | 1.15E+00 | 4.00E-02 | 1.53E-01 | 1.12E+00 | 1.56E-01 | 5.56E-01 | 1.22E+00 | 7.63E-02 | 2.15E-01 | -1.07E+00 | 2.50E-01 | 9.19E-01 | 1.22E+00 | 3.38E-01 | 9.30E-01 | 1.27E+00 |
| SOGA1 | 4.33E-05 | 2.26E-04 | -1.39E+00 | 8.34E-03 | 3.60E-02 | 1.08E+00 | 1.61E-06 | 6.83E-06 | -2.19E+00 | 1.27E-03 | 4.87E-03 | -1.29E+00 | 8.67E-03 | 3.09E-02 | -1.18E+00 | 4.42E-02 | 1.25E-01 | -1.00E+00 | 1.17E-02 | 4.28E-02 | -1.25E+00 | 2.70E-01 | 7.42E-01 | 1.15E+00 |
| ZC4H2 | 4.34E-05 | 2.26E-04 | -1.34E+00 | 2.21E-04 | 9.53E-04 | -1.13E+00 | 9.83E-03 | 4.18E-02 | -1.15E+00 | 1.17E-02 | 4.46E-02 | 1.06E+00 | 8.71E-04 | 3.10E-03 | -1.52E+00 | 9.11E-02 | 2.57E-01 | 1.12E+00 | 3.99E-03 | 1.47E-02 | -1.29E+00 | 5.85E-03 | 1.61E-02 | -1.26E+00 |
| NIPAL2 | 4.35E-05 | 2.27E-04 | -1.27E+00 | 1.28E-02 | 5.53E-02 | 1.17E+00 | 5.98E-05 | 2.54E-04 | -1.27E+00 | 2.98E-03 | 1.14E-02 | 1.05E+00 | 4.26E-02 | 1.52E-01 | -1.01E+00 | 2.95E-01 | 8.31E-01 | 1.31E+00 | 1.55E-02 | 5.71E-02 | -1.09E+00 | 4.95E-02 | 1.36E-01 | -1.02E+00 |
| FAM210B | 4.36E-05 | 2.27E-04 | -1.09E+00 | 2.08E-02 | 8.99E-02 | 1.25E+00 | 1.02E-02 | 4.34E-02 | -1.17E+00 | 7.70E-02 | 2.94E-01 | 1.16E+00 | 7.37E-03 | 2.63E-02 | -1.02E+00 | 2.11E-01 | 5.95E-01 | 1.28E+00 | 5.41E-04 | 1.99E-03 | -1.23E+00 | 4.20E-03 | 1.15E-02 | -1.14E+00 |
| MAP3K11 | 4.37E-05 | 2.28E-04 | -1.11E+00 | 1.20E-04 | 5.20E-04 | 1.00E+00 | 9.80E-08 | 4.17E-07 | -1.54E+00 | 4.06E-04 | 1.55E-03 | -1.05E+00 | 1.93E-05 | 6.87E-05 | -1.29E+00 | 3.18E-04 | 8.97E-04 | -1.10E+00 | 9.32E-05 | 3.42E-04 | -1.17E+00 | 5.92E-03 | 1.63E-02 | -1.00E+00 |
| TM7SF2 | 4.39E-05 | 2.29E-04 | -1.13E+00 | 6.00E-05 | 2.59E-04 | -1.01E+00 | 8.23E-03 | 3.50E-02 | -1.18E+00 | 3.73E-02 | 1.42E-01 | 1.05E+00 | 2.13E-01 | 7.59E-01 | 1.26E+00 | 1.55E-01 | 4.38E-01 | 1.33E+00 | 3.01E-02 | 1.11E-01 | 1.68E+00 | 1.26E-01 | 3.45E-01 | 1.57E+00 |
| PSMG1 | 4.39E-05 | 2.29E-04 | 1.01E+00 | 1.10E-02 | 4.75E-02 | 1.27E+00 | 2.44E-03 | 1.04E-02 | 1.00E+00 | 3.20E-02 | 1.22E-01 | 1.22E+00 | 5.05E-08 | 1.80E-07 | -1.37E+00 | 7.95E-03 | 2.24E-02 | 1.03E+00 | 3.50E-04 | 1.28E-03 | -1.10E+00 | 2.79E-01 | 7.65E-01 | 1.29E+00 |
| NXT2 | 4.40E-05 | 2.29E-04 | -1.41E+00 | 7.99E-04 | 3.45E-03 | -1.12E+00 | 1.18E-02 | 5.01E-02 | -1.06E+00 | 2.66E-05 | 1.02E-04 | -1.15E+00 | 1.78E-02 | 6.35E-02 | -1.13E+00 | 9.32E-02 | 2.63E-01 | 1.07E+00 | 1.94E-04 | 7.12E-04 | -1.19E+00 | 6.13E-02 | 1.68E-01 | 1.06E+00 |
| HYAL3 | 4.40E-05 | 2.29E-04 | -2.01E+00 | 2.20E-03 | 9.50E-03 | -1.22E+00 | 1.04E-04 | 4.41E-04 | -1.59E+00 | 2.59E-02 | 9.88E-02 | 1.08E+00 | 1.79E-16 | 6.39E-16 | -1.79E+00 | 9.20E-07 | 2.59E-06 | -1.09E+00 | 8.56E-06 | 3.14E-05 | -1.02E+00 | 7.44E-03 | 2.04E-02 | 1.17E+00 |
| CEBPZOS | 4.41E-05 | 2.30E-04 | -1.18E+00 | 2.73E-03 | 1.18E-02 | 1.11E+00 | 5.89E-04 | 2.50E-03 | -1.14E+00 | 5.57E-03 | 2.13E-02 | 1.11E+00 | 4.02E-05 | 1.43E-04 | -1.07E+00 | 9.85E-05 | 2.78E-04 | -1.01E+00 | 3.64E-02 | 1.34E-01 | -1.01E+00 | 4.27E-02 | 1.17E-01 | 1.04E+00 |
| CYB5D2 | 4.41E-05 | 2.30E-04 | -1.38E+00 | 1.29E-05 | 5.59E-05 | -1.42E+00 | 3.04E-03 | 1.29E-02 | -1.16E+00 | 2.05E-07 | 7.83E-07 | -1.57E+00 | 5.75E-03 | 2.05E-02 | -1.47E+00 | 1.15E-02 | 3.24E-02 | -1.34E+00 | 7.04E-02 | 2.58E-01 | 1.26E+00 | 1.57E-02 | 4.31E-02 | -1.12E+00 |
| FAM172A | 4.42E-05 | 2.30E-04 | -1.09E+00 | 2.48E-04 | 1.07E-03 | 1.03E+00 | 1.00E-04 | 4.26E-04 | -1.31E+00 | 5.14E-03 | 1.96E-02 | -1.00E+00 | 1.60E-02 | 5.71E-02 | 1.08E+00 | 1.16E-02 | 3.28E-02 | 1.08E+00 | 4.06E-04 | 1.49E-03 | -1.15E+00 | 7.48E-03 | 2.05E-02 | -1.06E+00 |
| CHIC1 | 4.42E-05 | 2.30E-04 | -1.49E+00 | 7.44E-05 | 3.21E-04 | -1.29E+00 | 1.71E-04 | 7.25E-04 | -1.51E+00 | 4.59E-03 | 1.75E-02 | -1.07E+00 | 4.64E-02 | 1.65E-01 | -1.32E+00 | 2.36E-01 | 6.64E-01 | 1.13E+00 | 1.79E-02 | 6.58E-02 | -1.18E+00 | 9.43E-02 | 2.59E-01 | -1.09E+00 |
| FAM86C2P | 4.44E-05 | 2.32E-04 | -1.96E+00 | 4.30E-04 | 1.86E-03 | -1.40E+00 | 5.04E-06 | 2.14E-05 | -1.52E+00 | 3.02E-07 | 1.16E-06 | -1.62E+00 | 7.88E-02 | 2.81E-01 | -1.28E+00 | 3.33E-01 | 9.39E-01 | 1.36E+00 | 2.41E-10 | 8.85E-10 | -2.27E+00 | 2.48E-07 | 6.82E-07 | -1.71E+00 |
| IDE | 4.45E-05 | 2.32E-04 | -1.79E+00 | 2.04E-02 | 8.83E-02 | 1.06E+00 | 1.90E-02 | 8.07E-02 | -1.10E+00 | 5.93E-02 | 2.27E-01 | 1.14E+00 | 1.73E-05 | 6.18E-05 | -1.46E+00 | 1.29E-02 | 3.63E-02 | 1.04E+00 | 2.81E-03 | 1.03E-02 | -1.05E+00 | 3.42E-02 | 9.39E-02 | 1.04E+00 |
| MRAS | 4.46E-05 | 2.32E-04 | -1.79E+00 | 7.06E-04 | 3.05E-03 | -1.28E+00 | 6.42E-02 | 2.73E-01 | -1.15E+00 | 3.03E-02 | 1.16E-01 | -1.18E+00 | 7.19E-04 | 2.56E-03 | -1.59E+00 | 4.69E-04 | 1.32E-03 | -1.63E+00 | 1.95E-01 | 7.16E-01 | 1.29E+00 | 3.36E-01 | 9.22E-01 | 1.01E+00 |
| GRK6 | 4.46E-05 | 2.32E-04 | -1.11E+00 | 1.01E-04 | 4.36E-04 | 1.05E+00 | 4.90E-04 | 2.08E-03 | -1.08E+00 | 7.48E-02 | 2.86E-01 | 1.25E+00 | 1.22E-05 | 4.35E-05 | -1.39E+00 | 3.05E-02 | 8.61E-02 | 1.12E+00 | 2.33E-01 | 8.53E-01 | 1.29E+00 | 2.73E-01 | 7.49E-01 | 1.26E+00 |
| AP006296.1 | 4.46E-05 | 2.33E-04 | -1.45E+00 | 7.96E-07 | 3.44E-06 | -1.65E+00 | 1.35E-07 | 5.74E-07 | -1.43E+00 | 2.65E-10 | 1.01E-09 | -1.49E+00 | 1.12E-06 | 4.00E-06 | -2.03E+00 | 2.39E-06 | 6.74E-06 | -1.92E+00 | 1.19E-14 | 4.36E-14 | -2.11E+00 | 1.19E-11 | 3.28E-11 | -1.83E+00 |
| EML3 | 4.48E-05 | 2.33E-04 | -1.18E+00 | 2.97E-04 | 1.28E-03 | -1.05E+00 | 8.48E-06 | 3.60E-05 | -1.52E+00 | 1.34E-03 | 5.13E-03 | -1.04E+00 | 1.73E-07 | 6.18E-07 | -1.65E+00 | 1.25E-04 | 3.53E-04 | -1.18E+00 | 4.17E-03 | 1.53E-02 | -1.07E+00 | 1.66E-01 | 4.56E-01 | 1.14E+00 |
| DNAJC25 | 4.48E-05 | 2.34E-04 | -1.51E+00 | 2.93E-05 | 1.26E-04 | -1.38E+00 | 1.38E-06 | 5.88E-06 | -1.49E+00 | 1.14E-06 | 4.34E-06 | -1.33E+00 | 5.79E-04 | 2.06E-03 | -1.28E+00 | 2.13E-05 | 6.01E-05 | -1.67E+00 | 7.68E-04 | 2.82E-03 | -1.18E+00 | 2.92E-04 | 8.02E-04 | -1.36E+00 |
| WARS2 | 4.48E-05 | 2.34E-04 | -1.32E+00 | 6.17E-05 | 2.67E-04 | -1.12E+00 | 3.31E-05 | 1.41E-04 | -1.25E+00 | 4.45E-05 | 1.70E-04 | -1.11E+00 | 4.36E-04 | 1.55E-03 | -1.28E+00 | 5.03E-03 | 1.42E-02 | -1.09E+00 | 1.37E-02 | 5.05E-02 | -1.13E+00 | 2.66E-02 | 7.30E-02 | -1.12E+00 |
| NFXL1 | 4.48E-05 | 2.34E-04 | -1.34E+00 | 2.14E-03 | 9.24E-03 | 1.01E+00 | 1.60E-03 | 6.79E-03 | -1.20E+00 | 2.26E-03 | 8.65E-03 | -1.01E+00 | 2.00E-02 | 7.14E-02 | -1.15E+00 | 1.77E-02 | 5.00E-02 | -1.14E+00 | 4.41E-02 | 1.62E-01 | -1.12E+00 | 3.14E-01 | 8.64E-01 | 1.18E+00 |
| SCNM1 | 4.50E-05 | 2.35E-04 | 1.00E+00 | 1.47E-03 | 6.35E-03 | 1.19E+00 | 2.10E-04 | 8.92E-04 | -1.11E+00 | 7.98E-04 | 3.05E-03 | 1.09E+00 | 1.05E-06 | 3.75E-06 | -1.22E+00 | 5.05E-06 | 1.42E-05 | -1.14E+00 | 1.55E-05 | 5.70E-05 | -1.10E+00 | 7.75E-06 | 2.13E-05 | -1.10E+00 |
| C12orf73 | 4.51E-05 | 2.35E-04 | -1.18E+00 | 2.18E-06 | 9.43E-06 | -1.25E+00 | 1.33E-03 | 5.63E-03 | -1.15E+00 | 6.09E-05 | 2.33E-04 | -1.17E+00 | 2.99E-07 | 1.06E-06 | -1.41E+00 | 2.62E-07 | 7.38E-07 | -1.47E+00 | 7.64E-06 | 2.80E-05 | -1.19E+00 | 2.71E-07 | 7.46E-07 | -1.38E+00 |
| LINC01341 | 4.51E-05 | 2.35E-04 | -1.48E+00 | 3.52E-02 | 1.52E-01 | 1.26E+00 | 1.11E-05 | 4.70E-05 | -2.00E+00 | 3.85E-03 | 1.47E-02 | -1.18E+00 | 6.34E-03 | 2.26E-02 | -1.93E+00 | 2.14E-02 | 6.03E-02 | -1.45E+00 | 2.53E-01 | 9.30E-01 | 1.11E+00 | 1.24E-01 | 3.39E-01 | 1.77E+00 |
| BEND5 | 4.52E-05 | 2.35E-04 | -1.20E+00 | 4.27E-03 | 1.85E-02 | 1.09E+00 | 1.15E-06 | 4.88E-06 | -1.37E+00 | 1.59E-05 | 6.07E-05 | -1.14E+00 | 4.20E-05 | 1.50E-04 | -1.29E+00 | 1.62E-06 | 4.58E-06 | -1.43E+00 | 1.49E-03 | 5.47E-03 | -1.32E+00 | 6.69E-04 | 1.84E-03 | -1.49E+00 |
| AP000568.1 | 4.52E-05 | 2.36E-04 | -1.27E+00 | 2.30E-09 | 9.92E-09 | -2.17E+00 | 1.68E-01 | 7.15E-01 | 1.15E+00 | 1.91E-03 | 7.29E-03 | -1.41E+00 | 2.00E-01 | 7.12E-01 | 1.33E+00 | 2.70E-03 | 7.61E-03 | -1.59E+00 | 1.41E-02 | 5.19E-02 | 1.10E+00 | 2.02E-09 | 5.55E-09 | -1.87E+00 |
| FAM174B | 4.52E-05 | 2.36E-04 | -1.78E+00 | 5.06E-02 | 2.18E-01 | 1.17E+00 | 1.10E-01 | 4.66E-01 | -1.11E+00 | 2.62E-01 | 1.00E+00 | 1.28E+00 | 9.45E-03 | 3.37E-02 | -1.67E+00 | 8.27E-02 | 2.33E-01 | -1.12E+00 | 4.48E-02 | 1.64E-01 | -1.14E+00 | 1.91E-01 | 5.25E-01 | -1.02E+00 |
| ANXA1 | 4.52E-05 | 2.36E-04 | 1.22E+00 | 9.45E-02 | 4.08E-01 | 1.11E+00 | 5.29E-02 | 2.25E-01 | 1.13E+00 | 2.33E-01 | 8.90E-01 | 1.07E+00 | 2.72E-02 | 9.70E-02 | 1.16E+00 | 5.12E-03 | 1.44E-02 | -1.28E+00 | 2.17E-18 | 7.96E-18 | 2.38E+00 | 2.04E-02 | 5.60E-02 | 1.34E+00 |
| DCAF15 | 4.54E-05 | 2.37E-04 | -1.17E+00 | 1.30E-03 | 5.61E-03 | 1.06E+00 | 2.08E-06 | 8.84E-06 | -1.30E+00 | 1.74E-05 | 6.64E-05 | -1.12E+00 | 1.52E-06 | 5.41E-06 | -1.43E+00 | 2.85E-04 | 8.04E-04 | -1.08E+00 | 1.86E-09 | 6.82E-09 | -1.43E+00 | 3.39E-04 | 9.30E-04 | -1.09E+00 |
| TRPT1 | 4.56E-05 | 2.38E-04 | -1.47E+00 | 3.32E-05 | 1.43E-04 | -1.31E+00 | 2.82E-06 | 1.20E-05 | -1.36E+00 | 3.07E-09 | 1.17E-08 | -1.50E+00 | 1.65E-03 | 5.87E-03 | -1.20E+00 | 7.23E-03 | 2.04E-02 | -1.10E+00 | 8.46E-02 | 3.10E-01 | 1.12E+00 | 8.28E-02 | 2.27E-01 | 1.02E+00 |
| NAXD | 4.58E-05 | 2.39E-04 | -1.43E+00 | 3.30E-03 | 1.43E-02 | 1.06E+00 | 3.17E-06 | 1.35E-05 | -1.66E+00 | 1.29E-04 | 4.93E-04 | -1.27E+00 | 3.19E-04 | 1.14E-03 | -1.34E+00 | 9.00E-02 | 2.54E-01 | 1.13E+00 | 2.59E-04 | 9.52E-04 | -1.22E+00 | 6.98E-03 | 1.92E-02 | -1.04E+00 |
| LY6G5C | 4.58E-05 | 2.39E-04 | -1.30E+00 | 9.91E-04 | 4.28E-03 | 1.03E+00 | 1.57E-05 | 6.69E-05 | -1.75E+00 | 1.06E-04 | 4.04E-04 | -1.48E+00 | 1.25E-01 | 4.47E-01 | 1.12E+00 | 1.02E-01 | 2.89E-01 | -1.15E+00 | 1.39E-03 | 5.09E-03 | -1.21E+00 | 1.03E-02 | 2.84E-02 | -1.16E+00 |
| URI1 | 4.58E-05 | 2.39E-04 | -1.01E+00 | 1.44E-06 | 6.21E-06 | 1.05E+00 | 7.79E-02 | 3.31E-01 | 1.12E+00 | 8.73E-04 | 3.34E-03 | 1.10E+00 | 5.16E-02 | 1.84E-01 | 1.04E+00 | 3.09E-02 | 8.70E-02 | 1.08E+00 | 2.30E-01 | 8.46E-01 | 1.24E+00 | 2.93E-01 | 8.06E-01 | 1.21E+00 |
| ZC3H7A | 4.59E-05 | 2.39E-04 | 1.01E+00 | 5.89E-07 | 2.55E-06 | 1.04E+00 | 3.85E-05 | 1.64E-04 | -1.20E+00 | 3.19E-07 | 1.22E-06 | -1.20E+00 | 1.37E-04 | 4.89E-04 | -1.12E+00 | 4.09E-04 | 1.15E-03 | -1.07E+00 | 2.78E-03 | 1.02E-02 | 1.02E+00 | 7.16E-05 | 1.97E-04 | -1.10E+00 |
| TARBP1 | 4.60E-05 | 2.40E-04 | -1.03E+00 | 4.53E-02 | 1.96E-01 | 1.30E+00 | 2.15E-05 | 9.12E-05 | -1.17E+00 | 4.41E-03 | 1.68E-02 | 1.14E+00 | 3.86E-05 | 1.38E-04 | -1.23E+00 | 2.60E-02 | 7.32E-02 | 1.12E+00 | 5.76E-05 | 2.11E-04 | -1.22E+00 | 3.52E-01 | 9.68E-01 | 1.29E+00 |
| BTN2A1 | 4.60E-05 | 2.40E-04 | -1.20E+00 | 2.21E-04 | 9.57E-04 | -1.06E+00 | 6.14E-02 | 2.61E-01 | 1.05E+00 | 1.02E-04 | 3.88E-04 | -1.36E+00 | 2.70E-03 | 9.62E-03 | -1.16E+00 | 1.78E-03 | 5.03E-03 | -1.15E+00 | 3.20E-04 | 1.18E-03 | -1.25E+00 | 3.40E-03 | 9.33E-03 | -1.22E+00 |
| RNASEH2A | 4.60E-05 | 2.40E-04 | -1.10E+00 | 1.90E-03 | 8.22E-03 | 1.15E+00 | 8.20E-02 | 3.48E-01 | 1.18E+00 | 1.63E-01 | 6.22E-01 | 1.17E+00 | 6.37E-06 | 2.27E-05 | -1.36E+00 | 4.95E-03 | 1.40E-02 | 1.02E+00 | 5.65E-02 | 2.07E-01 | 1.12E+00 | 2.51E-01 | 6.89E-01 | 1.39E+00 |
| C15orf40 | 4.60E-05 | 2.40E-04 | -1.11E+00 | 1.02E-02 | 4.40E-02 | 1.26E+00 | 2.14E-02 | 9.08E-02 | 1.00E+00 | 3.42E-02 | 1.31E-01 | 1.21E+00 | 2.07E-03 | 7.38E-03 | -1.10E+00 | 2.83E-01 | 7.98E-01 | 1.32E+00 | 8.49E-04 | 3.12E-03 | -1.02E+00 | 2.94E-02 | 8.09E-02 | 1.04E+00 |
| EIF3J-AS1 | 4.62E-05 | 2.41E-04 | -1.23E+00 | 2.28E-01 | 9.84E-01 | 1.41E+00 | 1.72E-03 | 7.30E-03 | -1.21E+00 | 1.63E-01 | 6.24E-01 | 1.31E+00 | 4.54E-08 | 1.62E-07 | -1.41E+00 | 1.16E-02 | 3.27E-02 | 1.16E+00 | 2.73E-03 | 1.00E-02 | -1.04E+00 | 2.20E-01 | 6.03E-01 | 1.11E+00 |
| PRSS2 | 4.62E-05 | 2.41E-04 | 2.93E+00 | 1.57E-02 | 6.79E-02 | 2.18E+00 | 1.11E-03 | 4.73E-03 | 4.59E+00 | 1.61E-02 | 6.16E-02 | 3.00E+00 | 2.80E-01 | 1.00E+00 | 1.00E+00 | 3.55E-01 | 1.00E+00 | -1.00E+00 | 2.53E-01 | 9.27E-01 | 1.09E+00 | 3.42E-01 | 9.40E-01 | 1.06E+00 |
| ZNF717 | 4.62E-05 | 2.41E-04 | -1.45E+00 | 4.21E-02 | 1.82E-01 | 1.25E+00 | 2.26E-05 | 9.61E-05 | -1.49E+00 | 3.60E-03 | 1.38E-02 | 1.00E+00 | 1.54E-02 | 5.51E-02 | -1.15E+00 | 1.18E-01 | 3.33E-01 | 1.13E+00 | 1.47E-04 | 5.40E-04 | -1.23E+00 | 1.58E-03 | 4.33E-03 | -1.18E+00 |
| EXOSC3 | 4.63E-05 | 2.41E-04 | -1.07E+00 | 8.52E-04 | 3.68E-03 | 1.21E+00 | 2.55E-08 | 1.08E-07 | -1.29E+00 | 9.45E-07 | 3.61E-06 | -1.05E+00 | 1.76E-06 | 6.29E-06 | -1.35E+00 | 1.55E-04 | 4.36E-04 | -1.12E+00 | 5.78E-12 | 2.12E-11 | -1.63E+00 | 3.34E-02 | 9.19E-02 | 1.08E+00 |
| ABCA5 | 4.64E-05 | 2.42E-04 | -1.18E+00 | 2.16E-02 | 9.34E-02 | 1.26E+00 | 4.38E-02 | 1.86E-01 | -1.01E+00 | 2.08E-01 | 7.96E-01 | 1.40E+00 | 7.12E-02 | 2.54E-01 | 1.05E+00 | 2.20E-01 | 6.21E-01 | 1.21E+00 | 1.87E-01 | 6.86E-01 | 1.08E+00 | 2.28E-01 | 6.25E-01 | 1.32E+00 |
| JHY | 4.66E-05 | 2.43E-04 | -1.14E+00 | 2.35E-04 | 1.02E-03 | -1.03E+00 | 7.35E-07 | 3.12E-06 | -1.70E+00 | 1.13E-06 | 4.31E-06 | -1.56E+00 | 1.11E-01 | 3.96E-01 | -1.04E+00 | 1.05E-01 | 2.95E-01 | -1.14E+00 | 5.50E-02 | 2.02E-01 | -1.12E+00 | 4.20E-03 | 1.15E-02 | -1.59E+00 |
| SP2 | 4.68E-05 | 2.44E-04 | -1.32E+00 | 2.58E-04 | 1.12E-03 | -1.13E+00 | 3.06E-04 | 1.30E-03 | -1.42E+00 | 1.35E-02 | 5.16E-02 | 1.05E+00 | 3.60E-03 | 1.28E-02 | -1.04E+00 | 7.38E-06 | 2.08E-05 | -1.58E+00 | 7.29E-05 | 2.68E-04 | -1.31E+00 | 1.80E-04 | 4.94E-04 | -1.31E+00 |
| ZNF726 | 4.68E-05 | 2.44E-04 | -1.24E+00 | 3.29E-05 | 1.42E-04 | -1.13E+00 | 6.76E-02 | 2.87E-01 | 1.07E+00 | 1.96E-01 | 7.49E-01 | 1.34E+00 | 1.11E-01 | 3.97E-01 | -1.00E+00 | 2.66E-01 | 7.51E-01 | 1.22E+00 | 3.63E-03 | 1.33E-02 | -1.06E+00 | 1.78E-01 | 4.88E-01 | 1.19E+00 |
| BCL10 | 4.68E-05 | 2.44E-04 | -1.11E+00 | 1.41E-10 | 6.11E-10 | -1.35E+00 | 5.54E-06 | 2.35E-05 | -1.37E+00 | 1.98E-11 | 7.55E-11 | -1.57E+00 | 2.91E-03 | 1.04E-02 | -1.22E+00 | 4.51E-04 | 1.27E-03 | -1.35E+00 | 1.79E-06 | 6.56E-06 | -1.22E+00 | 4.85E-10 | 1.33E-09 | -1.51E+00 |
| FTO | 4.68E-05 | 2.44E-04 | -1.18E+00 | 2.93E-04 | 1.27E-03 | 1.02E+00 | 4.13E-03 | 1.76E-02 | -1.19E+00 | 1.28E-02 | 4.88E-02 | 1.06E+00 | 9.86E-03 | 3.52E-02 | -1.10E+00 | 7.03E-03 | 1.98E-02 | -1.16E+00 | 1.01E-02 | 3.72E-02 | -1.09E+00 | 3.54E-02 | 9.72E-02 | 1.04E+00 |
| CINP | 4.69E-05 | 2.44E-04 | -1.15E+00 | 8.29E-05 | 3.58E-04 | -1.02E+00 | 6.60E-04 | 2.80E-03 | -1.08E+00 | 1.01E-03 | 3.86E-03 | 1.03E+00 | 1.26E-04 | 4.48E-04 | -1.21E+00 | 2.03E-02 | 5.73E-02 | 1.04E+00 | 1.65E-02 | 6.05E-02 | 1.05E+00 | 1.55E-01 | 4.25E-01 | 1.18E+00 |
| ATP5G2P4 | 4.69E-05 | 2.44E-04 | 1.55E+00 | 7.81E-08 | 3.37E-07 | 1.91E+00 | 1.65E-03 | 7.02E-03 | 1.53E+00 | 2.12E-04 | 8.12E-04 | 1.79E+00 | 1.33E-01 | 4.73E-01 | 1.33E+00 | 5.56E-02 | 1.57E-01 | 1.69E+00 | 8.84E-03 | 3.25E-02 | 1.48E+00 | 1.75E-05 | 4.81E-05 | 1.85E+00 |
| QTRT1 | 4.70E-05 | 2.45E-04 | -1.13E+00 | 2.31E-04 | 9.98E-04 | 1.08E+00 | 2.19E-03 | 9.32E-03 | -1.16E+00 | 3.62E-03 | 1.38E-02 | 1.01E+00 | 1.90E-03 | 6.78E-03 | -1.18E+00 | 2.15E-01 | 6.07E-01 | 1.34E+00 | 2.14E-02 | 7.85E-02 | 1.07E+00 | 2.68E-01 | 7.36E-01 | 1.19E+00 |
| ZNF660 | 4.71E-05 | 2.45E-04 | -1.33E+00 | 1.11E-01 | 4.81E-01 | 1.31E+00 | 9.88E-03 | 4.20E-02 | -1.18E+00 | 1.27E-01 | 4.84E-01 | 1.16E+00 | 9.92E-03 | 3.54E-02 | -1.03E+00 | 2.95E-01 | 8.33E-01 | 1.36E+00 | 4.74E-05 | 1.74E-04 | -1.35E+00 | 3.00E-03 | 8.25E-03 | -1.13E+00 |
| ACP5 | 4.71E-05 | 2.45E-04 | -1.73E+00 | 8.72E-04 | 3.77E-03 | -1.29E+00 | 1.89E-03 | 8.02E-03 | -1.63E+00 | 1.25E-03 | 4.77E-03 | -1.40E+00 | 7.88E-02 | 2.81E-01 | -2.01E+00 | 1.30E-01 | 3.67E-01 | -1.53E+00 | 9.11E-04 | 3.34E-03 | -1.44E+00 | 6.74E-02 | 1.85E-01 | -1.03E+00 |
| AC008957.1 | 4.71E-05 | 2.46E-04 | -1.71E+00 | 1.75E-04 | 7.58E-04 | -1.50E+00 | 1.22E-01 | 5.17E-01 | 1.37E+00 | 2.62E-01 | 1.00E+00 | 1.27E+00 | 2.71E-02 | 9.67E-02 | -1.45E+00 | 2.72E-02 | 7.65E-02 | -1.34E+00 | 9.17E-02 | 3.37E-01 | 1.56E+00 | 2.92E-01 | 8.01E-01 | 1.37E+00 |
| MCOLN2 | 4.74E-05 | 2.47E-04 | -1.57E+00 | 1.93E-02 | 8.34E-02 | 1.09E+00 | 1.59E-07 | 6.76E-07 | -1.79E+00 | 2.01E-05 | 7.67E-05 | -1.24E+00 | 3.08E-02 | 1.10E-01 | -1.41E+00 | 1.10E-01 | 3.09E-01 | -1.13E+00 | 2.29E-02 | 8.41E-02 | -1.48E+00 | 2.91E-02 | 8.01E-02 | -1.61E+00 |
| MORN2 | 4.75E-05 | 2.47E-04 | -1.17E+00 | 3.09E-05 | 1.33E-04 | -1.07E+00 | 2.05E-02 | 8.70E-02 | -1.00E+00 | 2.76E-06 | 1.05E-05 | -1.23E+00 | 8.79E-03 | 3.13E-02 | 1.05E+00 | 4.42E-03 | 1.25E-02 | 1.05E+00 | 2.02E-01 | 7.40E-01 | 1.35E+00 | 3.43E-01 | 9.42E-01 | 1.26E+00 |
| SH3TC1 | 4.77E-05 | 2.48E-04 | -1.21E+00 | 8.65E-02 | 3.74E-01 | 1.34E+00 | 4.94E-04 | 2.10E-03 | -1.66E+00 | 4.43E-03 | 1.69E-02 | -1.14E+00 | 2.71E-01 | 9.67E-01 | 1.45E+00 | 2.47E-01 | 6.97E-01 | -1.22E+00 | 2.63E-01 | 9.65E-01 | -1.02E+00 | 3.22E-01 | 8.84E-01 | -1.15E+00 |
| RPL7AP11 | 4.77E-05 | 2.49E-04 | -1.38E+00 | 5.73E-02 | 2.48E-01 | 1.86E+00 | 1.83E-02 | 7.78E-02 | -1.01E+00 | 1.24E-03 | 4.75E-03 | 2.05E+00 | 1.39E-02 | 4.94E-02 | -1.20E+00 | 2.10E-02 | 5.92E-02 | 1.96E+00 | 3.72E-03 | 1.37E-02 | -1.30E+00 | 2.64E-04 | 7.26E-04 | 1.86E+00 |
| UBA5 | 4.78E-05 | 2.49E-04 | -1.10E+00 | 1.45E-03 | 6.27E-03 | 1.15E+00 | 1.00E-02 | 4.25E-02 | -1.01E+00 | 1.51E-02 | 5.77E-02 | 1.11E+00 | 7.06E-04 | 2.52E-03 | -1.01E+00 | 6.92E-02 | 1.95E-01 | 1.26E+00 | 1.11E-01 | 4.09E-01 | 1.08E+00 | 3.41E-01 | 9.36E-01 | 1.26E+00 |
| ENPP5 | 4.80E-05 | 2.50E-04 | 2.54E+00 | 1.27E-04 | 5.47E-04 | 2.89E+00 | 1.02E-01 | 4.34E-01 | 1.46E+00 | 4.50E-03 | 1.72E-02 | 2.04E+00 | 7.14E-03 | 2.55E-02 | 4.07E+00 | 2.54E-03 | 7.15E-03 | 4.21E+00 | 1.24E-02 | 4.54E-02 | 2.56E+00 | 1.18E-01 | 3.25E-01 | 2.07E+00 |
| RNF145 | 4.83E-05 | 2.51E-04 | -1.20E+00 | 3.58E-10 | 1.55E-09 | -1.60E+00 | 3.77E-07 | 1.60E-06 | -1.50E+00 | 9.18E-11 | 3.51E-10 | -1.72E+00 | 2.26E-02 | 8.04E-02 | 1.11E+00 | 5.08E-07 | 1.43E-06 | -1.58E+00 | 8.25E-05 | 3.03E-04 | -1.02E+00 | 1.09E-08 | 2.99E-08 | -1.36E+00 |
| SMYD4 | 4.83E-05 | 2.52E-04 | -1.33E+00 | 1.24E-02 | 5.35E-02 | 1.21E+00 | 1.06E-05 | 4.49E-05 | -1.55E+00 | 2.42E-02 | 9.25E-02 | 1.09E+00 | 7.34E-03 | 2.62E-02 | -1.29E+00 | 7.25E-02 | 2.04E-01 | 1.04E+00 | 4.83E-06 | 1.77E-05 | -1.39E+00 | 7.76E-03 | 2.13E-02 | -1.03E+00 |
| CREB5 | 4.85E-05 | 2.53E-04 | -1.43E+00 | 3.68E-07 | 1.59E-06 | -2.07E+00 | 1.75E-01 | 7.43E-01 | 1.18E+00 | 2.01E-04 | 7.70E-04 | -1.85E+00 | 1.41E-01 | 5.04E-01 | 1.22E+00 | 1.37E-02 | 3.85E-02 | -1.68E+00 | 1.02E-01 | 3.74E-01 | -1.11E+00 | 1.42E-02 | 3.91E-02 | -1.36E+00 |
| TRAPPC9 | 4.85E-05 | 2.53E-04 | -1.55E+00 | 1.20E-02 | 5.19E-02 | 1.00E+00 | 2.08E-02 | 8.82E-02 | 1.04E+00 | 5.18E-03 | 1.98E-02 | 1.13E+00 | 1.78E-02 | 6.34E-02 | -1.19E+00 | 3.41E-02 | 9.60E-02 | -1.09E+00 | 1.15E-01 | 4.21E-01 | 1.47E+00 | 2.78E-01 | 7.63E-01 | 1.34E+00 |
| ITGA5 | 4.85E-05 | 2.53E-04 | -1.27E+00 | 1.61E-02 | 6.94E-02 | 1.18E+00 | 6.63E-07 | 2.82E-06 | -1.50E+00 | 1.50E-03 | 5.74E-03 | -1.02E+00 | 9.33E-02 | 3.33E-01 | 1.20E+00 | 1.71E-01 | 4.82E-01 | 1.72E+00 | 7.66E-02 | 2.81E-01 | 1.19E+00 | 2.00E-01 | 5.50E-01 | 1.24E+00 |
| AL132656.2 | 4.87E-05 | 2.54E-04 | -1.43E+00 | 4.64E-05 | 2.01E-04 | -1.38E+00 | 2.95E-05 | 1.25E-04 | -1.67E+00 | 1.52E-05 | 5.82E-05 | -1.61E+00 | 1.61E-06 | 5.74E-06 | -1.66E+00 | 5.43E-06 | 1.53E-05 | -1.59E+00 | 2.72E-04 | 1.00E-03 | -1.40E+00 | 7.87E-03 | 2.16E-02 | -1.15E+00 |
| MRPL11 | 4.87E-05 | 2.54E-04 | 1.02E+00 | 8.10E-11 | 3.50E-10 | 1.00E+00 | 1.72E-01 | 7.29E-01 | 1.20E+00 | 1.61E-01 | 6.14E-01 | 1.15E+00 | 6.63E-02 | 2.36E-01 | 1.16E+00 | 2.99E-01 | 8.42E-01 | 1.31E+00 | 1.12E-01 | 4.12E-01 | 1.11E+00 | 2.32E-01 | 6.36E-01 | 1.18E+00 |
| ELMOD3 | 4.87E-05 | 2.54E-04 | -1.40E+00 | 2.22E-01 | 9.57E-01 | 1.59E+00 | 3.59E-05 | 1.53E-04 | -1.42E+00 | 9.84E-03 | 3.76E-02 | 1.10E+00 | 2.51E-03 | 8.94E-03 | -1.22E+00 | 2.09E-01 | 5.90E-01 | 1.21E+00 | 5.12E-02 | 1.88E-01 | 1.07E+00 | 2.42E-01 | 6.65E-01 | 1.33E+00 |
| ETV3 | 4.88E-05 | 2.55E-04 | 2.02E+00 | 4.87E-04 | 2.11E-03 | -1.19E+00 | 2.32E-01 | 9.86E-01 | 1.32E+00 | 4.68E-04 | 1.79E-03 | -1.25E+00 | 2.36E-01 | 8.41E-01 | 1.41E+00 | 3.54E-04 | 9.98E-04 | -1.43E+00 | 1.62E-01 | 5.94E-01 | 1.34E+00 | 1.57E-02 | 4.31E-02 | -1.14E+00 |
| THNSL2 | 4.89E-05 | 2.55E-04 | -1.61E+00 | 1.61E-03 | 6.96E-03 | -1.15E+00 | 6.68E-03 | 2.84E-02 | -1.25E+00 | 1.13E-02 | 4.33E-02 | -1.06E+00 | 5.55E-05 | 1.98E-04 | -1.37E+00 | 2.08E-03 | 5.88E-03 | -1.06E+00 | 2.97E-03 | 1.09E-02 | -1.06E+00 | 1.83E-03 | 5.03E-03 | -1.20E+00 |
| C12orf65 | 4.89E-05 | 2.55E-04 | -1.08E+00 | 2.26E-04 | 9.76E-04 | 1.10E+00 | 7.53E-10 | 3.20E-09 | -1.61E+00 | 1.38E-04 | 5.25E-04 | -1.05E+00 | 9.39E-12 | 3.35E-11 | -1.89E+00 | 1.97E-06 | 5.54E-06 | -1.29E+00 | 8.51E-21 | 3.13E-20 | -1.93E+00 | 1.55E-05 | 4.27E-05 | -1.08E+00 |
| GSAP | 4.90E-05 | 2.55E-04 | -1.31E+00 | 8.27E-04 | 3.58E-03 | 1.01E+00 | 3.98E-05 | 1.69E-04 | -1.44E+00 | 1.20E-03 | 4.56E-03 | -1.17E+00 | 1.21E-02 | 4.33E-02 | -1.32E+00 | 3.00E-02 | 8.45E-02 | -1.12E+00 | 8.69E-05 | 3.19E-04 | -1.24E+00 | 4.12E-03 | 1.13E-02 | -1.16E+00 |
| SUSD3 | 4.92E-05 | 2.56E-04 | -1.17E+00 | 4.03E-03 | 1.74E-02 | 1.20E+00 | 1.51E-05 | 6.42E-05 | -1.35E+00 | 1.69E-06 | 6.47E-06 | -1.27E+00 | 5.04E-03 | 1.80E-02 | -1.66E+00 | 3.09E-02 | 8.71E-02 | -1.27E+00 | 1.80E-01 | 6.61E-01 | 1.10E+00 | 2.99E-01 | 8.22E-01 | 1.05E+00 |
| MINOS1 | 4.94E-05 | 2.58E-04 | -1.14E+00 | 9.96E-12 | 4.30E-11 | -1.61E+00 | 2.41E-04 | 1.02E-03 | -1.13E+00 | 7.69E-19 | 2.94E-18 | -1.93E+00 | 2.27E-04 | 8.10E-04 | -1.30E+00 | 1.34E-05 | 3.79E-05 | -1.45E+00 | 1.28E-08 | 4.69E-08 | -1.26E+00 | 5.98E-17 | 1.64E-16 | -1.61E+00 |
| POLR1B | 4.95E-05 | 2.58E-04 | -1.63E+00 | 3.59E-02 | 1.55E-01 | 1.20E+00 | 2.76E-09 | 1.17E-08 | -1.85E+00 | 2.42E-02 | 9.26E-02 | 1.21E+00 | 3.60E-02 | 1.28E-01 | -1.42E+00 | 3.50E-01 | 9.88E-01 | 1.25E+00 | 2.44E-05 | 8.95E-05 | -1.47E+00 | 1.94E-01 | 5.34E-01 | 1.20E+00 |
| SLC25A28 | 4.96E-05 | 2.59E-04 | -1.13E+00 | 6.79E-07 | 2.93E-06 | -1.10E+00 | 7.17E-09 | 3.05E-08 | -1.47E+00 | 2.75E-10 | 1.05E-09 | -1.45E+00 | 6.65E-02 | 2.37E-01 | 1.11E+00 | 3.37E-02 | 9.49E-02 | 1.06E+00 | 1.43E-05 | 5.26E-05 | -1.31E+00 | 2.21E-04 | 6.06E-04 | -1.21E+00 |
| CIZ1 | 4.98E-05 | 2.60E-04 | -1.09E+00 | 2.03E-05 | 8.76E-05 | 1.03E+00 | 4.91E-03 | 2.09E-02 | -1.14E+00 | 3.61E-03 | 1.38E-02 | -1.00E+00 | 3.34E-04 | 1.19E-03 | -1.09E+00 | 1.88E-03 | 5.31E-03 | 1.01E+00 | 9.35E-06 | 3.43E-05 | -1.15E+00 | 9.48E-05 | 2.60E-04 | -1.09E+00 |
| COMT | 4.99E-05 | 2.60E-04 | -1.03E+00 | 1.53E-05 | 6.61E-05 | 1.06E+00 | 7.43E-02 | 3.16E-01 | -1.04E+00 | 1.94E-01 | 7.41E-01 | 1.08E+00 | 8.22E-03 | 2.93E-02 | 1.04E+00 | 2.16E-01 | 6.09E-01 | 1.23E+00 | 1.73E-01 | 6.35E-01 | 1.13E+00 | 2.56E-01 | 7.04E-01 | 1.20E+00 |
| HBS1L | 4.99E-05 | 2.60E-04 | -1.06E+00 | 2.05E-10 | 8.88E-10 | -1.21E+00 | 1.33E-02 | 5.66E-02 | 1.02E+00 | 7.26E-06 | 2.77E-05 | -1.12E+00 | 1.69E-02 | 6.04E-02 | -1.01E+00 | 3.18E-01 | 8.96E-01 | 1.44E+00 | 2.58E-01 | 9.48E-01 | 1.17E+00 | 1.05E-01 | 2.89E-01 | 1.46E+00 |
| DET1 | 5.00E-05 | 2.61E-04 | -1.72E+00 | 3.09E-03 | 1.33E-02 | -1.09E+00 | 1.60E-04 | 6.78E-04 | -1.40E+00 | 1.52E-02 | 5.79E-02 | 1.10E+00 | 4.62E-04 | 1.65E-03 | -1.56E+00 | 1.81E-02 | 5.10E-02 | -1.12E+00 | 1.42E-08 | 5.21E-08 | -1.95E+00 | 2.94E-02 | 8.09E-02 | 1.01E+00 |
| AP000356.3 | 5.00E-05 | 2.61E-04 | -1.60E+00 | 7.19E-03 | 3.11E-02 | 1.03E+00 | 2.31E-01 | 9.80E-01 | 1.19E+00 | 1.63E-01 | 6.21E-01 | 1.18E+00 | 3.13E-03 | 1.12E-02 | -1.85E+00 | 1.78E-02 | 5.01E-02 | -1.32E+00 | 2.87E-06 | 1.05E-05 | -2.10E+00 | 1.43E-05 | 3.94E-05 | -1.85E+00 |
| FUT10 | 5.01E-05 | 2.61E-04 | -1.96E+00 | 1.02E-01 | 4.39E-01 | 1.19E+00 | 3.16E-07 | 1.34E-06 | -2.06E+00 | 3.16E-04 | 1.21E-03 | -1.28E+00 | 2.10E-04 | 7.48E-04 | -2.10E+00 | 4.41E-02 | 1.24E-01 | -1.09E+00 | 5.46E-04 | 2.01E-03 | -1.42E+00 | 3.20E-01 | 8.78E-01 | 1.38E+00 |
| CEP162 | 5.02E-05 | 2.62E-04 | -1.47E+00 | 1.35E-02 | 5.82E-02 | 1.10E+00 | 7.58E-06 | 3.22E-05 | -1.56E+00 | 7.86E-04 | 3.00E-03 | -1.13E+00 | 2.61E-03 | 9.31E-03 | -1.25E+00 | 1.19E-01 | 3.35E-01 | 1.13E+00 | 1.35E-04 | 4.96E-04 | -1.22E+00 | 9.12E-03 | 2.51E-02 | -1.06E+00 |
| CENPT | 5.02E-05 | 2.62E-04 | -1.23E+00 | 5.62E-03 | 2.43E-02 | 1.14E+00 | 1.68E-02 | 7.14E-02 | -1.01E+00 | 7.85E-02 | 3.00E-01 | 1.23E+00 | 4.68E-06 | 1.67E-05 | -1.48E+00 | 1.34E-02 | 3.77E-02 | 1.01E+00 | 1.05E-05 | 3.84E-05 | -1.13E+00 | 1.90E-03 | 5.23E-03 | -1.01E+00 |
| CHORDC1 | 5.02E-05 | 2.62E-04 | -1.06E+00 | 3.36E-11 | 1.45E-10 | -1.30E+00 | 4.70E-04 | 2.00E-03 | -1.15E+00 | 5.09E-06 | 1.94E-05 | -1.11E+00 | 2.26E-02 | 8.07E-02 | 1.11E+00 | 4.78E-04 | 1.35E-03 | -1.08E+00 | 9.62E-03 | 3.53E-02 | 1.08E+00 | 3.76E-02 | 1.03E-01 | 1.03E+00 |
| SELL | 5.02E-05 | 2.62E-04 | -1.13E+00 | 1.25E-01 | 5.41E-01 | 1.20E+00 | 7.39E-06 | 3.14E-05 | -1.42E+00 | 2.40E-03 | 9.17E-03 | -1.12E+00 | 2.19E-29 | 7.81E-29 | -2.52E+00 | 3.91E-31 | 1.10E-30 | -2.32E+00 | 6.04E-05 | 2.22E-04 | -2.27E+00 | 2.21E-02 | 6.07E-02 | -1.53E+00 |
| NSUN6 | 5.03E-05 | 2.62E-04 | -1.03E+00 | 3.64E-09 | 1.57E-08 | -1.16E+00 | 3.45E-02 | 1.47E-01 | 1.05E+00 | 1.75E-02 | 6.69E-02 | 1.11E+00 | 4.13E-02 | 1.47E-01 | 1.21E+00 | 8.49E-02 | 2.39E-01 | 1.21E+00 | 5.69E-05 | 2.09E-04 | -1.37E+00 | 1.29E-05 | 3.55E-05 | -1.48E+00 |
| RILPL2 | 5.03E-05 | 2.62E-04 | 1.06E+00 | 2.32E-09 | 1.00E-08 | 1.04E+00 | 2.49E-02 | 1.06E-01 | 1.11E+00 | 9.71E-04 | 3.71E-03 | 1.05E+00 | 3.59E-02 | 1.28E-01 | 1.17E+00 | 4.59E-04 | 1.29E-03 | 1.04E+00 | 1.99E-02 | 7.32E-02 | 1.05E+00 | 3.16E-02 | 8.67E-02 | 1.09E+00 |
| SLC1A5 | 5.03E-05 | 2.62E-04 | -1.46E+00 | 1.07E-04 | 4.60E-04 | -1.26E+00 | 3.57E-07 | 1.52E-06 | -1.59E+00 | 1.55E-03 | 5.92E-03 | -1.02E+00 | 3.20E-04 | 1.14E-03 | -1.84E+00 | 5.27E-03 | 1.49E-02 | -1.43E+00 | 1.20E-22 | 4.41E-22 | -2.14E+00 | 3.03E-07 | 8.31E-07 | -1.25E+00 |
| DNAJB4 | 5.04E-05 | 2.63E-04 | -1.22E+00 | 2.48E-06 | 1.07E-05 | -1.34E+00 | 1.82E-01 | 7.74E-01 | 1.06E+00 | 1.44E-01 | 5.49E-01 | 1.08E+00 | 9.33E-04 | 3.32E-03 | -1.27E+00 | 2.28E-03 | 6.43E-03 | -1.11E+00 | 8.16E-03 | 3.00E-02 | -1.08E+00 | 1.02E-01 | 2.81E-01 | 1.09E+00 |
| CCDC24 | 5.06E-05 | 2.64E-04 | -1.67E+00 | 2.11E-02 | 9.11E-02 | 1.05E+00 | 1.37E-01 | 5.83E-01 | -1.07E+00 | 6.71E-02 | 2.56E-01 | -1.26E+00 | 1.38E-01 | 4.93E-01 | 1.08E+00 | 8.99E-02 | 2.53E-01 | -1.08E+00 | 4.32E-02 | 1.58E-01 | 1.54E+00 | 7.38E-02 | 2.03E-01 | 1.68E+00 |
| NSRP1P1 | 5.09E-05 | 2.65E-04 | -1.05E+00 | 1.78E-09 | 7.70E-09 | -1.26E+00 | 2.98E-02 | 1.27E-01 | 1.02E+00 | 4.25E-03 | 1.62E-02 | 1.00E+00 | 2.61E-01 | 9.30E-01 | 1.26E+00 | 2.90E-01 | 8.17E-01 | 1.18E+00 | 3.28E-04 | 1.20E-03 | -1.20E+00 | 9.52E-05 | 2.62E-04 | -1.46E+00 |
| STX6 | 5.11E-05 | 2.66E-04 | -1.51E+00 | 7.53E-03 | 3.25E-02 | 1.03E+00 | 6.15E-03 | 2.61E-02 | -1.25E+00 | 7.85E-03 | 3.00E-02 | -1.05E+00 | 1.09E-03 | 3.89E-03 | -1.31E+00 | 2.98E-03 | 8.41E-03 | -1.13E+00 | 5.33E-03 | 1.96E-02 | -1.08E+00 | 2.03E-02 | 5.58E-02 | 1.12E+00 |
| CCL3 | 5.13E-05 | 2.67E-04 | 3.72E+00 | 9.53E-02 | 4.12E-01 | 1.79E+00 | 2.87E-03 | 1.22E-02 | 2.41E+00 | 2.01E-01 | 7.69E-01 | 1.43E+00 | 1.79E-01 | 6.38E-01 | -1.16E+00 | 1.86E-01 | 5.23E-01 | -1.33E+00 | 2.39E-01 | 8.77E-01 | 1.24E+00 | 3.64E-01 | 1.00E+00 | 1.00E+00 |
| MAPK6 | 5.15E-05 | 2.68E-04 | -1.13E+00 | 1.74E-11 | 7.53E-11 | -1.67E+00 | 2.72E-06 | 1.16E-05 | -1.37E+00 | 2.99E-13 | 1.14E-12 | -1.95E+00 | 7.58E-04 | 2.70E-03 | -1.16E+00 | 3.88E-06 | 1.09E-05 | -1.51E+00 | 1.03E-04 | 3.77E-04 | -1.24E+00 | 2.92E-06 | 8.01E-06 | -1.45E+00 |
| BMI1 | 5.17E-05 | 2.69E-04 | -1.10E+00 | 1.08E-01 | 4.66E-01 | 1.58E+00 | 2.77E-05 | 1.18E-04 | -1.24E+00 | 3.17E-02 | 1.21E-01 | 1.22E+00 | 2.28E-03 | 8.12E-03 | -1.10E+00 | 1.93E-01 | 5.44E-01 | 1.34E+00 | 4.75E-05 | 1.74E-04 | -1.26E+00 | 1.76E-01 | 4.85E-01 | 1.19E+00 |
| MEG3 | 5.17E-05 | 2.69E-04 | -1.09E+00 | 1.01E-01 | 4.38E-01 | 1.29E+00 | 7.80E-02 | 3.32E-01 | -1.04E+00 | 3.60E-04 | 1.37E-03 | 1.60E+00 | 1.95E-01 | 6.96E-01 | 1.62E+00 | 6.94E-02 | 1.96E-01 | 1.88E+00 | 1.61E-02 | 5.89E-02 | 2.76E+00 | 1.38E-04 | 3.79E-04 | 2.54E+00 |
| RSPH3 | 5.21E-05 | 2.72E-04 | -1.53E+00 | 9.85E-03 | 4.26E-02 | 1.03E+00 | 4.85E-03 | 2.06E-02 | -1.24E+00 | 3.22E-03 | 1.23E-02 | -1.07E+00 | 2.49E-06 | 8.88E-06 | -2.03E+00 | 8.37E-05 | 2.36E-04 | -1.47E+00 | 3.96E-02 | 1.45E-01 | -1.01E+00 | 5.10E-03 | 1.40E-02 | -1.28E+00 |
| STX8 | 5.21E-05 | 2.72E-04 | 1.02E+00 | 3.64E-08 | 1.57E-07 | -1.05E+00 | 3.87E-03 | 1.64E-02 | -1.06E+00 | 9.69E-05 | 3.70E-04 | 1.00E+00 | 6.40E-04 | 2.28E-03 | -1.02E+00 | 2.17E-03 | 6.12E-03 | 1.01E+00 | 2.39E-01 | 8.78E-01 | 1.21E+00 | 4.47E-02 | 1.23E-01 | 1.09E+00 |
| RHOQ | 5.21E-05 | 2.72E-04 | -1.12E+00 | 2.18E-05 | 9.42E-05 | -1.05E+00 | 1.56E-04 | 6.62E-04 | -1.24E+00 | 2.43E-04 | 9.27E-04 | -1.09E+00 | 1.12E-02 | 3.99E-02 | -1.26E+00 | 2.31E-02 | 6.52E-02 | -1.14E+00 | 2.66E-01 | 9.75E-01 | 1.19E+00 | 1.87E-01 | 5.15E-01 | 1.53E+00 |

| | | | | | | | | | | | | | | | | | | | | | | | | |
|---|---|---|---|---|---|---|---|---|---|---|---|---|---|---|---|---|---|---|---|---|---|---|---|---|
| FMO4 | 5.22E-05 | 2.72E-04 | -1.80E+00 | 1.58E-03 | 6.84E-03 | -1.22E+00 | 8.30E-03 | 3.53E-02 | -1.43E+00 | 1.24E-02 | 4.72E-02 | -1.22E+00 | 2.29E-04 | 8.17E-04 | -2.07E+00 | 1.43E-02 | 4.03E-02 | -1.37E+00 | 3.82E-04 | 1.40E-03 | -1.48E+00 | 1.25E-02 | 3.44E-02 | -1.15E+00 |
| UTP11 | 5.23E-05 | 2.72E-04 | -1.11E+00 | 1.25E-03 | 5.42E-03 | 1.12E+00 | 1.95E-04 | 8.27E-04 | -1.15E+00 | 2.45E-03 | 9.35E-03 | 1.06E+00 | 1.47E-07 | 5.25E-07 | -1.63E+00 | 3.99E-03 | 1.13E-02 | -1.00E+00 | 1.32E-06 | 4.86E-06 | -1.26E+00 | 6.07E-02 | 1.67E-01 | 1.17E+00 |
| RPP40 | 5.24E-05 | 2.73E-04 | -1.25E+00 | 1.51E-01 | 6.50E-01 | 1.64E+00 | 1.66E-07 | 7.07E-07 | -1.29E+00 | 2.75E-04 | 1.05E-03 | 1.08E+00 | 3.29E-04 | 1.17E-03 | -1.66E+00 | 2.72E-01 | 7.66E-01 | 1.44E+00 | 1.77E-04 | 6.48E-04 | -1.16E+00 | 3.57E-01 | 9.81E-01 | 1.32E+00 |
| LPIN2 | 5.24E-05 | 2.73E-04 | -1.08E+00 | 4.54E-07 | 1.96E-06 | -1.11E+00 | 1.25E-03 | 5.32E-03 | -1.23E+00 | 1.01E-03 | 3.86E-03 | -1.08E+00 | 4.68E-02 | 1.67E-01 | 1.15E+00 | 1.05E-01 | 2.97E-01 | 1.27E+00 | 9.71E-03 | 3.56E-02 | -1.05E+00 | 2.04E-03 | 5.61E-03 | -1.12E+00 |
| AC022107.1 | 5.24E-05 | 2.73E-04 | -1.51E+00 | 1.45E-04 | 6.26E-04 | -1.35E+00 | 3.49E-04 | 1.48E-03 | -1.62E+00 | 1.87E-03 | 7.16E-03 | -1.33E+00 | 1.00E-02 | 3.57E-02 | -1.05E+00 | 3.97E-03 | 1.12E-02 | -1.35E+00 | 5.40E-03 | 1.98E-02 | -1.23E+00 | 1.19E-01 | 3.27E-01 | 1.06E+00 |
| RASSF1 | 5.24E-05 | 2.73E-04 | 1.02E+00 | 1.52E-07 | 6.57E-07 | 1.02E+00 | 1.47E-02 | 6.24E-02 | 1.03E+00 | 1.59E-02 | 6.08E-02 | 1.08E+00 | 1.40E-02 | 5.01E-02 | 1.25E+00 | 2.83E-03 | 7.99E-03 | 1.15E+00 | 3.28E-03 | 1.20E-02 | 1.57E+00 | 3.36E-01 | 9.24E-01 | 1.28E+00 |
| AL133342.1 | 5.26E-05 | 2.74E-04 | 3.03E+00 | 1.13E-01 | 4.88E-01 | 1.73E+00 | 6.45E-05 | 2.74E-04 | 3.37E+00 | 3.60E-02 | 1.38E-01 | 2.30E+00 | 4.72E-03 | 1.68E-02 | 2.79E+00 | 2.91E-01 | 8.19E-01 | 1.53E+00 | 1.15E-01 | 4.23E-01 | 1.61E+00 | 6.66E-02 | 1.83E-01 | -1.10E+00 |
| CPSF7 | 5.26E-05 | 2.74E-04 | -1.15E+00 | 8.09E-03 | 3.50E-02 | 1.21E+00 | 5.51E-04 | 2.34E-03 | -1.16E+00 | 3.69E-02 | 1.41E-01 | 1.16E+00 | 1.33E-04 | 4.73E-04 | -1.29E+00 | 2.47E-02 | 6.95E-02 | 1.07E+00 | 1.14E-06 | 4.20E-06 | -1.32E+00 | 3.99E-02 | 1.10E-01 | 1.04E+00 |
| COG1 | 5.27E-05 | 2.75E-04 | -1.27E+00 | 6.61E-04 | 2.86E-03 | 1.01E+00 | 2.25E-03 | 9.55E-03 | -1.33E+00 | 6.12E-03 | 2.34E-02 | -1.15E+00 | 3.62E-02 | 1.29E-01 | -1.28E+00 | 6.48E-02 | 1.83E-01 | -1.12E+00 | 1.79E-06 | 6.58E-06 | -1.79E+00 | 1.65E-04 | 4.54E-04 | -1.54E+00 |
| C19orf38 | 5.28E-05 | 2.75E-04 | -1.02E+00 | 3.01E-06 | 1.30E-05 | 1.05E+00 | 2.19E-06 | 9.29E-06 | -1.27E+00 | 3.40E-08 | 1.30E-07 | -1.21E+00 | 1.52E-02 | 5.41E-02 | -1.17E+00 | 5.38E-03 | 1.52E-02 | -1.20E+00 | 1.99E-01 | 7.32E-01 | -1.07E+00 | 1.75E-01 | 4.81E-01 | -1.30E+00 |
| AC007383.1 | 5.29E-05 | 2.76E-04 | -1.48E+00 | 1.98E-06 | 8.54E-06 | -1.62E+00 | 2.42E-04 | 1.03E-03 | -1.36E+00 | 1.64E-06 | 6.26E-06 | -1.55E+00 | 1.71E-03 | 6.09E-03 | -1.27E+00 | 1.31E-03 | 3.68E-03 | -1.28E+00 | 2.35E-07 | 8.63E-07 | -1.62E+00 | 1.22E-07 | 3.35E-07 | -1.64E+00 |
| ZNF347 | 5.30E-05 | 2.76E-04 | -1.52E+00 | 2.93E-05 | 1.27E-04 | -1.34E+00 | 6.26E-05 | 2.66E-04 | -1.65E+00 | 1.22E-03 | 4.68E-03 | -1.28E+00 | 8.74E-03 | 3.12E-02 | -1.17E+00 | 1.13E-01 | 3.18E-01 | 1.11E+00 | 1.92E-07 | 7.04E-07 | -1.39E+00 | 1.71E-04 | 4.70E-04 | -1.11E+00 |
| HUS1 | 5.31E-05 | 2.77E-04 | -1.16E+00 | 5.73E-06 | 2.47E-05 | -1.20E+00 | 1.05E-06 | 4.48E-06 | -1.27E+00 | 5.82E-12 | 2.22E-11 | -1.50E+00 | 1.08E-03 | 3.86E-03 | -1.09E+00 | 7.25E-06 | 2.04E-05 | -1.46E+00 | 3.50E-03 | 1.28E-02 | -1.09E+00 | 4.45E-02 | 1.22E-01 | 1.02E+00 |
| ZC3H3 | 5.33E-05 | 2.78E-04 | -1.40E+00 | 1.84E-03 | 7.94E-03 | 1.01E+00 | 3.86E-05 | 1.64E-04 | -1.51E+00 | 8.99E-04 | 3.43E-03 | -1.17E+00 | 2.28E-06 | 8.13E-06 | -1.80E+00 | 1.09E-05 | 3.08E-05 | -1.75E+00 | 1.46E-06 | 5.35E-06 | -1.66E+00 | 2.75E-04 | 7.55E-04 | -1.34E+00 |
| FAM120AOS | 5.34E-05 | 2.78E-04 | -1.04E+00 | 1.04E-07 | 4.51E-07 | -1.02E+00 | 5.67E-05 | 2.41E-04 | -1.20E+00 | 2.55E-05 | 9.73E-05 | -1.11E+00 | 4.00E-02 | 1.42E-01 | 1.09E+00 | 1.37E-02 | 3.85E-02 | 1.00E+00 | 7.90E-02 | 2.90E-01 | 1.11E+00 | 8.54E-02 | 2.35E-01 | 1.13E+00 |
| ADARB1 | 5.34E-05 | 2.78E-04 | -1.41E+00 | 2.11E-03 | 9.10E-03 | -1.01E+00 | 5.71E-04 | 2.43E-03 | -1.39E+00 | 5.13E-02 | 1.96E-01 | 1.19E+00 | 5.85E-09 | 2.09E-08 | -1.91E+00 | 1.26E-04 | 3.54E-04 | -1.27E+00 | 2.45E-01 | 9.01E-01 | 1.27E+00 | 1.91E-01 | 5.23E-01 | 1.20E+00 |
| RFC3 | 5.36E-05 | 2.79E-04 | -1.69E+00 | 1.81E-03 | 7.81E-03 | -1.19E+00 | 3.22E-02 | 1.37E-01 | -1.01E+00 | 1.29E-01 | 4.94E-01 | 1.22E+00 | 9.77E-02 | 3.48E-01 | -1.05E+00 | 2.55E-01 | 7.20E-01 | 1.28E+00 | 2.73E-06 | 1.00E-05 | -1.29E+00 | 1.10E-02 | 3.02E-02 | 1.02E+00 |
| CCL4 | 5.37E-05 | 2.80E-04 | 8.56E+00 | 1.75E-01 | 7.58E-01 | 1.70E+00 | 1.98E-02 | 8.42E-02 | 1.62E+00 | 2.05E-01 | 7.84E-01 | -1.89E+00 | 1.44E-01 | 5.14E-01 | 8.11E+00 | 3.14E-01 | 8.84E-01 | 2.63E+00 | 1.22E-01 | 4.47E-01 | 5.46E+00 | 3.10E-01 | 8.51E-01 | -1.02E+00 |
| FBXO9 | 5.38E-05 | 2.81E-04 | -1.12E+00 | 3.26E-02 | 1.41E-01 | 1.31E+00 | 2.31E-04 | 9.80E-04 | -1.33E+00 | 5.74E-03 | 2.19E-02 | 1.01E+00 | 9.65E-05 | 3.44E-04 | -1.03E+00 | 7.76E-02 | 2.19E-01 | 1.25E+00 | 1.34E-02 | 4.93E-02 | 1.06E+00 | 4.86E-02 | 1.34E-01 | 1.10E+00 |
| RAD54L2 | 5.40E-05 | 2.81E-04 | -1.10E+00 | 3.81E-03 | 1.65E-02 | 1.23E+00 | 2.20E-02 | 9.33E-02 | -1.07E+00 | 1.69E-01 | 6.46E-01 | 1.26E+00 | 4.59E-02 | 1.64E-01 | 1.12E+00 | 3.00E-01 | 8.46E-01 | 1.41E+00 | 1.23E-01 | 4.53E-01 | 1.19E+00 | 3.48E-01 | 9.57E-01 | 1.41E+00 |
| HIST1H2AI | 5.40E-05 | 2.81E-04 | -1.37E+00 | 1.28E-08 | 5.54E-08 | -1.90E+00 | 2.13E-01 | 9.04E-01 | 2.08E+00 | 6.63E-04 | 2.53E-03 | -1.02E+00 | 4.68E-03 | 1.67E-02 | -1.71E+00 | 2.59E-04 | 7.29E-04 | -2.50E+00 | 6.45E-14 | 2.37E-13 | -1.19E+00 | 1.71E-24 | 4.68E-24 | -2.08E+00 |
| YEATS2 | 5.40E-05 | 2.82E-04 | -1.34E+00 | 2.21E-03 | 9.53E-03 | -1.01E+00 | 6.36E-05 | 2.70E-04 | -1.28E+00 | 2.93E-03 | 1.12E-02 | 1.05E+00 | 9.13E-05 | 3.25E-04 | -1.43E+00 | 1.68E-02 | 4.74E-02 | -1.03E+00 | 9.56E-05 | 3.51E-04 | -1.21E+00 | 7.35E-03 | 2.02E-02 | -1.00E+00 |
| MFSD5 | 5.42E-05 | 2.82E-04 | -1.78E+00 | 9.15E-04 | 3.95E-03 | -1.29E+00 | 7.57E-04 | 3.22E-03 | -1.39E+00 | 5.68E-03 | 2.17E-02 | -1.10E+00 | 1.88E-04 | 6.72E-04 | -1.74E+00 | 1.03E-02 | 2.90E-02 | -1.15E+00 | 1.12E-05 | 4.10E-05 | -1.51E+00 | 1.89E-03 | 5.19E-03 | -1.12E+00 |
| SLC2A13 | 5.42E-05 | 2.83E-04 | -1.53E+00 | 6.08E-03 | 2.63E-02 | 1.05E+00 | 3.26E-04 | 1.39E-03 | -1.43E+00 | 1.55E-03 | 5.92E-03 | -1.19E+00 | 3.36E-02 | 1.20E-01 | -1.05E+00 | 1.98E-01 | 5.59E-01 | 1.33E+00 | 9.68E-02 | 3.55E-01 | 1.05E+00 | 2.46E-01 | 6.76E-01 | 1.48E+00 |
| MCM5 | 5.42E-05 | 2.83E-04 | 1.07E+00 | 2.04E-06 | 8.83E-06 | 1.04E+00 | 2.63E-02 | 1.12E-01 | 1.33E+00 | 1.10E-01 | 4.19E-01 | 1.20E+00 | 2.28E-06 | 8.12E-06 | -1.37E+00 | 2.05E-04 | 5.78E-04 | -1.13E+00 | 1.83E-03 | 6.73E-03 | -1.12E+00 | 1.80E-01 | 4.95E-01 | 1.18E+00 |
| CBX4 | 5.42E-05 | 2.83E-04 | -1.23E+00 | 3.18E-07 | 1.37E-06 | -1.58E+00 | 9.15E-09 | 3.89E-08 | -1.67E+00 | 7.47E-13 | 2.85E-12 | -1.96E+00 | 2.85E-05 | 1.02E-04 | -1.83E+00 | 3.84E-05 | 1.08E-04 | -2.00E+00 | 1.53E-02 | 5.61E-02 | -1.20E+00 | 4.83E-04 | 1.33E-03 | -1.76E+00 |
| CLIC2 | 5.42E-05 | 2.83E-04 | -1.24E+00 | 9.99E-05 | 4.32E-04 | -1.09E+00 | 1.33E-02 | 5.64E-02 | -1.18E+00 | 3.71E-02 | 1.42E-01 | 1.07E+00 | 2.22E-01 | 7.90E-01 | 1.51E+00 | 6.30E-02 | 1.78E-01 | 2.03E+00 | 6.75E-09 | 2.48E-08 | -1.50E+00 | 1.91E-08 | 5.25E-08 | -1.49E+00 |
| ZNF891 | 5.43E-05 | 2.83E-04 | -1.36E+00 | 3.78E-02 | 1.63E-01 | 1.61E+00 | 2.53E-08 | 1.07E-07 | -1.60E+00 | 5.34E-03 | 2.04E-02 | 1.09E+00 | 1.20E-02 | 4.27E-02 | -1.32E+00 | 2.10E-01 | 5.92E-01 | 1.13E+00 | 5.94E-05 | 2.18E-04 | -1.52E+00 | 1.85E-01 | 5.09E-01 | 1.07E+00 |
| ZNF654 | 5.43E-05 | 2.83E-04 | -1.18E+00 | 1.82E-04 | 7.85E-04 | 1.02E+00 | 6.85E-06 | 2.91E-05 | -1.29E+00 | 1.04E-06 | 3.97E-06 | -1.18E+00 | 3.72E-05 | 1.33E-04 | -1.39E+00 | 4.62E-06 | 1.30E-05 | -1.46E+00 | 3.73E-05 | 1.37E-04 | -1.33E+00 | 2.73E-03 | 7.50E-03 | -1.04E+00 |
| MFSD10 | 5.44E-05 | 2.84E-04 | -1.10E+00 | 3.44E-06 | 1.48E-05 | 1.02E+00 | 1.64E-02 | 6.97E-02 | 1.07E+00 | 4.03E-02 | 1.54E-01 | 1.24E+00 | 2.39E-02 | 8.51E-02 | 1.08E+00 | 5.10E-02 | 1.44E-01 | 1.19E+00 | 1.47E-01 | 5.39E-01 | 1.35E+00 | 1.07E-01 | 2.93E-01 | 1.46E+00 |
| FSCN1 | 5.44E-05 | 2.84E-04 | -1.06E+00 | 2.72E-08 | 1.18E-07 | -1.12E+00 | 1.68E-02 | 7.13E-02 | 1.03E+00 | 1.29E-02 | 4.93E-02 | 1.19E+00 | 1.36E-06 | 4.86E-06 | -1.36E+00 | 3.16E-03 | 8.92E-03 | 1.05E+00 | 3.91E-02 | 1.43E-01 | 1.05E+00 | 2.53E-01 | 6.94E-01 | 1.18E+00 |
| TFE3 | 5.45E-05 | 2.84E-04 | -1.12E+00 | 6.88E-04 | 2.97E-03 | 1.11E+00 | 2.32E-05 | 9.86E-05 | -1.52E+00 | 1.80E-04 | 6.89E-04 | -1.27E+00 | 3.71E-03 | 1.32E-02 | -1.25E+00 | 6.49E-04 | 1.83E-03 | -1.52E+00 | 2.68E-05 | 9.84E-05 | -1.69E+00 | 4.74E-04 | 1.30E-03 | -1.43E+00 |
| LZTS3 | 5.46E-05 | 2.84E-04 | -1.11E+00 | 5.84E-12 | 2.52E-11 | -1.61E+00 | 3.14E-02 | 1.33E-01 | -1.08E+00 | 6.60E-04 | 2.52E-03 | -1.37E+00 | 3.54E-02 | 1.26E-01 | -1.70E+00 | 3.45E-02 | 9.71E-02 | -2.10E+00 | 1.40E-01 | 5.13E-01 | -1.10E+00 | 1.35E-01 | 3.71E-01 | -1.47E+00 |
| HSPA13 | 5.46E-05 | 2.85E-04 | -1.12E+00 | 5.00E-08 | 2.16E-07 | -1.33E+00 | 7.57E-04 | 3.22E-03 | -1.11E+00 | 1.07E-02 | 4.08E-02 | 1.11E+00 | 2.01E-06 | 7.16E-06 | -1.15E+00 | 1.07E-07 | 3.01E-07 | -1.39E+00 | 3.56E-05 | 1.31E-04 | -1.27E+00 | 4.69E-04 | 1.29E-03 | -1.19E+00 |
| COX17P1 | 5.47E-05 | 2.85E-04 | -1.38E+00 | 2.35E-09 | 1.02E-08 | -2.32E+00 | 1.53E-04 | 6.49E-04 | -1.17E+00 | 7.93E-15 | 3.03E-14 | -1.99E+00 | 1.74E-09 | 6.21E-09 | -1.81E+00 | 1.30E-15 | 3.67E-15 | -3.08E+00 | 8.64E-17 | 3.17E-16 | -1.66E+00 | 1.28E-29 | 3.52E-29 | -2.59E+00 |
| TTLL5 | 5.47E-05 | 2.85E-04 | -1.36E+00 | 5.31E-04 | 2.29E-03 | -1.06E+00 | 1.47E-05 | 6.26E-05 | -1.49E+00 | 3.97E-04 | 1.52E-03 | -1.13E+00 | 7.57E-05 | 2.70E-04 | -1.46E+00 | 4.01E-04 | 1.13E-03 | -1.34E+00 | 4.33E-03 | 1.59E-02 | -1.03E+00 | 6.01E-02 | 1.65E-01 | 1.06E+00 |
| FAM160A2 | 5.49E-05 | 2.86E-04 | -1.39E+00 | 2.37E-02 | 1.02E-01 | 1.20E+00 | 6.11E-03 | 2.60E-02 | -1.17E+00 | 4.84E-02 | 1.85E-01 | 1.09E+00 | 7.10E-02 | 2.53E-01 | -1.04E+00 | 2.17E-01 | 6.11E-01 | 1.29E+00 | 5.00E-04 | 1.84E-03 | -1.37E+00 | 4.67E-03 | 1.28E-02 | -1.31E+00 |
| FMNL3 | 5.51E-05 | 2.87E-04 | -1.40E+00 | 2.78E-02 | 1.20E-01 | 1.07E+00 | 4.04E-04 | 1.71E-03 | -1.38E+00 | 3.33E-02 | 1.27E-01 | 1.06E+00 | 4.75E-04 | 1.69E-03 | -1.45E+00 | 3.39E-03 | 9.54E-03 | -1.18E+00 | 2.70E-02 | 9.92E-02 | -1.02E+00 | 8.96E-02 | 2.46E-01 | 1.10E+00 |
| FBF1 | 5.52E-05 | 2.88E-04 | -1.70E+00 | 6.68E-04 | 2.89E-03 | -1.27E+00 | 1.15E-01 | 4.89E-01 | 1.09E+00 | 4.70E-02 | 1.80E-01 | 1.02E+00 | 1.44E-01 | 5.15E-01 | 1.14E+00 | 1.80E-01 | 5.08E-01 | 1.15E+00 | 2.26E-01 | 8.30E-01 | 1.26E+00 | 2.95E-01 | 8.10E-01 | 1.15E+00 |
| NUP54 | 5.52E-05 | 2.88E-04 | -1.07E+00 | 1.81E-04 | 7.81E-04 | 1.03E+00 | 2.41E-07 | 1.02E-06 | -1.30E+00 | 2.72E-07 | 1.04E-06 | -1.14E+00 | 4.29E-07 | 1.53E-06 | -1.33E+00 | 1.02E-04 | 2.86E-04 | -1.08E+00 | 3.86E-09 | 1.42E-08 | -1.39E+00 | 5.65E-05 | 1.55E-04 | -1.19E+00 |
| AC135507.2 | 5.55E-05 | 2.89E-04 | -2.24E+00 | 5.34E-05 | 2.31E-04 | -2.34E+00 | 9.54E-16 | 4.06E-15 | -2.81E+00 | 1.25E-10 | 4.77E-10 | -1.75E+00 | 6.90E-02 | 2.46E-01 | -1.10E+00 | 8.34E-03 | 2.35E-02 | -1.98E+00 | 5.16E-02 | 1.90E-01 | -1.30E+00 | 9.34E-02 | 2.57E-01 | -1.33E+00 |
| MYO5A | 5.55E-05 | 2.89E-04 | -1.25E+00 | 2.77E-04 | 1.19E-03 | 1.00E+00 | 3.89E-04 | 1.65E-03 | -1.23E+00 | 2.31E-04 | 8.81E-04 | -1.15E+00 | 1.44E-01 | 5.12E-01 | 1.37E+00 | 8.88E-02 | 2.50E-01 | 1.18E+00 | 1.23E-01 | 4.52E-01 | 1.04E+00 | 4.22E-02 | 1.16E-01 | 1.01E+00 |
| RPL34P1 | 5.56E-05 | 2.90E-04 | -1.27E+00 | 3.28E-09 | 1.42E-08 | -2.08E+00 | 6.09E-05 | 2.59E-04 | -1.38E+00 | 5.57E-10 | 2.13E-09 | -2.14E+00 | 2.15E-01 | 7.67E-01 | 1.42E+00 | 5.00E-03 | 1.41E-02 | -1.68E+00 | 1.85E-01 | 6.79E-01 | 1.14E+00 | 4.77E-02 | 1.31E-01 | -1.25E+00 |
| SLC35A5 | 5.56E-05 | 2.90E-04 | -1.38E+00 | 8.02E-04 | 3.47E-03 | -1.07E+00 | 2.72E-09 | 1.16E-08 | -2.09E+00 | 1.14E-06 | 4.35E-06 | -1.48E+00 | 4.37E-03 | 1.56E-02 | -1.14E+00 | 2.56E-02 | 7.23E-02 | -1.03E+00 | 1.61E-06 | 5.92E-06 | -1.39E+00 | 7.92E-05 | 2.17E-04 | -1.29E+00 |
| ZNF746 | 5.56E-05 | 2.90E-04 | -1.28E+00 | 4.40E-03 | 1.90E-02 | 1.06E+00 | 3.02E-04 | 1.28E-03 | -1.31E+00 | 1.51E-03 | 5.76E-03 | -1.09E+00 | 2.02E-02 | 7.21E-02 | -1.07E+00 | 2.81E-02 | 7.92E-02 | -1.02E+00 | 3.18E-03 | 1.17E-02 | -1.15E+00 | 1.03E-02 | 2.83E-02 | -1.18E+00 |
| ANXA3 | 5.56E-05 | 2.90E-04 | -1.41E+00 | 2.99E-02 | 1.29E-01 | 1.18E+00 | 1.60E-01 | 6.79E-01 | 1.08E+00 | 1.25E-01 | 4.78E-01 | 1.87E+00 | 1.99E-02 | 7.11E-02 | 1.02E+00 | 2.57E-01 | 7.25E-01 | 1.48E+00 | 1.89E-01 | 6.94E-01 | 1.49E+00 | 1.09E-01 | 3.00E-01 | 1.21E+00 |
| GSK3A | 5.56E-05 | 2.90E-04 | -1.11E+00 | 1.17E-04 | 5.05E-04 | 1.05E+00 | 2.18E-05 | 9.25E-05 | -1.19E+00 | 1.69E-04 | 6.44E-04 | 1.04E+00 | 1.84E-05 | 6.56E-05 | -1.22E+00 | 4.68E-05 | 1.32E-04 | -1.24E+00 | 7.86E-09 | 2.88E-08 | -1.28E+00 | 1.99E-05 | 5.46E-05 | -1.03E+00 |
| LCMT1 | 5.56E-05 | 2.90E-04 | -1.00E+00 | 1.79E-05 | 7.74E-05 | 1.09E+00 | 2.75E-02 | 1.17E-01 | 1.06E+00 | 5.77E-04 | 2.20E-03 | 1.05E+00 | 9.99E-07 | 3.56E-06 | -1.33E+00 | 1.10E-04 | 3.09E-04 | -1.11E+00 | 1.00E-02 | 3.67E-02 | -1.02E+00 | 9.07E-03 | 2.49E-02 | 1.03E+00 |
| CNOT10 | 5.58E-05 | 2.91E-04 | -1.24E+00 | 1.61E-02 | 6.96E-02 | 1.18E+00 | 5.02E-07 | 2.13E-06 | -1.36E+00 | 3.19E-08 | 1.22E-07 | -1.27E+00 | 7.39E-06 | 2.63E-05 | -1.36E+00 | 4.93E-03 | 1.39E-02 | 1.02E+00 | 2.34E-05 | 8.60E-05 | -1.09E+00 | 2.26E-03 | 6.21E-03 | -1.02E+00 |
| RCCD1 | 5.59E-05 | 2.91E-04 | -1.48E+00 | 1.47E-02 | 6.36E-02 | 1.16E+00 | 4.44E-06 | 1.89E-05 | -1.41E+00 | 1.97E-02 | 7.53E-02 | 1.19E+00 | 2.11E-06 | 7.54E-06 | -1.72E+00 | 4.97E-03 | 1.40E-02 | -1.09E+00 | 1.20E-09 | 4.40E-09 | -1.36E+00 | 5.11E-02 | 1.40E-01 | 1.13E+00 |
| SH3BGRL2 | 5.60E-05 | 2.92E-04 | -1.38E+00 | 5.54E-05 | 2.40E-04 | -1.21E+00 | 1.56E-01 | 6.64E-01 | 1.00E+00 | 7.30E-02 | 2.79E-01 | -1.00E+00 | 1.45E-02 | 5.16E-02 | -1.23E+00 | 2.36E-01 | 6.64E-01 | 1.23E+00 | 9.62E-03 | 3.53E-02 | 1.01E+00 | 5.77E-02 | 1.58E-01 | 1.13E+00 |
| CORO7 | 5.60E-05 | 2.92E-04 | -1.16E+00 | 1.71E-04 | 7.38E-04 | 1.01E+00 | 5.60E-02 | 2.38E-01 | 1.06E+00 | 7.86E-02 | 3.00E-01 | 1.22E+00 | 7.56E-04 | 2.70E-03 | -1.09E+00 | 4.57E-04 | 1.29E-03 | -1.08E+00 | 1.03E-04 | 3.79E-04 | -1.21E+00 | 1.91E-02 | 5.24E-02 | 1.05E+00 |
| NSMCE2 | 5.64E-05 | 2.94E-04 | -1.10E+00 | 7.29E-04 | 3.15E-03 | 1.08E+00 | 4.14E-02 | 1.76E-01 | 1.01E+00 | 5.72E-02 | 2.19E-01 | 1.16E+00 | 6.46E-05 | 2.30E-04 | -1.17E+00 | 1.99E-02 | 5.61E-02 | 1.12E+00 | 6.40E-02 | 2.35E-01 | 1.07E+00 | 1.37E-01 | 3.76E-01 | 1.12E+00 |
| PI4KA | 5.65E-05 | 2.94E-04 | -1.18E+00 | 2.27E-02 | 9.80E-02 | 1.19E+00 | 1.22E-01 | 5.17E-01 | 1.23E+00 | 2.29E-02 | 8.76E-02 | 1.25E+00 | 2.97E-02 | 1.06E-01 | -1.07E+00 | 8.89E-03 | 2.51E-02 | -1.12E+00 | 8.56E-03 | 3.14E-02 | -1.06E+00 | 1.02E-02 | 2.81E-02 | -1.05E+00 |
| AP003733.4 | 5.67E-05 | 2.96E-04 | 2.79E+00 | 2.06E-02 | 8.90E-02 | 2.24E+00 | 1.41E-03 | 6.00E-03 | 2.62E+00 | 7.59E-02 | 2.90E-01 | 2.05E+00 | 1.55E-02 | 5.52E-02 | 2.51E+00 | 2.81E-03 | 7.92E-03 | 2.95E+00 | 2.67E-01 | 9.80E-01 | 1.23E+00 | 3.21E-01 | 8.82E-01 | 1.27E+00 |
| HCG11 | 5.68E-05 | 2.96E-04 | -1.48E+00 | 1.39E-01 | 6.01E-01 | 1.42E+00 | 1.53E-07 | 6.49E-07 | -1.84E+00 | 7.05E-03 | 2.69E-02 | 1.08E+00 | 6.47E-03 | 2.31E-02 | -1.42E+00 | 2.98E-01 | 8.39E-01 | 1.35E+00 | 8.16E-07 | 3.00E-06 | -2.00E+00 | 4.43E-02 | 1.22E-01 | 1.03E+00 |
| DICER1-AS1 | 5.69E-05 | 2.96E-04 | -1.29E+00 | 5.14E-06 | 2.22E-05 | -1.42E+00 | 1.54E-01 | 6.55E-01 | 1.09E+00 | 5.28E-02 | 2.02E-01 | 1.05E+00 | 5.30E-02 | 1.89E-01 | 1.19E+00 | 8.15E-03 | 2.30E-02 | -1.19E+00 | 4.11E-03 | 1.51E-02 | -1.35E+00 | 7.75E-03 | 2.13E-02 | -1.30E+00 |
| PRKAR2B | 5.69E-05 | 2.96E-04 | -1.11E+00 | 8.06E-05 | 3.48E-04 | -1.01E+00 | 1.07E-01 | 4.54E-01 | 1.19E+00 | 2.07E-02 | 7.91E-02 | 1.11E+00 | 3.02E-04 | 1.08E-03 | 1.96E+00 | 4.28E-15 | 1.21E-14 | 3.19E+00 | 2.65E-03 | 9.71E-03 | 1.60E+00 | 1.72E-01 | 4.72E-01 | 1.24E+00 |
| LUC7L | 5.69E-05 | 2.97E-04 | -1.04E+00 | 9.69E-04 | 4.19E-03 | 1.11E+00 | 2.23E-06 | 9.46E-06 | -1.23E+00 | 2.82E-06 | 1.08E-05 | -1.04E+00 | 1.77E-06 | 6.31E-06 | -1.25E+00 | 3.11E-04 | 8.77E-04 | -1.06E+00 | 4.65E-04 | 1.71E-03 | -1.11E+00 | 2.45E-02 | 6.73E-02 | -1.02E+00 |
| ARHGEF10 | 5.71E-05 | 2.98E-04 | -1.77E+00 | 1.25E-02 | 5.39E-02 | 1.02E+00 | 7.98E-02 | 3.39E-01 | -1.05E+00 | 6.92E-02 | 2.64E-01 | 1.71E+00 | 1.39E-01 | 4.95E-01 | -1.04E+00 | 2.31E-01 | 6.51E-01 | 1.64E+00 | 1.45E-03 | 5.31E-03 | -1.46E+00 | 5.29E-02 | 1.45E-01 | 1.45E+00 |
| NAP1L2 | 5.74E-05 | 2.99E-04 | -1.65E+00 | 1.58E-03 | 6.82E-03 | -1.18E+00 | 3.32E-03 | 1.41E-02 | -1.33E+00 | 4.31E-02 | 1.65E-01 | -1.02E+00 | 1.48E-03 | 5.27E-03 | -1.87E+00 | 3.44E-02 | 9.69E-02 | -1.31E+00 | 9.44E-02 | 3.46E-01 | -1.08E+00 | 2.13E-01 | 5.84E-01 | 1.50E+00 |
| IGSF10 | 5.74E-05 | 2.99E-04 | -1.52E+00 | 1.06E-03 | 4.57E-03 | -1.11E+00 | 7.10E-04 | 3.02E-03 | -1.30E+00 | 1.40E-03 | 5.35E-03 | -1.12E+00 | 1.95E-03 | 6.94E-03 | -1.24E+00 | 2.16E-04 | 6.09E-04 | -1.47E+00 | 1.51E-02 | 5.53E-02 | -1.26E+00 | 1.27E-02 | 3.48E-02 | -1.29E+00 |
| LRRC75A-AS1 | 5.74E-05 | 2.99E-04 | -1.05E+00 | 1.87E-11 | 8.07E-11 | -1.11E+00 | 5.40E-03 | 2.29E-02 | -1.12E+00 | 2.89E-05 | 1.10E-04 | -1.14E+00 | 1.76E-03 | 6.26E-03 | -1.06E+00 | 1.24E-03 | 3.50E-03 | -1.10E+00 | 9.20E-02 | 3.38E-01 | 1.04E+00 | 1.95E-01 | 5.35E-01 | -1.02E+00 |
| PIP4K2C | 5.74E-05 | 2.99E-04 | -1.89E+00 | 4.41E-03 | 1.90E-02 | -1.09E+00 | 1.90E-02 | 8.08E-02 | -1.17E+00 | 4.06E-02 | 1.55E-01 | 1.02E+00 | 1.31E-03 | 4.68E-03 | -1.85E+00 | 1.12E-01 | 3.16E-01 | 1.17E+00 | 3.02E-02 | 1.11E-01 | -1.10E+00 | 3.36E-01 | 9.24E-01 | 1.23E+00 |
| AL359922.2 | 5.75E-05 | 2.99E-04 | -2.18E+00 | 5.18E-03 | 2.24E-02 | -1.27E+00 | 3.33E-02 | 1.41E-01 | -1.20E+00 | 7.31E-02 | 2.79E-01 | 1.06E+00 | 8.79E-02 | 3.14E-01 | -1.07E+00 | 3.17E-01 | 8.94E-01 | 1.17E+00 | 1.50E-02 | 5.51E-02 | -1.50E+00 | 3.55E-01 | 9.75E-01 | 1.12E+00 |
| AC133965.1 | 5.77E-05 | 3.00E-04 | -2.22E+00 | 4.72E-04 | 2.04E-03 | -1.62E+00 | 1.09E-02 | 4.65E-02 | -1.63E+00 | 5.70E-02 | 2.18E-01 | -1.14E+00 | 7.01E-08 | 2.50E-07 | -2.01E+00 | 9.35E-07 | 2.64E-06 | -1.62E+00 | 4.52E-07 | 1.66E-06 | -1.61E+00 | 5.97E-04 | 1.64E-03 | -1.23E+00 |
| PLEKHO1 | 5.77E-05 | 3.01E-04 | -1.08E+00 | 1.44E-06 | 6.21E-06 | -1.05E+00 | 1.18E-01 | 5.03E-01 | 1.14E+00 | 1.08E-03 | 4.13E-03 | 1.07E+00 | 6.32E-04 | 2.25E-03 | -1.17E+00 | 4.74E-06 | 1.34E-05 | -1.51E+00 | 2.53E-01 | 9.29E-01 | 3.35E+00 | 3.37E-01 | 9.26E-01 | 2.11E+00 |
| FAM226B | 5.78E-05 | 3.01E-04 | -1.49E+00 | 5.48E-02 | 2.37E-01 | 1.65E+00 | 7.44E-04 | 3.16E-03 | -1.60E+00 | 6.56E-02 | 2.51E-01 | 1.11E+00 | 1.11E-01 | 3.97E-01 | 1.97E+00 | 7.39E-02 | 2.08E-01 | 2.33E+00 | 2.58E-01 | 9.47E-01 | 1.25E+00 | 3.08E-01 | 8.47E-01 | 1.19E+00 |
| TXK | 5.78E-05 | 3.01E-04 | -1.75E+00 | 8.85E-02 | 3.82E-01 | 1.23E+00 | 7.03E-04 | 2.99E-03 | -1.57E+00 | 1.24E-02 | 4.73E-02 | -1.08E+00 | 3.74E-02 | 1.33E-01 | -1.09E+00 | 1.43E-01 | 4.02E-01 | 1.17E+00 | 1.83E-01 | 6.71E-01 | 1.17E+00 | 2.81E-01 | 7.72E-01 | 1.13E+00 |
| COPRS | 5.78E-05 | 3.01E-04 | -1.09E+00 | 2.25E-03 | 9.71E-03 | 1.14E+00 | 3.73E-02 | 1.59E-01 | 1.09E+00 | 2.77E-04 | 1.06E-03 | 1.04E+00 | 5.51E-06 | 1.96E-05 | -1.48E+00 | 1.82E-04 | 5.13E-04 | -1.27E+00 | 2.20E-02 | 8.09E-02 | 1.08E+00 | 1.77E-01 | 4.85E-01 | 1.43E+00 |
| PLCG1-AS1 | 5.81E-05 | 3.03E-04 | -1.64E+00 | 2.39E-07 | 1.03E-06 | -2.62E+00 | 3.07E-03 | 1.30E-02 | -1.27E+00 | 1.19E-04 | 4.55E-04 | -1.77E+00 | 7.97E-03 | 2.84E-02 | -1.29E+00 | 2.08E-05 | 5.87E-05 | -2.73E+00 | 2.50E-02 | 9.17E-02 | -1.34E+00 | 6.17E-04 | 1.69E-03 | -1.98E+00 |
| NCK1-AS1 | 5.81E-05 | 3.03E-04 | -1.41E+00 | 3.37E-05 | 1.46E-04 | -1.38E+00 | 5.79E-06 | 2.46E-05 | -1.74E+00 | 3.82E-06 | 1.46E-05 | -1.54E+00 | 3.41E-06 | 1.22E-05 | -1.61E+00 | 2.63E-05 | 7.41E-05 | -1.46E+00 | 7.68E-05 | 2.82E-04 | -1.25E+00 | 2.95E-04 | 8.11E-04 | -1.30E+00 |
| CENPE | 5.81E-05 | 3.03E-04 | -1.87E+00 | 3.92E-05 | 1.69E-04 | -1.73E+00 | 9.48E-04 | 4.03E-03 | -1.28E+00 | 1.12E-06 | 4.26E-06 | -1.27E+00 | 1.72E-08 | 6.13E-08 | -3.38E+00 | 2.52E-07 | 7.09E-07 | -2.28E+00 | 5.23E-07 | 1.92E-06 | -1.38E+00 | 6.33E-10 | 1.74E-09 | -1.38E+00 |
| ELOF1 | 5.81E-05 | 3.03E-04 | -1.09E+00 | 1.48E-03 | 6.39E-03 | 1.17E+00 | 7.81E-04 | 3.32E-03 | -1.17E+00 | 3.00E-03 | 1.15E-02 | 1.01E+00 | 7.24E-06 | 2.58E-05 | -1.21E+00 | 1.75E-02 | 4.93E-02 | 1.12E+00 | 1.89E-02 | 6.93E-02 | 1.05E+00 | 2.77E-01 | 7.62E-01 | 1.24E+00 |
| SLC25A5 | 5.82E-05 | 3.04E-04 | -1.10E+00 | 7.99E-08 | 3.45E-07 | -1.10E+00 | 7.90E-02 | 3.36E-01 | -1.06E+00 | 3.19E-02 | 1.22E-01 | -1.09E+00 | 5.87E-04 | 2.09E-03 | -1.14E+00 | 1.02E-01 | 2.88E-01 | -1.06E+00 | 1.96E-03 | 7.20E-03 | -1.09E+00 | 3.59E-01 | 9.86E-01 | 1.06E+00 |
| SMC6 | 5.83E-05 | 3.04E-04 | -1.39E+00 | 9.61E-03 | 4.15E-02 | 1.10E+00 | 1.18E-06 | 5.00E-06 | -1.25E+00 | 4.73E-03 | 1.81E-02 | 1.09E+00 | 2.25E-07 | 8.00E-07 | -1.71E+00 | 7.90E-03 | 2.23E-02 | 1.07E+00 | 1.13E-07 | 4.16E-07 | -1.29E+00 | 1.23E-03 | 3.37E-03 | -1.03E+00 |
| SHD | 5.86E-05 | 3.05E-04 | -1.95E+00 | 1.66E-05 | 7.16E-05 | -2.27E+00 | 3.10E-04 | 1.32E-03 | -1.47E+00 | 5.62E-03 | 2.15E-02 | -1.14E+00 | 2.31E-01 | 8.24E-01 | 1.46E+00 | 9.55E-02 | 2.69E-01 | -1.19E+00 | 2.72E-01 | 1.00E+00 | 1.00E+00 | 3.14E-01 | 8.63E-01 | 1.23E+00 |
| NAA35 | 5.87E-05 | 3.06E-04 | -1.11E+00 | 5.82E-06 | 2.51E-05 | -1.10E+00 | 3.05E-06 | 1.30E-05 | -1.30E+00 | 5.08E-04 | 1.94E-03 | 1.00E+00 | 1.10E-07 | 3.91E-07 | -1.43E+00 | 2.63E-04 | 7.41E-04 | 1.00E+00 | 1.66E-06 | 6.10E-06 | -1.22E+00 | 4.12E-05 | 1.13E-04 | -1.17E+00 |
| AC098592.2 | 5.88E-05 | 3.07E-04 | -1.68E+00 | 1.70E-04 | 7.34E-04 | -1.52E+00 | 5.45E-10 | 2.32E-09 | -1.99E+00 | 2.99E-09 | 1.14E-08 | -1.78E+00 | 8.62E-04 | 3.07E-03 | -1.75E+00 | 1.54E-03 | 4.35E-03 | -1.57E+00 | 1.00E-16 | 3.68E-16 | -2.58E+00 | 4.95E-12 | 1.36E-11 | -1.84E+00 |
| TOB1-AS1 | 5.91E-05 | 3.08E-04 | -1.72E+00 | 5.23E-03 | 2.26E-02 | -1.07E+00 | 2.22E-05 | 9.42E-05 | -1.93E+00 | 2.62E-05 | 1.00E-04 | -1.90E+00 | 1.31E-02 | 4.68E-02 | -1.82E+00 | 1.49E-01 | 4.21E-01 | -1.04E+00 | 2.48E-03 | 9.10E-03 | -1.94E+00 | 4.61E-02 | 1.27E-01 | -1.31E+00 |
| RAB18 | 5.92E-05 | 3.09E-04 | -1.06E+00 | 1.47E-03 | 6.34E-03 | 1.16E+00 | 1.06E-08 | 4.50E-08 | -1.40E+00 | 2.04E-04 | 7.79E-04 | -1.02E+00 | 4.61E-03 | 1.64E-02 | 1.06E+00 | 4.26E-04 | 1.20E-03 | -1.01E+00 | 8.56E-06 | 3.14E-05 | -1.15E+00 | 3.59E-03 | 9.87E-03 | -1.05E+00 |
| VMP1 | 5.94E-05 | 3.09E-04 | 1.82E+00 | 1.58E-02 | 6.83E-02 | 1.77E+00 | 5.37E-02 | 2.28E-01 | 1.50E+00 | 9.31E-04 | 3.56E-03 | 1.92E+00 | 4.87E-06 | 1.74E-05 | 2.18E+00 | 9.85E-11 | 2.78E-10 | 2.80E+00 | 2.35E-04 | 8.64E-04 | 2.25E+00 | 3.35E-05 | 9.20E-05 | 2.24E+00 |
| CBL | 5.94E-05 | 3.09E-04 | -1.05E+00 | 2.74E-04 | 1.18E-03 | 1.06E+00 | 3.30E-06 | 1.40E-05 | -1.24E+00 | 9.69E-05 | 3.70E-04 | 1.03E+00 | 6.49E-02 | 2.31E-01 | -1.00E+00 | 3.42E-02 | 9.65E-02 | -1.27E+00 | 1.54E-01 | 5.67E-01 | -1.01E+00 | 1.68E-01 | 4.62E-01 | -1.03E+00 |
| ABCD1 | 5.94E-05 | 3.10E-04 | -1.53E+00 | 9.01E-05 | 3.89E-04 | -1.34E+00 | 5.84E-06 | 2.48E-05 | -1.81E+00 | 2.67E-04 | 1.02E-03 | -1.31E+00 | 4.78E-03 | 1.71E-02 | -1.57E+00 | 5.12E-03 | 1.44E-02 | -1.48E+00 | 2.10E-02 | 7.72E-02 | -1.40E+00 | 1.36E-01 | 3.72E-01 | 1.04E+00 |
| MARK4 | 5.94E-05 | 3.10E-04 | -1.18E+00 | 3.57E-04 | 1.54E-03 | 1.01E+00 | 2.50E-03 | 1.06E-02 | -1.04E+00 | 1.65E-03 | 6.29E-03 | 1.10E+00 | 9.27E-03 | 3.31E-02 | -1.19E+00 | 6.38E-02 | 1.80E-01 | 1.00E+00 | 1.39E-02 | 5.11E-02 | -1.02E+00 | 4.67E-02 | 1.28E-01 | -1.00E+00 |
| FAM153B | 5.94E-05 | 3.10E-04 | -1.26E+00 | 3.67E-02 | 1.59E-01 | 1.21E+00 | 1.47E-04 | 6.23E-04 | -1.58E+00 | 1.35E-02 | 5.16E-02 | -1.02E+00 | 2.37E-01 | 8.45E-01 | 1.32E+00 | 1.65E-01 | 4.64E-01 | 1.81E+00 | 2.60E-01 | 9.56E-01 | 1.11E+00 | 3.62E-01 | 9.95E-01 | 1.16E+00 |
| SLC52A2 | 5.95E-05 | 3.10E-04 | -1.23E+00 | 9.55E-03 | 4.13E-02 | 1.16E+00 | 9.11E-04 | 3.87E-03 | -1.19E+00 | 5.78E-02 | 2.21E-01 | 1.12E+00 | 6.27E-08 | 2.23E-07 | -1.51E+00 | 4.05E-03 | 1.14E-02 | 1.00E+00 | 5.75E-04 | 2.11E-03 | -1.11E+00 | 3.50E-01 | 9.62E-01 | 1.24E+00 |
| RNLS | 5.97E-05 | 3.11E-04 | -1.26E+00 | 2.83E-03 | 1.22E-02 | 1.08E+00 | 1.68E-03 | 7.15E-03 | -1.41E+00 | 2.70E-02 | 1.03E-01 | 1.00E+00 | 4.76E-02 | 1.70E-01 | -1.11E+00 | 1.44E-01 | 4.06E-01 | 1.16E+00 | 1.16E-03 | 4.24E-03 | -1.29E+00 | 3.48E-01 | 9.55E-01 | 1.31E+00 |
| KATNBL1 | 5.97E-05 | 3.11E-04 | -1.01E+00 | 1.09E-01 | 4.70E-01 | 1.38E+00 | 1.99E-08 | 8.45E-08 | -1.38E+00 | 6.50E-04 | 2.48E-03 | 1.05E+00 | 1.06E-07 | 3.78E-07 | -1.31E+00 | 4.91E-03 | 1.39E-02 | 1.04E+00 | 3.48E-05 | 1.28E-04 | -1.20E+00 | 1.68E-03 | 4.63E-03 | -1.03E+00 |
| RPS15AP24 | 5.97E-05 | 3.11E-04 | 1.00E+00 | 7.30E-03 | 3.15E-02 | 1.32E+00 | 8.30E-03 | 3.53E-02 | -1.01E+00 | 2.15E-02 | 8.22E-02 | 1.18E+00 | 2.86E-04 | 1.02E-03 | -1.07E+00 | 6.82E-02 | 1.92E-01 | 1.26E+00 | 9.54E-04 | 3.50E-03 | -1.10E+00 | 1.89E-01 | 5.19E-01 | 1.27E+00 |
| EIF3D | 5.99E-05 | 3.12E-04 | -1.03E+00 | 4.96E-02 | 2.14E-01 | 1.10E+00 | 3.66E-02 | 1.56E-01 | -1.03E+00 | 1.58E-01 | 6.05E-01 | 1.11E+00 | 1.66E-02 | 5.93E-02 | -1.06E+00 | 1.90E-01 | 5.37E-01 | 1.11E+00 | 9.20E-02 | 3.38E-01 | -1.01E+00 | 3.61E-01 | 9.92E-01 | 1.05E+00 |
| SIPA1 | 6.00E-05 | 3.13E-04 | -1.01E+00 | 3.51E-04 | 1.51E-03 | 1.12E+00 | 5.62E-03 | 2.39E-02 | 1.00E+00 | 1.47E-03 | 5.61E-03 | 1.08E+00 | 3.10E-03 | 1.10E-02 | 1.02E+00 | 1.81E-01 | 5.11E-01 | 1.40E+00 | 1.62E-01 | 5.94E-01 | 1.14E+00 | 3.03E-01 | 8.32E-01 | 1.25E+00 |
| ADAT1 | 6.03E-05 | 3.14E-04 | -1.52E+00 | 2.38E-03 | 1.03E-02 | -1.03E+00 | 2.74E-07 | 1.16E-06 | -1.68E+00 | 1.23E-05 | 4.71E-05 | -1.25E+00 | 1.09E-02 | 3.89E-02 | -1.09E+00 | 5.32E-02 | 1.50E-01 | 1.03E+00 | 5.87E-04 | 2.16E-03 | -1.15E+00 | 3.90E-03 | 1.07E-02 | -1.13E+00 |
| AC005632.2 | 6.04E-05 | 3.15E-04 | -1.67E+00 | 1.55E-05 | 6.70E-05 | -1.71E+00 | 8.47E-02 | 3.60E-01 | 1.02E+00 | 3.56E-02 | 1.36E-01 | -1.01E+00 | 1.15E-02 | 4.08E-02 | -1.17E+00 | 1.15E-02 | 3.23E-02 | -1.13E+00 | 1.56E-03 | 5.71E-03 | -1.41E+00 | 6.93E-02 | 1.90E-01 | -1.02E+00 |
| DHRS1 | 6.05E-05 | 3.16E-04 | -1.19E+00 | 6.41E-02 | 2.77E-01 | 1.34E+00 | 2.50E-06 | 1.06E-05 | -1.35E+00 | 5.30E-04 | 2.02E-03 | -1.04E+00 | 8.97E-04 | 3.20E-03 | -1.19E+00 | 1.11E-01 | 3.13E-01 | 1.22E+00 | 1.48E-02 | 5.45E-02 | -1.04E+00 | 1.82E-03 | 5.01E-03 | -1.20E+00 |
| AC017071.1 | 6.06E-05 | 3.16E-04 | -2.04E+00 | 2.38E-03 | 1.03E-02 | -1.42E+00 | 1.03E-03 | 4.40E-03 | -1.73E+00 | 3.37E-02 | 1.29E-01 | -1.00E+00 | 5.08E-02 | 1.81E-01 | -1.36E+00 | 2.61E-01 | 7.36E-01 | 1.13E+00 | 1.60E-01 | 5.87E-01 | -1.17E+00 | 2.78E-01 | 7.64E-01 | 1.17E+00 |
| DPY30 | 6.06E-05 | 3.16E-04 | -1.01E+00 | 3.46E-04 | 1.49E-03 | 1.15E+00 | 1.70E-02 | 7.23E-02 | 1.06E+00 | 4.73E-02 | 1.81E-01 | 1.17E+00 | 2.44E-06 | 8.70E-06 | -1.20E+00 | 2.69E-03 | 7.58E-03 | 1.06E+00 | 4.80E-03 | 1.76E-02 | -1.00E+00 | 1.57E-01 | 4.31E-01 | 1.16E+00 |
| AC127391.1 | 6.08E-05 | 3.17E-04 | -1.77E+00 | 4.33E-04 | 1.87E-03 | -1.49E+00 | 1.78E-04 | 7.58E-04 | -1.72E+00 | 1.25E-04 | 4.77E-04 | -1.70E+00 | 4.16E-05 | 1.48E-04 | -2.27E+00 | 1.40E-03 | 3.94E-03 | -1.59E+00 | 6.04E-22 | 2.22E-21 | -2.36E+00 | 1.23E-15 | 3.38E-15 | -1.82E+00 |
| SLC38A6 | 6.09E-05 | 3.17E-04 | -1.18E+00 | 1.65E-03 | 7.12E-03 | 1.04E+00 | 7.42E-05 | 3.15E-04 | -1.46E+00 | 5.68E-03 | 2.17E-02 | 1.03E+00 | 2.46E-01 | 8.77E-01 | 1.42E+00 | 9.81E-02 | 2.76E-01 | 1.13E+00 | 2.40E-03 | 8.79E-03 | -1.30E+00 | 5.97E-03 | 1.64E-02 | -1.27E+00 |
| GDPGP1 | 6.10E-05 | 3.18E-04 | -1.52E+00 | 8.58E-03 | 3.71E-02 | 1.07E+00 | 4.68E-03 | 1.99E-02 | -1.59E+00 | 1.23E-01 | 4.70E-01 | 1.09E+00 | 5.56E-05 | 1.98E-04 | -1.80E+00 | 3.70E-03 | 1.04E-02 | -1.24E+00 | 2.34E-10 | 8.60E-10 | -2.51E+00 | 3.13E-06 | 8.60E-06 | -1.64E+00 |
| MED11 | 6.10E-05 | 3.18E-04 | -1.35E+00 | 2.10E-02 | 9.07E-02 | 1.15E+00 | 5.95E-06 | 2.53E-05 | -1.51E+00 | 5.80E-03 | 2.22E-02 | 1.07E+00 | 5.47E-04 | 1.95E-03 | -1.74E+00 | 2.02E-01 | 5.69E-01 | 1.21E+00 | 2.96E-07 | 1.09E-06 | -1.48E+00 | 6.41E-02 | 1.76E-01 | 1.11E+00 |
| FAM126B | 6.10E-05 | 3.18E-04 | -1.21E+00 | 1.40E-02 | 6.05E-02 | 1.15E+00 | 1.13E-02 | 4.80E-02 | -1.04E+00 | 8.21E-03 | 3.14E-02 | 1.04E+00 | 1.89E-05 | 6.74E-05 | -1.35E+00 | 7.37E-04 | 2.08E-03 | -1.15E+00 | 1.10E-02 | 4.05E-02 | -1.02E+00 | 3.50E-02 | 9.61E-02 | 1.05E+00 |
| AL136295.5 | 6.11E-05 | 3.19E-04 | -1.12E+00 | 1.95E-16 | 8.43E-16 | -2.37E+00 | 1.20E-03 | 5.12E-03 | -1.14E+00 | 5.49E-17 | 2.10E-16 | -2.90E+00 | 4.80E-07 | 1.71E-06 | -1.95E+00 | 2.08E-09 | 5.87E-09 | -2.58E+00 | 5.86E-06 | 2.15E-05 | -1.50E+00 | 6.53E-10 | 1.79E-09 | -2.45E+00 |
| MOCS3 | 6.12E-05 | 3.19E-04 | -1.30E+00 | 1.74E-05 | 7.50E-05 | -1.47E+00 | 2.90E-02 | 1.23E-01 | -1.16E+00 | 4.65E-02 | 1.78E-01 | -1.01E+00 | 1.95E-01 | 6.97E-01 | 1.22E+00 | 6.55E-02 | 1.85E-01 | -1.05E+00 | 3.45E-02 | 1.26E-01 | 1.05E+00 | 1.84E-02 | 5.05E-02 | -1.12E+00 |
| RASA4 | 6.12E-05 | 3.19E-04 | -1.41E+00 | 1.45E-02 | 6.27E-02 | 1.15E+00 | 3.98E-03 | 1.69E-02 | -1.27E+00 | 7.91E-03 | 3.02E-02 | -1.08E+00 | 2.62E-02 | 9.34E-02 | -1.47E+00 | 9.18E-02 | 2.59E-01 | -1.10E+00 | 4.23E-02 | 1.55E-01 | -1.31E+00 | 1.32E-01 | 3.63E-01 | -1.13E+00 |
| RPL4P5 | 6.12E-05 | 3.19E-04 | 1.51E+00 | 4.70E-21 | 2.03E-20 | 2.26E+00 | 1.31E-06 | 5.55E-06 | 1.68E+00 | 1.81E-17 | 6.92E-17 | 2.55E+00 | 2.49E-01 | 8.86E-01 | 1.34E+00 | 2.08E-08 | 5.86E-08 | 2.19E+00 | 8.50E-02 | 3.12E-01 | 1.41E+00 | 1.96E-11 | 5.38E-11 | 2.23E+00 |
| BCL6-AS1 | 6.13E-05 | 3.19E-04 | 2.43E+00 | 1.43E-01 | 6.18E-01 | -1.19E+00 | 1.22E-01 | 5.19E-01 | 1.37E+00 | 6.55E-02 | 2.50E-01 | -1.05E+00 | 5.38E-02 | 1.92E-01 | 2.80E+00 | 2.25E-01 | 6.33E-01 | 1.44E+00 | 8.40E-02 | 3.08E-01 | 1.66E+00 | 3.00E-01 | 8.23E-01 | 1.12E+00 |
| MFAP3L | 6.15E-05 | 3.20E-04 | -1.88E+00 | 2.06E-05 | 8.89E-05 | -2.12E+00 | 1.94E-02 | 8.23E-02 | -1.61E+00 | 1.89E-02 | 7.20E-02 | -1.68E+00 | 2.22E-01 | 7.91E-01 | -1.00E+00 | 3.17E-01 | 8.95E-01 | -1.02E+00 | 1.39E-03 | 5.09E-03 | 4.94E+00 | 6.89E-03 | 1.89E-02 | 3.45E+00 |
| USP40 | 6.16E-05 | 3.21E-04 | -1.25E+00 | 8.67E-03 | 3.75E-02 | 1.15E+00 | 7.72E-05 | 3.28E-04 | -1.53E+00 | 1.93E-02 | 7.36E-02 | 1.03E+00 | 1.03E-01 | 3.68E-01 | 1.10E+00 | 3.45E-01 | 9.73E-01 | 1.52E+00 | 2.44E-02 | 8.95E-02 | 1.01E+00 | 3.46E-01 | 9.51E-01 | 1.32E+00 |
| IGFLR1 | 6.16E-05 | 3.21E-04 | -1.70E+00 | 6.25E-04 | 2.70E-03 | -1.32E+00 | 1.19E-05 | 5.06E-05 | -1.38E+00 | 2.09E-05 | 7.99E-05 | -1.28E+00 | 2.88E-03 | 1.03E-02 | -1.51E+00 | 2.00E-03 | 5.63E-03 | -1.55E+00 | 2.56E-02 | 9.38E-02 | -1.26E+00 | 6.69E-02 | 1.84E-01 | -1.16E+00 |
| ZEB1-AS1 | 6.18E-05 | 3.22E-04 | -1.16E+00 | 2.38E-03 | 1.03E-02 | 1.12E+00 | 8.60E-04 | 3.65E-03 | -1.39E+00 | 4.55E-02 | 1.74E-01 | 1.10E+00 | 2.30E-01 | 8.20E-01 | 1.47E+00 | 1.09E-01 | 3.07E-01 | 2.23E+00 | 7.82E-02 | 2.87E-01 | -1.04E+00 | 2.09E-01 | 5.74E-01 | 1.08E+00 |
| SLC22A5 | 6.18E-05 | 3.22E-04 | -1.46E+00 | 3.23E-04 | 1.40E-03 | -1.22E+00 | 4.21E-12 | 1.79E-11 | -2.60E+00 | 3.55E-06 | 1.36E-05 | -1.39E+00 | 1.66E-03 | 5.90E-03 | -1.91E+00 | 7.44E-02 | 2.10E-01 | 1.09E+00 | 6.31E-04 | 2.32E-03 | -1.54E+00 | 2.11E-01 | 5.79E-01 | 1.15E+00 |
| EIF2S2P3 | 6.19E-05 | 3.23E-04 | -1.30E+00 | 3.23E-04 | 1.39E-03 | -1.02E+00 | 3.12E-08 | 1.32E-07 | -1.63E+00 | 1.31E-09 | 5.01E-09 | -1.60E+00 | 3.66E-02 | 1.31E-01 | -1.20E+00 | 1.04E-01 | 2.93E-01 | -1.07E+00 | 3.96E-03 | 1.45E-02 | -1.03E+00 | 4.39E-03 | 1.20E-02 | -1.07E+00 |
| RPL7P47 | 6.20E-05 | 3.23E-04 | -1.10E+00 | 8.32E-02 | 3.60E-01 | 1.20E+00 | 1.74E-02 | 7.41E-02 | -1.08E+00 | 4.60E-02 | 1.76E-01 | 1.20E+00 | 1.72E-03 | 6.12E-03 | -1.08E+00 | 8.60E-02 | 2.43E-01 | 1.24E+00 | 1.06E-07 | 3.89E-07 | -1.16E+00 | 1.37E-01 | 3.78E-01 | 1.19E+00 |
| ZNF439 | 6.20E-05 | 3.23E-04 | -1.07E+00 | 3.93E-02 | 1.70E-01 | 1.31E+00 | 4.19E-09 | 1.78E-08 | -1.52E+00 | 6.80E-03 | 2.60E-02 | 1.07E+00 | 2.58E-03 | 9.19E-03 | -1.06E+00 | 3.26E-01 | 9.19E-01 | 1.30E+00 | 3.23E-02 | 1.18E-01 | 1.08E+00 | 3.42E-01 | 9.39E-01 | 1.35E+00 |
| ABHD8 | 6.22E-05 | 3.24E-04 | -1.22E+00 | 2.73E-04 | 1.18E-03 | -1.05E+00 | 1.08E-04 | 4.59E-04 | -1.55E+00 | 1.29E-03 | 4.92E-03 | -1.25E+00 | 6.05E-05 | 2.16E-04 | -1.32E+00 | 3.90E-04 | 1.10E-03 | -1.19E+00 | 1.01E-01 | 3.72E-01 | 1.15E+00 | 2.44E-02 | 6.70E-02 | -1.06E+00 |
| CDAN1 | 6.22E-05 | 3.24E-04 | -1.43E+00 | 2.96E-03 | 1.28E-02 | -1.02E+00 | 9.25E-04 | 3.93E-03 | -1.59E+00 | 1.22E-02 | 4.65E-02 | -1.16E+00 | 1.49E-03 | 5.33E-03 | -1.41E+00 | 4.19E-02 | 1.18E-01 | 1.03E+00 | 8.59E-05 | 3.15E-04 | -1.55E+00 | 2.75E-02 | 7.54E-02 | -1.08E+00 |
| FRA10AC1 | 6.23E-05 | 3.25E-04 | -1.10E+00 | 1.99E-02 | 8.61E-02 | 1.11E+00 | 1.10E-03 | 4.69E-03 | -1.03E+00 | 8.63E-04 | 3.30E-03 | 1.03E+00 | 6.24E-03 | 2.22E-02 | -1.10E+00 | 2.06E-02 | 5.80E-02 | 1.02E+00 | 1.33E-02 | 4.88E-02 | -1.05E+00 | 1.56E-02 | 4.27E-02 | -1.01E+00 |
| KDELC1 | 6.24E-05 | 3.25E-04 | -1.63E+00 | 6.20E-03 | 2.68E-02 | -1.05E+00 | 1.02E-09 | 4.35E-09 | -1.87E+00 | 5.90E-08 | 2.25E-07 | -1.35E+00 | 3.38E-02 | 1.21E-01 | -1.50E+00 | 1.04E-01 | 2.94E-01 | -1.27E+00 | 1.89E-05 | 6.93E-05 | -1.73E+00 | 2.07E-02 | 5.70E-02 | -1.08E+00 |
| ATP6V1E2 | 6.24E-05 | 3.25E-04 | -1.24E+00 | 2.40E-08 | 1.04E-07 | -1.61E+00 | 5.75E-02 | 2.44E-01 | -1.05E+00 | 6.25E-02 | 2.39E-01 | 1.04E+00 | 2.36E-01 | 8.43E-01 | 1.12E+00 | 1.88E-01 | 5.30E-01 | -1.15E+00 | 7.00E-02 | 2.57E-01 | -1.30E+00 | 1.69E-02 | 4.63E-02 | -1.78E+00 |
| MRPL2 | 6.25E-05 | 3.26E-04 | -1.12E+00 | 9.07E-04 | 3.92E-03 | 1.13E+00 | 4.12E-03 | 1.75E-02 | -1.03E+00 | 2.49E-04 | 9.49E-04 | 1.05E+00 | 3.35E-06 | 1.19E-05 | -1.33E+00 | 5.99E-03 | 1.69E-02 | 1.05E+00 | 5.80E-05 | 2.13E-04 | -1.12E+00 | 6.61E-03 | 1.82E-02 | 1.03E+00 |
| AC005332.9 | 6.27E-05 | 3.27E-04 | -1.23E+00 | 5.72E-03 | 2.47E-02 | 1.10E+00 | 1.81E-02 | 7.67E-02 | -1.06E+00 | 1.56E-03 | 5.95E-03 | -1.05E+00 | 1.44E-03 | 5.15E-03 | -1.19E+00 | 5.10E-04 | 1.44E-03 | -1.27E+00 | 5.13E-02 | 1.88E-01 | 1.11E+00 | 5.34E-03 | 1.47E-02 | -1.06E+00 |
| MARS2 | 6.27E-05 | 3.27E-04 | -1.97E+00 | 2.78E-03 | 1.20E-02 | -1.28E+00 | 3.63E-09 | 1.54E-08 | -3.07E+00 | 1.13E-04 | 4.30E-04 | -1.48E+00 | 3.88E-05 | 1.39E-04 | -2.37E+00 | 5.99E-03 | 1.69E-02 | -1.21E+00 | 3.37E-06 | 1.24E-05 | -1.72E+00 | 4.05E-03 | 1.11E-02 | -1.15E+00 |
| HAUS2 | 6.27E-05 | 3.27E-04 | -1.12E+00 | 2.16E-08 | 9.33E-08 | -1.30E+00 | 1.13E-05 | 4.80E-05 | -1.23E+00 | 8.41E-06 | 3.21E-05 | -1.15E+00 | 8.21E-04 | 2.93E-03 | -1.04E+00 | 6.61E-05 | 1.86E-04 | -1.25E+00 | 2.96E-04 | 1.09E-03 | -1.16E+00 | 9.08E-04 | 2.49E-03 | -1.11E+00 |
| USP19 | 6.31E-05 | 3.29E-04 | -1.23E+00 | 8.53E-03 | 3.69E-02 | 1.18E+00 | 2.30E-03 | 9.76E-03 | -1.18E+00 | 3.86E-03 | 1.47E-02 | -1.04E+00 | 1.32E-03 | 4.70E-03 | -1.41E+00 | 7.92E-02 | 2.23E-01 | 1.02E+00 | 2.29E-05 | 8.41E-05 | -1.25E+00 | 1.31E-04 | 3.59E-04 | -1.22E+00 |
| ADIPOR2 | 6.34E-05 | 3.30E-04 | -1.15E+00 | 1.56E-04 | 6.75E-04 | 1.00E+00 | 2.73E-04 | 1.16E-03 | -1.20E+00 | 1.53E-03 | 5.83E-03 | -1.00E+00 | 1.76E-02 | 6.29E-02 | -1.13E+00 | 1.26E-01 | 3.56E-01 | 1.07E+00 | 1.64E-12 | 6.01E-12 | -1.62E+00 | 2.78E-06 | 7.64E-06 | -1.29E+00 |
| STRADA | 6.34E-05 | 3.30E-04 | -1.14E+00 | 2.30E-04 | 9.92E-04 | 1.01E+00 | 7.37E-05 | 3.13E-04 | -1.31E+00 | 2.34E-04 | 8.92E-04 | -1.12E+00 | 4.46E-04 | 1.59E-03 | -1.27E+00 | 9.44E-03 | 2.66E-02 | -1.03E+00 | 6.20E-03 | 2.28E-02 | -1.15E+00 | 1.50E-01 | 4.13E-01 | 1.11E+00 |
| IL18R1 | 6.37E-05 | 3.32E-04 | -1.68E+00 | 1.68E-03 | 7.28E-03 | -1.21E+00 | 2.73E-02 | 1.16E-01 | -1.25E+00 | 3.47E-02 | 1.33E-01 | -1.18E+00 | 8.66E-02 | 3.09E-01 | -1.14E+00 | 1.15E-03 | 3.25E-03 | -2.03E+00 | 6.10E-02 | 2.24E-01 | -1.29E+00 | 1.48E-01 | 4.08E-01 | -1.23E+00 |
| PRPF3 | 6.40E-05 | 3.33E-04 | -1.29E+00 | 5.28E-04 | 2.28E-03 | -1.09E+00 | 4.41E-06 | 1.87E-05 | -1.31E+00 | 9.77E-04 | 3.73E-03 | 1.03E+00 | 1.73E-04 | 6.18E-04 | -1.30E+00 | 1.24E-02 | 3.49E-02 | -1.02E+00 | 5.54E-11 | 2.03E-10 | -1.66E+00 | 2.44E-05 | 6.69E-05 | -1.18E+00 |
| MTRF1L | 6.43E-05 | 3.35E-04 | -1.20E+00 | 5.59E-05 | 2.41E-04 | -1.06E+00 | 3.07E-03 | 1.31E-02 | -1.26E+00 | 5.87E-05 | 2.24E-04 | -1.35E+00 | 2.33E-01 | 8.31E-01 | 1.41E+00 | 1.06E-01 | 2.99E-01 | 1.08E+00 | 2.56E-02 | 9.40E-02 | 1.13E+00 | 1.39E-02 | 3.80E-02 | 1.03E+00 |
| SLC2A8 | 6.43E-05 | 3.35E-04 | -1.79E+00 | 5.30E-04 | 2.29E-03 | -1.37E+00 | 3.10E-02 | 1.32E-01 | -1.15E+00 | 4.78E-02 | 1.83E-01 | 1.03E+00 | 4.14E-02 | 1.48E-01 | -1.35E+00 | 1.15E-02 | 3.25E-02 | -1.78E+00 | 2.48E-04 | 9.10E-04 | -1.72E+00 | 6.86E-02 | 1.88E-01 | -1.05E+00 |
| TAPT1 | 6.44E-05 | 3.35E-04 | -1.04E+00 | 6.50E-10 | 2.81E-09 | -1.16E+00 | 4.99E-05 | 2.12E-04 | -1.21E+00 | 1.38E-08 | 5.26E-08 | -1.28E+00 | 3.31E-02 | 1.18E-01 | 1.10E+00 | 1.71E-03 | 4.82E-03 | -1.09E+00 | 5.74E-05 | 2.11E-04 | -1.14E+00 | 2.75E-07 | 7.54E-07 | -1.41E+00 |
| ADAM15 | 6.44E-05 | 3.36E-04 | -1.21E+00 | 5.28E-05 | 2.28E-04 | -1.11E+00 | 3.31E-02 | 1.41E-01 | 1.07E+00 | 4.93E-02 | 1.88E-01 | 1.27E+00 | 1.55E-05 | 5.53E-05 | -2.08E+00 | 3.61E-04 | 1.02E-03 | -1.62E+00 | 7.59E-02 | 2.78E-01 | 1.08E+00 | 2.03E-01 | 5.57E-01 | 1.53E+00 |
| NSF | 6.44E-05 | 3.36E-04 | -1.54E+00 | 3.88E-02 | 1.67E-01 | 1.27E+00 | 1.59E-06 | 6.75E-06 | -1.45E+00 | 2.09E-03 | 7.97E-03 | 1.04E+00 | 6.34E-03 | 2.26E-02 | -1.14E+00 | 5.32E-02 | 1.50E-01 | 1.07E+00 | 7.58E-07 | 2.78E-06 | -1.41E+00 | 2.06E-02 | 5.66E-02 | -1.01E+00 |
| RPGR | 6.45E-05 | 3.36E-04 | -1.29E+00 | 5.14E-06 | 2.22E-05 | -1.38E+00 | 3.06E-06 | 1.30E-05 | -1.24E+00 | 1.89E-12 | 7.23E-12 | -1.58E+00 | 1.39E-04 | 4.95E-04 | -1.46E+00 | 1.75E-08 | 4.92E-08 | -2.34E+00 | 1.78E-01 | 6.53E-01 | 1.11E+00 | 1.42E-02 | 3.91E-02 | -1.26E+00 |
| CARS | 6.45E-05 | 3.36E-04 | -1.34E+00 | 3.37E-02 | 1.46E-01 | 1.26E+00 | 9.37E-08 | 3.98E-07 | -1.54E+00 | 2.41E-05 | 9.22E-05 | -1.11E+00 | 1.26E-11 | 4.48E-11 | -2.55E+00 | 4.54E-05 | 1.28E-04 | -1.17E+00 | 3.97E-04 | 1.46E-03 | -1.12E+00 | 9.06E-02 | 2.49E-01 | 1.09E+00 |
| L3MBTL2 | 6.47E-05 | 3.37E-04 | -1.41E+00 | 9.43E-02 | 4.07E-01 | 1.26E+00 | 1.47E-03 | 6.23E-03 | -1.30E+00 | 8.73E-02 | 3.34E-01 | 1.21E+00 | 2.37E-04 | 8.44E-04 | -1.27E+00 | 4.04E-02 | 1.14E-01 | 1.21E+00 | 1.95E-05 | 7.14E-05 | -1.35E+00 | 1.79E-01 | 4.91E-01 | 1.19E+00 |
| DDX39B | 6.47E-05 | 3.37E-04 | 1.01E+00 | 7.89E-05 | 3.41E-04 | 1.13E+00 | 3.75E-03 | 1.59E-02 | -1.08E+00 | 3.43E-04 | 1.31E-03 | 1.00E+00 | 1.15E-02 | 4.11E-02 | 1.10E+00 | 2.22E-02 | 6.25E-02 | 1.13E+00 | 2.62E-04 | 9.63E-04 | -1.12E+00 | 9.30E-03 | 2.55E-02 | -1.06E+00 |
| ZNF720 | 6.49E-05 | 3.38E-04 | -1.30E+00 | 1.96E-01 | 8.45E-01 | 1.50E+00 | 8.94E-04 | 3.80E-03 | -1.33E+00 | 5.14E-02 | 1.96E-01 | 1.14E+00 | 1.17E-04 | 4.18E-04 | -1.49E+00 | 1.10E-02 | 3.11E-02 | -1.09E+00 | 3.02E-03 | 1.11E-02 | -1.11E+00 | 3.08E-02 | 8.47E-02 | 1.02E+00 |
| ACAP1 | 6.50E-05 | 3.39E-04 | -1.03E+00 | 2.27E-02 | 9.80E-02 | 1.23E+00 | 3.63E-03 | 1.54E-02 | -1.16E+00 | 1.28E-03 | 4.88E-03 | -1.02E+00 | 1.10E-05 | 3.92E-05 | -1.17E+00 | 1.34E-04 | 3.77E-04 | -1.12E+00 | 2.86E-02 | 1.05E-01 | 1.08E+00 | 3.97E-03 | 1.09E-02 | -1.00E+00 |
| DPH1 | 6.50E-05 | 3.39E-04 | -1.94E+00 | 2.51E-03 | 1.08E-02 | -1.31E+00 | 1.05E-02 | 4.47E-02 | -1.30E+00 | 1.31E-02 | 5.01E-02 | -1.14E+00 | 1.59E-01 | 5.68E-01 | 1.02E+00 | 9.75E-02 | 2.75E-01 | -1.23E+00 | 2.10E-01 | 7.72E-01 | 1.18E+00 | 1.12E-01 | 3.08E-01 | -1.17E+00 |
| SP140L | 6.52E-05 | 3.40E-04 | -1.06E+00 | 5.56E-06 | 2.40E-05 | 1.03E+00 | 8.26E-05 | 3.51E-04 | -1.13E+00 | 8.33E-05 | 3.18E-04 | -1.06E+00 | 1.32E-02 | 4.71E-02 | -1.04E+00 | 7.99E-03 | 2.25E-02 | -1.04E+00 | 3.99E-08 | 1.46E-07 | -1.35E+00 | 1.55E-07 | 4.27E-07 | -1.36E+00 |
| IFT46 | 6.54E-05 | 3.41E-04 | -1.37E+00 | 1.63E-04 | 7.06E-04 | -1.22E+00 | 3.09E-02 | 1.31E-01 | -1.03E+00 | 6.06E-03 | 2.31E-02 | -1.04E+00 | 1.61E-03 | 5.76E-03 | -1.39E+00 | 6.34E-02 | 1.79E-01 | 1.03E+00 | 1.11E-03 | 4.07E-03 | -1.13E+00 | 7.91E-02 | 2.17E-01 | 1.17E+00 |
| SEM1 | 6.54E-05 | 3.41E-04 | -1.01E+00 | 5.70E-19 | 2.46E-18 | -1.22E+00 | 2.28E-01 | 9.68E-01 | 1.13E+00 | 1.65E-05 | 6.29E-05 | -1.14E+00 | 1.15E-03 | 4.11E-03 | -1.06E+00 | 4.92E-05 | 1.39E-04 | -1.12E+00 | 6.05E-02 | 2.22E-01 | 1.03E+00 | 6.12E-03 | 1.68E-02 | -1.14E+00 |
| RPL26 | 6.59E-05 | 3.43E-04 | 1.15E+00 | 8.60E-29 | 3.72E-28 | -1.40E+00 | 7.10E-04 | 3.02E-03 | 1.16E+00 | 1.01E-16 | 3.85E-16 | -1.38E+00 | 3.53E-03 | 1.26E-02 | 1.15E+00 | 1.30E-11 | 3.66E-11 | -1.36E+00 | 9.06E-02 | 3.32E-01 | 1.12E+00 | 1.64E-19 | 4.52E-19 | -1.41E+00 |
| SPR | 6.60E-05 | 3.44E-04 | -2.14E+00 | 2.91E-04 | 1.26E-03 | -1.84E+00 | 4.48E-04 | 1.90E-03 | -1.69E+00 | 5.29E-04 | 2.02E-03 | -1.47E+00 | 1.38E-01 | 4.94E-01 | -1.28E+00 | 2.67E-01 | 7.54E-01 | -1.01E+00 | 7.22E-06 | 2.65E-05 | -1.24E+00 | 8.00E-02 | 2.20E-01 | 1.26E+00 |
| AC009951.1 | 6.60E-05 | 3.44E-04 | -1.61E+00 | 2.68E-02 | 1.16E-01 | 1.14E+00 | 3.09E-07 | 1.31E-06 | -1.65E+00 | 1.43E-02 | 5.45E-02 | 1.13E+00 | 2.56E-04 | 9.12E-04 | -1.21E+00 | 3.43E-01 | 9.67E-01 | 1.40E+00 | 8.89E-06 | 3.26E-05 | -1.26E+00 | 3.49E-01 | 9.58E-01 | 1.35E+00 |
| SERGEF | 6.60E-05 | 3.44E-04 | -1.12E+00 | 8.97E-04 | 3.88E-03 | 1.09E+00 | 6.73E-03 | 2.86E-02 | -1.11E+00 | 7.79E-03 | 2.98E-02 | 1.04E+00 | 3.50E-05 | 1.25E-04 | -1.28E+00 | 5.57E-02 | 1.57E-01 | 1.12E+00 | 2.36E-01 | 8.66E-01 | 1.30E+00 | 2.82E-01 | 7.75E-01 | 1.42E+00 |
| MRPL47 | 6.61E-05 | 3.45E-04 | -1.01E+00 | 1.19E-08 | 5.16E-08 | -1.06E+00 | 3.03E-03 | 1.29E-02 | 1.08E+00 | 2.68E-05 | 1.02E-04 | 1.01E+00 | 1.88E-05 | 6.69E-05 | -1.11E+00 | 5.28E-03 | 1.49E-02 | 1.12E+00 | 3.55E-04 | 1.30E-03 | -1.12E+00 | 4.59E-02 | 1.26E-01 | 1.03E+00 |
| RASGRP1 | 6.66E-05 | 3.47E-04 | -1.39E+00 | 6.04E-05 | 2.61E-04 | -1.28E+00 | 4.68E-05 | 1.99E-04 | -1.50E+00 | 1.69E-05 | 6.44E-05 | -1.43E+00 | 8.31E-03 | 2.96E-02 | -1.40E+00 | 1.76E-02 | 4.97E-02 | -1.28E+00 | 2.41E-02 | 8.86E-02 | -1.22E+00 | 6.59E-02 | 1.81E-01 | -1.04E+00 |
| SWI5 | 6.66E-05 | 3.47E-04 | -1.15E+00 | 3.32E-04 | 1.43E-03 | 1.04E+00 | 2.15E-01 | 9.14E-01 | 1.12E+00 | 7.31E-02 | 2.79E-01 | 1.17E+00 | 1.52E-03 | 5.41E-03 | -1.08E+00 | 7.89E-03 | 2.22E-02 | 1.03E+00 | 1.23E-01 | 4.52E-01 | 1.26E+00 | 1.11E-02 | 3.04E-02 | 1.07E+00 |
| RASSF8-AS1 | 6.67E-05 | 3.48E-04 | -1.30E+00 | 7.19E-05 | 3.11E-04 | -1.23E+00 | 4.51E-02 | 1.91E-01 | -1.07E+00 | 7.13E-02 | 2.72E-01 | 1.03E+00 | 2.16E-01 | 7.69E-01 | 1.54E+00 | 3.17E-01 | 8.95E-01 | 1.08E+00 | 1.89E-01 | 6.92E-01 | 1.68E+00 | 3.36E-01 | 9.23E-01 | 1.31E+00 |
| PITPNA | 6.69E-05 | 3.49E-04 | -1.35E+00 | 3.19E-03 | 1.38E-02 | 1.02E+00 | 1.34E-04 | 5.71E-04 | -1.26E+00 | 5.13E-06 | 1.96E-05 | -1.25E+00 | 5.48E-03 | 1.95E-02 | -1.28E+00 | 3.82E-02 | 1.08E-01 | -1.02E+00 | 2.01E-03 | 7.38E-03 | -1.07E+00 | 2.06E-02 | 5.66E-02 | 1.03E+00 |
| CD48 | 6.71E-05 | 3.50E-04 | -1.94E+00 | 2.25E-05 | 9.73E-05 | -1.26E+00 | 1.80E-17 | 7.65E-17 | -2.06E+00 | 3.42E-39 | 1.31E-38 | -3.63E+00 | 2.20E-01 | 7.85E-01 | 1.00E+00 | 2.08E-01 | 5.87E-01 | -1.68E+00 | 8.29E-02 | 3.04E-01 | -1.73E+00 | 9.01E-02 | 2.48E-01 | -1.68E+00 |
| SLC41A1 | 6.72E-05 | 3.50E-04 | -1.47E+00 | 2.97E-04 | 1.29E-03 | -1.25E+00 | 1.27E-04 | 5.39E-04 | -1.59E+00 | 2.31E-04 | 8.83E-04 | -1.35E+00 | 4.74E-02 | 1.69E-01 | -1.41E+00 | 1.85E-01 | 5.22E-01 | 1.09E+00 | 1.47E-01 | 5.40E-01 | 1.07E+00 | 3.40E-01 | 9.35E-01 | 1.17E+00 |
| SGMS1-AS1 | 6.77E-05 | 3.53E-04 | -1.36E+00 | 5.31E-03 | 2.30E-02 | 1.05E+00 | 3.78E-02 | 1.61E-01 | -1.15E+00 | 1.38E-01 | 5.27E-01 | 1.20E+00 | 1.81E-02 | 6.46E-02 | -1.08E+00 | 1.31E-01 | 3.69E-01 | 1.22E+00 | 7.83E-02 | 2.87E-01 | -1.05E+00 | 7.20E-02 | 1.98E-01 | -1.14E+00 |
| CPSF3 | 6.77E-05 | 3.53E-04 | -1.10E+00 | 1.24E-04 | 5.35E-04 | 1.04E+00 | 1.25E-04 | 5.31E-04 | -1.14E+00 | 3.13E-04 | 1.19E-03 | 1.06E+00 | 5.48E-03 | 1.95E-02 | 1.09E+00 | 1.40E-01 | 3.96E-01 | 1.36E+00 | 1.56E-02 | 5.71E-02 | 1.04E+00 | 3.24E-01 | 8.91E-01 | 1.24E+00 |
| HIST3H2A | 6.78E-05 | 3.53E-04 | 1.02E+00 | 1.38E-07 | 5.96E-07 | 1.03E+00 | 3.30E-06 | 1.40E-05 | -1.29E+00 | 1.34E-05 | 5.11E-05 | -1.11E+00 | 3.30E-02 | 1.18E-01 | 1.02E+00 | 1.03E-01 | 2.91E-01 | 1.10E+00 | 4.38E-02 | 1.61E-01 | -1.19E+00 | 1.39E-05 | 3.82E-05 | -1.60E+00 |
| PPP1R35 | 6.80E-05 | 3.54E-04 | -1.06E+00 | 6.49E-04 | 2.81E-03 | 1.09E+00 | 3.66E-04 | 1.56E-03 | -1.16E+00 | 4.62E-05 | 1.76E-04 | -1.09E+00 | 1.73E-06 | 6.15E-06 | -1.21E+00 | 1.81E-02 | 5.11E-02 | 1.15E+00 | 4.40E-05 | 1.62E-04 | -1.12E+00 | 7.66E-02 | 2.11E-01 | 1.18E+00 |
| ZNF142 | 6.81E-05 | 3.55E-04 | -1.72E+00 | 7.56E-03 | 3.26E-02 | -1.08E+00 | 3.59E-03 | 1.53E-02 | -1.18E+00 | 4.43E-03 | 1.69E-02 | -1.06E+00 | 2.35E-03 | 8.38E-03 | -1.62E+00 | 9.91E-02 | 2.79E-01 | 1.15E+00 | 1.24E-01 | 4.54E-01 | 1.02E+00 | 3.63E-01 | 9.98E-01 | 1.24E+00 |
| EPS15L1 | 6.83E-05 | 3.56E-04 | -1.13E+00 | 1.38E-03 | 5.97E-03 | 1.05E+00 | 6.36E-05 | 2.70E-04 | -1.26E+00 | 2.02E-04 | 7.72E-04 | -1.05E+00 | 5.98E-12 | 2.13E-11 | -1.49E+00 | 2.36E-08 | 6.65E-08 | -1.25E+00 | 6.25E-03 | 2.29E-02 | 1.04E+00 | 9.20E-02 | 2.53E-01 | 1.13E+00 |
| ZNF627 | 6.83E-05 | 3.56E-04 | -1.51E+00 | 7.34E-05 | 3.17E-04 | -1.30E+00 | 1.67E-02 | 7.07E-02 | -1.38E+00 | 2.57E-02 | 9.83E-02 | -1.15E+00 | 1.26E-01 | 4.48E-01 | 1.80E+00 | 1.20E-01 | 3.39E-01 | 1.86E+00 | 6.82E-02 | 2.50E-01 | 1.02E+00 | 3.40E-01 | 9.33E-01 | 1.25E+00 |
| R3HCC1L | 6.84E-05 | 3.56E-04 | -1.34E+00 | 1.36E-03 | 5.90E-03 | -1.01E+00 | 6.24E-04 | 2.65E-03 | -1.23E+00 | 1.56E-03 | 5.97E-03 | -1.07E+00 | 9.52E-04 | 3.39E-03 | -1.15E+00 | 2.43E-04 | 6.85E-04 | -1.30E+00 | 6.63E-06 | 2.43E-05 | -1.31E+00 | 4.89E-06 | 1.34E-05 | -1.37E+00 |
| TRRAP | 6.84E-05 | 3.57E-04 | -1.30E+00 | 3.99E-02 | 1.73E-01 | 1.32E+00 | 1.99E-02 | 8.45E-02 | -1.04E+00 | 7.82E-02 | 2.99E-01 | 1.17E+00 | 1.03E-04 | 3.68E-04 | -1.45E+00 | 5.88E-03 | 1.66E-02 | -1.22E+00 | 3.07E-03 | 1.13E-02 | -1.09E+00 | 1.94E-02 | 5.32E-02 | 1.09E+00 |
| ANKLE1 | 6.85E-05 | 3.57E-04 | -2.17E+00 | 3.17E-02 | 1.37E-01 | 1.09E+00 | 2.70E-12 | 1.15E-11 | -2.15E+00 | 2.34E-03 | 8.94E-03 | 1.02E+00 | 2.79E-04 | 9.93E-04 | -2.42E+00 | 9.38E-02 | 2.64E-01 | 1.11E+00 | 4.15E-40 | 1.52E-39 | -2.78E+00 | 5.83E-11 | 1.60E-10 | -1.28E+00 |
| SACM1L | 6.86E-05 | 3.57E-04 | -1.08E+00 | 1.98E-03 | 8.55E-03 | 1.17E+00 | 1.22E-07 | 5.19E-07 | -1.46E+00 | 1.64E-06 | 6.25E-06 | -1.18E+00 | 9.88E-05 | 3.52E-04 | -1.15E+00 | 4.66E-03 | 1.31E-02 | 1.07E+00 | 5.63E-03 | 2.07E-02 | 1.02E+00 | 5.05E-03 | 1.39E-02 | -1.05E+00 |
| AL590762.1 | 6.90E-05 | 3.60E-04 | -1.35E+00 | 8.16E-05 | 3.53E-04 | -1.16E+00 | 2.16E-05 | 9.16E-05 | -1.46E+00 | 6.75E-03 | 2.58E-02 | -1.01E+00 | 2.28E-06 | 8.11E-06 | -1.49E+00 | 3.67E-03 | 1.03E-02 | 1.00E+00 | 1.14E-09 | 4.19E-09 | -1.62E+00 | 6.78E-06 | 1.86E-05 | -1.34E+00 |
| DNAJC24 | 6.92E-05 | 3.61E-04 | -1.59E+00 | 7.12E-02 | 3.08E-01 | 1.21E+00 | 1.42E-02 | 6.04E-02 | -1.25E+00 | 1.30E-01 | 4.95E-01 | 1.20E+00 | 8.23E-02 | 2.94E-01 | 1.06E+00 | 2.10E-01 | 5.92E-01 | 1.22E+00 | 1.33E-02 | 4.89E-02 | -1.10E+00 | 5.40E-02 | 1.48E-01 | 1.05E+00 |
| AP1S2 | 6.92E-05 | 3.61E-04 | 1.02E+00 | 1.49E-01 | 6.43E-01 | 1.42E+00 | 6.66E-03 | 2.83E-02 | -1.08E+00 | 1.42E-01 | 5.43E-01 | 1.29E+00 | 1.65E-06 | 5.88E-06 | -1.17E+00 | 1.45E-02 | 4.09E-02 | 1.15E+00 | 6.32E-07 | 2.32E-06 | -1.31E+00 | 1.13E-01 | 3.11E-01 | 1.10E+00 |
| IQCH-AS1 | 6.93E-05 | 3.61E-04 | -1.58E+00 | 2.65E-02 | 1.14E-01 | 1.07E+00 | 1.50E-04 | 6.38E-04 | -1.35E+00 | 7.00E-05 | 2.67E-04 | -1.21E+00 | 8.58E-06 | 3.06E-05 | -2.57E+00 | 1.86E-03 | 5.26E-03 | -1.49E+00 | 1.50E-03 | 5.51E-03 | -1.41E+00 | 1.09E-02 | 3.00E-02 | -1.23E+00 |
| RPL15P18 | 6.95E-05 | 3.62E-04 | -1.06E+00 | 5.94E-02 | 2.57E-01 | 1.11E+00 | 4.97E-03 | 2.11E-02 | -1.14E+00 | 2.33E-01 | 8.90E-01 | 1.07E+00 | 7.01E-05 | 2.50E-04 | -1.12E+00 | 3.31E-01 | 9.34E-01 | 1.15E+00 | 1.46E-03 | 5.36E-03 | -1.13E+00 | 3.41E-01 | 9.37E-01 | 1.04E+00 |
| ZNF93 | 6.96E-05 | 3.63E-04 | -1.21E+00 | 1.09E-05 | 4.69E-05 | -1.32E+00 | 1.80E-05 | 7.65E-05 | -1.32E+00 | 4.24E-07 | 1.62E-06 | -1.31E+00 | 8.96E-05 | 3.19E-04 | -1.55E+00 | 9.65E-07 | 2.72E-06 | -2.12E+00 | 2.17E-05 | 7.96E-05 | -1.26E+00 | 4.19E-06 | 1.15E-05 | -1.36E+00 |
| VPS11 | 6.97E-05 | 3.63E-04 | -1.28E+00 | 5.75E-03 | 2.49E-02 | 1.10E+00 | 1.03E-02 | 4.39E-02 | -1.12E+00 | 5.39E-02 | 2.06E-01 | 1.15E+00 | 1.36E-02 | 4.86E-02 | -1.01E+00 | 5.95E-02 | 1.68E-01 | 1.12E+00 | 5.43E-02 | 1.99E-01 | 1.06E+00 | 3.28E-02 | 9.01E-02 | 1.04E+00 |
| RBBP9 | 6.97E-05 | 3.63E-04 | -1.84E+00 | 3.62E-02 | 1.57E-01 | 1.12E+00 | 7.48E-03 | 3.18E-02 | -1.21E+00 | 1.50E-01 | 5.75E-01 | 1.26E+00 | 2.78E-05 | 9.92E-05 | -1.72E+00 | 3.83E-04 | 1.08E-03 | -1.39E+00 | 2.30E-06 | 8.44E-06 | -1.89E+00 | 2.07E-02 | 5.68E-02 | -1.13E+00 |
| ERGIC1 | 6.98E-05 | 3.64E-04 | 1.01E+00 | 5.50E-02 | 2.38E-01 | 1.33E+00 | 3.50E-03 | 1.49E-02 | -1.03E+00 | 8.77E-02 | 3.35E-01 | 1.22E+00 | 1.36E-04 | 4.86E-04 | -1.05E+00 | 1.69E-02 | 4.76E-02 | 1.11E+00 | 6.04E-05 | 2.22E-04 | -1.09E+00 | 1.03E-02 | 2.82E-02 | 1.01E+00 |
| PLAGL2 | 6.98E-05 | 3.64E-04 | -1.51E+00 | 4.59E-06 | 1.99E-05 | -1.68E+00 | 2.12E-05 | 9.00E-05 | -1.62E+00 | 1.02E-04 | 3.90E-04 | -1.34E+00 | 6.38E-04 | 2.28E-03 | -1.21E+00 | 1.36E-05 | 3.82E-05 | -1.55E+00 | 4.74E-06 | 1.74E-05 | -1.21E+00 | 6.94E-04 | 1.91E-03 | -1.08E+00 |
| AC138649.1 | 7.00E-05 | 3.65E-04 | -1.23E+00 | 2.86E-06 | 1.24E-05 | -1.24E+00 | 2.72E-03 | 1.16E-02 | -1.13E+00 | 3.26E-05 | 1.25E-04 | -1.27E+00 | 1.06E-01 | 3.79E-01 | 1.04E+00 | 1.09E-01 | 3.08E-01 | 1.00E+00 | 1.51E-03 | 5.53E-03 | -1.23E+00 | 1.06E-06 | 2.91E-06 | -1.67E+00 |
| CDCA7 | 7.01E-05 | 3.65E-04 | -1.04E+00 | 2.21E-01 | 9.55E-01 | 1.52E+00 | 5.19E-02 | 2.20E-01 | 1.03E+00 | 1.20E-01 | 4.58E-01 | 1.41E+00 | 8.81E-05 | 3.14E-04 | -1.43E+00 | 1.21E-02 | 3.40E-02 | 1.04E+00 | 5.41E-05 | 1.99E-04 | -1.19E+00 | 2.97E-01 | 8.16E-01 | 1.33E+00 |
| GP1BA | 7.01E-05 | 3.66E-04 | -2.62E+00 | 6.58E-03 | 2.84E-02 | -1.24E+00 | 1.19E-02 | 5.04E-02 | -1.71E+00 | 4.95E-02 | 1.89E-01 | -1.20E+00 | 1.04E-04 | 3.71E-04 | -2.63E+00 | 4.59E-04 | 1.29E-03 | -2.27E+00 | 2.03E-04 | 7.44E-04 | -1.01E+00 | 3.58E-02 | 9.84E-02 | -1.20E+00 |
| THOC6 | 7.02E-05 | 3.66E-04 | -1.13E+00 | 2.84E-03 | 1.23E-02 | 1.13E+00 | 4.89E-06 | 2.08E-05 | -1.20E+00 | 9.47E-05 | 3.62E-04 | -1.01E+00 | 1.45E-06 | 5.16E-06 | -1.29E+00 | 5.56E-02 | 1.57E-01 | 1.16E+00 | 2.20E-05 | 8.09E-05 | -1.06E+00 | 3.37E-02 | 9.26E-02 | 1.10E+00 |
| ATP6V1F | 7.06E-05 | 3.68E-04 | -1.10E+00 | 6.76E-06 | 2.92E-05 | -1.04E+00 | 1.14E-02 | 4.85E-02 | -1.10E+00 | 2.50E-02 | 9.55E-02 | -1.03E+00 | 1.63E-06 | 5.81E-06 | -1.13E+00 | 3.21E-04 | 9.05E-04 | -1.05E+00 | 6.78E-03 | 2.49E-02 | -1.04E+00 | 1.68E-01 | 4.62E-01 | 1.07E+00 |
| CLCN4 | 7.06E-05 | 3.68E-04 | -2.37E+00 | 9.51E-05 | 4.11E-04 | -2.31E+00 | 1.12E-01 | 4.74E-01 | -1.26E+00 | 9.98E-02 | 3.81E-01 | -1.38E+00 | 4.43E-02 | 1.58E-01 | -1.85E+00 | 6.36E-02 | 1.79E-01 | -1.71E+00 | 8.91E-13 | 3.27E-12 | -1.53E+00 | 1.01E-15 | 2.76E-15 | -1.87E+00 |
| AL591846.2 | 7.08E-05 | 3.69E-04 | 2.70E+00 | 1.75E-01 | 7.57E-01 | 1.40E+00 | 3.29E-02 | 1.40E-01 | 1.57E+00 | 1.28E-01 | 4.89E-01 | 1.48E+00 | 1.46E-01 | 5.21E-01 | -1.22E+00 | 2.07E-01 | 5.84E-01 | -1.18E+00 | 1.89E-01 | 6.93E-01 | -1.06E+00 | 3.43E-01 | 9.42E-01 | 1.10E+00 |
| MPDZ | 7.10E-05 | 3.70E-04 | -1.34E+00 | 2.12E-01 | 9.15E-01 | 1.55E+00 | 4.93E-03 | 2.09E-02 | -1.05E+00 | 2.26E-01 | 8.63E-01 | 1.48E+00 | 1.78E-02 | 6.35E-02 | -1.33E+00 | 1.10E-01 | 3.10E-01 | 1.10E+00 | 2.11E-03 | 7.75E-03 | -1.29E+00 | 3.57E-01 | 9.81E-01 | 1.27E+00 |
| CCDC86 | 7.12E-05 | 3.71E-04 | -2.01E+00 | 2.19E-03 | 9.47E-03 | -1.25E+00 | 8.40E-12 | 3.57E-11 | -2.54E+00 | 4.90E-08 | 1.87E-07 | -1.64E+00 | 4.39E-07 | 1.56E-06 | -3.53E+00 | 4.98E-03 | 1.40E-02 | -1.31E+00 | 1.44E-15 | 5.29E-15 | -2.18E+00 | 3.29E-05 | 9.03E-05 | -1.12E+00 |
| NEDD1 | 7.13E-05 | 3.71E-04 | -1.26E+00 | 6.88E-04 | 2.97E-03 | -1.01E+00 | 2.28E-07 | 9.68E-07 | -1.36E+00 | 1.51E-06 | 5.76E-06 | -1.17E+00 | 2.22E-04 | 7.93E-04 | -1.21E+00 | 3.53E-03 | 9.94E-03 | -1.04E+00 | 6.14E-11 | 2.25E-10 | -1.75E+00 | 1.21E-06 | 3.31E-06 | -1.40E+00 |
| ENSA | 7.15E-05 | 3.73E-04 | 1.02E+00 | 3.75E-07 | 1.62E-06 | 1.01E+00 | 3.66E-02 | 1.56E-01 | 1.01E+00 | 1.87E-02 | 7.15E-02 | 1.04E+00 | 3.04E-02 | 1.09E-01 | 1.08E+00 | 1.26E-01 | 3.56E-01 | 1.16E+00 | 1.84E-01 | 6.77E-01 | 1.02E+00 | 2.68E-01 | 7.36E-01 | 1.09E+00 |
| MAP3K8 | 7.16E-05 | 3.73E-04 | 2.08E+00 | 6.43E-02 | 2.78E-01 | 1.38E+00 | 1.76E-04 | 7.46E-04 | 1.99E+00 | 6.17E-02 | 2.36E-01 | 1.14E+00 | 1.39E-01 | 4.95E-01 | 1.62E+00 | 9.59E-07 | 2.70E-06 | -1.50E+00 | 2.41E-01 | 8.84E-01 | 1.53E+00 | 7.99E-05 | 2.19E-04 | -1.33E+00 |
| LDB1 | 7.19E-05 | 3.75E-04 | -1.22E+00 | 1.25E-03 | 5.38E-03 | 1.05E+00 | 1.21E-03 | 5.12E-03 | -1.14E+00 | 3.01E-04 | 1.15E-03 | -1.06E+00 | 2.71E-11 | 9.68E-11 | -1.57E+00 | 3.07E-03 | 8.66E-03 | -1.01E+00 | 4.57E-20 | 1.68E-19 | -1.55E+00 | 1.44E-08 | 3.94E-08 | -1.08E+00 |
| EMG1 | 7.21E-05 | 3.76E-04 | -1.15E+00 | 2.42E-03 | 1.05E-02 | 1.07E+00 | 4.25E-04 | 1.81E-03 | -1.10E+00 | 1.07E-03 | 4.08E-03 | 1.09E+00 | 3.00E-05 | 1.07E-04 | -1.43E+00 | 6.42E-02 | 1.81E-01 | 1.10E+00 | 3.29E-05 | 1.21E-04 | -1.20E+00 | 9.32E-02 | 2.56E-01 | 1.13E+00 |
| NDUFB11 | 7.25E-05 | 3.78E-04 | -1.05E+00 | 8.75E-11 | 3.78E-10 | -1.08E+00 | 1.73E-01 | 7.36E-01 | 1.02E+00 | 1.01E-02 | 3.86E-02 | -1.05E+00 | 1.41E-03 | 5.02E-03 | -1.08E+00 | 6.88E-02 | 1.94E-01 | 1.03E+00 | 2.57E-02 | 9.43E-02 | -1.04E+00 | 2.22E-01 | 6.11E-01 | 1.01E+00 |
| TRIM26 | 7.26E-05 | 3.79E-04 | -1.48E+00 | 1.38E-04 | 5.98E-04 | -1.29E+00 | 5.42E-03 | 2.30E-02 | -1.17E+00 | 2.63E-04 | 1.00E-03 | -1.19E+00 | 1.75E-02 | 6.23E-02 | -1.25E+00 | 1.41E-02 | 3.97E-02 | -1.32E+00 | 1.61E-11 | 5.90E-11 | -2.02E+00 | 1.60E-09 | 4.40E-09 | -1.77E+00 |
| WDR5B | 7.27E-05 | 3.79E-04 | -1.57E+00 | 9.00E-03 | 3.89E-02 | -1.02E+00 | 2.97E-05 | 1.26E-04 | -1.62E+00 | 1.08E-04 | 4.13E-04 | -1.36E+00 | 4.53E-04 | 1.62E-03 | -1.65E+00 | 5.23E-03 | 1.47E-02 | -1.32E+00 | 5.09E-02 | 1.87E-01 | -1.06E+00 | 1.51E-02 | 4.15E-02 | -1.18E+00 |
| MOB1B | 7.27E-05 | 3.79E-04 | -1.22E+00 | 2.85E-03 | 1.23E-02 | 1.04E+00 | 2.30E-02 | 9.77E-02 | -1.05E+00 | 2.75E-03 | 1.05E-02 | -1.05E+00 | 3.15E-02 | 1.12E-01 | 1.01E+00 | 1.68E-01 | 4.73E-01 | 1.17E+00 | 1.55E-08 | 5.69E-08 | -1.08E+00 | 6.33E-11 | 1.74E-10 | -1.27E+00 |
| MRPL28 | 7.27E-05 | 3.79E-04 | -1.02E+00 | 1.74E-05 | 7.52E-05 | 1.03E+00 | 3.96E-03 | 1.68E-02 | 1.00E+00 | 1.87E-05 | 7.14E-05 | -1.04E+00 | 1.69E-04 | 6.02E-04 | -1.00E+00 | 8.86E-03 | 2.50E-02 | 1.12E+00 | 1.39E-01 | 5.09E-01 | 1.15E+00 | 2.15E-01 | 5.90E-01 | 1.16E+00 |
| DDX3X | 7.30E-05 | 3.80E-04 | -1.04E+00 | 1.62E-14 | 7.01E-14 | -1.13E+00 | 1.28E-02 | 5.44E-02 | 1.01E+00 | 4.09E-03 | 1.56E-02 | 1.07E+00 | 2.65E-01 | 9.44E-01 | 1.11E+00 | 3.72E-02 | 1.05E-01 | 1.11E+00 | 2.07E-04 | 7.58E-04 | -1.08E+00 | 4.97E-03 | 1.37E-02 | -1.01E+00 |
| SIRT3 | 7.33E-05 | 3.82E-04 | -1.55E+00 | 4.97E-03 | 2.15E-02 | -1.09E+00 | 2.07E-01 | 8.81E-01 | 1.25E+00 | 2.40E-01 | 9.15E-01 | 1.36E+00 | 1.86E-03 | 6.63E-03 | -1.55E+00 | 2.42E-01 | 6.82E-01 | 1.26E+00 | 1.07E-02 | 3.93E-02 | 1.01E+00 | 3.70E-02 | 1.02E-01 | -1.05E+00 |
| PMS2CL | 7.33E-05 | 3.82E-04 | -1.35E+00 | 1.05E-04 | 4.54E-04 | -1.11E+00 | 1.47E-04 | 6.25E-04 | -1.36E+00 | 5.94E-06 | 2.27E-05 | -1.47E+00 | 8.93E-05 | 3.18E-04 | -1.42E+00 | 1.23E-06 | 3.46E-06 | -1.77E+00 | 5.28E-03 | 1.94E-02 | -1.15E+00 | 1.40E-02 | 3.85E-02 | -1.18E+00 |
| GTF2IRD2P1 | 7.35E-05 | 3.83E-04 | -1.30E+00 | 8.06E-06 | 3.48E-05 | -1.38E+00 | 1.43E-02 | 6.08E-02 | -1.12E+00 | 1.29E-04 | 4.93E-04 | -1.41E+00 | 2.24E-02 | 7.98E-02 | 1.00E+00 | 2.48E-02 | 6.99E-02 | -1.01E+00 | 1.28E-11 | 4.69E-11 | -1.51E+00 | 2.96E-11 | 8.14E-11 | -1.56E+00 |
| DPH2 | 7.36E-05 | 3.84E-04 | -1.41E+00 | 1.75E-03 | 7.58E-03 | -1.08E+00 | 9.38E-05 | 3.98E-04 | -1.39E+00 | 3.10E-02 | 1.18E-01 | 1.17E+00 | 4.52E-04 | 1.61E-03 | -1.45E+00 | 1.85E-01 | 5.21E-01 | 1.24E+00 | 1.90E-04 | 6.99E-04 | -1.12E+00 | 2.56E-01 | 7.04E-01 | 1.44E+00 |
| FN1 | 7.37E-05 | 3.84E-04 | -1.80E+00 | 5.52E-02 | 2.39E-01 | 1.05E+00 | 7.77E-08 | 3.30E-07 | -2.40E+00 | 8.30E-02 | 3.17E-01 | -1.00E+00 | 1.10E-02 | 3.94E-02 | -1.48E+00 | 3.14E-01 | 8.84E-01 | 1.52E+00 | 1.37E-03 | 5.04E-03 | -1.42E+00 | 3.04E-01 | 8.34E-01 | 1.52E+00 |
| SYNE3 | 7.40E-05 | 3.86E-04 | -1.06E+00 | 1.79E-01 | 7.73E-01 | 1.60E+00 | 3.63E-02 | 1.54E-01 | -1.10E+00 | 2.39E-01 | 9.12E-01 | 1.36E+00 | 2.26E-02 | 8.04E-02 | 1.10E+00 | 1.95E-01 | 5.51E-01 | 1.23E+00 | 1.50E-01 | 5.49E-01 | 1.53E+00 | 1.49E-01 | 4.10E-01 | 1.46E+00 |
| RABEP2 | 7.42E-05 | 3.87E-04 | -1.06E+00 | 4.04E-05 | 1.74E-04 | -1.03E+00 | 7.27E-05 | 3.09E-04 | -1.26E+00 | 6.56E-06 | 2.50E-05 | -1.24E+00 | 1.02E-03 | 3.64E-03 | -1.14E+00 | 7.03E-02 | 1.98E-01 | 1.17E+00 | 6.44E-05 | 2.36E-04 | -1.36E+00 | 1.53E-02 | 4.21E-02 | -1.07E+00 |
| SELENOI | 7.42E-05 | 3.87E-04 | -1.39E+00 | 3.58E-06 | 1.55E-05 | -1.48E+00 | 5.67E-02 | 2.41E-01 | -1.01E+00 | 5.69E-03 | 2.17E-02 | -1.04E+00 | 1.39E-02 | 4.96E-02 | -1.06E+00 | 3.17E-02 | 8.94E-02 | 1.02E+00 | 7.18E-03 | 2.63E-02 | 1.00E+00 | 6.08E-02 | 1.67E-01 | 1.05E+00 |
| TAF8 | 7.43E-05 | 3.87E-04 | -1.05E+00 | 2.00E-02 | 8.66E-02 | 1.34E+00 | 6.13E-05 | 2.61E-04 | -1.21E+00 | 1.57E-02 | 6.01E-02 | 1.19E+00 | 8.60E-03 | 3.07E-02 | -1.01E+00 | 2.41E-01 | 6.80E-01 | 1.30E+00 | 5.52E-05 | 2.03E-04 | -1.17E+00 | 2.48E-02 | 6.80E-02 | 1.02E+00 |
| MROH1 | 7.45E-05 | 3.88E-04 | -1.28E+00 | 2.13E-03 | 9.20E-03 | 1.01E+00 | 2.31E-05 | 9.81E-05 | -1.32E+00 | 2.61E-05 | 9.97E-05 | -1.16E+00 | 5.32E-04 | 1.90E-03 | -1.31E+00 | 6.01E-03 | 1.70E-02 | -1.15E+00 | 6.54E-03 | 2.40E-02 | -1.19E+00 | 1.92E-02 | 5.29E-02 | -1.11E+00 |
| POP4 | 7.46E-05 | 3.89E-04 | -1.14E+00 | 2.58E-02 | 1.12E-01 | 1.23E+00 | 2.65E-04 | 1.13E-03 | -1.11E+00 | 2.31E-02 | 8.82E-02 | 1.16E+00 | 8.27E-06 | 2.95E-05 | -1.37E+00 | 5.63E-02 | 1.59E-01 | 1.26E+00 | 8.54E-06 | 3.14E-05 | -1.20E+00 | 3.57E-02 | 9.82E-02 | 1.08E+00 |
| AP001172.1 | 7.47E-05 | 3.90E-04 | -1.55E+00 | 1.42E-04 | 6.16E-04 | -1.41E+00 | 3.84E-02 | 1.63E-01 | 1.74E+00 | 2.94E-02 | 1.12E-01 | 2.02E+00 | 2.42E-01 | 8.64E-01 | -1.11E+00 | 1.96E-01 | 5.53E-01 | -1.51E+00 | 1.22E-02 | 4.46E-02 | -1.58E+00 | 6.00E-02 | 1.65E-01 | -1.29E+00 |
| AP001830.1 | 7.49E-05 | 3.90E-04 | -1.44E+00 | 1.90E-05 | 8.20E-05 | -1.42E+00 | 4.78E-03 | 2.03E-02 | -1.35E+00 | 1.28E-03 | 4.88E-03 | -1.41E+00 | 2.18E-01 | 7.76E-01 | 1.13E+00 | 2.04E-01 | 5.74E-01 | -1.05E+00 | 1.32E-02 | 4.86E-02 | -1.70E+00 | 7.35E-02 | 2.02E-01 | -1.36E+00 |
| SSBP1 | 7.53E-05 | 3.92E-04 | 1.02E+00 | 1.02E-04 | 4.41E-04 | 1.10E+00 | 1.76E-01 | 7.48E-01 | 1.09E+00 | 2.37E-01 | 9.06E-01 | 1.16E+00 | 1.38E-05 | 4.94E-05 | -1.09E+00 | 1.04E-01 | 2.94E-01 | 1.15E+00 | 2.25E-01 | 8.27E-01 | 1.12E+00 | 6.81E-03 | 1.87E-02 | 1.36E+00 |
| NELFB | 7.53E-05 | 3.93E-04 | -1.12E+00 | 2.47E-03 | 1.07E-02 | 1.10E+00 | 5.99E-03 | 2.55E-02 | 1.01E+00 | 1.29E-02 | 4.93E-02 | 1.17E+00 | 2.04E-03 | 7.28E-03 | -1.04E+00 | 2.76E-01 | 7.77E-01 | 1.38E+00 | 2.36E-03 | 8.68E-03 | 1.03E+00 | 5.25E-02 | 1.44E-01 | 1.14E+00 |
| PEX11A | 7.54E-05 | 3.93E-04 | -1.52E+00 | 1.44E-01 | 6.24E-01 | 1.39E+00 | 6.95E-03 | 2.95E-02 | -1.42E+00 | 2.39E-01 | 9.15E-01 | 1.48E+00 | 1.36E-02 | 4.86E-02 | -1.13E+00 | 4.18E-02 | 1.18E-01 | 1.06E+00 | 1.85E-04 | 6.80E-04 | -1.35E+00 | 1.49E-01 | 4.08E-01 | 1.13E+00 |
| SUN1 | 7.55E-05 | 3.93E-04 | -1.30E+00 | 9.76E-04 | 4.22E-03 | -1.03E+00 | 1.07E-05 | 4.54E-05 | -1.39E+00 | 3.91E-05 | 1.49E-04 | -1.17E+00 | 1.00E-02 | 3.57E-02 | 1.12E+00 | 5.82E-02 | 1.64E-01 | 1.31E+00 | 3.79E-05 | 1.39E-04 | -1.19E+00 | 9.45E-02 | 2.60E-01 | 1.13E+00 |
| PPIP5K1 | 7.57E-05 | 3.95E-04 | -1.80E+00 | 1.70E-02 | 7.34E-02 | 1.05E+00 | 2.58E-03 | 1.10E-02 | -1.61E+00 | 3.97E-02 | 1.52E-01 | -1.10E+00 | 4.35E-02 | 1.55E-01 | -1.10E+00 | 2.75E-01 | 7.76E-01 | 1.37E+00 | 1.39E-03 | 5.09E-03 | -1.51E+00 | 3.38E-02 | 9.28E-02 | -1.19E+00 |
| NIF3L1 | 7.59E-05 | 3.96E-04 | -1.99E+00 | 7.23E-02 | 3.12E-01 | 1.20E+00 | 2.64E-05 | 1.12E-04 | -1.53E+00 | 2.14E-02 | 8.16E-02 | 1.13E+00 | 6.63E-03 | 2.36E-02 | -1.50E+00 | 1.43E-01 | 4.04E-01 | 1.07E+00 | 5.41E-10 | 1.99E-09 | -1.67E+00 | 1.60E-01 | 4.41E-01 | 1.24E+00 |
| PSMB4 | 7.61E-05 | 3.96E-04 | -1.06E+00 | 1.27E-04 | 5.50E-04 | 1.03E+00 | 1.51E-02 | 6.43E-02 | -1.15E+00 | 4.96E-03 | 1.89E-02 | -1.07E+00 | 3.54E-09 | 1.26E-08 | -1.31E+00 | 2.39E-04 | 6.73E-04 | -1.05E+00 | 2.97E-05 | 1.09E-04 | -1.26E+00 | 3.22E-02 | 8.84E-02 | -1.06E+00 |
| HESX1 | 7.61E-05 | 3.97E-04 | -1.20E+00 | 8.89E-03 | 3.84E-02 | 1.19E+00 | 2.48E-08 | 1.06E-07 | -1.75E+00 | 3.29E-08 | 1.26E-07 | -1.49E+00 | 2.22E-02 | 7.92E-02 | -1.97E+00 | 1.28E-01 | 3.61E-01 | -1.18E+00 | 3.06E-02 | 1.12E-01 | -1.38E+00 | 1.09E-01 | 3.00E-01 | -1.14E+00 |
| SYF2 | 7.62E-05 | 3.97E-04 | -1.00E+00 | 2.10E-05 | 9.05E-05 | 1.06E+00 | 8.44E-02 | 3.59E-01 | 1.02E+00 | 4.06E-02 | 1.55E-01 | 1.06E+00 | 2.46E-02 | 8.78E-02 | 1.10E+00 | 1.13E-03 | 3.18E-03 | -1.00E+00 | 9.06E-02 | 3.32E-01 | 1.10E+00 | 6.62E-03 | 1.82E-02 | -1.03E+00 |
| MSANTD2 | 7.64E-05 | 3.98E-04 | -1.79E+00 | 8.55E-03 | 3.70E-02 | -1.03E+00 | 4.73E-04 | 2.01E-03 | -1.72E+00 | 1.50E-02 | 5.75E-02 | -1.09E+00 | 3.30E-03 | 1.18E-02 | -1.73E+00 | 5.59E-02 | 1.58E-01 | -1.06E+00 | 1.16E-02 | 4.26E-02 | -1.29E+00 | 2.43E-01 | 6.68E-01 | 1.20E+00 |
| LRRC1 | 7.64E-05 | 3.98E-04 | -1.52E+00 | 3.40E-03 | 1.47E-02 | -1.02E+00 | 5.09E-05 | 2.16E-04 | -1.62E+00 | 5.37E-04 | 2.05E-03 | -1.25E+00 | 1.74E-02 | 6.20E-02 | -1.33E+00 | 1.47E-02 | 4.15E-02 | -1.42E+00 | 1.80E-04 | 6.61E-04 | -1.89E+00 | 1.57E-02 | 4.30E-02 | -1.38E+00 |
| ZNF358 | 7.65E-05 | 3.99E-04 | -1.30E+00 | 3.60E-02 | 1.56E-01 | 1.22E+00 | 3.31E-03 | 1.41E-02 | -1.19E+00 | 7.04E-03 | 2.69E-02 | -1.02E+00 | 3.37E-04 | 1.20E-03 | -1.18E+00 | 1.90E-03 | 5.37E-03 | -1.07E+00 | 6.00E-03 | 2.20E-02 | -1.05E+00 | 4.07E-03 | 1.12E-02 | -1.07E+00 |
| GPN3 | 7.67E-05 | 4.00E-04 | -1.34E+00 | 1.31E-03 | 5.68E-03 | -1.04E+00 | 1.89E-02 | 8.05E-02 | -1.01E+00 | 1.00E-02 | 3.84E-02 | 1.08E+00 | 8.04E-05 | 2.87E-04 | -1.52E+00 | 6.27E-03 | 1.77E-02 | -1.11E+00 | 1.38E-05 | 5.06E-05 | -1.17E+00 | 5.04E-03 | 1.39E-02 | 1.06E+00 |
| C11orf84 | 7.67E-05 | 4.00E-04 | -1.49E+00 | 8.16E-04 | 3.53E-03 | -1.21E+00 | 7.73E-04 | 3.28E-03 | -1.33E+00 | 3.63E-02 | 1.39E-01 | 1.15E+00 | 8.12E-04 | 2.89E-03 | -1.87E+00 | 5.09E-02 | 1.43E-01 | -1.09E+00 | 3.21E-06 | 1.18E-05 | -1.35E+00 | 4.64E-04 | 1.28E-03 | -1.16E+00 |
| ARHGAP25 | 7.68E-05 | 4.00E-04 | -1.12E+00 | 3.10E-04 | 1.34E-03 | 1.08E+00 | 1.01E-05 | 4.28E-05 | -1.36E+00 | 5.61E-04 | 2.14E-03 | -1.06E+00 | 1.37E-05 | 4.89E-05 | -1.84E+00 | 4.52E-06 | 1.27E-05 | -1.96E+00 | 1.04E-04 | 3.82E-04 | -1.61E+00 | 2.72E-05 | 7.48E-05 | -1.99E+00 |
| SCARB1 | 7.69E-05 | 4.01E-04 | -1.23E+00 | 1.13E-03 | 4.87E-03 | 1.02E+00 | 1.34E-02 | 5.67E-02 | -1.06E+00 | 9.99E-02 | 3.82E-01 | 1.27E+00 | 3.65E-03 | 1.30E-02 | -1.27E+00 | 2.17E-02 | 6.11E-02 | -1.05E+00 | 1.25E-03 | 4.58E-03 | -1.25E+00 | 2.56E-01 | 7.02E-01 | 1.20E+00 |
| GPR63 | 7.69E-05 | 4.01E-04 | -1.96E+00 | 2.86E-02 | 1.24E-01 | -1.10E+00 | 5.16E-02 | 2.19E-01 | -1.09E+00 | 1.74E-01 | 6.66E-01 | 1.55E+00 | 2.01E-02 | 7.17E-02 | -1.60E+00 | 1.29E-01 | 3.64E-01 | -1.02E+00 | 1.85E-07 | 6.78E-07 | -2.62E+00 | 6.84E-04 | 1.88E-03 | -1.55E+00 |
| RAF1 | 7.70E-05 | 4.01E-04 | -1.11E+00 | 5.05E-05 | 2.18E-04 | -1.01E+00 | 1.85E-06 | 7.85E-06 | -1.30E+00 | 3.05E-11 | 1.16E-10 | -1.37E+00 | 2.04E-08 | 7.28E-08 | -1.37E+00 | 1.04E-05 | 2.94E-05 | -1.18E+00 | 1.10E-04 | 4.02E-04 | -1.08E+00 | 2.27E-04 | 6.24E-04 | -1.08E+00 |
| PAQR6 | 7.71E-05 | 4.02E-04 | -1.65E+00 | 2.48E-05 | 1.07E-04 | -1.73E+00 | 1.59E-01 | 6.77E-01 | -1.06E+00 | 4.22E-02 | 1.61E-01 | -1.27E+00 | 7.45E-02 | 2.66E-01 | 1.84E+00 | 2.52E-01 | 7.10E-01 | -1.00E+00 | 1.53E-04 | 5.63E-04 | -1.69E+00 | 9.59E-05 | 2.63E-04 | -2.16E+00 |
| MYZAP | 7.72E-05 | 4.03E-04 | 2.79E+00 | 6.67E-08 | 2.88E-07 | 4.13E+00 | 2.82E-02 | 1.20E-01 | 2.42E+00 | 7.85E-03 | 3.00E-02 | 3.39E+00 | 1.11E-02 | 3.95E-02 | 3.38E+00 | 1.18E-01 | 3.32E-01 | 2.35E+00 | 2.32E-02 | 8.51E-02 | 5.17E+00 | 5.57E-02 | 1.53E-01 | 2.39E+00 |
| NUDT8 | 7.74E-05 | 4.04E-04 | 1.90E+00 | 1.72E-02 | 7.42E-02 | 1.83E+00 | 1.51E-09 | 6.44E-09 | 2.11E+00 | 1.69E-04 | 6.47E-04 | 1.87E+00 | 1.39E-01 | 4.97E-01 | 1.85E+00 | 2.25E-01 | 6.34E-01 | 1.87E+00 | 2.56E-01 | 9.41E-01 | 1.23E+00 | 3.16E-01 | 8.67E-01 | 1.34E+00 |
| WWP2 | 7.75E-05 | 4.04E-04 | -1.36E+00 | 3.01E-04 | 1.30E-03 | -1.12E+00 | 1.68E-02 | 7.15E-02 | -1.13E+00 | 2.34E-02 | 8.93E-02 | 1.01E+00 | 7.95E-04 | 2.83E-03 | -1.37E+00 | 5.15E-03 | 1.45E-02 | -1.16E+00 | 1.61E-02 | 5.90E-02 | -1.06E+00 | 1.21E-02 | 3.32E-02 | -1.16E+00 |
| COQ5 | 7.76E-05 | 4.04E-04 | -1.15E+00 | 1.31E-02 | 5.64E-02 | 1.22E+00 | 6.28E-07 | 2.67E-06 | -1.34E+00 | 1.10E-07 | 4.18E-07 | -1.26E+00 | 9.09E-05 | 3.24E-04 | -1.24E+00 | 2.76E-02 | 7.78E-02 | 1.11E+00 | 1.25E-06 | 4.57E-06 | -1.23E+00 | 8.95E-04 | 2.46E-03 | -1.05E+00 |

| | | | | | | | | | | | | | | | | | | | | | | | | |
|---|---|---|---|---|---|---|---|---|---|---|---|---|---|---|---|---|---|---|---|---|---|---|---|---|
| AC104563.1 | 7.76E-05 | 4.05E-04 | -1.10E+00 | 2.09E-97 | 9.05E-97 | -1.81E+00 | 5.27E-04 | 2.24E-03 | -1.12E+00 | 4.12E-61 | 1.57E-60 | -1.80E+00 | 6.21E-04 | 2.21E-03 | -1.13E+00 | 2.94E-45 | 8.28E-45 | -1.74E+00 | 5.68E-08 | 2.09E-07 | -1.17E+00 | 4.47E-65 | 1.23E-64 | -1.82E+00 |
| TBRG1 | 7.81E-05 | 4.07E-04 | -1.11E+00 | 5.65E-06 | 2.44E-05 | -1.09E+00 | 3.80E-03 | 1.62E-02 | -1.06E+00 | 9.50E-05 | 3.63E-04 | -1.08E+00 | 2.05E-02 | 7.31E-02 | -1.08E+00 | 6.61E-04 | 1.86E-03 | -1.28E+00 | 1.12E-03 | 4.12E-03 | 1.02E+00 | 4.72E-06 | 1.30E-05 | -1.25E+00 |
| TPI1P1 | 7.82E-05 | 4.07E-04 | -1.61E+00 | 2.65E-06 | 1.15E-05 | -1.91E+00 | 1.44E-05 | 6.10E-05 | -1.56E+00 | 1.38E-05 | 5.29E-05 | -1.38E+00 | 8.18E-02 | 2.92E-01 | 1.15E+00 | 2.85E-02 | 8.03E-02 | -1.03E+00 | 1.03E-01 | 3.77E-01 | 1.15E+00 | 4.38E-02 | 1.20E-01 | 1.03E+00 |
| MRRF | 7.82E-05 | 4.08E-04 | -1.10E+00 | 7.30E-05 | 3.16E-04 | -1.02E+00 | 2.95E-03 | 1.25E-02 | -1.20E+00 | 4.82E-03 | 1.84E-02 | -1.01E+00 | 4.99E-04 | 1.78E-03 | -1.46E+00 | 1.86E-01 | 5.25E-01 | 1.16E+00 | 2.59E-02 | 9.51E-02 | 1.05E+00 | 3.64E-02 | 9.99E-02 | 1.09E+00 |
| AC008982.2 | 7.83E-05 | 4.08E-04 | -1.24E+00 | 4.94E-05 | 2.14E-04 | -1.18E+00 | 6.42E-03 | 2.73E-02 | -1.14E+00 | 2.38E-03 | 9.10E-03 | -1.11E+00 | 1.43E-02 | 5.11E-02 | -1.09E+00 | 3.03E-02 | 8.55E-02 | -1.03E+00 | 1.11E-06 | 4.09E-06 | -1.73E+00 | 1.43E-03 | 3.93E-03 | -1.33E+00 |
| SERPINB6 | 7.87E-05 | 4.10E-04 | -1.10E+00 | 8.03E-02 | 3.47E-01 | 1.12E+00 | 6.69E-02 | 2.84E-01 | -1.06E+00 | 1.51E-01 | 5.75E-01 | 1.11E+00 | 1.08E-02 | 3.84E-02 | -1.00E+00 | 2.45E-01 | 6.91E-01 | 1.28E+00 | 3.06E-02 | 1.12E-01 | -1.01E+00 | 2.49E-01 | 6.84E-01 | 1.18E+00 |
| TBC1D8B | 7.89E-05 | 4.11E-04 | -1.40E+00 | 3.48E-04 | 1.50E-03 | -1.21E+00 | 1.15E-10 | 4.89E-10 | -2.72E+00 | 1.05E-05 | 4.01E-05 | -1.52E+00 | 1.29E-05 | 4.59E-05 | -1.69E+00 | 2.19E-04 | 6.16E-04 | -1.38E+00 | 2.44E-04 | 8.95E-04 | -1.42E+00 | 1.05E-02 | 2.89E-02 | -1.05E+00 |
| SLC35D1 | 7.90E-05 | 4.12E-04 | -1.44E+00 | 1.43E-04 | 6.17E-04 | -1.21E+00 | 2.98E-03 | 1.27E-02 | -1.32E+00 | 4.73E-03 | 1.81E-02 | -1.19E+00 | 2.43E-02 | 8.66E-02 | 1.00E+00 | 6.50E-03 | 1.83E-02 | -1.13E+00 | 5.44E-03 | 2.00E-02 | -1.09E+00 | 9.77E-04 | 2.68E-03 | -1.15E+00 |
| FBLN1 | 7.90E-05 | 4.12E-04 | -2.51E+00 | 1.16E-04 | 4.99E-04 | -2.45E+00 | 1.27E-10 | 5.40E-10 | -1.69E+00 | 1.18E-16 | 4.49E-16 | -1.86E+00 | 9.54E-03 | 3.40E-02 | -1.88E+00 | 5.64E-03 | 1.59E-02 | -2.30E+00 | 3.79E-02 | 1.39E-01 | -1.69E+00 | 6.28E-02 | 1.72E-01 | -1.70E+00 |
| CNEP1R1 | 7.90E-05 | 4.12E-04 | -1.58E+00 | 1.59E-03 | 6.87E-03 | -1.19E+00 | 1.18E-01 | 5.01E-01 | 1.09E+00 | 1.79E-01 | 6.83E-01 | 1.38E+00 | 3.70E-02 | 1.32E-01 | -1.03E+00 | 3.53E-03 | 9.95E-03 | -1.41E+00 | 2.72E-01 | 1.00E+00 | 1.32E+00 | 1.57E-01 | 4.30E-01 | 1.04E+00 |
| AC135983.2 | 7.91E-05 | 4.12E-04 | -1.15E+00 | 1.49E-01 | 6.46E-01 | 1.43E+00 | 8.48E-05 | 3.60E-04 | -1.32E+00 | 1.65E-03 | 6.32E-03 | 1.01E+00 | 2.03E-02 | 7.23E-02 | 1.00E+00 | 2.26E-01 | 6.37E-01 | 1.33E+00 | 2.10E-04 | 7.69E-04 | -1.14E+00 | 2.79E-02 | 7.67E-02 | 1.05E+00 |
| RSBN1L | 7.95E-05 | 4.15E-04 | -1.05E+00 | 3.60E-10 | 1.56E-09 | -1.16E+00 | 3.50E-04 | 1.49E-03 | -1.09E+00 | 1.79E-05 | 6.84E-05 | -1.06E+00 | 6.87E-09 | 2.45E-08 | -1.28E+00 | 4.50E-06 | 1.27E-05 | -1.12E+00 | 6.74E-07 | 2.47E-06 | -1.22E+00 | 2.51E-03 | 6.89E-03 | -1.00E+00 |
| ELMO1 | 7.96E-05 | 4.15E-04 | 1.05E+00 | 7.76E-04 | 3.35E-03 | 1.72E+00 | 3.99E-04 | 1.69E-03 | -1.12E+00 | 1.49E-01 | 5.68E-01 | 1.39E+00 | 1.11E-01 | 3.97E-01 | 1.41E+00 | 8.49E-02 | 2.39E-01 | 1.47E+00 | 3.14E-03 | 1.15E-02 | -1.06E+00 | 3.07E-01 | 8.44E-01 | 1.15E+00 |
| PTP4A3 | 7.96E-05 | 4.15E-04 | -1.19E+00 | 9.00E-02 | 3.89E-01 | 1.35E+00 | 1.10E-02 | 4.68E-02 | -1.19E+00 | 8.40E-03 | 3.21E-02 | -1.06E+00 | 1.90E-02 | 6.79E-02 | -1.29E+00 | 1.67E-01 | 4.70E-01 | 1.07E+00 | 2.02E-01 | 7.40E-01 | 1.16E+00 | 3.48E-01 | 9.56E-01 | 1.14E+00 |
| FLOT2 | 7.96E-05 | 4.15E-04 | -1.21E+00 | 2.03E-02 | 8.76E-02 | 1.26E+00 | 6.25E-05 | 2.65E-04 | -1.27E+00 | 1.83E-02 | 6.99E-02 | 1.18E+00 | 5.73E-08 | 2.04E-07 | -1.56E+00 | 2.17E-04 | 6.12E-04 | -1.04E+00 | 2.33E-08 | 8.55E-08 | -1.20E+00 | 2.36E-06 | 6.48E-06 | -1.13E+00 |
| SAMD15 | 7.98E-05 | 4.16E-04 | -1.91E+00 | 3.14E-06 | 1.35E-05 | -2.47E+00 | 7.60E-03 | 3.23E-02 | -1.16E+00 | 1.40E-04 | 5.36E-04 | -1.47E+00 | 7.37E-02 | 2.63E-01 | 1.89E+00 | 3.00E-01 | 8.47E-01 | 1.41E+00 | 2.40E-01 | 8.80E-01 | 1.15E+00 | 2.32E-01 | 6.36E-01 | -1.04E+00 |
| GPR3 | 7.98E-05 | 4.16E-04 | -1.57E+00 | 4.96E-07 | 2.14E-06 | -2.07E+00 | 2.13E-02 | 9.06E-02 | -1.12E+00 | 1.18E-03 | 4.52E-03 | -1.40E+00 | 1.72E-04 | 6.13E-04 | -1.44E+00 | 1.08E-06 | 3.03E-06 | -2.50E+00 | 1.13E-02 | 4.14E-02 | -1.70E+00 | 2.34E-02 | 6.44E-02 | -1.65E+00 |
| PCDHGA10 | 7.98E-05 | 4.16E-04 | -1.43E+00 | 1.02E-01 | 4.41E-01 | 1.23E+00 | 3.16E-06 | 1.34E-05 | -1.89E+00 | 1.02E-02 | 3.91E-02 | 1.02E+00 | 2.04E-03 | 7.28E-03 | -1.20E+00 | 8.43E-02 | 2.38E-01 | 1.20E+00 | 3.47E-02 | 1.27E-01 | -1.16E+00 | 2.65E-01 | 7.29E-01 | 1.10E+00 |
| AC034236.1 | 7.98E-05 | 4.16E-04 | 1.18E+00 | 2.52E-09 | 1.09E-08 | 1.29E+00 | 7.63E-03 | 3.24E-02 | 1.11E+00 | 1.78E-06 | 6.80E-06 | 1.26E+00 | 5.35E-02 | 1.91E-01 | 1.14E+00 | 2.12E-07 | 5.98E-07 | 1.34E+00 | 2.39E-01 | 8.78E-01 | 1.10E+00 | 1.34E-06 | 3.68E-06 | 1.26E+00 |
| COX20 | 8.00E-05 | 4.17E-04 | -1.36E+00 | 1.83E-06 | 7.91E-06 | -1.55E+00 | 8.20E-03 | 3.48E-02 | -1.34E+00 | 1.50E-02 | 5.75E-02 | -1.19E+00 | 1.99E-04 | 7.11E-04 | -1.57E+00 | 1.89E-05 | 5.32E-05 | -1.81E+00 | 6.70E-02 | 2.46E-01 | -1.09E+00 | 1.24E-02 | 3.42E-02 | -1.44E+00 |
| SLC4A8 | 8.01E-05 | 4.17E-04 | -1.62E+00 | 5.44E-03 | 2.35E-02 | -1.05E+00 | 1.54E-01 | 6.54E-01 | 1.04E+00 | 1.77E-01 | 6.77E-01 | 1.78E+00 | 5.09E-03 | 1.82E-02 | -1.16E+00 | 3.40E-02 | 9.58E-02 | 1.05E+00 | 4.32E-02 | 1.58E-01 | -1.10E+00 | 2.49E-01 | 6.85E-01 | 1.18E+00 |
| TIMM10 | 8.01E-05 | 4.17E-04 | -1.04E+00 | 6.27E-06 | 2.71E-05 | -1.07E+00 | 1.44E-01 | 6.13E-01 | 1.18E+00 | 1.00E-02 | 3.83E-02 | 1.12E+00 | 4.19E-03 | 1.49E-02 | -1.10E+00 | 1.76E-02 | 4.97E-02 | -1.06E+00 | 1.01E-01 | 3.71E-01 | 1.01E+00 | 6.13E-02 | 1.68E-01 | -1.05E+00 |
| PIGT | 8.01E-05 | 4.17E-04 | -1.01E+00 | 2.38E-05 | 1.03E-04 | 1.07E+00 | 4.40E-02 | 1.87E-01 | 1.03E+00 | 1.04E-02 | 3.95E-02 | 1.04E+00 | 5.15E-02 | 1.84E-01 | 1.17E+00 | 3.05E-02 | 8.59E-02 | 1.12E+00 | 4.11E-02 | 1.51E-01 | 1.07E+00 | 3.44E-01 | 9.44E-01 | 1.17E+00 |
| WDR49 | 8.02E-05 | 4.18E-04 | -1.29E+00 | 1.47E-01 | 6.35E-01 | 1.54E+00 | 1.79E-03 | 7.63E-03 | -1.38E+00 | 7.73E-02 | 2.95E-01 | 1.04E+00 | 2.42E-01 | 8.64E-01 | 1.53E+00 | 2.64E-01 | 7.45E-01 | 1.08E+00 | 1.25E-01 | 4.60E-01 | 1.73E+00 | 1.18E-01 | 3.25E-01 | 1.96E+00 |
| TMEM198B | 8.02E-05 | 4.18E-04 | -1.31E+00 | 1.98E-04 | 8.58E-04 | -1.16E+00 | 9.28E-04 | 3.94E-03 | -1.42E+00 | 2.30E-03 | 8.80E-03 | -1.29E+00 | 6.73E-02 | 2.40E-01 | -1.04E+00 | 1.82E-01 | 5.13E-01 | 1.08E+00 | 1.02E-02 | 3.76E-02 | -1.20E+00 | 1.77E-03 | 4.86E-03 | -1.40E+00 |
| PEX16 | 8.04E-05 | 4.19E-04 | 1.02E+00 | 1.73E-06 | 7.47E-06 | 1.03E+00 | 8.65E-03 | 3.68E-02 | -1.01E+00 | 1.36E-02 | 5.19E-02 | 1.08E+00 | 1.76E-03 | 6.28E-03 | 1.01E+00 | 4.98E-02 | 1.40E-01 | 1.17E+00 | 2.30E-01 | 8.44E-01 | 1.27E+00 | 1.43E-01 | 3.92E-01 | 1.16E+00 |
| CASKIN2 | 8.04E-05 | 4.19E-04 | -1.67E+00 | 1.95E-05 | 8.42E-05 | -1.79E+00 | 3.62E-03 | 1.54E-02 | -1.23E+00 | 1.40E-02 | 5.36E-02 | 1.01E+00 | 5.60E-02 | 2.00E-01 | -1.56E+00 | 1.79E-01 | 5.04E-01 | -1.29E+00 | 1.39E-02 | 5.11E-02 | -1.26E+00 | 9.21E-02 | 2.53E-01 | -1.03E+00 |
| STK25 | 8.04E-05 | 4.19E-04 | -1.02E+00 | 4.57E-04 | 1.97E-03 | 1.13E+00 | 4.02E-04 | 1.71E-03 | -1.18E+00 | 2.83E-03 | 1.08E-02 | 1.02E+00 | 5.34E-04 | 1.90E-03 | -1.02E+00 | 3.09E-02 | 8.72E-02 | 1.20E+00 | 8.46E-03 | 3.10E-02 | 1.00E+00 | 1.55E-01 | 4.25E-01 | 1.12E+00 |
| FBXW7 | 8.05E-05 | 4.19E-04 | -1.12E+00 | 3.09E-07 | 1.33E-06 | -1.09E+00 | 7.32E-05 | 3.11E-04 | -1.28E+00 | 3.12E-04 | 1.19E-03 | -1.07E+00 | 1.16E-04 | 4.15E-04 | -1.28E+00 | 2.54E-05 | 7.17E-05 | -1.31E+00 | 5.98E-08 | 2.20E-07 | -1.47E+00 | 9.23E-06 | 2.54E-05 | -1.35E+00 |
| LDHAP2 | 8.06E-05 | 4.20E-04 | -1.62E+00 | 2.26E-04 | 9.75E-04 | -1.46E+00 | 5.60E-02 | 2.38E-01 | -1.04E+00 | 5.28E-02 | 2.02E-01 | -1.04E+00 | 1.18E-03 | 4.22E-03 | -2.04E+00 | 6.58E-03 | 1.86E-02 | -1.62E+00 | 3.33E-03 | 1.22E-02 | -1.27E+00 | 1.00E-01 | 2.76E-01 | 1.07E+00 |
| LYST | 8.08E-05 | 4.21E-04 | -1.40E+00 | 5.08E-02 | 2.19E-01 | 1.16E+00 | 4.45E-07 | 1.89E-06 | -1.49E+00 | 9.87E-05 | 3.77E-04 | -1.20E+00 | 1.30E-03 | 4.65E-03 | -1.33E+00 | 7.71E-03 | 2.17E-02 | -1.16E+00 | 3.48E-03 | 1.28E-02 | -1.15E+00 | 6.60E-03 | 1.81E-02 | -1.12E+00 |
| KCNJ8 | 8.08E-05 | 4.21E-04 | -1.63E+00 | 9.47E-03 | 4.09E-02 | -1.02E+00 | 1.45E-02 | 6.18E-02 | -1.71E+00 | 1.66E-01 | 6.34E-01 | 1.16E+00 | 5.17E-05 | 1.84E-04 | -1.65E+00 | 4.66E-05 | 1.31E-04 | -1.67E+00 | 1.86E-01 | 6.84E-01 | -1.05E+00 | 3.24E-01 | 8.90E-01 | 1.04E+00 |
| RPL12P14 | 8.09E-05 | 4.21E-04 | -1.17E+00 | 4.21E-08 | 1.82E-07 | -1.48E+00 | 1.51E-01 | 6.41E-01 | 1.07E+00 | 3.68E-05 | 1.41E-04 | -1.50E+00 | 2.58E-03 | 9.20E-03 | -1.03E+00 | 9.49E-06 | 2.68E-05 | -1.48E+00 | 1.14E-03 | 4.19E-03 | -1.20E+00 | 6.20E-07 | 1.70E-06 | -1.67E+00 |
| KRT18P11 | 8.11E-05 | 4.23E-04 | 2.61E+00 | 2.82E-04 | 1.22E-03 | 3.01E+00 | 3.32E-03 | 1.41E-02 | 2.01E+00 | 3.18E-02 | 1.21E-01 | 1.97E+00 | 2.10E-01 | 7.49E-01 | 1.14E+00 | 3.19E-01 | 8.98E-01 | 1.23E+00 | 2.33E-01 | 8.56E-01 | 1.26E+00 | 2.28E-01 | 6.25E-01 | 1.46E+00 |
| COG8 | 8.13E-05 | 4.24E-04 | -1.62E+00 | 3.98E-02 | 1.72E-01 | 1.00E+00 | 3.40E-04 | 1.44E-03 | -1.43E+00 | 7.29E-03 | 2.78E-02 | -1.08E+00 | 3.17E-03 | 1.13E-02 | -1.39E+00 | 9.81E-02 | 2.76E-01 | 1.05E+00 | 1.15E-05 | 4.21E-05 | -1.49E+00 | 1.07E-01 | 2.95E-01 | 1.02E+00 |
| LINC01473 | 8.17E-05 | 4.26E-04 | -1.38E+00 | 3.37E-04 | 1.45E-03 | -1.13E+00 | 4.47E-04 | 1.90E-03 | -1.44E+00 | 1.04E-03 | 3.98E-03 | -1.19E+00 | 1.82E-01 | 6.50E-01 | 1.29E+00 | 2.16E-01 | 6.09E-01 | 1.77E+00 | 1.67E-01 | 6.14E-01 | 1.40E+00 | 2.20E-01 | 6.05E-01 | 1.51E+00 |
| POLI | 8.17E-05 | 4.26E-04 | -1.20E+00 | 1.30E-03 | 5.62E-03 | 1.08E+00 | 2.67E-04 | 1.13E-03 | -1.28E+00 | 9.43E-04 | 3.60E-03 | -1.02E+00 | 3.69E-03 | 1.31E-02 | -1.23E+00 | 1.01E-01 | 2.85E-01 | 1.22E+00 | 2.04E-04 | 7.50E-04 | -1.32E+00 | 1.04E-02 | 2.86E-02 | -1.11E+00 |
| AC146507.1 | 8.18E-05 | 4.26E-04 | -1.29E+00 | 2.74E-07 | 1.19E-06 | -1.60E+00 | 2.10E-02 | 8.93E-02 | -1.04E+00 | 1.04E-04 | 3.96E-04 | -1.36E+00 | 1.76E-03 | 6.27E-03 | -1.20E+00 | 2.04E-04 | 5.75E-04 | -1.56E+00 | 1.19E-03 | 4.37E-03 | -1.20E+00 | 2.28E-04 | 6.27E-04 | -1.30E+00 |
| NOP56P3 | 8.24E-05 | 4.30E-04 | -1.74E+00 | 2.69E-04 | 1.16E-03 | -1.47E+00 | 5.73E-04 | 2.44E-03 | -1.53E+00 | 4.85E-03 | 1.85E-02 | -1.09E+00 | 4.20E-02 | 1.50E-01 | -1.57E+00 | 2.67E-01 | 7.52E-01 | 1.20E+00 | 5.54E-06 | 2.03E-05 | -1.59E+00 | 2.35E-07 | 6.45E-07 | -1.78E+00 |
| SPATS2L | 8.26E-05 | 4.30E-04 | -1.18E+00 | 1.41E-02 | 6.08E-02 | 1.19E+00 | 2.31E-02 | 9.83E-02 | -1.12E+00 | 2.27E-01 | 8.66E-01 | 1.36E+00 | 2.95E-02 | 1.05E-01 | -1.50E+00 | 2.95E-02 | 8.31E-02 | -1.47E+00 | 6.61E-03 | 2.43E-02 | -1.91E+00 | 2.66E-02 | 7.32E-02 | -1.62E+00 |
| ZDHHC17 | 8.27E-05 | 4.31E-04 | -1.26E+00 | 1.11E-04 | 4.81E-04 | -1.11E+00 | 1.41E-03 | 5.99E-03 | -1.09E+00 | 4.68E-04 | 1.79E-03 | 1.02E+00 | 6.37E-05 | 2.27E-04 | -1.26E+00 | 3.35E-05 | 9.45E-05 | -1.27E+00 | 1.48E-04 | 5.45E-04 | -1.06E+00 | 2.02E-03 | 5.55E-03 | 1.05E+00 |
| NDUFB8 | 8.27E-05 | 4.31E-04 | -1.02E+00 | 2.24E-20 | 9.70E-20 | -1.35E+00 | 2.08E-02 | 8.82E-02 | -1.05E+00 | 1.41E-17 | 5.39E-17 | -1.57E+00 | 1.11E-04 | 3.96E-04 | -1.07E+00 | 4.26E-11 | 1.20E-10 | -1.44E+00 | 5.50E-02 | 2.02E-01 | -1.01E+00 | 3.32E-06 | 9.12E-06 | -1.41E+00 |
| AP000894.1 | 8.33E-05 | 4.34E-04 | -1.32E+00 | 1.03E-05 | 4.47E-05 | -1.30E+00 | 1.20E-06 | 5.11E-06 | -1.29E+00 | 5.68E-10 | 2.17E-09 | -1.34E+00 | 1.20E-06 | 4.29E-06 | -1.58E+00 | 2.92E-07 | 8.24E-07 | -1.67E+00 | 1.06E-02 | 3.90E-02 | -1.01E+00 | 4.29E-08 | 1.18E-07 | -1.45E+00 |
| NCOA2 | 8.35E-05 | 4.35E-04 | -1.14E+00 | 9.30E-04 | 4.02E-03 | 1.04E+00 | 7.66E-04 | 3.25E-03 | -1.22E+00 | 2.07E-04 | 7.92E-04 | -1.11E+00 | 2.69E-04 | 9.57E-04 | -1.16E+00 | 1.03E-04 | 2.90E-04 | -1.22E+00 | 4.85E-06 | 1.78E-05 | -1.30E+00 | 1.92E-03 | 5.28E-03 | -1.18E+00 |
| ELF4 | 8.35E-05 | 4.35E-04 | -1.37E+00 | 1.73E-04 | 7.45E-04 | -1.26E+00 | 2.90E-02 | 1.23E-01 | -1.10E+00 | 2.03E-03 | 7.77E-03 | -1.25E+00 | 2.30E-03 | 8.19E-03 | -1.17E+00 | 2.35E-02 | 6.63E-02 | 1.01E+00 | 1.46E-10 | 5.37E-10 | -1.72E+00 | 1.06E-06 | 2.90E-06 | -1.41E+00 |
| CHCHD5 | 8.38E-05 | 4.37E-04 | -1.02E+00 | 4.06E-10 | 1.76E-09 | -1.11E+00 | 1.41E-02 | 5.98E-02 | -1.11E+00 | 2.23E-07 | 8.53E-07 | -1.33E+00 | 2.96E-04 | 1.06E-03 | -1.02E+00 | 1.60E-03 | 4.51E-03 | 1.04E+00 | 7.68E-02 | 2.82E-01 | 1.04E+00 | 6.82E-04 | 1.87E-03 | -1.17E+00 |
| ME3 | 8.41E-05 | 4.38E-04 | -1.21E+00 | 4.49E-05 | 1.94E-04 | -1.06E+00 | 1.93E-02 | 8.19E-02 | 1.02E+00 | 2.45E-02 | 9.37E-02 | 1.14E+00 | 2.29E-03 | 8.16E-03 | -1.30E+00 | 5.65E-02 | 1.59E-01 | 1.15E+00 | 1.53E-04 | 5.62E-04 | -1.25E+00 | 1.50E-02 | 4.12E-02 | -1.05E+00 |
| HERC2P3 | 8.41E-05 | 4.38E-04 | -1.50E+00 | 1.14E-03 | 4.92E-03 | -1.07E+00 | 3.55E-04 | 1.51E-03 | -1.61E+00 | 1.32E-03 | 5.05E-03 | -1.18E+00 | 2.36E-01 | 8.43E-01 | 1.18E+00 | 1.95E-04 | 5.50E-04 | 2.41E+00 | 1.28E-01 | 4.70E-01 | 1.16E+00 | 9.90E-03 | 2.72E-02 | 1.02E+00 |
| DEAF1 | 8.44E-05 | 4.40E-04 | -1.12E+00 | 1.00E-02 | 4.33E-02 | 1.23E+00 | 5.24E-04 | 2.22E-03 | -1.06E+00 | 2.67E-02 | 1.02E-01 | 1.23E+00 | 1.78E-05 | 6.34E-05 | -1.28E+00 | 9.83E-03 | 2.77E-02 | 1.08E+00 | 1.76E-02 | 6.46E-02 | -1.05E+00 | 3.54E-01 | 9.73E-01 | 1.22E+00 |
| SMARCE1 | 8.46E-05 | 4.41E-04 | 1.05E+00 | 3.51E-03 | 1.51E-02 | 1.30E+00 | 9.53E-04 | 4.05E-03 | -1.09E+00 | 5.18E-03 | 1.98E-02 | 1.10E+00 | 1.20E-04 | 4.28E-04 | -1.07E+00 | 2.63E-03 | 7.40E-03 | 1.04E+00 | 2.79E-02 | 1.02E-01 | 1.07E+00 | 1.80E-01 | 4.94E-01 | 1.13E+00 |
| MIS12 | 8.52E-05 | 4.44E-04 | -1.07E+00 | 1.60E-05 | 6.90E-05 | -1.04E+00 | 6.91E-03 | 2.94E-02 | -1.07E+00 | 8.93E-03 | 3.41E-02 | 1.09E+00 | 5.14E-02 | 1.83E-01 | 1.21E+00 | 8.62E-03 | 2.43E-02 | 1.02E+00 | 1.97E-06 | 7.23E-06 | -1.32E+00 | 4.78E-04 | 1.31E-03 | -1.20E+00 |
| PLEKHA3 | 8.52E-05 | 4.44E-04 | -1.03E+00 | 5.65E-06 | 2.44E-05 | 1.03E+00 | 4.82E-02 | 2.05E-01 | 1.10E+00 | 1.69E-01 | 6.46E-01 | 1.30E+00 | 1.56E-01 | 5.57E-01 | 1.25E+00 | 8.87E-02 | 2.50E-01 | 1.21E+00 | 1.78E-01 | 6.53E-01 | 1.15E+00 | 1.17E-01 | 3.20E-01 | 1.13E+00 |
| BAD | 8.52E-05 | 4.44E-04 | 1.02E+00 | 7.01E-11 | 3.03E-10 | -1.24E+00 | 1.19E-01 | 5.07E-01 | 1.14E+00 | 3.06E-03 | 1.17E-02 | 1.04E+00 | 2.10E-03 | 7.49E-03 | -1.00E+00 | 2.15E-04 | 6.07E-04 | -1.14E+00 | 8.17E-02 | 3.00E-01 | 1.42E+00 | 1.65E-01 | 4.52E-01 | 1.17E+00 |
| SLC25A45 | 8.53E-05 | 4.44E-04 | -1.18E+00 | 2.72E-09 | 1.18E-08 | -1.48E+00 | 1.53E-02 | 6.52E-02 | -1.15E+00 | 3.01E-03 | 1.15E-02 | -1.11E+00 | 2.07E-04 | 7.38E-04 | -1.29E+00 | 2.58E-02 | 7.28E-02 | -1.04E+00 | 9.62E-05 | 3.53E-04 | -1.12E+00 | 1.68E-05 | 4.61E-05 | -1.16E+00 |
| PDE4B | 8.58E-05 | 4.47E-04 | 1.95E+00 | 1.68E-05 | 7.26E-05 | -1.70E+00 | 2.05E-04 | 8.71E-04 | 1.96E+00 | 7.60E-06 | 2.90E-05 | -1.83E+00 | 2.51E-01 | 8.97E-01 | 1.61E+00 | 1.22E-01 | 3.44E-01 | -1.92E+00 | 1.66E-01 | 6.08E-01 | 1.54E+00 | 2.57E-01 | 7.05E-01 | -1.61E+00 |
| CATSPER2 | 8.59E-05 | 4.48E-04 | -1.30E+00 | 1.14E-02 | 4.91E-02 | 1.15E+00 | 5.02E-04 | 2.13E-03 | -1.24E+00 | 4.03E-05 | 1.54E-04 | -1.26E+00 | 1.27E-04 | 4.51E-04 | -1.36E+00 | 3.45E-04 | 9.72E-04 | -1.30E+00 | 7.70E-08 | 2.83E-07 | -1.65E+00 | 7.12E-05 | 1.96E-04 | -1.28E+00 |
| OXCT1 | 8.61E-05 | 4.49E-04 | -1.20E+00 | 2.63E-05 | 1.13E-04 | -1.10E+00 | 1.29E-03 | 5.46E-03 | -1.04E+00 | 3.63E-04 | 1.39E-03 | 1.02E+00 | 1.37E-03 | 4.88E-03 | -1.18E+00 | 1.68E-02 | 4.73E-02 | -1.03E+00 | 2.47E-05 | 9.08E-05 | -1.06E+00 | 1.27E-03 | 3.50E-03 | 1.01E+00 |
| HMGB1P24 | 8.62E-05 | 4.49E-04 | -1.11E+00 | 2.14E-05 | 9.23E-05 | -1.08E+00 | 1.09E-01 | 4.62E-01 | 1.06E+00 | 2.76E-03 | 1.05E-02 | -1.01E+00 | 3.07E-05 | 1.10E-04 | -1.21E+00 | 1.71E-03 | 4.82E-03 | 1.02E+00 | 5.09E-09 | 1.87E-08 | -1.34E+00 | 3.05E-05 | 8.38E-05 | -1.21E+00 |
| ABL1 | 8.62E-05 | 4.49E-04 | -1.32E+00 | 2.19E-03 | 9.48E-03 | -1.01E+00 | 2.31E-01 | 9.83E-01 | 1.17E+00 | 1.98E-01 | 7.56E-01 | 1.21E+00 | 8.69E-04 | 3.10E-03 | -1.18E+00 | 7.94E-05 | 2.24E-04 | -1.34E+00 | 9.51E-07 | 3.49E-06 | -1.47E+00 | 6.66E-05 | 1.83E-04 | -1.31E+00 |
| MAP3K7CL | 8.64E-05 | 4.50E-04 | -1.54E+00 | 5.89E-05 | 2.55E-04 | -1.54E+00 | 4.51E-02 | 1.92E-01 | -1.29E+00 | 1.55E-01 | 5.93E-01 | 1.11E+00 | 1.63E-01 | 5.80E-01 | 1.28E+00 | 2.43E-01 | 6.86E-01 | 1.63E+00 | 9.14E-06 | 3.35E-05 | -1.87E+00 | 2.62E-04 | 7.20E-04 | -1.62E+00 |
| ASB6 | 8.68E-05 | 4.52E-04 | -1.34E+00 | 1.59E-03 | 6.87E-03 | -1.06E+00 | 3.62E-04 | 1.54E-03 | -1.33E+00 | 1.51E-03 | 5.78E-03 | -1.15E+00 | 1.70E-04 | 6.06E-04 | -1.67E+00 | 2.37E-02 | 6.69E-02 | -1.02E+00 | 8.35E-07 | 3.07E-06 | -1.43E+00 | 4.65E-02 | 1.28E-01 | 1.08E+00 |
| SURF1 | 8.70E-05 | 4.53E-04 | -1.01E+00 | 1.98E-04 | 8.56E-04 | 1.10E+00 | 4.50E-06 | 1.91E-05 | -1.19E+00 | 7.85E-07 | 3.00E-06 | -1.10E+00 | 2.44E-02 | 8.70E-02 | 1.07E+00 | 4.55E-02 | 1.28E-01 | 1.17E+00 | 2.36E-01 | 8.66E-01 | 1.23E+00 | 6.56E-02 | 1.80E-01 | 1.15E+00 |
| KAT2A | 8.70E-05 | 4.53E-04 | -1.14E+00 | 2.88E-03 | 1.25E-02 | 1.15E+00 | 1.98E-02 | 8.42E-02 | 1.02E+00 | 2.16E-01 | 8.25E-01 | 1.37E+00 | 6.13E-03 | 2.19E-02 | -1.24E+00 | 9.14E-02 | 2.58E-01 | 1.13E+00 | 1.88E-03 | 6.90E-03 | -1.14E+00 | 3.08E-01 | 8.45E-01 | 1.24E+00 |
| SMOX | 8.70E-05 | 4.53E-04 | 2.25E+00 | 2.25E-01 | 9.74E-01 | 1.56E+00 | 2.24E-01 | 9.50E-01 | 1.27E+00 | 4.58E-02 | 1.75E-01 | 1.07E+00 | 9.35E-11 | 3.33E-10 | 3.29E+00 | 1.37E-03 | 3.85E-03 | 2.16E+00 | 2.12E-07 | 7.77E-07 | 2.77E+00 | 4.36E-02 | 1.20E-01 | 1.65E+00 |
| GALNT6 | 8.70E-05 | 4.53E-04 | -1.42E+00 | 2.57E-02 | 1.11E-01 | 1.22E+00 | 1.55E-05 | 6.57E-05 | -1.67E+00 | 5.36E-03 | 2.05E-02 | -1.07E+00 | 4.78E-06 | 1.70E-05 | -1.46E+00 | 2.37E-03 | 6.68E-03 | -1.05E+00 | 4.64E-13 | 1.70E-12 | -1.97E+00 | 8.91E-06 | 2.45E-05 | -1.31E+00 |
| AC092821.1 | 8.72E-05 | 4.54E-04 | -1.38E+00 | 1.98E-03 | 8.57E-03 | 1.01E+00 | 3.75E-03 | 1.59E-02 | -1.32E+00 | 7.61E-03 | 2.91E-02 | -1.13E+00 | 3.94E-02 | 1.40E-01 | -1.23E+00 | 1.20E-01 | 3.37E-01 | 1.01E+00 | 2.65E-05 | 9.72E-05 | -1.72E+00 | 2.35E-02 | 6.45E-02 | -1.10E+00 |
| BCL2L1 | 8.74E-05 | 4.56E-04 | -1.35E+00 | 1.74E-02 | 7.50E-02 | 1.17E+00 | 9.40E-04 | 3.99E-03 | -1.36E+00 | 4.53E-02 | 1.73E-01 | 1.11E+00 | 6.69E-08 | 2.39E-07 | -1.35E+00 | 2.19E-04 | 6.19E-04 | -1.02E+00 | 1.41E-14 | 5.17E-14 | -1.20E+00 | 2.05E-08 | 5.63E-08 | -1.13E+00 |
| AC007406.5 | 8.77E-05 | 4.57E-04 | -1.37E+00 | 1.98E-02 | 8.54E-02 | 1.10E+00 | 2.96E-04 | 1.26E-03 | -1.51E+00 | 2.33E-03 | 8.89E-03 | -1.21E+00 | 1.87E-03 | 6.66E-03 | -1.74E+00 | 3.42E-02 | 9.64E-02 | -1.18E+00 | 1.10E-01 | 4.04E-01 | 1.14E+00 | 3.00E-02 | 8.24E-02 | -1.20E+00 |
| ZNF544 | 8.77E-05 | 4.57E-04 | -1.16E+00 | 6.14E-04 | 2.65E-03 | 1.03E+00 | 5.74E-08 | 2.44E-07 | -1.49E+00 | 5.13E-07 | 1.96E-06 | -1.23E+00 | 3.65E-04 | 1.30E-03 | -1.25E+00 | 6.94E-02 | 1.96E-01 | 1.12E+00 | 5.09E-09 | 1.87E-08 | -1.64E+00 | 3.69E-05 | 1.01E-04 | -1.24E+00 |
| MLH1 | 8.79E-05 | 4.58E-04 | -1.08E+00 | 1.61E-05 | 6.94E-05 | -1.06E+00 | 1.57E-02 | 6.67E-02 | -1.08E+00 | 5.31E-05 | 2.03E-04 | -1.21E+00 | 1.20E-04 | 4.28E-04 | -1.31E+00 | 3.43E-03 | 9.68E-03 | -1.05E+00 | 7.46E-05 | 2.74E-04 | -1.04E+00 | 1.22E-02 | 3.35E-02 | 1.11E+00 |
| TBL2 | 8.79E-05 | 4.58E-04 | -1.39E+00 | 2.28E-02 | 9.84E-02 | 1.17E+00 | 4.22E-06 | 1.79E-05 | -1.39E+00 | 6.27E-04 | 2.39E-03 | 1.02E+00 | 1.01E-05 | 3.61E-05 | -1.59E+00 | 2.35E-01 | 6.63E-01 | 1.31E+00 | 1.18E-07 | 4.34E-07 | -1.32E+00 | 5.37E-03 | 1.48E-02 | 1.02E+00 |
| GMNN | 8.82E-05 | 4.59E-04 | -1.54E+00 | 1.40E-03 | 6.06E-03 | -1.14E+00 | 1.69E-03 | 7.17E-03 | -1.17E+00 | 1.01E-03 | 3.86E-03 | -1.03E+00 | 2.01E-06 | 7.18E-06 | -2.11E+00 | 2.40E-05 | 6.77E-05 | -1.71E+00 | 7.16E-25 | 2.63E-24 | -1.90E+00 | 1.28E-05 | 3.53E-05 | -1.12E+00 |
| AC018738.1 | 8.82E-05 | 4.59E-04 | 1.12E+00 | 5.81E-39 | 2.51E-38 | -1.44E+00 | 8.89E-04 | 3.78E-03 | 1.12E+00 | 2.13E-24 | 8.14E-24 | -1.43E+00 | 7.88E-02 | 2.81E-01 | 1.07E+00 | 6.84E-17 | 1.93E-16 | -1.39E+00 | 2.39E-01 | 8.78E-01 | 1.06E+00 | 5.49E-30 | 1.51E-29 | -1.44E+00 |
| PSPH | 8.83E-05 | 4.60E-04 | -1.35E+00 | 4.26E-05 | 1.84E-04 | -1.19E+00 | 1.47E-03 | 6.24E-03 | -1.18E+00 | 1.77E-02 | 6.78E-02 | 1.12E+00 | 2.36E-01 | 8.42E-01 | 1.23E+00 | 7.45E-02 | 2.10E-01 | 2.07E+00 | 1.73E-04 | 6.37E-04 | -1.22E+00 | 5.29E-02 | 1.45E-01 | 1.11E+00 |
| AL355802.1 | 8.84E-05 | 4.60E-04 | 2.63E+00 | 1.69E-01 | 7.32E-01 | 1.73E+00 | 5.85E-04 | 2.49E-03 | 2.08E+00 | 1.58E-01 | 6.04E-01 | 1.61E+00 | 5.29E-02 | 1.89E-01 | 2.31E+00 | 1.06E-01 | 2.99E-01 | 1.99E+00 | 1.86E-05 | 6.81E-05 | 2.74E+00 | 4.20E-03 | 1.15E-02 | 2.36E+00 |
| AC011468.1 | 8.84E-05 | 4.60E-04 | -1.58E+00 | 5.89E-04 | 2.55E-03 | -1.23E+00 | 2.08E-02 | 8.83E-02 | -1.32E+00 | 1.18E-01 | 4.49E-01 | 1.03E+00 | 1.73E-02 | 6.15E-02 | -1.59E+00 | 7.65E-03 | 2.16E-02 | -1.86E+00 | 2.30E-01 | 8.46E-01 | 1.05E+00 | 1.20E-01 | 3.29E-01 | -1.16E+00 |
| AP3M2 | 8.85E-05 | 4.61E-04 | -1.15E+00 | 1.37E-03 | 5.94E-03 | 1.02E+00 | 7.28E-08 | 3.09E-07 | -1.87E+00 | 1.04E-02 | 3.98E-02 | -1.01E+00 | 3.47E-02 | 1.24E-01 | -1.10E+00 | 1.89E-02 | 5.33E-02 | -1.27E+00 | 5.60E-03 | 2.05E-02 | -1.19E+00 | 1.96E-01 | 5.37E-01 | 1.12E+00 |
| INPP5K | 8.89E-05 | 4.63E-04 | -1.28E+00 | 1.80E-05 | 7.78E-05 | -1.22E+00 | 2.20E-01 | 9.35E-01 | 1.24E+00 | 2.16E-01 | 8.27E-01 | 1.50E+00 | 1.43E-03 | 5.11E-03 | -1.45E+00 | 2.13E-02 | 6.00E-02 | -1.06E+00 | 1.08E-02 | 3.96E-02 | -1.09E+00 | 3.32E-03 | 9.12E-03 | -1.17E+00 |
| SAE1 | 8.89E-05 | 4.63E-04 | -1.16E+00 | 9.18E-04 | 3.97E-03 | 1.05E+00 | 9.13E-02 | 3.88E-01 | 1.10E+00 | 9.43E-02 | 3.60E-01 | 1.27E+00 | 5.71E-07 | 2.04E-06 | -1.45E+00 | 1.20E-02 | 3.38E-02 | 1.07E+00 | 1.25E-05 | 4.59E-05 | -1.16E+00 | 1.17E-01 | 3.22E-01 | 1.20E+00 |
| CEP57L1 | 8.90E-05 | 4.64E-04 | -1.14E+00 | 4.21E-03 | 1.82E-02 | 1.17E+00 | 1.46E-05 | 6.20E-05 | -1.34E+00 | 3.19E-02 | 1.22E-01 | 1.17E+00 | 5.31E-07 | 1.89E-06 | -1.58E+00 | 4.25E-03 | 1.20E-02 | -1.06E+00 | 2.01E-03 | 7.39E-03 | -1.07E+00 | 2.37E-02 | 6.50E-02 | 1.12E+00 |
| MIS18BP1 | 8.92E-05 | 4.65E-04 | -1.03E+00 | 1.78E-01 | 7.71E-01 | 1.33E+00 | 6.56E-02 | 2.79E-01 | -1.11E+00 | 2.46E-01 | 9.41E-01 | 1.16E+00 | 9.90E-03 | 3.53E-02 | 1.03E+00 | 1.56E-01 | 4.40E-01 | 1.14E+00 | 2.53E-04 | 9.30E-04 | -1.03E+00 | 9.11E-03 | 2.50E-02 | 1.02E+00 |
| USP22 | 8.94E-05 | 4.66E-04 | -1.01E+00 | 1.60E-13 | 6.92E-13 | -1.18E+00 | 1.67E-01 | 7.08E-01 | 1.14E+00 | 3.69E-05 | 1.41E-04 | -1.15E+00 | 1.30E-05 | 4.62E-05 | -1.13E+00 | 8.69E-09 | 2.45E-08 | -1.43E+00 | 7.99E-06 | 2.93E-05 | -1.15E+00 | 5.37E-09 | 1.48E-08 | -1.31E+00 |
| ZNF880 | 8.96E-05 | 4.67E-04 | -1.01E+00 | 6.33E-03 | 2.73E-02 | 1.23E+00 | 1.31E-02 | 5.56E-02 | -1.02E+00 | 1.41E-01 | 5.40E-01 | 1.37E+00 | 1.50E-01 | 5.36E-01 | 1.25E+00 | 1.82E-01 | 5.13E-01 | 1.29E+00 | 7.33E-02 | 2.69E-01 | 1.05E+00 | 2.17E-01 | 5.97E-01 | 1.15E+00 |
| ZNF680 | 8.96E-05 | 4.67E-04 | -1.24E+00 | 4.65E-02 | 2.01E-01 | 1.26E+00 | 3.64E-06 | 1.54E-05 | -1.46E+00 | 1.13E-05 | 4.32E-05 | -1.25E+00 | 1.55E-09 | 5.53E-09 | -1.55E+00 | 1.91E-06 | 5.38E-06 | -1.24E+00 | 5.00E-04 | 1.83E-03 | -1.09E+00 | 3.79E-03 | 1.04E-02 | -1.03E+00 |
| ATRN | 9.00E-05 | 4.69E-04 | -1.50E+00 | 4.37E-02 | 1.89E-01 | 1.19E+00 | 4.28E-07 | 1.82E-06 | -1.80E+00 | 2.73E-04 | 1.04E-03 | -1.15E+00 | 5.94E-03 | 2.12E-02 | -1.44E+00 | 3.32E-02 | 9.35E-02 | -1.15E+00 | 4.06E-06 | 1.49E-05 | -1.76E+00 | 1.89E-02 | 5.20E-02 | -1.07E+00 |
| HKR1 | 9.12E-05 | 4.75E-04 | -1.25E+00 | 4.17E-04 | 1.80E-03 | -1.06E+00 | 9.62E-08 | 4.09E-07 | -1.51E+00 | 2.59E-05 | 9.90E-05 | -1.15E+00 | 7.37E-05 | 2.63E-04 | -1.21E+00 | 1.01E-02 | 2.86E-02 | 1.07E+00 | 4.66E-05 | 1.71E-04 | -1.21E+00 | 7.62E-04 | 2.09E-03 | -1.07E+00 |
| ZNF471 | 9.14E-05 | 4.76E-04 | -1.91E+00 | 2.46E-03 | 1.07E-02 | -1.43E+00 | 1.61E-02 | 6.85E-02 | -1.40E+00 | 1.38E-01 | 5.27E-01 | 1.19E+00 | 2.08E-01 | 7.41E-01 | 1.68E+00 | 3.17E-01 | 8.93E-01 | 1.46E+00 | 3.97E-02 | 1.46E-01 | -1.29E+00 | 1.52E-01 | 4.19E-01 | -1.11E+00 |
| NAP1L1 | 9.14E-05 | 4.76E-04 | -1.19E+00 | 3.02E-02 | 1.30E-01 | -1.03E+00 | 6.24E-02 | 2.65E-01 | -1.10E+00 | 2.27E-01 | 8.66E-01 | 1.01E+00 | 2.31E-06 | 8.25E-06 | -1.07E+00 | 8.69E-04 | 2.45E-03 | 1.00E+00 | 2.19E-03 | 8.03E-03 | -1.09E+00 | 3.31E-01 | 9.10E-01 | 1.04E+00 |
| CSF2RB | 9.14E-05 | 4.76E-04 | -2.17E+00 | 6.46E-04 | 2.79E-03 | -1.56E+00 | 1.99E-12 | 8.48E-12 | -2.48E+00 | 1.71E-06 | 6.53E-06 | -1.22E+00 | 3.18E-29 | 1.13E-28 | -2.26E+00 | 2.06E-04 | 5.80E-04 | -1.18E+00 | 7.62E-22 | 2.80E-21 | -1.98E+00 | 1.56E-05 | 4.29E-05 | -1.22E+00 |
| SCML2 | 9.15E-05 | 4.77E-04 | -1.32E+00 | 3.46E-03 | 1.49E-02 | 1.08E+00 | 3.15E-06 | 1.34E-05 | -1.55E+00 | 2.14E-05 | 8.16E-05 | -1.26E+00 | 5.62E-03 | 2.00E-02 | -1.49E+00 | 1.89E-02 | 5.33E-02 | -1.21E+00 | 1.83E-04 | 6.70E-04 | -1.38E+00 | 1.51E-02 | 4.16E-02 | -1.03E+00 |
| WDR7 | 9.17E-05 | 4.78E-04 | -2.00E+00 | 5.63E-03 | 2.43E-02 | -1.24E+00 | 8.43E-04 | 3.58E-03 | -1.49E+00 | 3.81E-02 | 1.46E-01 | 1.09E+00 | 2.54E-02 | 9.07E-02 | -1.34E+00 | 1.87E-01 | 5.28E-01 | 1.09E+00 | 1.88E-05 | 6.91E-05 | -1.37E+00 | 1.25E-03 | 3.44E-03 | -1.13E+00 |
| TATDN1P1 | 9.22E-05 | 4.80E-04 | -1.05E+00 | 4.06E-06 | 1.75E-05 | 1.03E+00 | 1.85E-07 | 7.87E-07 | -1.37E+00 | 8.07E-08 | 3.08E-07 | -1.23E+00 | 1.34E-04 | 4.78E-04 | -1.16E+00 | 3.58E-05 | 1.01E-04 | -1.15E+00 | 4.36E-03 | 1.60E-02 | -1.04E+00 | 1.20E-02 | 3.30E-02 | 1.01E+00 |
| GBP3 | 9.23E-05 | 4.81E-04 | -1.32E+00 | 4.51E-02 | 1.95E-01 | 1.25E+00 | 2.69E-08 | 1.15E-07 | -1.93E+00 | 6.85E-05 | 2.62E-04 | -1.18E+00 | 1.49E-06 | 5.32E-06 | -1.85E+00 | 2.90E-03 | 8.17E-03 | -1.12E+00 | 2.65E-05 | 9.71E-05 | -1.50E+00 | 2.89E-03 | 7.94E-03 | -1.16E+00 |
| RICTOR | 9.23E-05 | 4.81E-04 | -1.03E+00 | 2.70E-04 | 1.17E-03 | 1.12E+00 | 1.63E-01 | 6.94E-01 | 1.15E+00 | 1.03E-02 | 3.93E-02 | 1.13E+00 | 4.75E-02 | 1.69E-01 | 1.18E+00 | 1.40E-04 | 3.95E-04 | -1.00E+00 | 5.75E-03 | 2.11E-02 | -1.04E+00 | 3.34E-03 | 9.18E-03 | -1.14E+00 |
| VDR | 9.23E-05 | 4.81E-04 | -2.06E+00 | 1.35E-03 | 5.85E-03 | -1.46E+00 | 1.62E-01 | 6.90E-01 | 1.20E+00 | 2.24E-01 | 8.55E-01 | 1.21E+00 | 1.36E-01 | 4.86E-01 | -1.28E+00 | 1.95E-01 | 5.49E-01 | -1.21E+00 | 1.41E-02 | 5.17E-02 | 1.02E+00 | 7.12E-03 | 1.96E-02 | 1.02E+00 |
| ITM2B | 9.23E-05 | 4.81E-04 | -1.09E+00 | 8.39E-03 | 3.63E-02 | -1.02E+00 | 3.97E-02 | 1.69E-01 | -1.03E+00 | 1.28E-02 | 4.90E-02 | -1.06E+00 | 3.60E-05 | 1.28E-04 | -1.16E+00 | 1.54E-04 | 4.35E-04 | -1.21E+00 | 7.67E-02 | 2.82E-01 | 1.02E+00 | 3.03E-03 | 8.32E-03 | -1.13E+00 |
| C15orf41 | 9.26E-05 | 4.82E-04 | -1.33E+00 | 4.44E-03 | 1.92E-02 | 1.06E+00 | 8.77E-08 | 3.73E-07 | -1.77E+00 | 3.43E-03 | 1.31E-02 | -1.02E+00 | 2.06E-05 | 7.35E-05 | -1.86E+00 | 2.52E-02 | 7.10E-02 | -1.08E+00 | 7.16E-07 | 2.63E-06 | -1.47E+00 | 1.24E-03 | 3.40E-03 | -1.13E+00 |
| SNRPGP15 | 9.27E-05 | 4.83E-04 | -1.20E+00 | 4.69E-06 | 2.03E-05 | -1.24E+00 | 5.60E-02 | 2.38E-01 | 1.08E+00 | 2.88E-04 | 1.10E-03 | -1.03E+00 | 1.25E-04 | 4.47E-04 | -1.30E+00 | 3.97E-03 | 1.12E-02 | -1.05E+00 | 1.98E-02 | 7.25E-02 | 1.04E+00 | 1.12E-02 | 3.07E-02 | 1.07E+00 |
| DPY19L2P3 | 9.30E-05 | 4.85E-04 | -1.50E+00 | 9.33E-04 | 4.03E-03 | -1.21E+00 | 1.44E-02 | 6.11E-02 | -1.28E+00 | 1.10E-01 | 4.20E-01 | 1.17E+00 | 8.26E-03 | 2.94E-02 | -1.20E+00 | 1.58E-02 | 4.44E-02 | -1.12E+00 | 5.73E-03 | 2.10E-02 | -1.13E+00 | 7.85E-02 | 2.16E-01 | 1.01E+00 |
| AL136380.1 | 9.30E-05 | 4.85E-04 | -1.14E+00 | 1.72E-44 | 7.41E-44 | -1.81E+00 | 1.56E-02 | 6.64E-02 | -1.13E+00 | 4.02E-18 | 1.54E-17 | -1.86E+00 | 1.07E-03 | 3.83E-03 | -1.08E+00 | 7.83E-18 | 2.21E-17 | -1.70E+00 | 1.35E-03 | 4.94E-03 | -1.13E+00 | 1.28E-13 | 3.52E-13 | -1.76E+00 |
| SLC25A26 | 9.31E-05 | 4.85E-04 | -1.01E+00 | 1.03E-04 | 4.44E-04 | 1.10E+00 | 8.24E-03 | 3.50E-02 | 1.03E+00 | 4.63E-03 | 1.77E-02 | 1.16E+00 | 6.07E-04 | 2.16E-03 | -1.10E+00 | 1.68E-02 | 4.74E-02 | 1.03E+00 | 3.62E-02 | 1.33E-01 | 1.08E+00 | 7.09E-02 | 1.95E-01 | 1.17E+00 |
| AC063947.2 | 9.31E-05 | 4.85E-04 | -2.03E+00 | 7.66E-05 | 3.31E-04 | -1.93E+00 | 1.57E-06 | 6.67E-06 | -2.23E+00 | 9.88E-06 | 3.77E-05 | -1.79E+00 | 4.55E-22 | 1.62E-21 | -3.07E+00 | 2.75E-18 | 7.76E-18 | -2.12E+00 | 2.15E-25 | 7.89E-25 | -2.59E+00 | 2.67E-23 | 7.33E-23 | -2.50E+00 |
| PCED1B-AS1 | 9.33E-05 | 4.86E-04 | -2.47E+00 | 7.17E-05 | 3.10E-04 | -2.56E+00 | 7.48E-02 | 3.18E-01 | -2.04E+00 | 3.81E-03 | 1.45E-02 | -3.52E+00 | 1.93E-01 | 6.88E-01 | 1.04E+00 | 1.70E-01 | 4.78E-01 | -1.25E+00 | 8.61E-02 | 3.16E-01 | 1.46E+00 | 2.75E-01 | 7.55E-01 | 1.18E+00 |
| MED12 | 9.39E-05 | 4.89E-04 | -1.11E+00 | 9.43E-03 | 4.08E-02 | 1.19E+00 | 8.36E-03 | 3.55E-02 | -1.09E+00 | 6.69E-02 | 2.55E-01 | 1.15E+00 | 1.08E-01 | 3.84E-01 | 1.10E+00 | 2.83E-01 | 7.97E-01 | 1.34E+00 | 1.59E-03 | 5.84E-03 | -1.09E+00 | 1.52E-02 | 4.17E-02 | 1.02E+00 |
| NR2C2 | 9.39E-05 | 4.89E-04 | -1.09E+00 | 2.10E-05 | 9.06E-05 | -1.10E+00 | 3.76E-03 | 1.60E-02 | -1.10E+00 | 1.61E-02 | 6.16E-02 | 1.11E+00 | 1.27E-02 | 4.53E-02 | -1.12E+00 | 1.20E-01 | 3.37E-01 | 1.13E+00 | 2.99E-03 | 1.10E-02 | -1.12E+00 | 5.68E-02 | 1.56E-01 | 1.06E+00 |
| PANK4 | 9.40E-05 | 4.90E-04 | -2.09E+00 | 7.14E-02 | 3.08E-01 | 1.18E+00 | 4.65E-04 | 1.97E-03 | -1.67E+00 | 1.03E-02 | 3.93E-02 | -1.14E+00 | 4.42E-04 | 1.58E-03 | -2.46E+00 | 8.59E-03 | 2.42E-02 | -1.37E+00 | 1.07E-04 | 3.94E-04 | -1.59E+00 | 9.84E-03 | 2.70E-02 | -1.24E+00 |
| THAP6 | 9.40E-05 | 4.90E-04 | -1.22E+00 | 3.25E-02 | 1.40E-01 | 1.21E+00 | 1.85E-05 | 7.87E-05 | -1.50E+00 | 5.59E-03 | 2.13E-02 | 1.04E+00 | 3.33E-04 | 1.19E-03 | -1.20E+00 | 3.37E-02 | 9.49E-02 | 1.09E+00 | 9.18E-03 | 3.37E-02 | -1.08E+00 | 1.42E-02 | 3.89E-02 | -1.12E+00 |
| CHD2 | 9.44E-05 | 4.92E-04 | 1.02E+00 | 6.30E-02 | 2.72E-01 | 1.36E+00 | 1.93E-06 | 8.18E-06 | -1.33E+00 | 2.71E-02 | 1.04E-01 | 1.12E+00 | 1.46E-04 | 5.22E-04 | 1.06E+00 | 1.46E-01 | 4.12E-01 | 1.32E+00 | 2.46E-04 | 9.03E-04 | -1.15E+00 | 1.16E-02 | 3.19E-02 | 1.03E+00 |
| MPRIP | 9.44E-05 | 4.92E-04 | -1.08E+00 | 6.07E-03 | 2.62E-02 | 1.16E+00 | 1.09E-01 | 4.62E-01 | -1.03E+00 | 2.25E-01 | 8.60E-01 | 1.23E+00 | 3.08E-03 | 1.10E-02 | -1.08E+00 | 2.89E-03 | 8.14E-03 | -1.11E+00 | 7.61E-03 | 2.79E-02 | 1.05E+00 | 8.74E-02 | 2.40E-01 | 1.16E+00 |
| TBL3 | 9.45E-05 | 4.93E-04 | -1.49E+00 | 2.71E-02 | 1.17E-01 | 1.10E+00 | 1.56E-02 | 6.61E-02 | -1.05E+00 | 2.00E-01 | 7.62E-01 | 1.45E+00 | 4.08E-02 | 1.46E-01 | -1.14E+00 | 2.44E-01 | 6.89E-01 | 1.47E+00 | 3.20E-04 | 1.18E-03 | -1.06E+00 | 3.40E-01 | 9.33E-01 | 1.28E+00 |
| MAK | 9.49E-05 | 4.95E-04 | -1.71E+00 | 1.18E-01 | 5.11E-01 | 1.37E+00 | 2.03E-01 | 8.63E-01 | 1.09E+00 | 1.19E-01 | 4.54E-01 | 1.57E+00 | 4.10E-03 | 1.46E-02 | -1.45E+00 | 7.81E-02 | 2.20E-01 | 1.15E+00 | 8.05E-07 | 2.95E-06 | -2.43E+00 | 2.67E-02 | 7.33E-02 | -1.18E+00 |
| GEMIN2 | 9.49E-05 | 4.95E-04 | -1.43E+00 | 1.75E-03 | 7.56E-03 | -1.12E+00 | 4.63E-02 | 1.97E-01 | 1.03E+00 | 2.43E-01 | 9.29E-01 | 1.41E+00 | 2.35E-06 | 8.37E-06 | -1.74E+00 | 1.47E-02 | 4.13E-02 | 1.14E+00 | 2.90E-02 | 1.07E-01 | 1.04E+00 | 3.08E-01 | 8.46E-01 | 1.33E+00 |
| RNF144B | 9.50E-05 | 4.95E-04 | -1.51E+00 | 1.71E-04 | 7.38E-04 | -1.23E+00 | 8.05E-03 | 3.42E-02 | -1.21E+00 | 8.19E-02 | 3.13E-01 | 1.26E+00 | 8.89E-03 | 3.17E-02 | -1.30E+00 | 1.72E-01 | 4.85E-01 | 1.53E+00 | 7.88E-04 | 2.89E-03 | -1.23E+00 | 2.78E-01 | 7.64E-01 | 1.22E+00 |
| DDX39A | 9.51E-05 | 4.95E-04 | -1.02E+00 | 3.77E-04 | 1.63E-03 | 1.10E+00 | 9.86E-03 | 4.19E-02 | -1.01E+00 | 1.51E-02 | 5.76E-02 | 1.09E+00 | 1.11E-08 | 3.96E-08 | -1.47E+00 | 1.26E-03 | 3.56E-03 | 1.01E+00 | 7.33E-09 | 2.69E-08 | -1.28E+00 | 7.12E-02 | 1.96E-01 | 1.06E+00 |
| NUP133 | 9.51E-05 | 4.95E-04 | -1.31E+00 | 2.21E-03 | 9.53E-03 | 1.02E+00 | 1.38E-03 | 5.86E-03 | -1.20E+00 | 7.81E-03 | 2.98E-02 | 1.05E+00 | 5.20E-05 | 1.85E-04 | -1.31E+00 | 8.32E-03 | 2.35E-02 | 1.00E+00 | 8.85E-04 | 3.25E-03 | -1.20E+00 | 3.06E-01 | 8.40E-01 | 1.23E+00 |
| YWHAZP3 | 9.52E-05 | 4.96E-04 | -1.08E+00 | 1.38E-14 | 5.94E-14 | -2.23E+00 | 9.41E-07 | 4.00E-06 | -1.29E+00 | 8.84E-15 | 3.38E-14 | -1.96E+00 | 2.93E-06 | 1.04E-05 | -1.13E+00 | 1.19E-14 | 3.36E-14 | -2.15E+00 | 2.47E-05 | 9.06E-05 | -1.09E+00 | 2.47E-15 | 6.80E-15 | -1.96E+00 |
| ARIH2 | 9.52E-05 | 4.96E-04 | -1.07E+00 | 2.24E-04 | 9.66E-04 | 1.03E+00 | 2.61E-05 | 1.11E-04 | -1.25E+00 | 2.64E-07 | 1.01E-06 | -1.19E+00 | 2.58E-08 | 9.19E-08 | -1.26E+00 | 3.42E-07 | 9.64E-07 | -1.12E+00 | 1.32E-01 | 4.83E-01 | 1.13E+00 | 1.91E-01 | 5.25E-01 | 1.17E+00 |
| MT-ND2 | 9.53E-05 | 4.96E-04 | 1.12E+00 | 4.22E-04 | 1.82E-03 | 1.10E+00 | 5.93E-07 | 2.52E-06 | 1.20E+00 | 1.38E-06 | 5.26E-06 | 1.19E+00 | 5.43E-08 | 1.94E-07 | 1.27E+00 | 5.66E-09 | 1.60E-08 | 1.28E+00 | 2.67E-17 | 9.80E-17 | 1.38E+00 | 5.92E-18 | 1.63E-17 | 1.36E+00 |
| BTBD1 | 9.53E-05 | 4.97E-04 | -1.16E+00 | 8.39E-03 | 3.63E-02 | 1.20E+00 | 7.44E-06 | 3.16E-05 | -1.27E+00 | 1.63E-04 | 6.22E-04 | -1.04E+00 | 4.21E-02 | 1.50E-01 | 1.14E+00 | 1.23E-01 | 3.47E-01 | 1.25E+00 | 7.77E-02 | 2.85E-01 | 1.29E+00 | 3.00E-01 | 8.24E-01 | 1.28E+00 |
| TMBIM1 | 9.55E-05 | 4.97E-04 | -1.14E+00 | 3.66E-02 | 1.58E-01 | 1.29E+00 | 3.69E-10 | 1.57E-09 | -1.78E+00 | 1.54E-05 | 5.87E-05 | -1.14E+00 | 1.72E-17 | 6.14E-17 | -1.80E+00 | 3.00E-07 | 8.46E-07 | -1.23E+00 | 8.73E-05 | 3.21E-04 | 1.02E+00 | 3.75E-02 | 1.03E-01 | -1.00E+00 |
| LMF1 | 9.57E-05 | 4.99E-04 | -1.25E+00 | 6.70E-06 | 2.89E-05 | -1.33E+00 | 1.28E-02 | 5.43E-02 | -1.10E+00 | 6.44E-03 | 2.46E-02 | -1.05E+00 | 2.56E-01 | 9.14E-01 | 1.50E+00 | 3.25E-01 | 9.15E-01 | 1.52E+00 | 4.22E-02 | 1.55E-01 | 1.03E+00 | 2.55E-01 | 6.99E-01 | 1.22E+00 |
| GAS2L3 | 9.59E-05 | 5.00E-04 | -2.16E+00 | 4.07E-04 | 1.76E-03 | -1.89E+00 | 1.08E-02 | 4.60E-02 | -1.51E+00 | 9.32E-04 | 3.56E-03 | -1.61E+00 | 2.00E-01 | 7.13E-01 | 1.50E+00 | 2.90E-01 | 8.16E-01 | 1.25E+00 | 7.14E-03 | 2.62E-02 | -1.31E+00 | 8.69E-02 | 2.39E-01 | -1.16E+00 |
| SGTB | 9.63E-05 | 5.02E-04 | -1.24E+00 | 1.07E-04 | 4.60E-04 | -1.10E+00 | 4.12E-05 | 1.75E-04 | -1.49E+00 | 1.03E-06 | 3.94E-06 | -1.45E+00 | 1.98E-01 | 7.07E-01 | 1.26E+00 | 2.06E-03 | 5.80E-03 | -1.37E+00 | 7.01E-05 | 2.57E-04 | -1.47E+00 | 1.58E-05 | 4.35E-05 | -1.66E+00 |
| THRB | 9.65E-05 | 5.03E-04 | -1.76E+00 | 2.76E-03 | 1.19E-02 | -1.09E+00 | 1.63E-01 | 6.91E-01 | -1.19E+00 | 1.86E-01 | 7.11E-01 | 1.48E+00 | 5.10E-02 | 1.82E-01 | 1.97E+00 | 2.78E-07 | 7.85E-07 | 3.52E+00 | 6.56E-24 | 2.41E-23 | 7.13E+00 | 2.77E-25 | 7.62E-25 | 7.48E+00 |
| FMO5 | 9.66E-05 | 5.03E-04 | -1.48E+00 | 5.45E-03 | 2.35E-02 | 1.00E+00 | 3.24E-02 | 1.38E-01 | -1.37E+00 | 2.47E-01 | 9.45E-01 | 1.27E+00 | 3.71E-04 | 1.32E-03 | -1.35E+00 | 5.29E-03 | 1.49E-02 | -1.14E+00 | 5.68E-06 | 2.08E-05 | -1.41E+00 | 2.13E-05 | 5.84E-05 | -1.50E+00 |
| SPAG16 | 9.66E-05 | 5.03E-04 | -1.17E+00 | 1.07E-02 | 4.64E-02 | 1.16E+00 | 1.53E-03 | 6.50E-03 | -1.09E+00 | 1.84E-04 | 7.02E-04 | -1.04E+00 | 1.76E-05 | 6.26E-05 | -1.56E+00 | 9.90E-05 | 2.79E-04 | -1.33E+00 | 3.28E-03 | 1.21E-02 | -1.04E+00 | 9.88E-03 | 2.71E-02 | 1.05E+00 |
| RPS6KB2 | 9.71E-05 | 5.06E-04 | -1.24E+00 | 3.24E-05 | 1.40E-04 | -1.18E+00 | 1.51E-02 | 6.40E-02 | -1.02E+00 | 2.50E-02 | 9.57E-02 | 1.09E+00 | 1.60E-02 | 5.69E-02 | -1.05E+00 | 1.45E-02 | 4.08E-02 | -1.12E+00 | 6.45E-03 | 2.37E-02 | -1.11E+00 | 2.44E-02 | 6.70E-02 | -1.09E+00 |
| SCYL3 | 9.73E-05 | 5.07E-04 | -1.50E+00 | 9.51E-02 | 4.11E-01 | 1.36E+00 | 6.49E-04 | 2.76E-03 | -1.43E+00 | 7.73E-02 | 2.95E-01 | 1.20E+00 | 3.06E-03 | 1.09E-02 | -1.55E+00 | 1.02E-01 | 2.88E-01 | 1.05E+00 | 4.57E-04 | 1.68E-03 | -1.69E+00 | 5.39E-02 | 1.48E-01 | -1.04E+00 |
| U2AF2 | 9.74E-05 | 5.07E-04 | -1.24E+00 | 6.63E-03 | 2.86E-02 | 1.13E+00 | 5.33E-06 | 2.27E-05 | -1.30E+00 | 1.27E-03 | 4.85E-03 | -1.03E+00 | 1.88E-03 | 6.71E-03 | -1.10E+00 | 4.05E-03 | 1.14E-02 | -1.07E+00 | 1.71E-07 | 6.28E-07 | -1.33E+00 | 1.67E-02 | 4.59E-02 | 1.06E+00 |
| RPL18A | 9.77E-05 | 5.09E-04 | 1.20E+00 | 3.38E-28 | 1.46E-27 | 1.71E+00 | 1.01E-02 | 4.31E-02 | 1.17E+00 | 1.02E-17 | 3.91E-17 | 1.72E+00 | 1.40E-02 | 4.99E-02 | 1.22E+00 | 4.84E-24 | 1.36E-23 | 1.87E+00 | 1.22E-01 | 4.49E-01 | 1.21E+00 | 3.97E-23 | 1.09E-22 | 1.81E+00 |
| SLC35F5 | 9.77E-05 | 5.09E-04 | -1.14E+00 | 6.77E-03 | 2.92E-02 | 1.17E+00 | 1.11E-07 | 4.72E-07 | -1.45E+00 | 1.67E-06 | 6.36E-06 | -1.16E+00 | 1.30E-02 | 4.65E-02 | 1.13E+00 | 1.88E-02 | 5.30E-02 | 1.19E+00 | 1.01E-01 | 3.72E-01 | 1.08E+00 | 2.47E-01 | 6.79E-01 | 1.19E+00 |
| SREK1 | 9.77E-05 | 5.09E-04 | 1.00E+00 | 4.94E-07 | 2.13E-06 | 1.01E+00 | 1.05E-01 | 4.45E-01 | 1.02E+00 | 1.13E-03 | 4.30E-03 | -1.02E+00 | 7.49E-02 | 2.67E-01 | 1.18E+00 | 1.55E-02 | 4.37E-02 | 1.11E+00 | 2.20E-01 | 8.09E-01 | 1.20E+00 | 2.66E-01 | 7.30E-01 | 1.11E+00 |
| ZNF583 | 9.78E-05 | 5.10E-04 | -1.37E+00 | 2.41E-02 | 1.04E-01 | 1.18E+00 | 5.87E-05 | 2.50E-04 | -1.61E+00 | 1.10E-03 | 4.21E-03 | -1.18E+00 | 5.20E-05 | 1.85E-04 | -1.84E+00 | 1.71E-03 | 4.81E-03 | -1.33E+00 | 9.49E-08 | 3.48E-07 | -1.79E+00 | 1.06E-05 | 2.91E-05 | -1.57E+00 |
| UBAP2L | 9.78E-05 | 5.10E-04 | -1.08E+00 | 4.89E-04 | 2.11E-03 | 1.07E+00 | 7.82E-06 | 3.32E-05 | -1.35E+00 | 5.58E-04 | 2.13E-03 | -1.02E+00 | 1.05E-04 | 3.73E-04 | -1.10E+00 | 2.25E-03 | 6.35E-03 | 1.03E+00 | 1.04E-06 | 3.80E-06 | -1.27E+00 | 6.85E-02 | 1.88E-01 | 1.13E+00 |
| AC022144.1 | 9.79E-05 | 5.10E-04 | -1.53E+00 | 1.70E-02 | 7.36E-02 | 1.09E+00 | 8.06E-05 | 3.42E-04 | -2.09E+00 | 4.28E-02 | 1.64E-01 | -1.02E+00 | 3.75E-10 | 1.34E-09 | -2.74E+00 | 6.22E-04 | 1.75E-03 | -1.26E+00 | 3.91E-05 | 1.44E-04 | -1.95E+00 | 2.11E-02 | 5.78E-02 | -1.08E+00 |
| MYPOP | 9.79E-05 | 5.10E-04 | -1.66E+00 | 4.33E-02 | 1.87E-01 | 1.10E+00 | 4.46E-06 | 1.89E-05 | -1.77E+00 | 4.46E-04 | 1.70E-03 | -1.24E+00 | 4.69E-04 | 1.67E-03 | -1.86E+00 | 1.65E-02 | 4.66E-02 | -1.11E+00 | 6.57E-03 | 2.41E-02 | -1.25E+00 | 3.60E-02 | 9.88E-02 | -1.26E+00 |
| CDC14B | 9.79E-05 | 5.10E-04 | -1.37E+00 | 2.42E-03 | 1.05E-02 | -1.07E+00 | 5.90E-03 | 2.51E-02 | -1.63E+00 | 1.15E-01 | 4.39E-01 | -1.02E+00 | 6.08E-03 | 2.17E-02 | -1.96E+00 | 1.08E-02 | 3.04E-02 | -1.81E+00 | 6.82E-02 | 2.50E-01 | -1.03E+00 | 1.34E-01 | 3.67E-01 | -1.12E+00 |
| DROSHA | 9.80E-05 | 5.11E-04 | -1.35E+00 | 3.77E-03 | 1.63E-02 | 1.01E+00 | 1.41E-03 | 5.98E-03 | -1.20E+00 | 1.70E-02 | 6.50E-02 | 1.03E+00 | 5.32E-05 | 1.90E-04 | -1.35E+00 | 3.29E-04 | 9.27E-04 | -1.17E+00 | 4.53E-04 | 1.66E-03 | -1.19E+00 | 1.33E-04 | 3.66E-04 | -1.18E+00 |
| SSR4 | 9.83E-05 | 5.12E-04 | -1.03E+00 | 3.30E-06 | 1.42E-05 | -1.01E+00 | 2.13E-01 | 9.06E-01 | 1.05E+00 | 2.59E-01 | 9.89E-01 | 1.12E+00 | 3.61E-03 | 1.29E-02 | -1.11E+00 | 1.87E-02 | 5.29E-02 | -1.10E+00 | 1.84E-01 | 6.74E-01 | 1.02E+00 | 2.79E-01 | 7.67E-01 | 1.04E+00 |
| SLAIN1 | 9.83E-05 | 5.12E-04 | -1.40E+00 | 1.37E-03 | 5.91E-03 | -1.04E+00 | 1.71E-08 | 7.27E-08 | -1.75E+00 | 2.16E-04 | 8.27E-04 | -1.12E+00 | 4.50E-04 | 1.61E-03 | -1.45E+00 | 1.14E-02 | 3.22E-02 | -1.05E+00 | 9.70E-06 | 3.56E-05 | -1.33E+00 | 1.32E-03 | 3.62E-03 | -1.10E+00 |
| SNORD3A | 9.84E-05 | 5.13E-04 | 2.20E+00 | 2.45E-05 | 1.06E-04 | -1.30E+00 | 1.75E-01 | 7.42E-01 | 1.33E+00 | 3.18E-08 | 1.21E-07 | -1.47E+00 | 6.37E-04 | 2.27E-03 | -1.34E+00 | 2.15E-03 | 6.06E-03 | -1.25E+00 | 1.33E-01 | 4.86E-01 | 1.08E+00 | 2.36E-01 | 6.47E-01 | 1.06E+00 |
| PPIAP72 | 9.84E-05 | 5.13E-04 | -1.55E+00 | 1.18E-04 | 5.09E-04 | -1.41E+00 | 1.75E-02 | 7.42E-02 | -1.26E+00 | 6.99E-02 | 2.67E-01 | 1.11E+00 | 2.76E-01 | 9.85E-01 | 1.34E+00 | 2.98E-01 | 8.39E-01 | 1.33E+00 | 3.56E-03 | 1.31E-02 | -1.15E+00 | 8.13E-03 | 2.23E-02 | -1.14E+00 |
| ZFYVE19 | 9.85E-05 | 5.13E-04 | -1.35E+00 | 1.72E-02 | 7.41E-02 | 1.12E+00 | 3.26E-04 | 1.38E-03 | -1.23E+00 | 1.14E-04 | 4.35E-04 | -1.14E+00 | 1.28E-03 | 4.55E-03 | -1.29E+00 | 2.31E-02 | 6.50E-02 | 1.01E+00 | 2.52E-04 | 9.25E-04 | -1.24E+00 | 2.10E-03 | 5.78E-03 | -1.14E+00 |
| ZNF77 | 9.89E-05 | 5.16E-04 | -1.45E+00 | 1.17E-03 | 5.06E-03 | -1.15E+00 | 2.63E-03 | 1.12E-02 | -1.36E+00 | 2.24E-03 | 8.54E-03 | -1.37E+00 | 1.15E-02 | 4.11E-02 | -1.09E+00 | 5.26E-03 | 1.48E-02 | -1.13E+00 | 1.19E-04 | 4.36E-04 | -1.40E+00 | 2.30E-03 | 6.32E-03 | -1.11E+00 |
| TMEM43 | 9.90E-05 | 5.16E-04 | -1.07E+00 | 2.07E-01 | 8.93E-01 | 1.46E+00 | 1.96E-04 | 8.34E-04 | -1.08E+00 | 1.24E-01 | 4.74E-01 | 1.31E+00 | 1.73E-02 | 6.19E-02 | 1.03E+00 | 1.71E-01 | 4.83E-01 | 1.25E+00 | 4.72E-03 | 1.73E-02 | -1.09E+00 | 3.37E-01 | 9.26E-01 | 1.19E+00 |
| LZIC | 9.91E-05 | 5.16E-04 | -1.11E+00 | 2.20E-03 | 9.50E-03 | 1.10E+00 | 1.07E-02 | 4.55E-02 | 1.04E+00 | 1.91E-03 | 7.31E-03 | 1.09E+00 | 9.54E-04 | 3.40E-03 | -1.11E+00 | 9.48E-03 | 2.67E-02 | 1.07E+00 | 6.79E-02 | 2.49E-01 | 1.11E+00 | 2.44E-02 | 6.70E-02 | 1.07E+00 |
| UBL7-AS1 | 9.91E-05 | 5.17E-04 | -1.06E+00 | 8.33E-02 | 3.60E-01 | 1.39E+00 | 2.43E-02 | 1.03E-01 | 1.08E+00 | 1.90E-01 | 7.27E-01 | 1.57E+00 | 3.45E-02 | 1.23E-01 | 1.00E+00 | 8.45E-02 | 2.38E-01 | 1.09E+00 | 7.39E-02 | 2.71E-01 | 1.07E+00 | 1.46E-01 | 4.01E-01 | 1.24E+00 |
| SLC39A9 | 9.92E-05 | 5.17E-04 | -1.28E+00 | 2.31E-03 | 9.96E-03 | 1.08E+00 | 2.69E-08 | 1.14E-07 | -1.48E+00 | 7.52E-08 | 2.87E-07 | -1.26E+00 | 1.71E-04 | 6.11E-04 | -1.50E+00 | 2.30E-04 | 6.49E-04 | -1.33E+00 | 2.97E-02 | 1.09E-01 | 1.01E+00 | 1.38E-01 | 3.80E-01 | 1.16E+00 |
| PEMT | 9.92E-05 | 5.17E-04 | -1.17E+00 | 1.83E-03 | 7.89E-03 | 1.02E+00 | 1.94E-03 | 8.22E-03 | -1.17E+00 | 4.28E-05 | 1.64E-04 | -1.15E+00 | 2.18E-02 | 7.79E-02 | -1.17E+00 | 2.27E-01 | 6.39E-01 | 1.15E+00 | 1.12E-03 | 4.11E-03 | -1.18E+00 | 2.25E-02 | 6.17E-02 | -1.07E+00 |
| SLC43A1 | 9.94E-05 | 5.18E-04 | -1.35E+00 | 1.05E-03 | 4.53E-03 | -1.05E+00 | 3.11E-04 | 1.32E-03 | -1.21E+00 | 3.85E-03 | 1.47E-02 | 1.03E+00 | 1.53E-07 | 5.46E-07 | -1.51E+00 | 1.11E-04 | 3.13E-04 | -1.08E+00 | 2.51E-04 | 9.20E-04 | -1.11E+00 | 1.14E-01 | 3.13E-01 | 1.18E+00 |
| CISD3 | 9.96E-05 | 5.19E-04 | -1.22E+00 | 7.62E-05 | 3.29E-04 | -1.08E+00 | 8.25E-02 | 3.50E-01 | 1.10E+00 | 3.27E-03 | 1.25E-02 | 1.10E+00 | 6.50E-02 | 2.32E-01 | 1.04E+00 | 2.28E-01 | 6.42E-01 | 1.25E+00 | 1.60E-01 | 5.88E-01 | 1.22E+00 | 1.90E-01 | 5.22E-01 | 1.42E+00 |
| FAM3C2 | 9.96E-05 | 5.19E-04 | -1.85E+00 | 3.58E-03 | 1.55E-02 | -1.28E+00 | 1.25E-01 | 5.32E-01 | 1.33E+00 | 1.33E-01 | 5.08E-01 | 1.63E+00 | 6.61E-02 | 2.36E-01 | -1.08E+00 | 5.32E-02 | 1.50E-01 | -1.22E+00 | 8.59E-02 | 3.15E-01 | 1.01E+00 | 1.44E-01 | 3.95E-01 | 1.01E+00 |
| FRG1EP | 1.00E-04 | 5.23E-04 | -1.04E+00 | 2.87E-06 | 1.24E-05 | 1.03E+00 | 8.94E-04 | 3.80E-03 | -1.17E+00 | 6.30E-05 | 2.41E-04 | -1.12E+00 | 7.40E-02 | 2.64E-01 | 1.11E+00 | 3.13E-02 | 8.84E-02 | 1.05E+00 | 2.50E-03 | 9.19E-03 | -1.06E+00 | 8.71E-04 | 2.39E-03 | -1.07E+00 |
| HAT1 | 1.01E-04 | 5.24E-04 | -1.02E+00 | 2.33E-04 | 1.01E-03 | 1.10E+00 | 3.96E-03 | 1.68E-02 | 1.03E+00 | 1.64E-05 | 6.26E-05 | 1.01E+00 | 3.07E-03 | 1.09E-02 | -1.06E+00 | 1.83E-03 | 5.17E-03 | -1.07E+00 | 1.08E-05 | 3.97E-05 | -1.18E+00 | 6.54E-03 | 1.80E-02 | -1.02E+00 |
| IQCG | 1.01E-04 | 5.24E-04 | -1.35E+00 | 1.03E-04 | 4.44E-04 | -1.28E+00 | 7.08E-03 | 3.01E-02 | -1.01E+00 | 2.84E-03 | 1.08E-02 | -1.08E+00 | 6.85E-03 | 2.44E-02 | -1.40E+00 | 5.24E-02 | 1.48E-01 | -1.07E+00 | 3.30E-02 | 1.21E-01 | -1.13E+00 | 3.56E-02 | 9.78E-02 | -1.14E+00 |
| ZNF669 | 1.01E-04 | 5.25E-04 | -1.09E+00 | 2.04E-08 | 8.80E-08 | -1.36E+00 | 2.18E-06 | 9.28E-06 | -1.33E+00 | 5.54E-08 | 2.12E-07 | -1.42E+00 | 4.44E-06 | 1.58E-05 | -1.34E+00 | 6.80E-07 | 1.92E-06 | -1.52E+00 | 2.07E-03 | 7.61E-03 | -1.22E+00 | 4.63E-04 | 1.27E-03 | -1.34E+00 |
| RAB2B | 1.01E-04 | 5.26E-04 | -1.04E+00 | 2.71E-04 | 1.17E-03 | 1.12E+00 | 3.80E-02 | 1.62E-01 | -1.01E+00 | 1.14E-03 | 4.36E-03 | -1.05E+00 | 2.92E-05 | 1.04E-04 | -1.16E+00 | 1.90E-04 | 5.36E-04 | -1.05E+00 | 2.86E-04 | 1.05E-03 | -1.16E+00 | 1.12E-03 | 3.08E-03 | -1.10E+00 |
| FAM102B | 1.01E-04 | 5.26E-04 | -1.28E+00 | 1.09E-02 | 4.70E-02 | 1.17E+00 | 4.18E-03 | 1.78E-02 | -1.17E+00 | 4.28E-02 | 1.64E-01 | 1.17E+00 | 5.33E-02 | 1.90E-01 | 1.14E+00 | 3.02E-03 | 8.51E-03 | -1.22E+00 | 4.13E-02 | 1.52E-01 | 1.00E+00 | 3.79E-03 | 1.04E-02 | -1.26E+00 |
| PCGF3 | 1.01E-04 | 5.26E-04 | -1.14E+00 | 1.09E-01 | 4.70E-01 | 1.41E+00 | 3.11E-02 | 1.32E-01 | -1.06E+00 | 2.22E-01 | 8.47E-01 | 1.35E+00 | 5.30E-03 | 1.89E-02 | -1.10E+00 | 2.10E-01 | 5.91E-01 | 1.34E+00 | 4.06E-03 | 1.49E-02 | -1.07E+00 | 3.49E-01 | 9.58E-01 | 1.23E+00 |
| ZNF827 | 1.01E-04 | 5.28E-04 | -1.20E+00 | 1.21E-03 | 5.24E-03 | 1.01E+00 | 1.91E-06 | 8.10E-06 | -1.41E+00 | 4.77E-05 | 1.82E-04 | -1.10E+00 | 2.47E-06 | 8.82E-06 | -1.62E+00 | 5.06E-04 | 1.43E-03 | -1.14E+00 | 2.70E-10 | 9.92E-10 | -1.87E+00 | 4.12E-06 | 1.13E-05 | -1.39E+00 |
| NT5C3B | 1.01E-04 | 5.28E-04 | -1.12E+00 | 2.26E-02 | 9.75E-02 | 1.15E+00 | 1.80E-03 | 7.63E-03 | -1.10E+00 | 8.87E-02 | 3.39E-01 | 1.25E+00 | 4.29E-02 | 1.53E-01 | 1.16E+00 | 2.37E-01 | 6.68E-01 | 1.31E+00 | 2.28E-01 | 8.37E-01 | 1.23E+00 | 3.83E-02 | 1.05E-01 | 1.55E+00 |
| C5orf22 | 1.01E-04 | 5.28E-04 | -1.26E+00 | 4.90E-05 | 2.12E-04 | -1.26E+00 | 1.78E-02 | 7.57E-02 | -1.12E+00 | 2.96E-03 | 1.13E-02 | -1.17E+00 | 2.83E-04 | 1.01E-03 | -1.61E+00 | 2.02E-03 | 5.70E-03 | -1.30E+00 | 1.11E-01 | 4.07E-01 | 1.07E+00 | 1.31E-01 | 3.59E-01 | 1.03E+00 |
| EDRF1 | 1.01E-04 | 5.28E-04 | -1.15E+00 | 2.45E-08 | 1.06E-07 | -1.40E+00 | 5.98E-03 | 2.54E-02 | -1.10E+00 | 4.73E-04 | 1.81E-03 | -1.14E+00 | 2.79E-02 | 9.96E-02 | 1.23E+00 | 3.09E-01 | 8.70E-01 | 1.84E+00 | 2.20E-01 | 8.07E-01 | 1.35E+00 | 2.48E-01 | 6.81E-01 | 1.43E+00 |
| CEP97 | 1.01E-04 | 5.29E-04 | -1.45E+00 | 1.53E-04 | 6.61E-04 | -1.28E+00 | 1.20E-04 | 5.09E-04 | -1.15E+00 | 2.96E-06 | 1.13E-05 | -1.20E+00 | 1.08E-02 | 3.87E-02 | -1.17E+00 | 5.82E-03 | 1.64E-02 | -1.14E+00 | 2.66E-04 | 9.77E-04 | -1.30E+00 | 1.40E-02 | 3.85E-02 | -1.08E+00 |
| CHPF2 | 1.01E-04 | 5.29E-04 | -1.04E+00 | 1.27E-06 | 5.49E-06 | -1.08E+00 | 3.82E-04 | 1.62E-03 | -1.18E+00 | 1.08E-02 | 4.13E-02 | 1.12E+00 | 2.83E-04 | 1.01E-03 | -1.08E+00 | 5.62E-05 | 1.59E-04 | -1.16E+00 | 8.58E-04 | 3.15E-03 | -1.09E+00 | 1.27E-03 | 3.49E-03 | -1.10E+00 |
| DDX56 | 1.02E-04 | 5.29E-04 | -1.12E+00 | 1.01E-01 | 4.37E-01 | 1.38E+00 | 3.88E-07 | 1.65E-06 | -1.45E+00 | 2.79E-03 | 1.07E-02 | -1.02E+00 | 5.49E-07 | 1.96E-06 | -1.63E+00 | 7.25E-02 | 2.04E-01 | 1.14E+00 | 3.24E-09 | 1.19E-08 | -1.38E+00 | 2.87E-02 | 7.88E-02 | 1.02E+00 |
| PRKCQ | 1.02E-04 | 5.30E-04 | -1.17E+00 | 2.89E-03 | 1.25E-02 | 1.14E+00 | 3.65E-04 | 1.55E-03 | -1.26E+00 | 3.53E-02 | 1.35E-01 | 1.11E+00 | 2.09E-04 | 7.44E-04 | -1.04E+00 | 1.49E-01 | 4.20E-01 | 1.20E+00 | 2.03E-11 | 7.46E-11 | -1.42E+00 | 2.15E-06 | 5.91E-06 | -1.22E+00 |
| SP3 | 1.02E-04 | 5.30E-04 | -1.05E+00 | 7.76E-04 | 3.35E-03 | 1.12E+00 | 4.11E-02 | 1.75E-01 | -1.02E+00 | 3.38E-04 | 1.29E-03 | -1.10E+00 | 1.01E-01 | 3.58E-01 | 1.22E+00 | 3.41E-02 | 9.60E-02 | 1.17E+00 | 1.41E-05 | 5.16E-05 | -1.16E+00 | 3.23E-06 | 8.86E-06 | -1.17E+00 |
| ZNF681 | 1.02E-04 | 5.31E-04 | -1.25E+00 | 2.24E-04 | 9.66E-04 | -1.15E+00 | 2.08E-09 | 8.83E-09 | -1.81E+00 | 4.86E-07 | 1.86E-06 | -1.37E+00 | 1.30E-07 | 4.63E-07 | -2.12E+00 | 3.44E-04 | 9.71E-04 | -1.24E+00 | 1.24E-03 | 4.54E-03 | -1.15E+00 | 2.26E-04 | 6.21E-04 | -1.30E+00 |
| ARMC8 | 1.02E-04 | 5.32E-04 | -1.04E+00 | 1.75E-04 | 7.57E-04 | 1.07E+00 | 9.94E-04 | 4.22E-03 | -1.09E+00 | 2.49E-03 | 9.52E-03 | 1.06E+00 | 7.97E-10 | 2.84E-09 | -1.42E+00 | 9.67E-05 | 2.72E-04 | -1.12E+00 | 1.60E-05 | 5.88E-05 | -1.27E+00 | 2.52E-02 | 6.92E-02 | -1.04E+00 |
| ZC3H18 | 1.02E-04 | 5.33E-04 | -1.21E+00 | 3.56E-03 | 1.54E-02 | 1.11E+00 | 1.28E-04 | 5.44E-04 | -1.19E+00 | 2.06E-03 | 7.88E-03 | 1.09E+00 | 1.99E-03 | 7.10E-03 | -1.09E+00 | 1.27E-02 | 3.59E-02 | 1.10E+00 | 7.22E-12 | 2.65E-11 | -1.70E+00 | 2.13E-05 | 5.84E-05 | -1.17E+00 |
| NDUFV2 | 1.03E-04 | 5.35E-04 | -1.36E+00 | 9.90E-05 | 4.28E-04 | -1.22E+00 | 2.94E-03 | 1.25E-02 | -1.28E+00 | 1.41E-03 | 5.40E-03 | -1.21E+00 | 1.56E-01 | 5.57E-01 | 1.22E+00 | 5.88E-02 | 1.66E-01 | 1.03E+00 | 4.79E-03 | 1.76E-02 | -1.04E+00 | 4.20E-04 | 1.15E-03 | -1.32E+00 |
| TMEM147 | 1.03E-04 | 5.36E-04 | -1.01E+00 | 4.30E-05 | 1.86E-04 | 1.09E+00 | 4.26E-04 | 1.81E-03 | -1.06E+00 | 3.01E-02 | 1.15E-01 | 1.18E+00 | 2.36E-03 | 8.41E-03 | -1.05E+00 | 2.35E-01 | 6.63E-01 | 1.25E+00 | 1.05E-02 | 3.84E-02 | -1.02E+00 | 2.51E-01 | 6.88E-01 | 1.24E+00 |
| LINC01138 | 1.03E-04 | 5.39E-04 | -1.14E+00 | 1.20E-01 | 5.17E-01 | 1.44E+00 | 1.42E-02 | 6.03E-02 | -1.11E+00 | 2.54E-01 | 9.70E-01 | 1.47E+00 | 1.92E-05 | 6.86E-05 | -1.44E+00 | 1.76E-03 | 4.96E-03 | 1.06E+00 | 2.63E-07 | 9.67E-07 | -1.89E+00 | 6.78E-05 | 1.86E-04 | -1.49E+00 |
| TXNDC11 | 1.04E-04 | 5.40E-04 | -1.20E+00 | 1.90E-05 | 8.20E-05 | -1.19E+00 | 7.43E-05 | 3.16E-04 | -1.27E+00 | 1.10E-06 | 4.18E-06 | -1.40E+00 | 1.38E-02 | 4.92E-02 | -1.01E+00 | 2.54E-03 | 7.15E-03 | -1.11E+00 | 2.49E-03 | 9.15E-03 | 1.03E+00 | 1.18E-02 | 3.25E-02 | -1.04E+00 |
| C6orf132 | 1.04E-04 | 5.41E-04 | -1.54E+00 | 4.46E-03 | 1.93E-02 | 1.00E+00 | 1.59E-03 | 6.74E-03 | -1.59E+00 | 1.13E-01 | 4.32E-01 | 1.11E+00 | 1.85E-02 | 6.61E-02 | -1.77E+00 | 7.42E-02 | 2.09E-01 | -1.30E+00 | 2.55E-02 | 9.35E-02 | -1.29E+00 | 8.65E-02 | 2.38E-01 | -1.16E+00 |
| ZSCAN30 | 1.04E-04 | 5.41E-04 | -1.12E+00 | 8.93E-03 | 3.86E-02 | 1.33E+00 | 1.37E-09 | 5.81E-09 | -1.84E+00 | 1.40E-06 | 5.35E-06 | -1.33E+00 | 6.06E-03 | 2.16E-02 | -1.10E+00 | 1.35E-02 | 3.81E-02 | -1.07E+00 | 3.59E-03 | 1.32E-02 | -1.16E+00 | 2.45E-01 | 6.73E-01 | 1.13E+00 |
| HSP90AB1 | 1.04E-04 | 5.41E-04 | -1.10E+00 | 2.84E-05 | 1.23E-04 | 1.18E+00 | 2.32E-02 | 9.85E-02 | -1.05E+00 | 3.99E-03 | 1.52E-02 | 1.14E+00 | 3.19E-04 | 1.14E-03 | -1.18E+00 | 1.33E-04 | 3.74E-04 | 1.23E+00 | 5.12E-04 | 1.88E-03 | -1.01E+00 | 5.39E-05 | 1.48E-04 | 1.25E+00 |
| MAK16 | 1.04E-04 | 5.41E-04 | -1.42E+00 | 1.03E-03 | 4.46E-03 | -1.10E+00 | 1.66E-04 | 7.05E-04 | -1.31E+00 | 2.03E-03 | 7.74E-03 | -1.02E+00 | 7.89E-02 | 2.81E-01 | 1.00E+00 | 1.10E-01 | 3.11E-01 | 1.26E+00 | 1.58E-05 | 5.80E-05 | -1.32E+00 | 2.53E-03 | 6.94E-03 | -1.15E+00 |
| VOPP1 | 1.04E-04 | 5.42E-04 | -1.48E+00 | 4.34E-03 | 1.88E-02 | 1.00E+00 | 5.90E-03 | 2.51E-02 | -1.21E+00 | 6.80E-02 | 2.60E-01 | 1.07E+00 | 3.19E-02 | 1.14E-01 | 1.00E+00 | 8.43E-02 | 2.38E-01 | 1.14E+00 | 1.85E-03 | 6.78E-03 | -1.01E+00 | 3.49E-01 | 9.58E-01 | 1.34E+00 |
| RABIF | 1.05E-04 | 5.47E-04 | -1.35E+00 | 1.07E-02 | 4.62E-02 | 1.14E+00 | 4.66E-05 | 1.98E-04 | -1.32E+00 | 3.18E-04 | 1.21E-03 | -1.06E+00 | 6.10E-05 | 2.17E-04 | -1.51E+00 | 1.49E-03 | 4.21E-03 | -1.13E+00 | 1.07E-03 | 3.93E-03 | -1.18E+00 | 3.19E-02 | 8.77E-02 | 1.07E+00 |
| MAP4K1 | 1.05E-04 | 5.47E-04 | -1.07E+00 | 8.32E-04 | 3.59E-03 | 1.12E+00 | 1.81E-04 | 7.67E-04 | -1.25E+00 | 7.44E-04 | 2.84E-03 | 1.00E+00 | 2.25E-03 | 8.01E-03 | -1.26E+00 | 1.11E-03 | 3.13E-03 | -1.27E+00 | 9.63E-04 | 3.53E-03 | -1.36E+00 | 2.03E-02 | 5.58E-02 | -1.03E+00 |
| RABGAP1L | 1.06E-04 | 5.52E-04 | -1.07E+00 | 1.28E-02 | 5.54E-02 | 1.16E+00 | 1.52E-02 | 6.46E-02 | -1.07E+00 | 4.51E-02 | 1.72E-01 | 1.09E+00 | 3.66E-03 | 1.30E-02 | 1.06E+00 | 7.21E-03 | 2.03E-02 | 1.16E+00 | 1.16E-01 | 4.24E-01 | 1.62E+00 | 2.95E-01 | 8.09E-01 | 1.34E+00 |
| RPS7P4 | 1.06E-04 | 5.53E-04 | -1.08E+00 | 4.86E-38 | 2.10E-37 | -1.70E+00 | 3.82E-02 | 1.62E-01 | -1.08E+00 | 2.82E-14 | 1.08E-13 | -1.64E+00 | 1.72E-04 | 6.13E-04 | -1.18E+00 | 2.33E-20 | 6.56E-20 | -1.82E+00 | 4.20E-04 | 1.54E-03 | -1.19E+00 | 1.63E-11 | 4.47E-11 | -1.67E+00 |
| NMNAT3 | 1.06E-04 | 5.54E-04 | -1.43E+00 | 1.66E-03 | 7.18E-03 | -1.04E+00 | 1.20E-06 | 5.09E-06 | -1.55E+00 | 2.05E-06 | 7.84E-06 | -1.32E+00 | 9.96E-09 | 3.55E-08 | -1.73E+00 | 1.47E-03 | 4.13E-03 | -1.05E+00 | 6.76E-13 | 2.48E-12 | -1.39E+00 | 3.74E-08 | 1.03E-07 | -1.15E+00 |
| EDA2R | 1.06E-04 | 5.54E-04 | -2.05E+00 | 4.15E-03 | 1.79E-02 | -1.28E+00 | 9.57E-15 | 4.07E-14 | -3.50E+00 | 3.55E-07 | 1.36E-06 | -1.61E+00 | 2.35E-01 | 8.37E-01 | 1.03E+00 | 3.22E-01 | 9.08E-01 | 1.26E+00 | 1.99E-04 | 7.29E-04 | -2.17E+00 | 2.06E-03 | 5.65E-03 | -1.76E+00 |
| TMEM167B | 1.06E-04 | 5.55E-04 | -1.03E+00 | 3.61E-05 | 1.56E-04 | -1.01E+00 | 3.87E-04 | 1.65E-03 | -1.20E+00 | 3.43E-03 | 1.31E-02 | -1.01E+00 | 7.51E-02 | 2.68E-01 | 1.11E+00 | 1.08E-02 | 3.06E-02 | 1.02E+00 | 2.72E-02 | 9.99E-02 | 1.05E+00 | 9.76E-03 | 2.68E-02 | 1.11E+00 |
| TMEM25 | 1.07E-04 | 5.56E-04 | -1.19E+00 | 1.57E-02 | 6.77E-02 | 1.16E+00 | 5.81E-03 | 2.47E-02 | -1.10E+00 | 3.78E-02 | 1.45E-01 | 1.14E+00 | 3.24E-02 | 1.16E-01 | -1.07E+00 | 5.89E-02 | 1.66E-01 | 1.01E+00 | 7.18E-02 | 2.63E-01 | 1.05E+00 | 1.83E-01 | 5.03E-01 | 1.21E+00 |
| PWAR6 | 1.07E-04 | 5.57E-04 | -1.56E+00 | 9.22E-02 | 3.98E-01 | 1.28E+00 | 7.15E-04 | 3.04E-03 | -1.91E+00 | 4.66E-02 | 1.78E-01 | -1.08E+00 | 3.74E-07 | 1.33E-06 | -2.14E+00 | 9.32E-02 | 2.63E-01 | 1.19E+00 | 5.07E-05 | 1.86E-04 | -1.55E+00 | 4.89E-02 | 1.34E-01 | 1.07E+00 |
| MRPS25 | 1.07E-04 | 5.57E-04 | -1.04E+00 | 1.44E-03 | 6.23E-03 | 1.15E+00 | 3.39E-05 | 1.44E-04 | -1.17E+00 | 2.40E-04 | 9.17E-04 | -1.02E+00 | 7.96E-04 | 2.84E-03 | -1.06E+00 | 7.86E-02 | 2.22E-01 | 1.21E+00 | 1.74E-03 | 6.40E-03 | -1.10E+00 | 3.42E-01 | 9.39E-01 | 1.14E+00 |
| PAK1 | 1.07E-04 | 5.58E-04 | -1.25E+00 | 1.53E-05 | 6.60E-05 | -1.21E+00 | 1.49E-08 | 6.35E-08 | -1.46E+00 | 7.72E-09 | 2.95E-08 | -1.39E+00 | 1.59E-02 | 5.66E-02 | -1.42E+00 | 1.24E-01 | 3.49E-01 | -1.02E+00 | 6.91E-05 | 2.53E-04 | -1.40E+00 | 5.79E-03 | 1.59E-02 | -1.10E+00 |
| WRAP53 | 1.07E-04 | 5.60E-04 | -1.26E+00 | 2.14E-04 | 9.25E-04 | -1.11E+00 | 2.27E-04 | 9.64E-04 | -1.27E+00 | 4.26E-03 | 1.63E-02 | 1.04E+00 | 1.77E-01 | 6.31E-01 | 1.27E+00 | 1.24E-01 | 3.48E-01 | 1.12E+00 | 1.03E-02 | 3.79E-02 | -1.05E+00 | 1.85E-01 | 5.09E-01 | 1.21E+00 |
| AC096642.1 | 1.08E-04 | 5.60E-04 | -2.32E+00 | 2.25E-01 | 9.70E-01 | 1.23E+00 | 2.03E-02 | 8.64E-02 | -1.73E+00 | 8.38E-02 | 3.20E-01 | 1.62E+00 | 1.94E-01 | 6.91E-01 | -1.07E+00 | 1.89E-01 | 5.34E-01 | 1.73E+00 | 5.63E-02 | 2.07E-01 | -1.56E+00 | 1.45E-01 | 3.99E-01 | -1.55E+00 |
| SDHAF4 | 1.08E-04 | 5.62E-04 | -1.32E+00 | 4.44E-05 | 1.92E-04 | -1.29E+00 | 5.44E-06 | 2.31E-05 | -1.54E+00 | 9.36E-07 | 3.57E-06 | -1.47E+00 | 8.32E-08 | 2.97E-07 | -2.15E+00 | 5.74E-07 | 1.62E-06 | -1.81E+00 | 1.19E-05 | 4.36E-05 | -1.34E+00 | 3.43E-06 | 9.43E-06 | -1.42E+00 |
| TMCO4 | 1.08E-04 | 5.63E-04 | -1.22E+00 | 4.18E-03 | 1.81E-02 | 1.12E+00 | 1.23E-02 | 5.24E-02 | 1.01E+00 | 1.02E-01 | 3.89E-01 | 1.34E+00 | 8.11E-03 | 2.89E-02 | -1.25E+00 | 4.04E-02 | 1.14E-01 | 1.02E+00 | 6.15E-02 | 2.26E-01 | -1.03E+00 | 2.03E-01 | 5.59E-01 | -1.01E+00 |
| CYCS | 1.09E-04 | 5.66E-04 | 1.03E+00 | 8.54E-37 | 3.69E-36 | -1.60E+00 | 9.95E-02 | 4.23E-01 | 1.11E+00 | 5.90E-06 | 2.25E-05 | -1.07E+00 | 1.10E-02 | 3.93E-02 | 1.10E+00 | 6.81E-08 | 1.92E-07 | -1.27E+00 | 6.60E-02 | 2.42E-01 | 1.01E+00 | 7.20E-02 | 1.98E-01 | -1.01E+00 |
| STRIP1 | 1.09E-04 | 5.66E-04 | -1.36E+00 | 1.32E-03 | 5.71E-03 | -1.09E+00 | 1.01E-04 | 4.30E-04 | -1.29E+00 | 1.39E-03 | 5.31E-03 | -1.02E+00 | 1.03E-04 | 3.66E-04 | -1.22E+00 | 2.03E-02 | 5.73E-02 | 1.24E+00 | 1.10E-10 | 4.04E-10 | -1.92E+00 | 2.62E-05 | 7.20E-05 | -1.39E+00 |
| AC018868.1 | 1.09E-04 | 5.69E-04 | 1.07E+00 | 8.16E-17 | 3.53E-16 | -1.24E+00 | 1.10E-01 | 4.69E-01 | 1.12E+00 | 1.35E-04 | 5.16E-04 | -1.06E+00 | 2.55E-03 | 9.11E-03 | 1.08E+00 | 3.79E-08 | 1.07E-07 | -1.21E+00 | 1.68E-02 | 6.16E-02 | 1.09E+00 | 1.31E-02 | 3.59E-02 | -1.02E+00 |
| PACSIN2 | 1.09E-04 | 5.69E-04 | -1.38E+00 | 1.07E-02 | 4.63E-02 | 1.06E+00 | 4.48E-04 | 1.90E-03 | -1.27E+00 | 4.59E-02 | 1.75E-01 | 1.11E+00 | 8.49E-05 | 3.03E-04 | -1.30E+00 | 2.91E-03 | 8.21E-03 | -1.05E+00 | 2.23E-03 | 8.19E-03 | 1.02E+00 | 1.76E-01 | 4.83E-01 | 1.24E+00 |
| ZMYM2 | 1.10E-04 | 5.71E-04 | -1.10E+00 | 5.30E-04 | 2.29E-03 | 1.03E+00 | 8.75E-06 | 3.72E-05 | -1.26E+00 | 4.86E-04 | 1.86E-03 | -1.04E+00 | 2.00E-03 | 7.12E-03 | -1.00E+00 | 4.27E-03 | 1.20E-02 | 1.04E+00 | 7.86E-08 | 2.88E-07 | -1.30E+00 | 5.59E-05 | 1.54E-04 | -1.22E+00 |
| IQCB1 | 1.10E-04 | 5.71E-04 | -1.09E+00 | 1.15E-06 | 4.95E-06 | -1.11E+00 | 8.74E-04 | 3.71E-03 | -1.01E+00 | 5.92E-05 | 2.26E-04 | -1.01E+00 | 1.54E-03 | 5.50E-03 | -1.01E+00 | 1.60E-04 | 4.51E-04 | -1.14E+00 | 1.74E-05 | 6.40E-05 | -1.16E+00 | 2.22E-06 | 6.10E-06 | -1.21E+00 |
| DNAJB2 | 1.10E-04 | 5.71E-04 | -1.14E+00 | 2.79E-04 | 1.21E-03 | -1.04E+00 | 6.24E-04 | 2.65E-03 | -1.18E+00 | 4.93E-03 | 1.88E-02 | -1.01E+00 | 2.58E-04 | 9.20E-04 | -1.03E+00 | 8.82E-04 | 2.49E-03 | -1.02E+00 | 7.76E-02 | 2.85E-01 | 1.14E+00 | 1.35E-02 | 3.70E-02 | -1.09E+00 |
| ZNF2 | 1.10E-04 | 5.71E-04 | -1.78E+00 | 2.36E-03 | 1.02E-02 | -1.34E+00 | 1.33E-04 | 5.66E-04 | -1.82E+00 | 9.38E-03 | 3.58E-02 | -1.14E+00 | 1.70E-03 | 6.07E-03 | -2.09E+00 | 2.89E-02 | 8.14E-02 | -1.38E+00 | 7.07E-09 | 2.59E-08 | -2.66E+00 | 1.23E-06 | 3.39E-06 | -2.14E+00 |
| ALDH16A1 | 1.10E-04 | 5.73E-04 | -1.19E+00 | 5.21E-04 | 2.25E-03 | -1.00E+00 | 6.94E-03 | 2.95E-02 | -1.01E+00 | 5.35E-03 | 2.04E-02 | 1.07E+00 | 9.85E-06 | 3.51E-05 | -1.27E+00 | 6.04E-04 | 1.70E-03 | -1.03E+00 | 8.47E-03 | 3.11E-02 | 1.05E+00 | 8.68E-02 | 2.38E-01 | 1.15E+00 |
| RAB8B | 1.10E-04 | 5.73E-04 | -1.14E+00 | 9.04E-09 | 3.91E-08 | -1.43E+00 | 6.48E-02 | 2.75E-01 | 1.11E+00 | 1.13E-03 | 4.31E-03 | -1.03E+00 | 6.06E-06 | 2.16E-05 | -1.34E+00 | 7.76E-07 | 2.19E-06 | -1.35E+00 | 5.51E-03 | 2.02E-02 | 1.09E+00 | 1.65E-03 | 4.54E-03 | -1.07E+00 |
| XXYLT1-AS2 | 1.11E-04 | 5.77E-04 | 1.00E+00 | 1.45E-01 | 6.28E-01 | 1.49E+00 | 1.46E-04 | 6.18E-04 | -1.70E+00 | 6.01E-03 | 2.30E-02 | -1.16E+00 | 2.19E-01 | 7.82E-01 | 1.02E+00 | 3.47E-01 | 9.79E-01 | 1.50E+00 | 2.71E-01 | 9.95E-01 | 1.24E+00 | 3.29E-01 | 9.03E-01 | 1.01E+00 |
| AC092053.3 | 1.11E-04 | 5.77E-04 | 3.01E+00 | 3.52E-02 | 1.52E-01 | 2.27E+00 | 2.61E-02 | 1.11E-01 | 1.93E+00 | 2.61E-01 | 9.97E-01 | 1.25E+00 | 2.19E-01 | 7.80E-01 | 1.42E+00 | 2.60E-01 | 7.32E-01 | 1.38E+00 | 1.39E-02 | 5.10E-02 | 2.59E+00 | 1.37E-01 | 3.77E-01 | 1.79E+00 |
| NMNAT1 | 1.11E-04 | 5.78E-04 | -1.61E+00 | 1.49E-02 | 6.44E-02 | 1.02E+00 | 7.72E-04 | 3.28E-03 | -1.62E+00 | 2.81E-02 | 1.07E-01 | -1.05E+00 | 1.16E-03 | 4.15E-03 | -1.14E+00 | 3.38E-02 | 9.54E-02 | 1.11E+00 | 2.44E-02 | 8.96E-02 | -1.02E+00 | 1.40E-01 | 3.84E-01 | 1.18E+00 |
| CFAP70 | 1.12E-04 | 5.83E-04 | -1.46E+00 | 1.10E-02 | 4.73E-02 | 1.05E+00 | 1.34E-03 | 5.70E-03 | -1.64E+00 | 1.20E-02 | 4.59E-02 | -1.12E+00 | 1.72E-01 | 6.15E-01 | 1.27E+00 | 2.21E-01 | 6.22E-01 | 1.75E+00 | 1.28E-01 | 4.69E-01 | 1.55E+00 | 2.77E-01 | 7.62E-01 | 1.23E+00 |
| SDCBP2-AS1 | 1.12E-04 | 5.83E-04 | -1.32E+00 | 9.01E-05 | 3.89E-04 | -1.22E+00 | 1.25E-03 | 5.29E-03 | -1.51E+00 | 6.87E-03 | 2.63E-02 | -1.08E+00 | 1.35E-02 | 4.81E-02 | -1.58E+00 | 5.85E-02 | 1.65E-01 | -1.08E+00 | 1.41E-01 | 5.17E-01 | -1.02E+00 | 2.82E-01 | 7.76E-01 | 1.07E+00 |
| APH1B | 1.12E-04 | 5.83E-04 | -1.13E+00 | 4.52E-03 | 1.96E-02 | 1.16E+00 | 6.79E-07 | 2.88E-06 | -1.50E+00 | 5.51E-03 | 2.10E-02 | 1.01E+00 | 7.17E-03 | 2.56E-02 | -1.04E+00 | 1.27E-02 | 3.59E-02 | 1.03E+00 | 3.16E-04 | 1.16E-03 | -1.14E+00 | 5.53E-04 | 1.52E-03 | -1.15E+00 |
| PROSER1 | 1.12E-04 | 5.84E-04 | -1.32E+00 | 7.17E-03 | 3.10E-02 | 1.05E+00 | 6.32E-03 | 2.69E-02 | -1.04E+00 | 6.21E-02 | 2.37E-01 | 1.25E+00 | 1.39E-04 | 4.97E-04 | -1.24E+00 | 9.41E-03 | 2.65E-02 | -1.01E+00 | 6.06E-10 | 2.23E-09 | -1.46E+00 | 1.02E-03 | 2.81E-03 | -1.04E+00 |
| PIK3R3 | 1.12E-04 | 5.84E-04 | -1.36E+00 | 3.91E-05 | 1.69E-04 | -1.43E+00 | 8.30E-12 | 3.53E-11 | -2.33E+00 | 6.53E-11 | 2.49E-10 | -1.95E+00 | 2.56E-08 | 9.11E-08 | -2.28E+00 | 2.15E-07 | 6.07E-07 | -1.82E+00 | 5.09E-04 | 1.87E-03 | -1.54E+00 | 7.24E-04 | 1.99E-03 | -1.64E+00 |
| ARCN1 | 1.12E-04 | 5.84E-04 | -1.07E+00 | 2.77E-03 | 1.20E-02 | 1.20E+00 | 4.04E-02 | 1.71E-01 | 1.04E+00 | 2.57E-02 | 9.82E-02 | 1.11E+00 | 7.42E-05 | 2.65E-04 | -1.18E+00 | 4.95E-03 | 1.39E-02 | 1.01E+00 | 9.27E-05 | 3.40E-04 | -1.20E+00 | 9.09E-04 | 2.50E-03 | -1.13E+00 |
| NAP1L5 | 1.12E-04 | 5.84E-04 | -1.44E+00 | 2.58E-06 | 1.11E-05 | -1.55E+00 | 8.75E-06 | 3.72E-05 | -1.43E+00 | 3.15E-08 | 1.20E-07 | -1.66E+00 | 2.57E-01 | 9.16E-01 | 1.54E+00 | 1.74E-01 | 4.91E-01 | 1.16E+00 | 3.18E-03 | 1.17E-02 | -1.27E+00 | 1.16E-02 | 3.18E-02 | -1.16E+00 |
| TPX2 | 1.12E-04 | 5.86E-04 | -1.58E+00 | 1.86E-03 | 8.03E-03 | -1.13E+00 | 2.44E-03 | 1.04E-02 | 1.57E+00 | 5.75E-02 | 2.19E-01 | 1.68E+00 | 2.61E-02 | 9.32E-02 | -1.40E+00 | 1.48E-01 | 4.16E-01 | 1.08E+00 | 9.94E-02 | 3.65E-01 | 1.03E+00 | 2.36E-01 | 6.47E-01 | 1.20E+00 |
| ANKRD20A5P | 1.13E-04 | 5.87E-04 | -1.28E+00 | 8.68E-02 | 3.75E-01 | 1.28E+00 | 2.89E-06 | 1.23E-05 | -1.61E+00 | 1.75E-03 | 6.67E-03 | -1.03E+00 | 2.20E-01 | 7.84E-01 | 1.05E+00 | 2.99E-01 | 8.42E-01 | 1.22E+00 | 1.96E-01 | 7.18E-01 | 1.15E+00 | 3.11E-01 | 8.55E-01 | 1.12E+00 |
| ANAPC7 | 1.13E-04 | 5.88E-04 | -1.15E+00 | 2.17E-01 | 9.39E-01 | 1.45E+00 | 4.83E-10 | 2.05E-09 | -1.48E+00 | 6.71E-05 | 2.56E-04 | -1.02E+00 | 1.95E-07 | 6.97E-07 | -1.38E+00 | 6.01E-03 | 1.69E-02 | 1.10E+00 | 1.20E-17 | 4.39E-17 | -1.78E+00 | 1.97E-03 | 5.40E-03 | -1.01E+00 |
| RAB6B | 1.13E-04 | 5.88E-04 | -2.38E+00 | 3.79E-04 | 1.64E-03 | -1.76E+00 | 2.85E-02 | 1.21E-01 | -1.63E+00 | 1.13E-01 | 4.30E-01 | -1.01E+00 | 2.34E-02 | 8.35E-02 | -1.20E+00 | 1.54E-02 | 4.33E-02 | 2.04E+00 | 2.71E-01 | 9.95E-01 | 1.15E+00 | 8.65E-02 | 2.38E-01 | 1.24E+00 |
| TBL1X | 1.13E-04 | 5.89E-04 | -1.49E+00 | 2.20E-03 | 9.52E-03 | -1.08E+00 | 1.40E-04 | 5.93E-04 | -1.59E+00 | 5.85E-03 | 2.24E-02 | -1.15E+00 | 4.09E-02 | 1.46E-01 | -1.05E+00 | 2.92E-01 | 8.24E-01 | 1.41E+00 | 1.30E-07 | 4.76E-07 | -1.36E+00 | 2.39E-03 | 6.56E-03 | -1.09E+00 |
| AC074327.1 | 1.13E-04 | 5.89E-04 | -1.03E+00 | 1.79E-06 | 7.73E-06 | -1.05E+00 | 2.03E-01 | 8.61E-01 | 1.13E+00 | 4.81E-03 | 1.84E-02 | -1.18E+00 | 1.59E-01 | 5.68E-01 | 1.81E+00 | 1.94E-01 | 5.47E-01 | 1.16E+00 | 9.93E-02 | 3.65E-01 | 1.83E+00 | 3.22E-02 | 8.84E-02 | -1.29E+00 |
| TOP3A | 1.13E-04 | 5.90E-04 | -1.41E+00 | 2.58E-02 | 1.11E-01 | 1.13E+00 | 6.67E-04 | 2.83E-03 | -1.30E+00 | 1.86E-02 | 7.10E-02 | 1.05E+00 | 1.33E-05 | 4.76E-05 | -1.70E+00 | 4.75E-04 | 1.34E-03 | -1.26E+00 | 3.60E-06 | 1.32E-05 | -1.29E+00 | 4.55E-03 | 1.25E-02 | -1.02E+00 |
| SDF2L1 | 1.13E-04 | 5.90E-04 | -1.11E+00 | 9.53E-03 | 4.12E-02 | 1.23E+00 | 2.17E-01 | 9.21E-01 | 1.24E+00 | 5.31E-02 | 2.03E-01 | 1.66E+00 | 2.33E-08 | 8.32E-08 | -1.52E+00 | 7.57E-04 | 2.13E-03 | 1.03E+00 | 1.13E-01 | 4.13E-01 | -1.05E+00 | 2.04E-04 | 5.62E-04 | 1.39E+00 |
| RYK | 1.13E-04 | 5.90E-04 | -1.13E+00 | 2.49E-04 | 1.07E-03 | 1.03E+00 | 5.50E-06 | 2.34E-05 | -1.43E+00 | 8.02E-05 | 3.06E-04 | -1.17E+00 | 2.30E-03 | 8.19E-03 | -1.05E+00 | 3.92E-02 | 1.10E-01 | 1.13E+00 | 5.15E-06 | 1.89E-05 | -1.26E+00 | 4.76E-04 | 1.31E-03 | -1.09E+00 |
| ARPC2 | 1.14E-04 | 5.94E-04 | -1.09E+00 | 6.67E-03 | 2.88E-02 | 1.02E+00 | 6.43E-02 | 2.73E-01 | -1.06E+00 | 8.47E-02 | 3.23E-01 | -1.04E+00 | 5.61E-02 | 2.00E-01 | -1.05E+00 | 4.36E-02 | 1.23E-01 | -1.06E+00 | 2.43E-01 | 8.92E-01 | 1.01E+00 | 3.04E-01 | 8.35E-01 | 1.02E+00 |
| TANGO6 | 1.14E-04 | 5.96E-04 | -2.04E+00 | 1.05E-03 | 4.52E-03 | -1.51E+00 | 4.63E-03 | 1.97E-02 | -1.38E+00 | 1.79E-02 | 6.86E-02 | -1.14E+00 | 3.10E-02 | 1.11E-01 | -1.34E+00 | 2.79E-01 | 7.88E-01 | 1.28E+00 | 6.61E-09 | 2.43E-08 | -1.81E+00 | 8.76E-07 | 2.41E-06 | -1.49E+00 |
| DNAJB14 | 1.14E-04 | 5.96E-04 | 1.04E+00 | 1.01E-04 | 4.36E-04 | 1.20E+00 | 2.03E-05 | 8.65E-05 | -1.17E+00 | 1.87E-04 | 7.14E-04 | -1.04E+00 | 6.03E-05 | 2.15E-04 | -1.07E+00 | 2.34E-04 | 6.60E-04 | 1.07E+00 | 1.93E-02 | 7.10E-02 | 1.06E+00 | 6.35E-02 | 1.74E-01 | 1.07E+00 |
| AC013399.1 | 1.14E-04 | 5.96E-04 | 2.91E+00 | 1.87E-01 | 8.06E-01 | 1.36E+00 | 1.56E-01 | 6.65E-01 | 1.31E+00 | 3.01E-02 | 1.15E-01 | -1.42E+00 | 5.19E-02 | 1.85E-01 | -1.13E+00 | 5.11E-02 | 1.44E-01 | -1.20E+00 | 2.44E-01 | 8.95E-01 | -1.05E+00 | 1.67E-01 | 4.58E-01 | -1.39E+00 |
| NPIPB8 | 1.14E-04 | 5.96E-04 | -1.57E+00 | 2.80E-03 | 1.21E-02 | -1.12E+00 | 1.58E-04 | 6.71E-04 | -1.56E+00 | 8.12E-06 | 3.10E-05 | -1.78E+00 | 2.52E-01 | 9.00E-01 | 1.28E+00 | 2.86E-01 | 8.05E-01 | 1.45E+00 | 1.57E-03 | 5.77E-03 | -1.31E+00 | 7.33E-04 | 2.01E-03 | -1.47E+00 |
| ITPRIPL1 | 1.15E-04 | 5.98E-04 | -1.66E+00 | 1.47E-02 | 6.37E-02 | -1.01E+00 | 8.29E-12 | 3.52E-11 | -1.91E+00 | 1.33E-06 | 5.10E-06 | -1.26E+00 | 2.55E-03 | 9.08E-03 | -2.67E+00 | 3.36E-02 | 9.47E-02 | -1.42E+00 | 6.17E-15 | 2.26E-14 | -2.57E+00 | 1.05E-07 | 2.89E-07 | -1.55E+00 |
| PMS2P3 | 1.15E-04 | 5.98E-04 | -1.27E+00 | 1.06E-05 | 4.57E-05 | -1.34E+00 | 3.51E-04 | 1.49E-03 | -1.41E+00 | 2.62E-05 | 1.00E-04 | -1.43E+00 | 1.07E-02 | 3.83E-02 | -1.02E+00 | 1.63E-02 | 4.60E-02 | -1.08E+00 | 1.80E-02 | 6.59E-02 | -1.02E+00 | 2.37E-02 | 6.50E-02 | -1.02E+00 |
| AC091132.1 | 1.15E-04 | 5.99E-04 | -1.43E+00 | 5.29E-03 | 2.29E-02 | -1.05E+00 | 8.12E-05 | 3.45E-04 | -1.30E+00 | 2.24E-05 | 8.57E-05 | -1.28E+00 | 2.52E-02 | 8.99E-02 | -1.14E+00 | 1.78E-02 | 5.03E-02 | -1.22E+00 | 3.08E-03 | 1.13E-02 | -1.68E+00 | 1.75E-03 | 4.80E-03 | -1.86E+00 |

| | | | | | | | | | | | | | | | | | | | | | | | | |
|---|---|---|---|---|---|---|---|---|---|---|---|---|---|---|---|---|---|---|---|---|---|---|---|---|
| FAM3A | 1.15E-04 | 6.02E-04 | -1.06E+00 | 7.32E-03 | 3.16E-02 | 1.19E+00 | 1.92E-04 | 8.14E-04 | -1.24E+00 | 1.18E-02 | 4.52E-02 | 1.07E+00 | 4.74E-03 | 1.69E-02 | 1.03E+00 | 5.72E-04 | 1.61E-03 | -1.00E+00 | 1.80E-01 | 6.59E-01 | 1.16E+00 | 6.71E-02 | 1.84E-01 | 1.08E+00 |
| FAM135A | 1.15E-04 | 6.02E-04 | -1.16E+00 | 2.44E-02 | 1.05E-01 | 1.24E+00 | 1.71E-04 | 7.25E-04 | -1.33E+00 | 3.55E-02 | 1.36E-01 | 1.18E+00 | 7.11E-03 | 2.54E-02 | -1.45E+00 | 8.53E-02 | 2.40E-01 | -1.10E+00 | 4.77E-03 | 1.75E-02 | -1.40E+00 | 2.32E-01 | 6.38E-01 | 1.08E+00 |
| PUS10 | 1.16E-04 | 6.03E-04 | -1.39E+00 | 3.30E-02 | 1.43E-01 | 1.21E+00 | 1.23E-07 | 5.25E-07 | -1.84E+00 | 7.10E-03 | 2.71E-02 | 1.04E+00 | 4.75E-04 | 1.69E-03 | -1.34E+00 | 2.75E-02 | 7.76E-02 | -1.01E+00 | 1.28E-05 | 4.69E-05 | -1.66E+00 | 7.92E-02 | 2.18E-01 | 1.03E+00 |
| ITCH | 1.16E-04 | 6.04E-04 | -1.06E+00 | 9.30E-04 | 4.02E-03 | 1.16E+00 | 8.54E-10 | 3.63E-09 | -1.63E+00 | 3.75E-08 | 1.43E-07 | -1.27E+00 | 2.61E-11 | 9.30E-11 | -1.61E+00 | 2.15E-10 | 6.06E-10 | -1.38E+00 | 1.88E-05 | 6.91E-05 | -1.21E+00 | 3.07E-07 | 8.44E-07 | -1.29E+00 |
| DHX16 | 1.16E-04 | 6.06E-04 | -1.51E+00 | 6.55E-04 | 2.83E-03 | -1.15E+00 | 2.50E-02 | 1.06E-01 | -1.06E+00 | 4.30E-02 | 1.64E-01 | 1.13E+00 | 2.97E-03 | 1.06E-02 | -1.40E+00 | 1.32E-01 | 3.72E-01 | 1.16E+00 | 4.30E-04 | 1.58E-03 | -1.24E+00 | 6.74E-05 | 1.85E-04 | -1.34E+00 |
| CPSF4 | 1.16E-04 | 6.06E-04 | -1.18E+00 | 9.66E-03 | 4.17E-02 | 1.19E+00 | 1.13E-03 | 4.82E-03 | -1.20E+00 | 1.85E-03 | 7.06E-03 | 1.00E+00 | 5.24E-05 | 1.87E-04 | -1.29E+00 | 5.49E-02 | 1.55E-01 | 1.18E+00 | 2.22E-03 | 8.14E-03 | -1.01E+00 | 2.94E-01 | 8.08E-01 | 1.38E+00 |
| ZNF296 | 1.16E-04 | 6.07E-04 | -1.71E+00 | 1.96E-03 | 8.48E-03 | -1.23E+00 | 3.03E-10 | 1.29E-09 | -2.36E+00 | 5.21E-07 | 1.99E-06 | -1.55E+00 | 4.46E-06 | 1.59E-05 | -1.84E+00 | 2.84E-06 | 7.99E-06 | -1.89E+00 | 1.23E-07 | 4.52E-07 | -1.87E+00 | 3.29E-04 | 9.03E-04 | -1.29E+00 |
| MED30 | 1.17E-04 | 6.08E-04 | -1.09E+00 | 2.20E-08 | 9.50E-08 | -1.21E+00 | 5.10E-03 | 2.17E-02 | -1.00E+00 | 1.75E-07 | 6.70E-07 | -1.15E+00 | 1.60E-03 | 5.70E-03 | 1.04E+00 | 2.68E-06 | 7.55E-06 | -1.09E+00 | 3.43E-06 | 1.26E-05 | -1.25E+00 | 4.54E-07 | 1.25E-06 | -1.36E+00 |
| YDJC | 1.17E-04 | 6.08E-04 | -1.04E+00 | 2.28E-04 | 9.86E-04 | 1.07E+00 | 4.36E-05 | 1.85E-04 | -1.11E+00 | 4.30E-02 | 1.64E-01 | 1.22E+00 | 2.23E-05 | 7.96E-05 | -1.19E+00 | 9.71E-03 | 2.74E-02 | 1.08E+00 | 1.45E-02 | 5.34E-02 | -1.06E+00 | 2.65E-01 | 7.29E-01 | 1.17E+00 |
| CAPS2 | 1.17E-04 | 6.10E-04 | -1.71E+00 | 3.28E-03 | 1.42E-02 | -1.11E+00 | 5.33E-04 | 2.26E-03 | -1.84E+00 | 5.73E-03 | 2.19E-02 | -1.41E+00 | 2.24E-01 | 7.99E-01 | 1.10E+00 | 2.86E-01 | 8.07E-01 | 1.15E+00 | 2.28E-04 | 8.36E-04 | -1.56E+00 | 3.58E-04 | 9.82E-04 | -1.80E+00 |
| UCHL3 | 1.17E-04 | 6.10E-04 | -1.00E+00 | 2.95E-06 | 1.27E-05 | -1.01E+00 | 8.14E-03 | 3.46E-02 | -1.01E+00 | 3.91E-07 | 1.49E-06 | -1.16E+00 | 4.77E-05 | 1.70E-04 | -1.28E+00 | 1.16E-03 | 3.28E-03 | -1.06E+00 | 2.80E-06 | 1.03E-05 | -1.13E+00 | 6.08E-03 | 1.67E-02 | 1.04E+00 |
| RNH1 | 1.17E-04 | 6.10E-04 | -1.04E+00 | 1.14E-02 | 4.95E-02 | 1.22E+00 | 7.60E-04 | 3.23E-03 | -1.10E+00 | 1.92E-01 | 7.33E-01 | 1.28E+00 | 3.61E-05 | 1.29E-04 | -1.12E+00 | 3.32E-01 | 9.37E-01 | 1.32E+00 | 1.34E-01 | 4.92E-01 | 1.10E+00 | 2.08E-01 | 5.70E-01 | 1.32E+00 |
| RRP9 | 1.17E-04 | 6.11E-04 | -1.39E+00 | 4.00E-03 | 1.73E-02 | -1.00E+00 | 5.41E-04 | 2.30E-03 | -1.13E+00 | 1.45E-01 | 5.53E-01 | 1.43E+00 | 8.18E-03 | 2.91E-02 | -1.30E+00 | 1.18E-01 | 3.33E-01 | 1.67E+00 | 9.13E-11 | 3.35E-10 | -1.47E+00 | 3.21E-02 | 8.83E-02 | 1.13E+00 |
| ISPD | 1.18E-04 | 6.13E-04 | -1.20E+00 | 1.66E-02 | 7.16E-02 | 1.10E+00 | 3.95E-07 | 1.68E-06 | -1.37E+00 | 2.92E-02 | 1.12E-01 | 1.17E+00 | 8.87E-05 | 3.16E-04 | -1.42E+00 | 5.87E-04 | 1.65E-03 | -1.16E+00 | 4.64E-07 | 1.70E-06 | -1.49E+00 | 5.44E-03 | 1.50E-02 | -1.10E+00 |
| FAM35DP | 1.18E-04 | 6.15E-04 | -1.60E+00 | 7.66E-04 | 3.31E-03 | -1.28E+00 | 3.06E-03 | 1.30E-02 | -1.46E+00 | 3.22E-03 | 1.23E-02 | -1.42E+00 | 9.02E-02 | 3.22E-01 | -1.45E+00 | 3.10E-01 | 8.75E-01 | 1.07E+00 | 1.94E-02 | 7.11E-02 | -1.17E+00 | 5.10E-02 | 1.40E-01 | -1.07E+00 |
| SETD4 | 1.18E-04 | 6.16E-04 | -1.29E+00 | 9.18E-03 | 3.97E-02 | 1.12E+00 | 3.90E-04 | 1.66E-03 | -1.24E+00 | 5.17E-03 | 1.98E-02 | 1.00E+00 | 1.19E-01 | 4.24E-01 | 1.16E+00 | 3.30E-01 | 9.29E-01 | 1.39E+00 | 2.47E-01 | 9.07E-01 | 1.16E+00 | 3.40E-01 | 9.34E-01 | 1.26E+00 |
| WDR13 | 1.18E-04 | 6.16E-04 | 1.02E+00 | 8.22E-05 | 3.55E-04 | 1.14E+00 | 4.31E-04 | 1.83E-03 | -1.11E+00 | 8.59E-06 | 3.28E-05 | -1.06E+00 | 1.02E-04 | 3.64E-04 | -1.12E+00 | 8.92E-05 | 2.51E-04 | -1.10E+00 | 3.38E-02 | 1.24E-01 | 1.06E+00 | 2.12E-01 | 5.83E-01 | 1.11E+00 |
| MYO9B | 1.18E-04 | 6.16E-04 | -1.00E+00 | 6.00E-05 | 2.59E-04 | 1.07E+00 | 4.16E-07 | 1.77E-06 | -1.26E+00 | 3.99E-06 | 1.52E-05 | -1.08E+00 | 2.90E-05 | 1.04E-04 | -1.17E+00 | 1.91E-06 | 5.38E-06 | -1.26E+00 | 7.02E-04 | 2.57E-03 | -1.14E+00 | 4.95E-02 | 1.36E-01 | 1.01E+00 |
| BICC1 | 1.18E-04 | 6.16E-04 | -1.47E+00 | 1.97E-03 | 8.53E-03 | -1.07E+00 | 7.91E-02 | 3.36E-01 | -1.14E+00 | 2.03E-01 | 7.77E-01 | 1.15E+00 | 2.50E-01 | 8.93E-01 | -1.06E+00 | 2.13E-01 | 6.01E-01 | 1.34E+00 | 2.52E-02 | 9.26E-02 | -1.44E+00 | 3.61E-01 | 9.93E-01 | 1.04E+00 |
| SWAP70 | 1.18E-04 | 6.17E-04 | -1.05E+00 | 1.35E-03 | 5.82E-03 | 1.12E+00 | 1.28E-03 | 5.46E-03 | -1.07E+00 | 2.09E-03 | 7.98E-03 | 1.04E+00 | 1.97E-10 | 7.01E-10 | -1.49E+00 | 1.13E-08 | 3.19E-08 | -1.31E+00 | 4.48E-04 | 1.64E-03 | -1.26E+00 | 3.95E-02 | 1.09E-01 | 1.01E+00 |
| ZNF623 | 1.20E-04 | 6.27E-04 | -1.47E+00 | 3.38E-03 | 1.46E-02 | -1.03E+00 | 2.40E-11 | 1.02E-10 | -2.59E+00 | 5.87E-06 | 2.24E-05 | -1.41E+00 | 6.67E-10 | 2.38E-09 | -3.21E+00 | 2.16E-05 | 6.08E-05 | -1.46E+00 | 1.01E-07 | 3.71E-07 | -1.83E+00 | 3.10E-04 | 8.53E-04 | -1.34E+00 |
| INTS4 | 1.21E-04 | 6.30E-04 | -1.28E+00 | 6.39E-03 | 2.76E-02 | 1.07E+00 | 4.56E-06 | 1.94E-05 | -1.47E+00 | 1.46E-03 | 5.57E-03 | 1.01E+00 | 1.31E-05 | 4.67E-05 | -1.67E+00 | 4.65E-03 | 1.31E-02 | -1.06E+00 | 5.65E-03 | 2.07E-02 | -1.01E+00 | 2.83E-03 | 7.78E-03 | -1.12E+00 |
| GFER | 1.21E-04 | 6.31E-04 | -1.11E+00 | 7.96E-05 | 3.44E-04 | -1.02E+00 | 4.69E-05 | 1.99E-04 | -1.26E+00 | 2.14E-07 | 8.16E-07 | -1.36E+00 | 1.57E-06 | 5.61E-06 | -1.43E+00 | 1.17E-08 | 3.30E-08 | -1.81E+00 | 1.39E-04 | 5.12E-04 | -1.18E+00 | 1.65E-04 | 4.53E-04 | -1.19E+00 |
| ADAMTS10 | 1.21E-04 | 6.32E-04 | -1.31E+00 | 7.46E-05 | 3.22E-04 | -1.29E+00 | 1.94E-04 | 8.22E-04 | -1.64E+00 | 2.05E-02 | 7.85E-02 | 1.04E+00 | 1.05E-01 | 3.74E-01 | -1.14E+00 | 9.41E-02 | 2.65E-01 | -1.25E+00 | 8.94E-02 | 3.28E-01 | 2.18E+00 | 1.01E-01 | 2.77E-01 | 1.99E+00 |
| MHENCR | 1.21E-04 | 6.32E-04 | 1.96E+00 | 1.03E-11 | 4.44E-11 | 3.02E+00 | 2.35E-01 | 9.97E-01 | 1.19E+00 | 2.92E-03 | 1.11E-02 | 1.71E+00 | 2.77E-01 | 9.88E-01 | 1.43E+00 | 5.69E-02 | 1.60E-01 | 2.12E+00 | 4.19E-02 | 1.54E-01 | 1.65E+00 | 7.06E-03 | 1.94E-02 | 2.11E+00 |
| AKTIP | 1.21E-04 | 6.32E-04 | -1.57E+00 | 3.89E-03 | 1.68E-02 | -1.14E+00 | 1.42E-03 | 6.03E-03 | -1.48E+00 | 2.27E-03 | 8.68E-03 | -1.34E+00 | 8.86E-02 | 3.16E-01 | -1.27E+00 | 1.62E-01 | 4.58E-01 | -1.02E+00 | 2.70E-01 | 9.92E-01 | 1.24E+00 | 3.48E-01 | 9.57E-01 | 1.15E+00 |
| UHRF1BP1 | 1.21E-04 | 6.32E-04 | -1.34E+00 | 6.41E-04 | 2.77E-03 | -1.08E+00 | 2.39E-06 | 1.01E-05 | -1.32E+00 | 1.23E-07 | 4.70E-07 | -1.25E+00 | 3.35E-02 | 1.19E-01 | -1.08E+00 | 7.11E-02 | 2.00E-01 | 1.03E+00 | 4.76E-03 | 1.75E-02 | -1.09E+00 | 5.82E-02 | 1.60E-01 | 1.16E+00 |
| GPC6 | 1.21E-04 | 6.33E-04 | -1.40E+00 | 9.02E-02 | 3.90E-01 | 1.31E+00 | 1.75E-07 | 7.42E-07 | -1.85E+00 | 8.98E-05 | 3.43E-04 | -1.20E+00 | 2.64E-02 | 9.42E-02 | -1.09E+00 | 2.68E-01 | 7.56E-01 | 1.44E+00 | 1.03E-01 | 3.78E-01 | 1.09E+00 | 1.52E-01 | 4.16E-01 | 1.16E+00 |
| PTGDR2 | 1.21E-04 | 6.33E-04 | -1.64E+00 | 4.66E-04 | 2.01E-03 | -1.34E+00 | 3.33E-03 | 1.42E-02 | -1.89E+00 | 3.41E-02 | 1.30E-01 | -1.19E+00 | 1.48E-12 | 5.29E-12 | -1.99E+00 | 1.49E-09 | 4.20E-09 | -1.46E+00 | 1.76E-03 | 6.46E-03 | -1.67E+00 | 1.67E-02 | 4.58E-02 | -1.35E+00 |
| DEPDC1B | 1.22E-04 | 6.34E-04 | -2.57E+00 | 3.62E-03 | 1.56E-02 | -1.36E+00 | 7.29E-03 | 3.10E-02 | -1.34E+00 | 3.91E-02 | 1.49E-01 | 1.13E+00 | 3.90E-02 | 1.39E-01 | -1.82E+00 | 3.13E-01 | 8.83E-01 | 1.24E+00 | 5.97E-10 | 2.19E-09 | -1.50E+00 | 2.31E-02 | 6.35E-02 | 1.15E+00 |
| COL9A3 | 1.22E-04 | 6.34E-04 | -1.65E+00 | 1.54E-02 | 6.64E-02 | -1.04E+00 | 6.52E-02 | 2.77E-01 | -1.14E+00 | 9.89E-02 | 3.78E-01 | 1.03E+00 | 5.96E-02 | 2.13E-01 | -1.39E+00 | 3.23E-01 | 9.09E-01 | 1.15E+00 | 1.12E-02 | 4.13E-02 | -1.23E+00 | 6.88E-03 | 1.89E-02 | -1.51E+00 |
| KAT6A | 1.22E-04 | 6.35E-04 | -1.08E+00 | 4.17E-05 | 1.80E-04 | 1.05E+00 | 8.36E-04 | 3.55E-03 | -1.08E+00 | 1.87E-05 | 7.15E-05 | -1.11E+00 | 8.59E-04 | 3.06E-03 | -1.07E+00 | 3.37E-05 | 9.49E-05 | -1.26E+00 | 9.47E-05 | 3.48E-04 | -1.25E+00 | 1.84E-03 | 5.05E-03 | -1.15E+00 |
| ANKRA2 | 1.22E-04 | 6.36E-04 | -1.14E+00 | 7.87E-03 | 3.40E-02 | 1.17E+00 | 6.73E-07 | 2.86E-06 | -1.54E+00 | 3.73E-03 | 1.43E-02 | 1.08E+00 | 2.18E-04 | 7.76E-04 | -1.21E+00 | 1.83E-02 | 5.15E-02 | 1.07E+00 | 4.37E-03 | 1.60E-02 | -1.10E+00 | 1.74E-02 | 4.78E-02 | 1.01E+00 |
| ZBED4 | 1.23E-04 | 6.39E-04 | -1.11E+00 | 2.67E-02 | 1.16E-01 | 1.27E+00 | 5.73E-05 | 2.43E-04 | -1.27E+00 | 4.21E-03 | 1.61E-02 | 1.07E+00 | 6.68E-05 | 2.38E-04 | -1.17E+00 | 6.95E-04 | 1.96E-03 | -1.03E+00 | 1.35E-05 | 4.95E-05 | -1.26E+00 | 1.60E-04 | 4.40E-04 | -1.16E+00 |
| ZER1 | 1.23E-04 | 6.41E-04 | -1.34E+00 | 4.97E-02 | 2.15E-01 | 1.20E+00 | 5.28E-04 | 2.24E-03 | -1.37E+00 | 1.18E-02 | 4.52E-02 | 1.01E+00 | 8.70E-04 | 3.10E-03 | -1.19E+00 | 1.59E-03 | 4.50E-03 | -1.10E+00 | 4.56E-03 | 1.67E-02 | -1.16E+00 | 6.98E-03 | 1.92E-02 | -1.11E+00 |
| PKN1 | 1.23E-04 | 6.41E-04 | 1.08E+00 | 8.82E-06 | 3.81E-05 | 1.12E+00 | 1.72E-01 | 7.33E-01 | 1.22E+00 | 3.44E-02 | 1.31E-01 | 1.21E+00 | 2.89E-03 | 1.03E-02 | 1.06E+00 | 1.61E-04 | 4.55E-04 | 1.07E+00 | 2.44E-01 | 8.94E-01 | 1.17E+00 | 1.80E-01 | 4.96E-01 | 1.12E+00 |
| COA1 | 1.23E-04 | 6.42E-04 | 1.05E+00 | 6.70E-05 | 2.89E-04 | 1.14E+00 | 6.22E-02 | 2.64E-01 | 1.17E+00 | 1.21E-01 | 4.61E-01 | 1.28E+00 | 6.27E-03 | 2.24E-02 | 1.07E+00 | 1.16E-01 | 3.27E-01 | 1.29E+00 | 1.59E-01 | 5.83E-01 | 1.10E+00 | 7.92E-02 | 2.17E-01 | 1.29E+00 |
| BUD13 | 1.23E-04 | 6.42E-04 | -1.55E+00 | 1.22E-03 | 5.25E-03 | -1.14E+00 | 2.15E-09 | 9.13E-09 | -1.74E+00 | 3.16E-07 | 1.21E-06 | -1.34E+00 | 1.05E-02 | 3.73E-02 | -1.41E+00 | 5.98E-02 | 1.69E-01 | -1.06E+00 | 2.98E-12 | 1.10E-11 | -1.88E+00 | 5.54E-05 | 1.52E-04 | -1.24E+00 |
| YIPF1 | 1.23E-04 | 6.42E-04 | -1.29E+00 | 2.19E-03 | 9.47E-03 | -1.01E+00 | 1.06E-04 | 4.49E-04 | -1.29E+00 | 8.30E-03 | 3.17E-02 | 1.06E+00 | 3.71E-08 | 1.32E-07 | -1.76E+00 | 4.08E-04 | 1.15E-03 | -1.17E+00 | 3.63E-07 | 1.33E-06 | -1.39E+00 | 3.74E-04 | 1.03E-03 | -1.15E+00 |
| NFYC-AS1 | 1.23E-04 | 6.43E-04 | 2.87E+00 | 4.27E-02 | 1.85E-01 | 2.19E+00 | 1.58E-02 | 6.73E-02 | 1.72E+00 | 1.88E-01 | 7.16E-01 | 1.46E+00 | 1.64E-01 | 5.85E-01 | 1.72E+00 | 1.91E-01 | 5.39E-01 | 1.81E+00 | 4.84E-02 | 1.78E-01 | 1.46E+00 | 3.43E-01 | 9.44E-01 | 1.08E+00 |
| TM9SF4 | 1.24E-04 | 6.44E-04 | -1.23E+00 | 1.25E-03 | 5.39E-03 | -1.03E+00 | 2.03E-03 | 8.64E-03 | -1.15E+00 | 1.61E-03 | 6.14E-03 | 1.01E+00 | 2.70E-02 | 9.64E-02 | 1.05E+00 | 1.82E-01 | 5.13E-01 | 1.22E+00 | 8.80E-03 | 3.23E-02 | -1.01E+00 | 8.94E-02 | 2.45E-01 | 1.12E+00 |
| CD79A | 1.24E-04 | 6.44E-04 | -1.28E+00 | 9.25E-03 | 4.00E-02 | 1.12E+00 | 4.28E-04 | 1.82E-03 | -1.56E+00 | 2.83E-02 | 1.08E-01 | 1.01E+00 | 7.79E-06 | 2.78E-05 | -1.66E+00 | 4.71E-04 | 1.33E-03 | -1.25E+00 | 2.67E-01 | 9.81E-01 | 1.24E+00 | 4.62E-02 | 1.27E-01 | 2.18E+00 |
| ATN1 | 1.24E-04 | 6.44E-04 | -1.04E+00 | 3.05E-07 | 1.32E-06 | -1.16E+00 | 8.78E-03 | 3.73E-02 | -1.06E+00 | 3.76E-03 | 1.43E-02 | 1.02E+00 | 6.50E-04 | 2.32E-03 | -1.17E+00 | 3.63E-04 | 1.02E-03 | -1.25E+00 | 9.53E-03 | 3.50E-02 | -1.07E+00 | 1.99E-02 | 5.47E-02 | -1.01E+00 |
| ARL8B | 1.24E-04 | 6.44E-04 | -1.04E+00 | 6.99E-07 | 3.02E-06 | -1.03E+00 | 1.33E-04 | 5.65E-04 | -1.12E+00 | 6.64E-07 | 2.54E-06 | -1.21E+00 | 1.47E-06 | 5.24E-06 | -1.17E+00 | 1.35E-08 | 3.81E-08 | -1.30E+00 | 2.69E-04 | 9.86E-04 | 1.01E+00 | 1.14E-03 | 3.13E-03 | 1.05E+00 |
| PRKY | 1.24E-04 | 6.45E-04 | -1.20E+00 | 1.32E-01 | 5.70E-01 | 1.37E+00 | 1.80E-06 | 7.64E-06 | -1.64E+00 | 1.22E-01 | 4.66E-01 | 1.14E+00 | 7.85E-03 | 2.80E-02 | -1.04E+00 | 3.41E-01 | 9.61E-01 | 1.55E+00 | 6.76E-07 | 2.48E-06 | -1.44E+00 | 1.05E-02 | 2.88E-02 | 1.01E+00 |
| PA2G4P6 | 1.24E-04 | 6.47E-04 | 2.46E+00 | 9.72E-04 | 4.20E-03 | 2.51E+00 | 5.85E-03 | 2.48E-02 | 1.77E+00 | 4.75E-02 | 1.82E-01 | 1.87E+00 | 1.71E-01 | 6.10E-01 | 1.55E+00 | 2.85E-01 | 8.03E-01 | 1.48E+00 | 9.92E-02 | 3.64E-01 | 1.55E+00 | 1.09E-01 | 2.99E-01 | 1.72E+00 |
| COQ10A | 1.24E-04 | 6.48E-04 | -1.18E+00 | 8.43E-06 | 3.64E-05 | -1.22E+00 | 7.10E-04 | 3.02E-03 | -1.17E+00 | 4.59E-03 | 1.75E-02 | 1.04E+00 | 5.82E-06 | 2.08E-05 | -1.47E+00 | 9.18E-04 | 2.59E-03 | -1.08E+00 | 4.95E-05 | 1.82E-04 | -1.23E+00 | 2.60E-04 | 7.15E-04 | -1.18E+00 |
| COX7BP2 | 1.25E-04 | 6.51E-04 | -1.35E+00 | 5.07E-03 | 2.19E-02 | 1.03E+00 | 4.78E-03 | 2.03E-02 | -1.11E+00 | 1.49E-04 | 5.68E-04 | -1.22E+00 | 5.44E-04 | 1.94E-03 | -1.33E+00 | 2.06E-04 | 5.81E-04 | -1.35E+00 | 1.94E-05 | 7.13E-05 | -1.37E+00 | 3.89E-07 | 1.07E-06 | -1.72E+00 |
| AC012615.1 | 1.25E-04 | 6.52E-04 | -1.47E+00 | 2.20E-05 | 9.51E-05 | -1.56E+00 | 2.53E-02 | 1.07E-01 | -1.21E+00 | 4.44E-03 | 1.70E-02 | -1.30E+00 | 1.70E-01 | 6.06E-01 | 1.24E+00 | 5.54E-02 | 1.56E-01 | -1.10E+00 | 2.62E-01 | 9.62E-01 | 1.26E+00 | 2.42E-01 | 6.65E-01 | 1.19E+00 |
| ZNF844 | 1.25E-04 | 6.52E-04 | -1.06E+00 | 4.78E-05 | 2.06E-04 | -1.03E+00 | 5.85E-02 | 2.49E-01 | 1.03E+00 | 1.13E-01 | 4.31E-01 | 1.21E+00 | 2.48E-03 | 8.86E-03 | -1.04E+00 | 1.72E-02 | 4.85E-02 | 1.04E+00 | 5.43E-06 | 1.99E-05 | -1.20E+00 | 5.17E-08 | 1.42E-07 | -1.50E+00 |
| C16orf70 | 1.26E-04 | 6.58E-04 | -1.96E+00 | 9.47E-02 | 4.09E-01 | 1.22E+00 | 6.83E-05 | 2.90E-04 | -1.94E+00 | 5.68E-03 | 2.17E-02 | -1.20E+00 | 2.55E-04 | 9.11E-04 | -1.73E+00 | 3.25E-03 | 9.17E-03 | -1.24E+00 | 2.96E-04 | 1.09E-03 | -1.60E+00 | 9.25E-03 | 2.54E-02 | -1.31E+00 |
| SECTM1 | 1.26E-04 | 6.58E-04 | -1.75E+00 | 1.72E-04 | 7.45E-04 | -1.61E+00 | 1.63E-01 | 6.93E-01 | -1.08E+00 | 9.14E-02 | 3.49E-01 | -1.27E+00 | 5.92E-02 | 2.11E-01 | -1.40E+00 | 5.45E-02 | 1.54E-01 | -1.57E+00 | 5.85E-02 | 2.15E-01 | -1.46E+00 | 5.13E-02 | 1.41E-01 | -1.62E+00 |
| PSD4 | 1.26E-04 | 6.59E-04 | -1.24E+00 | 4.40E-03 | 1.90E-02 | 1.07E+00 | 7.38E-05 | 3.14E-04 | -1.33E+00 | 5.08E-04 | 1.94E-03 | -1.07E+00 | 4.58E-06 | 1.63E-05 | -1.49E+00 | 8.17E-06 | 2.30E-05 | -1.34E+00 | 4.14E-05 | 1.52E-04 | -1.40E+00 | 2.33E-01 | 6.40E-01 | 1.21E+00 |
| ACAD8 | 1.27E-04 | 6.61E-04 | -1.41E+00 | 5.43E-02 | 2.35E-01 | 1.28E+00 | 8.25E-03 | 3.50E-02 | -1.13E+00 | 8.55E-02 | 3.26E-01 | 1.22E+00 | 2.46E-02 | 8.78E-02 | 1.03E+00 | 2.01E-01 | 5.66E-01 | 1.18E+00 | 2.31E-03 | 8.48E-03 | 1.15E+00 | 7.99E-02 | 2.20E-01 | 1.24E+00 |
| COX7A2 | 1.27E-04 | 6.64E-04 | -1.02E+00 | 2.14E-13 | 9.26E-13 | -1.09E+00 | 2.30E-01 | 9.79E-01 | 1.11E+00 | 4.66E-03 | 1.78E-02 | -1.04E+00 | 7.54E-02 | 2.69E-01 | 1.02E+00 | 8.22E-05 | 2.32E-04 | -1.13E+00 | 2.28E-01 | 8.35E-01 | 1.05E+00 | 5.18E-02 | 1.42E-01 | -1.09E+00 |
| UQCRH | 1.27E-04 | 6.64E-04 | 1.29E+00 | 1.69E-01 | 7.29E-01 | 1.20E+00 | 1.44E-06 | 6.13E-06 | 1.42E+00 | 1.42E-02 | 5.41E-02 | 1.24E+00 | 1.01E-01 | 3.59E-01 | 1.17E+00 | 2.35E-01 | 6.61E-01 | 1.16E+00 | 6.73E-02 | 2.47E-01 | 1.17E+00 | 6.64E-02 | 1.82E-01 | 1.14E+00 |
| PIGQ | 1.28E-04 | 6.65E-04 | -1.37E+00 | 1.60E-01 | 6.91E-01 | 1.39E+00 | 2.48E-03 | 1.05E-02 | -1.14E+00 | 2.85E-02 | 1.09E-01 | 1.12E+00 | 7.74E-03 | 2.76E-02 | -1.18E+00 | 8.08E-02 | 2.28E-01 | 1.12E+00 | 5.22E-03 | 1.91E-02 | -1.12E+00 | 2.11E-01 | 5.81E-01 | 1.27E+00 |
| BORCS5 | 1.28E-04 | 6.67E-04 | -1.21E+00 | 7.10E-04 | 3.07E-03 | -1.03E+00 | 4.63E-04 | 1.97E-03 | -1.26E+00 | 1.09E-02 | 4.17E-02 | 1.06E+00 | 1.02E-02 | 3.65E-02 | -1.15E+00 | 1.52E-02 | 4.28E-02 | -1.03E+00 | 2.10E-06 | 7.71E-06 | -1.48E+00 | 1.84E-02 | 5.05E-02 | -1.02E+00 |
| SLC30A9 | 1.28E-04 | 6.67E-04 | -1.12E+00 | 3.99E-04 | 1.73E-03 | 1.08E+00 | 3.81E-06 | 1.62E-05 | -1.25E+00 | 4.69E-04 | 1.79E-03 | 1.05E+00 | 8.66E-03 | 3.09E-02 | 1.02E+00 | 4.50E-02 | 1.27E-01 | 1.10E+00 | 1.12E-02 | 4.09E-02 | -1.05E+00 | 2.37E-02 | 6.51E-02 | -1.03E+00 |
| ZNF84 | 1.28E-04 | 6.67E-04 | -1.18E+00 | 2.36E-05 | 1.02E-04 | -1.14E+00 | 3.44E-08 | 1.46E-07 | -1.52E+00 | 8.04E-07 | 3.07E-06 | -1.25E+00 | 3.93E-03 | 1.40E-02 | -1.11E+00 | 4.04E-03 | 1.14E-02 | -1.09E+00 | 5.56E-06 | 2.04E-05 | -1.49E+00 | 1.79E-04 | 4.92E-04 | -1.27E+00 |
| CLOCK | 1.28E-04 | 6.68E-04 | -1.29E+00 | 1.26E-01 | 5.46E-01 | 1.36E+00 | 9.12E-07 | 3.87E-06 | -1.72E+00 | 2.26E-03 | 8.64E-03 | -1.04E+00 | 3.92E-04 | 1.40E-03 | -1.66E+00 | 2.25E-01 | 6.34E-01 | 1.17E+00 | 6.20E-05 | 2.28E-04 | -1.32E+00 | 3.32E-01 | 9.13E-01 | 1.38E+00 |
| RPL12P17 | 1.29E-04 | 6.70E-04 | -1.11E+00 | 8.61E-06 | 3.72E-05 | -1.13E+00 | 5.49E-02 | 2.33E-01 | -1.04E+00 | 4.45E-02 | 1.70E-01 | -1.05E+00 | 7.72E-02 | 2.75E-01 | -1.05E+00 | 2.05E-01 | 5.77E-01 | -1.02E+00 | 4.75E-06 | 1.74E-05 | -1.07E+00 | 2.75E-02 | 7.56E-02 | -1.04E+00 |
| ITGA2B | 1.29E-04 | 6.70E-04 | -1.92E+00 | 8.51E-04 | 3.68E-03 | -1.54E+00 | 1.85E-06 | 7.88E-06 | -1.64E+00 | 1.83E-05 | 6.98E-05 | -1.26E+00 | 1.68E-13 | 6.00E-13 | -1.34E+00 | 9.75E-04 | 2.75E-03 | -1.09E+00 | 4.12E-02 | 1.51E-01 | 1.04E+00 | 5.19E-02 | 1.43E-01 | 1.01E+00 |
| ALDH2 | 1.29E-04 | 6.71E-04 | -1.09E+00 | 3.20E-02 | 1.38E-01 | 1.18E+00 | 8.81E-04 | 3.74E-03 | -1.03E+00 | 8.27E-05 | 3.16E-04 | 1.01E+00 | 7.08E-07 | 2.52E-06 | -1.31E+00 | 4.25E-04 | 1.20E-03 | -1.00E+00 | 2.56E-03 | 9.40E-03 | 1.01E+00 | 2.94E-02 | 8.07E-02 | 1.01E+00 |
| SPAG7 | 1.29E-04 | 6.72E-04 | 1.02E+00 | 2.59E-04 | 1.12E-03 | 1.12E+00 | 1.91E-02 | 8.12E-02 | 1.01E+00 | 1.22E-04 | 4.68E-04 | -1.00E+00 | 1.05E-04 | 3.73E-04 | -1.02E+00 | 8.00E-03 | 2.25E-02 | 1.11E+00 | 1.13E-01 | 4.16E-01 | 1.17E+00 | 1.61E-01 | 4.42E-01 | 1.17E+00 |
| TTI1 | 1.29E-04 | 6.72E-04 | -1.30E+00 | 1.24E-03 | 5.36E-03 | 1.03E+00 | 6.25E-07 | 2.66E-06 | -1.55E+00 | 2.02E-03 | 7.71E-03 | -1.03E+00 | 1.98E-04 | 7.06E-04 | -1.74E+00 | 1.61E-02 | 4.54E-02 | -1.06E+00 | 2.48E-10 | 9.11E-10 | -1.58E+00 | 8.38E-04 | 2.30E-03 | -1.08E+00 |
| TMEM161A | 1.29E-04 | 6.72E-04 | -1.10E+00 | 1.95E-06 | 8.42E-06 | -1.16E+00 | 1.81E-04 | 7.67E-04 | -1.22E+00 | 3.57E-03 | 1.36E-02 | 1.07E+00 | 1.18E-02 | 4.20E-02 | -1.06E+00 | 2.07E-01 | 5.83E-01 | 1.20E+00 | 2.75E-02 | 1.01E-01 | 1.04E+00 | 7.17E-02 | 1.97E-01 | 1.12E+00 |
| ZNF704 | 1.29E-04 | 6.73E-04 | -1.37E+00 | 5.74E-02 | 2.48E-01 | 1.33E+00 | 2.31E-04 | 9.80E-04 | -1.14E+00 | 2.64E-04 | 1.01E-03 | 1.08E+00 | 7.78E-02 | 2.78E-01 | -1.05E+00 | 2.11E-01 | 5.94E-01 | 1.10E+00 | 4.05E-03 | 1.49E-02 | -1.29E+00 | 7.86E-02 | 2.16E-01 | 1.04E+00 |
| FAM76B | 1.29E-04 | 6.73E-04 | -1.25E+00 | 3.34E-05 | 1.44E-04 | -1.16E+00 | 8.89E-05 | 3.78E-04 | -1.17E+00 | 1.16E-03 | 4.41E-03 | 1.01E+00 | 4.55E-04 | 1.62E-03 | -1.13E+00 | 2.66E-04 | 7.49E-04 | -1.10E+00 | 1.80E-04 | 6.61E-04 | -1.03E+00 | 1.18E-03 | 3.25E-03 | -1.02E+00 |
| RPL35A | 1.29E-04 | 6.73E-04 | 1.07E+00 | 3.69E-10 | 1.59E-09 | -1.13E+00 | 2.55E-05 | 1.08E-04 | 1.11E+00 | 1.12E-04 | 4.28E-04 | -1.09E+00 | 3.70E-05 | 1.32E-04 | 1.13E+00 | 9.39E-05 | 2.65E-04 | -1.11E+00 | 8.87E-03 | 3.25E-02 | 1.10E+00 | 3.04E-06 | 8.34E-06 | -1.12E+00 |
| PSMC1P5 | 1.29E-04 | 6.73E-04 | -1.44E+00 | 6.95E-03 | 3.00E-02 | -1.02E+00 | 1.54E-04 | 6.56E-04 | -1.29E+00 | 2.33E-02 | 8.90E-02 | 1.16E+00 | 1.72E-02 | 6.15E-02 | -1.23E+00 | 3.29E-01 | 9.28E-01 | 1.35E+00 | 1.79E-09 | 6.58E-09 | -1.58E+00 | 6.44E-02 | 1.77E-01 | 1.13E+00 |
| AP3S1 | 1.29E-04 | 6.74E-04 | 1.07E+00 | 2.06E-03 | 8.89E-03 | 1.21E+00 | 6.58E-02 | 2.80E-01 | 1.02E+00 | 2.05E-01 | 7.84E-01 | 1.21E+00 | 9.69E-06 | 3.46E-05 | -1.34E+00 | 2.35E-02 | 6.62E-02 | -1.00E+00 | 5.28E-06 | 1.94E-05 | -1.12E+00 | 8.08E-03 | 2.22E-02 | -1.02E+00 |
| GOLGA4 | 1.30E-04 | 6.78E-04 | -1.21E+00 | 1.12E-06 | 4.83E-06 | -1.13E+00 | 2.81E-03 | 1.19E-02 | -1.10E+00 | 5.19E-06 | 1.98E-05 | -1.14E+00 | 1.24E-01 | 4.42E-01 | 1.12E+00 | 1.17E-04 | 3.31E-04 | -1.22E+00 | 3.18E-05 | 1.17E-04 | -1.08E+00 | 8.57E-05 | 2.35E-04 | -1.21E+00 |
| FSD1L | 1.30E-04 | 6.79E-04 | -1.48E+00 | 1.70E-02 | 7.33E-02 | 1.07E+00 | 1.52E-04 | 6.45E-04 | -1.34E+00 | 1.04E-02 | 3.98E-02 | 1.06E+00 | 3.98E-05 | 1.42E-04 | -1.55E+00 | 2.15E-03 | 6.05E-03 | -1.16E+00 | 1.38E-01 | 5.08E-01 | 1.06E+00 | 3.32E-01 | 9.11E-01 | 1.27E+00 |
| PTPA | 1.31E-04 | 6.80E-04 | -1.25E+00 | 8.57E-04 | 3.70E-03 | -1.02E+00 | 3.15E-04 | 1.34E-03 | -1.23E+00 | 2.84E-03 | 1.08E-02 | -1.01E+00 | 1.50E-03 | 5.34E-03 | -1.22E+00 | 5.63E-02 | 1.59E-01 | 1.07E+00 | 7.42E-05 | 2.72E-04 | -1.20E+00 | 2.45E-03 | 6.73E-03 | -1.07E+00 |
| HIST2H4A | 1.31E-04 | 6.80E-04 | -1.11E+00 | 1.44E-01 | 6.24E-01 | 1.40E+00 | 9.71E-04 | 4.12E-03 | -1.41E+00 | 3.01E-02 | 1.15E-01 | 1.05E+00 | 3.85E-02 | 1.37E-01 | -1.02E+00 | 3.38E-01 | 9.52E-01 | 1.33E+00 | 1.47E-02 | 5.39E-02 | -1.08E+00 | 1.18E-03 | 3.23E-03 | -1.44E+00 |
| DOPEY1 | 1.31E-04 | 6.81E-04 | -1.27E+00 | 7.52E-03 | 3.25E-02 | 1.10E+00 | 2.03E-04 | 8.65E-04 | -1.39E+00 | 9.43E-04 | 3.60E-03 | -1.14E+00 | 2.30E-04 | 8.21E-04 | -1.34E+00 | 7.55E-05 | 2.13E-04 | -1.55E+00 | 2.55E-04 | 9.34E-04 | -1.63E+00 | 2.25E-03 | 6.19E-03 | -1.35E+00 |
| MARCKSL1P2 | 1.31E-04 | 6.84E-04 | 2.49E+00 | 2.24E-01 | 9.66E-01 | 1.27E+00 | 5.85E-03 | 2.49E-02 | 2.17E+00 | 1.43E-01 | 5.48E-01 | 1.55E+00 | 1.18E-01 | 4.20E-01 | 1.84E+00 | 2.57E-01 | 7.24E-01 | 1.38E+00 | 2.26E-01 | 8.31E-01 | 1.16E+00 | 3.37E-01 | 9.25E-01 | -1.02E+00 |
| ZNF776 | 1.31E-04 | 6.84E-04 | -1.21E+00 | 2.14E-03 | 9.27E-03 | 1.03E+00 | 5.57E-03 | 2.37E-02 | -1.15E+00 | 4.53E-04 | 1.73E-03 | -1.18E+00 | 9.53E-02 | 3.40E-01 | 2.04E+00 | 1.57E-01 | 4.42E-01 | 1.95E+00 | 2.28E-01 | 8.36E-01 | 1.38E+00 | 1.96E-01 | 5.40E-01 | 1.14E+00 |
| LRRC6 | 1.31E-04 | 6.84E-04 | -1.58E+00 | 1.05E-04 | 4.55E-04 | -1.65E+00 | 3.99E-03 | 1.70E-02 | -1.34E+00 | 3.72E-03 | 1.42E-02 | -1.26E+00 | 8.00E-02 | 2.85E-01 | -1.26E+00 | 5.64E-02 | 1.59E-01 | -1.66E+00 | 1.75E-03 | 6.41E-03 | -1.27E+00 | 1.93E-04 | 5.30E-04 | -1.72E+00 |
| FNTB | 1.32E-04 | 6.89E-04 | -1.94E+00 | 2.66E-02 | 1.15E-01 | 1.04E+00 | 2.04E-03 | 8.68E-03 | -1.32E+00 | 3.06E-02 | 1.17E-01 | 1.15E+00 | 1.90E-03 | 6.77E-03 | -1.31E+00 | 1.81E-02 | 5.10E-02 | -1.04E+00 | 4.75E-07 | 1.74E-06 | -1.39E+00 | 1.65E-02 | 4.53E-02 | 1.02E+00 |
| SARNP | 1.32E-04 | 6.89E-04 | -1.33E+00 | 2.34E-03 | 1.01E-02 | -1.05E+00 | 6.47E-03 | 2.75E-02 | -1.28E+00 | 1.88E-04 | 7.17E-04 | -1.53E+00 | 2.04E-03 | 7.26E-03 | -1.26E+00 | 5.73E-03 | 1.62E-02 | -1.21E+00 | 9.21E-03 | 3.38E-02 | -1.07E+00 | 3.65E-02 | 1.00E-01 | -1.05E+00 |
| ZBTB33 | 1.32E-04 | 6.89E-04 | -1.28E+00 | 1.42E-03 | 6.15E-03 | -1.02E+00 | 3.04E-04 | 1.29E-03 | -1.49E+00 | 1.58E-02 | 6.02E-02 | -1.02E+00 | 1.04E-06 | 3.70E-06 | -1.59E+00 | 4.29E-05 | 1.21E-04 | -1.25E+00 | 5.71E-10 | 2.09E-09 | -1.59E+00 | 7.73E-04 | 2.12E-03 | 1.01E+00 |
| SLC9B2 | 1.33E-04 | 6.94E-04 | -1.21E+00 | 3.69E-03 | 1.60E-02 | 1.12E+00 | 3.04E-03 | 1.29E-02 | -1.11E+00 | 2.05E-02 | 7.85E-02 | 1.11E+00 | 1.38E-03 | 4.92E-03 | -1.10E+00 | 2.62E-03 | 7.38E-03 | 1.02E+00 | 3.07E-02 | 1.13E-01 | -1.03E+00 | 2.73E-01 | 7.50E-01 | 1.43E+00 |
| PCIF1 | 1.33E-04 | 6.94E-04 | -1.01E+00 | 4.35E-07 | 1.88E-06 | -1.04E+00 | 1.67E-02 | 7.12E-02 | -1.02E+00 | 5.89E-04 | 2.25E-03 | -1.04E+00 | 3.08E-02 | 1.10E-01 | 1.06E+00 | 4.10E-02 | 1.15E-01 | 1.10E+00 | 2.36E-01 | 8.68E-01 | 1.22E+00 | 1.94E-02 | 5.34E-02 | -1.02E+00 |
| BLM | 1.33E-04 | 6.95E-04 | -1.68E+00 | 4.28E-03 | 1.85E-02 | -1.15E+00 | 1.83E-01 | 7.76E-01 | 1.32E+00 | 1.17E-01 | 4.48E-01 | 1.33E+00 | 3.72E-04 | 1.33E-03 | -1.86E+00 | 8.49E-03 | 2.39E-02 | -1.25E+00 | 1.48E-06 | 5.44E-06 | -1.37E+00 | 3.41E-01 | 9.35E-01 | 1.30E+00 |
| TLR3 | 1.34E-04 | 7.00E-04 | -1.40E+00 | 4.19E-03 | 1.81E-02 | 1.04E+00 | 1.27E-01 | 5.41E-01 | 1.11E+00 | 2.48E-01 | 9.48E-01 | 1.48E+00 | 1.21E-01 | 4.30E-01 | -1.45E+00 | 1.78E-01 | 5.02E-01 | -1.33E+00 | 1.84E-02 | 6.76E-02 | -1.34E+00 | 3.96E-02 | 1.09E-01 | -1.21E+00 |
| ST7 | 1.34E-04 | 7.00E-04 | -1.27E+00 | 1.08E-03 | 4.67E-03 | 1.01E+00 | 2.93E-02 | 1.25E-01 | -1.11E+00 | 2.27E-02 | 8.67E-02 | -1.03E+00 | 1.30E-02 | 4.64E-02 | -1.15E+00 | 2.27E-02 | 6.41E-02 | -1.06E+00 | 7.29E-05 | 2.67E-04 | -1.15E+00 | 6.00E-02 | 1.65E-01 | 1.13E+00 |
| LIMK1 | 1.34E-04 | 7.00E-04 | -1.67E+00 | 1.25E-03 | 5.42E-03 | -1.29E+00 | 1.37E-03 | 5.80E-03 | -1.41E+00 | 4.36E-03 | 1.67E-02 | -1.17E+00 | 1.47E-02 | 5.24E-02 | -1.07E+00 | 5.38E-03 | 1.52E-02 | -1.26E+00 | 9.15E-06 | 3.36E-05 | -1.35E+00 | 1.39E-05 | 3.82E-05 | -1.54E+00 |
| AL162578.1 | 1.34E-04 | 7.00E-04 | -1.05E+00 | 1.86E-04 | 8.04E-04 | 1.06E+00 | 1.19E-02 | 5.07E-02 | -1.07E+00 | 8.99E-02 | 3.43E-01 | 1.22E+00 | 2.45E-04 | 8.73E-04 | -1.36E+00 | 4.46E-03 | 1.26E-02 | -1.13E+00 | 1.22E-05 | 4.49E-05 | -1.39E+00 | 5.05E-03 | 1.39E-02 | -1.02E+00 |
| FAM133B | 1.34E-04 | 7.00E-04 | -1.11E+00 | 1.62E-06 | 7.01E-06 | -1.21E+00 | 5.40E-03 | 2.29E-02 | -1.05E+00 | 1.68E-02 | 6.42E-02 | 1.16E+00 | 2.99E-04 | 1.07E-03 | -1.14E+00 | 2.34E-04 | 6.60E-04 | -1.21E+00 | 1.98E-01 | 7.28E-01 | 1.25E+00 | 4.41E-02 | 1.21E-01 | 1.60E+00 |
| FAM109A | 1.35E-04 | 7.01E-04 | -1.67E+00 | 1.26E-03 | 5.44E-03 | -1.33E+00 | 1.08E-03 | 4.60E-03 | -1.74E+00 | 4.26E-03 | 1.63E-02 | -1.39E+00 | 8.18E-02 | 2.92E-01 | -1.22E+00 | 1.80E-01 | 5.06E-01 | -1.03E+00 | 3.75E-03 | 1.38E-02 | -1.04E+00 | 2.41E-03 | 6.62E-03 | -1.22E+00 |
| ZNF234 | 1.35E-04 | 7.05E-04 | -1.63E+00 | 7.77E-02 | 3.36E-01 | 1.11E+00 | 3.56E-11 | 1.51E-10 | -2.58E+00 | 2.32E-04 | 8.88E-04 | -1.11E+00 | 6.65E-02 | 2.37E-01 | -1.10E+00 | 3.08E-01 | 8.67E-01 | 1.30E+00 | 4.32E-12 | 1.59E-11 | -2.57E+00 | 1.72E-06 | 4.72E-06 | -1.65E+00 |
| ZNF573 | 1.35E-04 | 7.05E-04 | -1.53E+00 | 9.59E-02 | 4.15E-01 | 1.24E+00 | 1.67E-04 | 7.08E-04 | -1.85E+00 | 9.58E-02 | 3.66E-01 | 1.17E+00 | 1.70E-03 | 6.07E-03 | -1.60E+00 | 2.50E-01 | 7.04E-01 | 1.31E+00 | 5.95E-05 | 2.19E-04 | -1.76E+00 | 2.47E-01 | 6.78E-01 | 1.20E+00 |
| RBMS1 | 1.35E-04 | 7.05E-04 | -1.07E+00 | 4.31E-03 | 1.86E-02 | 1.16E+00 | 2.68E-02 | 1.14E-01 | -1.03E+00 | 3.76E-03 | 1.43E-02 | 1.01E+00 | 9.91E-05 | 3.53E-04 | -1.17E+00 | 7.48E-06 | 2.11E-05 | -1.28E+00 | 3.48E-08 | 1.28E-07 | -1.52E+00 | 8.01E-04 | 2.20E-03 | -1.20E+00 |
| PPIL3 | 1.35E-04 | 7.05E-04 | -1.04E+00 | 5.46E-05 | 2.36E-04 | 1.02E+00 | 7.62E-07 | 3.24E-06 | -1.31E+00 | 3.12E-08 | 1.19E-07 | -1.25E+00 | 3.05E-07 | 1.09E-06 | -1.47E+00 | 1.42E-04 | 4.00E-04 | -1.09E+00 | 6.08E-09 | 2.23E-08 | -1.33E+00 | 7.03E-08 | 1.93E-07 | -1.29E+00 |
| PLPPR2 | 1.36E-04 | 7.07E-04 | -1.57E+00 | 3.78E-04 | 1.63E-03 | -1.34E+00 | 9.03E-03 | 3.83E-02 | -1.09E+00 | 6.52E-04 | 2.49E-03 | -1.18E+00 | 4.02E-02 | 1.43E-01 | -1.21E+00 | 6.84E-02 | 1.93E-01 | -1.30E+00 | 2.54E-01 | 9.32E-01 | 1.20E+00 | 3.20E-01 | 8.79E-01 | 1.29E+00 |
| CNNM1 | 1.36E-04 | 7.09E-04 | -1.83E+00 | 7.27E-02 | 3.14E-01 | 1.12E+00 | 8.72E-06 | 3.71E-05 | -1.75E+00 | 7.90E-03 | 3.02E-02 | 1.01E+00 | 2.77E-01 | 9.87E-01 | 1.18E+00 | 2.05E-01 | 5.79E-01 | 1.82E+00 | 8.18E-02 | 3.00E-01 | -1.37E+00 | 6.12E-02 | 1.68E-01 | 1.65E+00 |
| GLB1L | 1.36E-04 | 7.09E-04 | -1.47E+00 | 7.32E-03 | 3.16E-02 | 1.04E+00 | 3.62E-04 | 1.54E-03 | -1.47E+00 | 8.37E-04 | 3.20E-03 | -1.27E+00 | 3.82E-04 | 1.36E-03 | -1.57E+00 | 9.86E-03 | 2.78E-02 | -1.10E+00 | 1.54E-02 | 5.65E-02 | -1.19E+00 | 1.93E-01 | 5.31E-01 | 1.18E+00 |
| PNKP | 1.36E-04 | 7.09E-04 | 1.01E+00 | 4.18E-07 | 1.81E-06 | -1.08E+00 | 6.80E-04 | 2.89E-03 | -1.06E+00 | 1.56E-04 | 5.98E-04 | 1.04E+00 | 1.50E-03 | 5.34E-03 | 1.02E+00 | 2.34E-03 | 6.60E-03 | -1.02E+00 | 8.41E-03 | 3.09E-02 | 1.03E+00 | 2.69E-02 | 7.40E-02 | 1.02E+00 |
| TOP1MT | 1.37E-04 | 7.12E-04 | -1.05E+00 | 1.65E-04 | 7.12E-04 | 1.09E+00 | 1.19E-02 | 5.04E-02 | 1.02E+00 | 5.61E-03 | 2.14E-02 | 1.14E+00 | 4.68E-02 | 1.67E-01 | 1.00E+00 | 5.69E-02 | 1.60E-01 | 1.06E+00 | 1.64E-01 | 6.01E-01 | 1.00E+00 | 2.08E-01 | 5.71E-01 | 1.35E+00 |
| AC087359.1 | 1.37E-04 | 7.13E-04 | -1.57E+00 | 5.80E-04 | 2.51E-03 | -1.31E+00 | 1.72E-02 | 7.29E-02 | -1.28E+00 | 9.88E-03 | 3.78E-02 | -1.29E+00 | 9.26E-02 | 3.30E-01 | -1.31E+00 | 2.74E-01 | 7.72E-01 | 1.14E+00 | 7.23E-02 | 2.65E-01 | -1.20E+00 | 1.04E-01 | 2.87E-01 | -1.10E+00 |
| AC022558.1 | 1.37E-04 | 7.16E-04 | -1.66E+00 | 1.84E-03 | 7.95E-03 | -1.17E+00 | 1.66E-02 | 7.07E-02 | -1.18E+00 | 5.22E-03 | 2.00E-02 | -1.29E+00 | 3.92E-04 | 1.40E-03 | -1.95E+00 | 1.18E-03 | 3.32E-03 | -1.83E+00 | 8.18E-02 | 3.00E-01 | -1.15E+00 | 8.57E-02 | 2.35E-01 | -1.17E+00 |
| PTTG1IP | 1.38E-04 | 7.17E-04 | 1.03E+00 | 8.08E-03 | 3.49E-02 | 1.25E+00 | 2.16E-01 | 9.17E-01 | 1.23E+00 | 2.09E-01 | 7.99E-01 | 1.37E+00 | 5.91E-02 | 2.11E-01 | 1.27E+00 | 2.30E-02 | 6.47E-02 | 1.21E+00 | 8.55E-02 | 3.14E-01 | 1.63E+00 | 3.11E-01 | 8.54E-01 | 1.43E+00 |
| FDFT1 | 1.38E-04 | 7.18E-04 | 1.03E+00 | 1.22E-01 | 5.28E-01 | 1.48E+00 | 4.43E-03 | 1.88E-02 | -1.15E+00 | 2.33E-01 | 8.91E-01 | 1.28E+00 | 8.57E-08 | 3.06E-07 | -1.26E+00 | 2.94E-01 | 8.29E-01 | 1.52E+00 | 9.49E-04 | 3.48E-03 | -1.09E+00 | 3.47E-01 | 9.52E-01 | 1.27E+00 |
| DRC3 | 1.38E-04 | 7.19E-04 | -1.72E+00 | 4.44E-03 | 1.92E-02 | -1.16E+00 | 1.07E-05 | 4.56E-05 | -1.69E+00 | 7.91E-05 | 3.02E-04 | -1.38E+00 | 6.00E-03 | 2.14E-02 | -1.21E+00 | 1.03E-02 | 2.89E-02 | -1.09E+00 | 1.60E-02 | 5.88E-02 | -1.35E+00 | 5.06E-02 | 1.39E-01 | -1.14E+00 |
| AL009031.1 | 1.39E-04 | 7.22E-04 | -2.17E+00 | 5.06E-03 | 2.19E-02 | -1.22E+00 | 3.38E-11 | 1.43E-10 | -3.26E+00 | 9.71E-11 | 3.71E-10 | -2.64E+00 | 5.08E-13 | 1.81E-12 | -2.98E+00 | 1.17E-09 | 3.30E-09 | -2.04E+00 | 4.43E-12 | 1.63E-11 | -1.72E+00 | 2.89E-09 | 7.95E-09 | -1.42E+00 |
| PUS7 | 1.39E-04 | 7.25E-04 | -1.57E+00 | 3.16E-02 | 1.37E-01 | 1.12E+00 | 1.65E-02 | 7.03E-02 | -1.04E+00 | 1.99E-01 | 7.59E-01 | 1.52E+00 | 4.77E-03 | 1.70E-02 | -1.50E+00 | 2.95E-01 | 8.31E-01 | 1.59E+00 | 5.71E-04 | 2.09E-03 | -1.19E+00 | 3.13E-01 | 8.59E-01 | 1.15E+00 |
| JAM3 | 1.39E-04 | 7.26E-04 | -1.31E+00 | 9.69E-04 | 4.19E-03 | -1.05E+00 | 1.26E-05 | 5.36E-05 | -1.35E+00 | 1.42E-05 | 5.44E-05 | -1.15E+00 | 4.71E-03 | 1.68E-02 | -1.66E+00 | 1.09E-01 | 3.07E-01 | -1.01E+00 | 3.11E-05 | 1.14E-04 | -1.91E+00 | 1.62E-02 | 4.46E-02 | -1.20E+00 |
| PAF1 | 1.40E-04 | 7.28E-04 | -1.22E+00 | 1.06E-02 | 4.60E-02 | 1.18E+00 | 9.33E-05 | 3.96E-04 | -1.30E+00 | 2.43E-04 | 9.27E-04 | -1.12E+00 | 6.57E-05 | 2.34E-04 | -1.50E+00 | 3.66E-02 | 1.03E-01 | 1.03E+00 | 5.08E-05 | 1.86E-04 | -1.20E+00 | 2.80E-05 | 7.70E-05 | -1.18E+00 |
| ESD | 1.40E-04 | 7.28E-04 | 1.00E+00 | 2.06E-01 | 8.91E-01 | 1.25E+00 | 8.78E-03 | 3.73E-02 | -1.13E+00 | 7.27E-02 | 2.78E-01 | 1.02E+00 | 1.85E-04 | 6.59E-04 | -1.09E+00 | 2.35E-01 | 6.64E-01 | 1.15E+00 | 6.72E-03 | 2.47E-02 | -1.01E+00 | 2.69E-01 | 7.39E-01 | 1.14E+00 |
| AC097493.2 | 1.40E-04 | 7.28E-04 | -2.02E+00 | 1.15E-01 | 4.99E-01 | 1.16E+00 | 1.46E-04 | 6.19E-04 | -1.70E+00 | 2.80E-03 | 1.07E-02 | -1.21E+00 | 3.16E-05 | 1.13E-04 | -1.76E+00 | 2.03E-04 | 5.73E-04 | -1.55E+00 | 1.04E-02 | 3.81E-02 | -1.71E+00 | 2.55E-01 | 6.99E-01 | 1.05E+00 |
| ATXN7L3 | 1.42E-04 | 7.38E-04 | -1.18E+00 | 3.29E-06 | 1.42E-05 | -1.27E+00 | 5.00E-04 | 2.12E-03 | -1.25E+00 | 5.91E-04 | 2.26E-03 | -1.06E+00 | 3.88E-03 | 1.38E-02 | -1.14E+00 | 1.50E-03 | 4.24E-03 | -1.26E+00 | 7.83E-03 | 2.87E-02 | -1.05E+00 | 2.17E-03 | 5.96E-03 | -1.22E+00 |
| RUFY3 | 1.42E-04 | 7.38E-04 | -1.24E+00 | 1.23E-02 | 5.32E-02 | 1.12E+00 | 4.62E-07 | 1.96E-06 | -1.43E+00 | 1.40E-05 | 5.35E-05 | -1.19E+00 | 2.16E-04 | 7.71E-04 | -1.38E+00 | 2.66E-02 | 7.51E-02 | 1.05E+00 | 9.42E-07 | 3.46E-06 | -1.38E+00 | 3.21E-06 | 8.81E-06 | -1.35E+00 |
| DCAF5 | 1.42E-04 | 7.39E-04 | -1.04E+00 | 1.13E-04 | 4.89E-04 | 1.06E+00 | 1.17E-03 | 4.97E-03 | -1.13E+00 | 2.32E-02 | 8.86E-02 | 1.13E+00 | 5.76E-03 | 2.05E-02 | 1.04E+00 | 3.85E-02 | 1.09E-01 | 1.14E+00 | 9.62E-05 | 3.53E-04 | -1.17E+00 | 6.30E-04 | 1.73E-03 | -1.09E+00 |
| GLA | 1.42E-04 | 7.41E-04 | -1.11E+00 | 1.18E-04 | 5.09E-04 | -1.02E+00 | 5.19E-04 | 2.20E-03 | -1.05E+00 | 2.99E-06 | 1.14E-05 | -1.08E+00 | 9.30E-06 | 3.31E-05 | -1.31E+00 | 1.43E-04 | 4.02E-04 | -1.13E+00 | 2.22E-01 | 8.15E-01 | 1.31E+00 | 1.61E-01 | 4.43E-01 | 1.43E+00 |
| RPL24P4 | 1.42E-04 | 7.41E-04 | 1.22E+00 | 1.82E-30 | 7.86E-30 | 1.76E+00 | 2.38E-03 | 1.01E-02 | 1.22E+00 | 1.93E-19 | 7.37E-19 | 1.78E+00 | 9.88E-03 | 3.52E-02 | 1.22E+00 | 6.55E-16 | 1.85E-15 | 1.77E+00 | 2.05E-02 | 7.52E-02 | 1.26E+00 | 6.93E-20 | 1.90E-19 | 1.84E+00 |
| BRPF1 | 1.43E-04 | 7.44E-04 | -1.22E+00 | 2.23E-03 | 9.64E-03 | 1.01E+00 | 1.83E-13 | 7.76E-13 | -2.02E+00 | 3.92E-11 | 1.50E-10 | -1.62E+00 | 8.61E-03 | 3.07E-02 | -1.11E+00 | 1.67E-02 | 4.70E-02 | -1.17E+00 | 9.69E-05 | 3.56E-04 | -1.23E+00 | 6.28E-07 | 1.73E-06 | -1.52E+00 |
| FCGR1A | 1.43E-04 | 7.44E-04 | -2.61E+00 | 7.67E-05 | 3.31E-04 | -2.82E+00 | 1.17E-14 | 4.95E-14 | -1.61E+00 | 3.29E-15 | 1.26E-14 | -1.42E+00 | 1.09E-01 | 3.90E-01 | 2.90E+00 | 2.76E-01 | 7.78E-01 | 1.05E+00 | 2.55E-01 | 9.34E-01 | -1.02E+00 | 2.72E-01 | 7.48E-01 | -1.27E+00 |
| AL662907.2 | 1.43E-04 | 7.44E-04 | -1.35E+00 | 7.87E-05 | 3.40E-04 | -1.51E+00 | 2.25E-03 | 9.57E-03 | -1.55E+00 | 2.96E-04 | 1.13E-03 | -1.75E+00 | 2.24E-02 | 7.97E-02 | -1.19E+00 | 1.65E-03 | 4.65E-03 | -1.70E+00 | 1.44E-02 | 5.28E-02 | -1.43E+00 | 2.01E-03 | 5.53E-03 | -2.26E+00 |
| AC009120.1 | 1.43E-04 | 7.45E-04 | -1.78E+00 | 6.01E-03 | 2.60E-02 | -1.14E+00 | 1.33E-02 | 5.65E-02 | -1.33E+00 | 1.07E-01 | 4.10E-01 | 1.12E+00 | 2.41E-02 | 8.61E-02 | -1.68E+00 | 1.75E-01 | 4.94E-01 | -1.03E+00 | 4.22E-04 | 1.55E-03 | -2.02E+00 | 6.24E-02 | 1.71E-01 | -1.15E+00 |
| FTCDNL1 | 1.43E-04 | 7.45E-04 | -1.09E+00 | 1.00E-04 | 4.33E-04 | -1.01E+00 | 5.69E-04 | 2.42E-03 | -1.09E+00 | 1.46E-03 | 5.56E-03 | 1.06E+00 | 1.86E-02 | 6.62E-02 | -1.12E+00 | 6.94E-02 | 1.96E-01 | 1.06E+00 | 2.11E-01 | 7.76E-01 | 1.31E+00 | 1.21E-01 | 3.33E-01 | 1.48E+00 |
| RPS20P2 | 1.43E-04 | 7.46E-04 | 1.01E+00 | 1.58E-14 | 6.82E-14 | -1.33E+00 | 5.80E-02 | 2.46E-01 | 1.12E+00 | 7.81E-08 | 2.98E-07 | -1.17E+00 | 1.25E-03 | 4.47E-03 | 1.00E+00 | 5.37E-08 | 1.51E-07 | -1.27E+00 | 1.20E-06 | 4.40E-06 | -1.20E+00 | 1.16E-13 | 3.18E-13 | -1.61E+00 |
| COL6A1 | 1.43E-04 | 7.47E-04 | -1.33E+00 | 8.82E-03 | 3.81E-02 | 1.16E+00 | 3.38E-02 | 1.44E-01 | -1.24E+00 | 1.51E-01 | 5.76E-01 | 1.74E+00 | 1.98E-02 | 7.05E-02 | -1.10E+00 | 5.38E-02 | 1.52E-01 | 1.06E+00 | 1.42E-01 | 5.23E-01 | -1.13E+00 | 1.61E-01 | 4.43E-01 | -1.35E+00 |
| SLC35F2 | 1.43E-04 | 7.48E-04 | -1.22E+00 | 1.45E-03 | 6.25E-03 | 1.04E+00 | 3.18E-09 | 1.35E-08 | -1.48E+00 | 2.22E-06 | 8.49E-06 | -1.16E+00 | 3.84E-04 | 1.37E-03 | -1.56E+00 | 5.69E-02 | 1.61E-01 | 1.04E+00 | 3.96E-06 | 1.45E-05 | -1.47E+00 | 1.13E-01 | 3.12E-01 | 1.11E+00 |
| BRAT1 | 1.44E-04 | 7.49E-04 | -1.27E+00 | 1.58E-02 | 6.83E-02 | 1.16E+00 | 5.23E-03 | 2.22E-02 | -1.17E+00 | 8.34E-03 | 3.19E-02 | -1.02E+00 | 1.87E-03 | 6.65E-03 | -1.34E+00 | 1.12E-01 | 3.17E-01 | 1.12E+00 | 3.27E-05 | 1.20E-04 | -1.33E+00 | 1.41E-02 | 3.88E-02 | -1.06E+00 |
| ZNF684 | 1.44E-04 | 7.53E-04 | -1.41E+00 | 1.24E-04 | 5.37E-04 | -1.26E+00 | 7.65E-03 | 3.25E-02 | -1.23E+00 | 3.28E-03 | 1.25E-02 | -1.28E+00 | 9.90E-02 | 3.53E-01 | 1.08E+00 | 9.37E-02 | 2.64E-01 | 1.03E+00 | 7.88E-02 | 2.89E-01 | 1.10E+00 | 3.98E-02 | 1.09E-01 | -1.06E+00 |
| NAALAD2 | 1.45E-04 | 7.54E-04 | -1.50E+00 | 1.10E-03 | 4.77E-03 | -1.18E+00 | 1.78E-01 | 7.55E-01 | 1.06E+00 | 2.23E-01 | 8.50E-01 | 1.24E+00 | 2.51E-01 | 8.94E-01 | 1.30E+00 | 3.04E-01 | 8.57E-01 | 1.45E+00 | 1.75E-01 | 6.41E-01 | 1.01E+00 | 5.10E-02 | 1.40E-01 | -1.60E+00 |
| CORO1C | 1.46E-04 | 7.59E-04 | -1.07E+00 | 1.46E-03 | 6.31E-03 | 1.13E+00 | 1.56E-03 | 6.61E-03 | -1.15E+00 | 1.95E-02 | 7.46E-02 | 1.07E+00 | 7.61E-02 | 2.71E-01 | -1.02E+00 | 2.18E-01 | 6.15E-01 | 1.21E+00 | 1.00E-05 | 3.69E-05 | -1.13E+00 | 1.14E-04 | 3.12E-04 | -1.19E+00 |
| SFXN2 | 1.46E-04 | 7.62E-04 | -1.52E+00 | 4.67E-03 | 2.02E-02 | 1.01E+00 | 3.93E-03 | 1.67E-02 | -1.18E+00 | 3.63E-02 | 1.39E-01 | 1.14E+00 | 3.06E-02 | 1.09E-01 | -1.29E+00 | 1.76E-01 | 4.97E-01 | 1.03E+00 | 4.49E-05 | 1.65E-04 | -1.46E+00 | 7.89E-03 | 2.17E-02 | -1.12E+00 |
| AL133410.1 | 1.46E-04 | 7.62E-04 | -1.34E+00 | 1.32E-01 | 5.71E-01 | 1.40E+00 | 5.91E-02 | 2.51E-01 | -1.10E+00 | 1.27E-01 | 4.84E-01 | 1.17E+00 | 2.14E-01 | 7.64E-01 | 1.56E+00 | 2.33E-01 | 6.57E-01 | 1.71E+00 | 3.69E-02 | 1.35E-01 | -1.11E+00 | 2.33E-01 | 6.40E-01 | 1.11E+00 |
| SLC35B3 | 1.46E-04 | 7.63E-04 | -1.29E+00 | 2.85E-02 | 1.23E-01 | 1.22E+00 | 2.06E-07 | 8.77E-07 | -1.84E+00 | 2.34E-04 | 8.96E-04 | -1.18E+00 | 2.33E-06 | 8.29E-06 | -1.93E+00 | 1.65E-02 | 4.64E-02 | -1.03E+00 | 8.04E-04 | 2.95E-03 | -1.30E+00 | 6.11E-03 | 1.68E-02 | -1.13E+00 |
| SC5D | 1.47E-04 | 7.64E-04 | -1.10E+00 | 7.89E-03 | 3.41E-02 | 1.20E+00 | 1.28E-03 | 5.42E-03 | -1.29E+00 | 2.35E-01 | 8.97E-01 | 1.32E+00 | 3.50E-06 | 1.25E-05 | -1.63E+00 | 3.57E-03 | 1.01E-02 | -1.09E+00 | 9.78E-04 | 3.59E-03 | -1.12E+00 | 2.88E-03 | 7.92E-03 | -1.12E+00 |
| BAALC | 1.47E-04 | 7.64E-04 | 1.72E+00 | 8.30E-02 | 3.59E-01 | 1.57E+00 | 3.42E-02 | 1.45E-01 | 1.44E+00 | 9.44E-02 | 3.61E-01 | 1.34E+00 | 4.59E-04 | 1.64E-03 | 2.44E+00 | 5.08E-02 | 1.43E-01 | 1.91E+00 | 1.14E-04 | 4.20E-04 | 3.63E+00 | 1.97E-03 | 5.40E-03 | 3.34E+00 |
| CHN2 | 1.47E-04 | 7.66E-04 | -1.20E+00 | 2.82E-03 | 1.22E-02 | 1.09E+00 | 9.14E-12 | 3.88E-11 | -1.69E+00 | 9.22E-08 | 3.52E-07 | -1.22E+00 | 5.89E-05 | 2.10E-04 | -1.12E+00 | 1.32E-02 | 3.71E-02 | 1.13E+00 | 2.07E-04 | 7.61E-04 | -1.24E+00 | 9.90E-02 | 2.72E-01 | 1.09E+00 |
| LINC00957 | 1.48E-04 | 7.72E-04 | -1.61E+00 | 2.87E-02 | 1.24E-01 | 1.08E+00 | 3.37E-05 | 1.43E-04 | -1.81E+00 | 7.63E-04 | 2.92E-03 | -1.25E+00 | 5.50E-02 | 1.96E-01 | -1.50E+00 | 2.55E-01 | 7.18E-01 | 1.21E+00 | 1.39E-01 | 5.11E-01 | -1.02E+00 | 2.59E-01 | 7.10E-01 | 1.03E+00 |
| TRAPPC6A | 1.48E-04 | 7.72E-04 | 1.08E+00 | 1.02E-07 | 4.40E-07 | 1.03E+00 | 1.50E-03 | 6.36E-03 | -1.05E+00 | 2.90E-06 | 1.11E-05 | -1.15E+00 | 6.92E-03 | 2.47E-02 | 1.02E+00 | 2.62E-02 | 7.39E-02 | 1.05E+00 | 2.70E-01 | 9.91E-01 | 1.33E+00 | 8.07E-02 | 2.22E-01 | 1.15E+00 |
| PPP6R2 | 1.48E-04 | 7.73E-04 | -1.16E+00 | 5.04E-03 | 2.18E-02 | 1.17E+00 | 8.54E-05 | 3.63E-04 | -1.18E+00 | 2.85E-03 | 1.09E-02 | 1.08E+00 | 7.74E-04 | 2.76E-03 | -1.11E+00 | 1.80E-04 | 5.06E-04 | -1.15E+00 | 7.71E-06 | 2.83E-05 | -1.39E+00 | 4.12E-02 | 1.13E-01 | 1.04E+00 |
| DDX55 | 1.48E-04 | 7.73E-04 | -1.24E+00 | 1.98E-01 | 8.57E-01 | 1.47E+00 | 1.89E-05 | 8.03E-05 | -1.38E+00 | 3.47E-02 | 1.33E-01 | 1.16E+00 | 6.93E-03 | 2.47E-02 | -1.13E+00 | 3.06E-01 | 8.62E-01 | 1.39E+00 | 5.02E-08 | 1.84E-07 | -1.36E+00 | 1.89E-02 | 5.19E-02 | 1.10E+00 |
| ZNF410 | 1.49E-04 | 7.77E-04 | -1.10E+00 | 3.83E-06 | 1.65E-05 | -1.18E+00 | 2.33E-04 | 9.88E-04 | -1.34E+00 | 4.97E-07 | 1.90E-06 | -1.73E+00 | 1.78E-03 | 6.34E-03 | -1.29E+00 | 1.90E-03 | 5.35E-03 | -1.32E+00 | 1.06E-04 | 3.88E-04 | -1.38E+00 | 8.85E-06 | 2.43E-05 | -1.51E+00 |
| ANGPT2 | 1.49E-04 | 7.77E-04 | -1.60E+00 | 1.55E-02 | 6.69E-02 | 1.14E+00 | 7.22E-04 | 3.07E-03 | -1.65E+00 | 6.29E-02 | 2.40E-01 | 1.03E+00 | 9.61E-67 | 3.43E-66 | -6.63E+00 | 9.15E-55 | 2.58E-54 | -3.99E+00 | 1.50E-06 | 5.51E-06 | -1.88E+00 | 7.16E-07 | 1.97E-06 | -2.07E+00 |
| FKBP2 | 1.49E-04 | 7.78E-04 | -1.03E+00 | 2.88E-02 | 1.25E-01 | 1.23E+00 | 2.21E-01 | 9.39E-01 | 1.12E+00 | 2.22E-01 | 8.48E-01 | 1.22E+00 | 2.54E-05 | 9.07E-05 | -1.16E+00 | 1.54E-03 | 4.33E-03 | -1.02E+00 | 2.39E-01 | 8.75E-01 | 1.06E+00 | 1.15E-01 | 3.17E-01 | 1.14E+00 |
| TBCAP2 | 1.49E-04 | 7.78E-04 | -1.48E+00 | 8.46E-04 | 3.65E-03 | -1.24E+00 | 2.35E-09 | 9.97E-09 | -1.76E+00 | 3.09E-12 | 1.18E-11 | -2.12E+00 | 1.11E-03 | 3.95E-03 | -1.72E+00 | 1.46E-02 | 4.12E-02 | -1.15E+00 | 1.01E-14 | 3.72E-14 | -2.60E+00 | 1.68E-07 | 4.63E-07 | -1.62E+00 |
| FOXRED1 | 1.49E-04 | 7.79E-04 | -1.28E+00 | 2.63E-02 | 1.13E-01 | 1.26E+00 | 1.88E-02 | 7.98E-02 | -1.04E+00 | 2.12E-01 | 8.10E-01 | 1.34E+00 | 5.66E-02 | 2.02E-01 | 1.03E+00 | 3.09E-01 | 8.72E-01 | 1.54E+00 | 1.18E-04 | 4.33E-04 | -1.11E+00 | 1.49E-02 | 4.10E-02 | 1.07E+00 |
| TJAP1 | 1.50E-04 | 7.80E-04 | -1.45E+00 | 6.50E-02 | 2.81E-01 | 1.24E+00 | 5.83E-03 | 2.48E-02 | -1.13E+00 | 5.15E-02 | 1.97E-01 | 1.11E+00 | 6.87E-03 | 2.45E-02 | -1.26E+00 | 1.09E-01 | 3.07E-01 | 1.12E+00 | 5.26E-04 | 1.93E-03 | -1.23E+00 | 2.83E-01 | 7.77E-01 | 1.15E+00 |
| DCAF6 | 1.50E-04 | 7.81E-04 | -1.02E+00 | 7.87E-04 | 3.40E-03 | 1.14E+00 | 1.24E-06 | 5.26E-06 | -1.37E+00 | 1.42E-04 | 5.42E-04 | -1.02E+00 | 1.31E-04 | 4.68E-04 | -1.23E+00 | 1.06E-01 | 2.99E-01 | 1.35E+00 | 2.55E-07 | 9.37E-07 | -1.35E+00 | 1.50E-02 | 4.11E-02 | 1.03E+00 |
| AC025171.4 | 1.50E-04 | 7.82E-04 | -1.46E+00 | 1.82E-01 | 7.86E-01 | 1.51E+00 | 1.40E-02 | 5.93E-02 | -1.36E+00 | 1.78E-01 | 6.82E-01 | 1.18E+00 | 3.67E-02 | 1.31E-01 | -1.59E+00 | 8.64E-02 | 2.44E-01 | 1.60E+00 | 7.69E-03 | 2.82E-02 | -1.46E+00 | 2.74E-01 | 7.52E-01 | 1.05E+00 |
| COX18 | 1.50E-04 | 7.82E-04 | -1.26E+00 | 8.05E-03 | 3.48E-02 | 1.14E+00 | 2.24E-01 | 9.53E-01 | 1.21E+00 | 1.30E-01 | 4.96E-01 | 1.53E+00 | 7.89E-02 | 2.81E-01 | 1.06E+00 | 8.95E-02 | 2.52E-01 | 1.07E+00 | 1.76E-04 | 6.46E-04 | -1.48E+00 | 9.35E-02 | 2.57E-01 | -1.03E+00 |
| ZNF786 | 1.51E-04 | 7.85E-04 | -1.70E+00 | 2.50E-02 | 1.08E-01 | 1.07E+00 | 4.91E-06 | 2.09E-05 | -2.22E+00 | 2.15E-02 | 8.20E-02 | -1.08E+00 | 2.64E-02 | 9.41E-02 | -1.50E+00 | 2.63E-01 | 7.40E-01 | 1.22E+00 | 1.05E-02 | 3.87E-02 | -1.55E+00 | 3.52E-01 | 9.68E-01 | 1.16E+00 |
| TAB1 | 1.51E-04 | 7.87E-04 | -1.25E+00 | 9.30E-03 | 4.02E-02 | 1.16E+00 | 2.08E-04 | 8.83E-04 | -1.25E+00 | 8.29E-05 | 3.17E-04 | -1.15E+00 | 1.15E-02 | 4.11E-02 | -1.16E+00 | 8.22E-02 | 2.32E-01 | 1.07E+00 | 1.14E-06 | 4.18E-06 | -1.28E+00 | 1.14E-02 | 3.13E-02 | 1.05E+00 |
| RGPD3 | 1.51E-04 | 7.87E-04 | -1.10E+00 | 3.83E-08 | 1.66E-07 | -1.36E+00 | 2.36E-02 | 1.00E-01 | -1.04E+00 | 1.23E-04 | 4.70E-04 | -1.29E+00 | 2.43E-01 | 8.66E-01 | 1.22E+00 | 7.02E-03 | 1.98E-02 | -1.56E+00 | 2.26E-02 | 8.29E-02 | -1.09E+00 | 1.57E-03 | 4.31E-03 | -1.37E+00 |
| ALMS1 | 1.51E-04 | 7.87E-04 | -1.24E+00 | 9.29E-03 | 4.01E-02 | 1.11E+00 | 1.64E-04 | 6.95E-04 | -1.23E+00 | 4.41E-03 | 1.68E-02 | 1.03E+00 | 2.43E-01 | 8.68E-01 | 1.33E+00 | 2.98E-01 | 8.39E-01 | 1.45E+00 | 5.15E-05 | 1.89E-04 | -1.36E+00 | 3.33E-02 | 9.15E-02 | 1.01E+00 |
| HSD17B7 | 1.51E-04 | 7.89E-04 | -1.37E+00 | 3.39E-03 | 1.46E-02 | -1.04E+00 | 2.35E-03 | 9.98E-03 | -1.14E+00 | 1.91E-03 | 7.30E-03 | 1.02E+00 | 2.01E-03 | 7.18E-03 | -1.21E+00 | 7.09E-02 | 2.00E-01 | 1.13E+00 | 1.50E-03 | 5.52E-03 | -1.12E+00 | 7.24E-02 | 1.99E-01 | 1.11E+00 |
| MSTO1 | 1.52E-04 | 7.90E-04 | -1.61E+00 | 4.57E-04 | 1.97E-03 | -1.34E+00 | 1.34E-02 | 5.71E-02 | -1.23E+00 | 1.26E-02 | 4.83E-02 | -1.06E+00 | 1.23E-03 | 4.40E-03 | -1.47E+00 | 5.08E-02 | 1.43E-01 | 1.01E+00 | 6.51E-05 | 2.39E-04 | -1.27E+00 | 1.46E-01 | 4.00E-01 | 1.22E+00 |
| DUSP22 | 1.52E-04 | 7.90E-04 | -1.11E+00 | 1.13E-02 | 4.87E-02 | 1.18E+00 | 9.06E-06 | 3.85E-05 | -1.43E+00 | 8.04E-04 | 3.07E-03 | -1.08E+00 | 1.52E-03 | 5.42E-03 | -1.15E+00 | 3.82E-02 | 1.08E-01 | 1.02E+00 | 1.34E-02 | 4.92E-02 | -1.02E+00 | 9.34E-03 | 2.57E-02 | -1.08E+00 |
| AHI1 | 1.52E-04 | 7.91E-04 | -1.20E+00 | 9.13E-05 | 3.95E-04 | -1.07E+00 | 7.60E-07 | 3.23E-06 | -1.27E+00 | 2.21E-08 | 8.44E-08 | -1.23E+00 | 1.92E-05 | 6.83E-05 | -1.24E+00 | 2.53E-03 | 7.14E-03 | 1.05E+00 | 2.10E-04 | 7.71E-04 | -1.13E+00 | 1.57E-01 | 4.30E-01 | 1.13E+00 |
| EIF1AY | 1.52E-04 | 7.92E-04 | 1.03E+00 | 1.55E-04 | 6.68E-04 | 1.14E+00 | 8.08E-02 | 3.43E-01 | 1.02E+00 | 2.52E-02 | 9.61E-02 | 1.09E+00 | 1.87E-04 | 6.68E-04 | -1.07E+00 | 1.09E-02 | 3.07E-02 | 1.05E+00 | 1.95E-02 | 7.15E-02 | 1.01E+00 | 9.63E-02 | 2.65E-01 | 1.02E+00 |
| FAM3C | 1.52E-04 | 7.92E-04 | -1.41E+00 | 1.24E-01 | 5.38E-01 | 1.28E+00 | 1.52E-06 | 6.46E-06 | -1.54E+00 | 3.77E-04 | 1.44E-03 | -1.15E+00 | 8.64E-03 | 3.08E-02 | -1.12E+00 | 1.82E-02 | 5.13E-02 | -1.06E+00 | 1.98E-03 | 7.26E-03 | -1.14E+00 | 1.75E-01 | 4.81E-01 | 1.11E+00 |
| RAB32 | 1.52E-04 | 7.92E-04 | -1.09E+00 | 2.72E-09 | 1.18E-08 | -1.14E+00 | 6.03E-03 | 2.56E-02 | -1.18E+00 | 1.87E-02 | 7.14E-02 | -1.06E+00 | 1.78E-12 | 6.34E-12 | -1.51E+00 | 4.00E-11 | 1.13E-10 | -1.38E+00 | 3.93E-04 | 1.44E-03 | 1.25E+00 | 1.97E-05 | 5.40E-05 | -1.03E+00 |
| ZNF189 | 1.52E-04 | 7.94E-04 | -1.25E+00 | 1.02E-03 | 4.39E-03 | -1.03E+00 | 6.05E-06 | 2.57E-05 | -1.59E+00 | 6.06E-05 | 2.31E-04 | -1.22E+00 | 4.08E-06 | 1.45E-05 | -1.86E+00 | 7.30E-05 | 2.06E-04 | -1.53E+00 | 2.78E-04 | 1.02E-03 | -1.64E+00 | 3.97E-03 | 1.09E-02 | -1.41E+00 |
| APEH | 1.52E-04 | 7.94E-04 | -1.18E+00 | 3.61E-03 | 1.56E-02 | 1.06E+00 | 8.03E-07 | 3.41E-06 | -1.25E+00 | 6.03E-06 | 2.30E-05 | -1.05E+00 | 2.51E-08 | 8.95E-08 | -1.40E+00 | 2.21E-03 | 6.24E-03 | 1.01E+00 | 2.24E-03 | 8.23E-03 | -1.11E+00 | 8.58E-02 | 2.36E-01 | 1.06E+00 |
| TRIM6 | 1.53E-04 | 7.97E-04 | -1.82E+00 | 3.97E-03 | 1.72E-02 | -1.21E+00 | 5.07E-06 | 2.15E-05 | -1.81E+00 | 3.76E-05 | 1.44E-04 | -1.41E+00 | 5.33E-05 | 1.90E-04 | -2.87E+00 | 4.89E-04 | 1.38E-03 | -2.09E+00 | 5.41E-15 | 1.98E-14 | -3.18E+00 | 1.15E-10 | 3.17E-10 | -2.21E+00 |
| ABHD17B | 1.54E-04 | 8.02E-04 | -1.03E+00 | 1.38E-05 | 5.97E-05 | -1.02E+00 | 7.52E-05 | 3.20E-04 | -1.20E+00 | 4.04E-09 | 1.54E-08 | -1.35E+00 | 3.56E-02 | 1.27E-01 | 1.10E+00 | 1.16E-03 | 3.28E-03 | -1.26E+00 | 1.15E-03 | 4.22E-03 | -1.15E+00 | 7.33E-03 | 2.01E-02 | -1.14E+00 |
| MFSD14B | 1.54E-04 | 8.03E-04 | -1.32E+00 | 7.88E-02 | 3.40E-01 | 1.24E+00 | 5.76E-06 | 2.45E-05 | -1.37E+00 | 4.74E-03 | 1.81E-02 | 1.07E+00 | 1.00E-09 | 3.57E-09 | -2.38E+00 | 6.11E-05 | 1.72E-04 | -1.26E+00 | 3.01E-15 | 1.11E-14 | -1.92E+00 | 1.07E-05 | 2.94E-05 | -1.24E+00 |
| VMA21 | 1.54E-04 | 8.03E-04 | -1.03E+00 | 8.81E-07 | 3.81E-06 | -1.01E+00 | 1.38E-02 | 5.88E-02 | -1.00E+00 | 8.96E-03 | 3.42E-02 | 1.06E+00 | 1.46E-04 | 5.22E-04 | -1.13E+00 | 1.24E-03 | 3.51E-03 | -1.04E+00 | 6.16E-03 | 2.26E-02 | 1.02E+00 | 1.40E-01 | 3.84E-01 | 1.20E+00 |
| BORCS6 | 1.55E-04 | 8.06E-04 | -1.22E+00 | 5.44E-02 | 2.35E-01 | 1.31E+00 | 6.66E-03 | 2.83E-02 | -1.08E+00 | 2.13E-01 | 8.13E-01 | 1.41E+00 | 5.82E-12 | 2.07E-11 | -1.71E+00 | 5.05E-03 | 1.42E-02 | 1.12E+00 | 1.05E-05 | 3.86E-05 | -1.23E+00 | 7.22E-03 | 1.98E-02 | -1.00E+00 |
| HMGXB4 | 1.55E-04 | 8.07E-04 | -1.07E+00 | 1.16E-05 | 5.00E-05 | -1.10E+00 | 1.61E-01 | 6.85E-01 | 1.18E+00 | 8.43E-04 | 3.22E-03 | 1.05E+00 | 1.37E-05 | 4.88E-05 | -1.50E+00 | 2.69E-05 | 7.58E-05 | -1.32E+00 | 8.90E-09 | 3.27E-08 | -1.25E+00 | 1.35E-04 | 3.70E-04 | -1.03E+00 |
| OR2L2 | 1.55E-04 | 8.09E-04 | -1.56E+00 | 4.96E-02 | 2.14E-01 | 1.15E+00 | 6.50E-03 | 2.76E-02 | -1.59E+00 | 9.31E-02 | 3.56E-01 | 1.08E+00 | 2.13E-02 | 7.59E-02 | -1.44E+00 | 2.24E-01 | 6.32E-01 | 1.62E+00 | 4.82E-02 | 1.77E-01 | -1.54E+00 | 2.28E-01 | 6.27E-01 | 1.35E+00 |
| HEATR3 | 1.56E-04 | 8.12E-04 | -1.63E+00 | 1.52E-03 | 6.59E-03 | -1.19E+00 | 5.80E-06 | 2.46E-05 | -1.51E+00 | 5.61E-04 | 2.14E-03 | -1.13E+00 | 1.03E-03 | 3.66E-03 | -1.77E+00 | 2.01E-02 | 5.68E-02 | -1.22E+00 | 1.01E-05 | 3.70E-05 | -1.45E+00 | 9.22E-04 | 2.53E-03 | -1.21E+00 |
| SH3BP1 | 1.56E-04 | 8.12E-04 | -1.21E+00 | 8.07E-06 | 3.49E-05 | -1.26E+00 | 1.02E-06 | 4.32E-06 | -1.36E+00 | 8.80E-11 | 3.36E-10 | -1.55E+00 | 1.33E-04 | 4.75E-04 | -1.53E+00 | 6.66E-05 | 1.88E-04 | -1.59E+00 | 2.08E-01 | 7.64E-01 | 1.11E+00 | 1.65E-01 | 4.53E-01 | 1.02E+00 |
| LRRK2 | 1.56E-04 | 8.13E-04 | -2.07E+00 | 3.32E-03 | 1.43E-02 | -1.32E+00 | 3.84E-03 | 1.63E-02 | -1.84E+00 | 1.25E-02 | 4.76E-02 | -1.35E+00 | 5.16E-02 | 1.84E-01 | -2.16E+00 | 8.30E-02 | 2.34E-01 | -2.31E+00 | 2.02E-01 | 7.42E-01 | 1.08E+00 | 3.45E-01 | 9.48E-01 | 1.33E+00 |
| MEX3C | 1.56E-04 | 8.15E-04 | -1.16E+00 | 8.64E-06 | 3.73E-05 | -1.14E+00 | 7.83E-04 | 3.33E-03 | -1.20E+00 | 2.88E-08 | 1.10E-07 | -1.44E+00 | 1.76E-02 | 6.26E-02 | 1.05E+00 | 1.09E-04 | 3.07E-04 | -1.27E+00 | 7.50E-04 | 2.75E-03 | -1.26E+00 | 3.71E-05 | 1.02E-04 | -1.49E+00 |
| CHTOP | 1.57E-04 | 8.19E-04 | -1.04E+00 | 8.37E-04 | 3.62E-03 | 1.15E+00 | 7.28E-08 | 3.09E-07 | -1.41E+00 | 1.42E-04 | 5.41E-04 | -1.05E+00 | 4.41E-06 | 1.57E-05 | -1.38E+00 | 1.13E-01 | 3.20E-01 | 1.12E+00 | 3.12E-15 | 1.14E-14 | -1.63E+00 | 1.20E-04 | 3.31E-04 | -1.16E+00 |
| HLA-DRA | 1.57E-04 | 8.20E-04 | -1.38E+00 | 1.08E-01 | 4.65E-01 | -1.08E+00 | 1.83E-03 | 7.77E-03 | -1.34E+00 | 1.85E-01 | 7.06E-01 | -1.03E+00 | 7.25E-14 | 2.58E-13 | -1.47E+00 | 2.80E-09 | 7.89E-09 | -1.30E+00 | 2.60E-87 | 9.54E-87 | -2.35E+00 | 1.32E-12 | 3.62E-12 | -1.35E+00 |
| ATF7 | 1.57E-04 | 8.20E-04 | -1.04E+00 | 6.63E-04 | 2.86E-03 | 1.12E+00 | 2.83E-04 | 1.20E-03 | -1.15E+00 | 2.98E-03 | 1.14E-02 | 1.09E+00 | 3.31E-03 | 1.18E-02 | 1.04E+00 | 3.92E-02 | 1.11E-01 | 1.15E+00 | 1.14E-02 | 4.17E-02 | 1.04E+00 | 2.55E-02 | 7.01E-02 | 1.02E+00 |
| LINC01765 | 1.57E-04 | 8.20E-04 | -1.30E+00 | 2.50E-04 | 1.08E-03 | -1.19E+00 | 1.74E-03 | 7.40E-03 | -1.39E+00 | 7.56E-04 | 2.89E-03 | -1.41E+00 | 1.96E-01 | 6.99E-01 | 1.50E+00 | 2.07E-01 | 5.84E-01 | 1.65E+00 | 8.62E-02 | 3.16E-01 | 1.77E+00 | 3.30E-01 | 9.05E-01 | 1.25E+00 |
| IDH2 | 1.57E-04 | 8.20E-04 | 1.07E+00 | 3.91E-08 | 1.69E-07 | 1.07E+00 | 1.24E-01 | 5.26E-01 | 1.22E+00 | 1.66E-01 | 6.33E-01 | 1.14E+00 | 1.36E-05 | 4.83E-05 | -1.17E+00 | 1.86E-05 | 5.25E-05 | -1.10E+00 | 1.47E-01 | 5.40E-01 | 1.15E+00 | 5.99E-02 | 1.65E-01 | 1.26E+00 |
| SNRK | 1.58E-04 | 8.21E-04 | -1.11E+00 | 1.95E-07 | 8.44E-07 | -1.26E+00 | 3.24E-04 | 1.38E-03 | -1.16E+00 | 8.82E-08 | 3.37E-07 | -1.30E+00 | 8.20E-04 | 2.92E-03 | -1.10E+00 | 6.49E-05 | 1.83E-04 | -1.30E+00 | 9.89E-04 | 3.63E-03 | -1.23E+00 | 3.35E-05 | 9.21E-05 | -1.45E+00 |
| LMTK2 | 1.58E-04 | 8.24E-04 | -1.58E+00 | 1.05E-03 | 4.54E-03 | -1.19E+00 | 3.57E-07 | 1.52E-06 | -2.11E+00 | 8.40E-07 | 3.21E-06 | -1.85E+00 | 8.60E-02 | 3.06E-01 | 1.00E+00 | 8.11E-02 | 2.29E-01 | -1.18E+00 | 3.04E-04 | 1.11E-03 | -1.64E+00 | 1.11E-02 | 3.06E-02 | -1.23E+00 |
| PIGH | 1.58E-04 | 8.24E-04 | -1.22E+00 | 5.64E-02 | 2.44E-01 | 1.25E+00 | 2.45E-06 | 1.04E-05 | -1.56E+00 | 5.13E-06 | 1.96E-05 | -1.32E+00 | 1.68E-02 | 5.98E-02 | -1.07E+00 | 8.08E-02 | 2.28E-01 | 1.06E+00 | 1.69E-06 | 6.20E-06 | -1.29E+00 | 1.16E-04 | 3.17E-04 | -1.14E+00 |
| ROBO3 | 1.58E-04 | 8.26E-04 | -1.09E+00 | 3.49E-04 | 1.51E-03 | 1.05E+00 | 4.15E-02 | 1.76E-01 | 1.05E+00 | 3.61E-02 | 1.38E-01 | 1.17E+00 | 1.06E-01 | 3.78E-01 | 1.15E+00 | 1.85E-01 | 5.21E-01 | 1.28E+00 | 2.23E-01 | 8.18E-01 | 1.33E+00 | 1.03E-01 | 2.83E-01 | 1.59E+00 |
| AC064807.1 | 1.59E-04 | 8.27E-04 | -1.06E+00 | 1.38E-06 | 5.96E-06 | -1.15E+00 | 5.13E-02 | 2.18E-01 | -1.03E+00 | 6.53E-04 | 2.49E-03 | -1.21E+00 | 1.09E-01 | 3.89E-01 | 1.16E+00 | 3.10E-01 | 8.74E-01 | 1.54E+00 | 2.45E-01 | 8.99E-01 | 1.26E+00 | 2.26E-01 | 6.22E-01 | 1.27E+00 |
| WDR75 | 1.59E-04 | 8.27E-04 | -1.14E+00 | 7.76E-05 | 3.35E-04 | -1.09E+00 | 4.12E-07 | 1.75E-06 | -1.48E+00 | 3.04E-05 | 1.16E-04 | -1.17E+00 | 6.31E-04 | 2.25E-03 | -1.22E+00 | 4.21E-02 | 1.19E-01 | 1.10E+00 | 1.85E-16 | 6.80E-16 | -1.93E+00 | 8.36E-07 | 2.30E-06 | -1.21E+00 |
| SGK1 | 1.59E-04 | 8.29E-04 | 5.31E+00 | 2.06E-01 | 8.91E-01 | 1.33E+00 | 8.81E-10 | 3.74E-09 | 7.39E+00 | 1.93E-01 | 7.36E-01 | 1.41E+00 | 1.01E-19 | 3.59E-19 | 1.11E+01 | 3.45E-01 | 9.73E-01 | 2.32E+00 | 2.12E-03 | 7.78E-03 | 4.21E+00 | 2.01E-01 | 5.52E-01 | 1.61E+00 |
| MRPL24 | 1.60E-04 | 8.32E-04 | -1.22E+00 | 1.70E-02 | 7.34E-02 | 1.22E+00 | 2.13E-04 | 9.05E-04 | -1.12E+00 | 3.42E-03 | 1.31E-02 | 1.12E+00 | 5.25E-04 | 1.87E-03 | -1.21E+00 | 7.98E-02 | 2.25E-01 | 1.21E+00 | 1.47E-04 | 5.41E-04 | -1.09E+00 | 3.37E-01 | 9.27E-01 | 1.29E+00 |
| SLX1B | 1.60E-04 | 8.32E-04 | -1.50E+00 | 1.21E-05 | 5.25E-05 | -1.71E+00 | 1.25E-06 | 5.30E-06 | -1.34E+00 | 1.83E-11 | 6.99E-11 | -1.74E+00 | 3.57E-10 | 1.27E-09 | -2.40E+00 | 3.02E-09 | 8.52E-09 | -2.13E+00 | 3.32E-07 | 1.22E-06 | -1.19E+00 | 1.48E-12 | 4.05E-12 | -1.49E+00 |
| LINC00513 | 1.60E-04 | 8.35E-04 | -1.55E+00 | 5.45E-04 | 2.36E-03 | -1.33E+00 | 8.13E-03 | 3.45E-02 | -1.29E+00 | 2.79E-02 | 1.06E-01 | -1.03E+00 | 9.72E-02 | 3.47E-01 | 2.24E+00 | 1.25E-01 | 3.51E-01 | 2.17E+00 | 4.33E-02 | 1.59E-01 | -1.47E+00 | 2.93E-02 | 8.05E-02 | -1.73E+00 |
| LINC00937 | 1.60E-04 | 8.35E-04 | -1.98E+00 | 5.08E-05 | 2.20E-04 | -2.30E+00 | 1.44E-01 | 6.13E-01 | -1.21E+00 | 2.58E-01 | 9.86E-01 | 1.22E+00 | 1.99E-03 | 7.09E-03 | -1.55E+00 | 5.51E-03 | 1.55E-02 | -1.36E+00 | 5.59E-13 | 2.05E-12 | -1.78E+00 | 5.00E-11 | 1.37E-10 | -1.56E+00 |
| GNS | 1.60E-04 | 8.36E-04 | -1.39E+00 | 2.43E-02 | 1.05E-01 | 1.15E+00 | 2.47E-03 | 1.05E-02 | -1.32E+00 | 8.84E-03 | 3.38E-02 | -1.06E+00 | 2.14E-04 | 7.62E-04 | -1.61E+00 | 1.01E-02 | 2.86E-02 | -1.12E+00 | 2.67E-11 | 9.81E-11 | -1.69E+00 | 6.15E-07 | 1.69E-06 | -1.39E+00 |
| ZNF268 | 1.61E-04 | 8.36E-04 | -1.11E+00 | 4.45E-09 | 1.92E-08 | -1.46E+00 | 1.39E-02 | 5.89E-02 | -1.05E+00 | 2.55E-04 | 9.75E-04 | -1.18E+00 | 3.58E-03 | 1.28E-02 | 1.00E+00 | 1.39E-03 | 3.93E-03 | -1.15E+00 | 1.60E-02 | 5.87E-02 | -1.11E+00 | 9.31E-03 | 2.56E-02 | -1.10E+00 |
| PLEKHA8P1 | 1.61E-04 | 8.37E-04 | -1.52E+00 | 2.17E-02 | 9.39E-02 | 1.09E+00 | 4.29E-04 | 1.82E-03 | -1.35E+00 | 1.01E-03 | 3.86E-03 | -1.15E+00 | 5.02E-04 | 1.79E-03 | -1.58E+00 | 3.11E-02 | 8.77E-02 | -1.01E+00 | 1.85E-03 | 6.80E-03 | 1.09E+00 | 1.51E-02 | 4.15E-02 | -1.01E+00 |
| HSPE1 | 1.61E-04 | 8.37E-04 | -1.11E+00 | 5.65E-02 | 2.44E-01 | 1.26E+00 | 3.10E-02 | 1.32E-01 | 1.32E+00 | 4.80E-02 | 1.83E-01 | 1.50E+00 | 3.29E-02 | 1.17E-01 | 1.09E+00 | 3.08E-01 | 8.69E-01 | 1.44E+00 | 4.80E-02 | 1.76E-01 | 1.14E+00 | 4.99E-04 | 1.37E-03 | 1.73E+00 |
| SH3D21 | 1.61E-04 | 8.38E-04 | -1.06E+00 | 1.11E-07 | 4.81E-07 | -1.23E+00 | 2.55E-09 | 1.09E-08 | -1.93E+00 | 2.15E-05 | 8.21E-05 | -1.22E+00 | 1.91E-02 | 6.83E-02 | -1.14E+00 | 1.63E-02 | 4.59E-02 | -1.17E+00 | 1.30E-01 | 4.77E-01 | -1.07E+00 | 3.01E-01 | 8.26E-01 | 1.12E+00 |
| MAP2K6 | 1.61E-04 | 8.39E-04 | -1.35E+00 | 1.04E-02 | 4.51E-02 | -1.01E+00 | 1.31E-02 | 5.58E-02 | -1.26E+00 | 1.17E-01 | 4.47E-01 | 1.15E+00 | 8.16E-04 | 2.91E-03 | -1.64E+00 | 1.69E-02 | 4.75E-02 | -1.12E+00 | 4.25E-08 | 1.56E-07 | -2.56E+00 | 1.68E-05 | 4.62E-05 | -1.79E+00 |
| FAM199X | 1.61E-04 | 8.39E-04 | -1.20E+00 | 1.78E-02 | 7.69E-02 | 1.29E+00 | 1.83E-02 | 7.77E-02 | -1.04E+00 | 1.14E-02 | 4.35E-02 | 1.02E+00 | 2.72E-05 | 9.69E-05 | -1.40E+00 | 8.66E-04 | 2.44E-03 | -1.12E+00 | 7.58E-04 | 2.78E-03 | -1.13E+00 | 4.36E-03 | 1.20E-02 | -1.09E+00 |
| ZNF398 | 1.61E-04 | 8.39E-04 | -1.45E+00 | 1.02E-02 | 4.43E-02 | 1.09E+00 | 2.36E-05 | 1.00E-04 | -1.57E+00 | 1.30E-05 | 4.97E-05 | -1.51E+00 | 2.24E-03 | 8.00E-03 | -1.32E+00 | 4.37E-04 | 1.23E-03 | -1.57E+00 | 3.31E-03 | 1.21E-02 | -1.23E+00 | 4.66E-02 | 1.28E-01 | -1.05E+00 |
| COQ6 | 1.61E-04 | 8.40E-04 | -1.22E+00 | 4.53E-04 | 1.96E-03 | -1.11E+00 | 1.99E-02 | 8.44E-02 | -1.02E+00 | 1.72E-03 | 6.56E-03 | -1.08E+00 | 4.81E-03 | 1.72E-02 | -1.14E+00 | 4.31E-03 | 1.22E-02 | -1.14E+00 | 2.74E-03 | 1.01E-02 | -1.10E+00 | 2.53E-03 | 6.94E-03 | -1.11E+00 |
| TET1 | 1.62E-04 | 8.43E-04 | -1.47E+00 | 2.00E-02 | 8.66E-02 | 1.18E+00 | 3.18E-05 | 1.35E-04 | -1.48E+00 | 2.07E-04 | 7.92E-04 | -1.21E+00 | 5.92E-09 | 2.11E-08 | -1.86E+00 | 4.21E-05 | 1.19E-04 | -1.25E+00 | 1.37E-07 | 5.02E-07 | -1.88E+00 | 1.07E-04 | 2.93E-04 | -1.47E+00 |
| AL691432.1 | 1.62E-04 | 8.43E-04 | -1.52E+00 | 2.17E-02 | 9.39E-02 | 1.08E+00 | 3.20E-05 | 1.36E-04 | -1.79E+00 | 9.86E-04 | 3.76E-03 | -1.29E+00 | 5.24E-04 | 1.87E-03 | -1.59E+00 | 9.73E-03 | 2.74E-02 | -1.17E+00 | 3.27E-04 | 1.20E-03 | -1.55E+00 | 1.08E-02 | 2.96E-02 | -1.25E+00 |
| STK32B | 1.62E-04 | 8.44E-04 | -1.23E+00 | 5.31E-06 | 2.30E-05 | -1.36E+00 | 1.45E-04 | 6.16E-04 | -1.44E+00 | 1.38E-03 | 5.28E-03 | -1.13E+00 | 5.99E-02 | 2.14E-01 | -1.20E+00 | 8.16E-02 | 2.30E-01 | -1.12E+00 | 1.22E-01 | 4.50E-01 | -1.12E+00 | 2.19E-01 | 6.03E-01 | 1.52E+00 |
| TCN2 | 1.62E-04 | 8.44E-04 | -1.45E+00 | 9.28E-03 | 4.01E-02 | -1.00E+00 | 1.75E-01 | 7.43E-01 | 1.29E+00 | 1.80E-01 | 6.88E-01 | 1.14E+00 | 2.43E-01 | 8.65E-01 | 1.16E+00 | 1.11E-01 | 3.13E-01 | -1.47E+00 | 5.75E-02 | 2.11E-01 | -1.65E+00 | 4.26E-02 | 1.17E-01 | -1.99E+00 |
| RBM43 | 1.62E-04 | 8.46E-04 | -1.17E+00 | 6.16E-04 | 2.66E-03 | 1.06E+00 | 1.79E-03 | 7.60E-03 | -1.29E+00 | 1.46E-02 | 5.58E-02 | -1.01E+00 | 1.82E-01 | 6.50E-01 | 1.30E+00 | 3.01E-01 | 8.48E-01 | 1.47E+00 | 6.77E-04 | 2.49E-03 | -1.20E+00 | 1.52E-04 | 4.19E-04 | -1.21E+00 |
| FZD1 | 1.66E-04 | 8.63E-04 | -1.23E+00 | 4.38E-03 | 1.89E-02 | 1.05E+00 | 1.33E-01 | 5.66E-01 | 1.09E+00 | 1.10E-01 | 4.22E-01 | 1.59E+00 | 8.20E-03 | 2.93E-02 | -1.49E+00 | 1.60E-02 | 4.50E-02 | -1.56E+00 | 2.24E-01 | 8.22E-01 | 1.08E+00 | 3.53E-01 | 9.70E-01 | 1.22E+00 |
| RPUSD2 | 1.66E-04 | 8.65E-04 | -1.18E+00 | 2.06E-04 | 8.91E-04 | -1.04E+00 | 6.57E-02 | 2.79E-01 | 1.12E+00 | 3.44E-02 | 1.31E-01 | 1.19E+00 | 2.85E-02 | 1.02E-01 | -1.00E+00 | 7.65E-02 | 2.16E-01 | 1.04E+00 | 1.81E-03 | 6.65E-03 | -1.14E+00 | 6.34E-02 | 1.74E-01 | 1.11E+00 |
| EDC3 | 1.66E-04 | 8.66E-04 | -1.42E+00 | 5.15E-02 | 2.23E-01 | 1.20E+00 | 8.91E-03 | 3.79E-02 | -1.20E+00 | 5.52E-02 | 2.11E-01 | 1.08E+00 | 4.88E-04 | 1.74E-03 | -1.67E+00 | 1.12E-01 | 3.15E-01 | 1.22E+00 | 3.03E-05 | 1.11E-04 | -1.38E+00 | 4.94E-03 | 1.36E-02 | -1.13E+00 |
| PCNX3 | 1.66E-04 | 8.67E-04 | -1.53E+00 | 6.66E-03 | 2.88E-02 | 1.06E+00 | 2.02E-01 | 8.58E-01 | 1.10E+00 | 1.54E-01 | 5.88E-01 | 1.50E+00 | 9.47E-02 | 3.38E-01 | -1.10E+00 | 2.65E-01 | 7.48E-01 | 1.62E+00 | 4.66E-03 | 1.71E-02 | -1.35E+00 | 1.49E-01 | 4.10E-01 | 1.06E+00 |
| CCNC | 1.67E-04 | 8.68E-04 | 1.01E+00 | 1.54E-03 | 6.64E-03 | 1.23E+00 | 6.41E-04 | 2.72E-03 | -1.13E+00 | 2.95E-03 | 1.13E-02 | 1.02E+00 | 7.66E-03 | 2.73E-02 | 1.11E+00 | 8.32E-03 | 2.35E-02 | 1.14E+00 | 2.76E-02 | 1.01E-01 | 1.04E+00 | 1.40E-01 | 3.83E-01 | 1.11E+00 |
| TMEM202-AS1 | 1.67E-04 | 8.69E-04 | -1.16E+00 | 5.03E-08 | 2.17E-07 | -1.72E+00 | 9.51E-02 | 4.04E-01 | 1.40E+00 | 8.27E-03 | 3.16E-02 | -1.35E+00 | 2.49E-02 | 8.89E-02 | -1.19E+00 | 1.69E-02 | 4.75E-02 | -1.32E+00 | 2.39E-03 | 8.77E-03 | -1.42E+00 | 6.38E-03 | 1.75E-02 | -1.41E+00 |
| AC024075.2 | 1.67E-04 | 8.69E-04 | -1.73E+00 | 1.36E-01 | 5.86E-01 | 1.25E+00 | 2.77E-05 | 1.18E-04 | -1.67E+00 | 1.51E-02 | 5.77E-02 | -1.05E+00 | 1.76E-04 | 6.28E-04 | -1.97E+00 | 5.02E-02 | 1.41E-01 | -1.05E+00 | 1.94E-03 | 7.12E-03 | -1.41E+00 | 8.55E-02 | 2.35E-01 | 1.01E+00 |
| GPR141 | 1.67E-04 | 8.69E-04 | -1.28E+00 | 3.14E-05 | 1.36E-04 | -1.45E+00 | 7.35E-04 | 3.12E-03 | -1.50E+00 | 6.62E-04 | 2.53E-03 | -1.46E+00 | 7.04E-06 | 2.51E-05 | -1.09E+00 | 2.21E-11 | 6.22E-11 | -1.42E+00 | 1.72E-01 | 6.32E-01 | 1.13E+00 | 9.60E-02 | 2.64E-01 | -1.04E+00 |
| AC132192.2 | 1.67E-04 | 8.69E-04 | 2.40E+00 | 8.29E-02 | 3.58E-01 | 1.90E+00 | 2.50E-02 | 1.06E-01 | 1.58E+00 | 4.92E-02 | 1.88E-01 | 1.02E+00 | 1.44E-03 | 5.15E-03 | 3.13E+00 | 2.20E-01 | 6.19E-01 | 1.92E+00 | 2.21E-01 | 8.12E-01 | 1.37E+00 | 1.93E-01 | 5.30E-01 | 1.09E+00 |
| MEF2D | 1.67E-04 | 8.69E-04 | -1.18E+00 | 1.44E-06 | 6.21E-06 | -1.29E+00 | 1.55E-04 | 6.59E-04 | -1.29E+00 | 5.68E-07 | 2.17E-06 | -1.37E+00 | 1.26E-02 | 4.50E-02 | 1.07E+00 | 1.89E-04 | 5.32E-04 | -1.33E+00 | 8.33E-03 | 3.06E-02 | 1.03E+00 | 1.56E-04 | 4.30E-04 | -1.28E+00 |
| GM2A | 1.67E-04 | 8.69E-04 | -1.21E+00 | 1.21E-05 | 5.22E-05 | -1.27E+00 | 3.63E-04 | 1.54E-03 | -1.18E+00 | 6.07E-05 | 2.32E-04 | -1.12E+00 | 4.24E-03 | 1.51E-02 | -1.11E+00 | 4.75E-03 | 1.34E-02 | -1.08E+00 | 5.53E-05 | 2.03E-04 | -1.48E+00 | 6.51E-04 | 1.79E-03 | -1.33E+00 |
| PLAGL1 | 1.67E-04 | 8.71E-04 | -1.08E+00 | 1.62E-03 | 6.99E-03 | 1.10E+00 | 2.85E-06 | 1.21E-05 | -1.30E+00 | 3.97E-03 | 1.52E-02 | -1.03E+00 | 6.96E-07 | 2.48E-06 | -1.25E+00 | 5.03E-05 | 1.42E-04 | -1.07E+00 | 2.49E-02 | 9.15E-02 | -1.07E+00 | 3.54E-01 | 9.73E-01 | 1.20E+00 |
| SLC25A21-AS1 | 1.68E-04 | 8.73E-04 | -1.66E+00 | 9.49E-05 | 4.10E-04 | -1.60E+00 | 5.55E-02 | 2.36E-01 | -1.14E+00 | 7.17E-02 | 2.74E-01 | -1.02E+00 | 1.75E-01 | 6.23E-01 | 1.10E+00 | 1.81E-01 | 5.10E-01 | 1.72E+00 | 2.12E-01 | 7.79E-01 | 1.25E+00 | 4.95E-02 | 1.36E-01 | 1.70E+00 |
| KCNAB2 | 1.68E-04 | 8.73E-04 | 1.05E+00 | 7.40E-05 | 3.20E-04 | 1.11E+00 | 2.45E-03 | 1.04E-02 | -1.05E+00 | 3.92E-03 | 1.50E-02 | 1.09E+00 | 5.92E-03 | 2.11E-02 | -1.14E+00 | 2.30E-03 | 6.47E-03 | -1.19E+00 | 2.20E-01 | 8.07E-01 | 1.23E+00 | 3.27E-01 | 8.99E-01 | 1.43E+00 |
| PMM2 | 1.68E-04 | 8.75E-04 | -1.35E+00 | 4.72E-06 | 2.04E-05 | -1.48E+00 | 4.67E-03 | 1.98E-02 | -1.17E+00 | 3.61E-02 | 1.38E-01 | 1.15E+00 | 1.26E-03 | 4.48E-03 | -1.17E+00 | 5.30E-03 | 1.49E-02 | -1.08E+00 | 3.96E-05 | 1.45E-04 | -1.22E+00 | 1.57E-02 | 4.31E-02 | -1.01E+00 |
| TXNL4B | 1.68E-04 | 8.76E-04 | -1.38E+00 | 2.19E-04 | 9.44E-04 | -1.22E+00 | 1.41E-05 | 6.00E-05 | -1.42E+00 | 3.56E-07 | 1.36E-06 | -1.45E+00 | 2.35E-03 | 8.39E-03 | -1.41E+00 | 4.33E-02 | 1.22E-01 | -1.04E+00 | 6.64E-04 | 2.44E-03 | -1.17E+00 | 3.73E-02 | 1.02E-01 | 1.02E+00 |
| CXorf56 | 1.68E-04 | 8.77E-04 | -1.33E+00 | 7.28E-03 | 3.15E-02 | 1.05E+00 | 1.36E-05 | 5.76E-05 | -1.43E+00 | 2.68E-04 | 1.02E-03 | -1.19E+00 | 5.04E-04 | 1.80E-03 | -1.44E+00 | 2.05E-03 | 5.78E-03 | -1.22E+00 | 3.78E-06 | 1.39E-05 | -1.49E+00 | 5.98E-04 | 1.64E-03 | -1.21E+00 |
| MRPL45 | 1.68E-04 | 8.78E-04 | -1.01E+00 | 4.31E-02 | 1.86E-01 | 1.25E+00 | 9.60E-07 | 4.08E-06 | -1.26E+00 | 6.98E-05 | 2.67E-04 | -1.02E+00 | 4.47E-04 | 1.59E-03 | 1.01E+00 | 5.02E-02 | 1.41E-01 | 1.23E+00 | 5.90E-07 | 2.17E-06 | -1.18E+00 | 3.51E-06 | 9.63E-06 | -1.10E+00 |
| ZNF701 | 1.69E-04 | 8.79E-04 | -1.24E+00 | 1.81E-03 | 7.82E-03 | -1.02E+00 | 8.06E-04 | 3.42E-03 | -1.09E+00 | 2.14E-04 | 8.19E-04 | -1.06E+00 | 1.95E-03 | 6.95E-03 | -1.03E+00 | 2.67E-03 | 7.53E-03 | -1.02E+00 | 1.10E-02 | 4.03E-02 | 1.01E+00 | 6.41E-03 | 1.76E-02 | 1.01E+00 |
| ZNF611 | 1.69E-04 | 8.79E-04 | -1.37E+00 | 2.20E-04 | 9.50E-04 | -1.27E+00 | 2.92E-04 | 1.24E-03 | -1.31E+00 | 1.33E-02 | 5.08E-02 | 1.02E+00 | 3.86E-04 | 1.37E-03 | -1.39E+00 | 1.50E-05 | 4.22E-05 | -1.59E+00 | 3.46E-06 | 1.27E-05 | -1.63E+00 | 9.02E-05 | 2.48E-04 | -1.55E+00 |
| ZNF18 | 1.69E-04 | 8.79E-04 | -1.32E+00 | 5.36E-05 | 2.32E-04 | -1.30E+00 | 1.45E-03 | 6.16E-03 | -1.34E+00 | 6.95E-02 | 2.66E-01 | 1.14E+00 | 1.87E-02 | 6.67E-02 | -1.07E+00 | 1.61E-02 | 4.54E-02 | -1.16E+00 | 4.10E-03 | 1.50E-02 | -1.17E+00 | 5.47E-03 | 1.50E-02 | -1.08E+00 |
| MAP3K1 | 1.69E-04 | 8.80E-04 | -1.23E+00 | 1.22E-04 | 5.28E-04 | -1.11E+00 | 6.40E-07 | 2.72E-06 | -1.38E+00 | 7.39E-04 | 2.82E-03 | -1.04E+00 | 3.31E-06 | 1.18E-05 | -1.19E+00 | 6.70E-06 | 1.89E-05 | -1.25E+00 | 3.96E-05 | 1.45E-04 | -1.26E+00 | 6.12E-03 | 1.68E-02 | -1.07E+00 |
| AFAP1L2 | 1.69E-04 | 8.80E-04 | -1.38E+00 | 3.66E-02 | 1.58E-01 | 1.13E+00 | 1.23E-02 | 5.23E-02 | -1.27E+00 | 1.48E-02 | 5.64E-02 | -1.13E+00 | 9.86E-07 | 3.52E-06 | -1.75E+00 | 6.09E-06 | 1.72E-05 | -1.62E+00 | 7.58E-02 | 2.78E-01 | -1.31E+00 | 7.19E-02 | 1.98E-01 | -1.36E+00 |
| AL109755.1 | 1.69E-04 | 8.80E-04 | -1.13E+00 | 1.31E-06 | 5.66E-06 | -1.23E+00 | 2.22E-01 | 9.43E-01 | 1.16E+00 | 3.64E-02 | 1.39E-01 | 1.02E+00 | 9.29E-02 | 3.31E-01 | 1.14E+00 | 1.74E-02 | 4.92E-02 | -1.13E+00 | 2.68E-03 | 9.84E-03 | -1.82E+00 | 3.07E-03 | 8.42E-03 | -1.94E+00 |
| KARS | 1.69E-04 | 8.81E-04 | 1.00E+00 | 6.00E-06 | 2.59E-05 | 1.06E+00 | 2.29E-02 | 9.73E-02 | 1.03E+00 | 1.39E-03 | 5.32E-03 | 1.04E+00 | 1.38E-04 | 4.93E-04 | -1.04E+00 | 1.46E-02 | 4.13E-02 | 1.13E+00 | 2.21E-03 | 8.09E-03 | -1.06E+00 | 1.93E-01 | 5.31E-01 | 1.09E+00 |
| LMBR1L | 1.69E-04 | 8.82E-04 | -1.10E+00 | 7.57E-02 | 3.27E-01 | 1.32E+00 | 5.56E-04 | 2.36E-03 | -1.28E+00 | 4.18E-02 | 1.60E-01 | 1.11E+00 | 1.90E-04 | 6.78E-04 | -1.29E+00 | 3.44E-03 | 9.70E-03 | -1.13E+00 | 1.47E-06 | 5.38E-06 | -1.60E+00 | 1.76E-05 | 4.84E-05 | -1.51E+00 |
| MRPS30-DT | 1.69E-04 | 8.82E-04 | -1.58E+00 | 2.44E-04 | 1.06E-03 | -1.46E+00 | 4.16E-02 | 1.77E-01 | -1.34E+00 | 1.01E-01 | 3.87E-01 | -1.02E+00 | 8.58E-02 | 3.06E-01 | -1.06E+00 | 1.33E-01 | 3.75E-01 | -1.00E+00 | 1.58E-03 | 5.80E-03 | -1.18E+00 | 8.95E-04 | 2.46E-03 | -1.37E+00 |
| ZFAND4 | 1.69E-04 | 8.83E-04 | -1.27E+00 | 6.21E-05 | 2.69E-04 | -1.20E+00 | 1.19E-08 | 5.05E-08 | -1.72E+00 | 5.66E-10 | 2.16E-09 | -1.74E+00 | 2.21E-02 | 7.86E-02 | -1.11E+00 | 4.89E-02 | 1.38E-01 | 1.05E+00 | 7.58E-02 | 2.78E-01 | -1.07E+00 | 4.11E-02 | 1.13E-01 | -1.23E+00 |
| PELI2 | 1.70E-04 | 8.86E-04 | -1.15E+00 | 9.52E-04 | 4.12E-03 | 1.04E+00 | 2.24E-06 | 9.52E-06 | -1.46E+00 | 4.85E-06 | 1.85E-05 | -1.23E+00 | 3.49E-05 | 1.25E-04 | -1.30E+00 | 5.84E-05 | 1.65E-04 | -1.29E+00 | 3.47E-07 | 1.27E-06 | -1.53E+00 | 6.00E-08 | 1.65E-07 | -1.56E+00 |
| SSBP4 | 1.70E-04 | 8.88E-04 | 1.11E+00 | 2.78E-06 | 1.20E-05 | 1.15E+00 | 1.19E-02 | 5.06E-02 | -1.03E+00 | 2.14E-02 | 8.18E-02 | 1.10E+00 | 1.85E-02 | 6.61E-02 | 1.11E+00 | 2.70E-03 | 7.60E-03 | 1.08E+00 | 1.38E-01 | 5.07E-01 | 1.24E+00 | 9.20E-02 | 2.53E-01 | 1.35E+00 |

| | | | | | | | | | | | | | | | | | | | | | | | | |
|---|---|---|---|---|---|---|---|---|---|---|---|---|---|---|---|---|---|---|---|---|---|---|---|---|
| AL031320.2 | 1.71E-04 | 8.89E-04 | -1.70E+00 | 3.77E-02 | 1.63E-01 | 1.15E+00 | 3.69E-03 | 1.57E-02 | -1.58E+00 | 2.06E-01 | 7.86E-01 | 1.47E+00 | 3.52E-02 | 1.25E-01 | -1.05E+00 | 2.38E-01 | 6.70E-01 | 1.36E+00 | 1.92E-01 | 7.04E-01 | 1.19E+00 | 4.88E-03 | 1.34E-02 | 1.97E+00 |
| AAGAB | 1.71E-04 | 8.90E-04 | -1.39E+00 | 1.51E-04 | 6.54E-04 | -1.26E+00 | 2.00E-08 | 8.48E-08 | -1.56E+00 | 1.28E-08 | 4.87E-08 | -1.38E+00 | 2.37E-04 | 8.46E-04 | -1.28E+00 | 6.44E-03 | 1.82E-02 | 1.03E+00 | 4.50E-07 | 1.65E-06 | -1.29E+00 | 6.73E-03 | 1.85E-02 | 1.03E+00 |
| NORAD | 1.71E-04 | 8.91E-04 | -1.05E+00 | 1.03E-04 | 4.43E-04 | 1.08E+00 | 1.43E-04 | 6.09E-04 | -1.25E+00 | 4.40E-02 | 1.68E-01 | 1.01E+00 | 3.40E-05 | 1.21E-04 | -1.18E+00 | 9.23E-04 | 2.60E-03 | -1.07E+00 | 1.76E-05 | 6.47E-05 | 1.02E+00 | 2.88E-03 | 7.92E-03 | 1.04E+00 |
| UTP15 | 1.71E-04 | 8.91E-04 | -1.52E+00 | 7.27E-04 | 3.14E-03 | -1.28E+00 | 2.02E-04 | 8.56E-04 | -1.25E+00 | 5.32E-06 | 2.03E-05 | -1.28E+00 | 7.22E-03 | 2.57E-02 | -1.33E+00 | 1.17E-01 | 3.31E-01 | 1.10E+00 | 3.14E-07 | 1.15E-06 | -1.49E+00 | 1.21E-03 | 3.32E-03 | -1.14E+00 |
| KRT8P12 | 1.71E-04 | 8.91E-04 | -1.62E+00 | 8.09E-03 | 3.50E-02 | -1.16E+00 | 9.01E-03 | 3.83E-02 | -1.58E+00 | 1.24E-01 | 4.74E-01 | 1.08E+00 | 1.11E-02 | 3.97E-02 | -1.22E+00 | 1.13E-02 | 3.19E-02 | -1.29E+00 | 1.31E-03 | 4.81E-03 | -1.39E+00 | 7.64E-02 | 2.10E-01 | 1.04E+00 |
| ARRDC4 | 1.71E-04 | 8.92E-04 | -1.32E+00 | 5.59E-06 | 2.41E-05 | -1.53E+00 | 1.23E-07 | 5.23E-07 | -1.66E+00 | 4.61E-14 | 1.76E-13 | -2.49E+00 | 4.55E-04 | 1.62E-03 | -1.19E+00 | 1.09E-08 | 3.08E-08 | -1.95E+00 | 2.10E-02 | 7.69E-02 | 1.10E+00 | 5.63E-06 | 1.55E-05 | -1.32E+00 |
| EIF2S2 | 1.71E-04 | 8.93E-04 | 1.02E+00 | 3.10E-05 | 1.34E-04 | 1.11E+00 | 4.16E-03 | 1.77E-02 | -1.10E+00 | 9.25E-05 | 3.53E-04 | -1.06E+00 | 7.58E-07 | 2.70E-06 | -1.11E+00 | 4.37E-04 | 1.23E-03 | 1.05E+00 | 6.69E-03 | 2.46E-02 | -1.02E+00 | 3.09E-02 | 8.49E-02 | -1.00E+00 |
| PAM | 1.71E-04 | 8.94E-04 | -1.52E+00 | 2.81E-02 | 1.21E-01 | 1.02E+00 | 1.58E-02 | 6.71E-02 | -1.22E+00 | 1.32E-01 | 5.05E-01 | 1.18E+00 | 2.71E-01 | 9.65E-01 | 1.40E+00 | 7.78E-02 | 2.19E-01 | 2.29E+00 | 1.21E-04 | 4.45E-04 | -1.53E+00 | 4.23E-02 | 1.16E-01 | -1.05E+00 |
| C1orf162 | 1.72E-04 | 8.95E-04 | -1.01E+00 | 4.54E-09 | 1.96E-08 | -1.04E+00 | 1.65E-02 | 7.01E-02 | 1.04E+00 | 1.79E-04 | 6.85E-04 | 1.01E+00 | 1.92E-05 | 6.84E-05 | -1.14E+00 | 2.96E-07 | 8.34E-07 | -1.25E+00 | 1.43E-01 | 5.26E-01 | 1.01E+00 | 4.97E-02 | 1.37E-01 | -1.13E+00 |
| CYTH2 | 1.72E-04 | 8.97E-04 | -1.06E+00 | 6.32E-03 | 2.73E-02 | 1.23E+00 | 5.81E-05 | 2.47E-04 | -1.31E+00 | 2.76E-05 | 1.05E-04 | -1.15E+00 | 4.07E-03 | 1.45E-02 | 1.02E+00 | 3.89E-02 | 1.10E-01 | 1.13E+00 | 1.54E-02 | 5.64E-02 | 1.09E+00 | 1.21E-03 | 3.33E-03 | -1.05E+00 |
| RN7SL505P | 1.72E-04 | 8.97E-04 | -1.66E+00 | 1.48E-02 | 6.38E-02 | 1.04E+00 | 2.15E-02 | 9.15E-02 | -1.27E+00 | 3.25E-03 | 1.24E-02 | -1.52E+00 | 5.80E-02 | 2.07E-01 | -1.21E+00 | 3.02E-01 | 8.50E-01 | 1.20E+00 | 9.70E-03 | 3.56E-02 | -1.07E+00 | 5.44E-02 | 1.49E-01 | -1.07E+00 |
| ENO2 | 1.72E-04 | 8.98E-04 | -1.43E+00 | 7.49E-03 | 3.24E-02 | -1.01E+00 | 2.25E-06 | 9.56E-06 | -1.42E+00 | 2.23E-05 | 8.51E-05 | -1.15E+00 | 5.37E-07 | 1.91E-06 | -2.45E+00 | 3.26E-05 | 9.18E-05 | -1.63E+00 | 8.52E-04 | 3.13E-03 | -1.12E+00 | 3.81E-02 | 1.05E-01 | 1.02E+00 |
| RPS27AP19 | 1.73E-04 | 9.03E-04 | 1.93E+00 | 1.05E-02 | 4.52E-02 | 1.21E+00 | 8.42E-04 | 3.58E-03 | 1.71E+00 | 1.17E-01 | 4.46E-01 | 1.28E+00 | 1.57E-01 | 5.60E-01 | 1.51E+00 | 3.45E-02 | 9.73E-02 | 1.10E+00 | 9.40E-02 | 3.45E-01 | 1.39E+00 | 1.83E-01 | 5.03E-01 | 1.11E+00 |
| TEX2 | 1.73E-04 | 9.03E-04 | -1.37E+00 | 2.37E-03 | 1.02E-02 | -1.03E+00 | 1.61E-03 | 6.84E-03 | -1.36E+00 | 7.60E-02 | 2.90E-01 | 1.12E+00 | 9.67E-05 | 3.45E-04 | -1.22E+00 | 2.40E-02 | 6.77E-02 | 1.02E+00 | 2.29E-07 | 8.42E-07 | -1.35E+00 | 2.91E-03 | 7.99E-03 | -1.05E+00 |
| NET1 | 1.73E-04 | 9.04E-04 | -1.15E+00 | 3.96E-02 | 1.71E-01 | 1.22E+00 | 8.46E-04 | 3.59E-03 | -1.30E+00 | 1.55E-03 | 5.90E-03 | -1.07E+00 | 3.12E-03 | 1.11E-02 | -1.03E+00 | 2.30E-01 | 6.48E-01 | 1.35E+00 | 1.68E-09 | 6.16E-09 | -1.27E+00 | 3.94E-07 | 1.08E-06 | -1.24E+00 |
| AC007216.3 | 1.74E-04 | 9.04E-04 | -1.63E+00 | 4.77E-06 | 2.06E-05 | -2.10E+00 | 2.09E-02 | 8.86E-02 | -1.13E+00 | 7.15E-04 | 2.73E-03 | -1.59E+00 | 1.65E-11 | 5.90E-11 | -2.67E+00 | 4.22E-11 | 1.19E-10 | -2.39E+00 | 4.03E-02 | 1.48E-01 | -1.19E+00 | 7.57E-02 | 2.08E-01 | -1.04E+00 |
| ZNF275 | 1.74E-04 | 9.06E-04 | -1.18E+00 | 7.92E-06 | 3.42E-05 | -1.34E+00 | 3.55E-02 | 1.51E-01 | -1.05E+00 | 1.84E-02 | 7.02E-02 | -1.05E+00 | 2.20E-01 | 7.86E-01 | 1.58E+00 | 1.38E-01 | 3.88E-01 | 1.16E+00 | 3.68E-02 | 1.35E-01 | 1.04E+00 | 1.42E-03 | 3.90E-03 | -1.39E+00 |
| NFYA | 1.74E-04 | 9.08E-04 | -1.30E+00 | 1.55E-03 | 6.72E-03 | -1.05E+00 | 3.23E-05 | 1.37E-04 | -1.73E+00 | 3.11E-03 | 1.19E-02 | -1.20E+00 | 5.09E-04 | 1.82E-03 | -1.46E+00 | 3.41E-03 | 9.62E-03 | -1.25E+00 | 7.68E-06 | 2.82E-05 | -1.70E+00 | 2.29E-01 | 6.29E-01 | 1.12E+00 |
| AC007114.1 | 1.75E-04 | 9.11E-04 | -1.29E+00 | 9.96E-05 | 4.30E-04 | -1.37E+00 | 4.46E-02 | 1.90E-01 | -1.14E+00 | 3.14E-03 | 1.20E-02 | -1.60E+00 | 2.28E-01 | 8.11E-01 | 1.23E+00 | 1.79E-01 | 5.06E-01 | 1.07E+00 | 5.18E-02 | 1.90E-01 | 1.81E+00 | 3.16E-01 | 8.67E-01 | 1.21E+00 |
| PER3 | 1.75E-04 | 9.12E-04 | -1.66E+00 | 2.85E-03 | 1.23E-02 | -1.20E+00 | 2.43E-04 | 1.03E-03 | -1.58E+00 | 1.67E-03 | 6.37E-03 | -1.24E+00 | 2.67E-03 | 9.51E-03 | -1.34E+00 | 3.09E-02 | 8.71E-02 | 1.03E+00 | 1.82E-01 | 6.68E-01 | 1.11E+00 | 1.39E-01 | 3.83E-01 | 1.39E+00 |
| CDK5 | 1.76E-04 | 9.15E-04 | -1.30E+00 | 3.50E-04 | 1.51E-03 | -1.15E+00 | 1.29E-04 | 5.49E-04 | -1.14E+00 | 1.01E-07 | 3.86E-07 | -1.26E+00 | 1.26E-07 | 4.50E-07 | -1.74E+00 | 3.69E-05 | 1.04E-04 | -1.27E+00 | 9.62E-08 | 3.53E-07 | -1.39E+00 | 3.03E-11 | 8.31E-11 | -1.73E+00 |
| DPH3 | 1.77E-04 | 9.21E-04 | -1.13E+00 | 3.88E-10 | 1.67E-09 | -1.50E+00 | 6.47E-05 | 2.75E-04 | -1.22E+00 | 8.25E-09 | 3.15E-08 | -1.43E+00 | 2.11E-04 | 7.52E-04 | -1.29E+00 | 8.84E-06 | 2.49E-05 | -1.60E+00 | 6.57E-02 | 2.41E-01 | 1.08E+00 | 9.38E-05 | 2.58E-04 | -1.27E+00 |
| AC079328.1 | 1.77E-04 | 9.23E-04 | 3.24E+00 | 7.68E-02 | 3.32E-01 | 2.16E+00 | 5.96E-06 | 2.53E-05 | 2.29E+00 | 2.52E-01 | 9.63E-01 | 1.49E+00 | 1.64E-02 | 5.86E-02 | 2.22E+00 | 1.87E-01 | 5.28E-01 | 1.72E+00 | 4.68E-05 | 1.72E-04 | 2.03E+00 | 9.26E-02 | 2.54E-01 | 1.56E+00 |
| UBE2T | 1.77E-04 | 9.23E-04 | -2.52E+00 | 8.91E-03 | 3.85E-02 | -1.33E+00 | 2.17E-03 | 9.21E-03 | -1.12E+00 | 4.50E-02 | 1.72E-01 | 1.23E+00 | 2.93E-03 | 1.05E-02 | -3.04E+00 | 1.66E-01 | 4.69E-01 | 1.11E+00 | 2.16E-11 | 7.92E-11 | -1.58E+00 | 1.83E-02 | 5.02E-02 | -1.06E+00 |
| FUOM | 1.78E-04 | 9.25E-04 | -1.05E+00 | 5.39E-09 | 2.33E-08 | -1.27E+00 | 9.87E-02 | 4.19E-01 | 1.31E+00 | 2.78E-02 | 1.06E-01 | 1.09E+00 | 8.06E-02 | 2.88E-01 | 1.05E+00 | 1.45E-02 | 4.09E-02 | -1.12E+00 | 5.03E-02 | 1.85E-01 | 1.46E+00 | 1.77E-01 | 4.86E-01 | 1.28E+00 |
| MANSC1 | 1.78E-04 | 9.28E-04 | -1.42E+00 | 3.95E-02 | 1.71E-01 | 1.25E+00 | 3.87E-03 | 1.64E-02 | -1.59E+00 | 4.41E-02 | 1.69E-01 | -1.04E+00 | 7.33E-02 | 2.61E-01 | -1.18E+00 | 1.74E-01 | 4.92E-01 | -1.05E+00 | 4.08E-06 | 1.50E-05 | -1.33E+00 | 4.19E-04 | 1.15E-03 | -1.19E+00 |
| WASIR2 | 1.78E-04 | 9.29E-04 | -1.08E+00 | 1.67E-06 | 7.23E-06 | -1.07E+00 | 6.66E-02 | 2.83E-01 | 1.09E+00 | 1.44E-04 | 5.50E-04 | -1.04E+00 | 3.05E-03 | 1.09E-02 | -1.02E+00 | 1.09E-03 | 3.06E-03 | -1.08E+00 | 7.09E-03 | 2.60E-02 | -1.04E+00 | 5.76E-03 | 1.58E-02 | -1.07E+00 |
| CIPC | 1.79E-04 | 9.31E-04 | -1.63E+00 | 2.76E-03 | 1.19E-02 | -1.24E+00 | 3.27E-02 | 1.39E-01 | -1.16E+00 | 1.89E-01 | 7.21E-01 | 1.28E+00 | 9.63E-02 | 3.43E-01 | -1.06E+00 | 1.79E-01 | 5.04E-01 | -1.04E+00 | 2.29E-05 | 8.40E-05 | -1.48E+00 | 1.35E-03 | 3.70E-03 | -1.20E+00 |
| LONP1 | 1.79E-04 | 9.32E-04 | -1.18E+00 | 1.32E-03 | 5.71E-03 | 1.02E+00 | 6.38E-02 | 2.71E-01 | 1.05E+00 | 1.06E-02 | 4.04E-02 | 1.11E+00 | 5.39E-02 | 1.92E-01 | 1.13E+00 | 3.21E-01 | 9.04E-01 | 1.47E+00 | 1.18E-02 | 4.33E-02 | -1.06E+00 | 2.58E-01 | 7.09E-01 | 1.18E+00 |
| AL358472.2 | 1.79E-04 | 9.33E-04 | -1.82E+00 | 1.45E-02 | 6.28E-02 | -1.05E+00 | 5.83E-02 | 2.48E-01 | -1.24E+00 | 1.74E-01 | 6.63E-01 | 1.49E+00 | 3.35E-02 | 1.20E-01 | -1.36E+00 | 1.33E-01 | 3.75E-01 | 1.04E+00 | 3.86E-03 | 1.42E-02 | -1.22E+00 | 6.08E-02 | 1.67E-01 | 1.02E+00 |
| AC108488.1 | 1.79E-04 | 9.33E-04 | -1.82E+00 | 1.70E-02 | 7.34E-02 | -1.03E+00 | 9.37E-03 | 3.98E-02 | -1.26E+00 | 5.93E-04 | 2.27E-03 | -1.40E+00 | 1.07E-07 | 3.83E-07 | -2.26E+00 | 1.27E-04 | 3.58E-04 | -1.29E+00 | 1.86E-06 | 6.82E-06 | -1.79E+00 | 6.70E-07 | 1.84E-06 | -1.92E+00 |
| ZFP62 | 1.79E-04 | 9.33E-04 | -1.13E+00 | 2.39E-03 | 1.03E-02 | 1.11E+00 | 7.67E-04 | 3.26E-03 | -1.26E+00 | 3.17E-02 | 1.21E-01 | 1.08E+00 | 5.99E-02 | 2.14E-01 | -1.01E+00 | 2.33E-01 | 6.57E-01 | 1.19E+00 | 2.62E-04 | 9.62E-04 | -1.11E+00 | 2.36E-02 | 6.50E-02 | 1.12E+00 |
| EFNB2 | 1.80E-04 | 9.38E-04 | -1.38E+00 | 4.71E-07 | 2.04E-06 | -2.20E+00 | 6.78E-02 | 2.88E-01 | -1.27E+00 | 9.34E-02 | 3.57E-01 | -1.27E+00 | 2.76E-01 | 9.85E-01 | 1.12E+00 | 3.31E-01 | 9.32E-01 | 1.00E+00 | 1.89E-02 | 6.94E-02 | -1.09E+00 | 9.59E-03 | 2.63E-02 | -1.28E+00 |
| FARSB | 1.80E-04 | 9.38E-04 | -1.21E+00 | 1.50E-02 | 6.47E-02 | 1.15E+00 | 2.89E-03 | 1.23E-02 | -1.08E+00 | 7.73E-02 | 2.95E-01 | 1.28E+00 | 4.29E-03 | 1.53E-02 | -1.34E+00 | 2.35E-01 | 6.61E-01 | 1.28E+00 | 8.35E-02 | 3.07E-01 | 1.16E+00 | 1.05E-01 | 2.88E-01 | 1.44E+00 |
| PRDM2 | 1.80E-04 | 9.39E-04 | 1.01E+00 | 7.27E-05 | 3.14E-04 | 1.06E+00 | 1.08E-04 | 4.59E-04 | -1.07E+00 | 1.43E-07 | 5.48E-07 | -1.12E+00 | 3.18E-03 | 1.13E-02 | 1.02E+00 | 1.74E-04 | 4.91E-04 | -1.11E+00 | 4.51E-03 | 1.66E-02 | -1.06E+00 | 1.14E-03 | 3.14E-03 | -1.14E+00 |
| AL928654.5 | 1.81E-04 | 9.41E-04 | -1.93E+00 | 1.10E-02 | 4.74E-02 | -1.19E+00 | 1.26E-01 | 5.36E-01 | -1.00E+00 | 1.82E-01 | 6.96E-01 | 1.27E+00 | 2.30E-02 | 8.19E-02 | -1.38E+00 | 2.97E-02 | 8.37E-02 | -1.38E+00 | 1.75E-02 | 6.43E-02 | -1.70E+00 | 3.50E-01 | 9.61E-01 | 1.16E+00 |
| KLHL5 | 1.81E-04 | 9.43E-04 | -1.09E+00 | 2.94E-05 | 1.27E-04 | -1.03E+00 | 2.75E-04 | 1.17E-03 | -1.27E+00 | 6.57E-04 | 2.51E-03 | -1.12E+00 | 1.17E-01 | 4.19E-01 | 1.08E+00 | 2.58E-01 | 7.28E-01 | 1.54E+00 | 2.80E-07 | 1.03E-06 | -1.19E+00 | 3.17E-08 | 8.72E-08 | -1.29E+00 |
| CASC10 | 1.81E-04 | 9.43E-04 | -1.01E+00 | 2.12E-02 | 9.14E-02 | 1.35E+00 | 2.44E-06 | 1.04E-05 | -1.55E+00 | 3.11E-03 | 1.19E-02 | -1.07E+00 | 1.33E-01 | 4.73E-01 | -1.38E+00 | 2.91E-01 | 8.19E-01 | 1.19E+00 | 1.90E-01 | 6.99E-01 | -1.03E+00 | 2.82E-01 | 7.76E-01 | 1.12E+00 |
| SETP20 | 1.81E-04 | 9.45E-04 | -1.32E+00 | 1.45E-03 | 6.28E-03 | -1.05E+00 | 9.37E-04 | 3.98E-03 | -1.26E+00 | 4.17E-02 | 1.59E-01 | 1.11E+00 | 1.27E-01 | 4.54E-01 | 1.16E+00 | 7.50E-02 | 2.11E-01 | 1.17E+00 | 1.79E-02 | 6.58E-02 | 1.05E+00 | 1.98E-01 | 5.43E-01 | 1.23E+00 |
| NBPF26 | 1.81E-04 | 9.46E-04 | -1.08E+00 | 5.42E-06 | 2.34E-05 | -1.06E+00 | 4.54E-04 | 1.93E-03 | -1.12E+00 | 9.49E-07 | 3.62E-06 | -1.20E+00 | 5.60E-09 | 2.00E-08 | -1.37E+00 | 4.78E-07 | 1.35E-06 | -1.21E+00 | 4.59E-05 | 1.68E-04 | -1.06E+00 | 1.30E-08 | 3.57E-08 | -1.27E+00 |
| REPS2 | 1.82E-04 | 9.48E-04 | -1.46E+00 | 2.49E-03 | 1.08E-02 | -1.06E+00 | 2.88E-02 | 1.22E-01 | -1.10E+00 | 3.21E-02 | 1.23E-01 | -1.06E+00 | 1.89E-03 | 6.75E-03 | -1.48E+00 | 4.77E-04 | 1.34E-03 | -1.60E+00 | 4.63E-02 | 1.70E-01 | 1.11E+00 | 1.73E-03 | 4.75E-03 | -1.37E+00 |
| FAF1 | 1.82E-04 | 9.49E-04 | -1.18E+00 | 4.15E-05 | 1.79E-04 | -1.09E+00 | 1.71E-01 | 7.28E-01 | 1.14E+00 | 3.81E-03 | 1.45E-02 | -1.01E+00 | 3.71E-02 | 1.32E-01 | 1.03E+00 | 2.90E-02 | 8.17E-02 | 1.02E+00 | 5.52E-02 | 2.03E-01 | 1.00E+00 | 5.52E-02 | 1.52E-01 | 1.07E+00 |
| GK | 1.83E-04 | 9.54E-04 | -1.60E+00 | 3.60E-02 | 1.55E-01 | 1.06E+00 | 2.66E-04 | 1.13E-03 | -1.21E+00 | 3.11E-02 | 1.19E-01 | 1.17E+00 | 2.58E-01 | 9.22E-01 | 1.24E+00 | 1.55E-01 | 4.38E-01 | 1.73E+00 | 1.63E-03 | 5.98E-03 | -1.35E+00 | 3.33E-01 | 9.15E-01 | 1.15E+00 |
| IGSF8 | 1.83E-04 | 9.55E-04 | -1.08E+00 | 9.95E-03 | 4.30E-02 | 1.25E+00 | 4.91E-05 | 2.09E-04 | -1.26E+00 | 4.66E-04 | 1.78E-03 | -1.04E+00 | 4.56E-04 | 1.63E-03 | -1.11E+00 | 6.08E-03 | 1.71E-02 | 1.03E+00 | 1.32E-02 | 4.86E-02 | 1.05E+00 | 1.85E-01 | 5.09E-01 | 1.19E+00 |
| FBXO41 | 1.84E-04 | 9.57E-04 | -1.25E+00 | 1.43E-03 | 6.16E-03 | 1.03E+00 | 1.31E-03 | 5.56E-03 | -1.12E+00 | 2.52E-03 | 9.63E-03 | 1.03E+00 | 4.13E-04 | 1.47E-03 | -1.51E+00 | 2.63E-03 | 7.40E-03 | -1.34E+00 | 1.61E-03 | 5.89E-03 | -1.39E+00 | 6.85E-02 | 1.88E-01 | -1.02E+00 |
| RAD51B | 1.84E-04 | 9.57E-04 | -1.38E+00 | 1.11E-03 | 4.79E-03 | -1.15E+00 | 6.60E-02 | 2.80E-01 | -1.05E+00 | 2.05E-02 | 7.82E-02 | -1.06E+00 | 1.17E-02 | 4.18E-02 | -1.36E+00 | 1.81E-01 | 5.11E-01 | 1.23E+00 | 4.70E-03 | 1.72E-02 | -1.07E+00 | 6.01E-03 | 1.65E-02 | -1.10E+00 |
| EGLN2 | 1.84E-04 | 9.57E-04 | -1.10E+00 | 1.20E-04 | 5.18E-04 | -1.04E+00 | 2.59E-08 | 1.10E-07 | -1.53E+00 | 1.62E-09 | 6.18E-09 | -1.42E+00 | 4.24E-11 | 1.51E-10 | -1.91E+00 | 2.63E-11 | 7.41E-11 | -1.88E+00 | 1.61E-12 | 5.93E-12 | -1.37E+00 | 2.18E-10 | 5.99E-10 | -1.32E+00 |
| CKMT2-AS1 | 1.84E-04 | 9.58E-04 | -1.88E+00 | 1.44E-01 | 6.22E-01 | 1.36E+00 | 1.28E-05 | 5.43E-05 | -1.87E+00 | 7.07E-02 | 2.70E-01 | 1.20E+00 | 1.67E-03 | 5.97E-03 | -1.51E+00 | 3.71E-02 | 1.05E-01 | -1.01E+00 | 3.77E-06 | 1.39E-05 | -1.42E+00 | 1.33E-02 | 3.64E-02 | 1.02E+00 |
| RXRA | 1.84E-04 | 9.59E-04 | -1.09E+00 | 7.74E-03 | 3.35E-02 | 1.26E+00 | 2.42E-03 | 1.03E-02 | -1.19E+00 | 1.15E-03 | 4.40E-03 | -1.09E+00 | 1.83E-05 | 6.54E-05 | -1.62E+00 | 5.61E-04 | 1.58E-03 | -1.27E+00 | 7.50E-03 | 2.75E-02 | -1.32E+00 | 2.08E-01 | 5.71E-01 | 1.11E+00 |
| SKIL | 1.84E-04 | 9.59E-04 | 1.98E+00 | 1.04E-10 | 4.47E-10 | -2.12E+00 | 5.00E-04 | 2.12E-03 | 1.83E+00 | 1.79E-06 | 6.84E-06 | -2.62E+00 | 1.29E-01 | 4.59E-01 | 1.42E+00 | 9.34E-13 | 2.63E-12 | -2.34E+00 | 4.11E-02 | 1.51E-01 | 1.18E+00 | 1.06E-12 | 2.92E-12 | -2.43E+00 |
| MAB21L2 | 1.84E-04 | 9.61E-04 | -1.51E+00 | 1.60E-02 | 6.93E-02 | 1.00E+00 | 1.92E-01 | 8.16E-01 | 1.04E+00 | 1.77E-01 | 6.76E-01 | 1.07E+00 | 1.93E-01 | 6.89E-01 | 1.62E+00 | 1.27E-01 | 3.59E-01 | 2.02E+00 | 1.23E-02 | 4.53E-02 | -1.51E+00 | 4.65E-03 | 1.28E-02 | -1.84E+00 |
| C18orf25 | 1.85E-04 | 9.62E-04 | -1.00E+00 | 2.97E-09 | 1.28E-08 | -1.29E+00 | 7.43E-03 | 3.16E-02 | -1.00E+00 | 1.22E-09 | 4.66E-09 | -1.37E+00 | 2.58E-01 | 9.21E-01 | 1.36E+00 | 4.63E-03 | 1.31E-02 | 1.03E+00 | 6.83E-02 | 2.51E-01 | 1.20E+00 | 1.15E-04 | 3.16E-04 | -1.17E+00 |
| NR8F2 | 1.85E-04 | 9.63E-04 | -1.07E+00 | 3.34E-07 | 1.44E-06 | -1.21E+00 | 3.88E-03 | 1.65E-02 | -1.08E+00 | 5.27E-07 | 2.01E-06 | -1.34E+00 | 2.13E-04 | 7.59E-04 | -1.11E+00 | 5.62E-08 | 1.58E-07 | -1.57E+00 | 2.69E-05 | 9.86E-05 | -1.16E+00 | 1.53E-05 | 4.20E-05 | -1.34E+00 |
| PAXIP1 | 1.85E-04 | 9.64E-04 | -1.40E+00 | 3.00E-02 | 1.29E-01 | 1.20E+00 | 7.16E-04 | 3.04E-03 | -1.43E+00 | 1.76E-01 | 6.72E-01 | 1.21E+00 | 4.64E-03 | 1.65E-02 | -1.42E+00 | 3.17E-01 | 8.93E-01 | 1.44E+00 | 5.88E-08 | 2.16E-07 | -1.54E+00 | 7.72E-05 | 2.12E-04 | -1.21E+00 |
| AP2A2 | 1.86E-04 | 9.69E-04 | -1.15E+00 | 1.32E-04 | 5.72E-04 | -1.12E+00 | 2.20E-02 | 9.35E-02 | -1.00E+00 | 5.21E-02 | 1.99E-01 | 1.13E+00 | 1.62E-01 | 5.77E-01 | 1.30E+00 | 3.46E-01 | 9.74E-01 | 1.49E+00 | 2.09E-02 | 7.68E-02 | 1.05E+00 | 2.35E-01 | 6.45E-01 | 1.23E+00 |
| ABHD1 | 1.86E-04 | 9.72E-04 | -1.27E+00 | 1.18E-02 | 5.09E-02 | 1.16E+00 | 1.34E-02 | 5.69E-02 | -1.28E+00 | 2.64E-02 | 1.01E-01 | -1.04E+00 | 2.43E-03 | 8.66E-03 | -1.53E+00 | 1.58E-02 | 4.44E-02 | -1.17E+00 | 6.72E-02 | 2.47E-01 | 1.64E+00 | 2.69E-01 | 7.38E-01 | 1.41E+00 |
| CDH12 | 1.86E-04 | 9.72E-04 | -1.37E+00 | 8.27E-02 | 3.57E-01 | 1.33E+00 | 7.53E-04 | 3.20E-03 | -1.52E+00 | 3.88E-02 | 1.48E-01 | 1.02E+00 | 1.81E-06 | 6.44E-06 | -1.62E+00 | 1.74E-04 | 4.91E-04 | -1.20E+00 | 1.15E-03 | 4.23E-03 | -1.47E+00 | 1.18E-01 | 3.24E-01 | -1.03E+00 |
| COMMD4 | 1.87E-04 | 9.74E-04 | 1.01E+00 | 8.37E-05 | 3.62E-04 | 1.11E+00 | 2.04E-01 | 8.66E-01 | 1.12E+00 | 2.00E-02 | 7.65E-02 | 1.07E+00 | 2.12E-05 | 7.55E-05 | -1.14E+00 | 1.13E-04 | 3.19E-04 | -1.01E+00 | 4.44E-02 | 1.63E-01 | 1.04E+00 | 8.01E-02 | 2.20E-01 | 1.07E+00 |
| WDR46 | 1.87E-04 | 9.76E-04 | 1.03E+00 | 6.21E-04 | 2.68E-03 | 1.17E+00 | 7.43E-03 | 3.16E-02 | 1.00E+00 | 8.02E-04 | 3.07E-03 | 1.06E+00 | 5.51E-03 | 1.97E-02 | 1.03E+00 | 2.20E-01 | 6.22E-01 | 1.33E+00 | 1.08E-02 | 3.95E-02 | 1.15E+00 | 2.09E-01 | 5.74E-01 | 1.36E+00 |
| ABLIM1 | 1.87E-04 | 9.76E-04 | -1.05E+00 | 4.57E-03 | 1.98E-02 | 1.17E+00 | 5.26E-02 | 2.24E-01 | 1.02E+00 | 1.67E-02 | 6.38E-02 | 1.03E+00 | 6.69E-02 | 2.39E-01 | 1.05E+00 | 1.54E-01 | 4.35E-01 | 1.26E+00 | 2.57E-01 | 9.44E-01 | 1.50E+00 | 1.03E-01 | 2.84E-01 | 1.23E+00 |
| ARHGAP10 | 1.88E-04 | 9.79E-04 | -1.09E+00 | 5.96E-03 | 2.58E-02 | 1.18E+00 | 6.66E-03 | 2.83E-02 | -1.11E+00 | 1.52E-02 | 5.82E-02 | 1.08E+00 | 8.53E-03 | 3.04E-02 | -1.28E+00 | 1.26E-01 | 3.57E-01 | 1.03E+00 | 1.61E-01 | 5.92E-01 | 1.25E+00 | 2.65E-01 | 7.29E-01 | 1.34E+00 |
| TRAFD1 | 1.89E-04 | 9.83E-04 | -1.50E+00 | 6.78E-03 | 2.93E-02 | -1.03E+00 | 1.27E-07 | 5.40E-07 | -1.71E+00 | 1.10E-08 | 4.22E-08 | -1.61E+00 | 9.14E-07 | 3.26E-06 | -1.67E+00 | 1.08E-05 | 3.05E-05 | -1.57E+00 | 6.47E-10 | 2.37E-09 | -1.35E+00 | 4.98E-09 | 1.37E-08 | -1.32E+00 |
| FP565260.6 | 1.89E-04 | 9.84E-04 | -1.61E+00 | 1.68E-04 | 7.25E-04 | -1.65E+00 | 2.06E-01 | 8.73E-01 | 1.05E+00 | 5.09E-02 | 1.94E-01 | -1.29E+00 | 7.20E-03 | 2.57E-02 | -1.94E+00 | 3.03E-02 | 8.55E-02 | -1.49E+00 | 5.52E-04 | 2.03E-03 | -1.40E+00 | 3.61E-03 | 9.91E-03 | -1.25E+00 |
| RBM42 | 1.89E-04 | 9.84E-04 | -1.08E+00 | 5.72E-03 | 2.47E-02 | 1.16E+00 | 2.39E-02 | 1.02E-01 | 1.04E+00 | 1.32E-03 | 5.04E-03 | 1.08E+00 | 2.03E-05 | 7.23E-05 | -1.09E+00 | 3.07E-02 | 8.66E-02 | 1.20E+00 | 8.02E-04 | 2.94E-03 | -1.07E+00 | 7.02E-02 | 1.93E-01 | 1.12E+00 |
| SNX21 | 1.89E-04 | 9.85E-04 | -1.51E+00 | 2.82E-02 | 1.22E-01 | 1.07E+00 | 2.04E-01 | 8.66E-01 | 1.06E+00 | 2.57E-01 | 9.82E-01 | 1.26E+00 | 2.22E-01 | 7.92E-01 | 1.06E+00 | 3.07E-01 | 8.64E-01 | 1.15E+00 | 1.27E-01 | 4.66E-01 | 1.19E+00 | 8.00E-02 | 2.20E-01 | -1.06E+00 |
| EFHC1 | 1.89E-04 | 9.85E-04 | -1.09E+00 | 9.28E-04 | 4.01E-03 | 1.12E+00 | 1.52E-05 | 6.48E-05 | -1.37E+00 | 3.68E-04 | 1.41E-03 | -1.07E+00 | 2.44E-02 | 8.71E-02 | 1.12E+00 | 3.17E-01 | 8.94E-01 | 1.39E+00 | 3.74E-03 | 1.37E-02 | -1.11E+00 | 1.27E-02 | 3.47E-02 | 1.02E+00 |
| AREL1 | 1.91E-04 | 9.93E-04 | -1.36E+00 | 2.23E-05 | 9.64E-05 | -1.48E+00 | 1.93E-04 | 8.20E-04 | -1.69E+00 | 4.49E-03 | 1.72E-02 | -1.29E+00 | 4.32E-03 | 1.54E-02 | -1.33E+00 | 7.12E-04 | 2.01E-03 | -1.46E+00 | 2.13E-07 | 7.81E-07 | -1.56E+00 | 1.30E-07 | 3.56E-07 | -1.49E+00 |
| PRDM15 | 1.91E-04 | 9.95E-04 | -1.53E+00 | 3.48E-03 | 1.50E-02 | -1.08E+00 | 5.81E-02 | 2.47E-01 | -1.09E+00 | 4.81E-02 | 1.84E-01 | 1.11E+00 | 1.42E-01 | 5.06E-01 | -1.22E+00 | 3.21E-01 | 9.05E-01 | 1.15E+00 | 1.85E-02 | 6.79E-02 | -1.13E+00 | 4.87E-02 | 1.34E-01 | -1.09E+00 |
| MOK | 1.91E-04 | 9.98E-04 | -1.38E+00 | 1.61E-04 | 6.98E-04 | -1.21E+00 | 1.21E-04 | 5.14E-04 | -1.27E+00 | 1.05E-07 | 4.02E-07 | -1.47E+00 | 9.86E-02 | 3.51E-01 | 1.12E+00 | 1.29E-02 | 3.62E-02 | -1.35E+00 | 5.59E-02 | 2.05E-01 | 1.77E+00 | 2.43E-01 | 6.66E-01 | 1.52E+00 |
| TLK2P1 | 1.92E-04 | 9.98E-04 | -1.04E+00 | 3.89E-07 | 1.68E-06 | -1.23E+00 | 5.70E-08 | 2.42E-07 | -1.56E+00 | 3.19E-11 | 1.22E-10 | -1.80E+00 | 2.94E-03 | 1.05E-02 | -1.16E+00 | 6.50E-04 | 1.83E-03 | -1.30E+00 | 6.82E-05 | 2.50E-04 | -1.32E+00 | 9.18E-06 | 2.52E-05 | -1.43E+00 |
| CEP170P1 | 1.92E-04 | 1.00E-03 | -1.07E+00 | 2.32E-06 | 1.00E-05 | -1.14E+00 | 3.56E-06 | 1.51E-05 | -1.26E+00 | 1.41E-06 | 5.39E-06 | -1.16E+00 | 7.26E-11 | 2.59E-10 | -2.12E+00 | 2.50E-08 | 7.04E-08 | -1.54E+00 | 2.74E-06 | 1.00E-05 | -1.47E+00 | 1.87E-06 | 5.15E-06 | -1.58E+00 |
| MTF1 | 1.93E-04 | 1.00E-03 | -1.11E+00 | 7.89E-08 | 3.41E-07 | -1.51E+00 | 2.16E-03 | 9.17E-03 | -1.24E+00 | 9.63E-05 | 3.68E-04 | -1.33E+00 | 6.59E-03 | 2.35E-02 | -1.26E+00 | 1.79E-03 | 5.04E-03 | -1.55E+00 | 3.50E-08 | 1.28E-07 | -1.89E+00 | 8.50E-09 | 2.34E-08 | -2.04E+00 |
| ALG8 | 1.93E-04 | 1.01E-03 | -1.14E+00 | 4.25E-04 | 1.84E-03 | 1.01E+00 | 9.02E-06 | 3.83E-05 | -1.25E+00 | 2.01E-05 | 7.67E-05 | -1.09E+00 | 6.58E-05 | 2.35E-04 | -1.38E+00 | 8.41E-03 | 2.37E-02 | -1.07E+00 | 6.00E-05 | 2.20E-04 | -1.15E+00 | 3.18E-02 | 8.75E-02 | 1.06E+00 |
| RPL26P6 | 1.94E-04 | 1.01E-03 | -1.09E+00 | 1.71E-09 | 7.38E-09 | -1.64E+00 | 1.75E-03 | 7.44E-03 | -1.06E+00 | 4.48E-08 | 1.71E-07 | -1.55E+00 | 1.07E-06 | 3.80E-06 | -1.53E+00 | 1.39E-11 | 3.92E-11 | -2.57E+00 | 8.86E-03 | 3.25E-02 | -1.04E+00 | 1.09E-05 | 2.99E-05 | -1.60E+00 |
| PPP1R3E | 1.94E-04 | 1.01E-03 | -1.26E+00 | 9.15E-02 | 3.96E-01 | 1.75E+00 | 3.41E-04 | 1.45E-03 | -1.40E+00 | 2.43E-01 | 9.29E-01 | 1.47E+00 | 4.55E-03 | 1.62E-02 | -1.09E+00 | 2.14E-01 | 6.04E-01 | 1.49E+00 | 9.75E-02 | 3.58E-01 | 1.08E+00 | 3.14E-01 | 8.64E-01 | 1.28E+00 |
| KCTD10 | 1.94E-04 | 1.01E-03 | -1.46E+00 | 1.69E-04 | 7.29E-04 | -1.34E+00 | 7.32E-04 | 3.11E-03 | -1.23E+00 | 1.58E-05 | 6.03E-05 | -1.29E+00 | 1.82E-01 | 6.48E-01 | 1.23E+00 | 1.73E-01 | 4.88E-01 | 1.18E+00 | 1.41E-04 | 5.19E-04 | -1.27E+00 | 5.48E-03 | 1.51E-02 | -1.10E+00 |
| PMEPA1 | 1.95E-04 | 1.02E-03 | -1.33E+00 | 4.03E-05 | 1.74E-04 | -1.55E+00 | 2.70E-02 | 1.15E-01 | -1.37E+00 | 1.53E-02 | 5.85E-02 | -1.46E+00 | 1.56E-07 | 5.56E-07 | -2.77E+00 | 1.02E-07 | 2.88E-07 | -2.86E+00 | 2.59E-02 | 9.49E-02 | -1.35E+00 | 1.52E-02 | 4.17E-02 | -1.73E+00 |
| GRAMD2B | 1.95E-04 | 1.02E-03 | -1.38E+00 | 4.97E-04 | 2.15E-03 | -1.16E+00 | 1.42E-02 | 6.05E-02 | -1.13E+00 | 1.30E-01 | 4.96E-01 | 1.33E+00 | 1.38E-04 | 4.93E-04 | -2.21E+00 | 8.31E-03 | 2.34E-02 | -1.31E+00 | 2.69E-01 | 9.87E-01 | 1.28E+00 | 1.52E-01 | 4.16E-01 | -1.02E+00 |
| POC5 | 1.96E-04 | 1.02E-03 | -1.29E+00 | 2.76E-04 | 1.19E-03 | -1.14E+00 | 2.31E-05 | 9.81E-05 | -1.63E+00 | 1.74E-04 | 6.63E-04 | -1.26E+00 | 1.40E-02 | 4.98E-02 | -1.02E+00 | 5.34E-02 | 1.50E-01 | 1.07E+00 | 8.43E-08 | 3.10E-07 | -1.52E+00 | 1.03E-05 | 2.82E-05 | -1.25E+00 |
| ANKRD12 | 1.96E-04 | 1.02E-03 | -1.07E+00 | 1.76E-07 | 7.61E-07 | -1.12E+00 | 4.03E-02 | 1.71E-01 | 1.03E+00 | 8.83E-04 | 3.37E-03 | -1.02E+00 | 8.98E-02 | 3.20E-01 | 1.29E+00 | 2.69E-02 | 7.58E-02 | 1.18E+00 | 7.12E-04 | 2.61E-03 | -1.23E+00 | 4.23E-04 | 1.16E-03 | -1.19E+00 |
| KIAA2013 | 1.97E-04 | 1.03E-03 | -1.28E+00 | 4.30E-02 | 1.86E-01 | 1.30E+00 | 1.97E-05 | 8.38E-05 | -1.21E+00 | 2.52E-03 | 9.64E-03 | 1.14E+00 | 4.33E-07 | 1.54E-06 | -1.74E+00 | 2.42E-03 | 6.83E-03 | -1.06E+00 | 9.05E-02 | 3.32E-01 | 1.07E+00 | 1.91E-01 | 5.25E-01 | 1.37E+00 |
| SENP1 | 1.97E-04 | 1.03E-03 | -1.45E+00 | 2.07E-03 | 8.94E-03 | -1.08E+00 | 1.81E-02 | 7.71E-02 | -1.09E+00 | 8.46E-03 | 3.23E-02 | -1.02E+00 | 2.55E-04 | 9.10E-04 | -1.57E+00 | 2.36E-03 | 6.66E-03 | -1.21E+00 | 1.28E-07 | 4.69E-07 | -1.19E+00 | 5.81E-04 | 1.60E-03 | -1.11E+00 |
| PARGP1 | 1.98E-04 | 1.03E-03 | -1.40E+00 | 1.13E-02 | 4.87E-02 | 1.01E+00 | 5.37E-05 | 2.28E-04 | -1.42E+00 | 7.65E-04 | 2.92E-03 | -1.11E+00 | 7.81E-03 | 2.79E-02 | -1.16E+00 | 1.72E-02 | 4.84E-02 | -1.05E+00 | 1.48E-03 | 5.42E-03 | -1.09E+00 | 3.00E-03 | 8.24E-03 | -1.05E+00 |
| LSM8 | 1.98E-04 | 1.03E-03 | -1.03E+00 | 5.14E-03 | 2.22E-02 | 1.12E+00 | 1.47E-02 | 6.24E-02 | -1.01E+00 | 6.18E-02 | 2.36E-01 | 1.11E+00 | 9.54E-05 | 3.40E-04 | -1.01E+00 | 2.50E-03 | 7.05E-03 | 1.05E+00 | 6.74E-02 | 2.47E-01 | 1.06E+00 | 2.42E-01 | 6.64E-01 | 1.08E+00 |
| ZNF70 | 1.98E-04 | 1.03E-03 | -1.28E+00 | 9.56E-04 | 4.13E-03 | -1.03E+00 | 9.49E-05 | 4.03E-04 | -1.38E+00 | 6.63E-03 | 2.53E-02 | 1.06E+00 | 3.86E-03 | 1.38E-02 | -1.42E+00 | 2.51E-01 | 7.08E-01 | 1.28E+00 | 4.09E-02 | 1.50E-01 | -1.12E+00 | 5.56E-02 | 1.53E-01 | -1.15E+00 |
| PHKG2 | 1.99E-04 | 1.04E-03 | 1.01E+00 | 1.67E-02 | 7.20E-02 | 1.30E+00 | 1.74E-02 | 7.38E-02 | 1.00E+00 | 1.00E-03 | 3.82E-03 | -1.05E+00 | 1.05E-03 | 3.75E-03 | -1.09E+00 | 1.24E-03 | 3.50E-03 | -1.06E+00 | 9.70E-02 | 3.56E-01 | 1.08E+00 | 1.06E-01 | 2.91E-01 | 1.05E+00 |
| TNFRSF4 | 2.00E-04 | 1.04E-03 | -1.93E+00 | 2.33E-02 | 1.01E-01 | -1.05E+00 | 5.58E-02 | 2.37E-01 | 1.19E+00 | 5.40E-02 | 2.06E-01 | 1.01E+00 | 7.22E-02 | 2.57E-01 | -1.32E+00 | 7.75E-02 | 2.19E-01 | -1.44E+00 | 2.45E-01 | 8.98E-01 | 1.03E+00 | 1.35E-01 | 3.72E-01 | 1.38E+00 |
| RNF146 | 2.02E-04 | 1.05E-03 | -1.30E+00 | 1.82E-04 | 7.85E-04 | -1.20E+00 | 2.32E-07 | 9.88E-07 | -1.61E+00 | 8.74E-06 | 3.34E-05 | -1.25E+00 | 1.50E-07 | 5.35E-07 | -1.69E+00 | 6.61E-07 | 1.86E-06 | -1.49E+00 | 4.46E-11 | 1.64E-10 | -1.74E+00 | 1.57E-07 | 4.32E-07 | -1.43E+00 |
| TRAPPC2B | 2.02E-04 | 1.05E-03 | -1.00E+00 | 3.23E-06 | 1.40E-05 | -1.01E+00 | 3.37E-02 | 1.43E-01 | -1.05E+00 | 3.50E-03 | 1.34E-02 | -1.02E+00 | 8.77E-03 | 3.13E-02 | 1.04E+00 | 4.01E-03 | 1.13E-02 | -1.02E+00 | 1.09E-02 | 4.01E-02 | 1.01E+00 | 1.42E-02 | 3.90E-02 | -1.10E+00 |
| TAPBPL | 2.02E-04 | 1.05E-03 | -1.20E+00 | 1.97E-02 | 8.49E-02 | 1.21E+00 | 2.69E-02 | 1.14E-01 | -1.03E+00 | 2.43E-02 | 9.29E-02 | 1.08E+00 | 1.66E-04 | 5.93E-04 | -1.34E+00 | 1.91E-03 | 5.37E-03 | -1.14E+00 | 3.08E-04 | 1.13E-03 | -1.40E+00 | 4.80E-03 | 1.32E-02 | -1.34E+00 |
| WDR37 | 2.02E-04 | 1.05E-03 | -1.16E+00 | 6.60E-04 | 2.85E-03 | 1.04E+00 | 2.24E-03 | 9.53E-03 | -1.23E+00 | 2.37E-03 | 9.05E-03 | -1.12E+00 | 3.22E-03 | 1.15E-02 | -1.43E+00 | 1.72E-01 | 4.84E-01 | 1.16E+00 | 7.89E-04 | 2.89E-03 | -1.28E+00 | 1.64E-02 | 4.52E-02 | -1.14E+00 |
| IGF2BP3 | 2.02E-04 | 1.06E-03 | -1.23E+00 | 3.88E-08 | 1.68E-07 | -2.08E+00 | 3.07E-03 | 1.30E-02 | -1.47E+00 | 3.82E-05 | 1.46E-04 | -1.86E+00 | 5.75E-02 | 2.05E-01 | -1.31E+00 | 2.98E-02 | 8.41E-02 | -1.65E+00 | 2.62E-01 | 9.61E-01 | 1.11E+00 | 2.21E-01 | 6.08E-01 | -1.16E+00 |
| CHPF | 2.03E-04 | 1.06E-03 | -1.68E+00 | 6.83E-03 | 2.95E-02 | -1.15E+00 | 1.44E-01 | 6.10E-01 | 1.05E+00 | 2.54E-01 | 9.69E-01 | 1.33E+00 | 4.63E-03 | 1.65E-02 | -1.80E+00 | 1.10E-01 | 3.10E-01 | 1.03E+00 | 2.00E-01 | 7.35E-01 | 1.37E+00 | 2.96E-01 | 8.14E-01 | 1.24E+00 |
| IFNAR1 | 2.03E-04 | 1.06E-03 | -1.05E+00 | 4.62E-05 | 2.00E-04 | -1.05E+00 | 7.53E-09 | 3.20E-08 | -1.44E+00 | 2.13E-11 | 8.13E-11 | -1.46E+00 | 1.57E-04 | 5.59E-04 | -1.33E+00 | 4.00E-05 | 1.13E-04 | -1.29E+00 | 1.71E-08 | 6.26E-08 | -1.39E+00 | 2.57E-12 | 7.06E-12 | -1.71E+00 |
| C9orf85 | 2.03E-04 | 1.06E-03 | -1.05E+00 | 3.48E-03 | 1.50E-02 | 1.12E+00 | 3.53E-05 | 1.50E-04 | -1.30E+00 | 7.70E-06 | 2.94E-05 | -1.20E+00 | 1.05E-03 | 3.74E-03 | -1.12E+00 | 1.05E-03 | 2.95E-03 | -1.09E+00 | 2.27E-05 | 8.34E-05 | -1.21E+00 | 6.56E-05 | 1.80E-04 | -1.23E+00 |
| AC004890.1 | 2.04E-04 | 1.07E-03 | 1.01E+00 | 2.02E-08 | 8.72E-08 | -1.16E+00 | 9.22E-02 | 3.92E-01 | 1.16E+00 | 4.17E-04 | 1.59E-03 | 1.07E+00 | 2.72E-04 | 9.69E-04 | -1.09E+00 | 5.02E-05 | 1.41E-04 | -1.15E+00 | 5.91E-04 | 2.17E-03 | -1.06E+00 | 5.66E-05 | 1.56E-04 | -1.19E+00 |
| ZNF124 | 2.05E-04 | 1.07E-03 | -1.24E+00 | 4.95E-03 | 2.14E-02 | 1.08E+00 | 1.44E-04 | 6.14E-04 | -1.26E+00 | 2.02E-02 | 7.70E-02 | 1.14E+00 | 1.42E-05 | 5.06E-05 | -1.27E+00 | 3.16E-03 | 8.91E-03 | 1.01E+00 | 9.31E-04 | 3.42E-03 | -1.21E+00 | 2.37E-02 | 6.51E-02 | 1.03E+00 |
| HS2ST1 | 2.05E-04 | 1.07E-03 | -1.37E+00 | 1.07E-02 | 4.62E-02 | 1.09E+00 | 9.89E-04 | 4.20E-03 | -1.17E+00 | 2.26E-04 | 8.63E-04 | -1.08E+00 | 1.45E-09 | 5.17E-09 | -1.54E+00 | 6.35E-08 | 1.79E-07 | -1.35E+00 | 1.44E-04 | 5.27E-04 | -1.13E+00 | 1.82E-01 | 5.01E-01 | 1.24E+00 |
| RBSN | 2.05E-04 | 1.07E-03 | -1.32E+00 | 2.60E-04 | 1.12E-03 | -1.18E+00 | 3.92E-04 | 1.67E-03 | -1.41E+00 | 1.24E-03 | 4.73E-03 | -1.23E+00 | 5.49E-05 | 1.96E-04 | -1.43E+00 | 1.87E-05 | 5.26E-05 | -1.44E+00 | 2.14E-05 | 7.85E-05 | -1.36E+00 | 2.09E-03 | 5.73E-03 | -1.14E+00 |
| P3H3 | 2.05E-04 | 1.07E-03 | -1.54E+00 | 4.49E-02 | 1.94E-01 | 1.03E+00 | 2.49E-03 | 1.06E-02 | -1.44E+00 | 1.79E-01 | 6.85E-01 | 1.22E+00 | 2.70E-02 | 9.63E-02 | -1.65E+00 | 1.84E-01 | 5.19E-01 | 1.01E+00 | 2.31E-01 | 8.48E-01 | -1.12E+00 | 3.34E-01 | 9.17E-01 | -1.03E+00 |
| DDB2 | 2.05E-04 | 1.07E-03 | -1.12E+00 | 3.49E-06 | 1.51E-05 | -1.10E+00 | 2.21E-02 | 9.39E-02 | 1.06E+00 | 5.62E-04 | 2.15E-03 | -1.03E+00 | 1.77E-02 | 6.32E-02 | -1.05E+00 | 4.27E-04 | 1.20E-03 | -1.41E+00 | 2.14E-05 | 7.85E-05 | -1.12E+00 | 5.27E-06 | 1.45E-05 | -1.32E+00 |
| ZFHX2 | 2.06E-04 | 1.07E-03 | -1.58E+00 | 1.77E-03 | 7.65E-03 | -1.20E+00 | 3.16E-07 | 1.34E-06 | -2.45E+00 | 3.16E-04 | 1.21E-03 | -1.36E+00 | 2.19E-02 | 7.82E-02 | -1.58E+00 | 9.11E-02 | 2.57E-01 | -1.10E+00 | 4.57E-02 | 1.68E-01 | -1.35E+00 | 1.60E-01 | 4.39E-01 | -1.16E+00 |
| AEN | 2.06E-04 | 1.07E-03 | -1.60E+00 | 1.25E-03 | 5.39E-03 | -1.22E+00 | 3.50E-09 | 1.49E-08 | -1.92E+00 | 7.89E-08 | 3.01E-07 | -1.48E+00 | 8.25E-02 | 2.94E-01 | 1.03E+00 | 1.38E-01 | 3.88E-01 | 1.06E+00 | 6.97E-08 | 2.56E-07 | -1.36E+00 | 8.54E-05 | 2.35E-04 | -1.20E+00 |
| NCF4 | 2.06E-04 | 1.07E-03 | 1.01E+00 | 3.06E-03 | 1.32E-02 | 1.20E+00 | 9.98E-02 | 4.24E-01 | -1.05E+00 | 3.79E-02 | 1.45E-01 | 1.10E+00 | 5.41E-02 | 1.93E-01 | 1.10E+00 | 3.52E-01 | 9.92E-01 | 1.31E+00 | 2.52E-02 | 9.27E-02 | 1.06E+00 | 1.45E-01 | 3.97E-01 | 1.10E+00 |
| AC018755.3 | 2.06E-04 | 1.07E-03 | -1.47E+00 | 1.39E-02 | 6.01E-02 | 1.08E+00 | 3.71E-09 | 1.58E-08 | -1.95E+00 | 2.30E-06 | 8.77E-06 | -1.35E+00 | 5.98E-06 | 2.13E-05 | -1.79E+00 | 5.81E-03 | 1.64E-02 | -1.05E+00 | 3.90E-03 | 1.43E-02 | -1.03E+00 | 6.29E-02 | 1.73E-01 | 1.11E+00 |
| FAM105A | 2.07E-04 | 1.08E-03 | -2.58E+00 | 1.43E-03 | 6.16E-03 | -1.90E+00 | 2.17E-04 | 9.24E-04 | -1.74E+00 | 8.20E-07 | 3.13E-06 | -2.12E+00 | 7.42E-04 | 2.65E-03 | -1.99E+00 | 6.71E-05 | 1.89E-04 | -2.82E+00 | 5.00E-02 | 1.83E-01 | -1.30E+00 | 1.24E-01 | 3.40E-01 | -1.16E+00 |
| POLR1D | 2.07E-04 | 1.08E-03 | -1.07E+00 | 5.64E-04 | 2.44E-03 | -1.04E+00 | 2.48E-03 | 1.05E-02 | -1.13E+00 | 3.71E-03 | 1.42E-02 | -1.11E+00 | 3.39E-03 | 1.21E-02 | -1.05E+00 | 9.14E-02 | 2.58E-01 | -1.00E+00 | 1.07E-03 | 3.93E-03 | -1.06E+00 | 1.91E-02 | 5.26E-02 | -1.08E+00 |
| AL365217.1 | 2.07E-04 | 1.08E-03 | -1.25E+00 | 8.06E-06 | 3.48E-05 | -1.40E+00 | 5.06E-02 | 2.15E-01 | -1.00E+00 | 6.03E-02 | 2.30E-01 | 1.14E+00 | 9.90E-04 | 3.53E-03 | -1.35E+00 | 5.13E-04 | 1.45E-03 | -1.48E+00 | 7.66E-05 | 2.81E-04 | -1.35E+00 | 5.99E-06 | 1.64E-05 | -1.61E+00 |
| ZNF418 | 2.07E-04 | 1.08E-03 | -1.53E+00 | 9.72E-03 | 4.20E-02 | -1.03E+00 | 7.12E-03 | 3.02E-02 | -1.37E+00 | 6.60E-02 | 2.52E-01 | 1.04E+00 | 1.42E-01 | 5.05E-01 | -1.01E+00 | 2.38E-01 | 6.70E-01 | 1.17E+00 | 2.04E-01 | 7.48E-01 | 1.39E+00 | 5.39E-02 | 1.48E-01 | 1.85E+00 |
| PSMD9 | 2.08E-04 | 1.08E-03 | -1.43E+00 | 5.53E-04 | 2.39E-03 | -1.27E+00 | 5.48E-04 | 2.33E-03 | -1.49E+00 | 7.25E-04 | 2.77E-03 | -1.26E+00 | 4.32E-06 | 1.54E-05 | -2.09E+00 | 1.91E-04 | 5.38E-04 | -1.57E+00 | 3.76E-02 | 1.38E-01 | -1.11E+00 | 9.71E-04 | 2.67E-03 | -1.43E+00 |
| KPNA4 | 2.08E-04 | 1.08E-03 | -1.03E+00 | 3.35E-11 | 1.45E-10 | -1.16E+00 | 5.39E-03 | 2.29E-02 | -1.00E+00 | 1.97E-06 | 7.51E-06 | -1.14E+00 | 6.24E-04 | 2.23E-03 | -1.15E+00 | 5.10E-05 | 1.44E-04 | -1.25E+00 | 4.84E-03 | 1.78E-02 | -1.03E+00 | 3.45E-04 | 9.47E-04 | -1.14E+00 |
| LCP1 | 2.08E-04 | 1.08E-03 | 1.01E+00 | 2.71E-02 | 1.17E-01 | 1.20E+00 | 2.29E-01 | 9.75E-01 | 1.10E+00 | 2.49E-01 | 9.53E-01 | 1.18E+00 | 3.64E-04 | 1.30E-03 | -1.08E+00 | 6.18E-05 | 1.74E-04 | -1.12E+00 | 2.65E-02 | 9.71E-02 | 1.06E+00 | 1.68E-01 | 4.60E-01 | 1.14E+00 |
| PSMC5 | 2.08E-04 | 1.08E-03 | 1.02E+00 | 7.55E-05 | 3.26E-04 | 1.15E+00 | 1.30E-01 | 5.51E-01 | 1.09E+00 | 1.66E-01 | 6.33E-01 | 1.20E+00 | 2.88E-05 | 1.03E-04 | -1.00E+00 | 5.62E-02 | 1.58E-01 | 1.18E+00 | 1.97E-01 | 7.24E-01 | 1.15E+00 | 1.33E-01 | 3.67E-01 | 1.21E+00 |
| SNHG1 | 2.09E-04 | 1.09E-03 | 1.02E+00 | 2.07E-09 | 8.95E-09 | -1.04E+00 | 3.55E-03 | 1.51E-02 | -1.20E+00 | 2.82E-04 | 1.08E-03 | -1.19E+00 | 4.79E-04 | 1.71E-03 | -1.10E+00 | 3.70E-03 | 1.04E-02 | -1.07E+00 | 4.47E-03 | 1.64E-02 | -1.04E+00 | 3.28E-03 | 9.01E-03 | -1.12E+00 |
| AL096870.2 | 2.10E-04 | 1.09E-03 | -1.54E+00 | 3.30E-03 | 1.43E-02 | -1.10E+00 | 1.12E-01 | 4.77E-01 | 1.01E+00 | 1.11E-01 | 4.24E-01 | 1.19E+00 | 1.14E-06 | 4.06E-06 | -1.94E+00 | 1.04E-05 | 2.92E-05 | -1.50E+00 | 1.42E-03 | 5.23E-03 | -1.52E+00 | 4.07E-03 | 1.12E-02 | -1.47E+00 |
| METTL22 | 2.10E-04 | 1.09E-03 | -1.23E+00 | 4.68E-03 | 2.02E-02 | 1.10E+00 | 1.06E-03 | 4.49E-03 | -1.19E+00 | 4.52E-04 | 1.73E-03 | -1.14E+00 | 2.75E-03 | 9.82E-03 | -1.40E+00 | 5.37E-02 | 1.51E-01 | -1.01E+00 | 1.28E-03 | 4.70E-03 | -1.18E+00 | 1.49E-03 | 4.09E-03 | -1.10E+00 |
| GAS2 | 2.10E-04 | 1.10E-03 | 2.48E+00 | 1.80E-20 | 7.77E-20 | 6.25E+00 | 2.19E-01 | 9.30E-01 | 1.29E+00 | 3.53E-02 | 1.35E-01 | 2.54E+00 | 9.01E-02 | 3.21E-01 | 2.79E+00 | 5.70E-02 | 1.61E-01 | 4.17E+00 | 1.25E-01 | 4.59E-01 | 1.93E+00 | 7.65E-02 | 2.10E-01 | 2.75E+00 |
| KDM3B | 2.11E-04 | 1.10E-03 | -1.18E+00 | 3.09E-04 | 1.34E-03 | -1.03E+00 | 3.99E-02 | 1.70E-01 | 1.02E+00 | 1.44E-02 | 5.50E-02 | 1.11E+00 | 2.46E-04 | 8.77E-04 | -1.12E+00 | 3.12E-02 | 8.80E-02 | 1.08E+00 | 2.39E-03 | 8.76E-03 | -1.07E+00 | 5.08E-02 | 1.40E-01 | 1.12E+00 |
| ZNF277 | 2.11E-04 | 1.10E-03 | 1.01E+00 | 1.88E-04 | 8.14E-04 | 1.09E+00 | 5.13E-07 | 2.18E-06 | -1.36E+00 | 2.73E-06 | 1.04E-05 | -1.17E+00 | 2.67E-05 | 9.52E-05 | -1.24E+00 | 1.17E-05 | 3.28E-05 | -1.15E+00 | 4.08E-09 | 1.50E-08 | -1.40E+00 | 8.60E-10 | 2.36E-09 | -1.47E+00 |
| CST7 | 2.11E-04 | 1.10E-03 | -1.48E+00 | 4.98E-05 | 2.15E-04 | -1.60E+00 | 1.42E-04 | 6.03E-04 | -1.67E+00 | 9.21E-03 | 3.52E-02 | -1.30E+00 | 4.73E-07 | 1.69E-06 | -1.78E+00 | 1.71E-07 | 4.81E-07 | -1.89E+00 | 2.48E-02 | 9.10E-02 | 2.19E+00 | 1.81E-01 | 4.97E-01 | 1.49E+00 |
| TTYH3 | 2.12E-04 | 1.11E-03 | -1.37E+00 | 1.21E-03 | 5.25E-03 | -1.06E+00 | 1.60E-03 | 6.80E-03 | -1.22E+00 | 4.23E-03 | 1.62E-02 | -1.00E+00 | 3.89E-02 | 1.39E-01 | -1.11E+00 | 7.35E-02 | 2.07E-01 | 1.01E+00 | 1.18E-04 | 4.33E-04 | -1.27E+00 | 2.62E-05 | 7.19E-05 | -1.59E+00 |
| ELP3 | 2.13E-04 | 1.11E-03 | -1.22E+00 | 9.98E-03 | 4.31E-02 | 1.11E+00 | 1.68E-05 | 7.15E-05 | -1.29E+00 | 2.48E-03 | 9.49E-03 | 1.07E+00 | 1.93E-04 | 6.89E-04 | -1.24E+00 | 1.40E-02 | 3.96E-02 | 1.05E+00 | 1.11E-05 | 4.06E-05 | -1.24E+00 | 2.97E-02 | 8.16E-02 | 1.02E+00 |
| B9D2 | 2.13E-04 | 1.11E-03 | -1.25E+00 | 1.10E-04 | 4.76E-04 | -1.16E+00 | 7.45E-05 | 3.17E-04 | -1.23E+00 | 5.12E-05 | 1.96E-04 | -1.19E+00 | 6.75E-03 | 2.41E-02 | -1.23E+00 | 3.13E-03 | 8.82E-03 | -1.31E+00 | 2.16E-02 | 7.92E-02 | -1.23E+00 | 2.88E-02 | 7.90E-02 | -1.16E+00 |
| RCC1 | 2.14E-04 | 1.11E-03 | -1.19E+00 | 5.12E-04 | 2.21E-03 | -1.00E+00 | 1.01E-04 | 4.28E-04 | -1.17E+00 | 3.56E-03 | 1.36E-02 | 1.10E+00 | 1.46E-05 | 5.19E-05 | -1.81E+00 | 3.03E-03 | 8.55E-03 | -1.29E+00 | 7.99E-07 | 2.93E-06 | -1.19E+00 | 7.61E-02 | 2.09E-01 | 1.24E+00 |
| TERF1P2 | 2.14E-04 | 1.12E-03 | -1.44E+00 | 2.33E-02 | 1.01E-01 | 1.12E+00 | 2.27E-05 | 9.66E-05 | -1.54E+00 | 2.68E-06 | 1.02E-05 | -1.54E+00 | 1.76E-02 | 6.27E-02 | -1.40E+00 | 1.25E-01 | 3.54E-01 | 1.03E+00 | 2.87E-03 | 1.05E-02 | -1.15E+00 | 1.82E-01 | 5.01E-01 | 1.13E+00 |
| DLK1 | 2.14E-04 | 1.12E-03 | 1.57E+00 | 8.56E-06 | 3.70E-05 | 1.85E+00 | 1.26E-02 | 5.33E-02 | 2.18E+00 | 6.18E-09 | 2.36E-08 | 3.28E+00 | 9.23E-08 | 3.29E-07 | 6.14E+00 | 8.26E-07 | 2.33E-06 | 5.32E+00 | 3.12E-04 | 1.15E-03 | 7.88E+00 | 3.71E-06 | 1.02E-05 | 8.98E+00 |
| STAM2 | 2.14E-04 | 1.12E-03 | -1.17E+00 | 1.02E-02 | 4.42E-02 | 1.13E+00 | 9.94E-04 | 4.22E-03 | -1.20E+00 | 2.06E-02 | 7.87E-02 | 1.13E+00 | 3.36E-06 | 1.20E-05 | -1.49E+00 | 3.19E-03 | 8.99E-03 | -1.03E+00 | 2.76E-04 | 1.01E-03 | -1.36E+00 | 1.16E-02 | 3.20E-02 | -1.07E+00 |
| COX10 | 2.14E-04 | 1.12E-03 | -1.31E+00 | 2.70E-03 | 1.17E-02 | 1.02E+00 | 2.61E-02 | 1.11E-01 | 1.00E+00 | 2.31E-02 | 8.82E-02 | 1.07E+00 | 2.13E-04 | 7.61E-04 | -1.51E+00 | 1.43E-03 | 4.04E-03 | -1.23E+00 | 1.66E-02 | 6.10E-02 | 1.04E+00 | 1.69E-01 | 4.65E-01 | 1.21E+00 |
| IFT122 | 2.15E-04 | 1.12E-03 | -1.25E+00 | 5.00E-03 | 2.16E-02 | 1.07E+00 | 4.45E-04 | 1.89E-03 | -1.22E+00 | 7.74E-04 | 2.96E-03 | -1.08E+00 | 5.42E-03 | 1.93E-02 | -1.23E+00 | 3.19E-02 | 9.00E-02 | -1.02E+00 | 2.02E-01 | 7.43E-01 | 1.22E+00 | 1.46E-01 | 4.00E-01 | 1.02E+00 |
| ZNF417 | 2.16E-04 | 1.12E-03 | -1.44E+00 | 6.49E-04 | 2.80E-03 | -1.31E+00 | 1.39E-02 | 5.91E-02 | -1.26E+00 | 2.55E-02 | 9.73E-02 | -1.06E+00 | 7.74E-02 | 2.76E-01 | -1.12E+00 | 1.45E-01 | 4.09E-01 | 1.02E+00 | 1.35E-05 | 4.97E-05 | -1.73E+00 | 1.09E-03 | 2.99E-03 | -1.38E+00 |
| TOR4A | 2.16E-04 | 1.13E-03 | -1.51E+00 | 7.13E-03 | 3.08E-02 | -1.04E+00 | 5.42E-04 | 2.30E-03 | -1.43E+00 | 2.88E-04 | 1.10E-03 | -1.33E+00 | 1.12E-11 | 4.01E-11 | -1.92E+00 | 1.95E-09 | 5.50E-09 | -1.65E+00 | 1.24E-08 | 4.56E-08 | -1.25E+00 | 7.21E-09 | 1.98E-08 | -1.27E+00 |
| FAM21FP | 2.16E-04 | 1.13E-03 | -1.36E+00 | 1.15E-03 | 4.96E-03 | -1.11E+00 | 1.12E-03 | 4.76E-03 | -1.57E+00 | 2.39E-03 | 9.14E-03 | -1.29E+00 | 2.57E-02 | 9.16E-02 | -1.03E+00 | 7.90E-03 | 2.23E-02 | -1.23E+00 | 7.01E-02 | 2.57E-01 | -1.07E+00 | 1.51E-01 | 4.15E-01 | -1.02E+00 |
| SARAF | 2.17E-04 | 1.13E-03 | -1.07E+00 | 2.07E-02 | 8.92E-02 | 1.02E+00 | 2.48E-02 | 1.05E-01 | -1.09E+00 | 3.28E-02 | 1.25E-01 | -1.05E+00 | 1.22E-01 | 4.36E-01 | 1.05E+00 | 1.64E-01 | 4.62E-01 | 1.01E+00 | 4.09E-02 | 1.50E-01 | 1.01E+00 | 1.21E-02 | 3.32E-02 | -1.05E+00 |
| FANCL | 2.17E-04 | 1.13E-03 | -1.30E+00 | 5.73E-03 | 2.48E-02 | 1.06E+00 | 1.50E-06 | 6.38E-06 | -1.43E+00 | 2.63E-04 | 1.00E-03 | -1.06E+00 | 4.37E-04 | 1.56E-03 | -1.37E+00 | 4.28E-03 | 1.21E-02 | -1.15E+00 | 2.11E-11 | 7.76E-11 | -1.78E+00 | 7.69E-06 | 2.11E-05 | -1.36E+00 |
| NAXE | 2.17E-04 | 1.13E-03 | -1.05E+00 | 1.55E-03 | 6.69E-03 | 1.09E+00 | 5.10E-04 | 2.17E-03 | -1.12E+00 | 2.35E-05 | 8.98E-05 | -1.05E+00 | 4.58E-08 | 1.63E-07 | -1.24E+00 | 6.68E-04 | 1.88E-03 | 1.04E+00 | 2.19E-01 | 8.05E-01 | 1.15E+00 | 6.82E-02 | 1.87E-01 | 1.39E+00 |
| ZFP3 | 2.17E-04 | 1.13E-03 | -1.54E+00 | 1.05E-03 | 4.55E-03 | -1.26E+00 | 6.90E-02 | 2.93E-01 | -1.12E+00 | 8.88E-02 | 3.39E-01 | 1.02E+00 | 3.80E-03 | 1.36E-02 | -1.31E+00 | 9.59E-03 | 2.70E-02 | -1.23E+00 | 6.73E-02 | 2.47E-01 | -1.11E+00 | 2.25E-01 | 6.17E-01 | 1.09E+00 |
| TTLL12 | 2.17E-04 | 1.13E-03 | -1.45E+00 | 8.04E-03 | 3.48E-02 | 1.02E+00 | 2.61E-03 | 1.11E-02 | -1.24E+00 | 1.61E-01 | 6.14E-01 | 1.24E+00 | 9.79E-05 | 3.49E-04 | -1.50E+00 | 2.60E-01 | 7.32E-01 | 1.38E+00 | 9.76E-09 | 3.58E-08 | -1.33E+00 | 3.02E-01 | 8.29E-01 | 1.29E+00 |
| RHOU | 2.17E-04 | 1.13E-03 | -1.55E+00 | 3.86E-06 | 1.67E-05 | -2.23E+00 | 4.21E-16 | 1.79E-15 | -2.27E+00 | 6.23E-15 | 2.38E-14 | -1.92E+00 | 4.30E-04 | 1.53E-03 | -1.93E+00 | 7.45E-05 | 2.10E-04 | -2.52E+00 | 1.07E-02 | 3.92E-02 | -1.43E+00 | 4.23E-02 | 1.16E-01 | -1.39E+00 |
| AL512637.1 | 2.17E-04 | 1.13E-03 | -1.79E+00 | 2.18E-04 | 9.41E-04 | -1.77E+00 | 5.11E-02 | 2.17E-01 | 1.02E+00 | 3.25E-04 | 1.24E-03 | -1.47E+00 | 2.61E-01 | 9.31E-01 | 1.19E+00 | 3.36E-01 | 9.46E-01 | 1.25E+00 | 1.89E-03 | 6.95E-03 | -1.15E+00 | 7.04E-05 | 1.93E-04 | -1.33E+00 |
| APAF1 | 2.17E-04 | 1.13E-03 | -1.50E+00 | 1.39E-02 | 5.99E-02 | -1.02E+00 | 5.45E-07 | 2.32E-06 | -1.72E+00 | 8.81E-06 | 3.37E-05 | -1.37E+00 | 1.23E-01 | 4.37E-01 | 1.11E+00 | 5.17E-02 | 1.46E-01 | -1.09E+00 | 6.16E-05 | 2.26E-04 | -1.31E+00 | 2.34E-06 | 6.43E-06 | -1.58E+00 |
| PISD | 2.17E-04 | 1.13E-03 | -1.13E+00 | 3.26E-03 | 1.41E-02 | 1.11E+00 | 6.35E-06 | 2.70E-05 | -1.37E+00 | 3.91E-04 | 1.49E-03 | -1.08E+00 | 1.95E-05 | 6.95E-05 | -1.37E+00 | 4.78E-04 | 1.35E-03 | -1.13E+00 | 8.53E-08 | 3.13E-07 | -1.41E+00 | 3.12E-02 | 8.56E-02 | 1.04E+00 |
| E2F6 | 2.17E-04 | 1.13E-03 | -1.55E+00 | 2.48E-03 | 1.07E-02 | -1.16E+00 | 1.16E-04 | 4.92E-04 | -1.47E+00 | 9.30E-04 | 3.55E-03 | -1.17E+00 | 7.29E-07 | 2.60E-06 | -1.77E+00 | 9.71E-07 | 2.74E-06 | -1.74E+00 | 1.80E-09 | 6.61E-09 | -1.68E+00 | 1.31E-04 | 3.60E-04 | -1.15E+00 |
| GPR108 | 2.18E-04 | 1.13E-03 | -1.10E+00 | 4.25E-03 | 1.84E-02 | 1.12E+00 | 1.85E-03 | 7.84E-03 | -1.13E+00 | 4.39E-03 | 1.68E-02 | 1.03E+00 | 2.55E-01 | 9.09E-01 | 1.28E+00 | 1.77E-01 | 5.00E-01 | 1.24E+00 | 1.17E-01 | 4.31E-01 | 1.21E+00 | 2.79E-01 | 7.67E-01 | 1.22E+00 |
| PDLIM1 | 2.18E-04 | 1.13E-03 | -1.09E+00 | 1.43E-01 | 6.18E-01 | 1.13E+00 | 5.24E-03 | 2.23E-02 | -1.18E+00 | 5.96E-02 | 2.28E-01 | 1.01E+00 | 2.52E-02 | 8.99E-02 | -1.04E+00 | 1.91E-01 | 5.39E-01 | 1.16E+00 | 9.01E-05 | 3.31E-04 | 1.43E+00 | 1.37E-03 | 3.76E-03 | 1.30E+00 |
| COG6 | 2.18E-04 | 1.14E-03 | -1.33E+00 | 6.59E-03 | 2.85E-02 | 1.04E+00 | 2.25E-08 | 9.58E-08 | -1.85E+00 | 2.53E-06 | 9.68E-06 | -1.35E+00 | 1.31E-02 | 4.68E-02 | -1.08E+00 | 2.76E-01 | 7.78E-01 | 1.31E+00 | 2.13E-06 | 7.80E-06 | -1.16E+00 | 3.88E-05 | 1.06E-04 | -1.11E+00 |
| ATPAF2 | 2.19E-04 | 1.14E-03 | -1.08E+00 | 1.31E-02 | 5.66E-02 | 1.24E+00 | 4.30E-05 | 1.83E-04 | -1.22E+00 | 6.53E-05 | 2.50E-04 | -1.09E+00 | 1.35E-04 | 4.80E-04 | -1.36E+00 | 3.93E-02 | 1.11E-01 | 1.09E+00 | 5.94E-03 | 2.18E-02 | -1.03E+00 | 4.71E-02 | 1.29E-01 | 1.07E+00 |
| IFT22 | 2.20E-04 | 1.14E-03 | -1.14E+00 | 4.79E-06 | 2.07E-05 | -1.20E+00 | 1.99E-01 | 8.47E-01 | 1.13E+00 | 1.69E-03 | 6.46E-03 | -1.09E+00 | 2.32E-05 | 8.29E-05 | -1.38E+00 | 2.85E-05 | 8.04E-05 | -1.35E+00 | 2.19E-01 | 8.03E-01 | 1.33E+00 | 3.42E-01 | 9.40E-01 | 1.30E+00 |
| LINC02265 | 2.20E-04 | 1.15E-03 | 2.54E+00 | 1.37E-01 | 5.92E-01 | 1.19E+00 | 2.36E-02 | 1.00E-01 | 1.82E+00 | 6.06E-02 | 2.31E-01 | -1.11E+00 | 1.74E-01 | 6.22E-01 | 1.41E+00 | 3.27E-01 | 9.22E-01 | -1.11E+00 | 2.37E-01 | 8.71E-01 | 1.20E+00 | 2.85E-01 | 7.82E-01 | -1.15E+00 |
| AP001442.1 | 2.20E-04 | 1.15E-03 | -1.51E+00 | 1.60E-03 | 6.90E-03 | -1.17E+00 | 4.68E-07 | 1.99E-06 | -1.80E+00 | 1.64E-06 | 6.25E-06 | -1.52E+00 | 2.13E-01 | 7.59E-01 | -1.21E+00 | 2.73E-01 | 7.70E-01 | -1.19E+00 | 5.58E-02 | 2.05E-01 | -1.61E+00 | 1.80E-01 | 4.94E-01 | -1.17E+00 |
| GFOD2 | 2.21E-04 | 1.15E-03 | -1.35E+00 | 6.58E-02 | 2.84E-01 | 1.30E+00 | 2.33E-07 | 9.91E-07 | -1.59E+00 | 1.30E-03 | 4.97E-03 | -1.02E+00 | 2.22E-05 | 7.93E-05 | -1.66E+00 | 4.82E-02 | 1.36E-01 | 1.07E+00 | 1.47E-07 | 5.41E-07 | -1.40E+00 | 6.17E-03 | 1.69E-02 | 1.03E+00 |
| DLGAP1-AS2 | 2.21E-04 | 1.15E-03 | -1.83E+00 | 6.57E-05 | 2.84E-04 | -1.86E+00 | 7.12E-02 | 3.02E-01 | -1.11E+00 | 6.94E-02 | 2.65E-01 | -1.02E+00 | 2.43E-01 | 8.66E-01 | 1.35E+00 | 2.51E-01 | 7.08E-01 | 1.23E+00 | 1.28E-01 | 4.70E-01 | 1.36E+00 | 3.24E-01 | 8.89E-01 | 1.23E+00 |
| ZNF503 | 2.22E-04 | 1.16E-03 | -1.21E+00 | 8.44E-04 | 3.65E-03 | 1.02E+00 | 9.54E-08 | 4.05E-07 | -1.52E+00 | 4.42E-06 | 1.69E-05 | -1.27E+00 | 2.07E-02 | 7.37E-02 | -1.11E+00 | 9.59E-03 | 2.70E-02 | -1.28E+00 | 2.90E-04 | 1.06E-03 | -2.35E+00 | 6.67E-04 | 1.83E-03 | -2.30E+00 |
| Z98885.3 | 2.22E-04 | 1.16E-03 | -1.23E+00 | 1.29E-05 | 5.57E-05 | -1.31E+00 | 8.94E-03 | 3.80E-02 | -1.17E+00 | 9.09E-06 | 3.47E-05 | -1.66E+00 | 6.82E-03 | 2.43E-02 | -1.24E+00 | 3.04E-04 | 8.58E-04 | -1.67E+00 | 1.49E-04 | 5.47E-04 | -1.43E+00 | 3.12E-06 | 8.57E-06 | -1.87E+00 |
| AVP | 2.23E-04 | 1.16E-03 | -1.19E+00 | 1.16E-01 | 5.01E-01 | 1.07E+00 | 9.84E-04 | 4.18E-03 | 1.09E+00 | 2.30E-01 | 8.77E-01 | 1.38E+00 | 5.43E-02 | 1.93E-01 | -1.21E+00 | 1.14E-01 | 3.20E-01 | 1.05E+00 | 2.70E-01 | 9.92E-01 | 1.55E+00 | 1.88E-01 | 5.16E-01 | 2.73E+00 |
| NRIP3 | 2.23E-04 | 1.16E-03 | -1.29E+00 | 3.92E-04 | 1.69E-03 | -1.17E+00 | 1.33E-04 | 5.65E-04 | -1.27E+00 | 2.42E-06 | 9.25E-06 | -1.43E+00 | 1.67E-02 | 5.95E-02 | -1.43E+00 | 1.91E-02 | 5.37E-02 | -1.63E+00 | 1.52E-02 | 5.59E-02 | -1.51E+00 | 1.38E-01 | 3.79E-01 | -1.20E+00 |
| BRCC3P1 | 2.23E-04 | 1.16E-03 | -1.62E+00 | 8.92E-04 | 3.86E-03 | -1.35E+00 | 7.23E-06 | 3.07E-05 | -1.83E+00 | 7.81E-04 | 2.98E-03 | -1.16E+00 | 3.35E-03 | 1.19E-02 | -1.58E+00 | 3.68E-03 | 1.04E-02 | -1.51E+00 | 3.63E-03 | 1.33E-02 | -1.28E+00 | 1.34E-02 | 3.67E-02 | -1.20E+00 |
| TMEM229B | 2.23E-04 | 1.16E-03 | -1.90E+00 | 1.04E-02 | 4.50E-02 | -1.21E+00 | 8.63E-05 | 3.67E-04 | -1.60E+00 | 1.50E-02 | 5.72E-02 | -1.02E+00 | 1.77E-04 | 6.31E-04 | -2.30E+00 | 1.03E-03 | 2.91E-03 | -1.72E+00 | 3.66E-03 | 1.34E-02 | -1.54E+00 | 4.81E-02 | 1.32E-01 | -1.16E+00 |
| HIST1H2BD | 2.24E-04 | 1.17E-03 | -1.38E+00 | 1.12E-03 | 4.82E-03 | -1.18E+00 | 1.37E-04 | 5.82E-04 | -1.83E+00 | 1.30E-03 | 4.95E-03 | -1.31E+00 | 4.38E-04 | 1.56E-03 | -2.10E+00 | 3.06E-02 | 8.64E-02 | -1.14E+00 | 1.38E-06 | 5.06E-06 | -1.62E+00 | 4.83E-07 | 1.33E-06 | -1.72E+00 |
| ZFHX2-AS1 | 2.25E-04 | 1.17E-03 | -1.18E+00 | 7.09E-02 | 3.06E-01 | 1.35E+00 | 2.92E-03 | 1.24E-02 | -1.32E+00 | 7.69E-02 | 2.94E-01 | 1.17E+00 | 2.66E-01 | 9.50E-01 | 1.58E+00 | 1.06E-01 | 3.00E-01 | 2.03E+00 | 1.88E-01 | 6.91E-01 | 1.16E+00 | 1.52E-01 | 4.16E-01 | 1.58E+00 |
| ATP5G1P5 | 2.25E-04 | 1.17E-03 | -1.52E+00 | 2.87E-05 | 1.24E-04 | -1.73E+00 | 1.51E-02 | 6.40E-02 | -1.09E+00 | 1.17E-05 | 4.48E-05 | -1.59E+00 | 1.73E-05 | 6.17E-05 | -1.78E+00 | 9.27E-07 | 2.61E-06 | -2.39E+00 | 3.98E-02 | 1.46E-01 | 1.05E+00 | 7.22E-03 | 1.98E-02 | -1.06E+00 |
| AC010894.1 | 2.25E-04 | 1.17E-03 | -1.76E+00 | 9.77E-05 | 4.22E-04 | -1.81E+00 | 1.42E-04 | 6.02E-04 | -1.38E+00 | 2.54E-07 | 9.71E-07 | -1.59E+00 | 7.53E-03 | 2.69E-02 | -2.00E+00 | 8.38E-02 | 2.36E-01 | -1.18E+00 | 7.81E-12 | 2.86E-11 | -1.80E+00 | 4.91E-10 | 1.35E-09 | -1.62E+00 |
| NAF1 | 2.27E-04 | 1.18E-03 | -1.04E+00 | 7.68E-10 | 3.32E-09 | -1.37E+00 | 8.96E-03 | 3.81E-02 | -1.03E+00 | 4.33E-04 | 1.65E-03 | -1.04E+00 | 1.27E-02 | 4.51E-02 | -1.10E+00 | 5.92E-03 | 1.67E-02 | -1.17E+00 | 8.86E-03 | 3.25E-02 | 1.04E+00 | 3.67E-03 | 1.01E-02 | -1.03E+00 |
| GPR137B | 2.28E-04 | 1.19E-03 | -1.50E+00 | 7.46E-05 | 3.22E-04 | -1.41E+00 | 4.43E-02 | 1.88E-01 | -1.12E+00 | 1.52E-02 | 5.79E-02 | -1.06E+00 | 6.01E-02 | 2.14E-01 | -1.11E+00 | 7.02E-02 | 1.98E-01 | -1.13E+00 | 3.91E-04 | 1.44E-03 | -1.52E+00 | 5.88E-03 | 1.61E-02 | -1.24E+00 |
| DTX3 | 2.28E-04 | 1.19E-03 | -1.07E+00 | 8.51E-05 | 3.68E-04 | -1.05E+00 | 1.23E-01 | 5.23E-01 | 1.10E+00 | 1.85E-02 | 7.08E-02 | 1.02E+00 | 5.16E-02 | 1.84E-01 | 1.10E+00 | 4.33E-02 | 1.22E-01 | 1.14E+00 | 8.24E-03 | 3.03E-02 | -1.14E+00 | 1.08E-02 | 2.97E-02 | -1.20E+00 |
| TRIM13 | 2.29E-04 | 1.19E-03 | -1.09E+00 | 2.17E-02 | 9.38E-02 | 1.25E+00 | 1.82E-04 | 7.74E-04 | -1.26E+00 | 3.27E-03 | 1.25E-02 | 1.03E+00 | 9.53E-02 | 3.40E-01 | 1.09E+00 | 1.23E-01 | 3.47E-01 | 1.15E+00 | 1.51E-02 | 5.54E-02 | 1.09E+00 | 6.98E-02 | 1.92E-01 | 1.19E+00 |
| AC008993.2 | 2.30E-04 | 1.20E-03 | -1.08E+00 | 1.39E-01 | 6.02E-01 | 1.45E+00 | 1.32E-02 | 5.62E-02 | -1.06E+00 | 2.53E-01 | 9.68E-01 | 1.49E+00 | 1.15E-02 | 4.10E-02 | 1.09E+00 | 2.67E-01 | 7.54E-01 | 1.39E+00 | 4.63E-03 | 1.70E-02 | 1.05E+00 | 8.54E-02 | 2.34E-01 | 1.23E+00 |
| AC006064.2 | 2.30E-04 | 1.20E-03 | -1.54E+00 | 9.95E-06 | 4.30E-05 | -1.98E+00 | 3.00E-03 | 1.27E-02 | -1.45E+00 | 3.42E-03 | 1.31E-02 | -1.43E+00 | 2.27E-01 | 8.08E-01 | 1.15E+00 | 2.82E-01 | 7.94E-01 | 1.55E+00 | 2.34E-02 | 8.59E-02 | -1.35E+00 | 3.92E-04 | 1.08E-03 | -2.25E+00 |
| ZNF100 | 2.30E-04 | 1.20E-03 | -1.11E+00 | 7.30E-05 | 3.16E-04 | -1.05E+00 | 2.21E-03 | 9.40E-03 | -1.11E+00 | 4.03E-05 | 1.54E-04 | -1.29E+00 | 9.45E-02 | 3.37E-01 | 1.11E+00 | 3.42E-02 | 9.65E-02 | -1.00E+00 | 4.56E-03 | 1.67E-02 | -1.06E+00 | 6.01E-03 | 1.65E-02 | -1.08E+00 |
| CDC14A | 2.31E-04 | 1.20E-03 | -1.15E+00 | 3.42E-04 | 1.48E-03 | -1.02E+00 | 8.25E-05 | 3.51E-04 | -1.23E+00 | 2.46E-06 | 9.39E-06 | -1.23E+00 | 9.44E-05 | 3.36E-04 | -1.28E+00 | 1.20E-04 | 3.40E-04 | -1.22E+00 | 4.25E-03 | 1.56E-02 | -1.02E+00 | 9.94E-05 | 2.73E-04 | -1.24E+00 |
| TBC1D17 | 2.31E-04 | 1.21E-03 | -1.22E+00 | 2.70E-04 | 1.17E-03 | -1.11E+00 | 2.04E-04 | 8.69E-04 | -1.25E+00 | 8.45E-05 | 3.23E-04 | -1.18E+00 | 1.79E-02 | 6.37E-02 | -1.20E+00 | 1.18E-01 | 3.33E-01 | 1.16E+00 | 6.42E-04 | 2.35E-03 | -1.50E+00 | 6.59E-04 | 1.81E-03 | -1.53E+00 |
| NUDCD2 | 2.32E-04 | 1.21E-03 | 1.03E+00 | 5.78E-02 | 2.50E-01 | 1.32E+00 | 1.52E-02 | 6.46E-02 | 1.04E+00 | 1.27E-01 | 4.83E-01 | 1.28E+00 | 2.60E-03 | 9.28E-03 | -1.01E+00 | 2.86E-01 | 8.06E-01 | 1.33E+00 | 1.29E-02 | 4.74E-02 | 1.01E+00 | 1.77E-01 | 4.88E-01 | 1.12E+00 |
| GTF2IP12 | 2.32E-04 | 1.21E-03 | -1.37E+00 | 1.64E-04 | 7.11E-04 | -1.31E+00 | 1.23E-04 | 5.21E-04 | -1.49E+00 | 8.85E-05 | 3.38E-04 | -1.42E+00 | 5.10E-02 | 1.82E-01 | -1.03E+00 | 8.95E-02 | 2.52E-01 | 1.04E+00 | 1.13E-01 | 4.14E-01 | 1.11E+00 | 1.47E-02 | 4.05E-02 | -1.30E+00 |
| ABCD3 | 2.33E-04 | 1.21E-03 | -1.14E+00 | 2.27E-04 | 9.83E-04 | -1.03E+00 | 4.31E-04 | 1.83E-03 | -1.23E+00 | 5.14E-07 | 1.97E-06 | -1.37E+00 | 2.99E-03 | 1.07E-02 | -1.10E+00 | 3.53E-04 | 9.94E-04 | -1.20E+00 | 7.11E-08 | 2.61E-07 | -1.47E+00 | 3.49E-05 | 9.58E-05 | -1.21E+00 |
| POP5 | 2.33E-04 | 1.21E-03 | -1.04E+00 | 4.29E-04 | 1.85E-03 | 1.12E+00 | 4.82E-03 | 2.05E-02 | -1.08E+00 | 7.84E-03 | 2.99E-02 | 1.07E+00 | 1.09E-05 | 3.87E-05 | -1.14E+00 | 4.54E-03 | 1.28E-02 | 1.07E+00 | 3.14E-04 | 1.15E-03 | -1.06E+00 | 6.13E-03 | 1.68E-02 | 1.05E+00 |
| RBBP8 | 2.33E-04 | 1.21E-03 | -1.55E+00 | 1.29E-03 | 5.56E-03 | -1.21E+00 | 1.38E-06 | 5.85E-06 | -1.44E+00 | 4.19E-05 | 1.60E-04 | -1.13E+00 | 9.51E-07 | 3.39E-06 | -2.02E+00 | 1.70E-04 | 4.79E-04 | -1.36E+00 | 1.78E-16 | 6.54E-16 | -1.91E+00 | 2.01E-08 | 5.53E-08 | -1.30E+00 |
| TARBP2 | 2.33E-04 | 1.22E-03 | -1.37E+00 | 5.31E-04 | 2.29E-03 | -1.18E+00 | 2.24E-05 | 9.53E-05 | -1.39E+00 | 5.08E-04 | 1.94E-03 | -1.13E+00 | 8.49E-10 | 3.03E-09 | -2.32E+00 | 4.32E-05 | 1.22E-04 | -1.30E+00 | 4.37E-04 | 1.60E-03 | -1.15E+00 | 2.31E-01 | 6.33E-01 | 1.33E+00 |
| DENND6A | 2.34E-04 | 1.22E-03 | -1.06E+00 | 3.00E-08 | 1.30E-07 | -1.23E+00 | 5.99E-05 | 2.54E-04 | -1.13E+00 | 1.44E-06 | 5.50E-06 | -1.14E+00 | 3.75E-04 | 1.34E-03 | -1.11E+00 | 2.22E-05 | 6.25E-05 | -1.37E+00 | 3.31E-07 | 1.21E-06 | -1.37E+00 | 1.60E-03 | 4.40E-03 | -1.13E+00 |
| LEKR1 | 2.35E-04 | 1.23E-03 | 1.84E+00 | 2.14E-03 | 9.26E-03 | -1.73E+00 | 2.80E-04 | 1.19E-03 | 2.65E+00 | 1.74E-01 | 6.66E-01 | 1.42E+00 | 6.42E-02 | 2.29E-01 | 1.97E+00 | 3.19E-01 | 9.00E-01 | 1.18E+00 | 1.13E-01 | 4.15E-01 | 1.35E+00 | 1.62E-01 | 4.45E-01 | -1.22E+00 |
| FDX1 | 2.36E-04 | 1.23E-03 | -1.01E+00 | 2.81E-06 | 1.21E-05 | -1.03E+00 | 2.69E-02 | 1.14E-01 | 1.07E+00 | 1.55E-06 | 5.93E-06 | -1.13E+00 | 5.95E-02 | 2.12E-01 | 1.27E+00 | 2.14E-02 | 6.02E-02 | 1.13E+00 | 1.37E-03 | 5.04E-03 | -1.09E+00 | 6.24E-03 | 1.72E-02 | -1.05E+00 |
| HNRNPLL | 2.36E-04 | 1.23E-03 | -1.11E+00 | 1.78E-02 | 7.69E-02 | 1.25E+00 | 4.67E-03 | 1.98E-02 | -1.15E+00 | 1.61E-02 | 6.14E-02 | 1.08E+00 | 3.73E-03 | 1.33E-02 | 1.12E+00 | 4.83E-02 | 1.36E-01 | 1.23E+00 | 5.20E-08 | 1.91E-07 | -1.30E+00 | 5.31E-04 | 1.46E-03 | -1.04E+00 |
| PARP6 | 2.36E-04 | 1.23E-03 | -1.21E+00 | 1.03E-05 | 4.43E-05 | -1.25E+00 | 2.78E-05 | 1.18E-04 | -1.33E+00 | 6.72E-05 | 2.57E-04 | -1.17E+00 | 4.93E-02 | 1.76E-01 | 1.15E+00 | 1.48E-02 | 4.16E-02 | -1.05E+00 | 1.28E-01 | 4.69E-01 | 1.23E+00 | 3.68E-02 | 1.01E-01 | -1.06E+00 |
| DTWD2 | 2.38E-04 | 1.24E-03 | -1.43E+00 | 5.73E-04 | 2.48E-03 | -1.29E+00 | 1.72E-03 | 7.30E-03 | -1.46E+00 | 5.26E-02 | 2.01E-01 | 1.10E+00 | 4.74E-03 | 1.69E-02 | 1.03E+00 | 2.51E-03 | 7.08E-03 | -1.03E+00 | 2.67E-01 | 9.81E-01 | 1.46E+00 | 3.07E-01 | 8.45E-01 | 1.44E+00 |
| GNL1 | 2.38E-04 | 1.24E-03 | 1.05E+00 | 6.57E-13 | 2.84E-12 | -1.41E+00 | 1.10E-03 | 4.68E-03 | -1.08E+00 | 1.10E-09 | 4.19E-09 | -1.42E+00 | 1.63E-03 | 5.79E-03 | 1.05E+00 | 7.69E-06 | 2.17E-05 | -1.28E+00 | 3.94E-06 | 1.45E-05 | -1.25E+00 | 2.72E-06 | 7.46E-06 | -1.31E+00 |
| MITD1 | 2.38E-04 | 1.24E-03 | -1.06E+00 | 1.41E-02 | 6.09E-02 | 1.23E+00 | 1.35E-07 | 5.74E-07 | -1.35E+00 | 1.07E-07 | 4.10E-07 | -1.18E+00 | 5.94E-08 | 2.12E-07 | -1.39E+00 | 3.27E-03 | 9.22E-03 | 1.06E+00 | 1.87E-02 | 6.88E-02 | -1.06E+00 | 9.96E-02 | 2.74E-01 | 1.06E+00 |
| ATP5G1 | 2.39E-04 | 1.25E-03 | 1.06E+00 | 5.12E-07 | 2.21E-06 | 1.07E+00 | 1.47E-02 | 6.23E-02 | 1.36E+00 | 7.17E-02 | 2.74E-01 | 1.34E+00 | 8.57E-05 | 3.06E-04 | -1.08E+00 | 2.50E-02 | 7.04E-02 | 1.10E+00 | 1.11E-01 | 4.09E-01 | 1.12E+00 | 7.01E-02 | 1.93E-01 | 1.32E+00 |
| ANKS1A | 2.39E-04 | 1.25E-03 | -1.29E+00 | 3.23E-03 | 1.40E-02 | -1.02E+00 | 3.47E-03 | 1.47E-02 | -1.22E+00 | 9.41E-02 | 3.59E-01 | 1.20E+00 | 1.28E-03 | 4.58E-03 | -1.30E+00 | 2.54E-02 | 7.15E-02 | -1.02E+00 | 8.57E-03 | 3.15E-02 | -1.03E+00 | 6.59E-02 | 1.81E-01 | 1.04E+00 |
| RPL7P32 | 2.39E-04 | 1.25E-03 | -1.08E+00 | 4.68E-04 | 2.02E-03 | 2.12E+00 | 7.55E-03 | 3.21E-02 | -1.09E+00 | 1.81E-06 | 6.90E-06 | 2.31E+00 | 2.12E-02 | 7.57E-02 | -1.10E+00 | 4.51E-03 | 1.27E-02 | 2.08E+00 | 2.15E-02 | 7.90E-02 | -1.04E+00 | 3.89E-06 | 1.07E-05 | 2.37E+00 |
| EZH1 | 2.40E-04 | 1.25E-03 | -1.08E+00 | 6.33E-04 | 2.73E-03 | 1.03E+00 | 3.74E-04 | 1.59E-03 | -1.38E+00 | 3.77E-02 | 1.44E-01 | 1.13E+00 | 1.14E-01 | 4.08E-01 | 1.08E+00 | 2.03E-01 | 5.72E-01 | 1.17E+00 | 3.27E-03 | 1.20E-02 | -1.07E+00 | 1.93E-02 | 5.30E-02 | -1.02E+00 |
| QRSL1 | 2.41E-04 | 1.26E-03 | -1.11E+00 | 3.67E-04 | 1.58E-03 | 1.06E+00 | 1.00E-07 | 4.25E-07 | -1.29E+00 | 3.63E-08 | 1.39E-07 | -1.20E+00 | 6.60E-05 | 2.35E-04 | -1.10E+00 | 4.32E-03 | 1.22E-02 | 1.08E+00 | 2.25E-04 | 8.27E-04 | -1.12E+00 | 2.13E-02 | 5.84E-02 | 1.02E+00 |
| DNAJC13 | 2.42E-04 | 1.26E-03 | -1.20E+00 | 1.56E-02 | 6.72E-02 | 1.21E+00 | 1.32E-09 | 5.62E-09 | -1.73E+00 | 2.01E-05 | 7.66E-05 | -1.12E+00 | 3.63E-02 | 1.30E-01 | -1.07E+00 | 1.96E-01 | 5.52E-01 | 1.28E+00 | 1.33E-05 | 4.87E-05 | -1.36E+00 | 3.63E-02 | 9.96E-02 | 1.03E+00 |
| FAM234A | 2.42E-04 | 1.26E-03 | -1.18E+00 | 4.15E-03 | 1.79E-02 | 1.11E+00 | 9.32E-03 | 3.96E-02 | -1.09E+00 | 4.48E-03 | 1.71E-02 | -1.02E+00 | 1.84E-04 | 6.56E-04 | -1.20E+00 | 6.52E-03 | 1.84E-02 | 1.08E+00 | 6.57E-06 | 2.41E-05 | -1.15E+00 | 3.23E-04 | 8.88E-04 | -1.06E+00 |
| AL358794.1 | 2.43E-04 | 1.26E-03 | 1.01E+00 | 5.97E-05 | 2.58E-04 | 1.03E+00 | 3.83E-02 | 1.63E-01 | 1.01E+00 | 2.24E-02 | 8.55E-02 | 1.17E+00 | 1.93E-02 | 6.89E-02 | 1.07E+00 | 5.45E-02 | 1.54E-01 | 1.23E+00 | 1.08E-03 | 3.97E-03 | -1.12E+00 | 2.79E-02 | 7.66E-02 | 1.08E+00 |
| RBM33 | 2.43E-04 | 1.27E-03 | 1.02E+00 | 2.56E-08 | 1.11E-07 | -1.02E+00 | 9.32E-03 | 3.96E-02 | 1.04E+00 | 1.26E-04 | 4.83E-04 | -1.00E+00 | 7.15E-02 | 2.55E-01 | 1.21E+00 | 4.10E-04 | 1.16E-03 | -1.05E+00 | 4.91E-02 | 1.80E-01 | 1.14E+00 | 3.88E-03 | 1.07E-02 | 1.02E+00 |
| GUSBP1 | 2.43E-04 | 1.27E-03 | -1.40E+00 | 1.12E-02 | 4.82E-02 | 1.04E+00 | 1.37E-07 | 5.83E-07 | -1.65E+00 | 6.70E-06 | 2.56E-05 | -1.22E+00 | 1.88E-03 | 6.69E-03 | -1.27E+00 | 1.72E-02 | 4.85E-02 | -1.05E+00 | 6.03E-02 | 2.21E-01 | 1.15E+00 | 3.85E-03 | 1.06E-02 | -1.12E+00 |
| IFNGR1 | 2.43E-04 | 1.27E-03 | 1.97E+00 | 3.30E-02 | 1.43E-01 | 1.84E+00 | 2.93E-02 | 1.25E-01 | 1.49E+00 | 2.08E-01 | 7.94E-01 | 1.51E+00 | 2.67E-01 | 9.54E-01 | 1.45E+00 | 3.60E-02 | 1.02E-01 | 1.09E+00 | 8.38E-05 | 3.08E-04 | -1.15E+00 | 1.06E-03 | 2.93E-03 | -1.08E+00 |
| WDR45B | 2.44E-04 | 1.27E-03 | -1.10E+00 | 3.63E-04 | 1.57E-03 | 1.04E+00 | 1.05E-04 | 4.45E-04 | -1.28E+00 | 5.87E-05 | 2.24E-04 | -1.16E+00 | 3.38E-04 | 1.20E-03 | -1.11E+00 | 4.13E-04 | 1.16E-03 | -1.06E+00 | 1.96E-06 | 7.19E-06 | -1.20E+00 | 6.98E-04 | 1.92E-03 | -1.03E+00 |
| FTH1P11 | 2.45E-04 | 1.28E-03 | 1.01E+00 | 5.57E-06 | 2.41E-05 | 1.11E+00 | 3.28E-02 | 1.40E-01 | 1.03E+00 | 2.26E-01 | 8.63E-01 | 1.30E+00 | 4.35E-04 | 1.55E-03 | -1.00E+00 | 4.97E-02 | 1.40E-01 | 1.11E+00 | 9.19E-02 | 3.37E-01 | 1.06E+00 | 2.51E-01 | 6.90E-01 | 1.23E+00 |
| RPL13AP3 | 2.45E-04 | 1.28E-03 | 1.49E+00 | 2.23E-02 | 9.64E-02 | 1.48E+00 | 1.37E-02 | 5.80E-02 | 1.43E+00 | 8.41E-02 | 3.21E-01 | 1.45E+00 | 2.44E-01 | 8.71E-01 | 1.32E+00 | 1.56E-01 | 4.39E-01 | 1.44E+00 | 2.42E-01 | 8.87E-01 | 1.29E+00 | 5.18E-02 | 1.42E-01 | 1.42E+00 |
| ARL4AP5 | 2.46E-04 | 1.28E-03 | 1.96E+00 | 2.13E-01 | 9.20E-01 | 1.52E+00 | 3.18E-03 | 1.35E-02 | 1.71E+00 | 1.37E-01 | 5.25E-01 | 1.27E+00 | 4.74E-09 | 1.69E-08 | 3.33E+00 | 1.33E-03 | 3.75E-03 | 2.52E+00 | 1.60E-09 | 5.87E-09 | 2.32E+00 | 1.68E-05 | 4.61E-05 | 2.16E+00 |
| DNTTIP1 | 2.46E-04 | 1.28E-03 | -1.07E+00 | 9.63E-02 | 4.16E-01 | 1.37E+00 | 3.67E-03 | 1.56E-02 | -1.07E+00 | 2.61E-03 | 9.98E-03 | 1.01E+00 | 1.02E-05 | 3.62E-05 | -1.26E+00 | 1.17E-02 | 3.29E-02 | 1.09E+00 | 1.74E-03 | 6.40E-03 | -1.02E+00 | 9.53E-03 | 2.62E-02 | 1.04E+00 |
| SLC43A2 | 2.47E-04 | 1.29E-03 | -1.20E+00 | 1.85E-04 | 7.98E-04 | -1.14E+00 | 4.65E-09 | 1.97E-08 | -2.58E+00 | 5.86E-08 | 2.24E-07 | -2.17E+00 | 4.17E-03 | 1.49E-02 | -1.48E+00 | 1.32E-02 | 3.73E-02 | -1.31E+00 | 1.81E-06 | 6.63E-06 | -2.03E+00 | 1.17E-06 | 3.22E-06 | -2.41E+00 |
| RNF19B | 2.47E-04 | 1.29E-03 | -1.27E+00 | 1.96E-04 | 8.49E-04 | -1.22E+00 | 4.38E-02 | 1.86E-01 | -1.22E+00 | 2.07E-03 | 7.90E-03 | -1.54E+00 | 7.30E-02 | 2.60E-01 | -1.07E+00 | 1.90E-01 | 5.35E-01 | 1.10E+00 | 1.95E-04 | 7.15E-04 | -1.39E+00 | 1.25E-06 | 3.43E-06 | -2.08E+00 |
| BBS9 | 2.47E-04 | 1.29E-03 | -1.40E+00 | 3.94E-03 | 1.70E-02 | -1.02E+00 | 9.03E-02 | 3.84E-01 | -1.01E+00 | 2.14E-01 | 8.19E-01 | 1.30E+00 | 4.29E-02 | 1.53E-01 | -1.19E+00 | 6.27E-02 | 1.77E-01 | -1.09E+00 | 2.39E-01 | 8.76E-01 | 1.27E+00 | 3.51E-01 | 9.64E-01 | 1.27E+00 |
| PCGF6 | 2.47E-04 | 1.29E-03 | -1.44E+00 | 3.37E-03 | 1.46E-02 | -1.06E+00 | 9.30E-06 | 3.95E-05 | -1.42E+00 | 4.35E-04 | 1.66E-03 | -1.01E+00 | 1.32E-03 | 4.71E-03 | -2.36E+00 | 1.42E-01 | 4.01E-01 | -1.09E+00 | 1.69E-11 | 6.19E-11 | -1.65E+00 | 3.71E-04 | 1.02E-03 | -1.14E+00 |
| AC092718.7 | 2.47E-04 | 1.29E-03 | -1.38E+00 | 3.18E-05 | 1.37E-04 | -1.41E+00 | 1.78E-03 | 7.54E-03 | -1.34E+00 | 8.98E-06 | 3.43E-05 | -1.70E+00 | 4.48E-03 | 1.60E-02 | -1.44E+00 | 1.26E-03 | 3.55E-03 | -1.75E+00 | 3.04E-03 | 1.12E-02 | -1.39E+00 | 1.76E-04 | 4.83E-04 | -1.86E+00 |
| TMEM209 | 2.49E-04 | 1.30E-03 | -1.22E+00 | 2.13E-02 | 9.20E-02 | 1.21E+00 | 3.82E-02 | 1.62E-01 | -1.19E+00 | 1.24E-01 | 4.75E-01 | 1.14E+00 | 1.24E-02 | 4.43E-02 | -1.11E+00 | 2.76E-01 | 7.79E-01 | 1.57E+00 | 4.16E-03 | 1.53E-02 | -1.15E+00 | 3.54E-02 | 9.73E-02 | -1.04E+00 |
| PIP4K2B | 2.49E-04 | 1.30E-03 | -1.17E+00 | 3.23E-02 | 1.40E-01 | 1.27E+00 | 2.47E-03 | 1.05E-02 | -1.17E+00 | 4.24E-02 | 1.62E-01 | 1.21E+00 | 1.06E-02 | 3.77E-02 | -1.16E+00 | 2.51E-02 | 7.09E-02 | -1.04E+00 | 8.03E-04 | 2.95E-03 | -1.18E+00 | 3.88E-02 | 1.06E-01 | 1.08E+00 |
| PRRC2A | 2.49E-04 | 1.30E-03 | 1.01E+00 | 7.13E-03 | 3.08E-02 | 1.25E+00 | 1.26E-03 | 5.36E-03 | -1.06E+00 | 2.79E-02 | 1.07E-01 | 1.18E+00 | 1.82E-02 | 6.49E-02 | 1.13E+00 | 2.76E-02 | 7.78E-02 | 1.16E+00 | 5.88E-02 | 2.16E-01 | 1.13E+00 | 2.47E-01 | 6.77E-01 | 1.21E+00 |
| POMGNT2 | 2.49E-04 | 1.30E-03 | -2.09E+00 | 3.00E-02 | 1.30E-01 | -1.03E+00 | 9.20E-02 | 3.91E-01 | -1.10E+00 | 5.42E-02 | 2.07E-01 | -1.02E+00 | 2.31E-01 | 8.22E-01 | 1.08E+00 | 2.95E-01 | 8.32E-01 | 1.14E+00 | 9.71E-08 | 3.56E-07 | -1.93E+00 | 9.76E-03 | 2.68E-02 | -1.12E+00 |
| SETD9 | 2.50E-04 | 1.30E-03 | -1.15E+00 | 4.34E-04 | 1.87E-03 | 1.02E+00 | 2.76E-06 | 1.17E-05 | -1.32E+00 | 2.80E-06 | 1.07E-05 | -1.14E+00 | 6.63E-03 | 2.36E-02 | -1.10E+00 | 2.11E-03 | 5.94E-03 | -1.21E+00 | 6.76E-03 | 2.48E-02 | -1.10E+00 | 3.21E-02 | 8.81E-02 | -1.04E+00 |
| TRABD | 2.50E-04 | 1.30E-03 | 1.00E+00 | 1.10E-03 | 4.77E-03 | 1.13E+00 | 1.02E-05 | 4.34E-05 | -1.16E+00 | 2.35E-05 | 8.97E-05 | -1.01E+00 | 9.88E-05 | 3.52E-04 | -1.12E+00 | 2.14E-04 | 6.03E-04 | -1.03E+00 | 1.11E-03 | 4.06E-03 | 1.07E+00 | 1.13E-03 | 3.11E-03 | 1.05E+00 |
| PTPRF | 2.50E-04 | 1.30E-03 | -1.62E+00 | 9.61E-04 | 4.15E-03 | -1.31E+00 | 2.10E-02 | 8.91E-02 | -1.19E+00 | 6.45E-02 | 2.46E-01 | -1.02E+00 | 2.66E-01 | 9.47E-01 | 1.14E+00 | 3.03E-01 | 8.55E-01 | 1.12E+00 | 3.82E-02 | 1.40E-01 | -1.24E+00 | 2.16E-01 | 5.94E-01 | 1.05E+00 |
| NBPF15 | 2.50E-04 | 1.30E-03 | -1.24E+00 | 1.46E-03 | 6.32E-03 | -1.01E+00 | 5.70E-04 | 2.42E-03 | -1.45E+00 | 1.05E-03 | 4.01E-03 | -1.19E+00 | 7.12E-04 | 2.54E-03 | -1.23E+00 | 2.18E-03 | 6.16E-03 | -1.17E+00 | 1.17E-10 | 4.29E-10 | -1.58E+00 | 1.63E-08 | 4.47E-08 | -1.42E+00 |
| PATJ | 2.50E-04 | 1.31E-03 | -1.17E+00 | 2.06E-02 | 8.89E-02 | 1.17E+00 | 1.64E-02 | 6.96E-02 | -1.14E+00 | 1.02E-01 | 3.89E-01 | 1.14E+00 | 2.55E-01 | 9.09E-01 | 1.33E+00 | 2.69E-01 | 7.57E-01 | 1.73E+00 | 9.98E-02 | 3.66E-01 | -1.24E+00 | 1.73E-01 | 4.74E-01 | -1.09E+00 |
| MTPAP | 2.51E-04 | 1.31E-03 | -1.12E+00 | 3.15E-03 | 1.36E-02 | 1.13E+00 | 1.44E-05 | 6.11E-05 | -1.26E+00 | 4.30E-05 | 1.64E-04 | -1.01E+00 | 2.56E-02 | 9.12E-02 | 1.03E+00 | 3.87E-02 | 1.09E-01 | 1.12E+00 | 9.80E-05 | 3.60E-04 | -1.25E+00 | 1.34E-01 | 3.67E-01 | 1.11E+00 |
| CCDC51 | 2.51E-04 | 1.31E-03 | -1.62E+00 | 8.38E-02 | 3.62E-01 | 1.25E+00 | 1.72E-02 | 7.33E-02 | -1.12E+00 | 2.49E-01 | 9.52E-01 | 1.44E+00 | 1.71E-01 | 6.09E-01 | 1.14E+00 | 9.37E-02 | 2.64E-01 | 1.87E+00 | 6.20E-10 | 2.27E-09 | -1.77E+00 | 6.68E-02 | 1.84E-01 | 1.11E+00 |
| GUCY2D | 2.52E-04 | 1.31E-03 | -1.39E+00 | 1.92E-04 | 8.31E-04 | -1.37E+00 | 6.26E-02 | 2.66E-01 | -1.08E+00 | 2.12E-03 | 8.08E-03 | -1.47E+00 | 1.34E-01 | 4.77E-01 | -1.27E+00 | 1.11E-01 | 3.13E-01 | -1.65E+00 | 2.72E-01 | 1.00E+00 | 1.00E+00 | 3.37E-01 | 9.26E-01 | 1.06E+00 |
| CYSTM1 | 2.52E-04 | 1.31E-03 | -1.10E+00 | 3.98E-02 | 1.72E-01 | 1.23E+00 | 4.30E-02 | 1.83E-01 | 1.10E+00 | 4.17E-02 | 1.59E-01 | 1.22E+00 | 1.28E-02 | 4.57E-02 | -1.17E+00 | 3.71E-03 | 1.05E-02 | -1.40E+00 | 1.44E-01 | 5.28E-01 | 1.08E+00 | 2.57E-01 | 7.05E-01 | 1.12E+00 |
| TMEM68 | 2.52E-04 | 1.32E-03 | -1.19E+00 | 5.44E-04 | 2.35E-03 | -1.02E+00 | 2.67E-06 | 1.13E-05 | -1.32E+00 | 1.85E-03 | 7.07E-03 | 1.13E+00 | 3.76E-04 | 1.34E-03 | -1.27E+00 | 7.41E-02 | 2.09E-01 | 1.19E+00 | 3.18E-10 | 1.17E-09 | -1.35E+00 | 4.63E-07 | 1.27E-06 | -1.17E+00 |
| GASAL1 | 2.52E-04 | 1.32E-03 | 2.57E+00 | 8.22E-02 | 3.55E-01 | 1.02E+00 | 2.70E-03 | 1.15E-02 | 2.34E+00 | 2.21E-01 | 8.44E-01 | 1.19E+00 | 9.94E-02 | 3.54E-01 | -1.03E+00 | 1.52E-01 | 4.27E-01 | 1.01E+00 | 1.72E-01 | 6.32E-01 | 1.39E+00 | 3.08E-02 | 8.47E-02 | -1.45E+00 |
| TMEM185B | 2.53E-04 | 1.32E-03 | -1.22E+00 | 2.56E-04 | 1.11E-03 | -1.11E+00 | 1.78E-04 | 7.55E-04 | -1.31E+00 | 2.53E-03 | 9.66E-03 | 1.01E+00 | 4.99E-02 | 1.78E-01 | 1.09E+00 | 3.55E-01 | 1.00E+00 | 1.57E+00 | 1.66E-04 | 6.09E-04 | -1.33E+00 | 3.30E-02 | 9.05E-02 | 1.03E+00 |
| LINC00665 | 2.53E-04 | 1.32E-03 | -1.05E+00 | 6.93E-03 | 3.00E-02 | 1.21E+00 | 7.86E-10 | 3.34E-09 | -1.52E+00 | 5.97E-05 | 2.28E-04 | -1.04E+00 | 4.98E-06 | 1.77E-05 | -1.23E+00 | 2.00E-03 | 5.65E-03 | 1.02E+00 | 1.45E-06 | 5.33E-06 | -1.30E+00 | 1.24E-01 | 3.39E-01 | 1.11E+00 |
| AL121832.2 | 2.54E-04 | 1.32E-03 | -1.32E+00 | 2.30E-04 | 9.95E-04 | -1.23E+00 | 7.48E-02 | 3.18E-01 | 1.01E+00 | 4.09E-02 | 1.56E-01 | -1.01E+00 | 3.57E-02 | 1.27E-01 | -1.02E+00 | 4.70E-02 | 1.33E-01 | -1.16E+00 | 2.17E-01 | 7.98E-01 | 1.03E+00 | 2.24E-01 | 6.14E-01 | 1.02E+00 |
| RAD17 | 2.56E-04 | 1.33E-03 | -1.21E+00 | 5.82E-03 | 2.51E-02 | 1.09E+00 | 3.07E-04 | 1.30E-03 | -1.19E+00 | 6.85E-03 | 2.62E-02 | 1.07E+00 | 4.55E-06 | 1.62E-05 | -1.23E+00 | 1.98E-02 | 5.58E-02 | 1.24E+00 | 5.27E-13 | 1.93E-12 | -1.44E+00 | 3.67E-07 | 1.01E-06 | -1.19E+00 |
| DDIT3 | 2.56E-04 | 1.33E-03 | 1.08E+00 | 2.11E-28 | 9.11E-28 | -2.04E+00 | 3.60E-02 | 1.53E-01 | 1.03E+00 | 3.05E-11 | 1.16E-10 | -1.62E+00 | 3.87E-05 | 1.38E-04 | -1.12E+00 | 2.31E-16 | 6.50E-16 | -2.13E+00 | 3.72E-08 | 1.37E-07 | -1.24E+00 | 5.09E-16 | 1.40E-15 | -2.26E+00 |
| KIF11 | 2.57E-04 | 1.34E-03 | -1.58E+00 | 8.24E-02 | 3.56E-01 | 1.25E+00 | 9.32E-03 | 3.96E-02 | -1.14E+00 | 1.40E-01 | 5.36E-01 | 1.28E+00 | 2.07E-03 | 7.39E-03 | -1.98E+00 | 4.53E-02 | 1.28E-01 | -1.20E+00 | 1.91E-02 | 7.01E-02 | -1.13E+00 | 2.27E-01 | 6.24E-01 | 1.14E+00 |
| ZNF446 | 2.57E-04 | 1.34E-03 | -1.32E+00 | 2.50E-04 | 1.08E-03 | -1.17E+00 | 3.25E-03 | 1.38E-02 | -1.51E+00 | 3.72E-03 | 1.42E-02 | -1.34E+00 | 5.08E-02 | 1.81E-01 | -1.02E+00 | 2.24E-02 | 6.33E-02 | -1.20E+00 | 2.70E-01 | 9.92E-01 | 1.27E+00 | 2.03E-01 | 5.57E-01 | 1.09E+00 |
| CD4 | 2.58E-04 | 1.34E-03 | -2.36E+00 | 7.27E-04 | 3.14E-03 | -2.16E+00 | 8.94E-02 | 3.80E-01 | -1.51E+00 | 9.22E-02 | 3.52E-01 | -1.63E+00 | 8.22E-08 | 2.93E-07 | -2.11E+00 | 1.39E-06 | 3.91E-06 | -1.70E+00 | 1.11E-33 | 4.08E-33 | -2.66E+00 | 2.89E-24 | 7.94E-24 | -1.78E+00 |
| ELMSAN1 | 2.59E-04 | 1.35E-03 | -1.04E+00 | 4.39E-20 | 1.90E-19 | -1.58E+00 | 3.91E-03 | 1.66E-02 | -1.13E+00 | 6.95E-20 | 2.65E-19 | -2.02E+00 | 1.01E-03 | 3.61E-03 | -1.04E+00 | 3.38E-15 | 9.53E-15 | -1.85E+00 | 2.63E-09 | 9.67E-09 | -1.29E+00 | 2.81E-10 | 7.73E-10 | -1.53E+00 |
| AC087623.3 | 2.62E-04 | 1.37E-03 | 2.40E+00 | 6.83E-02 | 2.95E-01 | 1.11E+00 | 2.28E-04 | 9.69E-04 | 2.88E+00 | 1.86E-01 | 7.09E-01 | 1.71E+00 | 1.30E-01 | 4.64E-01 | 1.79E+00 | 3.33E-01 | 9.38E-01 | 1.34E+00 | 1.26E-02 | 4.64E-02 | 2.12E+00 | 1.43E-01 | 3.94E-01 | 1.54E+00 |
| LRRC42 | 2.63E-04 | 1.37E-03 | -1.48E+00 | 6.41E-02 | 2.77E-01 | 1.23E+00 | 3.38E-04 | 1.44E-03 | -1.22E+00 | 4.50E-02 | 1.72E-01 | 1.20E+00 | 3.16E-03 | 1.13E-02 | -1.42E+00 | 1.71E-02 | 4.82E-02 | -1.09E+00 | 3.37E-16 | 1.24E-15 | -1.83E+00 | 2.92E-08 | 8.01E-08 | -1.30E+00 |
| ASXL2 | 2.63E-04 | 1.37E-03 | -1.29E+00 | 9.70E-03 | 4.19E-02 | 1.08E+00 | 7.53E-07 | 3.20E-06 | -1.52E+00 | 2.67E-03 | 1.02E-02 | 1.01E+00 | 4.68E-04 | 1.67E-03 | -1.48E+00 | 5.12E-02 | 1.44E-01 | 1.02E+00 | 1.51E-16 | 5.53E-16 | -1.91E+00 | 9.67E-13 | 2.66E-12 | -1.58E+00 |

| | | | | | | | | | | | | | | | | | | | | | | | | |
|---|---|---|---|---|---|---|---|---|---|---|---|---|---|---|---|---|---|---|---|---|---|---|---|---|
| ZNF154 | 2.63E-04 | 1.37E-03 | -1.21E+00 | 7.60E-03 | 3.28E-02 | 1.03E+00 | 5.78E-05 | 2.46E-04 | -1.40E+00 | 1.29E-02 | 4.93E-02 | 1.06E+00 | 9.21E-03 | 3.28E-02 | 1.05E+00 | 4.61E-03 | 1.30E-02 | 1.00E+00 | 8.29E-02 | 3.04E-01 | 1.08E+00 | 4.00E-02 | 1.10E-01 | -1.03E+00 |
| SLC38A9 | 2.64E-04 | 1.38E-03 | -1.50E+00 | 6.35E-02 | 2.74E-01 | 1.20E+00 | 1.37E-07 | 5.84E-07 | -1.70E+00 | 4.07E-05 | 1.55E-04 | -1.19E+00 | 1.18E-03 | 4.22E-03 | -1.49E+00 | 8.54E-03 | 2.41E-02 | -1.12E+00 | 1.01E-06 | 3.70E-06 | -1.51E+00 | 1.52E-03 | 4.19E-03 | -1.17E+00 |
| VCL | 2.65E-04 | 1.38E-03 | 1.01E+00 | 6.09E-03 | 2.63E-02 | 1.20E+00 | 1.06E-01 | 4.49E-01 | 1.13E+00 | 2.30E-01 | 8.80E-01 | 1.34E+00 | 1.24E-02 | 4.43E-02 | 1.09E+00 | 3.05E-01 | 8.61E-01 | 1.26E+00 | 1.94E-06 | 7.13E-06 | 1.74E+00 | 1.21E-03 | 3.34E-03 | 1.36E+00 |
| CRACR2A | 2.65E-04 | 1.38E-03 | -1.31E+00 | 3.24E-03 | 1.40E-02 | 1.06E+00 | 3.88E-03 | 1.65E-02 | -1.15E+00 | 1.86E-02 | 7.10E-02 | 1.02E+00 | 9.28E-03 | 3.31E-02 | -1.22E+00 | 5.55E-02 | 1.56E-01 | -1.02E+00 | 1.48E-01 | 5.44E-01 | 1.08E+00 | 1.56E-01 | 4.28E-01 | 1.02E+00 |
| PLD1 | 2.66E-04 | 1.38E-03 | -1.52E+00 | 1.99E-04 | 8.58E-04 | -1.34E+00 | 4.73E-05 | 2.01E-04 | -1.33E+00 | 8.40E-03 | 3.21E-02 | 1.04E+00 | 7.31E-15 | 2.61E-14 | -1.67E+00 | 2.00E-14 | 5.63E-14 | -1.56E+00 | 1.59E-03 | 5.83E-03 | -1.41E+00 | 1.96E-01 | 5.38E-01 | 1.13E+00 |
| PRDX4 | 2.66E-04 | 1.39E-03 | -1.03E+00 | 3.59E-03 | 1.55E-02 | 1.14E+00 | 2.11E-01 | 8.95E-01 | 1.15E+00 | 2.10E-01 | 8.02E-01 | 1.28E+00 | 1.53E-02 | 5.46E-02 | 1.05E+00 | 3.44E-01 | 9.71E-01 | 1.28E+00 | 2.14E-02 | 7.86E-02 | 1.33E+00 | 1.59E-04 | 4.38E-04 | 1.58E+00 |
| APOBEC3F | 2.66E-04 | 1.39E-03 | -1.25E+00 | 2.13E-05 | 9.21E-05 | -1.41E+00 | 7.73E-04 | 3.28E-03 | -1.34E+00 | 3.19E-03 | 1.22E-02 | -1.20E+00 | 7.60E-04 | 2.71E-03 | -1.31E+00 | 2.05E-02 | 5.77E-02 | 1.19E+00 | 1.11E-04 | 4.08E-04 | -1.32E+00 | 1.74E-04 | 4.79E-04 | -1.47E+00 |
| SNRPGP2 | 2.66E-04 | 1.39E-03 | -1.15E+00 | 3.97E-05 | 1.72E-04 | -1.18E+00 | 1.80E-02 | 7.65E-02 | 1.06E+00 | 3.61E-06 | 1.38E-05 | -1.29E+00 | 8.52E-04 | 3.04E-03 | -1.38E+00 | 3.12E-03 | 8.80E-03 | -1.29E+00 | 1.50E-03 | 5.49E-03 | -1.07E+00 | 8.74E-02 | 2.40E-01 | 1.10E+00 |
| AQP3 | 2.66E-04 | 1.39E-03 | -1.98E+00 | 1.44E-03 | 6.22E-03 | -1.59E+00 | 1.66E-04 | 7.06E-04 | -2.58E+00 | 7.81E-03 | 2.98E-02 | -1.59E+00 | 3.49E-05 | 1.25E-04 | -1.41E+00 | 4.39E-04 | 1.24E-03 | -1.81E+00 | 1.14E-01 | 4.17E-01 | 1.89E+00 | 5.89E-02 | 1.62E-01 | 2.48E+00 |
| VPS33A | 2.67E-04 | 1.39E-03 | -1.30E+00 | 2.51E-02 | 1.09E-01 | 1.16E+00 | 1.22E-12 | 5.19E-12 | -1.98E+00 | 4.75E-08 | 1.81E-07 | -1.34E+00 | 5.30E-05 | 1.89E-04 | -1.59E+00 | 7.58E-03 | 2.14E-02 | -1.04E+00 | 1.96E-08 | 7.20E-08 | -1.53E+00 | 1.45E-05 | 3.98E-05 | -1.35E+00 |
| ZNF571-AS1 | 2.67E-04 | 1.39E-03 | -1.56E+00 | 1.06E-03 | 4.58E-03 | -1.29E+00 | 4.72E-02 | 2.01E-01 | -1.15E+00 | 2.08E-01 | 7.95E-01 | 1.36E+00 | 1.94E-01 | 6.93E-01 | 1.21E+00 | 1.24E-01 | 3.50E-01 | -1.15E+00 | 1.83E-03 | 6.73E-03 | -1.54E+00 | 2.11E-02 | 5.80E-02 | -1.12E+00 |
| TRMT1 | 2.68E-04 | 1.39E-03 | 1.04E+00 | 5.92E-04 | 2.56E-03 | 1.16E+00 | 7.17E-04 | 3.05E-03 | -1.11E+00 | 7.16E-03 | 2.73E-02 | 1.11E+00 | 1.59E-06 | 5.66E-06 | -1.25E+00 | 4.00E-05 | 1.13E-04 | -1.15E+00 | 9.27E-03 | 3.40E-02 | 1.02E+00 | 4.38E-02 | 1.20E-01 | 1.02E+00 |
| HIPK3 | 2.68E-04 | 1.40E-03 | -1.05E+00 | 1.57E-06 | 6.79E-06 | -1.09E+00 | 1.11E-01 | 4.72E-01 | 1.17E+00 | 1.95E-06 | 7.44E-06 | -1.16E+00 | 2.83E-02 | 1.01E-01 | 1.20E+00 | 8.58E-04 | 2.42E-03 | 1.01E+00 | 1.74E-01 | 6.39E-01 | 1.33E+00 | 2.92E-02 | 8.02E-02 | 1.11E+00 |
| EPB41L2 | 2.69E-04 | 1.40E-03 | -1.35E+00 | 1.04E-02 | 4.50E-02 | 1.04E+00 | 1.08E-04 | 4.57E-04 | -1.27E+00 | 9.68E-04 | 3.70E-03 | -1.03E+00 | 1.66E-03 | 5.92E-03 | -1.52E+00 | 1.40E-02 | 3.95E-02 | -1.17E+00 | 4.64E-04 | 1.70E-03 | -1.09E+00 | 1.76E-02 | 4.84E-02 | 1.01E+00 |
| EIF4A1P5 | 2.69E-04 | 1.40E-03 | -1.58E+00 | 1.41E-04 | 6.11E-04 | -1.53E+00 | 2.21E-03 | 9.40E-03 | -1.33E+00 | 4.65E-05 | 1.78E-04 | -1.46E+00 | 2.00E-02 | 7.13E-02 | -1.39E+00 | 7.32E-03 | 2.06E-02 | -2.02E+00 | 2.31E-01 | 8.48E-01 | 1.11E+00 | 7.07E-02 | 1.94E-01 | -1.25E+00 |
| MORN4 | 2.69E-04 | 1.40E-03 | -1.57E+00 | 9.56E-04 | 4.13E-03 | -1.37E+00 | 2.34E-02 | 9.94E-02 | -1.15E+00 | 9.39E-03 | 3.59E-02 | -1.17E+00 | 1.88E-01 | 6.70E-01 | -1.05E+00 | 2.53E-01 | 7.14E-01 | 1.02E+00 | 4.02E-03 | 1.47E-02 | -1.54E+00 | 4.55E-02 | 1.25E-01 | -1.20E+00 |
| HSDL1 | 2.70E-04 | 1.41E-03 | -1.47E+00 | 2.33E-03 | 1.01E-02 | -1.13E+00 | 1.80E-06 | 7.64E-06 | -1.84E+00 | 1.44E-04 | 5.48E-04 | -1.28E+00 | 1.10E-05 | 3.92E-05 | -1.63E+00 | 4.31E-03 | 1.22E-02 | -1.04E+00 | 1.21E-04 | 4.44E-04 | -1.34E+00 | 3.76E-02 | 1.03E-01 | 1.09E+00 |
| LRRC75B | 2.70E-04 | 1.41E-03 | -1.48E+00 | 1.05E-04 | 4.55E-04 | -1.43E+00 | 8.60E-05 | 3.65E-04 | -1.59E+00 | 7.34E-04 | 2.81E-03 | -1.23E+00 | 1.61E-04 | 5.75E-04 | -2.40E+00 | 1.89E-02 | 5.34E-02 | -1.20E+00 | 1.18E-05 | 4.32E-05 | -1.79E+00 | 2.78E-04 | 7.63E-04 | -1.45E+00 |
| AC027682.7 | 2.70E-04 | 1.41E-03 | -1.56E+00 | 7.86E-06 | 3.40E-05 | -2.06E+00 | 1.94E-02 | 8.23E-02 | -1.34E+00 | 3.63E-03 | 1.38E-02 | -1.55E+00 | 4.14E-04 | 1.48E-03 | -2.18E+00 | 8.00E-04 | 2.25E-03 | -1.93E+00 | 1.54E-01 | 5.65E-01 | 1.00E+00 | 2.11E-01 | 5.81E-01 | 1.05E+00 |
| CCDC152 | 2.71E-04 | 1.41E-03 | -1.16E+00 | 7.58E-04 | 3.27E-03 | 1.02E+00 | 1.59E-01 | 6.75E-01 | 1.14E+00 | 1.10E-01 | 4.21E-01 | 1.27E+00 | 1.07E-01 | 3.82E-01 | 1.10E+00 | 1.86E-01 | 5.25E-01 | 1.24E+00 | 4.46E-02 | 1.64E-01 | 1.49E+00 | 3.15E-01 | 8.66E-01 | 1.21E+00 |
| BCORL1 | 2.71E-04 | 1.41E-03 | -1.68E+00 | 2.04E-01 | 8.82E-01 | 1.39E+00 | 2.18E-03 | 9.28E-03 | -1.45E+00 | 1.91E-01 | 7.28E-01 | 1.26E+00 | 1.85E-05 | 6.59E-05 | -1.94E+00 | 9.94E-05 | 2.80E-04 | -1.69E+00 | 1.16E-05 | 4.26E-05 | -1.66E+00 | 8.15E-03 | 2.24E-02 | -1.08E+00 |
| GALNT10 | 2.71E-04 | 1.41E-03 | -1.09E+00 | 1.22E-03 | 5.25E-03 | 1.10E+00 | 8.19E-04 | 3.48E-03 | -1.30E+00 | 3.42E-02 | 1.31E-01 | 1.15E+00 | 6.20E-02 | 2.21E-01 | 1.34E+00 | 3.29E-01 | 9.27E-01 | 1.57E+00 | 1.80E-01 | 6.60E-01 | 1.28E+00 | 5.01E-02 | 1.38E-01 | 1.23E+00 |
| SRA1 | 2.72E-04 | 1.42E-03 | -1.09E+00 | 3.17E-05 | 1.37E-04 | -1.10E+00 | 3.78E-02 | 1.61E-01 | -1.00E+00 | 1.92E-04 | 7.34E-04 | -1.14E+00 | 1.06E-05 | 3.79E-05 | -1.17E+00 | 2.99E-07 | 8.42E-07 | -1.31E+00 | 1.38E-01 | 5.06E-01 | 1.15E+00 | 2.72E-01 | 7.48E-01 | 1.14E+00 |
| TMEM222 | 2.72E-04 | 1.42E-03 | -1.17E+00 | 7.89E-04 | 3.41E-03 | -1.00E+00 | 1.67E-02 | 7.11E-02 | -1.03E+00 | 3.06E-04 | 1.17E-03 | -1.07E+00 | 4.58E-04 | 1.63E-03 | -1.06E+00 | 2.76E-03 | 7.78E-03 | -1.02E+00 | 1.79E-01 | 6.57E-01 | 1.29E+00 | 2.82E-01 | 7.75E-01 | 1.35E+00 |
| MZF1 | 2.72E-04 | 1.42E-03 | -1.11E+00 | 9.90E-03 | 4.28E-02 | 1.15E+00 | 1.03E-02 | 4.38E-02 | -1.04E+00 | 1.25E-01 | 4.78E-01 | 1.30E+00 | 2.48E-02 | 8.85E-02 | 1.06E+00 | 2.31E-01 | 6.51E-01 | 1.28E+00 | 2.40E-01 | 8.80E-01 | 1.10E+00 | 1.30E-01 | 3.58E-01 | 1.39E+00 |
| MALSU1 | 2.72E-04 | 1.42E-03 | 1.02E+00 | 1.67E-02 | 7.21E-02 | 1.32E+00 | 8.64E-04 | 3.67E-03 | -1.02E+00 | 2.72E-02 | 1.04E-01 | 1.18E+00 | 3.32E-05 | 1.18E-04 | -1.19E+00 | 1.06E-01 | 2.99E-01 | 1.31E+00 | 6.02E-05 | 2.21E-04 | -1.03E+00 | 9.58E-02 | 2.63E-01 | 1.21E+00 |
| RPL17P40 | 2.74E-04 | 1.43E-03 | -1.37E+00 | 1.47E-01 | 6.36E-01 | 1.47E+00 | 8.94E-04 | 3.80E-03 | -1.58E+00 | 4.64E-02 | 1.77E-01 | 1.05E+00 | 1.35E-01 | 4.82E-01 | 1.11E+00 | 3.45E-01 | 9.71E-01 | 1.37E+00 | 3.84E-02 | 1.41E-01 | -1.48E+00 | 1.08E-01 | 2.97E-01 | -1.21E+00 |
| ILF3-AS1 | 2.74E-04 | 1.43E-03 | -1.07E+00 | 7.28E-04 | 3.15E-03 | 1.08E+00 | 5.40E-02 | 2.30E-01 | 1.04E+00 | 6.39E-02 | 2.44E-01 | 1.15E+00 | 1.04E-03 | 3.70E-03 | -1.00E+00 | 1.66E-01 | 4.67E-01 | 1.19E+00 | 7.20E-03 | 2.64E-02 | -1.10E+00 | 7.73E-02 | 2.12E-01 | 1.05E+00 |
| SNRNP25 | 2.75E-04 | 1.43E-03 | -1.11E+00 | 1.75E-05 | 7.56E-05 | -1.04E+00 | 8.89E-02 | 3.78E-01 | 1.24E+00 | 1.16E-01 | 4.42E-01 | 1.16E+00 | 8.54E-07 | 3.05E-06 | -1.57E+00 | 6.21E-04 | 1.75E-03 | -1.20E+00 | 1.64E-01 | 6.00E-01 | 1.14E+00 | 1.48E-01 | 4.05E-01 | 1.33E+00 |
| CYB5B | 2.76E-04 | 1.44E-03 | -1.06E+00 | 5.53E-02 | 2.39E-01 | 1.25E+00 | 9.16E-07 | 3.89E-06 | -1.28E+00 | 1.79E-05 | 6.84E-05 | -1.07E+00 | 7.03E-08 | 2.51E-07 | -1.31E+00 | 2.69E-05 | 7.58E-05 | -1.03E+00 | 4.29E-07 | 1.57E-06 | -1.29E+00 | 3.28E-03 | 9.01E-03 | -1.03E+00 |
| AL663070.1 | 2.76E-04 | 1.44E-03 | -1.20E+00 | 3.80E-40 | 1.64E-39 | -1.83E+00 | 1.85E-03 | 7.88E-03 | -1.26E+00 | 1.20E-20 | 4.58E-20 | -1.92E+00 | 3.12E-05 | 1.11E-04 | -1.23E+00 | 9.13E-19 | 2.57E-18 | -1.77E+00 | 8.64E-06 | 3.17E-05 | -1.25E+00 | 3.37E-15 | 9.26E-15 | -1.85E+00 |
| DHX32 | 2.77E-04 | 1.44E-03 | -1.14E+00 | 3.27E-03 | 1.41E-02 | 1.11E+00 | 3.88E-05 | 1.65E-04 | -1.25E+00 | 1.10E-02 | 4.19E-02 | 1.12E+00 | 1.68E-04 | 6.00E-04 | -1.27E+00 | 9.39E-03 | 2.65E-02 | -1.06E+00 | 2.74E-07 | 1.00E-06 | -1.38E+00 | 5.92E-04 | 1.63E-03 | -1.03E+00 |
| ASB7 | 2.78E-04 | 1.45E-03 | -1.47E+00 | 7.47E-03 | 3.23E-02 | 1.04E+00 | 6.40E-10 | 2.72E-09 | -1.89E+00 | 4.71E-07 | 1.80E-06 | -1.35E+00 | 5.34E-02 | 1.90E-01 | -1.00E+00 | 9.67E-02 | 2.73E-01 | 1.17E+00 | 1.85E-01 | 6.79E-01 | 1.19E+00 | 2.23E-01 | 6.13E-01 | 1.20E+00 |
| OCEL1 | 2.79E-04 | 1.45E-03 | -1.45E+00 | 1.47E-02 | 6.34E-02 | -1.01E+00 | 8.50E-10 | 3.61E-09 | -2.01E+00 | 5.42E-08 | 2.07E-07 | -1.64E+00 | 2.07E-07 | 7.39E-07 | -1.56E+00 | 6.81E-04 | 1.92E-03 | -1.13E+00 | 6.12E-03 | 2.25E-02 | -1.06E+00 | 4.46E-02 | 1.22E-01 | 1.06E+00 |
| GLRX2 | 2.79E-04 | 1.45E-03 | -1.06E+00 | 9.84E-04 | 4.25E-03 | 1.12E+00 | 9.89E-06 | 4.20E-05 | -1.26E+00 | 1.40E-04 | 5.34E-04 | 1.00E+00 | 2.07E-03 | 7.36E-03 | -1.10E+00 | 8.51E-03 | 2.40E-02 | 1.01E+00 | 1.81E-07 | 6.64E-07 | -1.42E+00 | 6.79E-03 | 1.87E-02 | -1.09E+00 |
| RASA1 | 2.79E-04 | 1.46E-03 | -1.12E+00 | 9.51E-05 | 4.11E-04 | -1.02E+00 | 2.28E-01 | 9.70E-01 | 1.22E+00 | 1.53E-01 | 5.83E-01 | 1.19E+00 | 5.82E-03 | 2.07E-02 | -1.12E+00 | 4.71E-03 | 1.33E-02 | -1.14E+00 | 4.60E-02 | 1.69E-01 | 1.23E+00 | 1.34E-03 | 3.67E-03 | -1.28E+00 |
| AL035404.1 | 2.80E-04 | 1.46E-03 | -1.65E+00 | 1.39E-04 | 6.01E-04 | -1.74E+00 | 1.43E-04 | 6.06E-04 | -1.56E+00 | 5.43E-07 | 2.07E-06 | -2.18E+00 | 3.94E-02 | 1.40E-01 | -1.20E+00 | 1.63E-02 | 4.60E-02 | -1.51E+00 | 2.05E-05 | 7.53E-05 | -1.36E+00 | 1.87E-05 | 5.14E-05 | -1.40E+00 |
| CAHM | 2.80E-04 | 1.46E-03 | 2.05E+00 | 8.75E-02 | 3.78E-01 | 1.66E+00 | 5.20E-02 | 2.21E-01 | 1.57E+00 | 9.23E-02 | 3.52E-01 | 1.24E+00 | 2.13E-01 | 7.59E-01 | 1.19E+00 | 3.32E-01 | 9.36E-01 | 1.34E+00 | 1.05E-01 | 3.85E-01 | -1.13E+00 | 3.40E-01 | 9.34E-01 | 1.19E+00 |
| MRPS33 | 2.80E-04 | 1.46E-03 | -1.02E+00 | 2.41E-09 | 1.04E-08 | -1.10E+00 | 2.01E-05 | 8.55E-05 | -1.22E+00 | 2.98E-07 | 1.14E-06 | -1.17E+00 | 1.63E-03 | 5.79E-03 | -1.09E+00 | 7.33E-02 | 2.07E-01 | 1.07E+00 | 4.23E-04 | 1.55E-03 | -1.10E+00 | 3.01E-03 | 8.27E-03 | -1.07E+00 |
| TBC1D32 | 2.81E-04 | 1.46E-03 | -1.34E+00 | 3.72E-03 | 1.61E-02 | -1.02E+00 | 5.24E-02 | 2.23E-01 | 1.01E+00 | 7.80E-02 | 2.98E-01 | 1.19E+00 | 2.36E-03 | 8.42E-03 | -1.42E+00 | 5.93E-02 | 1.67E-01 | 1.09E+00 | 4.98E-03 | 1.83E-02 | -1.30E+00 | 2.34E-02 | 6.42E-02 | -1.16E+00 |
| AL033543.1 | 2.82E-04 | 1.47E-03 | -1.41E+00 | 2.70E-02 | 1.17E-01 | 1.29E+00 | 1.80E-02 | 7.65E-02 | -1.44E+00 | 1.07E-01 | 4.10E-01 | 1.15E+00 | 2.18E-06 | 7.78E-06 | -2.05E+00 | 4.71E-02 | 1.33E-01 | 1.12E+00 | 4.24E-06 | 1.56E-05 | -1.47E+00 | 2.02E-02 | 5.54E-02 | -1.19E+00 |
| CD244 | 2.83E-04 | 1.47E-03 | -1.01E+00 | 2.22E-02 | 9.59E-02 | 1.32E+00 | 6.45E-09 | 2.74E-08 | -1.57E+00 | 3.53E-07 | 1.35E-06 | -1.29E+00 | 2.02E-02 | 7.22E-02 | 1.04E+00 | 1.08E-01 | 3.03E-01 | 1.14E+00 | 2.28E-01 | 8.38E-01 | 1.16E+00 | 3.11E-01 | 8.53E-01 | 1.24E+00 |
| AL121929.2 | 2.83E-04 | 1.47E-03 | -1.64E+00 | 6.72E-03 | 2.90E-02 | -1.13E+00 | 4.13E-06 | 1.76E-05 | -1.84E+00 | 1.48E-04 | 5.67E-04 | -1.36E+00 | 2.24E-02 | 7.98E-02 | -1.86E+00 | 9.84E-02 | 2.77E-01 | -1.23E+00 | 3.55E-02 | 1.30E-01 | -1.31E+00 | 2.45E-01 | 6.74E-01 | 1.04E+00 |
| LINC00624 | 2.84E-04 | 1.48E-03 | -1.50E+00 | 4.44E-04 | 1.92E-03 | -1.35E+00 | 3.18E-07 | 1.35E-06 | -1.81E+00 | 1.64E-05 | 6.26E-05 | -1.29E+00 | 1.13E-01 | 4.03E-01 | 1.03E+00 | 2.09E-01 | 5.89E-01 | 1.12E+00 | 1.08E-02 | 3.95E-02 | -1.19E+00 | 2.39E-02 | 6.57E-02 | -1.26E+00 |
| CHMP4BP1 | 2.85E-04 | 1.48E-03 | -1.40E+00 | 2.91E-04 | 1.26E-03 | -1.38E+00 | 7.79E-09 | 3.31E-08 | -2.02E+00 | 2.58E-09 | 9.84E-09 | -2.01E+00 | 9.94E-02 | 3.54E-01 | -1.16E+00 | 3.52E-02 | 9.93E-02 | -1.48E+00 | 1.12E-09 | 4.10E-09 | -2.00E+00 | 8.33E-10 | 2.29E-09 | -2.10E+00 |
| NIPA2 | 2.86E-04 | 1.49E-03 | -1.41E+00 | 7.24E-04 | 3.13E-03 | -1.21E+00 | 4.39E-04 | 1.86E-03 | -1.10E+00 | 1.49E-06 | 5.70E-06 | -1.21E+00 | 3.22E-06 | 1.15E-05 | -1.68E+00 | 3.32E-04 | 9.37E-04 | -1.33E+00 | 7.94E-06 | 2.91E-05 | -1.33E+00 | 4.60E-02 | 1.26E-01 | 1.04E+00 |
| RNVU1-15 | 2.86E-04 | 1.49E-03 | 2.71E+00 | 1.86E-02 | 8.02E-02 | -1.44E+00 | 3.98E-02 | 1.69E-01 | 1.61E+00 | 1.63E-03 | 6.21E-03 | -1.86E+00 | 1.02E-01 | 3.63E-01 | -1.15E+00 | 6.91E-02 | 1.95E-01 | -1.51E+00 | 4.04E-02 | 1.48E-01 | -1.55E+00 | 3.50E-01 | 9.62E-01 | 1.18E+00 |
| REX1BD | 2.87E-04 | 1.49E-03 | 1.00E+00 | 2.09E-05 | 9.04E-05 | 1.05E+00 | 1.44E-01 | 6.13E-01 | 1.25E+00 | 1.40E-01 | 5.34E-01 | 1.24E+00 | 1.09E-01 | 3.89E-01 | 1.20E+00 | 9.55E-03 | 2.69E-02 | 1.09E+00 | 2.40E-01 | 8.82E-01 | 1.29E+00 | 2.42E-01 | 6.66E-01 | 1.27E+00 |
| PGPEP1 | 2.87E-04 | 1.49E-03 | -1.02E+00 | 4.60E-05 | 1.99E-04 | 1.06E+00 | 4.03E-07 | 1.71E-06 | -1.39E+00 | 5.24E-06 | 2.00E-05 | -1.12E+00 | 1.68E-04 | 6.00E-04 | -1.21E+00 | 8.87E-05 | 2.50E-04 | -1.28E+00 | 8.78E-03 | 3.22E-02 | -1.09E+00 | 1.82E-03 | 5.01E-03 | -1.22E+00 |
| NDUFS4 | 2.88E-04 | 1.50E-03 | 1.08E+00 | 3.24E-05 | 1.40E-04 | 1.16E+00 | 6.34E-02 | 2.69E-01 | 1.12E+00 | 3.81E-03 | 1.45E-02 | 1.11E+00 | 2.00E-03 | 7.12E-03 | 1.07E+00 | 1.15E-02 | 3.23E-02 | 1.15E+00 | 2.99E-02 | 1.10E-01 | 1.06E+00 | 9.94E-02 | 2.73E-01 | 1.08E+00 |
| UBA3 | 2.88E-04 | 1.50E-03 | -1.11E+00 | 1.62E-01 | 7.00E-01 | 1.53E+00 | 1.24E-07 | 5.26E-07 | -1.52E+00 | 5.50E-04 | 2.10E-03 | -1.03E+00 | 1.56E-04 | 5.55E-04 | -1.12E+00 | 1.41E-01 | 3.96E-01 | 1.28E+00 | 1.81E-06 | 6.64E-06 | -1.20E+00 | 3.93E-02 | 1.08E-01 | 1.07E+00 |
| FNBP1L | 2.89E-04 | 1.51E-03 | -1.16E+00 | 1.04E-04 | 4.48E-04 | -1.11E+00 | 5.25E-04 | 2.23E-03 | -1.18E+00 | 2.19E-04 | 8.36E-04 | -1.14E+00 | 3.31E-02 | 1.18E-01 | 1.08E+00 | 2.69E-02 | 7.58E-02 | -1.01E+00 | 3.60E-02 | 1.32E-01 | 1.32E+00 | 1.21E-02 | 3.33E-02 | -1.00E+00 |
| SETD6 | 2.90E-04 | 1.51E-03 | -1.29E+00 | 9.44E-03 | 4.08E-02 | 1.13E+00 | 2.21E-01 | 9.39E-01 | 1.20E+00 | 2.00E-01 | 7.64E-01 | 1.40E+00 | 3.83E-02 | 1.37E-01 | -1.22E+00 | 2.34E-01 | 6.59E-01 | 1.20E+00 | 4.71E-03 | 1.73E-02 | -1.10E+00 | 1.84E-01 | 5.04E-01 | 1.23E+00 |
| AC013470.2 | 2.91E-04 | 1.52E-03 | -1.92E+00 | 2.69E-03 | 1.16E-02 | -1.51E+00 | 1.93E-01 | 8.21E-01 | 1.03E+00 | 2.25E-01 | 8.59E-01 | 1.07E+00 | 1.19E-01 | 4.25E-01 | -1.17E+00 | 3.30E-01 | 9.32E-01 | 1.37E+00 | 1.96E-01 | 7.19E-01 | -1.06E+00 | 2.12E-01 | 5.82E-01 | -1.13E+00 |
| ADGRG1 | 2.91E-04 | 1.52E-03 | -1.03E+00 | 2.28E-02 | 9.87E-02 | 1.51E+00 | 1.75E-08 | 7.45E-08 | -1.39E+00 | 3.51E-02 | 1.34E-01 | 1.16E+00 | 2.40E-06 | 8.55E-06 | -1.38E+00 | 4.98E-02 | 1.40E-01 | 1.13E+00 | 1.61E-02 | 5.90E-02 | 1.05E+00 | 2.56E-01 | 7.03E-01 | 1.18E+00 |
| UBAC2-AS1 | 2.92E-04 | 1.52E-03 | -1.81E+00 | 4.50E-04 | 1.94E-03 | -1.73E+00 | 2.65E-02 | 1.13E-01 | -1.20E+00 | 1.67E-03 | 6.37E-03 | -1.53E+00 | 3.67E-02 | 1.31E-01 | -1.32E+00 | 6.80E-02 | 1.92E-01 | -1.27E+00 | 2.59E-03 | 9.50E-03 | 1.03E+00 | 2.11E-03 | 5.78E-03 | -1.11E+00 |
| QSOX2 | 2.92E-04 | 1.52E-03 | -1.47E+00 | 1.52E-02 | 6.55E-02 | 1.10E+00 | 1.27E-04 | 5.38E-04 | -1.40E+00 | 5.99E-03 | 2.29E-02 | -1.04E+00 | 2.11E-04 | 7.51E-04 | -1.78E+00 | 1.13E-03 | 3.18E-03 | -1.40E+00 | 1.45E-03 | 5.32E-03 | -1.39E+00 | 1.88E-01 | 5.17E-01 | 1.12E+00 |
| FN3K | 2.92E-04 | 1.52E-03 | -1.50E+00 | 2.73E-02 | 1.18E-01 | 1.17E+00 | 3.71E-02 | 1.58E-01 | -1.31E+00 | 1.67E-01 | 6.36E-01 | 1.20E+00 | 6.80E-02 | 2.42E-01 | -1.05E+00 | 8.48E-02 | 2.39E-01 | 2.10E+00 | 8.48E-06 | 3.11E-05 | -1.12E+00 | 1.32E-04 | 3.64E-04 | -1.15E+00 |
| RAB40C | 2.92E-04 | 1.52E-03 | -1.38E+00 | 6.33E-02 | 2.73E-01 | 1.28E+00 | 7.16E-02 | 3.04E-01 | -1.05E+00 | 1.22E-01 | 4.66E-01 | 1.55E+00 | 2.12E-04 | 7.56E-04 | -1.64E+00 | 4.44E-02 | 1.25E-01 | 1.04E+00 | 1.38E-02 | 5.06E-02 | -1.18E+00 | 2.42E-01 | 6.65E-01 | 1.19E+00 |
| MICAL3 | 2.92E-04 | 1.52E-03 | -1.33E+00 | 6.15E-04 | 2.66E-03 | -1.08E+00 | 1.12E-01 | 4.74E-01 | -1.06E+00 | 2.22E-01 | 8.47E-01 | 1.40E+00 | 3.08E-03 | 1.10E-02 | -1.36E+00 | 5.70E-02 | 1.61E-01 | 1.04E+00 | 2.65E-05 | 9.72E-05 | -1.32E+00 | 3.38E-06 | 9.29E-06 | -1.44E+00 |
| ZNF592 | 2.93E-04 | 1.53E-03 | -1.63E+00 | 1.89E-01 | 8.15E-01 | 1.36E+00 | 4.20E-05 | 1.78E-04 | -1.52E+00 | 1.52E-02 | 5.80E-02 | 1.04E+00 | 1.36E-03 | 4.85E-03 | -1.39E+00 | 8.14E-04 | 2.29E-03 | -1.54E+00 | 4.88E-07 | 1.79E-06 | -2.03E+00 | 3.54E-03 | 9.71E-03 | -1.27E+00 |
| PIN1 | 2.93E-04 | 1.53E-03 | 1.08E+00 | 7.13E-04 | 3.08E-03 | 1.22E+00 | 1.49E-01 | 6.34E-01 | 1.17E+00 | 2.60E-02 | 9.92E-02 | 1.16E+00 | 3.33E-02 | 1.19E-01 | 1.15E+00 | 1.63E-01 | 4.59E-01 | 1.30E+00 | 1.17E-01 | 4.31E-01 | 1.06E+00 | 1.81E-01 | 4.97E-01 | 1.12E+00 |
| PXDC1 | 2.93E-04 | 1.53E-03 | -1.40E+00 | 1.04E-02 | 4.51E-02 | -1.03E+00 | 5.78E-03 | 2.46E-02 | -1.67E+00 | 1.41E-01 | 5.38E-01 | 1.02E+00 | 2.63E-01 | 9.36E-01 | 1.02E+00 | 3.20E-01 | 9.02E-01 | -1.02E+00 | 2.58E-01 | 9.47E-01 | 1.08E+00 | 2.55E-01 | 7.01E-01 | 1.25E+00 |
| EXOSC10 | 2.94E-04 | 1.53E-03 | -1.15E+00 | 2.04E-01 | 8.80E-01 | 1.47E+00 | 2.56E-10 | 1.09E-09 | -1.47E+00 | 3.98E-06 | 1.52E-05 | -1.06E+00 | 7.36E-04 | 2.63E-03 | -1.24E+00 | 4.35E-02 | 1.23E-01 | 1.21E+00 | 7.75E-08 | 2.84E-07 | -1.31E+00 | 9.75E-03 | 2.68E-02 | -1.04E+00 |
| CALCOCO2 | 2.95E-04 | 1.54E-03 | 1.48E+00 | 2.05E-01 | 8.85E-01 | 1.36E+00 | 4.26E-02 | 1.81E-01 | 1.18E+00 | 1.27E-01 | 4.85E-01 | 1.17E+00 | 6.21E-07 | 2.21E-06 | 1.02E+00 | 1.97E-10 | 5.56E-10 | -1.22E+00 | 1.19E-08 | 4.36E-08 | -1.35E+00 | 5.11E-08 | 1.40E-07 | -1.47E+00 |
| TBRG4 | 2.96E-04 | 1.54E-03 | -1.03E+00 | 2.49E-02 | 1.08E-01 | 1.36E+00 | 8.66E-06 | 3.68E-05 | -1.20E+00 | 5.51E-03 | 2.11E-02 | 1.14E+00 | 7.35E-05 | 2.62E-04 | -1.39E+00 | 8.67E-02 | 2.44E-01 | 1.17E+00 | 5.45E-05 | 2.00E-04 | -1.17E+00 | 3.40E-01 | 9.34E-01 | 1.21E+00 |
| BTBD11 | 2.96E-04 | 1.54E-03 | -1.22E+00 | 6.20E-04 | 2.68E-03 | -1.07E+00 | 1.55E-02 | 6.59E-02 | -1.11E+00 | 6.41E-03 | 2.45E-02 | -1.02E+00 | 2.52E-01 | 8.99E-01 | 1.51E+00 | 1.23E-01 | 3.46E-01 | 1.02E+00 | 1.76E-01 | 6.44E-01 | 1.48E+00 | 3.26E-01 | 8.96E-01 | 1.30E+00 |
| NADSYN1 | 2.97E-04 | 1.55E-03 | -1.03E+00 | 1.77E-04 | 7.64E-04 | 1.02E+00 | 1.53E-03 | 6.51E-03 | -1.11E+00 | 1.66E-02 | 6.34E-02 | 1.17E+00 | 4.56E-08 | 1.63E-07 | -1.30E+00 | 8.01E-05 | 2.26E-04 | -1.06E+00 | 7.91E-02 | 2.90E-01 | 1.19E+00 | 5.37E-02 | 1.47E-01 | 1.18E+00 |
| LETM2 | 2.97E-04 | 1.55E-03 | 2.50E+00 | 4.80E-02 | 2.07E-01 | 1.08E+00 | 7.74E-03 | 3.29E-02 | 1.74E+00 | 1.76E-01 | 6.74E-01 | 1.14E+00 | 4.99E-02 | 1.78E-01 | 1.88E+00 | 2.77E-01 | 7.82E-01 | 1.26E+00 | 2.16E-01 | 7.94E-01 | 1.35E+00 | 2.87E-02 | 7.88E-02 | -1.26E+00 |
| LINC00926 | 2.97E-04 | 1.55E-03 | -1.90E+00 | 8.34E-03 | 3.60E-02 | -1.12E+00 | 3.91E-02 | 1.66E-01 | -1.20E+00 | 2.88E-03 | 1.10E-02 | -1.44E+00 | 2.50E-01 | 8.91E-01 | 1.32E+00 | 3.45E-01 | 9.72E-01 | 1.20E+00 | 4.04E-02 | 1.48E-01 | -1.16E+00 | 1.02E-02 | 2.80E-02 | -1.47E+00 |
| AKAP8 | 2.98E-04 | 1.55E-03 | -1.10E+00 | 8.63E-09 | 3.73E-08 | -1.40E+00 | 2.80E-03 | 1.19E-02 | 1.02E+00 | 3.95E-10 | 1.51E-09 | -1.35E+00 | 3.28E-04 | 1.17E-03 | -1.06E+00 | 5.92E-06 | 1.67E-05 | -1.28E+00 | 9.49E-04 | 3.48E-03 | -1.13E+00 | 3.46E-04 | 9.51E-04 | -1.21E+00 |
| PYCR3 | 2.98E-04 | 1.55E-03 | -1.48E+00 | 1.99E-03 | 8.58E-03 | -1.16E+00 | 6.58E-02 | 2.80E-01 | 1.01E+00 | 1.07E-01 | 4.10E-01 | 1.15E+00 | 1.99E-02 | 7.08E-02 | -1.60E+00 | 9.78E-02 | 2.76E-01 | -1.04E+00 | 6.63E-02 | 2.43E-01 | -1.13E+00 | 2.70E-01 | 7.42E-01 | 1.25E+00 |
| RBMS1P1 | 2.99E-04 | 1.56E-03 | -1.43E+00 | 1.29E-02 | 5.56E-02 | 1.08E+00 | 2.65E-03 | 1.12E-02 | -1.25E+00 | 3.58E-03 | 1.37E-02 | -1.08E+00 | 4.04E-02 | 1.44E-01 | -1.04E+00 | 1.22E-01 | 3.43E-01 | 1.19E+00 | 8.17E-03 | 3.00E-02 | -1.28E+00 | 5.80E-02 | 1.59E-01 | -1.02E+00 |
| ZNF382 | 2.99E-04 | 1.56E-03 | -1.28E+00 | 6.42E-03 | 2.77E-02 | 1.10E+00 | 2.04E-07 | 8.67E-07 | -1.99E+00 | 2.45E-03 | 9.35E-03 | -1.02E+00 | 1.04E-03 | 3.72E-03 | -1.42E+00 | 9.78E-04 | 2.76E-03 | -1.52E+00 | 3.95E-03 | 1.45E-02 | -1.22E+00 | 4.41E-02 | 1.21E-01 | 1.02E+00 |
| FAM81B | 2.99E-04 | 1.56E-03 | -1.87E+00 | 5.03E-04 | 2.17E-03 | -1.71E+00 | 3.81E-02 | 1.62E-01 | -1.19E+00 | 1.46E-02 | 5.58E-02 | -1.33E+00 | 1.18E-03 | 4.22E-03 | -1.82E+00 | 4.22E-04 | 1.19E-03 | -1.96E+00 | 9.67E-02 | 3.55E-01 | 1.81E+00 | 2.57E-01 | 7.05E-01 | 1.13E+00 |
| LDLRAD4 | 3.00E-04 | 1.56E-03 | -1.36E+00 | 3.51E-04 | 1.52E-03 | -1.26E+00 | 6.80E-03 | 2.89E-02 | -1.18E+00 | 3.69E-03 | 1.41E-02 | -1.11E+00 | 2.50E-02 | 8.92E-02 | -1.17E+00 | 2.49E-01 | 7.02E-01 | 1.27E+00 | 4.91E-08 | 1.80E-07 | -1.36E+00 | 1.50E-06 | 4.13E-06 | -1.32E+00 |
| PURB | 3.00E-04 | 1.56E-03 | 1.67E+00 | 2.31E-03 | 9.97E-03 | 1.11E+00 | 5.55E-02 | 2.36E-01 | 1.36E+00 | 2.56E-02 | 9.79E-02 | 1.11E+00 | 1.76E-01 | 6.27E-01 | 1.39E+00 | 9.70E-04 | 2.73E-03 | -1.05E+00 | 8.25E-03 | 3.03E-02 | 1.02E+00 | 8.62E-03 | 2.37E-02 | -1.03E+00 |
| ZNF721 | 3.00E-04 | 1.56E-03 | -1.01E+00 | 6.26E-06 | 2.71E-05 | -1.07E+00 | 1.07E-03 | 4.54E-03 | -1.11E+00 | 6.41E-06 | 2.45E-05 | -1.17E+00 | 2.95E-02 | 1.05E-01 | 1.21E+00 | 1.80E-04 | 5.08E-04 | -1.12E+00 | 1.31E-02 | 4.82E-02 | -1.02E+00 | 7.29E-05 | 2.00E-04 | -1.21E+00 |
| AC027097.1 | 3.00E-04 | 1.56E-03 | -1.22E+00 | 5.01E-02 | 2.17E-01 | 1.29E+00 | 1.67E-07 | 7.10E-07 | -1.68E+00 | 1.33E-03 | 5.08E-03 | -1.04E+00 | 1.32E-02 | 4.71E-02 | -1.19E+00 | 2.01E-02 | 5.67E-02 | -1.12E+00 | 1.11E-03 | 4.07E-03 | -1.76E+00 | 1.24E-01 | 3.39E-01 | 1.01E+00 |
| AIP | 3.01E-04 | 1.57E-03 | 1.03E+00 | 3.69E-02 | 1.60E-01 | 1.24E+00 | 4.79E-04 | 2.04E-03 | -1.10E+00 | 3.56E-04 | 1.36E-03 | 1.05E+00 | 5.27E-04 | 1.88E-03 | -1.00E+00 | 1.58E-01 | 4.45E-01 | 1.23E+00 | 1.61E-03 | 5.90E-03 | -1.06E+00 | 3.63E-02 | 9.96E-02 | 1.08E+00 |
| AC018647.2 | 3.02E-04 | 1.57E-03 | -1.57E+00 | 6.75E-02 | 2.92E-01 | 1.17E+00 | 1.43E-05 | 6.09E-05 | -1.70E+00 | 1.69E-02 | 6.44E-02 | 1.05E+00 | 5.23E-05 | 1.87E-04 | -1.78E+00 | 6.43E-02 | 1.81E-01 | 1.07E+00 | 8.56E-03 | 3.14E-02 | -1.30E+00 | 5.86E-02 | 1.61E-01 | 1.01E+00 |
| DCPS | 3.03E-04 | 1.58E-03 | -1.12E+00 | 4.67E-04 | 2.02E-03 | 1.03E+00 | 2.07E-06 | 8.79E-06 | -1.21E+00 | 1.24E-07 | 4.72E-07 | -1.15E+00 | 2.72E-03 | 9.70E-03 | -1.22E+00 | 6.35E-02 | 1.79E-01 | 1.07E+00 | 1.13E-03 | 4.16E-03 | 1.02E+00 | 1.09E-01 | 3.00E-01 | 1.15E+00 |
| NDUFS5 | 3.03E-04 | 1.58E-03 | -1.04E+00 | 4.22E-25 | 1.82E-24 | -1.32E+00 | 1.59E-01 | 6.76E-01 | 1.12E+00 | 3.10E-05 | 1.18E-04 | -1.19E+00 | 2.14E-01 | 7.64E-01 | 1.18E+00 | 5.65E-04 | 1.59E-03 | -1.11E+00 | 2.42E-01 | 8.89E-01 | 1.06E+00 | 5.46E-02 | 1.50E-01 | -1.11E+00 |
| IFT80 | 3.03E-04 | 1.58E-03 | -1.23E+00 | 2.12E-02 | 9.18E-02 | 1.17E+00 | 1.97E-05 | 8.39E-05 | -1.48E+00 | 7.55E-03 | 2.88E-02 | 1.07E+00 | 5.51E-04 | 1.96E-03 | -1.23E+00 | 4.00E-02 | 1.13E-01 | 1.04E+00 | 6.00E-03 | 2.20E-02 | -1.03E+00 | 2.04E-01 | 5.60E-01 | 1.23E+00 |
| AC008440.1 | 3.04E-04 | 1.58E-03 | 2.12E+00 | 7.08E-02 | 3.06E-01 | 1.03E+00 | 3.53E-04 | 1.50E-03 | 2.27E+00 | 1.61E-01 | 6.15E-01 | 1.12E+00 | 8.26E-02 | 2.94E-01 | 2.13E+00 | 2.93E-01 | 8.27E-01 | 1.17E+00 | 3.73E-03 | 1.37E-02 | 3.30E+00 | 1.17E-01 | 3.22E-01 | 1.90E+00 |
| [illegible] | [illegible] | [illegible] | [illegible] | [illegible] | [illegible] | [illegible] | [illegible] | [illegible] | [illegible] | [illegible] | [illegible] | [illegible] | [illegible] | [illegible] | [illegible] | [illegible] | [illegible] | [illegible] | [illegible] | [illegible] | [illegible] | [illegible] | [illegible] | [illegible] |

| | | | | | | | | | | | | | | | | | | | | | | | | |
|---|---|---|---|---|---|---|---|---|---|---|---|---|---|---|---|---|---|---|---|---|---|---|---|---|
| TMEM14A | 3.78E-04 | 1.97E-03 | -1.07E+00 | 3.28E-03 | 1.42E-02 | 1.11E+00 | 4.38E-02 | 1.86E-01 | 1.07E+00 | 9.67E-02 | 3.69E-01 | 1.32E+00 | 1.22E-03 | 4.36E-03 | -1.24E+00 | 8.18E-03 | 2.30E-02 | -1.05E+00 | 7.17E-02 | 2.63E-01 | 1.09E+00 | 1.65E-01 | 4.53E-01 | 1.18E+00 |
| RNF14 | 3.78E-04 | 1.97E-03 | -1.16E+00 | 2.76E-03 | 1.19E-02 | 1.05E+00 | 1.06E-05 | 4.49E-05 | -1.52E+00 | 1.38E-05 | 5.26E-05 | -1.27E+00 | 4.04E-03 | 1.44E-02 | -1.17E+00 | 5.72E-02 | 1.61E-01 | 1.05E+00 | 6.37E-04 | 2.34E-03 | -1.11E+00 | 1.13E-02 | 3.10E-02 | 1.03E+00 |
| MAP3K21 | 3.79E-04 | 1.97E-03 | -1.59E+00 | 4.74E-03 | 2.05E-02 | -1.08E+00 | 2.65E-12 | 1.13E-11 | -2.11E+00 | 5.20E-10 | 1.99E-09 | -1.65E+00 | 1.54E-01 | 5.48E-01 | -1.06E+00 | 2.94E-01 | 8.28E-01 | 1.23E+00 | 1.21E-06 | 4.43E-06 | -2.50E+00 | 2.57E-04 | 7.05E-04 | -1.66E+00 |
| TUT1 | 3.79E-04 | 1.98E-03 | -1.57E+00 | 9.93E-03 | 4.29E-02 | -1.09E+00 | 3.82E-07 | 1.62E-06 | -1.85E+00 | 5.80E-04 | 2.21E-03 | -1.16E+00 | 1.13E-02 | 4.04E-02 | -1.20E+00 | 1.73E-01 | 4.89E-01 | 1.26E+00 | 9.07E-12 | 3.33E-11 | -2.12E+00 | 2.65E-05 | 7.29E-05 | -1.28E+00 |
| KIF21B | 3.80E-04 | 1.98E-03 | -1.34E+00 | 2.02E-05 | 8.74E-05 | -1.51E+00 | 2.11E-05 | 8.98E-05 | -1.70E+00 | 1.46E-03 | 5.57E-03 | -1.13E+00 | 1.35E-04 | 4.80E-04 | -1.52E+00 | 6.59E-05 | 1.86E-04 | -1.70E+00 | 1.03E-01 | 3.76E-01 | 1.03E+00 | 2.69E-01 | 7.38E-01 | 1.11E+00 |
| TRIM46 | 3.81E-04 | 1.99E-03 | -1.19E+00 | 5.32E-03 | 2.30E-02 | 1.02E+00 | 6.57E-03 | 2.79E-02 | -1.29E+00 | 7.85E-02 | 3.00E-01 | 1.14E+00 | 1.12E-02 | 4.01E-02 | -1.81E+00 | 1.39E-01 | 3.92E-01 | -1.01E+00 | 1.35E-01 | 4.96E-01 | -1.21E+00 | 3.08E-01 | 8.45E-01 | -1.02E+00 |
| AC007686.1 | 3.81E-04 | 1.99E-03 | -1.23E+00 | 2.64E-08 | 1.14E-07 | -1.94E+00 | 1.95E-01 | 8.27E-01 | 1.13E+00 | 4.55E-03 | 1.74E-02 | -1.37E+00 | 1.57E-01 | 5.60E-01 | 1.15E+00 | 1.46E-03 | 4.12E-03 | -2.11E+00 | 5.84E-02 | 2.14E-01 | 1.02E+00 | 7.46E-03 | 2.05E-02 | -1.39E+00 |
| ABHD14A-ACY1 | 3.83E-04 | 1.99E-03 | -1.48E+00 | 9.55E-05 | 4.13E-04 | -1.76E+00 | 1.84E-02 | 7.82E-02 | -1.21E+00 | 5.44E-02 | 2.08E-01 | 1.01E+00 | 5.94E-03 | 2.12E-02 | -1.40E+00 | 7.59E-03 | 2.14E-02 | -1.35E+00 | 2.95E-04 | 1.08E-03 | -1.90E+00 | 8.51E-02 | 2.34E-01 | -1.03E+00 |
| IL7 | 3.83E-04 | 2.00E-03 | -2.17E+00 | 1.84E-03 | 7.97E-03 | -1.67E+00 | 1.45E-02 | 6.17E-02 | -1.52E+00 | 9.44E-03 | 3.61E-02 | -1.75E+00 | 1.36E-21 | 4.84E-21 | -2.63E+00 | 2.32E-14 | 6.54E-14 | -1.65E+00 | 1.33E-41 | 4.88E-41 | -2.54E+00 | 4.04E-22 | 1.11E-21 | -1.71E+00 |
| ACOT4 | 3.85E-04 | 2.01E-03 | -1.60E+00 | 3.15E-02 | 1.36E-01 | 1.12E+00 | 1.99E-03 | 8.46E-03 | -1.77E+00 | 1.75E-01 | 6.68E-01 | 1.09E+00 | 9.27E-03 | 3.31E-02 | -1.44E+00 | 2.39E-01 | 6.75E-01 | 1.32E+00 | 2.83E-04 | 1.04E-03 | -1.00E+00 | 2.43E-01 | 6.67E-01 | 1.48E+00 |
| TMEM147-AS1 | 3.85E-04 | 2.01E-03 | -1.45E+00 | 1.55E-01 | 6.71E-01 | 1.53E+00 | 1.61E-04 | 6.83E-04 | -1.61E+00 | 1.80E-01 | 6.86E-01 | 1.30E+00 | 1.69E-01 | 6.04E-01 | 1.45E+00 | 3.97E-02 | 1.12E-01 | 1.83E+00 | 5.65E-02 | 2.07E-01 | -1.14E+00 | 1.24E-01 | 3.40E-01 | 1.37E+00 |
| HDHD2 | 3.86E-04 | 2.01E-03 | -1.70E+00 | 9.24E-04 | 3.99E-03 | -1.53E+00 | 1.40E-06 | 5.95E-06 | -1.94E+00 | 2.10E-04 | 8.02E-04 | -1.31E+00 | 1.03E-01 | 3.68E-01 | 1.86E+00 | 1.93E-01 | 5.44E-01 | 1.62E+00 | 4.74E-05 | 1.74E-04 | -1.87E+00 | 4.68E-03 | 1.29E-02 | -1.43E+00 |
| SEC11C | 3.87E-04 | 2.01E-03 | -1.02E+00 | 1.54E-03 | 6.65E-03 | 1.15E+00 | 1.10E-01 | 4.68E-01 | 1.10E+00 | 2.02E-02 | 7.73E-02 | 1.15E+00 | 4.55E-04 | 1.62E-03 | -1.09E+00 | 7.73E-03 | 2.18E-02 | 1.03E+00 | 1.72E-01 | 6.32E-01 | 1.12E+00 | 3.33E-01 | 9.14E-01 | 1.20E+00 |
| TLK2 | 3.88E-04 | 2.02E-03 | -1.01E+00 | 3.30E-04 | 1.43E-03 | 1.13E+00 | 6.06E-06 | 2.57E-05 | -1.19E+00 | 1.20E-07 | 4.57E-07 | -1.14E+00 | 1.47E-02 | 5.23E-02 | 1.04E+00 | 1.12E-02 | 3.14E-02 | 1.05E+00 | 2.36E-03 | 8.65E-03 | -1.03E+00 | 5.77E-04 | 1.58E-03 | -1.22E+00 |
| CFLAR | 3.90E-04 | 2.03E-03 | 1.04E+00 | 9.71E-03 | 4.19E-02 | 1.24E+00 | 1.78E-03 | 7.58E-03 | -1.14E+00 | 8.68E-03 | 3.32E-02 | 1.00E+00 | 5.37E-02 | 1.92E-01 | 1.10E+00 | 2.22E-01 | 6.25E-01 | 1.39E+00 | 2.98E-03 | 1.10E-02 | -1.07E+00 | 1.13E-03 | 3.12E-03 | -1.11E+00 |
| AC006277.1 | 3.91E-04 | 2.04E-03 | 2.28E+00 | 8.03E-02 | 3.47E-01 | 1.07E+00 | 1.37E-03 | 5.83E-03 | 2.12E+00 | 1.24E-01 | 4.73E-01 | 1.01E+00 | 1.84E-01 | 6.56E-01 | 1.60E+00 | 1.37E-01 | 3.87E-01 | -1.19E+00 | 1.83E-01 | 6.71E-01 | 1.47E+00 | 2.39E-01 | 6.58E-01 | -1.19E+00 |
| ZNF182 | 3.92E-04 | 2.04E-03 | -1.43E+00 | 2.60E-03 | 1.12E-02 | -1.09E+00 | 7.57E-04 | 3.22E-03 | -1.49E+00 | 3.20E-02 | 1.22E-01 | 1.03E+00 | 4.79E-06 | 1.71E-05 | -1.94E+00 | 1.22E-04 | 3.44E-04 | -1.55E+00 | 1.30E-04 | 4.76E-04 | -1.55E+00 | 3.41E-02 | 9.36E-02 | -1.08E+00 |
| EIF4G1 | 3.92E-04 | 2.04E-03 | -1.11E+00 | 2.18E-01 | 9.42E-01 | 1.38E+00 | 6.69E-02 | 2.84E-01 | 1.03E+00 | 9.95E-02 | 3.80E-01 | 1.23E+00 | 7.65E-04 | 2.73E-03 | -1.18E+00 | 2.20E-01 | 6.20E-01 | 1.22E+00 | 5.83E-05 | 2.14E-04 | -1.10E+00 | 1.68E-01 | 4.62E-01 | 1.19E+00 |
| RNF26 | 3.93E-04 | 2.05E-03 | -1.49E+00 | 4.93E-02 | 2.13E-01 | 1.23E+00 | 8.32E-04 | 3.53E-03 | -1.17E+00 | 2.88E-03 | 1.10E-02 | 1.01E+00 | 5.75E-04 | 2.05E-03 | -1.76E+00 | 4.12E-02 | 1.16E-01 | -1.10E+00 | 1.34E-05 | 4.91E-05 | -1.49E+00 | 1.14E-01 | 3.13E-01 | 1.17E+00 |
| AC005165.2 | 3.93E-04 | 2.05E-03 | 1.91E+00 | 8.41E-02 | 3.63E-01 | 1.81E+00 | 9.37E-04 | 3.98E-03 | 1.69E+00 | 1.75E-01 | 6.69E-01 | 1.42E+00 | 3.74E-04 | 1.33E-03 | 1.97E+00 | 6.84E-02 | 1.93E-01 | 1.68E+00 | 1.75E-02 | 6.42E-02 | 1.66E+00 | 2.97E-01 | 8.16E-01 | 1.37E+00 |
| TTPAL | 3.93E-04 | 2.05E-03 | -1.07E+00 | 3.76E-04 | 1.63E-03 | -1.02E+00 | 6.87E-02 | 2.92E-01 | 1.07E+00 | 7.79E-03 | 2.97E-02 | -1.03E+00 | 1.86E-01 | 6.64E-01 | 1.22E+00 | 7.82E-02 | 2.20E-01 | 1.11E+00 | 9.17E-02 | 3.37E-01 | 1.17E+00 | 8.16E-02 | 2.24E-01 | 1.13E+00 |
| ERICH6-AS1 | 3.93E-04 | 2.05E-03 | -1.06E+00 | 2.69E-04 | 1.16E-03 | -1.02E+00 | 7.07E-02 | 3.00E-01 | -1.03E+00 | 3.54E-02 | 1.35E-01 | 1.08E+00 | 2.65E-01 | 9.46E-01 | 1.34E+00 | 1.26E-01 | 3.55E-01 | 1.00E+00 | 9.00E-02 | 3.30E-01 | 1.59E+00 | 3.51E-01 | 9.64E-01 | 1.21E+00 |
| THAP2 | 3.93E-04 | 2.05E-03 | -1.23E+00 | 1.98E-03 | 8.54E-03 | -1.04E+00 | 7.40E-05 | 3.14E-04 | -1.51E+00 | 1.18E-05 | 4.50E-05 | -1.45E+00 | 2.66E-03 | 9.49E-03 | -1.20E+00 | 6.97E-03 | 1.97E-02 | -1.01E+00 | 8.99E-06 | 3.30E-05 | -1.52E+00 | 7.97E-04 | 2.19E-03 | -1.29E+00 |
| CDKL5 | 3.94E-04 | 2.05E-03 | -1.37E+00 | 6.71E-02 | 2.90E-01 | 1.24E+00 | 5.38E-04 | 2.28E-03 | -1.41E+00 | 5.05E-05 | 1.93E-04 | -1.50E+00 | 4.05E-02 | 1.44E-01 | -1.44E+00 | 5.85E-02 | 1.65E-01 | -1.34E+00 | 1.46E-01 | 5.37E-01 | -1.11E+00 | 2.66E-01 | 7.31E-01 | -1.01E+00 |
| ZNF641 | 3.96E-04 | 2.06E-03 | -1.38E+00 | 7.65E-03 | 3.31E-02 | -1.06E+00 | 9.41E-07 | 4.00E-06 | -2.01E+00 | 6.71E-06 | 2.56E-05 | -1.65E+00 | 1.19E-04 | 4.26E-04 | -1.90E+00 | 1.65E-03 | 4.64E-03 | -1.53E+00 | 4.43E-04 | 1.63E-03 | -1.18E+00 | 1.75E-06 | 4.82E-06 | -1.51E+00 |
| ECT2L | 3.96E-04 | 2.06E-03 | -1.63E+00 | 3.13E-03 | 1.35E-02 | -1.23E+00 | 9.13E-02 | 3.88E-01 | 1.02E+00 | 1.45E-01 | 5.53E-01 | 1.11E+00 | 2.47E-01 | 8.81E-01 | 1.37E+00 | 2.51E-01 | 7.09E-01 | 1.22E+00 | 2.02E-01 | 7.40E-01 | -1.05E+00 | 9.43E-02 | 2.59E-01 | -1.42E+00 |
| ZNF613 | 3.97E-04 | 2.07E-03 | -1.62E+00 | 1.73E-03 | 7.47E-03 | -1.47E+00 | 9.79E-03 | 4.16E-02 | -1.41E+00 | 5.50E-02 | 2.10E-01 | 1.00E+00 | 1.20E-01 | 4.27E-01 | -1.20E+00 | 3.24E-01 | 9.13E-01 | 1.18E+00 | 9.95E-02 | 3.65E-01 | -1.07E+00 | 1.06E-01 | 2.91E-01 | 1.04E+00 |
| SAMSN1 | 3.97E-04 | 2.07E-03 | -1.14E+00 | 6.71E-04 | 2.90E-03 | -1.00E+00 | 5.34E-04 | 2.27E-03 | -1.16E+00 | 7.25E-06 | 2.77E-05 | -1.19E+00 | 1.26E-08 | 4.49E-08 | -1.50E+00 | 1.85E-03 | 5.22E-03 | -1.07E+00 | 1.53E-09 | 5.61E-09 | -1.45E+00 | 8.13E-05 | 2.23E-04 | -1.14E+00 |
| PTGES2 | 3.97E-04 | 2.07E-03 | -1.21E+00 | 7.76E-03 | 3.35E-02 | 1.10E+00 | 9.49E-06 | 4.03E-05 | -1.41E+00 | 7.66E-03 | 2.93E-02 | 1.11E+00 | 4.15E-09 | 1.48E-08 | -1.91E+00 | 1.53E-02 | 4.30E-02 | 1.12E+00 | 6.70E-07 | 2.46E-06 | -1.08E+00 | 1.61E-01 | 4.43E-01 | 1.23E+00 |
| DOPEY2 | 3.98E-04 | 2.08E-03 | -1.25E+00 | 5.94E-02 | 2.57E-01 | 1.26E+00 | 7.70E-05 | 3.27E-04 | -1.27E+00 | 4.49E-04 | 1.72E-03 | -1.00E+00 | 1.50E-05 | 5.36E-05 | -1.38E+00 | 9.53E-06 | 2.69E-05 | -1.43E+00 | 3.48E-04 | 1.28E-03 | -1.62E+00 | 4.77E-02 | 1.31E-01 | -1.12E+00 |
| XPA | 3.98E-04 | 2.08E-03 | -1.00E+00 | 1.78E-03 | 7.69E-03 | 1.17E+00 | 5.07E-02 | 2.15E-01 | 1.04E+00 | 7.42E-03 | 2.83E-02 | 1.03E+00 | 1.05E-01 | 3.74E-01 | 1.05E+00 | 2.07E-01 | 5.84E-01 | 1.11E+00 | 9.99E-03 | 3.67E-02 | -1.19E+00 | 9.87E-02 | 2.71E-01 | -1.03E+00 |
| UBR4 | 3.99E-04 | 2.08E-03 | -1.11E+00 | 6.98E-03 | 3.02E-02 | 1.13E+00 | 7.57E-04 | 3.22E-03 | -1.16E+00 | 2.02E-04 | 7.70E-04 | -1.11E+00 | 9.20E-09 | 3.28E-08 | -1.31E+00 | 4.16E-05 | 1.17E-04 | -1.08E+00 | 2.99E-03 | 1.10E-02 | -1.00E+00 | 3.41E-03 | 9.37E-03 | -1.01E+00 |
| NOMO3 | 3.99E-04 | 2.08E-03 | -1.60E+00 | 2.64E-03 | 1.14E-02 | -1.19E+00 | 2.23E-02 | 9.49E-02 | -1.14E+00 | 4.59E-03 | 1.75E-02 | -1.27E+00 | 1.11E-01 | 3.95E-01 | -1.15E+00 | 1.03E-01 | 2.91E-01 | -1.14E+00 | 2.84E-03 | 1.04E-02 | -1.25E+00 | 3.77E-02 | 1.04E-01 | 1.01E+00 |
| CHEK1 | 3.99E-04 | 2.08E-03 | -1.06E+00 | 3.85E-04 | 1.67E-03 | 1.02E+00 | 4.01E-02 | 1.70E-01 | 1.33E+00 | 1.27E-01 | 4.83E-01 | 1.29E+00 | 2.23E-02 | 7.94E-02 | -1.00E+00 | 7.75E-03 | 2.18E-02 | -1.08E+00 | 3.14E-03 | 1.15E-02 | 1.00E+00 | 1.97E-01 | 5.40E-01 | 1.27E+00 |
| FNDC3B | 3.99E-04 | 2.08E-03 | -1.22E+00 | 1.78E-07 | 7.69E-07 | -1.40E+00 | 3.54E-03 | 1.50E-02 | -1.04E+00 | 5.52E-06 | 2.11E-05 | -1.16E+00 | 2.93E-03 | 1.04E-02 | -1.19E+00 | 8.56E-07 | 2.41E-06 | -1.81E+00 | 7.73E-08 | 2.84E-07 | -1.86E+00 | 6.41E-05 | 1.76E-04 | -1.37E+00 |
| RASAL3 | 3.99E-04 | 2.08E-03 | -1.06E+00 | 1.33E-03 | 5.75E-03 | 1.14E+00 | 1.67E-02 | 7.10E-02 | -1.03E+00 | 9.66E-03 | 3.69E-02 | 1.04E+00 | 1.04E-02 | 3.72E-02 | 1.07E+00 | 1.14E-01 | 3.22E-01 | 1.24E+00 | 2.08E-02 | 7.65E-02 | 1.04E+00 | 7.97E-05 | 2.19E-04 | -1.20E+00 |
| CDK16 | 4.00E-04 | 2.08E-03 | -1.15E+00 | 2.04E-03 | 8.81E-03 | -1.02E+00 | 5.45E-02 | 2.32E-01 | 1.03E+00 | 5.48E-03 | 2.09E-02 | -1.05E+00 | 1.10E-01 | 3.93E-01 | 1.18E+00 | 1.76E-01 | 4.97E-01 | 1.25E+00 | 4.27E-04 | 1.57E-03 | -1.15E+00 | 1.45E-02 | 3.98E-02 | 1.05E+00 |
| AC083798.2 | 4.00E-04 | 2.08E-03 | -1.12E+00 | 2.75E-08 | 1.19E-07 | -1.36E+00 | 6.32E-03 | 2.69E-02 | -1.16E+00 | 9.72E-05 | 3.71E-04 | -1.35E+00 | 1.31E-02 | 4.66E-02 | -1.16E+00 | 3.34E-03 | 9.43E-03 | -1.29E+00 | 1.70E-01 | 6.24E-01 | 1.10E+00 | 7.48E-04 | 2.06E-03 | -1.43E+00 |
| U2AF1L4 | 4.00E-04 | 2.09E-03 | 1.06E+00 | 1.11E-05 | 4.80E-05 | -1.01E+00 | 1.04E-02 | 4.41E-02 | -1.17E+00 | 6.34E-07 | 2.42E-06 | -1.48E+00 | 1.95E-04 | 6.95E-04 | -1.31E+00 | 1.85E-04 | 5.22E-04 | -1.29E+00 | 6.69E-04 | 2.45E-03 | -1.15E+00 | 2.16E-04 | 5.93E-04 | -1.26E+00 |
| NDUFB9 | 4.00E-04 | 2.09E-03 | 1.11E+00 | 2.12E-06 | 9.16E-06 | 1.10E+00 | 1.16E-01 | 4.94E-01 | 1.24E+00 | 7.13E-02 | 2.73E-01 | 1.18E+00 | 4.39E-02 | 1.56E-01 | 1.25E+00 | 1.53E-01 | 4.32E-01 | 1.32E+00 | 9.68E-02 | 3.55E-01 | 1.32E+00 | 2.50E-02 | 6.87E-02 | 1.40E+00 |
| ZBTB14 | 4.01E-04 | 2.09E-03 | -1.35E+00 | 4.08E-03 | 1.76E-02 | 1.01E+00 | 3.41E-08 | 1.45E-07 | -1.72E+00 | 6.12E-06 | 2.34E-05 | -1.29E+00 | 2.48E-05 | 8.85E-05 | -1.68E+00 | 3.59E-03 | 1.01E-02 | -1.20E+00 | 3.47E-04 | 1.27E-03 | -1.25E+00 | 2.04E-02 | 5.60E-02 | -1.03E+00 |
| RAB8A | 4.01E-04 | 2.09E-03 | 1.02E+00 | 6.75E-05 | 2.92E-04 | 1.05E+00 | 3.39E-04 | 1.44E-03 | -1.16E+00 | 1.94E-05 | 7.41E-05 | -1.14E+00 | 1.47E-02 | 5.24E-02 | 1.09E+00 | 3.89E-05 | 1.10E-04 | -1.21E+00 | 1.06E-01 | 3.89E-01 | 1.13E+00 | 1.72E-02 | 4.72E-02 | -1.06E+00 |
| GIPC2 | 4.01E-04 | 2.09E-03 | -1.12E+00 | 6.98E-02 | 3.02E-01 | 1.34E+00 | 9.77E-02 | 4.15E-01 | -1.13E+00 | 2.46E-01 | 9.39E-01 | 1.32E+00 | 2.06E-01 | 7.36E-01 | 1.75E+00 | 3.05E-02 | 8.59E-02 | 4.27E+00 | 1.94E-01 | 7.11E-01 | -1.05E+00 | 2.06E-01 | 5.67E-01 | -1.02E+00 |
| WDR44 | 4.03E-04 | 2.10E-03 | -1.47E+00 | 1.33E-03 | 5.77E-03 | -1.30E+00 | 2.29E-03 | 9.74E-03 | -1.42E+00 | 6.44E-03 | 2.46E-02 | -1.16E+00 | 3.60E-02 | 1.28E-01 | 1.03E+00 | 6.43E-02 | 1.81E-01 | 1.07E+00 | 3.94E-08 | 1.44E-07 | -1.47E+00 | 2.48E-08 | 6.80E-08 | -1.52E+00 |
| HSPBP1 | 4.03E-04 | 2.10E-03 | -1.21E+00 | 4.16E-03 | 1.80E-02 | 1.04E+00 | 9.45E-06 | 4.01E-05 | -1.31E+00 | 6.16E-06 | 2.35E-05 | -1.21E+00 | 4.97E-04 | 1.77E-03 | -1.21E+00 | 1.83E-02 | 5.15E-02 | 1.08E+00 | 4.17E-06 | 1.53E-05 | -1.22E+00 | 8.99E-03 | 2.47E-02 | 1.05E+00 |
| RAD51D | 4.03E-04 | 2.10E-03 | -1.08E+00 | 1.03E-01 | 4.43E-01 | 1.36E+00 | 1.46E-07 | 6.19E-07 | -1.45E+00 | 2.41E-04 | 9.21E-04 | 1.03E+00 | 1.13E-05 | 4.03E-05 | -1.60E+00 | 5.35E-03 | 1.51E-02 | -1.10E+00 | 5.36E-08 | 1.97E-07 | -1.45E+00 | 1.72E-01 | 4.73E-01 | 1.11E+00 |
| VAMP8 | 4.05E-04 | 2.11E-03 | -1.11E+00 | 2.20E-04 | 9.50E-04 | -1.08E+00 | 1.05E-02 | 4.44E-02 | -1.12E+00 | 2.17E-03 | 8.27E-03 | -1.18E+00 | 1.02E-08 | 3.64E-08 | -1.23E+00 | 3.84E-05 | 1.08E-04 | -1.16E+00 | 3.83E-03 | 1.41E-02 | -1.09E+00 | 2.00E-01 | 5.49E-01 | -1.02E+00 |
| KCTD9 | 4.05E-04 | 2.11E-03 | -1.28E+00 | 3.56E-04 | 1.54E-03 | -1.12E+00 | 8.81E-07 | 3.74E-06 | -1.57E+00 | 6.95E-06 | 2.65E-05 | -1.40E+00 | 5.38E-04 | 1.92E-03 | -1.69E+00 | 4.80E-04 | 1.35E-03 | -1.71E+00 | 1.13E-06 | 4.15E-06 | -1.39E+00 | 6.95E-07 | 1.91E-06 | -1.45E+00 |
| RNF217-AS1 | 4.05E-04 | 2.11E-03 | -1.61E+00 | 1.71E-03 | 7.39E-03 | -1.37E+00 | 4.30E-04 | 1.83E-03 | -1.87E+00 | 4.16E-04 | 1.59E-03 | -1.84E+00 | 2.01E-01 | 7.18E-01 | 1.21E+00 | 2.79E-01 | 7.86E-01 | 1.13E+00 | 2.68E-01 | 9.84E-01 | 1.40E+00 | 3.30E-01 | 9.06E-01 | 1.08E+00 |
| HIVEP3 | 4.06E-04 | 2.11E-03 | -1.01E+00 | 1.43E-01 | 6.19E-01 | 1.55E+00 | 1.55E-02 | 6.57E-02 | -1.02E+00 | 1.25E-01 | 4.78E-01 | 1.27E+00 | 1.89E-01 | 6.73E-01 | 1.87E+00 | 1.95E-01 | 5.49E-01 | 2.06E+00 | 1.98E-01 | 7.25E-01 | 1.02E+00 | 2.67E-01 | 7.34E-01 | 1.42E+00 |
| RPL7AP60 | 4.06E-04 | 2.11E-03 | -1.10E+00 | 8.62E-09 | 3.73E-08 | -1.49E+00 | 5.79E-03 | 2.46E-02 | -1.11E+00 | 3.55E-08 | 1.36E-07 | -1.60E+00 | 1.65E-01 | 5.88E-01 | 1.23E+00 | 1.47E-03 | 4.15E-03 | -1.28E+00 | 1.85E-01 | 6.80E-01 | 1.19E+00 | 1.30E-05 | 3.56E-05 | -1.52E+00 |
| FAM212A | 4.06E-04 | 2.12E-03 | -1.13E+00 | 3.05E-03 | 1.32E-02 | 1.85E+00 | 7.28E-10 | 3.09E-09 | -1.60E+00 | 7.11E-03 | 2.71E-02 | 1.14E+00 | 4.03E-09 | 1.44E-08 | -1.35E+00 | 7.75E-02 | 2.18E-01 | 1.26E+00 | 1.51E-05 | 5.56E-05 | 1.01E+00 | 8.87E-02 | 2.44E-01 | 1.19E+00 |
| LY75 | 4.06E-04 | 2.12E-03 | -1.31E+00 | 2.17E-01 | 9.38E-01 | 1.47E+00 | 3.42E-04 | 1.45E-03 | -1.29E+00 | 1.19E-01 | 4.53E-01 | 1.35E+00 | 2.13E-03 | 7.60E-03 | -1.45E+00 | 6.46E-02 | 1.82E-01 | 1.09E+00 | 2.25E-02 | 8.26E-02 | -1.17E+00 | 1.99E-01 | 5.47E-01 | 1.52E+00 |
| KLF10 | 4.06E-04 | 2.12E-03 | 1.07E+00 | 7.18E-26 | 3.10E-25 | -2.10E+00 | 7.94E-06 | 3.37E-05 | -1.15E+00 | 1.49E-12 | 5.69E-12 | -1.52E+00 | 5.61E-04 | 2.00E-03 | -1.27E+00 | 8.70E-10 | 2.45E-09 | -2.73E+00 | 3.66E-03 | 1.34E-02 | -1.01E+00 | 9.24E-10 | 2.54E-09 | -2.52E+00 |
| AL031282.2 | 4.07E-04 | 2.12E-03 | -1.30E+00 | 1.56E-02 | 6.73E-02 | 1.13E+00 | 4.00E-03 | 1.70E-02 | -1.32E+00 | 1.17E-02 | 4.45E-02 | -1.07E+00 | 1.52E-02 | 5.41E-02 | -1.22E+00 | 6.59E-02 | 1.86E-01 | 1.01E+00 | 5.75E-05 | 2.11E-04 | -1.70E+00 | 4.37E-02 | 1.20E-01 | -1.05E+00 |
| AC004951.1 | 4.09E-04 | 2.13E-03 | -1.68E+00 | 9.06E-03 | 3.91E-02 | -1.10E+00 | 3.63E-04 | 1.54E-03 | -1.45E+00 | 8.17E-03 | 3.12E-02 | -1.06E+00 | 1.90E-05 | 6.76E-05 | -2.16E+00 | 1.85E-04 | 5.23E-04 | -1.70E+00 | 5.37E-02 | 1.97E-01 | -1.23E+00 | 2.63E-01 | 7.22E-01 | 1.07E+00 |
| MARCH9 | 4.09E-04 | 2.13E-03 | -1.13E+00 | 9.05E-03 | 3.91E-02 | 1.19E+00 | 4.12E-03 | 1.75E-02 | -1.13E+00 | 9.47E-02 | 3.62E-01 | 1.22E+00 | 5.19E-03 | 1.85E-02 | -1.02E+00 | 2.51E-02 | 7.09E-02 | 1.03E+00 | 1.10E-01 | 4.05E-01 | 1.05E+00 | 3.37E-01 | 9.27E-01 | 1.27E+00 |
| AC099689.1 | 4.10E-04 | 2.14E-03 | -1.17E+00 | 1.04E-03 | 4.49E-03 | -1.09E+00 | 8.90E-02 | 3.78E-01 | -1.06E+00 | 2.60E-02 | 9.93E-02 | -1.09E+00 | 2.68E-01 | 9.56E-01 | 1.35E+00 | 3.34E-01 | 9.41E-01 | 1.26E+00 | 1.61E-03 | 5.89E-03 | -1.35E+00 | 3.82E-03 | 1.05E-02 | -1.32E+00 |
| ATP1B3 | 4.10E-04 | 2.14E-03 | 1.06E+00 | 1.76E-01 | 7.61E-01 | 1.45E+00 | 1.27E-01 | 5.41E-01 | 1.08E+00 | 2.56E-01 | 9.78E-01 | 1.34E+00 | 3.60E-05 | 1.28E-04 | -1.12E+00 | 2.53E-02 | 7.14E-02 | 1.09E+00 | 1.66E-02 | 6.10E-02 | 1.01E+00 | 3.91E-02 | 1.07E-01 | 1.43E+00 |
| ARFGEF2 | 4.11E-04 | 2.14E-03 | -1.33E+00 | 2.46E-03 | 1.06E-02 | -1.08E+00 | 3.40E-04 | 1.44E-03 | -1.24E+00 | 5.27E-05 | 2.01E-04 | -1.12E+00 | 4.38E-03 | 1.56E-02 | -1.12E+00 | 2.07E-03 | 5.82E-03 | -1.16E+00 | 9.74E-04 | 3.57E-03 | -1.24E+00 | 3.13E-02 | 8.59E-02 | -1.06E+00 |
| RMND5B | 4.11E-04 | 2.14E-03 | -1.11E+00 | 5.81E-04 | 2.51E-03 | 1.02E+00 | 1.81E-04 | 7.68E-04 | -1.37E+00 | 5.31E-05 | 2.03E-04 | -1.30E+00 | 1.52E-03 | 5.44E-03 | -1.11E+00 | 2.27E-02 | 6.41E-02 | 1.07E+00 | 4.29E-04 | 1.58E-03 | 1.02E+00 | 2.49E-02 | 6.85E-02 | 1.06E+00 |
| LDHB | 4.12E-04 | 2.15E-03 | -1.09E+00 | 1.63E-01 | 7.02E-01 | 1.03E+00 | 1.68E-01 | 7.13E-01 | -1.01E+00 | 2.13E-01 | 8.13E-01 | 1.04E+00 | 8.49E-04 | 3.03E-03 | -1.15E+00 | 3.00E-01 | 8.47E-01 | 1.04E+00 | 3.02E-02 | 1.11E-01 | -1.03E+00 | 6.43E-02 | 1.77E-01 | 1.13E+00 |
| TFCP2 | 4.12E-04 | 2.15E-03 | -1.19E+00 | 1.20E-01 | 5.20E-01 | 1.45E+00 | 4.30E-04 | 1.83E-03 | -1.26E+00 | 7.62E-02 | 2.91E-01 | 1.26E+00 | 1.78E-07 | 6.36E-07 | -1.56E+00 | 1.53E-03 | 4.33E-03 | -1.04E+00 | 2.03E-07 | 7.45E-07 | -1.27E+00 | 5.79E-02 | 1.59E-01 | 1.20E+00 |
| GCSHP5 | 4.14E-04 | 2.16E-03 | -1.11E+00 | 1.09E-06 | 4.69E-06 | -1.29E+00 | 5.35E-02 | 2.27E-01 | 1.15E+00 | 1.62E-02 | 6.19E-02 | 1.19E+00 | 2.71E-03 | 9.65E-03 | -1.21E+00 | 5.87E-02 | 1.66E-01 | 1.08E+00 | 2.13E-06 | 7.80E-06 | -1.25E+00 | 2.45E-03 | 6.74E-03 | -1.09E+00 |
| DAGLB | 4.15E-04 | 2.17E-03 | -1.23E+00 | 1.84E-03 | 7.95E-03 | -1.05E+00 | 1.87E-05 | 7.94E-05 | -1.19E+00 | 1.65E-06 | 6.32E-06 | -1.18E+00 | 5.39E-04 | 1.92E-03 | -1.80E+00 | 6.72E-03 | 1.89E-02 | -1.37E+00 | 3.83E-03 | 1.40E-02 | -1.55E+00 | 8.03E-03 | 2.21E-02 | -1.38E+00 |
| NAGA | 4.16E-04 | 2.17E-03 | -1.40E+00 | 2.58E-02 | 1.12E-01 | 1.15E+00 | 3.08E-03 | 1.31E-02 | -1.18E+00 | 2.11E-02 | 8.04E-02 | 1.07E+00 | 4.12E-05 | 1.47E-04 | -1.63E+00 | 5.28E-03 | 1.49E-02 | -1.09E+00 | 7.32E-02 | 2.69E-01 | 1.02E+00 | 7.02E-02 | 1.93E-01 | -1.11E+00 |
| DUSP18 | 4.16E-04 | 2.17E-03 | -1.27E+00 | 8.13E-02 | 3.51E-01 | 1.31E+00 | 6.66E-09 | 2.83E-08 | -2.38E+00 | 5.86E-05 | 2.24E-04 | -1.32E+00 | 1.38E-01 | 4.91E-01 | 1.09E+00 | 1.27E-01 | 3.59E-01 | -1.03E+00 | 1.90E-01 | 6.98E-01 | 1.01E+00 | 2.45E-01 | 6.73E-01 | -1.07E+00 |
| CNNM4 | 4.16E-04 | 2.17E-03 | -1.43E+00 | 2.91E-03 | 1.26E-02 | -1.15E+00 | 1.01E-04 | 4.30E-04 | -1.56E+00 | 8.53E-04 | 3.26E-03 | -1.19E+00 | 3.88E-02 | 1.38E-01 | -1.22E+00 | 6.97E-02 | 1.97E-01 | -1.07E+00 | 2.43E-02 | 8.93E-02 | -1.20E+00 | 2.15E-01 | 5.90E-01 | 1.03E+00 |
| NMT1 | 4.16E-04 | 2.17E-03 | -1.25E+00 | 7.30E-03 | 3.16E-02 | 1.07E+00 | 1.65E-01 | 7.01E-01 | 1.21E+00 | 2.25E-01 | 8.61E-01 | 1.44E+00 | 3.44E-06 | 1.23E-05 | -1.37E+00 | 7.93E-04 | 2.24E-03 | -1.05E+00 | 1.64E-04 | 6.00E-04 | -1.25E+00 | 5.75E-03 | 1.58E-02 | -1.07E+00 |
| AL035456.1 | 4.16E-04 | 2.17E-03 | -1.58E+00 | 1.85E-03 | 8.00E-03 | -1.27E+00 | 8.21E-02 | 3.49E-01 | -1.05E+00 | 1.99E-01 | 7.59E-01 | 1.29E+00 | 3.78E-02 | 1.35E-01 | -1.02E+00 | 2.36E-01 | 6.64E-01 | 1.29E+00 | 4.71E-09 | 1.73E-08 | -1.29E+00 | 1.11E-01 | 3.06E-01 | 1.21E+00 |
| ARL4AP2 | 4.17E-04 | 2.17E-03 | -1.13E+00 | 2.33E-08 | 1.01E-07 | -1.55E+00 | 1.39E-01 | 5.91E-01 | 1.14E+00 | 6.17E-04 | 2.36E-03 | -1.53E+00 | 4.11E-02 | 1.47E-01 | 1.86E+00 | 1.53E-01 | 4.32E-01 | 1.07E+00 | 4.06E-02 | 1.49E-01 | 1.53E+00 | 1.23E-01 | 3.38E-01 | 1.10E+00 |
| CPSF1 | 4.17E-04 | 2.17E-03 | -1.08E+00 | 7.15E-05 | 3.09E-04 | -1.03E+00 | 2.34E-01 | 9.95E-01 | 1.24E+00 | 7.59E-02 | 2.90E-01 | 1.31E+00 | 2.56E-01 | 9.14E-01 | 1.43E+00 | 1.74E-01 | 4.91E-01 | 1.48E+00 | 3.55E-04 | 1.30E-03 | -1.12E+00 | 1.45E-01 | 3.98E-01 | 1.14E+00 |
| SLC25A16 | 4.18E-04 | 2.18E-03 | -1.16E+00 | 5.06E-04 | 2.19E-03 | -1.09E+00 | 1.63E-02 | 6.92E-02 | -1.08E+00 | 4.33E-03 | 1.65E-02 | -1.05E+00 | 1.17E-03 | 4.18E-03 | -1.31E+00 | 8.48E-03 | 2.39E-02 | -1.12E+00 | 2.03E-03 | 7.44E-03 | -1.12E+00 | 5.74E-04 | 1.58E-03 | -1.14E+00 |
| AC011444.4 | 4.18E-04 | 2.18E-03 | 1.02E+00 | 4.01E-14 | 1.73E-13 | -1.55E+00 | 1.69E-03 | 7.16E-03 | -1.09E+00 | 7.31E-08 | 2.79E-07 | -1.39E+00 | 7.00E-05 | 2.50E-04 | -1.18E+00 | 1.03E-08 | 2.89E-08 | -1.64E+00 | 2.09E-06 | 7.66E-06 | -1.36E+00 | 3.03E-11 | 8.31E-11 | -2.04E+00 |
| MBTPS2 | 4.18E-04 | 2.18E-03 | -1.40E+00 | 5.87E-03 | 2.53E-02 | -1.03E+00 | 4.92E-05 | 2.09E-04 | -1.46E+00 | 4.04E-04 | 1.54E-03 | -1.21E+00 | 2.38E-01 | 8.48E-01 | 1.27E+00 | 1.87E-01 | 5.29E-01 | 1.89E+00 | 1.05E-02 | 3.87E-02 | -1.17E+00 | 2.35E-02 | 6.47E-02 | -1.13E+00 |
| AC022098.1 | 4.18E-04 | 2.18E-03 | -1.65E+00 | 5.13E-03 | 2.22E-02 | -1.14E+00 | 4.29E-03 | 1.82E-02 | -1.15E+00 | 1.16E-03 | 4.44E-03 | -1.16E+00 | 3.91E-02 | 1.39E-01 | -1.12E+00 | 8.48E-03 | 2.39E-02 | -1.41E+00 | 1.42E-03 | 5.22E-03 | -1.28E+00 | 5.99E-03 | 1.65E-02 | -1.18E+00 |
| STXBP1 | 4.18E-04 | 2.18E-03 | -1.70E+00 | 4.71E-02 | 2.03E-01 | 1.14E+00 | 2.04E-01 | 8.66E-01 | 1.26E+00 | 2.49E-01 | 9.51E-01 | 1.30E+00 | 2.35E-01 | 8.38E-01 | 1.53E+00 | 3.08E-01 | 8.69E-01 | 1.44E+00 | 1.12E-01 | 4.13E-01 | 1.21E+00 | 1.04E-01 | 2.86E-01 | 1.08E+00 |
| CASP7 | 4.19E-04 | 2.18E-03 | -1.14E+00 | 2.95E-03 | 1.28E-02 | 1.11E+00 | 1.63E-04 | 6.94E-04 | -1.17E+00 | 1.08E-03 | 4.13E-03 | -1.01E+00 | 2.91E-05 | 1.04E-04 | -2.02E+00 | 2.08E-03 | 5.85E-03 | -1.36E+00 | 4.58E-06 | 1.68E-05 | -1.37E+00 | 1.31E-03 | 3.60E-03 | -1.17E+00 |
| DNAJC14 | 4.19E-04 | 2.18E-03 | -1.45E+00 | 2.60E-05 | 1.12E-04 | -1.78E+00 | 1.72E-01 | 7.29E-01 | 1.06E+00 | 3.47E-02 | 1.33E-01 | -1.26E+00 | 1.58E-01 | 5.65E-01 | 1.03E+00 | 7.45E-02 | 2.10E-01 | -1.24E+00 | 7.46E-03 | 2.74E-02 | -1.24E+00 | 7.77E-03 | 2.14E-02 | -1.22E+00 |
| THUMPD3 | 4.21E-04 | 2.19E-03 | 1.01E+00 | 1.84E-08 | 7.94E-08 | -1.17E+00 | 5.81E-05 | 2.47E-04 | -1.16E+00 | 4.45E-04 | 1.70E-03 | 1.04E+00 | 1.24E-05 | 4.43E-05 | -1.16E+00 | 1.90E-07 | 5.35E-07 | -1.23E+00 | 2.05E-05 | 7.54E-05 | -1.19E+00 | 1.45E-06 | 3.99E-06 | -1.22E+00 |
| SLC9A8 | 4.21E-04 | 2.19E-03 | -1.39E+00 | 1.18E-02 | 5.12E-02 | 1.01E+00 | 2.21E-01 | 9.41E-01 | 1.14E+00 | 5.67E-02 | 2.17E-01 | -1.03E+00 | 3.38E-03 | 1.21E-02 | -1.47E+00 | 7.42E-03 | 2.09E-02 | -1.39E+00 | 2.77E-03 | 1.02E-02 | -1.23E+00 | 1.82E-02 | 5.00E-02 | -1.14E+00 |
| TNNI2 | 4.22E-04 | 2.20E-03 | -2.34E+00 | 1.10E-04 | 4.75E-04 | -2.75E+00 | 5.33E-02 | 2.26E-01 | -1.06E+00 | 1.13E-04 | 4.33E-04 | -1.43E+00 | 9.68E-03 | 3.45E-02 | -1.94E+00 | 8.63E-03 | 2.43E-02 | -2.16E+00 | 2.45E-01 | 8.98E-01 | -1.04E+00 | 3.06E-01 | 8.40E-01 | -1.09E+00 |
| NLRP1 | 4.22E-04 | 2.20E-03 | 1.00E+00 | 1.34E-08 | 5.81E-08 | -1.19E+00 | 1.71E-01 | 7.26E-01 | 1.33E+00 | 9.69E-02 | 3.70E-01 | 1.18E+00 | 8.74E-02 | 3.11E-01 | -1.03E+00 | 2.75E-02 | 7.76E-02 | -1.27E+00 | 9.41E-02 | 3.45E-01 | -1.15E+00 | 5.01E-02 | 1.38E-01 | -1.53E+00 |
| TRDMT1 | 4.22E-04 | 2.20E-03 | -1.13E+00 | 3.92E-03 | 1.70E-02 | 1.11E+00 | 5.78E-02 | 2.46E-01 | -1.06E+00 | 2.53E-01 | 9.66E-01 | 1.25E+00 | 6.50E-03 | 2.32E-02 | -1.31E+00 | 2.89E-02 | 8.14E-02 | -1.01E+00 | 4.32E-02 | 1.58E-01 | -1.18E+00 | 2.61E-01 | 7.16E-01 | 1.39E+00 |
| FBXO4 | 4.22E-04 | 2.20E-03 | -1.09E+00 | 1.05E-02 | 4.52E-02 | 1.20E+00 | 6.21E-03 | 2.64E-02 | -1.08E+00 | 3.83E-03 | 1.46E-02 | 1.04E+00 | 9.08E-04 | 3.24E-03 | -1.36E+00 | 1.35E-01 | 3.80E-01 | 1.14E+00 | 4.45E-04 | 1.63E-03 | -1.14E+00 | 4.35E-02 | 1.20E-01 | 1.03E+00 |
| LRP1 | 4.25E-04 | 2.21E-03 | -1.51E+00 | 1.55E-02 | 6.70E-02 | 1.04E+00 | 2.66E-02 | 1.13E-01 | -1.36E+00 | 3.29E-02 | 1.26E-01 | -1.28E+00 | 2.09E-01 | 7.44E-01 | 1.44E+00 | 3.55E-01 | 1.00E+00 | 1.09E+00 | 2.08E-01 | 7.64E-01 | 1.15E+00 | 3.24E-01 | 8.91E-01 | 1.05E+00 |
| CEP68 | 4.25E-04 | 2.21E-03 | -1.03E+00 | 3.72E-03 | 1.61E-02 | 1.24E+00 | 3.71E-03 | 1.58E-02 | -1.12E+00 | 3.42E-03 | 1.31E-02 | 1.02E+00 | 4.27E-04 | 1.52E-03 | -1.07E+00 | 1.22E-01 | 3.43E-01 | 1.22E+00 | 1.11E-03 | 4.06E-03 | -1.13E+00 | 6.14E-02 | 1.69E-01 | 1.07E+00 |
| ZNF682 | 4.26E-04 | 2.22E-03 | -1.63E+00 | 4.75E-03 | 2.05E-02 | -1.10E+00 | 1.58E-05 | 6.71E-05 | -1.70E+00 | 1.61E-04 | 6.14E-04 | -1.37E+00 | 9.07E-03 | 3.24E-02 | -1.19E+00 | 1.43E-02 | 4.04E-02 | -1.19E+00 | 1.11E-01 | 4.06E-01 | 1.13E+00 | 1.75E-01 | 4.81E-01 | 1.09E+00 |
| ZDHHC6 | 4.26E-04 | 2.22E-03 | -1.07E+00 | 2.35E-02 | 1.02E-01 | 1.28E+00 | 2.95E-05 | 1.25E-04 | -1.19E+00 | 3.42E-04 | 1.31E-03 | 1.03E+00 | 2.80E-04 | 9.97E-04 | -1.11E+00 | 1.14E-02 | 3.22E-02 | 1.15E+00 | 2.18E-03 | 8.01E-03 | -1.10E+00 | 1.61E-01 | 4.42E-01 | 1.14E+00 |
| NDUFB1P1 | 4.27E-04 | 2.22E-03 | -1.31E+00 | 2.06E-07 | 8.90E-07 | -2.13E+00 | 2.09E-01 | 8.89E-01 | 1.14E+00 | 1.73E-04 | 6.61E-04 | -1.71E+00 | 3.81E-02 | 1.36E-01 | 1.03E+00 | 2.89E-03 | 8.15E-03 | -1.34E+00 | 8.62E-02 | 3.16E-01 | 1.05E+00 | 2.39E-04 | 6.57E-04 | -1.44E+00 |
| PSPC1 | 4.29E-04 | 2.23E-03 | -1.08E+00 | 4.73E-05 | 2.04E-04 | -1.06E+00 | 1.09E-03 | 4.63E-03 | -1.05E+00 | 7.33E-06 | 2.80E-05 | -1.09E+00 | 5.57E-05 | 1.98E-04 | -1.19E+00 | 9.67E-05 | 2.72E-04 | -1.19E+00 | 2.19E-03 | 8.05E-03 | -1.02E+00 | 6.06E-05 | 1.66E-04 | -1.16E+00 |
| FAM89B | 4.29E-04 | 2.24E-03 | 1.03E+00 | 5.71E-05 | 2.47E-04 | 1.06E+00 | 1.34E-02 | 5.67E-02 | -1.02E+00 | 4.56E-04 | 1.74E-03 | -1.07E+00 | 2.30E-04 | 8.21E-04 | 1.04E+00 | 2.55E-05 | 7.19E-05 | -1.04E+00 | 5.07E-02 | 1.86E-01 | 1.18E+00 | 5.78E-04 | 1.59E-03 | -1.03E+00 |
| C16orf86 | 4.30E-04 | 2.24E-03 | -1.06E+00 | 1.71E-03 | 7.40E-03 | 1.11E+00 | 8.76E-02 | 3.72E-01 | 1.07E+00 | 2.40E-02 | 9.18E-02 | 1.09E+00 | 2.18E-01 | 7.78E-01 | 1.26E+00 | 2.52E-01 | 7.12E-01 | 1.32E+00 | 7.66E-02 | 2.81E-01 | 1.15E+00 | 5.29E-02 | 1.45E-01 | 1.09E+00 |
| LINC01252 | 4.31E-04 | 2.25E-03 | -1.48E+00 | 3.57E-02 | 1.54E-01 | 1.13E+00 | 1.08E-03 | 4.59E-03 | -1.60E+00 | 1.49E-02 | 5.68E-02 | -1.04E+00 | 8.79E-04 | 3.14E-03 | -1.78E+00 | 7.18E-03 | 2.02E-02 | -1.28E+00 | 6.23E-02 | 2.29E-01 | 1.02E+00 | 4.89E-02 | 1.34E-01 | -1.30E+00 |
| ZC3H7B | 4.31E-04 | 2.25E-03 | -1.23E+00 | 1.59E-03 | 6.87E-03 | -1.01E+00 | 1.11E-02 | 4.73E-02 | -1.07E+00 | 1.01E-01 | 3.87E-01 | 1.26E+00 | 4.66E-03 | 1.66E-02 | -1.19E+00 | 6.27E-02 | 1.77E-01 | 1.09E+00 | 6.57E-04 | 2.41E-03 | -1.28E+00 | 2.79E-02 | 7.67E-02 | -1.02E+00 |
| ASAP1 | 4.32E-04 | 2.25E-03 | -1.18E+00 | 1.13E-04 | 4.89E-04 | -1.10E+00 | 8.04E-04 | 3.41E-03 | -1.17E+00 | 2.50E-03 | 9.55E-03 | 1.00E+00 | 1.66E-07 | 5.90E-07 | -1.49E+00 | 1.61E-03 | 4.55E-03 | -1.01E+00 | 6.07E-16 | 2.23E-15 | -1.27E+00 | 1.35E-12 | 3.70E-12 | -1.27E+00 |
| BCAR3 | 4.32E-04 | 2.25E-03 | -1.49E+00 | 3.68E-03 | 1.59E-02 | -1.22E+00 | 8.69E-02 | 3.69E-01 | -1.15E+00 | 1.30E-01 | 4.98E-01 | 1.06E+00 | 1.69E-01 | 6.03E-01 | 1.03E+00 | 2.80E-01 | 7.91E-01 | 1.04E+00 | 2.70E-01 | 9.93E-01 | 1.18E+00 | 1.21E-01 | 3.33E-01 | -1.31E+00 |
| GSS | 4.32E-04 | 2.25E-03 | -1.06E+00 | 1.27E-04 | 5.51E-04 | -1.03E+00 | 3.06E-03 | 1.30E-02 | -1.08E+00 | 1.03E-05 | 3.94E-05 | -1.12E+00 | 8.14E-06 | 2.90E-05 | -1.36E+00 | 6.22E-03 | 1.75E-02 | 1.04E+00 | 2.13E-02 | 7.82E-02 | 1.07E+00 | 1.23E-01 | 3.38E-01 | 1.09E+00 |
| LINC00528 | 4.32E-04 | 2.25E-03 | -1.43E+00 | 3.97E-03 | 1.72E-02 | -1.06E+00 | 8.64E-03 | 3.67E-02 | -1.24E+00 | 1.36E-02 | 5.21E-02 | -1.07E+00 | 4.47E-03 | 1.60E-02 | -1.40E+00 | 1.55E-01 | 4.36E-01 | 1.18E+00 | 6.18E-02 | 2.27E-01 | 1.02E+00 | 3.40E-01 | 9.34E-01 | 1.33E+00 |
| ENPP2 | 4.35E-04 | 2.27E-03 | -1.97E+00 | 2.59E-04 | 1.12E-03 | -2.37E+00 | 1.59E-07 | 6.75E-07 | -2.76E+00 | 9.38E-06 | 3.58E-05 | -1.95E+00 | 5.77E-12 | 2.06E-11 | -2.31E+00 | 2.46E-11 | 6.93E-11 | -1.99E+00 | 1.72E-03 | 6.33E-03 | -1.44E+00 | 4.69E-04 | 1.29E-03 | -1.74E+00 |
| LRCH4 | 4.36E-04 | 2.27E-03 | -1.25E+00 | 2.56E-03 | 1.11E-02 | -1.10E+00 | 1.36E-01 | 5.78E-01 | 1.19E+00 | 1.37E-01 | 5.25E-01 | 1.24E+00 | 1.08E-01 | 3.87E-01 | 1.14E+00 | 8.53E-03 | 2.40E-02 | -1.29E+00 | 2.60E-01 | 9.56E-01 | 1.33E+00 | 3.64E-01 | 1.00E+00 | 1.34E+00 |
| TRIM65 | 4.37E-04 | 2.28E-03 | -1.29E+00 | 2.55E-02 | 1.10E-01 | 1.10E+00 | 6.19E-07 | 2.63E-06 | -1.42E+00 | 1.25E-03 | 4.77E-03 | 1.04E+00 | 7.69E-05 | 2.74E-04 | -1.65E+00 | 3.30E-03 | 9.31E-03 | -1.20E+00 | 2.33E-08 | 8.55E-08 | -1.53E+00 | 6.56E-04 | 1.80E-03 | -1.12E+00 |
| FAM86C1 | 4.37E-04 | 2.28E-03 | -1.48E+00 | 1.12E-02 | 4.83E-02 | 1.01E+00 | 1.09E-05 | 4.64E-05 | -1.67E+00 | 7.54E-04 | 2.88E-03 | -1.05E+00 | 2.49E-04 | 8.87E-04 | -1.90E+00 | 5.62E-02 | 1.58E-01 | 1.07E+00 | 7.96E-05 | 2.92E-04 | -1.66E+00 | 3.37E-01 | 9.25E-01 | 1.22E+00 |
| LDHAP5 | 4.37E-04 | 2.28E-03 | -1.51E+00 | 3.23E-05 | 1.40E-04 | -1.82E+00 | 9.02E-03 | 3.83E-02 | -1.21E+00 | 7.31E-05 | 2.79E-04 | -1.44E+00 | 5.28E-02 | 1.88E-01 | -1.28E+00 | 1.93E-01 | 5.43E-01 | 1.06E+00 | 1.45E-02 | 5.32E-02 | -1.08E+00 | 6.41E-03 | 1.76E-02 | -1.24E+00 |
| DNAJC19 | 4.38E-04 | 2.28E-03 | 1.02E+00 | 1.51E-03 | 6.53E-03 | 1.13E+00 | 6.00E-02 | 2.55E-01 | 1.16E+00 | 9.61E-02 | 3.67E-01 | 1.27E+00 | 4.14E-03 | 1.48E-02 | 1.03E+00 | 1.71E-01 | 4.81E-01 | 1.35E+00 | 1.11E-01 | 4.09E-01 | 1.12E+00 | 3.34E-01 | 9.17E-01 | 1.24E+00 |
| ANKRD44 | 4.38E-04 | 2.29E-03 | -1.01E+00 | 6.99E-03 | 3.02E-02 | 1.21E+00 | 1.87E-04 | 7.93E-04 | -1.27E+00 | 1.34E-05 | 5.11E-05 | -1.22E+00 | 1.24E-02 | 4.43E-02 | 1.13E+00 | 4.53E-03 | 1.28E-02 | 1.04E+00 | 1.53E-03 | 5.60E-03 | -1.50E+00 | 1.72E-02 | 4.73E-02 | -1.32E+00 |
| NUP155 | 4.38E-04 | 2.29E-03 | -1.43E+00 | 6.40E-02 | 2.76E-01 | 1.14E+00 | 7.91E-06 | 3.36E-05 | -1.44E+00 | 4.18E-03 | 1.60E-02 | 1.02E+00 | 3.57E-03 | 1.27E-02 | -1.34E+00 | 9.57E-02 | 2.70E-01 | 1.15E+00 | 7.10E-07 | 2.61E-06 | -1.51E+00 | 2.76E-01 | 7.58E-01 | 1.18E+00 |
| MRPL22 | 4.39E-04 | 2.29E-03 | 1.10E+00 | 1.40E-02 | 6.04E-02 | 1.35E+00 | 8.69E-04 | 3.69E-03 | -1.04E+00 | 3.46E-04 | 1.32E-03 | 1.01E+00 | 2.22E-04 | 7.93E-04 | -1.12E+00 | 1.60E-01 | 4.51E-01 | 1.24E+00 | 3.89E-03 | 1.43E-02 | 1.02E+00 | 6.17E-03 | 1.69E-02 | 1.01E+00 |
| VRK2 | 4.39E-04 | 2.29E-03 | -1.31E+00 | 6.93E-06 | 2.99E-05 | -1.45E+00 | 2.54E-08 | 1.08E-07 | -1.61E+00 | 4.97E-11 | 1.90E-10 | -1.75E+00 | 2.09E-02 | 7.47E-02 | -1.22E+00 | 3.81E-02 | 1.08E-01 | -1.06E+00 | 1.50E-06 | 5.50E-06 | -1.43E+00 | 2.67E-04 | 7.32E-04 | -1.20E+00 |
| ZFAS1 | 4.40E-04 | 2.29E-03 | 1.12E+00 | 1.53E-01 | 6.61E-01 | -1.03E+00 | 1.59E-01 | 6.74E-01 | -1.00E+00 | 7.45E-03 | 2.85E-02 | -1.07E+00 | 3.20E-06 | 1.14E-05 | 1.30E+00 | 3.29E-01 | 9.28E-01 | 1.02E+00 | 1.43E-12 | 5.23E-12 | 1.73E+00 | 2.64E-01 | 7.26E-01 | 1.12E+00 |
| TCF25 | 4.40E-04 | 2.29E-03 | -1.00E+00 | 2.74E-10 | 1.18E-09 | -1.08E+00 | 3.38E-02 | 1.43E-01 | -1.02E+00 | 2.65E-05 | 1.01E-04 | -1.15E+00 | 1.15E-03 | 4.10E-03 | -1.01E+00 | 3.79E-06 | 1.07E-05 | -1.21E+00 | 1.58E-01 | 5.80E-01 | 1.09E+00 | 5.72E-03 | 1.57E-02 | -1.09E+00 |
| PICALM | 4.40E-04 | 2.29E-03 | 1.06E+00 | 5.68E-06 | 2.45E-05 | 1.03E+00 | 4.45E-04 | 1.89E-03 | -1.10E+00 | 3.06E-06 | 1.17E-05 | -1.11E+00 | 1.36E-06 | 4.83E-06 | -1.31E+00 | 4.42E-05 | 1.25E-04 | -1.14E+00 | 1.12E-20 | 4.10E-20 | -1.97E+00 | 9.69E-16 | 2.66E-15 | -1.86E+00 |
| GPR180 | 4.41E-04 | 2.30E-03 | -1.37E+00 | 3.64E-02 | 1.57E-01 | 1.18E+00 | 1.08E-05 | 4.59E-05 | -1.74E+00 | 2.40E-02 | 9.16E-02 | -1.00E+00 | 1.78E-03 | 6.35E-03 | -1.35E+00 | 2.25E-02 | 6.35E-02 | -1.06E+00 | 1.45E-08 | 5.33E-08 | -1.52E+00 | 3.56E-04 | 9.79E-04 | -1.23E+00 |
| CENPJ | 4.43E-04 | 2.31E-03 | -1.26E+00 | 3.08E-03 | 1.33E-02 | -1.01E+00 | 4.96E-02 | 2.11E-01 | -1.03E+00 | 1.38E-01 | 5.25E-01 | 1.25E+00 | 2.37E-01 | 8.45E-01 | 1.40E+00 | 2.10E-01 | 5.93E-01 | 1.37E+00 | 1.87E-03 | 6.86E-03 | 1.54E+00 | 2.60E-07 | 7.14E-07 | 1.89E+00 |
| AC138392.1 | 4.45E-04 | 2.32E-03 | -1.39E+00 | 6.92E-03 | 2.99E-02 | 1.02E+00 | 1.37E-07 | 5.83E-07 | -1.97E+00 | 1.74E-06 | 6.64E-06 | -1.55E+00 | 6.16E-06 | 2.20E-05 | -2.04E+00 | 3.36E-04 | 9.48E-04 | -1.46E+00 | 1.77E-09 | 6.50E-09 | -1.48E+00 | 2.70E-09 | 7.42E-09 | -1.72E+00 |
| C12orf10 | 4.45E-04 | 2.32E-03 | 1.02E+00 | 1.37E-03 | 5.91E-03 | 1.20E+00 | 8.01E-04 | 3.40E-03 | -1.12E+00 | 2.23E-04 | 8.54E-04 | -1.04E+00 | 1.19E-05 | 4.26E-05 | -1.11E+00 | 5.42E-04 | 1.53E-03 | -1.02E+00 | 2.28E-04 | 8.36E-04 | -1.02E+00 | 2.67E-02 | 7.33E-02 | 1.08E+00 |
| TSPAN4 | 4.45E-04 | 2.32E-03 | -1.02E+00 | 2.25E-06 | 9.72E-06 | -1.09E+00 | 1.17E-04 | 4.95E-04 | -1.13E+00 | 5.21E-06 | 1.99E-05 | -1.06E+00 | 1.30E-14 | 4.62E-14 | -2.05E+00 | 1.52E-12 | 4.28E-12 | -1.85E+00 | 1.99E-05 | 7.31E-05 | -1.09E+00 | 2.33E-05 | 6.40E-05 | -1.26E+00 |
| CCDC57 | 4.46E-04 | 2.32E-03 | 1.03E+00 | 1.23E-11 | 5.31E-11 | -1.22E+00 | 2.02E-01 | 8.57E-01 | 1.25E+00 | 1.87E-04 | 7.14E-04 | -1.00E+00 | 1.85E-01 | 6.59E-01 | 1.19E+00 | 1.35E-03 | 3.80E-03 | -1.10E+00 | 2.10E-01 | 7.70E-01 | 1.31E+00 | 1.85E-01 | 5.07E-01 | 1.13E+00 |
| FAM129A | 4.46E-04 | 2.33E-03 | -1.09E+00 | 7.84E-08 | 3.39E-07 | -1.32E+00 | 1.53E-06 | 6.52E-06 | -1.35E+00 | 3.56E-06 | 1.36E-05 | -1.14E+00 | 3.64E-03 | 1.30E-02 | -1.24E+00 | 5.15E-03 | 1.45E-02 | -1.21E+00 | 2.41E-06 | 8.86E-06 | -1.33E+00 | 1.32E-05 | 3.62E-05 | -1.31E+00 |
| GPR135 | 4.47E-04 | 2.33E-03 | -1.19E+00 | 1.08E-02 | 4.68E-02 | 1.10E+00 | 4.18E-02 | 1.78E-01 | -1.27E+00 | 2.39E-01 | 9.14E-01 | 1.18E+00 | 2.02E-01 | 7.21E-01 | 1.33E+00 | 2.76E-01 | 7.79E-01 | 1.69E+00 | 4.35E-02 | 1.60E-01 | -1.17E+00 | 3.46E-01 | 9.51E-01 | 1.22E+00 |
| DXO | 4.48E-04 | 2.33E-03 | -1.01E+00 | 4.12E-05 | 1.78E-04 | -1.01E+00 | 2.53E-02 | 1.07E-01 | -1.07E+00 | 2.96E-03 | 1.13E-02 | -1.08E+00 | 1.44E-03 | 5.12E-03 | -1.08E+00 | 1.19E-02 | 3.35E-02 | 1.02E+00 | 2.01E-01 | 7.38E-01 | 1.26E+00 | 3.04E-01 | 8.36E-01 | 1.31E+00 |
| NCOA1 | 4.48E-04 | 2.34E-03 | 1.06E+00 | 9.98E-03 | 4.31E-02 | 1.25E+00 | 8.09E-06 | 3.44E-05 | -1.33E+00 | 1.00E-05 | 3.83E-05 | -1.07E+00 | 6.45E-05 | 2.30E-04 | -1.21E+00 | 3.39E-05 | 9.55E-05 | -1.26E+00 | 9.50E-03 | 3.49E-02 | -1.02E+00 | 6.07E-02 | 1.67E-01 | 1.04E+00 |
| ERCC6 | 4.49E-04 | 2.34E-03 | -1.19E+00 | 1.74E-02 | 7.53E-02 | 1.14E+00 | 6.55E-03 | 2.78E-02 | -1.35E+00 | 3.82E-02 | 1.46E-01 | -1.03E+00 | 4.96E-02 | 1.77E-01 | 1.13E+00 | 1.45E-01 | 4.07E-01 | 1.17E+00 | 6.90E-03 | 2.53E-02 | 1.01E+00 | 1.43E-03 | 3.93E-03 | -1.21E+00 |
| FAM168B | 4.49E-04 | 2.34E-03 | -1.09E+00 | 2.46E-04 | 1.06E-03 | 1.01E+00 | 5.48E-03 | 2.33E-02 | -1.12E+00 | 1.05E-02 | 4.01E-02 | 1.03E+00 | 3.68E-04 | 1.31E-03 | -1.11E+00 | 7.00E-05 | 1.97E-04 | -1.17E+00 | 4.30E-02 | 1.58E-01 | 1.05E+00 | 5.86E-02 | 1.61E-01 | 1.12E+00 |
| AC009961.1 | 4.50E-04 | 2.34E-03 | -1.32E+00 | 2.88E-03 | 1.24E-02 | -1.07E+00 | 8.34E-04 | 3.54E-03 | -1.62E+00 | 3.84E-02 | 1.46E-01 | 1.03E+00 | 2.10E-01 | 7.49E-01 | 1.55E+00 | 3.37E-01 | 9.49E-01 | 1.43E+00 | 2.71E-01 | 9.96E-01 | 1.19E+00 | 3.53E-01 | 9.68E-01 | 1.16E+00 |
| CCRL2 | 4.51E-04 | 2.35E-03 | -1.60E+00 | 4.62E-03 | 2.00E-02 | -1.13E+00 | 8.66E-03 | 3.68E-02 | -1.32E+00 | 9.45E-03 | 3.61E-02 | -1.25E+00 | 1.58E-02 | 5.64E-02 | -1.46E+00 | 8.75E-02 | 2.47E-01 | -1.15E+00 | 1.76E-01 | 6.45E-01 | 1.33E+00 | 1.33E-01 | 3.66E-01 | 1.15E+00 |
| AC005072.1 | 4.52E-04 | 2.35E-03 | -1.74E+00 | 2.15E-02 | 9.27E-02 | -1.12E+00 | 1.72E-03 | 7.31E-03 | -1.40E+00 | 5.63E-02 | 2.15E-01 | 1.03E+00 | 2.74E-01 | 9.77E-01 | 1.23E+00 | 3.02E-01 | 8.52E-01 | 1.44E+00 | 3.23E-06 | 1.18E-05 | -2.14E+00 | 2.18E-02 | 6.00E-02 | -1.14E+00 |
| AL390728.6 | 4.52E-04 | 2.36E-03 | -1.29E+00 | 1.77E-02 | 7.65E-02 | 1.09E+00 | 2.63E-15 | 1.12E-14 | -1.90E+00 | 2.67E-11 | 1.02E-10 | -1.36E+00 | 3.60E-09 | 1.28E-08 | -2.18E+00 | 6.74E-06 | 1.90E-05 | -1.44E+00 | 4.21E-07 | 1.54E-06 | -1.37E+00 | 7.47E-03 | 2.05E-02 | -1.02E+00 |
| IREB2 | 4.53E-04 | 2.36E-03 | 1.05E+00 | 1.60E-02 | 6.92E-02 | 1.29E+00 | 6.16E-02 | 2.62E-01 | 1.08E+00 | 2.62E-01 | 1.00E+00 | 1.36E+00 | 3.52E-02 | 1.25E-01 | 1.15E+00 | 7.71E-02 | 2.17E-01 | 1.15E+00 | 1.40E-01 | 5.12E-01 | 1.10E+00 | 2.16E-01 | 5.92E-01 | 1.26E+00 |
| FAM107B | 4.55E-04 | 2.37E-03 | -1.22E+00 | 2.16E-05 | 9.32E-05 | -1.27E+00 | 1.90E-01 | 8.07E-01 | 1.12E+00 | 6.34E-03 | 2.42E-02 | -1.10E+00 | 6.30E-02 | 2.25E-01 | -1.10E+00 | 7.83E-03 | 2.21E-02 | -1.72E+00 | 7.24E-02 | 2.66E-01 | 1.83E+00 | 8.92E-02 | 2.45E-01 | 1.67E+00 |
| NFATC3 | 4.55E-04 | 2.37E-03 | -1.34E+00 | 2.54E-02 | 1.10E-01 | 1.08E+00 | 4.83E-03 | 2.05E-02 | -1.21E+00 | 9.18E-02 | 3.51E-01 | 1.17E+00 | 1.76E-04 | 6.27E-04 | -1.42E+00 | 2.86E-04 | 8.06E-04 | -1.44E+00 | 2.36E-01 | 8.67E-01 | 1.08E+00 | 2.10E-01 | 5.76E-01 | -1.02E+00 |
| ZDHHC8 | 4.56E-04 | 2.37E-03 | -1.19E+00 | 5.99E-03 | 2.59E-02 | 1.03E+00 | 7.56E-02 | 3.21E-01 | 1.05E+00 | 1.50E-01 | 5.72E-01 | 1.25E+00 | 1.05E-01 | 3.75E-01 | 1.20E+00 | 2.81E-02 | 7.91E-02 | -1.07E+00 | 2.33E-01 | 8.57E-01 | 1.26E+00 | 1.09E-01 | 2.99E-01 | 1.60E+00 |
| SUV39H1 | 4.56E-04 | 2.38E-03 | -1.62E+00 | 8.81E-03 | 3.81E-02 | -1.09E+00 | 1.63E-03 | 6.93E-03 | -1.18E+00 | 5.15E-04 | 1.97E-03 | -1.12E+00 | 1.89E-02 | 6.73E-02 | -1.42E+00 | 1.91E-02 | 5.39E-02 | -1.63E+00 | 4.80E-05 | 1.76E-04 | -1.69E+00 | 1.80E-01 | 4.93E-01 | 1.06E+00 |
| AL080317.3 | 4.57E-04 | 2.38E-03 | -1.76E+00 | 7.37E-02 | 3.18E-01 | 1.17E+00 | 6.21E-03 | 2.64E-02 | -1.28E+00 | 2.20E-02 | 8.41E-02 | 1.03E+00 | 6.74E-05 | 2.40E-04 | -1.76E+00 | 1.06E-02 | 2.98E-02 | -1.07E+00 | 1.62E-03 | 5.93E-03 | -1.51E+00 | 1.37E-01 | 3.78E-01 | -1.02E+00 |
| AL392172.1 | 4.59E-04 | 2.39E-03 | -1.24E+00 | 7.07E-05 | 3.05E-04 | -1.24E+00 | 1.18E-01 | 5.02E-01 | 1.32E+00 | 8.44E-02 | 3.22E-01 | 1.02E+00 | 3.84E-04 | 1.37E-03 | -1.30E+00 | 9.47E-04 | 2.67E-03 | -1.25E+00 | 7.03E-02 | 2.58E-01 | 1.02E+00 | 1.80E-02 | 4.95E-02 | -1.09E+00 |
| LAMC1 | 4.59E-04 | 2.39E-03 | -1.26E+00 | 4.13E-02 | 1.78E-01 | 1.29E+00 | 8.24E-02 | 3.50E-01 | 1.07E+00 | 2.46E-01 | 9.41E-01 | 1.38E+00 | 2.28E-03 | 8.13E-03 | -1.20E+00 | 9.10E-03 | 2.57E-02 | -1.12E+00 | 5.79E-02 | 2.13E-01 | -1.06E+00 | 2.82E-01 | 7.75E-01 | 1.25E+00 |
| PI4KAP2 | 4.59E-04 | 2.39E-03 | -1.06E+00 | 1.35E-06 | 5.85E-06 | -1.27E+00 | 3.38E-02 | 1.44E-01 | -1.02E+00 | 3.57E-04 | 1.36E-03 | -1.18E+00 | 8.44E-02 | 3.01E-01 | 1.02E+00 | 9.59E-02 | 2.70E-01 | 1.04E+00 | 3.62E-05 | 1.33E-04 | -1.33E+00 | 3.86E-05 | 1.06E-04 | -1.27E+00 |
| RPS6KA3 | 4.60E-04 | 2.40E-03 | 1.00E+00 | 3.85E-02 | 1.66E-01 | 1.32E+00 | 3.51E-03 | 1.49E-02 | 1.02E+00 | 9.30E-03 | 3.55E-02 | 1.15E+00 | 3.51E-04 | 1.25E-03 | -1.15E+00 | 4.93E-04 | 1.39E-03 | -1.07E+00 | 8.66E-03 | 3.18E-02 | -1.01E+00 | 9.58E-02 | 2.63E-01 | 1.09E+00 |
| PAQR3 | 4.61E-04 | 2.40E-03 | -1.07E+00 | 1.14E-08 | 4.94E-08 | -1.46E+00 | 1.67E-03 | 7.09E-03 | -1.10E+00 | 7.75E-06 | 2.96E-05 | -1.23E+00 | 2.06E-01 | 7.35E-01 | 1.51E+00 | 3.46E-02 | 9.77E-02 | 1.02E+00 | 8.54E-02 | 3.13E-01 | 1.25E+00 | 3.93E-02 | 1.08E-01 | 1.16E+00 |
| NR2C1 | 4.61E-04 | 2.40E-03 | -1.03E+00 | 5.71E-03 | 2.47E-02 | 1.20E+00 | 2.33E-02 | 9.92E-02 | 1.03E+00 | 1.53E-03 | 5.83E-03 | 1.03E+00 | 7.02E-02 | 2.50E-01 | 1.15E+00 | 2.46E-02 | 6.94E-02 | -1.00E+00 | 1.50E-04 | 5.51E-04 | -1.14E+00 | 1.18E-03 | 3.24E-03 | -1.07E+00 |
| PEAR1 | 4.63E-04 | 2.41E-03 | -1.20E+00 | 5.28E-02 | 2.28E-01 | 1.31E+00 | 3.27E-02 | 1.39E-01 | -1.42E+00 | 2.53E-01 | 9.68E-01 | 1.31E+00 | 2.03E-01 | 7.25E-01 | 1.09E+00 | 1.98E-01 | 5.57E-01 | -1.08E+00 | 7.20E-02 | 2.64E-01 | 1.36E+00 | 8.33E-02 | 2.29E-01 | 1.17E+00 |
| RPSAP29 | 4.63E-04 | 2.41E-03 | 1.99E+00 | 1.70E-01 | 7.35E-01 | 1.68E+00 | 1.86E-02 | 7.92E-02 | 1.60E+00 | 2.83E-02 | 1.08E-01 | 1.19E+00 | 2.32E-01 | 8.26E-01 | 1.35E+00 | 2.32E-01 | 6.53E-01 | 1.30E+00 | 2.60E-02 | 9.56E-02 | 1.61E+00 | 2.83E-01 | 7.79E-01 | 1.46E+00 |
| ELP4 | 4.63E-04 | 2.42E-03 | -1.16E+00 | 5.04E-02 | 2.18E-01 | 1.25E+00 | 8.14E-08 | 3.46E-07 | -1.49E+00 | 1.21E-06 | 4.63E-06 | -1.22E+00 | 2.74E-08 | 9.78E-08 | -2.02E+00 | 8.18E-03 | 2.30E-02 | -1.04E+00 | 1.37E-07 | 5.04E-07 | -1.33E+00 | 1.77E-02 | 4.87E-02 | 1.06E+00 |
| YAE1D1 | 4.64E-04 | 2.42E-03 | -1.18E+00 | 6.18E-04 | 2.67E-03 | -1.03E+00 | 2.14E-06 | 9.10E-06 | -1.46E+00 | 5.82E-05 | 2.22E-04 | -1.17E+00 | 1.42E-02 | 5.05E-02 | -1.44E+00 | 2.00E-01 | 5.65E-01 | 1.20E+00 | 1.16E-07 | 4.25E-07 | -1.81E+00 | 3.33E-03 | 9.16E-03 | -1.19E+00 |
| JUP | 4.66E-04 | 2.43E-03 | -1.01E+00 | 2.10E-03 | 9.09E-03 | 1.21E+00 | 3.60E-02 | 1.53E-01 | -1.03E+00 | 3.53E-02 | 1.35E-01 | 1.09E+00 | 8.96E-03 | 3.20E-02 | -1.30E+00 | 3.63E-02 | 1.02E-01 | -1.06E+00 | 2.42E-03 | 8.88E-03 | -1.35E+00 | 1.61E-03 | 4.43E-03 | -1.50E+00 |
| AUTS2 | 4.68E-04 | 2.44E-03 | -1.16E+00 | 2.73E-02 | 1.18E-01 | 1.20E+00 | 2.41E-02 | 1.03E-01 | -1.06E+00 | 8.57E-02 | 3.27E-01 | 1.25E+00 | 3.45E-02 | 1.23E-01 | 1.00E+00 | 3.53E-01 | 9.95E-01 | 1.39E+00 | 4.75E-03 | 1.74E-02 | -1.06E+00 | 1.10E-01 | 3.03E-01 | 1.16E+00 |
| AL137792.1 | 4.68E-04 | 2.44E-03 | -1.33E+00 | 8.30E-05 | 3.58E-04 | -1.44E+00 | 7.50E-02 | 3.19E-01 | 1.01E+00 | 3.45E-05 | 1.32E-04 | -1.63E+00 | 1.42E-01 | 5.05E-01 | 1.02E+00 | 2.27E-03 | 6.41E-03 | -2.17E+00 | 2.34E-02 | 8.61E-02 | -1.08E+00 | 1.14E-04 | 3.12E-04 | -1.77E+00 |
| FAM104A | 4.69E-04 | 2.44E-03 | -1.62E+00 | 8.04E-03 | 3.47E-02 | -1.13E+00 | 1.25E-04 | 5.32E-04 | -1.79E+00 | 1.55E-03 | 5.90E-03 | -1.28E+00 | 2.63E-07 | 9.38E-07 | -2.21E+00 | 3.78E-05 | 1.07E-04 | -1.54E+00 | 9.18E-10 | 3.37E-09 | -1.37E+00 | 4.56E-09 | 1.25E-08 | -1.41E+00 |
| CHMP7 | 4.69E-04 | 2.44E-03 | -1.21E+00 | 2.24E-02 | 9.67E-02 | 1.17E+00 | 1.85E-03 | 7.85E-03 | -1.24E+00 | 5.30E-03 | 2.03E-02 | -1.02E+00 | 3.19E-06 | 1.14E-05 | -1.57E+00 | 9.46E-04 | 2.67E-03 | -1.09E+00 | 8.07E-09 | 2.96E-08 | -1.78E+00 | 1.18E-03 | 3.25E-03 | -1.16E+00 |
| CAT | 4.69E-04 | 2.44E-03 | -1.12E+00 | 8.37E-02 | 3.61E-01 | 1.05E+00 | 7.70E-05 | 3.27E-04 | -1.31E+00 | 4.73E-02 | 1.81E-01 | -1.07E+00 | 1.33E-03 | 4.76E-03 | 1.41E+00 | 4.28E-06 | 1.21E-05 | 1.53E+00 | 2.36E-01 | 8.67E-01 | 1.13E+00 | 2.09E-01 | 5.74E-01 | 1.10E+00 |
| YBX1P6 | 4.69E-04 | 2.44E-03 | 2.09E+00 | 1.48E-03 | 6.41E-03 | 2.29E+00 | 1.65E-09 | 7.02E-09 | 2.33E+00 | 5.26E-09 | 2.01E-08 | 2.66E+00 | 1.69E-02 | 6.04E-02 | 2.13E+00 | 8.51E-04 | 2.40E-03 | 2.52E+00 | 3.19E-03 | 1.17E-02 | 1.89E+00 | 4.18E-09 | 1.15E-08 | 2.41E+00 |
| COX5B | 4.69E-04 | 2.45E-03 | -1.01E+00 | 1.82E-12 | 7.86E-12 | -1.13E+00 | 1.30E-01 | 5.54E-01 | 1.15E+00 | 4.88E-02 | 1.87E-01 | 1.01E+00 | 5.37E-03 | 1.92E-02 | -1.07E+00 | 8.22E-03 | 2.32E-02 | -1.09E+00 | 2.33E-01 | 8.54E-01 | 1.08E+00 | 3.42E-01 | 9.39E-01 | 1.03E+00 |
| NFRKB | 4.71E-04 | 2.45E-03 | -1.22E+00 | 5.63E-02 | 2.43E-01 | 1.31E+00 | 9.70E-03 | 4.12E-02 | -1.14E+00 | 2.20E-01 | 8.39E-01 | 1.38E+00 | 1.95E-02 | 6.96E-02 | -1.12E+00 | 2.93E-01 | 8.26E-01 | 1.29E+00 | 4.35E-05 | 1.60E-04 | -1.22E+00 | 2.03E-02 | 5.57E-02 | -1.02E+00 |
| AC144652.1 | 4.71E-04 | 2.46E-03 | 2.25E+00 | 3.59E-02 | 1.55E-01 | 1.08E+00 | 1.43E-02 | 6.08E-02 | 2.02E+00 | 2.30E-01 | 8.79E-01 | 1.62E+00 | 2.12E-01 | 7.55E-01 | 1.31E+00 | 1.68E-01 | 4.73E-01 | 1.13E+00 | 1.87E-01 | 6.86E-01 | 1.50E+00 | 9.67E-02 | 2.66E-01 | 1.73E+00 |
| AC003681.1 | 4.71E-04 | 2.46E-03 | -1.06E+00 | 4.51E-05 | 1.95E-04 | -1.01E+00 | 2.96E-02 | 1.26E-01 | -1.11E+00 | 1.62E-04 | 6.20E-04 | -1.20E+00 | 7.10E-05 | 2.53E-04 | -1.11E+00 | 2.55E-10 | 7.20E-10 | -1.46E+00 | 1.54E-06 | 5.66E-06 | -1.50E+00 | 5.16E-11 | 1.42E-10 | -1.83E+00 |
| HOXB6 | 4.71E-04 | 2.46E-03 | -1.14E+00 | 7.38E-04 | 3.19E-03 | -1.08E+00 | 2.23E-04 | 9.45E-04 | -1.47E+00 | 7.49E-04 | 2.86E-03 | -1.27E+00 | 1.40E-02 | 4.98E-02 | -1.41E+00 | 1.41E-01 | 3.98E-01 | 1.04E+00 | 2.05E-02 | 7.53E-02 | -1.46E+00 | 1.71E-01 | 4.69E-01 | -1.09E+00 |
| AC004034.1 | 4.72E-04 | 2.46E-03 | -1.06E+00 | 1.67E-02 | 7.20E-02 | 1.24E+00 | 4.57E-02 | 1.94E-01 | -1.09E+00 | 3.74E-02 | 1.43E-01 | 1.02E+00 | 3.15E-03 | 1.12E-02 | -1.31E+00 | 2.43E-03 | 6.85E-03 | -1.30E+00 | 1.24E-01 | 4.56E-01 | -1.25E+00 | 1.86E-01 | 5.12E-01 | -1.26E+00 |
| C22orf34 | 4.73E-04 | 2.46E-03 | -1.46E+00 | 2.66E-03 | 1.15E-02 | -1.17E+00 | 1.12E-01 | 4.74E-01 | 1.09E+00 | 2.05E-01 | 7.81E-01 | 1.47E+00 | 1.26E-01 | 4.50E-01 | 1.86E+00 | 1.17E-01 | 3.29E-01 | 2.08E+00 | 7.22E-03 | 2.65E-02 | -1.43E+00 | 2.31E-01 | 6.35E-01 | 1.07E+00 |
| IRF8 | 4.74E-04 | 2.47E-03 | -1.09E+01 | 1.56E-03 | 6.73E-03 | -8.05E+00 | 5.22E-03 | 2.22E-02 | -1.51E+00 | 1.87E-06 | 7.15E-06 | -2.20E+00 | 2.80E-01 | 1.00E+00 | 1.00E+00 | 3.55E-01 | 1.00E+00 | 1.00E+00 | 2.45E-01 | 8.99E-01 | 1.08E+00 | 2.80E-01 | 7.69E-01 | -1.09E+00 |
| PRPF38B | 4.76E-04 | 2.48E-03 | -1.06E+00 | 5.96E-04 | 2.57E-03 | 1.03E+00 | 1.04E-03 | 4.43E-03 | -1.20E+00 | 4.73E-03 | 1.81E-02 | -1.08E+00 | 2.90E-04 | 1.03E-03 | -1.06E+00 | 1.15E-03 | 3.23E-03 | -1.03E+00 | 9.23E-05 | 3.39E-04 | -1.09E+00 | 1.55E-01 | 4.26E-01 | 1.05E+00 |
| GLRB | 4.77E-04 | 2.49E-03 | -2.03E+00 | 1.74E-02 | 7.51E-02 | -1.21E+00 | 8.99E-03 | 3.82E-02 | -1.63E+00 | 1.48E-01 | 5.66E-01 | 1.05E+00 | 2.75E-01 | 9.81E-01 | 1.00E+00 | 2.95E-01 | 8.31E-01 | 1.15E+00 | 1.56E-04 | 5.74E-04 | -1.40E+00 | 9.57E-03 | 2.63E-02 | -1.05E+00 |
| C19orf25 | 4.77E-04 | 2.49E-03 | -1.02E+00 | 4.93E-02 | 2.13E-01 | 1.28E+00 | 2.93E-04 | 1.25E-03 | -1.20E+00 | 4.54E-03 | 1.74E-02 | 1.07E+00 | 5.57E-02 | 1.99E-01 | 1.18E+00 | 2.47E-01 | 6.95E-01 | 1.35E+00 | 1.48E-04 | 5.43E-04 | -1.04E+00 | 6.65E-02 | 1.83E-01 | 1.23E+00 |
| SCRN1 | 4.77E-04 | 2.49E-03 | -1.08E+00 | 3.24E-02 | 1.40E-01 | 1.25E+00 | 2.24E-03 | 9.53E-03 | -1.19E+00 | 4.90E-02 | 1.87E-01 | 1.11E+00 | 2.05E-01 | 7.31E-01 | 1.25E+00 | 1.66E-01 | 4.68E-01 | 1.27E+00 | 3.98E-05 | 1.46E-04 | -1.19E+00 | 1.00E-03 | 2.75E-03 | -1.04E+00 |
| ACOT7 | 4.77E-04 | 2.49E-03 | -1.23E+00 | 1.04E-04 | 4.48E-04 | -1.24E+00 | 1.04E-02 | 4.41E-02 | 1.10E+00 | 4.12E-05 | 1.57E-04 | 1.03E+00 | 2.21E-04 | 7.88E-04 | -1.55E+00 | 5.24E-03 | 1.48E-02 | -1.18E+00 | 6.07E-04 | 2.23E-03 | -1.19E+00 | 2.85E-01 | 7.83E-01 | 1.24E+00 |
| WRN | 4.78E-04 | 2.49E-03 | -1.19E+00 | 1.25E-03 | 5.39E-03 | 1.00E+00 | 2.38E-02 | 1.01E-01 | -1.01E+00 | 2.21E-02 | 8.45E-02 | 1.13E+00 | 3.61E-02 | 1.29E-01 | 1.08E+00 | 2.36E-01 | 6.65E-01 | 1.33E+00 | 1.10E-09 | 4.05E-09 | -1.27E+00 | 4.29E-04 | 1.18E-03 | 1.04E+00 |
| LINC00205 | 4.80E-04 | 2.50E-03 | -1.37E+00 | 1.79E-01 | 7.72E-01 | 1.50E+00 | 4.78E-06 | 2.03E-05 | -2.02E+00 | 1.16E-03 | 4.44E-03 | -1.19E+00 | 3.32E-02 | 1.18E-01 | -1.52E+00 | 2.23E-01 | 6.29E-01 | 1.01E+00 | 1.86E-04 | 6.83E-04 | -1.96E+00 | 6.23E-02 | 1.71E-01 | -1.14E+00 |
| SCRN2 | 4.80E-04 | 2.50E-03 | -1.01E+00 | 5.80E-06 | 2.50E-05 | -1.07E+00 | 2.38E-02 | 1.01E-01 | -1.05E+00 | 2.82E-03 | 1.08E-02 | -1.03E+00 | 9.19E-04 | 3.28E-03 | -1.12E+00 | 2.45E-03 | 6.91E-03 | -1.05E+00 | 1.67E-01 | 6.11E-01 | 1.18E+00 | 1.75E-03 | 4.81E-03 | -1.17E+00 |
| JRK | 4.81E-04 | 2.50E-03 | -1.70E+00 | 4.92E-02 | 2.13E-01 | 1.95E+00 | 2.16E-06 | 9.18E-06 | -1.94E+00 | 1.53E-01 | 5.83E-01 | 1.25E+00 | 2.75E-02 | 9.80E-02 | -1.35E+00 | 1.47E-01 | 4.13E-01 | 1.73E+00 | 5.62E-02 | 2.06E-01 | 1.00E+00 | 1.13E-02 | 3.10E-02 | 1.79E+00 |
| FN3KRP | 4.81E-04 | 2.51E-03 | -1.17E+00 | 4.02E-02 | 1.74E-01 | 1.24E+00 | 5.04E-04 | 2.14E-03 | -1.19E+00 | 7.21E-02 | 2.75E-01 | 1.19E+00 | 3.15E-06 | 1.12E-05 | -1.68E+00 | 1.14E-02 | 3.21E-02 | 1.01E+00 | 4.69E-09 | 1.72E-08 | -1.39E+00 | 7.78E-02 | 2.14E-01 | 1.13E+00 |
| AC009506.1 | 4.81E-04 | 2.51E-03 | -1.27E+00 | 1.21E-02 | 5.24E-02 | 1.03E+00 | 1.27E-02 | 5.39E-02 | -1.17E+00 | 3.69E-02 | 1.41E-01 | 1.09E+00 | 3.21E-02 | 1.15E-01 | -1.06E+00 | 2.94E-01 | 8.30E-01 | 1.43E+00 | 3.44E-03 | 1.26E-02 | -1.12E+00 | 5.58E-02 | 1.53E-01 | 1.11E+00 |
| AL034397.3 | 4.82E-04 | 2.51E-03 | -1.10E+00 | 1.33E-01 | 5.74E-01 | 1.36E+00 | 6.06E-02 | 2.58E-01 | 1.05E+00 | 5.22E-04 | 1.99E-03 | 1.84E+00 | 3.27E-11 | 1.16E-10 | -1.57E+00 | 1.32E-01 | 3.72E-01 | 1.22E+00 | 1.81E-03 | 6.64E-03 | 1.08E+00 | 1.18E-01 | 3.24E-01 | 1.56E+00 |
| METTL15 | 4.82E-04 | 2.51E-03 | -1.21E+00 | 1.17E-05 | 5.08E-05 | -1.38E+00 | 4.88E-08 | 2.07E-07 | -1.75E+00 | 9.28E-10 | 3.54E-09 | -1.73E+00 | 2.62E-02 | 9.34E-02 | -1.11E+00 | 1.31E-02 | 3.70E-02 | -1.24E+00 | 1.38E-04 | 5.07E-04 | -1.18E+00 | 8.94E-05 | 2.45E-04 | -1.30E+00 |
| ZNF211 | 4.83E-04 | 2.51E-03 | -1.50E+00 | 3.79E-03 | 1.64E-02 | -1.16E+00 | 6.59E-02 | 2.80E-01 | -1.06E+00 | 3.19E-02 | 1.22E-01 | -1.11E+00 | 6.90E-02 | 2.46E-01 | 1.03E+00 | 2.98E-02 | 8.40E-02 | -1.20E+00 | 9.27E-04 | 3.40E-03 | -1.44E+00 | 5.14E-04 | 1.41E-03 | -1.48E+00 |
| TUBB6 | 4.83E-04 | 2.52E-03 | -1.07E+00 | 1.42E-02 | 6.13E-02 | 1.26E+00 | 1.10E-01 | 4.68E-01 | 1.19E+00 | 1.53E-01 | 5.84E-01 | 1.47E+00 | 4.50E-02 | 1.61E-01 | -1.05E+00 | 1.93E-01 | 5.45E-01 | 1.21E+00 | 4.37E-03 | 1.60E-02 | -1.19E+00 | 3.05E-01 | 8.37E-01 | 1.09E+00 |
| CRAT | 4.83E-04 | 2.52E-03 | -1.23E+00 | 2.31E-03 | 9.97E-03 | 1.05E+00 | 3.39E-03 | 1.44E-02 | -1.17E+00 | 3.55E-02 | 1.35E-01 | 1.14E+00 | 4.09E-04 | 1.46E-03 | -1.19E+00 | 1.39E-02 | 3.91E-02 | 1.07E+00 | 4.05E-03 | 1.49E-02 | 1.08E+00 | 1.65E-03 | 4.53E-03 | -1.04E+00 |
| SIRT1 | 4.83E-04 | 2.52E-03 | -1.13E+00 | 6.99E-09 | 3.02E-08 | -1.73E+00 | 1.89E-05 | 8.02E-05 | -1.40E+00 | 1.73E-06 | 6.61E-06 | -1.40E+00 | 2.63E-02 | 9.38E-02 | 1.06E+00 | 1.47E-02 | 4.15E-02 | -1.08E+00 | 4.69E-06 | 1.72E-05 | -1.25E+00 | 6.34E-06 | 1.74E-05 | -1.36E+00 |
| HARBI1 | 4.84E-04 | 2.52E-03 | -1.49E+00 | 8.94E-05 | 3.86E-04 | -1.62E+00 | 7.19E-03 | 3.05E-02 | -1.33E+00 | 1.91E-03 | 7.29E-03 | -1.33E+00 | 5.81E-04 | 2.07E-03 | -1.74E+00 | 5.76E-03 | 1.62E-02 | -1.21E+00 | 2.55E-05 | 9.38E-05 | -1.94E+00 | 1.75E-03 | 4.80E-03 | -1.52E+00 |
| P2RY11 | 4.84E-04 | 2.52E-03 | -1.07E+00 | 5.32E-06 | 2.30E-05 | -1.25E+00 | 1.61E-01 | 6.85E-01 | 1.14E+00 | 1.17E-02 | 4.48E-02 | 1.01E+00 | 1.44E-01 | 5.15E-01 | 1.99E+00 | 1.96E-01 | 5.53E-01 | 1.93E+00 | 2.38E-03 | 8.72E-03 | -1.20E+00 | 1.76E-03 | 4.84E-03 | -1.19E+00 |
| SPATA5 | 4.84E-04 | 2.52E-03 | -1.34E+00 | 1.22E-02 | 5.26E-02 | 1.02E+00 | 2.84E-03 | 1.21E-02 | -1.26E+00 | 6.82E-03 | 2.61E-02 | -1.08E+00 | 2.59E-02 | 9.23E-02 | -1.23E+00 | 2.51E-01 | 7.07E-01 | 1.13E+00 | 1.56E-08 | 5.71E-08 | -1.49E+00 | 6.40E-04 | 1.76E-03 | -1.07E+00 |
| PRRT3 | 4.85E-04 | 2.53E-03 | -1.39E+00 | 3.00E-03 | 1.29E-02 | -1.13E+00 | 2.15E-03 | 9.14E-03 | -1.29E+00 | 3.54E-02 | 1.35E-01 | 1.07E+00 | 3.07E-02 | 1.10E-01 | -1.22E+00 | 9.37E-02 | 2.64E-01 | -1.07E+00 | 1.09E-01 | 3.99E-01 | -1.02E+00 | 3.59E-01 | 9.86E-01 | 1.24E+00 |
| LINC00939 | 4.87E-04 | 2.54E-03 | -1.68E+00 | 1.41E-02 | 6.11E-02 | -1.16E+00 | 1.75E-01 | 7.43E-01 | -1.18E+00 | 1.70E-01 | 6.48E-01 | 1.25E+00 | 8.38E-02 | 2.99E-01 | -1.32E+00 | 2.74E-01 | 7.72E-01 | 1.47E+00 | 1.92E-01 | 7.06E-01 | 1.24E+00 | 3.42E-01 | 9.40E-01 | 1.28E+00 |
| AC011773.1 | 4.87E-04 | 2.54E-03 | 2.95E+00 | 1.53E-01 | 6.63E-01 | 1.62E+00 | 3.18E-03 | 1.35E-02 | 2.08E+00 | 2.31E-01 | 8.81E-01 | 1.21E+00 | 3.86E-04 | 1.37E-03 | 4.87E+00 | 1.33E-01 | 3.74E-01 | 2.20E+00 | 4.31E-03 | 1.58E-02 | 2.14E+00 | 2.39E-01 | 6.57E-01 | -1.05E+00 |
| ATP5L | 4.88E-04 | 2.54E-03 | -1.07E+00 | 6.48E-14 | 2.80E-13 | -1.20E+00 | 4.59E-02 | 1.95E-01 | -1.06E+00 | 1.32E-05 | 5.04E-05 | -1.21E+00 | 2.31E-02 | 8.25E-02 | -1.04E+00 | 3.50E-07 | 9.88E-07 | -1.25E+00 | 1.26E-02 | 4.63E-02 | -1.04E+00 | 5.73E-06 | 1.57E-05 | -1.22E+00 |
| ELOCP2 | 4.88E-04 | 2.54E-03 | -1.17E+00 | 5.52E-07 | 2.38E-06 | -1.51E+00 | 7.03E-02 | 2.99E-01 | 1.10E+00 | 6.63E-05 | 2.53E-04 | -1.18E+00 | 1.20E-03 | 4.26E-03 | -1.26E+00 | 2.37E-04 | 6.69E-04 | -1.27E+00 | 3.05E-15 | 1.12E-14 | -1.77E+00 | 8.61E-12 | 2.37E-11 | -1.50E+00 |
| TCEAL6 | 4.89E-04 | 2.55E-03 | -1.13E+00 | 4.17E-06 | 1.80E-05 | -1.35E+00 | 3.74E-03 | 1.59E-02 | -1.11E+00 | 2.47E-04 | 9.43E-04 | -1.24E+00 | 1.09E-03 | 3.88E-03 | -1.24E+00 | 9.55E-04 | 2.69E-03 | -1.31E+00 | 1.52E-02 | 5.58E-02 | -1.09E+00 | 2.32E-02 | 6.38E-02 | -1.10E+00 |
| RMDN1 | 4.90E-04 | 2.55E-03 | -1.07E+00 | 1.99E-04 | 8.58E-04 | 1.03E+00 | 4.57E-02 | 1.94E-01 | 1.01E+00 | 8.65E-04 | 3.30E-03 | 1.01E+00 | 4.26E-03 | 1.52E-02 | 1.05E+00 | 2.15E-02 | 6.07E-02 | 1.17E+00 | 7.03E-05 | 2.58E-04 | -1.11E+00 | 1.84E-03 | 5.06E-03 | -1.01E+00 |
| LIPT1 | 4.90E-04 | 2.56E-03 | -1.18E+00 | 1.36E-02 | 5.89E-02 | 1.18E+00 | 2.08E-04 | 8.83E-04 | -1.57E+00 | 1.70E-04 | 6.51E-04 | -1.37E+00 | 1.91E-04 | 6.81E-04 | -1.70E+00 | 1.08E-02 | 3.06E-02 | -1.10E+00 | 3.51E-03 | 1.29E-02 | -1.34E+00 | 8.65E-02 | 2.38E-01 | 1.03E+00 |
| FKBP1B | 4.92E-04 | 2.57E-03 | -1.39E+00 | 8.09E-02 | 3.50E-01 | 1.30E+00 | 1.49E-02 | 6.33E-02 | -1.31E+00 | 2.26E-01 | 8.65E-01 | 1.27E+00 | 9.18E-04 | 3.27E-03 | -1.14E+00 | 7.47E-02 | 2.11E-01 | 1.16E+00 | 2.62E-01 | 9.62E-01 | 1.42E+00 | 2.84E-01 | 7.79E-01 | 1.32E+00 |
| BRD7P2 | 4.92E-04 | 2.57E-03 | -1.20E+00 | 4.97E-02 | 2.15E-01 | 1.30E+00 | 6.02E-04 | 2.56E-03 | -1.23E+00 | 2.39E-01 | 9.13E-01 | 1.37E+00 | 4.05E-03 | 1.44E-02 | -1.09E+00 | 3.49E-01 | 9.85E-01 | 1.40E+00 | 5.85E-02 | 2.15E-01 | 1.02E+00 | 6.97E-02 | 1.92E-01 | 1.55E+00 |
| AC018445.4 | 4.94E-04 | 2.58E-03 | -1.37E+00 | 1.19E-03 | 5.15E-03 | -1.28E+00 | 1.23E-01 | 5.21E-01 | 1.05E+00 | 9.35E-02 | 3.57E-01 | 1.02E+00 | 7.42E-02 | 2.64E-01 | -1.14E+00 | 1.59E-01 | 4.49E-01 | -1.07E+00 | 4.00E-02 | 1.47E-01 | -1.30E+00 | 1.95E-01 | 5.36E-01 | -1.10E+00 |
| RNF4 | 4.95E-04 | 2.58E-03 | -1.10E+00 | 8.87E-05 | 3.83E-04 | -1.04E+00 | 3.04E-04 | 1.29E-03 | -1.10E+00 | 1.58E-04 | 6.05E-04 | -1.07E+00 | 1.34E-02 | 4.77E-02 | 1.07E+00 | 1.87E-04 | 5.26E-04 | -1.18E+00 | 3.83E-07 | 1.40E-06 | -1.31E+00 | 6.86E-04 | 1.88E-03 | -1.03E+00 |
| SNHG9 | 4.96E-04 | 2.59E-03 | -1.20E+00 | 2.00E-06 | 8.64E-06 | -1.56E+00 | 1.27E-04 | 5.38E-04 | -1.52E+00 | 9.56E-06 | 3.65E-05 | -1.70E+00 | 2.47E-01 | 8.80E-01 | 1.14E+00 | 5.92E-02 | 1.67E-01 | -1.52E+00 | 1.26E-01 | 4.63E-01 | 1.49E+00 | 8.21E-02 | 2.25E-01 | 1.01E+00 |
| AL645728.1 | 4.96E-04 | 2.59E-03 | 2.01E+00 | 9.15E-02 | 3.96E-01 | 1.18E+00 | 2.04E-03 | 8.68E-03 | 1.77E+00 | 2.57E-02 | 9.82E-02 | -1.12E+00 | 2.60E-01 | 9.29E-01 | 1.57E+00 | 1.07E-01 | 3.02E-01 | 1.07E+00 | 1.87E-01 | 6.85E-01 | 1.39E+00 | 2.15E-01 | 5.89E-01 | 1.09E+00 |
| TPST2 | 4.96E-04 | 2.59E-03 | 1.09E+00 | 2.10E-01 | 9.08E-01 | 1.51E+00 | 4.75E-02 | 2.02E-01 | 1.12E+00 | 1.55E-01 | 5.93E-01 | 1.55E+00 | 2.63E-02 | 9.39E-02 | 1.65E+00 | 1.85E-03 | 5.22E-03 | 1.69E+00 | 1.08E-02 | 3.98E-02 | 1.56E+00 | 2.07E-02 | 5.68E-02 | 1.45E+00 |
| FARS2 | 4.98E-04 | 2.59E-03 | -1.05E+00 | 1.95E-04 | 8.41E-04 | 1.01E+00 | 5.00E-03 | 2.13E-02 | -1.09E+00 | 1.08E-05 | 4.14E-05 | -1.18E+00 | 5.27E-05 | 1.88E-04 | -1.25E+00 | 2.49E-02 | 7.01E-02 | 1.12E+00 | 5.36E-03 | 1.97E-02 | -1.04E+00 | 6.95E-03 | 1.91E-02 | 1.02E+00 |
| PHKA2 | 4.98E-04 | 2.59E-03 | -1.24E+00 | 2.16E-03 | 9.33E-03 | -1.04E+00 | 2.14E-02 | 9.08E-02 | 1.00E+00 | 3.80E-03 | 1.45E-02 | 1.01E+00 | 1.27E-03 | 4.54E-03 | -1.14E+00 | 3.00E-03 | 8.46E-03 | -1.01E+00 | 4.71E-02 | 1.73E-01 | 1.14E+00 | 3.76E-02 | 1.03E-01 | 1.16E+00 |
| BCL2L2 | 4.98E-04 | 2.59E-03 | -1.27E+00 | 1.05E-03 | 4.54E-03 | -1.11E+00 | 7.72E-04 | 3.28E-03 | -1.25E+00 | 2.61E-03 | 9.97E-03 | -1.04E+00 | 6.26E-02 | 2.23E-01 | 1.07E+00 | 9.73E-03 | 2.74E-02 | -1.18E+00 | 2.57E-01 | 9.43E-01 | 1.35E+00 | 1.49E-01 | 4.10E-01 | 1.15E+00 |
| TIMM29 | 4.98E-04 | 2.60E-03 | -1.08E+00 | 4.50E-03 | 1.94E-02 | 1.08E+00 | 1.84E-02 | 7.83E-02 | -1.10E+00 | 7.11E-02 | 2.72E-01 | 1.13E+00 | 9.11E-06 | 3.25E-05 | -1.28E+00 | 4.05E-03 | 1.14E-02 | 1.06E+00 | 6.48E-04 | 2.38E-03 | -1.07E+00 | 4.11E-02 | 1.13E-01 | 1.04E+00 |
| ATP11A | 4.99E-04 | 2.60E-03 | -1.49E+00 | 1.46E-03 | 6.29E-03 | -1.23E+00 | 1.02E-02 | 4.34E-02 | -1.10E+00 | 3.59E-02 | 1.37E-01 | 1.10E+00 | 1.39E-02 | 4.97E-02 | -1.18E+00 | 1.86E-01 | 5.25E-01 | 1.25E+00 | 2.20E-04 | 8.09E-04 | -1.28E+00 | 7.42E-03 | 2.04E-02 | -1.06E+00 |
| RPL23AP3 | 4.99E-04 | 2.60E-03 | -1.28E+00 | 3.67E-03 | 1.59E-02 | -1.01E+00 | 1.37E-06 | 5.82E-06 | -1.76E+00 | 2.57E-04 | 9.81E-04 | -1.23E+00 | 5.79E-03 | 2.06E-02 | -1.41E+00 | 1.60E-02 | 4.50E-02 | -1.16E+00 | 1.90E-07 | 6.98E-07 | -1.76E+00 | 2.54E-04 | 6.97E-04 | -1.26E+00 |
| PYGO2 | 5.00E-04 | 2.61E-03 | -1.46E+00 | 1.44E-03 | 6.21E-03 | -1.21E+00 | 7.21E-04 | 3.06E-03 | -1.40E+00 | 4.48E-04 | 1.71E-03 | -1.26E+00 | 3.71E-06 | 1.32E-05 | -2.11E+00 | 4.85E-03 | 1.37E-02 | -1.17E+00 | 1.24E-21 | 4.54E-21 | -2.69E+00 | 3.16E-09 | 8.67E-09 | -1.46E+00 |
| PPAN | 5.01E-04 | 2.61E-03 | -1.13E+00 | 1.47E-04 | 6.37E-04 | -1.09E+00 | 3.54E-08 | 1.51E-07 | -1.40E+00 | 2.68E-07 | 1.02E-06 | -1.19E+00 | 8.81E-03 | 3.14E-02 | -1.34E+00 | 7.19E-02 | 2.03E-01 | 1.03E+00 | 1.38E-13 | 5.06E-13 | -1.75E+00 | 2.73E-10 | 7.50E-10 | -1.47E+00 |
| CARD9 | 5.01E-04 | 2.61E-03 | -1.16E+00 | 1.39E-04 | 6.01E-04 | -1.14E+00 | 1.69E-03 | 7.18E-03 | -1.28E+00 | 1.38E-02 | 5.25E-02 | -1.00E+00 | 1.48E-04 | 5.27E-04 | -1.53E+00 | 4.19E-03 | 1.18E-02 | -1.12E+00 | 1.48E-01 | 5.42E-01 | 1.08E+00 | 3.52E-01 | 9.66E-01 | 1.25E+00 |
| ILVBL | 5.02E-04 | 2.62E-03 | -1.16E+00 | 4.90E-03 | 2.12E-02 | 1.06E+00 | 8.67E-02 | 3.68E-01 | 1.06E+00 | 4.02E-02 | 1.53E-01 | 1.13E+00 | 1.05E-02 | 3.73E-02 | -1.08E+00 | 8.05E-02 | 2.27E-01 | 1.11E+00 | 1.75E-03 | 6.42E-03 | -1.14E+00 | 1.31E-01 | 3.59E-01 | 1.14E+00 |
| NOTCH2NL | 5.04E-04 | 2.63E-03 | -1.45E+00 | 4.13E-02 | 1.78E-01 | 1.18E+00 | 1.50E-02 | 6.36E-02 | -1.28E+00 | 1.27E-01 | 4.86E-01 | 1.65E+00 | 8.55E-06 | 3.05E-05 | -1.51E+00 | 3.21E-03 | 9.05E-03 | 1.02E+00 | 7.24E-03 | 2.66E-02 | -1.19E+00 | 2.15E-01 | 5.92E-01 | 1.12E+00 |
| AC090517.2 | 5.04E-04 | 2.63E-03 | -1.66E+00 | 1.65E-01 | 7.13E-01 | 1.32E+00 | 6.21E-03 | 2.64E-02 | -1.31E+00 | 1.65E-01 | 6.29E-01 | 1.25E+00 | 5.25E-02 | 1.87E-01 | -1.30E+00 | 2.02E-01 | 5.70E-01 | 1.04E+00 | 8.43E-02 | 3.09E-01 | 1.60E+00 | 2.67E-02 | 7.34E-02 | 2.02E+00 |
| LARP1B | 5.04E-04 | 2.63E-03 | 1.02E+00 | 3.58E-04 | 1.55E-03 | 1.10E+00 | 1.37E-02 | 5.83E-02 | -1.05E+00 | 7.52E-03 | 2.87E-02 | 1.04E+00 | 1.02E-04 | 3.64E-04 | -1.31E+00 | 6.07E-03 | 1.71E-02 | 1.01E+00 | 2.30E-02 | 8.43E-02 | 1.02E+00 | 1.63E-02 | 4.47E-02 | 1.00E+00 |
| SEC61B | 5.05E-04 | 2.63E-03 | 1.27E+00 | 5.83E-02 | 2.52E-01 | 1.13E+00 | 4.92E-04 | 2.09E-03 | 1.39E+00 | 3.06E-02 | 1.17E-01 | 1.30E+00 | 4.74E-03 | 1.69E-02 | -1.04E+00 | 2.34E-03 | 6.60E-03 | -1.08E+00 | 1.28E-02 | 4.68E-02 | 1.26E+00 | 1.57E-02 | 4.30E-02 | 1.24E+00 |
| FAM50A | 5.05E-04 | 2.63E-03 | -1.21E+00 | 1.13E-03 | 4.88E-03 | -1.02E+00 | 1.96E-01 | 8.35E-01 | 1.31E+00 | 1.13E-01 | 4.30E-01 | 1.35E+00 | 1.34E-03 | 4.77E-03 | -1.05E+00 | 5.12E-04 | 1.44E-03 | -1.12E+00 | 1.39E-01 | 5.10E-01 | 1.28E+00 | 2.63E-01 | 7.23E-01 | 1.32E+00 |
| HNRNPM | 5.06E-04 | 2.63E-03 | 1.01E+00 | 5.69E-02 | 2.46E-01 | 1.16E+00 | 1.18E-01 | 5.01E-01 | 1.10E+00 | 1.18E-01 | 4.49E-01 | 1.21E+00 | 2.94E-02 | 1.05E-01 | 1.04E+00 | 1.67E-01 | 4.70E-01 | 1.31E+00 | 3.66E-02 | 1.34E-01 | -1.04E+00 | 3.57E-01 | 9.80E-01 | 1.11E+00 |
| TPGS1 | 5.06E-04 | 2.63E-03 | 1.04E+00 | 8.59E-09 | 3.71E-08 | -1.07E+00 | 1.23E-02 | 5.21E-02 | 1.02E+00 | 1.10E-05 | 4.20E-05 | -1.08E+00 | 3.06E-06 | 1.09E-05 | -1.13E+00 | 2.22E-05 | 6.26E-05 | -1.11E+00 | 1.65E-03 | 6.07E-03 | -1.03E+00 | 2.44E-03 | 6.71E-03 | -1.06E+00 |
| AC090061.1 | 5.06E-04 | 2.64E-03 | -1.49E+00 | 6.45E-04 | 2.79E-03 | -1.36E+00 | 1.07E-03 | 4.57E-03 | -1.55E+00 | 5.35E-04 | 2.04E-03 | -1.44E+00 | 7.66E-02 | 2.73E-01 | -1.15E+00 | 2.02E-01 | 5.70E-01 | 1.16E+00 | 8.01E-02 | 2.94E-01 | -1.06E+00 | 1.25E-01 | 3.43E-01 | -1.15E+00 |
| NFAM1 | 5.06E-04 | 2.64E-03 | -1.58E+00 | 3.69E-03 | 1.59E-02 | -1.27E+00 | 1.49E-02 | 6.33E-02 | -1.49E+00 | 2.40E-02 | 9.18E-02 | -1.29E+00 | 5.14E-03 | 1.83E-02 | -1.45E+00 | 6.23E-03 | 1.76E-02 | -1.39E+00 | 5.32E-03 | 1.95E-02 | -1.57E+00 | 5.75E-02 | 1.58E-01 | -1.29E+00 |
| EHBP1 | 5.10E-04 | 2.66E-03 | -1.11E+00 | 9.73E-04 | 4.20E-03 | 1.02E+00 | 5.92E-03 | 2.52E-02 | -1.21E+00 | 2.03E-02 | 7.75E-02 | 1.02E+00 | 8.23E-03 | 2.94E-02 | -1.08E+00 | 7.32E-02 | 2.06E-01 | 1.16E+00 | 2.11E-07 | 7.73E-07 | -1.34E+00 | 2.16E-04 | 5.92E-04 | -1.10E+00 |
| SCHIP1 | 5.11E-04 | 2.66E-03 | -1.10E+00 | 1.98E-02 | 8.56E-02 | 1.84E+00 | 2.50E-02 | 1.06E-01 | -1.16E+00 | 3.68E-02 | 1.41E-01 | 1.94E+00 | 2.75E-01 | 9.80E-01 | 1.54E+00 | 3.02E-02 | 8.53E-02 | 2.46E+00 | 1.98E-01 | 7.28E-01 | 1.39E+00 | 3.37E-02 | 9.25E-02 | 2.36E+00 |
| MAML1 | 5.11E-04 | 2.66E-03 | -1.29E+00 | 6.97E-03 | 3.01E-02 | 1.04E+00 | 7.61E-05 | 3.23E-04 | -1.43E+00 | 4.35E-03 | 1.66E-02 | -1.06E+00 | 9.12E-04 | 3.25E-03 | -1.06E+00 | 6.22E-03 | 1.75E-02 | -1.01E+00 | 1.03E-10 | 3.77E-10 | -1.86E+00 | 4.14E-05 | 1.14E-04 | -1.22E+00 |
| HINFP | 5.11E-04 | 2.66E-03 | -1.16E+00 | 6.55E-02 | 2.83E-01 | 1.34E+00 | 3.22E-07 | 1.37E-06 | -1.80E+00 | 5.21E-03 | 1.99E-02 | 1.01E+00 | 3.59E-02 | 1.28E-01 | 1.00E+00 | 1.87E-01 | 5.27E-01 | 1.25E+00 | 2.66E-05 | 9.78E-05 | -1.32E+00 | 1.20E-03 | 3.29E-03 | -1.14E+00 |
| ZNHIT2 | 5.11E-04 | 2.67E-03 | -1.57E+00 | 1.30E-02 | 5.60E-02 | -1.05E+00 | 5.39E-02 | 2.29E-01 | -1.05E+00 | 2.23E-01 | 8.54E-01 | 1.43E+00 | 2.21E-02 | 7.89E-02 | -1.29E+00 | 1.64E-01 | 4.62E-01 | 1.10E+00 | 7.84E-05 | 2.88E-04 | -1.42E+00 | 1.15E-01 | 3.15E-01 | -1.03E+00 |
| TRO | 5.13E-04 | 2.67E-03 | -1.27E+00 | 2.45E-04 | 1.06E-03 | -1.24E+00 | 6.14E-03 | 2.61E-02 | -1.35E+00 | 1.63E-02 | 6.24E-02 | -1.09E+00 | 4.81E-03 | 1.72E-02 | -1.30E+00 | 1.58E-02 | 4.45E-02 | -1.19E+00 | 7.50E-02 | 2.75E-01 | -1.16E+00 | 3.37E-01 | 9.26E-01 | 1.24E+00 |
| ZNF527 | 5.14E-04 | 2.68E-03 | -1.27E+00 | 4.64E-04 | 2.00E-03 | -1.19E+00 | 1.39E-04 | 5.91E-04 | -1.34E+00 | 4.21E-06 | 1.61E-05 | -1.41E+00 | 9.69E-02 | 3.45E-01 | -1.12E+00 | 1.43E-01 | 4.02E-01 | -1.11E+00 | 2.67E-04 | 9.80E-04 | -1.55E+00 | 1.33E-03 | 3.65E-03 | -1.44E+00 |
| MTHFS | 5.15E-04 | 2.68E-03 | -1.55E+00 | 5.75E-03 | 2.48E-02 | -1.14E+00 | 3.31E-06 | 1.41E-05 | -1.86E+00 | 1.30E-05 | 4.97E-05 | -1.55E+00 | 1.24E-09 | 4.41E-09 | -3.53E+00 | 1.92E-06 | 5.42E-06 | -1.82E+00 | 2.49E-13 | 9.15E-13 | -2.06E+00 | 2.37E-09 | 6.52E-09 | -1.63E+00 |
| FAM49B | 5.15E-04 | 2.69E-03 | 1.03E+00 | 2.71E-04 | 1.17E-03 | 1.09E+00 | 1.20E-03 | 5.12E-03 | -1.07E+00 | 1.95E-04 | 7.46E-04 | -1.02E+00 | 6.25E-09 | 2.23E-08 | -1.30E+00 | 3.50E-06 | 9.88E-06 | -1.15E+00 | 6.10E-04 | 2.24E-03 | -1.07E+00 | 2.44E-02 | 6.70E-02 | 1.06E+00 |
| BCL9L | 5.16E-04 | 2.69E-03 | -1.28E+00 | 3.66E-02 | 1.58E-01 | 1.18E+00 | 1.68E-02 | 7.14E-02 | -1.33E+00 | 2.24E-01 | 8.57E-01 | 1.40E+00 | 1.41E-01 | 5.04E-01 | -1.39E+00 | 3.06E-01 | 8.62E-01 | 1.28E+00 | 1.15E-01 | 4.22E-01 | 1.04E+00 | 1.53E-01 | 4.20E-01 | -1.03E+00 |
| AC138058.1 | 5.16E-04 | 2.69E-03 | -1.34E+00 | 9.65E-04 | 4.17E-03 | -1.14E+00 | 7.51E-02 | 3.19E-01 | -1.11E+00 | 1.63E-01 | 6.22E-01 | 1.21E+00 | 1.13E-01 | 4.03E-01 | -1.24E+00 | 6.88E-02 | 1.94E-01 | -1.48E+00 | 6.94E-03 | 2.55E-02 | -1.44E+00 | 1.77E-02 | 4.85E-02 | -1.37E+00 |
| EEF2KMT | 5.18E-04 | 2.70E-03 | -2.22E+00 | 7.20E-03 | 3.11E-02 | -1.48E+00 | 6.68E-14 | 2.84E-13 | -2.93E+00 | 3.83E-09 | 1.46E-08 | -1.82E+00 | 2.29E-02 | 8.17E-02 | -1.71E+00 | 1.81E-01 | 5.10E-01 | 1.09E+00 | 1.57E-05 | 5.77E-05 | -1.80E+00 | 4.55E-02 | 1.25E-01 | 1.01E+00 |
| PINX1 | 5.18E-04 | 2.70E-03 | -1.41E+00 | 1.42E-02 | 6.15E-02 | 1.03E+00 | 4.62E-10 | 1.96E-09 | -1.78E+00 | 6.06E-09 | 2.31E-08 | -1.42E+00 | 1.90E-03 | 6.78E-03 | -1.45E+00 | 5.77E-02 | 1.63E-01 | 1.10E+00 | 2.31E-08 | 8.46E-08 | -1.54E+00 | 3.05E-02 | 8.38E-02 | 1.09E+00 |
| PLCB4 | 5.19E-04 | 2.71E-03 | -1.07E+00 | 8.95E-02 | 3.87E-01 | 1.34E+00 | 8.49E-02 | 3.61E-01 | 1.11E+00 | 9.31E-02 | 3.56E-01 | 1.25E+00 | 2.02E-01 | 7.20E-01 | 1.31E+00 | 2.27E-01 | 6.39E-01 | 1.20E+00 | 2.92E-03 | 1.07E-02 | -1.05E+00 | 3.91E-03 | 1.08E-02 | -1.14E+00 |
| SLC25A15 | 5.19E-04 | 2.71E-03 | -1.52E+00 | 3.66E-03 | 1.58E-02 | -1.16E+00 | 1.15E-01 | 4.87E-01 | 1.08E+00 | 6.23E-02 | 2.38E-01 | 1.07E+00 | 2.64E-01 | 9.43E-01 | 1.05E+00 | 2.90E-01 | 8.17E-01 | 1.19E+00 | 3.41E-02 | 1.25E-01 | -1.02E+00 | 8.46E-02 | 2.32E-01 | 1.06E+00 |
| ZNF197 | 5.21E-04 | 2.72E-03 | -1.28E+00 | 1.03E-03 | 4.46E-03 | -1.12E+00 | 1.74E-08 | 7.39E-08 | -1.72E+00 | 8.34E-07 | 3.19E-06 | -1.38E+00 | 1.34E-02 | 4.77E-02 | 1.04E+00 | 5.68E-02 | 1.60E-01 | 1.11E+00 | 2.70E-05 | 9.91E-05 | -1.25E+00 | 3.56E-06 | 9.78E-06 | -1.37E+00 |
| HNRNPCL2 | 5.21E-04 | 2.72E-03 | -1.56E+00 | 6.25E-03 | 2.70E-02 | -1.17E+00 | 2.68E-09 | 1.14E-08 | -2.05E+00 | 1.62E-09 | 6.20E-09 | -1.94E+00 | 1.94E-06 | 6.92E-06 | -2.22E+00 | 3.33E-06 | 9.39E-06 | -1.94E+00 | 1.21E-05 | 4.42E-05 | -1.61E+00 | 9.55E-04 | 2.62E-03 | -1.25E+00 |
| TACO1 | 5.22E-04 | 2.72E-03 | -1.24E+00 | 2.18E-02 | 9.44E-02 | 1.12E+00 | 3.88E-04 | 1.65E-03 | -1.14E+00 | 2.67E-02 | 1.02E-01 | 1.18E+00 | 7.93E-05 | 2.83E-04 | -1.38E+00 | 3.36E-02 | 9.47E-02 | 1.08E+00 | 6.27E-08 | 2.30E-07 | -1.20E+00 | 8.10E-06 | 2.23E-05 | -1.05E+00 |
| TP53I13 | 5.22E-04 | 2.72E-03 | -1.01E+00 | 5.36E-06 | 2.31E-05 | -1.02E+00 | 9.54E-02 | 4.05E-01 | 1.10E+00 | 3.70E-03 | 1.41E-02 | 1.04E+00 | 9.17E-03 | 3.27E-02 | 1.11E+00 | 5.22E-03 | 1.47E-02 | 1.08E+00 | 1.01E-02 | 3.72E-02 | 1.64E+00 | 1.08E-01 | 2.96E-01 | 1.48E+00 |

| | | | | | | | | | | | | | | | | | | | | | | | | |
|---|---|---|---|---|---|---|---|---|---|---|---|---|---|---|---|---|---|---|---|---|---|---|---|---|
| CLASP1 | 5.22E-04 | 2.72E-03 | -1.04E+00 | 9.33E-04 | 4.03E-03 | 1.05E+00 | 9.75E-02 | 4.14E-01 | 1.14E+00 | 7.61E-02 | 2.91E-01 | 1.22E+00 | 1.68E-03 | 5.99E-03 | 1.02E+00 | 3.19E-04 | 9.00E-04 | -1.08E+00 | 2.20E-04 | 8.08E-04 | -1.15E+00 | 3.11E-06 | 8.56E-06 | -1.35E+00 |
| AAAS | 5.22E-04 | 2.72E-03 | -1.16E+00 | 2.03E-02 | 8.77E-02 | 1.15E+00 | 1.15E-02 | 4.89E-02 | -1.06E+00 | 5.28E-02 | 2.02E-01 | 1.25E+00 | 1.08E-03 | 3.84E-03 | -1.36E+00 | 2.36E-01 | 6.66E-01 | 1.28E+00 | 2.64E-05 | 9.68E-05 | -1.17E+00 | 5.09E-02 | 1.40E-01 | 1.11E+00 |
| LMF2 | 5.23E-04 | 2.73E-03 | -1.11E+00 | 1.94E-03 | 8.39E-03 | 1.08E+00 | 1.03E-02 | 4.40E-02 | -1.06E+00 | 3.48E-02 | 1.33E-01 | 1.18E+00 | 6.47E-06 | 2.31E-05 | -1.41E+00 | 8.42E-05 | 2.37E-04 | -1.26E+00 | 1.04E-01 | 3.83E-01 | 1.03E+00 | 3.24E-01 | 8.89E-01 | 1.31E+00 |
| AC009403.1 | 5.26E-04 | 2.74E-03 | -1.26E+00 | 4.50E-02 | 1.94E-01 | 1.21E+00 | 6.78E-02 | 2.88E-01 | -1.06E+00 | 2.04E-01 | 7.79E-01 | 1.27E+00 | 1.53E-01 | 5.44E-01 | 1.03E+00 | 2.67E-01 | 7.52E-01 | 1.42E+00 | 2.10E-01 | 7.70E-01 | 1.44E+00 | 1.34E-01 | 3.69E-01 | 1.16E+00 |
| DRAM2 | 5.27E-04 | 2.75E-03 | 1.06E+00 | 2.06E-03 | 8.90E-03 | 1.18E+00 | 1.15E-03 | 4.87E-03 | -1.08E+00 | 5.78E-03 | 2.21E-02 | 1.06E+00 | 3.23E-05 | 1.15E-04 | -1.09E+00 | 6.01E-04 | 1.69E-03 | 1.01E+00 | 5.81E-04 | 2.13E-03 | -1.06E+00 | 2.42E-02 | 6.64E-02 | 1.08E+00 |
| ACD | 5.28E-04 | 2.75E-03 | -1.28E+00 | 6.67E-03 | 2.88E-02 | 1.05E+00 | 1.77E-03 | 7.51E-03 | -1.14E+00 | 5.29E-04 | 2.02E-03 | -1.03E+00 | 3.14E-04 | 1.12E-03 | -1.19E+00 | 1.37E-02 | 3.87E-02 | 1.09E+00 | 1.61E-03 | 5.92E-03 | -1.01E+00 | 1.85E-01 | 5.09E-01 | 1.20E+00 |
| LHPP | 5.28E-04 | 2.75E-03 | -1.49E+00 | 4.81E-04 | 2.08E-03 | -1.45E+00 | 9.96E-04 | 4.23E-03 | -1.35E+00 | 2.26E-04 | 8.62E-04 | -1.37E+00 | 1.13E-01 | 4.04E-01 | 1.36E+00 | 3.49E-01 | 9.84E-01 | 1.56E+00 | 1.00E-02 | 3.67E-02 | -1.01E+00 | 2.16E-01 | 5.94E-01 | 1.30E+00 |
| MSH3 | 5.28E-04 | 2.75E-03 | -1.15E+00 | 1.24E-02 | 5.34E-02 | 1.12E+00 | 3.88E-04 | 1.65E-03 | -1.31E+00 | 3.06E-04 | 1.17E-03 | -1.12E+00 | 4.10E-04 | 1.46E-03 | -1.33E+00 | 1.17E-03 | 3.31E-03 | -1.19E+00 | 2.91E-03 | 1.07E-02 | -1.10E+00 | 2.31E-03 | 6.33E-03 | -1.10E+00 |
| PIEZO1 | 5.28E-04 | 2.75E-03 | -1.07E+00 | 2.68E-03 | 1.16E-02 | 1.12E+00 | 1.58E-02 | 6.72E-02 | -1.04E+00 | 1.69E-02 | 6.47E-02 | 1.10E+00 | 2.90E-04 | 1.04E-03 | -1.34E+00 | 6.20E-02 | 1.75E-01 | 1.08E+00 | 7.39E-05 | 2.71E-04 | -1.14E+00 | 8.54E-02 | 2.35E-01 | 1.14E+00 |
| HIST1H4E | 5.29E-04 | 2.76E-03 | 2.06E+00 | 7.74E-03 | 3.35E-02 | -1.01E+00 | 3.24E-03 | 1.38E-02 | 2.06E+00 | 2.27E-01 | 8.66E-01 | 1.40E+00 | 2.28E-01 | 8.12E-01 | 1.58E+00 | 5.22E-02 | 1.47E-01 | -1.02E+00 | 7.77E-02 | 2.85E-01 | -1.08E+00 | 6.58E-03 | 1.81E-02 | -1.44E+00 |
| NOL7 | 5.30E-04 | 2.76E-03 | 1.06E+00 | 2.38E-10 | 1.03E-09 | 1.00E+00 | 9.75E-02 | 4.14E-01 | 1.08E+00 | 3.39E-03 | 1.29E-02 | 1.00E+00 | 7.31E-06 | 2.60E-05 | -1.08E+00 | 2.97E-05 | 8.37E-05 | -1.05E+00 | 1.12E-01 | 4.12E-01 | 1.11E+00 | 2.23E-01 | 6.13E-01 | 1.06E+00 |
| DCAF17 | 5.31E-04 | 2.76E-03 | -1.33E+00 | 3.66E-02 | 1.58E-01 | 1.13E+00 | 1.02E-04 | 4.32E-04 | -1.42E+00 | 7.80E-03 | 2.98E-02 | 1.01E+00 | 1.10E-02 | 3.94E-02 | -1.11E+00 | 9.83E-02 | 2.77E-01 | 1.21E+00 | 1.10E-05 | 4.03E-05 | -1.38E+00 | 1.92E-02 | 5.28E-02 | 1.01E+00 |
| ECD | 5.31E-04 | 2.77E-03 | -1.05E+00 | 6.38E-05 | 2.76E-04 | -1.05E+00 | 5.79E-03 | 2.46E-02 | -1.08E+00 | 7.00E-05 | 2.67E-04 | -1.11E+00 | 1.32E-03 | 4.69E-03 | -1.06E+00 | 2.01E-05 | 5.66E-05 | -1.27E+00 | 2.69E-05 | 9.87E-05 | -1.25E+00 | 6.59E-05 | 1.81E-04 | -1.17E+00 |
| CLCC1 | 5.31E-04 | 2.77E-03 | -1.41E+00 | 1.44E-02 | 6.22E-02 | 1.01E+00 | 2.10E-02 | 8.94E-02 | -1.09E+00 | 6.25E-03 | 2.39E-02 | -1.01E+00 | 1.88E-05 | 6.69E-05 | -1.86E+00 | 2.70E-04 | 7.61E-04 | -1.46E+00 | 4.88E-04 | 1.79E-03 | -1.19E+00 | 4.09E-05 | 1.12E-04 | -1.30E+00 |
| IFT57 | 5.32E-04 | 2.77E-03 | -1.01E+00 | 3.76E-06 | 1.63E-05 | -1.01E+00 | 3.01E-03 | 1.28E-02 | -1.08E+00 | 1.34E-05 | 5.11E-05 | -1.17E+00 | 1.80E-03 | 6.40E-03 | -1.07E+00 | 1.20E-02 | 3.38E-02 | 1.07E+00 | 2.52E-01 | 9.27E-01 | 1.46E+00 | 2.34E-01 | 6.41E-01 | 1.29E+00 |
| DARS2 | 5.33E-04 | 2.78E-03 | -1.45E+00 | 1.37E-01 | 5.94E-01 | 1.38E+00 | 2.09E-02 | 8.89E-02 | -1.06E+00 | 1.70E-01 | 6.49E-01 | 1.25E+00 | 2.22E-03 | 7.92E-03 | -1.60E+00 | 8.65E-02 | 2.44E-01 | 1.07E+00 | 1.01E-03 | 3.71E-03 | -1.06E+00 | 2.48E-01 | 6.80E-01 | 1.41E+00 |
| TNRC6C-AS1 | 5.33E-04 | 2.78E-03 | -1.27E+00 | 5.49E-03 | 2.37E-02 | 1.04E+00 | 5.19E-02 | 2.21E-01 | -1.09E+00 | 4.84E-02 | 1.85E-01 | 1.12E+00 | 2.13E-01 | 7.60E-01 | 1.26E+00 | 9.96E-02 | 2.81E-01 | -1.00E+00 | 3.68E-05 | 1.35E-04 | -1.78E+00 | 3.08E-04 | 8.46E-04 | -1.59E+00 |
| MLX | 5.33E-04 | 2.78E-03 | -1.12E+00 | 5.47E-03 | 2.36E-02 | 1.11E+00 | 1.46E-03 | 6.19E-03 | -1.16E+00 | 2.40E-03 | 9.16E-03 | 1.04E+00 | 2.00E-03 | 7.12E-03 | -1.02E+00 | 7.05E-03 | 1.99E-02 | 1.03E+00 | 1.42E-02 | 5.20E-02 | -1.06E+00 | 3.88E-02 | 1.07E-01 | 1.07E+00 |
| OXSM | 5.33E-04 | 2.78E-03 | -1.31E+00 | 6.72E-02 | 2.91E-01 | 1.25E+00 | 5.01E-03 | 2.13E-02 | -1.11E+00 | 4.68E-02 | 1.79E-01 | 1.16E+00 | 3.06E-04 | 1.09E-03 | -1.61E+00 | 9.74E-02 | 2.75E-01 | 1.20E+00 | 3.44E-06 | 1.26E-05 | -1.44E+00 | 1.69E-01 | 4.63E-01 | 1.10E+00 |
| PPP1R3F | 5.34E-04 | 2.78E-03 | -1.37E+00 | 1.24E-01 | 5.36E-01 | 1.37E+00 | 9.70E-09 | 4.12E-08 | -2.05E+00 | 4.03E-05 | 1.54E-04 | -1.30E+00 | 5.95E-02 | 2.12E-01 | -1.05E+00 | 2.96E-01 | 8.34E-01 | 1.33E+00 | 4.56E-02 | 1.68E-01 | -1.14E+00 | 1.80E-01 | 4.93E-01 | -1.02E+00 |
| INO80 | 5.34E-04 | 2.78E-03 | -1.13E+00 | 7.76E-03 | 3.35E-02 | 1.10E+00 | 5.87E-06 | 2.50E-05 | -1.18E+00 | 1.47E-04 | 5.63E-04 | 1.07E+00 | 1.08E-03 | 3.85E-03 | -1.03E+00 | 3.05E-03 | 8.61E-03 | 1.07E+00 | 4.24E-06 | 1.56E-05 | -1.22E+00 | 2.23E-05 | 6.13E-05 | -1.18E+00 |
| ZNF473 | 5.34E-04 | 2.78E-03 | -1.46E+00 | 2.70E-04 | 1.17E-03 | -1.50E+00 | 2.83E-03 | 1.20E-02 | -1.23E+00 | 6.36E-03 | 2.43E-02 | -1.04E+00 | 8.27E-02 | 2.95E-01 | -1.43E+00 | 1.51E-01 | 4.26E-01 | -1.17E+00 | 1.74E-05 | 6.37E-05 | -1.55E+00 | 9.07E-05 | 2.49E-04 | -1.49E+00 |
| AIF1 | 5.38E-04 | 2.80E-03 | 1.24E+00 | 1.73E-01 | 7.46E-01 | -1.01E+00 | 2.32E-03 | 9.85E-03 | 1.28E+00 | 1.64E-01 | 6.27E-01 | 1.06E+00 | 1.72E-01 | 6.14E-01 | 1.15E+00 | 2.65E-05 | 7.48E-05 | -1.18E+00 | 3.53E-06 | 1.29E-05 | 1.55E+00 | 1.19E-01 | 3.27E-01 | 1.21E+00 |
| EEF1A1P19 | 5.38E-04 | 2.80E-03 | -1.05E+00 | 5.64E-04 | 2.44E-03 | 1.10E+00 | 3.12E-02 | 1.32E-01 | 1.01E+00 | 3.86E-02 | 1.47E-01 | 1.14E+00 | 1.36E-01 | 4.86E-01 | 1.29E+00 | 1.27E-01 | 3.57E-01 | 1.27E+00 | 1.02E-01 | 3.75E-01 | 1.11E+00 | 1.14E-01 | 3.13E-01 | 1.11E+00 |
| MTND6P4 | 5.38E-04 | 2.80E-03 | -1.72E+00 | 1.19E-03 | 5.13E-03 | -1.48E+00 | 6.81E-03 | 2.89E-02 | -1.22E+00 | 7.88E-03 | 3.01E-02 | -1.12E+00 | 2.73E-01 | 9.75E-01 | 1.40E+00 | 3.13E-01 | 8.81E-01 | 1.50E+00 | 1.21E-05 | 4.42E-05 | -1.49E+00 | 5.01E-04 | 1.38E-03 | -1.28E+00 |
| NELFA | 5.38E-04 | 2.80E-03 | -1.05E+00 | 6.30E-02 | 2.72E-01 | 1.35E+00 | 1.82E-06 | 7.75E-06 | -1.32E+00 | 5.31E-06 | 2.03E-05 | -1.10E+00 | 2.68E-03 | 9.55E-03 | -1.25E+00 | 3.01E-01 | 8.48E-01 | 1.50E+00 | 2.26E-07 | 8.30E-07 | -1.31E+00 | 5.25E-04 | 1.44E-03 | -1.08E+00 |
| ATE1 | 5.41E-04 | 2.82E-03 | -1.06E+00 | 2.26E-03 | 9.77E-03 | 1.08E+00 | 1.34E-03 | 5.68E-03 | -1.12E+00 | 1.41E-03 | 5.40E-03 | -1.03E+00 | 1.23E-01 | 4.37E-01 | 1.19E+00 | 2.32E-01 | 6.55E-01 | 1.43E+00 | 2.91E-05 | 1.07E-04 | -1.10E+00 | 3.05E-03 | 8.37E-03 | -1.09E+00 |
| ZC3HAV1L | 5.42E-04 | 2.82E-03 | -1.56E+00 | 6.31E-02 | 2.73E-01 | 1.08E+00 | 5.60E-07 | 2.38E-06 | -2.12E+00 | 1.05E-02 | 3.99E-02 | 1.03E+00 | 1.36E-01 | 4.86E-01 | 1.02E+00 | 1.56E-01 | 4.41E-01 | 1.69E+00 | 7.28E-04 | 2.67E-03 | -1.25E+00 | 4.22E-02 | 1.16E-01 | 1.02E+00 |
| SNRPEP4 | 5.43E-04 | 2.83E-03 | 1.05E+00 | 1.11E-04 | 4.80E-04 | 1.12E+00 | 1.45E-01 | 6.15E-01 | 1.23E+00 | 8.61E-02 | 3.29E-01 | 1.20E+00 | 4.39E-03 | 1.56E-02 | 1.05E+00 | 1.03E-01 | 2.89E-01 | 1.21E+00 | 1.28E-02 | 4.69E-02 | 1.06E+00 | 3.38E-01 | 9.29E-01 | 1.22E+00 |
| SPRED1 | 5.44E-04 | 2.83E-03 | -1.43E+00 | 4.26E-03 | 1.84E-02 | -1.05E+00 | 4.24E-02 | 1.80E-01 | -1.13E+00 | 8.03E-02 | 3.07E-01 | 1.04E+00 | 1.76E-08 | 6.29E-08 | -2.12E+00 | 8.71E-07 | 2.45E-06 | -1.63E+00 | 9.79E-07 | 3.59E-06 | -1.59E+00 | 4.70E-07 | 1.29E-06 | -1.64E+00 |
| AC005332.6 | 5.44E-04 | 2.83E-03 | -1.32E+00 | 1.48E-02 | 6.38E-02 | 1.07E+00 | 2.83E-02 | 1.20E-01 | -1.43E+00 | 1.76E-01 | 6.74E-01 | -1.02E+00 | 1.02E-01 | 3.64E-01 | -1.16E+00 | 1.08E-01 | 3.04E-01 | -1.14E+00 | 1.90E-02 | 6.96E-02 | -1.32E+00 | 2.16E-01 | 5.93E-01 | 1.00E+00 |
| RPL21P11 | 5.44E-04 | 2.83E-03 | -1.45E+00 | 6.13E-05 | 2.65E-04 | -1.73E+00 | 5.32E-03 | 2.26E-02 | -1.31E+00 | 3.26E-05 | 1.25E-04 | -1.88E+00 | 2.31E-04 | 8.24E-04 | -1.54E+00 | 1.12E-03 | 3.15E-03 | -1.28E+00 | 1.80E-02 | 6.59E-02 | -1.15E+00 | 2.01E-03 | 5.52E-03 | -1.54E+00 |
| PLEKHF2 | 5.45E-04 | 2.84E-03 | 1.07E+00 | 2.57E-05 | 1.11E-04 | 1.03E+00 | 6.05E-06 | 2.57E-05 | -1.19E+00 | 8.15E-07 | 3.11E-06 | -1.18E+00 | 7.36E-05 | 2.62E-04 | -1.13E+00 | 1.39E-06 | 3.92E-06 | -1.30E+00 | 3.35E-08 | 1.23E-07 | 1.07E+00 | 1.07E-08 | 2.93E-08 | -1.21E+00 |
| AC008555.8 | 5.47E-04 | 2.85E-03 | -1.37E+00 | 6.87E-02 | 2.97E-01 | 1.20E+00 | 1.44E-04 | 6.14E-04 | -1.54E+00 | 1.01E-03 | 3.86E-03 | -1.26E+00 | 2.05E-05 | 7.30E-05 | -2.09E+00 | 1.67E-03 | 4.70E-03 | -1.36E+00 | 4.42E-02 | 1.62E-01 | -1.10E+00 | 2.30E-01 | 6.32E-01 | 1.17E+00 |
| MCM6 | 5.49E-04 | 2.86E-03 | -1.34E+00 | 3.11E-02 | 1.35E-01 | 1.17E+00 | 1.96E-01 | 8.33E-01 | 1.06E+00 | 8.29E-02 | 3.17E-01 | 1.38E+00 | 3.30E-02 | 1.18E-01 | -1.33E+00 | 1.80E-01 | 5.08E-01 | 1.02E+00 | 1.98E-09 | 7.26E-09 | -1.34E+00 | 1.93E-01 | 5.30E-01 | 1.16E+00 |
| DUSP7 | 5.50E-04 | 2.87E-03 | -1.60E+00 | 1.29E-01 | 5.59E-01 | 1.26E+00 | 2.50E-02 | 1.06E-01 | -1.19E+00 | 1.55E-01 | 5.93E-01 | 1.14E+00 | 2.04E-01 | 7.28E-01 | 1.17E+00 | 1.60E-01 | 4.50E-01 | 1.00E+00 | 1.07E-01 | 3.93E-01 | 1.14E+00 | 1.03E-01 | 2.84E-01 | -1.10E+00 |
| GAK | 5.50E-04 | 2.87E-03 | -1.08E+00 | 1.80E-04 | 7.80E-04 | 1.01E+00 | 2.78E-04 | 1.18E-03 | -1.22E+00 | 2.59E-04 | 9.91E-04 | -1.00E+00 | 7.07E-04 | 2.52E-03 | -1.19E+00 | 2.08E-04 | 5.86E-04 | -1.22E+00 | 3.04E-03 | 1.12E-02 | -1.05E+00 | 5.88E-02 | 1.62E-01 | 1.10E+00 |
| BLOC1S3 | 5.50E-04 | 2.87E-03 | -1.33E+00 | 2.43E-02 | 1.05E-01 | 1.12E+00 | 5.48E-03 | 2.33E-02 | -1.36E+00 | 3.55E-03 | 1.36E-02 | -1.22E+00 | 8.74E-03 | 3.12E-02 | -1.52E+00 | 9.50E-02 | 2.68E-01 | -1.04E+00 | 1.64E-04 | 6.03E-04 | -1.19E+00 | 2.54E-03 | 6.97E-03 | -1.17E+00 |
| QTRT2 | 5.50E-04 | 2.87E-03 | -1.24E+00 | 1.22E-02 | 5.29E-02 | 1.21E+00 | 5.76E-02 | 2.45E-01 | 1.08E+00 | 2.07E-01 | 7.90E-01 | 1.39E+00 | 4.23E-04 | 1.51E-03 | -1.36E+00 | 3.61E-02 | 1.02E-01 | 1.03E+00 | 5.87E-08 | 2.15E-07 | -1.50E+00 | 3.97E-03 | 1.09E-02 | -1.08E+00 |
| MDC1 | 5.51E-04 | 2.87E-03 | -1.35E+00 | 6.62E-02 | 2.86E-01 | 1.19E+00 | 2.72E-02 | 1.15E-01 | -1.07E+00 | 1.89E-01 | 7.24E-01 | 1.28E+00 | 1.01E-02 | 3.59E-02 | -1.14E+00 | 3.04E-01 | 8.57E-01 | 1.27E+00 | 2.81E-04 | 1.03E-03 | -1.14E+00 | 3.63E-02 | 9.98E-02 | 1.07E+00 |
| CCDC58 | 5.52E-04 | 2.88E-03 | -1.10E+00 | 4.02E-03 | 1.74E-02 | 1.07E+00 | 9.39E-02 | 3.99E-01 | 1.16E+00 | 1.71E-02 | 6.55E-02 | 1.16E+00 | 2.23E-05 | 7.95E-05 | -1.48E+00 | 6.74E-03 | 1.90E-02 | -1.00E+00 | 9.77E-04 | 3.58E-03 | -1.09E+00 | 8.15E-02 | 2.24E-01 | 1.09E+00 |
| MEA1 | 5.52E-04 | 2.88E-03 | -1.00E+00 | 1.54E-03 | 6.65E-03 | 1.12E+00 | 6.52E-02 | 2.77E-01 | 1.12E+00 | 9.06E-03 | 3.46E-02 | 1.17E+00 | 3.62E-08 | 1.29E-07 | -1.31E+00 | 6.70E-06 | 1.89E-05 | -1.12E+00 | 1.32E-01 | 4.86E-01 | 1.13E+00 | 1.92E-01 | 5.28E-01 | 1.12E+00 |
| DCAF13P3 | 5.53E-04 | 2.88E-03 | -1.32E+00 | 3.60E-04 | 1.56E-03 | -1.28E+00 | 8.30E-05 | 3.53E-04 | -1.45E+00 | 3.05E-06 | 1.16E-05 | -1.43E+00 | 2.13E-03 | 7.59E-03 | -1.43E+00 | 4.67E-03 | 1.32E-02 | -1.31E+00 | 1.28E-05 | 4.71E-05 | -1.24E+00 | 4.35E-07 | 1.20E-06 | -1.40E+00 |
| DOCK10 | 5.53E-04 | 2.88E-03 | -1.21E+00 | 1.63E-03 | 7.05E-03 | -1.01E+00 | 1.16E-03 | 4.95E-03 | -1.25E+00 | 4.73E-04 | 1.81E-03 | -1.13E+00 | 8.19E-03 | 2.92E-02 | -1.46E+00 | 2.94E-02 | 8.29E-02 | -1.19E+00 | 2.08E-02 | 7.64E-02 | -1.07E+00 | 2.71E-01 | 7.45E-01 | 1.35E+00 |
| TOB2P1 | 5.53E-04 | 2.88E-03 | -1.69E+00 | 5.80E-03 | 2.51E-02 | -1.22E+00 | 2.03E-01 | 8.64E-01 | 1.03E+00 | 7.93E-02 | 3.03E-01 | 1.76E+00 | 7.40E-02 | 2.64E-01 | 2.42E+00 | 3.23E-02 | 9.11E-02 | 2.69E+00 | 2.20E-01 | 8.09E-01 | 1.29E+00 | 2.09E-01 | 5.74E-01 | -1.05E+00 |
| DSE | 5.54E-04 | 2.89E-03 | -1.36E+00 | 3.03E-02 | 1.31E-01 | 1.06E+00 | 6.33E-05 | 2.69E-04 | -1.53E+00 | 1.45E-05 | 5.55E-05 | -1.46E+00 | 7.00E-04 | 2.50E-03 | -2.24E+00 | 1.19E-02 | 3.35E-02 | -1.32E+00 | 1.65E-02 | 6.07E-02 | -1.44E+00 | 1.51E-01 | 4.13E-01 | 1.04E+00 |
| DBR1 | 5.55E-04 | 2.89E-03 | -1.42E+00 | 1.89E-03 | 8.16E-03 | -1.14E+00 | 6.42E-04 | 2.73E-03 | -1.28E+00 | 1.06E-02 | 4.05E-02 | 1.02E+00 | 3.88E-02 | 1.38E-01 | -1.11E+00 | 7.80E-02 | 2.20E-01 | -1.02E+00 | 1.35E-04 | 4.96E-04 | -1.15E+00 | 6.18E-02 | 1.70E-01 | 1.21E+00 |
| PATZ1 | 5.56E-04 | 2.90E-03 | -1.07E+00 | 5.35E-02 | 2.31E-01 | 1.35E+00 | 7.22E-05 | 3.07E-04 | -1.20E+00 | 6.74E-03 | 2.58E-02 | 1.15E+00 | 7.67E-04 | 2.74E-03 | 1.00E+00 | 1.24E-03 | 3.51E-03 | 1.01E+00 | 6.71E-07 | 2.46E-06 | -1.49E+00 | 1.34E-02 | 3.68E-02 | -1.04E+00 |
| GABPA | 5.57E-04 | 2.90E-03 | -1.13E+00 | 6.57E-03 | 2.84E-02 | 1.11E+00 | 2.93E-05 | 1.24E-04 | -1.31E+00 | 2.11E-04 | 8.05E-04 | -1.15E+00 | 2.98E-05 | 1.06E-04 | -1.59E+00 | 1.46E-03 | 4.11E-03 | -1.20E+00 | 8.75E-05 | 3.21E-04 | -1.25E+00 | 3.43E-03 | 9.43E-03 | -1.12E+00 |
| RPL31P2 | 5.59E-04 | 2.91E-03 | -1.13E+00 | 2.34E-08 | 1.01E-07 | -1.86E+00 | 5.00E-02 | 2.12E-01 | 1.33E+00 | 6.88E-02 | 2.63E-01 | -1.09E+00 | 7.47E-02 | 2.66E-01 | 1.79E+00 | 1.04E-01 | 2.93E-01 | -1.00E+00 | 1.74E-01 | 6.39E-01 | 1.16E+00 | 8.27E-03 | 2.27E-02 | -1.24E+00 |
| ZFHX3 | 5.60E-04 | 2.92E-03 | -1.15E+00 | 1.34E-03 | 5.77E-03 | 1.02E+00 | 9.65E-03 | 4.10E-02 | -1.11E+00 | 5.05E-02 | 1.93E-01 | 1.11E+00 | 4.56E-03 | 1.63E-02 | -1.50E+00 | 1.05E-01 | 2.96E-01 | 1.06E+00 | 5.44E-03 | 1.99E-02 | -1.09E+00 | 2.72E-01 | 7.48E-01 | 1.18E+00 |
| MAIP1 | 5.60E-04 | 2.92E-03 | -1.05E+00 | 2.97E-04 | 1.28E-03 | 1.01E+00 | 1.79E-04 | 7.60E-04 | -1.14E+00 | 6.69E-06 | 2.56E-05 | -1.09E+00 | 7.22E-03 | 2.57E-02 | -1.06E+00 | 7.80E-02 | 2.20E-01 | 1.09E+00 | 5.27E-05 | 1.93E-04 | -1.11E+00 | 1.09E-03 | 2.99E-03 | -1.01E+00 |
| MLKL | 5.60E-04 | 2.92E-03 | -1.28E+00 | 2.00E-02 | 8.66E-02 | 1.16E+00 | 6.98E-08 | 2.96E-07 | -1.56E+00 | 3.77E-05 | 1.44E-04 | -1.09E+00 | 1.21E-04 | 4.30E-04 | -1.66E+00 | 2.22E-04 | 6.26E-04 | -1.58E+00 | 9.04E-07 | 3.32E-06 | -1.80E+00 | 3.94E-02 | 1.08E-01 | 1.03E+00 |
| CASP6 | 5.61E-04 | 2.93E-03 | -1.09E+00 | 5.07E-03 | 2.19E-02 | 1.09E+00 | 2.44E-06 | 1.03E-05 | -1.38E+00 | 3.43E-04 | 1.31E-03 | -1.09E+00 | 1.60E-04 | 5.72E-04 | -1.04E+00 | 2.00E-02 | 5.63E-02 | 1.11E+00 | 2.17E-04 | 7.95E-04 | -1.13E+00 | 1.06E-01 | 2.91E-01 | 1.05E+00 |
| TRAM2 | 5.62E-04 | 2.93E-03 | -1.36E+00 | 3.29E-02 | 1.42E-01 | 1.15E+00 | 5.53E-03 | 2.35E-02 | -1.35E+00 | 2.51E-02 | 9.58E-02 | 1.01E+00 | 1.25E-02 | 4.46E-02 | -1.22E+00 | 2.06E-01 | 5.80E-01 | 1.20E+00 | 9.96E-06 | 3.65E-05 | -1.46E+00 | 3.86E-04 | 1.06E-03 | -1.26E+00 |
| CYP1B1 | 5.64E-04 | 2.94E-03 | 3.34E+00 | 2.15E-01 | 9.31E-01 | 1.08E+00 | 2.85E-02 | 1.21E-01 | 2.24E+00 | 1.96E-01 | 7.49E-01 | -1.20E+00 | 8.68E-04 | 3.09E-03 | 4.31E+00 | 3.31E-01 | 9.34E-01 | 1.12E+00 | 4.60E-08 | 1.69E-07 | 4.97E+00 | 3.29E-01 | 9.04E-01 | 1.13E+00 |
| AGFG1 | 5.64E-04 | 2.94E-03 | -1.18E+00 | 9.65E-04 | 4.17E-03 | -1.01E+00 | 6.31E-03 | 2.68E-02 | -1.08E+00 | 6.96E-02 | 2.66E-01 | 1.20E+00 | 1.10E-01 | 3.90E-01 | 1.16E+00 | 5.69E-02 | 1.60E-01 | 1.13E+00 | 3.15E-02 | 1.15E-01 | 1.10E+00 | 2.63E-01 | 7.22E-01 | 1.28E+00 |
| SLFN13 | 5.65E-04 | 2.94E-03 | -1.41E+00 | 1.80E-03 | 7.78E-03 | -1.15E+00 | 7.91E-02 | 3.36E-01 | 1.05E+00 | 3.96E-03 | 1.51E-02 | -1.05E+00 | 1.21E-02 | 4.32E-02 | -1.47E+00 | 1.37E-02 | 3.86E-02 | -1.43E+00 | 1.63E-03 | 5.99E-03 | -1.10E+00 | 6.71E-04 | 1.84E-03 | -1.07E+00 |
| RRAGC | 5.65E-04 | 2.94E-03 | -1.29E+00 | 7.46E-02 | 3.22E-01 | 1.31E+00 | 2.17E-06 | 9.21E-06 | -1.61E+00 | 4.19E-05 | 1.60E-04 | -1.22E+00 | 8.12E-04 | 2.89E-03 | -1.49E+00 | 4.87E-02 | 1.37E-01 | 1.13E+00 | 4.26E-05 | 1.56E-04 | -1.58E+00 | 3.37E-03 | 9.27E-03 | -1.20E+00 |
| MAN2B2 | 5.65E-04 | 2.94E-03 | -1.06E+00 | 2.29E-02 | 9.88E-02 | 1.26E+00 | 8.79E-04 | 3.73E-03 | -1.30E+00 | 2.04E-02 | 7.80E-02 | 1.08E+00 | 1.56E-03 | 5.57E-03 | -1.37E+00 | 1.36E-02 | 3.84E-02 | -1.15E+00 | 2.32E-03 | 8.53E-03 | -1.17E+00 | 4.66E-04 | 1.28E-03 | -1.31E+00 |
| CENPW | 5.65E-04 | 2.94E-03 | -1.03E+00 | 2.48E-06 | 1.07E-05 | -1.11E+00 | 1.20E-03 | 5.08E-03 | 1.40E+00 | 1.54E-02 | 5.90E-02 | 1.01E+00 | 1.26E-03 | 4.49E-03 | 1.10E+00 | 2.43E-03 | 6.85E-03 | -1.00E+00 | 2.60E-01 | 9.53E-01 | 1.22E+00 | 2.66E-01 | 7.31E-01 | 1.24E+00 |
| NUDT9 | 5.66E-04 | 2.95E-03 | -1.25E+00 | 6.54E-03 | 2.83E-02 | 1.05E+00 | 5.12E-02 | 2.18E-01 | 1.03E+00 | 2.22E-01 | 8.48E-01 | 1.36E+00 | 2.33E-02 | 8.32E-02 | -1.03E+00 | 2.85E-01 | 8.04E-01 | 1.27E+00 | 1.35E-03 | 4.96E-03 | -1.04E+00 | 7.97E-02 | 2.19E-01 | 1.12E+00 |
| MRFAP1 | 5.66E-04 | 2.95E-03 | 1.05E+00 | 7.15E-08 | 3.09E-07 | 1.04E+00 | 1.31E-02 | 5.58E-02 | -1.12E+00 | 1.05E-02 | 4.00E-02 | -1.01E+00 | 1.01E-05 | 3.61E-05 | -1.16E+00 | 7.88E-03 | 2.22E-02 | -1.03E+00 | 8.28E-04 | 3.04E-03 | -1.07E+00 | 9.52E-02 | 2.61E-01 | 1.06E+00 |
| AURKAIP1 | 5.66E-04 | 2.95E-03 | 1.03E+00 | 1.05E-04 | 4.55E-04 | 1.11E+00 | 1.09E-01 | 4.61E-01 | 1.12E+00 | 1.99E-02 | 7.61E-02 | 1.08E+00 | 7.41E-05 | 2.64E-04 | -1.08E+00 | 7.83E-04 | 2.21E-03 | 1.05E+00 | 2.62E-01 | 9.62E-01 | 1.16E+00 | 6.58E-02 | 1.81E-01 | 1.30E+00 |
| AL451165.2 | 5.67E-04 | 2.95E-03 | -1.26E+00 | 2.91E-04 | 1.26E-03 | -1.19E+00 | 2.24E-02 | 9.51E-02 | -1.06E+00 | 1.39E-04 | 5.32E-04 | -1.40E+00 | 3.90E-02 | 1.39E-01 | 1.11E+00 | 5.05E-02 | 1.42E-01 | 1.05E+00 | 1.53E-01 | 5.60E-01 | 1.29E+00 | 3.74E-03 | 1.03E-02 | -1.41E+00 |
| PDCD6IPP2 | 5.67E-04 | 2.96E-03 | -1.06E+00 | 4.92E-03 | 2.13E-02 | 1.22E+00 | 2.86E-03 | 1.22E-02 | -1.11E+00 | 2.48E-04 | 9.46E-04 | -1.00E+00 | 5.47E-06 | 1.95E-05 | -1.38E+00 | 2.46E-05 | 6.94E-05 | -1.26E+00 | 1.66E-05 | 6.09E-05 | -1.25E+00 | 1.58E-04 | 4.33E-04 | -1.22E+00 |
| AL161457.2 | 5.68E-04 | 2.96E-03 | -1.38E+00 | 2.79E-03 | 1.20E-02 | -1.12E+00 | 1.63E-07 | 6.93E-07 | -2.23E+00 | 7.84E-05 | 2.99E-04 | -1.43E+00 | 1.99E-01 | 7.09E-01 | 1.26E+00 | 3.20E-01 | 9.02E-01 | 1.47E+00 | 2.53E-01 | 9.29E-01 | 1.09E+00 | 3.04E-01 | 8.35E-01 | 1.18E+00 |
| SLC25A13 | 5.68E-04 | 2.96E-03 | -1.13E+00 | 3.07E-03 | 1.33E-02 | 1.01E+00 | 3.72E-06 | 1.58E-05 | -1.43E+00 | 3.67E-06 | 1.40E-05 | -1.18E+00 | 2.57E-03 | 9.15E-03 | -1.07E+00 | 2.67E-02 | 7.53E-02 | 1.13E+00 | 3.92E-04 | 1.44E-03 | -1.07E+00 | 2.79E-03 | 7.67E-03 | -1.09E+00 |
| SIRT2 | 5.69E-04 | 2.97E-03 | -1.02E+00 | 2.03E-02 | 8.76E-02 | 1.28E+00 | 1.29E-02 | 5.48E-02 | -1.05E+00 | 9.08E-03 | 3.47E-02 | 1.06E+00 | 1.23E-03 | 4.37E-03 | -1.25E+00 | 1.58E-02 | 4.47E-02 | -1.04E+00 | 2.53E-02 | 9.29E-02 | 1.02E+00 | 3.06E-01 | 8.40E-01 | 1.24E+00 |
| ZNF354C | 5.70E-04 | 2.97E-03 | -1.43E+00 | 2.64E-03 | 1.14E-02 | -1.17E+00 | 8.03E-02 | 3.41E-01 | -1.02E+00 | 1.63E-01 | 6.21E-01 | 1.26E+00 | 9.66E-03 | 3.44E-02 | -1.20E+00 | 1.15E-02 | 3.24E-02 | -1.19E+00 | 5.39E-02 | 1.98E-01 | 1.04E+00 | 5.99E-03 | 1.65E-02 | -1.25E+00 |
| AC005839.1 | 5.72E-04 | 2.98E-03 | -1.33E+00 | 8.43E-03 | 3.64E-02 | 1.05E+00 | 4.11E-03 | 1.75E-02 | -1.30E+00 | 8.72E-03 | 3.33E-02 | -1.10E+00 | 1.39E-01 | 4.94E-01 | 1.02E+00 | 1.27E-01 | 3.59E-01 | -1.04E+00 | 1.11E-04 | 4.06E-04 | -1.56E+00 | 8.44E-02 | 2.32E-01 | -1.06E+00 |
| AC003102.1 | 5.73E-04 | 2.99E-03 | -1.52E+00 | 3.47E-03 | 1.50E-02 | -1.27E+00 | 3.66E-02 | 1.56E-01 | -1.13E+00 | 5.71E-03 | 2.18E-02 | -1.11E+00 | 4.46E-02 | 1.59E-01 | -1.19E+00 | 7.39E-02 | 2.08E-01 | -1.03E+00 | 9.90E-03 | 3.64E-02 | -1.24E+00 | 4.44E-02 | 1.22E-01 | -1.10E+00 |
| IRF3 | 5.74E-04 | 2.99E-03 | -1.04E+00 | 6.47E-03 | 2.79E-02 | 1.15E+00 | 5.83E-04 | 2.48E-03 | -1.22E+00 | 3.02E-03 | 1.15E-02 | -1.01E+00 | 3.13E-06 | 1.12E-05 | -1.37E+00 | 1.50E-02 | 4.22E-02 | 1.10E+00 | 1.16E-05 | 4.26E-05 | -1.14E+00 | 1.04E-02 | 2.86E-02 | 1.08E+00 |
| HNRNPA3P10 | 5.74E-04 | 2.99E-03 | -1.56E+00 | 2.68E-04 | 1.16E-03 | -1.53E+00 | 2.12E-03 | 8.99E-03 | -1.22E+00 | 1.34E-03 | 5.13E-03 | -1.12E+00 | 9.95E-03 | 3.55E-02 | -1.35E+00 | 6.56E-02 | 1.85E-01 | 1.07E+00 | 9.07E-05 | 3.33E-04 | -1.34E+00 | 6.44E-04 | 1.77E-03 | -1.17E+00 |
| FGD3 | 5.74E-04 | 2.99E-03 | -1.08E+00 | 1.33E-01 | 5.73E-01 | 1.42E+00 | 6.57E-06 | 2.79E-05 | -1.44E+00 | 8.72E-04 | 3.33E-03 | -1.07E+00 | 7.03E-06 | 2.51E-05 | -1.35E+00 | 6.70E-04 | 1.89E-03 | -1.10E+00 | 3.62E-06 | 1.33E-05 | -1.40E+00 | 3.60E-05 | 9.89E-05 | -1.30E+00 |
| AC021016.1 | 5.74E-04 | 2.99E-03 | 2.04E+00 | 8.47E-03 | 3.66E-02 | 2.08E+00 | 1.77E-03 | 7.51E-03 | 1.73E+00 | 9.98E-03 | 3.81E-02 | 1.96E+00 | 7.18E-02 | 2.56E-01 | 1.89E+00 | 5.17E-02 | 1.46E-01 | 2.06E+00 | 2.36E-01 | 8.65E-01 | 1.41E+00 | 5.88E-03 | 1.61E-02 | 2.00E+00 |
| LUZP1 | 5.75E-04 | 2.99E-03 | 1.08E+00 | 2.74E-11 | 1.18E-10 | -1.18E+00 | 2.84E-03 | 1.21E-02 | 1.01E+00 | 3.93E-10 | 1.50E-09 | -1.37E+00 | 1.22E-05 | 4.35E-05 | -1.22E+00 | 2.98E-10 | 8.39E-10 | -1.59E+00 | 4.61E-05 | 1.69E-04 | -1.16E+00 | 1.52E-09 | 4.17E-09 | -1.43E+00 |
| AC010095.1 | 5.75E-04 | 3.00E-03 | -1.33E+00 | 1.91E-06 | 8.25E-06 | -1.87E+00 | 3.83E-03 | 1.63E-02 | -1.10E+00 | 8.21E-13 | 3.14E-12 | -2.48E+00 | 1.61E-03 | 5.76E-03 | -1.10E+00 | 6.66E-07 | 1.88E-06 | -2.20E+00 | 5.78E-02 | 2.12E-01 | 1.08E+00 | 6.57E-06 | 1.80E-05 | -1.65E+00 |
| U1 | 5.75E-04 | 3.00E-03 | 2.02E+00 | 1.11E-04 | 4.78E-04 | -1.24E+00 | 1.10E-06 | 4.67E-06 | 2.14E+00 | 2.52E-02 | 9.61E-02 | -1.11E+00 | 9.79E-02 | 3.49E-01 | 1.27E+00 | 1.77E-02 | 4.99E-02 | -1.30E+00 | 7.93E-03 | 2.91E-02 | -1.28E+00 | 1.47E-02 | 4.05E-02 | -1.40E+00 |
| LINC00342 | 5.76E-04 | 3.00E-03 | 1.01E+00 | 4.84E-02 | 2.09E-01 | 1.40E+00 | 8.42E-03 | 3.58E-02 | 1.01E+00 | 1.52E-01 | 5.79E-01 | 1.33E+00 | 1.57E-01 | 5.60E-01 | 1.20E+00 | 3.44E-01 | 9.71E-01 | 1.43E+00 | 1.92E-01 | 7.06E-01 | 1.15E+00 | 2.33E-01 | 6.39E-01 | 1.37E+00 |
| SEMA6A | 5.77E-04 | 3.01E-03 | -1.62E+00 | 1.77E-03 | 7.63E-03 | -1.29E+00 | 1.66E-04 | 7.05E-04 | -1.54E+00 | 3.03E-03 | 1.16E-02 | -1.09E+00 | 7.38E-02 | 2.63E-01 | 1.04E+00 | 3.57E-02 | 1.01E-01 | -1.19E+00 | 7.60E-02 | 2.79E-01 | 1.12E+00 | 1.30E-01 | 3.57E-01 | -1.05E+00 |
| TMEM259 | 5.79E-04 | 3.02E-03 | 1.04E+00 | 8.88E-04 | 3.84E-03 | 1.18E+00 | 6.42E-04 | 2.73E-03 | -1.05E+00 | 1.01E-02 | 3.86E-02 | 1.23E+00 | 9.43E-05 | 3.36E-04 | -1.10E+00 | 9.71E-03 | 2.74E-02 | 1.06E+00 | 2.25E-02 | 8.26E-02 | 1.09E+00 | 1.78E-01 | 4.90E-01 | 1.20E+00 |
| C9orf40 | 5.79E-04 | 3.02E-03 | -1.46E+00 | 7.47E-03 | 3.23E-02 | -1.11E+00 | 3.86E-04 | 1.64E-03 | -1.50E+00 | 2.83E-05 | 1.08E-04 | -1.37E+00 | 4.79E-06 | 1.71E-05 | -2.38E+00 | 9.99E-04 | 2.82E-03 | -1.38E+00 | 7.95E-17 | 2.92E-16 | -1.60E+00 | 2.71E-09 | 7.45E-09 | -1.25E+00 |
| CES4A | 5.80E-04 | 3.02E-03 | -1.56E+00 | 1.02E-02 | 4.39E-02 | -1.15E+00 | 5.75E-02 | 2.44E-01 | -1.07E+00 | 7.24E-02 | 2.77E-01 | 1.07E+00 | 2.74E-01 | 9.77E-01 | 1.42E+00 | 2.66E-01 | 7.50E-01 | 1.65E+00 | 2.39E-01 | 8.77E-01 | 1.29E+00 | 2.32E-01 | 6.37E-01 | 1.44E+00 |
| PSMC6 | 5.81E-04 | 3.03E-03 | -1.01E+00 | 8.23E-02 | 3.56E-01 | 1.29E+00 | 4.68E-06 | 1.99E-05 | -1.22E+00 | 1.28E-04 | 4.89E-04 | -1.03E+00 | 3.34E-06 | 1.19E-05 | -1.10E+00 | 1.01E-02 | 2.85E-02 | 1.09E+00 | 7.96E-08 | 2.92E-07 | -1.30E+00 | 1.86E-02 | 5.10E-02 | 1.01E+00 |
| LAPTM5 | 5.81E-04 | 3.03E-03 | 1.03E+00 | 1.59E-01 | 6.86E-01 | 1.26E+00 | 2.98E-02 | 1.27E-01 | -1.02E+00 | 7.91E-02 | 3.02E-01 | 1.20E+00 | 1.71E-01 | 6.08E-01 | 1.00E+00 | 2.25E-01 | 6.34E-01 | -1.02E+00 | 2.30E-01 | 8.43E-01 | 1.13E+00 | 2.00E-01 | 5.50E-01 | 1.11E+00 |
| PDZD2 | 5.81E-04 | 3.03E-03 | 1.02E+00 | 2.20E-03 | 9.51E-03 | 1.17E+00 | 5.23E-02 | 2.22E-01 | 1.05E+00 | 4.88E-02 | 1.86E-01 | 1.19E+00 | 1.96E-02 | 6.98E-02 | -1.13E+00 | 8.13E-02 | 2.29E-01 | 1.10E+00 | 1.44E-02 | 5.29E-02 | -1.26E+00 | 7.44E-02 | 2.04E-01 | -1.11E+00 |
| ARL6IP6 | 5.81E-04 | 3.03E-03 | -1.05E+00 | 1.07E-03 | 4.63E-03 | 1.11E+00 | 1.58E-03 | 6.69E-03 | -1.15E+00 | 1.16E-04 | 4.45E-04 | -1.06E+00 | 1.30E-14 | 4.62E-14 | -2.01E+00 | 2.70E-07 | 7.62E-07 | -1.27E+00 | 1.86E-14 | 6.84E-14 | -1.48E+00 | 8.63E-05 | 2.37E-04 | -1.10E+00 |
| KIAA0100 | 5.82E-04 | 3.03E-03 | -1.13E+00 | 7.28E-04 | 3.14E-03 | -1.01E+00 | 3.38E-06 | 1.44E-05 | -1.20E+00 | 6.33E-05 | 2.42E-04 | 1.00E+00 | 2.82E-04 | 1.01E-03 | -1.11E+00 | 3.51E-04 | 9.90E-04 | -1.12E+00 | 6.23E-03 | 2.29E-02 | -1.14E+00 | 2.49E-01 | 6.85E-01 | 1.15E+00 |
| NDUFA12 | 5.82E-04 | 3.04E-03 | 1.09E+00 | 4.22E-04 | 1.82E-03 | 1.19E+00 | 2.06E-02 | 8.75E-02 | 1.26E+00 | 2.50E-01 | 9.55E-01 | 1.13E+00 | 4.25E-05 | 1.51E-04 | -1.07E+00 | 1.25E-02 | 3.53E-02 | 1.12E+00 | 1.78E-02 | 6.54E-02 | 1.04E+00 | 1.01E-01 | 2.77E-01 | 1.08E+00 |
| L2HGDH | 5.86E-04 | 3.06E-03 | -1.52E+00 | 1.04E-02 | 4.50E-02 | -1.06E+00 | 1.27E-02 | 5.42E-02 | -1.20E+00 | 4.24E-02 | 1.62E-01 | 1.12E+00 | 2.76E-02 | 9.83E-02 | -1.24E+00 | 1.02E-01 | 2.88E-01 | 1.10E+00 | 3.08E-03 | 1.13E-02 | -1.01E+00 | 1.64E-01 | 4.51E-01 | 1.21E+00 |
| CWC27 | 5.87E-04 | 3.06E-03 | 1.01E+00 | 3.77E-03 | 1.63E-02 | 1.17E+00 | 2.38E-03 | 1.01E-02 | 1.01E+00 | 5.65E-04 | 2.16E-03 | 1.07E+00 | 4.81E-04 | 1.71E-03 | -1.09E+00 | 1.06E-01 | 2.99E-01 | 1.22E+00 | 8.22E-02 | 3.02E-01 | 1.11E+00 | 3.46E-01 | 9.49E-01 | 1.15E+00 |
| RPL9P21 | 5.87E-04 | 3.06E-03 | -1.13E+00 | 3.90E-22 | 1.68E-21 | -1.41E+00 | 4.07E-02 | 1.73E-01 | -1.12E+00 | 1.80E-08 | 6.89E-08 | -1.39E+00 | 1.02E-03 | 3.62E-03 | -1.17E+00 | 1.81E-07 | 5.12E-07 | -1.37E+00 | 3.74E-05 | 1.37E-04 | -1.19E+00 | 8.11E-11 | 2.23E-10 | -1.45E+00 |
| CPT2 | 5.87E-04 | 3.06E-03 | -1.23E+00 | 3.20E-02 | 1.38E-01 | 1.17E+00 | 4.42E-05 | 1.88E-04 | -1.34E+00 | 4.93E-03 | 1.88E-02 | 1.05E+00 | 2.01E-05 | 7.17E-05 | -1.75E+00 | 9.19E-03 | 2.59E-02 | -1.03E+00 | 1.33E-08 | 4.90E-08 | -1.65E+00 | 6.52E-03 | 1.79E-02 | -1.04E+00 |
| MAPKAPK5 | 5.87E-04 | 3.06E-03 | -1.06E+00 | 3.49E-03 | 1.51E-02 | 1.11E+00 | 1.55E-06 | 6.58E-06 | -1.35E+00 | 3.37E-05 | 1.29E-04 | -1.09E+00 | 1.67E-01 | 5.95E-01 | 1.23E+00 | 1.04E-01 | 2.92E-01 | 1.13E+00 | 5.44E-04 | 2.00E-03 | -1.18E+00 | 1.10E-01 | 3.02E-01 | 1.12E+00 |
| SLC44A1 | 5.87E-04 | 3.06E-03 | -1.10E+00 | 1.03E-01 | 4.47E-01 | 1.36E+00 | 2.26E-04 | 9.59E-04 | -1.16E+00 | 3.07E-02 | 1.17E-01 | 1.19E+00 | 2.71E-03 | 9.67E-03 | 2.20E+00 | 2.40E-03 | 6.75E-03 | 2.34E+00 | 1.41E-05 | 5.17E-05 | 1.87E+00 | 5.90E-04 | 1.62E-03 | 1.96E+00 |
| OLA1P2 | 5.87E-04 | 3.06E-03 | -1.53E+00 | 1.70E-04 | 7.34E-04 | -1.68E+00 | 8.06E-06 | 3.43E-05 | -1.75E+00 | 3.84E-07 | 1.47E-06 | -1.94E+00 | 1.57E-03 | 5.61E-03 | -1.74E+00 | 6.64E-03 | 1.87E-02 | -1.31E+00 | 5.04E-07 | 1.85E-06 | -1.82E+00 | 2.85E-07 | 7.82E-07 | -1.83E+00 |
| CHAMP1 | 5.88E-04 | 3.06E-03 | -1.19E+00 | 9.53E-04 | 4.12E-03 | -1.08E+00 | 2.87E-03 | 1.22E-02 | -1.13E+00 | 9.03E-03 | 3.45E-02 | 1.11E+00 | 7.01E-03 | 2.50E-02 | -1.18E+00 | 8.70E-02 | 2.45E-01 | 1.02E+00 | 4.90E-06 | 1.80E-05 | -1.37E+00 | 1.39E-01 | 3.83E-01 | 1.08E+00 |
| SLC25A12 | 5.88E-04 | 3.06E-03 | -1.17E+00 | 7.40E-03 | 3.20E-02 | 1.13E+00 | 1.44E-07 | 6.12E-07 | -1.59E+00 | 1.25E-05 | 4.76E-05 | -1.16E+00 | 2.51E-02 | 8.97E-02 | -1.12E+00 | 2.14E-01 | 6.02E-01 | 1.23E+00 | 6.06E-07 | 2.22E-06 | -1.54E+00 | 1.33E-03 | 3.65E-03 | -1.13E+00 |
| CIR1 | 5.88E-04 | 3.06E-03 | 1.10E+00 | 1.45E-09 | 6.26E-09 | -1.09E+00 | 1.09E-02 | 4.64E-02 | -1.00E+00 | 3.83E-06 | 1.46E-05 | -1.20E+00 | 2.25E-02 | 8.04E-02 | 1.12E+00 | 2.04E-04 | 5.75E-04 | -1.03E+00 | 6.98E-04 | 2.56E-03 | -1.05E+00 | 1.14E-06 | 3.12E-06 | -1.25E+00 |
| UBBP1 | 5.88E-04 | 3.06E-03 | 1.44E+00 | 2.21E-01 | 9.53E-01 | 1.40E+00 | 2.31E-06 | 9.80E-06 | 1.49E+00 | 5.22E-03 | 2.00E-02 | 1.36E+00 | 1.47E-02 | 5.24E-02 | 1.58E+00 | 2.66E-01 | 7.51E-01 | 1.45E+00 | 6.68E-05 | 2.45E-04 | 1.58E+00 | 3.04E-03 | 8.36E-03 | 1.43E+00 |
| CSNK1A1 | 5.91E-04 | 3.08E-03 | 1.10E+00 | 1.27E-07 | 5.49E-07 | 1.08E+00 | 1.86E-01 | 7.92E-01 | 1.26E+00 | 6.50E-03 | 2.48E-02 | 1.17E+00 | 1.97E-02 | 7.02E-02 | 1.10E+00 | 2.30E-03 | 6.47E-03 | 1.12E+00 | 7.68E-02 | 2.82E-01 | 1.12E+00 | 2.00E-02 | 5.48E-02 | 1.10E+00 |
| AC107081.2 | 5.91E-04 | 3.08E-03 | -1.59E+00 | 4.50E-03 | 1.94E-02 | -1.23E+00 | 4.32E-05 | 1.84E-04 | -2.07E+00 | 2.25E-04 | 8.60E-04 | -1.66E+00 | 5.47E-02 | 1.95E-01 | -1.25E+00 | 1.48E-01 | 4.18E-01 | 1.02E+00 | 3.87E-04 | 1.42E-03 | -2.07E+00 | 2.63E-02 | 7.24E-02 | -1.34E+00 |
| FUS | 5.93E-04 | 3.09E-03 | -1.04E+00 | 2.79E-11 | 1.21E-10 | -1.12E+00 | 1.44E-01 | 6.13E-01 | 1.10E+00 | 1.76E-01 | 6.73E-01 | 1.06E+00 | 3.89E-05 | 1.39E-04 | -1.13E+00 | 2.39E-05 | 6.74E-05 | -1.19E+00 | 3.89E-03 | 1.43E-02 | -1.15E+00 | 7.65E-02 | 2.10E-01 | -1.02E+00 |
| LINC01515 | 5.93E-04 | 3.09E-03 | -1.25E+00 | 1.26E-02 | 5.43E-02 | 1.14E+00 | 9.38E-03 | 3.99E-02 | -1.29E+00 | 2.38E-01 | 9.10E-01 | 1.36E+00 | 1.94E-01 | 6.90E-01 | -1.11E+00 | 3.30E-01 | 9.31E-01 | 1.26E+00 | 1.30E-03 | 4.78E-03 | -1.57E+00 | 1.10E-01 | 3.03E-01 | 1.06E+00 |
| CSNK2A2 | 5.93E-04 | 3.09E-03 | -1.04E+00 | 2.87E-04 | 1.24E-03 | 1.10E+00 | 5.81E-06 | 2.47E-05 | -1.27E+00 | 2.65E-06 | 1.01E-05 | -1.14E+00 | 1.08E-06 | 3.86E-06 | -1.47E+00 | 4.57E-04 | 1.29E-03 | -1.12E+00 | 1.96E-02 | 7.20E-02 | 1.03E+00 | 1.01E-02 | 2.78E-02 | -1.03E+00 |
| TOGARAM1 | 5.93E-04 | 3.09E-03 | -1.02E+00 | 1.71E-04 | 7.40E-04 | 1.04E+00 | 3.27E-03 | 1.39E-02 | -1.06E+00 | 5.05E-05 | 1.93E-04 | -1.09E+00 | 1.54E-01 | 5.49E-01 | 1.17E+00 | 2.25E-02 | 6.34E-02 | -1.04E+00 | 1.53E-01 | 5.61E-01 | 1.27E+00 | 1.38E-01 | 3.79E-01 | 1.01E+00 |
| TRAF4 | 5.94E-04 | 3.10E-03 | -1.48E+00 | 2.26E-04 | 9.78E-04 | -1.53E+00 | 2.15E-07 | 9.14E-07 | -1.91E+00 | 1.02E-07 | 3.89E-07 | -1.80E+00 | 2.93E-03 | 1.04E-02 | -1.50E+00 | 3.65E-02 | 1.03E-01 | -1.13E+00 | 3.55E-07 | 1.30E-06 | -1.60E+00 | 1.04E-05 | 2.86E-05 | -1.63E+00 |
| TCTEX1D1 | 5.94E-04 | 3.10E-03 | 1.07E+00 | 1.01E-06 | 4.38E-06 | 1.06E+00 | 1.09E-03 | 4.65E-03 | -1.09E+00 | 1.38E-02 | 5.26E-02 | 1.09E+00 | 1.05E-02 | 3.74E-02 | 1.06E+00 | 2.23E-06 | 6.30E-06 | -1.48E+00 | 7.19E-02 | 2.64E-01 | -1.04E+00 | 1.15E-01 | 3.15E-01 | 1.02E+00 |
| FAM219A | 5.95E-04 | 3.10E-03 | -1.87E+00 | 4.82E-02 | 2.08E-01 | -1.11E+00 | 2.32E-01 | 9.87E-01 | 1.22E+00 | 1.04E-01 | 3.98E-01 | 1.57E+00 | 2.30E-02 | 8.19E-02 | -1.29E+00 | 5.70E-02 | 1.61E-01 | -1.12E+00 | 1.99E-02 | 7.31E-02 | -1.34E+00 | 1.12E-01 | 3.07E-01 | -1.06E+00 |
| ZNF71 | 5.96E-04 | 3.11E-03 | -1.38E+00 | 5.62E-02 | 2.43E-01 | 1.23E+00 | 1.00E-04 | 4.26E-04 | -1.48E+00 | 8.64E-03 | 3.30E-02 | 1.03E+00 | 5.22E-02 | 1.86E-01 | -1.28E+00 | 3.08E-01 | 8.69E-01 | 1.26E+00 | 2.87E-05 | 1.05E-04 | -1.53E+00 | 8.54E-02 | 2.34E-01 | 1.05E+00 |
| SPOUT1 | 5.99E-04 | 3.12E-03 | -1.22E+00 | 3.30E-03 | 1.43E-02 | 1.01E+00 | 1.19E-04 | 5.04E-04 | -1.24E+00 | 3.87E-05 | 1.48E-04 | -1.10E+00 | 1.80E-03 | 6.41E-03 | -1.27E+00 | 2.54E-02 | 7.15E-02 | -1.01E+00 | 8.14E-10 | 2.99E-09 | -1.43E+00 | 5.86E-03 | 1.61E-02 | 1.01E+00 |
| ARHGEF11 | 5.99E-04 | 3.12E-03 | -1.56E+00 | 8.50E-04 | 3.67E-03 | -1.53E+00 | 2.22E-03 | 9.42E-03 | -1.52E+00 | 9.51E-04 | 3.63E-03 | -1.60E+00 | 2.63E-01 | 9.38E-01 | 1.36E+00 | 2.57E-01 | 7.24E-01 | 1.49E+00 | 1.40E-02 | 5.14E-02 | -1.05E+00 | 5.35E-02 | 1.47E-01 | -1.05E+00 |
| BTRC | 6.00E-04 | 3.13E-03 | -1.20E+00 | 5.64E-03 | 2.44E-02 | 1.07E+00 | 6.11E-09 | 2.60E-08 | -2.14E+00 | 3.40E-04 | 1.30E-03 | -1.11E+00 | 1.98E-02 | 7.06E-02 | -1.12E+00 | 3.33E-02 | 9.39E-02 | -1.09E+00 | 3.21E-03 | 1.18E-02 | -1.28E+00 | 4.61E-02 | 1.27E-01 | -1.08E+00 |
| AC073263.2 | 6.02E-04 | 3.14E-03 | -1.48E+00 | 7.74E-04 | 3.34E-03 | -1.32E+00 | 1.53E-01 | 6.49E-01 | -1.18E+00 | 4.77E-02 | 1.82E-01 | -1.47E+00 | 2.98E-03 | 1.06E-02 | -1.62E+00 | 6.38E-04 | 1.80E-03 | -2.10E+00 | 1.13E-01 | 4.14E-01 | 1.70E+00 | 1.08E-01 | 2.96E-01 | 1.68E+00 |
| BEST1 | 6.02E-04 | 3.14E-03 | 3.52E+00 | 5.88E-02 | 2.54E-01 | 2.39E+00 | 1.45E-01 | 6.16E-01 | 1.32E+00 | 2.36E-01 | 9.03E-01 | 1.31E+00 | 2.11E-01 | 7.54E-01 | 1.38E+00 | 2.84E-01 | 7.99E-01 | 1.71E+00 | 1.98E-01 | 7.26E-01 | 1.11E+00 | 3.22E-01 | 8.85E-01 | 1.34E+00 |
| RPL19 | 6.03E-04 | 3.14E-03 | 1.08E+00 | 3.48E-02 | 1.50E-01 | 1.05E+00 | 1.07E-04 | 4.55E-04 | 1.12E+00 | 3.19E-03 | 1.22E-02 | 1.09E+00 | 7.66E-03 | 2.73E-02 | 1.09E+00 | 7.16E-03 | 2.02E-02 | 1.09E+00 | 1.50E-01 | 5.50E-01 | 1.09E+00 | 2.32E-02 | 6.39E-02 | 1.09E+00 |
| UXT | 6.03E-04 | 3.14E-03 | -1.05E+00 | 3.85E-07 | 1.66E-06 | -1.05E+00 | 1.91E-01 | 8.11E-01 | 1.02E+00 | 7.70E-02 | 2.94E-01 | -1.00E+00 | 2.22E-01 | 7.90E-01 | 1.10E+00 | 3.09E-01 | 8.72E-01 | 1.07E+00 | 1.06E-01 | 3.89E-01 | 1.02E+00 | 1.09E-01 | 3.00E-01 | -1.03E+00 |
| NEK9 | 6.03E-04 | 3.14E-03 | -1.14E+00 | 9.50E-02 | 4.11E-01 | 1.33E+00 | 2.05E-04 | 8.71E-04 | -1.27E+00 | 3.50E-02 | 1.34E-01 | 1.13E+00 | 8.01E-03 | 2.85E-02 | -1.01E+00 | 1.45E-01 | 4.08E-01 | 1.33E+00 | 2.84E-03 | 1.04E-02 | -1.08E+00 | 2.24E-01 | 6.14E-01 | 1.15E+00 |
| FAM174A | 6.04E-04 | 3.15E-03 | -1.26E+00 | 6.36E-04 | 2.75E-03 | -1.17E+00 | 2.92E-03 | 1.24E-02 | -1.22E+00 | 2.39E-05 | 9.12E-05 | -1.59E+00 | 2.00E-01 | 7.14E-01 | 1.23E+00 | 5.65E-02 | 1.59E-01 | -1.21E+00 | 1.20E-08 | 4.41E-08 | -1.50E+00 | 8.46E-08 | 2.32E-07 | -1.42E+00 |
| NEURL1 | 6.04E-04 | 3.15E-03 | -1.46E+00 | 1.92E-05 | 8.32E-05 | -1.93E+00 | 4.07E-07 | 1.73E-06 | -1.62E+00 | 9.31E-05 | 3.56E-04 | -1.20E+00 | 4.75E-03 | 1.69E-02 | -2.07E+00 | 7.80E-02 | 2.20E-01 | -1.14E+00 | 3.76E-06 | 1.38E-05 | -1.89E+00 | 3.44E-03 | 9.45E-03 | -1.20E+00 |
| TRIM59 | 6.06E-04 | 3.16E-03 | -1.22E+00 | 1.99E-01 | 8.62E-01 | 1.56E+00 | 1.36E-06 | 5.78E-06 | -1.62E+00 | 7.27E-04 | 2.78E-03 | -1.09E+00 | 1.63E-02 | 5.80E-02 | -1.12E+00 | 8.13E-02 | 2.29E-01 | 1.04E+00 | 4.25E-06 | 1.56E-05 | -1.40E+00 | 6.55E-02 | 1.80E-01 | 1.03E+00 |
| PAG1 | 6.06E-04 | 3.16E-03 | -1.23E+00 | 1.36E-03 | 5.88E-03 | 1.04E+00 | 3.88E-03 | 1.65E-02 | -1.24E+00 | 9.83E-04 | 3.75E-03 | -1.10E+00 | 2.66E-18 | 9.50E-18 | -2.11E+00 | 1.38E-23 | 3.89E-23 | -2.45E+00 | 2.00E-10 | 7.36E-10 | -2.67E+00 | 2.04E-05 | 5.61E-05 | -1.61E+00 |
| ZNF622 | 6.07E-04 | 3.16E-03 | -1.15E+00 | 2.68E-03 | 1.16E-02 | 1.07E+00 | 2.62E-03 | 1.11E-02 | -1.13E+00 | 2.07E-04 | 7.89E-04 | -1.09E+00 | 6.73E-05 | 2.40E-04 | -1.14E+00 | 5.29E-03 | 1.49E-02 | 1.02E+00 | 2.57E-07 | 9.44E-07 | -1.23E+00 | 1.26E-04 | 3.46E-04 | -1.04E+00 |
| CSGALNACT2 | 6.07E-04 | 3.16E-03 | -1.06E+00 | 4.28E-06 | 1.85E-05 | -1.06E+00 | 3.67E-03 | 1.56E-02 | -1.07E+00 | 3.42E-08 | 1.31E-07 | -1.31E+00 | 4.07E-04 | 1.45E-03 | -1.20E+00 | 3.10E-04 | 8.75E-04 | -1.31E+00 | 7.24E-04 | 2.66E-03 | -1.19E+00 | 1.17E-06 | 3.21E-06 | -1.59E+00 |
| ESRRA | 6.08E-04 | 3.17E-03 | -1.56E+00 | 7.18E-04 | 3.10E-03 | -1.46E+00 | 2.27E-16 | 9.64E-16 | -2.56E+00 | 2.01E-12 | 7.70E-12 | -1.73E+00 | 1.29E-09 | 4.58E-09 | -2.15E+00 | 2.05E-07 | 5.79E-07 | -1.65E+00 | 3.92E-07 | 1.44E-06 | -1.50E+00 | 5.47E-02 | 1.50E-01 | 1.10E+00 |
| ARHGAP4 | 6.08E-04 | 3.17E-03 | 1.03E+00 | 7.43E-03 | 3.21E-02 | 1.27E+00 | 1.78E-02 | 7.54E-02 | 1.10E+00 | 2.85E-02 | 1.09E-01 | 1.24E+00 | 4.13E-07 | 1.47E-06 | -1.38E+00 | 1.13E-05 | 3.19E-05 | -1.21E+00 | 4.61E-03 | 1.69E-02 | 1.06E+00 | 7.55E-03 | 2.07E-02 | 1.05E+00 |
| CHMP1A | 6.11E-04 | 3.18E-03 | -1.05E+00 | 8.13E-02 | 3.51E-01 | 1.32E+00 | 9.42E-03 | 4.00E-02 | 1.00E+00 | 4.29E-02 | 1.64E-01 | 1.19E+00 | 6.45E-09 | 2.30E-08 | -1.55E+00 | 8.07E-05 | 2.27E-04 | -1.14E+00 | 3.74E-03 | 1.37E-02 | 1.06E+00 | 2.91E-01 | 7.98E-01 | 1.30E+00 |
| ORC3 | 6.11E-04 | 3.18E-03 | -1.28E+00 | 2.12E-02 | 9.16E-02 | 1.07E+00 | 6.58E-07 | 2.79E-06 | -1.28E+00 | 1.73E-06 | 6.61E-06 | -1.15E+00 | 4.79E-07 | 1.71E-06 | -1.71E+00 | 8.66E-04 | 2.44E-03 | -1.09E+00 | 2.11E-04 | 7.76E-04 | -1.10E+00 | 3.27E-02 | 8.97E-02 | 1.12E+00 |
| MAX | 6.14E-04 | 3.20E-03 | 1.01E+00 | 1.37E-04 | 5.94E-04 | 1.05E+00 | 4.62E-07 | 1.96E-06 | -1.36E+00 | 2.71E-06 | 1.03E-05 | -1.23E+00 | 2.51E-01 | 8.93E-01 | 1.17E+00 | 7.02E-02 | 1.98E-01 | 1.12E+00 | 4.18E-02 | 1.53E-01 | 1.61E+00 | 9.22E-03 | 2.53E-02 | 1.22E+00 |
| INTS5 | 6.14E-04 | 3.20E-03 | -1.25E+00 | 6.77E-02 | 2.93E-01 | 1.27E+00 | 4.85E-06 | 2.06E-05 | -1.84E+00 | 6.69E-02 | 2.55E-01 | 1.12E+00 | 1.32E-04 | 4.72E-04 | -1.66E+00 | 3.55E-01 | 1.00E+00 | 1.43E+00 | 1.18E-09 | 4.32E-09 | -1.67E+00 | 3.57E-01 | 9.80E-01 | 1.42E+00 |
| KLHL13 | 6.17E-04 | 3.22E-03 | -1.39E+00 | 4.01E-03 | 1.73E-02 | -1.07E+00 | 7.35E-02 | 3.12E-01 | 1.12E+00 | 2.06E-01 | 7.85E-01 | 1.39E+00 | 1.81E-04 | 6.44E-04 | -1.60E+00 | 6.32E-02 | 1.78E-01 | 1.17E+00 | 1.80E-01 | 6.61E-01 | 1.32E+00 | 7.90E-02 | 2.17E-01 | 1.62E+00 |
| SUSD6 | 6.18E-04 | 3.22E-03 | -1.10E+00 | 1.33E-02 | 5.73E-02 | 1.24E+00 | 1.37E-03 | 5.82E-03 | -1.19E+00 | 1.54E-03 | 5.89E-03 | -1.03E+00 | 4.11E-04 | 1.47E-03 | -1.25E+00 | 1.16E-03 | 3.28E-03 | -1.10E+00 | 6.24E-05 | 2.29E-04 | -1.43E+00 | 3.21E-03 | 8.81E-03 | -1.12E+00 |
| ESYT3 | 6.18E-04 | 3.22E-03 | -1.48E+00 | 2.04E-02 | 8.84E-02 | 1.07E+00 | 4.33E-03 | 1.84E-02 | -1.43E+00 | 5.49E-03 | 2.10E-02 | -1.25E+00 | 1.36E-01 | 4.86E-01 | 1.16E+00 | 2.37E-01 | 6.69E-01 | 1.23E+00 | 2.01E-01 | 7.37E-01 | 1.32E+00 | 2.92E-01 | 8.02E-01 | 1.49E+00 |
| TPD52L1 | 6.20E-04 | 3.23E-03 | -1.35E+00 | 4.45E-02 | 1.92E-01 | 1.14E+00 | 1.92E-01 | 8.15E-01 | 1.15E+00 | 1.83E-01 | 7.00E-01 | 1.30E+00 | 1.06E-01 | 3.79E-01 | -1.66E+00 | 3.13E-01 | 8.82E-01 | 1.05E+00 | 2.15E-01 | 7.89E-01 | 1.13E+00 | 2.92E-01 | 8.03E-01 | 1.38E+00 |
| GZF1 | 6.20E-04 | 3.23E-03 | -1.16E+00 | 6.41E-06 | 2.77E-05 | -1.29E+00 | 5.56E-04 | 2.36E-03 | -1.31E+00 | 9.98E-06 | 3.81E-05 | -1.37E+00 | 2.76E-07 | 9.84E-07 | -1.67E+00 | 1.53E-08 | 4.32E-08 | -1.87E+00 | 6.05E-06 | 2.22E-05 | -1.72E+00 | 3.62E-05 | 9.94E-05 | -1.83E+00 |
| FEM1A | 6.20E-04 | 3.23E-03 | -1.60E+00 | 5.52E-05 | 2.38E-04 | -1.70E+00 | 1.87E-01 | 7.94E-01 | 1.23E+00 | 3.79E-02 | 1.45E-01 | 1.03E+00 | 2.79E-03 | 9.95E-03 | -1.88E+00 | 1.59E-01 | 4.49E-01 | 1.14E+00 | 1.38E-01 | 5.05E-01 | -1.01E+00 | 1.84E-02 | 5.06E-02 | -1.41E+00 |
| MTURN | 6.21E-04 | 3.24E-03 | 1.04E+00 | 1.87E-01 | 8.08E-01 | 1.41E+00 | 6.45E-02 | 2.74E-01 | -1.04E+00 | 8.36E-02 | 3.19E-01 | 1.37E+00 | 5.66E-02 | 2.02E-01 | 1.08E+00 | 2.99E-01 | 8.44E-01 | 1.17E+00 | 1.89E-01 | 6.95E-01 | 1.16E+00 | 3.03E-01 | 8.32E-01 | 1.15E+00 |
| AKT1S1 | 6.22E-04 | 3.24E-03 | -1.15E+00 | 2.36E-05 | 1.02E-04 | -1.27E+00 | 2.89E-04 | 1.23E-03 | -1.24E+00 | 1.36E-05 | 5.19E-05 | -1.19E+00 | 1.68E-07 | 5.98E-07 | -1.68E+00 | 2.50E-05 | 7.05E-05 | -1.25E+00 | 3.25E-05 | 1.19E-04 | -1.26E+00 | 1.58E-05 | 4.33E-05 | -1.29E+00 |
| ATG2A | 6.23E-04 | 3.25E-03 | -1.18E+00 | 8.92E-06 | 3.85E-05 | -1.35E+00 | 2.65E-02 | 1.13E-01 | -1.05E+00 | 4.11E-03 | 1.57E-02 | -1.09E+00 | 1.53E-01 | 5.44E-01 | 1.30E+00 | 1.56E-01 | 4.41E-01 | 1.16E+00 | 5.23E-02 | 1.92E-01 | -1.05E+00 | 1.85E-01 | 5.09E-01 | 1.01E+00 |
| RNF220 | 6.23E-04 | 3.25E-03 | 1.06E+00 | 2.84E-02 | 1.23E-01 | 1.33E+00 | 8.66E-03 | 3.68E-02 | 1.01E+00 | 1.81E-01 | 6.92E-01 | 1.37E+00 | 1.02E-01 | 3.64E-01 | 1.23E+00 | 3.50E-01 | 9.87E-01 | 1.41E+00 | 1.75E-01 | 6.41E-01 | 1.19E+00 | 3.27E-01 | 8.97E-01 | 1.35E+00 |
| TMEM208 | 6.24E-04 | 3.25E-03 | 1.06E+00 | 3.03E-05 | 1.31E-04 | 1.10E+00 | 1.67E-01 | 7.10E-01 | 1.14E+00 | 3.22E-02 | 1.23E-01 | 1.18E+00 | 1.81E-02 | 6.45E-02 | 1.12E+00 | 1.19E-02 | 3.35E-02 | 1.16E+00 | 1.48E-01 | 5.45E-01 | 1.17E+00 | 3.58E-01 | 9.83E-01 | 1.23E+00 |
| RBM19 | 6.24E-04 | 3.25E-03 | -1.16E+00 | 4.11E-02 | 1.78E-01 | 1.23E+00 | 4.73E-03 | 2.01E-02 | -1.10E+00 | 2.90E-03 | 1.11E-02 | 1.07E+00 | 2.42E-06 | 8.65E-06 | -1.45E+00 | 5.31E-04 | 1.50E-03 | -1.04E+00 | 7.14E-03 | 2.62E-02 | -1.15E+00 | 4.99E-02 | 1.37E-01 | 1.08E+00 |
| HDAC8 | 6.27E-04 | 3.27E-03 | -1.07E+00 | 1.55E-03 | 6.69E-03 | 1.11E+00 | 2.33E-01 | 9.90E-01 | 1.15E+00 | 1.90E-01 | 7.28E-01 | 1.42E+00 | 2.96E-03 | 1.05E-02 | 1.03E+00 | 5.13E-02 | 1.45E-01 | 1.24E+00 | 8.75E-04 | 3.21E-03 | -1.10E+00 | 2.73E-02 | 7.49E-02 | 1.01E+00 |
| TXNDC15 | 6.27E-04 | 3.27E-03 | -1.11E+00 | 2.02E-01 | 8.72E-01 | 1.37E+00 | 3.46E-08 | 1.47E-07 | -1.47E+00 | 2.59E-04 | 9.91E-04 | -1.04E+00 | 3.70E-06 | 1.32E-05 | -1.31E+00 | 1.55E-03 | 4.38E-03 | -1.00E+00 | 1.20E-07 | 4.42E-07 | -1.41E+00 | 1.47E-04 | 4.05E-04 | -1.09E+00 |
| NBEA | 6.28E-04 | 3.27E-03 | -1.14E+00 | 1.87E-01 | 8.10E-01 | 1.41E+00 | 3.58E-02 | 1.52E-01 | -1.09E+00 | 4.18E-02 | 1.60E-01 | 1.10E+00 | 2.53E-01 | 9.01E-01 | 1.23E+00 | 2.94E-01 | 8.29E-01 | 1.25E+00 | 1.04E-01 | 3.83E-01 | -1.07E+00 | 2.41E-01 | 6.63E-01 | 1.00E+00 |
| AC093157.1 | 6.28E-04 | 3.27E-03 | -1.40E+00 | 2.39E-02 | 1.03E-01 | 1.12E+00 | 1.37E-03 | 5.82E-03 | -1.40E+00 | 2.06E-02 | 7.88E-02 | 1.03E+00 | 3.38E-05 | 1.20E-04 | -1.77E+00 | 4.11E-02 | 1.16E-01 | 1.09E+00 | 8.23E-08 | 3.02E-07 | -1.74E+00 | 2.19E-02 | 6.02E-02 | -1.02E+00 |
| RHBDF2 | 6.29E-04 | 3.28E-03 | -1.53E+00 | 7.60E-03 | 3.28E-02 | 1.00E+00 | 4.14E-04 | 1.76E-03 | -1.40E+00 | 1.69E-04 | 6.47E-04 | -1.39E+00 | 2.79E-02 | 9.93E-02 | -2.20E+00 | 1.67E-01 | 4.72E-01 | 1.05E+00 | 4.80E-03 | 1.76E-02 | -1.41E+00 | 1.17E-01 | 3.20E-01 | -1.09E+00 |
| EHBP1L1 | 6.29E-04 | 3.28E-03 | -1.12E+00 | 5.54E-03 | 2.39E-02 | 1.07E+00 | 1.01E-02 | 4.31E-02 | -1.06E+00 | 5.67E-02 | 2.17E-01 | 1.18E+00 | 2.91E-03 | 1.04E-02 | -1.26E+00 | 2.38E-02 | 6.72E-02 | 1.05E+00 | 2.43E-01 | 8.92E-01 | 1.32E+00 | 2.58E-01 | 7.09E-01 | 1.30E+00 |
| PFKL | 6.30E-04 | 3.28E-03 | 1.11E+00 | 1.51E-03 | 6.54E-03 | 1.23E+00 | 7.19E-02 | 3.05E-01 | 1.17E+00 | 1.29E-01 | 4.93E-01 | 1.34E+00 | 9.18E-06 | 3.27E-05 | -1.16E+00 | 5.75E-04 | 1.62E-03 | 1.02E+00 | 2.33E-01 | 8.54E-01 | 1.27E+00 | 3.84E-02 | 1.05E-01 | 1.41E+00 |
| AL627308.2 | 6.32E-04 | 3.29E-03 | -1.39E+00 | 3.11E-06 | 1.34E-05 | -2.24E+00 | 9.88E-03 | 4.20E-02 | -1.20E+00 | 8.31E-05 | 3.17E-04 | -1.64E+00 | 8.38E-02 | 2.99E-01 | 1.00E+00 | 1.45E-01 | 4.08E-01 | 1.11E+00 | 3.31E-03 | 1.21E-02 | -1.06E+00 | 4.88E-05 | 1.34E-04 | -1.33E+00 |
| SLC35D2 | 6.32E-04 | 3.29E-03 | 1.08E+00 | 4.19E-05 | 1.81E-04 | 1.14E+00 | 9.36E-06 | 3.98E-05 | -1.22E+00 | 5.74E-04 | 2.19E-03 | 1.01E+00 | 1.65E-03 | 5.87E-03 | -1.02E+00 | 5.72E-02 | 1.61E-01 | 1.12E+00 | 7.35E-02 | 2.70E-01 | 1.27E+00 | 7.08E-04 | 1.95E-03 | -1.01E+00 |
| LINC01806 | 6.32E-04 | 3.29E-03 | -1.28E+00 | 1.76E-03 | 7.59E-03 | -1.12E+00 | 5.54E-02 | 2.35E-01 | -1.09E+00 | 5.63E-02 | 2.15E-01 | 1.08E+00 | 1.76E-03 | 6.26E-03 | -1.40E+00 | 6.49E-04 | 1.83E-03 | -1.51E+00 | 6.18E-02 | 2.27E-01 | -1.12E+00 | 1.27E-02 | 3.49E-02 | -1.53E+00 |
| SPDYE3 | 6.35E-04 | 3.31E-03 | -1.43E+00 | 1.06E-01 | 4.60E-01 | 1.32E+00 | 2.20E-02 | 9.34E-02 | -1.15E+00 | 4.70E-02 | 1.80E-01 | 1.02E+00 | 8.33E-06 | 2.97E-05 | -1.69E+00 | 2.27E-04 | 6.40E-04 | -1.40E+00 | 3.45E-05 | 1.27E-04 | -1.77E+00 | 1.16E-01 | 3.20E-01 | 1.01E+00 |
| AL513302.2 | 6.37E-04 | 3.32E-03 | -2.33E+00 | 1.68E-01 | 7.24E-01 | 1.01E+00 | 3.14E-03 | 1.34E-02 | -1.81E+00 | 1.58E-01 | 6.02E-01 | 1.45E+00 | 8.30E-02 | 2.96E-01 | -1.72E+00 | 1.88E-01 | 5.31E-01 | 1.46E+00 | 1.86E-01 | 6.84E-01 | -1.09E+00 | 7.23E-02 | 1.99E-01 | 1.52E+00 |
| DNM1L | 6.38E-04 | 3.32E-03 | 1.01E+00 | 1.31E-04 | 5.68E-04 | 1.08E+00 | 1.08E-02 | 4.58E-02 | -1.05E+00 | 1.11E-03 | 4.25E-03 | 1.01E+00 | 1.25E-01 | 4.45E-01 | 1.15E+00 | 9.37E-02 | 2.64E-01 | 1.10E+00 | 1.97E-01 | 7.25E-01 | 1.13E+00 | 1.53E-01 | 4.20E-01 | 1.06E+00 |
| TMEM91 | 6.38E-04 | 3.32E-03 | -1.01E+00 | 1.85E-02 | 8.00E-02 | 1.27E+00 | 3.62E-04 | 1.54E-03 | -1.25E+00 | 7.25E-03 | 2.77E-02 | 1.05E+00 | 5.67E-02 | 2.02E-01 | 1.17E+00 | 4.81E-02 | 1.36E-01 | 1.12E+00 | 1.47E-04 | 5.40E-04 | 2.31E+00 | 2.40E-01 | 6.59E-01 | 1.44E+00 |
| KLF5 | 6.38E-04 | 3.32E-03 | -1.04E+00 | 1.83E-05 | 7.93E-05 | -1.07E+00 | 5.67E-02 | 2.41E-01 | -1.06E+00 | 3.86E-02 | 1.48E-01 | -1.04E+00 | 9.38E-02 | 3.34E-01 | -1.00E+00 | 2.78E-02 | 7.83E-02 | -1.42E+00 | 7.54E-02 | 2.77E-01 | -1.66E+00 | 8.35E-02 | 2.29E-01 | -1.76E+00 |
| CCDC142 | 6.38E-04 | 3.32E-03 | -1.37E+00 | 1.02E-03 | 4.41E-03 | -1.27E+00 | 1.09E-02 | 4.65E-02 | -1.42E+00 | 1.95E-01 | 7.43E-01 | 1.26E+00 | 1.17E-02 | 4.16E-02 | -1.24E+00 | 1.69E-01 | 4.76E-01 | 1.20E+00 | 7.56E-02 | 2.77E-01 | -1.02E+00 | 2.96E-01 | 8.14E-01 | 1.25E+00 |
| RASSF8 | 6.38E-04 | 3.32E-03 | -1.17E+00 | 2.67E-03 | 1.15E-02 | 1.03E+00 | 9.29E-02 | 3.95E-01 | 1.02E+00 | 8.61E-02 | 3.29E-01 | 1.12E+00 | 2.28E-01 | 8.13E-01 | -1.12E+00 | 2.38E-01 | 6.70E-01 | -1.48E+00 | 1.97E-01 | 7.23E-01 | 1.39E+00 | 2.94E-01 | 8.08E-01 | -1.20E+00 |
| ZNF415 | 6.39E-04 | 3.33E-03 | -1.25E+00 | 2.60E-04 | 1.12E-03 | -1.35E+00 | 1.89E-01 | 8.01E-01 | 1.26E+00 | 1.20E-01 | 4.57E-01 | 1.11E+00 | 9.98E-04 | 3.56E-03 | -1.60E+00 | 9.92E-04 | 2.80E-03 | -1.53E+00 | 2.48E-01 | 9.11E-01 | 1.24E+00 | 1.82E-01 | 4.99E-01 | 1.07E+00 |
| AC090616.7 | 6.41E-04 | 3.34E-03 | -1.17E+00 | 2.45E-02 | 1.06E-01 | 1.20E+00 | 5.69E-07 | 2.42E-06 | -1.74E+00 | 9.42E-06 | 3.60E-05 | -1.43E+00 | 6.14E-04 | 2.19E-03 | -1.55E+00 | 6.77E-03 | 1.91E-02 | -1.22E+00 | 1.40E-04 | 5.13E-04 | -1.92E+00 | 1.03E-03 | 2.83E-03 | -1.51E+00 |
| ACBD5 | 6.41E-04 | 3.34E-03 | -1.10E+00 | 1.80E-03 | 7.76E-03 | 1.07E+00 | 2.72E-04 | 1.15E-03 | -1.15E+00 | 4.43E-05 | 1.69E-04 | -1.14E+00 | 1.81E-03 | 6.45E-03 | -1.30E+00 | 6.68E-03 | 1.88E-02 | -1.21E+00 | 5.39E-06 | 1.98E-05 | -1.36E+00 | 1.29E-05 | 3.54E-05 | -1.37E+00 |
| POLR1C | 6.43E-04 | 3.35E-03 | -1.24E+00 | 2.21E-03 | 9.53E-03 | -1.02E+00 | 1.49E-02 | 6.31E-02 | -1.00E+00 | 1.44E-01 | 5.52E-01 | 1.34E+00 | 9.98E-05 | 3.56E-04 | -1.47E+00 | 2.04E-02 | 5.76E-02 | 1.01E+00 | 2.48E-03 | 9.11E-03 | -1.17E+00 | 1.26E-01 | 3.46E-01 | 1.10E+00 |
| RTL5 | 6.43E-04 | 3.35E-03 | -1.54E+00 | 2.22E-03 | 9.59E-03 | -1.26E+00 | 5.70E-03 | 2.42E-02 | -1.67E+00 | 3.50E-02 | 1.34E-01 | -1.23E+00 | 3.97E-04 | 1.42E-03 | -1.61E+00 | 3.81E-02 | 1.07E-01 | 1.06E+00 | 1.23E-01 | 4.51E-01 | 1.59E+00 | 3.58E-01 | 9.82E-01 | 1.28E+00 |
| TET3 | 6.47E-04 | 3.37E-03 | -1.28E+00 | 7.70E-04 | 3.33E-03 | -1.16E+00 | 1.85E-02 | 7.88E-02 | -1.11E+00 | 6.26E-02 | 2.39E-01 | 1.10E+00 | 8.81E-07 | 3.14E-06 | -1.68E+00 | 1.59E-03 | 4.47E-03 | -1.08E+00 | 3.18E-08 | 1.17E-07 | -1.60E+00 | 4.26E-06 | 1.17E-05 | -1.36E+00 |
| PARP8 | 6.47E-04 | 3.37E-03 | -1.05E+00 | 1.16E-07 | 5.03E-07 | -1.29E+00 | 5.09E-02 | 2.16E-01 | 1.05E+00 | 1.12E-02 | 4.27E-02 | 1.04E+00 | 4.60E-03 | 1.64E-02 | -1.04E+00 | 3.27E-06 | 9.21E-06 | -1.65E+00 | 4.13E-03 | 1.51E-02 | -1.26E+00 | 4.53E-02 | 1.25E-01 | -1.04E+00 |
| VAC14 | 6.47E-04 | 3.37E-03 | -1.34E+00 | 5.91E-03 | 2.55E-02 | -1.03E+00 | 3.15E-06 | 1.34E-05 | -1.37E+00 | 8.15E-04 | 3.11E-03 | 1.05E+00 | 6.52E-06 | 2.33E-05 | -1.97E+00 | 1.18E-03 | 3.34E-03 | -1.13E+00 | 4.70E-07 | 1.72E-06 | -1.37E+00 | 7.09E-03 | 1.95E-02 | 1.03E+00 |
| RPS27P21 | 6.49E-04 | 3.38E-03 | 1.94E+00 | 6.41E-07 | 2.77E-06 | -1.34E+00 | 3.66E-05 | 1.56E-04 | 1.90E+00 | 2.64E-06 | 1.01E-05 | -1.45E+00 | 7.47E-03 | 2.66E-02 | 2.08E+00 | 3.79E-03 | 1.07E-02 | -1.04E+00 | 2.78E-04 | 1.02E-03 | 1.83E+00 | 3.35E-05 | 9.19E-05 | -1.55E+00 |
| LRFN4 | 6.51E-04 | 3.39E-03 | -1.27E+00 | 1.20E-02 | 5.17E-02 | 1.11E+00 | 4.55E-03 | 1.93E-02 | -1.17E+00 | 5.79E-02 | 2.21E-01 | 1.22E+00 | 2.33E-02 | 8.32E-02 | -1.33E+00 | 1.22E-01 | 3.44E-01 | 1.02E+00 | 6.08E-04 | 2.23E-03 | -1.18E+00 | 3.07E-02 | 8.43E-02 | 1.07E+00 |
| PPP6R3 | 6.51E-04 | 3.39E-03 | -1.05E+00 | 3.73E-03 | 1.61E-02 | 1.19E+00 | 1.72E-04 | 7.32E-04 | -1.29E+00 | 1.73E-02 | 6.63E-02 | 1.07E+00 | 2.53E-05 | 9.01E-05 | -1.24E+00 | 2.43E-03 | 6.84E-03 | -1.04E+00 | 1.31E-04 | 4.82E-04 | -1.15E+00 | 3.48E-02 | 9.55E-02 | -1.05E+00 |
| CCSER2 | 6.51E-04 | 3.39E-03 | -1.04E+00 | 1.32E-02 | 5.71E-02 | 1.19E+00 | 2.86E-02 | 1.21E-01 | -1.06E+00 | 6.33E-04 | 2.42E-03 | -1.15E+00 | 2.53E-01 | 9.01E-01 | 1.14E+00 | 2.54E-01 | 7.16E-01 | 1.05E+00 | 1.45E-01 | 5.32E-01 | 1.02E+00 | 1.29E-01 | 3.53E-01 | -1.08E+00 |
| AC012435.1 | 6.51E-04 | 3.40E-03 | -1.83E+00 | 3.18E-02 | 1.37E-01 | 1.08E+00 | 1.10E-01 | 4.67E-01 | -1.05E+00 | 9.71E-02 | 3.71E-01 | 1.74E+00 | 1.22E-02 | 4.34E-02 | -1.45E+00 | 2.13E-01 | 6.01E-01 | 1.21E+00 | 3.78E-02 | 1.39E-01 | -1.49E+00 | 1.03E-01 | 2.82E-01 | 1.48E+00 |
| C2orf42 | 6.52E-04 | 3.40E-03 | -1.66E+00 | 4.76E-03 | 2.06E-02 | -1.30E+00 | 1.40E-02 | 5.96E-02 | -1.28E+00 | 8.16E-02 | 3.12E-01 | 1.14E+00 | 1.20E-02 | 4.28E-02 | -1.42E+00 | 1.83E-01 | 5.15E-01 | 1.13E+00 | 1.92E-06 | 7.06E-06 | -1.75E+00 | 1.06E-04 | 2.91E-04 | -1.54E+00 |
| CTPS2 | 6.54E-04 | 3.41E-03 | -1.21E+00 | 7.18E-03 | 3.10E-02 | 1.11E+00 | 4.70E-03 | 1.99E-02 | -1.21E+00 | 3.99E-02 | 1.52E-01 | 1.08E+00 | 3.29E-02 | 1.17E-01 | -1.02E+00 | 6.27E-02 | 1.77E-01 | 1.03E+00 | 1.37E-08 | 5.03E-08 | -1.49E+00 | 1.74E-07 | 4.77E-07 | -1.43E+00 |
| CEP120 | 6.54E-04 | 3.41E-03 | -1.30E+00 | 2.73E-02 | 1.18E-01 | 1.15E+00 | 4.53E-07 | 1.93E-06 | -1.63E+00 | 3.80E-04 | 1.45E-03 | -1.11E+00 | 1.49E-01 | 5.32E-01 | 1.18E+00 | 2.25E-01 | 6.35E-01 | 1.23E+00 | 1.29E-03 | 4.75E-03 | -1.25E+00 | 7.40E-03 | 2.03E-02 | -1.09E+00 |
| RN7SL600P | 6.55E-04 | 3.42E-03 | 2.83E+00 | 2.81E-02 | 1.21E-01 | 2.34E+00 | 2.45E-04 | 1.04E-03 | 2.86E+00 | 2.86E-02 | 1.09E-01 | 2.36E+00 | 1.18E-02 | 4.21E-02 | 3.26E+00 | 1.99E-02 | 5.61E-02 | 3.17E+00 | 4.98E-02 | 1.83E-01 | 1.91E+00 | 2.23E-02 | 6.13E-02 | 2.54E+00 |
| DIRAS1 | 6.56E-04 | 3.42E-03 | -1.29E+00 | 1.15E-02 | 4.96E-02 | 1.05E+00 | 5.10E-03 | 2.17E-02 | -1.21E+00 | 4.11E-03 | 1.57E-02 | -1.12E+00 | 9.59E-03 | 3.42E-02 | -1.40E+00 | 8.42E-02 | 2.37E-01 | 1.02E+00 | 1.09E-01 | 3.99E-01 | -1.17E+00 | 3.11E-01 | 8.55E-01 | 1.29E+00 |
| TYW5 | 6.56E-04 | 3.42E-03 | -1.15E+00 | 6.70E-02 | 2.90E-01 | 1.31E+00 | 9.93E-04 | 4.22E-03 | -1.18E+00 | 1.33E-02 | 5.09E-02 | 1.12E+00 | 2.50E-02 | 8.92E-02 | -1.03E+00 | 1.38E-01 | 3.88E-01 | 1.18E+00 | 1.26E-02 | 4.63E-02 | 1.01E+00 | 5.13E-02 | 1.41E-01 | 1.05E+00 |
| AL136162.1 | 6.57E-04 | 3.42E-03 | -1.51E+00 | 7.34E-04 | 3.17E-03 | -1.38E+00 | 1.72E-02 | 7.30E-02 | -1.38E+00 | 1.35E-01 | 5.14E-01 | 1.16E+00 | 1.60E-01 | 5.72E-01 | 1.80E+00 | 1.64E-01 | 4.62E-01 | 1.82E+00 | 5.15E-02 | 1.89E-01 | -1.06E+00 | 2.17E-01 | 5.96E-01 | 1.06E+00 |
| UCK1 | 6.57E-04 | 3.42E-03 | -1.20E+00 | 5.62E-02 | 2.43E-01 | 1.25E+00 | 1.97E-01 | 8.35E-01 | 1.22E+00 | 8.90E-02 | 3.40E-01 | 1.27E+00 | 2.24E-01 | 8.00E-01 | 1.15E+00 | 3.16E-01 | 8.91E-01 | 1.17E+00 | 3.70E-02 | 1.36E-01 | 1.13E+00 | 3.48E-01 | 9.56E-01 | 1.35E+00 |
| TSPY26P | 6.58E-04 | 3.43E-03 | -1.51E+00 | 6.25E-02 | 2.70E-01 | 1.18E+00 | 4.79E-03 | 2.03E-02 | -1.28E+00 | 1.53E-02 | 5.85E-02 | -1.00E+00 | 1.12E-01 | 4.00E-01 | -1.02E+00 | 2.54E-01 | 7.16E-01 | 1.22E+00 | 4.04E-03 | 1.48E-02 | -1.41E+00 | 9.94E-02 | 2.73E-01 | -1.04E+00 |
| CSAG3 | 6.58E-04 | 3.43E-03 | -1.06E+00 | 1.47E-02 | 6.37E-02 | 1.14E+00 | 1.01E-02 | 4.30E-02 | -1.45E+00 | 8.71E-02 | 3.33E-01 | -1.03E+00 | 1.14E-01 | 4.05E-01 | -1.40E+00 | 1.01E-01 | 2.84E-01 | -1.72E+00 | 4.07E-02 | 1.49E-01 | -1.85E+00 | 1.38E-01 | 3.79E-01 | -1.38E+00 |
| GNG7 | 6.59E-04 | 3.43E-03 | -1.01E+00 | 1.15E-05 | 4.97E-05 | -1.04E+00 | 5.38E-02 | 2.29E-01 | 1.07E+00 | 7.93E-03 | 3.03E-02 | 1.10E+00 | 1.27E-05 | 4.54E-05 | -1.24E+00 | 3.00E-07 | 8.44E-07 | -1.43E+00 | 6.27E-02 | 2.30E-01 | 1.02E+00 | 8.19E-03 | 2.25E-02 | -1.20E+00 |
| EVI5 | 6.59E-04 | 3.44E-03 | -1.75E+00 | 4.72E-02 | 2.04E-01 | -1.03E+00 | 1.22E-07 | 5.17E-07 | -2.75E+00 | 9.43E-04 | 3.60E-03 | -1.29E+00 | 4.26E-07 | 1.52E-06 | -1.42E+00 | 8.07E-05 | 2.27E-04 | -1.15E+00 | 1.63E-06 | 5.98E-06 | -1.46E+00 | 2.89E-03 | 7.94E-03 | -1.15E+00 |
| RPL10P1 | 6.60E-04 | 3.44E-03 | -1.02E+00 | 5.42E-04 | 2.34E-03 | 1.06E+00 | 9.14E-02 | 3.88E-01 | 1.01E+00 | 1.77E-01 | 6.75E-01 | 1.47E+00 | 6.62E-02 | 2.36E-01 | 1.16E+00 | 5.81E-02 | 1.64E-01 | 1.16E+00 | 7.02E-03 | 2.58E-02 | -1.18E+00 | 1.86E-01 | 5.10E-01 | 1.19E+00 |
| HERC1 | 6.60E-04 | 3.44E-03 | -1.02E+00 | 1.45E-03 | 6.25E-03 | 1.14E+00 | 8.53E-04 | 3.63E-03 | -1.13E+00 | 1.85E-04 | 7.08E-04 | -1.04E+00 | 1.92E-04 | 6.84E-04 | -1.08E+00 | 3.54E-04 | 9.99E-04 | -1.12E+00 | 5.75E-04 | 2.11E-03 | 1.13E+00 | 1.40E-03 | 3.84E-03 | 1.00E+00 |
| RPL7AP34 | 6.60E-04 | 3.44E-03 | 1.00E+00 | 2.32E-06 | 1.00E-05 | 2.41E+00 | 1.53E-03 | 6.48E-03 | -1.07E+00 | 1.17E-04 | 4.47E-04 | 2.18E+00 | 3.02E-02 | 1.08E-01 | 1.01E+00 | 6.64E-08 | 1.87E-07 | 2.62E+00 | 2.38E-01 | 8.74E-01 | 1.32E+00 | 1.40E-18 | 3.84E-18 | 3.38E+00 |
| AC142472.1 | 6.61E-04 | 3.45E-03 | -1.21E+00 | 5.65E-02 | 2.44E-01 | 1.79E+00 | 1.29E-02 | 5.48E-02 | -1.25E+00 | 1.17E-01 | 4.48E-01 | 1.63E+00 | 3.74E-02 | 1.34E-01 | -1.03E+00 | 1.59E-02 | 4.48E-02 | 2.26E+00 | 2.24E-01 | 8.22E-01 | 1.54E+00 | 5.18E-02 | 1.42E-01 | 1.94E+00 |
| BOLA1 | 6.63E-04 | 3.45E-03 | -1.25E+00 | 8.29E-02 | 3.58E-01 | 1.32E+00 | 1.07E-02 | 4.54E-02 | -1.08E+00 | 1.24E-02 | 4.73E-02 | 1.06E+00 | 5.53E-05 | 1.97E-04 | -1.60E+00 | 6.26E-02 | 1.77E-01 | 1.11E+00 | 1.18E-06 | 4.35E-06 | -1.25E+00 | 3.64E-03 | 9.99E-03 | 1.03E+00 |
| ULK2 | 6.63E-04 | 3.45E-03 | -1.49E+00 | 1.46E-03 | 6.30E-03 | -1.28E+00 | 1.66E-02 | 7.05E-02 | -1.17E+00 | 9.21E-04 | 3.52E-03 | -1.43E+00 | 3.57E-03 | 1.27E-02 | 1.03E+00 | 2.74E-04 | 7.72E-04 | -1.18E+00 | 1.83E-01 | 6.73E-01 | 1.29E+00 | 3.35E-01 | 9.19E-01 | 1.14E+00 |
| CAMKMT | 6.63E-04 | 3.46E-03 | -1.31E+00 | 4.73E-03 | 2.04E-02 | -1.11E+00 | 1.77E-04 | 7.53E-04 | -1.39E+00 | 1.85E-03 | 7.08E-03 | -1.05E+00 | 2.30E-03 | 8.19E-03 | -1.31E+00 | 9.64E-02 | 2.72E-01 | 1.08E+00 | 2.15E-02 | 7.91E-02 | 1.04E+00 | 2.56E-01 | 7.04E-01 | 1.27E+00 |
| UTP4 | 6.66E-04 | 3.47E-03 | -1.25E+00 | 3.50E-02 | 1.51E-01 | 1.19E+00 | 6.42E-05 | 2.73E-04 | -1.33E+00 | 1.15E-02 | 4.40E-02 | 1.04E+00 | 1.79E-05 | 6.37E-05 | -1.52E+00 | 1.44E-01 | 4.06E-01 | 1.30E+00 | 4.16E-06 | 1.53E-05 | -1.48E+00 | 2.78E-01 | 7.65E-01 | 1.10E+00 |
| AC090543.2 | 6.67E-04 | 3.47E-03 | -1.11E+00 | 3.09E-12 | 1.33E-11 | -2.42E+00 | 2.35E-01 | 1.00E+00 | 1.17E+00 | 2.56E-07 | 9.79E-07 | -1.90E+00 | 9.25E-03 | 3.30E-02 | -1.04E+00 | 1.07E-09 | 3.01E-09 | -2.97E+00 | 1.85E-01 | 6.81E-01 | 1.20E+00 | 9.60E-05 | 2.64E-04 | -1.60E+00 |
| ARL16 | 6.67E-04 | 3.48E-03 | 1.06E+00 | 1.38E-02 | 5.95E-02 | 1.36E+00 | 7.80E-02 | 3.32E-01 | 1.13E+00 | 3.41E-02 | 1.30E-01 | 1.26E+00 | 7.18E-06 | 2.56E-05 | -1.10E+00 | 5.28E-03 | 1.49E-02 | 1.13E+00 | 3.33E-02 | 1.22E-01 | 1.14E+00 | 3.00E-01 | 8.23E-01 | 1.36E+00 |
| ZNF577 | 6.68E-04 | 3.48E-03 | -1.27E+00 | 2.40E-02 | 1.04E-01 | 1.14E+00 | 1.12E-03 | 4.75E-03 | -1.30E+00 | 2.30E-02 | 8.78E-02 | 1.03E+00 | 2.31E-01 | 8.25E-01 | 1.57E+00 | 2.16E-01 | 6.10E-01 | 1.28E+00 | 1.33E-01 | 4.90E-01 | 1.56E+00 | 1.77E-01 | 4.87E-01 | 1.54E+00 |
| ANKRD13D | 6.68E-04 | 3.48E-03 | 1.09E+00 | 1.19E-03 | 5.15E-03 | 1.21E+00 | 1.01E-04 | 4.29E-04 | -1.19E+00 | 1.25E-04 | 4.78E-04 | -1.10E+00 | 2.29E-02 | 8.18E-02 | 1.02E+00 | 1.26E-02 | 3.55E-02 | 1.00E+00 | 5.44E-05 | 2.00E-04 | -1.20E+00 | 2.10E-02 | 5.76E-02 | 1.09E+00 |
| PPRC2B | 6.69E-04 | 3.48E-03 | -1.10E+00 | 2.60E-04 | 1.12E-03 | -1.09E+00 | 8.31E-05 | 3.53E-04 | -1.21E+00 | 1.24E-05 | 4.72E-05 | -1.15E+00 | 1.95E-03 | 6.96E-03 | -1.07E+00 | 2.12E-05 | 5.98E-05 | -1.30E+00 | 2.62E-04 | 9.63E-04 | -1.11E+00 | 4.30E-03 | 1.18E-02 | -1.04E+00 |
| LINC00900 | 6.69E-04 | 3.48E-03 | -1.32E+00 | 8.70E-03 | 3.76E-02 | -1.04E+00 | 1.37E-02 | 5.81E-02 | -1.34E+00 | 1.98E-02 | 7.55E-02 | -1.16E+00 | 9.74E-02 | 3.47E-01 | -1.30E+00 | 2.13E-01 | 6.00E-01 | 1.06E+00 | 1.61E-01 | 5.92E-01 | -1.05E+00 | 2.64E-01 | 7.26E-01 | -1.07E+00 |
| RPL39P38 | 6.69E-04 | 3.49E-03 | -1.08E+00 | 1.00E-09 | 4.32E-09 | -1.88E+00 | 1.37E-01 | 5.82E-01 | 1.07E+00 | 7.70E-04 | 2.94E-03 | -1.40E+00 | 1.46E-02 | 5.22E-02 | -1.15E+00 | 4.45E-05 | 1.26E-04 | -2.06E+00 | 1.05E-02 | 3.86E-02 | -1.24E+00 | 2.20E-03 | 6.05E-03 | -1.55E+00 |
| RC3H2 | 6.70E-04 | 3.49E-03 | 1.01E+00 | 1.46E-03 | 6.33E-03 | 1.24E+00 | 2.98E-02 | 1.27E-01 | 1.04E+00 | 1.35E-02 | 5.16E-02 | 1.11E+00 | 1.18E-01 | 4.20E-01 | 1.15E+00 | 9.83E-03 | 2.77E-02 | -1.09E+00 | 6.00E-03 | 2.20E-02 | 1.00E+00 | 1.09E-02 | 3.00E-02 | -1.00E+00 |
| PITPNC1 | 6.71E-04 | 3.50E-03 | 1.03E+00 | 6.04E-04 | 2.61E-03 | 1.14E+00 | 2.72E-04 | 1.16E-03 | -1.26E+00 | 4.60E-06 | 1.76E-05 | -1.29E+00 | 4.03E-02 | 1.44E-01 | 1.12E+00 | 4.28E-03 | 1.21E-02 | -1.02E+00 | 2.53E-01 | 9.29E-01 | 1.00E+00 | 3.52E-01 | 9.66E-01 | 1.01E+00 |
| CDADC1 | 6.71E-04 | 3.50E-03 | 1.22E+00 | 1.22E-04 | 5.27E-04 | 1.24E+00 | 8.86E-03 | 3.76E-02 | -1.03E+00 | 1.52E-03 | 5.80E-03 | 1.02E+00 | 4.79E-02 | 1.71E-01 | 1.24E+00 | 1.40E-03 | 3.95E-03 | -1.09E+00 | 1.77E-03 | 6.51E-03 | -1.04E+00 | 7.84E-03 | 2.15E-02 | -1.02E+00 |
| SLC2A9 | 6.71E-04 | 3.50E-03 | -1.58E+00 | 8.40E-05 | 3.63E-04 | -2.01E+00 | 4.79E-03 | 2.03E-02 | -1.52E+00 | 2.58E-02 | 9.86E-02 | -1.19E+00 | 2.37E-01 | 8.45E-01 | 1.30E+00 | 1.51E-01 | 4.25E-01 | -1.07E+00 | 1.58E-01 | 5.78E-01 | 1.55E+00 | 1.13E-01 | 3.10E-01 | 1.51E+00 |
| ACTG1 | 6.71E-04 | 3.50E-03 | 1.13E+00 | 1.11E-07 | 4.79E-07 | 1.24E+00 | 1.16E-01 | 4.94E-01 | -1.03E+00 | 8.65E-02 | 3.30E-01 | 1.07E+00 | 1.11E-07 | 3.95E-07 | -1.26E+00 | 1.21E-03 | 3.41E-03 | -1.12E+00 | 2.89E-11 | 1.06E-10 | -1.23E+00 | 1.24E-03 | 3.39E-03 | -1.06E+00 |
| UBE2S | 6.72E-04 | 3.50E-03 | 1.99E+00 | 7.26E-03 | 3.14E-02 | 1.14E+00 | 2.16E-04 | 9.17E-04 | 1.82E+00 | 2.51E-01 | 9.57E-01 | 1.36E+00 | 5.57E-02 | 1.99E-01 | 1.69E+00 | 3.55E-01 | 1.00E+00 | 1.37E+00 | 2.30E-02 | 8.45E-02 | 1.58E+00 | 1.05E-01 | 2.89E-01 | 1.64E+00 |
| AC245100.8 | 6.72E-04 | 3.50E-03 | 1.02E+00 | 5.54E-06 | 2.39E-05 | -1.07E+00 | 5.70E-06 | 2.42E-05 | -1.34E+00 | 9.55E-10 | 3.65E-09 | -1.56E+00 | 8.08E-03 | 2.88E-02 | -1.15E+00 | 7.27E-02 | 2.05E-01 | 1.12E+00 | 7.19E-03 | 2.64E-02 | -1.20E+00 | 3.68E-04 | 1.01E-03 | -1.66E+00 |
| AC005831.1 | 6.74E-04 | 3.51E-03 | -1.65E+00 | 7.63E-03 | 3.30E-02 | -1.31E+00 | 6.21E-03 | 2.64E-02 | -1.35E+00 | 3.71E-03 | 1.42E-02 | -1.29E+00 | 1.69E-01 | 6.04E-01 | -1.10E+00 | 5.19E-02 | 1.46E-01 | -1.49E+00 | 1.56E-01 | 5.74E-01 | 1.04E+00 | 2.65E-01 | 7.29E-01 | 1.15E+00 |
| TSR3 | 6.75E-04 | 3.52E-03 | 1.02E+00 | 2.21E-04 | 9.55E-04 | 1.14E+00 | 8.61E-02 | 3.66E-01 | 1.12E+00 | 8.04E-03 | 3.07E-02 | 1.12E+00 | 2.32E-03 | 8.28E-03 | -1.05E+00 | 2.00E-02 | 5.63E-02 | 1.09E+00 | 4.24E-03 | 1.56E-02 | -1.00E+00 | 2.10E-01 | 5.77E-01 | 1.20E+00 |
| FAM173A | 6.75E-04 | 3.52E-03 | -1.19E+00 | 5.45E-06 | 2.36E-05 | -1.34E+00 | 7.32E-04 | 3.11E-03 | -1.12E+00 | 3.33E-08 | 1.27E-07 | -1.30E+00 | 1.80E-04 | 6.42E-04 | -1.10E+00 | 3.63E-04 | 1.02E-03 | -1.10E+00 | 2.66E-01 | 9.77E-01 | 1.21E+00 | 1.81E-01 | 4.96E-01 | 1.20E+00 |
| TMEM64 | 6.76E-04 | 3.52E-03 | -1.17E+00 | 4.26E-04 | 1.84E-03 | -1.12E+00 | 9.16E-03 | 3.89E-02 | -1.13E+00 | 2.61E-04 | 9.97E-04 | -1.30E+00 | 1.65E-01 | 5.90E-01 | 1.24E+00 | 8.08E-02 | 2.28E-01 | 1.14E+00 | 4.89E-03 | 1.80E-02 | 1.07E+00 | 5.61E-05 | 1.54E-04 | -1.34E+00 |
| BTF3 | 6.77E-04 | 3.53E-03 | 1.08E+00 | 1.79E-06 | 7.73E-06 | 1.12E+00 | 7.02E-05 | 2.98E-04 | 1.11E+00 | 2.70E-07 | 1.03E-06 | 1.16E+00 | 5.04E-02 | 1.80E-01 | 1.07E+00 | 9.20E-03 | 2.59E-02 | 1.09E+00 | 1.56E-01 | 5.74E-01 | 1.08E+00 | 1.86E-03 | 5.10E-03 | 1.12E+00 |
| TECPR1 | 6.78E-04 | 3.53E-03 | -1.27E+00 | 1.16E-02 | 5.03E-02 | 1.08E+00 | 4.60E-02 | 1.95E-01 | 1.02E+00 | 2.41E-02 | 9.20E-02 | -1.03E+00 | 2.17E-03 | 7.75E-03 | -1.30E+00 | 4.33E-02 | 1.22E-01 | 1.12E+00 | 4.64E-09 | 1.70E-08 | -1.76E+00 | 1.15E-04 | 3.16E-04 | -1.17E+00 |
| RRAGB | 6.79E-04 | 3.54E-03 | -1.04E+00 | 4.80E-03 | 2.07E-02 | 1.12E+00 | 7.63E-04 | 3.24E-03 | -1.23E+00 | 2.95E-03 | 1.13E-02 | 1.02E+00 | 6.55E-02 | 2.34E-01 | 1.16E+00 | 4.82E-02 | 1.36E-01 | 1.07E+00 | 4.57E-02 | 1.68E-01 | 1.12E+00 | 2.54E-01 | 6.99E-01 | 1.28E+00 |
| DTX2 | 6.80E-04 | 3.54E-03 | -1.44E+00 | 2.87E-02 | 1.24E-01 | 1.09E+00 | 2.06E-04 | 8.76E-04 | -1.64E+00 | 3.43E-03 | 1.31E-02 | -1.19E+00 | 9.39E-02 | 3.35E-01 | -1.09E+00 | 2.63E-01 | 7.42E-01 | 1.48E+00 | 5.56E-08 | 2.04E-07 | -2.15E+00 | 9.08E-04 | 2.49E-03 | -1.30E+00 |
| ZFP28 | 6.80E-04 | 3.54E-03 | -1.59E+00 | 5.19E-03 | 2.24E-02 | -1.21E+00 | 1.09E-04 | 4.62E-04 | -1.57E+00 | 6.81E-04 | 2.60E-03 | -1.28E+00 | 1.97E-01 | 7.02E-01 | 1.01E+00 | 1.41E-01 | 3.97E-01 | -1.14E+00 | 2.09E-04 | 7.66E-04 | -1.46E+00 | 2.40E-03 | 6.60E-03 | -1.20E+00 |
| AC092070.2 | 6.82E-04 | 3.55E-03 | -1.40E+00 | 1.47E-02 | 6.37E-02 | 1.06E+00 | 1.79E-02 | 7.62E-02 | -1.30E+00 | 8.61E-02 | 3.29E-01 | 1.04E+00 | 1.73E-01 | 6.15E-01 | 1.13E+00 | 1.08E-01 | 3.05E-01 | 1.04E+00 | 1.36E-01 | 4.99E-01 | -1.07E+00 | 3.40E-01 | 9.35E-01 | 1.23E+00 |
| SSX2IP | 6.82E-04 | 3.56E-03 | -1.24E+00 | 2.42E-03 | 1.04E-02 | -1.01E+00 | 1.29E-03 | 5.48E-03 | -1.33E+00 | 2.58E-02 | 9.86E-02 | 1.04E+00 | 2.13E-01 | 7.58E-01 | 1.43E+00 | 3.20E-01 | 9.02E-01 | 1.49E+00 | 1.77E-02 | 6.48E-02 | 1.44E+00 | 2.47E-01 | 6.79E-01 | 1.18E+00 |
| MED16 | 6.82E-04 | 3.56E-03 | -1.06E+00 | 3.39E-02 | 1.46E-01 | 1.30E+00 | 1.79E-04 | 7.60E-04 | -1.29E+00 | 4.42E-03 | 1.69E-02 | 1.01E+00 | 2.49E-03 | 8.87E-03 | -1.10E+00 | 2.91E-01 | 8.20E-01 | 1.35E+00 | 3.53E-05 | 1.30E-04 | -1.13E+00 | 1.15E-03 | 3.16E-03 | -1.08E+00 |
| HSPA14 | 6.83E-04 | 3.56E-03 | -1.14E+00 | 4.50E-04 | 1.94E-03 | -1.08E+00 | 7.48E-08 | 3.18E-07 | -1.34E+00 | 1.49E-11 | 5.67E-11 | -1.43E+00 | 1.05E-03 | 3.74E-03 | -1.38E+00 | 4.63E-03 | 1.31E-02 | -1.20E+00 | 8.87E-04 | 3.26E-03 | -1.08E+00 | 1.54E-03 | 4.23E-03 | -1.05E+00 |
| TMEM164 | 6.86E-04 | 3.57E-03 | -1.24E+00 | 8.17E-03 | 3.53E-02 | 1.08E+00 | 1.16E-01 | 4.91E-01 | 1.13E+00 | 1.78E-01 | 6.78E-01 | 1.30E+00 | 1.17E-01 | 4.16E-01 | 1.34E+00 | 3.46E-01 | 9.76E-01 | 1.48E+00 | 2.15E-02 | 7.88E-02 | -1.02E+00 | 2.63E-01 | 7.23E-01 | 1.08E+00 |
| BX088651.4 | 6.87E-04 | 3.58E-03 | -1.25E+00 | 1.63E-02 | 7.06E-02 | 1.19E+00 | 1.98E-04 | 8.43E-04 | -1.51E+00 | 2.69E-02 | 1.03E-01 | 1.00E+00 | 3.39E-03 | 1.21E-02 | -1.22E+00 | 2.72E-01 | 7.68E-01 | 1.36E+00 | 1.82E-04 | 6.68E-04 | -1.31E+00 | 1.22E-01 | 3.36E-01 | 1.13E+00 |
| LINC01684 | 6.88E-04 | 3.59E-03 | -1.18E+00 | 1.94E-01 | 8.37E-01 | 1.44E+00 | 3.97E-02 | 1.69E-01 | -1.03E+00 | 2.06E-01 | 7.86E-01 | 1.33E+00 | 2.74E-03 | 9.78E-03 | -1.17E+00 | 1.07E-01 | 3.00E-01 | 1.23E+00 | 5.55E-02 | 2.04E-01 | -1.01E+00 | 3.44E-01 | 9.45E-01 | 1.26E+00 |
| PLK3 | 6.88E-04 | 3.59E-03 | 2.03E+00 | 3.04E-04 | 1.31E-03 | -1.12E+00 | 1.18E-01 | 5.02E-01 | 1.36E+00 | 1.12E-04 | 4.30E-04 | -1.33E+00 | 1.84E-02 | 6.55E-02 | 1.76E+00 | 1.95E-03 | 5.50E-03 | -1.17E+00 | 1.47E-02 | 5.40E-02 | 1.85E+00 | 1.29E-02 | 3.55E-02 | -1.17E+00 |
| ACTG1P20 | 6.88E-04 | 3.59E-03 | -1.33E+00 | 5.71E-04 | 2.47E-03 | -1.27E+00 | 1.29E-12 | 5.48E-12 | -2.33E+00 | 8.08E-14 | 3.09E-13 | -2.31E+00 | 1.99E-04 | 7.09E-04 | -1.66E+00 | 4.98E-05 | 1.40E-04 | -1.92E+00 | 7.02E-02 | 2.58E-01 | -1.27E+00 | 6.03E-02 | 1.66E-01 | -1.57E+00 |
| HSF5 | 6.88E-04 | 3.59E-03 | -1.61E+00 | 8.20E-03 | 3.54E-02 | -1.15E+00 | 2.81E-04 | 1.20E-03 | -1.85E+00 | 5.50E-02 | 2.10E-01 | -1.04E+00 | 2.91E-02 | 1.04E-01 | -1.41E+00 | 2.44E-02 | 6.87E-02 | -1.46E+00 | 1.80E-01 | 6.62E-01 | -1.05E+00 | 2.24E-01 | 6.16E-01 | -1.00E+00 |
| WDR78 | 6.89E-04 | 3.59E-03 | -1.49E+00 | 5.86E-04 | 2.53E-03 | -1.45E+00 | 8.31E-03 | 3.53E-02 | -1.26E+00 | 1.30E-03 | 4.95E-03 | -1.35E+00 | 3.98E-05 | 1.42E-04 | -2.36E+00 | 3.70E-04 | 1.04E-03 | -1.63E+00 | 3.05E-02 | 1.12E-01 | -1.14E+00 | 3.26E-01 | 8.95E-01 | 1.23E+00 |
| RNF7 | 6.89E-04 | 3.59E-03 | 1.03E+00 | 1.54E-05 | 6.64E-05 | 1.05E+00 | 1.11E-02 | 4.71E-02 | -1.10E+00 | 4.59E-05 | 1.75E-04 | -1.15E+00 | 1.26E-03 | 4.48E-03 | -1.02E+00 | 1.08E-06 | 3.03E-06 | -1.14E+00 | 6.44E-03 | 2.37E-02 | 1.01E+00 | 1.82E-04 | 5.01E-04 | -1.06E+00 |
| MSANTD3 | 6.90E-04 | 3.59E-03 | -1.06E+00 | 2.55E-04 | 1.10E-03 | -1.03E+00 | 2.14E-04 | 9.09E-04 | -1.38E+00 | 8.45E-05 | 3.23E-04 | -1.21E+00 | 4.03E-04 | 1.44E-03 | -1.19E+00 | 5.14E-05 | 1.45E-04 | -1.28E+00 | 5.57E-03 | 2.04E-02 | 1.01E+00 | 1.81E-03 | 4.98E-03 | -1.02E+00 |
| HMBS | 6.92E-04 | 3.61E-03 | -1.14E+00 | 6.27E-04 | 2.71E-03 | -1.01E+00 | 1.73E-04 | 7.37E-04 | -1.15E+00 | 2.36E-07 | 9.00E-07 | -1.25E+00 | 6.94E-11 | 2.47E-10 | -1.54E+00 | 1.10E-02 | 3.09E-02 | 1.14E+00 | 1.91E-06 | 7.01E-06 | -1.18E+00 | 1.38E-03 | 3.80E-03 | -1.08E+00 |
| CDKN1C | 6.93E-04 | 3.61E-03 | -1.07E+00 | 3.29E-04 | 1.42E-03 | 1.02E+00 | 7.93E-05 | 3.37E-04 | -1.35E+00 | 1.16E-04 | 4.42E-04 | -1.21E+00 | 1.15E-02 | 4.11E-02 | 1.05E+00 | 4.80E-04 | 1.35E-03 | -1.18E+00 | 3.41E-02 | 1.25E-01 | 2.13E+00 | 1.49E-01 | 4.10E-01 | 1.88E+00 |
| STK36 | 6.94E-04 | 3.62E-03 | -1.70E+00 | 2.31E-02 | 9.98E-02 | -1.10E+00 | 5.05E-06 | 2.15E-05 | -1.74E+00 | 9.10E-05 | 3.48E-04 | -1.27E+00 | 2.51E-02 | 8.96E-02 | -1.50E+00 | 9.77E-02 | 2.75E-01 | -1.08E+00 | 2.50E-01 | 9.16E-01 | 1.20E+00 | 3.27E-01 | 8.98E-01 | 1.19E+00 |

| | | | | | | | | | | | | | | | | | | | | | | | | |
|---|---|---|---|---|---|---|---|---|---|---|---|---|---|---|---|---|---|---|---|---|---|---|---|---|
| GVINP1 | 6.97E-04 | 3.63E-03 | -1.34E+00 | 1.22E-01 | 5.26E-01 | 1.48E+00 | 6.38E-04 | 2.71E-03 | -1.58E+00 | 1.56E-02 | 5.97E-02 | -1.06E+00 | 2.61E-02 | 9.30E-02 | -2.15E+00 | 4.65E-02 | 1.31E-01 | -1.93E+00 | 6.24E-04 | 2.29E-03 | -1.99E+00 | 3.26E-03 | 8.95E-03 | -1.87E+00 |
| POLR1A | 6.97E-04 | 3.63E-03 | -1.33E+00 | 2.73E-02 | 1.18E-01 | 1.12E+00 | 3.54E-02 | 1.50E-01 | 1.05E+00 | 2.33E-01 | 8.90E-01 | 1.39E+00 | 1.25E-05 | 4.46E-05 | -1.70E+00 | 2.11E-03 | 5.94E-03 | -1.22E+00 | 9.89E-05 | 3.63E-04 | -1.23E+00 | 2.79E-01 | 7.66E-01 | 1.18E+00 |
| SAT1 | 6.98E-04 | 3.64E-03 | 1.75E+00 | 2.41E-05 | 1.04E-04 | 1.28E+00 | 2.87E-03 | 1.22E-02 | 1.76E+00 | 1.50E-02 | 5.75E-02 | 1.41E+00 | 4.40E-02 | 1.57E-01 | 1.57E+00 | 2.46E-01 | 6.94E-01 | 1.44E+00 | 5.43E-02 | 1.99E-01 | 1.58E+00 | 2.18E-01 | 5.98E-01 | 1.54E+00 |
| NPTN-IT1 | 7.02E-04 | 3.66E-03 | -1.39E+00 | 1.77E-03 | 7.64E-03 | -1.23E+00 | 1.01E-01 | 4.30E-01 | -1.06E+00 | 1.75E-01 | 6.68E-01 | 1.11E+00 | 2.60E-01 | 9.26E-01 | 1.27E+00 | 3.55E-01 | 1.00E+00 | 1.35E+00 | 1.23E-01 | 4.52E-01 | -1.06E+00 | 5.04E-02 | 1.38E-01 | -1.27E+00 |
| DHFR2 | 7.03E-04 | 3.66E-03 | -1.63E+00 | 1.37E-02 | 5.93E-02 | -1.08E+00 | 5.20E-06 | 2.21E-05 | -1.81E+00 | 4.32E-04 | 1.65E-03 | -1.32E+00 | 6.76E-04 | 2.41E-03 | -2.22E+00 | 6.97E-02 | 1.97E-01 | -1.08E+00 | 1.62E-03 | 5.95E-03 | -1.40E+00 | 4.18E-02 | 1.15E-01 | -1.06E+00 |
| CWC15 | 7.03E-04 | 3.67E-03 | 1.08E+00 | 2.45E-05 | 1.06E-04 | 1.06E+00 | 2.25E-03 | 9.56E-03 | -1.09E+00 | 4.21E-04 | 1.61E-03 | -1.06E+00 | 2.21E-03 | 7.88E-03 | 1.01E+00 | 1.18E-02 | 3.33E-02 | 1.12E+00 | 5.56E-02 | 2.04E-01 | 1.03E+00 | 9.53E-03 | 2.62E-02 | -1.05E+00 |
| RPL4P2 | 7.04E-04 | 3.67E-03 | -1.04E+00 | 4.10E-12 | 1.77E-11 | -1.73E+00 | 2.95E-03 | 1.25E-02 | -1.06E+00 | 1.18E-11 | 4.51E-11 | -1.63E+00 | 2.87E-04 | 1.02E-03 | -1.15E+00 | 1.13E-08 | 3.20E-08 | -1.73E+00 | 1.78E-08 | 6.52E-08 | -1.40E+00 | 3.81E-13 | 1.05E-12 | -1.66E+00 |
| FAM200A | 7.04E-04 | 3.67E-03 | -1.47E+00 | 1.11E-02 | 4.78E-02 | -1.03E+00 | 1.56E-03 | 6.64E-03 | -1.58E+00 | 4.09E-03 | 1.56E-02 | -1.45E+00 | 2.85E-04 | 1.02E-03 | -2.22E+00 | 3.07E-02 | 8.65E-02 | -1.03E+00 | 5.36E-06 | 1.97E-05 | -1.51E+00 | 2.36E-05 | 6.49E-05 | -1.40E+00 |
| LCA5 | 7.05E-04 | 3.67E-03 | -1.56E+00 | 1.66E-02 | 7.17E-02 | -1.00E+00 | 3.06E-02 | 1.30E-01 | -1.37E+00 | 4.06E-02 | 1.55E-01 | -1.27E+00 | 1.13E-01 | 4.03E-01 | -1.08E+00 | 3.55E-01 | 9.99E-01 | 1.18E+00 | 6.52E-03 | 2.39E-02 | 1.24E+00 | 8.47E-07 | 2.33E-06 | -1.28E+00 |
| NEDD9 | 7.05E-04 | 3.68E-03 | -1.96E+00 | 3.96E-03 | 1.71E-02 | -1.51E+00 | 9.37E-03 | 3.98E-02 | -1.40E+00 | 5.86E-03 | 2.24E-02 | -1.39E+00 | 3.18E-05 | 1.13E-04 | -1.40E+00 | 1.48E-04 | 4.16E-04 | -1.26E+00 | 2.60E-01 | 9.56E-01 | -1.05E+00 | 2.53E-01 | 6.94E-01 | -1.06E+00 |
| TOLLIP | 7.06E-04 | 3.68E-03 | -1.04E+00 | 4.17E-05 | 1.80E-04 | -1.06E+00 | 1.87E-05 | 7.97E-05 | -1.34E+00 | 1.34E-05 | 5.11E-05 | -1.12E+00 | 7.55E-06 | 2.69E-05 | -1.44E+00 | 7.15E-04 | 2.02E-03 | -1.08E+00 | 3.77E-03 | 1.38E-02 | 1.06E+00 | 5.92E-04 | 1.63E-03 | -1.00E+00 |
| ZAP70 | 7.07E-04 | 3.69E-03 | 2.66E+00 | 3.27E-02 | 1.41E-01 | 2.21E+00 | 1.63E-01 | 6.92E-01 | -1.30E+00 | 2.23E-01 | 8.50E-01 | -1.39E+00 | 2.20E-01 | 7.85E-01 | 1.32E+00 | 2.73E-01 | 7.70E-01 | 1.35E+00 | 1.72E-01 | 6.30E-01 | 1.21E+00 | 3.64E-01 | 1.00E+00 | -1.00E+00 |
| ST6GALNAC4 | 7.08E-04 | 3.69E-03 | -1.06E+00 | 1.04E-02 | 4.48E-02 | 1.19E+00 | 7.50E-04 | 3.19E-03 | -1.17E+00 | 2.26E-02 | 8.65E-02 | 1.15E+00 | 1.37E-08 | 4.89E-08 | -1.42E+00 | 8.86E-05 | 2.50E-04 | -1.09E+00 | 6.35E-06 | 2.33E-05 | -1.40E+00 | 1.12E-02 | 3.09E-02 | -1.09E+00 |
| AKAP8L | 7.09E-04 | 3.69E-03 | -1.15E+00 | 2.05E-05 | 8.84E-05 | -1.16E+00 | 1.08E-04 | 4.60E-04 | -1.25E+00 | 1.56E-07 | 5.96E-07 | -1.41E+00 | 9.69E-05 | 3.45E-04 | -1.19E+00 | 9.88E-08 | 2.78E-07 | -1.58E+00 | 4.59E-02 | 1.68E-01 | 1.09E+00 | 6.02E-03 | 1.65E-02 | -1.08E+00 |
| CETN2 | 7.09E-04 | 3.69E-03 | -1.05E+00 | 1.03E-03 | 4.44E-03 | 1.10E+00 | 7.24E-03 | 3.08E-02 | -1.05E+00 | 3.24E-04 | 1.24E-03 | 1.01E+00 | 2.95E-03 | 1.05E-02 | -1.18E+00 | 2.02E-01 | 5.70E-01 | 1.27E+00 | 1.30E-01 | 4.76E-01 | 1.27E+00 | 3.47E-01 | 9.53E-01 | 1.34E+00 |
| PABPC1L | 7.11E-04 | 3.70E-03 | -1.09E+00 | 2.37E-03 | 1.02E-02 | 1.09E+00 | 7.06E-04 | 3.00E-03 | -1.30E+00 | 6.37E-03 | 2.43E-02 | -1.05E+00 | 2.04E-01 | 7.28E-01 | 1.06E+00 | 2.17E-01 | 6.12E-01 | -1.09E+00 | 7.09E-02 | 2.60E-01 | -1.04E+00 | 1.57E-01 | 4.32E-01 | 1.14E+00 |
| CCDC58P1 | 7.11E-04 | 3.71E-03 | -1.56E+00 | 5.53E-03 | 2.39E-02 | -1.16E+00 | 1.12E-07 | 4.76E-07 | -1.82E+00 | 7.04E-09 | 2.69E-08 | -1.85E+00 | 2.27E-05 | 8.10E-05 | -3.13E+00 | 3.26E-04 | 9.19E-04 | -2.17E+00 | 1.03E-12 | 3.76E-12 | -2.09E+00 | 1.72E-08 | 4.72E-08 | -1.53E+00 |
| C6orf120 | 7.12E-04 | 3.71E-03 | -1.20E+00 | 1.30E-06 | 5.61E-06 | -1.42E+00 | 3.52E-07 | 1.50E-06 | -1.34E+00 | 4.37E-08 | 1.67E-07 | -1.39E+00 | 2.69E-02 | 9.57E-02 | -1.00E+00 | 7.06E-02 | 1.99E-01 | 1.15E+00 | 1.92E-04 | 7.04E-04 | -1.29E+00 | 4.38E-02 | 1.20E-01 | 1.01E+00 |
| AC215219.1 | 7.13E-04 | 3.71E-03 | -1.26E+00 | 2.22E-03 | 9.61E-03 | -1.17E+00 | 1.41E-03 | 5.98E-03 | -1.21E+00 | 4.84E-03 | 1.85E-02 | 1.03E+00 | 6.45E-03 | 2.30E-02 | -1.16E+00 | 3.95E-03 | 1.11E-02 | -1.25E+00 | 1.16E-02 | 4.26E-02 | -1.16E+00 | 9.48E-02 | 2.60E-01 | 1.13E+00 |
| PGAM1P6 | 7.13E-04 | 3.71E-03 | -1.66E+00 | 3.36E-04 | 1.45E-03 | -1.74E+00 | 1.79E-01 | 7.61E-01 | 1.16E+00 | 6.02E-02 | 2.30E-01 | -1.13E+00 | 1.30E-01 | 4.63E-01 | 1.06E+00 | 1.25E-03 | 3.53E-03 | -2.14E+00 | 2.38E-02 | 8.74E-02 | -1.24E+00 | 1.74E-02 | 4.79E-02 | -1.39E+00 |
| D2HGDH | 7.13E-04 | 3.71E-03 | -1.07E+00 | 1.57E-06 | 6.80E-06 | -1.25E+00 | 2.15E-01 | 9.15E-01 | 1.26E+00 | 1.34E-02 | 5.13E-02 | -1.03E+00 | 3.94E-02 | 1.40E-01 | 1.01E+00 | 1.23E-02 | 3.47E-02 | -1.15E+00 | 1.35E-01 | 4.95E-01 | 1.18E+00 | 9.88E-03 | 2.71E-02 | -1.07E+00 |
| PAPLN | 7.13E-04 | 3.72E-03 | -1.63E+00 | 1.32E-02 | 5.68E-02 | -1.08E+00 | 2.67E-03 | 1.13E-02 | -1.67E+00 | 1.15E-01 | 4.40E-01 | 1.11E+00 | 5.37E-02 | 1.92E-01 | -1.13E+00 | 1.81E-01 | 5.09E-01 | 1.10E+00 | 3.37E-04 | 1.24E-03 | -1.41E+00 | 5.96E-02 | 1.64E-01 | 1.01E+00 |
| CEP295 | 7.16E-04 | 3.73E-03 | -1.15E+00 | 1.74E-03 | 7.52E-03 | -1.00E+00 | 6.46E-04 | 2.75E-03 | -1.19E+00 | 5.60E-05 | 2.14E-04 | -1.21E+00 | 6.28E-03 | 2.24E-02 | -1.08E+00 | 9.96E-04 | 2.81E-03 | -1.30E+00 | 6.41E-03 | 2.35E-02 | 1.13E+00 | 1.34E-03 | 3.68E-03 | -1.03E+00 |
| AL096829.2 | 7.19E-04 | 3.75E-03 | -1.25E+00 | 1.06E-05 | 4.57E-05 | -1.82E+00 | 5.93E-05 | 2.52E-04 | -1.61E+00 | 8.73E-06 | 3.34E-05 | -1.57E+00 | 2.77E-02 | 9.88E-02 | -1.27E+00 | 1.29E-02 | 3.63E-02 | -1.37E+00 | 1.24E-04 | 4.57E-04 | -1.29E+00 | 2.66E-03 | 7.30E-03 | -1.17E+00 |
| AC011477.1 | 7.22E-04 | 3.76E-03 | -1.36E+00 | 3.30E-02 | 1.43E-01 | 1.06E+00 | 1.44E-04 | 6.10E-04 | -1.62E+00 | 9.42E-04 | 3.60E-03 | -1.30E+00 | 2.64E-01 | 9.42E-01 | 1.12E+00 | 1.11E-01 | 3.14E-01 | 1.82E+00 | 9.89E-02 | 3.63E-01 | -1.09E+00 | 2.53E-01 | 6.95E-01 | 1.44E+00 |
| ADAT2 | 7.23E-04 | 3.77E-03 | -1.15E+00 | 1.13E-04 | 4.88E-04 | -1.15E+00 | 1.13E-03 | 4.81E-03 | -1.26E+00 | 1.91E-03 | 7.29E-03 | -1.07E+00 | 2.67E-02 | 9.52E-02 | -1.02E+00 | 9.64E-03 | 2.72E-02 | -1.08E+00 | 8.89E-02 | 3.26E-01 | 1.09E+00 | 2.87E-01 | 7.88E-01 | 1.25E+00 |
| DHDDS | 7.24E-04 | 3.77E-03 | -1.15E+00 | 4.26E-03 | 1.84E-02 | 1.08E+00 | 1.07E-02 | 4.55E-02 | -1.09E+00 | 2.32E-01 | 8.88E-01 | 1.40E+00 | 7.46E-04 | 2.66E-03 | -1.23E+00 | 3.29E-02 | 9.28E-02 | 1.03E+00 | 3.46E-02 | 1.27E-01 | 1.07E+00 | 6.80E-02 | 1.87E-01 | 1.13E+00 |
| LINC00888 | 7.24E-04 | 3.77E-03 | -1.09E+00 | 7.12E-03 | 3.08E-02 | 1.16E+00 | 4.34E-06 | 1.84E-05 | -1.44E+00 | 1.88E-04 | 7.17E-04 | -1.11E+00 | 5.31E-03 | 1.89E-02 | -1.22E+00 | 4.06E-02 | 1.14E-01 | 1.06E+00 | 1.79E-02 | 6.56E-02 | 1.25E+00 | 3.15E-03 | 8.64E-03 | -1.18E+00 |
| AL031220.1 | 7.24E-04 | 3.77E-03 | -1.26E+00 | 5.24E-05 | 2.26E-04 | -1.48E+00 | 3.04E-02 | 1.29E-01 | -1.02E+00 | 7.90E-07 | 3.02E-06 | -1.89E+00 | 4.36E-03 | 1.55E-02 | -1.43E+00 | 8.37E-04 | 2.36E-03 | -1.62E+00 | 1.86E-01 | 6.83E-01 | 1.31E+00 | 1.63E-02 | 4.48E-02 | -1.12E+00 |
| PANK3 | 7.26E-04 | 3.78E-03 | -1.05E+00 | 1.41E-03 | 6.09E-03 | 1.09E+00 | 3.87E-04 | 1.64E-03 | -1.20E+00 | 1.94E-02 | 7.42E-02 | 1.14E+00 | 5.54E-05 | 1.98E-04 | -1.13E+00 | 3.55E-04 | 1.00E-03 | -1.13E+00 | 5.25E-07 | 1.93E-06 | -1.41E+00 | 4.28E-05 | 1.17E-04 | -1.16E+00 |
| GDN4L | 7.26E-04 | 3.78E-03 | 1.00E+00 | 2.09E-02 | 9.05E-02 | 1.31E+00 | 1.13E-02 | 4.80E-02 | -1.03E+00 | 3.72E-02 | 1.42E-01 | 1.19E+00 | 1.44E-02 | 5.13E-02 | 1.11E+00 | 1.32E-01 | 3.71E-01 | 1.26E+00 | 2.87E-05 | 1.05E-04 | -1.21E+00 | 4.30E-02 | 1.18E-01 | -1.03E+00 |
| FAM41C | 7.27E-04 | 3.79E-03 | -1.28E+00 | 9.17E-06 | 3.96E-05 | -1.57E+00 | 3.61E-02 | 1.53E-01 | -1.04E+00 | 7.96E-06 | 3.04E-05 | -1.60E+00 | 1.82E-03 | 6.48E-03 | -1.15E+00 | 7.60E-06 | 2.14E-05 | -2.03E+00 | 9.09E-03 | 3.34E-02 | -1.24E+00 | 1.46E-03 | 4.02E-03 | -1.55E+00 |
| AL050341.2 | 7.32E-04 | 3.82E-03 | -1.05E+00 | 2.42E-02 | 1.04E-01 | 1.22E+00 | 1.95E-03 | 8.28E-03 | -1.17E+00 | 5.58E-03 | 2.13E-02 | 1.04E+00 | 1.12E-03 | 4.00E-03 | -1.09E+00 | 6.11E-02 | 1.72E-01 | 1.23E+00 | 2.67E-02 | 9.81E-02 | 1.07E+00 | 1.34E-01 | 3.67E-01 | 1.10E+00 |
| DNAAF5 | 7.32E-04 | 3.82E-03 | -1.31E+00 | 3.37E-03 | 1.46E-02 | -1.05E+00 | 1.50E-02 | 6.36E-02 | -1.03E+00 | 2.39E-01 | 9.14E-01 | 1.52E+00 | 2.91E-02 | 1.04E-01 | -1.16E+00 | 3.48E-01 | 9.80E-01 | 1.51E+00 | 2.92E-04 | 1.07E-03 | -1.43E+00 | 3.56E-01 | 9.78E-01 | 1.20E+00 |
| MGST3 | 7.34E-04 | 3.83E-03 | -1.02E+00 | 3.22E-02 | 1.39E-01 | 1.19E+00 | 2.18E-01 | 9.28E-01 | 1.14E+00 | 2.26E-01 | 8.63E-01 | 1.19E+00 | 3.65E-09 | 1.30E-08 | -1.36E+00 | 1.47E-08 | 4.15E-08 | -1.23E+00 | 5.73E-07 | 2.10E-06 | -1.27E+00 | 9.59E-04 | 2.63E-03 | -1.02E+00 |
| MIO808 | 7.34E-04 | 3.83E-03 | 3.00E+00 | 2.03E-03 | 8.77E-03 | 3.22E+00 | 3.84E-02 | 1.63E-01 | 1.78E+00 | 1.72E-01 | 6.56E-01 | 1.70E+00 | 4.95E-03 | 1.77E-02 | 2.18E+00 | 2.67E-01 | 7.51E-01 | 1.65E+00 | 5.52E-03 | 2.02E-02 | 1.85E+00 | 2.80E-02 | 7.70E-02 | 2.00E+00 |
| CUL7 | 7.34E-04 | 3.83E-03 | -1.28E+00 | 4.17E-03 | 1.80E-02 | -1.03E+00 | 1.65E-01 | 7.03E-01 | 1.07E+00 | 1.02E-01 | 3.89E-01 | 1.09E+00 | 1.80E-01 | 6.42E-01 | 1.22E+00 | 2.34E-01 | 6.59E-01 | 1.28E+00 | 2.13E-02 | 7.81E-02 | -1.12E+00 | 1.10E-02 | 3.03E-02 | -1.35E+00 |
| FASTKD3 | 7.35E-04 | 3.83E-03 | -1.43E+00 | 1.73E-02 | 7.46E-02 | 1.00E+00 | 1.98E-08 | 8.41E-08 | -1.79E+00 | 3.08E-07 | 1.17E-06 | -1.43E+00 | 2.52E-01 | 8.98E-01 | 1.47E+00 | 3.54E-01 | 9.97E-01 | 1.48E+00 | 3.23E-06 | 1.18E-05 | -1.31E+00 | 3.38E-03 | 9.27E-03 | -1.00E+00 |
| PSMD10 | 7.35E-04 | 3.83E-03 | -1.20E+00 | 5.82E-03 | 2.52E-02 | 1.03E+00 | 7.99E-03 | 3.39E-02 | -1.06E+00 | 1.97E-02 | 7.54E-02 | 1.17E+00 | 2.43E-06 | 8.67E-06 | -1.50E+00 | 6.37E-02 | 1.80E-01 | 1.02E+00 | 2.63E-08 | 9.64E-08 | -1.38E+00 | 5.75E-04 | 1.58E-03 | -1.06E+00 |
| ZMIZ2 | 7.37E-04 | 3.84E-03 | -1.12E+00 | 7.15E-03 | 3.09E-02 | 1.16E+00 | 1.68E-02 | 7.12E-02 | -1.15E+00 | 1.27E-01 | 4.86E-01 | 1.24E+00 | 3.55E-03 | 1.26E-02 | -1.19E+00 | 1.71E-02 | 4.81E-02 | -1.06E+00 | 8.12E-04 | 2.98E-03 | -1.22E+00 | 6.37E-03 | 1.75E-02 | -1.17E+00 |
| ASCC2 | 7.40E-04 | 3.86E-03 | -1.02E+00 | 7.43E-02 | 3.21E-01 | 1.40E+00 | 1.34E-03 | 5.68E-03 | -1.16E+00 | 7.25E-02 | 2.77E-01 | 1.26E+00 | 1.30E-03 | 4.62E-03 | -1.12E+00 | 2.70E-01 | 7.62E-01 | 1.40E+00 | 2.61E-07 | 9.58E-07 | -1.33E+00 | 7.75E-03 | 2.13E-02 | -1.04E+00 |
| GPR160 | 7.42E-04 | 3.87E-03 | -1.25E+00 | 9.22E-04 | 3.98E-03 | -1.23E+00 | 4.48E-03 | 1.90E-02 | -1.09E+00 | 7.83E-05 | 2.99E-04 | -1.29E+00 | 8.27E-02 | 2.95E-01 | -1.14E+00 | 1.71E-01 | 4.81E-01 | -1.01E+00 | 3.17E-04 | 1.16E-03 | -1.37E+00 | 2.74E-04 | 7.51E-04 | -1.59E+00 |
| ZDHHC15 | 7.42E-04 | 3.87E-03 | -1.50E+00 | 3.77E-02 | 1.63E-01 | 1.17E+00 | 2.23E-01 | 9.50E-01 | 1.02E+00 | 1.70E-01 | 6.48E-01 | 1.39E+00 | 6.09E-03 | 2.17E-02 | -1.61E+00 | 4.61E-02 | 1.30E-01 | -1.12E+00 | 6.94E-02 | 2.55E-01 | 1.05E+00 | 3.03E-02 | 8.33E-02 | -1.03E+00 |
| SNX12 | 7.44E-04 | 3.88E-03 | -1.38E+00 | 1.05E-03 | 4.54E-03 | -1.22E+00 | 1.37E-01 | 5.80E-01 | 1.16E+00 | 1.35E-01 | 5.16E-01 | 1.30E+00 | 3.09E-02 | 1.10E-01 | -1.08E+00 | 1.34E-01 | 3.78E-01 | 1.15E+00 | 3.86E-04 | 1.42E-03 | -1.25E+00 | 1.52E-02 | 4.18E-02 | -1.04E+00 |
| PARP2 | 7.45E-04 | 3.89E-03 | -1.13E+00 | 3.22E-03 | 1.39E-02 | 1.09E+00 | 2.73E-02 | 1.16E-01 | -1.01E+00 | 1.32E-01 | 5.02E-01 | 1.28E+00 | 3.83E-02 | 1.36E-01 | -1.05E+00 | 1.48E-01 | 4.16E-01 | 1.13E+00 | 6.69E-04 | 2.46E-03 | -1.08E+00 | 5.85E-02 | 1.61E-01 | 1.17E+00 |
| STAG3L5P | 7.47E-04 | 3.89E-03 | 2.18E+00 | 1.50E-02 | 6.49E-02 | -1.11E+00 | 4.39E-03 | 1.87E-02 | 1.83E+00 | 3.31E-02 | 1.26E-01 | -1.07E+00 | 2.70E-02 | 9.62E-02 | 2.21E+00 | 2.70E-01 | 7.60E-01 | 1.33E+00 | 1.68E-01 | 6.17E-01 | 1.21E+00 | 1.61E-01 | 4.43E-01 | -1.13E+00 |
| ATG13 | 7.47E-04 | 3.89E-03 | -1.15E+00 | 1.16E-05 | 5.03E-05 | -1.23E+00 | 7.43E-02 | 3.16E-01 | 1.09E+00 | 2.07E-04 | 7.89E-04 | -1.10E+00 | 2.27E-04 | 8.09E-04 | -1.19E+00 | 5.33E-05 | 1.50E-04 | -1.29E+00 | 1.17E-04 | 4.30E-04 | -1.21E+00 | 3.13E-06 | 8.60E-06 | -1.38E+00 |
| ANKLE2 | 7.47E-04 | 3.89E-03 | 1.05E+00 | 9.74E-05 | 4.21E-04 | 1.11E+00 | 9.09E-02 | 3.86E-01 | 1.07E+00 | 2.06E-01 | 7.88E-01 | 1.23E+00 | 7.78E-02 | 2.78E-01 | 1.44E+00 | 2.91E-02 | 8.20E-02 | 1.37E+00 | 1.68E-03 | 6.17E-03 | -1.10E+00 | 9.41E-05 | 2.58E-04 | -1.18E+00 |
| CHFR | 7.47E-04 | 3.90E-03 | -1.07E+00 | 2.09E-03 | 9.01E-03 | 1.07E+00 | 3.12E-04 | 1.32E-03 | -1.28E+00 | 4.04E-04 | 1.54E-03 | -1.09E+00 | 2.94E-02 | 1.05E-01 | 1.10E+00 | 4.02E-02 | 1.13E-01 | 1.08E+00 | 7.51E-07 | 2.76E-06 | -1.42E+00 | 1.54E-05 | 4.23E-05 | -1.25E+00 |
| PFKP | 7.48E-04 | 3.90E-03 | 1.04E+00 | 1.05E-01 | 4.54E-01 | 1.42E+00 | 2.76E-02 | 1.17E-01 | 1.03E+00 | 2.56E-01 | 9.76E-01 | 1.40E+00 | 1.87E-03 | 6.67E-03 | 1.01E+00 | 1.27E-01 | 3.58E-01 | 1.26E+00 | 2.21E-03 | 8.13E-03 | -1.02E+00 | 3.35E-01 | 9.19E-01 | 1.26E+00 |
| AL353754.1 | 7.50E-04 | 3.91E-03 | -1.27E+00 | 4.28E-05 | 1.85E-04 | -1.37E+00 | 1.64E-03 | 6.97E-03 | -1.32E+00 | 1.22E-04 | 4.66E-04 | -1.47E+00 | 9.07E-04 | 3.24E-03 | -1.44E+00 | 3.76E-03 | 1.06E-02 | -1.34E+00 | 9.66E-04 | 3.55E-03 | -1.60E+00 | 5.75E-02 | 1.58E-01 | -1.14E+00 |
| ANAPC4 | 7.51E-04 | 3.91E-03 | -1.08E+00 | 2.05E-01 | 8.86E-01 | 1.53E+00 | 8.62E-03 | 3.66E-02 | -1.14E+00 | 6.28E-02 | 2.40E-01 | 1.17E+00 | 1.97E-03 | 7.02E-03 | -1.14E+00 | 8.46E-03 | 2.38E-02 | -1.09E+00 | 2.75E-03 | 1.01E-02 | -1.07E+00 | 9.75E-03 | 2.68E-02 | -1.04E+00 |
| AC002467.1 | 7.52E-04 | 3.92E-03 | -1.70E+00 | 7.84E-03 | 3.39E-02 | -1.21E+00 | 1.18E-03 | 5.03E-03 | -1.66E+00 | 1.73E-03 | 6.60E-03 | -1.50E+00 | 3.27E-06 | 1.17E-05 | -2.52E+00 | 5.37E-02 | 1.51E-01 | 1.07E+00 | 1.23E-08 | 4.50E-08 | -1.61E+00 | 3.36E-04 | 9.22E-04 | -1.17E+00 |
| YBX1P10 | 7.52E-04 | 3.92E-03 | 2.18E+00 | 1.48E-02 | 6.40E-02 | 2.12E+00 | 5.34E-04 | 2.27E-03 | 1.78E+00 | 2.17E-02 | 8.31E-02 | 1.79E+00 | 2.63E-01 | 9.38E-01 | 1.59E+00 | 3.27E-01 | 9.22E-01 | 1.54E+00 | 2.56E-04 | 9.41E-04 | 1.75E+00 | 3.18E-06 | 8.74E-06 | 2.03E+00 |
| PPP1R7 | 7.53E-04 | 3.92E-03 | 1.12E+00 | 1.87E-04 | 8.09E-04 | 1.21E+00 | 2.37E-03 | 1.01E-02 | -1.04E+00 | 9.50E-05 | 3.63E-04 | -1.03E+00 | 1.63E-05 | 5.82E-05 | -1.10E+00 | 1.47E-02 | 4.16E-02 | 1.14E+00 | 2.36E-01 | 8.67E-01 | 1.21E+00 | 3.61E-01 | 9.93E-01 | 1.23E+00 |
| MED25 | 7.53E-04 | 3.92E-03 | -1.18E+00 | 4.03E-04 | 1.74E-03 | -1.10E+00 | 1.43E-02 | 6.07E-02 | -1.02E+00 | 2.87E-03 | 1.09E-02 | 1.01E+00 | 5.98E-03 | 2.13E-02 | -1.13E+00 | 1.82E-01 | 5.13E-01 | 1.18E+00 | 5.69E-02 | 2.09E-01 | 1.15E+00 | 6.07E-02 | 1.67E-01 | 1.11E+00 |
| NRDE2 | 7.54E-04 | 3.93E-03 | -1.10E+00 | 2.75E-05 | 1.19E-04 | -1.12E+00 | 1.46E-02 | 6.19E-02 | -1.03E+00 | 2.62E-04 | 1.00E-03 | -1.06E+00 | 2.00E-01 | 7.14E-01 | 1.24E+00 | 8.03E-02 | 2.26E-01 | 1.07E+00 | 1.54E-01 | 5.65E-01 | 1.39E+00 | 7.35E-02 | 2.02E-01 | 1.11E+00 |
| AP001029.2 | 7.55E-04 | 3.93E-03 | -1.40E+00 | 4.91E-03 | 2.12E-02 | -1.11E+00 | 9.48E-07 | 4.03E-06 | -1.89E+00 | 3.94E-05 | 1.51E-04 | -1.42E+00 | 7.22E-05 | 2.57E-04 | -2.29E+00 | 1.55E-03 | 4.36E-03 | -1.54E+00 | 7.87E-02 | 2.89E-01 | -1.24E+00 | 2.08E-01 | 5.73E-01 | -1.06E+00 |
| L3MBTL4 | 7.56E-04 | 3.94E-03 | -1.03E+00 | 1.56E-02 | 6.76E-02 | 1.29E+00 | 4.29E-09 | 1.82E-08 | -1.58E+00 | 3.68E-06 | 1.41E-05 | -1.16E+00 | 9.63E-07 | 3.43E-06 | -1.79E+00 | 1.84E-04 | 5.18E-04 | -1.34E+00 | 1.15E-03 | 4.23E-03 | -1.61E+00 | 1.84E-01 | 5.05E-01 | 1.12E+00 |
| NEURL2 | 7.57E-04 | 3.94E-03 | -1.23E+00 | 4.41E-04 | 1.91E-03 | -1.25E+00 | 5.06E-04 | 2.15E-03 | -1.35E+00 | 8.07E-04 | 3.08E-03 | -1.19E+00 | 4.76E-02 | 1.70E-01 | -1.19E+00 | 8.28E-02 | 2.33E-01 | -1.14E+00 | 1.67E-01 | 6.14E-01 | 1.14E+00 | 1.72E-01 | 4.72E-01 | -1.01E+00 |
| ZNRF2 | 7.59E-04 | 3.96E-03 | -1.41E+00 | 1.71E-03 | 7.37E-03 | -1.18E+00 | 2.06E-04 | 8.75E-04 | -1.39E+00 | 3.21E-04 | 1.22E-03 | -1.24E+00 | 3.34E-02 | 1.19E-01 | -1.01E+00 | 4.82E-02 | 1.36E-01 | 1.02E+00 | 8.49E-03 | 3.12E-02 | 1.06E+00 | 5.75E-02 | 1.58E-01 | 1.10E+00 |
| TAX1BP3 | 7.60E-04 | 3.96E-03 | -1.35E+00 | 1.07E-06 | 4.63E-06 | -2.02E+00 | 3.63E-05 | 1.54E-04 | -1.59E+00 | 8.09E-08 | 3.09E-07 | -1.97E+00 | 9.83E-07 | 3.51E-06 | -1.40E+00 | 2.55E-07 | 7.18E-07 | -1.55E+00 | 1.11E-01 | 4.06E-01 | 1.12E+00 | 1.05E-03 | 2.88E-03 | -1.43E+00 |
| HAUS5 | 7.63E-04 | 3.98E-03 | -1.07E+00 | 1.12E-03 | 4.86E-03 | 1.07E+00 | 4.68E-02 | 1.99E-01 | 1.34E+00 | 1.13E-01 | 4.30E-01 | 1.51E+00 | 4.48E-03 | 1.60E-02 | -1.08E+00 | 3.38E-03 | 9.53E-03 | -1.12E+00 | 7.61E-02 | 2.79E-01 | 1.02E+00 | 7.07E-02 | 1.94E-01 | 1.04E+00 |
| INO80E | 7.64E-04 | 3.98E-03 | -1.27E+00 | 1.70E-02 | 7.32E-02 | 1.12E+00 | 1.89E-07 | 8.03E-07 | -1.58E+00 | 7.48E-08 | 2.86E-07 | -1.38E+00 | 7.30E-03 | 2.60E-02 | -1.25E+00 | 4.98E-02 | 1.40E-01 | 1.06E+00 | 4.27E-02 | 1.57E-01 | 1.01E+00 | 1.67E-01 | 4.58E-01 | 1.17E+00 |
| TMEM175 | 7.64E-04 | 3.98E-03 | -1.19E+00 | 1.57E-02 | 6.77E-02 | 1.14E+00 | 1.24E-02 | 5.26E-02 | -1.15E+00 | 5.19E-02 | 1.98E-01 | 1.05E+00 | 7.49E-02 | 2.67E-01 | 1.10E+00 | 2.61E-01 | 7.34E-01 | 1.23E+00 | 4.46E-03 | 1.64E-02 | -1.04E+00 | 1.33E-01 | 3.65E-01 | 1.20E+00 |
| SMKR1 | 7.65E-04 | 3.99E-03 | -1.47E+00 | 1.50E-03 | 6.47E-03 | -1.31E+00 | 6.54E-02 | 2.78E-01 | -1.03E+00 | 8.54E-02 | 3.26E-01 | 1.08E+00 | 6.74E-02 | 2.40E-01 | -1.73E+00 | 1.43E-01 | 4.02E-01 | -1.29E+00 | 6.17E-02 | 2.26E-01 | 1.65E+00 | 5.24E-03 | 1.44E-02 | 2.32E+00 |
| ZNF584 | 7.65E-04 | 3.99E-03 | -1.80E+00 | 8.67E-03 | 3.75E-02 | -1.22E+00 | 3.41E-05 | 1.45E-04 | -1.50E+00 | 6.37E-04 | 2.43E-03 | -1.16E+00 | 2.14E-03 | 7.62E-03 | -1.85E+00 | 4.56E-02 | 1.29E-01 | -1.20E+00 | 4.68E-03 | 1.72E-02 | -1.11E+00 | 2.53E-01 | 6.96E-01 | 1.20E+00 |
| LRRC37BP1 | 7.65E-04 | 3.99E-03 | -1.13E+00 | 2.29E-03 | 9.91E-03 | 1.06E+00 | 6.62E-04 | 2.81E-03 | -1.28E+00 | 1.87E-03 | 7.13E-03 | -1.10E+00 | 1.07E-04 | 3.82E-04 | -1.58E+00 | 4.42E-04 | 1.25E-03 | -1.40E+00 | 1.47E-02 | 5.41E-02 | -1.08E+00 | 4.77E-02 | 1.31E-01 | -1.12E+00 |
| TRPV2 | 7.66E-04 | 3.99E-03 | 1.03E+00 | 3.59E-02 | 1.55E-01 | 1.30E+00 | 3.59E-03 | 1.53E-02 | -1.08E+00 | 2.04E-01 | 7.80E-01 | 1.41E+00 | 7.20E-02 | 2.57E-01 | 1.09E+00 | 1.99E-01 | 5.61E-01 | 1.48E+00 | 8.76E-03 | 3.22E-02 | 1.01E+00 | 8.91E-02 | 2.45E-01 | 1.08E+00 |
| NECAP2 | 7.67E-04 | 4.00E-03 | 1.03E+00 | 1.14E-02 | 4.94E-02 | 1.26E+00 | 3.14E-02 | 1.33E-01 | 1.03E+00 | 7.56E-02 | 2.89E-01 | 1.23E+00 | 3.41E-05 | 1.22E-04 | -1.24E+00 | 2.56E-02 | 7.22E-02 | 1.14E+00 | 9.99E-04 | 3.67E-03 | -1.15E+00 | 3.00E-03 | 8.23E-03 | -1.06E+00 |
| FCHO1 | 7.69E-04 | 4.01E-03 | -1.25E+00 | 1.52E-02 | 6.58E-02 | 1.10E+00 | 3.68E-05 | 1.56E-04 | -1.37E+00 | 5.03E-03 | 1.92E-02 | 1.04E+00 | 5.32E-02 | 1.90E-01 | -1.14E+00 | 3.55E-01 | 1.00E+00 | 1.46E+00 | 5.04E-03 | 1.85E-02 | -1.26E+00 | 7.10E-03 | 1.95E-02 | -1.25E+00 |
| SNN | 7.69E-04 | 4.01E-03 | -1.30E+00 | 9.96E-05 | 4.30E-04 | -1.49E+00 | 5.43E-03 | 2.31E-02 | -1.31E+00 | 2.07E-02 | 7.90E-02 | -1.03E+00 | 1.86E-02 | 6.62E-02 | -1.34E+00 | 4.95E-03 | 1.40E-02 | -1.72E+00 | 4.40E-02 | 1.62E-01 | -1.49E+00 | 3.11E-02 | 8.56E-02 | -1.72E+00 |
| SNHG16 | 7.71E-04 | 4.02E-03 | 1.04E+00 | 9.25E-06 | 4.00E-05 | -1.07E+00 | 7.08E-05 | 3.01E-04 | -1.12E+00 | 8.64E-11 | 3.30E-10 | -1.36E+00 | 3.70E-05 | 1.32E-04 | -1.28E+00 | 4.94E-06 | 1.39E-05 | -1.39E+00 | 6.18E-07 | 2.27E-06 | -1.32E+00 | 1.98E-03 | 5.43E-03 | -1.13E+00 |
| RBM5-AS1 | 7.72E-04 | 4.02E-03 | -1.24E+00 | 4.23E-04 | 1.83E-03 | -1.24E+00 | 1.79E-01 | 7.60E-01 | 1.13E+00 | 1.96E-02 | 7.48E-02 | -1.03E+00 | 1.90E-01 | 6.78E-01 | 1.13E+00 | 9.03E-02 | 2.55E-01 | -1.05E+00 | 1.45E-04 | 5.30E-04 | -1.59E+00 | 1.97E-03 | 5.40E-03 | -1.35E+00 |
| TTC21B | 7.72E-04 | 4.02E-03 | -1.05E+00 | 2.86E-06 | 1.23E-05 | -1.20E+00 | 1.55E-01 | 6.60E-01 | 1.20E+00 | 1.26E-03 | 4.80E-03 | -1.05E+00 | 5.00E-04 | 1.80E-03 | -1.18E+00 | 1.24E-04 | 3.49E-04 | -1.27E+00 | 1.59E-02 | 5.83E-02 | -1.06E+00 | 6.86E-03 | 1.88E-02 | -1.11E+00 |
| AGK | 7.74E-04 | 4.03E-03 | -1.01E+00 | 9.53E-02 | 4.12E-01 | 1.65E+00 | 3.16E-03 | 1.34E-02 | -1.13E+00 | 1.02E-01 | 3.91E-01 | 1.24E+00 | 1.92E-03 | 6.85E-03 | -1.19E+00 | 1.97E-01 | 5.55E-01 | 1.23E+00 | 7.96E-05 | 2.92E-04 | -1.20E+00 | 2.06E-02 | 5.65E-02 | 1.06E+00 |
| TIGD1 | 7.75E-04 | 4.04E-03 | -1.18E+00 | 1.22E-01 | 5.29E-01 | 1.38E+00 | 3.19E-04 | 1.36E-03 | -1.32E+00 | 1.06E-01 | 4.04E-01 | 1.31E+00 | 2.32E-02 | 8.29E-02 | -1.05E+00 | 3.31E-01 | 9.32E-01 | 1.65E+00 | 2.05E-02 | 7.52E-02 | -1.01E+00 | 3.57E-01 | 9.80E-01 | 1.33E+00 |
| XPO4 | 7.75E-04 | 4.04E-03 | -1.08E+00 | 2.10E-02 | 9.09E-02 | 1.29E+00 | 7.47E-04 | 3.17E-03 | -1.18E+00 | 1.26E-03 | 4.83E-03 | 1.01E+00 | 2.33E-02 | 8.32E-02 | 1.02E+00 | 7.02E-02 | 1.98E-01 | 1.17E+00 | 8.24E-03 | 3.02E-02 | -1.06E+00 | 2.05E-01 | 5.64E-01 | 1.14E+00 |
| CCDC32 | 7.76E-04 | 4.05E-03 | -1.58E+00 | 5.48E-02 | 2.37E-01 | 1.15E+00 | 5.40E-03 | 2.29E-02 | -1.43E+00 | 4.07E-02 | 1.55E-01 | 1.02E+00 | 2.74E-05 | 9.78E-05 | -1.41E+00 | 2.01E-02 | 5.68E-02 | 1.08E+00 | 2.62E-05 | 9.63E-05 | -1.46E+00 | 4.59E-03 | 1.26E-02 | -1.12E+00 |
| RAP2A | 7.76E-04 | 4.05E-03 | -1.36E+00 | 2.37E-03 | 1.02E-02 | -1.15E+00 | 5.43E-04 | 2.31E-03 | -1.37E+00 | 5.76E-04 | 2.20E-03 | -1.22E+00 | 4.44E-04 | 1.58E-03 | -1.16E+00 | 1.85E-03 | 5.23E-03 | -1.04E+00 | 6.04E-04 | 2.22E-03 | -1.09E+00 | 7.38E-05 | 2.03E-04 | -1.27E+00 |
| MPZL2 | 7.80E-04 | 4.06E-03 | -1.24E+00 | 6.40E-02 | 2.76E-01 | 1.21E+00 | 1.33E-01 | 5.66E-01 | 1.01E+00 | 1.16E-01 | 4.44E-01 | 1.06E+00 | 7.69E-02 | 2.74E-01 | 2.51E+00 | 2.55E-01 | 7.18E-01 | 1.64E+00 | 1.84E-01 | 6.74E-01 | 1.41E+00 | 3.02E-01 | 8.30E-01 | -1.00E+00 |
| MOSPD3 | 7.80E-04 | 4.06E-03 | -1.19E+00 | 1.05E-02 | 4.56E-02 | 1.13E+00 | 6.58E-02 | 2.80E-01 | -1.06E+00 | 6.78E-02 | 2.59E-01 | 1.08E+00 | 1.05E-02 | 3.75E-02 | -1.11E+00 | 3.41E-01 | 9.62E-01 | 1.59E+00 | 2.49E-01 | 9.13E-01 | 1.23E+00 | 1.35E-01 | 3.72E-01 | 1.42E+00 |
| SLC25A53 | 7.80E-04 | 4.06E-03 | -1.38E+00 | 2.21E-03 | 9.54E-03 | -1.17E+00 | 6.57E-03 | 2.79E-02 | -1.30E+00 | 7.96E-03 | 3.04E-02 | -1.17E+00 | 5.11E-02 | 1.82E-01 | 1.13E+00 | 2.56E-01 | 7.22E-01 | 1.60E+00 | 1.66E-03 | 6.09E-03 | 1.98E+00 | 1.14E-01 | 3.14E-01 | 1.68E+00 |
| HMGN4 | 7.80E-04 | 4.07E-03 | -1.35E+00 | 2.67E-03 | 1.15E-02 | -1.15E+00 | 4.28E-02 | 1.82E-01 | -1.18E+00 | 1.08E-01 | 4.13E-01 | 1.03E+00 | 1.92E-05 | 6.84E-05 | -1.61E+00 | 5.78E-05 | 1.63E-04 | -1.53E+00 | 7.40E-06 | 2.72E-05 | -1.57E+00 | 5.49E-05 | 1.51E-04 | -1.57E+00 |
| WDFY4 | 7.81E-04 | 4.07E-03 | -1.29E+00 | 3.42E-02 | 1.48E-01 | 1.21E+00 | 6.75E-02 | 2.87E-01 | 1.01E+00 | 3.81E-03 | 1.45E-02 | -1.05E+00 | 5.82E-03 | 2.08E-02 | -1.39E+00 | 1.37E-02 | 3.86E-02 | -1.18E+00 | 8.98E-04 | 3.30E-03 | -1.67E+00 | 7.33E-02 | 2.01E-01 | -1.10E+00 |
| NEXN | 7.81E-04 | 4.07E-03 | 3.00E+00 | 9.75E-02 | 4.21E-01 | 1.94E+00 | 3.72E-04 | 1.58E-03 | 2.67E+00 | 1.59E-01 | 6.09E-01 | 1.56E+00 | 2.80E-01 | 9.99E-01 | 1.08E+00 | 9.71E-02 | 2.74E-01 | -1.38E+00 | 2.41E-02 | 8.86E-02 | 4.39E+00 | 3.53E-01 | 9.69E-01 | 2.88E+00 |
| PEX3 | 7.83E-04 | 4.08E-03 | -1.26E+00 | 2.00E-02 | 8.62E-02 | 1.08E+00 | 9.67E-05 | 4.11E-04 | -1.20E+00 | 8.94E-04 | 3.41E-03 | -1.05E+00 | 1.60E-04 | 5.71E-04 | -1.85E+00 | 9.56E-03 | 2.69E-02 | -1.15E+00 | 6.09E-06 | 2.23E-05 | -1.27E+00 | 3.51E-02 | 9.64E-02 | 1.09E+00 |
| PTK2 | 7.83E-04 | 4.08E-03 | -1.60E+00 | 1.17E-05 | 5.06E-05 | -2.44E+00 | 1.53E-01 | 6.51E-01 | -1.21E+00 | 8.40E-02 | 3.21E-01 | -1.55E+00 | 3.86E-05 | 1.38E-04 | -3.06E+00 | 6.16E-05 | 1.74E-04 | -3.06E+00 | 4.65E-13 | 1.71E-12 | -1.17E+00 | 3.05E-20 | 8.38E-20 | -1.49E+00 |
| AL035587.1 | 7.85E-04 | 4.09E-03 | -1.42E+00 | 4.15E-02 | 1.80E-01 | 1.20E+00 | 3.93E-03 | 1.67E-02 | -1.32E+00 | 4.59E-02 | 1.75E-01 | 1.17E+00 | 1.58E-01 | 5.62E-01 | 1.01E+00 | 2.47E-01 | 6.95E-01 | 1.14E+00 | 7.64E-02 | 2.80E-01 | -1.19E+00 | 3.31E-01 | 9.08E-01 | 1.22E+00 |
| PELI1 | 7.86E-04 | 4.09E-03 | 1.00E+00 | 2.93E-12 | 1.26E-11 | -1.76E+00 | 2.13E-01 | 9.04E-01 | 1.24E+00 | 7.26E-06 | 2.77E-05 | -1.47E+00 | 2.06E-01 | 7.35E-01 | 1.32E+00 | 2.31E-03 | 6.52E-03 | -1.38E+00 | 1.81E-01 | 6.64E-01 | 1.15E+00 | 5.84E-06 | 1.60E-05 | -1.80E+00 |
| GSTZ1 | 7.86E-04 | 4.09E-03 | -1.07E+00 | 7.28E-05 | 3.15E-04 | -1.12E+00 | 1.04E-03 | 4.41E-03 | -1.16E+00 | 1.39E-09 | 5.30E-09 | -1.47E+00 | 5.52E-06 | 1.97E-05 | -1.16E+00 | 8.68E-04 | 2.45E-03 | 1.02E+00 | 9.73E-02 | 3.57E-01 | 1.50E+00 | 9.54E-02 | 2.62E-01 | 1.07E+00 |
| FOXO4 | 7.86E-04 | 4.10E-03 | -1.26E+00 | 2.56E-02 | 1.11E-01 | 1.10E+00 | 2.20E-01 | 9.34E-01 | 1.24E+00 | 1.39E-01 | 5.30E-01 | 1.51E+00 | 1.87E-01 | 6.67E-01 | -1.09E+00 | 3.24E-01 | 9.14E-01 | 1.09E+00 | 2.41E-01 | 8.83E-01 | 1.33E+00 | 3.12E-01 | 8.57E-01 | 1.28E+00 |
| AC126389.1 | 7.89E-04 | 4.11E-03 | -1.16E+00 | 2.43E-04 | 1.05E-03 | -1.11E+00 | 4.60E-02 | 1.96E-01 | -1.11E+00 | 1.55E-02 | 5.92E-02 | -1.09E+00 | 2.60E-01 | 9.25E-01 | 1.29E+00 | 1.87E-01 | 5.27E-01 | 1.08E+00 | 5.18E-02 | 1.90E-01 | -1.05E+00 | 3.24E-02 | 8.90E-02 | -1.16E+00 |
| ITGB4 | 7.90E-04 | 4.12E-03 | -1.81E+00 | 2.44E-04 | 1.05E-03 | -2.38E+00 | 1.63E-02 | 6.92E-02 | -1.45E+00 | 7.03E-03 | 2.69E-02 | -1.59E+00 | 1.72E-01 | 6.15E-01 | -1.29E+00 | 2.80E-01 | 7.89E-01 | -1.06E+00 | 1.96E-05 | 7.19E-05 | -1.09E+00 | 3.81E-07 | 1.05E-06 | -1.25E+00 |
| FAM86EP | 7.92E-04 | 4.13E-03 | -1.63E+00 | 2.11E-02 | 9.12E-02 | -1.02E+00 | 1.14E-03 | 4.84E-03 | -1.49E+00 | 1.45E-02 | 5.55E-02 | -1.16E+00 | 2.17E-01 | 7.74E-01 | 1.11E+00 | 1.73E-01 | 4.89E-01 | -1.15E+00 | 2.12E-03 | 7.77E-03 | -1.66E+00 | 2.15E-01 | 5.91E-01 | 1.31E+00 |
| EPB41L3 | 7.93E-04 | 4.13E-03 | -1.08E+00 | 3.04E-03 | 1.31E-02 | 1.18E+00 | 1.34E-03 | 5.68E-03 | -1.27E+00 | 8.40E-03 | 3.21E-02 | 1.04E+00 | 2.37E-01 | 8.44E-01 | -1.12E+00 | 2.52E-01 | 7.09E-01 | -1.28E+00 | 2.55E-01 | 9.37E-01 | 1.06E+00 | 3.20E-01 | 8.80E-01 | 1.17E+00 |
| RHOBTB3 | 7.94E-04 | 4.14E-03 | 1.04E+00 | 3.70E-06 | 1.60E-05 | -1.04E+00 | 3.51E-02 | 1.49E-01 | -1.07E+00 | 3.68E-03 | 1.40E-02 | -1.04E+00 | 2.93E-03 | 1.04E-02 | -1.01E+00 | 1.99E-04 | 5.60E-04 | -1.08E+00 | 4.10E-09 | 1.50E-08 | -1.50E+00 | 3.28E-06 | 9.01E-06 | -1.28E+00 |
| RP2 | 7.95E-04 | 4.14E-03 | -1.13E+00 | 1.74E-03 | 7.54E-03 | 1.03E+00 | 3.38E-06 | 1.44E-05 | -1.44E+00 | 1.70E-08 | 6.51E-08 | -1.44E+00 | 1.34E-05 | 4.79E-05 | -1.42E+00 | 6.99E-07 | 1.97E-06 | -1.56E+00 | 5.50E-06 | 2.02E-05 | -1.45E+00 | 2.36E-05 | 6.49E-05 | -1.48E+00 |
| RPL12P42 | 7.95E-04 | 4.14E-03 | 1.04E+00 | 9.79E-07 | 4.23E-06 | -1.07E+00 | 3.28E-02 | 1.39E-01 | 1.06E+00 | 2.04E-04 | 7.79E-04 | -1.04E+00 | 1.51E-03 | 5.37E-03 | 1.01E+00 | 4.16E-02 | 1.17E-01 | 1.17E+00 | 4.75E-05 | 1.74E-04 | -1.17E+00 | 3.73E-07 | 1.02E-06 | -1.40E+00 |
| CHMP4C | 7.97E-04 | 4.15E-03 | -1.19E+00 | 8.12E-02 | 3.51E-01 | 1.29E+00 | 1.28E-05 | 5.42E-05 | -1.86E+00 | 3.16E-03 | 1.21E-02 | -1.14E+00 | 5.17E-03 | 1.84E-02 | -1.53E+00 | 9.14E-02 | 2.58E-01 | 1.02E+00 | 2.22E-03 | 8.13E-03 | -1.54E+00 | 1.03E-02 | 2.83E-02 | -1.28E+00 |
| PACRGL | 7.99E-04 | 4.16E-03 | -1.34E+00 | 1.38E-02 | 5.96E-02 | 1.05E+00 | 2.05E-04 | 8.73E-04 | -1.45E+00 | 1.10E-03 | 4.19E-03 | -1.11E+00 | 1.50E-04 | 5.35E-04 | -1.77E+00 | 1.01E-02 | 2.84E-02 | -1.15E+00 | 2.88E-05 | 1.06E-04 | -1.59E+00 | 2.19E-02 | 6.01E-02 | 1.00E+00 |
| ST3GAL3 | 7.99E-04 | 4.17E-03 | -1.36E+00 | 1.27E-02 | 5.50E-02 | 1.01E+00 | 3.99E-04 | 1.70E-03 | -1.43E+00 | 8.55E-04 | 3.27E-03 | -1.17E+00 | 5.73E-03 | 2.04E-02 | -1.12E+00 | 1.49E-01 | 4.19E-01 | 1.16E+00 | 1.75E-03 | 6.41E-03 | -1.17E+00 | 1.75E-01 | 4.81E-01 | 1.16E+00 |
| ZNF257 | 8.01E-04 | 4.18E-03 | -1.18E+00 | 2.19E-04 | 9.46E-04 | -1.28E+00 | 2.94E-06 | 1.25E-05 | -1.50E+00 | 3.66E-06 | 1.40E-05 | -1.30E+00 | 1.28E-01 | 4.55E-01 | 1.08E+00 | 9.20E-02 | 2.59E-01 | -1.00E+00 | 3.57E-02 | 1.31E-01 | 1.05E+00 | 4.39E-02 | 1.21E-01 | -1.00E+00 |
| LOXL1 | 8.02E-04 | 4.18E-03 | -1.25E+00 | 1.49E-03 | 6.42E-03 | -1.18E+00 | 2.52E-05 | 1.07E-04 | -1.53E+00 | 2.80E-04 | 1.07E-03 | -1.29E+00 | 2.80E-01 | 9.97E-01 | 1.09E+00 | 3.54E-01 | 9.98E-01 | 1.11E+00 | 2.06E-01 | 7.56E-01 | -1.15E+00 | 3.41E-01 | 9.37E-01 | 1.01E+00 |
| TBC1D1 | 8.02E-04 | 4.18E-03 | -1.02E+00 | 1.53E-03 | 6.61E-03 | 1.09E+00 | 7.10E-04 | 3.02E-03 | -1.04E+00 | 6.67E-06 | 2.55E-05 | -1.09E+00 | 1.42E-02 | 5.06E-02 | -1.00E+00 | 1.02E-01 | 2.89E-01 | 1.18E+00 | 4.26E-02 | 1.56E-01 | 1.07E+00 | 3.29E-01 | 9.02E-01 | 1.28E+00 |
| CCDC159 | 8.03E-04 | 4.18E-03 | -1.11E+00 | 1.04E-03 | 4.48E-03 | 1.00E+00 | 7.73E-03 | 3.28E-02 | -1.15E+00 | 1.17E-02 | 4.45E-02 | -1.02E+00 | 6.89E-03 | 2.46E-02 | -1.10E+00 | 7.81E-02 | 2.20E-01 | 1.11E+00 | 2.21E-01 | 8.13E-01 | 1.25E+00 | 2.16E-01 | 5.92E-01 | 1.13E+00 |
| EIF2B2 | 8.03E-04 | 4.19E-03 | 1.01E+00 | 2.80E-03 | 1.21E-02 | 1.16E+00 | 4.84E-06 | 2.06E-05 | -1.28E+00 | 6.33E-04 | 2.42E-03 | -1.01E+00 | 1.88E-04 | 6.72E-04 | -1.13E+00 | 7.73E-03 | 2.18E-02 | 1.13E+00 | 8.03E-03 | 2.95E-02 | 1.10E+00 | 6.15E-02 | 1.69E-01 | 1.20E+00 |
| GAS6 | 8.03E-04 | 4.19E-03 | -1.70E+00 | 1.21E-02 | 5.21E-02 | -1.34E+00 | 3.24E-04 | 1.38E-03 | -1.70E+00 | 1.76E-03 | 6.74E-03 | -1.37E+00 | 1.55E-01 | 5.53E-01 | 2.06E+00 | 2.02E-01 | 5.69E-01 | 2.00E+00 | 2.60E-01 | 9.56E-01 | 1.07E+00 | 1.95E-01 | 5.36E-01 | 1.41E+00 |
| PSTK | 8.04E-04 | 4.19E-03 | -1.22E+00 | 7.42E-03 | 3.21E-02 | 1.05E+00 | 2.01E-02 | 8.55E-02 | -1.13E+00 | 2.37E-02 | 9.07E-02 | 1.03E+00 | 2.28E-03 | 8.14E-03 | -1.55E+00 | 2.01E-02 | 5.67E-02 | -1.21E+00 | 4.21E-02 | 1.54E-01 | -1.01E+00 | 3.35E-01 | 9.19E-01 | 1.37E+00 |
| FOCAD | 8.06E-04 | 4.20E-03 | -1.03E+00 | 2.70E-02 | 1.17E-01 | 1.32E+00 | 2.05E-05 | 8.69E-05 | -1.31E+00 | 6.35E-03 | 2.42E-02 | 1.07E+00 | 1.90E-02 | 6.78E-02 | -1.04E+00 | 4.09E-02 | 1.15E-01 | 1.06E+00 | 6.32E-02 | 2.32E-01 | 1.01E+00 | 3.10E-01 | 8.51E-01 | 1.24E+00 |
| ZNF747 | 8.06E-04 | 4.20E-03 | -1.31E+00 | 4.81E-03 | 2.08E-02 | -1.05E+00 | 1.34E-03 | 5.69E-03 | -1.25E+00 | 1.56E-02 | 5.95E-02 | 1.06E+00 | 1.03E-04 | 3.67E-04 | -1.79E+00 | 7.67E-03 | 2.16E-02 | -1.19E+00 | 2.84E-03 | 1.04E-02 | -1.17E+00 | 7.61E-02 | 2.09E-01 | 1.02E+00 |
| NPRL3 | 8.06E-04 | 4.20E-03 | -1.06E+00 | 9.53E-03 | 4.12E-02 | 1.19E+00 | 1.13E-04 | 4.80E-04 | -1.32E+00 | 6.36E-04 | 2.43E-03 | -1.08E+00 | 1.71E-05 | 6.10E-05 | -1.08E+00 | 2.19E-02 | 6.18E-02 | 1.28E+00 | 1.97E-04 | 7.21E-04 | -1.17E+00 | 4.46E-03 | 1.23E-02 | -1.11E+00 |
| AC018371.2 | 8.09E-04 | 4.22E-03 | -1.41E+00 | 2.11E-03 | 9.12E-03 | -1.16E+00 | 1.06E-13 | 4.52E-13 | -2.17E+00 | 5.80E-10 | 2.22E-09 | -1.50E+00 | 4.44E-05 | 1.58E-04 | -1.48E+00 | 1.13E-06 | 3.20E-06 | -1.83E+00 | 4.83E-08 | 1.77E-07 | -1.54E+00 | 1.87E-07 | 5.14E-07 | -1.44E+00 |
| MELTF-AS1 | 8.09E-04 | 4.22E-03 | -1.26E+00 | 4.61E-02 | 1.99E-01 | 1.34E+00 | 6.24E-02 | 2.65E-01 | 1.11E+00 | 6.43E-02 | 2.45E-01 | 1.27E+00 | 9.17E-02 | 3.27E-01 | -1.09E+00 | 6.99E-02 | 1.97E-01 | -1.27E+00 | 1.78E-01 | 6.52E-01 | 1.35E+00 | 6.13E-02 | 1.68E-01 | 1.77E+00 |
| UPRT | 8.11E-04 | 4.23E-03 | -1.32E+00 | 1.40E-02 | 6.06E-02 | 1.00E+00 | 1.48E-05 | 6.27E-05 | -1.49E+00 | 5.46E-03 | 2.09E-02 | 1.05E+00 | 2.22E-04 | 7.93E-04 | -1.21E+00 | 2.72E-04 | 7.68E-04 | -1.29E+00 | 3.24E-05 | 1.19E-04 | -1.47E+00 | 6.03E-03 | 1.66E-02 | -1.20E+00 |
| ZNF682 | 8.11E-04 | 4.23E-03 | -1.16E+00 | 1.77E-01 | 7.65E-01 | 1.41E+00 | 5.78E-04 | 2.45E-03 | -1.39E+00 | 1.01E-01 | 3.86E-01 | 1.09E+00 | 1.72E-03 | 6.14E-03 | -1.53E+00 | 2.92E-02 | 8.24E-02 | -1.16E+00 | 6.99E-03 | 2.57E-02 | -1.52E+00 | 1.59E-01 | 4.37E-01 | -1.06E+00 |
| AC010173.1 | 8.12E-04 | 4.23E-03 | 2.42E+00 | 1.84E-01 | 7.96E-01 | 1.30E+00 | 3.66E-03 | 1.56E-02 | 2.12E+00 | 1.74E-01 | 6.63E-01 | 1.43E+00 | 2.21E-01 | 7.88E-01 | 1.36E+00 | 3.35E-01 | 9.45E-01 | 1.04E+00 | 1.93E-01 | 7.07E-01 | -1.17E+00 | 2.42E-01 | 6.66E-01 | -1.26E+00 |
| CD24 | 8.13E-04 | 4.24E-03 | -1.87E+00 | 4.18E-03 | 1.81E-02 | -1.49E+00 | 1.58E-01 | 6.69E-01 | 1.54E+00 | 6.08E-02 | 2.32E-01 | -1.43E+00 | 1.43E-01 | 5.10E-01 | -1.02E+00 | 1.34E-01 | 3.79E-01 | -1.14E+00 | 1.62E-01 | 5.94E-01 | -1.06E+00 | 3.32E-01 | 9.13E-01 | 1.21E+00 |
| GIT2 | 8.16E-04 | 4.25E-03 | -1.02E+00 | 4.68E-04 | 2.02E-03 | 1.10E+00 | 1.56E-05 | 6.63E-05 | -1.28E+00 | 1.48E-05 | 5.65E-05 | -1.11E+00 | 1.41E-02 | 5.04E-02 | -1.04E+00 | 6.46E-02 | 1.82E-01 | 1.09E+00 | 2.17E-04 | 7.97E-04 | -1.15E+00 | 1.43E-02 | 3.92E-02 | 1.02E+00 |
| PSKH1 | 8.17E-04 | 4.26E-03 | -1.39E+00 | 2.55E-02 | 1.10E-01 | 1.06E+00 | 1.17E-06 | 4.96E-06 | -1.91E+00 | 8.23E-03 | 3.15E-02 | -1.22E+00 | 1.77E-04 | 6.31E-04 | -1.43E+00 | 1.03E-02 | 2.92E-02 | -1.07E+00 | 9.09E-05 | 3.34E-04 | -1.32E+00 | 1.84E-02 | 5.05E-02 | -1.02E+00 |
| AL391121.1 | 8.17E-04 | 4.26E-03 | 1.99E+00 | 1.50E-02 | 6.48E-02 | 1.24E+00 | 9.64E-04 | 4.10E-03 | 1.85E+00 | 7.61E-02 | 2.91E-01 | 1.14E+00 | 9.02E-04 | 3.22E-03 | 2.26E+00 | 3.46E-01 | 9.76E-01 | 1.41E+00 | 1.49E-01 | 5.46E-01 | 1.62E+00 | 1.43E-02 | 3.92E-02 | -1.02E+00 |
| NBPF11 | 8.17E-04 | 4.26E-03 | -1.32E+00 | 9.20E-04 | 3.97E-03 | -1.23E+00 | 3.77E-02 | 1.60E-01 | -1.12E+00 | 7.14E-03 | 2.73E-02 | -1.31E+00 | 2.35E-03 | 8.37E-03 | -1.30E+00 | 3.81E-02 | 1.07E-01 | -1.10E+00 | 3.81E-02 | 1.40E-01 | -1.17E+00 | 1.00E-01 | 2.75E-01 | 1.01E+00 |
| HAGH | 8.18E-04 | 4.26E-03 | -1.20E+00 | 3.49E-02 | 1.51E-01 | 1.20E+00 | 2.62E-04 | 1.11E-03 | -1.27E+00 | 3.75E-03 | 1.43E-02 | 1.02E+00 | 6.40E-04 | 2.28E-03 | -1.23E+00 | 9.68E-02 | 2.73E-01 | 1.27E+00 | 3.00E-07 | 1.10E-06 | -1.19E+00 | 5.25E-06 | 1.44E-05 | -1.17E+00 |
| USMG5 | 8.18E-04 | 4.26E-03 | 1.02E+00 | 1.08E-31 | 4.68E-31 | -1.45E+00 | 5.74E-02 | 2.44E-01 | -1.01E+00 | 2.86E-13 | 1.09E-12 | -1.51E+00 | 8.93E-04 | 3.18E-03 | -1.12E+00 | 5.90E-16 | 1.66E-15 | -1.61E+00 | 1.37E-01 | 5.04E-01 | -1.00E+00 | 1.81E-05 | 4.96E-05 | -1.41E+00 |
| ZGLP1 | 8.19E-04 | 4.27E-03 | -1.39E+00 | 1.41E-01 | 6.09E-01 | 1.34E+00 | 1.27E-06 | 5.39E-06 | -2.09E+00 | 3.26E-05 | 1.25E-04 | -1.46E+00 | 6.23E-05 | 2.22E-04 | -2.49E+00 | 5.32E-03 | 1.50E-02 | -1.60E+00 | 6.34E-02 | 2.33E-01 | 1.99E+00 | 2.06E-01 | 5.66E-01 | 1.63E+00 |
| PNPLA6 | 8.19E-04 | 4.27E-03 | -1.32E+00 | 1.31E-02 | 5.68E-02 | 1.03E+00 | 1.43E-02 | 6.08E-02 | -1.09E+00 | 6.19E-02 | 2.36E-01 | 1.11E+00 | 6.01E-02 | 2.14E-01 | 1.14E+00 | 1.31E-01 | 3.70E-01 | 1.23E+00 | 2.04E-01 | 7.50E-01 | 1.35E+00 | 3.21E-01 | 8.82E-01 | 1.35E+00 |
| LINC01597 | 8.20E-04 | 4.27E-03 | -1.19E+00 | 4.28E-03 | 1.85E-02 | 1.05E+00 | 1.11E-01 | 4.70E-01 | 1.02E+00 | 5.75E-02 | 2.20E-01 | 1.02E+00 | 1.43E-01 | 5.10E-01 | -1.04E+00 | 3.31E-01 | 9.32E-01 | 1.30E+00 | 8.24E-02 | 3.03E-01 | -1.10E+00 | 1.21E-01 | 3.33E-01 | -1.15E+00 |
| ZNF202 | 8.20E-04 | 4.27E-03 | -1.27E+00 | 1.41E-03 | 6.11E-03 | -1.14E+00 | 4.55E-04 | 1.93E-03 | -1.34E+00 | 4.28E-03 | 1.63E-02 | 1.01E+00 | 1.73E-01 | 6.17E-01 | 1.16E+00 | 3.10E-01 | 8.73E-01 | 1.38E+00 | 2.46E-03 | 9.02E-03 | -1.29E+00 | 2.63E-03 | 7.22E-03 | -1.32E+00 |
| RPL10P8 | 8.21E-04 | 4.28E-03 | 2.63E+00 | 6.27E-04 | 2.71E-03 | 3.06E+00 | 9.35E-04 | 3.97E-03 | 2.43E+00 | 9.57E-03 | 3.66E-02 | 2.36E+00 | 1.03E-01 | 3.66E-01 | 2.00E+00 | 9.59E-03 | 2.70E-02 | 2.93E+00 | 7.48E-02 | 2.75E-01 | 1.69E+00 | 1.60E-01 | 4.39E-01 | 1.66E+00 |
| KIFC3 | 8.22E-04 | 4.28E-03 | -1.28E+00 | 1.25E-02 | 5.41E-02 | 1.03E+00 | 1.53E-02 | 6.48E-02 | -1.23E+00 | 1.09E-02 | 4.16E-02 | -1.19E+00 | 1.73E-01 | 6.16E-01 | 1.21E+00 | 7.16E-02 | 2.02E-01 | -1.08E+00 | 2.59E-03 | 9.49E-03 | 2.02E+00 | 1.35E-01 | 3.72E-01 | 1.18E+00 |
| METTL8 | 8.22E-04 | 4.28E-03 | -1.17E+00 | 1.65E-01 | 7.14E-01 | 1.41E+00 | 6.03E-07 | 2.56E-06 | -1.43E+00 | 9.52E-03 | 3.64E-02 | 1.16E+00 | 1.41E-05 | 5.02E-05 | -1.68E+00 | 6.79E-02 | 1.91E-01 | 1.16E+00 | 3.10E-03 | 1.14E-02 | -1.09E+00 | 1.18E-01 | 3.25E-01 | 1.13E+00 |
| TMTC4 | 8.23E-04 | 4.29E-03 | -1.19E+00 | 1.05E-01 | 4.53E-01 | 1.43E+00 | 1.80E-03 | 7.66E-03 | -1.27E+00 | 9.07E-02 | 3.46E-01 | 1.17E+00 | 3.78E-03 | 1.35E-02 | -1.41E+00 | 2.76E-02 | 7.77E-02 | -1.15E+00 | 1.29E-02 | 4.72E-02 | -1.25E+00 | 3.22E-01 | 8.84E-01 | 1.28E+00 |
| AC005261.2 | 8.24E-04 | 4.30E-03 | -1.10E+00 | 5.15E-04 | 2.22E-03 | -1.02E+00 | 4.71E-06 | 2.00E-05 | -1.29E+00 | 9.35E-07 | 3.57E-06 | -1.24E+00 | 8.82E-05 | 3.14E-04 | -1.11E+00 | 1.88E-05 | 5.29E-05 | -1.12E+00 | 1.93E-04 | 7.07E-04 | -1.17E+00 | 1.64E-03 | 4.50E-03 | -1.14E+00 |
| YLPM1 | 8.26E-04 | 4.31E-03 | -1.03E+00 | 4.06E-04 | 1.75E-03 | 1.05E+00 | 1.19E-01 | 5.04E-01 | 1.16E+00 | 9.45E-02 | 3.61E-01 | 1.19E+00 | 6.47E-04 | 2.31E-03 | -1.02E+00 | 1.08E-04 | 3.06E-04 | -1.08E+00 | 6.06E-04 | 2.22E-03 | 1.01E+00 | 1.50E-02 | 4.13E-02 | 1.09E+00 |
| ELOVL7 | 8.28E-04 | 4.32E-03 | -1.29E+00 | 3.25E-04 | 1.40E-03 | -1.23E+00 | 6.66E-03 | 2.83E-02 | -1.26E+00 | 3.09E-02 | 1.18E-01 | 1.08E+00 | 1.73E-03 | 6.18E-03 | 1.31E+00 | 2.04E-01 | 5.76E-01 | 1.31E+00 | 3.25E-02 | 1.19E-01 | 1.15E+00 | 1.08E-02 | 2.97E-02 | 1.06E+00 |
| GATA2-AS1 | 8.28E-04 | 4.32E-03 | -1.03E+00 | 1.88E-02 | 8.14E-02 | 1.22E+00 | 3.16E-03 | 1.34E-02 | -1.33E+00 | 1.99E-02 | 7.61E-02 | 1.04E+00 | 2.77E-03 | 9.87E-03 | -1.22E+00 | 2.66E-01 | 7.51E-01 | 1.22E+00 | 3.26E-05 | 1.20E-04 | -1.17E+00 | 1.15E-04 | 3.16E-04 | -1.17E+00 |
| ZFP69 | 8.30E-04 | 4.32E-03 | -1.67E+00 | 6.92E-02 | 2.99E-01 | 1.24E+00 | 2.30E-02 | 9.76E-02 | -1.40E+00 | 1.71E-01 | 6.52E-01 | 1.11E+00 | 1.41E-04 | 5.01E-04 | -2.06E+00 | 1.59E-02 | 4.49E-02 | -1.11E+00 | 6.57E-05 | 2.41E-04 | -1.61E+00 | 3.83E-02 | 1.05E-01 | -1.02E+00 |
| H2AFJ | 8.30E-04 | 4.32E-03 | 1.65E+00 | 2.02E-01 | 8.71E-01 | 1.48E+00 | 2.10E-02 | 8.91E-02 | 1.41E+00 | 1.14E-01 | 4.35E-01 | 1.30E+00 | 1.10E-01 | 3.92E-01 | 1.57E+00 | 3.47E-01 | 9.77E-01 | 1.43E+00 | 1.75E-01 | 6.41E-01 | 1.28E+00 | 3.10E-04 | 8.53E-04 | 1.04E+00 |
| ITPA | 8.31E-04 | 4.33E-03 | 1.02E+00 | 7.89E-05 | 3.41E-04 | 1.06E+00 | 5.00E-02 | 2.12E-01 | 1.09E+00 | 4.45E-02 | 1.70E-01 | 1.22E+00 | 3.25E-05 | 1.16E-04 | -1.14E+00 | 1.14E-02 | 3.21E-02 | 1.16E+00 | 2.60E-01 | 9.53E-01 | 1.22E+00 | 5.89E-02 | 1.62E-01 | 1.40E+00 |
| KCTD9P1 | 8.33E-04 | 4.34E-03 | -1.70E+00 | 1.26E-02 | 5.46E-02 | -1.21E+00 | 5.53E-05 | 2.35E-04 | -1.42E+00 | 1.60E-04 | 6.10E-04 | -1.36E+00 | 6.75E-05 | 2.41E-04 | -2.15E+00 | 2.72E-04 | 7.66E-04 | -1.92E+00 | 1.46E-07 | 5.37E-07 | -1.69E+00 | 3.50E-08 | 9.61E-08 | -1.87E+00 |
| UBR3 | 8.39E-04 | 4.37E-03 | -1.32E+00 | 2.80E-03 | 1.21E-02 | -1.14E+00 | 5.64E-02 | 2.40E-01 | 1.05E+00 | 7.23E-02 | 2.76E-01 | 1.22E+00 | 3.08E-02 | 1.10E-01 | 1.10E+00 | 1.30E-02 | 3.66E-02 | -1.04E+00 | 1.28E-09 | 4.68E-09 | -1.34E+00 | 6.78E-06 | 1.86E-05 | -1.18E+00 |
| VARS | 8.39E-04 | 4.37E-03 | -1.11E+00 | 5.93E-03 | 2.56E-02 | 1.11E+00 | 2.13E-03 | 9.05E-03 | -1.01E+00 | 1.66E-03 | 6.35E-03 | 1.10E+00 | 1.54E-01 | 5.50E-01 | 1.42E+00 | 3.04E-01 | 8.58E-01 | 1.74E+00 | 1.64E-02 | 6.01E-02 | 1.05E+00 | 1.88E-01 | 5.16E-01 | 1.33E+00 |
| CCNB1 | 8.40E-04 | 4.38E-03 | -1.45E+00 | 3.98E-02 | 1.72E-01 | 1.08E+00 | 3.98E-05 | 1.69E-04 | -1.12E+00 | 1.32E-04 | 5.06E-04 | 1.13E+00 | 2.05E-03 | 7.31E-03 | -1.76E+00 | 1.60E-01 | 4.50E-01 | 1.14E+00 | 2.23E-05 | 8.17E-05 | -1.21E+00 | 8.74E-02 | 2.40E-01 | 1.02E+00 |
| REEP3 | 8.41E-04 | 4.38E-03 | -1.28E+00 | 4.60E-03 | 1.99E-02 | 1.01E+00 | 1.65E-06 | 7.01E-06 | -1.53E+00 | 3.59E-07 | 1.37E-06 | -1.41E+00 | 2.02E-02 | 7.21E-02 | -1.56E+00 | 9.39E-02 | 2.65E-01 | -1.15E+00 | 2.29E-04 | 8.42E-04 | -1.29E+00 | 6.99E-03 | 1.92E-02 | -1.18E+00 |
| WDPCP | 8.42E-04 | 4.39E-03 | -1.15E+00 | 1.29E-02 | 5.57E-02 | 1.12E+00 | 1.44E-03 | 6.12E-03 | -1.14E+00 | 7.15E-04 | 2.73E-03 | -1.10E+00 | 1.08E-01 | 3.85E-01 | 1.14E+00 | 5.59E-02 | 1.58E-01 | -1.06E+00 | 4.23E-02 | 1.55E-01 | -1.20E+00 | 8.13E-02 | 2.23E-01 | -1.07E+00 |
| OSGEPL1 | 8.42E-04 | 4.39E-03 | -1.29E+00 | 1.05E-01 | 4.55E-01 | 1.59E+00 | 1.56E-09 | 6.62E-09 | -1.83E+00 | 7.76E-03 | 2.96E-02 | 1.08E+00 | 2.79E-02 | 9.96E-02 | 1.01E+00 | 2.48E-01 | 6.99E-01 | 1.58E+00 | 3.61E-08 | 1.33E-07 | -1.62E+00 | 1.45E-01 | 3.98E-01 | 1.14E+00 |
| RBM15B | 8.43E-04 | 4.39E-03 | 1.08E+00 | 6.27E-09 | 2.71E-08 | -1.11E+00 | 4.37E-03 | 1.86E-02 | -1.01E+00 | 2.56E-04 | 9.77E-04 | -1.01E+00 | 5.30E-08 | 1.89E-07 | -1.31E+00 | 1.62E-08 | 4.56E-08 | -1.30E+00 | 1.02E-05 | 3.75E-05 | -1.21E+00 | 2.49E-05 | 6.85E-05 | -1.13E+00 |
| AC095052.1 | 8.43E-04 | 4.39E-03 | -1.31E+00 | 9.60E-06 | 4.15E-05 | -2.00E+00 | 2.64E-03 | 1.12E-02 | -1.35E+00 | 6.64E-05 | 2.54E-04 | -1.78E+00 | 5.98E-02 | 2.13E-01 | -1.17E+00 | 6.20E-02 | 1.75E-01 | -1.15E+00 | 9.48E-02 | 3.48E-01 | 1.10E+00 | 1.72E-02 | 4.71E-02 | -1.18E+00 |
| VWA8 | 8.43E-04 | 4.39E-03 | -1.34E+00 | 1.88E-02 | 8.12E-02 | 1.08E+00 | 1.04E-04 | 4.44E-04 | -1.45E+00 | 1.61E-04 | 6.16E-04 | -1.33E+00 | 4.99E-03 | 1.78E-02 | -1.16E+00 | 1.91E-02 | 5.39E-02 | -1.09E+00 | 1.43E-06 | 5.27E-06 | -1.53E+00 | 8.50E-03 | 2.34E-02 | -1.11E+00 |
| PITRM1 | 8.43E-04 | 4.39E-03 | -1.17E+00 | 2.38E-03 | 1.03E-02 | 1.04E+00 | 2.92E-03 | 1.24E-02 | -1.18E+00 | 7.14E-03 | 2.73E-02 | 1.00E+00 | 7.09E-04 | 2.53E-03 | -1.29E+00 | 1.27E-02 | 3.59E-02 | 1.01E+00 | 2.14E-02 | 7.85E-02 | 1.02E+00 | 1.10E-01 | 3.03E-01 | 1.13E+00 |
| PLOD3 | 8.43E-04 | 4.40E-03 | -1.27E+00 | 7.28E-02 | 3.14E-01 | 1.21E+00 | 1.20E-02 | 5.09E-02 | 1.01E+00 | 6.80E-03 | 2.60E-02 | 1.13E+00 | 1.06E-03 | 3.77E-03 | -1.27E+00 | 6.42E-03 | 1.81E-02 | -1.15E+00 | 1.98E-03 | 7.28E-03 | -1.14E+00 | 3.82E-03 | 1.05E-02 | -1.16E+00 |
| AC108463.3 | 8.44E-04 | 4.40E-03 | -1.88E+00 | 7.21E-03 | 3.12E-02 | -1.23E+00 | 1.94E-03 | 8.26E-03 | -1.69E+00 | 2.80E-02 | 1.07E-01 | -1.06E+00 | 1.99E-06 | 7.09E-06 | -1.46E+00 | 1.24E-02 | 3.49E-02 | 1.06E+00 | 6.02E-10 | 2.21E-09 | -1.88E+00 | 7.26E-04 | 1.99E-03 | -1.16E+00 |
| AC091729.3 | 8.46E-04 | 4.41E-03 | -1.38E+00 | 3.11E-04 | 1.34E-03 | -1.38E+00 | 1.31E-01 | 5.57E-01 | 1.41E+00 | 2.50E-01 | 9.55E-01 | 1.22E+00 | 3.53E-02 | 1.26E-01 | -1.33E+00 | 2.18E-01 | 6.16E-01 | 1.08E+00 | 3.55E-02 | 1.30E-01 | 1.66E+00 | 1.36E-01 | 3.75E-01 | 1.13E+00 |
| RPRD2 | 8.48E-04 | 4.42E-03 | -1.05E+00 | 8.40E-04 | 3.63E-03 | 1.07E+00 | 1.45E-02 | 6.16E-02 | -1.12E+00 | 4.24E-03 | 1.62E-02 | -1.02E+00 | 5.19E-06 | 1.85E-05 | -1.34E+00 | 1.19E-04 | 3.36E-04 | -1.12E+00 | 9.51E-07 | 3.49E-06 | -1.28E+00 | 1.48E-04 | 4.08E-04 | -1.10E+00 |
| MCTS1 | 8.49E-04 | 4.42E-03 | 1.06E+00 | 2.79E-02 | 1.20E-01 | 1.32E+00 | 2.33E-02 | 9.90E-02 | 1.08E+00 | 4.35E-02 | 1.66E-01 | 1.29E+00 | 2.08E-02 | 7.42E-02 | 1.07E+00 | 3.42E-01 | 9.64E-01 | 1.38E+00 | 1.80E-02 | 6.59E-02 | 1.05E+00 | 2.47E-01 | 6.79E-01 | 1.35E+00 |
| AGMAT | 8.50E-04 | 4.43E-03 | -1.95E+00 | 1.16E-02 | 5.02E-02 | -1.34E+00 | 1.98E-03 | 8.39E-03 | -1.39E+00 | 1.14E-02 | 4.34E-02 | -1.06E+00 | 8.68E-02 | 3.10E-01 | -1.39E+00 | 1.68E-01 | 4.75E-01 | -1.14E+00 | 6.09E-03 | 2.24E-02 | -1.21E+00 | 1.48E-02 | 4.07E-02 | -1.23E+00 |
| GPAT4 | 8.51E-04 | 4.43E-03 | -1.17E+00 | 9.78E-03 | 4.23E-02 | 1.08E+00 | 1.18E-06 | 5.00E-06 | -1.56E+00 | 1.42E-05 | 5.44E-05 | -1.26E+00 | 6.20E-04 | 2.21E-03 | -1.24E+00 | 4.16E-04 | 1.17E-03 | -1.24E+00 | 7.31E-14 | 2.68E-13 | -1.89E+00 | 3.47E-08 | 9.54E-08 | -1.43E+00 |
| REG4 | 8.51E-04 | 4.44E-03 | 2.15E+00 | 2.20E-01 | 9.50E-01 | 1.22E+00 | 2.31E-02 | 9.80E-02 | 2.01E+00 | 1.64E-01 | 6.26E-01 | 1.42E+00 | 2.80E-01 | 1.00E+00 | 1.00E+00 | 3.28E-01 | 9.24E-01 | 1.09E+00 | 2.07E-01 | 7.61E-01 | 1.22E+00 | 3.28E-01 | 9.02E-01 | 1.07E+00 |
| PNPLA4 | 8.51E-04 | 4.44E-03 | -1.17E+00 | 1.07E-03 | 4.60E-03 | -1.04E+00 | 2.70E-02 | 1.15E-01 | -1.08E+00 | 6.72E-03 | 2.57E-02 | -1.03E+00 | 7.57E-03 | 2.70E-02 | -1.12E+00 | 1.62E-01 | 4.56E-01 | 1.18E+00 | 1.20E-01 | 4.39E-01 | 1.08E+00 | 2.37E-01 | 6.50E-01 | 1.17E+00 |
| SIA | 8.54E-04 | 4.45E-03 | 1.75E+00 | 1.97E-01 | 8.51E-01 | 1.58E+00 | 1.52E-03 | 6.44E-03 | 1.37E+00 | 1.64E-02 | 6.27E-02 | 1.49E+00 | 5.03E-14 | 1.79E-13 | 3.72E+00 | 6.06E-11 | 1.71E-10 | 2.94E+00 | 6.19E-21 | 2.27E-20 | 2.62E+00 | 3.86E-15 | 1.06E-14 | 2.42E+00 |
| ATP1A3 | 8.56E-04 | 4.46E-03 | -1.57E+00 | 1.09E-02 | 4.71E-02 | -1.12E+00 | 2.05E-03 | 8.73E-03 | -1.29E+00 | 4.85E-06 | 1.85E-05 | -1.55E+00 | 2.77E-01 | 9.88E-01 | 1.26E+00 | 2.17E-01 | 6.12E-01 | 1.02E+00 | 2.12E-01 | 7.77E-01 | 1.18E+00 | 2.79E-01 | 7.67E-01 | 1.36E+00 |
| CYB561D1 | 8.56E-04 | 4.46E-03 | -1.42E+00 | 4.26E-03 | 1.84E-02 | -1.16E+00 | 2.43E-04 | 1.02E-03 | -1.49E+00 | 1.52E-03 | 5.79E-03 | -1.11E+00 | 1.35E-01 | 4.82E-01 | 1.19E+00 | 2.82E-01 | 7.95E-01 | 1.65E+00 | 9.65E-03 | 3.54E-02 | -1.25E+00 | 1.42E-02 | 3.91E-02 | -1.26E+00 |
| NUDT16L1 | 8.56E-04 | 4.46E-03 | 1.00E+00 | 1.14E-02 | 4.95E-02 | 1.25E+00 | 1.89E-02 | 8.02E-02 | 1.07E+00 | 2.65E-02 | 1.01E-01 | 1.22E+00 | 9.98E-02 | 3.56E-01 | 1.18E+00 | 3.41E-01 | 9.63E-01 | 1.36E+00 | 1.10E-01 | 4.05E-01 | 1.07E+00 | 2.32E-01 | 6.37E-01 | 1.26E+00 |
| MAP1LC3A | 8.56E-04 | 4.46E-03 | -1.22E+00 | 4.19E-06 | 1.81E-05 | -1.53E+00 | 1.52E-02 | 6.47E-02 | -1.25E+00 | 4.07E-03 | 1.56E-02 | -1.37E+00 | 1.35E-04 | 4.80E-04 | -1.24E+00 | 1.21E-05 | 3.40E-05 | -1.52E+00 | 1.45E-01 | 5.33E-01 | 1.46E+00 | 1.22E-02 | 3.36E-02 | 1.07E+00 |
| EFCC1 | 8.57E-04 | 4.46E-03 | 1.82E+00 | 3.50E-04 | 1.51E-03 | 2.22E+00 | 4.57E-02 | 1.94E-01 | 1.98E+00 | 2.56E-02 | 9.79E-02 | 2.15E+00 | 2.49E-01 | 8.89E-01 | 1.43E+00 | 3.23E-01 | 9.11E-01 | 1.29E+00 | 1.07E-01 | 3.94E-01 | 1.55E+00 | 1.03E-01 | 2.83E-01 | 2.00E+00 |
| TRERF1 | 8.57E-04 | 4.47E-03 | -1.23E+00 | 6.30E-04 | 2.72E-03 | -1.11E+00 | 5.04E-04 | 2.14E-03 | -1.38E+00 | 2.46E-04 | 9.38E-04 | -1.21E+00 | 2.64E-06 | 9.41E-06 | -2.08E+00 | 3.92E-04 | 1.10E-03 | -1.43E+00 | 1.75E-02 | 6.42E-02 | -1.26E+00 | 6.40E-03 | 1.76E-02 | -1.44E+00 |
| ARHGAP5 | 8.59E-04 | 4.48E-03 | 1.05E+00 | 4.40E-03 | 1.90E-02 | 1.18E+00 | 2.43E-03 | 1.03E-02 | -1.13E+00 | 1.25E-03 | 4.76E-03 | -1.01E+00 | 2.13E-02 | 7.60E-02 | 1.07E+00 | 6.60E-02 | 7.32E-02 | 1.06E+00 | 6.53E-04 | 2.40E-03 | -1.05E+00 | 1.05E-05 | 2.89E-05 | -1.21E+00 |
| UBE2SP1 | 8.59E-04 | 4.48E-03 | -1.10E+00 | 4.45E-10 | 1.92E-09 | -1.97E+00 | 2.11E-01 | 8.94E-01 | 1.27E+00 | 9.70E-04 | 3.71E-03 | -1.15E+00 | 1.66E-03 | 5.93E-03 | -1.04E+00 | 2.47E-06 | 6.95E-06 | -1.66E+00 | 1.84E-04 | 6.76E-04 | -1.20E+00 | 2.39E-05 | 6.56E-05 | -1.36E+00 |
| MFSD9 | 8.60E-04 | 4.48E-03 | -1.57E+00 | 3.82E-03 | 1.65E-02 | -1.21E+00 | 8.15E-04 | 3.46E-03 | -1.57E+00 | 6.79E-04 | 2.59E-03 | -1.51E+00 | 9.45E-07 | 3.37E-06 | -1.98E+00 | 2.84E-07 | 8.00E-07 | -2.09E+00 | 1.01E-02 | 3.69E-02 | -1.19E+00 | 1.77E-02 | 4.85E-02 | -1.17E+00 |
| KPTN | 8.61E-04 | 4.48E-03 | -1.20E+00 | 2.75E-04 | 1.19E-03 | -1.21E+00 | 5.92E-02 | 2.52E-01 | 1.06E+00 | 1.51E-02 | 5.75E-02 | 1.06E+00 | 1.34E-04 | 4.77E-04 | -2.12E+00 | 2.52E-03 | 7.10E-03 | -1.52E+00 | 5.15E-02 | 1.89E-01 | 1.07E+00 | 9.30E-02 | 2.56E-01 | 1.12E+00 |
| NEU3 | 8.61E-04 | 4.49E-03 | -1.06E+00 | 6.23E-03 | 2.69E-02 | 1.30E+00 | 8.53E-06 | 3.62E-05 | -1.48E+00 | 3.03E-04 | 1.16E-03 | -1.10E+00 | 4.07E-03 | 1.45E-02 | -1.20E+00 | 1.59E-02 | 4.49E-02 | -1.03E+00 | 3.78E-04 | 1.39E-03 | -1.19E+00 | 5.08E-02 | 1.40E-01 | 1.05E+00 |
| ICAM5 | 8.62E-04 | 4.49E-03 | -1.29E+00 | 2.92E-02 | 1.26E-01 | 2.03E+00 | 4.66E-05 | 1.98E-04 | -1.82E+00 | 2.12E-01 | 8.09E-01 | 1.36E+00 | 2.22E-02 | 7.93E-02 | -1.01E+00 | 1.90E-01 | 5.36E-01 | 1.69E+00 | 4.38E-04 | 1.61E-03 | -1.15E+00 | 3.12E-01 | 8.57E-01 | 1.35E+00 |
| CXXC4 | 8.63E-04 | 4.50E-03 | -1.82E+00 | 6.99E-02 | 3.02E-01 | -1.01E+00 | 2.13E-01 | 9.07E-01 | 1.07E+00 | 6.76E-02 | 2.58E-01 | 1.79E+00 | 1.59E-01 | 5.66E-01 | -1.20E+00 | 2.94E-01 | 8.28E-01 | 1.05E+00 | 2.47E-01 | 9.05E-01 | 1.32E+00 | 2.42E-01 | 6.64E-01 | 1.50E+00 |
| YKT6 | 8.63E-04 | 4.50E-03 | -1.42E+00 | 1.33E-02 | 5.74E-02 | -1.07E+00 | 3.52E-03 | 1.50E-02 | -1.24E+00 | 4.28E-02 | 1.64E-01 | 1.12E+00 | 4.88E-02 | 1.74E-01 | 1.01E+00 | 1.17E-01 | 3.31E-01 | 1.14E+00 | 5.43E-02 | 1.99E-01 | -1.02E+00 | 3.56E-01 | 9.79E-01 | 1.31E+00 |
| LINC02328 | 8.64E-04 | 4.50E-03 | -1.13E+00 | 2.19E-04 | 9.46E-04 | -1.15E+00 | 7.66E-02 | 3.25E-01 | -1.01E+00 | 1.01E-02 | 3.84E-02 | -1.12E+00 | 2.73E-01 | 9.74E-01 | 1.26E+00 | 2.87E-01 | 8.09E-01 | 1.02E+00 | 1.18E-01 | 4.34E-01 | -1.35E+00 | 2.21E-01 | 6.08E-01 | -1.14E+00 |
| CLDN7 | 8.65E-04 | 4.51E-03 | 2.35E+00 | 2.59E-03 | 1.12E-02 | 2.30E+00 | 9.29E-05 | 3.95E-04 | 2.91E+00 | 3.47E-02 | 1.33E-01 | 2.37E+00 | 2.05E-01 | 7.30E-01 | 1.55E+00 | 3.14E-01 | 8.86E-01 | 1.42E+00 | 2.51E-01 | 9.22E-01 | 1.04E+00 | 3.30E-01 | 9.06E-01 | 1.02E+00 |
| SUPT20H | 8.65E-04 | 4.51E-03 | -1.10E+00 | 2.52E-02 | 1.09E-01 | 1.23E+00 | 1.89E-03 | 8.04E-03 | -1.10E+00 | 1.11E-01 | 4.22E-01 | 1.21E+00 | 7.47E-08 | 2.66E-07 | -1.42E+00 | 1.08E-04 | 3.04E-04 | -1.06E+00 | 3.19E-06 | 1.17E-05 | -1.15E+00 | 6.44E-04 | 1.77E-03 | -1.03E+00 |
| AC004076.2 | 8.65E-04 | 4.51E-03 | -1.39E+00 | 1.24E-01 | 5.35E-01 | 1.30E+00 | 1.28E-02 | 5.42E-02 | -1.25E+00 | 2.49E-01 | 9.52E-01 | 1.42E+00 | 1.86E-01 | 6.63E-01 | 1.44E+00 | 1.48E-01 | 4.17E-01 | 1.82E+00 | 2.09E-01 | 7.66E-01 | 1.29E+00 | 2.02E-01 | 5.55E-01 | 1.51E+00 |
| CDK5RAP1 | 8.66E-04 | 4.51E-03 | -1.09E+00 | 1.13E-02 | 4.88E-02 | 1.18E+00 | 1.56E-02 | 6.63E-02 | 1.01E+00 | 2.80E-04 | 1.07E-03 | -1.06E+00 | 1.08E-03 | 3.86E-03 | -1.21E+00 | 1.42E-02 | 4.01E-02 | -1.03E+00 | 8.29E-03 | 3.04E-02 | -1.01E+00 | 9.02E-02 | 2.48E-01 | 1.05E+00 |
| SESN2 | 8.68E-04 | 4.53E-03 | -1.15E+00 | 2.86E-05 | 1.24E-04 | -1.38E+00 | 1.50E-01 | 6.35E-01 | -1.03E+00 | 2.19E-05 | 8.37E-05 | -1.70E+00 | 4.24E-02 | 1.51E-01 | -1.21E+00 | 5.15E-03 | 1.45E-02 | -1.57E+00 | 2.69E-01 | 9.87E-01 | 1.17E+00 | 5.45E-02 | 1.50E-01 | -1.19E+00 |
| ZNF532 | 8.69E-04 | 4.53E-03 | -1.26E+00 | 1.65E-03 | 7.11E-03 | -1.18E+00 | 1.76E-02 | 7.46E-02 | -1.23E+00 | 7.85E-03 | 3.00E-02 | -1.20E+00 | 2.72E-02 | 9.71E-02 | -1.52E+00 | 6.72E-02 | 1.89E-01 | -1.17E+00 | 1.51E-01 | 5.55E-01 | -1.08E+00 | 2.81E-01 | 7.71E-01 | 1.01E+00 |
| C17orf58 | 8.70E-04 | 4.53E-03 | -1.58E+00 | 4.26E-03 | 1.84E-02 | -1.21E+00 | 9.66E-05 | 4.10E-04 | -1.71E+00 | 3.01E-04 | 1.15E-03 | -1.52E+00 | 1.49E-04 | 5.32E-04 | -1.14E+00 | 4.20E-04 | 1.18E-03 | -1.23E+00 | 1.79E-12 | 6.58E-12 | -1.38E+00 | 1.13E-08 | 3.10E-08 | -1.28E+00 |
| FICD | 8.70E-04 | 4.53E-03 | -1.44E+00 | 1.02E-04 | 4.42E-04 | -1.61E+00 | 5.23E-02 | 2.22E-01 | -1.27E+00 | 2.62E-02 | 1.00E-01 | -1.30E+00 | 3.43E-02 | 1.22E-01 | -1.13E+00 | 1.20E-02 | 3.39E-02 | -1.30E+00 | 6.61E-02 | 2.43E-01 | -1.24E+00 | 2.14E-01 | 5.88E-01 | -1.02E+00 |
| AC115837.1 | 8.70E-04 | 4.53E-03 | 1.05E+00 | 3.70E-02 | 1.60E-01 | 1.29E+00 | 4.25E-02 | 1.80E-01 | 1.04E+00 | 7.38E-02 | 2.82E-01 | 1.26E+00 | 2.21E-02 | 7.89E-02 | 1.06E+00 | 2.37E-01 | 6.68E-01 | 1.39E+00 | 1.05E-02 | 3.84E-02 | -1.01E+00 | 1.87E-01 | 5.13E-01 | 1.17E+00 |
| AC147067.1 | 8.70E-04 | 4.53E-03 | -1.13E+00 | 2.22E-03 | 9.60E-03 | 1.03E+00 | 6.15E-04 | 2.61E-03 | -1.58E+00 | 3.38E-03 | 1.29E-02 | -1.00E+00 | 5.16E-06 | 1.84E-05 | -1.34E+00 | 1.60E-03 | 4.52E-03 | 1.00E+00 | 1.70E-02 | 6.26E-02 | 1.07E+00 | 4.11E-03 | 1.13E-02 | -1.18E+00 |
| TMEM60 | 8.71E-04 | 4.54E-03 | -1.19E+00 | 1.56E-03 | 6.75E-03 | -1.07E+00 | 3.62E-07 | 1.54E-06 | -1.63E+00 | 5.46E-04 | 2.09E-03 | -1.09E+00 | 6.87E-07 | 2.45E-06 | -1.68E+00 | 1.47E-04 | 4.13E-04 | -1.27E+00 | 2.11E-06 | 7.74E-06 | -1.22E+00 | 2.45E-03 | 6.72E-03 | 1.01E+00 |
| AC024075.1 | 8.74E-04 | 4.56E-03 | -1.62E+00 | 2.50E-03 | 1.08E-02 | -1.47E+00 | 2.38E-05 | 1.01E-04 | -1.65E+00 | 1.69E-05 | 6.46E-05 | -1.60E+00 | 1.71E-05 | 6.09E-05 | -2.58E+00 | 8.86E-04 | 2.50E-03 | -1.56E+00 | 1.13E-02 | 4.14E-02 | -1.29E+00 | 2.14E-02 | 5.88E-02 | -1.28E+00 |
| ASS1P1 | 8.76E-04 | 4.56E-03 | -1.36E+00 | 5.67E-03 | 2.45E-02 | 2.19E+00 | 3.36E-02 | 1.43E-01 | -1.22E+00 | 5.56E-02 | 2.13E-01 | 1.94E+00 | 1.02E-01 | 3.65E-01 | -1.43E+00 | 3.16E-01 | 8.91E-01 | 1.31E+00 | 2.92E-02 | 1.07E-01 | -1.51E+00 | 3.42E-01 | 9.40E-01 | 1.18E+00 |
| AMD1 | 8.77E-04 | 4.57E-03 | 1.40E+00 | 6.66E-05 | 2.88E-04 | 1.54E+00 | 5.69E-04 | 2.42E-03 | 1.38E+00 | 3.47E-06 | 1.33E-05 | 1.62E+00 | 2.24E-01 | 7.99E-01 | 1.23E+00 | 5.76E-02 | 1.62E-01 | 1.35E+00 | 2.24E-01 | 8.23E-01 | 1.24E+00 | 4.72E-03 | 1.30E-02 | 1.40E+00 |
| POLR3GP1 | 8.78E-04 | 4.58E-03 | -1.73E+00 | 4.28E-04 | 1.85E-03 | -1.80E+00 | 1.46E-05 | 6.19E-05 | -1.92E+00 | 4.20E-05 | 1.61E-04 | -1.70E+00 | 6.09E-02 | 2.17E-01 | -1.24E+00 | 1.95E-01 | 5.49E-01 | -1.07E+00 | 5.11E-04 | 1.88E-03 | -1.38E+00 | 7.80E-02 | 2.14E-01 | 1.01E+00 |
| CHEK2 | 8.81E-04 | 4.59E-03 | -1.38E+00 | 7.57E-04 | 3.27E-03 | -1.23E+00 | 7.53E-02 | 3.20E-01 | -1.06E+00 | 2.20E-01 | 8.41E-01 | 1.34E+00 | 7.15E-02 | 2.55E-01 | -1.24E+00 | 3.41E-01 | 9.61E-01 | 1.23E+00 | 4.24E-05 | 1.56E-04 | -1.41E+00 | 2.61E-01 | 7.17E-01 | 1.27E+00 |
| ZDHHC24 | 8.82E-04 | 4.60E-03 | 1.00E+00 | 2.27E-03 | 9.82E-03 | 1.15E+00 | 5.65E-03 | 2.40E-02 | -1.07E+00 | 8.06E-03 | 3.08E-02 | 1.07E+00 | 7.01E-03 | 2.50E-02 | 1.04E+00 | 3.50E-03 | 9.87E-03 | 1.08E+00 | 9.63E-03 | 3.53E-02 | -1.00E+00 | 1.03E-01 | 2.83E-01 | 1.15E+00 |
| TBC1D31 | 8.86E-04 | 4.62E-03 | -1.10E+00 | 1.34E-01 | 5.81E-01 | 1.49E+00 | 6.23E-04 | 2.64E-03 | -1.27E+00 | 6.24E-02 | 2.38E-01 | 1.21E+00 | 1.53E-02 | 5.44E-02 | -1.14E+00 | 2.49E-01 | 7.02E-01 | 1.33E+00 | 1.02E-02 | 3.76E-02 | -1.17E+00 | 1.04E-01 | 2.85E-01 | -1.00E+00 |
| RASSF5 | 8.87E-04 | 4.62E-03 | -1.20E+00 | 1.28E-02 | 5.53E-02 | 1.07E+00 | 1.33E-03 | 5.65E-03 | -1.18E+00 | 2.33E-06 | 8.89E-06 | -1.39E+00 | 1.52E-04 | 5.43E-04 | -1.20E+00 | 1.25E-03 | 3.52E-03 | -1.13E+00 | 7.41E-10 | 2.72E-09 | -1.72E+00 | 5.20E-06 | 1.43E-05 | -1.25E+00 |
| ARFC3P1 | 8.87E-04 | 4.62E-03 | -1.22E+00 | 7.07E-02 | 3.06E-01 | 1.29E+00 | 3.82E-02 | 1.62E-01 | 1.56E+00 | 3.68E-03 | 1.41E-02 | 2.15E+00 | 1.81E-01 | 6.46E-01 | 1.17E+00 | 2.38E-01 | 6.70E-01 | 1.62E+00 | 2.21E-02 | 8.11E-02 | 1.64E+00 | 7.12E-04 | 1.95E-03 | 2.11E+00 |
| IPO13 | 8.88E-04 | 4.63E-03 | -1.54E+00 | 8.72E-02 | 3.77E-01 | 1.21E+00 | 3.73E-05 | 1.59E-04 | -1.82E+00 | 6.97E-03 | 2.66E-02 | -1.11E+00 | 9.73E-03 | 3.47E-02 | -1.26E+00 | 2.47E-01 | 6.98E-01 | 1.36E+00 | 1.13E-03 | 4.15E-03 | -1.23E+00 | 2.14E-01 | 5.89E-01 | 1.08E+00 |
| PSMD5 | 8.90E-04 | 4.64E-03 | -1.20E+00 | 1.09E-02 | 4.72E-02 | 1.12E+00 | 1.50E-04 | 6.37E-04 | -1.42E+00 | 4.09E-03 | 1.56E-02 | -1.05E+00 | 4.59E-04 | 1.64E-03 | -1.18E+00 | 7.09E-03 | 2.00E-02 | -1.00E+00 | 1.58E-06 | 5.81E-06 | -1.21E+00 | 3.79E-04 | 1.04E-03 | -1.09E+00 |
| WRAP73 | 8.90E-04 | 4.64E-03 | -1.28E+00 | 2.30E-01 | 9.95E-01 | 1.45E+00 | 7.03E-07 | 2.99E-06 | -1.55E+00 | 1.48E-02 | 5.65E-02 | 1.11E+00 | 3.86E-04 | 1.38E-03 | -1.33E+00 | 1.99E-02 | 5.61E-02 | 1.03E+00 | 2.94E-05 | 1.08E-04 | -1.26E+00 | 1.32E-01 | 3.63E-01 | 1.17E+00 |
| ODF2L | 8.90E-04 | 4.64E-03 | -1.00E+00 | 3.19E-04 | 1.38E-03 | 1.10E+00 | 2.40E-03 | 1.02E-02 | -1.10E+00 | 4.60E-05 | 1.76E-04 | -1.07E+00 | 1.60E-04 | 5.69E-04 | -1.07E+00 | 7.69E-06 | 2.17E-05 | -1.23E+00 | 2.86E-02 | 1.05E-01 | 1.02E+00 | 1.73E-03 | 4.75E-03 | -1.13E+00 |
| PJA1 | 8.95E-04 | 4.66E-03 | -1.28E+00 | 8.17E-02 | 3.53E-01 | 1.31E+00 | 3.18E-05 | 1.35E-04 | -1.47E+00 | 5.49E-03 | 2.10E-02 | -1.00E+00 | 2.21E-06 | 7.87E-06 | -1.70E+00 | 7.67E-05 | 2.16E-04 | -1.29E+00 | 2.28E-07 | 8.35E-07 | -1.67E+00 | 2.03E-03 | 5.57E-03 | -1.10E+00 |
| PHLDB3 | 8.96E-04 | 4.67E-03 | -1.21E+00 | 6.87E-04 | 2.97E-03 | -1.15E+00 | 1.40E-03 | 5.96E-03 | -1.37E+00 | 1.57E-03 | 6.00E-03 | -1.14E+00 | 5.00E-05 | 1.78E-04 | -1.51E+00 | 3.94E-03 | 1.11E-02 | -1.05E+00 | 3.58E-03 | 1.31E-02 | -1.21E+00 | 2.37E-01 | 6.51E-01 | 1.16E+00 |
| SRCAP | 8.99E-04 | 4.69E-03 | -1.22E+00 | 2.36E-04 | 1.02E-03 | -1.29E+00 | 1.50E-05 | 6.37E-05 | -1.57E+00 | 3.89E-05 | 1.49E-04 | -1.30E+00 | 2.42E-02 | 8.63E-02 | 1.06E+00 | 8.28E-03 | 2.33E-02 | -1.15E+00 | 4.80E-02 | 1.76E-01 | -1.07E+00 | 7.36E-03 | 2.02E-02 | -1.37E+00 |
| EXOC2 | 8.99E-04 | 4.69E-03 | -1.51E+00 | 8.69E-02 | 3.75E-01 | 1.16E+00 | 8.33E-04 | 3.54E-03 | -1.36E+00 | 3.99E-03 | 1.52E-02 | -1.08E+00 | 3.25E-03 | 1.16E-02 | -1.26E+00 | 3.52E-02 | 9.93E-02 | 1.04E+00 | 3.07E-05 | 1.13E-04 | -1.42E+00 | 1.84E-03 | 5.05E-03 | -1.14E+00 |
| SMARCA5-AS1 | 8.99E-04 | 4.69E-03 | -1.65E+00 | 8.69E-02 | 3.76E-01 | 1.16E+00 | 1.27E-10 | 5.40E-10 | -2.65E+00 | 7.91E-06 | 3.02E-05 | -1.40E+00 | 3.30E-03 | 1.18E-02 | -1.46E+00 | 1.13E-01 | 3.17E-01 | 1.09E+00 | 9.00E-03 | 3.30E-02 | -1.42E+00 | 1.56E-01 | 4.28E-01 | 1.01E+00 |
| PRMT1 | 9.00E-04 | 4.69E-03 | -1.00E+00 | 6.19E-03 | 2.67E-02 | 1.15E+00 | 2.25E-01 | 9.56E-01 | 1.14E+00 | 1.81E-01 | 6.93E-01 | 1.22E+00 | 2.09E-03 | 7.45E-03 | -1.01E+00 | 2.58E-01 | 7.28E-01 | 1.30E+00 | 2.70E-01 | 9.90E-01 | 1.14E+00 | 7.18E-03 | 1.97E-02 | 1.39E+00 |
| AC116913.1 | 9.00E-04 | 4.69E-03 | -1.80E+00 | 7.16E-02 | 3.09E-01 | 1.08E+00 | 1.05E-02 | 4.45E-02 | -1.32E+00 | 2.16E-02 | 8.26E-02 | -1.15E+00 | 1.83E-01 | 6.51E-01 | 1.64E+00 | 1.15E-01 | 3.26E-01 | 1.95E+00 | 3.72E-02 | 1.36E-01 | -1.18E+00 | 2.97E-01 | 8.16E-01 | 1.15E+00 |
| EEF1A1 | 9.01E-04 | 4.70E-03 | -1.01E+00 | 2.28E-01 | 9.84E-01 | 1.25E+00 | 9.60E-02 | 4.08E-01 | -1.02E+00 | 2.02E-02 | 7.70E-02 | 1.28E+00 | 3.40E-02 | 1.21E-01 | 1.02E+00 | 2.96E-01 | 8.36E-01 | 1.22E+00 | 5.32E-03 | 1.95E-02 | -1.02E+00 | 1.97E-01 | 5.41E-01 | 1.23E+00 |
| EAF1 | 9.03E-04 | 4.71E-03 | -1.21E+00 | 3.42E-03 | 1.48E-02 | 1.04E+00 | 1.54E-05 | 6.53E-05 | -1.47E+00 | 6.22E-06 | 2.38E-05 | -1.23E+00 | 9.01E-06 | 3.21E-05 | -1.87E+00 | 1.82E-04 | 5.12E-04 | -1.45E+00 | 8.75E-09 | 3.21E-08 | -1.75E+00 | 1.08E-07 | 2.97E-07 | -1.67E+00 |
| FBXL6 | 9.04E-04 | 4.71E-03 | -1.23E+00 | 1.46E-04 | 6.31E-04 | -1.24E+00 | 4.18E-05 | 1.78E-04 | -1.25E+00 | 8.67E-06 | 3.31E-05 | -1.22E+00 | 5.25E-02 | 1.87E-01 | 1.04E+00 | 6.31E-02 | 1.78E-01 | 1.12E+00 | 7.00E-02 | 2.57E-01 | 1.15E+00 | 2.87E-02 | 7.89E-02 | 1.03E+00 |
| MAPKAPK2 | 9.04E-04 | 4.71E-03 | 1.06E+00 | 2.05E-08 | 8.85E-08 | -1.28E+00 | 6.03E-02 | 2.56E-01 | 1.12E+00 | 3.25E-02 | 1.24E-01 | 1.20E+00 | 5.06E-05 | 1.80E-04 | -1.09E+00 | 2.19E-06 | 6.17E-06 | -1.22E+00 | 5.68E-05 | 2.08E-04 | -1.17E+00 | 8.44E-05 | 2.32E-04 | -1.14E+00 |
| RPL21 | 9.06E-04 | 4.72E-03 | 1.07E+00 | 3.00E-19 | 1.30E-18 | -1.65E+00 | 1.56E-01 | 6.64E-01 | 1.18E+00 | 6.11E-11 | 2.33E-10 | -1.41E+00 | 1.32E-02 | 4.69E-02 | 1.15E+00 | 5.37E-10 | 1.51E-09 | -1.50E+00 | 1.06E-01 | 3.91E-01 | 1.06E+00 | 2.19E-08 | 6.02E-08 | -1.49E+00 |
| LIPE | 9.07E-04 | 4.73E-03 | -1.38E+00 | 5.63E-04 | 2.43E-03 | -1.36E+00 | 2.55E-02 | 1.08E-01 | -1.19E+00 | 4.78E-02 | 1.83E-01 | -1.05E+00 | 2.16E-03 | 7.71E-03 | -1.61E+00 | 1.05E-02 | 2.97E-02 | -1.23E+00 | 4.52E-02 | 1.66E-01 | -1.16E+00 | 3.09E-01 | 8.49E-01 | 1.39E+00 |
| TGIF1 | 9.07E-04 | 4.73E-03 | 1.87E+00 | 3.11E-02 | 1.34E-01 | 1.94E+00 | 2.03E-01 | 8.62E-01 | 1.27E+00 | 1.74E-01 | 6.64E-01 | 1.25E+00 | 1.58E-01 | 5.57E-01 | 1.22E+00 | 2.87E-01 | 8.09E-01 | 1.23E+00 | 3.04E-02 | 1.11E-01 | 1.02E+00 | 3.14E-01 | 8.62E-01 | 1.26E+00 |
| SMIM1 | 9.09E-04 | 4.74E-03 | -1.14E+00 | 4.36E-06 | 1.88E-05 | -1.43E+00 | 2.07E-02 | 8.79E-02 | 1.08E+00 | 1.62E-03 | 6.18E-03 | -1.19E+00 | 1.24E-01 | 4.44E-01 | 1.72E+00 | 2.63E-01 | 7.42E-01 | 1.48E+00 | 3.46E-05 | 1.27E-04 | 1.70E+00 | 8.91E-02 | 2.45E-01 | 1.12E+00 |
| FAM43A | 9.10E-04 | 4.74E-03 | -1.21E+00 | 1.60E-02 | 6.90E-02 | 1.09E+00 | 9.02E-03 | 3.83E-02 | -1.16E+00 | 2.29E-01 | 8.73E-01 | 1.27E+00 | 1.79E-01 | 6.37E-01 | 1.22E+00 | 2.65E-01 | 7.47E-01 | 1.23E+00 | 2.37E-01 | 8.70E-01 | 1.11E+00 | 3.55E-01 | 9.74E-01 | 1.33E+00 |
| ERCC8 | 9.12E-04 | 4.75E-03 | -1.19E+00 | 6.47E-03 | 2.80E-02 | 1.10E+00 | 6.49E-03 | 2.76E-02 | -1.18E+00 | 1.22E-02 | 4.68E-02 | 1.03E+00 | 1.44E-02 | 5.15E-02 | -1.17E+00 | 7.04E-02 | 1.98E-01 | 1.08E+00 | 2.12E-04 | 7.77E-04 | -1.29E+00 | 4.58E-03 | 1.26E-02 | -1.14E+00 |
| TMEM120A | 9.13E-04 | 4.76E-03 | -1.42E+00 | 2.38E-03 | 1.03E-02 | -1.22E+00 | 2.37E-03 | 1.01E-02 | -1.33E+00 | 5.73E-03 | 2.19E-02 | -1.12E+00 | 3.15E-03 | 1.12E-02 | -1.24E+00 | 3.70E-02 | 1.04E-01 | 1.01E+00 | 1.06E-01 | 3.89E-01 | 1.50E+00 | 2.45E-01 | 6.74E-01 | 1.41E+00 |
| SORBS1 | 9.14E-04 | 4.77E-03 | -1.66E+00 | 2.14E-02 | 9.23E-02 | -1.09E+00 | 8.19E-03 | 3.48E-02 | -1.49E+00 | 6.35E-02 | 2.43E-01 | -1.03E+00 | 3.06E-18 | 1.09E-17 | -2.16E+00 | 5.15E-17 | 1.45E-16 | -2.01E+00 | 4.35E-16 | 1.60E-15 | -2.08E+00 | 2.50E-14 | 6.87E-14 | -2.05E+00 |
| EARS2 | 9.15E-04 | 4.77E-03 | -1.55E+00 | 1.24E-02 | 5.35E-02 | -1.07E+00 | 4.79E-03 | 2.04E-02 | -1.37E+00 | 1.37E-01 | 5.24E-01 | 1.28E+00 | 1.55E-02 | 5.53E-02 | -1.71E+00 | 2.85E-01 | 8.04E-01 | 1.21E+00 | 3.82E-04 | 1.40E-03 | -1.31E+00 | 3.50E-01 | 9.62E-01 | 1.30E+00 |
| SRP14-AS1 | 9.16E-04 | 4.77E-03 | -1.03E+00 | 1.70E-02 | 7.36E-02 | 1.24E+00 | 3.00E-02 | 1.28E-01 | -1.02E+00 | 2.63E-03 | 1.00E-02 | -1.03E+00 | 2.02E-02 | 7.20E-02 | -1.21E+00 | 1.13E-01 | 3.18E-01 | 1.07E+00 | 3.32E-02 | 1.22E-01 | -1.06E+00 | 1.29E-01 | 3.54E-01 | 1.07E+00 |
| RPL35AP21 | 9.17E-04 | 4.78E-03 | 1.92E+00 | 3.05E-02 | 1.32E-01 | 1.21E+00 | 8.79E-03 | 3.73E-02 | 1.61E+00 | 3.27E-02 | 1.25E-01 | 1.09E+00 | 2.11E-01 | 7.52E-01 | 1.56E+00 | 3.10E-01 | 8.75E-01 | 1.48E+00 | 6.91E-04 | 2.54E-03 | 2.09E+00 | 1.59E-01 | 4.37E-01 | 1.56E+00 |
| RGS5 | 9.19E-04 | 4.79E-03 | -1.44E+00 | 8.26E-03 | 3.57E-02 | -1.10E+00 | 7.13E-02 | 3.03E-01 | -1.09E+00 | 1.26E-01 | 4.80E-01 | 1.05E+00 | 3.57E-02 | 1.27E-01 | -1.24E+00 | 1.17E-01 | 3.30E-01 | -1.06E+00 | 1.85E-02 | 6.81E-02 | -1.08E+00 | 2.37E-02 | 6.50E-02 | -1.24E+00 |
| C15orf52 | 9.19E-04 | 4.79E-03 | -1.31E+00 | 4.39E-02 | 1.90E-01 | 1.72E+00 | 2.67E-02 | 1.13E-01 | -1.46E+00 | 1.61E-01 | 6.15E-01 | 1.86E+00 | 2.64E-01 | 9.42E-01 | 1.59E+00 | 2.97E-01 | 8.38E-01 | 1.76E+00 | 1.07E-01 | 3.94E-01 | -1.16E+00 | 2.27E-01 | 6.25E-01 | -1.04E+00 |
| LINC02019 | 9.19E-04 | 4.79E-03 | -1.42E+00 | 3.81E-03 | 1.65E-02 | -1.12E+00 | 1.83E-01 | 7.79E-01 | 1.13E+00 | 6.69E-02 | 2.55E-01 | -1.03E+00 | 1.82E-01 | 6.50E-01 | 1.69E+00 | 1.51E-01 | 4.25E-01 | 1.71E+00 | 2.70E-01 | 9.90E-01 | 1.31E+00 | 2.35E-01 | 6.47E-01 | 1.45E+00 |
| SPARC | 9.20E-04 | 4.79E-03 | 1.03E+00 | 7.74E-03 | 3.35E-02 | 1.27E+00 | 4.04E-02 | 1.72E-01 | -1.01E+00 | 7.12E-02 | 2.72E-01 | 1.21E+00 | 2.27E-02 | 8.09E-02 | -1.01E+00 | 9.90E-02 | 2.79E-01 | 1.20E+00 | 1.76E-01 | 6.46E-01 | 1.52E+00 | 4.73E-02 | 1.30E-01 | 1.71E+00 |
| CCDC69 | 9.20E-04 | 4.79E-03 | -1.10E+00 | 4.05E-02 | 1.75E-01 | 1.28E+00 | 2.00E-05 | 8.51E-05 | -1.34E+00 | 1.59E-04 | 6.06E-04 | -1.10E+00 | 1.18E-01 | 4.22E-01 | 1.28E+00 | 5.47E-02 | 1.54E-01 | 1.68E+00 | 4.97E-02 | 1.83E-01 | 1.04E+00 | 1.24E-01 | 3.42E-01 | 1.15E+00 |
| SNORA19 | 9.24E-04 | 4.82E-03 | -1.47E+00 | 3.07E-03 | 1.33E-02 | -1.19E+00 | 2.86E-03 | 1.21E-02 | -1.28E+00 | 2.71E-03 | 1.04E-02 | -1.22E+00 | 6.73E-03 | 2.40E-02 | -1.58E+00 | 3.27E-02 | 9.22E-02 | -1.17E+00 | 2.15E-03 | 7.87E-03 | -1.62E+00 | 4.63E-03 | 1.27E-02 | -1.57E+00 |
| GABBR1 | 9.25E-04 | 4.82E-03 | -1.10E+00 | 3.64E-02 | 1.57E-01 | 1.32E+00 | 5.44E-04 | 2.31E-03 | -1.28E+00 | 4.24E-04 | 1.62E-03 | -1.20E+00 | 2.55E-03 | 9.09E-03 | -1.03E+00 | 3.20E-02 | 9.03E-02 | 1.14E+00 | 2.82E-05 | 1.04E-04 | -1.22E+00 | 6.25E-04 | 1.72E-03 | -1.17E+00 |
| MAURD1 | 9.25E-04 | 4.82E-03 | -1.21E+00 | 4.53E-03 | 1.96E-02 | -1.03E+00 | 1.12E-02 | 4.74E-02 | -1.37E+00 | 8.38E-02 | 3.20E-01 | 1.00E+00 | 2.09E-03 | 7.46E-03 | -1.51E+00 | 3.19E-02 | 8.98E-02 | -1.06E+00 | 1.86E-03 | 6.84E-03 | -1.27E+00 | 7.19E-02 | 1.98E-01 | 1.02E+00 |
| UBXN7 | 9.27E-04 | 4.83E-03 | 1.04E+00 | 2.39E-05 | 1.03E-04 | 1.05E+00 | 1.86E-03 | 7.91E-03 | -1.16E+00 | 3.74E-04 | 1.43E-03 | -1.07E+00 | 9.88E-05 | 3.52E-04 | -1.07E+00 | 2.27E-05 | 6.40E-05 | -1.09E+00 | 5.15E-02 | 1.89E-01 | -1.05E+00 | 2.05E-02 | 5.62E-02 | -1.06E+00 |
| AC138969.1 | 9.27E-04 | 4.83E-03 | -1.55E+00 | 1.27E-04 | 5.51E-04 | -1.81E+00 | 1.85E-06 | 7.87E-06 | -2.03E+00 | 6.59E-06 | 2.52E-05 | -1.72E+00 | 4.66E-02 | 1.66E-01 | -1.04E+00 | 2.03E-02 | 5.73E-02 | -1.39E+00 | 3.61E-05 | 1.32E-04 | -1.47E+00 | 7.92E-08 | 2.18E-07 | -2.39E+00 |
| DPY19L1 | 9.28E-04 | 4.83E-03 | -1.70E+00 | 3.75E-03 | 1.64E-02 | -1.33E+00 | 3.33E-02 | 1.41E-01 | -1.43E+00 | 8.62E-02 | 3.29E-01 | -1.06E+00 | 2.22E-03 | 7.93E-03 | -1.39E+00 | 5.56E-03 | 1.57E-02 | -1.19E+00 | 2.71E-02 | 9.93E-02 | -1.15E+00 | 1.18E-01 | 3.25E-01 | -1.07E+00 |

| | | | | | | | | | | | | | | | | | | | | | | | | |
|---|---|---|---|---|---|---|---|---|---|---|---|---|---|---|---|---|---|---|---|---|---|---|---|---|
| WDR31 | 9.28E-04 | 4.84E-03 | -1.40E+00 | 3.11E-03 | 1.35E-02 | -1.18E+00 | 6.04E-05 | 2.56E-04 | -1.56E+00 | 2.48E-05 | 9.49E-05 | -1.43E+00 | 1.93E-01 | 6.87E-01 | 1.26E+00 | 1.13E-01 | 3.17E-01 | 2.09E+00 | 9.16E-02 | 3.36E-01 | 1.15E+00 | 2.87E-01 | 7.90E-01 | 1.28E+00 |
| POT1-AS1 | 9.30E-04 | 4.85E-03 | -1.65E+00 | 3.39E-02 | 1.46E-01 | -1.21E+00 | 1.97E-01 | 8.37E-01 | 1.11E+00 | 2.28E-01 | 8.72E-01 | 1.16E+00 | 2.28E-02 | 8.15E-02 | -1.80E+00 | 4.85E-02 | 1.37E-01 | -1.47E+00 | 2.69E-01 | 9.87E-01 | 1.05E+00 | 2.99E-01 | 8.22E-01 | 1.18E+00 |
| BTD | 9.34E-04 | 4.87E-03 | -1.32E+00 | 3.18E-02 | 1.37E-01 | 1.16E+00 | 2.31E-02 | 9.80E-02 | -1.17E+00 | 5.38E-02 | 2.05E-01 | 1.02E+00 | 5.52E-02 | 1.97E-01 | 1.04E+00 | 2.78E-01 | 7.83E-01 | 1.31E+00 | 6.44E-02 | 2.36E-01 | 1.10E+00 | 2.11E-02 | 5.80E-02 | -1.05E+00 |
| CXCL1 | 9.36E-04 | 4.88E-03 | -1.28E+00 | 2.31E-06 | 9.98E-06 | -2.70E+00 | 1.34E-02 | 5.70E-02 | 2.09E+00 | 9.46E-02 | 3.61E-01 | -1.27E+00 | 1.20E-01 | 4.28E-01 | -1.78E+00 | 1.77E-02 | 4.99E-02 | -3.70E+00 | 4.62E-02 | 1.69E-01 | 1.63E+00 | 9.61E-05 | 2.64E-04 | -2.31E+00 |
| AKAP13 | 9.37E-04 | 4.88E-03 | 1.10E+00 | 6.20E-04 | 2.68E-03 | 1.19E+00 | 9.14E-04 | 3.89E-03 | -1.09E+00 | 4.17E-04 | 1.59E-03 | 1.01E+00 | 6.43E-02 | 2.29E-01 | 1.15E+00 | 1.04E-03 | 2.93E-03 | -1.09E+00 | 1.01E-02 | 3.69E-02 | 1.00E+00 | 1.65E-04 | 4.53E-04 | -1.19E+00 |
| SYAP1 | 9.37E-04 | 4.88E-03 | 1.09E+00 | 3.34E-12 | 1.44E-11 | -1.31E+00 | 1.31E-01 | 5.57E-01 | 1.12E+00 | 2.46E-06 | 9.38E-06 | -1.25E+00 | 1.92E-01 | 6.84E-01 | 1.33E+00 | 1.97E-04 | 5.54E-04 | -1.11E+00 | 1.63E-03 | 5.99E-03 | 1.03E+00 | 3.88E-09 | 1.07E-08 | -1.37E+00 |
| GIMAP6 | 9.38E-04 | 4.89E-03 | 2.02E+00 | 6.78E-17 | 2.93E-16 | 3.61E+00 | 3.05E-02 | 1.30E-01 | -1.03E+00 | 2.08E-02 | 7.94E-02 | 1.86E+00 | 1.09E-01 | 3.89E-01 | 1.01E+00 | 2.54E-01 | 7.16E-01 | 1.23E+00 | 2.01E-02 | 7.37E-02 | -1.59E+00 | 2.10E-01 | 5.76E-01 | 1.01E+00 |
| C8orf88 | 9.38E-04 | 4.89E-03 | -1.24E+00 | 3.76E-07 | 1.63E-06 | -2.05E+00 | 2.84E-02 | 1.21E-01 | -1.21E+00 | 1.84E-02 | 7.02E-02 | -1.23E+00 | 2.00E-01 | 7.13E-01 | 1.23E+00 | 2.53E-01 | 7.13E-01 | 1.12E+00 | 2.35E-01 | 8.61E-01 | 1.07E+00 | 1.26E-01 | 3.47E-01 | -1.12E+00 |
| SHARPIN | 9.38E-04 | 4.89E-03 | -1.06E+00 | 5.68E-04 | 2.46E-03 | 1.06E+00 | 5.92E-04 | 2.52E-03 | -1.15E+00 | 6.72E-03 | 2.57E-02 | 1.10E+00 | 5.36E-04 | 1.91E-03 | -1.01E+00 | 5.17E-03 | 1.46E-02 | 1.10E+00 | 7.57E-02 | 2.78E-01 | 1.19E+00 | 2.94E-01 | 8.08E-01 | 1.27E+00 |
| ATP5G3 | 9.40E-04 | 4.90E-03 | -1.02E+00 | 1.03E-03 | 4.46E-03 | 1.03E+00 | 1.39E-01 | 5.92E-01 | 1.10E+00 | 1.74E-01 | 6.63E-01 | 1.02E+00 | 2.92E-05 | 1.04E-04 | -1.16E+00 | 9.88E-02 | 2.78E-01 | 1.01E+00 | 4.08E-02 | 1.50E-01 | -1.01E+00 | 1.42E-01 | 3.90E-01 | 1.13E+00 |
| OXNAD1 | 9.40E-04 | 4.90E-03 | -1.28E+00 | 8.26E-03 | 3.57E-02 | 1.02E+00 | 2.89E-02 | 1.23E-01 | -1.06E+00 | 5.31E-02 | 2.03E-01 | 1.08E+00 | 1.84E-01 | 6.56E-01 | 1.25E+00 | 1.81E-01 | 5.09E-01 | 1.09E+00 | 1.40E-03 | 5.13E-03 | -1.08E+00 | 1.28E-01 | 3.51E-01 | 1.17E+00 |
| MRPS10P1 | 9.41E-04 | 4.90E-03 | -1.27E+00 | 1.13E-05 | 4.87E-05 | -1.43E+00 | 6.88E-05 | 2.92E-04 | -1.26E+00 | 7.06E-07 | 2.70E-06 | -1.31E+00 | 4.39E-03 | 1.56E-02 | -1.08E+00 | 7.66E-03 | 2.16E-02 | -1.06E+00 | 5.31E-05 | 1.95E-04 | -1.22E+00 | 4.33E-08 | 1.19E-07 | -1.56E+00 |
| WDR86-AS1 | 9.42E-04 | 4.91E-03 | -1.29E+00 | 6.52E-03 | 2.82E-02 | -1.03E+00 | 5.03E-04 | 2.14E-03 | -1.46E+00 | 1.29E-04 | 4.93E-04 | -1.51E+00 | 8.61E-02 | 3.07E-01 | -1.28E+00 | 1.63E-01 | 4.61E-01 | -1.18E+00 | 2.23E-01 | 8.17E-01 | -1.08E+00 | 3.33E-01 | 9.15E-01 | 1.05E+00 |
| ZBTB42 | 9.42E-04 | 4.91E-03 | -1.33E+00 | 1.18E-02 | 5.10E-02 | -1.01E+00 | 1.59E-04 | 6.75E-04 | -1.62E+00 | 1.28E-02 | 4.89E-02 | -1.11E+00 | 7.22E-02 | 2.57E-01 | -1.05E+00 | 1.50E-01 | 4.22E-01 | 1.07E+00 | 1.17E-05 | 4.29E-05 | -1.79E+00 | 4.84E-03 | 1.33E-02 | -1.18E+00 |
| TLK2P2 | 9.42E-04 | 4.91E-03 | -1.49E+00 | 3.57E-02 | 1.54E-01 | 1.03E+00 | 1.13E-01 | 4.81E-01 | -1.03E+00 | 1.04E-01 | 3.98E-01 | 1.03E+00 | 2.41E-01 | 8.58E-01 | 1.21E+00 | 2.06E-01 | 5.80E-01 | 1.55E+00 | 6.37E-02 | 2.34E-01 | -1.30E+00 | 2.36E-01 | 6.47E-01 | 1.00E+00 |
| SDSL | 9.44E-04 | 4.92E-03 | 1.02E+00 | 6.70E-03 | 2.89E-02 | 1.22E+00 | 9.39E-05 | 3.99E-04 | -1.16E+00 | 1.56E-06 | 5.95E-06 | -1.14E+00 | 9.67E-05 | 3.45E-04 | -1.58E+00 | 6.87E-04 | 1.94E-03 | -1.41E+00 | 1.26E-04 | 4.61E-04 | -1.29E+00 | 2.10E-03 | 5.76E-03 | -1.08E+00 |
| VPS16 | 9.44E-04 | 4.92E-03 | -1.08E+00 | 6.37E-03 | 2.75E-02 | 1.12E+00 | 1.74E-04 | 7.37E-04 | -1.11E+00 | 1.30E-05 | 4.97E-05 | -1.06E+00 | 1.65E-02 | 5.88E-02 | 1.04E+00 | 2.21E-02 | 6.23E-02 | 1.07E+00 | 1.41E-01 | 5.16E-01 | 1.24E+00 | 2.35E-01 | 6.46E-01 | 1.29E+00 |
| VIM2P | 9.46E-04 | 4.93E-03 | 2.65E+00 | 8.22E-03 | 3.55E-02 | 2.72E+00 | 3.96E-06 | 1.68E-05 | 3.06E+00 | 2.00E-04 | 7.62E-04 | 3.04E+00 | 5.21E-02 | 1.86E-01 | 1.95E+00 | 8.95E-02 | 2.52E-01 | 1.79E+00 | 2.68E-01 | 9.85E-01 | 1.18E+00 | 2.66E-01 | 7.31E-01 | -1.02E+00 |
| ECSIT | 9.47E-04 | 4.94E-03 | -1.02E+00 | 2.09E-04 | 9.02E-04 | 1.04E+00 | 6.66E-07 | 2.83E-06 | -1.17E+00 | 4.85E-07 | 1.85E-06 | -1.07E+00 | 2.67E-11 | 9.51E-11 | -1.73E+00 | 9.12E-06 | 2.57E-05 | -1.19E+00 | 2.28E-07 | 8.35E-07 | -1.30E+00 | 8.56E-05 | 2.35E-04 | -1.14E+00 |
| AL645568.1 | 9.48E-04 | 4.94E-03 | -1.98E+00 | 4.24E-02 | 1.83E-01 | -1.08E+00 | 3.01E-03 | 1.28E-02 | -1.46E+00 | 4.43E-04 | 1.69E-03 | -1.61E+00 | 8.37E-02 | 2.98E-01 | -1.04E+00 | 3.55E-01 | 1.00E+00 | 1.54E+00 | 8.93E-02 | 3.28E-01 | -1.09E+00 | 2.66E-01 | 7.31E-01 | 1.04E+00 |
| DHRS11 | 9.49E-04 | 4.95E-03 | -1.11E+00 | 1.32E-03 | 5.69E-03 | -1.02E+00 | 4.45E-02 | 1.89E-01 | -1.03E+00 | 9.29E-02 | 3.55E-01 | 1.15E+00 | 6.52E-04 | 2.32E-03 | -1.18E+00 | 9.36E-04 | 2.64E-03 | -1.11E+00 | 8.66E-05 | 3.18E-04 | -1.08E+00 | 3.21E-05 | 8.83E-05 | -1.18E+00 |
| IRAK1BP1 | 9.49E-04 | 4.95E-03 | -1.20E+00 | 1.08E-02 | 4.66E-02 | 1.08E+00 | 3.89E-03 | 1.65E-02 | -1.12E+00 | 1.14E-02 | 4.34E-02 | 1.08E+00 | 1.01E-04 | 3.60E-04 | -1.51E+00 | 1.83E-03 | 5.15E-03 | -1.19E+00 | 1.28E-01 | 4.68E-01 | 1.03E+00 | 1.85E-01 | 5.08E-01 | 1.48E+00 |
| OTUD5 | 9.49E-04 | 4.95E-03 | 1.03E+00 | 2.43E-06 | 1.05E-05 | -1.07E+00 | 1.53E-01 | 6.49E-01 | 1.28E+00 | 2.90E-02 | 1.11E-01 | 1.11E+00 | 6.13E-03 | 2.18E-02 | -1.00E+00 | 4.10E-02 | 1.16E-01 | 1.13E+00 | 1.88E-02 | 6.89E-02 | 1.22E+00 | 2.27E-04 | 6.25E-04 | -1.11E+00 |
| MUS81 | 9.51E-04 | 4.96E-03 | 1.00E+00 | 4.70E-02 | 2.03E-01 | 1.32E+00 | 4.95E-04 | 2.10E-03 | -1.15E+00 | 5.58E-03 | 2.13E-02 | 1.08E+00 | 1.03E-03 | 3.66E-03 | -1.17E+00 | 1.04E-01 | 2.92E-01 | 1.20E+00 | 4.24E-04 | 1.55E-03 | -1.07E+00 | 2.92E-02 | 8.01E-02 | 1.08E+00 |
| NCR3 | 9.51E-04 | 4.96E-03 | 2.61E+00 | 1.30E-01 | 5.61E-01 | 1.53E+00 | 1.85E-01 | 7.85E-01 | 1.20E+00 | 2.32E-01 | 8.86E-01 | 1.23E+00 | 1.99E-01 | 7.11E-01 | 1.35E+00 | 1.17E-01 | 3.31E-01 | 1.86E+00 | 6.22E-02 | 2.28E-01 | 1.58E+00 | 8.88E-02 | 2.44E-01 | 1.68E+00 |
| TVP23B | 9.52E-04 | 4.96E-03 | -1.31E+00 | 2.13E-02 | 9.18E-02 | 1.13E+00 | 1.62E-01 | 6.90E-01 | 1.39E+00 | 1.76E-01 | 6.71E-01 | 1.35E+00 | 3.88E-02 | 1.38E-01 | -1.02E+00 | 2.55E-01 | 7.18E-01 | 1.35E+00 | 9.62E-02 | 3.53E-01 | 1.01E+00 | 1.67E-01 | 4.60E-01 | 1.09E+00 |
| TDP2 | 9.53E-04 | 4.97E-03 | -1.03E+00 | 1.41E-01 | 6.11E-01 | 1.50E+00 | 3.74E-04 | 1.59E-03 | -1.17E+00 | 2.27E-02 | 8.66E-02 | 1.20E+00 | 6.47E-05 | 2.31E-04 | -1.42E+00 | 2.28E-02 | 6.43E-02 | -1.01E+00 | 1.67E-04 | 6.14E-04 | -1.16E+00 | 5.46E-03 | 1.50E-02 | -1.05E+00 |
| SPACA9 | 9.55E-04 | 4.98E-03 | -1.06E+00 | 5.50E-05 | 2.38E-04 | -1.21E+00 | 8.19E-03 | 3.48E-02 | -1.10E+00 | 2.11E-03 | 8.05E-03 | -1.01E+00 | 2.09E-02 | 7.45E-02 | -1.31E+00 | 1.32E-01 | 3.73E-01 | 1.12E+00 | 9.27E-02 | 3.40E-01 | -1.02E+00 | 1.08E-02 | 2.97E-02 | -1.36E+00 |
| NAGLU | 9.59E-04 | 5.00E-03 | -1.17E+00 | 3.67E-03 | 1.59E-02 | -1.00E+00 | 4.28E-07 | 1.82E-06 | -1.40E+00 | 2.01E-07 | 7.67E-07 | -1.29E+00 | 3.56E-04 | 1.27E-03 | -1.50E+00 | 3.62E-02 | 1.02E-01 | -1.04E+00 | 5.11E-05 | 1.87E-04 | -1.22E+00 | 4.32E-03 | 1.19E-02 | -1.12E+00 |
| PARPBP | 9.61E-04 | 5.01E-03 | -1.45E+00 | 1.43E-03 | 6.19E-03 | -1.25E+00 | 1.15E-01 | 4.89E-01 | -1.01E+00 | 7.96E-03 | 3.04E-02 | -1.15E+00 | 7.54E-05 | 2.69E-04 | -1.79E+00 | 5.61E-05 | 1.58E-04 | -1.84E+00 | 8.61E-05 | 3.16E-04 | -1.18E+00 | 4.83E-02 | 1.33E-01 | 1.13E+00 |
| CLSTN1 | 9.61E-04 | 5.01E-03 | 1.04E+00 | 1.19E-01 | 5.15E-01 | 1.43E+00 | 7.54E-03 | 3.20E-02 | -1.02E+00 | 5.59E-02 | 2.13E-01 | 1.25E+00 | 6.73E-05 | 2.40E-04 | -1.30E+00 | 4.51E-04 | 1.27E-03 | -1.15E+00 | 2.16E-05 | 7.94E-05 | -1.14E+00 | 1.86E-02 | 5.11E-02 | 1.16E+00 |
| RPS15AP1 | 9.63E-04 | 5.02E-03 | 1.25E+00 | 1.85E-04 | 8.00E-04 | -1.02E+00 | 2.51E-04 | 1.07E-03 | 1.26E+00 | 5.85E-02 | 2.23E-01 | -1.02E+00 | 8.10E-03 | 2.89E-02 | 1.22E+00 | 3.51E-01 | 9.89E-01 | 1.04E+00 | 3.97E-02 | 1.46E-01 | 1.17E+00 | 2.94E-02 | 8.07E-02 | -1.04E+00 |
| UGDH-AS1 | 9.65E-04 | 5.03E-03 | -1.84E+00 | 7.31E-03 | 3.16E-02 | -1.32E+00 | 1.28E-03 | 5.43E-03 | -1.78E+00 | 6.04E-02 | 2.31E-01 | 1.02E+00 | 5.00E-08 | 1.78E-07 | -2.32E+00 | 2.69E-04 | 7.57E-04 | -1.34E+00 | 1.29E-04 | 4.74E-04 | -1.68E+00 | 8.34E-05 | 2.29E-04 | -1.72E+00 |
| GPN2 | 9.66E-04 | 5.03E-03 | -1.12E+00 | 2.20E-03 | 9.50E-03 | 1.01E+00 | 7.00E-05 | 2.97E-04 | -1.30E+00 | 1.59E-05 | 6.07E-05 | -1.21E+00 | 9.18E-03 | 3.27E-02 | -1.09E+00 | 5.68E-02 | 1.60E-01 | 1.03E+00 | 2.81E-04 | 1.03E-03 | -1.20E+00 | 7.80E-02 | 2.14E-01 | 1.05E+00 |
| CKLF | 9.68E-04 | 5.05E-03 | -1.00E+00 | 1.28E-08 | 5.55E-08 | -1.25E+00 | 4.11E-02 | 1.75E-01 | 1.04E+00 | 1.54E-12 | 5.88E-12 | -1.53E+00 | 7.41E-03 | 2.64E-02 | -1.48E+00 | 1.14E-02 | 3.21E-02 | -1.39E+00 | 1.36E-01 | 4.99E-01 | 1.34E+00 | 1.36E-02 | 3.73E-02 | -1.03E+00 |
| TNFRSF11A | 9.69E-04 | 5.05E-03 | -1.34E+00 | 1.09E-02 | 4.70E-02 | 1.05E+00 | 5.82E-03 | 2.47E-02 | -1.46E+00 | 5.32E-03 | 2.03E-02 | -1.50E+00 | 2.24E-01 | 7.97E-01 | 1.28E+00 | 3.03E-01 | 8.55E-01 | 1.25E+00 | 8.77E-02 | 3.22E-01 | -1.04E+00 | 1.11E-01 | 3.04E-01 | -1.22E+00 |
| LONRF1 | 9.69E-04 | 5.05E-03 | 1.06E+00 | 7.21E-09 | 3.12E-08 | -1.43E+00 | 1.93E-01 | 8.21E-01 | 1.24E+00 | 6.36E-03 | 2.43E-02 | -1.03E+00 | 5.48E-03 | 1.95E-02 | -1.06E+00 | 3.47E-03 | 9.79E-03 | -1.16E+00 | 1.91E-04 | 7.03E-04 | -1.67E+00 | 6.58E-03 | 1.81E-02 | -1.26E+00 |
| NCBP1 | 9.70E-04 | 5.05E-03 | -1.16E+00 | 3.29E-03 | 1.42E-02 | 1.09E+00 | 1.63E-03 | 6.93E-03 | -1.15E+00 | 4.20E-03 | 1.60E-02 | 1.04E+00 | 6.65E-05 | 2.37E-04 | -1.41E+00 | 2.48E-03 | 6.99E-03 | -1.09E+00 | 1.23E-06 | 4.50E-06 | -1.41E+00 | 3.19E-03 | 8.78E-03 | -1.13E+00 |
| TBC1D22A | 9.70E-04 | 5.06E-03 | -1.16E+00 | 8.82E-03 | 3.81E-02 | 1.07E+00 | 5.20E-06 | 2.21E-05 | -1.40E+00 | 2.33E-05 | 8.89E-05 | -1.17E+00 | 3.79E-06 | 1.35E-05 | -1.33E+00 | 1.42E-04 | 4.00E-04 | -1.13E+00 | 5.77E-05 | 2.12E-04 | -1.18E+00 | 4.70E-07 | 1.29E-06 | -1.41E+00 |
| RAB30 | 9.71E-04 | 5.06E-03 | 2.54E+00 | 1.70E-02 | 7.36E-02 | -1.25E+00 | 2.17E-06 | 9.23E-06 | 3.58E+00 | 1.49E-01 | 5.69E-01 | -1.20E+00 | 8.40E-02 | 3.00E-01 | 1.99E+00 | 3.20E-01 | 9.03E-01 | 1.38E+00 | 7.87E-02 | 2.89E-01 | 3.20E+00 | 1.70E-01 | 4.68E-01 | 1.55E+00 |
| AC079944.1 | 9.75E-04 | 5.08E-03 | -1.27E+00 | 4.59E-02 | 1.98E-01 | 1.79E+00 | 2.44E-02 | 1.04E-01 | -1.02E+00 | 1.75E-02 | 6.67E-02 | 1.89E+00 | 6.98E-02 | 2.49E-01 | 1.02E+00 | 7.03E-02 | 1.98E-01 | 2.00E+00 | 4.01E-02 | 1.47E-01 | -1.07E+00 | 3.75E-02 | 1.03E-01 | 1.66E+00 |
| Metazoa_SRP | 9.83E-04 | 5.12E-03 | 1.07E+00 | 1.46E-05 | 6.31E-05 | 1.04E+00 | 1.79E-05 | 7.61E-05 | -1.15E+00 | 4.31E-07 | 1.65E-06 | -1.12E+00 | 3.56E-02 | 1.27E-01 | 1.12E+00 | 2.36E-02 | 6.65E-02 | 1.08E+00 | 2.42E-02 | 8.90E-02 | 1.05E+00 | 2.72E-02 | 7.46E-02 | 1.02E+00 |
| DECR1 | 9.84E-04 | 5.13E-03 | 1.00E+00 | 1.08E-04 | 4.66E-04 | 1.05E+00 | 6.47E-03 | 2.75E-02 | -1.04E+00 | 1.92E-07 | 7.33E-07 | -1.14E+00 | 5.63E-06 | 2.01E-05 | -1.32E+00 | 4.35E-03 | 1.23E-02 | -1.02E+00 | 7.62E-05 | 2.80E-04 | -1.28E+00 | 8.68E-06 | 2.38E-05 | -1.36E+00 |
| CCNT2 | 9.85E-04 | 5.13E-03 | 1.05E+00 | 5.36E-03 | 2.32E-02 | 1.25E+00 | 2.24E-04 | 9.50E-04 | -1.24E+00 | 5.68E-03 | 2.17E-02 | 1.03E+00 | 2.60E-03 | 9.28E-03 | -1.11E+00 | 8.56E-02 | 2.41E-01 | 1.09E+00 | 1.86E-03 | 6.83E-03 | -1.11E+00 | 7.62E-02 | 2.09E-01 | 1.07E+00 |
| AC007066.1 | 9.89E-04 | 5.15E-03 | 1.60E+00 | 1.07E-02 | 4.64E-02 | 1.29E+00 | 9.72E-03 | 4.13E-02 | 1.50E+00 | 3.11E-03 | 1.19E-02 | 1.13E+00 | 7.39E-02 | 2.64E-01 | 1.23E+00 | 7.31E-04 | 2.06E-03 | 1.04E+00 | 9.39E-02 | 3.45E-01 | 1.40E+00 | 1.24E-02 | 3.41E-02 | 1.08E+00 |
| GPR18 | 9.89E-04 | 5.15E-03 | -1.37E+00 | 5.06E-04 | 2.19E-03 | -1.42E+00 | 5.99E-03 | 2.55E-02 | -1.25E+00 | 3.34E-04 | 1.28E-03 | -1.38E+00 | 2.70E-02 | 9.62E-02 | -1.17E+00 | 2.00E-02 | 5.64E-02 | -1.26E+00 | 9.20E-05 | 3.37E-04 | -1.45E+00 | 6.02E-05 | 1.65E-04 | -1.43E+00 |
| ZNF430 | 9.90E-04 | 5.16E-03 | 1.01E+00 | 2.15E-08 | 9.29E-08 | -1.14E+00 | 9.58E-02 | 4.07E-01 | 1.10E+00 | 2.18E-04 | 8.32E-04 | -1.06E+00 | 8.69E-03 | 3.10E-02 | -1.02E+00 | 1.05E-03 | 2.95E-03 | -1.03E+00 | 5.45E-02 | 2.00E-01 | 1.07E+00 | 3.09E-04 | 8.48E-04 | -1.19E+00 |
| RFESD | 9.92E-04 | 5.17E-03 | -2.00E+00 | 1.29E-02 | 5.55E-02 | -1.25E+00 | 2.40E-04 | 1.02E-03 | -2.16E+00 | 1.76E-03 | 6.73E-03 | -1.47E+00 | 1.70E-01 | 6.07E-01 | 1.71E+00 | 1.86E-01 | 5.24E-01 | 1.73E+00 | 7.14E-02 | 2.62E-01 | 1.05E+00 | 3.23E-01 | 8.87E-01 | 1.30E+00 |
| SPATA7 | 9.94E-04 | 5.18E-03 | -1.07E+00 | 5.23E-02 | 2.26E-01 | 1.28E+00 | 1.72E-04 | 7.29E-04 | -1.46E+00 | 7.47E-03 | 2.85E-02 | -1.04E+00 | 4.66E-03 | 1.66E-02 | -1.19E+00 | 3.68E-03 | 1.04E-02 | -1.24E+00 | 1.97E-02 | 7.24E-02 | -1.10E+00 | 4.76E-02 | 1.31E-01 | 1.07E+00 |
| AL592295.1 | 9.96E-04 | 5.19E-03 | -1.11E+00 | 8.20E-07 | 3.54E-06 | -1.59E+00 | 1.08E-01 | 4.60E-01 | 1.14E+00 | 1.21E-04 | 4.62E-04 | -1.29E+00 | 6.33E-03 | 2.26E-02 | -1.13E+00 | 7.65E-05 | 2.16E-04 | -1.57E+00 | 1.87E-01 | 6.87E-01 | 1.19E+00 | 2.84E-02 | 7.81E-02 | -1.09E+00 |
| KLHDC1 | 9.97E-04 | 5.19E-03 | -1.36E+00 | 2.27E-02 | 9.80E-02 | 1.13E+00 | 2.68E-02 | 1.14E-01 | -1.30E+00 | 1.73E-02 | 6.62E-02 | -1.29E+00 | 1.17E-01 | 4.17E-01 | 1.13E+00 | 1.63E-01 | 4.59E-01 | 1.99E+00 | 1.23E-02 | 4.50E-02 | -1.17E+00 | 2.76E-02 | 7.58E-02 | -1.23E+00 |
| TEFM | 9.97E-04 | 5.20E-03 | -1.17E+00 | 4.27E-02 | 1.84E-01 | 1.18E+00 | 1.91E-04 | 8.10E-04 | -1.25E+00 | 2.70E-02 | 1.03E-01 | 1.10E+00 | 7.42E-02 | 2.64E-01 | 1.13E+00 | 3.11E-01 | 8.78E-01 | 1.46E+00 | 5.50E-02 | 2.02E-01 | 1.07E+00 | 3.35E-02 | 9.19E-02 | -1.01E+00 |
| PAIP2B | 9.99E-04 | 5.20E-03 | -1.26E+00 | 1.23E-01 | 5.30E-01 | 1.35E+00 | 3.42E-04 | 1.45E-03 | -1.43E+00 | 4.95E-02 | 1.89E-01 | 1.21E+00 | 9.83E-07 | 3.51E-06 | -1.85E+00 | 6.52E-06 | 1.84E-05 | -1.75E+00 | 1.93E-02 | 7.10E-02 | 1.05E+00 | 7.00E-02 | 1.92E-01 | 1.05E+00 |
| KIF13B | 1.00E-03 | 5.24E-03 | -1.19E+00 | 3.78E-03 | 1.63E-02 | -1.01E+00 | 7.86E-02 | 3.34E-01 | 1.52E+00 | 2.11E-01 | 8.05E-01 | 1.52E+00 | 2.07E-01 | 7.37E-01 | 1.39E+00 | 3.09E-02 | 8.71E-02 | -1.10E+00 | 2.27E-03 | 8.34E-03 | -1.33E+00 | 1.40E-04 | 3.85E-04 | -1.61E+00 |
| SNHG15 | 1.01E-03 | 5.24E-03 | 1.07E+00 | 7.73E-06 | 3.34E-05 | 1.06E+00 | 1.02E-01 | 4.34E-01 | 1.04E+00 | 1.46E-02 | 5.57E-02 | 1.04E+00 | 3.32E-04 | 1.19E-03 | 1.03E+00 | 1.38E-05 | 3.89E-05 | -1.05E+00 | 2.59E-01 | 9.52E-01 | 1.33E+00 | 2.20E-01 | 6.05E-01 | 1.17E+00 |
| ZNF526 | 1.01E-03 | 5.24E-03 | -1.47E+00 | 3.67E-03 | 1.58E-02 | -1.35E+00 | 3.27E-05 | 1.39E-04 | -1.62E+00 | 3.61E-03 | 1.38E-02 | -1.11E+00 | 3.12E-03 | 1.11E-02 | -1.98E+00 | 1.33E-02 | 3.76E-02 | -1.50E+00 | 3.83E-07 | 1.41E-06 | -1.90E+00 | 2.27E-02 | 6.23E-02 | -1.03E+00 |
| CDKAL1 | 1.01E-03 | 5.25E-03 | -1.08E+00 | 7.56E-03 | 3.27E-02 | 1.16E+00 | 1.77E-01 | 7.53E-01 | 1.15E+00 | 2.38E-01 | 9.09E-01 | 1.26E+00 | 1.30E-05 | 4.65E-05 | -1.34E+00 | 1.38E-06 | 3.89E-06 | -1.51E+00 | 1.45E-03 | 5.33E-03 | -1.08E+00 | 1.85E-01 | 5.08E-01 | 1.19E+00 |
| ZBTB20-AS4 | 1.01E-03 | 5.25E-03 | -1.40E+00 | 6.46E-02 | 2.79E-01 | 1.21E+00 | 3.98E-02 | 1.69E-01 | -1.34E+00 | 4.79E-02 | 1.83E-01 | -1.17E+00 | 1.10E-05 | 3.90E-05 | -2.77E+00 | 1.11E-03 | 3.13E-03 | -1.60E+00 | 1.97E-01 | 7.25E-01 | 1.09E+00 | 2.06E-01 | 5.67E-01 | -1.17E+00 |
| TULP3 | 1.01E-03 | 5.27E-03 | -1.45E+00 | 1.28E-02 | 5.55E-02 | -1.02E+00 | 2.64E-04 | 1.12E-03 | -1.44E+00 | 2.88E-04 | 1.10E-03 | -1.31E+00 | 9.68E-02 | 3.45E-01 | -1.02E+00 | 3.30E-01 | 9.31E-01 | 1.43E+00 | 3.12E-05 | 1.14E-04 | -1.44E+00 | 8.28E-02 | 2.28E-01 | 1.09E+00 |
| PAIP2 | 1.01E-03 | 5.28E-03 | 1.13E+00 | 5.38E-12 | 2.33E-11 | -1.08E+00 | 1.44E-01 | 6.13E-01 | 1.09E+00 | 5.06E-05 | 1.93E-04 | -1.04E+00 | 1.34E-01 | 4.77E-01 | 1.34E+00 | 1.16E-02 | 3.27E-02 | 1.00E+00 | 1.71E-01 | 6.27E-01 | 1.27E+00 | 4.17E-03 | 1.15E-02 | -1.08E+00 |
| DCBLD1 | 1.01E-03 | 5.28E-03 | -1.82E+00 | 1.13E-02 | 4.87E-02 | -1.30E+00 | 9.43E-03 | 4.01E-02 | -1.32E+00 | 1.02E-03 | 3.90E-03 | -1.49E+00 | 2.53E-02 | 9.01E-02 | -1.52E+00 | 1.33E-01 | 3.74E-01 | 1.03E+00 | 5.89E-05 | 2.16E-04 | -1.67E+00 | 2.33E-02 | 6.41E-02 | -1.15E+00 |
| YWHAZP5 | 1.01E-03 | 5.29E-03 | -1.35E+00 | 3.91E-05 | 1.69E-04 | -1.91E+00 | 1.95E-02 | 8.27E-02 | -1.17E+00 | 2.25E-02 | 8.58E-02 | -1.05E+00 | 1.26E-02 | 4.50E-02 | -1.26E+00 | 5.44E-03 | 1.53E-02 | -1.45E+00 | 3.16E-03 | 1.16E-02 | -1.23E+00 | 4.41E-05 | 1.21E-04 | -1.78E+00 |
| AC023906.1 | 1.01E-03 | 5.29E-03 | 1.15E+00 | 9.15E-08 | 3.95E-07 | -1.00E+00 | 1.36E-01 | 5.79E-01 | 1.14E+00 | 3.04E-04 | 1.16E-03 | -1.03E+00 | 5.76E-03 | 2.05E-02 | 1.01E+00 | 2.33E-04 | 6.56E-04 | -1.12E+00 | 3.30E-02 | 1.21E-01 | 1.07E+00 | 2.09E-02 | 5.75E-02 | 1.03E+00 |
| FLYWCH1 | 1.02E-03 | 5.30E-03 | -1.01E+00 | 4.44E-02 | 1.92E-01 | 1.33E+00 | 1.46E-04 | 6.20E-04 | -1.22E+00 | 5.21E-03 | 1.99E-02 | 1.10E+00 | 3.31E-03 | 1.18E-02 | 1.02E+00 | 1.71E-01 | 4.81E-01 | 1.40E+00 | 1.32E-03 | 4.84E-03 | -1.12E+00 | 3.96E-02 | 1.09E-01 | 1.00E+00 |
| SNX13 | 1.02E-03 | 5.30E-03 | -1.02E+00 | 2.28E-03 | 9.84E-03 | 1.09E+00 | 1.05E-03 | 4.48E-03 | -1.14E+00 | 3.42E-04 | 1.31E-03 | -1.05E+00 | 2.65E-02 | 9.46E-02 | 1.09E+00 | 6.66E-02 | 1.88E-01 | 1.20E+00 | 1.85E-01 | 6.78E-01 | 1.26E+00 | 3.23E-01 | 8.87E-01 | 1.33E+00 |
| TTLL3 | 1.02E-03 | 5.31E-03 | -1.17E+00 | 2.92E-03 | 1.26E-02 | -1.02E+00 | 2.83E-02 | 1.20E-01 | 1.02E+00 | 7.71E-02 | 2.94E-01 | 1.26E+00 | 1.45E-01 | 5.15E-01 | 1.68E+00 | 2.88E-01 | 8.13E-01 | 1.53E+00 | 1.26E-01 | 4.61E-01 | 1.10E+00 | 1.26E-01 | 3.46E-01 | 1.06E+00 |
| RRM2B | 1.02E-03 | 5.31E-03 | -1.10E+00 | 1.16E-03 | 5.01E-03 | -1.02E+00 | 1.26E-02 | 5.35E-02 | -1.08E+00 | 1.87E-03 | 7.14E-03 | -1.10E+00 | 2.75E-01 | 9.82E-01 | 1.53E+00 | 3.43E-01 | 9.68E-01 | 1.49E+00 | 4.06E-02 | 1.49E-01 | 1.09E+00 | 1.72E-03 | 4.73E-03 | -1.23E+00 |
| LBH | 1.02E-03 | 5.31E-03 | -1.22E+00 | 1.57E-05 | 6.76E-05 | -1.37E+00 | 2.16E-03 | 9.19E-03 | -1.38E+00 | 1.04E-03 | 3.95E-03 | -1.25E+00 | 3.42E-04 | 1.22E-03 | -1.37E+00 | 1.35E-03 | 3.81E-03 | -1.22E+00 | 6.45E-10 | 2.37E-09 | -1.99E+00 | 5.31E-10 | 1.46E-09 | -2.06E+00 |
| MOSPD1 | 1.02E-03 | 5.31E-03 | -1.17E+00 | 2.69E-04 | 1.16E-03 | -1.15E+00 | 2.66E-03 | 1.13E-02 | -1.26E+00 | 5.09E-03 | 1.94E-02 | -1.08E+00 | 1.62E-03 | 5.77E-03 | -1.64E+00 | 1.76E-02 | 4.97E-02 | -1.20E+00 | 2.84E-03 | 1.04E-02 | -1.10E+00 | 3.50E-04 | 9.61E-04 | -1.31E+00 |
| AL645924.2 | 1.02E-03 | 5.31E-03 | -1.01E+00 | 2.83E-09 | 1.22E-08 | -1.35E+00 | 1.08E-02 | 4.60E-02 | -1.04E+00 | 9.92E-06 | 3.79E-05 | -1.30E+00 | 5.84E-02 | 2.08E-01 | 1.24E+00 | 6.20E-04 | 1.75E-03 | -1.07E+00 | 1.28E-01 | 4.70E-01 | 1.07E+00 | 1.94E-02 | 5.34E-02 | -1.08E+00 |
| ZNF793 | 1.02E-03 | 5.32E-03 | -1.08E+00 | 1.98E-01 | 8.57E-01 | 1.47E+00 | 2.02E-03 | 8.59E-03 | -1.36E+00 | 1.34E-01 | 5.11E-01 | 1.13E+00 | 2.74E-03 | 9.76E-03 | -1.29E+00 | 1.18E-01 | 3.33E-01 | 1.20E+00 | 5.66E-03 | 2.08E-02 | -1.16E+00 | 1.66E-01 | 4.56E-01 | 1.17E+00 |
| CCDC22 | 1.02E-03 | 5.32E-03 | -1.07E+00 | 9.02E-03 | 3.90E-02 | 1.21E+00 | 1.92E-01 | 8.14E-01 | 1.18E+00 | 2.56E-01 | 9.78E-01 | 1.42E+00 | 3.72E-03 | 1.32E-02 | 1.04E+00 | 1.07E-01 | 3.00E-01 | 1.28E+00 | 9.65E-02 | 3.54E-01 | 1.22E+00 | 3.35E-01 | 9.21E-01 | 1.38E+00 |
| GOSR2 | 1.02E-03 | 5.32E-03 | 1.05E+00 | 2.84E-03 | 1.23E-02 | 1.20E+00 | 2.28E-05 | 9.68E-05 | -1.30E+00 | 2.02E-03 | 7.72E-03 | 1.05E+00 | 9.46E-04 | 3.37E-03 | 1.02E+00 | 2.03E-02 | 5.73E-02 | 1.17E+00 | 1.63E-04 | 5.99E-04 | -1.08E+00 | 8.35E-02 | 2.29E-01 | 1.16E+00 |
| CDKN1B | 1.03E-03 | 5.34E-03 | 1.06E+00 | 5.21E-02 | 2.25E-01 | 1.33E+00 | 2.07E-01 | 8.79E-01 | 1.24E+00 | 2.61E-01 | 9.98E-01 | 1.42E+00 | 1.90E-01 | 6.76E-01 | 1.14E+00 | 2.90E-01 | 8.17E-01 | 1.31E+00 | 1.78E-03 | 6.53E-03 | -1.10E+00 | 4.21E-04 | 1.16E-03 | -1.15E+00 |
| DSCC1 | 1.03E-03 | 5.36E-03 | -1.59E+00 | 1.23E-03 | 5.33E-03 | -1.43E+00 | 1.90E-01 | 8.09E-01 | 1.14E+00 | 2.38E-02 | 9.11E-02 | 1.04E+00 | 1.88E-01 | 6.69E-01 | 1.03E+00 | 3.37E-01 | 9.49E-01 | 1.30E+00 | 3.99E-10 | 1.46E-09 | -1.60E+00 | 1.30E-01 | 3.57E-01 | 1.06E+00 |
| PDLIM5 | 1.03E-03 | 5.37E-03 | -1.22E+00 | 1.21E-02 | 5.21E-02 | 1.09E+00 | 3.70E-07 | 1.57E-06 | -1.66E+00 | 1.29E-04 | 4.93E-04 | -1.13E+00 | 9.22E-07 | 3.29E-06 | -1.48E+00 | 1.34E-03 | 3.79E-03 | -1.26E+00 | 2.23E-01 | 8.19E-01 | 1.14E+00 | 7.10E-02 | 1.95E-01 | 1.12E+00 |
| RN7SL288P | 1.03E-03 | 5.38E-03 | 3.23E+00 | 4.90E-02 | 2.12E-01 | 2.27E+00 | 6.43E-09 | 2.73E-08 | 4.33E+00 | 1.84E-02 | 7.02E-02 | 2.49E+00 | 5.75E-02 | 2.05E-01 | 2.35E+00 | 2.43E-01 | 6.85E-01 | 1.66E+00 | 1.08E-01 | 3.95E-01 | 1.30E+00 | 2.72E-01 | 7.46E-01 | 1.14E+00 |
| C16orf54 | 1.03E-03 | 5.39E-03 | 1.11E+00 | 1.25E-15 | 5.41E-15 | -1.29E+00 | 4.10E-04 | 1.74E-03 | -1.08E+00 | 3.30E-09 | 1.26E-08 | -1.23E+00 | 1.08E-01 | 3.85E-01 | 1.36E+00 | 2.91E-04 | 8.19E-04 | -1.18E+00 | 8.32E-04 | 3.05E-03 | 1.73E+00 | 6.13E-03 | 1.68E-02 | 1.00E+00 |
| CASC15 | 1.04E-03 | 5.40E-03 | 1.08E+00 | 7.68E-02 | 3.32E-01 | 1.67E+00 | 3.82E-07 | 1.62E-06 | -1.41E+00 | 5.02E-03 | 1.92E-02 | 1.05E+00 | 1.37E-06 | 4.88E-06 | -1.29E+00 | 1.31E-05 | 3.70E-05 | -1.25E+00 | 2.12E-01 | 7.79E-01 | 1.09E+00 | 2.86E-01 | 7.86E-01 | 1.11E+00 |
| EEPD1 | 1.04E-03 | 5.40E-03 | -1.48E+00 | 1.78E-01 | 7.68E-01 | 1.30E+00 | 1.83E-04 | 7.79E-04 | -1.55E+00 | 7.44E-02 | 2.84E-01 | 1.10E+00 | 7.79E-02 | 2.78E-01 | -1.31E+00 | 2.20E-01 | 6.19E-01 | 1.01E+00 | 8.66E-03 | 3.18E-02 | -1.56E+00 | 2.18E-01 | 5.99E-01 | 1.04E+00 |
| RPL7P56 | 1.04E-03 | 5.42E-03 | 1.82E+00 | 1.76E-01 | 7.59E-01 | 1.49E+00 | 1.21E-04 | 5.15E-04 | 1.77E+00 | 2.38E-01 | 9.11E-01 | 1.41E+00 | 3.44E-02 | 1.23E-01 | 1.79E+00 | 8.05E-02 | 2.27E-01 | 1.77E+00 | 1.02E-01 | 3.74E-01 | 1.48E+00 | 3.18E-01 | 8.74E-01 | 1.40E+00 |
| SPIDR | 1.04E-03 | 5.42E-03 | -1.05E+00 | 8.69E-04 | 3.76E-03 | 1.06E+00 | 8.43E-04 | 3.58E-03 | -1.07E+00 | 9.79E-05 | 3.74E-04 | 1.01E+00 | 1.08E-03 | 3.85E-03 | -1.13E+00 | 1.03E-02 | 2.91E-02 | -1.00E+00 | 1.29E-02 | 4.75E-02 | -1.08E+00 | 4.88E-02 | 1.34E-01 | -1.00E+00 |
| ZBED5 | 1.04E-03 | 5.42E-03 | 1.04E+00 | 5.68E-07 | 2.45E-06 | -1.02E+00 | 3.29E-02 | 1.40E-01 | 1.08E+00 | 7.11E-04 | 2.72E-03 | 1.05E+00 | 8.10E-02 | 2.89E-01 | 1.20E+00 | 1.48E-01 | 4.18E-01 | 1.22E+00 | 3.96E-02 | 1.45E-01 | 1.01E+00 | 4.47E-02 | 1.23E-01 | 1.03E+00 |
| AC002350.2 | 1.04E-03 | 5.44E-03 | -1.45E+00 | 1.48E-02 | 6.41E-02 | -1.09E+00 | 2.88E-02 | 1.22E-01 | -1.28E+00 | 2.52E-02 | 9.62E-02 | -1.18E+00 | 1.79E-01 | 6.38E-01 | 1.66E+00 | 2.35E-01 | 6.61E-01 | 1.47E+00 | 2.71E-01 | 9.95E-01 | 1.23E+00 | 2.58E-01 | 7.09E-01 | 1.21E+00 |
| IPO4 | 1.04E-03 | 5.44E-03 | -1.59E+00 | 5.66E-03 | 2.45E-02 | -1.25E+00 | 4.19E-03 | 1.78E-02 | -1.29E+00 | 5.89E-04 | 2.25E-03 | -1.35E+00 | 1.65E-01 | 5.89E-01 | -1.00E+00 | 2.12E-01 | 5.99E-01 | 1.09E+00 | 6.61E-03 | 2.43E-02 | -1.07E+00 | 2.64E-03 | 7.24E-03 | -1.18E+00 |
| AKAP7 | 1.04E-03 | 5.44E-03 | -1.10E+00 | 4.31E-03 | 1.86E-02 | 1.06E+00 | 8.95E-04 | 3.80E-03 | -1.18E+00 | 1.62E-05 | 6.19E-05 | -1.19E+00 | 1.25E-03 | 4.44E-03 | -1.35E+00 | 4.12E-04 | 1.16E-03 | -1.33E+00 | 1.13E-04 | 4.14E-04 | -1.23E+00 | 6.97E-07 | 1.91E-06 | -1.36E+00 |
| CRYBG3 | 1.05E-03 | 5.45E-03 | -1.84E+00 | 7.02E-03 | 3.03E-02 | -1.39E+00 | 8.66E-03 | 3.68E-02 | -1.25E+00 | 1.01E-02 | 3.86E-02 | -1.13E+00 | 2.45E-01 | 8.73E-01 | 1.09E+00 | 3.42E-01 | 9.64E-01 | 1.14E+00 | 4.27E-02 | 1.57E-01 | -1.12E+00 | 5.40E-02 | 1.48E-01 | -1.25E+00 |
| AC022415.1 | 1.05E-03 | 5.46E-03 | -1.71E+00 | 3.68E-02 | 1.59E-01 | 1.01E+00 | 3.20E-05 | 1.36E-04 | -1.73E+00 | 7.98E-04 | 3.05E-03 | -1.24E+00 | 7.28E-02 | 2.60E-01 | -1.09E+00 | 2.48E-01 | 7.00E-01 | 1.22E+00 | 2.29E-03 | 8.41E-03 | -1.43E+00 | 7.73E-03 | 2.12E-02 | -1.38E+00 |
| EEF1DP7 | 1.05E-03 | 5.46E-03 | -1.43E+00 | 2.52E-02 | 1.09E-01 | -1.02E+00 | 4.21E-02 | 1.79E-01 | -1.26E+00 | 3.41E-03 | 1.30E-02 | -1.49E+00 | 2.10E-01 | 7.51E-01 | 1.09E+00 | 6.87E-02 | 1.94E-01 | -1.32E+00 | 1.24E-03 | 4.55E-03 | -1.61E+00 | 2.61E-02 | 7.18E-02 | -1.15E+00 |
| RPP25 | 1.05E-03 | 5.46E-03 | -1.52E+00 | 1.80E-02 | 7.80E-02 | -1.07E+00 | 4.65E-05 | 1.98E-04 | -1.48E+00 | 1.88E-01 | 7.18E-01 | 1.37E+00 | 2.63E-01 | 9.39E-01 | 1.12E+00 | 2.73E-01 | 7.70E-01 | 1.50E+00 | 1.94E-02 | 7.13E-02 | -1.68E+00 | 1.98E-01 | 5.44E-01 | -1.06E+00 |
| CDPF1 | 1.05E-03 | 5.47E-03 | -1.32E+00 | 1.14E-01 | 4.92E-01 | 1.33E+00 | 2.61E-04 | 1.11E-03 | -1.52E+00 | 1.63E-02 | 6.21E-02 | -1.00E+00 | 8.18E-02 | 2.92E-01 | -1.02E+00 | 1.60E-01 | 4.51E-01 | 1.68E+00 | 2.01E-06 | 7.38E-06 | -1.57E+00 | 3.84E-03 | 1.05E-02 | -1.12E+00 |
| MED12L | 1.05E-03 | 5.47E-03 | 1.02E+00 | 3.22E-02 | 1.39E-01 | 1.28E+00 | 2.24E-02 | 9.54E-02 | 1.04E+00 | 2.14E-01 | 8.17E-01 | 1.55E+00 | 4.26E-02 | 1.52E-01 | 1.71E+00 | 1.70E-01 | 4.80E-01 | 1.52E+00 | 4.41E-04 | 1.62E-03 | 1.83E+00 | 6.30E-02 | 1.73E-01 | 1.48E+00 |
| PYM1 | 1.05E-03 | 5.47E-03 | 1.05E+00 | 1.87E-03 | 8.09E-03 | 1.25E+00 | 1.00E-02 | 4.25E-02 | -1.08E+00 | 3.78E-03 | 1.44E-02 | 1.03E+00 | 4.00E-04 | 1.43E-03 | -1.11E+00 | 4.21E-02 | 1.19E-01 | 1.19E+00 | 9.32E-03 | 3.42E-02 | -1.04E+00 | 5.27E-02 | 1.45E-01 | 1.03E+00 |
| PHF12 | 1.05E-03 | 5.48E-03 | -1.00E+00 | 1.47E-03 | 6.34E-03 | 1.14E+00 | 3.54E-02 | 1.50E-01 | -1.02E+00 | 6.07E-04 | 2.32E-03 | -1.12E+00 | 1.48E-04 | 5.29E-04 | -1.12E+00 | 1.23E-05 | 3.45E-05 | -1.40E+00 | 1.41E-02 | 5.17E-02 | 1.01E+00 | 4.21E-03 | 1.16E-02 | -1.20E+00 |
| FRS2 | 1.05E-03 | 5.49E-03 | -1.15E+00 | 2.93E-04 | 1.27E-03 | -1.10E+00 | 6.02E-02 | 2.56E-01 | 1.08E+00 | 2.05E-03 | 7.83E-03 | -1.04E+00 | 7.23E-04 | 2.58E-03 | -1.20E+00 | 7.90E-06 | 2.23E-05 | -1.61E+00 | 3.21E-02 | 1.18E-01 | -1.03E+00 | 5.22E-05 | 1.43E-04 | -1.65E+00 |
| AC011270.1 | 1.06E-03 | 5.50E-03 | -1.07E+00 | 1.06E-05 | 4.59E-05 | -1.22E+00 | 7.10E-05 | 3.02E-04 | -1.20E+00 | 1.06E-07 | 4.06E-07 | -1.36E+00 | 1.37E-05 | 4.90E-05 | -1.74E+00 | 2.79E-05 | 7.88E-05 | -1.56E+00 | 3.56E-05 | 1.31E-04 | -1.29E+00 | 6.92E-08 | 1.90E-07 | -1.59E+00 |
| AIFM1 | 1.06E-03 | 5.52E-03 | -1.21E+00 | 8.69E-03 | 3.75E-02 | 1.05E+00 | 1.12E-04 | 4.76E-04 | -1.26E+00 | 1.63E-05 | 6.21E-05 | -1.17E+00 | 9.43E-05 | 3.36E-04 | -1.25E+00 | 2.55E-02 | 7.19E-02 | 1.13E+00 | 2.21E-06 | 8.10E-06 | -1.21E+00 | 6.99E-05 | 1.92E-04 | -1.10E+00 |
| FANCM | 1.06E-03 | 5.52E-03 | -1.32E+00 | 5.28E-02 | 2.28E-01 | 1.28E+00 | 1.05E-04 | 4.46E-04 | -1.29E+00 | 3.34E-03 | 1.27E-02 | 1.01E+00 | 5.56E-06 | 1.98E-05 | -1.79E+00 | 5.10E-04 | 1.44E-03 | -1.26E+00 | 2.73E-05 | 1.00E-04 | -1.42E+00 | 4.95E-03 | 1.36E-02 | -1.09E+00 |
| KDM4B | 1.06E-03 | 5.53E-03 | -1.09E+00 | 6.01E-03 | 2.60E-02 | 1.12E+00 | 5.54E-07 | 2.35E-06 | -1.43E+00 | 1.77E-04 | 6.77E-04 | -1.08E+00 | 1.75E-02 | 6.23E-02 | 1.01E+00 | 2.75E-02 | 7.75E-02 | 1.07E+00 | 1.89E-03 | 6.92E-03 | -1.17E+00 | 1.07E-02 | 2.95E-02 | -1.13E+00 |
| RFX3 | 1.06E-03 | 5.54E-03 | -1.07E+00 | 5.84E-02 | 2.52E-01 | 1.28E+00 | 3.33E-05 | 1.41E-04 | -1.37E+00 | 5.65E-05 | 2.16E-04 | -1.17E+00 | 1.72E-03 | 6.13E-03 | -1.43E+00 | 3.68E-02 | 1.04E-01 | -1.07E+00 | 6.87E-03 | 2.52E-02 | -1.20E+00 | 6.52E-02 | 1.79E-01 | 1.09E+00 |
| PRPF31 | 1.06E-03 | 5.54E-03 | 1.09E+00 | 6.89E-03 | 2.98E-02 | 1.26E+00 | 1.09E-01 | 4.61E-01 | 1.13E+00 | 1.42E-01 | 5.43E-01 | 1.25E+00 | 2.43E-03 | 8.67E-03 | 1.00E+00 | 1.04E-01 | 2.92E-01 | 1.29E+00 | 7.03E-03 | 2.58E-02 | 1.10E+00 | 2.95E-01 | 8.10E-01 | 1.24E+00 |
| PA2G4P2 | 1.06E-03 | 5.54E-03 | -1.51E+00 | 5.94E-03 | 2.57E-02 | -1.11E+00 | 1.34E-02 | 5.67E-02 | -1.15E+00 | 4.92E-02 | 1.88E-01 | 1.08E+00 | 1.92E-01 | 6.84E-01 | 1.12E+00 | 2.24E-01 | 6.31E-01 | 1.66E+00 | 3.41E-02 | 1.25E-01 | 1.01E+00 | 2.81E-01 | 7.72E-01 | 1.23E+00 |
| PRKD3 | 1.06E-03 | 5.54E-03 | 1.01E+00 | 6.21E-04 | 2.68E-03 | 1.16E+00 | 2.96E-03 | 1.26E-02 | -1.05E+00 | 2.97E-02 | 1.13E-01 | 1.16E+00 | 5.52E-02 | 1.97E-01 | 1.62E+00 | 9.10E-02 | 2.56E-01 | 1.64E+00 | 1.70E-01 | 6.22E-01 | 1.18E+00 | 2.21E-01 | 6.06E-01 | 1.20E+00 |
| RNASEH1 | 1.06E-03 | 5.54E-03 | -1.09E+00 | 2.55E-04 | 1.10E-03 | -1.04E+00 | 1.33E-02 | 5.64E-02 | 1.04E+00 | 3.69E-04 | 1.41E-03 | 1.00E+00 | 6.27E-08 | 2.23E-07 | -1.45E+00 | 4.28E-06 | 1.21E-05 | -1.23E+00 | 6.06E-03 | 2.22E-02 | -1.02E+00 | 3.96E-02 | 1.09E-01 | 1.11E+00 |
| DMD | 1.06E-03 | 5.54E-03 | -1.34E+00 | 4.34E-03 | 1.87E-02 | -1.12E+00 | 5.75E-02 | 2.44E-01 | -1.18E+00 | 1.31E-01 | 4.99E-01 | 1.03E+00 | 1.73E-01 | 6.18E-01 | -1.00E+00 | 2.91E-01 | 8.22E-01 | -1.02E+00 | 2.22E-01 | 8.15E-01 | -1.03E+00 | 2.54E-01 | 6.97E-01 | 1.39E+00 |
| MRPL54 | 1.07E-03 | 5.55E-03 | 1.10E+00 | 4.77E-09 | 2.06E-08 | -1.03E+00 | 3.83E-02 | 1.63E-01 | 1.04E+00 | 9.54E-06 | 3.64E-05 | -1.07E+00 | 3.96E-02 | 1.41E-01 | 1.09E+00 | 1.79E-02 | 5.06E-02 | 1.02E+00 | 1.12E-01 | 4.10E-01 | 1.16E+00 | 4.88E-02 | 1.34E-01 | 1.03E+00 |
| NDUFB1 | 1.07E-03 | 5.56E-03 | 1.04E+00 | 1.71E-19 | 7.37E-19 | -1.70E+00 | 4.67E-04 | 1.98E-03 | -1.08E+00 | 5.69E-17 | 2.17E-16 | -1.71E+00 | 1.09E-04 | 3.88E-04 | -1.07E+00 | 4.18E-09 | 1.18E-08 | -1.40E+00 | 2.72E-03 | 1.00E-02 | -1.08E+00 | 1.22E-15 | 3.34E-15 | -1.84E+00 |
| JPX | 1.07E-03 | 5.57E-03 | -1.28E+00 | 2.46E-04 | 1.06E-03 | -1.23E+00 | 2.91E-03 | 1.24E-02 | -1.16E+00 | 2.83E-03 | 1.08E-02 | -1.04E+00 | 3.67E-02 | 1.31E-01 | 1.04E+00 | 4.27E-02 | 1.20E-01 | 1.07E+00 | 1.08E-03 | 3.98E-03 | -1.28E+00 | 5.83E-03 | 1.60E-02 | -1.16E+00 |
| KCTD17 | 1.07E-03 | 5.59E-03 | -1.13E+00 | 4.24E-02 | 1.83E-01 | 1.25E+00 | 4.65E-02 | 1.97E-01 | 1.02E+00 | 3.29E-02 | 1.26E-01 | 1.07E+00 | 7.85E-03 | 2.80E-02 | -1.47E+00 | 2.41E-02 | 6.79E-02 | -1.21E+00 | 4.93E-02 | 1.81E-01 | 1.71E+00 | 4.08E-03 | 1.12E-02 | 2.31E+00 |
| CEP192 | 1.08E-03 | 5.61E-03 | -1.20E+00 | 7.32E-03 | 3.16E-02 | 1.00E+00 | 4.22E-04 | 1.79E-03 | -1.30E+00 | 1.77E-02 | 6.76E-02 | 1.10E+00 | 5.29E-03 | 1.89E-02 | -1.13E+00 | 7.07E-02 | 1.99E-01 | 1.13E+00 | 7.46E-05 | 2.74E-04 | -1.17E+00 | 2.50E-03 | 6.87E-03 | -1.09E+00 |
| PTOV1 | 1.08E-03 | 5.62E-03 | -1.13E+00 | 1.71E-04 | 7.40E-04 | -1.10E+00 | 2.79E-03 | 1.19E-02 | -1.10E+00 | 4.13E-06 | 1.58E-05 | -1.24E+00 | 2.28E-05 | 8.13E-05 | -1.38E+00 | 4.59E-05 | 1.29E-04 | -1.28E+00 | 3.86E-03 | 1.42E-02 | -1.00E+00 | 6.75E-03 | 1.85E-02 | -1.01E+00 |
| SOCS7 | 1.08E-03 | 5.62E-03 | -1.27E+00 | 7.24E-04 | 3.13E-03 | -1.22E+00 | 1.64E-03 | 6.96E-03 | -1.32E+00 | 3.51E-02 | 1.34E-01 | 1.13E+00 | 8.11E-05 | 2.89E-04 | -1.73E+00 | 9.14E-03 | 2.58E-02 | -1.16E+00 | 9.12E-02 | 3.35E-01 | -1.06E+00 | 3.56E-01 | 9.77E-01 | 1.37E+00 |
| HYKK | 1.08E-03 | 5.62E-03 | -1.80E+00 | 3.66E-02 | 1.58E-01 | -1.19E+00 | 2.85E-02 | 1.21E-01 | -1.26E+00 | 1.98E-02 | 7.55E-02 | -1.22E+00 | 6.93E-02 | 2.47E-01 | -1.23E+00 | 7.34E-02 | 2.07E-01 | -1.21E+00 | 2.34E-02 | 8.58E-02 | -1.15E+00 | 1.28E-02 | 3.52E-02 | -1.28E+00 |
| FAM228B | 1.08E-03 | 5.62E-03 | -1.01E+00 | 8.91E-04 | 3.85E-03 | 1.06E+00 | 8.84E-06 | 3.76E-05 | -1.21E+00 | 6.04E-05 | 2.31E-04 | -1.04E+00 | 1.40E-01 | 5.00E-01 | 1.20E+00 | 1.89E-01 | 5.32E-01 | 1.27E+00 | 5.23E-03 | 1.92E-02 | -1.11E+00 | 3.35E-02 | 9.21E-02 | 1.04E+00 |
| ATP23 | 1.08E-03 | 5.62E-03 | -1.16E+00 | 9.37E-03 | 4.05E-02 | 1.14E+00 | 3.25E-07 | 1.38E-06 | -1.35E+00 | 1.17E-07 | 4.45E-07 | -1.23E+00 | 5.44E-07 | 1.94E-06 | -1.76E+00 | 2.84E-03 | 8.00E-03 | -1.02E+00 | 1.91E-06 | 7.00E-06 | -1.29E+00 | 7.66E-02 | 2.10E-01 | 1.10E+00 |
| PCK2 | 1.08E-03 | 5.63E-03 | -1.08E+00 | 4.23E-04 | 1.83E-03 | 1.01E+00 | 7.61E-06 | 3.23E-05 | -1.31E+00 | 2.50E-07 | 9.56E-07 | -1.34E+00 | 9.10E-02 | 3.25E-01 | -1.14E+00 | 1.44E-01 | 4.05E-01 | -1.10E+00 | 3.27E-06 | 1.20E-05 | -1.72E+00 | 1.01E-04 | 2.79E-04 | -1.45E+00 |
| SLC25A43 | 1.08E-03 | 5.64E-03 | -1.15E+00 | 8.03E-02 | 3.47E-01 | 1.32E+00 | 4.10E-04 | 1.74E-03 | -1.42E+00 | 9.08E-03 | 3.47E-02 | -1.04E+00 | 3.95E-03 | 1.41E-02 | -1.43E+00 | 9.27E-02 | 2.61E-01 | 1.11E+00 | 3.17E-02 | 1.16E-01 | -1.18E+00 | 3.45E-01 | 9.47E-01 | 1.24E+00 |
| IL13RA1 | 1.08E-03 | 5.65E-03 | -1.85E+00 | 1.90E-01 | 8.21E-01 | 1.28E+00 | 2.07E-03 | 8.79E-03 | -1.40E+00 | 2.28E-03 | 8.72E-03 | -1.31E+00 | 2.06E-01 | 7.34E-01 | 1.72E+00 | 2.99E-01 | 8.44E-01 | 1.25E+00 | 1.20E-01 | 4.42E-01 | -1.25E+00 | 1.42E-01 | 3.90E-01 | -1.35E+00 |
| PLK4 | 1.09E-03 | 5.66E-03 | -1.75E+00 | 2.30E-04 | 9.95E-04 | -2.10E+00 | 8.02E-02 | 3.41E-01 | 1.46E+00 | 1.03E-01 | 3.93E-01 | 1.18E+00 | 4.00E-02 | 1.43E-01 | -1.33E+00 | 3.86E-02 | 1.09E-01 | -1.48E+00 | 1.38E-02 | 5.07E-02 | -1.04E+00 | 2.00E-01 | 5.50E-01 | 1.42E+00 |
| TFPI | 1.09E-03 | 5.66E-03 | 1.65E+00 | 1.26E-02 | 5.44E-02 | 1.66E+00 | 4.57E-04 | 1.94E-03 | 1.64E+00 | 1.44E-01 | 5.50E-01 | 1.50E+00 | 3.64E-04 | 1.30E-03 | 2.48E+00 | 7.86E-02 | 2.21E-01 | 1.81E+00 | 3.34E-03 | 1.23E-02 | 1.75E+00 | 3.25E-01 | 8.92E-01 | 1.30E+00 |
| FFAR3 | 1.09E-03 | 5.66E-03 | 3.10E+00 | 1.66E-01 | 7.16E-01 | 1.22E+00 | 4.31E-04 | 1.83E-03 | 2.75E+00 | 2.15E-01 | 8.20E-01 | 1.11E+00 | 5.73E-02 | 2.04E-01 | 2.97E+00 | 2.87E-01 | 8.08E-01 | 1.28E+00 | 5.50E-02 | 2.02E-01 | 2.64E+00 | 3.64E-01 | 1.00E+00 | -1.00E+00 |
| BZW1 | 1.09E-03 | 5.68E-03 | 1.05E+00 | 7.69E-13 | 3.32E-12 | -1.17E+00 | 8.52E-03 | 3.62E-02 | -1.03E+00 | 7.19E-04 | 2.75E-03 | -1.02E+00 | 1.66E-02 | 5.90E-02 | 1.12E+00 | 8.22E-07 | 2.32E-06 | -1.13E+00 | 8.92E-02 | 3.27E-01 | 1.01E+00 | 5.63E-02 | 1.55E-01 | 1.00E+00 |
| STX11 | 1.09E-03 | 5.69E-03 | -1.21E+00 | 4.11E-07 | 1.78E-06 | -1.81E+00 | 1.00E-03 | 4.25E-03 | -1.40E+00 | 4.55E-05 | 1.74E-04 | -1.55E+00 | 1.74E-02 | 6.22E-02 | 1.08E+00 | 3.24E-06 | 9.13E-06 | -1.67E+00 | 4.15E-02 | 1.52E-01 | -1.11E+00 | 9.20E-04 | 2.53E-03 | -1.13E+00 |
| KLHL18 | 1.09E-03 | 5.70E-03 | -1.39E+00 | 3.73E-05 | 1.61E-04 | -1.75E+00 | 1.42E-02 | 6.05E-02 | -1.13E+00 | 5.75E-04 | 2.20E-03 | -1.26E+00 | 1.02E-04 | 3.63E-04 | -1.43E+00 | 1.39E-03 | 3.92E-03 | -1.29E+00 | 2.58E-13 | 9.48E-13 | -1.46E+00 | 3.78E-09 | 1.04E-08 | -1.30E+00 |
| MIER1 | 1.09E-03 | 5.70E-03 | 1.08E+00 | 6.82E-09 | 2.95E-08 | 1.01E+00 | 1.70E-02 | 7.22E-02 | 1.02E+00 | 1.77E-04 | 6.75E-04 | -1.03E+00 | 2.57E-02 | 9.15E-02 | 1.18E+00 | 1.33E-03 | 3.76E-03 | -1.01E+00 | 1.52E-01 | 5.59E-01 | 1.14E+00 | 2.51E-03 | 6.89E-03 | -1.07E+00 |
| TPSB2 | 1.09E-03 | 5.70E-03 | -1.63E+00 | 1.38E-04 | 5.95E-04 | -2.21E+00 | 4.23E-05 | 1.80E-04 | -1.65E+00 | 1.32E-06 | 5.06E-06 | -1.37E+00 | 7.95E-30 | 2.83E-29 | -2.88E+00 | 5.87E-51 | 1.66E-50 | -3.17E+00 | 4.19E-08 | 1.54E-07 | -2.31E+00 | 5.63E-06 | 1.55E-05 | -2.16E+00 |
| ZCWPW1 | 1.09E-03 | 5.70E-03 | -1.27E+00 | 2.08E-03 | 8.97E-03 | -1.10E+00 | 2.25E-02 | 9.55E-02 | -1.08E+00 | 8.86E-05 | 3.38E-04 | -1.46E+00 | 4.63E-02 | 1.65E-01 | -1.14E+00 | 5.15E-02 | 1.45E-01 | -1.19E+00 | 4.64E-02 | 1.70E-01 | -1.03E+00 | 7.42E-04 | 2.04E-03 | -1.61E+00 |
| GK5 | 1.10E-03 | 5.71E-03 | -1.06E+00 | 4.12E-02 | 1.78E-01 | 1.27E+00 | 2.02E-02 | 8.57E-02 | -1.01E+00 | 8.35E-02 | 3.19E-01 | 1.31E+00 | 5.71E-02 | 2.04E-01 | 1.05E+00 | 1.61E-01 | 4.53E-01 | 1.16E+00 | 3.26E-03 | 1.20E-02 | -1.26E+00 | 1.10E-01 | 3.02E-01 | 1.12E+00 |
| C10orf88 | 1.10E-03 | 5.71E-03 | -1.49E+00 | 9.59E-03 | 4.14E-02 | -1.16E+00 | 1.10E-03 | 4.68E-03 | -1.39E+00 | 4.42E-03 | 1.69E-02 | -1.12E+00 | 8.74E-05 | 3.12E-04 | -1.78E+00 | 6.87E-04 | 1.94E-03 | -1.52E+00 | 3.09E-07 | 1.13E-06 | -1.64E+00 | 3.03E-05 | 8.32E-05 | -1.42E+00 |
| SLC7A5 | 1.10E-03 | 5.71E-03 | -2.13E+00 | 4.03E-03 | 1.74E-02 | -1.56E+00 | 1.28E-01 | 5.43E-01 | 1.18E+00 | 1.01E-01 | 3.85E-01 | 1.29E+00 | 8.45E-03 | 3.01E-02 | -1.59E+00 | 3.17E-01 | 8.93E-01 | 1.39E+00 | 2.01E-07 | 7.39E-07 | -1.22E+00 | 3.13E-03 | 8.59E-03 | 1.02E+00 |
| ZNF66 | 1.10E-03 | 5.73E-03 | -1.10E+00 | 6.80E-03 | 2.94E-02 | 1.10E+00 | 1.59E-04 | 6.77E-04 | -1.50E+00 | 1.07E-04 | 4.07E-04 | -1.35E+00 | 1.65E-02 | 5.90E-02 | -1.09E+00 | 1.54E-02 | 4.33E-02 | -1.07E+00 | 7.12E-02 | 2.61E-01 | 1.18E+00 | 3.04E-01 | 8.34E-01 | 1.34E+00 |
| BTG3 | 1.10E-03 | 5.74E-03 | 1.08E+00 | 2.55E-26 | 1.10E-25 | -1.65E+00 | 7.03E-02 | 2.99E-01 | 1.08E+00 | 2.55E-13 | 9.74E-13 | -1.47E+00 | 2.12E-02 | 7.55E-02 | 1.24E+00 | 1.63E-08 | 4.59E-08 | -1.25E+00 | 4.72E-05 | 1.73E-04 | -1.23E+00 | 4.76E-12 | 1.31E-11 | -1.72E+00 |
| DGKA | 1.10E-03 | 5.74E-03 | -1.12E+00 | 4.45E-02 | 1.92E-01 | 1.22E+00 | 1.96E-03 | 8.32E-03 | -1.33E+00 | 1.16E-02 | 4.45E-02 | 1.01E+00 | 7.21E-02 | 2.57E-01 | 1.14E+00 | 2.46E-02 | 6.93E-02 | -1.05E+00 | 7.30E-02 | 2.68E-01 | -1.00E+00 | 6.98E-02 | 1.92E-01 | -1.06E+00 |
| RRN3 | 1.10E-03 | 5.75E-03 | -1.13E+00 | 3.25E-03 | 1.40E-02 | 1.05E+00 | 2.85E-05 | 1.21E-04 | -1.34E+00 | 3.37E-06 | 1.29E-05 | -1.23E+00 | 3.36E-05 | 1.20E-04 | -1.34E+00 | 6.10E-03 | 1.72E-02 | 1.04E+00 | 4.17E-26 | 1.53E-25 | -2.00E+00 | 5.67E-12 | 1.56E-11 | -1.33E+00 |
| PML | 1.10E-03 | 5.75E-03 | 1.11E+00 | 3.58E-02 | 1.55E-01 | 1.36E+00 | 3.16E-05 | 1.34E-04 | -1.19E+00 | 1.39E-02 | 5.29E-02 | 1.14E+00 | 1.65E-02 | 5.89E-02 | 1.01E+00 | 2.38E-01 | 6.70E-01 | 1.29E+00 | 1.41E-10 | 5.19E-10 | -1.57E+00 | 1.24E-06 | 3.39E-06 | -1.31E+00 |
| MESP1 | 1.10E-03 | 5.75E-03 | -1.16E+00 | 2.39E-02 | 1.03E-01 | 1.19E+00 | 2.30E-05 | 9.76E-05 | -1.26E+00 | 1.56E-04 | 5.97E-04 | -1.05E+00 | 1.63E-02 | 5.80E-02 | -1.22E+00 | 1.84E-01 | 5.20E-01 | 1.24E+00 | 1.99E-07 | 7.31E-07 | -1.44E+00 | 2.11E-02 | 5.80E-02 | 1.05E+00 |
| ATG16L1 | 1.11E-03 | 5.77E-03 | -1.51E+00 | 3.90E-03 | 1.69E-02 | -1.18E+00 | 2.12E-01 | 8.99E-01 | 1.25E+00 | 1.82E-01 | 6.93E-01 | 1.22E+00 | 1.54E-01 | 5.48E-01 | 1.02E+00 | 2.05E-01 | 5.77E-01 | 1.13E+00 | 2.10E-02 | 7.71E-02 | -1.11E+00 | 2.13E-02 | 5.86E-02 | -1.10E+00 |
| CPD | 1.11E-03 | 5.77E-03 | -1.19E+00 | 3.95E-02 | 1.71E-01 | 1.18E+00 | 2.51E-02 | 1.07E-01 | -1.04E+00 | 1.90E-02 | 7.26E-02 | 1.08E+00 | 2.36E-03 | 8.40E-03 | -1.16E+00 | 8.61E-04 | 2.43E-03 | -1.17E+00 | 4.49E-05 | 1.65E-04 | -1.16E+00 | 8.03E-03 | 2.21E-02 | 1.07E+00 |
| FAM91A1 | 1.11E-03 | 5.78E-03 | -1.00E+00 | 6.74E-05 | 2.91E-04 | -1.01E+00 | 7.89E-03 | 3.35E-02 | -1.02E+00 | 6.09E-04 | 2.33E-03 | -1.01E+00 | 2.59E-03 | 9.23E-03 | -1.04E+00 | 1.22E-03 | 3.45E-03 | -1.06E+00 | 5.66E-03 | 2.08E-02 | -1.08E+00 | 6.05E-03 | 1.66E-02 | -1.07E+00 |
| AC125807.1 | 1.11E-03 | 5.80E-03 | 1.65E+00 | 4.10E-09 | 1.77E-08 | -1.10E+00 | 1.57E-05 | 6.67E-05 | 1.86E+00 | 2.18E-02 | 8.32E-02 | 1.20E+00 | 3.24E-03 | 1.16E-02 | 1.89E+00 | 7.89E-02 | 2.22E-01 | 1.20E+00 | 6.98E-03 | 2.56E-02 | 1.58E+00 | 8.18E-03 | 2.25E-02 | 1.02E+00 |
| PPIL2 | 1.11E-03 | 5.81E-03 | -1.28E+00 | 2.03E-03 | 8.78E-03 | -1.24E+00 | 1.81E-02 | 7.69E-02 | -1.31E+00 | 7.22E-02 | 2.76E-01 | 1.02E+00 | 8.02E-02 | 2.86E-01 | 1.01E+00 | 5.20E-02 | 1.47E-01 | -1.14E+00 | 1.38E-04 | 5.06E-04 | -1.41E+00 | 1.19E-02 | 3.26E-02 | -1.04E+00 |
| DUSP28 | 1.11E-03 | 5.81E-03 | -1.06E+00 | 6.49E-06 | 2.80E-05 | -1.24E+00 | 2.24E-02 | 9.51E-02 | 1.01E+00 | 5.95E-03 | 2.27E-02 | -1.03E+00 | 2.33E-01 | 8.29E-01 | 1.42E+00 | 2.51E-01 | 7.08E-01 | 1.35E+00 | 3.27E-02 | 1.20E-01 | 1.12E+00 | 1.75E-02 | 4.80E-02 | -1.05E+00 |
| GHDC | 1.12E-03 | 5.83E-03 | -1.08E+00 | 4.56E-03 | 1.97E-02 | 1.11E+00 | 1.23E-02 | 5.21E-02 | -1.05E+00 | 2.58E-03 | 9.87E-03 | 1.02E+00 | 2.24E-04 | 7.98E-04 | -1.15E+00 | 2.47E-02 | 6.95E-02 | 1.15E+00 | 1.60E-03 | 5.88E-03 | 1.00E+00 | 3.54E-01 | 9.72E-01 | 1.37E+00 |
| KIAA1211 | 1.12E-03 | 5.84E-03 | -1.38E+00 | 6.38E-04 | 2.76E-03 | -1.34E+00 | 5.76E-03 | 2.45E-02 | -1.21E+00 | 1.66E-03 | 6.34E-03 | -1.21E+00 | 2.34E-03 | 8.33E-03 | -1.16E+00 | 4.82E-02 | 1.36E-01 | 1.10E+00 | 1.57E-02 | 5.78E-02 | -1.03E+00 | 5.25E-04 | 1.44E-03 | -1.29E+00 |
| FASTKD5 | 1.12E-03 | 5.84E-03 | -1.18E+00 | 1.75E-03 | 7.55E-03 | -1.06E+00 | 3.02E-02 | 1.28E-01 | 1.04E+00 | 7.64E-04 | 2.92E-03 | -1.12E+00 | 4.44E-02 | 1.58E-01 | 1.16E+00 | 2.99E-02 | 8.42E-02 | 1.06E+00 | 7.70E-02 | 2.83E-01 | 1.08E+00 | 3.07E-02 | 8.42E-02 | -1.05E+00 |
| PARP15 | 1.12E-03 | 5.84E-03 | -1.20E+00 | 4.03E-03 | 1.74E-02 | -1.05E+00 | 1.96E-02 | 8.32E-02 | -1.14E+00 | 2.36E-02 | 9.03E-02 | 1.06E+00 | 8.61E-02 | 3.07E-01 | -1.06E+00 | 8.05E-02 | 2.27E-01 | -1.08E+00 | 7.46E-02 | 2.74E-01 | 1.06E+00 | 5.18E-03 | 1.42E-02 | -1.12E+00 |
| AC008429.1 | 1.12E-03 | 5.85E-03 | -1.30E+00 | 1.08E-02 | 4.68E-02 | 1.03E+00 | 1.01E-03 | 4.29E-03 | -1.22E+00 | 2.09E-03 | 8.00E-03 | -1.15E+00 | 1.40E-01 | 5.01E-01 | -1.08E+00 | 1.42E-01 | 4.02E-01 | -1.09E+00 | 8.43E-03 | 3.09E-02 | -1.73E+00 | 1.82E-02 | 4.99E-02 | -1.65E+00 |
| H3F3AP6 | 1.12E-03 | 5.86E-03 | 2.28E+00 | 1.61E-01 | 6.95E-01 | 1.74E+00 | 9.19E-03 | 3.91E-02 | 1.67E+00 | 1.54E-01 | 5.89E-01 | 1.24E+00 | 1.17E-01 | 4.18E-01 | 1.75E+00 | 2.95E-01 | 8.32E-01 | 1.40E+00 | 6.75E-02 | 2.48E-01 | 1.51E+00 | 1.44E-01 | 3.96E-01 | 1.06E+00 |
| GATA3 | 1.13E-03 | 5.87E-03 | -1.17E+00 | 1.28E-06 | 5.52E-06 | -1.74E+00 | 1.93E-01 | 8.21E-01 | -1.56E+00 | 1.11E-01 | 4.23E-01 | -1.89E+00 | 2.80E-01 | 1.00E+00 | 1.00E+00 | 3.55E-01 | 1.00E+00 | -1.00E+00 | 2.72E-01 | 1.00E+00 | 1.00E+00 | 3.64E-01 | 1.00E+00 | -1.00E+00 |
| NOC4L | 1.13E-03 | 5.87E-03 | -1.21E+00 | 5.55E-04 | 2.40E-03 | -1.17E+00 | 1.78E-02 | 7.58E-02 | 1.01E+00 | 1.37E-02 | 5.25E-02 | 1.19E+00 | 2.54E-02 | 9.06E-02 | -1.05E+00 | 1.27E-01 | 3.59E-01 | 1.28E+00 | 3.64E-03 | 1.34E-02 | -1.04E+00 | 1.83E-01 | 5.01E-01 | 1.39E+00 |
| FBXL20 | 1.13E-03 | 5.87E-03 | -1.17E+00 | 1.07E-01 | 4.64E-01 | 1.26E+00 | 8.48E-05 | 3.60E-04 | -1.24E+00 | 1.05E-03 | 4.01E-03 | 1.01E+00 | 6.22E-04 | 2.22E-03 | -1.13E+00 | 2.83E-02 | 7.98E-02 | 1.01E+00 | 6.53E-03 | 2.40E-02 | -1.12E+00 | 9.88E-02 | 2.71E-01 | 1.01E+00 |
| SEC16A | 1.13E-03 | 5.91E-03 | -1.17E+00 | 1.06E-02 | 4.56E-02 | 1.07E+00 | 4.06E-06 | 1.73E-05 | -1.33E+00 | 3.58E-04 | 1.37E-03 | -1.02E+00 | 1.24E-01 | 4.40E-01 | 1.27E+00 | 1.53E-01 | 4.32E-01 | 1.30E+00 | 4.76E-02 | 1.75E-01 | -1.02E+00 | 1.52E-01 | 4.18E-01 | 1.14E+00 |
| RPS10P5 | 1.14E-03 | 5.92E-03 | -1.08E+00 | 1.06E-01 | 4.60E-01 | 1.77E+00 | 1.11E-04 | 4.73E-04 | -1.30E+00 | 6.72E-02 | 2.57E-01 | 1.70E+00 | 2.09E-03 | 7.44E-03 | -1.19E+00 | 2.43E-01 | 6.86E-01 | 1.66E+00 | 2.77E-02 | 1.02E-01 | -1.05E+00 | 3.96E-03 | 1.09E-02 | 1.73E+00 |
| BRMS1 | 1.14E-03 | 5.93E-03 | 1.03E+00 | 1.82E-03 | 7.88E-03 | 1.20E+00 | 3.84E-02 | 1.63E-01 | 1.07E+00 | 1.11E-01 | 4.23E-01 | 1.29E+00 | 1.39E-03 | 4.97E-03 | -1.01E+00 | 4.36E-02 | 1.23E-01 | 1.14E+00 | 1.99E-03 | 7.30E-03 | -1.01E+00 | 1.18E-01 | 3.25E-01 | 1.16E+00 |
| ZNF706 | 1.14E-03 | 5.94E-03 | 1.12E+00 | 1.96E-01 | 8.46E-01 | 1.49E+00 | 3.34E-03 | 1.42E-02 | 1.01E+00 | 9.93E-02 | 3.79E-01 | 1.24E+00 | 4.88E-03 | 1.74E-02 | 1.11E+00 | 6.28E-02 | 1.77E-01 | 1.19E+00 | 1.27E-05 | 4.65E-05 | -1.17E+00 | 9.05E-02 | 2.49E-01 | 1.09E+00 |
| DISC1 | 1.14E-03 | 5.94E-03 | -1.14E+00 | 4.33E-03 | 1.87E-02 | 1.13E+00 | 5.39E-05 | 2.29E-04 | -1.35E+00 | 6.02E-05 | 2.30E-04 | -1.25E+00 | 4.29E-02 | 1.53E-01 | -1.15E+00 | 1.85E-01 | 5.20E-01 | 1.09E+00 | 1.13E-02 | 4.15E-02 | -1.57E+00 | 1.94E-01 | 5.33E-01 | -1.05E+00 |
| RASSF4 | 1.14E-03 | 5.95E-03 | -1.83E+00 | 9.04E-03 | 3.91E-02 | -1.25E+00 | 1.05E-02 | 4.47E-02 | -1.47E+00 | 1.20E-02 | 4.60E-02 | -1.37E+00 | 4.73E-02 | 1.69E-01 | -1.40E+00 | 3.84E-02 | 1.08E-01 | -1.52E+00 | 2.07E-01 | 7.58E-01 | -1.11E+00 | 3.47E-01 | 9.54E-01 | 1.17E+00 |
| AC018529.2 | 1.14E-03 | 5.95E-03 | -1.38E+00 | 3.03E-02 | 1.31E-01 | 1.03E+00 | 3.69E-03 | 1.57E-02 | -1.50E+00 | 4.21E-02 | 1.61E-01 | -1.02E+00 | 4.44E-02 | 1.58E-01 | -1.36E+00 | 1.32E-01 | 3.73E-01 | -1.04E+00 | 2.62E-02 | 9.63E-02 | -1.41E+00 | 9.25E-02 | 2.54E-01 | -1.22E+00 |
| IDUA | 1.14E-03 | 5.96E-03 | -1.53E+00 | 3.69E-03 | 1.59E-02 | -1.26E+00 | 1.60E-02 | 6.80E-02 | -1.16E+00 | 2.47E-03 | 9.43E-03 | -1.27E+00 | 7.39E-02 | 2.64E-01 | -1.04E+00 | 7.71E-02 | 2.17E-01 | -1.18E+00 | 1.34E-02 | 4.92E-02 | -1.33E+00 | 2.54E-01 | 6.97E-01 | 1.09E+00 |
| EXOG | 1.15E-03 | 5.98E-03 | -1.12E+00 | 2.09E-03 | 9.03E-03 | 1.06E+00 | 6.02E-04 | 2.56E-03 | -1.19E+00 | 2.55E-03 | 9.73E-03 | -1.01E+00 | 2.62E-02 | 9.34E-02 | -1.01E+00 | 9.53E-02 | 2.69E-01 | 1.17E+00 | 5.04E-02 | 1.85E-01 | 1.04E+00 | 4.65E-02 | 1.28E-01 | 1.01E+00 |
| NEK3 | 1.15E-03 | 5.99E-03 | -1.03E+00 | 1.12E-05 | 4.85E-05 | -1.06E+00 | 6.78E-04 | 2.88E-03 | -1.32E+00 | 1.16E-04 | 4.41E-04 | -1.24E+00 | 8.56E-04 | 3.05E-03 | -1.29E+00 | 5.79E-04 | 1.63E-03 | -1.25E+00 | 6.90E-02 | 2.53E-01 | -1.03E+00 | 1.91E-02 | 5.24E-02 | -1.16E+00 |
| ZNF764 | 1.15E-03 | 6.01E-03 | -1.31E+00 | 7.92E-04 | 3.42E-03 | -1.24E+00 | 1.06E-02 | 4.52E-02 | -1.16E+00 | 2.69E-03 | 1.03E-02 | -1.10E+00 | 4.83E-03 | 1.72E-02 | -1.27E+00 | 4.08E-03 | 1.15E-02 | -1.30E+00 | 9.22E-03 | 3.39E-02 | -1.20E+00 | 2.22E-03 | 6.09E-03 | -1.25E+00 |
| NDUFB4 | 1.15E-03 | 6.02E-03 | -1.02E+00 | 8.71E-14 | 3.76E-13 | -1.22E+00 | 2.83E-02 | 1.20E-01 | -1.09E+00 | 7.32E-08 | 2.80E-07 | -1.35E+00 | 1.87E-06 | 6.65E-06 | -1.23E+00 | 6.10E-11 | 1.72E-10 | -1.36E+00 | 3.18E-02 | 1.17E-01 | -1.08E+00 | 9.76E-05 | 2.68E-04 | -1.30E+00 |
| UBE2SP2 | 1.16E-03 | 6.02E-03 | 1.04E+00 | 2.37E-11 | 1.02E-10 | -1.47E+00 | 6.32E-02 | 2.69E-01 | 1.15E+00 | 7.36E-05 | 2.81E-04 | -1.07E+00 | 1.20E-01 | 4.29E-01 | 1.25E+00 | 3.98E-04 | 1.12E-03 | -1.21E+00 | 2.71E-01 | 9.94E-01 | 1.16E+00 | 9.52E-02 | 2.61E-01 | 1.03E+00 |
| NAPA-AS1 | 1.16E-03 | 6.03E-03 | -1.25E+00 | 5.35E-05 | 2.31E-04 | -1.56E+00 | 2.10E-02 | 8.90E-02 | -1.28E+00 | 8.82E-03 | 3.37E-02 | -1.38E+00 | 1.82E-01 | 6.48E-01 | 1.17E+00 | 1.64E-01 | 4.63E-01 | 1.01E+00 | 5.86E-02 | 2.15E-01 | -1.28E+00 | 5.74E-02 | 1.58E-01 | -1.39E+00 |
| RINL | 1.16E-03 | 6.03E-03 | -1.19E+00 | 1.18E-02 | 5.10E-02 | 1.05E+00 | 4.88E-02 | 2.07E-01 | -1.18E+00 | 2.30E-01 | 8.77E-01 | 1.19E+00 | 4.51E-03 | 1.61E-02 | -1.10E+00 | 1.04E-01 | 2.92E-01 | 1.22E+00 | 1.71E-05 | 6.27E-05 | -1.36E+00 | 1.44E-01 | 3.95E-01 | 1.15E+00 |
| HEATR4 | 1.16E-03 | 6.05E-03 | -1.31E+00 | 2.69E-02 | 1.16E-01 | 1.10E+00 | 5.30E-04 | 2.25E-03 | -1.74E+00 | 2.49E-02 | 9.52E-02 | 1.01E+00 | 1.32E-01 | 4.72E-01 | -1.13E+00 | 2.49E-01 | 7.01E-01 | 1.61E+00 | 1.48E-01 | 5.42E-01 | 1.46E+00 | 2.08E-01 | 5.71E-01 | 1.31E+00 |
| ADAM9 | 1.16E-03 | 6.05E-03 | -1.16E+00 | 4.78E-04 | 2.06E-03 | -1.16E+00 | 1.31E-02 | 5.55E-02 | -1.13E+00 | 1.44E-03 | 5.50E-03 | -1.17E+00 | 4.24E-02 | 1.51E-01 | 1.09E+00 | 6.05E-03 | 1.71E-02 | -1.20E+00 | 9.56E-05 | 3.51E-04 | -1.25E+00 | 5.99E-07 | 1.65E-06 | -1.71E+00 |
| CNNM3 | 1.16E-03 | 6.06E-03 | -1.45E+00 | 3.08E-02 | 1.33E-01 | 1.04E+00 | 3.31E-05 | 1.41E-04 | -1.77E+00 | 1.02E-03 | 3.91E-03 | -1.21E+00 | 1.84E-02 | 6.55E-02 | -1.29E+00 | 1.46E-01 | 4.13E-01 | 1.15E+00 | 5.23E-13 | 1.92E-12 | -2.11E+00 | 3.33E-07 | 9.14E-07 | -1.51E+00 |
| PLCL2 | 1.16E-03 | 6.07E-03 | 1.03E+00 | 9.79E-03 | 4.23E-02 | 1.26E+00 | 4.17E-02 | 1.77E-01 | 1.05E+00 | 1.26E-01 | 4.81E-01 | 1.29E+00 | 7.69E-02 | 2.74E-01 | 1.05E+00 | 3.34E-01 | 9.40E-01 | 1.44E+00 | 1.98E-02 | 7.27E-02 | -1.20E+00 | 2.64E-01 | 7.25E-01 | 1.23E+00 |
| ISCA1P6 | 1.17E-03 | 6.08E-03 | -1.41E+00 | 4.68E-04 | 2.02E-03 | -1.46E+00 | 1.47E-02 | 6.26E-02 | -1.45E+00 | 9.14E-02 | 3.49E-01 | 1.01E+00 | 4.94E-03 | 1.76E-02 | -1.32E+00 | 8.04E-03 | 2.27E-02 | -1.26E+00 | 2.31E-03 | 8.49E-03 | -1.19E+00 | 7.88E-04 | 2.16E-03 | -1.44E+00 |
| AARS2 | 1.17E-03 | 6.09E-03 | -1.36E+00 | 1.29E-02 | 5.57E-02 | -1.05E+00 | 2.59E-03 | 1.10E-02 | -1.34E+00 | 2.54E-02 | 9.71E-02 | -1.05E+00 | 9.49E-03 | 3.39E-02 | -1.69E+00 | 2.33E-01 | 6.57E-01 | 1.24E+00 | 2.54E-04 | 9.30E-04 | -1.39E+00 | 3.78E-02 | 1.04E-01 | 1.07E+00 |
| ATG12 | 1.17E-03 | 6.09E-03 | -1.09E+00 | 9.69E-02 | 4.19E-01 | 1.29E+00 | 7.24E-03 | 3.08E-02 | -1.13E+00 | 1.31E-02 | 4.98E-02 | 1.07E+00 | 6.27E-02 | 2.24E-01 | 1.13E+00 | 6.57E-02 | 1.85E-01 | 1.05E+00 | 1.70E-02 | 6.23E-02 | 1.02E+00 | 1.13E-01 | 3.12E-01 | 1.14E+00 |
| MRM1 | 1.17E-03 | 6.11E-03 | -1.60E+00 | 4.03E-02 | 1.74E-01 | 1.10E+00 | 7.22E-03 | 3.07E-02 | -1.30E+00 | 2.26E-01 | 8.62E-01 | 1.31E+00 | 4.65E-02 | 1.66E-01 | -1.16E+00 | 1.48E-01 | 4.18E-01 | 1.82E+00 | 6.09E-11 | 2.24E-10 | -1.51E+00 | 6.36E-04 | 1.75E-03 | -1.02E+00 |
| C10orf25 | 1.17E-03 | 6.11E-03 | -1.47E+00 | 1.79E-01 | 7.73E-01 | 1.26E+00 | 2.85E-07 | 1.21E-06 | -2.03E+00 | 9.57E-04 | 3.66E-03 | -1.15E+00 | 9.89E-06 | 3.53E-05 | -2.81E+00 | 4.58E-02 | 1.29E-01 | -1.07E+00 | 5.85E-08 | 2.15E-07 | -1.86E+00 | 1.05E-03 | 2.88E-03 | -1.17E+00 |
| AC120114.2 | 1.17E-03 | 6.12E-03 | -1.50E+00 | 4.97E-03 | 2.15E-02 | -1.23E+00 | 1.22E-02 | 5.18E-02 | -1.14E+00 | 8.23E-03 | 3.14E-02 | -1.16E+00 | 8.54E-05 | 3.04E-04 | -2.10E+00 | 4.52E-04 | 1.27E-03 | -1.70E+00 | 2.28E-01 | 8.36E-01 | 1.10E+00 | 1.96E-01 | 5.38E-01 | -1.19E+00 |
| PILRA | 1.18E-03 | 6.13E-03 | -1.56E+00 | 1.03E-02 | 4.44E-02 | 1.01E+00 | 1.18E-02 | 5.00E-02 | -1.21E+00 | 1.84E-01 | 7.05E-01 | 1.25E+00 | 1.76E-01 | 6.27E-01 | 1.84E+00 | 3.49E-01 | 9.85E-01 | 1.38E+00 | 2.36E-01 | 8.67E-01 | -1.02E+00 | 3.18E-01 | 8.73E-01 | -1.04E+00 |
| TESK2 | 1.18E-03 | 6.14E-03 | -1.23E+00 | 3.92E-03 | 1.70E-02 | -1.06E+00 | 6.76E-02 | 2.87E-01 | 1.05E+00 | 1.17E-02 | 4.46E-02 | -1.13E+00 | 1.56E-01 | 5.55E-01 | 1.16E+00 | 1.13E-01 | 3.19E-01 | 1.04E+00 | 3.82E-02 | 1.40E-01 | 1.10E+00 | 9.61E-02 | 2.64E-01 | 1.12E+00 |
| ZNF318 | 1.18E-03 | 6.14E-03 | -1.46E+00 | 8.42E-03 | 3.64E-02 | -1.15E+00 | 3.45E-05 | 1.47E-04 | -1.87E+00 | 1.10E-02 | 4.22E-02 | -1.13E+00 | 5.09E-03 | 1.81E-02 | -1.31E+00 | 3.54E-02 | 9.98E-02 | -1.09E+00 | 2.00E-02 | 7.32E-02 | -1.02E+00 | 4.79E-02 | 1.31E-01 | -1.04E+00 |
| TMEM17 | 1.18E-03 | 6.16E-03 | -1.43E+00 | 1.35E-01 | 5.82E-01 | 1.66E+00 | 2.11E-06 | 8.97E-06 | -1.94E+00 | 2.61E-03 | 9.97E-03 | -1.07E+00 | 8.19E-05 | 2.92E-04 | -1.94E+00 | 1.83E-02 | 5.17E-02 | -1.06E+00 | 9.05E-07 | 3.32E-06 | -2.24E+00 | 3.46E-04 | 9.51E-04 | -1.54E+00 |
| AC005822.1 | 1.18E-03 | 6.16E-03 | -1.41E+00 | 1.44E-03 | 6.24E-03 | -1.31E+00 | 8.85E-02 | 3.76E-01 | 1.06E+00 | 2.22E-02 | 8.49E-02 | 1.01E+00 | 1.91E-04 | 6.82E-04 | -1.44E+00 | 1.54E-04 | 4.35E-04 | -1.47E+00 | 9.93E-02 | 3.65E-01 | 1.20E+00 | 6.53E-02 | 1.79E-01 | 1.09E+00 |
| AC105114.1 | 1.18E-03 | 6.16E-03 | -1.56E+00 | 8.44E-02 | 3.65E-01 | 1.18E+00 | 1.61E-01 | 6.85E-01 | -1.07E+00 | 4.84E-02 | 1.85E-01 | -1.42E+00 | 4.59E-04 | 1.64E-03 | -1.69E+00 | 2.27E-02 | 6.41E-02 | -1.06E+00 | 8.59E-02 | 3.15E-01 | -1.35E+00 | 1.67E-01 | 4.58E-01 | -1.13E+00 |
| TEC | 1.18E-03 | 6.17E-03 | 1.00E+00 | 8.94E-04 | 3.86E-03 | 1.12E+00 | 1.47E-02 | 6.23E-02 | -1.07E+00 | 1.12E-01 | 4.29E-01 | 1.23E+00 | 4.60E-02 | 1.64E-01 | 1.24E+00 | 4.59E-02 | 1.29E-01 | 1.19E+00 | 2.08E-01 | 7.63E-01 | 1.41E+00 | 2.84E-01 | 7.81E-01 | 1.26E+00 |
| SRF | 1.18E-03 | 6.17E-03 | -1.16E+00 | 8.85E-06 | 3.83E-05 | -1.61E+00 | 2.25E-02 | 9.58E-02 | -1.00E+00 | 3.82E-03 | 1.46E-02 | -1.01E+00 | 4.19E-02 | 1.50E-01 | -1.03E+00 | 4.19E-02 | 1.18E-01 | -1.00E+00 | 4.02E-03 | 1.48E-02 | -1.26E+00 | 2.71E-02 | 7.44E-02 | -1.17E+00 |
| LINC02232 | 1.19E-03 | 6.18E-03 | -2.02E+00 | 8.23E-02 | 3.56E-01 | -1.05E+00 | 1.84E-01 | 7.83E-01 | -1.01E+00 | 2.51E-01 | 9.58E-01 | 1.10E+00 | 5.38E-04 | 1.92E-03 | -2.04E+00 | 8.46E-03 | 2.39E-02 | -1.40E+00 | 2.03E-03 | 7.43E-03 | -1.78E+00 | 3.27E-02 | 8.97E-02 | -1.28E+00 |
| MUC12 | 1.19E-03 | 6.20E-03 | 2.24E+00 | 2.02E-01 | 8.75E-01 | 1.16E+00 | 2.24E-04 | 9.54E-04 | 2.50E+00 | 2.40E-01 | 9.16E-01 | 1.08E+00 | 1.94E-02 | 6.90E-02 | 2.19E+00 | 4.49E-02 | 1.27E-01 | -1.53E+00 | 4.08E-02 | 1.50E-01 | 1.70E+00 | 3.02E-01 | 8.29E-01 | -1.09E+00 |
| C9orf3 | 1.19E-03 | 6.22E-03 | -1.00E+00 | 9.60E-03 | 4.15E-02 | 1.22E+00 | 6.74E-02 | 2.86E-01 | 1.01E+00 | 1.18E-01 | 4.50E-01 | 1.22E+00 | 1.34E-01 | 4.79E-01 | 1.25E+00 | 3.07E-01 | 8.66E-01 | 1.53E+00 | 8.34E-04 | 3.06E-03 | -1.66E+00 | 4.43E-02 | 1.22E-01 | -1.20E+00 |
| SERPINH1 | 1.19E-03 | 6.22E-03 | -1.34E+00 | 2.26E-01 | 9.77E-01 | 1.52E+00 | 9.45E-06 | 4.01E-05 | -1.67E+00 | 1.05E-01 | 4.00E-01 | 1.28E+00 | 2.78E-05 | 9.92E-05 | -1.97E+00 | 3.59E-02 | 1.01E-01 | 1.01E+00 | 9.50E-02 | 3.49E-01 | -1.17E+00 | 7.49E-02 | 2.06E-01 | 1.44E+00 |
| MRPL3P1 | 1.19E-03 | 6.23E-03 | -1.27E+00 | 5.49E-05 | 2.37E-04 | -1.46E+00 | 2.04E-06 | 8.69E-06 | -1.35E+00 | 3.48E-07 | 1.33E-06 | -1.27E+00 | 1.07E-04 | 3.82E-04 | -1.60E+00 | 1.47E-05 | 4.16E-05 | -1.88E+00 | 1.26E-07 | 4.64E-07 | -1.28E+00 | 1.09E-11 | 2.98E-11 | -1.68E+00 |
| AL357055.3 | 1.20E-03 | 6.23E-03 | -1.92E+00 | 2.04E-01 | 8.81E-01 | 1.42E+00 | 3.20E-05 | 1.36E-04 | -1.96E+00 | 5.45E-02 | 2.08E-01 | 1.11E+00 | 2.63E-02 | 9.39E-02 | -1.48E+00 | 3.33E-01 | 9.39E-01 | 1.25E+00 | 5.66E-03 | 2.08E-02 | -1.71E+00 | 3.64E-01 | 9.99E-01 | 1.13E+00 |
| ZBTB4 | 1.20E-03 | 6.23E-03 | 1.01E+00 | 2.15E-07 | 9.30E-07 | -1.28E+00 | 1.43E-02 | 6.08E-02 | -1.03E+00 | 3.27E-03 | 1.25E-02 | -1.05E+00 | 7.00E-04 | 2.50E-03 | -1.16E+00 | 1.98E-03 | 5.58E-03 | -1.15E+00 | 1.66E-03 | 6.08E-03 | -1.20E+00 | 3.02E-04 | 8.30E-04 | -1.32E+00 |
| AC105052.4 | 1.20E-03 | 6.23E-03 | -1.32E+00 | 7.22E-03 | 3.12E-02 | -1.05E+00 | 8.89E-06 | 3.78E-05 | -1.91E+00 | 2.67E-02 | 1.02E-01 | 1.08E+00 | 2.00E-01 | 7.14E-01 | 1.17E+00 | 2.16E-01 | 6.08E-01 | 1.04E+00 | 6.09E-03 | 2.24E-02 | -1.61E+00 | 3.11E-02 | 8.55E-02 | -1.40E+00 |
| MIR3936HG | 1.20E-03 | 6.23E-03 | -1.38E+00 | 1.61E-02 | 6.97E-02 | 1.04E+00 | 5.91E-02 | 2.51E-01 | -1.18E+00 | 3.06E-02 | 1.17E-01 | -1.24E+00 | 1.44E-01 | 5.13E-01 | 1.01E+00 | 1.08E-01 | 3.04E-01 | -1.07E+00 | 2.59E-02 | 9.51E-02 | -1.38E+00 | 3.64E-01 | 1.00E+00 | 1.23E+00 |
| RINT1 | 1.20E-03 | 6.23E-03 | -1.39E+00 | 6.72E-03 | 2.90E-02 | -1.10E+00 | 4.79E-03 | 2.04E-02 | -1.26E+00 | 6.41E-03 | 2.45E-02 | -1.15E+00 | 5.24E-03 | 1.87E-02 | -1.23E+00 | 1.07E-02 | 3.01E-02 | -1.15E+00 | 3.97E-06 | 1.46E-05 | -1.54E+00 | 5.97E-04 | 1.64E-03 | -1.24E+00 |
| ELF1 | 1.20E-03 | 6.24E-03 | 1.08E+00 | 2.54E-15 | 1.10E-14 | -1.15E+00 | 1.53E-01 | 6.50E-01 | 1.08E+00 | 1.86E-07 | 7.09E-07 | -1.16E+00 | 7.84E-02 | 2.80E-01 | 1.08E+00 | 1.95E-05 | 5.50E-05 | -1.18E+00 | 1.86E-01 | 6.84E-01 | 1.37E+00 | 1.36E-02 | 3.73E-02 | -1.05E+00 |
| RARG | 1.20E-03 | 6.25E-03 | -1.50E+00 | 9.58E-04 | 4.14E-03 | -1.43E+00 | 5.29E-04 | 2.25E-03 | -2.22E+00 | 2.91E-02 | 1.11E-01 | -1.20E+00 | 2.33E-02 | 8.32E-02 | -1.57E+00 | 2.18E-01 | 6.15E-01 | 1.18E+00 | 1.62E-10 | 5.96E-10 | -2.03E+00 | 1.32E-05 | 3.62E-05 | -1.35E+00 |
| RARS2 | 1.20E-03 | 6.26E-03 | -1.15E+00 | 3.42E-02 | 1.48E-01 | 1.21E+00 | 1.94E-02 | 8.22E-02 | 1.01E+00 | 4.90E-02 | 1.87E-01 | 1.24E+00 | 7.11E-04 | 2.53E-03 | -1.20E+00 | 5.69E-02 | 1.60E-01 | 1.09E+00 | 2.38E-02 | 8.75E-02 | -1.04E+00 | 3.04E-01 | 8.35E-01 | 1.20E+00 |
| AC103988.2 | 1.20E-03 | 6.27E-03 | -1.32E+00 | 6.56E-04 | 2.84E-03 | -1.30E+00 | 2.10E-01 | 8.92E-01 | 1.15E+00 | 4.91E-02 | 1.87E-01 | 1.00E+00 | 2.84E-02 | 1.01E-01 | -1.06E+00 | 3.73E-03 | 1.05E-02 | -1.46E+00 | 2.04E-01 | 7.48E-01 | 1.22E+00 | 2.01E-03 | 5.52E-03 | -1.29E+00 |
| ALG2 | 1.20E-03 | 6.27E-03 | -1.12E+00 | 1.41E-04 | 6.09E-04 | -1.05E+00 | 6.21E-03 | 2.64E-02 | -1.08E+00 | 4.82E-03 | 1.84E-02 | -1.00E+00 | 1.22E-03 | 4.37E-03 | -1.14E+00 | 3.00E-06 | 8.45E-06 | -1.50E+00 | 1.65E-03 | 6.05E-03 | -1.24E+00 | 6.71E-03 | 1.84E-02 | -1.18E+00 |
| MFSD14C | 1.20E-03 | 6.27E-03 | -1.18E+00 | 1.95E-01 | 8.41E-01 | 1.53E+00 | 1.44E-06 | 6.13E-06 | -1.39E+00 | 2.99E-03 | 1.14E-02 | 1.05E+00 | 3.65E-02 | 1.30E-01 | 1.10E+00 | 3.42E-01 | 9.65E-01 | 1.49E+00 | 3.07E-08 | 1.13E-07 | -1.47E+00 | 1.45E-03 | 3.99E-03 | -1.08E+00 |
| DIAPH2 | 1.20E-03 | 6.27E-03 | -1.04E+00 | 6.53E-04 | 2.82E-03 | 1.02E+00 | 7.86E-02 | 3.34E-01 | 1.07E+00 | 7.06E-03 | 2.70E-02 | 1.06E+00 | 7.22E-02 | 2.58E-01 | 1.14E+00 | 4.55E-03 | 1.28E-02 | -1.26E+00 | 6.05E-02 | 2.22E-01 | -1.09E+00 | 1.17E-01 | 3.20E-01 | -1.03E+00 |
| ZNF319 | 1.20E-03 | 6.28E-03 | -1.53E+00 | 4.87E-03 | 2.10E-02 | -1.22E+00 | 1.44E-03 | 6.14E-03 | -1.42E+00 | 3.06E-02 | 1.17E-01 | 1.01E+00 | 1.31E-01 | 4.69E-01 | -1.06E+00 | 8.37E-02 | 2.36E-01 | -1.19E+00 | 2.69E-02 | 9.89E-02 | -1.32E+00 | 5.02E-02 | 1.38E-01 | -1.27E+00 |
| TOMM5 | 1.20E-03 | 6.28E-03 | -1.13E+00 | 3.13E-04 | 1.35E-03 | -1.15E+00 | 1.84E-02 | 7.81E-02 | -1.03E+00 | 3.03E-05 | 1.16E-04 | -1.20E+00 | 5.46E-04 | 1.95E-03 | -1.34E+00 | 4.90E-04 | 1.38E-03 | -1.34E+00 | 6.64E-02 | 2.44E-01 | 1.12E+00 | 4.62E-02 | 1.27E-01 | 1.04E+00 |
| MFSD4B | 1.21E-03 | 6.30E-03 | -1.20E+00 | 4.71E-02 | 2.04E-01 | 1.21E+00 | 1.39E-04 | 5.92E-04 | -1.37E+00 | 7.27E-04 | 2.78E-03 | -1.15E+00 | 7.27E-02 | 2.59E-01 | 1.01E+00 | 1.48E-01 | 4.17E-01 | 1.18E+00 | 7.37E-04 | 2.70E-03 | -1.10E+00 | 1.24E-01 | 3.41E-01 | 1.17E+00 |
| PACSIN3 | 1.21E-03 | 6.30E-03 | -1.38E+00 | 1.03E-02 | 4.44E-02 | 1.02E+00 | 5.93E-03 | 2.52E-02 | -1.13E+00 | 9.88E-02 | 3.77E-01 | 1.36E+00 | 2.24E-01 | 8.00E-01 | 1.32E+00 | 3.18E-01 | 8.97E-01 | 1.27E+00 | 8.15E-02 | 2.99E-01 | 1.58E+00 | 1.82E-01 | 5.00E-01 | 1.40E+00 |
| ERCC3 | 1.21E-03 | 6.32E-03 | -1.15E+00 | 1.02E-03 | 4.42E-03 | -1.08E+00 | 2.61E-02 | 1.11E-01 | -1.05E+00 | 1.54E-02 | 5.89E-02 | 1.08E+00 | 3.74E-04 | 1.33E-03 | -1.47E+00 | 3.35E-02 | 9.45E-02 | 1.05E+00 | 6.06E-09 | 2.23E-08 | -1.53E+00 | 5.33E-03 | 1.46E-02 | -1.02E+00 |
| ZNF619 | 1.22E-03 | 6.34E-03 | -1.40E+00 | 3.38E-02 | 1.46E-01 | 1.04E+00 | 7.19E-03 | 3.05E-02 | -1.18E+00 | 1.45E-02 | 5.55E-02 | -1.00E+00 | 8.94E-05 | 3.19E-04 | -2.02E+00 | 1.08E-03 | 3.05E-03 | -1.59E+00 | 2.39E-01 | 8.77E-01 | 1.19E+00 | 2.81E-01 | 7.71E-01 | 1.35E+00 |
| FXR2 | 1.22E-03 | 6.35E-03 | -1.46E+00 | 8.94E-02 | 3.86E-01 | 1.17E+00 | 1.44E-08 | 6.13E-08 | -1.82E+00 | 2.61E-05 | 9.98E-05 | -1.29E+00 | 4.06E-10 | 1.45E-09 | -2.34E+00 | 7.27E-06 | 2.05E-05 | -1.32E+00 | 9.13E-08 | 3.35E-07 | -1.69E+00 | 7.86E-05 | 2.16E-04 | -1.32E+00 |
| ING2 | 1.22E-03 | 6.37E-03 | 1.07E+00 | 4.68E-06 | 2.02E-05 | -1.08E+00 | 7.22E-03 | 3.07E-02 | 1.04E+00 | 1.54E-05 | 5.88E-05 | -1.13E+00 | 1.53E-01 | 5.45E-01 | 1.40E+00 | 2.06E-02 | 5.82E-02 | 1.15E+00 | 1.46E-01 | 5.36E-01 | 1.17E+00 | 6.83E-02 | 1.88E-01 | 1.03E+00 |
| DCLRE1A | 1.22E-03 | 6.37E-03 | -2.08E+00 | 5.58E-03 | 2.41E-02 | -1.51E+00 | 2.68E-02 | 1.14E-01 | -1.31E+00 | 2.50E-02 | 9.57E-02 | -1.31E+00 | 1.55E-01 | 5.53E-01 | 1.06E+00 | 2.90E-01 | 8.17E-01 | 1.24E+00 | 1.89E-01 | 6.95E-01 | 2.13E+00 | 2.47E-01 | 6.78E-01 | 1.52E+00 |
| PDK2 | 1.23E-03 | 6.39E-03 | -1.19E+00 | 1.13E-03 | 4.90E-03 | -1.16E+00 | 3.77E-04 | 1.60E-03 | -1.51E+00 | 3.54E-03 | 1.35E-02 | -1.13E+00 | 6.66E-02 | 2.38E-01 | -1.15E+00 | 1.24E-01 | 3.51E-01 | -1.01E+00 | 4.82E-03 | 1.77E-02 | -1.23E+00 | 3.33E-02 | 9.14E-02 | -1.07E+00 |
| ACADSB | 1.23E-03 | 6.40E-03 | -1.25E+00 | 1.14E-02 | 4.93E-02 | 1.08E+00 | 8.39E-02 | 3.56E-01 | 1.12E+00 | 4.92E-02 | 1.88E-01 | 1.12E+00 | 6.43E-02 | 2.29E-01 | 1.06E+00 | 1.09E-01 | 3.07E-01 | 1.19E+00 | 1.03E-01 | 3.79E-01 | 1.41E+00 | 1.59E-01 | 4.36E-01 | 1.42E+00 |
| SLC16A10 | 1.23E-03 | 6.43E-03 | -1.75E+00 | 2.66E-03 | 1.15E-02 | -1.50E+00 | 1.11E-01 | 4.72E-01 | 1.33E+00 | 9.62E-02 | 3.68E-01 | 1.62E+00 | 2.43E-03 | 8.66E-03 | -1.47E+00 | 4.55E-03 | 1.28E-02 | -1.58E+00 | 7.81E-13 | 2.87E-12 | -2.07E+00 | 6.83E-07 | 1.88E-06 | -1.40E+00 |
| MUTYH | 1.23E-03 | 6.43E-03 | -1.13E+00 | 3.90E-03 | 1.69E-02 | 1.04E+00 | 3.00E-03 | 1.27E-02 | -1.15E+00 | 1.47E-04 | 5.62E-04 | -1.15E+00 | 1.02E-01 | 3.63E-01 | 1.23E+00 | 2.18E-01 | 6.14E-01 | 1.44E+00 | 3.07E-03 | 1.13E-02 | -1.01E+00 | 1.85E-01 | 5.09E-01 | 1.22E+00 |
| RALGAPA1 | 1.23E-03 | 6.43E-03 | -1.09E+00 | 7.42E-03 | 3.21E-02 | 1.14E+00 | 1.88E-03 | 7.98E-03 | -1.19E+00 | 1.10E-02 | 4.19E-02 | 1.10E+00 | 1.07E-01 | 3.80E-01 | 1.31E+00 | 1.03E-01 | 2.90E-01 | 1.19E+00 | 2.76E-03 | 1.01E-02 | -1.13E+00 | 3.31E-03 | 9.08E-03 | -1.09E+00 |
| AC091132.5 | 1.23E-03 | 6.43E-03 | -1.28E+00 | 1.11E-03 | 4.78E-03 | -1.11E+00 | 2.01E-01 | 8.53E-01 | 1.19E+00 | 4.19E-02 | 1.60E-01 | 1.06E+00 | 7.73E-02 | 2.76E-01 | -1.07E+00 | 1.01E-01 | 2.84E-01 | 1.01E+00 | 1.93E-02 | 7.07E-02 | -1.38E+00 | 2.64E-02 | 7.24E-02 | -1.50E+00 |
| HSD17B1 | 1.23E-03 | 6.43E-03 | -1.28E+00 | 1.19E-04 | 5.15E-04 | -1.48E+00 | 1.20E-01 | 5.10E-01 | 1.07E+00 | 2.22E-01 | 8.49E-01 | 1.25E+00 | 9.92E-02 | 3.54E-01 | -1.08E+00 | 2.27E-01 | 6.39E-01 | 1.18E+00 | 4.03E-02 | 1.48E-01 | -1.25E+00 | 3.08E-01 | 8.46E-01 | 1.19E+00 |
| NLE1 | 1.23E-03 | 6.43E-03 | -1.08E+00 | 1.62E-04 | 6.99E-04 | -1.06E+00 | 2.40E-02 | 1.02E-01 | 1.04E+00 | 1.21E-02 | 4.61E-02 | 1.10E+00 | 2.19E-02 | 7.80E-02 | -1.10E+00 | 2.27E-01 | 6.41E-01 | 1.23E+00 | 6.99E-07 | 2.56E-06 | -1.33E+00 | 9.75E-04 | 2.68E-03 | -1.14E+00 |
| ARFIP2 | 1.23E-03 | 6.43E-03 | -1.05E+00 | 8.26E-03 | 3.57E-02 | 1.16E+00 | 1.53E-02 | 6.49E-02 | 1.01E+00 | 4.11E-03 | 1.57E-02 | 1.09E+00 | 1.60E-02 | 5.71E-02 | 1.02E+00 | 5.08E-02 | 1.43E-01 | 1.14E+00 | 1.32E-02 | 4.83E-02 | 1.00E+00 | 1.16E-01 | 3.19E-01 | 1.18E+00 |
| PIGP | 1.23E-03 | 6.44E-03 | 1.08E+00 | 1.27E-04 | 5.48E-04 | 1.14E+00 | 2.47E-05 | 1.05E-04 | -1.27E+00 | 3.03E-07 | 1.16E-06 | -1.18E+00 | 1.26E-05 | 4.48E-05 | -1.23E+00 | 9.53E-04 | 2.69E-03 | -1.02E+00 | 2.79E-06 | 1.02E-05 | -1.20E+00 | 1.51E-06 | 4.14E-06 | -1.25E+00 |
| LINC00663 | 1.24E-03 | 6.44E-03 | -1.64E+00 | 3.14E-02 | 1.36E-01 | -1.18E+00 | 1.49E-02 | 6.34E-02 | -1.25E+00 | 3.89E-02 | 1.49E-01 | -1.07E+00 | 5.13E-02 | 1.83E-01 | -1.12E+00 | 1.18E-01 | 3.33E-01 | -1.05E+00 | 4.25E-02 | 1.56E-01 | -1.46E+00 | 2.54E-01 | 6.99E-01 | 1.06E+00 |
| ZFPL1 | 1.24E-03 | 6.45E-03 | -1.14E+00 | 1.50E-05 | 6.50E-05 | -1.34E+00 | 2.34E-01 | 9.95E-01 | 1.10E+00 | 7.40E-02 | 2.83E-01 | -1.02E+00 | 4.06E-02 | 1.45E-01 | -1.10E+00 | 3.93E-02 | 1.11E-01 | -1.05E+00 | 4.79E-02 | 1.76E-01 | -1.18E+00 | 1.02E-01 | 2.79E-01 | -1.04E+00 |
| KLC1 | 1.24E-03 | 6.45E-03 | -1.04E+00 | 4.99E-02 | 2.15E-01 | 1.29E+00 | 5.42E-04 | 2.30E-03 | -1.17E+00 | 3.80E-03 | 1.45E-02 | 1.05E+00 | 7.36E-04 | 2.62E-03 | -1.08E+00 | 1.34E-02 | 3.77E-02 | 1.06E+00 | 2.22E-03 | 8.16E-03 | -1.05E+00 | 6.63E-03 | 1.82E-02 | 1.03E+00 |
| AADAT | 1.24E-03 | 6.45E-03 | -1.42E+00 | 3.34E-02 | 1.44E-01 | 1.04E+00 | 1.91E-04 | 8.10E-04 | -1.60E+00 | 3.54E-05 | 1.35E-04 | -1.65E+00 | 8.22E-02 | 2.93E-01 | -1.49E+00 | 7.02E-02 | 1.98E-01 | -1.62E+00 | 2.54E-02 | 9.34E-02 | -1.53E+00 | 1.86E-01 | 5.12E-01 | -1.05E+00 |
| RNF215 | 1.24E-03 | 6.46E-03 | -1.14E+00 | 2.61E-03 | 1.13E-02 | 1.02E+00 | 7.44E-02 | 3.16E-01 | -1.07E+00 | 1.44E-01 | 5.49E-01 | 1.18E+00 | 5.51E-03 | 1.96E-02 | -1.44E+00 | 9.62E-02 | 2.71E-01 | 1.08E+00 | 3.15E-02 | 1.16E-01 | -1.24E+00 | 2.62E-02 | 7.20E-02 | -1.27E+00 |
| IL1RAP | 1.24E-03 | 6.47E-03 | 1.83E+00 | 1.17E-01 | 5.05E-01 | 1.20E+00 | 2.85E-07 | 1.21E-06 | 1.81E+00 | 2.25E-01 | 8.58E-01 | 1.50E+00 | 4.65E-03 | 1.66E-02 | 3.12E+00 | 2.66E-01 | 7.51E-01 | 1.29E+00 | 1.21E-02 | 4.45E-02 | 1.71E+00 | 2.72E-01 | 7.46E-01 | 1.08E+00 |
| CHRNB1 | 1.24E-03 | 6.47E-03 | -1.09E+00 | 5.81E-03 | 2.51E-02 | 1.07E+00 | 1.11E-04 | 4.70E-04 | -1.16E+00 | 1.22E-05 | 4.65E-05 | -1.10E+00 | 1.41E-06 | 5.04E-06 | -1.65E+00 | 4.02E-06 | 1.13E-05 | -1.46E+00 | 1.28E-02 | 4.72E-02 | -1.06E+00 | 8.51E-02 | 2.34E-01 | 1.12E+00 |
| LINC01970 | 1.24E-03 | 6.47E-03 | 2.86E+00 | 9.48E-02 | 4.10E-01 | 1.69E+00 | 1.37E-01 | 5.83E-01 | 1.27E+00 | 1.18E-01 | 4.52E-01 | -1.12E+00 | 2.03E-01 | 7.24E-01 | 1.43E+00 | 2.89E-01 | 8.15E-01 | 1.35E+00 | 1.63E-02 | 5.98E-02 | 1.83E+00 | 2.00E-01 | 5.49E-01 | 1.31E+00 |
| TANC2 | 1.25E-03 | 6.49E-03 | -1.06E+00 | 1.26E-02 | 5.46E-02 | 1.18E+00 | 8.69E-02 | 3.69E-01 | 1.03E+00 | 3.85E-02 | 1.47E-01 | 1.05E+00 | 3.99E-02 | 1.42E-01 | 1.02E+00 | 3.16E-01 | 8.89E-01 | 1.44E+00 | 1.06E-04 | 3.88E-04 | -1.36E+00 | 3.20E-06 | 8.79E-06 | -1.49E+00 |
| RNVU1-7 | 1.25E-03 | 6.49E-03 | 2.20E+00 | 2.17E-03 | 9.37E-03 | -1.99E+00 | 1.29E-01 | 5.47E-01 | 1.37E+00 | 1.47E-02 | 5.61E-02 | -1.51E+00 | 1.01E-01 | 3.61E-01 | 1.01E+00 | 1.26E-02 | 3.57E-02 | -1.91E+00 | 2.45E-01 | 8.98E-01 | 1.25E+00 | 3.43E-01 | 9.43E-01 | 1.17E+00 |
| TMEM176B | 1.25E-03 | 6.50E-03 | -1.86E+00 | 1.10E-02 | 4.76E-02 | -1.06E+00 | 1.84E-07 | 7.82E-07 | -2.33E+00 | 1.93E-06 | 7.38E-06 | -2.34E+00 | 2.02E-02 | 7.19E-02 | -2.17E+00 | 6.53E-02 | 1.84E-01 | -1.65E+00 | 9.82E-02 | 3.61E-01 | -1.92E+00 | 1.76E-01 | 4.85E-01 | -1.50E+00 |
| TBKBP1 | 1.25E-03 | 6.50E-03 | -1.37E+00 | 2.71E-03 | 1.17E-02 | -1.23E+00 | 1.79E-01 | 7.62E-01 | 1.14E+00 | 2.40E-02 | 9.17E-02 | -1.14E+00 | 1.33E-01 | 4.75E-01 | -1.06E+00 | 2.79E-02 | 7.85E-02 | -1.70E+00 | 1.44E-01 | 5.29E-01 | 1.51E+00 | 3.35E-01 | 9.21E-01 | 1.25E+00 |

| | | | | | | | | | | | | | | | | | | | | | | | | |
|---|---|---|---|---|---|---|---|---|---|---|---|---|---|---|---|---|---|---|---|---|---|---|---|---|
| AL353662.2 | 1.25E-03 | 6.52E-03 | 1.74E+00 | 2.09E-01 | 9.03E-01 | 1.54E+00 | 9.31E-04 | 3.95E-03 | 1.65E+00 | 3.03E-02 | 1.16E-01 | 1.62E+00 | 1.35E-01 | 4.82E-01 | 1.49E+00 | 2.22E-01 | 6.27E-01 | 1.50E+00 | 2.92E-02 | 1.07E-01 | 1.53E+00 | 3.52E-01 | 9.67E-01 | 1.27E+00 |
| NPIPB6 | 1.25E-03 | 6.53E-03 | -1.30E+00 | 2.78E-03 | 1.20E-02 | -1.15E+00 | 1.11E-02 | 4.73E-02 | -1.25E+00 | 1.03E-02 | 3.93E-02 | -1.19E+00 | 4.81E-02 | 1.72E-01 | -1.32E+00 | 1.51E-01 | 4.25E-01 | 1.01E+00 | 3.95E-02 | 1.45E-01 | -1.26E+00 | 2.49E-01 | 6.85E-01 | 1.13E+00 |
| AP001486.2 | 1.25E-03 | 6.54E-03 | -1.36E+00 | 7.35E-02 | 3.17E-01 | 1.21E+00 | 6.81E-02 | 2.89E-01 | -1.04E+00 | 2.36E-02 | 9.01E-02 | 1.02E+00 | 2.44E-01 | 8.71E-01 | 1.36E+00 | 9.68E-02 | 2.73E-01 | 1.81E+00 | 1.61E-01 | 5.90E-01 | 1.09E+00 | 3.33E-01 | 9.14E-01 | 1.21E+00 |
| AC026316.1 | 1.25E-03 | 6.54E-03 | 2.61E+00 | 1.34E-01 | 5.78E-01 | 1.78E+00 | 4.21E-02 | 1.79E-01 | 1.63E+00 | 1.28E-01 | 4.88E-01 | 1.20E+00 | 2.52E-02 | 8.98E-02 | 2.66E+00 | 1.10E-01 | 3.11E-01 | 2.32E+00 | 4.38E-03 | 1.61E-02 | 2.18E+00 | 2.90E-01 | 7.97E-01 | 1.34E+00 |
| MT1X | 1.26E-03 | 6.57E-03 | -1.02E+00 | 5.93E-06 | 2.56E-05 | -1.27E+00 | 3.52E-04 | 1.49E-03 | -1.07E+00 | 2.16E-09 | 8.26E-09 | -1.46E+00 | 7.10E-02 | 2.53E-01 | 1.39E+00 | 5.85E-05 | 1.65E-04 | 1.18E+00 | 2.12E-01 | 7.77E-01 | 1.51E+00 | 4.41E-04 | 1.21E-03 | -1.04E+00 |
| TTC30B | 1.27E-03 | 6.60E-03 | -1.21E+00 | 1.63E-04 | 7.04E-04 | -1.42E+00 | 2.55E-03 | 1.08E-02 | -1.27E+00 | 3.98E-03 | 1.52E-02 | -1.16E+00 | 1.12E-01 | 3.99E-01 | 1.10E+00 | 1.91E-01 | 5.38E-01 | 1.22E+00 | 1.09E-01 | 4.01E-01 | 1.08E+00 | 1.13E-01 | 3.10E-01 | -1.09E+00 |
| TSNAX | 1.27E-03 | 6.60E-03 | 1.05E+00 | 8.53E-05 | 3.69E-04 | 1.06E+00 | 6.57E-03 | 2.79E-02 | -1.01E+00 | 7.14E-03 | 2.73E-02 | 1.10E+00 | 3.93E-05 | 1.40E-04 | -1.01E+00 | 7.72E-04 | 2.18E-03 | 1.11E+00 | 1.23E-01 | 4.51E-01 | 1.12E+00 | 7.44E-02 | 2.04E-01 | 1.08E+00 |
| AC002553.1 | 1.27E-03 | 6.61E-03 | -1.11E+00 | 1.29E-04 | 5.56E-04 | -1.14E+00 | 2.60E-03 | 1.10E-02 | -1.18E+00 | 9.33E-04 | 3.56E-03 | -1.09E+00 | 8.32E-04 | 2.97E-03 | -1.20E+00 | 5.97E-03 | 1.68E-02 | 1.02E+00 | 1.48E-01 | 5.44E-01 | 1.15E+00 | 3.86E-02 | 1.06E-01 | -1.04E+00 |
| CES2 | 1.28E-03 | 6.68E-03 | -1.21E+00 | 9.88E-03 | 4.27E-02 | 1.09E+00 | 1.11E-03 | 4.72E-03 | -1.31E+00 | 1.35E-02 | 5.17E-02 | -1.02E+00 | 6.61E-02 | 2.36E-01 | 1.14E+00 | 2.76E-02 | 7.78E-02 | 1.07E+00 | 2.47E-01 | 9.05E-01 | 1.26E+00 | 2.21E-01 | 6.07E-01 | 1.47E+00 |
| KCTD12 | 1.28E-03 | 6.69E-03 | -1.71E+00 | 1.16E-01 | 5.03E-01 | 1.26E+00 | 6.04E-03 | 2.57E-02 | -1.72E+00 | 1.14E-01 | 4.35E-01 | -1.03E+00 | 2.65E-01 | 9.46E-01 | 1.06E+00 | 3.48E-01 | 9.81E-01 | 1.21E+00 | 9.49E-03 | 3.48E-02 | -2.09E+00 | 1.07E-01 | 2.93E-01 | -1.48E+00 |
| IMMP2L | 1.28E-03 | 6.69E-03 | 1.06E+00 | 1.32E-02 | 5.71E-02 | 1.28E+00 | 1.88E-01 | 7.98E-01 | 1.21E+00 | 2.49E-01 | 9.50E-01 | 1.41E+00 | 3.36E-02 | 1.20E-01 | 1.17E+00 | 7.86E-02 | 2.22E-01 | 1.26E+00 | 2.29E-01 | 8.39E-01 | 1.19E+00 | 1.20E-01 | 3.31E-01 | 1.39E+00 |
| MTATP6P1 | 1.29E-03 | 6.71E-03 | 1.10E+00 | 2.06E-01 | 8.89E-01 | 1.01E+00 | 2.85E-04 | 1.21E-03 | 1.14E+00 | 5.38E-02 | 2.05E-01 | 1.08E+00 | 2.89E-04 | 1.03E-03 | 1.16E+00 | 9.02E-04 | 2.54E-03 | 1.15E+00 | 1.68E-03 | 6.16E-03 | 1.15E+00 | 8.28E-05 | 2.27E-04 | 1.16E+00 |
| MECOM | 1.29E-03 | 6.71E-03 | -1.04E+00 | 9.90E-03 | 4.28E-02 | 1.25E+00 | 6.84E-02 | 2.91E-01 | 1.04E+00 | 1.92E-02 | 7.33E-02 | 1.07E+00 | 2.05E-01 | 7.30E-01 | -1.00E+00 | 2.30E-01 | 6.49E-01 | 1.04E+00 | 7.18E-05 | 2.64E-04 | -1.42E+00 | 1.76E-03 | 4.85E-03 | -1.13E+00 |
| RAB33B | 1.29E-03 | 6.71E-03 | -1.01E+00 | 7.58E-12 | 3.27E-11 | -1.46E+00 | 4.70E-04 | 2.00E-03 | -1.42E+00 | 6.63E-07 | 2.53E-06 | -1.58E+00 | 8.50E-02 | 3.03E-01 | 1.13E+00 | 4.14E-03 | 1.17E-02 | -1.18E+00 | 1.76E-04 | 6.44E-04 | -1.48E+00 | 2.09E-04 | 5.75E-04 | -1.55E+00 |
| ATG4B | 1.29E-03 | 6.72E-03 | -1.16E+00 | 4.30E-02 | 1.86E-01 | 1.16E+00 | 7.15E-03 | 3.04E-02 | -1.01E+00 | 1.23E-04 | 4.71E-04 | -1.01E+00 | 3.74E-02 | 1.33E-01 | 1.08E+00 | 9.00E-02 | 2.54E-01 | 1.18E+00 | 9.78E-03 | 3.59E-02 | -1.01E+00 | 5.82E-03 | 1.60E-02 | -1.07E+00 |
| HERC2P10 | 1.29E-03 | 6.73E-03 | -1.28E+00 | 4.89E-04 | 2.11E-03 | -1.40E+00 | 2.06E-01 | 8.76E-01 | 1.37E+00 | 1.89E-01 | 7.21E-01 | 1.28E+00 | 1.26E-01 | 4.49E-01 | 1.03E+00 | 3.21E-02 | 9.05E-02 | -1.29E+00 | 6.39E-03 | 2.34E-02 | -1.07E+00 | 3.02E-03 | 8.29E-03 | -1.26E+00 |
| SRSF9 | 1.29E-03 | 6.74E-03 | -1.03E+00 | 2.72E-06 | 1.17E-05 | -1.17E+00 | 9.66E-02 | 4.10E-01 | 1.13E+00 | 5.45E-04 | 2.08E-03 | -1.03E+00 | 1.70E-04 | 6.06E-04 | -1.04E+00 | 3.14E-06 | 8.85E-06 | -1.19E+00 | 1.51E-03 | 5.56E-03 | -1.02E+00 | 1.42E-03 | 3.89E-03 | -1.01E+00 |
| FCER1A | 1.29E-03 | 6.74E-03 | -1.26E+00 | 1.11E-03 | 4.79E-03 | -1.31E+00 | 1.40E-01 | 5.94E-01 | 1.72E+00 | 1.53E-01 | 5.83E-01 | 2.98E+00 | 5.04E-02 | 1.80E-01 | 1.04E+00 | 9.47E-02 | 2.67E-01 | 1.17E+00 | 2.11E-01 | 7.73E-01 | 1.31E+00 | 3.23E-01 | 8.86E-01 | 1.24E+00 |
| AC068888.1 | 1.29E-03 | 6.75E-03 | -1.47E+00 | 2.51E-02 | 1.08E-01 | 1.04E+00 | 3.88E-04 | 1.65E-03 | -1.54E+00 | 1.01E-02 | 3.85E-02 | -1.09E+00 | 4.27E-03 | 1.52E-02 | -1.68E+00 | 2.09E-01 | 5.90E-01 | 1.19E+00 | 1.23E-04 | 4.50E-04 | -2.12E+00 | 3.42E-01 | 9.40E-01 | 1.31E+00 |
| FO393411.1 | 1.30E-03 | 6.77E-03 | -1.03E+00 | 2.10E-02 | 9.09E-02 | 1.05E+00 | 2.18E-01 | 9.28E-01 | 1.00E+00 | 2.53E-01 | 9.68E-01 | 1.06E+00 | 3.98E-03 | 1.42E-02 | -1.08E+00 | 1.13E-01 | 3.20E-01 | 1.03E+00 | 2.65E-05 | 9.72E-05 | -1.18E+00 | 3.25E-01 | 8.92E-01 | 1.03E+00 |
| HMGB1P51 | 1.30E-03 | 6.78E-03 | -1.13E+00 | 3.48E-04 | 1.50E-03 | -1.09E+00 | 1.14E-02 | 4.82E-02 | -1.32E+00 | 1.50E-02 | 5.73E-02 | -1.09E+00 | 1.17E-02 | 4.17E-02 | -1.44E+00 | 9.63E-02 | 2.72E-01 | 1.00E+00 | 1.29E-01 | 4.74E-01 | -1.01E+00 | 2.56E-01 | 7.04E-01 | 1.31E+00 |
| ITSN1 | 1.30E-03 | 6.79E-03 | -1.26E+00 | 3.41E-03 | 1.47E-02 | -1.06E+00 | 3.54E-03 | 1.51E-02 | -1.26E+00 | 3.70E-03 | 1.41E-02 | -1.18E+00 | 1.92E-01 | 6.83E-01 | 1.64E+00 | 3.35E-01 | 9.44E-01 | 1.17E+00 | 6.18E-03 | 2.27E-02 | -1.05E+00 | 1.59E-03 | 4.36E-03 | -1.41E+00 |
| SMPD2 | 1.31E-03 | 6.82E-03 | -1.01E+00 | 2.05E-04 | 8.85E-04 | -1.04E+00 | 2.34E-06 | 9.95E-06 | -1.34E+00 | 3.26E-06 | 1.25E-05 | -1.18E+00 | 5.09E-02 | 1.81E-01 | 1.05E+00 | 1.32E-01 | 3.71E-01 | 1.22E+00 | 1.06E-01 | 3.87E-01 | 1.07E+00 | 3.18E-01 | 8.73E-01 | 1.15E+00 |
| RAP2C-AS1 | 1.31E-03 | 6.82E-03 | -1.16E+00 | 7.11E-04 | 3.07E-03 | -1.20E+00 | 4.14E-02 | 1.76E-01 | -1.11E+00 | 3.85E-02 | 1.47E-01 | -1.03E+00 | 1.80E-01 | 6.42E-01 | 1.01E+00 | 3.44E-01 | 9.71E-01 | 1.36E+00 | 1.72E-01 | 6.30E-01 | 1.37E+00 | 3.43E-01 | 9.44E-01 | 1.23E+00 |
| ZNF44 | 1.31E-03 | 6.83E-03 | 1.07E+00 | 4.38E-09 | 1.89E-08 | -1.23E+00 | 4.04E-02 | 1.71E-01 | 1.01E+00 | 2.69E-04 | 1.03E-03 | -1.25E+00 | 1.11E-03 | 3.95E-03 | -1.13E+00 | 6.18E-04 | 1.74E-03 | -1.20E+00 | 6.73E-03 | 2.47E-02 | -1.34E+00 | 6.66E-03 | 1.83E-02 | -1.32E+00 |
| CRELD2 | 1.31E-03 | 6.84E-03 | -1.10E+00 | 5.45E-03 | 2.35E-02 | 1.09E+00 | 1.17E-01 | 4.99E-01 | 1.13E+00 | 2.91E-02 | 1.11E-01 | 1.13E+00 | 4.11E-08 | 1.47E-07 | -2.15E+00 | 2.51E-06 | 7.07E-06 | -1.67E+00 | 2.40E-02 | 8.79E-02 | -1.13E+00 | 3.42E-01 | 9.40E-01 | 1.15E+00 |
| AC055713.1 | 1.32E-03 | 6.87E-03 | -1.44E+00 | 1.13E-02 | 4.89E-02 | 1.01E+00 | 1.39E-01 | 5.90E-01 | 1.09E+00 | 2.52E-02 | 9.61E-02 | -1.05E+00 | 9.96E-02 | 3.55E-01 | -1.03E+00 | 3.51E-01 | 9.90E-01 | 1.48E+00 | 1.19E-02 | 4.38E-02 | -1.15E+00 | 1.31E-03 | 3.59E-03 | -1.45E+00 |
| CDT1 | 1.32E-03 | 6.88E-03 | -1.16E+00 | 3.09E-06 | 1.33E-05 | -1.33E+00 | 2.13E-01 | 9.07E-01 | 1.14E+00 | 2.54E-04 | 9.68E-04 | 1.05E+00 | 1.21E-04 | 4.32E-04 | -1.77E+00 | 5.97E-04 | 1.68E-03 | -1.39E+00 | 1.52E-04 | 5.58E-04 | -1.20E+00 | 3.62E-02 | 9.94E-02 | 1.00E+00 |
| TMSB4XP1 | 1.32E-03 | 6.88E-03 | -1.13E+00 | 3.56E-02 | 1.54E-01 | 1.23E+00 | 6.10E-07 | 2.59E-06 | -1.74E+00 | 1.56E-02 | 5.96E-02 | 1.08E+00 | 9.64E-06 | 3.44E-05 | -2.06E+00 | 1.13E-04 | 3.19E-04 | -1.40E+00 | 1.85E-03 | 6.78E-03 | -2.45E+00 | 5.33E-02 | 1.46E-01 | -1.86E+00 |
| GMIP | 1.32E-03 | 6.90E-03 | -1.23E+00 | 9.16E-03 | 3.96E-02 | 1.03E+00 | 2.06E-03 | 8.77E-03 | -1.24E+00 | 1.57E-03 | 6.01E-03 | -1.14E+00 | 6.14E-03 | 2.19E-02 | -1.25E+00 | 8.58E-02 | 2.42E-01 | 1.09E+00 | 3.31E-02 | 1.21E-01 | -1.03E+00 | 7.93E-02 | 2.18E-01 | 1.03E+00 |
| AGAP9 | 1.32E-03 | 6.90E-03 | -1.62E+00 | 6.24E-03 | 2.70E-02 | -1.25E+00 | 8.72E-02 | 3.71E-01 | 1.07E+00 | 1.39E-01 | 5.29E-01 | 1.19E+00 | 1.14E-01 | 4.07E-01 | 1.05E+00 | 2.07E-01 | 5.84E-01 | 1.04E+00 | 2.55E-01 | 9.36E-01 | 1.25E+00 | 2.36E-01 | 6.47E-01 | 1.38E+00 |
| LINC01128 | 1.33E-03 | 6.92E-03 | -1.59E+00 | 4.73E-02 | 2.05E-01 | 1.06E+00 | 5.71E-03 | 2.43E-02 | -1.45E+00 | 8.06E-02 | 3.08E-01 | 1.05E+00 | 3.58E-03 | 1.28E-02 | -1.49E+00 | 3.18E-02 | 8.96E-02 | -1.18E+00 | 3.88E-04 | 1.43E-03 | -1.40E+00 | 9.93E-03 | 2.73E-02 | -1.22E+00 |
| ZNF585B | 1.33E-03 | 6.94E-03 | -1.39E+00 | 3.39E-02 | 1.46E-01 | 1.00E+00 | 1.05E-07 | 4.48E-07 | -2.22E+00 | 9.78E-05 | 3.73E-04 | -1.37E+00 | 1.26E-02 | 4.49E-02 | -1.31E+00 | 4.98E-02 | 1.40E-01 | -1.05E+00 | 1.01E-01 | 3.72E-01 | 1.21E+00 | 8.79E-02 | 2.41E-01 | 1.00E+00 |
| ZRSR2 | 1.33E-03 | 6.94E-03 | -1.15E+00 | 1.87E-02 | 8.06E-02 | 1.11E+00 | 4.29E-04 | 1.82E-03 | -1.16E+00 | 2.52E-02 | 9.63E-02 | 1.24E+00 | 6.80E-03 | 2.43E-02 | -1.15E+00 | 9.20E-02 | 2.59E-01 | 1.10E+00 | 1.15E-02 | 4.21E-02 | -1.11E+00 | 9.53E-04 | 2.62E-03 | -1.26E+00 |
| SYT5 | 1.34E-03 | 6.96E-03 | 2.62E+00 | 2.07E-01 | 8.97E-01 | 1.45E+00 | 2.81E-02 | 1.19E-01 | 1.56E+00 | 1.26E-01 | 4.81E-01 | -1.23E+00 | 1.53E-05 | 5.44E-05 | 5.28E+00 | 3.20E-02 | 9.02E-02 | 2.79E+00 | 3.41E-04 | 1.25E-03 | 2.67E+00 | 3.11E-01 | 8.55E-01 | 1.21E+00 |
| RTN4IP1 | 1.34E-03 | 6.97E-03 | -1.54E+00 | 3.85E-03 | 1.67E-02 | -1.29E+00 | 8.31E-05 | 3.53E-04 | -1.54E+00 | 3.28E-04 | 1.25E-03 | -1.30E+00 | 9.91E-02 | 3.53E-01 | -1.15E+00 | 1.46E-01 | 4.13E-01 | -1.02E+00 | 9.15E-07 | 3.36E-06 | -1.57E+00 | 1.61E-04 | 4.41E-04 | -1.31E+00 |
| CENPL | 1.34E-03 | 6.99E-03 | -1.43E+00 | 7.75E-03 | 3.35E-02 | -1.08E+00 | 1.54E-02 | 6.55E-02 | -1.01E+00 | 1.09E-02 | 4.18E-02 | 1.06E+00 | 4.96E-02 | 1.77E-01 | -1.02E+00 | 1.45E-01 | 4.08E-01 | 1.14E+00 | 7.80E-03 | 2.86E-02 | -1.07E+00 | 9.16E-02 | 2.52E-01 | 1.10E+00 |
| ZNF10 | 1.34E-03 | 6.99E-03 | -1.20E+00 | 1.33E-04 | 5.76E-04 | -1.32E+00 | 2.30E-04 | 9.77E-04 | -1.44E+00 | 5.49E-07 | 2.10E-06 | -1.80E+00 | 2.59E-03 | 9.25E-03 | -1.19E+00 | 4.17E-04 | 1.17E-03 | -1.59E+00 | 3.30E-02 | 1.21E-01 | -1.04E+00 | 3.04E-02 | 8.34E-02 | -1.17E+00 |
| SEMA4A | 1.34E-03 | 7.00E-03 | -1.22E+00 | 1.26E-03 | 5.47E-03 | -1.11E+00 | 1.57E-02 | 6.67E-02 | -1.08E+00 | 8.83E-03 | 3.37E-02 | 1.02E+00 | 2.65E-02 | 9.44E-02 | -1.06E+00 | 1.88E-02 | 5.31E-02 | -1.12E+00 | 9.30E-02 | 3.41E-01 | -1.11E+00 | 2.18E-01 | 5.99E-01 | 1.06E+00 |
| AC084033.3 | 1.34E-03 | 7.01E-03 | -1.15E+00 | 6.92E-12 | 2.99E-11 | -1.35E+00 | 2.73E-03 | 1.16E-02 | -1.18E+00 | 6.52E-06 | 2.49E-05 | -1.27E+00 | 8.20E-02 | 2.92E-01 | -1.06E+00 | 2.57E-02 | 7.24E-02 | -1.20E+00 | 3.04E-02 | 1.12E-01 | -1.01E+00 | 1.47E-02 | 4.03E-02 | -1.22E+00 |
| MVB12A | 1.35E-03 | 7.04E-03 | 1.01E+00 | 6.70E-03 | 2.90E-02 | 1.22E+00 | 1.67E-01 | 7.10E-01 | 1.16E+00 | 1.53E-01 | 5.86E-01 | 1.34E+00 | 1.47E-03 | 5.24E-03 | -1.07E+00 | 1.25E-01 | 3.51E-01 | 1.37E+00 | 9.18E-02 | 3.37E-01 | 1.18E+00 | 3.08E-01 | 8.45E-01 | 1.42E+00 |
| THAP8 | 1.35E-03 | 7.04E-03 | -1.41E+00 | 3.75E-02 | 1.62E-01 | 1.06E+00 | 6.35E-04 | 2.70E-03 | -1.66E+00 | 2.74E-02 | 1.05E-01 | -1.06E+00 | 2.13E-04 | 7.58E-04 | -2.63E+00 | 1.06E-01 | 2.99E-01 | 1.01E+00 | 5.33E-07 | 1.96E-06 | -1.93E+00 | 6.04E-04 | 1.66E-03 | -1.28E+00 |
| NEK8 | 1.35E-03 | 7.05E-03 | -1.48E+00 | 1.99E-03 | 8.59E-03 | -1.28E+00 | 4.19E-03 | 1.78E-02 | -1.37E+00 | 6.01E-03 | 2.29E-02 | -1.20E+00 | 1.61E-02 | 5.73E-02 | -1.06E+00 | 2.09E-03 | 5.89E-03 | -1.36E+00 | 9.74E-03 | 3.57E-02 | -1.47E+00 | 8.65E-02 | 2.37E-01 | -1.09E+00 |
| HYOU1 | 1.35E-03 | 7.05E-03 | -1.59E+00 | 4.39E-03 | 1.90E-02 | -1.29E+00 | 1.25E-01 | 5.33E-01 | 1.12E+00 | 3.52E-02 | 1.35E-01 | -1.01E+00 | 2.06E-06 | 7.36E-06 | -1.77E+00 | 1.45E-06 | 4.09E-06 | -1.83E+00 | 7.07E-03 | 2.60E-02 | -1.17E+00 | 6.54E-03 | 1.80E-02 | -1.21E+00 |
| P2RY1 | 1.36E-03 | 7.09E-03 | -1.73E+00 | 4.13E-03 | 1.78E-02 | -1.38E+00 | 1.59E-02 | 6.76E-02 | -1.34E+00 | 2.48E-03 | 9.45E-03 | -1.54E+00 | 5.08E-02 | 1.81E-01 | 1.13E+00 | 1.86E-01 | 5.23E-01 | 1.28E+00 | 3.86E-03 | 1.42E-02 | 1.35E+00 | 1.68E-01 | 4.61E-01 | 1.25E+00 |
| AL137802.2 | 1.36E-03 | 7.09E-03 | -1.15E+00 | 5.72E-02 | 2.47E-01 | 1.24E+00 | 3.23E-02 | 1.37E-01 | -1.24E+00 | 1.11E-01 | 4.26E-01 | 1.02E+00 | 1.55E-03 | 5.52E-03 | -1.34E+00 | 2.31E-02 | 6.50E-02 | 1.04E+00 | 8.93E-02 | 3.28E-01 | 1.44E+00 | 1.73E-01 | 4.75E-01 | 1.41E+00 |
| CEP250 | 1.36E-03 | 7.10E-03 | -1.17E+00 | 3.07E-02 | 1.33E-01 | 1.16E+00 | 8.67E-02 | 3.68E-01 | 1.03E+00 | 2.52E-01 | 9.64E-01 | 1.39E+00 | 6.42E-04 | 2.29E-03 | -1.36E+00 | 1.59E-02 | 4.49E-02 | -1.06E+00 | 1.52E-01 | 5.59E-01 | 1.14E+00 | 2.03E-01 | 5.56E-01 | 1.40E+00 |
| ZBTB39 | 1.36E-03 | 7.11E-03 | -1.33E+00 | 5.16E-03 | 2.23E-02 | -1.14E+00 | 4.36E-07 | 1.85E-06 | -1.77E+00 | 2.67E-09 | 1.02E-08 | -1.92E+00 | 1.77E-01 | 6.32E-01 | 1.43E+00 | 2.34E-01 | 6.61E-01 | -1.05E+00 | 2.69E-01 | 9.86E-01 | 1.38E+00 | 2.65E-01 | 7.29E-01 | 1.15E+00 |
| CEP76 | 1.36E-03 | 7.11E-03 | -1.53E+00 | 6.75E-03 | 2.92E-02 | -1.21E+00 | 1.34E-02 | 5.68E-02 | -1.20E+00 | 4.50E-02 | 1.72E-01 | 1.03E+00 | 2.91E-03 | 1.04E-02 | -1.44E+00 | 3.29E-04 | 9.26E-04 | -1.70E+00 | 2.67E-04 | 9.80E-04 | -1.28E+00 | 1.34E-02 | 3.67E-02 | -1.14E+00 |
| SLC10A7 | 1.37E-03 | 7.13E-03 | -1.48E+00 | 1.39E-02 | 6.00E-02 | -1.09E+00 | 7.17E-03 | 3.05E-02 | -1.20E+00 | 1.63E-02 | 6.21E-02 | 1.00E+00 | 2.00E-05 | 7.14E-05 | -2.02E+00 | 4.08E-04 | 1.15E-03 | -1.39E+00 | 6.81E-03 | 2.50E-02 | -1.20E+00 | 1.36E-02 | 3.74E-02 | -1.16E+00 |
| AACS | 1.37E-03 | 7.14E-03 | -1.36E+00 | 5.42E-02 | 2.34E-01 | 1.16E+00 | 1.42E-01 | 6.01E-01 | 1.09E+00 | 2.51E-01 | 9.58E-01 | 1.38E+00 | 1.29E-02 | 4.62E-02 | -1.08E+00 | 9.82E-02 | 2.77E-01 | 1.21E+00 | 1.38E-02 | 5.05E-02 | -1.02E+00 | 4.14E-02 | 1.14E-01 | 1.00E+00 |
| AC068491.3 | 1.37E-03 | 7.14E-03 | -1.63E+00 | 2.62E-03 | 1.13E-02 | -1.50E+00 | 1.07E-02 | 4.54E-02 | -1.45E+00 | 1.81E-01 | 6.90E-01 | 1.27E+00 | 4.70E-04 | 1.68E-03 | -1.24E+00 | 1.94E-01 | 5.48E-01 | 1.51E+00 | 1.97E-11 | 7.24E-11 | -1.65E+00 | 3.98E-05 | 1.09E-04 | -1.22E+00 |
| DMAC2 | 1.37E-03 | 7.16E-03 | 1.07E+00 | 1.04E-03 | 4.49E-03 | 1.16E+00 | 6.81E-03 | 2.89E-02 | -1.06E+00 | 1.12E-02 | 4.29E-02 | 1.13E+00 | 1.08E-01 | 3.87E-01 | 1.38E+00 | 5.91E-02 | 1.67E-01 | 1.27E+00 | 2.28E-02 | 8.37E-02 | 1.08E+00 | 1.33E-01 | 3.65E-01 | 1.23E+00 |
| TMEM70 | 1.37E-03 | 7.16E-03 | -1.18E+00 | 6.13E-03 | 2.65E-02 | 1.06E+00 | 7.96E-07 | 3.38E-06 | -1.29E+00 | 9.92E-06 | 3.79E-05 | -1.08E+00 | 1.86E-03 | 6.63E-03 | -1.15E+00 | 3.74E-03 | 1.05E-02 | -1.03E+00 | 1.22E-06 | 4.48E-06 | -1.28E+00 | 5.52E-04 | 1.52E-03 | -1.10E+00 |
| AC009094.1 | 1.38E-03 | 7.21E-03 | -1.61E+00 | 2.70E-02 | 1.16E-01 | -1.04E+00 | 1.38E-06 | 5.88E-06 | -1.68E+00 | 3.01E-06 | 1.15E-05 | -1.34E+00 | 4.13E-09 | 1.47E-08 | -2.80E+00 | 2.47E-07 | 6.97E-07 | -1.85E+00 | 1.58E-16 | 5.80E-16 | -2.35E+00 | 6.82E-07 | 1.87E-06 | -1.38E+00 |
| TEX261 | 1.38E-03 | 7.21E-03 | -1.26E+00 | 2.67E-03 | 1.15E-02 | -1.09E+00 | 2.28E-03 | 9.69E-03 | -1.18E+00 | 1.98E-02 | 7.58E-02 | 1.06E+00 | 1.29E-02 | 4.58E-02 | 1.01E+00 | 5.87E-02 | 1.65E-01 | 1.12E+00 | 1.52E-05 | 5.59E-05 | -1.33E+00 | 2.16E-01 | 5.92E-01 | 1.13E+00 |
| KMT2B | 1.39E-03 | 7.23E-03 | -1.05E+00 | 2.98E-02 | 1.29E-01 | 1.25E+00 | 6.37E-02 | 2.71E-01 | 1.06E+00 | 4.53E-03 | 1.73E-02 | 1.09E+00 | 6.06E-02 | 2.16E-01 | 1.15E+00 | 7.41E-02 | 2.09E-01 | 1.14E+00 | 3.85E-04 | 1.41E-03 | -1.08E+00 | 6.80E-04 | 1.87E-03 | -1.08E+00 |
| STAMBPL1 | 1.39E-03 | 7.24E-03 | -1.24E+00 | 7.83E-03 | 3.38E-02 | 1.04E+00 | 4.45E-06 | 1.89E-05 | -1.57E+00 | 1.00E-05 | 3.82E-05 | -1.38E+00 | 7.69E-02 | 2.74E-01 | -1.24E+00 | 1.51E-01 | 4.27E-01 | 1.00E+00 | 7.34E-03 | 2.69E-02 | -1.42E+00 | 4.97E-02 | 1.37E-01 | -1.17E+00 |
| PCCA | 1.39E-03 | 7.25E-03 | -1.31E+00 | 3.70E-02 | 1.60E-01 | 1.13E+00 | 3.89E-05 | 1.65E-04 | -1.43E+00 | 9.45E-03 | 3.61E-02 | 1.09E+00 | 1.18E-02 | 4.20E-02 | -1.19E+00 | 3.51E-01 | 9.89E-01 | 1.54E+00 | 2.39E-05 | 8.78E-05 | -1.35E+00 | 5.03E-03 | 1.38E-02 | -1.14E+00 |
| CD99L2 | 1.39E-03 | 7.25E-03 | -1.08E+00 | 4.29E-03 | 1.85E-02 | 1.09E+00 | 6.99E-02 | 2.97E-01 | -1.05E+00 | 4.78E-02 | 1.82E-01 | 1.06E+00 | 2.43E-02 | 8.65E-02 | -1.07E+00 | 1.06E-01 | 2.99E-01 | 1.06E+00 | 3.17E-02 | 1.16E-01 | -1.02E+00 | 3.52E-01 | 9.66E-01 | 1.33E+00 |
| RPS15AP10 | 1.39E-03 | 7.26E-03 | -1.24E+00 | 1.52E-02 | 6.55E-02 | 1.08E+00 | 8.32E-04 | 3.54E-03 | -1.26E+00 | 2.95E-05 | 1.13E-04 | -1.27E+00 | 1.21E-02 | 4.32E-02 | -1.19E+00 | 3.10E-03 | 8.74E-03 | -1.34E+00 | 1.82E-08 | 6.67E-08 | -1.71E+00 | 1.25E-06 | 3.44E-06 | -1.45E+00 |
| TMUB1 | 1.40E-03 | 7.27E-03 | 1.04E+00 | 1.62E-04 | 7.01E-04 | 1.08E+00 | 1.43E-03 | 6.07E-03 | -1.06E+00 | 2.62E-05 | 1.00E-04 | -1.08E+00 | 8.29E-06 | 2.95E-05 | -1.25E+00 | 1.17E-05 | 3.30E-05 | -1.18E+00 | 1.39E-01 | 5.09E-01 | 1.20E+00 | 6.89E-03 | 1.89E-02 | 1.08E+00 |
| USP24 | 1.40E-03 | 7.27E-03 | -1.18E+00 | 4.30E-02 | 1.86E-01 | 1.15E+00 | 9.80E-03 | 4.16E-02 | -1.02E+00 | 9.32E-02 | 3.56E-01 | 1.28E+00 | 2.45E-04 | 8.73E-04 | -1.20E+00 | 1.64E-02 | 4.63E-02 | 1.08E+00 | 1.11E-05 | 4.06E-05 | -1.35E+00 | 3.18E-02 | 8.73E-02 | 1.04E+00 |
| AL365356.1 | 1.40E-03 | 7.29E-03 | -1.18E+00 | 4.26E-05 | 1.84E-04 | -1.26E+00 | 7.48E-06 | 3.18E-05 | -1.70E+00 | 8.31E-05 | 3.18E-04 | -1.32E+00 | 7.82E-04 | 2.79E-03 | -1.28E+00 | 2.99E-04 | 8.44E-04 | -1.41E+00 | 1.18E-05 | 4.31E-05 | -1.70E+00 | 9.73E-04 | 2.67E-03 | -1.34E+00 |
| PGBD1 | 1.40E-03 | 7.30E-03 | -1.24E+00 | 1.08E-01 | 4.67E-01 | 1.32E+00 | 4.23E-05 | 1.80E-04 | -1.47E+00 | 2.35E-02 | 8.96E-02 | 1.10E+00 | 5.11E-03 | 1.82E-02 | -1.14E+00 | 3.35E-01 | 9.44E-01 | 1.44E+00 | 4.58E-03 | 1.68E-02 | -1.13E+00 | 2.45E-01 | 6.74E-01 | 1.15E+00 |
| CCDC102A | 1.40E-03 | 7.32E-03 | -1.03E+00 | 6.33E-02 | 2.73E-01 | 1.34E+00 | 1.94E-03 | 8.23E-03 | -1.16E+00 | 6.55E-02 | 2.50E-01 | 1.21E+00 | 2.84E-06 | 1.01E-05 | -1.71E+00 | 9.63E-04 | 2.72E-03 | -1.22E+00 | 4.55E-05 | 1.67E-04 | -1.61E+00 | 2.75E-03 | 7.54E-03 | -1.33E+00 |
| TRMT2B | 1.41E-03 | 7.33E-03 | -1.23E+00 | 1.30E-02 | 5.60E-02 | 1.09E+00 | 1.03E-05 | 4.36E-05 | -1.52E+00 | 3.86E-04 | 1.48E-03 | -1.17E+00 | 3.14E-03 | 1.12E-02 | -1.07E+00 | 6.23E-02 | 1.76E-01 | 1.15E+00 | 7.76E-09 | 2.85E-08 | -1.42E+00 | 1.16E-08 | 3.18E-08 | -1.54E+00 |
| ZNF300 | 1.41E-03 | 7.33E-03 | -1.18E+00 | 8.54E-04 | 3.69E-03 | -1.21E+00 | 8.78E-05 | 3.73E-04 | -1.62E+00 | 6.23E-04 | 2.38E-03 | -1.35E+00 | 1.89E-01 | 6.73E-01 | 1.02E+00 | 3.35E-01 | 9.43E-01 | 1.14E+00 | 2.33E-01 | 8.54E-01 | 1.11E+00 | 3.09E-01 | 8.48E-01 | 1.21E+00 |
| ARNT2 | 1.41E-03 | 7.36E-03 | -1.34E+00 | 6.51E-03 | 2.81E-02 | -1.11E+00 | 2.26E-01 | 9.61E-01 | 1.10E+00 | 1.77E-01 | 6.76E-01 | 1.27E+00 | 5.56E-03 | 1.98E-02 | -1.65E+00 | 2.84E-02 | 8.00E-02 | -1.30E+00 | 4.70E-02 | 1.72E-01 | -1.39E+00 | 2.47E-01 | 6.78E-01 | 1.08E+00 |
| OSBPL11 | 1.42E-03 | 7.38E-03 | -1.30E+00 | 1.64E-04 | 7.11E-04 | -1.44E+00 | 3.19E-03 | 1.35E-02 | -1.27E+00 | 1.13E-04 | 4.30E-04 | -1.43E+00 | 3.52E-03 | 1.25E-02 | -1.70E+00 | 6.91E-03 | 1.95E-02 | -1.55E+00 | 8.44E-11 | 3.10E-10 | -1.99E+00 | 4.59E-09 | 1.26E-08 | -1.75E+00 |
| CTBP2P8 | 1.42E-03 | 7.40E-03 | -1.54E+00 | 1.85E-03 | 8.00E-03 | -1.45E+00 | 1.39E-04 | 5.92E-04 | -1.84E+00 | 8.17E-03 | 3.12E-02 | -1.18E+00 | 6.77E-03 | 2.41E-02 | -1.39E+00 | 3.30E-02 | 9.30E-02 | -1.21E+00 | 1.45E-05 | 5.30E-05 | -1.58E+00 | 3.26E-03 | 8.96E-03 | -1.24E+00 |
| SENP8 | 1.42E-03 | 7.41E-03 | -1.55E+00 | 2.50E-03 | 1.08E-02 | -1.40E+00 | 1.09E-04 | 4.63E-04 | -1.65E+00 | 9.64E-05 | 3.68E-04 | -1.55E+00 | 9.47E-02 | 3.38E-01 | -1.18E+00 | 3.33E-01 | 9.40E-01 | 1.47E+00 | 1.09E-02 | 4.00E-02 | -1.36E+00 | 7.61E-02 | 2.09E-01 | -1.12E+00 |
| SULF2 | 1.43E-03 | 7.47E-03 | -1.94E+00 | 1.04E-02 | 4.48E-02 | -1.45E+00 | 1.35E-02 | 5.76E-02 | -1.37E+00 | 1.84E-02 | 7.02E-02 | -1.30E+00 | 4.46E-02 | 1.59E-01 | -1.44E+00 | 8.60E-02 | 2.42E-01 | -1.17E+00 | 1.71E-01 | 6.28E-01 | 1.28E+00 | 1.29E-01 | 3.55E-01 | 1.59E+00 |
| CNOT3 | 1.43E-03 | 7.48E-03 | -1.09E+00 | 4.05E-04 | 1.75E-03 | -1.13E+00 | 1.82E-04 | 7.72E-04 | -1.39E+00 | 7.35E-04 | 2.81E-03 | -1.13E+00 | 1.12E-06 | 3.99E-06 | -1.64E+00 | 1.29E-06 | 3.62E-06 | -1.47E+00 | 1.53E-05 | 5.62E-05 | -1.36E+00 | 7.56E-05 | 2.08E-04 | -1.42E+00 |
| DENND1A | 1.44E-03 | 7.49E-03 | -1.35E+00 | 7.15E-03 | 3.09E-02 | -1.08E+00 | 3.77E-05 | 1.60E-04 | -1.31E+00 | 1.29E-06 | 4.92E-06 | -1.33E+00 | 2.00E-01 | 7.13E-01 | 1.22E+00 | 3.42E-01 | 9.64E-01 | 1.38E+00 | 1.53E-07 | 5.63E-07 | -1.53E+00 | 4.96E-05 | 1.36E-04 | -1.28E+00 |
| ST3GAL6 | 1.44E-03 | 7.49E-03 | -1.27E+00 | 3.09E-04 | 1.34E-03 | -1.26E+00 | 6.57E-05 | 2.79E-04 | -1.25E+00 | 2.43E-04 | 9.27E-04 | -1.04E+00 | 1.87E-01 | 6.66E-01 | 1.03E+00 | 1.83E-01 | 5.16E-01 | -1.10E+00 | 1.53E-01 | 5.60E-01 | -1.12E+00 | 1.87E-01 | 5.14E-01 | 1.48E+00 |
| RABL2B | 1.44E-03 | 7.49E-03 | -1.16E+00 | 1.61E-01 | 6.95E-01 | 1.47E+00 | 2.27E-04 | 9.64E-04 | -1.31E+00 | 1.53E-01 | 5.85E-01 | 1.34E+00 | 1.38E-04 | 4.91E-04 | -1.49E+00 | 1.43E-02 | 4.04E-02 | -1.04E+00 | 1.72E-01 | 6.31E-01 | 1.04E+00 | 2.49E-01 | 6.85E-01 | 1.39E+00 |
| PEX11G | 1.44E-03 | 7.50E-03 | -1.53E+00 | 1.40E-04 | 6.04E-04 | -2.03E+00 | 4.99E-05 | 2.12E-04 | -1.82E+00 | 2.57E-06 | 9.82E-06 | -2.17E+00 | 8.98E-04 | 3.20E-03 | -2.03E+00 | 1.08E-02 | 3.05E-02 | -1.47E+00 | 1.43E-01 | 5.23E-01 | 1.12E+00 | 1.31E-01 | 3.61E-01 | 1.11E+00 |
| ANKHD1 | 1.44E-03 | 7.51E-03 | -1.10E+00 | 1.14E-04 | 4.95E-04 | -1.13E+00 | 3.65E-04 | 1.55E-03 | -1.21E+00 | 9.82E-07 | 3.75E-06 | -1.32E+00 | 4.18E-02 | 1.49E-01 | 1.06E+00 | 1.54E-02 | 4.33E-02 | -1.09E+00 | 5.83E-03 | 2.14E-02 | -1.08E+00 | 3.86E-03 | 1.06E-02 | -1.10E+00 |
| ZNF385C | 1.44E-03 | 7.52E-03 | -1.23E+00 | 1.73E-02 | 7.49E-02 | 1.11E+00 | 2.80E-02 | 1.19E-01 | -1.16E+00 | 5.10E-02 | 1.95E-01 | 1.04E+00 | 2.26E-01 | 8.05E-01 | 1.54E+00 | 1.42E-01 | 4.00E-01 | 1.84E+00 | 9.63E-02 | 3.53E-01 | -1.17E+00 | 3.25E-01 | 8.94E-01 | 1.14E+00 |
| TRIM35 | 1.44E-03 | 7.52E-03 | -1.68E+00 | 1.21E-01 | 5.24E-01 | 1.09E+00 | 2.71E-05 | 1.15E-04 | -1.92E+00 | 3.67E-02 | 1.40E-01 | -1.04E+00 | 2.11E-06 | 7.52E-06 | -2.31E+00 | 4.35E-03 | 1.23E-02 | -1.30E+00 | 1.64E-04 | 6.02E-04 | -1.28E+00 | 2.47E-01 | 6.79E-01 | 1.30E+00 |
| P2RX1 | 1.44E-03 | 7.53E-03 | 1.02E+00 | 1.57E-01 | 6.79E-01 | 1.50E+00 | 3.39E-05 | 1.44E-04 | -1.23E+00 | 1.07E-02 | 4.10E-02 | 1.18E+00 | 8.73E-08 | 3.11E-07 | -1.47E+00 | 8.86E-04 | 2.50E-03 | -1.16E+00 | 1.35E-01 | 4.96E-01 | 1.31E+00 | 3.26E-01 | 8.96E-01 | 1.28E+00 |
| AFAP1L1 | 1.44E-03 | 7.53E-03 | -1.37E+00 | 2.78E-02 | 1.20E-01 | 1.14E+00 | 2.31E-01 | 9.82E-01 | -1.12E+00 | 1.58E-01 | 6.03E-01 | 1.26E+00 | 1.69E-01 | 6.01E-01 | -1.09E+00 | 2.52E-01 | 7.11E-01 | -1.01E+00 | 1.34E-01 | 4.93E-01 | 1.16E+00 | 1.24E-01 | 3.40E-01 | -1.19E+00 |
| ZNF324 | 1.45E-03 | 7.57E-03 | -1.32E+00 | 1.83E-01 | 7.92E-01 | 1.26E+00 | 1.96E-03 | 8.32E-03 | -1.15E+00 | 4.64E-02 | 1.77E-01 | 1.25E+00 | 5.98E-02 | 2.13E-01 | 1.02E+00 | 3.33E-01 | 9.39E-01 | 1.49E+00 | 2.72E-02 | 9.98E-02 | -1.08E+00 | 3.31E-01 | 9.08E-01 | 1.28E+00 |
| GYPC | 1.45E-03 | 7.57E-03 | -1.04E+00 | 6.58E-03 | 2.84E-02 | 1.06E+00 | 3.62E-02 | 1.54E-01 | -1.04E+00 | 1.40E-01 | 5.33E-01 | -1.02E+00 | 1.73E-03 | 6.17E-03 | -1.14E+00 | 8.62E-02 | 2.43E-01 | 1.22E+00 | 3.32E-04 | 1.22E-03 | -1.33E+00 | 1.48E-01 | 4.07E-01 | -1.12E+00 |
| POLR3C | 1.45E-03 | 7.57E-03 | -1.14E+00 | 6.64E-02 | 2.87E-01 | 1.30E+00 | 7.98E-05 | 3.39E-04 | -1.30E+00 | 1.16E-03 | 4.44E-03 | -1.05E+00 | 1.80E-04 | 6.41E-04 | -1.32E+00 | 1.30E-01 | 3.67E-01 | 1.35E+00 | 4.29E-04 | 1.58E-03 | -1.21E+00 | 1.73E-03 | 4.74E-03 | -1.15E+00 |
| DPYSL2 | 1.45E-03 | 7.58E-03 | -1.01E+00 | 5.73E-02 | 2.48E-01 | 1.40E+00 | 1.31E-01 | 5.59E-01 | 1.40E+00 | 2.13E-01 | 8.12E-01 | 1.51E+00 | 5.14E-02 | 1.83E-01 | 1.06E+00 | 8.70E-03 | 2.45E-02 | -1.10E+00 | 4.01E-02 | 1.47E-01 | -1.03E+00 | 5.13E-02 | 1.41E-01 | 1.02E+00 |
| SETD7 | 1.46E-03 | 7.60E-03 | 1.66E+00 | 3.07E-07 | 1.33E-06 | -1.24E+00 | 8.89E-03 | 3.78E-02 | 1.60E+00 | 3.35E-06 | 1.28E-05 | -1.20E+00 | 2.30E-01 | 8.18E-01 | 1.59E+00 | 5.46E-03 | 1.54E-02 | -1.24E+00 | 3.48E-04 | 1.28E-03 | -1.39E+00 | 2.65E-05 | 7.27E-05 | -1.79E+00 |
| GEMIN4 | 1.46E-03 | 7.61E-03 | -1.51E+00 | 2.16E-02 | 9.34E-02 | -1.02E+00 | 5.55E-09 | 2.36E-08 | -1.86E+00 | 8.36E-04 | 3.19E-03 | -1.06E+00 | 2.89E-03 | 1.03E-02 | -1.50E+00 | 2.79E-02 | 7.85E-02 | -1.14E+00 | 1.11E-12 | 4.09E-12 | -2.06E+00 | 1.12E-02 | 3.08E-02 | 1.02E+00 |
| AC012360.3 | 1.46E-03 | 7.61E-03 | -1.07E+00 | 1.60E-02 | 6.92E-02 | 1.18E+00 | 3.76E-03 | 1.60E-02 | -1.19E+00 | 1.02E-02 | 3.91E-02 | 1.06E+00 | 2.10E-01 | 7.50E-01 | 1.33E+00 | 3.49E-01 | 9.84E-01 | 1.49E+00 | 8.85E-03 | 3.25E-02 | -1.09E+00 | 1.59E-01 | 4.35E-01 | 1.18E+00 |
| ZNF579 | 1.46E-03 | 7.62E-03 | -1.12E+00 | 1.56E-02 | 6.75E-02 | 1.14E+00 | 6.37E-05 | 2.71E-04 | -1.44E+00 | 2.28E-03 | 8.72E-03 | -1.12E+00 | 2.89E-03 | 1.03E-02 | -1.15E+00 | 1.91E-02 | 5.38E-02 | 1.04E+00 | 3.12E-03 | 1.15E-02 | -1.25E+00 | 2.52E-01 | 6.92E-01 | 1.11E+00 |
| ULK4 | 1.46E-03 | 7.63E-03 | -1.15E+00 | 1.17E-03 | 5.05E-03 | -1.08E+00 | 6.24E-05 | 2.65E-04 | -1.21E+00 | 2.48E-06 | 9.47E-06 | -1.27E+00 | 1.90E-01 | 6.78E-01 | 1.15E+00 | 2.10E-01 | 5.93E-01 | 1.16E+00 | 3.37E-02 | 1.24E-01 | -1.10E+00 | 8.77E-03 | 2.41E-02 | -1.30E+00 |
| TBCD | 1.47E-03 | 7.64E-03 | -1.19E+00 | 3.25E-03 | 1.41E-02 | 1.00E+00 | 4.00E-02 | 1.70E-01 | -1.12E+00 | 4.07E-02 | 1.55E-01 | -1.03E+00 | 3.14E-04 | 1.12E-03 | -1.47E+00 | 1.15E-01 | 3.24E-01 | 1.07E+00 | 2.60E-03 | 9.53E-03 | -1.05E+00 | 1.59E-01 | 4.38E-01 | 1.30E+00 |
| ARFGAP1 | 1.47E-03 | 7.66E-03 | -1.18E+00 | 2.42E-02 | 1.05E-01 | 1.12E+00 | 4.84E-03 | 2.06E-02 | -1.21E+00 | 1.23E-02 | 4.68E-02 | 1.00E+00 | 2.15E-02 | 7.65E-02 | -1.16E+00 | 2.48E-01 | 6.99E-01 | 1.33E+00 | 1.32E-05 | 4.85E-05 | -1.33E+00 | 1.86E-03 | 5.11E-03 | -1.14E+00 |
| HLCS | 1.47E-03 | 7.67E-03 | -1.24E+00 | 1.33E-01 | 5.73E-01 | 1.36E+00 | 1.56E-02 | 6.64E-02 | -1.21E+00 | 4.98E-02 | 1.90E-01 | 1.09E+00 | 1.06E-01 | 3.77E-01 | 1.05E+00 | 2.80E-01 | 7.89E-01 | 1.30E+00 | 8.92E-02 | 3.27E-01 | -1.01E+00 | 2.58E-01 | 7.08E-01 | 1.09E+00 |
| PRR11 | 1.48E-03 | 7.70E-03 | -1.58E+00 | 3.28E-03 | 1.42E-02 | -1.43E+00 | 6.57E-10 | 2.79E-09 | -2.00E+00 | 2.88E-09 | 1.10E-08 | -1.77E+00 | 4.05E-02 | 1.44E-01 | -1.22E+00 | 1.63E-01 | 4.59E-01 | -1.00E+00 | 2.03E-12 | 7.46E-12 | -2.11E+00 | 4.59E-07 | 1.26E-06 | -1.43E+00 |
| SHF | 1.48E-03 | 7.71E-03 | -1.01E+00 | 5.04E-03 | 2.18E-02 | 1.17E+00 | 1.31E-02 | 5.58E-02 | -1.09E+00 | 4.51E-02 | 1.72E-01 | 1.15E+00 | 2.04E-03 | 7.28E-03 | -1.21E+00 | 1.17E-03 | 3.30E-03 | -1.27E+00 | 2.07E-01 | 7.59E-01 | 1.39E+00 | 3.49E-01 | 9.58E-01 | 1.29E+00 |
| PITPNM1 | 1.48E-03 | 7.73E-03 | -1.36E+00 | 1.70E-05 | 7.35E-05 | -1.74E+00 | 1.13E-01 | 4.79E-01 | 1.03E+00 | 1.12E-01 | 4.27E-01 | 1.14E+00 | 1.92E-01 | 6.84E-01 | 1.81E+00 | 3.48E-01 | 9.80E-01 | 1.20E+00 | 2.41E-01 | 8.84E-01 | 1.14E+00 | 1.75E-01 | 4.80E-01 | -1.41E+00 |
| KCTD21-AS1 | 1.48E-03 | 7.73E-03 | -1.33E+00 | 1.25E-03 | 5.39E-03 | -1.37E+00 | 9.19E-02 | 3.90E-01 | -1.05E+00 | 1.88E-01 | 7.18E-01 | 1.24E+00 | 2.46E-04 | 8.76E-04 | -1.66E+00 | 1.56E-03 | 4.39E-03 | -1.47E+00 | 5.25E-02 | 1.93E-01 | -1.02E+00 | 3.37E-01 | 9.27E-01 | 1.30E+00 |
| NGDN | 1.48E-03 | 7.73E-03 | -1.01E+00 | 2.81E-03 | 1.21E-02 | 1.06E+00 | 9.51E-02 | 4.04E-01 | 1.12E+00 | 1.61E-02 | 6.15E-02 | 1.14E+00 | 1.19E-01 | 4.26E-01 | 1.32E+00 | 2.35E-01 | 6.64E-01 | 1.38E+00 | 3.58E-05 | 1.31E-04 | -1.09E+00 | 8.33E-02 | 2.29E-01 | 1.13E+00 |
| AC226101.1 | 1.48E-03 | 7.74E-03 | 1.12E+00 | 8.12E-04 | 3.51E-03 | 1.22E+00 | 1.38E-02 | 5.86E-02 | 1.48E+00 | 2.15E-02 | 8.22E-02 | 1.67E+00 | 1.93E-01 | 6.89E-01 | 1.47E+00 | 3.23E-01 | 9.10E-01 | 1.58E+00 | 1.62E-01 | 5.93E-01 | 1.26E+00 | 9.23E-02 | 2.54E-01 | 1.55E+00 |
| ABCA1 | 1.49E-03 | 7.75E-03 | -1.18E+00 | 3.01E-09 | 1.30E-08 | -1.58E+00 | 2.23E-02 | 9.47E-02 | -1.11E+00 | 5.31E-06 | 2.03E-05 | -1.92E+00 | 3.88E-02 | 1.38E-01 | 1.14E+00 | 8.84E-06 | 2.49E-05 | -1.57E+00 | 8.59E-02 | 3.15E-01 | -1.02E+00 | 1.02E-02 | 2.79E-02 | -1.58E+00 |
| SEC24D | 1.49E-03 | 7.76E-03 | -1.29E+00 | 1.08E-01 | 4.68E-01 | 1.29E+00 | 1.42E-02 | 6.03E-02 | -1.09E+00 | 9.85E-02 | 3.76E-01 | 1.21E+00 | 3.62E-08 | 1.29E-07 | -1.53E+00 | 4.18E-07 | 1.18E-06 | -1.43E+00 | 7.37E-06 | 2.71E-05 | -1.29E+00 | 1.05E-03 | 2.89E-03 | -1.06E+00 |
| SEPT7P2 | 1.49E-03 | 7.77E-03 | -1.09E+00 | 1.32E-03 | 5.69E-03 | 1.00E+00 | 1.74E-03 | 7.41E-03 | -1.23E+00 | 4.65E-04 | 1.77E-03 | -1.23E+00 | 2.95E-02 | 1.05E-01 | 1.00E+00 | 1.05E-01 | 2.97E-01 | 1.09E+00 | 1.10E-01 | 4.03E-01 | 1.55E+00 | 2.59E-01 | 7.11E-01 | 1.43E+00 |
| DMXL1 | 1.49E-03 | 7.77E-03 | 1.01E+00 | 2.87E-06 | 1.24E-05 | -1.05E+00 | 4.13E-02 | 1.76E-01 | 1.09E+00 | 2.29E-02 | 8.75E-02 | 1.11E+00 | 2.29E-01 | 8.16E-01 | 1.24E+00 | 1.10E-01 | 3.11E-01 | 1.16E+00 | 3.14E-02 | 1.15E-01 | -1.00E+00 | 2.54E-02 | 6.97E-02 | -1.03E+00 |
| CACNB3 | 1.49E-03 | 7.79E-03 | -1.33E+00 | 7.31E-03 | 3.16E-02 | -1.11E+00 | 1.60E-01 | 6.79E-01 | 1.04E+00 | 1.80E-01 | 6.88E-01 | 1.54E+00 | 1.22E-01 | 4.34E-01 | -1.17E+00 | 2.48E-01 | 6.98E-01 | 1.02E+00 | 4.22E-02 | 1.55E-01 | 1.05E+00 | 9.87E-02 | 2.71E-01 | 1.08E+00 |
| AC133552.2 | 1.50E-03 | 7.80E-03 | -1.45E+00 | 1.04E-01 | 4.47E-01 | 1.24E+00 | 1.56E-02 | 6.64E-02 | -1.50E+00 | 1.84E-01 | 7.05E-01 | 1.53E+00 | 4.63E-02 | 1.65E-01 | 1.03E+00 | 2.19E-01 | 6.16E-01 | 1.79E+00 | 2.35E-03 | 8.62E-03 | -1.34E+00 | 3.80E-02 | 1.04E-01 | -1.09E+00 |
| SLC38A4 | 1.50E-03 | 7.81E-03 | -1.02E+00 | 1.30E-02 | 5.61E-02 | 1.21E+00 | 1.47E-04 | 6.24E-04 | -1.10E+00 | 6.31E-04 | 2.41E-03 | -1.04E+00 | 6.19E-05 | 2.21E-04 | -1.56E+00 | 1.19E-02 | 3.34E-02 | -1.08E+00 | 1.95E-01 | 7.17E-01 | 1.15E+00 | 3.06E-01 | 8.42E-01 | 1.29E+00 |
| RFC2 | 1.50E-03 | 7.82E-03 | 1.06E+00 | 2.85E-03 | 1.23E-02 | 1.17E+00 | 3.98E-02 | 1.69E-01 | 1.13E+00 | 1.04E-03 | 3.96E-03 | 1.08E+00 | 3.10E-07 | 1.11E-06 | -1.50E+00 | 1.28E-05 | 3.62E-05 | -1.30E+00 | 3.18E-05 | 1.17E-04 | -1.14E+00 | 3.45E-02 | 9.48E-02 | 1.11E+00 |
| RRP36 | 1.50E-03 | 7.83E-03 | -1.11E+00 | 5.15E-02 | 2.22E-01 | 1.29E+00 | 1.26E-03 | 5.36E-03 | -1.16E+00 | 8.67E-03 | 3.31E-02 | 1.07E+00 | 1.72E-03 | 6.15E-03 | -1.09E+00 | 5.19E-02 | 1.46E-01 | 1.20E+00 | 5.83E-04 | 2.14E-03 | -1.12E+00 | 2.73E-01 | 7.51E-01 | 1.19E+00 |
| AC062017.1 | 1.50E-03 | 7.84E-03 | 2.19E+00 | 8.58E-02 | 3.71E-01 | 1.72E+00 | 1.00E-02 | 4.26E-02 | 2.13E+00 | 4.76E-02 | 1.82E-01 | 1.93E+00 | 2.70E-01 | 9.62E-01 | 1.23E+00 | 2.61E-01 | 7.35E-01 | 1.58E+00 | 1.14E-03 | 4.18E-03 | 2.42E+00 | 1.32E-01 | 3.64E-01 | 1.56E+00 |
| ZNF772 | 1.51E-03 | 7.85E-03 | -1.39E+00 | 1.08E-02 | 4.67E-02 | 1.01E+00 | 2.28E-01 | 9.69E-01 | 1.14E+00 | 1.40E-01 | 5.36E-01 | 1.59E+00 | 2.55E-02 | 9.08E-02 | -1.19E+00 | 7.48E-02 | 2.11E-01 | -1.07E+00 | 9.00E-03 | 3.30E-02 | -1.27E+00 | 6.64E-03 | 1.82E-02 | -1.21E+00 |
| AL133492.1 | 1.51E-03 | 7.85E-03 | -1.41E+00 | 3.26E-02 | 1.41E-01 | -1.01E+00 | 1.71E-01 | 7.26E-01 | 1.16E+00 | 1.99E-01 | 7.62E-01 | 1.34E+00 | 9.14E-02 | 3.26E-01 | -1.80E+00 | 6.98E-02 | 1.97E-01 | -2.13E+00 | 1.51E-01 | 5.56E-01 | -1.25E+00 | 2.71E-01 | 7.44E-01 | -1.03E+00 |
| AC005288.1 | 1.51E-03 | 7.87E-03 | -1.52E+00 | 4.38E-03 | 1.89E-02 | -1.25E+00 | 1.94E-01 | 8.24E-01 | -1.01E+00 | 1.11E-01 | 4.24E-01 | -1.01E+00 | 1.52E-01 | 5.43E-01 | 1.13E+00 | 5.67E-02 | 1.60E-01 | -1.03E+00 | 1.24E-01 | 4.55E-01 | 1.10E+00 | 5.72E-02 | 1.57E-01 | -1.11E+00 |
| ADAMTS6 | 1.51E-03 | 7.88E-03 | -1.31E+00 | 9.90E-03 | 4.28E-02 | -1.01E+00 | 9.97E-06 | 4.24E-05 | -1.67E+00 | 7.88E-04 | 3.01E-03 | -1.24E+00 | 5.92E-02 | 2.11E-01 | -1.30E+00 | 3.30E-02 | 9.29E-02 | -1.65E+00 | 1.88E-05 | 6.91E-05 | -2.26E+00 | 6.30E-05 | 1.73E-04 | -1.80E+00 |
| LMBRD2 | 1.51E-03 | 7.88E-03 | -1.23E+00 | 9.75E-02 | 4.21E-01 | 1.32E+00 | 7.59E-06 | 3.22E-05 | -1.85E+00 | 4.17E-03 | 1.59E-02 | -1.10E+00 | 6.40E-06 | 2.28E-05 | -2.41E+00 | 7.48E-03 | 2.11E-02 | -1.27E+00 | 4.93E-02 | 1.81E-01 | -1.01E+00 | 1.96E-01 | 5.37E-01 | 1.44E+00 |
| MT1A | 1.51E-03 | 7.88E-03 | -1.68E+00 | 3.78E-03 | 1.63E-02 | -1.48E+00 | 1.51E-02 | 6.41E-02 | -1.33E+00 | 4.03E-04 | 1.54E-03 | -1.89E+00 | 6.90E-02 | 2.46E-01 | -1.38E+00 | 2.22E-01 | 6.25E-01 | 1.23E+00 | 2.20E-01 | 8.08E-01 | 1.15E+00 | 1.28E-01 | 3.51E-01 | 1.01E+00 |
| HECTD3 | 1.51E-03 | 7.88E-03 | -1.45E+00 | 1.86E-02 | 8.03E-02 | 1.03E+00 | 1.70E-04 | 7.24E-04 | -1.55E+00 | 1.86E-03 | 7.09E-03 | -1.06E+00 | 3.07E-07 | 1.10E-06 | -2.39E+00 | 1.39E-04 | 3.92E-04 | -1.42E+00 | 4.97E-07 | 1.83E-06 | -1.43E+00 | 4.89E-05 | 1.34E-04 | -1.25E+00 |
| NRGN | 1.51E-03 | 7.88E-03 | -1.41E+00 | 2.81E-04 | 1.22E-03 | -1.58E+00 | 3.86E-02 | 1.64E-01 | 1.17E+00 | 6.51E-04 | 2.48E-03 | -1.11E+00 | 1.27E-06 | 4.53E-06 | -2.30E+00 | 1.05E-06 | 2.95E-06 | -2.44E+00 | 2.16E-01 | 7.92E-01 | 1.87E+00 | 5.34E-02 | 1.47E-01 | 1.07E+00 |
| CMTR2 | 1.51E-03 | 7.89E-03 | -1.12E+00 | 7.75E-03 | 3.35E-02 | 1.12E+00 | 2.23E-05 | 9.48E-05 | -1.32E+00 | 3.08E-03 | 1.18E-02 | 1.00E+00 | 4.01E-05 | 1.43E-04 | -1.16E+00 | 1.67E-05 | 4.69E-05 | -1.31E+00 | 4.93E-03 | 1.81E-02 | -1.00E+00 | 7.36E-03 | 2.02E-02 | -1.00E+00 |
| SRM | 1.51E-03 | 7.89E-03 | 1.08E+00 | 3.43E-04 | 1.48E-03 | 1.16E+00 | 2.22E-01 | 9.44E-01 | 1.17E+00 | 2.48E-01 | 9.48E-01 | 1.25E+00 | 6.08E-07 | 2.17E-06 | -1.24E+00 | 2.15E-02 | 6.07E-02 | 1.14E+00 | 4.98E-03 | 1.83E-02 | 1.06E+00 | 3.57E-02 | 9.82E-02 | 1.45E+00 |
| PIGW | 1.52E-03 | 7.90E-03 | -1.45E+00 | 1.24E-02 | 5.36E-02 | -1.11E+00 | 9.59E-02 | 4.08E-01 | -1.16E+00 | 2.18E-01 | 8.32E-01 | 1.17E+00 | 1.06E-02 | 3.78E-02 | -1.64E+00 | 9.77E-02 | 2.76E-01 | -1.14E+00 | 6.69E-09 | 2.45E-08 | -2.05E+00 | 1.13E-02 | 3.11E-02 | 1.03E+00 |
| HSD17B10 | 1.52E-03 | 7.92E-03 | 1.02E+00 | 1.53E-02 | 6.61E-02 | 1.18E+00 | 1.42E-01 | 6.05E-01 | 1.19E+00 | 4.05E-02 | 1.55E-01 | 1.20E+00 | 2.10E-08 | 7.48E-08 | -1.26E+00 | 1.09E-03 | 3.06E-03 | 1.01E+00 | 4.34E-02 | 1.59E-01 | 1.08E+00 | 2.67E-01 | 7.34E-01 | 1.29E+00 |
| MAP3K7 | 1.52E-03 | 7.93E-03 | -1.07E+00 | 3.53E-02 | 1.53E-01 | 1.22E+00 | 6.23E-05 | 2.65E-04 | -1.32E+00 | 3.00E-03 | 1.15E-02 | -1.02E+00 | 9.20E-05 | 3.28E-04 | -1.31E+00 | 2.66E-03 | 7.49E-03 | -1.08E+00 | 6.21E-06 | 2.28E-05 | -1.48E+00 | 1.27E-02 | 3.48E-02 | -1.05E+00 |
| SAMD1 | 1.52E-03 | 7.94E-03 | -1.05E+00 | 3.58E-02 | 1.55E-01 | 1.31E+00 | 7.93E-02 | 3.37E-01 | 1.03E+00 | 1.78E-01 | 6.81E-01 | 1.38E+00 | 2.91E-02 | 1.04E-01 | -1.03E+00 | 2.14E-01 | 6.02E-01 | 1.25E+00 | 1.11E-01 | 4.09E-01 | 1.15E+00 | 9.56E-03 | 2.63E-02 | 1.61E+00 |
| C9orf84 | 1.53E-03 | 7.95E-03 | -1.74E+00 | 2.02E-01 | 8.72E-01 | 1.50E+00 | 4.82E-04 | 2.05E-03 | -1.69E+00 | 1.99E-01 | 7.60E-01 | 1.35E+00 | 8.77E-02 | 3.13E-01 | -1.44E+00 | 1.94E-01 | 5.47E-01 | 1.58E+00 | 1.55E-01 | 5.69E-01 | 1.54E+00 | 2.68E-02 | 7.35E-02 | 2.30E+00 |
| AC007319.1 | 1.53E-03 | 7.97E-03 | 1.82E+00 | 1.17E-03 | 5.06E-03 | 2.12E+00 | 9.18E-03 | 3.90E-02 | 1.82E+00 | 2.61E-02 | 9.96E-02 | 1.99E+00 | 1.36E-01 | 4.86E-01 | 1.66E+00 | 2.40E-01 | 6.75E-01 | 1.60E+00 | 1.47E-01 | 5.40E-01 | 1.58E+00 | 3.06E-01 | 8.39E-01 | 1.16E+00 |
| VWF | 1.53E-03 | 8.00E-03 | -1.07E+00 | 6.36E-02 | 2.75E-01 | 1.17E+00 | 8.17E-03 | 3.47E-02 | 1.94E+00 | 5.52E-05 | 2.11E-04 | 2.65E+00 | 1.86E-01 | 6.63E-01 | 1.38E+00 | 5.96E-02 | 1.68E-01 | -1.63E+00 | 1.85E-02 | 6.78E-02 | 1.82E+00 | 1.45E-01 | 4.00E-01 | 1.28E+00 |
| CDV3 | 1.54E-03 | 8.00E-03 | 1.13E+00 | 3.75E-06 | 1.62E-05 | 1.09E+00 | 1.15E-01 | 4.90E-01 | 1.07E+00 | 1.07E-01 | 4.09E-01 | 1.11E+00 | 5.99E-02 | 2.14E-01 | 1.17E+00 | 4.33E-03 | 1.22E-02 | 1.03E+00 | 1.68E-01 | 6.16E-01 | 1.14E+00 | 1.85E-01 | 5.09E-01 | 1.06E+00 |
| RN7SL449P | 1.54E-03 | 8.00E-03 | -1.38E+00 | 2.95E-02 | 1.27E-01 | 1.11E+00 | 2.27E-01 | 9.64E-01 | 1.21E+00 | 2.27E-01 | 8.68E-01 | 1.27E+00 | 2.39E-01 | 8.51E-01 | 1.15E+00 | 2.46E-01 | 6.94E-01 | 1.04E+00 | 6.73E-02 | 2.47E-01 | -1.23E+00 | 8.29E-02 | 2.28E-01 | -1.25E+00 |
| MATK | 1.54E-03 | 8.01E-03 | 1.07E+00 | 4.69E-04 | 2.02E-03 | 1.15E+00 | 1.43E-01 | 6.09E-01 | 1.07E+00 | 1.36E-01 | 5.21E-01 | 1.16E+00 | 1.30E-04 | 4.63E-04 | -1.05E+00 | 1.06E-02 | 2.98E-02 | 1.13E+00 | 2.59E-03 | 9.51E-03 | 1.02E+00 | 2.34E-01 | 6.42E-01 | 1.42E+00 |
| TAS2R4 | 1.54E-03 | 8.01E-03 | -1.79E+00 | 5.69E-03 | 2.46E-02 | 2.10E+00 | 1.48E-02 | 6.28E-02 | -1.48E+00 | 1.24E-01 | 4.74E-01 | 1.72E+00 | 1.17E-01 | 4.15E-01 | -1.59E+00 | 3.38E-01 | 9.53E-01 | 1.18E+00 | 2.50E-01 | 9.18E-01 | 1.09E+00 | 3.11E-01 | 8.55E-01 | 1.20E+00 |
| LRRC34 | 1.54E-03 | 8.02E-03 | -1.47E+00 | 3.02E-02 | 1.31E-01 | 1.03E+00 | 2.07E-04 | 8.81E-04 | -1.45E+00 | 2.14E-03 | 8.17E-03 | -1.14E+00 | 1.47E-02 | 5.26E-02 | -1.38E+00 | 3.15E-02 | 8.87E-02 | -1.19E+00 | 1.27E-02 | 4.67E-02 | -1.05E+00 | 3.40E-01 | 9.35E-01 | 1.37E+00 |
| AC245452.1 | 1.54E-03 | 8.03E-03 | -1.11E+00 | 1.57E-01 | 6.80E-01 | 1.45E+00 | 5.47E-03 | 2.32E-02 | -1.14E+00 | 5.33E-02 | 2.04E-01 | 1.22E+00 | 1.98E-03 | 7.05E-03 | -1.19E+00 | 2.17E-03 | 6.10E-03 | -1.23E+00 | 2.16E-01 | 7.94E-01 | 1.24E+00 | 2.64E-01 | 7.26E-01 | 1.21E+00 |
| TTC21B-AS1 | 1.54E-03 | 8.03E-03 | -1.71E+00 | 1.39E-01 | 6.00E-01 | 1.16E+00 | 1.95E-03 | 8.29E-03 | -1.60E+00 | 3.68E-02 | 1.41E-01 | -1.09E+00 | 1.62E-01 | 5.78E-01 | -1.03E+00 | 1.94E-01 | 5.47E-01 | 1.54E+00 | 2.69E-01 | 9.89E-01 | 1.12E+00 | 3.53E-01 | 9.71E-01 | 1.21E+00 |
| EIF3L | 1.55E-03 | 8.07E-03 | -1.01E+00 | 1.26E-04 | 5.46E-04 | -1.02E+00 | 2.18E-02 | 9.24E-02 | 1.05E+00 | 6.26E-04 | 2.39E-03 | -1.02E+00 | 3.87E-04 | 1.38E-03 | -1.29E+00 | 3.29E-05 | 9.27E-05 | -1.31E+00 | 6.62E-07 | 2.43E-06 | -1.31E+00 | 2.23E-06 | 6.13E-06 | -1.45E+00 |
| SPOCK3 | 1.55E-03 | 8.09E-03 | -1.08E+00 | 2.15E-01 | 9.28E-01 | 1.41E+00 | 2.17E-01 | 9.23E-01 | 1.23E+00 | 1.46E-02 | 5.58E-02 | 2.03E+00 | 2.36E-01 | 8.41E-01 | -1.07E+00 | 2.21E-01 | 6.23E-01 | 1.01E+00 | 2.14E-01 | 7.87E-01 | 1.46E+00 | 5.55E-02 | 1.52E-01 | 2.15E+00 |
| RANBP3 | 1.55E-03 | 8.10E-03 | -1.26E+00 | 3.43E-02 | 1.48E-01 | 1.14E+00 | 2.24E-03 | 9.53E-03 | -1.19E+00 | 4.16E-03 | 1.59E-02 | -1.02E+00 | 1.62E-02 | 5.78E-02 | -1.21E+00 | 1.91E-01 | 5.38E-01 | 1.12E+00 | 2.64E-05 | 9.69E-05 | -1.22E+00 | 6.19E-02 | 1.70E-01 | 1.13E+00 |
| AL354920.1 | 1.55E-03 | 8.10E-03 | -1.49E+00 | 2.69E-03 | 1.16E-02 | -1.32E+00 | 1.48E-01 | 6.28E-01 | -1.03E+00 | 8.43E-02 | 3.22E-01 | -1.01E+00 | 1.53E-01 | 5.44E-01 | -1.15E+00 | 2.45E-01 | 6.92E-01 | 1.03E+00 | 8.47E-02 | 3.11E-01 | 1.04E+00 | 1.39E-01 | 3.81E-01 | 1.11E+00 |
| BTBD2 | 1.55E-03 | 8.10E-03 | -1.03E+00 | 2.56E-02 | 1.11E-01 | 1.26E+00 | 1.72E-06 | 7.32E-06 | -1.40E+00 | 2.47E-05 | 9.45E-05 | -1.11E+00 | 2.34E-04 | 8.34E-04 | -1.34E+00 | 6.36E-02 | 1.79E-01 | 1.06E+00 | 1.38E-04 | 5.07E-04 | -1.09E+00 | 5.97E-05 | 1.64E-04 | -1.17E+00 |
| FTSJ1 | 1.55E-03 | 8.10E-03 | -1.35E+00 | 8.69E-02 | 3.75E-01 | 1.14E+00 | 9.52E-11 | 4.04E-10 | -1.77E+00 | 1.00E-08 | 3.82E-08 | -1.37E+00 | 2.12E-04 | 7.54E-04 | -1.56E+00 | 1.31E-02 | 3.68E-02 | -1.15E+00 | 1.28E-07 | 4.69E-07 | -1.57E+00 | 2.71E-02 | 7.45E-02 | 1.03E+00 |
| ZNF169 | 1.56E-03 | 8.12E-03 | -1.32E+00 | 1.80E-01 | 7.78E-01 | 1.45E+00 | 1.57E-03 | 6.69E-03 | -1.54E+00 | 2.03E-01 | 7.74E-01 | 1.22E+00 | 1.52E-02 | 5.44E-02 | -1.58E+00 | 3.21E-01 | 9.04E-01 | 1.20E+00 | 1.30E-02 | 4.78E-02 | -1.56E+00 | 3.12E-01 | 8.58E-01 | 1.07E+00 |
| ZNF561-AS1 | 1.56E-03 | 8.12E-03 | -1.15E+00 | 2.31E-02 | 9.97E-02 | 1.13E+00 | 2.25E-03 | 9.57E-03 | -1.33E+00 | 5.25E-02 | 2.01E-01 | 1.13E+00 | 6.74E-03 | 2.40E-02 | -1.05E+00 | 5.21E-02 | 1.47E-01 | 1.11E+00 | 6.67E-02 | 2.45E-01 | 1.01E+00 | 1.42E-01 | 3.90E-01 | 1.04E+00 |
| ARHGAP35 | 1.56E-03 | 8.15E-03 | -1.08E+00 | 1.02E-02 | 4.42E-02 | 1.15E+00 | 3.27E-05 | 1.39E-04 | -1.48E+00 | 1.07E-03 | 4.07E-03 | -1.11E+00 | 2.20E-06 | 7.86E-06 | -1.66E+00 | 2.59E-04 | 7.29E-04 | -1.26E+00 | 4.92E-08 | 1.81E-07 | -1.43E+00 | 7.31E-04 | 2.01E-03 | -1.10E+00 |
| ZNF529-AS1 | 1.57E-03 | 8.16E-03 | -1.19E+00 | 3.79E-03 | 1.64E-02 | -1.00E+00 | 1.13E-02 | 4.79E-02 | -1.25E+00 | 1.72E-02 | 6.58E-02 | -1.09E+00 | 1.87E-01 | 6.68E-01 | 1.22E+00 | 5.37E-02 | 1.51E-01 | -1.06E+00 | 5.70E-02 | 2.09E-01 | 1.04E+00 | 2.06E-04 | 5.65E-04 | -1.51E+00 |
| HECTD4 | 1.57E-03 | 8.17E-03 | 1.01E+00 | 5.72E-02 | 2.47E-01 | 1.33E+00 | 1.66E-03 | 7.07E-03 | -1.05E+00 | 5.01E-03 | 1.92E-02 | 1.07E+00 | 1.85E-02 | 6.60E-02 | -1.02E+00 | 1.01E-02 | 2.84E-02 | -1.01E+00 | 2.41E-01 | 8.85E-01 | 1.23E+00 | 7.61E-02 | 2.09E-01 | 1.07E+00 |
| RPS27AP12 | 1.57E-03 | 8.20E-03 | -1.25E+00 | 1.12E-02 | 4.84E-02 | 1.04E+00 | 5.42E-06 | 2.30E-05 | -1.37E+00 | 1.90E-04 | 7.24E-04 | -1.12E+00 | 5.01E-02 | 1.78E-01 | -1.04E+00 | 6.40E-02 | 1.80E-01 | -1.11E+00 | 1.32E-02 | 4.83E-02 | -1.09E+00 | 3.76E-02 | 1.03E-01 | -1.04E+00 |
| Z95331.1 | 1.57E-03 | 8.20E-03 | -1.30E+00 | 5.69E-02 | 2.46E-01 | 1.07E+00 | 8.41E-03 | 3.58E-02 | -1.30E+00 | 1.29E-02 | 4.91E-02 | -1.12E+00 | 1.05E-01 | 3.74E-01 | 1.10E+00 | 1.60E-01 | 4.50E-01 | 1.12E+00 | 7.51E-03 | 2.76E-02 | -1.29E+00 | 1.55E-01 | 4.24E-01 | 1.12E+00 |
| IMPAD1 | 1.58E-03 | 8.22E-03 | -1.16E+00 | 2.13E-06 | 9.22E-06 | -1.36E+00 | 1.84E-03 | 7.83E-03 | -1.10E+00 | 9.62E-05 | 3.67E-04 | -1.11E+00 | 1.93E-02 | 6.89E-02 | -1.06E+00 | 1.97E-02 | 5.55E-02 | -1.06E+00 | 7.99E-04 | 2.93E-03 | -1.04E+00 | 3.60E-02 | 9.90E-02 | 1.12E+00 |
| PLEKHA8 | 1.58E-03 | 8.24E-03 | -1.40E+00 | 1.57E-02 | 6.80E-02 | -1.03E+00 | 8.87E-03 | 3.77E-02 | -1.28E+00 | 4.11E-02 | 1.57E-01 | 1.02E+00 | 1.09E-05 | 3.89E-05 | -1.86E+00 | 2.35E-04 | 6.63E-04 | -1.41E+00 | 1.17E-07 | 4.29E-07 | -1.68E+00 | 1.47E-04 | 4.04E-04 | -1.33E+00 |
| NPDC1 | 1.58E-03 | 8.25E-03 | 1.08E+00 | 1.10E-01 | 4.76E-01 | 1.32E+00 | 1.10E-01 | 4.66E-01 | 1.09E+00 | 5.84E-02 | 2.23E-01 | 1.18E+00 | 4.73E-02 | 1.69E-01 | -1.22E+00 | 1.14E-01 | 3.20E-01 | 1.06E+00 | 1.45E-01 | 5.34E-01 | 1.75E+00 | 6.08E-02 | 1.67E-01 | 2.55E+00 |
| LINC01137 | 1.58E-03 | 8.26E-03 | -1.17E+00 | 3.14E-02 | 1.36E-01 | 1.13E+00 | 1.31E-04 | 5.55E-04 | -1.50E+00 | 2.01E-03 | 7.67E-03 | -1.13E+00 | 2.33E-01 | 8.32E-01 | 1.23E+00 | 2.72E-01 | 7.65E-01 | 1.29E+00 | 5.78E-02 | 2.12E-01 | -1.11E+00 | 3.14E-01 | 8.62E-01 | 1.39E+00 |
| IRF9 | 1.59E-03 | 8.26E-03 | 1.05E+00 | 5.43E-38 | 2.35E-37 | -2.03E+00 | 1.50E-02 | 6.36E-02 | 1.01E+00 | 5.59E-23 | 2.13E-22 | -2.17E+00 | 4.43E-05 | 1.58E-04 | -1.04E+00 | 5.00E-20 | 1.41E-19 | -2.42E+00 | 1.56E-06 | 5.74E-06 | -1.29E+00 | 2.84E-20 | 7.80E-20 | -2.60E+00 |
| RRNAD1 | 1.59E-03 | 8.27E-03 | -1.43E+00 | 8.98E-03 | 3.88E-02 | -1.09E+00 | 4.02E-02 | 1.71E-01 | -1.09E+00 | 1.15E-01 | 4.40E-01 | 1.18E+00 | 8.78E-04 | 3.13E-03 | -1.08E+00 | 9.37E-02 | 2.64E-01 | 1.31E+00 | 1.58E-01 | 5.79E-01 | 1.36E+00 | 2.47E-01 | 6.78E-01 | 1.41E+00 |
| RBM34 | 1.59E-03 | 8.31E-03 | -1.05E+00 | 1.13E-05 | 4.89E-05 | -1.23E+00 | 1.41E-02 | 6.00E-02 | -1.07E+00 | 1.87E-03 | 7.14E-03 | -1.11E+00 | 1.03E-01 | 3.68E-01 | 1.15E+00 | 9.32E-02 | 2.63E-01 | 1.10E+00 | 1.96E-02 | 7.19E-02 | 1.16E+00 | 3.94E-04 | 1.08E-03 | -1.15E+00 |
| LACC1 | 1.59E-03 | 8.31E-03 | -1.23E+00 | 1.78E-03 | 7.70E-03 | -1.06E+00 | 1.17E-03 | 4.98E-03 | -1.42E+00 | 1.70E-03 | 6.49E-03 | -1.22E+00 | 1.48E-02 | 5.26E-02 | -1.05E+00 | 2.40E-02 | 6.77E-02 | -1.11E+00 | 5.00E-02 | 1.83E-01 | -1.08E+00 | 2.16E-03 | 5.92E-03 | -1.56E+00 |
| FAM81A | 1.60E-03 | 8.33E-03 | -2.59E+00 | 8.82E-02 | 3.81E-01 | -1.22E+00 | 1.59E-01 | 6.76E-01 | 1.23E+00 | 4.47E-02 | 1.71E-01 | 1.61E+00 | 2.75E-01 | 9.80E-01 | 1.20E+00 | 3.29E-01 | 9.26E-01 | 1.16E+00 | 1.48E-01 | 5.42E-01 | 1.46E+00 | 8.16E-02 | 2.24E-01 | 1.82E+00 |
| AC025171.1 | 1.60E-03 | 8.33E-03 | 1.09E+00 | 2.38E-07 | 1.03E-06 | -1.07E+00 | 9.25E-02 | 3.93E-01 | 1.14E+00 | 8.50E-03 | 3.25E-02 | 1.06E+00 | 5.94E-02 | 2.12E-01 | 1.14E+00 | 1.64E-01 | 4.62E-01 | 1.26E+00 | 1.32E-02 | 4.83E-02 | 1.06E+00 | 1.23E-01 | 3.39E-01 | 1.17E+00 |
| FAM161B | 1.60E-03 | 8.33E-03 | -1.40E+00 | 1.18E-02 | 5.09E-02 | -1.04E+00 | 7.45E-03 | 3.16E-02 | -1.18E+00 | 3.31E-05 | 1.27E-04 | -1.82E+00 | 7.85E-02 | 2.80E-01 | -1.11E+00 | 4.75E-02 | 1.34E-01 | -1.32E+00 | 4.52E-02 | 1.66E-01 | -1.21E+00 | 2.56E-01 | 7.04E-01 | 1.08E+00 |
| AC034102.1 | 1.60E-03 | 8.33E-03 | -1.07E+00 | 2.82E-03 | 1.22E-02 | 1.12E+00 | 4.49E-03 | 1.91E-02 | -1.25E+00 | 4.12E-02 | 1.57E-01 | 1.05E+00 | 5.23E-03 | 1.87E-02 | -1.09E+00 | 6.92E-02 | 1.95E-01 | 1.17E+00 | 1.72E-01 | 6.32E-01 | 1.18E+00 | 2.48E-01 | 6.80E-01 | 1.35E+00 |
| LMBRD1 | 1.60E-03 | 8.34E-03 | 1.01E+00 | 4.31E-03 | 1.86E-02 | 1.11E+00 | 6.25E-06 | 2.66E-05 | -1.36E+00 | 1.14E-05 | 4.35E-05 | -1.20E+00 | 3.18E-03 | 1.14E-02 | -1.05E+00 | 4.06E-02 | 1.15E-01 | 1.15E+00 | 1.34E-03 | 4.92E-03 | -1.05E+00 | 7.23E-02 | 1.99E-01 | 1.14E+00 |
| TNFRSF10D | 1.60E-03 | 8.35E-03 | -1.35E+00 | 8.79E-04 | 3.80E-03 | -1.28E+00 | 5.55E-04 | 2.36E-03 | -1.69E+00 | 2.51E-05 | 9.57E-05 | -2.17E+00 | 5.57E-04 | 1.99E-03 | -1.90E+00 | 3.53E-04 | 9.96E-04 | -2.19E+00 | 2.07E-01 | 7.59E-01 | 1.13E+00 | 1.60E-01 | 4.38E-01 | -1.13E+00 |
| ERMARD | 1.60E-03 | 8.36E-03 | -1.39E+00 | 1.34E-02 | 5.80E-02 | 1.00E+00 | 3.22E-03 | 1.37E-02 | -1.25E+00 | 3.71E-03 | 1.42E-02 | -1.11E+00 | 2.49E-01 | 8.89E-01 | 1.28E+00 | 1.96E-01 | 5.51E-01 | 1.72E+00 | 1.77E-04 | 6.51E-04 | -1.41E+00 | 1.38E-03 | 3.80E-03 | -1.28E+00 |
| ZC3HAV1 | 1.60E-03 | 8.36E-03 | 1.11E+00 | 7.51E-05 | 3.25E-04 | 1.13E+00 | 4.80E-02 | 2.04E-01 | -1.08E+00 | 8.98E-04 | 3.43E-03 | -1.09E+00 | 2.14E-04 | 7.64E-04 | -1.10E+00 | 1.04E-03 | 2.93E-03 | -1.11E+00 | 6.44E-06 | 2.36E-05 | -1.32E+00 | 2.06E-04 | 5.65E-04 | -1.30E+00 |
| SPICE1 | 1.60E-03 | 8.36E-03 | -1.20E+00 | 3.84E-02 | 1.66E-01 | 1.24E+00 | 4.31E-02 | 1.83E-01 | 1.04E+00 | 8.80E-02 | 3.36E-01 | 1.24E+00 | 8.22E-03 | 2.93E-02 | -1.19E+00 | 3.11E-02 | 8.77E-02 | -1.07E+00 | 8.16E-04 | 2.99E-03 | -1.30E+00 | 9.81E-03 | 2.69E-02 | -1.16E+00 |
| STIM1 | 1.60E-03 | 8.36E-03 | 1.03E+00 | 3.41E-03 | 1.47E-02 | 1.18E+00 | 1.30E-04 | 5.53E-04 | -1.14E+00 | 3.71E-06 | 1.42E-05 | -1.15E+00 | 7.28E-03 | 2.60E-02 | -1.11E+00 | 2.66E-02 | 7.51E-02 | 1.04E+00 | 3.30E-04 | 1.21E-03 | 1.04E+00 | 2.07E-04 | 5.68E-04 | -1.11E+00 |
| CTPS1 | 1.61E-03 | 8.37E-03 | -1.12E+00 | 5.52E-04 | 2.38E-03 | -1.12E+00 | 1.08E-01 | 4.59E-01 | 1.07E+00 | 4.78E-02 | 1.83E-01 | 1.25E+00 | 2.55E-02 | 9.08E-02 | -1.40E+00 | 2.65E-01 | 7.46E-01 | 1.25E+00 | 2.60E-07 | 9.54E-07 | -1.20E+00 | 1.92E-01 | 5.27E-01 | 1.34E+00 |
| MBD3 | 1.61E-03 | 8.37E-03 | -1.06E+00 | 6.03E-05 | 2.61E-04 | -1.09E+00 | 1.54E-01 | 6.55E-01 | 1.15E+00 | 1.44E-02 | 5.50E-02 | 1.11E+00 | 1.05E-02 | 3.76E-02 | -1.05E+00 | 4.31E-02 | 1.21E-01 | 1.13E+00 | 2.47E-01 | 9.06E-01 | 1.28E+00 | 2.41E-01 | 6.62E-01 | 1.21E+00 |
| CEP83-AS1 | 1.61E-03 | 8.37E-03 | -1.19E+00 | 9.69E-04 | 4.19E-03 | -1.27E+00 | 2.96E-02 | 1.26E-01 | -1.21E+00 | 1.13E-01 | 4.33E-01 | 1.17E+00 | 1.20E-01 | 4.29E-01 | 1.96E+00 | 1.71E-01 | 4.81E-01 | 1.87E+00 | 4.12E-02 | 1.51E-01 | -1.28E+00 | 9.52E-02 | 2.62E-01 | -1.20E+00 |
| TNFRSF1B | 1.61E-03 | 8.38E-03 | -1.25E+00 | 1.63E-01 | 7.06E-01 | 1.60E+00 | 1.84E-07 | 7.80E-07 | -1.89E+00 | 6.88E-05 | 2.63E-04 | -1.35E+00 | 2.78E-01 | 9.91E-01 | 1.16E+00 | 3.17E-01 | 8.95E-01 | -1.15E+00 | 2.23E-01 | 8.18E-01 | -1.12E+00 | 2.79E-01 | 7.67E-01 | -1.15E+00 |
| LINC02018 | 1.61E-03 | 8.39E-03 | -1.26E+00 | 2.47E-03 | 1.07E-02 | -1.19E+00 | 5.71E-02 | 2.43E-01 | -1.24E+00 | 6.52E-02 | 2.49E-01 | -1.04E+00 | 2.71E-01 | 9.67E-01 | 1.31E+00 | 2.49E-01 | 7.01E-01 | 1.44E+00 | 4.71E-02 | 1.73E-01 | -1.58E+00 | 1.18E-01 | 3.24E-01 | -1.31E+00 |
| ZC3H8 | 1.61E-03 | 8.41E-03 | -1.07E+00 | 4.63E-02 | 2.00E-01 | 1.27E+00 | 4.46E-02 | 1.90E-01 | -1.04E+00 | 2.42E-01 | 9.24E-01 | 1.26E+00 | 4.80E-03 | 1.71E-02 | -1.14E+00 | 2.07E-01 | 5.85E-01 | 1.34E+00 | 1.24E-03 | 4.56E-03 | -1.08E+00 | 7.48E-02 | 2.05E-01 | 1.06E+00 |
| RNU6-10P | 1.61E-03 | 8.41E-03 | -1.47E+00 | 1.74E-04 | 7.52E-04 | -1.67E+00 | 1.31E-02 | 5.56E-02 | -1.15E+00 | 9.92E-05 | 3.79E-04 | -1.59E+00 | 6.09E-03 | 2.17E-02 | -1.38E+00 | 1.06E-02 | 2.97E-02 | -1.43E+00 | 1.83E-04 | 6.72E-04 | -1.64E+00 | 7.34E-04 | 2.02E-03 | -1.47E+00 |
| PDE3B | 1.62E-03 | 8.42E-03 | -1.57E+00 | 6.16E-04 | 2.66E-03 | -1.56E+00 | 2.38E-06 | 1.01E-05 | -1.63E+00 | 5.28E-07 | 2.02E-06 | -1.55E+00 | 4.13E-05 | 1.47E-04 | -2.06E+00 | 2.71E-09 | 7.65E-09 | -3.32E+00 | 6.50E-08 | 2.39E-07 | -2.13E+00 | 8.79E-08 | 2.41E-07 | -2.32E+00 |
| ZSCAN16 | 1.62E-03 | 8.44E-03 | -1.38E+00 | 9.15E-02 | 3.95E-01 | 1.25E+00 | 7.91E-06 | 3.36E-05 | -1.52E+00 | 3.79E-05 | 1.45E-04 | -1.29E+00 | 2.72E-03 | 9.68E-03 | -1.26E+00 | 7.52E-02 | 2.12E-01 | 1.15E+00 | 6.58E-05 | 2.42E-04 | -1.42E+00 | 2.71E-03 | 7.44E-03 | -1.18E+00 |
| RBM41 | 1.63E-03 | 8.47E-03 | -1.06E+00 | 2.42E-03 | 1.04E-02 | 1.95E+00 | 8.50E-04 | 3.61E-03 | -1.23E+00 | 1.13E-01 | 4.31E-01 | 1.23E+00 | 4.09E-02 | 1.46E-01 | 1.05E+00 | 3.08E-01 | 8.67E-01 | 1.38E+00 | 7.44E-03 | 2.73E-02 | -1.08E+00 | 2.67E-01 | 7.34E-01 | 1.21E+00 |
| AL355432.1 | 1.63E-03 | 8.48E-03 | -1.67E+00 | 9.00E-03 | 3.89E-02 | -1.34E+00 | 1.83E-01 | 7.80E-01 | 1.01E+00 | 1.76E-01 | 6.71E-01 | -1.05E+00 | 2.82E-05 | 1.01E-04 | -2.20E+00 | 2.00E-04 | 5.63E-04 | -1.80E+00 | 4.80E-19 | 1.76E-18 | -2.36E+00 | 1.72E-17 | 4.73E-17 | -2.15E+00 |
| HOXA6 | 1.63E-03 | 8.49E-03 | 1.00E+00 | 3.29E-02 | 1.42E-01 | 1.22E+00 | 1.05E-02 | 4.47E-02 | -1.07E+00 | 1.62E-01 | 6.19E-01 | 1.22E+00 | 7.23E-02 | 2.58E-01 | 1.10E+00 | 6.39E-02 | 1.80E-01 | 1.00E+00 | 7.23E-02 | 2.65E-01 | -1.22E+00 | 1.93E-01 | 5.29E-01 | 1.00E+00 |
| AP006623.1 | 1.63E-03 | 8.49E-03 | -1.22E+00 | 5.12E-03 | 2.21E-02 | -1.04E+00 | 5.45E-03 | 2.32E-02 | -1.30E+00 | 2.93E-03 | 1.12E-02 | -1.18E+00 | 1.78E-02 | 6.35E-02 | -1.80E+00 | 1.00E-01 | 2.82E-01 | -1.24E+00 | 1.06E-01 | 3.87E-01 | 1.09E+00 | 6.27E-02 | 1.72E-01 | -1.19E+00 |
| EPC1 | 1.63E-03 | 8.49E-03 | 1.09E+00 | 1.29E-12 | 5.56E-12 | -1.07E+00 | 1.15E-01 | 4.88E-01 | 1.08E+00 | 7.21E-04 | 2.75E-03 | -1.08E+00 | 1.18E-01 | 4.22E-01 | 1.25E+00 | 5.17E-04 | 1.46E-03 | 1.00E+00 | 2.48E-01 | 9.12E-01 | 1.17E+00 | 1.56E-03 | 4.29E-03 | -1.09E+00 |
| DGCR2 | 1.63E-03 | 8.51E-03 | -1.18E+00 | 1.50E-01 | 6.49E-01 | 1.40E+00 | 4.44E-05 | 1.88E-04 | -1.45E+00 | 1.81E-02 | 6.91E-02 | 1.13E+00 | 5.27E-04 | 1.88E-03 | -1.58E+00 | 2.01E-02 | 5.68E-02 | -1.07E+00 | 7.06E-05 | 2.59E-04 | -1.38E+00 | 1.20E-03 | 3.28E-03 | -1.26E+00 |
| RPL15P14 | 1.64E-03 | 8.54E-03 | -1.12E+00 | 6.23E-29 | 2.69E-28 | -1.50E+00 | 3.69E-04 | 1.57E-03 | -1.23E+00 | 8.65E-12 | 3.30E-11 | -1.53E+00 | 2.37E-03 | 8.46E-03 | -1.14E+00 | 2.73E-10 | 7.70E-10 | -1.46E+00 | 2.49E-04 | 9.15E-04 | -1.18E+00 | 4.67E-11 | 1.28E-10 | -1.53E+00 |
| AC093495.1 | 1.64E-03 | 8.54E-03 | 1.01E+00 | 2.05E-11 | 8.88E-11 | -1.97E+00 | 1.74E-03 | 7.39E-03 | -1.13E+00 | 7.57E-08 | 2.89E-07 | -1.98E+00 | 2.26E-01 | 8.06E-01 | 1.21E+00 | 1.61E-01 | 4.55E-01 | -1.12E+00 | 2.50E-01 | 9.16E-01 | 1.08E+00 | 2.92E-01 | 8.02E-01 | -1.12E+00 |
| TANGO2 | 1.64E-03 | 8.55E-03 | -1.39E+00 | 1.07E-02 | 4.63E-02 | -1.09E+00 | 1.62E-02 | 6.87E-02 | -1.37E+00 | 1.57E-02 | 6.00E-02 | -1.22E+00 | 6.89E-08 | 2.46E-07 | -2.42E+00 | 5.60E-04 | 1.58E-03 | -1.22E+00 | 2.21E-02 | 8.12E-02 | -1.12E+00 | 5.69E-02 | 1.56E-01 | -1.10E+00 |
| AC006238.1 | 1.64E-03 | 8.56E-03 | 2.56E+00 | 7.23E-02 | 3.12E-01 | 1.81E+00 | 4.56E-02 | 1.94E-01 | 1.68E+00 | 1.26E-01 | 4.82E-01 | -1.37E+00 | 9.26E-02 | 3.30E-01 | 2.39E+00 | 4.81E-03 | 1.36E-02 | 3.66E+00 | 1.39E-01 | 5.11E-01 | -1.05E+00 | 2.63E-01 | 7.22E-01 | 1.06E+00 |
| MDGA1 | 1.64E-03 | 8.56E-03 | -1.25E+00 | 1.13E-02 | 4.87E-02 | -1.02E+00 | 3.62E-09 | 1.54E-08 | -1.76E+00 | 2.12E-07 | 8.11E-07 | -1.46E+00 | 1.39E-01 | 4.96E-01 | -1.46E+00 | 1.90E-01 | 5.36E-01 | -1.40E+00 | 2.66E-01 | 9.77E-01 | -1.01E+00 | 3.50E-01 | 9.62E-01 | 1.03E+00 |
| AC015813.1 | 1.64E-03 | 8.56E-03 | -1.26E+00 | 3.61E-02 | 1.56E-01 | 1.08E+00 | 1.03E-02 | 4.39E-02 | -1.11E+00 | 1.97E-01 | 7.52E-01 | 1.35E+00 | 2.52E-02 | 8.97E-02 | -1.15E+00 | 2.07E-01 | 5.83E-01 | 1.23E+00 | 1.36E-02 | 5.00E-02 | -1.10E+00 | 3.33E-01 | 9.14E-01 | 1.26E+00 |
| PKIA | 1.64E-03 | 8.57E-03 | -1.33E+00 | 3.54E-02 | 1.53E-01 | 1.10E+00 | 2.24E-04 | 9.52E-04 | -1.57E+00 | 9.71E-03 | 3.71E-02 | -1.08E+00 | 3.12E-02 | 1.11E-01 | -1.09E+00 | 3.28E-01 | 9.25E-01 | 1.38E+00 | 1.26E-07 | 4.64E-07 | -1.12E+00 | 1.11E-09 | 3.06E-09 | -1.29E+00 |
| RPL21P122 | 1.64E-03 | 8.57E-03 | -1.01E+00 | 6.38E-06 | 2.76E-05 | -1.08E+00 | 1.60E-01 | 6.80E-01 | 1.26E+00 | 5.17E-03 | 1.98E-02 | 1.10E+00 | 2.48E-01 | 8.86E-01 | 1.39E+00 | 4.49E-05 | 1.26E-04 | -1.21E+00 | 1.06E-01 | 3.87E-01 | 1.21E+00 | 1.57E-03 | 4.31E-03 | -1.03E+00 |
| PPIAP16 | 1.65E-03 | 8.58E-03 | 1.11E+00 | 4.20E-02 | 1.81E-01 | 1.34E+00 | 1.12E-01 | 4.77E-01 | 1.21E+00 | 1.98E-01 | 7.55E-01 | 1.49E+00 | 5.84E-05 | 2.08E-04 | -1.09E+00 | 3.55E-01 | 1.00E+00 | 1.32E+00 | 3.60E-02 | 1.32E-01 | 1.02E+00 | 1.07E-01 | 2.93E-01 | 1.30E+00 |
| AL391425.1 | 1.65E-03 | 8.59E-03 | -1.35E+00 | 2.84E-03 | 1.23E-02 | -1.27E+00 | 4.18E-02 | 1.78E-01 | -1.30E+00 | 2.96E-02 | 1.13E-01 | -1.32E+00 | 3.12E-02 | 1.11E-01 | -1.75E+00 | 2.26E-01 | 6.38E-01 | -1.05E+00 | 4.94E-03 | 1.81E-02 | -1.77E+00 | 3.37E-02 | 9.25E-02 | -1.51E+00 |
| SLC46A1 | 1.65E-03 | 8.59E-03 | -1.38E+00 | 3.35E-03 | 1.45E-02 | -1.21E+00 | 5.39E-02 | 2.29E-01 | -1.10E+00 | 1.61E-01 | 6.14E-01 | 1.20E+00 | 8.04E-02 | 2.87E-01 | -1.11E+00 | 1.92E-01 | 5.42E-01 | 1.11E+00 | 7.17E-02 | 2.63E-01 | -1.06E+00 | 2.12E-01 | 5.83E-01 | 1.17E+00 |
| DDX3P3 | 1.65E-03 | 8.60E-03 | -1.14E+00 | 5.77E-06 | 2.49E-05 | -1.67E+00 | 1.98E-02 | 8.40E-02 | -1.23E+00 | 7.58E-06 | 2.90E-05 | -2.44E+00 | 9.61E-02 | 3.43E-01 | 1.16E+00 | 7.72E-05 | 2.18E-04 | -1.93E+00 | 2.63E-02 | 9.66E-02 | -1.04E+00 | 2.53E-05 | 6.94E-05 | -2.23E+00 |
| GOLGA2 | 1.65E-03 | 8.62E-03 | -1.08E+00 | 4.18E-02 | 1.81E-01 | 1.23E+00 | 1.10E-06 | 4.69E-06 | -1.52E+00 | 1.04E-04 | 3.98E-04 | -1.14E+00 | 2.50E-13 | 8.91E-13 | -2.00E+00 | 5.98E-07 | 1.69E-06 | -1.25E+00 | 1.96E-08 | 7.19E-08 | -1.47E+00 | 2.11E-07 | 5.79E-07 | -1.37E+00 |
| PRR12 | 1.66E-03 | 8.63E-03 | -1.45E+00 | 1.51E-02 | 6.50E-02 | -1.06E+00 | 5.65E-02 | 2.40E-01 | -1.09E+00 | 1.78E-01 | 6.81E-01 | 1.22E+00 | 8.96E-02 | 3.19E-01 | -1.12E+00 | 3.45E-01 | 9.73E-01 | 1.33E+00 | 4.30E-04 | 1.58E-03 | -1.32E+00 | 2.81E-03 | 7.72E-03 | -1.20E+00 |
| CBWD3 | 1.66E-03 | 8.66E-03 | 1.02E+00 | 1.15E-01 | 4.96E-01 | 1.51E+00 | 3.57E-02 | 1.52E-01 | -1.01E+00 | 5.29E-02 | 2.02E-01 | 1.57E+00 | 2.10E-01 | 7.47E-01 | 1.35E+00 | 3.19E-01 | 9.01E-01 | 1.56E+00 | 5.95E-03 | 2.18E-02 | -1.01E+00 | 1.57E-01 | 4.31E-01 | 1.26E+00 |
| SLC25A27 | 1.66E-03 | 8.67E-03 | -1.49E+00 | 1.36E-01 | 5.89E-01 | 1.18E+00 | 1.31E-03 | 5.56E-03 | -1.57E+00 | 8.67E-02 | 3.31E-01 | 1.07E+00 | 9.00E-02 | 3.21E-01 | 1.97E+00 | 2.58E-01 | 7.28E-01 | 1.51E+00 | 2.19E-01 | 8.04E-01 | 1.36E+00 | 1.46E-01 | 4.00E-01 | 1.67E+00 |
| ZNF175 | 1.66E-03 | 8.67E-03 | -1.45E+00 | 5.08E-02 | 2.20E-01 | 1.10E+00 | 3.87E-02 | 1.64E-01 | -1.25E+00 | 2.23E-01 | 8.51E-01 | 1.27E+00 | 5.36E-02 | 1.91E-01 | 1.13E+00 | 8.74E-02 | 2.46E-01 | 1.12E+00 | 1.01E-01 | 3.71E-01 | 1.66E+00 | 2.80E-02 | 7.69E-02 | 1.23E+00 |
| ANAPC2 | 1.66E-03 | 8.67E-03 | -1.18E+00 | 1.57E-02 | 6.77E-02 | 1.12E+00 | 3.40E-04 | 1.44E-03 | -1.46E+00 | 1.36E-04 | 5.19E-04 | -1.34E+00 | 1.81E-03 | 6.46E-03 | -1.51E+00 | 1.18E-01 | 3.33E-01 | 1.26E+00 | 3.88E-05 | 1.42E-04 | -1.62E+00 | 2.92E-02 | 8.01E-02 | -1.07E+00 |
| AD000090.1 | 1.66E-03 | 8.67E-03 | -1.22E+00 | 1.73E-05 | 7.47E-05 | -1.86E+00 | 1.77E-03 | 7.53E-03 | -1.34E+00 | 4.45E-04 | 1.70E-03 | -1.45E+00 | 2.02E-01 | 7.19E-01 | 1.78E+00 | 3.50E-02 | 9.86E-02 | -1.43E+00 | 6.54E-02 | 2.40E-01 | 1.00E+00 | 7.03E-03 | 1.93E-02 | -1.46E+00 |
| CCDC124 | 1.66E-03 | 8.67E-03 | 1.04E+00 | 2.88E-04 | 1.25E-03 | 1.09E+00 | 2.18E-02 | 9.25E-02 | 1.05E+00 | 9.12E-02 | 3.48E-01 | 1.23E+00 | 1.35E-04 | 4.80E-04 | -1.13E+00 | 1.16E-02 | 3.28E-02 | 1.08E+00 | 8.02E-05 | 2.94E-04 | -1.11E+00 | 1.09E-01 | 2.98E-01 | 1.17E+00 |
| ZNF711 | 1.67E-03 | 8.70E-03 | 1.04E+00 | 7.89E-03 | 3.41E-02 | 1.15E+00 | 3.90E-07 | 1.66E-06 | -1.30E+00 | 2.28E-06 | 8.71E-06 | -1.17E+00 | 4.50E-03 | 1.61E-02 | -1.08E+00 | 3.94E-04 | 1.11E-03 | -1.19E+00 | 1.50E-03 | 5.52E-03 | -1.05E+00 | 2.27E-02 | 6.23E-02 | -1.02E+00 |
| UVSSA | 1.67E-03 | 8.71E-03 | -1.12E+00 | 1.61E-03 | 6.94E-03 | -1.00E+00 | 2.51E-04 | 1.06E-03 | -1.38E+00 | 1.14E-04 | 4.35E-04 | -1.21E+00 | 7.24E-02 | 2.58E-01 | 1.13E+00 | 3.29E-02 | 9.29E-02 | -1.04E+00 | 1.62E-03 | 5.96E-03 | -1.32E+00 | 1.74E-02 | 4.78E-02 | -1.06E+00 |
| AP005060.1 | 1.67E-03 | 8.71E-03 | -1.32E+00 | 2.61E-03 | 1.13E-02 | -1.16E+00 | 6.38E-04 | 2.71E-03 | -1.47E+00 | 2.80E-03 | 1.07E-02 | -1.21E+00 | 1.06E-01 | 3.79E-01 | -1.22E+00 | 1.16E-01 | 3.28E-01 | -1.34E+00 | 1.38E-02 | 5.08E-02 | -1.48E+00 | 1.49E-02 | 4.08E-02 | -1.69E+00 |
| MICU3 | 1.68E-03 | 8.74E-03 | -1.01E+00 | 9.10E-02 | 3.93E-01 | 1.38E+00 | 7.61E-05 | 3.23E-04 | -1.45E+00 | 8.29E-03 | 3.17E-02 | 1.03E+00 | 8.62E-03 | 3.07E-02 | -1.31E+00 | 1.78E-01 | 5.02E-01 | 1.25E+00 | 3.34E-03 | 1.23E-02 | -1.41E+00 | 8.11E-02 | 2.23E-01 | -1.08E+00 |
| MMS19 | 1.68E-03 | 8.75E-03 | -1.26E+00 | 1.21E-01 | 5.24E-01 | 1.26E+00 | 6.95E-05 | 2.95E-04 | -1.29E+00 | 1.08E-01 | 4.13E-01 | 1.30E+00 | 7.79E-05 | 2.78E-04 | -1.42E+00 | 7.17E-02 | 2.02E-01 | 1.26E+00 | 1.79E-05 | 6.57E-05 | -1.20E+00 | 6.22E-02 | 1.71E-01 | 1.10E+00 |
| ZNF736 | 1.68E-03 | 8.75E-03 | 1.05E+00 | 1.81E-04 | 7.80E-04 | -1.01E+00 | 1.42E-01 | 6.05E-01 | 1.06E+00 | 2.25E-02 | 8.59E-02 | 1.02E+00 | 2.79E-01 | 9.94E-01 | 1.47E+00 | 3.53E-01 | 9.96E-01 | 1.47E+00 | 7.52E-04 | 2.76E-03 | -1.07E+00 | 5.17E-03 | 1.42E-02 | -1.09E+00 |
| TRMT6 | 1.68E-03 | 8.75E-03 | -1.19E+00 | 7.44E-03 | 3.21E-02 | -1.00E+00 | 3.21E-04 | 1.37E-03 | -1.18E+00 | 6.18E-03 | 2.36E-02 | 1.06E+00 | 2.63E-04 | 9.38E-04 | -1.38E+00 | 9.04E-03 | 2.55E-02 | -1.06E+00 | 4.30E-10 | 1.58E-09 | -1.48E+00 | 4.62E-07 | 1.27E-06 | -1.28E+00 |
| HMGB3P6 | 1.68E-03 | 8.75E-03 | -1.35E+00 | 4.10E-06 | 1.77E-05 | -2.09E+00 | 1.20E-02 | 5.10E-02 | -1.12E+00 | 7.60E-08 | 2.90E-07 | -1.68E+00 | 3.34E-06 | 1.19E-05 | -1.83E+00 | 1.56E-07 | 4.41E-07 | -1.95E+00 | 5.15E-11 | 1.89E-10 | -1.67E+00 | 3.49E-14 | 9.57E-14 | -1.85E+00 |
| DNAJB6 | 1.68E-03 | 8.77E-03 | 1.01E+00 | 1.50E-05 | 6.50E-05 | -1.01E+00 | 7.22E-03 | 3.07E-02 | -1.05E+00 | 3.24E-05 | 1.24E-04 | -1.12E+00 | 1.65E-01 | 5.87E-01 | 1.32E+00 | 5.28E-05 | 1.49E-04 | -1.04E+00 | 2.28E-03 | 8.38E-03 | -1.01E+00 | 1.86E-04 | 5.10E-04 | -1.18E+00 |
| ZKSCAN4 | 1.68E-03 | 8.78E-03 | -1.39E+00 | 1.89E-01 | 8.18E-01 | 1.29E+00 | 7.45E-05 | 3.17E-04 | -1.62E+00 | 3.72E-02 | 1.42E-01 | 1.06E+00 | 1.34E-04 | 4.78E-04 | -1.74E+00 | 1.10E-02 | 3.11E-02 | -1.04E+00 | 8.96E-02 | 3.29E-01 | -1.06E+00 | 2.00E-01 | 5.49E-01 | 1.15E+00 |
| EFCAB11 | 1.69E-03 | 8.78E-03 | -1.32E+00 | 1.76E-03 | 7.61E-03 | -1.21E+00 | 2.14E-01 | 9.09E-01 | 1.15E+00 | 2.50E-01 | 9.55E-01 | 1.42E+00 | 4.59E-04 | 1.64E-03 | -1.51E+00 | 8.01E-03 | 2.26E-02 | -1.09E+00 | 3.66E-03 | 1.34E-02 | -1.07E+00 | 5.08E-02 | 1.40E-01 | 1.06E+00 |
| PPP1R1B | 1.69E-03 | 8.80E-03 | 2.31E+00 | 8.05E-02 | 3.48E-01 | 1.61E+00 | 3.62E-02 | 1.54E-01 | 1.67E+00 | 1.57E-01 | 5.98E-01 | 1.30E+00 | 9.25E-03 | 3.30E-02 | 2.49E+00 | 2.53E-01 | 7.13E-01 | 1.35E+00 | 4.80E-03 | 1.76E-02 | 1.94E+00 | 8.87E-02 | 2.44E-01 | 1.67E+00 |
| DNAJC9 | 1.70E-03 | 8.84E-03 | -1.32E+00 | 2.65E-03 | 1.14E-02 | -1.06E+00 | 1.49E-01 | 6.33E-01 | 1.26E+00 | 2.51E-01 | 9.59E-01 | 1.36E+00 | 2.52E-10 | 8.98E-10 | -1.70E+00 | 1.44E-07 | 4.05E-07 | -1.30E+00 | 1.33E-09 | 4.89E-09 | -1.22E+00 | 2.77E-06 | 7.62E-06 | -1.12E+00 |
| AC006480.2 | 1.70E-03 | 8.88E-03 | -1.52E+00 | 4.96E-02 | 2.14E-01 | 1.85E+00 | 6.88E-05 | 2.92E-04 | -1.88E+00 | 2.40E-01 | 9.17E-01 | 1.34E+00 | 1.07E-02 | 3.82E-02 | -1.61E+00 | 3.46E-01 | 9.74E-01 | 1.33E+00 | 1.74E-05 | 6.40E-05 | -1.67E+00 | 1.12E-01 | 3.08E-01 | 1.06E+00 |
| TTC7B | 1.71E-03 | 8.92E-03 | -1.05E+00 | 2.24E-04 | 9.69E-04 | -1.09E+00 | 4.89E-03 | 2.08E-02 | -1.05E+00 | 1.00E-02 | 3.82E-02 | 1.08E+00 | 2.22E-04 | 7.91E-04 | -1.16E+00 | 7.58E-05 | 2.14E-04 | -1.19E+00 | 9.24E-08 | 3.39E-07 | -1.08E+00 | 3.16E-07 | 8.69E-07 | -1.16E+00 |
| MFGE8 | 1.71E-03 | 8.93E-03 | 1.02E+00 | 6.23E-02 | 2.69E-01 | 1.41E+00 | 9.56E-02 | 4.06E-01 | 1.16E+00 | 4.38E-02 | 1.67E-01 | 1.20E+00 | 5.89E-05 | 2.10E-04 | -1.24E+00 | 8.31E-07 | 2.34E-06 | -1.67E+00 | 2.39E-01 | 8.79E-01 | 1.34E+00 | 1.83E-01 | 5.03E-01 | 1.10E+00 |
| CROCCP2 | 1.72E-03 | 8.94E-03 | -1.04E+00 | 9.54E-02 | 4.12E-01 | 1.45E+00 | 1.64E-05 | 6.97E-05 | -1.32E+00 | 6.75E-03 | 2.58E-02 | 1.11E+00 | 3.40E-05 | 1.21E-04 | -1.46E+00 | 2.66E-01 | 7.51E-01 | 1.29E+00 | 4.67E-05 | 1.72E-04 | -1.09E+00 | 1.09E-01 | 3.00E-01 | 1.14E+00 |
| FAM160B1 | 1.72E-03 | 8.95E-03 | 1.66E+00 | 6.81E-02 | 2.94E-01 | 1.32E+00 | 4.23E-02 | 1.80E-01 | 1.13E+00 | 8.03E-07 | 3.07E-06 | -1.08E+00 | 2.74E-01 | 9.76E-01 | 1.42E+00 | 2.95E-02 | 8.32E-02 | 1.07E+00 | 2.26E-01 | 8.31E-01 | 1.25E+00 | 4.38E-04 | 1.20E-03 | -1.14E+00 |
| WDR35 | 1.72E-03 | 8.95E-03 | -1.26E+00 | 3.61E-03 | 1.56E-02 | -1.08E+00 | 1.67E-06 | 7.08E-06 | -1.58E+00 | 3.05E-05 | 1.17E-04 | -1.24E+00 | 1.76E-01 | 6.28E-01 | 1.28E+00 | 2.41E-01 | 6.80E-01 | 1.36E+00 | 1.99E-01 | 7.32E-01 | 1.39E+00 | 2.14E-01 | 5.89E-01 | 1.14E+00 |
| RARRES2 | 1.72E-03 | 8.96E-03 | -1.02E+00 | 6.83E-05 | 2.95E-04 | -1.17E+00 | 5.23E-02 | 2.22E-01 | 1.08E+00 | 1.93E-02 | 7.39E-02 | 1.02E+00 | 2.70E-01 | 9.63E-01 | -1.01E+00 | 3.24E-01 | 9.13E-01 | -1.11E+00 | 2.42E-01 | 8.89E-01 | 1.19E+00 | 3.23E-01 | 8.86E-01 | 1.18E+00 |
| MIOS | 1.72E-03 | 8.98E-03 | -1.08E+00 | 1.94E-05 | 8.36E-05 | -1.16E+00 | 6.39E-07 | 2.72E-06 | -1.39E+00 | 1.69E-05 | 6.47E-05 | -1.13E+00 | 1.05E-03 | 3.74E-03 | -1.13E+00 | 1.22E-02 | 3.44E-02 | 1.06E+00 | 6.43E-03 | 2.36E-02 | -1.10E+00 | 8.61E-02 | 2.37E-01 | 1.13E+00 |
| SNHG8 | 1.73E-03 | 9.02E-03 | 1.18E+00 | 1.83E-01 | 7.92E-01 | 1.09E+00 | 6.86E-02 | 2.91E-01 | 1.10E+00 | 2.33E-01 | 8.91E-01 | 1.04E+00 | 8.53E-03 | 3.04E-02 | 1.21E+00 | 2.47E-01 | 6.95E-01 | 1.07E+00 | 1.56E-03 | 5.74E-03 | 1.27E+00 | 1.23E-01 | 3.37E-01 | 1.09E+00 |
| AC145138.1 | 1.73E-03 | 9.02E-03 | -1.65E+00 | 7.66E-02 | 3.31E-01 | 1.12E+00 | 5.34E-05 | 2.27E-04 | -2.00E+00 | 2.44E-02 | 9.33E-02 | -1.06E+00 | 2.91E-03 | 1.04E-02 | -1.70E+00 | 4.33E-03 | 1.22E-02 | -1.44E+00 | 7.71E-06 | 2.83E-05 | -1.93E+00 | 2.72E-04 | 7.48E-04 | -1.60E+00 |
| LINC00484 | 1.73E-03 | 9.02E-03 | 2.87E+00 | 8.27E-02 | 3.58E-01 | 1.72E+00 | 4.04E-02 | 1.71E-01 | 1.87E+00 | 1.34E-01 | 5.13E-01 | 1.43E+00 | 2.31E-03 | 8.24E-03 | 3.84E+00 | 2.19E-01 | 6.17E-01 | 1.56E+00 | 1.99E-01 | 7.32E-01 | 1.07E+00 | 4.66E-02 | 1.28E-01 | -1.54E+00 |
| AL157938.2 | 1.74E-03 | 9.04E-03 | -1.18E+00 | 2.96E-03 | 1.28E-02 | -1.09E+00 | 4.16E-02 | 1.77E-01 | -1.15E+00 | 2.41E-02 | 9.22E-02 | -1.09E+00 | 1.15E-01 | 4.10E-01 | 1.05E+00 | 6.38E-02 | 1.80E-01 | -1.02E+00 | 6.89E-02 | 2.53E-01 | -1.18E+00 | 5.51E-02 | 1.51E-01 | -1.21E+00 |
| RPS4XP6 | 1.74E-03 | 9.05E-03 | -1.03E+00 | 8.28E-21 | 3.58E-20 | -1.34E+00 | 2.27E-01 | 9.66E-01 | 1.05E+00 | 6.73E-06 | 2.57E-05 | -1.24E+00 | 1.49E-02 | 5.30E-02 | 1.06E+00 | 4.83E-09 | 1.36E-08 | -1.24E+00 | 1.92E-01 | 7.06E-01 | 1.05E+00 | 3.82E-05 | 1.05E-04 | -1.27E+00 |
| TRIM23 | 1.74E-03 | 9.05E-03 | -1.12E+00 | 3.83E-03 | 1.65E-02 | 1.03E+00 | 1.91E-02 | 8.11E-02 | -1.13E+00 | 3.22E-02 | 1.23E-01 | 1.04E+00 | 1.69E-04 | 6.02E-04 | -1.45E+00 | 8.65E-03 | 2.44E-02 | -1.08E+00 | 1.41E-01 | 5.19E-01 | 1.16E+00 | 2.09E-01 | 5.74E-01 | 1.13E+00 |
| AL513497.1 | 1.74E-03 | 9.05E-03 | -1.65E+00 | 1.54E-03 | 6.67E-03 | -1.57E+00 | 4.26E-02 | 1.81E-01 | -1.16E+00 | 2.62E-02 | 1.00E-01 | -1.19E+00 | 7.20E-02 | 2.57E-01 | -1.14E+00 | 3.59E-02 | 1.01E-01 | -1.33E+00 | 1.67E-01 | 6.13E-01 | -1.12E+00 | 7.24E-02 | 1.99E-01 | -1.28E+00 |
| VPS26B | 1.74E-03 | 9.06E-03 | 1.01E+00 | 1.11E-03 | 4.82E-03 | 1.13E+00 | 4.14E-06 | 1.76E-05 | -1.27E+00 | 1.29E-04 | 4.92E-04 | 1.04E+00 | 6.18E-05 | 2.20E-04 | -1.34E+00 | 5.69E-03 | 1.60E-02 | -1.06E+00 | 7.02E-03 | 2.58E-02 | -1.05E+00 | 2.55E-01 | 7.00E-01 | 1.17E+00 |
| TRAF6 | 1.74E-03 | 9.08E-03 | -1.12E+00 | 4.06E-05 | 1.75E-04 | -1.23E+00 | 1.14E-01 | 4.83E-01 | 1.03E+00 | 1.53E-01 | 5.83E-01 | 1.19E+00 | 2.13E-01 | 7.60E-01 | 1.33E+00 | 3.22E-02 | 9.09E-02 | -1.01E+00 | 3.98E-04 | 1.46E-03 | -1.09E+00 | 9.94E-03 | 2.73E-02 | -1.03E+00 |
| ZBTB41 | 1.74E-03 | 9.09E-03 | -1.18E+00 | 5.26E-03 | 2.27E-02 | -1.03E+00 | 5.01E-04 | 2.13E-03 | -1.24E+00 | 6.45E-03 | 2.46E-02 | 1.05E+00 | 4.91E-03 | 1.75E-02 | -1.55E+00 | 6.70E-03 | 1.89E-02 | -1.53E+00 | 2.56E-02 | 9.41E-02 | -1.05E+00 | 1.97E-01 | 5.41E-01 | 1.16E+00 |

| | | | | | | | | | | | | | | | | | | | | | | | | |
|---|---|---|---|---|---|---|---|---|---|---|---|---|---|---|---|---|---|---|---|---|---|---|---|---|
| HDDC3 | 1.74E-03 | 9.09E-03 | -1.29E+00 | 2.86E-02 | 1.23E-01 | 1.15E+00 | 2.40E-03 | 1.02E-02 | -1.56E+00 | 1.36E-02 | 5.19E-02 | -1.18E+00 | 1.33E-02 | 4.75E-02 | -1.10E+00 | 6.63E-02 | 1.87E-01 | 1.05E+00 | 1.41E-04 | 5.16E-04 | -1.36E+00 | 2.10E-03 | 5.76E-03 | -1.17E+00 |
| PKN2 | 1.75E-03 | 9.10E-03 | 1.06E+00 | 4.90E-10 | 2.12E-09 | -1.07E+00 | 3.40E-04 | 1.44E-03 | -1.15E+00 | 1.68E-07 | 6.42E-07 | -1.25E+00 | 2.30E-03 | 8.20E-03 | -1.05E+00 | 4.18E-07 | 1.18E-06 | -1.31E+00 | 1.18E-02 | 4.34E-02 | 1.03E+00 | 9.48E-06 | 2.60E-05 | -1.25E+00 |
| AL109766.1 | 1.75E-03 | 9.10E-03 | 2.37E+00 | 5.77E-02 | 2.49E-01 | 2.03E+00 | 4.17E-04 | 1.77E-03 | 2.37E+00 | 2.09E-02 | 7.97E-02 | 2.21E+00 | 2.67E-01 | 9.51E-01 | 1.41E+00 | 2.24E-01 | 6.31E-01 | 1.69E+00 | 2.02E-03 | 7.42E-03 | 2.12E+00 | 9.84E-03 | 2.70E-02 | 2.00E+00 |
| AC025198.1 | 1.75E-03 | 9.11E-03 | -1.36E+00 | 3.13E-03 | 1.35E-02 | -1.21E+00 | 8.65E-03 | 3.67E-02 | -1.31E+00 | 1.14E-03 | 4.36E-03 | -1.39E+00 | 3.47E-03 | 1.24E-02 | -1.55E+00 | 5.46E-03 | 1.54E-02 | -1.52E+00 | 1.54E-05 | 5.64E-05 | -1.73E+00 | 4.10E-05 | 1.13E-04 | -1.68E+00 |
| AC092747.4 | 1.75E-03 | 9.11E-03 | -1.12E+00 | 2.00E-01 | 8.66E-01 | 1.54E+00 | 2.09E-04 | 8.88E-04 | -1.38E+00 | 8.26E-02 | 3.16E-01 | 1.17E+00 | 2.75E-01 | 9.79E-01 | 1.44E+00 | 8.97E-02 | 2.53E-01 | 1.96E+00 | 1.78E-05 | 6.54E-05 | -1.36E+00 | 1.34E-01 | 3.67E-01 | 1.13E+00 |
| MCPH1-AS1 | 1.75E-03 | 9.12E-03 | -1.57E+00 | 1.23E-01 | 5.33E-01 | 1.32E+00 | 1.97E-01 | 8.39E-01 | 1.08E+00 | 1.64E-01 | 6.25E-01 | 1.46E+00 | 1.48E-01 | 5.27E-01 | 1.09E+00 | 2.08E-01 | 5.86E-01 | 1.75E+00 | 2.90E-02 | 1.06E-01 | -1.31E+00 | 1.23E-01 | 3.37E-01 | -1.03E+00 |
| MND1 | 1.75E-03 | 9.12E-03 | -1.95E+00 | 1.46E-03 | 6.31E-03 | -1.86E+00 | 1.50E-01 | 6.36E-01 | 1.21E+00 | 8.36E-05 | 3.19E-04 | -1.11E+00 | 3.69E-02 | 1.31E-01 | -1.53E+00 | 3.78E-02 | 1.06E-01 | -1.65E+00 | 1.31E-01 | 4.81E-01 | 1.06E+00 | 5.41E-02 | 1.49E-01 | 1.49E+00 |
| AL365277.1 | 1.75E-03 | 9.12E-03 | -1.37E+00 | 2.49E-02 | 1.08E-01 | 1.06E+00 | 1.22E-03 | 5.19E-03 | -1.58E+00 | 7.39E-03 | 2.82E-02 | -1.21E+00 | 5.84E-02 | 2.08E-01 | 1.04E+00 | 1.84E-01 | 5.19E-01 | 1.24E+00 | 9.44E-02 | 3.46E-01 | -1.00E+00 | 4.34E-02 | 1.19E-01 | -1.27E+00 |
| AC008966.1 | 1.75E-03 | 9.12E-03 | -1.64E+00 | 9.15E-02 | 3.96E-01 | 1.14E+00 | 2.21E-02 | 9.38E-02 | -1.24E+00 | 1.98E-01 | 7.55E-01 | 1.28E+00 | 7.72E-02 | 2.75E-01 | -1.06E+00 | 2.07E-01 | 5.84E-01 | 1.24E+00 | 1.44E-04 | 5.29E-04 | -1.34E+00 | 4.56E-03 | 1.25E-02 | -1.10E+00 |
| EMC9 | 1.75E-03 | 9.13E-03 | -1.10E+00 | 5.92E-04 | 2.56E-03 | -1.07E+00 | 1.34E-01 | 5.71E-01 | 1.29E+00 | 9.32E-02 | 3.56E-01 | 1.17E+00 | 2.71E-04 | 9.64E-04 | -1.31E+00 | 3.47E-04 | 9.79E-04 | -1.26E+00 | 5.54E-02 | 2.03E-01 | -1.04E+00 | 6.22E-03 | 1.71E-02 | -1.31E+00 |
| RGL2 | 1.75E-03 | 9.14E-03 | 1.05E+00 | 1.51E-10 | 6.51E-10 | -1.31E+00 | 7.25E-03 | 3.08E-02 | 1.01E+00 | 5.48E-09 | 2.09E-08 | -1.45E+00 | 1.61E-05 | 5.75E-05 | -1.30E+00 | 9.35E-07 | 2.64E-06 | -1.43E+00 | 2.38E-02 | 8.75E-02 | 1.05E+00 | 3.57E-04 | 9.81E-04 | -1.29E+00 |
| CNRIP1 | 1.76E-03 | 9.16E-03 | -1.74E+00 | 1.25E-02 | 5.42E-02 | -1.24E+00 | 1.79E-01 | 7.60E-01 | 1.47E+00 | 2.93E-02 | 1.12E-01 | 2.27E+00 | 8.54E-06 | 3.04E-05 | -1.21E+00 | 2.77E-02 | 7.81E-02 | 1.24E+00 | 2.67E-01 | 9.80E-01 | -1.04E+00 | 1.78E-02 | 4.89E-02 | 1.21E+00 |
| AC074117.1 | 1.76E-03 | 9.17E-03 | -1.34E+00 | 9.38E-02 | 4.05E-01 | 1.18E+00 | 5.47E-05 | 2.33E-04 | -1.60E+00 | 9.20E-04 | 3.51E-03 | -1.19E+00 | 7.95E-03 | 2.83E-02 | -1.55E+00 | 1.26E-01 | 3.54E-01 | 1.04E+00 | 1.32E-02 | 4.86E-02 | -1.37E+00 | 1.67E-01 | 4.60E-01 | 1.01E+00 |
| POP7 | 1.76E-03 | 9.17E-03 | -1.19E+00 | 2.79E-02 | 1.20E-01 | 1.11E+00 | 7.42E-02 | 3.15E-01 | 1.03E+00 | 1.70E-01 | 6.48E-01 | 1.21E+00 | 2.52E-06 | 9.00E-06 | -1.55E+00 | 1.99E-02 | 5.61E-02 | 1.10E+00 | 8.65E-05 | 3.17E-04 | -1.03E+00 | 1.27E-01 | 3.50E-01 | 1.22E+00 |
| SPINT1-AS1 | 1.76E-03 | 9.17E-03 | -1.03E+00 | 1.70E-04 | 7.34E-04 | -1.08E+00 | 2.21E-01 | 9.41E-01 | 1.24E+00 | 1.27E-01 | 4.84E-01 | 1.08E+00 | 4.24E-02 | 1.51E-01 | -1.27E+00 | 5.65E-02 | 1.59E-01 | -1.24E+00 | 1.45E-01 | 5.33E-01 | -1.08E+00 | 3.27E-01 | 8.97E-01 | 1.13E+00 |
| SAP130 | 1.76E-03 | 9.19E-03 | -1.52E+00 | 2.32E-02 | 1.00E-01 | -1.07E+00 | 6.98E-03 | 2.97E-02 | -1.27E+00 | 2.28E-02 | 8.70E-02 | -1.08E+00 | 2.32E-06 | 8.27E-06 | -2.21E+00 | 3.02E-04 | 8.52E-04 | -1.43E+00 | 4.53E-08 | 1.66E-07 | -1.57E+00 | 2.61E-07 | 7.16E-07 | -1.46E+00 |
| CCDC30 | 1.77E-03 | 9.20E-03 | -1.34E+00 | 9.40E-02 | 4.06E-01 | 1.24E+00 | 3.26E-07 | 1.38E-06 | -1.91E+00 | 3.55E-06 | 1.35E-05 | -1.50E+00 | 7.95E-03 | 2.83E-02 | -1.13E+00 | 2.31E-02 | 6.52E-02 | -1.03E+00 | 1.18E-01 | 4.32E-01 | 1.12E+00 | 2.91E-01 | 7.99E-01 | 1.26E+00 |
| NKX2-3 | 1.77E-03 | 9.21E-03 | -1.31E+00 | 1.43E-04 | 6.18E-04 | 2.24E+00 | 7.87E-02 | 3.34E-01 | -1.19E+00 | 1.31E-01 | 4.99E-01 | 1.67E+00 | 1.56E-01 | 5.54E-01 | 1.90E+00 | 7.85E-02 | 2.21E-01 | 2.67E+00 | 1.04E-01 | 3.83E-01 | -1.40E+00 | 2.90E-01 | 7.97E-01 | 1.40E+00 |
| PPP1R8 | 1.77E-03 | 9.21E-03 | -1.16E+00 | 8.32E-02 | 3.60E-01 | 1.26E+00 | 6.84E-04 | 2.91E-03 | -1.14E+00 | 1.87E-02 | 7.13E-02 | 1.12E+00 | 1.70E-03 | 6.06E-03 | -1.22E+00 | 1.60E-01 | 4.50E-01 | 1.18E+00 | 4.94E-04 | 1.81E-03 | -1.07E+00 | 2.15E-01 | 5.90E-01 | 1.20E+00 |
| EIF4ENIF1 | 1.77E-03 | 9.21E-03 | -1.24E+00 | 3.46E-02 | 1.50E-01 | 1.08E+00 | 1.11E-03 | 4.72E-03 | -1.23E+00 | 1.04E-01 | 3.99E-01 | 1.25E+00 | 1.23E-04 | 4.38E-04 | -1.37E+00 | 7.94E-03 | 2.24E-02 | 1.02E+00 | 5.20E-07 | 1.91E-06 | -1.39E+00 | 2.35E-04 | 6.46E-04 | -1.12E+00 |
| PEX7 | 1.77E-03 | 9.21E-03 | -1.25E+00 | 2.13E-03 | 9.20E-03 | -1.04E+00 | 6.02E-02 | 2.56E-01 | -1.06E+00 | 3.05E-02 | 1.17E-01 | 1.05E+00 | 1.45E-02 | 5.16E-02 | -1.12E+00 | 9.16E-02 | 2.58E-01 | 1.09E+00 | 2.00E-01 | 7.32E-01 | 1.25E+00 | 2.42E-01 | 6.65E-01 | 1.30E+00 |
| ACAA1 | 1.77E-03 | 9.24E-03 | -1.25E+00 | 5.03E-03 | 2.17E-02 | -1.10E+00 | 1.81E-04 | 7.68E-04 | -1.42E+00 | 1.84E-04 | 7.04E-04 | -1.32E+00 | 1.84E-06 | 6.56E-06 | -2.49E+00 | 1.64E-02 | 4.63E-02 | -1.09E+00 | 4.06E-03 | 1.49E-02 | -1.27E+00 | 3.54E-02 | 9.72E-02 | -1.02E+00 |
| RAB35 | 1.77E-03 | 9.24E-03 | -1.24E+00 | 1.13E-02 | 4.90E-02 | 1.03E+00 | 1.55E-03 | 6.61E-03 | -1.19E+00 | 7.29E-03 | 2.79E-02 | 1.05E+00 | 5.99E-05 | 2.13E-04 | -1.27E+00 | 2.61E-05 | 7.36E-05 | -1.33E+00 | 9.49E-06 | 3.48E-05 | -1.32E+00 | 2.54E-03 | 6.98E-03 | 1.00E+00 |
| RRS1 | 1.78E-03 | 9.28E-03 | -1.15E+00 | 5.00E-02 | 2.16E-01 | 1.27E+00 | 9.47E-06 | 4.02E-05 | -1.43E+00 | 8.67E-02 | 3.31E-01 | 1.23E+00 | 1.18E-04 | 4.22E-04 | -1.47E+00 | 3.22E-01 | 9.07E-01 | 1.44E+00 | 3.99E-12 | 1.47E-11 | -1.69E+00 | 7.69E-02 | 2.11E-01 | 1.16E+00 |
| NEU1 | 1.78E-03 | 9.29E-03 | 2.07E+00 | 6.26E-10 | 2.71E-09 | -1.04E+00 | 2.14E-01 | 9.07E-01 | 1.43E+00 | 1.86E-05 | 7.11E-05 | -1.12E+00 | 1.76E-01 | 6.28E-01 | 1.56E+00 | 1.15E-03 | 3.25E-03 | -1.08E+00 | 1.53E-01 | 5.61E-01 | 1.54E+00 | 3.60E-03 | 9.89E-03 | -1.07E+00 |
| XIRP2 | 1.78E-03 | 9.30E-03 | 2.74E+00 | 1.41E-03 | 6.10E-03 | 3.40E+00 | 6.61E-02 | 2.81E-01 | 2.22E+00 | 2.13E-01 | 8.14E-01 | 1.61E+00 | 2.03E-01 | 7.22E-01 | -1.19E+00 | 2.86E-01 | 8.07E-01 | -1.18E+00 | 1.13E-01 | 4.15E-01 | 1.49E+00 | 1.92E-01 | 5.27E-01 | 1.29E+00 |
| WHRN | 1.79E-03 | 9.31E-03 | 2.16E+00 | 5.99E-03 | 2.59E-02 | -1.38E+00 | 6.87E-02 | 2.92E-01 | 1.46E+00 | 2.46E-02 | 9.39E-02 | -1.22E+00 | 5.88E-02 | 2.10E-01 | 1.96E+00 | 5.23E-02 | 1.47E-01 | -1.74E+00 | 4.04E-03 | 1.48E-02 | 2.86E+00 | 3.57E-01 | 9.80E-01 | 1.14E+00 |
| SMARCA4 | 1.79E-03 | 9.32E-03 | 1.05E+00 | 2.43E-02 | 1.05E-01 | 1.37E+00 | 4.51E-04 | 1.92E-03 | -1.03E+00 | 4.05E-04 | 1.55E-03 | 1.09E+00 | 1.83E-04 | 6.53E-04 | -1.15E+00 | 1.12E-01 | 3.16E-01 | 1.25E+00 | 8.66E-04 | 3.18E-03 | -1.04E+00 | 2.33E-01 | 6.39E-01 | 1.20E+00 |
| TAF1A | 1.79E-03 | 9.34E-03 | -1.56E+00 | 2.31E-02 | 9.98E-02 | 1.02E+00 | 4.39E-03 | 1.87E-02 | -1.30E+00 | 4.68E-02 | 1.79E-01 | 1.08E+00 | 4.61E-03 | 1.64E-02 | -1.55E+00 | 8.11E-02 | 2.29E-01 | 1.00E+00 | 6.05E-06 | 2.22E-05 | -1.85E+00 | 1.01E-03 | 2.78E-03 | -1.33E+00 |
| AC098934.1 | 1.80E-03 | 9.37E-03 | -1.07E+00 | 3.21E-03 | 1.39E-02 | 1.01E+00 | 1.04E-02 | 4.42E-02 | 1.75E+00 | 1.11E-01 | 4.22E-01 | 1.72E+00 | 1.77E-01 | 6.30E-01 | -1.13E+00 | 1.82E-01 | 5.13E-01 | -1.29E+00 | 1.03E-01 | 3.79E-01 | -1.08E+00 | 2.48E-01 | 6.81E-01 | 1.15E+00 |
| REV1 | 1.80E-03 | 9.37E-03 | 1.02E+00 | 1.60E-04 | 6.90E-04 | 1.05E+00 | 7.68E-03 | 3.26E-02 | 1.00E+00 | 1.20E-02 | 4.59E-02 | 1.14E+00 | 2.46E-01 | 8.76E-01 | 1.31E+00 | 1.16E-01 | 3.27E-01 | 1.22E+00 | 5.48E-04 | 2.01E-03 | -1.00E+00 | 1.92E-02 | 5.29E-02 | 1.04E+00 |
| SYNE2 | 1.80E-03 | 9.39E-03 | -1.92E+00 | 9.22E-02 | 3.98E-01 | 1.06E+00 | 1.18E-01 | 5.00E-01 | -1.95E+00 | 2.16E-01 | 8.26E-01 | -1.59E+00 | 4.94E-09 | 1.76E-08 | -2.18E+00 | 8.99E-05 | 2.53E-04 | -1.34E+00 | 3.17E-03 | 1.16E-02 | -1.29E+00 | 2.22E-01 | 6.11E-01 | 1.10E+00 |
| WDR66 | 1.81E-03 | 9.42E-03 | -1.61E+00 | 2.25E-02 | 9.73E-02 | -1.08E+00 | 2.40E-06 | 1.02E-05 | -1.96E+00 | 2.80E-04 | 1.07E-03 | -1.25E+00 | 1.24E-02 | 4.42E-02 | -1.80E+00 | 5.04E-02 | 1.42E-01 | -1.48E+00 | 4.17E-02 | 1.53E-01 | -1.39E+00 | 4.12E-02 | 1.13E-01 | -1.47E+00 |
| PTPDC1 | 1.81E-03 | 9.44E-03 | -1.39E+00 | 4.54E-02 | 1.96E-01 | 1.08E+00 | 1.09E-02 | 4.62E-02 | -1.18E+00 | 9.26E-02 | 3.54E-01 | 1.17E+00 | 9.27E-02 | 3.31E-01 | 1.01E+00 | 1.42E-01 | 4.00E-01 | -1.02E+00 | 2.16E-02 | 7.94E-02 | -1.37E+00 | 3.52E-01 | 9.66E-01 | 1.13E+00 |
| AC132008.2 | 1.81E-03 | 9.44E-03 | -1.49E+00 | 1.29E-02 | 5.56E-02 | -1.10E+00 | 8.81E-03 | 3.74E-02 | -1.29E+00 | 1.68E-02 | 6.41E-02 | -1.15E+00 | 1.22E-01 | 4.36E-01 | -1.08E+00 | 1.53E-01 | 4.32E-01 | -1.02E+00 | 1.10E-02 | 4.04E-02 | -1.03E+00 | 4.53E-02 | 1.25E-01 | -1.04E+00 |
| PHGDH | 1.81E-03 | 9.44E-03 | 1.08E+00 | 2.56E-02 | 1.10E-01 | 1.25E+00 | 1.44E-01 | 6.13E-01 | 1.10E+00 | 1.20E-01 | 4.57E-01 | 1.14E+00 | 2.35E-04 | 8.36E-04 | -1.42E+00 | 5.75E-05 | 1.62E-04 | -1.46E+00 | 2.23E-01 | 8.20E-01 | 1.53E+00 | 9.36E-02 | 2.57E-01 | 1.75E+00 |
| FTH1P23 | 1.81E-03 | 9.45E-03 | 1.78E+00 | 3.55E-03 | 1.53E-02 | 1.01E+00 | 4.85E-04 | 2.06E-03 | 1.87E+00 | 8.56E-02 | 3.27E-01 | 1.18E+00 | 3.70E-05 | 1.32E-04 | 2.25E+00 | 3.72E-02 | 1.05E-01 | 1.25E+00 | 2.46E-03 | 9.04E-03 | 1.65E+00 | 2.10E-02 | 5.76E-02 | 1.10E+00 |
| ALKBH2 | 1.81E-03 | 9.45E-03 | 1.07E+00 | 2.00E-03 | 8.62E-03 | 1.20E+00 | 7.90E-02 | 3.36E-01 | 1.05E+00 | 2.29E-01 | 8.74E-01 | 1.31E+00 | 8.08E-05 | 2.88E-04 | -1.30E+00 | 6.07E-03 | 1.71E-02 | -1.03E+00 | 1.89E-03 | 6.95E-03 | -1.17E+00 | 2.00E-01 | 5.48E-01 | 1.14E+00 |
| SPN53 | 1.82E-03 | 9.46E-03 | -1.02E+00 | 2.96E-09 | 1.28E-08 | -1.35E+00 | 1.90E-01 | 8.06E-01 | 1.18E+00 | 3.42E-05 | 1.31E-04 | -1.02E+00 | 2.52E-01 | 8.99E-01 | 1.01E+00 | 3.23E-01 | 9.09E-01 | -1.01E+00 | 1.84E-01 | 6.77E-01 | 1.51E+00 | 3.60E-01 | 9.89E-01 | -1.02E+00 |
| AP005899.1 | 1.82E-03 | 9.47E-03 | -1.18E+00 | 2.09E-04 | 9.02E-04 | -1.36E+00 | 3.58E-02 | 1.52E-01 | -1.05E+00 | 3.28E-03 | 1.25E-02 | -1.18E+00 | 2.12E-01 | 7.55E-01 | 1.37E+00 | 5.59E-02 | 1.57E-01 | -1.22E+00 | 9.85E-02 | 3.61E-01 | -1.03E+00 | 3.10E-01 | 8.51E-01 | 1.29E+00 |
| SLC35G1 | 1.82E-03 | 9.49E-03 | -1.18E+00 | 1.46E-01 | 6.32E-01 | 1.39E+00 | 1.35E-02 | 5.75E-02 | -1.13E+00 | 1.23E-01 | 4.69E-01 | 1.58E+00 | 9.02E-02 | 3.22E-01 | 1.05E+00 | 1.60E-01 | 4.51E-01 | 1.71E+00 | 2.17E-01 | 7.96E-01 | 1.19E+00 | 6.95E-04 | 1.91E-03 | 1.83E+00 |
| SELENON | 1.82E-03 | 9.50E-03 | -1.21E+00 | 3.68E-03 | 1.59E-02 | -1.05E+00 | 3.61E-02 | 1.53E-01 | -1.06E+00 | 1.64E-02 | 6.27E-02 | 1.03E+00 | 2.12E-04 | 7.56E-04 | -1.32E+00 | 2.75E-05 | 7.77E-05 | -1.56E+00 | 4.15E-03 | 1.52E-02 | -1.03E+00 | 7.90E-03 | 2.17E-02 | 1.06E+00 |
| C12orf43 | 1.82E-03 | 9.50E-03 | -1.09E+00 | 4.01E-02 | 1.73E-01 | 1.28E+00 | 8.71E-06 | 3.70E-05 | -1.24E+00 | 1.55E-03 | 5.94E-03 | 1.07E+00 | 1.05E-06 | 3.75E-06 | -1.48E+00 | 7.17E-03 | 2.02E-02 | 1.05E+00 | 8.77E-12 | 3.22E-11 | -1.55E+00 | 1.66E-06 | 4.55E-06 | -1.19E+00 |
| KCTD13 | 1.82E-03 | 9.50E-03 | -1.13E+00 | 9.27E-03 | 4.01E-02 | 1.13E+00 | 1.08E-04 | 4.59E-04 | -1.35E+00 | 1.50E-05 | 5.72E-05 | -1.29E+00 | 7.89E-04 | 2.81E-03 | -1.29E+00 | 2.15E-04 | 6.07E-04 | -1.40E+00 | 2.51E-05 | 9.23E-05 | -1.32E+00 | 6.67E-04 | 1.83E-03 | -1.19E+00 |
| OPRL1 | 1.82E-03 | 9.51E-03 | -1.33E+00 | 4.22E-02 | 1.82E-01 | 1.10E+00 | 1.45E-04 | 6.16E-04 | -1.52E+00 | 1.00E-02 | 3.82E-02 | -1.05E+00 | 2.08E-01 | 7.41E-01 | 1.12E+00 | 1.73E-01 | 4.88E-01 | -1.08E+00 | 1.84E-01 | 6.77E-01 | -1.21E+00 | 2.76E-01 | 7.58E-01 | -1.13E+00 |
| PBLD | 1.82E-03 | 9.51E-03 | -1.25E+00 | 2.33E-02 | 1.01E-01 | 1.05E+00 | 2.39E-02 | 1.01E-01 | -1.09E+00 | 2.74E-02 | 1.05E-01 | 1.10E+00 | 2.86E-03 | 1.02E-02 | -1.23E+00 | 2.38E-02 | 6.72E-02 | -1.02E+00 | 1.86E-03 | 6.83E-03 | -1.16E+00 | 5.27E-03 | 1.45E-02 | -1.06E+00 |
| NCOA6 | 1.83E-03 | 9.53E-03 | -1.16E+00 | 4.64E-03 | 2.01E-02 | -1.00E+00 | 3.68E-02 | 1.56E-01 | 1.01E+00 | 1.26E-01 | 4.82E-01 | 1.28E+00 | 6.43E-03 | 2.29E-02 | -1.08E+00 | 5.64E-03 | 1.59E-02 | -1.13E+00 | 1.32E-06 | 4.84E-06 | -1.36E+00 | 3.95E-03 | 1.09E-02 | -1.02E+00 |
| AC098818.2 | 1.83E-03 | 9.54E-03 | 2.15E+00 | 1.37E-01 | 5.94E-01 | 1.58E+00 | 4.52E-06 | 1.92E-05 | 3.05E+00 | 1.22E-01 | 4.66E-01 | 1.69E+00 | 1.30E-01 | 4.63E-01 | 1.55E+00 | 3.13E-01 | 8.84E-01 | 1.15E+00 | 1.49E-02 | 5.46E-02 | 2.08E+00 | 2.41E-02 | 6.62E-02 | 2.09E+00 |
| C3orf62 | 1.84E-03 | 9.58E-03 | -1.22E+00 | 2.48E-02 | 1.07E-01 | 1.12E+00 | 1.00E-05 | 4.25E-05 | -1.45E+00 | 7.77E-07 | 2.97E-06 | -1.42E+00 | 1.23E-03 | 4.39E-03 | -1.22E+00 | 8.92E-05 | 2.51E-04 | -1.46E+00 | 9.09E-08 | 3.34E-07 | -1.62E+00 | 7.87E-09 | 2.16E-08 | -1.73E+00 |
| MAST4 | 1.84E-03 | 9.61E-03 | 1.11E+00 | 9.27E-03 | 4.01E-02 | 1.28E+00 | 3.39E-03 | 1.44E-02 | -1.02E+00 | 1.36E-03 | 5.19E-03 | 1.09E+00 | 2.14E-01 | 7.64E-01 | 1.14E+00 | 1.10E-01 | 3.10E-01 | 1.45E+00 | 2.13E-02 | 7.83E-02 | 1.04E+00 | 4.49E-02 | 1.23E-01 | 1.02E+00 |
| ANGEL1 | 1.85E-03 | 9.65E-03 | -1.50E+00 | 1.14E-02 | 4.95E-02 | -1.15E+00 | 4.32E-06 | 1.83E-05 | -1.74E+00 | 8.36E-05 | 3.19E-04 | -1.28E+00 | 2.93E-03 | 1.05E-02 | -1.56E+00 | 1.20E-01 | 3.39E-01 | 1.10E+00 | 9.79E-03 | 3.59E-02 | -1.26E+00 | 1.87E-01 | 5.15E-01 | 1.46E+00 |
| BCLAF1P1 | 1.85E-03 | 9.65E-03 | -1.20E+00 | 6.34E-04 | 2.74E-03 | -1.12E+00 | 1.97E-02 | 8.37E-02 | -1.19E+00 | 1.54E-02 | 5.88E-02 | -1.07E+00 | 2.49E-02 | 8.87E-02 | -1.04E+00 | 6.16E-02 | 1.74E-01 | -1.01E+00 | 2.25E-03 | 8.27E-03 | -1.21E+00 | 1.75E-03 | 4.81E-03 | -1.31E+00 |
| ZFYVE21 | 1.85E-03 | 9.65E-03 | -1.22E+00 | 4.54E-02 | 1.96E-01 | 1.19E+00 | 1.22E-02 | 5.18E-02 | -1.33E+00 | 1.63E-01 | 6.23E-01 | 1.26E+00 | 4.99E-07 | 1.78E-06 | -1.95E+00 | 1.31E-03 | 3.70E-03 | -1.06E+00 | 1.56E-07 | 5.72E-07 | -1.07E+00 | 1.01E-03 | 2.78E-03 | -1.01E+00 |
| RPL31P7 | 1.85E-03 | 9.66E-03 | 1.06E+00 | 3.23E-19 | 1.40E-18 | -1.46E+00 | 1.01E-01 | 4.28E-01 | 1.15E+00 | 1.58E-09 | 6.04E-09 | -1.28E+00 | 1.43E-01 | 5.12E-01 | 1.28E+00 | 2.55E-07 | 7.18E-07 | -1.38E+00 | 5.99E-02 | 2.20E-01 | 1.12E+00 | 3.13E-06 | 8.61E-06 | -1.34E+00 |
| TRPS1 | 1.86E-03 | 9.69E-03 | 1.04E+00 | 3.59E-02 | 1.55E-01 | 1.74E+00 | 2.06E-01 | 8.77E-01 | 1.17E+00 | 2.56E-03 | 9.79E-03 | 1.86E+00 | 1.13E-02 | 4.03E-02 | 2.18E+00 | 3.15E-04 | 8.89E-04 | 3.04E+00 | 1.98E-01 | 7.26E-01 | 1.19E+00 | 2.68E-01 | 7.37E-01 | 1.39E+00 |
| TMCC1 | 1.86E-03 | 9.71E-03 | -1.24E+00 | 2.22E-03 | 9.57E-03 | -1.13E+00 | 5.02E-05 | 2.13E-04 | -1.51E+00 | 9.41E-04 | 3.59E-03 | -1.17E+00 | 8.04E-05 | 2.87E-04 | -1.50E+00 | 3.92E-04 | 1.10E-03 | -1.27E+00 | 8.29E-03 | 3.04E-02 | -1.21E+00 | 6.99E-02 | 1.92E-01 | -1.04E+00 |
| VSIG10L | 1.86E-03 | 9.72E-03 | -1.43E+00 | 4.15E-04 | 1.79E-03 | -1.62E+00 | 9.85E-02 | 4.18E-01 | -1.10E+00 | 1.71E-01 | 6.55E-01 | 1.10E+00 | 4.75E-02 | 1.69E-01 | -1.07E+00 | 9.24E-03 | 2.60E-02 | -1.39E+00 | 1.92E-02 | 7.03E-02 | -1.65E+00 | 1.15E-01 | 3.17E-01 | -1.22E+00 |
| KCTD18 | 1.87E-03 | 9.72E-03 | -1.06E+00 | 1.82E-02 | 7.88E-02 | 1.18E+00 | 5.46E-05 | 2.32E-04 | -1.24E+00 | 1.09E-03 | 4.16E-03 | 1.02E+00 | 6.20E-04 | 2.21E-03 | -1.11E+00 | 1.22E-02 | 3.44E-02 | 1.06E+00 | 8.86E-03 | 3.25E-02 | 1.00E+00 | 2.11E-01 | 5.80E-01 | 1.21E+00 |
| INTS12 | 1.87E-03 | 9.73E-03 | -1.02E+00 | 9.23E-04 | 3.99E-03 | 1.03E+00 | 4.28E-03 | 1.82E-02 | -1.15E+00 | 2.85E-03 | 1.09E-02 | -1.04E+00 | 1.59E-02 | 5.66E-02 | 1.14E+00 | 3.42E-02 | 9.64E-02 | 1.09E+00 | 5.94E-05 | 2.18E-04 | -1.10E+00 | 1.98E-06 | 5.43E-06 | -1.31E+00 |
| ALOX15B | 1.87E-03 | 9.75E-03 | -1.18E+00 | 2.46E-02 | 1.06E-01 | 1.10E+00 | 8.04E-02 | 3.42E-01 | 1.05E+00 | 4.78E-02 | 1.82E-01 | 1.06E+00 | 1.88E-01 | 6.71E-01 | -1.55E+00 | 2.90E-01 | 8.18E-01 | -1.09E+00 | 2.47E-01 | 9.08E-01 | 1.16E+00 | 3.49E-01 | 9.59E-01 | -1.00E+00 |
| YIPF2 | 1.87E-03 | 9.76E-03 | -1.26E+00 | 1.75E-02 | 7.55E-02 | 1.13E+00 | 3.04E-04 | 1.29E-03 | -1.37E+00 | 3.38E-03 | 1.29E-02 | -1.06E+00 | 2.22E-03 | 7.90E-03 | -1.62E+00 | 9.00E-02 | 2.54E-01 | -1.06E+00 | 1.67E-01 | 6.13E-01 | 1.10E+00 | 2.97E-01 | 8.15E-01 | 1.29E+00 |
| CUBN | 1.88E-03 | 9.78E-03 | -1.41E+00 | 1.81E-02 | 7.81E-02 | 1.02E+00 | 2.00E-01 | 8.49E-01 | 1.10E+00 | 1.57E-01 | 6.00E-01 | 1.46E+00 | 1.50E-01 | 5.34E-01 | 1.03E+00 | 2.81E-01 | 7.92E-01 | 1.15E+00 | 1.70E-01 | 6.25E-01 | 1.30E+00 | 1.99E-01 | 5.48E-01 | 1.43E+00 |
| FKBP15 | 1.88E-03 | 9.82E-03 | -1.46E+00 | 2.19E-01 | 9.44E-01 | 1.44E+00 | 5.24E-04 | 2.23E-03 | -1.46E+00 | 2.49E-02 | 9.52E-02 | 1.02E+00 | 1.79E-01 | 6.38E-01 | 1.05E+00 | 1.74E-01 | 4.91E-01 | 1.55E+00 | 5.29E-03 | 1.94E-02 | -1.17E+00 | 2.93E-01 | 8.04E-01 | 1.19E+00 |
| AC093525.4 | 1.89E-03 | 9.87E-03 | -1.45E+00 | 6.82E-03 | 2.95E-02 | -1.22E+00 | 7.99E-03 | 3.39E-02 | -1.35E+00 | 2.78E-02 | 1.06E-01 | -1.06E+00 | 1.33E-02 | 4.75E-02 | -1.52E+00 | 1.03E-01 | 2.91E-01 | 1.04E+00 | 2.36E-02 | 8.68E-02 | -1.23E+00 | 1.72E-01 | 4.73E-01 | 1.01E+00 |
| SLC26A2 | 1.89E-03 | 9.87E-03 | -1.15E+00 | 3.92E-03 | 1.69E-02 | -1.03E+00 | 4.28E-04 | 1.82E-03 | -1.35E+00 | 4.52E-02 | 1.73E-01 | 1.18E+00 | 1.47E-03 | 5.24E-03 | 2.44E+00 | 3.37E-03 | 9.49E-03 | 2.15E+00 | 2.04E-02 | 7.47E-02 | 1.03E+00 | 7.63E-02 | 2.10E-01 | 1.45E+00 |
| TXNDC16 | 1.91E-03 | 9.94E-03 | -1.27E+00 | 7.37E-03 | 3.19E-02 | -1.03E+00 | 3.79E-02 | 1.61E-01 | -1.06E+00 | 1.36E-02 | 5.20E-02 | -1.07E+00 | 1.39E-02 | 4.96E-02 | -1.25E+00 | 2.71E-02 | 7.64E-02 | -1.11E+00 | 1.73E-02 | 6.36E-02 | -1.07E+00 | 8.79E-03 | 2.42E-02 | -1.18E+00 |
| NRAV | 1.91E-03 | 9.97E-03 | -1.59E+00 | 1.39E-02 | 6.01E-02 | -1.16E+00 | 7.98E-08 | 3.39E-07 | -2.05E+00 | 1.12E-05 | 4.28E-05 | -1.55E+00 | 1.17E-02 | 4.19E-02 | -1.77E+00 | 3.33E-02 | 9.40E-02 | -1.42E+00 | 2.20E-02 | 8.07E-02 | -1.32E+00 | 4.85E-02 | 1.33E-01 | -1.24E+00 |
| C1QTNF3 | 1.92E-03 | 9.99E-03 | -1.45E+00 | 9.55E-02 | 4.13E-01 | 1.26E+00 | 2.17E-01 | 9.22E-01 | 1.20E+00 | 2.39E-01 | 9.13E-01 | 1.28E+00 | 6.06E-02 | 2.16E-01 | -1.19E+00 | 4.47E-02 | 1.26E-01 | -1.20E+00 | 9.50E-03 | 3.49E-02 | -1.40E+00 | 3.27E-01 | 8.98E-01 | 1.27E+00 |
| ZBTB48 | 1.92E-03 | 9.99E-03 | -1.08E+00 | 8.22E-02 | 3.55E-01 | 1.26E+00 | 2.52E-04 | 1.07E-03 | -1.23E+00 | 8.19E-03 | 3.13E-02 | 1.10E+00 | 1.01E-05 | 3.60E-05 | -1.40E+00 | 2.01E-03 | 5.68E-03 | -1.03E+00 | 3.19E-04 | 1.17E-03 | -1.34E+00 | 2.60E-01 | 7.14E-01 | 1.14E+00 |
| AC007191.1 | 1.92E-03 | 1.00E-02 | -1.26E+00 | 1.67E-02 | 7.20E-02 | 1.14E+00 | 1.93E-01 | 8.20E-01 | -1.02E+00 | 2.51E-01 | 9.58E-01 | 1.16E+00 | 3.61E-02 | 1.29E-01 | -1.42E+00 | 1.84E-01 | 5.19E-01 | -1.01E+00 | 9.76E-03 | 3.58E-02 | -1.29E+00 | 3.98E-02 | 1.09E-01 | -1.10E+00 |
| AMMECR1L | 1.92E-03 | 1.00E-02 | -1.08E+00 | 1.08E-04 | 4.66E-04 | -1.20E+00 | 2.32E-03 | 9.84E-03 | -1.21E+00 | 3.27E-05 | 1.25E-04 | -1.37E+00 | 2.66E-02 | 9.49E-02 | -1.08E+00 | 6.55E-04 | 1.85E-03 | -1.64E+00 | 2.50E-02 | 9.18E-02 | -1.07E+00 | 1.77E-02 | 4.87E-02 | -1.20E+00 |
| HPDL | 1.92E-03 | 1.00E-02 | -1.55E+00 | 6.56E-02 | 2.84E-01 | 1.13E+00 | 2.69E-04 | 1.14E-03 | -1.41E+00 | 5.01E-02 | 1.92E-01 | 1.55E+00 | 6.21E-03 | 2.22E-02 | -2.12E+00 | 3.05E-01 | 8.59E-01 | 1.29E+00 | 7.22E-06 | 2.65E-05 | -1.35E+00 | 1.13E-01 | 3.10E-01 | 1.53E+00 |
| SYNJ1 | 1.92E-03 | 1.00E-02 | -1.25E+00 | 2.47E-04 | 1.07E-03 | -1.45E+00 | 1.39E-02 | 5.93E-02 | -1.19E+00 | 7.44E-03 | 2.84E-02 | -1.21E+00 | 5.20E-02 | 1.85E-01 | -1.04E+00 | 3.15E-02 | 8.88E-02 | -1.05E+00 | 1.29E-05 | 4.74E-05 | -1.43E+00 | 3.06E-05 | 8.39E-05 | -1.57E+00 |
| ZNF550 | 1.92E-03 | 1.00E-02 | -1.15E+00 | 1.28E-02 | 5.55E-02 | 1.12E+00 | 7.79E-06 | 3.31E-05 | -1.34E+00 | 2.09E-04 | 7.98E-04 | -1.05E+00 | 2.31E-03 | 8.24E-03 | -1.26E+00 | 1.81E-02 | 5.09E-02 | -1.02E+00 | 1.41E-05 | 5.16E-05 | -1.50E+00 | 6.82E-02 | 1.87E-01 | 1.05E+00 |
| CPEB4 | 1.92E-03 | 1.00E-02 | -1.06E+00 | 6.91E-04 | 2.99E-03 | -1.02E+00 | 1.12E-03 | 4.75E-03 | -1.26E+00 | 1.49E-04 | 5.71E-04 | -1.20E+00 | 2.47E-01 | 8.81E-01 | 1.56E+00 | 1.34E-01 | 3.78E-01 | 1.14E+00 | 1.04E-06 | 3.80E-06 | 1.09E+00 | 1.40E-09 | 3.86E-09 | -1.19E+00 |
| SNRNP70 | 1.93E-03 | 1.00E-02 | 1.05E+00 | 6.93E-02 | 2.99E-01 | 1.27E+00 | 2.27E-01 | 9.64E-01 | 1.15E+00 | 2.35E-01 | 8.96E-01 | 1.27E+00 | 7.88E-02 | 2.81E-01 | 1.18E+00 | 1.70E-01 | 4.79E-01 | 1.43E+00 | 1.71E-01 | 6.28E-01 | 1.09E+00 | 6.93E-02 | 1.90E-01 | 1.32E+00 |
| PCSK1N | 1.93E-03 | 1.01E-02 | 2.42E+00 | 3.51E-04 | 1.52E-03 | 2.90E+00 | 8.82E-02 | 3.75E-01 | 1.91E+00 | 4.37E-02 | 1.67E-01 | 2.15E+00 | 2.43E-01 | 8.68E-01 | 1.13E+00 | 3.22E-01 | 9.07E-01 | 1.19E+00 | 1.39E-02 | 5.11E-02 | 2.70E+00 | 1.67E-02 | 4.58E-02 | 2.62E+00 |
| RNF121 | 1.93E-03 | 1.01E-02 | -1.21E+00 | 5.38E-03 | 2.32E-02 | -1.05E+00 | 1.36E-02 | 5.79E-02 | -1.21E+00 | 6.23E-02 | 2.38E-01 | 1.13E+00 | 9.94E-05 | 3.54E-04 | -1.46E+00 | 1.70E-02 | 4.79E-02 | 1.02E+00 | 1.72E-05 | 6.33E-05 | -1.31E+00 | 1.75E-03 | 4.80E-03 | -1.09E+00 |
| ADAM19 | 1.93E-03 | 1.01E-02 | -1.62E+00 | 1.46E-02 | 6.29E-02 | -1.21E+00 | 4.75E-04 | 2.02E-03 | -2.26E+00 | 4.82E-03 | 1.84E-02 | -1.69E+00 | 2.49E-01 | 8.88E-01 | 1.13E+00 | 1.72E-01 | 4.85E-01 | 1.69E+00 | 1.98E-01 | 7.27E-01 | -1.18E+00 | 3.63E-01 | 9.97E-01 | 1.08E+00 |
| STK39 | 1.93E-03 | 1.01E-02 | -1.37E+00 | 1.52E-02 | 6.58E-02 | -1.05E+00 | 7.05E-05 | 2.99E-04 | -1.31E+00 | 4.87E-04 | 1.86E-03 | -1.07E+00 | 3.77E-02 | 1.34E-01 | -1.16E+00 | 2.03E-01 | 5.73E-01 | 1.16E+00 | 1.28E-03 | 4.69E-03 | -1.32E+00 | 2.07E-01 | 5.69E-01 | 1.07E+00 |
| AGPAT3 | 1.93E-03 | 1.01E-02 | -1.28E+00 | 2.11E-03 | 9.10E-03 | -1.12E+00 | 1.05E-01 | 4.45E-01 | 1.04E+00 | 9.84E-02 | 3.76E-01 | 1.21E+00 | 8.38E-02 | 2.99E-01 | 1.17E+00 | 2.48E-01 | 6.98E-01 | 1.40E+00 | 1.25E-04 | 4.58E-04 | -1.19E+00 | 4.85E-02 | 1.33E-01 | 1.10E+00 |
| PLEKHM3 | 1.93E-03 | 1.01E-02 | -1.16E+00 | 3.51E-02 | 1.52E-01 | 1.18E+00 | 1.31E-04 | 5.58E-04 | -1.49E+00 | 1.59E-03 | 6.06E-03 | -1.10E+00 | 3.70E-03 | 1.32E-02 | -1.25E+00 | 5.32E-02 | 1.50E-01 | -1.01E+00 | 1.75E-09 | 6.43E-09 | -2.04E+00 | 4.77E-05 | 1.31E-04 | -1.36E+00 |
| AC104809.2 | 1.93E-03 | 1.01E-02 | -1.47E+00 | 6.83E-02 | 2.95E-01 | 1.06E+00 | 8.68E-03 | 3.69E-02 | -1.40E+00 | 2.00E-01 | 7.64E-01 | 1.28E+00 | 1.65E-01 | 5.87E-01 | 1.74E+00 | 3.47E-01 | 9.78E-01 | 1.19E+00 | 2.23E-01 | 8.17E-01 | 1.25E+00 | 1.44E-01 | 3.96E-01 | 1.57E+00 |
| TAF9B | 1.94E-03 | 1.01E-02 | -1.07E+00 | 2.36E-05 | 1.02E-04 | -1.14E+00 | 2.12E-03 | 9.00E-03 | -1.13E+00 | 1.62E-06 | 6.19E-06 | -1.21E+00 | 4.11E-03 | 1.46E-02 | -1.14E+00 | 5.30E-04 | 1.49E-03 | -1.27E+00 | 1.39E-02 | 5.09E-02 | -1.07E+00 | 1.05E-02 | 2.87E-02 | -1.07E+00 |
| KIAA0355 | 1.94E-03 | 1.01E-02 | -1.49E+00 | 2.23E-02 | 9.64E-02 | 1.01E+00 | 5.52E-04 | 2.35E-03 | -1.72E+00 | 2.24E-02 | 8.56E-02 | -1.10E+00 | 1.17E-02 | 4.18E-02 | -1.32E+00 | 1.09E-01 | 3.06E-01 | 1.16E+00 | 1.61E-03 | 5.90E-03 | -1.39E+00 | 3.11E-02 | 8.53E-02 | -1.11E+00 |
| COX19 | 1.94E-03 | 1.01E-02 | 1.02E+00 | 3.82E-09 | 1.65E-08 | -1.11E+00 | 4.28E-02 | 1.82E-01 | -1.02E+00 | 1.47E-02 | 5.60E-02 | 1.10E+00 | 5.68E-02 | 2.03E-01 | 1.14E+00 | 1.57E-03 | 4.43E-03 | -1.01E+00 | 1.66E-05 | 6.11E-05 | -1.28E+00 | 2.33E-05 | 6.41E-05 | -1.20E+00 |
| MTG1 | 1.94E-03 | 1.01E-02 | -1.38E+00 | 1.60E-01 | 6.91E-01 | 1.30E+00 | 2.20E-06 | 9.33E-06 | -1.71E+00 | 8.52E-04 | 3.26E-03 | -1.13E+00 | 1.77E-03 | 6.32E-03 | -1.62E+00 | 1.01E-01 | 2.84E-01 | -1.02E+00 | 2.16E-02 | 7.92E-02 | -1.12E+00 | 3.71E-02 | 1.02E-01 | -1.01E+00 |
| SERPINB8 | 1.94E-03 | 1.01E-02 | -2.20E+00 | 2.27E-03 | 9.81E-03 | -2.14E+00 | 2.07E-02 | 8.82E-02 | -1.05E+00 | 2.76E-09 | 1.05E-08 | -1.68E+00 | 1.06E-01 | 3.79E-01 | -1.44E+00 | 4.91E-02 | 1.38E-01 | -2.17E+00 | 1.94E-01 | 7.11E-01 | -1.26E+00 | 2.03E-01 | 5.58E-01 | -1.45E+00 |
| ATP9B | 1.94E-03 | 1.01E-02 | -1.13E+00 | 6.00E-03 | 2.59E-02 | 1.04E+00 | 4.57E-03 | 1.94E-02 | -1.27E+00 | 9.71E-03 | 3.71E-02 | -1.07E+00 | 1.07E-02 | 3.83E-02 | -1.05E+00 | 1.48E-01 | 4.18E-01 | 1.24E+00 | 1.02E-02 | 3.75E-02 | 1.03E+00 | 1.58E-02 | 4.33E-02 | -1.00E+00 |
| AC007541.1 | 1.94E-03 | 1.01E-02 | -1.29E+00 | 3.73E-02 | 1.61E-01 | 1.06E+00 | 5.01E-03 | 2.13E-02 | -1.64E+00 | 1.37E-01 | 5.23E-01 | 1.16E+00 | 1.53E-04 | 5.45E-04 | -1.68E+00 | 2.06E-03 | 5.82E-03 | -1.35E+00 | 1.60E-02 | 5.86E-02 | -1.05E+00 | 2.94E-01 | 8.08E-01 | 1.29E+00 |
| ZNF551 | 1.94E-03 | 1.01E-02 | -1.20E+00 | 6.09E-03 | 2.63E-02 | 1.02E+00 | 1.09E-03 | 4.61E-03 | -1.16E+00 | 3.66E-03 | 1.40E-02 | 1.06E+00 | 3.53E-02 | 1.26E-01 | 1.01E+00 | 2.44E-02 | 6.87E-02 | -1.03E+00 | 7.05E-05 | 2.59E-04 | -1.41E+00 | 6.49E-03 | 1.78E-02 | -1.03E+00 |
| CH17-340M24.3 | 1.95E-03 | 1.01E-02 | -1.09E+00 | 9.99E-05 | 4.32E-04 | -1.20E+00 | 1.72E-01 | 7.29E-01 | 1.11E+00 | 1.53E-02 | 5.85E-02 | -1.09E+00 | 1.54E-02 | 5.49E-02 | -1.01E+00 | 1.38E-03 | 3.89E-03 | -1.21E+00 | 2.29E-02 | 8.40E-02 | -1.06E+00 | 9.99E-03 | 2.74E-02 | -1.26E+00 |
| GPR155 | 1.95E-03 | 1.02E-02 | -1.47E+00 | 8.45E-02 | 3.65E-01 | 1.34E+00 | 4.21E-05 | 1.79E-04 | -2.09E+00 | 4.54E-03 | 1.73E-02 | -1.11E+00 | 2.07E-01 | 7.40E-01 | 1.02E+00 | 7.51E-02 | 2.12E-01 | 1.88E+00 | 2.61E-05 | 9.58E-05 | -1.35E+00 | 1.10E-02 | 3.02E-02 | 1.01E+00 |
| BCAS2P2 | 1.96E-03 | 1.02E-02 | -1.23E+00 | 5.08E-05 | 2.20E-04 | -1.60E+00 | 5.13E-03 | 2.18E-02 | -1.24E+00 | 1.14E-06 | 4.34E-06 | -2.30E+00 | 9.73E-03 | 3.47E-02 | -1.50E+00 | 1.45E-02 | 4.09E-02 | -1.36E+00 | 3.08E-02 | 1.13E-01 | -1.08E+00 | 1.44E-05 | 3.97E-05 | -1.91E+00 |
| RXFP1 | 1.96E-03 | 1.02E-02 | -1.81E+00 | 2.31E-01 | 9.98E-01 | 1.45E+00 | 3.23E-02 | 1.37E-01 | -1.47E+00 | 2.40E-01 | 9.18E-01 | 1.38E+00 | 2.57E-04 | 9.17E-04 | -3.14E+00 | 3.20E-01 | 9.02E-01 | 1.09E+00 | 3.15E-02 | 1.16E-01 | -1.37E+00 | 2.56E-01 | 7.04E-01 | 1.13E+00 |
| UFM1 | 1.96E-03 | 1.02E-02 | 1.07E+00 | 7.69E-04 | 3.32E-03 | 1.12E+00 | 2.89E-02 | 1.23E-01 | 1.02E+00 | 2.24E-04 | 8.57E-04 | 1.02E+00 | 3.72E-05 | 1.33E-04 | -1.11E+00 | 3.05E-03 | 8.59E-03 | -1.03E+00 | 2.77E-06 | 1.02E-05 | -1.15E+00 | 4.24E-03 | 1.16E-02 | -1.03E+00 |
| CCDC127 | 1.96E-03 | 1.02E-02 | -1.24E+00 | 1.22E-01 | 5.25E-01 | 1.31E+00 | 2.35E-03 | 9.96E-03 | -1.31E+00 | 1.09E-01 | 4.15E-01 | 1.19E+00 | 6.66E-04 | 2.37E-03 | -1.36E+00 | 1.93E-02 | 5.45E-02 | -1.00E+00 | 3.56E-10 | 1.31E-09 | -1.95E+00 | 7.83E-03 | 2.15E-02 | -1.07E+00 |
| NUDT6 | 1.96E-03 | 1.02E-02 | -1.52E+00 | 1.05E-01 | 4.54E-01 | 1.17E+00 | 2.28E-03 | 9.67E-03 | -1.50E+00 | 1.89E-03 | 7.21E-03 | -1.37E+00 | 4.25E-03 | 1.51E-02 | -1.65E+00 | 3.14E-01 | 8.86E-01 | 1.30E+00 | 2.22E-04 | 8.13E-04 | -1.31E+00 | 2.68E-03 | 7.36E-03 | -1.20E+00 |
| LINC01347 | 1.97E-03 | 1.03E-02 | -1.08E+00 | 3.65E-07 | 1.58E-06 | -1.76E+00 | 5.51E-02 | 2.34E-01 | -1.08E+00 | 1.75E-03 | 6.69E-03 | -1.49E+00 | 2.70E-01 | 9.63E-01 | 1.41E+00 | 1.12E-01 | 3.15E-01 | -1.00E+00 | 1.76E-01 | 6.45E-01 | 1.58E+00 | 4.78E-02 | 1.31E-01 | -1.15E+00 |
| RHOT2 | 1.97E-03 | 1.03E-02 | 1.02E+00 | 5.62E-03 | 2.43E-02 | 1.19E+00 | 4.25E-02 | 1.81E-01 | 1.03E+00 | 3.12E-02 | 1.19E-01 | 1.17E+00 | 5.87E-04 | 2.09E-03 | -1.14E+00 | 3.42E-02 | 9.65E-02 | 1.10E+00 | 1.58E-01 | 5.78E-01 | 1.21E+00 | 3.07E-01 | 8.44E-01 | 1.29E+00 |
| KDM2B | 1.97E-03 | 1.03E-02 | -1.12E+00 | 4.71E-02 | 2.03E-01 | 1.27E+00 | 3.88E-03 | 1.65E-02 | -1.14E+00 | 1.69E-02 | 6.44E-02 | 1.03E+00 | 2.29E-02 | 8.15E-02 | -1.15E+00 | 1.47E-01 | 4.15E-01 | 1.09E+00 | 1.21E-04 | 4.45E-04 | -1.21E+00 | 9.57E-02 | 2.63E-01 | 1.17E+00 |
| AC093726.1 | 1.97E-03 | 1.03E-02 | -1.45E+00 | 4.58E-02 | 1.98E-01 | 1.15E+00 | 1.09E-02 | 4.65E-02 | -1.43E+00 | 1.46E-01 | 5.59E-01 | 1.22E+00 | 1.76E-01 | 6.26E-01 | 1.01E+00 | 1.58E-01 | 4.45E-01 | 1.67E+00 | 1.66E-03 | 6.08E-03 | -1.69E+00 | 1.88E-02 | 5.15E-02 | -1.42E+00 |
| AC091167.5 | 1.98E-03 | 1.03E-02 | -1.53E+00 | 1.88E-02 | 8.13E-02 | -1.09E+00 | 1.26E-02 | 5.36E-02 | -1.26E+00 | 4.89E-03 | 1.87E-02 | -1.27E+00 | 2.44E-03 | 8.70E-03 | -1.61E+00 | 3.61E-03 | 1.02E-02 | -1.55E+00 | 3.09E-06 | 1.14E-05 | -1.77E+00 | 9.31E-05 | 2.56E-04 | -1.62E+00 |
| R3HCC1 | 1.98E-03 | 1.03E-02 | 1.09E+00 | 7.05E-05 | 3.05E-04 | 1.13E+00 | 5.75E-02 | 2.44E-01 | 1.06E+00 | 4.81E-03 | 1.84E-02 | 1.09E+00 | 1.77E-01 | 6.31E-01 | 1.31E+00 | 3.49E-01 | 9.85E-01 | 1.44E+00 | 2.51E-01 | 9.22E-01 | 1.23E+00 | 1.23E-01 | 3.38E-01 | 1.15E+00 |
| DNAJC9-AS1 | 1.98E-03 | 1.03E-02 | -1.41E+00 | 2.27E-01 | 9.79E-01 | 1.58E+00 | 2.10E-04 | 8.92E-04 | -1.68E+00 | 1.17E-01 | 4.46E-01 | 1.26E+00 | 6.88E-03 | 2.45E-02 | -1.60E+00 | 2.57E-01 | 7.26E-01 | 1.25E+00 | 3.84E-04 | 1.41E-03 | -1.47E+00 | 3.46E-01 | 9.51E-01 | 1.34E+00 |
| PKN3 | 1.98E-03 | 1.03E-02 | -1.12E+00 | 1.78E-02 | 7.67E-02 | 1.16E+00 | 1.94E-02 | 8.24E-02 | -1.05E+00 | 9.00E-02 | 3.44E-01 | 1.22E+00 | 1.00E-01 | 3.57E-01 | -1.05E+00 | 3.54E-01 | 9.97E-01 | 1.53E+00 | 9.80E-02 | 3.60E-01 | 1.07E+00 | 3.14E-01 | 8.64E-01 | 1.31E+00 |
| EHD1 | 1.98E-03 | 1.03E-02 | -1.23E+00 | 6.08E-02 | 2.63E-01 | 1.16E+00 | 6.41E-02 | 2.72E-01 | 1.05E+00 | 1.03E-01 | 3.94E-01 | 1.25E+00 | 5.34E-05 | 1.90E-04 | -1.49E+00 | 3.39E-03 | 9.54E-03 | -1.16E+00 | 4.95E-06 | 1.82E-05 | -1.05E+00 | 3.23E-05 | 8.88E-05 | -1.10E+00 |
| AL354872.1 | 1.99E-03 | 1.03E-02 | -1.53E+00 | 1.85E-03 | 7.99E-03 | -1.55E+00 | 1.08E-04 | 4.58E-04 | -1.43E+00 | 1.67E-06 | 6.38E-06 | -1.77E+00 | 6.21E-05 | 2.21E-04 | -1.82E+00 | 4.40E-06 | 1.24E-05 | -2.45E+00 | 1.85E-14 | 6.78E-14 | -2.12E+00 | 4.79E-13 | 1.31E-12 | -2.06E+00 |
| [illegible] | [illegible] | [illegible] | [illegible] | [illegible] | [illegible] | [illegible] | [illegible] | [illegible] | [illegible] | [illegible] | [illegible] | [illegible] | [illegible] | [illegible] | [illegible] | [illegible] | [illegible] | [illegible] | [illegible] | [illegible] | [illegible] | [illegible] | [illegible] | [illegible] |
| [illegible] | [illegible] | [illegible] | [illegible] | [illegible] | [illegible] | [illegible] | [illegible] | [illegible] | [illegible] | [illegible] | [illegible] | [illegible] | [illegible] | [illegible] | [illegible] | [illegible] | [illegible] | [illegible] | [illegible] | [illegible] | [illegible] | [illegible] | [illegible] | [illegible] |
| COG5 | 2.00E-03 | 1.04E-02 | -1.08E+00 | 9.40E-03 | 4.06E-02 | 1.14E+00 | 5.20E-06 | 2.21E-05 | -1.28E+00 | 8.67E-04 | 3.31E-03 | 1.04E+00 | 3.88E-04 | 1.38E-03 | -1.16E+00 | 2.81E-01 | 7.93E-01 | 1.01E+00 | 9.34E-04 | 3.43E-03 | -1.14E+00 | 6.18E-02 | 1.70E-01 | 1.08E+00 |
| AC104619.3 | 2.00E-03 | 1.04E-02 | -1.23E+00 | 2.17E-03 | 9.38E-03 | -1.04E+00 | 3.70E-02 | 1.57E-01 | -1.15E+00 | 4.52E-02 | 1.73E-01 | 1.02E+00 | 4.90E-03 | 1.75E-02 | -1.27E+00 | 7.17E-04 | 2.02E-03 | -1.52E+00 | 1.78E-02 | 6.55E-02 | -1.25E+00 | 3.35E-02 | 9.21E-02 | -1.24E+00 |
| CSTF3 | 2.01E-03 | 1.05E-02 | 1.03E+00 | 1.25E-03 | 5.40E-03 | 1.19E+00 | 2.75E-03 | 1.17E-02 | -1.08E+00 | 1.12E-03 | 4.26E-03 | 1.06E+00 | 8.51E-04 | 3.03E-03 | -1.07E+00 | 7.81E-03 | 2.20E-02 | 1.07E+00 | 4.77E-07 | 1.75E-06 | -1.22E+00 | 2.75E-02 | 7.56E-02 | 1.08E+00 |
| ZSCAN9 | 2.01E-03 | 1.05E-02 | -1.12E+00 | 7.16E-03 | 3.10E-02 | 1.11E+00 | 3.99E-05 | 1.70E-04 | -1.40E+00 | 1.23E-03 | 4.69E-03 | -1.06E+00 | 3.86E-05 | 1.38E-04 | -1.54E+00 | 1.40E-04 | 3.95E-04 | -1.37E+00 | 2.08E-07 | 7.62E-07 | -1.59E+00 | 3.40E-05 | 9.35E-05 | -1.31E+00 |
| FAM49A | 2.01E-03 | 1.05E-02 | -1.10E+00 | 4.49E-03 | 1.94E-02 | 1.00E+00 | 2.95E-05 | 1.25E-04 | -1.34E+00 | 2.99E-05 | 1.14E-04 | -1.17E+00 | 2.58E-02 | 9.20E-02 | -1.07E+00 | 3.89E-02 | 1.10E-01 | 1.04E+00 | 9.81E-03 | 3.60E-02 | -1.89E+00 | 7.74E-02 | 2.13E-01 | -1.18E+00 |
| MINCR | 2.02E-03 | 1.05E-02 | -1.10E+00 | 5.48E-03 | 2.37E-02 | 1.08E+00 | 1.37E-01 | 5.82E-01 | 1.01E+00 | 1.22E-03 | 4.67E-03 | -1.27E+00 | 9.37E-02 | 3.34E-01 | 1.77E+00 | 3.22E-01 | 9.08E-01 | 1.32E+00 | 2.23E-02 | 8.17E-02 | 1.62E+00 | 7.02E-02 | 1.93E-01 | 1.04E+00 |
| ADSL | 2.02E-03 | 1.05E-02 | -1.17E+00 | 4.45E-03 | 1.92E-02 | -1.03E+00 | 1.15E-03 | 4.87E-03 | -1.30E+00 | 3.51E-03 | 1.34E-02 | -1.03E+00 | 1.87E-05 | 6.68E-05 | -1.59E+00 | 1.44E-04 | 4.06E-04 | -1.33E+00 | 2.80E-06 | 1.03E-05 | -1.61E+00 | 1.60E-03 | 4.39E-03 | -1.33E+00 |
| AL356966.1 | 2.02E-03 | 1.05E-02 | -1.49E+00 | 3.02E-03 | 1.30E-02 | -1.39E+00 | 1.77E-01 | 7.52E-01 | -1.12E+00 | 2.34E-01 | 8.94E-01 | -1.02E+00 | 1.12E-02 | 3.98E-02 | -1.66E+00 | 2.13E-02 | 6.01E-02 | -1.54E+00 | 2.70E-01 | 9.92E-01 | 1.14E+00 | 2.67E-01 | 7.32E-01 | 1.17E+00 |
| SLC10A3 | 2.03E-03 | 1.06E-02 | -1.33E+00 | 6.32E-02 | 2.73E-01 | 1.18E+00 | 7.68E-04 | 3.26E-03 | -1.38E+00 | 1.91E-02 | 7.30E-02 | 1.08E+00 | 1.21E-05 | 4.31E-05 | -1.37E+00 | 3.86E-05 | 1.09E-04 | -1.25E+00 | 3.58E-02 | 1.31E-01 | 1.15E+00 | 2.37E-01 | 6.51E-01 | 1.18E+00 |
| KLHDC8B | 2.03E-03 | 1.06E-02 | -1.06E+00 | 6.16E-02 | 2.66E-01 | 1.24E+00 | 4.30E-05 | 1.83E-04 | -1.47E+00 | 6.88E-04 | 2.63E-03 | -1.05E+00 | 6.47E-03 | 2.31E-02 | -1.11E+00 | 9.20E-02 | 2.59E-01 | 1.15E+00 | 2.29E-05 | 8.40E-05 | -1.10E+00 | 1.13E-03 | 3.11E-03 | 1.04E+00 |
| TLCD2 | 2.03E-03 | 1.06E-02 | -1.52E+00 | 7.35E-02 | 3.17E-01 | 1.01E+00 | 4.32E-03 | 1.83E-02 | -1.81E+00 | 1.87E-02 | 7.16E-02 | -1.29E+00 | 1.43E-01 | 5.11E-01 | 1.89E+00 | 2.10E-01 | 5.93E-01 | 1.73E+00 | 8.92E-02 | 3.27E-01 | -1.27E+00 | 3.25E-01 | 8.92E-01 | 1.23E+00 |
| NOD2 | 2.03E-03 | 1.06E-02 | -1.47E+00 | 3.41E-02 | 1.47E-01 | -1.03E+00 | 1.80E-09 | 7.63E-09 | -2.38E+00 | 2.06E-09 | 7.88E-09 | -2.41E+00 | 9.32E-02 | 3.32E-01 | -1.41E+00 | 8.73E-02 | 2.46E-01 | -1.64E+00 | 5.94E-02 | 2.18E-01 | -1.37E+00 | 1.70E-01 | 4.67E-01 | -1.07E+00 |
| C17orf75 | 2.03E-03 | 1.06E-02 | -1.11E+00 | 1.24E-02 | 5.35E-02 | 1.05E+00 | 2.71E-05 | 1.15E-04 | -1.38E+00 | 7.01E-05 | 2.68E-04 | -1.23E+00 | 2.40E-04 | 8.57E-04 | -1.70E+00 | 3.36E-02 | 9.48E-02 | -1.00E+00 | 5.27E-05 | 1.93E-04 | -1.32E+00 | 2.34E-04 | 6.43E-04 | -1.30E+00 |
| AL513534.1 | 2.03E-03 | 1.06E-02 | -1.54E+00 | 3.01E-02 | 1.30E-01 | -1.07E+00 | 9.46E-03 | 4.02E-02 | -1.42E+00 | 3.71E-02 | 1.42E-01 | -1.01E+00 | 7.23E-05 | 2.58E-04 | -2.02E+00 | 1.80E-04 | 5.07E-04 | -1.81E+00 | 3.67E-06 | 1.35E-05 | -2.05E+00 | 4.85E-05 | 1.33E-04 | -2.00E+00 |
| ZDHHC4P1 | 2.04E-03 | 1.06E-02 | -1.72E+00 | 6.84E-02 | 2.95E-01 | 1.07E+00 | 1.15E-04 | 4.90E-04 | -1.83E+00 | 5.31E-04 | 2.03E-03 | -1.55E+00 | 1.22E-04 | 4.34E-04 | -1.59E+00 | 7.00E-05 | 1.97E-04 | -2.07E+00 | 3.15E-03 | 1.15E-02 | -1.21E+00 | 5.19E-04 | 1.42E-03 | -1.55E+00 |
| PPM1B | 2.04E-03 | 1.06E-02 | 1.05E+00 | 3.15E-05 | 1.36E-04 | 1.07E+00 | 7.40E-07 | 3.14E-06 | -1.31E+00 | 6.90E-08 | 2.64E-07 | -1.20E+00 | 1.37E-04 | 4.89E-04 | -1.17E+00 | 5.85E-04 | 1.65E-03 | -1.09E+00 | 1.66E-09 | 6.08E-09 | -1.43E+00 | 4.88E-06 | 1.34E-05 | -1.24E+00 |
| AC025161.1 | 2.04E-03 | 1.06E-02 | -1.44E+00 | 3.72E-03 | 1.61E-02 | -1.23E+00 | 1.20E-02 | 5.11E-02 | -1.10E+00 | 1.20E-03 | 4.58E-03 | -1.03E+00 | 5.55E-07 | 1.98E-06 | -2.17E+00 | 1.33E-06 | 3.74E-06 | -2.04E+00 | 2.32E-13 | 8.53E-13 | -1.75E+00 | 5.42E-06 | 1.49E-05 | -1.25E+00 |
| MCM4 | 2.04E-03 | 1.06E-02 | -1.33E+00 | 7.74E-03 | 3.35E-02 | -1.14E+00 | 5.00E-02 | 2.12E-01 | 1.11E+00 | 1.16E-01 | 4.42E-01 | 1.40E+00 | 1.05E-02 | 3.74E-02 | -1.67E+00 | 4.93E-02 | 1.39E-01 | -1.30E+00 | 2.23E-04 | 8.20E-04 | -1.35E+00 | 6.49E-02 | 1.78E-01 | 1.16E+00 |
| GBA | 2.04E-03 | 1.06E-02 | -1.27E+00 | 1.04E-02 | 4.47E-02 | -1.01E+00 | 1.59E-05 | 6.77E-05 | -1.35E+00 | 6.54E-04 | 2.50E-03 | -1.08E+00 | 3.81E-03 | 1.36E-02 | -1.36E+00 | 2.00E-01 | 5.63E-01 | 1.23E+00 | 1.48E-02 | 5.43E-02 | 1.04E+00 | 2.85E-01 | 7.84E-01 | 1.28E+00 |
| SPATA24 | 2.04E-03 | 1.06E-02 | -1.21E+00 | 1.05E-02 | 4.53E-02 | -1.03E+00 | 9.43E-04 | 4.01E-03 | -1.50E+00 | 9.44E-05 | 3.61E-04 | -1.80E+00 | 1.77E-01 | 6.30E-01 | 1.06E+00 | 1.69E-01 | 4.76E-01 | -1.06E+00 | 1.08E-02 | 3.96E-02 | 1.88E+00 | 3.00E-01 | 8.23E-01 | 1.38E+00 |
| MTRF1 | 2.05E-03 | 1.07E-02 | -1.18E+00 | 6.88E-03 | 2.97E-02 | -1.02E+00 | 2.78E-03 | 1.18E-02 | -1.31E+00 | 1.78E-03 | 6.81E-03 | -1.17E+00 | 5.97E-02 | 2.13E-01 | 1.16E+00 | 3.36E-02 | 9.48E-02 | -1.01E+00 | 1.37E-05 | 5.04E-05 | -1.31E+00 | 1.87E-04 | 5.14E-04 | -1.12E+00 |
| KMT2E | 2.06E-03 | 1.07E-02 | 1.03E+00 | 3.43E-06 | 1.48E-05 | -1.01E+00 | 6.16E-02 | 2.62E-01 | -1.04E+00 | 1.32E-02 | 5.06E-02 | -1.02E+00 | 2.67E-02 | 9.52E-02 | 1.06E+00 | 3.00E-03 | 8.46E-03 | -1.03E+00 | 6.74E-06 | 2.48E-05 | -1.25E+00 | 2.44E-04 | 6.69E-04 | -1.21E+00 |
| TSC1 | 2.06E-03 | 1.07E-02 | -1.21E+00 | 1.90E-02 | 8.21E-02 | 1.07E+00 | 5.19E-02 | 2.21E-01 | 1.03E+00 | 1.25E-01 | 4.79E-01 | 1.30E+00 | 2.91E-02 | 1.04E-01 | -1.01E+00 | 4.33E-02 | 1.22E-01 | -1.00E+00 | 3.51E-06 | 1.29E-05 | -1.43E+00 | 3.19E-05 | 8.76E-05 | -1.27E+00 |
| AC093627.4 | 2.07E-03 | 1.08E-02 | -1.49E+00 | 9.10E-02 | 3.93E-01 | 1.18E+00 | 2.66E-10 | 1.13E-09 | -2.18E+00 | 3.09E-07 | 1.18E-06 | -1.55E+00 | 9.81E-02 | 3.50E-01 | -1.25E+00 | 3.00E-01 | 8.45E-01 | 1.20E+00 | 4.48E-04 | 1.64E-03 | -2.36E+00 | 2.47E-02 | 6.78E-02 | -1.49E+00 |
| FLVCR1-AS1 | 2.07E-03 | 1.08E-02 | -1.16E+00 | 3.23E-02 | 1.40E-01 | 1.18E+00 | 2.95E-02 | 1.25E-01 | -1.10E+00 | 3.31E-02 | 1.26E-01 | 1.06E+00 | 4.85E-02 | 1.73E-01 | -1.18E+00 | 3.26E-01 | 9.20E-01 | 1.58E+00 | 3.52E-02 | 1.29E-01 | -1.11E+00 | 2.76E-01 | 7.58E-01 | 1.13E+00 |
| SPTY2D1 | 2.08E-03 | 1.08E-02 | -1.02E+00 | 5.66E-17 | 2.44E-16 | -2.02E+00 | 3.64E-02 | 1.55E-01 | 1.38E+00 | 3.35E-05 | 1.28E-04 | -1.45E+00 | 1.48E-05 | 5.28E-05 | -1.20E+00 | 9.18E-18 | 2.59E-17 | -2.29E+00 | 9.06E-09 | 3.33E-08 | -1.37E+00 | 3.16E-16 | 8.69E-16 | -1.78E+00 |
| KBTBD3 | 2.08E-03 | 1.09E-02 | -1.14E+00 | 4.06E-03 | 1.76E-02 | -1.06E+00 | 2.31E-01 | 9.83E-01 | 1.27E+00 | 1.38E-01 | 5.28E-01 | 1.19E+00 | 1.89E-02 | 6.74E-02 | -1.10E+00 | 8.48E-03 | 2.39E-02 | -1.20E+00 | 2.52E-02 | 9.23E-02 | -1.12E+00 | 1.37E-01 | 3.77E-01 | 1.00E+00 |
| CACNA1C | 2.08E-03 | 1.09E-02 | -1.38E+00 | 2.76E-03 | 1.19E-02 | -1.28E+00 | 3.02E-03 | 1.28E-02 | -1.34E+00 | 3.27E-02 | 1.25E-01 | 1.04E+00 | 2.68E-01 | 9.54E-01 | 1.15E+00 | 3.55E-01 | 1.00E+00 | 1.21E+00 | 1.63E-04 | 5.99E-04 | -2.13E+00 | 9.05E-03 | 2.49E-02 | -1.40E+00 |
| SUMO2P1 | 2.08E-03 | 1.09E-02 | -1.21E+00 | 1.03E-03 | 4.46E-03 | -1.19E+00 | 4.39E-04 | 1.87E-03 | -1.37E+00 | 6.01E-03 | 2.30E-02 | -1.03E+00 | 2.70E-03 | 9.62E-03 | -1.17E+00 | 3.74E-02 | 1.05E-01 | 1.08E+00 | 4.38E-03 | 1.61E-02 | -1.09E+00 | 2.79E-01 | 7.68E-01 | 1.38E+00 |
| ZNF813 | 2.08E-03 | 1.09E-02 | -1.15E+00 | 6.73E-02 | 2.91E-01 | 1.24E+00 | 3.39E-04 | 1.44E-03 | -1.43E+00 | 3.84E-04 | 1.47E-03 | -1.33E+00 | 2.34E-05 | 8.35E-05 | -2.06E+00 | 1.36E-03 | 3.84E-03 | -1.26E+00 | 3.61E-04 | 1.32E-03 | -1.36E+00 | 1.24E-02 | 3.40E-02 | -1.13E+00 |
| AC117513.1 | 2.08E-03 | 1.09E-02 | -1.36E+00 | 1.93E-01 | 8.36E-01 | 1.35E+00 | 3.18E-02 | 1.35E-01 | -1.33E+00 | 9.10E-02 | 3.47E-01 | 1.75E+00 | 3.84E-04 | 1.37E-03 | -1.59E+00 | 2.04E-03 | 5.76E-03 | -1.36E+00 | 8.96E-02 | 3.29E-01 | -1.13E+00 | 3.22E-01 | 8.85E-01 | 1.18E+00 |
| AL031595.2 | 2.09E-03 | 1.09E-02 | -1.23E+00 | 5.94E-03 | 2.57E-02 | -1.02E+00 | 9.46E-02 | 4.02E-01 | 1.10E+00 | 8.39E-02 | 3.20E-01 | 1.04E+00 | 2.32E-01 | 8.27E-01 | 1.26E+00 | 2.17E-01 | 6.11E-01 | -1.01E+00 | 1.64E-01 | 6.03E-01 | -1.00E+00 | 1.56E-01 | 4.28E-01 | -1.21E+00 |
| WASH4P | 2.09E-03 | 1.09E-02 | -1.35E+00 | 5.17E-03 | 2.23E-02 | -1.11E+00 | 2.81E-04 | 1.19E-03 | -1.60E+00 | 8.29E-02 | 3.17E-01 | 1.17E+00 | 1.21E-02 | 4.30E-02 | -1.24E+00 | 8.06E-02 | 2.27E-01 | -1.01E+00 | 1.07E-08 | 3.92E-08 | -1.99E+00 | 2.73E-03 | 7.50E-03 | -1.02E+00 |
| AC092683.1 | 2.09E-03 | 1.09E-02 | -1.45E+00 | 3.09E-02 | 1.33E-01 | 1.02E+00 | 7.87E-02 | 3.34E-01 | -1.08E+00 | 1.61E-01 | 6.15E-01 | 1.13E+00 | 2.07E-03 | 7.38E-03 | -1.72E+00 | 2.31E-02 | 6.50E-02 | -1.19E+00 | 3.23E-05 | 1.19E-04 | -1.75E+00 | 4.89E-05 | 1.34E-04 | -1.62E+00 |
| ANKRD13C | 2.09E-03 | 1.09E-02 | -1.10E+00 | 5.31E-03 | 2.30E-02 | 1.01E+00 | 1.75E-04 | 7.44E-04 | -1.45E+00 | 6.22E-05 | 2.37E-04 | -1.28E+00 | 1.49E-02 | 5.33E-02 | 1.00E+00 | 5.97E-03 | 1.68E-02 | -1.11E+00 | 3.90E-02 | 1.43E-01 | -1.02E+00 | 4.50E-02 | 1.24E-01 | -1.02E+00 |
| PTGER2 | 2.10E-03 | 1.09E-02 | 1.90E+00 | 9.05E-03 | 3.91E-02 | 2.02E+00 | 7.49E-03 | 3.18E-02 | -1.03E+00 | 6.20E-02 | 2.37E-01 | 1.22E+00 | 7.36E-03 | 2.62E-02 | 1.78E+00 | 3.22E-01 | 9.07E-01 | 1.33E+00 | 5.28E-17 | 1.94E-16 | 4.00E+00 | 2.23E-07 | 6.11E-07 | 2.57E+00 |
| SCYL1 | 2.10E-03 | 1.09E-02 | -1.17E+00 | 2.25E-01 | 9.73E-01 | 1.39E+00 | 8.14E-03 | 3.46E-02 | -1.08E+00 | 1.45E-01 | 5.56E-01 | 1.30E+00 | 3.44E-05 | 1.23E-04 | -1.39E+00 | 1.06E-01 | 3.00E-01 | 1.18E+00 | 1.98E-03 | 7.27E-03 | -1.09E+00 | 4.91E-02 | 1.35E-01 | 1.02E+00 |
| AC108673.3 | 2.10E-03 | 1.10E-02 | -1.53E+00 | 1.55E-01 | 6.68E-01 | 1.34E+00 | 1.11E-02 | 4.72E-02 | -1.42E+00 | 1.18E-01 | 4.51E-01 | 1.05E+00 | 3.94E-02 | 1.41E-01 | -1.40E+00 | 1.78E-01 | 5.01E-01 | -1.09E+00 | 1.48E-01 | 5.43E-01 | -1.04E+00 | 3.38E-01 | 9.28E-01 | 1.14E+00 |
| TSPAN32 | 2.10E-03 | 1.10E-02 | 1.08E+00 | 3.00E-02 | 1.30E-01 | 1.28E+00 | 6.15E-06 | 2.61E-05 | -1.26E+00 | 2.91E-04 | 1.11E-03 | 1.04E+00 | 1.59E-02 | 5.66E-02 | 1.05E+00 | 4.90E-02 | 1.38E-01 | 1.06E+00 | 1.52E-01 | 5.58E-01 | 1.37E+00 | 1.32E-01 | 3.62E-01 | 1.09E+00 |
| AC073111.5 | 2.10E-03 | 1.10E-02 | -1.24E+00 | 3.97E-02 | 1.71E-01 | 1.13E+00 | 1.35E-06 | 5.74E-06 | -1.57E+00 | 8.57E-06 | 3.27E-05 | -1.27E+00 | 1.76E-03 | 6.27E-03 | -1.42E+00 | 1.49E-02 | 4.21E-02 | -1.12E+00 | 2.55E-18 | 9.37E-18 | -2.40E+00 | 8.46E-16 | 2.32E-15 | -2.01E+00 |
| TAF13 | 2.11E-03 | 1.10E-02 | -1.17E+00 | 1.62E-05 | 7.01E-05 | -1.48E+00 | 7.12E-03 | 3.03E-02 | -1.25E+00 | 1.71E-02 | 6.53E-02 | -1.00E+00 | 1.43E-01 | 5.10E-01 | 1.30E+00 | 1.63E-03 | 4.58E-03 | -1.28E+00 | 1.92E-06 | 7.05E-06 | -1.57E+00 | 1.16E-02 | 3.18E-02 | -1.14E+00 |
| PARK7 | 2.11E-03 | 1.10E-02 | -1.01E+00 | 4.81E-02 | 2.08E-01 | 1.11E+00 | 1.55E-01 | 6.57E-01 | 1.12E+00 | 2.23E-01 | 8.53E-01 | 1.10E+00 | 5.69E-04 | 2.03E-03 | -1.07E+00 | 4.71E-02 | 1.33E-01 | 1.08E+00 | 1.62E-01 | 5.95E-01 | 1.12E+00 | 1.96E-02 | 5.38E-02 | 1.23E+00 |
| TAB3 | 2.11E-03 | 1.10E-02 | -1.12E+00 | 2.94E-03 | 1.27E-02 | 1.01E+00 | 1.07E-02 | 4.53E-02 | 1.02E+00 | 8.08E-04 | 3.09E-03 | -1.04E+00 | 1.41E-02 | 5.03E-02 | 1.06E+00 | 7.59E-02 | 2.14E-01 | 1.21E+00 | 1.30E-01 | 4.76E-01 | 1.18E+00 | 4.23E-02 | 1.16E-01 | 1.14E+00 |
| [illegible] | [illegible] | [illegible] | [illegible] | [illegible] | [illegible] | [illegible] | [illegible] | [illegible] | [illegible] | [illegible] | [illegible] | [illegible] | [illegible] | [illegible] | [illegible] | [illegible] | [illegible] | [illegible] | [illegible] | [illegible] | [illegible] | [illegible] | [illegible] | [illegible] |
| [illegible] | [illegible] | [illegible] | [illegible] | [illegible] | [illegible] | [illegible] | [illegible] | [illegible] | [illegible] | [illegible] | [illegible] | [illegible] | [illegible] | [illegible] | [illegible] | [illegible] | [illegible] | [illegible] | [illegible] | [illegible] | [illegible] | [illegible] | [illegible] | [illegible] |
| PFAS | 2.13E-03 | 1.11E-02 | -1.25E+00 | 9.45E-02 | 4.08E-01 | 1.31E+00 | 1.40E-02 | 5.96E-02 | -1.12E+00 | 9.72E-02 | 3.71E-01 | 1.55E+00 | 8.13E-03 | 2.90E-02 | -1.45E+00 | 3.02E-01 | 8.50E-01 | 1.34E+00 | 1.00E-02 | 3.67E-02 | -1.05E+00 | 1.04E-01 | 2.85E-01 | 1.45E+00 |
| MAP4K3 | 2.14E-03 | 1.11E-02 | 2.03E+00 | 2.11E-01 | 9.12E-01 | 1.58E+00 | 1.22E-01 | 5.19E-01 | 1.38E+00 | 1.36E-01 | 5.20E-01 | 1.22E+00 | 6.37E-02 | 2.27E-01 | 1.25E+00 | 1.06E-04 | 2.99E-04 | -1.28E+00 | 2.14E-01 | 7.86E-01 | 1.23E+00 | 4.66E-03 | 1.28E-02 | -1.24E+00 |
| GSTM4 | 2.14E-03 | 1.11E-02 | -1.12E+00 | 3.59E-03 | 1.55E-02 | 1.02E+00 | 2.31E-03 | 9.82E-03 | -1.20E+00 | 3.65E-03 | 1.40E-02 | -1.03E+00 | 2.59E-01 | 9.23E-01 | 1.44E+00 | 2.74E-01 | 7.72E-01 | 1.63E+00 | 2.03E-01 | 7.45E-01 | 1.14E+00 | 2.89E-01 | 7.94E-01 | 1.28E+00 |
| AF111169.1 | 2.14E-03 | 1.11E-02 | 1.04E+00 | 2.76E-04 | 1.19E-03 | 1.11E+00 | 3.77E-02 | 1.60E-01 | 1.13E+00 | 7.52E-03 | 2.87E-02 | 1.18E+00 | 2.34E-03 | 8.35E-03 | 1.09E+00 | 9.27E-04 | 2.61E-03 | 1.11E+00 | 4.64E-03 | 1.70E-02 | 1.01E+00 | 2.09E-01 | 5.74E-01 | 1.24E+00 |
| TMEM14EP | 2.14E-03 | 1.11E-02 | -1.31E+00 | 1.00E-02 | 4.33E-02 | -1.02E+00 | 2.35E-01 | 9.99E-01 | 1.28E+00 | 5.11E-03 | 1.95E-02 | -1.25E+00 | 1.97E-01 | 7.02E-01 | 1.04E+00 | 3.38E-01 | 9.53E-01 | 1.27E+00 | 9.42E-02 | 3.46E-01 | 1.02E+00 | 2.27E-01 | 6.23E-01 | 1.06E+00 |
| SIAE | 2.14E-03 | 1.12E-02 | -1.15E+00 | 3.76E-03 | 1.62E-02 | -1.01E+00 | 1.12E-03 | 4.76E-03 | -1.32E+00 | 4.37E-03 | 1.67E-02 | -1.05E+00 | 6.65E-05 | 2.37E-04 | -1.71E+00 | 3.57E-04 | 1.01E-03 | -1.48E+00 | 5.41E-02 | 1.99E-01 | 1.17E+00 | 1.26E-01 | 3.45E-01 | 1.01E+00 |
| SF3B6 | 2.14E-03 | 1.12E-02 | 1.04E+00 | 4.57E-05 | 1.98E-04 | 1.08E+00 | 1.51E-01 | 6.42E-01 | 1.15E+00 | 1.29E-01 | 4.92E-01 | 1.13E+00 | 2.78E-02 | 9.93E-02 | -1.03E+00 | 1.52E-01 | 4.27E-01 | 1.06E+00 | 1.01E-01 | 3.72E-01 | 1.16E+00 | 4.26E-02 | 1.17E-01 | 1.24E+00 |
| DLGAP3 | 2.14E-03 | 1.12E-02 | -1.20E+00 | 4.64E-04 | 2.00E-03 | -1.44E+00 | 1.91E-01 | 8.10E-01 | 1.06E+00 | 8.29E-02 | 3.17E-01 | 1.09E+00 | 2.64E-01 | 9.43E-01 | 1.05E+00 | 2.68E-01 | 7.57E-01 | 1.40E+00 | 1.27E-01 | 4.65E-01 | -1.18E+00 | 1.24E-01 | 3.40E-01 | -1.21E+00 |
| RNF38 | 2.15E-03 | 1.12E-02 | 1.10E+00 | 1.18E-04 | 5.10E-04 | 1.09E+00 | 1.94E-01 | 8.26E-01 | 1.16E+00 | 2.39E-03 | 9.13E-03 | -1.07E+00 | 3.57E-04 | 1.27E-03 | -1.08E+00 | 1.19E-05 | 3.35E-05 | -1.37E+00 | 7.96E-04 | 2.92E-03 | -1.30E+00 | 9.03E-04 | 2.48E-03 | -1.17E+00 |
| GLYCTK | 2.15E-03 | 1.12E-02 | -1.38E+00 | 1.65E-03 | 7.12E-03 | -1.32E+00 | 8.23E-04 | 3.50E-03 | -1.30E+00 | 2.89E-05 | 1.10E-04 | -1.50E+00 | 5.06E-05 | 1.80E-04 | -2.01E+00 | 3.15E-05 | 8.87E-05 | -2.07E+00 | 1.57E-04 | 5.77E-04 | -1.32E+00 | 1.95E-05 | 5.35E-05 | -1.63E+00 |
| MMD | 2.16E-03 | 1.13E-02 | -1.11E+00 | 2.86E-03 | 1.24E-02 | 1.03E+00 | 7.44E-04 | 3.16E-03 | -1.17E+00 | 3.50E-03 | 1.34E-02 | -1.04E+00 | 1.07E-01 | 3.81E-01 | 1.02E+00 | 3.11E-01 | 8.78E-01 | 1.25E+00 | 3.11E-05 | 1.14E-04 | 1.64E+00 | 6.67E-09 | 1.83E-08 | 1.38E+00 |
| KNSTRN | 2.16E-03 | 1.13E-02 | -1.15E+00 | 3.15E-03 | 1.36E-02 | -1.04E+00 | 5.83E-05 | 2.48E-04 | -1.40E+00 | 2.98E-05 | 1.14E-04 | -1.17E+00 | 1.84E-02 | 6.55E-02 | -1.06E+00 | 7.00E-02 | 1.97E-01 | 1.02E+00 | 1.18E-04 | 4.31E-04 | -1.18E+00 | 3.43E-02 | 9.43E-02 | 1.02E+00 |
| HIST1H2AH | 2.16E-03 | 1.13E-02 | -1.15E+00 | 5.14E-05 | 2.22E-04 | -1.59E+00 | 1.10E-01 | 4.66E-01 | 1.63E+00 | 2.76E-03 | 1.05E-02 | -1.21E+00 | 6.15E-03 | 2.19E-02 | -1.33E+00 | 4.22E-05 | 1.19E-04 | -2.35E+00 | 2.35E-07 | 8.62E-07 | -1.37E+00 | 1.13E-10 | 3.11E-10 | -1.93E+00 |
| PNP | 2.16E-03 | 1.13E-02 | -1.02E+00 | 4.75E-08 | 2.05E-07 | -1.39E+00 | 2.71E-09 | 1.15E-08 | -1.27E+00 | 7.67E-09 | 2.93E-08 | -1.18E+00 | 1.03E-07 | 3.68E-07 | -1.54E+00 | 4.03E-04 | 1.14E-03 | -1.10E+00 | 4.60E-17 | 1.69E-16 | -1.42E+00 | 5.10E-09 | 1.40E-08 | -1.29E+00 |
| TMEM201 | 2.17E-03 | 1.13E-02 | -1.42E+00 | 2.26E-02 | 9.77E-02 | 1.02E+00 | 3.23E-02 | 1.37E-01 | -1.06E+00 | 1.92E-02 | 7.35E-02 | 1.09E+00 | 7.07E-03 | 2.52E-02 | -1.49E+00 | 4.84E-03 | 1.37E-02 | -1.58E+00 | 1.89E-05 | 6.93E-05 | -1.54E+00 | 9.10E-02 | 2.50E-01 | 1.10E+00 |
| RNF126 | 2.17E-03 | 1.13E-02 | 1.11E+00 | 1.92E-07 | 8.29E-07 | -1.01E+00 | 1.90E-01 | 8.06E-01 | 1.18E+00 | 8.02E-02 | 3.06E-01 | 1.28E+00 | 2.78E-04 | 9.91E-04 | -1.03E+00 | 9.38E-03 | 2.65E-02 | 1.17E+00 | 1.09E-03 | 4.01E-03 | 1.02E+00 | 1.17E-02 | 3.22E-02 | 1.05E+00 |
| CTNND1 | 2.18E-03 | 1.14E-02 | -1.03E+00 | 6.59E-05 | 2.85E-04 | -1.05E+00 | 2.23E-02 | 9.46E-02 | 1.02E+00 | 2.31E-04 | 8.83E-04 | -1.16E+00 | 4.50E-03 | 1.60E-02 | -1.04E+00 | 4.46E-04 | 1.26E-03 | -1.31E+00 | 2.37E-10 | 8.70E-10 | -1.49E+00 | 6.95E-08 | 1.91E-07 | -1.38E+00 |
| SPAG5-AS1 | 2.18E-03 | 1.14E-02 | -1.65E+00 | 4.79E-02 | 2.07E-01 | -1.01E+00 | 1.44E-02 | 6.11E-02 | -1.49E+00 | 1.22E-02 | 4.65E-02 | -1.38E+00 | 7.47E-02 | 2.66E-01 | -1.06E+00 | 2.38E-01 | 6.72E-01 | 1.11E+00 | 2.26E-02 | 8.29E-02 | -1.50E+00 | 5.70E-02 | 1.57E-01 | -1.37E+00 |
| JDP2 | 2.18E-03 | 1.14E-02 | 2.26E+00 | 1.50E-01 | 6.47E-01 | 1.61E+00 | 3.56E-06 | 1.51E-05 | 2.76E+00 | 1.30E-02 | 4.98E-02 | 2.23E+00 | 4.73E-02 | 1.68E-01 | 2.88E+00 | 3.55E-01 | 1.00E+00 | 1.28E+00 | 2.71E-01 | 9.95E-01 | 1.04E+00 | 3.09E-01 | 8.49E-01 | -1.14E+00 |
| CCDC88B | 2.19E-03 | 1.14E-02 | -1.04E+00 | 3.17E-03 | 1.37E-02 | 1.09E+00 | 7.79E-02 | 3.31E-01 | 1.03E+00 | 6.35E-02 | 2.42E-01 | 1.17E+00 | 7.24E-03 | 2.58E-02 | 1.06E+00 | 2.42E-04 | 6.82E-04 | -1.06E+00 | 2.26E-01 | 8.30E-01 | 1.14E+00 | 2.31E-01 | 6.36E-01 | 1.38E+00 |
| SLC4A7 | 2.19E-03 | 1.14E-02 | -1.03E+00 | 4.07E-04 | 1.76E-03 | -1.05E+00 | 5.93E-07 | 2.52E-06 | -1.40E+00 | 1.39E-06 | 5.32E-06 | -1.19E+00 | 1.18E-03 | 4.21E-03 | -1.26E+00 | 2.79E-03 | 7.88E-03 | -1.27E+00 | 1.04E-06 | 3.83E-06 | -1.68E+00 | 5.31E-03 | 1.46E-02 | -1.18E+00 |
| LINC00954 | 2.19E-03 | 1.14E-02 | -1.25E+00 | 1.55E-01 | 6.71E-01 | 1.37E+00 | 8.19E-02 | 3.48E-01 | -1.16E+00 | 8.71E-02 | 3.33E-01 | 1.57E+00 | 4.42E-03 | 1.58E-02 | -1.11E+00 | 7.81E-02 | 2.20E-01 | 1.08E+00 | 2.41E-01 | 8.85E-01 | 1.33E+00 | 2.99E-01 | 8.22E-01 | 1.50E+00 |
| DPY19L2P1 | 2.19E-03 | 1.14E-02 | -1.99E+00 | 2.30E-02 | 9.93E-02 | -1.29E+00 | 1.28E-01 | 5.44E-01 | -1.05E+00 | 1.64E-01 | 6.28E-01 | 1.06E+00 | 3.55E-02 | 1.27E-01 | -1.13E+00 | 7.13E-02 | 2.01E-01 | 1.05E+00 | 5.72E-08 | 2.10E-07 | -1.41E+00 | 6.65E-06 | 1.83E-05 | -1.26E+00 |
| PTGR2 | 2.19E-03 | 1.14E-02 | -1.12E+00 | 2.02E-02 | 8.71E-02 | 1.11E+00 | 3.29E-04 | 1.40E-03 | -1.27E+00 | 1.72E-03 | 6.56E-03 | -1.05E+00 | 3.16E-02 | 1.13E-01 | -1.20E+00 | 3.32E-01 | 9.36E-01 | 1.38E+00 | 5.31E-04 | 1.95E-03 | -1.40E+00 | 4.85E-03 | 1.33E-02 | -1.23E+00 |
| KCNMB3P1 | 2.20E-03 | 1.14E-02 | -1.51E+00 | 9.25E-04 | 4.00E-03 | -1.58E+00 | 4.87E-03 | 2.07E-02 | -1.55E+00 | 2.98E-02 | 1.14E-01 | -1.12E+00 | 4.16E-03 | 1.48E-02 | -1.46E+00 | 3.19E-02 | 8.99E-02 | -1.13E+00 | 2.36E-02 | 8.67E-02 | -1.47E+00 | 1.58E-02 | 4.33E-02 | -1.32E+00 |
| DSTN | 2.20E-03 | 1.15E-02 | 1.09E+00 | 5.81E-03 | 2.51E-02 | 1.19E+00 | 1.89E-01 | 8.04E-01 | 1.16E+00 | 6.70E-03 | 2.56E-02 | 1.09E+00 | 1.61E-03 | 5.72E-03 | 1.05E+00 | 2.75E-02 | 7.76E-02 | 1.12E+00 | 1.23E-01 | 4.50E-01 | 1.34E+00 | 9.05E-02 | 2.49E-01 | 1.30E+00 |
| BMT2 | 2.21E-03 | 1.15E-02 | -1.06E+00 | 5.18E-03 | 2.24E-02 | 1.11E+00 | 1.36E-02 | 5.76E-02 | -1.16E+00 | 1.39E-02 | 5.33E-02 | 1.01E+00 | 9.57E-02 | 3.41E-01 | 1.15E+00 | 3.75E-02 | 1.06E-01 | -1.10E+00 | 2.14E-02 | 7.84E-02 | -1.02E+00 | 2.78E-03 | 7.64E-03 | -1.17E+00 |
| AP5Z1 | 2.21E-03 | 1.15E-02 | -1.07E+00 | 3.06E-02 | 1.32E-01 | 1.26E+00 | 1.42E-01 | 6.02E-01 | 1.12E+00 | 5.50E-02 | 2.10E-01 | 1.19E+00 | 8.30E-02 | 2.96E-01 | 1.04E+00 | 3.46E-01 | 9.76E-01 | 1.41E+00 | 3.04E-03 | 1.12E-02 | -1.02E+00 | 8.54E-03 | 2.35E-02 | -1.02E+00 |
| EP400 | 2.21E-03 | 1.15E-02 | -1.12E+00 | 1.87E-02 | 8.10E-02 | 1.12E+00 | 1.13E-03 | 4.82E-03 | -1.14E+00 | 1.69E-02 | 6.44E-02 | 1.13E+00 | 7.75E-03 | 2.76E-02 | 1.03E+00 | 1.11E-02 | 3.12E-02 | 1.02E+00 | 2.37E-06 | 8.69E-06 | -1.25E+00 | 6.55E-05 | 1.80E-04 | -1.14E+00 |
| CITED4 | 2.21E-03 | 1.15E-02 | -1.06E+00 | 2.19E-07 | 9.47E-07 | -1.68E+00 | 1.63E-05 | 6.91E-05 | 2.03E+00 | 5.31E-02 | 2.03E-01 | 1.25E+00 | 1.05E-01 | 3.75E-01 | 1.02E+00 | 3.26E-02 | 9.18E-02 | -1.61E+00 | 5.20E-02 | 1.91E-01 | -1.32E+00 | 8.15E-03 | 2.24E-02 | -2.05E+00 |
| CD5 | 2.21E-03 | 1.15E-02 | -1.07E+00 | 4.24E-02 | 1.83E-01 | 1.23E+00 | 1.34E-01 | 5.68E-01 | -1.17E+00 | 8.12E-02 | 3.10E-01 | -1.10E+00 | 1.28E-01 | 4.57E-01 | 2.07E+00 | 2.11E-01 | 5.94E-01 | 1.91E+00 | 2.47E-01 | 9.07E-01 | 1.18E+00 | 3.10E-01 | 8.51E-01 | 1.28E+00 |
| ZNF75A | 2.21E-03 | 1.15E-02 | -1.05E+00 | 1.17E-01 | 5.05E-01 | 1.36E+00 | 3.37E-03 | 1.43E-02 | -1.16E+00 | 1.92E-01 | 7.32E-01 | 1.29E+00 | 1.37E-02 | 4.89E-02 | 1.06E+00 | 9.72E-02 | 2.74E-01 | 1.19E+00 | 8.73E-02 | 3.20E-01 | 1.06E+00 | 1.09E-01 | 2.98E-01 | 1.47E+00 |
| PICK1 | 2.21E-03 | 1.15E-02 | -1.01E+00 | 3.98E-02 | 1.72E-01 | 1.28E+00 | 5.98E-02 | 2.54E-01 | 1.05E+00 | 6.62E-02 | 2.53E-01 | 1.21E+00 | 2.77E-02 | 9.89E-02 | 1.03E+00 | 6.86E-02 | 1.93E-01 | 1.07E+00 | 3.87E-02 | 1.42E-01 | 1.04E+00 | 1.47E-01 | 4.05E-01 | 1.14E+00 |
| HEBP2 | 2.21E-03 | 1.15E-02 | 1.11E+00 | 1.02E-09 | 4.40E-09 | -1.04E+00 | 1.61E-01 | 6.83E-01 | 1.07E+00 | 2.96E-04 | 1.13E-03 | -1.08E+00 | 5.43E-03 | 1.94E-02 | 1.02E+00 | 6.28E-04 | 1.77E-03 | -1.04E+00 | 1.84E-01 | 6.74E-01 | 1.14E+00 | 1.51E-02 | 4.15E-02 | 1.00E+00 |
| PCNX1 | 2.21E-03 | 1.15E-02 | 1.64E+00 | 2.16E-02 | 9.34E-02 | 1.73E+00 | 6.68E-02 | 2.84E-01 | 1.41E+00 | 2.54E-01 | 9.70E-01 | 1.44E+00 | 2.94E-05 | 1.05E-04 | 2.35E+00 | 1.79E-02 | 5.05E-02 | 1.85E+00 | 2.06E-02 | 7.57E-02 | 1.54E+00 | 7.44E-02 | 2.04E-01 | 1.49E+00 |
| GTF3C1 | 2.21E-03 | 1.15E-02 | -1.11E+00 | 2.01E-03 | 8.68E-03 | -1.07E+00 | 5.01E-04 | 2.13E-03 | -1.18E+00 | 3.61E-04 | 1.38E-03 | -1.08E+00 | 7.10E-05 | 2.53E-04 | -1.13E+00 | 2.99E-03 | 8.44E-03 | 1.03E+00 | 3.01E-02 | 1.10E-01 | 1.00E+00 | 1.12E-01 | 3.08E-01 | 1.14E+00 |
| ST20 | 2.22E-03 | 1.16E-02 | -1.23E+00 | 6.08E-03 | 2.63E-02 | -1.08E+00 | 5.06E-06 | 2.15E-05 | -1.37E+00 | 3.05E-06 | 1.16E-05 | -1.27E+00 | 9.51E-05 | 3.39E-04 | -1.42E+00 | 3.02E-03 | 8.52E-03 | -1.18E+00 | 1.21E-07 | 4.46E-07 | -1.44E+00 | 3.76E-09 | 1.03E-08 | -1.67E+00 |
| METTL14 | 2.22E-03 | 1.16E-02 | -1.11E+00 | 1.17E-03 | 5.06E-03 | -1.03E+00 | 6.28E-02 | 2.67E-01 | -1.05E+00 | 3.54E-02 | 1.35E-01 | 1.07E+00 | 2.79E-04 | 9.97E-04 | -1.09E+00 | 1.14E-03 | 3.22E-03 | -1.05E+00 | 4.39E-04 | 1.61E-03 | -1.21E+00 | 2.82E-03 | 7.75E-03 | -1.11E+00 |
| SMIM11B | 2.23E-03 | 1.16E-02 | -1.46E+00 | 1.67E-03 | 7.21E-03 | -1.44E+00 | 6.77E-04 | 2.88E-03 | -1.56E+00 | 5.57E-03 | 2.13E-02 | -1.20E+00 | 1.65E-01 | 5.89E-01 | 1.13E+00 | 2.76E-01 | 7.79E-01 | 1.26E+00 | 1.80E-01 | 6.62E-01 | 1.12E+00 | 4.95E-02 | 1.36E-01 | -1.26E+00 |
| ZNF710 | 2.24E-03 | 1.17E-02 | -1.48E+00 | 7.26E-02 | 3.14E-01 | 1.20E+00 | 4.78E-02 | 2.03E-01 | -1.23E+00 | 7.17E-02 | 2.74E-01 | -1.03E+00 | 5.08E-03 | 1.81E-02 | -1.37E+00 | 6.12E-03 | 1.72E-02 | -1.41E+00 | 2.21E-01 | 8.10E-01 | 1.16E+00 | 3.52E-01 | 9.66E-01 | 1.35E+00 |
| MAP10 | 2.24E-03 | 1.17E-02 | -1.14E+00 | 2.08E-01 | 9.00E-01 | 1.53E+00 | 1.02E-05 | 4.35E-05 | -1.71E+00 | 4.63E-04 | 1.77E-03 | -1.20E+00 | 1.91E-02 | 6.83E-02 | -1.22E+00 | 5.88E-02 | 1.66E-01 | -1.06E+00 | 8.68E-02 | 3.18E-01 | -1.03E+00 | 2.55E-01 | 7.02E-01 | 1.16E+00 |
| DPM1 | 2.25E-03 | 1.17E-02 | 1.01E+00 | 7.18E-02 | 3.10E-01 | 1.29E+00 | 1.23E-01 | 5.23E-01 | 1.06E+00 | 8.93E-02 | 3.41E-01 | 1.15E+00 | 1.53E-03 | 5.47E-03 | -1.14E+00 | 1.56E-01 | 4.40E-01 | 1.22E+00 | 8.14E-04 | 2.99E-03 | -1.02E+00 | 1.42E-01 | 3.90E-01 | 1.18E+00 |
| [illegible] | [illegible] | [illegible] | [illegible] | [illegible] | [illegible] | [illegible] | [illegible] | [illegible] | [illegible] | [illegible] | [illegible] | [illegible] | [illegible] | [illegible] | [illegible] | [illegible] | [illegible] | [illegible] | [illegible] | [illegible] | [illegible] | [illegible] | [illegible] | [illegible] |
| [illegible] | [illegible] | [illegible] | [illegible] | [illegible] | [illegible] | [illegible] | [illegible] | [illegible] | [illegible] | [illegible] | [illegible] | [illegible] | [illegible] | [illegible] | [illegible] | [illegible] | [illegible] | [illegible] | [illegible] | [illegible] | [illegible] | [illegible] | [illegible] | [illegible] |
| [illegible] | [illegible] | [illegible] | [illegible] | [illegible] | [illegible] | [illegible] | [illegible] | [illegible] | [illegible] | [illegible] | [illegible] | [illegible] | [illegible] | [illegible] | [illegible] | [illegible] | [illegible] | [illegible] | [illegible] | [illegible] | [illegible] | [illegible] | [illegible] | [illegible] |
| CCDC58P3 | 2.25E-03 | 1.17E-02 | -1.48E+00 | 2.27E-02 | 9.81E-02 | -1.01E+00 | 3.07E-08 | 1.30E-07 | -1.96E+00 | 8.52E-06 | 3.25E-05 | -1.29E+00 | 3.04E-02 | 1.08E-01 | -1.37E+00 | 1.19E-01 | 3.35E-01 | -1.07E+00 | 9.65E-12 | 3.54E-11 | -2.69E+00 | 3.40E-03 | 9.34E-03 | -1.12E+00 |
| AP000240.1 | 2.25E-03 | 1.17E-02 | -1.20E+00 | 8.06E-05 | 3.48E-04 | -1.62E+00 | 1.53E-04 | 6.48E-04 | -1.59E+00 | 6.27E-06 | 2.39E-05 | -1.61E+00 | 1.66E-03 | 5.94E-03 | -1.52E+00 | 4.19E-03 | 1.18E-02 | -1.38E+00 | 1.57E-03 | 5.74E-03 | -1.57E+00 | 6.07E-03 | 1.67E-02 | -1.38E+00 |
| GOLGA8M | 2.25E-03 | 1.17E-02 | -1.17E+00 | 1.05E-01 | 4.53E-01 | 1.35E+00 | 1.41E-04 | 5.98E-04 | -1.73E+00 | 1.17E-04 | 4.48E-04 | -1.68E+00 | 1.99E-01 | 7.11E-01 | 1.27E+00 | 2.75E-01 | 7.74E-01 | -1.08E+00 | 1.43E-01 | 5.25E-01 | 1.04E+00 | 2.35E-01 | 6.44E-01 | 1.09E+00 |
| C8orf44 | 2.26E-03 | 1.18E-02 | -1.11E+00 | 5.12E-02 | 2.21E-01 | 1.26E+00 | 1.33E-04 | 5.66E-04 | -1.55E+00 | 5.38E-02 | 2.06E-01 | 1.11E+00 | 6.88E-02 | 2.45E-01 | 1.06E+00 | 3.41E-01 | 9.62E-01 | 1.52E+00 | 1.29E-03 | 4.74E-03 | -1.18E+00 | 3.15E-02 | 8.65E-02 | -1.01E+00 |
| MTMR6 | 2.26E-03 | 1.18E-02 | -1.02E+00 | 4.16E-12 | 1.80E-11 | -1.91E+00 | 2.48E-02 | 1.05E-01 | -1.02E+00 | 1.78E-11 | 6.80E-11 | -1.89E+00 | 4.29E-03 | 1.53E-02 | -1.03E+00 | 1.18E-04 | 3.32E-04 | -1.31E+00 | 1.08E-04 | 3.96E-04 | -1.21E+00 | 2.29E-04 | 6.30E-04 | -1.27E+00 |
| TOM1L1 | 2.26E-03 | 1.18E-02 | -1.19E+00 | 1.36E-02 | 5.87E-02 | 1.04E+00 | 1.60E-02 | 6.78E-02 | -1.26E+00 | 1.27E-02 | 4.86E-02 | -1.17E+00 | 4.35E-02 | 1.55E-01 | -1.21E+00 | 1.09E-01 | 3.07E-01 | -1.04E+00 | 2.39E-01 | 8.79E-01 | 1.23E+00 | 3.22E-01 | 8.84E-01 | 1.30E+00 |
| C2CD2 | 2.26E-03 | 1.18E-02 | -1.16E+00 | 7.95E-02 | 3.44E-01 | 1.20E+00 | 1.14E-02 | 4.85E-02 | -1.18E+00 | 8.43E-02 | 3.22E-01 | 1.15E+00 | 2.24E-01 | 8.00E-01 | 1.05E+00 | 3.12E-01 | 8.80E-01 | 1.55E+00 | 1.36E-03 | 4.97E-03 | -1.52E+00 | 2.61E-03 | 7.18E-03 | -1.38E+00 |
| CHCHD6 | 2.27E-03 | 1.18E-02 | -1.06E+00 | 1.72E-04 | 7.45E-04 | -1.10E+00 | 1.20E-01 | 5.09E-01 | 1.14E+00 | 2.41E-02 | 9.22E-02 | 1.19E+00 | 2.22E-02 | 7.92E-02 | 1.05E+00 | 5.46E-04 | 1.54E-03 | -1.28E+00 | 2.58E-01 | 9.45E-01 | 1.27E+00 | 1.27E-01 | 3.48E-01 | 1.52E+00 |
| KPNA5 | 2.27E-03 | 1.18E-02 | -1.06E+00 | 2.80E-03 | 1.21E-02 | 1.05E+00 | 2.07E-02 | 8.80E-02 | 1.03E+00 | 1.85E-03 | 7.08E-03 | 1.01E+00 | 3.25E-02 | 1.16E-01 | -1.12E+00 | 3.24E-02 | 9.14E-02 | -1.05E+00 | 3.80E-03 | 1.39E-02 | -1.27E+00 | 3.01E-03 | 8.26E-03 | -1.21E+00 |
| MT-CO1 | 2.27E-03 | 1.18E-02 | -1.06E+00 | 1.87E-01 | 8.07E-01 | 1.00E+00 | 1.14E-03 | 4.84E-03 | -1.08E+00 | 1.54E-01 | 5.86E-01 | 1.01E+00 | 1.28E-06 | 4.57E-06 | -1.18E+00 | 2.61E-01 | 7.35E-01 | 1.03E+00 | 2.22E-05 | 8.13E-05 | -1.09E+00 | 1.46E-02 | 4.00E-02 | 1.09E+00 |
| ZNF286B | 2.27E-03 | 1.18E-02 | -1.13E+00 | 3.93E-02 | 1.70E-01 | 1.25E+00 | 1.42E-02 | 6.02E-02 | -1.02E+00 | 1.92E-02 | 7.35E-02 | 1.09E+00 | 2.81E-04 | 1.00E-03 | -1.40E+00 | 3.21E-03 | 9.04E-03 | -1.12E+00 | 2.57E-02 | 9.41E-02 | -1.02E+00 | 5.82E-02 | 1.60E-01 | 1.01E+00 |
| TXNRD2 | 2.27E-03 | 1.18E-02 | -1.08E+00 | 3.95E-03 | 1.70E-02 | 1.04E+00 | 2.65E-02 | 1.13E-01 | -1.03E+00 | 5.06E-02 | 1.93E-01 | 1.13E+00 | 2.10E-02 | 7.48E-02 | -1.11E+00 | 2.74E-01 | 7.72E-01 | 1.23E+00 | 4.33E-02 | 1.59E-01 | 1.13E+00 | 5.09E-02 | 1.40E-01 | 1.10E+00 |
| VAT1 | 2.27E-03 | 1.18E-02 | -1.01E+00 | 4.99E-03 | 2.16E-02 | 1.05E+00 | 3.04E-03 | 1.29E-02 | -1.11E+00 | 3.39E-03 | 1.29E-02 | 1.03E+00 | 8.16E-07 | 2.91E-06 | -1.31E+00 | 1.21E-05 | 3.40E-05 | -1.24E+00 | 5.46E-03 | 2.00E-02 | -1.16E+00 | 1.94E-01 | 5.34E-01 | 1.11E+00 |
| ZFP1 | 2.27E-03 | 1.18E-02 | -1.21E+00 | 7.22E-04 | 3.12E-03 | -1.26E+00 | 1.68E-02 | 7.14E-02 | -1.17E+00 | 3.55E-02 | 1.36E-01 | 1.05E+00 | 7.81E-05 | 2.79E-04 | -1.79E+00 | 1.24E-03 | 3.50E-03 | -1.29E+00 | 4.45E-04 | 1.64E-03 | -1.34E+00 | 3.46E-03 | 9.52E-03 | -1.28E+00 |
| GABPB2 | 2.28E-03 | 1.19E-02 | -1.27E+00 | 6.31E-02 | 2.73E-01 | 1.12E+00 | 3.63E-04 | 1.54E-03 | -1.36E+00 | 3.88E-03 | 1.48E-02 | -1.02E+00 | 5.32E-03 | 1.90E-02 | -1.51E+00 | 1.02E-01 | 2.89E-01 | 1.03E+00 | 4.13E-05 | 1.52E-04 | -1.32E+00 | 4.81E-02 | 1.32E-01 | 1.02E+00 |
| SLC25A35 | 2.28E-03 | 1.19E-02 | -1.28E+00 | 1.18E-02 | 5.10E-02 | -1.07E+00 | 1.35E-04 | 5.75E-04 | -1.57E+00 | 2.57E-03 | 9.83E-03 | -1.15E+00 | 3.02E-08 | 1.08E-07 | -1.97E+00 | 6.09E-06 | 1.72E-05 | -1.50E+00 | 5.31E-05 | 1.95E-04 | -1.53E+00 | 1.14E-02 | 3.14E-02 | -1.20E+00 |
| ARSK | 2.28E-03 | 1.19E-02 | -1.10E+00 | 1.96E-03 | 8.45E-03 | -1.03E+00 | 8.43E-06 | 3.58E-05 | -1.56E+00 | 7.88E-04 | 3.01E-03 | -1.14E+00 | 4.80E-06 | 1.71E-05 | -2.18E+00 | 2.92E-04 | 8.22E-04 | -1.58E+00 | 1.09E-03 | 4.02E-03 | -1.22E+00 | 1.27E-04 | 3.49E-04 | -1.38E+00 |
| CDC42P6 | 2.28E-03 | 1.19E-02 | 1.81E+00 | 1.27E-01 | 5.47E-01 | 1.45E+00 | 1.02E-02 | 4.35E-02 | 1.66E+00 | 2.20E-01 | 8.41E-01 | 1.48E+00 | 6.21E-02 | 2.21E-01 | 1.79E+00 | 2.45E-01 | 6.92E-01 | 1.37E+00 | 9.96E-04 | 3.66E-03 | 1.85E+00 | 1.24E-01 | 3.39E-01 | 1.61E+00 |
| RAB44 | 2.29E-03 | 1.19E-02 | -1.19E+00 | 1.84E-04 | 7.97E-04 | -1.29E+00 | 1.09E-05 | 4.62E-05 | -1.22E+00 | 2.38E-05 | 9.11E-05 | -1.06E+00 | 1.33E-04 | 4.74E-04 | -1.46E+00 | 3.23E-04 | 9.09E-04 | -1.28E+00 | 4.82E-02 | 1.77E-01 | -1.68E+00 | 2.23E-01 | 6.13E-01 | -1.08E+00 |
| AL353743.1 | 2.29E-03 | 1.19E-02 | -1.22E+00 | 1.06E-02 | 4.59E-02 | -1.03E+00 | 2.29E-03 | 9.73E-03 | -1.23E+00 | 1.64E-03 | 6.26E-03 | -1.06E+00 | 3.20E-03 | 1.14E-02 | -1.60E+00 | 7.84E-03 | 2.21E-02 | -1.32E+00 | 8.75E-03 | 3.21E-02 | -1.10E+00 | 6.74E-02 | 1.85E-01 | -1.07E+00 |
| TBC1D25 | 2.29E-03 | 1.19E-02 | -1.24E+00 | 9.67E-03 | 4.18E-02 | -1.02E+00 | 5.84E-04 | 2.48E-03 | -1.34E+00 | 3.57E-03 | 1.36E-02 | -1.07E+00 | 2.18E-02 | 7.76E-02 | -1.16E+00 | 1.05E-01 | 2.96E-01 | 1.11E+00 | 3.96E-06 | 1.45E-05 | -1.85E+00 | 1.32E-03 | 3.63E-03 | -1.42E+00 |
| MYL9 | 2.30E-03 | 1.20E-02 | 1.02E+00 | 1.63E-01 | 7.03E-01 | 1.69E+00 | 1.19E-01 | 5.07E-01 | 1.09E+00 | 1.05E-01 | 4.02E-01 | 1.74E+00 | 1.70E-01 | 6.08E-01 | 1.87E+00 | 1.63E-01 | 4.61E-01 | 1.92E+00 | 1.75E-01 | 6.41E-01 | 2.74E+00 | 3.60E-01 | 9.89E-01 | -1.25E+00 |
| PANK1 | 2.30E-03 | 1.20E-02 | -1.77E+00 | 1.14E-02 | 4.92E-02 | -1.36E+00 | 2.24E-02 | 9.51E-02 | -1.31E+00 | 5.99E-02 | 2.29E-01 | -1.05E+00 | 3.03E-03 | 1.08E-02 | -1.80E+00 | 6.32E-03 | 1.78E-02 | -1.80E+00 | 4.35E-13 | 1.59E-12 | -2.15E+00 | 8.41E-04 | 2.31E-03 | -1.07E+00 |
| SCAP | 2.30E-03 | 1.20E-02 | -1.08E+00 | 1.61E-01 | 6.97E-01 | 1.43E+00 | 2.17E-05 | 9.24E-05 | -1.20E+00 | 2.08E-02 | 7.96E-02 | 1.23E+00 | 3.62E-05 | 1.29E-04 | -1.37E+00 | 1.67E-03 | 4.72E-03 | -1.09E+00 | 1.04E-08 | 3.81E-08 | -1.42E+00 | 1.82E-02 | 5.00E-02 | 1.04E+00 |
| ZFAND2A | 2.30E-03 | 1.20E-02 | -1.02E+00 | 2.56E-05 | 1.11E-04 | -1.06E+00 | 2.25E-03 | 9.55E-03 | -1.02E+00 | 5.59E-03 | 2.14E-02 | 1.12E+00 | 8.91E-04 | 3.18E-03 | -1.13E+00 | 1.49E-04 | 4.20E-04 | -1.14E+00 | 2.70E-01 | 9.92E-01 | 1.20E+00 | 3.15E-01 | 8.66E-01 | 1.11E+00 |
| AC012358.3 | 2.30E-03 | 1.20E-02 | -1.14E+00 | 1.48E-03 | 6.40E-03 | -1.10E+00 | 3.02E-02 | 1.28E-01 | 1.04E+00 | 5.38E-05 | 2.06E-04 | -1.34E+00 | 7.65E-06 | 2.73E-05 | -1.50E+00 | 2.84E-06 | 8.00E-06 | -1.59E+00 | 6.76E-02 | 2.48E-01 | -1.03E+00 | 2.50E-03 | 6.85E-03 | -1.37E+00 |
| AC134407.3 | 2.30E-03 | 1.20E-02 | -1.59E+00 | 9.17E-02 | 3.96E-01 | 1.12E+00 | 1.37E-02 | 5.83E-02 | -1.35E+00 | 1.05E-01 | 4.00E-01 | 1.06E+00 | 2.90E-09 | 1.04E-08 | -4.03E+00 | 1.00E-06 | 2.82E-06 | -2.10E+00 | 1.20E-02 | 4.42E-02 | -1.42E+00 | 3.08E-01 | 8.46E-01 | 1.22E+00 |
| AL356273.3 | 2.30E-03 | 1.20E-02 | 3.19E+00 | 8.61E-02 | 3.72E-01 | 2.20E+00 | 4.19E-07 | 1.78E-06 | 3.31E+00 | 8.40E-02 | 3.21E-01 | -1.14E+00 | 5.34E-05 | 1.90E-04 | 3.84E+00 | 3.33E-01 | 9.39E-01 | 1.50E+00 | 1.13E-02 | 4.13E-02 | 1.90E+00 | 3.17E-01 | 8.71E-01 | 1.14E+00 |
| POLRMT | 2.30E-03 | 1.20E-02 | -1.19E+00 | 1.88E-01 | 8.11E-01 | 1.44E+00 | 1.40E-01 | 5.97E-01 | 1.11E+00 | 1.42E-02 | 5.41E-02 | 1.90E+00 | 3.60E-02 | 1.29E-01 | 1.00E+00 | 2.93E-01 | 8.27E-01 | 1.48E+00 | 4.82E-02 | 1.77E-01 | 1.04E+00 | 1.67E-01 | 4.60E-01 | 1.56E+00 |
| TATDN3 | 2.31E-03 | 1.20E-02 | -1.05E+00 | 1.16E-01 | 5.03E-01 | 1.38E+00 | 1.14E-03 | 4.86E-03 | -1.18E+00 | 1.40E-03 | 5.34E-03 | 1.00E+00 | 5.88E-04 | 2.10E-03 | -1.26E+00 | 4.42E-02 | 1.25E-01 | 1.14E+00 | 1.40E-04 | 5.14E-04 | -1.25E+00 | 5.01E-03 | 1.38E-02 | -1.10E+00 |
| KYAT1 | 2.31E-03 | 1.20E-02 | -1.48E+00 | 3.66E-02 | 1.58E-01 | 1.03E+00 | 2.69E-05 | 1.14E-04 | -1.55E+00 | 8.78E-03 | 3.35E-02 | 1.07E+00 | 7.94E-02 | 2.83E-01 | -1.15E+00 | 1.23E-01 | 3.47E-01 | -1.03E+00 | 1.90E-02 | 6.96E-02 | -1.08E+00 | 7.84E-02 | 2.15E-01 | 1.09E+00 |

| | | | | | | | | | | | | | | | | | | | | | | | | |
|---|---|---|---|---|---|---|---|---|---|---|---|---|---|---|---|---|---|---|---|---|---|---|---|---|
| MOB2 | 2.31E-03 | 1.21E-02 | 1.06E+00 | 5.19E-02 | 2.24E-01 | 1.36E+00 | 1.35E-03 | 5.74E-03 | -1.08E+00 | 2.08E-02 | 7.94E-02 | 1.25E+00 | 6.55E-02 | 2.34E-01 | 1.18E+00 | 1.71E-01 | 4.82E-01 | 1.53E+00 | 3.43E-03 | 1.26E-02 | 1.02E+00 | 5.25E-02 | 1.44E-01 | 1.12E+00 |
| PTMAP2 | 2.32E-03 | 1.21E-02 | 1.95E+00 | 2.30E-01 | 9.96E-01 | 1.61E+00 | 6.81E-06 | 2.89E-05 | 2.16E+00 | 7.55E-03 | 2.89E-02 | 1.86E+00 | 1.91E-01 | 6.80E-01 | 1.56E+00 | 2.46E-01 | 6.94E-01 | 1.31E+00 | 9.32E-06 | 3.42E-05 | 2.13E+00 | 3.56E-10 | 9.78E-10 | 2.58E+00 |
| RTL10 | 2.32E-03 | 1.21E-02 | -1.19E+00 | 4.32E-04 | 1.87E-03 | -1.14E+00 | 1.97E-07 | 8.37E-07 | -1.60E+00 | 2.18E-04 | 8.33E-04 | -1.13E+00 | 4.53E-02 | 1.62E-01 | -1.07E+00 | 2.14E-01 | 6.04E-01 | 1.23E+00 | 1.07E-08 | 3.93E-08 | -1.61E+00 | 2.71E-04 | 7.45E-04 | -1.28E+00 |
| ZNF334 | 2.32E-03 | 1.21E-02 | -1.05E+00 | 5.61E-02 | 2.42E-01 | 1.33E+00 | 2.04E-02 | 8.65E-02 | -1.38E+00 | 3.60E-02 | 1.37E-01 | -1.10E+00 | 7.10E-02 | 2.53E-01 | -1.11E+00 | 3.20E-01 | 9.01E-01 | 1.36E+00 | 1.13E-02 | 4.15E-02 | -1.15E+00 | 1.99E-01 | 5.46E-01 | 1.10E+00 |
| UCN | 2.32E-03 | 1.21E-02 | -1.32E+00 | 2.09E-01 | 9.03E-01 | 1.46E+00 | 2.73E-02 | 1.16E-01 | -1.24E+00 | 2.38E-01 | 9.10E-01 | 1.33E+00 | 1.97E-01 | 7.02E-01 | 1.08E+00 | 1.72E-01 | 4.86E-01 | 1.63E+00 | 2.07E-01 | 7.61E-01 | 1.13E+00 | 1.11E-01 | 3.05E-01 | 1.76E+00 |
| NACAP2 | 2.32E-03 | 1.21E-02 | -1.31E+00 | 5.49E-05 | 2.37E-04 | -1.63E+00 | 1.68E-03 | 7.15E-03 | -1.30E+00 | 6.00E-06 | 2.29E-05 | -1.91E+00 | 1.88E-03 | 6.71E-03 | -1.27E+00 | 3.85E-04 | 1.09E-03 | -1.71E+00 | 1.19E-06 | 4.37E-06 | -1.94E+00 | 1.65E-06 | 4.52E-06 | -1.94E+00 |
| SSBP2 | 2.32E-03 | 1.21E-02 | 1.05E+00 | 2.14E-01 | 9.26E-01 | 1.36E+00 | 9.58E-03 | 4.07E-02 | -1.08E+00 | 1.01E-02 | 3.87E-02 | 1.07E+00 | 9.34E-02 | 3.33E-01 | 1.27E+00 | 1.38E-01 | 3.89E-01 | 1.37E+00 | 6.94E-04 | 2.55E-03 | -1.05E+00 | 3.91E-03 | 1.07E-02 | -1.05E+00 |
| MORC2 | 2.33E-03 | 1.21E-02 | -1.32E+00 | 2.05E-04 | 8.88E-04 | -1.51E+00 | 1.28E-02 | 5.43E-02 | -1.20E+00 | 9.71E-02 | 3.71E-01 | 1.26E+00 | 2.04E-02 | 7.29E-02 | -1.08E+00 | 4.29E-02 | 1.21E-01 | 1.07E+00 | 2.76E-05 | 1.01E-04 | -1.66E+00 | 1.19E-03 | 3.28E-03 | -1.34E+00 |
| ADAM32 | 2.33E-03 | 1.21E-02 | -1.33E+00 | 7.79E-02 | 3.36E-01 | 1.25E+00 | 1.28E-03 | 5.43E-03 | -1.64E+00 | 3.86E-02 | 1.48E-01 | 1.01E+00 | 5.50E-02 | 1.96E-01 | -1.22E+00 | 3.31E-01 | 9.34E-01 | 1.40E+00 | 5.10E-02 | 1.87E-01 | 1.06E+00 | 2.10E-01 | 5.76E-01 | 1.27E+00 |
| CC2D1A | 2.33E-03 | 1.21E-02 | -1.14E+00 | 7.17E-02 | 3.10E-01 | 1.31E+00 | 4.73E-05 | 2.01E-04 | -1.45E+00 | 7.71E-03 | 2.94E-02 | -1.02E+00 | 8.77E-05 | 3.13E-04 | -1.67E+00 | 1.16E-02 | 3.26E-02 | -1.06E+00 | 2.25E-10 | 8.24E-10 | -1.54E+00 | 2.45E-03 | 6.73E-03 | -1.02E+00 |
| SLC2A11 | 2.33E-03 | 1.21E-02 | -1.12E+00 | 2.84E-02 | 1.23E-01 | 1.17E+00 | 1.17E-02 | 4.98E-02 | -1.23E+00 | 1.57E-01 | 5.99E-01 | 1.25E+00 | 3.32E-03 | 1.18E-02 | -1.37E+00 | 3.39E-02 | 9.57E-02 | -1.04E+00 | 5.97E-02 | 2.19E-01 | 1.03E+00 | 3.54E-01 | 9.72E-01 | 1.39E+00 |
| ALDH1L2 | 2.33E-03 | 1.22E-02 | -1.77E+00 | 6.70E-02 | 2.89E-01 | -1.03E+00 | 1.02E-04 | 4.34E-04 | -1.90E+00 | 1.05E-02 | 4.00E-02 | -1.22E+00 | 8.07E-02 | 2.88E-01 | -1.42E+00 | 8.62E-02 | 2.43E-01 | -1.42E+00 | 2.26E-01 | 8.30E-01 | -1.18E+00 | 2.40E-01 | 6.59E-01 | 1.05E+00 |
| BBS4 | 2.33E-03 | 1.22E-02 | -1.14E+00 | 9.36E-04 | 4.04E-03 | -1.15E+00 | 3.49E-04 | 1.48E-03 | -1.27E+00 | 1.57E-05 | 6.01E-05 | -1.35E+00 | 4.11E-03 | 1.46E-02 | -1.27E+00 | 1.20E-02 | 3.40E-02 | -1.06E+00 | 4.30E-02 | 1.58E-01 | 1.08E+00 | 2.02E-01 | 5.55E-01 | 1.18E+00 |
| AC018521.5 | 2.34E-03 | 1.22E-02 | -1.08E+00 | 6.10E-06 | 2.63E-05 | -1.31E+00 | 1.49E-02 | 6.33E-02 | -1.02E+00 | 2.32E-06 | 8.85E-06 | -1.41E+00 | 1.03E-02 | 3.68E-02 | 1.04E+00 | 1.25E-02 | 3.52E-02 | 1.11E+00 | 9.56E-02 | 3.51E-01 | 1.31E+00 | 2.49E-01 | 6.85E-01 | 1.10E+00 |
| TMEM108 | 2.35E-03 | 1.22E-02 | -1.54E+00 | 7.99E-03 | 3.45E-02 | -1.30E+00 | 6.74E-02 | 2.86E-01 | -1.36E+00 | 1.48E-01 | 5.66E-01 | -1.03E+00 | 5.64E-02 | 2.01E-01 | -1.24E+00 | 3.04E-02 | 8.56E-02 | -1.61E+00 | 2.76E-11 | 1.01E-10 | -2.15E+00 | 1.54E-06 | 4.22E-06 | -1.50E+00 |
| ZBTB2 | 2.35E-03 | 1.23E-02 | -1.33E+00 | 2.61E-03 | 1.13E-02 | -1.24E+00 | 5.14E-03 | 2.18E-02 | -1.17E+00 | 1.30E-03 | 4.97E-03 | -1.20E+00 | 7.33E-03 | 2.61E-02 | -1.14E+00 | 4.36E-04 | 1.23E-03 | -1.68E+00 | 7.76E-09 | 2.85E-08 | -1.65E+00 | 2.35E-07 | 6.46E-07 | -1.52E+00 |
| SRC | 2.35E-03 | 1.23E-02 | -1.34E+00 | 6.80E-02 | 2.94E-01 | 1.16E+00 | 1.16E-02 | 4.94E-02 | -1.45E+00 | 1.25E-01 | 4.79E-01 | 1.09E+00 | 1.50E-01 | 5.35E-01 | -1.03E+00 | 1.90E-01 | 5.35E-01 | -1.01E+00 | 2.77E-02 | 1.01E-01 | 1.45E+00 | 1.13E-01 | 3.09E-01 | 1.13E+00 |
| AC087650.1 | 2.35E-03 | 1.23E-02 | -1.24E+00 | 1.09E-05 | 4.69E-05 | -1.79E+00 | 2.07E-05 | 8.78E-05 | -1.53E+00 | 2.34E-04 | 8.96E-04 | -1.23E+00 | 5.77E-02 | 2.06E-01 | 1.05E+00 | 3.98E-03 | 1.12E-02 | -1.37E+00 | 5.68E-05 | 2.09E-04 | -1.85E+00 | 3.04E-03 | 8.36E-03 | -1.36E+00 |
| SPIN4 | 2.35E-03 | 1.23E-02 | -1.26E+00 | 9.48E-02 | 4.10E-01 | 1.22E+00 | 1.06E-02 | 4.51E-02 | -1.15E+00 | 2.42E-01 | 9.25E-01 | 1.39E+00 | 8.18E-03 | 2.92E-02 | -1.10E+00 | 1.14E-01 | 3.23E-01 | 1.23E+00 | 8.63E-09 | 3.17E-08 | -1.89E+00 | 2.16E-01 | 5.93E-01 | 1.11E+00 |
| ARHGEF25 | 2.36E-03 | 1.23E-02 | -1.34E+00 | 1.67E-02 | 7.23E-02 | -1.07E+00 | 1.94E-01 | 8.22E-01 | 1.13E+00 | 2.48E-01 | 9.48E-01 | 1.24E+00 | 7.00E-02 | 2.50E-01 | -1.13E+00 | 2.91E-02 | 8.20E-02 | -1.53E+00 | 1.20E-01 | 4.41E-01 | -1.15E+00 | 2.23E-01 | 6.11E-01 | -1.14E+00 |
| AL121658.1 | 2.36E-03 | 1.23E-02 | 2.40E+00 | 1.55E-01 | 6.71E-01 | 1.18E+00 | 1.42E-01 | 6.03E-01 | 1.35E+00 | 4.29E-03 | 1.64E-02 | -1.64E+00 | 2.62E-01 | 9.33E-01 | 1.28E+00 | 2.48E-01 | 6.99E-01 | 1.47E+00 | 2.15E-01 | 7.88E-01 | 1.27E+00 | 2.73E-01 | 7.51E-01 | 1.40E+00 |
| ZFAT | 2.36E-03 | 1.23E-02 | -1.35E+00 | 1.81E-02 | 7.82E-02 | -1.03E+00 | 4.17E-04 | 1.77E-03 | -1.47E+00 | 3.66E-04 | 1.40E-03 | -1.31E+00 | 1.58E-05 | 5.63E-05 | -1.71E+00 | 1.85E-04 | 5.23E-04 | -1.33E+00 | 1.39E-02 | 5.08E-02 | -1.04E+00 | 6.14E-02 | 1.69E-01 | 1.03E+00 |
| SGCB | 2.36E-03 | 1.23E-02 | -1.22E+00 | 8.48E-03 | 3.66E-02 | 1.04E+00 | 1.58E-01 | 6.72E-01 | 1.16E+00 | 2.37E-01 | 9.03E-01 | 1.47E+00 | 2.84E-02 | 1.01E-01 | -1.82E+00 | 7.79E-02 | 2.20E-01 | -1.35E+00 | 2.66E-01 | 9.77E-01 | 1.21E+00 | 1.70E-01 | 4.67E-01 | 1.79E+00 |
| TMSB10P1 | 2.36E-03 | 1.23E-02 | 1.73E+00 | 1.48E-06 | 6.39E-06 | 2.23E+00 | 6.52E-02 | 2.77E-01 | 1.47E+00 | 1.79E-03 | 6.85E-03 | 1.91E+00 | 2.09E-01 | 7.44E-01 | 1.43E+00 | 5.45E-03 | 1.54E-02 | 1.90E+00 | 3.99E-02 | 1.46E-01 | 1.47E+00 | 1.96E-04 | 5.40E-04 | 1.83E+00 |
| C19orf44 | 2.37E-03 | 1.23E-02 | -1.74E+00 | 3.91E-02 | 1.69E-01 | -1.06E+00 | 1.57E-04 | 6.67E-04 | -1.51E+00 | 5.95E-04 | 2.27E-03 | -1.41E+00 | 2.48E-01 | 8.83E-01 | 1.11E+00 | 2.60E-01 | 7.32E-01 | 1.00E+00 | 9.96E-03 | 3.65E-02 | -1.25E+00 | 7.12E-02 | 1.96E-01 | -1.11E+00 |
| DNAJB9 | 2.37E-03 | 1.23E-02 | 1.83E+00 | 3.33E-02 | 1.44E-01 | 1.20E+00 | 8.04E-02 | 3.42E-01 | 1.14E+00 | 4.24E-05 | 1.62E-04 | -1.21E+00 | 2.32E-02 | 8.28E-02 | 1.15E+00 | 2.23E-06 | 6.29E-06 | -1.38E+00 | 9.36E-02 | 3.43E-01 | 1.19E+00 | 2.48E-06 | 6.82E-06 | -1.60E+00 |
| TTC30A | 2.37E-03 | 1.24E-02 | -1.08E+00 | 4.97E-05 | 2.15E-04 | -1.44E+00 | 2.11E-01 | 8.98E-01 | 1.30E+00 | 1.69E-01 | 6.47E-01 | 1.24E+00 | 1.36E-01 | 4.84E-01 | 1.08E+00 | 2.89E-01 | 8.14E-01 | 1.32E+00 | 3.81E-02 | 1.40E-01 | -1.08E+00 | 1.11E-01 | 3.05E-01 | -1.00E+00 |
| SMARCA2 | 2.38E-03 | 1.24E-02 | 1.56E+00 | 8.89E-03 | 3.84E-02 | 1.65E+00 | 2.04E-01 | 8.66E-01 | 1.25E+00 | 2.02E-01 | 7.70E-01 | 1.37E+00 | 7.48E-02 | 2.67E-01 | 1.14E+00 | 2.09E-01 | 5.90E-01 | 1.16E+00 | 1.26E-02 | 4.63E-02 | 1.08E+00 | 2.47E-01 | 6.78E-01 | 1.21E+00 |
| NAA40 | 2.38E-03 | 1.24E-02 | -1.19E+00 | 2.28E-01 | 9.85E-01 | 1.40E+00 | 1.51E-07 | 6.41E-07 | -1.70E+00 | 6.73E-03 | 2.57E-02 | 1.05E+00 | 1.39E-04 | 4.95E-04 | -1.39E+00 | 5.73E-04 | 1.62E-03 | -1.26E+00 | 2.65E-08 | 9.73E-08 | -1.71E+00 | 2.23E-05 | 6.13E-05 | -1.41E+00 |
| TBC1D22B | 2.38E-03 | 1.24E-02 | -1.19E+00 | 3.31E-03 | 1.43E-02 | -1.06E+00 | 3.62E-02 | 1.54E-01 | 1.01E+00 | 5.57E-02 | 2.13E-01 | 1.19E+00 | 2.26E-01 | 8.07E-01 | 1.23E+00 | 1.63E-01 | 4.60E-01 | 2.08E+00 | 2.73E-05 | 1.00E-04 | -1.76E+00 | 6.77E-04 | 1.86E-03 | -1.42E+00 |
| RPS12P26 | 2.38E-03 | 1.24E-02 | 1.01E+00 | 2.55E-02 | 1.10E-01 | 1.13E+00 | 9.60E-02 | 4.08E-01 | 1.12E+00 | 3.72E-02 | 1.42E-01 | 1.28E+00 | 3.09E-02 | 1.10E-01 | 1.01E+00 | 1.97E-01 | 5.56E-01 | 1.14E+00 | 3.57E-02 | 1.31E-01 | 1.04E+00 | 1.46E-01 | 4.00E-01 | 1.22E+00 |
| FZD2 | 2.39E-03 | 1.24E-02 | -1.60E+00 | 5.81E-02 | 2.51E-01 | 1.07E+00 | 1.00E-04 | 4.25E-04 | -1.66E+00 | 1.33E-01 | 5.06E-01 | 1.30E+00 | 1.83E-07 | 6.52E-07 | -4.58E+00 | 9.93E-04 | 2.80E-03 | -1.57E+00 | 1.16E-06 | 4.26E-06 | -2.52E+00 | 2.41E-03 | 6.63E-03 | -1.39E+00 |
| ARHGAP32 | 2.39E-03 | 1.24E-02 | -1.24E+00 | 1.18E-01 | 5.09E-01 | 1.29E+00 | 2.99E-03 | 1.27E-02 | -1.35E+00 | 1.03E-02 | 3.92E-02 | 1.01E+00 | 1.95E-01 | 6.97E-01 | 1.65E+00 | 2.65E-01 | 7.47E-01 | 1.66E+00 | 2.45E-01 | 8.98E-01 | 1.24E+00 | 8.23E-02 | 2.26E-01 | 1.50E+00 |
| CD79B | 2.39E-03 | 1.24E-02 | 1.08E+00 | 4.36E-02 | 1.88E-01 | 1.36E+00 | 1.23E-03 | 5.23E-03 | -1.03E+00 | 8.71E-04 | 3.33E-03 | 1.12E+00 | 7.08E-03 | 2.52E-02 | -1.09E+00 | 2.63E-02 | 7.43E-02 | 1.01E+00 | 8.77E-02 | 3.22E-01 | 1.05E+00 | 1.06E-01 | 2.91E-01 | 1.07E+00 |
| ACO2 | 2.39E-03 | 1.25E-02 | -1.12E+00 | 1.35E-03 | 5.85E-03 | -1.12E+00 | 9.52E-02 | 4.05E-01 | 1.24E+00 | 8.49E-02 | 3.24E-01 | -1.03E+00 | 2.76E-01 | 9.85E-01 | 1.30E+00 | 1.60E-01 | 4.51E-01 | 1.07E+00 | 1.19E-01 | 4.38E-01 | 1.50E+00 | 1.89E-01 | 5.19E-01 | 1.55E+00 |
| RPL31P28 | 2.40E-03 | 1.25E-02 | -1.32E+00 | 1.04E-01 | 4.48E-01 | 1.34E+00 | 1.88E-01 | 7.99E-01 | 1.04E+00 | 2.44E-01 | 9.33E-01 | 1.27E+00 | 2.55E-02 | 9.08E-02 | -1.28E+00 | 1.56E-01 | 4.41E-01 | 1.14E+00 | 9.22E-02 | 3.38E-01 | -1.08E+00 | 2.95E-01 | 8.10E-01 | 1.12E+00 |
| ZNF226 | 2.40E-03 | 1.25E-02 | 1.02E+00 | 6.69E-02 | 2.89E-01 | 1.32E+00 | 3.09E-03 | 1.31E-02 | -1.09E+00 | 1.06E-02 | 4.06E-02 | 1.08E+00 | 6.91E-03 | 2.46E-02 | -1.03E+00 | 1.24E-01 | 3.51E-01 | 1.19E+00 | 4.09E-02 | 1.50E-01 | 1.11E+00 | 2.04E-02 | 5.61E-02 | 1.02E+00 |
| AL356055.1 | 2.40E-03 | 1.25E-02 | 2.85E+00 | 4.35E-06 | 1.88E-05 | 4.43E+00 | 6.08E-02 | 2.58E-01 | 1.96E+00 | 8.41E-04 | 3.21E-03 | 3.72E+00 | 1.94E-02 | 6.93E-02 | 3.13E+00 | 2.22E-04 | 6.27E-04 | 4.78E+00 | 3.71E-02 | 1.36E-01 | 2.20E+00 | 3.24E-02 | 8.91E-02 | 2.61E+00 |
| HSPE1P3 | 2.40E-03 | 1.25E-02 | -1.34E+00 | 2.21E-02 | 9.55E-02 | 1.10E+00 | 6.27E-02 | 2.66E-01 | 1.47E+00 | 1.12E-01 | 4.29E-01 | 1.57E+00 | 1.34E-01 | 4.78E-01 | 1.12E+00 | 2.33E-01 | 6.58E-01 | 1.67E+00 | 8.28E-02 | 3.04E-01 | 1.21E+00 | 1.80E-01 | 4.96E-01 | 1.48E+00 |
| ST7-OT4 | 2.41E-03 | 1.26E-02 | -1.21E+00 | 2.86E-04 | 1.23E-03 | -1.26E+00 | 6.75E-05 | 2.87E-04 | -1.43E+00 | 5.66E-06 | 2.16E-05 | -1.44E+00 | 1.21E-02 | 4.30E-02 | -1.17E+00 | 4.18E-04 | 1.18E-03 | -1.83E+00 | 2.24E-04 | 8.21E-04 | -1.54E+00 | 1.75E-04 | 4.82E-04 | -1.57E+00 |
| LNPK | 2.42E-03 | 1.26E-02 | -1.13E+00 | 6.33E-02 | 2.73E-01 | 1.26E+00 | 1.81E-02 | 7.69E-02 | -1.13E+00 | 7.68E-02 | 2.93E-01 | 1.14E+00 | 2.10E-07 | 7.50E-07 | -1.90E+00 | 1.96E-04 | 5.52E-04 | -1.29E+00 | 3.09E-04 | 1.13E-03 | -1.11E+00 | 1.29E-02 | 3.56E-02 | 1.06E+00 |
| PXMP2 | 2.42E-03 | 1.26E-02 | 1.05E+00 | 1.29E-05 | 5.56E-05 | -1.08E+00 | 2.07E-01 | 8.78E-01 | 1.23E+00 | 1.42E-06 | 5.41E-06 | -1.19E+00 | 4.52E-02 | 1.61E-01 | 1.00E+00 | 6.21E-03 | 1.75E-02 | -1.33E+00 | 2.19E-01 | 8.06E-01 | 1.33E+00 | 5.53E-02 | 1.52E-01 | -1.00E+00 |
| DVL3 | 2.42E-03 | 1.26E-02 | -1.14E+00 | 9.76E-03 | 4.22E-02 | 1.09E+00 | 1.40E-02 | 5.93E-02 | -1.08E+00 | 1.20E-02 | 4.57E-02 | 1.03E+00 | 5.10E-02 | 1.82E-01 | 1.14E+00 | 3.85E-02 | 1.08E-01 | 1.07E+00 | 4.84E-05 | 1.78E-04 | -1.10E+00 | 1.35E-02 | 3.72E-02 | 1.00E+00 |
| LIPT2 | 2.42E-03 | 1.26E-02 | -1.37E+00 | 5.70E-02 | 2.46E-01 | 1.15E+00 | 1.33E-02 | 5.66E-02 | -1.37E+00 | 1.39E-01 | 5.32E-01 | 1.44E+00 | 2.69E-01 | 9.60E-01 | 1.13E+00 | 2.03E-01 | 5.71E-01 | 1.52E+00 | 6.71E-04 | 2.46E-03 | -1.45E+00 | 6.99E-05 | 1.92E-04 | -1.80E+00 |
| SLC30A7 | 2.43E-03 | 1.27E-02 | -1.04E+00 | 3.79E-02 | 1.64E-01 | 1.27E+00 | 8.81E-05 | 3.74E-04 | -1.27E+00 | 2.34E-03 | 8.95E-03 | 1.01E+00 | 5.86E-05 | 2.09E-04 | -1.21E+00 | 1.88E-03 | 5.30E-03 | -1.06E+00 | 1.15E-03 | 4.22E-03 | -1.09E+00 | 1.61E-01 | 4.43E-01 | 1.19E+00 |
| RPL30P3 | 2.43E-03 | 1.27E-02 | -1.04E+00 | 1.30E-07 | 5.61E-07 | -1.45E+00 | 9.50E-02 | 4.04E-01 | 1.06E+00 | 2.74E-04 | 1.05E-03 | -1.34E+00 | 1.08E-01 | 3.86E-01 | 1.20E+00 | 7.29E-04 | 2.06E-03 | -1.62E+00 | 2.59E-02 | 9.49E-02 | -1.14E+00 | 2.59E-03 | 7.13E-03 | -1.35E+00 |
| SSBP3-AS1 | 2.44E-03 | 1.27E-02 | -1.48E+00 | 2.11E-01 | 9.11E-01 | 1.36E+00 | 9.17E-04 | 3.89E-03 | -1.32E+00 | 6.45E-02 | 2.46E-01 | 1.25E+00 | 2.24E-03 | 7.99E-03 | -1.64E+00 | 2.03E-01 | 5.72E-01 | 1.22E+00 | 4.04E-03 | 1.48E-02 | -1.61E+00 | 2.66E-01 | 7.30E-01 | 1.08E+00 |
| ZNF224 | 2.44E-03 | 1.27E-02 | 1.00E+00 | 1.43E-01 | 6.18E-01 | 1.49E+00 | 1.72E-02 | 7.30E-02 | 1.02E+00 | 1.14E-01 | 4.35E-01 | 1.35E+00 | 9.79E-02 | 3.49E-01 | 1.10E+00 | 2.90E-01 | 8.17E-01 | 1.28E+00 | 1.81E-01 | 6.65E-01 | 1.51E+00 | 3.37E-01 | 9.26E-01 | 1.37E+00 |
| UNC119B | 2.44E-03 | 1.27E-02 | 1.01E+00 | 5.81E-04 | 2.51E-03 | 1.07E+00 | 2.16E-01 | 9.16E-01 | 1.18E+00 | 4.11E-02 | 1.57E-01 | 1.16E+00 | 2.50E-01 | 8.93E-01 | 1.17E+00 | 3.15E-01 | 8.89E-01 | 1.37E+00 | 2.59E-01 | 9.50E-01 | 1.26E+00 | 3.33E-01 | 9.14E-01 | 1.23E+00 |
| HNRNPU | 2.44E-03 | 1.27E-02 | 1.02E+00 | 1.41E-03 | 6.09E-03 | -1.02E+00 | 2.20E-01 | 9.34E-01 | 1.07E+00 | 2.22E-01 | 8.50E-01 | 1.02E+00 | 5.36E-02 | 1.91E-01 | 1.02E+00 | 2.91E-01 | 8.21E-01 | 1.09E+00 | 6.62E-04 | 2.43E-03 | -1.16E+00 | 3.06E-01 | 8.41E-01 | 1.02E+00 |
| ZNF571 | 2.45E-03 | 1.27E-02 | -1.22E+00 | 1.21E-02 | 5.21E-02 | 1.04E+00 | 2.27E-03 | 9.66E-03 | -1.29E+00 | 8.49E-05 | 3.24E-04 | -1.36E+00 | 2.46E-02 | 8.78E-02 | -1.02E+00 | 7.21E-03 | 2.03E-02 | -1.18E+00 | 1.72E-01 | 6.32E-01 | 1.19E+00 | 1.39E-01 | 3.81E-01 | 1.08E+00 |
| UBXN1 | 2.45E-03 | 1.28E-02 | 1.05E+00 | 2.44E-03 | 1.06E-02 | 1.09E+00 | 2.84E-02 | 1.21E-01 | -1.06E+00 | 8.84E-04 | 3.38E-03 | -1.05E+00 | 1.05E-02 | 3.75E-02 | -1.03E+00 | 1.10E-02 | 3.10E-02 | -1.03E+00 | 2.25E-01 | 8.26E-01 | 1.09E+00 | 4.78E-02 | 1.31E-01 | -1.01E+00 |
| SEPT6 | 2.45E-03 | 1.28E-02 | -1.12E+00 | 2.65E-02 | 1.15E-01 | 1.21E+00 | 7.25E-04 | 3.08E-03 | -1.21E+00 | 2.02E-01 | 7.71E-01 | 1.03E+00 | 2.68E-03 | 9.54E-03 | 1.03E+00 | 1.30E-01 | 3.68E-01 | 1.16E+00 | 1.40E-02 | 5.14E-02 | -1.08E+00 | 2.77E-03 | 7.62E-03 | -1.14E+00 |
| AC004148.2 | 2.45E-03 | 1.28E-02 | -1.34E+00 | 6.19E-03 | 2.67E-02 | -1.03E+00 | 1.25E-01 | 5.30E-01 | -1.05E+00 | 7.36E-02 | 2.81E-01 | -1.02E+00 | 1.66E-01 | 5.93E-01 | 1.16E+00 | 3.49E-01 | 9.85E-01 | 1.49E+00 | 1.67E-01 | 6.14E-01 | 1.22E+00 | 1.35E-02 | 3.70E-02 | 1.83E+00 |
| AC073896.2 | 2.46E-03 | 1.28E-02 | -1.60E+00 | 6.41E-03 | 2.77E-02 | -1.34E+00 | 2.17E-02 | 9.21E-02 | -1.12E+00 | 3.43E-03 | 1.31E-02 | -1.29E+00 | 7.77E-02 | 2.77E-01 | -1.37E+00 | 3.35E-01 | 9.45E-01 | 1.33E+00 | 4.25E-03 | 1.56E-02 | -1.52E+00 | 5.40E-02 | 1.48E-01 | -1.13E+00 |
| ACTG1P9 | 2.46E-03 | 1.28E-02 | 2.17E+00 | 5.85E-02 | 2.53E-01 | 1.86E+00 | 3.73E-04 | 1.58E-03 | 2.11E+00 | 5.30E-02 | 2.03E-01 | 1.75E+00 | 1.48E-01 | 5.27E-01 | 1.69E+00 | 2.11E-01 | 5.94E-01 | 1.16E+00 | 8.84E-03 | 3.25E-02 | 2.71E+00 | 4.21E-02 | 1.16E-01 | 2.36E+00 |
| AC104794.2 | 2.46E-03 | 1.28E-02 | -1.25E+00 | 1.30E-03 | 5.61E-03 | -1.26E+00 | 3.31E-02 | 1.40E-01 | -1.28E+00 | 1.51E-01 | 5.78E-01 | 1.15E+00 | 1.20E-02 | 4.30E-02 | -1.16E+00 | 5.71E-04 | 1.61E-03 | -1.72E+00 | 4.12E-02 | 1.51E-01 | 1.09E+00 | 1.69E-02 | 4.64E-02 | 1.03E+00 |
| ARHGAP15 | 2.47E-03 | 1.29E-02 | 1.12E+00 | 2.88E-03 | 1.24E-02 | 1.31E+00 | 3.90E-03 | 1.66E-02 | -1.03E+00 | 1.90E-02 | 7.26E-02 | 1.18E+00 | 1.16E-02 | 4.14E-02 | 1.07E+00 | 1.44E-01 | 4.06E-01 | 1.22E+00 | 7.95E-03 | 2.92E-02 | -1.01E+00 | 7.83E-02 | 2.15E-01 | 1.08E+00 |
| CHIC2 | 2.47E-03 | 1.29E-02 | 1.11E+00 | 6.72E-10 | 2.91E-09 | -1.13E+00 | 4.63E-10 | 1.97E-09 | -1.42E+00 | 3.89E-14 | 1.49E-13 | -1.60E+00 | 7.44E-04 | 2.65E-03 | -1.03E+00 | 5.56E-07 | 1.57E-06 | -1.34E+00 | 3.60E-08 | 1.32E-07 | -1.38E+00 | 2.63E-06 | 7.23E-06 | -1.35E+00 |
| AC010186.3 | 2.47E-03 | 1.29E-02 | -1.14E+00 | 4.13E-03 | 1.78E-02 | -1.05E+00 | 2.27E-01 | 9.66E-01 | 1.16E+00 | 1.04E-01 | 3.99E-01 | 1.74E+00 | 2.53E-01 | 9.01E-01 | 1.19E+00 | 1.67E-01 | 4.70E-01 | 1.56E+00 | 3.06E-04 | 1.12E-03 | -1.70E+00 | 1.68E-03 | 4.61E-03 | -1.42E+00 |
| NAA60 | 2.47E-03 | 1.29E-02 | -1.01E+00 | 2.44E-02 | 1.05E-01 | 1.20E+00 | 2.55E-05 | 1.08E-04 | -1.19E+00 | 4.44E-07 | 1.69E-06 | -1.30E+00 | 2.77E-02 | 9.86E-02 | 1.06E+00 | 1.31E-02 | 3.70E-02 | -1.00E+00 | 1.64E-04 | 6.03E-04 | -1.13E+00 | 5.45E-04 | 1.50E-03 | -1.06E+00 |
| PHF10 | 2.47E-03 | 1.29E-02 | -1.18E+00 | 1.43E-01 | 6.17E-01 | 1.33E+00 | 5.39E-04 | 2.29E-03 | -1.15E+00 | 2.43E-02 | 9.28E-02 | 1.10E+00 | 9.02E-07 | 3.22E-06 | -1.60E+00 | 1.23E-04 | 3.46E-04 | -1.18E+00 | 7.65E-07 | 2.81E-06 | -1.41E+00 | 4.43E-04 | 1.22E-03 | -1.18E+00 |
| ZNF879 | 2.47E-03 | 1.29E-02 | -1.31E+00 | 1.02E-02 | 4.43E-02 | -1.11E+00 | 4.11E-02 | 1.75E-01 | -1.22E+00 | 2.04E-01 | 7.80E-01 | 1.27E+00 | 4.76E-02 | 1.70E-01 | 1.89E+00 | 4.65E-02 | 1.31E-01 | 2.29E+00 | 7.91E-02 | 2.90E-01 | -1.06E+00 | 3.46E-01 | 9.50E-01 | 1.36E+00 |
| FOSL1 | 2.48E-03 | 1.29E-02 | -1.05E+00 | 4.47E-10 | 1.93E-09 | -3.04E+00 | 7.08E-06 | 3.01E-05 | -1.42E+00 | 1.70E-14 | 6.48E-14 | -3.00E+00 | 6.64E-05 | 2.37E-04 | -2.58E+00 | 8.53E-05 | 2.41E-04 | -2.82E+00 | 7.52E-04 | 2.76E-03 | -1.47E+00 | 2.29E-03 | 6.28E-03 | -1.34E+00 |
| SPIN3 | 2.48E-03 | 1.29E-02 | -1.20E+00 | 4.51E-02 | 1.95E-01 | 1.27E+00 | 2.08E-04 | 8.76E-04 | -1.65E+00 | 2.27E-02 | 8.69E-02 | 1.06E+00 | 1.12E-01 | 4.00E-01 | 1.20E+00 | 2.23E-01 | 6.28E-01 | 1.86E+00 | 1.60E-05 | 5.88E-05 | -1.55E+00 | 5.68E-04 | 1.56E-03 | -1.33E+00 |
| HSF2 | 2.48E-03 | 1.29E-02 | -1.11E+00 | 6.32E-02 | 2.73E-01 | 1.24E+00 | 3.35E-02 | 1.42E-01 | -1.09E+00 | 7.78E-03 | 2.97E-02 | -1.07E+00 | 2.35E-04 | 8.40E-04 | -1.41E+00 | 1.06E-03 | 2.99E-03 | -1.31E+00 | 2.83E-03 | 1.04E-02 | -1.08E+00 | 6.03E-02 | 1.66E-01 | 1.04E+00 |
| AC010768.2 | 2.48E-03 | 1.29E-02 | 2.39E+00 | 3.88E-03 | 1.67E-02 | 2.71E+00 | 2.30E-02 | 9.76E-02 | 2.05E+00 | 4.44E-02 | 1.70E-01 | 2.14E+00 | 9.81E-02 | 3.50E-01 | 1.79E+00 | 5.90E-02 | 1.66E-01 | 1.87E+00 | 2.64E-01 | 9.69E-01 | 1.20E+00 | 5.49E-02 | 1.51E-01 | 1.83E+00 |
| ZNF117 | 2.48E-03 | 1.29E-02 | -1.06E+00 | 3.76E-03 | 1.63E-02 | 1.05E+00 | 2.66E-04 | 1.13E-03 | -1.44E+00 | 7.68E-04 | 2.93E-03 | -1.15E+00 | 5.31E-07 | 1.89E-06 | -1.55E+00 | 6.01E-04 | 1.69E-03 | -1.12E+00 | 2.86E-04 | 1.05E-03 | -1.21E+00 | 1.35E-02 | 3.70E-02 | -1.03E+00 |
| FEM1C | 2.48E-03 | 1.29E-02 | -1.08E+00 | 4.11E-11 | 1.78E-10 | -2.04E+00 | 2.03E-02 | 8.61E-02 | -1.19E+00 | 5.62E-07 | 2.15E-06 | -1.87E+00 | 1.36E-01 | 4.86E-01 | 1.69E+00 | 1.78E-02 | 5.01E-02 | -1.14E+00 | 2.47E-01 | 9.08E-01 | 1.19E+00 | 4.96E-02 | 1.36E-01 | -1.13E+00 |
| BBIP1 | 2.48E-03 | 1.29E-02 | 1.02E+00 | 2.28E-02 | 9.86E-02 | 1.24E+00 | 3.39E-02 | 1.44E-01 | 1.02E+00 | 1.02E-02 | 3.90E-02 | 1.10E+00 | 1.84E-02 | 6.58E-02 | -1.01E+00 | 6.52E-02 | 1.84E-01 | 1.13E+00 | 6.88E-03 | 2.52E-02 | -1.05E+00 | 5.62E-02 | 1.54E-01 | 1.07E+00 |
| HEATR5B | 2.48E-03 | 1.29E-02 | -1.07E+00 | 2.56E-02 | 1.10E-01 | 1.21E+00 | 2.34E-04 | 9.93E-04 | -1.21E+00 | 4.88E-02 | 1.87E-01 | 1.18E+00 | 1.60E-03 | 5.72E-03 | -1.08E+00 | 3.32E-03 | 9.35E-03 | -1.06E+00 | 2.29E-02 | 8.41E-02 | -1.01E+00 | 8.16E-02 | 2.24E-01 | 1.12E+00 |
| P2RY14 | 2.49E-03 | 1.30E-02 | 1.06E+00 | 1.63E-03 | 7.04E-03 | 1.13E+00 | 7.97E-02 | 3.39E-01 | 1.07E+00 | 4.18E-02 | 1.60E-01 | 1.02E+00 | 8.09E-03 | 2.88E-02 | 2.26E+00 | 8.01E-02 | 2.26E-01 | 1.41E+00 | 5.22E-05 | 1.92E-04 | 2.13E+00 | 3.62E-02 | 9.96E-02 | 1.68E+00 |
| TRMT12 | 2.50E-03 | 1.30E-02 | -1.03E+00 | 7.83E-04 | 3.38E-03 | -1.13E+00 | 5.43E-03 | 2.31E-02 | -1.15E+00 | 3.99E-04 | 1.52E-03 | -1.16E+00 | 2.64E-02 | 9.42E-02 | -1.01E+00 | 5.22E-02 | 1.47E-01 | 1.03E+00 | 1.26E-02 | 4.63E-02 | -1.05E+00 | 1.92E-01 | 5.28E-01 | 1.17E+00 |
| CD52 | 2.51E-03 | 1.31E-02 | 1.24E+00 | 4.36E-02 | 1.89E-01 | 1.15E+00 | 5.47E-02 | 2.33E-01 | 1.10E+00 | 2.57E-01 | 9.81E-01 | 1.13E+00 | 3.68E-19 | 1.31E-18 | -1.62E+00 | 3.14E-29 | 8.84E-29 | -1.74E+00 | 1.81E-09 | 6.63E-09 | -1.32E+00 | 1.21E-06 | 3.32E-06 | -1.18E+00 |
| HECTD1 | 2.51E-03 | 1.31E-02 | 1.03E+00 | 1.08E-01 | 4.66E-01 | 1.31E+00 | 3.40E-03 | 1.45E-02 | -1.08E+00 | 2.27E-01 | 8.67E-01 | 1.29E+00 | 1.94E-04 | 6.92E-04 | -1.02E+00 | 2.43E-02 | 6.84E-02 | 1.13E+00 | 6.07E-04 | 2.23E-03 | -1.01E+00 | 8.84E-02 | 2.43E-01 | 1.14E+00 |
| LHFPL2 | 2.51E-03 | 1.31E-02 | -1.26E+00 | 5.23E-02 | 2.26E-01 | 1.23E+00 | 8.34E-03 | 3.54E-02 | -1.29E+00 | 1.99E-01 | 7.61E-01 | 1.39E+00 | 1.84E-02 | 6.57E-02 | 1.05E+00 | 2.36E-01 | 6.65E-01 | 1.64E+00 | 1.75E-03 | 6.41E-03 | 1.04E+00 | 2.81E-01 | 7.71E-01 | 1.26E+00 |
| FOXO3 | 2.51E-03 | 1.31E-02 | 1.05E+00 | 6.41E-03 | 2.77E-02 | 1.13E+00 | 4.82E-03 | 2.05E-02 | 1.04E+00 | 9.49E-04 | 3.62E-03 | 1.06E+00 | 4.16E-02 | 1.48E-01 | 1.15E+00 | 8.07E-02 | 2.28E-01 | 1.14E+00 | 2.14E-16 | 7.86E-16 | -1.84E+00 | 5.61E-11 | 1.54E-10 | -1.54E+00 |
| FKBPL | 2.52E-03 | 1.31E-02 | -1.56E+00 | 8.54E-03 | 3.69E-02 | -1.24E+00 | 3.49E-05 | 1.48E-04 | -1.65E+00 | 8.78E-04 | 3.35E-03 | -1.21E+00 | 3.74E-04 | 1.33E-03 | -2.02E+00 | 5.14E-04 | 1.45E-03 | -1.69E+00 | 9.44E-02 | 3.46E-01 | 1.11E+00 | 2.93E-01 | 8.06E-01 | 1.29E+00 |
| NEDD4L | 2.52E-03 | 1.31E-02 | -1.19E+00 | 4.38E-03 | 1.89E-02 | -1.06E+00 | 2.66E-02 | 1.13E-01 | -1.22E+00 | 3.32E-02 | 1.27E-01 | -1.02E+00 | 5.79E-02 | 2.07E-01 | -1.03E+00 | 1.09E-01 | 3.08E-01 | 1.06E+00 | 3.52E-08 | 1.29E-07 | -1.28E+00 | 2.37E-09 | 6.52E-09 | -1.40E+00 |
| AKAP10 | 2.52E-03 | 1.32E-02 | -1.05E+00 | 2.47E-03 | 1.07E-02 | 1.06E+00 | 4.37E-07 | 1.86E-06 | -1.51E+00 | 3.89E-05 | 1.49E-04 | -1.17E+00 | 6.00E-03 | 2.14E-02 | -1.07E+00 | 1.19E-01 | 3.35E-01 | 1.22E+00 | 1.49E-03 | 5.47E-03 | -1.19E+00 | 9.20E-02 | 2.53E-01 | 1.12E+00 |
| LINC00861 | 2.53E-03 | 1.32E-02 | 3.83E+00 | 8.80E-03 | 3.80E-02 | 3.75E+00 | 1.97E-01 | 8.37E-01 | 1.07E+00 | 1.99E-01 | 7.62E-01 | 1.33E+00 | 1.42E-03 | 5.05E-03 | 6.30E+00 | 9.69E-02 | 2.73E-01 | 3.08E+00 | 2.50E-02 | 9.19E-02 | 3.02E+00 | 2.99E-01 | 8.22E-01 | 1.33E+00 |
| ZNF442 | 2.53E-03 | 1.32E-02 | -1.50E+00 | 7.94E-03 | 3.43E-02 | -1.25E+00 | 6.25E-04 | 2.66E-03 | -1.72E+00 | 2.22E-03 | 8.47E-03 | -1.48E+00 | 1.07E-01 | 3.82E-01 | 1.68E+00 | 1.63E-01 | 4.60E-01 | 1.78E+00 | 7.49E-04 | 2.75E-03 | -1.35E+00 | 5.06E-03 | 1.39E-02 | -1.34E+00 |
| HNRNPUL1 | 2.53E-03 | 1.32E-02 | 1.03E+00 | 2.60E-09 | 1.12E-08 | -1.10E+00 | 1.82E-02 | 7.73E-02 | -1.07E+00 | 1.51E-02 | 5.77E-02 | -1.02E+00 | 2.43E-08 | 8.66E-08 | -1.23E+00 | 1.01E-08 | 2.84E-08 | -1.30E+00 | 7.72E-06 | 2.83E-05 | -1.32E+00 | 4.86E-04 | 1.34E-03 | -1.26E+00 |
| BAHD1 | 2.53E-03 | 1.32E-02 | -1.29E+00 | 7.70E-03 | 3.33E-02 | -1.09E+00 | 1.73E-02 | 7.36E-02 | -1.13E+00 | 9.87E-02 | 3.77E-01 | 1.20E+00 | 1.43E-03 | 5.11E-03 | -1.30E+00 | 3.19E-02 | 8.99E-02 | 1.05E+00 | 1.11E-03 | 4.07E-03 | -1.41E+00 | 1.03E-01 | 2.83E-01 | 1.06E+00 |
| EMILIN1 | 2.53E-03 | 1.32E-02 | -1.24E+00 | 5.08E-02 | 2.19E-01 | 1.23E+00 | 1.50E-09 | 6.36E-09 | -1.55E+00 | 1.76E-04 | 6.72E-04 | 1.07E+00 | 1.37E-05 | 4.89E-05 | -1.72E+00 | 2.09E-03 | 5.89E-03 | -1.25E+00 | 5.16E-02 | 1.89E-01 | 1.08E+00 | 2.49E-01 | 6.84E-01 | 1.20E+00 |
| NOCT | 2.54E-03 | 1.32E-02 | -1.16E+00 | 1.46E-05 | 6.32E-05 | -1.51E+00 | 2.65E-03 | 1.13E-02 | -1.19E+00 | 2.72E-05 | 1.04E-04 | -1.35E+00 | 1.34E-02 | 4.79E-02 | -1.05E+00 | 7.44E-03 | 2.10E-02 | -1.09E+00 | 9.75E-05 | 3.58E-04 | -1.43E+00 | 1.03E-04 | 2.82E-04 | -1.45E+00 |
| MROH8 | 2.54E-03 | 1.32E-02 | -1.61E+00 | 2.67E-03 | 1.15E-02 | -1.70E+00 | 5.32E-02 | 2.26E-01 | -1.05E+00 | 3.81E-02 | 1.45E-01 | 1.02E+00 | 1.73E-01 | 6.16E-01 | -1.16E+00 | 1.11E-01 | 3.13E-01 | -1.40E+00 | 9.20E-05 | 3.37E-04 | -1.57E+00 | 4.91E-02 | 1.35E-01 | 1.05E+00 |
| NBPF9 | 2.54E-03 | 1.32E-02 | -1.39E+00 | 1.05E-03 | 4.54E-03 | -1.38E+00 | 7.41E-04 | 3.15E-03 | -1.54E+00 | 6.60E-04 | 2.52E-03 | -1.44E+00 | 1.30E-02 | 4.63E-02 | -1.27E+00 | 3.80E-02 | 1.07E-01 | -1.05E+00 | 5.60E-03 | 2.05E-02 | -1.40E+00 | 1.05E-01 | 2.88E-01 | -1.08E+00 |
| VPS33B | 2.55E-03 | 1.33E-02 | -1.34E+00 | 1.21E-02 | 5.24E-02 | -1.05E+00 | 8.09E-03 | 3.44E-02 | -1.13E+00 | 7.40E-02 | 2.83E-01 | 1.16E+00 | 2.21E-02 | 7.88E-02 | -1.05E+00 | 1.06E-01 | 2.98E-01 | 1.13E+00 | 4.12E-04 | 1.51E-03 | -1.21E+00 | 9.57E-02 | 2.63E-01 | 1.08E+00 |
| DBF4 | 2.55E-03 | 1.33E-02 | -1.17E+00 | 9.70E-03 | 4.19E-02 | 1.04E+00 | 1.59E-03 | 6.76E-03 | -1.19E+00 | 1.46E-04 | 5.58E-04 | -1.25E+00 | 1.42E-01 | 5.05E-01 | 1.13E+00 | 6.05E-02 | 1.70E-01 | -1.02E+00 | 6.89E-03 | 2.53E-02 | -1.06E+00 | 5.72E-02 | 1.57E-01 | 1.03E+00 |
| ACVR2B | 2.55E-03 | 1.33E-02 | -1.42E+00 | 4.75E-02 | 2.05E-01 | 1.05E+00 | 5.17E-02 | 2.19E-01 | -1.13E+00 | 2.50E-01 | 9.55E-01 | 1.30E+00 | 2.76E-03 | 9.83E-03 | -1.64E+00 | 4.18E-02 | 1.18E-01 | -1.21E+00 | 1.65E-03 | 6.05E-03 | -1.10E+00 | 3.77E-03 | 1.04E-02 | -1.05E+00 |
| SLC5A6 | 2.55E-03 | 1.33E-02 | -1.33E+00 | 2.77E-02 | 1.20E-01 | 1.08E+00 | 5.57E-07 | 2.37E-06 | -1.48E+00 | 1.67E-03 | 6.38E-03 | 1.04E+00 | 4.52E-09 | 1.61E-08 | -2.66E+00 | 6.66E-04 | 1.88E-03 | -1.19E+00 | 1.08E-04 | 3.95E-04 | -1.11E+00 | 2.57E-01 | 7.05E-01 | 1.35E+00 |
| SNRPFP1 | 2.56E-03 | 1.33E-02 | 1.65E+00 | 2.69E-02 | 1.16E-01 | 1.23E+00 | 1.20E-06 | 5.10E-06 | 1.87E+00 | 2.16E-01 | 8.25E-01 | 1.37E+00 | 2.07E-01 | 7.38E-01 | 1.53E+00 | 2.62E-01 | 7.39E-01 | 1.44E+00 | 2.18E-02 | 8.01E-02 | 1.55E+00 | 3.35E-01 | 9.20E-01 | 1.32E+00 |
| FBLN5 | 2.56E-03 | 1.34E-02 | 2.27E+00 | 4.58E-03 | 1.98E-02 | 2.50E+00 | 4.45E-02 | 1.89E-01 | -1.07E+00 | 4.58E-03 | 1.75E-02 | -1.22E+00 | 2.63E-01 | 9.36E-01 | 1.10E+00 | 3.21E-01 | 9.06E-01 | 1.19E+00 | 1.85E-01 | 6.81E-01 | -1.23E+00 | 3.36E-01 | 9.23E-01 | -1.02E+00 |
| PRR7 | 2.57E-03 | 1.34E-02 | 1.11E+00 | 7.05E-08 | 3.05E-07 | -1.05E+00 | 2.99E-05 | 1.27E-04 | -1.06E+00 | 1.10E-08 | 4.19E-08 | -1.19E+00 | 6.57E-02 | 2.34E-01 | -1.12E+00 | 1.10E-02 | 3.11E-02 | -1.55E+00 | 2.57E-02 | 9.45E-02 | 1.18E+00 | 3.96E-03 | 1.09E-02 | -1.37E+00 |
| ATF4P4 | 2.57E-03 | 1.34E-02 | 1.04E+00 | 1.80E-18 | 7.76E-18 | -1.37E+00 | 1.47E-03 | 6.24E-03 | -1.11E+00 | 6.66E-19 | 2.55E-18 | -1.63E+00 | 3.00E-02 | 1.07E-01 | 1.08E+00 | 6.23E-11 | 1.76E-10 | -1.45E+00 | 4.03E-03 | 1.48E-02 | 1.02E+00 | 8.92E-12 | 2.45E-11 | -1.46E+00 |
| RPL35P1 | 2.57E-03 | 1.34E-02 | 1.08E+00 | 2.91E-64 | 1.26E-63 | -1.54E+00 | 1.48E-03 | 6.30E-03 | 1.11E+00 | 2.52E-35 | 9.61E-35 | -1.50E+00 | 2.89E-02 | 1.03E-01 | 1.09E+00 | 4.52E-25 | 1.27E-24 | -1.44E+00 | 1.61E-01 | 5.92E-01 | 1.10E+00 | 1.69E-32 | 4.65E-32 | -1.43E+00 |
| PRICKLE3 | 2.58E-03 | 1.34E-02 | -1.24E+00 | 1.79E-02 | 7.74E-02 | 1.02E+00 | 1.44E-01 | 6.14E-01 | -1.10E+00 | 2.33E-01 | 8.90E-01 | 1.29E+00 | 7.62E-03 | 2.72E-02 | -1.96E+00 | 2.11E-01 | 5.95E-01 | 1.02E+00 | 8.10E-03 | 2.97E-02 | -1.31E+00 | 1.27E-02 | 3.48E-02 | -1.24E+00 |
| PSMA1 | 2.59E-03 | 1.35E-02 | 1.06E+00 | 2.90E-05 | 1.25E-04 | 1.07E+00 | 2.17E-01 | 9.24E-01 | 1.08E+00 | 1.02E-01 | 3.90E-01 | 1.05E+00 | 2.22E-04 | 7.90E-04 | -1.13E+00 | 5.99E-02 | 1.69E-01 | 1.00E+00 | 3.49E-02 | 1.28E-01 | -1.02E+00 | 3.54E-01 | 9.73E-01 | 1.06E+00 |
| AF117829.1 | 2.59E-03 | 1.35E-02 | -1.03E+00 | 7.61E-04 | 3.29E-03 | 1.03E+00 | 3.95E-04 | 1.68E-03 | -1.27E+00 | 2.33E-04 | 8.89E-04 | -1.13E+00 | 2.11E-01 | 7.53E-01 | 1.30E+00 | 1.92E-01 | 5.41E-01 | 1.19E+00 | 1.20E-04 | 4.39E-04 | -1.19E+00 | 7.88E-04 | 2.17E-03 | -1.17E+00 |
| ZNF140 | 2.60E-03 | 1.36E-02 | -1.30E+00 | 7.20E-04 | 3.11E-03 | -1.38E+00 | 5.82E-03 | 2.47E-02 | -1.30E+00 | 2.48E-02 | 9.46E-02 | -1.02E+00 | 3.82E-02 | 1.36E-01 | -1.12E+00 | 2.20E-03 | 6.21E-03 | -1.63E+00 | 5.72E-02 | 2.10E-01 | -1.04E+00 | 1.89E-02 | 5.18E-02 | -1.26E+00 |
| APTR | 2.60E-03 | 1.36E-02 | -1.10E+00 | 2.13E-01 | 9.20E-01 | 1.53E+00 | 1.40E-02 | 5.96E-02 | -1.07E+00 | 4.82E-02 | 1.84E-01 | 1.17E+00 | 9.01E-02 | 3.21E-01 | 1.01E+00 | 2.57E-01 | 7.25E-01 | 1.48E+00 | 2.79E-02 | 1.03E-01 | -1.15E+00 | 3.08E-01 | 8.46E-01 | 1.13E+00 |
| AC091132.4 | 2.60E-03 | 1.36E-02 | -1.23E+00 | 8.13E-04 | 3.51E-03 | -1.25E+00 | 5.30E-02 | 2.25E-01 | -1.01E+00 | 2.90E-02 | 1.11E-01 | -1.05E+00 | 2.59E-02 | 9.24E-02 | -1.48E+00 | 3.47E-02 | 9.77E-02 | -1.54E+00 | 1.02E-02 | 3.75E-02 | -1.43E+00 | 1.07E-01 | 2.93E-01 | -1.10E+00 |
| FGD2 | 2.60E-03 | 1.36E-02 | -1.81E+00 | 4.23E-03 | 1.83E-02 | -1.74E+00 | 2.97E-02 | 1.26E-01 | -1.38E+00 | 3.60E-02 | 1.38E-01 | -1.37E+00 | 2.80E-01 | 1.00E+00 | 1.00E+00 | 2.77E-01 | 7.82E-01 | 1.17E+00 | 2.56E-01 | 9.41E-01 | 1.01E+00 | 2.38E-01 | 6.54E-01 | -1.19E+00 |
| ABRAXAS2 | 2.61E-03 | 1.36E-02 | -1.13E+00 | 2.28E-02 | 9.84E-02 | 1.12E+00 | 1.86E-05 | 7.90E-05 | -1.29E+00 | 5.68E-04 | 2.17E-03 | -1.00E+00 | 1.19E-03 | 4.23E-03 | -1.09E+00 | 6.01E-02 | 1.70E-01 | 1.23E+00 | 7.16E-08 | 2.63E-07 | -1.44E+00 | 1.91E-04 | 5.24E-04 | -1.15E+00 |
| MELTF | 2.61E-03 | 1.36E-02 | -1.07E+00 | 2.47E-04 | 1.07E-03 | -1.07E+00 | 2.25E-02 | 9.54E-02 | 1.51E+00 | 3.54E-02 | 1.35E-01 | 1.15E+00 | 3.50E-02 | 1.25E-01 | -1.50E+00 | 5.66E-02 | 1.60E-01 | -1.27E+00 | 7.02E-02 | 2.58E-01 | -1.01E+00 | 3.18E-01 | 8.73E-01 | 1.32E+00 |
| ATAD3A | 2.61E-03 | 1.36E-02 | -1.11E+00 | 6.26E-03 | 2.71E-02 | 1.06E+00 | 6.43E-02 | 2.73E-01 | 1.32E+00 | 2.23E-01 | 8.52E-01 | 1.31E+00 | 7.16E-02 | 2.55E-01 | -1.00E+00 | 3.04E-01 | 8.57E-01 | 1.47E+00 | 5.10E-02 | 1.87E-01 | 1.09E+00 | 7.07E-03 | 1.94E-02 | 1.64E+00 |
| TECPR2 | 2.62E-03 | 1.36E-02 | -1.54E+00 | 7.35E-02 | 3.17E-01 | 1.03E+00 | 2.09E-03 | 8.88E-03 | -1.34E+00 | 2.10E-02 | 8.01E-02 | 1.01E+00 | 1.42E-01 | 5.05E-01 | 1.08E+00 | 2.40E-01 | 6.76E-01 | 1.25E+00 | 2.04E-01 | 7.49E-01 | 1.11E+00 | 1.69E-01 | 4.65E-01 | 1.07E+00 |
| RPL5P9 | 2.62E-03 | 1.36E-02 | 1.85E+00 | 2.78E-07 | 1.20E-06 | 2.67E+00 | 2.41E-05 | 1.03E-04 | 2.09E+00 | 6.84E-08 | 2.61E-07 | 2.82E+00 | 1.58E-02 | 5.64E-02 | 1.80E+00 | 4.79E-03 | 1.35E-02 | 1.96E+00 | 5.33E-02 | 1.96E-01 | 1.54E+00 | 5.82E-05 | 1.60E-04 | 2.13E+00 |
| IL12A-AS1 | 2.62E-03 | 1.37E-02 | -1.08E+00 | 1.17E-03 | 5.07E-03 | -1.01E+00 | 2.68E-02 | 1.14E-01 | -1.17E+00 | 1.31E-01 | 4.99E-01 | 1.21E+00 | 1.95E-01 | 6.96E-01 | 1.06E+00 | 1.71E-01 | 4.83E-01 | -1.17E+00 | 2.56E-01 | 9.40E-01 | 1.13E+00 | 3.42E-01 | 9.39E-01 | 1.14E+00 |
| CBR3-AS1 | 2.63E-03 | 1.37E-02 | -1.27E+00 | 7.21E-02 | 3.12E-01 | 1.22E+00 | 4.57E-04 | 1.94E-03 | -1.51E+00 | 6.05E-04 | 2.31E-03 | -1.36E+00 | 7.09E-03 | 2.53E-02 | -1.36E+00 | 3.63E-03 | 1.02E-02 | -1.55E+00 | 3.02E-02 | 1.11E-01 | -1.10E+00 | 5.38E-03 | 1.48E-02 | -1.33E+00 |
| MBIP | 2.63E-03 | 1.37E-02 | 1.04E+00 | 1.44E-02 | 6.23E-02 | 1.17E+00 | 3.99E-03 | 1.69E-02 | -1.09E+00 | 1.56E-04 | 5.94E-04 | -1.03E+00 | 7.61E-04 | 2.71E-03 | -1.17E+00 | 4.06E-02 | 1.14E-01 | 1.11E+00 | 4.63E-07 | 1.70E-06 | -1.24E+00 | 2.49E-05 | 6.85E-05 | -1.14E+00 |
| Z74021.1 | 2.63E-03 | 1.37E-02 | 1.43E+00 | 6.04E-05 | 2.61E-04 | 1.11E+00 | 1.43E-05 | 6.09E-05 | 1.65E+00 | 1.42E-01 | 5.43E-01 | 1.31E+00 | 8.33E-02 | 2.97E-01 | 1.39E+00 | 2.34E-01 | 6.59E-01 | 1.29E+00 | 1.11E-02 | 4.07E-02 | 1.50E+00 | 7.29E-02 | 2.00E-01 | 1.31E+00 |
| MAP2K4 | 2.63E-03 | 1.37E-02 | -1.19E+00 | 5.04E-02 | 2.18E-01 | 1.15E+00 | 6.26E-06 | 2.66E-05 | -1.57E+00 | 8.71E-04 | 3.33E-03 | -1.11E+00 | 8.75E-04 | 3.12E-03 | -1.28E+00 | 1.36E-03 | 3.84E-03 | -1.25E+00 | 1.05E-09 | 3.86E-09 | -1.93E+00 | 1.82E-05 | 4.99E-05 | -1.32E+00 |
| KCTD7 | 2.63E-03 | 1.37E-02 | -1.35E+00 | 6.18E-02 | 2.67E-01 | 1.76E+00 | 2.64E-02 | 1.12E-01 | -1.50E+00 | 1.32E-01 | 5.03E-01 | 1.55E+00 | 1.28E-01 | 4.55E-01 | 1.13E+00 | 2.68E-01 | 7.56E-01 | 1.87E+00 | 3.92E-05 | 1.44E-04 | -1.76E+00 | 1.82E-02 | 4.99E-02 | -1.16E+00 |
| AC010615.2 | 2.64E-03 | 1.38E-02 | -1.16E+00 | 1.00E-02 | 4.33E-02 | 1.04E+00 | 2.75E-05 | 1.17E-04 | -1.27E+00 | 1.25E-05 | 4.79E-05 | -1.19E+00 | 4.04E-06 | 1.44E-05 | -1.54E+00 | 2.02E-05 | 5.70E-05 | -1.30E+00 | 9.26E-13 | 3.40E-12 | -1.76E+00 | 2.40E-05 | 6.60E-05 | -1.28E+00 |
| MPL | 2.65E-03 | 1.38E-02 | -1.53E+00 | 3.14E-03 | 1.36E-02 | -1.40E+00 | 2.02E-01 | 8.59E-01 | 1.24E+00 | 2.03E-01 | 7.74E-01 | 1.09E+00 | 4.94E-02 | 1.76E-01 | -1.39E+00 | 2.31E-01 | 6.51E-01 | 1.15E+00 | 4.67E-02 | 1.71E-01 | 1.71E+00 | 3.31E-01 | 9.09E-01 | 1.29E+00 |
| ZNF567 | 2.65E-03 | 1.38E-02 | 1.02E+00 | 5.53E-05 | 2.39E-04 | -1.19E+00 | 5.71E-03 | 2.42E-02 | -1.05E+00 | 1.66E-04 | 6.33E-04 | -1.19E+00 | 1.76E-02 | 6.28E-02 | 1.07E+00 | 1.91E-05 | 5.39E-05 | -1.52E+00 | 6.25E-02 | 2.29E-01 | 1.14E+00 | 1.55E-05 | 4.26E-05 | -1.43E+00 |
| MCU | 2.66E-03 | 1.38E-02 | -1.30E+00 | 7.08E-02 | 3.06E-01 | 1.21E+00 | 2.11E-01 | 8.96E-01 | 1.19E+00 | 2.21E-01 | 8.46E-01 | 1.31E+00 | 7.83E-05 | 2.79E-04 | -1.36E+00 | 3.75E-02 | 1.06E-01 | 1.13E+00 | 1.23E-04 | 4.53E-04 | -1.06E+00 | 4.82E-03 | 1.32E-02 | -1.00E+00 |
| C4orf33 | 2.67E-03 | 1.39E-02 | -1.27E+00 | 1.13E-02 | 4.88E-02 | 1.02E+00 | 3.33E-03 | 1.41E-02 | -1.27E+00 | 7.51E-06 | 2.87E-05 | -1.75E+00 | 1.35E-06 | 4.82E-06 | -1.34E+00 | 1.59E-06 | 4.48E-06 | -1.35E+00 | 3.33E-11 | 1.22E-10 | -1.45E+00 | 2.52E-18 | 6.91E-18 | -2.32E+00 |
| CBLL1 | 2.68E-03 | 1.39E-02 | -1.00E+00 | 1.47E-03 | 6.37E-03 | 1.03E+00 | 2.03E-05 | 8.61E-05 | -1.22E+00 | 2.02E-05 | 7.71E-05 | -1.10E+00 | 1.56E-07 | 5.56E-07 | -1.33E+00 | 2.07E-05 | 5.84E-05 | -1.08E+00 | 1.69E-19 | 6.19E-19 | -1.99E+00 | 8.40E-10 | 2.31E-09 | -1.36E+00 |
| AC107068.1 | 2.68E-03 | 1.40E-02 | -1.55E+00 | 3.13E-02 | 1.35E-01 | -1.10E+00 | 2.40E-02 | 1.02E-01 | -1.24E+00 | 1.39E-02 | 5.31E-02 | -1.27E+00 | 7.20E-02 | 2.57E-01 | -1.25E+00 | 6.28E-02 | 1.77E-01 | -1.25E+00 | 3.26E-02 | 1.20E-01 | -1.46E+00 | 1.41E-01 | 3.86E-01 | -1.20E+00 |
| AC241585.2 | 2.68E-03 | 1.40E-02 | -1.25E+00 | 3.39E-02 | 1.47E-01 | 1.03E+00 | 2.62E-03 | 1.11E-02 | -1.40E+00 | 1.23E-03 | 4.69E-03 | -1.39E+00 | 1.71E-02 | 6.09E-02 | -1.32E+00 | 4.25E-02 | 1.20E-01 | -1.19E+00 | 1.01E-06 | 3.71E-06 | -1.58E+00 | 1.11E-05 | 3.04E-05 | -1.43E+00 |
| CU633967.1 | 2.68E-03 | 1.40E-02 | -1.90E+00 | 2.54E-02 | 1.10E-01 | -1.20E+00 | 7.38E-05 | 3.14E-04 | -1.93E+00 | 2.58E-02 | 9.86E-02 | -1.10E+00 | 2.51E-02 | 8.95E-02 | -2.09E+00 | 1.18E-01 | 3.32E-01 | -1.39E+00 | 2.89E-12 | 1.06E-11 | -2.06E+00 | 9.11E-04 | 2.50E-03 | -1.13E+00 |
| CCDC113 | 2.68E-03 | 1.40E-02 | -1.63E+00 | 1.43E-02 | 6.16E-02 | -1.25E+00 | 2.27E-02 | 9.63E-02 | -1.33E+00 | 1.37E-01 | 5.25E-01 | 1.09E+00 | 8.56E-02 | 3.05E-01 | -1.14E+00 | 1.38E-01 | 3.89E-01 | -1.18E+00 | 4.82E-02 | 1.77E-01 | -1.21E+00 | 3.14E-01 | 8.63E-01 | 1.25E+00 |
| UFD1 | 2.69E-03 | 1.40E-02 | 1.00E+00 | 5.13E-03 | 2.22E-02 | 1.12E+00 | 3.27E-02 | 1.39E-01 | 1.25E+00 | 2.44E-01 | 9.33E-01 | 1.27E+00 | 4.33E-02 | 1.55E-01 | 1.07E+00 | 4.34E-02 | 1.22E-01 | 1.15E+00 | 2.52E-01 | 9.26E-01 | 1.27E+00 | 2.73E-01 | 7.51E-01 | 1.20E+00 |
| MCPH1 | 2.70E-03 | 1.41E-02 | -1.02E+00 | 4.81E-04 | 2.08E-03 | 1.06E+00 | 5.87E-05 | 2.49E-04 | -1.34E+00 | 2.00E-05 | 7.64E-05 | -1.14E+00 | 1.03E-02 | 3.67E-02 | -1.02E+00 | 2.01E-02 | 5.67E-02 | -1.01E+00 | 2.09E-03 | 7.68E-03 | -1.07E+00 | 5.20E-06 | 1.43E-05 | -1.33E+00 |
| CD300LF | 2.70E-03 | 1.41E-02 | 1.05E+00 | 5.83E-02 | 2.52E-01 | 1.35E+00 | 2.34E-03 | 9.95E-03 | -1.14E+00 | 5.93E-03 | 2.27E-02 | 1.04E+00 | 3.60E-04 | 1.28E-03 | -1.45E+00 | 2.01E-02 | 5.68E-02 | -1.09E+00 | 1.78E-03 | 6.54E-03 | -1.16E+00 | 1.66E-02 | 4.57E-02 | -1.01E+00 |
| C6orf226 | 2.70E-03 | 1.41E-02 | 1.79E+00 | 3.89E-02 | 1.68E-01 | 1.23E+00 | 1.26E-02 | 5.35E-02 | 1.46E+00 | 7.82E-03 | 2.99E-02 | -1.07E+00 | 7.01E-04 | 2.50E-03 | 2.40E+00 | 3.52E-01 | 9.93E-01 | 1.54E+00 | 3.09E-11 | 1.13E-10 | 2.90E+00 | 9.16E-03 | 2.52E-02 | 1.80E+00 |
| ALDH7A1 | 2.70E-03 | 1.41E-02 | -1.31E+00 | 4.59E-02 | 1.99E-01 | 1.11E+00 | 2.34E-01 | 9.94E-01 | 1.18E+00 | 2.16E-01 | 8.24E-01 | 1.42E+00 | 2.33E-01 | 8.29E-01 | 1.26E+00 | 3.33E-01 | 9.40E-01 | 1.41E+00 | 1.27E-02 | 4.66E-02 | -1.23E+00 | 2.76E-02 | 7.58E-02 | -1.14E+00 |
| RUSC1-AS1 | 2.70E-03 | 1.41E-02 | -1.17E+00 | 2.72E-02 | 1.17E-01 | 2.11E+00 | 1.76E-03 | 7.49E-03 | -1.23E+00 | 2.30E-01 | 8.79E-01 | 1.38E+00 | 5.10E-02 | 1.82E-01 | -1.27E+00 | 1.19E-02 | 3.36E-02 | 2.17E+00 | 4.88E-05 | 1.79E-04 | -1.47E+00 | 1.04E-01 | 2.85E-01 | 1.60E+00 |
| CASC3 | 2.70E-03 | 1.41E-02 | -1.02E+00 | 6.55E-03 | 2.83E-02 | 1.10E+00 | 7.58E-06 | 3.22E-05 | -1.29E+00 | 2.84E-04 | 1.09E-03 | -1.01E+00 | 3.77E-03 | 1.35E-02 | -1.01E+00 | 1.32E-01 | 3.73E-01 | 1.24E+00 | 8.38E-06 | 3.08E-05 | -1.25E+00 | 1.39E-04 | 3.83E-04 | -1.15E+00 |
| LSM11 | 2.71E-03 | 1.41E-02 | -1.36E+00 | 4.60E-03 | 1.99E-02 | -1.24E+00 | 6.78E-03 | 2.88E-02 | -1.24E+00 | 1.20E-02 | 4.58E-02 | -1.06E+00 | 1.39E-02 | 4.96E-02 | -1.45E+00 | 8.84E-02 | 2.49E-01 | -1.12E+00 | 4.93E-03 | 1.81E-02 | -1.35E+00 | 2.55E-01 | 7.00E-01 | 1.18E+00 |
| GLB1L2 | 2.71E-03 | 1.41E-02 | -1.33E+00 | 1.18E-03 | 5.10E-03 | -1.48E+00 | 1.35E-03 | 5.75E-03 | -1.20E+00 | 8.80E-03 | 3.36E-02 | -1.07E+00 | 1.63E-01 | 5.81E-01 | 1.08E+00 | 6.93E-02 | 1.95E-01 | -1.61E+00 | 1.65E-01 | 6.04E-01 | 1.11E+00 | 5.97E-02 | 1.64E-01 | 1.82E+00 |
| GRAMD1A | 2.71E-03 | 1.41E-02 | 1.01E+00 | 2.32E-03 | 1.00E-02 | 1.13E+00 | 9.11E-06 | 3.87E-05 | -1.31E+00 | 3.02E-05 | 1.15E-04 | -1.14E+00 | 7.74E-07 | 2.76E-06 | -1.34E+00 | 8.53E-04 | 2.40E-03 | -1.02E+00 | 3.78E-03 | 1.39E-02 | -1.01E+00 | 5.59E-03 | 1.54E-02 | -1.07E+00 |
| CEP131 | 2.71E-03 | 1.41E-02 | -1.27E+00 | 9.15E-04 | 3.95E-03 | -1.31E+00 | 1.13E-02 | 4.80E-02 | 1.75E+00 | 1.08E-01 | 4.11E-01 | 1.62E+00 | 6.27E-02 | 2.24E-01 | 1.13E+00 | 4.26E-02 | 1.20E-01 | 1.03E+00 | 8.09E-02 | 2.97E-01 | -1.04E+00 | 1.50E-01 | 4.12E-01 | 1.12E+00 |
| PVT1 | 2.71E-03 | 1.41E-02 | -1.23E+00 | 1.60E-03 | 6.90E-03 | -1.23E+00 | 1.38E-05 | 5.85E-05 | -1.34E+00 | 4.63E-04 | 1.77E-03 | -1.08E+00 | 3.06E-02 | 1.09E-01 | 1.01E+00 | 3.46E-01 | 9.74E-01 | 1.43E+00 | 9.61E-08 | 3.53E-07 | -1.45E+00 | 4.76E-12 | 1.31E-11 | -1.56E+00 |
| MIPEP | 2.71E-03 | 1.41E-02 | -1.46E+00 | 9.71E-03 | 4.19E-02 | -1.15E+00 | 2.22E-07 | 9.41E-07 | -1.58E+00 | 3.18E-08 | 1.22E-07 | -1.55E+00 | 1.19E-09 | 4.26E-09 | -2.77E+00 | 2.32E-07 | 6.55E-07 | -1.92E+00 | 9.51E-09 | 3.49E-08 | -1.69E+00 | 6.10E-03 | 1.68E-02 | 1.05E+00 |
| RPS29P5 | 2.72E-03 | 1.42E-02 | 1.17E+00 | 6.01E-38 | 2.60E-37 | -1.52E+00 | 2.18E-01 | 9.24E-01 | 1.03E+00 | 3.43E-21 | 1.31E-20 | -1.68E+00 | 4.01E-03 | 1.43E-02 | 1.04E+00 | 7.87E-22 | 2.22E-21 | -1.84E+00 | 9.32E-03 | 3.42E-02 | -1.02E+00 | 3.55E-19 | 9.75E-19 | -1.89E+00 |
| ACYP1 | 2.72E-03 | 1.42E-02 | 1.05E+00 | 8.23E-02 | 3.56E-01 | 1.36E+00 | 1.30E-01 | 5.54E-01 | 1.31E+00 | 2.29E-01 | 8.76E-01 | 1.37E+00 | 3.47E-05 | 1.24E-04 | -1.12E+00 | 3.02E-02 | 8.51E-02 | 1.19E+00 | 1.33E-01 | 4.90E-01 | 1.13E+00 | 2.54E-01 | 6.97E-01 | 1.25E+00 |
| MPDU1 | 2.72E-03 | 1.42E-02 | 1.09E+00 | 6.29E-03 | 2.72E-02 | 1.20E+00 | 4.18E-05 | 1.77E-04 | -1.11E+00 | 1.97E-06 | 7.53E-06 | -1.07E+00 | 1.74E-02 | 6.20E-02 | 1.00E+00 | 3.10E-01 | 8.75E-01 | 1.39E+00 | 9.82E-03 | 3.60E-02 | 1.15E+00 | 1.74E-01 | 4.79E-01 | 1.23E+00 |
| PROCR | 2.72E-03 | 1.42E-02 | -1.53E+00 | 3.95E-02 | 1.71E-01 | -1.01E+00 | 1.98E-13 | 8.31E-13 | -2.89E+00 | 8.04E-08 | 3.07E-07 | -1.70E+00 | 2.76E-01 | 9.82E-01 | 1.18E+00 | 2.69E-01 | 7.57E-01 | 1.49E+00 | 1.73E-02 | 6.35E-02 | -1.62E+00 | 3.74E-02 | 1.03E-01 | -1.46E+00 |
| TXNP6 | 2.73E-03 | 1.42E-02 | 1.10E+00 | 2.87E-08 | 1.24E-07 | 1.05E+00 | 8.10E-02 | 3.44E-01 | 1.22E+00 | 2.84E-02 | 1.09E-01 | 1.07E+00 | 2.83E-03 | 1.01E-02 | 1.05E+00 | 1.89E-02 | 5.34E-02 | 1.07E+00 | 7.13E-02 | 2.62E-01 | 1.11E+00 | 1.31E-01 | 3.60E-01 | 1.09E+00 |
| TXLNGY | 2.73E-03 | 1.42E-02 | 1.09E+00 | 7.74E-03 | 3.34E-02 | 1.27E+00 | 1.30E-02 | 5.52E-02 | -1.08E+00 | 3.77E-02 | 1.44E-01 | 1.15E+00 | 3.20E-03 | 1.14E-02 | 1.08E+00 | 2.49E-02 | 7.01E-02 | 1.20E+00 | 7.13E-04 | 2.62E-03 | -1.05E+00 | 1.48E-01 | 4.07E-01 | 1.15E+00 |
| RPL26L1 | 2.73E-03 | 1.43E-02 | 1.03E+00 | 4.12E-03 | 1.78E-02 | 1.17E+00 | 1.36E-02 | 5.76E-02 | 1.47E+00 | 2.69E-02 | 1.03E-01 | 1.15E+00 | 1.24E-01 | 4.43E-01 | 1.22E+00 | 1.83E-01 | 5.17E-01 | 1.61E+00 | 4.79E-02 | 1.76E-01 | 1.15E+00 | 1.53E-01 | 4.19E-01 | 1.21E+00 |
| GPD1L | 2.74E-03 | 1.43E-02 | -1.59E+00 | 1.36E-02 | 5.86E-02 | -1.13E+00 | 4.28E-05 | 1.82E-04 | -1.54E+00 | 4.74E-03 | 1.81E-02 | -1.05E+00 | 4.02E-05 | 1.43E-04 | -2.59E+00 | 6.25E-03 | 1.76E-02 | -1.37E+00 | 5.58E-07 | 2.05E-06 | -1.47E+00 | 1.32E-05 | 3.62E-05 | -1.38E+00 |
| PDSS1 | 2.74E-03 | 1.43E-02 | -1.71E+00 | 3.28E-02 | 1.42E-01 | -1.09E+00 | 6.51E-02 | 2.76E-01 | 1.01E+00 | 3.43E-02 | 1.31E-01 | 1.10E+00 | 3.43E-02 | 1.22E-01 | -1.19E+00 | 1.69E-01 | 4.77E-01 | 1.26E+00 | 6.75E-13 | 2.48E-12 | -1.94E+00 | 1.63E-02 | 4.48E-02 | 1.03E+00 |
| TTTY14 | 2.74E-03 | 1.43E-02 | -1.15E+00 | [illegible] | 1.16E-01 | 1.10E+00 | 1.11E-02 | 4.73E-02 | -1.12E+00 | 6.69E-02 | 2.56E-01 | 1.20E+00 | 5.50E-02 | 1.96E-01 | -1.05E+00 | 1.25E-01 | 3.52E-01 | 1.16E+00 | 1.39E-01 | 5.10E-01 | 1.12E+00 | 2.86E-01 | 7.87E-01 | 1.33E+00 |
| ITGAX | 2.74E-03 | 1.43E-02 | -1.22E+00 | 2.48E-04 | 1.07E-03 | -1.23E+00 | 3.45E-09 | 1.47E-08 | -1.87E+00 | 1.71E-07 | 6.53E-07 | -1.39E+00 | 4.44E-05 | 1.58E-04 | -3.01E+00 | 1.04E-04 | 2.93E-04 | -2.90E+00 | 1.66E-01 | 6.08E-01 | -1.38E+00 | 2.99E-01 | 8.20E-01 | -1.07E+00 |
| IL15 | 2.75E-03 | 1.43E-02 | -1.14E+00 | 1.67E-01 | 7.21E-01 | 1.41E+00 | 2.55E-03 | 1.08E-02 | -1.54E+00 | 2.77E-02 | 1.06E-01 | -1.03E+00 | 1.63E-02 | 5.80E-02 | -1.27E+00 | 3.71E-02 | 1.05E-01 | -1.02E+00 | 1.80E-02 | 6.61E-02 | -1.10E+00 | 2.08E-01 | 5.72E-01 | 1.14E+00 |
| MFNG | 2.75E-03 | 1.43E-02 | 1.15E+00 | 1.38E-01 | 5.97E-01 | 1.47E+00 | 4.28E-02 | 1.82E-01 | -1.03E+00 | 1.02E-01 | 3.90E-01 | 1.40E+00 | 1.22E-05 | 4.33E-05 | -1.05E+00 | 4.13E-02 | 1.17E-01 | 1.25E+00 | 3.50E-02 | 1.28E-01 | 1.03E+00 | 3.34E-01 | 9.19E-01 | 1.22E+00 |
| STARD3 | 2.75E-03 | 1.43E-02 | 1.08E+00 | 5.95E-02 | 2.57E-01 | 1.41E+00 | 1.53E-03 | 6.49E-03 | -1.09E+00 | 4.83E-03 | 1.84E-02 | 1.11E+00 | 2.67E-01 | 9.53E-01 | 1.66E+00 | 2.21E-01 | 6.24E-01 | 1.76E+00 | 2.39E-01 | 8.78E-01 | 1.35E+00 | 2.77E-01 | 7.61E-01 | 1.34E+00 |
| AC244093.2 | 2.76E-03 | 1.44E-02 | -1.39E+00 | 2.91E-02 | 1.26E-01 | 1.01E+00 | 3.14E-02 | 1.33E-01 | -1.19E+00 | 1.56E-01 | 5.97E-01 | 1.16E+00 | 2.06E-02 | 7.35E-02 | -1.64E+00 | 3.25E-01 | 9.15E-01 | 1.28E+00 | 1.84E-01 | 6.74E-01 | 1.41E+00 | 2.73E-01 | 7.50E-01 | 1.39E+00 |
| PCNX2 | 2.76E-03 | 1.44E-02 | -1.13E+00 | 1.37E-02 | 5.90E-02 | 1.08E+00 | 1.61E-02 | 6.82E-02 | -1.18E+00 | 8.07E-03 | 3.08E-02 | -1.05E+00 | 3.16E-02 | 1.13E-01 | -1.16E+00 | 4.70E-02 | 1.32E-01 | -1.12E+00 | 8.77E-02 | 3.22E-01 | -1.23E+00 | 9.21E-02 | 2.53E-01 | -1.30E+00 |
| AL512643.1 | 2.76E-03 | 1.44E-02 | -1.57E+00 | 2.61E-02 | 1.13E-01 | -1.11E+00 | 2.77E-02 | 1.18E-01 | -1.58E+00 | 3.73E-02 | 1.42E-01 | -1.26E+00 | 2.49E-09 | 8.87E-09 | -2.18E+00 | 1.45E-03 | 4.08E-03 | -1.09E+00 | 3.80E-05 | 1.40E-04 | -1.66E+00 | 1.32E-01 | 3.64E-01 | 1.08E+00 |
| AL022328.4 | 2.76E-03 | 1.44E-02 | -1.19E+00 | 4.13E-03 | 1.79E-02 | -1.08E+00 | 3.03E-04 | 1.29E-03 | -1.50E+00 | 2.66E-05 | 1.02E-04 | -1.72E+00 | 2.56E-01 | 9.12E-01 | 1.22E+00 | 2.39E-01 | 6.73E-01 | 1.65E+00 | 8.14E-02 | 2.99E-01 | 1.91E+00 | 1.76E-01 | 4.84E-01 | 1.65E+00 |
| TBCCD1 | 2.76E-03 | 1.44E-02 | -1.19E+00 | 6.64E-02 | 2.87E-01 | 1.23E+00 | 7.14E-03 | 3.03E-02 | -1.27E+00 | 1.69E-02 | 6.44E-02 | -1.08E+00 | 7.13E-03 | 2.54E-02 | -1.64E+00 | 1.28E-01 | 3.62E-01 | 1.04E+00 | 3.01E-02 | 1.11E-01 | -1.21E+00 | 2.24E-01 | 6.14E-01 | 1.41E+00 |
| LIPC | 2.76E-03 | 1.44E-02 | -1.43E+00 | 1.04E-02 | 4.48E-02 | -1.08E+00 | 1.96E-01 | 8.32E-01 | 1.14E+00 | 2.23E-01 | 8.50E-01 | 1.51E+00 | 2.61E-02 | 9.29E-02 | 1.07E+00 | 7.41E-03 | 2.09E-02 | -1.18E+00 | 4.73E-02 | 1.74E-01 | 1.08E+00 | 2.36E-02 | 6.48E-02 | -1.22E+00 |
| MBLAC2 | 2.77E-03 | 1.44E-02 | -1.20E+00 | 4.32E-02 | 1.87E-01 | 1.17E+00 | 9.67E-05 | 4.11E-04 | -1.54E+00 | 1.97E-02 | 7.52E-02 | 1.00E+00 | 1.48E-01 | 5.28E-01 | 1.10E+00 | 3.48E-01 | 9.82E-01 | 1.52E+00 | 3.87E-03 | 1.42E-02 | -1.20E+00 | 3.01E-01 | 8.28E-01 | 1.22E+00 |
| ANK3 | 2.77E-03 | 1.44E-02 | 1.03E+00 | 4.96E-02 | 2.14E-01 | 1.26E+00 | 4.73E-03 | 2.01E-02 | -1.02E+00 | 4.54E-03 | 1.73E-02 | 1.01E+00 | 6.31E-02 | 2.25E-01 | -1.81E+00 | 2.27E-01 | 6.39E-01 | 1.03E+00 | 2.39E-01 | 8.78E-01 | 1.16E+00 | 3.12E-01 | 8.56E-01 | 1.16E+00 |
| AC068631.1 | 2.78E-03 | 1.45E-02 | -1.38E+00 | 1.48E-01 | 6.38E-01 | 1.37E+00 | 9.70E-03 | 4.12E-02 | -1.43E+00 | 4.06E-02 | 1.54E-01 | -1.24E+00 | 2.24E-01 | 8.00E-01 | 1.23E+00 | 2.73E-01 | 7.71E-01 | 1.37E+00 | 4.36E-02 | 1.60E-01 | -1.79E+00 | 2.57E-01 | 7.05E-01 | -1.12E+00 |
| FBXO18 | 2.78E-03 | 1.45E-02 | -1.25E+00 | 4.22E-03 | 1.82E-02 | -1.11E+00 | 1.78E-03 | 7.58E-03 | -1.37E+00 | 4.40E-03 | 1.68E-02 | -1.14E+00 | 1.25E-01 | 4.47E-01 | 1.13E+00 | 6.72E-02 | 1.89E-01 | -1.02E+00 | 5.67E-03 | 2.08E-02 | 1.01E+00 | 2.12E-03 | 5.82E-03 | -1.13E+00 |
| SGPP2 | 2.78E-03 | 1.45E-02 | -1.60E+00 | 1.90E-01 | 8.20E-01 | 1.19E+00 | 5.26E-03 | 2.23E-02 | -1.31E+00 | 1.47E-02 | 5.60E-02 | -1.18E+00 | 1.32E-01 | 4.71E-01 | 1.11E+00 | 2.76E-01 | 7.79E-01 | 1.63E+00 | 5.67E-05 | 2.08E-04 | -1.84E+00 | 8.61E-04 | 2.37E-03 | -1.57E+00 |
| METRN | 2.79E-03 | 1.45E-02 | -1.42E+00 | 2.98E-03 | 1.29E-02 | -1.35E+00 | 1.11E-01 | 4.71E-01 | 1.55E+00 | 9.76E-03 | 3.73E-02 | 2.36E+00 | 9.19E-02 | 3.28E-01 | 2.17E+00 | 1.63E-01 | 4.61E-01 | 1.94E+00 | 1.34E-03 | 4.93E-03 | 1.84E+00 | 3.75E-04 | 1.03E-03 | 2.09E+00 |
| ZNF768 | 2.79E-03 | 1.46E-02 | -1.00E+00 | 4.78E-03 | 2.07E-02 | 1.08E+00 | 8.10E-03 | 3.44E-02 | -1.09E+00 | 7.41E-04 | 2.83E-03 | -1.04E+00 | 1.57E-03 | 5.59E-03 | -1.22E+00 | 2.53E-02 | 7.14E-02 | 1.05E+00 | 3.76E-07 | 1.38E-06 | -1.39E+00 | 8.58E-05 | 2.36E-04 | -1.20E+00 |
| ING1 | 2.80E-03 | 1.46E-02 | -1.10E+00 | 6.26E-05 | 2.70E-04 | -1.32E+00 | 1.49E-04 | 6.34E-04 | -1.36E+00 | 3.89E-05 | 1.48E-04 | -1.27E+00 | 1.59E-03 | 5.68E-03 | -1.02E+00 | 1.67E-06 | 4.69E-06 | -1.45E+00 | 7.77E-03 | 2.85E-02 | 1.01E+00 | 4.20E-03 | 1.15E-02 | -1.12E+00 |
| FAM83D | 2.80E-03 | 1.46E-02 | -2.25E+00 | 4.53E-03 | 1.96E-02 | -2.06E+00 | 1.51E-01 | 6.43E-01 | -1.14E+00 | 2.52E-01 | 9.64E-01 | 1.16E+00 | 3.50E-08 | 1.25E-07 | -2.56E+00 | 4.14E-04 | 1.17E-03 | -1.40E+00 | 6.04E-38 | 2.22E-37 | -2.67E+00 | 4.43E-17 | 1.22E-16 | -1.61E+00 |
| ZNF876P | 2.80E-03 | 1.46E-02 | 2.73E+00 | 1.91E-02 | 8.25E-02 | 2.57E+00 | 1.13E-01 | 4.81E-01 | 1.96E+00 | 2.07E-01 | 7.90E-01 | 1.15E+00 | 2.79E-01 | 9.94E-01 | 1.16E+00 | 3.42E-01 | 9.64E-01 | 1.25E+00 | 2.52E-01 | 9.25E-01 | 1.25E+00 | 2.28E-01 | 6.26E-01 | 1.08E+00 |
| ATP7A | 2.81E-03 | 1.46E-02 | -1.29E+00 | 4.96E-02 | 2.14E-01 | 1.14E+00 | 8.09E-03 | 3.44E-02 | -1.38E+00 | 9.64E-02 | 3.68E-01 | 1.06E+00 | 2.58E-08 | 9.20E-08 | -1.94E+00 | 5.42E-04 | 1.53E-03 | -1.16E+00 | 3.86E-07 | 1.42E-06 | -1.48E+00 | 4.55E-03 | 1.25E-02 | -1.10E+00 |
| RPL34P27 | 2.81E-03 | 1.47E-02 | -1.11E+00 | 2.97E-43 | 1.28E-42 | -1.98E+00 | 5.49E-02 | 2.33E-01 | -1.06E+00 | 5.66E-21 | 2.16E-20 | -1.79E+00 | 4.90E-02 | 1.75E-01 | 1.01E+00 | 1.07E-20 | 3.00E-20 | -1.79E+00 | 6.51E-03 | 2.39E-02 | -1.08E+00 | 2.83E-21 | 7.78E-21 | -2.01E+00 |
| STXBP4 | 2.81E-03 | 1.47E-02 | -1.16E+00 | 1.04E-01 | 4.51E-01 | 1.66E+00 | 1.31E-07 | 5.58E-07 | -2.05E+00 | 3.30E-03 | 1.26E-02 | -1.05E+00 | 1.40E-07 | 5.00E-07 | -1.83E+00 | 5.62E-04 | 1.58E-03 | -1.20E+00 | 5.00E-05 | 1.83E-04 | -1.67E+00 | 2.04E-02 | 5.59E-02 | -1.12E+00 |
| ZNF16 | 2.81E-03 | 1.47E-02 | -1.19E+00 | 2.86E-02 | 1.24E-01 | 1.12E+00 | 4.66E-02 | 1.98E-01 | 1.03E+00 | 1.86E-01 | 7.11E-01 | 1.49E+00 | 1.28E-01 | 4.58E-01 | 1.08E+00 | 3.11E-01 | 8.76E-01 | 1.43E+00 | 1.43E-01 | 5.25E-01 | 1.12E+00 | 2.84E-01 | 7.80E-01 | 1.39E+00 |
| SMCR8 | 2.82E-03 | 1.47E-02 | -1.02E+00 | 8.02E-04 | 3.47E-03 | -1.09E+00 | 2.40E-03 | 1.02E-02 | -1.12E+00 | 3.09E-04 | 1.18E-03 | -1.16E+00 | 1.51E-02 | 5.38E-02 | -1.16E+00 | 2.82E-03 | 7.94E-03 | -1.53E+00 | 9.58E-03 | 3.51E-02 | -1.15E+00 | 1.89E-02 | 5.18E-02 | -1.16E+00 |
| ZNF665 | 2.82E-03 | 1.47E-02 | -1.22E+00 | 5.00E-02 | 2.16E-01 | 1.19E+00 | 9.82E-05 | 4.17E-04 | -1.38E+00 | 1.41E-03 | 5.37E-03 | -1.11E+00 | 1.78E-03 | 6.33E-03 | -1.51E+00 | 7.37E-02 | 2.08E-01 | 1.02E+00 | 4.87E-02 | 1.79E-01 | -1.05E+00 | 3.73E-02 | 1.03E-01 | -1.15E+00 |
| ZFAND2B | 2.82E-03 | 1.47E-02 | 1.08E+00 | 3.56E-02 | 1.54E-01 | 1.38E+00 | 3.85E-02 | 1.64E-01 | 1.09E+00 | 8.01E-02 | 3.06E-01 | 1.29E+00 | 8.38E-03 | 2.99E-02 | 1.08E+00 | 1.55E-01 | 4.38E-01 | 1.31E+00 | 2.26E-01 | 8.30E-01 | 1.53E+00 | 2.56E-01 | 7.04E-01 | 1.36E+00 |
| UBE2D2 | 2.83E-03 | 1.48E-02 | 1.08E+00 | 6.31E-03 | 2.73E-02 | 1.16E+00 | 1.12E-01 | 4.76E-01 | 1.07E+00 | 2.46E-01 | 9.41E-01 | 1.15E+00 | 1.51E-01 | 5.39E-01 | 1.13E+00 | 2.42E-01 | 6.83E-01 | 1.21E+00 | 8.49E-02 | 3.11E-01 | 1.02E+00 | 3.56E-01 | 9.77E-01 | 1.09E+00 |
| IFT52 | 2.84E-03 | 1.48E-02 | -1.02E+00 | 3.92E-03 | 1.70E-02 | 1.11E+00 | 1.05E-04 | 4.44E-04 | -1.17E+00 | 2.05E-06 | 7.84E-06 | -1.20E+00 | 5.92E-03 | 2.11E-02 | -1.07E+00 | 1.94E-01 | 5.47E-01 | 1.18E+00 | 7.57E-03 | 2.78E-02 | 1.01E+00 | 1.56E-02 | 4.28E-02 | 1.08E+00 |
| AC008677.2 | 2.85E-03 | 1.49E-02 | 1.62E+00 | 4.85E-02 | 2.10E-01 | 1.41E+00 | 1.12E-03 | 4.78E-03 | 1.61E+00 | 2.64E-02 | 1.01E-01 | 1.61E+00 | 1.05E-01 | 3.73E-01 | 1.59E+00 | 2.10E-01 | 5.91E-01 | 1.54E+00 | 1.63E-01 | 5.98E-01 | 1.36E+00 | 1.03E-01 | 2.82E-01 | 1.44E+00 |
| COL18A1 | 2.85E-03 | 1.49E-02 | -1.10E+00 | 5.33E-05 | 2.31E-04 | -1.33E+00 | 2.24E-01 | 9.51E-01 | 1.33E+00 | 6.78E-02 | 2.59E-01 | -1.00E+00 | 6.04E-02 | 2.16E-01 | 1.04E+00 | 2.91E-01 | 8.21E-01 | 1.37E+00 | 1.58E-01 | 5.79E-01 | 1.31E+00 | 6.71E-02 | 1.84E-01 | 1.06E+00 |
| ECT2 | 2.86E-03 | 1.49E-02 | -1.53E+00 | 5.49E-02 | 2.37E-01 | 1.07E+00 | 8.67E-04 | 3.68E-03 | -1.27E+00 | 2.50E-02 | 9.57E-02 | 1.09E+00 | 1.02E-01 | 3.64E-01 | -1.47E+00 | 2.71E-01 | 7.63E-01 | 1.16E+00 | 4.58E-04 | 1.68E-03 | -1.56E+00 | 1.21E-02 | 3.33E-02 | -1.20E+00 |
| CRAMP1 | 2.86E-03 | 1.49E-02 | -1.23E+00 | 1.48E-03 | 6.38E-03 | -1.21E+00 | 2.41E-04 | 1.02E-03 | -1.33E+00 | 1.21E-03 | 4.64E-03 | -1.12E+00 | 1.22E-01 | 4.37E-01 | 1.15E+00 | 8.49E-02 | 2.39E-01 | 1.04E+00 | 1.40E-03 | 5.14E-03 | -1.45E+00 | 1.85E-02 | 5.08E-02 | -1.10E+00 |
| PDZD8 | 2.86E-03 | 1.49E-02 | 1.02E+00 | 1.20E-01 | 5.16E-01 | 1.31E+00 | 8.42E-04 | 3.58E-03 | -1.22E+00 | 8.56E-02 | 3.27E-01 | 1.08E+00 | 1.05E-02 | 3.73E-02 | -1.07E+00 | 3.26E-01 | 9.20E-01 | 1.20E+00 | 9.17E-02 | 3.36E-01 | -1.04E+00 | 2.59E-01 | 7.12E-01 | 1.04E+00 |
| EPN2 | 2.86E-03 | 1.49E-02 | -1.19E+00 | 1.04E-03 | 4.51E-03 | -1.29E+00 | 3.83E-04 | 1.63E-03 | -1.82E+00 | 1.35E-03 | 5.14E-03 | -1.47E+00 | 2.14E-02 | 7.63E-02 | -1.07E+00 | 8.58E-03 | 2.42E-02 | -1.25E+00 | 8.08E-05 | 2.96E-04 | -1.40E+00 | 6.81E-06 | 1.87E-05 | -1.50E+00 |
| POP1 | 2.86E-03 | 1.49E-02 | -1.76E+00 | 5.48E-02 | 2.37E-01 | 1.03E+00 | 2.18E-05 | 9.24E-05 | -1.59E+00 | 6.61E-04 | 2.52E-03 | -1.18E+00 | 2.48E-03 | 8.84E-03 | -2.52E+00 | 2.66E-02 | 7.51E-02 | -1.54E+00 | 8.02E-06 | 2.94E-05 | -1.54E+00 | 2.19E-02 | 6.01E-02 | -1.08E+00 |
| SAP18P2 | 2.87E-03 | 1.50E-02 | 2.56E+00 | 4.65E-02 | 2.01E-01 | 2.15E+00 | 1.82E-02 | 7.72E-02 | 1.79E+00 | 2.20E-01 | 8.40E-01 | 1.40E+00 | 2.14E-01 | 7.62E-01 | 1.43E+00 | 3.55E-01 | 1.00E+00 | 1.18E+00 | 4.61E-02 | 1.69E-01 | 1.60E+00 | 3.04E-01 | 8.35E-01 | 1.15E+00 |
| FAS | 2.87E-03 | 1.50E-02 | -1.34E+00 | 1.96E-03 | 8.47E-03 | -1.33E+00 | 3.56E-09 | 1.51E-08 | -2.06E+00 | 2.04E-07 | 7.80E-07 | -1.68E+00 | 1.41E-01 | 5.02E-01 | -1.24E+00 | 2.10E-01 | 5.92E-01 | -1.07E+00 | 6.52E-07 | 2.39E-06 | -1.79E+00 | 5.34E-07 | 1.47E-06 | -1.86E+00 |
| FBXO46 | 2.88E-03 | 1.50E-02 | -1.27E+00 | 7.38E-02 | 3.19E-01 | 1.20E+00 | 4.30E-06 | 1.83E-05 | -1.82E+00 | 6.15E-03 | 2.35E-02 | -1.05E+00 | 1.06E-01 | 3.77E-01 | 1.01E+00 | 3.37E-01 | 9.49E-01 | 1.30E+00 | 6.42E-06 | 2.35E-05 | -1.50E+00 | 2.05E-02 | 5.63E-02 | 1.06E+00 |
| EIF4EBP1 | 2.88E-03 | 1.50E-02 | 1.04E+00 | 1.51E-03 | 6.53E-03 | 1.14E+00 | 1.47E-01 | 6.26E-01 | 1.05E+00 | 2.50E-01 | 9.55E-01 | 1.24E+00 | 4.69E-08 | 1.67E-07 | -1.25E+00 | 3.37E-03 | 9.49E-03 | 1.07E+00 | 2.60E-03 | 9.54E-03 | -1.04E+00 | 2.00E-01 | 5.49E-01 | 1.28E+00 |
| RPL21P1 | 2.88E-03 | 1.50E-02 | -1.01E+00 | 9.44E-09 | 4.08E-08 | -1.53E+00 | 1.53E-02 | 6.48E-02 | -1.08E+00 | 8.80E-07 | 3.36E-06 | -1.45E+00 | 1.08E-03 | 3.84E-03 | -1.16E+00 | 8.57E-07 | 2.41E-06 | -1.79E+00 | 2.90E-05 | 1.06E-04 | -1.21E+00 | 6.58E-08 | 1.81E-07 | -1.56E+00 |
| TCP11L1 | 2.89E-03 | 1.51E-02 | -1.07E+00 | 1.96E-02 | 8.49E-02 | 1.21E+00 | 1.49E-05 | 6.32E-05 | -1.39E+00 | 2.63E-04 | 1.01E-03 | -1.12E+00 | 1.33E-01 | 4.75E-01 | 1.04E+00 | 3.15E-02 | 8.89E-02 | -1.19E+00 | 1.35E-03 | 4.94E-03 | -1.43E+00 | 9.59E-03 | 2.64E-02 | -1.26E+00 |
| HNRNPLP2 | 2.90E-03 | 1.51E-02 | -1.16E+00 | 4.56E-03 | 1.97E-02 | -1.01E+00 | 3.60E-06 | 1.53E-05 | -1.40E+00 | 3.83E-06 | 1.46E-05 | -1.23E+00 | 2.49E-04 | 8.89E-04 | -1.27E+00 | 8.49E-04 | 2.39E-03 | -1.18E+00 | 2.11E-11 | 7.75E-11 | -1.45E+00 | 3.97E-12 | 1.09E-11 | -1.38E+00 |
| ZNF83 | 2.90E-03 | 1.51E-02 | 1.03E+00 | 5.67E-02 | 2.45E-01 | 1.35E+00 | 8.30E-03 | 3.53E-02 | -1.05E+00 | 4.67E-02 | 1.78E-01 | 1.22E+00 | 2.01E-02 | 7.18E-02 | 1.24E+00 | 1.69E-01 | 4.76E-01 | 1.31E+00 | 5.30E-02 | 1.94E-01 | 1.14E+00 | 1.18E-01 | 3.25E-01 | 1.13E+00 |
| RPS8 | 2.90E-03 | 1.51E-02 | -1.06E+00 | 8.35E-43 | 3.61E-42 | -1.37E+00 | 1.05E-01 | 4.45E-01 | -1.02E+00 | 6.04E-18 | 2.31E-17 | -1.28E+00 | 5.58E-02 | 1.99E-01 | -1.05E+00 | 7.38E-13 | 2.08E-12 | -1.26E+00 | 1.16E-05 | 4.24E-05 | -1.07E+00 | 1.39E-20 | 3.83E-20 | -1.31E+00 |
| MAGI2 | 2.90E-03 | 1.51E-02 | -1.21E+00 | 9.04E-03 | 3.91E-02 | 1.03E+00 | 7.28E-03 | 3.09E-02 | -1.32E+00 | 4.54E-03 | 1.74E-02 | -1.29E+00 | 2.27E-01 | 8.09E-01 | 1.29E+00 | 2.53E-01 | 7.14E-01 | 1.07E+00 | 1.72E-01 | 6.30E-01 | -1.03E+00 | 2.37E-01 | 6.51E-01 | -1.10E+00 |
| NDUFB7 | 2.90E-03 | 1.51E-02 | 1.07E+00 | 5.00E-09 | 2.16E-08 | -1.01E+00 | 1.74E-01 | 7.40E-01 | 1.17E+00 | 1.50E-02 | 5.71E-02 | 1.06E+00 | 3.72E-03 | 1.33E-02 | 1.09E+00 | 6.42E-04 | 1.81E-03 | 1.04E+00 | 1.06E-01 | 3.89E-01 | 1.22E+00 | 3.21E-01 | 8.81E-01 | 1.11E+00 |
| LINC00680 | 2.90E-03 | 1.51E-02 | -1.21E+00 | 7.16E-02 | 3.09E-01 | 1.30E+00 | 5.96E-03 | 2.53E-02 | -1.15E+00 | 3.43E-02 | 1.31E-01 | 1.10E+00 | 2.29E-01 | 8.16E-01 | 1.43E+00 | 2.88E-01 | 8.11E-01 | 1.56E+00 | 1.24E-03 | 4.56E-03 | -1.20E+00 | 3.10E-01 | 8.52E-01 | 1.30E+00 |
| PLAU | 2.90E-03 | 1.51E-02 | -1.10E+00 | 2.77E-06 | 1.20E-05 | -1.56E+00 | 1.01E-04 | 4.30E-04 | -1.22E+00 | 1.00E-14 | 3.84E-14 | -1.91E+00 | 8.24E-03 | 2.94E-02 | -1.57E+00 | 4.53E-04 | 1.28E-03 | -2.80E+00 | 9.99E-02 | 3.67E-01 | -1.27E+00 | 1.49E-01 | 4.08E-01 | -1.26E+00 |
| RRN3P3 | 2.90E-03 | 1.51E-02 | -1.33E+00 | 2.24E-01 | 9.69E-01 | 1.46E+00 | 8.60E-05 | 3.65E-04 | -1.61E+00 | 5.55E-03 | 2.12E-02 | -1.06E+00 | 2.04E-03 | 7.28E-03 | -1.39E+00 | 1.42E-01 | 4.01E-01 | 1.20E+00 | 6.49E-04 | 2.38E-03 | -1.40E+00 | 2.99E-01 | 8.22E-01 | 1.26E+00 |
| FADS3 | 2.90E-03 | 1.51E-02 | -1.28E+00 | 5.41E-02 | 2.34E-01 | 1.15E+00 | 7.10E-04 | 3.02E-03 | -1.34E+00 | 1.85E-03 | 7.06E-03 | -1.09E+00 | 1.06E-14 | 3.79E-14 | -1.80E+00 | 3.82E-04 | 1.08E-03 | 1.05E+00 | 7.70E-08 | 2.83E-07 | -1.47E+00 | 1.93E-02 | 5.29E-02 | 1.02E+00 |
| AL139288.1 | 2.91E-03 | 1.52E-02 | 5.54E+00 | 4.07E-02 | 1.76E-01 | 2.14E+00 | 9.67E-03 | 4.11E-02 | 3.16E+00 | 2.41E-01 | 9.22E-01 | 1.21E+00 | 9.89E-03 | 3.53E-02 | 5.89E+00 | 1.50E-01 | 4.23E-01 | 1.86E+00 | 2.48E-01 | 9.11E-01 | 1.27E+00 | 1.36E-01 | 3.73E-01 | -1.52E+00 |
| AL139384.1 | 2.91E-03 | 1.52E-02 | -1.48E+00 | 7.19E-03 | 3.11E-02 | -1.24E+00 | 7.32E-02 | 3.11E-01 | -1.03E+00 | 5.15E-03 | 1.97E-02 | -1.50E+00 | 1.81E-01 | 6.46E-01 | 1.04E+00 | 3.45E-01 | 9.74E-01 | 1.22E+00 | 2.46E-01 | 9.01E-01 | 1.43E+00 | 3.61E-01 | 9.91E-01 | 1.30E+00 |
| ZNF688 | 2.91E-03 | 1.52E-02 | 1.01E+00 | 8.29E-02 | 3.58E-01 | 1.40E+00 | 2.70E-02 | 1.15E-01 | -1.02E+00 | 6.02E-02 | 2.30E-01 | 1.18E+00 | 8.38E-03 | 2.99E-02 | 1.05E+00 | 2.10E-01 | 5.91E-01 | 1.34E+00 | 3.89E-02 | 1.43E-01 | 1.22E+00 | 3.98E-02 | 1.09E-01 | 1.17E+00 |
| STIL | 2.92E-03 | 1.52E-02 | -1.66E+00 | 6.22E-03 | 2.69E-02 | -1.40E+00 | 2.02E-01 | 8.58E-01 | 1.21E+00 | 1.95E-01 | 7.45E-01 | 1.19E+00 | 8.88E-03 | 3.17E-02 | -1.73E+00 | 9.03E-03 | 2.55E-02 | -1.69E+00 | 1.79E-05 | 6.58E-05 | -1.19E+00 | 3.15E-03 | 8.65E-03 | -1.09E+00 |
| ARHGEF34P | 2.92E-03 | 1.52E-02 | -1.06E+00 | 4.01E-03 | 1.73E-02 | 1.07E+00 | 2.98E-02 | 1.27E-01 | -1.08E+00 | 8.47E-03 | 3.23E-02 | -1.18E+00 | 3.05E-02 | 1.09E-01 | 2.64E+00 | 3.42E-01 | 9.64E-01 | 1.30E+00 | 5.56E-02 | 2.04E-01 | 2.14E+00 | 2.84E-01 | 7.81E-01 | 1.43E+00 |
| NUSAP1 | 2.92E-03 | 1.52E-02 | -1.45E+00 | 7.12E-03 | 3.08E-02 | -1.30E+00 | 5.60E-02 | 2.38E-01 | 1.74E+00 | 7.76E-02 | 2.96E-01 | 1.59E+00 | 1.17E-03 | 4.18E-03 | -2.18E+00 | 3.02E-03 | 8.51E-03 | -1.85E+00 | 9.22E-04 | 3.39E-03 | 1.01E+00 | 5.09E-02 | 1.40E-01 | 1.13E+00 |
| RABL3 | 2.92E-03 | 1.52E-02 | -1.44E+00 | 7.10E-03 | 3.07E-02 | -1.22E+00 | 4.53E-05 | 1.93E-04 | -1.48E+00 | 2.33E-05 | 8.90E-05 | -1.37E+00 | 6.40E-02 | 2.28E-01 | 1.01E+00 | 2.02E-01 | 5.69E-01 | 1.27E+00 | 6.15E-06 | 2.26E-05 | -1.28E+00 | 1.35E-04 | 3.70E-04 | -1.10E+00 |
| KRI1 | 2.93E-03 | 1.53E-02 | -1.18E+00 | 5.12E-02 | 2.21E-01 | 1.15E+00 | 1.22E-05 | 5.18E-05 | -1.29E+00 | 3.37E-02 | 1.29E-01 | 1.21E+00 | 6.01E-05 | 2.14E-04 | -1.20E+00 | 3.32E-03 | 9.35E-03 | -1.02E+00 | 1.16E-08 | 4.25E-08 | -1.43E+00 | 2.05E-03 | 5.64E-03 | -1.10E+00 |
| AC005776.2 | 2.93E-03 | 1.53E-02 | -1.21E+00 | 1.15E-06 | 4.96E-06 | -2.13E+00 | 3.91E-04 | 1.66E-03 | -1.51E+00 | 5.25E-07 | 2.01E-06 | -1.99E+00 | 5.52E-03 | 1.97E-02 | -1.24E+00 | 1.31E-04 | 3.70E-04 | -2.01E+00 | 4.27E-03 | 1.57E-02 | -1.47E+00 | 1.43E-03 | 3.93E-03 | -1.67E+00 |
| PXYLP1 | 2.94E-03 | 1.53E-02 | -1.51E+00 | 1.21E-01 | 5.21E-01 | 1.22E+00 | 1.83E-01 | 7.76E-01 | 1.14E+00 | 1.17E-01 | 4.47E-01 | 1.59E+00 | 2.78E-01 | 9.92E-01 | 1.01E+00 | 2.39E-01 | 6.73E-01 | 1.33E+00 | 8.41E-02 | 3.09E-01 | -1.22E+00 | 3.23E-01 | 8.88E-01 | 1.12E+00 |
| RPL4P3 | 2.94E-03 | 1.53E-02 | 1.01E+00 | 3.37E-09 | 1.46E-08 | -1.56E+00 | 2.98E-08 | 1.27E-07 | -1.38E+00 | 1.93E-15 | 7.35E-15 | -1.83E+00 | 4.67E-04 | 1.67E-03 | -1.14E+00 | 5.72E-05 | 1.61E-04 | -1.37E+00 | 2.98E-07 | 1.09E-06 | -1.28E+00 | 5.83E-11 | 1.60E-10 | -1.66E+00 |
| AC006330.1 | 2.95E-03 | 1.54E-02 | -1.42E+00 | 4.65E-04 | 2.01E-03 | -1.62E+00 | 3.91E-02 | 1.66E-01 | -1.12E+00 | 2.49E-03 | 9.49E-03 | -1.59E+00 | 1.31E-01 | 4.69E-01 | 1.09E+00 | 6.62E-02 | 1.87E-01 | -1.10E+00 | 1.90E-01 | 6.97E-01 | 1.02E+00 | 2.10E-01 | 5.77E-01 | 1.35E+00 |
| SH3GL1P1 | 2.95E-03 | 1.54E-02 | -1.36E+00 | 3.55E-03 | 1.53E-02 | -1.37E+00 | 4.61E-05 | 1.96E-04 | -1.84E+00 | 1.93E-04 | 7.37E-04 | -1.58E+00 | 1.58E-01 | 5.62E-01 | 1.59E+00 | 1.48E-01 | 4.16E-01 | -1.17E+00 | 4.48E-03 | 1.64E-02 | -1.43E+00 | 1.09E-03 | 2.99E-03 | -1.77E+00 |
| AC087752.3 | 2.95E-03 | 1.54E-02 | 2.94E+00 | 3.21E-02 | 1.39E-01 | 2.36E+00 | 2.01E-01 | 8.53E-01 | 1.31E+00 | 2.17E-01 | 8.28E-01 | 1.35E+00 | 2.01E-01 | 7.17E-01 | 1.27E+00 | 1.52E-01 | 4.29E-01 | 1.01E+00 | 5.13E-04 | 1.88E-03 | 2.96E+00 | 3.42E-01 | 9.39E-01 | 1.34E+00 |
| ZC3H12B | 2.95E-03 | 1.54E-02 | -1.23E+00 | 4.60E-03 | 1.99E-02 | -1.10E+00 | 1.93E-04 | 8.20E-04 | -1.68E+00 | 1.02E-02 | 3.88E-02 | -1.05E+00 | 8.30E-02 | 2.96E-01 | -1.35E+00 | 7.42E-02 | 2.09E-01 | -1.51E+00 | 1.06E-01 | 3.87E-01 | -1.12E+00 | 3.35E-01 | 9.21E-01 | 1.12E+00 |
| C17orf113 | 2.96E-03 | 1.54E-02 | -1.49E+00 | 4.44E-02 | 1.92E-01 | 1.02E+00 | 6.07E-04 | 2.58E-03 | -1.56E+00 | 7.37E-03 | 2.81E-02 | -1.11E+00 | 6.13E-02 | 2.18E-01 | -1.26E+00 | 2.73E-01 | 7.70E-01 | 1.50E+00 | 9.48E-04 | 3.48E-03 | -1.43E+00 | 1.58E-02 | 4.33E-02 | -1.13E+00 |
| GS1-124K5.4 | 2.96E-03 | 1.54E-02 | -1.11E+00 | 1.57E-03 | 6.80E-03 | -1.10E+00 | 1.05E-01 | 4.45E-01 | 1.16E+00 | 8.92E-03 | 3.41E-02 | 1.02E+00 | 7.52E-02 | 2.68E-01 | 2.00E+00 | 8.83E-02 | 2.49E-01 | 2.16E+00 | 1.66E-01 | 6.10E-01 | 1.11E+00 | 2.52E-01 | 6.93E-01 | 1.20E+00 |
| USP9X | 2.97E-03 | 1.55E-02 | 1.01E+00 | 2.48E-03 | 1.07E-02 | 1.17E+00 | 9.17E-03 | 3.90E-02 | -1.00E+00 | 2.05E-03 | 7.82E-03 | 1.03E+00 | 3.50E-03 | 1.25E-02 | -1.01E+00 | 8.47E-03 | 2.39E-02 | 1.05E+00 | 5.94E-06 | 2.18E-05 | -1.18E+00 | 2.72E-05 | 7.48E-05 | -1.18E+00 |
| IL12RB2 | 2.98E-03 | 1.55E-02 | 1.01E+00 | 1.76E-01 | 7.60E-01 | 1.56E+00 | 4.78E-02 | 2.03E-01 | 1.03E+00 | 2.90E-02 | 1.11E-01 | 1.10E+00 | 1.43E-01 | 5.10E-01 | 1.97E+00 | 2.83E-01 | 7.97E-01 | 1.64E+00 | 2.09E-01 | 7.67E-01 | 1.16E+00 | 5.84E-02 | 1.60E-01 | 2.14E+00 |
| AC126615.1 | 2.98E-03 | 1.55E-02 | -1.23E+00 | 3.62E-03 | 1.56E-02 | -1.10E+00 | 6.67E-04 | 2.83E-03 | -1.31E+00 | 1.36E-04 | 5.19E-04 | -1.40E+00 | 1.83E-03 | 6.52E-03 | -2.12E+00 | 1.15E-02 | 3.25E-02 | -1.56E+00 | 3.49E-03 | 1.28E-02 | -1.46E+00 | 2.31E-03 | 6.34E-03 | -1.54E+00 |
| RPL7P1 | 2.98E-03 | 1.55E-02 | 1.74E+00 | 4.96E-06 | 2.14E-05 | 2.30E+00 | 3.43E-03 | 1.46E-02 | 1.56E+00 | 3.80E-06 | 1.45E-05 | 2.15E+00 | 8.41E-02 | 3.00E-01 | 1.74E+00 | 3.55E-03 | 1.00E-02 | 2.12E+00 | 6.54E-03 | 2.40E-02 | 1.58E+00 | 1.39E-06 | 3.83E-06 | 2.10E+00 |
| RPS16 | 2.99E-03 | 1.56E-02 | 1.07E+00 | 4.05E-02 | 1.75E-01 | -1.03E+00 | 1.05E-02 | 4.44E-02 | 1.08E+00 | 1.58E-01 | 6.02E-01 | 1.00E+00 | 1.54E-01 | 5.50E-01 | 1.03E+00 | 2.32E-02 | 6.54E-02 | 1.08E+00 | 5.94E-02 | 2.18E-01 | 1.01E+00 | 3.13E-01 | 8.59E-01 | 1.03E+00 |
| POLR1E | 3.00E-03 | 1.56E-02 | 1.04E+00 | 1.02E-02 | 4.42E-02 | 1.25E+00 | 1.04E-02 | 4.43E-02 | -1.03E+00 | 2.59E-03 | 9.90E-03 | 1.09E+00 | 1.29E-04 | 4.61E-04 | -1.10E+00 | 2.90E-03 | 8.17E-03 | 1.13E+00 | 5.66E-06 | 2.08E-05 | -1.25E+00 | 1.75E-02 | 4.80E-02 | 1.02E+00 |
| SMPD1 | 3.00E-03 | 1.56E-02 | -1.24E+00 | 2.61E-02 | 1.13E-01 | 1.09E+00 | 3.48E-02 | 1.48E-01 | -1.08E+00 | 3.70E-02 | 1.41E-01 | -1.01E+00 | 1.60E-03 | 5.71E-03 | -1.18E+00 | 1.85E-02 | 5.22E-02 | -1.02E+00 | 4.21E-03 | 1.55E-02 | -1.12E+00 | 4.44E-02 | 1.22E-01 | -1.01E+00 |
| GADD45A | 3.01E-03 | 1.57E-02 | 1.25E+00 | 5.36E-05 | 2.32E-04 | 1.21E+00 | 1.97E-03 | 8.36E-03 | -1.05E+00 | 2.68E-06 | 1.02E-05 | -1.15E+00 | 4.17E-09 | 1.49E-08 | -1.42E+00 | 2.30E-09 | 6.48E-09 | -1.31E+00 | 9.05E-04 | 3.32E-03 | 1.17E+00 | 1.21E-08 | 3.32E-08 | -1.19E+00 |
| CAB39L | 3.01E-03 | 1.57E-02 | -1.33E+00 | 3.93E-04 | 1.70E-03 | -1.58E+00 | 1.91E-01 | 8.10E-01 | 1.18E+00 | 2.34E-01 | 8.92E-01 | 1.24E+00 | 1.80E-01 | 6.42E-01 | 1.53E+00 | 3.47E-01 | 9.77E-01 | 1.27E+00 | 1.27E-01 | 4.64E-01 | -1.07E+00 | 2.07E-01 | 5.69E-01 | 1.24E+00 |
| MBNL3 | 3.02E-03 | 1.58E-02 | -1.35E+00 | 5.52E-02 | 2.39E-01 | 1.10E+00 | 2.17E-03 | 9.21E-03 | -1.24E+00 | 6.63E-03 | 2.53E-02 | 1.03E+00 | 3.86E-04 | 1.38E-03 | -1.28E+00 | 1.75E-02 | 4.92E-02 | -1.00E+00 | 1.53E-02 | 5.61E-02 | -1.04E+00 | 2.18E-01 | 5.98E-01 | 1.13E+00 |
| AC105760.2 | 3.02E-03 | 1.58E-02 | -1.56E+00 | 2.21E-02 | 9.54E-02 | -1.11E+00 | 7.27E-05 | 3.09E-04 | -2.03E+00 | 4.66E-02 | 1.78E-01 | -1.02E+00 | 1.33E-01 | 4.75E-01 | 1.02E+00 | 2.61E-01 | 7.35E-01 | 1.17E+00 | 1.75E-01 | 6.41E-01 | 1.01E+00 | 1.12E-01 | 3.08E-01 | 1.55E+00 |
| RPL11P3 | 3.03E-03 | 1.58E-02 | 1.61E+00 | 2.21E-02 | 9.56E-02 | 1.34E+00 | 2.65E-03 | 1.12E-02 | 1.59E+00 | 2.61E-02 | 9.96E-02 | 1.15E+00 | 2.28E-04 | 8.11E-04 | 1.91E+00 | 3.16E-01 | 8.91E-01 | 1.38E+00 | 2.26E-02 | 8.31E-02 | 1.60E+00 | 1.27E-01 | 3.48E-01 | 1.18E+00 |
| MZF1-AS1 | 3.04E-03 | 1.58E-02 | 2.19E+00 | 2.44E-04 | 1.05E-03 | -1.20E+00 | 1.71E-06 | 7.26E-06 | 2.45E+00 | 1.23E-01 | 4.68E-01 | 1.15E+00 | 1.13E-01 | 4.03E-01 | 1.26E+00 | 7.02E-03 | 1.98E-02 | -1.46E+00 | 3.75E-02 | 1.38E-01 | 1.71E+00 | 2.10E-03 | 5.76E-03 | -1.45E+00 |
| BCS1L | 3.04E-03 | 1.58E-02 | -1.01E+00 | 1.57E-02 | 6.76E-02 | 1.19E+00 | 3.59E-02 | 1.53E-01 | 1.05E+00 | 6.71E-02 | 2.56E-01 | 1.24E+00 | 1.02E-03 | 3.63E-03 | -1.25E+00 | 7.64E-02 | 2.15E-01 | 1.13E+00 | 9.03E-04 | 3.32E-03 | -1.07E+00 | 1.22E-01 | 3.34E-01 | 1.23E+00 |
| HDAC9 | 3.04E-03 | 1.58E-02 | -1.72E+00 | 5.06E-04 | 2.19E-03 | -1.88E+00 | 3.83E-02 | 1.63E-01 | -1.33E+00 | 2.11E-02 | 8.07E-02 | -1.50E+00 | 7.79E-03 | 2.78E-02 | -2.30E+00 | 1.69E-02 | 4.76E-02 | -1.97E+00 | 2.57E-02 | 9.42E-02 | -1.05E+00 | 1.63E-02 | 4.47E-02 | -1.33E+00 |
| NOV | 3.04E-03 | 1.58E-02 | -1.32E+00 | 2.46E-05 | 1.06E-04 | -1.99E+00 | 9.83E-05 | 4.18E-04 | -1.62E+00 | 4.63E-06 | 1.77E-05 | -1.66E+00 | 2.23E-01 | 7.95E-01 | 1.05E+00 | 6.99E-02 | 1.97E-01 | -2.03E+00 | 9.54E-02 | 3.50E-01 | 2.04E+00 | 2.85E-01 | 7.84E-01 | 1.21E+00 |

| | | | | | | | | | | | | | | | | | | | | | | | | |
|---|---|---|---|---|---|---|---|---|---|---|---|---|---|---|---|---|---|---|---|---|---|---|---|---|
| AL034379.1 | 3.07E-03 | 1.60E-02 | -1.08E+00 | 7.90E-79 | 3.41E-78 | -1.87E+00 | 1.16E-01 | 4.94E-01 | -1.02E+00 | 1.69E-36 | 6.47E-36 | -1.75E+00 | 1.44E-01 | 5.14E-01 | -1.02E+00 | 5.71E-33 | 1.61E-32 | -1.79E+00 | 5.56E-02 | 2.04E-01 | -1.01E+00 | 3.98E-38 | 1.09E-37 | -1.74E+00 |
| AC002454.1 | 3.08E-03 | 1.61E-02 | 1.10E+00 | 9.64E-07 | 4.17E-06 | -1.19E+00 | 1.40E-02 | 5.94E-02 | -1.00E+00 | 2.17E-05 | 8.30E-05 | -1.10E+00 | 2.78E-01 | 9.91E-01 | 1.36E+00 | 4.86E-02 | 1.37E-01 | -1.00E+00 | 8.41E-02 | 3.09E-01 | 1.05E+00 | 1.09E-02 | 2.98E-02 | -1.16E+00 |
| ZNF607 | 3.09E-03 | 1.61E-02 | -1.55E+00 | 4.99E-02 | 2.15E-01 | 1.03E+00 | 1.77E-01 | 7.52E-01 | 1.08E+00 | 1.73E-01 | 6.61E-01 | 1.19E+00 | 2.33E-01 | 8.29E-01 | 1.27E+00 | 2.42E-01 | 6.82E-01 | 1.69E+00 | 3.17E-05 | 1.16E-04 | -1.93E+00 | 5.54E-02 | 1.52E-01 | 1.02E+00 |
| HNRNPA1P35 | 3.09E-03 | 1.61E-02 | -1.27E+00 | 3.63E-02 | 1.57E-01 | 1.11E+00 | 9.04E-05 | 3.84E-04 | -1.37E+00 | 1.01E-03 | 3.84E-03 | -1.11E+00 | 2.61E-02 | 9.29E-02 | -1.14E+00 | 1.95E-01 | 5.48E-01 | 1.20E+00 | 2.06E-05 | 7.55E-05 | -1.38E+00 | 4.46E-03 | 1.23E-02 | -1.10E+00 |
| RAPGEF3 | 3.10E-03 | 1.61E-02 | -1.16E+00 | 6.17E-03 | 2.67E-02 | -1.04E+00 | 2.25E-03 | 9.57E-03 | -1.51E+00 | 1.65E-02 | 6.31E-02 | -1.11E+00 | 5.65E-02 | 2.02E-01 | -1.32E+00 | 9.04E-02 | 2.55E-01 | -1.13E+00 | 2.35E-01 | 8.61E-01 | 1.15E+00 | 3.12E-01 | 8.56E-01 | -1.06E+00 |
| ZNF610 | 3.10E-03 | 1.61E-02 | -1.52E+00 | 1.38E-01 | 5.96E-01 | 1.24E+00 | 1.10E-02 | 4.69E-02 | -1.51E+00 | 2.66E-02 | 1.02E-01 | -1.20E+00 | 3.69E-04 | 1.32E-03 | -2.17E+00 | 3.83E-03 | 1.08E-02 | -1.51E+00 | 1.85E-02 | 6.80E-02 | -1.53E+00 | 1.80E-01 | 4.95E-01 | -1.01E+00 |
| ELF2 | 3.10E-03 | 1.62E-02 | 1.08E+00 | 2.23E-13 | 9.63E-13 | -1.21E+00 | 6.11E-03 | 2.60E-02 | 1.02E+00 | 3.62E-12 | 1.38E-11 | -1.33E+00 | 1.81E-02 | 6.47E-02 | 1.07E+00 | 1.73E-06 | 4.87E-06 | -1.26E+00 | 2.16E-05 | 7.93E-05 | -1.13E+00 | 1.30E-07 | 3.56E-07 | -1.34E+00 |
| TSEN54 | 3.10E-03 | 1.62E-02 | -1.21E+00 | 7.22E-02 | 3.12E-01 | 1.19E+00 | 1.05E-01 | 4.45E-01 | 1.00E+00 | 1.33E-01 | 5.07E-01 | 1.41E+00 | 2.30E-01 | 8.20E-01 | 1.31E+00 | 2.97E-02 | 8.37E-02 | 1.89E+00 | 1.51E-02 | 5.54E-02 | 1.06E+00 | 1.55E-01 | 4.25E-01 | 1.45E+00 |
| SEC14L1P1 | 3.10E-03 | 1.62E-02 | -1.64E+00 | 3.37E-02 | 1.45E-01 | -1.12E+00 | 1.59E-01 | 6.77E-01 | 1.03E+00 | 2.21E-01 | 8.43E-01 | 1.39E+00 | 1.73E-01 | 6.17E-01 | 1.01E+00 | 1.44E-01 | 4.07E-01 | 1.73E+00 | 1.49E-02 | 5.45E-02 | -1.20E+00 | 9.76E-03 | 2.68E-02 | -1.35E+00 |
| SFSWAP | 3.10E-03 | 1.62E-02 | 1.04E+00 | 3.85E-04 | 1.66E-03 | 1.12E+00 | 5.51E-03 | 2.34E-02 | -1.07E+00 | 9.54E-04 | 3.65E-03 | 1.01E+00 | 1.93E-06 | 6.89E-06 | -1.16E+00 | 2.06E-03 | 5.80E-03 | 1.06E+00 | 1.70E-04 | 6.24E-04 | -1.03E+00 | 4.36E-03 | 1.20E-02 | 1.04E+00 |
| AC131009.4 | 3.11E-03 | 1.62E-02 | -1.28E+00 | 7.42E-02 | 3.20E-01 | 1.20E+00 | 2.05E-01 | 8.73E-01 | 1.16E+00 | 2.32E-01 | 8.86E-01 | 1.27E+00 | 2.42E-01 | 8.61E-01 | 1.19E+00 | 1.82E-01 | 5.13E-01 | 1.08E+00 | 8.12E-02 | 2.98E-01 | 1.87E+00 | 1.70E-01 | 4.66E-01 | 1.66E+00 |
| CARD11 | 3.11E-03 | 1.62E-02 | -1.39E+00 | 8.67E-03 | 3.75E-02 | -1.13E+00 | 1.80E-01 | 7.66E-01 | 1.38E+00 | 6.82E-02 | 2.60E-01 | 1.73E+00 | 2.91E-02 | 1.04E-01 | -1.42E+00 | 1.64E-02 | 4.61E-02 | -1.63E+00 | 8.77E-03 | 3.22E-02 | -1.61E+00 | 8.29E-02 | 2.28E-01 | -1.23E+00 |
| TMEM160 | 3.11E-03 | 1.62E-02 | 1.16E+00 | 3.50E-03 | 1.51E-02 | 1.22E+00 | 6.73E-02 | 2.86E-01 | 1.27E+00 | 1.00E-01 | 3.82E-01 | 1.16E+00 | 1.20E-03 | 4.27E-03 | 1.02E+00 | 1.40E-05 | 3.95E-05 | -1.05E+00 | 2.34E-01 | 8.60E-01 | 1.21E+00 | 2.55E-01 | 7.00E-01 | 1.30E+00 |
| HINT1 | 3.13E-03 | 1.63E-02 | -1.07E+00 | 3.25E-02 | 1.41E-01 | -1.03E+00 | 1.21E-01 | 5.13E-01 | -1.02E+00 | 1.52E-01 | 5.82E-01 | -1.02E+00 | 2.52E-04 | 8.97E-04 | -1.12E+00 | 9.16E-03 | 2.58E-02 | -1.09E+00 | 9.44E-02 | 3.47E-01 | 1.02E+00 | 2.00E-01 | 5.48E-01 | 1.03E+00 |
| RBM4B | 3.13E-03 | 1.63E-02 | -1.03E+00 | 7.33E-02 | 3.17E-01 | 1.31E+00 | 2.18E-05 | 9.28E-05 | -1.34E+00 | 1.34E-03 | 5.14E-03 | -1.00E+00 | 3.94E-03 | 1.40E-02 | -1.14E+00 | 7.87E-03 | 2.22E-02 | -1.11E+00 | 5.93E-08 | 2.18E-07 | -1.46E+00 | 1.64E-03 | 4.49E-03 | -1.06E+00 |
| CCSER1 | 3.13E-03 | 1.63E-02 | -1.03E+00 | 8.88E-02 | 3.84E-01 | 1.31E+00 | 7.35E-03 | 3.12E-02 | -1.10E+00 | 3.19E-02 | 1.22E-01 | 1.15E+00 | 1.86E-01 | 6.63E-01 | 1.87E+00 | 1.47E-01 | 4.15E-01 | 2.07E+00 | 8.85E-03 | 3.25E-02 | -1.31E+00 | 1.23E-01 | 3.37E-01 | 1.02E+00 |
| AC135050.6 | 3.14E-03 | 1.64E-02 | -1.24E+00 | 1.01E-02 | 4.36E-02 | 1.00E+00 | 1.59E-03 | 6.77E-03 | -1.21E+00 | 2.82E-02 | 1.08E-01 | 1.06E+00 | 2.15E-03 | 7.66E-03 | -1.52E+00 | 1.96E-02 | 5.54E-02 | -1.08E+00 | 1.48E-03 | 5.43E-03 | -1.50E+00 | 2.74E-01 | 7.52E-01 | 1.07E+00 |
| TTF1 | 3.14E-03 | 1.64E-02 | -1.03E+00 | 1.07E-02 | 4.64E-02 | 1.11E+00 | 3.99E-02 | 1.70E-01 | 1.02E+00 | 1.74E-02 | 6.64E-02 | 1.03E+00 | 2.41E-04 | 8.58E-04 | -1.10E+00 | 2.88E-02 | 8.12E-02 | 1.11E+00 | 3.90E-03 | 1.43E-02 | -1.01E+00 | 5.28E-02 | 1.45E-01 | 1.12E+00 |
| AC027290.2 | 3.15E-03 | 1.64E-02 | 1.33E+00 | 2.96E-02 | 1.28E-01 | 1.17E+00 | 6.92E-03 | 2.94E-02 | 1.48E+00 | 1.52E-01 | 5.81E-01 | 1.22E+00 | 6.07E-04 | 2.16E-03 | 1.70E+00 | 1.43E-02 | 4.03E-02 | 1.65E+00 | 1.98E-01 | 7.28E-01 | 1.19E+00 | 8.02E-02 | 2.20E-01 | -1.01E+00 |
| AC127496.5 | 3.15E-03 | 1.64E-02 | -1.26E+00 | 7.82E-02 | 3.38E-01 | 1.30E+00 | 6.81E-06 | 2.89E-05 | -1.95E+00 | 2.31E-04 | 8.84E-04 | -1.35E+00 | 1.77E-01 | 6.32E-01 | -1.08E+00 | 2.83E-01 | 7.99E-01 | 1.06E+00 | 1.15E-01 | 4.22E-01 | -1.13E+00 | 3.01E-02 | 8.27E-02 | -1.52E+00 |
| SSR3 | 3.16E-03 | 1.65E-02 | 1.07E+00 | 4.40E-04 | 1.90E-03 | 1.15E+00 | 5.15E-02 | 2.19E-01 | 1.02E+00 | 7.17E-02 | 2.74E-01 | 1.12E+00 | 1.90E-08 | 6.76E-08 | -1.36E+00 | 9.64E-04 | 2.72E-03 | -1.14E+00 | 1.53E-03 | 5.63E-03 | -1.18E+00 | 9.23E-02 | 2.53E-01 | 1.00E+00 |
| FADS2 | 3.16E-03 | 1.65E-02 | 1.06E+00 | 1.33E-01 | 5.77E-01 | 1.35E+00 | 8.06E-02 | 3.43E-01 | 1.14E+00 | 2.16E-01 | 8.25E-01 | 1.52E+00 | 2.36E-03 | 8.43E-03 | -1.03E+00 | 1.89E-01 | 5.33E-01 | 1.29E+00 | 7.92E-03 | 2.91E-02 | -1.11E+00 | 3.15E-01 | 8.66E-01 | 1.13E+00 |
| IFT81 | 3.16E-03 | 1.65E-02 | -1.16E+00 | 8.94E-03 | 3.86E-02 | 1.02E+00 | 9.32E-05 | 3.96E-04 | -1.45E+00 | 1.29E-03 | 4.94E-03 | -1.21E+00 | 3.11E-09 | 1.11E-08 | -1.48E+00 | 1.01E-08 | 2.85E-08 | -1.46E+00 | 1.43E-01 | 5.24E-01 | 1.12E+00 | 8.14E-02 | 2.24E-01 | 1.07E+00 |
| ARHGAP21 | 3.17E-03 | 1.65E-02 | -1.07E+00 | 2.54E-02 | 1.10E-01 | 1.19E+00 | 1.20E-01 | 5.12E-01 | -1.02E+00 | 8.69E-02 | 3.32E-01 | 1.11E+00 | 1.32E-02 | 4.71E-02 | -1.02E+00 | 9.65E-03 | 2.72E-02 | -1.10E+00 | 1.73E-04 | 6.35E-04 | 1.09E+00 | 1.27E-05 | 3.50E-05 | -1.20E+00 |
| AC124248.1 | 3.17E-03 | 1.65E-02 | -1.15E+00 | 2.78E-03 | 1.20E-02 | -1.08E+00 | 5.19E-03 | 2.20E-02 | -1.45E+00 | 1.40E-02 | 5.34E-02 | -1.14E+00 | 1.24E-03 | 4.41E-03 | 1.02E+00 | 3.13E-04 | 8.81E-04 | -1.02E+00 | 4.94E-06 | 1.81E-05 | -1.31E+00 | 1.10E-11 | 3.01E-11 | -1.86E+00 |
| FAM35A | 3.17E-03 | 1.65E-02 | -1.15E+00 | 1.93E-02 | 8.33E-02 | 1.11E+00 | 5.97E-02 | 2.54E-01 | -1.06E+00 | 1.35E-01 | 5.16E-01 | 1.22E+00 | 6.31E-02 | 2.25E-01 | 1.03E+00 | 2.79E-01 | 7.88E-01 | 1.29E+00 | 9.47E-08 | 3.48E-07 | -1.42E+00 | 5.68E-03 | 1.56E-02 | 1.07E+00 |
| PDE6D | 3.17E-03 | 1.65E-02 | -1.09E+00 | 4.40E-03 | 1.90E-02 | 1.01E+00 | 8.25E-03 | 3.51E-02 | -1.04E+00 | 5.15E-03 | 1.97E-02 | 1.10E+00 | 4.87E-06 | 1.74E-05 | -1.40E+00 | 2.50E-05 | 7.05E-05 | -1.24E+00 | 1.99E-04 | 7.30E-04 | -1.18E+00 | 5.39E-02 | 1.48E-01 | 1.08E+00 |
| AC067852.6 | 3.18E-03 | 1.66E-02 | -1.25E+00 | 6.94E-05 | 3.00E-04 | -1.90E+00 | 7.32E-04 | 3.11E-03 | -1.53E+00 | 2.01E-06 | 7.69E-06 | -2.34E+00 | 8.17E-02 | 2.91E-01 | 1.18E+00 | 1.57E-02 | 4.43E-02 | -1.09E+00 | 2.67E-04 | 9.78E-04 | -1.22E+00 | 2.07E-04 | 5.68E-04 | -1.35E+00 |
| HIST2H2BE | 3.18E-03 | 1.66E-02 | 1.92E+00 | 1.65E-01 | 7.13E-01 | 1.63E+00 | 1.35E-01 | 5.75E-01 | 1.03E+00 | 1.25E-01 | 4.77E-01 | 1.75E+00 | 1.70E-01 | 6.07E-01 | 1.45E+00 | 1.55E-01 | 4.36E-01 | 1.63E+00 | 1.19E-01 | 4.37E-01 | 1.71E+00 | 3.08E-01 | 8.46E-01 | 1.35E+00 |
| BTAF1 | 3.19E-03 | 1.66E-02 | -1.01E+00 | 2.12E-06 | 9.16E-06 | -1.26E+00 | 1.11E-01 | 4.71E-01 | 1.15E+00 | 1.34E-04 | 5.11E-04 | -1.14E+00 | 2.29E-01 | 8.18E-01 | 1.64E+00 | 2.95E-03 | 8.31E-03 | -1.07E+00 | 2.62E-06 | 9.60E-06 | -1.15E+00 | 1.87E-11 | 5.13E-11 | -1.57E+00 |
| TRPV1 | 3.19E-03 | 1.66E-02 | -1.57E+00 | 7.06E-02 | 3.05E-01 | 1.07E+00 | 3.40E-02 | 1.45E-01 | -1.15E+00 | 2.40E-01 | 9.16E-01 | 1.43E+00 | 3.84E-02 | 1.37E-01 | -1.15E+00 | 5.40E-02 | 1.52E-01 | -1.07E+00 | 1.53E-01 | 5.61E-01 | -1.02E+00 | 3.77E-02 | 1.03E-01 | 1.74E+00 |
| ANKRD6 | 3.19E-03 | 1.66E-02 | -1.14E+00 | 1.44E-02 | 6.22E-02 | 1.11E+00 | 4.79E-03 | 2.03E-02 | -1.27E+00 | 1.44E-04 | 5.50E-04 | -1.51E+00 | 9.51E-02 | 3.39E-01 | 1.01E+00 | 1.58E-02 | 4.45E-02 | -1.57E+00 | 1.94E-01 | 7.12E-01 | -1.01E+00 | 1.55E-01 | 4.26E-01 | -1.22E+00 |
| CPNE7 | 3.20E-03 | 1.67E-02 | -1.32E+00 | 9.03E-04 | 3.90E-03 | -1.31E+00 | 1.42E-01 | 6.05E-01 | 1.08E+00 | 2.59E-02 | 9.90E-02 | 1.15E+00 | 8.66E-02 | 3.09E-01 | -1.71E+00 | 8.24E-02 | 2.32E-01 | -2.35E+00 | 5.59E-02 | 2.05E-01 | -1.23E+00 | 2.24E-01 | 6.14E-01 | 1.11E+00 |
| PLXND1 | 3.20E-03 | 1.67E-02 | 1.03E+00 | 1.29E-03 | 5.57E-03 | 1.12E+00 | 2.03E-04 | 8.62E-04 | -1.12E+00 | 7.07E-05 | 2.70E-04 | -1.04E+00 | 1.71E-02 | 6.08E-02 | -1.14E+00 | 1.04E-02 | 2.93E-02 | -1.15E+00 | 1.75E-03 | 6.41E-03 | -1.29E+00 | 1.90E-01 | 5.21E-01 | 1.23E+00 |
| SEC24B | 3.20E-03 | 1.67E-02 | -1.05E+00 | 3.10E-03 | 1.34E-02 | 1.05E+00 | 5.21E-02 | 2.21E-01 | 1.02E+00 | 2.92E-03 | 1.11E-02 | -1.03E+00 | 4.56E-02 | 1.63E-01 | 1.13E+00 | 2.18E-03 | 6.15E-03 | -1.23E+00 | 1.54E-06 | 5.64E-06 | -1.39E+00 | 4.49E-04 | 1.23E-03 | -1.22E+00 |
| BX679664.1 | 3.21E-03 | 1.67E-02 | 1.04E+00 | 1.71E-55 | 7.38E-55 | -1.82E+00 | 2.23E-01 | 9.49E-01 | 1.08E+00 | 1.50E-23 | 5.71E-23 | -1.85E+00 | 2.59E-01 | 9.24E-01 | 1.14E+00 | 5.48E-17 | 1.54E-16 | -1.70E+00 | 1.06E-01 | 3.88E-01 | 1.07E+00 | 2.99E-20 | 8.23E-20 | -1.99E+00 |
| FAM45BP | 3.21E-03 | 1.67E-02 | -1.67E+00 | 1.18E-02 | 5.09E-02 | -1.36E+00 | 2.92E-03 | 1.24E-02 | -1.43E+00 | 1.99E-02 | 7.62E-02 | -1.01E+00 | 3.22E-02 | 1.15E-01 | -1.23E+00 | 1.32E-01 | 3.71E-01 | -1.02E+00 | 9.98E-02 | 3.66E-01 | 1.27E+00 | 1.79E-01 | 4.91E-01 | 1.30E+00 |
| AC092139.3 | 3.21E-03 | 1.67E-02 | -1.59E+00 | 2.87E-02 | 1.24E-01 | 1.01E+00 | 3.09E-02 | 1.31E-01 | -1.13E+00 | 3.47E-02 | 1.33E-01 | 1.02E+00 | 2.79E-02 | 9.94E-02 | -1.37E+00 | 5.55E-02 | 1.56E-01 | -1.23E+00 | 3.01E-03 | 1.11E-02 | -1.51E+00 | 1.93E-02 | 5.29E-02 | -1.24E+00 |
| RSRC2 | 3.22E-03 | 1.68E-02 | 1.01E+00 | 2.30E-15 | 9.92E-15 | -1.20E+00 | 9.40E-03 | 4.00E-02 | -1.11E+00 | 3.14E-06 | 1.20E-05 | -1.28E+00 | 7.12E-04 | 2.54E-03 | 1.02E+00 | 2.30E-07 | 6.48E-07 | -1.21E+00 | 1.80E-04 | 6.59E-04 | -1.16E+00 | 1.59E-04 | 4.36E-04 | -1.27E+00 |
| CREM | 3.22E-03 | 1.68E-02 | -1.12E+00 | 2.63E-03 | 1.14E-02 | -1.07E+00 | 6.56E-02 | 2.79E-01 | -1.08E+00 | 5.08E-02 | 1.94E-01 | 1.02E+00 | 2.23E-01 | 7.96E-01 | 1.26E+00 | 6.52E-02 | 1.84E-01 | -1.02E+00 | 2.26E-02 | 8.31E-02 | -1.07E+00 | 1.69E-02 | 4.65E-02 | -1.20E+00 |
| EFL1 | 3.23E-03 | 1.68E-02 | -1.28E+00 | 3.37E-02 | 1.46E-01 | 1.02E+00 | 9.75E-02 | 4.14E-01 | 1.04E+00 | 1.26E-01 | 4.82E-01 | 1.50E+00 | 5.33E-04 | 1.90E-03 | -1.44E+00 | 1.73E-01 | 4.88E-01 | 1.34E+00 | 2.17E-03 | 7.97E-03 | -1.18E+00 | 2.27E-01 | 6.23E-01 | 1.17E+00 |
| AC073896.4 | 3.23E-03 | 1.69E-02 | -1.16E+00 | 1.45E-03 | 6.28E-03 | -1.18E+00 | 1.22E-06 | 5.16E-06 | -1.71E+00 | 4.52E-07 | 1.73E-06 | -1.67E+00 | 1.51E-05 | 5.37E-05 | -1.57E+00 | 1.97E-05 | 5.55E-05 | -1.53E+00 | 5.16E-06 | 1.89E-05 | -1.45E+00 | 1.74E-07 | 4.79E-07 | -1.79E+00 |
| AC006333.2 | 3.24E-03 | 1.69E-02 | -1.19E+00 | 3.51E-02 | 1.52E-01 | 1.13E+00 | 8.64E-03 | 3.67E-02 | -1.08E+00 | 2.00E-02 | 7.64E-02 | 1.12E+00 | 2.33E-01 | 8.31E-01 | 1.29E+00 | 1.22E-01 | 3.44E-01 | 1.06E+00 | 5.68E-04 | 2.09E-03 | -1.29E+00 | 4.22E-02 | 1.16E-01 | -1.04E+00 |
| GALNT14 | 3.25E-03 | 1.69E-02 | -1.10E+00 | 9.53E-02 | 4.12E-01 | 1.32E+00 | 7.12E-03 | 3.03E-02 | -1.05E+00 | 2.79E-03 | 1.07E-02 | 1.15E+00 | 8.01E-02 | 2.85E-01 | -1.35E+00 | 2.80E-01 | 7.90E-01 | 1.28E+00 | 5.94E-05 | 2.18E-04 | -1.49E+00 | 4.20E-02 | 1.15E-01 | 1.03E+00 |
| NFKB2 | 3.25E-03 | 1.70E-02 | 1.53E+00 | 3.70E-02 | 1.60E-01 | 1.26E+00 | 3.59E-02 | 1.53E-01 | 1.56E+00 | 6.60E-04 | 2.52E-03 | 1.12E+00 | 1.43E-03 | 5.11E-03 | 1.81E+00 | 1.62E-04 | 4.57E-04 | 1.01E+00 | 2.50E-01 | 9.17E-01 | 1.33E+00 | 4.76E-12 | 1.31E-11 | -1.51E+00 |
| TBC1D13 | 3.26E-03 | 1.70E-02 | -1.17E+00 | 1.30E-01 | 5.61E-01 | 1.34E+00 | 2.00E-02 | 8.49E-02 | 1.00E+00 | 1.14E-01 | 4.37E-01 | 1.26E+00 | 4.39E-02 | 1.57E-01 | 1.03E+00 | 1.61E-01 | 4.54E-01 | 1.12E+00 | 3.91E-02 | 1.44E-01 | 1.09E+00 | 8.14E-02 | 2.24E-01 | 1.16E+00 |
| DCTN4 | 3.26E-03 | 1.70E-02 | -1.02E+00 | 1.97E-04 | 8.50E-04 | -1.05E+00 | 4.31E-03 | 1.83E-02 | -1.07E+00 | 1.84E-03 | 7.03E-03 | 1.05E+00 | 3.47E-03 | 1.24E-02 | -1.02E+00 | 2.38E-03 | 6.70E-03 | -1.06E+00 | 2.13E-03 | 7.83E-03 | -1.14E+00 | 9.55E-02 | 2.62E-01 | 1.05E+00 |
| RAB7B | 3.26E-03 | 1.70E-02 | -1.62E+00 | 4.94E-02 | 2.14E-01 | -1.11E+00 | 3.87E-11 | 1.65E-10 | -1.78E+00 | 1.69E-01 | 6.46E-01 | 1.57E+00 | 2.03E-01 | 7.24E-01 | -1.26E+00 | 2.56E-01 | 7.22E-01 | -1.26E+00 | 2.03E-01 | 7.45E-01 | 1.22E+00 | 3.07E-01 | 8.43E-01 | 1.19E+00 |
| LINC01359 | 3.26E-03 | 1.70E-02 | -1.55E+00 | 1.07E-01 | 4.61E-01 | 1.12E+00 | 1.84E-04 | 7.83E-04 | -1.82E+00 | 5.95E-03 | 2.27E-02 | -1.23E+00 | 5.78E-03 | 2.06E-02 | -1.60E+00 | 6.15E-02 | 1.73E-01 | -1.11E+00 | 1.08E-01 | 3.95E-01 | -1.33E+00 | 2.30E-01 | 6.32E-01 | -1.18E+00 |
| SMO | 3.26E-03 | 1.70E-02 | -1.28E+00 | 1.49E-01 | 6.46E-01 | 1.25E+00 | 1.48E-02 | 6.30E-02 | -1.03E+00 | 1.19E-02 | 4.55E-02 | 1.08E+00 | 9.55E-04 | 3.41E-03 | -2.66E+00 | 4.34E-02 | 1.22E-01 | -1.12E+00 | 7.16E-04 | 2.63E-03 | -1.41E+00 | 2.46E-03 | 6.76E-03 | -1.25E+00 |
| PFN1P2 | 3.27E-03 | 1.70E-02 | -1.23E+00 | 9.92E-02 | 4.29E-01 | 1.28E+00 | 6.35E-06 | 2.70E-05 | -1.54E+00 | 2.21E-02 | 8.43E-02 | 1.16E+00 | 1.60E-04 | 5.71E-04 | -1.86E+00 | 1.15E-02 | 3.23E-02 | -1.16E+00 | 1.00E-05 | 3.67E-05 | -1.70E+00 | 7.37E-04 | 2.02E-03 | -1.38E+00 |
| UMAD1 | 3.27E-03 | 1.71E-02 | -1.11E+00 | 2.78E-02 | 1.20E-01 | 1.27E+00 | 3.02E-03 | 1.28E-02 | -1.37E+00 | 1.05E-01 | 4.01E-01 | 1.16E+00 | 1.64E-02 | 5.86E-02 | -1.22E+00 | 1.05E-01 | 2.97E-01 | 1.12E+00 | 1.45E-02 | 5.33E-02 | -1.10E+00 | 1.16E-01 | 3.19E-01 | 1.12E+00 |
| CELF2-AS1 | 3.27E-03 | 1.71E-02 | 1.98E+00 | 1.19E-01 | 5.15E-01 | 1.81E+00 | 1.76E-06 | 7.46E-06 | 2.19E+00 | 1.93E-02 | 7.39E-02 | 1.86E+00 | 1.13E-01 | 4.02E-01 | 1.25E+00 | 7.10E-02 | 2.00E-01 | 1.06E+00 | 1.53E-01 | 5.60E-01 | 1.21E+00 | 3.11E-02 | 8.54E-02 | -1.18E+00 |
| ZNF816 | 3.28E-03 | 1.71E-02 | -1.32E+00 | 2.09E-03 | 9.03E-03 | -1.30E+00 | 3.06E-02 | 1.30E-01 | -1.15E+00 | 1.53E-02 | 5.85E-02 | -1.16E+00 | 2.95E-03 | 1.05E-02 | -1.30E+00 | 2.93E-03 | 8.27E-03 | -1.43E+00 | 3.10E-04 | 1.14E-03 | -1.43E+00 | 5.64E-02 | 1.55E-01 | -1.14E+00 |
| PILRB | 3.29E-03 | 1.71E-02 | -1.08E+00 | 3.76E-02 | 1.62E-01 | 1.26E+00 | 1.08E-03 | 4.60E-03 | -1.28E+00 | 1.05E-02 | 4.00E-02 | -1.04E+00 | 6.81E-02 | 2.43E-01 | 1.13E+00 | 2.51E-01 | 7.08E-01 | 1.54E+00 | 1.29E-03 | 4.74E-03 | -1.20E+00 | 1.91E-01 | 5.25E-01 | 1.16E+00 |
| AC011447.3 | 3.29E-03 | 1.72E-02 | -1.20E+00 | 4.89E-06 | 2.11E-05 | -1.91E+00 | 2.16E-02 | 9.18E-02 | -1.03E+00 | 7.11E-04 | 2.71E-03 | -1.18E+00 | 1.54E-01 | 5.49E-01 | 1.12E+00 | 2.03E-02 | 5.72E-02 | -1.43E+00 | 1.23E-02 | 4.51E-02 | -1.35E+00 | 1.74E-01 | 4.78E-01 | -1.04E+00 |
| AC008731.1 | 3.29E-03 | 1.72E-02 | -1.22E+00 | 4.46E-03 | 1.93E-02 | -1.07E+00 | 2.87E-03 | 1.22E-02 | -1.38E+00 | 1.69E-03 | 6.45E-03 | -1.38E+00 | 1.34E-01 | 4.78E-01 | -1.02E+00 | 2.69E-02 | 7.58E-02 | -1.41E+00 | 2.08E-01 | 7.63E-01 | 1.12E+00 | 2.22E-01 | 6.10E-01 | 1.05E+00 |
| UBQLN4 | 3.29E-03 | 1.72E-02 | -1.33E+00 | 8.88E-03 | 3.84E-02 | -1.15E+00 | 5.03E-08 | 2.14E-07 | -1.79E+00 | 4.71E-05 | 1.80E-04 | -1.19E+00 | 4.01E-02 | 1.43E-01 | -1.15E+00 | 2.25E-01 | 6.34E-01 | 1.17E+00 | 2.19E-05 | 8.04E-05 | -1.37E+00 | 3.78E-05 | 1.04E-04 | -1.41E+00 |
| GTF3C2-AS1 | 3.31E-03 | 1.72E-02 | -1.17E+00 | 4.75E-03 | 2.05E-02 | -1.14E+00 | 6.10E-03 | 2.59E-02 | -1.57E+00 | 1.20E-01 | 4.58E-01 | 1.07E+00 | 4.06E-03 | 1.45E-02 | -2.01E+00 | 3.34E-02 | 9.41E-02 | -1.36E+00 | 4.02E-03 | 1.48E-02 | -1.73E+00 | 4.74E-02 | 1.30E-01 | -1.30E+00 |
| KCNG4 | 3.31E-03 | 1.73E-02 | -1.32E+00 | 3.55E-04 | 1.53E-03 | -1.60E+00 | 2.18E-01 | 9.25E-01 | 1.20E+00 | 2.31E-02 | 8.83E-02 | -1.18E+00 | 3.69E-02 | 1.31E-01 | -1.30E+00 | 5.51E-02 | 1.55E-01 | -1.39E+00 | 2.25E-01 | 8.26E-01 | 1.03E+00 | 3.34E-01 | 9.17E-01 | 1.05E+00 |
| PPP4R1L | 3.31E-03 | 1.73E-02 | -1.20E+00 | 3.54E-02 | 1.53E-01 | 1.14E+00 | 9.29E-02 | 3.95E-01 | -1.00E+00 | 1.33E-02 | 5.08E-02 | -1.32E+00 | 2.52E-01 | 8.98E-01 | 1.38E+00 | 3.22E-01 | 9.07E-01 | 1.28E+00 | 5.81E-02 | 2.13E-01 | -1.39E+00 | 1.02E-01 | 2.81E-01 | -1.30E+00 |
| ZSCAN26 | 3.32E-03 | 1.73E-02 | -1.15E+00 | 1.45E-01 | 6.26E-01 | 1.36E+00 | 7.52E-04 | 3.20E-03 | -1.34E+00 | 2.13E-02 | 8.12E-02 | 1.05E+00 | 7.51E-03 | 2.68E-02 | -1.16E+00 | 3.54E-02 | 9.97E-02 | 1.07E+00 | 1.17E-07 | 4.29E-07 | -1.75E+00 | 1.05E-03 | 2.88E-03 | -1.33E+00 |
| CEBPB | 3.33E-03 | 1.73E-02 | 1.88E+00 | 2.21E-01 | 9.53E-01 | 1.63E+00 | 3.71E-02 | 1.57E-01 | 1.50E+00 | 1.69E-01 | 6.47E-01 | 1.37E+00 | 1.31E-02 | 4.67E-02 | 1.24E+00 | 3.34E-04 | 9.41E-04 | -1.13E+00 | 1.31E-06 | 4.82E-06 | -1.23E+00 | 5.53E-03 | 1.52E-02 | -1.10E+00 |
| AGAP3 | 3.33E-03 | 1.73E-02 | 2.68E+00 | 3.52E-03 | 1.52E-02 | 3.11E+00 | 2.03E-01 | 8.63E-01 | 1.31E+00 | 2.14E-01 | 8.19E-01 | 1.52E+00 | 2.10E-01 | 7.47E-01 | 1.40E+00 | 2.02E-01 | 5.70E-01 | 1.52E+00 | 2.10E-01 | 7.72E-01 | 1.09E+00 | 7.70E-02 | 2.11E-01 | 1.60E+00 |
| PSMG4 | 3.33E-03 | 1.74E-02 | 1.02E+00 | 5.66E-03 | 2.44E-02 | 1.16E+00 | 1.03E-02 | 4.37E-02 | -1.04E+00 | 9.24E-03 | 3.53E-02 | 1.10E+00 | 4.26E-02 | 1.52E-01 | 1.04E+00 | 9.97E-02 | 2.81E-01 | 1.57E+00 | 6.53E-03 | 2.40E-02 | 1.00E+00 | 1.41E-01 | 3.86E-01 | 1.05E+00 |
| AMOTL1 | 3.34E-03 | 1.74E-02 | -1.31E+00 | 1.40E-02 | 6.04E-02 | -1.06E+00 | 4.24E-02 | 1.80E-01 | -1.05E+00 | 9.61E-02 | 3.67E-01 | 1.19E+00 | 2.30E-01 | 8.20E-01 | 1.54E+00 | 3.37E-01 | 9.49E-01 | 1.47E+00 | 1.80E-01 | 6.62E-01 | 1.18E+00 | 2.13E-01 | 5.84E-01 | 1.33E+00 |
| JPH4 | 3.34E-03 | 1.74E-02 | -1.66E+00 | 4.77E-02 | 2.06E-01 | -1.04E+00 | 7.63E-02 | 3.24E-01 | -1.23E+00 | 7.45E-02 | 2.84E-01 | -1.28E+00 | 1.37E-04 | 4.90E-04 | -1.97E+00 | 2.52E-04 | 7.10E-04 | -1.92E+00 | 2.09E-07 | 7.67E-07 | -1.68E+00 | 2.50E-06 | 6.85E-06 | -1.83E+00 |
| C2orf82 | 3.34E-03 | 1.74E-02 | 1.89E+00 | 1.02E-01 | 4.43E-01 | 1.45E+00 | 5.49E-02 | 2.33E-01 | 1.04E+00 | 6.17E-02 | 2.36E-01 | 1.13E+00 | 2.30E-01 | 8.20E-01 | -1.07E+00 | 3.38E-01 | 9.53E-01 | 1.28E+00 | 2.02E-01 | 7.40E-01 | 1.54E+00 | 2.11E-01 | 5.80E-01 | 1.67E+00 |
| SASS6 | 3.35E-03 | 1.74E-02 | -1.30E+00 | 1.65E-01 | 7.13E-01 | 1.71E+00 | 5.41E-02 | 2.30E-01 | -1.00E+00 | 1.35E-01 | 5.15E-01 | 1.26E+00 | 3.95E-05 | 1.41E-04 | -1.77E+00 | 1.87E-02 | 5.28E-02 | -1.04E+00 | 7.26E-03 | 2.67E-02 | -1.08E+00 | 3.09E-01 | 8.50E-01 | 1.24E+00 |
| TMTC2 | 3.35E-03 | 1.74E-02 | -1.23E+00 | 9.17E-02 | 3.96E-01 | 1.30E+00 | 4.51E-02 | 1.91E-01 | -1.02E+00 | 4.77E-02 | 1.82E-01 | 1.01E+00 | 2.97E-02 | 1.06E-01 | -1.49E+00 | 6.24E-02 | 1.76E-01 | -1.36E+00 | 2.26E-01 | 8.28E-01 | -1.12E+00 | 2.68E-01 | 7.37E-01 | 1.24E+00 |
| TPM3P9 | 3.35E-03 | 1.75E-02 | -1.42E+00 | 3.38E-02 | 1.46E-01 | -1.02E+00 | 2.21E-03 | 9.40E-03 | -1.28E+00 | 1.93E-02 | 7.38E-02 | 1.04E+00 | 4.99E-02 | 1.78E-01 | -1.01E+00 | 4.93E-02 | 1.39E-01 | -1.05E+00 | 1.18E-03 | 4.33E-03 | -1.34E+00 | 2.03E-01 | 5.58E-01 | 1.13E+00 |
| RALGAPA1P1 | 3.35E-03 | 1.75E-02 | -1.12E+00 | 2.39E-02 | 1.03E-01 | 1.11E+00 | 1.69E-01 | 7.17E-01 | 1.29E+00 | 1.66E-01 | 6.35E-01 | 1.52E+00 | 6.85E-02 | 2.44E-01 | -1.06E+00 | 1.86E-01 | 5.25E-01 | 1.23E+00 | 1.37E-01 | 5.01E-01 | 1.04E+00 | 2.42E-01 | 6.64E-01 | 1.14E+00 |
| ENO3 | 3.36E-03 | 1.75E-02 | -1.45E+00 | 4.13E-03 | 1.78E-02 | -1.41E+00 | 1.07E-03 | 4.57E-03 | -1.52E+00 | 9.23E-05 | 3.53E-04 | -1.56E+00 | 1.07E-05 | 3.81E-05 | -2.10E+00 | 2.04E-06 | 5.75E-06 | -2.37E+00 | 1.72E-01 | 6.30E-01 | 1.29E+00 | 1.82E-01 | 5.00E-01 | 1.12E+00 |
| CCDC136 | 3.36E-03 | 1.75E-02 | -1.10E+00 | 1.44E-02 | 6.23E-02 | 1.13E+00 | 3.56E-03 | 1.51E-02 | -1.33E+00 | 3.24E-02 | 1.24E-01 | 1.01E+00 | 2.72E-04 | 9.68E-04 | -1.44E+00 | 4.57E-04 | 1.29E-03 | -1.27E+00 | 1.50E-01 | 5.49E-01 | -1.12E+00 | 2.04E-01 | 5.61E-01 | -1.01E+00 |
| AL008729.1 | 3.36E-03 | 1.75E-02 | -1.22E+00 | 8.89E-03 | 3.84E-02 | 1.02E+00 | 1.99E-01 | 8.46E-01 | 1.07E+00 | 4.48E-02 | 1.71E-01 | -1.14E+00 | 1.09E-01 | 3.88E-01 | -1.00E+00 | 1.09E-01 | 3.08E-01 | -1.02E+00 | 1.53E-01 | 5.62E-01 | 1.01E+00 | 3.61E-01 | 9.91E-01 | 1.36E+00 |
| SLC25A25-AS1 | 3.36E-03 | 1.75E-02 | -1.15E+00 | 5.14E-04 | 2.22E-03 | -1.19E+00 | 1.53E-03 | 6.48E-03 | -1.39E+00 | 9.40E-04 | 3.59E-03 | -1.27E+00 | 1.60E-01 | 5.69E-01 | 1.10E+00 | 3.33E-01 | 9.37E-01 | 1.39E+00 | 8.97E-04 | 3.29E-03 | -1.33E+00 | 7.97E-02 | 2.19E-01 | 1.13E+00 |
| COPA | 3.36E-03 | 1.75E-02 | 1.11E+00 | 1.55E-03 | 6.68E-03 | 1.22E+00 | 3.00E-03 | 1.27E-02 | -1.11E+00 | 4.58E-02 | 1.75E-01 | 1.13E+00 | 7.07E-03 | 2.52E-02 | 1.02E+00 | 7.57E-02 | 2.13E-01 | 1.17E+00 | 2.45E-02 | 9.00E-02 | 1.07E+00 | 1.78E-01 | 4.88E-01 | 1.10E+00 |
| KLRB1 | 3.37E-03 | 1.75E-02 | -1.60E+00 | 1.32E-02 | 5.72E-02 | -1.27E+00 | 5.71E-03 | 2.43E-02 | -1.06E+00 | 5.56E-03 | 2.12E-02 | -1.44E+00 | 1.20E-02 | 4.28E-02 | -1.41E+00 | 1.18E-02 | 3.33E-02 | -1.51E+00 | 1.06E-01 | 3.91E-01 | 2.28E+00 | 2.82E-01 | 7.75E-01 | 1.05E+00 |
| AC093323.1 | 3.37E-03 | 1.75E-02 | -1.11E+00 | 2.11E-01 | 9.12E-01 | 1.56E+00 | 1.10E-05 | 4.65E-05 | -1.45E+00 | 8.07E-04 | 3.08E-03 | -1.04E+00 | 1.64E-02 | 5.84E-02 | -1.06E+00 | 2.77E-01 | 7.80E-01 | 1.59E+00 | 2.91E-03 | 1.07E-02 | -1.08E+00 | 2.88E-01 | 7.91E-01 | 1.26E+00 |
| NAIP | 3.37E-03 | 1.76E-02 | -1.20E+00 | 2.84E-02 | 1.23E-01 | 1.08E+00 | 2.29E-02 | 9.75E-02 | -1.08E+00 | 5.16E-02 | 1.97E-01 | 1.07E+00 | 2.18E-02 | 7.77E-02 | 1.14E+00 | 4.34E-02 | 1.22E-01 | 1.14E+00 | 2.93E-04 | 1.08E-03 | -1.44E+00 | 1.29E-01 | 3.55E-01 | 1.00E+00 |
| PPP1R10 | 3.37E-03 | 1.76E-02 | 1.53E+00 | 1.21E-03 | 5.22E-03 | 1.11E+00 | 1.65E-01 | 7.03E-01 | 1.11E+00 | 3.31E-03 | 1.26E-02 | 1.01E+00 | 3.23E-03 | 1.15E-02 | 1.10E+00 | 4.39E-03 | 1.24E-02 | 1.08E+00 | 1.48E-03 | 5.43E-03 | 1.04E+00 | 4.21E-03 | 1.16E-02 | -1.04E+00 |
| SLC23A2 | 3.37E-03 | 1.76E-02 | -1.27E+00 | 2.14E-02 | 9.23E-02 | 1.03E+00 | 5.15E-05 | 2.19E-04 | -1.32E+00 | 2.16E-03 | 8.27E-03 | 1.02E+00 | 2.20E-03 | 7.83E-03 | -1.63E+00 | 1.36E-02 | 3.84E-02 | -1.25E+00 | 8.04E-03 | 2.95E-02 | -1.30E+00 | 2.71E-01 | 7.44E-01 | 1.16E+00 |
| TXNRD3 | 3.37E-03 | 1.76E-02 | -1.31E+00 | 1.29E-02 | 5.58E-02 | -1.01E+00 | 1.19E-03 | 5.06E-03 | -1.38E+00 | 2.23E-02 | 8.51E-02 | 1.06E+00 | 3.79E-02 | 1.35E-01 | -1.31E+00 | 1.32E-01 | 3.73E-01 | 1.00E+00 | 4.09E-03 | 1.50E-02 | -1.37E+00 | 3.03E-03 | 8.32E-03 | -1.41E+00 |
| MRPL39 | 3.39E-03 | 1.77E-02 | -1.09E+00 | 1.40E-01 | 6.03E-01 | 1.36E+00 | 6.56E-03 | 2.79E-02 | -1.04E+00 | 5.97E-03 | 2.28E-02 | 1.14E+00 | 6.65E-03 | 2.37E-02 | -1.17E+00 | 9.70E-02 | 2.74E-01 | 1.29E+00 | 7.91E-04 | 2.90E-03 | -1.04E+00 | 2.76E-03 | 7.58E-03 | -1.02E+00 |
| ZSWIM4 | 3.39E-03 | 1.77E-02 | -1.27E+00 | 4.42E-04 | 1.91E-03 | -1.37E+00 | 2.31E-01 | 9.83E-01 | 1.19E+00 | 1.09E-01 | 4.15E-01 | 1.12E+00 | 5.65E-02 | 2.01E-01 | -1.02E+00 | 1.68E-02 | 4.73E-02 | -1.35E+00 | 1.84E-01 | 6.74E-01 | 1.13E+00 | 1.52E-01 | 4.17E-01 | -1.05E+00 |
| EEF1B2P6 | 3.39E-03 | 1.77E-02 | 1.30E+00 | 2.00E-06 | 8.64E-06 | -1.02E+00 | 5.43E-05 | 2.30E-04 | 1.52E+00 | 9.87E-02 | 3.77E-01 | 1.08E+00 | 2.00E-02 | 7.12E-02 | 1.45E+00 | 3.41E-01 | 9.62E-01 | 1.18E+00 | 2.60E-03 | 9.56E-03 | 1.47E+00 | 2.32E-01 | 6.36E-01 | 1.16E+00 |
| PSEN1 | 3.39E-03 | 1.77E-02 | -1.00E+00 | 1.48E-04 | 6.41E-04 | 1.00E+00 | 9.02E-02 | 3.83E-01 | 1.13E+00 | 6.68E-03 | 2.55E-02 | 1.02E+00 | 6.34E-03 | 2.26E-02 | -1.01E+00 | 4.95E-04 | 1.39E-03 | -1.16E+00 | 5.65E-03 | 2.08E-02 | 1.12E+00 | 1.04E-02 | 2.86E-02 | 1.03E+00 |
| BRWD3 | 3.40E-03 | 1.77E-02 | -1.14E+00 | 8.30E-02 | 3.59E-01 | 1.23E+00 | 1.38E-04 | 5.76E-04 | -1.39E+00 | 4.12E-03 | 1.57E-02 | -1.06E+00 | 5.93E-02 | 2.11E-01 | -1.01E+00 | 4.58E-02 | 1.29E-01 | -1.01E+00 | 1.85E-02 | 6.78E-02 | 1.68E+00 | 3.47E-01 | 9.52E-01 | 1.33E+00 |
| OR10A2 | 3.40E-03 | 1.77E-02 | -1.97E+00 | 1.48E-01 | 6.39E-01 | 1.06E+00 | 1.74E-01 | 7.41E-01 | 1.32E+00 | 1.05E-01 | 4.02E-01 | 1.62E+00 | 2.74E-01 | 9.76E-01 | 1.39E+00 | 3.22E-01 | 9.07E-01 | 1.23E+00 | 1.14E-02 | 4.17E-02 | -1.42E+00 | 3.47E-02 | 9.52E-02 | -1.27E+00 |
| PELP1 | 3.40E-03 | 1.77E-02 | -1.11E+00 | 1.20E-01 | 5.19E-01 | 1.42E+00 | 2.52E-02 | 1.07E-01 | 1.03E+00 | 5.36E-02 | 2.05E-01 | 1.22E+00 | 2.03E-04 | 7.23E-04 | -1.36E+00 | 1.89E-02 | 5.34E-02 | 1.06E+00 | 9.05E-09 | 3.32E-08 | -1.27E+00 | 1.35E-03 | 3.71E-03 | 1.04E+00 |
| TPRN | 3.41E-03 | 1.78E-02 | -1.78E+00 | 1.64E-02 | 7.08E-02 | -1.31E+00 | 1.59E-03 | 6.75E-03 | 2.09E+00 | 1.12E-02 | 4.27E-02 | 2.11E+00 | 4.00E-02 | 1.43E-01 | -1.12E+00 | 1.88E-02 | 5.30E-02 | -1.37E+00 | 2.59E-01 | 9.49E-01 | 1.26E+00 | 2.52E-01 | 6.93E-01 | 1.45E+00 |
| TNFAIP1 | 3.42E-03 | 1.78E-02 | -1.26E+00 | 2.06E-03 | 8.92E-03 | -1.26E+00 | 1.27E-05 | 5.42E-05 | -1.56E+00 | 3.07E-03 | 1.17E-02 | -1.03E+00 | 1.35E-05 | 4.80E-05 | -1.69E+00 | 6.00E-04 | 1.69E-03 | -1.20E+00 | 6.53E-04 | 2.40E-03 | -1.26E+00 | 5.53E-04 | 1.52E-03 | -1.29E+00 |
| SNHG5 | 3.43E-03 | 1.79E-02 | 1.14E+00 | 3.66E-10 | 1.58E-09 | -1.19E+00 | 1.40E-01 | 5.94E-01 | 1.07E+00 | 1.70E-04 | 6.50E-04 | -1.19E+00 | 2.00E-01 | 7.12E-01 | 1.07E+00 | 2.56E-04 | 7.21E-04 | -1.19E+00 | 1.45E-01 | 5.33E-01 | 1.06E+00 | 7.20E-04 | 1.98E-03 | -1.20E+00 |
| MARVELD1 | 3.43E-03 | 1.79E-02 | -1.12E+00 | 1.43E-01 | 6.17E-01 | 1.40E+00 | 3.01E-02 | 1.28E-01 | 1.00E+00 | 1.09E-01 | 4.17E-01 | 1.58E+00 | 5.43E-06 | 1.93E-05 | -1.85E+00 | 1.43E-02 | 4.02E-02 | 1.03E+00 | 9.35E-04 | 3.43E-03 | -1.11E+00 | 5.02E-02 | 1.38E-01 | 1.06E+00 |
| SLC1A4 | 3.43E-03 | 1.79E-02 | -1.06E+00 | 7.12E-02 | 3.08E-01 | 1.24E+00 | 5.72E-02 | 2.43E-01 | 1.11E+00 | 2.40E-01 | 9.17E-01 | 1.50E+00 | 1.56E-02 | 5.58E-02 | -1.24E+00 | 1.10E-01 | 3.11E-01 | 1.04E+00 | 4.30E-03 | 1.58E-02 | -1.36E+00 | 3.02E-01 | 8.29E-01 | 1.18E+00 |
| ADGRA3 | 3.44E-03 | 1.79E-02 | -1.19E+00 | 6.88E-02 | 2.97E-01 | 1.26E+00 | 2.15E-05 | 9.12E-05 | -1.35E+00 | 5.42E-03 | 2.07E-02 | 1.12E+00 | 2.85E-04 | 1.02E-03 | -1.61E+00 | 1.13E-01 | 3.19E-01 | 1.16E+00 | 6.68E-04 | 2.45E-03 | -1.28E+00 | 3.55E-01 | 9.74E-01 | 1.32E+00 |
| SNHG4 | 3.44E-03 | 1.79E-02 | -1.58E+00 | 1.54E-01 | 6.67E-01 | 1.34E+00 | 5.50E-04 | 2.34E-03 | -1.64E+00 | 6.73E-02 | 2.57E-01 | 1.16E+00 | 2.18E-01 | 7.76E-01 | -1.01E+00 | 3.63E-02 | 1.02E-01 | 2.43E+00 | 2.71E-03 | 9.93E-03 | -1.32E+00 | 2.73E-01 | 7.49E-01 | 1.24E+00 |
| LINC01399 | 3.44E-03 | 1.79E-02 | -1.26E+00 | 3.51E-04 | 1.52E-03 | -1.54E+00 | 1.12E-02 | 4.75E-02 | -1.19E+00 | 1.47E-03 | 5.61E-03 | -1.25E+00 | 3.18E-03 | 1.13E-02 | -1.40E+00 | 1.85E-03 | 5.22E-03 | -1.54E+00 | 3.62E-06 | 1.33E-05 | -1.68E+00 | 1.06E-04 | 2.92E-04 | -1.36E+00 |
| C2orf15 | 3.44E-03 | 1.79E-02 | -1.40E+00 | 4.61E-02 | 1.99E-01 | 1.02E+00 | 1.58E-02 | 6.71E-02 | -1.26E+00 | 2.96E-02 | 1.13E-01 | -1.03E+00 | 1.48E-01 | 5.27E-01 | 1.03E+00 | 2.07E-01 | 5.83E-01 | 1.15E+00 | 2.16E-01 | 7.93E-01 | 1.02E+00 | 2.79E-01 | 7.65E-01 | 1.03E+00 |
| SYNJ2 | 3.44E-03 | 1.79E-02 | 1.02E+00 | 3.19E-06 | 1.38E-05 | -1.23E+00 | 3.43E-02 | 1.46E-01 | 1.03E+00 | 2.09E-03 | 8.00E-03 | -1.00E+00 | 3.85E-06 | 1.37E-05 | 3.34E+00 | 1.80E-01 | 5.08E-01 | 1.87E+00 | 1.96E-01 | 7.20E-01 | 1.25E+00 | 5.65E-04 | 1.55E-03 | -1.34E+00 |
| BAP1 | 3.46E-03 | 1.80E-02 | -1.05E+00 | 1.01E-03 | 4.37E-03 | 1.03E+00 | 5.25E-03 | 2.23E-02 | -1.07E+00 | 2.39E-03 | 9.11E-03 | -1.03E+00 | 1.88E-01 | 6.69E-01 | 1.49E+00 | 2.11E-01 | 5.95E-01 | 1.29E+00 | 4.91E-02 | 1.80E-01 | 1.01E+00 | 1.11E-01 | 3.05E-01 | 1.09E+00 |
| RPSAP18 | 3.46E-03 | 1.80E-02 | -1.30E+00 | 2.15E-02 | 9.30E-02 | 1.04E+00 | 9.90E-02 | 4.20E-01 | -1.05E+00 | 1.20E-01 | 4.58E-01 | 1.09E+00 | 8.28E-03 | 2.95E-02 | -1.31E+00 | 3.09E-02 | 8.70E-02 | -1.12E+00 | 2.14E-02 | 7.86E-02 | -1.34E+00 | 5.56E-02 | 1.53E-01 | -1.21E+00 |
| ATAD5 | 3.46E-03 | 1.80E-02 | -1.03E+00 | 5.92E-02 | 2.56E-01 | 1.32E+00 | 2.17E-01 | 9.22E-01 | 1.12E+00 | 2.28E-01 | 8.69E-01 | 1.21E+00 | 7.29E-04 | 2.60E-03 | -1.14E+00 | 8.07E-03 | 2.27E-02 | 1.04E+00 | 9.08E-04 | 3.33E-03 | -1.19E+00 | 2.83E-01 | 7.76E-01 | 1.13E+00 |
| LINC00852 | 3.46E-03 | 1.80E-02 | -1.40E+00 | 5.26E-02 | 2.27E-01 | 1.10E+00 | 4.52E-02 | 1.92E-01 | -1.07E+00 | 4.20E-02 | 1.60E-01 | -1.05E+00 | 2.22E-01 | 7.90E-01 | -1.12E+00 | 2.24E-01 | 6.33E-01 | -1.27E+00 | 3.88E-02 | 1.42E-01 | -1.19E+00 | 1.53E-01 | 4.21E-01 | -1.05E+00 |
| GLMP | 3.46E-03 | 1.81E-02 | -1.39E+00 | 3.69E-02 | 1.59E-01 | -1.04E+00 | 2.18E-03 | 9.27E-03 | -1.71E+00 | 6.16E-03 | 2.35E-02 | -1.40E+00 | 1.29E-04 | 4.62E-04 | -1.77E+00 | 3.07E-04 | 8.64E-04 | -1.82E+00 | 8.31E-02 | 3.05E-01 | 1.02E+00 | 1.56E-01 | 4.28E-01 | 1.03E+00 |
| ELL | 3.47E-03 | 1.81E-02 | -1.27E+00 | 9.88E-05 | 4.27E-04 | -1.57E+00 | 2.86E-04 | 1.22E-03 | -1.30E+00 | 1.80E-04 | 6.86E-04 | -1.18E+00 | 7.08E-03 | 2.52E-02 | -1.26E+00 | 2.10E-02 | 5.93E-02 | -1.16E+00 | 2.07E-05 | 7.60E-05 | -1.54E+00 | 2.94E-03 | 8.07E-03 | -1.17E+00 |
| AP003086.1 | 3.48E-03 | 1.81E-02 | -1.22E+00 | 7.71E-03 | 3.33E-02 | -1.06E+00 | 2.19E-01 | 9.30E-01 | 1.17E+00 | 2.44E-01 | 9.31E-01 | 1.35E+00 | 1.65E-01 | 5.88E-01 | 1.60E+00 | 2.05E-01 | 5.79E-01 | 1.59E+00 | 3.75E-02 | 1.38E-01 | -1.27E+00 | 8.45E-02 | 2.32E-01 | -1.16E+00 |
| NUDT16P1 | 3.48E-03 | 1.82E-02 | -1.42E+00 | 1.63E-01 | 7.04E-01 | 1.34E+00 | 3.35E-02 | 1.42E-01 | -1.29E+00 | 1.59E-01 | 6.06E-01 | 1.07E+00 | 1.45E-01 | 5.15E-01 | -1.45E+00 | 2.99E-01 | 8.43E-01 | 1.42E+00 | 3.41E-03 | 1.25E-02 | -1.58E+00 | 2.91E-01 | 7.99E-01 | 1.18E+00 |
| CDK8 | 3.50E-03 | 1.82E-02 | -1.12E+00 | 2.75E-03 | 1.19E-02 | -1.01E+00 | 1.57E-02 | 6.68E-02 | -1.12E+00 | 4.83E-02 | 1.85E-01 | 1.07E+00 | 1.01E-05 | 3.60E-05 | -1.36E+00 | 1.57E-04 | 4.42E-04 | -1.13E+00 | 3.48E-07 | 1.28E-06 | -1.47E+00 | 1.20E-03 | 3.28E-03 | -1.12E+00 |
| VCPIP1 | 3.50E-03 | 1.82E-02 | 1.00E+00 | 9.48E-07 | 4.10E-06 | -1.21E+00 | 1.00E-03 | 4.25E-03 | -1.19E+00 | 5.17E-05 | 1.98E-04 | -1.24E+00 | 9.80E-04 | 3.50E-03 | -1.10E+00 | 3.96E-05 | 1.12E-04 | -1.17E+00 | 1.42E-03 | 5.22E-03 | -1.06E+00 | 5.90E-04 | 1.62E-03 | -1.15E+00 |
| GADD45B | 3.50E-03 | 1.83E-02 | 1.73E+00 | 5.58E-53 | 2.41E-52 | -5.81E+00 | 1.20E-04 | 5.10E-04 | -1.06E+00 | 3.65E-43 | 1.40E-42 | -5.57E+00 | 1.18E-01 | 4.20E-01 | 2.32E+00 | 2.19E-09 | 6.16E-09 | -3.69E+00 | 1.01E-01 | 3.72E-01 | 1.43E+00 | 5.81E-07 | 1.59E-06 | -4.04E+00 |
| TP73-AS1 | 3.51E-03 | 1.83E-02 | -1.40E+00 | 3.41E-02 | 1.47E-01 | 1.05E+00 | 2.86E-02 | 1.21E-01 | -1.15E+00 | 1.07E-01 | 4.07E-01 | 1.16E+00 | 1.79E-01 | 6.37E-01 | 1.78E+00 | 3.17E-01 | 8.94E-01 | 1.58E+00 | 1.23E-04 | 4.53E-04 | -1.29E+00 | 5.33E-03 | 1.47E-02 | -1.10E+00 |
| CBWD6 | 3.52E-03 | 1.84E-02 | 1.06E+00 | 7.82E-05 | 3.38E-04 | 1.07E+00 | 2.45E-03 | 1.04E-02 | -1.05E+00 | 1.73E-06 | 6.60E-06 | -1.09E+00 | 3.96E-06 | 1.41E-05 | -1.20E+00 | 1.29E-03 | 3.64E-03 | -1.05E+00 | 6.49E-02 | 2.38E-01 | 1.18E+00 | 2.91E-01 | 7.99E-01 | 1.33E+00 |
| CCNL2 | 3.53E-03 | 1.84E-02 | 1.04E+00 | 1.12E-01 | 4.84E-01 | 1.38E+00 | 6.43E-05 | 2.73E-04 | -1.23E+00 | 3.65E-03 | 1.40E-02 | 1.09E+00 | 4.22E-04 | 1.50E-03 | -1.08E+00 | 1.43E-02 | 4.04E-02 | 1.10E+00 | 9.20E-04 | 3.38E-03 | -1.09E+00 | 5.25E-02 | 1.44E-01 | 1.12E+00 |
| POLR2J4 | 3.53E-03 | 1.84E-02 | -1.12E+00 | 1.75E-03 | 7.58E-03 | -1.07E+00 | 8.52E-03 | 3.62E-02 | -1.14E+00 | 2.32E-02 | 8.87E-02 | 1.11E+00 | 4.67E-04 | 1.67E-03 | -1.39E+00 | 1.51E-04 | 4.25E-04 | -1.50E+00 | 6.30E-03 | 2.31E-02 | -1.04E+00 | 6.87E-04 | 1.89E-03 | -1.13E+00 |
| IL12A | 3.53E-03 | 1.84E-02 | -1.12E+00 | 8.06E-04 | 3.48E-03 | -1.10E+00 | 6.13E-05 | 2.60E-04 | -1.59E+00 | 1.20E-03 | 4.58E-03 | -1.20E+00 | 1.06E-02 | 3.79E-02 | -1.42E+00 | 2.76E-03 | 7.78E-03 | -1.80E+00 | 1.20E-02 | 4.39E-02 | -1.49E+00 | 3.82E-02 | 1.05E-01 | -1.28E+00 |
| ZNF343 | 3.53E-03 | 1.84E-02 | -1.18E+00 | 2.30E-04 | 9.95E-04 | -1.73E+00 | 1.12E-03 | 4.75E-03 | -1.46E+00 | 1.28E-03 | 4.90E-03 | -1.37E+00 | 2.71E-02 | 9.67E-02 | -1.42E+00 | 1.13E-01 | 3.19E-01 | -1.02E+00 | 2.33E-04 | 8.56E-04 | -2.06E+00 | 2.59E-01 | 7.10E-01 | 1.06E+00 |
| FKBP7 | 3.54E-03 | 1.85E-02 | 1.01E+00 | 6.09E-02 | 2.63E-01 | 1.34E+00 | 3.82E-02 | 1.62E-01 | -1.03E+00 | 1.18E-01 | 4.50E-01 | 1.31E+00 | 8.83E-04 | 3.15E-03 | -1.31E+00 | 6.96E-02 | 1.96E-01 | 1.13E+00 | 1.57E-04 | 5.77E-04 | -1.52E+00 | 4.14E-02 | 1.14E-01 | -1.07E+00 |
| BAHCC1 | 3.54E-03 | 1.85E-02 | -1.15E+00 | 6.61E-04 | 2.86E-03 | -1.22E+00 | 3.60E-12 | 1.53E-11 | -1.42E+00 | 2.24E-11 | 8.55E-11 | -1.20E+00 | 1.58E-02 | 5.62E-02 | -1.73E+00 | 3.05E-02 | 8.59E-02 | -1.58E+00 | 2.67E-01 | 9.81E-01 | 1.17E+00 | 3.63E-01 | 9.98E-01 | 1.31E+00 |
| PRUNE1 | 3.54E-03 | 1.85E-02 | -1.59E+00 | 2.11E-01 | 9.12E-01 | 1.33E+00 | 1.63E-03 | 6.91E-03 | -1.37E+00 | 1.25E-02 | 4.76E-02 | -1.07E+00 | 1.08E-05 | 3.86E-05 | -2.27E+00 | 1.53E-02 | 4.30E-02 | -1.16E+00 | 2.68E-08 | 9.84E-08 | -1.28E+00 | 1.96E-02 | 5.39E-02 | 1.14E+00 |
| DAZAP1 | 3.55E-03 | 1.85E-02 | 1.11E+00 | 5.45E-04 | 2.35E-03 | 1.17E+00 | 1.73E-04 | 7.36E-04 | -1.06E+00 | 1.23E-05 | 4.69E-05 | -1.05E+00 | 6.03E-05 | 2.15E-04 | -1.18E+00 | 3.82E-05 | 1.08E-04 | -1.06E+00 | 3.00E-05 | 1.10E-04 | -1.19E+00 | 6.50E-04 | 1.78E-03 | -1.11E+00 |
| AL022326.2 | 3.55E-03 | 1.85E-02 | -1.24E+00 | 2.99E-02 | 1.29E-01 | 1.09E+00 | 2.22E-02 | 9.43E-02 | -1.28E+00 | 9.34E-02 | 3.57E-01 | 1.05E+00 | 4.71E-02 | 1.68E-01 | -1.43E+00 | 1.13E-01 | 3.18E-01 | -1.19E+00 | 1.18E-01 | 4.31E-01 | -1.23E+00 | 2.36E-01 | 6.49E-01 | 1.49E+00 |
| NXPE3 | 3.56E-03 | 1.85E-02 | -1.07E+00 | 5.01E-03 | 2.17E-02 | 1.06E+00 | 2.71E-03 | 1.15E-02 | -1.24E+00 | 1.28E-02 | 4.88E-02 | 1.04E+00 | 1.67E-03 | 5.96E-03 | -1.02E+00 | 6.96E-04 | 1.96E-03 | -1.12E+00 | 1.71E-02 | 6.27E-02 | -1.01E+00 | 4.01E-02 | 1.10E-01 | 1.01E+00 |
| C8orf82 | 3.56E-03 | 1.85E-02 | -1.05E+00 | 2.66E-03 | 1.15E-02 | 1.04E+00 | 2.27E-02 | 9.66E-02 | -1.07E+00 | 1.16E-02 | 4.44E-02 | 1.02E+00 | 2.55E-01 | 9.09E-01 | 1.51E+00 | 3.04E-01 | 8.56E-01 | 1.45E+00 | 2.49E-01 | 9.13E-01 | 1.32E+00 | 7.80E-02 | 2.14E-01 | 1.14E+00 |
| LYRM9 | 3.57E-03 | 1.86E-02 | -1.14E+00 | 2.12E-02 | 9.14E-02 | 1.11E+00 | 1.82E-02 | 7.72E-02 | -1.20E+00 | 2.11E-02 | 8.07E-02 | -1.00E+00 | 3.91E-02 | 1.39E-01 | 1.10E+00 | 1.19E-01 | 3.36E-01 | 1.18E+00 | 1.73E-01 | 6.36E-01 | 1.20E+00 | 1.76E-02 | 4.84E-02 | -1.11E+00 |
| CUL9 | 3.57E-03 | 1.86E-02 | -1.16E+00 | 1.04E-02 | 4.51E-02 | 1.03E+00 | 1.49E-03 | 6.33E-03 | -1.11E+00 | 2.69E-02 | 1.03E-01 | 1.06E+00 | 4.68E-02 | 1.67E-01 | 1.15E+00 | 1.86E-01 | 5.24E-01 | 1.39E+00 | 1.50E-02 | 5.49E-02 | -1.30E+00 | 2.51E-01 | 6.88E-01 | 1.08E+00 |
| AP000769.1 | 3.57E-03 | 1.86E-02 | -1.20E+00 | 2.40E-02 | 1.04E-01 | 1.12E+00 | 1.91E-01 | 8.10E-01 | 1.23E+00 | 2.56E-01 | 9.78E-01 | 1.38E+00 | 2.59E-01 | 9.23E-01 | 1.10E+00 | 9.48E-02 | 2.67E-01 | -1.41E+00 | 1.51E-01 | 5.53E-01 | 1.56E+00 | 3.28E-01 | 9.00E-01 | 1.34E+00 |
| TBC1D10C | 3.58E-03 | 1.87E-02 | 1.11E+00 | 1.92E-01 | 8.29E-01 | 1.54E+00 | 4.63E-04 | 1.97E-03 | -1.09E+00 | 6.87E-04 | 2.62E-03 | 1.07E+00 | 6.92E-04 | 2.47E-03 | -1.10E+00 | 2.17E-03 | 6.11E-03 | 1.10E+00 | 6.03E-04 | 2.21E-03 | -1.16E+00 | 4.81E-04 | 1.32E-03 | -1.16E+00 |
| FBXL17 | 3.58E-03 | 1.87E-02 | -1.03E+00 | 6.40E-03 | 2.76E-02 | 1.10E+00 | 3.22E-02 | 1.37E-01 | -1.09E+00 | 1.16E-01 | 4.42E-01 | 1.18E+00 | 7.00E-03 | 2.49E-02 | -1.15E+00 | 6.62E-02 | 1.87E-01 | 1.14E+00 | 1.75E-01 | 6.42E-01 | 1.40E+00 | 2.11E-01 | 5.80E-01 | 1.45E+00 |
| PPP1R9B | 3.60E-03 | 1.88E-02 | -1.13E+00 | 8.42E-02 | 3.64E-01 | 1.31E+00 | 3.82E-02 | 1.62E-01 | -1.02E+00 | 2.06E-01 | 7.88E-01 | 1.37E+00 | 1.38E-04 | 4.93E-04 | -1.38E+00 | 8.92E-04 | 2.51E-03 | -1.22E+00 | 4.34E-04 | 1.59E-03 | -1.41E+00 | 4.87E-03 | 1.34E-02 | -1.24E+00 |
| PTH2R | 3.60E-03 | 1.88E-02 | -1.78E+00 | 4.69E-03 | 2.03E-02 | -1.75E+00 | 1.60E-01 | 6.79E-01 | 1.25E+00 | 1.87E-01 | 7.15E-01 | 1.40E+00 | 1.10E-01 | 3.92E-01 | 1.07E+00 | 6.71E-02 | 1.89E-01 | -1.08E+00 | 1.16E-04 | 4.25E-04 | -1.47E+00 | 7.65E-03 | 2.10E-02 | -1.25E+00 |
| AK9 | 3.61E-03 | 1.88E-02 | -1.02E+00 | 7.53E-02 | 3.26E-01 | 1.30E+00 | 8.78E-06 | 3.73E-05 | -1.53E+00 | 2.23E-02 | 8.54E-02 | 1.05E+00 | 1.54E-03 | 5.50E-03 | -1.27E+00 | 3.48E-01 | 9.80E-01 | 1.53E+00 | 2.70E-05 | 9.93E-05 | -1.24E+00 | 2.82E-04 | 7.73E-04 | -1.16E+00 |
| TAF9 | 3.61E-03 | 1.88E-02 | 1.15E+00 | 2.46E-07 | 1.06E-06 | 1.03E+00 | 8.74E-02 | 3.71E-01 | 1.12E+00 | 3.11E-03 | 1.19E-02 | 1.10E+00 | 4.08E-02 | 1.46E-01 | 1.11E+00 | 1.29E-01 | 3.63E-01 | 1.21E+00 | 1.56E-01 | 5.74E-01 | 1.06E+00 | 2.18E-01 | 5.99E-01 | 1.13E+00 |
| XPO5 | 3.61E-03 | 1.88E-02 | -1.25E+00 | 4.36E-02 | 1.88E-01 | 1.18E+00 | 2.75E-05 | 1.17E-04 | -1.30E+00 | 1.28E-02 | 4.88E-02 | 1.08E+00 | 2.75E-03 | 9.81E-03 | -1.39E+00 | 1.90E-01 | 5.36E-01 | 1.16E+00 | 3.11E-07 | 1.14E-06 | -1.25E+00 | 1.38E-01 | 3.80E-01 | 1.23E+00 |
| FCHO2 | 3.62E-03 | 1.88E-02 | 1.09E+00 | 3.16E-03 | 1.37E-02 | 1.13E+00 | 4.29E-05 | 1.82E-04 | -1.49E+00 | 1.01E-05 | 3.86E-05 | -1.43E+00 | 1.20E-01 | 4.27E-01 | 1.21E+00 | 4.41E-02 | 1.24E-01 | -1.07E+00 | 8.89E-04 | 3.26E-03 | -1.24E+00 | 3.20E-03 | 8.80E-03 | -1.12E+00 |
| PPP4R1 | 3.62E-03 | 1.89E-02 | 1.07E+00 | 1.14E-05 | 4.91E-05 | -1.02E+00 | 2.45E-02 | 1.04E-01 | 1.07E+00 | 7.58E-02 | 2.90E-01 | 1.23E+00 | 8.31E-02 | 2.96E-01 | 1.08E+00 | 7.38E-02 | 2.08E-01 | 1.08E+00 | 1.66E-02 | 6.10E-02 | -1.09E+00 | 1.83E-01 | 5.02E-01 | 1.05E+00 |
| KMT2E-AS1 | 3.62E-03 | 1.89E-02 | 1.80E+00 | 5.00E-03 | 2.16E-02 | 1.07E+00 | 9.83E-02 | 4.19E-01 | 1.69E+00 | 9.70E-03 | 3.71E-02 | -1.00E+00 | 2.14E-01 | 7.64E-01 | 1.42E+00 | 3.74E-02 | 1.06E-01 | 1.04E+00 | 1.82E-01 | 6.69E-01 | 1.44E+00 | 1.03E-01 | 2.84E-01 | -1.24E+00 |
| EFCAB7 | 3.62E-03 | 1.89E-02 | -1.21E+00 | 1.02E-01 | 4.40E-01 | 1.29E+00 | 1.18E-04 | 5.00E-04 | -1.38E+00 | 2.89E-02 | 1.10E-01 | 1.26E+00 | 6.51E-02 | 2.32E-01 | -1.05E+00 | 1.39E-01 | 3.93E-01 | 1.02E+00 | 4.02E-04 | 1.48E-03 | -1.52E+00 | 4.68E-02 | 1.28E-01 | -1.05E+00 |
| DCAKD | 3.62E-03 | 1.89E-02 | -1.17E+00 | 7.04E-02 | 3.04E-01 | 1.25E+00 | 8.03E-03 | 3.41E-02 | -1.15E+00 | 1.07E-02 | 4.09E-02 | 1.03E+00 | 6.44E-04 | 2.29E-03 | -1.41E+00 | 5.51E-02 | 1.55E-01 | 1.12E+00 | 2.70E-04 | 9.89E-04 | -1.41E+00 | 3.11E-01 | 8.55E-01 | 1.16E+00 |
| SMG5 | 3.62E-03 | 1.89E-02 | -1.15E+00 | 2.74E-02 | 1.19E-01 | 1.12E+00 | 4.41E-03 | 1.87E-02 | -1.17E+00 | 2.48E-03 | 9.48E-03 | -1.06E+00 | 1.87E-02 | 6.65E-02 | -1.08E+00 | 3.07E-02 | 8.66E-02 | -1.05E+00 | 7.08E-03 | 2.60E-02 | -1.13E+00 | 1.39E-01 | 3.83E-01 | 1.06E+00 |
| DHRS7B | 3.63E-03 | 1.89E-02 | 1.03E+00 | 1.10E-02 | 4.76E-02 | 1.19E+00 | 1.99E-01 | 8.48E-01 | 1.22E+00 | 3.47E-02 | 1.33E-01 | 1.04E+00 | 8.13E-03 | 2.90E-02 | -1.07E+00 | 1.57E-01 | 4.43E-01 | 1.22E+00 | 2.79E-05 | 1.02E-04 | -1.21E+00 | 2.34E-03 | 6.41E-03 | -1.08E+00 |
| WDR83 | 3.63E-03 | 1.89E-02 | -1.13E+00 | 1.79E-02 | 7.74E-02 | 1.08E+00 | 4.55E-02 | 1.93E-01 | 1.12E+00 | 2.50E-02 | 9.55E-02 | 1.23E+00 | 1.18E-06 | 4.21E-06 | -1.58E+00 | 3.90E-06 | 1.10E-05 | -1.45E+00 | 5.18E-01 | 1.16E+00 | 3.52E-01 | 9.66E-01 | 1.28E+00 | |
| SCAMP4 | 3.63E-03 | 1.89E-02 | -1.08E+00 | 5.89E-04 | 2.55E-03 | -1.12E+00 | 1.29E-03 | 5.49E-03 | -1.19E+00 | 1.85E-04 | 7.08E-04 | -1.12E+00 | 3.64E-13 | 1.30E-12 | -2.05E+00 | 1.91E-11 | 5.38E-11 | -1.87E+00 | 2.37E-03 | 8.70E-03 | -1.13E+00 | 2.62E-02 | 7.20E-02 | -1.06E+00 |
| PRRG2 | 3.63E-03 | 1.89E-02 | -1.22E+00 | 1.52E-04 | 6.57E-04 | -1.62E+00 | 8.29E-03 | 3.52E-02 | 1.77E+00 | 8.82E-02 | 3.37E-01 | -1.05E+00 | 2.00E-01 | 7.14E-01 | 1.33E+00 | 2.94E-01 | 8.28E-01 | 1.35E+00 | 2.22E-01 | 8.15E-01 | 1.26E+00 | 2.32E-01 | 6.36E-01 | -1.06E+00 |
| ABHD4 | 3.63E-03 | 1.89E-02 | -1.28E+00 | 1.28E-02 | 5.53E-02 | -1.01E+00 | 9.47E-03 | 4.02E-02 | -1.40E+00 | 1.77E-01 | 6.75E-01 | 1.25E+00 | 8.29E-03 | 2.96E-02 | -1.05E+00 | 1.01E-01 | 2.84E-01 | 1.15E+00 | 1.93E-01 | 7.09E-01 | 1.47E+00 | 1.67E-01 | 4.59E-01 | 1.17E+00 |
| H2AFX | 3.64E-03 | 1.90E-02 | -1.02E+00 | 6.20E-09 | 2.68E-08 | -1.54E+00 | 9.88E-02 | 4.20E-01 | 1.18E+00 | 8.91E-03 | 3.40E-02 | 1.23E+00 | 2.98E-08 | 1.06E-07 | -1.66E+00 | 5.52E-05 | 1.56E-04 | -1.16E+00 | 2.08E-10 | 7.64E-10 | -1.39E+00 | 2.17E-01 | 5.96E-01 | 1.17E+00 |
| GRIP1 | 3.65E-03 | 1.90E-02 | -1.34E+00 | 2.43E-02 | 1.05E-01 | -1.09E+00 | 1.88E-05 | 7.97E-05 | -1.59E+00 | 3.71E-03 | 1.42E-02 | 1.00E+00 | 2.58E-01 | 9.21E-01 | 1.20E+00 | 3.05E-01 | 8.59E-01 | 1.01E+00 | 1.85E-01 | 6.79E-01 | 1.45E+00 | 2.84E-01 | 7.80E-01 | 1.27E+00 |
| TUBG2 | 3.65E-03 | 1.90E-02 | -1.41E+00 | 1.57E-02 | 6.80E-02 | -1.12E+00 | 5.53E-02 | 2.35E-01 | -1.34E+00 | 1.43E-01 | 5.45E-01 | 1.06E+00 | 2.18E-01 | 7.76E-01 | 1.32E+00 | 1.33E-01 | 3.76E-01 | 2.13E+00 | 2.80E-04 | 1.03E-03 | 2.37E+00 | 4.16E-03 | 1.14E-02 | 2.27E+00 |
| SGO2 | 3.66E-03 | 1.91E-02 | -1.13E+00 | 2.45E-02 | 1.06E-01 | 1.10E+00 | 1.02E-05 | 4.35E-05 | -1.34E+00 | 1.28E-04 | 4.88E-04 | -1.08E+00 | 1.57E-04 | 5.61E-04 | -1.24E+00 | 2.19E-03 | 6.16E-03 | -1.06E+00 | 3.38E-04 | 1.24E-03 | -1.32E+00 | 1.16E-01 | 3.18E-01 | 1.01E+00 |
| AP000350.7 | 3.67E-03 | 1.91E-02 | -1.13E+00 | 1.02E-03 | 4.39E-03 | -1.43E+00 | 1.55E-03 | 6.59E-03 | -1.23E+00 | 2.87E-05 | 1.10E-04 | -1.76E+00 | 1.82E-01 | 6.51E-01 | -1.08E+00 | 3.23E-01 | 9.10E-01 | 1.16E+00 | 8.28E-02 | 3.04E-01 | -1.25E+00 | 1.22E-01 | 3.34E-01 | -1.25E+00 |
| URI5 | 3.67E-03 | 1.91E-02 | -1.35E+00 | 1.57E-01 | 6.80E-01 | 1.34E+00 | 6.99E-02 | 2.97E-01 | 1.01E+00 | 2.51E-01 | 9.58E-01 | 1.42E+00 | 1.13E-03 | 4.03E-03 | -1.34E+00 | 1.16E-02 | 3.28E-02 | -1.06E+00 | 7.97E-09 | 2.93E-08 | -1.77E+00 | 2.84E-04 | 7.79E-04 | -1.17E+00 |
| ACBD4 | 3.67E-03 | 1.91E-02 | -1.03E+00 | 2.56E-02 | 1.11E-01 | 1.22E+00 | 6.27E-03 | 2.66E-02 | -1.21E+00 | 3.88E-03 | 1.48E-02 | -1.04E+00 | 7.78E-03 | 2.77E-02 | -1.21E+00 | 2.47E-01 | 6.97E-01 | 1.50E+00 | 1.67E-02 | 6.11E-02 | -1.05E+00 | 5.42E-02 | 1.49E-01 | 1.04E+00 |
| MPST | 3.68E-03 | 1.92E-02 | -1.00E+00 | 1.78E-03 | 7.70E-03 | 1.08E+00 | 4.34E-02 | 1.85E-01 | 1.02E+00 | 1.55E-03 | 5.93E-03 | 1.02E+00 | 3.99E-03 | 1.42E-02 | 1.08E+00 | 3.79E-02 | 1.07E-01 | 1.67E+00 | 9.33E-03 | 3.43E-02 | -1.05E+00 | 1.13E-01 | 3.09E-01 | 1.00E+00 |
| SRD5A3-AS1 | 3.68E-03 | 1.92E-02 | -1.26E+00 | 1.09E-02 | 4.72E-02 | -1.11E+00 | 4.13E-03 | 1.76E-02 | -1.43E+00 | 7.69E-02 | 2.94E-01 | 1.07E+00 | 1.28E-01 | 4.56E-01 | -1.08E+00 | 1.57E-01 | 4.41E-01 | -1.10E+00 | 4.79E-04 | 1.76E-03 | -1.34E+00 | 1.04E-04 | 2.85E-04 | -1.57E+00 |
| PHC3 | 3.69E-03 | 1.92E-02 | 1.09E+00 | 2.85E-04 | 1.23E-03 | 1.11E+00 | 1.83E-02 | 7.77E-02 | 1.03E+00 | 1.77E-03 | 6.77E-03 | 1.03E+00 | 6.98E-02 | 2.49E-01 | 1.17E+00 | 1.83E-02 | 5.15E-02 | 1.10E+00 | 2.14E-02 | 7.86E-02 | 1.03E+00 | 9.16E-03 | 2.52E-02 | -1.06E+00 |
| ANP32B | 3.69E-03 | 1.92E-02 | -1.07E+00 | 1.93E-01 | 8.33E-01 | 1.04E+00 | 1.01E-03 | 4.30E-03 | -1.12E+00 | 4.75E-02 | 1.82E-01 | -1.05E+00 | 2.28E-03 | 8.12E-03 | -1.10E+00 | 3.17E-01 | 8.93E-01 | 1.05E+00 | 2.54E-02 | 9.32E-02 | -1.03E+00 | 3.45E-01 | 9.48E-01 | 1.04E+00 |
| USE1 | 3.69E-03 | 1.92E-02 | 1.07E+00 | 3.33E-06 | 1.44E-05 | 1.03E+00 | 8.15E-03 | 3.46E-02 | -1.01E+00 | 1.37E-03 | 5.24E-03 | -1.00E+00 | 1.35E-05 | 4.81E-05 | -1.18E+00 | 2.09E-03 | 5.88E-03 | 1.02E+00 | 2.31E-03 | 8.47E-03 | -1.01E+00 | 2.42E-04 | 6.64E-04 | -1.09E+00 |
| CAMTA1 | 3.69E-03 | 1.92E-02 | 1.12E+00 | 1.90E-06 | 8.22E-06 | 1.06E+00 | 1.98E-01 | 8.43E-01 | 1.21E+00 | 4.22E-02 | 1.61E-01 | 1.16E+00 | 5.97E-05 | 2.13E-04 | -1.14E+00 | 1.15E-03 | 3.24E-03 | -1.05E+00 | 1.51E-01 | 5.53E-01 | 1.17E+00 | 1.37E-01 | 3.75E-01 | 1.11E+00 |
| KCNMB4 | 3.69E-03 | 1.93E-02 | -1.02E+00 | 1.16E-01 | 5.01E-01 | 1.40E+00 | 9.84E-04 | 4.18E-03 | -1.25E+00 | 1.61E-02 | 6.16E-02 | 1.04E+00 | 2.09E-05 | 7.44E-05 | -1.48E+00 | 1.94E-04 | 5.46E-04 | -1.35E+00 | 2.67E-04 | 9.79E-04 | -1.47E+00 | 4.30E-04 | 1.18E-03 | -1.54E+00 |
| MARCH3 | 3.70E-03 | 1.93E-02 | -1.18E+00 | 5.56E-03 | 2.40E-02 | -1.08E+00 | 1.90E-01 | 8.07E-01 | 1.11E+00 | 2.40E-01 | 9.16E-01 | 1.28E+00 | 1.48E-02 | 5.27E-02 | -1.09E+00 | 1.25E-02 | 3.51E-02 | -1.13E+00 | 5.75E-03 | 2.11E-02 | -1.04E+00 | 7.66E-02 | 2.10E-01 | 1.08E+00 |
| MZT2B | 3.70E-03 | 1.93E-02 | 1.20E+00 | 4.45E-09 | 1.92E-08 | -1.13E+00 | 1.85E-04 | 7.88E-04 | 1.29E+00 | 1.76E-03 | 6.73E-03 | -1.11E+00 | 2.37E-01 | 8.44E-01 | 1.14E+00 | 2.67E-04 | 7.52E-04 | -1.10E+00 | 1.80E-02 | 6.61E-02 | 1.26E+00 | 1.08E-01 | 2.96E-01 | -1.04E+00 |
| KBTBD6 | 3.70E-03 | 1.93E-02 | -1.38E+00 | 1.93E-02 | 8.33E-02 | -1.04E+00 | 1.31E-10 | 5.59E-10 | -2.34E+00 | 7.52E-08 | 2.87E-07 | -1.67E+00 | 1.76E-01 | 6.27E-01 | 1.12E+00 | 2.21E-01 | 6.22E-01 | 1.09E+00 | 4.10E-04 | 1.50E-03 | -1.56E+00 | 6.03E-02 | 1.66E-01 | -1.00E+00 |
| ARL15 | 3.71E-03 | 1.93E-02 | -1.32E+00 | 2.63E-02 | 1.14E-01 | 1.04E+00 | 2.11E-05 | 8.98E-05 | -1.93E+00 | 7.14E-04 | 2.73E-03 | -1.28E+00 | 1.94E-14 | 6.92E-14 | -2.05E+00 | 8.64E-06 | 2.44E-05 | -1.19E+00 | 1.69E-04 | 6.21E-04 | -1.30E+00 | 8.52E-02 | 2.34E-01 | 1.09E+00 |
| ACSM3 | 3.73E-03 | 1.94E-02 | 1.03E+00 | 7.70E-03 | 3.33E-02 | 1.20E+00 | 4.11E-03 | 1.75E-02 | -1.28E+00 | 2.08E-01 | 7.96E-01 | 1.25E+00 | 1.32E-01 | 4.71E-01 | 1.31E+00 | 1.60E-02 | 4.51E-02 | 1.91E+00 | 1.53E-02 | 5.61E-02 | -1.14E+00 | 1.71E-01 | 4.69E-01 | 1.02E+00 |
| ABT1 | 3.73E-03 | 1.94E-02 | 1.05E+00 | 4.83E-02 | 2.09E-01 | 1.34E+00 | 1.32E-01 | 5.62E-01 | 1.18E+00 | 8.36E-02 | 3.19E-01 | 1.24E+00 | 2.05E-02 | 7.31E-02 | 1.08E+00 | 3.09E-01 | 8.70E-01 | 1.42E+00 | 1.94E-01 | 7.12E-01 | 1.19E+00 | 1.19E-01 | 3.27E-01 | 1.41E+00 |
| KDM4A-AS1 | 3.74E-03 | 1.95E-02 | -1.45E+00 | 5.00E-04 | 2.16E-03 | -1.92E+00 | 1.58E-01 | 6.70E-01 | 1.04E+00 | 2.48E-01 | 9.48E-01 | 1.29E+00 | 1.12E-01 | 3.99E-01 | 1.07E+00 | 2.10E-01 | 5.92E-01 | 1.08E+00 | 1.21E-01 | 4.45E-01 | -1.11E+00 | 1.64E-01 | 4.51E-01 | -1.13E+00 |
| UBXN8 | 3.74E-03 | 1.95E-02 | -1.33E+00 | 4.48E-02 | 1.94E-01 | 1.11E+00 | 5.34E-03 | 2.27E-02 | -1.19E+00 | 7.41E-03 | 2.83E-02 | -1.04E+00 | 5.85E-03 | 2.09E-02 | -1.31E+00 | 2.29E-01 | 6.45E-01 | 1.23E+00 | 1.20E-05 | 4.40E-05 | -1.41E+00 | 5.20E-05 | 1.43E-04 | -1.25E+00 |
| CDK20 | 3.74E-03 | 1.95E-02 | -1.51E+00 | 2.62E-02 | 1.13E-01 | -1.13E+00 | 6.18E-04 | 2.62E-03 | -1.66E+00 | 5.79E-02 | 2.21E-01 | 1.02E+00 | 1.11E-02 | 3.97E-02 | -1.83E+00 | 9.51E-02 | 2.68E-01 | -1.10E+00 | 2.37E-01 | 8.71E-01 | 1.06E+00 | 8.20E-02 | 2.25E-01 | 1.76E+00 |
| MED19 | 3.75E-03 | 1.95E-02 | -1.05E+00 | 7.14E-08 | 3.08E-07 | -1.40E+00 | 1.73E-02 | 7.35E-02 | -1.03E+00 | 1.17E-04 | 4.46E-04 | -1.20E+00 | 8.45E-09 | 3.01E-08 | -1.57E+00 | 4.46E-09 | 1.26E-08 | -1.52E+00 | 7.07E-05 | 2.60E-04 | -1.17E+00 | 8.49E-04 | 2.33E-03 | -1.10E+00 |
| GFOD1 | 3.75E-03 | 1.95E-02 | 1.82E+00 | 9.25E-03 | 4.00E-02 | 1.99E+00 | 1.06E-01 | 4.50E-01 | 1.32E+00 | 9.14E-02 | 3.49E-01 | 1.22E+00 | 1.34E-01 | 4.78E-01 | 1.27E+00 | 3.05E-02 | 8.59E-02 | 1.07E+00 | 4.29E-02 | 1.58E-01 | 1.07E+00 | 2.44E-01 | 6.71E-01 | 1.23E+00 |
| HELLPAR | 3.75E-03 | 1.95E-02 | -1.26E+00 | 8.98E-03 | 3.88E-02 | 1.09E+00 | 2.05E-01 | 8.73E-01 | 1.22E+00 | 1.62E-01 | 6.18E-01 | 1.59E+00 | 9.05E-03 | 3.23E-02 | -1.14E+00 | 1.47E-02 | 4.13E-02 | -1.20E+00 | 3.51E-03 | 1.29E-02 | -1.16E+00 | 3.33E-01 | 9.14E-01 | 1.50E+00 |
| WDR18 | 3.75E-03 | 1.96E-02 | 1.06E+00 | 2.30E-03 | 9.92E-03 | 1.16E+00 | 1.67E-01 | 7.11E-01 | 1.21E+00 | 9.85E-02 | 3.76E-01 | 1.25E+00 | 2.14E-03 | 7.64E-03 | -1.07E+00 | 9.64E-02 | 2.72E-01 | 1.18E+00 | 6.15E-02 | 2.26E-01 | 1.15E+00 | 2.74E-01 | 7.53E-01 | 1.23E+00 |
| C7orf31 | 3.76E-03 | 1.96E-02 | -1.04E+00 | 9.29E-03 | 4.01E-02 | 1.11E+00 | 5.69E-05 | 2.42E-04 | -1.36E+00 | 5.85E-04 | 2.23E-03 | -1.11E+00 | 2.68E-02 | 9.56E-02 | -1.05E+00 | 4.09E-02 | 1.15E-01 | -1.02E+00 | 8.35E-02 | 3.07E-01 | -1.00E+00 | 2.41E-01 | 6.61E-01 | 1.15E+00 |
| TP53I11 | 3.76E-03 | 1.96E-02 | -1.54E+00 | 6.01E-03 | 2.60E-02 | -1.35E+00 | 2.10E-03 | 8.90E-03 | -1.26E+00 | 3.30E-04 | 1.26E-03 | -1.18E+00 | 1.21E-01 | 4.32E-01 | -1.23E+00 | 1.94E-01 | 5.47E-01 | -1.03E+00 | 1.71E-01 | 6.26E-01 | -1.15E+00 | 3.46E-01 | 9.51E-01 | 1.10E+00 |
| PDIA5 | 3.77E-03 | 1.96E-02 | -1.05E+00 | 1.25E-02 | 5.38E-02 | 1.07E+00 | 1.90E-01 | 8.07E-01 | 1.27E+00 | 1.35E-01 | 5.15E-01 | 1.62E+00 | 1.49E-03 | 5.29E-03 | -1.21E+00 | 1.71E-02 | 4.83E-02 | -1.01E+00 | 9.72E-02 | 3.57E-01 | 1.26E+00 | 3.54E-01 | 9.72E-01 | 1.12E+00 |
| LINC00294 | 3.77E-03 | 1.96E-02 | -1.15E+00 | 2.37E-02 | 1.02E-01 | 1.08E+00 | 4.61E-05 | 1.96E-04 | -1.57E+00 | 3.05E-02 | 1.17E-01 | 1.13E+00 | 1.01E-02 | 3.59E-02 | -1.14E+00 | 7.29E-03 | 2.05E-02 | -1.18E+00 | 1.08E-06 | 3.97E-06 | -2.15E+00 | 1.11E-04 | 3.05E-04 | -1.63E+00 |
| AC007114.2 | 3.77E-03 | 1.96E-02 | -1.46E+00 | 2.67E-02 | 1.16E-01 | -1.07E+00 | 7.42E-02 | 3.15E-01 | -1.08E+00 | 2.49E-01 | 9.50E-01 | 1.36E+00 | 1.02E-01 | 3.64E-01 | -1.03E+00 | 9.39E-02 | 2.65E-01 | -1.15E+00 | 1.81E-01 | 6.66E-01 | 1.06E+00 | 2.53E-01 | 6.94E-01 | 1.04E+00 |
| SRP19 | 3.78E-03 | 1.97E-02 | 1.11E+00 | 3.67E-05 | 1.59E-04 | 1.04E+00 | 1.59E-02 | 6.76E-02 | 1.01E+00 | 1.50E-04 | 5.74E-04 | 1.01E+00 | 2.35E-06 | 8.40E-06 | -1.01E+00 | 1.77E-06 | 4.98E-06 | -1.02E+00 | 2.08E-03 | 7.62E-03 | -1.00E+00 | 6.83E-04 | 1.88E-03 | -1.07E+00 |
| PRIMPOL | 3.78E-03 | 1.97E-02 | -1.10E+00 | 6.36E-03 | 2.75E-02 | 1.05E+00 | 2.26E-05 | 9.61E-05 | -1.46E+00 | 1.60E-03 | 6.11E-03 | -1.06E+00 | 9.75E-06 | 3.48E-05 | -1.44E+00 | 4.60E-04 | 1.30E-03 | -1.19E+00 | 5.23E-03 | 1.92E-02 | -1.01E+00 | 7.11E-04 | 1.95E-03 | -1.09E+00 |
| ZSWIM9 | 3.79E-03 | 1.97E-02 | -1.11E+00 | 4.54E-02 | 1.96E-01 | 1.22E+00 | 1.62E-01 | 6.89E-01 | 1.04E+00 | 2.33E-01 | 8.91E-01 | 1.29E+00 | 2.00E-03 | 7.14E-03 | 1.19E+00 | 1.91E-01 | 5.38E-01 | 1.59E+00 | 2.83E-02 | 1.04E-01 | -1.08E+00 | 7.52E-02 | 2.07E-01 | 1.06E+00 |
| DIRC2 | 3.79E-03 | 1.98E-02 | -1.08E+00 | 4.88E-04 | 2.11E-03 | -1.19E+00 | 6.93E-02 | 2.94E-01 | 1.02E+00 | 2.35E-02 | 8.97E-02 | 1.03E+00 | 1.92E-01 | 6.86E-01 | 1.21E+00 | 1.01E-01 | 2.85E-01 | 1.03E+00 | 3.88E-03 | 1.42E-02 | -1.26E+00 | 3.40E-03 | 9.34E-03 | -1.38E+00 |
| MTM1 | 3.79E-03 | 1.98E-02 | -1.09E+00 | 3.40E-03 | 1.47E-02 | -1.01E+00 | 5.62E-04 | 2.39E-03 | -1.30E+00 | 3.08E-03 | 1.18E-02 | -1.03E+00 | 8.75E-04 | 3.12E-03 | -1.15E+00 | 2.32E-03 | 6.53E-03 | -1.11E+00 | 6.65E-02 | 2.44E-01 | 1.07E+00 | 2.09E-01 | 5.75E-01 | 1.09E+00 |
| SLC19A1 | 3.79E-03 | 1.98E-02 | -1.22E+00 | 1.48E-02 | 6.41E-02 | 1.05E+00 | 2.93E-05 | 1.24E-04 | -1.22E+00 | 2.46E-04 | 9.40E-04 | 1.00E+00 | 6.73E-03 | 2.40E-02 | -1.38E+00 | 1.92E-01 | 5.40E-01 | 1.25E+00 | 9.74E-03 | 3.57E-02 | -1.07E+00 | 9.19E-02 | 2.52E-01 | 1.56E+00 |
| AHCYL2 | 3.79E-03 | 1.98E-02 | -1.35E+00 | 1.80E-02 | 7.79E-02 | -1.03E+00 | 1.23E-02 | 5.22E-02 | -1.15E+00 | 1.92E-02 | 7.34E-02 | 1.03E+00 | 2.19E-03 | 7.80E-03 | -1.35E+00 | 5.20E-02 | 1.46E-01 | 1.05E+00 | 1.27E-03 | 4.66E-03 | -1.47E+00 | 6.27E-02 | 1.72E-01 | -1.00E+00 |
| STAC | 3.80E-03 | 1.98E-02 | -1.18E+00 | 9.66E-02 | 4.17E-01 | 1.34E+00 | 1.33E-03 | 5.67E-03 | -1.41E+00 | 4.77E-03 | 1.82E-02 | -1.10E+00 | 6.41E-03 | 2.29E-02 | -1.51E+00 | 2.35E-02 | 6.62E-02 | -1.20E+00 | 2.81E-05 | 1.03E-04 | -1.24E+00 | 2.22E-04 | 6.11E-04 | -1.13E+00 |
| TMEM187 | 3.80E-03 | 1.98E-02 | -1.34E+00 | 1.38E-01 | 5.98E-01 | 1.32E+00 | 3.22E-03 | 1.37E-02 | -1.54E+00 | 2.67E-03 | 1.02E-02 | -1.33E+00 | 1.50E-03 | 5.36E-03 | -1.82E+00 | 6.50E-03 | 1.83E-02 | -1.47E+00 | 1.80E-02 | 6.59E-02 | -1.12E+00 | 2.82E-01 | 7.74E-01 | 1.33E+00 |
| AC024940.2 | 3.80E-03 | 1.98E-02 | 1.14E+00 | 1.31E-03 | 5.68E-03 | 1.23E+00 | 1.14E-01 | 4.86E-01 | 1.13E+00 | 1.46E-01 | 5.59E-01 | 1.28E+00 | 1.37E-02 | 4.87E-02 | 1.17E+00 | 1.45E-01 | 4.09E-01 | 1.25E+00 | 1.97E-02 | 7.22E-02 | 1.06E+00 | 3.62E-01 | 9.95E-01 | 1.19E+00 |
| SMG1P2 | 3.80E-03 | 1.98E-02 | -1.03E+00 | 6.51E-02 | 2.81E-01 | 1.35E+00 | 1.27E-01 | 5.40E-01 | 1.14E+00 | 2.19E-01 | 8.37E-01 | 1.52E+00 | 8.50E-02 | 3.03E-01 | 1.26E+00 | 3.13E-01 | 8.82E-01 | 1.72E+00 | 2.29E-03 | 8.42E-03 | -1.01E+00 | 2.85E-01 | 7.82E-01 | 1.38E+00 |
| TLDC1 | 3.81E-03 | 1.98E-02 | -1.21E+00 | 1.54E-01 | 6.68E-01 | 1.33E+00 | 1.48E-03 | 6.28E-03 | -1.43E+00 | 2.83E-02 | 1.08E-01 | -1.04E+00 | 4.01E-04 | 1.43E-03 | -1.53E+00 | 1.43E-02 | 4.03E-02 | -1.07E+00 | 1.26E-04 | 4.63E-04 | -1.59E+00 | 1.80E-04 | 4.95E-04 | -1.67E+00 |
| AC105942.1 | 3.81E-03 | 1.99E-02 | -1.01E+00 | 1.11E-02 | 4.79E-02 | 1.12E+00 | 4.60E-02 | 1.96E-01 | -1.07E+00 | 1.86E-01 | 7.09E-01 | 1.23E+00 | 1.76E-01 | 6.28E-01 | 1.36E+00 | 2.87E-01 | 8.09E-01 | 1.27E+00 | 1.77E-01 | 6.49E-01 | -1.02E+00 | 3.57E-01 | 9.80E-01 | 1.24E+00 |
| AL391244.1 | 3.81E-03 | 1.99E-02 | -1.20E+00 | 1.57E-01 | 6.78E-01 | 1.33E+00 | 1.11E-04 | 4.70E-04 | -1.42E+00 | 1.98E-01 | 7.57E-01 | 1.29E+00 | 1.28E-03 | 4.57E-03 | -1.34E+00 | 2.50E-01 | 7.05E-01 | 1.50E+00 | 8.87E-03 | 3.26E-02 | 1.01E+00 | 6.55E-02 | 1.80E-01 | 1.55E+00 |
| SRMP2 | 3.82E-03 | 1.99E-02 | -1.52E+00 | 3.06E-02 | 1.32E-01 | -1.11E+00 | 9.59E-02 | 4.07E-01 | -1.02E+00 | 2.14E-01 | 8.16E-01 | 1.20E+00 | 1.76E-01 | 6.27E-01 | 1.01E+00 | 1.52E-01 | 4.29E-01 | -1.11E+00 | 3.76E-03 | 1.38E-02 | -1.63E+00 | 2.39E-01 | 6.57E-01 | 1.09E+00 |
| FP236383.1 | 3.82E-03 | 1.99E-02 | -1.02E+00 | 1.31E-01 | 5.67E-01 | 1.42E+00 | 1.82E-04 | 7.73E-04 | -1.38E+00 | 5.28E-03 | 2.02E-02 | -1.00E+00 | 2.20E-01 | 7.85E-01 | 1.43E+00 | 2.64E-01 | 7.43E-01 | 1.67E+00 | 2.77E-02 | 1.02E-01 | -1.10E+00 | 1.40E-02 | 3.84E-02 | -1.21E+00 |
| HIVEP1 | 3.82E-03 | 1.99E-02 | 1.01E+00 | 2.56E-06 | 1.11E-05 | -1.21E+00 | 2.27E-01 | 9.63E-01 | 1.18E+00 | 1.30E-03 | 4.97E-03 | -1.11E+00 | 8.83E-02 | 3.15E-01 | 1.03E+00 | 2.54E-02 | 7.16E-02 | -1.16E+00 | 2.03E-02 | 7.44E-02 | -1.25E+00 | 6.58E-02 | 1.81E-01 | -1.18E+00 |
| MED27 | 3.83E-03 | 1.99E-02 | -1.10E+00 | 1.72E-02 | 7.44E-02 | 1.07E+00 | 2.89E-03 | 1.23E-02 | -1.14E+00 | 4.15E-05 | 1.58E-04 | -1.18E+00 | 1.65E-06 | 5.89E-06 | -1.70E+00 | 1.57E-04 | 4.43E-04 | -1.23E+00 | 2.80E-06 | 1.03E-05 | -1.28E+00 | 1.94E-04 | 5.34E-04 | -1.10E+00 |
| RABGGTA | 3.83E-03 | 2.00E-02 | -1.17E+00 | 8.51E-03 | 3.68E-02 | 1.04E+00 | 1.20E-02 | 5.11E-02 | -1.13E+00 | 1.14E-03 | 4.36E-03 | -1.14E+00 | 2.10E-03 | 7.50E-03 | -1.08E+00 | 1.62E-02 | 4.56E-02 | 1.07E+00 | 2.79E-05 | 1.02E-04 | -1.37E+00 | 1.25E-04 | 3.43E-04 | -1.25E+00 |
| MIR181A2HG | 3.83E-03 | 2.00E-02 | -1.24E+00 | 2.55E-02 | 1.10E-01 | 1.01E+00 | 1.72E-03 | 7.30E-03 | -1.40E+00 | 3.30E-04 | 1.26E-03 | -1.51E+00 | 9.09E-03 | 3.24E-02 | -1.51E+00 | 1.25E-01 | 3.51E-01 | -1.04E+00 | 8.05E-05 | 2.96E-04 | -2.09E+00 | 2.65E-03 | 7.28E-03 | -1.60E+00 |
| PEBP1 | 3.84E-03 | 2.00E-02 | -1.03E+00 | 2.65E-02 | 1.14E-01 | 1.11E+00 | 1.52E-01 | 6.47E-01 | 1.04E+00 | 2.36E-01 | 9.00E-01 | 1.06E+00 | 4.37E-04 | 1.56E-03 | -1.10E+00 | 2.62E-01 | 7.38E-01 | 1.11E+00 | 2.07E-04 | 7.59E-04 | -1.13E+00 | 3.13E-01 | 8.61E-01 | 1.10E+00 |
| WDR55 | 3.84E-03 | 2.00E-02 | -1.07E+00 | 3.31E-03 | 1.43E-02 | -1.04E+00 | 9.39E-05 | 3.99E-04 | -1.28E+00 | 3.03E-04 | 1.16E-03 | -1.07E+00 | 3.58E-04 | 1.28E-03 | -1.16E+00 | 9.62E-04 | 2.71E-03 | -1.10E+00 | 6.48E-04 | 2.38E-03 | -1.16E+00 | 2.71E-03 | 7.44E-03 | -1.11E+00 |
| PIK3R4 | 3.84E-03 | 2.00E-02 | -1.26E+00 | 4.75E-03 | 2.05E-02 | -1.09E+00 | 3.71E-02 | 1.58E-01 | -1.07E+00 | 1.47E-01 | 5.60E-01 | 1.22E+00 | 2.59E-01 | 9.25E-01 | 1.30E+00 | 2.21E-01 | 6.23E-01 | 1.11E+00 | 2.54E-03 | 9.32E-03 | -1.32E+00 | 1.82E-01 | 5.01E-01 | 1.09E+00 |
| AC007485.2 | 3.85E-03 | 2.01E-02 | -1.55E+00 | 2.25E-01 | 9.72E-01 | 1.29E+00 | 1.12E-05 | 4.78E-05 | -2.59E+00 | 3.07E-03 | 1.17E-02 | -1.38E+00 | 1.03E-02 | 3.67E-02 | -1.75E+00 | 2.07E-01 | 5.84E-01 | 1.05E+00 | 8.75E-05 | 3.21E-04 | -1.90E+00 | 5.84E-03 | 1.60E-02 | -1.36E+00 |
| CTTNBP2NL | 3.85E-03 | 2.01E-02 | 1.85E+00 | 4.10E-02 | 1.77E-01 | 1.81E+00 | 8.27E-02 | 3.51E-01 | 1.37E+00 | 1.15E-01 | 4.40E-01 | 1.65E+00 | 1.92E-03 | 6.84E-03 | 1.38E+00 | 1.08E-01 | 3.05E-01 | 1.13E+00 | 1.07E-01 | 3.94E-01 | -1.08E+00 | 4.77E-02 | 1.31E-01 | -1.42E+00 |
| RERE | 3.86E-03 | 2.01E-02 | 1.09E+00 | 2.63E-04 | 1.14E-03 | 1.11E+00 | 9.59E-02 | 4.08E-01 | 1.20E+00 | 1.19E-02 | 4.53E-02 | 1.21E+00 | 6.81E-02 | 2.43E-01 | 1.24E+00 | 9.12E-03 | 2.57E-02 | 1.02E+00 | 4.80E-05 | 1.76E-04 | -1.16E+00 | 1.92E-03 | 5.29E-03 | -1.06E+00 |
| APLF | 3.88E-03 | 2.02E-02 | -1.59E+00 | 1.52E-02 | 6.57E-02 | -1.23E+00 | 1.32E-02 | 5.59E-02 | -1.35E+00 | 1.92E-02 | 7.34E-02 | -1.14E+00 | 4.22E-04 | 1.51E-03 | -1.55E+00 | 1.36E-02 | 3.83E-02 | -1.14E+00 | 3.55E-02 | 1.30E-01 | 1.11E+00 | 1.22E-01 | 3.36E-01 | 1.06E+00 |
| AL845472.1 | 3.88E-03 | 2.02E-02 | -1.22E+00 | 5.79E-02 | 2.50E-01 | 1.06E+00 | 9.63E-04 | 4.09E-03 | -1.56E+00 | 2.13E-02 | 8.14E-02 | -1.04E+00 | 2.13E-03 | 7.58E-03 | -1.24E+00 | 2.86E-01 | 8.07E-01 | -1.22E+00 | 5.23E-04 | 1.92E-03 | -1.88E+00 | 3.33E-02 | 9.16E-02 | -1.21E+00 |
| HSPE1P5 | 3.88E-03 | 2.02E-02 | -1.45E+00 | 1.13E-01 | 4.90E-01 | 1.24E+00 | 2.11E-01 | 8.97E-01 | 1.18E+00 | 6.63E-02 | 2.53E-01 | 1.74E+00 | 1.82E-01 | 6.50E-01 | 1.06E+00 | 1.09E-01 | 3.08E-01 | 1.83E+00 | 1.26E-01 | 4.63E-01 | 1.08E+00 | 6.41E-03 | 1.76E-02 | 1.73E+00 |
| ARID1A | 3.88E-03 | 2.02E-02 | 1.11E+00 | 3.07E-05 | 1.32E-04 | 1.09E+00 | 8.86E-02 | 3.76E-01 | 1.08E+00 | 6.07E-04 | 2.32E-03 | -1.01E+00 | 5.00E-04 | 1.78E-03 | 1.02E+00 | 5.84E-03 | 1.65E-02 | 1.08E+00 | 8.64E-06 | 3.17E-05 | -1.19E+00 | 1.54E-03 | 4.23E-03 | -1.09E+00 |
| IGHMBP2 | 3.90E-03 | 2.03E-02 | -1.37E+00 | 4.98E-02 | 2.15E-01 | 1.09E+00 | 1.01E-03 | 4.30E-03 | -1.36E+00 | 3.38E-03 | 1.29E-02 | -1.18E+00 | 2.42E-01 | 8.61E-01 | 1.25E+00 | 3.53E-01 | 9.96E-01 | 1.29E+00 | 7.32E-07 | 2.69E-06 | -1.50E+00 | 8.85E-02 | 2.43E-01 | 1.11E+00 |
| RPS7P11 | 3.90E-03 | 2.03E-02 | 1.08E+00 | 1.67E-09 | 7.22E-09 | -1.21E+00 | 2.09E-01 | 8.87E-01 | 1.19E+00 | 5.27E-06 | 2.01E-05 | -1.15E+00 | 1.31E-02 | 4.68E-02 | 1.06E+00 | 3.13E-05 | 8.84E-05 | -1.21E+00 | 1.80E-01 | 6.60E-01 | 1.22E+00 | 3.16E-06 | 8.69E-06 | -1.29E+00 |
| SIGLEC10 | 3.90E-03 | 2.03E-02 | -1.33E+00 | 8.92E-04 | 3.85E-03 | -1.37E+00 | 8.03E-03 | 3.41E-02 | -1.36E+00 | 8.11E-03 | 3.10E-02 | -1.24E+00 | 9.57E-33 | 3.41E-32 | -1.90E+00 | 8.98E-27 | 2.53E-26 | -1.67E+00 | 1.52E-02 | 5.59E-02 | -1.36E+00 | 1.93E-01 | 5.31E-01 | 1.02E+00 |
| AC010080.1 | 3.90E-03 | 2.03E-02 | -1.10E+00 | 8.66E-03 | 3.83E-02 | 2.21E+00 | 9.05E-03 | 3.84E-02 | -1.35E+00 | 9.62E-02 | 3.68E-01 | 1.56E+00 | 1.48E-04 | 5.27E-04 | -1.70E+00 | 2.56E-02 | 7.21E-02 | 1.09E+00 | 2.52E-03 | 9.24E-03 | -1.22E+00 | 3.44E-02 | 9.44E-02 | -1.22E+00 |
| SLC25A30 | 3.90E-03 | 2.03E-02 | -1.23E+00 | 1.09E-01 | 4.72E-01 | 1.56E+00 | 1.68E-02 | 7.13E-02 | -1.19E+00 | 7.42E-02 | 2.83E-01 | 1.23E+00 | 2.94E-02 | 1.05E-01 | -1.11E+00 | 8.62E-02 | 2.43E-01 | 1.16E+00 | 1.06E-01 | 3.87E-01 | -1.04E+00 | 2.82E-01 | 7.75E-01 | 1.06E+00 |
| ATP2A2 | 3.90E-03 | 2.04E-02 | -1.00E+00 | 1.61E-03 | 6.94E-03 | 1.07E+00 | 1.21E-02 | 5.16E-02 | 1.01E+00 | 2.78E-04 | 1.06E-03 | 1.03E+00 | 2.50E-01 | 8.91E-01 | 1.38E+00 | 3.06E-02 | 8.62E-02 | 1.09E+00 | 6.12E-10 | 2.24E-09 | -1.42E+00 | 4.27E-06 | 1.17E-05 | -1.25E+00 |
| SMN1 | 3.91E-03 | 2.04E-02 | 1.05E+00 | 1.18E-03 | 5.10E-03 | 1.06E+00 | 3.31E-03 | 1.41E-02 | -1.08E+00 | 3.06E-05 | 1.17E-04 | -1.09E+00 | 2.63E-03 | 9.37E-03 | 1.38E+00 | 3.62E-02 | 1.02E-01 | 1.05E+00 | 5.18E-02 | 1.90E-01 | 1.11E+00 | 2.18E-01 | 5.98E-01 | 1.15E+00 |
| BX323046.1 | 3.91E-03 | 2.04E-02 | -1.32E+00 | 1.22E-04 | 5.29E-04 | -1.79E+00 | 2.05E-01 | 8.72E-01 | 1.05E+00 | 2.33E-01 | 8.90E-01 | 1.19E+00 | 9.73E-02 | 3.47E-01 | -1.37E+00 | 7.02E-02 | 1.98E-01 | -1.58E+00 | 2.01E-01 | 7.37E-01 | 1.29E+00 | 2.94E-01 | 8.07E-01 | -1.04E+00 |
| GRB10 | 3.92E-03 | 2.04E-02 | -1.03E+00 | 8.39E-02 | 3.63E-01 | 1.34E+00 | 2.18E-03 | 9.27E-03 | -1.14E+00 | 4.82E-02 | 1.84E-01 | 1.19E+00 | 2.61E-02 | 9.32E-02 | -1.07E+00 | 7.45E-02 | 2.10E-01 | 1.11E+00 | 1.47E-03 | 5.40E-03 | 1.03E+00 | 7.85E-02 | 2.16E-01 | 1.20E+00 |
| LINC02044 | 3.92E-03 | 2.04E-02 | -1.33E+00 | 5.64E-02 | 2.43E-01 | 1.16E+00 | 1.71E-01 | 7.28E-01 | 1.17E+00 | 3.06E-02 | 1.17E-01 | -1.22E+00 | 9.72E-02 | 3.47E-01 | 1.58E+00 | 3.26E-01 | 9.20E-01 | 1.20E+00 | 1.89E-01 | 6.94E-01 | 1.04E+00 | 2.97E-01 | 8.16E-01 | 1.37E+00 |
| AF127577.6 | 3.92E-03 | 2.04E-02 | -1.15E+00 | 1.06E-07 | 4.60E-07 | 2.61E+00 | 2.59E-03 | 1.10E-02 | -1.36E+00 | 4.74E-02 | 1.81E-01 | 1.66E+00 | 1.05E-01 | 3.75E-01 | 1.91E+00 | 5.38E-01 | 1.87E+00 | 1.78E-01 | 6.54E-01 | 1.38E+00 | 1.50E-03 | 4.13E-03 | 2.26E+00 | |
| C2orf76 | 3.94E-03 | 2.05E-02 | -1.10E+00 | 1.19E-02 | 5.15E-02 | 1.05E+00 | 2.20E-01 | 9.33E-01 | 1.28E+00 | 2.34E-01 | 8.95E-01 | 1.54E+00 | 3.09E-04 | 1.10E-03 | -1.21E+00 | 1.45E-02 | 4.10E-02 | 1.14E+00 | 2.60E-05 | 9.56E-05 | -1.22E+00 | 5.95E-04 | 1.64E-03 | -1.09E+00 |
| FAM213A | 3.96E-03 | 2.06E-02 | 1.06E+00 | 2.53E-03 | 1.09E-02 | 1.17E+00 | 1.87E-01 | 7.96E-01 | 1.26E+00 | 2.58E-01 | 9.85E-01 | 1.41E+00 | 1.22E-02 | 4.36E-02 | 1.14E+00 | 3.13E-01 | 8.82E-01 | 1.53E+00 | 4.03E-04 | 1.48E-03 | -1.25E+00 | 3.07E-03 | 8.44E-03 | -1.26E+00 |
| AC027279.4 | 3.96E-03 | 2.07E-02 | -1.53E+00 | 2.14E-02 | 9.27E-02 | -1.14E+00 | 1.72E-01 | 7.33E-01 | 1.25E+00 | 1.01E-01 | 7.29E-01 | 1.34E+00 | 1.46E-01 | 5.20E-01 | 1.03E+00 | 3.16E-01 | 8.90E-01 | 1.41E+00 | 1.90E-01 | 6.96E-01 | 1.31E+00 | 3.13E-01 | 8.61E-01 | 1.16E+00 |
| SNRNP35 | 3.97E-03 | 2.07E-02 | -1.04E+00 | 5.43E-03 | 2.35E-02 | 1.11E+00 | 1.88E-04 | 8.00E-04 | -1.21E+00 | 6.04E-03 | 2.31E-02 | 1.07E+00 | 9.78E-03 | 3.49E-02 | -1.02E+00 | 4.08E-02 | 1.13E-01 | 1.08E+00 | 1.25E-05 | 4.59E-05 | -1.20E+00 | 1.93E-03 | 5.29E-03 | -1.01E+00 |
| C18orf32 | 3.97E-03 | 2.07E-02 | -1.23E+00 | 8.35E-04 | 3.61E-03 | -1.31E+00 | 2.94E-02 | 1.25E-01 | -1.16E+00 | 3.02E-02 | 1.15E-01 | -1.07E+00 | 5.48E-02 | 1.95E-01 | -1.04E+00 | 1.39E-02 | 3.92E-02 | -1.26E+00 | 2.30E-03 | 8.45E-03 | -1.52E+00 | 1.37E-02 | 3.77E-02 | -1.46E+00 |
| AP002414.4 | 3.97E-03 | 2.07E-02 | -1.28E+00 | 9.04E-04 | 3.90E-03 | -1.45E+00 | 9.23E-03 | 3.92E-02 | -1.23E+00 | 3.31E-03 | 1.26E-02 | -1.27E+00 | 2.19E-03 | 7.82E-03 | -1.65E+00 | 2.66E-03 | 7.50E-03 | -1.59E+00 | 2.19E-02 | 8.02E-02 | -1.24E+00 | 5.32E-02 | 1.46E-01 | -1.11E+00 |
| AL049776.1 | 3.98E-03 | 2.07E-02 | -1.24E+00 | 6.13E-05 | 2.65E-04 | -1.74E+00 | 1.56E-02 | 6.62E-02 | -1.28E+00 | 2.50E-02 | 9.53E-02 | -1.11E+00 | 1.44E-01 | 5.15E-01 | 1.14E+00 | 2.88E-01 | 8.13E-01 | 1.40E+00 | 2.52E-03 | 9.26E-03 | -1.39E+00 | 6.40E-03 | 1.76E-02 | -1.16E+00 |
| TSHR | 3.99E-03 | 2.08E-02 | -1.71E+00 | 1.10E-01 | 4.77E-01 | 1.08E+00 | 9.51E-02 | 4.04E-01 | -1.06E+00 | 1.85E-01 | 7.08E-01 | 1.13E+00 | 7.15E-03 | 2.55E-02 | -1.61E+00 | 2.24E-01 | 6.32E-01 | 1.15E+00 | 1.98E-02 | 7.27E-02 | -1.46E+00 | 1.71E-01 | 4.69E-01 | -1.02E+00 |
| CDC37L1 | 3.99E-03 | 2.08E-02 | -1.08E+00 | 7.90E-04 | 3.42E-03 | -1.08E+00 | 1.81E-01 | 7.67E-01 | 1.27E+00 | 1.56E-01 | 5.95E-01 | 1.28E+00 | 5.05E-03 | 1.80E-02 | -1.09E+00 | 3.23E-03 | 9.12E-03 | -1.09E+00 | 1.99E-01 | 7.32E-01 | 1.29E+00 | 1.51E-01 | 4.13E-01 | 1.18E+00 |
| FAN2 | 4.00E-03 | 2.09E-02 | -1.20E+00 | 2.13E-02 | 9.22E-02 | 1.06E+00 | 3.38E-02 | 1.44E-01 | -1.00E+00 | 5.89E-04 | 2.25E-03 | -1.23E+00 | 6.33E-02 | 2.26E-01 | 1.04E+00 | 1.25E-01 | 3.52E-01 | 1.10E+00 | 3.59E-03 | 1.32E-02 | -1.26E+00 | 5.81E-02 | 1.60E-01 | -1.11E+00 |
| NCAPH2 | 4.01E-03 | 2.09E-02 | -1.10E+00 | 4.78E-04 | 2.06E-03 | -1.16E+00 | 2.38E-02 | 1.01E-01 | 1.48E+00 | 8.47E-02 | 3.23E-01 | 1.17E+00 | 3.04E-02 | 1.08E-01 | -1.15E+00 | 3.82E-02 | 1.08E-01 | -1.10E+00 | 2.06E-01 | 7.56E-01 | 1.21E+00 | 2.34E-01 | 6.43E-01 | 1.45E+00 |
| PALLD | 4.01E-03 | 2.09E-02 | -1.18E+00 | 1.98E-01 | 8.57E-01 | 1.45E+00 | 4.58E-03 | 1.95E-02 | -1.24E+00 | 1.28E-02 | 4.90E-02 | 1.03E+00 | 8.61E-03 | 3.07E-02 | -1.45E+00 | 7.24E-02 | 2.04E-01 | -1.01E+00 | 1.26E-01 | 4.63E-01 | -1.23E+00 | 3.36E-01 | 9.22E-01 | 1.19E+00 |
| S100A9 | 4.02E-03 | 2.09E-02 | -1.76E+00 | 9.99E-04 | 4.32E-03 | 5.46E+00 | 8.57E-03 | 3.64E-02 | 1.18E+00 | 2.51E-05 | 9.58E-05 | -2.00E+00 | 1.05E-01 | 3.76E-01 | 5.06E+00 | 3.42E-02 | 9.65E-02 | -2.40E+00 | 2.27E-02 | 8.32E-02 | -2.52E+00 | 2.92E-02 | 8.03E-02 | -1.54E+00 |
| CABLES1 | 4.02E-03 | 2.09E-02 | -1.11E+00 | 5.71E-02 | 2.47E-01 | 1.25E+00 | 2.26E-02 | 9.59E-02 | -1.10E+00 | 8.90E-02 | 3.40E-01 | 1.21E+00 | 2.62E-01 | 9.33E-01 | 1.38E+00 | 1.13E-01 | 3.18E-01 | 1.96E+00 | 1.42E-04 | 5.21E-04 | 2.29E+00 | 1.08E-01 | 2.97E-01 | 1.53E+00 |

| | | | | | | | | | | | | | | | | | | | | | | | | |
|---|---|---|---|---|---|---|---|---|---|---|---|---|---|---|---|---|---|---|---|---|---|---|---|---|
| AC007382.1 | 4.02E-03 | 2.09E-02 | -1.39E+00 | 1.81E-01 | 7.84E-01 | 1.52E+00 | 6.97E-03 | 2.96E-02 | -1.38E+00 | 3.80E-02 | 1.45E-01 | -1.04E+00 | 2.35E-01 | 8.36E-01 | 1.26E+00 | 1.42E-01 | 4.00E-01 | 1.25E+00 | 7.97E-02 | 2.92E-01 | 1.36E+00 | 3.23E-01 | 8.86E-01 | 1.26E+00 |
| PEBP1P3 | 4.02E-03 | 2.09E-02 | 1.73E+00 | 5.49E-02 | 2.37E-01 | 1.69E+00 | 3.72E-03 | 1.58E-02 | 1.76E+00 | 3.27E-02 | 1.25E-01 | 1.82E+00 | 1.10E-03 | 3.94E-03 | 2.91E+00 | 1.04E-02 | 2.94E-02 | 2.45E+00 | 3.66E-04 | 1.34E-03 | 2.34E+00 | 2.58E-01 | 7.09E-01 | 1.18E+00 |
| SETDB1 | 4.03E-03 | 2.10E-02 | -1.02E+00 | 4.67E-05 | 2.02E-04 | -1.11E+00 | 2.42E-04 | 1.03E-03 | -1.22E+00 | 1.46E-05 | 5.56E-05 | -1.20E+00 | 1.33E-02 | 4.74E-02 | 1.11E+00 | 1.29E-03 | 3.65E-03 | -1.06E+00 | 6.79E-05 | 2.49E-04 | -1.17E+00 | 2.76E-05 | 7.59E-05 | -1.30E+00 |
| PTPN23 | 4.03E-03 | 2.10E-02 | -1.20E+00 | 1.95E-01 | 8.41E-01 | 1.54E+00 | 5.83E-03 | 2.48E-02 | -1.10E+00 | 3.99E-02 | 1.52E-01 | 1.22E+00 | 1.92E-02 | 6.85E-02 | -1.13E+00 | 6.88E-02 | 1.94E-01 | 1.05E+00 | 3.15E-02 | 1.16E-01 | -1.02E+00 | 2.11E-01 | 5.80E-01 | 1.11E+00 |
| PHYHD1 | 4.03E-03 | 2.10E-02 | -1.08E+00 | 9.15E-03 | 3.95E-02 | 1.07E+00 | 2.56E-04 | 1.09E-03 | -1.47E+00 | 1.51E-02 | 5.77E-02 | 1.00E+00 | 4.53E-02 | 1.62E-01 | -1.96E+00 | 1.02E-01 | 2.87E-01 | -1.47E+00 | 1.96E-02 | 7.20E-02 | -1.46E+00 | 1.82E-02 | 4.99E-02 | -1.63E+00 |
| AC078846.1 | 4.03E-03 | 2.10E-02 | -1.09E+00 | 8.77E-03 | 3.79E-02 | 1.04E+00 | 1.34E-01 | 5.68E-01 | 1.31E+00 | 1.30E-01 | 4.98E-01 | 1.21E+00 | 2.58E-01 | 9.18E-01 | 1.42E+00 | 1.41E-01 | 3.96E-01 | 1.67E+00 | 2.35E-01 | 8.62E-01 | 1.40E+00 | 2.14E-01 | 5.87E-01 | 1.61E+00 |
| MREG | 4.03E-03 | 2.10E-02 | 1.91E+00 | 1.21E-01 | 5.23E-01 | 1.25E+00 | 8.26E-02 | 3.51E-01 | 1.57E+00 | 1.28E-01 | 4.89E-01 | 1.75E+00 | 1.82E-01 | 6.50E-01 | -1.17E+00 | 3.36E-01 | 9.47E-01 | 1.31E+00 | 2.68E-02 | 9.82E-02 | -1.23E+00 | 6.66E-02 | 1.83E-01 | -1.19E+00 |
| PSEN2 | 4.04E-03 | 2.10E-02 | -1.16E+00 | 3.67E-03 | 1.58E-02 | -1.10E+00 | 3.19E-04 | 1.35E-03 | -1.51E+00 | 2.80E-03 | 1.07E-02 | -1.10E+00 | 1.81E-03 | 6.47E-03 | -1.37E+00 | 3.47E-02 | 9.79E-02 | 1.12E+00 | 6.49E-04 | 2.38E-03 | -1.30E+00 | 7.67E-02 | 2.11E-01 | 1.01E+00 |
| HEG1 | 4.04E-03 | 2.11E-02 | -1.20E+00 | 2.24E-02 | 9.68E-02 | -1.05E+00 | 5.37E-02 | 2.28E-01 | 1.01E+00 | 2.17E-02 | 8.30E-02 | -1.08E+00 | 2.66E-01 | 9.50E-01 | 1.33E+00 | 6.80E-02 | 1.92E-01 | -1.57E+00 | 2.62E-03 | 9.62E-03 | -1.67E+00 | 5.55E-03 | 1.52E-02 | -1.65E+00 |
| CLCN6 | 4.04E-03 | 2.11E-02 | -1.04E+00 | 7.48E-02 | 3.23E-01 | 1.36E+00 | 2.70E-02 | 1.15E-01 | 1.00E+00 | 1.60E-01 | 6.13E-01 | 1.36E+00 | 3.94E-02 | 1.41E-01 | -1.06E+00 | 1.33E-01 | 3.74E-01 | 1.15E+00 | 1.61E-01 | 5.90E-01 | 1.49E+00 | 2.41E-01 | 6.62E-01 | 1.43E+00 |
| SMURF2 | 4.04E-03 | 2.11E-02 | 1.05E+00 | 3.12E-04 | 1.35E-03 | 1.01E+00 | 4.23E-03 | 1.80E-02 | -1.03E+00 | 6.45E-07 | 2.46E-06 | -1.24E+00 | 1.37E-01 | 4.87E-01 | 1.61E+00 | 1.49E-01 | 4.20E-01 | 1.21E+00 | 2.40E-01 | 8.80E-01 | 1.28E+00 | 1.81E-01 | 4.98E-01 | 1.14E+00 |
| ATP5LP3 | 4.04E-03 | 2.11E-02 | 1.67E+00 | 1.83E-01 | 7.91E-01 | 1.46E+00 | 1.72E-05 | 7.30E-05 | 1.88E+00 | 9.10E-02 | 3.48E-01 | 1.59E+00 | 6.69E-02 | 2.39E-01 | 1.67E+00 | 3.35E-01 | 9.45E-01 | 1.47E+00 | 7.43E-04 | 2.73E-03 | 1.87E+00 | 1.72E-02 | 4.72E-02 | 1.77E+00 |
| INPP1 | 4.05E-03 | 2.11E-02 | -1.13E+00 | 1.09E-01 | 4.69E-01 | 1.24E+00 | 2.99E-03 | 1.27E-02 | -1.07E+00 | 1.45E-03 | 5.53E-03 | 1.08E+00 | 1.68E-04 | 5.99E-04 | -1.65E+00 | 1.20E-02 | 3.38E-02 | -1.11E+00 | 7.83E-05 | 2.87E-04 | -1.19E+00 | 1.61E-04 | 4.43E-04 | -1.16E+00 |
| SEPT7P9 | 4.06E-03 | 2.11E-02 | -1.22E+00 | 9.70E-04 | 4.19E-03 | -1.34E+00 | 7.83E-04 | 3.33E-03 | -1.31E+00 | 1.37E-04 | 5.23E-04 | -1.37E+00 | 1.01E-09 | 3.61E-09 | -1.82E+00 | 6.77E-14 | 1.91E-13 | -2.52E+00 | 4.42E-07 | 1.62E-06 | -1.52E+00 | 4.42E-06 | 1.21E-05 | -1.37E+00 |
| TRAF3 | 4.06E-03 | 2.11E-02 | -1.17E+00 | 1.08E-01 | 4.66E-01 | 1.25E+00 | 6.38E-06 | 2.71E-05 | -1.51E+00 | 6.47E-03 | 2.47E-02 | 1.04E+00 | 2.95E-02 | 1.05E-01 | -1.19E+00 | 2.89E-01 | 8.14E-01 | 1.27E+00 | 4.53E-02 | 1.66E-01 | -1.08E+00 | 7.52E-02 | 2.06E-01 | 1.44E+00 |
| LINC-PINT | 4.06E-03 | 2.12E-02 | 1.73E+00 | 1.20E-01 | 5.20E-01 | 1.43E+00 | 1.92E-01 | 8.14E-01 | 1.22E+00 | 2.95E-03 | 1.13E-02 | -1.13E+00 | 5.13E-03 | 1.83E-02 | 2.88E+00 | 3.00E-01 | 8.45E-01 | 1.63E+00 | 1.97E-01 | 7.23E-01 | 1.39E+00 | 1.01E-01 | 2.76E-01 | 1.01E+00 |
| RPS21 | 4.07E-03 | 2.12E-02 | 1.08E+00 | 1.77E-57 | 7.64E-57 | -1.62E+00 | 2.95E-02 | 1.25E-01 | 1.07E+00 | 6.58E-35 | 2.51E-34 | -1.58E+00 | 1.19E-01 | 4.24E-01 | 1.04E+00 | 6.54E-28 | 1.84E-27 | -1.58E+00 | 3.60E-03 | 1.32E-02 | 1.01E+00 | 9.51E-44 | 2.61E-43 | -1.65E+00 |
| AL162377.1 | 4.07E-03 | 2.12E-02 | 1.84E+00 | 1.57E-03 | 6.76E-03 | -1.18E+00 | 8.01E-02 | 3.40E-01 | 1.55E+00 | 1.42E-01 | 5.44E-01 | 1.14E+00 | 1.13E-01 | 4.04E-01 | 1.90E+00 | 2.46E-01 | 6.94E-01 | 1.54E+00 | 2.64E-01 | 9.68E-01 | 1.30E+00 | 3.61E-01 | 9.93E-01 | 1.29E+00 |
| CMTM7 | 4.08E-03 | 2.13E-02 | -1.04E+00 | 1.03E-02 | 4.46E-02 | 1.17E+00 | 1.69E-03 | 7.20E-03 | -1.10E+00 | 4.84E-04 | 1.85E-03 | 1.04E+00 | 9.48E-06 | 3.38E-05 | -1.47E+00 | 2.85E-05 | 8.03E-05 | -1.35E+00 | 8.65E-06 | 3.18E-05 | -1.27E+00 | 7.92E-03 | 2.18E-02 | -1.01E+00 |
| SMIM6 | 4.08E-03 | 2.13E-02 | 1.83E+00 | 3.76E-02 | 1.63E-01 | 1.15E+00 | 1.76E-01 | 7.46E-01 | 1.30E+00 | 1.49E-02 | 5.70E-02 | -1.15E+00 | 2.89E-02 | 1.03E-01 | -1.12E+00 | 8.13E-02 | 2.29E-01 | -1.00E+00 | 6.40E-02 | 2.35E-01 | 1.46E+00 | 7.20E-02 | 1.98E-01 | -1.14E+00 |
| RFC5 | 4.09E-03 | 2.13E-02 | -1.13E+00 | 1.93E-02 | 8.34E-02 | 1.06E+00 | 5.90E-02 | 2.51E-01 | 1.27E+00 | 1.28E-01 | 4.88E-01 | 1.35E+00 | 9.23E-02 | 3.29E-01 | 1.07E+00 | 3.42E-01 | 9.64E-01 | 1.61E+00 | 4.16E-02 | 1.53E-01 | 1.07E+00 | 3.26E-01 | 8.95E-01 | 1.28E+00 |
| ARMCX7P | 4.10E-03 | 2.14E-02 | 1.01E+00 | 2.18E-03 | 9.43E-03 | 1.11E+00 | 1.44E-06 | 6.10E-06 | -1.29E+00 | 3.67E-07 | 1.40E-06 | -1.19E+00 | 9.27E-04 | 3.30E-03 | -1.18E+00 | 1.69E-01 | 4.76E-01 | 1.25E+00 | 2.44E-03 | 8.95E-03 | -1.07E+00 | 5.08E-02 | 1.39E-01 | 1.04E+00 |
| ERV3-1 | 4.10E-03 | 2.14E-02 | 1.03E+00 | 1.16E-01 | 5.02E-01 | 1.59E+00 | 5.97E-02 | 2.54E-01 | 1.02E+00 | 1.54E-01 | 5.87E-01 | 1.47E+00 | 3.63E-02 | 1.29E-01 | 1.12E+00 | 3.17E-01 | 8.93E-01 | 1.41E+00 | 2.77E-02 | 1.02E-01 | 1.86E+00 | 1.62E-02 | 4.46E-02 | 1.71E+00 |
| APOO | 4.10E-03 | 2.14E-02 | -1.24E+00 | 4.97E-03 | 2.15E-02 | -1.12E+00 | 2.09E-03 | 8.88E-03 | -1.21E+00 | 2.69E-03 | 1.03E-02 | -1.01E+00 | 3.23E-03 | 1.15E-02 | -1.49E+00 | 6.05E-02 | 1.71E-01 | -1.08E+00 | 8.19E-02 | 3.01E-01 | 1.11E+00 | 2.57E-01 | 7.05E-01 | 1.20E+00 |
| KLHL15 | 4.10E-03 | 2.14E-02 | -1.06E+00 | 1.00E-06 | 4.33E-06 | -1.52E+00 | 7.19E-02 | 3.06E-01 | 1.47E+00 | 2.86E-03 | 1.09E-02 | -1.02E+00 | 1.49E-01 | 5.32E-01 | 1.27E+00 | 1.69E-03 | 4.77E-03 | -1.30E+00 | 2.63E-04 | 9.64E-04 | -1.15E+00 | 1.66E-07 | 4.55E-07 | -1.65E+00 |
| SAMD4A | 4.11E-03 | 2.14E-02 | -1.16E+00 | 1.69E-02 | 7.29E-02 | 1.15E+00 | 4.71E-09 | 2.00E-08 | -2.83E+00 | 1.69E-08 | 6.44E-08 | -2.56E+00 | 1.19E-01 | 4.26E-01 | -1.27E+00 | 6.18E-02 | 1.74E-01 | -2.00E+00 | 2.36E-01 | 8.65E-01 | 1.05E+00 | 2.95E-01 | 8.09E-01 | -1.12E+00 |
| MYBBP1A | 4.11E-03 | 2.14E-02 | -1.24E+00 | 9.39E-02 | 4.06E-01 | 1.22E+00 | 1.53E-05 | 6.49E-05 | -1.32E+00 | 2.26E-03 | 8.61E-03 | 1.02E+00 | 6.19E-04 | 2.21E-03 | -1.58E+00 | 1.29E-01 | 3.64E-01 | 1.14E+00 | 2.53E-06 | 9.30E-06 | -1.45E+00 | 2.91E-01 | 8.00E-01 | 1.25E+00 |
| FAM35BP | 4.11E-03 | 2.14E-02 | -1.28E+00 | 4.06E-03 | 1.76E-02 | -1.14E+00 | 9.57E-05 | 4.07E-04 | -1.50E+00 | 1.31E-03 | 5.02E-03 | -1.11E+00 | 7.60E-04 | 2.71E-03 | -1.40E+00 | 4.07E-03 | 1.15E-02 | -1.17E+00 | 4.44E-06 | 1.63E-05 | -1.37E+00 | 5.67E-05 | 1.56E-04 | -1.24E+00 |
| RCBTB2 | 4.12E-03 | 2.15E-02 | -1.24E+00 | 2.19E-02 | 9.47E-02 | 1.01E+00 | 7.92E-04 | 3.37E-03 | -1.35E+00 | 1.58E-03 | 6.02E-03 | -1.21E+00 | 2.59E-03 | 9.25E-03 | -1.47E+00 | 1.30E-01 | 3.66E-01 | 1.10E+00 | 1.43E-01 | 5.26E-01 | 1.16E+00 | 3.11E-01 | 8.54E-01 | 1.25E+00 |
| RAPGEF5 | 4.12E-03 | 2.15E-02 | -1.46E+00 | 8.03E-03 | 3.47E-02 | -1.29E+00 | 1.73E-08 | 7.35E-08 | -1.99E+00 | 8.27E-09 | 3.16E-08 | -1.72E+00 | 5.65E-02 | 2.01E-01 | -1.39E+00 | 4.20E-02 | 1.18E-01 | -1.56E+00 | 1.34E-06 | 4.92E-06 | -2.30E+00 | 2.10E-05 | 5.77E-05 | -2.00E+00 |
| JADE2 | 4.12E-03 | 2.15E-02 | 1.03E+00 | 2.13E-01 | 9.20E-01 | 1.59E+00 | 1.12E-03 | 4.74E-03 | -1.07E+00 | 5.82E-02 | 2.22E-01 | 1.33E+00 | 7.90E-02 | 2.82E-01 | 1.20E+00 | 1.55E-01 | 4.38E-01 | 1.32E+00 | 9.13E-03 | 3.35E-02 | -1.25E+00 | 2.16E-01 | 5.92E-01 | 1.09E+00 |
| AZU1 | 4.13E-03 | 2.15E-02 | -2.32E+00 | 7.88E-05 | 3.41E-04 | -5.28E+00 | 8.37E-24 | 3.56E-23 | -2.71E+00 | 1.11E-24 | 4.26E-24 | -2.88E+00 | 8.14E-02 | 2.90E-01 | -1.38E+00 | 8.43E-02 | 2.38E-01 | -1.57E+00 | 2.09E-01 | 7.69E-01 | -1.10E+00 | 1.18E-01 | 3.24E-01 | -1.91E+00 |
| AC067931.3 | 4.14E-03 | 2.16E-02 | -1.26E+00 | 6.38E-04 | 2.76E-03 | -1.50E+00 | 1.79E-01 | 7.62E-01 | 1.08E+00 | 9.37E-04 | 3.58E-03 | -1.76E+00 | 5.81E-02 | 2.07E-01 | -1.12E+00 | 1.61E-03 | 4.53E-03 | -2.26E+00 | 1.25E-03 | 4.60E-03 | -1.56E+00 | 6.23E-04 | 1.71E-03 | -1.80E+00 |
| LRRC63 | 4.14E-03 | 2.16E-02 | -1.26E+00 | 2.05E-02 | 8.85E-02 | -1.02E+00 | 1.65E-03 | 7.00E-03 | -1.62E+00 | 1.40E-02 | 5.34E-02 | -1.20E+00 | 2.07E-01 | 7.36E-01 | 1.44E+00 | 2.98E-01 | 8.39E-01 | 1.05E+00 | 2.52E-01 | 9.26E-01 | 1.10E+00 | 2.86E-01 | 7.85E-01 | 1.02E+00 |
| PPIAP2 | 4.14E-03 | 2.16E-02 | -1.29E+00 | 2.36E-02 | 1.02E-01 | 1.03E+00 | 2.21E-04 | 9.38E-04 | -1.38E+00 | 3.38E-02 | 1.29E-01 | 1.12E+00 | 3.34E-03 | 1.19E-02 | -2.16E+00 | 2.64E-01 | 7.44E-01 | 1.16E+00 | 1.54E-04 | 5.64E-04 | -1.69E+00 | 2.35E-01 | 6.44E-01 | 1.15E+00 |
| MNS1 | 4.14E-03 | 2.16E-02 | -1.45E+00 | 5.52E-04 | 2.38E-03 | -2.03E+00 | 5.76E-02 | 2.45E-01 | 1.29E+00 | 4.52E-02 | 1.73E-01 | -1.03E+00 | 5.63E-02 | 2.01E-01 | -1.18E+00 | 3.40E-02 | 9.58E-02 | -1.33E+00 | 5.81E-02 | 2.13E-01 | 1.44E+00 | 2.65E-01 | 7.29E-01 | 1.31E+00 |
| MSANTD4 | 4.15E-03 | 2.17E-02 | -1.09E+00 | 1.80E-03 | 7.76E-03 | -1.07E+00 | 1.11E-01 | 4.70E-01 | -1.02E+00 | 1.10E-01 | 4.18E-01 | 1.03E+00 | 1.52E-02 | 5.43E-02 | -1.02E+00 | 2.96E-02 | 8.36E-02 | 1.08E+00 | 1.43E-01 | 5.23E-01 | 1.18E+00 | 4.18E-02 | 1.15E-01 | -1.02E+00 |
| BCL2L11 | 4.15E-03 | 2.17E-02 | -1.20E+00 | 1.13E-02 | 4.86E-02 | -1.09E+00 | 3.65E-02 | 1.55E-01 | -1.17E+00 | 1.77E-02 | 6.75E-02 | -1.22E+00 | 4.41E-02 | 1.57E-01 | -1.20E+00 | 1.85E-01 | 5.23E-01 | 1.04E+00 | 2.21E-01 | 8.10E-01 | 1.28E+00 | 3.38E-01 | 9.29E-01 | 1.39E+00 |
| GLMN | 4.17E-03 | 2.18E-02 | -1.12E+00 | 9.94E-02 | 4.29E-01 | 1.23E+00 | 1.37E-02 | 5.81E-02 | -1.10E+00 | 1.53E-02 | 5.85E-02 | 1.07E+00 | 5.63E-02 | 2.01E-01 | -1.10E+00 | 3.14E-01 | 8.86E-01 | 1.37E+00 | 3.63E-03 | 1.33E-02 | -1.10E+00 | 1.19E-01 | 3.27E-01 | 1.19E+00 |
| MRPS28 | 4.17E-03 | 2.18E-02 | -1.59E+00 | 1.20E-02 | 5.17E-02 | -1.33E+00 | 1.09E-01 | 4.64E-01 | 1.06E+00 | 2.54E-01 | 9.70E-01 | 1.39E+00 | 2.70E-02 | 9.61E-02 | -1.42E+00 | 2.02E-02 | 5.69E-02 | -1.53E+00 | 4.89E-10 | 1.80E-09 | -1.74E+00 | 4.83E-08 | 1.33E-07 | -1.56E+00 |
| EIF1B | 4.18E-03 | 2.18E-02 | -1.02E+00 | 6.44E-20 | 2.78E-19 | -1.34E+00 | 6.73E-04 | 2.86E-03 | -1.23E+00 | 1.84E-12 | 7.05E-12 | -1.49E+00 | 2.56E-04 | 9.12E-04 | -1.23E+00 | 3.08E-13 | 8.68E-13 | -1.70E+00 | 1.98E-01 | 7.28E-01 | 1.17E+00 | 4.16E-03 | 1.14E-02 | -1.12E+00 |
| RPLP1 | 4.18E-03 | 2.18E-02 | -1.05E+00 | 1.42E-52 | 6.15E-52 | -1.40E+00 | 6.14E-02 | 2.61E-01 | -1.03E+00 | 4.93E-24 | 1.88E-23 | -1.32E+00 | 3.81E-03 | 1.36E-02 | -1.07E+00 | 5.03E-24 | 1.42E-23 | -1.36E+00 | 3.50E-08 | 1.28E-07 | -1.09E+00 | 4.35E-37 | 1.19E-36 | -1.38E+00 |
| RPL22L1 | 4.19E-03 | 2.18E-02 | 1.06E+00 | 1.22E-09 | 5.27E-09 | -1.07E+00 | 1.40E-01 | 5.97E-01 | 1.10E+00 | 1.44E-02 | 5.51E-02 | -1.07E+00 | 1.25E-02 | 4.47E-02 | 1.05E+00 | 8.17E-03 | 2.30E-02 | -1.01E+00 | 2.56E-01 | 9.38E-01 | 1.15E+00 | 2.02E-01 | 5.54E-01 | 1.15E+00 |
| WDTC1 | 4.19E-03 | 2.19E-02 | 1.05E+00 | 3.04E-02 | 1.32E-01 | 1.26E+00 | 2.83E-07 | 1.20E-06 | -1.44E+00 | 2.17E-05 | 8.29E-05 | -1.10E+00 | 1.68E-03 | 5.98E-03 | 1.02E+00 | 8.24E-03 | 2.32E-02 | 1.10E+00 | 4.13E-05 | 1.52E-04 | -1.43E+00 | 9.42E-02 | 2.59E-01 | 1.03E+00 |
| TM9SF1 | 4.20E-03 | 2.19E-02 | -1.01E+00 | 1.01E-04 | 4.37E-04 | -1.32E+00 | 1.33E-01 | 5.66E-01 | 1.12E+00 | 2.37E-01 | 9.06E-01 | 1.47E+00 | 1.84E-02 | 6.54E-02 | -1.09E+00 | 3.86E-03 | 1.09E-02 | -1.31E+00 | 3.14E-02 | 1.15E-01 | -1.01E+00 | 1.10E-02 | 3.04E-02 | -1.15E+00 |
| ZNF90 | 4.20E-03 | 2.19E-02 | -1.60E+00 | 7.46E-03 | 3.22E-02 | -1.46E+00 | 9.85E-05 | 4.18E-04 | -1.51E+00 | 3.80E-04 | 1.45E-03 | -1.30E+00 | 4.34E-02 | 1.55E-01 | -1.12E+00 | 3.66E-02 | 1.03E-01 | -1.31E+00 | 4.27E-02 | 1.57E-01 | -1.10E+00 | 6.82E-02 | 1.87E-01 | -1.05E+00 |
| BRPF3 | 4.21E-03 | 2.19E-02 | -1.29E+00 | 3.74E-03 | 1.62E-02 | -1.23E+00 | 1.14E-02 | 4.85E-02 | -1.17E+00 | 6.36E-03 | 2.43E-02 | -1.07E+00 | 2.61E-01 | 9.31E-01 | 1.28E+00 | 3.48E-01 | 9.81E-01 | 1.26E+00 | 1.99E-01 | 7.29E-01 | 1.26E+00 | 2.75E-01 | 7.55E-01 | 1.31E+00 |
| RAD54B | 4.22E-03 | 2.20E-02 | -1.60E+00 | 2.96E-02 | 1.28E-01 | -1.22E+00 | 1.61E-02 | 6.82E-02 | -1.27E+00 | 7.83E-02 | 2.99E-01 | 1.05E+00 | 1.12E-01 | 4.00E-01 | -1.43E+00 | 2.20E-01 | 6.21E-01 | -1.12E+00 | 3.22E-04 | 1.18E-03 | -1.30E+00 | 1.04E-01 | 2.86E-01 | 1.03E+00 |
| SAMD12 | 4.23E-03 | 2.20E-02 | -1.10E+00 | 2.95E-03 | 1.27E-02 | -1.11E+00 | 2.35E-01 | 9.97E-01 | 1.09E+00 | 1.31E-01 | 5.00E-01 | 1.48E+00 | 2.06E-01 | 7.35E-01 | 1.12E+00 | 3.12E-01 | 8.80E-01 | 1.21E+00 | 2.18E-01 | 8.01E-01 | 1.14E+00 | 1.99E-01 | 5.47E-01 | 1.08E+00 |
| FAT4 | 4.23E-03 | 2.21E-02 | -1.56E+00 | 1.18E-01 | 5.10E-01 | 1.08E+00 | 1.44E-01 | 6.11E-01 | -1.04E+00 | 1.35E-01 | 5.15E-01 | 1.02E+00 | 7.95E-03 | 2.83E-02 | -1.92E+00 | 5.18E-03 | 1.46E-02 | -2.27E+00 | 2.68E-01 | 9.84E-01 | 1.16E+00 | 2.67E-01 | 7.33E-01 | 1.25E+00 |
| GTPBP2 | 4.24E-03 | 2.21E-02 | -1.26E+00 | 6.43E-03 | 2.78E-02 | -1.16E+00 | 1.71E-02 | 7.27E-02 | -1.13E+00 | 3.79E-02 | 1.45E-01 | 1.07E+00 | 2.70E-02 | 9.62E-02 | -1.03E+00 | 1.41E-02 | 3.97E-02 | -1.14E+00 | 1.91E-02 | 7.02E-02 | 1.02E+00 | 1.10E-01 | 3.01E-01 | 1.18E+00 |
| PGLS | 4.26E-03 | 2.22E-02 | 1.07E+00 | 7.04E-04 | 3.04E-03 | 1.08E+00 | 1.28E-01 | 5.45E-01 | 1.02E+00 | 5.20E-02 | 1.99E-01 | 1.02E+00 | 1.10E-05 | 3.93E-05 | -1.13E+00 | 6.20E-04 | 1.75E-03 | -1.12E+00 | 2.85E-02 | 1.05E-01 | 1.05E+00 | 8.06E-02 | 2.21E-01 | 1.07E+00 |
| AC024568.1 | 4.26E-03 | 2.22E-02 | -1.13E+00 | 1.32E-02 | 5.72E-02 | 1.12E+00 | 2.89E-02 | 1.23E-01 | -1.22E+00 | 2.00E-01 | 7.63E-01 | 1.38E+00 | 3.98E-02 | 1.42E-01 | -1.15E+00 | 6.57E-02 | 1.85E-01 | 1.02E+00 | 1.34E-03 | 4.92E-03 | -1.35E+00 | 1.80E-02 | 4.95E-02 | -1.06E+00 |
| PI4K2B | 4.26E-03 | 2.22E-02 | -1.24E+00 | 3.65E-02 | 1.58E-01 | 1.10E+00 | 2.21E-03 | 9.39E-03 | -1.28E+00 | 5.50E-03 | 2.10E-02 | -1.07E+00 | 4.18E-03 | 1.49E-02 | -1.69E+00 | 1.16E-01 | 3.28E-01 | 1.14E+00 | 5.38E-10 | 1.97E-09 | -1.65E+00 | 1.19E-03 | 3.27E-03 | -1.07E+00 |
| DIP2B | 4.26E-03 | 2.22E-02 | -1.05E+00 | 1.30E-01 | 5.63E-01 | 1.35E+00 | 8.24E-03 | 3.50E-02 | -1.07E+00 | 1.99E-02 | 7.61E-02 | 1.17E+00 | 9.58E-02 | 3.42E-01 | 1.26E+00 | 3.49E-01 | 9.84E-01 | 1.46E+00 | 1.81E-03 | 6.65E-03 | -1.03E+00 | 3.88E-02 | 1.07E-01 | 1.12E+00 |
| AC093752.1 | 4.26E-03 | 2.22E-02 | 1.01E+00 | 3.23E-03 | 1.40E-02 | 1.15E+00 | 1.64E-04 | 6.95E-04 | -1.39E+00 | 3.86E-03 | 1.47E-02 | -1.02E+00 | 4.44E-03 | 1.58E-02 | -1.15E+00 | 5.08E-02 | 1.43E-01 | 1.02E+00 | 7.29E-04 | 2.68E-03 | -1.11E+00 | 9.42E-03 | 2.59E-02 | -1.04E+00 |
| THUMPD3-AS1 | 4.27E-03 | 2.22E-02 | 1.11E+00 | 1.18E-03 | 5.09E-03 | 1.16E+00 | 9.46E-03 | 4.02E-02 | -1.02E+00 | 2.83E-03 | 1.08E-02 | 1.11E+00 | 1.16E-01 | 4.12E-01 | 1.22E+00 | 1.79E-01 | 5.06E-01 | 1.29E+00 | 1.21E-01 | 4.43E-01 | 1.23E+00 | 2.61E-02 | 7.18E-02 | 1.20E+00 |
| AP1S3 | 4.27E-03 | 2.23E-02 | -1.75E+00 | 2.64E-02 | 1.14E-01 | -1.30E+00 | 1.36E-08 | 5.78E-08 | -2.16E+00 | 2.49E-05 | 9.51E-05 | -1.39E+00 | 5.93E-02 | 2.12E-01 | -1.38E+00 | 8.55E-02 | 2.41E-01 | -1.25E+00 | 1.53E-01 | 5.60E-01 | -1.14E+00 | 2.26E-01 | 6.20E-01 | 1.36E+00 |
| AC126474.2 | 4.27E-03 | 2.23E-02 | -1.32E+00 | 1.61E-03 | 6.95E-03 | -1.31E+00 | 1.18E-01 | 5.00E-01 | 1.07E+00 | 2.32E-02 | 8.88E-02 | 1.01E+00 | 3.15E-02 | 1.12E-01 | -1.02E+00 | 3.67E-02 | 1.03E-01 | -1.16E+00 | 9.66E-02 | 3.54E-01 | -1.02E+00 | 2.81E-01 | 7.71E-01 | 1.18E+00 |
| RFC4 | 4.28E-03 | 2.23E-02 | -1.07E+00 | 2.39E-02 | 1.03E-01 | 1.22E+00 | 2.12E-02 | 9.02E-02 | 1.42E+00 | 8.32E-02 | 3.18E-01 | 1.56E+00 | 2.02E-03 | 7.20E-03 | -1.13E+00 | 5.04E-02 | 1.42E-01 | 1.17E+00 | 1.12E-01 | 4.11E-01 | 1.10E+00 | 1.93E-03 | 5.30E-03 | 1.43E+00 |
| ACYP2 | 4.28E-03 | 2.23E-02 | 1.02E+00 | 6.03E-04 | 2.61E-03 | 1.08E+00 | 1.21E-03 | 5.12E-03 | -1.13E+00 | 1.42E-05 | 5.41E-05 | -1.13E+00 | 6.19E-04 | 2.21E-03 | -1.08E+00 | 3.37E-06 | 9.51E-06 | -1.28E+00 | 2.69E-01 | 9.87E-01 | 1.28E+00 | 2.73E-01 | 7.49E-01 | 1.18E+00 |
| NACA | 4.29E-03 | 2.23E-02 | 1.06E+00 | 4.70E-02 | 2.03E-01 | 1.03E+00 | 5.39E-02 | 2.29E-01 | 1.04E+00 | 1.46E-03 | 5.59E-03 | 1.10E+00 | 4.26E-02 | 1.52E-01 | 1.06E+00 | 9.00E-02 | 2.54E-01 | 1.04E+00 | 6.03E-02 | 2.21E-01 | 1.01E+00 | 3.39E-01 | 9.31E-01 | 1.02E+00 |
| IFFO2 | 4.30E-03 | 2.24E-02 | -1.32E+00 | 3.70E-04 | 1.60E-03 | -1.44E+00 | 5.27E-03 | 2.24E-02 | -1.23E+00 | 3.09E-04 | 1.18E-03 | -1.40E+00 | 2.88E-05 | 1.03E-04 | -1.78E+00 | 1.70E-05 | 4.79E-05 | -1.65E+00 | 2.00E-02 | 7.35E-02 | -1.29E+00 | 5.78E-02 | 1.59E-01 | -1.12E+00 |
| ZNF782 | 4.30E-03 | 2.24E-02 | -1.09E+00 | 1.59E-02 | 6.85E-02 | 1.04E+00 | 2.03E-02 | 8.61E-02 | -1.18E+00 | 1.60E-02 | 6.09E-02 | -1.04E+00 | 5.56E-03 | 1.98E-02 | -1.43E+00 | 1.17E-02 | 3.29E-02 | -1.30E+00 | 2.54E-02 | 9.32E-02 | -1.11E+00 | 3.04E-01 | 8.34E-01 | 1.25E+00 |
| RPL10P15 | 4.30E-03 | 2.24E-02 | 1.19E+00 | 4.62E-20 | 2.00E-19 | -1.40E+00 | 1.46E-01 | 6.19E-01 | 1.14E+00 | 2.17E-18 | 8.28E-18 | -1.68E+00 | 7.25E-02 | 2.59E-01 | 1.23E+00 | 7.46E-15 | 2.10E-14 | -1.67E+00 | 5.05E-02 | 1.85E-01 | 1.05E+00 | 1.73E-12 | 4.74E-12 | -1.76E+00 |
| LINC00921 | 4.30E-03 | 2.24E-02 | -1.11E+00 | 1.02E-01 | 4.39E-01 | 1.33E+00 | 2.15E-02 | 9.15E-02 | -1.23E+00 | 9.22E-02 | 3.52E-01 | 1.07E+00 | 1.92E-01 | 6.83E-01 | 1.06E+00 | 2.28E-01 | 6.43E-01 | 1.54E+00 | 9.62E-02 | 3.53E-01 | 1.10E+00 | 3.42E-01 | 9.39E-01 | 1.31E+00 |
| KLHL7 | 4.30E-03 | 2.24E-02 | -1.13E+00 | 4.12E-02 | 1.78E-01 | 1.16E+00 | 4.03E-04 | 1.71E-03 | -1.31E+00 | 5.17E-04 | 1.97E-03 | -1.13E+00 | 1.45E-07 | 5.18E-07 | -1.76E+00 | 5.17E-06 | 1.46E-05 | -1.33E+00 | 1.90E-04 | 6.98E-04 | -1.17E+00 | 8.80E-05 | 2.42E-04 | -1.15E+00 |
| TSPYL2 | 4.32E-03 | 2.25E-02 | 1.77E+00 | 9.43E-03 | 4.07E-02 | 1.19E+00 | 3.08E-02 | 1.31E-01 | 1.63E+00 | 2.01E-02 | 7.69E-02 | 1.10E+00 | 8.64E-04 | 3.08E-03 | 1.80E+00 | 3.65E-02 | 1.03E-01 | -1.01E+00 | 5.82E-05 | 2.14E-04 | 2.22E+00 | 1.03E-01 | 2.84E-01 | 1.00E+00 |
| KAT14 | 4.32E-03 | 2.25E-02 | -1.22E+00 | 2.26E-02 | 9.75E-02 | 1.08E+00 | 1.81E-03 | 7.68E-03 | -1.35E+00 | 2.43E-03 | 9.30E-03 | -1.29E+00 | 1.08E-03 | 3.86E-03 | -1.73E+00 | 9.27E-03 | 2.61E-02 | -1.32E+00 | 3.63E-03 | 1.33E-02 | -1.23E+00 | 3.79E-02 | 1.04E-01 | -1.08E+00 |
| PRELID3A | 4.32E-03 | 2.25E-02 | -1.23E+00 | 1.03E-02 | 4.44E-02 | -1.02E+00 | 7.44E-03 | 3.16E-02 | -1.22E+00 | 2.98E-02 | 1.14E-01 | 1.10E+00 | 1.00E-01 | 3.58E-01 | -1.21E+00 | 2.83E-01 | 7.97E-01 | 1.31E+00 | 4.86E-02 | 1.78E-01 | -1.30E+00 | 3.17E-01 | 8.70E-01 | 1.14E+00 |
| AC025279.1 | 4.33E-03 | 2.26E-02 | -1.21E+00 | 3.75E-03 | 1.62E-02 | -1.12E+00 | 1.82E-05 | 7.72E-05 | -2.01E+00 | 6.15E-05 | 2.35E-04 | -1.61E+00 | 1.86E-05 | 6.64E-05 | -1.63E+00 | 1.84E-05 | 5.19E-05 | -1.63E+00 | 7.50E-04 | 2.75E-03 | -1.71E+00 | 5.01E-02 | 1.38E-01 | -1.06E+00 |
| GNB1L | 4.35E-03 | 2.27E-02 | -1.32E+00 | 2.49E-03 | 1.08E-02 | -1.32E+00 | 1.48E-05 | 6.27E-05 | -1.39E+00 | 1.03E-03 | 3.94E-03 | -1.06E+00 | 2.22E-03 | 7.90E-03 | -2.18E+00 | 1.54E-01 | 4.33E-01 | 1.08E+00 | 7.98E-02 | 2.93E-01 | -1.04E+00 | 3.11E-01 | 8.53E-01 | 1.27E+00 |
| DOCK4 | 4.35E-03 | 2.27E-02 | 2.40E+00 | 7.51E-02 | 3.24E-01 | 1.77E+00 | 1.23E-01 | 5.23E-01 | 1.37E+00 | 1.76E-01 | 6.72E-01 | 1.40E+00 | 1.01E-04 | 3.59E-04 | 3.50E+00 | 2.66E-01 | 7.51E-01 | 1.26E+00 | 1.81E-01 | 6.65E-01 | 1.28E+00 | 2.62E-01 | 7.18E-01 | -1.17E+00 |
| HOMER3 | 4.35E-03 | 2.27E-02 | -2.36E+00 | 4.43E-03 | 1.91E-02 | -2.23E+00 | 1.24E-13 | 5.28E-13 | -2.48E+00 | 4.77E-12 | 1.82E-11 | -1.98E+00 | 2.67E-01 | 9.51E-01 | 1.28E+00 | 7.21E-02 | 2.03E-01 | -1.45E+00 | 6.88E-02 | 2.53E-01 | -1.70E+00 | 1.12E-01 | 3.08E-01 | -1.62E+00 |
| BACE1-AS | 4.35E-03 | 2.27E-02 | -1.66E+00 | 6.03E-02 | 2.60E-01 | 1.90E+00 | 1.50E-02 | 6.39E-02 | -1.37E+00 | 1.49E-01 | 5.70E-01 | 1.51E+00 | 4.76E-06 | 1.70E-05 | -2.77E+00 | 1.05E-02 | 2.97E-02 | -1.11E+00 | 1.17E-03 | 4.28E-03 | -1.63E+00 | 3.11E-01 | 8.55E-01 | 1.27E+00 |
| SPATA33 | 4.35E-03 | 2.27E-02 | -1.16E+00 | 1.83E-01 | 7.89E-01 | 1.37E+00 | 1.83E-02 | 7.78E-02 | -1.14E+00 | 3.65E-02 | 1.39E-01 | 1.11E+00 | 1.59E-01 | 5.66E-01 | 1.04E+00 | 2.11E-01 | 5.95E-01 | 1.10E+00 | 2.55E-01 | 9.37E-01 | 1.27E+00 | 3.52E-01 | 9.66E-01 | 1.32E+00 |
| ARHGEF37 | 4.36E-03 | 2.27E-02 | 1.90E+00 | 7.90E-02 | 3.42E-01 | 1.56E+00 | 1.75E-03 | 7.45E-03 | 2.25E+00 | 2.42E-01 | 9.23E-01 | 1.30E+00 | 1.03E-01 | 3.66E-01 | 1.98E+00 | 3.00E-01 | 8.47E-01 | 1.22E+00 | 4.94E-02 | 1.81E-01 | 1.52E+00 | 2.26E-01 | 6.20E-01 | 1.27E+00 |
| WDR47 | 4.36E-03 | 2.27E-02 | -1.17E+00 | 3.32E-03 | 1.44E-02 | -1.06E+00 | 4.87E-02 | 2.07E-01 | -1.14E+00 | 3.54E-02 | 1.35E-01 | -1.11E+00 | 7.61E-03 | 2.71E-02 | -1.19E+00 | 4.08E-03 | 1.15E-02 | -1.20E+00 | 4.89E-03 | 1.80E-02 | -1.22E+00 | 1.29E-03 | 3.55E-03 | -1.42E+00 |
| AL450384.2 | 4.36E-03 | 2.27E-02 | -1.08E+00 | 1.05E-01 | 4.52E-01 | 1.33E+00 | 1.56E-01 | 6.63E-01 | 1.12E+00 | 2.12E-01 | 8.10E-01 | 1.46E+00 | 1.56E-01 | 5.56E-01 | 1.72E+00 | 3.67E-02 | 1.03E-01 | 2.35E+00 | 1.39E-01 | 5.11E-01 | 1.03E+00 | 6.93E-02 | 1.90E-01 | 1.60E+00 |
| IDI1 | 4.37E-03 | 2.28E-02 | 1.12E+00 | 8.11E-05 | 3.51E-04 | 1.15E+00 | 1.84E-02 | 7.80E-02 | 1.03E+00 | 5.54E-02 | 2.12E-01 | 1.30E+00 | 2.03E-02 | 7.25E-02 | 1.36E+00 | 3.25E-01 | 9.16E-01 | 1.70E+00 | 4.93E-02 | 1.81E-01 | -1.00E+00 | 1.85E-01 | 5.09E-01 | 1.28E+00 |
| NPIPB2 | 4.37E-03 | 2.28E-02 | -1.21E+00 | 6.16E-03 | 2.66E-02 | -1.11E+00 | 3.13E-02 | 1.33E-01 | -1.14E+00 | 6.20E-02 | 2.37E-01 | 1.06E+00 | 6.63E-02 | 2.36E-01 | -1.01E+00 | 7.24E-02 | 2.04E-01 | -1.11E+00 | 1.11E-01 | 4.09E-01 | 1.19E+00 | 4.95E-03 | 1.36E-02 | -1.39E+00 |
| MRPS11 | 4.37E-03 | 2.28E-02 | -1.15E+00 | 4.23E-02 | 1.83E-01 | 1.16E+00 | 8.78E-03 | 3.73E-02 | -1.24E+00 | 1.28E-01 | 4.89E-01 | 1.27E+00 | 5.15E-03 | 1.84E-02 | -1.13E+00 | 1.41E-02 | 3.97E-02 | -1.10E+00 | 3.93E-05 | 1.44E-04 | -1.38E+00 | 1.18E-01 | 3.24E-01 | 1.12E+00 |
| AC006538.1 | 4.37E-03 | 2.28E-02 | -1.56E+00 | 1.61E-02 | 6.98E-02 | -1.28E+00 | 1.68E-04 | 7.16E-04 | -1.63E+00 | 1.26E-03 | 4.80E-03 | -1.33E+00 | 1.61E-02 | 5.76E-02 | -1.56E+00 | 1.09E-01 | 3.08E-01 | -1.05E+00 | 2.83E-04 | 1.04E-03 | -1.70E+00 | 4.09E-03 | 1.12E-02 | -1.38E+00 |
| GOLGA1 | 4.38E-03 | 2.28E-02 | -1.01E+00 | 5.57E-04 | 2.40E-03 | -1.15E+00 | 7.26E-02 | 3.08E-01 | -1.01E+00 | 7.02E-03 | 2.68E-02 | -1.09E+00 | 2.15E-02 | 7.67E-02 | -1.31E+00 | 2.74E-02 | 7.73E-02 | -1.28E+00 | 2.64E-03 | 9.68E-03 | -1.04E+00 | 5.67E-03 | 1.56E-02 | -1.19E+00 |
| AGPAT4-IT1 | 4.39E-03 | 2.29E-02 | -1.17E+00 | 1.90E-01 | 8.19E-01 | 1.34E+00 | 8.39E-03 | 3.56E-02 | -1.25E+00 | 1.55E-01 | 5.94E-01 | 1.22E+00 | 5.60E-02 | 2.00E-01 | 1.12E+00 | 1.67E-01 | 4.71E-01 | 1.88E+00 | 2.12E-02 | 7.77E-02 | -1.15E+00 | 2.82E-01 | 7.75E-01 | 1.40E+00 |
| ZNF444 | 4.39E-03 | 2.29E-02 | -1.02E+00 | 2.56E-02 | 1.11E-01 | 1.22E+00 | 3.67E-03 | 1.56E-02 | -1.26E+00 | 6.38E-03 | 2.44E-02 | -1.06E+00 | 8.11E-08 | 2.89E-07 | -1.76E+00 | 2.77E-05 | 7.82E-05 | -1.33E+00 | 3.39E-05 | 1.24E-04 | -1.29E+00 | 2.34E-04 | 6.44E-04 | -1.16E+00 |
| COASY | 4.41E-03 | 2.30E-02 | 1.02E+00 | 1.19E-02 | 5.14E-02 | 1.18E+00 | 4.97E-03 | 2.11E-02 | -1.05E+00 | 2.76E-03 | 1.06E-02 | 1.06E+00 | 2.58E-03 | 9.19E-03 | -1.09E+00 | 7.83E-02 | 2.21E-01 | 1.19E+00 | 9.14E-03 | 3.35E-02 | 1.08E+00 | 1.72E-01 | 4.73E-01 | 1.30E+00 |
| NSUN3 | 4.41E-03 | 2.30E-02 | -1.01E+00 | 3.39E-04 | 1.46E-03 | -1.02E+00 | 6.14E-04 | 2.61E-03 | -1.27E+00 | 1.48E-05 | 5.64E-05 | -1.33E+00 | 3.05E-02 | 1.09E-01 | 1.13E+00 | 2.04E-02 | 5.75E-02 | -1.05E+00 | 4.17E-04 | 1.53E-03 | -1.26E+00 | 7.00E-04 | 1.92E-03 | -1.26E+00 |
| TMEM217 | 4.41E-03 | 2.30E-02 | -1.32E+00 | 1.49E-03 | 6.46E-03 | -1.32E+00 | 7.34E-06 | 3.12E-05 | -1.55E+00 | 1.43E-05 | 5.47E-05 | -1.28E+00 | 3.22E-02 | 1.15E-01 | -1.46E+00 | 6.94E-02 | 1.96E-01 | -1.38E+00 | 1.67E-01 | 6.13E-01 | -1.02E+00 | 1.67E-01 | 4.59E-01 | -1.14E+00 |
| MMP25-AS1 | 4.42E-03 | 2.31E-02 | -1.36E+00 | 1.17E-01 | 5.06E-01 | 1.24E+00 | 7.28E-03 | 3.09E-02 | -1.25E+00 | 4.66E-02 | 1.78E-01 | 1.05E+00 | 2.75E-01 | 9.82E-01 | 1.38E+00 | 2.96E-01 | 8.36E-01 | 1.28E+00 | 2.32E-01 | 8.52E-01 | 1.34E+00 | 1.56E-01 | 4.29E-01 | 1.74E+00 |
| ELAC1 | 4.43E-03 | 2.31E-02 | -1.01E+00 | 7.75E-03 | 3.35E-02 | 1.08E+00 | 6.17E-05 | 2.62E-04 | -1.22E+00 | 1.15E-04 | 4.41E-04 | -1.06E+00 | 7.10E-03 | 2.53E-02 | -1.24E+00 | 1.28E-01 | 3.60E-01 | 1.07E+00 | 1.47E-04 | 5.40E-04 | -1.16E+00 | 1.43E-01 | 3.92E-01 | 1.18E+00 |
| AC096677.1 | 4.43E-03 | 2.31E-02 | -1.39E+00 | 2.11E-01 | 9.13E-01 | 1.33E+00 | 5.00E-04 | 2.12E-03 | -1.51E+00 | 6.02E-03 | 2.30E-02 | -1.12E+00 | 2.22E-02 | 7.91E-02 | -1.49E+00 | 1.10E-01 | 3.11E-01 | -1.02E+00 | 2.91E-04 | 1.07E-03 | -1.66E+00 | 5.57E-03 | 1.53E-02 | -1.29E+00 |
| ARMCX5-GPRASP2 | 4.43E-03 | 2.31E-02 | -1.45E+00 | 1.42E-02 | 6.15E-02 | -1.26E+00 | 6.84E-02 | 2.91E-01 | -1.15E+00 | 2.36E-02 | 9.03E-02 | -1.25E+00 | 2.29E-01 | 8.17E-01 | -1.03E+00 | 2.76E-01 | 7.78E-01 | 1.04E+00 | 1.58E-03 | 5.80E-03 | -1.49E+00 | 1.38E-02 | 3.80E-02 | -1.30E+00 |
| TSC22D1 | 4.44E-03 | 2.31E-02 | 1.82E+00 | 3.17E-04 | 1.37E-03 | 1.87E+00 | 2.17E-01 | 9.24E-01 | 1.37E+00 | 7.36E-02 | 2.81E-01 | 1.26E+00 | 1.84E-01 | 6.55E-01 | 1.67E+00 | 2.48E-05 | 7.00E-05 | 2.50E+00 | 1.50E-07 | 5.50E-07 | -1.33E+00 | 1.63E-01 | 4.47E-01 | 1.06E+00 |
| SMIM11A | 4.45E-03 | 2.32E-02 | 1.07E+00 | 2.57E-02 | 1.11E-01 | 1.33E+00 | 2.31E-01 | 9.83E-01 | 1.21E+00 | 2.09E-01 | 7.97E-01 | 1.29E+00 | 1.21E-01 | 4.33E-01 | 1.24E+00 | 2.18E-01 | 6.16E-01 | 1.48E+00 | 2.19E-01 | 8.02E-01 | 1.38E+00 | 3.03E-01 | 8.33E-01 | 1.33E+00 |
| KIF16B | 4.45E-03 | 2.32E-02 | -1.24E+00 | 1.39E-01 | 6.00E-01 | 1.30E+00 | 3.13E-04 | 1.33E-03 | -1.38E+00 | 2.21E-03 | 8.43E-03 | -1.09E+00 | 2.07E-01 | 7.36E-01 | 1.11E+00 | 1.36E-01 | 3.85E-01 | 1.77E+00 | 6.92E-03 | 2.54E-02 | -1.12E+00 | 2.39E-01 | 6.57E-01 | 1.12E+00 |
| ELK1 | 4.48E-03 | 2.33E-02 | -1.04E+00 | 4.29E-04 | 1.85E-03 | -1.05E+00 | 1.08E-02 | 4.58E-02 | -1.09E+00 | 3.11E-03 | 1.19E-02 | 1.02E+00 | 2.08E-02 | 7.43E-02 | -1.17E+00 | 2.91E-01 | 8.20E-01 | 1.50E+00 | 3.13E-05 | 1.15E-04 | -1.13E+00 | 1.07E-05 | 2.94E-05 | -1.20E+00 |
| MAP9 | 4.48E-03 | 2.33E-02 | -1.04E+00 | 2.74E-06 | 1.19E-05 | -1.37E+00 | 1.82E-02 | 7.71E-02 | -1.01E+00 | 1.46E-05 | 5.58E-05 | -1.38E+00 | 1.17E-01 | 4.17E-01 | 1.60E+00 | 1.96E-01 | 5.53E-01 | 1.16E+00 | 3.60E-03 | 1.32E-02 | 2.02E+00 | 2.64E-02 | 7.24E-02 | -1.16E+00 |
| MAPK8 | 4.48E-03 | 2.34E-02 | 1.09E+00 | 1.44E-03 | 6.22E-03 | 1.15E+00 | 3.68E-04 | 1.56E-03 | -1.05E+00 | 1.65E-02 | 6.31E-02 | 1.17E+00 | 4.59E-04 | 1.64E-03 | -1.08E+00 | 1.13E-04 | 3.19E-04 | -1.14E+00 | 1.24E-05 | 4.53E-05 | -1.29E+00 | 1.41E-03 | 3.86E-03 | -1.02E+00 |
| ARIH1 | 4.50E-03 | 2.34E-02 | 1.06E+00 | 4.42E-04 | 1.91E-03 | 1.15E+00 | 1.47E-01 | 6.23E-01 | 1.09E+00 | 3.10E-02 | 1.18E-01 | 1.03E+00 | 1.14E-02 | 4.06E-02 | 1.03E+00 | 1.79E-02 | 5.03E-02 | -1.08E+00 | 5.43E-04 | 1.99E-03 | -1.13E+00 | 1.49E-02 | 4.10E-02 | -1.05E+00 |
| WDR76 | 4.50E-03 | 2.34E-02 | -1.36E+00 | 1.14E-02 | 4.92E-02 | -1.14E+00 | 1.97E-02 | 8.38E-02 | 1.25E+00 | 6.37E-02 | 2.43E-01 | 1.39E+00 | 8.48E-02 | 3.02E-01 | -1.24E+00 | 3.38E-01 | 9.54E-01 | 1.36E+00 | 3.84E-04 | 1.41E-03 | -1.25E+00 | 2.69E-01 | 7.38E-01 | 1.19E+00 |
| TRIT1 | 4.50E-03 | 2.35E-02 | -1.13E+00 | 3.24E-02 | 1.40E-01 | 1.25E+00 | 2.30E-07 | 9.77E-07 | -1.48E+00 | 1.96E-04 | 7.47E-04 | -1.02E+00 | 4.23E-04 | 1.51E-03 | -1.25E+00 | 4.62E-02 | 1.30E-01 | 1.09E+00 | 1.81E-02 | 6.64E-02 | -1.06E+00 | 2.61E-01 | 7.16E-01 | 1.25E+00 |
| ARHGAP20 | 4.50E-03 | 2.35E-02 | -1.15E+00 | 1.97E-04 | 8.50E-04 | -1.27E+00 | 9.14E-02 | 3.88E-01 | 1.46E+00 | 1.75E-01 | 6.68E-01 | -1.15E+00 | 2.59E-01 | 9.22E-01 | 1.14E+00 | 1.52E-01 | 4.29E-01 | -1.83E+00 | 1.92E-01 | 7.05E-01 | 1.26E+00 | 3.31E-01 | 9.08E-01 | 1.09E+00 |
| CAPN15 | 4.51E-03 | 2.35E-02 | -1.15E+00 | 1.38E-01 | 5.95E-01 | 1.35E+00 | 2.08E-01 | 8.83E-01 | 1.12E+00 | 4.61E-02 | 1.76E-01 | 1.66E+00 | 8.68E-03 | 3.09E-02 | -1.31E+00 | 3.55E-01 | 1.00E+00 | 1.51E+00 | 8.67E-03 | 3.18E-02 | -1.14E+00 | 6.25E-02 | 1.72E-01 | 1.45E+00 |
| TRAK2 | 4.51E-03 | 2.35E-02 | -1.03E+00 | 9.40E-02 | 4.06E-01 | 1.33E+00 | 8.83E-03 | 3.75E-02 | -1.15E+00 | 6.00E-02 | 2.29E-01 | 1.19E+00 | 2.50E-03 | 8.93E-03 | -1.20E+00 | 1.84E-01 | 5.20E-01 | 1.28E+00 | 1.08E-16 | 3.98E-16 | -1.79E+00 | 1.95E-13 | 5.35E-13 | -1.64E+00 |
| CCDC163 | 4.52E-03 | 2.36E-02 | -1.66E+00 | 1.11E-01 | 4.81E-01 | 1.10E+00 | 7.41E-05 | 3.15E-04 | -2.05E+00 | 3.51E-03 | 1.34E-02 | -1.35E+00 | 4.76E-03 | 1.70E-02 | -1.79E+00 | 3.65E-02 | 1.03E-01 | -1.29E+00 | 4.19E-03 | 1.54E-02 | -1.25E+00 | 2.11E-02 | 5.80E-02 | -1.03E+00 |
| KIF7 | 4.53E-03 | 2.36E-02 | -1.21E+00 | 1.91E-02 | 8.27E-02 | -1.01E+00 | 1.85E-01 | 7.84E-01 | 1.23E+00 | 2.20E-01 | 8.42E-01 | 1.52E+00 | 7.56E-02 | 2.70E-01 | -1.29E+00 | 2.30E-01 | 6.49E-01 | 1.14E+00 | 2.33E-01 | 8.54E-01 | 1.06E+00 | 2.47E-01 | 6.78E-01 | -1.02E+00 |
| BCLAF1 | 4.53E-03 | 2.36E-02 | 1.12E+00 | 4.14E-12 | 1.79E-11 | -1.05E+00 | 5.34E-02 | 2.27E-01 | 1.32E+00 | 1.30E-01 | 4.98E-01 | 1.24E+00 | 2.23E-01 | 7.96E-01 | 1.27E+00 | 2.11E-02 | 5.95E-02 | 1.04E+00 | 5.13E-02 | 1.88E-01 | 1.11E+00 | 5.40E-02 | 1.48E-01 | 1.10E+00 |
| TSPAN10 | 4.53E-03 | 2.36E-02 | -1.36E+00 | 2.74E-02 | 1.18E-01 | -1.03E+00 | 7.68E-03 | 3.26E-02 | -1.33E+00 | 4.65E-02 | 1.78E-01 | 1.01E+00 | 1.00E-01 | 3.58E-01 | 1.14E+00 | 2.39E-01 | 6.74E-01 | 1.34E+00 | 5.73E-03 | 2.10E-02 | -1.46E+00 | 7.44E-02 | 2.04E-01 | -1.18E+00 |
| SLC22A15 | 4.54E-03 | 2.36E-02 | -1.31E+00 | 2.75E-03 | 1.19E-02 | -1.36E+00 | 1.81E-03 | 7.70E-03 | -1.34E+00 | 4.61E-04 | 1.76E-03 | -1.29E+00 | 4.27E-02 | 1.52E-01 | 1.01E+00 | 4.28E-03 | 1.21E-02 | -1.46E+00 | 7.84E-04 | 2.88E-03 | -1.40E+00 | 4.91E-05 | 1.35E-04 | -1.94E+00 |
| GLI4 | 4.59E-03 | 2.39E-02 | -1.04E+00 | 2.92E-02 | 1.26E-01 | 1.15E+00 | 1.04E-01 | 4.40E-01 | 1.15E+00 | 3.03E-02 | 1.16E-01 | 1.14E+00 | 1.94E-01 | 6.91E-01 | 1.16E+00 | 2.38E-01 | 6.72E-01 | 1.52E+00 | 7.66E-02 | 2.81E-01 | 1.16E+00 | 4.77E-02 | 1.31E-01 | 1.05E+00 |
| LDLRAD3 | 4.59E-03 | 2.39E-02 | 2.20E+00 | 4.98E-02 | 2.15E-01 | 1.93E+00 | 3.03E-03 | 1.29E-02 | 2.08E+00 | 1.89E-02 | 7.20E-02 | 2.12E+00 | 2.58E-04 | 9.21E-04 | -1.74E+00 | 1.28E-02 | 3.60E-02 | -1.19E+00 | 2.93E-10 | 1.08E-09 | -1.43E+00 | 3.89E-04 | 1.07E-03 | -1.05E+00 |
| NEURL1B | 4.61E-03 | 2.40E-02 | -1.22E+00 | 5.26E-02 | 2.27E-01 | 1.69E+00 | 1.07E-10 | 4.56E-10 | -2.12E+00 | 2.32E-02 | 8.86E-02 | 1.09E+00 | 2.02E-04 | 7.20E-04 | -1.75E+00 | 1.18E-01 | 3.32E-01 | 1.23E+00 | 7.00E-06 | 2.57E-05 | -1.71E+00 | 1.87E-04 | 5.14E-04 | -1.39E+00 |
| AC095055.1 | 4.62E-03 | 2.41E-02 | 1.94E+00 | 8.46E-02 | 3.65E-01 | 1.14E+00 | 5.21E-03 | 2.21E-02 | 1.92E+00 | 2.13E-01 | 8.14E-01 | 1.36E+00 | 1.76E-01 | 6.26E-01 | 1.62E+00 | 2.54E-01 | 7.15E-01 | 1.57E+00 | 2.13E-03 | 7.82E-03 | 1.83E+00 | 1.20E-01 | 3.29E-01 | -1.10E+00 |
| BLZF1 | 4.62E-03 | 2.41E-02 | -1.02E+00 | 1.08E-01 | 4.68E-01 | 1.28E+00 | 1.94E-06 | 8.23E-06 | -1.33E+00 | 5.24E-05 | 2.00E-04 | -1.08E+00 | 8.59E-03 | 3.06E-02 | -1.03E+00 | 1.46E-01 | 4.12E-01 | 1.15E+00 | 4.69E-07 | 1.72E-06 | -1.19E+00 | 4.29E-07 | 1.18E-06 | -1.26E+00 |
| ZNF606 | 4.63E-03 | 2.41E-02 | 1.02E+00 | 1.43E-01 | 6.18E-01 | 1.67E+00 | 1.30E-04 | 5.51E-04 | -1.48E+00 | 2.29E-01 | 8.73E-01 | 1.33E+00 | 1.45E-03 | 5.16E-03 | -1.04E+00 | 2.37E-01 | 6.68E-01 | 1.39E+00 | 1.11E-03 | 4.09E-03 | -1.20E+00 | 2.53E-01 | 6.95E-01 | 1.23E+00 |
| ABRACL | 4.64E-03 | 2.42E-02 | 1.12E+00 | 1.20E-05 | 5.20E-05 | 1.06E+00 | 3.54E-02 | 1.50E-01 | -1.05E+00 | 1.92E-02 | 7.34E-02 | -1.03E+00 | 9.36E-11 | 3.34E-10 | -1.39E+00 | 4.47E-08 | 1.26E-07 | -1.21E+00 | 9.20E-05 | 3.37E-04 | -1.25E+00 | 1.99E-02 | 5.46E-02 | -1.12E+00 |
| COBLL1 | 4.64E-03 | 2.42E-02 | -1.14E+00 | 8.04E-03 | 3.48E-02 | 1.01E+00 | 9.59E-07 | 4.07E-06 | -1.45E+00 | 1.22E-06 | 4.64E-06 | -1.25E+00 | 1.42E-01 | 5.07E-01 | -1.97E+00 | 2.96E-01 | 8.35E-01 | -1.18E+00 | 2.92E-02 | 1.07E-01 | -1.49E+00 | 1.34E-01 | 3.69E-01 | -1.24E+00 |
| MSH5 | 4.64E-03 | 2.42E-02 | -1.23E+00 | 6.42E-03 | 2.77E-02 | -1.07E+00 | 1.86E-02 | 7.91E-02 | -1.10E+00 | 5.95E-03 | 2.27E-02 | -1.11E+00 | 5.21E-04 | 1.86E-03 | -1.61E+00 | 2.07E-04 | 5.82E-04 | -1.76E+00 | 9.75E-02 | 3.58E-01 | -1.01E+00 | 4.24E-02 | 1.17E-01 | -1.09E+00 |
| DEPDC5 | 4.65E-03 | 2.42E-02 | -1.20E+00 | 3.90E-03 | 1.69E-02 | -1.14E+00 | 1.29E-02 | 5.50E-02 | -1.20E+00 | 7.04E-02 | 2.69E-01 | 1.13E+00 | 1.48E-03 | 5.29E-03 | -1.45E+00 | 4.69E-03 | 1.32E-02 | -1.30E+00 | 9.40E-03 | 3.45E-02 | -1.11E+00 | 1.83E-01 | 5.03E-01 | 1.19E+00 |
| MLF1 | 4.65E-03 | 2.42E-02 | -1.09E+00 | 2.56E-04 | 1.11E-03 | -1.25E+00 | 1.30E-01 | 5.53E-01 | 1.14E+00 | 1.89E-01 | 7.24E-01 | 1.30E+00 | 1.61E-01 | 5.74E-01 | 1.59E+00 | 3.00E-01 | 8.46E-01 | 1.37E+00 | 3.68E-02 | 1.35E-01 | 1.64E+00 | 2.30E-01 | 6.33E-01 | 1.42E+00 |
| LINC01637 | 4.66E-03 | 2.43E-02 | -1.15E+00 | 1.60E-03 | 6.92E-03 | -1.24E+00 | 6.57E-02 | 2.79E-01 | -1.02E+00 | 1.54E-01 | 5.90E-01 | 1.24E+00 | 2.89E-02 | 1.03E-01 | 1.08E+00 | 1.57E-02 | 4.42E-02 | 1.02E+00 | 2.05E-01 | 7.53E-01 | 1.34E+00 | 2.33E-01 | 6.40E-01 | 1.14E+00 |
| ZNF557 | 4.66E-03 | 2.43E-02 | -1.07E+00 | 7.47E-04 | 3.23E-03 | -1.04E+00 | 1.70E-06 | 7.22E-06 | -1.63E+00 | 1.27E-05 | 4.87E-05 | -1.38E+00 | 3.99E-10 | 1.42E-09 | -1.76E+00 | 7.13E-13 | 2.01E-12 | -2.04E+00 | 3.52E-04 | 1.29E-03 | -1.29E+00 | 3.78E-03 | 1.04E-02 | -1.18E+00 |
| GIMAP7 | 4.66E-03 | 2.43E-02 | 1.64E+00 | 3.33E-39 | 1.44E-38 | 3.95E+00 | 7.85E-02 | 3.34E-01 | -1.03E+00 | 5.80E-03 | 2.22E-02 | 1.81E+00 | 2.47E-01 | 8.79E-01 | 1.22E+00 | 2.82E-01 | 7.94E-01 | 1.64E+00 | 9.18E-03 | 3.37E-02 | -2.05E+00 | 2.89E-02 | 7.94E-02 | -1.64E+00 |
| PATL2 | 4.67E-03 | 2.43E-02 | -1.12E+00 | 3.39E-03 | 1.47E-02 | 1.03E+00 | 1.23E-02 | 5.23E-02 | -1.11E+00 | 1.47E-03 | 5.61E-03 | -1.19E+00 | 5.54E-04 | 1.98E-03 | -2.18E+00 | 8.76E-03 | 2.47E-02 | -1.53E+00 | 2.27E-03 | 8.34E-03 | -1.58E+00 | 1.38E-02 | 3.79E-02 | -1.42E+00 |
| RNF40 | 4.68E-03 | 2.44E-02 | -1.20E+00 | 6.38E-02 | 2.76E-01 | 1.25E+00 | 1.05E-03 | 4.44E-03 | -1.29E+00 | 4.88E-02 | 1.86E-01 | 1.18E+00 | 4.02E-02 | 1.43E-01 | -1.29E+00 | 2.29E-01 | 6.47E-01 | 1.11E+00 | 9.07E-03 | 3.33E-02 | -1.18E+00 | 1.80E-01 | 4.95E-01 | 1.10E+00 |
| XKR6 | 4.68E-03 | 2.44E-02 | -1.06E+00 | 1.41E-03 | 6.09E-03 | -1.08E+00 | 1.24E-03 | 5.28E-03 | -1.33E+00 | 1.07E-03 | 4.07E-03 | -1.12E+00 | 1.60E-01 | 5.70E-01 | 1.69E+00 | 3.50E-01 | 9.86E-01 | 1.52E+00 | 6.88E-02 | 2.53E-01 | 1.54E+00 | 1.06E-01 | 2.92E-01 | 1.07E+00 |
| CFAP69 | 4.69E-03 | 2.44E-02 | -1.37E+00 | 4.19E-02 | 1.81E-01 | -1.01E+00 | 1.70E-04 | 7.24E-04 | -1.68E+00 | 3.68E-04 | 1.41E-03 | -1.42E+00 | 1.08E-01 | 3.86E-01 | -1.13E+00 | 2.41E-01 | 6.79E-01 | 1.05E+00 | 2.34E-01 | 8.58E-01 | -1.12E+00 | 2.70E-01 | 7.42E-01 | -1.13E+00 |
| RPS10P16 | 4.69E-03 | 2.44E-02 | 2.63E+00 | 2.07E-02 | 8.92E-02 | 2.43E+00 | 7.98E-02 | 3.39E-01 | 1.01E+00 | 1.82E-01 | 6.94E-01 | 1.31E+00 | 2.65E-01 | 9.43E-01 | 1.15E+00 | 2.41E-01 | 6.79E-01 | 1.43E+00 | 6.73E-02 | 2.47E-01 | 1.59E+00 | 8.87E-02 | 2.44E-01 | 1.73E+00 |
| KIAA1522 | 4.69E-03 | 2.44E-02 | -1.09E+00 | 4.13E-02 | 1.78E-01 | 1.25E+00 | 1.01E-01 | 4.29E-01 | 1.13E+00 | 1.46E-01 | 5.56E-01 | 1.32E+00 | 2.93E-03 | 1.04E-02 | -1.14E+00 | 4.66E-03 | 1.31E-02 | -1.15E+00 | 6.86E-03 | 2.52E-02 | -1.10E+00 | 3.99E-02 | 1.10E-01 | -1.05E+00 |
| UGP2 | 4.69E-03 | 2.44E-02 | 1.18E+00 | 4.73E-03 | 2.04E-02 | 1.26E+00 | 1.08E-04 | 4.60E-04 | -1.13E+00 | 1.60E-02 | 6.09E-02 | 1.12E+00 | 6.73E-06 | 2.40E-05 | -1.06E+00 | 1.57E-02 | 4.42E-02 | 1.16E+00 | 2.34E-01 | 8.60E-01 | 1.12E+00 | 2.04E-01 | 5.61E-01 | 1.21E+00 |
| AC006548.3 | 4.70E-03 | 2.45E-02 | -1.41E+00 | 3.10E-02 | 1.34E-01 | -1.02E+00 | 6.97E-04 | 2.96E-03 | -1.51E+00 | 1.59E-02 | 6.08E-02 | -1.09E+00 | 7.69E-02 | 2.74E-01 | -1.15E+00 | 1.62E-01 | 4.56E-01 | 1.06E+00 | 6.11E-02 | 2.24E-01 | -1.14E+00 | 2.48E-01 | 6.81E-01 | 1.16E+00 |
| TIMM8A | 4.70E-03 | 2.45E-02 | -1.14E+00 | 5.41E-02 | 2.34E-01 | 1.13E+00 | 2.60E-03 | 1.10E-02 | -1.13E+00 | 1.64E-03 | 6.26E-03 | 1.02E+00 | 7.90E-03 | 2.82E-02 | -1.26E+00 | 6.99E-02 | 1.97E-01 | 1.03E+00 | 8.45E-03 | 3.10E-02 | -1.05E+00 | 3.50E-01 | 9.61E-01 | 1.27E+00 |
| GOLT1B | 4.72E-03 | 2.46E-02 | -1.13E+00 | 2.73E-03 | 1.18E-02 | -1.10E+00 | 9.04E-03 | 3.84E-02 | -1.12E+00 | 1.69E-03 | 6.47E-03 | -1.03E+00 | 1.36E-02 | 4.86E-02 | -1.00E+00 | 2.47E-03 | 6.97E-03 | -1.16E+00 | 9.25E-06 | 3.39E-05 | -1.21E+00 | 9.98E-05 | 2.74E-04 | -1.26E+00 |
| GIT1 | 4.72E-03 | 2.46E-02 | -1.10E+00 | 1.86E-01 | 8.03E-01 | 1.38E+00 | 8.54E-02 | 3.63E-01 | 1.14E+00 | 2.36E-01 | 9.02E-01 | 1.49E+00 | 1.14E-02 | 4.07E-02 | -1.13E+00 | 1.21E-01 | 3.41E-01 | 1.13E+00 | 3.64E-03 | 1.33E-02 | 1.00E+00 | 1.25E-02 | 3.43E-02 | 1.07E+00 |
| RHBDD3 | 4.73E-03 | 2.47E-02 | -1.13E+00 | 2.04E-02 | 8.83E-02 | 1.05E+00 | 3.89E-04 | 1.65E-03 | -1.10E+00 | 1.83E-02 | 6.99E-02 | 1.17E+00 | 1.11E-05 | 3.94E-05 | -1.46E+00 | 1.11E-04 | 3.13E-04 | -1.29E+00 | 1.35E-03 | 4.97E-03 | -1.03E+00 | 2.24E-01 | 6.14E-01 | 1.19E+00 |
| AIF1L | 4.74E-03 | 2.47E-02 | -1.39E+00 | 1.16E-02 | 5.00E-02 | -1.23E+00 | 1.91E-01 | 8.12E-01 | 1.25E+00 | 5.66E-02 | 2.16E-01 | 1.01E+00 | 2.31E-01 | 8.22E-01 | -1.22E+00 | 2.88E-01 | 8.13E-01 | -1.23E+00 | 1.35E-01 | 4.96E-01 | -1.23E+00 | 3.45E-01 | 9.48E-01 | 1.11E+00 |
| SNX8 | 4.74E-03 | 2.47E-02 | -1.45E+00 | 4.07E-03 | 1.76E-02 | -1.32E+00 | 8.12E-04 | 3.45E-03 | -1.35E+00 | 4.74E-04 | 1.81E-03 | -1.22E+00 | 1.56E-02 | 5.55E-02 | -1.24E+00 | 5.51E-02 | 1.55E-01 | -1.09E+00 | 2.81E-04 | 1.03E-03 | -1.03E+00 | 3.35E-05 | 9.21E-05 | -1.12E+00 |
| KIAA1468 | 4.74E-03 | 2.47E-02 | -1.03E+00 | 2.20E-02 | 9.49E-02 | 1.21E+00 | 2.79E-02 | 1.18E-01 | -1.08E+00 | 5.06E-03 | 1.93E-02 | -1.13E+00 | 5.17E-06 | 1.84E-05 | -1.43E+00 | 1.19E-04 | 3.36E-04 | -1.26E+00 | 3.56E-04 | 1.31E-03 | -1.13E+00 | 2.05E-03 | 5.62E-03 | -1.25E+00 |
| AC137499.1 | 4.74E-03 | 2.47E-02 | -1.46E+00 | 1.14E-02 | 4.95E-02 | -1.19E+00 | 1.60E-05 | 6.78E-05 | -1.61E+00 | 1.34E-05 | 5.11E-05 | -1.46E+00 | 1.63E-02 | 5.80E-02 | -1.44E+00 | 2.96E-01 | 8.35E-01 | 1.27E+00 | 2.75E-18 | 1.01E-17 | -1.98E+00 | 2.54E-15 | 6.97E-15 | -1.69E+00 |
| DOK1 | 4.75E-03 | 2.48E-02 | 1.07E+00 | 3.19E-02 | 1.38E-01 | 1.28E+00 | 1.66E-04 | 7.05E-04 | -1.18E+00 | 6.89E-04 | 2.63E-03 | -1.02E+00 | 3.04E-08 | 1.08E-07 | -1.66E+00 | 1.24E-03 | 3.51E-03 | -1.03E+00 | 2.74E-09 | 1.01E-08 | -1.36E+00 | 4.78E-06 | 1.31E-05 | -1.17E+00 |
| SWSAP1 | 4.77E-03 | 2.49E-02 | 1.00E+00 | 7.49E-02 | 3.24E-01 | 1.37E+00 | 2.05E-06 | 8.69E-06 | -1.51E+00 | 1.44E-03 | 5.52E-03 | -1.01E+00 | 1.80E-01 | 6.42E-01 | 1.30E+00 | 1.93E-01 | 5.43E-01 | 1.16E+00 | 5.71E-02 | 2.10E-01 | -1.12E+00 | 3.60E-01 | 9.89E-01 | 1.18E+00 |
| SCPEP1 | 4.77E-03 | 2.49E-02 | -2.68E+00 | 3.35E-02 | 1.45E-01 | -1.45E+00 | 7.82E-03 | 3.32E-02 | -1.65E+00 | 7.63E-03 | 2.91E-02 | -1.99E+00 | 6.02E-02 | 2.15E-01 | -1.33E+00 | 1.97E-01 | 5.56E-01 | 1.12E+00 | 2.82E-10 | 1.03E-09 | -1.30E+00 | 6.59E-13 | 1.81E-12 | -1.46E+00 |
| CD38 | 4.78E-03 | 2.49E-02 | 1.52E+00 | 2.28E-01 | 9.84E-01 | 1.43E+00 | 1.82E-01 | 7.73E-01 | 1.10E+00 | 1.93E-01 | 7.37E-01 | 1.30E+00 | 2.20E-04 | 7.86E-04 | 1.91E+00 | 2.70E-04 | 7.62E-04 | 1.94E+00 | 2.68E-02 | 9.82E-02 | 1.06E+00 | 3.80E-03 | 1.04E-02 | 1.02E+00 |
| AC007038.1 | 4.79E-03 | 2.50E-02 | -1.71E+00 | 1.87E-01 | 8.07E-01 | 1.35E+00 | 3.81E-02 | 1.62E-01 | -1.14E+00 | 2.42E-01 | 9.23E-01 | 1.38E+00 | 1.33E-01 | 4.75E-01 | 1.09E+00 | 1.71E-01 | 4.82E-01 | 1.71E+00 | 1.34E-01 | 4.93E-01 | -1.02E+00 | 3.04E-01 | 8.34E-01 | 1.29E+00 |
| PLXNB2 | 4.79E-03 | 2.50E-02 | -1.16E+00 | 2.28E-01 | 9.85E-01 | 1.46E+00 | 3.06E-03 | 1.30E-02 | -1.20E+00 | 6.00E-04 | 2.29E-03 | -1.13E+00 | 2.23E-01 | 7.95E-01 | 1.13E+00 | 3.53E-01 | 9.96E-01 | 1.47E+00 | 1.26E-01 | 4.64E-01 | 1.04E+00 | 1.50E-01 | 4.13E-01 | -1.04E+00 |
| HOMER1 | 4.79E-03 | 2.50E-02 | -1.05E+00 | 4.84E-02 | 2.09E-01 | 1.28E+00 | 1.98E-03 | 8.42E-03 | -1.29E+00 | 1.07E-01 | 4.10E-01 | 1.18E+00 | 8.21E-04 | 2.93E-03 | -1.48E+00 | 2.62E-02 | 7.37E-02 | -1.04E+00 | 5.13E-08 | 1.88E-07 | -2.23E+00 | 1.98E-04 | 5.45E-04 | -1.46E+00 |
| SARS2 | 4.80E-03 | 2.50E-02 | -1.31E+00 | 4.58E-03 | 1.98E-02 | -1.28E+00 | 4.39E-04 | 1.86E-03 | -1.27E+00 | 1.98E-06 | 7.58E-06 | -1.55E+00 | 1.07E-01 | 3.82E-01 | 1.01E+00 | 9.67E-02 | 2.73E-01 | -1.11E+00 | 3.38E-05 | 1.24E-04 | -1.35E+00 | 1.15E-05 | 3.15E-05 | -1.34E+00 |
| IFNAR2 | 4.80E-03 | 2.50E-02 | -1.07E+00 | 2.21E-03 | 9.53E-03 | 1.00E+00 | 9.62E-05 | 4.09E-04 | -1.17E+00 | 2.39E-06 | 9.13E-06 | -1.14E+00 | 1.13E-02 | 4.04E-02 | -1.12E+00 | 8.06E-02 | 2.27E-01 | 1.16E+00 | 5.68E-04 | 2.08E-03 | -1.09E+00 | 1.16E-02 | 3.19E-02 | 1.03E+00 |
| SCUBE3 | 4.81E-03 | 2.50E-02 | -1.38E+00 | 8.24E-02 | 3.56E-01 | 1.10E+00 | 1.54E-02 | 6.52E-02 | -1.18E+00 | 2.13E-01 | 8.14E-01 | 1.37E+00 | 2.21E-01 | 7.86E-01 | -1.01E+00 | 3.43E-01 | 9.66E-01 | 1.23E+00 | 3.26E-03 | 1.20E-02 | -1.60E+00 | 5.34E-03 | 1.47E-02 | -1.66E+00 |
| ZNF790-AS1 | 4.81E-03 | 2.51E-02 | -1.01E+00 | 3.15E-04 | 1.36E-03 | -1.02E+00 | 2.17E-01 | 9.22E-01 | 1.19E+00 | 5.05E-02 | 1.93E-01 | 1.09E+00 | 1.43E-02 | 5.10E-02 | 1.07E+00 | 1.16E-01 | 3.26E-01 | 1.26E+00 | 1.85E-01 | 6.79E-01 | 1.26E+00 | 2.76E-01 | 7.58E-01 | 1.16E+00 |
| C11orf95 | 4.82E-03 | 2.51E-02 | -1.08E+00 | 6.12E-02 | 2.65E-01 | 1.23E+00 | 2.96E-03 | 1.26E-02 | -1.36E+00 | 7.50E-02 | 2.87E-01 | 1.14E+00 | 2.74E-06 | 9.78E-06 | -2.07E+00 | 8.71E-03 | 2.45E-02 | -1.16E+00 | 6.73E-04 | 2.47E-03 | -1.45E+00 | 5.77E-02 | 1.58E-01 | -1.05E+00 |
| PITHD1 | 4.84E-03 | 2.52E-02 | 1.13E+00 | 7.82E-04 | 3.38E-03 | 1.20E+00 | 2.21E-03 | 9.38E-03 | 1.23E+00 | 9.57E-02 | 3.66E-01 | 1.42E+00 | 9.68E-04 | 3.45E-03 | -1.03E+00 | 6.48E-02 | 1.83E-01 | 1.21E+00 | 2.27E-01 | 8.32E-01 | 1.20E+00 | 2.98E-01 | 8.18E-01 | 1.26E+00 |
| DHTKD1 | 4.85E-03 | 2.53E-02 | -1.12E+00 | 9.62E-02 | 4.16E-01 | 1.26E+00 | 4.13E-07 | 1.75E-06 | -1.41E+00 | 3.68E-05 | 1.41E-04 | -1.07E+00 | 1.82E-04 | 6.50E-04 | -1.49E+00 | 1.47E-03 | 4.14E-03 | -1.24E+00 | 4.46E-08 | 1.64E-07 | -1.72E+00 | 1.62E-04 | 4.44E-04 | -1.33E+00 |
| AHSA2 | 4.85E-03 | 2.53E-02 | 1.11E+00 | 9.66E-02 | 4.17E-01 | 1.47E+00 | 1.48E-04 | 6.28E-04 | -1.11E+00 | 1.04E-02 | 3.99E-02 | 1.28E+00 | 1.09E-02 | 3.88E-02 | 1.01E+00 | 2.99E-01 | 8.44E-01 | 1.44E+00 | 2.15E-01 | 7.89E-01 | 1.37E+00 | 1.24E-01 | 3.40E-01 | 1.49E+00 |
| SLC16A4 | 4.85E-03 | 2.53E-02 | -1.55E+00 | 7.02E-03 | 3.03E-02 | 1.73E+00 | 2.74E-02 | 1.16E-01 | -1.13E+00 | 1.98E-03 | 7.58E-03 | 2.25E+00 | 2.65E-01 | 9.44E-01 | 1.54E+00 | 2.84E-04 | 8.00E-04 | 3.43E+00 | 6.04E-03 | 2.22E-02 | -1.20E+00 | 5.89E-03 | 1.62E-02 | 1.70E+00 |
| UBE2D4 | 4.86E-03 | 2.53E-02 | -1.12E+00 | 3.90E-02 | 1.68E-01 | 1.18E+00 | 6.87E-04 | 2.92E-03 | -1.37E+00 | 1.14E-01 | 4.36E-01 | 1.14E+00 | 8.38E-03 | 2.99E-02 | 1.01E+00 | 9.13E-02 | 2.57E-01 | 1.26E+00 | 1.39E-01 | 5.10E-01 | 1.04E+00 | 2.64E-01 | 7.25E-01 | 1.33E+00 |
| IL18 | 4.87E-03 | 2.54E-02 | 1.03E+00 | 1.21E-03 | 5.22E-03 | 1.02E+00 | 3.39E-03 | 1.44E-02 | -1.13E+00 | 3.27E-04 | 1.25E-03 | -1.07E+00 | 6.34E-03 | 2.26E-02 | -1.07E+00 | 1.62E-04 | 4.57E-04 | -1.13E+00 | 3.64E-02 | 1.34E-01 | -1.00E+00 | 2.08E-02 | 5.72E-02 | -1.07E+00 |
| H3F3AP5 | 4.87E-03 | 2.54E-02 | 1.66E+00 | 2.06E-01 | 8.88E-01 | 1.47E+00 | 2.28E-03 | 9.69E-03 | 1.57E+00 | 2.03E-01 | 7.75E-01 | 1.49E+00 | 9.74E-02 | 3.47E-01 | 1.30E+00 | 9.74E-02 | 2.74E-01 | 1.25E+00 | 5.44E-04 | 2.00E-03 | 1.67E+00 | 1.26E-01 | 3.46E-01 | 1.39E+00 |
| AC009061.2 | 4.87E-03 | 2.54E-02 | -1.31E+00 | 1.93E-01 | 8.34E-01 | 1.31E+00 | 5.39E-03 | 2.29E-02 | -1.28E+00 | 4.71E-02 | 1.80E-01 | 1.12E+00 | 1.44E-01 | 5.13E-01 | 1.08E+00 | 2.99E-01 | 8.43E-01 | 1.50E+00 | 9.64E-03 | 3.54E-02 | -1.17E+00 | 1.36E-01 | 3.75E-01 | 1.06E+00 |
| ZNF624 | 4.87E-03 | 2.54E-02 | -1.31E+00 | 4.34E-02 | 1.87E-01 | 1.07E+00 | 4.23E-02 | 1.80E-01 | -1.15E+00 | 2.22E-02 | 8.46E-02 | -1.14E+00 | 1.71E-01 | 6.08E-01 | 1.29E+00 | 1.27E-01 | 3.57E-01 | 1.15E+00 | 2.25E-01 | 8.26E-01 | 1.07E+00 | 1.25E-01 | 3.45E-01 | -1.08E+00 |
| ZNF222 | 4.87E-03 | 2.54E-02 | -1.18E+00 | 1.43E-06 | 6.16E-06 | -1.97E+00 | 1.85E-03 | 7.84E-03 | -1.23E+00 | 2.56E-08 | 9.77E-08 | -2.02E+00 | 1.78E-08 | 6.34E-08 | -1.88E+00 | 1.37E-11 | 3.86E-11 | -2.85E+00 | 4.65E-02 | 1.71E-01 | -1.11E+00 | 3.49E-03 | 9.58E-03 | -1.45E+00 |
| NDC1 | 4.89E-03 | 2.55E-02 | -1.33E+00 | 5.51E-02 | 2.38E-01 | 1.18E+00 | 3.46E-07 | 1.47E-06 | -1.41E+00 | 1.72E-03 | 6.55E-03 | 1.10E+00 | 7.91E-03 | 2.82E-02 | -1.47E+00 | 6.29E-02 | 1.77E-01 | -1.06E+00 | 7.07E-06 | 2.60E-05 | -1.51E+00 | 2.93E-02 | 8.05E-02 | 1.01E+00 |
| AC127496.7 | 4.89E-03 | 2.55E-02 | 1.76E+00 | 4.10E-02 | 1.77E-01 | 1.72E+00 | 1.69E-01 | 7.17E-01 | 1.17E+00 | 2.51E-01 | 9.58E-01 | 1.36E+00 | 1.76E-01 | 6.28E-01 | 1.92E+00 | 3.04E-01 | 8.57E-01 | 1.12E+00 | 1.34E-02 | 4.93E-02 | 2.94E+00 | 1.90E-01 | 5.21E-01 | 1.75E+00 |
| ACAD10 | 4.89E-03 | 2.55E-02 | -1.16E+00 | 1.13E-01 | 4.90E-01 | 1.32E+00 | 1.77E-06 | 7.52E-06 | -1.57E+00 | 5.56E-04 | 2.13E-03 | -1.10E+00 | 9.11E-04 | 3.25E-03 | -1.42E+00 | 5.40E-03 | 1.52E-02 | -1.17E+00 | 7.33E-03 | 2.69E-02 | -1.17E+00 | 1.14E-01 | 3.13E-01 | 1.01E+00 |
| PCDHGA8 | 4.89E-03 | 2.55E-02 | -1.13E+00 | 3.12E-02 | 1.35E-01 | 1.14E+00 | 7.89E-03 | 3.35E-02 | -1.23E+00 | 6.61E-02 | 2.53E-01 | 1.10E+00 | 1.28E-01 | 4.56E-01 | 1.71E+00 | 2.34E-01 | 6.60E-01 | 1.56E+00 | 1.75E-01 | 6.40E-01 | 1.36E+00 | 1.67E-01 | 4.57E-01 | 1.57E+00 |
| LEMD2 | 4.90E-03 | 2.56E-02 | 1.00E+00 | 5.25E-03 | 2.27E-02 | 1.12E+00 | 2.72E-05 | 1.15E-04 | -1.24E+00 | 3.76E-06 | 1.44E-05 | -1.20E+00 | 7.41E-03 | 2.64E-02 | -1.18E+00 | 1.81E-03 | 5.12E-03 | -1.38E+00 | 1.39E-04 | 5.09E-04 | -1.30E+00 | 2.26E-03 | 6.20E-03 | -1.20E+00 |
| SLC35B1 | 4.91E-03 | 2.56E-02 | 1.06E+00 | 3.80E-03 | 1.64E-02 | 1.17E+00 | 8.29E-02 | 3.52E-01 | 1.36E+00 | 1.58E-01 | 6.03E-01 | 1.24E+00 | 1.57E-03 | 5.58E-03 | -1.15E+00 | 1.05E-04 | 2.97E-04 | -1.23E+00 | 1.92E-01 | 7.06E-01 | 1.11E+00 | 2.00E-01 | 5.50E-01 | 1.10E+00 |
| CCNY | 4.92E-03 | 2.56E-02 | 1.11E+00 | 1.01E-01 | 4.37E-01 | 1.42E+00 | 2.46E-02 | 1.05E-01 | 1.03E+00 | 5.86E-02 | 2.24E-01 | 1.21E+00 | 1.03E-03 | 3.67E-03 | -1.03E+00 | 2.54E-03 | 7.17E-03 | 1.00E+00 | 7.59E-03 | 2.79E-02 | 1.07E+00 | 3.93E-02 | 1.08E-01 | 1.16E+00 |
| CREB3 | 4.92E-03 | 2.56E-02 | -1.03E+00 | 4.13E-02 | 1.78E-01 | 1.24E+00 | 9.43E-02 | 4.01E-01 | 1.10E+00 | 7.23E-02 | 2.76E-01 | 1.20E+00 | 4.00E-04 | 1.43E-03 | -1.15E+00 | 1.36E-02 | 3.84E-02 | 1.05E+00 | 2.03E-01 | 7.44E-01 | 1.26E+00 | 2.95E-01 | 8.10E-01 | 1.24E+00 |
| CROT | 4.92E-03 | 2.57E-02 | -1.25E+00 | 4.26E-02 | 1.84E-01 | 1.06E+00 | 3.07E-03 | 1.31E-02 | -1.27E+00 | 2.05E-02 | 7.82E-02 | 1.03E+00 | 1.76E-04 | 6.29E-04 | -1.37E+00 | 1.44E-03 | 4.07E-03 | -1.18E+00 | 2.32E-03 | 8.52E-03 | -1.12E+00 | 1.05E-02 | 2.89E-02 | -1.06E+00 |
| AP001033.2 | 4.93E-03 | 2.57E-02 | -1.55E+00 | 5.34E-02 | 2.31E-01 | -1.01E+00 | 3.78E-02 | 1.61E-01 | -1.27E+00 | 9.17E-02 | 3.50E-01 | -1.02E+00 | 2.21E-02 | 7.89E-02 | 3.03E+00 | 5.26E-02 | 1.48E-01 | 2.76E+00 | 1.44E-03 | 5.28E-03 | -1.18E+00 | 1.03E-01 | 2.83E-01 | 1.10E+00 |
| PCBD2 | 4.93E-03 | 2.57E-02 | -1.11E+00 | 1.62E-01 | 7.01E-01 | 1.49E+00 | 1.19E-02 | 5.07E-02 | -1.04E+00 | 1.78E-02 | 6.80E-02 | 1.16E+00 | 2.09E-04 | 7.45E-04 | -1.41E+00 | 4.44E-02 | 1.25E-01 | 1.12E+00 | 7.50E-04 | 2.75E-03 | -1.15E+00 | 1.24E-03 | 3.39E-03 | -1.14E+00 |
| UCHL5 | 4.93E-03 | 2.57E-02 | 1.00E+00 | 1.67E-02 | 7.20E-02 | 1.24E+00 | 2.37E-03 | 1.01E-02 | -1.07E+00 | 4.86E-04 | 1.85E-03 | 1.04E+00 | 9.07E-06 | 3.23E-05 | -1.18E+00 | 8.81E-03 | 2.48E-02 | 1.08E+00 | 4.86E-04 | 1.78E-03 | -1.15E+00 | 6.16E-02 | 1.69E-01 | 1.04E+00 |
| AC026979.3 | 4.93E-03 | 2.57E-02 | 1.94E+00 | 5.99E-03 | 2.59E-02 | 1.15E+00 | 4.65E-03 | 1.97E-02 | 1.74E+00 | 6.61E-02 | 2.53E-01 | 1.14E+00 | 3.04E-02 | 1.08E-01 | 1.98E+00 | 1.34E-01 | 3.78E-01 | 1.08E+00 | 1.81E-01 | 6.65E-01 | 1.47E+00 | 3.66E-03 | 1.01E-02 | -1.31E+00 |
| FBXL19-AS1 | 4.94E-03 | 2.57E-02 | -1.65E+00 | 2.10E-01 | 9.07E-01 | 1.36E+00 | 1.77E-06 | 7.50E-06 | -1.95E+00 | 7.55E-07 | 2.88E-06 | -1.96E+00 | 1.35E-02 | 4.80E-02 | -1.27E+00 | 2.30E-02 | 6.49E-02 | -1.19E+00 | 6.28E-02 | 2.31E-01 | -1.12E+00 | 1.44E-02 | 3.97E-02 | -1.49E+00 |
| SUV39H2 | 4.95E-03 | 2.58E-02 | -1.09E+00 | 1.36E-02 | 5.88E-02 | 1.04E+00 | 2.69E-04 | 1.14E-03 | -1.23E+00 | 1.03E-03 | 3.94E-03 | 1.01E+00 | 2.82E-03 | 1.01E-02 | -1.51E+00 | 1.74E-01 | 4.92E-01 | 1.14E+00 | 2.60E-07 | 9.54E-07 | -1.35E+00 | 5.84E-02 | 1.60E-01 | 1.10E+00 |
| ZHX2 | 4.96E-03 | 2.59E-02 | -1.08E+00 | 1.86E-02 | 8.04E-02 | 1.12E+00 | 1.95E-02 | 8.28E-02 | -1.19E+00 | 2.29E-03 | 8.73E-03 | -1.33E+00 | 1.05E-01 | 3.74E-01 | 1.03E+00 | 2.43E-01 | 6.84E-01 | 1.15E+00 | 1.83E-08 | 6.73E-08 | -2.46E+00 | 4.20E-04 | 1.15E-03 | -1.37E+00 |
| PBX3 | 4.96E-03 | 2.59E-02 | -1.01E+00 | 7.90E-02 | 3.41E-01 | 1.34E+00 | 3.08E-02 | 1.31E-01 | -1.03E+00 | 1.53E-02 | 5.85E-02 | 1.10E+00 | 2.33E-03 | 8.31E-03 | -1.13E+00 | 2.01E-02 | 5.68E-02 | 1.01E+00 | 1.48E-04 | 5.44E-04 | -1.22E+00 | 1.89E-02 | 5.20E-02 | 1.01E+00 |
| VPS51 | 4.97E-03 | 2.59E-02 | -1.09E+00 | 2.40E-02 | 1.04E-01 | 1.14E+00 | 5.88E-03 | 2.50E-02 | -1.29E+00 | 9.72E-03 | 3.71E-02 | -1.11E+00 | 1.86E-01 | 6.64E-01 | 1.47E+00 | 1.65E-01 | 4.65E-01 | 1.66E+00 | 3.18E-02 | 1.17E-01 | -1.06E+00 | 1.60E-02 | 4.39E-02 | -1.12E+00 |
| METTL25 | 4.99E-03 | 2.60E-02 | -1.14E+00 | 4.30E-02 | 1.86E-01 | 1.18E+00 | 4.78E-03 | 2.03E-02 | -1.19E+00 | 1.37E-02 | 5.22E-02 | -1.08E+00 | 8.35E-02 | 2.98E-01 | 1.01E+00 | 1.10E-01 | 3.10E-01 | 1.09E+00 | 9.50E-06 | 3.49E-05 | -1.66E+00 | 1.02E-03 | 2.79E-03 | -1.32E+00 |
| CDKN1A | 4.99E-03 | 2.60E-02 | 2.61E+00 | 1.71E-01 | 7.40E-01 | 1.52E+00 | 6.69E-02 | 2.84E-01 | 1.40E+00 | 6.39E-02 | 2.44E-01 | -1.47E+00 | 9.00E-02 | 3.21E-01 | 2.68E+00 | 3.29E-01 | 9.29E-01 | 1.12E+00 | 9.58E-02 | 3.52E-01 | 1.89E+00 | 7.66E-02 | 2.10E-01 | -1.57E+00 |
| VILL | 4.99E-03 | 2.60E-02 | -1.15E+00 | 4.79E-02 | 2.07E-01 | 1.18E+00 | 4.76E-02 | 2.02E-01 | -1.04E+00 | 2.63E-02 | 1.00E-01 | 1.06E+00 | 3.16E-03 | 1.13E-02 | -1.40E+00 | 6.38E-02 | 1.80E-01 | -1.04E+00 | 4.48E-02 | 1.64E-01 | 1.04E+00 | 4.40E-02 | 1.21E-01 | 1.07E+00 |
| KRT18 | 5.00E-03 | 2.60E-02 | 1.12E+00 | 6.44E-02 | 2.78E-01 | 1.29E+00 | 1.85E-01 | 7.87E-01 | 1.16E+00 | 8.34E-02 | 3.18E-01 | 1.16E+00 | 3.28E-06 | 1.17E-05 | -1.47E+00 | 3.90E-10 | 1.10E-09 | -1.86E+00 | 5.38E-03 | 1.98E-02 | -1.06E+00 | 6.13E-02 | 1.68E-01 | -1.02E+00 |
| ZNF850 | 5.00E-03 | 2.61E-02 | -1.51E+00 | 4.04E-02 | 1.75E-01 | 1.00E+00 | 1.76E-01 | 7.49E-01 | -1.02E+00 | 2.24E-01 | 8.55E-01 | 1.22E+00 | 1.47E-05 | 5.23E-05 | -1.69E+00 | 3.68E-05 | 1.04E-04 | -1.50E+00 | 5.37E-08 | 1.97E-07 | -1.41E+00 | 1.03E-08 | 2.83E-08 | -1.55E+00 |
| INAFM2 | 5.00E-03 | 2.61E-02 | -1.02E+00 | 4.97E-04 | 2.15E-03 | -1.11E+00 | 1.85E-02 | 7.84E-02 | -1.09E+00 | 7.21E-04 | 2.75E-03 | -1.16E+00 | 3.63E-02 | 1.29E-01 | 1.12E+00 | 9.37E-02 | 2.64E-01 | 1.10E+00 | 2.46E-02 | 9.02E-02 | 1.30E+00 | 3.82E-04 | 1.05E-03 | -1.14E+00 |
| SIRT7 | 5.00E-03 | 2.61E-02 | -1.05E+00 | 2.98E-02 | 1.29E-01 | 1.19E+00 | 8.27E-07 | 3.51E-06 | -1.45E+00 | 4.79E-05 | 1.83E-04 | -1.14E+00 | 2.58E-03 | 9.21E-03 | 1.01E+00 | 9.61E-04 | 2.71E-03 | -1.08E+00 | 4.93E-05 | 1.81E-04 | -1.22E+00 | 3.19E-02 | 8.75E-02 | 1.02E+00 |
| AL645933.2 | 5.01E-03 | 2.61E-02 | -1.37E+00 | 2.01E-01 | 8.70E-01 | 1.40E+00 | 1.20E-03 | 5.10E-03 | -1.33E+00 | 2.14E-02 | 8.18E-02 | 1.06E+00 | 6.87E-03 | 2.45E-02 | -2.21E+00 | 2.31E-01 | 6.50E-01 | 1.08E+00 | 5.57E-04 | 2.05E-03 | -1.87E+00 | 4.59E-02 | 1.26E-01 | -1.17E+00 |
| RGS3 | 5.02E-03 | 2.62E-02 | -1.10E+00 | 1.12E-03 | 4.83E-03 | -1.12E+00 | 8.61E-04 | 3.66E-03 | -1.23E+00 | 6.11E-05 | 2.33E-04 | -1.37E+00 | 6.34E-03 | 2.26E-02 | -1.16E+00 | 2.17E-03 | 6.11E-03 | -1.34E+00 | 6.80E-02 | 2.49E-01 | 1.95E+00 | 3.23E-01 | 8.89E-01 | 1.52E+00 |
| NKAPP1 | 5.02E-03 | 2.62E-02 | -1.06E+00 | 8.82E-03 | 3.81E-02 | 1.06E+00 | 8.71E-07 | 3.70E-06 | -1.75E+00 | 9.26E-06 | 3.54E-05 | -1.36E+00 | 2.38E-02 | 8.49E-02 | 1.06E+00 | 1.96E-01 | 5.53E-01 | 1.34E+00 | 2.03E-07 | 7.46E-07 | -1.38E+00 | 7.42E-05 | 2.04E-04 | -1.11E+00 |
| SKI | 5.03E-03 | 2.62E-02 | -1.13E+00 | 4.22E-04 | 1.82E-03 | -1.19E+00 | 4.10E-03 | 1.74E-02 | -1.12E+00 | 1.32E-04 | 5.03E-04 | -1.22E+00 | 2.62E-03 | 9.35E-03 | -1.11E+00 | 6.66E-05 | 1.88E-04 | -1.44E+00 | 7.94E-03 | 2.91E-02 | 1.01E+00 | 2.16E-04 | 5.93E-04 | -1.21E+00 |
| RIPK3 | 5.03E-03 | 2.62E-02 | -1.21E+00 | 1.55E-03 | 6.70E-03 | -1.20E+00 | 3.56E-02 | 1.51E-01 | -1.08E+00 | 1.25E-01 | 4.77E-01 | 1.25E+00 | 3.89E-03 | 1.39E-02 | -1.38E+00 | 4.51E-03 | 1.27E-02 | -1.46E+00 | 5.18E-02 | 1.90E-01 | -1.12E+00 | 1.75E-02 | 4.80E-02 | -1.49E+00 |
| SLC22A4 | 5.03E-03 | 2.62E-02 | -1.52E+00 | 2.60E-04 | 1.12E-03 | -2.38E+00 | 1.01E-04 | 4.28E-04 | -1.45E+00 | 1.26E-05 | 4.80E-05 | -1.40E+00 | 2.34E-01 | 8.35E-01 | -1.07E+00 | 3.11E-01 | 8.76E-01 | -1.05E+00 | 2.26E-01 | 8.29E-01 | 1.13E+00 | 3.64E-01 | 9.99E-01 | 1.11E+00 |
| AR | 5.03E-03 | 2.62E-02 | -1.43E+00 | 1.27E-01 | 5.49E-01 | 1.10E+00 | 5.67E-05 | 2.41E-04 | -1.72E+00 | 3.72E-03 | 1.42E-02 | -1.17E+00 | 2.51E-01 | 8.95E-01 | 1.00E+00 | 3.54E-01 | 9.97E-01 | 1.10E+00 | 1.72E-01 | 6.30E-01 | 1.34E+00 | 2.78E-01 | 7.65E-01 | 1.18E+00 |
| ACAP3 | 5.04E-03 | 2.62E-02 | -1.08E+00 | 2.80E-02 | 1.21E-01 | 1.12E+00 | 1.98E-01 | 8.41E-01 | 1.19E+00 | 2.62E-01 | 1.00E+00 | 1.41E+00 | 2.08E-01 | 7.42E-01 | 1.29E+00 | 1.98E-01 | 5.57E-01 | 1.27E+00 | 1.07E-01 | 3.93E-01 | -1.08E+00 | 2.47E-01 | 6.78E-01 | 1.29E+00 |
| AL136311.1 | 5.04E-03 | 2.62E-02 | -1.43E+00 | 2.82E-02 | 1.22E-01 | -1.06E+00 | 2.43E-02 | 1.03E-01 | -1.43E+00 | 2.85E-02 | 1.09E-01 | -1.19E+00 | 2.20E-01 | 7.83E-01 | 1.64E+00 | 1.25E-01 | 3.52E-01 | 1.01E+00 | 2.53E-01 | 9.30E-01 | 1.16E+00 | 2.29E-01 | 6.28E-01 | 1.10E+00 |
| DENND1C | 5.04E-03 | 2.63E-02 | -1.05E+00 | 2.50E-02 | 1.08E-01 | 1.17E+00 | 7.39E-03 | 3.14E-02 | -1.10E+00 | 4.27E-03 | 1.63E-02 | -1.02E+00 | 6.31E-02 | 2.25E-01 | 1.05E+00 | 4.30E-02 | 1.21E-01 | 1.02E+00 | 1.78E-01 | 6.54E-01 | 1.04E+00 | 2.06E-01 | 5.66E-01 | 1.01E+00 |
| AC012360.1 | 5.05E-03 | 2.63E-02 | -1.24E+00 | 8.89E-03 | 3.84E-02 | -1.13E+00 | 1.86E-01 | 7.91E-01 | 1.20E+00 | 1.53E-01 | 5.84E-01 | 1.16E+00 | 1.00E-01 | 3.58E-01 | -1.15E+00 | 2.31E-01 | 6.50E-01 | -1.01E+00 | 5.07E-02 | 1.86E-01 | -1.34E+00 | 2.96E-01 | 8.14E-01 | 1.09E+00 |
| SLC25A4 | 5.05E-03 | 2.63E-02 | -1.09E+00 | 1.18E-01 | 5.08E-01 | 1.35E+00 | 1.95E-01 | 8.28E-01 | 1.29E+00 | 1.56E-01 | 5.96E-01 | 1.53E+00 | 5.69E-07 | 2.03E-06 | -1.60E+00 | 1.52E-04 | 4.27E-04 | -1.19E+00 | 2.54E-02 | 9.31E-02 | -1.02E+00 | 3.08E-01 | 8.47E-01 | 1.28E+00 |
| PTPRD | 5.06E-03 | 2.64E-02 | 2.12E+00 | 1.58E-01 | 6.82E-01 | 1.42E+00 | 2.01E-04 | 8.53E-04 | 2.23E+00 | 2.53E-01 | 9.68E-01 | 1.34E+00 | 1.93E-01 | 6.90E-01 | 1.35E+00 | 3.55E-01 | 1.00E+00 | -1.00E+00 | 2.37E-01 | 8.71E-01 | -1.07E+00 | 3.57E-01 | 9.80E-01 | -1.02E+00 |
| CLIC1 | 5.08E-03 | 2.65E-02 | 1.11E+00 | 9.63E-04 | 4.16E-03 | 1.16E+00 | 7.20E-03 | 3.06E-02 | 1.11E+00 | 2.03E-03 | 7.74E-03 | 1.15E+00 | 1.36E-01 | 4.86E-01 | -1.02E+00 | 2.68E-01 | 7.55E-01 | 1.02E+00 | 7.51E-02 | 2.75E-01 | 1.08E+00 | 1.72E-02 | 4.71E-02 | 1.12E+00 |
| CYB5RL | 5.09E-03 | 2.65E-02 | -1.32E+00 | 1.73E-01 | 7.49E-01 | 1.40E+00 | 1.80E-02 | 7.66E-02 | -1.08E+00 | 9.89E-03 | 3.78E-02 | -1.15E+00 | 7.31E-03 | 2.61E-02 | -1.44E+00 | 1.40E-01 | 3.96E-01 | 1.17E+00 | 1.84E-01 | 6.74E-01 | 1.14E+00 | 2.37E-01 | 6.50E-01 | 1.39E+00 |
| UHRF2 | 5.09E-03 | 2.65E-02 | -1.01E+00 | 6.12E-05 | 2.65E-04 | -1.08E+00 | 1.90E-05 | 8.08E-05 | -1.16E+00 | 2.85E-06 | 1.09E-05 | -1.05E+00 | 4.97E-05 | 1.77E-04 | -1.20E+00 | 3.16E-04 | 8.91E-04 | -1.05E+00 | 3.79E-05 | 1.39E-04 | -1.23E+00 | 2.27E-04 | 6.23E-04 | -1.20E+00 |
| TCTEX1D2 | 5.10E-03 | 2.66E-02 | -1.10E+00 | 1.42E-04 | 6.15E-04 | -1.22E+00 | 3.24E-03 | 1.38E-02 | -1.05E+00 | 8.60E-10 | 3.28E-09 | -1.59E+00 | 1.61E-02 | 5.74E-02 | -1.15E+00 | 3.81E-03 | 1.07E-02 | -1.33E+00 | 5.16E-06 | 1.89E-05 | -1.17E+00 | 8.59E-08 | 2.36E-07 | -1.40E+00 |
| FAM3B | 5.10E-03 | 2.66E-02 | -1.58E+00 | 4.62E-02 | 2.00E-01 | -1.08E+00 | 5.17E-03 | 2.20E-02 | -1.34E+00 | 3.27E-02 | 1.25E-01 | -1.07E+00 | 5.02E-06 | 1.79E-05 | -2.08E+00 | 3.77E-06 | 1.06E-05 | -2.12E+00 | 2.42E-01 | 8.87E-01 | 1.10E+00 | 2.26E-01 | 6.20E-01 | 1.38E+00 |
| NAA10 | 5.10E-03 | 2.66E-02 | 1.09E+00 | 7.10E-05 | 3.07E-04 | 1.09E+00 | 1.20E-01 | 5.09E-01 | 1.05E+00 | 8.24E-03 | 3.15E-02 | 1.03E+00 | 4.90E-04 | 1.75E-03 | -1.07E+00 | 1.51E-01 | 4.26E-01 | 1.22E+00 | 2.90E-02 | 1.06E-01 | -1.06E+00 | 3.25E-01 | 8.92E-01 | 1.19E+00 |
| GIN1 | 5.11E-03 | 2.67E-02 | -1.36E+00 | 2.86E-02 | 1.24E-01 | 1.04E+00 | 3.94E-03 | 1.67E-02 | -1.51E+00 | 3.21E-02 | 1.23E-01 | -1.16E+00 | 6.29E-02 | 2.24E-01 | -1.13E+00 | 2.75E-01 | 7.75E-01 | 1.36E+00 | 1.10E-04 | 4.02E-04 | -1.75E+00 | 2.34E-02 | 6.42E-02 | -1.14E+00 |
| NRIP2 | 5.12E-03 | 2.67E-02 | -1.34E+00 | 4.01E-02 | 1.73E-01 | 1.07E+00 | 7.57E-03 | 3.22E-02 | -1.38E+00 | 1.14E-01 | 4.34E-01 | 1.17E+00 | 6.41E-02 | 2.29E-01 | -1.08E+00 | 2.23E-01 | 6.30E-01 | 1.23E+00 | 1.25E-02 | 4.59E-02 | -1.43E+00 | 2.81E-01 | 7.72E-01 | 1.10E+00 |
| FAM86DP | 5.12E-03 | 2.67E-02 | -1.02E+00 | 4.41E-03 | 1.91E-02 | 1.09E+00 | 2.10E-03 | 8.91E-03 | -1.20E+00 | 4.37E-02 | 1.67E-01 | 1.13E+00 | 1.95E-02 | 6.95E-02 | -1.07E+00 | 6.57E-02 | 1.85E-01 | 1.04E+00 | 1.25E-01 | 4.58E-01 | 1.06E+00 | 2.33E-01 | 6.41E-01 | 1.35E+00 |
| SLC39A10 | 5.12E-03 | 2.67E-02 | -1.03E+00 | 7.38E-02 | 3.19E-01 | 1.21E+00 | 1.05E-06 | 4.47E-06 | -1.23E+00 | 4.89E-05 | 1.87E-04 | -1.02E+00 | 1.74E-03 | 6.19E-03 | 1.10E+00 | 5.23E-03 | 1.48E-02 | 1.13E+00 | 7.41E-05 | 2.72E-04 | -1.22E+00 | 2.72E-02 | 7.48E-02 | 1.07E+00 |
| PRMT9 | 5.13E-03 | 2.67E-02 | 1.06E+00 | 1.63E-04 | 7.06E-04 | -1.11E+00 | 1.20E-01 | 5.09E-01 | 1.01E+00 | 1.26E-01 | 4.82E-01 | 1.12E+00 | 2.45E-01 | 8.75E-01 | 1.55E+00 | 2.15E-01 | 6.06E-01 | 1.29E+00 | 2.51E-01 | 9.20E-01 | 1.51E+00 | 2.32E-01 | 6.36E-01 | 1.26E+00 |
| NUDT14 | 5.13E-03 | 2.67E-02 | -1.19E+00 | 8.21E-03 | 3.55E-02 | -1.03E+00 | 1.90E-01 | 8.08E-01 | -1.09E+00 | 1.53E-01 | 5.83E-01 | -1.06E+00 | 1.17E-02 | 4.18E-02 | -1.65E+00 | 3.14E-02 | 8.86E-02 | -1.48E+00 | 2.57E-01 | 9.43E-01 | 1.20E+00 | 1.49E-01 | 4.10E-01 | 1.06E+00 |
| MPC2 | 5.13E-03 | 2.67E-02 | 1.31E+00 | 5.28E-06 | 2.28E-05 | 1.03E+00 | 1.01E-01 | 4.28E-01 | 1.18E+00 | 3.06E-04 | 1.17E-03 | -1.05E+00 | 1.11E-01 | 3.97E-01 | 1.45E+00 | 2.29E-01 | 6.45E-01 | 1.52E+00 | 7.42E-05 | 2.72E-04 | 1.64E+00 | 3.79E-02 | 1.04E-01 | 1.45E+00 |
| AC004520.1 | 5.14E-03 | 2.68E-02 | -1.21E+00 | 1.91E-01 | 8.27E-01 | 1.72E+00 | 7.55E-03 | 3.21E-02 | -1.20E+00 | 6.44E-03 | 2.46E-02 | -1.12E+00 | 3.12E-04 | 1.11E-03 | -1.39E+00 | 7.39E-05 | 2.08E-04 | -1.55E+00 | 2.20E-01 | 8.06E-01 | 1.22E+00 | 2.31E-01 | 6.33E-01 | 1.58E+00 |
| LINC00115 | 5.14E-03 | 2.68E-02 | -1.12E+00 | 9.62E-03 | 4.16E-02 | 1.02E+00 | 9.97E-02 | 4.24E-01 | -1.09E+00 | 2.20E-01 | 8.41E-01 | 1.21E+00 | 4.95E-02 | 1.76E-01 | -1.29E+00 | 3.08E-01 | 8.69E-01 | 1.28E+00 | 2.16E-01 | 7.93E-01 | 1.18E+00 | 1.98E-01 | 5.45E-01 | 1.51E+00 |
| ARL11 | 5.14E-03 | 2.68E-02 | 1.11E+00 | 2.25E-01 | 9.72E-01 | 1.67E+00 | 9.59E-10 | 4.08E-09 | -1.65E+00 | 1.05E-05 | 4.01E-05 | -1.15E+00 | 4.91E-04 | 1.75E-03 | -1.41E+00 | 7.21E-02 | 2.03E-01 | 1.11E+00 | 3.15E-03 | 1.15E-02 | -1.44E+00 | 9.28E-03 | 2.55E-02 | -1.29E+00 |
| ZNF585A | 5.15E-03 | 2.69E-02 | -1.33E+00 | 1.14E-01 | 4.94E-01 | 1.21E+00 | 3.90E-05 | 1.66E-04 | -1.69E+00 | 2.00E-04 | 7.62E-04 | -1.43E+00 | 1.39E-01 | 4.94E-01 | 1.02E+00 | 2.51E-01 | 7.09E-01 | 1.54E+00 | 1.91E-02 | 7.01E-02 | -1.20E+00 | 1.51E-01 | 4.14E-01 | 1.01E+00 |
| ROBO4 | 5.17E-03 | 2.70E-02 | 1.15E+00 | 2.03E-01 | 8.76E-01 | 1.58E+00 | 1.24E-01 | 5.27E-01 | 1.04E+00 | 2.32E-01 | 8.86E-01 | 1.32E+00 | 1.93E-01 | 6.89E-01 | 1.94E+00 | 2.32E-01 | 6.54E-01 | 1.84E+00 | 1.95E-01 | 7.14E-01 | -1.31E+00 | 3.47E-01 | 9.54E-01 | 1.02E+00 |
| AC008567.3 | 5.17E-03 | 2.70E-02 | -1.43E+00 | 9.84E-03 | 4.25E-02 | -1.28E+00 | 1.63E-01 | 6.92E-01 | 1.08E+00 | 1.12E-01 | 4.27E-01 | 1.03E+00 | 1.16E-02 | 4.15E-02 | -1.30E+00 | 6.43E-02 | 1.81E-01 | 1.03E+00 | 2.19E-01 | 8.04E-01 | 1.38E+00 | 3.46E-01 | 9.49E-01 | 1.22E+00 |
| KTI12 | 5.18E-03 | 2.70E-02 | -1.01E+00 | 1.12E-05 | 4.86E-05 | -1.20E+00 | 1.97E-06 | 8.37E-06 | -1.35E+00 | 2.95E-07 | 1.13E-06 | -1.26E+00 | 1.44E-04 | 5.13E-04 | -1.16E+00 | 7.59E-05 | 2.14E-04 | -1.28E+00 | 2.00E-04 | 7.34E-04 | -1.12E+00 | 3.02E-05 | 8.30E-05 | -1.17E+00 |
| DOK4 | 5.19E-03 | 2.71E-02 | -1.15E+00 | 1.44E-01 | 6.20E-01 | 1.32E+00 | 1.58E-01 | 6.70E-01 | 1.05E+00 | 2.38E-01 | 9.11E-01 | 1.37E+00 | 1.65E-01 | 5.88E-01 | 1.00E+00 | 1.56E-01 | 4.39E-01 | -1.17E+00 | 2.10E-01 | 7.72E-01 | 1.14E+00 | 3.34E-01 | 9.18E-01 | 1.23E+00 |
| HIVEP2 | 5.21E-03 | 2.72E-02 | 1.01E+00 | 4.64E-04 | 2.00E-03 | -1.10E+00 | 1.08E-04 | 4.60E-04 | -1.40E+00 | 7.23E-05 | 2.76E-04 | -1.29E+00 | 2.65E-02 | 9.44E-02 | -1.38E+00 | 1.44E-01 | 4.06E-01 | 1.02E+00 | 3.29E-04 | 1.21E-03 | -1.41E+00 | 1.11E-02 | 3.04E-02 | -1.08E+00 |
| SFPQ | 5.21E-03 | 2.72E-02 | 1.01E+00 | 5.48E-06 | 2.37E-05 | -1.03E+00 | 2.17E-01 | 9.22E-01 | 1.05E+00 | 1.24E-01 | 4.74E-01 | 1.02E+00 | 3.14E-03 | 1.12E-02 | -1.13E+00 | 2.82E-03 | 7.94E-03 | -1.09E+00 | 2.67E-04 | 9.78E-04 | -1.21E+00 | 8.80E-02 | 2.42E-01 | -1.09E+00 |
| AC090114.2 | 5.22E-03 | 2.72E-02 | 1.02E+00 | 4.04E-04 | 1.75E-03 | -1.03E+00 | 8.15E-02 | 3.46E-01 | -1.05E+00 | 4.93E-02 | 1.88E-01 | 1.01E+00 | 3.23E-02 | 1.15E-01 | 1.07E+00 | 9.25E-02 | 2.61E-01 | 1.16E+00 | 1.00E-01 | 3.68E-01 | 1.01E+00 | 9.37E-02 | 2.57E-01 | -1.04E+00 |
| ZNF775 | 5.22E-03 | 2.72E-02 | -1.20E+00 | 2.25E-01 | 9.74E-01 | 1.51E+00 | 2.39E-02 | 1.02E-01 | -1.04E+00 | 8.32E-02 | 3.18E-01 | 1.62E+00 | 1.49E-03 | 5.30E-03 | -1.47E+00 | 2.76E-01 | 7.79E-01 | 1.43E+00 | 3.90E-03 | 1.43E-02 | -1.17E+00 | 1.21E-01 | 3.31E-01 | 1.11E+00 |
| LRRC37A | 5.23E-03 | 2.73E-02 | -1.08E+00 | 6.49E-03 | 2.81E-02 | 1.06E+00 | 9.19E-07 | 3.90E-06 | -1.55E+00 | 2.15E-10 | 8.20E-10 | -1.92E+00 | 3.88E-04 | 1.38E-03 | -1.61E+00 | 5.06E-04 | 1.43E-03 | -1.56E+00 | 1.27E-02 | 4.68E-02 | -1.42E+00 | 9.39E-02 | 2.58E-01 | -1.26E+00 |
| TIPIN | 5.23E-03 | 2.73E-02 | -1.13E+00 | 5.29E-03 | 2.28E-02 | -1.03E+00 | 2.82E-04 | 1.20E-03 | -1.15E+00 | 1.64E-03 | 6.28E-03 | 1.00E+00 | 2.12E-02 | 7.55E-02 | -1.37E+00 | 2.51E-01 | 7.08E-01 | 1.19E+00 | 2.97E-07 | 1.09E-06 | -1.28E+00 | 2.60E-04 | 7.13E-04 | -1.08E+00 |
| PPP1R12B | 5.25E-03 | 2.73E-02 | -1.06E+00 | 1.39E-02 | 6.00E-02 | 1.09E+00 | 1.81E-02 | 7.69E-02 | -1.18E+00 | 2.32E-02 | 8.87E-02 | -1.00E+00 | 8.14E-02 | 2.90E-01 | -1.03E+00 | 1.25E-01 | 3.53E-01 | 1.07E+00 | 1.40E-05 | 5.13E-05 | -1.49E+00 | 5.55E-03 | 1.52E-02 | -1.16E+00 |
| AC046185.3 | 5.25E-03 | 2.74E-02 | 1.09E+00 | 4.50E-02 | 1.95E-01 | 1.33E+00 | 5.08E-06 | 2.16E-05 | -1.40E+00 | 2.43E-02 | 9.29E-02 | 1.15E+00 | 2.36E-07 | 8.43E-07 | -1.20E+00 | 8.66E-04 | 2.44E-03 | 1.05E+00 | 3.46E-05 | 1.27E-04 | -1.31E+00 | 1.26E-01 | 3.45E-01 | 1.17E+00 |
| ACER3 | 5.25E-03 | 2.74E-02 | -1.15E+00 | 7.10E-02 | 3.07E-01 | 1.25E+00 | 8.51E-04 | 3.62E-03 | -1.21E+00 | 9.86E-03 | 3.76E-02 | 1.09E+00 | 3.05E-03 | 1.09E-02 | -1.28E+00 | 8.49E-02 | 2.39E-01 | 1.09E+00 | 5.66E-04 | 2.08E-03 | -1.13E+00 | 3.49E-02 | 9.59E-02 | 1.03E+00 |
| PIBF1 | 5.25E-03 | 2.74E-02 | -1.02E+00 | 2.36E-02 | 1.02E-01 | 1.18E+00 | 6.19E-06 | 2.63E-05 | -1.28E+00 | 1.28E-04 | 4.88E-04 | -1.09E+00 | 1.03E-06 | 3.67E-06 | -1.35E+00 | 1.53E-04 | 4.31E-04 | -1.11E+00 | 8.93E-09 | 3.28E-08 | -1.48E+00 | 5.78E-05 | 1.59E-04 | -1.20E+00 |
| HSD11B1L | 5.27E-03 | 2.75E-02 | 1.05E+00 | 4.02E-04 | 1.74E-03 | -1.01E+00 | 1.24E-02 | 5.28E-02 | -1.25E+00 | 7.91E-03 | 3.02E-02 | -1.17E+00 | 2.26E-02 | 8.07E-02 | -1.17E+00 | 2.88E-02 | 8.11E-02 | -1.10E+00 | 1.96E-01 | 7.21E-01 | 1.08E+00 | 7.77E-02 | 2.13E-01 | -1.25E+00 |

| | | | | | | | | | | | | | | | | | | | | | | | | |
|---|---|---|---|---|---|---|---|---|---|---|---|---|---|---|---|---|---|---|---|---|---|---|---|---|
| ATP1B3P1 | 5.27E-03 | 2.75E-02 | -1.33E+00 | 4.34E-03 | 1.88E-02 | -1.38E+00 | 3.23E-03 | 1.37E-02 | -1.29E+00 | 4.93E-03 | 1.88E-02 | -1.17E+00 | 2.03E-01 | 7.26E-01 | 1.47E+00 | 2.84E-01 | 8.00E-01 | 1.48E+00 | 2.14E-01 | 7.87E-01 | 1.10E+00 | 2.02E-01 | 5.54E-01 | 1.49E+00 |
| AC106872.2 | 5.27E-03 | 2.75E-02 | -1.21E+00 | 4.34E-03 | 1.88E-02 | -1.21E+00 | 3.41E-02 | 1.45E-01 | -1.22E+00 | 7.44E-04 | 2.84E-03 | -1.64E+00 | 8.54E-02 | 3.04E-01 | -1.06E+00 | 8.27E-02 | 2.33E-01 | -1.14E+00 | 9.61E-03 | 3.53E-02 | 1.02E+00 | 2.08E-03 | 5.72E-03 | -1.24E+00 |
| SH2D3A | 5.28E-03 | 2.75E-02 | -1.37E+00 | 2.20E-01 | 9.50E-01 | 1.26E+00 | 2.50E-04 | 1.06E-03 | -2.25E+00 | 1.59E-02 | 6.06E-02 | -1.26E+00 | 2.04E-01 | 7.27E-01 | -1.17E+00 | 2.21E-01 | 6.24E-01 | 1.45E+00 | 2.64E-05 | 9.70E-05 | -2.05E+00 | 8.49E-04 | 2.33E-03 | -1.64E+00 |
| MAP3K10 | 5.28E-03 | 2.75E-02 | -1.25E+00 | 3.78E-02 | 1.63E-01 | 1.15E+00 | 1.65E-02 | 6.99E-02 | -1.21E+00 | 8.99E-02 | 3.43E-01 | 1.17E+00 | 5.91E-02 | 2.11E-01 | -1.49E+00 | 3.41E-01 | 9.62E-01 | 1.15E+00 | 2.70E-01 | 9.92E-01 | 1.04E+00 | 1.68E-01 | 4.61E-01 | 1.39E+00 |
| WWC2 | 5.30E-03 | 2.76E-02 | -1.17E+00 | 1.06E-01 | 4.58E-01 | 1.25E+00 | 9.76E-03 | 4.15E-02 | -1.22E+00 | 5.20E-02 | 1.99E-01 | 1.16E+00 | 2.03E-02 | 7.22E-02 | -1.12E+00 | 1.93E-01 | 5.45E-01 | 1.22E+00 | 4.64E-03 | 1.70E-02 | -1.14E+00 | 2.03E-02 | 5.58E-02 | -1.03E+00 |
| KIF27 | 5.30E-03 | 2.76E-02 | -1.13E+00 | 7.39E-03 | 3.19E-02 | -1.04E+00 | 7.54E-03 | 3.20E-02 | -1.14E+00 | 9.45E-04 | 3.61E-03 | -1.22E+00 | 1.05E-01 | 3.76E-01 | 1.13E+00 | 7.78E-02 | 2.19E-01 | 1.00E+00 | 3.41E-03 | 1.25E-02 | -1.35E+00 | 4.55E-02 | 1.25E-01 | -1.03E+00 |
| ACP1 | 5.30E-03 | 2.76E-02 | 1.11E+00 | 3.00E-05 | 1.30E-04 | 1.14E+00 | 1.52E-01 | 6.46E-01 | 1.02E+00 | 7.10E-02 | 2.71E-01 | 1.07E+00 | 1.22E-02 | 4.35E-02 | 1.15E+00 | 1.17E-01 | 3.30E-01 | 1.24E+00 | 2.67E-01 | 9.80E-01 | 1.19E+00 | 3.25E-01 | 8.92E-01 | 1.16E+00 |
| GAS8 | 5.30E-03 | 2.76E-02 | -1.22E+00 | 2.53E-03 | 1.09E-02 | -1.30E+00 | 9.06E-04 | 3.85E-03 | -1.56E+00 | 4.61E-03 | 1.76E-02 | -1.23E+00 | 4.83E-04 | 1.72E-03 | -2.01E+00 | 6.94E-04 | 1.96E-03 | -1.86E+00 | 9.14E-02 | 3.35E-01 | -1.07E+00 | 2.25E-02 | 6.19E-02 | -1.35E+00 |
| EEF1A1P4 | 5.30E-03 | 2.76E-02 | 1.01E+00 | 4.23E-03 | 1.83E-02 | 1.12E+00 | 3.79E-02 | 1.61E-01 | 1.01E+00 | 6.59E-03 | 2.52E-02 | 1.05E+00 | 6.09E-02 | 2.17E-01 | 1.12E+00 | 1.30E-02 | 3.67E-02 | 1.01E+00 | 7.77E-03 | 2.85E-02 | -1.01E+00 | 2.26E-03 | 6.21E-03 | -1.14E+00 |
| RPL39P3 | 5.30E-03 | 2.76E-02 | -1.07E+00 | 6.02E-35 | 2.60E-34 | -1.51E+00 | 1.31E-02 | 5.56E-02 | -1.08E+00 | 4.01E-24 | 1.53E-23 | -1.51E+00 | 3.80E-02 | 1.35E-01 | -1.07E+00 | 1.10E-21 | 3.09E-21 | -1.55E+00 | 3.63E-07 | 1.33E-06 | -1.15E+00 | 2.16E-26 | 5.94E-26 | -1.55E+00 |
| PRORSD1P | 5.32E-03 | 2.77E-02 | -1.18E+00 | 2.23E-01 | 9.65E-01 | 1.50E+00 | 9.03E-02 | 3.83E-01 | 1.05E+00 | 2.24E-01 | 8.54E-01 | 1.55E+00 | 2.57E-01 | 9.18E-01 | 1.49E+00 | 1.97E-01 | 5.55E-01 | 1.89E+00 | 2.16E-01 | 7.91E-01 | 1.29E+00 | 1.49E-01 | 4.09E-01 | 1.48E+00 |
| NSUN5P1 | 5.33E-03 | 2.78E-02 | 1.12E+00 | 5.99E-02 | 2.59E-01 | 1.75E+00 | 5.48E-03 | 2.33E-02 | 1.00E+00 | 1.89E-01 | 7.22E-01 | 1.53E+00 | 7.07E-02 | 2.52E-01 | 1.13E+00 | 3.61E-02 | 1.02E-01 | 1.80E+00 | 1.93E-01 | 7.07E-01 | 1.10E+00 | 1.71E-04 | 4.69E-04 | 1.82E+00 |
| SRGAP2 | 5.33E-03 | 2.78E-02 | -1.11E+00 | 5.91E-02 | 2.55E-01 | 1.15E+00 | 2.01E-02 | 8.56E-02 | -1.13E+00 | 5.82E-02 | 2.22E-01 | 1.12E+00 | 1.66E-01 | 5.90E-01 | 1.22E+00 | 1.31E-01 | 3.70E-01 | -1.00E+00 | 4.15E-04 | 1.52E-03 | -1.21E+00 | 1.25E-03 | 3.43E-03 | -1.18E+00 |
| VAV3-AS1 | 5.34E-03 | 2.78E-02 | -1.23E+00 | 6.28E-03 | 2.71E-02 | -1.15E+00 | 2.13E-01 | 9.07E-01 | 1.06E+00 | 2.57E-01 | 9.81E-01 | 1.20E+00 | 2.77E-01 | 9.88E-01 | 1.22E+00 | 1.96E-01 | 5.52E-01 | 1.43E+00 | 1.53E-01 | 5.60E-01 | -1.04E+00 | 1.35E-01 | 3.71E-01 | -1.08E+00 |
| GGACT | 5.34E-03 | 2.78E-02 | -1.22E+00 | 5.38E-05 | 2.33E-04 | -1.79E+00 | 2.95E-06 | 1.25E-05 | -1.92E+00 | 3.57E-06 | 1.36E-05 | -1.80E+00 | 1.68E-06 | 5.98E-06 | -3.07E+00 | 1.46E-04 | 4.12E-04 | -1.83E+00 | 1.52E-01 | 5.59E-01 | 1.17E+00 | 3.25E-02 | 8.94E-02 | -1.19E+00 |
| SLC38A10 | 5.35E-03 | 2.79E-02 | 1.03E+00 | 3.97E-02 | 1.71E-01 | 1.20E+00 | 8.70E-04 | 3.70E-03 | -1.05E+00 | 8.46E-03 | 3.23E-02 | 1.15E+00 | 1.05E-05 | 3.74E-05 | -1.29E+00 | 2.42E-03 | 6.83E-03 | -1.02E+00 | 2.95E-08 | 1.08E-07 | -1.54E+00 | 1.17E-01 | 3.22E-01 | 1.07E+00 |
| SLA2 | 5.35E-03 | 2.79E-02 | 1.02E+00 | 2.11E-03 | 9.12E-03 | 1.02E+00 | 4.19E-04 | 1.78E-03 | -1.29E+00 | 1.50E-05 | 5.73E-05 | -1.34E+00 | 4.25E-02 | 1.51E-01 | 1.07E+00 | 2.00E-03 | 5.62E-03 | -1.56E+00 | 3.10E-03 | 1.14E-02 | -1.03E+00 | 8.91E-04 | 2.45E-03 | -1.29E+00 |
| SEC61A2 | 5.35E-03 | 2.79E-02 | -1.20E+00 | 3.44E-02 | 1.48E-01 | 1.09E+00 | 1.39E-04 | 5.90E-04 | -1.46E+00 | 2.82E-05 | 1.08E-04 | -1.46E+00 | 3.50E-02 | 1.25E-01 | -1.26E+00 | 6.16E-02 | 1.74E-01 | -1.10E+00 | 3.89E-02 | 1.43E-01 | -1.08E+00 | 1.08E-02 | 2.97E-02 | -1.48E+00 |
| AC104763.1 | 5.36E-03 | 2.79E-02 | -1.22E+00 | 7.89E-02 | 3.41E-01 | 1.20E+00 | 1.35E-07 | 5.75E-07 | -1.94E+00 | 6.71E-05 | 2.56E-04 | -1.27E+00 | 8.25E-04 | 2.94E-03 | -2.19E+00 | 2.93E-02 | 8.26E-02 | -1.23E+00 | 8.49E-04 | 3.12E-03 | -1.71E+00 | 1.26E-01 | 3.46E-01 | -1.02E+00 |
| CMTM4 | 5.37E-03 | 2.80E-02 | -1.33E+00 | 2.24E-01 | 9.67E-01 | 1.37E+00 | 1.30E-02 | 5.51E-02 | -1.14E+00 | 2.46E-03 | 9.38E-03 | -1.24E+00 | 1.34E-01 | 4.78E-01 | -1.14E+00 | 7.12E-02 | 2.01E-01 | -1.58E+00 | 1.66E-01 | 6.09E-01 | -1.23E+00 | 2.94E-01 | 8.08E-01 | -1.18E+00 |
| ERAP2 | 5.38E-03 | 2.80E-02 | -1.16E+00 | 1.89E-01 | 8.16E-01 | 1.28E+00 | 6.37E-03 | 2.71E-02 | -1.03E+00 | 4.05E-03 | 1.55E-02 | 1.05E+00 | 1.12E-03 | 4.00E-03 | -1.70E+00 | 1.26E-02 | 3.56E-02 | -1.31E+00 | 2.58E-03 | 9.48E-03 | -1.43E+00 | 7.15E-02 | 1.96E-01 | -1.02E+00 |
| AC099677.1 | 5.39E-03 | 2.81E-02 | -1.30E+00 | 1.32E-03 | 5.72E-03 | -1.48E+00 | 7.00E-02 | 2.98E-01 | -1.09E+00 | 2.07E-02 | 7.90E-02 | -1.15E+00 | 2.33E-02 | 8.31E-02 | -1.14E+00 | 1.06E-02 | 2.98E-02 | -1.34E+00 | 2.04E-01 | 7.48E-01 | 1.09E+00 | 1.04E-01 | 2.85E-01 | -1.12E+00 |
| FANCD2 | 5.39E-03 | 2.81E-02 | -1.58E+00 | 1.41E-03 | 6.11E-03 | -1.68E+00 | 1.68E-01 | 7.12E-01 | 1.12E+00 | 5.64E-02 | 2.15E-01 | 1.05E+00 | 1.84E-02 | 6.58E-02 | -1.50E+00 | 1.04E-02 | 2.92E-02 | -1.89E+00 | 1.38E-01 | 5.05E-01 | 1.07E+00 | 1.72E-01 | 4.72E-01 | 1.34E+00 |
| PEA15 | 5.39E-03 | 2.81E-02 | -1.20E+00 | 1.54E-01 | 6.64E-01 | 1.28E+00 | 3.01E-02 | 1.28E-01 | -1.09E+00 | 1.44E-01 | 5.51E-01 | 1.24E+00 | 3.23E-02 | 1.15E-01 | 1.02E+00 | 9.54E-02 | 2.69E-01 | 1.23E+00 | 2.93E-03 | 1.08E-02 | 1.01E+00 | 8.83E-02 | 2.43E-01 | 1.19E+00 |
| ABHD17C | 5.40E-03 | 2.81E-02 | -1.48E+00 | 2.70E-02 | 1.16E-01 | -1.11E+00 | 7.64E-03 | 3.25E-02 | -1.30E+00 | 1.76E-02 | 6.74E-02 | -1.10E+00 | 1.68E-01 | 5.99E-01 | -1.16E+00 | 3.24E-01 | 9.13E-01 | 1.29E+00 | 4.88E-07 | 1.79E-06 | -1.93E+00 | 1.43E-03 | 3.93E-03 | -1.23E+00 |
| FAM110D | 5.41E-03 | 2.82E-02 | -1.56E+00 | 4.50E-03 | 1.95E-02 | -1.58E+00 | 2.60E-03 | 1.10E-02 | -1.46E+00 | 2.13E-03 | 8.14E-03 | -1.44E+00 | 1.03E-02 | 3.67E-02 | -1.99E+00 | 5.97E-02 | 1.68E-01 | -1.28E+00 | 8.87E-03 | 3.26E-02 | -1.36E+00 | 2.30E-01 | 6.33E-01 | 1.16E+00 |
| GALNTL6 | 5.42E-03 | 2.82E-02 | -1.17E+00 | 1.06E-01 | 4.57E-01 | 1.72E+00 | 8.30E-03 | 3.53E-02 | -1.43E+00 | 1.07E-01 | 4.11E-01 | 1.06E+00 | 1.75E-02 | 6.25E-02 | -2.62E+00 | 2.69E-02 | 7.58E-02 | -2.77E+00 | 2.40E-01 | 8.81E-01 | -1.09E+00 | 2.67E-01 | 7.35E-01 | 1.35E+00 |
| PPP1R26 | 5.42E-03 | 2.83E-02 | -1.47E+00 | 6.87E-02 | 2.97E-01 | 1.12E+00 | 1.76E-02 | 7.46E-02 | -1.16E+00 | 2.24E-01 | 8.55E-01 | 1.37E+00 | 1.69E-02 | 6.01E-02 | -1.32E+00 | 1.95E-01 | 5.49E-01 | 1.18E+00 | 2.82E-05 | 1.03E-04 | -1.63E+00 | 5.13E-03 | 1.41E-02 | -1.15E+00 |
| GIPC3 | 5.43E-03 | 2.83E-02 | -1.33E+00 | 1.27E-01 | 5.49E-01 | 1.14E+00 | 1.36E-01 | 5.78E-01 | -1.15E+00 | 2.08E-01 | 7.96E-01 | 1.25E+00 | 2.45E-01 | 8.75E-01 | 1.03E+00 | 2.76E-01 | 7.78E-01 | -1.15E+00 | 2.82E-06 | 1.03E-05 | -2.42E+00 | 3.41E-05 | 9.38E-05 | -1.97E+00 |
| CIC | 5.43E-03 | 2.83E-02 | 1.03E+00 | 6.88E-03 | 2.97E-02 | 1.19E+00 | 2.13E-02 | 9.03E-02 | -1.02E+00 | 7.52E-03 | 2.87E-02 | -1.01E+00 | 2.28E-03 | 8.14E-03 | -1.16E+00 | 1.24E-03 | 3.49E-03 | -1.32E+00 | 4.48E-04 | 1.64E-03 | -1.24E+00 | 1.95E-05 | 5.34E-05 | -1.55E+00 |
| NPRL2 | 5.46E-03 | 2.84E-02 | -1.05E+00 | 9.44E-02 | 4.08E-01 | 1.32E+00 | 2.23E-04 | 9.45E-04 | -1.22E+00 | 3.24E-02 | 1.24E-01 | 1.19E+00 | 8.51E-04 | 3.03E-03 | -1.40E+00 | 1.04E-01 | 2.94E-01 | 1.23E+00 | 8.47E-05 | 3.11E-04 | -1.16E+00 | 1.51E-01 | 4.16E-01 | 1.20E+00 |
| RANBP17 | 5.46E-03 | 2.84E-02 | -1.17E+00 | 3.20E-02 | 1.38E-01 | 1.11E+00 | 2.46E-05 | 1.05E-04 | -1.28E+00 | 1.55E-03 | 5.91E-03 | 1.03E+00 | 5.49E-04 | 1.96E-03 | -1.39E+00 | 1.05E-04 | 2.96E-04 | -1.59E+00 | 1.05E-04 | 3.84E-04 | -1.27E+00 | 4.19E-02 | 1.15E-01 | 1.04E+00 |
| TCEA3 | 5.46E-03 | 2.85E-02 | -1.19E+00 | 6.06E-03 | 2.62E-02 | -1.06E+00 | 2.69E-02 | 1.14E-01 | 1.62E+00 | 2.49E-01 | 9.50E-01 | 1.19E+00 | 1.17E-01 | 4.18E-01 | -1.54E+00 | 1.81E-01 | 5.10E-01 | -1.25E+00 | 2.62E-01 | 9.60E-01 | 1.02E+00 | 3.06E-01 | 8.40E-01 | 1.12E+00 |
| RAD17P2 | 5.46E-03 | 2.85E-02 | 2.29E+00 | 2.24E-02 | 9.67E-02 | 2.20E+00 | 1.48E-01 | 6.31E-01 | 1.09E+00 | 2.51E-01 | 9.58E-01 | 1.29E+00 | 2.72E-01 | 9.69E-01 | 1.25E+00 | 3.30E-01 | 9.31E-01 | 1.19E+00 | 2.27E-01 | 8.31E-01 | 1.17E+00 | 2.10E-01 | 5.78E-01 | 1.04E+00 |
| NDUFA4 | 5.46E-03 | 2.85E-02 | 1.44E+00 | 2.40E-02 | 1.04E-01 | 1.28E+00 | 1.77E-05 | 7.50E-05 | 1.75E+00 | 1.18E-01 | 4.50E-01 | 1.43E+00 | 2.75E-02 | 9.80E-02 | 1.47E+00 | 3.49E-01 | 9.85E-01 | 1.26E+00 | 7.95E-06 | 2.92E-05 | 1.63E+00 | 3.61E-02 | 9.92E-02 | 1.36E+00 |
| PPIAP46 | 5.46E-03 | 2.85E-02 | -1.09E+00 | 1.71E-05 | 7.40E-05 | -1.72E+00 | 1.00E-04 | 4.27E-04 | -1.30E+00 | 7.30E-09 | 2.79E-08 | -1.66E+00 | 3.76E-02 | 1.34E-01 | -1.08E+00 | 1.89E-02 | 5.33E-02 | -1.22E+00 | 1.33E-03 | 4.88E-03 | -1.06E+00 | 2.28E-06 | 6.27E-06 | -1.43E+00 |
| BARD1 | 5.46E-03 | 2.85E-02 | 1.64E+00 | 1.86E-01 | 8.02E-01 | 1.27E+00 | 1.70E-01 | 7.23E-01 | 1.25E+00 | 5.73E-03 | 2.19E-02 | 1.02E+00 | 1.05E-01 | 3.73E-01 | 1.11E+00 | 2.56E-02 | 7.23E-02 | -1.05E+00 | 3.44E-04 | 1.26E-03 | -1.25E+00 | 1.66E-04 | 4.55E-04 | -1.27E+00 |
| KDM7A | 5.47E-03 | 2.85E-02 | -1.22E+00 | 1.86E-02 | 8.05E-02 | 1.02E+00 | 8.29E-04 | 3.52E-03 | -1.69E+00 | 2.44E-03 | 9.33E-03 | -1.36E+00 | 1.93E-02 | 6.89E-02 | -1.13E+00 | 7.45E-02 | 2.10E-01 | 1.05E+00 | 1.45E-07 | 5.31E-07 | -1.85E+00 | 5.94E-04 | 1.63E-03 | -1.37E+00 |
| AL451007.1 | 5.47E-03 | 2.85E-02 | -1.09E+00 | 6.53E-06 | 2.82E-05 | -1.97E+00 | 1.82E-01 | 7.71E-01 | 1.01E+00 | 2.31E-02 | 8.81E-02 | -1.24E+00 | 1.28E-01 | 4.58E-01 | 1.13E+00 | 8.49E-03 | 2.39E-02 | -1.40E+00 | 4.52E-02 | 1.66E-01 | -1.15E+00 | 8.18E-04 | 2.25E-03 | -1.63E+00 |
| AC096915.1 | 5.48E-03 | 2.86E-02 | -1.08E+00 | 1.77E-02 | 7.65E-02 | 2.00E+00 | 3.92E-07 | 1.66E-06 | -1.69E+00 | 2.53E-01 | 9.68E-01 | 1.39E+00 | 1.13E-02 | 4.02E-02 | -1.48E+00 | 2.17E-02 | 6.13E-02 | 1.75E+00 | 6.34E-03 | 2.33E-02 | -1.19E+00 | 4.45E-03 | 1.22E-02 | 1.97E+00 |
| PWP2 | 5.48E-03 | 2.86E-02 | -1.19E+00 | 8.56E-02 | 3.70E-01 | 1.22E+00 | 6.70E-02 | 2.85E-01 | -1.02E+00 | 1.53E-01 | 5.85E-01 | 1.45E+00 | 3.86E-03 | 1.38E-02 | -1.29E+00 | 1.44E-01 | 4.05E-01 | 1.20E+00 | 9.24E-04 | 3.39E-03 | -1.16E+00 | 2.94E-01 | 8.07E-01 | 1.21E+00 |
| KMT5C | 5.49E-03 | 2.86E-02 | -1.09E+00 | 6.93E-03 | 2.99E-02 | -1.07E+00 | 1.54E-02 | 6.53E-02 | -1.08E+00 | 1.53E-03 | 5.84E-03 | -1.11E+00 | 8.09E-02 | 2.89E-01 | 2.37E+00 | 9.43E-02 | 2.66E-01 | 2.60E+00 | 1.45E-06 | 5.30E-06 | -1.62E+00 | 1.50E-02 | 4.12E-02 | -1.09E+00 |
| BOLA3-AS1 | 5.49E-03 | 2.86E-02 | -1.27E+00 | 4.32E-02 | 1.87E-01 | 1.20E+00 | 1.97E-02 | 8.38E-02 | -1.43E+00 | 3.60E-02 | 1.38E-01 | -1.14E+00 | 2.65E-30 | 9.46E-30 | -2.56E+00 | 1.33E-17 | 3.76E-17 | -1.47E+00 | 5.00E-20 | 1.84E-19 | -1.89E+00 | 2.02E-08 | 5.55E-08 | -1.28E+00 |
| GPR161 | 5.50E-03 | 2.87E-02 | -1.40E+00 | 2.32E-02 | 1.00E-01 | -1.10E+00 | 7.52E-04 | 3.20E-03 | -1.67E+00 | 1.14E-02 | 4.36E-02 | -1.16E+00 | 1.30E-02 | 4.65E-02 | -2.13E+00 | 3.88E-02 | 1.09E-01 | -1.71E+00 | 2.26E-08 | 8.28E-08 | -1.98E+00 | 1.14E-05 | 3.12E-05 | -1.53E+00 |
| BMF | 5.51E-03 | 2.87E-02 | 1.91E+00 | 9.76E-02 | 4.22E-01 | 1.76E+00 | 1.34E-01 | 5.67E-01 | 1.49E+00 | 2.23E-01 | 8.52E-01 | 1.44E+00 | 1.99E-01 | 7.08E-01 | -1.18E+00 | 3.37E-01 | 9.49E-01 | 1.33E+00 | 1.80E-01 | 6.60E-01 | 1.42E+00 | 3.64E-01 | 1.00E+00 | 1.00E+00 |
| DPY19L2P2 | 5.51E-03 | 2.87E-02 | -1.18E+00 | 7.30E-02 | 3.15E-01 | 1.23E+00 | 1.30E-02 | 5.54E-02 | -1.12E+00 | 1.46E-01 | 5.56E-01 | 1.31E+00 | 3.21E-02 | 1.14E-01 | -1.29E+00 | 1.47E-01 | 4.15E-01 | 1.08E+00 | 6.78E-04 | 2.49E-03 | -1.32E+00 | 1.77E-02 | 4.87E-02 | -1.06E+00 |
| LRRC19 | 5.52E-03 | 2.87E-02 | -1.29E+00 | 5.99E-02 | 2.59E-01 | 1.14E+00 | 1.31E-01 | 5.59E-01 | 1.03E+00 | 1.57E-01 | 6.00E-01 | 1.53E+00 | 1.47E-02 | 5.23E-02 | -1.52E+00 | 9.12E-02 | 2.57E-01 | -1.05E+00 | 1.39E-01 | 5.11E-01 | 1.08E+00 | 2.41E-01 | 6.61E-01 | 1.50E+00 |
| ARMC4 | 5.53E-03 | 2.88E-02 | -1.41E+00 | 1.92E-02 | 8.28E-02 | -1.16E+00 | 1.14E-02 | 4.83E-02 | -1.29E+00 | 1.63E-01 | 6.23E-01 | 1.16E+00 | 4.53E-02 | 1.62E-01 | -1.68E+00 | 1.55E-01 | 4.37E-01 | -1.13E+00 | 1.37E-04 | 5.02E-04 | -2.28E+00 | 4.20E-03 | 1.15E-02 | -1.61E+00 |
| ATP8A1 | 5.54E-03 | 2.89E-02 | 1.06E+00 | 2.06E-02 | 8.89E-02 | 1.22E+00 | 8.36E-03 | 3.55E-02 | -1.15E+00 | 1.73E-02 | 6.62E-02 | 1.03E+00 | 1.48E-06 | 5.26E-06 | -1.36E+00 | 1.48E-05 | 4.17E-05 | -1.26E+00 | 4.83E-04 | 1.77E-03 | -1.23E+00 | 1.90E-03 | 5.23E-03 | -1.37E+00 |
| TDG | 5.55E-03 | 2.89E-02 | -1.08E+00 | 3.54E-03 | 1.53E-02 | -1.04E+00 | 2.67E-04 | 1.13E-03 | -1.17E+00 | 1.12E-04 | 4.27E-04 | -1.15E+00 | 7.97E-05 | 2.84E-04 | -1.36E+00 | 1.90E-04 | 5.36E-04 | -1.27E+00 | 2.43E-05 | 8.90E-05 | -1.22E+00 | 1.04E-06 | 2.86E-06 | -1.39E+00 |
| PFKM | 5.56E-03 | 2.90E-02 | -1.12E+00 | 4.47E-02 | 1.93E-01 | 1.13E+00 | 4.46E-04 | 1.89E-03 | -1.37E+00 | 2.98E-02 | 1.14E-01 | 1.07E+00 | 5.09E-03 | 1.81E-02 | -1.39E+00 | 9.57E-02 | 2.70E-01 | 1.04E+00 | 3.74E-08 | 1.37E-07 | -1.58E+00 | 2.60E-05 | 7.15E-05 | -1.19E+00 |
| ZNF470 | 5.56E-03 | 2.90E-02 | -1.41E+00 | 4.79E-02 | 2.07E-01 | -1.00E+00 | 1.77E-02 | 7.50E-02 | -1.64E+00 | 1.93E-01 | 7.38E-01 | 1.09E+00 | 1.71E-04 | 6.09E-04 | -2.14E+00 | 9.00E-03 | 2.54E-02 | -1.25E+00 | 1.21E-02 | 4.44E-02 | -1.21E+00 | 8.94E-02 | 2.45E-01 | -1.05E+00 |
| SUGT1P3 | 5.57E-03 | 2.90E-02 | -1.31E+00 | 1.78E-01 | 7.71E-01 | 1.35E+00 | 7.94E-03 | 3.37E-02 | -1.34E+00 | 1.20E-01 | 4.58E-01 | 1.17E+00 | 1.82E-01 | 6.48E-01 | 1.07E+00 | 3.06E-01 | 8.61E-01 | 1.48E+00 | 1.24E-01 | 4.54E-01 | 1.04E+00 | 2.29E-01 | 6.28E-01 | 1.12E+00 |
| RSU1 | 5.57E-03 | 2.90E-02 | 1.09E+00 | 1.55E-02 | 6.70E-02 | 1.36E+00 | 2.49E-03 | 1.06E-02 | -1.06E+00 | 3.90E-02 | 1.49E-01 | 1.21E+00 | 1.19E-02 | 4.23E-02 | 1.04E+00 | 2.63E-01 | 7.42E-01 | 1.49E+00 | 5.30E-03 | 1.94E-02 | -1.08E+00 | 3.01E-01 | 8.26E-01 | 1.08E+00 |
| ARPIN | 5.58E-03 | 2.91E-02 | -1.29E+00 | 9.81E-03 | 4.24E-02 | -1.13E+00 | 4.46E-03 | 1.89E-02 | -1.65E+00 | 2.81E-02 | 1.07E-01 | -1.16E+00 | 3.23E-02 | 1.15E-01 | -1.50E+00 | 1.43E-01 | 4.02E-01 | -1.13E+00 | 4.98E-02 | 1.83E-01 | 1.52E+00 | 1.34E-01 | 3.67E-01 | 1.56E+00 |
| CCDC94 | 5.58E-03 | 2.91E-02 | 1.00E+00 | 7.39E-03 | 3.19E-02 | 1.17E+00 | 3.14E-03 | 1.34E-02 | -1.10E+00 | 5.86E-04 | 2.24E-03 | -1.00E+00 | 3.03E-03 | 1.08E-02 | 1.05E+00 | 3.07E-02 | 8.66E-02 | 1.15E+00 | 8.85E-07 | 3.25E-06 | -1.23E+00 | 2.53E-03 | 6.96E-03 | -1.02E+00 |
| INO80D | 5.58E-03 | 2.91E-02 | 1.08E+00 | 3.03E-03 | 1.31E-02 | 1.18E+00 | 4.65E-03 | 1.98E-02 | -1.04E+00 | 7.71E-05 | 2.95E-04 | -1.03E+00 | 4.92E-02 | 1.76E-01 | 1.22E+00 | 6.11E-03 | 1.72E-02 | 1.08E+00 | 4.62E-04 | 1.69E-03 | -1.19E+00 | 4.00E-04 | 1.10E-03 | -1.16E+00 |
| U2AF1L5 | 5.59E-03 | 2.91E-02 | 1.06E+00 | 1.52E-08 | 6.57E-08 | 1.02E+00 | 3.87E-03 | 1.65E-02 | 1.32E+00 | 2.04E-01 | 7.79E-01 | 1.17E+00 | 2.19E-02 | 7.80E-02 | 1.01E+00 | 1.36E-02 | 3.84E-02 | 1.04E+00 | 1.60E-01 | 5.87E-01 | 1.21E+00 | 1.19E-01 | 3.28E-01 | 1.19E+00 |
| WNK3 | 5.59E-03 | 2.91E-02 | -1.48E+00 | 1.91E-03 | 8.27E-03 | -1.59E+00 | 1.99E-01 | 8.46E-01 | 1.32E+00 | 2.53E-01 | 9.67E-01 | 1.19E+00 | 1.58E-01 | 5.62E-01 | 3.17E+00 | 4.34E-02 | 1.22E-01 | -1.45E+00 | 4.30E-06 | 1.58E-05 | 5.11E+00 | 2.84E-02 | 7.81E-02 | 2.57E+00 |
| YOD1 | 5.59E-03 | 2.92E-02 | 1.79E+00 | 1.11E-03 | 4.79E-03 | 2.17E+00 | 2.35E-01 | 1.00E+00 | 1.34E+00 | 1.81E-01 | 6.91E-01 | 1.36E+00 | 2.73E-01 | 9.75E-01 | 1.46E+00 | 2.32E-01 | 6.54E-01 | 1.29E+00 | 5.58E-02 | 2.05E-01 | 1.22E+00 | 4.45E-02 | 1.22E-01 | 1.13E+00 |
| CYTIP | 5.60E-03 | 2.92E-02 | 1.61E+00 | 8.18E-06 | 3.54E-05 | 1.04E+00 | 9.78E-05 | 4.16E-04 | 1.85E+00 | 2.14E-03 | 8.17E-03 | 1.02E+00 | 2.65E-01 | 9.46E-01 | 1.31E+00 | 2.70E-01 | 7.62E-01 | 1.21E+00 | 3.15E-02 | 1.16E-01 | -1.23E+00 | 1.18E-01 | 3.25E-01 | -1.06E+00 |
| ITGAL | 5.60E-03 | 2.92E-02 | -1.32E+00 | 2.58E-02 | 1.11E-01 | -1.03E+00 | 8.50E-03 | 3.61E-02 | -1.15E+00 | 1.69E-04 | 6.45E-04 | -1.27E+00 | 2.67E-01 | 9.52E-01 | 1.41E+00 | 2.52E-01 | 7.09E-01 | 1.25E+00 | 6.44E-02 | 2.36E-01 | -1.25E+00 | 1.14E-01 | 3.13E-01 | -1.13E+00 |
| GPSM2 | 5.61E-03 | 2.92E-02 | 1.09E+00 | 2.81E-04 | 1.21E-03 | 1.92E+00 | 3.86E-04 | 1.64E-03 | -1.10E+00 | 2.46E-01 | 9.39E-01 | 1.50E+00 | 1.97E-02 | 7.02E-02 | 1.09E+00 | 3.45E-01 | 9.73E-01 | 1.41E+00 | 9.42E-04 | 3.46E-03 | -1.17E+00 | 2.15E-02 | 5.91E-02 | 1.02E+00 |
| RPS7P3 | 5.61E-03 | 2.93E-02 | 1.11E+00 | 1.47E-10 | 6.37E-10 | -1.40E+00 | 3.25E-02 | 1.38E-01 | 1.02E+00 | 2.44E-07 | 9.32E-07 | -1.38E+00 | 1.87E-01 | 6.67E-01 | 1.64E+00 | 7.70E-04 | 2.17E-03 | -1.18E+00 | 1.73E-03 | 6.34E-03 | -1.07E+00 | 9.09E-11 | 2.50E-10 | -1.91E+00 |
| MTOR | 5.61E-03 | 2.93E-02 | -1.17E+00 | 1.16E-01 | 5.00E-01 | 1.31E+00 | 5.42E-05 | 2.30E-04 | -1.38E+00 | 4.77E-03 | 1.82E-02 | 1.00E+00 | 1.01E-02 | 3.60E-02 | -1.15E+00 | 7.57E-02 | 2.13E-01 | 1.05E+00 | 1.72E-05 | 6.31E-05 | -1.33E+00 | 2.50E-01 | 6.86E-01 | 1.18E+00 |
| CD6 | 5.62E-03 | 2.93E-02 | -1.05E+00 | 3.01E-02 | 1.30E-01 | 1.23E+00 | 3.40E-02 | 1.44E-01 | -1.28E+00 | 1.20E-01 | 4.60E-01 | -1.03E+00 | 9.77E-02 | 3.48E-01 | 1.13E+00 | 3.30E-01 | 9.29E-01 | 1.44E+00 | 9.68E-02 | 3.55E-01 | 1.18E+00 | 3.64E-01 | 1.00E+00 | 1.23E+00 |
| MPP7 | 5.63E-03 | 2.93E-02 | -1.08E+00 | 4.03E-04 | 1.74E-03 | -1.28E+00 | 9.97E-02 | 4.24E-01 | -1.10E+00 | 2.20E-01 | 8.42E-01 | 1.15E+00 | 1.78E-02 | 6.36E-02 | 2.48E+00 | 6.75E-03 | 1.90E-02 | 2.99E+00 | 1.90E-01 | 6.98E-01 | 1.19E+00 | 2.10E-02 | 5.77E-02 | -1.09E+00 |
| CYTH4 | 5.63E-03 | 2.93E-02 | -1.03E+00 | 5.84E-02 | 2.52E-01 | 1.56E+00 | 1.79E-02 | 7.62E-02 | -1.06E+00 | 2.50E-01 | 9.56E-01 | 1.31E+00 | 1.62E-01 | 5.78E-01 | 1.09E+00 | 1.79E-01 | 5.05E-01 | -1.01E+00 | 2.14E-01 | 7.84E-01 | 1.03E+00 | 3.25E-01 | 8.92E-01 | 1.05E+00 |
| CALM1 | 5.64E-03 | 2.94E-02 | 1.05E+00 | 1.56E-04 | 6.73E-04 | 1.11E+00 | 3.00E-02 | 1.27E-01 | 1.23E+00 | 1.78E-01 | 6.80E-01 | 1.21E+00 | 8.55E-02 | 3.05E-01 | 1.04E+00 | 9.74E-02 | 2.74E-01 | 1.05E+00 | 1.03E-01 | 3.78E-01 | 1.17E+00 | 1.64E-01 | 4.52E-01 | 1.10E+00 |
| RPL5 | 5.64E-03 | 2.94E-02 | 1.09E+00 | 1.86E-20 | 8.03E-20 | 1.32E+00 | 4.86E-04 | 2.06E-03 | 1.13E+00 | 3.55E-18 | 1.36E-17 | 1.37E+00 | 3.33E-02 | 1.19E-01 | 1.09E+00 | 1.67E-12 | 4.70E-12 | 1.33E+00 | 1.84E-01 | 6.75E-01 | 1.09E+00 | 2.31E-13 | 6.35E-13 | 1.30E+00 |
| PHTF1 | 5.66E-03 | 2.95E-02 | 1.10E+00 | 5.03E-04 | 2.17E-03 | 1.05E+00 | 1.06E-01 | 4.48E-01 | 1.03E+00 | 1.06E-02 | 4.05E-02 | -1.04E+00 | 2.97E-02 | 1.06E-01 | 1.57E+00 | 3.04E-01 | 8.57E-01 | 1.29E+00 | 1.88E-03 | 6.89E-03 | -1.17E+00 | 4.65E-08 | 1.28E-07 | -1.56E+00 |
| UBE2L3 | 5.67E-03 | 2.95E-02 | 1.07E+00 | 4.65E-02 | 2.01E-01 | 1.30E+00 | 9.98E-03 | 4.24E-02 | 1.01E+00 | 8.28E-02 | 3.16E-01 | 1.22E+00 | 1.53E-06 | 5.47E-06 | -1.33E+00 | 6.35E-02 | 1.79E-01 | 1.15E+00 | 3.73E-04 | 1.37E-03 | -1.07E+00 | 3.38E-01 | 9.29E-01 | 1.18E+00 |
| AC025171.2 | 5.67E-03 | 2.95E-02 | 1.77E+00 | 2.51E-03 | 1.08E-02 | -1.07E+00 | 5.83E-02 | 2.48E-01 | 1.48E+00 | 7.60E-03 | 2.90E-02 | -1.09E+00 | 1.42E-01 | 5.05E-01 | 1.55E+00 | 2.82E-01 | 7.95E-01 | 1.35E+00 | 2.66E-01 | 9.76E-01 | 1.18E+00 | 2.26E-01 | 6.22E-01 | 1.03E+00 |
| ZNF658 | 5.68E-03 | 2.96E-02 | -1.40E+00 | 3.02E-02 | 1.30E-01 | -1.09E+00 | 8.75E-03 | 3.72E-02 | -1.40E+00 | 5.41E-02 | 2.07E-01 | -1.09E+00 | 5.43E-02 | 1.93E-01 | -1.17E+00 | 1.83E-01 | 5.15E-01 | 1.13E+00 | 2.40E-01 | 8.82E-01 | 1.09E+00 | 1.87E-01 | 5.12E-01 | -1.04E+00 |
| ZFP82 | 5.69E-03 | 2.96E-02 | -1.02E+00 | 7.78E-02 | 3.36E-01 | 1.30E+00 | 6.81E-03 | 2.89E-02 | -1.28E+00 | 1.93E-01 | 7.36E-01 | 1.16E+00 | 1.06E-02 | 3.78E-02 | -1.18E+00 | 6.75E-02 | 1.90E-01 | 1.09E+00 | 3.18E-03 | 1.17E-02 | -1.27E+00 | 1.11E-01 | 3.04E-01 | 1.11E+00 |
| MCRIP2 | 5.69E-03 | 2.97E-02 | -1.00E+00 | 3.46E-03 | 1.50E-02 | 1.14E+00 | 2.18E-01 | 9.28E-01 | 1.25E+00 | 2.25E-01 | 8.59E-01 | 1.44E+00 | 2.05E-04 | 7.31E-04 | -1.20E+00 | 1.53E-01 | 4.32E-01 | 1.36E+00 | 2.07E-01 | 7.59E-01 | 1.23E+00 | 4.14E-02 | 1.14E-01 | 1.54E+00 |
| DNMT3B | 5.69E-03 | 2.97E-02 | -1.12E+00 | 1.80E-02 | 7.77E-02 | 1.10E+00 | 8.30E-04 | 3.53E-03 | -1.06E+00 | 1.52E-03 | 5.79E-03 | 1.03E+00 | 1.82E-03 | 6.47E-03 | -1.16E+00 | 9.12E-06 | 2.57E-05 | -1.61E+00 | 2.11E-03 | 7.75E-03 | -1.12E+00 | 1.45E-04 | 3.99E-04 | -1.34E+00 |
| RIC3 | 5.70E-03 | 2.97E-02 | 1.76E+00 | 1.84E-01 | 7.96E-01 | 1.58E+00 | 1.84E-02 | 7.83E-02 | 1.49E+00 | 2.38E-01 | 9.10E-01 | 1.23E+00 | 1.33E-02 | 4.74E-02 | 2.10E+00 | 6.16E-02 | 1.74E-01 | 1.81E+00 | 2.76E-02 | 1.01E-01 | 1.92E+00 | 2.30E-02 | 6.31E-02 | 2.29E+00 |
| ATP6V1H | 5.70E-03 | 2.97E-02 | 1.17E+00 | 4.01E-04 | 1.73E-03 | 1.16E+00 | 1.80E-02 | 7.65E-02 | 1.02E+00 | 1.69E-06 | 6.47E-06 | -1.13E+00 | 6.47E-04 | 2.31E-03 | -1.14E+00 | 3.47E-03 | 9.79E-03 | -1.07E+00 | 1.81E-08 | 6.66E-08 | -1.38E+00 | 2.16E-07 | 5.92E-07 | -1.33E+00 |
| PFDN4 | 5.70E-03 | 2.97E-02 | 1.14E+00 | 1.50E-12 | 6.50E-12 | -1.10E+00 | 8.10E-03 | 3.44E-02 | -1.09E+00 | 3.92E-08 | 1.50E-07 | -1.25E+00 | 1.28E-06 | 4.55E-06 | -1.10E+00 | 5.00E-06 | 1.41E-05 | -1.07E+00 | 6.26E-05 | 2.30E-04 | -1.18E+00 | 1.05E-07 | 2.88E-07 | -1.39E+00 |
| ATXN7L3B | 5.70E-03 | 2.97E-02 | 1.12E+00 | 1.69E-07 | 7.30E-07 | 1.03E+00 | 6.66E-03 | 2.83E-02 | -1.03E+00 | 4.60E-05 | 1.76E-04 | -1.10E+00 | 1.42E-05 | 5.07E-05 | -1.05E+00 | 9.63E-09 | 2.71E-08 | -1.21E+00 | 1.54E-03 | 5.65E-03 | -1.03E+00 | 2.05E-04 | 5.63E-04 | -1.12E+00 |
| GNPAT | 5.71E-03 | 2.98E-02 | 1.11E+00 | 3.00E-03 | 1.30E-02 | 1.24E+00 | 5.77E-04 | 2.45E-03 | -1.06E+00 | 1.64E-04 | 6.28E-04 | 1.03E+00 | 1.52E-03 | 5.41E-03 | 1.06E+00 | 2.06E-03 | 5.81E-03 | 1.11E+00 | 4.67E-04 | 1.71E-03 | -1.07E+00 | 1.37E-01 | 3.75E-01 | 1.15E+00 |
| RDH5 | 5.75E-03 | 2.99E-02 | -1.15E+00 | 1.22E-02 | 5.27E-02 | 1.09E+00 | 5.19E-02 | 2.21E-01 | -1.03E+00 | 1.32E-01 | 5.05E-01 | 1.24E+00 | 2.79E-01 | 9.95E-01 | 1.31E+00 | 3.45E-01 | 9.73E-01 | 1.40E+00 | 8.76E-02 | 3.22E-01 | 1.54E+00 | 1.10E-01 | 3.01E-01 | 1.65E+00 |
| RPL31 | 5.75E-03 | 3.00E-02 | 1.11E+00 | 1.84E-10 | 7.96E-10 | -1.17E+00 | 8.93E-03 | 3.79E-02 | 1.14E+00 | 1.16E-02 | 4.44E-02 | -1.07E+00 | 1.43E-02 | 5.09E-02 | 1.15E+00 | 2.86E-02 | 8.05E-02 | -1.11E+00 | 1.36E-01 | 5.00E-01 | 1.04E+00 | 1.71E-03 | 4.70E-03 | -1.13E+00 |
| AC005162.3 | 5.76E-03 | 3.00E-02 | -1.46E+00 | 4.20E-02 | 1.81E-01 | 1.04E+00 | 3.23E-02 | 1.37E-01 | -1.29E+00 | 2.11E-01 | 8.06E-01 | 1.45E+00 | 3.82E-02 | 1.36E-01 | -1.97E+00 | 3.42E-01 | 9.64E-01 | -1.02E+00 | 5.13E-05 | 1.88E-04 | -2.34E+00 | 4.96E-03 | 1.36E-02 | -1.55E+00 |
| PTPN9 | 5.77E-03 | 3.01E-02 | -1.23E+00 | 1.87E-01 | 8.06E-01 | 1.45E+00 | 2.33E-03 | 9.91E-03 | -1.20E+00 | 4.10E-02 | 1.57E-01 | 1.16E+00 | 2.79E-03 | 9.96E-03 | -1.04E+00 | 2.97E-02 | 8.36E-02 | 1.13E+00 | 4.28E-05 | 1.57E-04 | -1.23E+00 | 2.25E-02 | 6.19E-02 | 1.10E+00 |
| AC011466.4 | 5.77E-03 | 3.01E-02 | -1.65E+00 | 1.95E-01 | 8.41E-01 | 1.31E+00 | 1.15E-02 | 4.89E-02 | -1.55E+00 | 2.61E-01 | 9.96E-01 | 1.32E+00 | 9.27E-04 | 3.30E-03 | -2.47E+00 | 2.70E-02 | 7.61E-02 | -1.40E+00 | 5.63E-04 | 2.07E-03 | -2.08E+00 | 2.26E-01 | 6.20E-01 | 1.01E+00 |
| POC1B | 5.78E-03 | 3.01E-02 | -1.07E+00 | 1.46E-01 | 6.29E-01 | 1.34E+00 | 2.43E-02 | 1.03E-01 | 1.01E+00 | 9.01E-02 | 3.44E-01 | 1.27E+00 | 2.75E-03 | 9.82E-03 | -1.31E+00 | 7.58E-02 | 2.14E-01 | 1.03E+00 | 4.76E-03 | 1.75E-02 | -1.14E+00 | 3.94E-02 | 1.08E-01 | 1.03E+00 |
| MCC | 5.78E-03 | 3.01E-02 | -1.25E+00 | 2.70E-03 | 1.16E-02 | -1.38E+00 | 1.80E-01 | 7.65E-01 | -1.01E+00 | 3.81E-02 | 1.46E-01 | -1.30E+00 | 2.02E-01 | 7.19E-01 | 1.46E+00 | 3.29E-01 | 9.29E-01 | 1.28E+00 | 6.67E-02 | 2.45E-01 | -1.06E+00 | 1.04E-01 | 2.87E-01 | -1.18E+00 |
| AC016542.3 | 5.78E-03 | 3.01E-02 | -1.45E+00 | 1.67E-02 | 7.20E-02 | -1.20E+00 | 3.43E-02 | 1.46E-01 | -1.41E+00 | 1.38E-02 | 5.28E-02 | -1.58E+00 | 1.52E-02 | 5.40E-02 | -1.88E+00 | 2.36E-01 | 6.65E-01 | 1.06E+00 | 1.15E-02 | 4.23E-02 | -1.18E+00 | 6.42E-03 | 1.76E-02 | -1.37E+00 |
| NHSL1 | 5.78E-03 | 3.01E-02 | -1.39E+00 | 9.68E-05 | 4.18E-04 | -1.69E+00 | 7.30E-02 | 3.10E-01 | 1.42E+00 | 2.39E-01 | 9.12E-01 | 1.21E+00 | 2.62E-01 | 9.36E-01 | 1.61E+00 | 9.17E-02 | 2.59E-01 | -1.35E+00 | 1.16E-01 | 4.25E-01 | -1.24E+00 | 9.76E-02 | 2.68E-01 | -1.26E+00 |
| GSTP1 | 5.80E-03 | 3.02E-02 | 1.24E+00 | 1.14E-03 | 4.93E-03 | 1.31E+00 | 9.31E-05 | 3.96E-04 | 1.36E+00 | 4.51E-04 | 1.72E-03 | 1.32E+00 | 1.81E-08 | 6.45E-08 | 1.48E+00 | 1.39E-04 | 3.91E-04 | 1.32E+00 | 2.91E-27 | 1.07E-26 | 1.99E+00 | 4.55E-39 | 1.25E-38 | 2.11E+00 |
| AL442125.1 | 5.81E-03 | 3.03E-02 | -1.57E+00 | 2.26E-01 | 9.77E-01 | 1.43E+00 | 1.33E-03 | 5.65E-03 | -1.70E+00 | 2.55E-01 | 9.73E-01 | 1.38E+00 | 4.43E-03 | 1.58E-02 | -2.08E+00 | 1.55E-01 | 4.38E-01 | -1.16E+00 | 4.78E-02 | 1.75E-01 | -1.11E+00 | 1.32E-01 | 3.64E-01 | 1.07E+00 |
| METTL5 | 5.81E-03 | 3.03E-02 | 1.05E+00 | 1.63E-02 | 7.02E-02 | 1.24E+00 | 9.03E-03 | 3.84E-02 | 1.01E+00 | 2.66E-03 | 1.02E-02 | 1.08E+00 | 1.48E-04 | 5.29E-04 | -1.11E+00 | 7.18E-02 | 2.02E-01 | 1.15E+00 | 2.76E-02 | 1.01E-01 | 1.02E+00 | 1.56E-01 | 4.27E-01 | 1.09E+00 |
| TCF20 | 5.81E-03 | 3.03E-02 | -1.13E+00 | 1.44E-01 | 6.24E-01 | 1.38E+00 | 1.15E-02 | 4.89E-02 | -1.25E+00 | 2.47E-01 | 9.43E-01 | 1.40E+00 | 1.02E-01 | 3.65E-01 | -1.04E+00 | 2.69E-01 | 7.59E-01 | 1.19E+00 | 2.93E-03 | 1.08E-02 | -1.28E+00 | 1.57E-01 | 4.30E-01 | 1.15E+00 |
| INF2 | 5.81E-03 | 3.03E-02 | -1.08E+00 | 9.17E-03 | 3.96E-02 | 1.08E+00 | 1.71E-01 | 7.26E-01 | 1.25E+00 | 4.49E-02 | 1.71E-01 | 1.20E+00 | 1.89E-04 | 6.76E-04 | -1.65E+00 | 1.86E-03 | 5.24E-03 | -1.29E+00 | 1.64E-05 | 6.01E-05 | -1.16E+00 | 2.32E-03 | 6.38E-03 | -1.02E+00 |
| OSBPL2 | 5.81E-03 | 3.03E-02 | -1.04E+00 | 7.76E-02 | 3.35E-01 | 1.32E+00 | 6.99E-03 | 2.97E-02 | -1.17E+00 | 1.67E-02 | 6.39E-02 | 1.02E+00 | 4.30E-04 | 1.53E-03 | -1.49E+00 | 1.10E-02 | 3.10E-02 | -1.12E+00 | 7.74E-03 | 2.84E-02 | -1.13E+00 | 1.73E-02 | 4.75E-02 | -1.10E+00 |
| VRK1 | 5.81E-03 | 3.03E-02 | 1.09E+00 | 2.50E-03 | 1.08E-02 | 1.21E+00 | 1.51E-01 | 6.43E-01 | 1.26E+00 | 1.24E-01 | 4.75E-01 | 1.21E+00 | 2.55E-04 | 9.10E-04 | -1.12E+00 | 2.40E-02 | 6.77E-02 | 1.16E+00 | 7.23E-08 | 2.66E-07 | -1.19E+00 | 5.70E-04 | 1.57E-03 | -1.05E+00 |
| ATP5E | 5.82E-03 | 3.03E-02 | 1.19E+00 | 9.43E-11 | 4.08E-10 | -1.14E+00 | 3.02E-03 | 1.28E-02 | 1.28E+00 | 1.62E-04 | 6.20E-04 | -1.10E+00 | 2.39E-01 | 8.54E-01 | 1.15E+00 | 1.62E-07 | 4.58E-07 | -1.22E+00 | 4.41E-02 | 1.62E-01 | 1.19E+00 | 4.02E-03 | 1.10E-02 | -1.18E+00 |
| MTERF2 | 5.82E-03 | 3.03E-02 | -1.09E+00 | 3.74E-02 | 1.62E-01 | 1.22E+00 | 4.55E-05 | 1.93E-04 | -1.40E+00 | 4.14E-04 | 1.58E-03 | -1.10E+00 | 2.03E-02 | 7.23E-02 | 1.12E+00 | 8.96E-02 | 2.53E-01 | 1.24E+00 | 5.39E-05 | 1.98E-04 | -1.16E+00 | 3.68E-02 | 1.01E-01 | 1.08E+00 |
| SCFD2 | 5.82E-03 | 3.03E-02 | -1.18E+00 | 1.36E-01 | 5.89E-01 | 1.31E+00 | 1.89E-03 | 8.02E-03 | -1.33E+00 | 1.97E-02 | 7.53E-02 | 1.01E+00 | 8.60E-03 | 3.07E-02 | -1.25E+00 | 1.55E-01 | 4.36E-01 | 1.27E+00 | 4.00E-08 | 1.47E-07 | -1.54E+00 | 2.80E-02 | 7.68E-02 | 1.01E+00 |
| TTC39C | 5.85E-03 | 3.05E-02 | 1.11E+00 | 5.46E-07 | 2.36E-06 | -1.03E+00 | 1.69E-01 | 7.17E-01 | 1.21E+00 | 6.57E-04 | 2.51E-03 | 1.10E+00 | 1.07E-01 | 3.83E-01 | 1.64E+00 | 3.37E-01 | 9.50E-01 | 1.34E+00 | 2.22E-01 | 8.14E-01 | 1.17E+00 | 9.75E-02 | 2.68E-01 | 1.01E+00 |
| MED22 | 5.85E-03 | 3.05E-02 | -1.20E+00 | 1.62E-01 | 7.02E-01 | 1.37E+00 | 4.83E-02 | 2.05E-01 | -1.07E+00 | 1.49E-02 | 5.69E-02 | -1.09E+00 | 2.59E-02 | 9.22E-02 | -1.03E+00 | 3.87E-02 | 1.09E-01 | 1.02E+00 | 4.63E-02 | 1.70E-01 | 1.02E+00 | 2.09E-01 | 5.73E-01 | 1.25E+00 |
| PIGX | 5.85E-03 | 3.05E-02 | 1.01E+00 | 7.02E-03 | 3.03E-02 | 1.12E+00 | 3.49E-03 | 1.48E-02 | -1.04E+00 | 1.20E-03 | 4.57E-03 | 1.07E+00 | 7.96E-04 | 2.84E-03 | -1.17E+00 | 2.94E-02 | 8.28E-02 | 1.14E+00 | 1.28E-02 | 4.70E-02 | 1.05E+00 | 3.32E-01 | 9.12E-01 | 1.20E+00 |
| CSRNP1 | 5.86E-03 | 3.05E-02 | 1.67E+00 | 1.17E-10 | 5.07E-10 | -1.38E+00 | 1.83E-01 | 7.76E-01 | 1.34E+00 | 1.15E-07 | 4.39E-07 | -1.45E+00 | 3.23E-02 | 1.15E-01 | 1.78E+00 | 3.54E-05 | 9.99E-05 | -1.53E+00 | 2.09E-01 | 7.68E-01 | 1.38E+00 | 3.30E-09 | 9.06E-09 | -2.16E+00 |
| FCGR1B | 5.86E-03 | 3.05E-02 | -1.69E+00 | 1.98E-03 | 8.56E-03 | -2.43E+00 | 1.00E-32 | 4.27E-32 | -2.55E+00 | 3.73E-43 | 1.43E-42 | -3.20E+00 | 2.18E-01 | 7.76E-01 | -1.05E+00 | 2.98E-01 | 8.40E-01 | -1.01E+00 | 2.38E-01 | 8.75E-01 | 1.05E+00 | 2.63E-01 | 7.22E-01 | -1.32E+00 |
| TIPARP-AS1 | 5.86E-03 | 3.05E-02 | 1.10E+00 | 1.09E-07 | 4.72E-07 | -1.15E+00 | 1.46E-02 | 6.19E-02 | 1.01E+00 | 2.86E-06 | 1.09E-05 | -1.33E+00 | 2.77E-01 | 9.88E-01 | 1.39E+00 | 4.13E-02 | 1.16E-01 | 1.04E+00 | 5.57E-02 | 2.05E-01 | -1.05E+00 | 1.84E-03 | 5.05E-03 | -1.36E+00 |
| WDR4 | 5.86E-03 | 3.06E-02 | -1.22E+00 | 3.49E-02 | 1.51E-01 | 1.14E+00 | 9.78E-02 | 4.15E-01 | 1.09E+00 | 1.52E-01 | 5.79E-01 | 1.21E+00 | 1.09E-01 | 3.88E-01 | -1.21E+00 | 1.68E-01 | 4.75E-01 | 1.73E+00 | 3.69E-06 | 1.35E-05 | -1.39E+00 | 3.64E-01 | 1.00E+00 | 1.27E+00 |
| AP3S2 | 5.87E-03 | 3.06E-02 | -1.23E+00 | 3.21E-03 | 1.39E-02 | -1.17E+00 | 9.51E-02 | 4.04E-01 | -1.08E+00 | 5.31E-02 | 2.03E-01 | 1.02E+00 | 5.51E-03 | 1.96E-02 | -1.37E+00 | 1.52E-01 | 4.28E-01 | 1.09E+00 | 1.58E-01 | 5.78E-01 | 1.11E+00 | 3.40E-01 | 9.35E-01 | 1.17E+00 |
| NIPA1 | 5.88E-03 | 3.07E-02 | 1.26E+00 | 7.24E-03 | 3.13E-02 | 1.05E+00 | 3.94E-03 | 1.67E-02 | -1.05E+00 | 3.72E-04 | 1.42E-03 | -1.24E+00 | 4.57E-03 | 1.63E-02 | -1.19E+00 | 3.87E-04 | 1.09E-03 | -1.51E+00 | 1.47E-04 | 5.41E-04 | -1.38E+00 | 5.04E-03 | 1.39E-02 | -1.16E+00 |
| INVS | 5.89E-03 | 3.07E-02 | -1.16E+00 | 2.30E-02 | 9.93E-02 | 1.04E+00 | 2.24E-04 | 9.51E-04 | -1.48E+00 | 2.35E-03 | 8.99E-03 | -1.14E+00 | 3.82E-03 | 1.36E-02 | -1.23E+00 | 2.58E-02 | 7.28E-02 | -1.11E+00 | 5.91E-04 | 2.17E-03 | -1.25E+00 | 3.18E-02 | 8.72E-02 | 1.02E+00 |
| NATD1 | 5.89E-03 | 3.07E-02 | -1.12E+00 | 1.61E-02 | 6.96E-02 | 1.05E+00 | 1.47E-01 | 6.23E-01 | 1.08E+00 | 2.13E-01 | 8.14E-01 | 1.47E+00 | 1.14E-01 | 4.05E-01 | -1.06E+00 | 9.74E-02 | 2.74E-01 | -1.20E+00 | 1.15E-01 | 4.23E-01 | -1.03E+00 | 1.58E-01 | 4.33E-01 | -1.04E+00 |
| LINC02352 | 5.90E-03 | 3.07E-02 | -1.11E+00 | 1.46E-03 | 6.29E-03 | -1.09E+00 | 1.10E-01 | 4.69E-01 | 1.37E+00 | 2.37E-01 | 9.06E-01 | 1.19E+00 | 1.49E-01 | 5.30E-01 | 1.14E+00 | 9.49E-02 | 2.68E-01 | 1.03E+00 | 4.07E-02 | 1.49E-01 | -1.13E+00 | 3.79E-02 | 1.04E-01 | -1.38E+00 |
| NEK6 | 5.93E-03 | 3.09E-02 | -1.07E+00 | 1.12E-02 | 4.82E-02 | 1.08E+00 | 7.07E-02 | 3.00E-01 | 1.02E+00 | 7.77E-03 | 2.97E-02 | -1.02E+00 | 1.03E-02 | 3.67E-02 | -1.80E+00 | 9.35E-02 | 2.64E-01 | -1.25E+00 | 9.76E-02 | 3.58E-01 | 1.02E+00 | 2.26E-01 | 6.20E-01 | 1.13E+00 |
| AL031595.3 | 5.93E-03 | 3.09E-02 | -1.13E+00 | 9.31E-02 | 4.02E-01 | 1.26E+00 | 1.99E-01 | 8.48E-01 | 1.29E+00 | 1.41E-01 | 5.37E-01 | 1.66E+00 | 2.74E-03 | 9.76E-03 | -1.01E+00 | 2.97E-01 | 8.36E-01 | 1.50E+00 | 1.12E-05 | 4.11E-05 | -1.21E+00 | 2.82E-01 | 7.75E-01 | 1.22E+00 |
| RPL23AP2 | 5.95E-03 | 3.10E-02 | -1.07E+00 | 1.46E-22 | 6.31E-22 | -1.37E+00 | 1.98E-01 | 8.40E-01 | 1.01E+00 | 5.94E-05 | 2.27E-04 | -1.22E+00 | 3.16E-04 | 1.13E-03 | -1.15E+00 | 1.52E-08 | 4.29E-08 | -1.32E+00 | 7.00E-04 | 2.57E-03 | -1.12E+00 | 7.59E-08 | 2.09E-07 | -1.32E+00 |
| AC022497.1 | 5.95E-03 | 3.10E-02 | -1.34E+00 | 5.37E-02 | 2.32E-01 | 1.08E+00 | 5.14E-03 | 2.18E-02 | -1.75E+00 | 1.59E-02 | 6.07E-02 | -1.54E+00 | 7.13E-03 | 2.54E-02 | -1.31E+00 | 3.45E-02 | 9.72E-02 | -1.13E+00 | 1.48E-01 | 5.45E-01 | 1.39E+00 | 2.97E-01 | 8.16E-01 | 1.39E+00 |
| AL353763.2 | 5.95E-03 | 3.10E-02 | 1.03E+00 | 2.14E-01 | 9.25E-01 | 1.46E+00 | 4.47E-03 | 1.90E-02 | -1.08E+00 | 5.20E-02 | 1.99E-01 | 1.28E+00 | 1.54E-02 | 5.49E-02 | -1.05E+00 | 3.54E-01 | 9.98E-01 | 1.41E+00 | 1.49E-01 | 5.48E-01 | 1.19E+00 | 1.55E-01 | 4.27E-01 | 1.11E+00 |
| RPSAP36 | 5.95E-03 | 3.10E-02 | -1.28E+00 | 2.46E-03 | 1.06E-02 | -1.33E+00 | 5.57E-02 | 2.37E-01 | -1.20E+00 | 8.74E-02 | 3.34E-01 | -1.01E+00 | 1.85E-02 | 6.58E-02 | -1.36E+00 | 9.35E-03 | 2.64E-02 | -1.59E+00 | 1.48E-03 | 5.44E-03 | -1.31E+00 | 1.87E-03 | 5.14E-03 | -1.45E+00 |
| RANBP10 | 5.95E-03 | 3.10E-02 | -1.15E+00 | 7.34E-03 | 3.17E-02 | -1.04E+00 | 1.23E-01 | 5.23E-01 | 1.08E+00 | 7.79E-02 | 2.98E-01 | 1.19E+00 | 3.76E-03 | 1.34E-02 | -1.11E+00 | 2.14E-02 | 6.04E-02 | -1.03E+00 | 2.45E-02 | 9.00E-02 | -1.05E+00 | 3.57E-02 | 9.81E-02 | -1.04E+00 |
| ALG11 | 5.96E-03 | 3.10E-02 | -1.25E+00 | 1.20E-01 | 5.18E-01 | 1.31E+00 | 6.06E-05 | 2.58E-04 | -1.75E+00 | 1.20E-02 | 4.60E-02 | -1.05E+00 | 2.84E-04 | 1.01E-03 | -1.84E+00 | 9.29E-02 | 2.62E-01 | 1.08E+00 | 3.03E-06 | 1.11E-05 | -1.29E+00 | 2.19E-03 | 6.02E-03 | -1.08E+00 |
| IQSEC2 | 5.96E-03 | 3.11E-02 | -1.22E+00 | 3.50E-03 | 1.51E-02 | -1.29E+00 | 4.79E-02 | 2.04E-01 | -1.17E+00 | 4.68E-02 | 1.79E-01 | -1.13E+00 | 1.02E-01 | 3.63E-01 | -1.06E+00 | 2.68E-01 | 7.56E-01 | 1.62E+00 | 1.26E-01 | 4.63E-01 | 1.06E+00 | 1.74E-01 | 4.78E-01 | 1.04E+00 |
| RECQL5 | 5.97E-03 | 3.11E-02 | -1.07E+00 | 2.30E-03 | 9.93E-03 | -1.01E+00 | 4.41E-04 | 1.88E-03 | -1.29E+00 | 9.30E-04 | 3.55E-03 | -1.14E+00 | 6.81E-04 | 2.43E-03 | -1.61E+00 | 2.03E-03 | 5.71E-03 | -1.53E+00 | 4.68E-07 | 1.72E-06 | -1.23E+00 | 2.70E-05 | 7.41E-05 | -1.34E+00 |
| STMN3 | 5.97E-03 | 3.11E-02 | 1.11E+00 | 1.90E-01 | 8.22E-01 | 1.40E+00 | 5.47E-03 | 2.32E-02 | -1.07E+00 | 5.95E-03 | 2.27E-02 | 1.06E+00 | 1.08E-02 | 3.86E-02 | -1.04E+00 | 8.32E-03 | 2.35E-02 | 1.01E+00 | 7.45E-03 | 2.73E-02 | -1.00E+00 | 2.16E-01 | 5.92E-01 | 1.21E+00 |
| AC092798.1 | 5.97E-03 | 3.11E-02 | 1.94E+00 | 2.12E-01 | 9.16E-01 | 1.39E+00 | 2.89E-03 | 1.23E-02 | 2.01E+00 | 1.81E-01 | 6.90E-01 | 1.55E+00 | 9.39E-03 | 3.35E-02 | 2.35E+00 | 3.35E-01 | 9.43E-01 | 1.20E+00 | 1.22E-01 | 4.50E-01 | 1.39E+00 | 3.33E-01 | 9.14E-01 | 1.15E+00 |
| ZNF468 | 5.98E-03 | 3.12E-02 | -1.18E+00 | 8.49E-04 | 3.67E-03 | -1.32E+00 | 2.21E-02 | 9.39E-02 | -1.03E+00 | 7.74E-04 | 2.96E-03 | -1.27E+00 | 4.33E-02 | 1.54E-01 | 1.05E+00 | 1.29E-02 | 3.63E-02 | -1.11E+00 | 1.74E-04 | 6.38E-04 | -1.25E+00 | 2.46E-06 | 6.76E-06 | -1.57E+00 |
| CAVIN2 | 5.99E-03 | 3.12E-02 | 1.68E+00 | 1.81E-01 | 7.81E-01 | 1.38E+00 | 1.42E-04 | 6.01E-04 | 2.02E+00 | 2.09E-01 | 7.99E-01 | 1.44E+00 | 1.98E-13 | 7.06E-13 | -2.15E+00 | 5.23E-06 | 1.47E-05 | -1.54E+00 | 9.61E-09 | 3.53E-08 | -1.07E+00 | 6.60E-04 | 1.81E-03 | -1.12E+00 |
| PPHLN1 | 6.01E-03 | 3.13E-02 | 1.09E+00 | 3.66E-02 | 1.58E-01 | 1.28E+00 | 7.80E-02 | 3.31E-01 | 1.16E+00 | 1.57E-02 | 6.00E-02 | 1.15E+00 | 5.16E-04 | 1.84E-03 | 1.01E+00 | 4.29E-02 | 1.21E-01 | 1.19E+00 | 2.93E-03 | 1.08E-02 | -1.02E+00 | 1.46E-01 | 4.01E-01 | 1.14E+00 |
| HLA-DQA2 | 6.01E-03 | 3.13E-02 | 1.03E+00 | 6.66E-02 | 2.88E-01 | 1.61E+00 | 4.42E-02 | 1.88E-01 | 1.18E+00 | 1.13E-01 | 4.30E-01 | 1.71E+00 | 2.57E-03 | 9.16E-03 | -1.44E+00 | 1.25E-01 | 3.54E-01 | 1.13E+00 | 3.09E-03 | 1.13E-02 | -1.49E+00 | 2.01E-01 | 5.52E-01 | 1.24E+00 |
| SLAMF7 | 6.02E-03 | 3.14E-02 | -1.57E+00 | 5.82E-03 | 2.51E-02 | -1.50E+00 | 8.73E-03 | 3.71E-02 | -1.51E+00 | 1.14E-01 | 4.35E-01 | 1.06E+00 | 3.31E-02 | 1.18E-01 | -1.20E+00 | 2.92E-02 | 8.23E-02 | -1.45E+00 | 1.48E-01 | 5.42E-01 | -1.04E+00 | 7.73E-02 | 2.12E-01 | -1.33E+00 |
| RPL29P12 | 6.02E-03 | 3.14E-02 | 1.01E+00 | 7.20E-02 | 3.11E-01 | 1.74E+00 | 5.31E-03 | 2.26E-02 | -1.16E+00 | 1.48E-01 | 5.64E-01 | 1.56E+00 | 2.08E-01 | 7.40E-01 | 1.55E+00 | 5.30E-03 | 1.49E-02 | 2.59E+00 | 2.48E-01 | 9.12E-01 | 1.28E+00 | 1.32E-05 | 3.63E-05 | 2.51E+00 |
| ZNF790 | 6.03E-03 | 3.14E-02 | -1.06E+00 | 1.55E-02 | 6.68E-02 | 1.09E+00 | 2.05E-01 | 8.70E-01 | 1.24E+00 | 1.03E-01 | 3.95E-01 | 1.29E+00 | 2.10E-03 | 7.50E-03 | -1.07E+00 | 1.77E-03 | 4.99E-03 | -1.08E+00 | 6.28E-02 | 2.31E-01 | 1.02E+00 | 1.98E-02 | 5.43E-02 | -1.05E+00 |
| TXLNB | 6.04E-03 | 3.15E-02 | -1.00E+00 | 1.48E-02 | 6.41E-02 | 1.02E+00 | 4.75E-02 | 2.02E-01 | 1.05E+00 | 7.07E-04 | 2.70E-03 | -1.44E+00 | 2.33E-01 | 8.32E-01 | -1.05E+00 | 2.24E-01 | 6.33E-01 | -1.22E+00 | 2.17E-01 | 7.98E-01 | -1.02E+00 | 3.27E-01 | 8.98E-01 | 1.02E+00 |
| LTBP2 | 6.04E-03 | 3.15E-02 | -1.63E+00 | 8.39E-02 | 3.63E-01 | -1.08E+00 | 1.91E-01 | 8.12E-01 | -1.01E+00 | 1.04E-01 | 3.97E-01 | 1.58E+00 | 1.25E-01 | 4.46E-01 | -1.22E+00 | 3.25E-01 | 9.17E-01 | 1.19E+00 | 1.03E-06 | 3.78E-06 | -1.83E+00 | 8.56E-07 | 2.35E-06 | -1.75E+00 |
| AC004812.2 | 6.05E-03 | 3.15E-02 | -1.11E+00 | 2.88E-02 | 1.25E-01 | 1.14E+00 | 8.97E-04 | 3.81E-03 | -1.51E+00 | 8.21E-03 | 3.13E-02 | -1.15E+00 | 1.69E-01 | 6.03E-01 | 1.80E+00 | 5.16E-02 | 1.45E-01 | 2.20E+00 | 2.60E-01 | 9.53E-01 | 1.23E+00 | 1.77E-01 | 4.86E-01 | 1.34E+00 |
| NUTM2A-AS1 | 6.05E-03 | 3.15E-02 | -1.01E+00 | 3.58E-03 | 1.55E-02 | 1.07E+00 | 2.52E-04 | 1.07E-03 | -1.20E+00 | 5.32E-04 | 2.03E-03 | -1.04E+00 | 2.68E-05 | 9.57E-05 | -1.08E+00 | 1.58E-02 | 4.44E-02 | 1.15E+00 | 2.73E-02 | 1.00E-01 | 1.02E+00 | 2.69E-01 | 7.39E-01 | 1.24E+00 |
| PSD | 6.05E-03 | 3.15E-02 | -1.13E+00 | 1.75E-04 | 7.54E-04 | -1.36E+00 | 4.80E-02 | 2.04E-01 | 1.66E+00 | 5.41E-02 | 2.07E-01 | 1.96E+00 | 6.27E-02 | 2.24E-01 | -1.28E+00 | 1.35E-01 | 3.79E-01 | -1.13E+00 | 1.59E-04 | 5.82E-04 | -1.91E+00 | 2.49E-04 | 6.85E-04 | -1.86E+00 |
| RIC1 | 6.06E-03 | 3.16E-02 | 1.12E+00 | 4.15E-03 | 1.79E-02 | 1.20E+00 | 1.07E-02 | 4.54E-02 | -1.06E+00 | 7.94E-03 | 3.03E-02 | 1.02E+00 | 2.81E-02 | 1.00E-01 | 2.02E+00 | 1.38E-01 | 3.89E-01 | 1.64E+00 | 8.14E-03 | 2.99E-02 | -1.03E+00 | 2.43E-02 | 6.68E-02 | 1.03E+00 |
| NRG2 | 6.08E-03 | 3.17E-02 | -1.26E+00 | 2.31E-03 | 9.98E-03 | -1.41E+00 | 2.25E-01 | 9.56E-01 | 1.07E+00 | 1.92E-01 | 7.32E-01 | 1.06E+00 | 1.97E-01 | 7.02E-01 | 1.49E+00 | 2.56E-01 | 7.22E-01 | 1.47E+00 | 1.87E-01 | 6.86E-01 | 1.27E+00 | 1.14E-01 | 3.12E-01 | 1.68E+00 |
| SMAD9 | 6.08E-03 | 3.17E-02 | -1.69E+00 | 9.75E-02 | 4.21E-01 | 1.65E+00 | 1.69E-03 | 7.16E-03 | -1.69E+00 | 8.47E-02 | 3.23E-01 | 1.02E+00 | 2.59E-02 | 9.23E-02 | -1.61E+00 | 3.87E-02 | 1.09E-01 | -1.31E+00 | 2.29E-01 | 8.40E-01 | 1.18E+00 | 2.98E-03 | 8.19E-03 | 2.51E+00 |
| MRPL42P6 | 6.09E-03 | 3.17E-02 | -1.15E+00 | 9.94E-02 | 4.30E-01 | 1.32E+00 | 3.31E-04 | 1.41E-03 | -1.39E+00 | 7.70E-04 | 2.94E-03 | -1.18E+00 | 6.67E-02 | 2.38E-01 | -1.12E+00 | 6.32E-02 | 1.78E-01 | -1.22E+00 | 8.21E-02 | 3.01E-01 | -1.22E+00 | 3.23E-01 | 8.88E-01 | 1.16E+00 |
| ANKRD13B | 6.12E-03 | 3.19E-02 | -1.35E+00 | 2.07E-02 | 8.94E-02 | -1.06E+00 | 3.53E-04 | 1.50E-03 | -1.40E+00 | 9.74E-03 | 3.72E-02 | 1.01E+00 | 9.71E-04 | 3.46E-03 | -1.47E+00 | 4.52E-03 | 1.27E-02 | -1.23E+00 | 1.25E-04 | 4.59E-04 | -1.88E+00 | 3.96E-02 | 1.09E-01 | -1.16E+00 |
| KRT17P1 | 6.12E-03 | 3.19E-02 | 1.02E+00 | 6.90E-05 | 2.98E-04 | -1.25E+00 | 2.15E-04 | 9.14E-04 | -1.34E+00 | 1.87E-05 | 7.15E-05 | -1.42E+00 | 5.58E-03 | 1.99E-02 | -2.40E+00 | 1.25E-02 | 3.51E-02 | -1.85E+00 | 1.48E-01 | 5.42E-01 | 1.84E+00 | 3.37E-01 | 9.26E-01 | 1.24E+00 |
| HABP4 | 6.13E-03 | 3.19E-02 | -1.10E+00 | 7.93E-02 | 3.43E-01 | 1.38E+00 | 3.75E-02 | 1.59E-01 | -1.20E+00 | 1.53E-01 | 5.86E-01 | 1.13E+00 | 2.18E-01 | 7.78E-01 | 1.13E+00 | 2.92E-01 | 8.23E-01 | 1.54E+00 | 5.72E-02 | 2.10E-01 | -1.36E+00 | 3.52E-01 | 9.66E-01 | 1.28E+00 |
| AC009948.5 | 6.13E-03 | 3.20E-02 | -1.21E+00 | 1.37E-01 | 5.91E-01 | 1.65E+00 | 9.95E-04 | 4.23E-03 | -1.70E+00 | 1.16E-01 | 4.42E-01 | 1.14E+00 | 2.69E-03 | 9.59E-03 | -1.63E+00 | 1.55E-02 | 4.36E-02 | -1.26E+00 | 3.33E-02 | 1.22E-01 | -1.43E+00 | 2.85E-01 | 7.84E-01 | 1.42E+00 |
| KIAA0232 | 6.14E-03 | 3.20E-02 | -1.02E+00 | 2.76E-04 | 1.19E-03 | -1.11E+00 | 5.07E-06 | 2.15E-05 | -1.35E+00 | 3.54E-06 | 1.35E-05 | -1.25E+00 | 5.35E-06 | 1.91E-05 | -1.55E+00 | 1.88E-05 | 5.30E-05 | -1.36E+00 | 6.16E-05 | 2.26E-04 | -1.19E+00 | 6.38E-09 | 1.75E-08 | -1.36E+00 |
| IDH1 | 6.14E-03 | 3.20E-02 | -1.15E+00 | 8.98E-02 | 3.88E-01 | 1.63E+00 | 1.37E-07 | 5.81E-07 | -1.57E+00 | 1.09E-01 | 4.17E-01 | 1.24E+00 | 1.44E-06 | 5.14E-06 | -1.61E+00 | 2.61E-02 | 7.35E-02 | 1.03E+00 | 2.29E-09 | 8.40E-09 | -1.53E+00 | 7.43E-03 | 2.04E-02 | 1.03E+00 |
| AC010326.4 | 6.14E-03 | 3.20E-02 | -1.25E+00 | 4.57E-02 | 1.97E-01 | 1.05E+00 | 1.34E-02 | 5.70E-02 | -1.43E+00 | 7.05E-02 | 2.69E-01 | -1.07E+00 | 1.29E-01 | 4.59E-01 | 1.08E+00 | 1.10E-01 | 3.10E-01 | 1.00E+00 | 3.11E-04 | 1.14E-03 | -1.66E+00 | 4.27E-04 | 1.17E-03 | -1.69E+00 |
| AL080317.1 | 6.14E-03 | 3.20E-02 | -1.45E+00 | 7.24E-02 | 3.13E-01 | 1.07E+00 | 6.14E-02 | 2.61E-01 | -1.28E+00 | 2.38E-01 | 9.07E-01 | 1.19E+00 | 9.57E-02 | 3.41E-01 | -1.10E+00 | 3.22E-01 | 9.08E-01 | 1.20E+00 | 1.43E-03 | 5.24E-03 | -1.47E+00 | 9.36E-02 | 2.57E-01 | 1.06E+00 |
| ADCY6 | 6.14E-03 | 3.20E-02 | -1.21E+00 | 7.55E-02 | 3.26E-01 | 1.25E+00 | 1.92E-02 | 8.15E-02 | -1.47E+00 | 1.04E-01 | 3.98E-01 | -1.01E+00 | 9.38E-02 | 3.34E-01 | 1.10E+00 | 1.77E-01 | 4.99E-01 | 1.13E+00 | 9.02E-02 | 3.31E-01 | 1.19E+00 | 8.89E-03 | 2.44E-02 | -1.04E+00 |
| TMEM176A | 6.15E-03 | 3.20E-02 | -1.47E+00 | 7.36E-03 | 3.18E-02 | -1.28E+00 | 5.14E-04 | 2.19E-03 | -1.86E+00 | 2.56E-04 | 9.78E-04 | -2.21E+00 | 6.70E-02 | 2.39E-01 | -1.68E+00 | 8.24E-02 | 2.32E-01 | -1.69E+00 | 1.49E-01 | 5.48E-01 | -1.46E+00 | 2.27E-01 | 6.24E-01 | -1.21E+00 |
| MCMBP | 6.15E-03 | 3.20E-02 | -1.21E+00 | 1.21E-01 | 5.22E-01 | 1.28E+00 | 4.04E-02 | 1.72E-01 | 1.03E+00 | 6.83E-02 | 2.61E-01 | 1.21E+00 | 2.90E-05 | 1.03E-04 | -1.73E+00 | 1.52E-03 | 4.29E-03 | -1.28E+00 | 6.30E-09 | 2.31E-08 | -1.40E+00 | 1.40E-02 | 3.85E-02 | 1.05E+00 |
| CEBPD | 6.16E-03 | 3.21E-02 | 2.64E+00 | 2.22E-01 | 9.58E-01 | 1.45E+00 | 9.96E-02 | 4.23E-01 | 1.24E+00 | 9.86E-02 | 3.77E-01 | 1.22E+00 | 2.52E-01 | 8.99E-01 | 2.14E+00 | 3.12E-01 | 8.80E-01 | 1.26E+00 | 1.48E-01 | 5.44E-01 | 1.99E+00 | 3.56E-01 | 9.77E-01 | -1.01E+00 |
| GNG4 | 6.17E-03 | 3.21E-02 | 1.95E+00 | 3.78E-06 | 1.63E-05 | 2.75E+00 | 5.36E-05 | 2.28E-04 | 1.96E+00 | 2.95E-04 | 1.13E-03 | 2.17E+00 | 1.64E-01 | 5.86E-01 | 2.22E+00 | 2.39E-01 | 6.73E-01 | 1.99E+00 | 1.99E-01 | 7.30E-01 | -1.27E+00 | 3.27E-01 | 9.00E-01 | 1.25E+00 |
| NPAS1 | 6.17E-03 | 3.21E-02 | -1.40E+00 | 2.47E-02 | 1.07E-01 | -1.12E+00 | 2.31E-02 | 9.80E-02 | -1.52E+00 | 1.41E-01 | 5.40E-01 | -1.00E+00 | 2.35E-01 | 8.40E-01 | 1.07E+00 | 2.82E-01 | 7.95E-01 | 1.49E+00 | 2.70E-01 | 9.90E-01 | 1.02E+00 | 2.59E-01 | 7.12E-01 | 1.25E+00 |
| AC008549.3 | 6.17E-03 | 3.21E-02 | 1.82E+00 | 1.08E-02 | 4.68E-02 | 1.11E+00 | 7.05E-02 | 2.99E-01 | 1.40E+00 | 6.73E-04 | 2.57E-03 | -1.10E+00 | 2.71E-01 | 9.66E-01 | 1.51E+00 | 5.24E-03 | 1.48E-02 | -1.01E+00 | 5.14E-03 | 1.89E-02 | 1.97E+00 | 2.48E-02 | 6.82E-02 | 1.03E+00 |
| MYLK | 6.17E-03 | 3.21E-02 | -1.61E+00 | 2.05E-01 | 8.84E-01 | 1.41E+00 | 6.53E-04 | 2.78E-03 | -1.73E+00 | 1.57E-01 | 6.00E-01 | 1.24E+00 | 1.78E-02 | 6.34E-02 | -2.10E+00 | 2.72E-01 | 7.67E-01 | -1.08E+00 | 6.56E-03 | 2.41E-02 | 1.10E+00 | 6.36E-02 | 1.75E-01 | -1.26E+00 |
| JAML | 6.17E-03 | 3.22E-02 | -1.38E+00 | 5.29E-03 | 2.28E-02 | -1.20E+00 | 1.63E-03 | 6.93E-03 | 1.78E+00 | 2.50E-01 | 9.54E-01 | -1.00E+00 | 2.70E-01 | 9.63E-01 | 1.06E+00 | 3.03E-01 | 8.53E-01 | -1.21E+00 | 1.81E-01 | 6.65E-01 | -1.89E+00 | 2.75E-01 | 7.56E-01 | -1.25E+00 |
| RPL7P6 | 6.17E-03 | 3.22E-02 | 1.88E+00 | 1.34E-01 | 5.77E-01 | 1.39E+00 | 1.78E-01 | 7.57E-01 | 1.31E+00 | 1.38E-02 | 5.27E-02 | 1.05E+00 | 1.84E-01 | 6.55E-01 | 1.21E+00 | 1.71E-02 | 4.81E-02 | -1.15E+00 | 1.05E-01 | 3.86E-01 | 1.45E+00 | 4.09E-02 | 1.12E-01 | -1.04E+00 |
| AC092117.2 | 6.19E-03 | 3.23E-02 | -1.33E+00 | 3.92E-03 | 1.70E-02 | -1.28E+00 | 4.33E-03 | 1.84E-02 | -1.29E+00 | 3.19E-06 | 1.22E-05 | -1.83E+00 | 5.08E-02 | 1.81E-01 | -1.14E+00 | 5.04E-02 | 1.42E-01 | -1.24E+00 | 1.61E-01 | 5.91E-01 | -1.00E+00 | 1.11E-01 | 3.04E-01 | -1.10E+00 |
| SNHG10 | 6.19E-03 | 3.23E-02 | -1.02E+00 | 8.45E-03 | 3.65E-02 | 1.13E+00 | 3.29E-02 | 1.40E-01 | -1.05E+00 | 5.52E-02 | 2.11E-01 | 1.12E+00 | 1.40E-02 | 5.00E-02 | -1.07E+00 | 1.63E-01 | 4.59E-01 | 1.31E+00 | 1.13E-01 | 4.17E-01 | 1.15E+00 | 3.58E-01 | 9.83E-01 | 1.38E+00 |
| ADCK1 | 6.19E-03 | 3.23E-02 | -1.32E+00 | 2.82E-02 | 1.22E-01 | -1.05E+00 | 4.80E-03 | 2.04E-02 | -1.31E+00 | 1.06E-02 | 4.06E-02 | -1.13E+00 | 1.27E-03 | 4.52E-03 | -1.94E+00 | 9.64E-03 | 2.72E-02 | -1.26E+00 | 1.26E-05 | 4.64E-05 | -1.45E+00 | 1.55E-03 | 4.27E-03 | -1.16E+00 |
| ZCCHC14 | 6.19E-03 | 3.23E-02 | -1.38E+00 | 2.68E-02 | 1.16E-01 | -1.07E+00 | 2.23E-01 | 9.46E-01 | 1.11E+00 | 3.18E-02 | 1.22E-01 | -1.17E+00 | 4.97E-02 | 1.77E-01 | -1.08E+00 | 2.70E-02 | 7.61E-02 | -1.24E+00 | 1.46E-02 | 5.35E-02 | -1.14E+00 | 2.11E-03 | 5.80E-03 | -1.50E+00 |
| BEX2 | 6.20E-03 | 3.23E-02 | 1.10E+00 | 7.97E-06 | 3.45E-05 | 1.04E+00 | 3.14E-02 | 1.33E-01 | 1.23E+00 | 1.59E-01 | 6.06E-01 | 1.13E+00 | 6.23E-02 | 2.22E-01 | 1.17E+00 | 6.06E-02 | 1.71E-01 | 1.10E+00 | 1.47E-01 | 5.40E-01 | 1.31E+00 | 1.92E-01 | 5.28E-01 | 1.32E+00 |
| KRBOX4 | 6.20E-03 | 3.23E-02 | -1.00E+00 | 1.36E-01 | 5.89E-01 | 1.37E+00 | 2.24E-06 | 9.51E-06 | -1.49E+00 | 1.14E-04 | 4.35E-04 | -1.11E+00 | 1.01E-05 | 3.61E-05 | -1.45E+00 | 4.23E-03 | 1.19E-02 | -1.02E+00 | 7.15E-05 | 2.63E-04 | -1.17E+00 | 1.89E-03 | 5.19E-03 | -1.10E+00 |
| PCYT2 | 6.20E-03 | 3.23E-02 | -1.11E+00 | 8.47E-02 | 3.66E-01 | 1.25E+00 | 1.81E-02 | 7.68E-02 | -1.09E+00 | 5.27E-02 | 2.01E-01 | 1.13E+00 | 3.20E-13 | 1.14E-12 | -2.26E+00 | 1.80E-05 | 5.07E-05 | -1.32E+00 | 3.35E-05 | 1.23E-04 | -1.27E+00 | 2.80E-03 | 7.68E-03 | -1.07E+00 |
| LINC00426 | 6.21E-03 | 3.24E-02 | -1.15E+00 | 8.76E-03 | 3.79E-02 | -1.05E+00 | 3.61E-04 | 1.53E-03 | -1.47E+00 | 2.79E-02 | 1.06E-01 | 1.03E+00 | 2.12E-01 | 7.57E-01 | 1.71E+00 | 3.47E-01 | 9.78E-01 | 1.48E+00 | 1.97E-01 | 7.22E-01 | -1.01E+00 | 2.59E-01 | 7.12E-01 | -1.09E+00 |
| SRR | 6.21E-03 | 3.24E-02 | -1.42E+00 | 2.21E-02 | 9.55E-02 | -1.07E+00 | 2.31E-01 | 9.81E-01 | 1.18E+00 | 2.37E-01 | 9.07E-01 | 1.38E+00 | 1.29E-01 | 4.60E-01 | 1.15E+00 | 2.80E-01 | 7.90E-01 | 1.28E+00 | 1.91E-02 | 7.02E-02 | -1.13E+00 | 1.27E-01 | 3.48E-01 | -1.06E+00 |
| ERF | 6.22E-03 | 3.24E-02 | -1.04E+00 | 2.30E-04 | 9.93E-04 | -1.13E+00 | 2.58E-03 | 1.10E-02 | -1.08E+00 | 2.76E-08 | 1.05E-07 | -1.52E+00 | 6.83E-02 | 2.43E-01 | 1.03E+00 | 1.35E-01 | 3.82E-01 | 1.15E+00 | 2.85E-02 | 1.05E-01 | -1.03E+00 | 4.27E-03 | 1.17E-02 | -1.34E+00 |
| PIK3R6 | 6.22E-03 | 3.24E-02 | -1.13E+00 | 7.83E-03 | 3.38E-02 | -1.03E+00 | 6.58E-06 | 2.80E-05 | -1.69E+00 | 2.29E-04 | 8.76E-04 | -1.25E+00 | 1.05E-06 | 3.75E-06 | -1.87E+00 | 2.86E-03 | 8.06E-03 | -1.17E+00 | 9.83E-06 | 3.61E-05 | -1.38E+00 | 1.19E-04 | 3.27E-04 | -1.34E+00 |
| RPL22P13 | 6.23E-03 | 3.24E-02 | 1.88E+00 | 6.58E-04 | 2.84E-03 | 2.29E+00 | 1.98E-04 | 8.43E-04 | 1.83E+00 | 1.47E-03 | 5.60E-03 | 2.07E+00 | 1.56E-01 | 5.55E-01 | 1.65E+00 | 2.38E-03 | 6.70E-03 | 2.25E+00 | 2.66E-04 | 9.77E-04 | 1.88E+00 | 7.08E-08 | 1.94E-07 | 2.36E+00 |
| C2CD5 | 6.23E-03 | 3.25E-02 | -1.02E+00 | 3.14E-02 | 1.36E-01 | 1.18E+00 | 6.16E-02 | 2.62E-01 | 1.02E+00 | 9.79E-02 | 3.74E-01 | 1.19E+00 | 8.59E-02 | 3.06E-01 | -1.03E+00 | 1.53E-01 | 4.32E-01 | 1.02E+00 | 2.89E-05 | 1.06E-04 | -1.34E+00 | 9.51E-04 | 2.61E-03 | -1.18E+00 |
| PPP3CB-AS1 | 6.23E-03 | 3.25E-02 | -1.06E+00 | 1.06E-01 | 4.58E-01 | 1.36E+00 | 8.67E-03 | 3.68E-02 | -1.28E+00 | 4.82E-02 | 1.84E-01 | 1.08E+00 | 1.04E-01 | 3.70E-01 | 1.30E+00 | 8.72E-02 | 2.46E-01 | 1.95E+00 | 3.11E-04 | 1.14E-03 | -1.08E+00 | 8.77E-05 | 2.41E-04 | -1.21E+00 |
| MYOM2 | 6.26E-03 | 3.26E-02 | -1.41E+00 | 9.47E-02 | 4.09E-01 | 1.06E+00 | 1.74E-03 | 7.41E-03 | -1.34E+00 | 7.07E-03 | 2.70E-02 | -1.08E+00 | 5.66E-05 | 2.02E-04 | -1.85E+00 | 6.71E-04 | 1.89E-03 | -1.45E+00 | 4.74E-04 | 1.74E-03 | -1.52E+00 | 2.41E-03 | 6.62E-03 | -1.45E+00 |
| MON1A | 6.26E-03 | 3.26E-02 | -1.26E+00 | 6.56E-02 | 2.84E-01 | 1.16E+00 | 1.13E-02 | 4.79E-02 | -1.35E+00 | 2.28E-01 | 8.73E-01 | 1.21E+00 | 2.91E-02 | 1.04E-01 | -1.22E+00 | 2.22E-01 | 6.25E-01 | 1.26E+00 | 3.32E-04 | 1.22E-03 | -1.27E+00 | 1.30E-01 | 3.58E-01 | 1.48E+00 |
| CNN3 | 6.27E-03 | 3.27E-02 | 1.11E+00 | 1.62E-03 | 7.00E-03 | 1.18E+00 | 2.19E-02 | 9.30E-02 | 1.01E+00 | 1.13E-01 | 4.32E-01 | 1.29E+00 | 2.30E-04 | 8.18E-04 | -1.23E+00 | 2.10E-03 | 5.93E-03 | -1.06E+00 | 1.23E-02 | 4.53E-02 | -1.11E+00 | 7.12E-02 | 1.96E-01 | 1.58E+00 |
| RPL30P4 | 6.27E-03 | 3.27E-02 | -1.11E+00 | 3.26E-56 | 1.41E-55 | -2.02E+00 | 9.98E-02 | 4.24E-01 | -1.06E+00 | 1.32E-21 | 5.06E-21 | -1.91E+00 | 9.29E-02 | 3.31E-01 | -1.02E+00 | 1.90E-24 | 5.37E-24 | -1.87E+00 | 1.61E-02 | 5.90E-02 | -1.07E+00 | 1.05E-30 | 2.88E-30 | -2.01E+00 |
| AC148476.1 | 6.27E-03 | 3.27E-02 | 2.05E+00 | 7.46E-02 | 3.22E-01 | 1.86E+00 | 1.77E-01 | 7.54E-01 | 1.37E+00 | 1.48E-01 | 5.65E-01 | 1.63E+00 | 1.30E-01 | 4.65E-01 | 1.76E+00 | 1.55E-01 | 4.36E-01 | 1.95E+00 | 1.91E-01 | 7.02E-01 | 1.27E+00 | 1.83E-01 | 5.02E-01 | 1.64E+00 |
| ZNF528-AS1 | 6.28E-03 | 3.27E-02 | -1.35E+00 | 2.86E-02 | 1.24E-01 | -1.05E+00 | 1.61E-01 | 6.84E-01 | 1.34E+00 | 3.88E-02 | 1.48E-01 | 1.94E+00 | 1.11E-01 | 3.97E-01 | 1.18E+00 | 1.62E-01 | 4.56E-01 | 1.14E+00 | 6.11E-02 | 2.24E-01 | 1.53E+00 | 1.99E-01 | 5.47E-01 | 1.27E+00 |
| ZBTB45 | 6.29E-03 | 3.28E-02 | -1.21E+00 | 4.29E-02 | 1.85E-01 | 1.09E+00 | 1.66E-04 | 7.05E-04 | -1.39E+00 | 4.06E-03 | 1.55E-02 | -1.01E+00 | 1.31E-03 | 4.66E-03 | -1.54E+00 | 1.79E-03 | 5.04E-03 | -1.56E+00 | 3.58E-04 | 1.31E-03 | -1.47E+00 | 5.26E-04 | 1.45E-03 | -1.55E+00 |
| LRRC56 | 6.29E-03 | 3.28E-02 | -1.22E+00 | 1.19E-02 | 5.15E-02 | -1.05E+00 | 8.50E-03 | 3.61E-02 | -1.29E+00 | 5.62E-02 | 2.14E-01 | 1.02E+00 | 2.09E-01 | 7.46E-01 | 1.15E+00 | 3.41E-01 | 9.62E-01 | 1.33E+00 | 1.49E-02 | 5.46E-02 | 1.96E+00 | 3.41E-02 | 9.38E-02 | 2.11E+00 |
| ZNF385D-AS2 | 6.30E-03 | 3.28E-02 | 2.16E+00 | 5.22E-02 | 2.26E-01 | 1.88E+00 | 2.19E-01 | 9.32E-01 | 1.30E+00 | 1.02E-01 | 3.90E-01 | 1.71E+00 | 2.08E-01 | 7.43E-01 | 1.14E+00 | 2.93E-01 | 8.26E-01 | 1.22E+00 | 8.87E-02 | 3.26E-01 | -1.32E+00 | 2.34E-01 | 6.42E-01 | -1.07E+00 |
| MED20 | 6.30E-03 | 3.28E-02 | -1.24E+00 | 8.91E-03 | 3.85E-02 | -1.20E+00 | 1.26E-03 | 5.34E-03 | -1.24E+00 | 1.69E-02 | 6.45E-02 | 1.09E+00 | 5.27E-03 | 1.88E-02 | -1.68E+00 | 1.71E-01 | 4.83E-01 | 1.18E+00 | 1.84E-07 | 6.74E-07 | -1.61E+00 | 3.79E-03 | 1.04E-02 | -1.03E+00 |
| BZW1P2 | 6.32E-03 | 3.29E-02 | 1.07E+00 | 2.93E-14 | 1.27E-13 | -1.47E+00 | 2.31E-01 | 9.80E-01 | 1.24E+00 | 4.97E-04 | 1.90E-03 | -1.04E+00 | 7.41E-02 | 2.64E-01 | 1.25E+00 | 3.24E-05 | 9.13E-05 | -1.16E+00 | 2.00E-01 | 7.36E-01 | 1.32E+00 | 6.30E-02 | 1.73E-01 | 1.07E+00 |
| AC087284.1 | 6.32E-03 | 3.29E-02 | -1.50E+00 | 1.49E-01 | 6.42E-01 | 1.18E+00 | 1.20E-02 | 5.08E-02 | -1.38E+00 | 2.39E-01 | 9.14E-01 | 1.31E+00 | 1.64E-01 | 5.85E-01 | -1.21E+00 | 2.28E-01 | 6.42E-01 | 1.59E+00 | 3.25E-03 | 1.19E-02 | -1.60E+00 | 5.20E-02 | 1.43E-01 | -1.23E+00 |
| MYLIP | 6.32E-03 | 3.30E-02 | 1.03E+00 | 1.82E-04 | 7.85E-04 | -1.00E+00 | 4.04E-07 | 1.72E-06 | -1.41E+00 | 1.32E-06 | 5.05E-06 | -1.22E+00 | 1.45E-02 | 5.18E-02 | 1.18E+00 | 1.04E-03 | 2.93E-03 | -1.01E+00 | 7.02E-07 | 2.58E-06 | -1.33E+00 | 5.41E-06 | 1.49E-05 | -1.35E+00 |
| ZIK1 | 6.32E-03 | 3.30E-02 | 1.88E+00 | 2.01E-01 | 8.70E-01 | 1.45E+00 | 8.79E-02 | 3.73E-01 | 1.32E+00 | 1.44E-01 | 5.49E-01 | 1.11E+00 | 1.76E-01 | 6.27E-01 | 1.22E+00 | 2.03E-01 | 5.72E-01 | 1.24E+00 | 6.03E-02 | 2.21E-01 | -1.06E+00 | 4.03E-02 | 1.11E-01 | -1.07E+00 |
| FHOD1 | 6.33E-03 | 3.30E-02 | -1.14E+00 | 1.78E-01 | 7.67E-01 | 1.39E+00 | 3.73E-04 | 1.58E-03 | -1.32E+00 | 4.52E-03 | 1.73E-02 | 1.02E+00 | 1.18E-03 | 4.22E-03 | -1.31E+00 | 3.81E-02 | 1.07E-01 | 1.07E+00 | 1.43E-07 | 5.24E-07 | -1.55E+00 | 2.62E-03 | 7.19E-03 | -1.25E+00 |
| SP2-AS1 | 6.33E-03 | 3.30E-02 | -1.18E+00 | 1.23E-03 | 5.32E-03 | -1.35E+00 | 4.81E-02 | 2.04E-01 | -1.20E+00 | 1.16E-01 | 4.45E-01 | -1.09E+00 | 9.01E-02 | 3.21E-01 | -1.07E+00 | 3.27E-01 | 9.23E-01 | 1.34E+00 | 1.69E-02 | 6.19E-02 | -1.13E+00 | 5.46E-02 | 1.50E-01 | -1.15E+00 |
| WDR5 | 6.34E-03 | 3.30E-02 | -1.22E+00 | 2.32E-02 | 1.00E-01 | 1.03E+00 | 4.12E-04 | 1.75E-03 | -1.19E+00 | 2.38E-04 | 9.08E-04 | -1.07E+00 | 3.08E-03 | 1.10E-02 | -1.34E+00 | 6.45E-03 | 1.82E-02 | -1.20E+00 | 5.09E-05 | 1.87E-04 | -1.12E+00 | 5.53E-03 | 1.52E-02 | 1.09E+00 |
| GPLD1 | 6.34E-03 | 3.31E-02 | -1.45E+00 | 1.44E-01 | 6.22E-01 | 1.25E+00 | 2.17E-01 | 9.22E-01 | 1.18E+00 | 6.08E-02 | 2.32E-01 | 1.99E+00 | 1.92E-01 | 6.83E-01 | 1.22E+00 | 1.45E-01 | 4.08E-01 | 1.94E+00 | 2.12E-05 | 7.76E-05 | -1.60E+00 | 2.30E-02 | 6.33E-02 | -1.07E+00 |
| WHAMM | 6.35E-03 | 3.31E-02 | 1.09E+00 | 3.71E-10 | 1.61E-09 | -1.30E+00 | 4.60E-04 | 1.95E-03 | -1.17E+00 | 5.07E-08 | 1.94E-07 | -1.37E+00 | 6.06E-04 | 2.16E-03 | -1.21E+00 | 5.22E-05 | 1.47E-04 | -1.24E+00 | 3.30E-05 | 1.21E-04 | -1.16E+00 | 6.68E-12 | 1.84E-11 | -1.59E+00 |
| TRAPPC2L | 6.35E-03 | 3.31E-02 | 1.10E+00 | 7.41E-06 | 3.20E-05 | 1.04E+00 | 1.74E-01 | 7.41E-01 | 1.19E+00 | 9.48E-03 | 3.62E-02 | 1.06E+00 | 1.04E-03 | 3.71E-03 | -1.10E+00 | 3.13E-05 | 8.81E-05 | -1.16E+00 | 2.08E-01 | 7.62E-01 | 1.15E+00 | 2.03E-01 | 5.58E-01 | 1.21E+00 |
| AP003072.5 | 6.35E-03 | 3.31E-02 | -1.64E+00 | 1.51E-01 | 6.50E-01 | 1.19E+00 | 3.72E-02 | 1.58E-01 | -1.29E+00 | 2.39E-01 | 9.13E-01 | 1.19E+00 | 1.34E-01 | 4.79E-01 | -1.02E+00 | 2.26E-01 | 6.36E-01 | 1.12E+00 | 2.26E-01 | 8.30E-01 | 1.22E+00 | 3.31E-01 | 9.08E-01 | 1.09E+00 |
| WIPF3 | 6.35E-03 | 3.31E-02 | -1.50E+00 | 3.99E-02 | 1.73E-01 | -1.10E+00 | 2.61E-02 | 1.11E-01 | -1.35E+00 | 1.31E-02 | 4.99E-02 | -1.57E+00 | 1.68E-01 | 5.99E-01 | 1.32E+00 | 3.22E-01 | 9.08E-01 | 1.09E+00 | 3.51E-06 | 1.29E-05 | -2.56E+00 | 6.88E-03 | 1.89E-02 | -1.43E+00 |
| EIF3EP1 | 6.35E-03 | 3.31E-02 | 1.07E+00 | 3.31E-05 | 1.43E-04 | -1.01E+00 | 2.58E-02 | 1.10E-01 | -1.02E+00 | 9.58E-04 | 3.66E-03 | -1.04E+00 | 2.49E-01 | 8.88E-01 | 1.41E+00 | 7.21E-02 | 2.03E-01 | 1.11E+00 | 2.91E-02 | 1.07E-01 | 1.01E+00 | 1.44E-03 | 3.95E-03 | -1.21E+00 |
| ACRBP | 6.36E-03 | 3.31E-02 | -1.00E+00 | 2.62E-03 | 1.13E-02 | 1.02E+00 | 2.14E-05 | 9.10E-05 | -1.45E+00 | 4.89E-05 | 1.87E-04 | -1.20E+00 | 1.04E-04 | 3.71E-04 | -1.19E+00 | 1.44E-05 | 4.06E-05 | -1.29E+00 | 4.95E-03 | 1.82E-02 | 3.19E+00 | 3.59E-02 | 9.85E-02 | 2.64E+00 |
| AL132838.1 | 6.36E-03 | 3.31E-02 | -1.18E+00 | 1.18E-01 | 5.09E-01 | 1.72E+00 | 2.20E-01 | 9.33E-01 | 1.21E+00 | 6.98E-04 | 2.67E-03 | 2.50E+00 | 1.15E-01 | 4.11E-01 | 1.04E+00 | 5.96E-02 | 1.68E-01 | 2.06E+00 | 5.03E-02 | 1.84E-01 | -1.14E+00 | 9.17E-04 | 2.52E-03 | 2.17E+00 |
| HMGB2P1 | 6.37E-03 | 3.32E-02 | -1.02E+00 | 7.09E-07 | 3.06E-06 | -1.84E+00 | 1.67E-01 | 7.09E-01 | 1.25E+00 | 4.55E-04 | 1.74E-03 | -1.27E+00 | 2.85E-02 | 1.02E-01 | -1.07E+00 | 1.47E-03 | 4.13E-03 | -1.79E+00 | 4.84E-04 | 1.78E-03 | -1.25E+00 | 1.96E-09 | 5.38E-09 | -1.70E+00 |
| AC008770.1 | 6.38E-03 | 3.33E-02 | 1.05E+00 | 5.63E-05 | 2.43E-04 | 1.01E+00 | 1.10E-02 | 4.67E-02 | -1.02E+00 | 8.26E-05 | 3.16E-04 | -1.09E+00 | 3.30E-06 | 1.18E-05 | -1.11E+00 | 6.75E-07 | 1.90E-06 | -1.15E+00 | 1.13E-01 | 4.14E-01 | 1.08E+00 | 1.22E-02 | 3.34E-02 | -1.01E+00 |
| PLEKHO2 | 6.38E-03 | 3.33E-02 | -1.32E+00 | 1.84E-01 | 7.94E-01 | 1.31E+00 | 1.68E-02 | 7.15E-02 | -1.34E+00 | 1.66E-01 | 6.34E-01 | 1.24E+00 | 6.48E-04 | 2.31E-03 | -2.25E+00 | 2.33E-01 | 6.57E-01 | 1.14E+00 | 1.23E-05 | 4.50E-05 | -1.21E+00 | 7.36E-04 | 2.02E-03 | -1.26E+00 |
| DIS3L2 | 6.40E-03 | 3.34E-02 | 1.05E+00 | 8.86E-03 | 3.83E-02 | 1.11E+00 | 2.99E-04 | 1.27E-03 | -1.40E+00 | 3.66E-03 | 1.40E-02 | -1.11E+00 | 2.50E-03 | 8.93E-03 | -1.10E+00 | 4.99E-02 | 1.41E-01 | 1.23E+00 | 4.42E-02 | 1.62E-01 | 1.04E+00 | 1.03E-01 | 2.84E-01 | 1.05E+00 |
| AL158050.1 | 6.42E-03 | 3.35E-02 | -1.23E+00 | 5.85E-04 | 2.53E-03 | -1.47E+00 | 9.28E-02 | 3.94E-01 | 1.03E+00 | 8.91E-03 | 3.40E-02 | -1.27E+00 | 2.39E-02 | 8.53E-02 | -1.03E+00 | 3.54E-04 | 9.99E-04 | -1.76E+00 | 1.48E-02 | 5.42E-02 | 1.00E+00 | 3.86E-05 | 1.06E-04 | -1.51E+00 |
| NUP210L | 6.43E-03 | 3.35E-02 | -1.19E+00 | 2.57E-02 | 1.11E-01 | 1.02E+00 | 3.94E-02 | 1.67E-01 | -1.16E+00 | 3.29E-02 | 1.26E-01 | -1.13E+00 | 4.16E-04 | 1.48E-03 | -2.13E+00 | 1.88E-03 | 5.29E-03 | -1.60E+00 | 3.92E-02 | 1.44E-01 | -1.30E+00 | 8.61E-02 | 2.37E-01 | -1.21E+00 |
| NUTM2B-AS1 | 6.43E-03 | 3.35E-02 | 1.14E+00 | 6.05E-06 | 2.61E-05 | 1.12E+00 | 6.41E-02 | 2.72E-01 | 1.08E+00 | 3.85E-02 | 1.47E-01 | 1.17E+00 | 1.03E-01 | 3.66E-01 | 1.08E+00 | 7.32E-02 | 2.06E-01 | 1.12E+00 | 1.93E-02 | 7.08E-02 | 1.02E+00 | 9.86E-02 | 2.71E-01 | 1.07E+00 |
| AC036176.1 | 6.43E-03 | 3.35E-02 | -1.07E+00 | 7.34E-03 | 3.17E-02 | 1.07E+00 | 4.34E-02 | 1.84E-01 | 1.01E+00 | 3.47E-02 | 1.33E-01 | 1.07E+00 | 2.24E-01 | 7.97E-01 | 1.49E+00 | 1.76E-01 | 4.96E-01 | 1.15E+00 | 7.42E-02 | 2.72E-01 | -1.11E+00 | 1.71E-01 | 4.69E-01 | -1.00E+00 |
| SOCS6 | 6.45E-03 | 3.36E-02 | -1.05E+00 | 5.05E-06 | 2.18E-05 | -1.36E+00 | 5.28E-02 | 2.24E-01 | 1.04E+00 | 2.08E-04 | 7.93E-04 | -1.28E+00 | 2.54E-02 | 9.04E-02 | 1.09E+00 | 6.86E-05 | 1.93E-04 | -1.56E+00 | 1.43E-03 | 5.25E-03 | -1.19E+00 | 4.86E-05 | 1.34E-04 | -1.46E+00 |
| NPIPA1 | 6.45E-03 | 3.36E-02 | -1.25E+00 | 3.61E-03 | 1.56E-02 | -1.26E+00 | 3.87E-02 | 1.64E-01 | -1.12E+00 | 6.16E-02 | 2.35E-01 | 1.01E+00 | 3.76E-04 | 1.34E-03 | -1.79E+00 | 2.15E-03 | 6.06E-03 | -1.42E+00 | 3.60E-02 | 1.32E-01 | -1.19E+00 | 4.09E-02 | 1.12E-01 | -1.13E+00 |
| AC117382.1 | 6.45E-03 | 3.36E-02 | -1.06E+00 | 3.51E-05 | 1.52E-04 | -1.29E+00 | 5.17E-02 | 2.19E-01 | 1.48E+00 | 1.66E-01 | 6.36E-01 | 1.46E+00 | 4.29E-02 | 1.53E-01 | 1.02E+00 | 1.26E-02 | 3.56E-02 | -1.30E+00 | 2.02E-01 | 7.40E-01 | 1.46E+00 | 1.60E-01 | 4.40E-01 | 1.52E+00 |
| AC024995.1 | 6.46E-03 | 3.36E-02 | 1.12E+00 | 2.17E-17 | 9.39E-17 | -1.59E+00 | 3.95E-02 | 1.68E-01 | 1.30E+00 | 9.08E-05 | 3.47E-04 | -1.16E+00 | 1.75E-03 | 6.25E-03 | 1.01E+00 | 5.93E-07 | 1.67E-06 | -1.33E+00 | 1.12E-03 | 4.12E-03 | 1.01E+00 | 9.08E-11 | 2.49E-10 | -1.56E+00 |
| ZNF345 | 6.46E-03 | 3.36E-02 | -1.04E+00 | 1.66E-01 | 7.17E-01 | 1.52E+00 | 2.27E-03 | 9.65E-03 | -1.22E+00 | 1.38E-03 | 5.27E-03 | -1.16E+00 | 5.52E-02 | 1.97E-01 | -1.04E+00 | 3.22E-01 | 9.07E-01 | 1.39E+00 | 2.23E-03 | 8.17E-03 | -1.27E+00 | 6.93E-02 | 1.90E-01 | 1.01E+00 |
| ADA2 | 6.47E-03 | 3.37E-02 | -1.54E+00 | 8.74E-03 | 3.77E-02 | -1.35E+00 | 4.16E-06 | 1.77E-05 | -1.42E+00 | 1.79E-07 | 6.85E-07 | -1.41E+00 | 9.09E-02 | 3.24E-01 | -1.34E+00 | 9.13E-02 | 2.57E-01 | -1.43E+00 | 1.87E-01 | 6.87E-01 | -1.06E+00 | 2.82E-01 | 7.76E-01 | 1.01E+00 |
| MFSD4A | 6.47E-03 | 3.37E-02 | -1.35E+00 | 2.11E-02 | 9.12E-02 | -1.09E+00 | 5.23E-02 | 2.22E-01 | -1.13E+00 | 1.07E-01 | 4.10E-01 | 1.10E+00 | 2.79E-04 | 9.94E-04 | -2.58E+00 | 1.55E-03 | 4.38E-03 | -1.88E+00 | 5.71E-02 | 2.09E-01 | -1.33E+00 | 3.18E-01 | 8.75E-01 | 1.28E+00 |
| WDR19 | 6.47E-03 | 3.37E-02 | -1.16E+00 | 2.02E-01 | 8.72E-01 | 1.48E+00 | 2.44E-03 | 1.04E-02 | -1.24E+00 | 4.11E-03 | 1.57E-02 | -1.04E+00 | 1.22E-01 | 4.35E-01 | -1.07E+00 | 2.02E-01 | 5.69E-01 | 1.61E+00 | 1.35E-01 | 4.96E-01 | 1.02E+00 | 3.24E-01 | 8.91E-01 | 1.26E+00 |
| TATDN2 | 6.47E-03 | 3.37E-02 | -1.09E+00 | 6.45E-02 | 2.79E-01 | 1.17E+00 | 2.05E-01 | 8.70E-01 | 1.27E+00 | 2.39E-01 | 9.15E-01 | 1.44E+00 | 1.07E-01 | 3.83E-01 | -1.02E+00 | 1.53E-01 | 4.32E-01 | -1.04E+00 | 3.48E-04 | 1.28E-03 | -1.27E+00 | 1.01E-03 | 2.77E-03 | -1.24E+00 |
| INTS6 | 6.47E-03 | 3.37E-02 | 1.62E+00 | 1.95E-04 | 8.42E-04 | 1.19E+00 | 3.31E-06 | 1.41E-05 | 1.81E+00 | 2.39E-01 | 9.11E-01 | 1.26E+00 | 9.75E-02 | 3.48E-01 | 1.35E+00 | 1.65E-02 | 4.66E-02 | 1.17E+00 | 3.51E-02 | 1.29E-01 | 1.15E+00 | 4.21E-03 | 1.16E-02 | -1.01E+00 |
| AL449283.1 | 6.48E-03 | 3.38E-02 | -2.15E+00 | 2.91E-02 | 1.26E-01 | -1.49E+00 | 4.78E-02 | 2.03E-01 | -1.16E+00 | 3.13E-02 | 1.20E-01 | -1.18E+00 | 2.61E-01 | 9.32E-01 | 1.01E+00 | 3.29E-01 | 9.26E-01 | 1.23E+00 | 2.14E-03 | 7.86E-03 | -1.14E+00 | 4.10E-02 | 1.13E-01 | -1.07E+00 |
| LINC02453 | 6.49E-03 | 3.38E-02 | -1.43E+00 | 5.87E-03 | 2.54E-02 | -1.37E+00 | 2.23E-01 | 9.45E-01 | 1.08E+00 | 1.01E-01 | 3.84E-01 | -1.11E+00 | 1.98E-01 | 7.07E-01 | 1.63E+00 | 3.41E-01 | 9.63E-01 | 1.27E+00 | 4.53E-02 | 1.66E-01 | -1.35E+00 | 4.62E-02 | 1.27E-01 | -1.46E+00 |
| NUDT3 | 6.50E-03 | 3.39E-02 | -1.07E+00 | 2.49E-03 | 1.08E-02 | -1.06E+00 | 2.13E-01 | 9.03E-01 | 1.15E+00 | 1.28E-01 | 4.88E-01 | 1.18E+00 | 2.56E-01 | 9.13E-01 | 1.30E+00 | 2.21E-01 | 6.22E-01 | 1.19E+00 | 4.85E-05 | 1.78E-04 | -1.30E+00 | 1.31E-04 | 3.60E-04 | -1.27E+00 |
| ETV5 | 6.51E-03 | 3.39E-02 | -1.28E+00 | 4.33E-04 | 1.87E-03 | -1.71E+00 | 3.04E-03 | 1.29E-02 | 1.82E+00 | 1.04E-01 | 3.97E-01 | -1.18E+00 | 1.85E-02 | 6.58E-02 | 1.97E+00 | 3.28E-01 | 9.25E-01 | 1.03E+00 | 5.23E-03 | 1.92E-02 | -1.63E+00 | 1.84E-03 | 5.05E-03 | -1.76E+00 |
| SMC5-AS1 | 6.51E-03 | 3.39E-02 | -1.18E+00 | 2.10E-02 | 9.06E-02 | -1.02E+00 | 2.15E-01 | 9.12E-01 | 1.12E+00 | 1.74E-01 | 6.64E-01 | 1.08E+00 | 1.34E-01 | 4.76E-01 | 1.19E+00 | 5.01E-02 | 1.41E-01 | -1.14E+00 | 2.12E-01 | 7.77E-01 | 1.13E+00 | 2.69E-01 | 7.38E-01 | 1.42E+00 |
| ERCC2 | 6.52E-03 | 3.40E-02 | -1.11E+00 | 1.49E-02 | 6.42E-02 | 1.07E+00 | 9.64E-03 | 4.10E-02 | -1.19E+00 | 3.63E-03 | 1.39E-02 | -1.11E+00 | 1.27E-02 | 4.53E-02 | -1.14E+00 | 9.01E-02 | 2.54E-01 | 1.04E+00 | 3.90E-08 | 1.43E-07 | -1.54E+00 | 1.07E-03 | 2.95E-03 | -1.09E+00 |
| PLAUR | 6.52E-03 | 3.40E-02 | -1.28E+00 | 6.12E-04 | 2.65E-03 | -1.47E+00 | 1.02E-04 | 4.35E-04 | -1.36E+00 | 3.13E-06 | 1.20E-05 | -1.50E+00 | 3.56E-02 | 1.27E-01 | -1.82E+00 | 1.59E-02 | 4.48E-02 | -2.77E+00 | 2.66E-01 | 9.74E-01 | 1.15E+00 | 1.57E-01 | 4.30E-01 | 1.00E+00 |
| ENTPD7 | 6.53E-03 | 3.40E-02 | -1.42E+00 | 3.22E-02 | 1.39E-01 | -1.07E+00 | 1.40E-01 | 5.95E-01 | 1.34E+00 | 1.72E-01 | 6.56E-01 | 1.40E+00 | 4.89E-02 | 1.74E-01 | -1.15E+00 | 8.22E-02 | 2.32E-01 | -1.02E+00 | 1.50E-01 | 5.52E-01 | 1.02E+00 | 1.98E-01 | 5.45E-01 | 1.05E+00 |
| STK40 | 6.53E-03 | 3.40E-02 | -1.08E+00 | 5.00E-02 | 2.16E-01 | 1.15E+00 | 6.74E-03 | 2.86E-02 | -1.08E+00 | 5.68E-02 | 2.17E-01 | 1.18E+00 | 1.29E-03 | 4.61E-03 | -1.23E+00 | 1.17E-02 | 3.29E-02 | -1.05E+00 | 2.38E-02 | 8.74E-02 | -1.02E+00 | 6.84E-02 | 1.88E-01 | 1.18E+00 |
| HRAS | 6.55E-03 | 3.41E-02 | -1.13E+00 | 9.64E-03 | 4.17E-02 | -1.00E+00 | 2.21E-01 | 9.41E-01 | 1.24E+00 | 1.31E-01 | 5.00E-01 | 1.27E+00 | 8.70E-04 | 3.10E-03 | -1.28E+00 | 4.28E-03 | 1.21E-02 | -1.18E+00 | 7.76E-02 | 2.85E-01 | 1.17E+00 | 1.04E-02 | 2.85E-02 | 1.08E+00 |
| MS4A6A | 6.55E-03 | 3.41E-02 | 1.26E+00 | 3.30E-02 | 1.43E-01 | 1.18E+00 | 1.89E-02 | 8.04E-02 | 1.66E+00 | 1.26E-02 | 4.82E-02 | 2.00E+00 | 2.18E-08 | 7.77E-08 | 5.35E+00 | 5.06E-03 | 1.43E-02 | 3.42E+00 | 2.28E-01 | 8.36E-01 | 1.40E+00 | 3.55E-01 | 9.76E-01 | 1.01E+00 |
| STAT4 | 6.57E-03 | 3.42E-02 | 1.03E+00 | 8.92E-04 | 3.85E-03 | -1.07E+00 | 4.48E-05 | 1.90E-04 | -1.35E+00 | 3.45E-08 | 1.32E-07 | -1.86E+00 | 3.23E-03 | 1.15E-02 | -1.61E+00 | 2.13E-03 | 6.00E-03 | -1.67E+00 | 2.55E-03 | 9.35E-03 | -1.42E+00 | 6.79E-02 | 1.87E-01 | -1.06E+00 |
| ZDHHC5 | 6.57E-03 | 3.42E-02 | -1.08E+00 | 1.35E-04 | 5.82E-04 | -1.26E+00 | 3.62E-06 | 1.54E-05 | -1.32E+00 | 6.37E-08 | 2.43E-07 | -1.44E+00 | 1.83E-03 | 6.52E-03 | -1.14E+00 | 3.03E-05 | 8.53E-05 | -1.61E+00 | 1.12E-06 | 4.13E-06 | -1.32E+00 | 2.28E-06 | 6.27E-06 | -1.36E+00 |
| SEPSECS-AS1 | 6.57E-03 | 3.42E-02 | -1.12E+00 | 2.13E-01 | 9.20E-01 | 1.49E+00 | 5.43E-04 | 2.31E-03 | -1.44E+00 | 6.22E-02 | 2.38E-01 | 1.09E+00 | 9.55E-04 | 3.41E-03 | -1.19E+00 | 6.87E-03 | 1.94E-02 | -1.01E+00 | 9.00E-02 | 3.30E-01 | -1.04E+00 | 1.16E-01 | 3.19E-01 | -1.04E+00 |
| KCNMB1 | 6.58E-03 | 3.43E-02 | -1.20E+00 | 1.57E-03 | 6.79E-03 | -1.26E+00 | 7.06E-03 | 3.00E-02 | -1.12E+00 | 1.27E-03 | 4.84E-03 | -1.20E+00 | 1.64E-01 | 5.83E-01 | -1.01E+00 | 1.37E-01 | 3.85E-01 | -1.22E+00 | 1.98E-01 | 7.26E-01 | 1.03E+00 | 2.66E-01 | 7.32E-01 | -1.03E+00 |
| AP001437.1 | 6.59E-03 | 3.43E-02 | 2.26E+00 | 1.48E-01 | 6.41E-01 | 1.54E+00 | 8.03E-02 | 3.41E-01 | 1.33E+00 | 2.19E-01 | 8.37E-01 | 1.09E+00 | 1.68E-01 | 6.00E-01 | 1.66E+00 | 3.41E-01 | 9.61E-01 | 1.26E+00 | 1.15E-01 | 4.22E-01 | 1.58E+00 | 1.75E-01 | 4.81E-01 | -1.15E+00 |

| | | | | | | | | | | | | | | | | | | | | | | | | |
|---|---|---|---|---|---|---|---|---|---|---|---|---|---|---|---|---|---|---|---|---|---|---|---|---|
| TAOK2 | 6.61E-03 | 3.45E-02 | -1.14E+00 | 3.20E-02 | 1.38E-01 | 1.13E+00 | 6.42E-03 | 2.73E-02 | -1.13E+00 | 1.62E-02 | 6.19E-02 | 1.02E+00 | 3.66E-02 | 1.30E-01 | 1.03E+00 | 4.77E-02 | 1.34E-01 | 1.02E+00 | 1.35E-03 | 4.95E-03 | -1.12E+00 | 9.27E-03 | 2.55E-02 | -1.08E+00 |
| LLGL2 | 6.62E-03 | 3.45E-02 | -1.09E+00 | 4.30E-02 | 1.86E-01 | 1.21E+00 | 5.83E-02 | 2.48E-01 | -1.00E+00 | 2.20E-01 | 8.40E-01 | 1.31E+00 | 8.20E-03 | 2.93E-02 | -1.20E+00 | 6.49E-02 | 1.83E-01 | 1.12E+00 | 1.33E-01 | 4.90E-01 | -1.02E+00 | 3.36E-01 | 9.23E-01 | 1.18E+00 |
| FBXW5 | 6.62E-03 | 3.45E-02 | 1.05E+00 | 3.49E-02 | 1.51E-01 | 1.26E+00 | 3.80E-02 | 1.61E-01 | 1.06E+00 | 2.77E-02 | 1.06E-01 | 1.22E+00 | 1.75E-04 | 6.24E-04 | -1.11E+00 | 3.53E-02 | 9.94E-02 | 1.11E+00 | 9.50E-03 | 3.49E-02 | 1.02E+00 | 4.07E-02 | 1.12E-01 | 1.03E+00 |
| FER | 6.63E-03 | 3.46E-02 | -1.20E+00 | 1.40E-01 | 6.04E-01 | 1.24E+00 | 8.31E-05 | 3.53E-04 | -1.53E+00 | 2.40E-03 | 9.16E-03 | -1.14E+00 | 1.68E-03 | 5.99E-03 | 1.05E+00 | 1.99E-03 | 5.61E-03 | 1.05E+00 | 1.85E-01 | 6.79E-01 | 1.17E+00 | 2.89E-01 | 7.92E-01 | 1.25E+00 |
| DDX50P1 | 6.65E-03 | 3.47E-02 | -1.38E+00 | 1.55E-01 | 6.68E-01 | 1.34E+00 | 1.00E-04 | 4.25E-04 | -1.83E+00 | 9.73E-03 | 3.72E-02 | -1.15E+00 | 5.60E-03 | 2.00E-02 | -1.69E+00 | 1.47E-01 | 4.14E-01 | 1.12E+00 | 2.44E-05 | 8.95E-05 | -1.91E+00 | 6.71E-02 | 1.84E-01 | -1.15E+00 |
| SNRPD2 | 6.66E-03 | 3.47E-02 | -1.08E+00 | 1.60E-14 | 6.91E-14 | -1.25E+00 | 2.05E-01 | 8.70E-01 | 1.01E+00 | 2.07E-04 | 7.89E-04 | -1.18E+00 | 1.33E-04 | 4.74E-04 | -1.16E+00 | 8.64E-07 | 2.44E-06 | -1.24E+00 | 2.92E-02 | 1.07E-01 | -1.06E+00 | 2.71E-02 | 7.43E-02 | -1.12E+00 |
| VIM | 6.66E-03 | 3.47E-02 | 1.20E+00 | 1.33E-02 | 5.73E-02 | 1.16E+00 | 5.52E-04 | 2.34E-03 | 1.33E+00 | 1.17E-03 | 4.48E-03 | 1.33E+00 | 4.83E-02 | 1.72E-01 | 1.16E+00 | 2.72E-01 | 7.67E-01 | 1.07E+00 | 3.08E-02 | 1.13E-01 | 1.26E+00 | 3.83E-03 | 1.05E-02 | 1.24E+00 |
| XPR1 | 6.66E-03 | 3.47E-02 | -1.01E+00 | 1.05E-02 | 4.53E-02 | 1.09E+00 | 1.28E-06 | 5.44E-06 | -1.61E+00 | 8.11E-06 | 3.10E-05 | -1.33E+00 | 2.38E-01 | 8.48E-01 | 1.47E+00 | 1.54E-01 | 4.33E-01 | 1.85E+00 | 2.23E-01 | 8.20E-01 | 1.14E+00 | 2.45E-01 | 6.74E-01 | 1.06E+00 |
| C11orf1 | 6.66E-03 | 3.47E-02 | 1.13E+00 | 1.49E-07 | 6.44E-07 | 1.03E+00 | 2.28E-01 | 9.69E-01 | 1.22E+00 | 1.66E-02 | 6.34E-02 | 1.17E+00 | 1.83E-02 | 6.51E-02 | 1.06E+00 | 8.64E-02 | 2.44E-01 | 1.19E+00 | 1.21E-01 | 4.44E-01 | 1.35E+00 | 2.04E-01 | 5.61E-01 | 1.26E+00 |
| ABHD15 | 6.67E-03 | 3.47E-02 | -1.04E+00 | 8.04E-02 | 3.48E-01 | 1.39E+00 | 3.20E-11 | 1.36E-10 | -2.57E+00 | 1.69E-05 | 6.47E-05 | -1.23E+00 | 2.79E-02 | 9.93E-02 | -1.24E+00 | 2.28E-01 | 6.42E-01 | 1.17E+00 | 4.06E-03 | 1.49E-02 | -1.36E+00 | 7.73E-02 | 2.12E-01 | 1.05E+00 |
| ZNF582-AS1 | 6.68E-03 | 3.48E-02 | -1.17E+00 | 5.65E-02 | 2.44E-01 | 1.20E+00 | 6.70E-02 | 2.85E-01 | -1.18E+00 | 1.90E-01 | 7.27E-01 | 1.10E+00 | 1.39E-01 | 4.96E-01 | 1.01E+00 | 1.83E-01 | 5.16E-01 | 1.02E+00 | 2.63E-03 | 9.66E-03 | 2.18E+00 | 2.01E-01 | 5.53E-01 | 1.40E+00 |
| SGO1 | 6.69E-03 | 3.48E-02 | -1.76E+00 | 3.07E-02 | 1.33E-01 | -1.12E+00 | 3.22E-03 | 1.37E-02 | -1.21E+00 | 1.57E-03 | 5.99E-03 | -1.11E+00 | 5.86E-02 | 2.09E-01 | -1.64E+00 | 1.49E-01 | 4.19E-01 | -1.26E+00 | 2.26E-04 | 8.31E-04 | -1.40E+00 | 2.74E-01 | 7.52E-01 | 1.09E+00 |
| ASXL1 | 6.69E-03 | 3.49E-02 | 1.07E+00 | 2.56E-02 | 1.10E-01 | 1.24E+00 | 9.95E-02 | 4.23E-01 | 1.09E+00 | 4.25E-03 | 1.62E-02 | 1.05E+00 | 3.24E-03 | 1.15E-02 | -1.04E+00 | 1.07E-02 | 3.02E-02 | 1.02E+00 | 2.58E-04 | 9.46E-04 | -1.13E+00 | 1.34E-02 | 3.67E-02 | 1.03E+00 |
| ZNF253 | 6.69E-03 | 3.49E-02 | -1.09E+00 | 2.45E-04 | 1.06E-03 | -1.24E+00 | 1.02E-03 | 4.32E-03 | -1.19E+00 | 1.92E-03 | 7.35E-03 | -1.03E+00 | 2.46E-02 | 8.78E-02 | -1.01E+00 | 5.39E-02 | 1.52E-01 | 1.01E+00 | 1.09E-05 | 4.01E-05 | -1.56E+00 | 7.53E-03 | 2.07E-02 | -1.14E+00 |
| NUP107 | 6.71E-03 | 3.50E-02 | -1.04E+00 | 1.37E-01 | 5.90E-01 | 1.53E+00 | 3.11E-03 | 1.32E-02 | -1.13E+00 | 2.59E-01 | 9.89E-01 | 1.29E+00 | 1.06E-02 | 3.79E-02 | 1.01E+00 | 2.76E-01 | 7.79E-01 | 1.39E+00 | 1.34E-04 | 4.91E-04 | -1.09E+00 | 3.38E-01 | 9.27E-01 | 1.39E+00 |
| CLEC11A | 6.71E-03 | 3.50E-02 | -1.14E+00 | 7.51E-04 | 3.25E-03 | -1.08E+00 | 1.60E-03 | 6.81E-03 | -1.50E+00 | 2.83E-06 | 1.08E-05 | -1.60E+00 | 1.14E-01 | 4.05E-01 | 1.52E+00 | 1.79E-01 | 5.03E-01 | 1.55E+00 | 2.14E-01 | 7.87E-01 | 1.18E+00 | 2.13E-01 | 5.85E-01 | 1.10E+00 |
| GPHN | 6.71E-03 | 3.50E-02 | -1.01E+00 | 9.64E-03 | 4.17E-02 | 1.06E+00 | 8.21E-05 | 3.49E-04 | -1.20E+00 | 1.40E-04 | 5.36E-04 | -1.10E+00 | 2.24E-02 | 7.99E-02 | -1.23E+00 | 5.49E-02 | 1.55E-01 | -1.02E+00 | 3.83E-02 | 1.41E-01 | -1.04E+00 | 1.56E-01 | 4.27E-01 | 1.12E+00 |
| AC233968.1 | 6.72E-03 | 3.50E-02 | -1.23E+00 | 4.85E-02 | 2.10E-01 | 1.09E+00 | 1.73E-01 | 7.36E-01 | 1.03E+00 | 1.29E-01 | 4.94E-01 | 1.06E+00 | 2.74E-01 | 9.77E-01 | 1.24E+00 | 1.08E-01 | 3.06E-01 | 2.01E+00 | 1.90E-01 | 6.97E-01 | 1.08E+00 | 1.24E-01 | 3.40E-01 | -1.10E+00 |
| SHANK3 | 6.72E-03 | 3.50E-02 | 1.02E+00 | 1.08E-01 | 4.65E-01 | 1.41E+00 | 3.56E-03 | 1.51E-02 | -1.06E+00 | 2.87E-03 | 1.10E-02 | 1.01E+00 | 5.87E-02 | 2.09E-01 | -1.47E+00 | 1.67E-01 | 4.72E-01 | -1.01E+00 | 1.64E-03 | 6.02E-03 | -1.60E+00 | 8.86E-03 | 2.43E-02 | -1.36E+00 |
| LIMS3 | 6.73E-03 | 3.51E-02 | 2.35E+00 | 2.19E-01 | 9.47E-01 | 1.26E+00 | 2.28E-01 | 9.70E-01 | 1.24E+00 | 1.28E-01 | 4.89E-01 | -1.08E+00 | 8.38E-04 | 2.99E-03 | 4.11E+00 | 4.38E-02 | 1.23E-01 | 2.59E+00 | 1.99E-02 | 7.30E-02 | 1.69E+00 | 9.82E-02 | 2.70E-01 | 1.50E+00 |
| OGN | 6.75E-03 | 3.52E-02 | -1.77E+00 | 3.08E-02 | 1.33E-01 | -1.24E+00 | 5.80E-02 | 2.47E-01 | -1.11E+00 | 2.41E-01 | 9.19E-01 | 1.22E+00 | 3.47E-02 | 1.24E-01 | -1.34E+00 | 1.85E-01 | 5.20E-01 | 1.10E+00 | 6.59E-02 | 2.42E-01 | 1.10E+00 | 3.39E-01 | 9.31E-01 | 1.37E+00 |
| SMIM4 | 6.75E-03 | 3.52E-02 | -1.03E+00 | 7.68E-07 | 3.32E-06 | -1.60E+00 | 4.03E-02 | 1.71E-01 | 1.00E+00 | 8.99E-09 | 3.43E-08 | -1.95E+00 | 4.71E-02 | 1.68E-01 | 1.07E+00 | 1.10E-02 | 3.10E-02 | -1.19E+00 | 1.83E-01 | 6.70E-01 | 1.11E+00 | 1.12E-03 | 3.07E-03 | -1.30E+00 |
| ANKRD33B | 6.75E-03 | 3.52E-02 | 1.03E+00 | 1.92E-02 | 8.29E-02 | 1.26E+00 | 1.52E-02 | 6.44E-02 | -1.14E+00 | 9.40E-02 | 3.59E-01 | 1.12E+00 | 3.21E-02 | 1.15E-01 | -1.03E+00 | 3.55E-02 | 1.00E-01 | 1.02E+00 | 4.28E-02 | 1.57E-01 | 1.18E+00 | 8.98E-02 | 2.47E-01 | 1.16E+00 |
| POLA1 | 6.75E-03 | 3.52E-02 | -1.29E+00 | 6.41E-02 | 2.77E-01 | 1.09E+00 | 6.55E-02 | 2.78E-01 | 1.01E+00 | 2.28E-01 | 8.70E-01 | 1.38E+00 | 3.25E-02 | 1.16E-01 | -1.32E+00 | 1.03E-01 | 2.89E-01 | -1.01E+00 | 6.26E-06 | 2.30E-05 | -1.25E+00 | 4.82E-03 | 1.32E-02 | 1.05E+00 |
| SIDT1 | 6.75E-03 | 3.52E-02 | -1.18E+00 | 1.26E-01 | 5.46E-01 | 1.28E+00 | 3.55E-03 | 1.51E-02 | -1.20E+00 | 2.33E-03 | 8.91E-03 | -1.13E+00 | 1.14E-02 | 4.07E-02 | -1.38E+00 | 9.94E-03 | 2.80E-02 | -1.53E+00 | 2.19E-01 | 8.04E-01 | 1.45E+00 | 3.47E-01 | 9.53E-01 | 1.19E+00 |
| AL157895.1 | 6.76E-03 | 3.52E-02 | -1.09E+00 | 7.50E-03 | 3.24E-02 | -1.01E+00 | 2.24E-02 | 9.53E-02 | -1.01E+00 | 1.45E-01 | 5.54E-01 | 1.42E+00 | 9.78E-02 | 3.49E-01 | 1.18E+00 | 9.44E-05 | 2.66E-04 | 1.01E+00 | 1.79E-05 | 6.57E-05 | 1.63E+00 | 5.11E-03 | 1.40E-02 | 1.52E+00 |
| NCF1B | 6.77E-03 | 3.53E-02 | -1.93E+00 | 7.69E-03 | 3.32E-02 | -1.86E+00 | 1.36E-01 | 5.79E-01 | -1.14E+00 | 1.38E-01 | 5.27E-01 | -1.16E+00 | 2.25E-04 | 8.01E-04 | -1.36E+00 | 4.14E-06 | 1.17E-05 | -2.48E+00 | 8.14E-05 | 2.99E-04 | -1.87E+00 | 2.01E-04 | 5.52E-04 | -1.60E+00 |
| DUSP16 | 6.77E-03 | 3.53E-02 | -1.02E+00 | 2.85E-02 | 1.23E-01 | 1.25E+00 | 4.03E-02 | 1.71E-01 | -1.07E+00 | 1.55E-01 | 5.92E-01 | 1.30E+00 | 3.38E-02 | 1.20E-01 | -1.07E+00 | 9.58E-02 | 2.70E-01 | 1.01E+00 | 6.59E-02 | 2.42E-01 | -1.06E+00 | 2.99E-01 | 8.22E-01 | 1.19E+00 |
| PPM1M | 6.78E-03 | 3.53E-02 | -1.15E+00 | 1.22E-02 | 5.29E-02 | 1.00E+00 | 2.48E-03 | 1.05E-02 | -1.57E+00 | 4.71E-02 | 1.80E-01 | -1.04E+00 | 2.41E-02 | 8.60E-02 | -1.12E+00 | 6.03E-02 | 1.70E-01 | -1.11E+00 | 5.22E-02 | 1.91E-01 | 1.55E+00 | 3.01E-01 | 8.27E-01 | 1.27E+00 |
| SLC2A5 | 6.80E-03 | 3.54E-02 | 1.62E+00 | 1.87E-02 | 8.09E-02 | 1.79E+00 | 1.14E-04 | 4.85E-04 | 1.60E+00 | 1.89E-01 | 7.23E-01 | 1.40E+00 | 2.79E-01 | 9.95E-01 | 1.17E+00 | 3.51E-01 | 9.91E-01 | 1.30E+00 | 2.22E-01 | 8.15E-01 | -1.15E+00 | 3.27E-01 | 8.97E-01 | 1.03E+00 |
| CCDC59 | 6.82E-03 | 3.55E-02 | 1.05E+00 | 1.90E-23 | 8.21E-23 | -1.39E+00 | 1.60E-01 | 6.79E-01 | 1.13E+00 | 1.61E-07 | 6.16E-07 | -1.23E+00 | 4.72E-02 | 1.68E-01 | 1.06E+00 | 9.33E-08 | 2.63E-07 | -1.32E+00 | 3.85E-02 | 1.41E-01 | 1.01E+00 | 2.00E-04 | 5.48E-04 | -1.20E+00 |
| SNX7 | 6.82E-03 | 3.55E-02 | -1.22E+00 | 1.72E-01 | 7.45E-01 | 1.35E+00 | 7.87E-03 | 3.34E-02 | -1.27E+00 | 2.16E-02 | 8.24E-02 | 1.03E+00 | 4.93E-02 | 1.76E-01 | -1.42E+00 | 8.25E-02 | 2.33E-01 | -1.22E+00 | 5.86E-02 | 2.15E-01 | -1.42E+00 | 1.25E-01 | 3.43E-01 | -1.32E+00 |
| ZNF99 | 6.82E-03 | 3.55E-02 | -1.59E+00 | 1.85E-01 | 8.00E-01 | 1.14E+00 | 5.89E-05 | 2.50E-04 | -1.82E+00 | 1.38E-03 | 5.27E-03 | -1.33E+00 | 2.38E-01 | 8.49E-01 | 1.22E+00 | 2.85E-01 | 8.03E-01 | 1.25E+00 | 2.71E-01 | 9.93E-01 | 1.17E+00 | 1.39E-01 | 3.82E-01 | 1.52E+00 |
| EPS8L2 | 6.82E-03 | 3.56E-02 | 1.05E+00 | 4.73E-02 | 2.04E-01 | 1.34E+00 | 5.79E-03 | 2.46E-02 | -1.06E+00 | 6.19E-02 | 2.37E-01 | 1.32E+00 | 5.37E-03 | 1.91E-02 | -1.30E+00 | 7.06E-03 | 1.99E-02 | -1.17E+00 | 1.99E-02 | 7.31E-02 | -1.27E+00 | 1.75E-01 | 4.82E-01 | 1.04E+00 |
| ITGAV | 6.84E-03 | 3.57E-02 | 1.05E+00 | 1.06E-03 | 4.58E-03 | 1.05E+00 | 2.33E-01 | 9.90E-01 | 1.13E+00 | 1.72E-01 | 6.57E-01 | 1.43E+00 | 2.04E-01 | 7.27E-01 | 1.69E+00 | 1.40E-01 | 3.96E-01 | 1.04E+00 | 7.42E-02 | 2.72E-01 | 1.01E+00 | 1.81E-02 | 4.98E-02 | -1.14E+00 |
| PHYH | 6.84E-03 | 3.57E-02 | 1.08E+00 | 2.97E-05 | 1.28E-04 | 1.01E+00 | 8.54E-02 | 3.63E-01 | 1.04E+00 | 1.02E-02 | 3.88E-02 | 1.04E+00 | 2.09E-03 | 7.45E-03 | -1.00E+00 | 3.27E-02 | 9.22E-02 | 1.16E+00 | 9.06E-04 | 3.33E-03 | -1.14E+00 | 3.73E-03 | 1.03E-02 | -1.02E+00 |
| MRPS31 | 6.85E-03 | 3.57E-02 | 1.05E+00 | 4.77E-02 | 2.06E-01 | 1.27E+00 | 2.13E-03 | 9.03E-03 | -1.10E+00 | 1.07E-02 | 4.09E-02 | 1.06E+00 | 3.49E-08 | 1.24E-07 | -1.47E+00 | 1.36E-02 | 3.84E-02 | 1.03E+00 | 3.22E-06 | 1.18E-05 | -1.26E+00 | 3.58E-02 | 9.84E-02 | -1.01E+00 |
| RALGAPA2 | 6.85E-03 | 3.57E-02 | 1.07E+00 | 1.24E-04 | 5.36E-04 | 1.04E+00 | 1.73E-06 | 7.34E-06 | -1.40E+00 | 4.50E-04 | 1.72E-03 | 1.02E+00 | 2.95E-05 | 1.05E-04 | -1.37E+00 | 2.68E-03 | 7.56E-03 | -1.06E+00 | 1.08E-11 | 3.96E-11 | -1.67E+00 | 5.92E-06 | 1.63E-05 | -1.17E+00 |
| ADGRL1 | 6.86E-03 | 3.58E-02 | -1.04E+00 | 4.56E-03 | 1.97E-02 | 1.01E+00 | 1.87E-04 | 7.96E-04 | -1.25E+00 | 5.34E-05 | 2.04E-04 | -1.14E+00 | 1.12E-02 | 3.99E-02 | -1.41E+00 | 3.99E-02 | 1.12E-01 | -1.17E+00 | 1.86E-01 | 6.85E-01 | 1.05E+00 | 3.39E-01 | 9.31E-01 | 1.21E+00 |
| ZBED6CL | 6.87E-03 | 3.58E-02 | -1.30E+00 | 8.06E-02 | 3.48E-01 | 1.19E+00 | 1.96E-01 | 8.32E-01 | 1.12E+00 | 1.52E-01 | 5.82E-01 | 1.59E+00 | 2.57E-01 | 9.17E-01 | 1.41E+00 | 3.54E-01 | 9.98E-01 | 1.40E+00 | 9.80E-04 | 3.60E-03 | -1.40E+00 | 1.70E-02 | 4.66E-02 | -1.10E+00 |
| TRAF5 | 6.87E-03 | 3.58E-02 | 1.17E+00 | 7.47E-03 | 3.23E-02 | 1.27E+00 | 3.58E-02 | 1.52E-01 | 1.17E+00 | 8.44E-02 | 3.22E-01 | 1.31E+00 | 2.72E-01 | 9.72E-01 | 1.60E+00 | 8.15E-02 | 2.30E-01 | 1.13E+00 | 1.05E-01 | 3.85E-01 | 1.03E+00 | 7.50E-02 | 2.06E-01 | -1.12E+00 |
| LRP8 | 6.88E-03 | 3.59E-02 | -1.40E+00 | 3.92E-02 | 1.69E-01 | -1.05E+00 | 1.27E-02 | 5.39E-02 | -1.24E+00 | 2.08E-01 | 7.96E-01 | 1.26E+00 | 2.68E-02 | 9.56E-02 | -1.45E+00 | 1.71E-01 | 4.81E-01 | 1.00E+00 | 5.65E-03 | 2.07E-02 | -1.18E+00 | 1.61E-01 | 4.41E-01 | 1.17E+00 |
| AC008393.1 | 6.89E-03 | 3.59E-02 | -1.12E+00 | 2.15E-01 | 9.29E-01 | 1.57E+00 | 5.68E-06 | 2.42E-05 | -2.26E+00 | 7.95E-03 | 3.04E-02 | -1.09E+00 | 7.09E-02 | 2.53E-01 | -1.27E+00 | 3.31E-01 | 9.33E-01 | 1.24E+00 | 1.45E-03 | 5.32E-03 | -1.63E+00 | 3.09E-02 | 8.48E-02 | -1.19E+00 |
| MARCKSL1 | 6.89E-03 | 3.59E-02 | -1.03E+00 | 1.59E-01 | 6.89E-01 | 1.24E+00 | 3.55E-02 | 1.51E-01 | -1.08E+00 | 2.18E-01 | 8.33E-01 | 1.23E+00 | 1.82E-08 | 6.48E-08 | -1.38E+00 | 1.99E-06 | 5.62E-06 | -1.24E+00 | 3.78E-06 | 1.39E-05 | -1.25E+00 | 2.87E-01 | 7.87E-01 | 1.18E+00 |
| CD82 | 6.92E-03 | 3.61E-02 | 1.02E+00 | 7.60E-02 | 3.28E-01 | 1.36E+00 | 2.30E-07 | 9.79E-07 | -1.26E+00 | 5.24E-02 | 2.00E-01 | 1.17E+00 | 2.78E-02 | 9.93E-02 | -1.14E+00 | 3.42E-01 | 9.64E-01 | 1.08E+00 | 9.47E-04 | 3.47E-03 | -1.32E+00 | 9.61E-02 | 2.64E-01 | -1.01E+00 |
| AC012676.5 | 6.94E-03 | 3.62E-02 | -1.80E+00 | 1.27E-02 | 5.50E-02 | -1.59E+00 | 7.19E-02 | 3.05E-01 | -1.11E+00 | 1.09E-01 | 4.16E-01 | 1.01E+00 | 9.74E-04 | 3.47E-03 | -1.84E+00 | 6.03E-04 | 1.70E-03 | -1.71E+00 | 6.39E-07 | 2.35E-06 | -2.18E+00 | 1.16E-04 | 3.19E-04 | -1.69E+00 |
| AP4M1 | 6.94E-03 | 3.62E-02 | 1.02E+00 | 8.18E-03 | 3.53E-02 | 1.15E+00 | 3.90E-02 | 1.66E-01 | 1.01E+00 | 3.41E-02 | 1.30E-01 | 1.06E+00 | 6.45E-04 | 2.30E-03 | -1.25E+00 | 5.36E-03 | 1.51E-02 | -1.01E+00 | 4.02E-05 | 1.48E-04 | -1.30E+00 | 8.03E-03 | 2.21E-02 | -1.01E+00 |
| THEMIS2 | 6.95E-03 | 3.62E-02 | -1.05E+00 | 7.97E-02 | 3.44E-01 | 1.29E+00 | 1.03E-03 | 4.39E-03 | -1.29E+00 | 9.65E-03 | 3.68E-02 | 1.00E+00 | 6.16E-02 | 2.20E-01 | 1.97E+00 | 1.37E-01 | 3.86E-01 | 1.93E+00 | 6.45E-03 | 2.37E-02 | -1.05E+00 | 3.05E-02 | 8.39E-02 | 1.05E+00 |
| TTLL4 | 6.99E-03 | 3.64E-02 | -1.31E+00 | 9.64E-02 | 4.16E-01 | 1.29E+00 | 9.63E-07 | 4.09E-06 | -1.77E+00 | 8.45E-05 | 3.23E-04 | -1.26E+00 | 2.20E-01 | 7.85E-01 | 1.13E+00 | 1.07E-01 | 3.01E-01 | 2.30E+00 | 1.54E-06 | 5.65E-06 | -1.74E+00 | 1.74E-02 | 4.78E-02 | -1.03E+00 |
| LINC00216 | 7.00E-03 | 3.65E-02 | -1.19E+00 | 1.19E-02 | 5.14E-02 | -1.03E+00 | 6.14E-02 | 2.61E-01 | 1.04E+00 | 2.03E-01 | 7.75E-01 | 1.36E+00 | 1.97E-01 | 7.03E-01 | 1.06E+00 | 3.24E-01 | 9.14E-01 | 1.19E+00 | 1.57E-01 | 5.77E-01 | 1.32E+00 | 2.15E-01 | 5.90E-01 | 1.07E+00 |
| LLGL1 | 7.02E-03 | 3.66E-02 | -1.52E+00 | 4.50E-02 | 1.94E-01 | -1.06E+00 | 1.10E-01 | 4.67E-01 | -1.01E+00 | 2.82E-02 | 1.08E-01 | -1.13E+00 | 6.78E-03 | 2.42E-02 | -1.39E+00 | 2.78E-02 | 7.83E-02 | -1.11E+00 | 2.75E-02 | 1.01E-01 | -1.11E+00 | 2.54E-02 | 6.97E-02 | -1.18E+00 |
| CYP2R1 | 7.02E-03 | 3.66E-02 | -1.21E+00 | 1.94E-01 | 8.39E-01 | 1.48E+00 | 1.02E-06 | 4.32E-06 | -2.01E+00 | 4.58E-03 | 1.75E-02 | -1.05E+00 | 2.36E-02 | 8.42E-02 | 1.02E+00 | 2.69E-01 | 7.57E-01 | 1.38E+00 | 2.56E-05 | 9.39E-05 | -1.65E+00 | 1.46E-01 | 4.02E-01 | 1.08E+00 |
| LSM1 | 7.03E-03 | 3.66E-02 | -1.25E+00 | 2.21E-03 | 9.53E-03 | -1.32E+00 | 1.08E-01 | 4.58E-01 | -1.00E+00 | 2.02E-02 | 7.73E-02 | -1.19E+00 | 2.49E-01 | 8.89E-01 | 1.24E+00 | 2.94E-01 | 8.29E-01 | 1.48E+00 | 1.23E-01 | 4.53E-01 | 1.11E+00 | 1.20E-01 | 3.29E-01 | 1.03E+00 |
| AC007494.2 | 7.05E-03 | 3.67E-02 | -1.21E+00 | 1.03E-02 | 4.47E-02 | -1.10E+00 | 2.69E-02 | 1.14E-01 | -1.17E+00 | 1.02E-01 | 3.90E-01 | 1.11E+00 | 3.40E-03 | 1.21E-02 | -1.73E+00 | 2.30E-01 | 6.49E-01 | 1.21E+00 | 5.65E-02 | 2.07E-01 | -1.17E+00 | 1.40E-01 | 3.83E-01 | -1.03E+00 |
| KIAA0895 | 7.05E-03 | 3.68E-02 | -1.21E+00 | 1.48E-03 | 6.42E-03 | -1.28E+00 | 8.07E-02 | 3.43E-01 | 1.06E+00 | 5.47E-03 | 2.09E-02 | -1.09E+00 | 4.93E-02 | 1.76E-01 | -1.12E+00 | 2.23E-02 | 6.29E-02 | -1.38E+00 | 1.36E-01 | 4.98E-01 | 1.55E+00 | 3.07E-01 | 8.44E-01 | 1.30E+00 |
| CRIPT | 7.06E-03 | 3.68E-02 | 1.17E+00 | 1.07E-01 | 4.64E-01 | 1.62E+00 | 1.51E-03 | 6.40E-03 | -1.05E+00 | 2.68E-02 | 1.02E-01 | 1.16E+00 | 1.40E-01 | 4.98E-01 | 1.38E+00 | 1.75E-01 | 4.93E-01 | 1.39E+00 | 3.32E-02 | 1.22E-01 | 1.11E+00 | 3.55E-01 | 9.74E-01 | 1.18E+00 |
| RAB33A | 7.06E-03 | 3.68E-02 | 1.07E+00 | 1.73E-02 | 7.46E-02 | 1.21E+00 | 1.06E-01 | 4.50E-01 | 1.14E+00 | 2.56E-01 | 9.79E-01 | 1.48E+00 | 7.21E-02 | 2.57E-01 | 1.28E+00 | 3.19E-01 | 8.99E-01 | 1.49E+00 | 6.87E-06 | 2.52E-05 | -1.37E+00 | 2.30E-05 | 6.33E-05 | -1.28E+00 |
| CSRP2 | 7.07E-03 | 3.68E-02 | -1.50E+00 | 4.91E-03 | 2.12E-02 | -1.51E+00 | 6.19E-02 | 2.63E-01 | 1.36E+00 | 1.34E-01 | 5.12E-01 | 1.37E+00 | 3.14E-04 | 1.12E-03 | -2.35E+00 | 2.51E-03 | 7.09E-03 | -1.69E+00 | 5.12E-03 | 1.88E-02 | -1.69E+00 | 1.35E-02 | 3.71E-02 | -1.65E+00 |
| HMGB1P37 | 7.07E-03 | 3.68E-02 | 1.81E+00 | 1.07E-01 | 4.61E-01 | 1.70E+00 | 1.32E-05 | 5.62E-05 | 2.11E+00 | 5.27E-02 | 2.01E-01 | 1.88E+00 | 3.73E-02 | 1.33E-01 | 1.75E+00 | 7.39E-03 | 2.08E-02 | 2.15E+00 | 2.34E-04 | 8.58E-04 | 1.83E+00 | 2.09E-04 | 5.74E-04 | 1.93E+00 |
| HEMGN | 7.07E-03 | 3.69E-02 | 1.11E+00 | 1.28E-03 | 5.53E-03 | 1.79E+00 | 1.59E-01 | 6.74E-01 | 1.12E+00 | 4.68E-02 | 1.79E-01 | 1.75E+00 | 1.73E-01 | 6.16E-01 | 1.43E+00 | 4.13E-02 | 1.16E-01 | 1.73E+00 | 3.80E-07 | 1.40E-06 | 1.90E+00 | 2.14E-08 | 5.89E-08 | 2.12E+00 |
| AC008427.1 | 7.07E-03 | 3.69E-02 | 2.00E+00 | 2.00E-01 | 8.65E-01 | 1.38E+00 | 1.32E-01 | 5.61E-01 | 1.36E+00 | 2.37E-01 | 9.07E-01 | 1.42E+00 | 2.00E-01 | 7.13E-01 | 1.58E+00 | 2.46E-01 | 6.93E-01 | 1.61E+00 | 2.18E-02 | 8.00E-02 | 1.77E+00 | 6.34E-03 | 1.74E-02 | 2.13E+00 |
| AC006011.2 | 7.08E-03 | 3.69E-02 | -1.27E+00 | 2.74E-03 | 1.18E-02 | -1.41E+00 | 1.20E-04 | 5.10E-04 | -1.51E+00 | 6.64E-06 | 2.54E-05 | -1.65E+00 | 4.34E-02 | 1.55E-01 | -1.22E+00 | 2.31E-02 | 6.51E-02 | -1.45E+00 | 1.50E-04 | 5.51E-04 | -1.39E+00 | 6.50E-05 | 1.78E-04 | -1.36E+00 |
| MICA | 7.10E-03 | 3.70E-02 | 1.05E+00 | 2.49E-03 | 1.08E-02 | 1.07E+00 | 8.48E-02 | 3.60E-01 | -1.04E+00 | 1.55E-01 | 5.94E-01 | 1.20E+00 | 3.58E-02 | 1.28E-01 | -1.02E+00 | 1.46E-01 | 4.11E-01 | 1.20E+00 | 1.38E-03 | 5.06E-03 | -1.13E+00 | 3.15E-02 | 8.66E-02 | -1.00E+00 |
| CRTC2 | 7.10E-03 | 3.70E-02 | 1.02E+00 | 8.19E-03 | 3.54E-02 | 1.09E+00 | 6.25E-02 | 2.65E-01 | -1.03E+00 | 5.38E-02 | 2.05E-01 | 1.16E+00 | 1.08E-01 | 3.83E-01 | 1.07E+00 | 7.02E-02 | 1.98E-01 | -1.05E+00 | 9.57E-02 | 3.51E-01 | 1.13E+00 | 2.50E-01 | 6.87E-01 | 1.13E+00 |
| DHFR | 7.10E-03 | 3.70E-02 | -1.31E+00 | 2.09E-02 | 9.02E-02 | -1.09E+00 | 6.06E-02 | 2.57E-01 | 1.12E+00 | 4.18E-05 | 1.60E-04 | -1.01E+00 | 1.34E-02 | 4.80E-02 | -2.12E+00 | 8.29E-02 | 2.34E-01 | -1.25E+00 | 2.44E-03 | 8.95E-03 | -1.21E+00 | 3.35E-01 | 9.19E-01 | 1.13E+00 |
| AL354892.3 | 7.10E-03 | 3.70E-02 | -1.21E+00 | 4.62E-03 | 2.00E-02 | -1.23E+00 | 9.43E-03 | 4.01E-02 | -1.25E+00 | 3.02E-02 | 1.15E-01 | 1.03E+00 | 8.30E-03 | 2.96E-02 | -1.54E+00 | 1.06E-01 | 2.99E-01 | -1.00E+00 | 7.02E-03 | 2.58E-02 | -1.31E+00 | 2.28E-03 | 6.26E-03 | -1.61E+00 |
| PSMD12P | 7.12E-03 | 3.71E-02 | 2.85E+00 | 1.56E-02 | 6.73E-02 | 2.85E+00 | 9.78E-02 | 4.16E-01 | 1.45E+00 | 4.41E-02 | 1.68E-01 | 1.74E+00 | 1.34E-01 | 4.77E-01 | 1.97E+00 | 2.48E-02 | 7.00E-02 | 3.28E+00 | 5.15E-02 | 1.89E-01 | 1.70E+00 | 7.39E-04 | 2.03E-03 | 2.31E+00 |
| CENPQ | 7.12E-03 | 3.71E-02 | -1.42E+00 | 3.28E-02 | 1.42E-01 | -1.02E+00 | 8.48E-04 | 3.60E-03 | -1.39E+00 | 1.56E-02 | 5.98E-02 | -1.07E+00 | 1.21E-01 | 4.31E-01 | 1.06E+00 | 1.99E-01 | 5.62E-01 | 1.04E+00 | 1.54E-04 | 5.64E-04 | -1.17E+00 | 7.96E-04 | 2.19E-03 | -1.15E+00 |
| NUDT18 | 7.12E-03 | 3.71E-02 | -1.11E+00 | 1.42E-02 | 6.12E-02 | -1.01E+00 | 1.75E-05 | 7.44E-05 | -1.41E+00 | 3.44E-07 | 1.31E-06 | -1.49E+00 | 3.17E-05 | 1.13E-04 | -1.41E+00 | 3.28E-04 | 9.26E-04 | -1.29E+00 | 6.09E-03 | 2.23E-02 | -1.16E+00 | 5.27E-03 | 1.45E-02 | -1.23E+00 |
| NUDT13 | 7.14E-03 | 3.72E-02 | -1.15E+00 | 8.99E-03 | 3.88E-02 | -1.05E+00 | 1.74E-01 | 7.39E-01 | 1.27E+00 | 1.73E-01 | 6.59E-01 | 1.14E+00 | 8.83E-02 | 3.15E-01 | 1.10E+00 | 1.26E-01 | 3.54E-01 | 1.07E+00 | 1.38E-01 | 5.06E-01 | 1.09E+00 | 1.14E-01 | 3.13E-01 | 1.16E+00 |
| ZBTB10 | 7.16E-03 | 3.73E-02 | 1.51E+00 | 4.77E-04 | 2.06E-03 | 1.10E+00 | 8.89E-02 | 3.78E-01 | 1.43E+00 | 7.70E-05 | 2.94E-04 | -1.05E+00 | 5.92E-02 | 2.11E-01 | 1.72E+00 | 1.72E-04 | 4.86E-04 | -1.21E+00 | 9.56E-05 | 3.51E-04 | -1.25E+00 | 1.81E-09 | 4.97E-09 | -1.71E+00 |
| TMEM191A | 7.17E-03 | 3.74E-02 | 2.05E+00 | 2.98E-02 | 1.29E-01 | 2.15E+00 | 1.84E-01 | 7.81E-01 | 1.14E+00 | 2.69E-02 | 1.03E-01 | 1.02E+00 | 2.54E-01 | 9.05E-01 | 1.43E+00 | 3.27E-01 | 9.22E-01 | 1.35E+00 | 1.20E-01 | 4.39E-01 | -1.03E+00 | 1.92E-01 | 5.28E-01 | 1.01E+00 |
| TYMSOS | 7.17E-03 | 3.74E-02 | -1.01E+00 | 5.57E-06 | 2.41E-05 | -1.27E+00 | 1.16E-02 | 4.92E-02 | 1.01E+00 | 2.47E-10 | 9.43E-10 | -1.43E+00 | 1.24E-01 | 4.42E-01 | 1.23E+00 | 1.93E-02 | 5.43E-02 | -1.07E+00 | 4.02E-02 | 1.47E-01 | -1.08E+00 | 6.30E-03 | 1.73E-02 | -1.30E+00 |
| OXER1 | 7.17E-03 | 3.74E-02 | -1.42E+00 | 4.41E-02 | 1.90E-01 | -1.06E+00 | 1.30E-02 | 5.53E-02 | -1.27E+00 | 5.23E-02 | 2.00E-01 | -1.03E+00 | 7.27E-03 | 2.59E-02 | -1.40E+00 | 1.41E-02 | 3.98E-02 | -1.46E+00 | 8.40E-02 | 3.08E-01 | -1.27E+00 | 1.02E-01 | 2.80E-01 | -1.30E+00 |
| KSR1 | 7.18E-03 | 3.74E-02 | -1.03E+00 | 4.95E-03 | 2.14E-02 | 1.00E+00 | 3.22E-02 | 1.37E-01 | -1.25E+00 | 6.81E-02 | 2.60E-01 | -1.11E+00 | 1.19E-01 | 4.23E-01 | -1.14E+00 | 6.03E-02 | 1.70E-01 | -1.40E+00 | 1.02E-02 | 3.73E-02 | -1.18E+00 | 7.17E-02 | 1.97E-01 | -1.10E+00 |
| PPP1R3B | 7.18E-03 | 3.74E-02 | -1.10E+00 | 3.25E-04 | 1.41E-03 | -1.37E+00 | 3.75E-03 | 1.59E-02 | -1.51E+00 | 8.82E-04 | 3.37E-03 | -1.72E+00 | 3.07E-04 | 1.09E-03 | -1.68E+00 | 3.96E-03 | 1.12E-02 | -1.29E+00 | 8.80E-03 | 3.23E-02 | -1.20E+00 | 4.82E-02 | 1.32E-01 | -1.00E+00 |
| ARHGEF39 | 7.21E-03 | 3.76E-02 | -1.38E+00 | 1.10E-02 | 4.75E-02 | -1.30E+00 | 2.33E-02 | 9.91E-02 | -1.13E+00 | 2.35E-01 | 8.96E-01 | 1.42E+00 | 2.35E-01 | 8.39E-01 | 1.14E+00 | 2.10E-01 | 5.91E-01 | 1.04E+00 | 3.62E-04 | 1.33E-03 | -1.44E+00 | 4.11E-02 | 1.13E-01 | 1.05E+00 |
| NOL6 | 7.23E-03 | 3.77E-02 | -1.05E+00 | 1.32E-02 | 5.69E-02 | 1.17E+00 | 2.14E-02 | 9.09E-02 | -1.11E+00 | 1.42E-01 | 5.43E-01 | 1.27E+00 | 1.08E-01 | 3.85E-01 | 1.08E+00 | 1.78E-01 | 5.02E-01 | 1.76E+00 | 5.17E-02 | 1.90E-01 | -1.03E+00 | 2.17E-02 | 5.96E-02 | 1.67E+00 |
| PWWP2B | 7.23E-03 | 3.77E-02 | -1.02E+00 | 1.06E-01 | 4.56E-01 | 1.35E+00 | 2.76E-06 | 1.17E-05 | -1.47E+00 | 2.18E-05 | 8.34E-05 | -1.21E+00 | 3.91E-03 | 1.40E-02 | -1.15E+00 | 5.73E-02 | 1.62E-01 | 1.08E+00 | 6.00E-05 | 2.20E-04 | -1.48E+00 | 5.91E-03 | 1.62E-02 | -1.16E+00 |
| MYCN | 7.24E-03 | 3.77E-02 | 1.80E+00 | 1.09E-16 | 4.70E-16 | 3.69E+00 | 2.24E-01 | 9.53E-01 | 1.32E+00 | 4.18E-12 | 1.60E-11 | 3.22E+00 | 4.25E-02 | 1.52E-01 | 1.23E+00 | 1.12E-02 | 3.16E-02 | 2.36E+00 | 2.40E-01 | 8.80E-01 | 1.37E+00 | 1.84E-02 | 5.05E-02 | 1.96E+00 |
| SRGN | 7.25E-03 | 3.78E-02 | -1.03E+00 | 1.92E-08 | 8.32E-08 | -1.20E+00 | 4.84E-04 | 2.06E-03 | -1.16E+00 | 3.32E-04 | 1.27E-03 | -1.18E+00 | 9.22E-08 | 3.29E-07 | -1.51E+00 | 7.65E-12 | 2.16E-11 | -1.93E+00 | 3.00E-06 | 1.10E-05 | -1.36E+00 | 2.13E-05 | 5.85E-05 | -1.38E+00 |
| TMEM125 | 7.26E-03 | 3.78E-02 | -1.09E+00 | 2.59E-02 | 1.12E-01 | 1.08E+00 | 1.24E-02 | 5.25E-02 | -1.27E+00 | 1.81E-01 | 6.90E-01 | 1.23E+00 | 5.50E-03 | 1.96E-02 | -1.95E+00 | 8.37E-02 | 2.36E-01 | -1.04E+00 | 1.80E-01 | 6.61E-01 | -1.05E+00 | 2.79E-01 | 7.66E-01 | -1.10E+00 |
| PCNPP5 | 7.27E-03 | 3.79E-02 | -1.15E+00 | 2.05E-03 | 8.86E-03 | -1.24E+00 | 2.93E-02 | 1.25E-01 | -1.07E+00 | 9.85E-03 | 3.76E-02 | -1.14E+00 | 2.61E-01 | 9.32E-01 | 1.40E+00 | 1.86E-01 | 5.23E-01 | 1.82E+00 | 1.10E-02 | 4.02E-02 | 1.00E+00 | 1.66E-02 | 4.55E-02 | -1.06E+00 |
| ACAT2 | 7.27E-03 | 3.79E-02 | -1.07E+00 | 6.01E-02 | 2.60E-01 | 1.19E+00 | 7.22E-02 | 3.07E-01 | 1.02E+00 | 1.22E-01 | 4.66E-01 | 1.61E+00 | 2.26E-02 | 8.05E-02 | -1.09E+00 | 3.08E-01 | 8.67E-01 | 1.47E+00 | 2.14E-02 | 7.84E-02 | -1.19E+00 | 2.72E-01 | 7.46E-01 | 1.13E+00 |
| AP001148.1 | 7.27E-03 | 3.79E-02 | -1.17E+00 | 1.14E-01 | 4.93E-01 | 1.26E+00 | 4.15E-02 | 1.76E-01 | 1.03E+00 | 1.05E-01 | 4.00E-01 | 1.17E+00 | 9.45E-03 | 3.37E-02 | -1.37E+00 | 2.95E-02 | 8.32E-02 | -1.19E+00 | 2.26E-02 | 8.29E-02 | -1.40E+00 | 3.39E-01 | 9.31E-01 | 1.15E+00 |
| TSPAN6 | 7.27E-03 | 3.79E-02 | -1.19E+00 | 1.20E-01 | 5.19E-01 | 1.64E+00 | 3.80E-02 | 1.62E-01 | -1.31E+00 | 1.20E-01 | 4.60E-01 | 1.03E+00 | 1.83E-01 | 6.51E-01 | 1.54E+00 | 1.19E-01 | 3.34E-01 | 2.10E+00 | 5.41E-02 | 1.99E-01 | 1.68E+00 | 1.09E-01 | 2.99E-01 | 1.87E+00 |
| CWF19L1 | 7.28E-03 | 3.80E-02 | -1.10E+00 | 2.55E-03 | 1.10E-02 | -1.11E+00 | 9.60E-02 | 4.08E-01 | 1.12E+00 | 1.76E-02 | 6.71E-02 | 1.11E+00 | 1.88E-05 | 6.71E-05 | -1.38E+00 | 3.92E-05 | 1.10E-04 | -1.28E+00 | 2.17E-02 | 7.98E-02 | 1.01E+00 | 5.67E-02 | 1.56E-01 | 1.01E+00 |
| OR1C1 | 7.29E-03 | 3.80E-02 | -1.35E+00 | 1.35E-02 | 5.82E-02 | -1.07E+00 | 9.32E-02 | 3.96E-01 | -1.23E+00 | 2.38E-01 | 9.11E-01 | 1.03E+00 | 2.12E-01 | 7.57E-01 | -1.07E+00 | 2.39E-01 | 6.74E-01 | -1.03E+00 | 2.71E-01 | 9.95E-01 | 1.09E+00 | 3.33E-01 | 9.14E-01 | 1.14E+00 |
| ITGB3BP | 7.29E-03 | 3.80E-02 | 1.02E+00 | 6.32E-05 | 2.73E-04 | 1.01E+00 | 9.98E-03 | 4.24E-02 | -1.03E+00 | 1.52E-03 | 5.81E-03 | 1.06E+00 | 1.68E-03 | 5.99E-03 | -1.13E+00 | 9.95E-02 | 2.80E-01 | 1.19E+00 | 1.54E-02 | 5.64E-02 | 1.05E+00 | 1.51E-01 | 4.16E-01 | 1.18E+00 |
| LRWD1 | 7.30E-03 | 3.80E-02 | -1.14E+00 | 6.24E-02 | 2.70E-01 | 1.21E+00 | 1.39E-04 | 5.89E-04 | -1.36E+00 | 2.24E-04 | 8.56E-04 | -1.15E+00 | 2.04E-03 | 7.28E-03 | -1.14E+00 | 2.39E-02 | 6.73E-02 | 1.01E+00 | 8.31E-03 | 3.05E-02 | 1.03E+00 | 2.62E-01 | 7.21E-01 | 1.32E+00 |
| CDH1 | 7.30E-03 | 3.80E-02 | -2.41E+00 | 2.77E-02 | 1.20E-01 | -1.64E+00 | 1.82E-01 | 7.73E-01 | -1.05E+00 | 2.37E-01 | 9.07E-01 | 1.25E+00 | 2.12E-01 | 7.57E-01 | -1.19E+00 | 2.67E-01 | 7.52E-01 | 2.13E+00 | 3.16E-95 | 1.16E-94 | -3.91E+00 | 4.06E-41 | 1.12E-40 | -2.17E+00 |
| ACACA | 7.30E-03 | 3.81E-02 | 1.03E+00 | 8.95E-02 | 3.87E-01 | 1.38E+00 | 1.86E-01 | 7.91E-01 | 1.14E+00 | 1.03E-01 | 3.94E-01 | 1.19E+00 | 7.93E-02 | 2.83E-01 | 1.01E+00 | 1.22E-01 | 3.43E-01 | 1.12E+00 | 4.69E-02 | 1.72E-01 | 1.05E+00 | 2.35E-02 | 6.44E-02 | 1.02E+00 |
| LYRM4 | 7.30E-03 | 3.81E-02 | 1.05E+00 | 5.87E-03 | 2.54E-02 | 1.15E+00 | 5.67E-02 | 2.41E-01 | 1.13E+00 | 2.34E-02 | 8.94E-02 | 1.18E+00 | 6.53E-05 | 2.33E-04 | -1.00E+00 | 5.70E-02 | 1.61E-01 | 1.28E+00 | 1.93E-04 | 7.09E-04 | -1.02E+00 | 7.19E-02 | 1.98E-01 | 1.15E+00 |
| AC040977.1 | 7.31E-03 | 3.81E-02 | -1.27E+00 | 7.58E-02 | 3.28E-01 | 1.14E+00 | 4.53E-03 | 1.93E-02 | -1.42E+00 | 7.41E-02 | 2.83E-01 | 1.10E+00 | 1.85E-03 | 6.58E-03 | -1.59E+00 | 2.46E-01 | 6.94E-01 | 1.29E+00 | 1.54E-02 | 5.66E-02 | 1.87E+00 | 2.73E-01 | 7.51E-01 | 1.39E+00 |
| SH2D3C | 7.31E-03 | 3.81E-02 | -1.09E+00 | 4.15E-02 | 1.79E-01 | 1.94E+00 | 2.96E-06 | 1.26E-05 | -1.66E+00 | 1.29E-03 | 4.93E-03 | -1.07E+00 | 5.28E-02 | 1.88E-01 | -1.06E+00 | 3.10E-01 | 8.73E-01 | 1.25E+00 | 6.18E-02 | 2.27E-01 | -1.13E+00 | 2.93E-01 | 8.06E-01 | 1.19E+00 |
| C1orf74 | 7.32E-03 | 3.82E-02 | -1.60E+00 | 8.35E-02 | 3.61E-01 | 1.00E+00 | 5.66E-07 | 2.41E-06 | -2.15E+00 | 5.41E-05 | 2.07E-04 | -1.63E+00 | 2.13E-01 | 7.59E-01 | -1.00E+00 | 2.96E-01 | 8.34E-01 | 1.03E+00 | 2.09E-04 | 7.66E-04 | -1.30E+00 | 2.22E-04 | 6.09E-04 | -1.40E+00 |
| USP6NL | 7.34E-03 | 3.82E-02 | 1.05E+00 | 2.27E-03 | 9.82E-03 | 1.13E+00 | 2.09E-01 | 8.89E-01 | 1.22E+00 | 1.40E-02 | 5.36E-02 | -1.01E+00 | 1.14E-01 | 4.08E-01 | 1.16E+00 | 6.22E-02 | 1.75E-01 | 1.15E+00 | 5.20E-04 | 1.91E-03 | -1.12E+00 | 5.13E-05 | 1.41E-04 | -1.20E+00 |
| TEP1 | 7.34E-03 | 3.82E-02 | -1.05E+00 | 1.97E-01 | 8.51E-01 | 1.41E+00 | 2.24E-02 | 9.53E-02 | -1.19E+00 | 1.04E-01 | 3.96E-01 | 1.21E+00 | 5.21E-03 | 1.86E-02 | -1.62E+00 | 1.82E-02 | 5.13E-02 | -1.33E+00 | 8.24E-03 | 3.02E-02 | -1.27E+00 | 4.96E-02 | 1.36E-01 | -1.07E+00 |
| CCDC15 | 7.35E-03 | 3.83E-02 | -1.46E+00 | 3.88E-02 | 1.68E-01 | -1.06E+00 | 1.92E-03 | 8.18E-03 | -1.46E+00 | 3.64E-02 | 1.39E-01 | 1.03E+00 | 1.70E-01 | 6.06E-01 | 1.03E+00 | 2.92E-01 | 8.24E-01 | 1.18E+00 | 1.91E-05 | 6.99E-05 | -1.30E+00 | 8.25E-03 | 2.27E-02 | -1.03E+00 |
| RNF144A | 7.36E-03 | 3.83E-02 | -1.00E+00 | 1.25E-01 | 5.40E-01 | 1.42E+00 | 8.41E-05 | 3.57E-04 | -1.32E+00 | 8.24E-03 | 3.15E-02 | 1.05E+00 | 2.13E-02 | 7.60E-02 | -1.08E+00 | 5.98E-02 | 1.69E-01 | 1.03E+00 | 2.48E-03 | 9.11E-03 | -1.30E+00 | 3.65E-03 | 1.00E-02 | -1.24E+00 |
| TPTE2P1 | 7.36E-03 | 3.84E-02 | -1.85E+00 | 5.11E-02 | 2.21E-01 | -1.12E+00 | 1.85E-01 | 7.88E-01 | -1.01E+00 | 2.43E-01 | 9.29E-01 | 1.18E+00 | 2.28E-01 | 8.13E-01 | 1.39E+00 | 2.42E-01 | 6.81E-01 | 1.57E+00 | 6.95E-02 | 2.55E-01 | 1.06E+00 | 1.80E-01 | 4.94E-01 | 1.73E+00 |
| CCDC102B | 7.37E-03 | 3.84E-02 | -1.09E+00 | 1.47E-01 | 6.37E-01 | 1.60E+00 | 4.25E-02 | 1.81E-01 | -1.08E+00 | 1.40E-01 | 5.33E-01 | 1.27E+00 | 1.53E-01 | 5.47E-01 | 1.12E+00 | 1.82E-01 | 5.13E-01 | 1.12E+00 | 1.25E-01 | 4.60E-01 | 1.50E+00 | 2.42E-01 | 6.65E-01 | 1.52E+00 |
| PTK7 | 7.37E-03 | 3.84E-02 | 1.03E+00 | 9.12E-02 | 3.94E-01 | 1.40E+00 | 1.33E-04 | 5.66E-04 | -1.27E+00 | 1.51E-02 | 5.78E-02 | 1.11E+00 | 3.80E-05 | 1.35E-04 | -1.26E+00 | 5.01E-04 | 1.41E-03 | -1.13E+00 | 2.30E-05 | 8.45E-05 | -1.16E+00 | 9.69E-04 | 2.66E-03 | -1.04E+00 |
| CR545473.1 | 7.38E-03 | 3.85E-02 | -1.04E+00 | 7.58E-07 | 3.28E-06 | -1.76E+00 | 7.37E-06 | 3.13E-05 | -1.68E+00 | 2.51E-07 | 9.57E-07 | -2.03E+00 | 2.13E-06 | 7.58E-06 | -1.54E+00 | 6.80E-09 | 1.92E-08 | -2.18E+00 | 1.43E-01 | 5.23E-01 | 1.09E+00 | 5.59E-05 | 1.54E-04 | -1.96E+00 |
| RCAN1 | 7.38E-03 | 3.85E-02 | -1.40E+00 | 6.23E-02 | 2.69E-01 | 1.04E+00 | 1.19E-02 | 5.06E-02 | 1.01E+00 | 4.47E-02 | 1.71E-01 | 1.16E+00 | 2.77E-02 | 9.86E-02 | -1.21E+00 | 2.03E-01 | 5.73E-01 | 1.26E+00 | 4.69E-02 | 1.72E-01 | 1.07E+00 | 2.02E-01 | 5.54E-01 | 1.35E+00 |
| CDC7 | 7.38E-03 | 3.85E-02 | -1.53E+00 | 1.62E-02 | 7.01E-02 | -1.34E+00 | 4.64E-02 | 1.97E-01 | -1.06E+00 | 6.33E-02 | 2.42E-01 | 1.10E+00 | 1.63E-01 | 5.80E-01 | -1.14E+00 | 3.07E-01 | 8.66E-01 | 1.18E+00 | 5.13E-06 | 1.88E-05 | -1.31E+00 | 1.09E-02 | 3.01E-02 | 1.05E+00 |
| PRAF2 | 7.40E-03 | 3.85E-02 | -1.03E+00 | 6.88E-04 | 2.97E-03 | -1.08E+00 | 4.66E-04 | 1.98E-03 | -1.24E+00 | 9.23E-04 | 3.52E-03 | -1.09E+00 | 2.69E-05 | 9.58E-05 | -1.46E+00 | 2.30E-04 | 6.50E-04 | -1.23E+00 | 8.27E-03 | 3.04E-02 | -1.09E+00 | 1.86E-02 | 5.12E-02 | -1.05E+00 |
| PIGO | 7.40E-03 | 3.86E-02 | -1.11E+00 | 5.23E-02 | 2.26E-01 | 1.20E+00 | 6.21E-05 | 2.64E-04 | -1.28E+00 | 1.57E-05 | 6.01E-05 | -1.25E+00 | 5.94E-03 | 2.12E-02 | -1.22E+00 | 2.01E-02 | 5.66E-02 | -1.06E+00 | 2.88E-03 | 1.06E-02 | -1.20E+00 | 1.07E-02 | 2.95E-02 | -1.11E+00 |
| TEPSIN | 7.40E-03 | 3.86E-02 | -1.03E+00 | 6.11E-02 | 2.64E-01 | 1.23E+00 | 5.72E-08 | 2.43E-07 | -1.70E+00 | 5.94E-05 | 2.27E-04 | -1.11E+00 | 1.06E-04 | 3.78E-04 | -1.50E+00 | 1.07E-02 | 3.02E-02 | -1.05E+00 | 2.55E-12 | 9.35E-12 | -1.91E+00 | 2.34E-07 | 6.43E-07 | -1.46E+00 |
| SAP18 | 7.42E-03 | 3.87E-02 | 1.27E+00 | 4.54E-03 | 1.96E-02 | 1.10E+00 | 5.14E-03 | 2.19E-02 | 1.26E+00 | 2.19E-01 | 8.35E-01 | 1.07E+00 | 1.41E-01 | 5.03E-01 | 1.18E+00 | 3.03E-01 | 8.55E-01 | 1.15E+00 | 7.44E-02 | 2.73E-01 | 1.15E+00 | 3.13E-01 | 8.59E-01 | 1.05E+00 |
| HARS2 | 7.43E-03 | 3.87E-02 | -1.08E+00 | 3.23E-02 | 1.39E-01 | 1.19E+00 | 5.53E-04 | 2.35E-03 | -1.29E+00 | 8.18E-03 | 3.12E-02 | 1.04E+00 | 2.87E-05 | 1.02E-04 | -1.81E+00 | 2.90E-03 | 8.17E-03 | -1.25E+00 | 8.23E-06 | 3.02E-05 | -1.61E+00 | 2.27E-02 | 6.24E-02 | -1.04E+00 |
| KCNK6 | 7.43E-03 | 3.87E-02 | -1.67E+00 | 4.34E-02 | 1.88E-01 | -1.17E+00 | 2.68E-07 | 1.14E-06 | -2.51E+00 | 7.98E-05 | 3.05E-04 | -1.66E+00 | 6.75E-37 | 2.41E-36 | -3.60E+00 | 5.87E-29 | 1.65E-28 | -2.38E+00 | 7.39E-23 | 2.71E-22 | -1.32E+00 | 7.55E-18 | 2.07E-17 | -1.37E+00 |
| A2M-AS1 | 7.43E-03 | 3.87E-02 | -1.22E+00 | 1.56E-02 | 6.74E-02 | 1.01E+00 | 5.72E-03 | 2.43E-02 | -1.37E+00 | 1.15E-02 | 4.38E-02 | -1.21E+00 | 7.22E-05 | 2.57E-04 | -1.33E+00 | 5.11E-05 | 1.44E-04 | -1.45E+00 | 7.38E-02 | 2.71E-01 | -1.10E+00 | 3.35E-01 | 9.20E-01 | 1.21E+00 |
| PDCD6IPP1 | 7.44E-03 | 3.88E-02 | -1.15E+00 | 4.27E-02 | 1.84E-01 | 1.12E+00 | 4.73E-06 | 2.01E-05 | -1.99E+00 | 2.88E-03 | 1.10E-02 | -1.19E+00 | 2.06E-02 | 7.34E-02 | -1.34E+00 | 3.64E-02 | 1.03E-01 | -1.16E+00 | 2.26E-02 | 8.31E-02 | -1.15E+00 | 1.56E-02 | 4.30E-02 | -1.30E+00 |
| AL354696.2 | 7.46E-03 | 3.89E-02 | -1.32E+00 | 3.16E-02 | 1.37E-01 | -1.09E+00 | 1.69E-02 | 7.17E-02 | -1.39E+00 | 4.32E-03 | 1.65E-02 | -1.55E+00 | 2.57E-01 | 9.17E-01 | 1.36E+00 | 2.81E-01 | 7.92E-01 | 1.60E+00 | 1.64E-01 | 6.04E-01 | -1.10E+00 | 2.21E-01 | 6.06E-01 | 1.01E+00 |
| SLC7A7 | 7.46E-03 | 3.89E-02 | -1.91E+00 | 2.81E-02 | 1.22E-01 | -1.16E+00 | 1.44E-01 | 6.10E-01 | -1.06E+00 | 1.52E-02 | 5.79E-02 | -1.80E+00 | 2.77E-01 | 9.88E-01 | 1.13E+00 | 3.36E-01 | 9.48E-01 | 1.06E+00 | 6.81E-04 | 2.50E-03 | -1.54E+00 | 3.04E-04 | 8.36E-04 | -2.17E+00 |
| AC092068.2 | 7.46E-03 | 3.89E-02 | -1.32E+00 | 1.03E-02 | 4.46E-02 | -1.12E+00 | 3.82E-02 | 1.62E-01 | -1.24E+00 | 2.08E-01 | 7.94E-01 | 1.16E+00 | 6.98E-02 | 2.49E-01 | -1.51E+00 | 3.24E-01 | 9.14E-01 | 1.12E+00 | 6.49E-02 | 2.38E-01 | -1.52E+00 | 1.80E-01 | 4.95E-01 | -1.30E+00 |
| ATF5 | 7.47E-03 | 3.89E-02 | -1.10E+00 | 1.22E-03 | 5.29E-03 | -1.08E+00 | 3.52E-02 | 1.49E-01 | -1.04E+00 | 5.35E-03 | 2.04E-02 | -1.08E+00 | 7.99E-04 | 2.85E-03 | -1.28E+00 | 4.43E-03 | 1.25E-02 | -1.17E+00 | 1.65E-02 | 6.06E-02 | -1.13E+00 | 6.34E-05 | 1.74E-04 | -1.30E+00 |
| MITF | 7.48E-03 | 3.90E-02 | -1.78E+00 | 3.96E-02 | 1.71E-01 | -1.19E+00 | 9.00E-03 | 3.82E-02 | -1.33E+00 | 3.77E-02 | 1.44E-01 | 1.05E+00 | 1.43E-45 | 5.12E-45 | -3.83E+00 | 1.44E-35 | 4.07E-35 | -2.36E+00 | 8.59E-10 | 3.15E-09 | -1.75E+00 | 5.66E-04 | 1.56E-03 | -1.20E+00 |
| AC022098.4 | 7.48E-03 | 3.90E-02 | 1.91E+00 | 3.71E-02 | 1.60E-01 | -1.40E+00 | 7.98E-08 | 3.39E-07 | 3.70E+00 | 2.51E-01 | 9.59E-01 | 1.09E+00 | 2.29E-01 | 8.17E-01 | 1.32E+00 | 7.25E-02 | 2.04E-01 | -1.49E+00 | 1.85E-01 | 6.77E-01 | 1.01E+00 | 6.64E-02 | 1.83E-01 | -1.35E+00 |
| SLC37A4 | 7.49E-03 | 3.90E-02 | -1.10E+00 | 5.53E-03 | 2.39E-02 | -1.01E+00 | 8.89E-04 | 3.78E-03 | -1.02E+00 | 8.63E-03 | 3.30E-02 | 1.25E+00 | 9.81E-04 | 3.50E-03 | -1.52E+00 | 2.35E-02 | 6.62E-02 | -1.07E+00 | 4.35E-04 | 1.60E-03 | -1.15E+00 | 3.25E-01 | 8.92E-01 | 1.32E+00 |
| AC141586.1 | 7.50E-03 | 3.91E-02 | -1.09E+00 | 8.14E-03 | 3.52E-02 | 1.01E+00 | 1.99E-02 | 8.46E-02 | -1.11E+00 | 1.01E-01 | 3.84E-01 | 1.25E+00 | 4.94E-02 | 1.76E-01 | 1.03E+00 | 3.11E-02 | 8.77E-02 | 1.04E+00 | 4.76E-02 | 1.75E-01 | 1.02E+00 | 1.42E-02 | 3.89E-02 | -1.09E+00 |
| SCN3B | 7.51E-03 | 3.91E-02 | -1.50E+00 | 8.50E-02 | 3.67E-01 | -1.10E+00 | 4.67E-02 | 1.98E-01 | -1.28E+00 | 3.98E-02 | 1.52E-01 | -1.23E+00 | 1.97E-01 | 7.02E-01 | 1.39E+00 | 3.21E-01 | 9.05E-01 | 1.11E+00 | 2.66E-01 | 9.76E-01 | 1.02E+00 | 3.10E-01 | 8.52E-01 | 1.15E+00 |
| C16orf72 | 7.51E-03 | 3.91E-02 | 1.68E+00 | 1.97E-04 | 8.49E-04 | -1.06E+00 | 1.59E-02 | 6.75E-02 | 1.49E+00 | 1.97E-04 | 7.52E-04 | -1.26E+00 | 1.45E-01 | 5.18E-01 | 1.52E+00 | 3.12E-03 | 8.80E-03 | -1.14E+00 | 1.30E-01 | 4.77E-01 | 1.40E+00 | 2.70E-04 | 7.43E-04 | -1.35E+00 |
| FP671120.1 | 7.51E-03 | 3.91E-02 | -1.09E+00 | 3.62E-02 | 1.56E-01 | 1.15E+00 | 1.45E-04 | 6.18E-04 | -1.73E+00 | 1.13E-03 | 4.30E-03 | -1.37E+00 | 2.34E-01 | 8.34E-01 | 1.72E+00 | 1.14E-01 | 3.22E-01 | 2.02E+00 | 5.50E-02 | 2.02E-01 | -1.13E+00 | 1.60E-02 | 4.40E-02 | -1.44E+00 |
| DEDD | 7.52E-03 | 3.92E-02 | -1.00E+00 | 1.78E-03 | 7.70E-03 | 1.06E+00 | 1.55E-02 | 6.57E-02 | -1.03E+00 | 1.45E-03 | 5.53E-03 | 1.03E+00 | 3.97E-06 | 1.41E-05 | -1.50E+00 | 1.65E-05 | 4.66E-05 | -1.33E+00 | 1.80E-03 | 6.62E-03 | -1.13E+00 | 6.71E-04 | 1.84E-03 | -1.21E+00 |
| TSTD1 | 7.52E-03 | 3.92E-02 | 1.13E+00 | 1.29E-06 | 5.58E-06 | 1.05E+00 | 3.82E-02 | 1.62E-01 | 1.35E+00 | 3.92E-02 | 1.50E-01 | 1.15E+00 | 2.15E-01 | 7.65E-01 | 1.40E+00 | 2.34E-01 | 6.61E-01 | 1.37E+00 | 5.55E-08 | 2.04E-07 | 2.05E+00 | 1.27E-02 | 3.49E-02 | 1.47E+00 |
| ZSCAN2 | 7.54E-03 | 3.93E-02 | -1.03E+00 | 2.82E-02 | 1.22E-01 | 1.17E+00 | 3.62E-03 | 1.54E-02 | -1.06E+00 | 9.70E-05 | 3.70E-04 | -1.15E+00 | 2.86E-03 | 1.02E-02 | -1.08E+00 | 2.72E-05 | 7.66E-05 | -1.41E+00 | 1.26E-01 | 4.62E-01 | 1.03E+00 | 3.04E-03 | 8.35E-03 | -1.24E+00 |
| TBCE | 7.54E-03 | 3.93E-02 | -1.02E+00 | 3.17E-02 | 1.37E-01 | 1.26E+00 | 7.25E-03 | 3.08E-02 | -1.32E+00 | 8.08E-02 | 3.09E-01 | 1.07E+00 | 9.07E-04 | 3.23E-03 | -1.42E+00 | 1.46E-01 | 4.11E-01 | 1.19E+00 | 7.02E-04 | 2.57E-03 | -1.10E+00 | 5.34E-02 | 1.47E-01 | 1.06E+00 |
| RNF25 | 7.55E-03 | 3.94E-02 | -1.41E+00 | 3.20E-02 | 1.38E-01 | -1.09E+00 | 3.22E-02 | 1.37E-01 | -1.17E+00 | 9.98E-02 | 3.81E-01 | 1.11E+00 | 2.18E-03 | 7.79E-03 | -1.46E+00 | 2.66E-01 | 7.49E-01 | 1.35E+00 | 7.40E-02 | 2.71E-01 | 1.00E+00 | 2.11E-01 | 5.80E-01 | 1.07E+00 |
| ST3GAL5 | 7.58E-03 | 3.95E-02 | -1.50E+00 | 5.28E-03 | 2.28E-02 | -1.45E+00 | 2.72E-03 | 1.16E-02 | -1.48E+00 | 2.60E-05 | 9.92E-05 | -2.07E+00 | 8.93E-04 | 3.19E-03 | -1.41E+00 | 5.37E-03 | 1.51E-02 | -1.17E+00 | 6.40E-02 | 2.35E-01 | 1.67E+00 | 1.33E-01 | 3.64E-01 | 1.13E+00 |
| DOC2GP | 7.58E-03 | 3.95E-02 | 2.10E+00 | 1.50E-05 | 6.47E-05 | 3.37E+00 | 1.03E-01 | 4.38E-01 | 1.43E+00 | 1.75E-02 | 6.70E-02 | 1.86E+00 | 2.59E-01 | 9.22E-01 | 1.10E+00 | 2.90E-01 | 8.17E-01 | 1.33E+00 | 1.70E-01 | 6.23E-01 | 1.59E+00 | 3.37E-01 | 9.25E-01 | 1.09E+00 |
| HMGB1P31 | 7.59E-03 | 3.96E-02 | -1.22E+00 | 2.04E-01 | 8.79E-01 | 1.39E+00 | 7.57E-02 | 3.22E-01 | -1.21E+00 | 1.60E-01 | 6.10E-01 | 1.46E+00 | 1.02E-01 | 3.62E-01 | 1.86E+00 | 5.34E-02 | 1.50E-01 | 2.13E+00 | 1.74E-01 | 6.40E-01 | 1.06E+00 | 3.19E-01 | 8.77E-01 | 1.25E+00 |
| PLOD1 | 7.59E-03 | 3.96E-02 | -1.29E+00 | 6.71E-02 | 2.90E-01 | 1.12E+00 | 5.31E-05 | 2.26E-04 | -1.54E+00 | 1.49E-02 | 5.70E-02 | 1.07E+00 | 2.28E-01 | 8.12E-01 | -1.01E+00 | 1.51E-01 | 4.25E-01 | 1.52E+00 | 2.59E-05 | 9.52E-05 | -1.42E+00 | 3.44E-01 | 9.45E-01 | 1.32E+00 |
| ZNF519 | 7.59E-03 | 3.96E-02 | -1.13E+00 | 1.46E-01 | 6.32E-01 | 1.42E+00 | 2.47E-06 | 1.05E-05 | -1.51E+00 | 2.54E-02 | 9.69E-02 | 1.18E+00 | 1.84E-05 | 6.56E-05 | -1.83E+00 | 4.68E-03 | 1.32E-02 | -1.18E+00 | 4.28E-04 | 1.57E-03 | -1.39E+00 | 2.26E-01 | 6.21E-01 | 1.30E+00 |
| HIPK1-AS1 | 7.61E-03 | 3.97E-02 | 2.18E+00 | 1.35E-01 | 5.82E-01 | 1.32E+00 | 1.20E-02 | 5.09E-02 | 1.99E+00 | 3.60E-02 | 1.38E-01 | -1.32E+00 | 2.12E-01 | 7.55E-01 | 1.75E+00 | 2.86E-01 | 8.06E-01 | 1.30E+00 | 5.07E-02 | 1.86E-01 | 1.11E+00 | 1.20E-02 | 3.29E-02 | -1.22E+00 |
| STIM2 | 7.62E-03 | 3.97E-02 | 1.10E+00 | 1.62E-02 | 6.98E-02 | 1.23E+00 | 7.03E-02 | 2.99E-01 | 1.05E+00 | 1.67E-01 | 6.38E-01 | 1.31E+00 | 1.08E-03 | 3.84E-03 | -1.08E+00 | 2.01E-02 | 5.68E-02 | 1.07E+00 | 4.16E-02 | 1.53E-01 | 1.06E+00 | 1.71E-01 | 4.70E-01 | 1.16E+00 |
| STAG3L4 | 7.62E-03 | 3.97E-02 | -1.09E+00 | 2.99E-03 | 1.29E-02 | -1.12E+00 | 8.07E-06 | 3.43E-05 | -1.55E+00 | 4.49E-07 | 1.72E-06 | -1.63E+00 | 1.57E-02 | 5.61E-02 | -1.05E+00 | 2.47E-02 | 6.96E-02 | -1.00E+00 | 5.83E-03 | 2.14E-02 | -1.04E+00 | 2.71E-03 | 7.44E-03 | -1.11E+00 |
| KLF16 | 7.62E-03 | 3.97E-02 | -1.08E+00 | 8.58E-04 | 3.71E-03 | -1.15E+00 | 1.25E-02 | 5.29E-02 | -1.06E+00 | 1.14E-02 | 4.35E-02 | 1.05E+00 | 5.51E-02 | 1.97E-01 | -1.08E+00 | 4.29E-02 | 1.21E-01 | -1.04E+00 | 5.07E-02 | 1.86E-01 | 1.04E+00 | 6.71E-02 | 1.84E-01 | 1.08E+00 |
| EIF4EP2 | 7.63E-03 | 3.98E-02 | -1.39E+00 | 4.89E-03 | 2.11E-02 | -1.49E+00 | 5.12E-04 | 2.18E-03 | -1.45E+00 | 2.28E-04 | 8.72E-04 | -1.39E+00 | 3.71E-02 | 1.32E-01 | -1.48E+00 | 8.08E-02 | 2.28E-01 | -1.23E+00 | 1.86E-03 | 6.82E-03 | -1.21E+00 | 4.73E-03 | 1.30E-02 | -1.12E+00 |
| AL356599.1 | 7.63E-03 | 3.98E-02 | -1.21E+00 | 7.70E-02 | 3.33E-01 | 1.26E+00 | 6.85E-05 | 2.91E-04 | -1.90E+00 | 5.57E-03 | 2.13E-02 | -1.17E+00 | 6.86E-02 | 2.45E-01 | -1.31E+00 | 3.19E-01 | 8.99E-01 | 1.23E+00 | 3.52E-02 | 1.29E-01 | -1.20E+00 | 2.07E-01 | 5.68E-01 | 1.04E+00 |
| ACVR1B | 7.67E-03 | 4.00E-02 | -1.14E+00 | 3.54E-02 | 1.53E-01 | 1.08E+00 | 1.28E-01 | 5.42E-01 | 1.02E+00 | 1.45E-01 | 5.53E-01 | 1.18E+00 | 2.27E-01 | 8.10E-01 | 1.47E+00 | 3.62E-02 | 1.02E-01 | -1.06E+00 | 1.58E-02 | 5.81E-02 | -1.38E+00 | 6.22E-03 | 1.71E-02 | -1.44E+00 |
| SERINC5 | 7.69E-03 | 4.01E-02 | 1.56E+00 | 3.86E-02 | 1.67E-01 | 1.70E+00 | 1.92E-01 | 8.15E-01 | 1.10E+00 | 1.70E-02 | 6.51E-02 | 1.56E+00 | 1.49E-01 | 5.30E-01 | 1.28E+00 | 3.33E-01 | 9.37E-01 | 1.55E+00 | 1.56E-01 | 5.73E-01 | 1.00E+00 | 2.74E-01 | 7.53E-01 | 1.20E+00 |
| GTF2F2 | 7.70E-03 | 4.02E-02 | 1.05E+00 | 5.77E-03 | 2.49E-02 | 1.18E+00 | 7.13E-04 | 3.03E-03 | -1.07E+00 | 6.43E-03 | 2.45E-02 | 1.07E+00 | 4.68E-08 | 1.67E-07 | -1.24E+00 | 3.09E-03 | 8.71E-03 | 1.11E+00 | 4.45E-05 | 1.63E-04 | -1.17E+00 | 4.77E-02 | 1.31E-01 | 1.06E+00 |
| GPR22 | 7.71E-03 | 4.02E-02 | -1.27E+00 | 6.14E-04 | 2.65E-03 | -1.51E+00 | 9.47E-02 | 4.02E-01 | -1.12E+00 | 2.82E-04 | 1.08E-03 | -1.90E+00 | 2.43E-01 | 8.66E-01 | 1.65E+00 | 3.04E-01 | 8.58E-01 | 1.41E+00 | 3.97E-03 | 1.46E-02 | -1.15E+00 | 3.32E-03 | 9.12E-03 | -1.27E+00 |
| TAF1C | 7.72E-03 | 4.02E-02 | -1.36E+00 | 1.38E-01 | 5.98E-01 | 1.22E+00 | 1.32E-03 | 5.60E-03 | -1.38E+00 | 1.94E-02 | 7.41E-02 | 1.05E+00 | 7.16E-04 | 2.55E-03 | -1.60E+00 | 2.45E-02 | 6.91E-02 | -1.12E+00 | 3.70E-02 | 1.36E-01 | -1.09E+00 | 1.32E-01 | 3.62E-01 | 1.07E+00 |
| AC015987.1 | 7.75E-03 | 4.04E-02 | -1.18E+00 | 5.63E-02 | 2.43E-01 | 1.15E+00 | 5.81E-02 | 2.47E-01 | -1.23E+00 | 9.49E-02 | 3.62E-01 | -1.01E+00 | 1.34E-01 | 4.79E-01 | 1.09E+00 | 2.58E-01 | 7.28E-01 | 1.18E+00 | 9.76E-02 | 3.58E-01 | -1.13E+00 | 3.24E-01 | 8.89E-01 | 1.15E+00 |
| AC002310.1 | 7.76E-03 | 4.05E-02 | -1.05E+00 | 1.38E-01 | 5.97E-01 | 1.39E+00 | 5.20E-04 | 2.21E-03 | -1.45E+00 | 3.84E-03 | 1.47E-02 | -1.23E+00 | 3.31E-02 | 1.18E-01 | -1.23E+00 | 2.55E-01 | 7.19E-01 | 1.19E+00 | 3.45E-03 | 1.27E-02 | -1.37E+00 | 3.99E-04 | 1.10E-03 | -1.58E+00 |
| BAALC-AS1 | 7.77E-03 | 4.05E-02 | 1.84E+00 | 1.16E-02 | 5.00E-02 | 2.09E+00 | 7.49E-02 | 3.18E-01 | 1.60E+00 | 2.70E-02 | 1.03E-01 | 1.94E+00 | 4.77E-02 | 1.70E-01 | 2.53E+00 | 1.68E-01 | 4.73E-01 | 2.10E+00 | 1.54E-01 | 5.64E-01 | 1.54E+00 | 1.90E-01 | 5.21E-01 | 1.67E+00 |
| LMLN | 7.77E-03 | 4.05E-02 | -1.32E+00 | 3.23E-02 | 1.40E-01 | -1.11E+00 | 1.73E-06 | 7.35E-06 | -1.77E+00 | 1.12E-06 | 4.30E-06 | -1.73E+00 | 2.43E-01 | 8.68E-01 | 1.37E+00 | 1.43E-01 | 4.04E-01 | -1.20E+00 | 4.74E-07 | 1.74E-06 | -2.15E+00 | 3.39E-05 | 9.32E-05 | -1.96E+00 |
| AC122718.1 | 7.79E-03 | 4.06E-02 | 1.88E+00 | 2.36E-02 | 1.02E-01 | 1.10E+00 | 5.93E-04 | 2.52E-03 | 2.01E+00 | 1.36E-01 | 5.21E-01 | 1.31E+00 | 4.30E-03 | 1.53E-02 | 2.14E+00 | 1.35E-01 | 3.80E-01 | 1.12E+00 | 9.74E-03 | 3.58E-02 | 1.85E+00 | 2.23E-01 | 6.13E-01 | 1.44E+00 |
| FAM46C | 7.81E-03 | 4.07E-02 | -1.00E+00 | 2.83E-09 | 1.22E-08 | -2.24E+00 | 4.11E-03 | 1.75E-02 | -1.17E+00 | 7.51E-12 | 2.87E-11 | -2.53E+00 | 1.14E-04 | 4.05E-04 | -1.08E+00 | 7.96E-08 | 2.24E-07 | -1.60E+00 | 1.37E-09 | 5.02E-09 | -1.13E+00 | 8.78E-20 | 2.41E-19 | -1.73E+00 |
| LRRTM2 | 7.82E-03 | 4.08E-02 | -1.09E+00 | 9.47E-05 | 4.09E-04 | -1.37E+00 | 9.48E-02 | 4.03E-01 | -1.01E+00 | 5.46E-02 | 2.09E-01 | 1.02E+00 | 2.75E-01 | 9.79E-01 | 1.26E+00 | 1.55E-01 | 4.38E-01 | 1.04E+00 | 1.15E-01 | 4.21E-01 | -1.12E+00 | 1.41E-01 | 3.86E-01 | -1.04E+00 |
| AC067735.1 | 7.83E-03 | 4.08E-02 | -1.18E+00 | 5.82E-03 | 2.51E-02 | -1.16E+00 | 9.36E-03 | 3.98E-02 | -1.19E+00 | 1.68E-02 | 6.43E-02 | -1.03E+00 | 2.11E-01 | 7.51E-01 | -1.11E+00 | 2.69E-01 | 7.57E-01 | -1.08E+00 | 1.68E-01 | 6.16E-01 | -1.04E+00 | 1.43E-01 | 3.93E-01 | -1.19E+00 |
| IDH1-AS1 | 7.84E-03 | 4.09E-02 | -1.39E+00 | 2.29E-03 | 9.89E-03 | -1.69E+00 | 3.08E-02 | 1.31E-01 | -1.28E+00 | 3.20E-03 | 1.22E-02 | -1.45E+00 | 5.73E-03 | 2.04E-02 | -1.39E+00 | 1.91E-03 | 5.39E-03 | -1.61E+00 | 6.32E-04 | 2.32E-03 | -1.32E+00 | 3.27E-04 | 8.99E-04 | -1.49E+00 |
| AC234781.2 | 7.84E-03 | 4.09E-02 | -1.09E+00 | 8.09E-04 | 3.50E-03 | -1.45E+00 | 3.73E-03 | 1.59E-02 | -1.45E+00 | 6.45E-04 | 2.47E-03 | -1.64E+00 | 4.77E-02 | 1.70E-01 | -1.12E+00 | 8.20E-03 | 2.31E-02 | -1.49E+00 | 3.48E-04 | 1.28E-03 | -1.91E+00 | 5.07E-05 | 1.39E-04 | -2.28E+00 |
| CA5BP1 | 7.84E-03 | 4.09E-02 | -1.17E+00 | 8.52E-03 | 3.68E-02 | -1.10E+00 | 4.42E-06 | 1.88E-05 | -1.37E+00 | 1.09E-06 | 4.17E-06 | -1.28E+00 | 2.19E-02 | 7.81E-02 | -1.13E+00 | 2.15E-02 | 6.07E-02 | -1.19E+00 | 2.52E-02 | 9.27E-02 | -1.04E+00 | 1.25E-02 | 3.43E-02 | -1.09E+00 |
| BDH1 | 7.85E-03 | 4.09E-02 | -1.08E+00 | 4.81E-02 | 2.08E-01 | 1.22E+00 | 6.05E-06 | 2.57E-05 | -1.20E+00 | 1.33E-04 | 5.08E-04 | -1.02E+00 | 1.27E-02 | 4.53E-02 | -1.15E+00 | 1.47E-02 | 4.13E-02 | -1.18E+00 | 3.72E-02 | 1.36E-01 | 1.02E+00 | 2.35E-01 | 6.45E-01 | 1.13E+00 |
| TPT1P6 | 7.85E-03 | 4.09E-02 | 1.01E+00 | 2.83E-06 | 1.22E-05 | 2.19E+00 | 1.20E-01 | 5.10E-01 | 1.16E+00 | 1.38E-08 | 5.27E-08 | 2.67E+00 | 1.72E-01 | 6.14E-01 | 1.67E+00 | 2.13E-05 | 6.01E-05 | 3.18E+00 | 2.66E-01 | 9.75E-01 | 1.29E+00 | 2.92E-06 | 8.03E-06 | 2.40E+00 |
| SIAH2 | 7.85E-03 | 4.09E-02 | 1.04E+00 | 1.84E-03 | 7.95E-03 | 1.13E+00 | 2.16E-02 | 9.18E-02 | 1.02E+00 | 4.91E-02 | 1.87E-01 | 1.18E+00 | 2.46E-01 | 8.77E-01 | 1.85E+00 | 1.18E-01 | 3.32E-01 | 1.88E+00 | 5.61E-03 | 2.06E-02 | 1.17E+00 | 8.90E-02 | 2.44E-01 | 1.26E+00 |
| HAUS4 | 7.87E-03 | 4.10E-02 | -1.23E+00 | 4.01E-04 | 1.73E-03 | -1.44E+00 | 2.03E-02 | 8.62E-02 | -1.05E+00 | 4.98E-03 | 1.90E-02 | -1.06E+00 | 4.71E-02 | 1.68E-01 | -1.22E+00 | 3.26E-01 | 9.19E-01 | 1.27E+00 | 2.16E-01 | 7.94E-01 | 1.23E+00 | 6.75E-02 | 1.85E-01 | -1.06E+00 |
| GNA12 | 7.87E-03 | 4.10E-02 | -1.00E+00 | 7.41E-02 | 3.20E-01 | 1.27E+00 | 2.18E-01 | 9.26E-01 | 1.14E+00 | 6.79E-02 | 2.59E-01 | 1.10E+00 | 1.65E-01 | 5.89E-01 | 1.55E+00 | 1.17E-01 | 3.30E-01 | 1.70E+00 | 3.33E-05 | 1.22E-04 | -1.16E+00 | 5.08E-05 | 1.40E-04 | -1.18E+00 |
| ATP6V1G1 | 7.88E-03 | 4.11E-02 | 1.01E+00 | 2.74E-03 | 1.19E-02 | 1.09E+00 | 1.23E-01 | 5.22E-01 | -1.01E+00 | 9.69E-02 | 3.70E-01 | 1.02E+00 | 1.01E-03 | 3.61E-03 | -1.09E+00 | 1.10E-01 | 3.10E-01 | -1.01E+00 | 1.78E-02 | 6.53E-02 | -1.06E+00 | 7.33E-02 | 2.01E-01 | -1.04E+00 |
| XRCC4 | 7.88E-03 | 4.11E-02 | -1.06E+00 | 5.69E-02 | 2.46E-01 | 1.26E+00 | 9.76E-04 | 4.15E-03 | -1.20E+00 | 1.53E-03 | 5.83E-03 | -1.03E+00 | 7.71E-02 | 2.75E-01 | 1.05E+00 | 2.33E-01 | 6.56E-01 | 1.16E+00 | 2.42E-04 | 8.89E-04 | -1.19E+00 | 6.50E-03 | 1.79E-02 | -1.05E+00 |
| TES | 7.89E-03 | 4.11E-02 | 1.09E+00 | 5.77E-03 | 2.49E-02 | 1.15E+00 | 2.04E-02 | 8.67E-02 | -1.05E+00 | 1.35E-02 | 5.15E-02 | 1.03E+00 | 2.44E-04 | 8.70E-04 | -1.13E+00 | 2.14E-04 | 6.04E-04 | -1.15E+00 | 2.99E-05 | 1.10E-04 | -1.28E+00 | 1.44E-07 | 3.96E-07 | -1.36E+00 |
| LINC00506 | 7.89E-03 | 4.11E-02 | -1.29E+00 | 6.04E-02 | 2.61E-01 | 1.13E+00 | 2.70E-02 | 1.15E-01 | -1.31E+00 | 1.46E-01 | 5.56E-01 | 1.17E+00 | 1.89E-01 | 6.73E-01 | -1.04E+00 | 1.48E-01 | 4.18E-01 | -1.24E+00 | 2.41E-03 | 8.84E-03 | -1.76E+00 | 3.31E-01 | 9.08E-01 | 1.20E+00 |
| CACNA1C-AS1 | 7.89E-03 | 4.11E-02 | -1.13E+00 | 2.41E-03 | 1.04E-02 | -1.19E+00 | 3.57E-02 | 1.52E-01 | -1.20E+00 | 8.04E-03 | 3.07E-02 | -1.26E+00 | 8.64E-02 | 3.08E-01 | -1.89E+00 | 3.24E-01 | 9.13E-01 | 1.02E+00 | 9.17E-12 | 3.37E-11 | -2.81E+00 | 6.86E-07 | 1.89E-06 | -1.65E+00 |
| ACOX3 | 7.89E-03 | 4.11E-02 | -1.05E+00 | 6.04E-02 | 2.61E-01 | 1.23E+00 | 7.06E-02 | 3.00E-01 | 1.07E+00 | 3.05E-02 | 1.17E-01 | 1.12E+00 | 7.36E-03 | 2.62E-02 | -1.25E+00 | 4.70E-02 | 1.32E-01 | 1.04E+00 | 1.20E-01 | 4.41E-01 | 1.21E+00 | 1.82E-01 | 5.00E-01 | 1.13E+00 |
| PXN | 7.91E-03 | 4.12E-02 | -1.13E+00 | 1.15E-01 | 4.96E-01 | 1.20E+00 | 2.36E-04 | 1.00E-03 | -1.36E+00 | 4.92E-03 | 1.88E-02 | 1.02E+00 | 1.03E-02 | 3.67E-02 | -1.31E+00 | 3.93E-02 | 1.11E-01 | -1.09E+00 | 3.98E-02 | 1.46E-01 | -1.43E+00 | 2.32E-01 | 6.37E-01 | 1.38E+00 |
| RRAS2 | 7.93E-03 | 4.13E-02 | -1.21E+00 | 2.51E-03 | 1.09E-02 | -1.20E+00 | 9.73E-05 | 4.13E-04 | -1.51E+00 | 1.38E-07 | 5.25E-07 | -1.88E+00 | 1.25E-01 | 4.45E-01 | 1.03E+00 | 8.97E-02 | 2.53E-01 | -1.11E+00 | 6.57E-03 | 2.41E-02 | -1.11E+00 | 1.36E-04 | 3.73E-04 | -1.36E+00 |
| NUBP2 | 7.95E-03 | 4.14E-02 | 1.09E+00 | 1.14E-02 | 4.93E-02 | 1.25E+00 | 1.25E-01 | 5.32E-01 | 1.16E+00 | 1.40E-01 | 5.34E-01 | 1.26E+00 | 9.84E-05 | 3.51E-04 | -1.19E+00 | 8.55E-02 | 2.41E-01 | 1.13E+00 | 2.35E-02 | 8.62E-02 | 1.07E+00 | 3.42E-01 | 9.40E-01 | 1.33E+00 |
| MAPK8IP1 | 7.95E-03 | 4.15E-02 | -1.01E+00 | 2.18E-02 | 9.41E-02 | 1.18E+00 | 1.37E-05 | 5.81E-05 | -1.46E+00 | 3.00E-04 | 1.15E-03 | -1.14E+00 | 8.30E-02 | 2.96E-01 | -1.11E+00 | 2.74E-01 | 7.73E-01 | 1.34E+00 | 1.05E-02 | 3.85E-02 | -1.60E+00 | 9.22E-02 | 2.53E-01 | -1.14E+00 |
| GYS1 | 7.97E-03 | 4.15E-02 | -1.26E+00 | 1.67E-02 | 7.19E-02 | -1.04E+00 | 1.17E-03 | 4.99E-03 | -1.33E+00 | 4.56E-04 | 1.74E-03 | -1.31E+00 | 3.04E-03 | 1.08E-02 | -1.66E+00 | 1.01E-02 | 2.86E-02 | -1.33E+00 | 2.40E-02 | 8.80E-02 | -1.27E+00 | 2.32E-01 | 6.37E-01 | 1.08E+00 |
| RNF115 | 7.99E-03 | 4.16E-02 | 1.12E+00 | 4.15E-05 | 1.79E-04 | 1.09E+00 | 8.09E-06 | 3.44E-05 | -1.22E+00 | 5.74E-05 | 2.19E-04 | -1.10E+00 | 5.24E-03 | 1.87E-02 | 1.17E+00 | 2.62E-03 | 7.39E-03 | 1.15E+00 | 5.10E-02 | 1.87E-01 | 1.08E+00 | 1.03E-02 | 2.84E-02 | 1.02E+00 |
| ANLN | 7.99E-03 | 4.17E-02 | -1.70E+00 | 9.80E-03 | 4.23E-02 | -1.65E+00 | 1.82E-01 | 7.74E-01 | 1.46E+00 | 2.30E-01 | 8.77E-01 | 1.40E+00 | 1.93E-01 | 6.89E-01 | -1.19E+00 | 3.11E-01 | 8.76E-01 | 1.06E+00 | 5.85E-03 | 2.15E-02 | -1.27E+00 | 5.52E-03 | 1.52E-02 | -1.12E+00 |
| AC095057.3 | 8.01E-03 | 4.17E-02 | -1.34E+00 | 2.67E-02 | 1.15E-01 | 2.16E+00 | 2.27E-01 | 9.64E-01 | 1.20E+00 | 1.36E-03 | 5.21E-03 | 2.71E+00 | 2.58E-02 | 9.20E-02 | -1.42E+00 | 1.55E-01 | 4.36E-01 | -1.03E+00 | 1.11E-01 | 4.08E-01 | -1.11E+00 | 1.66E-01 | 4.56E-01 | -1.04E+00 |
| WWOX | 8.02E-03 | 4.18E-02 | -1.19E+00 | 6.49E-03 | 2.81E-02 | -1.10E+00 | 6.23E-02 | 2.65E-01 | 1.37E+00 | 7.84E-02 | 2.99E-01 | 1.21E+00 | 5.85E-03 | 2.09E-02 | -1.07E+00 | 3.78E-02 | 1.07E-01 | 1.09E+00 | 1.30E-01 | 4.77E-01 | 1.41E+00 | 2.57E-01 | 7.07E-01 | 1.16E+00 |
| AC113410.1 | 8.03E-03 | 4.19E-02 | 1.08E+00 | 3.44E-06 | 1.49E-05 | -1.11E+00 | 1.45E-01 | 6.18E-01 | 1.18E+00 | 2.46E-03 | 9.38E-03 | 1.06E+00 | 2.66E-01 | 9.47E-01 | 1.32E+00 | 1.19E-01 | 3.35E-01 | 1.17E+00 | 2.30E-01 | 8.46E-01 | 1.31E+00 | 7.32E-02 | 2.01E-01 | 1.18E+00 |
| NEDD8 | 8.05E-03 | 4.19E-02 | 1.43E+00 | 1.33E-01 | 5.77E-01 | 1.45E+00 | 2.87E-04 | 1.22E-03 | 1.52E+00 | 7.96E-02 | 3.04E-01 | 1.35E+00 | 2.59E-01 | 9.23E-01 | 1.31E+00 | 2.88E-01 | 8.11E-01 | 1.28E+00 | 6.54E-03 | 2.40E-02 | 1.49E+00 | 3.66E-02 | 1.01E-01 | 1.38E+00 |
| IL17RE | 8.06E-03 | 4.20E-02 | -1.10E+00 | 9.69E-03 | 4.19E-02 | 1.04E+00 | 2.84E-04 | 1.21E-03 | -1.47E+00 | 5.01E-07 | 1.92E-06 | -1.60E+00 | 1.11E-01 | 3.97E-01 | -1.39E+00 | 2.31E-01 | 6.50E-01 | -1.07E+00 | 7.73E-03 | 2.84E-02 | -1.16E+00 | 1.37E-01 | 3.75E-01 | 1.06E+00 |
| DPP9 | 8.08E-03 | 4.21E-02 | -1.00E+00 | 9.55E-03 | 4.13E-02 | 1.10E+00 | 1.34E-02 | 5.70E-02 | -1.07E+00 | 3.15E-02 | 1.20E-01 | 1.19E+00 | 1.94E-01 | 6.93E-01 | 1.27E+00 | 6.05E-02 | 1.71E-01 | 1.04E+00 | 2.63E-04 | 9.67E-04 | -1.22E+00 | 1.74E-02 | 4.79E-02 | -1.03E+00 |
| DNAJC1 | 8.08E-03 | 4.21E-02 | 1.03E+00 | 1.34E-03 | 5.80E-03 | 1.15E+00 | 7.24E-03 | 3.08E-02 | -1.03E+00 | 2.52E-03 | 9.61E-03 | 1.05E+00 | 3.32E-05 | 1.18E-04 | -1.10E+00 | 3.16E-04 | 8.92E-04 | -1.01E+00 | 1.49E-04 | 5.46E-04 | -1.18E+00 | 1.23E-01 | 3.39E-01 | 1.05E+00 |
| EIF5A2 | 8.09E-03 | 4.21E-02 | -1.07E+00 | 4.91E-04 | 2.12E-03 | -1.21E+00 | 2.07E-01 | 8.78E-01 | 1.18E+00 | 1.33E-01 | 5.09E-01 | 1.20E+00 | 1.55E-01 | 5.53E-01 | 1.29E+00 | 4.93E-02 | 1.39E-01 | -1.10E+00 | 9.11E-02 | 3.34E-01 | 1.03E+00 | 3.21E-02 | 8.81E-02 | -1.17E+00 |
| AL139089.1 | 8.10E-03 | 4.22E-02 | 1.96E+00 | 2.27E-02 | 9.81E-02 | 2.12E+00 | 1.44E-01 | 6.12E-01 | 1.39E+00 | 6.86E-02 | 2.62E-01 | 1.87E+00 | 1.79E-02 | 6.40E-02 | 2.54E+00 | 2.08E-01 | 5.87E-01 | 1.79E+00 | 7.48E-02 | 2.75E-01 | 1.73E+00 | 1.08E-01 | 2.97E-01 | 1.95E+00 |
| LINC01266 | 8.11E-03 | 4.23E-02 | -1.17E+00 | 8.13E-02 | 3.51E-01 | 1.15E+00 | 2.27E-01 | 9.67E-01 | 1.07E+00 | 8.79E-02 | 3.36E-01 | 1.78E+00 | 9.15E-02 | 3.26E-01 | 1.94E+00 | 6.07E-02 | 1.71E-01 | 2.41E+00 | 1.94E-01 | 7.11E-01 | -1.08E+00 | 3.46E-01 | 9.49E-01 | 1.05E+00 |
| LRRC20 | 8.12E-03 | 4.23E-02 | -1.20E+00 | 5.44E-03 | 2.35E-02 | -1.33E+00 | 6.98E-06 | 2.97E-05 | -1.69E+00 | 8.04E-04 | 3.07E-03 | -1.26E+00 | 1.33E-01 | 4.75E-01 | -1.62E+00 | 3.15E-01 | 8.89E-01 | 1.01E+00 | 1.25E-02 | 4.59E-02 | -1.15E+00 | 1.21E-01 | 3.32E-01 | -1.02E+00 |
| USP27X | 8.14E-03 | 4.24E-02 | -1.11E+00 | 3.35E-03 | 1.45E-02 | -1.10E+00 | 1.20E-03 | 5.10E-03 | -1.32E+00 | 3.79E-03 | 1.45E-02 | -1.15E+00 | 2.37E-03 | 8.46E-03 | -1.64E+00 | 7.91E-03 | 2.23E-02 | -1.38E+00 | 2.76E-02 | 1.01E-01 | -1.17E+00 | 4.19E-02 | 1.15E-01 | -1.21E+00 |
| STX18 | 8.14E-03 | 4.24E-02 | 1.03E+00 | 8.18E-03 | 3.53E-02 | 1.14E+00 | 8.28E-02 | 3.52E-01 | 1.08E+00 | 2.39E-02 | 9.12E-02 | 1.14E+00 | 3.82E-08 | 1.36E-07 | -1.88E+00 | 1.03E-03 | 2.90E-03 | -1.15E+00 | 2.57E-04 | 9.44E-04 | -1.12E+00 | 3.85E-02 | 1.06E-01 | 1.05E+00 |
| ZNF670 | 8.15E-03 | 4.25E-02 | -1.33E+00 | 2.26E-02 | 9.78E-02 | -1.14E+00 | 3.12E-02 | 1.33E-01 | -1.12E+00 | 1.37E-02 | 5.22E-02 | -1.16E+00 | 3.85E-02 | 1.37E-01 | -1.19E+00 | 4.06E-02 | 1.14E-01 | -1.29E+00 | 9.14E-12 | 3.35E-11 | -2.15E+00 | 2.64E-12 | 7.26E-12 | -2.26E+00 |
| AC106795.1 | 8.15E-03 | 4.25E-02 | -1.06E+00 | 1.88E-01 | 8.10E-01 | 1.40E+00 | 1.86E-01 | 7.92E-01 | 1.20E+00 | 1.08E-01 | 4.11E-01 | 1.31E+00 | 5.28E-02 | 1.88E-01 | 1.11E+00 | 3.17E-01 | 8.93E-01 | 1.69E+00 | 2.27E-01 | 8.32E-01 | 1.24E+00 | 7.76E-02 | 2.13E-01 | 1.46E+00 |
| MKL1 | 8.15E-03 | 4.25E-02 | -1.06E+00 | 1.05E-01 | 4.53E-01 | 1.30E+00 | 3.30E-02 | 1.40E-01 | 1.03E+00 | 4.90E-02 | 1.87E-01 | 1.13E+00 | 1.46E-02 | 5.21E-02 | -1.03E+00 | 1.67E-02 | 4.70E-02 | 1.02E+00 | 4.73E-03 | 1.74E-02 | -1.10E+00 | 1.26E-01 | 3.46E-01 | 1.16E+00 |
| SNRPC | 8.18E-03 | 4.26E-02 | 1.13E+00 | 1.32E-05 | 5.69E-05 | 1.15E+00 | 1.54E-02 | 6.54E-02 | 1.34E+00 | 2.54E-01 | 9.69E-01 | 1.24E+00 | 1.14E-01 | 4.06E-01 | 1.24E+00 | 2.28E-01 | 6.42E-01 | 1.29E+00 | 5.32E-02 | 1.95E-01 | 1.35E+00 | 4.22E-02 | 1.16E-01 | 1.36E+00 |
| DPCD | 8.18E-03 | 4.27E-02 | -1.04E+00 | 4.84E-03 | 2.09E-02 | 1.02E+00 | 4.70E-03 | 1.99E-02 | 1.07E+00 | 1.11E-06 | 4.24E-06 | -1.15E+00 | 2.84E-02 | 1.01E-01 | -1.09E+00 | 5.96E-02 | 1.68E-01 | 1.03E+00 | 2.23E-01 | 8.19E-01 | 1.36E+00 | 2.72E-01 | 7.46E-01 | 1.22E+00 |
| NOP14-AS1 | 8.20E-03 | 4.27E-02 | -1.15E+00 | 6.25E-03 | 2.70E-02 | -1.17E+00 | 6.13E-06 | 2.60E-05 | -1.70E+00 | 2.77E-06 | 1.06E-05 | -1.59E+00 | 2.86E-04 | 1.02E-03 | -1.73E+00 | 1.09E-03 | 3.08E-03 | -1.48E+00 | 2.24E-02 | 8.22E-02 | 1.06E+00 | 5.77E-02 | 1.58E-01 | 1.01E+00 |
| CDKN2AIP | 8.20E-03 | 4.28E-02 | 1.04E+00 | 6.46E-02 | 2.79E-01 | 1.26E+00 | 6.23E-02 | 2.65E-01 | 1.06E+00 | 6.36E-02 | 2.43E-01 | 1.20E+00 | 3.64E-02 | 1.30E-01 | 1.71E+00 | 2.37E-03 | 6.68E-03 | 2.03E+00 | 4.85E-02 | 1.78E-01 | 1.55E+00 | 1.23E-01 | 3.39E-01 | 1.48E+00 |
| RPL13AP20 | 8.20E-03 | 4.28E-02 | 1.82E+00 | 8.43E-02 | 3.64E-01 | 1.14E+00 | 2.54E-03 | 1.08E-02 | 1.71E+00 | 1.23E-01 | 4.71E-01 | 1.08E+00 | 2.25E-02 | 8.01E-02 | 2.32E+00 | 2.20E-01 | 6.21E-01 | 1.67E+00 | 2.00E-01 | 7.35E-01 | 1.23E+00 | 6.73E-02 | 1.85E-01 | -1.03E+00 |
| NUMBL | 8.22E-03 | 4.28E-02 | -1.26E+00 | 9.18E-02 | 3.97E-01 | 1.20E+00 | 5.45E-06 | 2.32E-05 | -1.79E+00 | 6.63E-03 | 2.53E-02 | -1.02E+00 | 5.47E-02 | 1.95E-01 | -1.17E+00 | 2.96E-01 | 8.36E-01 | 1.22E+00 | 7.18E-03 | 2.63E-02 | -1.50E+00 | 3.13E-01 | 8.58E-01 | 1.13E+00 |
| CCDC167 | 8.22E-03 | 4.29E-02 | 1.82E+00 | 1.16E-01 | 5.03E-01 | 1.42E+00 | 1.20E-04 | 5.09E-04 | 1.63E+00 | 9.07E-02 | 3.47E-01 | 1.25E+00 | 1.06E-03 | 3.79E-03 | -1.17E+00 | 3.96E-04 | 1.12E-03 | -1.19E+00 | 2.40E-01 | 8.80E-01 | 1.17E+00 | 3.03E-01 | 8.33E-01 | 1.25E+00 |
| AGBL5 | 8.24E-03 | 4.30E-02 | 1.05E+00 | 2.46E-02 | 1.06E-01 | 1.19E+00 | 1.04E-01 | 4.41E-01 | 1.07E+00 | 2.04E-02 | 7.80E-02 | 1.03E+00 | 1.09E-04 | 3.89E-04 | -1.29E+00 | 5.78E-03 | 1.63E-02 | 1.00E+00 | 2.49E-01 | 9.13E-01 | 1.35E+00 | 3.54E-01 | 9.73E-01 | 1.32E+00 |
| ADGRE2 | 8.25E-03 | 4.30E-02 | 1.06E+00 | 1.58E-02 | 6.81E-02 | 1.15E+00 | 1.06E-04 | 4.49E-04 | -1.10E+00 | 1.56E-02 | 5.96E-02 | 1.15E+00 | 1.51E-01 | 5.39E-01 | 1.19E+00 | 1.73E-01 | 4.86E-01 | 1.20E+00 | 1.99E-01 | 7.32E-01 | 1.29E+00 | 1.19E-01 | 3.27E-01 | 1.19E+00 |
| MTX3 | 8.26E-03 | 4.30E-02 | -1.32E+00 | 3.57E-02 | 1.54E-01 | -1.00E+00 | 4.99E-03 | 2.12E-02 | -1.17E+00 | 2.52E-03 | 9.61E-03 | -1.12E+00 | 2.46E-01 | 8.77E-01 | 1.23E+00 | 1.12E-01 | 3.16E-01 | 1.93E+00 | 1.35E-01 | 4.94E-01 | 1.02E+00 | 1.32E-01 | 3.63E-01 | -1.03E+00 |
| CIP2A | 8.27E-03 | 4.31E-02 | -1.22E+00 | 1.21E-02 | 5.25E-02 | -1.08E+00 | 7.96E-02 | 3.38E-01 | 1.01E+00 | 6.45E-03 | 2.46E-02 | -1.01E+00 | 1.30E-01 | 4.64E-01 | -1.23E+00 | 1.64E-01 | 4.63E-01 | -1.17E+00 | 3.48E-06 | 1.28E-05 | -1.49E+00 | 8.94E-02 | 2.45E-01 | 1.12E+00 |
| CCDC8 | 8.27E-03 | 4.31E-02 | -1.41E+00 | 1.42E-01 | 6.12E-01 | 1.20E+00 | 3.59E-07 | 1.53E-06 | -1.99E+00 | 2.79E-03 | 1.07E-02 | -1.10E+00 | 5.83E-02 | 2.08E-01 | -1.43E+00 | 1.76E-01 | 4.96E-01 | -1.00E+00 | 2.23E-02 | 8.18E-02 | -1.24E+00 | 3.31E-01 | 9.08E-01 | 1.29E+00 |
| NDRG1 | 8.28E-03 | 4.31E-02 | 1.08E+00 | 7.72E-03 | 3.33E-02 | 1.13E+00 | 1.04E-04 | 4.40E-04 | -1.18E+00 | 9.41E-04 | 3.59E-03 | 1.04E+00 | 1.88E-01 | 6.72E-01 | 1.33E+00 | 7.62E-02 | 2.15E-01 | 1.12E+00 | 4.93E-02 | 1.81E-01 | -1.02E+00 | 1.06E-01 | 2.92E-01 | 1.04E+00 |
| SLC9A6 | 8.28E-03 | 4.32E-02 | -1.41E+00 | 2.78E-02 | 1.20E-01 | -1.12E+00 | 5.38E-06 | 2.29E-05 | -1.69E+00 | 1.35E-05 | 5.17E-05 | -1.39E+00 | 1.20E-02 | 4.27E-02 | -1.40E+00 | 2.18E-02 | 6.15E-02 | -1.20E+00 | 6.93E-10 | 2.54E-09 | -2.20E+00 | 1.00E-02 | 2.76E-02 | -1.07E+00 |
| QPCTL | 8.29E-03 | 4.32E-02 | -1.15E+00 | 2.16E-02 | 9.34E-02 | -1.00E+00 | 5.92E-02 | 2.51E-01 | 1.57E+00 | 2.40E-01 | 9.18E-01 | 1.44E+00 | 6.79E-04 | 2.42E-03 | -1.72E+00 | 1.10E-02 | 3.11E-02 | -1.21E+00 | 1.70E-01 | 6.26E-01 | 1.41E+00 | 1.33E-01 | 3.66E-01 | 1.83E+00 |
| SRGAP1 | 8.30E-03 | 4.32E-02 | -1.06E+00 | 1.26E-01 | 5.46E-01 | 1.67E+00 | 2.98E-02 | 1.27E-01 | -1.15E+00 | 1.41E-01 | 5.38E-01 | 1.39E+00 | 1.19E-01 | 4.25E-01 | -1.07E+00 | 2.78E-01 | 7.84E-01 | 1.58E+00 | 6.03E-02 | 2.21E-01 | -1.37E+00 | 3.56E-01 | 9.77E-01 | 1.44E+00 |
| DPH6 | 8.31E-03 | 4.33E-02 | -1.05E+00 | 2.00E-02 | 8.64E-02 | 1.16E+00 | 1.11E-02 | 4.73E-02 | -1.21E+00 | 3.44E-02 | 1.31E-01 | 1.07E+00 | 7.63E-02 | 2.72E-01 | 1.19E+00 | 2.94E-01 | 8.27E-01 | 1.38E+00 | 1.90E-04 | 6.98E-04 | -1.18E+00 | 3.70E-03 | 1.02E-02 | 1.05E+00 |
| PEX13 | 8.31E-03 | 4.33E-02 | 1.10E+00 | 2.30E-04 | 9.94E-04 | -1.02E+00 | 7.24E-04 | 3.08E-03 | -1.09E+00 | 5.88E-05 | 2.25E-04 | -1.10E+00 | 2.65E-03 | 9.44E-03 | -1.14E+00 | 8.69E-04 | 2.45E-03 | -1.25E+00 | 4.38E-03 | 1.61E-02 | -1.01E+00 | 1.42E-04 | 3.90E-04 | -1.17E+00 |
| RREB1 | 8.31E-03 | 4.33E-02 | 1.03E+00 | 2.10E-04 | 9.09E-04 | 1.04E+00 | 1.35E-02 | 5.74E-02 | -1.07E+00 | 3.20E-05 | 1.22E-04 | -1.14E+00 | 4.04E-05 | 1.44E-04 | -1.14E+00 | 4.74E-07 | 1.34E-06 | -1.25E+00 | 2.25E-11 | 8.27E-11 | -1.75E+00 | 1.03E-04 | 2.83E-04 | -1.16E+00 |
| DFFB | 8.37E-03 | 4.36E-02 | 1.03E+00 | 3.15E-02 | 1.36E-01 | 1.24E+00 | 3.89E-06 | 1.65E-05 | -1.55E+00 | 2.03E-02 | 7.75E-02 | 1.09E+00 | 1.05E-03 | 3.73E-03 | -1.55E+00 | 1.87E-01 | 5.28E-01 | 1.26E+00 | 2.53E-05 | 9.29E-05 | -1.30E+00 | 1.34E-02 | 3.68E-02 | 1.07E+00 |
| AC107375.1 | 8.37E-03 | 4.36E-02 | -1.13E+00 | 1.30E-02 | 5.61E-02 | 1.01E+00 | 1.69E-01 | 7.17E-01 | 1.13E+00 | 9.59E-02 | 3.66E-01 | 1.05E+00 | 2.05E-01 | 7.29E-01 | 1.28E+00 | 3.29E-01 | 9.26E-01 | 1.45E+00 | 1.61E-01 | 5.90E-01 | 1.37E+00 | 2.57E-01 | 7.05E-01 | 1.10E+00 |
| NUDT16 | 8.37E-03 | 4.36E-02 | 1.20E+00 | 2.65E-03 | 1.14E-02 | 1.20E+00 | 4.85E-05 | 2.06E-04 | -1.12E+00 | 1.62E-04 | 6.19E-04 | 1.04E+00 | 1.85E-08 | 6.58E-08 | -1.71E+00 | 1.19E-08 | 3.36E-08 | -1.63E+00 | 8.39E-11 | 3.08E-10 | -2.00E+00 | 1.48E-03 | 4.06E-03 | -1.22E+00 |
| CCSAP | 8.38E-03 | 4.37E-02 | -1.04E+00 | 3.48E-03 | 1.50E-02 | -1.01E+00 | 2.24E-04 | 9.52E-04 | -1.30E+00 | 1.03E-03 | 3.94E-03 | -1.07E+00 | 1.37E-04 | 4.88E-04 | -1.22E+00 | 2.72E-05 | 7.66E-05 | -1.33E+00 | 1.95E-05 | 7.14E-05 | -1.33E+00 | 3.99E-05 | 1.10E-04 | -1.50E+00 |
| HMGB1P5 | 8.40E-03 | 4.38E-02 | 1.23E+00 | 4.90E-06 | 2.12E-05 | 1.13E+00 | 1.17E-02 | 4.95E-02 | 1.51E+00 | 1.79E-01 | 6.84E-01 | 1.34E+00 | 2.44E-01 | 8.72E-01 | 1.26E+00 | 1.08E-01 | 3.05E-01 | 1.24E+00 | 2.20E-01 | 8.09E-01 | 1.35E+00 | 3.32E-01 | 9.13E-01 | 1.35E+00 |
| AF131215.2 | 8.40E-03 | 4.38E-02 | -1.43E+00 | 1.74E-01 | 7.50E-01 | 1.29E+00 | 3.11E-02 | 1.32E-01 | -1.33E+00 | 2.57E-01 | 9.83E-01 | 1.31E+00 | 1.71E-01 | 6.09E-01 | -1.11E+00 | 3.52E-01 | 9.92E-01 | 1.24E+00 | 1.29E-02 | 4.73E-02 | -1.51E+00 | 3.14E-01 | 8.63E-01 | 1.09E+00 |
| ABCG2 | 8.40E-03 | 4.38E-02 | -1.33E+00 | 1.22E-01 | 5.26E-01 | 1.15E+00 | 4.51E-02 | 1.92E-01 | -1.34E+00 | 6.22E-02 | 2.38E-01 | -1.27E+00 | 2.76E-01 | 9.85E-01 | -1.01E+00 | 2.16E-01 | 6.10E-01 | 1.51E+00 | 1.07E-07 | 3.94E-07 | -1.93E+00 | 3.12E-04 | 8.57E-04 | -1.45E+00 |
| LPP-AS2 | 8.41E-03 | 4.38E-02 | -1.09E+00 | 1.21E-04 | 5.24E-04 | -1.52E+00 | 7.24E-02 | 3.08E-01 | 1.03E+00 | 1.36E-01 | 5.20E-01 | 1.18E+00 | 7.17E-02 | 2.56E-01 | -1.32E+00 | 8.45E-02 | 2.38E-01 | -1.27E+00 | 1.14E-01 | 4.20E-01 | -1.08E+00 | 5.54E-02 | 1.52E-01 | -1.25E+00 |
| IDNK | 8.44E-03 | 4.40E-02 | 1.03E+00 | 1.68E-02 | 7.24E-02 | 1.20E+00 | 5.65E-04 | 2.40E-03 | -1.38E+00 | 2.23E-03 | 8.53E-03 | -1.14E+00 | 1.56E-03 | 5.57E-03 | -1.07E+00 | 6.94E-02 | 1.96E-01 | 1.16E+00 | 2.14E-01 | 7.86E-01 | 1.24E+00 | 2.64E-01 | 7.26E-01 | 1.17E+00 |
| TMEM94 | 8.44E-03 | 4.40E-02 | -1.05E+00 | 1.25E-01 | 5.42E-01 | 1.30E+00 | 1.10E-06 | 4.67E-06 | -1.39E+00 | 9.86E-06 | 3.77E-05 | -1.18E+00 | 5.26E-04 | 1.88E-03 | -1.34E+00 | 7.54E-03 | 2.12E-02 | -1.06E+00 | 6.85E-03 | 2.51E-02 | -1.10E+00 | 3.54E-02 | 9.72E-02 | 1.03E+00 |
| PRMT5 | 8.44E-03 | 4.40E-02 | -1.39E+00 | 2.02E-02 | 8.74E-02 | -1.16E+00 | 1.27E-01 | 5.40E-01 | 1.02E+00 | 2.18E-01 | 8.33E-01 | 1.22E+00 | 2.49E-04 | 8.88E-04 | -2.11E+00 | 3.84E-03 | 1.08E-02 | -1.38E+00 | 3.75E-03 | 1.37E-02 | -1.21E+00 | 4.63E-03 | 1.27E-02 | -1.34E+00 |
| CCL4L2 | 8.44E-03 | 4.40E-02 | 3.37E+00 | 1.48E-01 | 6.37E-01 | 1.25E+00 | 3.72E-02 | 1.58E-01 | 2.41E+00 | 2.62E-01 | 1.00E+00 | 1.09E+00 | 1.41E-01 | 5.02E-01 | 1.46E+00 | 3.15E-01 | 8.87E-01 | 1.12E+00 | 2.23E-01 | 8.19E-01 | -1.02E+00 | 2.47E-01 | 6.78E-01 | -1.15E+00 |
| TLR4 | 8.44E-03 | 4.40E-02 | -1.31E+00 | 7.74E-03 | 3.35E-02 | -1.39E+00 | 1.92E-04 | 8.14E-04 | -1.30E+00 | 4.97E-03 | 1.90E-02 | 1.03E+00 | 8.51E-02 | 3.03E-01 | -1.83E+00 | 1.44E-01 | 4.07E-01 | -1.57E+00 | 1.07E-01 | 3.91E-01 | -1.61E+00 | 2.17E-01 | 5.97E-01 | -1.42E+00 |
| ANKRD20A3 | 8.44E-03 | 4.40E-02 | 2.32E+00 | 2.15E-01 | 9.27E-01 | 1.36E+00 | 3.99E-02 | 1.69E-01 | 1.77E+00 | 1.75E-01 | 6.69E-01 | 1.42E+00 | 8.52E-02 | 3.04E-01 | 1.79E+00 | 3.48E-01 | 9.80E-01 | 1.22E+00 | 1.93E-07 | 7.07E-07 | 3.48E+00 | 2.61E-02 | 7.18E-02 | 2.08E+00 |
| DNM1 | 8.49E-03 | 4.42E-02 | 1.04E+00 | 8.95E-02 | 3.87E-01 | 1.35E+00 | 1.39E-03 | 5.92E-03 | -1.28E+00 | 5.01E-02 | 1.92E-01 | 1.18E+00 | 2.18E-01 | 7.77E-01 | 1.06E+00 | 2.00E-01 | 5.64E-01 | -1.00E+00 | 5.75E-02 | 2.11E-01 | -1.10E+00 | 9.35E-02 | 2.57E-01 | -1.04E+00 |
| ACY3 | 8.49E-03 | 4.42E-02 | 1.13E+00 | 1.30E-02 | 5.62E-02 | 1.28E+00 | 7.52E-02 | 3.20E-01 | 1.30E+00 | 2.40E-01 | 9.18E-01 | -1.12E+00 | 2.67E-01 | 9.53E-01 | 1.23E+00 | 3.51E-01 | 9.89E-01 | 1.18E+00 | 2.52E-01 | 9.24E-01 | 1.25E+00 | 3.36E-01 | 9.22E-01 | 1.19E+00 |
| NAALADL2 | 8.51E-03 | 4.43E-02 | -1.13E+00 | 5.06E-02 | 2.19E-01 | 1.24E+00 | 1.17E-01 | 4.97E-01 | 1.12E+00 | 2.39E-01 | 9.12E-01 | 1.47E+00 | 2.28E-01 | 8.12E-01 | 1.31E+00 | 2.87E-01 | 8.09E-01 | 1.60E+00 | 2.24E-01 | 8.23E-01 | 1.25E+00 | 1.78E-01 | 4.88E-01 | 1.58E+00 |
| TCHH | 8.51E-03 | 4.43E-02 | 1.99E+00 | 2.31E-01 | 1.00E+00 | -1.00E+00 | 2.92E-02 | 1.24E-01 | 1.87E+00 | 9.79E-02 | 3.74E-01 | 1.34E+00 | 1.65E-01 | 5.90E-01 | 1.55E+00 | 3.55E-01 | 1.00E+00 | -1.00E+00 | 6.37E-02 | 2.34E-01 | 1.39E+00 | 4.15E-02 | 1.14E-01 | 1.79E+00 |
| NANOS3 | 8.51E-03 | 4.43E-02 | -1.10E+00 | 1.80E-05 | 7.78E-05 | -1.70E+00 | 3.24E-02 | 1.38E-01 | -1.19E+00 | 9.86E-03 | 3.76E-02 | -1.13E+00 | 4.61E-02 | 1.64E-01 | -1.15E+00 | 2.70E-03 | 7.61E-03 | -2.09E+00 | 1.97E-01 | 7.22E-01 | -1.03E+00 | 1.85E-01 | 5.09E-01 | -1.30E+00 |
| AC073320.1 | 8.51E-03 | 4.43E-02 | 2.24E+00 | 1.68E-01 | 7.24E-01 | 1.18E+00 | 1.07E-01 | 4.56E-01 | 1.40E+00 | 1.31E-01 | 5.00E-01 | 1.11E+00 | 3.90E-03 | 1.39E-02 | -1.21E+00 | 1.81E-03 | 5.11E-03 | -1.36E+00 | 4.43E-02 | 1.62E-01 | -1.54E+00 | 1.71E-01 | 4.69E-01 | -1.20E+00 |
| ADAMTS1 | 8.51E-03 | 4.44E-02 | -1.47E+00 | 2.44E-02 | 1.05E-01 | -1.20E+00 | 1.08E-01 | 4.59E-01 | -1.09E+00 | 2.32E-01 | 8.87E-01 | 1.20E+00 | 7.22E-02 | 2.57E-01 | 2.96E+00 | 1.70E-01 | 4.80E-01 | 1.96E+00 | 2.84E-02 | 1.04E-01 | 2.67E+00 | 2.01E-01 | 5.52E-01 | -1.05E+00 |
| LINC01012 | 8.51E-03 | 4.44E-02 | -1.28E+00 | 1.27E-01 | 5.49E-01 | 1.57E+00 | 1.45E-01 | 6.18E-01 | -1.02E+00 | 9.08E-03 | 3.47E-02 | 2.43E+00 | 1.72E-01 | 6.15E-01 | 1.09E+00 | 8.45E-02 | 2.38E-01 | 2.14E+00 | 2.57E-01 | 9.43E-01 | 1.21E+00 | 1.88E-01 | 5.16E-01 | 1.61E+00 |

| | | | | | | | | | | | | | | | | | | | | | | | | |
|---|---|---|---|---|---|---|---|---|---|---|---|---|---|---|---|---|---|---|---|---|---|---|---|---|
| PKP2 | 8.53E-03 | 4.44E-02 | -1.08E+00 | 6.88E-02 | 2.97E-01 | 1.21E+00 | 3.19E-03 | 1.36E-02 | -1.18E+00 | 1.23E-02 | 4.70E-02 | 1.07E+00 | 7.52E-02 | 2.68E-01 | 1.02E+00 | 3.20E-01 | 9.02E-01 | 1.40E+00 | 2.36E-02 | 8.66E-02 | -1.08E+00 | 2.73E-01 | 7.49E-01 | 1.22E+00 |
| TRMT10B | 8.53E-03 | 4.44E-02 | -1.01E+00 | 2.30E-01 | 9.92E-01 | 1.45E+00 | 4.56E-07 | 1.94E-06 | -1.64E+00 | 3.20E-04 | 1.22E-03 | -1.14E+00 | 6.96E-06 | 2.48E-05 | -1.52E+00 | 5.60E-02 | 1.58E-01 | 1.05E+00 | 1.06E-03 | 3.90E-03 | -1.14E+00 | 1.78E-02 | 4.89E-02 | 1.02E+00 |
| GPRASP2 | 8.53E-03 | 4.44E-02 | -1.06E+00 | 1.09E-01 | 4.72E-01 | 1.74E+00 | 5.10E-04 | 2.17E-03 | -1.57E+00 | 5.57E-03 | 2.13E-02 | -1.11E+00 | 1.96E-01 | 6.98E-01 | 1.68E+00 | 1.91E-01 | 5.38E-01 | 2.00E+00 | 2.35E-01 | 8.61E-01 | 1.14E+00 | 8.99E-02 | 2.47E-01 | 1.80E+00 |
| BRI3 | 8.53E-03 | 4.44E-02 | 1.03E+00 | 1.18E-05 | 5.08E-05 | 1.03E+00 | 1.63E-02 | 6.93E-02 | -1.01E+00 | 1.14E-03 | 4.37E-03 | -1.04E+00 | 5.14E-04 | 1.83E-03 | -1.03E+00 | 9.27E-04 | 2.61E-03 | -1.04E+00 | 1.12E-03 | 4.12E-03 | -1.07E+00 | 9.53E-03 | 2.62E-02 | -1.01E+00 |
| ORMDL3 | 8.54E-03 | 4.45E-02 | -1.19E+00 | 9.75E-02 | 4.21E-01 | 1.19E+00 | 4.94E-03 | 2.10E-02 | -1.30E+00 | 1.33E-01 | 5.08E-01 | 1.24E+00 | 1.70E-03 | 6.07E-03 | -1.05E+00 | 2.06E-02 | 5.80E-02 | 1.11E+00 | 4.04E-05 | 1.48E-04 | -1.19E+00 | 5.85E-03 | 1.61E-02 | -1.08E+00 |
| DNAJA4 | 8.54E-03 | 4.45E-02 | -1.47E+00 | 7.97E-03 | 3.45E-02 | -1.31E+00 | 2.02E-02 | 8.57E-02 | -1.12E+00 | 2.16E-02 | 8.24E-02 | 1.00E+00 | 4.91E-02 | 1.75E-01 | -1.27E+00 | 1.42E-01 | 4.01E-01 | -1.02E+00 | 2.57E-04 | 9.44E-04 | -1.41E+00 | 3.87E-02 | 1.06E-01 | 1.05E+00 |
| NFE2L1 | 8.54E-03 | 4.45E-02 | -1.09E+00 | 4.16E-02 | 1.80E-01 | 1.18E+00 | 1.14E-02 | 4.83E-02 | -1.08E+00 | 2.25E-02 | 8.59E-02 | 1.04E+00 | 5.30E-02 | 1.89E-01 | 1.09E+00 | 1.88E-01 | 5.30E-01 | 1.28E+00 | 1.16E-01 | 4.26E-01 | 1.11E+00 | 2.18E-02 | 5.99E-02 | 1.01E+00 |
| CRY2 | 8.55E-03 | 4.45E-02 | -1.21E+00 | 5.01E-03 | 2.17E-02 | -1.23E+00 | 4.50E-03 | 1.91E-02 | -1.29E+00 | 1.63E-02 | 6.23E-02 | -1.03E+00 | 1.17E-02 | 4.19E-02 | -1.24E+00 | 1.84E-02 | 5.20E-02 | -1.20E+00 | 4.80E-04 | 1.76E-03 | -1.49E+00 | 9.17E-03 | 2.52E-02 | -1.18E+00 |
| LINC02175 | 8.55E-03 | 4.46E-02 | -1.30E+00 | 7.08E-02 | 3.06E-01 | 1.04E+00 | 1.54E-03 | 6.55E-03 | -1.59E+00 | 2.66E-02 | 1.02E-01 | -1.14E+00 | 1.06E-01 | 3.76E-01 | -1.03E+00 | 3.08E-01 | 8.68E-01 | 1.26E+00 | 2.44E-01 | 8.97E-01 | 1.34E+00 | 2.94E-01 | 8.07E-01 | 1.09E+00 |
| DOHH | 8.55E-03 | 4.46E-02 | -1.17E+00 | 1.28E-01 | 5.55E-01 | 1.33E+00 | 6.50E-02 | 2.76E-01 | 1.08E+00 | 1.12E-01 | 4.29E-01 | 1.28E+00 | 8.38E-03 | 2.99E-02 | -1.19E+00 | 3.40E-01 | 9.57E-01 | 1.44E+00 | 5.22E-05 | 1.91E-04 | -1.13E+00 | 4.46E-03 | 1.23E-02 | 1.06E+00 |
| JOSD2 | 8.56E-03 | 4.46E-02 | -1.01E+00 | 1.86E-03 | 8.02E-03 | -1.06E+00 | 1.42E-01 | 6.02E-01 | 1.08E+00 | 7.54E-02 | 2.88E-01 | 1.14E+00 | 4.26E-02 | 1.52E-01 | 1.11E+00 | 3.52E-02 | 9.92E-02 | 1.08E+00 | 1.05E-01 | 3.85E-01 | 1.47E+00 | 1.86E-01 | 5.10E-01 | 1.27E+00 |
| AC069224.1 | 8.56E-03 | 4.46E-02 | -1.21E+00 | 1.00E-01 | 4.34E-01 | 1.17E+00 | 1.56E-02 | 6.63E-02 | -1.51E+00 | 8.43E-02 | 3.22E-01 | -1.01E+00 | 2.35E-01 | 8.36E-01 | 1.12E+00 | 1.31E-01 | 3.68E-01 | 1.89E+00 | 3.80E-03 | 1.39E-02 | -1.48E+00 | 1.23E-02 | 3.38E-02 | -1.22E+00 |
| GOLGA6L10 | 8.56E-03 | 4.46E-02 | -1.36E+00 | 2.17E-02 | 9.37E-02 | -1.13E+00 | 5.97E-04 | 2.54E-03 | -1.54E+00 | 1.49E-05 | 5.71E-05 | -1.91E+00 | 2.19E-01 | 7.81E-01 | 1.39E+00 | 3.10E-02 | 8.75E-02 | -1.25E+00 | 7.33E-03 | 2.69E-02 | -1.22E+00 | 3.00E-02 | 8.23E-02 | -1.06E+00 |
| FAM133CP | 8.56E-03 | 4.46E-02 | -1.03E+00 | 1.45E-06 | 6.27E-06 | -1.49E+00 | 1.75E-02 | 7.44E-02 | 1.05E+00 | 2.54E-08 | 9.69E-08 | -1.56E+00 | 4.67E-02 | 1.67E-01 | 1.22E+00 | 2.14E-05 | 6.02E-05 | -1.51E+00 | 1.79E-02 | 6.58E-02 | -1.00E+00 | 1.54E-07 | 4.22E-07 | -1.49E+00 |
| PRSS57 | 8.57E-03 | 4.47E-02 | -1.13E+00 | 1.64E-01 | 7.11E-01 | -1.01E+00 | 1.23E-02 | 5.23E-02 | -1.17E+00 | 1.22E-01 | 4.65E-01 | -1.06E+00 | 9.11E-02 | 3.25E-01 | -1.01E+00 | 3.27E-01 | 9.22E-01 | 1.03E+00 | 4.09E-04 | 1.50E-03 | 1.30E+00 | 4.00E-06 | 1.10E-05 | 1.31E+00 |
| RGPD5 | 8.58E-03 | 4.47E-02 | 1.09E+00 | 2.45E-08 | 1.06E-07 | -1.28E+00 | 1.61E-03 | 6.85E-03 | -1.10E+00 | 2.39E-07 | 9.15E-07 | -1.40E+00 | 2.48E-01 | 8.83E-01 | 1.51E+00 | 2.77E-02 | 7.82E-02 | -1.02E+00 | 6.10E-04 | 2.24E-03 | -1.09E+00 | 1.81E-06 | 4.98E-06 | -1.42E+00 |
| AC008781.2 | 8.60E-03 | 4.48E-02 | -1.28E+00 | 7.40E-03 | 3.20E-02 | -1.30E+00 | 1.50E-01 | 6.39E-01 | 1.36E+00 | 1.02E-01 | 3.89E-01 | -1.04E+00 | 2.15E-01 | 7.65E-01 | 1.50E+00 | 2.75E-01 | 7.75E-01 | 1.48E+00 | 1.57E-01 | 5.76E-01 | -1.02E+00 | 1.37E-02 | 3.76E-02 | -1.71E+00 |
| ORC5 | 8.60E-03 | 4.48E-02 | -1.07E+00 | 7.60E-02 | 3.29E-01 | 1.27E+00 | 1.58E-02 | 6.70E-02 | -1.08E+00 | 2.38E-01 | 9.10E-01 | 1.37E+00 | 2.89E-02 | 1.03E-01 | -1.00E+00 | 1.39E-01 | 3.90E-01 | 1.27E+00 | 3.61E-07 | 1.32E-06 | -1.62E+00 | 1.56E-01 | 4.27E-01 | 1.12E+00 |
| JADE3 | 8.61E-03 | 4.49E-02 | -1.25E+00 | 1.59E-01 | 6.88E-01 | 1.44E+00 | 1.99E-02 | 8.47E-02 | -1.17E+00 | 1.32E-01 | 5.06E-01 | 1.17E+00 | 1.34E-01 | 4.79E-01 | -1.03E+00 | 2.30E-01 | 6.48E-01 | 1.73E+00 | 2.47E-05 | 9.08E-05 | -1.61E+00 | 9.58E-03 | 2.63E-02 | -1.21E+00 |
| PSMA8 | 8.62E-03 | 4.49E-02 | -1.40E+00 | 9.27E-03 | 4.01E-02 | -1.26E+00 | 1.09E-01 | 4.64E-01 | 1.36E+00 | 2.62E-01 | 1.00E+00 | 1.26E+00 | 2.34E-02 | 8.35E-02 | -1.68E+00 | 1.28E-01 | 3.61E-01 | 1.01E+00 | 1.80E-02 | 6.60E-02 | -1.34E+00 | 1.88E-02 | 5.17E-02 | -1.41E+00 |
| EGFL7 | 8.64E-03 | 4.50E-02 | -1.05E+00 | 1.43E-01 | 6.19E-01 | 1.20E+00 | 4.25E-03 | 1.81E-02 | -1.12E+00 | 2.22E-01 | 8.49E-01 | 1.20E+00 | 6.88E-05 | 2.45E-04 | 1.02E+00 | 1.77E-02 | 5.00E-02 | 1.15E+00 | 1.05E-01 | 3.84E-01 | 1.55E+00 | 9.62E-02 | 2.64E-01 | 1.10E+00 |
| RPS6KA2 | 8.64E-03 | 4.51E-02 | -1.23E+00 | 2.87E-02 | 1.24E-01 | 1.05E+00 | 7.81E-03 | 3.32E-02 | -1.38E+00 | 3.13E-04 | 1.20E-03 | -1.81E+00 | 2.65E-01 | 9.46E-01 | 1.31E+00 | 2.75E-01 | 7.75E-01 | 1.53E+00 | 1.47E-01 | 5.41E-01 | -1.32E+00 | 2.15E-01 | 5.90E-01 | -1.36E+00 |
| ITGAE | 8.64E-03 | 4.51E-02 | 1.51E+00 | 1.56E-01 | 6.73E-01 | 1.39E+00 | 1.10E-01 | 4.68E-01 | 1.27E+00 | 2.45E-01 | 9.34E-01 | 1.30E+00 | 9.09E-04 | 3.24E-03 | 1.76E+00 | 5.21E-03 | 1.47E-02 | 1.60E+00 | 1.49E-05 | 5.48E-05 | 1.90E+00 | 1.69E-03 | 4.65E-03 | 1.73E+00 |
| AC079035.1 | 8.66E-03 | 4.51E-02 | -1.27E+00 | 4.34E-04 | 1.87E-03 | -1.54E+00 | 1.07E-02 | 4.57E-02 | -1.26E+00 | 1.93E-03 | 7.38E-03 | -1.42E+00 | 1.42E-02 | 5.08E-02 | -1.34E+00 | 9.63E-02 | 2.71E-01 | -1.01E+00 | 7.44E-02 | 2.73E-01 | -1.00E+00 | 1.68E-01 | 4.63E-01 | 1.14E+00 |
| IGKC | 8.67E-03 | 4.52E-02 | -7.30E+00 | 1.96E-02 | 8.46E-02 | 1.58E+00 | 2.08E-01 | 8.82E-01 | 1.15E+00 | 2.09E-01 | 7.97E-01 | -1.01E+00 | 9.86E-02 | 3.51E-01 | -1.35E+00 | 3.05E-01 | 8.61E-01 | 1.22E+00 | 9.31E-02 | 3.42E-01 | 1.44E+00 | 3.64E-01 | 1.00E+00 | 1.00E+00 |
| SCRN3 | 8.69E-03 | 4.53E-02 | -1.05E+00 | 4.35E-02 | 1.88E-01 | 1.26E+00 | 2.79E-04 | 1.18E-03 | -1.29E+00 | 1.09E-02 | 4.16E-02 | 1.08E+00 | 9.82E-03 | 3.50E-02 | -1.20E+00 | 3.34E-01 | 9.41E-01 | 1.35E+00 | 4.84E-09 | 1.78E-08 | -1.42E+00 | 1.64E-04 | 4.49E-04 | -1.09E+00 |
| SNORD3B-2 | 8.70E-03 | 4.54E-02 | 1.32E+00 | 5.01E-58 | 2.17E-57 | -2.73E+00 | 7.44E-02 | 3.16E-01 | 1.13E+00 | 1.65E-25 | 6.31E-25 | -2.30E+00 | 1.50E-19 | 5.36E-19 | -1.59E+00 | 5.55E-19 | 1.57E-18 | -1.91E+00 | 4.26E-07 | 1.56E-06 | -1.18E+00 | 1.09E-08 | 2.99E-08 | -1.62E+00 |
| ATG101 | 8.70E-03 | 4.54E-02 | 2.40E+00 | 8.14E-02 | 3.52E-01 | 2.00E+00 | 4.13E-03 | 1.76E-02 | 2.43E+00 | 1.62E-01 | 6.20E-01 | 1.66E+00 | 1.46E-02 | 5.21E-02 | 2.79E+00 | 1.29E-01 | 3.63E-01 | 2.08E+00 | 6.81E-02 | 2.50E-01 | 1.71E+00 | 7.29E-02 | 2.00E-01 | -1.29E+00 |
| USP9Y | 8.71E-03 | 4.54E-02 | -1.04E+00 | 7.58E-02 | 3.27E-01 | 1.24E+00 | 1.48E-03 | 6.30E-03 | -1.23E+00 | 4.37E-03 | 1.67E-02 | -1.01E+00 | 2.20E-01 | 7.85E-01 | 1.26E+00 | 3.26E-01 | 9.18E-01 | 1.34E+00 | 1.85E-02 | 6.78E-02 | -1.05E+00 | 3.81E-02 | 1.05E-01 | -1.02E+00 |
| FRMD8 | 8.71E-03 | 4.54E-02 | -1.10E+00 | 3.01E-02 | 1.30E-01 | 1.15E+00 | 1.20E-03 | 5.08E-03 | -1.27E+00 | 1.96E-03 | 7.50E-03 | -1.11E+00 | 4.68E-02 | 1.67E-01 | -1.09E+00 | 3.31E-01 | 9.33E-01 | 1.29E+00 | 1.90E-04 | 6.99E-04 | -1.00E+00 | 2.18E-01 | 5.99E-01 | 1.27E+00 |
| CHMP4A | 8.71E-03 | 4.54E-02 | -1.06E+00 | 1.11E-03 | 4.80E-03 | -1.15E+00 | 7.62E-03 | 3.24E-02 | -1.11E+00 | 1.84E-03 | 7.01E-03 | -1.08E+00 | 5.59E-03 | 1.99E-02 | -1.30E+00 | 5.63E-03 | 1.59E-02 | -1.33E+00 | 1.03E-02 | 3.78E-02 | -1.10E+00 | 9.58E-03 | 2.63E-02 | -1.19E+00 |
| ZW10 | 8.74E-03 | 4.55E-02 | -1.10E+00 | 8.62E-02 | 3.73E-01 | 1.32E+00 | 1.08E-01 | 4.57E-01 | -1.03E+00 | 5.10E-02 | 1.95E-01 | 1.03E+00 | 3.01E-02 | 1.07E-01 | -1.19E+00 | 2.50E-01 | 7.05E-01 | 1.13E+00 | 4.49E-04 | 1.65E-03 | -1.08E+00 | 3.26E-03 | 8.95E-03 | 1.02E+00 |
| TMEM154 | 8.75E-03 | 4.56E-02 | 1.10E+00 | 8.21E-02 | 3.55E-01 | 1.52E+00 | 2.98E-03 | 1.27E-02 | -1.12E+00 | 2.75E-02 | 1.05E-01 | 1.14E+00 | 1.11E-05 | 3.95E-05 | -1.85E+00 | 7.28E-06 | 2.05E-05 | -1.73E+00 | 7.72E-06 | 2.83E-05 | -1.39E+00 | 7.31E-05 | 2.01E-04 | -1.42E+00 |
| FDXR | 8.75E-03 | 4.56E-02 | -1.06E+00 | 2.22E-02 | 9.60E-02 | 1.10E+00 | 4.14E-07 | 1.76E-06 | -1.52E+00 | 9.45E-10 | 3.61E-09 | -1.59E+00 | 2.17E-04 | 7.72E-04 | -1.61E+00 | 7.11E-03 | 2.01E-02 | -1.11E+00 | 1.25E-03 | 4.58E-03 | -1.04E+00 | 1.12E-05 | 3.09E-05 | -1.32E+00 |
| ARHGEF1 | 8.75E-03 | 4.56E-02 | 1.20E+00 | 5.67E-02 | 2.45E-01 | 1.43E+00 | 2.74E-02 | 1.17E-01 | 1.11E+00 | 1.03E-02 | 3.93E-02 | 1.16E+00 | 1.82E-04 | 6.47E-04 | -1.08E+00 | 6.17E-03 | 1.74E-02 | 1.06E+00 | 9.12E-02 | 3.35E-01 | 1.13E+00 | 2.05E-01 | 5.63E-01 | 1.25E+00 |
| DNAJC27-AS1 | 8.75E-03 | 4.56E-02 | -1.02E+00 | 2.11E-04 | 9.11E-04 | -1.29E+00 | 1.53E-01 | 6.51E-01 | 1.41E+00 | 5.45E-02 | 2.08E-01 | -1.15E+00 | 1.85E-02 | 6.61E-02 | -1.46E+00 | 8.06E-02 | 2.27E-01 | -1.09E+00 | 7.20E-02 | 2.64E-01 | -1.11E+00 | 7.26E-02 | 1.99E-01 | -1.18E+00 |
| KDM5D | 8.75E-03 | 4.56E-02 | 1.12E+00 | 1.83E-02 | 7.93E-02 | 1.29E+00 | 4.39E-03 | 1.86E-02 | -1.23E+00 | 1.64E-02 | 6.26E-02 | 1.02E+00 | 6.52E-02 | 2.32E-01 | 1.18E+00 | 1.51E-01 | 4.27E-01 | 1.30E+00 | 7.78E-02 | 2.86E-01 | 1.09E+00 | 1.94E-01 | 5.33E-01 | 1.13E+00 |
| ARHGAP11A | 8.78E-03 | 4.58E-02 | -1.78E+00 | 1.38E-01 | 5.94E-01 | 1.10E+00 | 1.67E-01 | 7.07E-01 | 1.02E+00 | 1.72E-01 | 6.55E-01 | 1.27E+00 | 4.92E-03 | 1.76E-02 | -1.75E+00 | 2.59E-02 | 7.29E-02 | -1.42E+00 | 4.12E-04 | 1.51E-03 | -1.29E+00 | 3.12E-01 | 8.58E-01 | 1.13E+00 |
| SART1 | 8.78E-03 | 4.58E-02 | 1.03E+00 | 6.93E-02 | 2.99E-01 | 1.30E+00 | 7.09E-03 | 3.01E-02 | -1.07E+00 | 6.41E-03 | 2.45E-02 | 1.03E+00 | 6.86E-02 | 2.45E-01 | 1.10E+00 | 2.68E-01 | 7.56E-01 | 1.39E+00 | 5.58E-04 | 2.05E-03 | -1.13E+00 | 2.67E-01 | 7.32E-01 | 1.15E+00 |
| AC087257.2 | 8.80E-03 | 4.59E-02 | -1.10E+00 | 3.26E-04 | 1.41E-03 | -1.44E+00 | 1.18E-01 | 5.01E-01 | 1.49E+00 | 1.52E-01 | 5.79E-01 | 1.25E+00 | 1.78E-01 | 6.36E-01 | -1.02E+00 | 1.08E-01 | 3.06E-01 | -1.21E+00 | 1.97E-01 | 7.25E-01 | 1.03E+00 | 2.45E-01 | 6.72E-01 | 1.01E+00 |
| ARL8A | 8.81E-03 | 4.59E-02 | 1.05E+00 | 9.30E-03 | 4.02E-02 | 1.20E+00 | 1.62E-06 | 6.89E-06 | -1.39E+00 | 1.52E-05 | 5.80E-05 | -1.10E+00 | 1.06E-08 | 3.77E-08 | -1.41E+00 | 6.64E-10 | 1.87E-09 | -1.42E+00 | 1.85E-07 | 6.81E-07 | -1.49E+00 | 6.15E-06 | 1.69E-05 | -1.34E+00 |
| SCUBE1 | 8.82E-03 | 4.59E-02 | -1.51E+00 | 1.78E-03 | 7.71E-03 | -1.90E+00 | 1.87E-01 | 7.94E-01 | 1.02E+00 | 1.59E-01 | 6.07E-01 | 1.00E+00 | 1.35E-13 | 4.80E-13 | -2.10E+00 | 1.81E-19 | 5.11E-19 | -3.85E+00 | 7.80E-02 | 2.86E-01 | -1.28E+00 | 4.07E-02 | 1.12E-01 | -1.64E+00 |
| MOB3C | 8.83E-03 | 4.60E-02 | -1.27E+00 | 8.23E-02 | 3.56E-01 | 1.08E+00 | 7.34E-07 | 3.12E-06 | -2.14E+00 | 1.99E-04 | 7.61E-04 | -1.41E+00 | 8.31E-02 | 2.96E-01 | -1.20E+00 | 3.21E-01 | 9.04E-01 | 1.24E+00 | 8.10E-08 | 2.97E-07 | 1.54E+00 | 1.30E-03 | 3.57E-03 | 1.71E+00 |
| C18orf15 | 8.83E-03 | 4.60E-02 | -1.39E+00 | 2.58E-02 | 1.12E-01 | -1.16E+00 | 1.98E-01 | 8.41E-01 | 1.23E+00 | 7.98E-02 | 3.05E-01 | -1.18E+00 | 1.97E-01 | 7.02E-01 | -1.10E+00 | 3.39E-01 | 9.57E-01 | 1.17E+00 | 5.93E-02 | 2.18E-01 | -1.06E+00 | 3.62E-01 | 9.94E-01 | 1.11E+00 |
| MED9 | 8.83E-03 | 4.60E-02 | 1.09E+00 | 3.42E-04 | 1.48E-03 | -1.04E+00 | 1.43E-02 | 6.09E-02 | -1.13E+00 | 4.61E-03 | 1.76E-02 | -1.07E+00 | 7.23E-04 | 2.58E-03 | -1.36E+00 | 5.12E-02 | 1.44E-01 | 1.01E+00 | 1.36E-03 | 4.99E-03 | -1.12E+00 | 2.50E-02 | 6.87E-02 | 1.02E+00 |
| NUDT1 | 8.83E-03 | 4.60E-02 | 1.14E+00 | 1.63E-03 | 7.03E-03 | 1.17E+00 | 1.96E-03 | 8.32E-03 | 1.49E+00 | 1.39E-01 | 5.31E-01 | 1.36E+00 | 1.82E-02 | 6.47E-02 | 1.18E+00 | 4.08E-04 | 1.15E-03 | 1.04E+00 | 1.83E-03 | 6.73E-03 | 1.57E+00 | 7.28E-03 | 2.00E-02 | 1.46E+00 |
| ERICH1 | 8.84E-03 | 4.61E-02 | 1.20E+00 | 8.02E-07 | 3.46E-06 | 1.12E+00 | 2.77E-03 | 1.18E-02 | -1.06E+00 | 4.86E-05 | 1.86E-04 | -1.03E+00 | 2.06E-03 | 7.33E-03 | -1.00E+00 | 1.75E-02 | 4.93E-02 | 1.09E+00 | 7.92E-06 | 2.91E-05 | -1.17E+00 | 3.88E-07 | 1.06E-06 | -1.27E+00 |
| PGP | 8.85E-03 | 4.61E-02 | -1.06E+00 | 1.61E-02 | 6.98E-02 | 1.07E+00 | 3.74E-02 | 1.59E-01 | 1.33E+00 | 2.14E-01 | 8.16E-01 | 1.25E+00 | 2.12E-05 | 7.55E-05 | -1.21E+00 | 3.63E-05 | 1.02E-04 | -1.13E+00 | 1.08E-01 | 3.96E-01 | 1.19E+00 | 1.41E-02 | 3.88E-02 | 1.40E+00 |
| MNAT1 | 8.85E-03 | 4.61E-02 | 1.06E+00 | 1.27E-06 | 5.49E-06 | -1.14E+00 | 5.03E-04 | 2.14E-03 | -1.13E+00 | 1.98E-10 | 7.57E-10 | -1.44E+00 | 1.34E-03 | 4.79E-03 | -1.05E+00 | 2.02E-06 | 5.70E-06 | -1.32E+00 | 1.27E-05 | 4.66E-05 | -1.02E+00 | 3.03E-09 | 8.34E-09 | -1.31E+00 |
| SERTAD3 | 8.85E-03 | 4.61E-02 | -1.13E+00 | 6.18E-05 | 2.67E-04 | -1.49E+00 | 6.23E-03 | 2.65E-02 | -1.23E+00 | 4.18E-05 | 1.60E-04 | -1.63E+00 | 2.63E-01 | 9.37E-01 | 1.28E+00 | 8.93E-03 | 2.52E-02 | -1.23E+00 | 6.30E-03 | 2.31E-02 | 1.04E+00 | 1.70E-05 | 4.68E-05 | -1.56E+00 |
| LINC01003 | 8.86E-03 | 4.62E-02 | 1.13E+00 | 4.02E-04 | 1.74E-03 | 1.18E+00 | 1.14E-02 | 4.85E-02 | -1.07E+00 | 4.74E-05 | 1.81E-04 | -1.18E+00 | 3.59E-02 | 1.28E-01 | 1.13E+00 | 3.40E-01 | 9.57E-01 | 1.42E+00 | 1.94E-01 | 7.11E-01 | 1.31E+00 | 2.44E-01 | 6.70E-01 | 1.32E+00 |
| ZNF599 | 8.87E-03 | 4.62E-02 | -1.18E+00 | 6.28E-02 | 2.71E-01 | 1.09E+00 | 9.54E-02 | 4.05E-01 | -1.06E+00 | 8.36E-02 | 3.19E-01 | 1.09E+00 | 1.47E-02 | 5.24E-02 | -1.24E+00 | 2.06E-02 | 5.80E-02 | -1.14E+00 | 1.92E-02 | 7.03E-02 | -1.15E+00 | 2.59E-02 | 7.13E-02 | -1.09E+00 |
| LAMB1 | 8.87E-03 | 4.62E-02 | -1.12E+00 | 3.87E-03 | 1.67E-02 | -1.10E+00 | 3.82E-02 | 1.63E-01 | -1.12E+00 | 1.30E-02 | 4.97E-02 | -1.29E+00 | 7.84E-02 | 2.79E-01 | -1.13E+00 | 2.31E-01 | 6.50E-01 | 1.29E+00 | 1.22E-02 | 4.48E-02 | -1.13E+00 | 3.45E-02 | 9.48E-02 | -1.23E+00 |
| AC079598.4 | 8.88E-03 | 4.63E-02 | -1.27E+00 | 1.51E-01 | 6.53E-01 | 1.31E+00 | 2.14E-06 | 9.09E-06 | -1.74E+00 | 1.29E-02 | 4.92E-02 | 1.11E+00 | 2.34E-04 | 8.34E-04 | -2.06E+00 | 5.71E-02 | 1.61E-01 | 1.04E+00 | 7.15E-11 | 2.63E-10 | -1.90E+00 | 2.33E-02 | 6.41E-02 | 1.08E+00 |
| MIR17HG | 8.88E-03 | 4.63E-02 | -1.23E+00 | 3.37E-02 | 1.46E-01 | 1.86E+00 | 4.31E-02 | 1.83E-01 | -1.15E+00 | 1.58E-01 | 6.03E-01 | 1.48E+00 | 4.64E-02 | 1.66E-01 | -1.29E+00 | 1.49E-01 | 4.20E-01 | -1.01E+00 | 2.23E-01 | 8.18E-01 | 1.13E+00 | 1.46E-01 | 4.01E-01 | 1.56E+00 |
| ATP2A1 | 8.91E-03 | 4.64E-02 | 1.03E+00 | 1.04E-02 | 4.51E-02 | 1.18E+00 | 5.83E-02 | 2.48E-01 | 1.13E+00 | 3.00E-02 | 1.15E-01 | 1.13E+00 | 2.84E-02 | 1.01E-01 | -1.10E+00 | 1.69E-02 | 4.76E-02 | -1.28E+00 | 2.60E-01 | 9.54E-01 | 1.14E+00 | 3.48E-01 | 9.57E-01 | 1.23E+00 |
| PTTG1 | 8.92E-03 | 4.65E-02 | -1.10E+00 | 5.59E-05 | 2.41E-04 | -1.21E+00 | 3.59E-02 | 1.52E-01 | 1.44E+00 | 2.71E-08 | 1.04E-07 | 1.09E+00 | 3.37E-02 | 1.20E-01 | 1.04E+00 | 1.03E-01 | 2.90E-01 | 1.25E+00 | 1.05E-01 | 3.86E-01 | 1.04E+00 | 3.15E-01 | 8.66E-01 | 1.22E+00 |
| GLIF1 | 8.92E-03 | 4.65E-02 | 1.05E+00 | 5.82E-03 | 2.52E-02 | 1.15E+00 | 1.36E-04 | 5.78E-04 | -1.33E+00 | 3.80E-04 | 1.45E-03 | -1.07E+00 | 4.89E-04 | 1.75E-03 | -1.30E+00 | 4.83E-02 | 1.36E-01 | 1.10E+00 | 9.98E-04 | 3.66E-03 | -1.06E+00 | 4.24E-03 | 1.17E-02 | 1.05E+00 |
| ZNF205 | 8.94E-03 | 4.66E-02 | -1.34E+00 | 1.12E-01 | 4.86E-01 | 1.24E+00 | 1.58E-03 | 6.69E-03 | -1.61E+00 | 4.36E-02 | 1.66E-01 | 1.07E+00 | 1.03E-01 | 3.65E-01 | 1.05E+00 | 2.21E-01 | 6.22E-01 | 1.76E+00 | 2.10E-01 | 7.71E-01 | 1.16E+00 | 9.31E-02 | 2.56E-01 | -1.13E+00 |
| LINC02210 | 8.95E-03 | 4.66E-02 | 1.00E+00 | 1.70E-01 | 7.36E-01 | 1.45E+00 | 1.14E-04 | 4.84E-04 | -1.55E+00 | 4.22E-02 | 1.61E-01 | 1.12E+00 | 3.56E-03 | 1.27E-02 | -1.10E+00 | 4.16E-02 | 1.17E-01 | 1.20E+00 | 5.29E-03 | 1.94E-02 | -1.16E+00 | 1.53E-02 | 4.21E-02 | -1.07E+00 |
| AC107464.3 | 8.95E-03 | 4.66E-02 | -1.07E+00 | 1.14E-01 | 4.94E-01 | 1.34E+00 | 6.13E-03 | 2.61E-02 | -1.29E+00 | 1.03E-02 | 3.92E-02 | -1.10E+00 | 1.89E-02 | 6.75E-02 | -1.28E+00 | 2.88E-01 | 8.11E-01 | 1.32E+00 | 1.47E-01 | 5.40E-01 | 1.03E+00 | 2.56E-01 | 7.04E-01 | 1.15E+00 |
| PTK2B | 8.95E-03 | 4.66E-02 | 1.02E+00 | 1.44E-01 | 6.21E-01 | 1.32E+00 | 3.07E-03 | 1.30E-02 | -1.08E+00 | 7.74E-02 | 2.96E-01 | 1.26E+00 | 4.01E-02 | 1.43E-01 | 1.02E+00 | 1.92E-01 | 5.42E-01 | 1.21E+00 | 7.36E-02 | 2.70E-01 | 1.09E+00 | 3.19E-02 | 8.75E-02 | 1.09E+00 |
| ARHGAP27 | 8.96E-03 | 4.67E-02 | -1.04E+00 | 8.89E-04 | 3.84E-03 | -1.03E+00 | 1.09E-02 | 4.65E-02 | -1.05E+00 | 8.82E-03 | 3.37E-02 | 1.09E+00 | 2.35E-02 | 8.39E-02 | -1.60E+00 | 8.49E-02 | 2.39E-01 | -1.27E+00 | 8.62E-02 | 3.16E-01 | -1.05E+00 | 1.68E-01 | 4.63E-01 | 1.02E+00 |
| RPS6KL1 | 8.96E-03 | 4.67E-02 | -1.20E+00 | 4.21E-04 | 1.82E-03 | -1.54E+00 | 3.86E-03 | 1.64E-02 | -1.10E+00 | 2.59E-06 | 9.90E-06 | -1.75E+00 | 1.30E-02 | 4.63E-02 | -1.55E+00 | 1.68E-02 | 4.73E-02 | -1.51E+00 | 2.68E-04 | 9.85E-04 | -1.27E+00 | 3.28E-03 | 9.00E-03 | -1.11E+00 |
| PDK3 | 8.97E-03 | 4.67E-02 | -1.14E+00 | 1.24E-02 | 5.36E-02 | -1.04E+00 | 3.72E-02 | 1.58E-01 | -1.12E+00 | 2.49E-02 | 9.51E-02 | -1.01E+00 | 1.18E-02 | 4.21E-02 | 1.05E+00 | 1.10E-01 | 3.11E-01 | 1.30E+00 | 2.57E-01 | 9.44E-01 | 1.23E+00 | 3.56E-03 | 9.77E-03 | -1.04E+00 |
| ZNF587B | 8.98E-03 | 4.68E-02 | -1.33E+00 | 9.56E-02 | 4.13E-01 | 1.11E+00 | 1.68E-01 | 7.16E-01 | 1.11E+00 | 1.49E-01 | 5.70E-01 | 1.60E+00 | 3.18E-02 | 1.13E-01 | 1.01E+00 | 1.47E-01 | 4.13E-01 | 1.16E+00 | 1.14E-02 | 4.20E-02 | -1.34E+00 | 9.31E-02 | 2.56E-01 | -1.04E+00 |
| SPRY1 | 8.99E-03 | 4.69E-02 | 1.84E+00 | 6.34E-02 | 2.74E-01 | 1.29E+00 | 4.62E-03 | 1.96E-02 | 1.94E+00 | 6.68E-02 | 2.55E-01 | 1.18E+00 | 2.00E-01 | 7.13E-01 | 1.69E+00 | 2.97E-01 | 8.36E-01 | 1.35E+00 | 2.61E-01 | 9.56E-01 | 1.48E+00 | 2.42E-03 | 6.66E-03 | -1.21E+00 |
| AC019181.1 | 9.01E-03 | 4.69E-02 | -1.07E+00 | 1.52E-03 | 6.57E-03 | -1.17E+00 | 1.21E-01 | 5.15E-01 | 1.37E+00 | 5.10E-02 | 1.95E-01 | -1.05E+00 | 6.40E-02 | 2.28E-01 | -1.23E+00 | 3.59E-02 | 1.01E-01 | -1.39E+00 | 5.21E-02 | 1.91E-01 | -1.11E+00 | 7.96E-02 | 2.19E-01 | -1.13E+00 |
| POLRMTP1 | 9.02E-03 | 4.70E-02 | -1.27E+00 | 1.66E-02 | 7.15E-02 | -1.11E+00 | 2.40E-03 | 1.02E-02 | -1.34E+00 | 5.00E-03 | 1.91E-02 | -1.08E+00 | 5.76E-02 | 2.05E-01 | -1.09E+00 | 2.97E-01 | 8.38E-01 | 1.39E+00 | 4.94E-03 | 1.81E-02 | -1.13E+00 | 5.85E-02 | 1.61E-01 | 1.01E+00 |
| RPS15A | 9.03E-03 | 4.70E-02 | 1.05E+00 | 1.74E-41 | 7.54E-41 | -1.35E+00 | 1.27E-03 | 5.41E-03 | 1.09E+00 | 1.91E-22 | 7.30E-22 | -1.29E+00 | 1.34E-01 | 4.77E-01 | 1.03E+00 | 9.95E-19 | 2.80E-18 | -1.31E+00 | 1.55E-01 | 5.68E-01 | 1.07E+00 | 2.94E-26 | 8.06E-26 | -1.33E+00 |
| AC104986.2 | 9.03E-03 | 4.71E-02 | -1.08E+00 | 9.03E-04 | 3.90E-03 | -1.30E+00 | 4.97E-05 | 2.11E-04 | -1.62E+00 | 2.65E-07 | 1.01E-06 | -1.95E+00 | 2.07E-01 | 7.40E-01 | 1.44E+00 | 3.32E-02 | 9.37E-02 | -1.19E+00 | 1.19E-02 | 4.37E-02 | -1.12E+00 | 1.05E-02 | 2.90E-02 | -1.21E+00 |
| SEPT9 | 9.04E-03 | 4.71E-02 | 1.14E+00 | 3.39E-03 | 1.47E-02 | 1.14E+00 | 1.48E-01 | 6.30E-01 | 1.26E+00 | 6.67E-02 | 2.55E-01 | 1.20E+00 | 3.22E-02 | 1.15E-01 | 1.08E+00 | 9.10E-03 | 2.56E-02 | 1.07E+00 | 9.18E-03 | 3.37E-02 | -1.10E+00 | 1.52E-03 | 4.18E-03 | -1.25E+00 |
| RFFL | 9.04E-03 | 4.71E-02 | 1.07E+00 | 8.15E-04 | 3.52E-03 | -1.00E+00 | 1.08E-01 | 4.61E-01 | 1.14E+00 | 3.91E-05 | 1.49E-04 | -1.28E+00 | 2.67E-03 | 9.52E-03 | -1.15E+00 | 9.90E-05 | 2.79E-04 | -1.45E+00 | 2.01E-03 | 7.36E-03 | -1.05E+00 | 4.12E-08 | 1.13E-07 | -1.74E+00 |
| NOD1 | 9.04E-03 | 4.71E-02 | -1.30E+00 | 1.12E-02 | 4.85E-02 | -1.16E+00 | 3.26E-02 | 1.39E-01 | -1.13E+00 | 2.87E-02 | 1.09E-01 | -1.07E+00 | 1.30E-01 | 4.64E-01 | -1.10E+00 | 7.02E-02 | 1.98E-01 | -1.33E+00 | 9.33E-02 | 3.42E-01 | -1.07E+00 | 3.29E-01 | 9.03E-01 | 1.21E+00 |
| OMG | 9.05E-03 | 4.72E-02 | -1.28E+00 | 1.16E-01 | 5.01E-01 | 1.21E+00 | 3.41E-02 | 1.45E-01 | -1.09E+00 | 7.60E-02 | 2.90E-01 | 1.23E+00 | 2.40E-01 | 8.55E-01 | 1.21E+00 | 2.98E-01 | 8.39E-01 | 1.59E+00 | 1.19E-04 | 4.38E-04 | -1.75E+00 | 7.97E-04 | 2.19E-03 | -1.64E+00 |
| CELF2 | 9.08E-03 | 4.73E-02 | 1.26E+00 | 6.04E-02 | 2.61E-01 | 1.28E+00 | 1.14E-01 | 4.83E-01 | 1.15E+00 | 2.07E-01 | 7.90E-01 | 1.13E+00 | 1.32E-02 | 4.69E-02 | 1.05E+00 | 9.36E-04 | 2.64E-03 | -1.01E+00 | 8.59E-03 | 3.15E-02 | -1.06E+00 | 3.38E-04 | 9.29E-04 | -1.11E+00 |
| RACGAP1 | 9.09E-03 | 4.74E-02 | -1.65E+00 | 6.93E-02 | 2.99E-01 | -1.05E+00 | 7.82E-05 | 3.32E-04 | -1.43E+00 | 5.01E-03 | 1.92E-02 | 1.08E+00 | 5.30E-03 | 1.89E-02 | -1.84E+00 | 7.45E-02 | 2.10E-01 | -1.13E+00 | 1.36E-09 | 5.00E-09 | -1.57E+00 | 3.40E-03 | 9.35E-03 | -1.01E+00 |
| GPKOW | 9.10E-03 | 4.74E-02 | -1.23E+00 | 1.18E-01 | 5.11E-01 | 1.24E+00 | 1.05E-01 | 4.44E-01 | 1.05E+00 | 6.86E-02 | 2.62E-01 | 1.16E+00 | 6.02E-05 | 2.15E-04 | -1.42E+00 | 5.52E-03 | 1.55E-02 | -1.02E+00 | 1.76E-02 | 6.45E-02 | 1.02E+00 | 1.56E-01 | 4.28E-01 | 1.13E+00 |
| ADORA2A-AS1 | 9.10E-03 | 4.74E-02 | -1.45E+00 | 1.22E-01 | 5.28E-01 | 1.10E+00 | 7.68E-02 | 3.26E-01 | -1.13E+00 | 3.46E-02 | 1.32E-01 | -1.31E+00 | 1.11E-01 | 3.96E-01 | -1.15E+00 | 5.19E-02 | 1.46E-01 | -1.67E+00 | 2.62E-01 | 9.61E-01 | 1.22E+00 | 1.27E-01 | 3.49E-01 | -1.20E+00 |
| AC104825.2 | 9.10E-03 | 4.74E-02 | -1.69E+00 | 1.54E-01 | 6.67E-01 | 1.50E+00 | 4.03E-05 | 1.71E-04 | -1.76E+00 | 4.69E-03 | 1.79E-02 | -1.17E+00 | 1.98E-03 | 7.05E-03 | -1.65E+00 | 2.76E-03 | 7.77E-03 | -1.73E+00 | 8.23E-03 | 3.02E-02 | -1.57E+00 | 1.62E-02 | 4.46E-02 | -1.45E+00 |
| HIRIP3 | 9.12E-03 | 4.75E-02 | -1.14E+00 | 3.87E-02 | 1.67E-01 | 1.06E+00 | 1.35E-02 | 5.72E-02 | 1.48E+00 | 2.19E-01 | 8.36E-01 | 1.49E+00 | 2.94E-02 | 1.05E-01 | -1.19E+00 | 9.88E-02 | 2.79E-01 | 1.06E+00 | 1.90E-01 | 6.96E-01 | 1.15E+00 | 2.19E-03 | 6.02E-03 | 1.44E+00 |
| LSP1 | 9.12E-03 | 4.75E-02 | -1.07E+00 | 1.50E-04 | 6.47E-04 | -1.07E+00 | 6.42E-03 | 2.73E-02 | -1.17E+00 | 3.97E-03 | 1.51E-02 | -1.14E+00 | 2.67E-08 | 9.53E-08 | -1.29E+00 | 4.55E-09 | 1.28E-08 | -1.30E+00 | 5.31E-02 | 1.95E-01 | 1.06E+00 | 2.31E-03 | 6.34E-03 | -1.08E+00 |
| NEURL4 | 9.12E-03 | 4.76E-02 | -1.20E+00 | 5.71E-02 | 2.47E-01 | 1.15E+00 | 2.09E-01 | 8.90E-01 | 1.20E+00 | 2.40E-01 | 9.18E-01 | 1.38E+00 | 5.06E-02 | 1.80E-01 | -1.50E+00 | 3.22E-01 | 9.07E-01 | 1.27E+00 | 9.69E-02 | 3.56E-01 | -1.09E+00 | 3.11E-01 | 8.55E-01 | 1.22E+00 |
| PIAS2 | 9.13E-03 | 4.76E-02 | 1.00E+00 | 2.93E-02 | 1.27E-01 | 1.19E+00 | 9.27E-02 | 3.94E-01 | 1.11E+00 | 2.10E-01 | 8.03E-01 | 1.34E+00 | 1.02E-01 | 3.63E-01 | 1.13E+00 | 1.64E-01 | 4.63E-01 | 1.14E+00 | 1.49E-02 | 5.45E-02 | -1.06E+00 | 9.25E-02 | 2.54E-01 | 1.06E+00 |
| USP6 | 9.13E-03 | 4.76E-02 | -1.42E+00 | 5.65E-02 | 2.44E-01 | -1.02E+00 | 7.18E-02 | 3.05E-01 | -1.05E+00 | 2.04E-01 | 7.78E-01 | 1.46E+00 | 1.08E-02 | 3.85E-02 | -1.24E+00 | 5.60E-02 | 1.58E-01 | -1.03E+00 | 1.94E-01 | 7.12E-01 | -1.01E+00 | 4.92E-03 | 1.35E-02 | 2.14E+00 |
| E2F3 | 9.13E-03 | 4.76E-02 | -1.01E+00 | 3.85E-03 | 1.66E-02 | 1.02E+00 | 3.90E-02 | 1.66E-01 | 1.08E+00 | 2.68E-04 | 1.02E-03 | -1.11E+00 | 8.27E-04 | 2.95E-03 | -1.11E+00 | 1.60E-02 | 4.51E-02 | 1.01E+00 | 8.27E-07 | 3.04E-06 | -1.28E+00 | 4.10E-04 | 1.13E-03 | -1.13E+00 |
| TMEM143 | 9.14E-03 | 4.77E-02 | -1.13E+00 | 3.07E-03 | 1.33E-02 | -1.17E+00 | 8.63E-02 | 3.66E-01 | 1.06E+00 | 2.58E-01 | 9.84E-01 | 1.45E+00 | 1.28E-01 | 4.56E-01 | 1.04E+00 | 3.53E-01 | 9.96E-01 | 1.44E+00 | 6.91E-02 | 2.54E-01 | -1.11E+00 | 3.42E-01 | 9.40E-01 | 1.18E+00 |
| USP30 | 9.14E-03 | 4.77E-02 | -1.17E+00 | 2.40E-02 | 1.04E-01 | -1.01E+00 | 3.45E-02 | 1.47E-01 | -1.20E+00 | 2.06E-02 | 7.87E-02 | -1.11E+00 | 2.20E-02 | 7.85E-02 | -1.25E+00 | 7.45E-02 | 2.10E-01 | -1.02E+00 | 4.42E-03 | 1.62E-02 | -1.29E+00 | 3.87E-02 | 1.06E-01 | -1.03E+00 |
| PARP16 | 9.16E-03 | 4.77E-02 | 1.03E+00 | 6.82E-03 | 2.95E-02 | 1.11E+00 | 9.45E-03 | 4.02E-02 | -1.07E+00 | 1.66E-02 | 6.34E-02 | 1.12E+00 | 1.73E-01 | 6.18E-01 | 1.18E+00 | 2.15E-01 | 6.06E-01 | 1.64E+00 | 8.07E-04 | 2.96E-03 | -1.31E+00 | 1.65E-01 | 4.53E-01 | 1.13E+00 |
| EED | 9.16E-03 | 4.77E-02 | -1.11E+00 | 2.37E-02 | 1.02E-01 | 1.04E+00 | 3.51E-05 | 1.49E-04 | -1.27E+00 | 3.48E-06 | 1.33E-05 | -1.23E+00 | 4.94E-02 | 1.76E-01 | 1.21E+00 | 4.78E-02 | 1.35E-01 | 1.08E+00 | 1.63E-06 | 5.98E-06 | -1.31E+00 | 5.55E-03 | 1.52E-02 | -1.02E+00 |
| H3F3C | 9.16E-03 | 4.78E-02 | 1.03E+00 | 1.14E-10 | 4.94E-10 | -1.18E+00 | 2.45E-02 | 1.04E-01 | -1.05E+00 | 1.61E-03 | 6.14E-03 | -1.15E+00 | 2.02E-02 | 7.20E-02 | -1.00E+00 | 5.46E-04 | 1.54E-03 | -1.09E+00 | 2.55E-06 | 9.37E-06 | -1.30E+00 | 5.03E-09 | 1.38E-08 | -1.49E+00 |
| AC018413.1 | 9.17E-03 | 4.78E-02 | -1.32E+00 | 3.97E-03 | 1.71E-02 | -1.40E+00 | 1.74E-01 | 7.41E-01 | 1.15E+00 | 1.65E-01 | 6.28E-01 | 1.21E+00 | 9.76E-02 | 3.48E-01 | 1.60E+00 | 2.58E-01 | 7.29E-01 | 1.29E+00 | 1.13E-01 | 4.16E-01 | -1.16E+00 | 6.29E-02 | 1.73E-01 | -1.50E+00 |
| EOGT | 9.17E-03 | 4.78E-02 | -1.15E+00 | 3.66E-02 | 1.58E-01 | 1.10E+00 | 1.04E-01 | 4.43E-01 | 1.08E+00 | 3.14E-03 | 1.20E-02 | -1.19E+00 | 7.57E-02 | 2.70E-01 | 1.06E+00 | 1.23E-01 | 3.47E-01 | 1.03E+00 | 8.87E-02 | 3.25E-01 | 1.13E+00 | 1.16E-01 | 3.20E-01 | 1.06E+00 |
| RPL39L | 9.17E-03 | 4.78E-02 | -1.07E+00 | 3.33E-90 | 1.44E-89 | -1.92E+00 | 5.97E-02 | 2.54E-01 | -1.05E+00 | 8.10E-58 | 3.09E-57 | -1.89E+00 | 2.04E-01 | 7.27E-01 | -1.01E+00 | 3.19E-49 | 8.99E-49 | -1.91E+00 | 5.02E-07 | 1.84E-06 | -1.11E+00 | 5.11E-65 | 1.40E-64 | -1.93E+00 |
| HNRNPA3P6 | 9.18E-03 | 4.78E-02 | 1.06E+00 | 2.54E-06 | 1.10E-05 | -1.07E+00 | 1.45E-01 | 6.18E-01 | 1.32E+00 | 4.25E-02 | 1.62E-01 | 1.21E+00 | 2.59E-01 | 9.23E-01 | 1.35E+00 | 7.97E-03 | 2.25E-02 | 1.04E+00 | 1.85E-01 | 6.79E-01 | 1.22E+00 | 5.51E-02 | 1.51E-01 | 1.09E+00 |
| RPL37P2 | 9.19E-03 | 4.79E-02 | -1.07E+00 | 9.16E-67 | 3.96E-66 | -2.14E+00 | 5.70E-02 | 2.42E-01 | -1.09E+00 | 1.28E-26 | 4.87E-26 | -2.11E+00 | 1.74E-02 | 6.19E-02 | -1.09E+00 | 6.37E-30 | 1.80E-29 | -2.10E+00 | 4.87E-04 | 1.79E-03 | -1.13E+00 | 1.52E-27 | 4.18E-27 | -2.13E+00 |
| ZNF138 | 9.21E-03 | 4.80E-02 | 1.01E+00 | 1.25E-01 | 5.41E-01 | 1.40E+00 | 2.48E-03 | 1.05E-02 | -1.13E+00 | 1.34E-03 | 5.11E-03 | -1.03E+00 | 3.37E-04 | 1.20E-03 | -1.27E+00 | 4.67E-03 | 1.32E-02 | -1.15E+00 | 3.19E-04 | 1.17E-03 | -1.07E+00 | 2.18E-04 | 5.98E-04 | -1.18E+00 |
| SNAPC4 | 9.21E-03 | 4.80E-02 | -1.04E+00 | 8.65E-04 | 3.74E-03 | -1.18E+00 | 2.10E-01 | 8.93E-01 | 1.13E+00 | 1.19E-01 | 4.54E-01 | 1.14E+00 | 1.25E-01 | 4.46E-01 | 1.10E+00 | 2.98E-01 | 8.40E-01 | 1.33E+00 | 3.46E-02 | 1.27E-01 | 1.00E+00 | 1.12E-01 | 3.09E-01 | 1.01E+00 |
| NSA2 | 9.21E-03 | 4.80E-02 | 1.06E+00 | 4.68E-06 | 2.02E-05 | 1.00E+00 | 5.81E-02 | 2.47E-01 | 1.05E+00 | 3.47E-02 | 1.32E-01 | 1.07E+00 | 1.31E-02 | 4.68E-02 | 1.03E+00 | 1.48E-02 | 4.17E-02 | 1.02E+00 | 5.07E-02 | 1.86E-01 | 1.00E+00 | 2.89E-02 | 7.95E-02 | -1.05E+00 |
| THRB-AS1 | 9.22E-03 | 4.81E-02 | -1.46E+00 | 5.37E-03 | 2.32E-02 | -1.54E+00 | 1.42E-01 | 6.03E-01 | -1.25E+00 | 1.96E-01 | 7.50E-01 | -1.08E+00 | 6.32E-02 | 2.26E-01 | 2.40E+00 | 7.73E-02 | 2.18E-01 | 2.07E+00 | 1.12E-01 | 4.09E-01 | 1.41E+00 | 2.49E-01 | 6.85E-01 | 1.09E+00 |
| TMEM14DP | 9.22E-03 | 4.81E-02 | 1.06E+00 | 9.33E-07 | 4.03E-06 | -1.31E+00 | 1.49E-01 | 6.34E-01 | 1.22E+00 | 7.99E-06 | 3.05E-05 | -1.30E+00 | 2.36E-01 | 8.41E-01 | 1.42E+00 | 1.09E-01 | 3.07E-01 | 1.08E+00 | 3.48E-02 | 1.28E-01 | 1.09E+00 | 9.77E-12 | 2.68E-11 | -1.66E+00 |
| SEMA4F | 9.23E-03 | 4.81E-02 | -1.15E+00 | 1.07E-01 | 4.63E-01 | 1.30E+00 | 3.52E-02 | 1.50E-01 | -1.30E+00 | 2.58E-01 | 9.86E-01 | 1.30E+00 | 7.15E-02 | 2.55E-01 | -1.17E+00 | 2.27E-01 | 6.41E-01 | 1.19E+00 | 3.41E-02 | 1.25E-01 | -1.26E+00 | 2.86E-01 | 7.87E-01 | 1.08E+00 |
| ZNF552 | 9.24E-03 | 4.81E-02 | -1.01E+00 | 9.22E-02 | 3.98E-01 | 1.43E+00 | 1.97E-01 | 8.37E-01 | 1.19E+00 | 1.04E-01 | 3.98E-01 | 1.14E+00 | 7.20E-02 | 2.57E-01 | 1.15E+00 | 1.84E-01 | 5.19E-01 | 1.27E+00 | 3.80E-03 | 1.40E-02 | -1.27E+00 | 3.08E-02 | 8.46E-02 | -1.08E+00 |
| DCUN1D2 | 9.24E-03 | 4.81E-02 | -1.25E+00 | 2.23E-02 | 9.64E-02 | -1.11E+00 | 1.86E-01 | 7.92E-01 | 1.32E+00 | 9.15E-02 | 3.50E-01 | 1.16E+00 | 1.07E-02 | 3.81E-02 | -1.58E+00 | 7.79E-02 | 2.20E-01 | -1.09E+00 | 2.08E-02 | 7.62E-02 | -1.16E+00 | 2.55E-01 | 7.01E-01 | 1.30E+00 |
| MKS1 | 9.26E-03 | 4.82E-02 | -1.14E+00 | 9.73E-02 | 4.21E-01 | 1.33E+00 | 2.84E-06 | 1.21E-05 | -1.49E+00 | 2.63E-04 | 1.01E-03 | -1.13E+00 | 1.82E-03 | 6.47E-03 | -1.37E+00 | 1.90E-01 | 5.36E-01 | 1.36E+00 | 2.10E-09 | 7.70E-09 | -1.69E+00 | 6.86E-04 | 1.88E-03 | -1.16E+00 |
| CLEC1B | 9.26E-03 | 4.82E-02 | -2.02E+00 | 1.03E-02 | 4.43E-02 | -2.05E+00 | 1.90E-01 | 8.07E-01 | 1.26E+00 | 1.69E-01 | 6.46E-01 | 1.33E+00 | 9.02E-02 | 3.21E-01 | -1.82E+00 | 9.55E-02 | 2.69E-01 | -2.67E+00 | 1.27E-01 | 4.67E-01 | 2.00E+00 | 1.20E-01 | 3.30E-01 | 1.09E+00 |
| POLR2J | 9.27E-03 | 4.83E-02 | 1.12E+00 | 1.62E-08 | 7.01E-08 | -1.01E+00 | 2.23E-01 | 9.49E-01 | 1.20E+00 | 6.05E-06 | 2.31E-05 | -1.08E+00 | 4.65E-04 | 1.66E-03 | 1.06E+00 | 1.29E-05 | 3.64E-05 | -1.11E+00 | 2.09E-01 | 7.66E-01 | 1.20E+00 | 2.08E-01 | 5.70E-01 | 1.13E+00 |
| STX17-AS1 | 9.28E-03 | 4.83E-02 | 1.02E+00 | 2.28E-04 | 9.86E-04 | -1.15E+00 | 4.75E-02 | 2.02E-01 | -1.16E+00 | 3.09E-02 | 1.18E-01 | -1.14E+00 | 2.33E-01 | 8.30E-01 | 1.34E+00 | 1.50E-01 | 4.23E-01 | 1.09E+00 | 7.42E-02 | 2.72E-01 | 1.74E+00 | 2.74E-01 | 7.52E-01 | 1.53E+00 |
| UBR7 | 9.29E-03 | 4.84E-02 | -1.18E+00 | 1.71E-01 | 7.38E-01 | 1.47E+00 | 1.52E-02 | 6.47E-02 | -1.01E+00 | 1.39E-02 | 5.32E-02 | 1.14E+00 | 8.25E-05 | 2.94E-04 | -1.70E+00 | 2.40E-03 | 6.76E-03 | -1.26E+00 | 1.32E-09 | 4.86E-09 | -1.35E+00 | 2.11E-04 | 5.81E-04 | -1.04E+00 |
| ALDH4A1 | 9.31E-03 | 4.85E-02 | -1.26E+00 | 8.64E-03 | 3.73E-02 | -1.28E+00 | 4.29E-02 | 1.82E-01 | -1.11E+00 | 5.90E-02 | 2.25E-01 | 1.02E+00 | 1.49E-08 | 5.30E-08 | -2.26E+00 | 1.12E-05 | 3.17E-05 | -1.53E+00 | 2.92E-02 | 1.07E-01 | -1.08E+00 | 1.14E-01 | 3.12E-01 | 1.05E+00 |
| FRG1BP | 9.31E-03 | 4.85E-02 | -1.04E+00 | 3.99E-03 | 1.72E-02 | -1.02E+00 | 1.36E-02 | 5.76E-02 | -1.08E+00 | 6.07E-03 | 2.32E-02 | -1.06E+00 | 1.91E-04 | 6.81E-04 | -1.29E+00 | 8.42E-04 | 2.37E-03 | -1.19E+00 | 1.37E-01 | 5.04E-01 | 1.08E+00 | 4.61E-02 | 1.27E-01 | -1.06E+00 |
| OTUD3 | 9.31E-03 | 4.85E-02 | 1.03E+00 | 1.78E-03 | 7.71E-03 | 1.03E+00 | 1.02E-03 | 4.33E-03 | -1.10E+00 | 1.25E-03 | 4.76E-03 | -1.04E+00 | 1.19E-03 | 4.23E-03 | -1.50E+00 | 2.03E-02 | 5.74E-02 | -1.10E+00 | 3.72E-03 | 1.37E-02 | -1.28E+00 | 8.46E-03 | 2.32E-02 | -1.16E+00 |
| ALDOC | 9.33E-03 | 4.86E-02 | -1.13E+00 | 4.85E-02 | 2.09E-01 | 1.15E+00 | 1.29E-01 | 5.49E-01 | 1.06E+00 | 3.92E-02 | 1.50E-01 | -1.05E+00 | 7.39E-04 | 2.64E-03 | -1.68E+00 | 1.65E-01 | 4.66E-01 | 1.19E+00 | 1.13E-01 | 4.14E-01 | -1.04E+00 | 1.71E-01 | 4.71E-01 | 1.63E+00 |
| AC009118.3 | 9.33E-03 | 4.86E-02 | -1.37E+00 | 6.13E-02 | 2.65E-01 | 1.01E+00 | 1.05E-02 | 4.45E-02 | -1.53E+00 | 9.68E-02 | 3.70E-01 | 1.00E+00 | 4.04E-05 | 1.44E-04 | -2.31E+00 | 1.20E-04 | 3.38E-04 | -2.31E+00 | 2.60E-01 | 9.54E-01 | 1.22E+00 | 3.35E-01 | 9.20E-01 | 1.22E+00 |
| ADAP2 | 9.34E-03 | 4.87E-02 | -2.31E+00 | 1.00E-02 | 4.33E-02 | -2.29E+00 | 9.47E-02 | 4.03E-01 | -1.24E+00 | 1.73E-01 | 6.59E-01 | -1.09E+00 | 1.95E-01 | 6.95E-01 | 1.24E+00 | 3.20E-01 | 9.02E-01 | 1.07E+00 | 1.45E-01 | 5.31E-01 | -1.03E+00 | 2.21E-01 | 6.06E-01 | -1.03E+00 |
| NT5DC3 | 9.34E-03 | 4.87E-02 | 1.76E+00 | 1.94E-01 | 8.40E-01 | 1.53E+00 | 6.85E-05 | 2.91E-04 | 1.88E+00 | 2.42E-01 | 9.25E-01 | 1.51E+00 | 2.53E-01 | 9.01E-01 | 1.35E+00 | 2.91E-01 | 8.20E-01 | 1.21E+00 | 1.78E-02 | 6.53E-02 | -1.37E+00 | 1.37E-01 | 3.76E-01 | -1.01E+00 |
| EVI5L | 9.35E-03 | 4.87E-02 | -1.30E+00 | 6.30E-02 | 2.72E-01 | 1.06E+00 | 2.13E-01 | 9.07E-01 | 1.08E+00 | 2.51E-01 | 9.57E-01 | 1.21E+00 | 4.08E-02 | 1.45E-01 | -1.32E+00 | 3.53E-01 | 9.96E-01 | 1.48E+00 | 4.50E-02 | 1.65E-01 | -1.01E+00 | 1.50E-01 | 4.12E-01 | 1.11E+00 |
| KCNA3 | 9.36E-03 | 4.88E-02 | 2.29E+00 | 1.19E-01 | 5.12E-01 | 1.57E+00 | 1.19E-01 | 5.04E-01 | 1.37E+00 | 9.89E-02 | 3.78E-01 | -1.14E+00 | 8.93E-03 | 3.18E-02 | 3.82E+00 | 3.04E-02 | 8.56E-02 | 3.09E+00 | 1.44E-02 | 5.27E-02 | 3.25E+00 | 1.89E-01 | 5.18E-01 | 1.92E+00 |
| CALN1 | 9.38E-03 | 4.89E-02 | 1.09E+00 | 8.47E-02 | 3.66E-01 | 1.36E+00 | 7.39E-02 | 3.14E-01 | 1.23E+00 | 1.61E-01 | 6.15E-01 | 1.45E+00 | 1.72E-01 | 6.15E-01 | 1.89E+00 | 3.31E-01 | 9.34E-01 | 1.45E+00 | 2.38E-01 | 8.72E-01 | 1.13E+00 | 2.99E-01 | 8.22E-01 | 1.59E+00 |
| ZNF846 | 9.39E-03 | 4.89E-02 | -1.12E+00 | 2.85E-03 | 1.23E-02 | -1.23E+00 | 1.52E-01 | 6.47E-01 | 1.04E+00 | 1.50E-02 | 5.71E-02 | -1.18E+00 | 2.32E-02 | 8.28E-02 | -1.02E+00 | 1.44E-03 | 4.07E-03 | -1.38E+00 | 1.80E-03 | 6.61E-03 | -1.24E+00 | 1.20E-02 | 3.30E-02 | -1.23E+00 |
| ZNF337-AS1 | 9.39E-03 | 4.89E-02 | -1.21E+00 | 1.12E-02 | 4.85E-02 | -1.09E+00 | 9.17E-05 | 3.90E-04 | -1.51E+00 | 5.09E-04 | 1.94E-03 | -1.23E+00 | 5.17E-03 | 1.84E-02 | -1.76E+00 | 2.61E-02 | 7.37E-02 | -1.34E+00 | 6.88E-04 | 2.53E-03 | -1.29E+00 | 6.17E-04 | 1.69E-03 | -1.25E+00 |
| RASSF7 | 9.43E-03 | 4.91E-02 | -1.04E+00 | 3.75E-02 | 1.62E-01 | 1.15E+00 | 6.80E-02 | 2.89E-01 | 1.05E+00 | 1.51E-01 | 5.76E-01 | 1.29E+00 | 1.59E-03 | 5.66E-03 | -1.49E+00 | 1.19E-02 | 3.35E-02 | -1.21E+00 | 5.20E-04 | 1.91E-03 | -1.17E+00 | 2.45E-02 | 6.74E-02 | 1.05E+00 |
| LSM3 | 9.43E-03 | 4.92E-02 | 1.17E+00 | 4.19E-08 | 1.81E-07 | 1.07E+00 | 1.53E-03 | 6.51E-03 | 1.49E+00 | 2.00E-01 | 7.64E-01 | 1.23E+00 | 1.91E-01 | 6.81E-01 | 1.35E+00 | 5.40E-02 | 1.52E-01 | 1.15E+00 | 5.77E-04 | 2.12E-03 | 1.53E+00 | 2.48E-03 | 6.82E-03 | 1.41E+00 |
| ZNF510 | 9.44E-03 | 4.92E-02 | -1.10E+00 | 2.94E-02 | 1.27E-01 | 1.10E+00 | 1.78E-02 | 7.54E-02 | -1.03E+00 | 4.16E-02 | 1.59E-01 | 1.11E+00 | 1.59E-02 | 5.66E-02 | -1.07E+00 | 2.58E-02 | 7.27E-02 | -1.10E+00 | 8.57E-02 | 3.15E-01 | 1.12E+00 | 1.05E-01 | 2.88E-01 | 1.11E+00 |
| ZDHHC23 | 9.44E-03 | 4.92E-02 | -1.13E+00 | 1.20E-02 | 5.18E-02 | -1.04E+00 | 4.63E-05 | 1.97E-04 | -1.46E+00 | 3.65E-04 | 1.40E-03 | -1.16E+00 | 2.85E-02 | 1.02E-01 | -1.40E+00 | 9.33E-02 | 2.63E-01 | -1.06E+00 | 1.76E-02 | 6.44E-02 | -1.30E+00 | 2.90E-01 | 7.95E-01 | 1.19E+00 |
| AC083880.1 | 9.46E-03 | 4.93E-02 | 2.13E+00 | 1.97E-01 | 8.50E-01 | 1.35E+00 | 1.20E-01 | 5.10E-01 | 1.36E+00 | 1.47E-01 | 5.63E-01 | 1.09E+00 | 6.64E-03 | 2.37E-02 | 2.91E+00 | 2.79E-01 | 7.87E-01 | 1.35E+00 | 1.65E-01 | 6.05E-01 | 1.33E+00 | 3.57E-01 | 9.80E-01 | 1.26E+00 |
| PRDM4 | 9.50E-03 | 4.95E-02 | -1.23E+00 | 8.70E-03 | 3.76E-02 | -1.11E+00 | 5.48E-03 | 2.33E-02 | -1.26E+00 | 2.82E-02 | 1.08E-01 | 1.07E+00 | 5.15E-05 | 1.84E-04 | -1.45E+00 | 4.80E-04 | 1.35E-03 | -1.16E+00 | 5.98E-07 | 2.19E-06 | -1.34E+00 | 1.63E-06 | 4.48E-06 | -1.47E+00 |
| NEK5 | 9.50E-03 | 4.95E-02 | -1.26E+00 | 9.22E-02 | 3.98E-01 | 1.15E+00 | 1.21E-02 | 5.14E-02 | -1.43E+00 | 5.10E-02 | 1.95E-01 | -1.09E+00 | 3.08E-02 | 1.10E-01 | -1.11E+00 | 2.59E-01 | 7.29E-01 | 1.28E+00 | 2.35E-01 | 8.64E-01 | 1.06E+00 | 1.29E-01 | 3.56E-01 | 1.40E+00 |
| UNC119 | 9.50E-03 | 4.95E-02 | -1.05E+00 | 2.69E-02 | 1.16E-01 | 1.21E+00 | 8.67E-03 | 3.68E-02 | -1.05E+00 | 1.23E-01 | 4.69E-01 | 1.37E+00 | 1.80E-03 | 6.43E-03 | -1.10E+00 | 2.60E-01 | 7.34E-01 | 1.30E+00 | 2.16E-01 | 7.91E-01 | 1.27E+00 | 4.53E-02 | 1.25E-01 | 1.47E+00 |
| TRMT10C | 9.51E-03 | 4.96E-02 | 1.13E+00 | 2.66E-08 | 1.15E-07 | -1.03E+00 | 1.28E-01 | 5.43E-01 | 1.02E+00 | 1.20E-01 | 4.60E-01 | 1.14E+00 | 1.54E-01 | 5.48E-01 | 1.26E+00 | 1.16E-01 | 3.26E-01 | 1.27E+00 | 1.69E-06 | 6.20E-06 | -1.22E+00 | 8.99E-02 | 2.47E-01 | 1.08E+00 |
| CCDC173 | 9.52E-03 | 4.96E-02 | -1.27E+00 | 6.59E-03 | 2.85E-02 | -1.27E+00 | 1.81E-01 | 7.71E-01 | 1.19E+00 | 2.30E-01 | 8.80E-01 | 1.04E+00 | 1.39E-02 | 4.94E-02 | -1.22E+00 | 4.01E-02 | 1.13E-01 | 1.00E+00 | 4.27E-03 | 1.57E-02 | -1.29E+00 | 7.53E-03 | 2.07E-02 | -1.33E+00 |
| AC104248.1 | 9.52E-03 | 4.96E-02 | -1.04E+00 | 1.03E-04 | 4.44E-04 | -1.70E+00 | 7.43E-02 | 3.16E-01 | 1.58E+00 | 1.28E-01 | 4.89E-01 | 1.52E+00 | 2.54E-01 | 9.06E-01 | 1.33E+00 | 6.67E-02 | 1.88E-01 | -1.55E+00 | 1.42E-01 | 5.23E-01 | 1.25E+00 | 2.36E-01 | 6.49E-01 | -1.37E+00 |
| KDM6B | 9.52E-03 | 4.96E-02 | 1.06E+00 | 7.49E-13 | 3.24E-12 | -1.45E+00 | 2.74E-02 | 1.16E-01 | 1.11E+00 | 3.10E-10 | 1.18E-09 | -1.48E+00 | 3.44E-05 | 1.23E-04 | -1.26E+00 | 2.30E-07 | 6.48E-07 | -1.50E+00 | 6.14E-11 | 2.25E-10 | -1.78E+00 | 6.38E-12 | 1.75E-11 | -2.21E+00 |
| PRSS23 | 9.53E-03 | 4.97E-02 | -1.45E+00 | 2.31E-01 | 1.00E+00 | 1.65E+00 | 1.36E-01 | 5.78E-01 | -1.29E+00 | 1.88E-01 | 7.18E-01 | 1.48E+00 | 1.83E-01 | 6.52E-01 | 1.75E+00 | 1.06E-01 | 2.99E-01 | 1.91E+00 | 2.66E-01 | 9.76E-01 | 1.35E+00 | 1.93E-01 | 5.31E-01 | 1.10E+00 |
| TCAF2 | 9.54E-03 | 4.97E-02 | -1.06E+00 | 2.62E-02 | 1.13E-01 | 1.15E+00 | 4.17E-02 | 1.77E-01 | -1.24E+00 | 3.37E-02 | 1.29E-01 | -1.10E+00 | 4.11E-02 | 1.46E-01 | -1.22E+00 | 3.10E-02 | 8.73E-02 | -1.40E+00 | 2.10E-02 | 7.72E-02 | -1.66E+00 | 3.76E-02 | 1.03E-01 | -1.56E+00 |
| MAST4-AS1 | 9.56E-03 | 4.98E-02 | -1.31E+00 | 2.33E-03 | 1.01E-02 | -1.46E+00 | 1.97E-01 | 8.37E-01 | 1.11E+00 | 2.46E-01 | 9.41E-01 | 1.27E+00 | 1.84E-02 | 6.57E-02 | -1.34E+00 | 2.27E-01 | 6.39E-01 | 1.17E+00 | 1.61E-01 | 5.91E-01 | 1.05E+00 | 5.57E-03 | 1.53E-02 | -1.38E+00 |
| TNF | 9.57E-03 | 4.99E-02 | 2.14E+00 | 9.73E-02 | 4.20E-01 | -1.19E+00 | 3.82E-02 | 1.62E-01 | 1.75E+00 | 2.36E-01 | 9.02E-01 | -1.01E+00 | 2.68E-01 | 9.56E-01 | -1.06E+00 | 2.17E-01 | 6.13E-01 | -1.73E+00 | 1.96E-01 | 7.20E-01 | 2.63E+00 | 1.46E-01 | 4.02E-01 | 1.54E+00 |
| PIGB | 9.58E-03 | 4.99E-02 | -1.17E+00 | 7.65E-02 | 3.30E-01 | 1.21E+00 | 3.43E-04 | 1.46E-03 | -1.40E+00 | 1.18E-03 | 4.50E-03 | -1.16E+00 | 2.61E-02 | 9.31E-02 | -1.12E+00 | 2.27E-01 | 6.40E-01 | 1.24E+00 | 2.03E-02 | 7.44E-02 | -1.10E+00 | 3.28E-01 | 9.00E-01 | 1.36E+00 |
| CLYBL | 9.58E-03 | 4.99E-02 | -1.22E+00 | 8.90E-03 | 3.85E-02 | -1.11E+00 | 1.17E-01 | 4.97E-01 | 1.40E+00 | 1.38E-01 | 5.27E-01 | 1.62E+00 | 1.44E-01 | 5.14E-01 | 1.09E+00 | 2.51E-01 | 7.06E-01 | 1.63E+00 | 1.50E-02 | 5.52E-02 | -1.04E+00 | 2.64E-03 | 7.26E-03 | -1.23E+00 |
| RSRC1 | 9.59E-03 | 5.00E-02 | 1.14E+00 | 1.43E-04 | 6.19E-04 | 1.13E+00 | 1.71E-02 | 7.28E-02 | 1.01E+00 | 1.01E-03 | 3.88E-03 | 1.07E+00 | 1.46E-05 | 5.20E-05 | -1.15E+00 | 5.36E-04 | 1.51E-03 | 1.00E+00 | 1.29E-02 | 4.75E-02 | -1.03E+00 | 4.34E-02 | 1.19E-01 | 1.06E+00 |
| C1orf112 | 9.60E-03 | 5.00E-02 | -1.41E+00 | 2.69E-02 | 1.16E-01 | -1.20E+00 | 1.66E-02 | 7.06E-02 | 1.99E+00 | 2.36E-02 | 9.02E-02 | 2.19E+00 | 1.33E-01 | 4.75E-01 | -1.12E+00 | 2.84E-02 | 8.01E-02 | -1.88E+00 | 1.27E-01 | 4.65E-01 | 1.17E+00 | 2.51E-01 | 6.89E-01 | 1.23E+00 |
| AL162274.1 | 9.61E-03 | 5.01E-02 | 2.28E+00 | 2.30E-01 | 9.96E-01 | 1.47E+00 | 2.07E-01 | 8.77E-01 | 1.24E+00 | 1.70E-01 | 6.49E-01 | 1.10E+00 | 1.59E-01 | 5.66E-01 | 1.47E+00 | 3.32E-01 | 9.35E-01 | 1.63E+00 | 7.51E-02 | 2.76E-01 | 1.82E+00 | 3.47E-01 | 9.53E-01 | 1.50E+00 |
| SEPT7-AS1 | 9.62E-03 | 5.01E-02 | -1.30E+00 | 1.18E-01 | 5.12E-01 | 1.16E+00 | 3.07E-04 | 1.30E-03 | -1.87E+00 | 1.71E-04 | 6.52E-04 | -1.98E+00 | 1.27E-01 | 4.54E-01 | -1.12E+00 | 3.55E-01 | 1.00E+00 | 1.27E+00 | 2.52E-01 | 9.24E-01 | 1.18E+00 | 3.15E-01 | 8.66E-01 | -1.01E+00 |
| HOXB2 | 9.62E-03 | 5.02E-02 | 1.14E+00 | 8.45E-04 | 3.65E-03 | 1.06E+00 | 5.65E-02 | 2.40E-01 | 1.18E+00 | 8.61E-03 | 3.29E-02 | 1.05E+00 | 7.00E-02 | 2.50E-01 | 1.32E+00 | 6.20E-02 | 1.75E-01 | 1.17E+00 | 7.43E-03 | 2.73E-02 | -1.01E+00 | 1.44E-02 | 3.95E-02 | 1.05E+00 |
| AC011447.7 | 9.62E-03 | 5.02E-02 | -1.60E+00 | 1.20E-02 | 5.17E-02 | -1.53E+00 | 2.55E-03 | 1.09E-02 | -1.29E+00 | 4.79E-03 | 1.83E-02 | -1.11E+00 | 1.65E-01 | 5.87E-01 | -1.23E+00 | 2.63E-01 | 7.42E-01 | 1.08E+00 | 2.44E-01 | 8.96E-01 | 1.20E+00 | 2.84E-02 | 7.79E-02 | 1.80E+00 |
| SPINK4 | 9.63E-03 | 5.02E-02 | -1.36E+00 | 7.04E-03 | 3.04E-02 | -1.49E+00 | 2.04E-03 | 8.68E-03 | -1.59E+00 | 1.66E-04 | 6.33E-04 | -2.14E+00 | 1.18E-01 | 4.20E-01 | 1.04E+00 | 4.35E-03 | 1.23E-02 | -1.27E+00 | 1.27E-01 | 4.66E-01 | 1.54E+00 | 1.82E-02 | 5.00E-02 | 2.44E+00 |
| RC3H1-IT1 | 9.65E-03 | 5.03E-02 | -1.26E+00 | 6.86E-03 | 2.97E-02 | -1.23E+00 | 1.35E-01 | 5.72E-01 | 1.12E+00 | 9.89E-03 | 3.78E-02 | -1.18E+00 | 1.88E-01 | 6.69E-01 | 1.11E+00 | 6.53E-02 | 1.84E-01 | -1.31E+00 | 2.47E-01 | 9.08E-01 | 1.16E+00 | 6.19E-02 | 1.70E-01 | -1.35E+00 |
| TOLLIP-AS1 | 9.66E-03 | 5.03E-02 | -1.38E+00 | 1.21E-02 | 5.22E-02 | -1.26E+00 | 1.09E-01 | 4.64E-01 | -1.02E+00 | 6.77E-03 | 2.58E-02 | -1.50E+00 | 1.39E-03 | 4.97E-03 | -1.38E+00 | 5.39E-02 | 1.52E-01 | -1.01E+00 | 2.38E-01 | 8.73E-01 | 1.28E+00 | 2.84E-01 | 7.81E-01 | 1.31E+00 |
| FBXO42 | 9.67E-03 | 5.04E-02 | 1.08E+00 | 4.64E-05 | 2.01E-04 | -1.06E+00 | 1.91E-01 | 8.11E-01 | 1.11E+00 | 4.45E-04 | 1.70E-03 | -1.21E+00 | 7.29E-02 | 2.60E-01 | 1.21E+00 | 3.26E-03 | 9.20E-03 | -1.16E+00 | 6.64E-06 | 2.44E-05 | -1.58E+00 | 1.83E-04 | 5.03E-04 | -1.41E+00 |
| AL139424.3 | 9.68E-03 | 5.05E-02 | -1.22E+00 | 1.40E-02 | 6.03E-02 | -1.05E+00 | 1.36E-01 | 5.76E-01 | 1.00E+00 | 2.00E-01 | 7.63E-01 | 1.15E+00 | 6.13E-02 | 2.19E-01 | -1.38E+00 | 2.65E-01 | 7.47E-01 | 1.17E+00 | 2.57E-02 | 9.42E-02 | -1.34E+00 | 1.61E-01 | 4.42E-01 | -1.04E+00 |
| RPL36A | 9.68E-03 | 5.05E-02 | 1.69E+00 | 3.70E-04 | 1.60E-03 | 2.13E+00 | 1.04E-01 | 4.43E-01 | 1.32E+00 | 1.58E-03 | 6.05E-03 | 1.98E+00 | 2.32E-01 | 8.26E-01 | 1.62E+00 | 1.44E-02 | 4.07E-02 | 2.16E+00 | 9.52E-03 | 3.49E-02 | 1.74E+00 | 4.43E-05 | 1.22E-04 | 2.31E+00 |
| PTPRJ | 9.71E-03 | 5.06E-02 | -1.70E+00 | 2.07E-02 | 8.93E-02 | -1.41E+00 | 2.09E-01 | 8.89E-01 | -1.01E+00 | 1.32E-01 | 5.04E-01 | -1.06E+00 | 1.39E-01 | 4.97E-01 | 1.59E+00 | 3.55E-01 | 1.00E+00 | -1.00E+00 | 1.23E-01 | 4.52E-01 | 2.88E+00 | 2.25E-01 | 6.18E-01 | 1.62E+00 |
| CDPG1 | 9.72E-03 | 5.07E-02 | -1.02E+00 | 7.09E-02 | 3.06E-01 | 1.29E+00 | 1.99E-05 | 8.45E-05 | -1.24E+00 | 1.84E-03 | 7.04E-03 | 1.06E+00 | 3.71E-05 | 1.32E-04 | -1.33E+00 | 3.13E-02 | 8.82E-02 | 1.13E+00 | 1.45E-08 | 5.33E-08 | -1.36E+00 | 1.41E-03 | 3.88E-03 | -1.01E+00 |
| AL583805.1 | 9.72E-03 | 5.07E-02 | 2.32E+00 | 3.53E-02 | 1.53E-01 | 2.24E+00 | 1.63E-01 | 6.93E-01 | 1.27E+00 | 2.43E-01 | 9.29E-01 | 1.39E+00 | 2.16E-01 | 7.71E-01 | 1.61E+00 | 3.55E-01 | 1.00E+00 | 1.36E+00 | 1.61E-01 | 5.92E-01 | 1.47E+00 | 1.80E-01 | 4.94E-01 | 1.54E+00 |
| OPHN1 | 9.72E-03 | 5.07E-02 | -1.25E+00 | 7.24E-02 | 3.13E-01 | 1.20E+00 | 3.72E-02 | 1.58E-01 | -1.05E+00 | 2.99E-02 | 1.14E-01 | -1.04E+00 | 1.74E-01 | 6.21E-01 | -1.05E+00 | 1.63E-01 | 4.59E-01 | -1.14E+00 | 1.23E-01 | 4.52E-01 | 1.29E+00 | 3.73E-02 | 1.02E-01 | -1.17E+00 |
| SNX33 | 9.72E-03 | 5.07E-02 | -1.39E+00 | 7.08E-02 | 3.06E-01 | 1.02E+00 | 8.85E-07 | 3.76E-06 | -2.20E+00 | 6.26E-05 | 2.39E-04 | -1.49E+00 | 1.72E-03 | 6.14E-03 | -1.73E+00 | 7.02E-02 | 1.98E-01 | -1.06E+00 | 6.56E-02 | 2.41E-01 | -1.19E+00 | 3.46E-01 | 9.51E-01 | 1.26E+00 |
| AL162456.1 | 9.73E-03 | 5.07E-02 | -1.19E+00 | 6.65E-02 | 2.87E-01 | 1.05E+00 | 1.87E-01 | 7.93E-01 | 1.10E+00 | 6.67E-02 | 2.55E-01 | 1.72E+00 | 2.69E-01 | 9.58E-01 | 1.27E+00 | 2.15E-01 | 6.06E-01 | -1.08E+00 | 4.26E-02 | 1.56E-01 | -1.36E+00 | 2.30E-01 | 6.32E-01 | -1.08E+00 |
| LAYN | 9.74E-03 | 5.08E-02 | 1.92E+00 | 1.98E-04 | 8.57E-04 | 2.54E+00 | 4.47E-02 | 1.90E-01 | 1.63E+00 | 1.90E-01 | 7.25E-01 | 1.32E+00 | 2.32E-01 | 8.26E-01 | -1.27E+00 | 2.54E-01 | 7.15E-01 | -1.58E+00 | 1.61E-01 | 5.90E-01 | 1.62E+00 | 1.97E-01 | 5.40E-01 | 1.78E+00 |
| OSM | 9.74E-03 | 5.08E-02 | 2.29E+00 | 2.57E-04 | 1.11E-03 | -1.23E+00 | 1.41E-01 | 5.97E-01 | 1.57E+00 | 3.24E-03 | 1.24E-02 | -1.21E+00 | 2.08E-01 | 7.43E-01 | 1.98E+00 | 2.49E-02 | 7.03E-02 | -1.26E+00 | 7.92E-03 | 2.91E-02 | -1.12E+00 | 6.66E-03 | 1.83E-02 | -1.25E+00 |
| PPP2R2D | 9.76E-03 | 5.08E-02 | 1.06E+00 | 2.75E-05 | 1.19E-04 | -1.10E+00 | 4.22E-03 | 1.79E-02 | -1.05E+00 | 1.83E-07 | 7.00E-07 | -1.21E+00 | 9.77E-05 | 3.48E-04 | -1.06E+00 | 2.61E-04 | 7.37E-04 | 1.01E+00 | 5.21E-03 | 1.91E-02 | 1.03E+00 | 6.98E-05 | 1.92E-04 | -1.18E+00 |
| SLC19A2 | 9.77E-03 | 5.09E-02 | -1.29E+00 | 1.02E-03 | 4.40E-03 | -1.61E+00 | 1.08E-03 | 4.59E-03 | -1.34E+00 | 5.93E-04 | 2.27E-03 | -1.20E+00 | 2.76E-01 | 9.85E-01 | 1.40E+00 | 1.34E-01 | 3.79E-01 | 1.11E+00 | 1.66E-01 | 6.08E-01 | 1.47E+00 | 6.16E-02 | 1.69E-01 | -1.07E+00 |
| VPS50 | 9.77E-03 | 5.09E-02 | -1.09E+00 | 2.51E-02 | 1.09E-01 | 1.07E+00 | 2.95E-03 | 1.25E-02 | -1.38E+00 | 4.30E-02 | 1.64E-01 | -1.03E+00 | 4.07E-03 | 1.45E-02 | -1.22E+00 | 1.21E-01 | 3.40E-01 | 1.16E+00 | 2.90E-02 | 1.06E-01 | 1.02E+00 | 1.50E-01 | 4.12E-01 | 1.05E+00 |
| NT5C2 | 9.78E-03 | 5.09E-02 | -1.05E+00 | 2.27E-03 | 9.80E-03 | -1.07E+00 | 1.53E-03 | 6.51E-03 | -1.28E+00 | 1.76E-04 | 6.74E-04 | -1.30E+00 | 1.08E-02 | 3.84E-02 | -1.09E+00 | 3.49E-04 | 9.83E-04 | -1.32E+00 | 1.38E-01 | 5.06E-01 | 1.05E+00 | 1.71E-02 | 4.70E-02 | -1.18E+00 |
| RIPK2 | 9.78E-03 | 5.10E-02 | 1.22E+00 | 1.38E-22 | 5.98E-22 | -1.42E+00 | 7.81E-02 | 3.32E-01 | 1.16E+00 | 1.55E-12 | 5.93E-12 | -1.41E+00 | 1.70E-01 | 6.05E-01 | 1.39E+00 | 6.25E-05 | 1.76E-04 | -1.11E+00 | 3.13E-07 | 1.15E-06 | -1.24E+00 | 2.65E-12 | 7.28E-12 | -1.63E+00 |
| RGP1 | 9.78E-03 | 5.10E-02 | 1.04E+00 | 1.69E-01 | 7.29E-01 | 1.36E+00 | 2.99E-02 | 1.27E-01 | -1.06E+00 | 4.39E-02 | 1.68E-01 | 1.08E+00 | 4.89E-03 | 1.74E-02 | -1.24E+00 | 7.32E-03 | 2.06E-02 | -1.20E+00 | 5.68E-03 | 2.09E-02 | -1.22E+00 | 8.23E-02 | 2.26E-01 | -1.04E+00 |
| AL163051.1 | 9.81E-03 | 5.11E-02 | 2.39E+00 | 9.38E-02 | 4.05E-01 | 1.96E+00 | 4.36E-03 | 1.85E-02 | 1.91E+00 | 1.36E-01 | 5.19E-01 | 1.07E+00 | 1.67E-01 | 5.94E-01 | 1.22E+00 | 3.49E-01 | 9.83E-01 | 1.41E+00 | 1.79E-01 | 6.58E-01 | 1.24E+00 | 1.13E-01 | 3.12E-01 | 1.06E+00 |
| AC127070.2 | 9.82E-03 | 5.12E-02 | 2.18E+00 | 1.93E-02 | 8.35E-02 | 2.17E+00 | 1.73E-01 | 7.33E-01 | 1.29E+00 | 1.77E-01 | 6.77E-01 | 1.09E+00 | 9.44E-02 | 3.37E-01 | -1.20E+00 | 1.56E-01 | 4.40E-01 | -1.08E+00 | 2.25E-01 | 8.25E-01 | 1.22E+00 | 3.25E-01 | 8.92E-01 | 1.09E+00 |
| NANP | 9.86E-03 | 5.14E-02 | -1.18E+00 | 1.32E-02 | 5.70E-02 | -1.08E+00 | 3.70E-03 | 1.57E-02 | -1.28E+00 | 3.73E-03 | 1.43E-02 | -1.17E+00 | 5.14E-03 | 1.83E-02 | -1.47E+00 | 1.67E-02 | 4.70E-02 | -1.26E+00 | 8.99E-10 | 3.30E-09 | -1.58E+00 | 8.90E-06 | 2.44E-05 | -1.22E+00 |
| ATPIF1 | 9.86E-03 | 5.14E-02 | 1.08E+00 | 5.84E-07 | 2.52E-06 | 1.04E+00 | 5.78E-02 | 2.46E-01 | 1.20E+00 | 1.72E-03 | 6.57E-03 | -1.06E+00 | 9.40E-03 | 3.35E-02 | 1.02E+00 | 2.70E-02 | 7.62E-02 | 1.04E+00 | 2.69E-01 | 9.87E-01 | 1.09E+00 | 1.03E-01 | 2.84E-01 | -1.08E+00 |
| TCF3 | 9.89E-03 | 5.15E-02 | 1.04E+00 | 3.38E-02 | 1.46E-01 | 1.15E+00 | 5.16E-02 | 2.19E-01 | 1.11E+00 | 2.19E-01 | 8.38E-01 | 1.43E+00 | 2.46E-01 | 8.78E-01 | 1.36E+00 | 1.31E-01 | 3.70E-01 | 1.23E+00 | 1.28E-02 | 4.71E-02 | -1.02E+00 | 3.12E-01 | 8.57E-01 | 1.27E+00 |
| RPL17P34 | 9.89E-03 | 5.16E-02 | 1.68E+00 | 5.73E-02 | 2.48E-01 | 1.26E+00 | 2.56E-02 | 1.09E-01 | 1.49E+00 | 3.49E-03 | 1.33E-02 | 1.07E+00 | 2.35E-01 | 8.39E-01 | 1.34E+00 | 7.00E-03 | 1.97E-02 | 1.03E+00 | 1.04E-01 | 3.82E-01 | 1.45E+00 | 7.92E-02 | 2.18E-01 | 1.06E+00 |
| PRKCA | 9.91E-03 | 5.16E-02 | -1.08E+00 | 9.35E-03 | 4.04E-02 | 1.07E+00 | 1.30E-02 | 5.52E-02 | -1.10E+00 | 2.56E-02 | 9.79E-02 | 1.06E+00 | 2.72E-02 | 9.71E-02 | -1.10E+00 | 4.02E-03 | 1.13E-02 | -1.35E+00 | 5.42E-02 | 1.99E-01 | 1.24E+00 | 1.80E-01 | 4.95E-01 | 1.31E+00 |
| AC093677.2 | 9.91E-03 | 5.16E-02 | -1.18E+00 | 1.32E-02 | 5.70E-02 | -1.09E+00 | 1.65E-01 | 7.01E-01 | 1.47E+00 | 2.13E-01 | 8.14E-01 | 1.29E+00 | 6.99E-02 | 2.49E-01 | -1.01E+00 | 2.28E-01 | 6.42E-01 | 1.13E+00 | 2.45E-01 | 8.99E-01 | 1.32E+00 | 3.99E-02 | 1.10E-01 | -1.29E+00 |
| GSDMB | 9.92E-03 | 5.17E-02 | -1.35E+00 | 2.30E-01 | 9.95E-01 | 1.31E+00 | 2.38E-02 | 1.01E-01 | -1.30E+00 | 8.45E-02 | 3.23E-01 | -1.02E+00 | 2.29E-01 | 8.15E-01 | 1.26E+00 | 2.50E-02 | 7.04E-02 | 2.53E+00 | 8.51E-02 | 3.12E-01 | -1.03E+00 | 1.16E-01 | 3.19E-01 | -1.01E+00 |
| LINC00309 | 9.92E-03 | 5.17E-02 | 2.07E+00 | 2.15E-01 | 9.29E-01 | 1.28E+00 | 1.62E-01 | 6.88E-01 | 1.01E+00 | 1.67E-02 | 6.37E-02 | -1.59E+00 | 1.72E-01 | 6.15E-01 | 1.45E+00 | 3.29E-01 | 9.27E-01 | 1.06E+00 | 2.14E-01 | 7.85E-01 | -1.12E+00 | 1.77E-01 | 4.87E-01 | -1.35E+00 |
| CLEC7A | 9.93E-03 | 5.18E-02 | 1.05E+00 | 4.59E-02 | 1.98E-01 | 1.34E+00 | 8.49E-02 | 3.61E-01 | -1.32E+00 | 1.03E-02 | 3.93E-02 | -1.55E+00 | 2.69E-01 | 9.61E-01 | 1.35E+00 | 2.15E-01 | 6.07E-01 | 1.04E+00 | 5.56E-02 | 2.04E-01 | -1.77E+00 | 7.18E-02 | 1.97E-01 | -1.85E+00 |
| URB2 | 9.94E-03 | 5.18E-02 | -1.21E+00 | 1.19E-02 | 5.15E-02 | -1.14E+00 | 2.52E-05 | 1.07E-04 | -1.62E+00 | 6.22E-05 | 2.38E-04 | -1.40E+00 | 1.46E-02 | 5.20E-02 | -1.65E+00 | 1.96E-02 | 5.52E-02 | -1.45E+00 | 2.72E-02 | 1.00E-01 | -1.09E+00 | 6.70E-02 | 1.84E-01 | -1.06E+00 |
| COQ7 | 9.95E-03 | 5.18E-02 | 1.08E+00 | 1.38E-02 | 5.97E-02 | 1.27E+00 | 9.85E-02 | 4.19E-01 | 1.12E+00 | 2.13E-01 | 8.14E-01 | 1.46E+00 | 7.12E-04 | 2.54E-03 | -1.07E+00 | 6.92E-02 | 1.95E-01 | 1.28E+00 | 1.66E-03 | 6.07E-03 | -1.17E+00 | 5.10E-02 | 1.40E-01 | 1.06E+00 |
| SENP3 | 9.96E-03 | 5.19E-02 | -1.34E+00 | 1.28E-02 | 5.51E-02 | -1.24E+00 | 1.97E-03 | 8.37E-03 | -1.38E+00 | 1.19E-03 | 4.55E-03 | -1.26E+00 | 1.40E-01 | 4.98E-01 | -1.01E+00 | 2.54E-01 | 7.16E-01 | 1.19E+00 | 3.04E-04 | 1.11E-03 | -1.43E+00 | 1.66E-02 | 4.56E-02 | -1.14E+00 |
| RYR3 | 9.97E-03 | 5.19E-02 | -1.83E+00 | 5.11E-02 | 2.21E-01 | -1.19E+00 | 1.69E-01 | 7.16E-01 | 1.10E+00 | 2.47E-01 | 9.44E-01 | 1.47E+00 | 1.14E-03 | 4.06E-03 | -1.09E+00 | 1.44E-01 | 4.05E-01 | 1.52E+00 | 5.38E-03 | 1.98E-02 | -1.02E+00 | 3.82E-03 | 1.05E-02 | -1.06E+00 |
| PNRC1 | 9.98E-03 | 5.20E-02 | -1.06E+00 | 1.51E-03 | 6.53E-03 | -1.10E+00 | 9.73E-05 | 4.13E-04 | -1.25E+00 | 1.01E-05 | 3.84E-05 | -1.27E+00 | 1.73E-01 | 6.17E-01 | 1.10E+00 | 2.93E-02 | 8.26E-02 | -1.01E+00 | 4.98E-07 | 1.83E-06 | -1.19E+00 | 5.94E-12 | 1.63E-11 | -1.50E+00 |
| AC105339.2 | 1.00E-02 | 5.21E-02 | -1.34E+00 | 8.84E-02 | 3.82E-01 | 1.69E+00 | 2.50E-03 | 1.06E-02 | -1.55E+00 | 1.12E-01 | 4.29E-01 | 1.15E+00 | 1.73E-01 | 6.18E-01 | 1.19E+00 | 3.16E-01 | 8.92E-01 | 1.33E+00 | 2.67E-02 | 9.82E-02 | -1.20E+00 | 3.46E-01 | 9.51E-01 | 1.22E+00 |
| AC008026.3 | 1.00E-02 | 5.22E-02 | -1.06E+00 | 2.12E-03 | 9.18E-03 | -1.10E+00 | 1.96E-01 | 8.31E-01 | 1.16E+00 | 8.09E-02 | 3.09E-01 | 1.06E+00 | 1.73E-01 | 6.17E-01 | 1.09E+00 | 2.03E-02 | 5.71E-02 | -1.32E+00 | 7.71E-03 | 2.83E-02 | -1.25E+00 | 1.29E-03 | 3.53E-03 | -1.62E+00 |
| AC040169.1 | 1.00E-02 | 5.22E-02 | -1.33E+00 | 9.37E-04 | 4.05E-03 | -1.73E+00 | 2.31E-01 | 9.81E-01 | 1.17E+00 | 1.50E-01 | 5.74E-01 | 1.12E+00 | 1.61E-01 | 5.74E-01 | 1.05E+00 | 2.90E-01 | 8.18E-01 | 1.18E+00 | 1.22E-01 | 4.46E-01 | 1.56E+00 | 2.83E-01 | 7.76E-01 | 1.28E+00 |
| CLPB | 1.00E-02 | 5.22E-02 | -1.32E+00 | 7.31E-02 | 3.16E-01 | 1.14E+00 | 1.56E-02 | 6.63E-02 | -1.01E+00 | 1.35E-04 | 5.16E-04 | -1.10E+00 | 2.93E-03 | 1.05E-02 | -1.63E+00 | 1.30E-02 | 3.67E-02 | -1.34E+00 | 3.22E-02 | 1.18E-01 | -1.04E+00 | 3.58E-01 | 9.83E-01 | 1.22E+00 |
| RPTOR | 1.00E-02 | 5.22E-02 | -1.25E+00 | 5.01E-02 | 2.16E-01 | 1.05E+00 | 7.52E-02 | 3.20E-01 | -1.10E+00 | 2.21E-01 | 8.43E-01 | 1.33E+00 | 2.06E-02 | 7.36E-02 | -1.03E+00 | 1.41E-01 | 3.98E-01 | 1.26E+00 | 1.57E-02 | 5.76E-02 | -1.08E+00 | 1.14E-01 | 3.13E-01 | 1.19E+00 |
| TRPC1 | 1.00E-02 | 5.22E-02 | -1.17E+00 | 5.90E-02 | 2.55E-01 | 1.15E+00 | 1.99E-05 | 8.47E-05 | -1.60E+00 | 6.94E-06 | 2.65E-05 | -1.59E+00 | 1.41E-02 | 5.04E-02 | -1.30E+00 | 6.59E-02 | 1.86E-01 | -1.02E+00 | 9.70E-02 | 3.56E-01 | -1.08E+00 | 1.27E-01 | 3.48E-01 | -1.00E+00 |
| PWARS | 1.00E-02 | 5.22E-02 | -1.27E+00 | 1.82E-02 | 7.85E-02 | -1.11E+00 | 2.41E-02 | 1.03E-01 | -1.30E+00 | 6.23E-02 | 2.38E-01 | -1.04E+00 | 2.09E-01 | 7.46E-01 | 1.33E+00 | 3.15E-01 | 8.87E-01 | 1.57E+00 | 5.90E-04 | 2.17E-03 | -1.45E+00 | 1.27E-02 | 3.49E-02 | -1.21E+00 |
| RPS4XP7 | 1.00E-02 | 5.23E-02 | 1.10E+00 | 2.73E-08 | 1.18E-07 | -1.53E+00 | 4.53E-02 | 1.93E-01 | 1.36E+00 | 1.20E-01 | 4.58E-01 | 1.14E+00 | 1.57E-02 | 5.59E-02 | -1.07E+00 | 8.89E-03 | 2.51E-02 | -1.13E+00 | 1.82E-01 | 6.69E-01 | 1.10E+00 | 1.47E-01 | 4.03E-01 | 1.02E+00 |
| AC105399.1 | 1.00E-02 | 5.23E-02 | -1.04E+00 | 6.82E-05 | 2.95E-04 | -1.68E+00 | 1.72E-01 | 7.32E-01 | 1.13E+00 | 3.35E-02 | 1.28E-01 | -1.00E+00 | 1.02E-01 | 3.64E-01 | 1.08E+00 | 3.12E-02 | 8.79E-02 | -1.16E+00 | 2.42E-01 | 8.87E-01 | 1.22E+00 | 5.61E-02 | 1.54E-01 | -1.14E+00 |
| CUL3 | 1.00E-02 | 5.23E-02 | 1.05E+00 | 9.29E-09 | 4.01E-08 | -1.18E+00 | 2.33E-01 | 9.90E-01 | 1.17E+00 | 1.28E-05 | 4.90E-05 | -1.16E+00 | 9.85E-02 | 3.51E-01 | 1.42E+00 | 7.79E-03 | 2.19E-02 | -1.09E+00 | 3.59E-02 | 1.32E-01 | 1.12E+00 | 2.56E-05 | 7.03E-05 | -1.20E+00 |
| AP003108.4 | 1.00E-02 | 5.23E-02 | -1.35E+00 | 5.51E-02 | 2.38E-01 | -1.05E+00 | 1.74E-03 | 7.38E-03 | -1.45E+00 | 4.86E-02 | 1.86E-01 | 1.03E+00 | 1.50E-02 | 5.34E-02 | -1.20E+00 | 2.92E-02 | 8.22E-02 | -1.19E+00 | 7.77E-02 | 2.85E-01 | -1.00E+00 | 2.22E-01 | 6.10E-01 | 1.71E+00 |
| NARFL | 1.00E-02 | 5.23E-02 | -1.11E+00 | 5.54E-02 | 2.39E-01 | 1.20E+00 | 4.07E-04 | 1.73E-03 | -1.19E+00 | 1.84E-03 | 7.02E-03 | 1.03E+00 | 2.13E-02 | 7.61E-02 | -1.08E+00 | 3.40E-01 | 9.59E-01 | 1.50E+00 | 1.76E-04 | 6.45E-04 | -1.30E+00 | 9.71E-03 | 2.67E-02 | -1.03E+00 |
| DGKG | 1.01E-02 | 5.25E-02 | -1.39E+00 | 3.51E-02 | 1.52E-01 | -1.09E+00 | 8.59E-04 | 3.65E-03 | -1.48E+00 | 1.23E-03 | 4.71E-03 | -1.21E+00 | 1.85E-10 | 6.60E-10 | -2.79E+00 | 1.59E-11 | 4.47E-11 | -2.91E+00 | 2.13E-03 | 7.82E-03 | -1.23E+00 | 1.47E-03 | 4.05E-03 | -1.73E+00 |
| AL138762.1 | 1.01E-02 | 5.27E-02 | -1.24E+00 | 1.41E-01 | 6.10E-01 | 1.24E+00 | 1.41E-02 | 5.99E-02 | -1.44E+00 | 3.47E-02 | 1.32E-01 | -1.14E+00 | 9.16E-12 | 3.27E-11 | -3.27E+00 | 2.48E-11 | 7.00E-11 | -2.88E+00 | 1.28E-01 | 4.70E-01 | 1.10E+00 | 3.42E-02 | 9.41E-02 | -1.18E+00 |
| TFPT | 1.01E-02 | 5.27E-02 | 1.04E+00 | 1.31E-02 | 5.68E-02 | 1.15E+00 | 2.18E-02 | 9.25E-02 | -1.03E+00 | 9.12E-02 | 3.48E-01 | 1.20E+00 | 9.87E-04 | 3.52E-03 | -1.10E+00 | 1.15E-02 | 3.23E-02 | 1.02E+00 | 4.17E-03 | 1.53E-02 | 1.01E+00 | 1.53E-02 | 4.20E-02 | 1.05E+00 |
| DIEXF | 1.01E-02 | 5.27E-02 | -1.09E+00 | 9.27E-02 | 4.01E-01 | 1.35E+00 | 2.19E-02 | 9.29E-02 | -1.43E+00 | 1.87E-01 | 7.16E-01 | 1.31E+00 | 2.62E-02 | 9.34E-02 | -1.11E+00 | 2.22E-01 | 6.25E-01 | 1.61E+00 | 1.34E-10 | 4.93E-10 | -1.65E+00 | 2.06E-06 | 5.67E-06 | -1.28E+00 |
| ZBTB34 | 1.01E-02 | 5.27E-02 | -1.17E+00 | 1.18E-02 | 5.08E-02 | -1.14E+00 | 3.08E-02 | 1.31E-01 | -1.16E+00 | 1.20E-02 | 4.59E-02 | -1.18E+00 | 4.27E-02 | 1.52E-01 | -1.10E+00 | 4.84E-02 | 1.37E-01 | -1.09E+00 | 1.58E-01 | 5.81E-01 | -1.02E+00 | 1.05E-02 | 2.88E-02 | -1.67E+00 |
| AC100810.1 | 1.01E-02 | 5.27E-02 | 1.13E+00 | 2.28E-02 | 9.86E-02 | 1.26E+00 | 6.76E-02 | 2.87E-01 | 1.02E+00 | 1.12E-02 | 4.27E-02 | 1.10E+00 | 1.01E-01 | 3.61E-01 | 1.28E+00 | 3.12E-01 | 8.78E-01 | 1.51E+00 | 2.45E-01 | 8.97E-01 | 1.31E+00 | 2.21E-01 | 6.08E-01 | 1.18E+00 |
| ELOB | 1.01E-02 | 5.27E-02 | 1.01E+00 | 8.53E-20 | 3.68E-19 | -1.26E+00 | 4.46E-02 | 1.90E-01 | 1.18E+00 | 4.18E-05 | 1.60E-04 | -1.16E+00 | 2.83E-03 | 1.01E-02 | -1.01E+00 | 1.33E-07 | 3.76E-07 | -1.18E+00 | 2.70E-01 | 9.91E-01 | 1.09E+00 | 2.93E-02 | 8.04E-02 | -1.12E+00 |
| NECAB1 | 1.01E-02 | 5.28E-02 | -1.03E+00 | 1.43E-02 | 6.18E-02 | 1.11E+00 | 1.88E-01 | 7.98E-01 | 1.19E+00 | 1.39E-01 | 5.30E-01 | 1.22E+00 | 1.72E-01 | 6.12E-01 | 1.73E+00 | 3.51E-02 | 9.89E-02 | 2.11E+00 | 2.04E-01 | 7.48E-01 | 1.18E+00 | 2.28E-01 | 6.26E-01 | 1.26E+00 |
| AC023355.1 | 1.01E-02 | 5.28E-02 | 2.12E+00 | 4.07E-02 | 1.76E-01 | 1.96E+00 | 2.21E-01 | 9.40E-01 | 1.19E+00 | 2.21E-01 | 8.45E-01 | 1.25E+00 | 2.58E-01 | 9.19E-01 | 1.24E+00 | 2.31E-01 | 6.51E-01 | 1.70E+00 | 5.83E-02 | 2.14E-01 | -1.25E+00 | 3.33E-01 | 9.14E-01 | 1.30E+00 |
| KCTD2 | 1.01E-02 | 5.28E-02 | -1.12E+00 | 1.03E-01 | 4.46E-01 | 1.26E+00 | 2.83E-03 | 1.20E-02 | -1.17E+00 | 8.02E-03 | 3.06E-02 | 1.02E+00 | 3.66E-04 | 1.31E-03 | -1.72E+00 | 1.38E-02 | 3.89E-02 | -1.19E+00 | 4.21E-04 | 1.54E-03 | -1.19E+00 | 7.12E-05 | 1.96E-04 | -1.31E+00 |
| FBXW4 | 1.02E-02 | 5.30E-02 | -1.05E+00 | 1.22E-02 | 5.29E-02 | 1.05E+00 | 3.43E-02 | 1.46E-01 | -1.08E+00 | 8.16E-02 | 3.12E-01 | 1.21E+00 | 1.18E-01 | 4.20E-01 | 1.19E+00 | 2.23E-01 | 6.29E-01 | 1.30E+00 | 2.15E-03 | 7.91E-03 | -1.07E+00 | 3.73E-03 | 1.02E-02 | -1.05E+00 |
| NRSN2 | 1.02E-02 | 5.30E-02 | -1.22E+00 | 1.48E-02 | 6.41E-02 | -1.11E+00 | 3.41E-03 | 1.45E-02 | -1.29E+00 | 2.26E-02 | 8.64E-02 | 1.05E+00 | 7.32E-02 | 2.61E-01 | -1.07E+00 | 8.30E-02 | 2.34E-01 | -1.11E+00 | 1.56E-01 | 5.74E-01 | 1.46E+00 | 1.11E-01 | 3.05E-01 | 1.72E+00 |
| ZDHHC14 | 1.02E-02 | 5.30E-02 | -1.13E+00 | 1.17E-01 | 5.04E-01 | 1.25E+00 | 6.17E-03 | 2.62E-02 | -1.12E+00 | 5.63E-02 | 2.15E-01 | 1.18E+00 | 3.64E-03 | 1.30E-02 | -1.13E+00 | 1.09E-01 | 3.08E-01 | 1.25E+00 | 4.65E-02 | 1.71E-01 | 1.00E+00 | 1.15E-01 | 3.16E-01 | 1.49E+00 |
| CRIP1 | 1.02E-02 | 5.30E-02 | 1.97E+00 | 1.82E-02 | 7.85E-02 | -1.06E+00 | 2.24E-01 | 9.50E-01 | 1.21E+00 | 5.53E-04 | 2.11E-03 | -1.38E+00 | 2.38E-01 | 8.47E-01 | 1.61E+00 | 2.79E-01 | 7.87E-01 | 1.30E+00 | 1.42E-01 | 5.20E-01 | 1.66E+00 | 3.34E-01 | 9.17E-01 | 1.03E+00 |
| POLR3B | 1.02E-02 | 5.30E-02 | -1.41E+00 | 1.44E-02 | 6.24E-02 | -1.32E+00 | 1.41E-04 | 6.00E-04 | -1.46E+00 | 5.89E-04 | 2.25E-03 | -1.13E+00 | 1.86E-02 | 6.64E-02 | -1.32E+00 | 2.49E-01 | 7.01E-01 | 1.32E+00 | 3.12E-06 | 1.14E-05 | -1.85E+00 | 2.26E-03 | 6.20E-03 | -1.19E+00 |
| KLHL22 | 1.02E-02 | 5.30E-02 | -1.24E+00 | 4.26E-02 | 1.84E-01 | 1.05E+00 | 1.84E-02 | 7.82E-02 | -1.30E+00 | 9.26E-02 | 3.54E-01 | 1.03E+00 | 1.11E-01 | 3.95E-01 | -1.08E+00 | 2.34E-01 | 6.59E-01 | 1.04E+00 | 3.47E-02 | 1.27E-01 | 1.04E+00 | 3.31E-03 | 9.08E-03 | -1.18E+00 |
| AC046158.4 | 1.02E-02 | 5.31E-02 | -1.25E+00 | 9.64E-02 | 4.17E-01 | 1.22E+00 | 1.72E-01 | 7.29E-01 | -1.07E+00 | 1.27E-01 | 4.86E-01 | 1.43E+00 | 7.73E-02 | 2.75E-01 | -1.21E+00 | 7.82E-02 | 2.20E-01 | -1.16E+00 | 3.02E-02 | 1.11E-01 | -1.29E+00 | 2.14E-01 | 5.86E-01 | -1.06E+00 |
| FAM213B | 1.02E-02 | 5.33E-02 | -1.20E+00 | 7.73E-02 | 3.34E-01 | 1.25E+00 | 9.26E-02 | 3.94E-01 | -1.02E+00 | 2.15E-01 | 8.20E-01 | 1.35E+00 | 9.70E-03 | 3.46E-02 | -1.06E+00 | 1.17E-01 | 3.31E-01 | 1.21E+00 | 2.49E-01 | 9.13E-01 | 1.36E+00 | 2.18E-02 | 5.99E-02 | 1.69E+00 |
| RPL3P7 | 1.02E-02 | 5.33E-02 | 1.72E+00 | 2.65E-03 | 1.15E-02 | 2.14E+00 | 7.34E-02 | 3.12E-01 | 1.50E+00 | 6.92E-03 | 2.64E-02 | 2.05E+00 | 1.47E-01 | 5.23E-01 | 1.63E+00 | 2.08E-02 | 5.88E-02 | 2.25E+00 | 8.59E-02 | 3.15E-01 | 1.65E+00 | 2.37E-03 | 6.50E-03 | 2.18E+00 |
| AC119396.1 | 1.02E-02 | 5.34E-02 | 2.06E+00 | 1.86E-01 | 8.05E-01 | 1.76E+00 | 2.04E-01 | 8.68E-01 | 1.26E+00 | 5.96E-02 | 2.28E-01 | 1.16E+00 | 2.08E-01 | 7.41E-01 | 1.32E+00 | 1.85E-01 | 5.23E-01 | 1.07E+00 | 1.70E-01 | 6.25E-01 | -1.01E+00 | 1.11E-01 | 3.04E-01 | -1.33E+00 |
| CAMSAP1 | 1.03E-02 | 5.34E-02 | -1.27E+00 | 1.86E-01 | 8.04E-01 | 1.36E+00 | 8.07E-05 | 3.43E-04 | -1.34E+00 | 8.73E-05 | 3.33E-04 | -1.21E+00 | 1.59E-02 | 5.66E-02 | -1.03E+00 | 2.60E-02 | 7.34E-02 | 1.03E+00 | 1.32E-03 | 4.83E-03 | -1.25E+00 | 2.47E-02 | 6.77E-02 | -1.07E+00 |
| SRRM2-AS1 | 1.03E-02 | 5.35E-02 | -1.39E+00 | 8.96E-04 | 3.87E-03 | -1.94E+00 | 2.03E-02 | 8.64E-02 | 1.91E+00 | 1.56E-01 | 5.95E-01 | 1.43E+00 | 2.47E-01 | 8.80E-01 | 1.30E+00 | 3.54E-01 | 9.98E-01 | 1.32E+00 | 6.89E-02 | 2.53E-01 | 1.72E+00 | 3.08E-01 | 8.46E-01 | 1.08E+00 |
| TMEM8A | 1.03E-02 | 5.36E-02 | -1.14E+00 | 2.30E-02 | 9.95E-02 | 1.10E+00 | 2.32E-01 | 9.85E-01 | 1.09E+00 | 1.61E-01 | 6.15E-01 | 1.17E+00 | 1.09E-01 | 3.89E-01 | 1.06E+00 | 1.60E-01 | 4.50E-01 | 1.15E+00 | 1.37E-01 | 5.01E-01 | 1.09E+00 | 3.62E-01 | 9.94E-01 | 1.32E+00 |
| FTH1P5 | 1.03E-02 | 5.36E-02 | 1.10E+00 | 7.85E-04 | 3.39E-03 | 1.16E+00 | 6.26E-02 | 2.66E-01 | 1.07E+00 | 2.63E-02 | 1.00E-01 | 1.17E+00 | 6.01E-03 | 2.14E-02 | -1.07E+00 | 1.06E-02 | 2.98E-02 | -1.02E+00 | 1.12E-01 | 4.10E-01 | 1.04E+00 | 3.15E-01 | 8.65E-01 | 1.17E+00 |
| SIRT5 | 1.03E-02 | 5.37E-02 | 1.01E+00 | 6.22E-02 | 2.69E-01 | 1.25E+00 | 9.90E-05 | 4.21E-04 | -1.36E+00 | 3.75E-04 | 1.43E-03 | -1.12E+00 | 5.17E-05 | 1.84E-04 | -1.55E+00 | 3.85E-03 | 1.08E-02 | -1.18E+00 | 2.76E-03 | 1.01E-02 | -1.05E+00 | 4.21E-02 | 1.16E-01 | 1.06E+00 |
| HEATR5A | 1.03E-02 | 5.37E-02 | -1.34E+00 | 1.45E-01 | 6.26E-01 | 1.78E+00 | 1.56E-05 | 6.62E-05 | -1.81E+00 | 3.51E-03 | 1.34E-02 | -1.19E+00 | 4.02E-02 | 1.44E-01 | -1.42E+00 | 2.89E-01 | 8.15E-01 | 1.30E+00 | 3.20E-05 | 1.18E-04 | -2.11E+00 | 1.86E-03 | 5.11E-03 | -1.69E+00 |
| TRIM9 | 1.03E-02 | 5.38E-02 | -1.31E+00 | 1.83E-01 | 7.92E-01 | 1.45E+00 | 2.56E-05 | 1.09E-04 | -1.44E+00 | 3.07E-02 | 1.17E-01 | 1.10E+00 | 7.31E-02 | 2.61E-01 | -1.94E+00 | 1.36E-01 | 3.84E-01 | -1.52E+00 | 1.39E-01 | 5.10E-01 | -1.35E+00 | 2.98E-01 | 8.18E-01 | 1.30E+00 |
| FBXO25 | 1.03E-02 | 5.38E-02 | -1.08E+00 | 5.47E-02 | 2.36E-01 | 1.21E+00 | 5.14E-03 | 2.19E-02 | -1.12E+00 | 2.64E-02 | 1.01E-01 | 1.09E+00 | 2.15E-03 | 7.67E-03 | -1.19E+00 | 3.41E-02 | 9.61E-02 | 1.07E+00 | 9.79E-04 | 3.59E-03 | -1.19E+00 | 9.62E-02 | 2.64E-01 | 1.04E+00 |
| YBX3 | 1.03E-02 | 5.38E-02 | 1.14E+00 | 7.38E-02 | 3.19E-01 | -1.01E+00 | 6.97E-03 | 2.96E-02 | 1.21E+00 | 5.18E-02 | 1.98E-01 | 1.13E+00 | 2.37E-02 | 8.46E-02 | 1.16E+00 | 7.20E-02 | 2.03E-01 | 1.09E+00 | 5.50E-04 | 2.02E-03 | -1.05E+00 | 1.27E-01 | 3.48E-01 | 1.02E+00 |
| GATAD2B | 1.03E-02 | 5.38E-02 | 1.08E+00 | 3.06E-02 | 1.32E-01 | 1.29E+00 | 1.76E-03 | 7.49E-03 | -1.15E+00 | 1.35E-02 | 5.16E-02 | 1.08E+00 | 1.24E-02 | 4.41E-02 | 1.08E+00 | 1.34E-02 | 3.77E-02 | 1.05E+00 | 1.24E-06 | 4.53E-06 | -1.32E+00 | 4.15E-06 | 1.14E-05 | -1.20E+00 |
| IFT43 | 1.03E-02 | 5.38E-02 | 1.12E+00 | 6.81E-05 | 2.94E-04 | -1.06E+00 | 1.04E-01 | 4.41E-01 | 1.34E+00 | 1.20E-01 | 4.58E-01 | 1.20E+00 | 3.36E-02 | 1.20E-01 | 1.13E+00 | 1.92E-01 | 5.40E-01 | 1.36E+00 | 2.54E-01 | 9.32E-01 | 1.28E+00 | 1.54E-01 | 4.24E-01 | 1.13E+00 |
| RAPGEF1 | 1.03E-02 | 5.39E-02 | -1.12E+00 | 6.34E-02 | 2.74E-01 | 1.14E+00 | 1.21E-01 | 5.16E-01 | 1.12E+00 | 6.05E-02 | 2.31E-01 | 1.13E+00 | 1.24E-01 | 4.41E-01 | -1.09E+00 | 8.39E-02 | 2.36E-01 | -1.21E+00 | 7.42E-02 | 2.72E-01 | -1.10E+00 | 2.79E-01 | 7.67E-01 | 1.15E+00 |
| BNIP1 | 1.03E-02 | 5.39E-02 | -1.04E+00 | 5.01E-03 | 2.17E-02 | -1.02E+00 | 2.63E-04 | 1.12E-03 | -1.15E+00 | 5.06E-06 | 1.93E-05 | -1.18E+00 | 1.17E-09 | 4.16E-09 | -1.81E+00 | 2.87E-08 | 8.09E-08 | -1.55E+00 | 6.63E-05 | 2.43E-04 | -1.13E+00 | 7.22E-05 | 1.98E-04 | -1.17E+00 |
| FAM227A | 1.03E-02 | 5.39E-02 | -1.56E+00 | 8.55E-02 | 3.70E-01 | 1.02E+00 | 1.35E-04 | 5.72E-04 | -1.41E+00 | 1.75E-03 | 6.67E-03 | -1.14E+00 | 3.30E-03 | 1.18E-02 | -2.04E+00 | 3.10E-02 | 8.73E-02 | -1.31E+00 | 3.45E-02 | 1.27E-01 | -1.46E+00 | 2.93E-01 | 8.06E-01 | 1.12E+00 |
| AC002464.1 | 1.04E-02 | 5.40E-02 | -1.19E+00 | 3.14E-02 | 1.36E-01 | 1.06E+00 | 6.79E-04 | 2.89E-03 | -1.46E+00 | 1.50E-03 | 5.73E-03 | -1.17E+00 | 7.73E-02 | 2.76E-01 | -1.45E+00 | 8.72E-02 | 2.46E-01 | -1.58E+00 | 2.59E-01 | 9.50E-01 | -1.13E+00 | 3.62E-01 | 9.93E-01 | -1.12E+00 |
| ZNF649 | 1.04E-02 | 5.41E-02 | -1.06E+00 | 1.97E-03 | 8.53E-03 | -1.16E+00 | 7.70E-05 | 3.27E-04 | -1.43E+00 | 1.96E-05 | 7.50E-05 | -1.44E+00 | 3.14E-02 | 1.12E-01 | -1.27E+00 | 1.08E-01 | 3.03E-01 | 1.04E+00 | 2.25E-04 | 8.25E-04 | -1.09E+00 | 9.37E-03 | 2.57E-02 | -1.03E+00 |
| GAB1 | 1.04E-02 | 5.42E-02 | -1.03E+00 | 1.05E-01 | 4.53E-01 | 1.28E+00 | 2.49E-02 | 1.06E-01 | -1.17E+00 | 1.95E-02 | 7.45E-02 | -1.14E+00 | 9.50E-02 | 3.39E-01 | 1.14E+00 | 2.90E-01 | 8.18E-01 | 1.27E+00 | 1.33E-08 | 4.87E-08 | -1.45E+00 | 5.20E-04 | 1.43E-03 | 1.01E+00 |
| MROH7 | 1.04E-02 | 5.43E-02 | -1.08E+00 | 1.99E-02 | 8.58E-02 | 1.05E+00 | 2.19E-01 | 9.30E-01 | 1.15E+00 | 1.79E-01 | 6.84E-01 | 1.10E+00 | 1.98E-01 | 7.07E-01 | 1.16E+00 | 2.12E-01 | 5.97E-01 | -1.05E+00 | 1.09E-01 | 3.99E-01 | -1.22E+00 | 1.67E-01 | 4.60E-01 | -1.11E+00 |
| BACE2 | 1.04E-02 | 5.44E-02 | -1.46E+00 | 6.15E-02 | 2.66E-01 | -1.03E+00 | 3.95E-02 | 1.68E-01 | -1.44E+00 | 2.36E-01 | 9.00E-01 | 1.20E+00 | 9.02E-30 | 3.22E-29 | -3.44E+00 | 2.11E-27 | 5.93E-27 | -3.26E+00 | 1.74E-05 | 6.40E-05 | -1.60E+00 | 8.31E-02 | 2.28E-01 | -1.01E+00 |

| | | | | | | | | | | | | | | | | | | | | | | | | |
|---|---|---|---|---|---|---|---|---|---|---|---|---|---|---|---|---|---|---|---|---|---|---|---|---|
| ERVK3-1 | 1.04E-02 | 5.44E-02 | 1.01E+00 | 3.56E-02 | 1.54E-01 | 1.16E+00 | 6.18E-02 | 2.63E-01 | 1.05E+00 | 1.66E-01 | 6.34E-01 | 1.28E+00 | 8.32E-02 | 2.97E-01 | 1.18E+00 | 1.20E-01 | 3.38E-01 | 1.17E+00 | 5.13E-03 | 1.88E-02 | -1.09E+00 | 1.48E-01 | 4.06E-01 | 1.14E+00 |
| AL442128.2 | 1.05E-02 | 5.45E-02 | 2.07E+00 | 1.88E-01 | 8.10E-01 | 1.36E+00 | 2.27E-01 | 9.63E-01 | 1.14E+00 | 5.65E-02 | 2.16E-01 | -1.13E+00 | 1.64E-01 | 5.86E-01 | 1.76E+00 | 1.10E-01 | 3.11E-01 | 2.12E+00 | 1.35E-01 | 4.94E-01 | -1.07E+00 | 1.32E-01 | 3.63E-01 | -1.19E+00 |
| EXOC6B | 1.05E-02 | 5.46E-02 | -1.10E+00 | 2.24E-02 | 9.68E-02 | 1.04E+00 | 1.39E-03 | 5.92E-03 | -1.39E+00 | 9.24E-03 | 3.53E-02 | -1.06E+00 | 2.01E-01 | 7.16E-01 | 1.60E+00 | 1.71E-01 | 4.82E-01 | 1.80E+00 | 4.67E-04 | 1.71E-03 | -1.33E+00 | 5.58E-03 | 1.53E-02 | -1.12E+00 |
| PPP1R37 | 1.05E-02 | 5.46E-02 | -1.12E+00 | 3.80E-02 | 1.64E-01 | 1.05E+00 | 1.76E-03 | 7.49E-03 | -1.26E+00 | 2.18E-02 | 8.34E-02 | 1.07E+00 | 6.23E-03 | 2.22E-02 | -1.22E+00 | 5.74E-03 | 1.62E-02 | -1.30E+00 | 2.57E-05 | 9.45E-05 | -1.35E+00 | 2.84E-04 | 7.81E-04 | -1.30E+00 |
| CBWD5 | 1.05E-02 | 5.46E-02 | 1.18E+00 | 8.03E-02 | 3.47E-01 | 1.38E+00 | 5.21E-02 | 2.21E-01 | -1.02E+00 | 2.33E-01 | 8.91E-01 | 1.26E+00 | 2.97E-03 | 1.06E-02 | 1.08E+00 | 4.69E-02 | 1.32E-01 | 1.28E+00 | 5.19E-02 | 1.90E-01 | 1.41E+00 | 1.18E-02 | 3.23E-02 | 1.50E+00 |
| GHRL | 1.05E-02 | 5.46E-02 | -1.18E+00 | 4.59E-03 | 1.98E-02 | -1.27E+00 | 1.07E-01 | 4.56E-01 | -1.12E+00 | 2.38E-01 | 9.10E-01 | 1.30E+00 | 1.98E-01 | 7.07E-01 | 1.06E+00 | 1.59E-01 | 4.49E-01 | -1.06E+00 | 7.60E-02 | 2.79E-01 | -1.13E+00 | 1.50E-01 | 4.12E-01 | -1.08E+00 |
| SMURF1 | 1.05E-02 | 5.47E-02 | -1.05E+00 | 9.96E-05 | 4.31E-04 | -1.40E+00 | 3.47E-02 | 1.48E-01 | 1.01E+00 | 3.67E-06 | 1.40E-05 | -1.56E+00 | 1.90E-01 | 6.79E-01 | 1.43E+00 | 2.07E-01 | 5.83E-01 | 1.04E+00 | 2.17E-01 | 7.98E-01 | 1.14E+00 | 9.49E-03 | 2.61E-02 | -1.35E+00 |
| FSIP2 | 1.05E-02 | 5.47E-02 | -1.56E+00 | 8.69E-02 | 3.75E-01 | -1.07E+00 | 1.79E-01 | 7.59E-01 | -1.10E+00 | 4.60E-02 | 1.76E-01 | 1.68E+00 | 6.93E-02 | 2.47E-01 | -1.28E+00 | 3.45E-01 | 9.71E-01 | 1.28E+00 | 2.70E-01 | 9.89E-01 | 1.03E+00 | 3.51E-01 | 9.63E-01 | 1.09E+00 |
| SOD2 | 1.05E-02 | 5.48E-02 | 1.59E+00 | 9.45E-02 | 4.08E-01 | 1.57E+00 | 1.98E-01 | 8.41E-01 | 1.34E+00 | 4.54E-02 | 1.73E-01 | 1.33E+00 | 4.00E-07 | 1.43E-06 | 2.98E+00 | 2.53E-09 | 7.13E-09 | 2.77E+00 | 2.18E-17 | 8.01E-17 | 2.62E+00 | 4.46E-13 | 1.22E-12 | 2.19E+00 |
| SLC11A1 | 1.05E-02 | 5.48E-02 | -1.02E+00 | 9.62E-03 | 4.16E-02 | 1.02E+00 | 1.26E-01 | 5.37E-01 | -1.07E+00 | 1.33E-01 | 5.08E-01 | 1.04E+00 | 4.24E-02 | 1.51E-01 | -1.03E+00 | 9.01E-02 | 2.54E-01 | 1.00E+00 | 1.18E-05 | 4.33E-05 | -1.12E+00 | 3.19E-06 | 8.78E-06 | -1.37E+00 |
| C21orf2 | 1.05E-02 | 5.48E-02 | -1.12E+00 | 3.32E-03 | 1.44E-02 | -1.12E+00 | 2.07E-01 | 8.81E-01 | 1.29E+00 | 3.85E-02 | 1.47E-01 | 1.09E+00 | 2.25E-01 | 8.03E-01 | 1.26E+00 | 1.99E-01 | 5.62E-01 | 1.24E+00 | 6.09E-02 | 2.23E-01 | 1.05E+00 | 2.35E-02 | 6.44E-02 | 1.04E+00 |
| IFT172 | 1.05E-02 | 5.49E-02 | -1.03E+00 | 1.06E-01 | 4.58E-01 | 1.35E+00 | 3.88E-03 | 1.65E-02 | -1.30E+00 | 1.71E-01 | 6.51E-01 | 1.22E+00 | 1.62E-01 | 5.78E-01 | 1.19E+00 | 2.87E-01 | 8.10E-01 | 1.48E+00 | 1.78E-01 | 6.52E-01 | 1.12E+00 | 1.78E-01 | 4.88E-01 | 1.54E+00 |
| AC068152.1 | 1.05E-02 | 5.49E-02 | -1.02E+00 | 4.16E-02 | 1.80E-01 | 1.15E+00 | 7.96E-04 | 3.38E-03 | -1.44E+00 | 1.41E-02 | 5.37E-02 | -1.07E+00 | 1.76E-02 | 6.28E-02 | -1.25E+00 | 1.41E-02 | 3.97E-02 | -1.28E+00 | 1.06E-01 | 3.87E-01 | 1.02E+00 | 3.40E-01 | 9.33E-01 | 1.27E+00 |
| AL021453.1 | 1.05E-02 | 5.49E-02 | 1.82E+00 | 3.30E-03 | 1.43E-02 | 1.18E+00 | 6.35E-02 | 2.70E-01 | 1.06E+00 | 7.50E-03 | 2.87E-02 | -1.00E+00 | 1.40E-01 | 4.99E-01 | 1.23E+00 | 3.33E-01 | 9.39E-01 | 1.31E+00 | 1.27E-01 | 4.66E-01 | 1.21E+00 | 1.70E-01 | 4.66E-01 | 1.20E+00 |
| PGAP1 | 1.06E-02 | 5.51E-02 | -1.05E+00 | 2.00E-01 | 8.63E-01 | 1.52E+00 | 1.80E-02 | 7.64E-02 | -1.06E+00 | 5.70E-02 | 2.18E-01 | 1.08E+00 | 9.78E-02 | 3.49E-01 | -1.38E+00 | 1.94E-01 | 5.46E-01 | -1.08E+00 | 2.15E-01 | 7.90E-01 | 1.10E+00 | 1.99E-01 | 5.47E-01 | 1.37E+00 |
| FOXP1-AS1 | 1.06E-02 | 5.51E-02 | -1.08E+00 | 5.91E-03 | 2.55E-02 | -1.00E+00 | 6.17E-02 | 2.62E-01 | -1.09E+00 | 9.05E-02 | 3.46E-01 | 1.10E+00 | 8.76E-03 | 3.12E-02 | -1.38E+00 | 1.03E-02 | 2.89E-02 | -1.27E+00 | 6.67E-03 | 2.45E-02 | -1.31E+00 | 1.77E-02 | 4.87E-02 | -1.15E+00 |
| ATP6V1C1 | 1.06E-02 | 5.52E-02 | -1.07E+00 | 2.25E-02 | 9.73E-02 | 1.08E+00 | 3.12E-08 | 1.33E-07 | -1.83E+00 | 2.51E-05 | 9.58E-05 | -1.33E+00 | 8.57E-03 | 3.06E-02 | -1.14E+00 | 3.95E-03 | 1.11E-02 | -1.26E+00 | 1.31E-06 | 4.82E-06 | -1.55E+00 | 6.35E-06 | 1.75E-05 | -1.55E+00 |
| SOS2 | 1.06E-02 | 5.54E-02 | 1.15E+00 | 1.00E-02 | 4.34E-02 | 1.20E+00 | 1.70E-05 | 7.23E-05 | -1.37E+00 | 5.23E-04 | 2.00E-03 | -1.06E+00 | 2.42E-02 | 8.63E-02 | 1.03E+00 | 5.96E-04 | 1.68E-03 | -1.14E+00 | 1.09E-03 | 3.99E-03 | -1.09E+00 | 1.70E-04 | 4.68E-04 | -1.22E+00 |
| SATB1 | 1.06E-02 | 5.54E-02 | 1.12E+00 | 1.20E-03 | 5.18E-03 | 1.18E+00 | 1.96E-02 | 8.33E-02 | -1.02E+00 | 4.42E-03 | 1.69E-02 | 1.07E+00 | 5.09E-03 | 1.81E-02 | -1.00E+00 | 1.40E-02 | 3.95E-02 | -1.02E+00 | 6.72E-10 | 2.47E-09 | -1.55E+00 | 1.78E-06 | 4.89E-06 | -1.36E+00 |
| AL359232.1 | 1.06E-02 | 5.54E-02 | -1.12E+00 | 1.16E-03 | 5.02E-03 | -1.22E+00 | 1.56E-02 | 6.64E-02 | 1.87E+00 | 1.66E-01 | 6.33E-01 | 1.41E+00 | 2.43E-01 | 8.65E-01 | 1.41E+00 | 1.21E-01 | 3.42E-01 | -1.13E+00 | 1.79E-01 | 6.58E-01 | 1.10E+00 | 3.49E-02 | 9.59E-02 | -1.29E+00 |
| CEACAM21 | 1.07E-02 | 5.56E-02 | 1.86E+00 | 1.17E-01 | 5.04E-01 | 1.64E+00 | 7.63E-02 | 3.24E-01 | 1.04E+00 | 1.19E-03 | 4.53E-03 | -1.16E+00 | 2.50E-01 | 8.93E-01 | 1.49E+00 | 2.35E-01 | 6.64E-01 | 1.82E+00 | 9.76E-02 | 3.58E-01 | 1.63E+00 | 2.72E-01 | 7.47E-01 | 1.51E+00 |
| CTHRC1 | 1.07E-02 | 5.56E-02 | 2.13E+00 | 2.07E-02 | 8.95E-02 | 2.28E+00 | 1.32E-01 | 5.60E-01 | 1.39E+00 | 9.33E-02 | 3.57E-01 | 1.73E+00 | 7.78E-02 | 2.77E-01 | 2.69E+00 | 6.86E-02 | 1.93E-01 | 3.14E+00 | 2.58E-03 | 9.46E-03 | 1.87E+00 | 6.42E-03 | 1.76E-02 | 2.05E+00 |
| B9D1 | 1.07E-02 | 5.56E-02 | -1.68E+00 | 6.37E-03 | 2.75E-02 | -1.99E+00 | 7.42E-02 | 3.15E-01 | 1.44E+00 | 2.21E-01 | 8.45E-01 | 1.16E+00 | 2.41E-03 | 8.58E-03 | -1.69E+00 | 2.94E-02 | 8.29E-02 | -1.17E+00 | 2.41E-02 | 8.83E-02 | -1.05E+00 | 2.08E-02 | 5.71E-02 | 1.04E+00 |
| AC016355.1 | 1.07E-02 | 5.56E-02 | -1.18E+00 | 2.55E-02 | 1.10E-01 | -1.11E+00 | 3.58E-02 | 1.52E-01 | -1.11E+00 | 7.76E-02 | 2.97E-01 | 1.10E+00 | 2.63E-01 | 9.37E-01 | 1.26E+00 | 2.77E-01 | 7.80E-01 | 1.49E+00 | 2.40E-01 | 8.80E-01 | 1.09E+00 | 3.52E-01 | 9.66E-01 | 1.17E+00 |
| AL353898.1 | 1.07E-02 | 5.58E-02 | -1.03E+00 | 1.48E-06 | 6.38E-06 | -1.97E+00 | 6.79E-02 | 2.89E-01 | 1.06E+00 | 1.03E-06 | 3.94E-06 | -1.74E+00 | 1.92E-02 | 6.84E-02 | -1.20E+00 | 1.90E-03 | 5.35E-03 | -1.78E+00 | 2.11E-02 | 7.76E-02 | -1.06E+00 | 6.93E-06 | 1.90E-05 | -1.67E+00 |
| ZNF496 | 1.08E-02 | 5.61E-02 | 1.04E+00 | 2.44E-02 | 1.06E-01 | 1.17E+00 | 2.19E-02 | 9.29E-02 | -1.02E+00 | 2.10E-02 | 8.02E-02 | 1.12E+00 | 8.25E-04 | 2.94E-03 | -1.09E+00 | 5.41E-03 | 1.52E-02 | 1.12E+00 | 5.31E-03 | 1.95E-02 | -1.12E+00 | 2.91E-02 | 8.00E-02 | -1.02E+00 |
| OXLD1 | 1.08E-02 | 5.62E-02 | 1.08E+00 | 7.64E-03 | 3.30E-02 | 1.26E+00 | 7.65E-06 | 3.25E-05 | -1.22E+00 | 1.06E-03 | 4.05E-03 | 1.03E+00 | 4.75E-04 | 1.69E-03 | 1.02E+00 | 3.13E-01 | 8.82E-01 | 1.44E+00 | 1.25E-01 | 4.60E-01 | 1.21E+00 | 1.41E-01 | 3.87E-01 | 1.41E+00 |
| RPL12P37 | 1.08E-02 | 5.62E-02 | -1.03E+00 | 2.10E-04 | 9.06E-04 | -1.39E+00 | 6.08E-02 | 2.58E-01 | -1.01E+00 | 2.15E-02 | 8.22E-02 | -1.10E+00 | 1.86E-01 | 6.63E-01 | 1.12E+00 | 3.98E-02 | 1.12E-01 | -1.34E+00 | 3.09E-02 | 1.14E-01 | -1.14E+00 | 7.22E-02 | 1.98E-01 | -1.05E+00 |
| ZNF710-AS1 | 1.08E-02 | 5.62E-02 | -1.22E+00 | 5.46E-02 | 2.36E-01 | 1.13E+00 | 1.09E-01 | 4.65E-01 | 1.40E+00 | 6.75E-02 | 2.58E-01 | 1.67E+00 | 6.16E-02 | 2.20E-01 | 2.11E+00 | 1.55E-01 | 4.36E-01 | 1.95E+00 | 9.65E-02 | 3.54E-01 | 1.53E+00 | 2.91E-01 | 7.99E-01 | 1.14E+00 |
| MIR762HG | 1.08E-02 | 5.62E-02 | 1.85E+00 | 1.35E-01 | 5.81E-01 | 1.13E+00 | 6.52E-03 | 2.77E-02 | -1.28E+00 | 5.23E-05 | 2.00E-04 | -1.64E+00 | 1.14E-01 | 4.08E-01 | -1.03E+00 | 2.19E-02 | 6.18E-02 | -1.37E+00 | 2.36E-01 | 8.66E-01 | 1.15E+00 | 1.69E-01 | 4.63E-01 | -1.22E+00 |
| MAN2C1 | 1.08E-02 | 5.62E-02 | -1.06E+00 | 2.30E-02 | 9.92E-02 | 1.09E+00 | 1.94E-01 | 8.25E-01 | 1.12E+00 | 1.56E-01 | 5.95E-01 | 1.23E+00 | 1.91E-01 | 6.82E-01 | 1.39E+00 | 3.72E-02 | 1.05E-01 | 1.03E+00 | 1.10E-02 | 4.04E-02 | -1.10E+00 | 1.07E-01 | 2.94E-01 | 1.13E+00 |
| AC002558.1 | 1.08E-02 | 5.63E-02 | -1.27E+00 | 6.65E-03 | 2.87E-02 | -1.23E+00 | 1.73E-01 | 7.34E-01 | 1.08E+00 | 2.26E-01 | 8.61E-01 | 1.38E+00 | 2.09E-01 | 7.45E-01 | 1.16E+00 | 2.56E-01 | 7.20E-01 | 1.56E+00 | 8.97E-02 | 3.29E-01 | 1.07E+00 | 2.13E-01 | 5.84E-01 | 1.47E+00 |
| HBEGF | 1.08E-02 | 5.64E-02 | 2.10E+00 | 8.29E-02 | 3.58E-01 | -1.54E+00 | 1.04E-02 | 4.42E-02 | 1.90E+00 | 4.66E-02 | 1.78E-01 | -1.67E+00 | 1.93E-01 | 6.87E-01 | 1.56E+00 | 1.87E-01 | 5.28E-01 | -1.36E+00 | 2.13E-01 | 7.80E-01 | 1.39E+00 | 3.21E-03 | 8.81E-03 | -1.71E+00 |
| AC133065.2 | 1.08E-02 | 5.64E-02 | -1.04E+00 | 3.96E-02 | 1.71E-01 | 1.23E+00 | 6.78E-02 | 2.88E-01 | -1.23E+00 | 2.57E-01 | 9.83E-01 | 1.27E+00 | 5.87E-02 | 2.09E-01 | -1.38E+00 | 7.69E-02 | 2.17E-01 | -1.43E+00 | 3.50E-03 | 1.29E-02 | -2.23E+00 | 1.31E-02 | 3.60E-02 | -2.05E+00 |
| ZNF528 | 1.08E-02 | 5.64E-02 | -1.15E+00 | 3.72E-02 | 1.61E-01 | 1.05E+00 | 1.28E-02 | 5.44E-02 | -1.38E+00 | 4.48E-02 | 1.71E-01 | -1.01E+00 | 4.55E-03 | 1.62E-02 | -1.42E+00 | 1.52E-02 | 4.29E-02 | -1.27E+00 | 8.85E-04 | 3.25E-03 | -1.12E+00 | 6.36E-02 | 1.75E-01 | 1.01E+00 |
| GORAB | 1.08E-02 | 5.65E-02 | -1.20E+00 | 9.76E-02 | 4.22E-01 | 1.15E+00 | 1.10E-04 | 4.67E-04 | -1.70E+00 | 7.19E-04 | 2.75E-03 | -1.35E+00 | 2.04E-02 | 7.28E-02 | -1.40E+00 | 2.47E-01 | 6.95E-01 | 1.18E+00 | 8.69E-09 | 3.19E-08 | -1.80E+00 | 3.22E-06 | 8.85E-06 | -1.52E+00 |
| CNNM2 | 1.09E-02 | 5.66E-02 | 1.05E+00 | 4.31E-03 | 1.86E-02 | 1.13E+00 | 7.88E-02 | 3.35E-01 | 1.58E+00 | 2.42E-01 | 9.26E-01 | 1.58E+00 | 7.87E-02 | 2.81E-01 | 1.04E+00 | 6.57E-02 | 1.85E-01 | 1.06E+00 | 4.00E-03 | 1.47E-02 | -1.11E+00 | 2.48E-03 | 6.81E-03 | -1.07E+00 |
| HIST1H3H | 1.09E-02 | 5.67E-02 | -1.54E+00 | 7.25E-03 | 3.13E-02 | -1.55E+00 | 1.42E-01 | 6.01E-01 | -1.04E+00 | 2.28E-03 | 8.72E-03 | -1.67E+00 | 3.07E-03 | 1.09E-02 | -1.58E+00 | 7.88E-03 | 2.22E-02 | -1.38E+00 | 2.38E-02 | 8.72E-02 | 1.21E+00 | 1.46E-03 | 4.01E-03 | 1.04E+00 |
| C5 | 1.09E-02 | 5.67E-02 | -1.30E+00 | 9.38E-02 | 4.05E-01 | 1.11E+00 | 6.03E-02 | 2.56E-01 | -1.04E+00 | 1.25E-01 | 4.77E-01 | 1.16E+00 | 4.05E-03 | 1.44E-02 | -2.04E+00 | 1.46E-02 | 4.11E-02 | -1.49E+00 | 2.10E-02 | 7.71E-02 | 1.04E+00 | 6.91E-02 | 1.90E-01 | 1.01E+00 |
| RELT | 1.09E-02 | 5.67E-02 | -1.05E+00 | 9.23E-03 | 3.99E-02 | 1.00E+00 | 2.80E-04 | 1.19E-03 | -1.28E+00 | 8.93E-07 | 3.41E-06 | -1.46E+00 | 3.57E-05 | 1.27E-04 | -1.60E+00 | 9.98E-05 | 2.81E-04 | -1.44E+00 | 9.59E-04 | 3.52E-03 | -1.16E+00 | 2.81E-03 | 7.73E-03 | -1.14E+00 |
| BDH2P1 | 1.09E-02 | 5.67E-02 | -1.43E+00 | 2.01E-01 | 8.67E-01 | 1.29E+00 | 1.71E-01 | 7.26E-01 | 1.33E+00 | 1.79E-01 | 6.85E-01 | 1.51E+00 | 2.24E-02 | 7.99E-02 | -1.51E+00 | 3.37E-01 | 9.49E-01 | 1.26E+00 | 4.20E-02 | 1.54E-01 | -1.15E+00 | 6.51E-02 | 1.79E-01 | -1.18E+00 |
| DPEP2 | 1.09E-02 | 5.69E-02 | 1.21E+00 | 4.10E-02 | 1.77E-01 | 1.38E+00 | 2.89E-04 | 1.23E-03 | -1.19E+00 | 1.86E-04 | 7.11E-04 | -1.03E+00 | 2.02E-02 | 7.19E-02 | 1.03E+00 | 1.33E-01 | 3.74E-01 | 1.31E+00 | 2.11E-01 | 7.76E-01 | 1.25E+00 | 2.36E-01 | 6.49E-01 | 1.16E+00 |
| RETSAT | 1.09E-02 | 5.70E-02 | -1.19E+00 | 1.26E-01 | 5.43E-01 | 1.24E+00 | 5.01E-07 | 2.13E-06 | -1.62E+00 | 4.48E-05 | 1.71E-04 | -1.19E+00 | 5.96E-02 | 2.13E-01 | -1.05E+00 | 2.14E-01 | 6.04E-01 | 1.19E+00 | 4.01E-02 | 1.47E-01 | -1.07E+00 | 8.92E-02 | 2.45E-01 | 1.02E+00 |
| RMI2 | 1.09E-02 | 5.70E-02 | -1.23E+00 | 3.38E-02 | 1.46E-01 | 1.06E+00 | 1.72E-02 | 7.31E-02 | -1.11E+00 | 1.23E-01 | 4.70E-01 | 1.21E+00 | 5.63E-02 | 2.01E-01 | -1.43E+00 | 1.40E-01 | 3.94E-01 | -1.26E+00 | 3.26E-07 | 1.20E-06 | -1.33E+00 | 1.11E-01 | 3.04E-01 | 1.08E+00 |
| DSTNP2 | 1.09E-02 | 5.70E-02 | -1.22E+00 | 8.89E-02 | 3.84E-01 | 1.25E+00 | 2.76E-06 | 1.17E-05 | -1.70E+00 | 6.25E-06 | 2.39E-05 | -1.38E+00 | 1.08E-02 | 3.86E-02 | -1.22E+00 | 1.65E-02 | 4.66E-02 | -1.23E+00 | 4.90E-09 | 1.80E-08 | -1.72E+00 | 4.91E-06 | 1.35E-05 | -1.36E+00 |
| AC103876.1 | 1.10E-02 | 5.72E-02 | -1.08E+00 | 2.30E-02 | 9.93E-02 | 1.13E+00 | 3.62E-02 | 1.54E-01 | -1.18E+00 | 1.61E-01 | 6.16E-01 | 1.18E+00 | 6.46E-02 | 2.30E-01 | -1.35E+00 | 1.55E-01 | 4.37E-01 | -1.10E+00 | 3.75E-03 | 1.38E-02 | -1.67E+00 | 5.17E-02 | 1.42E-01 | -1.20E+00 |
| JPT2 | 1.10E-02 | 5.72E-02 | -1.09E+00 | 1.16E-02 | 4.99E-02 | -1.04E+00 | 4.57E-02 | 1.94E-01 | -1.01E+00 | 2.04E-01 | 7.80E-01 | 1.30E+00 | 8.99E-04 | 3.21E-03 | -1.70E+00 | 3.44E-02 | 9.70E-02 | -1.06E+00 | 1.97E-07 | 7.24E-07 | -1.28E+00 | 1.57E-01 | 4.32E-01 | 1.22E+00 |
| SEPSECS | 1.10E-02 | 5.73E-02 | -1.04E+00 | 5.13E-03 | 2.22E-02 | -1.07E+00 | 2.69E-03 | 1.14E-02 | -1.36E+00 | 2.09E-02 | 7.99E-02 | -1.03E+00 | 4.06E-03 | 1.45E-02 | -1.22E+00 | 4.79E-02 | 1.35E-01 | 1.07E+00 | 8.66E-04 | 3.18E-03 | -1.20E+00 | 1.77E-02 | 4.87E-02 | -1.06E+00 |
| LRP12 | 1.10E-02 | 5.73E-02 | -1.03E+00 | 3.53E-05 | 1.52E-04 | -1.35E+00 | 4.69E-03 | 1.99E-02 | -1.22E+00 | 8.93E-07 | 3.41E-06 | -1.77E+00 | 1.51E-01 | 5.39E-01 | 1.13E+00 | 6.10E-03 | 1.72E-02 | -1.32E+00 | 1.04E-02 | 3.83E-02 | 1.10E+00 | 6.99E-08 | 1.92E-07 | -1.33E+00 |
| DLEU2 | 1.10E-02 | 5.73E-02 | 1.85E+00 | 1.48E-01 | 6.40E-01 | 1.65E+00 | 2.33E-01 | 9.92E-01 | 1.28E+00 | 1.97E-01 | 7.51E-01 | 1.59E+00 | 2.58E-01 | 9.21E-01 | 1.41E+00 | 1.80E-01 | 5.08E-01 | -1.00E+00 | 9.86E-02 | 3.62E-01 | -1.06E+00 | 1.86E-01 | 5.11E-01 | 1.01E+00 |
| SRGAP2B | 1.10E-02 | 5.73E-02 | -1.18E+00 | 2.48E-02 | 1.07E-01 | 1.01E+00 | 1.13E-03 | 4.82E-03 | -1.25E+00 | 2.56E-04 | 9.79E-04 | -1.30E+00 | 1.24E-01 | 4.40E-01 | 1.12E+00 | 2.20E-01 | 6.20E-01 | 1.22E+00 | 4.01E-03 | 1.47E-02 | -1.20E+00 | 3.59E-02 | 9.87E-02 | -1.04E+00 |
| SLC10A5 | 1.10E-02 | 5.73E-02 | -1.08E+00 | 6.78E-03 | 2.93E-02 | -1.02E+00 | 1.42E-02 | 6.03E-02 | -1.31E+00 | 2.54E-02 | 9.69E-02 | -1.13E+00 | 1.83E-15 | 6.53E-15 | -2.17E+00 | 4.95E-19 | 1.40E-18 | -2.48E+00 | 1.79E-01 | 6.55E-01 | -1.38E+00 | 2.79E-01 | 7.68E-01 | -1.20E+00 |
| ZNF737 | 1.10E-02 | 5.74E-02 | 1.03E+00 | 5.28E-04 | 2.28E-03 | -1.09E+00 | 7.20E-04 | 3.06E-03 | -1.14E+00 | 3.19E-05 | 1.22E-04 | -1.26E+00 | 1.79E-05 | 6.36E-05 | -1.56E+00 | 9.70E-05 | 2.73E-04 | -1.38E+00 | 8.99E-03 | 3.30E-02 | -1.08E+00 | 1.15E-03 | 3.16E-03 | -1.34E+00 |
| LINC00909 | 1.10E-02 | 5.74E-02 | 1.10E+00 | 5.10E-03 | 2.21E-02 | 1.10E+00 | 5.11E-02 | 2.17E-01 | 1.02E+00 | 2.30E-02 | 8.78E-02 | 1.09E+00 | 1.04E-01 | 3.69E-01 | 1.36E+00 | 3.18E-02 | 8.98E-02 | 1.22E+00 | 1.03E-01 | 3.76E-01 | 1.16E+00 | 3.07E-01 | 8.44E-01 | 1.25E+00 |
| TACC1 | 1.10E-02 | 5.74E-02 | 1.51E+00 | 5.64E-10 | 2.44E-09 | -1.19E+00 | 3.65E-02 | 1.55E-01 | 1.48E+00 | 3.14E-07 | 1.20E-06 | -1.23E+00 | 7.14E-07 | 2.54E-06 | 2.70E+00 | 8.27E-03 | 2.33E-02 | 1.07E+00 | 2.27E-02 | 8.33E-02 | 1.43E+00 | 7.02E-06 | 1.93E-05 | -1.29E+00 |
| CCDC149 | 1.10E-02 | 5.75E-02 | -1.10E+00 | 5.83E-02 | 2.52E-01 | 1.19E+00 | 1.83E-01 | 7.79E-01 | 1.15E+00 | 7.50E-02 | 2.86E-01 | 1.08E+00 | 9.34E-02 | 3.33E-01 | -1.21E+00 | 8.00E-02 | 2.26E-01 | -1.54E+00 | 5.27E-02 | 1.93E-01 | -1.32E+00 | 1.46E-01 | 4.02E-01 | -1.19E+00 |
| ZBTB43 | 1.10E-02 | 5.75E-02 | 1.08E+00 | 4.37E-07 | 1.89E-06 | -1.13E+00 | 1.01E-03 | 4.27E-03 | -1.22E+00 | 9.68E-10 | 3.70E-09 | -1.64E+00 | 1.03E-03 | 3.67E-03 | -1.04E+00 | 1.02E-07 | 2.88E-07 | -1.44E+00 | 1.14E-02 | 4.18E-02 | -1.11E+00 | 2.38E-06 | 6.54E-06 | -1.67E+00 |
| ZNF57 | 1.10E-02 | 5.76E-02 | -1.14E+00 | 4.59E-02 | 1.98E-01 | 1.10E+00 | 4.30E-04 | 1.83E-03 | -1.20E+00 | 6.18E-05 | 2.36E-04 | -1.31E+00 | 1.65E-01 | 5.88E-01 | 1.16E+00 | 2.38E-01 | 6.70E-01 | 1.23E+00 | 1.09E-03 | 3.98E-03 | -1.53E+00 | 1.26E-02 | 3.47E-02 | -1.29E+00 |
| DAPK2 | 1.10E-02 | 5.76E-02 | -1.37E+00 | 3.20E-02 | 1.38E-01 | -1.11E+00 | 6.98E-03 | 2.97E-02 | -1.30E+00 | 7.34E-03 | 2.80E-02 | -1.16E+00 | 7.81E-02 | 2.78E-01 | -1.07E+00 | 1.07E-02 | 3.01E-02 | -1.60E+00 | 1.08E-01 | 3.95E-01 | 1.15E+00 | 1.91E-01 | 5.23E-01 | 1.04E+00 |
| MT-ND4L | 1.10E-02 | 5.76E-02 | 1.04E+00 | 3.84E-22 | 1.66E-21 | -1.33E+00 | 3.43E-02 | 1.46E-01 | 1.23E+00 | 9.29E-03 | 3.55E-02 | -1.07E+00 | 1.97E-01 | 7.03E-01 | 1.17E+00 | 1.11E-05 | 3.13E-05 | -1.20E+00 | 6.85E-02 | 2.51E-01 | 1.26E+00 | 1.60E-02 | 4.40E-02 | -1.06E+00 |
| SCAND1 | 1.11E-02 | 5.76E-02 | 1.11E+00 | 1.48E-06 | 6.39E-06 | 1.03E+00 | 5.81E-02 | 2.47E-01 | 1.18E+00 | 6.71E-02 | 2.56E-01 | 1.01E+00 | 3.60E-02 | 1.28E-01 | 1.12E+00 | 8.77E-03 | 2.47E-02 | 1.03E+00 | 7.22E-02 | 2.65E-01 | 1.26E+00 | 2.99E-01 | 8.22E-01 | 1.17E+00 |
| RPL34 | 1.11E-02 | 5.76E-02 | -1.06E+00 | 1.08E-35 | 4.68E-35 | -1.38E+00 | 2.21E-01 | 9.39E-01 | 1.01E+00 | 3.49E-15 | 1.33E-14 | -1.27E+00 | 1.78E-01 | 6.33E-01 | -1.02E+00 | 2.32E-19 | 6.54E-19 | -1.36E+00 | 2.96E-06 | 1.09E-05 | -1.08E+00 | 9.11E-30 | 2.50E-29 | -1.41E+00 |
| SKAP1-AS1 | 1.11E-02 | 5.76E-02 | 1.01E+00 | 8.87E-03 | 3.83E-02 | 1.09E+00 | 6.69E-03 | 2.84E-02 | -1.48E+00 | 3.11E-03 | 1.19E-02 | -1.49E+00 | 1.45E-01 | 5.19E-01 | 1.00E+00 | 2.04E-01 | 5.74E-01 | 1.03E+00 | 8.44E-02 | 3.10E-01 | 1.63E+00 | 1.04E-01 | 2.86E-01 | 1.84E+00 |
| AL590617.2 | 1.11E-02 | 5.77E-02 | -1.15E+00 | 3.57E-02 | 1.54E-01 | 1.10E+00 | 4.14E-03 | 1.76E-02 | -1.37E+00 | 4.64E-03 | 1.77E-02 | -1.21E+00 | 9.36E-02 | 3.34E-01 | -1.35E+00 | 8.02E-02 | 2.26E-01 | -1.43E+00 | 2.46E-02 | 9.04E-02 | -1.56E+00 | 6.89E-02 | 1.89E-01 | -1.29E+00 |
| CXorf40B | 1.11E-02 | 5.78E-02 | -1.29E+00 | 1.15E-01 | 4.96E-01 | 1.25E+00 | 3.65E-02 | 1.55E-01 | -1.22E+00 | 1.60E-01 | 6.12E-01 | 1.47E+00 | 1.96E-01 | 6.98E-01 | 1.16E+00 | 6.99E-02 | 1.97E-01 | 2.35E+00 | 8.69E-02 | 3.19E-01 | 1.57E+00 | 5.77E-05 | 1.59E-04 | 2.49E+00 |
| FAAP100 | 1.11E-02 | 5.78E-02 | -1.13E+00 | 1.88E-02 | 8.12E-02 | 1.03E+00 | 1.35E-01 | 5.72E-01 | 1.35E+00 | 5.67E-02 | 2.17E-01 | 1.72E+00 | 9.48E-05 | 3.38E-04 | -1.54E+00 | 6.99E-03 | 1.97E-02 | -1.06E+00 | 1.56E-04 | 5.74E-04 | -1.25E+00 | 1.49E-01 | 4.08E-01 | 1.13E+00 |
| KATNAL2 | 1.11E-02 | 5.78E-02 | -1.25E+00 | 9.63E-02 | 4.16E-01 | 1.20E+00 | 9.47E-03 | 4.02E-02 | -1.39E+00 | 1.56E-01 | 5.97E-01 | 1.25E+00 | 2.28E-01 | 8.12E-01 | 1.14E+00 | 2.52E-01 | 7.10E-01 | 1.58E+00 | 5.71E-02 | 2.10E-01 | -1.10E+00 | 2.04E-01 | 5.59E-01 | 1.03E+00 |
| KIAA0391 | 1.11E-02 | 5.79E-02 | -1.33E+00 | 1.21E-01 | 5.21E-01 | 1.14E+00 | 6.22E-02 | 2.64E-01 | -1.16E+00 | 2.42E-02 | 9.24E-02 | -1.08E+00 | 1.11E-02 | 3.97E-02 | -1.32E+00 | 9.49E-02 | 2.67E-01 | 1.16E+00 | 1.09E-01 | 4.02E-01 | 1.01E+00 | 1.92E-01 | 5.28E-01 | 1.08E+00 |
| APOC1 | 1.12E-02 | 5.81E-02 | -2.00E+00 | 1.91E-02 | 8.24E-02 | -1.70E+00 | 5.34E-02 | 2.27E-01 | -1.43E+00 | 5.87E-02 | 2.24E-01 | -1.33E+00 | 1.96E-07 | 6.97E-07 | -2.39E+00 | 2.60E-01 | 7.32E-01 | 1.02E+00 | 1.27E-15 | 4.65E-15 | -1.85E+00 | 2.72E-14 | 7.47E-14 | -1.86E+00 |
| SSSCA1-AS1 | 1.12E-02 | 5.82E-02 | -1.16E+00 | 3.58E-02 | 1.55E-01 | 1.03E+00 | 1.86E-01 | 7.89E-01 | 1.15E+00 | 8.16E-02 | 3.12E-01 | 1.02E+00 | 2.36E-01 | 8.41E-01 | 1.35E+00 | 2.74E-01 | 7.71E-01 | 1.68E+00 | 2.68E-01 | 9.84E-01 | 1.20E+00 | 9.47E-02 | 2.60E-01 | -1.21E+00 |
| TMEM11 | 1.12E-02 | 5.83E-02 | 1.09E+00 | 7.52E-02 | 3.25E-01 | 1.33E+00 | 6.13E-02 | 2.60E-01 | 1.05E+00 | 1.93E-01 | 7.38E-01 | 1.42E+00 | 4.97E-07 | 1.77E-06 | -1.26E+00 | 4.58E-06 | 1.29E-05 | -1.17E+00 | 5.35E-10 | 1.96E-09 | -1.46E+00 | 7.03E-04 | 1.93E-03 | -1.12E+00 |
| SMYD5 | 1.12E-02 | 5.83E-02 | -1.26E+00 | 3.70E-02 | 1.60E-01 | -1.01E+00 | 2.19E-04 | 9.29E-04 | -1.36E+00 | 7.70E-04 | 2.94E-03 | -1.13E+00 | 1.57E-01 | 5.60E-01 | 1.13E+00 | 1.32E-01 | 3.72E-01 | 1.01E+00 | 5.50E-05 | 2.02E-04 | -1.27E+00 | 1.83E-02 | 5.02E-02 | -1.01E+00 |
| AC137630.3 | 1.12E-02 | 5.83E-02 | -1.15E+00 | 1.01E-04 | 4.35E-04 | -1.52E+00 | 8.52E-02 | 3.62E-01 | 1.00E+00 | 2.47E-03 | 9.43E-03 | -1.34E+00 | 1.09E-02 | 3.90E-02 | -1.45E+00 | 2.55E-02 | 7.19E-02 | -1.20E+00 | 7.35E-02 | 2.70E-01 | -1.18E+00 | 4.55E-02 | 1.25E-01 | -1.38E+00 |
| SERPINB9 | 1.12E-02 | 5.83E-02 | -1.01E+00 | 8.60E-02 | 3.72E-01 | 1.57E+00 | 1.17E-03 | 4.95E-03 | -1.36E+00 | 6.83E-02 | 2.61E-01 | 1.15E+00 | 1.24E-03 | 4.41E-03 | -1.31E+00 | 1.98E-01 | 5.59E-01 | 1.21E+00 | 6.50E-08 | 2.39E-07 | -1.45E+00 | 3.95E-04 | 1.08E-03 | -1.13E+00 |
| AC016894.1 | 1.12E-02 | 5.83E-02 | 1.06E+00 | 9.57E-04 | 4.13E-03 | -1.05E+00 | 1.19E-01 | 5.06E-01 | 1.08E+00 | 1.78E-01 | 6.79E-01 | 1.28E+00 | 1.19E-04 | 4.26E-04 | -1.46E+00 | 5.23E-04 | 1.47E-03 | -1.28E+00 | 1.29E-02 | 4.73E-02 | -1.11E+00 | 2.59E-02 | 7.11E-02 | -1.03E+00 |
| AL049872.1 | 1.12E-02 | 5.84E-02 | 1.08E+00 | 4.18E-02 | 1.80E-01 | 1.19E+00 | 1.28E-01 | 5.44E-01 | 1.04E+00 | 1.47E-01 | 5.63E-01 | 1.22E+00 | 1.12E-04 | 3.99E-04 | -1.20E+00 | 2.03E-02 | 5.73E-02 | 1.00E+00 | 2.46E-05 | 9.04E-05 | -1.19E+00 | 3.58E-01 | 9.84E-01 | 1.13E+00 |
| ATP8B2 | 1.12E-02 | 5.84E-02 | -1.00E+00 | 2.29E-01 | 9.91E-01 | 1.62E+00 | 1.20E-02 | 5.11E-02 | -1.21E+00 | 8.21E-03 | 3.14E-02 | -1.06E+00 | 1.42E-01 | 5.06E-01 | 1.32E+00 | 1.21E-02 | 3.42E-02 | 1.04E+00 | 1.95E-03 | 7.15E-03 | -1.35E+00 | 2.66E-03 | 7.32E-03 | -1.36E+00 |
| NBPF19 | 1.12E-02 | 5.84E-02 | 1.10E+00 | 2.40E-04 | 1.04E-03 | 1.03E+00 | 9.90E-02 | 4.20E-01 | 1.09E+00 | 7.67E-02 | 2.93E-01 | 1.23E+00 | 3.32E-02 | 1.18E-01 | 1.13E+00 | 1.22E-01 | 3.45E-01 | 1.19E+00 | 1.35E-01 | 4.95E-01 | 1.26E+00 | 3.48E-03 | 9.57E-03 | -1.08E+00 |
| NEK4 | 1.13E-02 | 5.87E-02 | -1.35E+00 | 3.68E-02 | 1.59E-01 | -1.03E+00 | 3.60E-04 | 1.53E-03 | -1.50E+00 | 1.68E-03 | 6.42E-03 | -1.25E+00 | 2.96E-06 | 1.05E-05 | -2.13E+00 | 3.60E-05 | 1.02E-04 | -1.60E+00 | 2.54E-04 | 9.32E-04 | -1.60E+00 | 4.99E-03 | 1.37E-02 | -1.32E+00 |
| GLUD2 | 1.13E-02 | 5.88E-02 | -1.15E+00 | 1.78E-03 | 7.70E-03 | -1.26E+00 | 2.19E-01 | 9.32E-01 | 1.11E+00 | 1.03E-01 | 3.92E-01 | 1.13E+00 | 1.85E-01 | 6.61E-01 | 1.57E+00 | 1.87E-01 | 5.28E-01 | 1.63E+00 | 5.07E-02 | 1.86E-01 | 1.62E+00 | 3.59E-01 | 9.87E-01 | 1.17E+00 |
| SNIP1 | 1.13E-02 | 5.88E-02 | -1.04E+00 | 2.98E-04 | 1.29E-03 | -1.24E+00 | 1.07E-01 | 4.55E-01 | 1.11E+00 | 5.65E-03 | 2.16E-02 | -1.07E+00 | 2.79E-01 | 9.94E-01 | 1.39E+00 | 1.51E-01 | 4.25E-01 | 1.14E+00 | 1.25E-03 | 4.59E-03 | -1.13E+00 | 3.64E-04 | 1.00E-03 | -1.20E+00 |
| AL928970.1 | 1.13E-02 | 5.89E-02 | -1.10E+00 | 1.30E-02 | 5.63E-02 | 1.05E+00 | 1.55E-01 | 6.60E-01 | 1.37E+00 | 1.32E-01 | 5.05E-01 | 1.16E+00 | 1.69E-01 | 6.02E-01 | 1.60E+00 | 7.27E-02 | 2.05E-01 | -1.14E+00 | 1.74E-01 | 6.40E-01 | 1.45E+00 | 5.45E-02 | 1.50E-01 | -1.14E+00 |
| MMAB | 1.13E-02 | 5.89E-02 | 1.17E+00 | 6.96E-03 | 3.01E-02 | 1.26E+00 | 1.03E-01 | 4.36E-01 | 1.15E+00 | 6.07E-02 | 2.32E-01 | 1.25E+00 | 3.63E-02 | 1.30E-01 | 1.21E+00 | 3.54E-01 | 9.98E-01 | 1.59E+00 | 4.64E-02 | 1.70E-01 | 1.38E+00 | 1.39E-02 | 3.82E-02 | 1.59E+00 |
| PRSS36 | 1.13E-02 | 5.89E-02 | -1.36E+00 | 2.44E-02 | 1.05E-01 | -1.20E+00 | 1.65E-01 | 7.01E-01 | 1.04E+00 | 2.32E-01 | 8.87E-01 | 1.27E+00 | 1.23E-02 | 4.40E-02 | -1.69E+00 | 4.23E-02 | 1.19E-01 | -1.35E+00 | 2.46E-01 | 9.03E-01 | 1.07E+00 | 1.87E-01 | 5.15E-01 | 1.01E+00 |
| S100B | 1.13E-02 | 5.89E-02 | -1.08E+00 | 7.94E-03 | 3.43E-02 | -1.02E+00 | 1.00E-02 | 4.27E-02 | -1.04E+00 | 9.93E-05 | 3.79E-04 | -1.24E+00 | 7.17E-02 | 2.56E-01 | -1.24E+00 | 1.50E-01 | 4.24E-01 | -1.05E+00 | 1.60E-01 | 5.86E-01 | 1.61E+00 | 3.50E-01 | 9.61E-01 | 1.19E+00 |
| DDX11 | 1.13E-02 | 5.90E-02 | -1.13E+00 | 6.67E-02 | 2.88E-01 | 1.15E+00 | 1.80E-01 | 7.66E-01 | 1.19E+00 | 1.30E-01 | 4.96E-01 | 1.54E+00 | 2.14E-01 | 7.65E-01 | 1.30E+00 | 2.60E-01 | 7.34E-01 | 1.72E+00 | 3.63E-04 | 1.33E-03 | -1.24E+00 | 2.21E-01 | 6.07E-01 | 1.32E+00 |
| NOXA1 | 1.13E-02 | 5.90E-02 | -1.14E+00 | 8.66E-02 | 3.74E-01 | 1.17E+00 | 2.02E-01 | 8.60E-01 | 1.14E+00 | 2.18E-01 | 8.34E-01 | 1.43E+00 | 1.80E-03 | 6.42E-03 | -1.62E+00 | 9.40E-03 | 2.65E-02 | -1.31E+00 | 1.04E-01 | 3.80E-01 | 1.53E+00 | 3.25E-01 | 8.93E-01 | 1.06E+00 |
| HMGN3-AS1 | 1.13E-02 | 5.90E-02 | 2.20E+00 | 2.20E-02 | 9.49E-02 | 2.32E+00 | 1.67E-01 | 7.10E-01 | 1.05E+00 | 2.09E-01 | 7.99E-01 | 1.30E+00 | 7.47E-03 | 2.66E-02 | 2.05E+00 | 2.72E-01 | 7.66E-01 | 1.49E+00 | 7.07E-04 | 2.59E-03 | 2.07E+00 | 1.28E-02 | 3.51E-02 | 1.82E+00 |
| NMT2 | 1.13E-02 | 5.91E-02 | -1.27E+00 | 2.25E-01 | 9.71E-01 | 1.44E+00 | 5.15E-03 | 2.19E-02 | -1.67E+00 | 7.62E-02 | 2.91E-01 | 1.06E+00 | 8.57E-09 | 3.06E-08 | -1.62E+00 | 1.04E-03 | 2.93E-03 | -1.06E+00 | 6.69E-09 | 2.45E-08 | -1.75E+00 | 4.53E-05 | 1.24E-04 | -1.31E+00 |
| FGR | 1.14E-02 | 5.92E-02 | -1.47E+00 | 2.98E-02 | 1.29E-01 | -1.01E+00 | 4.12E-02 | 1.75E-01 | -1.21E+00 | 1.75E-02 | 6.68E-02 | -1.27E+00 | 9.43E-02 | 3.36E-01 | -1.59E+00 | 1.22E-01 | 3.44E-01 | -1.65E+00 | 1.38E-01 | 5.05E-01 | -1.11E+00 | 3.35E-01 | 9.19E-01 | 1.17E+00 |
| RNASET2 | 1.14E-02 | 5.92E-02 | 1.00E+00 | 5.81E-04 | 2.51E-03 | 1.07E+00 | 1.60E-01 | 6.79E-01 | 1.12E+00 | 2.04E-01 | 7.81E-01 | 1.12E+00 | 7.04E-06 | 2.51E-05 | -1.02E+00 | 1.16E-05 | 3.28E-05 | -1.01E+00 | 1.05E-03 | 3.84E-03 | 1.02E+00 | 1.33E-02 | 3.65E-02 | 1.07E+00 |
| ZBED3-AS1 | 1.14E-02 | 5.93E-02 | -1.13E+00 | 1.96E-01 | 8.45E-01 | 1.45E+00 | 7.76E-03 | 3.30E-02 | -1.22E+00 | 2.26E-01 | 8.64E-01 | 1.26E+00 | 8.51E-02 | 3.03E-01 | 1.19E+00 | 9.71E-02 | 2.74E-01 | 1.13E+00 | 1.15E-03 | 4.21E-03 | -1.55E+00 | 2.54E-01 | 6.98E-01 | 1.21E+00 |
| WDR27 | 1.14E-02 | 5.95E-02 | -1.07E+00 | 6.62E-02 | 2.86E-01 | 1.21E+00 | 5.41E-04 | 2.30E-03 | -1.38E+00 | 1.85E-02 | 7.06E-02 | -1.00E+00 | 5.43E-03 | 1.94E-02 | -1.21E+00 | 1.42E-01 | 4.01E-01 | 1.25E+00 | 3.89E-02 | 1.43E-01 | -1.06E+00 | 1.68E-01 | 4.61E-01 | 1.13E+00 |
| YWHAZ | 1.14E-02 | 5.96E-02 | 1.35E+00 | 7.70E-13 | 3.33E-12 | -1.08E+00 | 4.82E-02 | 2.05E-01 | 1.29E+00 | 3.65E-03 | 1.40E-02 | -1.03E+00 | 2.51E-01 | 8.95E-01 | 1.22E+00 | 2.03E-04 | 5.73E-04 | -1.11E+00 | 2.46E-01 | 9.04E-01 | 1.11E+00 | 1.07E-03 | 2.93E-03 | -1.20E+00 |
| LINC00476 | 1.14E-02 | 5.96E-02 | -1.06E+00 | 1.07E-01 | 4.64E-01 | 1.25E+00 | 5.73E-03 | 2.44E-02 | -1.21E+00 | 1.69E-02 | 6.46E-02 | 1.03E+00 | 2.05E-03 | 7.29E-03 | -1.07E+00 | 2.64E-02 | 7.44E-02 | 1.11E+00 | 3.56E-03 | 1.31E-02 | -1.14E+00 | 3.55E-02 | 9.74E-02 | -1.05E+00 |
| EIF2B3 | 1.14E-02 | 5.96E-02 | -1.03E+00 | 6.09E-02 | 2.63E-01 | 1.16E+00 | 3.21E-04 | 1.36E-03 | -1.16E+00 | 5.57E-04 | 2.13E-03 | -1.02E+00 | 4.02E-02 | 1.44E-01 | -1.10E+00 | 2.16E-01 | 6.09E-01 | 1.58E+00 | 1.91E-05 | 7.01E-05 | -1.14E+00 | 5.95E-02 | 1.63E-01 | 1.19E+00 |
| BIN1 | 1.14E-02 | 5.97E-02 | 1.44E+00 | 1.74E-02 | 7.52E-02 | 1.60E+00 | 1.95E-01 | 8.27E-01 | 1.27E+00 | 1.63E-01 | 6.23E-01 | 1.44E+00 | 1.90E-01 | 6.77E-01 | 1.49E+00 | 3.27E-01 | 9.22E-01 | 1.51E+00 | 4.51E-03 | 1.66E-02 | -1.11E+00 | 4.33E-02 | 1.19E-01 | -1.12E+00 |
| FP565260.1 | 1.15E-02 | 5.97E-02 | -1.38E+00 | 2.44E-02 | 1.06E-01 | -1.24E+00 | 1.73E-04 | 7.37E-04 | -1.47E+00 | 3.60E-04 | 1.37E-03 | -1.34E+00 | 1.53E-02 | 5.46E-02 | -1.53E+00 | 1.21E-01 | 3.41E-01 | 1.01E+00 | 3.75E-08 | 1.38E-07 | -1.70E+00 | 7.46E-08 | 2.05E-07 | -1.77E+00 |
| SLC18B1 | 1.15E-02 | 5.98E-02 | -1.31E+00 | 6.46E-02 | 2.79E-01 | 1.04E+00 | 4.97E-02 | 2.11E-01 | -1.25E+00 | 9.86E-02 | 3.76E-01 | 1.04E+00 | 3.97E-02 | 1.42E-01 | -1.27E+00 | 2.55E-01 | 7.19E-01 | 1.29E+00 | 1.54E-01 | 5.66E-01 | 1.14E+00 | 3.50E-01 | 9.61E-01 | 1.28E+00 |
| AP000936.3 | 1.15E-02 | 5.98E-02 | 1.18E+00 | 3.72E-45 | 1.61E-44 | -1.72E+00 | 2.00E-02 | 8.48E-02 | 1.16E+00 | 1.08E-18 | 4.12E-18 | -1.71E+00 | 4.37E-02 | 1.56E-01 | 1.21E+00 | 9.94E-18 | 2.80E-17 | -1.67E+00 | 6.13E-02 | 2.25E-01 | 1.24E+00 | 6.84E-16 | 1.88E-15 | -1.63E+00 |
| RPL21P134 | 1.15E-02 | 5.99E-02 | 1.09E+00 | 1.04E-02 | 4.48E-02 | 1.11E+00 | 2.08E-02 | 8.85E-02 | 1.19E+00 | 1.36E-01 | 5.20E-01 | 1.14E+00 | 1.38E-01 | 4.93E-01 | 1.13E+00 | 3.34E-01 | 9.42E-01 | 1.10E+00 | 2.29E-01 | 8.41E-01 | 1.15E+00 | 3.46E-01 | 9.51E-01 | 1.10E+00 |
| RNY3P8 | 1.15E-02 | 5.99E-02 | -1.69E+00 | 2.68E-02 | 1.16E-01 | -1.37E+00 | 3.70E-02 | 1.57E-01 | -1.43E+00 | 7.80E-03 | 2.98E-02 | -1.79E+00 | 2.68E-02 | 9.56E-02 | 1.11E+00 | 2.92E-03 | 8.23E-03 | -1.16E+00 | 8.25E-02 | 3.03E-01 | 1.12E+00 | 4.03E-02 | 1.11E-01 | 1.06E+00 |
| DAPL1 | 1.15E-02 | 6.00E-02 | 2.08E+00 | 1.53E-01 | 6.61E-01 | 1.40E+00 | 1.43E-01 | 6.06E-01 | 1.21E+00 | 1.25E-01 | 4.77E-01 | -1.10E+00 | 4.22E-02 | 1.50E-01 | -1.30E+00 | 4.53E-03 | 1.28E-02 | -2.02E+00 | 1.62E-01 | 5.96E-01 | 1.26E+00 | 3.25E-01 | 8.93E-01 | 1.12E+00 |
| SEC22A | 1.15E-02 | 6.00E-02 | -1.14E+00 | 1.91E-01 | 8.25E-01 | 1.36E+00 | 2.51E-05 | 1.07E-04 | -1.54E+00 | 4.75E-03 | 1.81E-02 | -1.05E+00 | 2.73E-03 | 9.72E-03 | -1.61E+00 | 7.24E-02 | 2.04E-01 | -1.06E+00 | 1.36E-03 | 4.99E-03 | -1.09E+00 | 1.20E-02 | 3.30E-02 | -1.05E+00 |
| ZNF17 | 1.15E-02 | 6.01E-02 | -1.17E+00 | 3.91E-02 | 1.69E-01 | 1.04E+00 | 3.70E-03 | 1.57E-02 | -1.42E+00 | 7.56E-02 | 2.89E-01 | 1.12E+00 | 1.87E-02 | 6.67E-02 | -1.02E+00 | 1.44E-02 | 4.07E-02 | -1.09E+00 | 1.51E-07 | 5.54E-07 | -1.85E+00 | 3.96E-05 | 1.09E-04 | -1.39E+00 |
| GSTCD | 1.15E-02 | 6.01E-02 | -1.22E+00 | 1.50E-01 | 6.47E-01 | 1.40E+00 | 2.56E-02 | 1.09E-01 | -1.07E+00 | 1.71E-02 | 6.55E-02 | 1.05E+00 | 1.04E-03 | 3.71E-03 | -1.71E+00 | 2.92E-02 | 8.23E-02 | -1.06E+00 | 3.46E-04 | 1.27E-03 | -1.20E+00 | 2.81E-03 | 7.73E-03 | -1.09E+00 |
| PDE12 | 1.15E-02 | 6.01E-02 | 1.10E+00 | 1.16E-06 | 5.03E-06 | -1.22E+00 | 1.12E-02 | 4.76E-02 | -1.07E+00 | 7.05E-04 | 2.69E-03 | -1.10E+00 | 2.21E-01 | 7.88E-01 | 1.51E+00 | 8.49E-02 | 2.39E-01 | 1.10E+00 | 1.52E-03 | 5.56E-03 | -1.04E+00 | 3.86E-02 | 1.06E-01 | 1.07E+00 |
| VNN2 | 1.15E-02 | 6.01E-02 | -1.58E+00 | 3.69E-02 | 1.59E-01 | -1.29E+00 | 1.44E-01 | 6.13E-01 | -1.02E+00 | 3.97E-02 | 1.52E-01 | -1.33E+00 | 8.50E-02 | 3.03E-01 | 2.14E+00 | 6.88E-02 | 1.94E-01 | 1.98E+00 | 1.69E-01 | 6.21E-01 | 1.37E+00 | 1.83E-01 | 5.04E-01 | -1.06E+00 |
| NUP50-AS1 | 1.15E-02 | 6.02E-02 | -1.26E+00 | 1.57E-01 | 6.77E-01 | 1.28E+00 | 2.29E-04 | 9.73E-04 | -1.69E+00 | 9.52E-03 | 3.63E-02 | -1.07E+00 | 2.82E-04 | 1.00E-03 | -2.00E+00 | 1.37E-02 | 3.85E-02 | -1.22E+00 | 6.06E-09 | 2.23E-08 | -2.36E+00 | 5.16E-03 | 1.42E-02 | -1.08E+00 |
| AP4E1 | 1.16E-02 | 6.04E-02 | -1.30E+00 | 8.94E-02 | 3.86E-01 | 1.14E+00 | 6.49E-02 | 2.76E-01 | -1.06E+00 | 1.01E-01 | 3.87E-01 | 1.15E+00 | 9.82E-02 | 3.50E-01 | 2.05E+00 | 3.31E-01 | 9.34E-01 | 1.35E+00 | 1.17E-01 | 4.29E-01 | 1.05E+00 | 3.46E-01 | 9.50E-01 | 1.22E+00 |
| CAMK2D | 1.16E-02 | 6.05E-02 | -1.32E+00 | 2.29E-02 | 9.90E-02 | -1.32E+00 | 6.06E-05 | 2.57E-04 | -1.75E+00 | 3.19E-05 | 1.22E-04 | -1.72E+00 | 2.79E-01 | 9.94E-01 | 1.14E+00 | 3.29E-01 | 9.27E-01 | 1.16E+00 | 3.77E-02 | 1.38E-01 | -1.19E+00 | 1.19E-01 | 3.26E-01 | -1.29E+00 |
| CCDC62 | 1.16E-02 | 6.06E-02 | -1.06E+00 | 1.76E-01 | 7.59E-01 | 1.45E+00 | 7.95E-03 | 3.38E-02 | -1.18E+00 | 3.73E-02 | 1.42E-01 | 1.14E+00 | 7.98E-02 | 2.85E-01 | 1.06E+00 | 1.22E-01 | 3.45E-01 | 1.02E+00 | 2.23E-01 | 8.18E-01 | 1.38E+00 | 2.71E-01 | 7.44E-01 | 1.40E+00 |
| AC022007.1 | 1.16E-02 | 6.06E-02 | -1.47E+00 | 3.93E-02 | 1.70E-01 | -1.14E+00 | 1.25E-01 | 5.32E-01 | 1.34E+00 | 2.54E-01 | 9.69E-01 | 1.39E+00 | 2.97E-02 | 1.06E-01 | -1.53E+00 | 5.45E-02 | 1.54E-01 | -1.39E+00 | 1.61E-01 | 5.90E-01 | 1.13E+00 | 3.29E-01 | 9.02E-01 | 1.31E+00 |
| CLMN | 1.16E-02 | 6.07E-02 | -1.06E+00 | 3.42E-03 | 1.48E-02 | -1.06E+00 | 2.01E-09 | 8.53E-09 | -1.98E+00 | 5.06E-07 | 1.93E-06 | -1.38E+00 | 1.50E-03 | 5.36E-03 | -1.46E+00 | 7.63E-03 | 2.15E-02 | -1.23E+00 | 2.24E-01 | 8.22E-01 | 1.65E+00 | 2.47E-01 | 6.79E-01 | 1.13E+00 |
| AC020658.5 | 1.16E-02 | 6.07E-02 | -1.36E+00 | 2.29E-01 | 9.90E-01 | 1.39E+00 | 1.67E-01 | 7.08E-01 | 1.37E+00 | 6.80E-03 | 2.60E-02 | 2.41E+00 | 1.55E-01 | 5.53E-01 | -1.16E+00 | 3.08E-01 | 8.67E-01 | 1.29E+00 | 3.78E-02 | 1.39E-01 | -1.60E+00 | 3.11E-01 | 8.53E-01 | 1.18E+00 |
| ADCY7 | 1.17E-02 | 6.08E-02 | -1.10E+00 | 1.21E-01 | 5.22E-01 | 1.32E+00 | 1.54E-01 | 6.55E-01 | 1.07E+00 | 1.79E-01 | 6.83E-01 | 1.21E+00 | 1.26E-01 | 4.50E-01 | 1.10E+00 | 9.15E-02 | 2.58E-01 | -1.09E+00 | 1.32E-01 | 4.84E-01 | -1.01E+00 | 1.41E-02 | 3.87E-02 | -1.36E+00 |
| TPT1P4 | 1.17E-02 | 6.08E-02 | 1.71E+00 | 2.74E-05 | 1.18E-04 | 2.35E+00 | 5.91E-03 | 2.51E-02 | 1.69E+00 | 1.23E-04 | 4.69E-04 | 2.20E+00 | 2.01E-01 | 7.18E-01 | 1.45E+00 | 1.50E-01 | 4.24E-01 | 1.70E+00 | 4.76E-02 | 1.75E-01 | 1.58E+00 | 2.96E-02 | 8.14E-02 | 1.85E+00 |
| ZNF34 | 1.17E-02 | 6.08E-02 | -1.19E+00 | 1.95E-02 | 8.42E-02 | -1.03E+00 | 7.37E-05 | 3.13E-04 | -1.58E+00 | 1.04E-03 | 3.98E-03 | -1.24E+00 | 2.60E-01 | 9.27E-01 | 1.34E+00 | 1.45E-01 | 4.09E-01 | 1.81E+00 | 3.26E-02 | 1.20E-01 | -1.15E+00 | 1.11E-01 | 3.06E-01 | -1.02E+00 |
| CCDC77 | 1.17E-02 | 6.09E-02 | -1.16E+00 | 8.59E-02 | 3.71E-01 | 1.23E+00 | 4.92E-07 | 2.09E-06 | -1.88E+00 | 1.34E-02 | 5.11E-02 | -1.02E+00 | 5.39E-04 | 1.92E-03 | -1.79E+00 | 2.35E-03 | 6.63E-03 | -1.40E+00 | 6.72E-06 | 2.47E-05 | -1.41E+00 | 7.86E-03 | 2.16E-02 | -1.08E+00 |
| CYTH1 | 1.17E-02 | 6.10E-02 | 1.64E+00 | 2.28E-01 | 9.87E-01 | 1.57E+00 | 2.15E-01 | 9.13E-01 | 1.29E+00 | 1.57E-02 | 5.99E-02 | 1.18E+00 | 1.87E-01 | 6.67E-01 | 1.40E+00 | 4.75E-02 | 1.34E-01 | 1.09E+00 | 2.24E-01 | 8.23E-01 | 1.22E+00 | 1.12E-01 | 3.09E-01 | -1.05E+00 |
| DNAJB12 | 1.17E-02 | 6.10E-02 | 1.13E+00 | 2.13E-05 | 9.21E-05 | 1.03E+00 | 5.02E-02 | 2.13E-01 | 1.09E+00 | 2.35E-04 | 8.96E-04 | -1.02E+00 | 1.75E-03 | 6.23E-03 | 1.10E+00 | 4.39E-05 | 1.24E-04 | -1.04E+00 | 1.87E-02 | 6.86E-02 | 1.01E+00 | 1.88E-03 | 5.16E-03 | -1.04E+00 |
| BACH1 | 1.17E-02 | 6.10E-02 | 1.12E+00 | 5.70E-08 | 2.46E-07 | -1.12E+00 | 3.35E-03 | 1.42E-02 | -1.07E+00 | 1.50E-07 | 5.74E-07 | -1.24E+00 | 6.89E-02 | 2.46E-01 | 1.19E+00 | 4.39E-05 | 1.24E-04 | -1.21E+00 | 5.34E-02 | 1.96E-01 | 1.20E+00 | 2.47E-05 | 6.80E-05 | -1.30E+00 |
| NAT8B | 1.17E-02 | 6.11E-02 | -1.34E+00 | 5.93E-02 | 2.56E-01 | 1.01E+00 | 1.03E-02 | 4.38E-02 | -1.58E+00 | 1.48E-01 | 5.67E-01 | 1.09E+00 | 2.04E-01 | 7.28E-01 | 1.33E+00 | 1.24E-01 | 3.49E-01 | -1.02E+00 | 2.42E-02 | 8.89E-02 | 2.18E+00 | 2.17E-02 | 5.95E-02 | 1.08E+00 |
| FCMR | 1.17E-02 | 6.12E-02 | 1.78E+00 | 5.48E-07 | 2.37E-06 | 2.89E+00 | 3.96E-02 | 1.68E-01 | 1.64E+00 | 8.26E-02 | 3.15E-01 | 1.60E+00 | 2.62E-01 | 9.34E-01 | 1.19E+00 | 2.99E-01 | 8.43E-01 | -1.20E+00 | 1.99E-01 | 7.32E-01 | 1.26E+00 | 2.84E-01 | 7.80E-01 | 1.28E+00 |
| INTS9 | 1.18E-02 | 6.13E-02 | -1.23E+00 | 1.12E-01 | 4.82E-01 | 1.23E+00 | 5.04E-05 | 2.14E-04 | -1.67E+00 | 9.73E-03 | 3.72E-02 | -1.08E+00 | 3.47E-04 | 1.24E-03 | -1.67E+00 | 3.52E-02 | 9.93E-02 | 1.06E+00 | 1.28E-03 | 4.69E-03 | -1.16E+00 | 3.54E-01 | 9.72E-01 | 1.35E+00 |
| TSTD3 | 1.18E-02 | 6.13E-02 | -1.12E+00 | 6.19E-02 | 2.67E-01 | 1.16E+00 | 6.09E-02 | 2.59E-01 | -1.29E+00 | 7.74E-02 | 2.96E-01 | -1.19E+00 | 1.65E-02 | 5.88E-02 | 1.00E+00 | 2.66E-02 | 7.49E-02 | 1.07E+00 | 1.60E-01 | 5.89E-01 | 1.10E+00 | 9.30E-02 | 2.55E-01 | 1.05E+00 |
| ZNF845 | 1.18E-02 | 6.13E-02 | -1.14E+00 | 2.16E-04 | 9.35E-04 | -1.54E+00 | 1.58E-02 | 6.72E-02 | -1.10E+00 | 1.56E-05 | 5.96E-05 | -1.81E+00 | 2.62E-01 | 9.35E-01 | 1.52E+00 | 7.32E-02 | 2.06E-01 | 1.11E+00 | 5.30E-02 | 1.94E-01 | 1.04E+00 | 2.42E-03 | 6.64E-03 | -1.37E+00 |
| RPL13P6 | 1.18E-02 | 6.14E-02 | -1.03E+00 | 1.85E-07 | 7.99E-07 | -1.66E+00 | 2.28E-01 | 9.67E-01 | 1.23E+00 | 1.97E-05 | 7.53E-05 | -1.41E+00 | 2.66E-01 | 9.50E-01 | 1.49E+00 | 1.69E-03 | 4.76E-03 | -1.37E+00 | 2.02E-01 | 7.42E-01 | 1.16E+00 | 3.46E-06 | 9.50E-06 | -1.84E+00 |
| SLC12A7 | 1.18E-02 | 6.15E-02 | -1.12E+00 | 1.03E-01 | 4.44E-01 | 1.19E+00 | 8.63E-04 | 3.67E-03 | -1.27E+00 | 6.11E-03 | 2.33E-02 | 1.02E+00 | 5.65E-05 | 2.01E-04 | -1.66E+00 | 1.97E-03 | 5.55E-03 | -1.31E+00 | 6.31E-10 | 2.32E-09 | -2.34E+00 | 9.38E-06 | 2.58E-05 | -1.52E+00 |
| ZNF616 | 1.18E-02 | 6.16E-02 | -1.20E+00 | 1.20E-01 | 5.20E-01 | 1.32E+00 | 7.38E-06 | 3.13E-05 | -2.90E+00 | 4.75E-03 | 1.81E-02 | -1.46E+00 | 6.96E-05 | 2.48E-04 | -2.34E+00 | 1.62E-03 | 4.56E-03 | -1.51E+00 | 4.34E-03 | 1.59E-02 | -1.10E+00 | 1.54E-03 | 4.24E-03 | -1.35E+00 |
| ATG5 | 1.18E-02 | 6.17E-02 | 1.06E+00 | 6.93E-03 | 3.00E-02 | 1.12E+00 | 6.66E-05 | 2.83E-04 | -1.20E+00 | 1.32E-05 | 5.06E-05 | -1.08E+00 | 1.38E-07 | 4.91E-07 | -1.45E+00 | 1.05E-03 | 2.97E-03 | -1.03E+00 | 8.17E-07 | 3.00E-06 | -1.22E+00 | 2.35E-02 | 6.46E-02 | 1.08E+00 |
| PMS2P7 | 1.19E-02 | 6.18E-02 | -1.20E+00 | 2.13E-02 | 9.19E-02 | -1.03E+00 | 3.68E-07 | 1.57E-06 | -1.96E+00 | 2.81E-05 | 1.07E-04 | -1.38E+00 | 4.48E-05 | 1.60E-04 | -2.15E+00 | 3.47E-04 | 9.78E-04 | -1.80E+00 | 1.70E-01 | 6.25E-01 | 1.19E+00 | 2.61E-01 | 7.16E-01 | 1.16E+00 |
| ALKBH7 | 1.19E-02 | 6.18E-02 | 1.14E+00 | 9.70E-05 | 4.19E-04 | 1.16E+00 | 1.91E-01 | 8.11E-01 | 1.06E+00 | 8.26E-02 | 3.16E-01 | 1.12E+00 | 1.78E-03 | 6.34E-03 | 1.00E+00 | 1.11E-02 | 3.13E-02 | 1.05E+00 | 1.17E-01 | 4.28E-01 | 1.10E+00 | 2.23E-01 | 6.13E-01 | 1.06E+00 |
| CASC19 | 1.19E-02 | 6.19E-02 | 2.12E+00 | 6.33E-02 | 2.73E-01 | 2.30E+00 | 2.93E-02 | 1.25E-01 | 2.07E+00 | 1.20E-01 | 4.57E-01 | 2.00E+00 | 2.66E-01 | 9.50E-01 | 1.28E+00 | 2.82E-01 | 7.95E-01 | -1.12E+00 | 2.24E-01 | 8.21E-01 | -1.16E+00 | 2.68E-01 | 7.36E-01 | 1.28E+00 |
| TOMM40L | 1.19E-02 | 6.20E-02 | -1.25E+00 | 3.91E-02 | 1.69E-01 | -1.03E+00 | 1.12E-01 | 4.74E-01 | 1.00E+00 | 1.09E-01 | 4.16E-01 | 1.74E+00 | 5.38E-02 | 1.92E-01 | -1.23E+00 | 1.62E-01 | 4.58E-01 | 1.02E+00 | 2.35E-09 | 8.64E-09 | -1.89E+00 | 9.41E-05 | 2.58E-04 | -1.25E+00 |
| ZNF558 | 1.20E-02 | 6.23E-02 | -1.22E+00 | 9.45E-02 | 4.08E-01 | 1.12E+00 | 2.40E-04 | 1.02E-03 | -1.53E+00 | 7.59E-03 | 2.90E-02 | -1.06E+00 | 1.92E-01 | 6.85E-01 | 1.17E+00 | 3.06E-01 | 8.62E-01 | 1.35E+00 | 2.03E-04 | 7.44E-04 | -1.56E+00 | 1.60E-01 | 4.39E-01 | 1.14E+00 |
| GPR107 | 1.20E-02 | 6.23E-02 | 1.00E+00 | 1.58E-02 | 6.83E-02 | 1.13E+00 | 9.48E-02 | 4.03E-01 | 1.05E+00 | 1.05E-02 | 4.00E-02 | -1.05E+00 | 2.33E-01 | 8.30E-01 | 1.28E+00 | 2.23E-01 | 6.28E-01 | 1.18E+00 | 8.43E-02 | 3.09E-01 | 1.11E+00 | 2.31E-02 | 6.34E-02 | -1.06E+00 |
| HERC2P9 | 1.20E-02 | 6.24E-02 | -1.14E+00 | 2.64E-02 | 1.14E-01 | 1.05E+00 | 1.27E-05 | 5.38E-05 | -1.59E+00 | 7.71E-06 | 2.95E-05 | -1.34E+00 | 6.66E-02 | 2.38E-01 | 1.04E+00 | 2.31E-02 | 6.53E-02 | 1.95E+00 | 4.67E-02 | 1.71E-01 | 1.24E+00 | 1.01E-01 | 2.78E-01 | 1.17E+00 |
| AC008074.3 | 1.20E-02 | 6.24E-02 | -1.05E+00 | 2.03E-03 | 8.75E-03 | -1.04E+00 | 1.22E-01 | 5.19E-01 | -1.07E+00 | 2.32E-02 | 8.86E-02 | -1.23E+00 | 1.82E-01 | 6.50E-01 | 1.22E+00 | 2.12E-01 | 5.98E-01 | 1.21E+00 | 2.40E-01 | 8.82E-01 | 1.57E+00 | 1.00E-01 | 2.75E-01 | -1.26E+00 |
| SH3D19 | 1.20E-02 | 6.25E-02 | -1.07E+00 | 4.70E-02 | 2.03E-01 | 1.14E+00 | 1.24E-04 | 5.27E-04 | -1.38E+00 | 2.09E-03 | 8.00E-03 | -1.04E+00 | 2.76E-02 | 9.85E-02 | -1.14E+00 | 1.81E-01 | 5.09E-01 | 1.11E+00 | 1.56E-01 | 5.72E-01 | 1.13E+00 | 3.08E-01 | 8.47E-01 | 1.19E+00 |
| DRAXIN | 1.20E-02 | 6.26E-02 | -1.37E+00 | 6.69E-02 | 2.89E-01 | -1.02E+00 | 5.03E-06 | 2.14E-05 | -1.82E+00 | 4.06E-04 | 1.55E-03 | -1.36E+00 | 1.41E-01 | 5.01E-01 | -1.08E+00 | 2.90E-01 | 8.17E-01 | 1.10E+00 | 2.54E-01 | 9.32E-01 | 1.04E+00 | 3.04E-01 | 8.35E-01 | 1.04E+00 |
| AC090950.2 | 1.20E-02 | 6.26E-02 | 1.98E+00 | 1.56E-01 | 6.74E-01 | 1.70E+00 | 6.72E-02 | 2.85E-01 | 1.49E+00 | 1.13E-01 | 4.31E-01 | 1.67E+00 | 2.26E-04 | 8.07E-04 | 3.26E+00 | 2.25E-02 | 6.35E-02 | 2.60E+00 | 2.39E-03 | 8.77E-03 | 2.54E+00 | 1.37E-02 | 3.76E-02 | 2.56E+00 |
| BCOR | 1.21E-02 | 6.29E-02 | -1.02E+00 | 1.85E-01 | 7.99E-01 | 1.42E+00 | 1.81E-04 | 7.67E-04 | -1.27E+00 | 1.09E-01 | 4.17E-01 | 1.30E+00 | 2.03E-02 | 7.25E-02 | -1.11E+00 | 2.50E-01 | 7.04E-01 | 1.27E+00 | 7.99E-07 | 2.93E-06 | -1.40E+00 | 1.03E-02 | 2.84E-02 | 1.04E+00 |
| MRPL1 | 1.21E-02 | 6.31E-02 | 1.16E+00 | 2.44E-02 | 1.05E-01 | 1.32E+00 | 1.85E-03 | 7.84E-03 | -1.03E+00 | 8.80E-03 | 3.36E-02 | 1.10E+00 | 6.28E-05 | 2.24E-04 | -1.14E+00 | 3.30E-02 | 9.30E-02 | 1.15E+00 | 3.62E-04 | 1.33E-03 | -1.10E+00 | 1.54E-01 | 4.23E-01 | 1.09E+00 |
| AC114490.1 | 1.21E-02 | 6.33E-02 | -1.26E+00 | 1.21E-01 | 5.24E-01 | 1.16E+00 | 1.23E-01 | 5.22E-01 | -1.06E+00 | 1.05E-01 | 4.01E-01 | 1.62E+00 | 2.88E-02 | 1.03E-01 | -1.52E+00 | 2.98E-01 | 8.40E-01 | 1.29E+00 | 2.34E-01 | 8.60E-01 | 1.31E+00 | 2.27E-01 | 6.23E-01 | 1.38E+00 |
| PLEC | 1.22E-02 | 6.35E-02 | 1.82E+00 | 5.04E-03 | 2.18E-02 | 2.24E+00 | 1.96E-01 | 8.31E-01 | 1.32E+00 | 2.29E-01 | 8.77E-01 | 1.10E+00 | 2.63E-01 | 9.39E-01 | 1.36E+00 | 3.30E-01 | 9.31E-01 | 1.20E+00 | 1.96E-02 | 7.19E-02 | 3.57E+00 | 1.64E-01 | 4.50E-01 | 2.38E+00 |
| ARHGEF26 | 1.22E-02 | 6.35E-02 | -1.38E+00 | 1.01E-01 | 4.35E-01 | 1.12E+00 | 4.40E-09 | 1.87E-08 | -2.30E+00 | 5.00E-06 | 1.91E-05 | -1.43E+00 | 2.57E-07 | 9.16E-07 | -2.86E+00 | 1.20E-06 | 3.38E-06 | -2.48E+00 | 8.15E-14 | 2.99E-13 | -3.26E+00 | 4.83E-07 | 1.33E-06 | -1.69E+00 |
| RHOBTB2 | 1.22E-02 | 6.35E-02 | -1.49E+00 | 1.89E-02 | 8.16E-02 | -1.29E+00 | 9.67E-02 | 4.11E-01 | -1.21E+00 | 1.59E-01 | 6.06E-01 | 1.32E+00 | 2.77E-02 | 9.88E-02 | -1.56E+00 | 1.63E-01 | 4.59E-01 | -1.04E+00 | 4.65E-04 | 1.71E-03 | -1.46E+00 | 5.34E-03 | 1.47E-02 | -1.39E+00 |
| ACLY | 1.22E-02 | 6.36E-02 | 1.07E+00 | 2.14E-01 | 9.23E-01 | 1.54E+00 | 1.67E-04 | 7.09E-04 | -1.14E+00 | 2.28E-01 | 8.71E-01 | 1.46E+00 | 8.63E-07 | 3.08E-06 | -1.31E+00 | 4.07E-02 | 1.15E-01 | 1.22E+00 | 4.93E-07 | 1.81E-06 | -1.30E+00 | 3.23E-01 | 8.89E-01 | 1.20E+00 |
| ZNF41 | 1.22E-02 | 6.37E-02 | -1.18E+00 | 5.35E-02 | 2.31E-01 | 1.05E+00 | 5.34E-03 | 2.27E-02 | -1.33E+00 | 3.69E-03 | 1.41E-02 | -1.24E+00 | 3.38E-02 | 1.21E-01 | -1.43E+00 | 2.62E-02 | 7.38E-02 | -1.50E+00 | 9.22E-03 | 3.38E-02 | -1.24E+00 | 1.18E-01 | 3.25E-01 | -1.08E+00 |
| PRKCB | 1.22E-02 | 6.37E-02 | 1.09E+00 | 1.64E-01 | 7.08E-01 | 1.42E+00 | 1.95E-02 | 8.29E-02 | -1.02E+00 | 7.54E-02 | 2.88E-01 | 1.21E+00 | 9.75E-03 | 3.48E-02 | 1.09E+00 | 1.38E-01 | 3.89E-01 | 1.33E+00 | 3.14E-02 | 1.15E-01 | -1.03E+00 | 8.51E-03 | 2.34E-02 | -1.15E+00 |
| RABGEF1 | 1.22E-02 | 6.38E-02 | -1.02E+00 | 2.93E-05 | 1.26E-04 | -1.64E+00 | 4.12E-03 | 1.75E-02 | -1.14E+00 | 1.84E-06 | 7.02E-06 | -1.56E+00 | 8.02E-02 | 2.86E-01 | 1.21E+00 | 1.62E-03 | 4.57E-03 | -1.32E+00 | 1.05E-03 | 3.85E-03 | -1.14E+00 | 5.48E-06 | 1.50E-05 | -1.44E+00 |
| SPNS1 | 1.22E-02 | 6.38E-02 | -1.24E+00 | 9.33E-02 | 4.03E-01 | 1.15E+00 | 2.42E-02 | 1.03E-01 | -1.29E+00 | 8.18E-02 | 3.12E-01 | 1.01E+00 | 1.23E-06 | 4.39E-06 | -2.24E+00 | 4.35E-06 | 1.23E-05 | -2.09E+00 | 9.37E-04 | 3.44E-03 | -1.51E+00 | 8.58E-03 | 2.36E-02 | -1.35E+00 |
| SEPT8 | 1.23E-02 | 6.39E-02 | -1.19E+00 | 4.65E-02 | 2.01E-01 | 1.09E+00 | 2.36E-02 | 1.00E-01 | -1.14E+00 | 2.58E-01 | 9.84E-01 | 1.46E+00 | 5.03E-04 | 1.79E-03 | -1.76E+00 | 3.39E-03 | 9.56E-03 | -1.39E+00 | 1.15E-07 | 4.22E-07 | -1.68E+00 | 1.36E-05 | 3.73E-05 | -1.34E+00 |
| AC027309.2 | 1.23E-02 | 6.39E-02 | 1.04E+00 | 1.49E-04 | 6.43E-04 | -1.06E+00 | 4.67E-02 | 1.98E-01 | 1.29E+00 | 3.68E-03 | 1.40E-02 | -1.05E+00 | 4.00E-02 | 1.42E-01 | 1.09E+00 | 1.36E-02 | 3.84E-02 | 1.05E+00 | 3.50E-02 | 1.29E-01 | 1.08E+00 | 7.05E-03 | 1.94E-02 | -1.01E+00 |
| STT3B | 1.23E-02 | 6.39E-02 | 1.19E+00 | 1.17E-03 | 5.05E-03 | 1.27E+00 | 2.04E-01 | 8.66E-01 | 1.25E+00 | 4.82E-02 | 1.84E-01 | 1.21E+00 | 1.36E-04 | 4.84E-04 | -1.11E+00 | 6.78E-05 | 1.91E-04 | -1.11E+00 | 1.25E-02 | 4.59E-02 | 1.05E+00 | 1.79E-02 | 4.93E-02 | 1.04E+00 |
| RPL5P18 | 1.23E-02 | 6.39E-02 | 1.18E+00 | 6.02E-03 | 2.60E-02 | 1.98E+00 | 1.37E-01 | 5.83E-01 | 1.32E+00 | 7.29E-08 | 2.78E-07 | 2.33E+00 | 8.31E-02 | 2.96E-01 | 1.72E+00 | 3.85E-05 | 1.09E-04 | 2.54E+00 | 1.50E-01 | 5.51E-01 | 1.40E+00 | 1.32E-04 | 3.62E-04 | 2.03E+00 |
| TMEM134 | 1.23E-02 | 6.39E-02 | 1.16E+00 | 2.09E-03 | 9.04E-03 | 1.22E+00 | 2.04E-01 | 8.66E-01 | 1.25E+00 | 1.24E-01 | 4.75E-01 | 1.26E+00 | 2.01E-01 | 7.16E-01 | 1.42E+00 | 2.60E-01 | 7.32E-01 | 1.25E+00 | 2.33E-01 | 8.54E-01 | 1.39E+00 | 1.72E-01 | 4.73E-01 | 1.16E+00 |
| KLHDC2 | 1.23E-02 | 6.40E-02 | 1.11E+00 | 2.73E-03 | 1.18E-02 | 1.11E+00 | 5.80E-02 | 2.46E-01 | 1.03E+00 | 8.44E-03 | 3.23E-02 | -1.01E+00 | 2.49E-02 | 8.87E-02 | 1.09E+00 | 1.89E-02 | 5.33E-02 | -1.02E+00 | 6.93E-04 | 2.54E-03 | -1.19E+00 | 1.33E-04 | 3.66E-04 | -1.28E+00 |
| ZNF174 | 1.23E-02 | 6.40E-02 | -1.14E+00 | 5.10E-02 | 2.20E-01 | 1.12E+00 | 5.85E-05 | 2.49E-04 | -1.43E+00 | 6.85E-03 | 2.62E-02 | 1.02E+00 | 6.75E-05 | 2.40E-04 | -1.45E+00 | 6.28E-05 | 1.77E-04 | -1.46E+00 | 1.02E-10 | 3.73E-10 | -1.85E+00 | 1.34E-10 | 3.67E-10 | -1.79E+00 |
| PAPSS2 | 1.23E-02 | 6.41E-02 | -1.58E+00 | 3.38E-02 | 1.46E-01 | -1.33E+00 | 9.92E-03 | 4.22E-02 | -1.52E+00 | 7.76E-02 | 2.96E-01 | -1.13E+00 | 1.54E-01 | 5.51E-01 | -1.17E+00 | 2.65E-01 | 7.48E-01 | 1.03E+00 | 2.16E-01 | 7.92E-01 | 1.31E+00 | 2.77E-01 | 7.61E-01 | 1.08E+00 |
| GMFG | 1.24E-02 | 6.44E-02 | 1.12E+00 | 4.64E-02 | 2.00E-01 | 1.01E+00 | 5.54E-03 | 2.35E-02 | 1.11E+00 | 2.07E-01 | 7.90E-01 | 1.02E+00 | 9.31E-02 | 3.32E-01 | 1.11E+00 | 3.02E-01 | 8.51E-01 | 1.06E+00 | 1.48E-05 | 5.41E-05 | 1.28E+00 | 2.58E-02 | 7.09E-02 | -1.02E+00 |
| AC008026.2 | 1.24E-02 | 6.46E-02 | 1.47E+00 | 1.37E-02 | 5.94E-02 | 1.32E+00 | 2.09E-03 | 8.90E-03 | 1.46E+00 | 1.01E-01 | 3.86E-01 | 1.44E+00 | 1.76E-01 | 6.28E-01 | 1.51E+00 | 2.29E-01 | 6.46E-01 | 1.42E+00 | 9.17E-03 | 3.36E-02 | 1.45E+00 | 1.40E-02 | 3.84E-02 | 1.44E+00 |
| C17orf100 | 1.24E-02 | 6.46E-02 | -1.13E+00 | 1.18E-01 | 5.11E-01 | 1.29E+00 | 6.67E-02 | 2.83E-01 | -1.20E+00 | 2.27E-01 | 8.68E-01 | 1.45E+00 | 1.26E-05 | 4.49E-05 | -1.93E+00 | 7.87E-03 | 2.22E-02 | -1.11E+00 | 1.64E-02 | 6.02E-02 | -1.29E+00 | 2.80E-01 | 7.70E-01 | 1.43E+00 |
| CIB2 | 1.24E-02 | 6.47E-02 | -1.27E+00 | 3.27E-03 | 1.41E-02 | -1.40E+00 | 7.07E-03 | 3.00E-02 | -1.25E+00 | 2.50E-02 | 9.55E-02 | -1.03E+00 | 1.01E-04 | 3.62E-04 | -2.02E+00 | 7.59E-04 | 2.14E-03 | -1.56E+00 | 1.71E-01 | 6.28E-01 | 1.17E+00 | 2.45E-01 | 6.73E-01 | 1.27E+00 |
| FKBP11 | 1.24E-02 | 6.48E-02 | 1.16E+00 | 5.32E-03 | 2.30E-02 | 1.24E+00 | 3.27E-02 | 1.39E-01 | 1.09E+00 | 3.03E-02 | 1.16E-01 | 1.21E+00 | 2.38E-11 | 8.50E-11 | -1.80E+00 | 3.64E-05 | 1.03E-04 | -1.13E+00 | 5.75E-03 | 2.11E-02 | 1.09E+00 | 3.54E-01 | 9.72E-01 | 1.34E+00 |
| BANP | 1.24E-02 | 6.48E-02 | 1.03E+00 | 5.74E-02 | 2.48E-01 | 1.25E+00 | 1.57E-03 | 6.69E-03 | -1.25E+00 | 6.98E-03 | 2.67E-02 | 1.01E+00 | 1.19E-01 | 4.24E-01 | 1.20E+00 | 1.47E-01 | 4.14E-01 | 1.23E+00 | 5.24E-02 | 1.92E-01 | 1.04E+00 | 9.11E-02 | 2.50E-01 | 1.08E+00 |
| ANKRD36B | 1.25E-02 | 6.49E-02 | 1.02E+00 | 1.25E-01 | 5.42E-01 | 1.34E+00 | 3.74E-02 | 1.59E-01 | 1.05E+00 | 3.50E-02 | 1.34E-01 | 1.26E+00 | 1.45E-03 | 5.16E-03 | -1.08E+00 | 8.36E-03 | 2.36E-02 | 1.05E+00 | 2.68E-02 | 9.83E-02 | 1.02E+00 | 2.47E-01 | 6.78E-01 | 1.22E+00 |
| BCL2L12 | 1.25E-02 | 6.50E-02 | -1.03E+00 | 3.44E-02 | 1.48E-01 | 1.20E+00 | 9.77E-02 | 4.15E-01 | 1.05E+00 | 5.69E-02 | 2.17E-01 | 1.16E+00 | 1.40E-03 | 4.97E-03 | -1.32E+00 | 1.42E-01 | 4.02E-01 | 1.17E+00 | 1.48E-03 | 5.43E-03 | 1.00E+00 | 2.45E-01 | 6.73E-01 | 1.23E+00 |
| HIST1H1B | 1.25E-02 | 6.50E-02 | -1.20E+00 | 1.45E-03 | 6.29E-03 | -1.35E+00 | 1.24E-01 | 5.27E-01 | 1.82E+00 | 1.42E-01 | 5.41E-01 | 1.58E+00 | 2.32E-02 | 8.28E-02 | -1.04E+00 | 2.13E-02 | 6.01E-02 | -1.12E+00 | 1.12E-03 | 4.10E-03 | 1.51E+00 | 4.15E-04 | 1.14E-03 | 1.26E+00 |
| LINC02384 | 1.25E-02 | 6.51E-02 | 1.03E+00 | 9.34E-02 | 4.04E-01 | 1.33E+00 | 2.30E-02 | 9.77E-02 | -1.10E+00 | 9.74E-02 | 3.72E-01 | 1.19E+00 | 2.47E-01 | 8.80E-01 | 1.57E+00 | 3.89E-02 | 1.10E-01 | 2.46E+00 | 1.32E-04 | 4.86E-04 | 2.17E+00 | 2.48E-01 | 6.80E-01 | 1.43E+00 |
| C2orf92 | 1.25E-02 | 6.51E-02 | 1.04E+00 | 7.27E-04 | 3.14E-03 | -1.12E+00 | 6.39E-02 | 2.71E-01 | -1.06E+00 | 8.90E-02 | 3.40E-01 | 1.10E+00 | 2.07E-01 | 7.39E-01 | 1.31E+00 | 1.46E-01 | 4.13E-01 | 1.12E+00 | 1.85E-01 | 6.79E-01 | 1.19E+00 | 1.39E-01 | 3.81E-01 | -1.01E+00 |
| POLK | 1.25E-02 | 6.52E-02 | 1.12E+00 | 3.01E-04 | 1.30E-03 | 1.17E+00 | 7.64E-03 | 3.24E-02 | 1.12E+00 | 1.56E-03 | 5.96E-03 | 1.16E+00 | 1.04E-03 | 3.69E-03 | -1.05E+00 | 1.99E-03 | 5.62E-03 | -1.05E+00 | 5.41E-02 | 1.99E-01 | 1.15E+00 | 1.27E-03 | 3.49E-03 | -1.06E+00 |
| AD000864.1 | 1.25E-02 | 6.52E-02 | 1.77E+00 | 7.43E-03 | 3.21E-02 | -1.19E+00 | 8.84E-02 | 3.76E-01 | 1.05E+00 | 1.09E-04 | 4.17E-04 | -1.62E+00 | 2.52E-02 | 8.99E-02 | -1.56E+00 | 5.62E-02 | 1.58E-01 | -1.32E+00 | 1.98E-01 | 7.25E-01 | 1.03E+00 | 3.43E-01 | 9.42E-01 | 1.18E+00 |
| LYZ | 1.26E-02 | 6.57E-02 | -1.65E+00 | 5.87E-03 | 2.54E-02 | -1.38E+00 | 1.61E-01 | 6.86E-01 | 1.19E+00 | 3.44E-02 | 1.32E-01 | -1.37E+00 | 1.07E-04 | 3.80E-04 | -1.44E+00 | 1.50E-04 | 4.22E-04 | -1.44E+00 | 5.67E-02 | 2.08E-01 | 1.37E+00 | 3.04E-01 | 8.35E-01 | 1.03E+00 |
| THRA1/BTR | 1.26E-02 | 6.57E-02 | -1.21E+00 | 2.02E-01 | 8.71E-01 | 1.43E+00 | 8.08E-02 | 3.43E-01 | -1.06E+00 | 5.91E-02 | 2.26E-01 | -1.08E+00 | 1.90E-01 | 6.77E-01 | 1.93E+00 | 3.44E-01 | 9.71E-01 | 1.27E+00 | 1.59E-01 | 5.82E-01 | 1.22E+00 | 1.76E-01 | 4.83E-01 | 1.50E+00 |
| C7orf49 | 1.26E-02 | 6.57E-02 | -1.17E+00 | 2.98E-02 | 1.29E-01 | 1.01E+00 | 6.56E-06 | 2.79E-05 | -1.81E+00 | 3.84E-03 | 1.47E-02 | -1.13E+00 | 6.73E-03 | 2.40E-02 | -1.35E+00 | 4.23E-02 | 1.19E-01 | -1.06E+00 | 9.17E-02 | 3.36E-01 | 1.07E+00 | 1.91E-02 | 5.26E-02 | -1.15E+00 |
| PLAG1 | 1.26E-02 | 6.58E-02 | 1.69E+00 | 1.11E-01 | 4.78E-01 | 1.69E+00 | 1.34E-01 | 5.71E-01 | 1.03E+00 | 1.07E-01 | 4.09E-01 | 1.15E+00 | 2.20E-01 | 7.86E-01 | 1.37E+00 | 3.26E-01 | 9.19E-01 | 1.37E+00 | 1.13E-01 | 4.16E-01 | 1.09E+00 | 2.26E-01 | 6.20E-01 | 1.21E+00 |
| WDR25 | 1.26E-02 | 6.58E-02 | -1.21E+00 | 3.24E-02 | 1.40E-01 | 1.05E+00 | 3.91E-04 | 1.66E-03 | -1.46E+00 | 4.69E-03 | 1.79E-02 | -1.09E+00 | 1.67E-01 | 5.95E-01 | 1.23E+00 | 1.79E-01 | 5.05E-01 | 1.81E+00 | 2.38E-05 | 8.74E-05 | -1.39E+00 | 1.44E-05 | 3.95E-05 | -1.34E+00 |
| AC099548.2 | 1.27E-02 | 6.60E-02 | -1.07E+00 | 7.05E-02 | 3.05E-01 | 1.24E+00 | 1.18E-02 | 5.03E-02 | -1.20E+00 | 8.73E-02 | 3.34E-01 | 1.10E+00 | 7.20E-02 | 2.57E-01 | -1.30E+00 | 1.47E-01 | 4.14E-01 | -1.16E+00 | 5.02E-03 | 1.84E-02 | -1.92E+00 | 1.95E-02 | 5.37E-02 | -1.69E+00 |
| NUP35 | 1.27E-02 | 6.61E-02 | -1.06E+00 | 2.49E-02 | 1.07E-01 | 1.08E+00 | 2.19E-03 | 9.30E-03 | -1.32E+00 | 1.66E-02 | 6.34E-02 | 1.00E+00 | 7.27E-02 | 2.59E-01 | -1.12E+00 | 1.77E-01 | 4.98E-01 | 1.04E+00 | 8.24E-09 | 3.02E-08 | -1.63E+00 | 2.31E-06 | 6.34E-06 | -1.41E+00 |
| SLC16A2 | 1.27E-02 | 6.62E-02 | 1.03E+00 | 1.42E-01 | 6.14E-01 | 1.38E+00 | 6.17E-02 | 2.62E-01 | -1.07E+00 | 6.84E-02 | 2.61E-01 | 1.08E+00 | 1.31E-01 | 4.67E-01 | -1.79E+00 | 3.51E-01 | 9.89E-01 | -1.02E+00 | 2.40E-01 | 8.80E-01 | -1.04E+00 | 2.77E-01 | 7.61E-01 | -1.27E+00 |
| AC011773.4 | 1.27E-02 | 6.63E-02 | 2.00E+00 | 1.03E-01 | 4.44E-01 | 1.76E+00 | 8.11E-02 | 3.45E-01 | 1.62E+00 | 1.36E-01 | 5.20E-01 | 1.60E+00 | 7.02E-02 | 2.50E-01 | 2.42E+00 | 1.94E-01 | 5.47E-01 | 1.86E+00 | 2.47E-01 | 9.07E-01 | 1.19E+00 | 1.55E-01 | 4.27E-01 | 1.55E+00 |
| MZT2A | 1.27E-02 | 6.63E-02 | 1.32E+00 | 1.44E-02 | 6.21E-02 | 1.37E+00 | 2.54E-04 | 1.08E-03 | 1.45E+00 | 1.58E-03 | 6.02E-03 | 1.44E+00 | 9.76E-02 | 3.48E-01 | 1.18E+00 | 1.24E-01 | 3.49E-01 | 1.38E+00 | 1.98E-01 | 7.26E-01 | 1.26E+00 | 3.05E-03 | 8.37E-03 | 1.43E+00 |
| AC006122.1 | 1.27E-02 | 6.63E-02 | -1.09E+00 | 6.05E-04 | 2.61E-03 | -1.47E+00 | 2.39E-02 | 1.02E-01 | -1.14E+00 | 5.33E-05 | 2.04E-04 | -1.73E+00 | 7.57E-03 | 2.70E-02 | -1.43E+00 | 6.05E-04 | 1.71E-03 | -2.18E+00 | 7.80E-03 | 2.86E-02 | -1.18E+00 | 1.07E-04 | 2.93E-04 | -2.05E+00 |
| JOSD1 | 1.27E-02 | 6.64E-02 | 1.11E+00 | 6.80E-08 | 2.94E-07 | -1.19E+00 | 2.32E-01 | 9.86E-01 | 1.27E+00 | 4.24E-04 | 1.62E-03 | 1.04E+00 | 1.02E-01 | 3.64E-01 | 1.27E+00 | 4.67E-04 | 1.32E-03 | -1.14E+00 | 5.62E-04 | 2.06E-03 | -1.03E+00 | 3.03E-06 | 8.32E-06 | -1.23E+00 |
| AL133243.2 | 1.28E-02 | 6.65E-02 | -1.04E+00 | 5.74E-02 | 2.48E-01 | 1.28E+00 | 8.45E-02 | 3.59E-01 | -1.08E+00 | 1.45E-01 | 5.53E-01 | 1.11E+00 | 2.76E-02 | 9.83E-02 | -1.20E+00 | 6.02E-02 | 1.70E-01 | -1.04E+00 | 1.39E-03 | 5.12E-03 | -1.35E+00 | 1.61E-01 | 4.43E-01 | 1.17E+00 |
| TBC1D14 | 1.28E-02 | 6.65E-02 | -1.17E+00 | 3.63E-02 | 1.57E-01 | 1.10E+00 | 2.34E-02 | 9.94E-02 | -1.11E+00 | 1.04E-01 | 3.97E-01 | 1.27E+00 | 5.05E-02 | 1.80E-01 | -1.03E+00 | 2.49E-02 | 7.01E-02 | -1.16E+00 | 6.04E-02 | 2.22E-01 | 1.06E+00 | 1.17E-01 | 3.20E-01 | 1.07E+00 |
| CHAF1A | 1.28E-02 | 6.67E-02 | -1.05E+00 | 1.75E-02 | 7.55E-02 | 1.03E+00 | 1.31E-01 | 5.56E-01 | 1.22E+00 | 2.04E-01 | 7.79E-01 | 1.38E+00 | 2.30E-03 | 8.20E-03 | -1.63E+00 | 1.95E-02 | 5.50E-02 | -1.28E+00 | 5.48E-02 | 2.01E-01 | 1.00E+00 | 2.18E-02 | 5.99E-02 | 1.36E+00 |
| ZNF251 | 1.28E-02 | 6.68E-02 | -1.11E+00 | 2.05E-01 | 8.87E-01 | 1.47E+00 | 6.92E-02 | 2.94E-01 | -1.04E+00 | 2.43E-01 | 9.26E-01 | 1.28E+00 | 7.17E-02 | 2.56E-01 | 1.10E+00 | 2.67E-01 | 7.53E-01 | 1.35E+00 | 3.06E-02 | 1.12E-01 | 1.02E+00 | 3.52E-01 | 9.67E-01 | 1.37E+00 |
| OSBPL5 | 1.28E-02 | 6.69E-02 | -1.39E+00 | 2.30E-02 | 9.93E-02 | -1.18E+00 | 8.02E-05 | 3.41E-04 | -1.53E+00 | 7.52E-03 | 2.87E-02 | -1.09E+00 | 6.15E-02 | 2.19E-01 | 1.01E+00 | 1.69E-01 | 4.76E-01 | 1.33E+00 | 8.06E-02 | 2.96E-01 | -1.06E+00 | 3.33E-01 | 9.14E-01 | 1.23E+00 |
| ZNF561 | 1.28E-02 | 6.70E-02 | 1.10E+00 | 2.71E-05 | 1.17E-04 | -1.16E+00 | 1.94E-04 | 8.26E-04 | -1.11E+00 | 5.47E-06 | 2.09E-05 | -1.16E+00 | 8.85E-05 | 3.15E-04 | -1.29E+00 | 1.40E-03 | 3.96E-03 | -1.11E+00 | 3.19E-05 | 1.17E-04 | -1.25E+00 | 1.85E-04 | 5.08E-04 | -1.15E+00 |
| GPATCH2 | 1.29E-02 | 6.70E-02 | 1.10E+00 | 2.75E-02 | 1.19E-01 | 1.26E+00 | 2.36E-07 | 1.00E-06 | -1.34E+00 | 3.94E-06 | 1.50E-05 | -1.09E+00 | 1.28E-02 | 4.56E-02 | -1.08E+00 | 2.66E-01 | 7.51E-01 | 1.24E+00 | 2.48E-01 | 9.10E-01 | 1.25E+00 | 8.09E-02 | 2.22E-01 | 1.10E+00 |
| LIPH | 1.29E-02 | 6.71E-02 | -1.33E+00 | 1.73E-01 | 7.48E-01 | 1.25E+00 | 8.92E-02 | 3.79E-01 | 1.42E+00 | 6.40E-02 | 2.44E-01 | 1.79E+00 | 3.26E-02 | 1.16E-01 | -1.46E+00 | 9.88E-02 | 2.78E-01 | -1.11E+00 | 8.03E-02 | 2.95E-01 | 1.66E+00 | 2.28E-01 | 6.27E-01 | 1.89E+00 |
| FBXO30 | 1.29E-02 | 6.73E-02 | -1.09E+00 | 6.57E-03 | 2.84E-02 | -1.11E+00 | 7.79E-02 | 3.31E-01 | 1.03E+00 | 5.79E-02 | 2.21E-01 | 1.08E+00 | 9.70E-03 | 3.46E-02 | -1.30E+00 | 1.41E-03 | 3.98E-03 | -1.53E+00 | 1.06E-06 | 3.87E-06 | -1.69E+00 | 1.25E-07 | 3.43E-07 | -2.02E+00 |
| ZNF75D | 1.29E-02 | 6.73E-02 | -1.29E+00 | 2.27E-01 | 9.82E-01 | 1.37E+00 | 3.95E-04 | 1.68E-03 | -1.64E+00 | 1.29E-03 | 4.92E-03 | -1.29E+00 | 8.56E-02 | 3.05E-01 | -1.23E+00 | 1.86E-01 | 5.25E-01 | 1.64E+00 | 1.65E-01 | 6.06E-01 | -1.03E+00 | 1.49E-01 | 4.09E-01 | 1.48E+00 |
| PPOX | 1.29E-02 | 6.74E-02 | -1.03E+00 | 9.72E-02 | 4.20E-01 | 1.36E+00 | 1.88E-02 | 8.00E-02 | -1.15E+00 | 1.45E-01 | 5.52E-01 | 1.19E+00 | 4.04E-06 | 1.44E-05 | -1.47E+00 | 2.08E-01 | 5.87E-01 | 1.32E+00 | 3.00E-04 | 1.10E-03 | -1.04E+00 | 1.41E-03 | 3.88E-03 | 1.02E+00 |
| CD83 | 1.29E-02 | 6.74E-02 | 1.05E+00 | 3.67E-13 | 1.58E-12 | -2.31E+00 | 1.94E-02 | 8.24E-02 | 1.01E+00 | 1.87E-07 | 7.14E-07 | -1.90E+00 | 3.00E-03 | 1.07E-02 | -1.47E+00 | 5.53E-05 | 1.56E-04 | -2.45E+00 | 2.79E-04 | 1.02E-03 | -1.36E+00 | 3.98E-05 | 1.09E-04 | -2.17E+00 |
| FNIP1 | 1.29E-02 | 6.74E-02 | 1.10E+00 | 1.11E-01 | 4.81E-01 | 1.43E+00 | 3.67E-02 | 1.56E-01 | -1.09E+00 | 2.54E-01 | 9.72E-01 | 1.20E+00 | 1.36E-03 | 4.86E-03 | -1.06E+00 | 7.30E-03 | 2.06E-02 | -1.02E+00 | 6.73E-04 | 2.47E-03 | -1.14E+00 | 6.22E-03 | 1.71E-02 | -1.07E+00 |
| AFG3L1P | 1.29E-02 | 6.74E-02 | -1.24E+00 | 1.01E-01 | 4.37E-01 | 1.25E+00 | 3.22E-02 | 1.37E-01 | -1.12E+00 | 1.14E-01 | 4.35E-01 | 1.15E+00 | 6.24E-02 | 2.22E-01 | -1.18E+00 | 3.51E-01 | 9.90E-01 | 1.46E+00 | 6.54E-04 | 2.40E-03 | -1.15E+00 | 6.74E-02 | 1.85E-01 | 1.16E+00 |
| RPL37P23 | 1.29E-02 | 6.74E-02 | -1.02E+00 | 4.02E-62 | 1.74E-61 | -2.02E+00 | 2.11E-01 | 8.95E-01 | 1.02E+00 | 7.70E-20 | 2.94E-19 | -1.92E+00 | 1.24E-02 | 4.42E-02 | -1.04E+00 | 4.56E-30 | 1.28E-29 | -2.13E+00 | 1.07E-03 | 3.91E-03 | -1.13E+00 | 1.64E-24 | 4.52E-24 | -2.16E+00 |
| CENPV | 1.29E-02 | 6.75E-02 | 1.06E+00 | 5.12E-02 | 2.21E-01 | 1.29E+00 | 1.01E-02 | 4.28E-02 | -1.06E+00 | 9.86E-02 | 3.76E-01 | 1.26E+00 | 1.49E-04 | 5.32E-04 | -1.27E+00 | 3.44E-02 | 9.71E-02 | -1.01E+00 | 2.33E-01 | 8.54E-01 | 1.13E+00 | 1.32E-03 | 3.63E-03 | 1.49E+00 |
| CYP3A5 | 1.30E-02 | 6.76E-02 | -1.15E+00 | 1.47E-01 | 6.36E-01 | 1.70E+00 | 2.84E-03 | 1.21E-02 | -1.40E+00 | 1.45E-01 | 5.54E-01 | 1.30E+00 | 1.44E-02 | 5.13E-02 | -1.36E+00 | 1.15E-01 | 3.24E-01 | 1.07E+00 | 1.26E-01 | 4.61E-01 | 1.42E+00 | 1.70E-01 | 4.68E-01 | 1.56E+00 |
| CCNK | 1.30E-02 | 6.76E-02 | 1.03E+00 | 3.47E-08 | 1.50E-07 | -1.53E+00 | 4.26E-03 | 1.81E-02 | 1.03E+00 | 3.27E-07 | 1.25E-06 | -1.26E+00 | 3.47E-02 | 1.24E-01 | 1.12E+00 | 1.96E-04 | 5.52E-04 | -1.28E+00 | 2.70E-03 | 9.92E-03 | -1.04E+00 | 7.73E-07 | 2.12E-06 | -1.40E+00 |
| ZHX3 | 1.30E-02 | 6.76E-02 | -1.15E+00 | 4.25E-02 | 1.84E-01 | 1.09E+00 | 2.78E-02 | 1.18E-01 | -1.24E+00 | 1.20E-01 | 4.60E-01 | 1.20E+00 | 2.18E-01 | 7.76E-01 | 1.23E+00 | 2.26E-01 | 6.36E-01 | 1.53E+00 | 2.15E-02 | 7.88E-02 | 1.02E+00 | 2.81E-01 | 7.73E-01 | 1.31E+00 |
| EIF3KP1 | 1.30E-02 | 6.78E-02 | 2.11E+00 | 6.02E-03 | 2.60E-02 | 2.63E+00 | 7.02E-02 | 2.98E-01 | 1.39E+00 | 3.42E-03 | 1.31E-02 | 2.20E+00 | 5.79E-02 | 2.07E-01 | 2.21E+00 | 5.33E-03 | 1.50E-02 | 2.88E+00 | 1.83E-01 | 6.70E-01 | 1.36E+00 | 1.14E-01 | 3.13E-01 | 1.66E+00 |
| KTN1-AS1 | 1.30E-02 | 6.79E-02 | -1.00E+00 | 4.65E-03 | 2.01E-02 | 1.00E+00 | 6.21E-06 | 2.64E-05 | -1.68E+00 | 4.04E-07 | 1.54E-06 | -1.61E+00 | 2.79E-01 | 9.94E-01 | 1.35E+00 | 2.15E-01 | 6.06E-01 | 1.80E+00 | 1.76E-04 | 6.46E-04 | -1.35E+00 | 5.49E-03 | 1.51E-02 | -1.12E+00 |
| KATNAL1 | 1.30E-02 | 6.80E-02 | 1.05E+00 | 1.62E-02 | 7.02E-02 | 1.17E+00 | 1.72E-01 | 7.31E-01 | 1.31E+00 | 2.35E-01 | 8.98E-01 | 1.36E+00 | 1.15E-03 | 4.09E-03 | -1.24E+00 | 2.70E-02 | 7.60E-02 | 1.03E+00 | 8.49E-02 | 3.12E-01 | 1.09E+00 | 1.42E-02 | 3.89E-02 | -1.14E+00 |
| UBALD2 | 1.30E-02 | 6.80E-02 | 1.12E+00 | 1.84E-10 | 7.94E-10 | -1.21E+00 | 1.04E-01 | 4.44E-01 | 1.26E+00 | 2.66E-09 | 1.01E-08 | -1.25E+00 | 6.87E-03 | 2.45E-02 | -1.01E+00 | 3.61E-05 | 1.02E-04 | -1.20E+00 | 3.00E-02 | 1.10E-01 | 1.16E+00 | 2.63E-06 | 7.23E-06 | -1.20E+00 |
| RPSAP11 | 1.31E-02 | 6.81E-02 | -1.11E+00 | 1.76E-01 | 7.61E-01 | 1.74E+00 | 3.01E-04 | 1.28E-03 | -1.52E+00 | 5.14E-02 | 1.96E-01 | 1.69E+00 | 4.88E-02 | 1.74E-01 | -1.07E+00 | 3.14E-02 | 8.86E-02 | 2.21E+00 | 1.88E-03 | 6.91E-03 | -1.42E+00 | 6.28E-02 | 1.72E-01 | 1.48E+00 |
| CHERP | 1.31E-02 | 6.83E-02 | -1.04E+00 | 5.69E-05 | 2.46E-04 | -1.33E+00 | 2.71E-03 | 1.15E-02 | -1.08E+00 | 4.04E-08 | 1.54E-07 | -1.53E+00 | 1.40E-01 | 4.98E-01 | 1.14E+00 | 5.24E-02 | 1.48E-01 | -1.04E+00 | 1.22E-01 | 4.47E-01 | 1.12E+00 | 5.09E-02 | 1.40E-01 | -1.00E+00 |
| PUDP | 1.31E-02 | 6.83E-02 | -1.51E+00 | 1.21E-01 | 5.22E-01 | 1.11E+00 | 2.12E-02 | 9.03E-02 | -1.26E+00 | 8.06E-02 | 3.08E-01 | 1.07E+00 | 1.30E-02 | 4.65E-02 | -1.70E+00 | 3.17E-02 | 8.94E-02 | -1.50E+00 | 1.49E-05 | 5.48E-05 | -1.66E+00 | 1.18E-01 | 3.25E-01 | 1.09E+00 |
| LRRIQ3 | 1.31E-02 | 6.83E-02 | -1.10E+00 | 4.02E-02 | 1.74E-01 | 1.13E+00 | 1.34E-08 | 5.70E-08 | -2.31E+00 | 2.30E-05 | 8.79E-05 | -1.48E+00 | 3.09E-02 | 1.10E-01 | -1.43E+00 | 3.19E-01 | 8.99E-01 | 1.55E+00 | 3.91E-02 | 1.43E-01 | -1.27E+00 | 1.80E-01 | 4.95E-01 | 1.02E+00 |
| HDHD5 | 1.31E-02 | 6.84E-02 | 1.06E+00 | 1.22E-02 | 5.26E-02 | 1.23E+00 | 9.22E-06 | 3.92E-05 | -1.13E+00 | 8.38E-05 | 3.20E-04 | 1.06E+00 | 2.40E-03 | 8.55E-03 | -1.07E+00 | 1.30E-01 | 3.68E-01 | 1.38E+00 | 3.12E-03 | 1.15E-02 | -1.02E+00 | 2.54E-01 | 6.99E-01 | 1.32E+00 |
| NCK2 | 1.31E-02 | 6.85E-02 | 1.71E+00 | 1.88E-01 | 8.12E-01 | 1.54E+00 | 1.98E-01 | 8.40E-01 | 1.36E+00 | 8.48E-02 | 3.24E-01 | 1.30E+00 | 3.43E-02 | 1.22E-01 | 1.12E+00 | 2.54E-03 | 7.17E-03 | -1.17E+00 | 1.81E-01 | 6.66E-01 | 1.97E+00 | 2.60E-01 | 7.14E-01 | 1.67E+00 |
| FAM214B | 1.31E-02 | 6.85E-02 | 1.03E+00 | 1.66E-02 | 7.19E-02 | 1.15E+00 | 1.17E-01 | 4.95E-01 | 1.38E+00 | 1.66E-01 | 6.34E-01 | 1.18E+00 | 1.17E-03 | 4.18E-03 | -1.11E+00 | 4.58E-02 | 1.29E-01 | 1.08E+00 | 7.82E-02 | 2.87E-01 | 1.82E+00 | 3.13E-04 | 8.59E-04 | 1.10E+00 |
| PTGES3P2 | 1.32E-02 | 6.86E-02 | -1.27E+00 | 1.94E-02 | 8.37E-02 | -1.13E+00 | 1.64E-01 | 6.97E-01 | 1.02E+00 | 1.44E-01 | 5.52E-01 | -1.07E+00 | 2.40E-01 | 8.57E-01 | 1.50E+00 | 1.49E-01 | 4.19E-01 | -1.09E+00 | 2.30E-03 | 8.44E-03 | 2.34E+00 | 2.36E-01 | 6.47E-01 | 1.31E+00 |
| LINC01659 | 1.32E-02 | 6.87E-02 | -1.04E+00 | 1.73E-02 | 7.49E-02 | 1.05E+00 | 9.50E-04 | 4.04E-03 | -1.51E+00 | 1.27E-01 | 4.84E-01 | 1.18E+00 | 2.74E-01 | 9.77E-01 | 1.25E+00 | 3.17E-01 | 8.93E-01 | 1.38E+00 | 5.93E-02 | 2.18E-01 | -1.57E+00 | 1.41E-01 | 3.86E-01 | -1.36E+00 |
| INTS6L | 1.32E-02 | 6.87E-02 | 1.02E+00 | 6.09E-03 | 2.63E-02 | 1.04E+00 | 5.24E-07 | 2.23E-06 | -1.74E+00 | 1.41E-05 | 5.37E-05 | -1.35E+00 | 9.01E-02 | 3.21E-01 | 1.15E+00 | 2.26E-01 | 6.37E-01 | 1.50E+00 | 4.43E-05 | 1.62E-04 | -1.24E+00 | 9.82E-04 | 2.70E-03 | -1.15E+00 |
| AMBRA1 | 1.32E-02 | 6.87E-02 | -1.21E+00 | 3.95E-02 | 1.71E-01 | -1.03E+00 | 8.69E-03 | 3.69E-02 | -1.19E+00 | 1.38E-02 | 5.26E-02 | -1.03E+00 | 4.97E-03 | 1.77E-02 | -1.55E+00 | 1.79E-02 | 5.04E-02 | -1.37E+00 | 1.16E-04 | 4.27E-04 | -1.63E+00 | 4.08E-03 | 1.12E-02 | -1.37E+00 |
| VDAC1P8 | 1.32E-02 | 6.90E-02 | -1.17E+00 | 1.19E-02 | 5.16E-02 | -1.08E+00 | 9.74E-02 | 4.14E-01 | -1.01E+00 | 1.19E-02 | 4.55E-02 | -1.03E+00 | 2.24E-05 | 7.97E-05 | -1.95E+00 | 1.36E-03 | 3.83E-03 | -1.39E+00 | 5.22E-02 | 1.92E-01 | 1.07E+00 | 3.03E-01 | 8.32E-01 | 1.23E+00 |
| CDK18 | 1.33E-02 | 6.91E-02 | -1.03E+00 | 1.96E-01 | 8.46E-01 | 1.46E+00 | 1.87E-06 | 7.95E-06 | -1.86E+00 | 1.72E-05 | 6.56E-05 | -1.51E+00 | 3.44E-03 | 1.23E-02 | -1.46E+00 | 5.74E-03 | 1.62E-02 | -1.34E+00 | 4.93E-02 | 1.81E-01 | -1.22E+00 | 2.21E-01 | 6.08E-01 | -1.01E+00 |
| TAMM41 | 1.33E-02 | 6.91E-02 | 1.01E+00 | 3.75E-02 | 1.62E-01 | 1.23E+00 | 1.51E-05 | 6.42E-05 | -1.40E+00 | 3.04E-03 | 1.16E-02 | 1.05E+00 | 6.98E-04 | 2.49E-03 | -1.54E+00 | 2.85E-01 | 8.04E-01 | 1.34E+00 | 1.17E-06 | 4.30E-06 | -1.33E+00 | 4.02E-03 | 1.11E-02 | -1.02E+00 |
| ATXN7 | 1.33E-02 | 6.92E-02 | 1.03E+00 | 8.94E-02 | 3.86E-01 | 1.25E+00 | 8.01E-02 | 3.40E-01 | 1.11E+00 | 4.22E-03 | 1.61E-02 | 1.08E+00 | 2.29E-01 | 8.17E-01 | 1.29E+00 | 1.07E-01 | 3.03E-01 | 1.22E+00 | 1.59E-02 | 5.85E-02 | 1.03E+00 | 4.39E-02 | 1.21E-01 | 1.04E+00 |
| RPS9 | 1.33E-02 | 6.93E-02 | 1.05E+00 | 3.70E-02 | 1.60E-01 | 1.04E+00 | 1.14E-04 | 4.83E-04 | 1.11E+00 | 2.66E-03 | 1.02E-02 | 1.09E+00 | 8.55E-06 | 3.05E-05 | 1.15E+00 | 7.29E-04 | 2.06E-03 | 1.11E+00 | 7.80E-02 | 2.86E-01 | 1.11E+00 | 1.18E-01 | 3.25E-01 | 1.04E+00 |
| PPIL6 | 1.33E-02 | 6.93E-02 | -1.20E+00 | 1.28E-01 | 5.55E-01 | 1.29E+00 | 7.57E-03 | 3.22E-02 | -1.39E+00 | 5.06E-02 | 1.93E-01 | -1.01E+00 | 1.17E-01 | 4.17E-01 | -1.21E+00 | 1.66E-01 | 4.67E-01 | -1.11E+00 | 4.98E-02 | 1.83E-01 | -1.31E+00 | 1.25E-01 | 3.43E-01 | -1.10E+00 |
| LINC01602 | 1.33E-02 | 6.93E-02 | 2.43E+00 | 2.09E-01 | 9.02E-01 | 1.59E+00 | 8.12E-03 | 3.45E-02 | 2.00E+00 | 3.15E-02 | 1.20E-01 | 1.99E+00 | 6.24E-02 | 2.22E-01 | 2.20E+00 | 1.98E-01 | 5.58E-01 | 2.02E+00 | 5.05E-03 | 1.85E-02 | 1.73E+00 | 5.67E-03 | 1.56E-02 | 1.83E+00 |
| PIK3CA | 1.33E-02 | 6.94E-02 | 1.16E+00 | 1.59E-06 | 6.85E-06 | -1.16E+00 | 2.09E-02 | 8.86E-02 | -1.04E+00 | 1.76E-05 | 6.72E-05 | -1.32E+00 | 4.42E-02 | 1.58E-01 | 1.08E+00 | 4.47E-04 | 1.26E-03 | -1.19E+00 | 2.25E-02 | 8.27E-02 | 1.03E+00 | 1.08E-04 | 2.96E-04 | -1.35E+00 |
| ZNF282 | 1.33E-02 | 6.94E-02 | -1.09E+00 | 6.87E-02 | 2.97E-01 | 1.25E+00 | 8.95E-02 | 3.80E-01 | 1.03E+00 | 9.04E-02 | 3.45E-01 | 1.15E+00 | 7.82E-02 | 2.79E-01 | 1.13E+00 | 1.99E-01 | 5.61E-01 | 1.34E+00 | 2.52E-01 | 9.23E-01 | 1.23E+00 | 3.39E-01 | 9.30E-01 | 1.36E+00 |
| MPPE1 | 1.33E-02 | 6.95E-02 | -1.12E+00 | 1.28E-01 | 5.53E-01 | 1.29E+00 | 4.38E-02 | 1.86E-01 | -1.16E+00 | 2.24E-01 | 8.55E-01 | 1.33E+00 | 1.90E-01 | 6.78E-01 | 1.35E+00 | 1.20E-01 | 3.38E-01 | 1.83E+00 | 1.45E-01 | 5.33E-01 | 1.11E+00 | 8.74E-02 | 2.40E-01 | 1.58E+00 |
| GPSM1 | 1.33E-02 | 6.95E-02 | 1.06E+00 | 6.68E-03 | 2.89E-02 | 1.17E+00 | 3.75E-02 | 1.59E-01 | 1.07E+00 | 4.40E-03 | 1.68E-02 | 1.15E+00 | 1.24E-01 | 4.43E-01 | 1.05E+00 | 3.10E-01 | 8.75E-01 | 1.37E+00 | 2.42E-01 | 8.88E-01 | 1.28E+00 | 2.23E-01 | 6.12E-01 | 1.55E+00 |
| AL353194.1 | 1.34E-02 | 6.97E-02 | -1.06E+00 | 6.53E-02 | 2.82E-01 | 1.24E+00 | 7.39E-05 | 3.14E-04 | -1.64E+00 | 3.27E-03 | 1.25E-02 | -1.09E+00 | 1.26E-01 | 4.49E-01 | 1.18E+00 | 2.23E-01 | 6.28E-01 | 1.28E+00 | 1.47E-01 | 5.39E-01 | 1.13E+00 | 2.61E-01 | 7.18E-01 | 1.24E+00 |

| | | | | | | | | | | | | | | | | | | | | | | | | |
|---|---|---|---|---|---|---|---|---|---|---|---|---|---|---|---|---|---|---|---|---|---|---|---|---|
| SMG7-AS1 | 1.34E-02 | 6.97E-02 | 1.84E+00 | 1.79E-01 | 7.73E-01 | 1.07E+00 | 8.18E-02 | 3.48E-01 | 1.47E+00 | 1.24E-01 | 4.74E-01 | 1.03E+00 | 1.26E-01 | 4.48E-01 | 2.00E+00 | 1.75E-01 | 4.94E-01 | -1.02E+00 | 7.19E-03 | 2.64E-02 | 2.29E+00 | 9.87E-02 | 2.71E-01 | 1.76E+00 |
| LRRFIP1 | 1.34E-02 | 6.97E-02 | 1.09E+00 | 1.70E-01 | 7.32E-01 | 1.28E+00 | 1.78E-01 | 7.58E-01 | 1.05E+00 | 2.02E-01 | 7.72E-01 | 1.07E+00 | 6.35E-04 | 2.27E-03 | -1.01E+00 | 1.02E-03 | 2.88E-03 | -1.07E+00 | 1.48E-04 | 5.44E-04 | -1.09E+00 | 4.01E-02 | 1.10E-01 | 1.01E+00 |
| ZNF133 | 1.34E-02 | 6.98E-02 | -1.16E+00 | 6.81E-02 | 2.94E-01 | 1.18E+00 | 7.35E-02 | 3.12E-01 | -1.03E+00 | 5.68E-02 | 2.17E-01 | 1.01E+00 | 4.10E-02 | 1.46E-01 | -1.20E+00 | 1.23E-01 | 3.47E-01 | 1.01E+00 | 2.28E-02 | 8.38E-02 | -1.27E+00 | 1.93E-01 | 5.29E-01 | 1.03E+00 |
| AC127070.5 | 1.34E-02 | 6.98E-02 | -1.28E+00 | 1.55E-02 | 6.72E-02 | -1.24E+00 | 6.66E-02 | 2.83E-01 | -1.15E+00 | 1.78E-01 | 6.79E-01 | 1.16E+00 | 2.54E-01 | 9.07E-01 | 1.32E+00 | 1.67E-01 | 4.72E-01 | 1.07E+00 | 1.73E-01 | 6.35E-01 | 1.04E+00 | 3.57E-01 | 9.81E-01 | 1.30E+00 |
| ZZEF1 | 1.34E-02 | 6.98E-02 | 1.07E+00 | 1.67E-02 | 7.23E-02 | 1.16E+00 | 3.56E-03 | 1.51E-02 | -1.16E+00 | 1.42E-02 | 5.44E-02 | 1.02E+00 | 1.01E-01 | 3.61E-01 | 1.17E+00 | 1.06E-01 | 2.98E-01 | 1.19E+00 | 1.29E-01 | 4.74E-01 | 1.31E+00 | 2.56E-02 | 7.02E-02 | 1.05E+00 |
| ZNF653 | 1.34E-02 | 6.99E-02 | -1.34E+00 | 2.12E-02 | 9.17E-02 | -1.20E+00 | 3.41E-06 | 1.45E-05 | -1.99E+00 | 2.24E-04 | 8.56E-04 | -1.32E+00 | 4.12E-08 | 1.47E-07 | -3.61E+00 | 4.23E-05 | 1.19E-04 | -1.83E+00 | 9.29E-02 | 3.41E-01 | -1.17E+00 | 8.31E-02 | 2.28E-01 | -1.31E+00 |
| GFPT1 | 1.34E-02 | 7.00E-02 | -1.09E+00 | 2.25E-03 | 9.70E-03 | -1.17E+00 | 1.89E-04 | 8.02E-04 | -1.28E+00 | 5.75E-04 | 2.20E-03 | -1.12E+00 | 4.97E-02 | 1.77E-01 | 1.01E+00 | 2.73E-03 | 7.70E-03 | -1.21E+00 | 2.14E-02 | 7.86E-02 | 1.01E+00 | 1.06E-01 | 2.91E-01 | 1.03E+00 |
| DLGAP4-AS1 | 1.34E-02 | 7.01E-02 | -1.17E+00 | 1.49E-01 | 6.45E-01 | 1.37E+00 | 1.61E-02 | 6.85E-02 | -1.42E+00 | 2.54E-02 | 9.72E-02 | -1.17E+00 | 1.78E-01 | 6.36E-01 | -1.14E+00 | 1.47E-01 | 4.15E-01 | -1.33E+00 | 1.98E-01 | 7.27E-01 | 1.36E+00 | 2.67E-01 | 7.35E-01 | 1.30E+00 |
| NTHL1 | 1.34E-02 | 7.01E-02 | 1.11E+00 | 5.00E-02 | 2.16E-01 | 1.32E+00 | 6.42E-02 | 2.73E-01 | 1.12E+00 | 4.97E-02 | 1.90E-01 | 1.24E+00 | 4.83E-04 | 1.72E-03 | -1.04E+00 | 4.00E-02 | 1.13E-01 | 1.20E+00 | 5.09E-03 | 1.87E-02 | -1.00E+00 | 8.37E-02 | 2.30E-01 | 1.22E+00 |
| GPS2 | 1.35E-02 | 7.01E-02 | 1.08E+00 | 4.17E-04 | 1.80E-03 | -1.10E+00 | 1.78E-02 | 7.58E-02 | -1.08E+00 | 3.46E-03 | 1.32E-02 | -1.04E+00 | 3.78E-04 | 1.35E-03 | -1.12E+00 | 4.25E-04 | 1.20E-03 | -1.23E+00 | 9.06E-04 | 3.33E-03 | -1.28E+00 | 2.64E-05 | 7.24E-05 | -1.70E+00 |
| RCBTB1 | 1.35E-02 | 7.01E-02 | -1.05E+00 | 8.03E-02 | 3.47E-01 | 1.18E+00 | 4.82E-02 | 2.05E-01 | 1.11E+00 | 8.40E-02 | 3.21E-01 | 1.27E+00 | 1.64E-01 | 5.86E-01 | 1.24E+00 | 2.27E-01 | 6.39E-01 | 1.32E+00 | 5.68E-03 | 2.09E-02 | -1.11E+00 | 2.68E-02 | 7.35E-02 | 1.01E+00 |
| CD63 | 1.35E-02 | 7.02E-02 | 1.02E+00 | 1.35E-01 | 5.82E-01 | 1.14E+00 | 5.73E-02 | 2.44E-01 | 1.16E+00 | 2.64E-03 | 1.01E-02 | 1.39E+00 | 9.34E-02 | 3.33E-01 | -1.10E+00 | 1.40E-02 | 3.95E-02 | -1.15E+00 | 2.85E-06 | 1.05E-05 | 1.35E+00 | 2.51E-07 | 6.90E-07 | 1.35E+00 |
| RPS4XP22 | 1.35E-02 | 7.03E-02 | -1.03E+00 | 1.26E-03 | 5.44E-03 | -1.15E+00 | 2.04E-01 | 8.65E-01 | 1.24E+00 | 3.68E-02 | 1.41E-01 | -1.04E+00 | 8.86E-02 | 3.16E-01 | 1.08E+00 | 9.05E-02 | 2.55E-01 | 1.10E+00 | 1.28E-01 | 4.69E-01 | 1.06E+00 | 6.92E-03 | 1.90E-02 | -1.60E+00 |
| CNKSR2 | 1.35E-02 | 7.03E-02 | 2.32E+00 | 4.47E-02 | 1.93E-01 | 2.22E+00 | 4.53E-02 | 1.92E-01 | -1.00E+00 | 4.50E-02 | 1.72E-01 | 1.13E+00 | 2.02E-01 | 7.21E-01 | 1.09E+00 | 3.22E-01 | 9.07E-01 | 1.31E+00 | 1.24E-01 | 4.57E-01 | -1.20E+00 | 2.58E-01 | 7.09E-01 | 1.24E+00 |
| PHC2 | 1.35E-02 | 7.04E-02 | 1.06E+00 | 2.02E-02 | 8.74E-02 | 1.20E+00 | 4.65E-02 | 1.97E-01 | 1.09E+00 | 2.60E-03 | 9.94E-03 | 1.04E+00 | 1.58E-01 | 5.65E-01 | 1.20E+00 | 1.07E-01 | 3.01E-01 | 1.15E+00 | 2.26E-01 | 8.28E-01 | 1.39E+00 | 3.23E-01 | 8.86E-01 | 1.38E+00 |
| ZNF578 | 1.35E-02 | 7.05E-02 | -1.25E+00 | 3.46E-02 | 1.49E-01 | -1.04E+00 | 3.40E-02 | 1.44E-01 | -1.34E+00 | 1.74E-01 | 6.66E-01 | 1.11E+00 | 5.18E-02 | 1.85E-01 | -1.04E+00 | 5.16E-02 | 1.46E-01 | -1.10E+00 | 3.31E-02 | 1.21E-01 | -1.21E+00 | 6.75E-02 | 1.86E-01 | -1.18E+00 |
| RTKN | 1.35E-02 | 7.05E-02 | -1.07E+00 | 1.62E-01 | 7.00E-01 | 1.39E+00 | 5.32E-06 | 2.26E-05 | -1.69E+00 | 3.23E-03 | 1.23E-02 | -1.05E+00 | 8.60E-07 | 3.07E-06 | -2.13E+00 | 1.37E-02 | 3.87E-02 | 1.01E+00 | 3.60E-05 | 1.32E-04 | -1.48E+00 | 1.00E-03 | 2.76E-03 | -1.46E+00 |
| LYPLAL1 | 1.35E-02 | 7.06E-02 | 1.12E+00 | 2.11E-02 | 9.13E-02 | 1.98E+00 | 6.69E-04 | 2.84E-03 | -1.27E+00 | 2.90E-02 | 1.11E-01 | 1.09E+00 | 6.28E-02 | 2.24E-01 | 1.03E+00 | 3.17E-01 | 8.93E-01 | 1.33E+00 | 6.12E-08 | 2.25E-07 | -1.37E+00 | 3.30E-03 | 9.06E-03 | -1.11E+00 |
| DDX5 | 1.36E-02 | 7.06E-02 | -1.02E+00 | 1.76E-09 | 7.62E-09 | -1.15E+00 | 3.32E-03 | 1.41E-02 | -1.11E+00 | 4.13E-03 | 1.58E-02 | -1.10E+00 | 2.97E-03 | 1.06E-02 | -1.10E+00 | 3.95E-05 | 1.11E-04 | -1.17E+00 | 8.25E-06 | 3.03E-05 | -1.16E+00 | 2.94E-03 | 8.08E-03 | -1.14E+00 |
| AL135925.1 | 1.36E-02 | 7.08E-02 | 1.18E+00 | 2.35E-03 | 1.01E-02 | 1.19E+00 | 7.97E-02 | 3.39E-01 | 1.11E+00 | 9.96E-04 | 3.80E-03 | -1.00E+00 | 2.12E-01 | 7.56E-01 | 1.49E+00 | 2.39E-02 | 6.74E-02 | 1.02E+00 | 1.66E-01 | 6.10E-01 | 1.52E+00 | 1.92E-03 | 5.28E-03 | 1.01E+00 |
| CPEB2 | 1.36E-02 | 7.08E-02 | -1.15E+00 | 2.61E-03 | 1.13E-02 | -1.27E+00 | 1.67E-01 | 7.10E-01 | 1.16E+00 | 2.32E-02 | 8.85E-02 | -1.19E+00 | 2.63E-01 | 9.38E-01 | 1.27E+00 | 2.93E-01 | 8.27E-01 | 1.22E+00 | 3.27E-02 | 1.20E-01 | 1.25E+00 | 1.58E-02 | 4.35E-02 | -1.09E+00 |
| PDE3A | 1.36E-02 | 7.08E-02 | -1.87E+00 | 1.56E-02 | 6.76E-02 | -1.95E+00 | 5.08E-02 | 2.16E-01 | -1.79E+00 | 9.40E-02 | 3.59E-01 | -1.47E+00 | 7.49E-02 | 2.67E-01 | -1.01E+00 | 1.18E-01 | 3.33E-01 | -1.10E+00 | 1.39E-01 | 5.11E-01 | 1.29E+00 | 1.38E-03 | 3.80E-03 | -1.04E+00 |
| FBXO32 | 1.36E-02 | 7.08E-02 | 1.07E+00 | 1.79E-01 | 7.75E-01 | 1.48E+00 | 2.41E-04 | 1.02E-03 | -1.31E+00 | 8.02E-03 | 3.06E-02 | 1.12E+00 | 2.01E-02 | 7.16E-02 | -1.20E+00 | 3.80E-02 | 1.07E-01 | -1.25E+00 | 1.59E-01 | 5.85E-01 | -1.06E+00 | 1.53E-01 | 4.19E-01 | -1.24E+00 |
| AC083904.1 | 1.36E-02 | 7.09E-02 | -1.16E+00 | 7.15E-04 | 3.09E-03 | -1.43E+00 | 6.34E-04 | 2.69E-03 | -1.33E+00 | 4.98E-06 | 1.90E-05 | -1.54E+00 | 1.97E-03 | 7.04E-03 | -1.18E+00 | 1.34E-04 | 3.78E-04 | -1.56E+00 | 8.52E-04 | 3.13E-03 | -1.26E+00 | 1.02E-03 | 2.79E-03 | -1.21E+00 |
| AL355001.2 | 1.37E-02 | 7.11E-02 | 1.02E+00 | 1.38E-02 | 5.98E-02 | 1.09E+00 | 4.75E-06 | 2.02E-05 | -1.45E+00 | 2.53E-06 | 9.67E-06 | -1.32E+00 | 5.92E-04 | 2.11E-03 | -1.34E+00 | 2.57E-03 | 7.24E-03 | -1.27E+00 | 1.72E-02 | 6.33E-02 | -1.03E+00 | 9.98E-04 | 2.74E-03 | -1.22E+00 |
| TPCN2 | 1.37E-02 | 7.12E-02 | -1.04E+00 | 2.20E-01 | 9.51E-01 | 1.58E+00 | 4.15E-02 | 1.76E-01 | -1.09E+00 | 2.28E-01 | 8.71E-01 | 1.25E+00 | 6.20E-05 | 2.21E-04 | -1.35E+00 | 4.84E-04 | 1.36E-03 | -1.20E+00 | 1.68E-03 | 6.17E-03 | -1.15E+00 | 7.88E-02 | 2.17E-01 | 1.08E+00 |
| AC015871.3 | 1.37E-02 | 7.13E-02 | -1.04E+00 | 5.06E-02 | 2.19E-01 | 1.89E+00 | 1.87E-03 | 7.93E-03 | -1.28E+00 | 1.17E-01 | 4.47E-01 | 1.22E+00 | 2.29E-02 | 8.16E-02 | -1.12E+00 | 1.69E-01 | 4.76E-01 | 1.19E+00 | 2.05E-02 | 7.52E-02 | -1.20E+00 | 3.64E-01 | 9.99E-01 | 1.30E+00 |
| CPM | 1.37E-02 | 7.13E-02 | -1.16E+00 | 4.02E-02 | 1.74E-01 | -1.04E+00 | 6.54E-02 | 2.78E-01 | -1.20E+00 | 2.20E-02 | 8.42E-02 | -1.53E+00 | 2.22E-01 | 7.92E-01 | 1.67E+00 | 8.72E-02 | 2.46E-01 | -1.28E+00 | 4.99E-03 | 1.83E-02 | -1.04E+00 | 4.38E-04 | 1.20E-03 | -1.84E+00 |
| AC016735.1 | 1.37E-02 | 7.15E-02 | 1.75E+00 | 2.25E-01 | 9.71E-01 | 1.37E+00 | 9.85E-02 | 4.19E-01 | 1.37E+00 | 1.94E-01 | 7.42E-01 | 1.16E+00 | 2.15E-01 | 7.66E-01 | 1.39E+00 | 3.18E-01 | 8.97E-01 | 1.18E+00 | 2.72E-01 | 1.00E+00 | 1.00E+00 | 3.41E-01 | 9.35E-01 | 1.06E+00 |
| ADCY3 | 1.38E-02 | 7.18E-02 | -1.12E+00 | 1.70E-02 | 7.36E-02 | -1.00E+00 | 2.23E-02 | 9.49E-02 | -1.07E+00 | 7.90E-04 | 3.02E-03 | -1.23E+00 | 2.15E-01 | 7.68E-01 | 1.03E+00 | 3.03E-01 | 8.55E-01 | 1.08E+00 | 4.48E-02 | 1.64E-01 | -1.27E+00 | 1.70E-01 | 4.68E-01 | 1.03E+00 |
| AC134407.2 | 1.38E-02 | 7.18E-02 | -1.29E+00 | 1.83E-01 | 7.90E-01 | 1.25E+00 | 1.64E-07 | 6.97E-07 | -2.08E+00 | 1.07E-05 | 4.08E-05 | -1.40E+00 | 1.44E-02 | 5.12E-02 | -1.42E+00 | 2.41E-01 | 6.78E-01 | 1.30E+00 | 4.16E-10 | 1.53E-09 | -2.65E+00 | 1.34E-05 | 3.69E-05 | -1.55E+00 |
| ESR2 | 1.38E-02 | 7.19E-02 | -1.17E+00 | 2.10E-01 | 9.08E-01 | 1.34E+00 | 6.92E-02 | 2.94E-01 | -1.63E+00 | 7.41E-02 | 2.83E-01 | -1.51E+00 | 2.06E-06 | 7.34E-06 | -1.64E+00 | 7.34E-02 | 2.07E-01 | 1.24E+00 | 1.51E-02 | 5.54E-02 | -1.40E+00 | 2.79E-01 | 7.65E-01 | 1.29E+00 |
| IRS2 | 1.38E-02 | 7.19E-02 | 1.65E+00 | 2.42E-02 | 1.05E-01 | 1.28E+00 | 1.31E-01 | 5.57E-01 | 1.24E+00 | 3.22E-03 | 1.23E-02 | 1.04E+00 | 3.95E-02 | 1.41E-01 | 1.72E+00 | 2.91E-02 | 8.20E-02 | 1.22E+00 | 7.71E-02 | 2.83E-01 | 1.38E+00 | 2.56E-03 | 7.04E-03 | -1.09E+00 |
| ZSCAN16-AS1 | 1.38E-02 | 7.22E-02 | 1.16E+00 | 1.65E-03 | 7.11E-03 | 1.13E+00 | 9.84E-02 | 4.18E-01 | 1.14E+00 | 1.82E-05 | 6.95E-05 | -1.15E+00 | 4.66E-04 | 1.66E-03 | -1.24E+00 | 3.13E-05 | 8.84E-05 | -1.40E+00 | 1.41E-01 | 5.17E-01 | 1.09E+00 | 2.12E-05 | 5.83E-05 | -1.34E+00 |
| MGAT4B | 1.39E-02 | 7.22E-02 | 1.11E+00 | 8.91E-02 | 3.85E-01 | 1.31E+00 | 1.69E-01 | 7.16E-01 | 1.16E+00 | 2.43E-01 | 9.27E-01 | 1.33E+00 | 3.90E-02 | 1.39E-01 | 1.19E+00 | 2.40E-01 | 6.75E-01 | 1.37E+00 | 2.63E-02 | 9.67E-02 | 1.73E+00 | 2.17E-02 | 5.96E-02 | 1.57E+00 |
| ACTR3B | 1.39E-02 | 7.23E-02 | -1.39E+00 | 9.58E-02 | 4.14E-01 | 1.11E+00 | 1.18E-02 | 5.01E-02 | -1.08E+00 | 1.12E-02 | 4.27E-02 | -1.02E+00 | 5.42E-03 | 1.93E-02 | -1.32E+00 | 1.54E-01 | 4.33E-01 | 1.22E+00 | 5.69E-03 | 2.09E-02 | -1.09E+00 | 2.49E-01 | 6.85E-01 | 1.47E+00 |
| SPAG8 | 1.39E-02 | 7.24E-02 | -1.43E+00 | 4.18E-02 | 1.81E-01 | -1.10E+00 | 6.67E-04 | 2.83E-03 | -1.69E+00 | 4.71E-03 | 1.80E-02 | -1.35E+00 | 2.38E-01 | 8.48E-01 | 1.09E+00 | 2.98E-01 | 8.40E-01 | 1.43E+00 | 2.11E-01 | 7.73E-01 | 1.21E+00 | 1.08E-01 | 2.97E-01 | 1.59E+00 |
| C9orf152 | 1.39E-02 | 7.25E-02 | -1.22E+00 | 6.68E-02 | 2.88E-01 | 1.86E+00 | 2.10E-01 | 8.94E-01 | 1.27E+00 | 1.42E-01 | 5.44E-01 | 1.60E+00 | 1.75E-01 | 6.25E-01 | -1.33E+00 | 2.76E-01 | 7.77E-01 | -1.17E+00 | 1.61E-01 | 5.90E-01 | 1.40E+00 | 1.31E-01 | 3.61E-01 | 1.66E+00 |
| DCTN6 | 1.39E-02 | 7.25E-02 | 1.10E+00 | 1.43E-02 | 6.18E-02 | 1.21E+00 | 7.37E-04 | 3.13E-03 | -1.20E+00 | 1.27E-04 | 4.83E-04 | -1.11E+00 | 1.78E-02 | 6.33E-02 | 1.07E+00 | 2.06E-02 | 5.80E-02 | 1.10E+00 | 7.57E-05 | 2.78E-04 | -1.16E+00 | 3.94E-03 | 1.08E-02 | -1.03E+00 |
| HES4 | 1.39E-02 | 7.25E-02 | 2.00E+00 | 2.20E-01 | 9.52E-01 | 1.50E+00 | 1.54E-02 | 6.55E-02 | 1.02E+00 | 1.53E-01 | 5.86E-01 | -1.99E+00 | 2.10E-01 | 7.47E-01 | 1.31E+00 | 3.55E-01 | 1.00E+00 | 1.00E+00 | 8.21E-02 | 3.01E-01 | -1.35E+00 | 8.45E-02 | 2.32E-01 | -1.39E+00 |
| TMC6 | 1.40E-02 | 7.28E-02 | 1.01E+00 | 6.85E-02 | 2.96E-01 | 1.27E+00 | 1.48E-03 | 6.29E-03 | -1.17E+00 | 2.67E-02 | 1.02E-01 | 1.11E+00 | 2.09E-03 | 7.46E-03 | -1.14E+00 | 1.77E-02 | 4.99E-02 | 1.03E+00 | 5.39E-03 | 1.98E-02 | -1.08E+00 | 5.97E-02 | 1.64E-01 | 1.05E+00 |
| GDAP1 | 1.40E-02 | 7.28E-02 | -1.15E+00 | 3.68E-03 | 1.59E-02 | -1.18E+00 | 9.83E-03 | 4.18E-02 | -1.24E+00 | 4.53E-03 | 1.73E-02 | -1.24E+00 | 3.60E-03 | 1.28E-02 | -1.21E+00 | 2.33E-03 | 6.56E-03 | -1.30E+00 | 6.21E-03 | 2.28E-02 | -1.07E+00 | 3.22E-04 | 8.85E-04 | -1.26E+00 |
| AC092279.1 | 1.40E-02 | 7.28E-02 | 1.06E+00 | 8.07E-03 | 3.49E-02 | 1.17E+00 | 1.18E-02 | 5.00E-02 | -1.05E+00 | 5.86E-03 | 2.24E-02 | -1.02E+00 | 4.16E-02 | 1.48E-01 | -1.05E+00 | 1.53E-01 | 4.32E-01 | 1.06E+00 | 1.43E-02 | 5.24E-02 | -1.09E+00 | 5.82E-03 | 1.60E-02 | -1.17E+00 |
| RPL32P26 | 1.40E-02 | 7.29E-02 | -1.29E+00 | 4.59E-02 | 1.98E-01 | -1.00E+00 | 2.60E-02 | 1.10E-01 | -1.33E+00 | 6.43E-03 | 2.45E-02 | -1.47E+00 | 4.44E-09 | 1.58E-08 | -2.04E+00 | 3.35E-07 | 9.46E-07 | -1.71E+00 | 3.67E-02 | 1.35E-01 | -1.24E+00 | 9.76E-02 | 2.68E-01 | -1.09E+00 |
| TEX14 | 1.40E-02 | 7.31E-02 | 2.10E+00 | 9.02E-02 | 3.90E-01 | -1.22E+00 | 1.53E-01 | 6.49E-01 | 1.37E+00 | 1.65E-02 | 6.29E-02 | -1.56E+00 | 1.44E-02 | 5.13E-02 | 2.34E+00 | 2.15E-01 | 6.07E-01 | -1.05E+00 | 1.08E-02 | 3.95E-02 | 2.16E+00 | 2.68E-01 | 7.35E-01 | 1.23E+00 |
| ZBTB7B | 1.40E-02 | 7.31E-02 | -1.23E+00 | 4.56E-02 | 1.97E-01 | 1.08E+00 | 2.16E-02 | 9.18E-02 | -1.38E+00 | 1.04E-01 | 3.98E-01 | 1.09E+00 | 1.96E-01 | 7.00E-01 | 1.19E+00 | 3.42E-01 | 9.64E-01 | 1.37E+00 | 2.16E-01 | 7.92E-01 | 1.32E+00 | 2.42E-01 | 6.66E-01 | 1.17E+00 |
| KALRN | 1.41E-02 | 7.33E-02 | -1.73E+00 | 2.14E-02 | 9.26E-02 | -1.56E+00 | 1.56E-01 | 6.63E-01 | -1.14E+00 | 2.59E-01 | 9.88E-01 | 1.15E+00 | 1.77E-01 | 6.30E-01 | -1.24E+00 | 3.06E-01 | 8.64E-01 | 1.06E+00 | 2.67E-03 | 9.82E-03 | -1.14E+00 | 1.89E-03 | 5.20E-03 | -1.45E+00 |
| AC023590.1 | 1.41E-02 | 7.34E-02 | -1.11E+00 | 4.71E-02 | 2.04E-01 | 1.04E+00 | 9.61E-02 | 4.08E-01 | -1.13E+00 | 1.81E-01 | 6.92E-01 | 1.10E+00 | 4.45E-02 | 1.59E-01 | -1.19E+00 | 5.25E-02 | 1.48E-01 | -1.25E+00 | 1.35E-01 | 4.95E-01 | -1.13E+00 | 8.29E-02 | 2.28E-01 | -1.30E+00 |
| ARHGAP42 | 1.41E-02 | 7.34E-02 | -1.83E+00 | 2.23E-02 | 9.64E-02 | -1.64E+00 | 1.93E-01 | 8.18E-01 | -1.01E+00 | 2.26E-01 | 8.65E-01 | 1.03E+00 | 1.65E-01 | 5.88E-01 | 1.17E+00 | 3.09E-01 | 8.71E-01 | 1.35E+00 | 9.64E-02 | 3.54E-01 | 1.18E+00 | 1.11E-02 | 3.06E-02 | -1.06E+00 |
| PEX14 | 1.41E-02 | 7.36E-02 | -1.13E+00 | 1.02E-02 | 4.39E-02 | -1.08E+00 | 2.49E-02 | 1.06E-01 | -1.03E+00 | 2.07E-01 | 7.89E-01 | 1.29E+00 | 1.60E-04 | 5.71E-04 | -1.84E+00 | 1.44E-01 | 4.05E-01 | 1.26E+00 | 1.34E-07 | 4.92E-07 | -1.73E+00 | 8.18E-05 | 2.25E-04 | -1.32E+00 |
| DNASE1 | 1.41E-02 | 7.37E-02 | -1.00E+00 | 2.30E-01 | 9.96E-01 | 1.49E+00 | 2.77E-04 | 1.18E-03 | -1.43E+00 | 2.34E-03 | 8.92E-03 | -1.07E+00 | 7.42E-03 | 2.65E-02 | 1.02E+00 | 2.53E-01 | 7.13E-01 | 1.38E+00 | 5.68E-04 | 2.08E-03 | -1.30E+00 | 8.87E-02 | 2.44E-01 | 1.07E+00 |
| AL596202.1 | 1.42E-02 | 7.38E-02 | -1.24E+00 | 1.72E-01 | 7.42E-01 | 1.54E+00 | 2.40E-03 | 1.02E-02 | -1.54E+00 | 1.68E-01 | 6.41E-01 | 1.37E+00 | 7.96E-04 | 2.84E-03 | -1.90E+00 | 1.67E-01 | 4.72E-01 | 1.15E+00 | 5.35E-05 | 1.96E-04 | -1.93E+00 | 9.48E-02 | 2.61E-01 | 1.03E+00 |
| LRRFIP1P1 | 1.42E-02 | 7.39E-02 | 1.02E+00 | 1.39E-02 | 6.02E-02 | 1.12E+00 | 1.83E-03 | 7.76E-03 | -1.21E+00 | 1.39E-02 | 5.33E-02 | 1.07E+00 | 1.34E-02 | 4.76E-02 | 1.07E+00 | 1.07E-01 | 3.03E-01 | 1.19E+00 | 2.30E-01 | 8.43E-01 | 1.29E+00 | 1.79E-01 | 4.93E-01 | 1.43E+00 |
| PROCA1 | 1.42E-02 | 7.40E-02 | -1.10E+00 | 8.13E-02 | 3.51E-01 | 1.20E+00 | 1.47E-03 | 6.27E-03 | -1.42E+00 | 3.50E-03 | 1.34E-02 | -1.14E+00 | 1.28E-02 | 4.56E-02 | -1.26E+00 | 4.07E-02 | 1.15E-01 | -1.17E+00 | 1.50E-01 | 5.49E-01 | 1.38E+00 | 2.07E-01 | 5.68E-01 | 1.17E+00 |
| ABAT | 1.42E-02 | 7.41E-02 | -1.10E+00 | 3.46E-02 | 1.50E-01 | 1.10E+00 | 1.18E-04 | 5.02E-04 | -1.41E+00 | 5.32E-03 | 2.03E-02 | -1.07E+00 | 1.99E-02 | 7.08E-02 | -1.17E+00 | 5.87E-02 | 1.66E-01 | -1.03E+00 | 6.91E-02 | 2.54E-01 | 1.01E+00 | 1.76E-01 | 4.84E-01 | 1.02E+00 |
| AC138207.8 | 1.42E-02 | 7.41E-02 | -1.11E+00 | 8.84E-03 | 3.82E-02 | -1.12E+00 | 2.16E-01 | 9.17E-01 | 1.32E+00 | 1.17E-02 | 4.47E-02 | -1.19E+00 | 1.94E-01 | 6.92E-01 | -1.16E+00 | 2.05E-01 | 5.79E-01 | -1.17E+00 | 2.63E-01 | 9.64E-01 | 1.01E+00 | 3.46E-01 | 9.51E-01 | 1.08E+00 |
| PMS2P1 | 1.42E-02 | 7.42E-02 | 1.11E+00 | 6.81E-02 | 2.94E-01 | 1.28E+00 | 7.70E-03 | 3.27E-02 | -1.09E+00 | 5.87E-02 | 2.24E-01 | 1.17E+00 | 4.12E-03 | 1.47E-02 | -1.10E+00 | 8.67E-02 | 2.44E-01 | 1.27E+00 | 7.11E-04 | 2.61E-03 | -1.04E+00 | 2.11E-01 | 5.79E-01 | 1.27E+00 |
| AL391807.1 | 1.43E-02 | 7.43E-02 | 1.71E+00 | 3.82E-02 | 1.65E-01 | 1.78E+00 | 9.91E-02 | 4.21E-01 | 1.54E+00 | 1.57E-01 | 6.01E-01 | 1.55E+00 | 2.80E-01 | 1.00E+00 | 1.00E+00 | 3.55E-01 | 1.00E+00 | 1.00E+00 | 2.04E-01 | 7.49E-01 | 1.26E+00 | 3.37E-01 | 9.26E-01 | 1.08E+00 |
| AMPD3 | 1.43E-02 | 7.45E-02 | -1.03E+00 | 3.22E-02 | 1.39E-01 | 1.05E+00 | 3.87E-04 | 1.64E-03 | -1.54E+00 | 4.42E-04 | 1.69E-03 | -1.35E+00 | 1.47E-08 | 5.24E-08 | -2.20E+00 | 2.76E-05 | 7.78E-05 | -1.43E+00 | 4.79E-08 | 1.76E-07 | -1.89E+00 | 4.52E-05 | 1.24E-04 | -1.41E+00 |
| AC073335.2 | 1.43E-02 | 7.46E-02 | -1.35E+00 | 2.70E-02 | 1.17E-01 | -1.08E+00 | 7.72E-03 | 3.28E-02 | -1.40E+00 | 2.72E-03 | 1.04E-02 | -1.50E+00 | 6.57E-12 | 2.34E-11 | -2.43E+00 | 3.24E-06 | 9.12E-06 | -1.37E+00 | 2.97E-03 | 1.09E-02 | -1.13E+00 | 9.28E-03 | 2.55E-02 | -1.12E+00 |
| TDRKH | 1.44E-02 | 7.48E-02 | -1.62E+00 | 1.18E-01 | 5.12E-01 | 1.02E+00 | 1.11E-01 | 4.73E-01 | -1.12E+00 | 1.89E-01 | 7.21E-01 | 1.09E+00 | 5.86E-02 | 2.09E-01 | 1.00E+00 | 2.10E-01 | 5.92E-01 | 1.24E+00 | 2.51E-02 | 9.23E-02 | -1.01E+00 | 3.39E-01 | 9.30E-01 | 1.31E+00 |
| TIMM9 | 1.44E-02 | 7.49E-02 | 1.13E+00 | 3.28E-02 | 1.42E-01 | 1.30E+00 | 1.02E-03 | 4.32E-03 | -1.04E+00 | 1.02E-02 | 3.89E-02 | 1.14E+00 | 5.40E-02 | 1.93E-01 | 1.10E+00 | 3.38E-01 | 9.54E-01 | 1.29E+00 | 5.36E-07 | 1.97E-06 | -1.27E+00 | 2.07E-02 | 5.68E-02 | 1.01E+00 |
| ANAPC1 | 1.44E-02 | 7.49E-02 | -1.01E+00 | 6.46E-02 | 2.79E-01 | 1.25E+00 | 1.92E-02 | 8.15E-02 | 1.03E+00 | 2.21E-02 | 8.43E-02 | 1.21E+00 | 9.80E-02 | 3.50E-01 | 1.13E+00 | 1.77E-01 | 4.99E-01 | 1.19E+00 | 1.20E-01 | 4.40E-01 | 1.13E+00 | 1.23E-01 | 3.38E-01 | 1.15E+00 |
| KIAA0895L | 1.44E-02 | 7.50E-02 | -1.51E+00 | 6.55E-02 | 2.83E-01 | -1.03E+00 | 5.64E-03 | 2.40E-02 | -1.38E+00 | 7.58E-02 | 2.90E-01 | 1.04E+00 | 3.64E-03 | 1.30E-02 | -1.36E+00 | 7.61E-03 | 2.14E-02 | -1.28E+00 | 1.82E-01 | 6.68E-01 | 1.13E+00 | 3.63E-01 | 9.98E-01 | 1.24E+00 |
| VANGL2 | 1.44E-02 | 7.51E-02 | -1.23E+00 | 1.80E-01 | 7.78E-01 | 1.44E+00 | 1.11E-04 | 4.72E-04 | -1.95E+00 | 7.84E-04 | 3.00E-03 | -1.57E+00 | 2.70E-02 | 9.62E-02 | -1.30E+00 | 2.62E-01 | 7.38E-01 | 1.31E+00 | 2.42E-05 | 8.88E-05 | -1.54E+00 | 2.24E-03 | 6.16E-03 | -1.25E+00 |
| TATDN1 | 1.45E-02 | 7.54E-02 | 1.50E+00 | 5.05E-02 | 2.18E-01 | 1.67E+00 | 1.48E-02 | 6.28E-02 | 1.54E+00 | 2.16E-03 | 8.25E-03 | 1.88E+00 | 2.69E-01 | 9.58E-01 | 1.54E+00 | 1.22E-01 | 3.43E-01 | 1.83E+00 | 1.52E-01 | 5.56E-01 | 1.27E+00 | 1.76E-02 | 4.84E-02 | 1.63E+00 |
| RPL9P3 | 1.45E-02 | 7.55E-02 | 1.09E+00 | 1.20E-08 | 5.17E-08 | -1.51E+00 | 5.57E-02 | 2.37E-01 | 1.11E+00 | 1.38E-08 | 5.28E-08 | -1.56E+00 | 1.20E-01 | 4.27E-01 | 1.26E+00 | 5.08E-05 | 1.43E-04 | -1.41E+00 | 5.98E-02 | 2.20E-01 | -1.10E+00 | 8.31E-05 | 2.28E-04 | -1.78E+00 |
| AC104461.1 | 1.45E-02 | 7.55E-02 | -1.18E+00 | 5.10E-02 | 2.20E-01 | 1.05E+00 | 4.99E-04 | 2.12E-03 | -1.99E+00 | 5.05E-03 | 1.93E-02 | -1.49E+00 | 4.56E-02 | 1.63E-01 | -1.08E+00 | 1.96E-01 | 5.52E-01 | 1.09E+00 | 1.91E-01 | 7.02E-01 | 1.41E+00 | 9.54E-03 | 2.62E-02 | -1.51E+00 |
| FAM13A-AS1 | 1.45E-02 | 7.57E-02 | 1.00E+00 | 1.81E-01 | 7.82E-01 | 1.53E+00 | 4.58E-02 | 1.95E-01 | 1.01E+00 | 7.05E-02 | 2.69E-01 | 1.15E+00 | 1.80E-01 | 6.41E-01 | 1.73E+00 | 2.92E-01 | 8.22E-01 | 1.69E+00 | 1.55E-01 | 5.68E-01 | -1.05E+00 | 1.20E-01 | 3.31E-01 | -1.13E+00 |
| MYCT1 | 1.45E-02 | 7.57E-02 | 1.18E+00 | 1.86E-01 | 8.05E-01 | 1.53E+00 | 1.89E-01 | 8.03E-01 | 1.29E+00 | 4.44E-02 | 1.70E-01 | 1.81E+00 | 2.04E-01 | 7.28E-01 | 1.40E+00 | 2.83E-01 | 7.97E-01 | 1.35E+00 | 2.16E-02 | 7.93E-02 | -1.01E+00 | 1.49E-02 | 4.09E-02 | -1.02E+00 |
| AC093142.1 | 1.45E-02 | 7.57E-02 | 1.05E+00 | 2.33E-08 | 1.01E-07 | -1.81E+00 | 2.32E-02 | 9.87E-02 | 1.50E+00 | 1.21E-02 | 4.62E-02 | -1.17E+00 | 2.91E-02 | 1.04E-01 | -1.01E+00 | 4.68E-05 | 1.32E-04 | -1.90E+00 | 1.06E-01 | 3.88E-01 | 1.03E+00 | 5.62E-04 | 1.54E-03 | -1.54E+00 |
| SPTLC1 | 1.45E-02 | 7.57E-02 | 1.05E+00 | 2.65E-04 | 1.15E-03 | -1.17E+00 | 5.99E-03 | 2.55E-02 | -1.11E+00 | 1.24E-03 | 4.72E-03 | -1.10E+00 | 6.20E-03 | 2.21E-02 | -1.00E+00 | 7.91E-03 | 2.23E-02 | -1.00E+00 | 2.31E-01 | 8.49E-01 | 1.08E+00 | 6.75E-03 | 1.86E-02 | -1.15E+00 |
| SUGT1P2 | 1.45E-02 | 7.58E-02 | -1.27E+00 | 5.47E-02 | 2.37E-01 | 1.02E+00 | 6.05E-02 | 2.57E-01 | -1.03E+00 | 1.48E-01 | 5.64E-01 | 1.16E+00 | 2.15E-01 | 7.66E-01 | 1.29E+00 | 3.19E-01 | 9.00E-01 | 1.36E+00 | 6.08E-02 | 2.23E-01 | -1.01E+00 | 3.10E-01 | 8.51E-01 | 1.18E+00 |
| AC005972.1 | 1.46E-02 | 7.58E-02 | -1.05E+00 | 1.26E-01 | 5.46E-01 | 1.59E+00 | 1.34E-02 | 5.70E-02 | -1.27E+00 | 2.54E-01 | 9.69E-01 | 1.44E+00 | 1.29E-01 | 4.61E-01 | 1.15E+00 | 1.88E-02 | 5.30E-02 | 2.41E+00 | 1.77E-03 | 6.48E-03 | -1.11E+00 | 2.65E-01 | 7.29E-01 | 1.28E+00 |
| AC009962.1 | 1.46E-02 | 7.59E-02 | -1.34E+00 | 5.32E-02 | 2.30E-01 | -1.03E+00 | 1.68E-01 | 7.15E-01 | -1.10E+00 | 7.01E-02 | 2.68E-01 | -1.29E+00 | 1.90E-01 | 6.78E-01 | 1.43E+00 | 3.09E-01 | 8.72E-01 | 1.29E+00 | 1.03E-01 | 3.79E-01 | -1.03E+00 | 1.92E-01 | 5.28E-01 | 1.08E+00 |
| ASH1L-AS1 | 1.46E-02 | 7.59E-02 | -1.11E+00 | 7.78E-03 | 3.36E-02 | -1.05E+00 | 1.84E-01 | 7.83E-01 | 1.13E+00 | 2.09E-02 | 7.99E-02 | 1.01E+00 | 8.31E-02 | 2.96E-01 | 1.91E+00 | 1.48E-01 | 4.18E-01 | 1.82E+00 | 2.55E-01 | 9.37E-01 | 1.27E+00 | 4.36E-02 | 1.20E-01 | 1.10E+00 |
| NTMT1 | 1.46E-02 | 7.60E-02 | -1.29E+00 | 1.75E-02 | 7.54E-02 | -1.25E+00 | 3.90E-05 | 1.66E-04 | -1.76E+00 | 6.23E-04 | 2.38E-03 | -1.38E+00 | 2.27E-02 | 8.09E-02 | -1.34E+00 | 5.67E-02 | 1.60E-01 | -1.18E+00 | 2.94E-06 | 1.08E-05 | -1.84E+00 | 2.71E-03 | 7.44E-03 | -1.16E+00 |
| NUDT2 | 1.46E-02 | 7.60E-02 | 1.10E+00 | 2.45E-03 | 1.06E-02 | 1.11E+00 | 1.79E-02 | 7.59E-02 | -1.08E+00 | 3.09E-04 | 1.18E-03 | -1.13E+00 | 2.35E-03 | 8.39E-03 | -1.04E+00 | 4.82E-04 | 1.36E-03 | -1.02E+00 | 7.97E-02 | 2.92E-01 | 1.12E+00 | 2.54E-02 | 6.97E-02 | 1.04E+00 |
| SMAD1 | 1.46E-02 | 7.60E-02 | -1.02E+00 | 7.92E-02 | 3.42E-01 | 1.26E+00 | 7.62E-05 | 3.24E-04 | -1.32E+00 | 7.12E-05 | 2.72E-04 | -1.18E+00 | 7.94E-05 | 2.83E-04 | -2.01E+00 | 7.81E-03 | 2.20E-02 | -1.13E+00 | 3.86E-06 | 1.42E-05 | -1.60E+00 | 1.71E-03 | 4.69E-03 | -1.21E+00 |
| SGMS1 | 1.46E-02 | 7.61E-02 | -1.14E+00 | 1.14E-02 | 4.93E-02 | -1.11E+00 | 8.60E-03 | 3.65E-02 | -1.44E+00 | 1.52E-02 | 5.82E-02 | -1.24E+00 | 4.25E-06 | 1.51E-05 | -1.34E+00 | 7.29E-06 | 2.05E-05 | -1.19E+00 | 3.07E-11 | 1.13E-10 | -1.23E+00 | 1.01E-12 | 2.79E-12 | -1.41E+00 |
| AC022364.1 | 1.46E-02 | 7.61E-02 | -1.12E+00 | 6.41E-02 | 2.77E-01 | 1.12E+00 | 1.54E-01 | 6.55E-01 | 1.26E+00 | 2.38E-01 | 9.09E-01 | 1.11E+00 | 1.78E-01 | 6.36E-01 | 1.70E+00 | 1.55E-01 | 4.38E-01 | 2.03E+00 | 3.04E-02 | 1.11E-01 | -1.50E+00 | 1.31E-01 | 3.60E-01 | -1.16E+00 |
| DEPDC4 | 1.47E-02 | 7.66E-02 | -1.52E+00 | 7.49E-02 | 3.24E-01 | -1.05E+00 | 7.99E-03 | 3.40E-02 | -1.41E+00 | 1.56E-02 | 5.94E-02 | -1.28E+00 | 2.17E-01 | 7.73E-01 | 1.17E+00 | 9.88E-02 | 2.79E-01 | -1.09E+00 | 9.56E-05 | 3.51E-04 | -1.43E+00 | 1.29E-02 | 3.55E-02 | -1.20E+00 |
| AC083899.1 | 1.47E-02 | 7.67E-02 | -1.27E+00 | 6.53E-02 | 2.82E-01 | 1.07E+00 | 9.58E-05 | 4.07E-04 | -1.57E+00 | 2.70E-02 | 1.03E-01 | 1.04E+00 | 5.95E-02 | 2.12E-01 | -1.24E+00 | 1.82E-01 | 5.13E-01 | 1.08E+00 | 5.55E-03 | 2.04E-02 | -1.33E+00 | 1.91E-01 | 5.25E-01 | 1.04E+00 |
| PIP5K1A | 1.48E-02 | 7.69E-02 | -1.07E+00 | 2.57E-02 | 1.11E-01 | 1.16E+00 | 2.63E-02 | 1.12E-01 | -1.11E+00 | 2.21E-02 | 8.44E-02 | 1.00E+00 | 5.30E-04 | 1.89E-03 | -1.12E+00 | 7.88E-03 | 2.22E-02 | 1.18E+00 | 4.08E-05 | 1.50E-04 | -1.30E+00 | 5.25E-04 | 1.44E-03 | -1.21E+00 |
| TGFB3 | 1.48E-02 | 7.70E-02 | -1.30E+00 | 4.41E-02 | 1.91E-01 | -1.08E+00 | 1.01E-01 | 4.30E-01 | -1.14E+00 | 7.67E-03 | 2.93E-02 | -1.48E+00 | 2.23E-02 | 7.94E-02 | -1.31E+00 | 7.90E-03 | 2.23E-02 | -1.74E+00 | 2.38E-01 | 8.72E-01 | 1.26E+00 | 2.12E-01 | 5.82E-01 | -1.05E+00 |
| LCMT1-AS2 | 1.48E-02 | 7.70E-02 | -1.94E+00 | 7.35E-02 | 3.18E-01 | -1.31E+00 | 2.22E-01 | 9.44E-01 | 1.19E+00 | 1.24E-01 | 4.74E-01 | 1.44E+00 | 2.63E-01 | 9.39E-01 | 1.09E+00 | 3.33E-01 | 9.39E-01 | 1.11E+00 | 1.97E-01 | 7.24E-01 | 1.56E+00 | 6.51E-02 | 1.79E-01 | 1.85E+00 |
| CSAD | 1.48E-02 | 7.71E-02 | 1.13E+00 | 3.69E-02 | 1.60E-01 | 1.28E+00 | 4.36E-04 | 1.85E-03 | -1.30E+00 | 1.17E-04 | 4.47E-04 | -1.20E+00 | 2.43E-01 | 8.65E-01 | 1.45E+00 | 9.93E-02 | 2.80E-01 | 1.01E+00 | 7.14E-03 | 2.62E-02 | -1.19E+00 | 9.24E-02 | 2.54E-01 | -1.08E+00 |
| KDM4D | 1.48E-02 | 7.71E-02 | -1.46E+00 | 1.59E-01 | 6.87E-01 | 1.23E+00 | 2.94E-02 | 1.25E-01 | -1.28E+00 | 4.47E-02 | 1.71E-01 | -1.09E+00 | 8.37E-02 | 2.98E-01 | -1.20E+00 | 2.21E-01 | 6.22E-01 | 1.11E+00 | 1.76E-02 | 6.47E-02 | -1.43E+00 | 5.49E-02 | 1.51E-01 | -1.25E+00 |
| ROR2 | 1.48E-02 | 7.72E-02 | -1.16E+00 | 1.95E-03 | 8.44E-03 | -1.63E+00 | 1.47E-01 | 6.24E-01 | 1.24E+00 | 2.57E-01 | 9.83E-01 | 1.17E+00 | 1.04E-01 | 3.72E-01 | -1.08E+00 | 4.50E-02 | 1.27E-01 | -1.32E+00 | 9.96E-02 | 3.65E-01 | -1.24E+00 | 2.75E-01 | 7.54E-01 | -1.15E+00 |
| AC073389.1 | 1.48E-02 | 7.73E-02 | 1.09E+00 | 6.69E-03 | 2.89E-02 | 1.06E+00 | 8.15E-03 | 3.46E-02 | -1.17E+00 | 8.72E-04 | 3.33E-03 | -1.27E+00 | 1.51E-01 | 5.40E-01 | 1.05E+00 | 2.76E-01 | 7.79E-01 | 1.16E+00 | 7.03E-02 | 2.58E-01 | -1.07E+00 | 2.05E-01 | 5.63E-01 | 1.03E+00 |
| ZNF501 | 1.49E-02 | 7.77E-02 | -1.12E+00 | 4.59E-02 | 1.98E-01 | 1.12E+00 | 2.31E-03 | 9.80E-03 | -1.46E+00 | 7.10E-02 | 2.71E-01 | 1.12E+00 | 4.53E-02 | 1.61E-01 | -1.03E+00 | 5.82E-02 | 1.64E-01 | -1.03E+00 | 3.71E-05 | 1.36E-04 | -1.77E+00 | 5.20E-04 | 1.43E-03 | -1.38E+00 |
| CDC42SE2 | 1.49E-02 | 7.78E-02 | 1.06E+00 | 1.19E-04 | 5.13E-04 | 1.06E+00 | 5.60E-02 | 2.38E-01 | 1.02E+00 | 1.66E-02 | 6.34E-02 | 1.06E+00 | 1.13E-03 | 4.04E-03 | -1.07E+00 | 2.26E-03 | 6.38E-03 | -1.06E+00 | 5.83E-06 | 2.14E-05 | -1.25E+00 | 1.85E-04 | 5.07E-04 | -1.14E+00 |
| AC099811.4 | 1.50E-02 | 7.79E-02 | -1.41E+00 | 7.31E-02 | 3.16E-01 | -1.07E+00 | 2.29E-01 | 9.71E-01 | 1.10E+00 | 1.41E-02 | 5.40E-02 | 2.38E+00 | 2.74E-01 | 9.77E-01 | 1.09E+00 | 9.80E-02 | 2.76E-01 | 1.56E+00 | 2.14E-01 | 7.86E-01 | 1.06E+00 | 3.64E-01 | 9.99E-01 | 1.17E+00 |
| ZNF549 | 1.50E-02 | 7.81E-02 | -1.08E+00 | 7.84E-02 | 3.39E-01 | 1.16E+00 | 7.14E-05 | 3.03E-04 | -1.65E+00 | 1.25E-02 | 4.76E-02 | 1.02E+00 | 2.15E-01 | 7.66E-01 | 1.17E+00 | 3.43E-01 | 9.66E-01 | 1.31E+00 | 9.11E-02 | 3.34E-01 | 1.04E+00 | 2.73E-01 | 7.51E-01 | 1.38E+00 |
| ST13P15 | 1.50E-02 | 7.82E-02 | -1.23E+00 | 8.85E-02 | 3.82E-01 | 1.15E+00 | 5.99E-03 | 2.55E-02 | -1.41E+00 | 1.00E-01 | 3.82E-01 | 1.14E+00 | 2.27E-01 | 8.08E-01 | 1.27E+00 | 7.20E-02 | 2.03E-01 | -1.09E+00 | 2.54E-01 | 9.34E-01 | 1.22E+00 | 7.40E-02 | 2.03E-01 | -1.10E+00 |
| TMEM161B-AS1 | 1.50E-02 | 7.82E-02 | -1.06E+00 | 1.06E-01 | 4.57E-01 | 1.67E+00 | 1.69E-03 | 7.17E-03 | -1.32E+00 | 1.25E-01 | 4.76E-01 | 1.26E+00 | 1.17E-01 | 4.16E-01 | 1.86E+00 | 4.64E-05 | 1.31E-04 | 3.25E+00 | 1.28E-01 | 4.71E-01 | 1.16E+00 | 2.08E-02 | 5.71E-02 | 1.75E+00 |
| URB1 | 1.50E-02 | 7.82E-02 | -1.08E+00 | 1.83E-01 | 7.92E-01 | 1.39E+00 | 2.68E-02 | 1.14E-01 | -1.08E+00 | 1.83E-01 | 7.00E-01 | 1.44E+00 | 1.87E-05 | 6.67E-05 | -1.73E+00 | 1.13E-02 | 3.19E-02 | 1.01E+00 | 1.89E-02 | 6.92E-02 | 1.01E+00 | 1.99E-01 | 5.47E-01 | 1.43E+00 |
| RPL24P2 | 1.50E-02 | 7.83E-02 | -1.05E+00 | 1.90E-52 | 8.21E-52 | -1.76E+00 | 1.06E-01 | 4.52E-01 | -1.03E+00 | 2.15E-22 | 8.22E-22 | -1.75E+00 | 1.25E-01 | 4.44E-01 | -1.00E+00 | 1.94E-18 | 5.46E-18 | -1.66E+00 | 6.11E-02 | 2.24E-01 | -1.06E+00 | 2.96E-16 | 8.12E-16 | -1.64E+00 |
| AC093536.1 | 1.50E-02 | 7.83E-02 | -1.10E+00 | 4.26E-02 | 1.84E-01 | 1.14E+00 | 1.13E-01 | 4.82E-01 | 1.52E+00 | 1.96E-02 | 7.49E-02 | 2.11E+00 | 8.72E-02 | 3.11E-01 | 1.70E+00 | 1.35E-01 | 3.81E-01 | 1.82E+00 | 2.33E-01 | 8.56E-01 | 1.23E+00 | 7.70E-02 | 2.11E-01 | 1.65E+00 |
| RAB20 | 1.50E-02 | 7.83E-02 | -1.24E+00 | 1.08E-01 | 4.67E-01 | 1.26E+00 | 1.14E-04 | 4.86E-04 | -1.68E+00 | 3.32E-04 | 1.27E-03 | -1.53E+00 | 6.45E-06 | 2.30E-05 | -3.77E+00 | 2.00E-04 | 5.65E-04 | -2.12E+00 | 5.94E-04 | 2.18E-03 | -1.81E+00 | 1.37E-02 | 3.77E-02 | -1.38E+00 |
| PUSL1 | 1.50E-02 | 7.83E-02 | -1.19E+00 | 2.49E-02 | 1.08E-01 | -1.02E+00 | 4.98E-02 | 2.12E-01 | 1.05E+00 | 2.70E-03 | 1.03E-02 | -1.04E+00 | 1.73E-01 | 6.15E-01 | -1.04E+00 | 3.31E-01 | 9.33E-01 | 1.33E+00 | 7.01E-05 | 2.57E-04 | -1.24E+00 | 1.37E-01 | 3.76E-01 | 1.16E+00 |
| ERMAP | 1.50E-02 | 7.84E-02 | -1.24E+00 | 4.59E-02 | 1.98E-01 | 1.09E+00 | 6.98E-03 | 2.96E-02 | -1.28E+00 | 1.26E-01 | 4.82E-01 | 1.25E+00 | 9.42E-06 | 3.36E-05 | -1.96E+00 | 3.06E-02 | 8.63E-02 | 1.06E+00 | 6.28E-20 | 2.31E-19 | -1.80E+00 | 1.33E-17 | 3.66E-17 | -1.81E+00 |
| YBX1 | 1.50E-02 | 7.84E-02 | -1.03E+00 | 2.65E-05 | 1.15E-04 | -1.07E+00 | 8.83E-02 | 3.75E-01 | 1.05E+00 | 7.83E-02 | 2.99E-01 | -1.03E+00 | 2.59E-01 | 9.25E-01 | 1.03E+00 | 2.65E-01 | 7.48E-01 | 1.04E+00 | 3.00E-03 | 1.10E-02 | -1.07E+00 | 2.39E-01 | 6.56E-01 | -1.01E+00 |
| AC125611.2 | 1.51E-02 | 7.85E-02 | -1.30E+00 | 2.14E-02 | 9.23E-02 | -1.19E+00 | 1.72E-03 | 7.30E-03 | -1.75E+00 | 4.13E-02 | 1.58E-01 | -1.11E+00 | 1.70E-01 | 6.06E-01 | 1.06E+00 | 7.50E-02 | 2.11E-01 | 2.06E+00 | 2.48E-02 | 9.11E-02 | -1.29E+00 | 3.43E-01 | 9.44E-01 | 1.31E+00 |
| POMT2 | 1.51E-02 | 7.85E-02 | -1.35E+00 | 1.77E-02 | 7.66E-02 | -1.28E+00 | 3.19E-02 | 1.35E-01 | -1.10E+00 | 2.69E-02 | 1.03E-01 | 1.02E+00 | 5.12E-02 | 1.83E-01 | -1.30E+00 | 1.34E-01 | 3.78E-01 | -1.10E+00 | 2.45E-01 | 9.00E-01 | 1.12E+00 | 1.23E-01 | 3.38E-01 | 1.61E+00 |
| AC006012.1 | 1.51E-02 | 7.88E-02 | -1.21E+00 | 3.07E-02 | 1.32E-01 | -1.01E+00 | 2.00E-01 | 8.51E-01 | 1.13E+00 | 2.14E-01 | 8.17E-01 | 1.20E+00 | 2.27E-01 | 8.08E-01 | 1.27E+00 | 1.16E-01 | 3.28E-01 | 1.96E+00 | 1.61E-01 | 5.90E-01 | -1.08E+00 | 1.92E-01 | 5.27E-01 | -1.06E+00 |
| SCAND2P | 1.51E-02 | 7.88E-02 | -1.34E+00 | 7.15E-02 | 3.09E-01 | 1.00E+00 | 8.51E-02 | 3.62E-01 | 1.00E+00 | 2.09E-01 | 7.99E-01 | 1.50E+00 | 2.10E-01 | 7.49E-01 | 1.18E+00 | 1.79E-01 | 5.05E-01 | 1.06E+00 | 3.44E-02 | 1.26E-01 | 1.85E+00 | 1.94E-01 | 5.34E-01 | 1.49E+00 |
| AL355336.1 | 1.51E-02 | 7.88E-02 | -1.05E+00 | 5.61E-04 | 2.42E-03 | -1.34E+00 | 1.70E-02 | 7.23E-02 | -1.11E+00 | 1.38E-02 | 5.26E-02 | -1.05E+00 | 6.06E-02 | 2.16E-01 | 2.05E+00 | 3.11E-01 | 8.77E-01 | 1.25E+00 | 5.98E-05 | 2.20E-04 | -1.68E+00 | 1.52E-04 | 4.18E-04 | -1.57E+00 |
| SKIDA1 | 1.51E-02 | 7.89E-02 | -1.25E+00 | 8.12E-02 | 3.51E-01 | 1.67E+00 | 1.92E-02 | 8.17E-02 | -1.26E+00 | 6.60E-02 | 2.52E-01 | 1.10E+00 | 2.10E-01 | 7.48E-01 | 1.02E+00 | 2.39E-01 | 6.74E-01 | 1.01E+00 | 4.60E-03 | 1.69E-02 | -2.16E+00 | 1.36E-01 | 3.73E-01 | -1.14E+00 |
| BRD3OS | 1.52E-02 | 7.90E-02 | -1.21E+00 | 3.50E-02 | 1.51E-01 | 1.01E+00 | 1.04E-07 | 4.44E-07 | -1.82E+00 | 1.00E-05 | 3.84E-05 | -1.26E+00 | 1.67E-03 | 5.95E-03 | -1.74E+00 | 5.98E-03 | 1.69E-02 | -1.45E+00 | 1.08E-03 | 3.97E-03 | -1.63E+00 | 4.94E-02 | 1.36E-01 | -1.19E+00 |
| AC245100.4 | 1.52E-02 | 7.91E-02 | -1.06E+00 | 6.25E-02 | 2.70E-01 | 1.22E+00 | 2.74E-03 | 1.16E-02 | -1.45E+00 | 9.38E-03 | 3.58E-02 | -1.20E+00 | 1.69E-01 | 6.03E-01 | 1.02E+00 | 3.40E-01 | 9.59E-01 | 1.31E+00 | 8.96E-05 | 3.29E-04 | -1.70E+00 | 3.29E-02 | 9.03E-02 | -1.08E+00 |
| RNASEH1-AS1 | 1.52E-02 | 7.91E-02 | 1.06E+00 | 8.40E-02 | 3.63E-01 | 1.36E+00 | 4.96E-02 | 2.11E-01 | 1.15E+00 | 2.94E-02 | 1.12E-01 | 1.25E+00 | 2.00E-01 | 7.14E-01 | 1.24E+00 | 3.49E-01 | 9.84E-01 | 1.33E+00 | 5.76E-02 | 2.12E-01 | 1.06E+00 | 3.40E-01 | 9.34E-01 | 1.31E+00 |
| LINC00467 | 1.52E-02 | 7.91E-02 | -1.09E+00 | 2.44E-03 | 1.05E-02 | -1.16E+00 | 3.25E-02 | 1.38E-01 | -1.11E+00 | 2.25E-03 | 8.60E-03 | -1.25E+00 | 1.02E-03 | 3.63E-03 | -1.25E+00 | 9.92E-03 | 2.80E-02 | -1.07E+00 | 1.99E-01 | 7.32E-01 | 1.42E+00 | 2.45E-03 | 6.74E-03 | -1.15E+00 |
| UNKL | 1.52E-02 | 7.92E-02 | -1.07E+00 | 2.21E-02 | 9.54E-02 | 1.10E+00 | 1.71E-01 | 7.29E-01 | 1.25E+00 | 5.27E-02 | 2.01E-01 | 1.16E+00 | 2.35E-01 | 8.39E-01 | 1.24E+00 | 3.42E-01 | 9.65E-01 | 1.45E+00 | 1.14E-03 | 4.17E-03 | -1.24E+00 | 8.76E-04 | 2.41E-03 | -1.31E+00 |
| RPS4XP14 | 1.52E-02 | 7.93E-02 | 1.96E+00 | 5.89E-02 | 2.54E-01 | 1.24E+00 | 2.24E-01 | 9.51E-01 | 1.25E+00 | 1.04E-03 | 3.98E-03 | -1.15E+00 | 2.35E-01 | 8.37E-01 | 1.44E+00 | 1.23E-01 | 3.47E-01 | 1.07E+00 | 1.60E-01 | 5.88E-01 | 1.40E+00 | 8.07E-03 | 2.22E-02 | -1.34E+00 |
| ZG16B | 1.52E-02 | 7.94E-02 | 1.79E+00 | 1.70E-01 | 7.35E-01 | 1.56E+00 | 1.48E-01 | 6.30E-01 | 1.43E+00 | 1.85E-01 | 7.06E-01 | 1.60E+00 | 6.78E-02 | 2.42E-01 | 1.84E+00 | 1.60E-01 | 4.51E-01 | 1.90E+00 | 4.58E-02 | 1.68E-01 | 1.58E+00 | 2.18E-01 | 6.00E-01 | 1.37E+00 |
| AL162430.1 | 1.52E-02 | 7.94E-02 | 1.07E+00 | 3.24E-10 | 1.40E-09 | -1.19E+00 | 2.78E-03 | 1.18E-02 | 1.11E+00 | 2.84E-06 | 1.08E-05 | -1.17E+00 | 2.51E-02 | 8.94E-02 | 1.08E+00 | 1.41E-03 | 3.98E-03 | -1.13E+00 | 2.25E-01 | 8.27E-01 | 1.11E+00 | 3.04E-07 | 8.34E-07 | -1.17E+00 |
| POLR3G | 1.53E-02 | 7.95E-02 | -1.20E+00 | 1.36E-01 | 5.87E-01 | 1.24E+00 | 5.91E-02 | 2.51E-01 | -1.19E+00 | 1.84E-01 | 7.03E-01 | 1.17E+00 | 2.59E-02 | 9.23E-02 | -1.20E+00 | 2.28E-01 | 6.42E-01 | 1.17E+00 | 5.04E-03 | 1.85E-02 | -1.13E+00 | 3.39E-01 | 9.30E-01 | 1.38E+00 |
| DDO | 1.53E-02 | 7.96E-02 | 1.06E+00 | 2.96E-02 | 1.28E-01 | 1.20E+00 | 5.21E-04 | 2.21E-03 | -1.31E+00 | 2.37E-03 | 9.04E-03 | -1.09E+00 | 3.96E-02 | 1.41E-01 | -1.37E+00 | 2.09E-01 | 5.90E-01 | 1.11E+00 | 6.80E-04 | 2.50E-03 | -1.62E+00 | 1.66E-02 | 4.56E-02 | -1.39E+00 |
| ZNF217 | 1.53E-02 | 7.97E-02 | 1.08E+00 | 7.49E-05 | 3.24E-04 | -1.08E+00 | 9.02E-02 | 3.83E-01 | 1.16E+00 | 6.15E-03 | 2.35E-02 | 1.05E+00 | 4.24E-02 | 1.51E-01 | 1.09E+00 | 1.90E-02 | 5.36E-02 | -1.08E+00 | 3.88E-02 | 1.42E-01 | 1.16E+00 | 3.58E-05 | 9.84E-05 | -1.31E+00 |
| SNRPB | 1.53E-02 | 7.97E-02 | 1.13E+00 | 7.99E-04 | 3.45E-03 | 1.10E+00 | 9.05E-03 | 3.84E-02 | 1.30E+00 | 9.98E-02 | 3.81E-01 | 1.23E+00 | 4.58E-02 | 1.63E-01 | 1.15E+00 | 1.38E-01 | 3.89E-01 | 1.17E+00 | 1.55E-01 | 5.68E-01 | 1.12E+00 | 6.29E-02 | 1.73E-01 | 1.18E+00 |
| MAPK8IP3 | 1.53E-02 | 7.98E-02 | -1.09E+00 | 2.56E-02 | 1.10E-01 | 1.06E+00 | 1.19E-01 | 5.07E-01 | 1.32E+00 | 1.94E-01 | 7.41E-01 | 1.48E+00 | 1.65E-01 | 5.87E-01 | 1.07E+00 | 1.02E-01 | 2.88E-01 | -1.11E+00 | 2.72E-01 | 9.99E-01 | 1.45E+00 | 2.61E-01 | 7.17E-01 | 1.23E+00 |
| RAB21 | 1.53E-02 | 7.99E-02 | 1.07E+00 | 3.33E-14 | 1.44E-13 | -1.45E+00 | 1.21E-03 | 5.15E-03 | -1.05E+00 | 8.37E-13 | 3.20E-12 | -1.49E+00 | 2.16E-03 | 7.72E-03 | -1.04E+00 | 1.50E-06 | 4.22E-06 | -1.34E+00 | 1.65E-03 | 6.07E-03 | -1.02E+00 | 1.31E-04 | 3.60E-04 | -1.14E+00 |
| AL133330.1 | 1.53E-02 | 7.99E-02 | -1.15E+00 | 5.24E-04 | 2.27E-03 | -1.30E+00 | 9.61E-03 | 4.08E-02 | -1.16E+00 | 2.03E-04 | 7.77E-04 | -1.22E+00 | 1.64E-01 | 5.83E-01 | 1.13E+00 | 1.50E-02 | 4.22E-02 | -1.92E+00 | 2.03E-01 | 7.43E-01 | 1.33E+00 | 3.01E-01 | 8.27E-01 | 1.11E+00 |
| LHFPL6 | 1.54E-02 | 8.01E-02 | -1.05E+00 | 2.06E-01 | 8.90E-01 | 1.61E+00 | 2.77E-03 | 1.18E-02 | -1.31E+00 | 2.19E-01 | 8.35E-01 | 1.38E+00 | 2.60E-02 | 9.27E-02 | -1.40E+00 | 2.04E-01 | 5.75E-01 | 2.02E+00 | 2.55E-06 | 9.35E-06 | -1.25E+00 | 1.81E-06 | 4.98E-06 | -1.20E+00 |
| APRT | 1.54E-02 | 8.02E-02 | 1.02E+00 | 2.39E-02 | 1.03E-01 | 1.07E+00 | 1.62E-01 | 6.87E-01 | 1.09E+00 | 7.27E-02 | 2.78E-01 | 1.17E+00 | 1.93E-01 | 6.89E-01 | 1.16E+00 | 1.65E-01 | 4.65E-01 | 1.22E+00 | 6.98E-02 | 2.56E-01 | 1.21E+00 | 8.98E-02 | 2.47E-01 | 1.19E+00 |
| BCKDHB | 1.54E-02 | 8.04E-02 | 1.05E+00 | 6.85E-02 | 2.96E-01 | 1.36E+00 | 2.08E-03 | 8.86E-03 | -1.05E+00 | 1.79E-03 | 6.84E-03 | 1.10E+00 | 1.22E-03 | 4.35E-03 | -1.11E+00 | 8.40E-03 | 2.37E-02 | 1.09E+00 | 5.22E-04 | 1.92E-03 | -1.05E+00 | 7.93E-03 | 2.18E-02 | 1.06E+00 |
| AC108676.1 | 1.54E-02 | 8.04E-02 | -1.23E+00 | 1.59E-02 | 6.87E-02 | -1.18E+00 | 1.11E-03 | 4.73E-03 | -1.44E+00 | 8.55E-03 | 3.27E-02 | -1.17E+00 | 2.75E-01 | 9.82E-01 | 1.19E+00 | 3.25E-01 | 9.17E-01 | 1.13E+00 | 8.71E-03 | 3.20E-02 | -1.42E+00 | 8.54E-02 | 2.35E-01 | -1.12E+00 |
| FOXP1 | 1.54E-02 | 8.04E-02 | 2.21E+00 | 3.71E-02 | 1.60E-01 | 2.29E+00 | 3.42E-03 | 1.45E-02 | 2.10E+00 | 2.44E-04 | 9.30E-04 | 2.81E+00 | 1.86E-01 | 6.62E-01 | 1.14E+00 | 2.62E-01 | 7.39E-01 | 1.80E+00 | 1.07E-02 | 3.92E-02 | 2.29E+00 | 2.05E-02 | 5.64E-02 | 2.24E+00 |
| TRAPPC13 | 1.55E-02 | 8.06E-02 | -1.03E+00 | 2.30E-02 | 9.95E-02 | 1.05E+00 | 4.01E-07 | 1.70E-06 | -1.51E+00 | 2.79E-07 | 1.07E-06 | -1.41E+00 | 8.73E-03 | 3.11E-02 | -1.36E+00 | 3.49E-01 | 9.84E-01 | 1.39E+00 | 6.64E-05 | 2.44E-04 | -1.34E+00 | 8.95E-03 | 2.46E-02 | -1.09E+00 |
| INPP4B | 1.55E-02 | 8.06E-02 | 1.14E+00 | 3.55E-03 | 1.53E-02 | 1.24E+00 | 3.80E-05 | 1.62E-04 | -1.36E+00 | 1.86E-02 | 7.10E-02 | 1.10E+00 | 1.70E-01 | 6.07E-01 | 1.31E+00 | 2.15E-01 | 6.07E-01 | 1.30E+00 | 1.91E-01 | 7.01E-01 | 1.27E+00 | 1.34E-01 | 3.67E-01 | 1.41E+00 |
| AL121672.2 | 1.55E-02 | 8.08E-02 | 1.01E+00 | 4.02E-02 | 1.74E-01 | 1.14E+00 | 1.20E-01 | 5.11E-01 | 1.47E+00 | 2.54E-01 | 9.69E-01 | 1.36E+00 | 2.44E-01 | 8.71E-01 | 1.20E+00 | 2.75E-01 | 7.75E-01 | 1.56E+00 | 1.93E-01 | 7.08E-01 | -1.01E+00 | 2.91E-01 | 7.98E-01 | 1.11E+00 |
| AC099811.1 | 1.55E-02 | 8.10E-02 | -1.50E+00 | 1.69E-02 | 7.32E-02 | -1.38E+00 | 2.02E-02 | 8.59E-02 | -1.47E+00 | 1.24E-01 | 4.72E-01 | -1.02E+00 | 2.11E-01 | 7.51E-01 | 1.06E+00 | 2.68E-01 | 7.56E-01 | 1.43E+00 | 1.78E-01 | 6.54E-01 | 1.47E+00 | 3.29E-01 | 9.04E-01 | 1.40E+00 |
| AC092164.1 | 1.55E-02 | 8.10E-02 | -1.27E+00 | 3.05E-02 | 1.32E-01 | -1.10E+00 | 1.75E-01 | 7.44E-01 | 1.06E+00 | 1.16E-01 | 4.45E-01 | -1.05E+00 | 2.26E-01 | 8.07E-01 | 1.29E+00 | 3.03E-01 | 8.55E-01 | 1.27E+00 | 2.03E-01 | 7.44E-01 | 1.11E+00 | 2.62E-01 | 7.19E-01 | 1.15E+00 |
| PHRF1 | 1.56E-02 | 8.11E-02 | -1.05E+00 | 4.45E-02 | 1.92E-01 | 1.15E+00 | 5.13E-04 | 2.18E-03 | -1.23E+00 | 2.39E-02 | 9.12E-02 | 1.18E+00 | 9.33E-02 | 3.32E-01 | 1.07E+00 | 1.11E-01 | 3.12E-01 | 1.05E+00 | 1.23E-05 | 4.50E-05 | -1.38E+00 | 8.72E-02 | 2.40E-01 | 1.03E+00 |
| AC018638.7 | 1.56E-02 | 8.11E-02 | -1.19E+00 | 3.61E-02 | 1.56E-01 | -1.11E+00 | 1.31E-03 | 5.57E-03 | -1.81E+00 | 1.04E-02 | 3.97E-02 | -1.29E+00 | 1.04E-01 | 3.70E-01 | -1.09E+00 | 4.98E-02 | 1.40E-01 | -1.41E+00 | 3.81E-02 | 1.40E-01 | -1.80E+00 | 8.36E-02 | 2.30E-01 | -1.75E+00 |
| PPP1R12A-AS1 | 1.56E-02 | 8.13E-02 | -1.09E+00 | 1.23E-01 | 5.33E-01 | 1.36E+00 | 1.99E-02 | 8.45E-02 | -1.40E+00 | 1.11E-02 | 4.24E-02 | -1.46E+00 | 2.01E-01 | 7.15E-01 | -1.05E+00 | 2.03E-01 | 5.73E-01 | 1.50E+00 | 2.26E-01 | 8.29E-01 | 1.30E+00 | 1.58E-02 | 4.33E-02 | 1.79E+00 |
| HSP90AA4P | 1.56E-02 | 8.13E-02 | 2.22E+00 | 2.64E-02 | 1.14E-01 | 2.44E+00 | 4.88E-02 | 2.07E-01 | 1.47E+00 | 1.87E-01 | 7.15E-01 | 1.61E+00 | 2.15E-01 | 7.66E-01 | 1.55E+00 | 7.06E-02 | 1.99E-01 | 2.07E+00 | 6.67E-03 | 2.45E-02 | 1.85E+00 | 7.66E-08 | 2.11E-07 | 2.80E+00 |
| JMJD6 | 1.56E-02 | 8.13E-02 | 1.10E+00 | 5.40E-08 | 2.33E-07 | -1.53E+00 | 3.92E-03 | 1.67E-02 | 1.01E+00 | 1.29E-04 | 4.92E-04 | -1.13E+00 | 1.19E-04 | 4.24E-04 | -1.15E+00 | 5.21E-08 | 1.47E-07 | -1.56E+00 | 1.22E-04 | 4.46E-04 | -1.18E+00 | 1.05E-06 | 2.87E-06 | -1.41E+00 |
| PMM1 | 1.57E-02 | 8.16E-02 | -1.20E+00 | 8.33E-02 | 3.60E-01 | 1.13E+00 | 1.14E-01 | 4.86E-01 | 1.33E+00 | 1.05E-01 | 4.00E-01 | 1.22E+00 | 2.49E-02 | 8.88E-02 | -1.10E+00 | 2.78E-02 | 7.85E-02 | -1.19E+00 | 1.14E-04 | 4.19E-04 | -1.42E+00 | 2.89E-02 | 7.94E-02 | 1.01E+00 |
| INPP5E | 1.57E-02 | 8.17E-02 | -1.21E+00 | 2.37E-02 | 1.02E-01 | -1.04E+00 | 5.75E-03 | 2.44E-02 | -1.38E+00 | 2.65E-02 | 1.01E-01 | -1.04E+00 | 6.03E-02 | 2.15E-01 | 1.05E+00 | 1.67E-01 | 4.70E-01 | 1.21E+00 | 2.56E-02 | 9.39E-02 | 1.00E+00 | 3.09E-01 | 8.49E-01 | 1.26E+00 |
| LDLRAP1 | 1.57E-02 | 8.18E-02 | -1.11E+00 | 1.62E-01 | 7.02E-01 | 1.31E+00 | 1.40E-01 | 5.97E-01 | 1.06E+00 | 1.97E-01 | 7.54E-01 | 1.19E+00 | 1.41E-01 | 5.04E-01 | 1.19E+00 | 3.09E-01 | 8.71E-01 | 1.73E+00 | 3.76E-03 | 1.38E-02 | 1.23E+00 | 3.39E-01 | 9.31E-01 | 1.45E+00 |
| PCYT1B | 1.57E-02 | 8.20E-02 | -1.24E+00 | 1.73E-02 | 7.49E-02 | -1.10E+00 | 9.86E-05 | 4.19E-04 | -1.81E+00 | 7.70E-05 | 2.94E-04 | -1.77E+00 | 2.53E-02 | 9.03E-02 | 1.03E+00 | 7.06E-02 | 1.99E-01 | 1.11E+00 | 2.17E-06 | 7.96E-06 | -1.41E+00 | 4.25E-05 | 1.17E-04 | -1.29E+00 |
| FGFBP3 | 1.58E-02 | 8.21E-02 | 1.04E+00 | 1.91E-02 | 8.26E-02 | 1.10E+00 | 1.76E-01 | 7.47E-01 | 1.28E+00 | 5.08E-02 | 1.94E-01 | 1.65E+00 | 3.86E-02 | 1.37E-01 | -1.17E+00 | 1.98E-01 | 5.58E-01 | 1.18E+00 | 2.11E-01 | 7.75E-01 | 1.27E+00 | 2.31E-01 | 6.34E-01 | 1.39E+00 |
| KLHL32 | 1.58E-02 | 8.23E-02 | 1.89E+00 | 1.15E-01 | 4.96E-01 | 1.61E+00 | 1.73E-01 | 7.34E-01 | 1.33E+00 | 1.04E-01 | 3.96E-01 | 1.71E+00 | 2.24E-01 | 7.99E-01 | 1.29E+00 | 1.71E-01 | 4.82E-01 | 1.74E+00 | 2.70E-01 | 9.90E-01 | 1.19E+00 | 1.42E-01 | 3.91E-01 | 1.58E+00 |
| CLSTN3 | 1.58E-02 | 8.23E-02 | -1.08E+00 | 1.16E-01 | 5.01E-01 | 1.32E+00 | 3.54E-04 | 1.50E-03 | -1.60E+00 | 6.15E-04 | 2.35E-03 | -1.39E+00 | 4.56E-07 | 1.63E-06 | -1.59E+00 | 8.87E-07 | 2.50E-06 | -1.41E+00 | 2.82E-05 | 1.03E-04 | -2.11E+00 | 2.04E-03 | 5.59E-03 | -1.56E+00 |
| ZNF317 | 1.58E-02 | 8.25E-02 | -1.02E+00 | 4.75E-03 | 2.05E-02 | -1.03E+00 | 1.21E-01 | 5.12E-01 | 1.08E+00 | 4.38E-04 | 1.67E-03 | -1.33E+00 | 2.46E-05 | 8.78E-05 | -1.29E+00 | 4.73E-05 | 1.33E-04 | -1.26E+00 | 2.11E-06 | 7.73E-06 | -1.25E+00 | 7.68E-07 | 2.11E-06 | -1.26E+00 |
| RDH14 | 1.58E-02 | 8.26E-02 | 1.20E+00 | 4.55E-03 | 1.97E-02 | 1.22E+00 | 1.53E-04 | 6.48E-04 | -1.12E+00 | 1.50E-06 | 5.74E-06 | -1.17E+00 | 1.51E-03 | 5.37E-03 | -1.02E+00 | 4.19E-02 | 1.18E-01 | 1.16E+00 | 2.75E-05 | 1.01E-04 | -1.20E+00 | 3.41E-06 | 9.38E-06 | -1.22E+00 |
| ARRB2 | 1.58E-02 | 8.26E-02 | 1.11E+00 | 9.26E-03 | 4.00E-02 | 1.24E+00 | 1.24E-02 | 5.25E-02 | 1.01E+00 | 8.82E-03 | 3.37E-02 | 1.10E+00 | 2.52E-06 | 8.97E-06 | -1.23E+00 | 2.70E-04 | 7.62E-04 | -1.04E+00 | 2.70E-01 | 9.90E-01 | 1.26E+00 | 3.64E-01 | 9.99E-01 | 1.24E+00 |
| CFAP44 | 1.59E-02 | 8.28E-02 | -1.15E+00 | 1.11E-01 | 4.78E-01 | 1.30E+00 | 3.33E-06 | 1.41E-05 | -1.52E+00 | 5.96E-06 | 2.28E-05 | -1.36E+00 | 1.57E-01 | 5.60E-01 | 1.16E+00 | 6.80E-02 | 1.92E-01 | -1.18E+00 | 2.57E-02 | 9.43E-02 | -1.06E+00 | 1.33E-01 | 3.65E-01 | 1.11E+00 |
| AL162390.1 | 1.59E-02 | 8.30E-02 | 1.06E+00 | 1.03E-05 | 4.46E-05 | -1.35E+00 | 3.48E-02 | 1.48E-01 | -1.01E+00 | 6.05E-06 | 2.31E-05 | -1.46E+00 | 5.66E-02 | 2.02E-01 | 1.12E+00 | 9.12E-03 | 2.57E-02 | -1.16E+00 | 1.21E-01 | 4.42E-01 | 1.33E+00 | 1.29E-01 | 3.56E-01 | -1.09E+00 |
| AL157815.1 | 1.59E-02 | 8.30E-02 | -1.09E+00 | 1.89E-01 | 8.17E-01 | 1.45E+00 | 2.16E-02 | 9.16E-02 | -1.25E+00 | 1.72E-01 | 6.58E-01 | 1.24E+00 | 6.79E-02 | 2.42E-01 | -1.30E+00 | 3.09E-01 | 8.70E-01 | 1.17E+00 | 2.44E-01 | 8.94E-01 | 1.08E+00 | 3.65E-02 | 1.00E-01 | 1.71E+00 |
| AC016757.1 | 1.59E-02 | 8.30E-02 | -1.11E+00 | 4.39E-02 | 1.90E-01 | 1.13E+00 | 2.13E-02 | 9.03E-02 | -1.32E+00 | 6.63E-02 | 2.53E-01 | -1.07E+00 | 2.10E-02 | 7.50E-02 | -1.05E+00 | 7.52E-02 | 2.12E-01 | 1.14E+00 | 2.52E-01 | 9.25E-01 | 1.37E+00 | 2.18E-02 | 6.00E-02 | -1.03E+00 |
| FASN | 1.59E-02 | 8.31E-02 | -1.03E+00 | 1.86E-01 | 8.04E-01 | 1.40E+00 | 8.85E-02 | 3.76E-01 | -1.02E+00 | 1.21E-01 | 4.61E-01 | 1.45E+00 | 1.61E-02 | 5.76E-02 | -1.21E+00 | 1.07E-01 | 3.03E-01 | 1.63E+00 | 6.22E-03 | 2.28E-02 | -1.02E+00 | 7.73E-02 | 2.12E-01 | 1.49E+00 |
| CDK5RAP3 | 1.59E-02 | 8.31E-02 | -1.20E+00 | 2.61E-02 | 1.13E-01 | -1.07E+00 | 2.26E-02 | 9.62E-02 | -1.19E+00 | 1.53E-01 | 5.83E-01 | 1.19E+00 | 2.43E-01 | 8.66E-01 | 1.45E+00 | 3.22E-01 | 9.09E-01 | 1.42E+00 | 1.61E-01 | 5.90E-01 | 1.19E+00 | 9.59E-02 | 2.63E-01 | 1.46E+00 |
| AC004943.2 | 1.59E-02 | 8.31E-02 | -1.28E+00 | 2.58E-02 | 1.12E-01 | -1.15E+00 | 2.26E-01 | 9.62E-01 | 1.21E+00 | 5.30E-02 | 2.02E-01 | -1.01E+00 | 3.74E-03 | 1.33E-02 | -1.58E+00 | 2.35E-02 | 6.64E-02 | -1.18E+00 | 3.54E-02 | 1.30E-01 | -1.10E+00 | 1.97E-01 | 5.42E-01 | 1.21E+00 |
| HSPA8P1 | 1.60E-02 | 8.32E-02 | -1.30E+00 | 1.61E-01 | 6.97E-01 | 1.31E+00 | 2.25E-01 | 9.56E-01 | 1.18E+00 | 2.32E-03 | 8.87E-03 | 2.33E+00 | 1.43E-01 | 5.10E-01 | -1.15E+00 | 2.94E-01 | 8.29E-01 | 1.43E+00 | 1.71E-01 | 6.26E-01 | 1.03E+00 | 1.42E-02 | 3.89E-02 | 1.67E+00 |
| LSR | 1.60E-02 | 8.33E-02 | -1.22E+00 | 2.94E-03 | 1.27E-02 | -1.44E+00 | 1.73E-01 | 7.36E-01 | 1.06E+00 | 1.94E-01 | 7.40E-01 | 1.13E+00 | 1.06E-01 | 3.77E-01 | -1.08E+00 | 2.76E-02 | 7.78E-02 | -1.71E+00 | 4.08E-02 | 1.50E-01 | 2.11E+00 | 3.02E-01 | 8.29E-01 | 1.23E+00 |
| AP001324.1 | 1.60E-02 | 8.34E-02 | -1.05E+00 | 7.38E-55 | 3.19E-54 | -1.47E+00 | 1.63E-01 | 6.92E-01 | -1.01E+00 | 8.21E-26 | 3.14E-25 | -1.37E+00 | 1.62E-01 | 5.78E-01 | -1.02E+00 | 1.94E-22 | 5.47E-22 | -1.39E+00 | 5.72E-05 | 2.10E-04 | -1.03E+00 | 4.32E-34 | 1.19E-33 | -1.40E+00 |
| DNAJC17 | 1.60E-02 | 8.35E-02 | -1.06E+00 | 1.01E-01 | 4.38E-01 | 1.21E+00 | 1.14E-02 | 4.85E-02 | 1.01E+00 | 2.79E-03 | 1.07E-02 | 1.00E+00 | 6.27E-03 | 2.23E-02 | -1.10E+00 | 4.21E-02 | 1.19E-01 | 1.12E+00 | 9.89E-06 | 3.63E-05 | -1.22E+00 | 1.82E-02 | 5.00E-02 | 1.03E+00 |
| SGTA | 1.60E-02 | 8.35E-02 | 1.11E+00 | 9.34E-04 | 4.03E-03 | 1.11E+00 | 6.62E-04 | 2.81E-03 | -1.05E+00 | 8.26E-05 | 3.16E-04 | 1.03E+00 | 8.77E-08 | 3.13E-07 | -1.45E+00 | 1.60E-03 | 4.51E-03 | -1.00E+00 | 3.01E-02 | 1.11E-01 | 1.07E+00 | 1.90E-01 | 5.23E-01 | 1.18E+00 |
| CRIP1P4 | 1.60E-02 | 8.35E-02 | 1.56E+00 | 3.81E-02 | 1.65E-01 | -1.06E+00 | 6.11E-03 | 2.60E-02 | 1.51E+00 | 6.86E-02 | 2.62E-01 | 1.01E+00 | 2.57E-01 | 9.16E-01 | 1.33E+00 | 8.23E-02 | 2.32E-01 | -1.49E+00 | 1.53E-01 | 5.61E-01 | 1.62E+00 | 3.59E-01 | 9.87E-01 | 1.05E+00 |
| PTDSS2 | 1.60E-02 | 8.36E-02 | 1.02E+00 | 6.29E-02 | 2.72E-01 | 1.24E+00 | 1.91E-07 | 8.13E-07 | -1.43E+00 | 7.95E-06 | 3.04E-05 | -1.08E+00 | 2.31E-05 | 8.23E-05 | -1.40E+00 | 1.46E-02 | 4.13E-02 | 1.10E+00 | 2.64E-02 | 9.69E-02 | 1.01E+00 | 7.20E-02 | 1.98E-01 | 1.53E+00 |
| SNHG11 | 1.61E-02 | 8.37E-02 | -1.03E+00 | 1.17E-02 | 5.07E-02 | 1.04E+00 | 1.08E-02 | 4.61E-02 | -1.05E+00 | 5.00E-03 | 1.91E-02 | 1.00E+00 | 4.94E-02 | 1.76E-01 | 1.24E+00 | 1.24E-02 | 3.49E-02 | 1.04E+00 | 6.66E-03 | 2.45E-02 | 1.03E+00 | 7.88E-06 | 2.16E-05 | -1.28E+00 |
| MYOM1 | 1.61E-02 | 8.38E-02 | -1.31E+00 | 1.60E-03 | 6.90E-03 | -1.91E+00 | 5.76E-02 | 2.45E-01 | -1.34E+00 | 1.42E-01 | 5.41E-01 | -1.09E+00 | 2.08E-01 | 7.41E-01 | 1.18E+00 | 2.00E-01 | 5.64E-01 | -1.00E+00 | 2.26E-01 | 8.31E-01 | 1.28E+00 | 3.27E-01 | 8.97E-01 | 1.19E+00 |
| PHPT1 | 1.61E-02 | 8.39E-02 | 1.20E+00 | 2.92E-04 | 1.26E-03 | 1.19E+00 | 1.87E-01 | 7.93E-01 | 1.21E+00 | 1.27E-01 | 4.85E-01 | 1.18E+00 | 7.10E-03 | 2.53E-02 | 1.10E+00 | 1.06E-01 | 2.98E-01 | 1.19E+00 | 7.96E-02 | 2.92E-01 | 1.39E+00 | 1.95E-01 | 5.36E-01 | 1.29E+00 |
| AC093227.1 | 1.61E-02 | 8.40E-02 | -1.38E+00 | 1.82E-01 | 7.85E-01 | 1.48E+00 | 1.27E-03 | 5.38E-03 | -1.99E+00 | 2.21E-01 | 8.44E-01 | 1.24E+00 | 7.40E-04 | 2.64E-03 | -2.32E+00 | 2.31E-01 | 6.50E-01 | 1.31E+00 | 3.25E-03 | 1.19E-02 | -1.54E+00 | 1.72E-01 | 4.73E-01 | 1.06E+00 |
| AC010378.1 | 1.61E-02 | 8.41E-02 | 1.80E+00 | 2.76E-02 | 1.19E-01 | -1.17E+00 | 3.50E-04 | 1.49E-03 | 2.77E+00 | 1.87E-01 | 7.15E-01 | 1.45E+00 | 1.11E-02 | 3.95E-02 | 2.44E+00 | 3.35E-01 | 9.43E-01 | 1.37E+00 | 1.96E-01 | 7.20E-01 | 1.38E+00 | 9.31E-02 | 2.56E-01 | -1.08E+00 |
| TIFA | 1.62E-02 | 8.42E-02 | 1.19E+00 | 1.82E-04 | 7.87E-04 | 1.01E+00 | 9.89E-05 | 4.20E-04 | -1.27E+00 | 8.66E-07 | 3.31E-06 | -1.36E+00 | 1.37E-02 | 4.89E-02 | -1.02E+00 | 9.73E-03 | 2.74E-02 | -1.09E+00 | 2.75E-03 | 1.01E-02 | -1.20E+00 | 4.00E-04 | 1.10E-03 | -1.35E+00 |
| ZNF212 | 1.62E-02 | 8.42E-02 | -1.33E+00 | 9.73E-02 | 4.20E-01 | 1.13E+00 | 2.24E-06 | 9.52E-06 | -2.09E+00 | 3.11E-03 | 1.19E-02 | -1.15E+00 | 6.01E-05 | 2.14E-04 | -1.77E+00 | 3.75E-04 | 1.06E-03 | -1.47E+00 | 7.58E-12 | 2.78E-11 | -2.31E+00 | 2.77E-07 | 7.61E-07 | -1.63E+00 |
| AC147651.3 | 1.62E-02 | 8.43E-02 | 1.01E+00 | 1.00E-02 | 4.33E-02 | 1.06E+00 | 2.22E-01 | 9.43E-01 | 1.22E+00 | 9.23E-02 | 3.52E-01 | 1.15E+00 | 2.21E-01 | 7.89E-01 | 1.41E+00 | 2.05E-01 | 5.78E-01 | 1.21E+00 | 2.40E-01 | 8.80E-01 | 1.30E+00 | 3.34E-01 | 9.18E-01 | 1.47E+00 |
| IGF1R | 1.62E-02 | 8.43E-02 | 1.15E+00 | 2.15E-01 | 9.31E-01 | 1.55E+00 | 1.06E-03 | 4.48E-03 | -1.16E+00 | 3.46E-02 | 1.32E-01 | 1.14E+00 | 8.90E-02 | 3.17E-01 | 1.25E+00 | 2.81E-01 | 7.93E-01 | 1.34E+00 | 2.75E-02 | 1.01E-01 | 1.22E+00 | 6.98E-02 | 1.92E-01 | 1.19E+00 |
| HMGB1P10 | 1.62E-02 | 8.45E-02 | 1.45E+00 | 3.16E-02 | 1.37E-01 | 1.63E+00 | 3.36E-04 | 1.43E-03 | 1.80E+00 | 1.16E-03 | 4.41E-03 | 1.86E+00 | 1.44E-02 | 5.13E-02 | 1.16E+00 | 2.93E-01 | 8.27E-01 | 1.41E+00 | 8.93E-02 | 3.28E-01 | 1.50E+00 | 1.19E-03 | 3.27E-03 | 1.74E+00 |
| C11orf24 | 1.62E-02 | 8.46E-02 | -1.40E+00 | 5.15E-02 | 2.23E-01 | -1.01E+00 | 1.93E-01 | 8.21E-01 | 1.30E+00 | 1.83E-01 | 6.98E-01 | 1.25E+00 | 2.16E-06 | 7.69E-06 | -2.17E+00 | 1.27E-04 | 3.58E-04 | -1.68E+00 | 1.15E-03 | 4.22E-03 | -1.06E+00 | 4.31E-03 | 1.18E-02 | -1.23E+00 |
| FLNA | 1.62E-02 | 8.46E-02 | 1.48E+00 | 1.12E-02 | 4.82E-02 | 1.71E+00 | 1.46E-01 | 6.21E-01 | 1.48E+00 | 5.49E-02 | 2.10E-01 | 1.61E+00 | 1.88E-01 | 6.70E-01 | 1.23E+00 | 3.22E-01 | 9.08E-01 | 1.35E+00 | 6.19E-02 | 2.27E-01 | 1.61E+00 | 5.64E-02 | 1.55E-01 | 1.38E+00 |
| VPS13D | 1.63E-02 | 8.47E-02 | -1.07E+00 | 6.34E-02 | 2.74E-01 | 1.22E+00 | 6.49E-02 | 2.76E-01 | 1.02E+00 | 2.50E-01 | 9.55E-01 | 1.37E+00 | 5.92E-02 | 2.11E-01 | 1.25E+00 | 1.51E-01 | 4.25E-01 | 1.39E+00 | 3.77E-06 | 1.39E-05 | -1.42E+00 | 6.27E-03 | 1.72E-02 | -1.17E+00 |
| BABAM2 | 1.63E-02 | 8.48E-02 | 1.22E+00 | 1.29E-03 | 5.56E-03 | 1.16E+00 | 1.21E-02 | 5.16E-02 | -1.08E+00 | 5.51E-04 | 2.10E-03 | -1.08E+00 | 2.62E-02 | 9.33E-02 | 1.30E+00 | 2.75E-03 | 7.76E-03 | 1.02E+00 | 2.47E-04 | 9.05E-04 | -1.00E+00 | 4.16E-03 | 1.14E-02 | 1.06E+00 |
| ATAD3B | 1.63E-02 | 8.48E-02 | -1.06E+00 | 4.26E-02 | 1.84E-01 | 1.15E+00 | 4.69E-02 | 1.99E-01 | 1.01E+00 | 1.58E-01 | 6.05E-01 | 1.34E+00 | 1.49E-01 | 5.30E-01 | 1.15E+00 | 1.41E-01 | 3.98E-01 | 1.78E+00 | 2.76E-03 | 1.01E-02 | 1.00E+00 | 1.39E-01 | 3.81E-01 | 1.48E+00 |
| COMMD6 | 1.63E-02 | 8.49E-02 | -1.06E+00 | 2.10E-04 | 9.09E-04 | -1.10E+00 | 2.30E-02 | 9.78E-02 | -1.08E+00 | 1.23E-02 | 4.68E-02 | -1.06E+00 | 2.28E-02 | 8.12E-02 | -1.04E+00 | 4.26E-05 | 1.20E-04 | -1.15E+00 | 2.42E-02 | 8.89E-02 | 1.03E+00 | 8.88E-05 | 2.44E-04 | -1.14E+00 |
| AC122129.1 | 1.63E-02 | 8.50E-02 | -1.36E+00 | 2.04E-01 | 8.81E-01 | 1.37E+00 | 3.00E-03 | 1.28E-02 | -1.69E+00 | 1.34E-02 | 5.11E-02 | -1.32E+00 | 1.36E-01 | 4.86E-01 | -1.30E+00 | 2.76E-01 | 7.79E-01 | -1.11E+00 | 8.90E-04 | 3.27E-03 | -1.43E+00 | 5.47E-03 | 1.50E-02 | -1.27E+00 |
| NPIPA7 | 1.63E-02 | 8.50E-02 | 1.74E+00 | 1.38E-01 | 5.94E-01 | 1.31E+00 | 2.16E-01 | 9.17E-01 | 1.26E+00 | 2.55E-01 | 9.75E-01 | 1.49E+00 | 5.09E-02 | 1.81E-01 | 1.09E+00 | 1.75E-02 | 4.93E-02 | -1.04E+00 | 1.57E-01 | 5.76E-01 | 1.11E+00 | 4.56E-02 | 1.25E-01 | 1.07E+00 |
| C16orf58 | 1.63E-02 | 8.51E-02 | -1.18E+00 | 2.16E-02 | 9.34E-02 | -1.08E+00 | 1.48E-02 | 6.27E-02 | 2.12E+00 | 2.18E-02 | 8.33E-02 | 2.15E+00 | 7.43E-02 | 2.65E-01 | -1.00E+00 | 2.97E-02 | 8.39E-02 | -1.29E+00 | 5.68E-02 | 2.09E-01 | -1.01E+00 | 1.43E-01 | 3.93E-01 | 1.04E+00 |
| FAM171A2 | 1.63E-02 | 8.52E-02 | 1.90E+00 | 1.59E-01 | 6.89E-01 | 1.46E+00 | 1.05E-01 | 4.47E-01 | 1.36E+00 | 2.30E-01 | 8.79E-01 | 1.02E+00 | 2.22E-01 | 7.91E-01 | 1.41E+00 | 2.15E-01 | 6.06E-01 | 1.09E+00 | 2.13E-01 | 7.82E-01 | 1.07E+00 | 2.43E-01 | 6.67E-01 | -1.06E+00 |
| MFAP4 | 1.64E-02 | 8.53E-02 | -1.03E+00 | 3.00E-03 | 1.30E-02 | -1.00E+00 | 3.10E-15 | 1.32E-14 | -2.07E+00 | 1.44E-15 | 5.48E-15 | -1.78E+00 | 2.59E-01 | 9.23E-01 | 1.22E+00 | 2.97E-01 | 8.36E-01 | -1.18E+00 | 2.58E-01 | 9.47E-01 | 1.04E+00 | 3.46E-01 | 9.50E-01 | 1.05E+00 |
| TMEM44 | 1.64E-02 | 8.53E-02 | 1.73E+00 | 2.28E-02 | 9.87E-02 | 2.02E+00 | 1.86E-02 | 7.92E-02 | 1.60E+00 | 5.12E-02 | 1.95E-01 | 1.70E+00 | 2.63E-03 | 9.38E-03 | 2.35E+00 | 1.65E-01 | 4.65E-01 | 1.68E+00 | 1.92E-01 | 7.06E-01 | 1.41E+00 | 1.98E-01 | 5.44E-01 | 1.41E+00 |
| FCGR1CP | 1.64E-02 | 8.54E-02 | -1.80E+00 | 3.17E-02 | 1.37E-01 | -1.51E+00 | 9.01E-25 | 3.83E-24 | -2.88E+00 | 8.56E-21 | 3.27E-20 | -1.93E+00 | 2.53E-01 | 9.01E-01 | 1.13E+00 | 2.35E-01 | 6.63E-01 | -1.29E+00 | 2.18E-01 | 8.00E-01 | 1.16E+00 | 2.91E-01 | 8.00E-01 | -1.09E+00 |
| GALR3 | 1.64E-02 | 8.54E-02 | 2.17E+00 | 6.21E-02 | 2.68E-01 | 2.00E+00 | 8.35E-02 | 3.55E-01 | 1.58E+00 | 2.47E-01 | 9.45E-01 | 1.37E+00 | 1.61E-01 | 5.75E-01 | 1.59E+00 | 2.36E-01 | 6.66E-01 | 1.44E+00 | 1.33E-01 | 4.89E-01 | 1.55E+00 | 2.07E-01 | 5.68E-01 | 1.57E+00 |
| TMIGD2 | 1.65E-02 | 8.60E-02 | 1.14E+00 | 1.56E-02 | 6.73E-02 | 1.24E+00 | 1.92E-05 | 8.17E-05 | -1.16E+00 | 1.70E-06 | 6.49E-06 | -1.13E+00 | 1.84E-05 | 6.55E-05 | -1.40E+00 | 2.49E-04 | 7.02E-04 | -1.14E+00 | 2.56E-01 | 9.38E-01 | 1.34E+00 | 2.04E-01 | 5.62E-01 | 1.51E+00 |
| PIK3C2A | 1.65E-02 | 8.62E-02 | 1.08E+00 | 3.68E-02 | 1.59E-01 | 1.33E+00 | 1.02E-01 | 4.35E-01 | 1.04E+00 | 4.15E-02 | 1.59E-01 | 1.10E+00 | 1.07E-02 | 3.83E-02 | 1.08E+00 | 1.83E-02 | 5.15E-02 | 1.14E+00 | 3.37E-03 | 1.24E-02 | -1.08E+00 | 1.99E-03 | 5.46E-03 | -1.12E+00 |
| MTMR9 | 1.65E-02 | 8.62E-02 | 1.01E+00 | 1.59E-01 | 6.87E-01 | 1.39E+00 | 7.29E-03 | 3.10E-02 | -1.08E+00 | 2.02E-02 | 7.72E-02 | 1.12E+00 | 2.98E-04 | 1.06E-03 | -1.21E+00 | 1.61E-03 | 4.55E-03 | -1.11E+00 | 2.29E-05 | 8.39E-05 | -1.28E+00 | 2.94E-04 | 8.07E-04 | -1.21E+00 |
| SLC39A13 | 1.66E-02 | 8.63E-02 | -1.07E+00 | 9.97E-03 | 4.31E-02 | -1.02E+00 | 2.01E-02 | 8.55E-02 | -1.04E+00 | 3.58E-02 | 1.37E-01 | 1.11E+00 | 6.14E-04 | 2.19E-03 | -1.37E+00 | 1.39E-02 | 3.91E-02 | -1.11E+00 | 1.79E-02 | 6.55E-02 | -1.02E+00 | 2.29E-02 | 6.29E-02 | -1.04E+00 |
| RNF169 | 1.66E-02 | 8.63E-02 | 1.75E+00 | 1.31E-02 | 5.65E-02 | 1.22E+00 | 8.00E-02 | 3.40E-01 | 1.43E+00 | 1.56E-01 | 5.97E-01 | 1.19E+00 | 1.93E-02 | 6.89E-02 | 1.84E+00 | 2.15E-01 | 6.05E-01 | 1.20E+00 | 1.20E-02 | 4.42E-02 | 1.65E+00 | 6.19E-02 | 1.70E-01 | 1.09E+00 |
| RPS27AP16 | 1.66E-02 | 8.65E-02 | -1.04E+00 | 6.76E-17 | 2.92E-16 | -1.20E+00 | 9.64E-03 | 4.09E-02 | -1.06E+00 | 7.76E-14 | 2.96E-13 | -1.22E+00 | 8.03E-02 | 2.86E-01 | -1.04E+00 | 1.08E-09 | 3.04E-09 | -1.20E+00 | 2.77E-06 | 1.02E-05 | -1.09E+00 | 3.29E-17 | 9.05E-17 | -1.24E+00 |
| SURF2 | 1.66E-02 | 8.65E-02 | 1.13E+00 | 2.83E-03 | 1.22E-02 | 1.09E+00 | 1.14E-02 | 4.86E-02 | 1.08E+00 | 2.37E-02 | 9.06E-02 | 1.25E+00 | 5.06E-05 | 1.80E-04 | -1.28E+00 | 4.58E-03 | 1.29E-02 | 1.05E+00 | 8.91E-02 | 3.27E-01 | 1.03E+00 | 3.44E-01 | 9.46E-01 | 1.19E+00 |
| BMP1 | 1.66E-02 | 8.65E-02 | -1.00E+00 | 6.04E-02 | 2.61E-01 | 1.24E+00 | 3.28E-02 | 1.39E-01 | 1.01E+00 | 1.50E-02 | 5.74E-02 | 1.06E+00 | 1.00E-01 | 3.57E-01 | 1.01E+00 | 9.81E-02 | 2.76E-01 | -1.07E+00 | 2.96E-03 | 1.09E-02 | -1.31E+00 | 9.22E-02 | 2.53E-01 | 1.04E+00 |
| AC115618.3 | 1.66E-02 | 8.66E-02 | -1.26E+00 | 2.22E-01 | 9.61E-01 | 1.45E+00 | 9.37E-02 | 3.98E-01 | -1.08E+00 | 1.73E-01 | 6.62E-01 | 1.16E+00 | 3.19E-02 | 1.14E-01 | -1.41E+00 | 3.89E-02 | 1.10E-01 | -1.45E+00 | 2.72E-01 | 9.97E-01 | 1.17E+00 | 2.31E-01 | 6.33E-01 | 1.55E+00 |
| AC009831.4 | 1.66E-02 | 8.66E-02 | 1.95E+00 | 7.15E-02 | 3.09E-01 | 1.84E+00 | 8.81E-03 | 3.74E-02 | 1.65E+00 | 1.32E-01 | 5.06E-01 | 1.04E+00 | 2.15E-01 | 7.65E-01 | 1.61E+00 | 3.40E-01 | 9.57E-01 | 1.26E+00 | 2.47E-01 | 9.06E-01 | 1.14E+00 | 3.58E-01 | 9.82E-01 | 1.06E+00 |
| AC125257.1 | 1.67E-02 | 8.69E-02 | -1.15E+00 | 2.85E-02 | 1.23E-01 | 1.02E+00 | 6.02E-03 | 2.56E-02 | -1.40E+00 | 2.70E-02 | 1.03E-01 | 1.00E+00 | 1.47E-02 | 5.24E-02 | -1.36E+00 | 4.48E-02 | 1.26E-01 | -1.12E+00 | 1.78E-03 | 6.54E-03 | -1.56E+00 | 3.29E-02 | 9.04E-02 | -1.15E+00 |
| TCF7 | 1.67E-02 | 8.70E-02 | 1.60E+00 | 3.20E-04 | 1.38E-03 | -1.11E+00 | 7.42E-07 | 3.15E-06 | 2.24E+00 | 1.15E-01 | 4.40E-01 | 1.10E+00 | 2.52E-01 | 8.99E-01 | 1.44E+00 | 2.74E-02 | 7.72E-02 | 1.02E+00 | 7.66E-02 | 2.81E-01 | 1.63E+00 | 1.15E-01 | 3.17E-01 | 1.02E+00 |
| AC025164.2 | 1.67E-02 | 8.71E-02 | 1.85E+00 | 1.35E-01 | 5.82E-01 | 1.68E+00 | 2.14E-01 | 9.10E-01 | 1.26E+00 | 1.89E-01 | 7.24E-01 | 1.47E+00 | 2.48E-01 | 8.83E-01 | 1.53E+00 | 2.68E-01 | 7.55E-01 | 1.62E+00 | 7.40E-02 | 2.71E-01 | 1.77E+00 | 3.52E-01 | 9.66E-01 | 1.22E+00 |
| ZNF287 | 1.67E-02 | 8.71E-02 | -1.25E+00 | 5.56E-02 | 2.40E-01 | 1.04E+00 | 1.60E-01 | 6.80E-01 | 1.10E+00 | 2.33E-01 | 8.90E-01 | 1.25E+00 | 2.17E-02 | 7.72E-02 | -1.37E+00 | 2.90E-02 | 8.19E-02 | -1.28E+00 | 9.19E-03 | 3.37E-02 | -1.23E+00 | 5.59E-02 | 1.53E-01 | -1.04E+00 |
| PLEKHM1P1 | 1.67E-02 | 8.72E-02 | -1.05E+00 | 2.78E-04 | 1.20E-03 | -1.28E+00 | 4.42E-02 | 1.88E-01 | 1.08E+00 | 4.33E-04 | 1.65E-03 | -1.28E+00 | 2.69E-01 | 9.58E-01 | 1.51E+00 | 2.13E-01 | 5.99E-01 | 1.27E+00 | 9.54E-05 | 3.50E-04 | -1.43E+00 | 1.13E-06 | 3.11E-06 | -1.73E+00 |
| DAG1 | 1.67E-02 | 8.72E-02 | -1.24E+00 | 6.15E-02 | 2.66E-01 | 1.05E+00 | 4.84E-02 | 2.06E-01 | -1.16E+00 | 6.16E-02 | 2.35E-01 | 1.06E+00 | 2.12E-01 | 7.56E-01 | 1.06E+00 | 1.21E-02 | 3.42E-02 | 2.26E+00 | 4.82E-07 | 1.77E-06 | -1.10E+00 | 3.04E-03 | 8.36E-03 | 1.17E+00 |
| AL139289.2 | 1.68E-02 | 8.74E-02 | -1.18E+00 | 2.04E-02 | 8.80E-02 | -1.19E+00 | 8.53E-02 | 3.62E-01 | -1.21E+00 | 2.08E-01 | 7.93E-01 | 1.10E+00 | 1.34E-01 | 4.78E-01 | 1.05E+00 | 2.83E-01 | 7.97E-01 | 1.26E+00 | 9.52E-02 | 3.49E-01 | 1.50E+00 | 3.51E-01 | 9.63E-01 | 1.25E+00 |
| AL391001.1 | 1.68E-02 | 8.74E-02 | -1.61E+00 | 2.52E-02 | 1.09E-01 | -1.51E+00 | 3.43E-02 | 1.46E-01 | -1.56E+00 | 8.00E-02 | 3.05E-01 | -1.25E+00 | 6.46E-09 | 2.30E-08 | -3.07E+00 | 1.92E-03 | 5.42E-03 | -1.13E+00 | 4.90E-02 | 1.80E-01 | -1.24E+00 | 2.05E-01 | 5.64E-01 | -1.06E+00 |
| STAG3 | 1.68E-02 | 8.74E-02 | -1.10E+00 | 1.36E-03 | 5.89E-03 | -1.36E+00 | 3.38E-02 | 1.44E-01 | -1.05E+00 | 7.71E-03 | 2.95E-02 | -1.18E+00 | 2.58E-01 | 9.21E-01 | 1.42E+00 | 3.50E-01 | 9.88E-01 | 1.39E+00 | 8.49E-02 | 3.12E-01 | -1.02E+00 | 3.38E-02 | 9.27E-02 | -1.25E+00 |
| CCT6P1 | 1.68E-02 | 8.75E-02 | -1.25E+00 | 5.63E-02 | 2.43E-01 | 1.02E+00 | 3.08E-02 | 1.31E-01 | -1.36E+00 | 1.37E-01 | 5.24E-01 | 1.11E+00 | 2.08E-01 | 7.41E-01 | 1.56E+00 | 1.96E-01 | 5.53E-01 | 1.78E+00 | 3.69E-02 | 1.35E-01 | 1.66E+00 | 2.79E-01 | 7.67E-01 | 1.37E+00 |
| PLEKHG1 | 1.68E-02 | 8.76E-02 | -1.87E+00 | 2.37E-02 | 1.03E-01 | -1.79E+00 | 2.21E-01 | 9.40E-01 | 1.05E+00 | 2.62E-01 | 1.00E+00 | -1.00E+00 | 4.93E-04 | 1.76E-03 | -3.01E+00 | 4.87E-02 | 1.37E-01 | -1.16E+00 | 1.52E-46 | 5.57E-46 | -2.49E+00 | 7.25E-43 | 1.99E-42 | -2.36E+00 |
| ZNF524 | 1.68E-02 | 8.76E-02 | 1.01E+00 | 2.82E-02 | 1.22E-01 | 1.20E+00 | 6.31E-09 | 2.68E-08 | -1.53E+00 | 5.80E-05 | 2.22E-04 | -1.07E+00 | 9.27E-06 | 3.31E-05 | -1.25E+00 | 5.06E-04 | 1.43E-03 | 1.02E+00 | 7.67E-05 | 2.82E-04 | -1.31E+00 | 4.28E-03 | 1.18E-02 | -1.11E+00 |
| CSF1 | 1.68E-02 | 8.77E-02 | -1.15E+00 | 9.30E-03 | 4.02E-02 | -1.27E+00 | 2.62E-03 | 1.11E-02 | -1.10E+00 | 1.53E-04 | 5.84E-04 | -1.40E+00 | 1.12E-04 | 3.98E-04 | 2.06E+00 | 2.77E-01 | 7.80E-01 | 1.17E+00 | 2.41E-01 | 8.85E-01 | 1.41E+00 | 6.86E-04 | 1.88E-03 | -1.08E+00 |
| HMGB1P20 | 1.69E-02 | 8.79E-02 | 2.06E+00 | 1.23E-01 | 5.30E-01 | 1.72E+00 | 4.91E-03 | 2.09E-02 | 1.95E+00 | 1.43E-01 | 5.48E-01 | 1.59E+00 | 1.56E-01 | 5.55E-01 | 1.56E+00 | 3.15E-01 | 8.89E-01 | 1.38E+00 | 1.82E-02 | 6.67E-02 | 1.85E+00 | 5.51E-02 | 1.51E-01 | 1.70E+00 |
| AC233266.2 | 1.69E-02 | 8.80E-02 | -1.23E+00 | 7.58E-02 | 3.27E-01 | 1.13E+00 | 5.05E-04 | 2.15E-03 | -1.66E+00 | 2.12E-02 | 8.10E-02 | -1.02E+00 | 7.13E-03 | 2.54E-02 | -1.43E+00 | 9.21E-02 | 2.60E-01 | 1.07E+00 | 9.34E-09 | 3.43E-08 | -1.55E+00 | 1.63E-09 | 4.47E-09 | -1.70E+00 |
| UBXN2B | 1.69E-02 | 8.81E-02 | -1.03E+00 | 1.24E-01 | 5.37E-01 | 1.36E+00 | 1.57E-04 | 6.67E-04 | -1.37E+00 | 1.93E-03 | 7.36E-03 | -1.09E+00 | 3.58E-02 | 1.28E-01 | -1.05E+00 | 3.19E-01 | 9.00E-01 | 1.40E+00 | 1.16E-01 | 4.27E-01 | 1.17E+00 | 7.92E-02 | 2.17E-01 | 1.10E+00 |
| GDPD5 | 1.69E-02 | 8.81E-02 | -1.26E+00 | 2.29E-01 | 9.88E-01 | 1.47E+00 | 1.50E-01 | 6.37E-01 | 1.13E+00 | 1.86E-01 | 7.11E-01 | 1.32E+00 | 2.75E-01 | 9.81E-01 | 1.24E+00 | 8.15E-02 | 2.30E-01 | 2.33E+00 | 2.61E-01 | 9.60E-01 | 1.31E+00 | 4.53E-03 | 1.24E-02 | 2.30E+00 |
| IGF2BP2-AS1 | 1.69E-02 | 8.82E-02 | 1.60E+00 | 1.15E-01 | 4.95E-01 | 1.20E+00 | 1.56E-01 | 6.61E-01 | 1.22E+00 | 1.41E-02 | 5.40E-02 | -1.13E+00 | 1.39E-02 | 4.95E-02 | 2.43E+00 | 1.27E-01 | 3.57E-01 | -1.08E+00 | 2.53E-01 | 9.27E-01 | 1.30E+00 | 3.73E-02 | 1.02E-01 | -1.28E+00 |
| CLUH | 1.69E-02 | 8.82E-02 | -1.21E+00 | 8.39E-02 | 3.62E-01 | 1.10E+00 | 8.23E-02 | 3.50E-01 | 1.06E+00 | 7.10E-02 | 2.71E-01 | 1.24E+00 | 2.69E-03 | 9.58E-03 | -1.53E+00 | 8.49E-02 | 2.39E-01 | -1.01E+00 | 3.28E-02 | 1.21E-01 | -1.06E+00 | 2.35E-02 | 6.45E-02 | 1.54E+00 |
| INPP5F | 1.69E-02 | 8.82E-02 | -1.08E+00 | 3.96E-02 | 1.71E-01 | 1.16E+00 | 1.11E-02 | 4.73E-02 | -1.06E+00 | 4.83E-03 | 1.84E-02 | 1.02E+00 | 2.07E-01 | 7.40E-01 | 1.34E+00 | 2.16E-01 | 6.09E-01 | 1.16E+00 | 1.44E-01 | 5.29E-01 | -1.03E+00 | 2.66E-01 | 7.31E-01 | 1.11E+00 |
| TMEM199 | 1.69E-02 | 8.82E-02 | -1.10E+00 | 3.76E-02 | 1.62E-01 | 1.06E+00 | 8.39E-02 | 3.56E-01 | 1.05E+00 | 5.11E-02 | 1.95E-01 | 1.12E+00 | 1.43E-03 | 5.10E-03 | -1.24E+00 | 8.56E-04 | 2.41E-03 | -1.32E+00 | 7.36E-03 | 2.70E-02 | -1.13E+00 | 3.38E-01 | 9.29E-01 | 1.28E+00 |
| STX4 | 1.69E-02 | 8.82E-02 | 1.19E+00 | 2.17E-02 | 9.36E-02 | 1.30E+00 | 2.27E-02 | 9.66E-02 | 1.01E+00 | 2.22E-01 | 8.48E-01 | 1.36E+00 | 1.63E-03 | 5.81E-03 | 1.07E+00 | 1.16E-01 | 3.26E-01 | 1.34E+00 | 5.51E-02 | 2.02E-01 | 1.20E+00 | 3.07E-02 | 8.43E-02 | 1.15E+00 |
| LNX2 | 1.69E-02 | 8.83E-02 | -1.17E+00 | 4.03E-02 | 1.74E-01 | 1.06E+00 | 2.19E-02 | 9.30E-02 | -1.03E+00 | 5.48E-02 | 2.09E-01 | 1.10E+00 | 1.22E-01 | 4.36E-01 | 1.12E+00 | 3.19E-01 | 8.98E-01 | 1.38E+00 | 2.44E-02 | 8.95E-02 | -1.02E+00 | 1.36E-01 | 3.75E-01 | 1.12E+00 |
| FKRP | 1.69E-02 | 8.83E-02 | -1.09E+00 | 1.10E-03 | 4.76E-03 | -1.22E+00 | 4.03E-02 | 1.71E-01 | -1.03E+00 | 6.40E-02 | 2.44E-01 | 1.15E+00 | 1.27E-01 | 4.54E-01 | 1.12E+00 | 1.17E-01 | 3.31E-01 | 1.10E+00 | 2.58E-01 | 9.47E-01 | 1.24E+00 | 3.33E-01 | 9.14E-01 | 1.18E+00 |
| KCNQ5 | 1.69E-02 | 8.83E-02 | 1.07E+00 | 4.20E-03 | 1.81E-02 | 1.08E+00 | 7.09E-03 | 3.01E-02 | -1.09E+00 | 2.54E-01 | 9.72E-01 | 1.41E+00 | 2.13E-02 | 7.60E-02 | -1.22E+00 | 8.65E-02 | 2.44E-01 | 1.02E+00 | 5.13E-07 | 1.88E-06 | -1.48E+00 | 8.75E-03 | 2.40E-02 | 1.01E+00 |
| AC109460.1 | 1.70E-02 | 8.83E-02 | -1.12E+00 | 1.92E-01 | 8.31E-01 | 1.42E+00 | 2.29E-07 | 9.73E-07 | -1.79E+00 | 1.09E-06 | 4.15E-06 | -1.48E+00 | 1.42E-04 | 5.08E-04 | -1.66E+00 | 9.81E-05 | 2.77E-04 | -1.72E+00 | 4.38E-04 | 1.61E-03 | -1.81E+00 | 1.58E-02 | 4.33E-02 | -1.19E+00 |
| SH3BGR | 1.70E-02 | 8.85E-02 | -1.11E+00 | 2.17E-02 | 9.37E-02 | 1.03E+00 | 3.02E-02 | 1.29E-01 | -1.04E+00 | 2.63E-03 | 1.00E-02 | -1.21E+00 | 1.03E-01 | 3.68E-01 | -1.04E+00 | 3.46E-01 | 9.74E-01 | 1.41E+00 | 1.79E-03 | 6.58E-03 | 1.91E+00 | 3.51E-01 | 9.63E-01 | 1.01E+00 |
| AL604028.2 | 1.70E-02 | 8.85E-02 | -1.23E+00 | 5.71E-02 | 2.47E-01 | 1.04E+00 | 1.90E-01 | 8.09E-01 | 1.12E+00 | 1.02E-01 | 3.89E-01 | 1.06E+00 | 1.39E-01 | 4.96E-01 | 1.10E+00 | 2.01E-01 | 5.67E-01 | 1.11E+00 | 2.07E-02 | 7.61E-02 | -1.14E+00 | 7.61E-02 | 2.09E-01 | -1.12E+00 |
| AC007878.1 | 1.70E-02 | 8.85E-02 | -1.17E+00 | 3.53E-02 | 1.52E-01 | -1.03E+00 | 4.57E-04 | 1.94E-03 | -1.72E+00 | 1.96E-02 | 7.49E-02 | -1.22E+00 | 9.03E-02 | 3.22E-01 | -1.05E+00 | 3.17E-01 | 8.93E-01 | 1.62E+00 | 2.18E-03 | 8.00E-03 | -1.35E+00 | 7.39E-02 | 2.03E-01 | 1.02E+00 |
| AC006994.2 | 1.70E-02 | 8.86E-02 | 1.44E+00 | 1.56E-01 | 6.75E-01 | -1.07E+00 | 1.32E-01 | 5.60E-01 | 1.44E+00 | 2.16E-01 | 8.26E-01 | 1.21E+00 | 7.89E-02 | 2.81E-01 | 2.35E+00 | 2.07E-01 | 5.83E-01 | 1.71E+00 | 2.28E-01 | 8.38E-01 | 1.52E+00 | 2.96E-01 | 8.13E-01 | 1.35E+00 |
| GALM | 1.70E-02 | 8.86E-02 | 1.06E+00 | 2.94E-03 | 1.27E-02 | -1.04E+00 | 7.65E-03 | 3.25E-02 | -1.25E+00 | 1.16E-03 | 4.44E-03 | -1.35E+00 | 6.04E-07 | 2.15E-06 | -1.65E+00 | 8.74E-04 | 2.46E-03 | -1.17E+00 | 1.19E-09 | 4.38E-09 | -1.71E+00 | 1.20E-05 | 3.30E-05 | -1.30E+00 |
| ANKRD16 | 1.70E-02 | 8.86E-02 | -1.02E+00 | 3.54E-02 | 1.53E-01 | 1.14E+00 | 1.83E-03 | 7.76E-03 | -1.19E+00 | 3.58E-03 | 1.37E-02 | 1.00E+00 | 7.71E-04 | 2.75E-03 | -1.68E+00 | 1.30E-01 | 3.67E-01 | 1.15E+00 | 1.70E-03 | 6.24E-03 | -1.13E+00 | 6.49E-02 | 1.78E-01 | 1.11E+00 |
| OXT | 1.70E-02 | 8.88E-02 | -1.01E+00 | 9.04E-06 | 3.91E-05 | -1.26E+00 | 2.09E-01 | 8.88E-01 | 1.34E+00 | 2.02E-01 | 7.73E-01 | 1.23E+00 | 1.53E-01 | 5.46E-01 | 1.05E+00 | 1.52E-01 | 4.30E-01 | -1.22E+00 | 2.66E-01 | 9.77E-01 | 1.03E+00 | 3.45E-01 | 9.49E-01 | 1.14E+00 |
| TMEM62 | 1.70E-02 | 8.88E-02 | -1.01E+00 | 4.58E-02 | 1.98E-01 | 1.19E+00 | 6.04E-05 | 2.57E-04 | -1.43E+00 | 2.78E-05 | 1.06E-04 | -1.42E+00 | 9.24E-02 | 3.30E-01 | 1.31E+00 | 1.68E-01 | 4.75E-01 | 1.25E+00 | 2.62E-02 | 9.63E-02 | 1.19E+00 | 1.68E-02 | 4.61E-02 | 1.02E+00 |
| CRHBP | 1.70E-02 | 8.88E-02 | -1.03E+00 | 3.44E-04 | 1.49E-03 | 1.45E+00 | 1.03E-02 | 4.39E-02 | -1.02E+00 | 9.05E-02 | 3.46E-01 | 1.37E+00 | 2.79E-08 | 9.96E-08 | 2.10E+00 | 1.69E-03 | 4.76E-03 | 1.77E+00 | 5.20E-02 | 1.91E-01 | 2.85E+00 | 3.16E-02 | 8.67E-02 | 3.10E+00 |

| | | | | | | | | | | | | | | | | | | | | | | | | |
|---|---|---|---|---|---|---|---|---|---|---|---|---|---|---|---|---|---|---|---|---|---|---|---|---|
| PMS2P4 | 1.70E-02 | 8.88E-02 | -1.02E+00 | 1.84E-02 | 7.97E-02 | -1.00E+00 | 1.71E-06 | 7.26E-06 | -1.85E+00 | 2.94E-06 | 1.12E-05 | -1.67E+00 | 5.88E-03 | 2.09E-02 | -1.24E+00 | 6.99E-03 | 1.97E-02 | -1.33E+00 | 9.75E-03 | 3.58E-02 | -1.15E+00 | 1.26E-02 | 3.46E-02 | -1.12E+00 |
| HIC1 | 1.70E-02 | 8.88E-02 | -1.09E+00 | 1.01E-02 | 4.35E-02 | -1.13E+00 | 8.07E-02 | 3.43E-01 | -1.04E+00 | 9.70E-02 | 3.71E-01 | 1.08E+00 | 2.60E-02 | 9.26E-02 | -1.42E+00 | 1.37E-02 | 3.87E-02 | -1.68E+00 | 2.38E-02 | 8.75E-02 | -1.25E+00 | 8.28E-02 | 2.27E-01 | -1.16E+00 |
| MCRIP1 | 1.71E-02 | 8.89E-02 | 1.19E+00 | 9.99E-03 | 4.32E-02 | 1.30E+00 | 2.11E-01 | 8.96E-01 | 1.21E+00 | 6.02E-02 | 2.30E-01 | 1.48E+00 | 2.68E-03 | 9.56E-03 | 1.09E+00 | 3.12E-03 | 8.80E-03 | 1.09E+00 | 1.90E-01 | 6.98E-01 | 1.24E+00 | 1.53E-01 | 4.20E-01 | 1.34E+00 |
| RBAK | 1.71E-02 | 8.90E-02 | 1.04E+00 | 4.13E-02 | 1.78E-01 | 1.24E+00 | 4.04E-06 | 1.72E-05 | -1.48E+00 | 3.81E-06 | 1.46E-05 | -1.31E+00 | 2.74E-01 | 9.76E-01 | 1.27E+00 | 5.96E-02 | 1.68E-01 | -1.25E+00 | 1.85E-01 | 6.79E-01 | 1.13E+00 | 2.63E-01 | 7.23E-01 | 1.42E+00 |
| AC004540.1 | 1.71E-02 | 8.91E-02 | 1.09E+00 | 2.92E-03 | 1.26E-02 | 1.04E+00 | 1.62E-01 | 6.90E-01 | 1.18E+00 | 3.95E-02 | 1.51E-01 | 1.04E+00 | 2.62E-01 | 9.36E-01 | 1.59E+00 | 2.72E-01 | 7.67E-01 | 1.36E+00 | 5.41E-02 | 1.99E-01 | 1.67E+00 | 1.35E-01 | 3.71E-01 | 1.69E+00 |
| ARHGEF19 | 1.71E-02 | 8.91E-02 | -1.22E+00 | 1.13E-01 | 4.88E-01 | 1.18E+00 | 1.19E-02 | 5.06E-02 | -1.19E+00 | 1.77E-01 | 6.77E-01 | 1.30E+00 | 1.70E-01 | 6.06E-01 | 1.04E+00 | 2.42E-01 | 6.82E-01 | 1.56E+00 | 2.57E-02 | 9.42E-02 | -1.09E+00 | 1.31E-01 | 3.59E-01 | 1.04E+00 |
| ZSCAN31 | 1.71E-02 | 8.92E-02 | 1.25E+00 | 8.56E-04 | 3.70E-03 | 1.10E+00 | 1.98E-02 | 8.40E-02 | 1.51E+00 | 4.51E-02 | 1.72E-01 | 1.63E+00 | 2.50E-02 | 8.90E-02 | 1.16E+00 | 1.42E-01 | 4.02E-01 | 1.43E+00 | 1.11E-01 | 4.07E-01 | 1.11E+00 | 1.92E-01 | 5.28E-01 | 1.09E+00 |
| ESAM | 1.71E-02 | 8.93E-02 | -1.08E+00 | 2.24E-01 | 9.69E-01 | 1.63E+00 | 1.93E-01 | 8.20E-01 | 1.21E+00 | 2.08E-01 | 7.96E-01 | 1.21E+00 | 2.27E-01 | 8.10E-01 | 1.09E+00 | 2.06E-01 | 5.80E-01 | -1.19E+00 | 1.35E-01 | 4.97E-01 | 2.10E+00 | 1.22E-01 | 3.34E-01 | 1.23E+00 |
| ZNF81 | 1.71E-02 | 8.93E-02 | -1.15E+00 | 4.40E-02 | 1.90E-01 | 1.04E+00 | 8.79E-02 | 3.73E-01 | -1.06E+00 | 2.57E-01 | 9.81E-01 | 1.26E+00 | 1.00E-01 | 3.58E-01 | 1.01E+00 | 2.46E-01 | 6.92E-01 | 1.28E+00 | 1.52E-01 | 5.56E-01 | 1.17E+00 | 2.63E-01 | 7.23E-01 | 1.08E+00 |
| CPA3 | 1.72E-02 | 8.95E-02 | -1.24E+00 | 2.45E-05 | 1.06E-04 | -1.47E+00 | 7.35E-05 | 3.12E-04 | -1.38E+00 | 2.59E-03 | 9.87E-03 | -1.12E+00 | 2.32E-02 | 8.28E-02 | -1.24E+00 | 1.93E-04 | 5.43E-04 | -1.42E+00 | 1.68E-01 | 6.17E-01 | 1.10E+00 | 8.33E-02 | 2.29E-01 | 1.14E+00 |
| AL592295.2 | 1.72E-02 | 8.96E-02 | -1.04E+00 | 1.00E-03 | 4.32E-03 | -1.29E+00 | 4.92E-03 | 2.09E-02 | -1.24E+00 | 4.54E-04 | 1.73E-03 | -1.46E+00 | 7.14E-02 | 2.54E-01 | -1.09E+00 | 2.37E-02 | 6.67E-02 | -1.48E+00 | 8.79E-02 | 3.23E-01 | 1.55E+00 | 2.54E-01 | 6.97E-01 | 1.06E+00 |
| PPP5D1 | 1.72E-02 | 8.98E-02 | -1.42E+00 | 3.87E-02 | 1.67E-01 | -1.12E+00 | 2.23E-01 | 9.46E-01 | 1.41E+00 | 1.64E-01 | 6.28E-01 | 1.15E+00 | 4.09E-02 | 1.46E-01 | -1.36E+00 | 9.13E-02 | 2.57E-01 | -1.13E+00 | 2.39E-01 | 8.75E-01 | 1.16E+00 | 2.34E-01 | 6.43E-01 | -1.10E+00 |
| AC135178.3 | 1.72E-02 | 8.98E-02 | -1.15E+00 | 1.10E-02 | 4.74E-02 | -1.15E+00 | 1.33E-01 | 5.64E-01 | 1.07E+00 | 5.81E-02 | 2.22E-01 | 1.77E+00 | 7.92E-02 | 2.82E-01 | -1.13E+00 | 1.70E-01 | 4.80E-01 | 1.06E+00 | 1.18E-01 | 4.34E-01 | -1.05E+00 | 1.18E-02 | 3.24E-02 | -1.55E+00 |
| MLLT11 | 1.72E-02 | 8.99E-02 | 1.09E+00 | 1.02E-01 | 4.40E-01 | 1.39E+00 | 1.09E-01 | 4.65E-01 | 1.09E+00 | 1.59E-01 | 6.08E-01 | 1.28E+00 | 4.44E-07 | 1.58E-06 | -1.23E+00 | 2.21E-06 | 6.24E-06 | -1.12E+00 | 2.01E-08 | 7.39E-08 | -1.31E+00 | 7.41E-05 | 2.03E-04 | -1.21E+00 |
| MEGF6 | 1.73E-02 | 9.01E-02 | 1.69E+00 | 2.13E-01 | 9.20E-01 | 1.59E+00 | 1.73E-01 | 7.36E-01 | 1.19E+00 | 8.73E-02 | 3.33E-01 | 1.24E+00 | 1.07E-01 | 3.82E-01 | 1.88E+00 | 2.92E-01 | 8.24E-01 | 1.48E+00 | 1.73E-01 | 6.34E-01 | 1.55E+00 | 3.39E-01 | 9.32E-01 | 1.24E+00 |
| DPY19L3 | 1.73E-02 | 9.02E-02 | -1.04E+00 | 5.39E-02 | 2.33E-01 | 1.22E+00 | 1.06E-02 | 4.50E-02 | -1.15E+00 | 3.45E-02 | 1.32E-01 | 1.09E+00 | 2.35E-01 | 8.38E-01 | 1.30E+00 | 3.32E-01 | 9.36E-01 | 1.37E+00 | 2.80E-02 | 1.03E-01 | 1.03E+00 | 3.14E-01 | 8.64E-01 | 1.38E+00 |
| RPL30P2 | 1.73E-02 | 9.02E-02 | 1.06E+00 | 5.78E-06 | 2.50E-05 | -1.41E+00 | 7.29E-02 | 3.10E-01 | 1.08E+00 | 6.34E-04 | 2.42E-03 | -1.14E+00 | 1.00E-01 | 3.57E-01 | 1.71E+00 | 4.13E-02 | 1.17E-01 | -1.06E+00 | 2.72E-01 | 9.99E-01 | 1.23E+00 | 1.13E-02 | 3.11E-02 | -1.18E+00 |
| AC021321.1 | 1.73E-02 | 9.02E-02 | -1.14E+00 | 1.02E-01 | 4.42E-01 | 1.27E+00 | 1.90E-01 | 8.07E-01 | 1.19E+00 | 1.16E-01 | 4.44E-01 | 1.22E+00 | 1.34E-01 | 4.78E-01 | 1.56E+00 | 6.90E-02 | 1.94E-01 | 1.96E+00 | 2.49E-01 | 9.15E-01 | 1.20E+00 | 3.24E-01 | 8.90E-01 | 1.27E+00 |
| PTBP2 | 1.73E-02 | 9.03E-02 | 1.08E+00 | 2.01E-03 | 8.69E-03 | 1.10E+00 | 2.12E-01 | 9.00E-01 | 1.22E+00 | 1.24E-02 | 4.72E-02 | 1.15E+00 | 1.79E-01 | 6.37E-01 | 1.40E+00 | 1.37E-01 | 3.86E-01 | 1.14E+00 | 1.19E-02 | 4.37E-02 | 1.08E+00 | 6.99E-03 | 1.92E-02 | 1.00E+00 |
| AC010889.1 | 1.73E-02 | 9.04E-02 | -1.40E+00 | 8.31E-02 | 3.59E-01 | -1.03E+00 | 9.65E-02 | 4.10E-01 | 1.44E+00 | 4.29E-02 | 1.64E-01 | 1.90E+00 | 8.94E-02 | 3.19E-01 | -1.14E+00 | 3.50E-01 | 9.86E-01 | 1.15E+00 | 2.53E-01 | 9.27E-01 | 1.08E+00 | 3.08E-01 | 8.47E-01 | 1.17E+00 |
| CASS4 | 1.74E-02 | 9.05E-02 | -1.33E+00 | 1.84E-02 | 7.93E-02 | -1.32E+00 | 1.80E-01 | 7.63E-01 | 1.07E+00 | 1.72E-01 | 6.57E-01 | 1.08E+00 | 2.73E-04 | 9.74E-04 | -1.65E+00 | 4.57E-04 | 1.29E-03 | -1.69E+00 | 3.04E-04 | 1.11E-03 | -1.09E+00 | 6.77E-06 | 1.86E-05 | -1.55E+00 |
| KRT18P67 | 1.74E-02 | 9.05E-02 | 1.07E+00 | 1.27E-03 | 5.47E-03 | -1.10E+00 | 1.88E-03 | 7.97E-03 | -1.53E+00 | 1.73E-02 | 6.62E-02 | -1.15E+00 | 6.67E-03 | 2.38E-02 | -1.28E+00 | 9.22E-03 | 2.60E-02 | -1.28E+00 | 2.53E-02 | 9.29E-02 | -1.32E+00 | 2.56E-02 | 7.02E-02 | -1.39E+00 |
| GVQW2 | 1.74E-02 | 9.06E-02 | -1.39E+00 | 5.08E-02 | 2.20E-01 | -1.12E+00 | 1.36E-02 | 5.80E-02 | 1.80E+00 | 5.09E-02 | 1.95E-01 | 1.70E+00 | 1.79E-01 | 6.37E-01 | -1.08E+00 | 2.60E-01 | 7.33E-01 | -1.03E+00 | 2.27E-01 | 8.32E-01 | -1.07E+00 | 1.73E-01 | 4.74E-01 | -1.41E+00 |
| AL445645.1 | 1.74E-02 | 9.08E-02 | -1.07E+00 | 4.04E-05 | 1.75E-04 | -1.53E+00 | 1.29E-01 | 5.50E-01 | 1.07E+00 | 3.06E-02 | 1.17E-01 | -1.09E+00 | 3.61E-02 | 1.29E-01 | 2.89E+00 | 1.74E-01 | 4.91E-01 | 1.97E+00 | 7.72E-02 | 2.83E-01 | 1.74E+00 | 3.40E-01 | 9.34E-01 | 1.20E+00 |
| MFSD12 | 1.74E-02 | 9.08E-02 | -1.10E+00 | 1.60E-01 | 6.91E-01 | 1.34E+00 | 2.69E-04 | 1.14E-03 | -1.42E+00 | 7.95E-03 | 3.04E-02 | -1.02E+00 | 5.26E-02 | 1.88E-01 | -1.16E+00 | 1.58E-01 | 4.45E-01 | 1.10E+00 | 1.83E-02 | 6.73E-02 | -1.53E+00 | 2.44E-01 | 6.71E-01 | -1.02E+00 |
| TNFRSF19 | 1.75E-02 | 9.10E-02 | -1.64E+00 | 2.34E-02 | 1.01E-01 | -1.64E+00 | 5.67E-02 | 2.41E-01 | -1.35E+00 | 2.28E-01 | 8.71E-01 | 1.25E+00 | 2.20E-03 | 7.84E-03 | -1.32E+00 | 1.42E-03 | 3.99E-03 | -1.42E+00 | 2.35E-14 | 8.62E-14 | -3.20E+00 | 1.47E-08 | 4.04E-08 | -2.12E+00 |
| FBN1 | 1.75E-02 | 9.11E-02 | -1.03E+00 | 5.07E-02 | 2.19E-01 | 1.18E+00 | 6.90E-02 | 2.93E-01 | 1.05E+00 | 1.87E-01 | 7.16E-01 | 1.31E+00 | 7.43E-02 | 2.65E-01 | 1.76E+00 | 1.40E-08 | 3.94E-08 | 2.73E+00 | 4.48E-03 | 1.64E-02 | 1.66E+00 | 5.02E-06 | 1.38E-05 | 1.92E+00 |
| AC010969.2 | 1.75E-02 | 9.11E-02 | -1.53E+00 | 8.08E-02 | 3.49E-01 | 1.74E+00 | 7.35E-02 | 3.12E-01 | -1.07E+00 | 6.14E-02 | 2.34E-01 | -1.05E+00 | 3.28E-04 | 1.17E-03 | -1.84E+00 | 4.44E-03 | 1.25E-02 | -1.30E+00 | 1.00E-03 | 3.68E-03 | -1.39E+00 | 1.92E-02 | 5.27E-02 | -1.20E+00 |
| SMG1P4 | 1.75E-02 | 9.12E-02 | -1.04E+00 | 6.18E-02 | 2.67E-01 | 1.20E+00 | 5.33E-02 | 2.27E-01 | -1.02E+00 | 1.16E-02 | 4.45E-02 | 1.03E+00 | 3.12E-02 | 1.11E-01 | -1.03E+00 | 1.48E-02 | 4.18E-02 | -1.16E+00 | 2.81E-05 | 1.03E-04 | -1.18E+00 | 8.05E-04 | 2.21E-03 | -1.13E+00 |
| RPL6P25 | 1.76E-02 | 9.15E-02 | -1.14E+00 | 4.12E-02 | 1.78E-01 | 1.07E+00 | 2.21E-05 | 9.38E-05 | -1.54E+00 | 7.54E-08 | 2.88E-07 | -1.89E+00 | 1.87E-01 | 6.65E-01 | 1.61E+00 | 1.97E-01 | 5.56E-01 | 1.71E+00 | 1.33E-03 | 4.89E-03 | -1.47E+00 | 4.77E-03 | 1.31E-02 | -1.40E+00 |
| AC027031.2 | 1.76E-02 | 9.16E-02 | -1.07E+00 | 6.56E-02 | 2.84E-01 | 1.20E+00 | 8.54E-02 | 3.63E-01 | -1.14E+00 | 8.22E-02 | 3.14E-01 | 1.50E+00 | 2.00E-02 | 7.14E-02 | -1.26E+00 | 2.19E-01 | 6.16E-01 | 1.19E+00 | 2.00E-01 | 7.35E-01 | 1.46E+00 | 2.74E-02 | 7.52E-02 | 1.99E+00 |
| NACC2 | 1.76E-02 | 9.17E-02 | -1.10E+00 | 1.14E-01 | 4.94E-01 | 1.22E+00 | 4.03E-03 | 1.71E-02 | -1.31E+00 | 7.03E-02 | 2.69E-01 | 1.08E+00 | 1.15E-02 | 4.11E-02 | -1.42E+00 | 7.28E-02 | 2.05E-01 | -1.10E+00 | 4.80E-05 | 1.76E-04 | -1.66E+00 | 3.44E-03 | 9.45E-03 | -1.23E+00 |
| TTC12 | 1.76E-02 | 9.17E-02 | -1.16E+00 | 7.01E-02 | 3.03E-01 | 1.15E+00 | 3.02E-04 | 1.28E-03 | -1.60E+00 | 3.34E-03 | 1.27E-02 | -1.18E+00 | 1.65E-01 | 5.90E-01 | 1.00E+00 | 2.82E-01 | 7.95E-01 | 1.10E+00 | 2.06E-02 | 7.54E-02 | -1.32E+00 | 2.68E-01 | 7.36E-01 | 1.09E+00 |
| STEAP3 | 1.76E-02 | 9.19E-02 | -1.65E+00 | 9.63E-02 | 4.16E-01 | 1.08E+00 | 8.69E-05 | 3.69E-04 | -1.59E+00 | 1.54E-04 | 5.89E-04 | -1.37E+00 | 9.95E-03 | 3.55E-02 | -1.53E+00 | 2.57E-01 | 7.24E-01 | 1.59E+00 | 1.03E-08 | 3.79E-08 | -1.23E+00 | 2.72E-02 | 7.48E-02 | 1.13E+00 |
| AC087071.1 | 1.77E-02 | 9.20E-02 | -1.06E+00 | 9.37E-06 | 4.05E-05 | -1.78E+00 | 7.69E-02 | 3.27E-01 | -1.06E+00 | 5.88E-03 | 2.24E-02 | -1.64E+00 | 1.54E-01 | 5.50E-01 | 1.04E+00 | 2.73E-01 | 7.70E-01 | 1.16E+00 | 1.80E-01 | 6.61E-01 | 1.28E+00 | 1.32E-01 | 3.64E-01 | -1.26E+00 |
| C1orf186 | 1.77E-02 | 9.22E-02 | -1.03E+00 | 1.47E-01 | 6.36E-01 | 1.38E+00 | 2.12E-03 | 8.99E-03 | -1.28E+00 | 6.21E-03 | 2.37E-02 | 1.45E+00 | 3.88E-06 | 1.38E-05 | -1.39E+00 | 1.44E-05 | 4.07E-05 | -1.10E+00 | 8.86E-03 | 3.25E-02 | -1.47E+00 | 1.53E-01 | 4.20E-01 | 1.32E+00 |
| EIF3G | 1.77E-02 | 9.22E-02 | 1.05E+00 | 1.43E-01 | 6.19E-01 | 1.18E+00 | 1.46E-01 | 6.19E-01 | 1.01E+00 | 2.11E-01 | 8.05E-01 | 1.11E+00 | 5.93E-02 | 2.11E-01 | 1.06E+00 | 2.42E-01 | 6.81E-01 | 1.19E+00 | 1.54E-01 | 5.66E-01 | 1.09E+00 | 2.38E-01 | 6.53E-01 | 1.17E+00 |
| ORADV1 | 1.77E-02 | 9.22E-02 | -1.01E+00 | 2.03E-02 | 8.77E-02 | 1.09E+00 | 4.77E-03 | 2.03E-02 | -1.11E+00 | 2.25E-02 | 8.61E-02 | 1.12E+00 | 9.67E-03 | 3.45E-02 | -1.16E+00 | 4.32E-02 | 1.22E-01 | -1.01E+00 | 1.20E-01 | 4.41E-01 | 1.16E+00 | 1.17E-01 | 3.22E-01 | 1.09E+00 |
| DYNLL1P7 | 1.77E-02 | 9.23E-02 | -1.29E+00 | 8.04E-02 | 3.48E-01 | 1.72E+00 | 1.06E-01 | 4.52E-01 | -1.04E+00 | 2.91E-03 | 1.11E-02 | 2.21E+00 | 8.64E-02 | 3.08E-01 | -1.20E+00 | 2.42E-02 | 6.83E-02 | 2.37E+00 | 2.71E-01 | 9.95E-01 | 1.17E+00 | 3.04E-06 | 8.34E-06 | 2.75E+00 |
| CTSH | 1.77E-02 | 9.23E-02 | -1.69E+00 | 6.98E-02 | 3.02E-01 | -1.06E+00 | 2.10E-01 | 8.92E-01 | -1.06E+00 | 9.00E-02 | 3.44E-01 | -1.51E+00 | 1.56E-03 | 5.58E-03 | -2.96E+00 | 7.23E-02 | 2.04E-01 | -1.25E+00 | 1.56E-06 | 5.73E-06 | -1.16E+00 | 1.45E-12 | 3.99E-12 | -1.50E+00 |
| ADCY10P1 | 1.77E-02 | 9.24E-02 | -1.29E+00 | 8.26E-02 | 3.57E-01 | 1.85E+00 | 6.34E-02 | 2.69E-01 | -1.23E+00 | 1.13E-01 | 4.33E-01 | 1.67E+00 | 1.65E-01 | 5.87E-01 | -1.21E+00 | 1.44E-01 | 4.06E-01 | 1.90E+00 | 6.87E-02 | 2.52E-01 | -1.41E+00 | 3.24E-01 | 8.91E-01 | -1.06E+00 |
| MMP17 | 1.78E-02 | 9.26E-02 | 1.76E+00 | 2.14E-01 | 9.24E-01 | 1.22E+00 | 1.90E-01 | 8.08E-01 | 1.24E+00 | 5.57E-02 | 2.13E-01 | -1.25E+00 | 1.49E-01 | 5.32E-01 | -1.49E+00 | 2.25E-01 | 6.35E-01 | -1.42E+00 | 1.98E-01 | 7.26E-01 | 1.23E+00 | 3.18E-02 | 8.72E-02 | 1.81E+00 |
| POLR2J2 | 1.78E-02 | 9.26E-02 | -1.12E+00 | 9.68E-02 | 4.18E-01 | 1.21E+00 | 8.41E-02 | 3.57E-01 | -1.06E+00 | 1.53E-01 | 5.85E-01 | 1.19E+00 | 1.21E-01 | 4.31E-01 | 1.11E+00 | 2.89E-01 | 8.14E-01 | 1.43E+00 | 2.52E-01 | 9.26E-01 | 1.24E+00 | 3.54E-01 | 9.72E-01 | 1.28E+00 |
| AC017083.1 | 1.78E-02 | 9.29E-02 | 1.69E+00 | 1.65E-01 | 7.11E-01 | 1.39E+00 | 4.47E-02 | 1.90E-01 | 1.57E+00 | 2.35E-01 | 8.98E-01 | 1.25E+00 | 6.83E-02 | 2.44E-01 | 2.25E+00 | 3.03E-01 | 8.53E-01 | 1.45E+00 | 7.18E-02 | 2.64E-01 | 1.79E+00 | 3.59E-01 | 9.87E-01 | 1.23E+00 |
| HSPA1B | 1.78E-02 | 9.29E-02 | -1.05E+00 | 1.91E-06 | 8.25E-06 | -1.40E+00 | 3.83E-03 | 1.63E-02 | -1.29E+00 | 1.07E-04 | 4.07E-04 | -1.41E+00 | 3.29E-05 | 1.17E-04 | -1.48E+00 | 1.17E-04 | 3.30E-04 | -1.44E+00 | 8.40E-03 | 3.08E-02 | -1.20E+00 | 8.33E-03 | 2.29E-02 | -1.28E+00 |
| SH3BGRL3 | 1.78E-02 | 9.29E-02 | -1.02E+00 | 1.18E-01 | 5.10E-01 | 1.15E+00 | 3.87E-05 | 1.64E-04 | -1.28E+00 | 2.09E-01 | 8.00E-01 | 1.03E+00 | 1.05E-07 | 3.75E-07 | -1.49E+00 | 5.76E-08 | 1.62E-07 | -1.43E+00 | 9.84E-04 | 3.61E-03 | -1.08E+00 | 1.44E-01 | 3.96E-01 | -1.10E+00 |
| ZNF283 | 1.78E-02 | 9.30E-02 | -1.20E+00 | 1.92E-01 | 8.30E-01 | 1.47E+00 | 5.46E-03 | 2.32E-02 | -1.53E+00 | 1.39E-01 | 5.31E-01 | 1.20E+00 | 2.42E-04 | 8.61E-04 | -1.68E+00 | 2.00E-01 | 5.63E-01 | 1.42E+00 | 4.57E-03 | 1.68E-02 | -1.31E+00 | 9.96E-02 | 2.74E-01 | -1.05E+00 |
| CALHM6 | 1.78E-02 | 9.30E-02 | 1.11E+00 | 1.76E-01 | 7.62E-01 | 1.25E+00 | 1.96E-01 | 8.34E-01 | 1.01E+00 | 3.68E-02 | 1.40E-01 | 1.01E+00 | 6.55E-06 | 2.34E-05 | 1.03E+00 | 1.31E-02 | 3.69E-02 | 1.21E+00 | 4.69E-08 | 1.72E-07 | -1.24E+00 | 2.74E-02 | 7.54E-02 | -1.01E+00 |
| MOV10 | 1.79E-02 | 9.32E-02 | -1.05E+00 | 1.80E-01 | 7.78E-01 | 1.34E+00 | 7.33E-04 | 3.12E-03 | -1.04E+00 | 1.03E-03 | 3.93E-03 | 1.12E+00 | 1.44E-05 | 5.14E-05 | -2.15E+00 | 1.11E-03 | 3.13E-03 | -1.35E+00 | 1.45E-05 | 5.33E-05 | -1.49E+00 | 7.75E-03 | 2.13E-02 | -1.18E+00 |
| GNAS | 1.79E-02 | 9.34E-02 | 1.23E+00 | 5.70E-03 | 2.46E-02 | 1.09E+00 | 9.62E-04 | 4.09E-03 | 1.26E+00 | 9.33E-02 | 3.56E-01 | 1.17E+00 | 1.33E-01 | 4.73E-01 | 1.13E+00 | 2.16E-01 | 6.08E-01 | 1.04E+00 | 7.57E-02 | 2.78E-01 | 1.21E+00 | 1.58E-01 | 4.34E-01 | 1.17E+00 |
| KRAS | 1.79E-02 | 9.34E-02 | 1.16E+00 | 2.59E-02 | 1.12E-01 | 1.32E+00 | 1.05E-02 | 4.47E-02 | 1.02E+00 | 2.29E-03 | 8.76E-03 | 1.07E+00 | 1.84E-02 | 6.56E-02 | 1.14E+00 | 2.27E-02 | 6.40E-02 | 1.16E+00 | 3.41E-03 | 1.25E-02 | 1.07E+00 | 5.34E-03 | 1.47E-02 | 1.06E+00 |
| CCNE2 | 1.79E-02 | 9.35E-02 | -1.33E+00 | 3.67E-02 | 1.59E-01 | -1.16E+00 | 9.98E-02 | 4.24E-01 | 1.45E+00 | 2.52E-01 | 9.61E-01 | 1.30E+00 | 9.31E-03 | 3.32E-02 | -1.71E+00 | 3.19E-03 | 9.00E-03 | -2.23E+00 | 2.66E-01 | 9.76E-01 | 1.35E+00 | 8.72E-02 | 2.40E-01 | 1.00E+00 |
| ARHGAP6 | 1.80E-02 | 9.37E-02 | -1.77E+00 | 1.91E-02 | 8.27E-02 | -1.79E+00 | 1.06E-01 | 4.49E-01 | -1.30E+00 | 8.16E-02 | 3.12E-01 | -1.42E+00 | 5.90E-02 | 2.10E-01 | 1.44E+00 | 8.38E-02 | 2.36E-01 | 1.28E+00 | 4.57E-07 | 1.68E-06 | 1.83E+00 | 1.66E-02 | 4.55E-02 | 1.37E+00 |
| AL451074.5 | 1.80E-02 | 9.39E-02 | -1.01E+00 | 6.10E-04 | 2.64E-03 | -1.21E+00 | 1.98E-03 | 8.39E-03 | -1.30E+00 | 3.67E-05 | 1.40E-04 | -1.54E+00 | 1.26E-02 | 4.51E-02 | -1.23E+00 | 1.83E-02 | 5.15E-02 | -1.17E+00 | 8.27E-02 | 3.03E-01 | 1.05E+00 | 5.92E-04 | 1.63E-03 | -1.52E+00 |
| GTF2B | 1.80E-02 | 9.40E-02 | 1.18E+00 | 1.80E-06 | 7.79E-06 | -1.08E+00 | 7.87E-02 | 3.34E-01 | 1.08E+00 | 6.66E-04 | 2.55E-03 | -1.01E+00 | 1.85E-01 | 6.59E-01 | 1.42E+00 | 9.62E-02 | 2.71E-01 | 1.13E+00 | 6.58E-02 | 2.42E-01 | 1.20E+00 | 4.45E-04 | 1.22E-03 | -1.03E+00 |
| ZBTB8A | 1.80E-02 | 9.41E-02 | 1.07E+00 | 1.79E-02 | 7.72E-02 | 1.20E+00 | 1.18E-02 | 5.03E-02 | -1.09E+00 | 9.58E-04 | 3.66E-03 | -1.07E+00 | 7.99E-02 | 2.85E-01 | 1.05E+00 | 1.15E-02 | 3.23E-02 | -1.04E+00 | 4.29E-02 | 1.57E-01 | 1.02E+00 | 7.37E-03 | 2.02E-02 | -1.15E+00 |
| OMD | 1.81E-02 | 9.41E-02 | -1.41E+00 | 3.35E-02 | 1.45E-01 | -1.21E+00 | 1.50E-01 | 6.36E-01 | 1.26E+00 | 2.01E-01 | 7.68E-01 | 1.21E+00 | 1.60E-01 | 5.69E-01 | -1.03E+00 | 2.86E-01 | 8.05E-01 | 1.17E+00 | 3.30E-02 | 1.21E-01 | -1.21E+00 | 5.67E-02 | 1.56E-01 | -1.07E+00 |
| AL669831.1 | 1.81E-02 | 9.42E-02 | -1.11E+00 | 2.44E-03 | 1.05E-02 | -1.26E+00 | 2.25E-03 | 9.55E-03 | -1.43E+00 | 6.82E-05 | 2.60E-04 | -1.75E+00 | 3.67E-02 | 1.31E-01 | -1.17E+00 | 4.53E-02 | 1.28E-01 | -1.19E+00 | 6.36E-02 | 2.33E-01 | -1.00E+00 | 1.95E-01 | 5.37E-01 | 1.13E+00 |
| AC092868.3 | 1.81E-02 | 9.43E-02 | 1.13E+00 | 7.47E-07 | 3.23E-06 | -1.28E+00 | 6.56E-06 | 2.79E-05 | -1.44E+00 | 3.62E-10 | 1.38E-09 | -1.90E+00 | 1.39E-02 | 4.95E-02 | -1.07E+00 | 8.02E-03 | 2.26E-02 | -1.11E+00 | 6.22E-05 | 2.28E-04 | -1.38E+00 | 7.65E-07 | 2.10E-06 | -1.70E+00 |
| FGD4 | 1.81E-02 | 9.43E-02 | -1.01E+00 | 6.25E-02 | 2.70E-01 | 1.21E+00 | 1.38E-01 | 5.85E-01 | -1.06E+00 | 2.27E-01 | 8.66E-01 | 1.13E+00 | 1.53E-01 | 5.44E-01 | 1.17E+00 | 8.55E-02 | 2.41E-01 | -1.17E+00 | 2.19E-01 | 8.06E-01 | 1.16E+00 | 1.08E-01 | 2.97E-01 | -1.08E+00 |
| CCNF | 1.81E-02 | 9.44E-02 | -1.45E+00 | 2.97E-02 | 1.28E-01 | -1.27E+00 | 1.38E-01 | 5.86E-01 | 1.12E+00 | 2.74E-02 | 1.05E-01 | 1.83E+00 | 1.56E-02 | 5.55E-02 | -1.89E+00 | 4.16E-02 | 1.17E-01 | -1.58E+00 | 7.02E-08 | 2.58E-07 | -1.68E+00 | 6.86E-04 | 1.88E-03 | -1.31E+00 |
| ZNF362 | 1.81E-02 | 9.44E-02 | -1.02E+00 | 8.13E-04 | 3.51E-03 | -1.06E+00 | 1.32E-03 | 5.63E-03 | -1.22E+00 | 9.55E-03 | 3.65E-02 | 1.04E+00 | 6.83E-03 | 2.44E-02 | -1.06E+00 | 2.26E-02 | 6.36E-02 | 1.12E+00 | 2.80E-04 | 1.03E-03 | -1.12E+00 | 1.76E-02 | 4.83E-02 | -1.03E+00 |
| AC116049.1 | 1.81E-02 | 9.46E-02 | -1.12E+00 | 1.18E-01 | 5.11E-01 | 1.62E+00 | 1.90E-03 | 8.09E-03 | -1.43E+00 | 2.03E-01 | 7.75E-01 | 1.30E+00 | 1.14E-02 | 4.06E-02 | -1.21E+00 | 3.06E-01 | 8.63E-01 | 1.45E+00 | 1.04E-01 | 3.82E-01 | 1.03E+00 | 4.41E-02 | 1.21E-01 | 1.79E+00 |
| ARHGDIB | 1.82E-02 | 9.46E-02 | -1.04E+00 | 3.53E-02 | 1.53E-01 | 1.12E+00 | 1.53E-01 | 6.50E-01 | -1.02E+00 | 1.47E-01 | 5.60E-01 | 1.06E+00 | 6.17E-05 | 2.20E-04 | -1.16E+00 | 5.48E-02 | 1.54E-01 | -1.07E+00 | 1.63E-08 | 5.97E-08 | -1.16E+00 | 2.41E-02 | 6.61E-02 | -1.03E+00 |
| STK11IP | 1.82E-02 | 9.46E-02 | -1.09E+00 | 4.72E-02 | 2.04E-01 | 1.17E+00 | 1.13E-01 | 4.79E-01 | 1.01E+00 | 5.03E-02 | 1.92E-01 | 1.07E+00 | 6.20E-02 | 2.21E-01 | -1.13E+00 | 1.97E-01 | 5.56E-01 | 1.13E+00 | 4.05E-02 | 1.49E-01 | -1.02E+00 | 1.61E-01 | 4.42E-01 | 1.15E+00 |
| AP001347.1 | 1.82E-02 | 9.48E-02 | -1.07E+00 | 2.04E-02 | 8.81E-02 | 1.05E+00 | 8.37E-02 | 3.55E-01 | 1.44E+00 | 2.14E-01 | 8.17E-01 | 1.27E+00 | 2.24E-01 | 7.98E-01 | 1.25E+00 | 2.33E-01 | 6.56E-01 | 1.22E+00 | 7.13E-02 | 2.62E-01 | 1.50E+00 | 1.44E-01 | 3.96E-01 | 1.66E+00 |
| DNAJB1 | 1.82E-02 | 9.48E-02 | 1.19E+00 | 3.33E-11 | 1.44E-10 | -1.29E+00 | 3.25E-02 | 1.38E-01 | 1.04E+00 | 9.08E-09 | 3.47E-08 | -1.35E+00 | 7.36E-04 | 2.62E-03 | -1.08E+00 | 2.07E-07 | 5.83E-07 | -1.49E+00 | 4.06E-03 | 1.49E-02 | -1.02E+00 | 6.83E-04 | 1.88E-03 | -1.21E+00 |
| TSPAN14 | 1.82E-02 | 9.49E-02 | 1.18E+00 | 1.55E-04 | 6.69E-04 | 1.03E+00 | 2.80E-04 | 1.19E-03 | -1.16E+00 | 1.46E-02 | 5.59E-02 | 1.15E+00 | 5.24E-06 | 1.87E-05 | -1.28E+00 | 3.17E-03 | 8.93E-03 | 1.03E+00 | 8.39E-13 | 3.08E-12 | -1.46E+00 | 8.01E-07 | 2.20E-06 | -1.22E+00 |
| VPS9D1-AS1 | 1.82E-02 | 9.49E-02 | -1.39E+00 | 8.91E-03 | 3.85E-02 | -1.51E+00 | 5.31E-03 | 2.26E-02 | -1.40E+00 | 2.21E-02 | 8.43E-02 | -1.14E+00 | 2.02E-03 | 7.21E-03 | -2.24E+00 | 4.29E-02 | 1.21E-01 | -1.27E+00 | 6.99E-07 | 2.57E-06 | -1.71E+00 | 1.96E-03 | 5.37E-03 | -1.20E+00 |
| HAUS6P1 | 1.82E-02 | 9.50E-02 | -1.22E+00 | 2.99E-03 | 1.29E-02 | -1.51E+00 | 6.51E-04 | 2.77E-03 | -1.27E+00 | 1.97E-05 | 7.53E-05 | -1.46E+00 | 5.02E-04 | 1.79E-03 | -1.78E+00 | 2.43E-03 | 6.85E-03 | -1.60E+00 | 5.42E-06 | 1.99E-05 | -1.56E+00 | 1.27E-07 | 3.49E-07 | -1.84E+00 |
| CCDC157 | 1.83E-02 | 9.52E-02 | -1.28E+00 | 9.21E-02 | 3.98E-01 | 1.12E+00 | 4.72E-04 | 2.00E-03 | -1.60E+00 | 5.39E-04 | 2.06E-03 | -1.51E+00 | 2.02E-01 | 7.19E-01 | 1.49E+00 | 1.23E-01 | 3.47E-01 | 1.72E+00 | 1.64E-01 | 6.00E-01 | -1.02E+00 | 1.14E-01 | 3.14E-01 | -1.17E+00 |
| TMEM44-AS1 | 1.83E-02 | 9.52E-02 | 1.04E+00 | 2.00E-02 | 8.62E-02 | 1.09E+00 | 2.96E-02 | 1.26E-01 | -1.22E+00 | 2.53E-02 | 9.65E-02 | -1.20E+00 | 4.32E-02 | 1.54E-01 | -1.26E+00 | 1.04E-01 | 2.93E-01 | -1.11E+00 | 1.00E-01 | 3.67E-01 | -1.20E+00 | 2.83E-01 | 7.76E-01 | -1.04E+00 |
| ZNF101 | 1.83E-02 | 9.55E-02 | 1.04E+00 | 2.19E-01 | 9.47E-01 | 1.60E+00 | 1.86E-04 | 7.92E-04 | -1.25E+00 | 7.32E-06 | 2.80E-05 | -1.40E+00 | 5.92E-03 | 2.11E-02 | -1.11E+00 | 3.85E-02 | 1.09E-01 | 1.05E+00 | 6.44E-04 | 2.36E-03 | -1.05E+00 | 3.24E-03 | 8.90E-03 | -1.05E+00 |
| IGHM | 1.84E-02 | 9.61E-02 | 1.43E+00 | 2.59E-11 | 1.12E-10 | 2.16E+00 | 1.32E-01 | 5.62E-01 | 1.32E+00 | 6.31E-02 | 2.41E-01 | 1.48E+00 | 2.13E-01 | 7.60E-01 | 1.27E+00 | 3.52E-01 | 9.91E-01 | 1.56E+00 | 2.02E-01 | 7.43E-01 | 1.70E+00 | 1.07E-01 | 2.94E-01 | 2.82E+00 |
| MATN1-AS1 | 1.84E-02 | 9.61E-02 | -1.13E+00 | 6.61E-05 | 2.86E-04 | -1.89E+00 | 1.76E-01 | 7.47E-01 | 1.29E+00 | 1.47E-01 | 5.60E-01 | 1.00E+00 | 1.79E-01 | 6.40E-01 | 1.24E+00 | 9.37E-02 | 2.64E-01 | -1.03E+00 | 8.47E-04 | 3.11E-03 | -1.63E+00 | 4.00E-03 | 1.10E-02 | -1.38E+00 |
| UQCRB | 1.85E-02 | 9.62E-02 | 1.17E+00 | 1.50E-01 | 6.48E-01 | 1.10E+00 | 6.75E-02 | 2.87E-01 | 1.12E+00 | 1.89E-01 | 7.23E-01 | 1.07E+00 | 9.30E-02 | 3.31E-01 | 1.03E+00 | 1.17E-02 | 3.29E-02 | -1.03E+00 | 5.66E-02 | 2.08E-01 | 1.02E+00 | 8.07E-02 | 2.22E-01 | -1.03E+00 |
| PRDM11 | 1.85E-02 | 9.63E-02 | -1.09E+00 | 2.11E-01 | 9.11E-01 | 1.39E+00 | 8.10E-06 | 3.44E-05 | -1.83E+00 | 3.65E-04 | 1.39E-03 | -1.36E+00 | 3.12E-02 | 1.11E-01 | -1.21E+00 | 2.81E-01 | 7.92E-01 | 1.29E+00 | 3.27E-04 | 1.20E-03 | -1.54E+00 | 1.55E-02 | 4.27E-02 | -1.14E+00 |
| ATG16L2 | 1.85E-02 | 9.65E-02 | 1.13E+00 | 2.27E-05 | 9.83E-05 | 1.05E+00 | 3.35E-02 | 1.42E-01 | 1.04E+00 | 6.98E-03 | 2.67E-02 | 1.04E+00 | 6.83E-04 | 2.44E-03 | -1.13E+00 | 1.33E-02 | 3.76E-02 | -1.00E+00 | 3.07E-02 | 1.13E-01 | -1.00E+00 | 2.75E-02 | 7.55E-02 | 1.00E+00 |
| ZSWIM1 | 1.85E-02 | 9.67E-02 | -1.17E+00 | 1.72E-01 | 7.43E-01 | 1.67E+00 | 2.40E-06 | 1.02E-05 | -2.06E+00 | 1.07E-03 | 4.09E-03 | -1.22E+00 | 1.40E-02 | 4.97E-02 | -1.13E+00 | 5.97E-02 | 1.68E-01 | 1.00E+00 | 6.65E-03 | 2.44E-02 | -1.47E+00 | 2.23E-01 | 6.13E-01 | 1.46E+00 |
| MIPOL1 | 1.86E-02 | 9.67E-02 | -1.11E+00 | 1.24E-01 | 5.34E-01 | 1.27E+00 | 5.25E-02 | 2.23E-01 | -1.18E+00 | 1.33E-01 | 5.09E-01 | 1.14E+00 | 2.57E-01 | 9.18E-01 | 1.40E+00 | 1.48E-01 | 4.18E-01 | 1.84E+00 | 6.36E-03 | 2.34E-02 | -1.30E+00 | 3.40E-01 | 9.35E-01 | 1.26E+00 |
| VPS18 | 1.86E-02 | 9.67E-02 | -1.01E+00 | 7.95E-02 | 3.43E-01 | 1.22E+00 | 4.67E-05 | 1.98E-04 | -1.47E+00 | 4.40E-04 | 1.68E-03 | -1.19E+00 | 1.50E-02 | 5.36E-02 | -1.12E+00 | 7.75E-02 | 2.19E-01 | 1.08E+00 | 9.90E-05 | 3.64E-04 | -1.45E+00 | 2.56E-03 | 7.03E-03 | -1.23E+00 |
| MPV17L2 | 1.86E-02 | 9.68E-02 | -1.10E+00 | 1.08E-01 | 4.66E-01 | 1.35E+00 | 6.97E-04 | 2.96E-03 | -1.47E+00 | 8.96E-04 | 3.42E-03 | -1.32E+00 | 1.64E-03 | 5.84E-03 | -1.67E+00 | 6.75E-03 | 1.90E-02 | -1.31E+00 | 1.27E-11 | 4.66E-11 | -2.03E+00 | 4.11E-04 | 1.13E-03 | -1.25E+00 |
| ZNF621 | 1.86E-02 | 9.69E-02 | 1.10E+00 | 2.65E-02 | 1.15E-01 | 1.31E+00 | 2.29E-04 | 9.71E-04 | -1.18E+00 | 3.00E-03 | 1.15E-02 | 1.05E+00 | 6.09E-02 | 2.17E-01 | 1.16E+00 | 3.36E-02 | 9.48E-02 | 1.06E+00 | 8.06E-02 | 2.96E-01 | 1.09E+00 | 2.61E-01 | 7.17E-01 | 1.15E+00 |
| KLF1 | 1.86E-02 | 9.71E-02 | -1.64E+00 | 5.52E-02 | 2.38E-01 | -1.17E+00 | 3.90E-04 | 1.66E-03 | -2.28E+00 | 7.38E-02 | 2.82E-01 | -1.06E+00 | 5.77E-56 | 2.06E-55 | -3.62E+00 | 2.99E-01 | 8.42E-01 | 1.19E+00 | 1.28E-37 | 4.70E-37 | -2.35E+00 | 1.00E-03 | 2.75E-03 | -1.15E+00 |
| TMEM63A | 1.86E-02 | 9.71E-02 | 1.01E+00 | 1.18E-01 | 5.11E-01 | 1.65E+00 | 1.29E-02 | 5.50E-02 | -1.09E+00 | 1.76E-02 | 6.72E-02 | 1.04E+00 | 5.64E-04 | 2.01E-03 | -1.77E+00 | 8.97E-03 | 2.53E-02 | -1.18E+00 | 2.81E-09 | 1.03E-08 | -1.19E+00 | 9.79E-05 | 2.69E-04 | -1.03E+00 |
| SNX11 | 1.86E-02 | 9.71E-02 | -1.11E+00 | 1.43E-01 | 6.20E-01 | 1.34E+00 | 2.81E-04 | 1.19E-03 | -1.41E+00 | 1.21E-03 | 4.63E-03 | 1.08E+00 | 6.92E-03 | 2.47E-02 | -1.15E+00 | 1.53E-01 | 4.32E-01 | 1.21E+00 | 1.57E-04 | 5.77E-04 | -1.20E+00 | 1.08E-03 | 2.97E-03 | -1.09E+00 |
| FAAP24 | 1.86E-02 | 9.71E-02 | -1.21E+00 | 1.96E-02 | 8.49E-02 | -1.10E+00 | 2.48E-04 | 1.05E-03 | -1.37E+00 | 6.03E-03 | 2.30E-02 | 1.02E+00 | 2.21E-01 | 7.88E-01 | 1.40E+00 | 2.89E-01 | 8.15E-01 | 1.22E+00 | 3.03E-02 | 1.11E-01 | 1.08E+00 | 1.24E-03 | 3.41E-03 | -1.08E+00 |
| EIF4A1 | 1.86E-02 | 9.71E-02 | 1.12E+00 | 8.90E-03 | 3.85E-02 | 1.26E+00 | 1.76E-02 | 7.48E-02 | 1.09E+00 | 1.56E-01 | 5.94E-01 | 1.68E+00 | 1.04E-02 | 3.69E-02 | 1.15E+00 | 2.25E-01 | 6.33E-01 | 1.60E+00 | 2.45E-01 | 8.98E-01 | 1.09E+00 | 1.70E-03 | 4.67E-03 | 1.71E+00 |
| AK7 | 1.87E-02 | 9.73E-02 | -1.47E+00 | 5.75E-02 | 2.49E-01 | -1.19E+00 | 1.55E-01 | 6.57E-01 | -1.10E+00 | 1.88E-01 | 7.19E-01 | -1.07E+00 | 2.50E-01 | 8.93E-01 | 1.12E+00 | 1.76E-01 | 4.97E-01 | 1.62E+00 | 1.05E-09 | 3.86E-09 | -1.59E+00 | 8.39E-11 | 2.31E-10 | -1.76E+00 |
| PRKD2 | 1.87E-02 | 9.73E-02 | 1.02E+00 | 2.40E-02 | 1.04E-01 | 1.13E+00 | 9.27E-04 | 3.94E-03 | -1.20E+00 | 2.69E-04 | 1.03E-03 | -1.33E+00 | 1.46E-02 | 5.20E-02 | -1.22E+00 | 7.57E-03 | 2.13E-02 | -1.36E+00 | 7.44E-06 | 2.73E-05 | -1.14E+00 | 7.52E-09 | 2.07E-08 | -1.69E+00 |
| TUBB4B | 1.87E-02 | 9.75E-02 | 1.71E+00 | 1.01E-01 | 4.34E-01 | 1.36E+00 | 8.08E-05 | 3.43E-04 | 2.05E+00 | 1.67E-01 | 6.40E-01 | 1.92E+00 | 1.47E-01 | 5.24E-01 | 1.28E+00 | 3.90E-02 | 1.10E-01 | 1.03E+00 | 1.78E-03 | 6.52E-03 | 1.70E+00 | 1.80E-01 | 4.96E-01 | 1.68E+00 |
| RPS4Y1 | 1.87E-02 | 9.75E-02 | -1.02E+00 | 1.27E-01 | 5.51E-01 | 1.04E+00 | 2.12E-01 | 9.03E-01 | 1.03E+00 | 8.72E-02 | 3.33E-01 | 1.01E+00 | 2.09E-01 | 7.47E-01 | -1.01E+00 | 3.04E-01 | 8.57E-01 | -1.00E+00 | 1.28E-03 | 4.70E-03 | 1.03E+00 | 1.65E-02 | 4.54E-02 | -1.01E+00 |
| PROSER2 | 1.87E-02 | 9.75E-02 | -1.01E+00 | 1.05E-02 | 4.55E-02 | 1.04E+00 | 4.27E-03 | 1.81E-02 | -1.13E+00 | 3.33E-04 | 1.27E-03 | -1.12E+00 | 2.63E-01 | 9.36E-01 | 1.36E+00 | 1.75E-01 | 4.92E-01 | 1.62E+00 | 4.05E-03 | 1.49E-02 | 1.06E+00 | 2.57E-04 | 7.07E-04 | -1.18E+00 |
| B3GALT6 | 1.87E-02 | 9.76E-02 | -1.01E+00 | 2.37E-02 | 1.02E-01 | 1.08E+00 | 2.30E-02 | 9.78E-02 | 1.01E+00 | 4.62E-02 | 1.77E-01 | 1.26E+00 | 3.36E-02 | 1.20E-01 | -1.22E+00 | 3.10E-01 | 8.75E-01 | 1.22E+00 | 1.80E-01 | 6.61E-01 | 1.34E+00 | 7.23E-02 | 1.99E-01 | 1.47E+00 |
| ST13P6 | 1.88E-02 | 9.78E-02 | -1.20E+00 | 2.29E-01 | 9.89E-01 | 1.45E+00 | 2.07E-01 | 8.82E-01 | 1.12E+00 | 1.59E-02 | 6.06E-02 | 1.98E+00 | 8.40E-02 | 2.99E-01 | 1.86E+00 | 1.99E-02 | 5.60E-02 | 2.43E+00 | 2.06E-01 | 7.58E-01 | 1.17E+00 | 1.17E-04 | 3.23E-04 | 2.45E+00 |
| AC021087.3 | 1.88E-02 | 9.80E-02 | -1.33E+00 | 1.71E-01 | 7.38E-01 | 1.22E+00 | 9.63E-03 | 4.09E-02 | -1.39E+00 | 1.20E-02 | 4.59E-02 | -1.33E+00 | 1.10E-01 | 3.90E-01 | 1.09E+00 | 1.56E-01 | 4.40E-01 | 1.03E+00 | 2.18E-01 | 7.99E-01 | 1.18E+00 | 2.06E-01 | 5.66E-01 | 1.47E+00 |
| SETD1A | 1.88E-02 | 9.81E-02 | -1.01E+00 | 2.00E-01 | 8.66E-01 | 1.42E+00 | 5.83E-02 | 2.48E-01 | 1.00E+00 | 1.07E-01 | 4.08E-01 | 1.16E+00 | 2.15E-01 | 7.67E-01 | 1.25E+00 | 2.94E-01 | 8.29E-01 | 1.33E+00 | 7.89E-02 | 2.90E-01 | -1.01E+00 | 1.34E-01 | 3.68E-01 | 1.06E+00 |
| IL11RA | 1.88E-02 | 9.81E-02 | 1.06E+00 | 9.53E-02 | 4.12E-01 | 1.36E+00 | 1.56E-01 | 6.64E-01 | 1.13E+00 | 1.32E-01 | 5.05E-01 | 1.54E+00 | 1.59E-01 | 5.65E-01 | 1.98E+00 | 2.14E-01 | 6.04E-01 | 1.88E+00 | 2.26E-01 | 8.28E-01 | 1.10E+00 | 2.84E-01 | 7.79E-01 | 1.07E+00 |
| AC136475.1 | 1.89E-02 | 9.83E-02 | 1.03E+00 | 1.80E-02 | 7.79E-02 | 1.16E+00 | 4.77E-02 | 2.03E-01 | -1.23E+00 | 1.21E-01 | 4.60E-01 | 1.04E+00 | 1.88E-01 | 6.69E-01 | 1.27E+00 | 1.28E-01 | 3.60E-01 | 1.15E+00 | 2.49E-01 | 9.14E-01 | 1.08E+00 | 2.99E-01 | 8.20E-01 | 1.29E+00 |
| HSP90AA6P | 1.89E-02 | 9.84E-02 | -1.05E+00 | 1.98E-01 | 8.54E-01 | 1.52E+00 | 5.72E-02 | 2.43E-01 | 1.32E+00 | 1.11E-02 | 4.23E-02 | 1.81E+00 | 9.35E-02 | 3.33E-01 | 1.17E+00 | 1.46E-01 | 4.12E-01 | 1.82E+00 | 5.36E-02 | 1.97E-01 | 1.39E+00 | 9.41E-12 | 2.59E-11 | 2.73E+00 |
| NDUFAF2 | 1.89E-02 | 9.84E-02 | 1.18E+00 | 3.52E-04 | 1.52E-03 | 1.12E+00 | 3.17E-02 | 1.35E-01 | 1.36E+00 | 1.02E-02 | 3.89E-02 | 1.14E+00 | 1.22E-01 | 4.35E-01 | 1.21E+00 | 2.68E-01 | 7.55E-01 | 1.45E+00 | 7.44E-03 | 2.73E-02 | 1.00E+00 | 4.23E-02 | 1.16E-01 | 1.06E+00 |
| FAM156B | 1.89E-02 | 9.85E-02 | 1.09E+00 | 4.75E-03 | 2.05E-02 | 1.10E+00 | 5.73E-02 | 2.43E-01 | 1.40E+00 | 1.54E-01 | 5.89E-01 | 1.46E+00 | 2.35E-01 | 8.39E-01 | 1.43E+00 | 1.27E-01 | 3.57E-01 | 1.78E+00 | 4.48E-03 | 1.64E-02 | -1.08E+00 | 2.21E-01 | 6.06E-01 | 1.18E+00 |
| MTMR1 | 1.89E-02 | 9.86E-02 | -1.11E+00 | 1.92E-01 | 8.30E-01 | 1.50E+00 | 3.08E-05 | 1.31E-04 | -1.42E+00 | 4.54E-04 | 1.74E-03 | -1.11E+00 | 7.15E-03 | 2.55E-02 | -1.69E+00 | 1.60E-01 | 4.51E-01 | 1.07E+00 | 9.99E-05 | 3.67E-04 | -1.44E+00 | 3.40E-03 | 9.33E-03 | -1.21E+00 |
| MTX1 | 1.89E-02 | 9.87E-02 | 1.09E+00 | 1.88E-01 | 8.13E-01 | 1.24E+00 | 1.09E-01 | 4.64E-01 | -1.04E+00 | 7.86E-03 | 3.00E-02 | -1.34E+00 | 7.43E-02 | 2.65E-01 | -1.24E+00 | 1.82E-01 | 5.13E-01 | 1.08E+00 | 7.19E-02 | 2.64E-01 | -1.08E+00 | 3.57E-02 | 9.80E-02 | -1.28E+00 |
| C15orf39 | 1.90E-02 | 9.88E-02 | 1.16E+00 | 8.66E-02 | 3.74E-01 | 1.38E+00 | 1.22E-02 | 5.19E-02 | -1.06E+00 | 2.48E-01 | 9.49E-01 | 1.38E+00 | 2.40E-01 | 8.55E-01 | 1.41E+00 | 2.70E-01 | 7.61E-01 | 1.60E+00 | 4.81E-03 | 1.76E-02 | 1.02E+00 | 2.76E-01 | 7.59E-01 | 1.24E+00 |
| ZNF788 | 1.90E-02 | 9.89E-02 | -1.14E+00 | 1.34E-01 | 5.78E-01 | 1.35E+00 | 3.98E-02 | 1.69E-01 | -1.13E+00 | 2.66E-02 | 1.02E-01 | -1.05E+00 | 2.02E-01 | 7.21E-01 | 1.10E+00 | 3.06E-01 | 8.62E-01 | 1.54E+00 | 5.68E-06 | 2.08E-05 | -1.40E+00 | 4.95E-04 | 1.36E-03 | -1.12E+00 |
| AL031727.1 | 1.90E-02 | 9.89E-02 | -1.05E+00 | 7.29E-93 | 3.15E-92 | -1.79E+00 | 3.54E-02 | 1.50E-01 | -1.05E+00 | 1.09E-57 | 4.17E-57 | -1.78E+00 | 5.29E-02 | 1.89E-01 | -1.06E+00 | 2.85E-46 | 8.04E-46 | -1.77E+00 | 1.38E-05 | 5.08E-05 | -1.09E+00 | 2.37E-59 | 6.50E-59 | -1.77E+00 |
| ZBED3 | 1.91E-02 | 9.93E-02 | 1.15E+00 | 3.78E-04 | 1.63E-03 | 1.10E+00 | 3.08E-02 | 1.31E-01 | 1.02E+00 | 1.41E-02 | 5.40E-02 | 1.10E+00 | 1.06E-04 | 3.78E-04 | -1.11E+00 | 2.01E-03 | 5.66E-03 | 1.01E+00 | 3.95E-04 | 1.45E-03 | -1.17E+00 | 7.57E-03 | 2.08E-02 | -1.04E+00 |
| AP001462.1 | 1.91E-02 | 9.94E-02 | 2.07E+00 | 5.65E-02 | 2.44E-01 | 1.97E+00 | 3.77E-02 | 1.60E-01 | 1.58E+00 | 6.94E-02 | 2.65E-01 | 1.64E+00 | 1.07E-01 | 3.82E-01 | 1.95E+00 | 1.55E-01 | 4.38E-01 | 1.94E+00 | 4.28E-02 | 1.57E-01 | 1.81E+00 | 3.33E-01 | 9.14E-01 | 1.21E+00 |
| CLP1 | 1.91E-02 | 9.94E-02 | 1.03E+00 | 8.58E-04 | 3.71E-03 | -1.17E+00 | 2.04E-02 | 8.65E-02 | -1.03E+00 | 4.70E-04 | 1.79E-03 | -1.23E+00 | 1.76E-02 | 6.26E-02 | -1.04E+00 | 7.28E-03 | 2.05E-02 | -1.16E+00 | 8.12E-04 | 2.98E-03 | -1.24E+00 | 2.45E-04 | 6.73E-04 | -1.34E+00 |
| RPLP1P6 | 1.91E-02 | 9.94E-02 | 1.82E+00 | 1.91E-01 | 8.25E-01 | 1.56E+00 | 7.20E-02 | 3.06E-01 | 1.45E+00 | 1.53E-01 | 5.83E-01 | 1.16E+00 | 2.76E-01 | 9.83E-01 | 1.51E+00 | 4.38E-02 | 1.23E-01 | -1.03E+00 | 5.61E-02 | 2.06E-01 | 1.02E+00 | 2.97E-02 | 8.17E-02 | -1.11E+00 |
| SMARCA1 | 1.91E-02 | 9.96E-02 | -1.05E+00 | 1.83E-01 | 7.90E-01 | 1.35E+00 | 3.94E-03 | 1.68E-02 | -1.23E+00 | 2.45E-02 | 9.36E-02 | 1.01E+00 | 5.08E-02 | 1.81E-01 | 1.11E+00 | 2.98E-02 | 8.41E-02 | 1.04E+00 | 1.69E-01 | 6.21E-01 | -1.01E+00 | 8.37E-02 | 2.30E-01 | -1.17E+00 |
| DYRK1A | 1.92E-02 | 9.99E-02 | 1.11E+00 | 3.16E-03 | 1.36E-02 | 1.14E+00 | 1.41E-01 | 5.98E-01 | 1.08E+00 | 7.68E-05 | 2.93E-04 | -1.14E+00 | 1.67E-02 | 5.94E-02 | -1.00E+00 | 1.21E-02 | 3.40E-02 | -1.05E+00 | 2.07E-06 | 7.59E-06 | -1.09E+00 | 4.65E-09 | 1.28E-08 | -1.31E+00 |
| ZNF461 | 1.92E-02 | 1.00E-01 | -1.05E+00 | 9.61E-03 | 4.15E-02 | -1.07E+00 | 9.59E-02 | 4.08E-01 | -1.03E+00 | 5.69E-02 | 2.17E-01 | -1.07E+00 | 8.09E-02 | 2.89E-01 | 1.15E+00 | 6.97E-03 | 1.97E-02 | -1.43E+00 | 2.10E-01 | 7.71E-01 | 1.23E+00 | 2.02E-01 | 5.56E-01 | -1.00E+00 |
| S100A8 | 1.92E-02 | 1.00E-01 | -1.40E+00 | 1.49E-02 | 6.45E-02 | 2.72E+00 | 7.50E-02 | 3.19E-01 | -1.17E+00 | 2.76E-03 | 1.06E-02 | -2.99E+00 | 1.45E-01 | 5.16E-01 | 5.81E+00 | 8.30E-02 | 2.34E-01 | -1.74E+00 | 2.21E-02 | 8.12E-02 | -4.36E+00 | 1.22E-02 | 3.35E-02 | -2.19E+00 |
| EEF1AKMT3 | 1.92E-02 | 1.00E-01 | -1.03E+00 | 6.40E-02 | 2.77E-01 | 1.14E+00 | 2.01E-02 | 8.53E-02 | -1.14E+00 | 5.50E-02 | 2.10E-01 | 1.17E+00 | 2.76E-01 | 9.84E-01 | 1.27E+00 | 8.52E-02 | 2.40E-01 | 1.91E+00 | 2.32E-02 | 8.52E-02 | -1.16E+00 | 3.32E-01 | 9.13E-01 | 1.15E+00 |
| RAB39B | 1.93E-02 | 1.00E-01 | 1.03E+00 | 6.20E-02 | 2.68E-01 | 1.23E+00 | 2.31E-02 | 9.81E-02 | -1.08E+00 | 8.02E-03 | 3.06E-02 | -1.09E+00 | 1.11E-03 | 3.96E-03 | -1.45E+00 | 3.13E-03 | 8.84E-03 | -1.31E+00 | 1.43E-03 | 5.26E-03 | -1.25E+00 | 1.68E-03 | 4.62E-03 | -1.24E+00 |
| SLC29A2 | 1.93E-02 | 1.01E-01 | -1.04E+00 | 2.28E-01 | 9.84E-01 | 1.54E+00 | 6.13E-05 | 2.60E-04 | -1.56E+00 | 1.00E-01 | 3.83E-01 | 1.28E+00 | 1.76E-02 | 6.26E-02 | -1.69E+00 | 2.82E-01 | 7.95E-01 | 1.41E+00 | 2.36E-01 | 8.68E-01 | 1.17E+00 | 1.65E-01 | 4.53E-01 | 1.48E+00 |
| TRIP10 | 1.93E-02 | 1.01E-01 | -1.11E+00 | 1.64E-01 | 7.09E-01 | 1.37E+00 | 1.19E-02 | 5.05E-02 | -1.68E+00 | 6.50E-02 | 2.48E-01 | -1.22E+00 | 3.61E-02 | 1.29E-01 | -1.39E+00 | 2.19E-01 | 6.18E-01 | 1.25E+00 | 1.95E-02 | 7.17E-02 | 1.25E+00 | 1.98E-03 | 5.45E-03 | -1.07E+00 |
| AC096577.1 | 1.93E-02 | 1.01E-01 | -1.18E+00 | 5.02E-02 | 2.17E-01 | -1.01E+00 | 1.55E-01 | 6.58E-01 | -1.06E+00 | 2.62E-01 | 1.00E+00 | 1.24E+00 | 4.01E-08 | 1.43E-07 | -1.66E+00 | 1.04E-05 | 2.92E-05 | -1.29E+00 | 1.60E-01 | 5.87E-01 | 1.12E+00 | 1.27E-02 | 3.49E-02 | -1.34E+00 |
| ATP11B | 1.94E-02 | 1.01E-01 | 1.08E+00 | 8.44E-03 | 3.65E-02 | 1.12E+00 | 1.29E-02 | 5.47E-02 | -1.03E+00 | 6.37E-05 | 2.43E-04 | -1.12E+00 | 2.00E-01 | 7.14E-01 | 1.50E+00 | 1.12E-01 | 3.15E-01 | 1.15E+00 | 1.72E-01 | 6.33E-01 | 1.34E+00 | 3.45E-02 | 9.49E-02 | 1.18E+00 |
| PPP5C | 1.94E-02 | 1.01E-01 | 1.07E+00 | 2.11E-02 | 9.13E-02 | 1.20E+00 | 3.10E-06 | 1.32E-05 | -1.26E+00 | 5.78E-08 | 2.21E-07 | -1.20E+00 | 2.25E-03 | 8.01E-03 | -1.51E+00 | 2.66E-02 | 7.50E-02 | -1.17E+00 | 7.15E-05 | 2.63E-04 | -1.15E+00 | 7.82E-02 | 2.15E-01 | 1.13E+00 |
| CD9 | 1.94E-02 | 1.01E-01 | 1.38E+00 | 8.36E-04 | 3.61E-03 | 1.64E+00 | 8.74E-02 | 3.71E-01 | 1.28E+00 | 1.83E-02 | 6.97E-02 | 1.58E+00 | 5.61E-02 | 2.00E-01 | 1.06E+00 | 5.31E-03 | 1.50E-02 | -1.12E+00 | 3.20E-04 | 1.17E-03 | 1.45E+00 | 4.97E-02 | 1.37E-01 | 1.05E+00 |
| TMEM150C | 1.95E-02 | 1.02E-01 | -1.19E+00 | 1.30E-01 | 5.64E-01 | 1.07E+00 | 2.07E-02 | 8.80E-02 | -1.10E+00 | 1.28E-01 | 4.90E-01 | 1.32E+00 | 1.28E-03 | 4.57E-03 | -2.06E+00 | 1.46E-02 | 4.12E-02 | -1.30E+00 | 2.35E-06 | 8.63E-06 | -1.51E+00 | 3.83E-04 | 1.05E-03 | -1.22E+00 |
| AMZ2 | 1.95E-02 | 1.02E-01 | 1.11E+00 | 1.82E-02 | 7.85E-02 | 1.28E+00 | 4.57E-03 | 1.94E-02 | -1.11E+00 | 3.98E-04 | 1.52E-03 | -1.06E+00 | 4.08E-04 | 1.45E-03 | -1.58E+00 | 5.98E-04 | 1.69E-03 | -1.39E+00 | 3.17E-03 | 1.16E-02 | -1.17E+00 | 2.89E-03 | 7.93E-03 | -1.15E+00 |
| CAPG | 1.95E-02 | 1.02E-01 | 1.14E+00 | 1.03E-05 | 4.45E-05 | 1.02E+00 | 4.75E-02 | 2.02E-01 | 1.34E+00 | 1.55E-02 | 5.94E-02 | 1.09E+00 | 4.85E-08 | 1.73E-07 | -1.72E+00 | 1.72E-11 | 4.86E-11 | -1.88E+00 | 3.91E-02 | 1.43E-01 | -1.26E+00 | 1.38E-01 | 3.79E-01 | -1.10E+00 |
| AC124045.1 | 1.95E-02 | 1.02E-01 | -1.10E+00 | 3.05E-02 | 1.32E-01 | -1.00E+00 | 1.45E-02 | 6.17E-02 | -1.43E+00 | 5.42E-02 | 2.07E-01 | -1.09E+00 | 1.61E-01 | 5.74E-01 | 1.00E+00 | 2.94E-01 | 8.28E-01 | 1.17E+00 | 1.00E-01 | 3.67E-01 | 1.02E+00 | 1.05E-01 | 2.87E-01 | -1.07E+00 |
| PYGB | 1.96E-02 | 1.02E-01 | -1.02E+00 | 2.90E-02 | 1.25E-01 | 1.11E+00 | 1.20E-04 | 5.09E-04 | -1.38E+00 | 2.07E-02 | 7.92E-02 | 1.12E+00 | 6.18E-03 | 2.21E-02 | -1.31E+00 | 9.94E-03 | 2.80E-02 | -1.22E+00 | 1.13E-02 | 4.14E-02 | -1.10E+00 | 2.99E-01 | 8.20E-01 | 1.33E+00 |
| FIRRE | 1.96E-02 | 1.02E-01 | -1.00E+00 | 1.07E-01 | 4.60E-01 | 1.41E+00 | 1.47E-03 | 6.26E-03 | -1.27E+00 | 1.42E-01 | 5.43E-01 | 1.34E+00 | 1.14E-03 | 4.07E-03 | -1.42E+00 | 1.84E-02 | 5.20E-02 | -1.05E+00 | 8.99E-03 | 3.30E-02 | -1.11E+00 | 2.88E-01 | 7.91E-01 | 1.29E+00 |
| ABCB9 | 1.96E-02 | 1.02E-01 | -1.25E+00 | 1.46E-01 | 6.32E-01 | 1.20E+00 | 5.71E-03 | 2.42E-02 | -1.29E+00 | 1.34E-03 | 5.13E-03 | -1.38E+00 | 6.74E-02 | 2.40E-01 | -1.14E+00 | 1.68E-01 | 4.75E-01 | 1.13E+00 | 2.08E-01 | 7.65E-01 | 1.15E+00 | 3.25E-01 | 8.92E-01 | 1.13E+00 |
| TOP2A | 1.97E-02 | 1.02E-01 | -2.24E+00 | 2.61E-02 | 1.13E-01 | -2.12E+00 | 2.25E-01 | 9.56E-01 | 1.14E+00 | 1.60E-01 | 6.10E-01 | 2.21E+00 | 2.76E-01 | 9.85E-01 | 1.20E+00 | 3.42E-01 | 9.64E-01 | 1.61E+00 | 5.10E-02 | 1.87E-01 | 1.16E+00 | 4.30E-02 | 1.18E-01 | 2.05E+00 |
| AC024257.3 | 1.97E-02 | 1.02E-01 | 1.08E+00 | 2.73E-02 | 1.18E-01 | 1.12E+00 | 2.07E-01 | 8.81E-01 | 1.15E+00 | 1.21E-01 | 4.63E-01 | 1.09E+00 | 6.19E-02 | 2.21E-01 | -1.41E+00 | 1.87E-01 | 5.28E-01 | -1.10E+00 | 1.15E-01 | 4.23E-01 | -1.14E+00 | 3.62E-01 | 9.94E-01 | 1.20E+00 |
| TBC1D10A | 1.97E-02 | 1.03E-01 | 1.79E+00 | 6.75E-03 | 2.92E-02 | 2.21E+00 | 2.23E-01 | 9.49E-01 | 1.35E+00 | 1.92E-01 | 7.34E-01 | 1.72E+00 | 3.72E-04 | 1.32E-03 | -1.31E+00 | 2.76E-02 | 7.78E-02 | 1.07E+00 | 1.53E-01 | 5.62E-01 | 1.02E+00 | 2.67E-01 | 7.33E-01 | 1.30E+00 |
| GTF2E1 | 1.98E-02 | 1.03E-01 | -1.45E+00 | 1.81E-01 | 7.82E-01 | 1.24E+00 | 1.84E-05 | 7.82E-05 | -1.69E+00 | 3.27E-03 | 1.25E-02 | -1.12E+00 | 4.39E-02 | 1.57E-01 | -1.21E+00 | 6.18E-02 | 1.74E-01 | -1.32E+00 | 5.76E-11 | 2.12E-10 | -1.81E+00 | 1.05E-06 | 2.88E-06 | -1.39E+00 |
| VAMP1 | 1.98E-02 | 1.03E-01 | -1.17E+00 | 1.15E-01 | 4.96E-01 | 1.26E+00 | 7.25E-04 | 3.08E-03 | -1.56E+00 | 1.56E-02 | 5.96E-02 | -1.11E+00 | 2.63E-01 | 9.38E-01 | 1.39E+00 | 2.93E-01 | 8.25E-01 | 1.25E+00 | 1.25E-02 | 4.57E-02 | -1.08E+00 | 5.21E-03 | 1.43E-02 | -1.28E+00 |
| PGBD4 | 1.98E-02 | 1.03E-01 | -1.11E+00 | 2.56E-02 | 1.11E-01 | -1.12E+00 | 2.47E-07 | 1.05E-06 | -2.01E+00 | 5.73E-05 | 2.19E-04 | -1.35E+00 | 1.03E-01 | 3.67E-01 | -1.05E+00 | 1.53E-01 | 4.31E-01 | -1.02E+00 | 5.71E-03 | 2.10E-02 | -1.13E+00 | 4.70E-02 | 1.29E-01 | -1.04E+00 |
| CABLES2 | 1.98E-02 | 1.03E-01 | -1.33E+00 | 9.68E-02 | 4.18E-01 | 1.12E+00 | 6.86E-02 | 2.91E-01 | -1.07E+00 | 1.22E-02 | 4.65E-02 | -1.22E+00 | 7.45E-02 | 2.66E-01 | -1.34E+00 | 1.96E-01 | 5.53E-01 | 1.02E+00 | 4.69E-03 | 1.72E-02 | -1.28E+00 | 9.06E-02 | 2.49E-01 | -1.01E+00 |
| MYH3 | 1.98E-02 | 1.03E-01 | -1.32E+00 | 9.13E-02 | 3.94E-01 | 1.03E+00 | 9.99E-02 | 4.24E-01 | -1.15E+00 | 2.50E-01 | 9.56E-01 | 1.21E+00 | 2.33E-01 | 8.32E-01 | 1.07E+00 | 2.69E-01 | 7.58E-01 | 1.11E+00 | 1.69E-01 | 6.19E-01 | -1.06E+00 | 2.05E-01 | 5.62E-01 | -1.04E+00 |
| ZNF862 | 1.98E-02 | 1.03E-01 | -1.25E+00 | 1.64E-01 | 7.11E-01 | 1.44E+00 | 2.62E-02 | 1.12E-01 | -1.22E+00 | 1.69E-01 | 6.46E-01 | 1.23E+00 | 1.30E-01 | 4.65E-01 | -1.04E+00 | 3.47E-01 | 9.77E-01 | 1.39E+00 | 1.20E-01 | 4.41E-01 | -1.03E+00 | 1.62E-01 | 4.46E-01 | -1.03E+00 |
| ZNF687 | 1.99E-02 | 1.03E-01 | -1.12E+00 | 2.43E-02 | 1.05E-01 | 1.01E+00 | 9.94E-05 | 4.22E-04 | -1.68E+00 | 6.47E-04 | 2.47E-03 | -1.39E+00 | 7.31E-05 | 2.61E-04 | -1.56E+00 | 2.68E-03 | 7.55E-03 | -1.27E+00 | 1.92E-05 | 7.05E-05 | -1.21E+00 | 2.38E-05 | 6.54E-05 | -1.35E+00 |
| FAR1 | 1.99E-02 | 1.04E-01 | 1.06E+00 | 5.24E-05 | 2.26E-04 | -1.15E+00 | 3.09E-04 | 1.31E-03 | -1.14E+00 | 7.20E-05 | 2.75E-04 | -1.07E+00 | 4.66E-02 | 1.66E-01 | 1.12E+00 | 1.18E-02 | 3.32E-02 | 1.01E+00 | 5.74E-05 | 2.11E-04 | -1.21E+00 | 1.70E-02 | 4.68E-02 | -1.03E+00 |
| KIAA0556 | 1.99E-02 | 1.04E-01 | -1.23E+00 | 1.10E-01 | 4.77E-01 | 1.15E+00 | 6.54E-02 | 2.78E-01 | -1.01E+00 | 1.01E-01 | 3.87E-01 | 1.19E+00 | 5.23E-02 | 1.86E-01 | -1.26E+00 | 2.04E-01 | 5.76E-01 | 1.22E+00 | 1.87E-01 | 6.87E-01 | 1.18E+00 | 3.26E-01 | 8.96E-01 | 1.34E+00 |
| AC024060.1 | 1.99E-02 | 1.04E-01 | -1.35E+00 | 5.52E-02 | 2.38E-01 | 1.77E+00 | 1.37E-03 | 5.82E-03 | -1.50E+00 | 1.66E-01 | 6.35E-01 | 1.38E+00 | 1.21E-01 | 4.32E-01 | 1.02E+00 | 2.98E-01 | 8.41E-01 | 1.37E+00 | 7.73E-03 | 2.84E-02 | -1.38E+00 | 3.15E-01 | 8.66E-01 | 1.36E+00 |
| LGALSL | 1.99E-02 | 1.04E-01 | -1.17E+00 | 2.25E-03 | 9.71E-03 | -1.49E+00 | 1.17E-01 | 4.97E-01 | 1.54E+00 | 2.52E-01 | 9.64E-01 | 1.16E+00 | 3.94E-02 | 1.40E-01 | 3.46E+00 | 2.98E-01 | 8.40E-01 | -1.12E+00 | 5.98E-15 | 2.20E-14 | 6.75E+00 | 2.21E-01 | 6.06E-01 | 2.28E+00 |
| DBN1 | 1.99E-02 | 1.04E-01 | 1.10E+00 | 1.16E-01 | 5.02E-01 | 1.32E+00 | 2.97E-05 | 1.26E-04 | -1.32E+00 | 1.11E-03 | 4.24E-03 | 1.01E+00 | 1.42E-13 | 5.08E-13 | -1.88E+00 | 5.16E-11 | 1.45E-10 | -1.49E+00 | 9.30E-02 | 3.41E-01 | 1.13E+00 | 2.78E-01 | 7.64E-01 | 1.39E+00 |
| GPATCH3 | 1.99E-02 | 1.04E-01 | -1.02E+00 | 5.34E-02 | 2.31E-01 | 1.20E+00 | 1.83E-02 | 7.77E-02 | -1.12E+00 | 9.78E-03 | 3.73E-02 | -1.00E+00 | 5.07E-03 | 1.81E-02 | -1.33E+00 | 1.35E-01 | 3.80E-01 | 1.14E+00 | 7.29E-05 | 2.68E-04 | -1.25E+00 | 1.35E-02 | 3.70E-02 | -1.03E+00 |
| DNAJB5 | 2.00E-02 | 1.04E-01 | 1.77E+00 | 1.19E-01 | 5.15E-01 | 1.07E+00 | 5.56E-05 | 2.36E-04 | 2.53E+00 | 2.48E-01 | 9.46E-01 | 1.19E+00 | 1.29E-01 | 4.62E-01 | -1.16E+00 | 1.18E-01 | 3.32E-01 | -1.27E+00 | 2.45E-01 | 9.00E-01 | 1.07E+00 | 3.56E-01 | 9.78E-01 | 1.07E+00 |
| PINLYP | 2.00E-02 | 1.04E-01 | -1.18E+00 | 3.35E-02 | 1.45E-01 | -1.04E+00 | 1.53E-02 | 6.48E-02 | -1.18E+00 | 6.11E-03 | 2.34E-02 | -1.11E+00 | 1.83E-01 | 6.52E-01 | 1.01E+00 | 2.78E-01 | 7.84E-01 | 1.42E+00 | 1.11E-02 | 4.09E-02 | -1.08E+00 | 3.40E-02 | 9.34E-02 | -1.06E+00 |
| ZBTB11-AS1 | 2.00E-02 | 1.04E-01 | 1.03E+00 | 2.10E-02 | 9.06E-02 | 1.11E+00 | 3.20E-02 | 1.36E-01 | -1.05E+00 | 9.76E-04 | 3.73E-03 | -1.32E+00 | 9.99E-02 | 3.56E-01 | 1.06E+00 | 2.49E-01 | 7.02E-01 | 1.23E+00 | 2.29E-03 | 8.39E-03 | -1.48E+00 | 1.11E-02 | 3.06E-02 | -1.28E+00 |
| COX16 | 2.00E-02 | 1.04E-01 | -1.00E+00 | 1.29E-02 | 5.56E-02 | 1.09E+00 | 1.03E-01 | 4.36E-01 | 1.07E+00 | 2.01E-01 | 7.69E-01 | 1.33E+00 | 8.65E-03 | 3.08E-02 | -1.29E+00 | 2.25E-02 | 6.35E-02 | -1.13E+00 | 1.00E-03 | 3.68E-03 | -1.13E+00 | 2.97E-05 | 8.17E-05 | -1.35E+00 |
| CRDCC | 2.00E-02 | 1.04E-01 | 1.01E+00 | 3.71E-02 | 1.60E-01 | 1.16E+00 | 2.07E-01 | 8.80E-01 | 1.16E+00 | 1.41E-01 | 5.37E-01 | 1.14E+00 | 2.03E-03 | 7.23E-03 | -1.61E+00 | 6.11E-02 | 1.72E-01 | -1.06E+00 | 1.35E-02 | 4.95E-02 | -1.22E+00 | 1.63E-01 | 4.47E-01 | 1.13E+00 |
| GLUD1P3 | 2.00E-02 | 1.04E-01 | 2.10E+00 | 6.33E-02 | 2.74E-01 | 2.00E+00 | 1.28E-01 | 5.44E-01 | 1.33E+00 | 1.81E-01 | 6.91E-01 | 1.09E+00 | 1.76E-01 | 6.26E-01 | 1.24E+00 | 1.69E-01 | 4.77E-01 | 1.16E+00 | 2.27E-01 | 8.32E-01 | 1.32E+00 | 1.28E-02 | 3.51E-02 | -1.55E+00 |
| RPL12P15 | 2.00E-02 | 1.04E-01 | 1.01E+00 | 4.39E-05 | 1.90E-04 | -1.53E+00 | 1.79E-01 | 7.61E-01 | 1.16E+00 | 2.49E-02 | 9.51E-02 | -1.09E+00 | 2.01E-01 | 7.16E-01 | 1.29E+00 | 2.25E-02 | 6.34E-02 | -1.29E+00 | 9.73E-02 | 3.57E-01 | 1.02E+00 | 1.16E-02 | 3.18E-02 | -1.27E+00 |
| GLCCI1 | 2.00E-02 | 1.04E-01 | 1.10E+00 | 2.03E-01 | 8.79E-01 | 1.57E+00 | 6.98E-04 | 2.96E-03 | -1.16E+00 | 1.97E-03 | 7.53E-03 | 1.05E+00 | 5.32E-03 | 1.90E-02 | -1.07E+00 | 3.35E-02 | 9.45E-02 | 1.14E+00 | 1.61E-03 | 5.92E-03 | -1.34E+00 | 7.97E-02 | 2.19E-01 | 1.02E+00 |
| KCP | 2.01E-02 | 1.05E-01 | -1.27E+00 | 2.03E-03 | 8.75E-03 | -1.78E+00 | 7.46E-03 | 3.17E-02 | -1.37E+00 | 1.05E-02 | 4.02E-02 | -1.23E+00 | 4.52E-02 | 1.61E-01 | -1.37E+00 | 1.44E-01 | 4.05E-01 | 1.02E+00 | 3.92E-03 | 1.44E-02 | -1.44E+00 | 3.10E-02 | 8.51E-02 | -1.07E+00 |
| CEP112 | 2.02E-02 | 1.05E-01 | -1.07E+00 | 2.26E-01 | 9.77E-01 | 1.52E+00 | 2.23E-02 | 9.49E-02 | -1.26E+00 | 4.38E-02 | 1.67E-01 | -1.05E+00 | 7.10E-02 | 2.53E-01 | -1.56E+00 | 1.22E-01 | 3.44E-01 | -1.30E+00 | 1.53E-01 | 5.62E-01 | -1.22E+00 | 1.80E-01 | 4.96E-01 | -1.28E+00 |
| FAR1-IT1 | 2.02E-02 | 1.05E-01 | -1.32E+00 | 3.33E-02 | 1.44E-01 | -1.08E+00 | 1.76E-02 | 7.46E-02 | -1.33E+00 | 4.17E-02 | 1.59E-01 | -1.12E+00 | 7.98E-02 | 2.84E-01 | -1.51E+00 | 2.39E-01 | 6.75E-01 | 1.18E+00 | 2.37E-01 | 8.72E-01 | 1.24E+00 | 2.72E-01 | 7.47E-01 | 1.17E+00 |
| CENPX | 2.02E-02 | 1.05E-01 | 1.20E+00 | 5.46E-03 | 2.36E-02 | 1.31E+00 | 9.21E-02 | 3.91E-01 | 1.35E+00 | 1.11E-01 | 4.24E-01 | 1.42E+00 | 5.62E-02 | 2.00E-01 | 1.21E+00 | 1.62E-01 | 4.56E-01 | 1.49E+00 | 2.52E-02 | 9.23E-02 | 1.29E+00 | 3.91E-05 | 1.08E-04 | 1.60E+00 |
| NUDT7 | 2.02E-02 | 1.05E-01 | -1.16E+00 | 3.45E-02 | 1.49E-01 | 1.01E+00 | 5.12E-03 | 2.18E-02 | -1.17E+00 | 1.52E-04 | 5.82E-04 | -1.39E+00 | 2.40E-01 | 8.57E-01 | -1.02E+00 | 3.13E-01 | 8.82E-01 | 1.35E+00 | 9.18E-02 | 3.37E-01 | -1.19E+00 | 5.70E-02 | 1.57E-01 | -1.32E+00 |
| IL6STP1 | 2.02E-02 | 1.05E-01 | -1.23E+00 | 1.98E-02 | 8.57E-02 | -1.22E+00 | 2.23E-02 | 9.47E-02 | -1.46E+00 | 2.77E-02 | 1.06E-01 | -1.35E+00 | 9.35E-02 | 3.33E-01 | 1.06E+00 | 3.36E-03 | 9.46E-03 | -2.42E+00 | 2.37E-03 | 8.70E-03 | 1.37E+00 | 3.35E-07 | 9.20E-07 | -1.78E+00 |
| USP30-AS1 | 2.03E-02 | 1.06E-01 | 1.01E+00 | 3.79E-04 | 1.64E-03 | -1.19E+00 | 1.98E-04 | 8.43E-04 | -1.27E+00 | 2.89E-05 | 1.11E-04 | -1.27E+00 | 1.66E-02 | 5.92E-02 | -1.94E+00 | 1.34E-02 | 3.77E-02 | -2.26E+00 | 4.43E-02 | 1.63E-01 | -1.66E+00 | 8.97E-02 | 2.46E-01 | -1.43E+00 |
| AC109322.1 | 2.03E-02 | 1.06E-01 | 1.04E+00 | 3.99E-05 | 1.72E-04 | -2.03E+00 | 5.50E-02 | 2.34E-01 | 1.07E+00 | 5.81E-05 | 2.22E-04 | -1.47E+00 | 9.83E-03 | 3.50E-02 | -1.36E+00 | 1.69E-02 | 4.77E-02 | -1.30E+00 | 5.37E-02 | 1.97E-01 | -1.08E+00 | 7.34E-02 | 2.02E-01 | -1.15E+00 |
| KCNQ1 | 2.03E-02 | 1.06E-01 | -1.35E+00 | 1.70E-02 | 7.36E-02 | -1.28E+00 | 1.90E-01 | 8.09E-01 | -1.03E+00 | 2.21E-01 | 8.44E-01 | 1.13E+00 | 2.51E-01 | 8.95E-01 | 1.19E+00 | 2.52E-01 | 7.09E-01 | 1.44E+00 | 2.28E-02 | 8.38E-02 | 1.96E+00 | 2.02E-01 | 5.54E-01 | 1.53E+00 |
| AC004656.1 | 2.03E-02 | 1.06E-01 | -1.17E+00 | 7.13E-02 | 3.08E-01 | 1.14E+00 | 3.97E-03 | 1.69E-02 | -1.52E+00 | 1.29E-01 | 4.93E-01 | 1.08E+00 | 2.80E-01 | 9.99E-01 | 1.23E+00 | 5.40E-02 | 1.52E-01 | 1.94E+00 | 2.23E-01 | 8.18E-01 | 1.31E+00 | 4.44E-02 | 1.22E-01 | 1.73E+00 |
| MAP2K1 | 2.03E-02 | 1.06E-01 | -1.04E+00 | 2.07E-01 | 8.95E-01 | 1.47E+00 | 2.34E-03 | 9.94E-03 | -1.11E+00 | 1.73E-04 | 6.60E-04 | -1.03E+00 | 2.84E-02 | 1.01E-01 | -1.05E+00 | 1.77E-01 | 4.99E-01 | 1.23E+00 | 2.24E-03 | 8.23E-03 | -1.13E+00 | 3.93E-02 | 1.08E-01 | 1.06E+00 |
| ABCB10 | 2.03E-02 | 1.06E-01 | -1.14E+00 | 5.69E-02 | 2.46E-01 | 1.09E+00 | 2.22E-02 | 9.42E-02 | -1.08E+00 | 1.34E-02 | 5.13E-02 | 1.03E+00 | 5.49E-02 | 1.96E-01 | 1.00E+00 | 2.39E-01 | 6.74E-01 | 1.57E+00 | 7.65E-06 | 2.81E-05 | -1.20E+00 | 5.66E-02 | 1.56E-01 | 1.07E+00 |
| RLF | 2.03E-02 | 1.06E-01 | 1.08E+00 | 4.22E-07 | 1.82E-06 | -1.27E+00 | 7.96E-02 | 3.38E-01 | 1.12E+00 | 1.01E-05 | 3.87E-05 | -1.19E+00 | 1.58E-01 | 5.63E-01 | 1.27E+00 | 9.35E-07 | 2.64E-06 | -1.58E+00 | 5.56E-05 | 2.04E-04 | 1.01E+00 | 2.56E-14 | 7.02E-14 | -1.64E+00 |
| AC064836.3 | 2.03E-02 | 1.06E-01 | 1.88E+00 | 1.98E-01 | 8.55E-01 | 1.65E+00 | 1.42E-01 | 6.04E-01 | 1.29E+00 | 1.05E-01 | 4.00E-01 | 1.11E+00 | 2.09E-01 | 7.46E-01 | 1.06E+00 | 1.71E-01 | 4.82E-01 | -1.14E+00 | 2.26E-01 | 8.30E-01 | 1.25E+00 | 3.28E-01 | 9.01E-01 | 1.05E+00 |
| KCNC1 | 2.03E-02 | 1.06E-01 | -1.14E+00 | 1.99E-01 | 8.62E-01 | 1.58E+00 | 1.85E-01 | 7.84E-01 | 1.23E+00 | 6.04E-03 | 2.31E-02 | 2.42E+00 | 2.35E-01 | 8.38E-01 | 1.40E+00 | 5.58E-02 | 1.57E-01 | 2.38E+00 | 7.79E-02 | 2.86E-01 | -1.27E+00 | 2.77E-01 | 7.60E-01 | 1.19E+00 |
| ZNF787 | 2.03E-02 | 1.06E-01 | 1.61E+00 | 2.13E-01 | 9.19E-01 | 1.51E+00 | 6.98E-03 | 2.96E-02 | 1.69E+00 | 2.50E-01 | 9.53E-01 | 1.45E+00 | 5.47E-03 | 1.95E-02 | 2.25E+00 | 2.24E-01 | 6.32E-01 | 1.65E+00 | 5.22E-03 | 1.91E-02 | 1.71E+00 | 3.05E-01 | 8.38E-01 | 1.48E+00 |
| LRRC45 | 2.04E-02 | 1.06E-01 | -1.31E+00 | 5.55E-02 | 2.40E-01 | -1.06E+00 | 2.60E-05 | 1.11E-04 | -1.41E+00 | 6.67E-02 | 2.55E-01 | 1.22E+00 | 1.01E-02 | 3.59E-02 | -1.63E+00 | 1.29E-01 | 3.65E-01 | 1.03E+00 | 4.65E-02 | 1.71E-01 | 1.03E+00 | 1.20E-01 | 3.29E-01 | 1.09E+00 |
| SLC7A8 | 2.04E-02 | 1.06E-01 | -1.19E+00 | 1.43E-01 | 6.20E-01 | 1.14E+00 | 4.40E-02 | 1.87E-01 | -1.02E+00 | 7.45E-02 | 2.84E-01 | 1.23E+00 | 2.31E-01 | 8.24E-01 | 1.07E+00 | 2.12E-01 | 5.97E-01 | -1.29E+00 | 2.68E-01 | 9.84E-01 | 1.14E+00 | 2.71E-01 | 7.45E-01 | -1.18E+00 |
| DHFRP1 | 2.04E-02 | 1.06E-01 | -1.78E+00 | 8.58E-02 | 3.71E-01 | -1.12E+00 | 1.50E-01 | 6.37E-01 | 1.31E+00 | 2.49E-01 | 9.53E-01 | 1.42E+00 | 2.30E-01 | 8.19E-01 | 1.12E+00 | 2.06E-01 | 5.80E-01 | 1.00E+00 | 1.71E-01 | 6.27E-01 | 1.08E+00 | 2.96E-01 | 8.13E-01 | 1.32E+00 |
| UBA52P5 | 2.04E-02 | 1.07E-01 | 1.16E+00 | 2.03E-01 | 8.76E-01 | 1.54E+00 | 1.09E-01 | 4.65E-01 | 1.16E+00 | 3.73E-02 | 1.43E-01 | 1.71E+00 | 9.02E-02 | 3.21E-01 | 1.16E+00 | 1.96E-02 | 5.53E-02 | 1.82E+00 | 1.98E-01 | 7.25E-01 | 1.38E+00 | 2.84E-03 | 7.81E-03 | 1.87E+00 |
| VARS2 | 2.05E-02 | 1.07E-01 | -1.04E+00 | 1.51E-02 | 6.52E-02 | 1.03E+00 | 3.18E-05 | 1.35E-04 | -1.40E+00 | 2.29E-06 | 8.73E-06 | -1.42E+00 | 1.84E-01 | 6.55E-01 | 1.49E+00 | 1.47E-01 | 4.14E-01 | 1.78E+00 | 2.68E-04 | 9.84E-04 | -1.19E+00 | 2.38E-02 | 6.54E-02 | 1.04E+00 |
| LPAR6 | 2.05E-02 | 1.07E-01 | 1.95E+00 | 2.18E-01 | 9.44E-01 | 1.73E+00 | 1.58E-01 | 6.73E-01 | 1.41E+00 | 7.99E-02 | 3.05E-01 | 1.28E+00 | 1.73E-01 | 6.15E-01 | 1.84E+00 | 1.12E-01 | 3.15E-01 | 2.14E+00 | 3.06E-02 | 1.12E-01 | -1.10E+00 | 2.03E-02 | 5.56E-02 | -1.19E+00 |
| RBL1 | 2.05E-02 | 1.07E-01 | -1.10E+00 | 1.06E-01 | 4.60E-01 | 1.34E+00 | 1.30E-02 | 5.52E-02 | -1.09E+00 | 1.40E-01 | 5.33E-01 | 1.22E+00 | 4.58E-02 | 1.63E-01 | -1.27E+00 | 8.03E-02 | 2.26E-01 | -1.14E+00 | 5.63E-08 | 2.07E-07 | -1.48E+00 | 1.04E-01 | 2.86E-01 | 1.11E+00 |
| RPLP2 | 2.05E-02 | 1.07E-01 | -1.04E+00 | 4.05E-46 | 1.75E-45 | -1.39E+00 | 1.61E-02 | 6.84E-02 | -1.05E+00 | 3.34E-27 | 1.28E-26 | -1.37E+00 | 7.10E-04 | 2.53E-03 | -1.10E+00 | 7.03E-20 | 1.98E-19 | -1.34E+00 | 1.87E-12 | 6.85E-12 | -1.16E+00 | 3.24E-37 | 8.91E-37 | -1.43E+00 |
| CD320 | 2.05E-02 | 1.07E-01 | -1.13E+00 | 3.09E-03 | 1.33E-02 | -1.27E+00 | 3.15E-02 | 1.34E-01 | 1.05E+00 | 2.05E-03 | 7.82E-03 | 1.07E+00 | 9.72E-02 | 3.46E-01 | 1.03E+00 | 2.32E-01 | 6.55E-01 | 1.70E+00 | 1.24E-02 | 4.57E-02 | -1.03E+00 | 5.21E-02 | 1.43E-01 | 1.53E+00 |
| SLC12A9 | 2.05E-02 | 1.07E-01 | 1.06E+00 | 1.46E-01 | 6.29E-01 | 1.42E+00 | 4.98E-02 | 2.12E-01 | -1.05E+00 | 1.18E-01 | 4.49E-01 | 1.15E+00 | 1.07E-01 | 3.83E-01 | -1.16E+00 | 2.87E-01 | 8.08E-01 | 1.54E+00 | 5.05E-02 | 1.85E-01 | -1.10E+00 | 3.11E-01 | 8.55E-01 | 1.12E+00 |
| C19orf47 | 2.06E-02 | 1.07E-01 | -1.15E+00 | 1.03E-02 | 4.46E-02 | -1.16E+00 | 7.04E-05 | 2.99E-04 | -1.44E+00 | 1.06E-04 | 4.03E-04 | -1.36E+00 | 1.10E-01 | 3.93E-01 | -1.02E+00 | 2.97E-01 | 8.38E-01 | 1.31E+00 | 9.67E-03 | 3.55E-02 | -1.06E+00 | 1.09E-02 | 3.00E-02 | -1.17E+00 |
| ABR | 2.06E-02 | 1.07E-01 | 1.03E+00 | 1.86E-01 | 8.04E-01 | 1.60E+00 | 1.43E-01 | 6.08E-01 | 1.19E+00 | 1.10E-01 | 4.18E-01 | 1.45E+00 | 1.67E-02 | 5.96E-02 | -1.32E+00 | 1.85E-02 | 5.21E-02 | -1.38E+00 | 7.72E-02 | 2.84E-01 | -1.15E+00 | 2.03E-01 | 5.57E-01 | 1.07E+00 |
| B3GALT4 | 2.06E-02 | 1.07E-01 | -1.21E+00 | 2.09E-01 | 9.02E-01 | 1.29E+00 | 8.21E-02 | 3.49E-01 | -1.12E+00 | 2.18E-01 | 8.31E-01 | 1.30E+00 | 1.73E-06 | 6.15E-06 | -1.80E+00 | 2.26E-04 | 6.38E-04 | -1.38E+00 | 5.74E-02 | 2.11E-01 | 1.07E+00 | 5.71E-02 | 1.57E-01 | -1.15E+00 |
| FUT4 | 2.06E-02 | 1.07E-01 | 1.07E+00 | 1.47E-02 | 6.33E-02 | 1.12E+00 | 3.91E-04 | 1.66E-03 | -1.18E+00 | 1.89E-04 | 7.21E-04 | -1.12E+00 | 9.76E-03 | 3.48E-02 | -1.45E+00 | 8.04E-02 | 2.27E-01 | -1.04E+00 | 4.31E-04 | 1.58E-03 | -1.51E+00 | 2.08E-03 | 5.71E-03 | -1.37E+00 |
| ENKD1 | 2.07E-02 | 1.08E-01 | 1.00E+00 | 1.04E-02 | 4.50E-02 | 1.03E+00 | 2.57E-02 | 1.09E-01 | -1.05E+00 | 1.49E-03 | 5.68E-03 | -1.06E+00 | 2.09E-01 | 7.46E-01 | 1.13E+00 | 2.20E-01 | 6.20E-01 | 1.15E+00 | 1.70E-01 | 6.25E-01 | 1.22E+00 | 3.44E-01 | 9.45E-01 | 1.36E+00 |
| RPL17P50 | 2.07E-02 | 1.08E-01 | 1.08E+00 | 2.10E-04 | 9.08E-04 | -1.10E+00 | 4.56E-02 | 1.94E-01 | 1.06E+00 | 1.05E-04 | 4.02E-04 | -1.17E+00 | 2.65E-01 | 9.46E-01 | 1.40E+00 | 9.78E-02 | 2.76E-01 | 1.14E+00 | 8.01E-02 | 2.94E-01 | 1.11E+00 | 1.70E-03 | 4.67E-03 | -1.22E+00 |
| TADA1 | 2.07E-02 | 1.08E-01 | 1.01E+00 | 7.48E-02 | 3.23E-01 | 1.27E+00 | 3.17E-04 | 1.35E-03 | -1.46E+00 | 6.72E-03 | 2.57E-02 | -1.06E+00 | 9.01E-02 | 3.21E-01 | -1.14E+00 | 3.31E-01 | 9.33E-01 | 1.31E+00 | 4.99E-08 | 1.83E-07 | -1.58E+00 | 1.58E-03 | 4.35E-03 | -1.12E+00 |
| FADS1 | 2.07E-02 | 1.08E-01 | 1.03E+00 | 2.11E-01 | 9.10E-01 | 1.40E+00 | 3.12E-02 | 1.32E-01 | 1.06E+00 | 2.36E-02 | 9.00E-02 | 1.68E+00 | 9.87E-06 | 3.52E-05 | -1.08E+00 | 3.20E-01 | 9.03E-01 | 1.39E+00 | 2.57E-02 | 9.43E-02 | 1.09E+00 | 2.26E-03 | 6.21E-03 | 1.63E+00 |
| AL080317.2 | 2.07E-02 | 1.08E-01 | -1.30E+00 | 1.55E-01 | 6.68E-01 | 1.67E+00 | 1.24E-01 | 5.27E-01 | -1.17E+00 | 2.12E-01 | 8.12E-01 | 1.31E+00 | 1.83E-01 | 6.51E-01 | 1.20E+00 | 1.42E-01 | 4.01E-01 | 1.83E+00 | 4.40E-02 | 1.61E-01 | -1.39E+00 | 2.98E-01 | 8.19E-01 | 1.02E+00 |
| GDF11 | 2.07E-02 | 1.08E-01 | 1.02E+00 | 1.17E-03 | 5.08E-03 | -1.12E+00 | 1.07E-02 | 4.55E-02 | -1.13E+00 | 1.05E-03 | 4.00E-03 | -1.09E+00 | 5.73E-07 | 2.04E-06 | -1.03E+00 | 3.62E-05 | 1.02E-04 | -1.00E+00 | 2.31E-01 | 8.48E-01 | 1.32E+00 | 8.20E-03 | 2.25E-02 | 1.09E+00 |
| RGPD1 | 2.07E-02 | 1.08E-01 | 1.10E+00 | 7.64E-05 | 3.30E-04 | -1.06E+00 | 1.79E-02 | 7.60E-02 | -1.04E+00 | 2.07E-07 | 7.92E-07 | -1.64E+00 | 2.18E-01 | 7.79E-01 | 1.46E+00 | 2.21E-03 | 6.23E-03 | -1.39E+00 | 7.17E-02 | 2.63E-01 | 1.11E+00 | 2.10E-05 | 5.77E-05 | -1.79E+00 |
| TMEM63B | 2.08E-02 | 1.08E-01 | -1.32E+00 | 9.18E-02 | 3.97E-01 | 1.15E+00 | 5.87E-04 | 2.49E-03 | -1.67E+00 | 1.89E-04 | 7.20E-04 | -1.76E+00 | 8.14E-05 | 2.90E-04 | -2.19E+00 | 2.47E-02 | 6.95E-02 | -1.13E+00 | 1.49E-08 | 5.47E-08 | -1.57E+00 | 3.22E-06 | 8.85E-06 | -1.42E+00 |
| NPC1 | 2.08E-02 | 1.08E-01 | -1.12E+00 | 8.86E-03 | 3.83E-02 | -1.11E+00 | 7.09E-02 | 3.01E-01 | -1.10E+00 | 2.81E-02 | 1.07E-01 | -1.14E+00 | 6.55E-02 | 2.33E-01 | 1.06E+00 | 2.62E-02 | 7.38E-02 | -1.22E+00 | 4.48E-03 | 1.65E-02 | -1.22E+00 | 1.64E-02 | 4.51E-02 | -1.14E+00 |
| FCHSD1 | 2.08E-02 | 1.09E-01 | -1.09E+00 | 3.62E-02 | 1.56E-01 | 1.04E+00 | 9.84E-04 | 4.18E-03 | -1.61E+00 | 3.15E-03 | 1.20E-02 | -1.29E+00 | 6.83E-02 | 2.44E-01 | -1.03E+00 | 4.59E-02 | 1.29E-01 | -1.13E+00 | 2.97E-02 | 1.09E-01 | -1.24E+00 | 2.22E-01 | 6.11E-01 | 1.11E+00 |
| SCAF8 | 2.08E-02 | 1.09E-01 | 1.09E+00 | 2.74E-04 | 1.18E-03 | -1.03E+00 | 1.00E-02 | 4.27E-02 | -1.03E+00 | 3.31E-04 | 1.26E-03 | -1.05E+00 | 3.14E-02 | 1.12E-01 | 1.14E+00 | 1.87E-02 | 5.28E-02 | 1.13E+00 | 5.04E-03 | 1.85E-02 | 1.02E+00 | 1.31E-04 | 3.59E-04 | -1.14E+00 |
| RPL7AP8 | 2.08E-02 | 1.09E-01 | -1.07E+00 | 5.77E-03 | 2.49E-02 | -1.20E+00 | 1.96E-01 | 8.33E-01 | 1.10E+00 | 5.94E-03 | 2.27E-02 | -1.26E+00 | 2.74E-01 | 9.77E-01 | 1.41E+00 | 1.50E-01 | 4.22E-01 | -1.09E+00 | 2.40E-02 | 8.81E-02 | 1.78E+00 | 3.42E-01 | 9.40E-01 | 1.34E+00 |
| SLC17A9 | 2.08E-02 | 1.09E-01 | 1.19E+00 | 3.96E-02 | 1.71E-01 | 1.39E+00 | 2.13E-01 | 9.03E-01 | 1.01E+00 | 9.28E-02 | 3.55E-01 | 1.28E+00 | 6.75E-03 | 2.41E-02 | -1.67E+00 | 1.09E-01 | 3.08E-01 | 1.04E+00 | 1.99E-01 | 7.32E-01 | 1.50E+00 | 1.35E-02 | 3.70E-02 | 2.49E+00 |
| ZNF525 | 2.08E-02 | 1.09E-01 | -1.06E+00 | 5.88E-03 | 2.54E-02 | -1.08E+00 | 1.82E-02 | 7.72E-02 | -1.15E+00 | 7.60E-02 | 2.90E-01 | 1.11E+00 | 2.58E-01 | 9.19E-01 | 1.39E+00 | 3.33E-01 | 9.38E-01 | 1.34E+00 | 2.05E-01 | 7.51E-01 | 1.27E+00 | 4.80E-02 | 1.32E-01 | 1.02E+00 |
| NDUFB10 | 2.08E-02 | 1.09E-01 | 1.09E+00 | 2.70E-04 | 1.17E-03 | 1.08E+00 | 1.07E-01 | 4.56E-01 | 1.11E+00 | 2.15E-01 | 8.19E-01 | 1.13E+00 | 3.69E-03 | 1.32E-02 | -1.00E+00 | 8.05E-02 | 2.27E-01 | 1.09E+00 | 2.09E-01 | 7.67E-01 | 1.16E+00 | 5.84E-02 | 1.61E-01 | 1.21E+00 |
| ZNF114 | 2.08E-02 | 1.09E-01 | -1.30E+00 | 6.95E-02 | 3.00E-01 | -1.02E+00 | 2.15E-01 | 9.14E-01 | 1.11E+00 | 1.35E-01 | 5.16E-01 | 1.08E+00 | 3.81E-03 | 1.36E-02 | -1.57E+00 | 2.79E-02 | 7.87E-02 | -1.15E+00 | 3.33E-02 | 1.22E-01 | -1.35E+00 | 1.57E-01 | 4.31E-01 | -1.02E+00 |
| AC004980.1 | 2.08E-02 | 1.09E-01 | -1.15E+00 | 9.43E-02 | 4.07E-01 | 1.15E+00 | 4.17E-02 | 1.77E-01 | -1.02E+00 | 5.29E-02 | 2.02E-01 | 1.17E+00 | 1.54E-02 | 5.49E-02 | -1.03E+00 | 9.97E-03 | 2.81E-02 | -1.15E+00 | 2.12E-02 | 7.79E-02 | -1.12E+00 | 1.19E-01 | 3.27E-01 | 1.04E+00 |
| SENCR | 2.09E-02 | 1.09E-01 | 1.06E+00 | 1.26E-01 | 5.45E-01 | 1.47E+00 | 1.13E-01 | 4.81E-01 | -1.02E+00 | 1.20E-01 | 4.57E-01 | 1.13E+00 | 1.17E-01 | 4.18E-01 | -1.14E+00 | 2.71E-01 | 7.63E-01 | 1.07E+00 | 2.33E-01 | 8.56E-01 | 1.91E+00 | 3.25E-01 | 8.92E-01 | 1.25E+00 |
| GALNT3 | 2.09E-02 | 1.09E-01 | -1.35E+00 | 1.66E-03 | 7.16E-03 | -1.79E+00 | 4.13E-13 | 1.76E-12 | -1.66E+00 | 5.09E-26 | 1.94E-25 | -2.37E+00 | 2.71E-01 | 9.67E-01 | 1.34E+00 | 2.35E-01 | 6.62E-01 | -1.25E+00 | 2.25E-01 | 8.26E-01 | 1.22E+00 | 3.24E-01 | 8.89E-01 | 1.15E+00 |
| ANKS3 | 2.09E-02 | 1.09E-01 | 1.05E+00 | 5.89E-02 | 2.54E-01 | 1.27E+00 | 1.93E-02 | 8.20E-02 | -1.12E+00 | 1.34E-01 | 5.13E-01 | 1.25E+00 | 5.37E-02 | 1.92E-01 | 1.08E+00 | 3.15E-01 | 8.89E-01 | 1.41E+00 | 4.41E-02 | 1.62E-01 | 1.02E+00 | 2.67E-01 | 7.33E-01 | 1.26E+00 |
| SLC25A14 | 2.09E-02 | 1.09E-01 | -1.03E+00 | 3.18E-02 | 1.37E-01 | 1.09E+00 | 4.25E-02 | 1.80E-01 | 1.02E+00 | 5.18E-02 | 1.98E-01 | 1.23E+00 | 1.11E-01 | 3.96E-01 | -1.00E+00 | 7.81E-02 | 2.20E-01 | -1.15E+00 | 1.94E-02 | 7.11E-02 | -1.20E+00 | 3.18E-01 | 8.72E-01 | 1.19E+00 |
| AC009245.1 | 2.09E-02 | 1.09E-01 | 1.80E+00 | 4.79E-03 | 2.07E-02 | 2.30E+00 | 9.00E-02 | 3.83E-01 | 1.48E+00 | 3.79E-03 | 1.45E-02 | 2.04E+00 | 1.55E-01 | 5.52E-01 | 1.31E+00 | 2.57E-02 | 7.25E-02 | 1.99E+00 | 1.82E-01 | 6.70E-01 | 1.34E+00 | 8.70E-04 | 2.39E-03 | 2.09E+00 |
| NEMP2 | 2.09E-02 | 1.09E-01 | 1.03E+00 | 1.45E-02 | 6.28E-02 | 1.16E+00 | 4.32E-05 | 1.84E-04 | -1.24E+00 | 9.11E-04 | 3.48E-03 | -1.02E+00 | 7.57E-02 | 2.70E-01 | 1.11E+00 | 5.86E-02 | 1.65E-01 | 1.05E+00 | 4.07E-03 | 1.49E-02 | -1.20E+00 | 1.41E-02 | 3.87E-02 | -1.08E+00 |
| BRWD1-AS2 | 2.10E-02 | 1.09E-01 | -1.24E+00 | 1.53E-01 | 6.61E-01 | 1.15E+00 | 1.53E-01 | 6.48E-01 | 1.03E+00 | 1.46E-01 | 5.58E-01 | 1.15E+00 | 2.51E-01 | 8.95E-01 | 1.21E+00 | 3.44E-01 | 9.70E-01 | 1.24E+00 | 2.62E-02 | 9.63E-02 | -1.24E+00 | 1.75E-01 | 4.80E-01 | 1.01E+00 |
| SGSM2 | 2.10E-02 | 1.10E-01 | 1.12E+00 | 8.81E-02 | 3.80E-01 | 1.38E+00 | 3.42E-03 | 1.45E-02 | -1.12E+00 | 1.05E-02 | 4.02E-02 | 1.04E+00 | 2.48E-01 | 8.83E-01 | 1.36E+00 | 2.74E-01 | 7.73E-01 | 1.28E+00 | 2.29E-01 | 8.42E-01 | 1.27E+00 | 2.49E-01 | 6.83E-01 | 1.45E+00 |
| RTN3P1 | 2.11E-02 | 1.10E-01 | 2.23E+00 | 9.74E-03 | 4.21E-02 | 2.83E+00 | 1.55E-01 | 6.58E-01 | 1.43E+00 | 1.09E-01 | 4.14E-01 | 1.69E+00 | 1.02E-01 | 3.64E-01 | 1.82E+00 | 1.38E-01 | 3.89E-01 | 1.91E+00 | 4.65E-02 | 1.71E-01 | 1.68E+00 | 2.91E-03 | 8.00E-03 | 2.39E+00 |
| NPM1P9 | 2.11E-02 | 1.10E-01 | 1.94E+00 | 6.06E-02 | 2.62E-01 | 1.92E+00 | 9.09E-02 | 3.86E-01 | 1.53E+00 | 2.11E-01 | 8.05E-01 | 1.47E+00 | 1.10E-01 | 3.93E-01 | 2.03E+00 | 7.65E-02 | 2.16E-01 | 2.30E+00 | 5.74E-02 | 2.11E-01 | 1.51E+00 | 8.72E-02 | 2.40E-01 | 1.61E+00 |
| PRADC1 | 2.11E-02 | 1.10E-01 | 1.00E+00 | 1.60E-02 | 6.89E-02 | 1.05E+00 | 1.58E-01 | 6.71E-01 | 1.23E+00 | 1.48E-01 | 5.66E-01 | 1.50E+00 | 3.15E-02 | 1.12E-01 | -1.01E+00 | 8.53E-02 | 2.40E-01 | 1.07E+00 | 2.41E-02 | 8.85E-02 | 1.03E+00 | 5.42E-02 | 1.49E-01 | 1.50E+00 |
| FSTL3 | 2.11E-02 | 1.10E-01 | -1.07E+00 | 6.84E-02 | 2.95E-01 | 1.09E+00 | 2.97E-02 | 1.26E-01 | -1.19E+00 | 1.68E-01 | 6.42E-01 | 1.20E+00 | 1.13E-01 | 4.03E-01 | -1.01E+00 | 1.67E-01 | 4.70E-01 | 1.05E+00 | 2.13E-01 | 7.83E-01 | 1.11E+00 | 3.61E-01 | 9.91E-01 | 1.28E+00 |
| MDFIC | 2.11E-02 | 1.10E-01 | 1.00E+00 | 2.14E-04 | 9.27E-04 | -1.11E+00 | 4.68E-02 | 1.99E-01 | 1.01E+00 | 5.81E-09 | 2.22E-08 | -1.52E+00 | 2.36E-01 | 8.41E-01 | 1.39E+00 | 3.37E-01 | 9.49E-01 | 1.25E+00 | 1.17E-01 | 4.30E-01 | -1.62E+00 | 1.80E-01 | 4.94E-01 | -1.61E+00 |
| RPL15P13 | 2.11E-02 | 1.10E-01 | -1.01E+00 | 2.78E-02 | 1.20E-01 | 1.03E+00 | 5.39E-02 | 2.29E-01 | 1.00E+00 | 1.18E-03 | 4.50E-03 | -1.23E+00 | 2.16E-01 | 7.69E-01 | 1.21E+00 | 1.44E-01 | 4.07E-01 | 1.07E+00 | 2.57E-01 | 9.42E-01 | 1.23E+00 | 3.00E-02 | 8.24E-02 | -1.02E+00 |
| GOLGA2P5 | 2.12E-02 | 1.10E-01 | -1.20E+00 | 1.42E-02 | 6.13E-02 | -1.14E+00 | 8.10E-02 | 3.44E-01 | 1.28E+00 | 1.45E-01 | 5.53E-01 | 1.07E+00 | 8.73E-02 | 3.11E-01 | -1.45E+00 | 1.54E-01 | 4.34E-01 | -1.23E+00 | 1.07E-01 | 3.91E-01 | -1.10E+00 | 3.40E-01 | 9.33E-01 | 1.29E+00 |
| AC079331.2 | 2.12E-02 | 1.10E-01 | -1.24E+00 | 1.12E-01 | 4.86E-01 | 1.06E+00 | 8.49E-02 | 3.61E-01 | -1.08E+00 | 1.78E-01 | 6.79E-01 | 1.23E+00 | 5.21E-04 | 1.86E-03 | -2.62E+00 | 3.54E-03 | 9.98E-03 | -1.98E+00 | 1.90E-01 | 6.97E-01 | 1.01E+00 | 2.33E-01 | 6.40E-01 | 1.00E+00 |
| YES1 | 2.12E-02 | 1.10E-01 | 1.12E+00 | 8.73E-06 | 3.77E-05 | -1.12E+00 | 1.40E-02 | 5.95E-02 | -1.02E+00 | 4.82E-05 | 1.84E-04 | -1.12E+00 | 1.86E-03 | 6.63E-03 | -1.32E+00 | 1.07E-05 | 3.02E-05 | -1.98E+00 | 1.00E-04 | 3.67E-04 | -1.21E+00 | 4.55E-09 | 1.25E-08 | -2.27E+00 |
| GSKIP | 2.13E-02 | 1.11E-01 | 1.15E+00 | 6.62E-03 | 2.86E-02 | 1.09E+00 | 2.15E-03 | 9.12E-03 | -1.13E+00 | 1.46E-03 | 5.59E-03 | -1.12E+00 | 3.58E-06 | 1.27E-05 | -1.38E+00 | 1.97E-05 | 5.55E-05 | -1.35E+00 | 1.17E-17 | 4.29E-17 | -1.90E+00 | 9.51E-12 | 2.61E-11 | -1.52E+00 |
| AC097376.2 | 2.14E-02 | 1.11E-01 | 1.10E+00 | 1.55E-02 | 6.70E-02 | 1.20E+00 | 3.08E-02 | 1.31E-01 | 1.07E+00 | 2.60E-03 | 9.95E-03 | 1.05E+00 | 1.19E-01 | 4.26E-01 | 1.29E+00 | 1.11E-01 | 3.14E-01 | 1.20E+00 | 3.79E-02 | 1.39E-01 | -1.04E+00 | 9.18E-02 | 2.52E-01 | 1.08E+00 |
| STX2 | 2.14E-02 | 1.11E-01 | -1.04E+00 | 9.76E-02 | 4.22E-01 | 1.22E+00 | 6.73E-03 | 2.86E-02 | -1.12E+00 | 1.87E-03 | 7.14E-03 | -1.10E+00 | 2.84E-02 | 1.01E-01 | -1.19E+00 | 3.27E-01 | 9.22E-01 | 1.32E+00 | 2.67E-01 | 9.82E-01 | 1.31E+00 | 2.43E-01 | 6.67E-01 | 1.16E+00 |
| CDC42P5 | 2.14E-02 | 1.11E-01 | -1.16E+00 | 5.29E-02 | 2.29E-01 | 1.07E+00 | 1.56E-01 | 6.64E-01 | 1.40E+00 | 4.74E-02 | 1.81E-01 | 1.87E+00 | 7.28E-02 | 2.60E-01 | 1.95E+00 | 2.60E-01 | 7.32E-01 | 1.50E+00 | 2.22E-01 | 8.15E-01 | 1.29E+00 | 6.93E-02 | 1.90E-01 | 1.70E+00 |
| Z97832.2 | 2.15E-02 | 1.12E-01 | -1.34E+00 | 1.52E-01 | 6.56E-01 | 1.13E+00 | 4.29E-04 | 1.82E-03 | -1.68E+00 | 5.67E-02 | 2.17E-01 | 1.03E+00 | 1.51E-01 | 5.40E-01 | 1.67E+00 | 6.51E-02 | 1.84E-01 | 2.29E+00 | 1.12E-01 | 4.12E-01 | -1.35E+00 | 2.86E-01 | 7.87E-01 | 1.04E+00 |
| AL589843.2 | 2.15E-02 | 1.12E-01 | -1.19E+00 | 1.98E-01 | 8.55E-01 | 1.30E+00 | 7.81E-02 | 3.32E-01 | -1.18E+00 | 2.60E-01 | 9.94E-01 | 1.22E+00 | 1.20E-02 | 4.30E-02 | -1.52E+00 | 2.13E-02 | 5.99E-02 | -1.45E+00 | 1.50E-01 | 5.49E-01 | 1.28E+00 | 3.23E-01 | 8.86E-01 | 1.15E+00 |
| RPL31P63 | 2.15E-02 | 1.12E-01 | 2.12E+00 | 8.94E-02 | 3.86E-01 | 1.99E+00 | 4.62E-02 | 1.96E-01 | -1.11E+00 | 4.30E-03 | 1.64E-02 | -1.23E+00 | 1.02E-01 | 3.64E-01 | -1.04E+00 | 2.04E-01 | 5.75E-01 | 1.21E+00 | 2.72E-01 | 9.97E-01 | 1.38E+00 | 3.17E-01 | 8.71E-01 | 1.27E+00 |
| MVD | 2.15E-02 | 1.12E-01 | 1.64E+00 | 3.91E-02 | 1.69E-01 | 1.78E+00 | 3.02E-02 | 1.28E-01 | 1.55E+00 | 1.64E-02 | 6.25E-02 | 1.90E+00 | 1.75E-01 | 6.25E-01 | 1.60E+00 | 7.13E-02 | 2.01E-01 | 1.76E+00 | 4.12E-02 | 1.51E-01 | 1.63E+00 | 4.45E-03 | 1.22E-02 | 2.00E+00 |
| AC011978.2 | 2.15E-02 | 1.12E-01 | 1.88E+00 | 1.51E-01 | 6.51E-01 | 1.10E+00 | 1.39E-02 | 5.91E-02 | 1.74E+00 | 1.57E-01 | 5.98E-01 | 1.28E+00 | 1.36E-01 | 4.86E-01 | 1.36E+00 | 2.12E-01 | 5.99E-01 | -1.12E+00 | 2.28E-01 | 8.37E-01 | -1.06E+00 | 2.71E-01 | 7.45E-01 | -1.01E+00 |
| FAM173B | 2.15E-02 | 1.12E-01 | -1.04E+00 | 5.49E-02 | 2.37E-01 | 1.16E+00 | 9.12E-07 | 3.87E-06 | -1.62E+00 | 7.67E-06 | 2.93E-05 | -1.29E+00 | 3.29E-05 | 1.17E-04 | -1.67E+00 | 4.04E-03 | 1.14E-02 | -1.18E+00 | 9.34E-05 | 3.43E-04 | -1.19E+00 | 1.08E-02 | 2.97E-02 | -1.03E+00 |
| FAM120B | 2.16E-02 | 1.12E-01 | -1.05E+00 | 6.49E-02 | 2.80E-01 | 1.23E+00 | 1.10E-03 | 4.67E-03 | -1.48E+00 | 3.69E-02 | 1.41E-01 | 1.01E+00 | 1.37E-03 | 4.89E-03 | -1.72E+00 | 1.89E-01 | 5.33E-01 | 1.25E+00 | 3.12E-04 | 1.15E-03 | -1.31E+00 | 9.01E-02 | 2.47E-01 | 1.08E+00 |
| CSNK1E | 2.16E-02 | 1.12E-01 | 1.15E+00 | 2.34E-03 | 1.01E-02 | 1.17E+00 | 1.06E-01 | 4.49E-01 | 1.22E+00 | 1.17E-01 | 4.46E-01 | 1.35E+00 | 1.39E-03 | 4.97E-03 | -1.03E+00 | 8.76E-03 | 2.47E-02 | 1.10E+00 | 1.95E-03 | 7.14E-03 | -1.06E+00 | 2.00E-02 | 5.49E-02 | 1.06E+00 |
| ST7L | 2.16E-02 | 1.12E-01 | 1.10E+00 | 1.90E-02 | 8.23E-02 | 1.15E+00 | 1.16E-01 | 4.92E-01 | 1.31E+00 | 1.26E-01 | 4.80E-01 | 1.18E+00 | 1.98E-03 | 7.07E-03 | -1.04E+00 | 4.98E-05 | 1.40E-04 | -1.28E+00 | 2.85E-05 | 1.05E-04 | -1.19E+00 | 3.57E-03 | 9.82E-03 | -1.06E+00 |
| B3GLCT | 2.16E-02 | 1.13E-01 | -1.23E+00 | 1.12E-01 | 4.82E-01 | 1.19E+00 | 1.88E-01 | 8.00E-01 | 1.04E+00 | 2.49E-01 | 9.51E-01 | 1.33E+00 | 1.33E-01 | 4.72E-01 | -1.42E+00 | 2.33E-01 | 6.56E-01 | -1.17E+00 | 9.81E-03 | 3.60E-02 | -1.23E+00 | 2.59E-02 | 7.13E-02 | -1.16E+00 |

| | | | | | | | | | | | | | | | | | | | | | | | | |
|---|---|---|---|---|---|---|---|---|---|---|---|---|---|---|---|---|---|---|---|---|---|---|---|---|
| ZFY | 2.16E-02 | 1.13E-01 | 1.18E+00 | 7.68E-10 | 3.32E-09 | -1.42E+00 | 8.69E-02 | 3.69E-01 | 1.08E+00 | 2.03E-05 | 7.74E-05 | -1.24E+00 | 2.39E-01 | 8.51E-01 | 1.51E+00 | 1.17E-03 | 3.30E-03 | -1.14E+00 | 3.17E-03 | 1.16E-02 | -1.09E+00 | 1.16E-04 | 3.18E-04 | -1.31E+00 |
| PGGHG | 2.16E-02 | 1.13E-01 | 1.14E+00 | 6.04E-04 | 2.61E-03 | -1.03E+00 | 8.67E-02 | 3.68E-01 | 1.00E+00 | 5.98E-02 | 2.29E-01 | 1.02E+00 | 7.56E-04 | 2.70E-03 | -1.64E+00 | 7.84E-03 | 2.21E-02 | -1.33E+00 | 1.32E-01 | 4.84E-01 | -1.03E+00 | 1.60E-01 | 4.40E-01 | 1.00E+00 |
| ZNF235 | 2.16E-02 | 1.13E-01 | -1.19E+00 | 8.09E-02 | 3.50E-01 | 1.15E+00 | 7.60E-02 | 3.23E-01 | -1.12E+00 | 7.21E-02 | 2.75E-01 | -1.05E+00 | 8.01E-02 | 2.86E-01 | 1.96E+00 | 5.54E-02 | 1.56E-01 | 2.20E+00 | 4.28E-03 | 1.57E-02 | -1.62E+00 | 1.87E-01 | 5.15E-01 | 1.01E+00 |
| SCAF1 | 2.17E-02 | 1.13E-01 | -1.03E+00 | 1.25E-01 | 5.41E-01 | 1.36E+00 | 1.93E-03 | 8.20E-03 | -1.31E+00 | 9.90E-02 | 3.78E-01 | 1.25E+00 | 1.53E-03 | 5.46E-03 | -1.36E+00 | 2.15E-02 | 6.07E-02 | -1.16E+00 | 5.45E-06 | 2.00E-05 | -1.43E+00 | 7.57E-02 | 2.08E-01 | 1.04E+00 |
| PLLP | 2.17E-02 | 1.13E-01 | -1.19E+00 | 2.31E-01 | 1.00E+00 | 1.44E+00 | 2.27E-03 | 9.63E-03 | -1.64E+00 | 3.36E-02 | 1.28E-01 | 1.00E+00 | 1.94E-01 | 6.91E-01 | 1.65E+00 | 4.00E-03 | 1.13E-02 | 2.90E+00 | 7.66E-02 | 2.81E-01 | 1.59E+00 | 1.30E-01 | 3.57E-01 | 1.76E+00 |
| VASH1 | 2.17E-02 | 1.13E-01 | -1.11E+00 | 2.52E-02 | 1.09E-01 | -1.12E+00 | 1.78E-01 | 7.57E-01 | 1.15E+00 | 1.79E-01 | 6.86E-01 | 1.29E+00 | 1.44E-01 | 5.13E-01 | 1.80E+00 | 2.44E-01 | 6.88E-01 | 1.64E+00 | 6.26E-02 | 2.30E-01 | -1.17E+00 | 6.09E-02 | 1.67E-01 | -1.32E+00 |
| MCUB | 2.17E-02 | 1.13E-01 | 1.57E+00 | 4.20E-02 | 1.81E-01 | 1.74E+00 | 1.19E-02 | 5.07E-02 | -1.00E+00 | 2.67E-02 | 1.02E-01 | 1.14E+00 | 1.01E-01 | 3.61E-01 | 1.23E+00 | 3.23E-01 | 9.11E-01 | 1.40E+00 | 6.31E-04 | 2.31E-03 | -1.08E+00 | 5.25E-02 | 1.44E-01 | 1.05E+00 |
| RCOR1 | 2.17E-02 | 1.13E-01 | 1.17E+00 | 8.72E-04 | 3.77E-03 | 1.10E+00 | 3.50E-05 | 1.49E-04 | -1.19E+00 | 3.22E-03 | 1.23E-02 | 1.11E+00 | 1.09E-02 | 3.87E-02 | 1.03E+00 | 1.04E-02 | 2.94E-02 | 1.01E+00 | 4.63E-02 | 1.70E-01 | 1.17E+00 | 3.23E-02 | 8.86E-02 | 1.11E+00 |
| ENC1 | 2.17E-02 | 1.13E-01 | -1.29E+00 | 8.91E-03 | 3.85E-02 | -1.51E+00 | 4.21E-06 | 1.79E-05 | -1.83E+00 | 5.22E-06 | 2.00E-05 | -1.74E+00 | 3.79E-03 | 1.35E-02 | -1.58E+00 | 3.45E-04 | 9.72E-04 | -2.17E+00 | 7.75E-03 | 2.85E-02 | -1.72E+00 | 1.09E-02 | 2.98E-02 | -1.71E+00 |
| TAF6L | 2.17E-02 | 1.13E-01 | -1.07E+00 | 5.63E-02 | 2.43E-01 | 1.20E+00 | 4.91E-05 | 2.09E-04 | -1.54E+00 | 3.00E-03 | 1.14E-02 | -1.12E+00 | 6.42E-03 | 2.29E-02 | -1.33E+00 | 7.05E-02 | 1.99E-01 | 1.03E+00 | 1.91E-02 | 7.01E-02 | -1.20E+00 | 7.90E-02 | 2.17E-01 | -1.15E+00 |
| AL603756.1 | 2.18E-02 | 1.13E-01 | -1.08E+00 | 2.75E-03 | 1.19E-02 | -1.33E+00 | 2.29E-01 | 9.73E-01 | 1.18E+00 | 1.44E-01 | 5.49E-01 | 1.13E+00 | 5.67E-02 | 2.02E-01 | -1.01E+00 | 1.22E-01 | 3.43E-01 | -1.03E+00 | 2.10E-01 | 7.70E-01 | 1.22E+00 | 3.22E-02 | 8.84E-02 | -1.65E+00 |
| DENND5A | 2.18E-02 | 1.14E-01 | 1.32E+00 | 8.90E-06 | 3.85E-05 | 1.16E+00 | 1.21E-01 | 5.14E-01 | 1.34E+00 | 3.12E-06 | 1.19E-05 | -1.17E+00 | 7.00E-03 | 2.50E-02 | -1.03E+00 | 8.82E-05 | 2.49E-04 | -1.46E+00 | 1.06E-01 | 3.88E-01 | 1.03E+00 | 1.31E-02 | 3.61E-02 | -1.24E+00 |
| SYP | 2.18E-02 | 1.14E-01 | -1.12E+00 | 4.63E-02 | 2.00E-01 | 1.09E+00 | 1.92E-01 | 8.14E-01 | -1.07E+00 | 2.49E-01 | 9.52E-01 | 1.11E+00 | 1.20E-01 | 4.26E-01 | -1.03E+00 | 1.13E-01 | 3.20E-01 | -1.15E+00 | 2.69E-01 | 9.86E-01 | 1.21E+00 | 2.98E-01 | 8.18E-01 | -1.02E+00 |
| ARMC9 | 2.18E-02 | 1.14E-01 | -1.07E+00 | 5.10E-02 | 2.20E-01 | 1.10E+00 | 9.49E-04 | 4.03E-03 | -1.43E+00 | 2.09E-02 | 7.98E-02 | -1.07E+00 | 3.34E-02 | 1.19E-01 | -1.29E+00 | 4.42E-02 | 1.25E-01 | -1.28E+00 | 1.61E-01 | 5.89E-01 | 1.36E+00 | 1.58E-01 | 4.33E-01 | 1.60E+00 |
| ZNF250 | 2.18E-02 | 1.14E-01 | -1.05E+00 | 2.60E-02 | 1.12E-01 | 1.08E+00 | 6.23E-09 | 2.65E-08 | -1.93E+00 | 3.10E-07 | 1.18E-06 | -1.48E+00 | 4.21E-05 | 1.50E-04 | -2.33E+00 | 6.48E-03 | 1.83E-02 | -1.27E+00 | 2.64E-05 | 9.69E-05 | -1.61E+00 | 2.67E-04 | 7.34E-04 | -1.53E+00 |
| ZNF727 | 2.18E-02 | 1.14E-01 | 1.04E+00 | 1.87E-03 | 8.07E-03 | -1.20E+00 | 8.84E-02 | 3.76E-01 | 1.02E+00 | 3.30E-02 | 1.26E-01 | -1.11E+00 | 1.30E-02 | 4.64E-02 | -1.25E+00 | 8.93E-03 | 2.52E-02 | -1.42E+00 | 1.21E-01 | 4.43E-01 | 1.63E+00 | 2.47E-01 | 6.78E-01 | 1.48E+00 |
| WBP2NL | 2.19E-02 | 1.14E-01 | -1.09E+00 | 1.56E-01 | 6.75E-01 | 1.48E+00 | 1.01E-01 | 4.28E-01 | -1.15E+00 | 1.60E-01 | 6.09E-01 | 1.44E+00 | 2.33E-01 | 8.29E-01 | 1.56E+00 | 5.86E-02 | 1.65E-01 | 2.08E+00 | 6.74E-03 | 2.47E-02 | -1.43E+00 | 2.75E-02 | 7.55E-02 | -1.36E+00 |
| DHRS3 | 2.19E-02 | 1.14E-01 | 3.00E+00 | 1.66E-01 | 7.17E-01 | 1.63E+00 | 3.24E-02 | 1.38E-01 | 2.56E+00 | 1.66E-01 | 6.34E-01 | 2.10E+00 | 5.65E-07 | 2.01E-06 | 2.23E+00 | 3.54E-24 | 9.98E-24 | 3.59E+00 | 2.72E-05 | 9.97E-05 | 1.52E+00 | 1.01E-02 | 2.78E-02 | 1.31E+00 |
| SLC9B1 | 2.19E-02 | 1.14E-01 | -1.26E+00 | 6.93E-02 | 2.99E-01 | 1.02E+00 | 1.27E-01 | 5.40E-01 | -1.03E+00 | 1.99E-01 | 7.61E-01 | 1.07E+00 | 1.58E-01 | 5.63E-01 | 1.06E+00 | 2.88E-01 | 8.13E-01 | 1.48E+00 | 1.15E-01 | 4.21E-01 | 1.46E+00 | 2.43E-01 | 6.67E-01 | 1.35E+00 |
| SDCCAG3 | 2.20E-02 | 1.14E-01 | 1.01E+00 | 3.71E-02 | 1.60E-01 | 1.22E+00 | 7.36E-03 | 3.13E-02 | -1.10E+00 | 1.27E-01 | 4.84E-01 | 1.24E+00 | 2.61E-05 | 9.29E-05 | -1.31E+00 | 8.19E-02 | 2.31E-01 | 1.27E+00 | 1.72E-01 | 6.32E-01 | 1.32E+00 | 9.98E-02 | 2.74E-01 | 1.55E+00 |
| PPM1N | 2.20E-02 | 1.15E-01 | -1.02E+00 | 2.71E-02 | 1.17E-01 | 1.09E+00 | 1.27E-01 | 5.41E-01 | 1.14E+00 | 2.55E-01 | 9.74E-01 | 1.44E+00 | 2.30E-01 | 8.20E-01 | 1.16E+00 | 1.32E-01 | 3.71E-01 | -1.09E+00 | 9.87E-03 | 3.62E-02 | 1.03E+00 | 2.00E-03 | 5.51E-03 | -1.23E+00 |
| LRCH2 | 2.20E-02 | 1.15E-01 | -1.04E+00 | 8.36E-02 | 3.61E-01 | 1.22E+00 | 1.16E-02 | 4.92E-02 | -1.47E+00 | 1.01E-02 | 3.87E-02 | -1.39E+00 | 5.89E-02 | 2.10E-01 | -1.32E+00 | 1.54E-01 | 4.35E-01 | -1.04E+00 | 2.47E-01 | 9.07E-01 | 1.29E+00 | 2.00E-01 | 5.49E-01 | -1.06E+00 |
| AMZ2P1 | 2.20E-02 | 1.15E-01 | 1.15E+00 | 3.60E-03 | 1.56E-02 | 1.16E+00 | 1.34E-02 | 5.69E-02 | -1.02E+00 | 1.79E-03 | 6.83E-03 | -1.06E+00 | 2.73E-02 | 9.75E-02 | 1.06E+00 | 1.28E-01 | 3.62E-01 | 1.21E+00 | 7.75E-03 | 2.85E-02 | -1.01E+00 | 4.36E-04 | 1.20E-03 | -1.22E+00 |
| AC106820.4 | 2.21E-02 | 1.15E-01 | -1.28E+00 | 6.12E-03 | 2.64E-02 | -1.49E+00 | 1.81E-01 | 7.70E-01 | 1.31E+00 | 1.89E-02 | 7.20E-02 | -1.30E+00 | 1.22E-01 | 4.35E-01 | 1.64E+00 | 2.38E-01 | 6.70E-01 | 1.40E+00 | 4.42E-03 | 1.62E-02 | -1.83E+00 | 1.10E-02 | 3.02E-02 | -1.71E+00 |
| AK5 | 2.21E-02 | 1.15E-01 | -1.29E+00 | 1.69E-02 | 7.28E-02 | -1.39E+00 | 2.10E-03 | 8.91E-03 | -1.07E+00 | 1.48E-07 | 5.65E-07 | -1.46E+00 | 1.37E-01 | 4.88E-01 | -1.04E+00 | 2.19E-01 | 6.17E-01 | -1.08E+00 | 2.00E-01 | 7.33E-01 | 1.14E+00 | 2.53E-01 | 6.95E-01 | 1.02E+00 |
| STAG1 | 2.22E-02 | 1.15E-01 | 1.08E+00 | 4.41E-02 | 1.91E-01 | 1.22E+00 | 7.98E-02 | 3.39E-01 | 1.07E+00 | 6.91E-02 | 2.64E-01 | 1.20E+00 | 1.31E-02 | 4.68E-02 | 1.03E+00 | 6.36E-02 | 1.79E-01 | 1.13E+00 | 2.98E-02 | 1.09E-01 | 1.00E+00 | 2.67E-04 | 7.34E-04 | -1.22E+00 |
| AC017104.3 | 2.22E-02 | 1.15E-01 | 2.25E+00 | 3.53E-02 | 1.52E-01 | 2.45E+00 | 2.30E-03 | 9.79E-03 | 2.09E+00 | 1.19E-02 | 4.55E-02 | 2.04E+00 | 6.57E-02 | 2.34E-01 | -1.14E+00 | 1.45E-01 | 4.09E-01 | 1.03E+00 | 2.38E-01 | 8.74E-01 | 1.17E+00 | 2.80E-01 | 7.68E-01 | 1.29E+00 |
| ZNF761 | 2.22E-02 | 1.15E-01 | -1.11E+00 | 3.91E-02 | 1.69E-01 | -1.02E+00 | 1.28E-02 | 5.43E-02 | -1.21E+00 | 3.20E-02 | 1.22E-01 | 1.10E+00 | 3.07E-02 | 1.10E-01 | -1.08E+00 | 6.69E-02 | 1.89E-01 | 1.06E+00 | 7.42E-04 | 2.72E-03 | -1.13E+00 | 4.78E-05 | 1.31E-04 | -1.35E+00 |
| FAM206A | 2.22E-02 | 1.16E-01 | -1.28E+00 | 8.11E-02 | 3.50E-01 | 1.06E+00 | 8.66E-02 | 3.68E-01 | -1.00E+00 | 8.49E-02 | 3.24E-01 | 1.14E+00 | 4.13E-02 | 1.47E-01 | -1.03E+00 | 2.01E-01 | 5.68E-01 | 1.27E+00 | 3.93E-07 | 1.44E-06 | -1.67E+00 | 3.26E-04 | 8.97E-04 | -1.27E+00 |
| CENPO | 2.22E-02 | 1.16E-01 | -1.56E+00 | 4.57E-02 | 1.98E-01 | -1.37E+00 | 5.02E-02 | 2.13E-01 | -1.05E+00 | 2.00E-01 | 7.64E-01 | 1.32E+00 | 8.75E-02 | 3.12E-01 | -1.05E+00 | 1.42E-01 | 4.00E-01 | 1.00E+00 | 5.32E-04 | 1.95E-03 | -1.28E+00 | 2.30E-01 | 6.31E-01 | 1.12E+00 |
| TRIM39 | 2.22E-02 | 1.16E-01 | -1.09E+00 | 5.19E-03 | 2.24E-02 | -1.19E+00 | 5.38E-04 | 2.28E-03 | -1.56E+00 | 2.59E-03 | 9.90E-03 | -1.27E+00 | 8.54E-02 | 3.05E-01 | -1.02E+00 | 9.71E-02 | 2.74E-01 | -1.10E+00 | 6.17E-03 | 2.27E-02 | -1.27E+00 | 5.07E-02 | 1.39E-01 | -1.02E+00 |
| APBB3 | 2.22E-02 | 1.16E-01 | -1.06E+00 | 2.41E-02 | 1.04E-01 | 1.02E+00 | 1.55E-01 | 6.57E-01 | 1.07E+00 | 2.24E-01 | 8.55E-01 | 1.27E+00 | 1.27E-01 | 4.53E-01 | 1.03E+00 | 2.54E-01 | 7.16E-01 | 1.18E+00 | 1.90E-02 | 6.98E-02 | -1.23E+00 | 2.31E-02 | 6.35E-02 | -1.24E+00 |
| MXD3 | 2.23E-02 | 1.16E-01 | -1.09E+00 | 1.61E-01 | 6.95E-01 | 1.40E+00 | 8.67E-02 | 3.68E-01 | 1.57E+00 | 6.33E-02 | 2.42E-01 | 1.89E+00 | 1.15E-01 | 4.09E-01 | -1.02E+00 | 3.38E-01 | 9.52E-01 | 1.38E+00 | 7.17E-02 | 2.63E-01 | 1.18E+00 | 1.47E-02 | 4.03E-02 | 1.41E+00 |
| CASZ1 | 2.23E-02 | 1.16E-01 | -1.15E+00 | 2.32E-05 | 1.00E-04 | -1.67E+00 | 1.57E-02 | 6.66E-02 | -1.29E+00 | 3.96E-03 | 1.51E-02 | -1.45E+00 | 2.95E-03 | 1.05E-02 | -1.38E+00 | 3.70E-04 | 1.04E-03 | -1.78E+00 | 2.17E-02 | 7.96E-02 | -1.44E+00 | 1.04E-02 | 2.85E-02 | -1.97E+00 |
| SEC14L1 | 2.23E-02 | 1.16E-01 | 1.21E+00 | 5.73E-06 | 2.48E-05 | -1.04E+00 | 1.93E-05 | 8.19E-05 | -1.22E+00 | 6.35E-09 | 2.43E-08 | -1.32E+00 | 1.61E-05 | 5.74E-05 | -1.17E+00 | 5.56E-08 | 1.57E-07 | -1.44E+00 | 1.42E-02 | 5.19E-02 | 1.26E+00 | 3.19E-04 | 8.75E-04 | -1.14E+00 |
| GYG2 | 2.23E-02 | 1.16E-01 | -1.19E+00 | 9.32E-03 | 4.03E-02 | -1.37E+00 | 2.24E-04 | 9.50E-04 | -1.81E+00 | 7.32E-04 | 2.79E-03 | -1.51E+00 | 6.85E-02 | 2.44E-01 | -1.14E+00 | 2.20E-01 | 6.21E-01 | 1.16E+00 | 1.29E-02 | 4.74E-02 | -1.35E+00 | 3.31E-02 | 9.08E-02 | -1.15E+00 |
| CDK17 | 2.23E-02 | 1.16E-01 | 1.09E+00 | 5.90E-06 | 2.55E-05 | -1.21E+00 | 1.21E-02 | 5.14E-02 | -1.11E+00 | 7.12E-05 | 2.72E-04 | -1.29E+00 | 2.56E-01 | 9.14E-01 | 1.32E+00 | 6.70E-02 | 1.89E-01 | 1.07E+00 | 1.42E-07 | 5.22E-07 | -1.33E+00 | 2.95E-07 | 8.11E-07 | -1.26E+00 |
| CMSS1 | 2.23E-02 | 1.16E-01 | 1.07E+00 | 1.70E-02 | 7.33E-02 | 1.21E+00 | 1.15E-01 | 4.90E-01 | 1.11E+00 | 4.50E-02 | 1.72E-01 | 1.21E+00 | 1.95E-03 | 6.96E-03 | -1.14E+00 | 3.80E-02 | 1.07E-01 | 1.07E+00 | 2.15E-03 | 7.87E-03 | 1.03E+00 | 4.83E-02 | 1.33E-01 | 1.08E+00 |
| EVPL | 2.23E-02 | 1.16E-01 | 1.01E+00 | 2.28E-01 | 9.85E-01 | 1.49E+00 | 2.96E-02 | 1.26E-01 | -1.11E+00 | 1.73E-01 | 6.60E-01 | 1.26E+00 | 5.18E-02 | 1.85E-01 | -1.69E+00 | 2.39E-01 | 6.72E-01 | -1.10E+00 | 1.61E-01 | 5.92E-01 | -1.05E+00 | 1.22E-01 | 3.34E-01 | -1.21E+00 |
| CDR2L | 2.24E-02 | 1.16E-01 | -1.34E+00 | 4.77E-02 | 2.06E-01 | -1.15E+00 | 2.95E-02 | 1.25E-01 | -1.27E+00 | 4.99E-03 | 1.90E-02 | -1.42E+00 | 1.15E-01 | 4.09E-01 | -1.67E+00 | 1.49E-01 | 4.19E-01 | -1.67E+00 | 1.80E-01 | 6.60E-01 | 1.42E+00 | 3.04E-01 | 8.36E-01 | -1.06E+00 |
| AC018529.3 | 2.24E-02 | 1.17E-01 | -1.16E+00 | 2.28E-01 | 9.86E-01 | 1.44E+00 | 9.43E-03 | 4.01E-02 | -1.41E+00 | 6.50E-02 | 2.48E-01 | -1.05E+00 | 1.78E-02 | 6.34E-02 | -1.93E+00 | 7.30E-02 | 2.06E-01 | -1.37E+00 | 2.10E-02 | 7.71E-02 | -1.04E+00 | 4.93E-03 | 1.35E-02 | -1.60E+00 |
| INPPL1 | 2.24E-02 | 1.17E-01 | -1.06E+00 | 2.13E-02 | 9.20E-02 | 1.03E+00 | 2.00E-02 | 8.50E-02 | -1.02E+00 | 3.85E-04 | 1.47E-03 | -1.15E+00 | 5.55E-06 | 1.98E-05 | -1.32E+00 | 4.11E-07 | 1.16E-06 | -1.40E+00 | 3.49E-02 | 1.28E-01 | 1.05E+00 | 3.73E-02 | 1.03E-01 | -1.01E+00 |
| NSMF | 2.24E-02 | 1.17E-01 | -1.01E+00 | 1.55E-02 | 6.71E-02 | 1.03E+00 | 2.09E-01 | 8.86E-01 | 1.31E+00 | 2.03E-01 | 7.74E-01 | 1.32E+00 | 1.69E-02 | 6.04E-02 | -1.15E+00 | 1.29E-01 | 3.64E-01 | 1.10E+00 | 2.29E-01 | 8.39E-01 | 1.17E+00 | 1.14E-01 | 3.12E-01 | 1.51E+00 |
| MAMDC2-AS1 | 2.24E-02 | 1.17E-01 | 1.63E+00 | 8.36E-02 | 3.61E-01 | 1.70E+00 | 2.30E-01 | 9.77E-01 | 1.02E+00 | 1.89E-01 | 7.23E-01 | 1.30E+00 | 2.99E-02 | 1.06E-01 | 4.38E+00 | 2.65E-01 | 7.48E-01 | 1.90E+00 | 8.47E-02 | 3.11E-01 | 2.78E+00 | 1.37E-01 | 3.76E-01 | 2.45E+00 |
| EID1 | 2.24E-02 | 1.17E-01 | -1.01E+00 | 3.73E-02 | 1.61E-01 | 1.18E+00 | 1.18E-01 | 5.01E-01 | 1.08E+00 | 7.05E-04 | 2.69E-03 | 1.26E+00 | 1.60E-01 | 5.72E-01 | 1.17E+00 | 7.64E-04 | 2.15E-03 | 1.32E+00 | 2.15E-02 | 7.91E-02 | 1.16E+00 | 1.01E-03 | 2.77E-03 | 1.25E+00 |
| RPL12P10 | 2.25E-02 | 1.17E-01 | 1.96E+00 | 2.14E-01 | 9.23E-01 | 1.55E+00 | 3.78E-02 | 1.60E-01 | 1.70E+00 | 2.36E-01 | 9.03E-01 | 1.44E+00 | 7.75E-02 | 2.76E-01 | 1.87E+00 | 2.81E-01 | 7.92E-01 | 1.51E+00 | 3.12E-03 | 1.15E-02 | 2.46E+00 | 2.47E-01 | 6.77E-01 | 1.55E+00 |
| CLEC2D | 2.25E-02 | 1.17E-01 | 1.06E+00 | 1.47E-01 | 6.37E-01 | 1.55E+00 | 1.90E-01 | 8.07E-01 | 1.06E+00 | 2.47E-01 | 9.43E-01 | 1.22E+00 | 2.84E-03 | 1.01E-02 | -1.36E+00 | 2.02E-01 | 5.68E-01 | 1.31E+00 | 1.08E-01 | 3.98E-01 | -1.02E+00 | 1.92E-01 | 5.28E-01 | 1.01E+00 |
| ZNF394 | 2.25E-02 | 1.17E-01 | 1.17E+00 | 4.25E-12 | 1.84E-11 | -1.28E+00 | 1.33E-01 | 5.64E-01 | 1.17E+00 | 1.20E-05 | 4.57E-05 | -1.12E+00 | 2.00E-01 | 7.13E-01 | 1.46E+00 | 7.67E-05 | 2.16E-04 | -1.14E+00 | 4.61E-02 | 1.69E-01 | 1.36E+00 | 7.16E-03 | 1.97E-02 | -1.08E+00 |
| MORN1 | 2.25E-02 | 1.17E-01 | -1.19E+00 | 8.62E-02 | 3.73E-01 | 1.12E+00 | 7.10E-02 | 3.02E-01 | -1.05E+00 | 1.56E-02 | 5.98E-02 | -1.10E+00 | 1.01E-02 | 3.60E-02 | -1.31E+00 | 7.02E-02 | 1.98E-01 | -1.05E+00 | 8.28E-02 | 3.04E-01 | -1.02E+00 | 2.34E-01 | 6.43E-01 | 1.16E+00 |
| CCDC85C | 2.26E-02 | 1.18E-01 | -1.12E+00 | 3.48E-03 | 1.50E-02 | -1.32E+00 | 1.07E-01 | 4.56E-01 | -1.01E+00 | 2.36E-04 | 9.00E-04 | -1.49E+00 | 2.55E-01 | 9.09E-01 | 1.52E+00 | 1.69E-01 | 4.76E-01 | 1.16E+00 | 1.77E-03 | 6.50E-03 | -1.24E+00 | 8.42E-06 | 2.31E-05 | -1.82E+00 |
| RBM26-AS1 | 2.26E-02 | 1.18E-01 | -1.16E+00 | 3.84E-02 | 1.66E-01 | -1.12E+00 | 1.83E-01 | 7.79E-01 | 1.06E+00 | 1.44E-01 | 5.49E-01 | 1.47E+00 | 8.85E-04 | 3.15E-03 | -2.15E+00 | 7.79E-03 | 2.19E-02 | -1.46E+00 | 9.47E-02 | 3.48E-01 | 1.62E+00 | 2.79E-01 | 7.68E-01 | 1.42E+00 |
| SLC22A18 | 2.26E-02 | 1.18E-01 | -1.14E+00 | 1.08E-02 | 4.67E-02 | -1.20E+00 | 8.71E-02 | 3.70E-01 | 1.39E+00 | 1.62E-01 | 6.20E-01 | 1.45E+00 | 1.12E-01 | 3.99E-01 | 1.08E+00 | 1.46E-01 | 4.11E-01 | 1.22E+00 | 4.81E-03 | 1.76E-02 | -1.40E+00 | 1.90E-01 | 5.21E-01 | 1.06E+00 |
| CR392039.4 | 2.26E-02 | 1.18E-01 | -1.12E+00 | 2.20E-01 | 9.51E-01 | 1.34E+00 | 1.93E-01 | 8.19E-01 | 1.15E+00 | 2.04E-01 | 7.80E-01 | 1.61E+00 | 1.52E-01 | 5.43E-01 | 1.08E+00 | 1.91E-01 | 5.38E-01 | 1.75E+00 | 1.26E-01 | 4.63E-01 | 1.06E+00 | 3.57E-01 | 9.81E-01 | 1.31E+00 |
| AL390728.4 | 2.27E-02 | 1.18E-01 | 1.04E+00 | 1.66E-02 | 7.18E-02 | 1.04E+00 | 1.79E-02 | 7.62E-02 | -1.08E+00 | 9.09E-03 | 3.47E-02 | -1.00E+00 | 2.43E-04 | 8.68E-04 | -1.45E+00 | 3.33E-04 | 9.39E-04 | -1.47E+00 | 1.86E-03 | 6.83E-03 | -1.30E+00 | 3.86E-03 | 1.06E-02 | -1.25E+00 |
| ZNF730 | 2.27E-02 | 1.18E-01 | -1.33E+00 | 7.51E-02 | 3.24E-01 | -1.05E+00 | 2.13E-01 | 9.04E-01 | 1.28E+00 | 1.59E-01 | 6.06E-01 | 1.63E+00 | 8.40E-02 | 2.99E-01 | -1.35E+00 | 1.24E-01 | 3.50E-01 | -1.28E+00 | 2.97E-06 | 1.09E-05 | -1.81E+00 | 1.51E-03 | 4.15E-03 | -1.34E+00 |
| SMAGP | 2.27E-02 | 1.18E-01 | -1.01E+00 | 2.01E-02 | 8.68E-02 | 1.09E+00 | 1.27E-01 | 5.38E-01 | 1.06E+00 | 6.90E-02 | 2.63E-01 | 1.05E+00 | 2.61E-01 | 9.32E-01 | 1.19E+00 | 3.18E-01 | 8.95E-01 | 1.12E+00 | 7.31E-03 | 2.68E-02 | -1.17E+00 | 3.59E-01 | 9.87E-01 | 1.36E+00 |
| AL450405.1 | 2.27E-02 | 1.18E-01 | 1.35E+00 | 4.40E-32 | 1.90E-31 | 2.51E+00 | 1.19E-02 | 5.07E-02 | 1.42E+00 | 7.81E-15 | 2.98E-14 | 2.47E+00 | 5.01E-02 | 1.79E-01 | 1.49E+00 | 1.23E-21 | 3.47E-21 | 2.79E+00 | 2.54E-03 | 9.31E-03 | 1.52E+00 | 7.73E-22 | 2.12E-21 | 2.70E+00 |
| GTF2IP20 | 2.27E-02 | 1.18E-01 | 1.03E+00 | 3.03E-02 | 1.31E-01 | 1.20E+00 | 5.28E-02 | 2.24E-01 | -1.03E+00 | 1.93E-02 | 7.36E-02 | -1.03E+00 | 7.00E-02 | 2.49E-01 | 1.07E+00 | 1.30E-01 | 3.66E-01 | 1.09E+00 | 7.70E-03 | 2.82E-02 | 1.02E+00 | 1.67E-02 | 4.59E-02 | -1.04E+00 |
| ETS1 | 2.27E-02 | 1.18E-01 | 1.53E+00 | 1.41E-01 | 6.09E-01 | 1.52E+00 | 1.52E-01 | 6.45E-01 | 1.06E+00 | 1.61E-01 | 6.16E-01 | 1.15E+00 | 2.10E-01 | 7.50E-01 | 1.58E+00 | 2.27E-01 | 6.40E-01 | 1.16E+00 | 5.45E-03 | 2.00E-02 | -1.17E+00 | 2.11E-02 | 5.80E-02 | -1.08E+00 |
| UBL5 | 2.28E-02 | 1.19E-01 | 1.10E+00 | 1.82E-15 | 7.84E-15 | -1.15E+00 | 5.40E-02 | 2.30E-01 | 1.21E+00 | 3.62E-03 | 1.38E-02 | -1.08E+00 | 5.16E-06 | 1.84E-05 | -1.19E+00 | 2.61E-10 | 7.35E-10 | -1.36E+00 | 2.42E-01 | 8.89E-01 | 1.09E+00 | 6.41E-03 | 1.76E-02 | -1.24E+00 |
| STARD9 | 2.28E-02 | 1.19E-01 | 1.05E+00 | 1.17E-01 | 5.04E-01 | 1.59E+00 | 1.38E-01 | 5.85E-01 | 1.17E+00 | 4.20E-02 | 1.60E-01 | 1.63E+00 | 2.28E-02 | 8.11E-02 | -1.49E+00 | 8.54E-02 | 2.41E-01 | -1.11E+00 | 3.12E-03 | 1.15E-02 | -1.71E+00 | 1.45E-02 | 3.97E-02 | -1.37E+00 |
| AC009159.4 | 2.28E-02 | 1.19E-01 | -1.35E+00 | 1.65E-01 | 7.15E-01 | 1.06E+00 | 1.41E-02 | 6.00E-02 | -1.53E+00 | 1.77E-02 | 6.78E-02 | -1.45E+00 | 7.79E-02 | 2.78E-01 | -1.26E+00 | 2.88E-01 | 8.12E-01 | 1.07E+00 | 1.08E-01 | 3.96E-01 | -1.16E+00 | 2.42E-01 | 6.65E-01 | 1.05E+00 |
| RANBP1 | 2.28E-02 | 1.19E-01 | 1.15E+00 | 1.25E-06 | 5.41E-06 | 1.12E+00 | 9.43E-03 | 4.01E-02 | 1.42E+00 | 1.36E-01 | 5.20E-01 | 1.37E+00 | 3.37E-05 | 1.20E-04 | -1.14E+00 | 9.24E-02 | 2.61E-01 | 1.17E+00 | 1.55E-01 | 5.67E-01 | 1.17E+00 | 5.35E-03 | 1.47E-02 | 1.40E+00 |
| FOXO1 | 2.29E-02 | 1.19E-01 | 1.53E+00 | 2.30E-02 | 9.95E-02 | 1.76E+00 | 1.76E-02 | 7.46E-02 | 1.83E+00 | 2.34E-01 | 8.95E-01 | 1.37E+00 | 2.94E-02 | 1.05E-01 | 2.36E+00 | 1.84E-01 | 5.19E-01 | 1.79E+00 | 2.32E-01 | 8.53E-01 | 1.34E+00 | 1.06E-01 | 2.91E-01 | 1.23E+00 |
| RN7SL308P | 2.29E-02 | 1.20E-01 | 1.74E+00 | 2.02E-01 | 8.72E-01 | 1.28E+00 | 3.65E-02 | 1.55E-01 | 1.56E+00 | 1.12E-01 | 4.27E-01 | 1.01E+00 | 4.32E-02 | 1.54E-01 | 1.86E+00 | 2.98E-01 | 8.39E-01 | -1.11E+00 | 1.30E-01 | 4.78E-01 | 1.30E+00 | 2.09E-01 | 5.74E-01 | -1.27E+00 |
| MAN2A2 | 2.30E-02 | 1.20E-01 | -1.09E+00 | 1.66E-01 | 7.16E-01 | 1.38E+00 | 2.45E-02 | 1.04E-01 | -1.29E+00 | 5.69E-02 | 2.17E-01 | -1.05E+00 | 9.38E-02 | 3.35E-01 | 1.22E+00 | 1.85E-01 | 5.22E-01 | 1.27E+00 | 1.51E-02 | 5.55E-02 | 1.10E+00 | 9.70E-03 | 2.66E-02 | 1.10E+00 |
| ZNF134 | 2.30E-02 | 1.20E-01 | 1.12E+00 | 3.63E-04 | 1.57E-03 | 1.03E+00 | 7.31E-02 | 3.11E-01 | 1.03E+00 | 5.44E-04 | 2.08E-03 | -1.11E+00 | 6.88E-02 | 2.45E-01 | 1.13E+00 | 3.77E-03 | 1.06E-02 | -1.06E+00 | 6.16E-03 | 2.26E-02 | -1.09E+00 | 6.18E-04 | 1.70E-03 | -1.25E+00 |
| AC211476.2 | 2.30E-02 | 1.20E-01 | -1.33E+00 | 1.02E-01 | 4.39E-01 | 1.05E+00 | 1.92E-01 | 8.14E-01 | 1.38E+00 | 7.25E-02 | 2.77E-01 | -1.04E+00 | 2.38E-01 | 8.49E-01 | -1.02E+00 | 3.32E-01 | 9.37E-01 | 1.07E+00 | 6.10E-02 | 2.24E-01 | -1.17E+00 | 8.11E-02 | 2.23E-01 | -1.20E+00 |
| PPP1R26-AS1 | 2.30E-02 | 1.20E-01 | -1.24E+00 | 1.53E-01 | 6.60E-01 | 1.62E+00 | 1.11E-02 | 4.70E-02 | -1.51E+00 | 2.99E-02 | 1.14E-01 | -1.19E+00 | 8.99E-07 | 3.21E-06 | -2.56E+00 | 1.82E-05 | 5.13E-05 | -1.81E+00 | 2.74E-03 | 1.00E-02 | -1.48E+00 | 3.42E-02 | 9.40E-02 | -1.15E+00 |
| ESYT2 | 2.30E-02 | 1.20E-01 | 1.49E+00 | 1.24E-01 | 5.38E-01 | 1.55E+00 | 1.67E-01 | 7.10E-01 | 1.21E+00 | 1.12E-01 | 4.27E-01 | 1.20E+00 | 6.06E-02 | 2.16E-01 | 1.70E+00 | 2.95E-01 | 8.31E-01 | 1.44E+00 | 1.25E-01 | 4.60E-01 | 1.44E+00 | 3.15E-01 | 8.67E-01 | 1.31E+00 |
| CCDC106 | 2.31E-02 | 1.20E-01 | 1.02E+00 | 3.47E-02 | 1.50E-01 | 1.13E+00 | 1.88E-04 | 8.00E-04 | -1.22E+00 | 4.94E-04 | 1.89E-03 | -1.04E+00 | 2.66E-03 | 9.49E-03 | -1.08E+00 | 2.67E-02 | 7.52E-02 | 1.15E+00 | 1.02E-02 | 3.75E-02 | 1.05E+00 | 1.91E-01 | 5.25E-01 | 1.22E+00 |
| ARL13B | 2.31E-02 | 1.20E-01 | -1.19E+00 | 2.80E-02 | 1.21E-01 | -1.05E+00 | 2.41E-04 | 1.02E-03 | -1.45E+00 | 1.27E-03 | 4.86E-03 | -1.23E+00 | 1.22E-02 | 4.34E-02 | -1.25E+00 | 2.64E-02 | 7.43E-02 | -1.30E+00 | 9.26E-08 | 3.40E-07 | -1.55E+00 | 1.71E-05 | 4.69E-05 | -1.27E+00 |
| HBP1 | 2.31E-02 | 1.20E-01 | 1.12E+00 | 4.16E-04 | 1.80E-03 | 1.01E+00 | 1.44E-01 | 6.14E-01 | 1.22E+00 | 3.42E-02 | 1.31E-01 | 1.12E+00 | 9.62E-02 | 3.43E-01 | 1.11E+00 | 1.18E-01 | 3.33E-01 | 1.08E+00 | 5.31E-04 | 1.95E-03 | -1.11E+00 | 3.86E-05 | 1.06E-04 | -1.15E+00 |
| FAAP20 | 2.31E-02 | 1.20E-01 | 1.21E+00 | 7.06E-07 | 3.05E-06 | 1.03E+00 | 1.94E-01 | 8.26E-01 | 1.10E+00 | 5.93E-02 | 2.26E-01 | 1.15E+00 | 8.61E-03 | 3.07E-02 | 1.07E+00 | 1.59E-02 | 4.49E-02 | 1.11E+00 | 1.36E-02 | 4.97E-02 | -1.04E+00 | 2.12E-02 | 5.83E-02 | 1.00E+00 |
| DUSP5 | 2.31E-02 | 1.20E-01 | 1.01E+00 | 1.64E-07 | 7.09E-07 | -3.05E+00 | 4.45E-03 | 1.89E-02 | -1.15E+00 | 8.31E-07 | 3.17E-06 | -2.43E+00 | 7.61E-03 | 2.71E-02 | -1.46E+00 | 1.06E-04 | 2.98E-04 | -2.91E+00 | 2.17E-01 | 7.98E-01 | 1.47E+00 | 1.51E-02 | 4.15E-02 | -1.25E+00 |
| PHTF2 | 2.31E-02 | 1.21E-01 | -1.01E+00 | 3.34E-02 | 1.44E-01 | 1.12E+00 | 3.87E-02 | 1.65E-01 | -1.03E+00 | 7.61E-04 | 2.91E-03 | -1.21E+00 | 5.12E-03 | 1.83E-02 | 1.03E+00 | 9.23E-05 | 2.60E-04 | -1.25E+00 | 2.16E-01 | 7.92E-01 | 1.24E+00 | 1.21E-01 | 3.32E-01 | 1.13E+00 |
| ENTPD3-AS1 | 2.32E-02 | 1.21E-01 | -1.03E+00 | 1.78E-03 | 7.71E-03 | -1.23E+00 | 5.00E-03 | 2.12E-02 | -1.44E+00 | 5.66E-04 | 2.16E-03 | -1.51E+00 | 2.46E-01 | 8.77E-01 | 1.11E+00 | 3.48E-01 | 9.80E-01 | 1.28E+00 | 1.78E-02 | 6.52E-02 | -1.08E+00 | 1.99E-02 | 5.47E-02 | -1.15E+00 |
| DGKQ | 2.32E-02 | 1.21E-01 | -1.07E+00 | 4.61E-02 | 1.99E-01 | 1.07E+00 | 1.62E-01 | 6.87E-01 | 1.15E+00 | 1.58E-01 | 6.05E-01 | 1.25E+00 | 1.87E-04 | 6.68E-04 | -1.78E+00 | 8.61E-04 | 2.43E-03 | -1.57E+00 | 7.50E-02 | 2.75E-01 | 1.03E+00 | 2.67E-01 | 7.33E-01 | 1.17E+00 |
| ZNF384 | 2.32E-02 | 1.21E-01 | -1.17E+00 | 1.02E-02 | 4.40E-02 | -1.20E+00 | 1.21E-01 | 5.14E-01 | 1.03E+00 | 9.71E-02 | 3.71E-01 | 1.05E+00 | 1.80E-01 | 6.41E-01 | 1.22E+00 | 6.27E-02 | 1.77E-01 | -1.35E+00 | 1.63E-04 | 5.99E-04 | -1.83E+00 | 2.07E-03 | 5.69E-03 | -1.50E+00 |
| PKNOX1 | 2.33E-02 | 1.21E-01 | 1.05E+00 | 1.01E-01 | 4.38E-01 | 1.35E+00 | 6.57E-06 | 2.79E-05 | -1.44E+00 | 3.53E-03 | 1.35E-02 | 1.03E+00 | 9.26E-07 | 3.30E-06 | -1.48E+00 | 7.15E-05 | 2.02E-04 | -1.28E+00 | 9.06E-04 | 3.33E-03 | -1.32E+00 | 6.76E-02 | 1.86E-01 | 1.04E+00 |
| YY2 | 2.33E-02 | 1.21E-01 | -1.23E+00 | 1.73E-02 | 7.49E-02 | -1.21E+00 | 7.81E-02 | 3.32E-01 | -1.10E+00 | 1.84E-01 | 7.04E-01 | 1.60E+00 | 7.26E-02 | 2.59E-01 | -1.26E+00 | 3.12E-01 | 8.79E-01 | 1.16E+00 | 6.79E-03 | 2.49E-02 | -1.19E+00 | 2.24E-02 | 6.14E-02 | -1.12E+00 |
| BACH2 | 2.33E-02 | 1.22E-01 | -1.01E+00 | 1.94E-01 | 8.40E-01 | 1.52E+00 | 1.15E-01 | 4.87E-01 | -1.09E+00 | 1.78E-01 | 6.80E-01 | 1.16E+00 | 2.71E-01 | 9.66E-01 | 1.15E+00 | 2.82E-01 | 7.95E-01 | 1.38E+00 | 7.56E-02 | 2.78E-01 | -1.13E+00 | 2.62E-01 | 7.19E-01 | 1.08E+00 |
| PRKG1 | 2.33E-02 | 1.22E-01 | 1.18E+00 | 3.99E-02 | 1.72E-01 | 1.75E+00 | 1.76E-01 | 7.49E-01 | 1.26E+00 | 3.39E-02 | 1.30E-01 | 1.89E+00 | 9.01E-02 | 3.21E-01 | 1.66E+00 | 3.53E-02 | 9.96E-02 | 1.82E+00 | 3.04E-03 | 1.12E-02 | 1.80E+00 | 2.85E-02 | 7.83E-02 | 1.62E+00 |
| AK1 | 2.34E-02 | 1.22E-01 | -1.44E+00 | 1.43E-01 | 6.17E-01 | 1.09E+00 | 2.25E-01 | 9.56E-01 | 1.04E+00 | 2.19E-01 | 8.37E-01 | 1.21E+00 | 4.78E-02 | 1.71E-01 | -1.31E+00 | 1.09E-01 | 3.08E-01 | -1.03E+00 | 8.58E-03 | 3.15E-02 | -1.24E+00 | 1.28E-02 | 3.51E-02 | -1.22E+00 |
| RPL18AP16 | 2.34E-02 | 1.22E-01 | -1.01E+00 | 1.16E-02 | 5.00E-02 | 1.05E+00 | 2.23E-02 | 9.49E-02 | 1.79E+00 | 2.22E-01 | 8.46E-01 | 1.41E+00 | 5.42E-02 | 1.93E-01 | 2.05E+00 | 3.11E-01 | 8.77E-01 | 1.31E+00 | 2.30E-01 | 8.46E-01 | 1.12E+00 | 2.69E-01 | 7.39E-01 | 1.08E+00 |
| NFKB1 | 2.34E-02 | 1.22E-01 | 1.04E+00 | 9.38E-03 | 4.05E-02 | 1.13E+00 | 6.00E-03 | 2.55E-02 | -1.02E+00 | 6.23E-04 | 2.38E-03 | 1.00E+00 | 9.54E-04 | 3.40E-03 | -1.41E+00 | 1.46E-02 | 4.12E-02 | -1.20E+00 | 1.35E-05 | 4.97E-05 | -1.21E+00 | 5.40E-02 | 1.48E-01 | 1.09E+00 |
| TNFSF4 | 2.34E-02 | 1.22E-01 | 1.10E+00 | 1.25E-01 | 5.41E-01 | 1.50E+00 | 3.31E-03 | 1.41E-02 | -1.32E+00 | 1.67E-02 | 6.36E-02 | -1.07E+00 | 6.92E-02 | 2.47E-01 | -1.43E+00 | 1.72E-01 | 4.84E-01 | -1.01E+00 | 1.23E-01 | 4.50E-01 | 2.13E+00 | 1.68E-01 | 4.62E-01 | 1.79E+00 |
| EIF5 | 2.34E-02 | 1.22E-01 | 1.04E+00 | 6.58E-06 | 2.84E-05 | 1.01E+00 | 9.78E-03 | 4.16E-02 | -1.08E+00 | 3.70E-02 | 1.41E-01 | 1.07E+00 | 1.70E-06 | 6.06E-06 | -1.06E+00 | 3.41E-03 | 9.62E-03 | -1.02E+00 | 2.95E-02 | 1.08E-01 | 1.03E+00 | 1.10E-01 | 3.03E-01 | 1.28E+00 |
| ARHGEF40 | 2.34E-02 | 1.22E-01 | 1.80E+00 | 2.20E-02 | 9.50E-02 | 1.25E+00 | 7.62E-02 | 3.24E-01 | 1.25E+00 | 2.95E-05 | 1.13E-04 | -1.09E+00 | 2.07E-02 | 7.39E-02 | -1.07E+00 | 2.01E-03 | 5.66E-03 | -1.52E+00 | 7.59E-02 | 2.79E-01 | -1.11E+00 | 5.01E-02 | 1.38E-01 | -1.52E+00 |
| COL9A2 | 2.34E-02 | 1.22E-01 | 1.09E+00 | 1.88E-03 | 8.14E-03 | -1.05E+00 | 1.09E-01 | 4.64E-01 | 1.11E+00 | 2.56E-01 | 9.79E-01 | 1.42E+00 | 1.35E-02 | 4.80E-02 | -1.06E+00 | 6.02E-02 | 1.70E-01 | 1.08E+00 | 2.64E-01 | 9.68E-01 | 1.18E+00 | 3.31E-01 | 9.09E-01 | 1.27E+00 |
| DPY19L1P1 | 2.35E-02 | 1.22E-01 | -1.23E+00 | 1.74E-02 | 7.52E-02 | -1.22E+00 | 1.03E-04 | 4.40E-04 | -1.70E+00 | 8.29E-04 | 3.17E-03 | -1.26E+00 | 1.68E-02 | 6.00E-02 | -1.11E+00 | 7.84E-03 | 2.21E-02 | -1.27E+00 | 7.41E-02 | 2.72E-01 | 1.04E+00 | 4.21E-02 | 1.16E-01 | -1.17E+00 |
| PRPF18 | 2.36E-02 | 1.23E-01 | 1.17E+00 | 7.10E-04 | 3.07E-03 | 1.08E+00 | 3.41E-02 | 1.45E-01 | 1.09E+00 | 7.12E-05 | 2.72E-04 | -1.10E+00 | 1.85E-01 | 6.60E-01 | 1.33E+00 | 1.48E-01 | 4.18E-01 | 1.29E+00 | 1.09E-04 | 4.01E-04 | -1.17E+00 | 2.51E-05 | 6.91E-05 | -1.24E+00 |
| MERTK | 2.36E-02 | 1.23E-01 | -1.04E+00 | 2.25E-01 | 9.74E-01 | 1.54E+00 | 2.31E-02 | 9.80E-02 | -1.23E+00 | 8.44E-02 | 3.22E-01 | 1.08E+00 | 1.31E-01 | 4.67E-01 | 1.15E+00 | 1.36E-01 | 3.83E-01 | -1.02E+00 | 1.95E-01 | 7.14E-01 | 1.35E+00 | 3.47E-01 | 9.52E-01 | 1.22E+00 |
| SLF1 | 2.36E-02 | 1.23E-01 | 1.03E+00 | 4.09E-02 | 1.77E-01 | 1.15E+00 | 2.22E-02 | 9.41E-02 | -1.05E+00 | 2.81E-02 | 1.07E-01 | 1.10E+00 | 7.09E-04 | 2.53E-03 | -1.19E+00 | 2.34E-03 | 6.61E-03 | -1.11E+00 | 5.04E-03 | 1.85E-02 | 1.01E+00 | 1.99E-01 | 5.46E-01 | 1.24E+00 |
| CSNK1G2 | 2.36E-02 | 1.23E-01 | 1.17E+00 | 4.41E-02 | 1.90E-01 | 1.35E+00 | 2.58E-02 | 1.10E-01 | 1.07E+00 | 9.98E-02 | 3.81E-01 | 1.36E+00 | 2.20E-03 | 7.84E-03 | 1.09E+00 | 2.28E-02 | 6.44E-02 | 1.21E+00 | 1.83E-02 | 6.73E-02 | 1.08E+00 | 6.16E-02 | 1.69E-01 | 1.15E+00 |
| CXCL3 | 2.36E-02 | 1.23E-01 | 1.94E+00 | 1.73E-02 | 7.49E-02 | -1.04E+00 | 4.71E-04 | 2.00E-03 | 2.91E+00 | 1.92E-01 | 7.33E-01 | 1.29E+00 | 2.09E-01 | 7.46E-01 | -1.32E+00 | 8.05E-08 | 2.27E-07 | -7.00E+00 | 1.00E-01 | 3.69E-01 | -1.10E+00 | 1.73E-03 | 4.75E-03 | -2.09E+00 |
| AC007066.2 | 2.36E-02 | 1.23E-01 | -1.16E+00 | 3.36E-02 | 1.45E-01 | 1.85E+00 | 2.30E-01 | 9.79E-01 | 1.19E+00 | 8.30E-02 | 3.17E-01 | 1.83E+00 | 2.41E-01 | 8.59E-01 | -1.01E+00 | 2.78E-01 | 7.83E-01 | 1.07E+00 | 2.64E-01 | 9.69E-01 | 1.11E+00 | 1.35E-01 | 3.71E-01 | 1.50E+00 |
| FRRS1 | 2.36E-02 | 1.23E-01 | -1.29E+00 | 2.70E-02 | 1.17E-01 | -1.27E+00 | 5.03E-02 | 2.14E-01 | -1.25E+00 | 8.21E-02 | 3.14E-01 | -1.04E+00 | 4.22E-05 | 1.51E-04 | -1.35E+00 | 1.27E-05 | 3.58E-05 | -1.48E+00 | 4.15E-03 | 1.52E-02 | -1.12E+00 | 1.98E-04 | 5.45E-04 | -1.27E+00 |
| MT-TT | 2.37E-02 | 1.24E-01 | 2.29E+00 | 2.21E-01 | 9.53E-01 | 1.39E+00 | 1.30E-01 | 5.53E-01 | 1.26E+00 | 4.49E-03 | 1.72E-02 | -1.47E+00 | 1.00E-01 | 3.58E-01 | 1.78E+00 | 1.75E-01 | 4.94E-01 | 1.08E+00 | 4.03E-02 | 1.48E-01 | 1.55E+00 | 1.42E-01 | 3.90E-01 | 1.13E+00 |
| AL691432.2 | 2.37E-02 | 1.24E-01 | 1.01E+00 | 1.98E-02 | 8.55E-02 | 1.09E+00 | 1.30E-02 | 5.50E-02 | -1.14E+00 | 3.59E-03 | 1.37E-02 | -1.05E+00 | 4.89E-02 | 1.74E-01 | 1.13E+00 | 1.57E-02 | 4.42E-02 | -1.09E+00 | 2.72E-02 | 9.99E-02 | -1.13E+00 | 2.76E-03 | 7.58E-03 | -1.39E+00 |
| RTRAF | 2.37E-02 | 1.24E-01 | 1.03E+00 | 3.62E-03 | 1.56E-02 | 1.07E+00 | 1.91E-01 | 8.11E-01 | 1.07E+00 | 2.53E-01 | 9.65E-01 | 1.07E+00 | 6.79E-03 | 2.42E-02 | -1.01E+00 | 2.61E-01 | 7.37E-01 | 1.09E+00 | 1.34E-01 | 4.92E-01 | 1.11E+00 | 1.27E-01 | 3.48E-01 | 1.12E+00 |
| ATG4D | 2.37E-02 | 1.24E-01 | -1.22E+00 | 1.07E-01 | 4.63E-01 | 1.15E+00 | 4.11E-03 | 1.75E-02 | -1.38E+00 | 4.31E-02 | 1.65E-01 | -1.01E+00 | 3.57E-05 | 1.27E-04 | -2.18E+00 | 3.23E-03 | 9.11E-03 | -1.36E+00 | 1.25E-03 | 4.60E-03 | -1.16E+00 | 5.02E-02 | 1.38E-01 | -1.04E+00 |
| RALY-AS1 | 2.38E-02 | 1.24E-01 | -1.12E+00 | 1.55E-03 | 6.72E-03 | -1.41E+00 | 9.41E-03 | 4.00E-02 | -1.22E+00 | 4.56E-04 | 1.74E-03 | -1.49E+00 | 1.64E-01 | 5.86E-01 | 1.09E+00 | 3.50E-01 | 9.86E-01 | 1.39E+00 | 9.81E-03 | 3.60E-02 | -1.25E+00 | 6.42E-03 | 1.76E-02 | -1.39E+00 |
| CRTC1 | 2.38E-02 | 1.24E-01 | 1.07E+00 | 9.97E-02 | 4.31E-01 | 1.31E+00 | 1.46E-02 | 6.20E-02 | -1.02E+00 | 1.06E-01 | 4.05E-01 | 1.31E+00 | 2.46E-03 | 8.78E-03 | -1.24E+00 | 2.68E-02 | 7.55E-02 | 1.07E+00 | 2.71E-01 | 9.95E-01 | 1.32E+00 | 3.04E-01 | 8.36E-01 | 1.23E+00 |
| SPDYA | 2.38E-02 | 1.24E-01 | 1.77E+00 | 5.19E-02 | 2.24E-01 | -1.14E+00 | 7.01E-02 | 2.98E-01 | 1.64E+00 | 2.57E-01 | 9.83E-01 | 1.18E+00 | 1.95E-01 | 6.95E-01 | 1.05E+00 | 5.95E-02 | 1.68E-01 | -1.56E+00 | 4.76E-02 | 1.75E-01 | 1.67E+00 | 2.53E-01 | 6.94E-01 | 1.02E+00 |
| AC009093.2 | 2.38E-02 | 1.24E-01 | -1.18E+00 | 1.59E-01 | 6.85E-01 | 1.30E+00 | 1.54E-02 | 6.53E-02 | -1.35E+00 | 1.95E-01 | 7.46E-01 | 1.24E+00 | 1.78E-05 | 6.35E-05 | -1.79E+00 | 1.72E-03 | 4.86E-03 | -1.24E+00 | 4.86E-08 | 1.79E-07 | -2.13E+00 | 2.69E-03 | 7.40E-03 | -1.31E+00 |
| MYO1F | 2.39E-02 | 1.25E-01 | 1.14E+00 | 2.31E-01 | 1.00E+00 | 1.47E+00 | 7.19E-04 | 3.05E-03 | -1.10E+00 | 1.28E-02 | 4.88E-02 | 1.14E+00 | 4.12E-09 | 1.47E-08 | -1.31E+00 | 5.03E-07 | 1.42E-06 | -1.17E+00 | 1.65E-04 | 6.07E-04 | -1.43E+00 | 2.06E-03 | 5.65E-03 | -1.26E+00 |
| SCAMP5 | 2.40E-02 | 1.25E-01 | -1.11E+00 | 1.14E-02 | 4.93E-02 | -1.13E+00 | 1.77E-03 | 7.51E-03 | -1.50E+00 | 9.50E-03 | 3.63E-02 | -1.19E+00 | 7.63E-02 | 2.72E-01 | -1.30E+00 | 5.62E-02 | 1.58E-01 | -1.56E+00 | 2.39E-01 | 8.75E-01 | 1.21E+00 | 3.30E-01 | 9.07E-01 | 1.19E+00 |
| RHOA | 2.40E-02 | 1.25E-01 | 1.09E+00 | 2.19E-01 | 9.47E-01 | 1.17E+00 | 4.46E-02 | 1.90E-01 | -1.07E+00 | 1.88E-01 | 7.19E-01 | 1.03E+00 | 2.50E-05 | 8.91E-05 | -1.14E+00 | 7.53E-02 | 2.12E-01 | -1.02E+00 | 2.25E-01 | 8.27E-01 | 1.03E+00 | 2.09E-01 | 5.75E-01 | 1.08E+00 |
| RBM47 | 2.40E-02 | 1.25E-01 | -1.41E+00 | 2.83E-02 | 1.22E-01 | -1.36E+00 | 1.13E-01 | 4.81E-01 | -1.41E+00 | 1.61E-02 | 6.16E-02 | -2.00E+00 | 1.77E-01 | 6.32E-01 | -1.20E+00 | 2.63E-01 | 7.42E-01 | -1.10E+00 | 8.14E-02 | 2.99E-01 | -1.31E+00 | 2.19E-01 | 6.02E-01 | -1.10E+00 |
| AC067817.1 | 2.40E-02 | 1.25E-01 | 1.85E+00 | 7.45E-02 | 3.22E-01 | -1.20E+00 | 2.36E-04 | 1.00E-03 | 2.68E+00 | 1.96E-01 | 7.50E-01 | -1.03E+00 | 4.90E-06 | 1.75E-05 | 5.70E+00 | 1.71E-01 | 4.81E-01 | 2.01E+00 | 8.69E-09 | 3.19E-08 | 4.16E+00 | 2.99E-01 | 8.21E-01 | 1.24E+00 |
| AL354714.2 | 2.40E-02 | 1.25E-01 | -1.22E+00 | 7.38E-03 | 3.19E-02 | -1.35E+00 | 4.26E-02 | 1.81E-01 | -1.04E+00 | 1.40E-03 | 5.33E-03 | -1.20E+00 | 2.72E-02 | 9.69E-02 | -1.13E+00 | 8.72E-03 | 2.46E-02 | -1.29E+00 | 6.84E-08 | 2.51E-07 | -1.62E+00 | 1.95E-07 | 5.36E-07 | -1.57E+00 |
| LCMT2 | 2.40E-02 | 1.25E-01 | -1.18E+00 | 6.22E-03 | 2.69E-02 | -1.35E+00 | 2.74E-03 | 1.16E-02 | -1.37E+00 | 6.55E-03 | 2.50E-02 | -1.21E+00 | 8.54E-02 | 3.05E-01 | -1.12E+00 | 8.08E-02 | 2.28E-01 | -1.18E+00 | 6.76E-03 | 2.48E-02 | -1.21E+00 | 1.02E-01 | 2.80E-01 | 1.03E+00 |
| GINS2 | 2.40E-02 | 1.25E-01 | 2.50E+00 | 1.61E-01 | 6.94E-01 | 1.82E+00 | 2.41E-07 | 1.02E-06 | 1.59E+00 | 7.12E-03 | 2.72E-02 | 1.50E+00 | 2.14E-01 | 7.63E-01 | -1.12E+00 | 2.09E-01 | 5.88E-01 | -1.35E+00 | 2.52E-01 | 9.27E-01 | 1.07E+00 | 1.29E-05 | 3.53E-05 | 1.61E+00 |
| MAFTRR | 2.40E-02 | 1.25E-01 | -1.26E+00 | 1.93E-01 | 8.33E-01 | 1.18E+00 | 1.86E-07 | 7.91E-07 | -2.28E+00 | 2.85E-05 | 1.09E-04 | -1.48E+00 | 3.73E-02 | 1.33E-01 | -1.24E+00 | 2.33E-01 | 6.56E-01 | 1.31E+00 | 2.25E-01 | 8.24E-01 | 1.35E+00 | 1.02E-01 | 2.81E-01 | 1.03E+00 |
| CHRAC1 | 2.40E-02 | 1.25E-01 | 1.13E+00 | 1.32E-06 | 5.69E-06 | -1.11E+00 | 1.29E-01 | 5.46E-01 | 1.15E+00 | 7.14E-03 | 2.73E-02 | 1.13E+00 | 7.95E-03 | 2.83E-02 | -1.12E+00 | 1.95E-03 | 5.50E-03 | -1.23E+00 | 4.89E-06 | 1.80E-05 | -1.18E+00 | 1.49E-04 | 4.09E-04 | -1.13E+00 |
| AC069544.1 | 2.41E-02 | 1.26E-01 | -1.17E+00 | 5.34E-02 | 2.31E-01 | 1.04E+00 | 8.64E-04 | 3.67E-03 | -1.42E+00 | 7.80E-04 | 2.98E-03 | -1.33E+00 | 1.42E-01 | 5.05E-01 | 1.87E+00 | 8.07E-02 | 2.27E-01 | 2.32E+00 | 6.60E-02 | 2.42E-01 | -1.37E+00 | 1.38E-01 | 3.78E-01 | -1.26E+00 |
| FBXO5 | 2.41E-02 | 1.26E-01 | -1.31E+00 | 1.11E-01 | 4.79E-01 | 1.11E+00 | 9.62E-02 | 4.09E-01 | 1.03E+00 | 2.51E-01 | 9.59E-01 | 1.27E+00 | 2.34E-01 | 8.34E-01 | 1.27E+00 | 3.50E-01 | 9.85E-01 | 1.52E+00 | 1.51E-08 | 5.55E-08 | -1.70E+00 | 3.32E-03 | 9.11E-03 | -1.16E+00 |
| RANBP9 | 2.41E-02 | 1.26E-01 | 1.09E+00 | 5.41E-04 | 2.34E-03 | 1.01E+00 | 3.64E-02 | 1.55E-01 | 1.12E+00 | 4.93E-06 | 1.88E-05 | -1.18E+00 | 5.63E-03 | 2.01E-02 | -1.08E+00 | 2.67E-03 | 7.53E-03 | -1.14E+00 | 1.19E-05 | 4.37E-05 | -1.20E+00 | 7.28E-05 | 2.00E-04 | -1.17E+00 |
| AC022413.1 | 2.42E-02 | 1.26E-01 | -1.02E+00 | 1.54E-02 | 6.65E-02 | -1.02E+00 | 5.87E-02 | 2.49E-01 | -1.01E+00 | 5.39E-02 | 2.06E-01 | 1.04E+00 | 4.24E-02 | 1.51E-01 | -1.02E+00 | 1.13E-01 | 3.20E-01 | 1.14E+00 | 1.53E-01 | 5.62E-01 | 1.15E+00 | 2.67E-01 | 7.32E-01 | 1.27E+00 |
| SHMT1 | 2.42E-02 | 1.26E-01 | -1.09E+00 | 3.25E-02 | 1.40E-01 | 1.08E+00 | 8.78E-04 | 3.73E-03 | -1.11E+00 | 1.50E-03 | 5.74E-03 | 1.02E+00 | 1.31E-02 | 4.66E-02 | -1.39E+00 | 4.50E-02 | 1.27E-01 | -1.16E+00 | 4.49E-03 | 1.65E-02 | -1.03E+00 | 3.25E-01 | 8.93E-01 | 1.30E+00 |
| LARS2 | 2.42E-02 | 1.26E-01 | -1.01E+00 | 2.15E-01 | 9.30E-01 | 1.52E+00 | 1.25E-01 | 5.32E-01 | 1.13E+00 | 1.47E-01 | 5.63E-01 | 1.27E+00 | 2.97E-02 | 1.06E-01 | -1.30E+00 | 2.17E-01 | 6.12E-01 | 1.12E+00 | 1.19E-02 | 4.37E-02 | -1.04E+00 | 2.86E-01 | 7.87E-01 | 1.29E+00 |
| NACAP1 | 2.42E-02 | 1.26E-01 | 1.76E+00 | 1.97E-01 | 8.50E-01 | 1.57E+00 | 2.11E-01 | 8.98E-01 | 1.29E+00 | 2.12E-01 | 8.11E-01 | 1.40E+00 | 2.55E-01 | 9.09E-01 | 1.26E+00 | 1.71E-01 | 4.81E-01 | 1.56E+00 | 1.97E-01 | 7.24E-01 | 1.20E+00 | 2.94E-01 | 8.09E-01 | 1.22E+00 |
| NRF1 | 2.42E-02 | 1.26E-01 | -1.20E+00 | 1.16E-01 | 5.03E-01 | 1.21E+00 | 8.60E-02 | 3.65E-01 | 1.09E+00 | 1.97E-01 | 7.52E-01 | 1.36E+00 | 1.78E-03 | 6.35E-03 | -1.50E+00 | 2.28E-03 | 6.44E-03 | -1.41E+00 | 1.65E-04 | 6.07E-04 | -1.34E+00 | 1.36E-01 | 3.73E-01 | 1.22E+00 |
| PARD3 | 2.42E-02 | 1.26E-01 | 1.07E+00 | 1.47E-01 | 6.35E-01 | 1.38E+00 | 6.60E-05 | 2.80E-04 | -1.37E+00 | 4.99E-03 | 1.91E-02 | 1.07E+00 | 1.47E-01 | 5.25E-01 | -1.63E+00 | 3.16E-01 | 8.90E-01 | -1.09E+00 | 9.31E-02 | 3.42E-01 | 1.57E+00 | 3.34E-01 | 9.19E-01 | 1.35E+00 |
| KIAA1958 | 2.43E-02 | 1.26E-01 | -1.03E+00 | 5.33E-02 | 2.30E-01 | 1.23E+00 | 2.90E-04 | 1.23E-03 | -1.22E+00 | 1.83E-03 | 6.97E-03 | 1.01E+00 | 1.97E-03 | 7.02E-03 | -1.30E+00 | 7.09E-03 | 2.00E-02 | -1.15E+00 | 3.49E-02 | 1.28E-01 | -1.29E+00 | 1.51E-01 | 4.14E-01 | 1.04E+00 |
| TREML2 | 2.43E-02 | 1.27E-01 | -1.45E+00 | 1.86E-02 | 8.06E-02 | -1.57E+00 | 2.28E-01 | 9.68E-01 | 1.13E+00 | 2.59E-01 | 9.88E-01 | 1.19E+00 | 2.00E-07 | 7.13E-07 | -2.33E+00 | 4.57E-06 | 1.29E-05 | -1.83E+00 | 3.13E-09 | 1.15E-08 | -1.38E+00 | 1.48E-09 | 4.08E-09 | -1.68E+00 |
| VCPKMT | 2.43E-02 | 1.27E-01 | -1.02E+00 | 1.34E-02 | 5.77E-02 | 1.02E+00 | 6.22E-04 | 2.64E-03 | -1.15E+00 | 2.79E-04 | 1.06E-03 | -1.11E+00 | 8.98E-02 | 3.20E-01 | 1.17E+00 | 3.51E-02 | 9.91E-02 | 1.03E+00 | 2.05E-02 | 7.52E-02 | 1.01E+00 | 3.50E-03 | 9.61E-03 | -1.15E+00 |
| RPL21AP69 | 2.43E-02 | 1.27E-01 | 1.19E+00 | 9.78E-22 | 4.23E-21 | -1.47E+00 | 3.16E-02 | 1.34E-01 | 1.10E+00 | 5.68E-18 | 2.17E-17 | -1.57E+00 | 6.66E-02 | 2.37E-01 | 1.25E+00 | 8.07E-10 | 2.28E-09 | -1.42E+00 | 2.53E-01 | 9.28E-01 | 1.22E+00 | 1.65E-05 | 4.52E-05 | -1.26E+00 |
| AC027702.1 | 2.43E-02 | 1.27E-01 | -1.16E+00 | 7.55E-02 | 3.26E-01 | 1.15E+00 | 4.82E-02 | 2.05E-01 | -1.29E+00 | 1.84E-01 | 7.02E-01 | 1.15E+00 | 7.47E-02 | 2.66E-01 | -1.44E+00 | 1.70E-01 | 4.80E-01 | -1.14E+00 | 4.35E-06 | 1.60E-05 | -1.60E+00 | 3.32E-03 | 9.12E-03 | -1.20E+00 |
| WASH2P | 2.43E-02 | 1.27E-01 | 1.15E+00 | 2.56E-11 | 1.11E-10 | -1.08E+00 | 2.29E-01 | 9.71E-01 | 1.24E+00 | 1.04E-03 | 3.96E-03 | 1.02E+00 | 1.27E-03 | 4.54E-03 | 1.61E+00 | 2.52E-01 | 7.10E-01 | 1.18E+00 | 1.94E-03 | 7.12E-03 | 1.55E+00 | 3.05E-02 | 8.39E-02 | 1.04E+00 |
| CDC25B | 2.43E-02 | 1.27E-01 | -1.27E+00 | 2.92E-02 | 1.26E-01 | -1.22E+00 | 1.93E-01 | 8.19E-01 | 1.10E+00 | 1.52E-01 | 5.81E-01 | 1.19E+00 | 1.90E-01 | 6.78E-01 | -1.21E+00 | 3.49E-01 | 9.85E-01 | 1.44E+00 | 4.52E-02 | 1.66E-01 | -1.07E+00 | 3.39E-01 | 9.31E-01 | 1.23E+00 |
| HIST1H4A | 2.43E-02 | 1.27E-01 | 2.10E+00 | 2.23E-01 | 9.64E-01 | 1.35E+00 | 3.49E-02 | 1.48E-01 | 1.67E+00 | 1.37E-01 | 5.22E-01 | 1.52E+00 | 2.71E-01 | 9.68E-01 | 1.22E+00 | 2.95E-01 | 8.31E-01 | 1.42E+00 | 1.84E-01 | 6.77E-01 | 1.46E+00 | 2.35E-01 | 6.45E-01 | 1.32E+00 |
| EAPP | 2.44E-02 | 1.27E-01 | 1.12E+00 | 1.09E-04 | 4.73E-04 | 1.07E+00 | 1.62E-01 | 6.89E-01 | 1.20E+00 | 1.73E-02 | 6.59E-02 | 1.15E+00 | 2.05E-03 | 7.31E-03 | 1.08E+00 | 4.18E-03 | 1.18E-02 | 1.09E+00 | 2.26E-01 | 8.29E-01 | 1.13E+00 | 1.34E-02 | 3.67E-02 | -1.06E+00 |
| OPA3 | 2.44E-02 | 1.27E-01 | -1.06E+00 | 5.87E-02 | 2.54E-01 | 1.11E+00 | 3.95E-04 | 1.68E-03 | -1.51E+00 | 8.86E-03 | 3.38E-02 | -1.12E+00 | 1.94E-06 | 6.92E-06 | -1.91E+00 | 2.45E-04 | 6.90E-04 | -1.28E+00 | 1.81E-08 | 6.65E-08 | -1.77E+00 | 3.05E-03 | 8.39E-03 | -1.13E+00 |
| SSH3 | 2.44E-02 | 1.27E-01 | 1.08E+00 | 8.47E-02 | 3.66E-01 | 1.44E+00 | 5.53E-03 | 2.35E-02 | -1.17E+00 | 2.56E-02 | 9.77E-02 | 1.08E+00 | 2.59E-02 | 9.23E-02 | -1.21E+00 | 1.81E-01 | 5.11E-01 | 1.13E+00 | 1.46E-01 | 5.35E-01 | 1.18E+00 | 9.26E-02 | 2.54E-01 | -1.04E+00 |
| MGAT5 | 2.45E-02 | 1.28E-01 | -1.07E+00 | 6.06E-02 | 2.62E-01 | 1.12E+00 | 7.91E-02 | 3.36E-01 | -1.12E+00 | 2.36E-01 | 9.01E-01 | 1.30E+00 | 1.48E-01 | 5.28E-01 | 1.19E+00 | 1.35E-01 | 3.79E-01 | 1.20E+00 | 9.90E-02 | 3.64E-01 | 1.10E+00 | 1.98E-01 | 5.43E-01 | 1.50E+00 |
| PLCB3 | 2.45E-02 | 1.28E-01 | -1.10E+00 | 1.80E-01 | 7.77E-01 | 1.38E+00 | 2.21E-01 | 9.39E-01 | 1.14E+00 | 2.34E-01 | 8.96E-01 | 1.36E+00 | 1.31E-02 | 4.66E-02 | -1.13E+00 | 5.40E-03 | 1.52E-02 | -1.32E+00 | 1.09E-05 | 4.00E-05 | -1.54E+00 | 6.77E-02 | 1.86E-01 | 1.10E+00 |
| AC090058.1 | 2.45E-02 | 1.28E-01 | 1.05E+00 | 9.67E-06 | 4.18E-05 | -1.56E+00 | 7.42E-02 | 3.15E-01 | 1.51E+00 | 5.40E-03 | 2.06E-02 | -1.19E+00 | 1.71E-01 | 6.09E-01 | 1.43E+00 | 8.05E-03 | 2.27E-02 | -1.74E+00 | 9.08E-02 | 3.33E-01 | 1.41E+00 | 4.75E-02 | 1.31E-01 | -1.29E+00 |
| AC023449.2 | 2.46E-02 | 1.28E-01 | -1.43E+00 | 6.56E-02 | 2.84E-01 | -1.11E+00 | 3.24E-02 | 1.38E-01 | -1.23E+00 | 2.02E-01 | 7.71E-01 | 1.24E+00 | 8.91E-02 | 3.18E-01 | 1.77E+00 | 1.70E-01 | 4.80E-01 | 1.66E+00 | 2.50E-01 | 9.17E-01 | 1.20E+00 | 3.11E-01 | 8.54E-01 | 1.06E+00 |
| L3MBTL1 | 2.47E-02 | 1.29E-01 | -1.02E+00 | 1.55E-01 | 6.69E-01 | 1.43E+00 | 3.42E-03 | 1.45E-02 | -1.48E+00 | 2.00E-02 | 7.64E-02 | -1.15E+00 | 1.81E-01 | 6.45E-01 | 1.48E+00 | 1.51E-01 | 4.26E-01 | 1.75E+00 | 1.94E-01 | 7.13E-01 | 1.15E+00 | 2.61E-01 | 7.18E-01 | 1.06E+00 |
| ZNF215 | 2.47E-02 | 1.29E-01 | -1.22E+00 | 4.26E-02 | 1.84E-01 | -1.02E+00 | 1.25E-01 | 5.33E-01 | -1.05E+00 | 1.67E-01 | 6.37E-01 | 1.09E+00 | 3.22E-02 | 1.15E-01 | -1.94E+00 | 1.08E-01 | 3.03E-01 | -1.32E+00 | 2.29E-03 | 8.39E-03 | -1.22E+00 | 2.43E-01 | 6.68E-01 | 1.42E+00 |
| ZNF440 | 2.47E-02 | 1.29E-01 | 1.02E+00 | 1.30E-02 | 5.62E-02 | 1.06E+00 | 1.38E-04 | 5.84E-04 | -1.49E+00 | 2.26E-04 | 8.65E-04 | -1.24E+00 | 2.62E-01 | 9.36E-01 | 1.42E+00 | 1.32E-01 | 3.73E-01 | 1.16E+00 | 7.05E-04 | 2.59E-03 | -1.21E+00 | 7.10E-04 | 1.95E-03 | -1.25E+00 |
| AC092368.3 | 2.48E-02 | 1.29E-01 | -1.04E+00 | 1.86E-02 | 8.03E-02 | -1.05E+00 | 1.44E-01 | 6.12E-01 | 1.09E+00 | 3.81E-02 | 1.45E-01 | -1.07E+00 | 1.74E-01 | 6.20E-01 | 1.17E+00 | 1.22E-01 | 3.45E-01 | -1.18E+00 | 4.15E-02 | 1.52E-01 | -1.30E+00 | 6.73E-02 | 1.85E-01 | -1.26E+00 |
| TMC3-AS1 | 2.48E-02 | 1.29E-01 | -1.30E+00 | 1.20E-01 | 5.18E-01 | 1.15E+00 | 1.11E-01 | 4.71E-01 | -1.05E+00 | 1.89E-01 | 7.21E-01 | 1.20E+00 | 7.71E-03 | 2.75E-02 | -1.70E+00 | 1.28E-02 | 3.59E-02 | -1.64E+00 | 6.69E-04 | 2.45E-03 | -1.57E+00 | 6.65E-03 | 1.83E-02 | -1.38E+00 |
| NBAT1 | 2.48E-02 | 1.29E-01 | -1.11E+00 | 9.22E-03 | 3.99E-02 | -1.23E+00 | 9.22E-02 | 3.92E-01 | -1.06E+00 | 2.53E-01 | 9.67E-01 | 1.36E+00 | 1.42E-02 | 5.05E-02 | -1.24E+00 | 4.61E-03 | 1.30E-02 | -1.49E+00 | 2.19E-01 | 8.03E-01 | 1.30E+00 | 2.74E-01 | 7.52E-01 | -1.19E+00 |
| PRSS27 | 2.49E-02 | 1.30E-01 | -1.29E+00 | 1.85E-02 | 8.00E-02 | -1.35E+00 | 1.29E-02 | 5.48E-02 | 2.00E+00 | 8.66E-02 | 3.31E-01 | 1.67E+00 | 2.46E-02 | 8.76E-02 | -1.28E+00 | 1.19E-02 | 3.35E-02 | -1.58E+00 | 5.26E-04 | 1.93E-03 | -1.92E+00 | 6.98E-03 | 1.92E-02 | -1.52E+00 |
| PKIG | 2.49E-02 | 1.30E-01 | 1.17E+00 | 3.74E-03 | 1.61E-02 | 1.14E+00 | 2.69E-03 | 1.14E-02 | -1.06E+00 | 3.59E-03 | 1.37E-02 | 1.08E+00 | 1.12E-02 | 3.99E-02 | 1.58E+00 | 5.75E-04 | 1.62E-03 | 1.82E+00 | 2.67E-01 | 9.81E-01 | 1.16E+00 | 3.02E-01 | 8.30E-01 | 1.22E+00 |
| NCAPGP1 | 2.49E-02 | 1.30E-01 | -1.21E+00 | 1.79E-01 | 7.73E-01 | 1.58E+00 | 6.49E-04 | 2.76E-03 | -1.68E+00 | 9.95E-03 | 3.80E-02 | -1.15E+00 | 6.05E-03 | 2.16E-02 | -1.61E+00 | 1.01E-02 | 2.85E-02 | -1.40E+00 | 5.53E-02 | 2.03E-01 | -1.27E+00 | 2.90E-01 | 7.97E-01 | 1.21E+00 |
| AL445189.2 | 2.49E-02 | 1.30E-01 | 1.07E+00 | 2.43E-04 | 1.05E-03 | -1.20E+00 | 5.43E-02 | 2.31E-01 | 1.03E+00 | 6.01E-09 | 2.29E-08 | -1.75E+00 | 3.87E-02 | 1.38E-01 | -1.01E+00 | 6.82E-03 | 1.92E-02 | -1.19E+00 | 1.02E-02 | 3.74E-02 | -1.00E+00 | 1.21E-10 | 3.32E-10 | -1.68E+00 |
| HELQ | 2.50E-02 | 1.30E-01 | 1.06E+00 | 6.01E-02 | 2.60E-01 | 1.32E+00 | 6.15E-03 | 2.62E-02 | -1.13E+00 | 2.78E-02 | 1.06E-01 | 1.13E+00 | 2.23E-01 | 7.96E-01 | 1.39E+00 | 2.60E-01 | 7.32E-01 | 1.49E+00 | 7.35E-02 | 2.70E-01 | 1.17E+00 | 7.20E-02 | 1.98E-01 | 1.11E+00 |
| LINC02021 | 2.50E-02 | 1.30E-01 | 1.02E+00 | 2.36E-03 | 1.02E-02 | -1.09E+00 | 6.12E-03 | 2.60E-02 | -1.32E+00 | 1.81E-02 | 6.92E-02 | -1.11E+00 | 7.57E-02 | 2.70E-01 | -1.18E+00 | 1.78E-01 | 5.00E-01 | 1.01E+00 | 1.53E-01 | 5.62E-01 | 1.56E+00 | 3.51E-01 | 9.64E-01 | 1.33E+00 |
| EGOT | 2.50E-02 | 1.31E-01 | -1.00E+00 | 2.73E-02 | 1.18E-01 | 1.05E+00 | 4.71E-02 | 2.00E-01 | -1.05E+00 | 1.67E-02 | 6.37E-02 | -1.06E+00 | 1.38E-01 | 4.91E-01 | 1.85E+00 | 2.84E-01 | 8.01E-01 | 1.57E+00 | 2.70E-01 | 9.89E-01 | 1.23E+00 | 3.11E-01 | 8.55E-01 | 1.19E+00 |
| AC073136.2 | 2.51E-02 | 1.31E-01 | -1.07E+00 | 7.09E-02 | 3.06E-01 | 1.24E+00 | 1.09E-01 | 4.64E-01 | 1.49E+00 | 1.66E-01 | 6.35E-01 | 1.58E+00 | 6.24E-02 | 2.22E-01 | -1.27E+00 | 8.37E-02 | 2.36E-01 | -1.17E+00 | 2.56E-01 | 9.39E-01 | 1.24E+00 | 3.60E-01 | 9.88E-01 | 1.32E+00 |
| ALMS1-IT1 | 2.51E-02 | 1.31E-01 | -1.36E+00 | 4.80E-02 | 2.07E-01 | -1.24E+00 | 3.21E-04 | 1.37E-03 | -1.57E+00 | 5.22E-05 | 1.99E-04 | -1.65E+00 | 4.77E-02 | 1.70E-01 | -1.27E+00 | 3.14E-02 | 8.84E-02 | -1.53E+00 | 2.69E-01 | 9.86E-01 | 1.13E+00 | 1.93E-01 | 5.31E-01 | -1.11E+00 |
| DBI | 2.52E-02 | 1.31E-01 | 1.42E+00 | 4.78E-02 | 2.06E-01 | 1.28E+00 | 7.45E-03 | 3.17E-02 | 1.33E+00 | 1.30E-01 | 4.97E-01 | 1.18E+00 | 1.04E-01 | 3.72E-01 | 1.71E+00 | 1.39E-02 | 3.91E-02 | 1.10E+00 | 7.12E-02 | 2.61E-01 | 1.43E+00 | 3.58E-01 | 9.85E-01 | 1.29E+00 |
| EBP | 2.52E-02 | 1.31E-01 | 1.18E+00 | 1.34E-02 | 5.78E-02 | 1.31E+00 | 6.69E-02 | 2.84E-01 | 1.29E+00 | 1.53E-01 | 5.83E-01 | 1.45E+00 | 8.70E-03 | 3.10E-02 | 1.11E+00 | 2.39E-01 | 6.75E-01 | 1.43E+00 | 2.06E-01 | 7.56E-01 | 1.15E+00 | 5.39E-02 | 1.48E-01 | 1.37E+00 |
| AC020663.3 | 2.52E-02 | 1.31E-01 | 1.01E+00 | 6.88E-04 | 2.97E-03 | -1.45E+00 | 1.63E-01 | 6.95E-01 | 1.08E+00 | 3.06E-02 | 1.17E-01 | -1.09E+00 | 1.48E-01 | 5.26E-01 | 1.54E+00 | 2.41E-01 | 6.80E-01 | 1.44E+00 | 8.19E-03 | 3.00E-02 | -1.26E+00 | 1.12E-03 | 3.06E-03 | -1.74E+00 |
| IPP | 2.52E-02 | 1.31E-01 | -1.36E+00 | 6.19E-02 | 2.67E-01 | -1.02E+00 | 1.66E-01 | 7.05E-01 | -1.04E+00 | 2.51E-01 | 9.59E-01 | 1.25E+00 | 2.26E-01 | 8.05E-01 | -1.01E+00 | 5.36E-02 | 1.51E-01 | 2.02E+00 | 4.47E-04 | 1.64E-03 | -1.36E+00 | 2.27E-04 | 6.24E-04 | -1.50E+00 |
| HIST1H2BB | 2.52E-02 | 1.31E-01 | -1.19E+00 | 6.33E-03 | 2.73E-02 | -1.49E+00 | 3.36E-02 | 1.43E-01 | -1.27E+00 | 9.06E-03 | 3.46E-02 | -1.29E+00 | 2.74E-01 | 9.76E-01 | 1.25E+00 | 3.53E-01 | 9.96E-01 | 1.27E+00 | 1.54E-02 | 5.66E-02 | -1.19E+00 | 5.33E-04 | 1.47E-03 | -1.70E+00 |
| ANKRD9 | 2.52E-02 | 1.32E-01 | -1.26E+00 | 9.12E-03 | 3.94E-02 | -1.58E+00 | 8.78E-05 | 3.73E-04 | -2.11E+00 | 6.58E-04 | 2.51E-03 | -1.56E+00 | 3.81E-02 | 1.36E-01 | -1.15E+00 | 3.67E-03 | 1.03E-02 | -1.61E+00 | 2.25E-01 | 8.24E-01 | 2.06E+00 | 3.95E-03 | 1.09E-02 | 1.13E+00 |
| EEF1A1P25 | 2.52E-02 | 1.32E-01 | -1.04E+00 | 6.44E-03 | 2.78E-02 | -1.11E+00 | 8.99E-02 | 3.82E-01 | 1.09E+00 | 1.53E-02 | 5.83E-02 | 1.00E+00 | 5.57E-02 | 1.99E-01 | -1.02E+00 | 3.52E-02 | 9.93E-02 | -1.07E+00 | 2.41E-01 | 8.84E-01 | 1.33E+00 | 3.81E-02 | 1.05E-01 | -1.08E+00 |
| AC074032.1 | 2.52E-02 | 1.32E-01 | 1.06E+00 | 1.29E-03 | 5.56E-03 | -1.24E+00 | 1.01E-01 | 4.27E-01 | 1.03E+00 | 1.18E-02 | 4.50E-02 | -1.13E+00 | 2.09E-01 | 7.45E-01 | 1.44E+00 | 1.01E-01 | 2.85E-01 | -1.11E+00 | 6.70E-03 | 2.46E-02 | -1.27E+00 | 2.45E-04 | 6.73E-04 | -1.66E+00 |
| AP003392.4 | 2.53E-02 | 1.32E-01 | -1.29E+00 | 1.21E-01 | 5.22E-01 | 1.13E+00 | 1.20E-02 | 5.10E-02 | -1.19E+00 | 7.08E-02 | 2.70E-01 | 1.04E+00 | 6.76E-02 | 2.41E-01 | -1.77E+00 | 3.51E-01 | 9.90E-01 | 1.23E+00 | 6.63E-02 | 2.43E-01 | -1.21E+00 | 5.68E-02 | 1.56E-01 | -1.23E+00 |
| GMPPA | 2.53E-02 | 1.32E-01 | -1.08E+00 | 7.15E-02 | 3.09E-01 | 1.18E+00 | 1.73E-02 | 7.36E-02 | -1.05E+00 | 4.08E-02 | 1.56E-01 | 1.16E+00 | 1.53E-04 | 5.45E-04 | -1.23E+00 | 5.54E-03 | 1.56E-02 | -1.01E+00 | 4.46E-09 | 1.64E-08 | -1.37E+00 | 7.43E-03 | 2.04E-02 | 1.06E+00 |
| AC092803.2 | 2.53E-02 | 1.32E-01 | 1.08E+00 | 1.53E-02 | 6.61E-02 | 1.15E+00 | 2.35E-01 | 1.00E+00 | 1.20E+00 | 2.62E-01 | 1.00E+00 | 1.37E+00 | 4.79E-02 | 1.71E-01 | -1.26E+00 | 1.96E-01 | 5.51E-01 | 1.18E+00 | 2.06E-01 | 7.54E-01 | 1.16E+00 | 1.95E-01 | 5.37E-01 | 1.75E+00 |
| HPSE | 2.53E-02 | 1.32E-01 | -1.76E+00 | 3.06E-02 | 1.32E-01 | -1.73E+00 | 5.63E-03 | 2.39E-02 | -1.87E+00 | 4.89E-03 | 1.87E-02 | -2.03E+00 | 1.06E-01 | 3.78E-01 | -1.88E+00 | 1.41E-01 | 3.98E-01 | -1.31E+00 | 1.26E-02 | 4.63E-02 | 1.52E+00 | 1.83E-05 | 5.04E-05 | -1.35E+00 |
| CD226 | 2.53E-02 | 1.32E-01 | -1.62E+00 | 5.07E-02 | 2.19E-01 | -1.45E+00 | 1.49E-02 | 6.34E-02 | -1.59E+00 | 1.74E-02 | 6.66E-02 | -1.43E+00 | 2.25E-01 | 8.01E-01 | 1.37E+00 | 2.27E-01 | 6.40E-01 | 1.59E+00 | 8.82E-02 | 3.24E-01 | 2.93E+00 | 2.41E-03 | 6.63E-03 | 1.31E+00 |
| AL359636.1 | 2.53E-02 | 1.32E-01 | 1.10E+00 | 7.92E-03 | 3.42E-02 | 1.12E+00 | 2.30E-01 | 9.79E-01 | 1.38E+00 | 1.84E-01 | 7.03E-01 | 1.46E+00 | 8.17E-02 | 2.91E-01 | 1.16E+00 | 1.57E-02 | 4.42E-02 | -1.01E+00 | 1.83E-01 | 6.73E-01 | 1.46E+00 | 3.14E-01 | 8.62E-01 | 1.46E+00 |
| MAGIX | 2.54E-02 | 1.32E-01 | 1.69E+00 | 8.36E-02 | 3.61E-01 | -1.12E+00 | 6.08E-02 | 2.58E-01 | 1.57E+00 | 1.63E-01 | 6.22E-01 | -1.09E+00 | 9.18E-02 | 3.27E-01 | 2.35E+00 | 9.09E-02 | 2.56E-01 | 1.06E+00 | 2.15E-01 | 7.88E-01 | 1.21E+00 | 1.13E-02 | 3.11E-02 | -1.22E+00 |
| MFN2 | 2.54E-02 | 1.32E-01 | -1.06E+00 | 7.67E-02 | 3.32E-01 | 1.16E+00 | 2.56E-02 | 1.09E-01 | -1.17E+00 | 1.12E-01 | 4.27E-01 | 1.13E+00 | 1.21E-03 | 4.33E-03 | -1.61E+00 | 9.40E-03 | 2.65E-02 | -1.23E+00 | 2.81E-06 | 1.03E-05 | -1.40E+00 | 6.48E-04 | 1.78E-03 | -1.18E+00 |
| ANKDD1A | 2.54E-02 | 1.32E-01 | 1.00E+00 | 2.03E-01 | 8.78E-01 | 1.62E+00 | 5.24E-02 | 2.23E-01 | -1.17E+00 | 2.17E-01 | 8.30E-01 | 1.41E+00 | 1.78E-01 | 6.36E-01 | 1.14E+00 | 2.88E-01 | 8.12E-01 | 1.35E+00 | 7.42E-02 | 2.72E-01 | -1.20E+00 | 2.33E-01 | 6.41E-01 | 1.03E+00 |
| AC020928.1 | 2.54E-02 | 1.32E-01 | -1.18E+00 | 1.56E-01 | 6.75E-01 | 1.21E+00 | 6.11E-05 | 2.60E-04 | -1.58E+00 | 1.84E-02 | 7.03E-02 | 1.10E+00 | 2.74E-02 | 9.77E-02 | -1.31E+00 | 2.98E-01 | 8.41E-01 | 1.23E+00 | 2.42E-01 | 8.87E-01 | 1.14E+00 | 9.16E-03 | 2.52E-02 | 1.74E+00 |
| TTC33 | 2.54E-02 | 1.33E-01 | -1.12E+00 | 5.36E-02 | 2.32E-01 | 1.03E+00 | 3.64E-02 | 1.55E-01 | -1.06E+00 | 1.37E-01 | 5.23E-01 | 1.21E+00 | 1.09E-03 | 3.88E-03 | -1.58E+00 | 2.84E-01 | 8.02E-01 | 1.25E+00 | 3.76E-06 | 1.38E-05 | 1.06E+00 | 2.68E-02 | 7.35E-02 | 1.12E+00 |
| AL354710.1 | 2.55E-02 | 1.33E-01 | -1.40E+00 | 1.34E-01 | 5.78E-01 | 1.51E+00 | 9.21E-02 | 3.91E-01 | 1.50E+00 | 2.59E-02 | 9.88E-02 | 2.05E+00 | 9.31E-02 | 3.32E-01 | -1.13E+00 | 3.51E-01 | 9.91E-01 | 1.35E+00 | 1.94E-01 | 7.12E-01 | 1.05E+00 | 3.73E-02 | 1.02E-01 | 1.82E+00 |
| TNK2 | 2.55E-02 | 1.33E-01 | 1.11E+00 | 2.82E-02 | 1.22E-01 | 1.20E+00 | 4.48E-04 | 1.90E-03 | -1.26E+00 | 9.43E-03 | 3.60E-02 | 1.08E+00 | 3.71E-02 | 1.32E-01 | 1.17E+00 | 7.86E-02 | 2.22E-01 | 1.21E+00 | 1.21E-03 | 4.46E-03 | -1.23E+00 | 6.06E-03 | 1.66E-02 | -1.14E+00 |
| JHDM1D-AS1 | 2.55E-02 | 1.33E-01 | -1.18E+00 | 2.12E-01 | 9.17E-01 | 1.41E+00 | 4.55E-02 | 1.93E-01 | -1.37E+00 | 2.33E-01 | 8.91E-01 | 1.21E+00 | 2.58E-01 | 9.18E-01 | 1.41E+00 | 1.31E-01 | 3.70E-01 | 1.83E+00 | 1.34E-01 | 4.90E-01 | 1.20E+00 | 7.68E-02 | 2.11E-01 | -1.07E+00 |
| FTH1P1 | 2.55E-02 | 1.33E-01 | -1.01E+00 | 3.49E-05 | 1.51E-04 | -1.83E+00 | 4.22E-02 | 1.79E-01 | -1.01E+00 | 1.72E-04 | 6.58E-04 | -1.75E+00 | 7.99E-04 | 2.85E-03 | -1.04E+00 | 3.49E-13 | 9.85E-13 | -2.62E+00 | 1.89E-12 | 6.95E-12 | -1.64E+00 | 1.38E-22 | 3.80E-22 | -2.84E+00 |
| ALCAM | 2.55E-02 | 1.33E-01 | 1.57E+00 | 1.13E-02 | 4.88E-02 | 1.95E+00 | 1.76E-01 | 7.47E-01 | 1.19E+00 | 2.12E-01 | 8.09E-01 | 1.47E+00 | 7.91E-03 | 2.82E-02 | -1.09E+00 | 2.73E-03 | 7.70E-03 | -1.26E+00 | 4.45E-02 | 1.63E-01 | -1.10E+00 | 1.71E-01 | 4.70E-01 | 1.16E+00 |
| AP003392.1 | 2.55E-02 | 1.33E-01 | -1.11E+00 | 9.44E-02 | 4.08E-01 | 1.11E+00 | 4.72E-02 | 2.00E-01 | -1.18E+00 | 2.47E-01 | 9.45E-01 | 1.30E+00 | 2.87E-04 | 1.02E-03 | -2.38E+00 | 9.84E-02 | 2.77E-01 | -1.10E+00 | 7.42E-02 | 2.72E-01 | -1.21E+00 | 1.02E-01 | 2.80E-01 | -1.20E+00 |
| CDK5RAP2 | 2.56E-02 | 1.33E-01 | -1.08E+00 | 1.13E-01 | 4.87E-01 | 1.20E+00 | 1.34E-02 | 5.69E-02 | 1.00E+00 | 7.14E-03 | 2.73E-02 | 1.07E+00 | 1.58E-02 | 5.64E-02 | -1.05E+00 | 1.01E-02 | 2.84E-02 | -1.02E+00 | 2.69E-02 | 9.88E-02 | -1.06E+00 | 2.09E-03 | 5.74E-03 | -1.15E+00 |
| TFCP2L1 | 2.56E-02 | 1.33E-01 | -1.40E+00 | 3.97E-02 | 1.72E-01 | -1.34E+00 | 1.67E-01 | 7.09E-01 | 1.08E+00 | 1.95E-02 | 7.46E-02 | -1.19E+00 | 2.02E-01 | 7.22E-01 | 1.21E+00 | 3.20E-01 | 9.02E-01 | 1.10E+00 | 2.36E-01 | 8.65E-01 | 1.07E+00 | 3.64E-01 | 1.00E+00 | 1.00E+00 |
| AC096667.1 | 2.56E-02 | 1.33E-01 | 1.04E+00 | 2.02E-01 | 8.72E-01 | 1.56E+00 | 6.76E-02 | 2.87E-01 | -1.12E+00 | 8.15E-02 | 3.11E-01 | 1.05E+00 | 1.26E-02 | 4.50E-02 | -1.45E+00 | 2.01E-02 | 5.67E-02 | -1.31E+00 | 2.12E-02 | 7.77E-02 | -1.12E+00 | 1.97E-02 | 5.40E-02 | -1.38E+00 |
| ZNF441 | 2.56E-02 | 1.34E-01 | -1.08E+00 | 5.44E-02 | 2.35E-01 | 1.06E+00 | 1.44E-01 | 6.10E-01 | -1.00E+00 | 7.12E-02 | 2.72E-01 | -1.01E+00 | 2.00E-01 | 7.15E-01 | 1.23E+00 | 2.77E-01 | 7.81E-01 | 1.53E+00 | 3.33E-02 | 1.22E-01 | -1.22E+00 | 2.15E-01 | 5.91E-01 | 1.07E+00 |
| BCL2 | 2.56E-02 | 1.34E-01 | 1.09E+00 | 1.28E-04 | 5.53E-04 | 1.03E+00 | 1.19E-05 | 5.05E-05 | -1.27E+00 | 1.86E-05 | 7.09E-05 | -1.04E+00 | 2.12E-01 | 7.56E-01 | 1.43E+00 | 1.45E-02 | 4.10E-02 | 1.10E+00 | 8.91E-02 | 3.27E-01 | 1.46E+00 | 6.63E-02 | 1.82E-01 | 1.06E+00 |
| ENO1-IT1 | 2.56E-02 | 1.34E-01 | -1.28E+00 | 1.90E-01 | 8.20E-01 | 1.24E+00 | 6.69E-04 | 2.84E-03 | -1.46E+00 | 5.40E-02 | 2.06E-01 | 1.15E+00 | 1.31E-01 | 4.65E-01 | -1.11E+00 | 5.61E-02 | 1.58E-01 | -1.64E+00 | 2.14E-01 | 7.87E-01 | 1.07E+00 | 2.32E-01 | 6.37E-01 | -1.09E+00 |
| HIST1H2AG | 2.57E-02 | 1.34E-01 | 1.00E+00 | 5.71E-02 | 2.47E-01 | 1.23E+00 | 5.57E-02 | 2.37E-01 | 1.54E+00 | 1.96E-01 | 7.50E-01 | 1.26E+00 | 1.99E-01 | 7.09E-01 | 1.14E+00 | 3.00E-01 | 8.46E-01 | 1.23E+00 | 6.73E-02 | 2.47E-01 | 1.17E+00 | 5.03E-03 | 1.38E-02 | -1.13E+00 |
| CRY1 | 2.57E-02 | 1.34E-01 | 1.10E+00 | 4.33E-02 | 1.87E-01 | 1.23E+00 | 1.16E-02 | 4.91E-02 | -1.13E+00 | 2.20E-01 | 8.38E-01 | 1.36E+00 | 1.32E-01 | 4.69E-01 | 1.23E+00 | 3.47E-01 | 9.77E-01 | 1.55E+00 | 3.00E-05 | 1.10E-04 | -1.18E+00 | 1.10E-04 | 3.02E-04 | -1.25E+00 |
| SOCS5 | 2.57E-02 | 1.34E-01 | 1.18E+00 | 2.14E-02 | 9.23E-02 | 1.30E+00 | 5.81E-02 | 2.47E-01 | 1.02E+00 | 9.90E-03 | 3.78E-02 | 1.03E+00 | 8.68E-03 | 3.09E-02 | -1.15E+00 | 8.97E-03 | 2.53E-02 | -1.19E+00 | 1.39E-02 | 5.09E-02 | -1.07E+00 | 9.71E-02 | 2.67E-01 | 1.05E+00 |
| MFSD7 | 2.57E-02 | 1.34E-01 | -1.09E+00 | 3.40E-02 | 1.47E-01 | 1.01E+00 | 1.31E-02 | 5.56E-02 | -1.28E+00 | 3.66E-02 | 1.40E-01 | -1.07E+00 | 1.83E-02 | 6.51E-02 | -1.30E+00 | 3.65E-02 | 1.03E-01 | -1.20E+00 | 1.29E-01 | 4.72E-01 | -1.06E+00 | 3.51E-01 | 9.65E-01 | 1.22E+00 |
| RELL1 | 2.57E-02 | 1.34E-01 | 1.12E+00 | 2.30E-04 | 9.95E-04 | -1.23E+00 | 1.52E-01 | 6.47E-01 | 1.34E+00 | 1.86E-04 | 7.10E-04 | -1.15E+00 | 1.70E-01 | 6.07E-01 | 1.94E+00 | 3.49E-01 | 9.83E-01 | 1.28E+00 | 2.36E-01 | 8.66E-01 | 1.11E+00 | 1.50E-01 | 4.13E-01 | -1.35E+00 |
| BFAR | 2.57E-02 | 1.34E-01 | 1.09E+00 | 1.45E-02 | 6.26E-02 | 1.20E+00 | 9.96E-03 | 4.23E-02 | -1.03E+00 | 2.57E-03 | 9.83E-03 | -1.00E+00 | 6.96E-07 | 2.48E-06 | -1.15E+00 | 4.29E-05 | 1.21E-04 | -1.07E+00 | 2.30E-03 | 8.43E-03 | -1.04E+00 | 5.10E-02 | 1.40E-01 | 1.03E+00 |
| SCN2A | 2.57E-02 | 1.34E-01 | -1.35E+00 | 2.16E-01 | 9.33E-01 | 1.30E+00 | 1.07E-03 | 4.56E-03 | -1.72E+00 | 2.57E-03 | 9.81E-03 | -1.56E+00 | 2.79E-03 | 9.93E-03 | -1.57E+00 | 1.43E-03 | 4.02E-03 | -1.82E+00 | 3.24E-02 | 1.19E-01 | 1.63E+00 | 2.65E-01 | 7.29E-01 | 1.27E+00 |
| RPL23AP43 | 2.58E-02 | 1.34E-01 | 1.18E+00 | 1.10E-11 | 4.75E-11 | -1.15E+00 | 3.40E-02 | 1.44E-01 | 1.18E+00 | 6.38E-04 | 2.44E-03 | -1.14E+00 | 6.13E-02 | 2.19E-01 | 1.24E+00 | 1.02E-02 | 2.89E-02 | -1.07E+00 | 2.70E-01 | 9.92E-01 | 1.13E+00 | 1.31E-02 | 3.61E-02 | -1.10E+00 |
| Z97989.1 | 2.58E-02 | 1.34E-01 | -1.16E+00 | 9.55E-02 | 4.13E-01 | 1.08E+00 | 1.08E-01 | 4.60E-01 | -1.04E+00 | 7.00E-02 | 2.68E-01 | -1.12E+00 | 6.33E-02 | 2.26E-01 | -1.54E+00 | 1.60E-01 | 4.51E-01 | -1.17E+00 | 9.18E-02 | 3.37E-01 | -1.39E+00 | 3.35E-01 | 9.20E-01 | -1.03E+00 |
| DARS-AS1 | 2.58E-02 | 1.34E-01 | -1.22E+00 | 3.94E-02 | 1.70E-01 | -1.03E+00 | 6.37E-02 | 2.71E-01 | -1.11E+00 | 1.27E-01 | 4.84E-01 | 1.03E+00 | 1.87E-01 | 6.66E-01 | 1.55E+00 | 3.37E-01 | 9.49E-01 | 1.17E+00 | 1.20E-01 | 4.40E-01 | -1.46E+00 | 2.96E-01 | 8.14E-01 | -1.01E+00 |
| SMIM24 | 2.58E-02 | 1.35E-01 | 1.13E+00 | 6.86E-06 | 2.97E-05 | 1.04E+00 | 1.10E-01 | 4.68E-01 | -1.05E+00 | 3.10E-04 | 1.18E-03 | -1.20E+00 | 3.69E-06 | 1.32E-05 | -1.11E+00 | 6.39E-10 | 1.80E-09 | -1.28E+00 | 5.55E-02 | 2.04E-01 | 1.37E+00 | 1.82E-01 | 5.00E-01 | 1.28E+00 |
| KCNAB1 | 2.59E-02 | 1.35E-01 | -1.23E+00 | 2.01E-01 | 8.68E-01 | 1.22E+00 | 1.06E-02 | 4.51E-02 | 2.12E+00 | 1.22E-01 | 4.65E-01 | 1.83E+00 | 8.23E-02 | 2.93E-01 | -1.14E+00 | 4.12E-02 | 1.16E-01 | -1.37E+00 | 1.77E-02 | 6.49E-02 | 1.81E+00 | 2.07E-01 | 5.69E-01 | 1.47E+00 |
| SLC15A4 | 2.59E-02 | 1.35E-01 | -1.04E+00 | 1.92E-01 | 8.29E-01 | 1.46E+00 | 1.43E-02 | 6.09E-02 | -1.04E+00 | 2.78E-02 | 1.06E-01 | 1.13E+00 | 1.54E-01 | 5.49E-01 | 1.31E+00 | 2.12E-01 | 5.98E-01 | 1.25E+00 | 1.86E-03 | 6.83E-03 | -1.45E+00 | 7.27E-02 | 2.00E-01 | -1.07E+00 |
| HIST2H2AA3 | 2.59E-02 | 1.35E-01 | 1.69E+00 | 1.93E-04 | 8.33E-04 | 1.01E+00 | 1.64E-02 | 6.97E-02 | 1.65E+00 | 1.65E-01 | 6.31E-01 | 1.31E+00 | 1.99E-03 | 7.08E-03 | 2.00E+00 | 2.67E-01 | 7.53E-01 | 1.18E+00 | 9.90E-02 | 3.63E-01 | 1.70E+00 | 6.18E-02 | 1.70E-01 | -1.30E+00 |
| AC008764.10 | 2.60E-02 | 1.35E-01 | 1.85E+00 | 1.50E-01 | 6.50E-01 | 1.03E+00 | 1.54E-01 | 6.54E-01 | 1.19E+00 | 5.01E-03 | 1.92E-02 | -1.57E+00 | 2.02E-02 | 7.22E-02 | 2.28E+00 | 2.95E-01 | 8.33E-01 | 1.15E+00 | 2.12E-01 | 7.79E-01 | 1.12E+00 | 1.70E-02 | 4.68E-02 | -1.36E+00 |
| PSMC1 | 2.60E-02 | 1.36E-01 | -1.20E+00 | 8.40E-03 | 3.63E-02 | -1.42E+00 | 2.02E-02 | 8.57E-02 | 1.74E+00 | 2.34E-01 | 8.94E-01 | 1.34E+00 | 1.33E-01 | 4.76E-01 | -1.00E+00 | 3.18E-02 | 8.98E-02 | -1.42E+00 | 1.22E-01 | 4.47E-01 | 1.03E+00 | 2.75E-02 | 7.55E-02 | -1.26E+00 |
| SOS1-IT1 | 2.60E-02 | 1.36E-01 | -1.41E+00 | 4.45E-02 | 1.92E-01 | -1.17E+00 | 1.13E-02 | 4.82E-02 | -1.43E+00 | 1.18E-01 | 4.52E-01 | 1.16E+00 | 6.17E-04 | 2.20E-03 | -2.99E+00 | 1.85E-02 | 5.22E-02 | -1.60E+00 | 3.24E-08 | 1.19E-07 | -1.84E+00 | 1.62E-03 | 4.45E-03 | -1.18E+00 |
| WEE1 | 2.61E-02 | 1.36E-01 | 1.15E+00 | 1.76E-01 | 7.59E-01 | 1.55E+00 | 4.92E-04 | 2.09E-03 | 1.88E+00 | 3.80E-04 | 1.45E-03 | 2.18E+00 | 2.59E-01 | 9.24E-01 | 1.68E+00 | 2.93E-01 | 8.27E-01 | 1.68E+00 | 2.18E-05 | 8.02E-05 | -1.22E+00 | 2.96E-01 | 8.14E-01 | 1.28E+00 |
| RAB15 | 2.61E-02 | 1.36E-01 | -1.16E+00 | 1.89E-01 | 8.15E-01 | 1.34E+00 | 1.69E-01 | 7.17E-01 | 1.07E+00 | 2.08E-01 | 7.96E-01 | 1.40E+00 | 9.95E-03 | 3.55E-02 | -1.27E+00 | 1.45E-02 | 4.10E-02 | -1.39E+00 | 8.30E-02 | 3.05E-01 | -1.32E+00 | 2.66E-01 | 7.31E-01 | 1.05E+00 |
| ZNF19 | 2.61E-02 | 1.36E-01 | -1.19E+00 | 6.04E-02 | 2.61E-01 | -1.00E+00 | 9.96E-03 | 4.23E-02 | -1.38E+00 | 2.61E-01 | 9.95E-01 | 1.39E+00 | 5.55E-02 | 1.98E-01 | -1.22E+00 | 7.93E-02 | 2.23E-01 | -1.11E+00 | 1.73E-01 | 6.36E-01 | 1.38E+00 | 3.06E-01 | 8.41E-01 | 1.31E+00 |
| ENOX2 | 2.61E-02 | 1.36E-01 | -1.15E+00 | 1.79E-01 | 7.72E-01 | 1.32E+00 | 1.56E-02 | 6.63E-02 | -1.19E+00 | 1.61E-01 | 6.14E-01 | 1.20E+00 | 5.40E-05 | 1.93E-04 | -1.44E+00 | 6.65E-03 | 1.87E-02 | -1.06E+00 | 2.38E-04 | 8.74E-04 | -1.18E+00 | 1.63E-03 | 4.48E-03 | -1.10E+00 |
| ARHGAP5-AS1 | 2.61E-02 | 1.36E-01 | 1.07E+00 | 1.96E-01 | 8.49E-01 | 1.80E+00 | 1.87E-01 | 7.94E-01 | 1.29E+00 | 2.11E-01 | 8.06E-01 | 1.63E+00 | 1.86E-01 | 6.62E-01 | 1.86E+00 | 2.17E-01 | 6.11E-01 | 1.85E+00 | 3.46E-02 | 1.27E-01 | -1.13E+00 | 7.02E-02 | 1.93E-01 | -1.01E+00 |
| NAPG | 2.61E-02 | 1.36E-01 | 1.08E+00 | 3.07E-06 | 1.33E-05 | -1.19E+00 | 2.60E-03 | 1.10E-02 | -1.12E+00 | 5.13E-06 | 1.96E-05 | -1.21E+00 | 5.87E-02 | 2.09E-01 | 1.07E+00 | 1.46E-03 | 4.12E-03 | -1.24E+00 | 8.75E-05 | 3.21E-04 | -1.32E+00 | 5.55E-03 | 1.53E-02 | -1.10E+00 |
| CD200 | 2.61E-02 | 1.36E-01 | 1.08E+00 | 1.84E-01 | 7.93E-01 | 1.50E+00 | 1.57E-02 | 6.69E-02 | -1.16E+00 | 1.36E-01 | 5.21E-01 | 1.20E+00 | 2.61E-01 | 9.30E-01 | 1.04E+00 | 3.24E-01 | 9.15E-01 | 1.26E+00 | 2.68E-01 | 9.84E-01 | 1.04E+00 | 3.23E-01 | 8.87E-01 | 1.20E+00 |
| AC135050.7 | 2.63E-02 | 1.37E-01 | -1.26E+00 | 6.25E-02 | 2.70E-01 | 1.02E+00 | 2.32E-01 | 9.84E-01 | 1.31E+00 | 2.57E-01 | 9.83E-01 | 1.41E+00 | 9.11E-02 | 3.25E-01 | -1.12E+00 | 1.13E-01 | 3.20E-01 | -1.06E+00 | 1.77E-01 | 6.51E-01 | -1.02E+00 | 2.81E-01 | 7.72E-01 | 1.06E+00 |
| AL121839.2 | 2.63E-02 | 1.37E-01 | -1.13E+00 | 5.83E-02 | 2.52E-01 | 1.77E+00 | 3.52E-02 | 1.50E-01 | -1.18E+00 | 2.09E-01 | 7.96E-01 | 1.30E+00 | 2.61E-01 | 9.31E-01 | -1.08E+00 | 1.13E-01 | 3.19E-01 | 1.98E+00 | 6.35E-02 | 2.33E-01 | 1.05E+00 | 3.18E-01 | 8.75E-01 | 1.35E+00 |
| MIR503HG | 2.64E-02 | 1.38E-01 | 1.08E+00 | 3.33E-02 | 1.44E-01 | 2.10E+00 | 6.74E-03 | 2.87E-02 | -1.29E+00 | 2.44E-01 | 9.32E-01 | 1.53E+00 | 3.87E-02 | 1.38E-01 | -1.50E+00 | 1.93E-01 | 5.43E-01 | 1.17E+00 | 2.58E-01 | 9.46E-01 | 1.36E+00 | 3.44E-01 | 9.45E-01 | 1.31E+00 |
| ZNF784 | 2.64E-02 | 1.38E-01 | 1.06E+00 | 2.71E-04 | 1.17E-03 | -1.28E+00 | 1.57E-05 | 6.67E-05 | -1.46E+00 | 2.26E-07 | 8.63E-07 | -1.74E+00 | 6.86E-02 | 2.44E-01 | 1.02E+00 | 3.31E-02 | 9.34E-02 | -1.20E+00 | 9.61E-02 | 3.53E-01 | -1.08E+00 | 1.52E-01 | 4.17E-01 | -1.01E+00 |
| RASSF1-AS1 | 2.65E-02 | 1.38E-01 | 1.87E+00 | 2.61E-02 | 1.13E-01 | -1.18E+00 | 2.64E-03 | 1.12E-02 | 2.23E+00 | 2.11E-01 | 8.06E-01 | 1.24E+00 | 1.98E-01 | 7.07E-01 | 1.56E+00 | 3.11E-01 | 8.75E-01 | 1.12E+00 | 2.18E-01 | 8.01E-01 | 1.11E+00 | 1.05E-01 | 2.88E-01 | -1.15E+00 |
| RNF32 | 2.65E-02 | 1.38E-01 | -1.46E+00 | 2.07E-02 | 8.96E-02 | -1.40E+00 | 2.10E-01 | 8.93E-01 | 1.02E+00 | 1.57E-01 | 5.98E-01 | 1.10E+00 | 2.25E-02 | 8.01E-02 | -1.07E+00 | 7.90E-02 | 2.23E-01 | -1.01E+00 | 2.51E-01 | 9.23E-01 | 1.06E+00 | 3.64E-01 | 1.00E+00 | 1.10E+00 |
| IGFBP7-AS1 | 2.66E-02 | 1.38E-01 | 2.03E+00 | 4.74E-02 | 2.05E-01 | 2.13E+00 | 8.45E-02 | 3.59E-01 | 1.44E+00 | 1.58E-01 | 6.02E-01 | 1.46E+00 | 2.36E-01 | 8.42E-01 | 1.54E+00 | 2.87E-01 | 8.09E-01 | 1.43E+00 | 2.49E-01 | 9.13E-01 | 1.22E+00 | 2.26E-01 | 6.20E-01 | 1.09E+00 |
| KBTBD12 | 2.66E-02 | 1.39E-01 | -1.36E+00 | 7.34E-02 | 3.17E-01 | -1.05E+00 | 1.29E-01 | 5.50E-01 | -1.19E+00 | 8.92E-02 | 3.41E-01 | -1.30E+00 | 2.61E-01 | 9.31E-01 | 1.07E+00 | 2.53E-01 | 7.13E-01 | 1.43E+00 | 9.82E-02 | 3.60E-01 | -1.02E+00 | 3.61E-01 | 9.91E-01 | 1.14E+00 |
| AC021078.1 | 2.66E-02 | 1.39E-01 | 1.15E+00 | 1.17E-04 | 5.04E-04 | 1.04E+00 | 5.72E-02 | 2.43E-01 | 1.01E+00 | 2.36E-06 | 9.01E-06 | -1.17E+00 | 1.17E-02 | 4.16E-02 | 1.11E+00 | 3.36E-04 | 9.48E-04 | -1.05E+00 | 2.09E-02 | 7.68E-02 | 1.06E+00 | 1.07E-03 | 2.93E-03 | -1.11E+00 |
| DHX38 | 2.67E-02 | 1.39E-01 | 1.04E+00 | 1.44E-01 | 6.24E-01 | 1.37E+00 | 7.81E-02 | 3.32E-01 | 1.34E+00 | 2.00E-01 | 7.63E-01 | 1.25E+00 | 1.57E-01 | 5.60E-01 | 1.18E+00 | 8.81E-02 | 2.48E-01 | 1.06E+00 | 1.40E-01 | 5.15E-01 | 1.08E+00 | 2.19E-01 | 6.02E-01 | 1.08E+00 |
| AMOT | 2.67E-02 | 1.39E-01 | 1.00E+00 | 2.17E-01 | 9.39E-01 | 1.50E+00 | 2.10E-01 | 8.92E-01 | 1.18E+00 | 1.34E-01 | 5.12E-01 | 1.15E+00 | 5.08E-02 | 1.81E-01 | -1.20E+00 | 1.02E-01 | 2.88E-01 | -1.22E+00 | 1.95E-01 | 7.16E-01 | 1.01E+00 | 3.10E-01 | 8.52E-01 | 1.31E+00 |
| MECR | 2.67E-02 | 1.39E-01 | 1.04E+00 | 3.54E-02 | 1.53E-01 | 1.14E+00 | 3.33E-03 | 1.42E-02 | -1.01E+00 | 3.27E-03 | 1.25E-02 | 1.08E+00 | 5.96E-02 | 2.13E-01 | 1.06E+00 | 2.13E-01 | 6.02E-01 | 1.54E+00 | 1.86E-03 | 6.81E-03 | -1.06E+00 | 1.45E-01 | 4.00E-01 | 1.13E+00 |
| LINC00511 | 2.68E-02 | 1.39E-01 | -1.44E+00 | 4.91E-02 | 2.12E-01 | -1.22E+00 | 1.90E-01 | 8.07E-01 | -1.03E+00 | 2.34E-01 | 8.93E-01 | 1.07E+00 | 8.67E-02 | 3.09E-01 | -1.05E+00 | 1.13E-01 | 3.19E-01 | 1.02E+00 | 5.20E-02 | 1.91E-01 | 1.61E+00 | 1.94E-01 | 5.32E-01 | 1.16E+00 |
| AL355075.2 | 2.68E-02 | 1.40E-01 | -1.00E+00 | 1.02E-01 | 4.41E-01 | 1.32E+00 | 1.92E-02 | 8.17E-02 | -1.13E+00 | 3.13E-02 | 1.20E-01 | 1.04E+00 | 4.37E-04 | 1.56E-03 | -1.71E+00 | 6.35E-02 | 1.79E-01 | -1.01E+00 | 5.60E-02 | 2.06E-01 | 1.06E+00 | 3.30E-01 | 9.07E-01 | 1.39E+00 |
| TST | 2.68E-02 | 1.40E-01 | -1.29E+00 | 4.96E-02 | 2.14E-01 | -1.03E+00 | 7.36E-06 | 3.13E-05 | -1.49E+00 | 6.26E-07 | 2.39E-06 | -1.33E+00 | 3.90E-03 | 1.39E-02 | -1.30E+00 | 1.16E-03 | 3.28E-03 | 2.15E+00 | 2.00E-24 | 7.32E-24 | -1.98E+00 | 1.07E-09 | 2.95E-09 | -1.46E+00 |
| MTA1 | 2.68E-02 | 1.40E-01 | -1.01E+00 | 8.43E-03 | 3.64E-02 | -1.03E+00 | 7.56E-02 | 3.21E-01 | 1.03E+00 | 8.01E-02 | 3.06E-01 | 1.24E+00 | 1.27E-01 | 4.51E-01 | 1.68E+00 | 2.43E-01 | 6.85E-01 | 1.56E+00 | 1.91E-01 | 7.00E-01 | 1.44E+00 | 3.09E-01 | 8.48E-01 | 1.40E+00 |
| ARHGEF12 | 2.69E-02 | 1.40E-01 | -1.50E+00 | 6.71E-02 | 2.90E-01 | -1.02E+00 | 4.93E-02 | 2.09E-01 | -1.40E+00 | 3.63E-02 | 1.39E-01 | -1.71E+00 | 5.35E-03 | 1.91E-02 | -1.32E+00 | 1.21E-01 | 3.40E-01 | 1.10E+00 | 1.01E-01 | 3.70E-01 | 1.37E+00 | 8.62E-11 | 2.37E-10 | -1.29E+00 |
| DSN1 | 2.69E-02 | 1.40E-01 | -1.20E+00 | 5.00E-02 | 2.16E-01 | 1.00E+00 | 5.34E-02 | 2.27E-01 | 1.02E+00 | 9.97E-02 | 3.81E-01 | 1.54E+00 | 1.19E-02 | 4.23E-02 | -1.83E+00 | 6.94E-02 | 1.96E-01 | -1.22E+00 | 9.82E-14 | 3.60E-13 | -1.51E+00 | 2.65E-01 | 7.29E-01 | 1.20E+00 |
| MEN1 | 2.69E-02 | 1.40E-01 | 1.06E+00 | 7.41E-02 | 3.20E-01 | 1.85E+00 | 1.70E-04 | 7.20E-04 | -1.32E+00 | 1.95E-03 | 7.44E-03 | -1.03E+00 | 1.56E-04 | 5.55E-04 | -1.66E+00 | 2.87E-01 | 8.09E-01 | 1.41E+00 | 3.77E-08 | 1.38E-07 | -1.45E+00 | 1.57E-02 | 4.30E-02 | 1.07E+00 |
| PDE7B | 2.69E-02 | 1.40E-01 | -1.22E+00 | 7.14E-02 | 3.09E-01 | -1.03E+00 | 2.35E-06 | 1.00E-05 | -1.92E+00 | 2.07E-03 | 7.90E-03 | -1.16E+00 | 5.92E-02 | 2.11E-01 | -1.36E+00 | 1.37E-01 | 3.86E-01 | -1.08E+00 | 5.42E-02 | 1.99E-01 | -1.45E+00 | 1.81E-01 | 4.98E-01 | -1.14E+00 |
| TRIB2 | 2.69E-02 | 1.40E-01 | 2.88E+00 | 4.38E-02 | 1.89E-01 | 3.16E+00 | 1.38E-01 | 5.85E-01 | 1.70E+00 | 1.51E-01 | 5.78E-01 | 2.02E+00 | 1.08E-03 | 3.84E-03 | 2.05E+00 | 3.79E-27 | 1.07E-26 | 4.15E+00 | 8.34E-05 | 3.06E-04 | -1.06E+00 | 3.40E-01 | 9.34E-01 | 1.19E+00 |
| GATA1 | 2.69E-02 | 1.40E-01 | -2.23E+00 | 4.57E-02 | 1.97E-01 | -1.74E+00 | 2.96E-02 | 1.26E-01 | -1.63E+00 | 9.15E-02 | 3.49E-01 | -1.07E+00 | 9.37E-25 | 3.34E-24 | -1.78E+00 | 4.31E-02 | 1.21E-01 | 1.09E+00 | 1.42E-21 | 5.20E-21 | -1.46E+00 | 1.08E-07 | 2.97E-07 | -1.19E+00 |

| | | | | | | | | | | | | | | | | | | | | | | | | |
|---|---|---|---|---|---|---|---|---|---|---|---|---|---|---|---|---|---|---|---|---|---|---|---|---|
| SYTL3 | 2.69E-02 | 1.40E-01 | -1.23E+00 | 5.21E-03 | 2.25E-02 | -1.43E+00 | 6.49E-04 | 2.76E-03 | -1.65E+00 | 9.30E-04 | 3.55E-03 | -1.55E+00 | 1.90E-01 | 6.76E-01 | 1.18E+00 | 8.91E-04 | 2.51E-03 | -1.86E+00 | 2.33E-01 | 8.54E-01 | 1.06E+00 | 1.23E-01 | 3.39E-01 | -1.42E+00 |
| BCO2 | 2.69E-02 | 1.40E-01 | -1.06E+00 | 2.04E-02 | 8.83E-02 | -1.17E+00 | 9.83E-03 | 4.18E-02 | -1.43E+00 | 3.23E-02 | 1.23E-01 | -1.09E+00 | 2.31E-01 | 8.24E-01 | 1.23E+00 | 3.55E-01 | 1.00E+00 | 1.46E+00 | 1.79E-01 | 6.58E-01 | -1.06E+00 | 1.71E-01 | 4.69E-01 | -1.13E+00 |
| ARL6 | 2.69E-02 | 1.40E-01 | -1.22E+00 | 2.27E-01 | 9.79E-01 | 1.39E+00 | 2.22E-04 | 9.43E-04 | -1.79E+00 | 6.96E-02 | 2.66E-01 | 1.05E+00 | 2.72E-01 | 9.69E-01 | 1.51E+00 | 3.50E-01 | 9.86E-01 | 1.41E+00 | 4.73E-03 | 1.74E-02 | -1.67E+00 | 1.40E-01 | 3.85E-01 | -1.05E+00 |
| LINK | 2.69E-02 | 1.40E-01 | 1.78E+00 | 1.47E-01 | 6.35E-01 | 1.64E+00 | 1.38E-01 | 5.87E-01 | 1.17E+00 | 4.36E-02 | 1.67E-01 | 1.13E+00 | 6.39E-02 | 2.28E-01 | 1.09E+00 | 1.28E-01 | 3.61E-01 | 1.17E+00 | 2.46E-02 | 9.04E-02 | -1.09E+00 | 2.22E-02 | 6.09E-02 | -1.03E+00 |
| CSF3R | 2.70E-02 | 1.41E-01 | 1.02E+00 | 4.54E-02 | 1.96E-01 | 1.12E+00 | 7.03E-03 | 2.99E-02 | -1.08E+00 | 2.26E-01 | 8.63E-01 | 1.16E+00 | 2.20E-06 | 7.85E-06 | -1.34E+00 | 9.62E-09 | 2.71E-08 | -1.63E+00 | 1.66E-01 | 6.10E-01 | 1.14E+00 | 3.07E-01 | 8.44E-01 | 1.16E+00 |
| AC104843.1 | 2.71E-02 | 1.41E-01 | 1.36E+00 | 6.00E-25 | 2.59E-24 | -1.30E+00 | 1.06E-02 | 4.51E-02 | 1.35E+00 | 5.38E-06 | 2.05E-05 | -1.14E+00 | 1.73E-01 | 6.18E-01 | 1.47E+00 | 5.15E-06 | 1.45E-05 | -1.09E+00 | 1.07E-01 | 3.92E-01 | 1.32E+00 | 2.94E-04 | 8.07E-04 | -1.22E+00 |
| RBMX2 | 2.71E-02 | 1.41E-01 | 1.13E+00 | 3.08E-02 | 1.33E-01 | 1.25E+00 | 8.77E-03 | 3.73E-02 | -1.07E+00 | 1.68E-01 | 6.44E-01 | 1.27E+00 | 9.16E-02 | 3.26E-01 | 1.16E+00 | 9.87E-02 | 2.78E-01 | 1.73E+00 | 7.32E-03 | 2.69E-02 | 1.01E+00 | 2.22E-01 | 6.09E-01 | 1.27E+00 |
| MYO1D | 2.72E-02 | 1.42E-01 | -1.07E+00 | 1.64E-02 | 7.10E-02 | -1.05E+00 | 1.80E-01 | 7.64E-01 | 1.32E+00 | 2.53E-01 | 9.67E-01 | 1.30E+00 | 1.33E-01 | 4.75E-01 | -1.17E+00 | 2.64E-01 | 7.43E-01 | 1.16E+00 | 8.54E-02 | 3.13E-01 | -1.07E+00 | 7.02E-02 | 1.93E-01 | -1.17E+00 |
| DTYMK | 2.72E-02 | 1.42E-01 | 1.03E+00 | 4.76E-02 | 2.06E-01 | 1.22E+00 | 5.86E-02 | 2.49E-01 | 1.32E+00 | 2.01E-01 | 7.68E-01 | 1.40E+00 | 6.25E-07 | 2.23E-06 | -1.46E+00 | 6.61E-03 | 1.86E-02 | -1.04E+00 | 7.85E-02 | 2.88E-01 | 1.34E+00 | 4.25E-03 | 1.17E-02 | 1.45E+00 |
| SLC4A4 | 2.72E-02 | 1.42E-01 | -1.23E+00 | 3.54E-02 | 1.53E-01 | -1.05E+00 | 2.62E-02 | 1.11E-01 | -1.14E+00 | 8.76E-03 | 3.35E-02 | -1.16E+00 | 4.61E-02 | 1.64E-01 | -1.17E+00 | 8.13E-02 | 2.29E-01 | -1.02E+00 | 7.49E-05 | 2.75E-04 | -1.34E+00 | 3.21E-04 | 8.81E-04 | -1.32E+00 |
| ITPK1-AS1 | 2.72E-02 | 1.42E-01 | 1.81E+00 | 2.94E-02 | 1.27E-01 | 2.11E+00 | 6.81E-02 | 2.90E-01 | -1.02E+00 | 2.03E-02 | 7.76E-02 | -1.06E+00 | 1.49E-01 | 5.31E-01 | 1.00E+00 | 3.20E-01 | 9.02E-01 | 1.43E+00 | 5.67E-02 | 2.08E-01 | -1.13E+00 | 6.00E-02 | 1.65E-01 | -1.21E+00 |
| RPL13AP6 | 2.73E-02 | 1.42E-01 | 1.16E+00 | 1.31E-04 | 5.65E-04 | 1.03E+00 | 1.93E-02 | 8.22E-02 | 1.04E+00 | 8.69E-07 | 3.32E-06 | -1.22E+00 | 1.06E-02 | 3.78E-02 | 1.07E+00 | 9.50E-05 | 2.68E-04 | -1.15E+00 | 8.03E-02 | 2.95E-01 | 1.51E+00 | 2.95E-01 | 8.09E-01 | 1.36E+00 |
| MOB3A | 2.74E-02 | 1.43E-01 | 1.08E+00 | 7.45E-02 | 3.22E-01 | 1.32E+00 | 6.85E-02 | 2.91E-01 | 1.01E+00 | 3.03E-03 | 1.16E-02 | -1.20E+00 | 6.76E-03 | 2.41E-02 | 1.06E+00 | 6.04E-05 | 1.70E-04 | -1.37E+00 | 7.07E-02 | 2.59E-01 | 1.18E+00 | 6.11E-03 | 1.68E-02 | -1.13E+00 |
| VWCE | 2.74E-02 | 1.43E-01 | -1.16E+00 | 6.69E-02 | 2.89E-01 | 1.06E+00 | 1.11E-02 | 4.73E-02 | -1.13E+00 | 1.71E-01 | 6.51E-01 | 1.34E+00 | 3.56E-02 | 1.27E-01 | -1.80E+00 | 2.33E-01 | 6.58E-01 | 1.18E+00 | 1.66E-03 | 6.10E-03 | -1.43E+00 | 2.71E-02 | 7.44E-02 | -1.03E+00 |
| LY96 | 2.74E-02 | 1.43E-01 | -1.43E+00 | 2.70E-02 | 1.16E-01 | -1.53E+00 | 2.12E-01 | 9.01E-01 | -1.05E+00 | 2.11E-01 | 8.05E-01 | -1.05E+00 | 2.50E-02 | 8.91E-02 | -1.14E+00 | 6.92E-03 | 1.95E-02 | -1.49E+00 | 8.10E-08 | 2.97E-07 | -2.15E+00 | 3.21E-08 | 8.81E-08 | -2.45E+00 |
| IQCC | 2.74E-02 | 1.43E-01 | -1.18E+00 | 3.95E-03 | 1.71E-02 | -1.43E+00 | 4.59E-02 | 1.95E-01 | -1.16E+00 | 1.09E-01 | 4.17E-01 | 1.08E+00 | 1.38E-01 | 4.92E-01 | 1.77E+00 | 2.76E-01 | 7.79E-01 | 1.58E+00 | 5.65E-02 | 2.07E-01 | -1.14E+00 | 8.64E-02 | 2.37E-01 | -1.10E+00 |
| LINC02581 | 2.74E-02 | 1.43E-01 | -1.10E+00 | 1.22E-01 | 5.25E-01 | 1.24E+00 | 7.57E-02 | 3.22E-01 | -1.12E+00 | 2.11E-01 | 8.05E-01 | 1.37E+00 | 6.55E-02 | 2.34E-01 | -1.11E+00 | 1.44E-01 | 4.06E-01 | 1.05E+00 | 9.68E-02 | 3.55E-01 | -1.13E+00 | 1.96E-01 | 5.37E-01 | 1.01E+00 |
| PTPRM | 2.74E-02 | 1.43E-01 | -1.15E+00 | 8.27E-02 | 3.57E-01 | 1.11E+00 | 4.13E-02 | 1.75E-01 | 1.01E+00 | 5.36E-02 | 2.05E-01 | 1.08E+00 | 1.08E-01 | 3.83E-01 | -1.30E+00 | 1.80E-01 | 5.06E-01 | -1.11E+00 | 2.23E-01 | 8.17E-01 | -1.13E+00 | 3.60E-01 | 9.90E-01 | 1.04E+00 |
| KIF18A | 2.74E-02 | 1.43E-01 | -1.04E+00 | 1.30E-03 | 5.61E-03 | -1.23E+00 | 2.18E-02 | 9.28E-02 | -1.13E+00 | 2.67E-03 | 1.02E-02 | -1.13E+00 | 1.73E-01 | 6.17E-01 | 1.12E+00 | 7.95E-02 | 2.24E-01 | -1.37E+00 | 1.52E-01 | 5.58E-01 | -1.00E+00 | 1.77E-02 | 4.87E-02 | -1.11E+00 |
| CTBP1-AS2 | 2.74E-02 | 1.43E-01 | 1.10E+00 | 7.50E-02 | 3.24E-01 | 1.31E+00 | 1.21E-03 | 5.14E-03 | -1.24E+00 | 6.28E-02 | 2.40E-01 | 1.20E+00 | 3.96E-02 | 1.41E-01 | -1.20E+00 | 2.52E-02 | 7.11E-02 | -1.25E+00 | 1.03E-01 | 3.78E-01 | 1.06E+00 | 2.04E-01 | 5.59E-01 | 1.14E+00 |
| MTMR7 | 2.75E-02 | 1.44E-01 | -1.12E+00 | 1.18E-01 | 5.11E-01 | 1.27E+00 | 9.34E-03 | 3.97E-02 | -1.40E+00 | 4.49E-02 | 1.71E-01 | -1.07E+00 | 1.58E-01 | 5.64E-01 | 1.65E+00 | 3.49E-01 | 9.84E-01 | 1.30E+00 | 2.02E-01 | 7.41E-01 | -1.07E+00 | 3.23E-01 | 8.88E-01 | 1.14E+00 |
| CSF1R | 2.76E-02 | 1.44E-01 | -1.32E+00 | 2.75E-02 | 1.19E-01 | -1.36E+00 | 8.43E-09 | 3.58E-08 | -2.11E+00 | 2.72E-09 | 1.04E-08 | -2.17E+00 | 1.55E-01 | 5.53E-01 | -1.41E+00 | 6.20E-02 | 1.75E-01 | -2.53E+00 | 1.89E-01 | 6.94E-01 | 1.04E+00 | 1.57E-01 | 4.31E-01 | -1.25E+00 |
| SUFU | 2.76E-02 | 1.44E-01 | -1.01E+00 | 1.70E-01 | 7.35E-01 | 1.41E+00 | 2.18E-05 | 9.27E-05 | -1.57E+00 | 1.64E-04 | 6.27E-04 | -1.33E+00 | 5.95E-02 | 2.12E-01 | -1.02E+00 | 3.29E-01 | 9.27E-01 | 1.38E+00 | 2.48E-03 | 9.12E-03 | -1.32E+00 | 6.00E-02 | 1.65E-01 | 1.00E+00 |
| AC079331.1 | 2.76E-02 | 1.44E-01 | -1.43E+00 | 1.00E-01 | 4.34E-01 | -1.00E+00 | 6.57E-02 | 2.79E-01 | -1.34E+00 | 1.96E-01 | 7.49E-01 | 1.08E+00 | 6.57E-02 | 2.34E-01 | -1.06E+00 | 6.23E-02 | 1.76E-01 | -1.19E+00 | 1.95E-01 | 7.17E-01 | 1.33E+00 | 3.02E-01 | 8.29E-01 | 1.13E+00 |
| CAPRIN2 | 2.76E-02 | 1.44E-01 | -1.02E+00 | 1.11E-01 | 4.79E-01 | 1.27E+00 | 7.31E-04 | 3.11E-03 | -1.29E+00 | 9.61E-04 | 3.67E-03 | -1.17E+00 | 2.20E-02 | 7.84E-02 | -1.12E+00 | 2.81E-01 | 7.92E-01 | 1.60E+00 | 1.18E-04 | 4.32E-04 | -1.10E+00 | 2.03E-03 | 5.58E-03 | 1.03E+00 |
| SLC2A1 | 2.76E-02 | 1.44E-01 | 1.04E+00 | 2.96E-04 | 1.28E-03 | -1.66E+00 | 2.66E-03 | 1.13E-02 | -1.33E+00 | 3.05E-04 | 1.17E-03 | -1.40E+00 | 2.16E-01 | 7.69E-01 | 1.51E+00 | 1.88E-02 | 5.31E-02 | 1.14E+00 | 7.44E-09 | 2.73E-08 | -1.26E+00 | 1.21E-08 | 3.32E-08 | -1.27E+00 |
| FAM193B | 2.76E-02 | 1.44E-01 | -1.07E+00 | 8.45E-03 | 3.65E-02 | -1.10E+00 | 1.74E-02 | 7.40E-02 | 1.02E+00 | 1.16E-01 | 4.42E-01 | 1.38E+00 | 2.35E-01 | 8.37E-01 | 1.42E+00 | 1.25E-01 | 3.52E-01 | 1.81E+00 | 1.83E-01 | 6.72E-01 | 1.40E+00 | 3.48E-01 | 9.56E-01 | 1.24E+00 |
| AC097263.1 | 2.76E-02 | 1.44E-01 | 1.91E+00 | 2.14E-03 | 9.27E-03 | 2.55E+00 | 1.43E-09 | 6.10E-09 | 2.28E+00 | 1.36E-14 | 5.20E-14 | 3.17E+00 | 1.85E-01 | 6.61E-01 | 1.12E+00 | 6.79E-03 | 1.91E-02 | 2.30E+00 | 6.71E-04 | 2.46E-03 | 1.97E+00 | 4.33E-18 | 1.19E-17 | 3.21E+00 |
| AP000648.3 | 2.77E-02 | 1.44E-01 | 1.12E+00 | 1.99E-02 | 8.58E-02 | 1.10E+00 | 1.05E-01 | 4.45E-01 | -1.03E+00 | 7.40E-02 | 2.83E-01 | 1.12E+00 | 2.73E-01 | 9.74E-01 | 1.43E+00 | 3.89E-02 | 1.10E-01 | -1.23E+00 | 1.47E-01 | 5.41E-01 | 1.33E+00 | 2.31E-01 | 6.35E-01 | 1.38E+00 |
| HVCN1 | 2.77E-02 | 1.44E-01 | -1.04E+00 | 1.21E-01 | 5.24E-01 | 1.74E+00 | 2.94E-03 | 1.25E-02 | -1.33E+00 | 8.49E-04 | 3.24E-03 | -1.33E+00 | 1.80E-01 | 6.41E-01 | -1.41E+00 | 3.14E-01 | 8.85E-01 | 1.41E+00 | 2.27E-01 | 8.32E-01 | 1.33E+00 | 2.52E-01 | 6.91E-01 | 1.39E+00 |
| [illegible] | | | | | | | | | | | | | | | | | | | | | | | | |
| INTS1 | 2.88E-02 | 1.50E-01 | 1.02E+00 | 5.95E-02 | 2.57E-01 | 1.24E+00 | 1.30E-01 | 5.53E-01 | 1.38E+00 | 1.97E-01 | 7.53E-01 | 1.58E+00 | 1.41E-01 | 5.04E-01 | 1.17E+00 | 3.53E-01 | 9.95E-01 | 1.37E+00 | 1.25E-01 | 4.59E-01 | 1.19E+00 | 2.11E-01 | 5.80E-01 | 1.45E+00 |
| ZNF678 | 2.88E-02 | 1.50E-01 | 1.10E+00 | 1.66E-01 | 7.15E-01 | 1.44E+00 | 5.49E-02 | 2.33E-01 | -1.10E+00 | 2.47E-01 | 9.45E-01 | 1.34E+00 | 7.82E-04 | 2.79E-03 | -1.07E+00 | 4.58E-04 | 1.29E-03 | -1.09E+00 | 7.72E-02 | 2.83E-01 | 1.07E+00 | 5.90E-02 | 1.62E-01 | -1.01E+00 |
| EXTL3 | 2.88E-02 | 1.50E-01 | -1.17E+00 | 9.43E-02 | 4.08E-01 | 1.16E+00 | 6.16E-04 | 2.62E-03 | -1.48E+00 | 3.71E-02 | 1.42E-01 | 1.06E+00 | 5.37E-03 | 1.92E-02 | -1.49E+00 | 6.15E-02 | 1.73E-01 | 1.00E+00 | 1.69E-03 | 6.19E-03 | -1.63E+00 | 2.54E-03 | 6.98E-03 | -1.70E+00 |
| GADD45G | 2.88E-02 | 1.50E-01 | 2.45E+00 | 1.23E-01 | 5.32E-01 | 2.17E+00 | 2.18E-01 | 9.25E-01 | 1.11E+00 | 2.50E-01 | 9.56E-01 | 1.33E+00 | 2.59E-01 | 9.22E-01 | 1.54E+00 | 3.49E-02 | 9.84E-02 | 2.39E+00 | 5.85E-03 | 2.15E-02 | 1.71E+00 | 9.35E-04 | 2.57E-03 | -1.02E+00 |
| AC092620.2 | 2.88E-02 | 1.50E-01 | 1.03E+00 | 9.65E-02 | 4.17E-01 | 1.34E+00 | 1.99E-02 | 8.47E-02 | -1.07E+00 | 1.94E-02 | 7.43E-02 | 1.09E+00 | 1.08E-01 | 3.84E-01 | 1.17E+00 | 1.14E-06 | 3.22E-06 | 2.93E+00 | 2.27E-01 | 8.32E-01 | 1.14E+00 | 4.31E-04 | 1.18E-03 | 2.22E+00 |
| NLRX1 | 2.89E-02 | 1.50E-01 | -1.15E+00 | 1.49E-01 | 6.42E-01 | 1.27E+00 | 6.86E-02 | 2.91E-01 | -1.05E+00 | 4.24E-02 | 1.62E-01 | 1.15E+00 | 3.02E-03 | 1.08E-02 | -1.46E+00 | 5.38E-02 | 1.52E-01 | -1.04E+00 | 4.24E-06 | 1.56E-05 | -1.48E+00 | 5.85E-03 | 1.61E-02 | -1.00E+00 |
| ZNF165 | 2.90E-02 | 1.51E-01 | 2.09E+00 | 2.07E-01 | 8.96E-01 | 1.39E+00 | 5.67E-03 | 2.41E-02 | -1.31E+00 | 4.34E-03 | 1.66E-02 | -1.23E+00 | 7.17E-02 | 2.56E-01 | -1.31E+00 | 8.45E-02 | 2.38E-01 | -1.23E+00 | 1.14E-01 | 4.17E-01 | -1.22E+00 | 1.99E-02 | 5.45E-02 | -1.84E+00 |
| SIGLEC17P | 2.90E-02 | 1.51E-01 | 1.77E+00 | 8.87E-02 | 3.83E-01 | -1.08E+00 | 2.94E-02 | 1.25E-01 | 1.55E+00 | 1.47E-02 | 5.62E-02 | -1.28E+00 | 2.63E-03 | 9.39E-03 | -3.01E+00 | 1.60E-06 | 4.51E-06 | -4.76E+00 | 2.38E-01 | 8.73E-01 | 1.18E+00 | 3.18E-01 | 8.73E-01 | 1.20E+00 |
| AC131212.2 | 2.90E-02 | 1.51E-01 | 1.07E+00 | 1.44E-01 | 6.22E-01 | 1.46E+00 | 7.07E-03 | 3.01E-02 | -1.16E+00 | 5.00E-02 | 1.91E-01 | 1.20E+00 | 4.00E-02 | 1.42E-01 | 1.10E+00 | 9.29E-02 | 2.62E-01 | 1.17E+00 | 1.27E-01 | 4.66E-01 | -1.00E+00 | 1.05E-01 | 2.89E-01 | 1.51E+00 |
| OR7E128P | 2.90E-02 | 1.51E-01 | -1.31E+00 | 4.74E-02 | 2.05E-01 | -1.21E+00 | 4.39E-02 | 1.87E-01 | -1.17E+00 | 7.72E-02 | 2.95E-01 | 1.04E+00 | 1.52E-01 | 5.42E-01 | 1.64E+00 | 2.91E-01 | 8.19E-01 | 1.26E+00 | 5.37E-02 | 1.97E-01 | -1.08E+00 | 3.25E-01 | 8.92E-01 | 1.13E+00 |
| TAF15 | 2.91E-02 | 1.51E-01 | 1.08E+00 | 1.99E-01 | 8.61E-01 | 1.28E+00 | 3.14E-02 | 1.34E-01 | 1.04E+00 | 2.29E-01 | 8.77E-01 | 1.29E+00 | 3.31E-04 | 1.18E-03 | -1.08E+00 | 7.72E-02 | 2.18E-01 | 1.21E+00 | 3.23E-02 | 1.18E-01 | 1.06E+00 | 3.51E-01 | 9.64E-01 | 1.27E+00 |
| AL357055.1 | 2.91E-02 | 1.52E-01 | 1.02E+00 | 7.67E-04 | 3.31E-03 | -1.32E+00 | 1.79E-01 | 7.59E-01 | 1.11E+00 | 4.33E-04 | 1.65E-03 | -1.40E+00 | 2.43E-03 | 8.65E-03 | -1.54E+00 | 5.60E-03 | 1.58E-02 | -1.29E+00 | 5.21E-02 | 1.91E-01 | -1.07E+00 | 3.49E-03 | 9.58E-03 | -1.41E+00 |
| SLC7A11 | 2.91E-02 | 1.52E-01 | -1.38E+00 | 2.29E-02 | 9.90E-02 | -1.46E+00 | 1.53E-01 | 6.50E-01 | -1.23E+00 | 9.42E-02 | 3.60E-01 | -1.38E+00 | 1.42E-01 | 5.08E-01 | 2.05E+00 | 8.22E-02 | 2.32E-01 | -1.03E+00 | 2.47E-01 | 9.06E-01 | -1.01E+00 | 7.95E-02 | 2.18E-01 | -1.44E+00 |
| HAGHL | 2.91E-02 | 1.52E-01 | -1.19E+00 | 5.91E-02 | 2.55E-01 | -1.19E+00 | 3.83E-02 | 1.63E-01 | -1.24E+00 | 1.59E-01 | 6.09E-01 | 1.10E+00 | 2.37E-01 | 8.45E-01 | 1.01E+00 | 3.22E-01 | 9.08E-01 | 1.12E+00 | 1.01E-01 | 3.70E-01 | 1.67E+00 | 2.70E-01 | 7.41E-01 | 1.56E+00 |
| TAGAP | 2.91E-02 | 1.52E-01 | 1.22E+00 | 1.99E-04 | 8.60E-04 | 1.14E+00 | 2.09E-01 | 8.89E-01 | 1.32E+00 | 1.11E-02 | 4.24E-02 | 1.18E+00 | 2.71E-03 | 9.67E-03 | 2.65E+00 | 1.93E-01 | 5.45E-01 | 1.22E+00 | 5.65E-05 | 2.07E-04 | 2.60E+00 | 1.71E-01 | 4.69E-01 | 1.56E+00 |
| ATP5HP1 | 2.92E-02 | 1.52E-01 | -1.19E+00 | 1.26E-01 | 5.44E-01 | 1.14E+00 | 4.88E-03 | 2.07E-02 | -1.35E+00 | 3.95E-03 | 1.51E-02 | -1.28E+00 | 4.20E-02 | 1.50E-01 | -1.14E+00 | 2.08E-01 | 5.88E-01 | 1.21E+00 | 1.61E-03 | 5.89E-03 | -1.24E+00 | 1.73E-02 | 4.76E-02 | -1.11E+00 |
| MED14OS | 2.92E-02 | 1.52E-01 | -1.01E+00 | 6.47E-02 | 2.80E-01 | 1.18E+00 | 1.18E-02 | 4.99E-02 | -1.30E+00 | 6.69E-02 | 2.55E-01 | 1.04E+00 | 3.92E-04 | 1.40E-03 | -1.42E+00 | 1.07E-03 | 3.01E-03 | -1.32E+00 | 2.71E-02 | 9.95E-02 | 1.77E+00 | 2.67E-01 | 7.33E-01 | 1.36E+00 |
| FOXC1 | 2.92E-02 | 1.52E-01 | 2.00E+00 | 1.31E-01 | 5.68E-01 | -1.19E+00 | 1.30E-01 | 5.51E-01 | 1.62E+00 | 1.82E-01 | 6.93E-01 | 1.11E+00 | 3.30E-03 | 1.18E-02 | -1.68E+00 | 2.59E-03 | 7.31E-03 | -1.81E+00 | 1.00E-01 | 3.68E-01 | -1.04E+00 | 2.17E-02 | 5.95E-02 | -1.49E+00 |
| NUDT12 | 2.92E-02 | 1.52E-01 | -1.08E+00 | 1.80E-01 | 7.79E-01 | 1.33E+00 | 8.83E-03 | 3.75E-02 | -1.26E+00 | 1.50E-02 | 5.74E-02 | -1.06E+00 | 2.62E-01 | 9.35E-01 | 1.41E+00 | 2.72E-02 | 7.68E-02 | 2.34E+00 | 2.26E-02 | 8.31E-02 | -1.06E+00 | 3.01E-01 | 8.26E-01 | 1.36E+00 |
| PHF13 | 2.92E-02 | 1.52E-01 | -1.22E+00 | 4.22E-02 | 1.82E-01 | -1.09E+00 | 9.94E-04 | 4.22E-03 | -1.92E+00 | 1.41E-02 | 5.38E-02 | -1.27E+00 | 3.44E-04 | 1.22E-03 | -1.84E+00 | 6.05E-04 | 1.70E-03 | -1.78E+00 | 5.36E-14 | 1.97E-13 | -1.90E+00 | 7.46E-13 | 2.05E-12 | -1.86E+00 |
| ZRANB1 | 2.92E-02 | 1.52E-01 | 1.05E+00 | 2.18E-05 | 9.40E-05 | -1.38E+00 | 7.83E-04 | 3.33E-03 | -1.18E+00 | 2.15E-07 | 8.22E-07 | -1.50E+00 | 1.51E-01 | 5.38E-01 | 1.15E+00 | 7.35E-03 | 2.07E-02 | -1.21E+00 | 2.11E-04 | 7.74E-04 | -1.22E+00 | 1.24E-06 | 3.40E-06 | -1.60E+00 |
| PHF19 | 2.92E-02 | 1.52E-01 | 1.09E+00 | 7.79E-04 | 3.37E-03 | 1.05E+00 | 4.41E-04 | 1.87E-03 | -1.03E+00 | 7.64E-05 | 2.92E-04 | 1.07E+00 | 1.90E-02 | 6.77E-02 | 1.01E+00 | 1.30E-03 | 3.67E-03 | -1.26E+00 | 3.62E-02 | 1.33E-01 | 1.06E+00 | 1.85E-01 | 5.09E-01 | 1.24E+00 |
| CYP2B7P | 2.93E-02 | 1.52E-01 | -1.04E+00 | 1.63E-03 | 7.02E-03 | -1.26E+00 | 1.59E-04 | 6.77E-04 | -1.43E+00 | 2.13E-05 | 8.14E-05 | -1.63E+00 | 1.67E-02 | 5.97E-02 | -1.78E+00 | 3.23E-02 | 9.10E-02 | -1.63E+00 | 5.61E-03 | 2.06E-02 | -1.60E+00 | 1.20E-02 | 3.29E-02 | -1.64E+00 |
| MED6 | 2.93E-02 | 1.53E-01 | 1.18E+00 | 6.95E-07 | 3.00E-06 | -1.06E+00 | 3.58E-03 | 1.52E-02 | 1.02E+00 | 1.28E-03 | 4.89E-03 | 1.11E+00 | 1.54E-01 | 5.48E-01 | 1.38E+00 | 6.82E-02 | 1.92E-01 | 1.22E+00 | 1.80E-03 | 6.61E-03 | -1.04E+00 | 4.15E-04 | 1.14E-03 | -1.08E+00 |
| TCAP | 2.93E-02 | 1.53E-01 | 1.84E+00 | 1.12E-02 | 4.84E-02 | 2.36E+00 | 9.06E-02 | 3.85E-01 | 1.58E+00 | 1.20E-01 | 4.57E-01 | 1.61E+00 | 6.02E-06 | 2.15E-05 | 5.07E+00 | 4.76E-02 | 1.34E-01 | 2.77E+00 | 7.26E-02 | 2.66E-01 | 2.01E+00 | 1.29E-02 | 3.55E-02 | 2.53E+00 |
| AL355922.1 | 2.94E-02 | 1.53E-01 | -1.32E+00 | 1.37E-02 | 5.93E-02 | -1.51E+00 | 1.05E-03 | 4.47E-03 | -1.63E+00 | 9.33E-06 | 3.56E-05 | -1.64E+00 | 2.78E-01 | 9.89E-01 | 1.19E+00 | 3.19E-01 | 8.98E-01 | -1.11E+00 | 2.00E-01 | 7.33E-01 | 1.34E+00 | 3.64E-01 | 1.00E+00 | -1.00E+00 |
| EFCAB2 | 2.94E-02 | 1.53E-01 | 1.03E+00 | 1.31E-01 | 5.67E-01 | 1.27E+00 | 1.18E-03 | 5.03E-03 | -1.20E+00 | 4.09E-04 | 1.56E-03 | -1.01E+00 | 6.43E-04 | 2.29E-03 | -1.24E+00 | 8.89E-03 | 2.51E-02 | 1.04E+00 | 7.21E-02 | 2.65E-01 | 1.01E+00 | 2.76E-01 | 7.58E-01 | 1.14E+00 |
| PIDD1 | 2.95E-02 | 1.54E-01 | -1.02E+00 | 1.40E-02 | 6.04E-02 | -1.05E+00 | 1.75E-01 | 7.42E-01 | 1.20E+00 | 2.48E-01 | 9.46E-01 | 1.57E+00 | 2.40E-01 | 8.57E-01 | 1.27E+00 | 2.65E-01 | 7.47E-01 | 1.22E+00 | 1.24E-01 | 4.56E-01 | 1.15E+00 | 9.54E-02 | 2.62E-01 | 1.52E+00 |
| PTENP1-AS | 2.95E-02 | 1.54E-01 | 1.72E+00 | 1.47E-01 | 6.34E-01 | 1.55E+00 | 1.91E-01 | 8.11E-01 | 1.31E+00 | 8.14E-02 | 3.11E-01 | 1.83E+00 | 2.28E-03 | 8.13E-03 | -1.75E+00 | 1.50E-03 | 4.23E-03 | -1.88E+00 | 4.39E-03 | 1.61E-02 | -1.40E+00 | 2.36E-03 | 6.49E-03 | -1.54E+00 |
| AP001330.4 | 2.95E-02 | 1.54E-01 | 1.80E+00 | 2.11E-02 | 9.10E-02 | -1.20E+00 | 1.40E-01 | 5.96E-01 | 1.39E+00 | 5.73E-04 | 2.19E-03 | -1.61E+00 | 4.68E-02 | 1.67E-01 | 2.18E+00 | 2.74E-01 | 7.73E-01 | 1.16E+00 | 3.28E-03 | 1.21E-02 | 1.95E+00 | 1.47E-01 | 4.04E-01 | -1.04E+00 |
| SLC27A4 | 2.95E-02 | 1.54E-01 | -1.25E+00 | 1.40E-01 | 6.04E-01 | 1.27E+00 | 6.57E-02 | 2.79E-01 | -1.06E+00 | 1.05E-01 | 4.02E-01 | 1.13E+00 | 1.32E-02 | 4.72E-02 | -1.28E+00 | 1.03E-01 | 2.90E-01 | 1.06E+00 | 5.80E-02 | 2.13E-01 | -1.07E+00 | 6.42E-02 | 1.76E-01 | 1.53E+00 |
| PSMG3-AS1 | 2.95E-02 | 1.54E-01 | -1.02E+00 | 1.62E-02 | 7.02E-02 | -1.07E+00 | 1.53E-02 | 6.49E-02 | -1.43E+00 | 1.35E-02 | 5.16E-02 | -1.33E+00 | 2.85E-02 | 1.02E-01 | -1.41E+00 | 2.48E-01 | 7.00E-01 | 1.25E+00 | 3.83E-04 | 1.41E-03 | -1.58E+00 | 8.92E-04 | 2.45E-03 | -1.44E+00 |
| DKFZP434A062 | 2.96E-02 | 1.54E-01 | 1.01E+00 | 1.57E-02 | 6.76E-02 | -1.00E+00 | 3.38E-03 | 1.44E-02 | -1.48E+00 | 2.59E-04 | 9.88E-04 | -1.57E+00 | 1.79E-01 | 6.37E-01 | -1.10E+00 | 2.45E-01 | 6.92E-01 | -1.02E+00 | 2.26E-01 | 8.29E-01 | -1.20E+00 | 3.13E-01 | 8.59E-01 | -1.15E+00 |
| TRAP1 | 2.96E-02 | 1.54E-01 | -1.16E+00 | 1.61E-01 | 6.97E-01 | 1.22E+00 | 2.24E-01 | 9.53E-01 | 1.18E+00 | 1.97E-01 | 7.51E-01 | 1.20E+00 | 2.00E-01 | 7.13E-01 | 1.04E+00 | 3.13E-01 | 8.82E-01 | 1.17E+00 | 3.90E-02 | 1.43E-01 | 1.69E+00 | 9.24E-02 | 2.54E-01 | 1.64E+00 |
| GRM3 | 2.96E-02 | 1.54E-01 | -1.14E+00 | 1.29E-01 | 5.57E-01 | 1.65E+00 | 1.43E-01 | 6.09E-01 | 1.13E+00 | 1.01E-01 | 3.87E-01 | 1.00E+00 | 1.31E-02 | 4.67E-02 | -1.45E+00 | 9.91E-02 | 2.79E-01 | -1.01E+00 | 6.19E-02 | 2.27E-01 | -1.48E+00 | 2.88E-01 | 7.90E-01 | -1.01E+00 |
| PANO1 | 2.96E-02 | 1.54E-01 | -1.29E+00 | 1.49E-01 | 6.45E-01 | 1.16E+00 | 6.73E-02 | 2.86E-01 | -1.09E+00 | 3.55E-02 | 1.36E-01 | -1.14E+00 | 1.78E-03 | 6.33E-03 | -1.92E+00 | 1.60E-02 | 4.52E-02 | -1.35E+00 | 2.63E-02 | 9.66E-02 | -1.37E+00 | 1.00E-01 | 2.75E-01 | -1.14E+00 |
| NUPR2 | 2.96E-02 | 1.54E-01 | -1.31E+00 | 1.32E-01 | 5.72E-01 | 1.26E+00 | 1.78E-01 | 7.57E-01 | 1.04E+00 | 2.31E-01 | 8.82E-01 | 1.32E+00 | 1.49E-01 | 5.30E-01 | -1.33E+00 | 2.55E-01 | 7.18E-01 | -1.16E+00 | 1.74E-01 | 6.37E-01 | 1.31E+00 | 3.38E-01 | 9.28E-01 | -1.01E+00 |
| IQCE | 2.96E-02 | 1.55E-01 | -1.14E+00 | 7.60E-02 | 3.29E-01 | 1.09E+00 | 5.08E-04 | 2.16E-03 | -1.57E+00 | 1.71E-02 | 6.53E-02 | -1.07E+00 | 1.45E-02 | 5.18E-02 | -1.63E+00 | 3.77E-02 | 1.06E-01 | -1.22E+00 | 6.77E-03 | 2.48E-02 | -1.23E+00 | 2.85E-02 | 7.82E-02 | -1.07E+00 |
| AC140076.1 | 2.97E-02 | 1.55E-01 | 1.72E+00 | 5.47E-03 | 2.36E-02 | -1.00E+00 | 1.94E-02 | 8.26E-02 | 1.59E+00 | 1.29E-01 | 4.92E-01 | 1.12E+00 | 1.39E-01 | 4.94E-01 | 1.31E+00 | 6.27E-03 | 1.77E-02 | -1.17E+00 | 2.29E-01 | 8.42E-01 | 1.39E+00 | 2.89E-02 | 7.93E-02 | -1.02E+00 |
| CDK2AP1 | 2.97E-02 | 1.55E-01 | 1.88E+00 | 1.91E-01 | 8.27E-01 | 1.20E+00 | 8.60E-02 | 3.65E-01 | 1.44E+00 | 5.13E-02 | 1.96E-01 | -1.19E+00 | 1.76E-01 | 6.26E-01 | 1.12E+00 | 1.35E-01 | 3.81E-01 | -1.24E+00 | 2.63E-01 | 9.66E-01 | 1.17E+00 | 3.35E-01 | 9.21E-01 | 1.35E+00 |
| [illegible] | | | | | | | | | | | | | | | | | | | | | | | | |

| | | | | | | | | | | | | | | | | | | | | | | | | |
|---|---|---|---|---|---|---|---|---|---|---|---|---|---|---|---|---|---|---|---|---|---|---|---|---|
| CYFIP2 | 3.26E-02 | 1.70E-01 | 1.17E+00 | 1.95E-01 | 8.42E-01 | 1.62E+00 | 6.10E-03 | 2.59E-02 | -1.13E+00 | 9.87E-02 | 3.77E-01 | 1.18E+00 | 3.63E-02 | 1.29E-01 | 1.16E+00 | 8.02E-02 | 2.26E-01 | 1.16E+00 | 8.44E-02 | 3.10E-01 | 1.12E+00 | 1.78E-01 | 4.90E-01 | 1.17E+00 |
| PLPP6 | 3.26E-02 | 1.70E-01 | -1.44E+00 | 1.78E-01 | 7.68E-01 | 1.08E+00 | 1.30E-02 | 5.52E-02 | -1.34E+00 | 1.81E-01 | 6.92E-01 | 1.25E+00 | 2.33E-05 | 8.31E-05 | -2.12E+00 | 4.85E-03 | 1.37E-02 | -1.33E+00 | 6.32E-04 | 2.32E-03 | -1.47E+00 | 1.12E-02 | 3.06E-02 | -1.12E+00 |
| HOXA10-AS | 3.27E-02 | 1.71E-01 | -1.06E+00 | 2.17E-02 | 9.39E-02 | -1.05E+00 | 4.03E-02 | 1.71E-01 | -1.20E+00 | 1.04E-02 | 3.97E-02 | -1.39E+00 | 5.45E-03 | 1.94E-02 | -1.41E+00 | 1.19E-02 | 3.35E-02 | -1.38E+00 | 1.27E-02 | 4.66E-02 | -1.32E+00 | 1.18E-02 | 3.23E-02 | -1.30E+00 |
| ZNF180 | 3.27E-02 | 1.71E-01 | -1.06E+00 | 4.42E-02 | 1.91E-01 | 1.02E+00 | 1.26E-01 | 5.37E-01 | 1.43E+00 | 2.08E-01 | 7.94E-01 | 1.37E+00 | 4.00E-02 | 1.43E-01 | -1.00E+00 | 3.57E-03 | 1.01E-02 | -1.48E+00 | 6.36E-02 | 2.33E-01 | -1.02E+00 | 1.08E-01 | 2.97E-01 | 1.09E+00 |
| WASH6P | 3.28E-02 | 1.71E-01 | 1.42E+00 | 1.31E-03 | 5.65E-03 | 1.80E+00 | 1.04E-02 | 4.40E-02 | 1.43E+00 | 1.81E-06 | 6.90E-06 | 2.05E+00 | 9.00E-02 | 3.21E-01 | 1.62E+00 | 3.59E-04 | 1.01E-03 | 2.02E+00 | 7.15E-04 | 2.62E-03 | 1.64E+00 | 3.77E-04 | 1.04E-03 | 1.74E+00 |
| PLGLB1 | 3.28E-02 | 1.71E-01 | 1.04E+00 | 2.40E-02 | 1.04E-01 | 1.07E+00 | 2.20E-01 | 9.36E-01 | 1.31E+00 | 2.45E-01 | 9.38E-01 | 1.44E+00 | 1.27E-02 | 4.54E-02 | -1.20E+00 | 2.27E-02 | 6.41E-02 | -1.19E+00 | 8.62E-02 | 3.16E-01 | 1.05E+00 | 8.58E-02 | 2.36E-01 | -1.08E+00 |
| AC002558.3 | 3.28E-02 | 1.71E-01 | -1.25E+00 | 6.87E-02 | 2.97E-01 | 1.00E+00 | 4.31E-04 | 1.83E-03 | -1.58E+00 | 9.05E-04 | 3.46E-03 | -1.37E+00 | 2.46E-03 | 8.78E-03 | -1.22E+00 | 3.00E-02 | 8.45E-02 | -1.00E+00 | 7.53E-03 | 2.76E-02 | -1.16E+00 | 9.13E-02 | 2.51E-01 | 1.11E+00 |
| TTC24 | 3.28E-02 | 1.71E-01 | 1.01E+00 | 8.67E-03 | 3.75E-02 | -1.07E+00 | 5.42E-05 | 2.30E-04 | -1.87E+00 | 1.94E-04 | 7.42E-04 | -1.60E+00 | 1.37E-02 | 4.87E-02 | -1.70E+00 | 2.87E-02 | 8.10E-02 | -1.47E+00 | 9.17E-02 | 3.36E-01 | 1.75E+00 | 2.58E-01 | 7.08E-01 | 1.41E+00 |
| FBXO36 | 3.28E-02 | 1.71E-01 | -1.16E+00 | 7.83E-02 | 3.38E-01 | 1.10E+00 | 2.06E-01 | 8.76E-01 | 1.23E+00 | 2.95E-02 | 1.13E-01 | -1.02E+00 | 1.34E-01 | 4.77E-01 | 1.84E+00 | 2.65E-01 | 7.48E-01 | 1.51E+00 | 1.38E-01 | 5.07E-01 | 1.55E+00 | 5.78E-02 | 1.59E-01 | 1.77E+00 |
| KLF8 | 3.28E-02 | 1.71E-01 | -1.15E+00 | 1.44E-01 | 6.24E-01 | 1.20E+00 | 1.09E-02 | 4.63E-02 | -1.51E+00 | 1.12E-01 | 4.26E-01 | 1.08E+00 | 7.31E-02 | 2.61E-01 | -1.06E+00 | 2.27E-01 | 6.39E-01 | 1.53E+00 | 7.79E-02 | 2.86E-01 | -1.22E+00 | 2.74E-01 | 7.51E-01 | 1.22E+00 |
| AL031432.1 | 3.29E-02 | 1.71E-01 | -1.07E+00 | 1.49E-02 | 6.46E-02 | -1.29E+00 | 8.07E-03 | 3.43E-02 | -1.21E+00 | 2.60E-03 | 9.94E-03 | -1.31E+00 | 1.92E-02 | 6.83E-02 | -1.56E+00 | 5.01E-03 | 1.41E-02 | -2.09E+00 | 5.03E-03 | 1.85E-02 | -1.27E+00 | 1.04E-04 | 2.87E-04 | -1.85E+00 |
| NINJ2 | 3.29E-02 | 1.71E-01 | -1.02E+00 | 4.26E-02 | 1.84E-01 | 1.10E+00 | 7.61E-02 | 3.23E-01 | -1.13E+00 | 1.25E-03 | 4.77E-03 | -1.38E+00 | 6.38E-03 | 2.28E-02 | -1.09E+00 | 6.64E-03 | 1.87E-02 | -1.16E+00 | 7.16E-02 | 2.63E-01 | 1.16E+00 | 2.81E-03 | 7.73E-03 | -1.14E+00 |
| C3orf33 | 3.29E-02 | 1.72E-01 | 1.11E+00 | 4.43E-03 | 1.91E-02 | -1.01E+00 | 2.07E-01 | 8.79E-01 | 1.23E+00 | 1.33E-02 | 5.08E-02 | -1.00E+00 | 1.79E-02 | 6.40E-02 | -1.10E+00 | 1.11E-02 | 3.12E-02 | -1.15E+00 | 3.46E-03 | 1.27E-02 | -1.08E+00 | 1.83E-05 | 5.02E-05 | -1.56E+00 |
| FBXO38 | 3.29E-02 | 1.72E-01 | 1.01E+00 | 1.54E-01 | 6.66E-01 | 1.32E+00 | 8.70E-05 | 3.70E-04 | -1.32E+00 | 3.30E-04 | 1.26E-03 | -1.13E+00 | 1.06E-01 | 3.77E-01 | 1.05E+00 | 2.34E-01 | 6.59E-01 | 1.23E+00 | 1.95E-06 | 7.17E-06 | -1.76E+00 | 1.94E-02 | 5.32E-02 | -1.08E+00 |
| TRIM7 | 3.29E-02 | 1.72E-01 | -1.38E+00 | 6.56E-02 | 2.83E-01 | -1.28E+00 | 1.29E-01 | 5.48E-01 | 1.39E+00 | 4.37E-02 | 1.67E-01 | 2.03E+00 | 2.38E-01 | 8.47E-01 | 1.08E+00 | 3.41E-01 | 9.62E-01 | 1.13E+00 | 1.22E-01 | 4.49E-01 | -1.03E+00 | 1.13E-01 | 3.12E-01 | 1.50E+00 |
| ZMYND19 | 3.29E-02 | 1.72E-01 | -1.12E+00 | 6.75E-02 | 2.91E-01 | 1.08E+00 | 4.59E-04 | 1.95E-03 | -1.25E+00 | 3.67E-02 | 1.40E-01 | 1.20E+00 | 2.51E-05 | 8.96E-05 | -1.87E+00 | 2.58E-04 | 7.28E-04 | -1.37E+00 | 8.69E-06 | 3.19E-05 | -1.24E+00 | 4.18E-02 | 1.15E-01 | 1.10E+00 |
| TONSL | 3.30E-02 | 1.72E-01 | -1.14E+00 | 2.09E-02 | 9.02E-02 | -1.19E+00 | 1.83E-01 | 7.76E-01 | 1.35E+00 | 2.51E-01 | 9.58E-01 | 1.44E+00 | 5.13E-02 | 1.83E-01 | -1.09E+00 | 1.59E-02 | 4.48E-02 | -1.37E+00 | 1.59E-05 | 5.84E-05 | -1.12E+00 | 1.62E-03 | 4.45E-03 | -1.03E+00 |
| CATSPERG | 3.30E-02 | 1.72E-01 | -1.13E+00 | 4.77E-02 | 2.06E-01 | 1.02E+00 | 2.01E-01 | 8.55E-01 | 1.10E+00 | 1.75E-01 | 6.69E-01 | 1.12E+00 | 2.42E-01 | 8.64E-01 | 1.23E+00 | 2.13E-01 | 5.99E-01 | 1.08E+00 | 6.88E-02 | 2.53E-01 | 1.74E+00 | 1.71E-01 | 4.69E-01 | 1.47E+00 |
| AL050331.3 | 3.30E-02 | 1.72E-01 | 1.17E+00 | 9.48E-07 | 4.10E-06 | -1.54E+00 | 4.16E-02 | 1.77E-01 | -1.08E+00 | 4.59E-04 | 1.75E-03 | -1.57E+00 | 6.85E-04 | 2.44E-03 | -1.58E+00 | 3.89E-04 | 1.10E-03 | -1.72E+00 | 2.09E-01 | 7.68E-01 | 1.04E+00 | 3.07E-01 | 8.43E-01 | 1.13E+00 |
| RPS14P4 | 3.31E-02 | 1.73E-01 | -1.23E+00 | 5.89E-02 | 2.54E-01 | -1.08E+00 | 2.25E-01 | 9.56E-01 | 1.17E+00 | 1.89E-01 | 7.23E-01 | 1.10E+00 | 1.42E-02 | 5.06E-02 | -1.27E+00 | 2.30E-01 | 6.48E-01 | 1.25E+00 | 6.06E-04 | 2.22E-03 | -1.26E+00 | 1.40E-02 | 3.84E-02 | -1.07E+00 |
| AC018754.1 | 3.31E-02 | 1.73E-01 | -1.19E+00 | 9.65E-02 | 4.17E-01 | 1.08E+00 | 1.00E-02 | 4.26E-02 | -1.40E+00 | 1.53E-02 | 5.86E-02 | -1.24E+00 | 1.30E-01 | 4.62E-01 | 1.04E+00 | 1.18E-01 | 3.32E-01 | 1.04E+00 | 1.06E-04 | 3.90E-04 | -1.95E+00 | 6.95E-04 | 1.91E-03 | -1.91E+00 |
| PGAM5 | 3.32E-02 | 1.73E-01 | 1.12E+00 | 1.11E-01 | 4.78E-01 | 1.40E+00 | 3.69E-02 | 1.57E-01 | -1.01E+00 | 3.36E-02 | 1.29E-01 | 1.09E+00 | 1.43E-02 | 5.09E-02 | -1.31E+00 | 2.72E-01 | 7.68E-01 | 1.29E+00 | 3.92E-04 | 1.44E-03 | -1.18E+00 | 8.82E-02 | 2.42E-01 | 1.12E+00 |
| UAP1L1 | 3.33E-02 | 1.73E-01 | -1.01E+00 | 8.51E-02 | 3.68E-01 | 1.28E+00 | 1.11E-01 | 4.72E-01 | 1.07E+00 | 1.76E-01 | 6.74E-01 | 1.25E+00 | 1.93E-03 | 6.89E-03 | -2.50E+00 | 6.81E-02 | 1.92E-01 | -1.04E+00 | 2.24E-02 | 8.21E-02 | -1.29E+00 | 1.50E-01 | 4.13E-01 | -1.03E+00 |
| ZNF227 | 3.33E-02 | 1.73E-01 | 1.18E+00 | 1.13E-01 | 4.88E-01 | 1.49E+00 | 9.80E-03 | 4.16E-02 | -1.09E+00 | 5.16E-02 | 1.97E-01 | 1.14E+00 | 1.10E-01 | 3.94E-01 | 1.14E+00 | 2.06E-01 | 5.80E-01 | 1.30E+00 | 5.04E-03 | 1.85E-02 | 1.00E+00 | 8.05E-03 | 2.21E-02 | -1.06E+00 |
| AC009113.1 | 3.33E-02 | 1.73E-01 | -1.21E+00 | 2.18E-01 | 9.44E-01 | 1.32E+00 | 3.30E-02 | 1.40E-01 | -1.13E+00 | 1.51E-01 | 5.77E-01 | 1.22E+00 | 2.74E-03 | 9.76E-03 | -1.36E+00 | 2.24E-02 | 6.33E-02 | -1.12E+00 | 6.64E-02 | 2.44E-01 | -1.02E+00 | 2.16E-01 | 5.93E-01 | 1.14E+00 |
| NDUFA7 | 3.34E-02 | 1.74E-01 | -1.01E+00 | 8.67E-04 | 3.75E-03 | -1.37E+00 | 2.19E-02 | 9.32E-02 | -1.13E+00 | 9.64E-03 | 3.68E-02 | -1.20E+00 | 2.44E-01 | 8.69E-01 | 1.30E+00 | 2.91E-01 | 8.21E-01 | 1.25E+00 | 4.71E-02 | 1.73E-01 | -1.03E+00 | 1.13E-03 | 3.12E-03 | -1.60E+00 |
| ZNF438 | 3.34E-02 | 1.74E-01 | -1.28E+00 | 2.27E-01 | 9.82E-01 | 1.52E+00 | 1.80E-06 | 7.64E-06 | -1.79E+00 | 5.97E-05 | 2.28E-04 | -1.39E+00 | 1.27E-05 | 4.54E-05 | -2.21E+00 | 1.20E-04 | 3.38E-04 | -1.69E+00 | 9.07E-03 | 3.33E-02 | 1.01E+00 | 1.26E-02 | 3.47E-02 | -1.27E+00 |
| SEPT4 | 3.34E-02 | 1.74E-01 | -1.60E+00 | 4.59E-02 | 1.98E-01 | -1.55E+00 | 4.58E-02 | 1.95E-01 | -1.34E+00 | 4.58E-02 | 1.75E-01 | -1.37E+00 | 2.38E-01 | 8.48E-01 | 1.34E+00 | 2.52E-01 | 7.09E-01 | 1.45E+00 | 9.93E-02 | 3.65E-01 | 1.69E+00 | 9.06E-02 | 2.49E-01 | -1.07E+00 |
| PHF8 | 3.34E-02 | 1.74E-01 | 1.06E+00 | 7.61E-03 | 3.29E-02 | 1.01E+00 | 2.42E-03 | 1.03E-02 | -1.23E+00 | 3.03E-02 | 1.16E-01 | 1.11E+00 | 6.06E-02 | 2.16E-01 | -1.01E+00 | 1.32E-01 | 3.72E-01 | 1.09E+00 | 1.90E-03 | 6.97E-03 | -1.14E+00 | 1.08E-01 | 2.96E-01 | 1.07E+00 |
| XYLB | 3.34E-02 | 1.74E-01 | -1.91E+00 | 1.73E-01 | 7.48E-01 | -1.05E+00 | 4.62E-04 | 1.96E-03 | -1.73E+00 | 2.63E-03 | 1.00E-02 | -1.34E+00 | 2.72E-02 | 9.69E-02 | -1.75E+00 | 8.71E-02 | 2.46E-01 | -1.22E+00 | 3.01E-13 | 1.10E-12 | -2.19E+00 | 1.96E-03 | 5.37E-03 | -1.05E+00 |
| PTCD1 | 3.34E-02 | 1.74E-01 | 1.75E+00 | 1.36E-01 | 5.88E-01 | 1.29E+00 | 2.34E-01 | 9.96E-01 | 1.21E+00 | 2.21E-03 | 8.44E-03 | -1.30E+00 | 4.30E-02 | 1.53E-01 | -1.07E+00 | 1.19E-01 | 3.35E-01 | 1.05E+00 | 2.71E-01 | 9.95E-01 | 1.23E+00 | 1.23E-01 | 3.39E-01 | 1.09E+00 |
| CBX7 | 3.34E-02 | 1.74E-01 | -1.00E+00 | 2.00E-01 | 8.64E-01 | 1.47E+00 | 3.17E-03 | 1.35E-02 | -1.48E+00 | 5.08E-02 | 1.94E-01 | 1.02E+00 | 9.40E-02 | 3.35E-01 | -1.10E+00 | 2.51E-01 | 7.09E-01 | 1.14E+00 | 3.74E-03 | 1.37E-02 | -1.40E+00 | 4.92E-02 | 1.35E-01 | -1.14E+00 |
| CLIC4 | 3.35E-02 | 1.74E-01 | -1.03E+00 | 5.43E-02 | 2.34E-01 | 1.10E+00 | 2.81E-02 | 1.19E-01 | -1.01E+00 | 1.12E-03 | 4.26E-03 | -1.10E+00 | 1.07E-02 | 3.81E-02 | -1.16E+00 | 1.19E-01 | 3.35E-01 | 1.14E+00 | 1.60E-03 | 5.86E-03 | -1.01E+00 | 1.57E-03 | 4.32E-03 | -1.07E+00 |
| CTU2 | 3.35E-02 | 1.74E-01 | -1.11E+00 | 6.31E-02 | 2.73E-01 | 1.13E+00 | 3.56E-02 | 1.51E-01 | -1.15E+00 | 1.77E-01 | 6.76E-01 | 1.44E+00 | 1.68E-01 | 5.99E-01 | -1.05E+00 | 1.59E-01 | 4.48E-01 | 1.63E+00 | 9.21E-03 | 3.38E-02 | 1.05E+00 | 5.15E-02 | 1.41E-01 | 1.66E+00 |
| ZNF696 | 3.36E-02 | 1.75E-01 | -1.29E+00 | 1.96E-01 | 8.48E-01 | 1.29E+00 | 1.66E-06 | 7.05E-06 | -2.14E+00 | 4.02E-03 | 1.54E-02 | -1.07E+00 | 3.57E-02 | 1.27E-01 | -1.59E+00 | 2.21E-01 | 6.22E-01 | 1.05E+00 | 2.74E-06 | 1.00E-05 | -1.77E+00 | 2.46E-02 | 6.77E-02 | -1.07E+00 |
| SLC26A1 | 3.36E-02 | 1.75E-01 | -1.25E+00 | 1.61E-02 | 6.95E-02 | -1.31E+00 | 6.36E-02 | 2.70E-01 | 1.51E+00 | 1.34E-01 | 5.10E-01 | 1.40E+00 | 1.22E-01 | 4.35E-01 | -1.08E+00 | 3.22E-01 | 9.07E-01 | 1.52E+00 | 1.48E-01 | 5.44E-01 | -1.10E+00 | 2.03E-01 | 5.58E-01 | -1.02E+00 |
| RPS10P2 | 3.36E-02 | 1.75E-01 | 1.78E+00 | 1.44E-01 | 6.22E-01 | 1.60E+00 | 4.34E-02 | 1.84E-01 | 1.62E+00 | 1.91E-01 | 7.31E-01 | 1.50E+00 | 2.70E-01 | 9.62E-01 | 1.34E+00 | 2.43E-01 | 6.85E-01 | 1.73E+00 | 2.08E-02 | 7.63E-02 | 1.96E+00 | 1.37E-01 | 3.77E-01 | 1.69E+00 |
| TMSB15B | 3.36E-02 | 1.75E-01 | 1.03E+00 | 1.46E-02 | 6.31E-02 | 1.05E+00 | 7.36E-02 | 3.13E-01 | 1.06E+00 | 2.08E-02 | 7.94E-02 | 1.15E+00 | 1.17E-05 | 4.16E-05 | -1.28E+00 | 9.98E-05 | 2.81E-04 | -1.17E+00 | 2.68E-01 | 9.83E-01 | 1.26E+00 | 3.10E-01 | 8.52E-01 | 1.32E+00 |
| AC092828.1 | 3.36E-02 | 1.75E-01 | -1.06E+00 | 2.08E-01 | 8.99E-01 | 1.32E+00 | 2.31E-01 | 9.83E-01 | 1.23E+00 | 1.69E-01 | 6.47E-01 | 1.54E+00 | 9.28E-02 | 3.31E-01 | -1.12E+00 | 3.32E-01 | 9.36E-01 | 1.21E+00 | 2.13E-01 | 7.83E-01 | 1.37E+00 | 8.38E-03 | 2.30E-02 | -1.10E+00 |
| AC067945.1 | 3.36E-02 | 1.75E-01 | -1.02E+00 | 3.19E-06 | 1.38E-05 | -2.63E+00 | 7.29E-02 | 3.10E-01 | 1.06E+00 | 2.48E-05 | 9.48E-05 | -2.03E+00 | 2.63E-01 | 9.39E-01 | 1.33E+00 | 2.06E-01 | 5.81E-01 | -1.18E+00 | 8.61E-02 | 3.16E-01 | -1.23E+00 | 1.28E-02 | 3.50E-02 | -1.80E+00 |
| CU633906.2 | 3.36E-02 | 1.75E-01 | -1.02E+00 | 2.89E-03 | 1.25E-02 | -1.31E+00 | 1.33E-01 | 5.65E-01 | 1.40E+00 | 3.40E-02 | 1.30E-01 | 1.05E+00 | 1.80E-03 | 6.43E-03 | -1.42E+00 | 3.85E-03 | 1.08E-02 | -1.33E+00 | 1.71E-02 | 6.29E-02 | 1.07E+00 | 2.35E-03 | 6.45E-03 | -1.07E+00 |
| MAGOH2P | 3.37E-02 | 1.76E-01 | 1.00E+00 | 1.62E-01 | 7.01E-01 | 1.68E+00 | 2.84E-02 | 1.21E-01 | 1.79E+00 | 8.00E-03 | 3.05E-02 | 2.33E+00 | 1.43E-01 | 5.09E-01 | 1.74E+00 | 1.48E-02 | 4.17E-02 | 2.57E+00 | 1.55E-01 | 5.71E-01 | 1.49E+00 | 7.66E-04 | 2.10E-03 | 2.23E+00 |
| AL391294.1 | 3.37E-02 | 1.76E-01 | 1.80E+00 | 1.08E-01 | 4.65E-01 | 1.76E+00 | 1.29E-01 | 5.46E-01 | 1.42E+00 | 1.50E-01 | 5.71E-01 | 1.51E+00 | 1.12E-01 | 3.98E-01 | 1.71E+00 | 1.72E-01 | 4.84E-01 | 1.80E+00 | 3.42E-03 | 1.26E-02 | 2.31E+00 | 2.13E-01 | 5.86E-01 | 1.59E+00 |
| AL133245.1 | 3.37E-02 | 1.76E-01 | -1.11E+00 | 5.19E-02 | 2.24E-01 | 1.05E+00 | 1.43E-02 | 6.07E-02 | -1.35E+00 | 6.89E-02 | 2.63E-01 | -1.00E+00 | 1.28E-01 | 4.58E-01 | 1.12E+00 | 2.52E-01 | 7.11E-01 | 1.56E+00 | 9.85E-02 | 3.62E-01 | -1.19E+00 | 1.51E-01 | 4.13E-01 | -1.13E+00 |
| ART5 | 3.38E-02 | 1.76E-01 | -1.60E+00 | 1.73E-02 | 7.47E-02 | -1.78E+00 | 2.64E-02 | 1.12E-01 | 1.69E+00 | 2.05E-01 | 7.81E-01 | 1.16E+00 | 8.19E-02 | 2.92E-01 | 1.50E+00 | 2.93E-01 | 8.25E-01 | 1.12E+00 | 2.42E-01 | 8.89E-01 | -1.03E+00 | 2.99E-01 | 8.21E-01 | -1.23E+00 |
| DMKN | 3.38E-02 | 1.76E-01 | -1.04E+00 | 1.57E-01 | 6.79E-01 | 1.20E+00 | 5.42E-02 | 2.30E-01 | 1.54E+00 | 1.34E-01 | 5.13E-01 | 1.40E+00 | 2.14E-01 | 7.61E-01 | 1.22E+00 | 3.22E-01 | 9.07E-01 | 1.16E+00 | 2.44E-01 | 8.97E-01 | 1.05E+00 | 2.33E-01 | 6.41E-01 | 1.24E+00 |
| GATAD2A | 3.38E-02 | 1.76E-01 | 1.09E+00 | 8.89E-03 | 3.84E-02 | 1.11E+00 | 1.27E-01 | 5.41E-01 | 1.11E+00 | 2.10E-02 | 8.02E-02 | 1.10E+00 | 2.83E-02 | 1.01E-01 | 1.05E+00 | 8.74E-04 | 2.47E-03 | -1.16E+00 | 1.53E-02 | 5.61E-02 | 1.04E+00 | 5.45E-02 | 1.50E-01 | 1.10E+00 |
| KHDC1 | 3.38E-02 | 1.76E-01 | 1.01E+00 | 1.47E-01 | 6.37E-01 | 1.69E+00 | 1.79E-03 | 7.61E-03 | -1.25E+00 | 2.51E-02 | 9.60E-02 | 1.08E+00 | 1.24E-02 | 4.42E-02 | -1.48E+00 | 1.74E-01 | 4.91E-01 | 1.12E+00 | 1.37E-01 | 5.03E-01 | 1.26E+00 | 3.54E-01 | 9.73E-01 | 1.39E+00 |
| CENPK | 3.39E-02 | 1.76E-01 | 1.03E+00 | 7.37E-03 | 3.18E-02 | -1.03E+00 | 8.61E-02 | 3.66E-01 | 1.29E+00 | 6.61E-02 | 2.53E-01 | 1.25E+00 | 5.78E-06 | 2.06E-05 | -1.89E+00 | 2.86E-04 | 8.06E-04 | -1.49E+00 | 3.58E-04 | 1.31E-03 | -1.05E+00 | 1.85E-01 | 5.09E-01 | 1.15E+00 |
| DGKZP1 | 3.39E-02 | 1.77E-01 | 1.01E+00 | 7.21E-02 | 3.11E-01 | 1.16E+00 | 1.03E-01 | 4.36E-01 | 1.02E+00 | 2.15E-01 | 8.20E-01 | 1.27E+00 | 7.74E-02 | 2.76E-01 | 1.98E+00 | 2.81E-01 | 7.91E-01 | 1.51E+00 | 4.04E-02 | 1.48E-01 | -1.10E+00 | 1.62E-01 | 4.45E-01 | 1.07E+00 |
| MYL12A | 3.39E-02 | 1.77E-01 | 1.19E+00 | 2.77E-02 | 1.20E-01 | 1.31E+00 | 2.12E-01 | 9.01E-01 | 1.09E+00 | 2.01E-01 | 7.68E-01 | 1.14E+00 | 1.05E-02 | 3.75E-02 | -1.06E+00 | 2.15E-01 | 6.05E-01 | 1.03E+00 | 2.49E-01 | 9.13E-01 | 1.09E+00 | 2.73E-01 | 7.50E-01 | -1.06E+00 |
| RAB11FIP3 | 3.39E-02 | 1.77E-01 | -1.11E+00 | 3.80E-02 | 1.64E-01 | -1.10E+00 | 1.21E-01 | 5.13E-01 | -1.08E+00 | 1.93E-01 | 7.38E-01 | 1.10E+00 | 1.25E-01 | 4.44E-01 | 1.03E+00 | 9.32E-02 | 2.63E-01 | -1.07E+00 | 5.41E-02 | 1.99E-01 | 1.72E+00 | 2.33E-01 | 6.41E-01 | 1.44E+00 |
| LINC01725 | 3.39E-02 | 1.77E-01 | -1.33E+00 | 1.93E-01 | 8.35E-01 | 1.09E+00 | 1.01E-03 | 4.27E-03 | -1.78E+00 | 7.89E-03 | 3.01E-02 | -1.29E+00 | 3.25E-02 | 1.16E-01 | -1.90E+00 | 8.54E-02 | 2.41E-01 | -1.31E+00 | 1.52E-01 | 5.58E-01 | 1.23E+00 | 1.45E-01 | 3.99E-01 | 1.62E+00 |
| ZNF652 | 3.39E-02 | 1.77E-01 | 1.47E+00 | 1.17E-01 | 5.03E-01 | 1.63E+00 | 1.21E-02 | 5.13E-02 | 1.03E+00 | 1.91E-03 | 7.30E-03 | 1.08E+00 | 2.20E-07 | 7.84E-07 | -1.41E+00 | 6.06E-07 | 1.71E-06 | -1.43E+00 | 3.27E-08 | 1.20E-07 | -1.61E+00 | 2.16E-05 | 5.94E-05 | -1.20E+00 |
| AC244394.1 | 3.39E-02 | 1.77E-01 | 1.08E+00 | 1.22E-04 | 5.27E-04 | -1.31E+00 | 1.42E-02 | 6.05E-02 | -1.16E+00 | 9.81E-05 | 3.75E-04 | -1.55E+00 | 6.87E-02 | 2.45E-01 | 1.09E+00 | 7.16E-03 | 2.02E-02 | -1.35E+00 | 3.38E-02 | 1.24E-01 | -1.08E+00 | 1.04E-02 | 2.85E-02 | -1.49E+00 |
| TEPP | 3.40E-02 | 1.77E-01 | 1.01E+00 | 1.03E-01 | 4.45E-01 | 1.26E+00 | 1.42E-01 | 6.05E-01 | 1.10E+00 | 1.16E-01 | 4.43E-01 | 1.16E+00 | 7.01E-02 | 2.50E-01 | 2.77E+00 | 8.98E-02 | 2.53E-01 | 2.55E+00 | 3.92E-02 | 1.44E-01 | 2.13E+00 | 2.28E-01 | 6.26E-01 | 1.48E+00 |
| AC091271.1 | 3.40E-02 | 1.77E-01 | 1.60E+00 | 3.20E-05 | 1.38E-04 | -1.08E+00 | 1.68E-02 | 7.12E-02 | 1.58E+00 | 6.44E-02 | 2.46E-01 | 1.07E+00 | 5.73E-02 | 2.04E-01 | 1.12E+00 | 1.67E-01 | 4.70E-01 | 1.13E+00 | 1.74E-01 | 6.39E-01 | 1.28E+00 | 1.71E-02 | 4.71E-02 | -1.14E+00 |
| C19orf81 | 3.40E-02 | 1.77E-01 | 1.04E+00 | 1.78E-07 | 7.70E-07 | -2.44E+00 | 4.47E-05 | 1.90E-04 | 2.47E+00 | 1.53E-01 | 5.84E-01 | 1.02E+00 | 1.43E-02 | 5.09E-02 | 2.99E+00 | 2.68E-01 | 7.56E-01 | 1.53E+00 | 2.71E-01 | 9.95E-01 | 1.18E+00 | 2.76E-01 | 7.57E-01 | 1.04E+00 |
| SLC8A1 | 3.40E-02 | 1.77E-01 | -1.57E+00 | 1.39E-01 | 6.00E-01 | -1.02E+00 | 5.32E-02 | 2.26E-01 | 1.29E+00 | 1.37E-01 | 5.23E-01 | -1.18E+00 | 2.45E-01 | 8.74E-01 | 1.25E+00 | 3.55E-01 | 1.00E+00 | 1.00E+00 | 2.65E-01 | 9.72E-01 | -1.01E+00 | 2.78E-01 | 7.62E-01 | -1.19E+00 |
| PFKFB3 | 3.41E-02 | 1.78E-01 | -1.14E+00 | 1.24E-02 | 5.36E-02 | -1.22E+00 | 3.05E-03 | 1.30E-02 | -1.35E+00 | 2.61E-03 | 9.98E-03 | -1.30E+00 | 4.74E-02 | 1.69E-01 | -1.13E+00 | 6.38E-02 | 1.80E-01 | -1.09E+00 | 9.16E-04 | 3.36E-03 | -1.28E+00 | 4.55E-02 | 1.25E-01 | -1.04E+00 |
| ARHGAP33 | 3.41E-02 | 1.78E-01 | 1.01E+00 | 2.14E-02 | 9.23E-02 | 1.11E+00 | 1.81E-01 | 7.69E-01 | 1.16E+00 | 2.27E-02 | 8.69E-02 | 1.03E+00 | 2.43E-01 | 8.66E-01 | 1.24E+00 | 2.23E-01 | 6.28E-01 | 1.05E+00 | 1.92E-02 | 7.04E-02 | -1.09E+00 | 7.22E-03 | 1.98E-02 | -1.17E+00 |
| PYROXD2 | 3.42E-02 | 1.78E-01 | -1.27E+00 | 7.04E-02 | 3.04E-01 | 1.01E+00 | 8.98E-03 | 3.82E-02 | -1.29E+00 | 3.06E-02 | 1.17E-01 | 1.01E+00 | 6.03E-04 | 2.15E-03 | -1.33E+00 | 7.44E-03 | 2.10E-02 | -1.19E+00 | 7.40E-04 | 2.72E-03 | -1.29E+00 | 9.16E-03 | 2.52E-02 | -1.12E+00 |
| CYCSP45 | 3.42E-02 | 1.78E-01 | 1.83E+00 | 1.36E-01 | 5.89E-01 | 1.15E+00 | 3.86E-03 | 1.64E-02 | 1.94E+00 | 1.44E-01 | 5.51E-01 | 1.05E+00 | 2.46E-01 | 8.77E-01 | 1.34E+00 | 5.86E-02 | 1.65E-01 | -1.32E+00 | 9.76E-02 | 3.58E-01 | 1.42E+00 | 7.51E-02 | 2.06E-01 | -1.10E+00 |
| NUP85 | 3.42E-02 | 1.78E-01 | -1.03E+00 | 2.17E-01 | 9.38E-01 | 1.44E+00 | 7.98E-02 | 3.39E-01 | 1.06E+00 | 2.09E-01 | 7.97E-01 | 1.44E+00 | 6.88E-03 | 2.45E-02 | -1.30E+00 | 2.90E-01 | 8.17E-01 | 1.44E+00 | 7.73E-08 | 2.84E-07 | -1.40E+00 | 9.25E-02 | 2.54E-01 | 1.15E+00 |
| AP1G2 | 3.42E-02 | 1.78E-01 | -1.19E+00 | 1.07E-01 | 4.63E-01 | 1.58E+00 | 9.80E-04 | 4.16E-03 | -1.59E+00 | 1.67E-02 | 6.37E-02 | -1.09E+00 | 1.84E-01 | 6.55E-01 | 1.10E+00 | 2.98E-01 | 8.40E-01 | 1.37E+00 | 3.86E-03 | 1.42E-02 | -1.52E+00 | 5.37E-03 | 1.47E-02 | -1.61E+00 |
| CHD7 | 3.43E-02 | 1.79E-01 | -2.12E+00 | 3.70E-02 | 1.60E-01 | -2.31E+00 | 2.05E-01 | 8.73E-01 | -1.09E+00 | 2.09E-01 | 8.00E-01 | -1.09E+00 | 9.68E-05 | 3.45E-04 | -2.17E+00 | 3.50E-06 | 9.88E-06 | -3.31E+00 | 2.12E-30 | 7.78E-30 | -2.99E+00 | 5.59E-21 | 1.54E-20 | -2.15E+00 |
| WWC3 | 3.43E-02 | 1.79E-01 | -1.07E+00 | 2.30E-01 | 9.92E-01 | 1.39E+00 | 2.95E-04 | 1.25E-03 | -1.39E+00 | 6.49E-03 | 2.48E-02 | -1.03E+00 | 2.60E-01 | 9.27E-01 | 1.36E+00 | 2.62E-01 | 7.38E-01 | 1.41E+00 | 1.47E-03 | 5.41E-03 | -1.26E+00 | 2.22E-02 | 6.09E-02 | 1.00E+00 |
| HELLS | 3.43E-02 | 1.79E-01 | 1.18E+00 | 5.42E-03 | 2.34E-02 | 1.06E+00 | 3.45E-02 | 1.47E-01 | 1.36E+00 | 1.80E-01 | 6.86E-01 | 1.33E+00 | 2.72E-02 | 9.71E-02 | -1.07E+00 | 1.34E-02 | 3.78E-02 | -1.17E+00 | 2.15E-04 | 7.87E-04 | -1.17E+00 | 1.38E-01 | 3.79E-01 | 1.05E+00 |
| AIMP1P1 | 3.44E-02 | 1.79E-01 | -1.18E+00 | 1.01E-01 | 4.35E-01 | 1.10E+00 | 1.46E-01 | 6.21E-01 | 1.34E+00 | 7.68E-02 | 2.93E-01 | 1.80E+00 | 1.25E-01 | 4.47E-01 | -1.29E+00 | 1.88E-01 | 5.29E-01 | -1.19E+00 | 1.28E-01 | 4.71E-01 | 1.12E+00 | 3.64E-01 | 1.00E+00 | 1.23E+00 |
| AHDC1 | 3.44E-02 | 1.79E-01 | 1.14E+00 | 3.58E-02 | 1.55E-01 | 1.22E+00 | 2.11E-04 | 8.96E-04 | -1.39E+00 | 1.25E-03 | 4.77E-03 | -1.15E+00 | 1.11E-01 | 3.97E-01 | -1.14E+00 | 2.06E-01 | 5.80E-01 | -1.06E+00 | 1.11E-01 | 4.09E-01 | 1.11E+00 | 1.25E-01 | 3.42E-01 | -1.09E+00 |
| PLCB1 | 3.44E-02 | 1.79E-01 | 1.41E+00 | 1.53E-01 | 6.59E-01 | 1.41E+00 | 6.39E-02 | 2.72E-01 | 1.02E+00 | 1.37E-02 | 5.24E-02 | 1.07E+00 | 1.82E-01 | 6.47E-01 | 1.27E+00 | 6.40E-02 | 1.80E-01 | 1.03E+00 | 2.36E-01 | 8.65E-01 | 1.12E+00 | 2.03E-01 | 5.59E-01 | -1.14E+00 |
| AFG1L | 3.44E-02 | 1.79E-01 | -1.07E+00 | 9.44E-02 | 4.08E-01 | 1.19E+00 | 4.33E-03 | 1.84E-02 | -1.33E+00 | 3.51E-04 | 1.34E-03 | -1.37E+00 | 7.12E-03 | 2.54E-02 | -1.13E+00 | 2.71E-02 | 7.65E-02 | -1.08E+00 | 2.02E-04 | 7.43E-04 | -1.44E+00 | 6.31E-02 | 1.73E-01 | -1.04E+00 |
| CFL1 | 3.44E-02 | 1.79E-01 | -1.04E+00 | 1.81E-01 | 7.82E-01 | 1.05E+00 | 2.31E-01 | 9.80E-01 | -1.00E+00 | 8.32E-02 | 3.18E-01 | 1.06E+00 | 9.74E-06 | 3.47E-05 | -1.20E+00 | 2.78E-03 | 7.85E-03 | -1.12E+00 | 5.75E-02 | 2.11E-01 | -1.07E+00 | 3.11E-01 | 8.53E-01 | -1.01E+00 |
| COL14A1 | 3.44E-02 | 1.79E-01 | -1.89E+00 | 2.27E-02 | 9.80E-02 | -2.02E+00 | 1.72E-01 | 7.31E-01 | 1.17E+00 | 2.39E-01 | 9.12E-01 | 1.06E+00 | 1.73E-01 | 6.15E-01 | 1.67E+00 | 2.11E-01 | 5.95E-01 | 1.04E+00 | 8.09E-11 | 2.97E-10 | -2.42E+00 | 8.14E-04 | 2.24E-03 | -1.28E+00 |
| AC093642.2 | 3.44E-02 | 1.79E-01 | -1.19E+00 | 6.46E-02 | 2.79E-01 | -1.02E+00 | 1.98E-01 | 8.41E-01 | 1.10E+00 | 1.63E-01 | 6.24E-01 | 1.18E+00 | 1.32E-01 | 4.71E-01 | -1.47E+00 | 2.21E-01 | 6.24E-01 | -1.17E+00 | 2.08E-01 | 7.63E-01 | 1.03E+00 | 3.62E-01 | 9.94E-01 | 1.10E+00 |
| LINC01521 | 3.45E-02 | 1.80E-01 | 1.03E+00 | 2.12E-01 | 9.17E-01 | 1.54E+00 | 2.17E-02 | 9.24E-02 | -1.10E+00 | 6.08E-02 | 2.32E-01 | 1.12E+00 | 6.01E-02 | 2.14E-01 | -1.18E+00 | 1.51E-01 | 4.27E-01 | 1.04E+00 | 2.82E-02 | 1.04E-01 | -1.29E+00 | 5.39E-02 | 1.48E-01 | -1.27E+00 |
| SMIM26 | 3.45E-02 | 1.80E-01 | 1.05E+00 | 9.15E-05 | 3.95E-04 | 1.00E+00 | 2.63E-02 | 1.12E-01 | -1.07E+00 | 2.02E-04 | 7.72E-04 | -1.13E+00 | 8.15E-03 | 2.91E-02 | -1.01E+00 | 4.71E-04 | 1.33E-03 | -1.09E+00 | 2.66E-03 | 9.78E-03 | -1.06E+00 | 4.64E-04 | 1.28E-03 | -1.16E+00 |
| IGHD | 3.45E-02 | 1.80E-01 | 1.41E+00 | 1.26E-01 | 5.44E-01 | 1.44E+00 | 1.57E-01 | 6.67E-01 | 1.12E+00 | 8.09E-02 | 3.09E-01 | 1.16E+00 | 2.23E-01 | 7.94E-01 | -1.06E+00 | 3.30E-01 | 9.30E-01 | 1.16E+00 | 2.28E-01 | 8.35E-01 | -1.12E+00 | 3.20E-01 | 8.80E-01 | -1.02E+00 |
| ZNF416 | 3.46E-02 | 1.80E-01 | -1.15E+00 | 5.95E-02 | 2.57E-01 | -1.01E+00 | 2.08E-01 | 8.85E-01 | 1.15E+00 | 4.23E-02 | 1.62E-01 | -1.08E+00 | 5.84E-02 | 2.08E-01 | 1.04E+00 | 2.50E-03 | 7.06E-03 | -1.74E+00 | 2.50E-03 | 9.19E-03 | -1.35E+00 | 8.05E-03 | 2.21E-02 | -1.30E+00 |
| TTC28-AS1 | 3.47E-02 | 1.81E-01 | 1.22E+00 | 2.50E-02 | 1.08E-01 | 1.33E+00 | 7.62E-04 | 3.24E-03 | -1.07E+00 | 7.07E-04 | 2.70E-03 | 1.02E+00 | 1.13E-01 | 4.02E-01 | 1.15E+00 | 3.22E-01 | 9.08E-01 | 1.36E+00 | 2.10E-01 | 7.71E-01 | 1.33E+00 | 7.18E-02 | 1.97E-01 | 1.12E+00 |
| RPS17P2 | 3.47E-02 | 1.81E-01 | 1.50E+00 | 1.03E-03 | 4.46E-03 | 1.19E+00 | 3.35E-03 | 1.42E-02 | 1.62E+00 | 5.43E-03 | 2.08E-02 | 1.18E+00 | 2.25E-01 | 8.02E-01 | 1.40E+00 | 3.86E-02 | 1.09E-01 | 1.11E+00 | 1.77E-01 | 6.49E-01 | 1.46E+00 | 1.65E-01 | 4.54E-01 | 1.21E+00 |
| LZTR1 | 3.47E-02 | 1.81E-01 | -1.01E+00 | 2.06E-01 | 8.90E-01 | 1.51E+00 | 7.63E-02 | 3.24E-01 | 1.11E+00 | 2.00E-01 | 7.65E-01 | 1.63E+00 | 5.58E-03 | 1.99E-02 | -1.32E+00 | 3.58E-02 | 1.01E-01 | -1.05E+00 | 3.97E-03 | 1.46E-02 | -1.22E+00 | 9.39E-02 | 2.58E-01 | 1.12E+00 |
| DLG3 | 3.47E-02 | 1.81E-01 | -1.15E+00 | 2.30E-02 | 9.96E-02 | -1.11E+00 | 1.45E-01 | 6.15E-01 | 1.08E+00 | 5.26E-02 | 2.01E-01 | -1.05E+00 | 2.77E-01 | 9.88E-01 | 1.10E+00 | 3.25E-01 | 9.15E-01 | 1.27E+00 | 8.48E-02 | 3.11E-01 | -1.02E+00 | 3.62E-01 | 9.94E-01 | 1.13E+00 |
| TKT | 3.47E-02 | 1.81E-01 | 1.07E+00 | 4.88E-05 | 2.11E-04 | 1.18E+00 | 1.24E-01 | 5.26E-01 | -1.02E+00 | 1.26E-01 | 4.81E-01 | 1.08E+00 | 2.57E-02 | 9.15E-02 | -1.11E+00 | 1.15E-01 | 3.24E-01 | 1.03E+00 | 1.63E-07 | 6.00E-07 | -1.18E+00 | 1.00E-01 | 2.75E-01 | 1.01E+00 |
| PHOSPHO2 | 3.47E-02 | 1.81E-01 | 1.07E+00 | 4.65E-02 | 2.01E-01 | 1.17E+00 | 1.99E-05 | 8.44E-05 | -1.51E+00 | 1.02E-05 | 3.90E-05 | -1.37E+00 | 6.66E-04 | 2.37E-03 | -1.25E+00 | 3.88E-02 | 1.09E-01 | 1.20E+00 | 6.12E-05 | 2.25E-04 | -1.41E+00 | 1.10E-02 | 3.02E-02 | -1.00E+00 |
| RASGRP2 | 3.47E-02 | 1.81E-01 | 1.40E+00 | 1.83E-01 | 7.92E-01 | 1.46E+00 | 2.34E-01 | 9.94E-01 | 1.18E+00 | 5.11E-02 | 1.95E-01 | 1.17E+00 | 1.91E-03 | 6.82E-03 | 1.04E+00 | 1.66E-03 | 4.68E-03 | 1.01E+00 | 2.19E-01 | 8.05E-01 | 1.33E+00 | 3.19E-01 | 8.76E-01 | 1.18E+00 |
| HMGB3P24 | 3.47E-02 | 1.81E-01 | -1.16E+00 | 2.53E-02 | 1.09E-01 | -1.21E+00 | 1.36E-04 | 5.77E-04 | -1.44E+00 | 7.04E-07 | 2.69E-06 | -1.71E+00 | 1.72E-01 | 6.14E-01 | 1.65E+00 | 2.10E-01 | 5.92E-01 | 1.69E+00 | 2.95E-03 | 1.08E-02 | -1.32E+00 | 1.15E-03 | 3.15E-03 | -1.36E+00 |
| ARHGAP26 | 3.48E-02 | 1.81E-01 | 1.60E+00 | 1.85E-01 | 7.97E-01 | 1.39E+00 | 1.68E-02 | 7.12E-02 | 1.59E+00 | 2.56E-01 | 9.79E-01 | 1.42E+00 | 2.86E-02 | 1.02E-01 | 1.00E+00 | 2.47E-03 | 6.96E-03 | -1.52E+00 | 2.28E-01 | 8.38E-01 | 1.16E+00 | 3.28E-01 | 9.02E-01 | 1.22E+00 |
| NLRP3 | 3.49E-02 | 1.82E-01 | 1.66E+00 | 3.04E-03 | 1.31E-02 | -1.18E+00 | 3.91E-02 | 1.66E-01 | 1.08E+00 | 1.09E-04 | 4.14E-04 | -1.37E+00 | 1.47E-01 | 5.23E-01 | 1.94E+00 | 3.04E-02 | 8.56E-02 | -1.32E+00 | 6.10E-02 | 2.24E-01 | 1.90E+00 | 2.30E-01 | 6.32E-01 | -1.25E+00 |
| C10orf10 | 3.49E-02 | 1.82E-01 | 1.16E+00 | 5.81E-02 | 2.51E-01 | 1.80E+00 | 4.82E-02 | 2.05E-01 | 1.05E+00 | 2.55E-01 | 9.74E-01 | 1.45E+00 | 2.65E-01 | 9.44E-01 | 1.82E+00 | 1.96E-01 | 5.53E-01 | -1.59E+00 | 5.74E-02 | 2.11E-01 | 3.47E+00 | 2.56E-01 | 7.04E-01 | 1.61E+00 |
| PIGV | 3.49E-02 | 1.82E-01 | 1.06E+00 | 1.88E-01 | 8.12E-01 | 1.52E+00 | 9.55E-05 | 4.06E-04 | -1.34E+00 | 3.57E-04 | 1.36E-03 | -1.11E+00 | 1.55E-03 | 5.52E-03 | -1.50E+00 | 3.27E-01 | 9.22E-01 | 1.33E+00 | 4.42E-05 | 1.62E-04 | -1.42E+00 | 1.89E-02 | 5.20E-02 | -1.08E+00 |
| LILRB2 | 3.49E-02 | 1.82E-01 | -1.05E+00 | 1.59E-01 | 6.86E-01 | 1.49E+00 | 4.28E-04 | 1.82E-03 | -1.87E+00 | 5.61E-04 | 2.14E-03 | -1.69E+00 | 2.78E-01 | 9.91E-01 | 1.20E+00 | 3.42E-01 | 9.64E-01 | 1.17E+00 | 1.52E-01 | 5.59E-01 | -1.42E+00 | 1.48E-01 | 4.08E-01 | -1.85E+00 |
| ZC3H12C | 3.50E-02 | 1.82E-01 | 1.27E+00 | 7.07E-06 | 3.06E-05 | -1.04E+00 | 1.82E-02 | 7.74E-02 | -1.14E+00 | 2.46E-04 | 9.40E-04 | -1.43E+00 | 7.69E-02 | 2.74E-01 | 1.02E+00 | 4.52E-02 | 1.28E-01 | -1.21E+00 | 2.38E-03 | 8.73E-03 | -1.20E+00 | 2.92E-02 | 8.02E-02 | 1.04E+00 |
| HIP1R | 3.50E-02 | 1.82E-01 | 1.09E+00 | 1.70E-02 | 7.34E-02 | 1.06E+00 | 4.74E-02 | 2.01E-01 | -1.00E+00 | 9.30E-04 | 3.55E-03 | -1.40E+00 | 2.66E-01 | 9.50E-01 | 1.31E+00 | 2.79E-01 | 7.87E-01 | 1.15E+00 | 1.40E-01 | 5.13E-01 | -1.10E+00 | 2.17E-01 | 5.97E-01 | -1.12E+00 |
| AL117332.1 | 3.50E-02 | 1.82E-01 | 1.09E+00 | 3.97E-05 | 1.72E-04 | -1.43E+00 | 1.46E-01 | 6.19E-01 | 1.09E+00 | 3.09E-02 | 1.18E-01 | 1.04E+00 | 3.77E-02 | 1.35E-01 | -1.12E+00 | 8.75E-03 | 2.47E-02 | -1.61E+00 | 2.03E-01 | 7.47E-01 | 1.35E+00 | 2.59E-01 | 7.10E-01 | 1.15E+00 |
| ACAD9 | 3.50E-02 | 1.82E-01 | 1.01E+00 | 6.15E-02 | 2.66E-01 | 1.16E+00 | 1.85E-02 | 7.86E-02 | -1.05E+00 | 6.55E-04 | 2.50E-03 | -1.08E+00 | 9.94E-04 | 3.54E-03 | -1.35E+00 | 8.50E-03 | 2.40E-02 | -1.05E+00 | 6.01E-05 | 2.21E-04 | -1.17E+00 | 1.72E-02 | 4.73E-02 | 1.01E+00 |
| CHMP1B | 3.50E-02 | 1.82E-01 | 1.13E+00 | 2.10E-06 | 9.09E-06 | -1.18E+00 | 1.64E-02 | 6.99E-02 | 1.01E+00 | 2.14E-05 | 8.19E-05 | -1.15E+00 | 1.28E-04 | 4.58E-04 | 1.07E+00 | 2.57E-06 | 7.24E-06 | -1.06E+00 | 4.59E-06 | 1.69E-05 | -1.26E+00 | 7.42E-06 | 2.04E-05 | -1.31E+00 |
| KNOP1P4 | 3.50E-02 | 1.82E-01 | -1.13E+00 | 8.46E-02 | 3.65E-01 | 1.11E+00 | 1.40E-01 | 5.94E-01 | 1.11E+00 | 1.63E-01 | 6.22E-01 | 1.22E+00 | 2.44E-02 | 8.68E-02 | -1.85E+00 | 1.65E-01 | 4.65E-01 | -1.14E+00 | 2.00E-01 | 7.34E-01 | 1.05E+00 | 1.42E-01 | 3.89E-01 | 1.75E+00 |
| KIAA0087 | 3.50E-02 | 1.82E-01 | 2.43E+00 | 2.06E-01 | 8.88E-01 | 1.44E+00 | 1.07E-01 | 4.56E-01 | 1.60E+00 | 1.94E-01 | 7.40E-01 | -1.03E+00 | 1.47E-01 | 5.24E-01 | 1.68E+00 | 2.44E-01 | 6.87E-01 | 1.35E+00 | 2.38E-01 | 8.74E-01 | -1.05E+00 | 3.59E-01 | 9.87E-01 | 1.07E+00 |
| AC002429.2 | 3.50E-02 | 1.83E-01 | -1.12E+00 | 1.56E-01 | 6.73E-01 | 1.21E+00 | 9.69E-02 | 4.12E-01 | -1.16E+00 | 5.01E-02 | 1.92E-01 | -1.27E+00 | 2.17E-01 | 7.74E-01 | 1.19E+00 | 1.42E-01 | 3.99E-01 | -1.28E+00 | 6.48E-02 | 2.38E-01 | 1.48E+00 | 2.70E-01 | 7.42E-01 | 1.19E+00 |
| AC002094.5 | 3.51E-02 | 1.83E-01 | 1.01E+00 | 1.79E-01 | 7.72E-01 | 1.43E+00 | 5.64E-02 | 2.39E-01 | -1.11E+00 | 1.00E-01 | 3.83E-01 | 1.07E+00 | 6.37E-02 | 2.27E-01 | 1.04E+00 | 1.91E-01 | 5.38E-01 | 1.22E+00 | 5.53E-02 | 2.03E-01 | -1.12E+00 | 1.04E-01 | 2.86E-01 | -1.11E+00 |
| TSKS | 3.51E-02 | 1.83E-01 | 1.70E+00 | 9.40E-02 | 4.06E-01 | 1.74E+00 | 5.36E-02 | 2.28E-01 | 1.56E+00 | 9.94E-02 | 3.80E-01 | 1.63E+00 | 2.49E-01 | 8.88E-01 | 1.22E+00 | 2.37E-01 | 6.69E-01 | 1.61E+00 | 8.48E-02 | 3.11E-01 | 1.33E+00 | 1.50E-02 | 4.12E-02 | 1.78E+00 |
| CKAP2L | 3.51E-02 | 1.83E-01 | -1.42E+00 | 4.69E-02 | 2.02E-01 | -1.33E+00 | 2.27E-01 | 9.64E-01 | 1.17E+00 | 9.60E-02 | 3.67E-01 | 1.59E+00 | 1.55E-01 | 5.52E-01 | -1.46E+00 | 2.48E-01 | 6.99E-01 | -1.27E+00 | 2.49E-02 | 9.12E-02 | -1.31E+00 | 3.23E-01 | 8.88E-01 | 1.00E+00 |
| ZSCAN12 | 3.51E-02 | 1.83E-01 | -1.08E+00 | 3.34E-02 | 1.44E-01 | -1.02E+00 | 1.35E-01 | 5.72E-01 | 1.02E+00 | 8.61E-02 | 3.29E-01 | 1.57E+00 | 2.04E-01 | 7.28E-01 | 1.35E+00 | 2.40E-01 | 6.75E-01 | 1.37E+00 | 1.45E-01 | 5.33E-01 | 1.12E+00 | 3.16E-01 | 8.69E-01 | 1.16E+00 |
| AC008735.1 | 3.52E-02 | 1.84E-01 | -1.16E+00 | 6.41E-02 | 2.77E-01 | -1.04E+00 | 1.14E-01 | 4.85E-01 | 1.47E+00 | 2.49E-01 | 9.51E-01 | 1.30E+00 | 1.78E-01 | 6.33E-01 | 1.20E+00 | 1.43E-01 | 4.02E-01 | -1.06E+00 | 6.16E-02 | 2.26E-01 | -1.20E+00 | 1.35E-01 | 3.72E-01 | -1.02E+00 |
| RUFY2 | 3.53E-02 | 1.84E-01 | 1.10E+00 | 4.24E-02 | 1.83E-01 | 1.19E+00 | 2.00E-01 | 8.48E-01 | 1.32E+00 | 3.87E-02 | 1.48E-01 | 1.28E+00 | 1.72E-01 | 6.13E-01 | 1.49E+00 | 1.52E-01 | 4.28E-01 | 1.32E+00 | 1.11E-01 | 4.07E-01 | 1.25E+00 | 1.03E-01 | 2.84E-01 | 1.19E+00 |
| ARHGEF7 | 3.53E-02 | 1.84E-01 | -1.00E+00 | 2.19E-01 | 9.46E-01 | 1.39E+00 | 2.10E-01 | 8.92E-01 | 1.21E+00 | 1.48E-01 | 5.66E-01 | 1.21E+00 | 2.49E-01 | 8.87E-01 | 1.30E+00 | 3.17E-01 | 8.94E-01 | 1.41E+00 | 1.80E-03 | 6.62E-03 | -1.07E+00 | 4.20E-03 | 1.15E-02 | -1.08E+00 |
| DLC1 | 3.54E-02 | 1.84E-01 | -2.80E+00 | 5.00E-02 | 2.16E-01 | -2.58E+00 | 3.15E-02 | 1.34E-01 | -2.25E+00 | 3.16E-02 | 1.21E-01 | -2.42E+00 | 3.84E-31 | 1.37E-30 | -2.24E+00 | 1.90E-11 | 5.36E-11 | -1.16E+00 | 1.82E-39 | 6.67E-39 | -1.73E+00 | 2.52E-21 | 6.92E-21 | -1.29E+00 |
| RECK | 3.54E-02 | 1.84E-01 | 1.09E+00 | 6.91E-03 | 2.99E-02 | 1.05E+00 | 5.54E-02 | 2.36E-01 | 1.37E+00 | 1.10E-01 | 4.18E-01 | 1.09E+00 | 3.69E-02 | 1.31E-01 | -1.13E+00 | 1.21E-01 | 3.41E-01 | 1.01E+00 | 9.11E-02 | 3.34E-01 | 1.12E+00 | 3.20E-02 | 8.79E-02 | -1.14E+00 |
| SLC25A23 | 3.54E-02 | 1.84E-01 | 1.02E+00 | 1.76E-01 | 7.62E-01 | 1.37E+00 | 1.12E-03 | 4.74E-03 | -1.28E+00 | 1.72E-02 | 6.59E-02 | -1.02E+00 | 1.67E-02 | 5.97E-02 | -1.50E+00 | 3.55E-01 | 1.00E+00 | 1.26E+00 | 2.27E-03 | 8.34E-03 | -1.29E+00 | 2.19E-02 | 6.02E-02 | -1.18E+00 |
| AL157392.3 | 3.54E-02 | 1.84E-01 | -1.12E+00 | 1.08E-01 | 4.66E-01 | 1.81E+00 | 1.16E-01 | 4.95E-01 | -1.06E+00 | 2.54E-01 | 9.69E-01 | 1.26E+00 | 1.48E-01 | 5.28E-01 | 1.03E+00 | 2.66E-01 | 7.49E-01 | 1.49E+00 | 2.89E-02 | 1.06E-01 | -1.12E+00 | 3.34E-01 | 9.17E-01 | 1.25E+00 |
| TTL | 3.55E-02 | 1.85E-01 | 1.08E+00 | 3.35E-02 | 1.45E-01 | 1.18E+00 | 1.01E-02 | 4.29E-02 | -1.06E+00 | 5.15E-04 | 1.97E-03 | -1.12E+00 | 8.07E-04 | 2.88E-03 | -1.66E+00 | 6.87E-02 | 1.94E-01 | -1.07E+00 | 2.70E-07 | 9.91E-07 | -1.35E+00 | 4.07E-04 | 1.12E-03 | -1.15E+00 |
| DAPK3 | 3.55E-02 | 1.85E-01 | 1.13E+00 | 3.42E-02 | 1.48E-01 | 1.18E+00 | 1.07E-01 | 4.53E-01 | 1.08E+00 | 1.13E-01 | 4.33E-01 | 1.19E+00 | 2.23E-02 | 7.96E-02 | 1.14E+00 | 1.80E-02 | 5.08E-02 | 1.11E+00 | 2.15E-03 | 7.89E-03 | -1.17E+00 | 1.62E-02 | 4.46E-02 | -1.14E+00 |
| VWA1 | 3.55E-02 | 1.85E-01 | -1.00E+00 | 1.69E-01 | 7.28E-01 | 1.33E+00 | 3.14E-02 | 1.33E-01 | 1.64E+00 | 7.61E-02 | 2.91E-01 | 1.61E+00 | 1.93E-01 | 6.87E-01 | 1.53E+00 | 2.92E-01 | 8.22E-01 | 1.34E+00 | 1.98E-01 | 7.29E-01 | 1.21E+00 | 2.95E-01 | 8.11E-01 | 1.28E+00 |
| AC144530.1 | 3.55E-02 | 1.85E-01 | 1.47E+00 | 1.89E-01 | 8.15E-01 | 1.43E+00 | 4.11E-03 | 1.75E-02 | 1.68E+00 | 1.36E-01 | 5.18E-01 | 1.55E+00 | 1.06E-01 | 3.77E-01 | 1.60E+00 | 2.79E-01 | 7.88E-01 | 1.58E+00 | 4.94E-02 | 1.81E-01 | 1.13E+00 | 2.63E-01 | 7.22E-01 | 1.24E+00 |
| LINC02447 | 3.55E-02 | 1.85E-01 | -1.00E+00 | 2.96E-03 | 1.28E-02 | -1.15E+00 | 1.51E-01 | 6.43E-01 | 1.28E+00 | 1.69E-01 | 6.46E-01 | 1.01E+00 | 3.29E-04 | 1.17E-03 | -1.66E+00 | 6.65E-04 | 1.88E-03 | -1.57E+00 | 1.14E-01 | 4.20E-01 | 1.46E+00 | 2.80E-01 | 7.69E-01 | -1.06E+00 |
| P2RY12 | 3.56E-02 | 1.85E-01 | 1.01E+00 | 1.01E-01 | 4.34E-01 | 1.30E+00 | 1.91E-01 | 8.11E-01 | 1.28E+00 | 2.37E-01 | 9.06E-01 | 1.30E+00 | 2.35E-01 | 8.39E-01 | 1.68E+00 | 3.02E-01 | 8.51E-01 | 1.33E+00 | 5.41E-02 | 1.99E-01 | 1.68E+00 | 3.00E-01 | 8.25E-01 | 1.31E+00 |
| ZNF564 | 3.56E-02 | 1.85E-01 | 1.01E+00 | 3.46E-03 | 1.49E-02 | -1.17E+00 | 2.69E-02 | 1.14E-01 | -1.25E+00 | 5.42E-02 | 2.07E-01 | 1.02E+00 | 2.15E-01 | 7.66E-01 | 1.28E+00 | 3.28E-01 | 9.26E-01 | 1.29E+00 | 1.57E-01 | 5.77E-01 | -1.12E+00 | 2.93E-01 | 8.05E-01 | 1.07E+00 |
| VMAC | 3.56E-02 | 1.85E-01 | 1.06E+00 | 2.63E-02 | 1.13E-01 | 1.11E+00 | 6.82E-03 | 2.90E-02 | -1.20E+00 | 1.57E-02 | 6.01E-02 | 1.06E+00 | 1.27E-01 | 4.52E-01 | 1.31E+00 | 2.06E-01 | 5.81E-01 | 1.36E+00 | 6.22E-03 | 2.28E-02 | -1.23E+00 | 6.86E-02 | 1.88E-01 | -1.01E+00 |
| RPL21P93 | 3.56E-02 | 1.86E-01 | 1.07E+00 | 2.92E-04 | 1.26E-03 | -1.37E+00 | 1.97E-02 | 8.38E-02 | 1.03E+00 | 1.54E-09 | 5.89E-09 | -1.86E+00 | 8.85E-02 | 3.15E-01 | 1.19E+00 | 4.58E-05 | 1.29E-04 | -1.84E+00 | 2.62E-02 | 9.61E-02 | 1.06E+00 | 2.35E-06 | 6.45E-06 | -1.68E+00 |
| TTC3P1 | 3.56E-02 | 1.86E-01 | 1.16E+00 | 1.06E-01 | 4.60E-01 | 1.61E+00 | 1.74E-01 | 7.38E-01 | 1.13E+00 | 8.11E-02 | 3.10E-01 | 1.52E+00 | 1.56E-01 | 5.58E-01 | 1.20E+00 | 1.52E-01 | 4.28E-01 | 1.47E+00 | 9.01E-02 | 3.31E-01 | 1.45E+00 | 1.18E-02 | 3.23E-02 | 1.67E+00 |
| PRR7-AS1 | 3.57E-02 | 1.86E-01 | 1.02E+00 | 3.45E-02 | 1.49E-01 | 1.02E+00 | 1.76E-01 | 7.49E-01 | 1.00E+00 | 1.87E-01 | 7.14E-01 | 1.46E+00 | 2.73E-01 | 9.75E-01 | 1.38E+00 | 1.35E-01 | 3.81E-01 | -1.07E+00 | 4.21E-04 | 1.54E-03 | -1.20E+00 | 6.50E-05 | 1.78E-04 | -1.73E+00 |
| CTNNA3 | 3.57E-02 | 1.86E-01 | 1.15E+00 | 1.38E-01 | 5.97E-01 | 1.73E+00 | 3.24E-02 | 1.38E-01 | -1.08E+00 | 9.56E-02 | 3.65E-01 | 1.85E+00 | 9.94E-03 | 3.54E-02 | -1.22E+00 | 1.31E-01 | 3.69E-01 | 1.26E+00 | 1.65E-01 | 6.05E-01 | 1.53E+00 | 1.81E-01 | 4.96E-01 | 1.53E+00 |
| TMEM255B | 3.58E-02 | 1.86E-01 | 1.95E+00 | 3.98E-02 | 1.72E-01 | 2.23E+00 | 1.33E-01 | 5.66E-01 | 1.36E+00 | 8.55E-03 | 3.27E-02 | 2.22E+00 | 4.74E-02 | 1.69E-01 | 2.30E+00 | 4.86E-02 | 1.37E-01 | 2.52E+00 | 1.56E-01 | 5.74E-01 | 1.37E+00 | 1.06E-01 | 2.91E-01 | 1.55E+00 |
| RPL12P46 | 3.58E-02 | 1.86E-01 | 1.12E+00 | 7.25E-05 | 3.13E-04 | -1.44E+00 | 9.33E-02 | 3.96E-01 | 1.10E+00 | 4.55E-05 | 1.74E-04 | -1.53E+00 | 1.23E-03 | 4.38E-03 | -1.38E+00 | 1.58E-06 | 4.44E-06 | -2.56E+00 | 9.43E-04 | 3.46E-03 | -1.33E+00 | 5.77E-07 | 1.58E-06 | -2.20E+00 |
| ZFAND3 | 3.58E-02 | 1.87E-01 | 1.11E+00 | 6.62E-02 | 2.86E-01 | 1.24E+00 | 2.48E-02 | 1.05E-01 | 1.01E+00 | 4.98E-03 | 1.90E-02 | 1.07E+00 | 1.24E-04 | 4.41E-04 | -1.08E+00 | 2.38E-05 | 6.71E-05 | -1.08E+00 | 5.47E-03 | 2.01E-02 | 1.00E+00 | 7.63E-03 | 2.10E-02 | -1.02E+00 |
| RPL23AP74 | 3.58E-02 | 1.87E-01 | 1.74E+00 | 1.93E-01 | 8.34E-01 | 1.27E+00 | 1.41E-01 | 5.97E-01 | 1.15E+00 | 1.46E-03 | 5.56E-03 | -1.35E+00 | 1.30E-03 | 4.65E-03 | 3.85E+00 | 1.07E-01 | 3.01E-01 | 2.24E+00 | 1.93E-02 | 7.07E-02 | 1.92E+00 | 9.53E-02 | 2.62E-01 | -1.04E+00 |
| AL031428.1 | 3.59E-02 | 1.87E-01 | -1.03E+00 | 2.11E-01 | 9.12E-01 | 1.40E+00 | 1.19E-02 | 5.07E-02 | -1.44E+00 | 1.05E-01 | 4.02E-01 | 1.04E+00 | 3.94E-02 | 1.40E-01 | -1.15E+00 | 2.03E-01 | 5.73E-01 | 1.14E+00 | 6.51E-02 | 2.39E-01 | -1.03E+00 | 2.72E-01 | 7.47E-01 | 1.26E+00 |
| ECM2 | 3.59E-02 | 1.87E-01 | -1.49E+00 | 2.15E-01 | 9.28E-01 | 1.17E+00 | 5.57E-02 | 2.37E-01 | -1.12E+00 | 2.83E-02 | 1.08E-01 | -1.12E+00 | 8.13E-03 | 2.90E-02 | -2.18E+00 | 8.67E-02 | 2.44E-01 | -1.15E+00 | 1.38E-01 | 5.05E-01 | 1.11E+00 | 1.50E-01 | 4.13E-01 | 1.64E+00 |
| MICALL1 | 3.59E-02 | 1.87E-01 | 1.02E+00 | 1.24E-01 | 5.37E-01 | 1.34E+00 | 5.53E-02 | 2.35E-01 | -1.02E+00 | 5.95E-02 | 2.27E-01 | 1.13E+00 | 7.81E-04 | 2.78E-03 | -1.53E+00 | 3.20E-03 | 9.03E-03 | -1.36E+00 | 2.66E-01 | 9.77E-01 | 1.21E+00 | 2.42E-01 | 6.65E-01 | 1.18E+00 |
| KRT18P55 | 3.59E-02 | 1.87E-01 | 1.78E+00 | 4.24E-02 | 1.83E-01 | 2.05E+00 | 3.36E-04 | 1.43E-03 | 2.51E+00 | 2.73E-03 | 1.04E-02 | 2.66E+00 | 1.12E-01 | 4.01E-01 | -1.19E+00 | 1.03E-01 | 2.91E-01 | -1.35E+00 | 1.41E-01 | 5.18E-01 | 1.49E+00 | 3.07E-01 | 8.44E-01 | 1.26E+00 |
| VPS26BP1 | 3.60E-02 | 1.87E-01 | -1.21E+00 | 7.67E-02 | 3.32E-01 | -1.09E+00 | 2.16E-01 | 9.16E-01 | 1.22E+00 | 2.26E-01 | 8.63E-01 | 1.27E+00 | 3.80E-03 | 1.35E-02 | -2.18E+00 | 1.12E-01 | 3.16E-01 | -1.10E+00 | 1.82E-01 | 6.69E-01 | -1.09E+00 | 1.26E-01 | 3.46E-01 | 1.60E+00 |
| SARM1 | 3.60E-02 | 1.87E-01 | -1.02E+00 | 6.75E-02 | 2.92E-01 | 1.11E+00 | 1.98E-02 | 8.41E-02 | -1.27E+00 | 2.06E-01 | 7.87E-01 | 1.28E+00 | 3.69E-02 | 1.32E-01 | -1.27E+00 | 2.95E-01 | 8.31E-01 | 1.43E+00 | 6.30E-02 | 2.31E-01 | -1.10E+00 | 2.39E-01 | 6.57E-01 | 1.28E+00 |
| AC074141.1 | 3.60E-02 | 1.88E-01 | -1.11E+00 | 1.47E-01 | 6.37E-01 | 1.37E+00 | 9.85E-02 | 4.19E-01 | -1.11E+00 | 1.13E-01 | 4.33E-01 | 1.01E+00 | 1.78E-01 | 6.35E-01 | 1.02E+00 | 3.06E-01 | 8.62E-01 | 1.29E+00 | 2.48E-01 | 9.11E-01 | 1.42E+00 | 3.02E-01 | 8.29E-01 | 1.31E+00 |
| AC008555.5 | 3.60E-02 | 1.88E-01 | 1.03E+00 | 2.27E-01 | 9.83E-01 | 1.54E+00 | 1.39E-01 | 5.90E-01 | 1.04E+00 | 2.09E-01 | 7.99E-01 | 1.54E+00 | 4.03E-02 | 1.44E-01 | -1.14E+00 | 1.43E-01 | 4.02E-01 | 1.08E+00 | 8.86E-04 | 3.25E-03 | -1.30E+00 | 1.12E-02 | 3.07E-02 | -1.11E+00 |
| AL359643.3 | 3.61E-02 | 1.88E-01 | -1.04E+00 | 2.36E-02 | 1.02E-01 | -1.03E+00 | 2.89E-04 | 1.23E-03 | -1.61E+00 | 3.94E-03 | 1.51E-02 | -1.22E+00 | 1.30E-01 | 4.64E-01 | -1.26E+00 | 2.00E-01 | 5.63E-01 | -1.08E+00 | 2.19E-02 | 8.02E-02 | -1.53E+00 | 1.37E-01 | 3.76E-01 | -1.18E+00 |
| AC021945.1 | 3.61E-02 | 1.88E-01 | -1.31E+00 | 3.01E-02 | 1.30E-01 | -1.38E+00 | 6.02E-02 | 2.56E-01 | 1.59E+00 | 1.27E-01 | 4.83E-01 | 1.51E+00 | 1.23E-01 | 4.37E-01 | -1.94E+00 | 2.27E-01 | 6.40E-01 | -1.43E+00 | 1.77E-01 | 6.50E-01 | 1.31E+00 | 1.51E-02 | 4.15E-02 | 1.80E+00 |
| MAD2L1BP | 3.61E-02 | 1.88E-01 | 1.03E+00 | 2.56E-03 | 1.10E-02 | -1.14E+00 | 1.62E-03 | 6.89E-03 | -1.19E+00 | 7.24E-04 | 2.76E-03 | -1.10E+00 | 3.96E-02 | 1.41E-01 | 1.15E+00 | 1.19E-01 | 3.35E-01 | 1.22E+00 | 1.07E-04 | 3.93E-04 | -1.09E+00 | 7.83E-05 | 2.15E-04 | -1.03E+00 |
| VSTM1 | 3.61E-02 | 1.88E-01 | -1.40E+00 | 2.26E-02 | 9.75E-02 | -1.59E+00 | 1.85E-01 | 7.85E-01 | 1.13E+00 | 7.86E-02 | 3.00E-01 | -1.12E+00 | 2.21E-04 | 7.89E-04 | -1.33E+00 | 4.35E-07 | 1.23E-06 | -2.17E+00 | 5.38E-06 | 1.98E-05 | -1.96E+00 | 3.22E-06 | 8.83E-06 | -2.26E+00 |
| IL1RL1 | 3.61E-02 | 1.88E-01 | -1.44E+00 | 2.80E-02 | 1.21E-01 | -2.57E+00 | 1.96E-01 | 8.32E-01 | 1.16E+00 | 2.60E-01 | 9.95E-01 | 1.23E+00 | 3.32E-08 | 1.18E-07 | 1.22E+00 | 8.85E-30 | 2.49E-29 | -2.49E+00 | 3.30E-02 | 1.21E-01 | -1.62E+00 | 2.54E-01 | 6.97E-01 | -1.15E+00 |
| CDCA8 | 3.61E-02 | 1.88E-01 | -1.58E+00 | 6.04E-02 | 2.61E-01 | -1.48E+00 | 1.23E-01 | 5.23E-01 | 1.05E+00 | 6.84E-02 | 2.61E-01 | 1.21E+00 | 3.73E-02 | 1.33E-01 | -2.26E+00 | 1.40E-01 | 3.96E-01 | -1.36E+00 | 7.01E-02 | 2.57E-01 | -1.15E+00 | 3.46E-02 | 9.50E-02 | 1.34E+00 |
| MALT1 | 3.62E-02 | 1.89E-01 | 1.01E+00 | 5.83E-03 | 2.52E-02 | -1.02E+00 | 1.99E-01 | 8.45E-01 | 1.22E+00 | 2.09E-02 | 8.00E-02 | 1.08E+00 | 1.01E-02 | 3.61E-02 | -1.40E+00 | 1.20E-01 | 3.37E-01 | 1.00E+00 | 7.81E-04 | 2.87E-03 | -1.36E+00 | 3.34E-02 | 9.16E-02 | -1.04E+00 |
| AC004898.1 | 3.62E-02 | 1.89E-01 | 1.02E+00 | 2.84E-02 | 1.23E-01 | 1.07E+00 | 2.12E-04 | 8.99E-04 | -1.51E+00 | 2.13E-04 | 8.15E-04 | -1.38E+00 | 1.70E-01 | 6.07E-01 | -1.18E+00 | 3.11E-01 | 8.76E-01 | 1.09E+00 | 2.10E-03 | 7.72E-03 | -1.47E+00 | 1.98E-02 | 5.44E-02 | -1.24E+00 |
| HHAT | 3.63E-02 | 1.89E-01 | 1.04E+00 | 8.31E-03 | 3.59E-02 | -1.01E+00 | 3.68E-02 | 1.56E-01 | -1.09E+00 | 9.67E-02 | 3.69E-01 | 1.19E+00 | 1.08E-01 | 3.85E-01 | -1.08E+00 | 5.36E-02 | 1.51E-01 | -1.21E+00 | 1.97E-01 | 7.24E-01 | -1.04E+00 | 2.28E-01 | 6.26E-01 | -1.16E+00 |
| AGBL3 | 3.63E-02 | 1.89E-01 | -1.12E+00 | 2.16E-01 | 9.35E-01 | 1.49E+00 | 1.08E-01 | 4.58E-01 | -1.15E+00 | 2.16E-01 | 8.24E-01 | 1.36E+00 | 8.42E-02 | 3.00E-01 | -1.15E+00 | 2.25E-01 | 6.33E-01 | 1.21E+00 | 1.80E-02 | 6.60E-02 | 1.70E+00 | 9.28E-02 | 2.55E-01 | 1.55E+00 |
| UQCRFS1P1 | 3.64E-02 | 1.89E-01 | 1.71E+00 | 4.25E-03 | 1.84E-02 | 2.37E+00 | 4.91E-02 | 2.09E-01 | 1.58E+00 | 1.69E-02 | 6.47E-02 | 1.97E+00 | 2.06E-01 | 7.36E-01 | 1.26E+00 | 2.86E-01 | 8.07E-01 | 1.54E+00 | 5.97E-02 | 2.19E-01 | 1.52E+00 | 7.96E-03 | 2.19E-02 | 1.78E+00 |
| C16orf74 | 3.64E-02 | 1.90E-01 | 2.01E+00 | 4.54E-02 | 1.96E-01 | 1.06E+00 | 2.27E-02 | 9.67E-02 | 1.29E+00 | 9.23E-02 | 3.52E-01 | 1.37E+00 | 2.71E-01 | 9.67E-01 | 1.09E+00 | 3.46E-01 | 9.75E-01 | 1.17E+00 | 2.42E-01 | 8.88E-01 | 1.13E+00 | 2.08E-01 | 5.71E-01 | 1.75E+00 |
| SLFN14 | 3.64E-02 | 1.90E-01 | -1.41E+00 | 4.20E-02 | 1.81E-01 | -1.42E+00 | 1.85E-01 | 7.84E-01 | 1.17E+00 | 2.04E-01 | 7.80E-01 | 1.17E+00 | 3.31E-05 | 1.18E-04 | -1.75E+00 | 1.60E-04 | 4.51E-04 | -1.67E+00 | 1.11E-03 | 4.06E-03 | 1.35E+00 | 6.44E-07 | 1.77E-06 | -1.61E+00 |
| ITM2C | 3.64E-02 | 1.90E-01 | 1.28E+00 | 2.06E-04 | 8.90E-04 | 1.53E+00 | 2.51E-02 | 1.07E-01 | 1.19E+00 | 2.08E-02 | 7.93E-02 | 1.25E+00 | 1.30E-03 | 4.63E-03 | 1.18E+00 | 2.50E-04 | 7.05E-04 | 1.16E+00 | 1.16E-01 | 4.25E-01 | 1.45E+00 | 8.30E-03 | 2.28E-02 | 1.56E+00 |
| AL445472.1 | 3.64E-02 | 1.90E-01 | 1.08E+00 | 3.06E-02 | 1.32E-01 | 1.16E+00 | 1.08E-02 | 4.58E-02 | -1.39E+00 | 9.85E-03 | 3.76E-02 | -1.19E+00 | 4.13E-05 | 1.47E-04 | -1.30E+00 | 4.37E-06 | 1.23E-05 | -1.47E+00 | 1.32E-02 | 4.86E-02 | -1.29E+00 | 1.87E-03 | 5.14E-03 | -1.46E+00 |
| UBE2D3P2 | 3.64E-02 | 1.90E-01 | -1.04E+00 | 6.13E-02 | 2.65E-01 | 1.18E+00 | 2.04E-01 | 8.69E-01 | 1.23E+00 | 9.42E-02 | 3.60E-01 | 1.09E+00 | 1.03E-01 | 3.68E-01 | 1.07E+00 | 1.67E-01 | 4.71E-01 | 1.10E+00 | 1.84E-01 | 6.77E-01 | 1.26E+00 | 3.10E-01 | 8.53E-01 | 1.14E+00 |
| AC015799.1 | 3.64E-02 | 1.90E-01 | 1.05E+00 | 3.27E-20 | 1.41E-19 | -2.69E+00 | 1.10E-04 | 4.69E-04 | -1.18E+00 | 2.58E-21 | 9.86E-21 | -3.24E+00 | 1.16E-02 | 4.13E-02 | 1.05E+00 | 5.66E-10 | 1.60E-09 | -2.32E+00 | 2.63E-02 | 9.67E-02 | -1.11E+00 | 1.24E-09 | 3.41E-09 | -2.76E+00 |
| MED29 | 3.65E-02 | 1.90E-01 | 1.16E+00 | 3.86E-02 | 1.67E-01 | 1.29E+00 | 6.99E-03 | 2.97E-02 | -1.04E+00 | 1.65E-04 | 6.32E-04 | -1.10E+00 | 4.67E-03 | 1.67E-02 | -1.03E+00 | 1.84E-02 | 5.20E-02 | 1.12E+00 | 2.64E-02 | 9.70E-02 | 1.01E+00 | 9.59E-02 | 2.63E-01 | 1.06E+00 |
| AL450326.1 | 3.65E-02 | 1.90E-01 | -1.04E+00 | 1.05E-02 | 4.53E-02 | -1.15E+00 | 1.25E-02 | 5.33E-02 | -1.21E+00 | 7.46E-02 | 2.85E-01 | 1.13E+00 | 1.39E-03 | 4.97E-03 | -1.45E+00 | 5.78E-03 | 1.63E-02 | -1.29E+00 | 3.81E-02 | 1.40E-01 | 1.01E+00 | 2.25E-01 | 6.17E-01 | 1.14E+00 |
| AC005070.3 | 3.65E-02 | 1.90E-01 | 1.52E+00 | 1.62E-01 | 7.00E-01 | 1.11E+00 | 3.40E-02 | 1.45E-01 | 1.60E+00 | 1.75E-01 | 6.67E-01 | 1.13E+00 | 2.47E-01 | 8.82E-01 | 1.34E+00 | 9.46E-03 | 2.67E-02 | -1.46E+00 | 1.13E-01 | 4.16E-01 | 1.44E+00 | 9.79E-02 | 2.69E-01 | -1.05E+00 |
| SOX6 | 3.65E-02 | 1.90E-01 | 1.89E+00 | 6.71E-02 | 2.90E-01 | 2.04E+00 | 8.23E-02 | 3.50E-01 | -1.11E+00 | 6.74E-02 | 2.57E-01 | -1.12E+00 | 9.87E-02 | 3.52E-01 | 1.83E+00 | 8.04E-02 | 2.27E-01 | 2.18E+00 | 5.04E-02 | 1.85E-01 | -1.06E+00 | 2.83E-03 | 7.78E-03 | -1.48E+00 |
| AC074085.2 | 3.65E-02 | 1.90E-01 | 1.64E+00 | 1.43E-01 | 6.19E-01 | 1.53E+00 | 8.76E-02 | 3.72E-01 | 1.48E+00 | 1.22E-01 | 4.65E-01 | 1.20E+00 | 1.91E-02 | 6.82E-02 | 2.17E+00 | 2.12E-01 | 5.97E-01 | 1.72E+00 | 2.42E-02 | 8.89E-02 | 1.88E+00 | 7.15E-02 | 1.96E-01 | 1.79E+00 |
| ZNF407 | 3.66E-02 | 1.91E-01 | 1.02E+00 | 2.60E-02 | 1.12E-01 | 1.07E+00 | 3.84E-02 | 1.63E-01 | -1.09E+00 | 2.54E-02 | 9.71E-02 | 1.02E+00 | 2.35E-01 | 8.36E-01 | 1.27E+00 | 2.37E-01 | 6.69E-01 | 1.19E+00 | 6.57E-03 | 2.41E-02 | -1.18E+00 | 5.78E-03 | 1.59E-02 | -1.19E+00 |
| AC040162.1 | 3.66E-02 | 1.91E-01 | 1.06E+00 | 1.15E-02 | 4.96E-02 | -1.03E+00 | 1.72E-03 | 7.32E-03 | -1.40E+00 | 1.01E-02 | 3.85E-02 | -1.10E+00 | 5.45E-03 | 1.94E-02 | -1.26E+00 | 6.59E-02 | 1.86E-01 | 1.09E+00 | 1.18E-03 | 4.32E-03 | -1.46E+00 | 7.65E-03 | 2.10E-02 | -1.34E+00 |
| AC012184.3 | 3.66E-02 | 1.91E-01 | 1.05E+00 | 6.72E-02 | 2.90E-01 | 1.07E+00 | 1.13E-01 | 4.81E-01 | 1.15E+00 | 5.40E-02 | 2.06E-01 | 1.03E+00 | 1.64E-01 | 5.86E-01 | 1.08E+00 | 3.30E-01 | 9.29E-01 | 1.46E+00 | 1.13E-01 | 4.15E-01 | 1.18E+00 | 6.44E-02 | 1.77E-01 | -1.06E+00 |
| ZNF708 | 3.67E-02 | 1.91E-01 | 1.26E+00 | 9.27E-03 | 4.01E-02 | 1.32E+00 | 1.51E-03 | 6.40E-03 | -1.15E+00 | 6.27E-03 | 2.39E-02 | -1.01E+00 | 1.96E-02 | 6.99E-02 | 1.75E+00 | 1.41E-02 | 3.98E-02 | 1.83E+00 | 2.15E-01 | 7.88E-01 | 1.26E+00 | 2.84E-01 | 7.80E-01 | 1.22E+00 |
| VDAC1P4 | 3.67E-02 | 1.91E-01 | 1.93E+00 | 1.08E-01 | 4.68E-01 | 1.66E+00 | 1.02E-01 | 4.32E-01 | 1.41E+00 | 1.47E-01 | 5.61E-01 | 1.02E+00 | 2.64E-01 | 9.41E-01 | 1.15E+00 | 2.63E-01 | 7.42E-01 | 1.09E+00 | 8.49E-02 | 3.12E-01 | 1.62E+00 | 2.10E-01 | 5.76E-01 | 1.46E+00 |
| C5orf34 | 3.67E-02 | 1.91E-01 | -1.08E+00 | 5.93E-02 | 2.56E-01 | 1.03E+00 | 9.18E-02 | 3.90E-01 | 1.04E+00 | 2.58E-01 | 9.87E-01 | 1.36E+00 | 5.00E-02 | 1.78E-01 | -1.77E+00 | 2.61E-01 | 7.36E-01 | -1.00E+00 | 5.42E-02 | 1.99E-01 | -1.11E+00 | 1.88E-01 | 5.17E-01 | 1.00E+00 |
| AC068620.2 | 3.67E-02 | 1.92E-01 | 1.61E+00 | 1.60E-01 | 6.92E-01 | 1.13E+00 | 1.86E-01 | 7.90E-01 | 1.21E+00 | 3.13E-02 | 1.19E-01 | -1.08E+00 | 3.22E-02 | 1.15E-01 | 2.15E+00 | 2.65E-01 | 7.46E-01 | 1.44E+00 | 2.33E-01 | 8.57E-01 | 1.14E+00 | 2.68E-01 | 7.37E-01 | -1.05E+00 |
| CHAC2 | 3.68E-02 | 1.92E-01 | -1.11E+00 | 8.82E-02 | 3.81E-01 | 1.04E+00 | 3.79E-03 | 1.61E-02 | -1.21E+00 | 1.54E-03 | 5.88E-03 | -1.14E+00 | 9.94E-02 | 3.54E-01 | 1.05E+00 | 1.09E-01 | 3.08E-01 | -1.09E+00 | 7.52E-08 | 2.76E-07 | -1.42E+00 | 2.82E-01 | 7.76E-01 | 1.15E+00 |
| LCK | 3.68E-02 | 1.92E-01 | 1.74E+00 | 1.25E-02 | 5.41E-02 | 2.21E+00 | 1.15E-01 | 4.90E-01 | -1.69E+00 | 1.88E-01 | 7.17E-01 | -1.53E+00 | 2.57E-01 | 9.16E-01 | 1.13E+00 | 3.06E-01 | 8.62E-01 | -1.09E+00 | 2.21E-01 | 8.11E-01 | 1.16E+00 | 1.27E-01 | 3.48E-01 | 1.61E+00 |
| CCDC97 | 3.68E-02 | 1.92E-01 | 1.11E+00 | 7.12E-04 | 3.07E-03 | 1.02E+00 | 1.05E-03 | 4.48E-03 | -1.15E+00 | 4.87E-03 | 1.86E-02 | 1.06E+00 | 2.03E-02 | 7.26E-02 | 1.15E+00 | 2.16E-02 | 6.08E-02 | 1.11E+00 | 2.65E-02 | 9.72E-02 | -1.02E+00 | 1.94E-01 | 5.32E-01 | 1.14E+00 |
| KAT2B | 3.69E-02 | 1.92E-01 | 2.01E+00 | 1.13E-01 | 4.90E-01 | 1.82E+00 | 3.19E-02 | 1.36E-01 | -1.09E+00 | 1.54E-02 | 5.89E-02 | -1.18E+00 | 1.24E-02 | 4.41E-02 | -1.81E+00 | 6.75E-02 | 1.90E-01 | -1.11E+00 | 7.90E-08 | 2.90E-07 | -1.42E+00 | 2.83E-09 | 7.78E-09 | -1.65E+00 |
| TMEM184B | 3.69E-02 | 1.92E-01 | -1.05E+00 | 1.03E-01 | 4.46E-01 | 1.28E+00 | 7.08E-02 | 3.01E-01 | -1.13E+00 | 1.97E-01 | 7.51E-01 | 1.41E+00 | 2.19E-03 | 7.80E-03 | -1.25E+00 | 4.96E-03 | 1.40E-02 | -1.25E+00 | 1.89E-05 | 6.93E-05 | -1.31E+00 | 4.45E-02 | 1.22E-01 | 1.12E+00 |
| RPL34P34 | 3.69E-02 | 1.92E-01 | -1.02E+00 | 6.61E-48 | 2.86E-47 | -1.88E+00 | 2.28E-01 | 9.71E-01 | 1.03E+00 | 2.94E-22 | 1.12E-21 | -1.87E+00 | 1.92E-01 | 6.85E-01 | 1.06E+00 | 3.95E-21 | 1.11E-20 | -1.84E+00 | 1.38E-03 | 5.06E-03 | -1.03E+00 | 1.61E-24 | 4.42E-24 | -1.92E+00 |
| AC087388.1 | 3.69E-02 | 1.92E-01 | -1.01E+00 | 1.24E-02 | 5.37E-02 | -1.20E+00 | 2.31E-05 | 9.79E-05 | -1.59E+00 | 1.60E-04 | 6.12E-04 | -1.29E+00 | 7.10E-02 | 2.53E-01 | -1.17E+00 | 2.30E-01 | 6.50E-01 | 1.05E+00 | 2.09E-02 | 7.69E-02 | -1.21E+00 | 1.72E-01 | 4.72E-01 | 1.11E+00 |
| DMTN | 3.69E-02 | 1.92E-01 | -1.80E+00 | 4.70E-02 | 2.03E-01 | -1.78E+00 | 1.33E-01 | 5.64E-01 | 1.65E+00 | 1.88E-01 | 7.17E-01 | 1.50E+00 | 1.10E-02 | 3.94E-02 | -1.12E+00 | 9.59E-02 | 2.70E-01 | 1.08E+00 | 4.08E-05 | 1.50E-04 | 1.03E+00 | 6.08E-17 | 1.67E-16 | -1.44E+00 |
| RPS23P8 | 3.69E-02 | 1.93E-01 | 1.04E+00 | 1.10E-06 | 4.75E-06 | -1.13E+00 | 1.00E-02 | 4.27E-02 | 1.07E+00 | 8.37E-03 | 3.20E-02 | -1.08E+00 | 4.89E-03 | 1.74E-02 | 1.10E+00 | 4.97E-02 | 1.40E-01 | -1.06E+00 | 9.55E-02 | 3.51E-01 | 1.04E+00 | 2.26E-04 | 6.20E-04 | -1.10E+00 |
| AC083843.2 | 3.70E-02 | 1.93E-01 | 1.78E+00 | 2.19E-01 | 9.45E-01 | 1.27E+00 | 1.85E-02 | 7.84E-02 | 1.94E+00 | 1.14E-01 | 4.36E-01 | 1.04E+00 | 1.07E-01 | 3.81E-01 | 1.91E+00 | 1.62E-01 | 4.57E-01 | 2.04E+00 | 4.78E-03 | 1.75E-02 | 2.24E+00 | 2.23E-01 | 6.12E-01 | 1.46E+00 |
| SLC8B1 | 3.70E-02 | 1.93E-01 | 1.14E+00 | 3.97E-02 | 1.71E-01 | 1.22E+00 | 1.70E-02 | 7.24E-02 | 1.01E+00 | 1.71E-03 | 6.53E-03 | 1.01E+00 | 2.23E-02 | 7.96E-02 | -1.16E+00 | 1.21E-01 | 3.40E-01 | 1.08E+00 | 1.74E-02 | 6.38E-02 | -1.18E+00 | 3.87E-02 | 1.06E-01 | -1.17E+00 |
| KIF23 | 3.70E-02 | 1.93E-01 | -1.48E+00 | 8.90E-02 | 3.85E-01 | 1.01E+00 | 1.34E-01 | 5.70E-01 | -1.03E+00 | 2.36E-01 | 9.02E-01 | 1.42E+00 | 2.97E-02 | 1.06E-01 | -2.35E+00 | 2.48E-02 | 6.98E-02 | -2.74E+00 | 1.06E-06 | 3.89E-06 | -1.46E+00 | 1.57E-04 | 4.31E-04 | -1.14E+00 |
| HSP90AA5P | 3.71E-02 | 1.93E-01 | 2.04E+00 | 3.63E-05 | 1.57E-04 | 4.09E+00 | 8.21E-02 | 3.49E-01 | 1.48E+00 | 7.52E-03 | 2.87E-02 | 2.13E+00 | 2.63E-01 | 9.38E-01 | 1.31E+00 | 1.43E-02 | 4.04E-02 | 2.62E+00 | 4.55E-02 | 1.67E-01 | 1.68E+00 | 9.22E-13 | 2.53E-12 | 3.63E+00 |
| ADRB2 | 3.71E-02 | 1.93E-01 | 1.39E+00 | 7.80E-02 | 3.37E-01 | -1.96E+00 | 6.46E-02 | 2.75E-01 | 1.71E+00 | 3.26E-02 | 1.24E-01 | 1.02E+00 | 1.61E-01 | 5.75E-01 | 1.06E+00 | 8.12E-02 | 2.29E-01 | -1.57E+00 | 1.82E-01 | 6.67E-01 | 1.35E+00 | 2.41E-01 | 6.63E-01 | 1.04E+00 |
| LILRB1 | 3.71E-02 | 1.93E-01 | -1.40E+00 | 5.22E-02 | 2.26E-01 | -1.32E+00 | 4.02E-02 | 1.71E-01 | -1.56E+00 | 3.42E-02 | 1.31E-01 | -1.61E+00 | 2.80E-01 | 1.00E+00 | 1.00E+00 | 3.55E-01 | 1.00E+00 | -1.00E+00 | 2.72E-01 | 1.00E+00 | 1.00E+00 | 3.64E-01 | 1.00E+00 | -1.00E+00 |
| ATG7 | 3.72E-02 | 1.94E-01 | -1.08E+00 | 7.62E-02 | 3.29E-01 | 1.11E+00 | 1.90E-08 | 8.07E-08 | -1.59E+00 | 1.91E-08 | 7.31E-08 | -1.32E+00 | 7.77E-02 | 2.77E-01 | -1.03E+00 | 1.43E-01 | 4.02E-01 | 1.03E+00 | 2.43E-05 | 8.92E-05 | -1.76E+00 | 4.63E-03 | 1.27E-02 | -1.23E+00 |
| CNTLN | 3.72E-02 | 1.94E-01 | 1.16E+00 | 1.92E-02 | 8.30E-02 | 1.34E+00 | 6.66E-02 | 2.83E-01 | 1.03E+00 | 6.42E-02 | 2.45E-01 | 1.15E+00 | 1.91E-04 | 6.81E-04 | -1.33E+00 | 5.23E-03 | 1.48E-02 | -1.08E+00 | 3.56E-05 | 1.31E-04 | -1.29E+00 | 7.19E-02 | 1.98E-01 | -1.07E+00 |
| FIG4 | 3.72E-02 | 1.94E-01 | 1.02E+00 | 2.72E-02 | 1.17E-01 | 1.10E+00 | 3.48E-07 | 1.48E-06 | -1.63E+00 | 4.10E-07 | 1.57E-06 | -1.49E+00 | 2.72E-05 | 9.72E-05 | -1.61E+00 | 1.38E-04 | 3.89E-04 | -1.50E+00 | 3.06E-02 | 1.12E-01 | 1.02E+00 | 6.28E-02 | 1.73E-01 | 1.00E+00 |
| THAP9 | 3.72E-02 | 1.94E-01 | -1.15E+00 | 3.37E-02 | 1.45E-01 | -1.16E+00 | 6.01E-02 | 2.55E-01 | 1.00E+00 | 1.61E-02 | 6.16E-02 | -1.12E+00 | 1.90E-02 | 6.76E-02 | -1.41E+00 | 6.16E-02 | 1.74E-01 | -1.07E+00 | 4.98E-02 | 1.83E-01 | -1.16E+00 | 1.07E-01 | 2.95E-01 | -1.08E+00 |
| RPL10P13 | 3.73E-02 | 1.94E-01 | 1.19E+00 | 4.50E-07 | 1.94E-06 | -1.23E+00 | 2.33E-01 | 9.90E-01 | 1.26E+00 | 1.37E-06 | 5.23E-06 | -1.43E+00 | 4.74E-04 | 1.69E-03 | -1.30E+00 | 3.83E-08 | 1.08E-07 | -1.91E+00 | 2.90E-04 | 1.07E-03 | -1.13E+00 | 2.00E-08 | 5.49E-08 | -1.59E+00 |
| FCN1 | 3.73E-02 | 1.94E-01 | -1.98E+00 | 6.93E-02 | 2.99E-01 | 2.70E+00 | 1.75E-01 | 7.42E-01 | -1.78E+00 | 3.78E-02 | 1.44E-01 | -4.90E+00 | 1.25E-01 | 4.47E-01 | -1.37E+00 | 1.25E-01 | 3.53E-01 | -1.94E+00 | 9.80E-02 | 3.60E-01 | -3.24E+00 | 1.11E-01 | 3.04E-01 | -2.84E+00 |
| TEDC2 | 3.73E-02 | 1.94E-01 | -1.71E+00 | 7.92E-02 | 3.42E-01 | -1.28E+00 | 8.20E-02 | 3.49E-01 | 1.42E+00 | 1.80E-01 | 6.88E-01 | 1.44E+00 | 1.11E-01 | 3.94E-01 | -1.26E+00 | 1.01E-01 | 2.84E-01 | -1.59E+00 | 6.52E-03 | 2.39E-02 | -1.07E+00 | 2.45E-01 | 6.73E-01 | 1.42E+00 |
| AC130448.1 | 3.73E-02 | 1.94E-01 | -1.30E+00 | 1.45E-01 | 6.26E-01 | 1.12E+00 | 1.01E-03 | 4.27E-03 | -1.66E+00 | 2.26E-03 | 8.64E-03 | -1.39E+00 | 2.69E-02 | 9.60E-02 | -1.64E+00 | 2.09E-01 | 5.89E-01 | 1.04E+00 | 6.85E-02 | 2.51E-01 | -1.27E+00 | 1.48E-01 | 4.08E-01 | -1.00E+00 |
| DCLK2 | 3.73E-02 | 1.94E-01 | 1.03E+00 | 5.02E-02 | 2.17E-01 | 1.14E+00 | 1.67E-01 | 7.09E-01 | 1.40E+00 | 1.75E-01 | 6.68E-01 | 1.63E+00 | 1.13E-03 | 4.03E-03 | -2.11E+00 | 3.93E-03 | 1.11E-02 | -1.86E+00 | 2.01E-01 | 7.39E-01 | 1.09E+00 | 2.34E-01 | 6.42E-01 | 1.35E+00 |
| YES1P1 | 3.73E-02 | 1.94E-01 | 1.77E+00 | 1.25E-01 | 5.41E-01 | 1.61E+00 | 1.51E-01 | 6.40E-01 | 1.14E+00 | 1.97E-01 | 7.54E-01 | 1.20E+00 | 1.82E-01 | 6.50E-01 | 1.32E+00 | 1.25E-01 | 3.52E-01 | 1.15E+00 | 9.89E-02 | 3.63E-01 | -1.15E+00 | 8.89E-02 | 2.44E-01 | -1.24E+00 |
| ZNF419 | 3.73E-02 | 1.94E-01 | 1.07E+00 | 4.80E-02 | 2.08E-01 | 1.23E+00 | 1.87E-01 | 7.93E-01 | 1.12E+00 | 1.35E-01 | 5.17E-01 | 1.22E+00 | 1.94E-01 | 6.93E-01 | 1.38E+00 | 1.86E-01 | 5.23E-01 | 1.23E+00 | 1.16E-01 | 4.27E-01 | 1.05E+00 | 1.01E-01 | 2.76E-01 | 1.04E+00 |
| AC026462.1 | 3.74E-02 | 1.95E-01 | 1.13E+00 | 2.45E-04 | 1.06E-03 | -1.28E+00 | 1.45E-02 | 6.15E-02 | 1.60E+00 | 2.28E-01 | 8.70E-01 | 1.37E+00 | 1.78E-01 | 6.35E-01 | 1.47E+00 | 3.43E-01 | 9.68E-01 | 1.26E+00 | 2.63E-02 | 9.67E-02 | 1.65E+00 | 2.97E-01 | 8.15E-01 | 1.17E+00 |
| TCTA | 3.74E-02 | 1.95E-01 | 1.11E+00 | 1.25E-01 | 5.38E-01 | 1.36E+00 | 1.62E-01 | 6.87E-01 | 1.20E+00 | 1.62E-01 | 6.17E-01 | 1.36E+00 | 1.32E-01 | 4.70E-01 | 1.26E+00 | 2.34E-01 | 6.59E-01 | 1.31E+00 | 1.37E-02 | 5.03E-02 | 1.09E+00 | 2.45E-02 | 6.74E-02 | 1.05E+00 |
| DDIAS | 3.74E-02 | 1.95E-01 | -1.43E+00 | 1.13E-01 | 4.89E-01 | -1.00E+00 | 5.95E-02 | 2.53E-01 | -1.20E+00 | 7.61E-02 | 2.91E-01 | -1.06E+00 | 1.92E-01 | 6.84E-01 | -1.05E+00 | 2.87E-01 | 8.10E-01 | 1.50E+00 | 6.17E-05 | 2.27E-04 | -1.40E+00 | 2.86E-01 | 7.85E-01 | 1.21E+00 |
| VN1R1 | 3.75E-02 | 1.95E-01 | -1.11E+00 | 2.08E-01 | 9.00E-01 | 1.44E+00 | 2.23E-01 | 9.48E-01 | 1.25E+00 | 2.03E-01 | 7.75E-01 | 1.51E+00 | 3.50E-02 | 1.25E-01 | 2.08E+00 | 8.54E-02 | 2.41E-01 | 1.96E+00 | 2.17E-01 | 7.98E-01 | 1.03E+00 | 3.64E-01 | 1.00E+00 | 1.24E+00 |
| BTBD9 | 3.75E-02 | 1.95E-01 | -1.06E+00 | 5.89E-03 | 2.54E-02 | -1.16E+00 | 9.24E-02 | 3.93E-01 | 1.35E+00 | 1.72E-01 | 6.56E-01 | 1.11E+00 | 3.04E-06 | 1.08E-05 | -1.66E+00 | 1.99E-06 | 5.62E-06 | -1.58E+00 | 7.76E-02 | 2.85E-01 | -1.00E+00 | 1.92E-01 | 5.27E-01 | 1.07E+00 |
| RGL4 | 3.75E-02 | 1.95E-01 | 1.44E+00 | 9.28E-03 | 4.01E-02 | 1.24E+00 | 2.67E-03 | 1.13E-02 | 1.58E+00 | 2.24E-02 | 8.54E-02 | 1.16E+00 | 1.93E-01 | 6.90E-01 | 1.82E+00 | 2.71E-01 | 7.63E-01 | 1.28E+00 | 2.27E-01 | 8.32E-01 | -1.03E+00 | 3.26E-01 | 8.96E-01 | 1.09E+00 |
| AC090181.2 | 3.76E-02 | 1.96E-01 | 1.70E+00 | 2.01E-01 | 8.70E-01 | 1.45E+00 | 1.30E-03 | 5.53E-03 | -1.36E+00 | 1.18E-05 | 4.51E-05 | -1.70E+00 | 2.64E-02 | 9.41E-02 | -1.14E+00 | 9.57E-02 | 2.70E-01 | 1.08E+00 | 3.17E-03 | 1.16E-02 | -1.57E+00 | 4.29E-02 | 1.18E-01 | -1.25E+00 |
| ISPD-AS1 | 3.76E-02 | 1.96E-01 | -1.23E+00 | 1.74E-01 | 7.52E-01 | 1.29E+00 | 8.11E-02 | 3.44E-01 | -1.04E+00 | 1.24E-01 | 4.73E-01 | 1.10E+00 | 2.34E-05 | 8.35E-05 | -1.75E+00 | 5.89E-02 | 1.66E-01 | 1.11E+00 | 5.02E-03 | 1.84E-02 | -1.26E+00 | 7.22E-02 | 1.98E-01 | -1.04E+00 |
| RPS24P7 | 3.77E-02 | 1.96E-01 | 1.46E+00 | 3.49E-06 | 1.51E-05 | 1.09E+00 | 4.17E-03 | 1.77E-02 | 1.51E+00 | 6.97E-02 | 2.66E-01 | 1.20E+00 | 1.11E-01 | 3.95E-01 | 1.46E+00 | 2.81E-02 | 7.92E-02 | 1.20E+00 | 4.98E-02 | 1.83E-01 | 1.50E+00 | 7.21E-02 | 1.98E-01 | 1.13E+00 |
| RPL21P75 | 3.77E-02 | 1.96E-01 | -1.00E+00 | 3.82E-21 | 1.65E-20 | -1.26E+00 | 3.71E-02 | 1.58E-01 | 1.08E+00 | 1.33E-09 | 5.08E-09 | -1.23E+00 | 1.92E-01 | 6.85E-01 | 1.03E+00 | 9.94E-08 | 2.80E-07 | -1.22E+00 | 4.62E-02 | 1.70E-01 | 1.02E+00 | 4.71E-11 | 1.29E-10 | -1.27E+00 |
| EDDM13 | 3.77E-02 | 1.96E-01 | -1.20E+00 | 2.83E-02 | 1.22E-01 | -1.20E+00 | 7.49E-02 | 3.18E-01 | -1.15E+00 | 1.60E-01 | 6.11E-01 | 1.57E+00 | 1.11E-01 | 3.96E-01 | -1.21E+00 | 2.76E-01 | 7.77E-01 | 1.15E+00 | 1.43E-01 | 5.25E-01 | 1.06E+00 | 2.08E-01 | 5.72E-01 | 1.06E+00 |
| SNX9 | 3.77E-02 | 1.96E-01 | 1.15E+00 | 1.24E-03 | 5.37E-03 | 1.07E+00 | 1.92E-01 | 8.15E-01 | 1.17E+00 | 4.71E-03 | 1.80E-02 | -1.11E+00 | 8.55E-03 | 3.05E-02 | -1.12E+00 | 7.00E-05 | 1.97E-04 | -1.98E+00 | 1.12E-11 | 4.10E-11 | -2.23E+00 | 3.77E-11 | 1.04E-10 | -2.07E+00 |
| C1orf159 | 3.77E-02 | 1.97E-01 | -1.20E+00 | 6.68E-02 | 2.89E-01 | -1.01E+00 | 2.00E-01 | 8.52E-01 | 1.17E+00 | 1.78E-01 | 6.79E-01 | 1.43E+00 | 1.83E-01 | 6.51E-01 | 1.07E+00 | 1.28E-01 | 3.60E-01 | -1.16E+00 | 8.30E-03 | 3.05E-02 | -1.34E+00 | 1.54E-01 | 4.22E-01 | 1.08E+00 |
| RPS10 | 3.79E-02 | 1.97E-01 | 1.17E+00 | 4.24E-05 | 1.83E-04 | -1.08E+00 | 6.74E-04 | 2.86E-03 | -1.09E+00 | 2.44E-05 | 9.32E-05 | -1.15E+00 | 1.44E-02 | 5.14E-02 | -1.18E+00 | 1.03E-02 | 2.89E-02 | -1.13E+00 | 3.69E-03 | 1.36E-02 | -1.02E+00 | 4.36E-05 | 1.20E-04 | -1.24E+00 |
| NAT8L | 3.79E-02 | 1.97E-01 | -1.28E+00 | 8.32E-02 | 3.60E-01 | -1.11E+00 | 1.63E-04 | 6.92E-04 | -1.65E+00 | 2.64E-02 | 1.01E-01 | -1.01E+00 | 2.57E-01 | 9.18E-01 | 1.04E+00 | 3.04E-01 | 8.57E-01 | -1.06E+00 | 2.41E-01 | 8.86E-01 | -1.04E+00 | 3.14E-01 | 8.63E-01 | 1.25E+00 |
| E4F1 | 3.79E-02 | 1.98E-01 | -1.10E+00 | 1.18E-01 | 5.09E-01 | 1.21E+00 | 3.44E-03 | 1.46E-02 | -1.14E+00 | 4.85E-02 | 1.85E-01 | 1.11E+00 | 5.24E-02 | 1.87E-01 | -1.13E+00 | 2.89E-01 | 8.16E-01 | 1.28E+00 | 8.25E-02 | 3.03E-01 | -1.01E+00 | 1.86E-01 | 5.11E-01 | 1.34E+00 |
| N6AMT1 | 3.79E-02 | 1.98E-01 | -1.14E+00 | 2.28E-01 | 9.84E-01 | 1.41E+00 | 4.70E-03 | 2.00E-02 | -1.59E+00 | 1.81E-02 | 6.93E-02 | -1.21E+00 | 1.64E-01 | 5.86E-01 | 1.07E+00 | 1.48E-01 | 4.18E-01 | 1.62E+00 | 4.59E-04 | 1.68E-03 | -1.13E+00 | 7.02E-02 | 1.93E-01 | 1.07E+00 |
| GPX1P1 | 3.79E-02 | 1.98E-01 | 1.23E+00 | 1.88E-03 | 8.14E-03 | 1.88E+00 | 2.34E-01 | 9.95E-01 | 1.13E+00 | 1.70E-04 | 6.51E-04 | 1.79E+00 | 5.31E-02 | 1.89E-01 | 1.16E+00 | 7.62E-03 | 2.15E-02 | 1.63E+00 | 1.41E-01 | 5.19E-01 | 1.37E+00 | 3.59E-05 | 9.87E-05 | 1.78E+00 |
| LINC00174 | 3.80E-02 | 1.98E-01 | 1.54E+00 | 1.25E-02 | 5.38E-02 | 1.78E+00 | 1.41E-01 | 5.99E-01 | 1.28E+00 | 2.28E-01 | 8.70E-01 | 1.46E+00 | 1.67E-02 | 5.97E-02 | 1.13E+00 | 2.65E-01 | 7.47E-01 | 1.40E+00 | 3.42E-02 | 1.25E-01 | 1.58E+00 | 8.70E-04 | 2.39E-03 | 1.81E+00 |
| HLA-J | 3.80E-02 | 1.98E-01 | -1.00E+00 | 2.03E-01 | 8.76E-01 | 1.47E+00 | 6.69E-02 | 2.84E-01 | -1.05E+00 | 1.37E-01 | 5.25E-01 | 1.64E+00 | 3.96E-02 | 1.41E-01 | -1.27E+00 | 3.34E-01 | 9.41E-01 | 1.36E+00 | 2.68E-02 | 9.84E-02 | -1.15E+00 | 3.52E-01 | 9.66E-01 | 1.24E+00 |
| WDR90 | 3.81E-02 | 1.98E-01 | -1.02E+00 | 3.05E-02 | 1.32E-01 | -1.04E+00 | 1.39E-01 | 5.92E-01 | 1.21E+00 | 4.05E-02 | 1.55E-01 | 1.11E+00 | 2.34E-01 | 8.35E-01 | 1.24E+00 | 2.51E-01 | 7.07E-01 | 1.10E+00 | 1.35E-01 | 4.95E-01 | 1.23E+00 | 3.00E-01 | 8.23E-01 | 1.50E+00 |
| ZBTB26 | 3.81E-02 | 1.98E-01 | 1.00E+00 | 1.83E-01 | 7.89E-01 | 1.35E+00 | 8.47E-04 | 3.60E-03 | -1.38E+00 | 1.48E-02 | 5.67E-02 | 1.03E+00 | 2.24E-03 | 8.00E-03 | -1.19E+00 | 2.15E-03 | 6.07E-03 | -1.21E+00 | 8.09E-02 | 2.97E-01 | -1.01E+00 | 5.51E-02 | 1.51E-01 | -1.16E+00 |
| AC017100.1 | 3.81E-02 | 1.99E-01 | 1.69E+00 | 3.22E-02 | 1.39E-01 | 1.93E+00 | 5.22E-02 | 2.22E-01 | 1.55E+00 | 8.10E-02 | 3.09E-01 | 1.67E+00 | 1.49E-01 | 5.30E-01 | 1.18E+00 | 5.51E-02 | 1.55E-01 | -1.05E+00 | 1.14E-01 | 4.18E-01 | 1.10E+00 | 3.64E-01 | 1.00E+00 | 1.32E+00 |
| AC005062.1 | 3.81E-02 | 1.99E-01 | -1.12E+00 | 1.95E-02 | 8.45E-02 | -1.22E+00 | 5.88E-03 | 2.50E-02 | -1.40E+00 | 1.99E-02 | 7.61E-02 | -1.07E+00 | 1.58E-02 | 5.62E-02 | -1.60E+00 | 1.07E-02 | 3.03E-02 | -1.96E+00 | 2.28E-01 | 8.37E-01 | 1.25E+00 | 1.25E-01 | 3.43E-01 | 1.70E+00 |
| ADCK5 | 3.81E-02 | 1.99E-01 | -1.24E+00 | 3.95E-02 | 1.71E-01 | -1.16E+00 | 2.88E-04 | 1.22E-03 | -1.72E+00 | 2.06E-03 | 7.88E-03 | -1.36E+00 | 1.66E-02 | 5.92E-02 | -1.34E+00 | 3.82E-02 | 1.08E-01 | -1.26E+00 | 1.50E-01 | 5.50E-01 | 1.16E+00 | 4.84E-02 | 1.33E-01 | 1.04E+00 |
| AC116050.1 | 3.81E-02 | 1.99E-01 | -1.01E+00 | 8.83E-02 | 3.81E-01 | 1.21E+00 | 1.14E-02 | 4.86E-02 | -1.29E+00 | 2.19E-01 | 8.38E-01 | 1.58E+00 | 2.97E-02 | 1.06E-01 | -1.20E+00 | 2.27E-01 | 6.40E-01 | 1.33E+00 | 1.25E-01 | 4.58E-01 | -1.02E+00 | 2.55E-02 | 7.00E-02 | 1.72E+00 |
| LINC01426 | 3.82E-02 | 1.99E-01 | -1.28E+00 | 4.59E-02 | 1.98E-01 | -1.17E+00 | 2.30E-01 | 9.77E-01 | 1.12E+00 | 1.22E-01 | 4.66E-01 | -1.05E+00 | 2.15E-01 | 7.66E-01 | 1.39E+00 | 1.26E-01 | 3.55E-01 | 1.76E+00 | 2.07E-06 | 7.59E-06 | -2.16E+00 | 6.46E-03 | 1.77E-02 | -1.28E+00 |
| NAT1 | 3.82E-02 | 1.99E-01 | -1.17E+00 | 4.56E-02 | 1.97E-01 | -1.10E+00 | 2.94E-02 | 1.25E-01 | -1.30E+00 | 1.80E-01 | 6.89E-01 | 1.13E+00 | 5.94E-02 | 2.12E-01 | 1.07E+00 | 1.93E-02 | 5.44E-02 | -1.33E+00 | 3.59E-04 | 1.32E-03 | -1.42E+00 | 3.72E-02 | 1.02E-01 | -1.04E+00 |
| KIF14 | 3.82E-02 | 1.99E-01 | -1.34E+00 | 9.44E-02 | 4.08E-01 | -1.02E+00 | 3.66E-02 | 1.56E-01 | -1.02E+00 | 5.16E-02 | 1.97E-01 | 1.12E+00 | 1.38E-02 | 4.92E-02 | -2.06E+00 | 2.94E-02 | 8.29E-02 | -1.79E+00 | 1.49E-07 | 5.46E-07 | -1.73E+00 | 1.61E-04 | 4.41E-04 | -1.35E+00 |
| MIB2 | 3.83E-02 | 2.00E-01 | 1.18E+00 | 7.04E-02 | 3.04E-01 | 1.37E+00 | 2.09E-01 | 8.86E-01 | 1.32E+00 | 2.36E-01 | 9.01E-01 | 1.50E+00 | 1.29E-01 | 4.60E-01 | 1.55E+00 | 3.45E-01 | 9.72E-01 | 1.33E+00 | 4.45E-02 | 1.63E-01 | 1.36E+00 | 1.22E-02 | 3.36E-02 | 1.54E+00 |
| AC009487.2 | 3.83E-02 | 2.00E-01 | 1.06E+00 | 3.17E-02 | 1.37E-01 | 1.10E+00 | 1.31E-01 | 5.55E-01 | 1.33E+00 | 4.60E-02 | 1.76E-01 | -1.02E+00 | 2.26E-01 | 8.06E-01 | 1.24E+00 | 2.32E-01 | 6.53E-01 | 1.08E+00 | 4.10E-04 | 1.51E-03 | -1.24E+00 | 5.07E-03 | 1.39E-02 | -1.27E+00 |
| CCNB2 | 3.83E-02 | 2.00E-01 | -1.33E+00 | 7.75E-02 | 3.35E-01 | -1.18E+00 | 4.88E-02 | 2.08E-01 | 1.19E+00 | 8.02E-06 | 3.06E-05 | 1.01E+00 | 1.79E-02 | 6.37E-02 | -1.76E+00 | 2.80E-02 | 7.88E-02 | -1.34E+00 | 2.19E-05 | 8.04E-05 | -1.12E+00 | 7.95E-03 | 2.19E-02 | 1.07E+00 |
| RPL10P3 | 3.83E-02 | 2.00E-01 | 1.16E+00 | 4.18E-03 | 1.80E-02 | 1.10E+00 | 1.08E-01 | 4.59E-01 | 1.43E+00 | 1.37E-01 | 5.23E-01 | 1.33E+00 | 5.28E-03 | 1.88E-02 | -1.08E+00 | 3.20E-03 | 9.03E-03 | -1.08E+00 | 2.20E-01 | 8.08E-01 | 1.30E+00 | 3.16E-01 | 8.69E-01 | 1.22E+00 |
| AC009318.3 | 3.84E-02 | 2.00E-01 | -1.10E+00 | 1.20E-01 | 5.19E-01 | 1.17E+00 | 8.24E-03 | 3.50E-02 | -1.18E+00 | 4.30E-02 | 1.64E-01 | 1.11E+00 | 3.13E-02 | 1.11E-01 | -1.43E+00 | 7.25E-02 | 2.04E-01 | -1.17E+00 | 3.23E-02 | 1.19E-01 | -1.23E+00 | 5.94E-02 | 1.63E-01 | -1.17E+00 |
| SPA17 | 3.84E-02 | 2.00E-01 | 1.16E+00 | 3.93E-09 | 1.70E-08 | -1.85E+00 | 1.15E-01 | 4.87E-01 | 1.39E+00 | 1.52E-03 | 5.79E-03 | -1.33E+00 | 1.10E-02 | 3.92E-02 | -1.22E+00 | 2.80E-03 | 7.88E-03 | -1.52E+00 | 5.66E-02 | 2.08E-01 | -1.02E+00 | 1.94E-02 | 5.32E-02 | -1.18E+00 |
| LOXL3 | 3.85E-02 | 2.01E-01 | 1.03E+00 | 7.02E-02 | 3.03E-01 | 1.23E+00 | 7.26E-05 | 3.08E-04 | -1.36E+00 | 7.01E-03 | 2.68E-02 | 1.05E+00 | 7.78E-03 | 2.77E-02 | -1.71E+00 | 8.62E-02 | 2.43E-01 | -1.02E+00 | 4.49E-04 | 1.65E-03 | -1.41E+00 | 5.37E-04 | 1.47E-03 | -1.49E+00 |
| AP4S1 | 3.86E-02 | 2.01E-01 | 1.14E+00 | 2.56E-02 | 1.11E-01 | 1.26E+00 | 1.51E-01 | 6.43E-01 | 1.13E+00 | 5.73E-02 | 2.19E-01 | 1.13E+00 | 2.53E-03 | 9.01E-03 | 1.06E+00 | 9.24E-03 | 2.60E-02 | 1.13E+00 | 1.07E-01 | 3.94E-01 | 1.10E+00 | 2.10E-01 | 5.77E-01 | 1.16E+00 |
| LST1 | 3.87E-02 | 2.02E-01 | 1.19E+00 | 1.32E-01 | 5.69E-01 | 1.20E+00 | 1.25E-03 | 5.30E-03 | 1.39E+00 | 1.64E-02 | 6.28E-02 | 1.37E+00 | 1.13E-08 | 4.04E-08 | -1.26E+00 | 5.69E-15 | 1.60E-14 | -1.45E+00 | 8.80E-03 | 3.23E-02 | -1.05E+00 | 1.02E-08 | 2.81E-08 | -1.26E+00 |
| LSM12P1 | 3.88E-02 | 2.02E-01 | -1.02E+00 | 2.12E-01 | 9.16E-01 | 1.41E+00 | 7.39E-02 | 3.14E-01 | 1.04E+00 | 1.71E-01 | 6.54E-01 | 1.29E+00 | 1.70E-01 | 6.07E-01 | 1.23E+00 | 6.13E-02 | 1.73E-01 | 1.04E+00 | 2.29E-01 | 8.39E-01 | 1.36E+00 | 3.47E-01 | 9.54E-01 | 1.30E+00 |
| TMEM53 | 3.88E-02 | 2.02E-01 | -1.07E+00 | 8.58E-02 | 3.71E-01 | 1.10E+00 | 1.06E-04 | 4.49E-04 | -1.44E+00 | 3.17E-05 | 1.21E-04 | -1.35E+00 | 4.96E-05 | 1.77E-04 | -1.64E+00 | 2.91E-03 | 8.22E-03 | -1.17E+00 | 1.14E-04 | 4.18E-04 | -1.27E+00 | 1.82E-03 | 4.99E-03 | -1.12E+00 |
| GAPDHP63 | 3.88E-02 | 2.02E-01 | 1.71E+00 | 9.45E-02 | 4.08E-01 | 1.74E+00 | 1.58E-01 | 6.70E-01 | 1.26E+00 | 7.65E-02 | 2.92E-01 | 1.17E+00 | 1.27E-01 | 4.54E-01 | 1.66E+00 | 3.53E-01 | 9.96E-01 | 1.24E+00 | 2.42E-02 | 8.89E-02 | 1.94E+00 | 5.52E-02 | 1.52E-01 | 1.91E+00 |
| LCA5L | 3.88E-02 | 2.02E-01 | -1.17E+00 | 2.14E-01 | 9.26E-01 | 1.43E+00 | 9.11E-05 | 3.87E-04 | -1.82E+00 | 2.63E-03 | 1.00E-02 | -1.27E+00 | 2.37E-02 | 8.45E-02 | -1.48E+00 | 2.75E-01 | 7.76E-01 | 1.31E+00 | 1.54E-01 | 5.67E-01 | 1.05E+00 | 3.12E-01 | 8.57E-01 | 1.22E+00 |

| | | | | | | | | | | | | | | | | | | | | | | | | |
|---|---|---|---|---|---|---|---|---|---|---|---|---|---|---|---|---|---|---|---|---|---|---|---|---|
| AL122035.2 | 3.89E-02 | 2.03E-01 | -1.27E+00 | 1.64E-01 | 7.11E-01 | 1.04E+00 | 1.27E-01 | 5.42E-01 | 1.11E+00 | 2.60E-01 | 9.94E-01 | 1.38E+00 | 1.47E-03 | 5.23E-03 | -1.86E+00 | 2.82E-02 | 7.95E-02 | -1.18E+00 | 5.40E-05 | 1.98E-04 | -1.58E+00 | 5.46E-03 | 1.50E-02 | -1.20E+00 |
| TRMT10A | 3.89E-02 | 2.03E-01 | -1.31E+00 | 6.67E-03 | 2.88E-02 | -1.75E+00 | 9.38E-05 | 3.98E-04 | -1.44E+00 | 3.62E-06 | 1.38E-05 | -1.60E+00 | 1.80E-02 | 6.40E-02 | -1.59E+00 | 8.19E-02 | 2.31E-01 | -1.06E+00 | 6.74E-02 | 2.47E-01 | -1.07E+00 | 5.11E-02 | 1.40E-01 | -1.19E+00 |
| GPR183 | 3.89E-02 | 2.03E-01 | 1.85E+00 | 1.48E-01 | 6.38E-01 | 1.02E+00 | 1.00E-01 | 4.27E-01 | 1.78E+00 | 6.41E-02 | 2.45E-01 | -1.14E+00 | 7.64E-08 | 2.72E-07 | 5.12E+00 | 2.18E-01 | 6.15E-01 | 2.18E+00 | 4.47E-03 | 1.64E-02 | 2.81E+00 | 3.44E-01 | 9.46E-01 | 1.37E+00 |
| BRCA1 | 3.90E-02 | 2.03E-01 | -1.18E+00 | 4.09E-02 | 1.77E-01 | -1.05E+00 | 3.91E-02 | 1.66E-01 | 1.12E+00 | 5.20E-03 | 1.99E-02 | 1.44E+00 | 8.12E-02 | 2.89E-01 | -1.06E+00 | 1.60E-01 | 4.50E-01 | 1.00E+00 | 3.79E-03 | 1.39E-02 | -1.05E+00 | 1.80E-01 | 4.93E-01 | 1.32E+00 |
| CHCHD4 | 3.90E-02 | 2.03E-01 | -1.05E+00 | 1.38E-01 | 5.98E-01 | 1.38E+00 | 3.14E-04 | 1.34E-03 | -1.31E+00 | 8.42E-03 | 3.21E-02 | 1.05E+00 | 2.31E-03 | 8.25E-03 | -1.48E+00 | 3.19E-01 | 8.99E-01 | 1.30E+00 | 3.21E-05 | 1.18E-04 | -1.24E+00 | 1.93E-01 | 5.31E-01 | 1.18E+00 |
| RPS23P1 | 3.90E-02 | 2.03E-01 | 1.10E+00 | 3.70E-04 | 1.60E-03 | -1.20E+00 | 2.17E-01 | 9.24E-01 | 1.19E+00 | 2.97E-03 | 1.13E-02 | -1.12E+00 | 9.72E-03 | 3.47E-02 | -1.32E+00 | 8.84E-03 | 2.49E-02 | -1.29E+00 | 2.57E-01 | 9.44E-01 | 1.12E+00 | 1.77E-03 | 4.87E-03 | -1.65E+00 |
| AC091390.3 | 3.90E-02 | 2.03E-01 | 1.00E+00 | 4.34E-03 | 1.87E-02 | -1.26E+00 | 2.35E-02 | 9.98E-02 | -1.16E+00 | 3.69E-03 | 1.41E-02 | -1.39E+00 | 2.34E-03 | 8.36E-03 | -1.82E+00 | 1.04E-02 | 2.92E-02 | -1.47E+00 | 1.98E-02 | 7.28E-02 | -1.15E+00 | 8.61E-02 | 2.36E-01 | 1.06E+00 |
| UBXN11 | 3.90E-02 | 2.03E-01 | 1.05E+00 | 2.46E-02 | 1.06E-01 | 1.10E+00 | 1.24E-05 | 5.26E-05 | -1.41E+00 | 4.01E-05 | 1.53E-04 | -1.22E+00 | 2.70E-01 | 9.64E-01 | 1.34E+00 | 1.04E-01 | 2.93E-01 | -1.25E+00 | 2.72E-01 | 9.98E-01 | 1.44E+00 | 2.26E-01 | 6.22E-01 | 1.17E+00 |
| MFSD6 | 3.91E-02 | 2.04E-01 | 1.17E+00 | 7.93E-02 | 3.43E-01 | 1.39E+00 | 6.95E-02 | 2.95E-01 | 1.02E+00 | 2.22E-02 | 8.49E-02 | 1.06E+00 | 6.51E-03 | 2.32E-02 | -1.20E+00 | 1.92E-03 | 5.42E-03 | -1.31E+00 | 2.65E-01 | 9.74E-01 | 1.44E+00 | 5.23E-02 | 1.44E-01 | -1.02E+00 |
| RPL23AP57 | 3.91E-02 | 2.04E-01 | -1.12E+00 | 1.46E-02 | 6.31E-02 | -1.23E+00 | 2.24E-04 | 9.52E-04 | -1.49E+00 | 3.21E-06 | 1.23E-05 | -1.87E+00 | 2.14E-01 | 7.62E-01 | 1.50E+00 | 2.52E-01 | 7.09E-01 | 1.01E+00 | 3.26E-04 | 1.20E-03 | -1.37E+00 | 9.60E-07 | 2.64E-06 | -1.98E+00 |
| AC018628.1 | 3.91E-02 | 2.04E-01 | 1.07E+00 | 9.93E-03 | 4.29E-02 | 1.09E+00 | 6.79E-04 | 2.89E-03 | -1.11E+00 | 9.14E-05 | 3.49E-04 | -1.11E+00 | 2.43E-03 | 8.67E-03 | -1.09E+00 | 1.37E-04 | 3.87E-04 | -1.15E+00 | 2.64E-04 | 9.70E-04 | -1.19E+00 | 3.15E-05 | 8.65E-05 | -1.32E+00 |
| AC083799.1 | 3.91E-02 | 2.04E-01 | 1.68E+00 | 8.22E-02 | 3.55E-01 | 1.78E+00 | 2.72E-02 | 1.16E-01 | 1.85E+00 | 2.44E-01 | 9.33E-01 | 1.41E+00 | 8.30E-02 | 2.96E-01 | -1.01E+00 | 6.06E-02 | 1.71E-01 | -1.13E+00 | 5.15E-02 | 1.89E-01 | 1.56E+00 | 2.59E-01 | 7.11E-01 | 1.61E+00 |
| SIRPB1 | 3.92E-02 | 2.04E-01 | -1.81E+00 | 5.66E-02 | 2.44E-01 | -1.66E+00 | 3.44E-03 | 1.46E-02 | -1.45E+00 | 7.27E-04 | 2.78E-03 | -1.62E+00 | 2.45E-01 | 8.74E-01 | 1.21E+00 | 3.20E-01 | 9.02E-01 | 1.20E+00 | 1.14E-01 | 4.19E-01 | -1.17E+00 | 1.66E-01 | 4.55E-01 | -1.22E+00 |
| AL355338.1 | 3.92E-02 | 2.04E-01 | 1.08E+00 | 3.73E-02 | 1.61E-01 | 1.18E+00 | 4.45E-04 | 1.89E-03 | -1.40E+00 | 5.39E-02 | 2.06E-01 | 1.19E+00 | 1.08E-02 | 3.86E-02 | -1.11E+00 | 6.41E-02 | 1.81E-01 | 1.13E+00 | 1.83E-02 | 6.71E-02 | 1.05E+00 | 3.31E-01 | 9.08E-01 | 1.29E+00 |
| CDC42EP2 | 3.93E-02 | 2.05E-01 | -1.13E+00 | 1.87E-02 | 8.10E-02 | -1.33E+00 | 1.64E-01 | 6.95E-01 | 1.05E+00 | 2.41E-01 | 9.22E-01 | 1.24E+00 | 2.43E-01 | 8.66E-01 | 1.31E+00 | 3.18E-01 | 8.96E-01 | 1.08E+00 | 2.34E-03 | 8.60E-03 | -1.82E+00 | 1.83E-01 | 5.04E-01 | 1.01E+00 |
| AC079610.2 | 3.93E-02 | 2.05E-01 | 1.02E+00 | 1.54E-01 | 6.64E-01 | 1.38E+00 | 4.46E-02 | 1.90E-01 | -1.01E+00 | 1.80E-02 | 6.86E-02 | 1.03E+00 | 1.52E-01 | 5.43E-01 | 1.12E+00 | 2.17E-01 | 6.13E-01 | 1.29E+00 | 6.12E-04 | 2.25E-03 | -1.81E+00 | 8.54E-03 | 2.35E-02 | -1.49E+00 |
| CLK2 | 3.93E-02 | 2.05E-01 | -1.03E+00 | 2.07E-01 | 8.97E-01 | 1.36E+00 | 1.06E-02 | 4.48E-02 | -1.11E+00 | 1.60E-01 | 6.11E-01 | 1.35E+00 | 5.23E-02 | 1.87E-01 | -1.09E+00 | 1.43E-01 | 4.02E-01 | 1.07E+00 | 3.61E-05 | 1.33E-04 | -1.37E+00 | 4.10E-02 | 1.13E-01 | 1.05E+00 |
| PACS2 | 3.93E-02 | 2.05E-01 | 1.08E+00 | 1.33E-01 | 5.73E-01 | 1.34E+00 | 4.58E-02 | 1.95E-01 | -1.03E+00 | 1.06E-01 | 4.04E-01 | 1.24E+00 | 1.61E-03 | 5.76E-03 | -1.26E+00 | 9.86E-03 | 2.78E-02 | -1.14E+00 | 6.72E-02 | 2.47E-01 | 1.09E+00 | 3.38E-01 | 9.29E-01 | 1.22E+00 |
| XPO6 | 3.95E-02 | 2.06E-01 | 1.07E+00 | 9.03E-02 | 3.90E-01 | 1.29E+00 | 5.89E-03 | 2.50E-02 | -1.10E+00 | 9.61E-03 | 3.67E-02 | 1.10E+00 | 2.43E-03 | 8.67E-03 | -1.32E+00 | 7.37E-02 | 2.08E-01 | 1.12E+00 | 4.77E-04 | 1.75E-03 | -1.18E+00 | 3.51E-02 | 9.63E-02 | 1.02E+00 |
| RPL17P26 | 3.95E-02 | 2.06E-01 | 1.05E+00 | 2.02E-02 | 8.72E-02 | 1.92E+00 | 1.06E-01 | 4.50E-01 | 1.12E+00 | 1.23E-03 | 4.70E-03 | 2.33E+00 | 1.94E-01 | 6.93E-01 | 1.30E+00 | 1.55E-03 | 4.37E-03 | 2.70E+00 | 5.51E-02 | 2.02E-01 | -1.03E+00 | 1.39E-03 | 3.81E-03 | 1.97E+00 |
| WNT2B | 3.95E-02 | 2.06E-01 | -1.12E+00 | 1.14E-01 | 4.93E-01 | 1.65E+00 | 1.15E-01 | 4.89E-01 | -1.23E+00 | 3.42E-02 | 1.31E-01 | 1.73E+00 | 8.13E-03 | 2.90E-02 | -1.50E+00 | 1.51E-01 | 4.27E-01 | 1.06E+00 | 1.22E-01 | 4.48E-01 | -1.10E+00 | 2.01E-01 | 5.51E-01 | 1.63E+00 |
| DPY19L2P4 | 3.95E-02 | 2.06E-01 | -1.17E+00 | 1.49E-01 | 6.43E-01 | 1.21E+00 | 2.10E-01 | 8.92E-01 | 1.12E+00 | 1.74E-01 | 6.64E-01 | 1.49E+00 | 7.41E-02 | 2.64E-01 | 1.99E+00 | 9.09E-02 | 2.56E-01 | 2.21E+00 | 2.09E-01 | 7.67E-01 | 1.14E+00 | 2.96E-01 | 8.14E-01 | 1.32E+00 |
| SLC11A2 | 3.96E-02 | 2.06E-01 | -1.08E+00 | 2.09E-01 | 9.04E-01 | 1.29E+00 | 2.43E-04 | 1.03E-03 | -1.35E+00 | 5.68E-02 | 2.17E-01 | 1.13E+00 | 1.18E-07 | 4.20E-07 | -1.74E+00 | 2.74E-03 | 7.71E-03 | -1.02E+00 | 2.33E-05 | 8.55E-05 | -1.29E+00 | 1.93E-01 | 5.29E-01 | 1.14E+00 |
| IGLL3P | 3.96E-02 | 2.06E-01 | -1.20E+00 | 4.22E-02 | 1.82E-01 | -1.18E+00 | 1.26E-01 | 5.36E-01 | 1.03E+00 | 1.97E-01 | 7.52E-01 | 1.34E+00 | 8.05E-02 | 2.87E-01 | -1.32E+00 | 1.45E-01 | 4.08E-01 | -1.12E+00 | 2.40E-01 | 8.79E-01 | 1.05E+00 | 2.53E-01 | 6.95E-01 | 1.47E+00 |
| ZNF815P | 3.96E-02 | 2.06E-01 | 1.04E+00 | 5.95E-02 | 2.57E-01 | 1.15E+00 | 2.05E-01 | 8.71E-01 | 1.17E+00 | 2.00E-01 | 7.62E-01 | 1.17E+00 | 1.23E-02 | 4.39E-02 | -1.31E+00 | 1.45E-01 | 4.08E-01 | 1.15E+00 | 1.22E-01 | 4.48E-01 | -1.06E+00 | 1.91E-01 | 5.23E-01 | -1.08E+00 |
| AL591034.2 | 3.96E-02 | 2.06E-01 | -1.19E+00 | 3.40E-02 | 1.47E-01 | -1.23E+00 | 2.81E-02 | 1.19E-01 | -1.02E+00 | 4.03E-05 | 1.54E-04 | -1.44E+00 | 6.87E-03 | 2.45E-02 | -1.81E+00 | 1.01E-02 | 2.86E-02 | -1.62E+00 | 1.74E-06 | 6.39E-06 | -1.49E+00 | 1.39E-04 | 3.82E-04 | -1.30E+00 |
| RPEL1 | 3.96E-02 | 2.07E-01 | -1.25E+00 | 8.04E-02 | 3.47E-01 | -1.03E+00 | 4.56E-11 | 1.94E-10 | -2.57E+00 | 5.43E-07 | 2.07E-06 | -1.56E+00 | 1.10E-02 | 3.91E-02 | -1.47E+00 | 3.16E-02 | 8.90E-02 | -1.21E+00 | 2.61E-04 | 9.59E-04 | -1.64E+00 | 6.93E-02 | 1.90E-01 | -1.01E+00 |
| SUPT3H | 3.97E-02 | 2.07E-01 | 1.15E+00 | 2.70E-02 | 1.16E-01 | 1.25E+00 | 3.39E-02 | 1.44E-01 | 1.01E+00 | 1.85E-02 | 7.07E-02 | 1.04E+00 | 1.21E-05 | 4.32E-05 | -1.46E+00 | 6.02E-04 | 1.70E-03 | -1.14E+00 | 3.77E-02 | 1.38E-01 | -1.07E+00 | 7.76E-02 | 2.13E-01 | 1.07E+00 |
| TTK | 3.97E-02 | 2.07E-01 | -1.49E+00 | 4.53E-02 | 1.96E-01 | -1.52E+00 | 1.54E-01 | 6.52E-01 | 1.10E+00 | 2.46E-01 | 9.40E-01 | 1.43E+00 | 1.09E-01 | 3.87E-01 | -1.83E+00 | 2.44E-01 | 6.87E-01 | -1.29E+00 | 5.90E-02 | 2.17E-01 | 1.19E+00 | 2.95E-01 | 8.10E-01 | 1.16E+00 |
| AL353759.1 | 3.97E-02 | 2.07E-01 | -1.28E+00 | 1.38E-01 | 5.97E-01 | 1.00E+00 | 2.27E-01 | 9.65E-01 | 1.02E+00 | 2.51E-01 | 9.60E-01 | 1.14E+00 | 7.13E-02 | 2.54E-01 | -1.12E+00 | 2.43E-01 | 6.86E-01 | 1.39E+00 | 2.49E-01 | 9.13E-01 | 1.08E+00 | 3.02E-01 | 8.29E-01 | 1.21E+00 |
| PLCH1 | 3.97E-02 | 2.07E-01 | -1.34E+00 | 1.25E-01 | 5.40E-01 | -1.09E+00 | 1.45E-01 | 6.18E-01 | 1.14E+00 | 2.62E-01 | 1.00E+00 | -1.00E+00 | 2.80E-01 | 1.00E+00 | 1.00E+00 | 3.05E-01 | 8.60E-01 | 1.13E+00 | 1.13E-01 | 4.15E-01 | 2.58E+00 | 1.22E-03 | 3.36E-03 | -1.31E+00 |
| PAXX | 3.97E-02 | 2.07E-01 | 1.19E+00 | 5.07E-03 | 2.19E-02 | 1.24E+00 | 6.84E-02 | 2.91E-01 | 1.30E+00 | 1.92E-01 | 7.33E-01 | 1.29E+00 | 7.67E-05 | 2.73E-04 | -1.15E+00 | 3.95E-02 | 1.11E-01 | 1.21E+00 | 3.41E-03 | 1.25E-02 | 1.04E+00 | 2.43E-01 | 6.67E-01 | 1.34E+00 |
| SNX16 | 3.98E-02 | 2.07E-01 | -1.01E+00 | 1.37E-01 | 5.92E-01 | 1.27E+00 | 6.90E-06 | 2.93E-05 | -1.58E+00 | 1.43E-06 | 5.48E-06 | -1.48E+00 | 1.95E-02 | 6.94E-02 | -1.11E+00 | 1.15E-02 | 3.24E-02 | -1.11E+00 | 6.87E-06 | 2.52E-05 | -1.73E+00 | 3.84E-04 | 1.06E-03 | -1.40E+00 |
| PCSK6 | 3.98E-02 | 2.08E-01 | -1.13E+00 | 2.00E-01 | 8.64E-01 | 1.21E+00 | 1.22E-01 | 5.19E-01 | 1.16E+00 | 7.83E-02 | 2.99E-01 | -1.15E+00 | 1.95E-01 | 6.97E-01 | 1.06E+00 | 3.49E-01 | 9.84E-01 | 1.35E+00 | 2.28E-01 | 8.38E-01 | 1.02E+00 | 2.98E-01 | 8.19E-01 | -1.14E+00 |
| RTTN | 3.98E-02 | 2.08E-01 | 1.04E+00 | 1.80E-02 | 7.77E-02 | 1.04E+00 | 2.23E-01 | 9.47E-01 | 1.16E+00 | 9.15E-02 | 3.50E-01 | 1.13E+00 | 6.55E-02 | 2.33E-01 | -1.11E+00 | 1.15E-01 | 3.24E-01 | -1.03E+00 | 1.27E-02 | 4.67E-02 | -1.00E+00 | 1.43E-01 | 3.93E-01 | 1.17E+00 |
| CEBPE | 3.99E-02 | 2.08E-01 | -2.19E+00 | 1.79E-01 | 7.75E-01 | -1.43E+00 | 8.81E-08 | 3.74E-07 | -1.88E+00 | 2.21E-06 | 8.42E-06 | -1.49E+00 | 1.48E-01 | 5.27E-01 | -1.27E+00 | 1.75E-01 | 4.93E-01 | -1.42E+00 | 2.30E-01 | 8.43E-01 | -1.06E+00 | 3.27E-01 | 8.98E-01 | 1.05E+00 |
| SLFNL1-AS1 | 3.99E-02 | 2.08E-01 | 1.02E+00 | 2.17E-02 | 9.37E-02 | 1.06E+00 | 1.79E-01 | 7.62E-01 | 1.19E+00 | 4.47E-02 | 1.71E-01 | 1.12E+00 | 1.34E-01 | 4.78E-01 | 1.00E+00 | 1.84E-01 | 5.20E-01 | 1.05E+00 | 2.95E-02 | 1.08E-01 | -1.15E+00 | 1.35E-01 | 3.71E-01 | -1.00E+00 |
| AL358942.1 | 4.00E-02 | 2.08E-01 | -1.17E+00 | 6.99E-02 | 3.02E-01 | -1.08E+00 | 1.02E-01 | 4.33E-01 | -1.01E+00 | 5.13E-02 | 1.96E-01 | -1.01E+00 | 4.12E-02 | 1.47E-01 | -1.81E+00 | 3.12E-01 | 8.79E-01 | 1.41E+00 | 2.46E-01 | 9.05E-01 | 1.29E+00 | 1.72E-01 | 4.71E-01 | 1.45E+00 |
| RPS26 | 4.00E-02 | 2.09E-01 | 1.20E+00 | 6.03E-03 | 2.61E-02 | 1.20E+00 | 2.23E-01 | 9.46E-01 | 1.12E+00 | 2.62E-01 | 1.00E+00 | 1.35E+00 | 1.40E-03 | 4.99E-03 | -1.18E+00 | 2.27E-03 | 6.41E-03 | -1.08E+00 | 1.97E-04 | 7.23E-04 | -1.15E+00 | 2.20E-03 | 6.04E-03 | -1.04E+00 |
| AP002026.1 | 4.01E-02 | 2.09E-01 | -1.06E+00 | 3.40E-02 | 1.47E-01 | -1.10E+00 | 3.15E-03 | 1.34E-02 | -1.75E+00 | 7.14E-03 | 2.73E-02 | -1.55E+00 | 1.78E-01 | 6.36E-01 | 1.66E+00 | 2.59E-01 | 7.29E-01 | 1.60E+00 | 9.26E-02 | 3.40E-01 | -1.25E+00 | 1.94E-01 | 5.34E-01 | -1.12E+00 |
| POLR2B | 4.01E-02 | 2.09E-01 | 1.19E+00 | 4.27E-02 | 1.85E-01 | 1.36E+00 | 2.69E-02 | 1.14E-01 | 1.06E+00 | 8.67E-02 | 3.31E-01 | 1.26E+00 | 9.09E-02 | 3.24E-01 | 1.13E+00 | 1.56E-01 | 4.41E-01 | 1.21E+00 | 6.41E-02 | 2.35E-01 | 1.18E+00 | 2.10E-01 | 5.77E-01 | 1.32E+00 |
| DNAAF2 | 4.01E-02 | 2.09E-01 | 1.24E+00 | 2.06E-07 | 8.88E-07 | -1.21E+00 | 1.34E-01 | 5.70E-01 | 1.40E+00 | 7.88E-04 | 3.01E-03 | 1.01E+00 | 5.09E-02 | 1.81E-01 | 1.69E+00 | 3.03E-01 | 8.55E-01 | 1.29E+00 | 1.55E-01 | 5.71E-01 | 1.29E+00 | 1.20E-02 | 3.30E-02 | 1.01E+00 |
| SEMA3F-AS1 | 4.01E-02 | 2.09E-01 | 1.13E+00 | 1.18E-02 | 5.11E-02 | 1.09E+00 | 3.61E-03 | 1.53E-02 | -1.24E+00 | 1.24E-04 | 4.75E-04 | -1.35E+00 | 5.28E-02 | 1.88E-01 | 1.02E+00 | 5.96E-02 | 1.68E-01 | 1.01E+00 | 2.09E-03 | 7.68E-03 | -1.28E+00 | 3.15E-02 | 8.65E-02 | -1.04E+00 |
| CARMIL2 | 4.02E-02 | 2.09E-01 | 1.07E+00 | 1.27E-01 | 5.47E-01 | 1.26E+00 | 1.87E-01 | 7.93E-01 | 1.23E+00 | 2.50E-01 | 9.54E-01 | 1.44E+00 | 9.63E-02 | 3.43E-01 | 1.04E+00 | 1.91E-01 | 5.38E-01 | 1.18E+00 | 1.59E-01 | 5.82E-01 | 1.01E+00 | 3.22E-01 | 8.86E-01 | 1.15E+00 |
| MKI67 | 4.02E-02 | 2.09E-01 | -1.80E+00 | 4.10E-02 | 1.77E-01 | -1.94E+00 | 1.33E-01 | 5.67E-01 | 1.31E+00 | 6.59E-02 | 2.52E-01 | 1.41E+00 | 1.11E-01 | 3.94E-01 | -2.31E+00 | 2.30E-01 | 6.48E-01 | -1.21E+00 | 1.15E-05 | 4.22E-05 | -1.42E+00 | 3.05E-02 | 8.38E-02 | 1.02E+00 |
| WDHD1 | 4.02E-02 | 2.09E-01 | -1.50E+00 | 1.14E-01 | 4.94E-01 | -1.10E+00 | 1.72E-03 | 7.32E-03 | -1.07E+00 | 2.22E-02 | 8.49E-02 | 1.20E+00 | 2.31E-02 | 8.22E-02 | -2.32E+00 | 1.83E-01 | 5.16E-01 | -1.22E+00 | 3.14E-10 | 1.15E-09 | -1.60E+00 | 2.98E-02 | 8.18E-02 | 1.03E+00 |
| POFUT1 | 4.02E-02 | 2.09E-01 | -1.11E+00 | 1.87E-01 | 8.06E-01 | 1.42E+00 | 4.11E-03 | 1.75E-02 | -1.42E+00 | 7.69E-02 | 2.94E-01 | 1.04E+00 | 1.43E-01 | 5.11E-01 | 1.15E+00 | 2.22E-01 | 6.25E-01 | 1.80E+00 | 1.21E-02 | 4.45E-02 | -1.04E+00 | 1.14E-02 | 3.13E-02 | -1.02E+00 |
| AC018926.2 | 4.02E-02 | 2.09E-01 | 1.78E+00 | 1.83E-01 | 7.92E-01 | 1.36E+00 | 2.15E-01 | 9.14E-01 | 1.34E+00 | 8.91E-02 | 3.41E-01 | 1.17E+00 | 4.54E-02 | 1.62E-01 | 3.25E+00 | 7.45E-02 | 2.10E-01 | 3.10E+00 | 2.19E-01 | 8.05E-01 | -1.14E+00 | 2.53E-01 | 6.94E-01 | -1.26E+00 |
| RN7SKP255 | 4.03E-02 | 2.10E-01 | 1.13E+00 | 9.22E-11 | 3.98E-10 | -2.92E+00 | 2.64E-04 | 1.12E-03 | -1.27E+00 | 9.48E-14 | 3.62E-13 | -3.25E+00 | 1.29E-01 | 4.62E-01 | 1.56E+00 | 7.84E-03 | 2.21E-02 | -1.93E+00 | 2.28E-01 | 8.37E-01 | 1.12E+00 | 6.93E-03 | 1.90E-02 | -1.77E+00 |
| UEVLD | 4.03E-02 | 2.10E-01 | -1.16E+00 | 8.13E-02 | 3.51E-01 | 1.05E+00 | 2.89E-02 | 1.23E-01 | -1.36E+00 | 1.77E-01 | 6.77E-01 | 1.04E+00 | 5.32E-02 | 1.90E-01 | -1.15E+00 | 3.38E-01 | 9.51E-01 | 1.44E+00 | 1.77E-06 | 6.48E-06 | -1.85E+00 | 2.54E-04 | 6.98E-04 | -1.51E+00 |
| MTFP1 | 4.05E-02 | 2.11E-01 | 1.85E+00 | 6.42E-02 | 2.77E-01 | 1.91E+00 | 3.04E-02 | 1.29E-01 | 1.81E+00 | 1.25E-01 | 4.79E-01 | 1.56E+00 | 1.48E-01 | 5.27E-01 | 1.67E+00 | 3.00E-01 | 8.47E-01 | 1.18E+00 | 1.24E-01 | 4.55E-01 | 1.07E+00 | 3.83E-02 | 1.05E-01 | -1.08E+00 |
| DENND5B | 4.05E-02 | 2.11E-01 | -1.01E+00 | 3.05E-03 | 1.32E-02 | -1.08E+00 | 7.86E-02 | 3.34E-01 | 1.03E+00 | 1.05E-01 | 4.03E-01 | 1.26E+00 | 3.23E-03 | 1.15E-02 | -1.38E+00 | 2.55E-03 | 7.18E-03 | -1.52E+00 | 7.71E-03 | 2.83E-02 | -1.28E+00 | 5.13E-02 | 1.41E-01 | -1.05E+00 |
| ASPM | 4.06E-02 | 2.11E-01 | -1.00E+00 | 7.94E-03 | 3.43E-02 | -1.56E+00 | 6.41E-02 | 2.73E-01 | 1.01E+00 | 2.26E-03 | 8.62E-03 | 1.27E+00 | 2.71E-01 | 9.65E-01 | 1.06E+00 | 2.57E-01 | 7.24E-01 | 1.93E+00 | 9.08E-03 | 3.33E-02 | -1.40E+00 | 3.39E-01 | 9.32E-01 | 1.09E+00 |
| GCSH | 4.06E-02 | 2.11E-01 | 1.01E+00 | 5.78E-02 | 2.50E-01 | 1.17E+00 | 3.28E-02 | 1.39E-01 | -1.14E+00 | 1.25E-01 | 4.78E-01 | 1.17E+00 | 3.36E-02 | 1.20E-01 | -1.21E+00 | 3.51E-01 | 9.90E-01 | 1.62E+00 | 6.43E-04 | 2.36E-03 | -1.35E+00 | 3.98E-02 | 1.09E-01 | 1.00E+00 |
| FOXJ3 | 4.06E-02 | 2.12E-01 | 1.16E+00 | 2.65E-06 | 1.14E-05 | -1.16E+00 | 4.95E-02 | 2.10E-01 | 1.11E+00 | 5.78E-04 | 2.21E-03 | -1.02E+00 | 1.06E-02 | 3.77E-02 | 1.06E+00 | 2.11E-03 | 5.94E-03 | -1.04E+00 | 4.57E-03 | 1.68E-02 | -1.06E+00 | 3.77E-03 | 1.04E-02 | -1.07E+00 |
| C11orf96 | 4.06E-02 | 2.12E-01 | 1.95E+00 | 1.68E-02 | 7.25E-02 | -1.91E+00 | 8.10E-02 | 3.44E-01 | -1.01E+00 | 1.58E-03 | 6.03E-03 | -1.68E+00 | 2.25E-01 | 8.02E-01 | 1.51E+00 | 8.49E-02 | 2.39E-01 | -1.18E+00 | 5.65E-02 | 2.07E-01 | 2.59E+00 | 3.76E-03 | 1.03E-02 | -1.07E+00 |
| OTUD1 | 4.07E-02 | 2.12E-01 | 1.13E+00 | 4.24E-02 | 1.83E-01 | 1.21E+00 | 5.79E-02 | 2.46E-01 | 1.00E+00 | 1.51E-02 | 5.77E-02 | 1.02E+00 | 6.30E-02 | 2.25E-01 | -1.17E+00 | 6.33E-02 | 1.79E-01 | -1.08E+00 | 4.69E-06 | 1.72E-05 | -1.63E+00 | 3.84E-06 | 1.06E-05 | -1.72E+00 |
| BIRC5 | 4.07E-02 | 2.12E-01 | -2.01E+00 | 4.26E-02 | 1.84E-01 | -1.88E+00 | 2.03E-02 | 8.61E-02 | 1.57E+00 | 2.37E-01 | 9.06E-01 | 1.51E+00 | 2.71E-01 | 9.66E-01 | 1.75E+00 | 3.55E-01 | 1.00E+00 | 1.64E+00 | 7.78E-02 | 2.85E-01 | -1.12E+00 | 5.72E-02 | 1.57E-01 | 1.01E+00 |
| RASL10B | 4.07E-02 | 2.12E-01 | -1.22E+00 | 4.67E-02 | 2.02E-01 | -1.17E+00 | 5.22E-06 | 2.22E-05 | -1.69E+00 | 8.54E-04 | 3.26E-03 | -1.15E+00 | 1.24E-01 | 4.43E-01 | -1.48E+00 | 2.34E-01 | 6.59E-01 | -1.18E+00 | 2.68E-01 | 9.83E-01 | 1.08E+00 | 1.48E-02 | 4.06E-02 | 1.94E+00 |
| AC004982.2 | 4.07E-02 | 2.12E-01 | 1.01E+00 | 1.09E-01 | 4.73E-01 | 1.25E+00 | 1.14E-01 | 4.85E-01 | 1.37E+00 | 2.28E-01 | 8.69E-01 | 1.05E+00 | 7.49E-02 | 2.67E-01 | 1.88E+00 | 2.39E-01 | 6.74E-01 | 1.53E+00 | 1.77E-01 | 6.50E-01 | 1.46E+00 | 2.06E-01 | 5.66E-01 | -1.08E+00 |
| HNRNPKP4 | 4.07E-02 | 2.12E-01 | 1.10E+00 | 2.91E-03 | 1.26E-02 | -1.03E+00 | 1.34E-01 | 5.71E-01 | 1.08E+00 | 1.23E-02 | 4.69E-02 | 1.02E+00 | 1.25E-01 | 4.46E-01 | 1.29E+00 | 1.30E-01 | 3.67E-01 | 1.26E+00 | 2.67E-01 | 9.81E-01 | 1.32E+00 | 2.21E-01 | 6.06E-01 | 1.51E+00 |
| LRRC49 | 4.08E-02 | 2.13E-01 | -1.14E+00 | 4.23E-03 | 1.83E-02 | -1.53E+00 | 2.64E-02 | 1.12E-01 | -1.20E+00 | 2.99E-02 | 1.14E-01 | -1.09E+00 | 5.00E-02 | 1.78E-01 | -1.15E+00 | 4.11E-02 | 1.16E-01 | -1.20E+00 | 1.18E-02 | 4.32E-02 | -1.37E+00 | 4.49E-02 | 1.23E-01 | -1.20E+00 |
| GGH | 4.09E-02 | 2.13E-01 | -1.08E+00 | 4.15E-02 | 1.79E-01 | 1.10E+00 | 1.28E-01 | 5.42E-01 | 1.34E+00 | 1.25E-01 | 4.78E-01 | 1.36E+00 | 1.92E-01 | 6.83E-01 | 1.04E+00 | 2.55E-01 | 7.19E-01 | 1.06E+00 | 3.47E-02 | 1.27E-01 | -1.04E+00 | 3.58E-01 | 9.83E-01 | 1.13E+00 |
| ZNF107 | 4.09E-02 | 2.13E-01 | 1.64E+00 | 4.59E-02 | 1.98E-01 | 1.24E+00 | 1.41E-01 | 5.98E-01 | 1.11E+00 | 1.99E-03 | 7.59E-03 | -1.12E+00 | 7.24E-03 | 2.58E-02 | -1.04E+00 | 5.09E-03 | 1.43E-02 | -1.12E+00 | 1.37E-01 | 5.01E-01 | 1.20E+00 | 7.94E-04 | 2.18E-03 | -1.25E+00 |
| AL359644.1 | 4.11E-02 | 2.14E-01 | 1.73E+00 | 1.24E-01 | 5.35E-01 | 1.57E+00 | 1.89E-01 | 8.02E-01 | 1.16E+00 | 6.89E-02 | 2.63E-01 | -1.06E+00 | 3.83E-02 | 1.36E-01 | 2.29E+00 | 1.92E-01 | 5.42E-01 | 1.79E+00 | 9.48E-02 | 3.48E-01 | 1.78E+00 | 3.17E-01 | 8.70E-01 | 1.36E+00 |
| AC005912.1 | 4.11E-02 | 2.14E-01 | 1.07E+00 | 3.19E-40 | 1.38E-39 | -1.48E+00 | 5.45E-02 | 2.32E-01 | 1.07E+00 | 1.47E-23 | 5.60E-23 | -1.47E+00 | 4.77E-02 | 1.70E-01 | 1.10E+00 | 3.06E-20 | 8.63E-20 | -1.48E+00 | 7.35E-02 | 2.70E-01 | 1.14E+00 | 1.83E-23 | 5.02E-23 | -1.45E+00 |
| DENND4A | 4.11E-02 | 2.14E-01 | 1.22E+00 | 4.18E-03 | 1.81E-02 | 1.23E+00 | 1.47E-01 | 6.24E-01 | 1.13E+00 | 6.15E-02 | 2.35E-01 | 1.17E+00 | 6.46E-02 | 2.30E-01 | 1.27E+00 | 8.29E-02 | 2.34E-01 | 1.19E+00 | 2.95E-02 | 1.08E-01 | 1.12E+00 | 2.88E-03 | 7.90E-03 | -1.01E+00 |
| AC010864.1 | 4.11E-02 | 2.14E-01 | 1.89E+00 | 4.15E-02 | 1.79E-01 | 2.14E+00 | 1.96E-01 | 8.31E-01 | 1.20E+00 | 2.26E-01 | 8.63E-01 | 1.43E+00 | 2.63E-01 | 9.36E-01 | 1.27E+00 | 1.04E-01 | 2.94E-01 | 1.91E+00 | 1.70E-01 | 6.23E-01 | 1.44E+00 | 3.06E-01 | 8.40E-01 | 1.35E+00 |
| CCNA2 | 4.11E-02 | 2.14E-01 | -1.23E+00 | 6.62E-02 | 2.86E-01 | -1.06E+00 | 7.45E-02 | 3.16E-01 | -1.26E+00 | 2.62E-01 | 1.00E+00 | 1.22E+00 | 3.79E-02 | 1.35E-01 | -2.13E+00 | 9.07E-02 | 2.56E-01 | -1.81E+00 | 1.12E-03 | 4.10E-03 | -1.72E+00 | 2.78E-01 | 7.63E-01 | -1.13E+00 |
| DPPA4 | 4.11E-02 | 2.14E-01 | 1.27E+00 | 1.49E-03 | 6.44E-03 | 1.24E+00 | 3.04E-05 | 1.29E-04 | -1.15E+00 | 4.46E-04 | 1.70E-03 | 1.00E+00 | 2.22E-04 | 7.91E-04 | -1.12E+00 | 9.71E-04 | 2.74E-03 | -1.02E+00 | 2.24E-08 | 8.22E-08 | -1.37E+00 | 2.51E-03 | 6.89E-03 | -1.09E+00 |
| C17orf53 | 4.12E-02 | 2.15E-01 | -1.28E+00 | 7.20E-02 | 3.11E-01 | -1.10E+00 | 1.79E-01 | 7.62E-01 | 1.16E+00 | 2.39E-01 | 9.13E-01 | 1.29E+00 | 1.18E-02 | 4.22E-02 | -2.04E+00 | 2.92E-02 | 8.22E-02 | -1.75E+00 | 1.66E-05 | 6.09E-05 | -1.60E+00 | 8.86E-02 | 2.43E-01 | 1.00E+00 |
| HIST1H2BF | 4.13E-02 | 2.15E-01 | 1.04E+00 | 7.40E-03 | 3.20E-02 | -1.17E+00 | 1.36E-02 | 5.76E-02 | -1.21E+00 | 4.53E-03 | 1.73E-02 | -1.31E+00 | 2.40E-01 | 8.54E-01 | 1.45E+00 | 2.37E-01 | 6.69E-01 | 1.72E+00 | 3.69E-02 | 1.35E-01 | 1.02E+00 | 9.79E-04 | 2.69E-03 | -1.30E+00 |
| ATG14 | 4.13E-02 | 2.15E-01 | 1.11E+00 | 2.10E-03 | 9.06E-03 | 1.06E+00 | 3.25E-02 | 1.38E-01 | 1.05E+00 | 1.95E-06 | 7.45E-06 | -1.37E+00 | 1.87E-03 | 6.68E-03 | -1.10E+00 | 2.83E-04 | 7.98E-04 | -1.23E+00 | 2.76E-02 | 1.01E-01 | 1.01E+00 | 1.21E-03 | 3.33E-03 | -1.17E+00 |
| HERC3 | 4.13E-02 | 2.15E-01 | 1.04E+00 | 1.84E-02 | 7.96E-02 | 1.06E+00 | 2.15E-01 | 9.14E-01 | 1.21E+00 | 2.82E-02 | 1.08E-01 | -1.01E+00 | 7.86E-04 | 2.80E-03 | -1.42E+00 | 2.38E-03 | 6.72E-03 | -1.26E+00 | 2.58E-05 | 9.47E-05 | -1.48E+00 | 4.36E-05 | 1.20E-04 | -1.50E+00 |
| TRMT61B | 4.14E-02 | 2.16E-01 | 1.09E+00 | 1.25E-01 | 5.38E-01 | 1.40E+00 | 1.24E-02 | 5.27E-02 | -1.08E+00 | 3.65E-02 | 1.40E-01 | 1.11E+00 | 1.93E-02 | 6.88E-02 | 1.01E+00 | 2.16E-02 | 6.08E-02 | -1.06E+00 | 1.95E-01 | 7.17E-01 | 1.16E+00 | 3.62E-01 | 9.94E-01 | 1.24E+00 |
| NCAPD3 | 4.14E-02 | 2.16E-01 | -1.02E+00 | 1.24E-01 | 5.34E-01 | 1.35E+00 | 5.81E-04 | 2.47E-03 | -1.27E+00 | 2.84E-03 | 1.08E-02 | 1.02E+00 | 7.18E-02 | 2.56E-01 | 1.05E+00 | 2.72E-01 | 7.66E-01 | 1.27E+00 | 2.61E-05 | 9.58E-05 | -1.25E+00 | 9.19E-02 | 2.52E-01 | 1.10E+00 |
| ACSF2 | 4.15E-02 | 2.16E-01 | 1.16E+00 | 1.08E-01 | 4.68E-01 | 1.46E+00 | 3.46E-03 | 1.47E-02 | -1.17E+00 | 2.70E-02 | 1.03E-01 | 1.11E+00 | 6.80E-03 | 2.42E-02 | -1.55E+00 | 3.01E-02 | 8.49E-02 | -1.19E+00 | 8.55E-03 | 3.14E-02 | -1.17E+00 | 1.10E-02 | 3.03E-02 | -1.11E+00 |
| ZKSCAN5 | 4.15E-02 | 2.16E-01 | 1.11E+00 | 1.54E-02 | 6.64E-02 | 1.02E+00 | 4.99E-02 | 2.12E-01 | -1.09E+00 | 1.74E-01 | 6.65E-01 | 1.27E+00 | 9.71E-02 | 3.46E-01 | 1.02E+00 | 6.27E-02 | 1.77E-01 | -1.10E+00 | 3.08E-02 | 1.13E-01 | -1.00E+00 | 1.05E-02 | 2.88E-02 | -1.14E+00 |
| MED13L | 4.16E-02 | 2.17E-01 | 1.32E+00 | 4.87E-02 | 2.11E-01 | 1.24E+00 | 2.38E-02 | 1.01E-01 | -1.07E+00 | 9.40E-04 | 3.59E-03 | -1.04E+00 | 6.02E-03 | 2.15E-02 | -1.08E+00 | 3.42E-05 | 9.65E-05 | -1.24E+00 | 1.52E-05 | 5.57E-05 | -1.09E+00 | 1.51E-03 | 4.15E-03 | -1.06E+00 |
| AC092171.2 | 4.16E-02 | 2.17E-01 | -1.35E+00 | 1.25E-01 | 5.40E-01 | 1.01E+00 | 1.62E-09 | 6.87E-09 | -2.73E+00 | 3.88E-05 | 1.48E-04 | -1.50E+00 | 1.40E-03 | 4.99E-03 | -1.68E+00 | 5.47E-03 | 1.54E-02 | -1.38E+00 | 2.96E-02 | 1.09E-01 | -1.32E+00 | 2.67E-01 | 7.33E-01 | 1.27E+00 |
| AC008536.3 | 4.17E-02 | 2.17E-01 | -1.24E+00 | 2.07E-01 | 8.96E-01 | 1.39E+00 | 2.26E-02 | 9.58E-02 | -1.43E+00 | 1.58E-01 | 6.02E-01 | 1.17E+00 | 4.76E-02 | 1.70E-01 | -1.22E+00 | 2.43E-01 | 6.85E-01 | 1.12E+00 | 1.96E-02 | 7.21E-02 | -1.19E+00 | 1.13E-01 | 3.11E-01 | -1.06E+00 |
| SUCNR1 | 4.17E-02 | 2.17E-01 | -1.62E+00 | 1.64E-02 | 7.10E-02 | -2.28E+00 | 5.74E-08 | 2.44E-07 | -1.42E+00 | 4.21E-11 | 1.61E-10 | -1.84E+00 | 2.57E-01 | 9.16E-01 | 1.12E+00 | 2.82E-01 | 7.95E-01 | 1.49E+00 | 6.24E-02 | 2.29E-01 | -1.15E+00 | 6.15E-02 | 1.69E-01 | 1.30E+00 |
| PPIB | 4.17E-02 | 2.17E-01 | 1.07E+00 | 6.99E-02 | 3.02E-01 | 1.21E+00 | 1.07E-03 | 4.56E-03 | -1.21E+00 | 8.24E-06 | 3.15E-05 | -1.38E+00 | 6.55E-05 | 2.34E-04 | -1.45E+00 | 3.56E-06 | 1.00E-05 | -1.82E+00 | 2.58E-03 | 9.48E-03 | -1.04E+00 | 1.79E-03 | 4.92E-03 | -1.19E+00 |
| RGS17 | 4.17E-02 | 2.17E-01 | -1.04E+00 | 2.11E-01 | 9.12E-01 | 1.56E+00 | 1.45E-01 | 6.17E-01 | -1.02E+00 | 2.24E-01 | 8.57E-01 | 1.45E+00 | 2.75E-01 | 9.81E-01 | 1.32E+00 | 3.36E-01 | 9.48E-01 | 1.35E+00 | 1.73E-01 | 6.34E-01 | 1.18E+00 | 3.13E-01 | 8.59E-01 | 1.28E+00 |
| CLDND2 | 4.17E-02 | 2.17E-01 | 1.56E+00 | 1.81E-01 | 7.82E-01 | 1.47E+00 | 3.52E-02 | 1.50E-01 | -1.04E+00 | 4.37E-03 | 1.67E-02 | -1.09E+00 | 1.46E-01 | 5.21E-01 | 1.76E+00 | 3.01E-01 | 8.50E-01 | 1.50E+00 | 1.96E-01 | 7.19E-01 | 1.09E+00 | 3.09E-01 | 8.47E-01 | 1.22E+00 |
| MKL2 | 4.17E-02 | 2.18E-01 | 1.16E+00 | 1.25E-01 | 5.40E-01 | 1.41E+00 | 1.03E-03 | 4.38E-03 | -1.29E+00 | 1.06E-02 | 4.05E-02 | -1.01E+00 | 7.33E-04 | 2.61E-03 | -1.24E+00 | 3.59E-03 | 1.01E-02 | -1.12E+00 | 5.48E-04 | 2.01E-03 | -1.20E+00 | 7.14E-04 | 1.96E-03 | -1.17E+00 |
| CENPS | 4.17E-02 | 2.18E-01 | -1.21E+00 | 1.41E-01 | 6.11E-01 | 1.29E+00 | 6.24E-02 | 2.65E-01 | 1.35E+00 | 7.18E-02 | 2.74E-01 | 1.16E+00 | 3.69E-02 | 1.31E-01 | -1.43E+00 | 1.17E-01 | 3.29E-01 | -1.07E+00 | 1.47E-01 | 5.38E-01 | 1.11E+00 | 1.46E-01 | 4.02E-01 | 1.44E+00 |
| MYO19 | 4.19E-02 | 2.18E-01 | 1.10E+00 | 3.82E-02 | 1.65E-01 | 1.19E+00 | 1.85E-01 | 7.85E-01 | 1.18E+00 | 9.71E-02 | 3.71E-01 | 1.23E+00 | 7.97E-02 | 2.84E-01 | 1.17E+00 | 2.81E-01 | 7.93E-01 | 1.47E+00 | 2.69E-02 | 9.89E-02 | -1.03E+00 | 2.86E-02 | 7.86E-02 | 1.06E+00 |
| AL731569.1 | 4.20E-02 | 2.19E-01 | -1.15E+00 | 1.49E-01 | 6.46E-01 | 1.13E+00 | 2.28E-01 | 9.71E-01 | 1.03E+00 | 1.02E-01 | 3.88E-01 | -1.20E+00 | 1.83E-01 | 6.53E-01 | 1.22E+00 | 3.27E-01 | 9.22E-01 | -1.01E+00 | 1.94E-01 | 7.12E-01 | 1.10E+00 | 1.25E-01 | 3.42E-01 | 1.53E+00 |
| PIGG | 4.20E-02 | 2.19E-01 | 1.14E+00 | 6.01E-03 | 2.60E-02 | 1.08E+00 | 7.28E-03 | 3.09E-02 | -1.10E+00 | 4.27E-03 | 1.63E-02 | -1.02E+00 | 3.26E-02 | 1.16E-01 | 1.03E+00 | 1.07E-02 | 3.01E-02 | -1.02E+00 | 1.77E-02 | 6.50E-02 | -1.10E+00 | 9.52E-03 | 2.61E-02 | -1.12E+00 |
| VPS37C | 4.21E-02 | 2.19E-01 | -1.22E+00 | 6.48E-02 | 2.80E-01 | -1.02E+00 | 2.92E-03 | 1.24E-02 | -1.42E+00 | 5.17E-04 | 1.97E-03 | -1.42E+00 | 4.93E-04 | 1.76E-03 | -2.13E+00 | 7.84E-03 | 2.21E-02 | -1.42E+00 | 6.47E-08 | 2.38E-07 | -1.66E+00 | 1.45E-04 | 3.98E-04 | -1.21E+00 |
| CA5B | 4.21E-02 | 2.20E-01 | 1.43E+00 | 1.29E-01 | 5.57E-01 | 1.35E+00 | 2.32E-01 | 9.85E-01 | 1.28E+00 | 7.82E-02 | 2.99E-01 | 1.25E+00 | 2.21E-03 | 7.89E-03 | 2.59E+00 | 2.91E-01 | 8.21E-01 | 1.69E+00 | 8.24E-06 | 3.03E-05 | 2.65E+00 | 1.45E-01 | 3.98E-01 | 1.63E+00 |
| AC013394.1 | 4.21E-02 | 2.20E-01 | 1.18E+00 | 4.71E-02 | 2.03E-01 | 1.29E+00 | 2.32E-01 | 9.86E-01 | 1.13E+00 | 2.09E-01 | 7.99E-01 | 1.25E+00 | 2.68E-01 | 9.55E-01 | 1.30E+00 | 2.41E-01 | 6.80E-01 | 1.51E+00 | 8.00E-02 | 2.94E-01 | 1.01E+00 | 1.80E-01 | 4.95E-01 | 1.02E+00 |
| STAU2 | 4.21E-02 | 2.20E-01 | 1.14E+00 | 5.79E-02 | 2.50E-01 | 1.34E+00 | 3.02E-02 | 1.28E-01 | -1.10E+00 | 7.97E-02 | 3.04E-01 | 1.09E+00 | 4.30E-04 | 1.53E-03 | -1.20E+00 | 1.92E-02 | 5.42E-02 | -1.02E+00 | 7.93E-02 | 2.91E-01 | 1.08E+00 | 1.82E-02 | 5.01E-02 | -1.00E+00 |
| HYPK | 4.22E-02 | 2.20E-01 | -1.10E+00 | 1.34E-01 | 5.79E-01 | 1.26E+00 | 5.28E-02 | 2.24E-01 | -1.06E+00 | 7.34E-02 | 2.80E-01 | 1.05E+00 | 4.23E-03 | 1.51E-02 | -1.60E+00 | 6.50E-02 | 1.83E-01 | -1.16E+00 | 3.08E-07 | 1.13E-06 | -2.23E+00 | 4.92E-04 | 1.35E-03 | -1.44E+00 |
| FZR1 | 4.22E-02 | 2.20E-01 | 1.16E+00 | 1.32E-02 | 5.71E-02 | 1.07E+00 | 2.26E-01 | 9.62E-01 | 1.17E+00 | 7.47E-02 | 2.85E-01 | 1.05E+00 | 1.88E-02 | 6.70E-02 | 1.07E+00 | 1.42E-02 | 3.99E-02 | -1.04E+00 | 2.57E-03 | 9.42E-03 | -1.19E+00 | 1.23E-02 | 3.37E-02 | -1.08E+00 |
| TTC32 | 4.22E-02 | 2.20E-01 | 1.18E+00 | 3.28E-03 | 1.42E-02 | 1.17E+00 | 3.85E-02 | 1.63E-01 | 1.04E+00 | 2.09E-03 | 8.00E-03 | 1.02E+00 | 6.78E-02 | 2.42E-01 | 1.22E+00 | 6.47E-03 | 1.82E-02 | 1.08E+00 | 8.39E-04 | 3.08E-03 | -1.10E+00 | 8.70E-05 | 2.39E-04 | -1.26E+00 |
| PRDM8 | 4.22E-02 | 2.20E-01 | 1.74E+00 | 2.24E-01 | 9.67E-01 | 1.46E+00 | 1.48E-01 | 6.31E-01 | 1.36E+00 | 3.00E-02 | 1.14E-01 | 1.14E+00 | 1.67E-01 | 5.95E-01 | 1.47E+00 | 4.65E-02 | 1.31E-01 | -1.02E+00 | 1.90E-01 | 6.98E-01 | 1.53E+00 | 1.78E-01 | 4.90E-01 | -1.04E+00 |
| ZNF707 | 4.23E-02 | 2.20E-01 | -1.11E+00 | 2.16E-01 | 9.31E-01 | 1.49E+00 | 8.84E-02 | 3.76E-01 | 1.05E+00 | 9.21E-02 | 3.52E-01 | 1.10E+00 | 7.40E-02 | 2.64E-01 | 2.02E+00 | 2.74E-01 | 7.72E-01 | 1.48E+00 | 2.13E-02 | 7.81E-02 | 1.91E+00 | 2.68E-01 | 7.37E-01 | 1.15E+00 |
| RPS11 | 4.23E-02 | 2.21E-01 | 1.04E+00 | 2.28E-04 | 9.86E-04 | 1.10E+00 | 1.49E-01 | 6.35E-01 | 1.02E+00 | 1.60E-07 | 6.10E-07 | 1.17E+00 | 3.88E-02 | 1.38E-01 | -1.05E+00 | 2.50E-02 | 7.04E-02 | 1.07E+00 | 2.11E-08 | 7.75E-08 | -1.12E+00 | 3.70E-02 | 1.02E-01 | 1.08E+00 |
| UFC1 | 4.24E-02 | 2.21E-01 | -1.01E+00 | 3.77E-03 | 1.63E-02 | 1.02E+00 | 1.85E-01 | 7.87E-01 | 1.03E+00 | 2.07E-01 | 7.91E-01 | 1.10E+00 | 7.75E-02 | 2.76E-01 | 1.03E+00 | 1.02E-01 | 2.87E-01 | 1.07E+00 | 1.85E-01 | 6.80E-01 | 1.10E+00 | 2.53E-01 | 6.96E-01 | 1.06E+00 |
| AC005726.4 | 4.25E-02 | 2.21E-01 | -1.09E+00 | 1.99E-01 | 8.62E-01 | 1.33E+00 | 3.77E-03 | 1.60E-02 | -1.31E+00 | 3.22E-03 | 1.23E-02 | -1.18E+00 | 2.16E-01 | 7.70E-01 | 1.28E+00 | 2.30E-01 | 6.49E-01 | 1.05E+00 | 8.05E-02 | 2.95E-01 | -1.13E+00 | 2.73E-01 | 7.51E-01 | 1.03E+00 |
| TMEM67 | 4.26E-02 | 2.22E-01 | 1.20E+00 | 9.69E-04 | 4.19E-03 | 1.03E+00 | 6.90E-02 | 2.93E-01 | 1.08E+00 | 2.30E-04 | 8.77E-04 | -1.19E+00 | 5.73E-02 | 2.04E-01 | 1.69E+00 | 2.77E-01 | 7.82E-01 | 1.42E+00 | 4.11E-03 | 1.51E-02 | 1.57E+00 | 1.72E-01 | 4.71E-01 | 1.27E+00 |
| SBDSP1 | 4.26E-02 | 2.22E-01 | 1.09E+00 | 1.97E-06 | 8.50E-06 | -1.72E+00 | 5.97E-02 | 2.54E-01 | -1.04E+00 | 2.37E-03 | 9.06E-03 | -1.37E+00 | 2.13E-02 | 7.58E-02 | -1.08E+00 | 1.90E-04 | 5.34E-04 | -1.71E+00 | 4.58E-03 | 1.68E-02 | -1.10E+00 | 1.35E-04 | 3.71E-04 | -1.59E+00 |
| GTF2IP1 | 4.26E-02 | 2.22E-01 | -1.03E+00 | 1.82E-01 | 7.86E-01 | 1.52E+00 | 4.75E-03 | 2.02E-02 | -1.30E+00 | 6.31E-03 | 2.41E-02 | -1.09E+00 | 2.36E-03 | 8.40E-03 | -1.36E+00 | 5.99E-02 | 1.69E-01 | 1.05E+00 | 1.88E-04 | 6.91E-04 | -1.40E+00 | 6.61E-03 | 1.82E-02 | -1.26E+00 |
| PRR29 | 4.27E-02 | 2.22E-01 | -1.22E+00 | 2.02E-02 | 8.75E-02 | -1.33E+00 | 2.25E-02 | 9.55E-02 | -1.26E+00 | 6.61E-02 | 2.53E-01 | -1.05E+00 | 1.11E-02 | 3.94E-02 | -1.33E+00 | 8.64E-02 | 2.44E-01 | -1.05E+00 | 4.93E-02 | 1.81E-01 | 1.14E+00 | 1.44E-01 | 3.97E-01 | 1.36E+00 |
| MXD1 | 4.27E-02 | 2.22E-01 | 1.12E+00 | 1.68E-09 | 7.24E-09 | -1.22E+00 | 1.34E-01 | 5.71E-01 | 1.28E+00 | 1.77E-04 | 6.74E-04 | -1.06E+00 | 1.73E-01 | 6.17E-01 | 1.41E+00 | 7.16E-03 | 2.02E-02 | -1.02E+00 | 6.68E-02 | 2.45E-01 | 1.15E+00 | 5.15E-07 | 1.42E-06 | -1.26E+00 |
| C17orf67 | 4.27E-02 | 2.22E-01 | 1.10E+00 | 6.62E-02 | 2.86E-01 | 1.23E+00 | 1.56E-02 | 6.64E-02 | -1.07E+00 | 1.05E-01 | 4.02E-01 | 1.27E+00 | 8.79E-02 | 3.14E-01 | 2.08E+00 | 2.01E-01 | 5.67E-01 | 1.69E+00 | 1.52E-01 | 5.57E-01 | 1.37E+00 | 3.24E-01 | 8.91E-01 | 1.11E+00 |
| AL161911.1 | 4.27E-02 | 2.23E-01 | -1.02E+00 | 9.30E-03 | 4.02E-02 | -1.20E+00 | 2.02E-02 | 8.59E-02 | -1.25E+00 | 1.68E-02 | 6.43E-02 | -1.20E+00 | 1.96E-01 | 6.98E-01 | 1.62E+00 | 2.21E-01 | 6.24E-01 | 1.58E+00 | 1.57E-01 | 5.77E-01 | -1.02E+00 | 3.57E-02 | 9.79E-02 | -1.32E+00 |
| ZKSCAN2 | 4.28E-02 | 2.23E-01 | -1.01E+00 | 5.38E-02 | 2.32E-01 | 1.12E+00 | 6.19E-02 | 2.63E-01 | -1.13E+00 | 3.96E-02 | 1.51E-01 | -1.06E+00 | 1.23E-01 | 4.39E-01 | 1.45E+00 | 9.96E-02 | 2.81E-01 | -1.09E+00 | 5.66E-02 | 2.08E-01 | 1.82E+00 | 2.09E-02 | 5.74E-02 | 1.81E+00 |
| PHF2 | 4.28E-02 | 2.23E-01 | -1.05E+00 | 1.76E-01 | 7.59E-01 | 1.38E+00 | 8.44E-02 | 3.59E-01 | 1.15E+00 | 6.35E-02 | 2.43E-01 | 1.16E+00 | 2.61E-02 | 9.32E-02 | -1.00E+00 | 2.66E-02 | 7.48E-02 | -1.01E+00 | 8.07E-07 | 2.96E-06 | -1.59E+00 | 3.28E-03 | 9.02E-03 | -1.07E+00 |
| AL035448.1 | 4.28E-02 | 2.23E-01 | -1.05E+00 | 1.55E-01 | 6.70E-01 | 1.28E+00 | 4.19E-03 | 1.78E-02 | -1.39E+00 | 3.58E-02 | 1.37E-01 | 1.01E+00 | 1.29E-01 | 4.60E-01 | 1.10E+00 | 2.91E-01 | 8.21E-01 | 1.55E+00 | 7.79E-02 | 2.86E-01 | 1.69E+00 | 3.59E-01 | 9.86E-01 | 1.25E+00 |
| C1RL-AS1 | 4.28E-02 | 2.23E-01 | 1.11E+00 | 1.02E-02 | 4.42E-02 | 1.04E+00 | 3.05E-05 | 1.29E-04 | -1.71E+00 | 1.13E-04 | 4.30E-04 | -1.48E+00 | 9.46E-03 | 3.37E-02 | -1.42E+00 | 8.84E-03 | 2.49E-02 | -1.54E+00 | 9.21E-03 | 3.38E-02 | -1.26E+00 | 6.84E-02 | 1.88E-01 | -1.02E+00 |
| NRBP2 | 4.28E-02 | 2.23E-01 | 1.60E+00 | 8.24E-03 | 3.56E-02 | 2.03E+00 | 2.36E-02 | 1.00E-01 | 2.18E+00 | 7.13E-02 | 2.72E-01 | 2.10E+00 | 2.23E-01 | 7.96E-01 | 1.46E+00 | 2.97E-01 | 8.36E-01 | 1.53E+00 | 2.70E-01 | 9.92E-01 | -1.07E+00 | 1.91E-01 | 5.26E-01 | 1.25E+00 |
| RPL38 | 4.29E-02 | 2.24E-01 | 1.06E+00 | 2.63E-69 | 1.14E-68 | -1.57E+00 | 5.34E-03 | 2.27E-02 | 1.10E+00 | 2.97E-32 | 1.14E-31 | -1.50E+00 | 6.93E-02 | 2.47E-01 | 1.08E+00 | 7.23E-27 | 2.04E-26 | -1.48E+00 | 2.10E-01 | 7.69E-01 | 1.01E+00 | 9.07E-42 | 2.49E-41 | -1.58E+00 |
| CCNA1 | 4.30E-02 | 2.24E-01 | -1.21E+00 | 2.43E-02 | 1.05E-01 | -1.73E+00 | 5.62E-08 | 2.39E-07 | -2.08E+00 | 1.08E-03 | 4.11E-03 | -1.01E+00 | 5.48E-09 | 1.95E-08 | -1.46E+00 | 4.17E-11 | 1.18E-10 | -2.30E+00 | 1.01E-01 | 3.72E-01 | -1.37E+00 | 1.24E-01 | 3.42E-01 | -1.50E+00 |
| OSBPL3 | 4.30E-02 | 2.24E-01 | 1.07E+00 | 1.59E-02 | 6.88E-02 | 1.10E+00 | 1.31E-02 | 5.56E-02 | -1.06E+00 | 5.66E-02 | 2.16E-01 | 1.20E+00 | 2.50E-02 | 8.93E-02 | 1.05E+00 | 4.58E-03 | 1.29E-02 | -1.09E+00 | 1.78E-01 | 6.53E-01 | 1.25E+00 | 2.45E-01 | 6.74E-01 | 1.18E+00 |
| DUSP11 | 4.30E-02 | 2.24E-01 | 1.24E+00 | 6.47E-02 | 2.79E-01 | 1.37E+00 | 9.08E-06 | 3.86E-05 | -1.34E+00 | 1.09E-03 | 4.16E-03 | -1.00E+00 | 2.20E-02 | 7.86E-02 | 1.08E+00 | 6.27E-02 | 1.77E-01 | 1.16E+00 | 4.19E-03 | 1.54E-02 | -1.04E+00 | 6.81E-03 | 1.87E-02 | 1.01E+00 |
| TRMU | 4.31E-02 | 2.24E-01 | 1.11E+00 | 2.17E-01 | 9.39E-01 | 1.68E+00 | 3.93E-02 | 1.67E-01 | -1.05E+00 | 2.40E-01 | 9.18E-01 | 1.34E+00 | 9.22E-03 | 3.29E-02 | -1.10E+00 | 1.19E-01 | 3.36E-01 | 1.24E+00 | 3.35E-02 | 1.23E-01 | -1.05E+00 | 1.77E-01 | 4.85E-01 | 1.33E+00 |
| LATS2 | 4.31E-02 | 2.25E-01 | 1.19E+00 | 8.27E-07 | 3.57E-06 | -1.11E+00 | 1.80E-01 | 7.63E-01 | 1.23E+00 | 4.39E-04 | 1.68E-03 | -1.08E+00 | 2.27E-01 | 8.08E-01 | 1.31E+00 | 8.48E-02 | 2.39E-01 | -1.12E+00 | 1.08E-01 | 3.98E-01 | 1.08E+00 | 2.30E-02 | 6.31E-02 | -1.17E+00 |
| MPZL1 | 4.31E-02 | 2.25E-01 | 1.16E+00 | 3.76E-03 | 1.63E-02 | 1.12E+00 | 5.02E-02 | 2.13E-01 | 1.07E+00 | 1.29E-03 | 4.94E-03 | 1.02E+00 | 1.39E-03 | 4.96E-03 | -1.15E+00 | 2.34E-02 | 6.58E-02 | 1.06E+00 | 1.72E-01 | 6.32E-01 | 1.24E+00 | 2.95E-01 | 8.12E-01 | 1.18E+00 |
| PTCH2 | 4.31E-02 | 2.25E-01 | -1.03E+00 | 1.01E-04 | 4.38E-04 | -1.89E+00 | 1.68E-03 | 7.14E-03 | -1.38E+00 | 1.52E-04 | 5.80E-04 | -1.69E+00 | 3.50E-02 | 1.25E-01 | -1.40E+00 | 7.94E-02 | 2.24E-01 | -1.29E+00 | 5.65E-03 | 2.07E-02 | -1.58E+00 | 2.32E-02 | 6.38E-02 | -1.44E+00 |
| AP5B1 | 4.31E-02 | 2.25E-01 | -1.05E+00 | 6.33E-02 | 2.74E-01 | 1.07E+00 | 3.82E-06 | 1.62E-05 | -1.57E+00 | 2.52E-02 | 9.63E-02 | 1.14E+00 | 1.20E-03 | 4.30E-03 | -2.45E+00 | 1.05E-02 | 2.97E-02 | -1.45E+00 | 1.62E-02 | 5.94E-02 | -1.32E+00 | 2.37E-01 | 6.50E-01 | 1.12E+00 |
| C18orf8 | 4.31E-02 | 2.25E-01 | 1.10E+00 | 1.81E-01 | 7.84E-01 | 1.71E+00 | 3.55E-04 | 1.51E-03 | -1.31E+00 | 1.92E-03 | 7.32E-03 | -1.10E+00 | 2.22E-01 | 7.90E-01 | 1.37E+00 | 2.32E-01 | 6.53E-01 | 1.35E+00 | 2.25E-02 | 8.25E-02 | -1.07E+00 | 5.54E-02 | 1.52E-01 | -1.03E+00 |
| ZNF219 | 4.31E-02 | 2.25E-01 | 1.18E+00 | 4.66E-03 | 2.01E-02 | 1.17E+00 | 7.13E-03 | 3.03E-02 | -1.13E+00 | 3.04E-03 | 1.16E-02 | -1.03E+00 | 1.28E-03 | 4.55E-03 | -1.22E+00 | 4.74E-03 | 1.34E-02 | -1.09E+00 | 1.97E-01 | 7.23E-01 | 1.09E+00 | 3.60E-01 | 9.90E-01 | 1.23E+00 |
| P2RY13 | 4.31E-02 | 2.25E-01 | 1.05E+00 | 6.66E-02 | 2.88E-01 | 1.16E+00 | 9.14E-02 | 3.88E-01 | 1.02E+00 | 1.79E-01 | 6.84E-01 | 1.25E+00 | 2.69E-02 | 9.58E-02 | 1.18E+00 | 1.07E-01 | 3.02E-01 | 1.39E+00 | 3.77E-02 | 1.38E-01 | 1.55E+00 | 3.46E-01 | 9.50E-01 | 1.33E+00 |
| CREBRF | 4.32E-02 | 2.25E-01 | 1.25E+00 | 1.94E-07 | 8.37E-07 | 1.07E+00 | 1.62E-01 | 6.90E-01 | 1.11E+00 | 4.41E-06 | 1.69E-05 | -1.16E+00 | 3.91E-03 | 1.39E-02 | 1.09E+00 | 3.95E-05 | 1.11E-04 | -1.01E+00 | 7.59E-03 | 2.79E-02 | -1.02E+00 | 2.93E-07 | 8.04E-07 | -1.39E+00 |
| ADGRE1 | 4.32E-02 | 2.25E-01 | 1.26E+00 | 1.81E-01 | 7.83E-01 | 1.64E+00 | 2.38E-02 | 1.01E-01 | 1.03E+00 | 1.81E-01 | 6.91E-01 | 1.27E+00 | 2.50E-01 | 8.91E-01 | 1.47E+00 | 2.58E-01 | 7.27E-01 | 1.04E+00 | 2.42E-01 | 8.90E-01 | 1.35E+00 | 3.56E-01 | 9.77E-01 | 1.07E+00 |
| AC079228.1 | 4.32E-02 | 2.25E-01 | -1.02E+00 | 4.27E-02 | 1.85E-01 | 1.89E+00 | 2.09E-01 | 8.87E-01 | 1.21E+00 | 1.22E-02 | 4.65E-02 | 2.02E+00 | 1.31E-01 | 4.66E-01 | 1.14E+00 | 6.82E-02 | 1.92E-01 | 1.74E+00 | 1.16E-01 | 4.27E-01 | 1.10E+00 | 1.13E-03 | 3.11E-03 | 1.80E+00 |
| AFF2 | 4.32E-02 | 2.25E-01 | -1.42E+00 | 4.10E-02 | 1.77E-01 | -1.54E+00 | 1.59E-02 | 6.74E-02 | -1.21E+00 | 6.27E-03 | 2.40E-02 | -1.10E+00 | 5.15E-03 | 1.84E-02 | -1.25E+00 | 3.90E-02 | 1.10E-01 | -1.02E+00 | 1.72E-01 | 6.33E-01 | 1.02E+00 | 1.63E-01 | 4.49E-01 | -1.17E+00 |
| PCNPP1 | 4.32E-02 | 2.25E-01 | -1.03E+00 | 3.29E-02 | 1.42E-01 | -1.09E+00 | 3.56E-11 | 1.51E-10 | -2.38E+00 | 1.05E-08 | 3.99E-08 | -1.68E+00 | 1.58E-04 | 5.64E-04 | -2.02E+00 | 1.31E-04 | 3.70E-04 | -2.12E+00 | 4.48E-05 | 1.64E-04 | -1.74E+00 | 1.10E-03 | 3.03E-03 | -1.52E+00 |
| LAMTOR4 | 4.33E-02 | 2.26E-01 | 1.09E+00 | 2.66E-06 | 1.15E-05 | 1.05E+00 | 1.67E-01 | 7.09E-01 | 1.06E+00 | 3.16E-03 | 1.21E-02 | -1.00E+00 | 2.78E-01 | 9.91E-01 | 1.13E+00 | 3.01E-02 | 8.50E-02 | -1.00E+00 | 1.32E-01 | 4.85E-01 | 1.20E+00 | 2.31E-01 | 6.35E-01 | 1.11E+00 |
| AC117395.1 | 4.33E-02 | 2.26E-01 | -1.09E+00 | 6.20E-02 | 2.68E-01 | 1.07E+00 | 4.29E-03 | 1.82E-02 | -1.32E+00 | 1.18E-04 | 4.49E-04 | -1.52E+00 | 2.66E-01 | 9.48E-01 | 1.23E+00 | 2.79E-01 | 7.87E-01 | 1.14E+00 | 1.16E-02 | 4.27E-02 | -1.22E+00 | 4.58E-02 | 1.26E-01 | -1.06E+00 |
| AC023157.3 | 4.33E-02 | 2.26E-01 | 1.18E+00 | 2.44E-04 | 1.05E-03 | 2.30E+00 | 3.62E-04 | 1.54E-03 | -1.41E+00 | 2.59E-01 | 9.89E-01 | 1.38E+00 | 5.22E-02 | 1.86E-01 | 1.09E+00 | 7.22E-02 | 2.03E-01 | 2.18E+00 | 2.60E-01 | 9.56E-01 | 1.44E+00 | 7.96E-02 | 2.19E-01 | 1.92E+00 |
| SOCS2-AS1 | 4.33E-02 | 2.26E-01 | 1.06E+00 | 9.32E-02 | 4.03E-01 | 1.30E+00 | 7.49E-02 | 3.18E-01 | 1.07E+00 | 1.79E-01 | 6.84E-01 | 1.32E+00 | 2.02E-01 | 7.21E-01 | 1.37E+00 | 2.05E-01 | 5.78E-01 | 1.35E+00 | 1.73E-01 | 6.35E-01 | 1.61E+00 | 2.57E-01 | 7.06E-01 | 1.64E+00 |
| CLCN7 | 4.33E-02 | 2.26E-01 | -1.14E+00 | 1.84E-02 | 7.93E-02 | -1.14E+00 | 1.49E-01 | 6.35E-01 | 1.04E+00 | 1.35E-01 | 5.16E-01 | 1.13E+00 | 1.56E-02 | 5.56E-02 | -1.40E+00 | 3.28E-02 | 9.24E-02 | -1.25E+00 | 1.99E-01 | 7.30E-01 | -1.10E+00 | 2.50E-01 | 6.87E-01 | 1.24E+00 |
| PNMA3 | 4.34E-02 | 2.26E-01 | 1.03E+00 | 1.74E-01 | 7.53E-01 | 1.47E+00 | 6.45E-03 | 2.74E-02 | -1.29E+00 | 1.61E-01 | 6.14E-01 | 1.21E+00 | 7.31E-02 | 2.61E-01 | -1.42E+00 | 1.69E-01 | 4.77E-01 | -1.04E+00 | 2.71E-01 | 9.94E-01 | 1.16E+00 | 1.43E-01 | 3.92E-01 | 1.64E+00 |
| CSTF2 | 4.34E-02 | 2.26E-01 | -1.26E+00 | 1.90E-01 | 8.23E-01 | 1.51E+00 | 2.59E-06 | 1.10E-05 | -1.65E+00 | 4.28E-05 | 1.64E-04 | -1.29E+00 | 3.47E-03 | 1.24E-02 | -2.19E+00 | 7.92E-02 | 2.23E-01 | -1.11E+00 | 5.47E-12 | 2.01E-11 | -1.68E+00 | 1.37E-03 | 3.76E-03 | 1.00E+00 |
| AC105074.1 | 4.34E-02 | 2.26E-01 | 1.52E+00 | 1.13E-01 | 4.89E-01 | 1.51E+00 | 8.56E-02 | 3.64E-01 | -1.48E+00 | 2.31E-01 | 8.84E-01 | -1.07E+00 | 2.00E-01 | 7.12E-01 | -1.17E+00 | 2.84E-01 | 8.00E-01 | -1.09E+00 | 2.72E-01 | 1.00E+00 | 1.00E+00 | 3.64E-01 | 1.00E+00 | 1.00E+00 |
| OBSCN | 4.34E-02 | 2.26E-01 | 1.08E+00 | 9.90E-02 | 4.28E-01 | 1.70E+00 | 3.62E-02 | 1.54E-01 | -1.16E+00 | 1.18E-01 | 4.52E-01 | 1.63E+00 | 1.78E-01 | 6.34E-01 | 1.26E+00 | 2.99E-01 | 8.42E-01 | 1.51E+00 | 2.31E-01 | 8.48E-01 | 1.21E+00 | 7.06E-02 | 1.94E-01 | 1.65E+00 |
| ALG14 | 4.35E-02 | 2.27E-01 | -1.12E+00 | 1.40E-01 | 6.07E-01 | 1.22E+00 | 1.58E-01 | 6.71E-01 | 1.06E+00 | 8.24E-02 | 3.15E-01 | 1.18E+00 | 6.53E-03 | 2.33E-02 | -1.42E+00 | 9.94E-03 | 2.80E-02 | -1.44E+00 | 1.72E-03 | 6.33E-03 | -1.24E+00 | 3.80E-02 | 1.04E-01 | -1.06E+00 |
| RRAS | 4.35E-02 | 2.27E-01 | 1.16E+00 | 1.05E-01 | 4.52E-01 | 1.35E+00 | 2.20E-01 | 9.36E-01 | 1.24E+00 | 2.17E-01 | 8.27E-01 | 1.50E+00 | 2.24E-01 | 8.00E-01 | 1.14E+00 | 1.96E-01 | 5.51E-01 | 1.06E+00 | 2.97E-02 | 1.09E-01 | 2.27E+00 | 3.16E-01 | 8.69E-01 | 1.48E+00 |
| AL138847.2 | 4.36E-02 | 2.27E-01 | -1.14E+00 | 2.52E-02 | 1.09E-01 | -1.21E+00 | 8.91E-02 | 3.79E-01 | -1.01E+00 | 7.17E-02 | 2.74E-01 | -1.13E+00 | 2.60E-02 | 9.28E-02 | -1.35E+00 | 6.28E-02 | 1.77E-01 | -1.18E+00 | 4.76E-04 | 1.75E-03 | -1.75E+00 | 3.33E-04 | 9.14E-04 | -1.97E+00 |
| KATNBL1P6 | 4.36E-02 | 2.27E-01 | 1.75E+00 | 7.80E-02 | 3.37E-01 | 1.82E+00 | 8.70E-02 | 3.70E-01 | 1.56E+00 | 6.69E-02 | 2.55E-01 | 1.79E+00 | 2.54E-01 | 9.06E-01 | 1.55E+00 | 2.56E-01 | 7.22E-01 | 1.60E+00 | 6.76E-02 | 2.48E-01 | 1.58E+00 | 2.60E-02 | 7.14E-02 | 1.78E+00 |
| RRN3P2 | 4.37E-02 | 2.28E-01 | -1.20E+00 | 7.70E-02 | 3.33E-01 | -1.05E+00 | 1.29E-03 | 5.49E-03 | -1.61E+00 | 5.23E-03 | 2.00E-02 | -1.24E+00 | 1.06E-04 | 3.78E-04 | -1.80E+00 | 2.82E-03 | 7.95E-03 | -1.19E+00 | 1.82E-06 | 6.70E-06 | -2.06E+00 | 1.30E-02 | 3.56E-02 | -1.20E+00 |
| ITFG2-AS1 | 4.37E-02 | 2.28E-01 | 1.11E+00 | 3.55E-02 | 1.53E-01 | 1.19E+00 | 8.44E-05 | 3.58E-04 | -1.63E+00 | 3.00E-06 | 1.15E-05 | -1.79E+00 | 1.98E-01 | 7.08E-01 | 1.53E+00 | 2.85E-01 | 8.04E-01 | 1.17E+00 | 2.45E-01 | 9.00E-01 | 1.24E+00 | 1.77E-01 | 4.86E-01 | 1.05E+00 |
| DCAF1 | 4.38E-02 | 2.28E-01 | -1.01E+00 | 9.08E-02 | 3.92E-01 | 1.18E+00 | 2.42E-03 | 1.03E-02 | -1.18E+00 | 6.13E-03 | 2.34E-02 | 1.07E+00 | 7.79E-02 | 2.78E-01 | -1.02E+00 | 1.59E-01 | 4.47E-01 | 1.11E+00 | 2.70E-05 | 9.90E-05 | -1.37E+00 | 7.57E-04 | 2.08E-03 | -1.15E+00 |
| AL356535.1 | 4.38E-02 | 2.28E-01 | 1.24E+00 | 1.22E-02 | 5.29E-02 | 1.31E+00 | 2.28E-01 | 9.70E-01 | 1.24E+00 | 2.33E-01 | 8.91E-01 | 1.42E+00 | 2.63E-01 | 9.37E-01 | 1.33E+00 | 1.80E-01 | 5.06E-01 | 1.56E+00 | 1.39E-01 | 5.10E-01 | 1.37E+00 | 2.13E-01 | 5.85E-01 | 1.38E+00 |
| MORC4 | 4.38E-02 | 2.28E-01 | -1.00E+00 | 1.05E-01 | 4.53E-01 | 1.27E+00 | 3.13E-02 | 1.33E-01 | -1.27E+00 | 4.56E-02 | 1.74E-01 | -1.04E+00 | 1.16E-01 | 4.12E-01 | 2.06E+00 | 2.32E-01 | 6.54E-01 | 1.75E+00 | 1.41E-01 | 5.18E-01 | 1.60E+00 | 2.21E-01 | 6.08E-01 | 1.50E+00 |
| MTND5P2 | 4.39E-02 | 2.29E-01 | -1.19E+00 | 1.80E-01 | 7.77E-01 | 1.28E+00 | 1.64E-01 | 6.97E-01 | 1.10E+00 | 9.90E-02 | 3.78E-01 | -1.01E+00 | 1.08E-01 | 3.83E-01 | -1.06E+00 | 6.58E-02 | 1.86E-01 | -1.13E+00 | 5.58E-04 | 2.05E-03 | -1.41E+00 | 2.78E-03 | 7.65E-03 | -1.48E+00 |
| SLC6A16 | 4.39E-02 | 2.29E-01 | -1.06E+00 | 1.67E-01 | 7.20E-01 | 1.53E+00 | 1.59E-02 | 6.74E-02 | -1.22E+00 | 1.36E-02 | 5.21E-02 | -1.21E+00 | 1.54E-01 | 5.51E-01 | 1.03E+00 | 2.61E-01 | 7.36E-01 | 1.07E+00 | 2.33E-01 | 8.56E-01 | 1.18E+00 | 3.41E-01 | 9.36E-01 | 1.07E+00 |
| TBC1D12 | 4.39E-02 | 2.29E-01 | -1.06E+00 | 5.00E-02 | 2.16E-01 | 1.02E+00 | 1.47E-03 | 6.23E-03 | -1.49E+00 | 1.39E-02 | 5.33E-02 | -1.04E+00 | 1.81E-01 | 6.46E-01 | 1.04E+00 | 3.20E-01 | 9.02E-01 | 1.66E+00 | 8.44E-02 | 3.10E-01 | -1.15E+00 | 1.36E-02 | 3.73E-02 | -1.66E+00 |
| RPL15P20 | 4.39E-02 | 2.29E-01 | 1.05E+00 | 8.46E-03 | 3.65E-02 | -1.01E+00 | 9.01E-02 | 3.83E-01 | 1.01E+00 | 3.57E-04 | 1.36E-03 | -1.50E+00 | 1.07E-01 | 3.80E-01 | 1.18E+00 | 5.30E-03 | 1.49E-02 | -1.43E+00 | 1.39E-02 | 5.11E-02 | -1.13E+00 | 1.77E-04 | 4.87E-04 | -1.56E+00 |
| NAIF1 | 4.40E-02 | 2.29E-01 | -1.10E+00 | 1.46E-02 | 6.32E-02 | -1.25E+00 | 6.67E-02 | 2.83E-01 | -1.05E+00 | 1.47E-01 | 5.61E-01 | 1.27E+00 | 2.82E-05 | 1.01E-04 | -1.61E+00 | 1.25E-04 | 3.53E-04 | -1.41E+00 | 6.35E-05 | 2.33E-04 | -1.40E+00 | 1.27E-03 | 3.48E-03 | -1.36E+00 |
| NUDT4P1 | 4.40E-02 | 2.29E-01 | -1.03E+00 | 4.39E-02 | 1.90E-01 | 1.06E+00 | 1.41E-02 | 5.98E-02 | -1.31E+00 | 7.66E-03 | 2.92E-02 | -1.29E+00 | 8.82E-03 | 3.14E-02 | -1.23E+00 | 5.46E-03 | 1.54E-02 | -1.22E+00 | 5.64E-05 | 2.07E-04 | -1.20E+00 | 8.07E-07 | 2.22E-06 | -1.63E+00 |
| TDP1 | 4.40E-02 | 2.29E-01 | 1.06E+00 | 7.07E-02 | 3.05E-01 | 1.20E+00 | 1.15E-01 | 4.90E-01 | 1.10E+00 | 1.74E-01 | 6.63E-01 | 1.28E+00 | 2.74E-04 | 9.77E-04 | -1.50E+00 | 8.42E-03 | 2.37E-02 | -1.15E+00 | 5.60E-03 | 2.06E-02 | -1.14E+00 | 3.44E-01 | 9.45E-01 | 1.24E+00 |
| FRMD3 | 4.41E-02 | 2.30E-01 | 1.92E+00 | 1.51E-01 | 6.54E-01 | 1.55E+00 | 2.19E-01 | 9.29E-01 | 1.16E+00 | 1.26E-01 | 4.81E-01 | 1.66E+00 | 1.44E-01 | 5.14E-01 | 1.72E+00 | 4.49E-02 | 1.27E-01 | 2.89E+00 | 2.12E-03 | 7.80E-03 | 2.39E+00 | 2.57E-05 | 7.06E-05 | 3.26E+00 |
| RN7SKP71 | 4.41E-02 | 2.30E-01 | 1.12E+00 | 2.29E-06 | 9.90E-06 | -2.00E+00 | 7.37E-04 | 3.13E-03 | -1.36E+00 | 9.68E-06 | 3.70E-05 | -2.00E+00 | 1.46E-03 | 5.20E-03 | -1.24E+00 | 6.71E-08 | 1.89E-07 | -2.94E+00 | 1.95E-01 | 7.16E-01 | 1.21E+00 | 2.79E-02 | 7.66E-02 | -1.55E+00 |
| RPA4 | 4.41E-02 | 2.30E-01 | -1.18E+00 | 1.62E-01 | 7.02E-01 | 1.19E+00 | 1.02E-02 | 4.33E-02 | -1.22E+00 | 2.18E-01 | 8.33E-01 | 1.27E+00 | 2.14E-01 | 7.64E-01 | 1.24E+00 | 3.46E-01 | 9.77E-01 | 1.02E+00 | 2.52E-01 | 9.23E-01 | 1.03E+00 | 2.42E-01 | 6.65E-01 | 1.35E+00 |
| FBRS | 4.41E-02 | 2.30E-01 | 1.17E+00 | 2.66E-08 | 1.15E-07 | -1.16E+00 | 8.83E-02 | 3.75E-01 | 1.11E+00 | 3.45E-05 | 1.32E-04 | -1.08E+00 | 7.17E-03 | 2.56E-02 | 1.04E+00 | 1.42E-04 | 4.00E-04 | -1.09E+00 | 2.45E-01 | 9.00E-01 | 1.36E+00 | 2.24E-03 | 6.15E-03 | -1.06E+00 |
| ABHD11 | 4.41E-02 | 2.30E-01 | -1.34E+00 | 7.46E-02 | 3.22E-01 | -1.14E+00 | 1.05E-01 | 4.46E-01 | 1.02E+00 | 4.13E-02 | 1.58E-01 | -1.07E+00 | 2.14E-01 | 7.62E-01 | 1.35E+00 | 8.71E-02 | 2.45E-01 | 1.24E+00 | 9.32E-10 | 3.42E-09 | 2.43E+00 | 6.96E-03 | 1.91E-02 | 1.52E+00 |
| ASB1 | 4.41E-02 | 2.30E-01 | 1.00E+00 | 1.30E-02 | 5.60E-02 | -1.10E+00 | 1.96E-03 | 8.32E-03 | -1.34E+00 | 2.21E-03 | 8.44E-03 | -1.22E+00 | 4.76E-03 | 1.70E-02 | -1.26E+00 | 5.41E-02 | 1.53E-01 | 1.09E+00 | 1.11E-02 | 4.06E-02 | 1.02E+00 | 9.30E-02 | 2.55E-01 | 1.17E+00 |
| TAF4 | 4.41E-02 | 2.30E-01 | -1.20E+00 | 2.15E-01 | 9.28E-01 | 1.26E+00 | 1.92E-03 | 8.15E-03 | -1.39E+00 | 4.54E-02 | 1.73E-01 | 1.05E+00 | 4.16E-02 | 1.48E-01 | -1.22E+00 | 2.60E-01 | 7.34E-01 | 1.30E+00 | 1.85E-01 | 6.78E-01 | 1.12E+00 | 2.07E-01 | 5.68E-01 | 1.35E+00 |
| AC005336.2 | 4.42E-02 | 2.30E-01 | -1.01E+00 | 5.19E-02 | 2.24E-01 | 2.04E+00 | 2.12E-01 | 8.99E-01 | 1.27E+00 | 6.91E-06 | 2.64E-05 | 3.37E+00 | 4.57E-02 | 1.63E-01 | 3.48E+00 | 4.07E-04 | 1.15E-03 | 5.95E+00 | 1.45E-01 | 5.32E-01 | 1.47E+00 | 1.37E-01 | 3.76E-01 | 1.87E+00 |
| VTI1A | 4.43E-02 | 2.31E-01 | 1.09E+00 | 1.33E-01 | 5.74E-01 | 1.33E+00 | 2.47E-03 | 1.05E-02 | -1.06E+00 | 5.76E-03 | 2.20E-02 | 1.11E+00 | 2.00E-01 | 7.14E-01 | 1.24E+00 | 1.23E-01 | 3.46E-01 | 1.17E+00 | 2.53E-02 | 9.30E-02 | 1.03E+00 | 1.61E-01 | 4.42E-01 | 1.20E+00 |
| AC233289.1 | 4.43E-02 | 2.31E-01 | -1.17E+00 | 6.82E-02 | 2.95E-01 | -1.11E+00 | 3.56E-02 | 1.51E-01 | -1.37E+00 | 1.21E-01 | 4.63E-01 | 1.02E+00 | 4.91E-03 | 1.75E-02 | -1.51E+00 | 2.02E-02 | 5.69E-02 | -1.19E+00 | 2.03E-02 | 7.44E-02 | -1.48E+00 | 3.39E-01 | 9.31E-01 | 1.13E+00 |
| TCEANC | 4.43E-02 | 2.31E-01 | -1.12E+00 | 1.90E-01 | 8.19E-01 | 1.53E+00 | 1.55E-02 | 6.58E-02 | -1.48E+00 | 8.63E-03 | 3.30E-02 | -1.54E+00 | 2.14E-03 | 7.63E-03 | -1.60E+00 | 1.36E-04 | 3.82E-04 | -2.03E+00 | 1.78E-03 | 6.52E-03 | -1.38E+00 | 2.76E-02 | 7.59E-02 | -1.16E+00 |
| SHC1 | 4.44E-02 | 2.31E-01 | 1.11E+00 | 8.84E-04 | 3.82E-03 | -1.21E+00 | 7.62E-02 | 3.24E-01 | 1.12E+00 | 2.07E-04 | 7.92E-04 | -1.32E+00 | 1.69E-03 | 6.04E-03 | -1.31E+00 | 1.64E-03 | 4.61E-03 | -1.33E+00 | 3.45E-08 | 1.26E-07 | -1.64E+00 | 1.99E-06 | 5.46E-06 | -1.43E+00 |
| CLK1 | 4.45E-02 | 2.32E-01 | 1.47E+00 | 4.75E-02 | 2.05E-01 | 1.33E+00 | 1.72E-02 | 7.30E-02 | -1.07E+00 | 5.67E-05 | 2.17E-04 | -1.11E+00 | 3.30E-03 | 1.18E-02 | 1.04E+00 | 4.07E-06 | 1.15E-05 | -1.18E+00 | 7.69E-05 | 2.82E-04 | -1.18E+00 | 9.97E-08 | 2.74E-07 | -1.48E+00 |
| SDK2 | 4.45E-02 | 2.32E-01 | 1.35E+00 | 1.61E-01 | 6.97E-01 | 1.19E+00 | 7.80E-02 | 3.32E-01 | 1.02E+00 | 5.79E-06 | 2.21E-05 | -1.37E+00 | 2.65E-01 | 9.43E-01 | -1.14E+00 | 3.18E-01 | 8.96E-01 | -1.24E+00 | 2.15E-01 | 7.88E-01 | 1.31E+00 | 3.02E-01 | 8.30E-01 | 1.30E+00 |
| GPM6B | 4.45E-02 | 2.32E-01 | 1.04E+00 | 1.83E-01 | 7.92E-01 | 1.43E+00 | 5.65E-04 | 2.40E-03 | -1.22E+00 | 4.99E-02 | 1.90E-01 | 1.18E+00 | 1.76E-01 | 6.29E-01 | 1.38E+00 | 3.27E-01 | 9.22E-01 | 1.50E+00 | 1.16E-01 | 4.25E-01 | -1.23E+00 | 1.21E-02 | 3.32E-02 | 1.76E+00 |
| CROCCP3 | 4.45E-02 | 2.32E-01 | -1.00E+00 | 2.11E-01 | 9.13E-01 | 1.43E+00 | 9.31E-03 | 3.95E-02 | -1.50E+00 | 1.40E-01 | 5.36E-01 | 1.10E+00 | 1.23E-01 | 4.39E-01 | 1.04E+00 | 3.35E-01 | 9.45E-01 | 1.33E+00 | 5.17E-02 | 1.90E-01 | -1.14E+00 | 1.21E-01 | 3.34E-01 | -1.01E+00 |
| ZBTB20-AS5 | 4.46E-02 | 2.32E-01 | -1.02E+00 | 3.75E-02 | 1.62E-01 | -1.01E+00 | 2.23E-01 | 9.48E-01 | 1.22E+00 | 2.28E-01 | 8.71E-01 | 1.18E+00 | 2.63E-01 | 9.37E-01 | 1.31E+00 | 1.67E-01 | 4.71E-01 | -1.02E+00 | 1.11E-01 | 4.08E-01 | -1.06E+00 | 6.60E-02 | 1.81E-01 | -1.31E+00 |
| ZNF646 | 4.46E-02 | 2.32E-01 | -1.07E+00 | 1.38E-01 | 5.95E-01 | 1.26E+00 | 4.50E-04 | 1.91E-03 | -1.42E+00 | 4.19E-04 | 1.60E-03 | -1.32E+00 | 6.14E-03 | 2.19E-02 | -1.51E+00 | 1.42E-01 | 4.02E-01 | 1.06E+00 | 4.46E-02 | 1.64E-01 | -1.17E+00 | 5.74E-02 | 1.58E-01 | -1.09E+00 |
| ABCA2 | 4.46E-02 | 2.32E-01 | 1.11E+00 | 4.99E-02 | 2.16E-01 | 1.19E+00 | 7.55E-03 | 3.21E-02 | -1.01E+00 | 1.33E-01 | 5.07E-01 | 1.36E+00 | 1.58E-01 | 5.63E-01 | -1.03E+00 | 9.02E-02 | 2.54E-01 | -1.35E+00 | 2.16E-01 | 7.92E-01 | 1.10E+00 | 2.42E-01 | 6.64E-01 | 1.53E+00 |
| GPR171 | 4.46E-02 | 2.33E-01 | 1.59E+00 | 3.18E-02 | 1.37E-01 | 1.90E+00 | 2.26E-01 | 9.62E-01 | 1.14E+00 | 2.48E-01 | 9.49E-01 | 1.30E+00 | 1.52E-02 | 5.41E-02 | 1.66E+00 | 1.89E-01 | 5.33E-01 | 1.39E+00 | 2.87E-08 | 1.05E-07 | 2.45E+00 | 3.62E-03 | 9.95E-03 | 1.85E+00 |
| AC107308.1 | 4.47E-02 | 2.33E-01 | -1.03E+00 | 5.46E-03 | 2.36E-02 | 2.28E+00 | 4.66E-03 | 1.98E-02 | -1.67E+00 | 4.31E-02 | 1.65E-01 | -1.12E+00 | 5.79E-02 | 2.06E-01 | -1.56E+00 | 9.52E-02 | 2.68E-01 | -1.40E+00 | 1.81E-01 | 6.66E-01 | -1.08E+00 | 3.64E-01 | 1.00E+00 | 1.13E+00 |
| ZNF582 | 4.47E-02 | 2.33E-01 | -1.10E+00 | 4.99E-02 | 2.15E-01 | -1.03E+00 | 9.33E-03 | 3.96E-02 | -1.34E+00 | 1.27E-02 | 4.85E-02 | -1.09E+00 | 5.13E-03 | 1.83E-02 | -1.20E+00 | 4.59E-03 | 1.29E-02 | -1.36E+00 | 7.93E-02 | 2.91E-01 | 1.09E+00 | 7.86E-03 | 2.16E-02 | -1.18E+00 |
| TNFRSF14-AS1 | 4.47E-02 | 2.33E-01 | -1.52E+00 | 6.87E-02 | 2.97E-01 | -1.41E+00 | 1.69E-01 | 7.18E-01 | 1.01E+00 | 1.79E-01 | 6.82E-01 | -1.02E+00 | 6.12E-06 | 2.18E-05 | -1.48E+00 | 1.66E-13 | 4.67E-13 | -2.90E+00 | 4.33E-02 | 1.59E-01 | 1.03E+00 | 7.58E-03 | 2.08E-02 | -1.32E+00 |
| LSS | 4.47E-02 | 2.33E-01 | 1.71E+00 | 1.57E-02 | 6.77E-02 | 2.14E+00 | 1.21E-01 | 5.14E-01 | 1.50E+00 | 7.82E-02 | 2.99E-01 | 1.85E+00 | 9.58E-02 | 3.42E-01 | -1.02E+00 | 3.24E-01 | 9.14E-01 | 1.33E+00 | 2.03E-01 | 7.47E-01 | 1.31E+00 | 2.00E-01 | 5.50E-01 | 1.61E+00 |
| AC108704.2 | 4.48E-02 | 2.33E-01 | -1.08E+00 | 1.50E-01 | 6.46E-01 | 1.42E+00 | 5.52E-02 | 2.35E-01 | -1.10E+00 | 6.27E-02 | 2.40E-01 | 1.79E+00 | 2.60E-01 | 9.26E-01 | 1.21E+00 | 3.29E-01 | 9.26E-01 | 1.01E+00 | 1.98E-01 | 7.27E-01 | 1.37E+00 | 1.32E-01 | 3.62E-01 | 1.64E+00 |
| AL590426.1 | 4.48E-02 | 2.34E-01 | -1.17E+00 | 1.31E-01 | 5.65E-01 | 1.48E+00 | 1.04E-01 | 4.43E-01 | -1.18E+00 | 2.23E-01 | 8.50E-01 | 1.39E+00 | 9.15E-02 | 3.26E-01 | -1.04E+00 | 3.38E-01 | 9.52E-01 | 1.25E+00 | 2.52E-01 | 9.25E-01 | 1.06E+00 | 1.05E-01 | 2.89E-01 | 1.70E+00 |
| TUBGCP3 | 4.49E-02 | 2.34E-01 | 1.06E+00 | 7.27E-02 | 3.14E-01 | 1.18E+00 | 1.93E-05 | 8.20E-05 | -1.21E+00 | 4.91E-04 | 1.88E-03 | 1.02E+00 | 9.49E-02 | 3.38E-01 | 1.25E+00 | 2.17E-01 | 6.13E-01 | 1.35E+00 | 2.47E-04 | 9.05E-04 | -1.19E+00 | 2.16E-02 | 5.95E-02 | 1.06E+00 |
| EXD5 | 4.49E-02 | 2.34E-01 | -1.16E+00 | 9.99E-02 | 4.32E-01 | 1.03E+00 | 7.55E-02 | 3.21E-01 | -1.17E+00 | 2.14E-01 | 8.16E-01 | 1.18E+00 | 2.12E-01 | 7.55E-01 | -1.06E+00 | 2.79E-01 | 7.86E-01 | 1.47E+00 | 1.26E-03 | 4.61E-03 | -1.38E+00 | 5.64E-02 | 1.55E-01 | 1.03E+00 |
| MACROD2 | 4.49E-02 | 2.34E-01 | 1.97E+00 | 5.91E-02 | 2.55E-01 | 2.25E+00 | 8.94E-02 | 3.80E-01 | 1.07E+00 | 1.23E-01 | 4.71E-01 | 1.28E+00 | 1.88E-01 | 6.70E-01 | 1.73E+00 | 6.81E-02 | 1.92E-01 | 2.17E+00 | 4.93E-02 | 1.81E-01 | 1.53E+00 | 1.64E-01 | 4.51E-01 | 1.48E+00 |
| PA2G4P4 | 4.49E-02 | 2.34E-01 | 1.66E+00 | 1.35E-01 | 5.83E-01 | 1.74E+00 | 1.00E-02 | 4.25E-02 | 1.67E+00 | 3.04E-02 | 1.16E-01 | 1.81E+00 | 2.12E-01 | 7.56E-01 | 1.52E+00 | 7.80E-02 | 2.20E-01 | 1.94E+00 | 2.50E-01 | 9.17E-01 | 1.35E+00 | 5.43E-04 | 1.49E-03 | 1.95E+00 |
| AC005540.1 | 4.49E-02 | 2.34E-01 | 1.79E+00 | 1.26E-01 | 5.46E-01 | 1.23E+00 | 2.44E-02 | 1.04E-01 | 1.90E+00 | 2.57E-01 | 9.83E-01 | 1.37E+00 | 1.96E-01 | 6.97E-01 | 1.04E+00 | 1.75E-01 | 4.94E-01 | -1.25E+00 | 2.57E-01 | 9.45E-01 | 1.25E+00 | 1.09E-01 | 2.99E-01 | -1.00E+00 |
| PLXNA3 | 4.50E-02 | 2.34E-01 | -1.06E+00 | 1.03E-01 | 4.47E-01 | 1.15E+00 | 3.75E-03 | 1.59E-02 | -1.66E+00 | 8.63E-03 | 3.30E-02 | -1.38E+00 | 1.12E-05 | 4.00E-05 | -1.55E+00 | 3.05E-05 | 8.60E-05 | -1.47E+00 | 1.78E-01 | 6.52E-01 | 1.60E+00 | 1.80E-01 | 4.93E-01 | 1.73E+00 |
| SLC25A42 | 4.50E-02 | 2.34E-01 | 1.08E+00 | 1.20E-01 | 5.17E-01 | 1.28E+00 | 8.21E-02 | 3.49E-01 | 1.02E+00 | 1.98E-01 | 7.56E-01 | 1.26E+00 | 1.66E-01 | 5.92E-01 | 1.19E+00 | 1.63E-01 | 4.61E-01 | 1.73E+00 | 5.95E-06 | 2.19E-05 | -1.33E+00 | 3.26E-04 | 8.94E-04 | -1.21E+00 |
| AL445524.1 | 4.50E-02 | 2.34E-01 | -1.35E+00 | 4.65E-02 | 2.01E-01 | -1.27E+00 | 1.23E-01 | 5.24E-01 | 1.51E+00 | 2.13E-01 | 8.15E-01 | 1.25E+00 | 1.43E-01 | 5.11E-01 | 1.13E+00 | 2.01E-02 | 5.66E-02 | -1.81E+00 | 1.03E-03 | 3.79E-03 | -1.33E+00 | 2.64E-02 | 7.26E-02 | -1.00E+00 |
| BTF3P7 | 4.50E-02 | 2.34E-01 | 1.54E+00 | 2.15E-09 | 9.31E-09 | 2.54E+00 | 2.13E-04 | 9.05E-04 | 1.69E+00 | 3.44E-16 | 1.31E-15 | 2.75E+00 | 1.45E-01 | 5.16E-01 | 1.29E+00 | 1.99E-04 | 5.61E-04 | 2.18E+00 | 5.37E-02 | 1.97E-01 | 1.47E+00 | 9.23E-06 | 2.54E-05 | 2.23E+00 |
| PTGES3P3 | 4.50E-02 | 2.35E-01 | 1.83E+00 | 2.58E-02 | 1.11E-01 | 2.31E+00 | 2.32E-03 | 9.88E-03 | 1.78E+00 | 1.44E-04 | 5.51E-04 | 2.42E+00 | 1.45E-01 | 5.17E-01 | 1.61E+00 | 2.68E-02 | 7.55E-02 | 2.34E+00 | 2.03E-07 | 7.45E-07 | 3.17E+00 | 3.04E-11 | 8.35E-11 | 4.16E+00 |
| AVEN | 4.50E-02 | 2.35E-01 | 1.21E+00 | 2.48E-02 | 1.07E-01 | 1.30E+00 | 1.98E-01 | 8.41E-01 | 1.26E+00 | 1.54E-01 | 5.87E-01 | 1.27E+00 | 1.79E-02 | 6.37E-02 | -1.01E+00 | 1.58E-02 | 4.44E-02 | 1.03E+00 | 1.45E-01 | 5.34E-01 | 1.34E+00 | 1.68E-01 | 4.61E-01 | 1.16E+00 |
| SLC20A2 | 4.50E-02 | 2.35E-01 | 1.14E+00 | 2.41E-02 | 1.04E-01 | 1.20E+00 | 3.50E-02 | 1.49E-01 | 1.03E+00 | 1.71E-01 | 6.52E-01 | 1.34E+00 | 4.70E-03 | 1.68E-02 | -1.06E+00 | 9.43E-02 | 2.66E-01 | 1.18E+00 | 6.96E-03 | 2.56E-02 | -1.07E+00 | 6.31E-02 | 1.73E-01 | 1.05E+00 |
| AC007365.1 | 4.50E-02 | 2.35E-01 | 1.59E+00 | 1.38E-01 | 5.94E-01 | 1.12E+00 | 1.80E-01 | 7.64E-01 | 1.02E+00 | 4.15E-02 | 1.59E-01 | -1.25E+00 | 2.41E-01 | 8.60E-01 | 1.25E+00 | 2.31E-01 | 6.52E-01 | -1.07E+00 | 2.35E-01 | 8.64E-01 | 1.01E+00 | 1.26E-01 | 3.47E-01 | -1.39E+00 |
| SFMBT2 | 4.50E-02 | 2.35E-01 | 1.01E+00 | 3.56E-03 | 1.54E-02 | -1.11E+00 | 9.95E-02 | 4.23E-01 | 1.07E+00 | 3.10E-03 | 1.18E-02 | -1.14E+00 | 2.33E-01 | 8.29E-01 | 1.69E+00 | 2.80E-01 | 7.90E-01 | 1.70E+00 | 8.76E-02 | 3.21E-01 | 1.54E+00 | 3.56E-01 | 9.78E-01 | 1.30E+00 |
| CETP | 4.51E-02 | 2.35E-01 | -1.26E+00 | 2.17E-02 | 9.36E-02 | -1.60E+00 | 8.47E-02 | 3.60E-01 | -1.26E+00 | 5.44E-02 | 2.08E-01 | -1.44E+00 | 8.41E-03 | 3.00E-02 | -1.58E+00 | 8.66E-04 | 2.44E-03 | -2.30E+00 | 1.85E-01 | 6.81E-01 | 1.12E+00 | 2.64E-03 | 7.24E-03 | -1.74E+00 |
| TOMM7 | 4.51E-02 | 2.35E-01 | 1.12E+00 | 1.30E-12 | 5.62E-12 | -1.18E+00 | 2.35E-02 | 9.97E-02 | 1.16E+00 | 1.33E-05 | 5.10E-05 | -1.19E+00 | 1.13E-02 | 4.03E-02 | 1.25E+00 | 7.04E-05 | 1.99E-04 | -1.13E+00 | 1.63E-01 | 5.97E-01 | 1.17E+00 | 4.63E-06 | 1.27E-05 | -1.23E+00 |
| NDUFV2-AS1 | 4.51E-02 | 2.35E-01 | 1.60E+00 | 1.44E-01 | 6.24E-01 | 1.40E+00 | 3.28E-02 | 1.39E-01 | 1.57E+00 | 2.61E-01 | 9.97E-01 | 1.25E+00 | 3.27E-03 | 1.17E-02 | 2.65E+00 | 2.01E-01 | 5.67E-01 | 1.65E+00 | 1.75E-01 | 6.42E-01 | 1.40E+00 | 2.33E-01 | 6.41E-01 | 1.15E+00 |
| XRRA1 | 4.51E-02 | 2.35E-01 | -1.11E+00 | 7.64E-02 | 3.30E-01 | 1.06E+00 | 1.02E-02 | 4.33E-02 | -1.27E+00 | 4.41E-02 | 1.68E-01 | 1.00E+00 | 1.15E-01 | 4.11E-01 | 1.04E+00 | 1.82E-01 | 5.13E-01 | 1.21E+00 | 2.28E-01 | 8.35E-01 | 1.24E+00 | 8.23E-02 | 2.26E-01 | -1.31E+00 |
| PLEKHG4 | 4.51E-02 | 2.35E-01 | 1.01E+00 | 7.02E-02 | 3.03E-01 | 1.13E+00 | 6.17E-08 | 2.62E-07 | -1.62E+00 | 1.24E-06 | 4.74E-06 | -1.25E+00 | 3.48E-02 | 1.24E-01 | -1.12E+00 | 6.16E-02 | 1.74E-01 | -1.04E+00 | 1.09E-01 | 4.02E-01 | -1.00E+00 | 3.36E-01 | 9.22E-01 | 1.36E+00 |
| TMEM131L | 4.52E-02 | 2.35E-01 | 1.07E+00 | 3.03E-02 | 1.31E-01 | 1.11E+00 | 4.77E-03 | 2.03E-02 | -1.16E+00 | 5.25E-03 | 2.01E-02 | 1.00E+00 | 1.48E-01 | 5.26E-01 | 1.06E+00 | 2.58E-01 | 7.26E-01 | 1.14E+00 | 9.40E-05 | 3.45E-04 | -1.39E+00 | 4.43E-02 | 1.22E-01 | 1.04E+00 |
| AL356489.2 | 4.52E-02 | 2.36E-01 | -1.09E+00 | 8.69E-02 | 3.75E-01 | 1.09E+00 | 1.59E-01 | 6.77E-01 | 1.20E+00 | 1.27E-01 | 4.85E-01 | 1.68E+00 | 2.37E-01 | 8.46E-01 | 1.47E+00 | 9.12E-02 | 2.57E-01 | 2.04E+00 | 1.14E-01 | 4.17E-01 | -1.02E+00 | 3.43E-01 | 9.43E-01 | 1.24E+00 |
| MED7 | 4.52E-02 | 2.36E-01 | 1.11E+00 | 4.63E-02 | 2.00E-01 | 1.69E+00 | 2.87E-03 | 1.22E-02 | -1.18E+00 | 1.22E-01 | 4.66E-01 | 1.23E+00 | 4.97E-02 | 1.77E-01 | 1.05E+00 | 3.55E-01 | 1.00E+00 | 1.38E+00 | 2.17E-02 | 7.95E-02 | 1.04E+00 | 2.72E-01 | 7.48E-01 | 1.18E+00 |
| AC105429.1 | 4.53E-02 | 2.36E-01 | -1.22E+00 | 1.23E-01 | 5.33E-01 | 1.07E+00 | 8.51E-03 | 3.62E-02 | -1.77E+00 | 3.42E-03 | 1.31E-02 | -2.01E+00 | 2.68E-01 | 9.56E-01 | 1.20E+00 | 3.29E-01 | 9.27E-01 | 1.44E+00 | 2.46E-01 | 9.04E-01 | 1.22E+00 | 3.35E-01 | 9.20E-01 | -1.00E+00 |
| TSPAN15 | 4.55E-02 | 2.37E-01 | -1.06E+00 | 2.20E-01 | 9.52E-01 | 1.51E+00 | 1.58E-01 | 6.70E-01 | -1.04E+00 | 2.20E-01 | 8.39E-01 | 1.34E+00 | 1.65E-01 | 5.87E-01 | -1.09E+00 | 2.35E-01 | 6.62E-01 | 1.03E+00 | 2.05E-01 | 7.53E-01 | 1.69E+00 | 3.06E-01 | 8.40E-01 | 1.13E+00 |
| FANCG | 4.56E-02 | 2.38E-01 | -1.06E+00 | 7.55E-02 | 3.26E-01 | 1.12E+00 | 1.65E-02 | 7.01E-02 | -1.01E+00 | 2.15E-03 | 8.20E-03 | 1.01E+00 | 2.81E-02 | 1.00E-01 | -1.10E+00 | 8.39E-02 | 2.36E-01 | 1.00E+00 | 2.70E-01 | 9.90E-01 | 1.24E+00 | 8.37E-02 | 2.30E-01 | 1.46E+00 |
| ZNF221 | 4.57E-02 | 2.38E-01 | -1.17E+00 | 9.91E-02 | 4.28E-01 | 1.03E+00 | 3.42E-04 | 1.45E-03 | -1.85E+00 | 7.78E-03 | 2.97E-02 | -1.23E+00 | 2.56E-01 | 9.14E-01 | 1.17E+00 | 2.85E-01 | 8.04E-01 | 1.36E+00 | 1.62E-02 | 5.96E-02 | -1.34E+00 | 1.56E-01 | 4.27E-01 | 1.08E+00 |
| KLLN | 4.57E-02 | 2.38E-01 | -1.11E+00 | 2.91E-02 | 1.26E-01 | -1.19E+00 | 4.64E-03 | 1.97E-02 | -1.17E+00 | 2.17E-03 | 8.28E-03 | -1.36E+00 | 1.01E-01 | 3.62E-01 | -1.38E+00 | 7.95E-02 | 2.24E-01 | -1.46E+00 | 4.42E-03 | 1.62E-02 | -1.54E+00 | 1.65E-02 | 4.54E-02 | -1.52E+00 |
| GORASP1 | 4.57E-02 | 2.38E-01 | 1.01E+00 | 8.57E-03 | 3.70E-02 | -1.17E+00 | 6.12E-03 | 2.60E-02 | -1.18E+00 | 2.69E-03 | 1.03E-02 | -1.15E+00 | 5.45E-02 | 1.94E-01 | 1.02E+00 | 6.69E-02 | 1.89E-01 | -1.01E+00 | 1.62E-01 | 5.93E-01 | 1.22E+00 | 1.93E-01 | 5.29E-01 | 1.21E+00 |
| AL359711.2 | 4.57E-02 | 2.38E-01 | 1.06E+00 | 4.15E-02 | 1.79E-01 | 1.16E+00 | 8.66E-02 | 3.68E-01 | 1.43E+00 | 7.71E-02 | 2.94E-01 | 1.03E+00 | 1.29E-03 | 4.60E-03 | -1.67E+00 | 7.72E-04 | 2.18E-03 | -1.65E+00 | 2.45E-01 | 9.01E-01 | 1.25E+00 | 3.51E-01 | 9.65E-01 | 1.16E+00 |
| AC008467.1 | 4.58E-02 | 2.38E-01 | -1.08E+00 | 1.89E-01 | 8.18E-01 | 1.26E+00 | 7.22E-02 | 3.07E-01 | -1.05E+00 | 9.48E-02 | 3.62E-01 | 1.18E+00 | 2.71E-01 | 9.68E-01 | 1.26E+00 | 2.68E-01 | 7.57E-01 | 1.10E+00 | 2.26E-01 | 8.28E-01 | 1.02E+00 | 2.26E-01 | 6.20E-01 | 1.44E+00 |
| SLC35E2 | 4.58E-02 | 2.39E-01 | -1.09E+00 | 2.55E-02 | 1.10E-01 | -1.10E+00 | 1.67E-03 | 7.08E-03 | -1.51E+00 | 2.25E-02 | 8.58E-02 | -1.07E+00 | 1.50E-01 | 5.35E-01 | 1.25E+00 | 2.06E-01 | 5.82E-01 | 1.13E+00 | 1.60E-01 | 5.87E-01 | 1.17E+00 | 2.24E-01 | 6.14E-01 | 1.50E+00 |
| AC078789.1 | 4.58E-02 | 2.39E-01 | 1.60E+00 | 1.73E-01 | 7.50E-01 | 1.25E+00 | 6.87E-02 | 2.92E-01 | 1.69E+00 | 1.04E-01 | 3.98E-01 | 1.70E+00 | 2.66E-01 | 9.48E-01 | 1.19E+00 | 2.87E-01 | 8.10E-01 | 1.00E+00 | 5.09E-02 | 1.87E-01 | 1.67E+00 | 4.18E-02 | 1.15E-01 | 2.07E+00 |
| RPL7P52 | 4.58E-02 | 2.39E-01 | 1.65E+00 | 9.51E-02 | 4.11E-01 | 1.13E+00 | 7.44E-02 | 3.16E-01 | 1.62E+00 | 1.65E-01 | 6.28E-01 | 1.61E+00 | 2.49E-01 | 8.88E-01 | 1.28E+00 | 1.11E-01 | 3.14E-01 | -1.06E+00 | 1.17E-01 | 4.28E-01 | 1.63E+00 | 2.65E-01 | 7.29E-01 | 1.37E+00 |
| NOVA1 | 4.58E-02 | 2.39E-01 | 1.10E+00 | 5.73E-02 | 2.47E-01 | 1.26E+00 | 1.15E-02 | 4.91E-02 | -1.26E+00 | 5.96E-02 | 2.28E-01 | 1.02E+00 | 2.34E-01 | 8.35E-01 | 1.57E+00 | 3.45E-01 | 9.72E-01 | 1.48E+00 | 4.21E-02 | 1.55E-01 | -1.11E+00 | 7.27E-02 | 2.00E-01 | -1.06E+00 |
| NPM1P46 | 4.60E-02 | 2.40E-01 | 1.12E+00 | 9.94E-04 | 4.29E-03 | -1.13E+00 | 3.68E-02 | 1.56E-01 | -1.05E+00 | 3.37E-05 | 1.29E-04 | -1.31E+00 | 6.38E-02 | 2.27E-01 | -1.07E+00 | 4.96E-02 | 1.40E-01 | -1.08E+00 | 9.29E-02 | 3.41E-01 | 1.08E+00 | 1.16E-01 | 3.19E-01 | 1.07E+00 |
| HIST1H2BL | 4.61E-02 | 2.40E-01 | -1.25E+00 | 3.71E-02 | 1.60E-01 | -1.34E+00 | 5.05E-02 | 2.15E-01 | 1.54E+00 | 1.36E-01 | 5.19E-01 | 1.46E+00 | 1.88E-01 | 6.69E-01 | 1.23E+00 | 4.07E-02 | 1.15E-01 | -1.29E+00 | 1.84E-01 | 6.77E-01 | 1.03E+00 | 9.12E-02 | 2.50E-01 | -1.25E+00 |
| ZNF793-AS1 | 4.61E-02 | 2.40E-01 | 1.04E+00 | 3.51E-04 | 1.51E-03 | -1.38E+00 | 1.51E-01 | 6.42E-01 | 1.34E+00 | 2.43E-01 | 9.30E-01 | 1.48E+00 | 2.19E-01 | 7.81E-01 | 1.51E+00 | 2.84E-01 | 8.00E-01 | 1.64E+00 | 7.06E-03 | 2.59E-02 | 1.97E+00 | 1.19E-01 | 3.27E-01 | 1.66E+00 |
| IQSEC1 | 4.62E-02 | 2.41E-01 | 1.00E+00 | 6.46E-02 | 2.79E-01 | 1.20E+00 | 7.25E-02 | 3.08E-01 | -1.14E+00 | 1.23E-01 | 4.70E-01 | 1.03E+00 | 2.80E-01 | 9.99E-01 | 1.28E+00 | 2.35E-01 | 6.64E-01 | 1.65E+00 | 2.15E-02 | 7.90E-02 | -1.03E+00 | 2.45E-01 | 6.73E-01 | 1.24E+00 |
| SETP11 | 4.63E-02 | 2.41E-01 | 1.09E+00 | 1.70E-01 | 7.36E-01 | 1.38E+00 | 1.59E-02 | 6.77E-02 | -1.15E+00 | 3.22E-02 | 1.23E-01 | 1.08E+00 | 1.72E-01 | 6.14E-01 | 1.13E+00 | 1.95E-01 | 5.49E-01 | 1.15E+00 | 8.26E-02 | 3.03E-01 | 1.34E+00 | 5.66E-02 | 1.55E-01 | 1.62E+00 |
| PCDHGB6 | 4.63E-02 | 2.41E-01 | -1.04E+00 | 7.58E-02 | 3.27E-01 | 1.17E+00 | 2.75E-05 | 1.17E-04 | -2.09E+00 | 9.08E-04 | 3.47E-03 | -1.30E+00 | 2.44E-02 | 8.72E-02 | -1.42E+00 | 6.70E-02 | 1.89E-01 | -1.19E+00 | 1.10E-01 | 4.02E-01 | -1.16E+00 | 1.03E-01 | 2.84E-01 | -1.16E+00 |

| | | | | | | | | | | | | | | | | | | | | | | | | |
|---|---|---|---|---|---|---|---|---|---|---|---|---|---|---|---|---|---|---|---|---|---|---|---|---|
| TNS3 | 4.68E-02 | 2.44E-01 | 1.14E+00 | 4.99E-02 | 2.16E-01 | 1.58E+00 | 2.14E-01 | 9.09E-01 | 1.18E+00 | 6.77E-02 | 2.58E-01 | 1.54E+00 | 2.42E-01 | 8.62E-01 | 1.75E+00 | 7.27E-02 | 2.05E-01 | 2.07E+00 | 1.44E-02 | 5.30E-02 | 1.93E+00 | 1.90E-04 | 5.23E-04 | 2.66E+00 |
| C8orf37 | 4.68E-02 | 2.44E-01 | -1.26E+00 | 3.98E-02 | 1.72E-01 | -1.24E+00 | 9.99E-04 | 4.25E-03 | -1.79E+00 | 2.04E-02 | 7.80E-02 | -1.27E+00 | 2.11E-01 | 7.51E-01 | -1.10E+00 | 3.43E-01 | 9.68E-01 | 1.10E+00 | 3.04E-04 | 1.11E-03 | -1.61E+00 | 9.57E-02 | 2.63E-01 | 1.00E+00 |
| AL627308.1 | 4.68E-02 | 2.44E-01 | -1.07E+00 | 1.92E-01 | 8.29E-01 | 1.60E+00 | 1.63E-01 | 6.93E-01 | 1.05E+00 | 2.19E-01 | 8.36E-01 | 1.24E+00 | 2.54E-01 | 9.07E-01 | 1.39E+00 | 3.50E-01 | 9.86E-01 | 1.43E+00 | 1.05E-01 | 3.84E-01 | 1.09E+00 | 3.15E-01 | 8.66E-01 | 1.32E+00 |
| ZNF213 | 4.70E-02 | 2.45E-01 | -1.36E+00 | 3.07E-02 | 1.33E-01 | -1.42E+00 | 2.96E-02 | 1.26E-01 | 1.01E+00 | 4.00E-02 | 1.53E-01 | 1.04E+00 | 7.79E-04 | 2.78E-03 | -1.42E+00 | 1.83E-02 | 5.15E-02 | -1.07E+00 | 1.17E-02 | 4.28E-02 | 1.40E+00 | 5.09E-02 | 1.40E-01 | 1.09E+00 |
| MMAA | 4.71E-02 | 2.45E-01 | 1.65E+00 | 1.09E-01 | 4.71E-01 | 1.68E+00 | 1.22E-01 | 5.18E-01 | 1.07E+00 | 4.60E-02 | 1.76E-01 | 1.08E+00 | 1.37E-01 | 4.88E-01 | -1.26E+00 | 1.04E-01 | 2.92E-01 | 1.91E+00 | 2.02E-01 | 7.40E-01 | 1.04E+00 | 1.63E-01 | 4.49E-01 | 1.49E+00 |
| PTGER4 | 4.71E-02 | 2.45E-01 | 1.49E+00 | 1.88E-05 | 8.11E-05 | -1.30E+00 | 2.16E-01 | 9.17E-01 | 1.26E+00 | 1.85E-06 | 7.06E-06 | -1.38E+00 | 2.02E-02 | 7.22E-02 | 2.17E+00 | 1.03E-01 | 2.90E-01 | -1.06E+00 | 1.19E-01 | 4.38E-01 | 1.60E+00 | 8.83E-02 | 2.43E-01 | -1.29E+00 |
| ADD2 | 4.71E-02 | 2.46E-01 | -1.83E+00 | 5.84E-02 | 2.52E-01 | -1.74E+00 | 2.22E-01 | 9.43E-01 | 1.13E+00 | 2.61E-01 | 9.96E-01 | 1.06E+00 | 2.53E-01 | 9.02E-01 | 1.43E+00 | 1.04E-03 | 2.92E-03 | 3.83E+00 | 1.51E-05 | 5.54E-05 | -1.10E+00 | 8.43E-03 | 2.32E-02 | 1.12E+00 |
| RETREG3 | 4.72E-02 | 2.46E-01 | 1.04E+00 | 7.31E-02 | 3.16E-01 | 1.23E+00 | 1.89E-05 | 8.05E-05 | -1.46E+00 | 9.66E-04 | 3.69E-03 | -1.03E+00 | 5.43E-03 | 1.94E-02 | -1.06E+00 | 3.38E-01 | 9.52E-01 | 1.33E+00 | 3.76E-02 | 1.38E-01 | 1.10E+00 | 7.70E-02 | 2.12E-01 | 1.15E+00 |
| E2F5 | 4.72E-02 | 2.46E-01 | 1.07E+00 | 2.25E-01 | 9.73E-01 | 1.55E+00 | 8.09E-02 | 3.44E-01 | 1.52E+00 | 6.86E-02 | 2.62E-01 | 1.81E+00 | 9.58E-02 | 3.42E-01 | 1.06E+00 | 2.89E-01 | 8.15E-01 | 1.26E+00 | 8.39E-02 | 3.08E-01 | 1.68E+00 | 1.96E-02 | 5.38E-02 | 2.04E+00 |
| HNRNPA1P39 | 4.73E-02 | 2.46E-01 | 1.11E+00 | 8.49E-02 | 3.67E-01 | 1.29E+00 | 1.29E-01 | 5.50E-01 | 1.15E+00 | 5.45E-02 | 2.08E-01 | 1.25E+00 | 1.97E-01 | 7.03E-01 | 1.27E+00 | 2.96E-01 | 8.36E-01 | 1.49E+00 | 1.57E-01 | 5.77E-01 | 1.21E+00 | 2.26E-01 | 6.20E-01 | 1.20E+00 |
| RIOK3 | 4.73E-02 | 2.46E-01 | 1.10E+00 | 2.24E-03 | 9.66E-03 | 1.16E+00 | 5.91E-03 | 2.51E-02 | -1.03E+00 | 2.00E-03 | 7.62E-03 | 1.03E+00 | 9.28E-04 | 3.31E-03 | -1.05E+00 | 3.91E-05 | 1.10E-04 | -1.11E+00 | 7.05E-02 | 2.59E-01 | 1.28E+00 | 4.66E-04 | 1.28E-03 | -1.04E+00 |
| AC005229.4 | 4.73E-02 | 2.46E-01 | 1.04E+00 | 1.44E-01 | 6.22E-01 | 1.32E+00 | 9.49E-02 | 4.03E-01 | 1.10E+00 | 1.82E-01 | 6.93E-01 | 1.23E+00 | 3.50E-03 | 1.25E-02 | -1.44E+00 | 1.14E-01 | 3.21E-01 | 1.21E+00 | 1.82E-05 | 6.69E-05 | -1.43E+00 | 5.08E-05 | 1.40E-04 | -1.54E+00 |
| AL096865.1 | 4.73E-02 | 2.46E-01 | 1.95E+00 | 3.99E-06 | 1.72E-05 | 3.83E+00 | 2.89E-02 | 1.23E-01 | 1.29E+00 | 7.90E-02 | 3.02E-01 | 1.34E+00 | 2.24E-01 | 7.97E-01 | 1.71E+00 | 2.94E-01 | 8.28E-01 | 1.50E+00 | 2.67E-01 | 9.79E-01 | 1.08E+00 | 2.29E-01 | 6.30E-01 | 1.47E+00 |
| AP000350.6 | 4.73E-02 | 2.46E-01 | -1.27E+00 | 2.08E-01 | 9.00E-01 | 1.16E+00 | 1.26E-01 | 5.35E-01 | 1.39E+00 | 1.77E-02 | 6.78E-02 | 2.30E+00 | 2.57E-01 | 9.15E-01 | 1.25E+00 | 1.42E-01 | 4.01E-01 | 1.76E+00 | 6.71E-02 | 2.46E-01 | -1.29E+00 | 2.41E-01 | 6.62E-01 | 1.31E+00 |
| UBXN6 | 4.74E-02 | 2.47E-01 | 1.15E+00 | 2.30E-03 | 9.93E-03 | 1.15E+00 | 1.22E-01 | 5.20E-01 | 1.01E+00 | 8.83E-02 | 3.37E-01 | 1.09E+00 | 3.88E-02 | 1.38E-01 | 1.20E+00 | 5.37E-02 | 1.51E-01 | 1.22E+00 | 3.65E-02 | 1.34E-01 | 1.43E+00 | 3.28E-01 | 9.01E-01 | 1.15E+00 |
| AC011939.1 | 4.74E-02 | 2.47E-01 | 1.23E+00 | 6.52E-11 | 2.82E-10 | -1.43E+00 | 1.11E-04 | 4.73E-04 | -1.21E+00 | 1.09E-09 | 4.15E-09 | -1.68E+00 | 8.55E-02 | 3.05E-01 | 1.84E+00 | 7.45E-03 | 2.10E-02 | -1.01E+00 | 3.66E-03 | 1.34E-02 | 1.10E+00 | 4.55E-09 | 1.25E-08 | -1.33E+00 |
| FRMD4B | 4.75E-02 | 2.48E-01 | 1.04E+00 | 4.02E-02 | 1.74E-01 | 1.49E+00 | 2.67E-04 | 1.13E-03 | -1.23E+00 | 2.24E-03 | 8.57E-03 | -1.02E+00 | 7.18E-06 | 2.56E-05 | -1.34E+00 | 2.18E-06 | 6.16E-06 | -1.34E+00 | 1.26E-05 | 4.62E-05 | 1.20E+00 | 7.98E-03 | 2.19E-02 | -1.04E+00 |
| DENND4B | 4.75E-02 | 2.48E-01 | 1.11E+00 | 6.55E-02 | 2.83E-01 | 1.34E+00 | 1.67E-03 | 7.12E-03 | -1.21E+00 | 1.07E-01 | 4.07E-01 | 1.25E+00 | 4.37E-04 | 1.56E-03 | -1.21E+00 | 9.26E-03 | 2.61E-02 | 1.05E+00 | 5.89E-04 | 2.16E-03 | -1.17E+00 | 4.64E-02 | 1.28E-01 | 1.04E+00 |
| HCN3 | 4.75E-02 | 2.48E-01 | -1.26E+00 | 2.06E-01 | 8.89E-01 | 1.42E+00 | 1.16E-01 | 4.91E-01 | -1.13E+00 | 2.41E-01 | 9.22E-01 | 1.38E+00 | 2.72E-01 | 9.69E-01 | 1.19E+00 | 2.97E-01 | 8.36E-01 | 1.62E+00 | 9.70E-06 | 3.56E-05 | -1.44E+00 | 4.67E-02 | 1.28E-01 | 1.11E+00 |
| SLC3A1 | 4.75E-02 | 2.48E-01 | -1.05E+00 | 4.86E-03 | 2.10E-02 | -1.34E+00 | 8.56E-05 | 3.64E-04 | -1.48E+00 | 2.31E-06 | 8.82E-06 | -2.01E+00 | 2.72E-01 | 9.71E-01 | 1.44E+00 | 1.69E-01 | 4.76E-01 | 1.03E+00 | 1.88E-01 | 6.92E-01 | 1.00E+00 | 2.42E-01 | 6.65E-01 | 1.53E+00 |
| GLRX3P2 | 4.76E-02 | 2.48E-01 | -1.09E+00 | 2.06E-01 | 8.91E-01 | 1.54E+00 | 1.30E-01 | 5.54E-01 | 1.37E+00 | 7.14E-03 | 2.73E-02 | 2.11E+00 | 2.28E-01 | 8.13E-01 | -1.09E+00 | 2.42E-02 | 6.83E-02 | 2.00E+00 | 1.09E-01 | 4.00E-01 | 1.49E+00 | 4.37E-05 | 1.20E-04 | 2.15E+00 |
| FGFRL1 | 4.76E-02 | 2.48E-01 | -1.05E+00 | 1.05E-01 | 4.52E-01 | 1.18E+00 | 1.55E-01 | 6.59E-01 | 1.16E+00 | 1.85E-01 | 7.07E-01 | 1.57E+00 | 8.77E-02 | 3.13E-01 | -1.32E+00 | 1.10E-01 | 3.10E-01 | -1.51E+00 | 8.07E-02 | 2.96E-01 | 2.03E+00 | 3.35E-01 | 9.19E-01 | -1.01E+00 |
| PXN-AS1 | 4.76E-02 | 2.48E-01 | -1.04E+00 | 2.15E-01 | 9.28E-01 | 1.38E+00 | 1.83E-01 | 7.78E-01 | 1.14E+00 | 1.63E-01 | 6.21E-01 | 1.51E+00 | 5.90E-02 | 2.11E-01 | 1.15E+00 | 7.41E-02 | 2.09E-01 | 1.12E+00 | 9.96E-02 | 3.65E-01 | 1.45E+00 | 1.94E-01 | 5.32E-01 | 1.47E+00 |
| CYBB | 4.76E-02 | 2.48E-01 | -2.41E+00 | 7.54E-02 | 3.26E-01 | -1.56E+00 | 1.50E-01 | 6.37E-01 | 1.20E+00 | 4.92E-02 | 1.88E-01 | -1.72E+00 | 1.71E-01 | 6.08E-01 | 1.53E+00 | 1.98E-01 | 5.57E-01 | 1.59E+00 | 2.69E-01 | 9.88E-01 | -1.15E+00 | 7.78E-02 | 2.14E-01 | 1.19E+00 |
| SSR4P1 | 4.76E-02 | 2.48E-01 | -1.03E+00 | 4.27E-02 | 1.84E-01 | -1.05E+00 | 1.81E-01 | 7.69E-01 | 1.25E+00 | 2.38E-01 | 9.11E-01 | 1.21E+00 | 2.02E-01 | 7.19E-01 | 1.80E+00 | 2.99E-01 | 8.44E-01 | 1.43E+00 | 1.32E-02 | 4.84E-02 | 1.84E+00 | 2.89E-01 | 7.94E-01 | 1.21E+00 |
| DUS1L | 4.76E-02 | 2.48E-01 | 1.17E+00 | 1.36E-01 | 5.89E-01 | 1.44E+00 | 1.96E-01 | 8.32E-01 | 1.16E+00 | 9.29E-02 | 3.55E-01 | 1.23E+00 | 2.97E-03 | 1.06E-02 | 1.03E+00 | 1.25E-02 | 3.52E-02 | 1.09E+00 | 1.91E-02 | 7.03E-02 | 1.11E+00 | 2.72E-01 | 7.48E-01 | 1.32E+00 |
| HMGN2P6 | 4.77E-02 | 2.49E-01 | 1.89E+00 | 2.31E-01 | 9.96E-01 | 1.28E+00 | 6.78E-04 | 2.88E-03 | 1.96E+00 | 1.07E-01 | 4.10E-01 | 1.03E+00 | 2.61E-01 | 9.31E-01 | 1.40E+00 | 1.78E-01 | 5.01E-01 | -1.01E+00 | 3.57E-02 | 1.31E-01 | 1.03E+00 | 2.52E-04 | 6.92E-04 | -1.42E+00 |
| MYOT | 4.78E-02 | 2.49E-01 | -1.09E+00 | 9.61E-02 | 4.15E-01 | 1.61E+00 | 4.88E-03 | 2.07E-02 | -1.38E+00 | 2.62E-01 | 9.99E-01 | 1.38E+00 | 8.66E-02 | 3.09E-01 | -1.17E+00 | 2.58E-01 | 7.28E-01 | 1.17E+00 | 8.74E-02 | 3.21E-01 | -1.15E+00 | 2.09E-01 | 5.74E-01 | 1.05E+00 |
| CGREF1 | 4.78E-02 | 2.49E-01 | 1.05E+00 | 9.19E-02 | 3.97E-01 | 1.29E+00 | 1.92E-02 | 8.17E-02 | 1.02E+00 | 1.63E-01 | 6.22E-01 | 1.33E+00 | 1.03E-01 | 3.66E-01 | 1.06E+00 | 1.65E-01 | 4.64E-01 | 1.80E+00 | 1.44E-01 | 5.28E-01 | 1.23E+00 | 1.92E-01 | 5.27E-01 | 1.44E+00 |
| TMEM99 | 4.78E-02 | 2.49E-01 | 1.06E+00 | 3.01E-02 | 1.30E-01 | 1.04E+00 | 1.13E-01 | 4.82E-01 | 1.34E+00 | 2.18E-01 | 8.33E-01 | 1.32E+00 | 5.75E-02 | 2.05E-01 | -1.17E+00 | 9.84E-02 | 2.77E-01 | -1.05E+00 | 1.11E-01 | 4.08E-01 | -1.13E+00 | 2.73E-01 | 7.51E-01 | 1.08E+00 |
| RTKN2 | 4.78E-02 | 2.49E-01 | -1.20E+00 | 6.56E-02 | 2.83E-01 | -1.09E+00 | 5.76E-03 | 2.45E-02 | -1.16E+00 | 1.62E-04 | 6.19E-04 | -1.23E+00 | 5.73E-02 | 2.04E-01 | -1.28E+00 | 5.35E-02 | 1.51E-01 | -1.58E+00 | 9.76E-02 | 3.58E-01 | 1.04E+00 | 2.59E-01 | 7.11E-01 | 1.29E+00 |
| TPM2 | 4.78E-02 | 2.49E-01 | 1.36E+00 | 7.41E-04 | 3.20E-03 | 1.67E+00 | 1.54E-01 | 6.54E-01 | 1.42E+00 | 1.20E-03 | 4.60E-03 | 1.98E+00 | 2.30E-02 | 8.20E-02 | -1.13E+00 | 4.57E-02 | 1.29E-01 | 1.14E+00 | 2.44E-01 | 8.94E-01 | 2.00E+00 | 2.04E-01 | 5.60E-01 | 1.55E+00 |
| RPS4XP20 | 4.78E-02 | 2.49E-01 | 1.53E+00 | 7.91E-03 | 3.42E-02 | -1.17E+00 | 7.41E-02 | 3.15E-01 | 1.51E+00 | 4.33E-03 | 1.66E-02 | -1.09E+00 | 2.74E-01 | 9.76E-01 | 1.48E+00 | 4.55E-02 | 1.28E-01 | 1.02E+00 | 9.96E-03 | 3.65E-02 | 1.67E+00 | 1.89E-01 | 5.20E-01 | -1.00E+00 |
| LINGO3 | 4.79E-02 | 2.50E-01 | 1.23E+00 | 4.56E-02 | 1.97E-01 | 1.32E+00 | 4.41E-04 | 1.87E-03 | -1.29E+00 | 1.64E-01 | 6.27E-01 | 1.27E+00 | 2.10E-01 | 7.50E-01 | 1.27E+00 | 2.39E-01 | 6.75E-01 | 1.24E+00 | 2.29E-01 | 8.41E-01 | 1.23E+00 | 2.86E-01 | 7.85E-01 | 1.40E+00 |
| TDRD7 | 4.79E-02 | 2.50E-01 | -1.02E+00 | 1.07E-01 | 4.62E-01 | 1.18E+00 | 1.36E-03 | 5.78E-03 | -1.48E+00 | 2.49E-03 | 9.52E-03 | -1.26E+00 | 1.03E-02 | 3.68E-02 | -1.52E+00 | 1.48E-02 | 4.16E-02 | -1.36E+00 | 9.28E-06 | 3.40E-05 | -1.92E+00 | 2.50E-06 | 6.85E-06 | -2.20E+00 |
| AL118506.1 | 4.81E-02 | 2.51E-01 | 1.00E+00 | 1.22E-01 | 5.28E-01 | 1.20E+00 | 5.63E-02 | 2.39E-01 | -1.22E+00 | 2.39E-01 | 9.13E-01 | 1.31E+00 | 1.29E-01 | 4.60E-01 | 1.11E+00 | 8.47E-02 | 2.39E-01 | -1.03E+00 | 1.09E-01 | 3.99E-01 | 1.54E+00 | 3.50E-01 | 9.61E-01 | 1.20E+00 |
| GNG2 | 4.81E-02 | 2.51E-01 | 2.12E+00 | 2.06E-01 | 8.91E-01 | 1.65E+00 | 2.78E-03 | 1.18E-02 | 2.27E+00 | 1.77E-01 | 6.75E-01 | 1.54E+00 | 2.19E-02 | 7.80E-02 | -1.20E+00 | 2.87E-01 | 8.09E-01 | 1.35E+00 | 7.26E-11 | 2.66E-10 | -1.44E+00 | 8.52E-09 | 2.34E-08 | -1.23E+00 |
| SPTAN1 | 4.81E-02 | 2.51E-01 | 1.29E+00 | 2.05E-04 | 8.86E-04 | 1.15E+00 | 4.57E-03 | 1.94E-02 | -1.10E+00 | 2.91E-04 | 1.11E-03 | -1.05E+00 | 7.96E-03 | 2.84E-02 | -1.09E+00 | 2.45E-02 | 6.91E-02 | -1.11E+00 | 7.95E-02 | 2.92E-01 | -1.13E+00 | 1.15E-02 | 3.15E-02 | -1.45E+00 |
| THADA | 4.81E-02 | 2.51E-01 | 1.03E+00 | 1.15E-01 | 4.97E-01 | 1.55E+00 | 1.67E-02 | 7.08E-02 | -1.09E+00 | 2.20E-01 | 8.42E-01 | 1.36E+00 | 1.37E-01 | 4.90E-01 | 1.30E+00 | 2.58E-01 | 7.26E-01 | 1.36E+00 | 1.65E-01 | 6.07E-01 | 1.22E+00 | 3.51E-01 | 9.65E-01 | 1.35E+00 |
| HJURP | 4.82E-02 | 2.51E-01 | -1.55E+00 | 8.40E-02 | 3.63E-01 | -1.32E+00 | 2.26E-01 | 9.59E-01 | 1.21E+00 | 2.58E-01 | 9.86E-01 | 1.30E+00 | 2.37E-01 | 8.44E-01 | -1.08E+00 | 2.93E-01 | 8.26E-01 | -1.13E+00 | 3.34E-02 | 1.22E-01 | -1.06E+00 | 2.02E-01 | 5.54E-01 | 1.25E+00 |
| F8 | 4.82E-02 | 2.51E-01 | -1.04E+00 | 5.36E-02 | 2.32E-01 | 1.04E+00 | 1.91E-01 | 8.10E-01 | -1.01E+00 | 2.43E-01 | 9.29E-01 | 1.19E+00 | 9.36E-02 | 3.34E-01 | -1.21E+00 | 2.33E-01 | 6.57E-01 | 1.01E+00 | 1.26E-01 | 4.62E-01 | -1.05E+00 | 2.28E-01 | 6.27E-01 | 1.07E+00 |
| LKAAEAR1 | 4.82E-02 | 2.51E-01 | 1.34E+00 | 1.85E-01 | 7.99E-01 | 1.32E+00 | 1.21E-06 | 5.16E-06 | 1.94E+00 | 3.63E-03 | 1.39E-02 | 1.89E+00 | 1.87E-01 | 6.66E-01 | -1.19E+00 | 1.69E-01 | 4.76E-01 | -1.52E+00 | 2.30E-01 | 8.46E-01 | -1.04E+00 | 2.41E-01 | 6.62E-01 | 1.73E+00 |
| PLGRKT | 4.83E-02 | 2.52E-01 | 1.61E+00 | 1.77E-01 | 7.66E-01 | 1.62E+00 | 7.16E-02 | 3.04E-01 | 1.36E+00 | 2.55E-01 | 9.74E-01 | 1.40E+00 | 1.56E-01 | 5.55E-01 | 1.46E+00 | 1.64E-01 | 4.63E-01 | 1.28E+00 | 1.66E-04 | 6.08E-04 | 1.64E+00 | 3.38E-04 | 9.30E-04 | 1.66E+00 |
| AP005403.1 | 4.83E-02 | 2.52E-01 | -1.00E+00 | 1.03E-01 | 4.44E-01 | 1.23E+00 | 2.36E-02 | 1.00E-01 | -1.03E+00 | 7.25E-04 | 2.77E-03 | -1.17E+00 | 1.37E-01 | 4.89E-01 | 1.09E+00 | 2.26E-01 | 6.37E-01 | 1.30E+00 | 1.75E-01 | 6.42E-01 | 1.29E+00 | 5.84E-02 | 1.60E-01 | 1.10E+00 |
| AL390039.1 | 4.83E-02 | 2.52E-01 | 1.71E+00 | 4.70E-02 | 2.03E-01 | 2.02E+00 | 4.27E-03 | 1.82E-02 | 1.72E+00 | 4.14E-02 | 1.58E-01 | 1.80E+00 | 2.34E-01 | 8.36E-01 | 1.62E+00 | 4.17E-02 | 1.18E-01 | 2.06E+00 | 4.29E-03 | 1.57E-02 | 1.84E+00 | 6.37E-04 | 1.75E-03 | 1.93E+00 |
| LINC00649 | 4.83E-02 | 2.52E-01 | -1.11E+00 | 8.04E-02 | 3.47E-01 | 1.03E+00 | 4.08E-02 | 1.73E-01 | -1.35E+00 | 4.10E-02 | 1.57E-01 | -1.44E+00 | 7.16E-02 | 2.55E-01 | -1.48E+00 | 1.78E-01 | 5.01E-01 | -1.14E+00 | 3.09E-08 | 1.13E-07 | -2.24E+00 | 1.18E-07 | 3.25E-07 | -2.21E+00 |
| AL357568.2 | 4.85E-02 | 2.53E-01 | -1.14E+00 | 1.01E-01 | 4.37E-01 | -1.04E+00 | 6.58E-02 | 2.80E-01 | -1.37E+00 | 6.01E-02 | 2.30E-01 | -1.40E+00 | 3.02E-05 | 1.08E-04 | -1.60E+00 | 3.23E-03 | 9.10E-03 | -1.14E+00 | 1.88E-01 | 6.89E-01 | 1.16E+00 | 7.71E-02 | 2.12E-01 | 1.59E+00 |
| AC092902.5 | 4.85E-02 | 2.53E-01 | -1.14E+00 | 1.27E-01 | 5.47E-01 | 1.71E+00 | 7.62E-02 | 3.24E-01 | -1.17E+00 | 2.49E-01 | 9.53E-01 | 1.31E+00 | 2.52E-01 | 8.98E-01 | 1.07E+00 | 3.09E-01 | 8.70E-01 | 1.21E+00 | 1.70E-01 | 6.23E-01 | -1.11E+00 | 2.42E-01 | 6.65E-01 | -1.08E+00 |
| RPL37AP8 | 4.85E-02 | 2.53E-01 | 1.07E+00 | 2.14E-56 | 9.27E-56 | -1.89E+00 | 1.92E-01 | 8.15E-01 | 1.03E+00 | 1.30E-24 | 4.95E-24 | -2.07E+00 | 7.00E-02 | 2.49E-01 | 1.02E+00 | 1.12E-17 | 3.17E-17 | -1.76E+00 | 4.65E-02 | 1.71E-01 | -1.03E+00 | 3.89E-17 | 1.07E-16 | -1.91E+00 |
| BRD1 | 4.85E-02 | 2.53E-01 | 1.12E+00 | 4.13E-02 | 1.79E-01 | 1.21E+00 | 1.23E-01 | 5.24E-01 | 1.09E+00 | 1.31E-02 | 5.01E-02 | -1.00E+00 | 1.12E-02 | 4.01E-02 | -1.07E+00 | 3.25E-03 | 9.17E-03 | -1.30E+00 | 2.73E-06 | 1.00E-05 | -1.31E+00 | 1.27E-04 | 3.50E-04 | -1.29E+00 |
| SYCP3 | 4.85E-02 | 2.53E-01 | 1.04E+00 | 2.31E-01 | 9.99E-01 | 1.54E+00 | 1.13E-01 | 4.80E-01 | 1.42E+00 | 3.23E-02 | 1.23E-01 | 2.21E+00 | 2.64E-02 | 9.42E-02 | -1.34E+00 | 3.49E-01 | 9.84E-01 | 1.37E+00 | 1.44E-01 | 5.27E-01 | 1.02E+00 | 1.63E-01 | 4.48E-01 | 1.06E+00 |
| RPL23AP49 | 4.86E-02 | 2.53E-01 | 1.21E+00 | 8.85E-03 | 3.82E-02 | 1.14E+00 | 7.67E-02 | 3.26E-01 | -1.00E+00 | 8.18E-02 | 3.13E-01 | 1.13E+00 | 2.60E-01 | 9.28E-01 | 1.19E+00 | 1.05E-01 | 2.96E-01 | 1.06E+00 | 2.30E-01 | 8.44E-01 | 1.25E+00 | 1.14E-01 | 3.14E-01 | -1.10E+00 |
| RPL12P12 | 4.86E-02 | 2.54E-01 | 1.34E+00 | 1.98E-03 | 8.55E-03 | 1.14E+00 | 3.71E-02 | 1.57E-01 | 1.33E+00 | 4.79E-02 | 1.83E-01 | 1.12E+00 | 1.96E-01 | 7.00E-01 | 1.30E+00 | 1.85E-01 | 5.22E-01 | 1.20E+00 | 1.96E-01 | 7.19E-01 | 1.30E+00 | 2.26E-01 | 6.20E-01 | 1.13E+00 |
| AC073073.1 | 4.87E-02 | 2.54E-01 | -1.00E+00 | 1.16E-01 | 5.01E-01 | 1.27E+00 | 9.29E-02 | 3.95E-01 | 1.42E+00 | 1.02E-01 | 3.89E-01 | 1.10E+00 | 2.35E-01 | 8.38E-01 | 1.35E+00 | 1.90E-01 | 5.35E-01 | 1.78E+00 | 1.97E-01 | 7.23E-01 | 1.45E+00 | 2.52E-01 | 6.91E-01 | 1.31E+00 |
| PTP4A2P1 | 4.87E-02 | 2.54E-01 | -1.06E+00 | 2.29E-01 | 9.88E-01 | 1.46E+00 | 4.32E-02 | 1.83E-01 | -1.12E+00 | 1.31E-01 | 5.02E-01 | 1.25E+00 | 1.11E-01 | 3.97E-01 | 1.01E+00 | 2.35E-01 | 6.64E-01 | 1.60E+00 | 4.08E-02 | 1.50E-01 | -1.05E+00 | 3.48E-01 | 9.57E-01 | 1.24E+00 |
| CBX3P3 | 4.88E-02 | 2.54E-01 | -1.48E+00 | 1.72E-01 | 7.43E-01 | 1.25E+00 | 1.89E-01 | 8.03E-01 | 1.01E+00 | 1.18E-01 | 4.49E-01 | -1.05E+00 | 2.11E-02 | 7.51E-02 | -1.61E+00 | 2.35E-01 | 6.64E-01 | 1.21E+00 | 5.66E-03 | 2.08E-02 | -1.22E+00 | 4.09E-04 | 1.12E-03 | -1.65E+00 |
| TOM1L2 | 4.88E-02 | 2.54E-01 | 1.10E+00 | 1.70E-01 | 7.34E-01 | 1.45E+00 | 5.49E-02 | 2.33E-01 | -1.04E+00 | 1.60E-01 | 6.12E-01 | 1.29E+00 | 1.19E-01 | 4.26E-01 | 1.24E+00 | 2.33E-01 | 6.57E-01 | 1.33E+00 | 1.16E-01 | 4.26E-01 | 1.33E+00 | 1.03E-01 | 2.83E-01 | 1.13E+00 |
| SLC2A4RG | 4.88E-02 | 2.54E-01 | 1.12E+00 | 1.42E-04 | 6.15E-04 | -1.13E+00 | 1.87E-02 | 7.95E-02 | 1.44E+00 | 1.84E-01 | 7.04E-01 | 1.34E+00 | 6.83E-02 | 2.44E-01 | -1.19E+00 | 4.61E-02 | 1.30E-01 | -1.30E+00 | 7.41E-02 | 2.72E-01 | 1.46E+00 | 2.77E-01 | 7.62E-01 | 1.33E+00 |
| TLN2 | 4.88E-02 | 2.55E-01 | -1.16E+00 | 3.74E-02 | 1.62E-01 | -1.23E+00 | 1.88E-04 | 8.01E-04 | -1.51E+00 | 3.58E-05 | 1.37E-04 | -1.48E+00 | 2.67E-04 | 9.52E-04 | -1.76E+00 | 9.74E-05 | 2.75E-04 | -1.92E+00 | 4.30E-02 | 1.58E-01 | -1.20E+00 | 1.25E-01 | 3.42E-01 | -1.02E+00 |
| RUSC2 | 4.89E-02 | 2.55E-01 | -1.04E+00 | 1.15E-02 | 4.96E-02 | -1.17E+00 | 1.77E-02 | 7.50E-02 | -1.21E+00 | 1.01E-02 | 3.88E-02 | -1.20E+00 | 5.57E-02 | 1.99E-01 | 2.09E+00 | 1.19E-01 | 3.34E-01 | 2.09E+00 | 2.45E-01 | 8.99E-01 | 1.04E+00 | 7.40E-02 | 2.03E-01 | -1.47E+00 |
| BCDIN3D-AS1 | 4.89E-02 | 2.55E-01 | -1.12E+00 | 1.64E-02 | 7.10E-02 | -1.23E+00 | 1.88E-05 | 7.99E-05 | -1.90E+00 | 2.51E-05 | 9.57E-05 | -1.63E+00 | 1.42E-02 | 5.07E-02 | -1.26E+00 | 1.55E-02 | 4.36E-02 | -1.39E+00 | 1.04E-01 | 3.82E-01 | -1.02E+00 | 1.93E-02 | 5.30E-02 | -1.37E+00 |
| ZCRB1 | 4.89E-02 | 2.55E-01 | 1.15E+00 | 5.49E-05 | 2.37E-04 | 1.08E+00 | 1.15E-01 | 4.88E-01 | 1.06E+00 | 5.46E-03 | 2.09E-02 | 1.04E+00 | 1.79E-01 | 6.39E-01 | 1.23E+00 | 1.61E-01 | 4.54E-01 | 1.26E+00 | 1.48E-01 | 5.42E-01 | 1.27E+00 | 2.75E-01 | 7.55E-01 | 1.14E+00 |
| EAF2 | 4.89E-02 | 2.55E-01 | 1.20E+00 | 1.84E-04 | 7.97E-04 | 1.14E+00 | 2.31E-01 | 9.83E-01 | 1.20E+00 | 3.35E-03 | 1.28E-02 | 1.04E+00 | 1.60E-01 | 5.72E-01 | 1.14E+00 | 3.26E-02 | 9.20E-02 | 1.03E+00 | 1.32E-01 | 4.83E-01 | 1.12E+00 | 2.73E-01 | 7.49E-01 | 1.13E+00 |
| EBF3 | 4.89E-02 | 2.55E-01 | -1.10E+00 | 9.67E-02 | 4.18E-01 | 1.12E+00 | 1.25E-01 | 5.29E-01 | 1.31E+00 | 1.88E-02 | 7.17E-02 | 2.18E+00 | 4.44E-03 | 1.58E-02 | -2.21E+00 | 1.76E-02 | 4.96E-02 | -1.79E+00 | 6.53E-03 | 2.40E-02 | -1.73E+00 | 7.85E-02 | 2.16E-01 | -1.28E+00 |
| AKT3-IT1 | 4.89E-02 | 2.55E-01 | 1.01E+00 | 1.49E-02 | 6.44E-02 | 1.00E+00 | 2.93E-03 | 1.24E-02 | -1.39E+00 | 4.50E-03 | 1.72E-02 | -1.24E+00 | 1.08E-01 | 3.87E-01 | -1.16E+00 | 3.68E-02 | 1.04E-01 | -1.70E+00 | 4.06E-02 | 1.49E-01 | -1.46E+00 | 7.40E-02 | 2.03E-01 | -1.40E+00 |
| TUBA1C | 4.90E-02 | 2.56E-01 | 1.80E+00 | 1.42E-01 | 6.12E-01 | 1.66E+00 | 1.12E-01 | 4.74E-01 | 1.09E+00 | 1.42E-01 | 5.43E-01 | 1.24E+00 | 2.33E-01 | 8.31E-01 | 1.22E+00 | 7.70E-02 | 2.17E-01 | -1.13E+00 | 8.22E-03 | 3.02E-02 | -1.09E+00 | 1.87E-01 | 5.14E-01 | 1.13E+00 |
| NPW | 4.90E-02 | 2.56E-01 | 1.06E+00 | 2.11E-02 | 9.12E-02 | 1.09E+00 | 1.69E-01 | 7.18E-01 | 1.06E+00 | 6.69E-02 | 2.55E-01 | 1.29E+00 | 1.36E-02 | 4.84E-02 | -1.31E+00 | 1.73E-01 | 4.89E-01 | 1.04E+00 | 4.37E-02 | 1.60E-01 | 1.06E+00 | 1.34E-03 | 3.68E-03 | 1.96E+00 |
| AL035701.1 | 4.91E-02 | 2.56E-01 | 1.51E+00 | 2.04E-01 | 8.81E-01 | 1.19E+00 | 1.52E-01 | 6.47E-01 | 1.29E+00 | 2.31E-01 | 8.81E-01 | 1.31E+00 | 2.17E-01 | 7.73E-01 | 1.48E+00 | 2.83E-01 | 7.97E-01 | 1.12E+00 | 4.84E-03 | 1.78E-02 | -1.42E+00 | 3.85E-02 | 1.06E-01 | -1.20E+00 |
| RHD | 4.91E-02 | 2.56E-01 | -1.15E+00 | 8.58E-02 | 3.71E-01 | 1.05E+00 | 1.13E-01 | 4.80E-01 | -1.16E+00 | 2.40E-01 | 9.16E-01 | 1.41E+00 | 2.23E-02 | 7.95E-02 | -1.50E+00 | 2.89E-02 | 8.14E-02 | -1.43E+00 | 8.56E-03 | 3.14E-02 | 1.20E+00 | 1.39E-06 | 3.83E-06 | -1.21E+00 |
| RPS2P45 | 4.91E-02 | 2.56E-01 | -1.28E+00 | 1.71E-01 | 7.37E-01 | 1.08E+00 | 1.97E-01 | 8.37E-01 | 1.12E+00 | 1.30E-01 | 4.97E-01 | 1.38E+00 | 8.19E-02 | 2.92E-01 | -1.30E+00 | 3.32E-01 | 9.36E-01 | 1.15E+00 | 7.28E-04 | 2.67E-03 | -1.96E+00 | 1.77E-02 | 4.87E-02 | -1.47E+00 |
| DHX35 | 4.92E-02 | 2.56E-01 | 1.06E+00 | 2.93E-03 | 1.27E-02 | -1.19E+00 | 1.08E-01 | 4.58E-01 | 1.37E+00 | 1.53E-01 | 5.86E-01 | 1.19E+00 | 4.26E-03 | 1.52E-02 | -1.33E+00 | 7.39E-02 | 2.08E-01 | 1.03E+00 | 4.75E-03 | 1.74E-02 | -1.15E+00 | 9.91E-04 | 2.72E-03 | -1.22E+00 |
| UBASH3B | 4.92E-02 | 2.56E-01 | 1.95E+00 | 1.09E-01 | 4.72E-01 | 1.42E+00 | 1.74E-01 | 7.40E-01 | 1.31E+00 | 4.42E-03 | 1.69E-02 | -1.08E+00 | 2.53E-01 | 9.02E-01 | 1.12E+00 | 1.36E-01 | 3.83E-01 | -1.38E+00 | 6.72E-03 | 2.47E-02 | -1.08E+00 | 2.43E-03 | 6.68E-03 | -1.65E+00 |
| CLEC9A | 4.93E-02 | 2.57E-01 | 1.14E+00 | 9.87E-06 | 4.26E-05 | 2.17E+00 | 1.16E-01 | 4.91E-01 | 1.40E+00 | 8.29E-03 | 3.17E-02 | 2.03E+00 | 2.30E-01 | 8.20E-01 | 1.53E+00 | 2.22E-01 | 6.26E-01 | 1.78E+00 | 1.81E-01 | 6.63E-01 | 1.53E+00 | 1.00E-01 | 2.75E-01 | 2.11E+00 |
| SMYD2 | 4.93E-02 | 2.57E-01 | 1.05E+00 | 3.90E-03 | 1.69E-02 | -1.12E+00 | 2.02E-02 | 8.60E-02 | -1.08E+00 | 9.88E-03 | 3.78E-02 | -1.02E+00 | 6.64E-03 | 2.37E-02 | -1.24E+00 | 3.62E-02 | 1.02E-01 | -1.06E+00 | 1.82E-04 | 6.69E-04 | -1.25E+00 | 1.04E-02 | 2.86E-02 | -1.09E+00 |
| ZNF185 | 4.93E-02 | 2.57E-01 | 1.09E+00 | 1.89E-01 | 8.18E-01 | 1.41E+00 | 2.65E-04 | 1.13E-03 | -1.34E+00 | 2.87E-02 | 1.10E-01 | 1.11E+00 | 1.01E-01 | 3.61E-01 | -1.01E+00 | 2.42E-02 | 6.81E-02 | -1.34E+00 | 1.75E-01 | 6.41E-01 | 1.53E+00 | 3.24E-01 | 8.91E-01 | 1.26E+00 |
| AC005261.6 | 4.93E-02 | 2.57E-01 | 1.13E+00 | 2.73E-02 | 1.18E-01 | 1.02E+00 | 6.70E-02 | 2.85E-01 | 1.01E+00 | 5.37E-02 | 2.05E-01 | 1.09E+00 | 3.40E-03 | 1.21E-02 | -1.14E+00 | 4.13E-05 | 1.17E-04 | -2.11E+00 | 6.73E-02 | 2.47E-01 | -1.15E+00 | 1.16E-01 | 3.18E-01 | -1.07E+00 |
| KRT18P20 | 4.95E-02 | 2.58E-01 | 1.57E+00 | 2.08E-01 | 9.00E-01 | 1.44E+00 | 2.99E-03 | 1.27E-02 | 1.89E+00 | 2.53E-01 | 9.68E-01 | 1.19E+00 | 1.80E-01 | 6.41E-01 | -1.25E+00 | 2.39E-01 | 6.74E-01 | -1.24E+00 | 8.16E-02 | 2.99E-01 | 1.82E+00 | 3.03E-01 | 8.32E-01 | 1.38E+00 |
| VPS9D1 | 4.95E-02 | 2.58E-01 | 1.12E+00 | 1.39E-02 | 6.01E-02 | 1.02E+00 | 8.84E-02 | 3.76E-01 | 1.11E+00 | 2.85E-02 | 1.09E-01 | 1.06E+00 | 1.94E-02 | 6.91E-02 | -1.23E+00 | 1.79E-01 | 5.05E-01 | 1.25E+00 | 9.54E-02 | 3.50E-01 | 1.09E+00 | 2.75E-02 | 7.55E-02 | -1.06E+00 |
| AC006449.6 | 4.97E-02 | 2.59E-01 | -1.02E+00 | 2.21E-01 | 9.54E-01 | 1.47E+00 | 4.01E-03 | 1.70E-02 | -1.39E+00 | 1.21E-03 | 4.63E-03 | -1.31E+00 | 2.09E-01 | 7.47E-01 | 1.09E+00 | 8.18E-02 | 2.31E-01 | 2.14E+00 | 7.24E-03 | 2.66E-02 | -1.06E+00 | 9.16E-02 | 2.52E-01 | 1.13E+00 |
| GPC2 | 4.97E-02 | 2.59E-01 | 1.01E+00 | 1.43E-01 | 6.19E-01 | 1.28E+00 | 1.59E-05 | 6.75E-05 | -1.77E+00 | 3.54E-04 | 1.35E-03 | -1.22E+00 | 3.28E-03 | 1.17E-02 | -1.25E+00 | 7.17E-04 | 2.02E-03 | -1.29E+00 | 3.27E-02 | 1.20E-01 | -1.51E+00 | 1.70E-01 | 4.66E-01 | -1.15E+00 |
| OCRL | 4.97E-02 | 2.59E-01 | -1.08E+00 | 1.98E-01 | 8.57E-01 | 1.49E+00 | 2.71E-06 | 1.15E-05 | -1.77E+00 | 2.08E-05 | 7.93E-05 | -1.34E+00 | 2.15E-01 | 7.65E-01 | 1.04E+00 | 2.43E-01 | 6.84E-01 | 1.39E+00 | 1.21E-02 | 4.44E-02 | -1.28E+00 | 1.14E-01 | 3.12E-01 | -1.13E+00 |
| FRMD6-AS1 | 4.97E-02 | 2.59E-01 | -1.03E+00 | 1.85E-01 | 8.00E-01 | 1.63E+00 | 9.84E-02 | 4.18E-01 | -1.08E+00 | 2.20E-01 | 8.40E-01 | 1.51E+00 | 1.64E-01 | 5.85E-01 | 1.30E+00 | 6.22E-03 | 1.75E-02 | 2.53E+00 | 8.80E-02 | 3.23E-01 | -1.03E+00 | 2.04E-05 | 5.59E-05 | 2.25E+00 |
| POLE4 | 4.97E-02 | 2.59E-01 | 1.07E+00 | 1.32E-02 | 5.70E-02 | 1.17E+00 | 5.41E-02 | 2.30E-01 | 1.32E+00 | 1.09E-01 | 4.15E-01 | 1.24E+00 | 8.43E-02 | 3.00E-01 | 1.19E+00 | 2.46E-02 | 6.92E-02 | 1.11E+00 | 2.37E-02 | 8.71E-02 | 1.50E+00 | 4.22E-02 | 1.16E-01 | 1.45E+00 |
| CDH12P2 | 4.98E-02 | 2.59E-01 | 1.01E+00 | 5.19E-02 | 2.24E-01 | 2.04E+00 | 2.41E-02 | 1.02E-01 | -1.31E+00 | 7.42E-02 | 2.84E-01 | 1.72E+00 | 2.14E-01 | 7.64E-01 | 1.23E+00 | 1.01E-01 | 2.84E-01 | 1.81E+00 | 5.15E-02 | 1.89E-01 | -1.11E+00 | 9.06E-02 | 2.49E-01 | 1.69E+00 |
| HIST1H2AL | 4.98E-02 | 2.60E-01 | -1.10E+00 | 2.14E-03 | 9.24E-03 | -1.62E+00 | 1.46E-01 | 6.19E-01 | 1.55E+00 | 3.97E-03 | 1.52E-02 | -1.28E+00 | 7.72E-02 | 2.75E-01 | -1.23E+00 | 1.96E-01 | 5.54E-01 | 1.06E+00 | 7.77E-03 | 2.85E-02 | 1.10E+00 | 1.33E-06 | 3.66E-06 | -1.44E+00 |
| AKT1 | 4.98E-02 | 2.60E-01 | 1.15E+00 | 4.59E-02 | 1.98E-01 | 1.23E+00 | 2.11E-01 | 8.95E-01 | 1.23E+00 | 2.08E-01 | 7.94E-01 | 1.36E+00 | 2.70E-02 | 9.64E-02 | 1.15E+00 | 2.15E-03 | 6.05E-03 | -1.01E+00 | 1.72E-01 | 6.32E-01 | 1.30E+00 | 2.77E-01 | 7.60E-01 | 1.31E+00 |
| AC110285.7 | 4.98E-02 | 2.60E-01 | 1.01E+00 | 2.22E-01 | 9.59E-01 | 1.56E+00 | 3.82E-02 | 1.62E-01 | -1.22E+00 | 2.25E-01 | 8.60E-01 | 1.26E+00 | 1.57E-02 | 5.58E-02 | -1.09E+00 | 3.99E-03 | 1.12E-02 | -1.24E+00 | 1.61E-01 | 5.91E-01 | -1.52E+00 | 3.49E-01 | 9.58E-01 | -1.03E+00 |
| YPEL4 | 4.99E-02 | 2.60E-01 | 2.06E+00 | 5.61E-02 | 2.42E-01 | 2.22E+00 | 1.90E-01 | 8.06E-01 | -1.03E+00 | 2.28E-01 | 8.70E-01 | 1.07E+00 | 1.45E-01 | 5.16E-01 | 1.79E+00 | 6.88E-02 | 1.94E-01 | 2.33E+00 | 7.35E-05 | 2.70E-04 | -1.13E+00 | 2.83E-05 | 7.77E-05 | -1.15E+00 |
| TRG-AS1 | 4.99E-02 | 2.60E-01 | -1.45E+00 | 9.70E-02 | 4.19E-01 | -1.15E+00 | 3.17E-02 | 1.35E-01 | -1.06E+00 | 2.21E-02 | 8.45E-02 | -1.10E+00 | 6.77E-02 | 2.41E-01 | -1.38E+00 | 6.70E-02 | 1.89E-01 | -1.53E+00 | 9.71E-02 | 3.56E-01 | -1.08E+00 | 3.68E-02 | 1.01E-01 | -1.18E+00 |
| CNN2 | 4.99E-02 | 2.60E-01 | 1.25E+00 | 7.83E-03 | 3.38E-02 | 1.29E+00 | 8.25E-07 | 3.50E-06 | -1.19E+00 | 1.96E-09 | 7.48E-09 | -1.20E+00 | 1.57E-04 | 5.58E-04 | -1.16E+00 | 1.56E-06 | 4.40E-06 | -1.47E+00 | 5.30E-05 | 1.94E-04 | -2.12E+00 | 8.89E-03 | 2.44E-02 | -1.52E+00 |
| FAM227B | 4.99E-02 | 2.60E-01 | 1.02E+00 | 1.79E-02 | 7.72E-02 | -1.02E+00 | 4.26E-02 | 1.81E-01 | -1.19E+00 | 8.02E-03 | 3.06E-02 | -1.27E+00 | 4.53E-02 | 1.62E-01 | 1.05E+00 | 2.54E-01 | 7.15E-01 | 1.37E+00 | 1.00E-01 | 3.68E-01 | -1.01E+00 | 6.64E-02 | 1.82E-01 | -1.11E+00 |
| NCALD | 5.00E-02 | 2.60E-01 | 1.70E+00 | 4.09E-02 | 1.77E-01 | 2.06E+00 | 2.33E-01 | 9.90E-01 | 1.22E+00 | 4.61E-02 | 1.76E-01 | -1.11E+00 | 2.20E-01 | 7.86E-01 | 1.32E+00 | 3.31E-01 | 9.32E-01 | 1.05E+00 | 2.07E-01 | 7.59E-01 | 1.22E+00 | 3.20E-01 | 8.79E-01 | -1.02E+00 |
| PHC1P1 | 5.00E-02 | 2.60E-01 | 1.63E+00 | 4.36E-02 | 1.88E-01 | 1.98E+00 | 2.20E-01 | 9.35E-01 | 1.13E+00 | 2.51E-01 | 9.58E-01 | 1.27E+00 | 1.61E-01 | 5.74E-01 | 1.18E+00 | 1.11E-01 | 3.13E-01 | 1.02E+00 | 6.60E-02 | 2.42E-01 | -1.14E+00 | 1.67E-02 | 4.58E-02 | -1.52E+00 |
| NDRG2 | 5.00E-02 | 2.61E-01 | 1.10E+00 | 1.40E-01 | 6.07E-01 | 1.55E+00 | 1.95E-01 | 8.28E-01 | 1.25E+00 | 6.93E-02 | 2.65E-01 | 1.69E+00 | 1.60E-01 | 5.69E-01 | -1.07E+00 | 3.19E-01 | 8.99E-01 | 1.39E+00 | 2.37E-01 | 8.71E-01 | 1.15E+00 | 3.35E-01 | 9.19E-01 | 1.34E+00 |
| PLP2 | 5.01E-02 | 2.61E-01 | 1.29E+00 | 7.91E-02 | 3.42E-01 | 1.26E+00 | 4.85E-02 | 2.06E-01 | -1.02E+00 | 6.22E-02 | 2.38E-01 | 1.05E+00 | 3.24E-05 | 1.15E-04 | -1.20E+00 | 7.86E-04 | 2.22E-03 | -1.07E+00 | 1.10E-01 | 4.02E-01 | -1.08E+00 | 1.34E-01 | 3.67E-01 | -1.04E+00 |
| AL596094.1 | 5.01E-02 | 2.61E-01 | -1.17E+00 | 1.09E-01 | 4.69E-01 | 1.01E+00 | 1.69E-01 | 7.20E-01 | 1.07E+00 | 2.92E-02 | 1.11E-01 | -1.13E+00 | 1.41E-01 | 5.01E-01 | 1.83E+00 | 1.47E-01 | 4.15E-01 | 1.97E+00 | 2.64E-01 | 9.68E-01 | 1.06E+00 | 3.14E-01 | 8.63E-01 | 1.21E+00 |
| MIR99AHG | 5.01E-02 | 2.61E-01 | 1.59E+00 | 1.18E-01 | 5.09E-01 | 1.67E+00 | 8.78E-03 | 3.73E-02 | 1.98E+00 | 1.76E-02 | 6.71E-02 | 2.16E+00 | 5.52E-02 | 1.97E-01 | 2.52E+00 | 7.65E-02 | 2.16E-01 | 2.51E+00 | 1.73E-01 | 6.37E-01 | 1.43E+00 | 2.36E-02 | 6.50E-02 | -1.40E+00 |
| ZNF540 | 5.02E-02 | 2.62E-01 | -1.25E+00 | 1.58E-01 | 6.83E-01 | 1.43E+00 | 4.74E-02 | 2.02E-01 | -1.41E+00 | 5.97E-02 | 2.28E-01 | -1.25E+00 | 9.05E-02 | 3.23E-01 | -1.28E+00 | 2.57E-01 | 7.25E-01 | 1.14E+00 | 2.66E-01 | 9.74E-01 | 1.05E+00 | 3.52E-01 | 9.68E-01 | 1.24E+00 |
| SMTN | 5.02E-02 | 2.62E-01 | -1.14E+00 | 7.57E-02 | 3.27E-01 | -1.04E+00 | 1.32E-01 | 5.59E-01 | -1.13E+00 | 1.64E-01 | 6.28E-01 | 1.47E+00 | 4.43E-04 | 1.58E-03 | -2.59E+00 | 6.23E-03 | 1.76E-02 | -1.74E+00 | 2.08E-01 | 7.63E-01 | 2.64E+00 | 3.23E-01 | 8.89E-01 | 1.33E+00 |
| EHD3 | 5.02E-02 | 2.62E-01 | -1.05E+00 | 1.86E-01 | 8.03E-01 | 1.37E+00 | 3.54E-02 | 1.50E-01 | -1.32E+00 | 2.61E-01 | 9.97E-01 | 1.40E+00 | 1.31E-03 | 4.68E-03 | -1.22E+00 | 2.29E-02 | 6.47E-02 | 1.09E+00 | 3.78E-08 | 1.39E-07 | -1.32E+00 | 4.52E-05 | 1.24E-04 | -1.14E+00 |
| GALNT12 | 5.04E-02 | 2.62E-01 | 1.01E+00 | 4.63E-02 | 2.00E-01 | 1.12E+00 | 2.21E-01 | 9.40E-01 | 1.28E+00 | 1.25E-01 | 4.79E-01 | 1.26E+00 | 2.63E-01 | 9.38E-01 | 1.33E+00 | 2.07E-01 | 5.82E-01 | 1.15E+00 | 2.46E-01 | 9.03E-01 | 1.32E+00 | 1.65E-01 | 4.54E-01 | 1.69E+00 |
| KANSL3 | 5.04E-02 | 2.63E-01 | 1.10E+00 | 7.92E-02 | 3.42E-01 | 1.24E+00 | 3.41E-03 | 1.45E-02 | -1.15E+00 | 1.25E-03 | 4.76E-03 | -1.13E+00 | 8.98E-03 | 3.20E-02 | 1.09E+00 | 5.41E-04 | 1.53E-03 | -1.12E+00 | 1.60E-03 | 5.87E-03 | -1.17E+00 | 2.67E-02 | 7.33E-02 | -1.08E+00 |
| AC130895.1 | 5.04E-02 | 2.63E-01 | 1.72E+00 | 1.44E-01 | 6.24E-01 | 1.65E+00 | 4.72E-06 | 2.00E-05 | -1.56E+00 | 1.89E-05 | 7.23E-05 | -1.31E+00 | 2.46E-02 | 8.79E-02 | -1.02E+00 | 4.15E-02 | 1.17E-01 | -1.01E+00 | 1.77E-01 | 6.50E-01 | 1.09E+00 | 1.37E-01 | 3.77E-01 | -1.07E+00 |
| LINC01550 | 5.04E-02 | 2.63E-01 | 1.11E+00 | 2.12E-01 | 9.15E-01 | 1.56E+00 | 2.45E-02 | 1.04E-01 | -1.20E+00 | 6.93E-02 | 2.65E-01 | 1.06E+00 | 2.57E-01 | 9.15E-01 | 1.17E+00 | 3.37E-01 | 9.49E-01 | 1.20E+00 | 7.87E-02 | 2.89E-01 | 1.66E+00 | 2.15E-01 | 5.91E-01 | 1.45E+00 |
| LINC02285 | 5.04E-02 | 2.63E-01 | 1.01E+00 | 9.88E-02 | 4.27E-01 | 1.22E+00 | 1.09E-02 | 4.64E-02 | -1.26E+00 | 4.69E-03 | 1.79E-02 | -1.35E+00 | 1.19E-05 | 4.23E-05 | -1.81E+00 | 7.45E-03 | 2.10E-02 | -1.07E+00 | 1.68E-01 | 6.17E-01 | 1.10E+00 | 3.39E-01 | 9.31E-01 | 1.33E+00 |
| RNF103 | 5.05E-02 | 2.63E-01 | 1.07E+00 | 1.28E-01 | 5.52E-01 | 1.31E+00 | 1.51E-01 | 6.42E-01 | 1.12E+00 | 7.67E-02 | 2.93E-01 | 1.24E+00 | 1.70E-01 | 6.07E-01 | 1.37E+00 | 5.96E-02 | 1.68E-01 | 1.07E+00 | 1.41E-02 | 5.19E-02 | -1.09E+00 | 1.13E-01 | 3.11E-01 | 1.05E+00 |
| ANKAR | 5.05E-02 | 2.63E-01 | 1.22E+00 | 1.41E-01 | 6.09E-01 | 1.77E+00 | 2.24E-05 | 9.52E-05 | -1.50E+00 | 5.13E-03 | 1.96E-02 | 1.05E+00 | 2.08E-01 | 7.41E-01 | 1.50E+00 | 8.18E-02 | 2.31E-01 | 1.93E+00 | 2.21E-02 | 8.11E-02 | -1.02E+00 | 9.63E-02 | 2.65E-01 | 1.12E+00 |
| EEFSEC | 5.05E-02 | 2.63E-01 | 1.01E+00 | 4.59E-02 | 1.98E-01 | 1.09E+00 | 8.37E-03 | 3.56E-02 | -1.19E+00 | 2.98E-03 | 1.14E-02 | -1.11E+00 | 1.56E-02 | 5.56E-02 | -1.01E+00 | 7.99E-02 | 2.25E-01 | 1.16E+00 | 5.28E-03 | 1.94E-02 | -1.06E+00 | 2.10E-01 | 5.76E-01 | 1.18E+00 |
| LINC01023 | 5.06E-02 | 2.64E-01 | 1.67E+00 | 4.07E-02 | 1.76E-01 | 1.27E+00 | 1.81E-01 | 7.67E-01 | 1.10E+00 | 2.48E-03 | 9.46E-03 | -1.16E+00 | 7.81E-02 | 2.78E-01 | 1.79E+00 | 3.43E-01 | 9.68E-01 | 1.44E+00 | 1.82E-01 | 6.69E-01 | 1.47E+00 | 8.27E-02 | 2.27E-01 | 1.06E+00 |
| AC132192.1 | 5.06E-02 | 2.64E-01 | -1.05E+00 | 3.40E-02 | 1.47E-01 | -1.03E+00 | 1.23E-03 | 5.20E-03 | -1.28E+00 | 2.97E-04 | 1.13E-03 | -1.29E+00 | 4.98E-02 | 1.77E-01 | -1.19E+00 | 1.26E-02 | 3.54E-02 | -1.47E+00 | 1.73E-02 | 6.34E-02 | -1.09E+00 | 4.14E-02 | 1.14E-01 | -1.03E+00 |
| MRPL30 | 5.06E-02 | 2.64E-01 | 1.09E+00 | 7.47E-02 | 3.23E-01 | 1.26E+00 | 4.79E-04 | 2.04E-03 | -1.19E+00 | 5.82E-04 | 2.22E-03 | -1.07E+00 | 6.73E-03 | 2.40E-02 | -1.07E+00 | 2.49E-02 | 7.01E-02 | 1.05E+00 | 1.65E-05 | 6.04E-05 | -1.33E+00 | 7.85E-04 | 2.16E-03 | -1.16E+00 |
| CD109 | 5.07E-02 | 2.64E-01 | 1.12E+00 | 2.12E-02 | 9.17E-02 | 1.20E+00 | 4.62E-02 | 1.96E-01 | 1.08E+00 | 1.47E-05 | 5.62E-05 | -1.19E+00 | 2.13E-01 | 7.61E-01 | 1.30E+00 | 2.28E-01 | 6.42E-01 | 1.24E+00 | 6.29E-03 | 2.31E-02 | -1.07E+00 | 1.19E-03 | 3.27E-03 | -1.15E+00 |
| NRXN3 | 5.07E-02 | 2.64E-01 | -1.14E+00 | 2.01E-01 | 8.67E-01 | 1.33E+00 | 4.75E-02 | 2.02E-01 | -1.32E+00 | 2.28E-01 | 8.70E-01 | 1.26E+00 | 2.64E-02 | 9.42E-02 | -1.90E+00 | 3.25E-02 | 9.16E-02 | -1.87E+00 | 2.18E-01 | 8.01E-01 | -1.15E+00 | 2.75E-01 | 7.55E-01 | -1.17E+00 |
| BX537318.1 | 5.08E-02 | 2.65E-01 | 1.05E+00 | 9.40E-03 | 4.06E-02 | 1.01E+00 | 6.56E-04 | 2.79E-03 | -1.56E+00 | 2.98E-03 | 1.14E-02 | -1.35E+00 | 2.76E-02 | 9.83E-02 | -1.29E+00 | 4.48E-02 | 1.26E-01 | -1.16E+00 | 5.20E-02 | 1.91E-01 | -1.17E+00 | 1.41E-01 | 3.88E-01 | 1.00E+00 |
| AL662844.4 | 5.08E-02 | 2.65E-01 | 1.24E+00 | 1.01E-13 | 4.37E-13 | 2.84E+00 | 6.64E-03 | 2.82E-02 | -1.13E+00 | 5.73E-03 | 2.19E-02 | 1.86E+00 | 1.70E-01 | 6.07E-01 | 1.60E+00 | 8.07E-06 | 2.27E-05 | 2.98E+00 | 2.27E-01 | 8.35E-01 | 1.25E+00 | 4.53E-02 | 1.24E-01 | 1.74E+00 |
| ZNF554 | 5.08E-02 | 2.65E-01 | 1.09E+00 | 6.06E-03 | 2.62E-02 | 1.04E+00 | 1.98E-02 | 8.40E-02 | -1.17E+00 | 4.10E-02 | 1.57E-01 | 1.04E+00 | 5.94E-02 | 2.12E-01 | 1.02E+00 | 2.76E-01 | 7.77E-01 | 1.34E+00 | 7.66E-05 | 2.81E-04 | -1.28E+00 | 7.51E-04 | 2.06E-03 | -1.22E+00 |
| ZNF829 | 5.08E-02 | 2.65E-01 | 1.03E+00 | 3.55E-03 | 1.53E-02 | -1.20E+00 | 1.57E-01 | 6.69E-01 | 1.21E+00 | 1.53E-02 | 5.85E-02 | 1.05E+00 | 1.17E-01 | 4.17E-01 | 1.60E+00 | 2.76E-01 | 7.77E-01 | 1.70E+00 | 9.43E-02 | 3.46E-01 | 1.67E+00 | 3.00E-01 | 8.24E-01 | 1.36E+00 |
| AC117503.5 | 5.09E-02 | 2.65E-01 | -1.12E+00 | 1.10E-01 | 4.75E-01 | 1.14E+00 | 1.46E-01 | 6.21E-01 | 1.41E+00 | 2.44E-01 | 9.33E-01 | 1.29E+00 | 2.95E-04 | 1.05E-03 | -1.28E+00 | 1.10E-05 | 3.09E-05 | -1.52E+00 | 1.15E-01 | 4.23E-01 | 1.07E+00 | 1.44E-01 | 3.96E-01 | 1.09E+00 |
| NECTIN1 | 5.09E-02 | 2.65E-01 | -1.28E+00 | 2.86E-02 | 1.24E-01 | -1.39E+00 | 2.45E-02 | 1.04E-01 | -1.20E+00 | 1.65E-02 | 6.31E-02 | -1.08E+00 | 1.76E-03 | 6.27E-03 | -1.12E+00 | 3.18E-03 | 8.96E-03 | -1.00E+00 | 9.51E-07 | 3.49E-06 | -1.31E+00 | 2.38E-02 | 6.55E-02 | 1.00E+00 |
| TCEAL3 | 5.09E-02 | 2.65E-01 | 1.25E+00 | 1.36E-02 | 5.87E-02 | 1.26E+00 | 1.24E-01 | 5.26E-01 | 1.39E+00 | 1.27E-01 | 4.86E-01 | 1.35E+00 | 2.26E-01 | 8.07E-01 | 1.35E+00 | 5.24E-02 | 1.48E-01 | 1.10E+00 | 6.62E-02 | 2.43E-01 | 1.43E+00 | 3.33E-01 | 9.15E-01 | 1.26E+00 |
| ANKS6 | 5.09E-02 | 2.65E-01 | 1.05E+00 | 1.79E-01 | 7.72E-01 | 1.34E+00 | 4.83E-02 | 2.05E-01 | -1.02E+00 | 1.16E-01 | 4.41E-01 | 1.29E+00 | 2.00E-01 | 7.13E-01 | 1.26E+00 | 1.97E-01 | 5.56E-01 | 1.94E+00 | 1.00E-01 | 3.68E-01 | 1.16E+00 | 3.40E-01 | 9.33E-01 | 1.35E+00 |
| AC107956.1 | 5.10E-02 | 2.66E-01 | 1.65E+00 | 5.22E-02 | 2.26E-01 | 1.23E+00 | 3.41E-02 | 1.45E-01 | 1.51E+00 | 7.56E-02 | 2.89E-01 | 1.06E+00 | 9.02E-02 | 3.22E-01 | 1.77E+00 | 1.23E-01 | 3.46E-01 | 1.14E+00 | 9.42E-04 | 3.46E-03 | 2.00E+00 | 5.66E-02 | 1.56E-01 | 1.02E+00 |
| PHLPP1 | 5.11E-02 | 2.66E-01 | -1.03E+00 | 1.76E-01 | 7.58E-01 | 1.29E+00 | 8.98E-04 | 3.81E-03 | -1.32E+00 | 1.69E-04 | 6.44E-04 | -1.32E+00 | 2.33E-01 | 8.32E-01 | 1.46E+00 | 2.29E-01 | 6.46E-01 | 1.31E+00 | 4.48E-02 | 1.64E-01 | 1.03E+00 | 6.21E-03 | 1.71E-02 | -1.16E+00 |
| GOLGA6L9 | 5.11E-02 | 2.66E-01 | -1.01E+00 | 5.41E-02 | 2.34E-01 | 1.08E+00 | 8.31E-05 | 3.53E-04 | -1.48E+00 | 2.08E-06 | 7.95E-06 | -1.60E+00 | 5.48E-03 | 1.95E-02 | -1.46E+00 | 3.22E-03 | 9.08E-03 | -1.59E+00 | 1.89E-06 | 6.95E-06 | -1.59E+00 | 2.92E-08 | 8.03E-08 | -1.92E+00 |
| MTSS1 | 5.11E-02 | 2.66E-01 | -1.01E+00 | 4.70E-02 | 2.03E-01 | 1.01E+00 | 1.65E-01 | 7.02E-01 | 1.36E+00 | 1.41E-01 | 5.37E-01 | -1.07E+00 | 5.11E-03 | 1.82E-02 | -1.27E+00 | 4.86E-02 | 1.37E-01 | 1.01E+00 | 4.36E-25 | 1.60E-24 | -1.37E+00 | 4.27E-30 | 1.17E-29 | -1.65E+00 |
| AC020658.1 | 5.12E-02 | 2.67E-01 | 1.74E+00 | 1.18E-01 | 5.11E-01 | 1.24E+00 | 1.99E-01 | 8.47E-01 | 1.29E+00 | 8.84E-02 | 3.38E-01 | 1.15E+00 | 2.28E-01 | 8.13E-01 | 1.29E+00 | 2.92E-02 | 8.23E-02 | -1.38E+00 | 2.39E-01 | 8.75E-01 | 1.34E+00 | 2.91E-01 | 7.99E-01 | 1.23E+00 |
| NRIP1 | 5.12E-02 | 2.67E-01 | 1.03E+00 | 7.12E-02 | 3.08E-01 | 1.14E+00 | 1.88E-01 | 7.98E-01 | 1.15E+00 | 7.71E-02 | 2.95E-01 | 1.13E+00 | 2.24E-02 | 7.97E-02 | 1.90E+00 | 7.01E-02 | 1.98E-01 | 1.31E+00 | 6.24E-05 | 2.29E-04 | 1.59E+00 | 1.09E-01 | 3.00E-01 | 1.39E+00 |
| SNHG12 | 5.12E-02 | 2.67E-01 | 1.35E+00 | 1.22E-01 | 5.25E-01 | 1.42E+00 | 2.87E-03 | 1.22E-02 | -1.10E+00 | 2.22E-02 | 8.49E-02 | 1.18E+00 | 2.26E-01 | 8.05E-01 | 1.42E+00 | 2.19E-01 | 6.19E-01 | 1.60E+00 | 1.85E-01 | 6.78E-01 | 1.95E+00 | 2.66E-01 | 7.30E-01 | 1.50E+00 |
| SPOPL | 5.13E-02 | 2.67E-01 | 1.11E+00 | 2.50E-03 | 1.08E-02 | -1.03E+00 | 4.77E-03 | 2.02E-02 | -1.13E+00 | 1.74E-04 | 6.66E-04 | -1.15E+00 | 4.96E-02 | 1.77E-01 | 1.82E+00 | 2.98E-01 | 8.40E-01 | 1.47E+00 | 5.10E-02 | 1.87E-01 | 1.41E+00 | 1.19E-01 | 3.28E-01 | 1.33E+00 |
| AC104232.1 | 5.13E-02 | 2.67E-01 | -1.41E+00 | 5.37E-02 | 2.32E-01 | -1.46E+00 | 4.92E-03 | 2.09E-02 | -1.99E+00 | 4.48E-03 | 1.71E-02 | -1.98E+00 | 8.96E-26 | 3.20E-25 | -4.28E+00 | 2.13E-17 | 5.99E-17 | -2.02E+00 | 1.18E-03 | 4.34E-03 | -1.43E+00 | 8.73E-03 | 2.40E-02 | -1.29E+00 |
| NXF1 | 5.13E-02 | 2.68E-01 | 1.09E+00 | 2.96E-02 | 1.28E-01 | 1.16E+00 | 9.76E-07 | 4.15E-06 | -1.28E+00 | 3.05E-08 | 1.16E-07 | -1.30E+00 | 2.96E-03 | 1.06E-02 | -1.12E+00 | 1.02E-03 | 2.88E-03 | -1.20E+00 | 1.83E-10 | 6.72E-10 | -1.53E+00 | 7.10E-06 | 1.95E-05 | -1.28E+00 |
| ZGRF1 | 5.14E-02 | 2.68E-01 | -1.06E+00 | 1.83E-02 | 7.89E-02 | -1.31E+00 | 1.88E-01 | 7.97E-01 | 1.05E+00 | 1.66E-01 | 6.35E-01 | 1.17E+00 | 7.05E-02 | 2.51E-01 | -1.33E+00 | 8.03E-02 | 2.26E-01 | -1.40E+00 | 1.89E-01 | 6.93E-01 | 1.08E+00 | 5.73E-02 | 1.57E-01 | 1.46E+00 |
| LTBP1 | 5.14E-02 | 2.68E-01 | -2.39E+00 | 7.77E-02 | 3.36E-01 | -2.03E+00 | 1.27E-03 | 5.38E-03 | -3.91E+00 | 2.02E-02 | 7.73E-02 | -1.73E+00 | 4.43E-16 | 1.58E-15 | -4.54E+00 | 3.41E-16 | 9.61E-16 | -3.58E+00 | 1.26E-01 | 4.63E-01 | -1.14E+00 | 1.17E-02 | 3.23E-02 | -1.14E+00 |
| TTLL10 | 5.14E-02 | 2.68E-01 | -1.04E+00 | 5.39E-02 | 2.33E-01 | 1.04E+00 | 1.47E-01 | 6.27E-01 | -1.09E+00 | 2.61E-01 | 9.98E-01 | 1.24E+00 | 5.59E-04 | 1.99E-03 | -1.12E+00 | 3.89E-05 | 1.10E-04 | -1.37E+00 | 2.79E-02 | 1.02E-01 | -1.10E+00 | 6.03E-03 | 1.66E-02 | -1.26E+00 |
| PIK3R5 | 5.14E-02 | 2.68E-01 | 1.03E+00 | 7.54E-02 | 3.26E-01 | 1.72E+00 | 2.89E-02 | 1.23E-01 | -1.14E+00 | 6.02E-02 | 2.30E-01 | 1.10E+00 | 1.05E-02 | 3.73E-02 | -1.49E+00 | 5.73E-03 | 1.61E-02 | -1.68E+00 | 1.82E-01 | 6.70E-01 | -1.03E+00 | 1.69E-01 | 4.63E-01 | -1.18E+00 |
| SAAL1 | 5.14E-02 | 2.68E-01 | 1.11E+00 | 6.36E-02 | 2.75E-01 | 1.28E+00 | 1.03E-04 | 4.39E-04 | -1.14E+00 | 4.61E-06 | 1.76E-05 | -1.10E+00 | 9.84E-05 | 3.51E-04 | -1.59E+00 | 6.81E-02 | 1.92E-01 | 1.05E+00 | 3.72E-04 | 1.37E-03 | -1.08E+00 | 1.99E-01 | 5.48E-01 | 1.21E+00 |
| DRG2 | 5.15E-02 | 2.68E-01 | 1.05E+00 | 2.79E-02 | 1.20E-01 | 1.15E+00 | 7.55E-02 | 3.21E-01 | 1.45E+00 | 1.74E-01 | 6.63E-01 | 1.32E+00 | 1.03E-01 | 3.69E-01 | 1.17E+00 | 2.53E-01 | 7.12E-01 | 1.57E+00 | 2.13E-02 | 7.81E-02 | -1.05E+00 | 2.30E-02 | 6.31E-02 | -1.04E+00 |
| NRP2 | 5.15E-02 | 2.68E-01 | 1.77E+00 | 1.35E-03 | 5.84E-03 | 3.11E+00 | 1.04E-02 | 4.42E-02 | 1.93E+00 | 2.36E-02 | 9.01E-02 | 2.22E+00 | 1.50E-01 | 5.36E-01 | 2.03E+00 | 1.73E-01 | 4.88E-01 | -1.34E+00 | 4.12E-02 | 1.51E-01 | 2.41E+00 | 2.55E-01 | 7.01E-01 | 1.47E+00 |
| DYRK3 | 5.15E-02 | 2.68E-01 | 1.09E+00 | 5.48E-04 | 2.37E-03 | -1.34E+00 | 1.70E-01 | 7.22E-01 | 1.05E+00 | 3.42E-02 | 1.31E-01 | -1.08E+00 | 2.70E-01 | 9.63E-01 | 1.33E+00 | 1.86E-01 | 5.23E-01 | -1.05E+00 | 2.25E-01 | 8.26E-01 | 1.18E+00 | 2.39E-01 | 6.56E-01 | 1.11E+00 |
| METRNL | 5.17E-02 | 2.69E-01 | -1.01E+00 | 2.34E-03 | 1.01E-02 | -1.52E+00 | 1.01E-02 | 4.27E-02 | -1.14E+00 | 1.51E-04 | 5.78E-04 | -1.54E+00 | 2.47E-01 | 8.79E-01 | 1.46E+00 | 2.71E-02 | 7.65E-02 | -1.81E+00 | 7.29E-02 | 2.68E-01 | -1.50E+00 | 6.92E-02 | 1.90E-01 | -1.64E+00 |
| TRMT112P6 | 5.17E-02 | 2.69E-01 | 1.04E+00 | 5.80E-03 | 2.50E-02 | 1.93E+00 | 1.03E-01 | 4.36E-01 | 1.42E+00 | 8.47E-06 | 3.24E-05 | 2.47E+00 | 2.12E-01 | 7.57E-01 | 1.21E+00 | 1.76E-03 | 4.95E-03 | 2.31E+00 | 2.67E-01 | 9.78E-01 | 1.28E+00 | 4.09E-06 | 1.12E-05 | 2.27E+00 |
| RDM1 | 5.17E-02 | 2.69E-01 | -1.40E+00 | 7.09E-02 | 3.06E-01 | -1.31E+00 | 1.88E-01 | 7.99E-01 | 1.10E+00 | 2.07E-01 | 7.91E-01 | 1.08E+00 | 2.27E-01 | 8.08E-01 | -1.11E+00 | 2.54E-01 | 7.16E-01 | -1.20E+00 | 2.46E-01 | 9.01E-01 | 1.27E+00 | 1.30E-02 | 3.57E-02 | -1.33E+00 |
| AC011939.2 | 5.17E-02 | 2.69E-01 | 1.56E+00 | 9.45E-05 | 4.08E-04 | -1.24E+00 | 2.26E-02 | 9.62E-02 | 1.59E+00 | 3.65E-04 | 1.39E-03 | -1.36E+00 | 4.04E-05 | 1.44E-04 | 2.55E+00 | 9.07E-02 | 2.56E-01 | 1.07E+00 | 1.65E-02 | 6.07E-02 | 1.41E+00 | 4.10E-04 | 1.13E-03 | -1.39E+00 |
| NRM | 5.17E-02 | 2.70E-01 | 1.16E+00 | 3.12E-02 | 1.35E-01 | 1.20E+00 | 1.24E-02 | 5.29E-02 | -1.18E+00 | 4.82E-02 | 1.84E-01 | 1.08E+00 | 4.55E-02 | 1.62E-01 | -1.04E+00 | 1.96E-01 | 5.54E-01 | 1.28E+00 | 1.69E-02 | 6.19E-02 | -1.01E+00 | 7.87E-02 | 2.16E-01 | 1.06E+00 |
| ARID3B | 5.19E-02 | 2.70E-01 | -1.07E+00 | 6.85E-02 | 2.96E-01 | 1.04E+00 | 1.09E-01 | 4.65E-01 | 1.03E+00 | 6.31E-03 | 2.41E-02 | -1.24E+00 | 4.81E-08 | 1.72E-07 | -1.71E+00 | 2.55E-08 | 7.19E-08 | -1.73E+00 | 2.37E-01 | 8.70E-01 | 1.35E+00 | 1.33E-01 | 3.64E-01 | -1.04E+00 |
| ZNF689 | 5.19E-02 | 2.70E-01 | 1.21E+00 | 7.46E-05 | 3.22E-04 | -1.12E+00 | 1.86E-03 | 7.91E-03 | -1.04E+00 | 1.47E-06 | 5.60E-06 | -1.32E+00 | 1.54E-01 | 5.49E-01 | 1.31E+00 | 7.30E-03 | 2.06E-02 | -1.15E+00 | 7.90E-02 | 2.90E-01 | 1.26E+00 | 1.89E-04 | 5.20E-04 | -1.22E+00 |
| KIF2C | 5.19E-02 | 2.70E-01 | -1.31E+00 | 4.70E-02 | 2.03E-01 | -1.38E+00 | 2.66E-02 | 1.13E-01 | 1.64E+00 | 2.59E-02 | 9.90E-02 | 1.96E+00 | 1.07E-01 | 3.83E-01 | -1.73E+00 | 1.66E-01 | 4.69E-01 | -1.36E+00 | 9.01E-02 | 3.31E-01 | 1.29E+00 | 3.52E-03 | 9.68E-03 | 1.68E+00 |
| KHNYN | 5.19E-02 | 2.71E-01 | 1.01E+00 | 1.10E-02 | 4.74E-02 | -1.09E+00 | 1.83E-03 | 7.77E-03 | -1.19E+00 | 3.39E-04 | 1.30E-03 | -1.14E+00 | 4.18E-03 | 1.49E-02 | -1.40E+00 | 2.29E-02 | 6.46E-02 | -1.10E+00 | 8.52E-04 | 3.13E-03 | -1.25E+00 | 3.76E-07 | 1.03E-06 | -2.05E+00 |
| SRP54-AS1 | 5.20E-02 | 2.71E-01 | -1.00E+00 | 8.62E-02 | 3.72E-01 | 1.23E+00 | 7.47E-02 | 3.17E-01 | -1.00E+00 | 2.79E-02 | 1.06E-01 | -1.09E+00 | 1.31E-02 | 4.67E-02 | -1.12E+00 | 6.50E-02 | 1.83E-01 | 1.02E+00 | 1.70E-01 | 6.25E-01 | 1.20E+00 | 1.85E-01 | 5.08E-01 | 1.16E+00 |
| E2F8 | 5.20E-02 | 2.71E-01 | -1.51E+00 | 9.59E-02 | 4.14E-01 | -1.20E+00 | 2.70E-02 | 1.15E-01 | 1.72E+00 | 2.62E-01 | 1.00E+00 | 1.15E+00 | 2.53E-02 | 9.01E-02 | -1.80E+00 | 8.10E-02 | 2.28E-01 | -1.34E+00 | 1.28E-01 | 4.68E-01 | 1.13E+00 | 1.48E-01 | 4.06E-01 | 1.13E+00 |
| THOP1 | 5.20E-02 | 2.71E-01 | 1.07E+00 | 1.08E-01 | 4.66E-01 | 1.30E+00 | 1.66E-01 | 7.05E-01 | 1.33E+00 | 1.40E-01 | 5.35E-01 | 1.36E+00 | 1.56E-01 | 5.56E-01 | 1.65E+00 | 2.20E-01 | 6.21E-01 | 1.62E+00 | 4.50E-02 | 1.65E-01 | 1.09E+00 | 3.59E-03 | 9.87E-03 | 1.71E+00 |
| ANXA2P2 | 5.20E-02 | 2.71E-01 | 1.75E+00 | 1.24E-04 | 5.37E-04 | 3.53E+00 | 1.35E-03 | 5.76E-03 | 2.09E+00 | 2.04E-08 | 7.78E-08 | 3.60E+00 | 4.94E-02 | 1.76E-01 | 3.17E+00 | 9.17E-03 | 2.59E-02 | 4.39E+00 | 6.27E-02 | 2.30E-01 | 2.10E+00 | 6.28E-03 | 1.73E-02 | 2.89E+00 |
| AL138756.1 | 5.21E-02 | 2.71E-01 | -1.32E+00 | 1.71E-01 | 7.38E-01 | 1.24E+00 | 6.59E-02 | 2.80E-01 | -1.40E+00 | 1.55E-01 | 5.93E-01 | 1.41E+00 | 2.35E-03 | 8.37E-03 | -1.49E+00 | 3.18E-03 | 8.97E-03 | -1.35E+00 | 2.51E-01 | 9.20E-01 | 1.37E+00 | 2.86E-01 | 7.86E-01 | 1.26E+00 |
| LYSMD4 | 5.21E-02 | 2.71E-01 | -1.10E+00 | 2.48E-02 | 1.07E-01 | -1.12E+00 | 1.83E-02 | 7.78E-02 | -1.17E+00 | 7.75E-03 | 2.96E-02 | -1.19E+00 | 1.47E-02 | 5.24E-02 | -1.25E+00 | 7.09E-03 | 2.00E-02 | -1.38E+00 | 1.78E-03 | 6.55E-03 | -1.21E+00 | 2.40E-02 | 6.60E-02 | -1.06E+00 |
| AC007228.2 | 5.21E-02 | 2.72E-01 | 1.00E+00 | 1.87E-03 | 8.08E-03 | -1.48E+00 | 4.32E-13 | 1.83E-12 | -2.07E+00 | 4.15E-15 | 1.58E-14 | -2.44E+00 | 3.38E-02 | 1.21E-01 | -1.29E+00 | 1.88E-02 | 5.30E-02 | -1.63E+00 | 1.18E-05 | 4.34E-05 | -1.63E+00 | 1.11E-05 | 3.05E-05 | -1.66E+00 |
| RPS3AP36 | 5.22E-02 | 2.72E-01 | 1.14E+00 | 1.25E-03 | 5.40E-03 | -1.05E+00 | 3.85E-02 | 1.64E-01 | 1.01E+00 | 1.36E-02 | 5.19E-02 | 1.03E+00 | 4.62E-02 | 1.65E-01 | 1.06E+00 | 3.24E-03 | 9.12E-03 | -1.23E+00 | 3.54E-02 | 1.30E-01 | -1.09E+00 | 3.36E-02 | 9.24E-02 | -1.13E+00 |
| OR7E38P | 5.24E-02 | 2.73E-01 | -1.15E+00 | 9.44E-02 | 4.08E-01 | 1.10E+00 | 1.03E-03 | 4.37E-03 | -1.60E+00 | 5.66E-04 | 2.16E-03 | -1.52E+00 | 1.78E-02 | 6.36E-02 | -1.52E+00 | 9.72E-02 | 2.74E-01 | -1.05E+00 | 5.18E-03 | 1.90E-02 | -1.21E+00 | 3.91E-05 | 1.07E-04 | -1.58E+00 |
| MED18 | 5.24E-02 | 2.73E-01 | -1.01E+00 | 1.73E-01 | 7.49E-01 | 1.33E+00 | 1.32E-07 | 5.59E-07 | -1.84E+00 | 6.50E-06 | 2.48E-05 | -1.39E+00 | 5.92E-03 | 2.11E-02 | -1.41E+00 | 1.88E-02 | 5.30E-02 | -1.22E+00 | 1.76E-07 | 6.46E-07 | -1.59E+00 | 6.96E-05 | 1.91E-04 | -1.27E+00 |
| HSPBAP1 | 5.24E-02 | 2.73E-01 | 1.01E+00 | 2.20E-01 | 9.50E-01 | 1.44E+00 | 5.06E-03 | 2.15E-02 | -1.28E+00 | 1.01E-01 | 3.85E-01 | 1.18E+00 | 2.33E-01 | 8.29E-01 | 1.23E+00 | 2.56E-01 | 7.23E-01 | 1.65E+00 | 1.58E-03 | 5.78E-03 | -1.22E+00 | 3.67E-03 | 1.01E-02 | -1.15E+00 |
| PPBP | 5.24E-02 | 2.73E-01 | -1.34E+00 | 4.95E-02 | 2.14E-01 | 1.28E+00 | 2.25E-01 | 9.57E-01 | 8.35E+00 | 1.23E-01 | 4.70E-01 | -1.39E+00 | 2.80E-01 | 9.99E-01 | 8.58E+00 | 3.09E-01 | 8.71E-01 | 4.81E+00 | 9.51E-02 | 3.49E-01 | 1.51E+00 | 2.56E-02 | 7.02E-02 | -1.22E+00 |
| AC026979.2 | 5.24E-02 | 2.73E-01 | -1.06E+00 | 1.07E-02 | 4.64E-02 | 2.35E+00 | 1.06E-01 | 4.52E-01 | -1.05E+00 | 2.08E-01 | 7.96E-01 | 1.22E+00 | 1.64E-01 | 5.85E-01 | 1.14E+00 | 3.38E-01 | 9.53E-01 | 1.31E+00 | 2.67E-01 | 9.78E-01 | 1.33E+00 | 1.60E-01 | 4.39E-01 | 1.66E+00 |
| LINC00672 | 5.25E-02 | 2.73E-01 | -1.03E+00 | 2.11E-01 | 9.13E-01 | 1.35E+00 | 4.96E-03 | 2.11E-02 | -1.48E+00 | 1.91E-02 | 7.29E-02 | -1.13E+00 | 2.70E-02 | 9.64E-02 | -1.24E+00 | 7.52E-02 | 2.12E-01 | 1.10E+00 | 9.21E-02 | 3.38E-01 | -1.33E+00 | 2.76E-01 | 7.57E-01 | 1.02E+00 |
| RPL23AP53 | 5.25E-02 | 2.73E-01 | 1.84E+00 | 6.09E-02 | 2.63E-01 | 1.93E+00 | 1.70E-01 | 7.24E-01 | -1.04E+00 | 9.16E-02 | 3.50E-01 | -1.19E+00 | 3.00E-02 | 1.07E-01 | -1.09E+00 | 4.84E-03 | 1.37E-02 | -1.60E+00 | 2.10E-01 | 7.70E-01 | 1.04E+00 | 1.53E-01 | 4.20E-01 | 1.03E+00 |
| DGUOK-AS1 | 5.26E-02 | 2.74E-01 | 1.02E+00 | 6.59E-02 | 2.85E-01 | 1.11E+00 | 2.13E-01 | 9.04E-01 | 1.19E+00 | 1.80E-01 | 6.88E-01 | 1.14E+00 | 4.78E-03 | 1.70E-02 | -1.45E+00 | 3.52E-02 | 9.93E-02 | -1.07E+00 | 4.94E-07 | 1.81E-06 | -1.46E+00 | 3.21E-04 | 8.83E-04 | -1.21E+00 |
| TAF1D | 5.26E-02 | 2.74E-01 | 1.16E+00 | 6.49E-02 | 2.80E-01 | 1.08E+00 | 2.35E-01 | 1.00E+00 | 1.02E+00 | 2.30E-02 | 8.78E-02 | -1.02E+00 | 5.81E-02 | 2.07E-01 | 1.21E+00 | 2.38E-01 | 6.70E-01 | 1.10E+00 | 1.14E-01 | 4.17E-01 | 1.28E+00 | 2.78E-01 | 7.65E-01 | 1.08E+00 |
| SLMAP | 5.27E-02 | 2.75E-01 | 1.10E+00 | 2.25E-02 | 9.73E-02 | 1.16E+00 | 1.66E-05 | 7.05E-05 | -1.34E+00 | 2.90E-05 | 1.11E-04 | -1.10E+00 | 3.02E-04 | 1.08E-03 | -1.07E+00 | 8.06E-03 | 2.27E-02 | 1.01E+00 | 4.77E-04 | 1.75E-03 | -1.15E+00 | 1.90E-02 | 5.21E-02 | -1.08E+00 |
| N4BP2L2-IT2 | 5.27E-02 | 2.75E-01 | 1.07E+00 | 1.12E-01 | 4.83E-01 | 1.77E+00 | 1.19E-01 | 5.04E-01 | 1.06E+00 | 2.52E-01 | 9.62E-01 | 1.30E+00 | 6.10E-02 | 2.17E-01 | -1.10E+00 | 2.95E-01 | 8.31E-01 | 1.50E+00 | 1.99E-01 | 7.32E-01 | 1.15E+00 | 2.98E-01 | 8.19E-01 | 1.34E+00 |
| TOX | 5.28E-02 | 2.75E-01 | 1.01E+00 | 1.66E-01 | 7.17E-01 | 1.32E+00 | 1.86E-01 | 7.91E-01 | 1.20E+00 | 1.51E-01 | 5.78E-01 | 1.30E+00 | 9.90E-02 | 3.53E-01 | 1.24E+00 | 3.08E-01 | 8.68E-01 | 1.47E+00 | 1.43E-01 | 5.24E-01 | 1.38E+00 | 2.45E-02 | 6.73E-02 | 1.71E+00 |
| MAP3K12 | 5.28E-02 | 2.75E-01 | 1.07E+00 | 1.50E-02 | 6.48E-02 | -1.11E+00 | 4.33E-03 | 1.84E-02 | -1.27E+00 | 2.58E-03 | 9.86E-03 | -1.18E+00 | 3.02E-02 | 1.08E-01 | -1.31E+00 | 6.06E-02 | 1.71E-01 | -1.14E+00 | 2.39E-01 | 8.76E-01 | 1.02E+00 | 3.27E-01 | 8.98E-01 | 1.01E+00 |
| IRX3 | 5.28E-02 | 2.75E-01 | -1.59E+00 | 5.69E-02 | 2.46E-01 | -1.97E+00 | 5.03E-07 | 2.14E-06 | -2.20E+00 | 1.61E-09 | 6.14E-09 | -2.64E+00 | 2.72E-01 | 9.69E-01 | 1.16E+00 | 2.63E-01 | 7.41E-01 | -1.12E+00 | 1.78E-01 | 6.52E-01 | 2.20E+00 | 3.53E-01 | 9.70E-01 | 1.21E+00 |
| FRG1 | 5.28E-02 | 2.75E-01 | 1.27E+00 | 1.91E-01 | 8.24E-01 | 1.50E+00 | 1.68E-01 | 7.15E-01 | 1.32E+00 | 2.35E-01 | 8.96E-01 | 1.42E+00 | 2.15E-02 | 7.67E-02 | -1.02E+00 | 1.24E-01 | 3.50E-01 | 1.20E+00 | 5.56E-02 | 2.04E-01 | 1.15E+00 | 1.73E-01 | 4.75E-01 | 1.22E+00 |
| LGMN | 5.28E-02 | 2.75E-01 | -2.62E+00 | 1.97E-01 | 8.50E-01 | -1.57E+00 | 2.42E-02 | 1.03E-01 | -1.72E+00 | 5.68E-02 | 2.17E-01 | -1.33E+00 | 2.78E-01 | 9.90E-01 | 1.18E+00 | 1.29E-01 | 3.64E-01 | 1.04E+00 | 1.36E-04 | 4.97E-04 | 2.96E+00 | 5.65E-02 | 1.55E-01 | 4.41E+00 |
| AC004918.1 | 5.29E-02 | 2.75E-01 | 1.15E+00 | 1.92E-01 | 8.28E-01 | 1.58E+00 | 9.49E-02 | 4.03E-01 | 1.05E+00 | 1.23E-01 | 4.68E-01 | 1.71E+00 | 2.00E-01 | 7.14E-01 | 1.20E+00 | 2.76E-01 | 7.79E-01 | 1.19E+00 | 9.74E-02 | 3.57E-01 | -1.12E+00 | 3.22E-01 | 8.84E-01 | 1.15E+00 |
| SFI1 | 5.29E-02 | 2.76E-01 | 1.55E+00 | 1.93E-01 | 8.34E-01 | 1.50E+00 | 4.11E-02 | 1.74E-01 | 1.50E+00 | 2.13E-01 | 8.12E-01 | 1.59E+00 | 4.97E-03 | 1.77E-02 | -1.25E+00 | 2.53E-02 | 7.13E-02 | -1.02E+00 | 1.52E-01 | 5.57E-01 | 1.06E+00 | 1.32E-01 | 3.62E-01 | 1.52E+00 |
| RPL13P2 | 5.30E-02 | 2.76E-01 | 1.21E+00 | 1.07E-06 | 4.64E-06 | 2.45E+00 | 1.29E-01 | 5.49E-01 | 1.36E+00 | 2.22E-06 | 8.49E-06 | 2.45E+00 | 2.75E-01 | 9.81E-01 | 1.40E+00 | 2.80E-04 | 7.90E-04 | 2.37E+00 | 2.54E-01 | 9.34E-01 | 1.29E+00 | 8.05E-09 | 2.21E-08 | 2.43E+00 |
| PALB2 | 5.30E-02 | 2.76E-01 | -1.15E+00 | 2.08E-01 | 8.98E-01 | 1.40E+00 | 2.29E-07 | 9.72E-07 | -1.75E+00 | 1.27E-03 | 4.84E-03 | -1.02E+00 | 9.70E-03 | 3.46E-02 | -1.15E+00 | 2.50E-02 | 7.05E-02 | -1.07E+00 | 2.96E-03 | 1.09E-02 | -1.36E+00 | 3.41E-02 | 9.38E-02 | -1.07E+00 |
| KCMF1 | 5.30E-02 | 2.76E-01 | 1.22E+00 | 1.66E-04 | 7.16E-04 | 1.10E+00 | 9.47E-02 | 4.02E-01 | 1.15E+00 | 6.84E-04 | 2.61E-03 | 1.04E+00 | 1.40E-01 | 4.98E-01 | 1.27E+00 | 5.45E-02 | 1.54E-01 | 1.12E+00 | 8.73E-02 | 3.20E-01 | 1.12E+00 | 8.77E-04 | 2.41E-03 | -1.03E+00 |
| ZNF841 | 5.30E-02 | 2.76E-01 | 1.06E+00 | 1.11E-02 | 4.81E-02 | -1.13E+00 | 1.91E-01 | 8.10E-01 | 1.18E+00 | 5.76E-02 | 2.20E-01 | 1.07E+00 | 8.46E-02 | 3.02E-01 | -1.01E+00 | 7.10E-02 | 2.00E-01 | -1.15E+00 | 3.59E-02 | 1.32E-01 | 1.08E+00 | 1.21E-02 | 3.33E-02 | -1.11E+00 |
| GPAT2P2 | 5.30E-02 | 2.76E-01 | 1.62E+00 | 1.41E-01 | 6.08E-01 | 1.68E+00 | 1.15E-01 | 4.89E-01 | 1.36E+00 | 1.93E-01 | 7.38E-01 | 1.55E+00 | 1.42E-01 | 5.06E-01 | 1.66E+00 | 1.23E-01 | 3.47E-01 | 1.82E+00 | 1.60E-01 | 5.86E-01 | 1.12E+00 | 2.01E-01 | 5.52E-01 | 1.43E+00 |
| RALGAPB | 5.31E-02 | 2.77E-01 | 1.07E+00 | 1.80E-01 | 7.76E-01 | 1.39E+00 | 7.03E-05 | 2.99E-04 | -1.41E+00 | 2.85E-03 | 1.09E-02 | -1.07E+00 | 3.05E-02 | 1.09E-01 | -1.01E+00 | 1.34E-01 | 3.78E-01 | 1.13E+00 | 6.43E-05 | 2.36E-04 | -1.30E+00 | 1.69E-01 | 4.63E-01 | 1.17E+00 |
| NCF1 | 5.31E-02 | 2.77E-01 | -1.55E+00 | 2.30E-02 | 9.93E-02 | -1.94E+00 | 1.15E-01 | 4.91E-01 | 1.52E+00 | 1.81E-01 | 6.90E-01 | -1.17E+00 | 5.23E-04 | 1.86E-03 | -1.45E+00 | 4.41E-04 | 1.24E-03 | -1.53E+00 | 3.75E-08 | 1.38E-07 | -2.54E+00 | 1.16E-07 | 3.19E-07 | -2.22E+00 |
| SIPA1L1 | 5.32E-02 | 2.77E-01 | 1.01E+00 | 1.83E-01 | 7.89E-01 | 1.39E+00 | 3.08E-02 | 1.31E-01 | -1.01E+00 | 2.36E-02 | 9.00E-02 | 1.09E+00 | 1.30E-01 | 4.64E-01 | 1.68E+00 | 1.17E-01 | 3.31E-01 | -1.13E+00 | 9.20E-02 | 3.38E-01 | 1.64E+00 | 1.39E-01 | 3.82E-01 | 1.65E+00 |
| SGMS2 | 5.32E-02 | 2.77E-01 | -1.04E+00 | 7.63E-02 | 3.30E-01 | 1.01E+00 | 1.69E-01 | 7.17E-01 | 1.26E+00 | 1.63E-02 | 6.21E-02 | -1.22E+00 | 1.82E-01 | 6.50E-01 | 1.48E+00 | 3.29E-01 | 9.26E-01 | 1.24E+00 | 2.22E-01 | 8.13E-01 | 1.18E+00 | 3.39E-01 | 9.31E-01 | 1.11E+00 |
| IL18BP | 5.32E-02 | 2.77E-01 | -1.64E+00 | 1.36E-01 | 5.87E-01 | 1.29E+00 | 8.05E-02 | 3.42E-01 | -1.12E+00 | 1.46E-01 | 5.58E-01 | 1.15E+00 | 1.76E-01 | 6.28E-01 | -1.09E+00 | 3.24E-01 | 9.13E-01 | 1.10E+00 | 5.52E-02 | 2.03E-01 | -1.12E+00 | 2.55E-01 | 7.00E-01 | 1.08E+00 |
| HIST2H2BD | 5.32E-02 | 2.78E-01 | 1.12E+00 | 3.30E-04 | 1.43E-03 | -1.24E+00 | 6.30E-02 | 2.68E-01 | 1.05E+00 | 5.99E-02 | 2.29E-01 | 1.20E+00 | 1.13E-02 | 4.04E-02 | -1.34E+00 | 3.45E-03 | 9.73E-03 | -1.54E+00 | 3.73E-02 | 1.37E-01 | -1.11E+00 | 5.23E-02 | 1.44E-01 | -1.12E+00 |
| AC055764.1 | 5.33E-02 | 2.78E-01 | 1.12E+00 | 8.44E-02 | 3.65E-01 | 1.30E+00 | 1.88E-01 | 7.99E-01 | 1.09E+00 | 2.28E-01 | 8.70E-01 | 1.23E+00 | 2.01E-01 | 7.16E-01 | 1.14E+00 | 2.46E-01 | 6.92E-01 | 1.59E+00 | 2.42E-01 | 8.90E-01 | 1.27E+00 | 3.03E-01 | 8.32E-01 | 1.09E+00 |
| FMN1 | 5.34E-02 | 2.78E-01 | -1.42E+00 | 7.50E-02 | 3.24E-01 | -1.29E+00 | 1.83E-02 | 7.77E-02 | -1.43E+00 | 4.68E-02 | 1.79E-01 | -1.07E+00 | 3.68E-06 | 1.31E-05 | -1.71E+00 | 1.95E-06 | 5.50E-06 | -1.73E+00 | 1.25E-02 | 4.59E-02 | -1.27E+00 | 1.32E-03 | 3.63E-03 | -1.56E+00 |
| AL359220.1 | 5.35E-02 | 2.79E-01 | 1.18E+00 | 3.45E-03 | 1.49E-02 | 2.21E+00 | 6.37E-02 | 2.70E-01 | -1.31E+00 | 2.38E-01 | 9.10E-01 | 1.27E+00 | 1.93E-01 | 6.89E-01 | 1.72E+00 | 2.87E-01 | 8.08E-01 | 1.72E+00 | 1.83E-01 | 6.72E-01 | -1.04E+00 | 1.11E-01 | 3.04E-01 | -1.72E+00 |
| AP001885.1 | 5.35E-02 | 2.79E-01 | 1.05E+00 | 4.31E-04 | 1.86E-03 | -1.59E+00 | 1.05E-01 | 4.46E-01 | 1.02E+00 | 4.90E-02 | 1.87E-01 | -1.04E+00 | 1.74E-01 | 6.20E-01 | 1.10E+00 | 6.12E-02 | 1.73E-01 | -1.15E+00 | 2.26E-01 | 8.31E-01 | 1.20E+00 | 8.32E-02 | 2.28E-01 | -1.21E+00 |
| PARD6G | 5.35E-02 | 2.79E-01 | -1.03E+00 | 3.22E-02 | 1.39E-01 | -1.09E+00 | 8.60E-03 | 3.66E-02 | -1.42E+00 | 2.40E-01 | 9.16E-01 | 1.28E+00 | 1.03E-01 | 3.68E-01 | -1.39E+00 | 1.70E-01 | 4.79E-01 | -1.16E+00 | 1.89E-03 | 6.94E-03 | -1.73E+00 | 6.03E-03 | 1.66E-02 | -1.57E+00 |
| HYMAI | 5.35E-02 | 2.79E-01 | 1.16E+00 | 2.04E-01 | 8.82E-01 | 1.50E+00 | 9.31E-02 | 3.95E-01 | 1.02E+00 | 2.28E-01 | 8.70E-01 | 1.39E+00 | 2.74E-01 | 9.77E-01 | 1.55E+00 | 1.72E-01 | 4.85E-01 | 1.24E+00 | 1.65E-01 | 6.06E-01 | 1.40E+00 | 3.35E-01 | 9.20E-01 | 1.45E+00 |
| AL034370.1 | 5.35E-02 | 2.79E-01 | -1.01E+00 | 8.97E-02 | 3.87E-01 | 1.15E+00 | 9.38E-02 | 3.99E-01 | 1.39E+00 | 1.73E-01 | 6.61E-01 | 1.48E+00 | 5.77E-02 | 2.06E-01 | 1.14E+00 | 3.17E-02 | 8.94E-02 | 1.01E+00 | 7.26E-02 | 2.66E-01 | 1.10E+00 | 1.80E-01 | 4.93E-01 | 1.25E+00 |
| RPS20P10 | 5.35E-02 | 2.79E-01 | 1.08E+00 | 1.88E-18 | 8.13E-18 | -1.25E+00 | 1.88E-01 | 7.99E-01 | 1.05E+00 | 1.56E-05 | 5.95E-05 | -1.23E+00 | 8.49E-02 | 3.03E-01 | 1.05E+00 | 4.02E-06 | 1.13E-05 | -1.21E+00 | 2.13E-02 | 7.82E-02 | -1.01E+00 | 1.09E-05 | 2.99E-05 | -1.24E+00 |
| CAD | 5.36E-02 | 2.79E-01 | 1.07E+00 | 2.28E-01 | 9.86E-01 | 1.44E+00 | 2.07E-01 | 8.82E-01 | 1.27E+00 | 4.60E-02 | 1.76E-01 | 1.84E+00 | 2.41E-01 | 8.60E-01 | 1.47E+00 | 2.68E-01 | 7.55E-01 | 1.64E+00 | 2.67E-01 | 9.80E-01 | 1.16E+00 | 1.26E-02 | 3.47E-02 | 1.67E+00 |
| CDC42-IT1 | 5.36E-02 | 2.79E-01 | 1.62E+00 | 2.10E-01 | 9.09E-01 | 1.33E+00 | 1.50E-02 | 6.36E-02 | 1.75E+00 | 1.02E-01 | 3.91E-01 | 1.60E+00 | 2.73E-01 | 9.72E-01 | 1.55E+00 | 3.52E-01 | 9.92E-01 | 1.40E+00 | 9.09E-02 | 3.34E-01 | 1.63E+00 | 3.26E-01 | 8.96E-01 | 1.32E+00 |
| TC2N | 5.36E-02 | 2.79E-01 | 1.57E+00 | 1.27E-01 | 5.51E-01 | 1.36E+00 | 1.93E-01 | 8.20E-01 | 1.08E+00 | 1.64E-01 | 6.27E-01 | 1.22E+00 | 3.23E-03 | 1.15E-02 | -1.43E+00 | 3.96E-03 | 1.12E-02 | -1.37E+00 | 2.57E-01 | 9.44E-01 | 1.00E+00 | 2.12E-01 | 5.82E-01 | -1.27E+00 |
| AL157871.3 | 5.36E-02 | 2.79E-01 | -1.02E+00 | 1.52E-02 | 6.58E-02 | -1.23E+00 | 3.41E-02 | 1.45E-01 | -1.01E+00 | 1.61E-03 | 6.15E-03 | -1.20E+00 | 8.62E-03 | 3.07E-02 | -1.78E+00 | 3.04E-02 | 8.56E-02 | -1.42E+00 | 1.58E-05 | 5.80E-05 | -1.32E+00 | 1.05E-06 | 2.87E-06 | -1.47E+00 |
| AC098617.1 | 5.37E-02 | 2.80E-01 | -1.18E+00 | 8.52E-02 | 3.68E-01 | -1.12E+00 | 1.36E-01 | 5.78E-01 | -1.11E+00 | 1.92E-01 | 7.35E-01 | 1.04E+00 | 2.12E-08 | 7.57E-08 | -2.40E+00 | 1.56E-05 | 4.39E-05 | -1.50E+00 | 1.44E-02 | 5.29E-02 | 1.03E+00 | 4.71E-02 | 1.29E-01 | -1.10E+00 |
| ARHGAP9 | 5.37E-02 | 2.80E-01 | 1.24E+00 | 1.84E-04 | 7.96E-04 | 1.16E+00 | 7.15E-04 | 3.04E-03 | -1.07E+00 | 5.18E-09 | 1.98E-08 | -1.24E+00 | 2.30E-02 | 8.19E-02 | 1.19E+00 | 5.62E-04 | 1.58E-03 | -1.03E+00 | 1.90E-02 | 6.99E-02 | 1.09E+00 | 1.71E-02 | 4.70E-02 | -1.01E+00 |
| AC009220.1 | 5.37E-02 | 2.80E-01 | 1.43E+00 | 7.85E-10 | 3.39E-09 | -1.32E+00 | 8.49E-03 | 3.61E-02 | 1.59E+00 | 1.50E-06 | 5.73E-06 | -1.19E+00 | 5.23E-03 | 1.87E-02 | 1.73E+00 | 7.16E-02 | 2.02E-01 | 1.04E+00 | 1.69E-01 | 6.20E-01 | 1.38E+00 | 2.12E-06 | 5.83E-06 | -1.31E+00 |
| IFI27 | 5.37E-02 | 2.80E-01 | -1.31E+00 | 1.18E-01 | 5.08E-01 | -1.08E+00 | 1.40E-02 | 5.94E-02 | -1.57E+00 | 6.92E-03 | 2.64E-02 | -1.78E+00 | 2.72E-01 | 9.68E-01 | 1.00E+00 | 2.48E-01 | 6.98E-01 | -1.04E+00 | 4.51E-97 | 1.66E-96 | -1.22E+01 | 4.59E-107 | 1.26E-106 | -2.13E+01 |
| KLHL21 | 5.38E-02 | 2.80E-01 | -1.00E+00 | 2.02E-01 | 8.72E-01 | 1.31E+00 | 4.19E-03 | 1.78E-02 | -1.19E+00 | 1.09E-01 | 4.16E-01 | 1.19E+00 | 2.41E-02 | 8.61E-02 | -1.39E+00 | 2.34E-01 | 6.60E-01 | 1.21E+00 | 4.15E-07 | 1.52E-06 | -1.58E+00 | 4.67E-03 | 1.28E-02 | -1.07E+00 |
| RPS24P17 | 5.39E-02 | 2.81E-01 | 1.64E+00 | 2.88E-05 | 1.25E-04 | -1.05E+00 | 7.29E-04 | 3.10E-03 | 1.82E+00 | 1.03E-01 | 3.95E-01 | 1.26E+00 | 1.67E-02 | 5.97E-02 | 2.14E+00 | 1.56E-01 | 4.39E-01 | 1.26E+00 | 1.83E-02 | 6.73E-02 | 1.74E+00 | 2.09E-01 | 5.73E-01 | 1.09E+00 |
| TRAIP | 5.39E-02 | 2.81E-01 | -1.18E+00 | 5.01E-02 | 2.16E-01 | -1.25E+00 | 1.96E-01 | 8.34E-01 | 1.10E+00 | 1.38E-01 | 5.27E-01 | 1.51E+00 | 2.30E-01 | 8.20E-01 | 1.11E+00 | 3.48E-01 | 9.80E-01 | 1.39E+00 | 1.93E-01 | 7.09E-01 | 1.34E+00 | 9.08E-02 | 2.49E-01 | 1.51E+00 |
| SLC25A51 | 5.39E-02 | 2.81E-01 | 1.68E+00 | 2.22E-01 | 9.58E-01 | 1.44E+00 | 6.52E-02 | 2.77E-01 | -1.04E+00 | 5.31E-02 | 2.03E-01 | 1.10E+00 | 9.37E-02 | 3.34E-01 | 1.93E+00 | 2.34E-01 | 6.61E-01 | 1.81E+00 | 3.74E-03 | 1.37E-02 | -1.23E+00 | 6.91E-03 | 1.90E-02 | -1.15E+00 |
| ZNF3 | 5.39E-02 | 2.81E-01 | 1.71E+00 | 3.38E-09 | 1.46E-08 | -1.27E+00 | 1.69E-01 | 7.17E-01 | 1.34E+00 | 2.19E-06 | 8.35E-06 | -1.21E+00 | 1.96E-03 | 6.99E-03 | 1.04E+00 | 4.04E-06 | 1.14E-05 | -1.27E+00 | 7.53E-08 | 2.76E-07 | -1.30E+00 | 1.37E-09 | 3.75E-09 | -1.50E+00 |
| AL022311.1 | 5.39E-02 | 2.81E-01 | 1.10E+00 | 6.19E-03 | 2.68E-02 | 1.08E+00 | 1.27E-01 | 5.41E-01 | 1.21E+00 | 5.74E-03 | 2.19E-02 | -1.01E+00 | 6.32E-02 | 2.25E-01 | 1.08E+00 | 1.19E-02 | 3.36E-02 | -1.09E+00 | 4.44E-02 | 1.63E-01 | -1.11E+00 | 5.47E-02 | 1.50E-01 | -1.06E+00 |
| AC015971.1 | 5.40E-02 | 2.81E-01 | 1.10E+00 | 1.67E-01 | 7.23E-01 | 1.67E+00 | 4.36E-02 | 1.85E-01 | -1.03E+00 | 3.65E-02 | 1.39E-01 | 1.11E+00 | 2.12E-02 | 7.55E-02 | -1.14E+00 | 6.02E-02 | 1.70E-01 | -1.00E+00 | 1.34E-01 | 4.92E-01 | 1.49E+00 | 3.13E-01 | 8.61E-01 | 1.39E+00 |
| PLPPR3 | 5.41E-02 | 2.82E-01 | -1.79E+00 | 4.99E-02 | 2.16E-01 | -2.08E+00 | 7.37E-14 | 3.13E-13 | -1.82E+00 | 2.10E-16 | 8.02E-16 | -1.97E+00 | 1.98E-01 | 7.04E-01 | -1.17E+00 | 2.03E-01 | 5.71E-01 | -1.39E+00 | 2.24E-01 | 8.21E-01 | -1.01E+00 | 2.75E-01 | 7.55E-01 | -1.10E+00 |
| SNRPD1 | 5.41E-02 | 2.82E-01 | 1.19E+00 | 3.93E-02 | 1.70E-01 | 1.38E+00 | 3.65E-03 | 1.55E-02 | 1.38E+00 | 2.09E-03 | 7.99E-03 | 1.51E+00 | 3.26E-04 | 1.16E-03 | 1.07E+00 | 2.40E-01 | 6.77E-01 | 1.40E+00 | 2.03E-01 | 7.45E-01 | 1.22E+00 | 4.56E-04 | 1.25E-03 | 1.58E+00 |
| AC005000.1 | 5.42E-02 | 2.82E-01 | 1.81E+00 | 2.10E-01 | 9.08E-01 | 1.27E+00 | 4.28E-02 | 1.82E-01 | 1.68E+00 | 1.76E-01 | 6.72E-01 | 1.47E+00 | 1.67E-01 | 5.94E-01 | 1.60E+00 | 1.00E-01 | 2.82E-01 | 2.10E+00 | 1.07E-01 | 3.92E-01 | -1.02E+00 | 1.22E-01 | 3.35E-01 | 1.55E+00 |
| DTNB | 5.42E-02 | 2.82E-01 | 1.19E+00 | 7.15E-02 | 3.09E-01 | 1.33E+00 | 4.52E-03 | 1.92E-02 | -1.09E+00 | 1.19E-02 | 4.56E-02 | 1.11E+00 | 1.18E-01 | 4.21E-01 | 1.13E+00 | 1.63E-01 | 4.59E-01 | 1.16E+00 | 7.44E-02 | 2.73E-01 | 1.00E+00 | 1.79E-01 | 4.90E-01 | 1.11E+00 |
| BTBD6 | 5.42E-02 | 2.82E-01 | 2.00E+00 | 1.52E-01 | 6.56E-01 | 1.64E+00 | 1.73E-01 | 7.34E-01 | 1.30E+00 | 1.50E-01 | 5.71E-01 | 1.08E+00 | 1.98E-01 | 7.04E-01 | 1.47E+00 | 2.56E-01 | 7.22E-01 | 1.12E+00 | 8.35E-04 | 3.06E-03 | 2.18E+00 | 2.00E-01 | 5.49E-01 | 1.54E+00 |
| AC009812.1 | 5.42E-02 | 2.83E-01 | 1.13E+00 | 1.74E-01 | 7.50E-01 | 1.53E+00 | 1.21E-02 | 5.14E-02 | -1.18E+00 | 7.24E-02 | 2.77E-01 | 1.11E+00 | 1.52E-01 | 5.40E-01 | 1.88E+00 | 2.43E-01 | 6.84E-01 | 1.86E+00 | 8.67E-03 | 3.18E-02 | -1.57E+00 | 1.24E-01 | 3.41E-01 | 1.01E+00 |
| EXOC7 | 5.42E-02 | 2.83E-01 | 1.12E+00 | 1.87E-01 | 8.10E-01 | 1.37E+00 | 1.17E-01 | 4.98E-01 | 1.23E+00 | 2.41E-01 | 9.21E-01 | 1.27E+00 | 5.25E-03 | 1.87E-02 | -1.04E+00 | 3.00E-02 | 8.47E-02 | 1.12E+00 | 1.24E-01 | 4.55E-01 | 1.20E+00 | 1.62E-01 | 4.45E-01 | 1.40E+00 |
| AC122688.3 | 5.42E-02 | 2.83E-01 | 1.05E+00 | 7.94E-02 | 3.43E-01 | 1.17E+00 | 1.77E-01 | 7.52E-01 | 1.20E+00 | 2.08E-01 | 7.94E-01 | 1.31E+00 | 1.84E-02 | 6.57E-02 | -1.32E+00 | 8.62E-02 | 2.43E-01 | -1.11E+00 | 1.54E-01 | 5.65E-01 | -1.08E+00 | 1.75E-01 | 4.80E-01 | 1.46E+00 |
| CKB | 5.43E-02 | 2.83E-01 | 1.11E+00 | 8.28E-02 | 3.58E-01 | 1.24E+00 | 6.84E-02 | 2.91E-01 | 1.68E+00 | 1.19E-01 | 4.56E-01 | 1.77E+00 | 1.77E-04 | 6.30E-04 | -1.45E+00 | 2.86E-02 | 8.06E-02 | -1.01E+00 | 4.75E-05 | 1.74E-04 | -1.42E+00 | 6.28E-06 | 1.72E-05 | -1.51E+00 |
| FBXO34 | 5.43E-02 | 2.83E-01 | 1.24E+00 | 3.32E-04 | 1.43E-03 | 1.07E+00 | 1.45E-01 | 6.15E-01 | 1.42E+00 | 9.81E-03 | 3.75E-02 | 1.12E+00 | 9.09E-02 | 3.24E-01 | 1.79E+00 | 1.37E-01 | 3.86E-01 | 1.36E+00 | 1.36E-03 | 4.97E-03 | -1.09E+00 | 9.01E-04 | 2.48E-03 | -1.11E+00 |
| CDK1 | 5.43E-02 | 2.83E-01 | -1.64E+00 | 1.60E-01 | 6.91E-01 | -1.03E+00 | 1.24E-02 | 5.26E-02 | 1.69E+00 | 6.87E-03 | 2.63E-02 | 2.35E+00 | 5.84E-02 | 2.08E-01 | -2.62E+00 | 1.21E-01 | 3.42E-01 | -2.06E+00 | 2.50E-05 | 9.16E-05 | -1.72E+00 | 1.89E-02 | 5.18E-02 | 1.54E+00 |
| TINAGL1 | 5.44E-02 | 2.83E-01 | 1.00E+00 | 2.87E-02 | 1.24E-01 | 1.00E+00 | 2.77E-08 | 1.18E-07 | -1.84E+00 | 9.66E-12 | 3.69E-11 | -2.25E+00 | 1.17E-01 | 4.18E-01 | -1.39E+00 | 1.00E-01 | 2.82E-01 | -1.68E+00 | 2.16E-01 | 7.92E-01 | -1.04E+00 | 2.68E-01 | 7.37E-01 | -1.17E+00 |
| SNORA33 | 5.44E-02 | 2.84E-01 | 1.70E+00 | 5.12E-02 | 2.21E-01 | 2.07E+00 | 2.14E-01 | 9.10E-01 | 1.21E+00 | 9.84E-02 | 3.76E-01 | 1.75E+00 | 2.41E-01 | 8.60E-01 | 1.15E+00 | 3.27E-01 | 9.21E-01 | 1.33E+00 | 4.86E-02 | 1.78E-01 | -1.10E+00 | 2.82E-01 | 7.75E-01 | 1.36E+00 |
| GSTO3P | 5.45E-02 | 2.84E-01 | 1.73E+00 | 1.19E-01 | 5.14E-01 | 1.68E+00 | 2.77E-02 | 1.18E-01 | 1.68E+00 | 1.79E-01 | 6.84E-01 | 1.55E+00 | 3.64E-02 | 1.30E-01 | 2.60E+00 | 3.10E-02 | 8.75E-02 | 2.78E+00 | 2.20E-08 | 8.08E-08 | 3.28E+00 | 5.64E-03 | 1.55E-02 | 2.21E+00 |
| COX7B | 5.45E-02 | 2.84E-01 | 1.16E+00 | 3.55E-14 | 1.53E-13 | -1.11E+00 | 1.99E-01 | 8.44E-01 | 1.14E+00 | 2.31E-04 | 8.84E-04 | -1.15E+00 | 3.29E-06 | 1.17E-05 | -1.13E+00 | 3.50E-09 | 9.87E-09 | -1.26E+00 | 2.46E-01 | 9.01E-01 | 1.09E+00 | 1.07E-01 | 2.93E-01 | -1.01E+00 |
| FAM157A | 5.45E-02 | 2.84E-01 | 1.03E+00 | 5.46E-03 | 2.36E-02 | -1.19E+00 | 2.28E-01 | 9.69E-01 | 1.09E+00 | 2.26E-01 | 8.65E-01 | 1.12E+00 | 2.47E-01 | 8.80E-01 | 1.32E+00 | 3.46E-01 | 9.76E-01 | 1.35E+00 | 2.27E-01 | 8.34E-01 | 1.41E+00 | 2.24E-01 | 6.14E-01 | 1.34E+00 |
| HCG27 | 5.46E-02 | 2.85E-01 | -1.35E+00 | 3.17E-02 | 1.37E-01 | 1.72E+00 | 1.59E-01 | 6.74E-01 | 1.38E+00 | 2.66E-03 | 1.01E-02 | 2.54E+00 | 2.72E-01 | 9.71E-01 | -1.08E+00 | 3.79E-02 | 1.07E-01 | 2.27E+00 | 2.61E-01 | 9.60E-01 | 1.13E+00 | 1.39E-01 | 3.81E-01 | 1.53E+00 |
| AC008429.2 | 5.47E-02 | 2.85E-01 | 1.01E+00 | 6.26E-02 | 2.70E-01 | 1.97E+00 | 9.56E-02 | 4.06E-01 | 1.04E+00 | 3.66E-02 | 1.40E-01 | 1.86E+00 | 9.35E-02 | 3.33E-01 | -1.27E+00 | 2.58E-01 | 7.28E-01 | 1.72E+00 | 2.33E-01 | 8.55E-01 | 1.32E+00 | 5.56E-05 | 1.53E-04 | 3.13E+00 |
| MXRA7 | 5.48E-02 | 2.86E-01 | 1.32E+00 | 2.17E-01 | 9.39E-01 | 1.22E+00 | 7.05E-02 | 2.99E-01 | 1.59E+00 | 2.49E-01 | 9.52E-01 | 1.36E+00 | 2.05E-01 | 7.31E-01 | 1.35E+00 | 1.01E-01 | 2.84E-01 | -1.34E+00 | 1.81E-01 | 6.65E-01 | 1.40E+00 | 3.55E-01 | 9.76E-01 | 1.37E+00 |
| CDYL2 | 5.49E-02 | 2.86E-01 | -1.12E+00 | 1.38E-01 | 5.96E-01 | 1.11E+00 | 3.71E-02 | 1.58E-01 | -1.15E+00 | 1.23E-01 | 4.69E-01 | 1.15E+00 | 3.31E-03 | 1.18E-02 | -1.57E+00 | 2.71E-02 | 7.64E-02 | -1.10E+00 | 8.06E-02 | 2.96E-01 | -1.14E+00 | 2.36E-01 | 6.48E-01 | 1.43E+00 |
| RDH10 | 5.50E-02 | 2.86E-01 | -1.44E+00 | 2.11E-01 | 9.11E-01 | 1.20E+00 | 1.85E-01 | 7.84E-01 | 1.02E+00 | 2.60E-01 | 9.94E-01 | 1.26E+00 | 8.35E-02 | 2.98E-01 | -1.01E+00 | 7.20E-02 | 2.03E-01 | -1.34E+00 | 1.96E-01 | 7.20E-01 | 1.02E+00 | 2.20E-01 | 6.05E-01 | -1.12E+00 |
| DYNC2H1 | 5.52E-02 | 2.88E-01 | -1.21E+00 | 7.84E-02 | 3.39E-01 | 1.68E+00 | 8.27E-05 | 3.51E-04 | -1.49E+00 | 7.71E-05 | 2.95E-04 | -1.41E+00 | 3.90E-02 | 1.39E-01 | -1.03E+00 | 7.01E-02 | 1.98E-01 | -1.10E+00 | 2.01E-01 | 7.39E-01 | 1.21E+00 | 3.04E-01 | 8.35E-01 | -1.01E+00 |
| CATSPER2P1 | 5.52E-02 | 2.88E-01 | -1.01E+00 | 6.94E-02 | 3.00E-01 | 1.14E+00 | 5.23E-03 | 2.22E-02 | -1.38E+00 | 1.08E-02 | 4.11E-02 | -1.14E+00 | 5.26E-03 | 1.88E-02 | -1.29E+00 | 3.39E-03 | 9.55E-03 | -1.40E+00 | 1.22E-01 | 4.48E-01 | 1.18E+00 | 1.97E-01 | 5.40E-01 | 1.24E+00 |
| PPP6R1 | 5.52E-02 | 2.88E-01 | 1.06E+00 | 1.43E-01 | 6.18E-01 | 1.29E+00 | 1.33E-03 | 5.63E-03 | -1.12E+00 | 1.85E-03 | 7.08E-03 | 1.01E+00 | 1.98E-02 | 7.07E-02 | -1.09E+00 | 1.61E-01 | 4.55E-01 | 1.25E+00 | 2.56E-01 | 9.41E-01 | 1.30E+00 | 1.14E-01 | 3.14E-01 | 1.18E+00 |

| | | | | | | | | | | | | | | | | | | | | | | | | |
|---|---|---|---|---|---|---|---|---|---|---|---|---|---|---|---|---|---|---|---|---|---|---|---|---|
| MTCO2P2 | 5.52E-02 | 2.88E-01 | 1.13E+00 | 4.17E-05 | 1.80E-04 | -1.86E+00 | 1.80E-01 | 7.63E-01 | 1.30E+00 | 4.77E-06 | 1.82E-05 | -2.03E+00 | 1.87E-03 | 6.66E-03 | -1.30E+00 | 2.38E-05 | 6.72E-05 | -1.96E+00 | 6.72E-03 | 2.47E-02 | -1.15E+00 | 1.17E-05 | 3.23E-05 | -1.60E+00 |
| CARD19 | 5.52E-02 | 2.88E-01 | 1.32E+00 | 1.35E-01 | 5.85E-01 | 1.46E+00 | 2.00E-01 | 8.52E-01 | 1.26E+00 | 2.13E-01 | 8.14E-01 | 1.33E+00 | 2.73E-01 | 9.72E-01 | 1.33E+00 | 3.65E-02 | 1.03E-01 | 1.07E+00 | 5.80E-02 | 2.13E-01 | 1.64E+00 | 2.90E-01 | 7.97E-01 | 1.20E+00 |
| FNIP2 | 5.52E-02 | 2.88E-01 | 1.12E+00 | 1.58E-02 | 6.83E-02 | 1.27E+00 | 3.59E-05 | 1.52E-04 | -1.29E+00 | 9.26E-06 | 3.54E-05 | -1.19E+00 | 8.74E-03 | 3.12E-02 | -1.19E+00 | 2.94E-02 | 8.29E-02 | -1.08E+00 | 2.98E-06 | 1.09E-05 | -1.56E+00 | 4.11E-06 | 1.13E-05 | -1.60E+00 |
| STPG1 | 5.53E-02 | 2.88E-01 | -1.16E+00 | 1.83E-01 | 7.89E-01 | 1.20E+00 | 2.93E-02 | 1.24E-01 | -1.25E+00 | 1.16E-02 | 4.44E-02 | -1.22E+00 | 1.51E-02 | 5.37E-02 | -1.30E+00 | 6.11E-02 | 1.72E-01 | -1.03E+00 | 1.86E-03 | 6.83E-03 | -1.49E+00 | 1.59E-02 | 4.37E-02 | -1.31E+00 |
| EIF3LP1 | 5.53E-02 | 2.88E-01 | 1.04E+00 | 3.22E-03 | 1.39E-02 | -1.24E+00 | 1.29E-02 | 5.48E-02 | 1.67E+00 | 8.51E-02 | 3.25E-01 | -1.07E+00 | 2.80E-01 | 1.00E+00 | 1.33E+00 | 8.05E-02 | 2.27E-01 | -1.09E+00 | 6.25E-02 | 2.29E-01 | -1.10E+00 | 2.01E-02 | 5.52E-02 | -1.19E+00 |
| AC012306.2 | 5.53E-02 | 2.88E-01 | 1.68E+00 | 8.52E-03 | 3.68E-02 | 2.42E+00 | 4.08E-02 | 1.73E-01 | 1.79E+00 | 2.04E-03 | 7.80E-03 | 2.73E+00 | 1.41E-01 | 5.04E-01 | 1.77E+00 | 6.54E-02 | 1.84E-01 | 2.10E+00 | 5.36E-03 | 1.97E-02 | 1.99E+00 | 1.51E-02 | 4.15E-02 | 2.15E+00 |
| XRCC2 | 5.54E-02 | 2.88E-01 | -1.37E+00 | 4.50E-02 | 1.94E-01 | -1.38E+00 | 5.36E-02 | 2.28E-01 | 1.01E+00 | 2.39E-01 | 9.14E-01 | 1.37E+00 | 2.01E-02 | 7.17E-02 | -1.49E+00 | 1.71E-01 | 4.83E-01 | -1.01E+00 | 8.49E-02 | 3.12E-01 | 1.19E+00 | 1.33E-03 | 3.65E-03 | 1.69E+00 |
| RN7SL172P | 5.54E-02 | 2.89E-01 | 1.04E+00 | 1.35E-01 | 5.84E-01 | 1.27E+00 | 1.56E-01 | 6.62E-01 | 1.37E+00 | 1.88E-01 | 7.18E-01 | 1.51E+00 | 2.29E-01 | 8.17E-01 | 1.20E+00 | 1.93E-01 | 5.43E-01 | -1.15E+00 | 1.34E-02 | 4.94E-02 | -1.38E+00 | 2.46E-02 | 6.75E-02 | -1.29E+00 |
| MEIS1 | 5.55E-02 | 2.89E-01 | 1.10E+00 | 2.46E-02 | 1.07E-01 | 1.50E+00 | 1.05E-01 | 4.45E-01 | 1.30E+00 | 5.19E-02 | 1.98E-01 | 1.67E+00 | 2.15E-02 | 7.66E-02 | 1.15E+00 | 1.06E-02 | 2.98E-02 | 1.14E+00 | 4.52E-03 | 1.66E-02 | 1.53E+00 | 7.48E-02 | 2.05E-01 | 1.34E+00 |
| FGFR1 | 5.55E-02 | 2.89E-01 | -1.15E+00 | 3.86E-03 | 1.67E-02 | -1.50E+00 | 2.32E-01 | 9.86E-01 | 1.29E+00 | 4.66E-02 | 1.78E-01 | -1.05E+00 | 2.00E-01 | 7.14E-01 | -1.23E+00 | 2.83E-01 | 7.99E-01 | -1.10E+00 | 9.29E-02 | 3.41E-01 | -1.31E+00 | 2.34E-01 | 6.42E-01 | -1.00E+00 |
| DOT1L | 5.55E-02 | 2.89E-01 | 1.08E+00 | 2.30E-02 | 9.95E-02 | 1.06E+00 | 7.67E-02 | 3.26E-01 | 1.02E+00 | 9.57E-04 | 3.66E-03 | -1.07E+00 | 2.69E-01 | 9.58E-01 | 1.31E+00 | 3.43E-02 | 9.68E-02 | -1.15E+00 | 3.23E-04 | 1.19E-03 | -1.13E+00 | 1.14E-06 | 3.14E-06 | -1.35E+00 |
| AC087283.1 | 5.56E-02 | 2.90E-01 | -1.09E+00 | 1.87E-02 | 8.08E-02 | -1.35E+00 | 2.99E-04 | 1.27E-03 | -1.57E+00 | 8.15E-05 | 3.11E-04 | -1.51E+00 | 3.79E-02 | 1.35E-01 | -1.07E+00 | 6.11E-03 | 1.72E-02 | -1.66E+00 | 2.14E-03 | 7.86E-03 | -1.36E+00 | 1.14E-01 | 3.13E-01 | 1.06E+00 |
| AC025594.2 | 5.56E-02 | 2.90E-01 | 1.65E+00 | 1.79E-01 | 7.75E-01 | 1.44E+00 | 1.79E-03 | 7.61E-03 | -1.35E+00 | 1.93E-03 | 7.38E-03 | -1.29E+00 | 5.23E-02 | 1.86E-01 | -1.30E+00 | 6.81E-02 | 1.92E-01 | -1.39E+00 | 1.27E-01 | 4.66E-01 | 1.52E+00 | 2.97E-01 | 8.15E-01 | 1.35E+00 |
| AP4B1 | 5.57E-02 | 2.90E-01 | 1.09E+00 | 3.33E-02 | 1.44E-01 | 1.18E+00 | 2.15E-01 | 9.14E-01 | 1.15E+00 | 2.48E-01 | 9.47E-01 | 1.26E+00 | 5.44E-02 | 1.94E-01 | 1.10E+00 | 9.96E-02 | 2.81E-01 | 1.20E+00 | 7.59E-03 | 2.79E-02 | -1.12E+00 | 3.12E-02 | 8.57E-02 | -1.08E+00 |
| VAC14-AS1 | 5.57E-02 | 2.90E-01 | -1.09E+00 | 1.03E-02 | 4.44E-02 | -1.41E+00 | 1.37E-01 | 5.81E-01 | 1.06E+00 | 2.34E-02 | 8.93E-02 | -1.16E+00 | 5.83E-02 | 2.08E-01 | -1.29E+00 | 4.29E-02 | 1.21E-01 | -1.38E+00 | 1.94E-01 | 7.11E-01 | 1.27E+00 | 3.47E-01 | 9.52E-01 | 1.17E+00 |
| TPCN1 | 5.57E-02 | 2.90E-01 | 1.10E+00 | 5.46E-02 | 2.36E-01 | 1.16E+00 | 2.14E-03 | 9.09E-03 | -1.44E+00 | 8.78E-02 | 3.35E-01 | 1.09E+00 | 2.75E-11 | 9.81E-11 | -1.41E+00 | 1.06E-11 | 2.98E-11 | -1.47E+00 | 1.34E-12 | 4.91E-12 | -1.52E+00 | 2.35E-03 | 6.45E-03 | 1.00E+00 |
| LINC02323 | 5.57E-02 | 2.90E-01 | 1.05E+00 | 3.74E-03 | 1.62E-02 | -1.26E+00 | 1.28E-01 | 5.42E-01 | 1.06E+00 | 1.51E-01 | 5.77E-01 | 1.16E+00 | 1.54E-01 | 5.51E-01 | 1.76E+00 | 2.37E-01 | 6.69E-01 | 1.65E+00 | 1.05E-01 | 3.84E-01 | 1.00E+00 | 2.82E-01 | 7.76E-01 | 1.18E+00 |
| AC026356.2 | 5.57E-02 | 2.90E-01 | -1.09E+00 | 1.22E-02 | 5.26E-02 | 1.78E+00 | 4.94E-05 | 2.10E-04 | -1.78E+00 | 1.81E-01 | 6.90E-01 | 1.57E+00 | 6.83E-02 | 2.44E-01 | -1.42E+00 | 1.02E-01 | 2.86E-01 | 2.06E+00 | 6.69E-03 | 2.46E-02 | -1.93E+00 | 1.57E-01 | 4.31E-01 | 1.42E+00 |
| AL022322.1 | 5.57E-02 | 2.90E-01 | -1.07E+00 | 1.36E-01 | 5.88E-01 | 1.23E+00 | 1.83E-01 | 7.77E-01 | 1.25E+00 | 1.79E-01 | 6.85E-01 | 1.07E+00 | 1.65E-01 | 5.90E-01 | 1.00E+00 | 6.03E-02 | 1.70E-01 | -1.35E+00 | 9.45E-02 | 3.47E-01 | 1.38E+00 | 3.58E-01 | 9.83E-01 | 1.17E+00 |
| RPS15AP40 | 5.58E-02 | 2.91E-01 | -1.06E+00 | 6.54E-02 | 2.82E-01 | 1.10E+00 | 9.29E-02 | 3.95E-01 | 1.02E+00 | 5.11E-02 | 1.95E-01 | 1.05E+00 | 2.88E-02 | 1.03E-01 | -1.09E+00 | 7.55E-03 | 2.13E-02 | -1.26E+00 | 6.81E-02 | 2.50E-01 | 1.57E+00 | 3.21E-01 | 8.81E-01 | 1.31E+00 |
| HAUS8 | 5.58E-02 | 2.91E-01 | -1.06E+00 | 6.68E-02 | 2.89E-01 | -1.03E+00 | 1.01E-03 | 4.30E-03 | -1.35E+00 | 1.81E-02 | 6.90E-02 | -1.03E+00 | 1.91E-01 | 6.82E-01 | -1.04E+00 | 3.02E-01 | 8.50E-01 | 1.12E+00 | 8.70E-12 | 3.19E-11 | -1.80E+00 | 6.92E-04 | 1.90E-03 | -1.18E+00 |
| PLEKHA4 | 5.58E-02 | 2.91E-01 | 1.03E+00 | 1.67E-01 | 7.22E-01 | 1.33E+00 | 1.42E-01 | 6.04E-01 | 1.11E+00 | 1.45E-01 | 5.52E-01 | 1.18E+00 | 1.50E-02 | 5.35E-02 | -1.33E+00 | 1.54E-02 | 4.34E-02 | -1.38E+00 | 1.69E-01 | 6.22E-01 | 1.41E+00 | 2.86E-01 | 7.86E-01 | 1.32E+00 |
| MSMO1 | 5.58E-02 | 2.91E-01 | 1.17E+00 | 1.80E-01 | 7.79E-01 | 1.66E+00 | 2.07E-01 | 8.80E-01 | -1.01E+00 | 1.59E-03 | 6.08E-03 | 1.89E+00 | 1.22E-01 | 4.37E-01 | 1.10E+00 | 5.93E-02 | 1.67E-01 | 1.78E+00 | 3.43E-02 | 1.26E-01 | -1.03E+00 | 1.03E-01 | 2.83E-01 | 1.66E+00 |
| AC026401.3 | 5.58E-02 | 2.91E-01 | 1.01E+00 | 1.52E-02 | 6.58E-02 | -1.10E+00 | 6.60E-02 | 2.80E-01 | 1.52E+00 | 2.03E-01 | 7.76E-01 | 1.30E+00 | 9.99E-02 | 3.56E-01 | -1.03E+00 | 9.58E-02 | 2.70E-01 | -1.17E+00 | 8.08E-05 | 2.97E-04 | -1.74E+00 | 1.67E-03 | 4.59E-03 | -1.44E+00 |
| TSPAN9 | 5.58E-02 | 2.91E-01 | -1.31E+00 | 7.11E-02 | 3.07E-01 | -1.31E+00 | 2.35E-01 | 1.00E+00 | 1.00E+00 | 2.62E-01 | 1.00E+00 | -1.00E+00 | 1.50E-01 | 5.35E-01 | -1.24E+00 | 2.41E-01 | 6.78E-01 | -1.08E+00 | 1.41E-02 | 5.18E-02 | 2.01E+00 | 2.73E-01 | 7.49E-01 | 1.62E+00 |
| LHX4 | 5.59E-02 | 2.91E-01 | 1.01E+00 | 2.29E-01 | 9.89E-01 | 1.48E+00 | 1.68E-01 | 7.14E-01 | 1.12E+00 | 2.60E-01 | 9.92E-01 | 1.40E+00 | 2.82E-02 | 1.01E-01 | -1.38E+00 | 3.37E-01 | 9.49E-01 | 1.35E+00 | 1.78E-04 | 6.53E-04 | -1.73E+00 | 1.46E-01 | 4.01E-01 | 1.06E+00 |
| AC046176.1 | 5.59E-02 | 2.91E-01 | 1.03E+00 | 1.39E-01 | 6.03E-01 | 1.33E+00 | 4.94E-02 | 2.10E-01 | -1.08E+00 | 1.35E-01 | 5.16E-01 | 1.19E+00 | 1.15E-01 | 4.10E-01 | 1.06E+00 | 2.42E-01 | 6.81E-01 | 1.53E+00 | 2.18E-01 | 7.99E-01 | 1.28E+00 | 1.40E-01 | 3.85E-01 | 1.16E+00 |
| CDC20 | 5.59E-02 | 2.92E-01 | -1.54E+00 | 1.02E-01 | 4.41E-01 | -1.10E+00 | 3.52E-03 | 1.50E-02 | 1.71E+00 | 8.73E-02 | 3.33E-01 | 1.34E+00 | 1.35E-01 | 4.82E-01 | -1.38E+00 | 2.03E-01 | 5.73E-01 | -1.02E+00 | 3.45E-02 | 1.27E-01 | -1.32E+00 | 3.32E-01 | 9.13E-01 | -1.10E+00 |
| MTCL1 | 5.60E-02 | 2.92E-01 | -1.56E+00 | 7.00E-02 | 3.02E-01 | -1.44E+00 | 1.07E-05 | 4.56E-05 | -1.99E+00 | 1.26E-03 | 4.80E-03 | -1.38E+00 | 2.38E-01 | 8.48E-01 | 1.16E+00 | 3.26E-01 | 9.19E-01 | 1.50E+00 | 3.22E-06 | 1.18E-05 | -1.60E+00 | 1.01E-01 | 2.76E-01 | 1.01E+00 |
| PEAK3 | 5.60E-02 | 2.92E-01 | 1.76E+00 | 9.45E-02 | 4.08E-01 | 1.94E+00 | 2.12E-01 | 9.03E-01 | 1.05E+00 | 1.41E-01 | 5.39E-01 | 1.15E+00 | 9.56E-03 | 3.41E-02 | -1.47E+00 | 1.77E-02 | 4.98E-02 | -1.37E+00 | 5.34E-02 | 1.96E-01 | -1.25E+00 | 8.44E-02 | 2.32E-01 | -1.22E+00 |
| CABP4 | 5.60E-02 | 2.92E-01 | 1.73E+00 | 5.01E-02 | 2.17E-01 | 1.96E+00 | 1.61E-01 | 6.85E-01 | 1.29E+00 | 2.23E-01 | 8.53E-01 | 1.21E+00 | 2.05E-01 | 7.32E-01 | 1.55E+00 | 2.87E-01 | 8.08E-01 | 1.50E+00 | 2.53E-01 | 9.28E-01 | 1.10E+00 | 2.59E-01 | 7.12E-01 | -1.04E+00 |
| AC106791.1 | 5.60E-02 | 2.92E-01 | 1.01E+00 | 1.40E-02 | 6.04E-02 | -1.05E+00 | 9.14E-02 | 3.88E-01 | -1.03E+00 | 1.61E-02 | 6.14E-02 | -1.09E+00 | 1.52E-02 | 5.41E-02 | -1.04E+00 | 2.03E-02 | 5.74E-02 | -1.07E+00 | 1.30E-01 | 4.77E-01 | 1.01E+00 | 2.68E-01 | 7.36E-01 | 1.11E+00 |
| CDCA5 | 5.61E-02 | 2.92E-01 | -1.44E+00 | 4.03E-02 | 1.74E-01 | -1.74E+00 | 1.16E-02 | 4.94E-02 | 1.39E+00 | 1.73E-01 | 6.63E-01 | 1.31E+00 | 1.75E-01 | 6.23E-01 | -1.52E+00 | 2.83E-01 | 7.99E-01 | -1.11E+00 | 1.61E-01 | 5.90E-01 | 1.05E+00 | 1.01E-05 | 2.76E-05 | 1.55E+00 |
| AC096751.1 | 5.61E-02 | 2.92E-01 | 1.66E+00 | 4.65E-04 | 2.01E-03 | 3.42E+00 | 1.98E-02 | 8.39E-02 | -1.32E+00 | 1.30E-01 | 4.98E-01 | 1.15E+00 | 1.85E-02 | 6.59E-02 | 2.41E+00 | 1.86E-01 | 5.24E-01 | 1.69E+00 | 2.57E-01 | 9.43E-01 | -1.06E+00 | 2.83E-01 | 7.77E-01 | 1.39E+00 |
| ANKRD49 | 5.61E-02 | 2.93E-01 | 1.29E+00 | 5.84E-05 | 2.52E-04 | 1.01E+00 | 8.46E-05 | 3.60E-04 | -1.25E+00 | 1.22E-07 | 4.66E-07 | -1.37E+00 | 1.87E-03 | 6.66E-03 | -1.01E+00 | 1.37E-04 | 3.86E-04 | -1.19E+00 | 5.43E-02 | 1.99E-01 | 1.11E+00 | 5.55E-03 | 1.53E-02 | -1.06E+00 |
| NAT14 | 5.62E-02 | 2.93E-01 | 1.21E+00 | 1.27E-02 | 5.49E-02 | 1.27E+00 | 3.45E-05 | 1.47E-04 | -1.16E+00 | 6.30E-04 | 2.41E-03 | 1.01E+00 | 4.12E-02 | 1.47E-01 | 1.15E+00 | 2.60E-01 | 7.32E-01 | 1.28E+00 | 2.58E-01 | 9.47E-01 | 1.22E+00 | 1.39E-01 | 3.81E-01 | 1.35E+00 |
| AC132872.1 | 5.62E-02 | 2.93E-01 | 1.61E+00 | 6.35E-02 | 2.74E-01 | 1.81E+00 | 7.33E-02 | 3.11E-01 | -1.08E+00 | 1.58E-01 | 6.04E-01 | 1.10E+00 | 1.72E-01 | 6.12E-01 | 1.71E+00 | 6.14E-02 | 1.73E-01 | 2.47E+00 | 6.74E-02 | 2.47E-01 | -1.36E+00 | 3.15E-01 | 8.66E-01 | 1.08E+00 |
| SRRD | 5.62E-02 | 2.93E-01 | 1.01E+00 | 1.45E-02 | 6.28E-02 | -1.13E+00 | 2.26E-02 | 9.59E-02 | -1.21E+00 | 1.33E-01 | 5.09E-01 | 1.18E+00 | 1.17E-02 | 4.16E-02 | -1.30E+00 | 5.37E-02 | 1.51E-01 | -1.07E+00 | 2.61E-01 | 9.60E-01 | 1.21E+00 | 2.21E-01 | 6.06E-01 | 1.37E+00 |
| LINC01465 | 5.62E-02 | 2.93E-01 | 1.50E+00 | 2.62E-02 | 1.13E-01 | -1.01E+00 | 7.69E-02 | 3.27E-01 | 1.13E+00 | 5.02E-04 | 1.92E-03 | -1.22E+00 | 1.73E-01 | 6.18E-01 | 1.06E+00 | 1.31E-01 | 3.69E-01 | -1.07E+00 | 1.65E-01 | 6.04E-01 | 1.44E+00 | 2.99E-01 | 8.20E-01 | 1.23E+00 |
| C10orf111 | 5.62E-02 | 2.93E-01 | 1.03E+00 | 7.50E-05 | 3.24E-04 | -1.77E+00 | 3.70E-02 | 1.57E-01 | -1.22E+00 | 4.33E-04 | 1.65E-03 | -1.76E+00 | 2.64E-01 | 9.40E-01 | 1.14E+00 | 1.96E-01 | 5.52E-01 | 1.01E+00 | 3.59E-02 | 1.32E-01 | -1.19E+00 | 4.56E-02 | 1.25E-01 | -1.22E+00 |
| SERP1 | 5.62E-02 | 2.93E-01 | -1.00E+00 | 1.58E-01 | 6.84E-01 | 1.05E+00 | 1.06E-01 | 4.52E-01 | 1.00E+00 | 1.99E-01 | 7.58E-01 | 1.06E+00 | 1.44E-01 | 5.13E-01 | 1.01E+00 | 2.80E-01 | 7.91E-01 | 1.03E+00 | 8.57E-03 | 3.14E-02 | -1.03E+00 | 2.47E-01 | 6.78E-01 | 1.01E+00 |
| TGFBI | 5.62E-02 | 2.93E-01 | -1.46E+00 | 1.22E-01 | 5.29E-01 | 1.04E+00 | 1.31E-01 | 5.58E-01 | -1.29E+00 | 1.49E-01 | 5.71E-01 | -1.29E+00 | 2.80E-01 | 1.00E+00 | 1.00E+00 | 3.55E-01 | 1.00E+00 | -1.00E+00 | 2.06E-01 | 7.57E-01 | 1.56E+00 | 3.20E-01 | 8.79E-01 | -1.01E+00 |
| AC099670.1 | 5.64E-02 | 2.94E-01 | 1.21E+00 | 5.43E-04 | 2.35E-03 | -1.26E+00 | 1.42E-02 | 6.05E-02 | -1.04E+00 | 1.52E-07 | 5.80E-07 | -1.65E+00 | 5.03E-02 | 1.79E-01 | 1.03E+00 | 2.09E-03 | 5.89E-03 | -1.48E+00 | 1.88E-03 | 6.90E-03 | -1.31E+00 | 2.69E-04 | 7.38E-04 | -1.36E+00 |
| SNTB2 | 5.65E-02 | 2.94E-01 | -1.38E+00 | 1.41E-01 | 6.09E-01 | 1.02E+00 | 1.63E-01 | 6.93E-01 | -1.04E+00 | 2.21E-01 | 8.43E-01 | 1.13E+00 | 2.23E-03 | 7.95E-03 | -1.55E+00 | 6.28E-02 | 1.77E-01 | -1.00E+00 | 1.89E-07 | 6.95E-07 | -1.53E+00 | 7.97E-04 | 2.19E-03 | -1.13E+00 |
| C10orf143 | 5.66E-02 | 2.95E-01 | 1.04E+00 | 5.13E-02 | 2.22E-01 | 1.13E+00 | 7.43E-03 | 3.16E-02 | -1.19E+00 | 4.24E-04 | 1.62E-03 | -1.31E+00 | 2.89E-02 | 1.03E-01 | -1.15E+00 | 1.48E-01 | 4.16E-01 | 1.25E+00 | 2.82E-02 | 1.04E-01 | -1.00E+00 | 2.31E-02 | 6.34E-02 | -1.09E+00 |
| DNMBP | 5.66E-02 | 2.95E-01 | -1.00E+00 | 6.24E-02 | 2.70E-01 | 1.07E+00 | 1.37E-01 | 5.82E-01 | 1.05E+00 | 1.03E-01 | 3.93E-01 | 1.05E+00 | 1.57E-02 | 5.59E-02 | -1.26E+00 | 1.18E-02 | 3.34E-02 | -1.41E+00 | 9.88E-03 | 3.63E-02 | -1.19E+00 | 2.87E-03 | 7.88E-03 | -1.43E+00 |
| FYCO1 | 5.66E-02 | 2.95E-01 | -1.16E+00 | 5.78E-02 | 2.50E-01 | -1.12E+00 | 6.27E-04 | 2.66E-03 | -1.38E+00 | 1.27E-02 | 4.83E-02 | -1.01E+00 | 2.18E-03 | 7.78E-03 | -1.61E+00 | 3.61E-02 | 1.02E-01 | -1.03E+00 | 2.93E-04 | 1.08E-03 | -1.56E+00 | 4.42E-03 | 1.21E-02 | -1.38E+00 |
| NAMPT | 5.66E-02 | 2.95E-01 | 1.17E+00 | 8.13E-16 | 3.51E-15 | -1.99E+00 | 1.42E-05 | 6.05E-05 | 2.00E+00 | 1.76E-02 | 6.73E-02 | 1.00E+00 | 9.01E-02 | 3.21E-01 | 2.11E+00 | 6.94E-02 | 1.96E-01 | 1.00E+00 | 4.44E-02 | 1.63E-01 | -1.00E+00 | 1.61E-02 | 4.41E-02 | -1.22E+00 |
| OSER1 | 5.66E-02 | 2.95E-01 | 1.17E+00 | 1.85E-09 | 7.97E-09 | -1.27E+00 | 3.26E-02 | 1.38E-01 | 1.39E+00 | 1.59E-05 | 6.09E-05 | -1.15E+00 | 5.59E-03 | 1.99E-02 | -1.03E+00 | 3.05E-06 | 8.60E-06 | -1.40E+00 | 6.75E-10 | 2.48E-09 | -1.29E+00 | 2.99E-13 | 8.21E-13 | -1.65E+00 |
| CACYBPP2 | 5.66E-02 | 2.95E-01 | 1.68E+00 | 1.40E-01 | 6.05E-01 | 1.77E+00 | 9.62E-03 | 4.09E-02 | 1.63E+00 | 1.76E-03 | 6.73E-03 | 2.02E+00 | 2.38E-01 | 8.48E-01 | 1.32E+00 | 2.39E-01 | 6.73E-01 | 1.53E+00 | 4.06E-02 | 1.49E-01 | 1.65E+00 | 9.59E-05 | 2.63E-04 | 2.21E+00 |
| AC025171.5 | 5.67E-02 | 2.96E-01 | 1.75E+00 | 1.07E-01 | 4.62E-01 | 1.72E+00 | 1.73E-01 | 7.34E-01 | -1.01E+00 | 9.85E-02 | 3.76E-01 | -1.06E+00 | 2.45E-01 | 8.73E-01 | 1.13E+00 | 3.50E-01 | 9.88E-01 | 1.08E+00 | 4.45E-02 | 1.63E-01 | -1.32E+00 | 1.09E-02 | 3.00E-02 | -1.58E+00 |
| ZCCHC18 | 5.67E-02 | 2.96E-01 | -1.07E+00 | 7.89E-02 | 3.41E-01 | -1.03E+00 | 1.74E-01 | 7.40E-01 | 1.02E+00 | 2.09E-01 | 8.00E-01 | 1.45E+00 | 1.50E-01 | 5.35E-01 | 1.44E+00 | 3.13E-01 | 8.83E-01 | 1.51E+00 | 2.30E-01 | 8.45E-01 | 1.33E+00 | 2.18E-01 | 6.00E-01 | 1.42E+00 |
| STK35 | 5.67E-02 | 2.96E-01 | 1.06E+00 | 1.49E-02 | 6.45E-02 | -1.03E+00 | 2.27E-01 | 9.65E-01 | 1.26E+00 | 2.39E-01 | 9.12E-01 | 1.32E+00 | 2.01E-01 | 7.18E-01 | 1.38E+00 | 3.23E-01 | 9.10E-01 | 1.45E+00 | 2.13E-01 | 7.80E-01 | 1.24E+00 | 3.05E-01 | 8.37E-01 | 1.30E+00 |
| NUP62CL | 5.67E-02 | 2.96E-01 | -1.31E+00 | 1.43E-01 | 6.17E-01 | 1.05E+00 | 2.12E-03 | 9.01E-03 | -1.46E+00 | 2.25E-03 | 8.60E-03 | -1.24E+00 | 1.79E-01 | 6.38E-01 | 1.74E+00 | 3.31E-01 | 9.33E-01 | 1.20E+00 | 1.16E-05 | 4.26E-05 | -1.35E+00 | 2.61E-02 | 7.18E-02 | 1.04E+00 |
| AC139713.2 | 5.68E-02 | 2.96E-01 | -1.11E+00 | 1.30E-01 | 5.61E-01 | 1.20E+00 | 1.55E-01 | 6.60E-01 | 1.05E+00 | 1.84E-01 | 7.02E-01 | 1.03E+00 | 4.96E-02 | 1.77E-01 | -1.24E+00 | 2.90E-01 | 8.17E-01 | 1.51E+00 | 1.45E-06 | 5.32E-06 | -1.22E+00 | 3.78E-03 | 1.04E-02 | -1.01E+00 |
| CR1 | 5.68E-02 | 2.96E-01 | 1.03E+00 | 2.31E-01 | 1.00E+00 | 1.52E+00 | 1.77E-01 | 7.51E-01 | -1.12E+00 | 1.78E-01 | 6.79E-01 | 1.48E+00 | 3.31E-03 | 1.18E-02 | 2.24E+00 | 1.19E-06 | 3.36E-06 | 3.43E+00 | 2.85E-03 | 1.05E-02 | 1.65E+00 | 6.15E-02 | 1.69E-01 | 1.04E+00 |
| AL136968.2 | 5.68E-02 | 2.96E-01 | 1.27E+00 | 2.32E-07 | 1.00E-06 | 1.01E+00 | 5.17E-02 | 2.20E-01 | 1.34E+00 | 1.38E-03 | 5.28E-03 | 1.03E+00 | 2.14E-01 | 7.62E-01 | 1.30E+00 | 1.00E-04 | 2.82E-04 | -1.02E+00 | 2.08E-01 | 7.64E-01 | 1.20E+00 | 5.46E-04 | 1.50E-03 | -1.13E+00 |
| AC118658.2 | 5.69E-02 | 2.97E-01 | 1.69E+00 | 2.21E-01 | 9.53E-01 | 1.30E+00 | 3.26E-02 | 1.38E-01 | -1.19E+00 | 1.17E-01 | 4.47E-01 | 1.06E+00 | 2.57E-01 | 9.17E-01 | 1.24E+00 | 2.78E-01 | 7.83E-01 | -1.04E+00 | 9.12E-02 | 3.35E-01 | 1.51E+00 | 9.53E-02 | 2.62E-01 | 1.61E+00 |
| AC112191.2 | 5.70E-02 | 2.97E-01 | 1.03E+00 | 1.42E-36 | 6.15E-36 | -1.53E+00 | 1.09E-01 | 4.64E-01 | -1.03E+00 | 2.39E-14 | 9.14E-14 | -1.58E+00 | 2.12E-04 | 7.56E-04 | -1.14E+00 | 1.20E-19 | 3.39E-19 | -1.78E+00 | 1.54E-04 | 5.66E-04 | -1.24E+00 | 1.28E-17 | 3.53E-17 | -1.84E+00 |
| AC002470.1 | 5.70E-02 | 2.97E-01 | -1.05E+00 | 8.95E-02 | 3.87E-01 | 1.05E+00 | 7.14E-02 | 3.04E-01 | -1.03E+00 | 4.61E-03 | 1.76E-02 | -1.37E+00 | 1.94E-01 | 6.92E-01 | 1.05E+00 | 2.48E-01 | 7.00E-01 | 1.12E+00 | 2.02E-03 | 7.40E-03 | -1.25E+00 | 6.56E-05 | 1.80E-04 | -1.76E+00 |
| DNAJC6 | 5.70E-02 | 2.97E-01 | -1.37E+00 | 1.39E-01 | 6.01E-01 | -1.00E+00 | 1.10E-03 | 4.66E-03 | -2.00E+00 | 9.29E-03 | 3.55E-02 | -1.22E+00 | 4.55E-05 | 1.62E-04 | -1.35E+00 | 1.94E-01 | 5.47E-01 | 1.39E+00 | 3.63E-22 | 1.33E-21 | -1.61E+00 | 2.92E-08 | 8.03E-08 | -1.11E+00 |
| GLT1D1 | 5.71E-02 | 2.97E-01 | 1.45E+00 | 2.31E-01 | 1.00E+00 | -1.00E+00 | 1.34E-02 | 5.71E-02 | 1.58E+00 | 1.96E-01 | 7.48E-01 | 1.11E+00 | 2.80E-01 | 1.00E+00 | -1.00E+00 | 3.55E-01 | 1.00E+00 | -1.00E+00 | 2.05E-01 | 7.52E-01 | -1.10E+00 | 2.24E-01 | 6.15E-01 | -1.19E+00 |
| PCDHB13 | 5.71E-02 | 2.98E-01 | 1.03E+00 | 1.64E-01 | 7.10E-01 | 1.27E+00 | 4.37E-02 | 1.86E-01 | -1.20E+00 | 1.23E-02 | 4.72E-02 | -1.34E+00 | 4.08E-02 | 1.46E-01 | -1.31E+00 | 6.23E-02 | 1.76E-01 | -1.17E+00 | 1.37E-01 | 5.01E-01 | 1.42E+00 | 3.29E-01 | 9.05E-01 | 1.17E+00 |
| NCAPG | 5.71E-02 | 2.98E-01 | -1.50E+00 | 8.72E-02 | 3.77E-01 | -1.28E+00 | 9.37E-02 | 3.98E-01 | 1.28E+00 | 1.76E-01 | 6.74E-01 | 1.38E+00 | 1.64E-01 | 5.86E-01 | -1.33E+00 | 2.16E-01 | 6.08E-01 | -1.33E+00 | 1.95E-03 | 7.15E-03 | -1.36E+00 | 1.52E-01 | 4.18E-01 | -1.09E+00 |
| RPS27AP1 | 5.72E-02 | 2.98E-01 | 1.69E+00 | 7.61E-02 | 3.29E-01 | 1.34E+00 | 2.23E-02 | 9.46E-02 | 1.68E+00 | 1.65E-01 | 6.32E-01 | 1.26E+00 | 6.24E-02 | 2.22E-01 | 1.92E+00 | 3.06E-01 | 8.63E-01 | 1.38E+00 | 1.54E-02 | 5.64E-02 | 1.68E+00 | 3.10E-01 | 8.51E-01 | 1.11E+00 |
| FBXO8 | 5.72E-02 | 2.98E-01 | 1.77E+00 | 2.03E-01 | 8.75E-01 | 1.42E+00 | 8.00E-02 | 3.40E-01 | 1.54E+00 | 2.13E-01 | 8.13E-01 | 1.28E+00 | 5.62E-02 | 2.00E-01 | 1.88E+00 | 2.27E-01 | 6.40E-01 | 1.50E+00 | 1.72E-01 | 6.32E-01 | 1.36E+00 | 7.36E-02 | 2.02E-01 | -1.08E+00 |
| ZNF225 | 5.72E-02 | 2.98E-01 | -1.01E+00 | 9.24E-02 | 3.99E-01 | 1.09E+00 | 1.84E-01 | 7.80E-01 | 1.19E+00 | 3.91E-02 | 1.49E-01 | -1.19E+00 | 1.19E-02 | 4.23E-02 | -1.46E+00 | 1.64E-01 | 4.63E-01 | 1.07E+00 | 2.54E-01 | 9.32E-01 | 1.21E+00 | 3.51E-01 | 9.65E-01 | 1.35E+00 |
| NPIPB13 | 5.72E-02 | 2.98E-01 | 1.06E+00 | 1.47E-01 | 6.35E-01 | 1.41E+00 | 7.03E-02 | 2.99E-01 | 1.48E+00 | 1.78E-01 | 6.80E-01 | 1.57E+00 | 7.64E-02 | 2.72E-01 | 1.86E+00 | 6.53E-02 | 1.84E-01 | 1.87E+00 | 8.25E-02 | 3.03E-01 | 1.14E+00 | 1.42E-01 | 3.91E-01 | 1.50E+00 |
| GCNT4 | 5.74E-02 | 2.99E-01 | -1.48E+00 | 4.88E-02 | 2.11E-01 | -1.66E+00 | 2.19E-01 | 9.30E-01 | -1.02E+00 | 2.05E-01 | 7.84E-01 | -1.07E+00 | 1.49E-03 | 5.32E-03 | -1.96E+00 | 3.18E-04 | 8.95E-04 | -2.25E+00 | 1.94E-04 | 7.12E-04 | -1.44E+00 | 3.09E-04 | 8.49E-04 | -1.30E+00 |
| ADAMTS14 | 5.74E-02 | 2.99E-01 | -1.27E+00 | 7.92E-02 | 3.42E-01 | -1.20E+00 | 7.27E-02 | 3.09E-01 | -1.24E+00 | 2.15E-01 | 8.21E-01 | 1.13E+00 | 2.70E-02 | 9.61E-02 | -1.40E+00 | 3.23E-01 | 9.10E-01 | 1.26E+00 | 1.15E-06 | 4.20E-06 | -1.54E+00 | 3.20E-03 | 8.79E-03 | -1.10E+00 |
| AL449106.1 | 5.74E-02 | 2.99E-01 | -1.00E+00 | 7.49E-02 | 3.24E-01 | 1.07E+00 | 6.24E-02 | 2.65E-01 | -1.14E+00 | 8.26E-02 | 3.16E-01 | -1.03E+00 | 9.50E-04 | 3.39E-03 | -1.72E+00 | 1.03E-03 | 2.91E-03 | -1.89E+00 | 5.89E-02 | 2.16E-01 | -1.28E+00 | 5.61E-02 | 1.54E-01 | -1.31E+00 |
| AC007683.1 | 5.75E-02 | 3.00E-01 | 1.14E+00 | 2.09E-02 | 9.03E-02 | 2.07E+00 | 6.08E-02 | 2.58E-01 | 1.04E+00 | 2.68E-03 | 1.02E-02 | 2.16E+00 | 5.40E-03 | 1.93E-02 | -1.15E+00 | 1.84E-01 | 5.19E-01 | 1.77E+00 | 3.31E-02 | 1.21E-01 | -1.03E+00 | 1.91E-02 | 5.26E-02 | 1.88E+00 |
| SYCE1L | 5.75E-02 | 3.00E-01 | 1.12E+00 | 1.84E-02 | 7.97E-02 | 1.10E+00 | 1.77E-01 | 7.53E-01 | 1.21E+00 | 2.81E-03 | 1.07E-02 | -1.14E+00 | 1.72E-01 | 6.12E-01 | 1.24E+00 | 6.71E-02 | 1.89E-01 | -1.05E+00 | 1.45E-01 | 5.32E-01 | 1.55E+00 | 2.83E-01 | 7.77E-01 | 1.16E+00 |
| B4GALT4 | 5.75E-02 | 3.00E-01 | 1.14E+00 | 5.44E-04 | 2.35E-03 | -1.04E+00 | 1.65E-01 | 7.00E-01 | 1.21E+00 | 2.04E-03 | 7.78E-03 | -1.05E+00 | 8.30E-02 | 2.96E-01 | 1.18E+00 | 5.96E-03 | 1.68E-02 | -1.01E+00 | 2.06E-02 | 7.57E-02 | 1.15E+00 | 1.52E-04 | 4.18E-04 | -1.20E+00 |
| SUB1 | 5.76E-02 | 3.00E-01 | 1.11E+00 | 1.39E-02 | 6.01E-02 | 1.14E+00 | 2.27E-01 | 9.63E-01 | 1.04E+00 | 9.04E-02 | 3.45E-01 | 1.01E+00 | 8.87E-03 | 3.16E-02 | -1.06E+00 | 1.55E-01 | 4.38E-01 | 1.15E+00 | 8.25E-02 | 3.03E-01 | -1.02E+00 | 3.08E-01 | 8.45E-01 | 1.10E+00 |
| EMP3 | 5.77E-02 | 3.01E-01 | 1.34E+00 | 2.17E-01 | 9.39E-01 | 1.37E+00 | 4.98E-02 | 2.11E-01 | 1.09E+00 | 2.18E-01 | 8.31E-01 | 1.18E+00 | 5.32E-02 | 1.90E-01 | 1.04E+00 | 3.49E-01 | 9.85E-01 | 1.13E+00 | 1.63E-01 | 5.98E-01 | 1.32E+00 | 3.54E-01 | 9.72E-01 | 1.07E+00 |
| C1orf132 | 5.77E-02 | 3.01E-01 | 1.44E+00 | 1.80E-03 | 7.76E-03 | 2.06E+00 | 1.06E-01 | 4.52E-01 | 1.62E+00 | 1.48E-01 | 5.66E-01 | 1.81E+00 | 1.36E-01 | 4.86E-01 | 1.72E+00 | 1.24E-01 | 3.50E-01 | 1.75E+00 | 1.46E-01 | 5.36E-01 | 1.72E+00 | 3.36E-01 | 9.23E-01 | 1.21E+00 |
| CRISPLD2 | 5.77E-02 | 3.01E-01 | -1.27E+00 | 2.15E-01 | 9.31E-01 | 1.26E+00 | 7.66E-02 | 3.26E-01 | -1.11E+00 | 8.69E-02 | 3.32E-01 | -1.14E+00 | 2.72E-01 | 9.71E-01 | -1.02E+00 | 2.80E-01 | 7.89E-01 | 1.19E+00 | 2.29E-01 | 8.39E-01 | -1.03E+00 | 1.56E-01 | 4.27E-01 | 1.52E+00 |
| AGA | 5.77E-02 | 3.01E-01 | 1.15E+00 | 5.29E-02 | 2.28E-01 | 1.25E+00 | 2.76E-02 | 1.17E-01 | 1.06E+00 | 6.74E-03 | 2.58E-02 | 1.09E+00 | 7.98E-02 | 2.85E-01 | 1.25E+00 | 8.09E-02 | 2.28E-01 | 1.27E+00 | 1.51E-01 | 5.56E-01 | 1.22E+00 | 1.09E-01 | 3.00E-01 | 1.08E+00 |
| CEBPB-AS1 | 5.78E-02 | 3.01E-01 | 1.05E+00 | 2.83E-02 | 1.22E-01 | -1.07E+00 | 2.19E-03 | 9.29E-03 | -1.40E+00 | 6.26E-03 | 2.39E-02 | -1.14E+00 | 2.50E-01 | 8.90E-01 | 1.21E+00 | 2.88E-01 | 8.11E-01 | 1.15E+00 | 1.73E-01 | 6.34E-01 | -1.02E+00 | 3.09E-01 | 8.49E-01 | 1.20E+00 |
| AC007229.1 | 5.78E-02 | 3.01E-01 | -1.01E+00 | 8.30E-02 | 3.59E-01 | 1.16E+00 | 2.00E-01 | 8.52E-01 | 1.15E+00 | 1.89E-01 | 7.22E-01 | 1.44E+00 | 1.05E-01 | 3.74E-01 | 1.08E+00 | 3.46E-01 | 9.74E-01 | 1.36E+00 | 1.40E-01 | 5.12E-01 | 1.17E+00 | 5.27E-03 | 1.45E-02 | 1.79E+00 |
| RFX1 | 5.78E-02 | 3.01E-01 | 1.58E+00 | 3.97E-03 | 1.72E-02 | -1.10E+00 | 1.33E-01 | 5.64E-01 | 1.23E+00 | 1.60E-05 | 6.10E-05 | -1.31E+00 | 1.29E-02 | 4.59E-02 | -1.15E+00 | 1.07E-02 | 3.01E-02 | -1.19E+00 | 3.22E-02 | 1.18E-01 | 1.05E+00 | 3.16E-02 | 8.69E-02 | 1.02E+00 |
| BCL6 | 5.79E-02 | 3.02E-01 | 1.01E+00 | 1.26E-02 | 5.43E-02 | -1.15E+00 | 2.32E-03 | 9.88E-03 | -1.36E+00 | 1.96E-04 | 7.49E-04 | -1.54E+00 | 2.72E-01 | 9.70E-01 | 1.31E+00 | 3.35E-01 | 9.45E-01 | 1.00E+00 | 2.48E-01 | 9.10E-01 | 1.11E+00 | 1.56E-01 | 4.27E-01 | -1.38E+00 |
| AC025040.2 | 5.80E-02 | 3.02E-01 | 1.01E+00 | 1.32E-02 | 5.70E-02 | -1.20E+00 | 1.20E-02 | 5.09E-02 | -1.21E+00 | 6.11E-04 | 2.33E-03 | -1.64E+00 | 2.54E-01 | 9.07E-01 | 1.30E+00 | 6.07E-02 | 1.71E-01 | -1.49E+00 | 7.62E-02 | 2.80E-01 | -1.01E+00 | 2.01E-02 | 5.52E-02 | -1.33E+00 |
| GARNL3 | 5.80E-02 | 3.02E-01 | -1.13E+00 | 1.30E-01 | 5.62E-01 | 1.06E+00 | 2.28E-01 | 9.68E-01 | 1.05E+00 | 1.77E-01 | 6.76E-01 | 1.35E+00 | 8.66E-02 | 3.09E-01 | -1.31E+00 | 1.14E-01 | 3.22E-01 | -1.27E+00 | 6.56E-02 | 2.41E-01 | -1.31E+00 | 2.48E-01 | 6.81E-01 | 1.04E+00 |
| CYP4F2 | 5.80E-02 | 3.02E-01 | -1.05E+00 | 3.79E-02 | 1.64E-01 | -1.10E+00 | 8.60E-02 | 3.65E-01 | 1.02E+00 | 2.37E-01 | 9.06E-01 | 1.30E+00 | 2.68E-01 | 9.57E-01 | 1.58E+00 | 2.58E-01 | 7.28E-01 | 1.57E+00 | 2.66E-01 | 9.74E-01 | 1.06E+00 | 3.44E-01 | 9.45E-01 | 1.13E+00 |
| AC012485.3 | 5.81E-02 | 3.03E-01 | 1.31E+00 | 2.26E-01 | 9.76E-01 | -1.12E+00 | 2.18E-02 | 9.28E-02 | 1.55E+00 | 2.01E-01 | 7.69E-01 | -1.08E+00 | 1.89E-01 | 6.73E-01 | 1.32E+00 | 1.30E-01 | 3.65E-01 | 1.63E+00 | 1.32E-01 | 4.84E-01 | -1.25E+00 | 3.42E-01 | 9.40E-01 | 1.09E+00 |
| CNOT7P1 | 5.81E-02 | 3.03E-01 | 1.07E+00 | 9.15E-02 | 3.96E-01 | 1.28E+00 | 3.51E-02 | 1.49E-01 | -1.13E+00 | 1.54E-01 | 5.88E-01 | 1.22E+00 | 2.07E-01 | 7.37E-01 | 1.27E+00 | 1.55E-01 | 4.38E-01 | 1.85E+00 | 8.84E-02 | 3.24E-01 | -1.00E+00 | 3.35E-01 | 9.20E-01 | 1.22E+00 |
| AC090686.1 | 5.81E-02 | 3.03E-01 | 1.05E+00 | 5.87E-02 | 2.54E-01 | 1.17E+00 | 7.34E-03 | 3.12E-02 | -1.15E+00 | 6.85E-05 | 2.62E-04 | -1.38E+00 | 1.16E-01 | 4.14E-01 | 1.15E+00 | 4.13E-02 | 1.16E-01 | -1.07E+00 | 1.57E-01 | 5.76E-01 | 1.14E+00 | 8.72E-02 | 2.39E-01 | 1.00E+00 |
| CD2 | 5.81E-02 | 3.03E-01 | -2.23E+00 | 5.32E-02 | 2.30E-01 | -2.44E+00 | 1.24E-01 | 5.27E-01 | -3.00E+00 | 4.10E-02 | 1.57E-01 | -3.83E+00 | 7.32E-02 | 2.61E-01 | -1.35E+00 | 4.82E-02 | 1.36E-01 | -1.71E+00 | 4.25E-02 | 1.56E-01 | 2.69E+00 | 2.37E-01 | 6.52E-01 | 1.05E+00 |
| TMEM241 | 5.81E-02 | 3.03E-01 | 1.03E+00 | 2.26E-01 | 9.77E-01 | 1.37E+00 | 1.15E-01 | 4.91E-01 | 1.04E+00 | 7.22E-02 | 2.76E-01 | 1.16E+00 | 1.53E-01 | 5.45E-01 | 1.15E+00 | 2.87E-01 | 8.09E-01 | 1.27E+00 | 1.88E-01 | 6.90E-01 | 1.18E+00 | 3.63E-01 | 9.97E-01 | 1.18E+00 |
| USMG5P1 | 5.82E-02 | 3.03E-01 | 1.13E+00 | 1.31E-02 | 5.65E-02 | -1.04E+00 | 9.37E-02 | 3.98E-01 | 1.03E+00 | 2.98E-03 | 1.14E-02 | -1.20E+00 | 3.56E-03 | 1.27E-02 | -1.29E+00 | 1.15E-03 | 3.25E-03 | -1.37E+00 | 1.84E-04 | 6.75E-04 | -1.39E+00 | 7.99E-05 | 2.20E-04 | -1.45E+00 |
| NABP1 | 5.82E-02 | 3.03E-01 | 1.42E+00 | 8.59E-03 | 3.71E-02 | 1.03E+00 | 1.24E-01 | 5.27E-01 | 1.15E+00 | 2.16E-04 | 8.24E-04 | -1.12E+00 | 2.54E-01 | 9.06E-01 | 1.34E+00 | 3.55E-01 | 1.00E+00 | 1.51E+00 | 1.79E-01 | 6.55E-01 | 1.39E+00 | 2.34E-01 | 6.42E-01 | 1.24E+00 |
| FCGR2A | 5.82E-02 | 3.03E-01 | -2.23E+00 | 7.58E-02 | 3.27E-01 | -2.13E+00 | 1.70E-01 | 7.23E-01 | -1.08E+00 | 3.63E-02 | 1.39E-01 | -1.87E+00 | 1.32E-01 | 4.71E-01 | 2.11E+00 | 2.58E-02 | 7.29E-02 | 2.34E+00 | 6.13E-05 | 2.25E-04 | 1.33E+00 | 1.91E-04 | 5.24E-04 | 1.20E+00 |
| RPS19BP1 | 5.82E-02 | 3.03E-01 | 1.33E+00 | 1.07E-01 | 4.62E-01 | 1.31E+00 | 4.76E-02 | 2.02E-01 | 1.31E+00 | 2.33E-01 | 8.92E-01 | 1.33E+00 | 2.31E-01 | 8.25E-01 | 1.39E+00 | 2.70E-01 | 7.62E-01 | 1.30E+00 | 1.47E-03 | 5.40E-03 | 1.51E+00 | 7.19E-02 | 1.98E-01 | 1.29E+00 |
| HBD | 5.83E-02 | 3.04E-01 | 2.13E+00 | 2.50E-02 | 1.08E-01 | 1.58E+00 | 6.86E-02 | 2.91E-01 | 1.38E+00 | 1.04E-01 | 3.98E-01 | 1.61E+00 | 9.40E-04 | 3.35E-03 | -1.99E+00 | 2.90E-01 | 8.17E-01 | -1.13E+00 | 8.38E-03 | 3.07E-02 | -1.02E+00 | 1.47E-01 | 4.04E-01 | -1.08E+00 |
| TRIM45 | 5.83E-02 | 3.04E-01 | 1.03E+00 | 3.97E-02 | 1.71E-01 | 1.02E+00 | 1.47E-02 | 6.25E-02 | -1.21E+00 | 2.35E-03 | 8.98E-03 | -1.28E+00 | 3.02E-02 | 1.08E-01 | -1.17E+00 | 4.38E-02 | 1.24E-01 | -1.21E+00 | 3.45E-02 | 1.26E-01 | -1.11E+00 | 1.03E-01 | 2.82E-01 | -1.06E+00 |
| RPS3AP42 | 5.83E-02 | 3.04E-01 | 1.78E+00 | 4.13E-02 | 1.79E-01 | 1.04E+00 | 7.56E-02 | 3.21E-01 | 1.42E+00 | 4.60E-02 | 1.76E-01 | -1.02E+00 | 7.20E-02 | 2.57E-01 | 1.09E+00 | 1.77E-02 | 5.00E-02 | -1.13E+00 | 2.52E-01 | 9.24E-01 | 1.10E+00 | 2.70E-03 | 7.40E-03 | -1.90E+00 |
| LENG8-AS1 | 5.84E-02 | 3.04E-01 | 1.67E+00 | 1.11E-01 | 4.81E-01 | 1.74E+00 | 2.43E-02 | 1.03E-01 | -1.18E+00 | 2.30E-01 | 8.80E-01 | 1.30E+00 | 2.48E-01 | 8.83E-01 | 1.12E+00 | 1.76E-01 | 4.96E-01 | 1.07E+00 | 2.38E-01 | 8.75E-01 | 1.30E+00 | 1.26E-01 | 3.45E-01 | 1.42E+00 |
| AC078795.1 | 5.84E-02 | 3.04E-01 | -1.02E+00 | 1.33E-01 | 5.74E-01 | 1.20E+00 | 1.81E-01 | 7.69E-01 | 1.11E+00 | 1.15E-01 | 4.41E-01 | 1.79E+00 | 7.27E-02 | 2.59E-01 | -1.18E+00 | 1.90E-01 | 5.35E-01 | 1.07E+00 | 8.57E-02 | 3.14E-01 | -1.22E+00 | 1.37E-01 | 3.76E-01 | -1.12E+00 |
| SLC9A1 | 5.85E-02 | 3.05E-01 | -1.01E+00 | 1.74E-02 | 7.53E-02 | -1.08E+00 | 7.73E-03 | 3.28E-02 | -1.18E+00 | 7.83E-03 | 2.99E-02 | -1.09E+00 | 1.04E-02 | 3.73E-02 | -1.40E+00 | 8.18E-02 | 2.31E-01 | 1.01E+00 | 9.56E-04 | 3.51E-03 | 1.02E+00 | 1.15E-02 | 3.16E-02 | 1.12E+00 |
| SRD5A1 | 5.85E-02 | 3.05E-01 | -1.07E+00 | 1.56E-01 | 6.75E-01 | 1.29E+00 | 5.18E-02 | 2.20E-01 | -1.01E+00 | 1.26E-01 | 4.81E-01 | 1.67E+00 | 4.37E-03 | 1.56E-02 | -1.71E+00 | 7.87E-02 | 2.22E-01 | -1.09E+00 | 1.72E-02 | 6.30E-02 | -1.18E+00 | 2.51E-01 | 6.91E-01 | 1.35E+00 |
| ORC1 | 5.86E-02 | 3.05E-01 | -1.39E+00 | 6.41E-02 | 2.77E-01 | -1.40E+00 | 6.38E-02 | 2.71E-01 | 1.08E+00 | 8.51E-05 | 3.25E-04 | -1.19E+00 | 2.26E-02 | 8.07E-02 | -1.65E+00 | 1.62E-02 | 4.57E-02 | -1.82E+00 | 2.82E-03 | 1.04E-02 | -1.21E+00 | 2.24E-01 | 6.15E-01 | 1.26E+00 |
| C3orf70 | 5.86E-02 | 3.06E-01 | -1.21E+00 | 2.03E-01 | 8.77E-01 | 1.42E+00 | 6.24E-02 | 2.65E-01 | -1.13E+00 | 1.89E-01 | 7.22E-01 | 1.60E+00 | 2.11E-01 | 7.51E-01 | 1.31E+00 | 8.00E-02 | 2.26E-01 | 2.08E+00 | 3.00E-07 | 1.10E-06 | -1.53E+00 | 2.68E-03 | 7.36E-03 | 1.01E+00 |
| ZNF276 | 5.86E-02 | 3.06E-01 | 1.17E+00 | 3.67E-05 | 1.59E-04 | -1.12E+00 | 4.44E-03 | 1.89E-02 | -1.04E+00 | 3.29E-08 | 1.26E-07 | -1.35E+00 | 1.00E-01 | 3.57E-01 | 1.26E+00 | 2.53E-02 | 7.14E-02 | -1.01E+00 | 1.53E-04 | 5.62E-04 | -1.16E+00 | 3.60E-07 | 9.90E-07 | -1.41E+00 |
| FIS1 | 5.87E-02 | 3.06E-01 | 1.10E+00 | 2.74E-02 | 1.18E-01 | 1.13E+00 | 1.62E-01 | 6.88E-01 | 1.14E+00 | 6.59E-02 | 2.52E-01 | 1.15E+00 | 8.66E-03 | 3.09E-02 | 1.05E+00 | 1.33E-01 | 3.75E-01 | 1.09E+00 | 1.57E-01 | 5.75E-01 | 1.22E+00 | 1.80E-01 | 4.93E-01 | 1.23E+00 |
| ITGB1P1 | 5.87E-02 | 3.06E-01 | 1.07E+00 | 1.73E-01 | 7.48E-01 | 1.45E+00 | 2.37E-03 | 1.01E-02 | -1.36E+00 | 3.98E-03 | 1.52E-02 | -1.14E+00 | 2.63E-02 | 9.37E-02 | -1.03E+00 | 2.33E-01 | 6.56E-01 | 1.43E+00 | 3.15E-04 | 1.15E-03 | -1.18E+00 | 1.82E-02 | 5.01E-02 | -1.02E+00 |
| TMEM254-AS1 | 5.87E-02 | 3.06E-01 | -1.04E+00 | 4.55E-02 | 1.97E-01 | -1.04E+00 | 1.53E-01 | 6.49E-01 | 1.05E+00 | 1.98E-01 | 7.57E-01 | 1.23E+00 | 5.24E-02 | 1.87E-01 | -1.52E+00 | 3.31E-02 | 9.32E-02 | -1.60E+00 | 1.75E-01 | 6.44E-01 | -1.25E+00 | 2.97E-01 | 8.17E-01 | -1.17E+00 |
| TMEM128 | 5.89E-02 | 3.07E-01 | 1.05E+00 | 1.65E-01 | 7.15E-01 | 1.62E+00 | 8.69E-04 | 3.69E-03 | -1.40E+00 | 1.26E-01 | 4.81E-01 | 1.16E+00 | 5.11E-03 | 1.82E-02 | -1.29E+00 | 2.51E-01 | 7.08E-01 | 1.34E+00 | 2.77E-03 | 1.02E-02 | -1.15E+00 | 8.41E-02 | 2.31E-01 | 1.19E+00 |
| USP12 | 5.89E-02 | 3.07E-01 | 1.06E+00 | 1.29E-04 | 5.59E-04 | -1.20E+00 | 1.30E-02 | 5.51E-02 | -1.12E+00 | 8.56E-05 | 3.27E-04 | -1.25E+00 | 6.53E-04 | 2.33E-03 | -1.18E+00 | 5.28E-06 | 1.49E-05 | -1.39E+00 | 7.43E-07 | 2.73E-06 | -1.11E+00 | 4.27E-13 | 1.17E-12 | -1.51E+00 |
| AC027559.1 | 5.90E-02 | 3.07E-01 | 1.25E+00 | 1.17E-01 | 5.05E-01 | 1.45E+00 | 2.70E-02 | 1.15E-01 | -1.01E+00 | 3.87E-02 | 1.48E-01 | 1.21E+00 | 1.14E-02 | 4.06E-02 | -1.00E+00 | 4.25E-02 | 1.20E-01 | 1.12E+00 | 2.57E-01 | 9.44E-01 | 1.24E+00 | 3.54E-01 | 9.73E-01 | 1.25E+00 |
| APOM | 5.91E-02 | 3.08E-01 | 1.04E+00 | 3.10E-03 | 1.34E-02 | -1.34E+00 | 2.12E-01 | 8.99E-01 | 1.08E+00 | 3.33E-02 | 1.27E-01 | -1.13E+00 | 1.53E-01 | 5.47E-01 | 1.61E+00 | 3.25E-01 | 9.15E-01 | 1.28E+00 | 6.33E-02 | 2.32E-01 | 1.40E+00 | 6.94E-02 | 1.91E-01 | -1.02E+00 |
| AC093635.1 | 5.93E-02 | 3.09E-01 | 1.57E+00 | 6.78E-02 | 2.93E-01 | 1.15E+00 | 1.49E-01 | 6.32E-01 | 1.26E+00 | 1.57E-01 | 6.00E-01 | -1.01E+00 | 2.40E-01 | 8.54E-01 | 1.52E+00 | 3.32E-01 | 9.36E-01 | 1.36E+00 | 1.15E-01 | 4.23E-01 | 1.49E+00 | 1.46E-01 | 4.01E-01 | 1.38E+00 |
| AC092902.2 | 5.93E-02 | 3.09E-01 | 1.13E+00 | 2.29E-01 | 9.91E-01 | 1.45E+00 | 5.96E-02 | 2.53E-01 | -1.06E+00 | 1.10E-01 | 4.18E-01 | 1.17E+00 | 6.22E-02 | 2.22E-01 | -1.08E+00 | 8.94E-02 | 2.52E-01 | -1.01E+00 | 2.68E-01 | 9.83E-01 | 1.14E+00 | 3.26E-01 | 8.95E-01 | 1.09E+00 |
| MTMR3 | 5.94E-02 | 3.10E-01 | -1.06E+00 | 1.43E-01 | 6.20E-01 | 1.25E+00 | 1.47E-01 | 6.25E-01 | 1.05E+00 | 1.90E-01 | 7.27E-01 | 1.19E+00 | 1.59E-03 | 5.66E-03 | -1.45E+00 | 3.85E-04 | 1.09E-03 | -1.57E+00 | 5.13E-03 | 1.88E-02 | -1.25E+00 | 4.87E-03 | 1.34E-02 | -1.35E+00 |
| AC040970.1 | 5.95E-02 | 3.10E-01 | 1.14E+00 | 1.13E-04 | 4.88E-04 | -1.70E+00 | 8.19E-02 | 3.48E-01 | 1.33E+00 | 1.70E-01 | 6.51E-01 | 1.04E+00 | 7.71E-02 | 2.75E-01 | -1.10E+00 | 4.63E-02 | 1.31E-01 | -1.26E+00 | 5.74E-02 | 2.11E-01 | -1.01E+00 | 1.15E-02 | 3.17E-02 | -1.32E+00 |
| C3orf58 | 5.95E-02 | 3.10E-01 | 1.17E+00 | 6.74E-02 | 2.91E-01 | 1.24E+00 | 5.43E-04 | 2.31E-03 | -1.27E+00 | 4.69E-03 | 1.79E-02 | -1.04E+00 | 2.28E-03 | 8.12E-03 | -1.29E+00 | 4.75E-03 | 1.34E-02 | -1.38E+00 | 2.25E-01 | 8.26E-01 | 1.29E+00 | 1.93E-03 | 5.29E-03 | -1.14E+00 |
| AC006042.3 | 5.96E-02 | 3.11E-01 | 1.54E+00 | 3.58E-04 | 1.55E-03 | 2.71E+00 | 9.22E-02 | 3.92E-01 | 1.07E+00 | 2.29E-01 | 8.75E-01 | 1.39E+00 | 2.21E-01 | 7.89E-01 | 1.21E+00 | 2.86E-01 | 8.07E-01 | 1.22E+00 | 2.61E-01 | 9.60E-01 | 1.21E+00 | 2.67E-01 | 7.33E-01 | 1.44E+00 |
| ZNF35 | 5.96E-02 | 3.11E-01 | 1.64E+00 | 5.35E-02 | 2.31E-01 | 1.15E+00 | 2.23E-01 | 9.46E-01 | 1.25E+00 | 2.19E-02 | 8.38E-02 | -1.00E+00 | 1.60E-01 | 5.69E-01 | 1.47E+00 | 1.64E-01 | 4.62E-01 | 1.13E+00 | 2.50E-01 | 9.17E-01 | 1.37E+00 | 1.30E-02 | 3.57E-02 | -1.05E+00 |
| TIGD2 | 5.96E-02 | 3.11E-01 | -1.00E+00 | 4.30E-02 | 1.86E-01 | -1.02E+00 | 1.47E-04 | 6.23E-04 | -1.64E+00 | 1.43E-03 | 5.46E-03 | -1.23E+00 | 4.91E-07 | 1.75E-06 | -2.65E+00 | 7.09E-04 | 2.00E-03 | -1.33E+00 | 8.09E-05 | 2.97E-04 | -1.31E+00 | 1.17E-04 | 3.23E-04 | -1.38E+00 |
| HNRNPA3P5 | 5.97E-02 | 3.11E-01 | -1.16E+00 | 9.65E-02 | 4.17E-01 | 1.02E+00 | 9.40E-02 | 3.99E-01 | -1.03E+00 | 2.01E-01 | 7.66E-01 | 1.20E+00 | 1.98E-01 | 7.07E-01 | 1.08E+00 | 2.42E-01 | 6.83E-01 | 1.07E+00 | 6.10E-02 | 2.24E-01 | -1.06E+00 | 1.99E-01 | 5.47E-01 | 1.10E+00 |
| AC007182.2 | 5.97E-02 | 3.11E-01 | 1.50E+00 | 5.56E-05 | 2.40E-04 | 1.07E+00 | 9.75E-02 | 4.14E-01 | 1.43E+00 | 9.14E-07 | 3.49E-06 | -1.12E+00 | 1.36E-01 | 4.84E-01 | 1.39E+00 | 2.02E-04 | 5.68E-04 | -1.08E+00 | 1.61E-01 | 5.91E-01 | 1.21E+00 | 2.09E-04 | 5.74E-04 | -1.25E+00 |
| LINC01529 | 5.98E-02 | 3.12E-01 | -1.17E+00 | 2.29E-01 | 9.88E-01 | 1.28E+00 | 2.29E-01 | 9.73E-01 | 1.11E+00 | 6.16E-02 | 2.35E-01 | 1.66E+00 | 1.80E-01 | 6.41E-01 | -1.27E+00 | 2.82E-01 | 7.95E-01 | 1.21E+00 | 3.01E-02 | 1.10E-01 | -1.29E+00 | 3.61E-01 | 9.92E-01 | 1.17E+00 |
| AC127070.3 | 5.98E-02 | 3.12E-01 | -1.07E+00 | 7.12E-02 | 3.07E-01 | 1.09E+00 | 2.28E-01 | 9.69E-01 | 1.25E+00 | 2.57E-01 | 9.83E-01 | 1.30E+00 | 1.78E-01 | 6.34E-01 | 1.15E+00 | 7.25E-02 | 2.04E-01 | -1.19E+00 | 1.74E-01 | 6.39E-01 | 1.32E+00 | 2.85E-01 | 7.84E-01 | 1.15E+00 |
| HDAC4 | 5.99E-02 | 3.12E-01 | 1.00E+00 | 5.88E-02 | 2.54E-01 | 1.08E+00 | 3.02E-03 | 1.28E-02 | -1.11E+00 | 5.19E-02 | 1.98E-01 | 1.21E+00 | 2.72E-01 | 9.68E-01 | 1.53E+00 | 2.48E-01 | 6.99E-01 | 1.36E+00 | 9.56E-03 | 3.51E-02 | -1.29E+00 | 2.63E-01 | 7.23E-01 | 1.10E+00 |
| IGFBP6 | 5.99E-02 | 3.12E-01 | 1.53E+00 | 1.85E-02 | 7.99E-02 | 1.18E+00 | 1.37E-01 | 5.81E-01 | 1.50E+00 | 1.24E-01 | 4.75E-01 | 1.41E+00 | 1.39E-02 | 4.95E-02 | -1.08E+00 | 9.24E-03 | 2.60E-02 | -1.23E+00 | 2.39E-01 | 8.78E-01 | 1.02E+00 | 2.70E-01 | 7.42E-01 | -1.04E+00 |
| GIMAP8 | 5.99E-02 | 3.12E-01 | 1.61E+00 | 5.97E-05 | 2.58E-04 | 2.98E+00 | 2.15E-01 | 9.12E-01 | 1.32E+00 | 2.50E-01 | 9.54E-01 | 1.33E+00 | 2.63E-01 | 9.39E-01 | 1.04E+00 | 3.40E-01 | 9.58E-01 | 1.04E+00 | 2.53E-01 | 9.28E-01 | 1.00E+00 | 3.42E-01 | 9.40E-01 | -1.01E+00 |
| ZNF449 | 6.01E-02 | 3.13E-01 | 1.72E+00 | 1.97E-01 | 8.50E-01 | 1.48E+00 | 1.19E-02 | 5.04E-02 | -1.26E+00 | 4.52E-02 | 1.73E-01 | 1.06E+00 | 9.87E-03 | 3.52E-02 | -1.51E+00 | 2.39E-02 | 6.72E-02 | -1.35E+00 | 9.26E-02 | 3.40E-01 | -1.09E+00 | 1.72E-01 | 4.73E-01 | -1.18E+00 |
| AC016773.1 | 6.01E-02 | 3.13E-01 | 1.00E+00 | 3.86E-04 | 1.67E-03 | 2.82E+00 | 2.38E-02 | 1.01E-01 | -1.58E+00 | 3.54E-03 | 1.35E-02 | 2.46E+00 | 1.60E-01 | 5.72E-01 | -1.19E+00 | 4.96E-02 | 1.40E-01 | 1.90E+00 | 2.67E-01 | 9.79E-01 | 1.22E+00 | 2.08E-03 | 5.73E-03 | 2.31E+00 |
| GAS2L1 | 6.01E-02 | 3.13E-01 | 1.99E+00 | 1.44E-01 | 6.23E-01 | 1.73E+00 | 6.89E-02 | 2.93E-01 | 1.99E+00 | 2.14E-01 | 8.18E-01 | 1.52E+00 | 3.32E-16 | 1.18E-15 | 4.94E+00 | 8.20E-11 | 2.31E-10 | 3.84E+00 | 9.31E-11 | 3.42E-10 | 2.54E+00 | 1.70E-04 | 4.67E-04 | 1.79E+00 |
| GUSBP5 | 6.01E-02 | 3.13E-01 | -1.19E+00 | 1.44E-01 | 6.23E-01 | 1.14E+00 | 1.43E-01 | 6.10E-01 | 1.40E+00 | 3.59E-02 | 1.37E-01 | 2.21E+00 | 2.17E-01 | 7.73E-01 | 1.17E+00 | 1.90E-01 | 5.35E-01 | 1.34E+00 | 1.77E-03 | 6.48E-03 | -1.22E+00 | 3.30E-01 | 9.05E-01 | 1.29E+00 |
| FBXL12 | 6.01E-02 | 3.13E-01 | 1.24E+00 | 3.91E-02 | 1.69E-01 | 1.27E+00 | 2.01E-04 | 8.56E-04 | -1.19E+00 | 1.59E-04 | 6.07E-04 | -1.05E+00 | 7.75E-03 | 2.76E-02 | 1.11E+00 | 1.26E-03 | 3.56E-03 | -1.09E+00 | 3.74E-05 | 1.37E-04 | -1.19E+00 | 3.97E-05 | 1.09E-04 | -1.22E+00 |
| RAB11FIP4 | 6.02E-02 | 3.14E-01 | 1.18E+00 | 2.44E-03 | 1.06E-02 | -1.03E+00 | 2.24E-01 | 9.51E-01 | 1.24E+00 | 3.66E-02 | 1.40E-01 | -1.05E+00 | 2.05E-01 | 7.33E-01 | 1.11E+00 | 6.81E-02 | 1.92E-01 | -1.64E+00 | 2.19E-01 | 8.03E-01 | -1.11E+00 | 3.49E-01 | 9.59E-01 | 1.11E+00 |
| ZFP69B | 6.02E-02 | 3.14E-01 | -1.15E+00 | 6.18E-03 | 2.67E-02 | -1.41E+00 | 8.60E-02 | 3.65E-01 | -1.01E+00 | 5.36E-02 | 2.05E-01 | -1.09E+00 | 1.06E-01 | 3.78E-01 | 1.01E+00 | 1.84E-02 | 5.18E-02 | -1.50E+00 | 9.02E-04 | 3.31E-03 | -1.44E+00 | 3.36E-04 | 9.22E-04 | -1.69E+00 |
| RBM45 | 6.03E-02 | 3.14E-01 | -1.05E+00 | 1.36E-01 | 5.90E-01 | 1.54E+00 | 1.08E-06 | 4.61E-06 | -1.56E+00 | 1.55E-04 | 5.93E-04 | -1.15E+00 | 2.38E-03 | 8.48E-03 | -1.51E+00 | 1.63E-01 | 4.60E-01 | 1.06E+00 | 9.08E-03 | 3.33E-02 | -1.08E+00 | 1.19E-02 | 3.28E-02 | -1.10E+00 |
| LINC01678 | 6.03E-02 | 3.14E-01 | 1.05E+00 | 1.56E-01 | 6.72E-01 | 1.31E+00 | 1.78E-01 | 7.57E-01 | 1.22E+00 | 1.74E-01 | 6.63E-01 | 1.22E+00 | 1.27E-01 | 4.53E-01 | 1.76E+00 | 1.32E-01 | 3.73E-01 | 1.08E+00 | 5.96E-02 | 2.19E-01 | 2.08E+00 | 1.85E-01 | 5.09E-01 | 1.63E+00 |
| HSPE1P8 | 6.03E-02 | 3.14E-01 | -1.01E+00 | 2.32E-02 | 1.00E-01 | -1.23E+00 | 2.29E-02 | 9.74E-02 | -1.13E+00 | 8.17E-04 | 3.12E-03 | -1.46E+00 | 2.15E-01 | 7.67E-01 | -1.01E+00 | 3.19E-01 | 9.00E-01 | 1.11E+00 | 5.65E-03 | 2.07E-02 | -1.09E+00 | 2.00E-04 | 5.50E-04 | -1.42E+00 |
| VASH2 | 6.03E-02 | 3.14E-01 | -1.07E+00 | 4.15E-02 | 1.79E-01 | -1.15E+00 | 3.82E-02 | 1.62E-01 | -1.13E+00 | 1.14E-01 | 4.34E-01 | 1.11E+00 | 4.17E-07 | 1.49E-06 | -3.20E+00 | 1.25E-04 | 3.54E-04 | -1.95E+00 | 1.13E-04 | 4.14E-04 | -1.86E+00 | 5.75E-02 | 1.58E-01 | 1.01E+00 |
| NCK1 | 6.03E-02 | 3.14E-01 | 1.23E+00 | 1.79E-05 | 7.71E-05 | 1.00E+00 | 1.62E-02 | 6.88E-02 | 1.04E+00 | 3.44E-05 | 1.31E-04 | -1.03E+00 | 6.62E-02 | 2.36E-01 | 1.17E+00 | 1.19E-02 | 3.35E-02 | 1.03E+00 | 4.68E-05 | 1.72E-04 | -1.16E+00 | 6.97E-07 | 1.91E-06 | -1.30E+00 |
| AC090950.1 | 6.04E-02 | 3.15E-01 | 1.49E+00 | 1.17E-01 | 5.06E-01 | 1.56E+00 | 3.52E-02 | 1.49E-01 | 1.80E+00 | 2.50E-01 | 9.57E-01 | 1.38E+00 | 2.86E-03 | 1.02E-02 | 3.59E+00 | 2.48E-02 | 6.99E-02 | 2.80E+00 | 1.36E-03 | 4.98E-03 | 3.31E+00 | 6.11E-02 | 1.68E-01 | 2.27E+00 |
| MTPN | 6.04E-02 | 3.15E-01 | 1.07E+00 | 2.18E-02 | 9.41E-02 | 1.12E+00 | 1.27E-02 | 5.39E-02 | -1.02E+00 | 9.67E-04 | 3.69E-03 | -1.06E+00 | 5.72E-02 | 2.04E-01 | 1.05E+00 | 1.12E-03 | 3.16E-03 | -1.08E+00 | 2.06E-01 | 7.56E-01 | 1.04E+00 | 2.48E-02 | 6.81E-02 | -1.08E+00 |
| CYSLTR2 | 6.05E-02 | 3.15E-01 | 1.54E+00 | 1.92E-01 | 8.31E-01 | 1.44E+00 | 7.98E-02 | 3.39E-01 | 1.19E+00 | 1.02E-01 | 3.89E-01 | 1.61E+00 | 1.53E-02 | 5.45E-02 | 3.76E+00 | 4.01E-02 | 1.13E-01 | 2.90E+00 | 7.44E-02 | 2.73E-01 | 1.76E+00 | 3.56E-01 | 9.77E-01 | 1.20E+00 |
| SEMA7A | 6.05E-02 | 3.15E-01 | 1.03E+00 | 8.24E-04 | 3.56E-03 | -1.64E+00 | 3.21E-02 | 1.36E-01 | -1.27E+00 | 4.85E-02 | 1.85E-01 | -1.15E+00 | 7.12E-12 | 2.54E-11 | -1.94E+00 | 1.24E-13 | 3.50E-13 | -2.68E+00 | 4.90E-03 | 1.80E-02 | -1.79E+00 | 2.35E-02 | 6.47E-02 | -1.49E+00 |
| ARMCX4 | 6.05E-02 | 3.15E-01 | -1.09E+00 | 2.10E-01 | 9.06E-01 | 1.21E+00 | 7.08E-02 | 3.01E-01 | 1.01E+00 | 4.18E-02 | 1.60E-01 | 1.02E+00 | 9.73E-02 | 3.47E-01 | 1.11E+00 | 1.39E-01 | 3.90E-01 | 1.13E+00 | 1.46E-01 | 5.36E-01 | 1.15E+00 | 1.71E-01 | 4.71E-01 | 1.04E+00 |
| AL355472.1 | 6.06E-02 | 3.16E-01 | 1.18E+00 | 1.01E-01 | 4.37E-01 | 1.35E+00 | 1.88E-01 | 8.00E-01 | 1.33E+00 | 1.21E-01 | 4.64E-01 | 1.21E+00 | 1.56E-01 | 5.56E-01 | 1.75E+00 | 3.08E-01 | 8.68E-01 | 1.38E+00 | 3.49E-02 | 1.28E-01 | 1.51E+00 | 2.48E-01 | 6.82E-01 | 1.36E+00 |
| ETFB | 6.06E-02 | 3.16E-01 | 1.11E+00 | 2.84E-03 | 1.23E-02 | 1.12E+00 | 2.04E-01 | 8.66E-01 | 1.24E+00 | 2.59E-02 | 9.89E-02 | 1.14E+00 | 2.82E-03 | 1.01E-02 | -1.01E+00 | 5.01E-03 | 1.41E-02 | 1.03E+00 | 1.95E-01 | 7.16E-01 | 1.30E+00 | 3.07E-01 | 8.43E-01 | 1.31E+00 |
| CCDC134 | 6.06E-02 | 3.16E-01 | -1.12E+00 | 4.55E-02 | 1.97E-01 | -1.19E+00 | 1.42E-01 | 6.05E-01 | -1.00E+00 | 2.27E-02 | 8.65E-02 | -1.33E+00 | 8.22E-02 | 2.93E-01 | -1.38E+00 | 3.52E-01 | 9.93E-01 | 1.18E+00 | 1.54E-01 | 5.65E-01 | 1.11E+00 | 2.64E-01 | 7.25E-01 | 1.21E+00 |
| MTHFD1L | 6.07E-02 | 3.16E-01 | 1.15E+00 | 1.31E-02 | 5.64E-02 | 1.12E+00 | 1.88E-01 | 7.98E-01 | 1.28E+00 | 4.47E-02 | 1.71E-01 | 1.34E+00 | 2.08E-01 | 7.41E-01 | 1.47E+00 | 3.41E-01 | 9.62E-01 | 1.29E+00 | 2.68E-01 | 9.84E-01 | 1.29E+00 | 1.06E-01 | 2.90E-01 | 1.56E+00 |
| MIATNB | 6.07E-02 | 3.16E-01 | -1.01E+00 | 1.29E-01 | 5.56E-01 | 1.19E+00 | 3.91E-02 | 1.66E-01 | -1.04E+00 | 2.03E-02 | 7.75E-02 | 1.03E+00 | 1.60E-01 | 5.71E-01 | 1.20E+00 | 3.07E-01 | 8.67E-01 | 1.31E+00 | 1.42E-01 | 5.20E-01 | 1.46E+00 | 2.71E-01 | 7.45E-01 | 1.28E+00 |
| PSAT1 | 6.08E-02 | 3.17E-01 | -1.67E+00 | 1.95E-01 | 8.44E-01 | -1.16E+00 | 6.12E-05 | 2.60E-04 | -2.68E+00 | 1.09E-03 | 4.18E-03 | -1.85E+00 | 7.49E-02 | 2.67E-01 | -1.64E+00 | 8.80E-02 | 2.48E-01 | -1.70E+00 | 5.63E-03 | 2.07E-02 | -1.43E+00 | 1.88E-01 | 5.15E-01 | 1.07E+00 |
| BRIP1 | 6.10E-02 | 3.18E-01 | -1.15E+00 | 5.20E-02 | 2.25E-01 | -1.24E+00 | 1.61E-01 | 6.84E-01 | 1.35E+00 | 2.50E-01 | 9.57E-01 | 1.45E+00 | 3.02E-02 | 1.08E-01 | -1.61E+00 | 6.04E-03 | 1.70E-02 | -2.50E+00 | 7.86E-09 | 2.88E-08 | -1.47E+00 | 1.38E-02 | 3.79E-02 | 1.01E+00 |
| ATF4P3 | 6.11E-02 | 3.18E-01 | 1.17E+00 | 2.31E-01 | 1.00E+00 | 1.38E+00 | 9.63E-04 | 4.09E-03 | -1.06E+00 | 3.95E-03 | 1.51E-02 | 1.12E+00 | 2.53E-03 | 9.03E-03 | -1.03E+00 | 3.41E-01 | 9.62E-01 | 1.22E+00 | 7.41E-02 | 2.72E-01 | 1.13E+00 | 3.17E-01 | 8.69E-01 | 1.27E+00 |
| COX20P2 | 6.11E-02 | 3.18E-01 | 1.11E+00 | 3.55E-03 | 1.53E-02 | -1.26E+00 | 1.51E-01 | 6.43E-01 | 1.17E+00 | 7.13E-04 | 2.72E-03 | -1.34E+00 | 7.23E-03 | 2.58E-02 | -1.11E+00 | 4.11E-03 | 1.16E-02 | -1.25E+00 | 1.62E-04 | 5.94E-04 | -1.11E+00 | 2.05E-04 | 5.62E-04 | -1.08E+00 |
| LRR1 | 6.11E-02 | 3.18E-01 | 1.08E+00 | 1.25E-01 | 5.41E-01 | 1.29E+00 | 5.62E-02 | 2.39E-01 | 1.07E+00 | 4.74E-02 | 1.81E-01 | 1.22E+00 | 6.11E-02 | 2.18E-01 | -1.04E+00 | 2.00E-01 | 5.64E-01 | 1.11E+00 | 6.90E-05 | 2.53E-04 | -1.26E+00 | 1.88E-01 | 5.16E-01 | 1.19E+00 |
| PJVK | 6.14E-02 | 3.20E-01 | 1.04E+00 | 1.16E-02 | 5.02E-02 | -1.20E+00 | 1.41E-01 | 5.99E-01 | 1.01E+00 | 2.00E-01 | 7.63E-01 | 1.15E+00 | 1.76E-01 | 6.27E-01 | -1.02E+00 | 3.43E-01 | 9.67E-01 | 1.23E+00 | 1.41E-01 | 5.17E-01 | 1.04E+00 | 9.12E-02 | 2.50E-01 | -1.19E+00 |
| AL117379.1 | 6.14E-02 | 3.20E-01 | 1.09E+00 | 3.73E-02 | 1.61E-01 | 2.08E+00 | 6.95E-02 | 2.95E-01 | -1.16E+00 | 1.21E-01 | 4.61E-01 | 1.74E+00 | 2.46E-01 | 8.76E-01 | 1.44E+00 | 1.79E-01 | 5.04E-01 | 1.89E+00 | 1.76E-02 | 6.44E-02 | 2.34E+00 | 2.16E-02 | 5.93E-02 | 2.30E+00 |
| CCDC71L | 6.14E-02 | 3.20E-01 | 1.27E+00 | 1.36E-03 | 5.87E-03 | 1.17E+00 | 7.25E-02 | 3.08E-01 | 1.39E+00 | 1.73E-01 | 6.60E-01 | 1.47E+00 | 6.75E-02 | 2.41E-01 | 1.67E+00 | 2.00E-01 | 5.63E-01 | 1.22E+00 | 1.50E-01 | 5.51E-01 | 1.46E+00 | 1.15E-01 | 3.17E-01 | 1.19E+00 |
| NTAN1 | 6.14E-02 | 3.20E-01 | -1.02E+00 | 1.39E-01 | 6.01E-01 | 1.25E+00 | 1.35E-04 | 5.72E-04 | -1.51E+00 | 1.76E-02 | 6.74E-02 | -1.01E+00 | 1.19E-06 | 4.23E-06 | -2.19E+00 | 1.32E-06 | 3.71E-06 | -2.14E+00 | 1.37E-06 | 5.04E-06 | -1.50E+00 | 1.86E-06 | 5.12E-06 | -1.56E+00 |
| ARNTL | 6.15E-02 | 3.21E-01 | -1.14E+00 | 8.34E-02 | 3.61E-01 | -1.01E+00 | 6.47E-02 | 2.75E-01 | -1.15E+00 | 5.10E-02 | 1.95E-01 | -1.15E+00 | 5.48E-02 | 1.95E-01 | 1.19E+00 | 4.90E-02 | 1.38E-01 | 1.01E+00 | 3.61E-03 | 1.32E-02 | -1.30E+00 | 2.13E-03 | 5.85E-03 | -1.49E+00 |
| AC007613.1 | 6.15E-02 | 3.21E-01 | 1.47E+00 | 2.13E-01 | 9.19E-01 | 1.19E+00 | 1.63E-02 | 6.91E-02 | 1.82E+00 | 2.32E-01 | 8.87E-01 | 1.40E+00 | 4.40E-02 | 1.57E-01 | -1.54E+00 | 6.74E-02 | 1.90E-01 | -1.51E+00 | 2.42E-01 | 8.89E-01 | 1.21E+00 | 2.44E-01 | 6.70E-01 | 1.11E+00 |
| TRIM28 | 6.16E-02 | 3.21E-01 | 1.23E+00 | 2.88E-02 | 1.25E-01 | 1.27E+00 | 7.71E-03 | 3.27E-02 | 1.40E+00 | 9.68E-02 | 3.70E-01 | 1.46E+00 | 6.42E-02 | 2.29E-01 | 1.29E+00 | 4.08E-02 | 1.15E-01 | 1.28E+00 | 1.40E-01 | 5.13E-01 | 1.36E+00 | 1.23E-02 | 3.37E-02 | 1.53E+00 |
| AC135626.1 | 6.16E-02 | 3.21E-01 | 1.71E+00 | 2.13E-01 | 9.20E-01 | 1.32E+00 | 1.10E-01 | 4.68E-01 | -1.00E+00 | 2.42E-01 | 9.23E-01 | 1.24E+00 | 9.65E-03 | 3.44E-02 | 3.58E+00 | 2.61E-02 | 7.34E-02 | 2.89E+00 | 2.42E-01 | 8.87E-01 | 1.10E+00 | 1.66E-01 | 4.55E-01 | -1.06E+00 |
| RNU4-2 | 6.16E-02 | 3.21E-01 | 1.55E+00 | 9.47E-05 | 4.09E-04 | -2.45E+00 | 2.12E-02 | 9.02E-02 | -1.10E+00 | 1.23E-05 | 4.68E-05 | -2.13E+00 | 7.04E-02 | 2.51E-01 | 2.47E+00 | 2.60E-01 | 7.33E-01 | 1.37E+00 | 1.31E-01 | 4.82E-01 | -1.20E+00 | 1.57E-01 | 4.32E-01 | -1.26E+00 |
| AL356968.2 | 6.17E-02 | 3.22E-01 | -1.02E+00 | 8.94E-02 | 3.86E-01 | 1.05E+00 | 1.27E-01 | 5.40E-01 | 1.34E+00 | 1.75E-01 | 6.70E-01 | 1.02E+00 | 1.19E-01 | 4.25E-01 | -1.06E+00 | 1.87E-01 | 5.28E-01 | 1.02E+00 | 3.97E-04 | 1.46E-03 | -1.06E+00 | 1.21E-04 | 3.33E-04 | -1.26E+00 |
| AL354707.1 | 6.18E-02 | 3.22E-01 | 1.14E+00 | 2.24E-01 | 9.70E-01 | 1.41E+00 | 1.69E-02 | 7.17E-02 | -1.20E+00 | 1.37E-01 | 5.25E-01 | 1.12E+00 | 2.20E-02 | 7.84E-02 | -1.00E+00 | 2.42E-01 | 6.82E-01 | 1.22E+00 | 1.20E-01 | 4.41E-01 | 1.32E+00 | 2.58E-01 | 7.09E-01 | 1.15E+00 |
| MT1L | 6.18E-02 | 3.22E-01 | -1.33E+00 | 5.28E-02 | 2.28E-01 | -1.50E+00 | 2.18E-01 | 9.25E-01 | -1.03E+00 | 2.30E-01 | 8.79E-01 | -1.03E+00 | 1.46E-01 | 5.21E-01 | -1.23E+00 | 1.95E-01 | 5.51E-01 | -1.23E+00 | 6.13E-07 | 2.25E-06 | -1.58E+00 | 1.73E-09 | 4.74E-09 | -1.88E+00 |
| ZNF569 | 6.18E-02 | 3.22E-01 | 1.04E+00 | 1.35E-01 | 5.85E-01 | 1.70E+00 | 2.37E-02 | 1.01E-01 | -1.16E+00 | 3.86E-02 | 1.47E-01 | 1.04E+00 | 3.76E-03 | 1.34E-02 | -1.31E+00 | 3.06E-02 | 8.63E-02 | -1.00E+00 | 1.99E-03 | 7.30E-03 | -1.24E+00 | 5.25E-03 | 1.44E-02 | -1.25E+00 |
| SLC2A3 | 6.18E-02 | 3.22E-01 | 1.98E+00 | 1.83E-01 | 7.92E-01 | 1.01E+00 | 9.40E-02 | 3.99E-01 | 1.77E+00 | 1.02E-01 | 3.90E-01 | -1.31E+00 | 1.55E-01 | 5.54E-01 | -1.20E+00 | 5.32E-03 | 1.50E-02 | -2.10E+00 | 1.16E-04 | 4.25E-04 | 1.88E+00 | 3.80E-02 | 1.04E-01 | 1.11E+00 |
| PRKCH | 6.19E-02 | 3.22E-01 | 1.19E+00 | 4.70E-02 | 2.03E-01 | 1.62E+00 | 1.57E-01 | 6.69E-01 | 1.37E+00 | 5.47E-02 | 2.09E-01 | 1.69E+00 | 2.44E-01 | 8.71E-01 | 1.34E+00 | 1.31E-01 | 3.68E-01 | 1.04E+00 | 1.86E-01 | 6.84E-01 | 1.15E+00 | 2.81E-01 | 7.73E-01 | 1.20E+00 |
| AP001363.1 | 6.19E-02 | 3.22E-01 | 1.56E+00 | 6.09E-02 | 2.63E-01 | -1.11E+00 | 4.55E-03 | 1.93E-02 | 1.86E+00 | 1.71E-01 | 6.53E-01 | 1.06E+00 | 2.88E-02 | 1.03E-01 | 2.50E+00 | 9.73E-02 | 2.74E-01 | -1.10E+00 | 1.74E-01 | 6.40E-01 | 1.30E+00 | 3.33E-01 | 9.14E-01 | 1.16E+00 |
| AC025048.2 | 6.19E-02 | 3.23E-01 | -1.04E+00 | 2.02E-01 | 8.71E-01 | 1.26E+00 | 1.22E-01 | 5.18E-01 | -1.08E+00 | 2.58E-01 | 9.84E-01 | 1.23E+00 | 4.59E-02 | 1.64E-01 | -1.43E+00 | 6.39E-02 | 1.80E-01 | -1.45E+00 | 1.02E-01 | 3.75E-01 | -1.14E+00 | 3.01E-01 | 8.26E-01 | 1.10E+00 |
| AL033527.5 | 6.19E-02 | 3.23E-01 | -1.13E+00 | 2.26E-01 | 9.77E-01 | 1.26E+00 | 7.06E-02 | 3.00E-01 | -1.09E+00 | 2.19E-01 | 8.38E-01 | 1.45E+00 | 1.20E-01 | 4.28E-01 | -1.08E+00 | 2.50E-01 | 7.04E-01 | 1.63E+00 | 1.23E-01 | 4.52E-01 | 1.03E+00 | 1.81E-01 | 4.96E-01 | -1.05E+00 |
| AC005261.4 | 6.19E-02 | 3.23E-01 | 1.53E+00 | 1.41E-01 | 6.10E-01 | 1.56E+00 | 1.53E-01 | 6.49E-01 | 1.40E+00 | 2.08E-01 | 7.96E-01 | 1.34E+00 | 2.60E-01 | 9.27E-01 | 1.36E+00 | 2.63E-01 | 7.41E-01 | 1.14E+00 | 1.72E-01 | 6.32E-01 | 1.10E+00 | 2.19E-01 | 6.01E-01 | 1.04E+00 |
| CENPU | 6.20E-02 | 3.23E-01 | 1.39E+00 | 2.28E-01 | 9.85E-01 | 1.38E+00 | 1.75E-02 | 7.43E-02 | 1.41E+00 | 5.38E-02 | 2.06E-01 | 1.18E+00 | 1.43E-01 | 5.11E-01 | 1.26E+00 | 2.12E-01 | 5.99E-01 | 1.32E+00 | 2.71E-01 | 9.96E-01 | 1.21E+00 | 1.63E-01 | 4.47E-01 | 1.14E+00 |
| C19orf73 | 6.20E-02 | 3.23E-01 | -1.02E+00 | 1.05E-03 | 4.52E-03 | -1.50E+00 | 8.69E-02 | 3.69E-01 | 1.06E+00 | 1.30E-02 | 4.95E-02 | -1.09E+00 | 2.21E-01 | 7.88E-01 | 1.09E+00 | 2.41E-01 | 6.78E-01 | 1.54E+00 | 1.23E-03 | 4.53E-03 | -1.41E+00 | 1.93E-02 | 5.29E-02 | -1.08E+00 |
| UBR1 | 6.20E-02 | 3.23E-01 | 1.17E+00 | 1.74E-03 | 7.52E-03 | -1.03E+00 | 2.07E-03 | 8.79E-03 | -1.24E+00 | 3.80E-03 | 1.45E-02 | -1.10E+00 | 2.55E-01 | 9.11E-01 | 1.47E+00 | 3.10E-01 | 8.75E-01 | 1.44E+00 | 1.44E-01 | 5.29E-01 | 1.18E+00 | 3.41E-01 | 9.37E-01 | 1.22E+00 |
| AOX2P | 6.20E-02 | 3.23E-01 | 1.68E+00 | 1.99E-03 | 8.59E-03 | 2.57E+00 | 1.39E-02 | 5.90E-02 | -1.32E+00 | 1.66E-01 | 6.34E-01 | 1.43E+00 | 1.74E-01 | 6.19E-01 | 1.82E+00 | 3.37E-01 | 9.50E-01 | 1.35E+00 | 2.67E-01 | 9.81E-01 | 1.01E+00 | 2.45E-01 | 6.73E-01 | 1.35E+00 |
| ZNF773 | 6.21E-02 | 3.24E-01 | 1.11E+00 | 5.04E-04 | 2.18E-03 | -1.25E+00 | 3.68E-02 | 1.56E-01 | -1.05E+00 | 1.98E-02 | 7.55E-02 | 1.02E+00 | 3.25E-03 | 1.16E-02 | -1.22E+00 | 1.56E-04 | 4.41E-04 | -1.65E+00 | 2.36E-01 | 8.65E-01 | 1.18E+00 | 5.98E-02 | 1.64E-01 | -1.16E+00 |
| FLJ21408 | 6.22E-02 | 3.24E-01 | 1.49E+00 | 7.97E-02 | 3.44E-01 | -1.13E+00 | 8.02E-02 | 3.41E-01 | 1.60E+00 | 2.12E-01 | 8.09E-01 | 1.31E+00 | 6.78E-02 | 2.42E-01 | 1.71E+00 | 2.11E-01 | 5.94E-01 | 1.04E+00 | 7.91E-02 | 2.90E-01 | 1.33E+00 | 3.17E-01 | 8.71E-01 | -1.02E+00 |
| TIGD6 | 6.22E-02 | 3.24E-01 | -1.04E+00 | 2.15E-01 | 9.31E-01 | 1.29E+00 | 1.30E-01 | 5.54E-01 | 1.06E+00 | 2.52E-01 | 9.62E-01 | 1.49E+00 | 2.21E-01 | 7.88E-01 | 1.19E+00 | 6.04E-02 | 1.70E-01 | 1.93E+00 | 2.61E-01 | 9.59E-01 | 1.39E+00 | 2.26E-01 | 6.20E-01 | 1.45E+00 |
| FAM207A | 6.23E-02 | 3.25E-01 | 1.57E+00 | 1.18E-01 | 5.09E-01 | 1.70E+00 | 4.68E-03 | 1.99E-02 | 1.69E+00 | 2.50E-01 | 9.54E-01 | 1.38E+00 | 4.04E-02 | 1.44E-01 | 2.16E+00 | 8.51E-03 | 2.40E-02 | 2.61E+00 | 2.32E-05 | 8.51E-05 | 2.01E+00 | 2.21E-02 | 6.06E-02 | 1.61E+00 |
| AC092171.3 | 6.23E-02 | 3.25E-01 | 1.10E+00 | 1.22E-02 | 5.26E-02 | 1.03E+00 | 8.92E-03 | 3.79E-02 | -1.23E+00 | 1.13E-01 | 4.32E-01 | 1.18E+00 | 1.48E-01 | 5.28E-01 | -1.02E+00 | 8.68E-02 | 2.45E-01 | -1.23E+00 | 3.18E-02 | 1.17E-01 | -1.83E+00 | 1.25E-01 | 3.42E-01 | -1.34E+00 |
| GNAI3 | 6.23E-02 | 3.25E-01 | 1.31E+00 | 2.80E-06 | 1.21E-05 | 1.08E+00 | 1.33E-01 | 5.67E-01 | 1.25E+00 | 2.40E-02 | 9.18E-02 | 1.08E+00 | 3.59E-02 | 1.28E-01 | 1.19E+00 | 4.11E-03 | 1.16E-02 | 1.04E+00 | 6.09E-02 | 2.24E-01 | 1.07E+00 | 5.77E-04 | 1.58E-03 | -1.10E+00 |
| ARMCX5 | 6.23E-02 | 3.25E-01 | 1.09E+00 | 1.75E-02 | 7.58E-02 | 1.08E+00 | 1.07E-04 | 4.54E-04 | -1.40E+00 | 5.05E-03 | 1.93E-02 | -1.06E+00 | 4.03E-02 | 1.44E-01 | -1.06E+00 | 1.90E-01 | 5.37E-01 | 1.21E+00 | 1.77E-03 | 6.50E-03 | -1.14E+00 | 2.04E-02 | 5.62E-02 | -1.03E+00 |
| AC021188.1 | 6.23E-02 | 3.25E-01 | -1.00E+00 | 8.25E-02 | 3.56E-01 | 1.04E+00 | 1.07E-03 | 4.56E-03 | -1.56E+00 | 3.46E-03 | 1.32E-02 | -1.24E+00 | 9.70E-02 | 3.46E-01 | -1.04E+00 | 2.70E-01 | 7.62E-01 | 1.30E+00 | 2.65E-02 | 9.72E-02 | -1.35E+00 | 6.17E-02 | 1.70E-01 | -1.15E+00 |
| FZD6 | 6.23E-02 | 3.25E-01 | 1.20E+00 | 2.54E-02 | 1.10E-01 | 1.19E+00 | 1.29E-01 | 5.50E-01 | 1.49E+00 | 7.15E-02 | 2.73E-01 | 1.17E+00 | 1.97E-01 | 7.02E-01 | 1.32E+00 | 2.19E-01 | 6.17E-01 | 1.34E+00 | 4.74E-02 | 1.74E-01 | 1.47E+00 | 3.33E-01 | 9.14E-01 | 1.23E+00 |
| CTU1 | 6.24E-02 | 3.25E-01 | 1.63E+00 | 1.52E-01 | 6.59E-01 | 1.75E+00 | 2.34E-01 | 9.95E-01 | 1.27E+00 | 1.74E-01 | 6.66E-01 | 1.54E+00 | 3.63E-03 | 1.29E-02 | 2.97E+00 | 2.08E-03 | 5.85E-03 | 3.28E+00 | 2.16E-01 | 7.92E-01 | 1.19E+00 | 2.29E-01 | 6.30E-01 | 1.40E+00 |
| SELENOP | 6.24E-02 | 3.25E-01 | 1.31E+00 | 2.27E-02 | 9.80E-02 | 1.23E+00 | 1.19E-01 | 5.08E-01 | 1.35E+00 | 1.53E-01 | 5.84E-01 | 1.21E+00 | 1.63E-01 | 5.80E-01 | 1.14E+00 | 1.59E-01 | 4.47E-01 | -1.09E+00 | 5.47E-02 | 2.01E-01 | 1.81E+00 | 7.57E-02 | 2.08E-01 | 1.84E+00 |
| AC092652.1 | 6.24E-02 | 3.25E-01 | 1.03E+00 | 2.19E-01 | 9.48E-01 | 1.41E+00 | 1.49E-02 | 6.34E-02 | -1.18E+00 | 3.42E-02 | 1.31E-01 | -1.05E+00 | 1.49E-02 | 5.30E-02 | -1.37E+00 | 7.41E-02 | 2.09E-01 | -1.20E+00 | 3.75E-04 | 1.38E-03 | -1.65E+00 | 2.09E-02 | 5.74E-02 | -1.19E+00 |
| MYL5 | 6.24E-02 | 3.25E-01 | 1.07E+00 | 4.20E-02 | 1.81E-01 | 1.08E+00 | 1.64E-01 | 6.96E-01 | 1.18E+00 | 7.69E-02 | 2.94E-01 | 1.14E+00 | 2.58E-01 | 9.22E-01 | 1.30E+00 | 9.74E-02 | 2.75E-01 | 1.08E+00 | 2.02E-01 | 7.41E-01 | 1.47E+00 | 3.52E-01 | 9.66E-01 | 1.30E+00 |
| PCDHGA5 | 6.24E-02 | 3.25E-01 | 1.13E+00 | 1.76E-01 | 7.62E-01 | 1.39E+00 | 1.56E-01 | 6.65E-01 | -1.08E+00 | 1.79E-01 | 6.83E-01 | 1.47E+00 | 2.47E-01 | 8.81E-01 | 1.29E+00 | 3.15E-01 | 8.89E-01 | 1.23E+00 | 2.47E-01 | 9.06E-01 | 1.17E+00 | 3.60E-01 | 9.88E-01 | 1.37E+00 |
| ZNF674 | 6.24E-02 | 3.25E-01 | 1.00E+00 | 1.67E-01 | 7.22E-01 | 1.25E+00 | 7.32E-04 | 3.11E-03 | -1.77E+00 | 1.24E-02 | 4.73E-02 | -1.25E+00 | 1.58E-01 | 5.64E-01 | 1.07E+00 | 1.50E-01 | 4.22E-01 | 1.05E+00 | 9.33E-02 | 3.42E-01 | -1.35E+00 | 1.80E-01 | 4.94E-01 | -1.16E+00 |
| AC005520.2 | 6.25E-02 | 3.25E-01 | 1.69E+00 | 1.70E-01 | 7.33E-01 | 1.65E+00 | 1.17E-02 | 4.98E-02 | -1.38E+00 | 7.62E-03 | 2.91E-02 | -1.39E+00 | 2.51E-01 | 8.96E-01 | 1.17E+00 | 1.81E-01 | 5.10E-01 | -1.07E+00 | 1.05E-01 | 3.87E-01 | 1.50E+00 | 1.74E-01 | 4.77E-01 | 1.05E+00 |
| C2CD2L | 6.26E-02 | 3.26E-01 | 1.10E+00 | 9.05E-02 | 3.91E-01 | 1.23E+00 | 6.28E-02 | 2.67E-01 | -1.25E+00 | 1.48E-01 | 5.64E-01 | -1.01E+00 | 9.63E-02 | 3.43E-01 | -1.22E+00 | 6.02E-02 | 1.70E-01 | -1.59E+00 | 1.75E-01 | 6.43E-01 | 1.16E+00 | 3.04E-01 | 8.35E-01 | 1.16E+00 |
| DLGAP1-AS1 | 6.26E-02 | 3.26E-01 | 1.12E+00 | 1.19E-01 | 5.15E-01 | 1.26E+00 | 1.50E-03 | 6.37E-03 | -1.67E+00 | 1.73E-03 | 6.62E-03 | -1.53E+00 | 9.31E-06 | 3.32E-05 | -2.01E+00 | 1.09E-02 | 3.08E-02 | -1.11E+00 | 2.50E-02 | 9.16E-02 | -1.11E+00 | 5.80E-04 | 1.59E-03 | -1.57E+00 |
| FAM239A | 6.26E-02 | 3.26E-01 | 1.06E+00 | 2.24E-03 | 9.67E-03 | -1.39E+00 | 4.80E-02 | 2.04E-01 | -1.11E+00 | 3.13E-02 | 1.19E-01 | -1.11E+00 | 1.13E-01 | 4.03E-01 | -1.14E+00 | 1.72E-01 | 4.84E-01 | 1.05E+00 | 9.96E-02 | 3.65E-01 | -1.17E+00 | 1.81E-01 | 4.96E-01 | 1.34E+00 |
| AC005498.3 | 6.26E-02 | 3.27E-01 | -1.03E+00 | 1.81E-01 | 7.80E-01 | 1.27E+00 | 2.13E-04 | 9.04E-04 | -1.78E+00 | 1.45E-03 | 5.54E-03 | -1.51E+00 | 4.33E-03 | 1.54E-02 | -2.08E+00 | 9.48E-02 | 2.67E-01 | -1.18E+00 | 2.15E-01 | 7.88E-01 | 1.24E+00 | 3.02E-01 | 8.29E-01 | 1.09E+00 |
| BBC3 | 6.27E-02 | 3.27E-01 | 1.18E+00 | 5.51E-04 | 2.38E-03 | -1.40E+00 | 2.76E-03 | 1.17E-02 | -1.42E+00 | 1.86E-04 | 7.09E-04 | -1.71E+00 | 7.50E-04 | 2.67E-03 | -1.15E+00 | 2.65E-05 | 7.47E-05 | -1.56E+00 | 1.01E-26 | 3.71E-26 | -1.08E+00 | 4.09E-39 | 1.12E-38 | -2.20E+00 |
| PLGLB2 | 6.27E-02 | 3.27E-01 | 1.64E+00 | 9.78E-02 | 4.23E-01 | 1.18E+00 | 1.50E-01 | 6.37E-01 | 1.32E+00 | 6.03E-03 | 2.30E-02 | -1.34E+00 | 2.48E-01 | 8.86E-01 | 1.19E+00 | 6.84E-02 | 1.93E-01 | -1.48E+00 | 1.01E-01 | 3.71E-01 | 1.03E+00 | 4.02E-03 | 1.10E-02 | -1.94E+00 |
| CCL23 | 6.29E-02 | 3.28E-01 | -1.19E+00 | 2.81E-02 | 1.22E-01 | -1.53E+00 | 3.22E-02 | 1.37E-01 | -1.06E+00 | 3.81E-04 | 1.46E-03 | -1.56E+00 | 5.07E-07 | 1.81E-06 | -1.25E+00 | 7.21E-09 | 2.03E-08 | -1.47E+00 | 5.49E-03 | 2.02E-02 | -1.21E+00 | 1.44E-01 | 3.96E-01 | 1.05E+00 |
| ARHGAP19 | 6.29E-02 | 3.28E-01 | -1.07E+00 | 8.58E-02 | 3.71E-01 | 1.03E+00 | 1.21E-01 | 5.13E-01 | 1.06E+00 | 1.27E-01 | 4.86E-01 | 1.08E+00 | 1.96E-01 | 7.00E-01 | 1.17E+00 | 3.42E-01 | 9.64E-01 | 1.34E+00 | 9.87E-02 | 3.62E-01 | 1.12E+00 | 2.04E-01 | 5.59E-01 | 1.27E+00 |
| CDKN2AIPNL | 6.29E-02 | 3.28E-01 | 1.27E+00 | 4.40E-03 | 1.90E-02 | 1.24E+00 | 1.86E-01 | 7.92E-01 | 1.19E+00 | 6.83E-02 | 2.61E-01 | 1.16E+00 | 1.79E-02 | 6.39E-02 | -1.01E+00 | 1.60E-01 | 4.52E-01 | 1.14E+00 | 1.15E-03 | 4.24E-03 | -1.04E+00 | 6.61E-03 | 1.82E-02 | 1.04E+00 |
| AL356000.1 | 6.29E-02 | 3.28E-01 | 1.71E+00 | 1.57E-01 | 6.78E-01 | 1.50E+00 | 7.52E-02 | 3.19E-01 | 1.43E+00 | 2.52E-01 | 9.63E-01 | 1.23E+00 | 2.08E-01 | 7.41E-01 | -1.10E+00 | 2.16E-01 | 6.09E-01 | 1.49E+00 | 1.14E-02 | 4.17E-02 | 1.79E+00 | 1.85E-01 | 5.07E-01 | 1.45E+00 |
| SRSF3 | 6.29E-02 | 3.28E-01 | -1.01E+00 | 7.56E-02 | 3.27E-01 | 1.17E+00 | 1.47E-02 | 6.26E-02 | -1.09E+00 | 2.54E-01 | 9.70E-01 | 1.04E+00 | 2.16E-13 | 7.72E-13 | -1.42E+00 | 6.74E-02 | 1.90E-01 | -1.00E+00 | 1.03E-06 | 3.78E-06 | -1.25E+00 | 3.33E-01 | 9.15E-01 | 1.04E+00 |
| GUK1 | 6.30E-02 | 3.28E-01 | 1.14E+00 | 1.85E-04 | 7.98E-04 | 1.13E+00 | 2.11E-01 | 8.96E-01 | 1.13E+00 | 3.68E-02 | 1.41E-01 | 1.11E+00 | 8.62E-03 | 3.07E-02 | 1.08E+00 | 6.27E-03 | 1.77E-02 | 1.05E+00 | 1.96E-01 | 7.19E-01 | 1.16E+00 | 3.56E-01 | 9.77E-01 | 1.08E+00 |
| C1orf198 | 6.30E-02 | 3.28E-01 | -1.03E+00 | 2.89E-02 | 1.25E-01 | -1.08E+00 | 3.86E-02 | 1.64E-01 | -1.08E+00 | 7.82E-03 | 2.99E-02 | -1.09E+00 | 1.24E-01 | 4.42E-01 | -1.11E+00 | 2.77E-01 | 7.82E-01 | 1.27E+00 | 6.21E-04 | 2.28E-03 | -1.20E+00 | 1.39E-05 | 3.81E-05 | -1.60E+00 |
| ALOX5 | 6.30E-02 | 3.28E-01 | 1.69E+00 | 7.41E-02 | 3.20E-01 | 1.77E+00 | 1.84E-01 | 7.83E-01 | 1.34E+00 | 9.52E-02 | 3.64E-01 | 1.73E+00 | 2.76E-01 | 9.86E-01 | 1.49E+00 | 3.55E-01 | 1.00E+00 | 1.43E+00 | 2.64E-01 | 9.68E-01 | 1.08E+00 | 1.62E-01 | 4.46E-01 | 2.47E+00 |
| SZRD1 | 6.30E-02 | 3.29E-01 | 1.17E+00 | 8.70E-03 | 3.76E-02 | 1.14E+00 | 1.09E-01 | 4.63E-01 | 1.16E+00 | 2.39E-02 | 9.12E-02 | 1.18E+00 | 1.66E-03 | 5.92E-03 | 1.04E+00 | 4.05E-05 | 1.14E-04 | -1.12E+00 | 2.42E-02 | 8.89E-02 | 1.12E+00 | 3.41E-02 | 9.36E-02 | 1.15E+00 |
| BRMS1L | 6.31E-02 | 3.29E-01 | 1.06E+00 | 1.29E-01 | 5.55E-01 | 1.26E+00 | 5.86E-03 | 2.49E-02 | -1.42E+00 | 7.27E-02 | 2.78E-01 | 1.03E+00 | 2.18E-01 | 7.79E-01 | 1.73E+00 | 3.14E-01 | 8.85E-01 | 1.48E+00 | 6.92E-04 | 2.54E-03 | -1.07E+00 | 5.46E-02 | 1.50E-01 | 1.08E+00 |
| LINC00173 | 6.31E-02 | 3.29E-01 | 1.46E+00 | 2.15E-03 | 9.28E-03 | 1.84E+00 | 1.35E-03 | 5.75E-03 | -1.19E+00 | 2.61E-01 | 9.96E-01 | 1.27E+00 | 6.32E-02 | 2.25E-01 | 1.25E+00 | 3.45E-01 | 9.72E-01 | 1.41E+00 | 9.54E-03 | 3.50E-02 | 1.61E+00 | 2.88E-01 | 7.91E-01 | 1.26E+00 |
| FAM160B2 | 6.31E-02 | 3.29E-01 | 1.03E+00 | 1.48E-01 | 6.38E-01 | 1.48E+00 | 9.23E-03 | 3.92E-02 | -1.26E+00 | 1.15E-01 | 4.41E-01 | 1.13E+00 | 2.39E-04 | 8.51E-04 | -1.38E+00 | 3.16E-02 | 8.91E-02 | 1.11E+00 | 6.56E-02 | 2.41E-01 | 1.03E+00 | 2.69E-01 | 7.38E-01 | 1.26E+00 |
| KLHL35 | 6.33E-02 | 3.30E-01 | 1.72E+00 | 1.70E-01 | 7.36E-01 | 1.67E+00 | 4.72E-02 | 2.01E-01 | 1.46E+00 | 1.50E-02 | 5.71E-02 | 1.97E+00 | 2.11E-01 | 7.51E-01 | 1.56E+00 | 1.61E-01 | 4.55E-01 | 1.97E+00 | 6.53E-02 | 2.40E-01 | 1.59E+00 | 1.80E-01 | 4.95E-01 | 1.52E+00 |
| CD7 | 6.33E-02 | 3.30E-01 | 1.42E+00 | 2.27E-04 | 9.83E-04 | -1.11E+00 | 2.75E-03 | 1.17E-02 | -1.24E+00 | 3.05E-06 | 1.16E-05 | -1.58E+00 | 1.48E-05 | 5.29E-05 | -1.03E+00 | 1.90E-08 | 5.35E-08 | -1.20E+00 | 7.53E-04 | 2.76E-03 | -1.02E+00 | 2.35E-05 | 6.46E-05 | -1.30E+00 |
| EIF3E | 6.35E-02 | 3.31E-01 | 1.30E+00 | 7.22E-02 | 3.12E-01 | 1.47E+00 | 3.63E-02 | 1.54E-01 | 1.25E+00 | 2.78E-03 | 1.06E-02 | 1.48E+00 | 1.26E-01 | 4.48E-01 | 1.35E+00 | 6.38E-03 | 1.80E-02 | 1.46E+00 | 1.61E-01 | 5.92E-01 | 1.27E+00 | 3.41E-02 | 9.37E-02 | 1.38E+00 |
| RIMKLA | 6.35E-02 | 3.31E-01 | -1.06E+00 | 2.15E-02 | 9.31E-02 | -1.26E+00 | 3.33E-05 | 1.42E-04 | -1.63E+00 | 3.58E-03 | 1.37E-02 | -1.06E+00 | 7.67E-02 | 2.73E-01 | -1.38E+00 | 1.28E-01 | 3.61E-01 | -1.13E+00 | 1.34E-01 | 4.91E-01 | -1.03E+00 | 2.59E-01 | 7.12E-01 | 1.08E+00 |
| STBD1 | 6.36E-02 | 3.31E-01 | 1.21E+00 | 4.71E-02 | 2.04E-01 | 1.27E+00 | 1.24E-01 | 5.26E-01 | 1.06E+00 | 3.84E-02 | 1.47E-01 | -1.02E+00 | 1.14E-01 | 4.05E-01 | 2.11E+00 | 1.99E-01 | 5.60E-01 | 2.14E+00 | 2.60E-01 | 9.53E-01 | 1.17E+00 | 3.49E-01 | 9.59E-01 | 1.26E+00 |
| C3orf38 | 6.36E-02 | 3.32E-01 | 1.22E+00 | 2.44E-02 | 1.05E-01 | 1.27E+00 | 4.66E-06 | 1.98E-05 | -1.27E+00 | 5.03E-08 | 1.92E-07 | -1.28E+00 | 6.37E-02 | 2.27E-01 | 1.23E+00 | 7.09E-03 | 2.00E-02 | 1.06E+00 | 6.06E-04 | 2.22E-03 | -1.03E+00 | 3.00E-07 | 8.25E-07 | -1.27E+00 |
| FLJ31104 | 6.36E-02 | 3.32E-01 | 1.54E+00 | 2.17E-01 | 9.37E-01 | 1.41E+00 | 1.99E-01 | 8.47E-01 | 1.25E+00 | 1.17E-01 | 4.45E-01 | -1.11E+00 | 1.38E-01 | 4.93E-01 | -1.21E+00 | 1.80E-01 | 5.08E-01 | -1.17E+00 | 1.81E-01 | 6.64E-01 | 1.44E+00 | 1.62E-01 | 4.45E-01 | -1.04E+00 |
| DPP10-AS1 | 6.36E-02 | 3.32E-01 | -1.16E+00 | 1.81E-01 | 7.81E-01 | 1.14E+00 | 1.06E-02 | 4.52E-02 | -1.32E+00 | 9.35E-03 | 3.57E-02 | -1.42E+00 | 2.48E-01 | 8.86E-01 | 1.26E+00 | 3.39E-01 | 9.56E-01 | 1.33E+00 | 1.75E-01 | 6.41E-01 | 1.35E+00 | 3.28E-01 | 9.00E-01 | 1.22E+00 |
| ARHGAP31 | 6.37E-02 | 3.32E-01 | -1.06E+00 | 1.09E-01 | 4.72E-01 | 1.15E+00 | 6.45E-03 | 2.74E-02 | -1.42E+00 | 3.03E-02 | 1.16E-01 | -1.03E+00 | 6.27E-03 | 2.24E-02 | -1.59E+00 | 5.22E-02 | 1.47E-01 | -1.17E+00 | 2.90E-02 | 1.07E-01 | -1.17E+00 | 2.60E-01 | 7.14E-01 | 1.22E+00 |
| TBCEL | 6.37E-02 | 3.32E-01 | -1.04E+00 | 3.11E-02 | 1.34E-01 | -1.03E+00 | 1.63E-02 | 6.91E-02 | -1.27E+00 | 2.77E-02 | 1.06E-01 | -1.10E+00 | 3.14E-03 | 1.12E-02 | -1.36E+00 | 2.60E-03 | 7.34E-03 | -1.49E+00 | 5.78E-03 | 2.12E-02 | 1.01E+00 | 5.32E-06 | 1.46E-05 | -1.58E+00 |
| FO393415.2 | 6.38E-02 | 3.32E-01 | 1.11E+00 | 3.10E-04 | 1.34E-03 | -1.59E+00 | 3.47E-02 | 1.47E-01 | -1.05E+00 | 2.53E-04 | 9.67E-04 | -1.55E+00 | 1.54E-01 | 5.48E-01 | 1.11E+00 | 4.55E-02 | 1.28E-01 | -1.24E+00 | 5.71E-02 | 2.10E-01 | 1.01E+00 | 5.03E-04 | 1.38E-03 | -1.96E+00 |
| KIF18P | 6.38E-02 | 3.32E-01 | -1.01E+00 | 1.08E-01 | 4.67E-01 | 1.19E+00 | 5.21E-06 | 2.21E-05 | -1.81E+00 | 1.45E-04 | 5.54E-04 | -1.29E+00 | 3.74E-06 | 1.33E-05 | -1.89E+00 | 2.12E-06 | 5.98E-06 | -2.06E+00 | 4.18E-11 | 1.53E-10 | -1.75E+00 | 1.05E-08 | 2.88E-08 | -1.76E+00 |
| RNASE6 | 6.38E-02 | 3.32E-01 | -4.02E+00 | 6.89E-02 | 2.98E-01 | -4.67E+00 | 1.75E-01 | 7.45E-01 | 1.00E+00 | 8.09E-02 | 3.09E-01 | -1.49E+00 | 7.61E-04 | 2.71E-03 | -1.42E+00 | 3.14E-01 | 8.84E-01 | 1.44E+00 | 3.28E-48 | 1.20E-47 | -2.56E+00 | 3.61E-38 | 9.91E-38 | -2.10E+00 |
| MIR133A1HG | 6.38E-02 | 3.33E-01 | -1.19E+00 | 1.91E-01 | 8.27E-01 | 1.21E+00 | 1.91E-04 | 8.13E-04 | -1.64E+00 | 1.01E-03 | 3.86E-03 | -1.41E+00 | 7.22E-02 | 2.57E-01 | -1.07E+00 | 1.90E-02 | 5.36E-02 | -1.38E+00 | 4.79E-03 | 1.76E-02 | -1.34E+00 | 2.45E-03 | 6.73E-03 | -1.52E+00 |
| GDF10 | 6.39E-02 | 3.33E-01 | 1.56E+00 | 4.03E-02 | 1.74E-01 | 2.10E+00 | 1.01E-02 | 4.29E-02 | 2.42E+00 | 6.81E-03 | 2.60E-02 | 2.87E+00 | 1.73E-05 | 6.17E-05 | 5.13E+00 | 7.92E-08 | 2.23E-07 | 7.45E+00 | 2.34E-01 | 8.58E-01 | 1.21E+00 | 1.67E-01 | 4.59E-01 | 1.61E+00 |
| FRMD6 | 6.39E-02 | 3.33E-01 | -1.15E+00 | 1.08E-01 | 4.66E-01 | 1.05E+00 | 1.43E-01 | 6.06E-01 | -1.11E+00 | 2.00E-01 | 7.65E-01 | 1.06E+00 | 6.29E-03 | 2.24E-02 | -1.44E+00 | 1.88E-01 | 5.30E-01 | 1.07E+00 | 1.46E-07 | 5.38E-07 | -1.84E+00 | 3.09E-04 | 8.49E-04 | -1.33E+00 |
| AC091042.1 | 6.39E-02 | 3.33E-01 | 1.05E+00 | 3.35E-05 | 1.45E-04 | -1.84E+00 | 2.30E-01 | 9.78E-01 | 1.23E+00 | 1.26E-04 | 4.83E-04 | -1.60E+00 | 2.93E-03 | 1.05E-02 | -1.27E+00 | 2.03E-06 | 5.72E-06 | -2.28E+00 | 2.46E-03 | 9.04E-03 | -1.22E+00 | 3.69E-08 | 1.01E-07 | -2.19E+00 |
| RAB31 | 6.41E-02 | 3.34E-01 | 1.08E+00 | 1.90E-01 | 8.20E-01 | -1.08E+00 | 4.76E-03 | 2.02E-02 | -1.32E+00 | 5.17E-03 | 1.97E-02 | -1.24E+00 | 2.32E-01 | 8.25E-01 | 1.29E+00 | 2.90E-01 | 8.17E-01 | 1.31E+00 | 1.91E-01 | 7.02E-01 | -1.26E+00 | 1.70E-01 | 4.67E-01 | -1.51E+00 |
| FAM53B | 6.41E-02 | 3.34E-01 | 1.04E+00 | 8.73E-02 | 3.77E-01 | 1.17E+00 | 3.09E-03 | 1.31E-02 | -1.11E+00 | 6.60E-03 | 2.52E-02 | 1.09E+00 | 1.90E-03 | 6.79E-03 | -1.14E+00 | 1.06E-02 | 2.99E-02 | 1.03E+00 | 1.16E-06 | 4.26E-06 | -1.54E+00 | 2.41E-04 | 6.61E-04 | -1.22E+00 |
| SMIM13 | 6.41E-02 | 3.34E-01 | 1.05E+00 | 2.20E-02 | 9.50E-02 | -1.15E+00 | 1.64E-02 | 6.96E-02 | -1.02E+00 | 3.04E-03 | 1.16E-02 | -1.05E+00 | 1.20E-02 | 4.29E-02 | -1.31E+00 | 1.06E-03 | 3.00E-03 | -1.77E+00 | 3.82E-02 | 1.40E-01 | -1.12E+00 | 1.50E-02 | 4.11E-02 | -1.25E+00 |
| PLEKHG2 | 6.42E-02 | 3.34E-01 | 1.21E+00 | 3.40E-03 | 1.47E-02 | -1.07E+00 | 2.31E-01 | 9.83E-01 | 1.30E+00 | 1.93E-02 | 7.38E-02 | 1.01E+00 | 8.64E-02 | 3.08E-01 | -1.02E+00 | 1.35E-01 | 3.80E-01 | 1.05E+00 | 9.23E-02 | 3.39E-01 | -1.05E+00 | 6.68E-02 | 1.84E-01 | -1.15E+00 |

| | | | | | | | | | | | | | | | | | | | | | | | | |
|---|---|---|---|---|---|---|---|---|---|---|---|---|---|---|---|---|---|---|---|---|---|---|---|---|
| AUH | 6.42E-02 | 3.35E-01 | 1.23E+00 | 9.49E-02 | 4.10E-01 | 1.36E+00 | 1.21E-05 | 5.13E-05 | -1.41E+00 | 2.50E-03 | 9.54E-03 | 1.00E+00 | 1.18E-01 | 4.20E-01 | 1.26E+00 | 1.18E-02 | 3.33E-02 | 1.04E+00 | 3.67E-08 | 1.35E-07 | -1.41E+00 | 4.40E-04 | 1.21E-03 | -1.12E+00 |
| AP002884.3 | 6.42E-02 | 3.35E-01 | 1.04E+00 | 1.67E-01 | 7.23E-01 | 1.31E+00 | 1.08E-01 | 4.57E-01 | 1.42E+00 | 1.37E-01 | 5.22E-01 | 1.51E+00 | 2.56E-01 | 9.13E-01 | 1.23E+00 | 3.17E-01 | 8.93E-01 | 1.21E+00 | 2.26E-01 | 8.28E-01 | -1.09E+00 | 3.35E-01 | 9.21E-01 | 1.09E+00 |
| CORO2A | 6.43E-02 | 3.35E-01 | 1.11E+00 | 1.93E-02 | 8.33E-02 | 1.13E+00 | 3.95E-02 | 1.68E-01 | 1.00E+00 | 5.56E-03 | 2.12E-02 | 1.06E+00 | 5.73E-03 | 2.04E-02 | -1.31E+00 | 3.94E-02 | 1.11E-01 | -1.03E+00 | 3.41E-06 | 1.25E-05 | -2.36E+00 | 2.54E-06 | 6.96E-06 | -2.33E+00 |
| NCLN | 6.43E-02 | 3.35E-01 | 1.12E+00 | 1.19E-02 | 5.14E-02 | 1.06E+00 | 8.46E-02 | 3.60E-01 | 1.10E+00 | 1.89E-01 | 7.20E-01 | 1.38E+00 | 4.34E-02 | 1.55E-01 | -1.00E+00 | 1.56E-01 | 4.40E-01 | 1.21E+00 | 4.22E-03 | 1.55E-02 | -1.09E+00 | 9.87E-02 | 2.71E-01 | 1.08E+00 |
| UBB | 6.43E-02 | 3.35E-01 | -1.02E+00 | 1.17E-14 | 5.05E-14 | -1.29E+00 | 2.24E-01 | 9.52E-01 | -1.01E+00 | 6.53E-05 | 2.50E-04 | -1.26E+00 | 2.22E-01 | 7.92E-01 | 1.07E+00 | 3.76E-05 | 1.06E-04 | -1.24E+00 | 2.18E-01 | 8.01E-01 | 1.12E+00 | 6.28E-03 | 1.73E-02 | -1.19E+00 |
| CXorf57 | 6.44E-02 | 3.35E-01 | 1.64E+00 | 2.09E-01 | 9.04E-01 | 1.21E+00 | 6.80E-02 | 2.89E-01 | 1.55E+00 | 1.04E-01 | 3.96E-01 | 1.56E+00 | 6.81E-04 | 2.43E-03 | 5.08E+00 | 2.67E-04 | 7.52E-04 | 5.83E+00 | 3.27E-12 | 1.20E-11 | 5.00E+00 | 3.13E-06 | 8.59E-06 | 3.88E+00 |
| PROS1 | 6.44E-02 | 3.36E-01 | -1.48E+00 | 9.34E-02 | 4.04E-01 | -1.41E+00 | 1.96E-01 | 8.31E-01 | 1.33E+00 | 2.26E-01 | 8.64E-01 | -1.02E+00 | 1.04E-01 | 3.72E-01 | -1.83E+00 | 1.13E-01 | 3.18E-01 | -1.68E+00 | 9.79E-06 | 3.59E-05 | -1.51E+00 | 2.69E-04 | 7.38E-04 | -1.55E+00 |
| SLC25A44 | 6.44E-02 | 3.36E-01 | 1.16E+00 | 1.12E-02 | 4.82E-02 | 1.01E+00 | 1.43E-01 | 6.09E-01 | 1.35E+00 | 1.87E-01 | 7.16E-01 | 1.21E+00 | 6.64E-03 | 2.37E-02 | -1.06E+00 | 3.70E-05 | 1.04E-04 | -1.75E+00 | 1.27E-03 | 4.65E-03 | -1.34E+00 | 8.75E-03 | 2.40E-02 | -1.31E+00 |
| DPRXP4 | 6.46E-02 | 3.37E-01 | 1.63E+00 | 2.88E-02 | 1.24E-01 | 2.13E+00 | 1.80E-01 | 7.66E-01 | -1.03E+00 | 3.01E-02 | 1.15E-01 | 1.74E+00 | 2.50E-01 | 8.93E-01 | 1.33E+00 | 1.09E-01 | 3.07E-01 | 2.07E+00 | 1.72E-01 | 6.33E-01 | -1.02E+00 | 1.86E-01 | 5.11E-01 | 1.47E+00 |
| TEKT2 | 6.47E-02 | 3.37E-01 | 1.57E+00 | 7.27E-02 | 3.14E-01 | 1.06E+00 | 1.24E-01 | 5.28E-01 | 1.51E+00 | 1.97E-01 | 7.51E-01 | 1.23E+00 | 2.43E-01 | 8.66E-01 | 1.25E+00 | 1.02E-01 | 2.89E-01 | -1.17E+00 | 3.05E-02 | 1.12E-01 | 2.05E+00 | 3.59E-01 | 9.87E-01 | 1.13E+00 |
| ATP6V1G2 | 6.47E-02 | 3.37E-01 | 1.10E+00 | 3.87E-03 | 1.67E-02 | -1.27E+00 | 9.77E-02 | 4.15E-01 | 1.05E+00 | 3.18E-03 | 1.21E-02 | -1.26E+00 | 1.17E-02 | 4.16E-02 | -1.80E+00 | 1.71E-02 | 4.82E-02 | -1.71E+00 | 6.93E-02 | 2.54E-01 | -1.33E+00 | 5.12E-02 | 1.41E-01 | -1.59E+00 |
| SSH2 | 6.47E-02 | 3.37E-01 | 1.19E+00 | 5.00E-04 | 2.16E-03 | 1.08E+00 | 1.63E-01 | 6.93E-01 | 1.09E+00 | 4.66E-03 | 1.78E-02 | -1.03E+00 | 2.32E-01 | 8.28E-01 | 1.35E+00 | 1.63E-03 | 4.58E-03 | -1.15E+00 | 5.42E-05 | 1.99E-04 | -1.25E+00 | 6.40E-08 | 1.76E-07 | -1.55E+00 |
| USP3 | 6.47E-02 | 3.37E-01 | 1.20E+00 | 2.03E-02 | 8.77E-02 | 1.23E+00 | 2.99E-04 | 1.27E-03 | -1.25E+00 | 5.40E-07 | 2.06E-06 | -1.39E+00 | 1.29E-04 | 4.60E-04 | -1.21E+00 | 2.09E-07 | 5.89E-07 | -1.58E+00 | 2.64E-12 | 9.70E-12 | -1.57E+00 | 2.13E-15 | 5.85E-15 | -2.03E+00 |
| PTGFRN | 6.49E-02 | 3.38E-01 | -1.33E+00 | 8.27E-02 | 3.58E-01 | -1.34E+00 | 1.25E-06 | 5.32E-06 | -2.03E+00 | 1.93E-06 | 7.39E-06 | -1.93E+00 | 2.80E-01 | 1.00E+00 | -1.00E+00 | 2.66E-01 | 7.49E-01 | 1.34E+00 | 3.62E-08 | 1.33E-07 | -1.98E+00 | 4.70E-05 | 1.29E-04 | -1.50E+00 |
| FOXJ2 | 6.49E-02 | 3.38E-01 | 1.19E+00 | 4.34E-02 | 1.88E-01 | 1.20E+00 | 2.14E-01 | 9.08E-01 | 1.22E+00 | 1.75E-01 | 6.68E-01 | 1.47E+00 | 8.07E-02 | 2.88E-01 | -1.00E+00 | 3.05E-01 | 8.59E-01 | 1.46E+00 | 4.12E-04 | 1.51E-03 | -1.20E+00 | 2.97E-02 | 8.16E-02 | 1.00E+00 |
| AC131971.1 | 6.49E-02 | 3.38E-01 | -1.15E+00 | 1.34E-01 | 5.79E-01 | 1.04E+00 | 1.44E-01 | 6.12E-01 | 1.01E+00 | 2.14E-01 | 8.17E-01 | 1.18E+00 | 2.60E-01 | 9.28E-01 | 1.12E+00 | 1.97E-01 | 5.54E-01 | -1.08E+00 | 1.79E-01 | 6.56E-01 | 1.41E+00 | 2.47E-01 | 6.79E-01 | 1.30E+00 |
| TROAP | 6.50E-02 | 3.39E-01 | -1.46E+00 | 4.70E-02 | 2.03E-01 | -1.88E+00 | 7.67E-03 | 3.26E-02 | 1.90E+00 | 1.54E-01 | 5.87E-01 | 1.50E+00 | 1.95E-01 | 6.95E-01 | -1.31E+00 | 2.29E-01 | 6.46E-01 | -1.39E+00 | 1.94E-01 | 7.12E-01 | 1.01E+00 | 2.80E-01 | 7.70E-01 | 1.06E+00 |
| AC022150.4 | 6.50E-02 | 3.39E-01 | 1.03E+00 | 1.90E-01 | 8.22E-01 | 1.54E+00 | 9.24E-02 | 3.93E-01 | -1.06E+00 | 1.35E-01 | 5.16E-01 | 1.12E+00 | 2.46E-02 | 8.78E-02 | -1.53E+00 | 8.28E-02 | 2.34E-01 | -1.33E+00 | 8.07E-04 | 2.96E-03 | -1.63E+00 | 6.10E-02 | 1.68E-01 | -1.15E+00 |
| AL136295.6 | 6.51E-02 | 3.39E-01 | -1.06E+00 | 2.20E-01 | 9.50E-01 | 1.36E+00 | 9.83E-02 | 4.18E-01 | -1.20E+00 | 1.55E-01 | 5.91E-01 | -1.02E+00 | 1.41E-01 | 5.01E-01 | -1.09E+00 | 1.97E-01 | 5.55E-01 | 1.58E+00 | 2.72E-02 | 9.99E-02 | -1.60E+00 | 5.56E-02 | 1.53E-01 | -1.46E+00 |
| AC069185.1 | 6.52E-02 | 3.40E-01 | 1.01E+00 | 2.91E-02 | 1.26E-01 | -1.16E+00 | 1.25E-01 | 5.33E-01 | 1.00E+00 | 2.54E-01 | 9.70E-01 | 1.28E+00 | 1.45E-01 | 5.15E-01 | 1.15E+00 | 2.90E-01 | 8.17E-01 | 1.31E+00 | 5.08E-02 | 1.86E-01 | 1.78E+00 | 5.43E-02 | 1.49E-01 | 1.92E+00 |
| RPL36AL | 6.53E-02 | 3.40E-01 | 1.01E+00 | 5.25E-03 | 2.27E-02 | -1.02E+00 | 1.90E-02 | 8.07E-02 | -1.06E+00 | 7.72E-03 | 2.95E-02 | -1.07E+00 | 2.56E-03 | 9.14E-03 | -1.11E+00 | 1.26E-04 | 3.54E-04 | -1.16E+00 | 6.60E-03 | 2.42E-02 | -1.05E+00 | 1.99E-02 | 5.46E-02 | -1.05E+00 |
| EPB41L4A-AS1 | 6.53E-02 | 3.40E-01 | 1.23E+00 | 9.07E-02 | 3.92E-01 | 1.16E+00 | 2.05E-01 | 8.73E-01 | 1.05E+00 | 1.71E-01 | 6.52E-01 | 1.09E+00 | 1.27E-03 | 4.51E-03 | 1.49E+00 | 3.11E-02 | 8.77E-02 | 1.37E+00 | 1.90E-03 | 6.99E-03 | 1.62E+00 | 3.47E-01 | 9.54E-01 | 1.18E+00 |
| BAALC-AS2 | 6.54E-02 | 3.41E-01 | 1.25E+00 | 1.99E-03 | 8.60E-03 | 1.12E+00 | 1.19E-04 | 5.07E-04 | -1.27E+00 | 6.36E-08 | 2.43E-07 | -1.62E+00 | 1.24E-01 | 4.43E-01 | 2.18E+00 | 3.38E-01 | 9.51E-01 | 1.51E+00 | 1.86E-01 | 6.81E-01 | 1.41E+00 | 3.36E-01 | 9.22E-01 | 1.21E+00 |
| AAR2 | 6.56E-02 | 3.42E-01 | 1.04E+00 | 1.44E-01 | 6.24E-01 | 1.36E+00 | 3.19E-05 | 1.35E-04 | -1.23E+00 | 3.59E-03 | 1.37E-02 | 1.19E+00 | 2.26E-02 | 8.04E-02 | -1.04E+00 | 2.81E-01 | 7.92E-01 | 1.40E+00 | 2.29E-04 | 8.41E-04 | -1.18E+00 | 8.58E-02 | 2.36E-01 | 1.14E+00 |
| BUB1 | 6.56E-02 | 3.42E-01 | -1.07E+00 | 1.46E-01 | 6.29E-01 | 1.17E+00 | 8.04E-02 | 3.42E-01 | -1.05E+00 | 1.18E-01 | 4.49E-01 | 1.17E+00 | 1.80E-01 | 6.43E-01 | -1.38E+00 | 3.28E-01 | 9.25E-01 | 1.32E+00 | 1.49E-01 | 5.47E-01 | 1.03E+00 | 3.22E-01 | 8.84E-01 | 1.14E+00 |
| TELO2 | 6.57E-02 | 3.42E-01 | 1.81E+00 | 2.27E-01 | 9.82E-01 | 1.60E+00 | 2.96E-03 | 1.26E-02 | -1.06E+00 | 3.12E-08 | 1.19E-07 | -1.33E+00 | 2.31E-02 | 8.22E-02 | -1.39E+00 | 2.17E-02 | 6.11E-02 | -1.44E+00 | 4.37E-02 | 1.60E-01 | -1.12E+00 | 7.74E-02 | 2.13E-01 | 1.03E+00 |
| AC099811.6 | 6.57E-02 | 3.42E-01 | -1.14E+00 | 1.41E-01 | 6.08E-01 | 1.66E+00 | 2.99E-10 | 1.27E-09 | -2.63E+00 | 4.64E-09 | 1.77E-08 | -2.20E+00 | 8.20E-04 | 2.92E-03 | -1.98E+00 | 3.52E-02 | 9.93E-02 | -1.04E+00 | 2.57E-04 | 9.42E-04 | -1.52E+00 | 5.46E-03 | 1.50E-02 | -1.40E+00 |
| AC008868.1 | 6.57E-02 | 3.43E-01 | 1.02E+00 | 1.56E-01 | 6.75E-01 | 1.49E+00 | 6.00E-02 | 2.55E-01 | 1.49E+00 | 2.54E-02 | 9.69E-02 | 1.83E+00 | 1.29E-01 | 4.60E-01 | 1.81E+00 | 3.07E-04 | 8.65E-04 | 3.12E+00 | 1.35E-01 | 4.96E-01 | 1.50E+00 | 6.80E-04 | 1.87E-03 | 2.23E+00 |
| AC092171.1 | 6.57E-02 | 3.43E-01 | 1.47E+00 | 1.37E-01 | 5.91E-01 | 1.25E+00 | 1.06E-01 | 4.52E-01 | 1.43E+00 | 1.48E-01 | 5.64E-01 | 1.31E+00 | 1.35E-01 | 4.82E-01 | 1.21E+00 | 1.69E-01 | 4.76E-01 | 1.21E+00 | 8.55E-02 | 3.14E-01 | -1.08E+00 | 5.40E-02 | 1.48E-01 | -1.12E+00 |
| RPL21P44 | 6.57E-02 | 3.43E-01 | 1.09E+00 | 1.50E-01 | 6.50E-01 | 1.56E+00 | 5.14E-05 | 2.18E-04 | -1.51E+00 | 1.47E-04 | 5.61E-04 | -1.36E+00 | 2.47E-01 | 8.82E-01 | 1.33E+00 | 2.64E-01 | 7.44E-01 | 1.54E+00 | 1.52E-01 | 5.59E-01 | -1.13E+00 | 3.34E-01 | 9.16E-01 | 1.19E+00 |
| CACNA2D4 | 6.57E-02 | 3.43E-01 | 1.09E+00 | 1.99E-02 | 8.60E-02 | 1.04E+00 | 2.10E-02 | 8.91E-02 | -1.03E+00 | 2.22E-02 | 8.49E-02 | 1.14E+00 | 2.81E-07 | 1.00E-06 | -1.79E+00 | 5.85E-07 | 1.65E-06 | -1.62E+00 | 1.66E-01 | 6.11E-01 | 1.00E+00 | 4.81E-02 | 1.32E-01 | -1.43E+00 |
| AC022395.1 | 6.58E-02 | 3.43E-01 | 1.73E+00 | 1.13E-01 | 4.86E-01 | 1.68E+00 | 1.41E-01 | 5.97E-01 | 1.14E+00 | 6.99E-02 | 2.67E-01 | 1.07E+00 | 2.11E-01 | 7.53E-01 | 1.64E+00 | 1.82E-01 | 5.12E-01 | 1.15E+00 | 1.08E-01 | 3.95E-01 | -1.06E+00 | 3.51E-01 | 9.64E-01 | 1.19E+00 |
| RPL17P22 | 6.59E-02 | 3.43E-01 | 1.68E+00 | 2.15E-01 | 9.27E-01 | 1.56E+00 | 3.31E-04 | 1.41E-03 | 2.08E+00 | 2.19E-02 | 8.35E-02 | 1.97E+00 | 2.47E-03 | 8.79E-03 | 2.39E+00 | 5.24E-02 | 1.48E-01 | 2.10E+00 | 2.01E-04 | 7.36E-04 | 2.31E+00 | 5.41E-02 | 1.49E-01 | 1.75E+00 |
| PLAC9 | 6.59E-02 | 3.44E-01 | 1.66E+00 | 2.22E-01 | 9.61E-01 | 1.39E+00 | 1.61E-02 | 6.86E-02 | 1.93E+00 | 1.59E-01 | 6.08E-01 | 1.48E+00 | 2.28E-01 | 8.13E-01 | 1.51E+00 | 1.39E-01 | 3.93E-01 | -1.11E+00 | 2.52E-01 | 9.25E-01 | -1.06E+00 | 2.78E-01 | 7.63E-01 | -1.16E+00 |
| PCSK4 | 6.59E-02 | 3.44E-01 | 1.77E+00 | 2.09E-01 | 9.01E-01 | 1.61E+00 | 6.41E-03 | 2.73E-02 | -1.28E+00 | 3.24E-03 | 1.24E-02 | -1.39E+00 | 1.87E-01 | 6.66E-01 | 1.02E+00 | 1.68E-01 | 4.75E-01 | -1.06E+00 | 3.08E-03 | 1.13E-02 | -1.76E+00 | 5.59E-02 | 1.54E-01 | -1.26E+00 |
| LDLR | 6.60E-02 | 3.44E-01 | 1.17E+00 | 4.55E-10 | 1.97E-09 | -1.32E+00 | 1.84E-02 | 7.81E-02 | 1.29E+00 | 1.22E-02 | 4.67E-02 | -1.06E+00 | 8.24E-02 | 2.94E-01 | 1.46E+00 | 1.22E-01 | 3.43E-01 | 1.46E+00 | 2.44E-01 | 8.96E-01 | 1.08E+00 | 2.56E-01 | 7.04E-01 | 1.22E+00 |
| GRIK5 | 6.61E-02 | 3.44E-01 | 1.19E+00 | 4.51E-03 | 1.95E-02 | 1.06E+00 | 3.86E-02 | 1.64E-01 | -1.01E+00 | 1.06E-02 | 4.04E-02 | 1.05E+00 | 4.96E-02 | 1.77E-01 | 1.11E+00 | 2.72E-03 | 7.67E-03 | -1.19E+00 | 2.66E-01 | 9.77E-01 | 1.24E+00 | 2.86E-01 | 7.85E-01 | 1.48E+00 |
| PET100 | 6.61E-02 | 3.45E-01 | 1.26E+00 | 5.44E-02 | 2.35E-01 | 1.34E+00 | 1.69E-02 | 7.19E-02 | -1.07E+00 | 1.03E-04 | 3.95E-04 | -1.22E+00 | 6.62E-02 | 2.36E-01 | 1.17E+00 | 1.07E-02 | 3.01E-02 | -1.01E+00 | 1.11E-01 | 4.07E-01 | 1.28E+00 | 9.79E-05 | 2.69E-04 | -1.26E+00 |
| ATP5IP1 | 6.62E-02 | 3.45E-01 | 1.67E+00 | 3.53E-03 | 1.53E-02 | 2.49E+00 | 4.53E-02 | 1.92E-01 | 1.60E+00 | 2.85E-03 | 1.09E-02 | 2.27E+00 | 1.07E-01 | 3.80E-01 | 1.08E+00 | 3.15E-01 | 8.89E-01 | 1.50E+00 | 1.30E-02 | 4.77E-02 | 1.86E+00 | 2.22E-06 | 6.11E-06 | 2.78E+00 |
| UCKL1 | 6.63E-02 | 3.46E-01 | 1.14E+00 | 5.65E-02 | 2.44E-01 | 1.27E+00 | 1.51E-03 | 6.44E-03 | -1.24E+00 | 1.35E-03 | 5.15E-03 | -1.08E+00 | 2.96E-05 | 1.06E-04 | -1.41E+00 | 3.84E-04 | 1.08E-03 | -1.15E+00 | 1.62E-01 | 5.94E-01 | 1.10E+00 | 2.15E-02 | 5.92E-02 | -1.01E+00 |
| CGNL1 | 6.63E-02 | 3.46E-01 | -1.87E+00 | 1.26E-01 | 5.45E-01 | -1.36E+00 | 2.30E-01 | 9.77E-01 | 1.12E+00 | 2.44E-01 | 9.34E-01 | -1.03E+00 | 6.37E-06 | 2.27E-05 | 3.15E+00 | 7.11E-06 | 2.00E-05 | 3.45E+00 | 1.58E-08 | 5.78E-08 | 2.62E+00 | 4.91E-02 | 1.35E-01 | 1.66E+00 |
| ALG5 | 6.64E-02 | 3.46E-01 | 1.26E+00 | 9.34E-02 | 4.03E-01 | 1.36E+00 | 1.58E-01 | 6.70E-01 | 1.32E+00 | 2.38E-01 | 9.08E-01 | 1.44E+00 | 1.34E-01 | 4.78E-01 | 1.30E+00 | 1.80E-01 | 5.08E-01 | 1.26E+00 | 1.99E-01 | 7.30E-01 | 1.15E+00 | 2.48E-01 | 6.82E-01 | 1.19E+00 |
| FANCI | 6.65E-02 | 3.46E-01 | -1.00E+00 | 2.99E-02 | 1.29E-01 | -1.11E+00 | 5.85E-03 | 2.48E-02 | 1.42E+00 | 1.41E-01 | 5.37E-01 | 1.50E+00 | 1.37E-02 | 4.88E-02 | -1.28E+00 | 7.73E-02 | 2.18E-01 | 1.04E+00 | 3.57E-02 | 1.31E-01 | 1.09E+00 | 2.60E-01 | 7.13E-01 | 1.24E+00 |
| AC090181.3 | 6.65E-02 | 3.47E-01 | 1.12E+00 | 3.59E-02 | 1.55E-01 | 1.15E+00 | 2.23E-01 | 9.48E-01 | 1.18E+00 | 2.36E-01 | 9.00E-01 | 1.32E+00 | 6.53E-02 | 2.33E-01 | 1.09E+00 | 4.19E-02 | 1.18E-01 | 1.05E+00 | 6.66E-02 | 2.44E-01 | -1.06E+00 | 2.81E-01 | 7.71E-01 | 1.34E+00 |
| USP21 | 6.65E-02 | 3.47E-01 | 1.08E+00 | 2.04E-02 | 8.80E-02 | 1.03E+00 | 1.04E-01 | 4.44E-01 | -1.04E+00 | 4.45E-02 | 1.70E-01 | -1.11E+00 | 2.04E-02 | 7.29E-02 | -1.28E+00 | 3.24E-01 | 9.15E-01 | 1.35E+00 | 7.33E-02 | 2.69E-01 | 1.24E+00 | 1.21E-01 | 3.32E-01 | 1.19E+00 |
| HNRNPRP1 | 6.67E-02 | 3.47E-01 | 1.69E+00 | 1.24E-01 | 5.37E-01 | 1.73E+00 | 3.72E-02 | 1.58E-01 | 1.76E+00 | 4.76E-02 | 1.82E-01 | 1.98E+00 | 2.65E-01 | 9.43E-01 | 1.27E+00 | 2.91E-01 | 8.21E-01 | 1.16E+00 | 3.39E-03 | 1.24E-02 | 2.15E+00 | 8.38E-03 | 2.30E-02 | 2.12E+00 |
| MGMT | 6.67E-02 | 3.48E-01 | 1.17E+00 | 7.67E-02 | 3.32E-01 | 1.32E+00 | 3.92E-02 | 1.67E-01 | 1.38E+00 | 2.20E-01 | 8.40E-01 | 1.35E+00 | 2.04E-01 | 7.29E-01 | 1.26E+00 | 2.85E-01 | 8.04E-01 | 1.33E+00 | 2.09E-01 | 7.67E-01 | 1.29E+00 | 2.69E-01 | 7.39E-01 | 1.21E+00 |
| AC131212.3 | 6.68E-02 | 3.48E-01 | 1.75E+00 | 8.12E-02 | 3.51E-01 | 1.83E+00 | 1.75E-01 | 7.43E-01 | 1.28E+00 | 7.54E-02 | 2.88E-01 | 1.53E+00 | 1.95E-01 | 6.95E-01 | 1.55E+00 | 3.38E-01 | 9.52E-01 | 1.38E+00 | 6.91E-02 | 2.54E-01 | -1.26E+00 | 3.16E-01 | 8.69E-01 | 1.02E+00 |
| AP006621.3 | 6.68E-02 | 3.48E-01 | 1.06E+00 | 2.08E-02 | 8.99E-02 | 1.91E+00 | 1.97E-02 | 8.35E-02 | -1.16E+00 | 1.53E-01 | 5.83E-01 | 1.27E+00 | 2.52E-01 | 8.97E-01 | 1.46E+00 | 1.66E-01 | 4.69E-01 | 1.84E+00 | 9.34E-02 | 3.43E-01 | 1.43E+00 | 1.68E-01 | 4.61E-01 | 1.45E+00 |
| KIF4A | 6.68E-02 | 3.48E-01 | -1.50E+00 | 9.56E-02 | 4.13E-01 | -1.36E+00 | 1.60E-02 | 6.79E-02 | 1.93E+00 | 1.24E-01 | 4.74E-01 | 1.98E+00 | 2.54E-01 | 9.06E-01 | 1.41E+00 | 3.55E-01 | 1.00E+00 | 1.00E+00 | 2.20E-01 | 8.09E-01 | 1.06E+00 | 8.53E-02 | 2.34E-01 | 1.23E+00 |
| NDC80 | 6.69E-02 | 3.49E-01 | 1.08E+00 | 5.43E-02 | 2.34E-01 | 1.03E+00 | 1.80E-01 | 7.64E-01 | 1.41E+00 | 1.30E-01 | 4.98E-01 | 1.36E+00 | 5.40E-02 | 1.92E-01 | -1.11E+00 | 2.49E-02 | 7.03E-02 | -1.84E+00 | 8.41E-03 | 3.09E-02 | -1.09E+00 | 1.31E-01 | 3.59E-01 | 1.10E+00 |
| PTOV1-AS1 | 6.69E-02 | 3.49E-01 | 1.11E+00 | 1.20E-01 | 5.17E-01 | 1.21E+00 | 1.63E-01 | 6.93E-01 | 1.14E+00 | 1.02E-01 | 3.90E-01 | 1.41E+00 | 5.84E-02 | 2.08E-01 | 2.27E+00 | 3.44E-01 | 9.71E-01 | 1.30E+00 | 1.04E-02 | 3.81E-02 | 1.97E+00 | 2.77E-01 | 7.62E-01 | 1.30E+00 |
| HBBP1 | 6.69E-02 | 3.49E-01 | -1.75E+00 | 8.54E-02 | 3.69E-01 | -1.73E+00 | 2.13E-01 | 9.07E-01 | 1.08E+00 | 2.62E-01 | 1.00E+00 | 1.00E+00 | 2.48E-01 | 8.85E-01 | 2.74E+00 | 3.35E-01 | 9.44E-01 | 1.21E+00 | 4.74E-08 | 1.74E-07 | -1.65E+00 | 3.69E-18 | 1.01E-17 | -4.67E+00 |
| ARRDC1-AS1 | 6.69E-02 | 3.49E-01 | 1.08E+00 | 1.08E-01 | 4.68E-01 | 1.85E+00 | 5.00E-05 | 2.13E-04 | -1.37E+00 | 4.53E-03 | 1.73E-02 | 1.04E+00 | 1.53E-02 | 5.45E-02 | -1.30E+00 | 3.02E-01 | 8.50E-01 | 1.31E+00 | 3.91E-03 | 1.44E-02 | -1.06E+00 | 4.76E-02 | 1.31E-01 | -1.01E+00 |
| ESCO2 | 6.69E-02 | 3.49E-01 | -1.67E+00 | 8.61E-02 | 3.72E-01 | -1.63E+00 | 5.99E-02 | 2.55E-01 | 1.53E+00 | 5.95E-02 | 2.27E-01 | 1.69E+00 | 1.88E-01 | 6.70E-01 | -2.39E+00 | 1.97E-01 | 5.54E-01 | -2.91E+00 | 9.63E-02 | 3.53E-01 | -1.23E+00 | 6.63E-03 | 1.82E-02 | 1.23E+00 |
| ADAP1 | 6.69E-02 | 3.49E-01 | 1.09E+00 | 5.94E-03 | 2.57E-02 | -1.09E+00 | 5.70E-03 | 2.42E-02 | 1.04E+00 | 1.12E-05 | 4.28E-05 | -1.13E+00 | 4.83E-03 | 1.72E-02 | -1.60E+00 | 8.27E-04 | 2.33E-03 | -2.17E+00 | 8.86E-02 | 3.25E-01 | -1.19E+00 | 9.72E-02 | 2.67E-01 | -1.21E+00 |
| PLPP7 | 6.69E-02 | 3.49E-01 | -1.11E+00 | 1.01E-01 | 4.37E-01 | 1.01E+00 | 9.98E-03 | 4.24E-02 | -1.51E+00 | 1.16E-02 | 4.44E-02 | -1.35E+00 | 1.08E-02 | 3.87E-02 | -1.78E+00 | 9.74E-02 | 2.74E-01 | -1.18E+00 | 5.11E-02 | 1.88E-01 | -1.02E+00 | 1.03E-01 | 2.83E-01 | 1.09E+00 |
| ULK1 | 6.70E-02 | 3.49E-01 | 1.08E+00 | 1.72E-01 | 7.44E-01 | 1.39E+00 | 6.32E-04 | 2.68E-03 | -1.31E+00 | 2.09E-05 | 7.98E-05 | -1.39E+00 | 1.26E-03 | 4.50E-03 | -1.31E+00 | 3.07E-04 | 8.66E-04 | -1.40E+00 | 6.00E-02 | 2.20E-01 | 1.01E+00 | 1.45E-02 | 3.98E-02 | -1.21E+00 |
| TPPP3 | 6.71E-02 | 3.50E-01 | 1.86E+00 | 2.30E-01 | 9.94E-01 | 1.28E+00 | 1.38E-01 | 5.87E-01 | 1.58E+00 | 8.44E-02 | 3.22E-01 | 1.02E+00 | 2.08E-01 | 7.43E-01 | -1.07E+00 | 1.95E-01 | 5.50E-01 | -1.20E+00 | 1.23E-01 | 4.53E-01 | 1.41E+00 | 3.64E-01 | 1.00E+00 | 1.00E+00 |
| CCDC180 | 6.72E-02 | 3.50E-01 | -1.05E+00 | 1.86E-01 | 8.02E-01 | 1.44E+00 | 4.28E-02 | 1.82E-01 | -1.26E+00 | 1.37E-01 | 5.23E-01 | 1.09E+00 | 2.61E-02 | 9.29E-02 | -1.48E+00 | 1.30E-01 | 3.68E-01 | -1.08E+00 | 9.01E-02 | 3.31E-01 | -1.18E+00 | 2.12E-01 | 5.81E-01 | 1.01E+00 |
| AL109918.1 | 6.72E-02 | 3.50E-01 | -1.14E+00 | 2.00E-01 | 8.66E-01 | 1.43E+00 | 1.85E-01 | 7.86E-01 | 1.16E+00 | 5.01E-02 | 1.91E-01 | 1.80E+00 | 1.17E-01 | 4.19E-01 | -1.08E+00 | 2.16E-01 | 6.10E-01 | 1.00E+00 | 1.22E-04 | 4.47E-04 | -1.43E+00 | 2.29E-02 | 6.29E-02 | -1.17E+00 |
| HSP90AB4P | 6.72E-02 | 3.50E-01 | 1.04E+00 | 3.90E-03 | 1.69E-02 | -1.33E+00 | 3.69E-02 | 1.57E-01 | -1.24E+00 | 3.12E-03 | 1.19E-02 | -1.38E+00 | 1.87E-01 | 6.68E-01 | 1.83E+00 | 3.15E-01 | 8.87E-01 | 1.28E+00 | 1.67E-01 | 6.11E-01 | 1.02E+00 | 3.06E-01 | 8.42E-01 | 1.41E+00 |
| IFI27L1 | 6.73E-02 | 3.51E-01 | 1.16E+00 | 2.83E-02 | 1.22E-01 | 1.13E+00 | 9.15E-02 | 3.89E-01 | 1.31E+00 | 1.40E-02 | 5.36E-02 | 1.04E+00 | 2.27E-01 | 8.08E-01 | 1.40E+00 | 1.11E-01 | 3.13E-01 | 1.20E+00 | 1.61E-01 | 5.90E-01 | 1.26E+00 | 1.55E-01 | 4.25E-01 | 1.12E+00 |
| SLC15A3 | 6.73E-02 | 3.51E-01 | -1.26E+00 | 1.33E-01 | 5.74E-01 | 1.01E+00 | 2.06E-01 | 8.76E-01 | 1.05E+00 | 1.81E-01 | 6.92E-01 | -1.02E+00 | 8.19E-03 | 2.92E-02 | -1.63E+00 | 1.48E-01 | 4.17E-01 | 1.15E+00 | 6.48E-02 | 2.38E-01 | -1.16E+00 | 2.29E-02 | 6.29E-02 | -1.50E+00 |
| AL031282.1 | 6.73E-02 | 3.51E-01 | 1.06E+00 | 1.72E-01 | 7.43E-01 | 1.55E+00 | 5.34E-06 | 2.27E-05 | -1.88E+00 | 1.08E-03 | 4.11E-03 | -1.19E+00 | 4.46E-02 | 1.59E-01 | -1.12E+00 | 2.96E-01 | 8.35E-01 | 1.43E+00 | 1.63E-01 | 5.98E-01 | 1.16E+00 | 1.26E-01 | 3.47E-01 | -1.02E+00 |
| COL5A1 | 6.74E-02 | 3.51E-01 | -1.07E+00 | 1.44E-02 | 6.20E-02 | -1.24E+00 | 3.95E-02 | 1.68E-01 | -1.18E+00 | 7.04E-03 | 2.69E-02 | -1.41E+00 | 1.43E-07 | 5.09E-07 | -2.66E+00 | 4.32E-08 | 1.22E-07 | -3.00E+00 | 1.18E-01 | 4.32E-01 | -1.17E+00 | 1.61E-01 | 4.43E-01 | -1.39E+00 |
| FAM83A-AS1 | 6.74E-02 | 3.51E-01 | 1.06E+00 | 2.13E-02 | 9.19E-02 | -1.12E+00 | 6.67E-02 | 2.83E-01 | -1.43E+00 | 6.17E-02 | 2.36E-01 | -1.43E+00 | 1.77E-04 | 6.33E-04 | -1.68E+00 | 6.41E-04 | 1.81E-03 | -1.48E+00 | 1.67E-01 | 6.11E-01 | 1.02E+00 | 6.82E-02 | 1.87E-01 | -1.35E+00 |
| ENGASE | 6.74E-02 | 3.51E-01 | 1.07E+00 | 9.84E-02 | 4.25E-01 | 1.64E+00 | 3.67E-03 | 1.56E-02 | -1.14E+00 | 1.63E-01 | 6.23E-01 | 1.41E+00 | 2.43E-02 | 8.65E-02 | -1.09E+00 | 1.53E-01 | 4.31E-01 | 1.21E+00 | 1.81E-01 | 6.66E-01 | 1.23E+00 | 3.24E-01 | 8.89E-01 | 1.47E+00 |
| CMTR1 | 6.74E-02 | 3.51E-01 | 1.13E+00 | 1.11E-01 | 4.79E-01 | 1.29E+00 | 1.81E-02 | 7.69E-02 | -1.04E+00 | 1.45E-03 | 5.54E-03 | -1.07E+00 | 2.12E-02 | 7.57E-02 | -1.07E+00 | 4.39E-03 | 1.24E-02 | -1.36E+00 | 1.32E-03 | 4.86E-03 | -1.21E+00 | 9.87E-04 | 2.71E-03 | -1.21E+00 |
| FAM229A | 6.74E-02 | 3.51E-01 | 1.57E+00 | 4.96E-02 | 2.14E-01 | -1.03E+00 | 7.43E-02 | 3.16E-01 | 1.41E+00 | 1.47E-02 | 5.61E-02 | -1.20E+00 | 1.10E-01 | 3.90E-01 | 1.86E+00 | 3.13E-01 | 8.83E-01 | 1.47E+00 | 8.01E-03 | 2.94E-02 | -1.19E+00 | 1.06E-03 | 2.91E-03 | -1.55E+00 |
| GPR82 | 6.75E-02 | 3.52E-01 | 1.07E+00 | 5.05E-02 | 2.18E-01 | 1.97E+00 | 2.07E-03 | 8.78E-03 | -1.64E+00 | 6.42E-02 | 2.45E-01 | 1.01E+00 | 2.00E-01 | 7.14E-01 | 1.07E+00 | 3.00E-01 | 8.46E-01 | 1.51E+00 | 1.95E-01 | 7.15E-01 | 1.08E+00 | 1.77E-01 | 4.87E-01 | -1.12E+00 |
| AL133477.1 | 6.75E-02 | 3.52E-01 | 1.30E+00 | 4.35E-05 | 1.88E-04 | -1.04E+00 | 9.73E-02 | 4.13E-01 | 1.12E+00 | 7.23E-05 | 2.76E-04 | -1.14E+00 | 1.10E-01 | 3.94E-01 | 1.35E+00 | 3.47E-03 | 9.77E-03 | -1.03E+00 | 2.57E-01 | 9.43E-01 | 1.34E+00 | 1.15E-02 | 3.16E-02 | -1.06E+00 |
| KHDRBS2 | 6.77E-02 | 3.53E-01 | 1.18E+00 | 1.85E-01 | 8.00E-01 | 1.66E+00 | 1.58E-01 | 6.71E-01 | 1.33E+00 | 1.15E-01 | 4.39E-01 | 1.55E+00 | 2.62E-01 | 9.35E-01 | 1.28E+00 | 2.87E-01 | 8.09E-01 | 1.42E+00 | 1.32E-01 | 4.86E-01 | 1.62E+00 | 1.35E-01 | 3.71E-01 | 1.61E+00 |
| PIH1D2 | 6.78E-02 | 3.53E-01 | -1.27E+00 | 5.74E-02 | 2.48E-01 | -1.31E+00 | 2.16E-01 | 9.17E-01 | 1.10E+00 | 8.78E-02 | 3.35E-01 | -1.18E+00 | 8.61E-03 | 3.07E-02 | -1.53E+00 | 1.84E-02 | 5.18E-02 | -1.33E+00 | 1.93E-01 | 7.10E-01 | 1.37E+00 | 1.60E-01 | 4.40E-01 | -1.12E+00 |
| AC145207.5 | 6.78E-02 | 3.53E-01 | -1.01E+00 | 5.55E-02 | 2.40E-01 | 1.04E+00 | 1.34E-02 | 5.70E-02 | 1.61E+00 | 2.62E-02 | 1.00E-01 | 1.83E+00 | 2.31E-01 | 8.25E-01 | 1.63E+00 | 1.41E-01 | 3.97E-01 | 1.10E+00 | 1.96E-01 | 7.20E-01 | 1.07E+00 | 1.90E-01 | 5.23E-01 | 1.29E+00 |
| BTBD3 | 6.78E-02 | 3.53E-01 | 1.22E+00 | 4.64E-03 | 2.01E-02 | 1.06E+00 | 4.08E-02 | 1.74E-01 | 1.06E+00 | 2.79E-03 | 1.07E-02 | 1.01E+00 | 2.37E-01 | 8.47E-01 | 1.31E+00 | 4.20E-02 | 1.18E-01 | -1.04E+00 | 4.19E-05 | 1.54E-04 | 2.24E+00 | 1.73E-01 | 4.74E-01 | 1.50E+00 |
| AC097376.1 | 6.79E-02 | 3.54E-01 | 1.59E+00 | 1.22E-01 | 5.26E-01 | 1.64E+00 | 1.50E-01 | 6.37E-01 | 1.38E+00 | 2.21E-01 | 8.44E-01 | 1.37E+00 | 1.03E-01 | 3.69E-01 | 2.03E+00 | 1.21E-01 | 3.40E-01 | 2.03E+00 | 2.68E-01 | 9.83E-01 | 1.42E+00 | 3.07E-01 | 8.43E-01 | 1.30E+00 |
| MIER2 | 6.79E-02 | 3.54E-01 | -1.10E+00 | 2.31E-01 | 1.00E+00 | 1.35E+00 | 1.89E-01 | 8.02E-01 | 1.10E+00 | 1.92E-01 | 7.33E-01 | 1.41E+00 | 6.71E-02 | 2.39E-01 | -1.52E+00 | 3.01E-01 | 8.47E-01 | 1.20E+00 | 1.03E-07 | 3.78E-07 | -1.82E+00 | 1.65E-04 | 4.53E-04 | -1.36E+00 |
| NDUFS6 | 6.79E-02 | 3.54E-01 | 1.23E+00 | 4.50E-07 | 1.94E-06 | 1.09E+00 | 6.61E-03 | 2.81E-02 | 1.46E+00 | 1.42E-01 | 5.43E-01 | 1.18E+00 | 2.63E-02 | 9.39E-02 | 1.09E+00 | 7.09E-03 | 2.00E-02 | 1.01E+00 | 2.44E-01 | 8.97E-01 | 1.18E+00 | 3.59E-01 | 9.86E-01 | 1.17E+00 |
| DENND1B | 6.79E-02 | 3.54E-01 | 1.14E+00 | 6.80E-02 | 2.94E-01 | 1.22E+00 | 1.24E-05 | 5.28E-05 | -1.23E+00 | 6.96E-07 | 2.66E-06 | -1.15E+00 | 2.76E-03 | 9.83E-03 | -1.09E+00 | 1.40E-05 | 3.96E-05 | -1.31E+00 | 5.38E-03 | 1.98E-02 | -1.12E+00 | 1.87E-02 | 5.15E-02 | -1.06E+00 |
| ABHD17A | 6.80E-02 | 3.54E-01 | 1.61E+00 | 1.32E-01 | 5.72E-01 | 1.68E+00 | 6.93E-02 | 2.95E-01 | 1.48E+00 | 1.20E-01 | 4.57E-01 | 1.34E+00 | 1.19E-01 | 4.23E-01 | 1.80E+00 | 1.13E-01 | 3.18E-01 | 1.98E+00 | 5.62E-02 | 2.06E-01 | 1.81E+00 | 4.99E-02 | 1.37E-01 | 1.83E+00 |
| AL137786.1 | 6.80E-02 | 3.54E-01 | 1.01E+00 | 4.77E-02 | 2.06E-01 | -1.06E+00 | 5.83E-02 | 2.48E-01 | -1.15E+00 | 1.05E-01 | 4.02E-01 | -1.01E+00 | 4.49E-07 | 1.60E-06 | -1.68E+00 | 2.56E-04 | 7.22E-04 | -1.19E+00 | 3.91E-05 | 1.43E-04 | -1.60E+00 | 4.46E-04 | 1.22E-03 | -1.45E+00 |
| AC092865.4 | 6.80E-02 | 3.54E-01 | 1.53E+00 | 8.87E-04 | 3.83E-03 | -1.02E+00 | 6.60E-03 | 2.81E-02 | 1.55E+00 | 1.36E-03 | 5.21E-03 | -1.05E+00 | 7.50E-02 | 2.68E-01 | 1.23E+00 | 7.01E-03 | 1.98E-02 | 1.04E+00 | 9.33E-02 | 3.42E-01 | 1.45E+00 | 1.09E-04 | 3.00E-04 | -1.30E+00 |
| JAZF1 | 6.80E-02 | 3.54E-01 | -1.32E+00 | 7.07E-02 | 3.06E-01 | -1.27E+00 | 2.20E-02 | 9.37E-02 | -1.59E+00 | 2.79E-02 | 1.06E-01 | -1.48E+00 | 1.33E-01 | 4.75E-01 | -1.16E+00 | 3.30E-01 | 9.29E-01 | 1.51E+00 | 9.34E-08 | 3.43E-07 | -1.36E+00 | 1.05E-04 | 2.87E-04 | -1.14E+00 |
| DHX37 | 6.80E-02 | 3.55E-01 | 1.04E+00 | 1.08E-01 | 4.68E-01 | 1.25E+00 | 1.11E-01 | 4.72E-01 | 1.08E+00 | 1.95E-01 | 7.46E-01 | 1.27E+00 | 6.82E-03 | 2.43E-02 | -1.36E+00 | 1.01E-01 | 2.85E-01 | 1.06E+00 | 3.04E-02 | 1.12E-01 | 1.08E+00 | 1.55E-01 | 4.27E-01 | 1.47E+00 |
| EIF4A1P2 | 6.81E-02 | 3.55E-01 | 1.24E+00 | 5.40E-02 | 2.33E-01 | 1.34E+00 | 9.71E-02 | 4.12E-01 | 1.16E+00 | 1.14E-01 | 4.35E-01 | 1.24E+00 | 1.31E-03 | 4.67E-03 | 1.01E+00 | 2.94E-01 | 8.29E-01 | 1.21E+00 | 1.38E-01 | 5.08E-01 | 1.27E+00 | 1.92E-01 | 5.27E-01 | 1.30E+00 |
| SBK1 | 6.81E-02 | 3.55E-01 | -1.03E+00 | 7.60E-02 | 3.28E-01 | -1.02E+00 | 2.24E-01 | 9.50E-01 | 1.14E+00 | 1.38E-01 | 5.26E-01 | -1.04E+00 | 2.72E-01 | 9.70E-01 | 1.24E+00 | 2.87E-01 | 8.08E-01 | 1.16E+00 | 2.42E-01 | 8.90E-01 | 1.31E+00 | 2.35E-01 | 6.44E-01 | 1.01E+00 |
| YWHAH | 6.81E-02 | 3.55E-01 | 1.49E+00 | 6.50E-04 | 2.81E-03 | 1.23E+00 | 1.31E-01 | 5.56E-01 | 1.22E+00 | 6.19E-02 | 2.37E-01 | 1.10E+00 | 1.43E-04 | 5.10E-04 | 1.09E+00 | 3.76E-03 | 1.06E-02 | 1.09E+00 | 4.36E-05 | 1.60E-04 | -1.04E+00 | 5.68E-04 | 1.56E-03 | -1.02E+00 |
| HCCS | 6.81E-02 | 3.55E-01 | 1.22E+00 | 2.04E-04 | 8.80E-04 | -1.25E+00 | 1.56E-01 | 6.63E-01 | 1.11E+00 | 2.04E-03 | 7.78E-03 | -1.13E+00 | 7.19E-04 | 2.56E-03 | -1.28E+00 | 1.60E-03 | 4.51E-03 | -1.15E+00 | 2.81E-02 | 1.03E-01 | 1.05E+00 | 1.72E-01 | 4.71E-01 | 1.22E+00 |
| AL162424.1 | 6.82E-02 | 3.55E-01 | -1.10E+00 | 9.17E-02 | 3.96E-01 | -1.11E+00 | 2.34E-01 | 9.96E-01 | 1.11E+00 | 2.23E-01 | 8.51E-01 | 1.02E+00 | 1.38E-04 | 4.94E-04 | -1.59E+00 | 5.16E-04 | 1.45E-03 | -1.46E+00 | 2.07E-01 | 7.61E-01 | 1.95E+00 | 5.09E-03 | 1.40E-02 | -1.15E+00 |
| PRRG1 | 6.82E-02 | 3.56E-01 | 1.01E+00 | 1.32E-01 | 5.72E-01 | 1.23E+00 | 5.74E-02 | 2.44E-01 | -1.23E+00 | 6.73E-02 | 2.57E-01 | -1.19E+00 | 2.15E-01 | 7.66E-01 | -1.21E+00 | 3.34E-01 | 9.41E-01 | -1.08E+00 | 1.10E-03 | 4.04E-03 | -1.68E+00 | 4.18E-02 | 1.15E-01 | -1.16E+00 |
| OSGIN2 | 6.82E-02 | 3.56E-01 | 1.21E+00 | 1.75E-04 | 7.54E-04 | 1.06E+00 | 2.15E-01 | 9.15E-01 | 1.21E+00 | 2.62E-03 | 1.00E-02 | 1.02E+00 | 1.52E-01 | 5.42E-01 | 1.35E+00 | 1.64E-03 | 4.61E-03 | -1.26E+00 | 1.51E-02 | 5.55E-02 | 1.02E+00 | 3.22E-04 | 8.85E-04 | -1.19E+00 |
| AC098487.1 | 6.84E-02 | 3.56E-01 | 1.09E+00 | 5.56E-03 | 2.40E-02 | -1.16E+00 | 7.23E-02 | 3.07E-01 | 1.02E+00 | 6.94E-03 | 2.65E-02 | -1.16E+00 | 1.65E-01 | 5.87E-01 | 1.05E+00 | 3.48E-01 | 9.82E-01 | 1.33E+00 | 2.16E-04 | 7.92E-04 | -1.28E+00 | 2.23E-04 | 6.13E-04 | -1.24E+00 |
| CERS2 | 6.84E-02 | 3.56E-01 | -1.13E+00 | 1.72E-01 | 7.44E-01 | 1.20E+00 | 1.74E-04 | 7.39E-04 | -1.75E+00 | 7.66E-03 | 2.93E-02 | -1.16E+00 | 1.07E-01 | 3.80E-01 | 1.02E+00 | 2.03E-01 | 5.73E-01 | 1.12E+00 | 3.67E-03 | 1.35E-02 | -1.19E+00 | 6.61E-03 | 1.82E-02 | -1.17E+00 |
| ALG10B | 6.86E-02 | 3.58E-01 | 1.20E+00 | 7.27E-02 | 3.14E-01 | 1.28E+00 | 6.19E-04 | 2.63E-03 | -1.28E+00 | 1.73E-02 | 6.60E-02 | 1.08E+00 | 1.18E-02 | 4.21E-02 | -1.11E+00 | 1.56E-01 | 4.40E-01 | 1.23E+00 | 5.80E-03 | 2.13E-02 | -1.18E+00 | 1.52E-01 | 4.18E-01 | 1.10E+00 |
| ZNF45 | 6.86E-02 | 3.58E-01 | -1.00E+00 | 1.92E-01 | 8.30E-01 | 1.42E+00 | 6.43E-06 | 2.73E-05 | -1.89E+00 | 5.73E-03 | 2.19E-02 | -1.05E+00 | 3.04E-04 | 1.08E-03 | -1.55E+00 | 1.86E-03 | 5.25E-03 | -1.32E+00 | 2.54E-03 | 9.34E-03 | -1.31E+00 | 5.19E-03 | 1.43E-02 | -1.25E+00 |
| AC032044.2 | 6.87E-02 | 3.58E-01 | 1.01E+00 | 3.08E-02 | 1.33E-01 | 1.02E+00 | 2.00E-03 | 8.52E-03 | -1.25E+00 | 1.36E-05 | 5.19E-05 | -1.52E+00 | 8.19E-02 | 2.92E-01 | -1.03E+00 | 8.74E-02 | 2.46E-01 | -1.06E+00 | 2.27E-01 | 8.33E-01 | 1.28E+00 | 1.04E-01 | 2.86E-01 | -1.04E+00 |
| AC004241.3 | 6.87E-02 | 3.58E-01 | -1.03E+00 | 4.79E-02 | 2.07E-01 | -1.04E+00 | 4.45E-02 | 1.89E-01 | -1.14E+00 | 2.98E-02 | 1.14E-01 | -1.10E+00 | 7.34E-04 | 2.62E-03 | -1.73E+00 | 3.73E-04 | 1.05E-03 | -1.78E+00 | 2.62E-02 | 9.60E-02 | -1.30E+00 | 4.66E-03 | 1.28E-02 | -1.52E+00 |
| AL591845.1 | 6.87E-02 | 3.58E-01 | 1.10E+00 | 3.44E-02 | 1.49E-01 | 2.03E+00 | 1.52E-03 | 6.44E-03 | -1.33E+00 | 1.94E-01 | 7.42E-01 | 1.33E+00 | 5.56E-03 | 1.98E-02 | -1.48E+00 | 2.20E-03 | 6.21E-03 | -1.55E+00 | 2.58E-01 | 9.46E-01 | 1.22E+00 | 2.62E-01 | 7.20E-01 | 1.57E+00 |
| EEF1DP3 | 6.88E-02 | 3.58E-01 | -1.21E+00 | 8.55E-02 | 3.70E-01 | -1.17E+00 | 5.85E-02 | 2.48E-01 | 1.59E+00 | 2.36E-01 | 9.03E-01 | 1.36E+00 | 2.70E-01 | 9.62E-01 | 1.21E+00 | 3.49E-01 | 9.83E-01 | 1.21E+00 | 2.41E-01 | 8.84E-01 | 1.25E+00 | 2.34E-01 | 6.43E-01 | 1.16E+00 |
| MAST2 | 6.88E-02 | 3.58E-01 | 1.07E+00 | 2.04E-01 | 8.82E-01 | 1.44E+00 | 8.97E-02 | 3.81E-01 | -1.00E+00 | 1.33E-01 | 5.07E-01 | 1.56E+00 | 2.41E-03 | 8.61E-03 | -1.64E+00 | 1.17E-02 | 3.29E-02 | -1.48E+00 | 5.24E-02 | 1.92E-01 | -1.11E+00 | 1.97E-02 | 5.40E-02 | 1.68E+00 |
| GREB1L | 6.88E-02 | 3.58E-01 | -1.13E+00 | 1.22E-01 | 5.26E-01 | 1.02E+00 | 1.30E-01 | 5.50E-01 | 1.04E+00 | 9.14E-02 | 3.49E-01 | 1.03E+00 | 1.92E-01 | 6.85E-01 | 1.35E+00 | 3.38E-01 | 9.52E-01 | 1.18E+00 | 5.29E-03 | 1.94E-02 | -1.17E+00 | 9.18E-03 | 2.52E-02 | -1.30E+00 |
| SLC16A14 | 6.89E-02 | 3.59E-01 | -1.08E+00 | 6.08E-02 | 2.63E-01 | -1.07E+00 | 1.74E-02 | 7.40E-02 | 1.45E+00 | 7.17E-02 | 2.74E-01 | -1.20E+00 | 2.61E-01 | 9.31E-01 | 1.15E+00 | 1.84E-01 | 5.19E-01 | -1.23E+00 | 1.45E-01 | 5.31E-01 | 1.01E+00 | 1.79E-01 | 4.92E-01 | -1.10E+00 |
| PLEKHB1 | 6.89E-02 | 3.59E-01 | 1.03E+00 | 1.68E-01 | 7.26E-01 | 1.49E+00 | 1.55E-01 | 6.57E-01 | 1.16E+00 | 8.18E-02 | 3.13E-01 | 1.11E+00 | 4.22E-03 | 1.50E-02 | -1.66E+00 | 4.52E-02 | 1.27E-01 | -1.13E+00 | 6.33E-02 | 2.32E-01 | -1.34E+00 | 1.54E-01 | 4.22E-01 | -1.13E+00 |
| YBX1P1 | 6.90E-02 | 3.60E-01 | 1.32E+00 | 8.26E-02 | 3.57E-01 | 1.29E+00 | 1.49E-02 | 6.33E-02 | 1.47E+00 | 1.55E-01 | 5.91E-01 | 1.37E+00 | 1.08E-01 | 3.86E-01 | 1.41E+00 | 3.87E-02 | 1.09E-01 | 1.56E+00 | 2.25E-01 | 8.26E-01 | 1.25E+00 | 2.61E-02 | 7.16E-02 | 1.42E+00 |
| PRIM1 | 6.90E-02 | 3.60E-01 | 1.25E+00 | 5.51E-02 | 2.38E-01 | 1.37E+00 | 7.16E-02 | 3.04E-01 | 1.17E+00 | 3.40E-03 | 1.30E-02 | 1.17E+00 | 2.97E-02 | 1.06E-01 | -1.14E+00 | 2.23E-01 | 6.28E-01 | 1.26E+00 | 1.27E-01 | 4.68E-01 | 1.15E+00 | 2.63E-01 | 7.23E-01 | 1.29E+00 |
| SRP14 | 6.90E-02 | 3.60E-01 | 1.00E+00 | 6.82E-03 | 2.95E-02 | -1.02E+00 | 5.54E-02 | 2.35E-01 | 1.07E+00 | 1.10E-01 | 4.20E-01 | -1.00E+00 | 1.03E-03 | 3.66E-03 | -1.08E+00 | 1.27E-01 | 3.59E-01 | -1.02E+00 | 8.55E-02 | 3.14E-01 | -1.01E+00 | 1.20E-01 | 3.31E-01 | -1.04E+00 |
| COL6A4P2 | 6.91E-02 | 3.60E-01 | 1.01E+00 | 1.92E-01 | 8.29E-01 | 1.48E+00 | 1.58E-04 | 6.72E-04 | -2.07E+00 | 4.70E-03 | 1.80E-02 | -1.26E+00 | 2.52E-01 | 8.99E-01 | 1.36E+00 | 2.77E-01 | 7.82E-01 | 1.67E+00 | 5.94E-02 | 2.18E-01 | -1.32E+00 | 7.46E-02 | 2.05E-01 | -1.44E+00 |
| DDAH2 | 6.91E-02 | 3.60E-01 | 1.15E+00 | 9.40E-02 | 4.06E-01 | 1.22E+00 | 5.48E-02 | 2.33E-01 | 1.01E+00 | 1.36E-01 | 5.19E-01 | 1.14E+00 | 3.65E-02 | 1.30E-01 | -1.02E+00 | 3.21E-06 | 9.05E-06 | -1.18E+00 | 1.18E-02 | 4.32E-02 | 1.13E+00 | 1.34E-02 | 3.67E-02 | 1.09E+00 |
| INE1 | 6.91E-02 | 3.60E-01 | 1.19E+00 | 3.39E-02 | 1.46E-01 | 1.03E+00 | 4.06E-04 | 1.73E-03 | -1.46E+00 | 1.31E-04 | 5.00E-04 | -1.46E+00 | 2.29E-01 | 8.18E-01 | 1.43E+00 | 3.48E-01 | 9.81E-01 | 1.31E+00 | 5.74E-02 | 2.11E-01 | -1.17E+00 | 8.08E-02 | 2.22E-01 | -1.15E+00 |
| AL162231.1 | 6.91E-02 | 3.60E-01 | 1.07E+00 | 1.36E-02 | 5.88E-02 | -1.29E+00 | 4.41E-02 | 1.87E-01 | 1.67E+00 | 2.41E-01 | 9.22E-01 | 1.14E+00 | 7.07E-02 | 2.52E-01 | -1.54E+00 | 9.05E-02 | 2.55E-01 | -1.40E+00 | 1.67E-02 | 6.15E-02 | 1.79E+00 | 2.98E-01 | 8.19E-01 | 1.19E+00 |
| CNR2 | 6.91E-02 | 3.60E-01 | -1.32E+00 | 1.60E-01 | 6.90E-01 | 1.02E+00 | 8.15E-02 | 3.46E-01 | -1.32E+00 | 1.54E-01 | 5.89E-01 | -1.12E+00 | 1.18E-28 | 4.19E-28 | -4.09E+00 | 4.33E-16 | 1.22E-15 | -1.87E+00 | 1.04E-01 | 3.81E-01 | 1.08E+00 | 3.09E-02 | 8.49E-02 | -1.35E+00 |
| CCDC84 | 6.91E-02 | 3.60E-01 | 1.21E+00 | 3.68E-02 | 1.59E-01 | 1.28E+00 | 7.66E-02 | 3.26E-01 | -1.03E+00 | 3.69E-02 | 1.41E-01 | 1.07E+00 | 1.28E-01 | 4.58E-01 | 1.30E+00 | 3.03E-01 | 8.55E-01 | 1.50E+00 | 1.70E-01 | 6.22E-01 | 1.34E+00 | 3.46E-01 | 9.49E-01 | 1.25E+00 |
| TMPO-AS1 | 6.93E-02 | 3.61E-01 | 1.01E+00 | 7.74E-02 | 3.34E-01 | 1.12E+00 | 2.35E-01 | 1.00E+00 | 1.13E+00 | 1.36E-02 | 5.21E-02 | -1.04E+00 | 6.95E-02 | 2.48E-01 | -1.18E+00 | 1.51E-01 | 4.26E-01 | 1.03E+00 | 7.50E-06 | 2.75E-05 | -1.58E+00 | 3.26E-01 | 8.96E-01 | 1.23E+00 |
| ACSM1 | 6.93E-02 | 3.61E-01 | 1.90E+00 | 2.04E-01 | 8.79E-01 | 1.46E+00 | 3.60E-02 | 1.53E-01 | -1.37E+00 | 8.24E-02 | 3.15E-01 | -1.09E+00 | 2.51E-01 | 8.93E-01 | 1.46E+00 | 1.70E-01 | 4.78E-01 | 1.89E+00 | 2.86E-02 | 1.05E-01 | 1.03E+00 | 8.90E-02 | 2.44E-01 | 1.10E+00 |
| SOSTDC1 | 6.93E-02 | 3.61E-01 | -1.46E+00 | 1.59E-01 | 6.87E-01 | 1.53E+00 | 1.72E-05 | 7.32E-05 | -2.59E+00 | 9.56E-02 | 3.65E-01 | 1.84E+00 | 1.25E-05 | 4.47E-05 | -2.51E+00 | 3.08E-01 | 8.69E-01 | 1.48E+00 | 1.18E-02 | 4.32E-02 | -1.34E+00 | 9.09E-02 | 2.50E-01 | 1.33E+00 |
| ZNF570 | 6.93E-02 | 3.61E-01 | -1.05E+00 | 8.22E-03 | 3.55E-02 | -1.32E+00 | 3.95E-02 | 1.68E-01 | -1.01E+00 | 1.94E-02 | 7.42E-02 | 1.06E+00 | 4.28E-03 | 1.53E-02 | -1.40E+00 | 9.34E-04 | 2.63E-03 | -1.75E+00 | 1.81E-01 | 6.66E-01 | 1.12E+00 | 8.23E-02 | 2.26E-01 | -1.11E+00 |
| AC005086.1 | 6.94E-02 | 3.61E-01 | 1.16E+00 | 2.64E-02 | 1.14E-01 | 1.07E+00 | 5.10E-02 | 2.17E-01 | -1.14E+00 | 2.70E-04 | 1.03E-03 | -1.55E+00 | 1.02E-01 | 3.62E-01 | 1.06E+00 | 1.65E-01 | 4.66E-01 | 1.09E+00 | 1.14E-03 | 4.19E-03 | -1.62E+00 | 6.37E-04 | 1.75E-03 | -1.76E+00 |
| FPGT-TNNI3K | 6.94E-02 | 3.62E-01 | 1.77E+00 | 1.67E-02 | 7.20E-02 | 2.38E+00 | 6.18E-02 | 2.63E-01 | -1.22E+00 | 1.90E-01 | 7.24E-01 | 1.14E+00 | 1.11E-01 | 3.96E-01 | -1.08E+00 | 1.16E-01 | 3.28E-01 | -1.12E+00 | 4.94E-02 | 1.81E-01 | 1.52E+00 | 2.20E-01 | 6.05E-01 | 1.39E+00 |
| SLC31A2 | 6.95E-02 | 3.62E-01 | 1.71E+00 | 4.13E-02 | 1.78E-01 | 2.10E+00 | 1.63E-02 | 6.94E-02 | 1.66E+00 | 1.37E-01 | 5.25E-01 | 1.62E+00 | 1.71E-01 | 6.08E-01 | 1.07E+00 | 2.12E-01 | 5.98E-01 | 1.04E+00 | 2.11E-01 | 7.74E-01 | -1.23E+00 | 3.36E-01 | 9.22E-01 | -1.08E+00 |
| NFKBIA | 6.95E-02 | 3.62E-01 | 1.21E+00 | 8.54E-09 | 3.69E-08 | -1.19E+00 | 3.82E-03 | 1.62E-02 | -1.05E+00 | 1.37E-06 | 5.23E-06 | -1.32E+00 | 2.28E-18 | 8.14E-18 | -1.69E+00 | 1.20E-24 | 3.38E-24 | -2.09E+00 | 7.82E-45 | 2.87E-44 | -2.42E+00 | 2.14E-42 | 5.89E-42 | -3.22E+00 |
| AC099494.1 | 6.96E-02 | 3.63E-01 | 1.51E+00 | 1.99E-01 | 8.61E-01 | 1.36E+00 | 1.14E-01 | 4.85E-01 | 1.36E+00 | 8.39E-02 | 3.21E-01 | 1.77E+00 | 1.19E-01 | 4.24E-01 | -1.31E+00 | 2.76E-01 | 7.79E-01 | 1.04E+00 | 1.35E-01 | 4.96E-01 | -1.17E+00 | 2.59E-01 | 7.12E-01 | 1.38E+00 |
| RASL10A | 6.96E-02 | 3.63E-01 | 1.70E+00 | 5.89E-02 | 2.55E-01 | 1.01E+00 | 3.32E-02 | 1.41E-01 | 1.99E+00 | 2.50E-02 | 9.55E-02 | 2.24E+00 | 1.36E-03 | 4.85E-03 | 3.16E+00 | 2.86E-01 | 8.07E-01 | 1.38E+00 | 2.14E-01 | 7.84E-01 | 1.28E+00 | 3.10E-01 | 8.50E-01 | 1.30E+00 |
| CR381670.1 | 6.96E-02 | 3.63E-01 | 1.13E+00 | 3.10E-02 | 1.34E-01 | 1.02E+00 | 2.18E-01 | 9.25E-01 | 1.23E+00 | 2.53E-01 | 9.68E-01 | 1.40E+00 | 2.51E-01 | 8.94E-01 | 1.17E+00 | 3.06E-01 | 8.62E-01 | 1.34E+00 | 2.23E-01 | 8.19E-01 | 1.24E+00 | 2.12E-01 | 5.82E-01 | 1.27E+00 |
| MRC2 | 6.96E-02 | 3.63E-01 | 1.23E+00 | 2.83E-03 | 1.22E-02 | -1.02E+00 | 5.70E-02 | 2.42E-01 | -1.00E+00 | 2.43E-02 | 9.29E-02 | 1.04E+00 | 4.83E-02 | 1.72E-01 | -1.49E+00 | 4.96E-02 | 1.40E-01 | -1.52E+00 | 2.36E-01 | 8.65E-01 | 1.28E+00 | 3.16E-01 | 8.67E-01 | 1.25E+00 |
| SKA3 | 6.96E-02 | 3.63E-01 | -1.17E+00 | 2.40E-02 | 1.04E-01 | -1.62E+00 | 5.88E-02 | 2.50E-01 | 1.51E+00 | 1.86E-02 | 7.10E-02 | 1.17E+00 | 2.31E-01 | 8.22E-01 | 1.53E+00 | 2.90E-01 | 8.17E-01 | -1.03E+00 | 2.20E-01 | 8.09E-01 | 1.50E+00 | 2.52E-01 | 6.91E-01 | 1.59E+00 |
| RBM44 | 6.97E-02 | 3.63E-01 | 1.47E+00 | 2.31E-01 | 9.98E-01 | 1.13E+00 | 8.25E-02 | 3.50E-01 | 1.31E+00 | 1.10E-01 | 4.19E-01 | 1.46E+00 | 2.75E-01 | 9.81E-01 | 1.20E+00 | 3.07E-01 | 8.67E-01 | 1.53E+00 | 8.61E-02 | 3.16E-01 | 1.60E+00 | 1.65E-01 | 4.53E-01 | 1.47E+00 |
| TRDC | 6.97E-02 | 3.63E-01 | -2.22E+00 | 5.95E-02 | 2.57E-01 | -2.45E+00 | 7.67E-03 | 3.26E-02 | -2.80E+00 | 6.60E-03 | 2.52E-02 | -2.97E+00 | 2.13E-01 | 7.60E-01 | 1.46E+00 | 9.21E-02 | 2.60E-01 | 1.09E+00 | 7.60E-06 | 2.79E-05 | -1.70E+00 | 3.31E-05 | 9.10E-05 | -1.80E+00 |
| AC034236.2 | 6.98E-02 | 3.64E-01 | 1.06E+00 | 1.84E-01 | 7.94E-01 | 1.40E+00 | 5.59E-02 | 2.38E-01 | -1.12E+00 | 8.46E-03 | 3.23E-02 | -1.31E+00 | 1.34E-01 | 4.78E-01 | -1.03E+00 | 2.44E-01 | 6.88E-01 | 1.18E+00 | 1.77E-01 | 6.48E-01 | 1.12E+00 | 2.09E-01 | 5.73E-01 | 1.61E+00 |
| AC091133.4 | 6.99E-02 | 3.64E-01 | 1.12E+00 | 2.88E-02 | 1.24E-01 | 1.04E+00 | 3.80E-05 | 1.61E-04 | -1.30E+00 | 6.15E-07 | 2.35E-06 | -1.42E+00 | 8.24E-05 | 2.94E-04 | -1.89E+00 | 1.07E-03 | 3.02E-03 | -1.42E+00 | 4.93E-09 | 1.81E-08 | -1.58E+00 | 4.70E-07 | 1.29E-06 | -1.35E+00 |
| AL136295.7 | 7.02E-02 | 3.66E-01 | -1.07E+00 | 1.85E-01 | 8.00E-01 | 1.32E+00 | 2.22E-01 | 9.43E-01 | 1.20E+00 | 2.14E-01 | 8.16E-01 | 1.40E+00 | 2.69E-01 | 9.61E-01 | 1.34E+00 | 1.47E-01 | 4.14E-01 | 1.80E+00 | 2.26E-01 | 8.31E-01 | 1.17E+00 | 1.94E-01 | 5.33E-01 | 1.54E+00 |
| AC016831.1 | 7.02E-02 | 3.66E-01 | 1.13E+00 | 6.78E-05 | 2.93E-04 | -1.22E+00 | 1.89E-01 | 8.03E-01 | 1.29E+00 | 8.23E-04 | 3.14E-03 | -1.47E+00 | 7.49E-02 | 2.67E-01 | 2.01E+00 | 2.08E-01 | 5.86E-01 | 1.81E+00 | 1.14E-01 | 4.18E-01 | 1.11E+00 | 8.42E-03 | 2.31E-02 | -1.61E+00 |
| NCOR2 | 7.02E-02 | 3.66E-01 | 1.12E+00 | 1.33E-01 | 5.73E-01 | 1.35E+00 | 4.51E-02 | 1.92E-01 | 1.39E+00 | 1.51E-01 | 5.77E-01 | 1.50E+00 | 8.64E-02 | 3.08E-01 | 1.12E+00 | 3.17E-01 | 8.94E-01 | 1.36E+00 | 4.32E-02 | 1.58E-01 | 1.09E+00 | 4.27E-02 | 1.17E-01 | 1.03E+00 |
| AC090152.1 | 7.04E-02 | 3.67E-01 | 1.19E+00 | 6.00E-02 | 2.59E-01 | 1.38E+00 | 5.94E-02 | 2.53E-01 | 1.43E+00 | 2.55E-01 | 9.75E-01 | 1.47E+00 | 2.62E-01 | 9.33E-01 | 1.45E+00 | 3.04E-01 | 8.57E-01 | 1.44E+00 | 7.22E-02 | 2.65E-01 | 1.45E+00 | 6.00E-03 | 1.65E-02 | 1.79E+00 |
| AL603914.1 | 7.04E-02 | 3.67E-01 | 1.50E+00 | 7.55E-02 | 3.26E-01 | 1.66E+00 | 2.28E-02 | 9.69E-02 | -1.26E+00 | 4.53E-02 | 1.73E-01 | -1.09E+00 | 4.90E-08 | 1.75E-07 | -2.74E+00 | 1.16E-05 | 3.27E-05 | -1.84E+00 | 1.37E-05 | 5.01E-05 | -2.39E+00 | 9.34E-03 | 2.57E-02 | -1.28E+00 |
| ASB14 | 7.04E-02 | 3.67E-01 | 1.15E+00 | 1.48E-01 | 6.38E-01 | 1.43E+00 | 1.93E-04 | 8.18E-04 | -1.21E+00 | 2.20E-03 | 8.40E-03 | 1.02E+00 | 6.17E-03 | 2.20E-02 | -1.45E+00 | 1.55E-02 | 4.37E-02 | -1.29E+00 | 1.57E-01 | 5.77E-01 | 1.02E+00 | 1.36E-01 | 3.74E-01 | -1.07E+00 |
| KANSL1-AS1 | 7.05E-02 | 3.67E-01 | 1.06E+00 | 7.55E-02 | 3.26E-01 | 1.17E+00 | 1.35E-01 | 5.76E-01 | 1.44E+00 | 6.35E-02 | 2.43E-01 | 1.03E+00 | 2.47E-01 | 8.79E-01 | 1.10E+00 | 9.90E-02 | 2.79E-01 | -1.09E+00 | 2.58E-01 | 9.45E-01 | 1.17E+00 | 6.82E-02 | 1.87E-01 | -1.28E+00 |
| VEGFB | 7.05E-02 | 3.67E-01 | 1.18E+00 | 4.59E-02 | 1.98E-01 | 1.28E+00 | 9.84E-02 | 4.18E-01 | 1.42E+00 | 1.12E-01 | 4.27E-01 | 1.30E+00 | 1.04E-01 | 3.70E-01 | 1.12E+00 | 1.27E-01 | 3.59E-01 | 1.13E+00 | 1.92E-01 | 7.04E-01 | 1.19E+00 | 2.27E-01 | 6.23E-01 | 1.25E+00 |
| FAM219B | 7.05E-02 | 3.68E-01 | 1.13E+00 | 8.69E-02 | 3.75E-01 | 1.26E+00 | 8.56E-02 | 3.64E-01 | 1.04E+00 | 2.56E-01 | 9.79E-01 | 1.37E+00 | 7.22E-03 | 2.57E-02 | -1.13E+00 | 2.16E-02 | 6.09E-02 | -1.03E+00 | 9.53E-03 | 3.50E-02 | -1.09E+00 | 3.50E-02 | 9.61E-02 | -1.11E+00 |
| PGM2L1 | 7.05E-02 | 3.68E-01 | 1.53E+00 | 8.04E-02 | 3.47E-01 | 1.50E+00 | 9.35E-02 | 3.97E-01 | 1.17E+00 | 4.56E-02 | 1.74E-01 | 1.22E+00 | 1.30E-01 | 4.64E-01 | 1.70E+00 | 1.63E-01 | 4.58E-01 | 1.26E+00 | 2.46E-01 | 9.04E-01 | 1.34E+00 | 1.46E-01 | 4.00E-01 | 1.10E+00 |
| ZUFSP | 7.06E-02 | 3.68E-01 | 1.21E+00 | 1.95E-02 | 8.41E-02 | 1.11E+00 | 4.89E-02 | 2.08E-01 | 1.02E+00 | 2.69E-02 | 1.03E-01 | 1.05E+00 | 6.60E-02 | 2.35E-01 | 1.14E+00 | 2.86E-01 | 8.05E-01 | 1.41E+00 | 1.82E-03 | 6.68E-03 | -1.17E+00 | 1.42E-02 | 3.89E-02 | -1.06E+00 |
| RPS15AP29 | 7.09E-02 | 3.70E-01 | 1.09E+00 | 2.43E-03 | 1.05E-02 | -1.39E+00 | 1.69E-01 | 7.16E-01 | 1.36E+00 | 1.85E-02 | 7.07E-02 | -1.22E+00 | 1.94E-01 | 6.91E-01 | 1.57E+00 | 3.90E-02 | 1.10E-01 | -1.25E+00 | 2.39E-01 | 8.78E-01 | 1.30E+00 | 1.13E-02 | 3.11E-02 | -1.61E+00 |
| HNRNPLP1 | 7.09E-02 | 3.70E-01 | 1.60E+00 | 2.23E-01 | 9.64E-01 | 1.34E+00 | 3.94E-02 | 1.68E-01 | 1.69E+00 | 1.82E-01 | 6.94E-01 | 1.62E+00 | 2.51E-01 | 8.96E-01 | 1.30E+00 | 2.06E-01 | 5.81E-01 | 1.06E+00 | 6.62E-02 | 2.43E-01 | 1.02E+00 | 3.43E-01 | 9.42E-01 | 1.21E+00 |
| INCENP | 7.10E-02 | 3.70E-01 | -1.29E+00 | 1.95E-01 | 8.43E-01 | 1.15E+00 | 2.25E-01 | 9.56E-01 | 1.19E+00 | 1.84E-01 | 7.01E-01 | 1.41E+00 | 6.72E-02 | 2.40E-01 | -1.33E+00 | 1.58E-01 | 4.47E-01 | -1.07E+00 | 2.60E-01 | 9.53E-01 | 1.16E+00 | 2.60E-01 | 7.15E-01 | 1.26E+00 |
| STAR | 7.10E-02 | 3.70E-01 | 1.58E+00 | 4.72E-02 | 2.04E-01 | 2.03E+00 | 1.26E-01 | 5.37E-01 | -1.05E+00 | 2.51E-01 | 9.58E-01 | 1.25E+00 | 1.84E-01 | 6.55E-01 | -1.22E+00 | 3.45E-01 | 9.71E-01 | 1.19E+00 | 4.02E-02 | 1.48E-01 | -1.41E+00 | 2.76E-01 | 7.59E-01 | -1.01E+00 |
| GLRX | 7.10E-02 | 3.70E-01 | 1.18E+00 | 6.08E-02 | 2.63E-01 | 1.28E+00 | 5.89E-02 | 2.50E-01 | 1.00E+00 | 5.99E-05 | 2.29E-04 | -1.14E+00 | 6.54E-05 | 2.33E-04 | -1.28E+00 | 2.61E-03 | 7.36E-03 | -1.05E+00 | 7.77E-07 | 2.85E-06 | -1.23E+00 | 5.55E-03 | 1.52E-02 | -1.04E+00 |
| AC103702.1 | 7.10E-02 | 3.70E-01 | 1.01E+00 | 1.49E-01 | 6.43E-01 | 1.48E+00 | 1.75E-01 | 7.42E-01 | 1.38E+00 | 9.54E-02 | 3.65E-01 | 1.76E+00 | 2.54E-01 | 9.04E-01 | 1.25E+00 | 1.13E-01 | 3.20E-01 | -1.26E+00 | 1.55E-01 | 5.67E-01 | 1.66E+00 | 2.99E-01 | 8.20E-01 | 1.41E+00 |
| AC234775.3 | 7.10E-02 | 3.70E-01 | 1.00E+00 | 5.30E-02 | 2.29E-01 | -1.04E+00 | 1.18E-01 | 5.01E-01 | 1.31E+00 | 2.14E-01 | 8.17E-01 | 1.06E+00 | 1.94E-01 | 6.93E-01 | 1.10E+00 | 2.99E-01 | 8.43E-01 | 1.48E+00 | 1.47E-01 | 5.39E-01 | 1.06E+00 | 1.53E-01 | 4.21E-01 | 1.16E+00 |
| PVR | 7.10E-02 | 3.70E-01 | 1.77E+00 | 5.70E-02 | 2.46E-01 | 2.07E+00 | 6.50E-04 | 2.76E-03 | -1.43E+00 | 5.19E-03 | 1.98E-02 | -1.12E+00 | 8.91E-02 | 3.18E-01 | -1.32E+00 | 2.59E-01 | 7.29E-01 | 1.24E+00 | 1.18E-05 | 4.34E-05 | -1.96E+00 | 9.38E-04 | 2.58E-03 | -1.44E+00 |
| C2orf27A | 7.11E-02 | 3.70E-01 | -1.15E+00 | 2.60E-02 | 1.12E-01 | 2.02E+00 | 7.82E-02 | 3.32E-01 | -1.08E+00 | 2.13E-01 | 8.15E-01 | 1.37E+00 | 2.55E-01 | 9.10E-01 | 1.17E+00 | 1.39E-01 | 3.93E-01 | 1.57E+00 | 6.65E-02 | 2.44E-01 | -1.09E+00 | 3.19E-01 | 8.75E-01 | 1.33E+00 |
| NRSN2-AS1 | 7.11E-02 | 3.71E-01 | 1.18E+00 | 7.56E-03 | 3.27E-02 | -1.02E+00 | 1.73E-06 | 7.36E-06 | -1.63E+00 | 1.44E-06 | 5.49E-06 | -1.50E+00 | 1.10E-02 | 3.92E-02 | -1.07E+00 | 2.96E-03 | 8.33E-03 | -1.15E+00 | 1.54E-02 | 5.67E-02 | -1.33E+00 | 1.06E-01 | 2.92E-01 | -1.07E+00 |
| DPP4 | 7.11E-02 | 3.71E-01 | 1.31E+00 | 5.76E-02 | 2.49E-01 | -1.19E+00 | 1.77E-01 | 7.53E-01 | 1.25E+00 | 2.21E-01 | 8.44E-01 | -1.02E+00 | 2.14E-01 | 7.62E-01 | 1.27E+00 | 2.95E-01 | 8.33E-01 | 1.21E+00 | 2.19E-01 | 8.05E-01 | 1.01E+00 | 2.90E-01 | 7.98E-01 | 1.17E+00 |
| ADCY9 | 7.11E-02 | 3.71E-01 | 1.07E+00 | 1.84E-01 | 7.97E-01 | 1.66E+00 | 1.41E-03 | 6.00E-03 | -1.43E+00 | 5.89E-02 | 2.25E-01 | 1.10E+00 | 1.26E-01 | 4.50E-01 | -1.26E+00 | 2.83E-01 | 7.99E-01 | 1.21E+00 | 6.87E-02 | 2.52E-01 | -1.56E+00 | 3.60E-01 | 9.90E-01 | 1.11E+00 |
| AC068491.4 | 7.12E-02 | 3.71E-01 | -1.22E+00 | 1.14E-01 | 4.94E-01 | -1.07E+00 | 6.18E-03 | 2.63E-02 | -1.83E+00 | 1.18E-01 | 4.51E-01 | 1.14E+00 | 7.69E-07 | 2.74E-06 | -2.00E+00 | 4.53E-02 | 1.28E-01 | 1.07E+00 | 7.93E-14 | 2.91E-13 | -1.76E+00 | 6.09E-06 | 1.67E-05 | -1.23E+00 |
| USP45 | 7.12E-02 | 3.71E-01 | 1.09E+00 | 1.06E-01 | 4.58E-01 | 1.80E+00 | 2.52E-05 | 1.07E-04 | -1.68E+00 | 2.99E-03 | 1.14E-02 | -1.07E+00 | 2.75E-03 | 9.79E-03 | -1.25E+00 | 6.88E-02 | 1.94E-01 | 1.15E+00 | 5.89E-04 | 2.16E-03 | -1.22E+00 | 5.71E-03 | 1.57E-02 | -1.09E+00 |
| AL021707.6 | 7.12E-02 | 3.71E-01 | 1.04E+00 | 3.02E-02 | 1.31E-01 | -1.37E+00 | 8.67E-02 | 3.68E-01 | -1.10E+00 | 1.65E-01 | 6.31E-01 | 1.15E+00 | 8.28E-02 | 2.95E-01 | -1.15E+00 | 1.97E-01 | 5.55E-01 | 1.17E+00 | 1.39E-08 | 5.11E-08 | -1.80E+00 | 5.59E-07 | 1.54E-06 | -1.53E+00 |
| MCM10 | 7.13E-02 | 3.72E-01 | -1.62E+00 | 5.95E-02 | 2.57E-01 | -2.06E+00 | 1.16E-01 | 4.95E-01 | -1.01E+00 | 1.67E-01 | 6.39E-01 | 1.22E+00 | 1.78E-01 | 6.34E-01 | -1.82E+00 | 2.47E-01 | 6.96E-01 | -1.53E+00 | 2.40E-02 | 8.82E-02 | -1.01E+00 | 2.93E-04 | 8.06E-04 | 1.73E+00 |
| AL590764.1 | 7.13E-02 | 3.72E-01 | -1.10E+00 | 2.21E-01 | 9.53E-01 | 1.33E+00 | 4.02E-02 | 1.71E-01 | -1.07E+00 | 6.04E-02 | 2.31E-01 | 1.11E+00 | 4.28E-04 | 1.53E-03 | -1.33E+00 | 8.94E-03 | 2.52E-02 | -1.11E+00 | 2.13E-03 | 7.82E-03 | -1.47E+00 | 2.99E-01 | 8.21E-01 | 1.17E+00 |
| SETBP1 | 7.14E-02 | 3.72E-01 | 1.40E+00 | 1.32E-04 | 5.71E-04 | 2.07E+00 | 1.85E-02 | 7.87E-02 | 1.92E+00 | 3.75E-03 | 1.43E-02 | 2.50E+00 | 2.37E-01 | 8.46E-01 | 1.54E+00 | 3.01E-01 | 8.48E-01 | 1.47E+00 | 2.47E-01 | 9.06E-01 | 1.15E+00 | 3.30E-01 | 9.08E-01 | 1.23E+00 |
| TMEM104 | 7.15E-02 | 3.73E-01 | 1.71E+00 | 4.28E-02 | 1.85E-01 | 2.04E+00 | 4.47E-03 | 1.90E-02 | -1.45E+00 | 1.22E-01 | 4.66E-01 | 1.18E+00 | 1.86E-01 | 6.64E-01 | 1.68E+00 | 2.52E-01 | 7.09E-01 | 1.62E+00 | 9.73E-02 | 3.57E-01 | 1.20E+00 | 3.33E-02 | 9.14E-02 | -1.11E+00 |
| TRPM2 | 7.16E-02 | 3.73E-01 | -1.35E+00 | 9.22E-02 | 3.98E-01 | -1.20E+00 | 2.87E-03 | 1.22E-02 | 1.70E+00 | 1.03E-01 | 3.94E-01 | 1.40E+00 | 2.48E-01 | 8.86E-01 | 1.23E+00 | 3.55E-01 | 1.00E+00 | 1.00E+00 | 2.44E-01 | 8.95E-01 | -1.02E+00 | 2.99E-01 | 8.21E-01 | -1.10E+00 |
| ARMC2 | 7.16E-02 | 3.73E-01 | 1.00E+00 | 1.25E-02 | 5.38E-02 | -1.20E+00 | 3.75E-03 | 1.59E-02 | -1.26E+00 | 2.25E-03 | 8.60E-03 | -1.18E+00 | 1.54E-01 | 5.48E-01 | 1.11E+00 | 1.76E-01 | 4.96E-01 | 1.04E+00 | 4.92E-03 | 1.81E-02 | -1.54E+00 | 8.60E-03 | 2.36E-02 | -1.42E+00 |
| SH3RF1 | 7.17E-02 | 3.74E-01 | 1.21E+00 | 3.82E-04 | 1.65E-03 | 1.05E+00 | 6.61E-02 | 2.81E-01 | 1.03E+00 | 1.59E-02 | 6.06E-02 | 1.06E+00 | 4.56E-02 | 1.63E-01 | -1.01E+00 | 2.80E-02 | 7.91E-02 | -1.12E+00 | 4.52E-02 | 1.66E-01 | -1.21E+00 | 2.04E-01 | 5.59E-01 | 1.02E+00 |
| MADD | 7.17E-02 | 3.74E-01 | 1.17E+00 | 5.91E-02 | 2.55E-01 | 1.84E+00 | 6.69E-04 | 2.84E-03 | -1.20E+00 | 5.98E-02 | 2.28E-01 | 1.27E+00 | 2.69E-03 | 9.59E-03 | -1.01E+00 | 1.65E-01 | 4.64E-01 | 1.27E+00 | 1.07E-02 | 3.94E-02 | 1.10E+00 | 1.05E-01 | 2.89E-01 | 1.25E+00 |
| AC011379.1 | 7.19E-02 | 3.74E-01 | 1.09E+00 | 2.10E-11 | 9.07E-11 | -2.35E+00 | 4.87E-02 | 2.07E-01 | -1.19E+00 | 1.96E-06 | 7.48E-06 | -3.66E+00 | 1.74E-01 | 6.19E-01 | 1.68E+00 | 5.12E-02 | 1.44E-01 | -2.03E+00 | 1.35E-01 | 4.95E-01 | 1.42E+00 | 2.69E-01 | 7.38E-01 | -1.24E+00 |
| U62317.3 | 7.19E-02 | 3.74E-01 | 1.67E+00 | 1.30E-01 | 5.61E-01 | 1.63E+00 | 2.10E-01 | 8.94E-01 | 1.18E+00 | 2.30E-01 | 8.78E-01 | 1.22E+00 | 1.07E-01 | 3.82E-01 | -1.26E+00 | 2.49E-01 | 7.02E-01 | 1.13E+00 | 7.81E-03 | 2.87E-02 | -1.27E+00 | 2.33E-01 | 6.41E-01 | 1.13E+00 |
| SLC17A5 | 7.19E-02 | 3.75E-01 | 1.11E+00 | 1.32E-01 | 5.72E-01 | 1.33E+00 | 8.41E-03 | 3.57E-02 | -1.18E+00 | 3.16E-02 | 1.21E-01 | 1.15E+00 | 6.44E-03 | 2.30E-02 | -1.42E+00 | 1.25E-02 | 3.52E-02 | -1.32E+00 | 3.23E-02 | 1.19E-01 | -1.05E+00 | 1.46E-02 | 4.02E-02 | -1.15E+00 |
| AC004803.1 | 7.20E-02 | 3.75E-01 | -1.01E+00 | 1.35E-01 | 5.82E-01 | 1.68E+00 | 5.49E-02 | 2.33E-01 | -1.09E+00 | 9.98E-02 | 3.81E-01 | 1.11E+00 | 8.28E-02 | 2.95E-01 | -1.13E+00 | 4.45E-02 | 1.26E-01 | -1.45E+00 | 1.17E-01 | 4.28E-01 | -1.05E+00 | 3.43E-01 | 9.41E-01 | 1.17E+00 |
| AL137145.2 | 7.20E-02 | 3.75E-01 | -1.03E+00 | 9.15E-02 | 3.96E-01 | 1.11E+00 | 1.95E-02 | 8.29E-02 | -1.29E+00 | 4.67E-03 | 1.78E-02 | -1.51E+00 | 6.78E-02 | 2.42E-01 | -1.32E+00 | 1.54E-01 | 4.33E-01 | -1.19E+00 | 1.56E-01 | 5.73E-01 | 1.38E+00 | 1.60E-01 | 4.40E-01 | 1.61E+00 |
| AP002495.1 | 7.22E-02 | 3.76E-01 | -1.14E+00 | 2.31E-01 | 1.00E+00 | 1.35E+00 | 1.95E-01 | 8.27E-01 | 1.13E+00 | 2.47E-01 | 9.45E-01 | 1.21E+00 | 2.33E-01 | 8.29E-01 | 1.49E+00 | 1.53E-01 | 4.32E-01 | 1.80E+00 | 2.68E-01 | 9.84E-01 | 1.41E+00 | 3.26E-01 | 8.96E-01 | 1.42E+00 |
| ZSCAN5A | 7.22E-02 | 3.76E-01 | 1.03E+00 | 9.47E-04 | 4.09E-03 | -1.44E+00 | 9.99E-04 | 4.24E-03 | -1.36E+00 | 6.18E-04 | 2.36E-03 | -1.27E+00 | 2.13E-02 | 7.59E-02 | -1.37E+00 | 8.61E-02 | 2.43E-01 | 1.04E+00 | 3.10E-02 | 1.14E-01 | -1.12E+00 | 7.50E-02 | 2.06E-01 | 1.00E+00 |
| BCR | 7.22E-02 | 3.76E-01 | 1.25E+00 | 7.33E-02 | 3.17E-01 | 1.34E+00 | 5.02E-02 | 2.13E-01 | 1.40E+00 | 6.39E-02 | 2.44E-01 | 1.20E+00 | 1.75E-01 | 6.23E-01 | 1.57E+00 | 3.24E-01 | 9.15E-01 | 1.50E+00 | 1.86E-03 | 6.84E-03 | -1.04E+00 | 1.79E-02 | 4.92E-02 | -1.01E+00 |
| KAZN | 7.22E-02 | 3.76E-01 | -1.10E+00 | 1.36E-01 | 5.86E-01 | 1.19E+00 | 7.14E-02 | 3.03E-01 | -1.09E+00 | 1.63E-01 | 6.21E-01 | 1.27E+00 | 5.40E-10 | 1.92E-09 | -2.47E+00 | 3.85E-04 | 1.09E-03 | -1.37E+00 | 1.00E-04 | 3.67E-04 | 1.05E+00 | 3.06E-02 | 8.42E-02 | 1.17E+00 |
| APOBEC2 | 7.22E-02 | 3.76E-01 | 1.73E+00 | 2.08E-02 | 8.98E-02 | 2.37E+00 | 1.54E-01 | 6.56E-01 | 1.00E+00 | 1.73E-01 | 6.60E-01 | 1.45E+00 | 8.74E-02 | 3.12E-01 | 2.05E+00 | 3.16E-03 | 8.90E-03 | 4.16E+00 | 1.72E-01 | 6.31E-01 | 1.45E+00 | 6.53E-02 | 1.79E-01 | 1.88E+00 |
| CCDC121 | 7.23E-02 | 3.77E-01 | -1.01E+00 | 1.47E-01 | 6.33E-01 | 1.51E+00 | 1.68E-03 | 7.15E-03 | -1.72E+00 | 1.40E-02 | 5.36E-02 | -1.27E+00 | 1.48E-01 | 5.28E-01 | -1.06E+00 | 2.57E-01 | 7.25E-01 | 1.10E+00 | 7.88E-03 | 2.89E-02 | -1.48E+00 | 2.17E-02 | 5.96E-02 | -1.38E+00 |
| RAB3C | 7.24E-02 | 3.77E-01 | -1.26E+00 | 1.33E-01 | 5.75E-01 | -1.11E+00 | 1.94E-01 | 8.24E-01 | 1.19E+00 | 2.16E-01 | 8.26E-01 | -1.04E+00 | 1.57E-03 | 5.61E-03 | -1.39E+00 | 2.88E-02 | 8.12E-02 | 1.04E+00 | 1.42E-01 | 5.22E-01 | 1.71E+00 | 1.37E-02 | 3.77E-02 | -1.18E+00 |
| GINS3 | 7.24E-02 | 3.77E-01 | -1.34E+00 | 1.29E-01 | 5.59E-01 | 1.03E+00 | 7.59E-03 | 3.23E-02 | -1.21E+00 | 9.05E-02 | 3.46E-01 | 1.17E+00 | 8.80E-02 | 3.14E-01 | -1.50E+00 | 1.63E-01 | 4.61E-01 | -1.30E+00 | 3.11E-10 | 1.14E-09 | -1.83E+00 | 1.38E-01 | 3.79E-01 | 1.12E+00 |
| MYL12B | 7.25E-02 | 3.78E-01 | 1.02E+00 | 2.19E-02 | 9.48E-02 | 1.21E+00 | 1.96E-01 | 8.32E-01 | 1.01E+00 | 1.27E-01 | 4.84E-01 | 1.09E+00 | 5.84E-03 | 2.08E-02 | -1.06E+00 | 1.48E-01 | 4.18E-01 | 1.10E+00 | 2.16E-01 | 7.92E-01 | 1.01E+00 | 1.22E-01 | 3.36E-01 | 1.09E+00 |
| CCDC162P | 7.25E-02 | 3.78E-01 | 1.58E+00 | 1.62E-01 | 6.99E-01 | 1.52E+00 | 2.05E-01 | 8.71E-01 | 1.14E+00 | 1.61E-01 | 6.16E-01 | 1.49E+00 | 2.69E-02 | 9.61E-02 | -1.56E+00 | 2.79E-01 | 7.88E-01 | 1.14E+00 | 2.55E-01 | 9.35E-01 | -1.08E+00 | 1.66E-01 | 4.55E-01 | 1.31E+00 |
| SLC44A4 | 7.25E-02 | 3.78E-01 | 1.48E+00 | 8.59E-02 | 3.71E-01 | -1.00E+00 | 1.06E-01 | 4.49E-01 | -1.01E+00 | 5.63E-02 | 2.15E-01 | -1.07E+00 | 4.30E-03 | 1.53E-02 | -1.35E+00 | 1.00E-03 | 2.82E-03 | -1.70E+00 | 1.60E-01 | 5.86E-01 | 1.23E+00 | 2.59E-01 | 7.10E-01 | 1.23E+00 |
| ZXDA | 7.25E-02 | 3.78E-01 | 1.58E+00 | 1.95E-02 | 8.42E-02 | -1.10E+00 | 7.16E-02 | 3.04E-01 | 1.62E+00 | 1.62E-01 | 6.20E-01 | 1.57E+00 | 2.71E-01 | 9.67E-01 | 1.47E+00 | 2.76E-01 | 7.77E-01 | 1.61E+00 | 9.83E-02 | 3.61E-01 | -1.07E+00 | 1.90E-01 | 5.22E-01 | 1.03E+00 |
| GSN | 7.27E-02 | 3.79E-01 | 1.22E+00 | 7.39E-03 | 3.19E-02 | 1.72E+00 | 1.74E-02 | 7.38E-02 | 1.36E+00 | 8.87E-02 | 3.39E-01 | 1.50E+00 | 2.02E-01 | 7.19E-01 | 1.59E+00 | 2.72E-02 | 7.67E-02 | 1.10E+00 | 4.16E-04 | 1.53E-03 | 1.78E+00 | 3.56E-02 | 9.79E-02 | 1.31E+00 |
| FO393401.1 | 7.27E-02 | 3.79E-01 | 1.20E+00 | 2.03E-04 | 8.77E-04 | -1.28E+00 | 1.09E-01 | 4.62E-01 | 1.00E+00 | 4.50E-04 | 1.72E-03 | -1.53E+00 | 4.43E-03 | 1.58E-02 | -1.08E+00 | 2.40E-03 | 6.76E-03 | -1.18E+00 | 2.62E-01 | 9.62E-01 | 1.21E+00 | 1.65E-01 | 4.53E-01 | 1.12E+00 |
| RAB3D | 7.28E-02 | 3.79E-01 | 1.21E+00 | 2.05E-02 | 8.85E-02 | 1.20E+00 | 1.46E-02 | 6.19E-02 | -1.00E+00 | 6.67E-03 | 2.55E-02 | 1.12E+00 | 6.76E-03 | 2.41E-02 | -1.19E+00 | 1.81E-03 | 5.10E-03 | -1.29E+00 | 7.83E-02 | 2.88E-01 | -1.19E+00 | 3.54E-01 | 9.73E-01 | 1.30E+00 |
| ANKMY1 | 7.28E-02 | 3.80E-01 | 1.05E+00 | 2.25E-02 | 9.74E-02 | -1.02E+00 | 2.14E-01 | 9.11E-01 | 1.18E+00 | 2.47E-01 | 9.44E-01 | 1.33E+00 | 6.90E-02 | 2.46E-01 | 1.02E+00 | 1.98E-01 | 5.58E-01 | 1.17E+00 | 1.14E-03 | 4.20E-03 | -1.12E+00 | 7.52E-05 | 2.06E-04 | -1.30E+00 |
| KRBA1 | 7.29E-02 | 3.80E-01 | 1.06E+00 | 1.26E-01 | 5.43E-01 | 1.28E+00 | 1.92E-01 | 8.17E-01 | 1.17E+00 | 1.17E-01 | 4.46E-01 | 1.14E+00 | 7.20E-02 | 2.57E-01 | -1.03E+00 | 1.47E-01 | 4.14E-01 | 1.04E+00 | 1.30E-01 | 4.77E-01 | 1.04E+00 | 3.55E-01 | 9.76E-01 | 1.28E+00 |
| PSMB7 | 7.31E-02 | 3.81E-01 | 1.25E+00 | 2.08E-03 | 8.98E-03 | 1.22E+00 | 1.03E-01 | 4.37E-01 | 1.28E+00 | 2.17E-01 | 8.29E-01 | 1.25E+00 | 6.90E-02 | 2.46E-01 | 1.16E+00 | 8.40E-02 | 2.37E-01 | 1.40E+00 | 1.86E-01 | 6.81E-01 | 1.22E+00 | 1.64E-01 | 4.50E-01 | 1.28E+00 |
| AL606834.3 | 7.31E-02 | 3.81E-01 | 1.58E+00 | 1.45E-01 | 6.27E-01 | 1.57E+00 | 2.35E-01 | 9.98E-01 | 1.17E+00 | 1.74E-01 | 6.63E-01 | 1.41E+00 | 2.51E-01 | 8.95E-01 | 1.35E+00 | 2.52E-01 | 7.12E-01 | 1.10E+00 | 1.17E-01 | 4.29E-01 | 1.62E+00 | 3.34E-01 | 9.17E-01 | 1.16E+00 |
| FANCA | 7.31E-02 | 3.81E-01 | 1.02E+00 | 8.23E-02 | 3.56E-01 | 1.14E+00 | 1.24E-01 | 5.25E-01 | 1.39E+00 | 2.23E-01 | 8.52E-01 | 1.52E+00 | 7.48E-02 | 2.67E-01 | -1.37E+00 | 1.39E-01 | 3.93E-01 | -1.19E+00 | 9.95E-04 | 3.65E-03 | -1.08E+00 | 3.12E-01 | 8.56E-01 | 1.42E+00 |
| STK24 | 7.31E-02 | 3.81E-01 | 1.58E+00 | 3.63E-02 | 1.57E-01 | -1.05E+00 | 5.28E-03 | 2.24E-02 | 1.91E+00 | 2.11E-01 | 8.05E-01 | 1.24E+00 | 6.56E-02 | 2.34E-01 | 1.87E+00 | 2.79E-01 | 7.85E-01 | 1.17E+00 | 4.12E-02 | 1.51E-01 | 1.45E+00 | 3.28E-02 | 9.00E-02 | -1.22E+00 |
| AP003108.2 | 7.32E-02 | 3.81E-01 | 1.09E+00 | 9.30E-03 | 4.02E-02 | -1.01E+00 | 1.07E-01 | 4.54E-01 | 1.36E+00 | 5.80E-02 | 2.21E-01 | -1.08E+00 | 7.77E-02 | 2.77E-01 | 2.01E+00 | 2.57E-01 | 7.25E-01 | 1.68E+00 | 2.18E-01 | 8.00E-01 | 1.18E+00 | 3.54E-01 | 9.73E-01 | 1.05E+00 |
| C16orf91 | 7.32E-02 | 3.81E-01 | 1.20E+00 | 2.13E-05 | 9.20E-05 | -1.25E+00 | 4.64E-04 | 1.97E-03 | -1.14E+00 | 4.69E-06 | 1.79E-05 | -1.15E+00 | 1.29E-01 | 4.60E-01 | 1.32E+00 | 3.29E-03 | 9.26E-03 | -1.09E+00 | 5.91E-02 | 2.17E-01 | 1.22E+00 | 1.12E-02 | 3.09E-02 | 1.06E+00 |
| LINC01004 | 7.32E-02 | 3.82E-01 | 1.44E+00 | 1.04E-01 | 4.51E-01 | 1.04E+00 | 1.70E-01 | 7.24E-01 | 1.33E+00 | 3.33E-02 | 1.27E-01 | -1.19E+00 | 2.28E-01 | 8.12E-01 | 1.23E+00 | 3.19E-01 | 9.00E-01 | 1.25E+00 | 1.63E-01 | 5.97E-01 | 1.34E+00 | 2.77E-01 | 7.60E-01 | 1.30E+00 |
| RFX8 | 7.33E-02 | 3.82E-01 | -1.02E+00 | 1.50E-02 | 6.48E-02 | -1.40E+00 | 6.92E-09 | 2.94E-08 | -1.63E+00 | 1.07E-08 | 4.09E-08 | -1.35E+00 | 1.12E-03 | 4.00E-03 | -2.65E+00 | 5.08E-04 | 1.43E-03 | -3.09E+00 | 2.42E-01 | 8.90E-01 | 1.15E+00 | 2.59E-01 | 7.10E-01 | -1.21E+00 |
| OR4D1 | 7.33E-02 | 3.82E-01 | -1.07E+00 | 5.15E-02 | 2.23E-01 | -1.21E+00 | 1.84E-01 | 7.80E-01 | -1.10E+00 | 1.58E-01 | 6.05E-01 | -1.11E+00 | 1.94E-01 | 6.93E-01 | 1.06E+00 | 2.23E-01 | 6.28E-01 | 1.73E+00 | 1.05E-04 | 3.87E-04 | -1.07E+00 | 2.70E-05 | 7.43E-05 | -1.34E+00 |
| CTH | 7.33E-02 | 3.82E-01 | 1.05E+00 | 1.61E-01 | 6.94E-01 | 1.30E+00 | 2.12E-02 | 9.02E-02 | -1.20E+00 | 7.62E-02 | 2.91E-01 | 1.11E+00 | 1.44E-01 | 5.14E-01 | 1.23E+00 | 3.07E-01 | 8.64E-01 | 1.32E+00 | 7.51E-04 | 2.76E-03 | -1.28E+00 | 1.16E-03 | 3.19E-03 | -1.29E+00 |
| DMC1 | 7.34E-02 | 3.83E-01 | -1.19E+00 | 2.24E-02 | 9.70E-02 | -1.80E+00 | 1.93E-01 | 8.22E-01 | 1.09E+00 | 1.80E-01 | 6.86E-01 | 1.10E+00 | 2.51E-01 | 8.96E-01 | -1.11E+00 | 2.82E-01 | 7.96E-01 | -1.32E+00 | 1.15E-04 | 4.21E-04 | -1.30E+00 | 7.85E-06 | 2.16E-05 | -1.35E+00 |
| LRRC37B | 7.35E-02 | 3.83E-01 | 1.26E+00 | 2.54E-03 | 1.10E-02 | 1.13E+00 | 1.22E-02 | 5.18E-02 | 1.04E+00 | 3.04E-05 | 1.16E-04 | -1.05E+00 | 1.78E-02 | 6.33E-02 | 1.02E+00 | 6.37E-04 | 1.80E-03 | -1.21E+00 | 1.66E-01 | 6.09E-01 | 1.05E+00 | 6.59E-04 | 1.81E-03 | -1.32E+00 |
| POLE | 7.35E-02 | 3.83E-01 | 1.63E+00 | 1.99E-01 | 8.61E-01 | 1.62E+00 | 1.61E-01 | 6.83E-01 | 1.47E+00 | 5.86E-02 | 2.24E-01 | 1.34E+00 | 1.87E-01 | 6.68E-01 | 1.15E+00 | 1.37E-01 | 3.88E-01 | 1.02E+00 | 1.93E-01 | 7.08E-01 | 1.30E+00 | 2.34E-01 | 6.42E-01 | 1.33E+00 |
| AC004893.2 | 7.35E-02 | 3.83E-01 | -1.03E+00 | 2.47E-02 | 1.07E-01 | -1.09E+00 | 2.49E-03 | 1.06E-02 | 2.18E+00 | 2.39E-01 | 9.12E-01 | 1.32E+00 | 4.23E-02 | 1.51E-01 | -1.10E+00 | 9.51E-03 | 2.68E-02 | -1.33E+00 | 2.69E-01 | 9.87E-01 | 1.25E+00 | 3.58E-01 | 9.84E-01 | 1.36E+00 |
| PARG | 7.35E-02 | 3.83E-01 | 1.16E+00 | 2.19E-01 | 9.47E-01 | 1.50E+00 | 1.40E-02 | 5.93E-02 | -1.11E+00 | 9.14E-02 | 3.49E-01 | 1.19E+00 | 5.48E-02 | 1.95E-01 | -1.03E+00 | 2.98E-01 | 8.40E-01 | 1.68E+00 | 2.01E-07 | 7.37E-07 | -1.81E+00 | 8.90E-02 | 2.44E-01 | 1.09E+00 |
| YJEFN3 | 7.36E-02 | 3.83E-01 | -1.01E+00 | 2.26E-02 | 9.75E-02 | -1.24E+00 | 6.70E-02 | 2.85E-01 | 1.09E+00 | 2.50E-02 | 9.55E-02 | 1.01E+00 | 2.00E-01 | 7.14E-01 | 1.11E+00 | 1.87E-01 | 5.27E-01 | 1.07E+00 | 1.82E-01 | 6.69E-01 | 1.04E+00 | 1.00E-01 | 2.75E-01 | -1.11E+00 |
| SH2B1 | 7.36E-02 | 3.83E-01 | 1.23E+00 | 4.90E-03 | 2.12E-02 | 1.18E+00 | 6.90E-04 | 2.93E-03 | -1.15E+00 | 1.30E-03 | 4.98E-03 | -1.03E+00 | 1.80E-02 | 6.41E-02 | -1.00E+00 | 2.55E-02 | 7.20E-02 | 1.02E+00 | 6.62E-03 | 2.43E-02 | -1.13E+00 | 4.55E-02 | 1.25E-01 | -1.02E+00 |
| SMIM5 | 7.36E-02 | 3.84E-01 | 1.62E+00 | 1.35E-01 | 5.82E-01 | 1.49E+00 | 3.06E-02 | 1.30E-01 | -1.46E+00 | 2.51E-02 | 9.58E-02 | -1.60E+00 | 2.57E-01 | 9.16E-01 | 1.25E+00 | 2.16E-01 | 6.09E-01 | 1.07E+00 | 4.07E-02 | 1.49E-01 | 1.51E+00 | 3.96E-02 | 1.09E-01 | 1.00E+00 |
| GCN1 | 7.37E-02 | 3.84E-01 | 1.07E+00 | 1.90E-01 | 8.23E-01 | 1.34E+00 | 1.28E-01 | 5.42E-01 | 1.11E+00 | 1.83E-01 | 6.99E-01 | 1.49E+00 | 1.10E-02 | 3.92E-02 | -1.04E+00 | 1.12E-01 | 3.16E-01 | 1.32E+00 | 2.53E-01 | 9.30E-01 | 1.21E+00 | 4.68E-03 | 1.29E-02 | 1.69E+00 |
| LGALS3 | 7.37E-02 | 3.84E-01 | 1.61E+00 | 3.54E-02 | 1.53E-01 | 1.97E+00 | 3.20E-02 | 1.36E-01 | 1.97E+00 | 2.00E-03 | 7.62E-03 | 2.70E+00 | 2.35E-01 | 8.37E-01 | 1.43E+00 | 3.17E-01 | 8.95E-01 | 1.01E+00 | 8.92E-02 | 3.27E-01 | 1.40E+00 | 1.53E-01 | 4.20E-01 | 1.30E+00 |
| EPGN | 7.37E-02 | 3.84E-01 | 1.09E+00 | 3.92E-19 | 1.70E-18 | 3.23E+00 | 1.90E-01 | 8.09E-01 | 1.34E+00 | 4.46E-09 | 1.70E-08 | 3.59E+00 | 2.52E-01 | 9.00E-01 | 1.39E+00 | 9.22E-12 | 2.60E-11 | 3.20E+00 | 1.34E-03 | 4.91E-03 | -1.08E+00 | 4.57E-03 | 1.25E-02 | 1.83E+00 |
| IRF6 | 7.37E-02 | 3.84E-01 | -1.10E+00 | 5.90E-02 | 2.55E-01 | -1.22E+00 | 2.01E-01 | 8.56E-01 | 1.05E+00 | 2.24E-01 | 8.55E-01 | 1.07E+00 | 8.85E-02 | 3.15E-01 | 2.06E+00 | 2.93E-02 | 8.27E-02 | 2.69E+00 | 1.74E-05 | 6.37E-05 | -1.57E+00 | 1.78E-04 | 4.90E-04 | -1.43E+00 |
| AF131216.1 | 7.38E-02 | 3.85E-01 | -1.15E+00 | 7.15E-02 | 3.09E-01 | -1.08E+00 | 2.24E-01 | 9.51E-01 | 1.24E+00 | 2.41E-01 | 9.21E-01 | 1.38E+00 | 1.88E-01 | 6.71E-01 | -1.17E+00 | 2.73E-01 | 7.71E-01 | -1.02E+00 | 2.02E-01 | 7.43E-01 | 1.10E+00 | 5.20E-02 | 1.43E-01 | -1.15E+00 |
| AC013470.3 | 7.39E-02 | 3.85E-01 | 1.03E+00 | 1.19E-01 | 5.12E-01 | 1.22E+00 | 1.70E-02 | 7.24E-02 | 1.89E+00 | 1.25E-01 | 4.77E-01 | 1.76E+00 | 2.95E-02 | 1.05E-01 | -1.37E+00 | 2.80E-01 | 7.90E-01 | 1.31E+00 | 2.48E-01 | 9.12E-01 | 1.30E+00 | 1.36E-01 | 3.73E-01 | 1.53E+00 |
| GNE | 7.40E-02 | 3.86E-01 | -1.02E+00 | 4.31E-02 | 1.86E-01 | -1.01E+00 | 5.57E-02 | 2.37E-01 | -1.10E+00 | 2.12E-02 | 8.08E-02 | -1.01E+00 | 1.77E-01 | 6.32E-01 | 1.58E+00 | 1.83E-01 | 5.15E-01 | 1.60E+00 | 1.67E-01 | 6.15E-01 | -1.03E+00 | 1.56E-02 | 4.30E-02 | -1.36E+00 |
| ZNF90P1 | 7.41E-02 | 3.86E-01 | 1.07E+00 | 2.05E-01 | 8.84E-01 | 1.51E+00 | 1.89E-02 | 8.03E-02 | -1.28E+00 | 2.43E-02 | 9.30E-02 | -1.21E+00 | 1.87E-01 | 6.67E-01 | 1.02E+00 | 2.19E-01 | 6.17E-01 | 1.64E+00 | 1.57E-03 | 5.75E-03 | -1.40E+00 | 6.31E-02 | 1.73E-01 | -1.10E+00 |
| AC139792.1 | 7.41E-02 | 3.86E-01 | 1.15E+00 | 1.15E-01 | 4.99E-01 | 1.32E+00 | 4.42E-05 | 1.88E-04 | -1.45E+00 | 3.73E-04 | 1.43E-03 | -1.14E+00 | 1.18E-01 | 4.22E-01 | -1.04E+00 | 1.82E-01 | 5.13E-01 | 1.08E+00 | 1.15E-02 | 4.23E-02 | -1.14E+00 | 1.21E-02 | 3.32E-02 | -1.17E+00 |
| PLS1 | 7.41E-02 | 3.86E-01 | -1.08E+00 | 1.09E-01 | 4.69E-01 | 1.09E+00 | 1.55E-02 | 6.60E-02 | -1.15E+00 | 1.12E-01 | 4.29E-01 | 1.27E+00 | 2.68E-01 | 9.54E-01 | 1.30E+00 | 1.33E-01 | 3.75E-01 | 1.83E+00 | 5.98E-11 | 2.19E-10 | -1.59E+00 | 5.58E-10 | 1.53E-09 | -1.57E+00 |
| PCAT6 | 7.42E-02 | 3.87E-01 | 1.18E+00 | 4.72E-02 | 2.04E-01 | 1.19E+00 | 5.90E-03 | 2.51E-02 | -1.48E+00 | 9.17E-03 | 3.50E-02 | -1.33E+00 | 1.19E-01 | 4.23E-01 | -1.05E+00 | 2.53E-01 | 7.13E-01 | 1.15E+00 | 2.20E-02 | 8.07E-02 | -1.44E+00 | 2.19E-02 | 6.01E-02 | -1.51E+00 |
| HIST1H3I | 7.42E-02 | 3.87E-01 | -1.21E+00 | 1.30E-02 | 5.60E-02 | -1.83E+00 | 2.80E-02 | 1.19E-01 | 1.53E+00 | 1.14E-01 | 4.37E-01 | -1.15E+00 | 1.33E-01 | 4.76E-01 | -1.14E+00 | 3.83E-02 | 1.08E-01 | -1.70E+00 | 3.82E-03 | 1.40E-02 | -1.06E+00 | 9.96E-05 | 2.74E-04 | -1.39E+00 |
| STARD4 | 7.42E-02 | 3.87E-01 | 1.20E+00 | 2.14E-01 | 9.25E-01 | 1.48E+00 | 7.67E-02 | 3.26E-01 | 1.07E+00 | 1.80E-01 | 6.88E-01 | 1.42E+00 | 7.85E-02 | 2.80E-01 | 1.15E+00 | 4.75E-02 | 1.34E-01 | -1.02E+00 | 3.23E-03 | 1.18E-02 | -1.16E+00 | 1.73E-01 | 4.75E-01 | 1.10E+00 |
| DNHD1 | 7.43E-02 | 3.87E-01 | 1.51E+00 | 7.38E-02 | 3.19E-01 | 1.68E+00 | 2.75E-02 | 1.17E-01 | -1.10E+00 | 5.69E-02 | 2.18E-01 | 1.17E+00 | 1.41E-03 | 5.05E-03 | -1.67E+00 | 5.50E-02 | 1.55E-01 | -1.04E+00 | 2.54E-01 | 9.32E-01 | 1.11E+00 | 1.76E-01 | 4.84E-01 | 1.58E+00 |
| CXCL12 | 7.43E-02 | 3.87E-01 | -1.25E+00 | 1.26E-01 | 5.44E-01 | -1.02E+00 | 1.70E-01 | 7.23E-01 | -1.08E+00 | 1.62E-01 | 6.20E-01 | -1.15E+00 | 1.08E-01 | 3.85E-01 | 1.57E+00 | 8.35E-02 | 2.35E-01 | 2.04E+00 | 1.34E-02 | 4.93E-02 | 1.89E+00 | 1.80E-01 | 4.94E-01 | -1.32E+00 |
| PODXL2 | 7.43E-02 | 3.87E-01 | 1.26E+00 | 2.86E-02 | 1.24E-01 | 1.28E+00 | 1.22E-01 | 5.19E-01 | 1.22E+00 | 4.73E-02 | 1.81E-01 | 1.46E+00 | 3.25E-02 | 1.16E-01 | -1.05E+00 | 3.21E-01 | 9.06E-01 | 1.41E+00 | 8.81E-02 | 3.23E-01 | 1.07E+00 | 3.09E-02 | 8.48E-02 | 1.60E+00 |
| AL590326.2 | 7.44E-02 | 3.88E-01 | 1.64E+00 | 1.05E-01 | 4.55E-01 | 1.70E+00 | 6.45E-02 | 2.74E-01 | 1.57E+00 | 1.27E-01 | 4.85E-01 | 1.65E+00 | 5.81E-02 | 2.07E-01 | 2.81E+00 | 4.67E-02 | 1.32E-01 | 2.88E+00 | 1.58E-01 | 5.81E-01 | 1.27E+00 | 2.49E-01 | 6.84E-01 | 1.33E+00 |
| AC108134.3 | 7.44E-02 | 3.88E-01 | -1.18E+00 | 2.07E-01 | 8.95E-01 | 1.36E+00 | 1.04E-01 | 4.40E-01 | 1.01E+00 | 2.51E-01 | 9.58E-01 | 1.43E+00 | 1.12E-01 | 3.99E-01 | -1.13E+00 | 3.46E-02 | 9.76E-02 | -1.31E+00 | 8.47E-02 | 3.11E-01 | 1.68E+00 | 1.22E-01 | 3.35E-01 | 1.74E+00 |
| LINC02076 | 7.45E-02 | 3.88E-01 | 1.05E+00 | 1.35E-01 | 5.82E-01 | 1.19E+00 | 8.81E-02 | 3.74E-01 | 1.59E+00 | 1.08E-01 | 4.13E-01 | 1.63E+00 | 1.12E-01 | 3.98E-01 | 1.13E+00 | 9.92E-02 | 2.80E-01 | -1.10E+00 | 7.07E-02 | 2.59E-01 | 1.81E+00 | 2.27E-01 | 6.23E-01 | 1.22E+00 |
| APOBR | 7.45E-02 | 3.88E-01 | 1.65E+00 | 1.48E-01 | 6.40E-01 | 1.25E+00 | 2.21E-01 | 9.40E-01 | 1.25E+00 | 2.08E-01 | 7.95E-01 | 1.21E+00 | 4.63E-02 | 1.65E-01 | 1.06E+00 | 3.88E-02 | 1.09E-01 | -1.01E+00 | 2.67E-01 | 9.78E-01 | 1.23E+00 | 1.50E-01 | 4.13E-01 | 1.78E+00 |
| TAS2R64P | 7.46E-02 | 3.89E-01 | -1.07E+00 | 9.49E-02 | 4.10E-01 | 1.01E+00 | 2.78E-04 | 1.18E-03 | -1.76E+00 | 1.02E-02 | 3.91E-02 | -1.18E+00 | 2.25E-01 | 8.01E-01 | 1.43E+00 | 3.29E-01 | 9.26E-01 | 1.22E+00 | 1.09E-01 | 4.01E-01 | 1.39E+00 | 2.00E-01 | 5.51E-01 | -1.15E+00 |
| PRKCE | 7.46E-02 | 3.89E-01 | 1.62E+00 | 1.99E-01 | 8.58E-01 | 1.60E+00 | 6.69E-03 | 2.84E-02 | 1.02E+00 | 7.34E-04 | 2.81E-03 | -1.01E+00 | 6.31E-02 | 2.25E-01 | 1.99E+00 | 2.70E-01 | 7.63E-01 | 1.58E+00 | 2.24E-01 | 8.21E-01 | 1.17E+00 | 2.97E-01 | 8.15E-01 | 1.14E+00 |
| KBTBD8 | 7.46E-02 | 3.89E-01 | 1.75E+00 | 4.62E-02 | 2.00E-01 | -1.20E+00 | 2.26E-01 | 9.59E-01 | 1.13E+00 | 2.44E-02 | 9.32E-02 | -1.61E+00 | 2.28E-01 | 8.13E-01 | 1.24E+00 | 1.67E-02 | 4.70E-02 | -1.62E+00 | 1.17E-01 | 4.28E-01 | 1.50E+00 | 1.65E-01 | 4.53E-01 | 1.26E+00 |
| AC011498.7 | 7.47E-02 | 3.89E-01 | 1.12E+00 | 1.60E-01 | 6.92E-01 | 1.39E+00 | 2.00E-01 | 8.51E-01 | 1.19E+00 | 1.36E-01 | 5.20E-01 | 1.19E+00 | 7.92E-02 | 2.82E-01 | -1.03E+00 | 1.31E-01 | 3.69E-01 | 1.11E+00 | 2.04E-01 | 7.50E-01 | 1.49E+00 | 3.12E-01 | 8.56E-01 | 1.39E+00 |
| TRIM73 | 7.48E-02 | 3.90E-01 | -1.02E+00 | 4.73E-02 | 2.04E-01 | 1.04E+00 | 8.06E-02 | 3.43E-01 | 1.48E+00 | 4.87E-02 | 1.86E-01 | 1.86E+00 | 1.73E-01 | 6.17E-01 | 1.03E+00 | 7.34E-02 | 2.07E-01 | -1.34E+00 | 2.26E-01 | 8.30E-01 | 1.15E+00 | 6.58E-02 | 1.81E-01 | -1.32E+00 |
| AP001024.1 | 7.52E-02 | 3.92E-01 | 1.54E+00 | 1.56E-06 | 6.73E-06 | 1.06E+00 | 6.58E-04 | 2.79E-03 | 1.58E+00 | 1.82E-02 | 6.96E-02 | 1.11E+00 | 7.70E-03 | 2.74E-02 | 1.90E+00 | 1.95E-01 | 5.49E-01 | 1.56E+00 | 4.02E-02 | 1.47E-01 | 1.36E+00 | 1.99E-01 | 5.46E-01 | 1.10E+00 |
| STAT3 | 7.52E-02 | 3.92E-01 | 1.33E+00 | 8.01E-07 | 3.46E-06 | 1.02E+00 | 6.32E-02 | 2.68E-01 | 1.04E+00 | 3.43E-03 | 1.31E-02 | -1.04E+00 | 6.29E-02 | 2.24E-01 | 1.26E+00 | 7.10E-02 | 2.00E-01 | 1.21E+00 | 8.91E-02 | 3.27E-01 | 1.17E+00 | 9.42E-03 | 2.59E-02 | -1.06E+00 |
| ATXN1L | 7.52E-02 | 3.92E-01 | 1.21E+00 | 2.22E-01 | 9.59E-01 | 1.58E+00 | 9.42E-02 | 4.00E-01 | 1.07E+00 | 3.12E-02 | 1.19E-01 | 1.03E+00 | 2.45E-01 | 8.74E-01 | 1.41E+00 | 1.69E-01 | 4.76E-01 | 1.03E+00 | 2.33E-01 | 8.56E-01 | 1.12E+00 | 1.32E-01 | 3.61E-01 | -1.02E+00 |

| | | | | | | | | | | | | | | | | | | | | | | | | |
|---|---|---|---|---|---|---|---|---|---|---|---|---|---|---|---|---|---|---|---|---|---|---|---|---|
| PRRT4 | 7.52E-02 | 3.92E-01 | 1.43E+00 | 1.43E-01 | 6.19E-01 | 1.29E+00 | 2.37E-02 | 1.01E-01 | 1.50E+00 | 2.10E-01 | 8.02E-01 | 1.32E+00 | 2.80E-01 | 1.00E+00 | -1.00E+00 | 3.55E-01 | 1.00E+00 | -1.00E+00 | 2.67E-01 | 9.81E-01 | -1.07E+00 | 2.53E-01 | 6.95E-01 | 1.08E+00 |
| MAP2K2 | 7.53E-02 | 3.92E-01 | 1.42E+00 | 8.24E-02 | 3.56E-01 | 1.37E+00 | 1.07E-01 | 4.53E-01 | 1.32E+00 | 1.38E-01 | 5.27E-01 | 1.22E+00 | 5.50E-02 | 1.96E-01 | 1.55E+00 | 9.72E-02 | 2.74E-01 | 1.44E+00 | 5.57E-03 | 2.04E-02 | 1.57E+00 | 1.82E-02 | 4.99E-02 | 1.43E+00 |
| AC120053.1 | 7.53E-02 | 3.92E-01 | -1.18E+00 | 6.98E-02 | 3.02E-01 | 1.95E+00 | 1.06E-02 | 4.50E-02 | -1.45E+00 | 4.86E-02 | 1.86E-01 | 1.00E+00 | 4.57E-05 | 1.63E-04 | -2.03E+00 | 2.07E-03 | 5.85E-03 | -1.35E+00 | 2.29E-01 | 8.40E-01 | 1.14E+00 | 1.22E-01 | 3.34E-01 | 1.50E+00 |
| AC104996.2 | 7.54E-02 | 3.93E-01 | 1.01E+00 | 3.16E-02 | 1.37E-01 | -1.20E+00 | 1.87E-01 | 7.94E-01 | 1.07E+00 | 5.06E-02 | 1.93E-01 | -1.07E+00 | 2.17E-04 | 7.75E-04 | -2.07E+00 | 3.51E-04 | 9.91E-04 | -1.97E+00 | 5.75E-05 | 2.11E-04 | -1.48E+00 | 6.04E-05 | 1.66E-04 | -1.53E+00 |
| LINC00910 | 7.54E-02 | 3.93E-01 | 1.05E+00 | 5.45E-02 | 2.36E-01 | 1.06E+00 | 7.36E-04 | 3.13E-03 | -1.54E+00 | 3.42E-03 | 1.31E-02 | -1.14E+00 | 1.34E-01 | 4.77E-01 | 1.05E+00 | 2.63E-01 | 7.40E-01 | 1.73E+00 | 3.09E-02 | 1.13E-01 | -1.09E+00 | 5.69E-02 | 1.56E-01 | -1.12E+00 |
| METTL18 | 7.54E-02 | 3.93E-01 | 1.11E+00 | 1.47E-01 | 6.34E-01 | 1.34E+00 | 8.23E-02 | 3.50E-01 | -1.02E+00 | 1.51E-01 | 5.76E-01 | 1.30E+00 | 1.36E-03 | 4.87E-03 | -1.25E+00 | 9.43E-02 | 2.66E-01 | 1.17E+00 | 2.66E-01 | 9.75E-01 | 1.28E+00 | 3.63E-01 | 9.98E-01 | 1.22E+00 |
| C7orf43 | 7.56E-02 | 3.94E-01 | 1.05E+00 | 1.38E-01 | 5.98E-01 | 1.18E+00 | 7.81E-03 | 3.32E-02 | -1.33E+00 | 1.56E-02 | 5.96E-02 | -1.06E+00 | 3.31E-02 | 1.18E-01 | -1.37E+00 | 3.06E-01 | 8.62E-01 | 1.47E+00 | 4.82E-04 | 1.77E-03 | -1.26E+00 | 6.83E-05 | 1.88E-04 | -1.58E+00 |
| TYMS | 7.57E-02 | 3.94E-01 | 1.22E+00 | 1.02E-02 | 4.41E-02 | 1.13E+00 | 1.76E-02 | 7.48E-02 | 1.45E+00 | 1.92E-01 | 7.35E-01 | 1.35E+00 | 1.13E-03 | 4.03E-03 | -1.38E+00 | 2.02E-02 | 5.68E-02 | -1.22E+00 | 2.14E-01 | 7.87E-01 | 1.11E+00 | 7.97E-05 | 2.19E-04 | 1.45E+00 |
| CCT6B | 7.58E-02 | 3.95E-01 | 1.51E+00 | 6.72E-02 | 2.90E-01 | 1.05E+00 | 1.63E-01 | 6.92E-01 | 1.05E+00 | 4.04E-03 | 1.54E-02 | -1.46E+00 | 7.73E-02 | 2.75E-01 | 1.74E+00 | 3.06E-01 | 8.63E-01 | 1.31E+00 | 9.00E-02 | 3.30E-01 | 1.52E+00 | 2.38E-01 | 6.54E-01 | 1.12E+00 |
| PKD2L2 | 7.58E-02 | 3.95E-01 | 1.16E+00 | 1.07E-01 | 4.63E-01 | 1.33E+00 | 7.24E-02 | 3.08E-01 | -1.01E+00 | 1.90E-01 | 7.24E-01 | 1.33E+00 | 8.36E-02 | 2.98E-01 | -1.05E+00 | 1.69E-01 | 4.76E-01 | 1.15E+00 | 1.84E-01 | 6.77E-01 | 1.08E+00 | 2.94E-01 | 8.07E-01 | 1.30E+00 |
| SUMF2 | 7.58E-02 | 3.95E-01 | 1.15E+00 | 7.70E-02 | 3.33E-01 | 1.33E+00 | 9.46E-03 | 4.02E-02 | -1.14E+00 | 4.00E-03 | 1.53E-02 | -1.01E+00 | 2.41E-04 | 8.60E-04 | -1.03E+00 | 1.82E-02 | 5.13E-02 | 1.20E+00 | 2.58E-01 | 9.49E-01 | 1.29E+00 | 1.37E-01 | 3.78E-01 | 1.26E+00 |
| RARRES1 | 7.59E-02 | 3.96E-01 | -1.26E+00 | 1.05E-01 | 4.52E-01 | -1.19E+00 | 5.44E-02 | 2.31E-01 | -1.36E+00 | 5.40E-02 | 2.06E-01 | -1.33E+00 | 4.56E-02 | 1.63E-01 | -1.30E+00 | 8.42E-02 | 2.37E-01 | -1.15E+00 | 2.63E-03 | 9.65E-03 | -1.29E+00 | 6.35E-02 | 1.74E-01 | 1.02E+00 |
| AL021807.1 | 7.61E-02 | 3.96E-01 | 1.40E+00 | 6.81E-02 | 2.94E-01 | 1.55E+00 | 2.07E-02 | 8.80E-02 | -1.40E+00 | 5.39E-02 | 2.06E-01 | -1.20E+00 | 1.08E-01 | 3.87E-01 | 1.71E+00 | 7.70E-02 | 2.17E-01 | 1.80E+00 | 3.66E-02 | 1.34E-01 | -1.58E+00 | 2.72E-01 | 7.48E-01 | 1.49E+00 |
| NUP188 | 7.61E-02 | 3.97E-01 | -1.01E+00 | 2.26E-01 | 9.75E-01 | 1.37E+00 | 4.39E-02 | 1.86E-01 | 1.01E+00 | 2.13E-01 | 8.13E-01 | 1.44E+00 | 1.15E-02 | 4.11E-02 | -1.16E+00 | 2.72E-01 | 7.67E-01 | 1.31E+00 | 6.59E-04 | 2.42E-03 | -1.20E+00 | 3.04E-01 | 8.35E-01 | 1.21E+00 |
| EXOC8 | 7.61E-02 | 3.97E-01 | 1.18E+00 | 7.25E-04 | 3.13E-03 | -1.05E+00 | 1.34E-01 | 5.71E-01 | 1.14E+00 | 4.25E-03 | 1.62E-02 | -1.11E+00 | 2.22E-01 | 7.92E-01 | 1.37E+00 | 1.06E-01 | 2.98E-01 | 1.01E+00 | 2.00E-01 | 7.34E-01 | 1.04E+00 | 2.69E-01 | 7.38E-01 | 1.10E+00 |
| ZNF316 | 7.61E-02 | 3.97E-01 | 1.17E+00 | 1.17E-01 | 5.07E-01 | 1.37E+00 | 6.08E-02 | 2.58E-01 | 1.06E+00 | 1.33E-01 | 5.09E-01 | 1.29E+00 | 1.61E-01 | 5.75E-01 | 1.55E+00 | 3.27E-01 | 9.21E-01 | 1.25E+00 | 2.05E-01 | 7.53E-01 | 1.07E+00 | 1.93E-01 | 5.30E-01 | 1.03E+00 |
| MEX3B | 7.62E-02 | 3.97E-01 | 1.24E+00 | 1.87E-01 | 8.07E-01 | 1.61E+00 | 1.44E-05 | 6.10E-05 | -1.23E+00 | 3.82E-02 | 1.46E-01 | 1.19E+00 | 2.75E-05 | 9.81E-05 | -1.29E+00 | 8.21E-02 | 2.31E-01 | 1.23E+00 | 7.39E-10 | 2.71E-09 | -1.45E+00 | 1.72E-04 | 4.72E-04 | -1.12E+00 |
| AC241585.1 | 7.62E-02 | 3.97E-01 | -1.25E+00 | 1.35E-01 | 5.84E-01 | -1.07E+00 | 6.63E-02 | 2.82E-01 | -1.24E+00 | 8.22E-02 | 3.14E-01 | -1.09E+00 | 1.68E-01 | 5.98E-01 | -1.09E+00 | 3.17E-01 | 8.95E-01 | 1.39E+00 | 4.51E-03 | 1.66E-02 | 1.84E+00 | 1.49E-01 | 4.09E-01 | 1.31E+00 |
| FAM19A2 | 7.63E-02 | 3.97E-01 | 1.25E+00 | 1.97E-02 | 8.51E-02 | 1.33E+00 | 8.19E-02 | 3.48E-01 | 1.03E+00 | 1.45E-01 | 5.56E-01 | 1.28E+00 | 1.94E-03 | 6.90E-03 | 1.06E+00 | 3.64E-02 | 1.03E-01 | 1.10E+00 | 1.62E-01 | 5.96E-01 | 1.05E+00 | 6.56E-02 | 1.80E-01 | -1.03E+00 |
| ELFN1-AS1 | 7.63E-02 | 3.98E-01 | -1.26E+00 | 8.01E-02 | 3.46E-01 | -1.27E+00 | 1.75E-01 | 7.42E-01 | 1.01E+00 | 5.67E-02 | 2.17E-01 | -1.33E+00 | 4.27E-02 | 1.52E-01 | -1.32E+00 | 1.44E-01 | 4.06E-01 | -1.07E+00 | 2.52E-01 | 9.26E-01 | 1.14E+00 | 1.67E-01 | 4.58E-01 | -1.20E+00 |
| AC011592.1 | 7.63E-02 | 3.98E-01 | 1.45E+00 | 1.55E-01 | 6.69E-01 | 1.38E+00 | 1.72E-01 | 7.32E-01 | 1.16E+00 | 1.55E-01 | 5.94E-01 | -1.08E+00 | 1.12E-01 | 4.00E-01 | 1.53E+00 | 1.57E-01 | 4.42E-01 | 1.55E+00 | 1.49E-01 | 5.47E-01 | 1.42E+00 | 8.53E-02 | 2.34E-01 | 1.36E+00 |
| AGPAT4 | 7.64E-02 | 3.98E-01 | 1.25E+00 | 1.37E-01 | 5.93E-01 | 1.68E+00 | 6.08E-04 | 2.58E-03 | -1.17E+00 | 3.53E-02 | 1.35E-01 | 1.23E+00 | 6.73E-02 | 2.40E-01 | 1.20E+00 | 2.60E-01 | 7.32E-01 | 1.35E+00 | 3.48E-03 | 1.28E-02 | -1.02E+00 | 3.47E-02 | 9.53E-02 | -1.01E+00 |
| HSP90B2P | 7.65E-02 | 3.98E-01 | 1.01E+00 | 1.66E-01 | 7.16E-01 | 1.34E+00 | 6.53E-02 | 2.77E-01 | 1.57E+00 | 5.67E-03 | 2.17E-02 | 2.11E+00 | 6.41E-02 | 2.28E-01 | 1.02E+00 | 1.68E-01 | 4.75E-01 | 1.22E+00 | 1.10E-01 | 4.04E-01 | 1.52E+00 | 6.13E-03 | 1.68E-02 | 1.96E+00 |
| AC145285.2 | 7.66E-02 | 3.99E-01 | -1.01E+00 | 7.39E-02 | 3.19E-01 | 1.07E+00 | 2.08E-01 | 8.83E-01 | 1.22E+00 | 1.75E-01 | 6.69E-01 | 1.11E+00 | 2.52E-01 | 8.98E-01 | 1.04E+00 | 1.41E-01 | 3.98E-01 | 1.56E+00 | 1.43E-01 | 5.24E-01 | -1.17E+00 | 7.52E-02 | 2.07E-01 | -1.36E+00 |
| AC018809.2 | 7.66E-02 | 3.99E-01 | -1.19E+00 | 1.78E-01 | 7.70E-01 | 1.13E+00 | 1.73E-02 | 7.35E-02 | -1.22E+00 | 5.20E-04 | 1.99E-03 | -1.45E+00 | 8.68E-03 | 3.10E-02 | -1.17E+00 | 6.37E-04 | 1.80E-03 | -1.64E+00 | 2.18E-01 | 8.01E-01 | 1.14E+00 | 1.12E-01 | 3.09E-01 | -1.13E+00 |
| CD93 | 7.66E-02 | 3.99E-01 | -1.15E+00 | 1.46E-01 | 6.32E-01 | 1.12E+00 | 1.48E-03 | 6.29E-03 | -1.44E+00 | 2.61E-06 | 9.97E-06 | -1.94E+00 | 6.59E-02 | 2.35E-01 | -1.67E+00 | 9.43E-02 | 2.66E-01 | -1.66E+00 | 2.27E-01 | 8.33E-01 | -1.07E+00 | 2.18E-01 | 5.99E-01 | -1.24E+00 |
| HIST1H2AE | 7.66E-02 | 3.99E-01 | 1.60E+00 | 1.28E-02 | 5.55E-02 | -1.02E+00 | 1.22E-01 | 5.19E-01 | 1.07E+00 | 1.60E-02 | 6.10E-02 | -1.12E+00 | 7.97E-02 | 2.84E-01 | -1.04E+00 | 7.66E-02 | 2.16E-01 | -1.00E+00 | 2.17E-02 | 7.96E-02 | -1.12E+00 | 9.25E-03 | 2.54E-02 | -1.02E+00 |
| PPP2R5D | 7.67E-02 | 4.00E-01 | 1.08E+00 | 1.71E-01 | 7.39E-01 | 1.58E+00 | 1.78E-03 | 7.58E-03 | -1.36E+00 | 3.49E-02 | 1.33E-01 | -1.00E+00 | 7.55E-03 | 2.69E-02 | -1.16E+00 | 4.11E-02 | 1.16E-01 | -1.01E+00 | 1.05E-05 | 3.85E-05 | -1.29E+00 | 2.87E-03 | 7.89E-03 | -1.07E+00 |
| SMG1P5 | 7.67E-02 | 4.00E-01 | 1.15E+00 | 1.47E-01 | 6.37E-01 | 1.34E+00 | 1.76E-01 | 7.47E-01 | 1.08E+00 | 1.31E-01 | 5.02E-01 | 1.06E+00 | 2.78E-01 | 9.91E-01 | 1.38E+00 | 3.46E-01 | 9.74E-01 | 1.42E+00 | 4.13E-02 | 1.52E-01 | 1.78E+00 | 2.22E-01 | 6.10E-01 | 1.37E+00 |
| AC099778.1 | 7.67E-02 | 4.00E-01 | 1.65E+00 | 5.34E-02 | 2.31E-01 | 1.20E+00 | 3.35E-02 | 1.42E-01 | 1.51E+00 | 2.48E-01 | 9.45E-01 | 1.33E+00 | 8.29E-03 | 2.96E-02 | -1.37E+00 | 1.01E-01 | 2.85E-01 | -1.06E+00 | 9.44E-02 | 3.46E-01 | -1.11E+00 | 3.59E-01 | 9.85E-01 | 1.21E+00 |
| ZFYVE1 | 7.67E-02 | 4.00E-01 | 1.15E+00 | 1.26E-01 | 5.44E-01 | 1.34E+00 | 4.03E-02 | 1.71E-01 | -1.09E+00 | 3.94E-02 | 1.50E-01 | 1.07E+00 | 5.67E-05 | 2.02E-04 | -1.40E+00 | 2.45E-04 | 6.91E-04 | -1.26E+00 | 5.65E-04 | 2.07E-03 | -1.17E+00 | 1.71E-04 | 4.70E-04 | -1.32E+00 |
| PTPRS | 7.68E-02 | 4.00E-01 | 1.05E+00 | 6.08E-02 | 2.63E-01 | 1.08E+00 | 7.84E-02 | 3.33E-01 | 1.48E+00 | 1.47E-01 | 5.63E-01 | 1.09E+00 | 1.87E-01 | 6.68E-01 | 1.09E+00 | 2.27E-01 | 6.41E-01 | 1.15E+00 | 4.01E-02 | 1.47E-01 | -1.57E+00 | 1.38E-01 | 3.80E-01 | -1.25E+00 |
| AC023509.4 | 7.69E-02 | 4.01E-01 | 1.11E+00 | 4.35E-03 | 1.88E-02 | -1.31E+00 | 6.03E-02 | 2.56E-01 | 1.53E+00 | 5.40E-02 | 2.06E-01 | -1.15E+00 | 2.30E-01 | 8.20E-01 | 1.64E+00 | 3.03E-01 | 8.56E-01 | 1.41E+00 | 1.60E-01 | 5.86E-01 | -1.14E+00 | 2.43E-01 | 6.66E-01 | 1.35E+00 |
| AP001000.1 | 7.70E-02 | 4.01E-01 | 1.20E+00 | 4.22E-04 | 1.82E-03 | -1.05E+00 | 9.51E-02 | 4.04E-01 | 1.05E+00 | 4.95E-04 | 1.89E-03 | -1.18E+00 | 1.31E-01 | 4.67E-01 | 1.19E+00 | 1.03E-04 | 2.91E-04 | -1.50E+00 | 1.04E-01 | 3.80E-01 | 1.22E+00 | 3.71E-02 | 1.02E-01 | -1.05E+00 |
| PALM | 7.70E-02 | 4.01E-01 | 1.08E+00 | 4.82E-02 | 2.08E-01 | 1.17E+00 | 2.15E-02 | 9.15E-02 | 1.04E+00 | 5.58E-02 | 2.13E-01 | 1.28E+00 | 1.05E-02 | 3.73E-02 | -1.07E+00 | 7.39E-02 | 2.08E-01 | 1.10E+00 | 1.29E-02 | 4.74E-02 | -1.32E+00 | 2.34E-01 | 6.42E-01 | 1.09E+00 |
| DHRS4L2 | 7.70E-02 | 4.01E-01 | 1.28E+00 | 7.10E-02 | 3.07E-01 | 1.72E+00 | 1.08E-01 | 4.59E-01 | 1.34E+00 | 2.89E-03 | 1.10E-02 | 1.78E+00 | 1.05E-01 | 3.74E-01 | 1.19E+00 | 3.15E-01 | 8.89E-01 | 1.53E+00 | 1.39E-01 | 5.10E-01 | 1.45E+00 | 5.30E-03 | 1.46E-02 | 1.85E+00 |
| LCN2 | 7.70E-02 | 4.01E-01 | -1.47E+00 | 8.39E-02 | 3.63E-01 | -1.57E+00 | 1.87E-01 | 7.96E-01 | 1.25E+00 | 2.14E-01 | 8.17E-01 | 1.29E+00 | 1.90E-04 | 6.78E-04 | -1.27E+00 | 8.61E-04 | 2.43E-03 | -1.18E+00 | 6.34E-04 | 2.33E-03 | 1.70E+00 | 8.61E-09 | 2.36E-08 | -1.25E+00 |
| SLC25A5-AS1 | 7.71E-02 | 4.02E-01 | 1.06E+00 | 5.10E-02 | 2.21E-01 | 1.04E+00 | 2.20E-01 | 9.34E-01 | 1.14E+00 | 2.03E-01 | 7.75E-01 | 1.16E+00 | 1.93E-03 | 6.88E-03 | -1.63E+00 | 4.51E-02 | 1.27E-01 | -1.08E+00 | 1.75E-01 | 6.44E-01 | 1.38E+00 | 2.77E-01 | 7.60E-01 | 1.40E+00 |
| TMSB4XP8 | 7.71E-02 | 4.02E-01 | -1.14E+00 | 2.01E-01 | 8.68E-01 | 1.38E+00 | 1.57E-02 | 6.67E-02 | -1.40E+00 | 5.25E-02 | 2.00E-01 | -1.07E+00 | 5.23E-03 | 1.87E-02 | -1.72E+00 | 4.33E-03 | 1.22E-02 | -1.73E+00 | 9.73E-08 | 3.57E-07 | -1.76E+00 | 2.25E-07 | 6.19E-07 | -1.92E+00 |
| AL353621.1 | 7.71E-02 | 4.02E-01 | 1.58E+00 | 2.11E-01 | 9.11E-01 | 1.44E+00 | 8.06E-02 | 3.43E-01 | 1.42E+00 | 3.77E-02 | 1.44E-01 | 1.90E+00 | 1.30E-01 | 4.62E-01 | 1.86E+00 | 2.35E-01 | 6.61E-01 | 1.65E+00 | 6.62E-03 | 2.43E-02 | 2.16E+00 | 2.83E-03 | 7.77E-03 | 2.62E+00 |
| AMZ1 | 7.71E-02 | 4.02E-01 | 1.15E+00 | 1.10E-02 | 4.74E-02 | -1.07E+00 | 1.50E-01 | 6.37E-01 | 1.03E+00 | 3.71E-02 | 1.42E-01 | -1.20E+00 | 6.50E-02 | 2.32E-01 | -1.30E+00 | 1.41E-01 | 3.99E-01 | -1.09E+00 | 6.44E-04 | 2.36E-03 | -1.51E+00 | 9.31E-03 | 2.56E-02 | -1.19E+00 |
| FAN1 | 7.72E-02 | 4.02E-01 | 1.03E+00 | 5.43E-02 | 2.34E-01 | 1.06E+00 | 1.42E-02 | 6.04E-02 | -1.09E+00 | 4.13E-02 | 1.58E-01 | 1.14E+00 | 9.53E-02 | 3.40E-01 | 1.01E+00 | 2.56E-01 | 7.22E-01 | 1.35E+00 | 2.54E-01 | 9.32E-01 | 1.17E+00 | 1.37E-01 | 3.75E-01 | 1.48E+00 |
| CYCSP39 | 7.73E-02 | 4.03E-01 | 1.07E+00 | 6.28E-05 | 2.71E-04 | -2.48E+00 | 1.42E-01 | 6.03E-01 | 1.01E+00 | 3.02E-06 | 1.15E-05 | -2.46E+00 | 3.18E-02 | 1.13E-01 | 2.29E+00 | 1.36E-01 | 3.84E-01 | -1.35E+00 | 3.59E-02 | 1.32E-01 | -1.00E+00 | 4.51E-04 | 1.24E-03 | -1.46E+00 |
| FAM169A | 7.73E-02 | 4.03E-01 | 1.54E+00 | 1.45E-02 | 6.28E-02 | 1.12E+00 | 2.09E-01 | 8.88E-01 | 1.12E+00 | 2.01E-02 | 7.70E-02 | -1.02E+00 | 2.57E-01 | 9.17E-01 | 1.44E+00 | 3.46E-02 | 9.75E-02 | -1.18E+00 | 2.62E-01 | 9.63E-01 | 1.27E+00 | 1.72E-01 | 4.74E-01 | 1.11E+00 |
| AC093616.1 | 7.74E-02 | 4.04E-01 | -1.03E+00 | 1.89E-01 | 8.17E-01 | 1.27E+00 | 1.58E-03 | 6.71E-03 | -1.36E+00 | 5.46E-04 | 2.08E-03 | -1.39E+00 | 2.39E-01 | 8.51E-01 | 1.49E+00 | 2.93E-01 | 8.26E-01 | 1.34E+00 | 2.71E-01 | 9.96E-01 | 1.21E+00 | 3.08E-01 | 8.45E-01 | 1.25E+00 |
| AC242376.2 | 7.75E-02 | 4.04E-01 | -1.19E+00 | 1.84E-02 | 7.95E-02 | 1.86E+00 | 7.99E-03 | 3.40E-02 | -1.38E+00 | 2.51E-01 | 9.58E-01 | 1.40E+00 | 2.75E-01 | 9.81E-01 | 1.25E+00 | 2.89E-01 | 8.15E-01 | 1.17E+00 | 1.25E-02 | 4.60E-02 | -1.19E+00 | 1.46E-01 | 4.01E-01 | 1.16E+00 |
| TCTEX1D4 | 7.75E-02 | 4.04E-01 | 1.69E+00 | 6.75E-02 | 2.92E-01 | 1.83E+00 | 2.40E-02 | 1.02E-01 | 1.89E+00 | 1.46E-01 | 5.58E-01 | 1.51E+00 | 6.92E-02 | 2.47E-01 | 2.02E+00 | 2.61E-01 | 7.35E-01 | 1.03E+00 | 1.56E-03 | 5.72E-03 | 2.77E+00 | 1.46E-01 | 4.02E-01 | 1.65E+00 |
| U2AF1 | 7.75E-02 | 4.04E-01 | 1.25E+00 | 8.07E-02 | 3.49E-01 | 1.68E+00 | 5.01E-03 | 2.13E-02 | 1.56E+00 | 1.57E-07 | 5.99E-07 | 2.26E+00 | 2.65E-03 | 9.44E-03 | 1.03E+00 | 1.13E-01 | 3.18E-01 | 1.62E+00 | 2.14E-01 | 7.85E-01 | 1.23E+00 | 2.74E-06 | 7.52E-06 | 1.89E+00 |
| ATP2A1-AS1 | 7.77E-02 | 4.05E-01 | -1.24E+00 | 3.73E-02 | 1.61E-01 | -1.52E+00 | 7.22E-02 | 3.07E-01 | 1.31E+00 | 3.21E-02 | 1.23E-01 | -1.05E+00 | 4.22E-02 | 1.50E-01 | -1.92E+00 | 9.37E-02 | 2.64E-01 | -1.52E+00 | 4.81E-02 | 1.76E-01 | -1.10E+00 | 2.56E-03 | 7.03E-03 | -1.46E+00 |
| AC091153.1 | 7.77E-02 | 4.05E-01 | 1.06E+00 | 7.14E-03 | 3.08E-02 | 1.14E+00 | 3.82E-02 | 1.62E-01 | 1.09E+00 | 3.70E-03 | 1.41E-02 | 1.16E+00 | 1.28E-01 | 4.55E-01 | 1.11E+00 | 1.98E-03 | 5.59E-03 | 1.21E+00 | 2.01E-02 | 7.36E-02 | 1.05E+00 | 5.47E-02 | 1.50E-01 | 1.16E+00 |
| CHTF18 | 7.77E-02 | 4.05E-01 | -1.05E+00 | 6.78E-02 | 2.93E-01 | -1.13E+00 | 2.10E-01 | 8.91E-01 | 1.17E+00 | 1.52E-02 | 5.80E-02 | -1.04E+00 | 1.82E-01 | 6.48E-01 | -1.20E+00 | 2.87E-01 | 8.09E-01 | 1.08E+00 | 6.06E-02 | 2.23E-01 | 1.02E+00 | 2.55E-01 | 6.99E-01 | 1.26E+00 |
| DHRS12 | 7.77E-02 | 4.05E-01 | 1.18E+00 | 2.02E-01 | 8.71E-01 | 1.50E+00 | 9.96E-02 | 4.23E-01 | 1.25E+00 | 6.55E-02 | 2.50E-01 | 1.49E+00 | 7.05E-02 | 2.51E-01 | 1.01E+00 | 2.30E-01 | 6.48E-01 | 1.30E+00 | 2.42E-01 | 8.87E-01 | 1.47E+00 | 2.66E-01 | 7.31E-01 | 1.31E+00 |
| AFF3 | 7.77E-02 | 4.05E-01 | 1.23E+00 | 6.20E-02 | 2.68E-01 | 1.32E+00 | 1.99E-02 | 8.45E-02 | 1.54E+00 | 1.12E-01 | 4.29E-01 | 1.56E+00 | 9.26E-02 | 3.30E-01 | 1.88E+00 | 1.55E-01 | 4.36E-01 | 1.76E+00 | 4.95E-03 | 1.82E-02 | -1.35E+00 | 7.78E-03 | 2.14E-02 | -1.20E+00 |
| [illegible] | [illegible] | [illegible] | [illegible] | [illegible] | [illegible] | [illegible] | [illegible] | [illegible] | [illegible] | [illegible] | [illegible] | [illegible] | [illegible] | [illegible] | [illegible] | [illegible] | [illegible] | [illegible] | [illegible] | [illegible] | [illegible] | [illegible] | [illegible] | [illegible] |

| | | | | | | | | | | | | | | | | | | | | | | | | |
|---|---|---|---|---|---|---|---|---|---|---|---|---|---|---|---|---|---|---|---|---|---|---|---|---|
| TUBA1B | 8.58E-02 | 4.47E-01 | 1.59E+00 | 2.36E-02 | 1.02E-01 | 2.02E+00 | 1.10E-01 | 4.66E-01 | 1.58E+00 | 5.28E-03 | 2.02E-02 | 2.44E+00 | 2.36E-01 | 8.42E-01 | 1.34E+00 | 3.20E-01 | 9.02E-01 | 1.83E+00 | 2.31E-01 | 8.48E-01 | 1.34E+00 | 4.36E-02 | 1.20E-01 | 1.90E+00 |
| EIF5AP4 | 8.59E-02 | 4.48E-01 | 1.59E+00 | 2.26E-01 | 9.77E-01 | 1.39E+00 | 4.31E-02 | 1.83E-01 | 1.50E+00 | 1.25E-01 | 4.79E-01 | 1.67E+00 | 6.95E-02 | 2.48E-01 | 2.10E+00 | 8.66E-02 | 2.44E-01 | 2.16E+00 | 7.34E-04 | 2.69E-03 | 1.83E+00 | 5.72E-06 | 1.57E-05 | 2.26E+00 |
| AL139393.2 | 8.59E-02 | 4.48E-01 | 1.53E+00 | 4.94E-02 | 2.14E-01 | -1.06E+00 | 1.75E-01 | 7.42E-01 | 1.22E+00 | 1.83E-02 | 6.98E-02 | -1.55E+00 | 2.16E-01 | 7.71E-01 | 1.53E+00 | 3.53E-01 | 9.94E-01 | 1.20E+00 | 1.59E-01 | 5.84E-01 | 1.05E+00 | 3.09E-01 | 8.49E-01 | 1.27E+00 |
| ABCA7 | 8.59E-02 | 4.48E-01 | -1.01E+00 | 6.04E-02 | 2.61E-01 | -1.03E+00 | 4.27E-02 | 1.82E-01 | -1.23E+00 | 1.13E-01 | 4.31E-01 | 1.07E+00 | 2.66E-01 | 9.50E-01 | 1.61E+00 | 3.08E-01 | 8.69E-01 | 1.54E+00 | 1.60E-03 | 5.89E-03 | 1.01E+00 | 4.21E-03 | 1.16E-02 | -1.15E+00 |
| TUBBP1 | 8.59E-02 | 4.48E-01 | 1.07E+00 | 1.87E-01 | 8.08E-01 | 1.40E+00 | 3.46E-03 | 1.47E-02 | 1.73E+00 | 1.64E-04 | 6.27E-04 | 2.21E+00 | 2.00E-02 | 7.12E-02 | -1.00E+00 | 9.09E-02 | 2.56E-01 | 1.88E+00 | 1.77E-01 | 6.51E-01 | 1.45E+00 | 8.28E-08 | 2.28E-07 | 2.41E+00 |
| C9orf72 | 8.60E-02 | 4.48E-01 | 1.38E+00 | 2.22E-01 | 9.59E-01 | 1.46E+00 | 1.25E-01 | 5.31E-01 | 1.06E+00 | 1.26E-01 | 4.83E-01 | 1.22E+00 | 2.28E-01 | 8.13E-01 | 1.23E+00 | 3.44E-01 | 9.70E-01 | 1.23E+00 | 8.20E-03 | 3.01E-02 | -1.14E+00 | 5.35E-02 | 1.47E-01 | -1.03E+00 |
| ADPGK-AS1 | 8.60E-02 | 4.48E-01 | -1.01E+00 | 5.47E-04 | 2.36E-03 | -1.95E+00 | 2.02E-01 | 8.59E-01 | 1.33E+00 | 1.29E-02 | 4.94E-02 | -1.09E+00 | 3.82E-02 | 1.36E-01 | -1.14E+00 | 6.09E-03 | 1.72E-02 | -2.38E+00 | 1.44E-01 | 5.27E-01 | -1.04E+00 | 2.26E-01 | 6.21E-01 | 1.02E+00 |
| IQGAP3 | 8.60E-02 | 4.48E-01 | 1.46E+00 | 2.31E-01 | 1.00E+00 | 1.00E+00 | 2.69E-02 | 1.14E-01 | 1.66E+00 | 1.68E-01 | 6.42E-01 | 1.33E+00 | 2.50E-01 | 8.92E-01 | 1.12E+00 | 2.98E-01 | 8.41E-01 | 1.20E+00 | 1.50E-06 | 5.51E-06 | -1.90E+00 | 4.17E-05 | 1.15E-04 | -1.76E+00 |
| SPIN2A | 8.61E-02 | 4.49E-01 | 1.15E+00 | 1.97E-01 | 8.50E-01 | 1.46E+00 | 7.11E-02 | 3.02E-01 | -1.06E+00 | 1.52E-01 | 5.80E-01 | 1.45E+00 | 1.73E-01 | 6.16E-01 | 1.19E+00 | 1.44E-01 | 4.07E-01 | 1.89E+00 | 6.16E-02 | 2.26E-01 | 1.67E+00 | 4.80E-02 | 1.32E-01 | 1.83E+00 |
| RP9 | 8.62E-02 | 4.49E-01 | 1.18E+00 | 7.56E-04 | 3.27E-03 | -1.11E+00 | 7.11E-02 | 3.02E-01 | 1.38E+00 | 2.17E-01 | 8.30E-01 | 1.30E+00 | 8.34E-06 | 2.98E-05 | -1.39E+00 | 1.29E-05 | 3.65E-05 | -1.33E+00 | 1.32E-01 | 4.83E-01 | 1.07E+00 | 8.45E-03 | 2.32E-02 | -1.08E+00 |
| AL117328.2 | 8.62E-02 | 4.49E-01 | 1.16E+00 | 2.37E-02 | 1.02E-01 | -1.70E+00 | 1.10E-02 | 4.68E-02 | 1.75E+00 | 1.26E-01 | 4.81E-01 | 1.41E+00 | 9.36E-02 | 3.34E-01 | 1.99E+00 | 3.34E-01 | 9.41E-01 | 1.23E+00 | 1.90E-01 | 6.97E-01 | 1.23E+00 | 3.21E-01 | 8.83E-01 | 1.20E+00 |
| SH3TC2 | 8.63E-02 | 4.50E-01 | -1.18E+00 | 1.28E-01 | 5.53E-01 | -1.05E+00 | 3.60E-02 | 1.53E-01 | -1.31E+00 | 1.96E-01 | 7.48E-01 | 1.35E+00 | 2.39E-01 | 8.52E-01 | 1.26E+00 | 3.04E-01 | 8.58E-01 | -1.01E+00 | 1.21E-02 | 4.45E-02 | 1.99E+00 | 1.91E-01 | 5.25E-01 | 1.39E+00 |
| FAM212B | 8.63E-02 | 4.50E-01 | 1.46E+00 | 4.83E-02 | 2.09E-01 | -1.02E+00 | 1.34E-01 | 5.70E-01 | 1.08E+00 | 1.32E-02 | 5.06E-02 | -1.22E+00 | 1.17E-01 | 4.19E-01 | 1.81E+00 | 2.39E-01 | 6.73E-01 | 1.33E+00 | 4.01E-05 | 1.47E-04 | -1.12E+00 | 1.11E-03 | 3.04E-03 | -1.09E+00 |
| AC025034.1 | 8.63E-02 | 4.50E-01 | 1.07E+00 | 2.13E-01 | 9.20E-01 | 1.40E+00 | 1.17E-02 | 4.98E-02 | -1.03E+00 | 1.47E-03 | 5.63E-03 | -1.09E+00 | 7.42E-02 | 2.65E-01 | 2.35E+00 | 1.37E-01 | 3.86E-01 | 2.09E+00 | 2.07E-01 | 7.61E-01 | 1.29E+00 | 1.88E-01 | 5.17E-01 | 1.55E+00 |
| PCBP4 | 8.64E-02 | 4.50E-01 | -1.03E+00 | 2.29E-01 | 9.92E-01 | 1.40E+00 | 2.50E-02 | 1.06E-01 | -1.21E+00 | 1.68E-01 | 6.43E-01 | 1.18E+00 | 1.07E-01 | 3.81E-01 | -1.14E+00 | 2.95E-01 | 8.33E-01 | 1.27E+00 | 1.58E-01 | 5.81E-01 | 1.37E+00 | 1.75E-01 | 4.80E-01 | 1.48E+00 |
| AC234772.3 | 8.64E-02 | 4.50E-01 | 1.16E+00 | 1.12E-01 | 4.84E-01 | 1.78E+00 | 1.97E-01 | 8.38E-01 | 1.15E+00 | 5.59E-02 | 2.14E-01 | 2.00E+00 | 2.72E-01 | 9.69E-01 | 1.15E+00 | 1.26E-01 | 3.55E-01 | 1.70E+00 | 1.67E-01 | 6.13E-01 | 1.36E+00 | 3.45E-01 | 9.47E-01 | 1.29E+00 |
| TMEM39A | 8.64E-02 | 4.50E-01 | 1.18E+00 | 5.12E-02 | 2.21E-01 | 1.18E+00 | 1.07E-03 | 4.56E-03 | -1.12E+00 | 2.76E-08 | 1.05E-07 | -1.64E+00 | 5.74E-04 | 2.05E-03 | -1.28E+00 | 5.12E-05 | 1.44E-04 | -1.50E+00 | 3.97E-02 | 1.46E-01 | -1.07E+00 | 6.08E-03 | 1.67E-02 | -1.26E+00 |
| TTI2 | 8.65E-02 | 4.51E-01 | 1.55E+00 | 1.68E-01 | 7.24E-01 | 1.49E+00 | 5.22E-03 | 2.22E-02 | -1.35E+00 | 2.10E-02 | 8.01E-02 | -1.09E+00 | 2.36E-01 | 8.43E-01 | 1.26E+00 | 2.38E-01 | 6.72E-01 | 1.16E+00 | 2.48E-01 | 9.08E-01 | 1.36E+00 | 2.06E-01 | 5.66E-01 | 1.51E+00 |
| MANEA | 8.65E-02 | 4.51E-01 | 1.13E+00 | 1.96E-01 | 8.46E-01 | 1.40E+00 | 8.01E-04 | 3.40E-03 | -1.24E+00 | 3.81E-03 | 1.45E-02 | -1.02E+00 | 4.24E-03 | 1.51E-02 | -1.18E+00 | 3.91E-02 | 1.10E-01 | 1.07E+00 | 9.52E-05 | 3.49E-04 | -1.48E+00 | 9.04E-02 | 2.48E-01 | -1.02E+00 |
| H1FX-AS1 | 8.66E-02 | 4.51E-01 | 1.70E+00 | 2.01E-01 | 8.68E-01 | 1.39E+00 | 1.38E-01 | 5.87E-01 | 1.10E+00 | 1.96E-01 | 7.48E-01 | 1.23E+00 | 1.47E-01 | 5.25E-01 | 1.08E+00 | 2.77E-01 | 7.82E-01 | 1.07E+00 | 2.19E-01 | 8.05E-01 | 1.03E+00 | 2.13E-01 | 5.85E-01 | -1.10E+00 |
| WDR6 | 8.67E-02 | 4.52E-01 | 1.10E+00 | 5.81E-02 | 2.51E-01 | 1.05E+00 | 3.64E-03 | 1.55E-02 | -1.29E+00 | 9.23E-04 | 3.52E-03 | -1.26E+00 | 3.72E-04 | 1.33E-03 | -1.48E+00 | 5.44E-04 | 1.53E-03 | -1.42E+00 | 5.64E-02 | 2.07E-01 | -1.03E+00 | 7.66E-02 | 2.10E-01 | -1.04E+00 |
| CIRBP-AS1 | 8.67E-02 | 4.52E-01 | 1.61E+00 | 4.60E-02 | 1.99E-01 | 1.00E+00 | 1.32E-02 | 5.60E-02 | 1.66E+00 | 2.59E-01 | 9.89E-01 | 1.24E+00 | 1.02E-01 | 3.63E-01 | -1.17E+00 | 1.43E-01 | 4.03E-01 | -1.22E+00 | 8.10E-02 | 2.97E-01 | -1.23E+00 | 1.81E-01 | 4.96E-01 | 1.00E+00 |
| AP002748.3 | 8.68E-02 | 4.52E-01 | 1.11E+00 | 5.54E-02 | 2.39E-01 | 1.13E+00 | 1.63E-02 | 6.91E-02 | -1.12E+00 | 6.74E-03 | 2.58E-02 | -1.02E+00 | 2.24E-01 | 8.00E-01 | 1.27E+00 | 2.49E-01 | 7.01E-01 | 1.61E+00 | 2.42E-01 | 8.87E-01 | 1.21E+00 | 2.85E-01 | 7.84E-01 | 1.21E+00 |
| RPS27P27 | 8.69E-02 | 4.53E-01 | 1.04E+00 | 4.69E-02 | 2.03E-01 | 2.17E+00 | 1.86E-01 | 7.91E-01 | 1.08E+00 | 1.33E-01 | 5.09E-01 | 1.55E+00 | 2.13E-01 | 7.59E-01 | 1.61E+00 | 2.39E-02 | 6.73E-02 | 2.57E+00 | 2.58E-01 | 9.46E-01 | 1.36E+00 | 1.65E-01 | 4.54E-01 | 1.39E+00 |
| ATP7B | 8.70E-02 | 4.53E-01 | 1.07E+00 | 2.30E-01 | 9.95E-01 | 1.57E+00 | 3.54E-02 | 1.50E-01 | -1.23E+00 | 5.63E-02 | 2.15E-01 | -1.02E+00 | 2.54E-01 | 9.05E-01 | 1.43E+00 | 1.42E-02 | 4.00E-02 | 2.25E+00 | 2.76E-03 | 1.01E-02 | -1.01E+00 | 3.36E-04 | 9.23E-04 | -1.10E+00 |
| GJA4 | 8.71E-02 | 4.54E-01 | -1.29E+00 | 1.26E-01 | 5.44E-01 | -1.07E+00 | 2.34E-01 | 9.96E-01 | 1.23E+00 | 2.04E-01 | 7.81E-01 | 1.86E+00 | 1.91E-02 | 6.81E-02 | 2.00E+00 | 9.71E-06 | 2.74E-05 | 2.86E+00 | 1.19E-01 | 4.38E-01 | 1.58E+00 | 2.80E-03 | 7.68E-03 | 1.61E+00 |
| AC141586.4 | 8.71E-02 | 4.54E-01 | 1.08E+00 | 1.29E-01 | 5.55E-01 | 1.30E+00 | 1.56E-01 | 6.63E-01 | 1.09E+00 | 1.69E-01 | 6.45E-01 | 1.20E+00 | 7.15E-03 | 2.55E-02 | -1.55E+00 | 2.72E-02 | 7.66E-02 | -1.24E+00 | 1.06E-02 | 3.90E-02 | -1.44E+00 | 8.93E-03 | 2.45E-02 | -1.61E+00 |
| TGFB1I1 | 8.72E-02 | 4.54E-01 | 1.23E+00 | 5.25E-02 | 2.27E-01 | 1.77E+00 | 2.24E-01 | 9.53E-01 | 1.40E+00 | 2.61E-02 | 9.95E-02 | 1.92E+00 | 1.55E-01 | 5.51E-01 | 1.14E+00 | 2.72E-01 | 7.66E-01 | 1.49E+00 | 7.76E-02 | 2.85E-01 | 1.56E+00 | 2.92E-01 | 8.01E-01 | 1.16E+00 |
| ZBTB40 | 8.73E-02 | 4.55E-01 | 1.13E+00 | 1.45E-01 | 6.25E-01 | 1.72E+00 | 1.46E-07 | 6.19E-07 | -1.60E+00 | 5.31E-07 | 2.03E-06 | -1.35E+00 | 9.45E-02 | 3.37E-01 | 1.13E+00 | 1.81E-01 | 5.09E-01 | 1.26E+00 | 1.00E-02 | 3.67E-02 | -1.07E+00 | 2.83E-02 | 7.78E-02 | 1.01E+00 |
| SCOC-AS1 | 8.74E-02 | 4.56E-01 | 1.03E+00 | 6.09E-03 | 2.63E-02 | -1.34E+00 | 1.41E-01 | 5.98E-01 | 1.14E+00 | 6.33E-03 | 2.42E-02 | -1.22E+00 | 1.80E-01 | 6.43E-01 | 1.09E+00 | 1.75E-01 | 4.94E-01 | 1.01E+00 | 1.12E-01 | 4.12E-01 | 1.09E+00 | 1.22E-03 | 3.35E-03 | -1.41E+00 |
| RPL19P21 | 8.74E-02 | 4.56E-01 | 1.57E+00 | 1.90E-01 | 8.21E-01 | 1.51E+00 | 1.09E-01 | 4.64E-01 | 1.40E+00 | 2.52E-01 | 9.64E-01 | 1.40E+00 | 1.38E-01 | 4.92E-01 | 1.73E+00 | 2.88E-01 | 8.12E-01 | 1.20E+00 | 2.01E-01 | 7.37E-01 | 1.43E+00 | 2.89E-01 | 7.93E-01 | 1.37E+00 |
| IL17RA | 8.74E-02 | 4.56E-01 | 1.16E+00 | 2.05E-02 | 8.87E-02 | 1.09E+00 | 2.60E-05 | 1.10E-04 | -1.30E+00 | 1.39E-03 | 5.30E-03 | 1.03E+00 | 4.11E-05 | 1.47E-04 | -1.64E+00 | 2.76E-05 | 7.78E-05 | -1.83E+00 | 5.10E-02 | 1.87E-01 | -1.02E+00 | 1.32E-01 | 3.62E-01 | -1.00E+00 |
| UBE2C | 8.74E-02 | 4.56E-01 | -1.26E+00 | 1.16E-01 | 5.00E-01 | -1.20E+00 | 2.10E-01 | 8.94E-01 | 1.71E+00 | 2.07E-01 | 7.91E-01 | 2.49E+00 | 1.34E-01 | 4.78E-01 | -2.16E+00 | 2.66E-01 | 7.51E-01 | -1.46E+00 | 3.31E-03 | 1.21E-02 | 1.09E+00 | 7.42E-05 | 2.04E-04 | 1.43E+00 |
| S100A11 | 8.75E-02 | 4.56E-01 | 1.20E+00 | 1.88E-02 | 8.14E-02 | 1.95E+00 | 1.03E-01 | 4.36E-01 | -1.07E+00 | 2.74E-02 | 1.05E-01 | -1.10E+00 | 2.09E-05 | 7.45E-05 | -1.77E+00 | 1.30E-04 | 3.67E-04 | -1.57E+00 | 2.94E-03 | 1.08E-02 | -1.62E+00 | 2.97E-02 | 8.16E-02 | -1.48E+00 |
| AL391069.2 | 8.78E-02 | 4.57E-01 | -1.24E+00 | 1.85E-01 | 8.00E-01 | 1.05E+00 | 1.07E-01 | 4.53E-01 | 1.37E+00 | 1.43E-02 | 5.47E-02 | -1.55E+00 | 4.12E-02 | 1.47E-01 | -1.30E+00 | 6.87E-02 | 1.94E-01 | -1.16E+00 | 2.65E-01 | 9.73E-01 | 1.19E+00 | 5.85E-02 | 1.61E-01 | -1.17E+00 |
| COX7C | 8.79E-02 | 4.58E-01 | 1.12E+00 | 2.85E-16 | 1.23E-15 | -1.22E+00 | 7.82E-02 | 3.32E-01 | 1.10E+00 | 2.79E-06 | 1.07E-05 | -1.23E+00 | 5.92E-02 | 2.11E-01 | -1.00E+00 | 1.34E-06 | 3.78E-06 | -1.24E+00 | 2.19E-01 | 8.03E-01 | 1.09E+00 | 1.02E-03 | 2.80E-03 | -1.17E+00 |
| SLC25A29 | 8.79E-02 | 4.58E-01 | 1.10E+00 | 3.61E-02 | 1.56E-01 | 1.05E+00 | 6.45E-04 | 2.74E-03 | -1.37E+00 | 1.21E-03 | 4.63E-03 | -1.24E+00 | 1.51E-02 | 5.39E-02 | -1.45E+00 | 4.16E-02 | 1.17E-01 | -1.24E+00 | 3.57E-04 | 1.31E-03 | -1.58E+00 | 1.10E-03 | 3.02E-03 | -1.46E+00 |
| SYVN1 | 8.82E-02 | 4.60E-01 | 1.20E+00 | 1.23E-01 | 5.30E-01 | 1.91E+00 | 4.18E-03 | 1.78E-02 | -1.26E+00 | 1.40E-01 | 5.34E-01 | 1.26E+00 | 8.54E-04 | 3.05E-03 | -1.50E+00 | 4.26E-02 | 1.20E-01 | -1.03E+00 | 4.55E-04 | 1.67E-03 | -1.37E+00 | 9.38E-02 | 2.58E-01 | 1.02E+00 |
| LTB4R | 8.83E-02 | 4.60E-01 | 1.12E+00 | 8.31E-02 | 3.59E-01 | 1.25E+00 | 5.45E-02 | 2.32E-01 | 1.02E+00 | 2.05E-02 | 7.84E-02 | 1.16E+00 | 2.46E-01 | 8.78E-01 | 1.44E+00 | 1.49E-01 | 4.21E-01 | 1.08E+00 | 1.62E-01 | 5.93E-01 | 1.08E+00 | 5.95E-02 | 1.64E-01 | -1.10E+00 |
| NUP58 | 8.83E-02 | 4.60E-01 | 1.11E+00 | 2.75E-09 | 1.19E-08 | -1.65E+00 | 2.41E-03 | 1.03E-02 | -1.04E+00 | 5.97E-12 | 2.28E-11 | -1.39E+00 | 7.78E-05 | 2.77E-04 | -1.27E+00 | 6.82E-09 | 1.92E-08 | -1.61E+00 | 1.17E-10 | 4.28E-10 | -1.41E+00 | 1.03E-11 | 2.82E-11 | -1.48E+00 |
| AC008764.6 | 8.84E-02 | 4.60E-01 | -1.00E+00 | 2.17E-02 | 9.38E-02 | -1.20E+00 | 1.90E-01 | 8.06E-01 | 1.06E+00 | 1.75E-01 | 6.70E-01 | 1.08E+00 | 2.24E-03 | 8.00E-03 | -1.73E+00 | 5.21E-04 | 1.47E-03 | -2.20E+00 | 1.83E-01 | 6.70E-01 | 1.02E+00 | 2.48E-01 | 6.80E-01 | 1.03E+00 |
| RETREG1 | 8.84E-02 | 4.61E-01 | 1.52E+00 | 2.17E-01 | 9.40E-01 | 1.60E+00 | 1.05E-01 | 4.46E-01 | 1.32E+00 | 2.28E-01 | 8.72E-01 | 1.43E+00 | 2.44E-01 | 8.70E-01 | 1.23E+00 | 2.42E-01 | 6.83E-01 | 1.24E+00 | 2.53E-01 | 9.29E-01 | 1.16E+00 | 3.46E-01 | 9.50E-01 | 1.30E+00 |
| WASF2 | 8.85E-02 | 4.61E-01 | 1.33E+00 | 1.20E-01 | 5.19E-01 | 1.34E+00 | 2.32E-01 | 9.86E-01 | 1.16E+00 | 9.36E-02 | 3.58E-01 | 1.17E+00 | 4.11E-03 | 1.46E-02 | -1.04E+00 | 4.85E-05 | 1.37E-04 | -1.09E+00 | 1.32E-02 | 4.86E-02 | 1.02E+00 | 2.29E-04 | 6.30E-04 | -1.16E+00 |
| ZNF674-AS1 | 8.85E-02 | 4.61E-01 | 1.21E+00 | 1.93E-02 | 8.33E-02 | 1.27E+00 | 1.71E-03 | 7.25E-03 | -1.25E+00 | 1.83E-03 | 6.98E-03 | -1.09E+00 | 1.85E-02 | 6.61E-02 | -1.07E+00 | 8.65E-02 | 2.44E-01 | 1.09E+00 | 7.94E-04 | 2.91E-03 | -1.41E+00 | 4.52E-02 | 1.24E-01 | -1.08E+00 |
| CES3 | 8.85E-02 | 4.61E-01 | -1.21E+00 | 1.49E-01 | 6.43E-01 | 1.02E+00 | 2.11E-02 | 8.95E-02 | -1.29E+00 | 3.91E-02 | 1.49E-01 | -1.24E+00 | 9.74E-03 | 3.47E-02 | -1.87E+00 | 7.79E-02 | 2.20E-01 | -1.27E+00 | 9.41E-02 | 3.45E-01 | 1.45E+00 | 2.14E-01 | 5.87E-01 | 1.28E+00 |
| BNIP3P1 | 8.86E-02 | 4.62E-01 | 1.62E+00 | 1.65E-01 | 7.13E-01 | 1.56E+00 | 1.96E-02 | 8.31E-02 | 1.62E+00 | 2.37E-02 | 9.06E-02 | 2.00E+00 | 2.04E-01 | 7.29E-01 | 1.59E+00 | 1.23E-01 | 3.47E-01 | 2.04E+00 | 9.25E-02 | 3.40E-01 | 1.38E+00 | 1.45E-01 | 3.99E-01 | 1.38E+00 |
| BX248409.1 | 8.87E-02 | 4.62E-01 | 1.13E+00 | 2.69E-47 | 1.16E-46 | -1.74E+00 | 6.13E-02 | 2.60E-01 | 1.22E+00 | 1.10E-16 | 4.21E-16 | -1.60E+00 | 2.50E-01 | 8.90E-01 | 1.19E+00 | 2.88E-15 | 8.12E-15 | -1.61E+00 | 2.28E-01 | 8.37E-01 | 1.16E+00 | 1.43E-16 | 3.92E-16 | -1.82E+00 |
| ANP32D | 8.88E-02 | 4.63E-01 | -1.03E+00 | 4.90E-02 | 2.12E-01 | -1.18E+00 | 2.38E-03 | 1.01E-02 | -1.44E+00 | 5.02E-05 | 1.92E-04 | -1.88E+00 | 7.26E-05 | 2.59E-04 | -2.25E+00 | 4.47E-04 | 1.26E-03 | -1.80E+00 | 2.49E-12 | 9.12E-12 | -1.96E+00 | 9.73E-15 | 2.67E-14 | -2.19E+00 |
| C8orf76 | 8.88E-02 | 4.63E-01 | 1.21E+00 | 1.72E-02 | 7.44E-02 | 1.09E+00 | 1.22E-03 | 5.19E-03 | -1.20E+00 | 5.57E-06 | 2.13E-05 | -1.31E+00 | 4.12E-03 | 1.47E-02 | -1.10E+00 | 9.57E-04 | 2.70E-03 | -1.23E+00 | 1.48E-09 | 5.43E-09 | -1.65E+00 | 1.37E-08 | 3.77E-08 | -1.51E+00 |
| AC010761.1 | 8.88E-02 | 4.63E-01 | 1.02E+00 | 1.32E-01 | 5.71E-01 | 1.20E+00 | 2.00E-02 | 8.48E-02 | -1.17E+00 | 1.92E-01 | 7.32E-01 | 1.31E+00 | 2.37E-01 | 8.46E-01 | 1.43E+00 | 3.04E-01 | 8.58E-01 | 1.21E+00 | 1.46E-01 | 5.36E-01 | -1.01E+00 | 2.49E-01 | 6.85E-01 | 1.02E+00 |
| MEG8 | 8.89E-02 | 4.63E-01 | 1.19E+00 | 4.59E-02 | 1.98E-01 | 1.90E+00 | 1.70E-02 | 7.21E-02 | -1.56E+00 | 1.56E-01 | 5.97E-01 | 1.41E+00 | 2.30E-01 | 8.21E-01 | 1.45E+00 | 1.34E-01 | 3.76E-01 | 2.02E+00 | 1.65E-01 | 6.06E-01 | 1.05E+00 | 3.26E-01 | 8.96E-01 | 1.34E+00 |
| NKG7 | 8.89E-02 | 4.64E-01 | 1.25E+00 | 2.02E-03 | 8.71E-03 | 1.13E+00 | 1.97E-01 | 8.37E-01 | 1.06E+00 | 2.43E-01 | 9.27E-01 | 1.11E+00 | 3.61E-03 | 1.29E-02 | -1.55E+00 | 4.72E-04 | 1.33E-03 | -1.95E+00 | 2.23E-01 | 8.20E-01 | 1.17E+00 | 2.59E-01 | 7.12E-01 | 1.09E+00 |
| NNAT | 8.91E-02 | 4.64E-01 | -1.05E+00 | 8.73E-02 | 3.77E-01 | -1.11E+00 | 2.12E-01 | 9.03E-01 | 1.27E+00 | 1.11E-01 | 4.23E-01 | 1.12E+00 | 1.51E-01 | 5.40E-01 | 1.20E+00 | 1.87E-01 | 5.28E-01 | 1.30E+00 | 1.58E-01 | 5.79E-01 | 1.42E+00 | 3.61E-01 | 9.91E-01 | 1.29E+00 |
| INHBA | 8.92E-02 | 4.65E-01 | -1.05E+00 | 2.73E-02 | 1.18E-01 | -1.36E+00 | 9.83E-14 | 4.18E-13 | -2.23E+00 | 1.36E-16 | 5.19E-16 | -2.81E+00 | 1.23E-01 | 4.38E-01 | -1.17E+00 | 7.63E-02 | 2.15E-01 | -1.54E+00 | 2.08E-01 | 7.62E-01 | -1.15E+00 | 3.44E-01 | 9.44E-01 | -1.06E+00 |
| FOSL2 | 8.92E-02 | 4.65E-01 | 1.35E+00 | 5.02E-08 | 2.17E-07 | -1.57E+00 | 4.89E-02 | 2.08E-01 | -1.00E+00 | 2.76E-11 | 1.06E-10 | -1.51E+00 | 7.00E-02 | 2.50E-01 | -1.03E+00 | 6.65E-09 | 1.88E-08 | -3.14E+00 | 2.72E-01 | 9.97E-01 | 1.13E+00 | 5.95E-02 | 1.64E-01 | -1.73E+00 |
| WWC1 | 8.92E-02 | 4.65E-01 | 1.00E+00 | 1.39E-01 | 5.98E-01 | 1.12E+00 | 1.89E-02 | 8.01E-02 | -1.19E+00 | 5.87E-02 | 2.24E-01 | 1.05E+00 | 2.09E-01 | 7.46E-01 | -1.03E+00 | 2.55E-01 | 7.19E-01 | -1.05E+00 | 2.98E-02 | 1.09E-01 | -1.43E+00 | 1.97E-01 | 5.41E-01 | 1.47E+00 |
| UCA1 | 8.92E-02 | 4.65E-01 | 1.66E+00 | 8.31E-02 | 3.59E-01 | 1.86E+00 | 9.85E-02 | 4.19E-01 | 1.53E+00 | 1.91E-01 | 7.29E-01 | 1.32E+00 | 1.33E-01 | 4.74E-01 | 1.51E+00 | 5.48E-02 | 1.55E-01 | 2.12E+00 | 8.37E-02 | 3.07E-01 | 1.56E+00 | 3.64E-01 | 9.99E-01 | 1.33E+00 |
| AP000873.2 | 8.94E-02 | 4.66E-01 | 1.50E+00 | 1.76E-01 | 7.61E-01 | 1.55E+00 | 1.45E-01 | 6.17E-01 | 1.27E+00 | 4.74E-02 | 1.81E-01 | 1.01E+00 | 2.46E-01 | 8.76E-01 | 1.18E+00 | 1.58E-01 | 4.45E-01 | 1.70E+00 | 1.78E-01 | 6.53E-01 | 1.42E+00 | 2.67E-01 | 7.34E-01 | 1.40E+00 |
| SLC37A3 | 8.95E-02 | 4.66E-01 | 1.17E+00 | 2.16E-02 | 9.33E-02 | 1.13E+00 | 1.07E-03 | 4.53E-03 | -1.39E+00 | 2.43E-03 | 9.28E-03 | -1.17E+00 | 1.59E-01 | 5.67E-01 | 1.16E+00 | 7.23E-02 | 2.04E-01 | 1.00E+00 | 2.96E-03 | 1.09E-02 | -1.22E+00 | 6.04E-05 | 1.66E-04 | -1.52E+00 |
| AC068473.5 | 8.95E-02 | 4.66E-01 | 1.13E+00 | 1.26E-01 | 5.44E-01 | 1.23E+00 | 3.52E-03 | 1.50E-02 | -1.39E+00 | 4.22E-02 | 1.61E-01 | 1.03E+00 | 6.50E-02 | 2.32E-01 | 2.05E+00 | 2.28E-01 | 6.43E-01 | 1.70E+00 | 2.26E-01 | 8.30E-01 | 1.27E+00 | 2.97E-01 | 8.17E-01 | 1.07E+00 |
| LINC00894 | 8.95E-02 | 4.66E-01 | -1.04E+00 | 2.02E-01 | 8.75E-01 | 1.45E+00 | 1.06E-01 | 4.52E-01 | -1.05E+00 | 1.04E-01 | 3.98E-01 | 1.66E+00 | 2.06E-01 | 7.33E-01 | 1.39E+00 | 1.02E-01 | 2.87E-01 | 1.81E+00 | 2.72E-01 | 1.00E+00 | 1.10E+00 | 3.17E-01 | 8.70E-01 | 1.01E+00 |
| ZNF778 | 8.96E-02 | 4.67E-01 | 1.03E+00 | 3.85E-02 | 1.67E-01 | -1.12E+00 | 9.72E-02 | 4.13E-01 | -1.03E+00 | 8.96E-02 | 3.42E-01 | 1.09E+00 | 2.07E-02 | 7.37E-02 | -1.60E+00 | 4.70E-02 | 1.32E-01 | -1.44E+00 | 1.88E-01 | 6.91E-01 | 1.67E+00 | 1.85E-01 | 5.09E-01 | -1.09E+00 |
| MAST3 | 8.97E-02 | 4.67E-01 | 1.60E+00 | 4.31E-02 | 1.86E-01 | 1.99E+00 | 2.09E-01 | 8.89E-01 | 1.08E+00 | 1.47E-01 | 5.62E-01 | 1.47E+00 | 1.26E-03 | 4.50E-03 | -1.58E+00 | 2.41E-03 | 6.80E-03 | -1.47E+00 | 1.96E-01 | 7.18E-01 | 1.03E+00 | 3.02E-01 | 8.29E-01 | 1.15E+00 |
| ZFYVE28 | 8.98E-02 | 4.68E-01 | 1.53E+00 | 2.00E-01 | 8.63E-01 | 1.33E+00 | 8.46E-02 | 3.60E-01 | -1.01E+00 | 2.93E-02 | 1.12E-01 | -1.06E+00 | 9.56E-04 | 3.41E-03 | -1.74E+00 | 4.15E-03 | 1.17E-02 | -1.35E+00 | 2.50E-05 | 9.17E-05 | -1.71E+00 | 4.86E-06 | 1.34E-05 | -2.04E+00 |
| AC079316.2 | 8.98E-02 | 4.68E-01 | 1.14E+00 | 1.23E-03 | 5.32E-03 | -1.44E+00 | 1.87E-01 | 7.96E-01 | 1.34E+00 | 3.44E-02 | 1.32E-01 | -1.16E+00 | 5.87E-02 | 2.09E-01 | -1.01E+00 | 4.85E-02 | 1.37E-01 | -1.10E+00 | 6.96E-02 | 2.56E-01 | -1.36E+00 | 1.45E-01 | 3.99E-01 | -1.23E+00 |
| PKD1 | 8.98E-02 | 4.68E-01 | 1.15E+00 | 1.05E-01 | 4.52E-01 | 1.37E+00 | 6.17E-02 | 2.62E-01 | 1.37E+00 | 8.34E-02 | 3.19E-01 | 1.60E+00 | 3.61E-02 | 1.29E-01 | -1.06E+00 | 1.88E-01 | 5.30E-01 | 1.26E+00 | 5.47E-02 | 2.01E-01 | 1.04E+00 | 3.38E-01 | 9.29E-01 | 1.24E+00 |
| HIRA | 8.98E-02 | 4.68E-01 | 1.11E+00 | 1.82E-01 | 7.85E-01 | 1.33E+00 | 1.01E-01 | 4.29E-01 | 1.02E+00 | 2.51E-01 | 9.58E-01 | 1.38E+00 | 5.33E-03 | 1.90E-02 | -1.53E+00 | 2.31E-01 | 6.51E-01 | 1.22E+00 | 6.06E-04 | 2.22E-03 | -1.36E+00 | 1.36E-01 | 3.74E-01 | 1.07E+00 |
| MYH9 | 8.99E-02 | 4.68E-01 | 1.18E+00 | 4.59E-02 | 1.98E-01 | 1.55E+00 | 3.24E-02 | 1.38E-01 | 1.08E+00 | 2.17E-01 | 8.28E-01 | 1.30E+00 | 2.37E-01 | 8.46E-01 | 1.46E+00 | 1.92E-01 | 5.42E-01 | 1.33E+00 | 4.82E-02 | 1.77E-01 | 1.41E+00 | 5.98E-02 | 1.64E-01 | 1.37E+00 |
| AL357093.2 | 8.99E-02 | 4.68E-01 | 1.08E+00 | 1.54E-01 | 6.65E-01 | 1.25E+00 | 5.83E-02 | 2.48E-01 | -1.02E+00 | 2.68E-02 | 1.02E-01 | -1.03E+00 | 2.42E-01 | 8.64E-01 | 1.19E+00 | 2.06E-01 | 5.80E-01 | 1.59E+00 | 1.50E-01 | 5.50E-01 | -1.13E+00 | 9.88E-02 | 2.71E-01 | -1.32E+00 |
| AC003071.1 | 8.99E-02 | 4.69E-01 | 1.14E+00 | 1.08E-01 | 4.66E-01 | 1.65E+00 | 7.77E-03 | 3.30E-02 | -1.05E+00 | 1.50E-01 | 5.74E-01 | 1.42E+00 | 1.28E-01 | 4.55E-01 | 1.18E+00 | 1.48E-01 | 4.18E-01 | 1.79E+00 | 1.82E-01 | 6.68E-01 | 1.16E+00 | 8.01E-02 | 2.20E-01 | 1.60E+00 |
| AL031708.1 | 9.00E-02 | 4.69E-01 | 1.09E+00 | 4.95E-02 | 2.14E-01 | -1.01E+00 | 2.25E-02 | 9.56E-02 | -1.24E+00 | 6.75E-02 | 2.58E-01 | -1.06E+00 | 2.64E-01 | 9.40E-01 | 1.33E+00 | 2.04E-01 | 5.74E-01 | 1.64E+00 | 1.02E-01 | 3.74E-01 | -1.19E+00 | 2.44E-01 | 6.71E-01 | 1.32E+00 |
| GCOM2 | 9.00E-02 | 4.69E-01 | 1.00E+00 | 1.86E-01 | 8.03E-01 | 1.29E+00 | 1.96E-02 | 8.34E-02 | -1.24E+00 | 1.02E-01 | 3.90E-01 | 1.11E+00 | 3.87E-04 | 1.38E-03 | -1.62E+00 | 1.47E-03 | 4.14E-03 | -1.42E+00 | 2.42E-02 | 8.88E-02 | -1.13E+00 | 1.69E-02 | 4.65E-02 | -1.25E+00 |
| KLC4 | 9.01E-02 | 4.70E-01 | 1.14E+00 | 1.49E-01 | 6.42E-01 | 1.31E+00 | 7.76E-04 | 3.30E-03 | -1.34E+00 | 1.44E-04 | 5.51E-04 | -1.40E+00 | 4.88E-02 | 1.74E-01 | -1.08E+00 | 1.74E-01 | 4.91E-01 | 1.20E+00 | 1.42E-01 | 5.23E-01 | 1.07E+00 | 2.06E-01 | 5.65E-01 | 1.04E+00 |
| CXCL2 | 9.03E-02 | 4.70E-01 | 1.31E+00 | 1.63E-03 | 7.02E-03 | -1.27E+00 | 2.08E-02 | 8.85E-02 | 3.36E+00 | 1.63E-01 | 6.23E-01 | 1.27E+00 | 2.09E-01 | 7.46E-01 | -1.21E+00 | 1.38E-03 | 3.89E-03 | -2.78E+00 | 5.70E-02 | 2.09E-01 | -1.19E+00 | 1.04E-02 | 2.86E-02 | -2.16E+00 |
| DGKH | 9.03E-02 | 4.71E-01 | 1.15E+00 | 1.32E-01 | 5.69E-01 | 1.67E+00 | 4.89E-05 | 2.08E-04 | -1.75E+00 | 2.88E-03 | 1.10E-02 | -1.13E+00 | 6.79E-04 | 2.42E-03 | -2.49E+00 | 4.12E-03 | 1.16E-02 | -1.82E+00 | 9.19E-03 | 3.37E-02 | 1.10E+00 | 6.11E-02 | 1.68E-01 | 1.19E+00 |
| RN7SL233P | 9.04E-02 | 4.71E-01 | -1.06E+00 | 5.47E-03 | 2.36E-02 | -1.57E+00 | 3.49E-03 | 1.48E-02 | -1.29E+00 | 2.05E-06 | 7.85E-06 | -2.11E+00 | 2.47E-01 | 8.80E-01 | 1.39E+00 | 8.79E-03 | 2.48E-02 | -2.05E+00 | 2.10E-02 | 7.71E-02 | 1.79E+00 | 1.20E-01 | 3.30E-01 | -1.37E+00 |
| KLHL2 | 9.04E-02 | 4.71E-01 | 1.14E+00 | 2.19E-01 | 9.45E-01 | 1.46E+00 | 4.85E-02 | 2.06E-01 | -1.04E+00 | 4.07E-02 | 1.55E-01 | 1.07E+00 | 4.97E-03 | 1.77E-02 | -1.68E+00 | 3.89E-02 | 1.10E-01 | -1.21E+00 | 2.21E-07 | 8.10E-07 | -1.75E+00 | 2.35E-04 | 6.45E-04 | -1.24E+00 |
| AP001107.1 | 9.04E-02 | 4.71E-01 | 1.08E+00 | 4.93E-03 | 2.13E-02 | -1.36E+00 | 1.76E-01 | 7.49E-01 | 1.07E+00 | 7.10E-02 | 2.71E-01 | -1.02E+00 | 1.25E-01 | 4.45E-01 | 1.65E+00 | 2.41E-01 | 6.80E-01 | 1.38E+00 | 6.28E-02 | 2.30E-01 | -1.05E+00 | 3.00E-02 | 8.23E-02 | -1.21E+00 |
| EVI2B | 9.04E-02 | 4.71E-01 | 1.33E+00 | 9.24E-02 | 3.99E-01 | 1.46E+00 | 1.44E-01 | 6.12E-01 | 1.12E+00 | 2.43E-01 | 9.30E-01 | 1.21E+00 | 2.33E-01 | 8.29E-01 | 1.28E+00 | 1.28E-01 | 3.59E-01 | 1.08E+00 | 7.47E-02 | 2.74E-01 | 1.83E+00 | 2.11E-01 | 5.80E-01 | 1.68E+00 |
| TPK1 | 9.05E-02 | 4.72E-01 | 1.11E+00 | 3.86E-02 | 1.67E-01 | 1.10E+00 | 8.35E-02 | 3.55E-01 | 1.45E+00 | 9.16E-02 | 3.50E-01 | 1.22E+00 | 2.07E-01 | 7.38E-01 | 1.29E+00 | 1.51E-02 | 4.26E-02 | -1.39E+00 | 2.47E-01 | 9.06E-01 | 1.34E+00 | 2.24E-01 | 6.15E-01 | 1.10E+00 |
| SGPL1 | 9.05E-02 | 4.72E-01 | 1.09E+00 | 4.56E-02 | 1.97E-01 | 1.23E+00 | 7.66E-04 | 3.26E-03 | -1.15E+00 | 6.80E-03 | 2.60E-02 | 1.09E+00 | 7.54E-03 | 2.69E-02 | -1.49E+00 | 3.08E-02 | 8.69E-02 | -1.12E+00 | 3.31E-03 | 1.21E-02 | -1.53E+00 | 1.05E-03 | 2.88E-03 | -1.98E+00 |
| TRAC | 9.05E-02 | 4.72E-01 | 1.32E+00 | 2.06E-01 | 8.89E-01 | 1.08E+00 | 1.05E-02 | 4.44E-02 | -4.13E+00 | 2.26E-03 | 8.62E-03 | -4.64E+00 | 1.05E-01 | 3.73E-01 | 1.71E+00 | 2.78E-01 | 7.85E-01 | 1.20E+00 | 2.72E-01 | 1.00E+00 | 1.00E+00 | 3.64E-01 | 1.00E+00 | 1.00E+00 |
| BTG3-AS1 | 9.06E-02 | 4.72E-01 | 1.50E+00 | 1.63E-01 | 7.04E-01 | 1.17E+00 | 1.42E-01 | 6.04E-01 | 1.54E+00 | 2.34E-01 | 8.95E-01 | 1.51E+00 | 2.78E-01 | 9.91E-01 | 1.22E+00 | 8.83E-02 | 2.49E-01 | 1.76E+00 | 4.34E-02 | 1.59E-01 | -1.22E+00 | 1.87E-01 | 5.13E-01 | -1.04E+00 |
| CTSG | 9.06E-02 | 4.72E-01 | 1.83E+00 | 1.62E-01 | 7.02E-01 | -1.11E+00 | 2.54E-02 | 1.08E-01 | 1.33E+00 | 5.51E-03 | 2.11E-02 | -1.05E+00 | 1.84E-01 | 6.57E-01 | -3.59E+00 | 2.45E-01 | 6.92E-01 | -3.42E+00 | 2.16E-01 | 7.92E-01 | 1.24E+00 | 3.51E-01 | 9.65E-01 | 1.04E+00 |
| AC011815.2 | 9.06E-02 | 4.72E-01 | 1.57E+00 | 3.35E-02 | 1.45E-01 | 2.07E+00 | 1.75E-01 | 7.43E-01 | -1.03E+00 | 1.71E-01 | 6.55E-01 | 1.53E+00 | 5.88E-02 | 2.10E-01 | 2.27E+00 | 2.64E-02 | 7.44E-02 | 3.58E+00 | 1.43E-01 | 5.26E-01 | 1.47E+00 | 1.00E-01 | 2.75E-01 | 1.77E+00 |
| DNAJC27 | 9.07E-02 | 4.73E-01 | 1.36E+00 | 3.66E-02 | 1.58E-01 | -1.06E+00 | 2.39E-03 | 1.02E-02 | -1.23E+00 | 6.45E-06 | 2.46E-05 | -1.54E+00 | 1.66E-01 | 5.91E-01 | 1.51E+00 | 7.13E-02 | 2.01E-01 | -1.09E+00 | 6.45E-02 | 2.37E-01 | 1.10E+00 | 2.79E-03 | 7.67E-03 | -1.31E+00 |
| LINC02367 | 9.07E-02 | 4.73E-01 | 1.54E+00 | 1.35E-01 | 5.84E-01 | 1.64E+00 | 1.31E-02 | 5.57E-02 | -1.16E+00 | 4.35E-02 | 1.66E-01 | 1.04E+00 | 2.02E-01 | 7.22E-01 | -1.04E+00 | 3.26E-01 | 9.18E-01 | 1.41E+00 | 1.74E-01 | 6.39E-01 | -1.01E+00 | 1.94E-01 | 5.32E-01 | 1.03E+00 |
| ZDHHC18 | 9.08E-02 | 4.73E-01 | 1.17E+00 | 1.42E-01 | 6.14E-01 | 1.87E+00 | 1.07E-01 | 4.56E-01 | 1.10E+00 | 1.98E-01 | 7.55E-01 | 1.39E+00 | 4.42E-02 | 1.57E-01 | -1.32E+00 | 2.84E-01 | 8.02E-01 | 1.24E+00 | 1.28E-01 | 4.70E-01 | 1.04E+00 | 1.81E-01 | 4.97E-01 | 1.50E+00 |
| ZNF700 | 9.09E-02 | 4.74E-01 | 1.10E+00 | 3.45E-02 | 1.49E-01 | 1.03E+00 | 3.91E-02 | 1.66E-01 | -1.02E+00 | 5.36E-03 | 2.05E-02 | -1.26E+00 | 1.71E-01 | 6.11E-01 | 1.28E+00 | 2.06E-01 | 5.80E-01 | 1.17E+00 | 3.44E-02 | 1.26E-01 | -1.17E+00 | 1.55E-02 | 4.25E-02 | -1.34E+00 |
| PYY2 | 9.10E-02 | 4.74E-01 | -1.36E+00 | 1.08E-01 | 4.67E-01 | -1.32E+00 | 1.96E-01 | 8.31E-01 | -1.11E+00 | 2.42E-01 | 9.25E-01 | -1.05E+00 | 1.15E-03 | 4.09E-03 | -1.05E+00 | 2.08E-07 | 5.86E-07 | -1.50E+00 | 8.09E-05 | 2.97E-04 | -1.87E+00 | 5.09E-03 | 1.40E-02 | -1.31E+00 |
| FAM60CP | 9.10E-02 | 4.74E-01 | 1.57E+00 | 5.50E-02 | 2.38E-01 | 1.90E+00 | 4.76E-02 | 2.02E-01 | 1.60E+00 | 8.28E-03 | 3.16E-02 | 2.12E+00 | 1.25E-01 | 4.44E-01 | 1.70E+00 | 4.55E-02 | 1.28E-01 | 1.96E+00 | 2.19E-01 | 8.03E-01 | 1.21E+00 | 2.10E-01 | 5.77E-01 | 1.39E+00 |
| MT-TY | 9.10E-02 | 4.74E-01 | 1.07E+00 | 2.65E-02 | 1.14E-01 | -1.20E+00 | 2.12E-01 | 9.03E-01 | 1.16E+00 | 8.47E-02 | 3.24E-01 | -1.05E+00 | 2.05E-01 | 7.32E-01 | 1.07E+00 | 2.18E-01 | 6.15E-01 | -1.08E+00 | 9.61E-02 | 3.53E-01 | 1.44E+00 | 1.42E-01 | 3.89E-01 | 1.05E+00 |
| GEMIN8 | 9.11E-02 | 4.75E-01 | 1.13E+00 | 8.92E-02 | 3.86E-01 | 1.24E+00 | 1.17E-06 | 4.96E-06 | -1.45E+00 | 7.96E-05 | 3.04E-04 | -1.12E+00 | 2.61E-04 | 9.29E-04 | -1.34E+00 | 1.36E-02 | 3.83E-02 | -1.02E+00 | 1.22E-05 | 4.48E-05 | -1.32E+00 | 2.34E-03 | 6.42E-03 | -1.03E+00 |
| AC020978.8 | 9.11E-02 | 4.75E-01 | 1.17E+00 | 1.88E-01 | 8.10E-01 | 1.58E+00 | 1.95E-03 | 8.28E-03 | -1.36E+00 | 9.80E-02 | 3.74E-01 | 1.16E+00 | 6.35E-03 | 2.26E-02 | -1.13E+00 | 1.16E-01 | 3.28E-01 | 1.25E+00 | 4.68E-02 | 1.72E-01 | -1.16E+00 | 3.20E-01 | 8.78E-01 | 1.24E+00 |
| AC005083.1 | 9.12E-02 | 4.75E-01 | 1.10E+00 | 3.51E-02 | 1.52E-01 | -1.04E+00 | 1.79E-01 | 7.61E-01 | 1.02E+00 | 9.89E-02 | 3.78E-01 | -1.11E+00 | 2.61E-01 | 9.29E-01 | 1.23E+00 | 2.73E-01 | 7.70E-01 | 1.45E+00 | 2.54E-01 | 9.31E-01 | 1.15E+00 | 1.38E-02 | 3.79E-02 | -1.54E+00 |
| NAPA | 9.12E-02 | 4.75E-01 | 1.07E+00 | 3.96E-02 | 1.71E-01 | -1.06E+00 | 8.59E-02 | 3.65E-01 | 1.00E+00 | 5.58E-03 | 2.13E-02 | -1.30E+00 | 1.06E-01 | 3.79E-01 | -1.08E+00 | 1.43E-01 | 4.04E-01 | -1.01E+00 | 8.90E-02 | 3.27E-01 | 1.04E+00 | 5.16E-02 | 1.42E-01 | -1.09E+00 |
| NCOA7 | 9.12E-02 | 4.75E-01 | 1.32E+00 | 3.85E-02 | 1.66E-01 | 1.27E+00 | 9.98E-02 | 4.24E-01 | 1.26E+00 | 4.13E-02 | 1.58E-01 | 1.16E+00 | 3.27E-02 | 1.17E-01 | 1.14E+00 | 7.29E-05 | 2.06E-04 | -1.22E+00 | 3.78E-04 | 1.39E-03 | -1.03E+00 | 4.64E-07 | 1.27E-06 | -1.25E+00 |
| NETO2 | 9.14E-02 | 4.76E-01 | -1.25E+00 | 1.86E-01 | 8.05E-01 | -1.03E+00 | 3.97E-13 | 1.69E-12 | -2.08E+00 | 1.84E-10 | 7.03E-10 | -1.66E+00 | 2.32E-01 | 8.29E-01 | 1.26E+00 | 3.34E-01 | 9.42E-01 | 1.10E+00 | 1.69E-01 | 6.21E-01 | -1.04E+00 | 3.08E-01 | 8.46E-01 | 1.05E+00 |
| RNVU1-6 | 9.15E-02 | 4.77E-01 | 1.51E+00 | 2.63E-02 | 1.14E-01 | -1.22E+00 | 2.19E-01 | 9.31E-01 | 1.16E+00 | 2.10E-02 | 8.02E-02 | -1.23E+00 | 9.58E-05 | 3.42E-04 | -1.92E+00 | 1.98E-05 | 5.59E-05 | -2.84E+00 | 1.12E-05 | 4.11E-05 | -2.11E+00 | 7.30E-06 | 2.00E-05 | -2.34E+00 |
| AL162151.2 | 9.18E-02 | 4.78E-01 | 1.20E+00 | 1.20E-02 | 5.19E-02 | 1.03E+00 | 5.80E-03 | 2.46E-02 | -1.11E+00 | 6.07E-05 | 2.32E-04 | -1.36E+00 | 1.29E-03 | 4.61E-03 | -1.34E+00 | 3.78E-05 | 1.07E-04 | -1.68E+00 | 1.31E-01 | 4.82E-01 | 1.12E+00 | 3.04E-03 | 8.36E-03 | -1.58E+00 |
| RPSAP6 | 9.19E-02 | 4.79E-01 | 1.06E+00 | 5.10E-02 | 2.20E-01 | 1.10E+00 | 5.39E-02 | 2.29E-01 | -1.07E+00 | 8.89E-04 | 3.39E-03 | -1.25E+00 | 1.72E-01 | 6.15E-01 | 1.04E+00 | 2.93E-01 | 8.27E-01 | 1.25E+00 | 1.94E-02 | 7.13E-02 | -1.20E+00 | 3.21E-03 | 8.81E-03 | -1.47E+00 |
| ASGR2 | 9.20E-02 | 4.79E-01 | -1.32E+00 | 1.15E-01 | 4.97E-01 | -1.32E+00 | 5.40E-02 | 2.29E-01 | -1.13E+00 | 2.06E-02 | 7.86E-02 | -1.37E+00 | 2.80E-01 | 1.00E+00 | 1.00E+00 | 3.55E-01 | 1.00E+00 | -1.00E+00 | 2.07E-01 | 7.61E-01 | -1.05E+00 | 3.30E-01 | 9.07E-01 | -1.09E+00 |
| MPEG1 | 9.20E-02 | 4.79E-01 | -2.60E+00 | 1.69E-01 | 7.32E-01 | -1.92E+00 | 2.11E-01 | 8.97E-01 | -1.01E+00 | 1.84E-01 | 7.03E-01 | -1.20E+00 | 2.80E-01 | 1.00E+00 | 1.00E+00 | 3.55E-01 | 1.00E+00 | 1.00E+00 | 1.26E-01 | 4.61E-01 | -1.70E+00 | 1.53E-01 | 4.19E-01 | -1.67E+00 |
| FRAT1 | 9.20E-02 | 4.79E-01 | 1.34E+00 | 1.43E-03 | 6.17E-03 | -1.16E+00 | 4.81E-03 | 2.04E-02 | 1.68E+00 | 1.36E-01 | 5.21E-01 | 1.19E+00 | 1.15E-01 | 4.10E-01 | 1.24E+00 | 1.38E-01 | 3.88E-01 | 1.19E+00 | 1.90E-02 | 6.97E-02 | 1.10E+00 | 2.60E-01 | 7.14E-01 | 1.30E+00 |
| TNNC2 | 9.20E-02 | 4.79E-01 | 1.37E+00 | 1.23E-01 | 5.32E-01 | 1.12E+00 | 2.25E-01 | 9.56E-01 | 1.15E+00 | 1.47E-02 | 5.63E-02 | -1.51E+00 | 2.04E-01 | 7.27E-01 | 1.52E+00 | 1.70E-01 | 4.78E-01 | -1.13E+00 | 3.34E-02 | 1.22E-01 | 2.26E+00 | 2.09E-01 | 5.74E-01 | 1.57E+00 |
| TTN-AS1 | 9.22E-02 | 4.80E-01 | 1.20E+00 | 8.81E-02 | 3.80E-01 | 1.25E+00 | 1.26E-01 | 5.35E-01 | 1.39E+00 | 4.12E-02 | 1.57E-01 | 1.82E+00 | 4.21E-02 | 1.50E-01 | 1.06E+00 | 7.63E-02 | 2.15E-01 | 1.13E+00 | 1.80E-01 | 6.61E-01 | 1.16E+00 | 8.32E-02 | 2.29E-01 | 1.00E+00 |
| HOXB4 | 9.22E-02 | 4.80E-01 | 1.44E+00 | 9.56E-03 | 4.13E-02 | 1.89E+00 | 1.35E-03 | 5.74E-03 | -1.18E+00 | 5.27E-02 | 2.01E-01 | 1.24E+00 | 2.91E-02 | 1.04E-01 | -1.01E+00 | 5.06E-02 | 1.43E-01 | 1.06E+00 | 5.76E-08 | 2.11E-07 | -1.64E+00 | 7.39E-04 | 2.03E-03 | -1.16E+00 |
| AC132872.4 | 9.22E-02 | 4.80E-01 | 1.49E+00 | 2.42E-02 | 1.05E-01 | -1.48E+00 | 1.98E-01 | 8.41E-01 | 1.09E+00 | 3.43E-02 | 1.31E-01 | -1.37E+00 | 2.71E-01 | 9.68E-01 | 1.28E+00 | 4.70E-02 | 1.32E-01 | -1.49E+00 | 8.62E-02 | 3.16E-01 | 1.74E+00 | 5.18E-02 | 1.42E-01 | -1.61E+00 |
| AURKA | 9.22E-02 | 4.81E-01 | -1.33E+00 | 1.48E-01 | 6.39E-01 | -1.11E+00 | 2.00E-02 | 8.49E-02 | -1.64E+00 | 3.38E-02 | 1.29E-01 | -1.25E+00 | 3.59E-02 | 1.28E-01 | -1.52E+00 | 8.45E-02 | 2.38E-01 | -1.35E+00 | 1.12E-08 | 4.13E-08 | -1.73E+00 | 7.15E-05 | 1.97E-04 | -1.47E+00 |
| AC007431.3 | 9.23E-02 | 4.81E-01 | 1.01E+00 | 1.49E-01 | 6.44E-01 | 1.05E+00 | 5.36E-04 | 2.28E-03 | -1.45E+00 | 2.03E-02 | 7.74E-02 | -1.03E+00 | 1.83E-02 | 6.51E-02 | -1.69E+00 | 7.74E-02 | 2.18E-01 | -1.23E+00 | 1.17E-01 | 4.29E-01 | -1.11E+00 | 1.46E-01 | 4.00E-01 | -1.14E+00 |
| Z73417.1 | 9.24E-02 | 4.81E-01 | 1.56E+00 | 2.24E-01 | 9.67E-01 | 1.48E+00 | 2.31E-01 | 9.81E-01 | 1.21E+00 | 2.19E-01 | 8.37E-01 | 1.33E+00 | 2.52E-01 | 8.98E-01 | 1.37E+00 | 1.71E-01 | 4.83E-01 | 1.84E+00 | 2.33E-01 | 8.56E-01 | 1.13E+00 | 2.37E-01 | 6.51E-01 | 1.45E+00 |
| ZDHHC1 | 9.24E-02 | 4.81E-01 | 1.75E+00 | 2.16E-01 | 9.31E-01 | 1.39E+00 | 5.12E-02 | 2.18E-01 | 1.79E+00 | 5.69E-02 | 2.17E-01 | -1.18E+00 | 1.78E-01 | 6.36E-01 | -1.17E+00 | 2.66E-01 | 7.51E-01 | 1.01E+00 | 8.69E-02 | 3.19E-01 | 1.61E+00 | 3.00E-01 | 8.23E-01 | -1.03E+00 |
| PAN3-AS1 | 9.26E-02 | 4.83E-01 | 1.42E+00 | 5.75E-02 | 2.48E-01 | 1.73E+00 | 1.84E-01 | 7.81E-01 | 1.13E+00 | 9.00E-02 | 3.44E-01 | 1.15E+00 | 1.51E-01 | 5.40E-01 | 1.92E+00 | 3.12E-01 | 8.80E-01 | 1.36E+00 | 3.81E-02 | 1.40E-01 | 2.34E+00 | 1.63E-01 | 4.48E-01 | 1.75E+00 |
| AL118556.1 | 9.26E-02 | 4.83E-01 | 1.63E+00 | 1.23E-01 | 5.30E-01 | 1.72E+00 | 2.18E-01 | 9.26E-01 | 1.18E+00 | 3.48E-02 | 1.33E-01 | -1.13E+00 | 1.70E-01 | 6.06E-01 | 1.21E+00 | 1.72E-01 | 4.85E-01 | 1.19E+00 | 8.49E-02 | 3.12E-01 | 1.04E+00 | 1.89E-02 | 5.20E-02 | -1.23E+00 |
| ZFPM1 | 9.27E-02 | 4.83E-01 | 1.69E+00 | 9.82E-02 | 4.24E-01 | 1.93E+00 | 1.92E-01 | 8.14E-01 | 1.28E+00 | 1.39E-01 | 5.30E-01 | 1.76E+00 | 2.64E-01 | 9.40E-01 | 1.37E+00 | 7.33E-02 | 2.07E-01 | 2.10E+00 | 5.01E-02 | 1.84E-01 | 1.42E+00 | 2.53E-01 | 6.96E-01 | 1.33E+00 |
| RPS28 | 9.28E-02 | 4.84E-01 | 1.09E+00 | 5.44E-33 | 2.35E-32 | -1.49E+00 | 2.10E-01 | 8.92E-01 | 1.11E+00 | 6.01E-09 | 2.29E-08 | -1.38E+00 | 9.75E-02 | 3.48E-01 | 1.06E+00 | 1.43E-10 | 4.04E-10 | -1.46E+00 | 1.16E-02 | 4.27E-02 | -1.07E+00 | 4.73E-11 | 1.30E-10 | -1.63E+00 |
| COMMD3 | 9.28E-02 | 4.84E-01 | 1.30E+00 | 1.35E-01 | 5.82E-01 | 1.56E+00 | 8.38E-02 | 3.56E-01 | 1.09E+00 | 1.73E-01 | 6.61E-01 | 1.40E+00 | 2.75E-07 | 9.80E-07 | -1.10E+00 | 9.07E-03 | 2.56E-02 | 1.17E+00 | 6.11E-04 | 2.24E-03 | -1.11E+00 | 1.63E-01 | 4.48E-01 | 1.10E+00 |
| C10orf76 | 9.28E-02 | 4.84E-01 | 1.24E+00 | 4.93E-02 | 2.13E-01 | 1.22E+00 | 3.39E-03 | 1.44E-02 | -1.12E+00 | 1.04E-03 | 3.98E-03 | -1.08E+00 | 4.99E-02 | 1.78E-01 | 1.05E+00 | 2.41E-01 | 6.79E-01 | 1.33E+00 | 1.80E-02 | 6.60E-02 | 1.64E+00 | 5.36E-02 | 1.47E-01 | 1.57E+00 |
| AC036176.2 | 9.30E-02 | 4.85E-01 | -1.03E+00 | 2.21E-01 | 9.53E-01 | 1.38E+00 | 1.10E-01 | 4.67E-01 | 1.51E+00 | 8.91E-02 | 3.40E-01 | 1.64E+00 | 8.26E-02 | 2.94E-01 | -1.15E+00 | 1.46E-01 | 4.11E-01 | -1.04E+00 | 1.91E-01 | 7.02E-01 | 1.05E+00 | 1.67E-01 | 4.57E-01 | -1.13E+00 |
| LINC01215 | 9.32E-02 | 4.86E-01 | 1.47E+00 | 6.82E-02 | 2.95E-01 | -1.15E+00 | 3.66E-02 | 1.56E-01 | 1.62E+00 | 4.94E-02 | 1.89E-01 | -1.10E+00 | 2.20E-01 | 7.86E-01 | 1.09E+00 | 2.60E-01 | 7.32E-01 | -1.18E+00 | 1.02E-01 | 3.75E-01 | -1.17E+00 | 1.25E-02 | 3.43E-02 | -1.80E+00 |
| PREX1 | 9.32E-02 | 4.86E-01 | 1.40E+00 | 1.50E-01 | 6.49E-01 | 1.38E+00 | 1.73E-01 | 7.34E-01 | 1.39E+00 | 7.38E-02 | 2.82E-01 | 1.32E+00 | 9.05E-02 | 3.23E-01 | 1.26E+00 | 1.55E-01 | 4.37E-01 | 1.35E+00 | 3.34E-02 | 1.23E-01 | 1.11E+00 | 1.96E-02 | 5.38E-02 | 1.08E+00 |
| CDCA3 | 9.34E-02 | 4.87E-01 | -1.21E+00 | 1.44E-01 | 6.24E-01 | -1.02E+00 | 3.52E-02 | 1.50E-01 | 1.56E+00 | 4.36E-02 | 1.67E-01 | 2.09E+00 | 2.51E-01 | 8.95E-01 | 1.12E+00 | 2.23E-01 | 6.29E-01 | -1.24E+00 | 2.71E-01 | 9.93E-01 | 1.01E+00 | 9.19E-02 | 2.53E-01 | 1.22E+00 |
| GLOD5 | 9.34E-02 | 4.87E-01 | -1.67E+00 | 1.56E-01 | 6.73E-01 | -1.25E+00 | 1.35E-01 | 5.76E-01 | -1.34E+00 | 2.24E-01 | 8.55E-01 | 1.01E+00 | 2.12E-67 | 7.55E-67 | -3.27E+00 | 1.78E-36 | 5.03E-36 | -1.66E+00 | 1.01E-17 | 3.72E-17 | -2.52E+00 | 4.35E-02 | 1.20E-01 | 1.02E+00 |
| MVK | 9.36E-02 | 4.88E-01 | 1.15E+00 | 1.37E-01 | 5.92E-01 | 1.84E+00 | 2.24E-01 | 9.51E-01 | 1.31E+00 | 1.40E-01 | 5.35E-01 | 1.61E+00 | 1.90E-02 | 6.78E-02 | -1.25E+00 | 1.40E-01 | 3.96E-01 | 1.05E+00 | 1.26E-01 | 4.63E-01 | 1.08E+00 | 6.88E-02 | 1.89E-01 | 1.59E+00 |
| AL096701.1 | 9.36E-02 | 4.88E-01 | 1.20E+00 | 2.25E-02 | 9.73E-02 | -1.00E+00 | 1.15E-01 | 4.90E-01 | 1.35E+00 | 8.53E-02 | 3.26E-01 | 1.05E+00 | 1.96E-01 | 7.00E-01 | 1.16E+00 | 6.04E-02 | 1.70E-01 | -1.23E+00 | 2.39E-02 | 8.79E-02 | -1.15E+00 | 2.33E-03 | 6.40E-03 | -1.45E+00 |
| PRKAR1B | 9.37E-02 | 4.88E-01 | 1.05E+00 | 1.44E-01 | 6.23E-01 | 1.25E+00 | 9.90E-02 | 4.21E-01 | -1.01E+00 | 4.90E-02 | 1.87E-01 | -1.01E+00 | 5.00E-02 | 1.78E-01 | -1.17E+00 | 1.94E-01 | 5.47E-01 | 1.20E+00 | 2.13E-03 | 7.84E-03 | -1.08E+00 | 8.03E-03 | 2.21E-02 | 1.01E+00 |
| C2orf40 | 9.38E-02 | 4.89E-01 | 1.19E+00 | 1.16E-01 | 5.01E-01 | 1.35E+00 | 6.35E-02 | 2.70E-01 | 1.57E+00 | 2.43E-01 | 9.28E-01 | 1.22E+00 | 2.53E-01 | 9.01E-01 | 1.11E+00 | 3.55E-01 | 1.00E+00 | -1.00E+00 | 2.23E-01 | 8.18E-01 | 1.11E+00 | 2.97E-01 | 8.15E-01 | 1.13E+00 |
| AL137003.2 | 9.39E-02 | 4.90E-01 | 1.47E+00 | 2.24E-01 | 9.70E-01 | 1.34E+00 | 1.39E-01 | 5.90E-01 | -1.13E+00 | 1.39E-01 | 5.32E-01 | 1.53E+00 | 1.19E-01 | 4.26E-01 | 1.93E+00 | 3.12E-03 | 8.79E-03 | 3.66E+00 | 1.34E-01 | 4.92E-01 | 1.46E+00 | 8.66E-02 | 2.38E-01 | 1.50E+00 |
| COHCR1 | 9.39E-02 | 4.90E-01 | 1.21E+00 | 5.95E-02 | 2.57E-01 | 1.26E+00 | 2.75E-03 | 1.17E-02 | -1.07E+00 | 1.27E-03 | 4.83E-03 | -1.00E+00 | 8.50E-02 | 3.03E-01 | -1.00E+00 | 6.15E-02 | 1.73E-01 | 1.06E+00 | 1.84E-01 | 6.74E-01 | 1.19E+00 | 6.20E-02 | 1.70E-01 | 1.02E+00 |
| BICRA | 9.40E-02 | 4.90E-01 | 1.22E+00 | 8.35E-02 | 3.61E-01 | 1.23E+00 | 2.79E-02 | 1.19E-01 | -1.25E+00 | 2.54E-02 | 9.71E-02 | -1.13E+00 | 4.25E-02 | 1.51E-01 | -1.18E+00 | 8.42E-02 | 2.37E-01 | -1.11E+00 | 4.58E-02 | 1.68E-01 | -1.20E+00 | 1.14E-01 | 3.13E-01 | -1.00E+00 |
| SNORD3C | 9.40E-02 | 4.90E-01 | 1.25E+00 | 1.19E-04 | 5.13E-04 | -1.34E+00 | 1.20E-01 | 5.10E-01 | 1.12E+00 | 5.13E-05 | 1.96E-04 | -2.01E+00 | 1.60E-02 | 5.69E-02 | -1.05E+00 | 4.62E-03 | 1.30E-02 | -1.45E+00 | 2.30E-02 | 8.43E-02 | -1.32E+00 | 2.32E-02 | 6.37E-02 | -1.31E+00 |
| NPIPB5 | 9.40E-02 | 4.90E-01 | 1.53E+00 | 4.14E-02 | 1.79E-01 | 1.17E+00 | 1.67E-01 | 7.10E-01 | 1.12E+00 | 1.97E-02 | 7.53E-02 | 1.11E+00 | 2.54E-01 | 9.06E-01 | 1.25E+00 | 3.53E-01 | 9.95E-01 | 1.33E+00 | 9.28E-03 | 3.41E-02 | 1.07E+00 | 1.03E-01 | 2.82E-01 | 1.09E+00 |
| EXOSC6 | 9.43E-02 | 4.92E-01 | 1.17E+00 | 9.71E-03 | 4.20E-02 | 1.23E+00 | 2.20E-01 | 9.34E-01 | 1.21E+00 | 8.89E-04 | 3.40E-03 | 1.05E+00 | 7.27E-02 | 2.59E-01 | 1.07E+00 | 1.30E-01 | 3.66E-01 | 1.15E+00 | 2.53E-01 | 9.28E-01 | 1.15E+00 | 1.42E-01 | 3.90E-01 | 1.11E+00 |
| ZNF777 | 9.44E-02 | 4.92E-01 | 1.19E+00 | 1.65E-01 | 7.14E-01 | 1.40E+00 | 2.29E-01 | 9.72E-01 | 1.23E+00 | 9.00E-02 | 3.44E-01 | 1.18E+00 | 2.36E-01 | 8.43E-01 | 1.30E+00 | 2.27E-01 | 6.41E-01 | 1.80E+00 | 2.09E-02 | 7.66E-02 | -1.02E+00 | 1.03E-01 | 2.83E-01 | 1.01E+00 |
| IVNS1ABP | 9.44E-02 | 4.92E-01 | 1.49E+00 | 5.21E-05 | 2.25E-04 | -1.14E+00 | 8.89E-03 | 3.78E-02 | 1.55E+00 | 9.33E-03 | 3.56E-02 | -1.00E+00 | 9.07E-03 | 3.23E-02 | 2.06E+00 | 1.76E-01 | 4.97E-01 | 1.33E+00 | 5.10E-02 | 1.87E-01 | 1.67E+00 | 3.34E-01 | 9.18E-01 | 1.40E+00 |
| AF213884.3 | 9.44E-02 | 4.92E-01 | 1.48E+00 | 3.60E-02 | 1.56E-01 | -1.08E+00 | 1.10E-02 | 4.68E-02 | 1.81E+00 | 1.62E-01 | 6.20E-01 | 1.00E+00 | 2.15E-01 | 7.67E-01 | 1.53E+00 | 3.40E-01 | 9.57E-01 | 1.32E+00 | 2.54E-01 | 9.32E-01 | 1.08E+00 | 1.46E-01 | 4.00E-01 | -1.27E+00 |
| AL591623.1 | 9.45E-02 | 4.92E-01 | 1.06E+00 | 5.80E-03 | 2.51E-02 | 2.67E+00 | 1.52E-01 | 6.45E-01 | 1.04E+00 | 9.25E-02 | 3.54E-01 | 1.68E+00 | 9.93E-03 | 3.54E-02 | 2.53E+00 | 7.81E-04 | 2.20E-03 | 3.35E+00 | 1.83E-02 | 6.71E-02 | 1.99E+00 | 1.76E-08 | 4.83E-08 | 3.76E+00 |
| LINC00562 | 9.45E-02 | 4.92E-01 | 1.00E+00 | 1.77E-01 | 7.63E-01 | 1.54E+00 | 2.26E-02 | 9.62E-02 | -1.13E+00 | 5.71E-03 | 2.18E-02 | -1.24E+00 | 9.36E-03 | 3.34E-02 | -1.59E+00 | 4.45E-02 | 1.25E-01 | -1.22E+00 | 7.77E-03 | 2.85E-02 | -1.58E+00 | 6.05E-02 | 1.66E-01 | -1.28E+00 |
| EMP2 | 9.45E-02 | 4.92E-01 | 1.38E+00 | 1.15E-01 | 4.95E-01 | 1.60E+00 | 1.49E-01 | 6.32E-01 | 1.03E+00 | 6.05E-02 | 2.31E-01 | -1.21E+00 | 2.16E-01 | 7.71E-01 | -1.10E+00 | 2.70E-01 | 7.60E-01 | -1.09E+00 | 1.01E-02 | 3.72E-02 | -1.64E+00 | 2.88E-01 | 7.91E-01 | -1.13E+00 |
| ARHGAP44 | 9.45E-02 | 4.92E-01 | -1.01E+00 | 1.03E-01 | 4.44E-01 | 1.66E+00 | 1.43E-01 | 6.09E-01 | -1.12E+00 | 9.31E-03 | 3.56E-02 | 2.22E+00 | 2.68E-01 | 9.54E-01 | 1.38E+00 | 2.27E-01 | 6.41E-01 | 1.71E+00 | 4.01E-02 | 1.47E-01 | -1.42E+00 | 2.55E-01 | 6.99E-01 | 1.06E+00 |
| COMTD1 | 9.46E-02 | 4.93E-01 | 1.22E+00 | 1.30E-01 | 5.62E-01 | 1.37E+00 | 9.75E-02 | 4.14E-01 | 1.36E+00 | 6.17E-02 | 2.36E-01 | 1.19E+00 | 2.77E-01 | 9.89E-01 | 1.53E+00 | 2.97E-01 | 8.37E-01 | 1.62E+00 | 8.54E-02 | 3.13E-01 | 1.12E+00 | 7.59E-02 | 2.08E-01 | 1.06E+00 |
| MYLPF | 9.46E-02 | 4.93E-01 | 1.54E+00 | 2.69E-02 | 1.16E-01 | -1.32E+00 | 9.17E-02 | 3.90E-01 | 1.30E+00 | 6.82E-03 | 2.61E-02 | -1.26E+00 | 2.44E-02 | 8.70E-02 | -1.05E+00 | 4.35E-05 | 1.23E-04 | -2.26E+00 | 5.97E-02 | 2.19E-01 | 1.49E+00 | 3.06E-01 | 8.40E-01 | -1.03E+00 |
| IGFBP7 | 9.47E-02 | 4.93E-01 | 1.22E+00 | 2.03E-01 | 8.77E-01 | 1.30E+00 | 1.84E-01 | 7.80E-01 | 1.08E+00 | 1.87E-01 | 7.13E-01 | 1.10E+00 | 1.02E-03 | 3.64E-03 | 1.76E+00 | 2.10E-03 | 5.93E-03 | 1.64E+00 | 7.40E-02 | 2.72E-01 | 1.38E+00 | 4.32E-02 | 1.19E-01 | 1.37E+00 |
| TCFL5 | 9.48E-02 | 4.94E-01 | -1.01E+00 | 1.55E-01 | 6.69E-01 | 1.15E+00 | 1.28E-01 | 5.42E-01 | 1.09E+00 | 9.23E-02 | 3.53E-01 | 1.16E+00 | 2.47E-01 | 8.79E-01 | 1.18E+00 | 3.10E-01 | 8.75E-01 | 1.59E+00 | 4.02E-03 | 1.47E-02 | -1.21E+00 | 1.30E-02 | 3.58E-02 | -1.10E+00 |
| EXT1 | 9.49E-02 | 4.94E-01 | -1.09E+00 | 1.12E-01 | 4.84E-01 | -1.05E+00 | 1.91E-01 | 8.13E-01 | 1.21E+00 | 2.55E-01 | 9.74E-01 | 1.18E+00 | 6.39E-02 | 2.28E-01 | -1.06E+00 | 9.83E-03 | 2.77E-02 | -1.41E+00 | 9.19E-03 | 3.37E-02 | -1.23E+00 | 2.00E-01 | 5.48E-01 | 1.12E+00 |
| CD40LG | 9.49E-02 | 4.94E-01 | -1.37E+00 | 1.30E-01 | 5.61E-01 | -1.28E+00 | 1.33E-01 | 5.66E-01 | -1.93E+00 | 1.57E-01 | 5.99E-01 | -1.82E+00 | 1.74E-01 | 6.22E-01 | 1.84E+00 | 1.25E-03 | 3.53E-03 | 4.05E+00 | 4.81E-17 | 1.76E-16 | -1.66E+00 | 1.59E-11 | 4.37E-11 | -1.26E+00 |
| PSD3 | 9.49E-02 | 4.94E-01 | 1.07E+00 | 1.84E-01 | 7.97E-01 | 1.37E+00 | 2.44E-02 | 1.04E-01 | -1.03E+00 | 9.79E-03 | 3.74E-02 | 1.00E+00 | 1.40E-01 | 4.98E-01 | -1.07E+00 | 1.45E-01 | 4.08E-01 | -1.05E+00 | 6.61E-03 | 2.43E-02 | -1.30E+00 | 6.29E-02 | 1.73E-01 | -1.04E+00 |
| SNTA1 | 9.50E-02 | 4.95E-01 | 1.04E+00 | 1.64E-01 | 7.10E-01 | 1.28E+00 | 3.64E-02 | 1.55E-01 | 1.56E+00 | 1.98E-01 | 7.56E-01 | 1.38E+00 | 7.64E-02 | 2.72E-01 | 2.30E+00 | 1.56E-02 | 4.40E-02 | 2.99E+00 | 7.22E-06 | 2.65E-05 | 4.18E+00 | 2.30E-03 | 6.33E-03 | 3.60E+00 |
| RN7SL382P | 9.50E-02 | 4.95E-01 | 1.17E+00 | 6.39E-03 | 2.76E-02 | -1.17E+00 | 2.40E-02 | 1.02E-01 | -1.11E+00 | 8.81E-07 | 3.37E-06 | -2.12E+00 | 1.34E-01 | 4.78E-01 | 1.90E+00 | 2.72E-01 | 7.67E-01 | 1.62E+00 | 1.93E-02 | 7.10E-02 | -1.25E+00 | 1.08E-02 | 2.97E-02 | -1.52E+00 |
| AMN1 | 9.51E-02 | 4.96E-01 | 1.02E+00 | 9.18E-02 | 3.97E-01 | 1.03E+00 | 4.55E-02 | 1.93E-01 | -1.16E+00 | 1.39E-02 | 5.30E-02 | -1.23E+00 | 1.83E-01 | 6.51E-01 | 1.27E+00 | 5.34E-02 | 1.50E-01 | -1.27E+00 | 3.34E-02 | 1.23E-01 | -1.28E+00 | 3.34E-01 | 9.17E-01 | 1.26E+00 |
| AC009145.4 | 9.53E-02 | 4.96E-01 | -1.00E+00 | 1.20E-01 | 5.16E-01 | 1.15E+00 | 3.79E-02 | 1.61E-01 | -1.26E+00 | 1.01E-01 | 3.87E-01 | -1.00E+00 | 2.22E-01 | 7.91E-01 | 1.15E+00 | 2.30E-01 | 6.47E-01 | 1.23E+00 | 8.43E-02 | 3.09E-01 | 1.32E+00 | 3.12E-01 | 8.56E-01 | -1.12E+00 |
| MTFR2 | 9.53E-02 | 4.97E-01 | -1.18E+00 | 2.03E-01 | 8.77E-01 | 1.25E+00 | 8.83E-02 | 3.75E-01 | 1.31E+00 | 1.46E-01 | 5.56E-01 | 1.39E+00 | 1.97E-01 | 7.04E-01 | -1.20E+00 | 3.49E-01 | 9.84E-01 | 1.21E+00 | 1.18E-01 | 4.31E-01 | 1.06E+00 | 1.67E-01 | 4.60E-01 | 1.29E+00 |
| AC123912.4 | 9.55E-02 | 4.98E-01 | 1.05E+00 | 7.04E-02 | 3.04E-01 | 1.91E+00 | 7.47E-02 | 3.17E-01 | 1.03E+00 | 3.98E-02 | 1.52E-01 | 2.12E+00 | 1.90E-01 | 6.78E-01 | 1.69E+00 | 1.22E-04 | 3.44E-04 | 2.81E+00 | 5.25E-02 | 1.93E-01 | 1.82E+00 | 2.45E-02 | 6.73E-02 | 2.01E+00 |
| RBM4 | 9.55E-02 | 4.98E-01 | 1.38E+00 | 8.54E-07 | 3.69E-06 | 1.06E+00 | 1.56E-01 | 6.61E-01 | 1.28E+00 | 1.21E-04 | 4.61E-04 | -1.02E+00 | 5.22E-02 | 1.86E-01 | 1.38E+00 | 4.96E-04 | 1.40E-03 | 1.06E+00 | 3.28E-02 | 1.21E-01 | 1.26E+00 | 2.61E-03 | 7.16E-03 | 1.00E+00 |
| AC026124.2 | 9.55E-02 | 4.98E-01 | 1.27E+00 | 2.03E-04 | 8.79E-04 | 1.05E+00 | 2.02E-01 | 8.58E-01 | 1.25E+00 | 1.92E-02 | 7.34E-02 | 1.08E+00 | 8.54E-02 | 3.05E-01 | 1.33E+00 | 3.29E-03 | 9.26E-03 | -1.00E+00 | 4.67E-02 | 1.71E-01 | 1.08E+00 | 8.41E-03 | 2.31E-02 | -1.10E+00 |
| HMGXB3 | 9.56E-02 | 4.98E-01 | 1.23E+00 | 6.50E-02 | 2.81E-01 | 1.23E+00 | 5.34E-04 | 2.27E-03 | -1.34E+00 | 8.02E-05 | 3.06E-04 | -1.28E+00 | 3.12E-03 | 1.11E-02 | -1.37E+00 | 4.76E-03 | 1.34E-02 | -1.47E+00 | 2.83E-02 | 1.04E-01 | -1.07E+00 | 7.21E-02 | 1.98E-01 | 1.03E+00 |
| GTF3A | 9.56E-02 | 4.98E-01 | 1.13E+00 | 3.83E-02 | 1.66E-01 | 1.20E+00 | 8.31E-02 | 3.53E-01 | 1.20E+00 | 4.71E-02 | 1.80E-01 | 1.25E+00 | 1.50E-02 | 5.36E-02 | 1.06E+00 | 3.15E-01 | 8.89E-01 | 1.20E+00 | 2.50E-01 | 9.19E-01 | 1.13E+00 | 1.42E-02 | 3.91E-02 | 1.33E+00 |
| FANCE | 9.57E-02 | 4.99E-01 | 1.01E+00 | 1.91E-01 | 8.24E-01 | 1.55E+00 | 2.44E-05 | 1.04E-04 | -1.35E+00 | 1.81E-05 | 6.90E-05 | -1.22E+00 | 1.26E-02 | 4.50E-02 | -1.44E+00 | 9.05E-02 | 2.55E-01 | 1.02E+00 | 1.37E-11 | 5.04E-11 | -1.52E+00 | 3.27E-07 | 8.98E-07 | -1.32E+00 |
| STON1 | 9.58E-02 | 4.99E-01 | 1.18E+00 | 2.13E-01 | 9.21E-01 | 1.50E+00 | 2.20E-01 | 9.36E-01 | 1.16E+00 | 1.38E-01 | 5.27E-01 | 1.07E+00 | 2.38E-01 | 8.48E-01 | 1.39E+00 | 3.55E-01 | 1.00E+00 | 1.36E+00 | 2.25E-01 | 8.26E-01 | 1.20E+00 | 2.56E-01 | 7.02E-01 | 1.29E+00 |
| TSC22D2 | 9.60E-02 | 5.00E-01 | 1.33E+00 | 2.61E-11 | 1.13E-10 | -1.60E+00 | 7.63E-02 | 3.24E-01 | 1.57E+00 | 3.00E-03 | 1.15E-02 | 1.04E+00 | 1.24E-01 | 4.41E-01 | 1.57E+00 | 2.94E-04 | 8.28E-04 | -1.35E+00 | 2.70E-05 | 9.92E-05 | -1.09E+00 | 1.08E-14 | 2.96E-14 | -1.77E+00 |
| STAM-AS1 | 9.60E-02 | 5.00E-01 | 1.50E+00 | 9.32E-02 | 4.03E-01 | 1.14E+00 | 3.14E-02 | 1.33E-01 | 1.72E+00 | 9.35E-02 | 3.57E-01 | 1.85E+00 | 3.37E-02 | 1.20E-01 | 2.66E+00 | 1.94E-01 | 5.48E-01 | 1.96E+00 | 1.70E-01 | 6.22E-01 | 1.28E+00 | 3.59E-01 | 9.85E-01 | 1.27E+00 |
| FTH1P8 | 9.60E-02 | 5.00E-01 | 1.26E+00 | 3.07E-02 | 1.33E-01 | 1.25E+00 | 7.78E-02 | 3.30E-01 | 1.25E+00 | 2.33E-01 | 8.90E-01 | 1.32E+00 | 1.55E-01 | 5.51E-01 | 1.19E+00 | 1.79E-01 | 5.06E-01 | 1.17E+00 | 2.09E-01 | 7.66E-01 | 1.15E+00 | 1.73E-01 | 4.75E-01 | 1.19E+00 |
| LPXN | 9.60E-02 | 5.00E-01 | 1.09E+00 | 1.32E-03 | 5.69E-03 | -1.59E+00 | 2.22E-01 | 9.43E-01 | 1.23E+00 | 7.08E-04 | 2.71E-03 | -1.59E+00 | 4.42E-03 | 1.58E-02 | -1.36E+00 | 1.52E-02 | 4.29E-02 | -1.13E+00 | 2.17E-04 | 7.96E-04 | -1.18E+00 | 1.01E-05 | 2.77E-05 | -1.63E+00 |
| TREM1 | 9.60E-02 | 5.01E-01 | 1.32E+00 | 2.31E-01 | 1.00E+00 | 1.00E+00 | 1.60E-01 | 6.80E-01 | 1.33E+00 | 1.59E-01 | 6.08E-01 | -1.03E+00 | 2.80E-01 | 1.00E+00 | 1.00E+00 | 3.55E-01 | 1.00E+00 | 1.00E+00 | 2.33E-01 | 8.57E-01 | 1.10E+00 | 3.10E-01 | 8.52E-01 | -1.03E+00 |
| TEAD4 | 9.61E-02 | 5.01E-01 | -1.11E+00 | 1.47E-01 | 6.36E-01 | 1.00E+00 | 1.70E-03 | 7.23E-03 | -1.56E+00 | 2.40E-01 | 9.19E-01 | 1.33E+00 | 2.80E-01 | 1.00E+00 | 1.00E+00 | 3.27E-01 | 9.21E-01 | 1.08E+00 | 5.37E-04 | 1.97E-03 | -1.45E+00 | 7.62E-02 | 2.09E-01 | -1.03E+00 |
| AP000866.6 | 9.61E-02 | 5.01E-01 | -1.11E+00 | 1.20E-01 | 5.20E-01 | -1.08E+00 | 1.28E-02 | 5.43E-02 | -1.69E+00 | 3.49E-02 | 1.33E-01 | -1.37E+00 | 6.80E-02 | 2.43E-01 | -1.36E+00 | 1.81E-01 | 5.10E-01 | 1.01E+00 | 1.55E-01 | 5.67E-01 | 1.37E+00 | 1.06E-01 | 2.92E-01 | -1.29E+00 |
| HMGN1P36 | 9.61E-02 | 5.01E-01 | 1.09E+00 | 3.88E-03 | 1.67E-02 | -1.60E+00 | 1.55E-02 | 6.59E-02 | -1.19E+00 | 4.43E-05 | 1.69E-04 | -1.94E+00 | 4.11E-02 | 1.47E-01 | -1.15E+00 | 1.37E-02 | 3.87E-02 | -1.28E+00 | 1.50E-01 | 5.50E-01 | 1.21E+00 | 3.18E-04 | 8.74E-04 | -1.67E+00 |
| BX679664.3 | 9.62E-02 | 5.01E-01 | 1.44E+00 | 2.31E-01 | 1.00E+00 | 1.47E+00 | 5.89E-02 | 2.50E-01 | 1.40E+00 | 1.84E-01 | 7.02E-01 | 1.34E+00 | 3.70E-02 | 1.32E-01 | 1.14E+00 | 2.86E-02 | 8.07E-02 | 1.16E+00 | 2.30E-01 | 8.43E-01 | 1.22E+00 | 1.97E-01 | 5.41E-01 | 1.10E+00 |
| HAUS1 | 9.62E-02 | 5.01E-01 | 1.17E+00 | 8.93E-03 | 3.86E-02 | 1.13E+00 | 5.26E-03 | 2.24E-02 | -1.07E+00 | 1.62E-05 | 6.20E-05 | -1.12E+00 | 1.16E-04 | 4.12E-04 | -1.17E+00 | 2.19E-03 | 6.17E-03 | 1.00E+00 | 1.13E-02 | 4.13E-02 | 1.03E+00 | 1.13E-02 | 3.11E-02 | 1.02E+00 |
| DEPDC1 | 9.64E-02 | 5.02E-01 | 1.59E+00 | 2.31E-01 | 9.97E-01 | 1.13E+00 | 1.37E-02 | 5.83E-02 | -1.54E+00 | 1.15E-01 | 4.39E-01 | -1.05E+00 | 1.72E-01 | 6.14E-01 | -1.15E+00 | 2.00E-01 | 5.64E-01 | -1.23E+00 | 1.08E-06 | 3.95E-06 | -1.90E+00 | 5.56E-05 | 1.53E-04 | -1.56E+00 |
| ANAPC15 | 9.64E-02 | 5.02E-01 | 1.46E+00 | 1.83E-01 | 7.91E-01 | 1.43E+00 | 2.22E-01 | 9.43E-01 | 1.14E+00 | 2.35E-02 | 8.97E-02 | -1.41E+00 | 3.04E-02 | 1.09E-01 | -1.18E+00 | 6.26E-03 | 1.76E-02 | -1.57E+00 | 2.05E-01 | 7.53E-01 | 1.33E+00 | 2.60E-01 | 7.14E-01 | 1.20E+00 |
| AC027279.1 | 9.64E-02 | 5.03E-01 | 1.03E+00 | 1.42E-01 | 6.15E-01 | 1.08E+00 | 2.22E-01 | 9.41E-01 | 1.22E+00 | 1.58E-01 | 6.04E-01 | 1.22E+00 | 6.32E-02 | 2.25E-01 | 1.01E+00 | 1.00E-01 | 2.83E-01 | 1.01E+00 | 2.48E-01 | 9.12E-01 | 1.32E+00 | 3.57E-01 | 9.81E-01 | 1.32E+00 |
| SAP30L-AS1 | 9.66E-02 | 5.03E-01 | -1.01E+00 | 1.94E-01 | 8.40E-01 | 1.60E+00 | 1.96E-01 | 8.33E-01 | 1.10E+00 | 1.60E-01 | 6.13E-01 | 1.12E+00 | 4.46E-03 | 1.59E-02 | 2.25E+00 | 2.34E-03 | 6.60E-03 | 2.46E+00 | 8.66E-02 | 3.18E-01 | 1.10E+00 | 2.83E-01 | 7.77E-01 | 1.21E+00 |
| AC004381.2 | 9.66E-02 | 5.03E-01 | 1.44E+00 | 1.54E-01 | 6.67E-01 | -1.14E+00 | 5.95E-02 | 2.53E-01 | 1.79E+00 | 2.56E-01 | 9.79E-01 | 1.36E+00 | 2.30E-03 | 8.21E-03 | 2.84E+00 | 7.58E-03 | 2.14E-02 | 2.73E+00 | 8.01E-04 | 2.94E-03 | -1.24E+00 | 3.37E-05 | 9.25E-05 | -1.33E+00 |
| CTTNBP2 | 9.66E-02 | 5.04E-01 | 1.55E+00 | 1.68E-01 | 7.28E-01 | 1.12E+00 | 1.01E-01 | 4.27E-01 | -1.10E+00 | 1.43E-02 | 5.47E-02 | -1.57E+00 | 1.11E-05 | 3.95E-05 | -2.14E+00 | 9.97E-09 | 2.81E-08 | -3.08E+00 | 4.02E-08 | 1.48E-07 | -1.98E+00 | 5.26E-14 | 1.44E-13 | -4.02E+00 |
| MAN1A1 | 9.67E-02 | 5.04E-01 | 1.24E+00 | 1.54E-01 | 6.64E-01 | 1.47E+00 | 2.39E-05 | 1.02E-04 | -1.15E+00 | 4.21E-03 | 1.61E-02 | 1.14E+00 | 1.32E-03 | 4.71E-03 | -1.16E+00 | 9.95E-02 | 2.81E-01 | 1.15E+00 | 9.02E-04 | 3.31E-03 | -1.06E+00 | 1.39E-01 | 3.82E-01 | 1.17E+00 |
| ADPRM | 9.68E-02 | 5.04E-01 | 1.58E+00 | 5.63E-04 | 2.43E-03 | 2.24E+00 | 1.72E-01 | 7.29E-01 | 1.11E+00 | 1.69E-02 | 6.46E-02 | 1.62E+00 | 2.42E-01 | 8.62E-01 | 1.50E+00 | 8.76E-03 | 2.47E-02 | 2.05E+00 | 3.10E-04 | 1.14E-03 | 1.94E+00 | 9.72E-03 | 2.67E-02 | 1.81E+00 |
| GBF1 | 9.69E-02 | 5.05E-01 | 1.12E+00 | 1.83E-01 | 7.91E-01 | 1.59E+00 | 4.29E-05 | 1.82E-04 | -1.42E+00 | 1.67E-02 | 6.39E-02 | 1.08E+00 | 4.72E-03 | 1.68E-02 | -1.11E+00 | 1.32E-02 | 3.73E-02 | -1.04E+00 | 8.63E-05 | 3.17E-04 | -1.31E+00 | 5.77E-04 | 1.59E-03 | -1.09E+00 |
| LYPD8 | 9.70E-02 | 5.06E-01 | -1.17E+00 | 1.02E-01 | 4.40E-01 | -1.25E+00 | 7.50E-02 | 3.19E-01 | -1.25E+00 | 3.04E-02 | 1.16E-01 | -1.49E+00 | 2.03E-01 | 7.23E-01 | 1.30E+00 | 3.27E-01 | 9.22E-01 | 1.03E+00 | 4.97E-84 | 1.82E-83 | -4.77E+00 | 4.53E-77 | 1.24E-76 | -4.32E+00 |
| AC008969.1 | 9.70E-02 | 5.06E-01 | 1.20E+00 | 1.22E-01 | 5.26E-01 | 1.39E+00 | 6.74E-02 | 2.87E-01 | 1.02E+00 | 1.08E-02 | 4.11E-02 | -1.03E+00 | 8.57E-02 | 3.06E-01 | 1.72E+00 | 8.31E-04 | 2.34E-03 | 2.39E+00 | 2.25E-01 | 8.27E-01 | 1.21E+00 | 2.81E-01 | 7.72E-01 | 1.33E+00 |
| SMG8 | 9.71E-02 | 5.06E-01 | 1.07E+00 | 7.14E-02 | 3.09E-01 | 1.00E+00 | 2.35E-01 | 9.96E-01 | 1.00E+00 | 2.44E-01 | 9.31E-01 | 1.12E+00 | 1.48E-01 | 5.28E-01 | 1.07E+00 | 6.03E-02 | 1.70E-01 | -1.32E+00 | 2.66E-02 | 9.77E-02 | -1.25E+00 | 1.71E-01 | 4.69E-01 | 1.06E+00 |
| AC023886.2 | 9.72E-02 | 5.06E-01 | 1.25E+00 | 2.04E-04 | 8.83E-04 | -1.03E+00 | 9.58E-03 | 4.07E-02 | 1.62E+00 | 9.47E-02 | 3.62E-01 | 1.27E+00 | 2.69E-01 | 9.60E-01 | 1.49E+00 | 2.84E-01 | 7.99E-01 | 1.38E+00 | 1.24E-02 | 4.55E-02 | 1.76E+00 | 2.11E-01 | 5.80E-01 | 1.25E+00 |
| WDR53 | 9.72E-02 | 5.06E-01 | 1.08E+00 | 6.73E-02 | 2.91E-01 | 1.12E+00 | 1.91E-01 | 8.13E-01 | 1.29E+00 | 1.69E-01 | 6.45E-01 | 1.21E+00 | 3.88E-02 | 1.38E-01 | 1.00E+00 | 7.86E-02 | 2.22E-01 | -1.02E+00 | 1.56E-06 | 5.74E-06 | -1.54E+00 | 3.54E-06 | 9.72E-06 | -1.44E+00 |
| AL731563.3 | 9.73E-02 | 5.07E-01 | 1.02E+00 | 1.84E-01 | 7.94E-01 | 1.49E+00 | 9.49E-04 | 4.03E-03 | -1.94E+00 | 1.06E-02 | 4.04E-02 | -1.31E+00 | 4.53E-02 | 1.62E-01 | -1.23E+00 | 6.66E-02 | 1.88E-01 | -1.15E+00 | 7.09E-02 | 2.60E-01 | -1.33E+00 | 2.09E-01 | 5.74E-01 | -1.08E+00 |
| AC116618.1 | 9.73E-02 | 5.07E-01 | 1.13E+00 | 7.91E-02 | 3.42E-01 | 1.28E+00 | 1.99E-01 | 8.45E-01 | 1.31E+00 | 1.96E-01 | 7.49E-01 | 1.34E+00 | 2.70E-01 | 9.63E-01 | 1.28E+00 | 3.55E-01 | 1.00E+00 | 1.41E+00 | 1.83E-01 | 6.72E-01 | 1.14E+00 | 9.60E-02 | 2.64E-01 | 1.01E+00 |
| GAPDHP44 | 9.73E-02 | 5.07E-01 | 1.51E+00 | 1.04E-01 | 4.51E-01 | 1.01E+00 | 1.20E-01 | 5.10E-01 | 1.08E+00 | 9.08E-03 | 3.47E-02 | -1.14E+00 | 2.75E-01 | 9.80E-01 | 1.40E+00 | 8.80E-02 | 2.48E-01 | -1.21E+00 | 5.41E-03 | 1.98E-02 | -1.31E+00 | 2.46E-02 | 6.75E-02 | -1.15E+00 |
| TPT1P1 | 9.74E-02 | 5.07E-01 | 1.45E+00 | 3.34E-08 | 1.44E-07 | 2.92E+00 | 1.38E-01 | 5.87E-01 | 1.48E+00 | 4.77E-07 | 1.82E-06 | 3.26E+00 | 2.55E-01 | 9.09E-01 | 1.39E+00 | 5.74E-03 | 1.62E-02 | 2.62E+00 | 1.03E-01 | 3.78E-01 | 1.75E+00 | 5.43E-04 | 1.49E-03 | 2.77E+00 |
| LINC02486 | 9.74E-02 | 5.07E-01 | 1.58E+00 | 1.34E-02 | 5.79E-02 | 2.42E+00 | 1.13E-01 | 4.79E-01 | -1.03E+00 | 9.39E-02 | 3.59E-01 | 1.55E+00 | 5.18E-02 | 1.85E-01 | 2.56E+00 | 9.30E-03 | 2.62E-02 | 3.56E+00 | 1.59E-01 | 5.82E-01 | 1.33E+00 | 6.82E-02 | 1.87E-01 | 1.79E+00 |
| UHRF1 | 9.74E-02 | 5.07E-01 | -1.23E+00 | 1.90E-01 | 8.23E-01 | 1.10E+00 | 1.32E-02 | 5.62E-02 | 1.30E+00 | 1.95E-01 | 7.46E-01 | 1.29E+00 | 1.19E-02 | 4.25E-02 | -3.12E+00 | 4.90E-02 | 1.38E-01 | -1.89E+00 | 1.92E-03 | 7.04E-03 | -1.23E+00 | 5.98E-02 | 1.64E-01 | 1.50E+00 |
| MYCNOS | 9.74E-02 | 5.08E-01 | 1.13E+00 | 7.11E-02 | 3.07E-01 | 1.15E+00 | 7.74E-03 | 3.29E-02 | 2.08E+00 | 4.43E-02 | 1.69E-01 | 2.11E+00 | 2.36E-01 | 8.41E-01 | 1.26E+00 | 1.95E-02 | 5.50E-02 | -1.23E+00 | 1.56E-01 | 5.74E-01 | 1.23E+00 | 2.26E-01 | 6.20E-01 | 1.33E+00 |
| AC092881.1 | 9.74E-02 | 5.08E-01 | 1.06E+00 | 6.07E-02 | 2.62E-01 | 1.01E+00 | 2.13E-01 | 9.05E-01 | 1.37E+00 | 1.67E-01 | 6.39E-01 | 1.33E+00 | 2.16E-01 | 7.71E-01 | 1.05E+00 | 2.86E-01 | 8.07E-01 | 1.23E+00 | 3.85E-04 | 1.41E-03 | -1.24E+00 | 5.35E-05 | 1.47E-04 | -1.49E+00 |
| AC118553.2 | 9.75E-02 | 5.08E-01 | -1.02E+00 | 7.81E-02 | 3.38E-01 | -1.05E+00 | 1.10E-01 | 4.67E-01 | -1.11E+00 | 1.26E-01 | 4.80E-01 | 1.02E+00 | 2.60E-01 | 9.26E-01 | 1.11E+00 | 3.34E-01 | 9.42E-01 | 1.14E+00 | 1.17E-01 | 4.30E-01 | -1.10E+00 | 9.61E-02 | 2.64E-01 | -1.13E+00 |
| AC025181.2 | 9.75E-02 | 5.08E-01 | 1.28E+00 | 1.20E-03 | 5.19E-03 | 1.07E+00 | 1.34E-01 | 5.71E-01 | 1.36E+00 | 1.67E-03 | 6.38E-03 | -1.04E+00 | 3.11E-02 | 1.11E-01 | 1.75E+00 | 1.66E-01 | 4.68E-01 | -1.00E+00 | 2.47E-01 | 9.07E-01 | 1.47E+00 | 1.08E-01 | 2.98E-01 | 1.04E+00 |
| SYCE2 | 9.77E-02 | 5.09E-01 | 1.68E+00 | 1.65E-01 | 7.12E-01 | 1.63E+00 | 4.02E-03 | 1.71E-02 | -1.14E+00 | 3.52E-05 | 1.35E-04 | -1.36E+00 | 2.54E-01 | 9.06E-01 | 1.33E+00 | 2.30E-01 | 6.49E-01 | 1.73E+00 | 1.27E-01 | 4.67E-01 | 1.40E+00 | 2.72E-01 | 7.46E-01 | 1.09E+00 |
| RPS4XP9 | 9.77E-02 | 5.09E-01 | 1.23E+00 | 6.02E-03 | 2.60E-02 | -1.02E+00 | 2.58E-02 | 1.10E-01 | 1.61E+00 | 9.05E-02 | 3.46E-01 | 1.01E+00 | 3.53E-02 | 1.26E-01 | -1.08E+00 | 1.92E-04 | 5.41E-04 | -1.65E+00 | 2.02E-01 | 7.43E-01 | 1.14E+00 | 5.15E-02 | 1.41E-01 | -1.04E+00 |
| AL592437.2 | 9.77E-02 | 5.09E-01 | 1.46E+00 | 8.52E-02 | 3.68E-01 | -1.06E+00 | 9.10E-02 | 3.87E-01 | -1.04E+00 | 1.79E-03 | 6.82E-03 | -1.72E+00 | 1.62E-01 | 5.78E-01 | 1.68E+00 | 3.00E-01 | 8.45E-01 | 1.32E+00 | 1.63E-01 | 5.99E-01 | -1.16E+00 | 2.07E-01 | 5.68E-01 | -1.11E+00 |
| LTBP4 | 9.78E-02 | 5.09E-01 | 1.25E+00 | 1.95E-01 | 8.41E-01 | 1.56E+00 | 6.33E-02 | 2.69E-01 | 1.48E+00 | 4.20E-03 | 1.60E-02 | 1.90E+00 | 9.42E-04 | 3.36E-03 | -1.08E+00 | 1.79E-01 | 5.04E-01 | 1.39E+00 | 2.61E-04 | 9.57E-04 | 2.04E+00 | 5.53E-04 | 1.52E-03 | 2.12E+00 |
| ACTA1 | 9.79E-02 | 5.10E-01 | 1.05E+00 | 1.11E-02 | 4.80E-02 | -1.46E+00 | 7.40E-02 | 3.15E-01 | 1.03E+00 | 1.89E-04 | 7.22E-04 | -1.60E+00 | 7.74E-03 | 2.76E-02 | -1.39E+00 | 1.92E-03 | 5.42E-03 | -1.77E+00 | 1.64E-03 | 6.02E-03 | -1.38E+00 | 1.14E-04 | 3.14E-04 | -1.51E+00 |
| KIAA0513 | 9.79E-02 | 5.10E-01 | -1.15E+00 | 1.44E-01 | 6.24E-01 | -1.02E+00 | 1.60E-01 | 6.79E-01 | -1.11E+00 | 5.47E-02 | 2.09E-01 | -1.72E+00 | 1.74E-01 | 6.20E-01 | -1.05E+00 | 2.98E-01 | 8.40E-01 | 1.10E+00 | 6.43E-03 | 2.36E-02 | 1.43E+00 | 4.12E-04 | 1.13E-03 | 1.05E+00 |
| AL159152.1 | 9.80E-02 | 5.11E-01 | 1.38E+00 | 8.89E-02 | 3.84E-01 | 1.59E+00 | 2.34E-01 | 9.96E-01 | 1.17E+00 | 3.33E-03 | 1.27E-02 | 2.66E+00 | 2.51E-01 | 8.95E-01 | 1.45E+00 | 7.31E-02 | 2.06E-01 | 2.31E+00 | 9.80E-02 | 3.60E-01 | 1.64E+00 | 2.06E-03 | 5.66E-03 | 2.68E+00 |
| BFSP2 | 9.81E-02 | 5.11E-01 | -1.27E+00 | 5.51E-02 | 2.38E-01 | -1.78E+00 | 1.17E-01 | 4.98E-01 | -1.08E+00 | 1.07E-01 | 4.10E-01 | -1.07E+00 | 1.20E-04 | 4.29E-04 | -1.24E+00 | 6.27E-10 | 1.77E-09 | -1.78E+00 | 2.62E-01 | 9.63E-01 | 1.14E+00 | 1.89E-01 | 5.18E-01 | 1.45E+00 |
| AC010618.3 | 9.83E-02 | 5.12E-01 | 1.02E+00 | 5.15E-02 | 2.22E-01 | 1.99E+00 | 2.21E-02 | 9.41E-02 | -1.08E+00 | 7.12E-02 | 2.72E-01 | 1.18E+00 | 1.31E-02 | 4.65E-02 | -1.56E+00 | 8.73E-02 | 2.46E-01 | -1.19E+00 | 2.47E-01 | 9.06E-01 | 1.27E+00 | 1.07E-01 | 2.93E-01 | 1.80E+00 |
| DOCK3 | 9.83E-02 | 5.12E-01 | 1.17E+00 | 1.35E-03 | 5.85E-03 | -1.45E+00 | 3.70E-03 | 1.57E-02 | -1.52E+00 | 1.85E-02 | 7.07E-02 | -1.15E+00 | 2.51E-01 | 8.95E-01 | 1.28E+00 | 3.08E-01 | 8.68E-01 | 1.25E+00 | 4.02E-02 | 1.48E-01 | -1.23E+00 | 8.80E-02 | 2.42E-01 | -1.14E+00 |
| CDYL | 9.83E-02 | 5.12E-01 | 1.39E+00 | 1.39E-02 | 6.01E-02 | 1.04E+00 | 8.33E-02 | 3.54E-01 | 1.12E+00 | 1.99E-02 | 7.62E-02 | 1.13E+00 | 1.03E-01 | 3.67E-01 | 1.13E+00 | 3.08E-01 | 8.67E-01 | 1.45E+00 | 2.85E-02 | 1.05E-01 | 1.19E+00 | 5.72E-02 | 1.57E-01 | 1.15E+00 |
| IGF2R | 9.83E-02 | 5.12E-01 | -1.06E+00 | 1.08E-01 | 4.68E-01 | 1.02E+00 | 1.68E-01 | 7.14E-01 | 1.20E+00 | 1.28E-01 | 4.88E-01 | -1.16E+00 | 2.43E-01 | 8.66E-01 | 1.31E+00 | 3.34E-01 | 9.41E-01 | 1.03E+00 | 1.42E-01 | 5.23E-01 | 1.25E+00 | 2.90E-01 | 7.97E-01 | 1.13E+00 |
| AC091390.2 | 9.83E-02 | 5.12E-01 | 1.16E+00 | 6.62E-02 | 2.86E-01 | 2.03E+00 | 1.36E-01 | 5.77E-01 | 1.37E+00 | 1.63E-01 | 6.23E-01 | 1.54E+00 | 1.67E-01 | 5.97E-01 | 1.14E+00 | 3.38E-01 | 9.52E-01 | 1.34E+00 | 2.47E-01 | 9.07E-01 | 1.35E+00 | 5.69E-02 | 1.56E-01 | 1.76E+00 |
| RAD51AP1 | 9.84E-02 | 5.13E-01 | 1.82E+00 | 2.13E-01 | 9.18E-01 | 1.21E+00 | 1.11E-04 | 4.73E-04 | 1.71E+00 | 1.38E-01 | 5.26E-01 | 1.41E+00 | 2.70E-01 | 9.63E-01 | -1.15E+00 | 2.51E-01 | 7.07E-01 | -1.60E+00 | 2.37E-01 | 8.70E-01 | -1.02E+00 | 1.28E-02 | 3.52E-02 | 1.31E+00 |
| CD96 | 9.84E-02 | 5.13E-01 | -1.89E+00 | 1.47E-01 | 6.33E-01 | -1.74E+00 | 1.92E-01 | 8.17E-01 | -1.12E+00 | 1.34E-02 | 5.13E-02 | -1.72E+00 | 2.28E-01 | 8.12E-01 | 1.20E+00 | 2.25E-01 | 6.35E-01 | -1.17E+00 | 8.39E-05 | 3.08E-04 | 4.02E+00 | 9.94E-02 | 2.73E-01 | 1.63E+00 |
| STARD7-AS1 | 9.85E-02 | 5.13E-01 | 1.06E+00 | 1.02E-01 | 4.40E-01 | 1.12E+00 | 3.54E-02 | 1.50E-01 | -1.14E+00 | 2.57E-02 | 9.83E-02 | -1.04E+00 | 3.03E-02 | 1.08E-01 | -1.22E+00 | 6.42E-02 | 1.81E-01 | -1.36E+00 | 2.60E-01 | 9.53E-01 | 1.31E+00 | 3.01E-01 | 8.28E-01 | 1.28E+00 |
| SNX32 | 9.85E-02 | 5.13E-01 | 1.21E+00 | 2.84E-02 | 1.23E-01 | 1.05E+00 | 3.16E-03 | 1.34E-02 | -1.31E+00 | 8.22E-04 | 3.14E-03 | -1.32E+00 | 2.74E-01 | 9.78E-01 | 1.37E+00 | 3.47E-01 | 9.77E-01 | 1.43E+00 | 1.67E-01 | 6.13E-01 | 1.04E+00 | 2.63E-01 | 7.22E-01 | 1.13E+00 |
| MANEA-AS1 | 9.85E-02 | 5.13E-01 | -1.10E+00 | 6.66E-03 | 2.88E-02 | 1.95E+00 | 1.34E-02 | 5.70E-02 | -1.46E+00 | 1.89E-01 | 7.23E-01 | 1.57E+00 | 4.10E-03 | 1.46E-02 | -1.22E+00 | 2.76E-01 | 7.79E-01 | 1.31E+00 | 1.82E-01 | 6.66E-01 | 1.32E+00 | 2.40E-03 | 6.58E-03 | 1.85E+00 |
| AC125437.2 | 9.85E-02 | 5.14E-01 | 1.52E+00 | 1.63E-01 | 7.05E-01 | 1.58E+00 | 1.88E-01 | 7.97E-01 | 1.24E+00 | 1.97E-02 | 7.53E-02 | -1.29E+00 | 2.79E-01 | 9.93E-01 | 1.23E+00 | 3.25E-01 | 9.17E-01 | 1.16E+00 | 2.09E-01 | 7.66E-01 | 1.27E+00 | 2.80E-01 | 7.69E-01 | 1.34E+00 |
| AC003075.1 | 9.85E-02 | 5.14E-01 | 1.41E+00 | 3.38E-02 | 1.46E-01 | -1.26E+00 | 3.79E-02 | 1.61E-01 | 1.76E+00 | 6.30E-02 | 2.41E-01 | 1.91E+00 | 2.74E-01 | 9.77E-01 | 1.49E+00 | 3.39E-01 | 9.55E-01 | 1.31E+00 | 2.16E-01 | 7.92E-01 | 1.24E+00 | 2.30E-01 | 6.32E-01 | -1.05E+00 |
| ABTB2 | 9.86E-02 | 5.14E-01 | 1.41E+00 | 2.13E-01 | 9.22E-01 | 1.26E+00 | 2.26E-01 | 9.59E-01 | 1.01E+00 | 2.59E-01 | 9.90E-01 | 1.09E+00 | 2.72E-01 | 9.71E-01 | 1.10E+00 | 2.90E-01 | 8.17E-01 | 1.03E+00 | 2.18E-02 | 8.00E-02 | 1.63E+00 | 7.67E-02 | 2.11E-01 | 1.57E+00 |
| NPM1P6 | 9.87E-02 | 5.14E-01 | 1.26E+00 | 2.09E-01 | 9.01E-01 | 1.58E+00 | 3.28E-02 | 1.40E-01 | 1.49E+00 | 6.71E-04 | 2.56E-03 | 1.95E+00 | 2.73E-01 | 9.73E-01 | 1.54E+00 | 1.87E-02 | 5.26E-02 | 2.31E+00 | 1.34E-02 | 4.91E-02 | 1.59E+00 | 1.97E-05 | 5.40E-05 | 2.07E+00 |
| HAL | 9.88E-02 | 5.15E-01 | 1.57E+00 | 1.37E-01 | 5.91E-01 | -1.09E+00 | 2.91E-06 | 1.24E-05 | -1.47E+00 | 1.99E-03 | 7.59E-03 | -1.02E+00 | 2.31E-01 | 8.25E-01 | 1.13E+00 | 3.05E-01 | 8.59E-01 | -1.41E+00 | 2.22E-01 | 8.17E-01 | 1.24E+00 | 2.71E-01 | 7.44E-01 | 1.30E+00 |
| ZXDB | 9.88E-02 | 5.15E-01 | 1.34E+00 | 3.37E-02 | 1.45E-01 | 1.02E+00 | 7.65E-02 | 3.25E-01 | 1.00E+00 | 2.06E-03 | 7.89E-03 | -1.40E+00 | 5.44E-03 | 1.94E-02 | -1.25E+00 | 2.51E-03 | 7.09E-03 | -1.56E+00 | 1.98E-02 | 7.28E-02 | -1.17E+00 | 4.56E-02 | 1.25E-01 | -1.10E+00 |
| AC079203.1 | 9.90E-02 | 5.16E-01 | 1.06E+00 | 8.90E-02 | 3.84E-01 | 1.11E+00 | 2.17E-01 | 9.23E-01 | 1.31E+00 | 2.57E-01 | 9.81E-01 | 1.44E+00 | 1.73E-01 | 6.17E-01 | 1.55E+00 | 4.34E-02 | 1.22E-01 | -1.59E+00 | 2.70E-01 | 9.90E-01 | 1.28E+00 | 9.19E-02 | 2.52E-01 | -1.22E+00 |
| REXO1 | 9.90E-02 | 5.16E-01 | 1.16E+00 | 9.01E-03 | 3.89E-02 | -1.04E+00 | 1.44E-01 | 6.13E-01 | 1.14E+00 | 2.19E-02 | 8.35E-02 | -1.03E+00 | 4.87E-02 | 1.74E-01 | 1.01E+00 | 4.88E-02 | 1.37E-01 | -1.01E+00 | 1.04E-04 | 3.81E-04 | -1.28E+00 | 2.44E-04 | 6.70E-04 | -1.27E+00 |
| RPS15AP17 | 9.91E-02 | 5.16E-01 | 1.51E+00 | 2.00E-01 | 8.62E-01 | 1.48E+00 | 1.72E-01 | 7.32E-01 | 1.34E+00 | 1.60E-01 | 6.13E-01 | 1.19E+00 | 1.72E-01 | 6.15E-01 | 1.60E+00 | 2.72E-01 | 7.67E-01 | 1.53E+00 | 1.99E-01 | 7.30E-01 | 1.02E+00 | 1.24E-01 | 3.39E-01 | -1.14E+00 |
| NME9 | 9.92E-02 | 5.17E-01 | 1.03E+00 | 2.31E-01 | 9.98E-01 | 1.45E+00 | 1.81E-01 | 7.69E-01 | 1.09E+00 | 4.84E-02 | 1.85E-01 | 1.71E+00 | 1.08E-03 | 3.84E-03 | -1.38E+00 | 5.20E-02 | 1.46E-01 | 1.02E+00 | 8.26E-02 | 3.03E-01 | 1.12E+00 | 4.76E-02 | 1.31E-01 | 1.76E+00 |
| KNTC1 | 9.92E-02 | 5.17E-01 | 1.49E+00 | 2.10E-01 | 9.08E-01 | 1.52E+00 | 2.57E-04 | 1.09E-03 | 1.78E+00 | 8.39E-03 | 3.21E-02 | 1.79E+00 | 8.03E-02 | 2.86E-01 | 1.01E+00 | 3.39E-01 | 9.57E-01 | 1.47E+00 | 2.58E-02 | 9.46E-02 | 1.06E+00 | 3.31E-01 | 9.10E-01 | 1.25E+00 |
| AHR | 9.93E-02 | 5.17E-01 | 1.37E+00 | 5.54E-02 | 2.40E-01 | 1.31E+00 | 2.63E-02 | 1.12E-01 | 1.53E+00 | 2.48E-01 | 9.48E-01 | 1.42E+00 | 5.72E-10 | 2.04E-09 | 3.64E+00 | 3.92E-05 | 1.10E-04 | 2.91E+00 | 7.83E-09 | 2.87E-08 | 2.77E+00 | 1.34E-01 | 3.68E-01 | 1.67E+00 |
| AL360270.1 | 9.93E-02 | 5.17E-01 | 1.56E+00 | 9.51E-02 | 4.11E-01 | 1.78E+00 | 2.08E-01 | 8.85E-01 | 1.24E+00 | 2.55E-01 | 9.74E-01 | 1.51E+00 | 9.58E-02 | 3.42E-01 | 1.77E+00 | 8.87E-02 | 2.50E-01 | 1.98E+00 | 2.65E-01 | 9.73E-01 | 1.14E+00 | 2.04E-01 | 5.61E-01 | -1.09E+00 |
| ZNF487 | 9.93E-02 | 5.18E-01 | 1.22E+00 | 3.58E-02 | 1.55E-01 | 1.06E+00 | 2.81E-02 | 1.19E-01 | -1.00E+00 | 1.04E-02 | 3.99E-02 | 1.05E+00 | 1.24E-04 | 4.41E-04 | -1.31E+00 | 2.29E-06 | 6.47E-06 | -1.70E+00 | 1.06E-02 | 3.90E-02 | 2.22E+00 | 2.61E-01 | 7.17E-01 | 1.26E+00 |
| RIN3 | 9.94E-02 | 5.18E-01 | 1.23E+00 | 1.38E-01 | 5.97E-01 | 1.42E+00 | 4.46E-02 | 1.90E-01 | 1.15E+00 | 4.62E-02 | 1.77E-01 | 1.24E+00 | 9.85E-05 | 3.51E-04 | -1.15E+00 | 2.22E-03 | 6.26E-03 | -1.03E+00 | 5.32E-03 | 1.95E-02 | -1.03E+00 | 2.26E-02 | 6.21E-02 | 1.05E+00 |
| ADORA2B | 9.95E-02 | 5.19E-01 | 1.11E+00 | 1.36E-01 | 5.86E-01 | 1.28E+00 | 1.81E-02 | 7.69E-02 | -1.29E+00 | 1.05E-01 | 4.02E-01 | 1.09E+00 | 1.27E-05 | 4.53E-05 | -1.75E+00 | 2.58E-04 | 7.28E-04 | -1.39E+00 | 3.49E-19 | 1.28E-18 | -2.32E+00 | 1.56E-07 | 4.30E-07 | -1.28E+00 |
| GFI1 | 9.95E-02 | 5.19E-01 | -1.18E+00 | 4.40E-02 | 1.90E-01 | -1.43E+00 | 9.54E-02 | 4.06E-01 | 1.12E+00 | 2.56E-03 | 9.79E-03 | -1.05E+00 | 2.80E-01 | 1.00E+00 | 1.00E+00 | 3.36E-01 | 9.47E-01 | 1.07E+00 | 2.60E-01 | 9.53E-01 | -1.05E+00 | 3.62E-01 | 9.93E-01 | 1.01E+00 |
| AL139246.5 | 9.99E-02 | 5.20E-01 | 1.63E+00 | 1.13E-01 | 4.90E-01 | 1.23E+00 | 8.86E-02 | 3.76E-01 | 1.04E+00 | 1.54E-02 | 5.87E-02 | -1.18E+00 | 4.98E-02 | 1.77E-01 | -1.01E+00 | 1.68E-02 | 4.73E-02 | -1.19E+00 | 2.71E-01 | 9.95E-01 | 1.43E+00 | 2.21E-01 | 6.07E-01 | 1.10E+00 |
| NHLRC1 | 1.00E-01 | 5.21E-01 | -1.14E+00 | 1.79E-01 | 7.71E-01 | 1.43E+00 | 8.58E-07 | 3.65E-06 | -2.23E+00 | 3.07E-04 | 1.17E-03 | -1.36E+00 | 3.15E-04 | 1.12E-03 | -2.61E+00 | 1.10E-01 | 3.09E-01 | 1.04E+00 | 2.31E-15 | 8.47E-15 | -2.73E+00 | 2.36E-02 | 6.48E-02 | -1.01E+00 |
| IL2RA | 1.00E-01 | 5.22E-01 | 1.31E+00 | 1.31E-01 | 5.67E-01 | 1.59E+00 | 3.46E-03 | 1.47E-02 | -1.66E+00 | 6.20E-02 | 2.37E-01 | -1.02E+00 | 1.22E-01 | 4.37E-01 | -1.48E+00 | 1.98E-01 | 5.59E-01 | -1.17E+00 | 1.56E-01 | 5.73E-01 | -1.22E+00 | 2.57E-01 | 7.05E-01 | -1.12E+00 |
| KNL1 | 1.00E-01 | 5.22E-01 | -1.04E+00 | 1.48E-02 | 6.38E-02 | -1.71E+00 | 1.33E-01 | 5.64E-01 | 1.09E+00 | 4.94E-03 | 1.89E-02 | 1.03E+00 | 2.09E-02 | 7.44E-02 | -1.66E+00 | 1.10E-01 | 3.11E-01 | -1.03E+00 | 1.98E-02 | 7.28E-02 | -1.17E+00 | 6.98E-02 | 1.92E-01 | 1.03E+00 |

| | | | | | | | | | | | | | | | | | | | | | | | | |
|---|---|---|---|---|---|---|---|---|---|---|---|---|---|---|---|---|---|---|---|---|---|---|---|---|
| TBCAP1 | 1.00E-01 | 5.23E-01 | 1.46E+00 | 7.24E-02 | 3.13E-01 | 1.32E+00 | 5.46E-03 | 2.32E-02 | 1.67E+00 | 2.38E-01 | 9.09E-01 | 1.56E+00 | 2.45E-01 | 8.72E-01 | 1.39E+00 | 2.94E-01 | 8.28E-01 | 1.45E+00 | 1.23E-01 | 4.50E-01 | 1.40E+00 | 3.08E-01 | 8.45E-01 | 1.28E+00 |
| RPS8P4 | 1.00E-01 | 5.24E-01 | 1.06E+00 | 5.93E-02 | 2.56E-01 | 1.11E+00 | 1.12E-01 | 4.75E-01 | 1.13E+00 | 2.21E-01 | 8.44E-01 | 1.49E+00 | 1.74E-01 | 6.22E-01 | 1.66E+00 | 2.91E-01 | 8.20E-01 | 1.56E+00 | 2.48E-01 | 9.12E-01 | 1.26E+00 | 3.36E-02 | 9.24E-02 | 1.80E+00 |
| Z97192.1 | 1.01E-01 | 5.24E-01 | 1.06E+00 | 2.19E-01 | 9.46E-01 | 1.46E+00 | 1.28E-02 | 5.46E-02 | -1.30E+00 | 4.44E-02 | 1.70E-01 | -1.02E+00 | 2.27E-01 | 8.10E-01 | 1.10E+00 | 3.55E-01 | 1.00E+00 | 1.25E+00 | 3.10E-02 | 1.14E-01 | -1.48E+00 | 1.99E-01 | 5.47E-01 | 1.12E+00 |
| IL12RB1 | 1.01E-01 | 5.25E-01 | 1.17E+00 | 7.93E-02 | 3.42E-01 | 1.16E+00 | 1.40E-02 | 5.93E-02 | -1.21E+00 | 1.69E-02 | 6.47E-02 | -1.09E+00 | 1.72E-01 | 6.14E-01 | 1.05E+00 | 2.68E-01 | 7.55E-01 | 1.24E+00 | 6.22E-02 | 2.28E-01 | -1.18E+00 | 5.26E-02 | 1.44E-01 | -1.32E+00 |
| AC007390.2 | 1.01E-01 | 5.25E-01 | 1.19E+00 | 2.31E-01 | 1.00E+00 | 1.41E+00 | 9.79E-02 | 4.16E-01 | -1.04E+00 | 7.55E-02 | 2.88E-01 | 1.04E+00 | 2.57E-01 | 9.18E-01 | 1.30E+00 | 3.52E-01 | 9.93E-01 | 1.34E+00 | 9.98E-02 | 3.66E-01 | 1.64E+00 | 3.47E-01 | 9.54E-01 | 1.29E+00 |
| BX322784.1 | 1.01E-01 | 5.25E-01 | 1.65E+00 | 1.46E-03 | 6.31E-03 | 3.07E+00 | 5.02E-03 | 2.13E-02 | 1.99E+00 | 3.05E-05 | 1.17E-04 | 3.06E+00 | 2.20E-01 | 7.86E-01 | 1.23E+00 | 3.13E-02 | 8.84E-02 | 2.41E+00 | 9.49E-03 | 3.48E-02 | 1.72E+00 | 4.01E-08 | 1.10E-07 | 2.83E+00 |
| AQP10 | 1.01E-01 | 5.25E-01 | 1.55E+00 | 2.18E-01 | 9.43E-01 | 1.06E+00 | 1.47E-01 | 6.27E-01 | 1.47E+00 | 2.29E-01 | 8.73E-01 | 1.15E+00 | 1.43E-02 | 5.11E-02 | 1.08E+00 | 7.99E-03 | 2.25E-02 | -1.13E+00 | 9.83E-02 | 3.61E-01 | 3.26E+00 | 3.34E-01 | 9.17E-01 | 1.74E+00 |
| S100A6 | 1.01E-01 | 5.25E-01 | 1.10E+00 | 1.92E-04 | 8.28E-04 | 1.05E+00 | 1.21E-01 | 5.16E-01 | 1.13E+00 | 1.78E-01 | 6.79E-01 | 1.11E+00 | 1.71E-03 | 6.10E-03 | -1.21E+00 | 7.32E-03 | 2.06E-02 | -1.27E+00 | 1.30E-01 | 4.76E-01 | -1.03E+00 | 5.91E-03 | 1.62E-02 | -1.25E+00 |
| MEIS2 | 1.01E-01 | 5.25E-01 | -1.12E+00 | 1.31E-01 | 5.65E-01 | -1.05E+00 | 1.96E-01 | 8.32E-01 | -1.06E+00 | 2.02E-01 | 7.72E-01 | -1.10E+00 | 2.64E-07 | 9.41E-07 | -2.11E+00 | 2.84E-06 | 8.00E-06 | -1.72E+00 | 5.03E-03 | 1.85E-02 | -1.15E+00 | 4.71E-03 | 1.29E-02 | -1.12E+00 |
| NEB | 1.01E-01 | 5.25E-01 | -1.09E+00 | 2.22E-01 | 9.59E-01 | 1.28E+00 | 7.57E-02 | 3.22E-01 | -1.19E+00 | 1.46E-01 | 5.57E-01 | 1.47E+00 | 1.10E-01 | 3.94E-01 | -1.14E+00 | 2.24E-01 | 6.32E-01 | 1.67E+00 | 4.79E-02 | 1.76E-01 | -1.18E+00 | 2.95E-01 | 8.11E-01 | 1.17E+00 |
| CRYM | 1.01E-01 | 5.25E-01 | -1.44E+00 | 1.85E-01 | 8.01E-01 | 1.09E+00 | 1.98E-01 | 8.42E-01 | -1.08E+00 | 2.48E-01 | 9.48E-01 | 1.16E+00 | 4.81E-05 | 1.72E-04 | -1.16E+00 | 1.85E-01 | 5.20E-01 | 1.61E+00 | 2.74E-11 | 1.00E-10 | -1.21E+00 | 7.89E-14 | 2.17E-13 | -1.39E+00 |
| LTK | 1.01E-01 | 5.25E-01 | -1.21E+00 | 1.37E-01 | 5.90E-01 | -1.11E+00 | 1.16E-02 | 4.91E-02 | -1.71E+00 | 7.88E-02 | 3.01E-01 | -1.13E+00 | 2.45E-01 | 8.74E-01 | 1.34E+00 | 3.27E-01 | 9.22E-01 | 1.29E+00 | 4.02E-03 | 1.47E-02 | 1.93E+00 | 3.38E-01 | 9.28E-01 | 1.12E+00 |
| CLSPN | 1.01E-01 | 5.25E-01 | -1.12E+00 | 6.95E-02 | 3.00E-01 | -1.56E+00 | 4.98E-02 | 2.12E-01 | 1.06E+00 | 1.84E-01 | 7.02E-01 | -1.07E+00 | 1.47E-01 | 5.25E-01 | -1.96E+00 | 2.34E-01 | 6.59E-01 | -1.33E+00 | 6.06E-04 | 2.22E-03 | -1.25E+00 | 1.69E-01 | 4.64E-01 | 1.19E+00 |
| C2orf27AP3 | 1.01E-01 | 5.26E-01 | 1.09E+00 | 9.89E-02 | 4.27E-01 | 1.15E+00 | 3.56E-09 | 1.51E-08 | -2.16E+00 | 3.78E-07 | 1.44E-06 | -1.51E+00 | 7.62E-02 | 2.72E-01 | -1.14E+00 | 5.49E-02 | 1.55E-01 | -1.25E+00 | 2.80E-02 | 1.03E-01 | 2.17E+00 | 1.20E-01 | 3.29E-01 | 1.70E+00 |
| AL035563.1 | 1.01E-01 | 5.26E-01 | 1.37E+00 | 1.22E-02 | 5.27E-02 | 1.90E+00 | 7.33E-02 | 3.11E-01 | 1.01E+00 | 6.65E-02 | 2.54E-01 | 1.10E+00 | 2.22E-01 | 7.92E-01 | 1.17E+00 | 2.22E-01 | 6.25E-01 | 1.56E+00 | 1.75E-01 | 6.42E-01 | 1.12E+00 | 3.62E-01 | 9.95E-01 | 1.29E+00 |
| HIST1H2BC | 1.01E-01 | 5.26E-01 | 1.37E+00 | 1.09E-01 | 4.72E-01 | 1.09E+00 | 1.74E-01 | 7.39E-01 | 1.22E+00 | 1.07E-01 | 4.10E-01 | 1.13E+00 | 3.01E-02 | 1.07E-01 | -1.01E+00 | 3.70E-02 | 1.04E-01 | -1.00E+00 | 5.08E-02 | 1.86E-01 | 1.48E+00 | 1.87E-01 | 5.15E-01 | 1.09E+00 |
| HNRNPABP1 | 1.01E-01 | 5.26E-01 | 1.03E+00 | 6.71E-02 | 2.90E-01 | -1.12E+00 | 6.51E-03 | 2.77E-02 | -1.26E+00 | 9.76E-04 | 3.73E-03 | -1.35E+00 | 7.86E-04 | 2.80E-03 | -1.86E+00 | 1.88E-03 | 5.30E-03 | -1.60E+00 | 3.86E-02 | 1.42E-01 | 1.17E+00 | 1.76E-05 | 4.84E-05 | -1.46E+00 |
| MVB12B | 1.01E-01 | 5.26E-01 | 1.07E+00 | 1.09E-01 | 4.69E-01 | 1.17E+00 | 5.41E-02 | 2.30E-01 | -1.02E+00 | 2.54E-02 | 9.72E-02 | 1.09E+00 | 8.64E-03 | 3.08E-02 | -1.38E+00 | 2.09E-03 | 5.89E-03 | -1.62E+00 | 2.84E-02 | 1.04E-01 | -1.38E+00 | 6.25E-02 | 1.72E-01 | -1.24E+00 |
| AC245060.5 | 1.01E-01 | 5.26E-01 | 1.57E+00 | 6.41E-02 | 2.77E-01 | 1.74E+00 | 1.90E-01 | 8.07E-01 | 1.31E+00 | 1.68E-01 | 6.42E-01 | 1.23E+00 | 2.79E-01 | 9.96E-01 | 1.47E+00 | 1.95E-01 | 5.50E-01 | 1.60E+00 | 2.94E-02 | 1.08E-01 | 1.98E+00 | 1.15E-01 | 3.17E-01 | 1.70E+00 |
| AL031594.1 | 1.01E-01 | 5.26E-01 | 1.10E+00 | 1.36E-03 | 5.89E-03 | -1.41E+00 | 2.00E-01 | 8.51E-01 | 1.16E+00 | 5.74E-02 | 2.19E-01 | -1.08E+00 | 1.18E-03 | 4.19E-03 | -1.56E+00 | 2.57E-03 | 7.24E-03 | -1.44E+00 | 3.84E-02 | 1.41E-01 | -1.26E+00 | 6.78E-02 | 1.86E-01 | -1.11E+00 |
| CA1 | 1.01E-01 | 5.26E-01 | 2.32E+00 | 1.35E-01 | 5.83E-01 | 2.11E+00 | 2.10E-01 | 8.92E-01 | 1.01E+00 | 1.96E-01 | 7.50E-01 | 1.62E+00 | 1.59E-02 | 5.66E-02 | 1.27E+01 | 3.07E-04 | 8.64E-04 | 2.37E+01 | 3.21E-86 | 1.18E-85 | 8.88E+00 | 3.65E-25 | 1.00E-24 | 4.88E+00 |
| TUBB2A | 1.01E-01 | 5.27E-01 | 1.06E+00 | 7.61E-03 | 3.29E-02 | -2.11E+00 | 4.01E-02 | 1.70E-01 | -1.24E+00 | 7.09E-04 | 2.71E-03 | -2.90E+00 | 7.17E-07 | 2.56E-06 | -1.52E+00 | 5.57E-07 | 1.57E-06 | -1.92E+00 | 1.14E-01 | 4.18E-01 | 1.46E+00 | 9.03E-08 | 2.48E-07 | -1.59E+00 |
| S1PR1 | 1.01E-01 | 5.27E-01 | 1.02E+00 | 8.93E-02 | 3.86E-01 | 1.00E+00 | 7.05E-02 | 2.99E-01 | -1.38E+00 | 4.21E-02 | 1.61E-01 | -1.68E+00 | 2.61E-07 | 9.32E-07 | -1.79E+00 | 7.37E-12 | 2.08E-11 | -3.98E+00 | 9.12E-08 | 3.35E-07 | -2.38E+00 | 1.88E-07 | 5.16E-07 | -2.41E+00 |
| SPATA1 | 1.01E-01 | 5.27E-01 | 1.49E+00 | 9.94E-02 | 4.29E-01 | 1.78E+00 | 1.40E-01 | 5.94E-01 | -1.00E+00 | 7.95E-02 | 3.04E-01 | -1.04E+00 | 3.91E-02 | 1.39E-01 | -1.24E+00 | 3.48E-01 | 9.81E-01 | 1.32E+00 | 2.62E-01 | 9.62E-01 | 1.20E+00 | 2.17E-01 | 5.95E-01 | 1.03E+00 |
| PACS1 | 1.01E-01 | 5.28E-01 | 1.22E+00 | 1.80E-01 | 7.76E-01 | 1.68E+00 | 6.29E-04 | 2.67E-03 | -1.25E+00 | 8.37E-02 | 3.20E-01 | 1.20E+00 | 6.23E-02 | 2.22E-01 | 1.12E+00 | 1.60E-01 | 4.51E-01 | 1.25E+00 | 2.10E-02 | 7.72E-02 | 1.10E+00 | 3.57E-02 | 9.81E-02 | 1.10E+00 |
| HEXIM1 | 1.01E-01 | 5.28E-01 | 1.31E+00 | 6.81E-02 | 2.94E-01 | 1.77E+00 | 5.45E-02 | 2.31E-01 | 1.42E+00 | 7.98E-02 | 3.05E-01 | 1.60E+00 | 2.69E-01 | 9.60E-01 | 1.35E+00 | 8.77E-02 | 2.47E-01 | 1.59E+00 | 1.91E-01 | 7.01E-01 | 1.15E+00 | 1.53E-01 | 4.21E-01 | 1.22E+00 |
| ZNF350 | 1.01E-01 | 5.29E-01 | 1.29E+00 | 5.83E-03 | 2.52E-02 | 1.09E+00 | 1.33E-01 | 5.65E-01 | 1.10E+00 | 2.57E-02 | 9.81E-02 | 1.12E+00 | 1.51E-02 | 5.37E-02 | 1.19E+00 | 4.56E-05 | 1.29E-04 | -1.18E+00 | 8.75E-02 | 3.21E-01 | 1.09E+00 | 9.26E-03 | 2.54E-02 | -1.04E+00 |
| JAG1 | 1.01E-01 | 5.29E-01 | 1.46E+00 | 8.34E-02 | 3.60E-01 | 1.77E+00 | 1.45E-03 | 6.16E-03 | -1.42E+00 | 8.58E-03 | 3.28E-02 | -1.26E+00 | 1.74E-01 | 6.19E-01 | -1.51E+00 | 3.42E-01 | 9.63E-01 | 1.23E+00 | 5.20E-02 | 1.91E-01 | -1.49E+00 | 9.23E-02 | 2.54E-01 | -1.50E+00 |
| USP46-AS1 | 1.02E-01 | 5.30E-01 | 1.19E+00 | 1.09E-01 | 4.73E-01 | 1.18E+00 | 2.27E-01 | 9.66E-01 | 1.13E+00 | 2.18E-01 | 8.33E-01 | 1.16E+00 | 7.94E-02 | 2.83E-01 | 2.40E+00 | 9.83E-02 | 2.77E-01 | 2.40E+00 | 7.14E-02 | 2.62E-01 | 1.68E+00 | 1.68E-01 | 4.61E-01 | 1.60E+00 |
| Z93930.2 | 1.02E-01 | 5.30E-01 | 1.50E+00 | 3.51E-02 | 1.52E-01 | 1.20E+00 | 1.65E-02 | 7.01E-02 | 1.66E+00 | 2.20E-01 | 8.39E-01 | 1.42E+00 | 2.52E-01 | 8.97E-01 | 1.26E+00 | 1.40E-01 | 3.95E-01 | 1.02E+00 | 1.75E-01 | 6.41E-01 | 1.15E+00 | 1.44E-01 | 3.94E-01 | 1.02E+00 |
| HDX | 1.02E-01 | 5.30E-01 | 1.00E+00 | 2.05E-01 | 8.87E-01 | 1.30E+00 | 1.42E-02 | 6.04E-02 | -1.35E+00 | 7.27E-02 | 2.78E-01 | 1.02E+00 | 1.57E-04 | 5.58E-04 | -1.55E+00 | 4.92E-04 | 1.39E-03 | -1.48E+00 | 4.68E-05 | 1.72E-04 | -2.12E+00 | 3.77E-03 | 1.04E-02 | -1.37E+00 |
| LY86 | 1.02E-01 | 5.30E-01 | -1.33E+00 | 6.37E-02 | 2.75E-01 | -1.56E+00 | 9.59E-03 | 4.07E-02 | -2.65E+00 | 2.13E-03 | 8.12E-03 | -3.22E+00 | 2.34E-01 | 8.34E-01 | 1.12E+00 | 3.55E-01 | 1.00E+00 | 1.00E+00 | 2.69E-01 | 9.88E-01 | 1.08E+00 | 2.55E-01 | 7.00E-01 | -1.09E+00 |
| NPIPA2 | 1.02E-01 | 5.30E-01 | 1.43E+00 | 1.99E-01 | 8.61E-01 | 1.43E+00 | 1.41E-01 | 5.98E-01 | 1.38E+00 | 6.02E-02 | 2.30E-01 | 1.07E+00 | 7.13E-04 | 2.54E-03 | -1.33E+00 | 1.32E-04 | 3.73E-04 | -1.59E+00 | 1.15E-01 | 4.21E-01 | 1.25E+00 | 2.46E-01 | 6.76E-01 | 1.29E+00 |
| PRKRIP1 | 1.02E-01 | 5.30E-01 | 1.18E+00 | 1.21E-02 | 5.23E-02 | 1.04E+00 | 2.23E-04 | 9.49E-04 | -1.23E+00 | 1.64E-05 | 6.28E-05 | -1.28E+00 | 1.16E-02 | 4.14E-02 | -1.07E+00 | 6.16E-02 | 1.74E-01 | 1.12E+00 | 5.34E-03 | 1.96E-02 | 1.07E+00 | 2.87E-04 | 7.87E-04 | -1.11E+00 |
| TRMO | 1.02E-01 | 5.31E-01 | 1.25E+00 | 6.98E-02 | 3.02E-01 | 1.28E+00 | 1.84E-01 | 7.80E-01 | 1.10E+00 | 9.46E-02 | 3.61E-01 | 1.11E+00 | 2.41E-02 | 8.61E-02 | -1.03E+00 | 2.99E-02 | 8.43E-02 | 1.02E+00 | 4.80E-02 | 1.76E-01 | 1.02E+00 | 5.53E-02 | 1.52E-01 | 1.06E+00 |
| DCP1A | 1.02E-01 | 5.31E-01 | 1.28E+00 | 3.18E-04 | 1.38E-03 | 1.09E+00 | 3.16E-03 | 1.34E-02 | 1.01E+00 | 1.69E-06 | 6.46E-06 | -1.19E+00 | 1.99E-02 | 7.08E-02 | 1.06E+00 | 3.77E-04 | 1.06E-03 | -1.16E+00 | 1.89E-06 | 6.95E-06 | -1.40E+00 | 9.68E-08 | 2.66E-07 | -1.48E+00 |
| GIMAP2 | 1.02E-01 | 5.31E-01 | 1.48E+00 | 1.39E-10 | 5.99E-10 | 2.42E+00 | 6.49E-03 | 2.76E-02 | -1.01E+00 | 1.41E-01 | 5.40E-01 | 1.54E+00 | 2.23E-02 | 7.96E-02 | -1.18E+00 | 3.37E-01 | 9.49E-01 | 1.30E+00 | 7.38E-07 | 2.71E-06 | -1.82E+00 | 3.19E-03 | 8.75E-03 | -1.15E+00 |
| HIST1H3C | 1.02E-01 | 5.31E-01 | -1.08E+00 | 3.56E-02 | 1.54E-01 | -1.50E+00 | 1.62E-02 | 6.90E-02 | -1.07E+00 | 7.35E-04 | 2.81E-03 | -1.46E+00 | 2.73E-01 | 9.72E-01 | 1.18E+00 | 3.38E-01 | 9.53E-01 | 1.09E+00 | 1.71E-02 | 6.27E-02 | 1.45E+00 | 5.89E-02 | 1.62E-01 | -1.35E+00 |
| FANCB | 1.02E-01 | 5.31E-01 | 1.11E+00 | 3.01E-02 | 1.30E-01 | -1.21E+00 | 8.81E-02 | 3.74E-01 | 1.34E+00 | 7.73E-03 | 2.95E-02 | -1.17E+00 | 6.75E-03 | 2.41E-02 | -1.56E+00 | 1.37E-03 | 3.85E-03 | -2.01E+00 | 4.21E-02 | 1.54E-01 | 1.00E+00 | 6.69E-02 | 1.84E-01 | 1.01E+00 |
| AC007546.2 | 1.02E-01 | 5.32E-01 | -1.09E+00 | 1.12E-01 | 4.82E-01 | -1.06E+00 | 2.06E-01 | 8.76E-01 | 1.06E+00 | 2.30E-01 | 8.77E-01 | 1.22E+00 | 9.20E-02 | 3.28E-01 | -1.11E+00 | 1.72E-01 | 4.84E-01 | 1.00E+00 | 1.92E-01 | 7.06E-01 | 1.46E+00 | 3.55E-01 | 9.75E-01 | 1.33E+00 |
| ARID5A | 1.02E-01 | 5.32E-01 | 1.22E+00 | 1.48E-06 | 6.39E-06 | -1.28E+00 | 1.09E-01 | 4.65E-01 | 1.17E+00 | 1.17E-03 | 4.48E-03 | -1.04E+00 | 1.26E-02 | 4.48E-02 | 1.03E+00 | 4.63E-04 | 1.30E-03 | -1.58E+00 | 1.49E-02 | 5.47E-02 | -1.01E+00 | 1.56E-03 | 4.30E-03 | -1.28E+00 |
| RUNDC3B | 1.02E-01 | 5.32E-01 | -1.02E+00 | 9.42E-03 | 4.07E-02 | 2.08E+00 | 1.73E-01 | 7.34E-01 | -1.08E+00 | 1.03E-01 | 3.93E-01 | 1.48E+00 | 1.93E-03 | 6.89E-03 | -3.19E+00 | 7.62E-02 | 2.15E-01 | -1.21E+00 | 1.37E-01 | 5.01E-01 | -1.29E+00 | 5.59E-02 | 1.54E-01 | 1.89E+00 |
| ZWINT | 1.02E-01 | 5.32E-01 | 1.08E+00 | 5.31E-02 | 2.29E-01 | -1.12E+00 | 2.04E-04 | 8.65E-04 | 1.57E+00 | 6.19E-02 | 2.36E-01 | 1.51E+00 | 1.04E-01 | 3.69E-01 | -1.30E+00 | 1.61E-01 | 4.55E-01 | -1.24E+00 | 1.75E-01 | 6.42E-01 | 1.02E+00 | 8.71E-05 | 2.39E-04 | 1.29E+00 |
| FOXM1 | 1.02E-01 | 5.33E-01 | -1.11E+00 | 1.79E-01 | 7.72E-01 | 1.04E+00 | 1.92E-02 | 8.14E-02 | 1.74E+00 | 7.06E-02 | 2.70E-01 | 1.79E+00 | 1.24E-02 | 4.41E-02 | -2.27E+00 | 8.29E-02 | 2.34E-01 | -1.41E+00 | 1.80E-01 | 6.61E-01 | -1.02E+00 | 2.34E-01 | 6.42E-01 | -1.06E+00 |
| AC090373.1 | 1.02E-01 | 5.33E-01 | -1.07E+00 | 1.62E-01 | 7.00E-01 | 1.14E+00 | 2.84E-02 | 1.21E-01 | -1.41E+00 | 1.45E-01 | 5.54E-01 | 1.11E+00 | 2.79E-01 | 9.94E-01 | 1.27E+00 | 3.52E-01 | 9.91E-01 | 1.19E+00 | 2.09E-01 | 7.68E-01 | 1.07E+00 | 1.32E-01 | 3.62E-01 | 1.63E+00 |
| CR391992.1 | 1.02E-01 | 5.33E-01 | 1.11E+00 | 5.35E-02 | 2.31E-01 | 1.07E+00 | 1.21E-02 | 5.16E-02 | -1.04E+00 | 3.17E-05 | 1.21E-04 | -1.27E+00 | 4.06E-03 | 1.45E-02 | -1.18E+00 | 1.99E-02 | 5.61E-02 | -1.01E+00 | 2.46E-01 | 9.01E-01 | 1.21E+00 | 1.22E-01 | 3.36E-01 | 1.09E+00 |
| IL2RG | 1.02E-01 | 5.33E-01 | 1.14E+00 | 7.51E-04 | 3.24E-03 | 1.07E+00 | 4.05E-02 | 1.72E-01 | -1.14E+00 | 2.61E-04 | 9.98E-04 | -1.21E+00 | 5.90E-02 | 2.10E-01 | 1.02E+00 | 1.01E-01 | 2.85E-01 | 1.09E+00 | 1.11E-01 | 4.09E-01 | 1.16E+00 | 1.28E-01 | 3.52E-01 | -1.02E+00 |
| SH2D4A | 1.02E-01 | 5.33E-01 | 1.28E+00 | 2.01E-01 | 8.69E-01 | 1.27E+00 | 2.27E-01 | 9.63E-01 | 1.26E+00 | 1.17E-01 | 4.46E-01 | 1.20E+00 | 2.68E-01 | 9.56E-01 | 1.03E+00 | 3.14E-01 | 8.85E-01 | -1.11E+00 | 4.38E-02 | 1.61E-01 | -1.42E+00 | 1.07E-01 | 2.93E-01 | -1.21E+00 |
| AP000344.1 | 1.02E-01 | 5.33E-01 | 1.20E+00 | 4.07E-03 | 1.76E-02 | -1.17E+00 | 2.87E-03 | 1.22E-02 | -1.07E+00 | 3.03E-03 | 1.16E-02 | -1.02E+00 | 2.70E-01 | 9.64E-01 | 1.23E+00 | 3.02E-01 | 8.52E-01 | 1.05E+00 | 1.96E-01 | 7.20E-01 | 1.02E+00 | 1.99E-01 | 5.46E-01 | -1.03E+00 |
| TFEB | 1.02E-01 | 5.33E-01 | 1.06E+00 | 1.10E-01 | 4.75E-01 | 1.70E+00 | 1.21E-01 | 5.14E-01 | 1.08E+00 | 2.22E-01 | 8.48E-01 | 1.29E+00 | 2.75E-01 | 9.79E-01 | 1.04E+00 | 2.88E-01 | 8.12E-01 | 1.27E+00 | 1.04E-01 | 3.81E-01 | 1.74E+00 | 1.91E-01 | 5.24E-01 | 1.58E+00 |
| SLC44A2 | 1.02E-01 | 5.33E-01 | 1.13E+00 | 5.07E-03 | 2.19E-02 | 1.02E+00 | 8.10E-02 | 3.44E-01 | 1.48E+00 | 1.24E-01 | 4.75E-01 | 1.24E+00 | 1.43E-02 | 5.10E-02 | 1.05E+00 | 1.05E-02 | 2.97E-02 | -1.01E+00 | 6.53E-03 | 2.40E-02 | 1.06E+00 | 4.73E-04 | 1.30E-03 | -1.08E+00 |
| GOLGA2P7 | 1.02E-01 | 5.33E-01 | 1.04E+00 | 3.20E-02 | 1.38E-01 | -1.11E+00 | 1.55E-03 | 6.60E-03 | -1.31E+00 | 3.57E-04 | 1.36E-03 | -1.22E+00 | 4.12E-02 | 1.47E-01 | -1.10E+00 | 7.77E-02 | 2.19E-01 | -1.03E+00 | 3.04E-04 | 1.12E-03 | -1.59E+00 | 3.52E-04 | 9.66E-04 | -1.52E+00 |
| MAP3K3 | 1.02E-01 | 5.33E-01 | 1.24E+00 | 8.93E-02 | 3.86E-01 | 1.35E+00 | 1.40E-02 | 5.96E-02 | -1.09E+00 | 7.17E-04 | 2.74E-03 | -1.21E+00 | 2.55E-01 | 9.11E-01 | 1.43E+00 | 2.60E-01 | 7.32E-01 | 1.18E+00 | 2.18E-04 | 8.01E-04 | -1.25E+00 | 1.12E-03 | 3.09E-03 | -1.28E+00 |
| ACVR2A | 1.02E-01 | 5.33E-01 | 1.21E+00 | 1.78E-01 | 7.71E-01 | 1.39E+00 | 2.27E-01 | 9.64E-01 | 1.26E+00 | 1.88E-01 | 7.18E-01 | 1.36E+00 | 2.12E-01 | 7.56E-01 | 1.08E+00 | 2.63E-01 | 7.42E-01 | 1.15E+00 | 1.53E-01 | 5.63E-01 | 1.52E+00 | 2.68E-01 | 7.37E-01 | 1.22E+00 |
| MEF2A | 1.03E-01 | 5.35E-01 | 1.26E+00 | 4.17E-05 | 1.80E-04 | -1.12E+00 | 3.48E-04 | 1.48E-03 | -1.17E+00 | 2.84E-08 | 1.09E-07 | -1.33E+00 | 8.13E-03 | 2.90E-02 | 1.06E+00 | 1.19E-05 | 3.34E-05 | -1.47E+00 | 1.95E-03 | 7.15E-03 | -1.13E+00 | 1.37E-05 | 3.78E-05 | -1.52E+00 |
| AC007342.7 | 1.03E-01 | 5.35E-01 | 1.60E+00 | 6.81E-02 | 2.94E-01 | 1.85E+00 | 1.98E-02 | 8.42E-02 | -1.40E+00 | 1.92E-02 | 7.35E-02 | -1.24E+00 | 2.33E-01 | 8.29E-01 | 1.28E+00 | 3.10E-01 | 8.74E-01 | 1.32E+00 | 3.27E-02 | 1.20E-01 | 1.78E+00 | 3.50E-01 | 9.61E-01 | 1.15E+00 |
| MS4A7 | 1.03E-01 | 5.36E-01 | 1.52E+00 | 2.12E-01 | 9.14E-01 | 1.50E+00 | 1.21E-01 | 5.16E-01 | -1.08E+00 | 4.93E-02 | 1.88E-01 | 1.03E+00 | 1.11E-01 | 3.96E-01 | -1.11E+00 | 1.17E-01 | 3.29E-01 | -1.18E+00 | 2.28E-01 | 8.38E-01 | 1.12E+00 | 2.07E-01 | 5.69E-01 | -1.11E+00 |
| RPL7P18 | 1.03E-01 | 5.36E-01 | -1.14E+00 | 2.26E-01 | 9.77E-01 | 1.20E+00 | 1.04E-01 | 4.42E-01 | -1.08E+00 | 9.09E-02 | 3.47E-01 | 1.59E+00 | 1.41E-01 | 5.04E-01 | 1.53E+00 | 2.49E-01 | 7.03E-01 | 1.38E+00 | 2.26E-01 | 8.28E-01 | 1.20E+00 | 3.62E-01 | 9.94E-01 | 1.08E+00 |
| MICAL2 | 1.03E-01 | 5.36E-01 | 1.06E+00 | 8.84E-02 | 3.82E-01 | 1.03E+00 | 1.95E-01 | 8.27E-01 | 1.06E+00 | 1.67E-01 | 6.38E-01 | 1.19E+00 | 7.81E-03 | 2.79E-02 | 1.18E+00 | 2.14E-01 | 6.03E-01 | 1.57E+00 | 1.31E-07 | 4.81E-07 | -1.23E+00 | 4.98E-08 | 1.37E-07 | -1.25E+00 |
| GPR89B | 1.03E-01 | 5.36E-01 | 1.05E+00 | 1.35E-01 | 5.82E-01 | 1.48E+00 | 1.40E-01 | 5.96E-01 | 1.45E+00 | 2.61E-02 | 9.95E-02 | 1.99E+00 | 1.65E-01 | 5.90E-01 | 1.24E+00 | 1.97E-01 | 5.56E-01 | 1.29E+00 | 6.01E-04 | 2.21E-03 | -1.21E+00 | 9.94E-02 | 2.73E-01 | 1.17E+00 |
| RPL12P38 | 1.03E-01 | 5.36E-01 | -1.03E+00 | 3.80E-38 | 1.64E-37 | -1.67E+00 | 1.56E-01 | 6.63E-01 | -1.00E+00 | 5.02E-16 | 1.92E-15 | -1.56E+00 | 1.15E-01 | 4.09E-01 | -1.02E+00 | 2.20E-14 | 6.21E-14 | -1.49E+00 | 1.24E-01 | 4.54E-01 | 1.02E+00 | 7.13E-14 | 1.96E-13 | -1.56E+00 |
| TCP1P1 | 1.03E-01 | 5.36E-01 | 1.14E+00 | 1.44E-01 | 6.22E-01 | 1.71E+00 | 1.98E-01 | 8.40E-01 | 1.17E+00 | 2.88E-02 | 1.10E-01 | 1.83E+00 | 1.48E-01 | 5.28E-01 | 1.12E+00 | 1.47E-01 | 4.14E-01 | 1.76E+00 | 1.26E-01 | 4.62E-01 | 1.16E+00 | 4.72E-04 | 1.30E-03 | 2.18E+00 |
| GABPB1 | 1.03E-01 | 5.36E-01 | 1.21E+00 | 1.02E-01 | 4.41E-01 | 1.45E+00 | 2.03E-01 | 8.62E-01 | 1.06E+00 | 2.16E-01 | 8.25E-01 | 1.16E+00 | 1.79E-02 | 6.37E-02 | 1.15E+00 | 3.55E-01 | 1.00E+00 | 1.29E+00 | 4.34E-03 | 1.59E-02 | -1.13E+00 | 1.00E-02 | 2.75E-02 | -1.07E+00 |
| BECN1 | 1.03E-01 | 5.36E-01 | 1.07E+00 | 2.03E-01 | 8.77E-01 | 1.37E+00 | 1.79E-04 | 7.61E-04 | -1.41E+00 | 2.52E-03 | 9.64E-03 | -1.06E+00 | 5.08E-05 | 1.81E-04 | -1.33E+00 | 8.35E-05 | 2.35E-04 | -1.25E+00 | 1.13E-03 | 4.17E-03 | -1.17E+00 | 1.16E-01 | 3.20E-01 | 1.11E+00 |
| SLC24A3 | 1.03E-01 | 5.37E-01 | -1.01E+00 | 1.30E-01 | 5.61E-01 | 1.06E+00 | 1.98E-01 | 8.41E-01 | 1.28E+00 | 5.63E-02 | 2.15E-01 | 1.89E+00 | 3.26E-02 | 1.16E-01 | -1.14E+00 | 9.10E-03 | 2.57E-02 | -1.26E+00 | 7.01E-02 | 2.57E-01 | 1.65E+00 | 1.11E-01 | 3.05E-01 | 1.44E+00 |
| RPL37AP1 | 1.03E-01 | 5.37E-01 | 1.29E+00 | 7.23E-22 | 3.13E-21 | -1.30E+00 | 1.30E-01 | 5.54E-01 | 1.26E+00 | 4.52E-10 | 1.73E-09 | -1.24E+00 | 1.26E-01 | 4.50E-01 | 1.22E+00 | 1.69E-07 | 4.77E-07 | -1.26E+00 | 1.11E-01 | 4.09E-01 | 1.11E+00 | 1.56E-09 | 4.27E-09 | -1.38E+00 |
| CIT | 1.03E-01 | 5.38E-01 | -1.16E+00 | 1.17E-01 | 5.04E-01 | -1.16E+00 | 3.31E-02 | 1.40E-01 | 1.46E+00 | 1.56E-01 | 5.94E-01 | 1.56E+00 | 2.73E-01 | 9.75E-01 | 1.17E+00 | 3.09E-01 | 8.70E-01 | 1.50E+00 | 1.51E-02 | 5.56E-02 | 1.33E+00 | 4.75E-04 | 1.30E-03 | 1.61E+00 |
| AC006387.1 | 1.03E-01 | 5.38E-01 | 1.06E+00 | 5.24E-02 | 2.26E-01 | -1.11E+00 | 1.67E-01 | 7.11E-01 | 1.04E+00 | 2.22E-01 | 8.49E-01 | 1.24E+00 | 1.80E-01 | 6.41E-01 | 1.51E+00 | 1.24E-07 | 3.50E-07 | 4.16E+00 | 6.61E-03 | 2.43E-02 | 1.71E+00 | 2.13E-07 | 5.84E-07 | 2.14E+00 |
| NAP1L4P3 | 1.03E-01 | 5.38E-01 | 1.22E+00 | 1.29E-01 | 5.59E-01 | 1.77E+00 | 6.78E-02 | 2.88E-01 | 1.05E+00 | 4.98E-02 | 1.90E-01 | 1.60E+00 | 2.48E-01 | 8.85E-01 | 1.42E+00 | 2.14E-01 | 6.02E-01 | 1.92E+00 | 2.44E-01 | 8.96E-01 | 1.25E+00 | 8.67E-03 | 2.38E-02 | 2.10E+00 |
| ABHD12 | 1.04E-01 | 5.40E-01 | 1.21E+00 | 5.82E-03 | 2.51E-02 | 1.02E+00 | 1.76E-01 | 7.49E-01 | 1.14E+00 | 3.25E-03 | 1.24E-02 | 1.07E+00 | 1.35E-01 | 4.80E-01 | 1.03E+00 | 4.71E-02 | 1.33E-01 | -1.17E+00 | 2.77E-03 | 1.02E-02 | -1.10E+00 | 1.30E-01 | 3.56E-01 | 1.05E+00 |
| BBOF1 | 1.04E-01 | 5.40E-01 | 1.43E+00 | 8.59E-02 | 3.71E-01 | 1.77E+00 | 4.88E-02 | 2.07E-01 | -1.09E+00 | 7.15E-04 | 2.73E-03 | -1.54E+00 | 2.12E-01 | 7.55E-01 | 1.11E+00 | 3.49E-01 | 9.84E-01 | 1.44E+00 | 1.55E-01 | 5.68E-01 | 1.35E+00 | 2.66E-01 | 7.32E-01 | 1.05E+00 |
| HINT1P1 | 1.04E-01 | 5.40E-01 | 1.31E+00 | 1.42E-06 | 6.16E-06 | 2.62E+00 | 5.64E-02 | 2.39E-01 | 1.53E+00 | 7.32E-10 | 2.80E-09 | 3.00E+00 | 2.44E-01 | 8.70E-01 | 1.30E+00 | 3.36E-04 | 9.48E-04 | 2.62E+00 | 1.43E-01 | 5.24E-01 | 1.58E+00 | 2.61E-06 | 7.16E-06 | 2.82E+00 |
| CD86 | 1.04E-01 | 5.40E-01 | -1.11E+00 | 1.34E-01 | 5.81E-01 | -1.06E+00 | 3.34E-04 | 1.42E-03 | -1.60E+00 | 6.67E-06 | 2.55E-05 | -1.92E+00 | 2.80E-01 | 1.00E+00 | 1.00E+00 | 3.55E-01 | 1.00E+00 | 1.00E+00 | 1.43E-01 | 5.23E-01 | -1.15E+00 | 1.68E-01 | 4.61E-01 | -1.19E+00 |
| AC100788.2 | 1.04E-01 | 5.41E-01 | 1.13E+00 | 4.48E-02 | 1.94E-01 | -1.03E+00 | 6.60E-02 | 2.81E-01 | -1.01E+00 | 2.39E-02 | 9.13E-02 | -1.08E+00 | 1.07E-01 | 3.82E-01 | 1.23E+00 | 2.04E-02 | 5.75E-02 | -1.29E+00 | 1.40E-01 | 5.13E-01 | -1.20E+00 | 1.36E-01 | 3.73E-01 | -1.40E+00 |
| HNRNPCP2 | 1.04E-01 | 5.41E-01 | 1.22E+00 | 1.56E-01 | 6.72E-01 | 1.44E+00 | 1.47E-02 | 6.25E-02 | 1.50E+00 | 3.38E-03 | 1.29E-02 | 1.78E+00 | 2.17E-01 | 7.75E-01 | 1.35E+00 | 2.96E-01 | 8.36E-01 | 1.48E+00 | 5.50E-02 | 2.02E-01 | 1.48E+00 | 5.48E-03 | 1.51E-02 | 1.70E+00 |
| AC002075.2 | 1.04E-01 | 5.41E-01 | 1.10E+00 | 4.27E-18 | 1.85E-17 | -1.31E+00 | 5.09E-02 | 2.16E-01 | 1.12E+00 | 2.15E-08 | 8.20E-08 | -1.29E+00 | 6.31E-02 | 2.25E-01 | 1.15E+00 | 8.14E-06 | 2.30E-05 | -1.21E+00 | 2.11E-01 | 7.76E-01 | 1.15E+00 | 1.76E-06 | 4.84E-06 | -1.25E+00 |
| ZNF230 | 1.04E-01 | 5.42E-01 | 1.03E+00 | 2.31E-01 | 1.00E+00 | 1.30E+00 | 8.22E-02 | 3.49E-01 | 1.05E+00 | 1.72E-03 | 6.57E-03 | -1.08E+00 | 2.71E-01 | 9.65E-01 | 1.47E+00 | 1.59E-01 | 4.49E-01 | -1.10E+00 | 5.54E-04 | 2.03E-03 | -1.85E+00 | 1.25E-02 | 3.43E-02 | -1.40E+00 |
| GSTM2 | 1.04E-01 | 5.42E-01 | 1.55E+00 | 2.07E-01 | 8.96E-01 | 1.60E+00 | 1.61E-02 | 6.86E-02 | 1.76E+00 | 3.07E-02 | 1.17E-01 | 2.02E+00 | 6.68E-05 | 2.38E-04 | 2.61E+00 | 4.66E-03 | 1.31E-02 | 2.27E+00 | 2.29E-03 | 8.40E-03 | 2.12E+00 | 1.63E-02 | 4.47E-02 | 1.95E+00 |
| AC112496.1 | 1.04E-01 | 5.42E-01 | -1.04E+00 | 1.11E-01 | 4.80E-01 | 1.17E+00 | 4.27E-03 | 1.82E-02 | -1.57E+00 | 1.68E-02 | 6.42E-02 | -1.24E+00 | 1.65E-01 | 5.87E-01 | 1.50E+00 | 1.95E-01 | 5.49E-01 | 1.59E+00 | 1.04E-02 | 3.83E-02 | -1.35E+00 | 5.54E-02 | 1.52E-01 | -1.04E+00 |
| AC020915.4 | 1.04E-01 | 5.42E-01 | 1.05E+00 | 1.14E-01 | 4.92E-01 | 1.11E+00 | 4.33E-02 | 1.84E-01 | -1.13E+00 | 3.23E-03 | 1.23E-02 | -1.40E+00 | 2.41E-01 | 8.58E-01 | 1.50E+00 | 1.26E-01 | 3.56E-01 | 1.97E+00 | 2.64E-02 | 9.68E-02 | -1.07E+00 | 5.63E-04 | 1.55E-03 | -1.58E+00 |
| TBXA2R | 1.04E-01 | 5.42E-01 | 1.17E+00 | 1.58E-01 | 6.84E-01 | 1.67E+00 | 1.16E-02 | 4.95E-02 | -1.43E+00 | 2.18E-01 | 8.34E-01 | 1.32E+00 | 8.16E-02 | 2.91E-01 | 3.37E+00 | 1.09E-01 | 3.08E-01 | 3.26E+00 | 1.96E-02 | 7.19E-02 | 1.08E+00 | 1.33E-01 | 3.67E-01 | -1.01E+00 |
| RHEBP1 | 1.04E-01 | 5.42E-01 | 1.05E+00 | 1.31E-01 | 5.67E-01 | 1.15E+00 | 1.36E-01 | 5.76E-01 | -1.02E+00 | 1.13E-02 | 4.32E-02 | -1.21E+00 | 1.57E-03 | 5.61E-03 | -1.31E+00 | 1.18E-04 | 3.32E-04 | -1.69E+00 | 1.35E-05 | 4.97E-05 | -1.65E+00 | 9.24E-05 | 2.54E-04 | -1.73E+00 |
| GCAT | 1.04E-01 | 5.43E-01 | 1.44E+00 | 9.82E-02 | 4.24E-01 | 1.30E+00 | 1.64E-01 | 6.97E-01 | 1.33E+00 | 5.19E-02 | 1.98E-01 | 1.22E+00 | 2.08E-01 | 7.40E-01 | 1.55E+00 | 4.59E-02 | 1.29E-01 | 2.03E+00 | 4.04E-02 | 1.48E-01 | 1.59E+00 | 1.47E-02 | 4.05E-02 | 1.72E+00 |
| PIMREG | 1.04E-01 | 5.43E-01 | -1.22E+00 | 1.52E-01 | 6.59E-01 | -1.08E+00 | 4.96E-03 | 2.11E-02 | 1.56E+00 | 1.87E-01 | 7.16E-01 | 1.02E+00 | 2.14E-01 | 7.64E-01 | -1.15E+00 | 2.61E-01 | 7.37E-01 | -1.23E+00 | 5.28E-04 | 1.94E-03 | -1.75E+00 | 7.88E-03 | 2.16E-02 | -1.61E+00 |
| DCP1B | 1.04E-01 | 5.44E-01 | 1.07E+00 | 1.14E-01 | 4.92E-01 | 1.58E+00 | 2.69E-11 | 1.14E-10 | -2.02E+00 | 5.65E-06 | 2.16E-05 | -1.19E+00 | 8.48E-05 | 3.02E-04 | -2.05E+00 | 5.49E-02 | 1.55E-01 | 1.09E+00 | 1.21E-02 | 4.43E-02 | -1.31E+00 | 2.54E-01 | 6.99E-01 | 1.32E+00 |
| YWHAZP4 | 1.04E-01 | 5.44E-01 | 1.22E+00 | 8.23E-04 | 3.56E-03 | -1.42E+00 | 2.17E-01 | 9.23E-01 | 1.16E+00 | 5.84E-02 | 2.23E-01 | -1.02E+00 | 2.07E-02 | 7.39E-02 | 1.29E+00 | 4.64E-05 | 1.31E-04 | -1.36E+00 | 1.06E-02 | 3.90E-02 | 1.56E+00 | 4.65E-02 | 1.28E-01 | -1.21E+00 |
| AL359922.3 | 1.04E-01 | 5.44E-01 | 1.16E+00 | 1.44E-01 | 6.24E-01 | 1.65E+00 | 1.56E-01 | 6.63E-01 | 1.06E+00 | 1.74E-03 | 6.63E-03 | 2.22E+00 | 6.69E-02 | 2.38E-01 | 1.15E+00 | 2.26E-01 | 6.38E-01 | 1.36E+00 | 2.70E-01 | 9.92E-01 | 1.29E+00 | 4.00E-03 | 1.10E-02 | 1.98E+00 |
| SRP72P2 | 1.04E-01 | 5.44E-01 | 1.55E+00 | 1.62E-01 | 7.02E-01 | 1.67E+00 | 7.86E-03 | 3.34E-02 | 2.11E+00 | 2.13E-02 | 8.15E-02 | 2.20E+00 | 4.99E-02 | 1.78E-01 | 2.29E+00 | 9.19E-02 | 2.59E-01 | 2.14E+00 | 8.07E-02 | 2.96E-01 | 1.40E+00 | 6.44E-02 | 1.77E-01 | 1.53E+00 |
| ZNF274 | 1.04E-01 | 5.44E-01 | 1.21E+00 | 1.78E-01 | 7.68E-01 | 1.41E+00 | 1.01E-02 | 4.29E-02 | -1.06E+00 | 5.07E-02 | 1.94E-01 | 1.15E+00 | 8.15E-03 | 2.91E-02 | 1.17E+00 | 1.70E-01 | 4.78E-01 | 1.30E+00 | 1.31E-03 | 4.81E-03 | -1.11E+00 | 1.07E-01 | 2.94E-01 | 1.13E+00 |
| AC128707.1 | 1.04E-01 | 5.44E-01 | 1.50E+00 | 7.30E-02 | 3.15E-01 | 1.80E+00 | 3.51E-02 | 1.49E-01 | -1.16E+00 | 1.62E-02 | 6.18E-02 | -1.21E+00 | 3.73E-02 | 1.33E-01 | -2.22E+00 | 6.69E-02 | 1.89E-01 | -2.02E+00 | 9.46E-02 | 3.47E-01 | -1.16E+00 | 2.07E-02 | 5.68E-02 | -1.59E+00 |
| EIF2AK3 | 1.04E-01 | 5.44E-01 | 1.02E+00 | 1.96E-01 | 8.46E-01 | 1.22E+00 | 4.33E-02 | 1.84E-01 | -1.09E+00 | 2.22E-02 | 8.47E-02 | -1.02E+00 | 1.82E-01 | 6.47E-01 | 1.13E+00 | 1.66E-01 | 4.68E-01 | -1.02E+00 | 3.33E-03 | 1.22E-02 | -1.58E+00 | 3.41E-02 | 9.38E-02 | -1.26E+00 |
| MS4A4E | 1.04E-01 | 5.44E-01 | 1.64E+00 | 2.25E-01 | 9.74E-01 | 1.26E+00 | 1.68E-01 | 7.13E-01 | 1.44E+00 | 1.04E-01 | 3.97E-01 | 2.11E+00 | 1.41E-15 | 5.03E-15 | 3.76E+00 | 6.90E-12 | 1.94E-11 | 3.23E+00 | 7.06E-14 | 2.59E-13 | 6.76E+00 | 2.61E-05 | 7.16E-05 | 4.30E+00 |
| KIAA0825 | 1.04E-01 | 5.44E-01 | -1.03E+00 | 9.51E-02 | 4.11E-01 | 1.01E+00 | 1.42E-02 | 6.01E-02 | -1.27E+00 | 4.50E-02 | 1.72E-01 | -1.04E+00 | 2.56E-01 | 9.14E-01 | 1.47E+00 | 2.92E-01 | 8.23E-01 | 1.52E+00 | 1.78E-03 | 6.53E-03 | -1.69E+00 | 1.71E-03 | 4.71E-03 | -1.88E+00 |
| UBXN10-AS1 | 1.05E-01 | 5.45E-01 | -1.01E+00 | 6.92E-02 | 2.99E-01 | -1.16E+00 | 5.08E-02 | 2.16E-01 | 1.50E+00 | 2.01E-01 | 7.69E-01 | -1.01E+00 | 1.03E-01 | 3.67E-01 | -1.41E+00 | 1.54E-01 | 4.34E-01 | -1.31E+00 | 9.48E-04 | 3.48E-03 | 1.77E+00 | 3.76E-03 | 1.03E-02 | 1.90E+00 |
| RUSC1 | 1.05E-01 | 5.45E-01 | 1.20E+00 | 5.35E-02 | 2.31E-01 | 1.86E+00 | 8.40E-04 | 3.57E-03 | -1.17E+00 | 5.88E-02 | 2.24E-01 | 1.21E+00 | 7.63E-03 | 2.72E-02 | -1.19E+00 | 3.67E-02 | 1.03E-01 | -1.01E+00 | 2.66E-03 | 9.77E-03 | -1.14E+00 | 5.95E-02 | 1.63E-01 | 1.03E+00 |
| OFD1 | 1.05E-01 | 5.45E-01 | 1.25E+00 | 2.31E-01 | 1.00E+00 | 1.54E+00 | 1.33E-01 | 5.66E-01 | 1.14E+00 | 2.24E-01 | 8.56E-01 | 1.40E+00 | 5.57E-03 | 1.98E-02 | 1.07E+00 | 1.30E-02 | 3.67E-02 | 1.12E+00 | 1.47E-01 | 5.41E-01 | 1.16E+00 | 1.08E-01 | 2.98E-01 | 1.13E+00 |
| HNRNPA1P28 | 1.05E-01 | 5.45E-01 | 1.42E+00 | 2.19E-01 | 9.47E-01 | 1.34E+00 | 8.32E-02 | 3.54E-01 | -1.08E+00 | 5.69E-02 | 2.18E-01 | -1.06E+00 | 1.63E-01 | 5.81E-01 | -1.01E+00 | 3.53E-01 | 9.96E-01 | 1.40E+00 | 9.34E-02 | 3.43E-01 | -1.04E+00 | 1.48E-01 | 4.07E-01 | -1.06E+00 |
| LDHBP2 | 1.05E-01 | 5.46E-01 | 1.08E+00 | 1.22E-03 | 5.28E-03 | 2.56E+00 | 3.22E-02 | 1.37E-01 | -1.07E+00 | 4.34E-02 | 1.66E-01 | 1.85E+00 | 1.70E-02 | 6.07E-02 | -1.42E+00 | 9.08E-02 | 2.56E-01 | 1.87E+00 | 2.07E-01 | 7.60E-01 | 1.21E+00 | 2.48E-09 | 6.81E-09 | 2.74E+00 |
| METTL17 | 1.05E-01 | 5.47E-01 | 1.25E+00 | 2.06E-01 | 8.90E-01 | 1.52E+00 | 1.21E-02 | 5.12E-02 | 1.04E+00 | 2.09E-01 | 7.98E-01 | 1.45E+00 | 6.66E-02 | 2.37E-01 | 1.15E+00 | 2.42E-01 | 6.84E-01 | 1.63E+00 | 3.59E-04 | 1.32E-03 | -1.07E+00 | 1.19E-01 | 3.26E-01 | 1.17E+00 |
| IKZF2 | 1.05E-01 | 5.47E-01 | 1.15E+00 | 2.19E-01 | 9.44E-01 | 1.37E+00 | 6.81E-06 | 2.89E-05 | -1.47E+00 | 1.89E-02 | 7.20E-02 | 1.03E+00 | 7.01E-06 | 2.50E-05 | -1.27E+00 | 3.96E-05 | 1.12E-04 | -1.26E+00 | 2.32E-04 | 8.52E-04 | -1.89E+00 | 1.28E-02 | 3.52E-02 | -1.27E+00 |
| MAP3K20-AS1 | 1.05E-01 | 5.48E-01 | 1.02E+00 | 1.45E-01 | 6.28E-01 | 1.22E+00 | 1.15E-01 | 4.88E-01 | -1.11E+00 | 9.63E-02 | 3.68E-01 | -1.02E+00 | 1.98E-02 | 7.07E-02 | -1.61E+00 | 5.10E-02 | 1.44E-01 | -1.29E+00 | 1.14E-01 | 4.19E-01 | -1.23E+00 | 2.99E-01 | 8.21E-01 | 1.06E+00 |
| KIF9-AS1 | 1.05E-01 | 5.48E-01 | 1.06E+00 | 1.77E-01 | 7.65E-01 | 1.32E+00 | 4.30E-02 | 1.83E-01 | -1.20E+00 | 2.88E-02 | 1.10E-01 | -1.13E+00 | 1.88E-04 | 6.72E-04 | -1.61E+00 | 2.33E-04 | 6.57E-04 | -1.62E+00 | 1.71E-03 | 6.28E-03 | -1.52E+00 | 1.24E-03 | 3.39E-03 | -1.58E+00 |
| IFRD1 | 1.05E-01 | 5.49E-01 | 1.26E+00 | 1.22E-11 | 5.27E-11 | -1.28E+00 | 1.48E-01 | 6.29E-01 | 1.09E+00 | 3.91E-07 | 1.49E-06 | -1.35E+00 | 1.17E-03 | 4.19E-03 | 1.12E+00 | 7.29E-09 | 2.06E-08 | -1.34E+00 | 1.30E-02 | 4.77E-02 | 1.12E+00 | 2.74E-07 | 7.51E-07 | -1.36E+00 |
| AKAP5 | 1.05E-01 | 5.49E-01 | 1.08E+00 | 9.30E-02 | 4.02E-01 | 1.02E+00 | 2.62E-02 | 1.11E-01 | -1.24E+00 | 1.50E-01 | 5.71E-01 | 1.18E+00 | 2.15E-03 | 7.68E-03 | -1.59E+00 | 8.81E-03 | 2.48E-02 | -1.40E+00 | 6.20E-02 | 2.27E-01 | -1.12E+00 | 2.79E-01 | 7.66E-01 | 1.13E+00 |
| AC072061.1 | 1.05E-01 | 5.49E-01 | 1.18E+00 | 2.29E-02 | 9.89E-02 | -1.05E+00 | 1.81E-01 | 7.70E-01 | 1.27E+00 | 1.40E-02 | 5.33E-02 | -1.35E+00 | 1.56E-01 | 5.57E-01 | 1.02E+00 | 1.79E-01 | 5.06E-01 | -1.03E+00 | 3.56E-02 | 1.31E-01 | 2.32E+00 | 1.67E-01 | 4.60E-01 | 1.73E+00 |
| RPL23AP18 | 1.05E-01 | 5.49E-01 | 1.25E+00 | 2.29E-08 | 9.88E-08 | -1.72E+00 | 2.86E-03 | 1.21E-02 | 1.63E+00 | 8.97E-06 | 3.43E-05 | -1.44E+00 | 2.19E-02 | 7.82E-02 | 2.22E+00 | 2.46E-01 | 6.93E-01 | 1.26E+00 | 1.57E-01 | 5.76E-01 | 1.22E+00 | 6.19E-04 | 1.70E-03 | -1.40E+00 |
| OLA1P1 | 1.06E-01 | 5.50E-01 | 1.54E+00 | 1.20E-01 | 5.19E-01 | 1.75E+00 | 2.32E-03 | 9.86E-03 | 1.97E+00 | 9.17E-04 | 3.50E-03 | 2.24E+00 | 7.00E-03 | 2.50E-02 | 2.97E+00 | 1.38E-02 | 3.88E-02 | 2.81E+00 | 2.45E-01 | 9.00E-01 | 1.39E+00 | 3.01E-01 | 8.26E-01 | 1.41E+00 |
| C9orf64 | 1.06E-01 | 5.50E-01 | 1.07E+00 | 9.31E-02 | 4.02E-01 | 1.81E+00 | 3.58E-03 | 1.52E-02 | -1.38E+00 | 2.22E-02 | 8.47E-02 | -1.01E+00 | 2.44E-04 | 8.71E-04 | -1.98E+00 | 8.22E-02 | 2.32E-01 | 1.01E+00 | 1.36E-04 | 5.00E-04 | -1.28E+00 | 5.75E-02 | 1.58E-01 | 1.03E+00 |
| AC108161.1 | 1.06E-01 | 5.50E-01 | 1.51E+00 | 1.93E-02 | 8.34E-02 | -1.02E+00 | 1.45E-01 | 6.16E-01 | 1.18E+00 | 5.54E-03 | 2.12E-02 | -1.08E+00 | 4.94E-02 | 1.76E-01 | -1.12E+00 | 4.73E-02 | 1.33E-01 | -1.15E+00 | 7.83E-02 | 2.88E-01 | 1.03E+00 | 1.65E-04 | 4.54E-04 | -1.76E+00 |
| WDR74 | 1.06E-01 | 5.50E-01 | 1.38E+00 | 2.53E-03 | 1.09E-02 | -1.26E+00 | 2.04E-04 | 8.66E-04 | -1.42E+00 | 5.44E-08 | 2.08E-07 | -2.01E+00 | 4.62E-02 | 1.65E-01 | -1.11E+00 | 5.59E-02 | 1.58E-01 | -1.20E+00 | 2.84E-04 | 1.04E-03 | -1.58E+00 | 2.81E-04 | 7.71E-04 | -1.59E+00 |
| LRRC8B | 1.06E-01 | 5.50E-01 | 1.23E+00 | 6.12E-03 | 2.64E-02 | 1.06E+00 | 1.28E-02 | 5.43E-02 | -1.04E+00 | 8.11E-04 | 3.10E-03 | -1.01E+00 | 5.39E-02 | 1.92E-01 | 1.18E+00 | 1.01E-04 | 2.86E-04 | -1.49E+00 | 9.96E-05 | 3.66E-04 | 1.01E+00 | 1.57E-06 | 4.32E-06 | -1.25E+00 |
| CCDC120 | 1.06E-01 | 5.52E-01 | 1.48E+00 | 1.64E-01 | 7.08E-01 | 1.51E+00 | 1.58E-01 | 6.69E-01 | 1.13E+00 | 1.33E-01 | 5.07E-01 | 1.29E+00 | 1.68E-01 | 5.99E-01 | 1.70E+00 | 4.18E-02 | 1.18E-01 | 2.42E+00 | 5.32E-02 | 1.95E-01 | -1.08E+00 | 1.31E-01 | 3.59E-01 | 1.04E+00 |
| LINC01126 | 1.06E-01 | 5.52E-01 | 1.32E+00 | 6.09E-03 | 2.63E-02 | 2.04E+00 | 2.35E-01 | 9.97E-01 | 1.28E+00 | 7.89E-03 | 3.01E-02 | 2.24E+00 | 1.36E-01 | 4.84E-01 | 1.94E+00 | 4.02E-02 | 1.13E-01 | 3.19E+00 | 1.97E-01 | 7.22E-01 | 1.35E+00 | 3.79E-02 | 1.04E-01 | 2.21E+00 |
| LRRC75A | 1.06E-01 | 5.52E-01 | -1.02E+00 | 2.31E-01 | 9.98E-01 | 1.44E+00 | 1.01E-01 | 4.28E-01 | -1.07E+00 | 1.52E-01 | 5.81E-01 | 1.17E+00 | 1.01E-01 | 3.59E-01 | 1.05E+00 | 2.33E-01 | 6.57E-01 | 1.66E+00 | 1.97E-01 | 7.25E-01 | 1.34E+00 | 8.88E-02 | 2.44E-01 | -1.01E+00 |
| FBXL8 | 1.06E-01 | 5.53E-01 | 1.11E+00 | 4.97E-02 | 2.15E-01 | 1.92E+00 | 4.38E-02 | 1.86E-01 | -1.18E+00 | 2.11E-01 | 8.07E-01 | 1.29E+00 | 2.25E-01 | 8.01E-01 | 1.15E+00 | 6.77E-02 | 1.91E-01 | 2.14E+00 | 4.77E-02 | 1.75E-01 | -1.24E+00 | 1.61E-01 | 4.43E-01 | 1.00E+00 |
| AP000919.1 | 1.06E-01 | 5.54E-01 | 1.14E+00 | 8.43E-02 | 3.64E-01 | 1.08E+00 | 2.73E-02 | 1.16E-01 | -1.11E+00 | 2.24E-01 | 8.57E-01 | 1.37E+00 | 1.02E-01 | 3.63E-01 | -1.03E+00 | 3.51E-01 | 9.89E-01 | 1.40E+00 | 1.91E-01 | 7.00E-01 | 1.40E+00 | 9.52E-02 | 2.62E-01 | 1.60E+00 |
| LILRA2 | 1.06E-01 | 5.55E-01 | 1.45E+00 | 2.71E-02 | 1.17E-01 | -1.12E+00 | 1.37E-01 | 5.83E-01 | 1.19E+00 | 9.50E-03 | 3.63E-02 | 1.02E+00 | 6.74E-02 | 2.40E-01 | -1.03E+00 | 2.77E-02 | 7.82E-02 | -1.16E+00 | 2.00E-01 | 7.35E-01 | -1.49E+00 | 2.72E-01 | 7.48E-01 | -1.48E+00 |
| AGAP6 | 1.06E-01 | 5.55E-01 | 1.05E+00 | 1.60E-01 | 6.92E-01 | 1.63E+00 | 1.74E-01 | 7.39E-01 | 1.08E+00 | 1.31E-01 | 5.00E-01 | 1.61E+00 | 1.97E-01 | 7.04E-01 | 1.11E+00 | 2.55E-01 | 7.18E-01 | 1.61E+00 | 1.63E-01 | 6.00E-01 | 1.14E+00 | 3.61E-01 | 9.91E-01 | 1.27E+00 |
| SREBF2 | 1.06E-01 | 5.55E-01 | 1.14E+00 | 1.24E-01 | 5.36E-01 | 1.58E+00 | 7.36E-02 | 3.13E-01 | 1.15E+00 | 2.55E-01 | 9.74E-01 | 1.47E+00 | 2.03E-01 | 7.25E-01 | 1.31E+00 | 1.40E-01 | 3.95E-01 | 1.17E+00 | 2.59E-01 | 9.52E-01 | 1.29E+00 | 1.41E-01 | 3.88E-01 | 1.45E+00 |
| ZNF853 | 1.06E-01 | 5.55E-01 | 1.11E+00 | 2.06E-01 | 8.89E-01 | 1.37E+00 | 2.17E-01 | 9.22E-01 | 1.15E+00 | 6.76E-02 | 2.58E-01 | 1.73E+00 | 2.10E-01 | 7.48E-01 | 1.64E+00 | 6.11E-02 | 1.72E-01 | 2.44E+00 | 2.16E-01 | 7.94E-01 | 1.07E+00 | 2.41E-01 | 6.61E-01 | 1.47E+00 |
| AL353898.3 | 1.06E-01 | 5.55E-01 | 1.09E+00 | 1.63E-01 | 7.03E-01 | 1.52E+00 | 5.33E-03 | 2.26E-02 | -1.41E+00 | 1.64E-02 | 6.28E-02 | -1.23E+00 | 4.03E-03 | 1.44E-02 | -1.87E+00 | 9.97E-02 | 2.81E-01 | -1.10E+00 | 9.45E-02 | 3.47E-01 | -1.02E+00 | 3.38E-01 | 9.29E-01 | 1.39E+00 |
| RNF139-AS1 | 1.06E-01 | 5.55E-01 | 1.13E+00 | 1.40E-01 | 6.05E-01 | 1.73E+00 | 1.76E-01 | 7.48E-01 | 1.54E+00 | 4.86E-02 | 1.86E-01 | 2.12E+00 | 1.13E-01 | 4.04E-01 | 2.13E+00 | 4.71E-03 | 1.33E-02 | 3.29E+00 | 5.41E-03 | 1.98E-02 | 1.82E+00 | 4.01E-02 | 1.10E-01 | 1.89E+00 |
| HMMR | 1.07E-01 | 5.55E-01 | -1.34E+00 | 1.23E-01 | 5.32E-01 | -1.10E+00 | 2.03E-01 | 8.63E-01 | 1.26E+00 | 2.54E-01 | 9.70E-01 | 1.69E+00 | 2.65E-02 | 9.43E-02 | -1.92E+00 | 6.53E-02 | 1.84E-01 | -1.48E+00 | 5.55E-02 | 2.04E-01 | -1.39E+00 | 2.19E-01 | 6.00E-01 | -1.19E+00 |
| RNF111 | 1.07E-01 | 5.55E-01 | 1.66E+00 | 3.77E-02 | 1.63E-01 | 1.17E+00 | 2.25E-01 | 9.54E-01 | 1.32E+00 | 3.59E-03 | 1.37E-02 | -1.09E+00 | 4.11E-06 | 1.46E-05 | 3.12E+00 | 1.22E-01 | 3.44E-01 | 1.91E+00 | 2.09E-01 | 7.66E-01 | 1.33E+00 | 6.07E-03 | 1.67E-02 | -1.14E+00 |
| AURKB | 1.07E-01 | 5.56E-01 | -1.03E+00 | 9.99E-02 | 4.31E-01 | -1.08E+00 | 9.91E-02 | 4.21E-01 | 1.70E+00 | 1.82E-01 | 6.96E-01 | 1.69E+00 | 4.30E-02 | 1.53E-01 | -1.97E+00 | 8.31E-02 | 2.34E-01 | -1.56E+00 | 1.20E-01 | 4.42E-01 | 1.55E+00 | 5.44E-02 | 1.49E-01 | 1.73E+00 |
| CDKL1 | 1.07E-01 | 5.56E-01 | 1.03E+00 | 1.59E-01 | 6.86E-01 | 1.56E+00 | 3.59E-02 | 1.52E-01 | -1.21E+00 | 2.28E-01 | 8.73E-01 | 1.37E+00 | 2.56E-01 | 9.12E-01 | 1.29E+00 | 3.31E-01 | 9.33E-01 | 1.39E+00 | 1.96E-05 | 7.20E-05 | -1.21E+00 | 3.77E-02 | 1.04E-01 | 1.21E+00 |
| AL118516.1 | 1.07E-01 | 5.57E-01 | 1.41E+00 | 6.97E-04 | 3.01E-03 | -1.07E+00 | 5.80E-02 | 2.47E-01 | 1.19E+00 | 2.07E-03 | 7.90E-03 | -1.00E+00 | 2.62E-01 | 9.33E-01 | 1.46E+00 | 3.23E-01 | 9.10E-01 | 1.40E+00 | 1.10E-01 | 4.05E-01 | 1.43E+00 | 5.83E-02 | 1.60E-01 | 1.73E+00 |
| AC106795.2 | 1.07E-01 | 5.57E-01 | -1.04E+00 | 2.08E-01 | 8.99E-01 | 1.27E+00 | 2.34E-01 | 9.93E-01 | 1.11E+00 | 1.36E-01 | 5.19E-01 | 1.42E+00 | 3.13E-02 | 1.12E-01 | -1.22E+00 | 7.36E-02 | 2.08E-01 | -1.14E+00 | 1.96E-02 | 7.19E-02 | -1.12E+00 | 3.74E-02 | 1.03E-01 | -1.05E+00 |
| C17orf97 | 1.07E-01 | 5.58E-01 | 1.10E+00 | 3.90E-02 | 1.69E-01 | -1.02E+00 | 1.36E-01 | 5.79E-01 | 1.01E+00 | 1.10E-01 | 4.21E-01 | 1.09E+00 | 1.06E-01 | 3.79E-01 | 1.02E+00 | 2.54E-01 | 7.16E-01 | 1.18E+00 | 1.86E-03 | 6.82E-03 | -1.29E+00 | 3.73E-04 | 1.02E-03 | -1.54E+00 |
| WDFY2 | 1.07E-01 | 5.58E-01 | 1.24E+00 | 2.11E-04 | 9.14E-04 | 1.06E+00 | 1.32E-01 | 5.63E-01 | 1.48E+00 | 4.47E-03 | 1.71E-02 | 1.23E+00 | 2.55E-01 | 9.08E-01 | 1.57E+00 | 4.84E-02 | 1.36E-01 | 1.25E+00 | 1.07E-01 | 3.91E-01 | 1.50E+00 | 2.52E-01 | 6.92E-01 | 1.32E+00 |
| TP53INP2 | 1.07E-01 | 5.59E-01 | 1.22E+00 | 1.84E-03 | 7.95E-03 | -1.61E+00 | 4.64E-02 | 1.97E-01 | -1.06E+00 | 3.22E-06 | 1.23E-05 | -2.51E+00 | 8.36E-03 | 2.98E-02 | -1.00E+00 | 1.42E-05 | 4.00E-05 | -1.87E+00 | 2.96E-02 | 1.09E-01 | -1.08E+00 | 1.50E-05 | 4.13E-05 | -2.50E+00 |
| LINC02035 | 1.07E-01 | 5.59E-01 | 1.09E+00 | 2.25E-01 | 9.72E-01 | 1.40E+00 | 3.46E-02 | 1.47E-01 | -1.29E+00 | 2.50E-01 | 9.55E-01 | 1.32E+00 | 3.62E-02 | 1.29E-01 | -1.43E+00 | 1.47E-01 | 4.15E-01 | 1.00E+00 | 1.25E-01 | 4.60E-01 | -1.18E+00 | 2.30E-01 | 6.32E-01 | -1.00E+00 |
| TRIB1 | 1.07E-01 | 5.60E-01 | 1.53E+00 | 8.16E-04 | 3.52E-03 | -1.62E+00 | 1.87E-01 | 7.92E-01 | 1.48E+00 | 1.55E-08 | 5.91E-08 | -1.87E+00 | 7.17E-02 | 2.56E-01 | 2.70E+00 | 3.31E-01 | 9.33E-01 | 1.48E+00 | 6.34E-02 | 2.33E-01 | -1.06E+00 | 5.69E-02 | 1.56E-01 | -1.34E+00 |
| MPO | 1.08E-01 | 5.61E-01 | -8.95E+00 | 2.14E-02 | 9.24E-02 | -2.97E+01 | 8.33E-12 | 3.54E-11 | -1.67E+00 | 1.85E-22 | 7.07E-22 | -2.30E+00 | 2.36E-01 | 8.43E-01 | 1.49E+00 | 2.72E-01 | 7.66E-01 | -1.44E+00 | 2.57E-01 | 9.44E-01 | 1.96E+00 | 2.20E-01 | 6.03E-01 | -1.37E+00 |
| VNN1 | 1.08E-01 | 5.62E-01 | -1.05E+00 | 1.69E-01 | 7.32E-01 | 1.41E+00 | 1.93E-01 | 8.22E-01 | 1.13E+00 | 1.78E-01 | 6.81E-01 | 1.28E+00 | 3.57E-02 | 1.27E-01 | -1.48E+00 | 3.47E-01 | 9.78E-01 | 1.22E+00 | 2.65E-01 | 9.72E-01 | 1.34E+00 | 1.97E-01 | 5.42E-01 | 1.38E+00 |
| AQP11 | 1.08E-01 | 5.62E-01 | 1.18E+00 | 8.01E-02 | 3.46E-01 | 1.89E+00 | 1.58E-01 | 6.70E-01 | 1.05E+00 | 2.33E-01 | 8.90E-01 | 1.28E+00 | 2.25E-02 | 8.03E-02 | -1.50E+00 | 2.62E-01 | 7.39E-01 | 1.22E+00 | 1.80E-01 | 6.60E-01 | 1.19E+00 | 3.37E-01 | 9.25E-01 | 1.23E+00 |
| WBP4 | 1.08E-01 | 5.63E-01 | 1.21E+00 | 2.86E-02 | 1.24E-01 | 1.19E+00 | 1.12E-01 | 4.77E-01 | 1.02E+00 | 9.98E-04 | 3.81E-03 | -1.12E+00 | 1.31E-04 | 4.67E-04 | -1.11E+00 | 4.79E-06 | 1.35E-05 | -1.34E+00 | 1.14E-08 | 4.19E-08 | -1.21E+00 | 1.40E-07 | 3.85E-07 | -1.24E+00 |
| SERPINB10 | 1.08E-01 | 5.63E-01 | -1.34E+00 | 1.23E-01 | 5.31E-01 | -1.27E+00 | 1.36E-02 | 5.76E-02 | 1.96E+00 | 1.23E-01 | 4.70E-01 | 1.33E+00 | 2.50E-01 | 8.93E-01 | 1.13E+00 | 3.55E-01 | 1.00E+00 | 1.00E+00 | 2.49E-01 | 9.13E-01 | 1.04E+00 | 3.64E-01 | 1.00E+00 | -1.00E+00 |
| HGS | 1.08E-01 | 5.63E-01 | 1.24E+00 | 1.15E-01 | 4.99E-01 | 1.38E+00 | 1.49E-01 | 6.33E-01 | 1.39E+00 | 2.52E-01 | 9.62E-01 | 1.34E+00 | 1.18E-01 | 4.22E-01 | 1.05E+00 | 2.05E-01 | 5.77E-01 | 1.70E+00 | 2.23E-01 | 8.17E-01 | 1.13E+00 | 3.22E-01 | 8.85E-01 | 1.26E+00 |
| RPL17P33 | 1.08E-01 | 5.63E-01 | 1.53E+00 | 1.39E-01 | 5.99E-01 | 1.58E+00 | 8.12E-03 | 3.45E-02 | 1.89E+00 | 1.59E-01 | 6.06E-01 | 1.52E+00 | 2.36E-01 | 8.42E-01 | 1.13E+00 | 3.55E-01 | 1.00E+00 | 1.36E+00 | 1.16E-02 | 4.27E-02 | 2.04E+00 | 7.51E-02 | 2.06E-01 | 1.84E+00 |
| AC002367.1 | 1.08E-01 | 5.63E-01 | 1.54E+00 | 6.30E-02 | 2.72E-01 | 1.22E+00 | 2.55E-02 | 1.08E-01 | 1.64E+00 | 1.03E-01 | 3.95E-01 | 1.18E+00 | 1.02E-01 | 3.63E-01 | 1.80E+00 | 2.18E-01 | 6.13E-01 | 1.20E+00 | 6.89E-02 | 2.53E-01 | 1.55E+00 | 1.61E-01 | 4.43E-01 | 1.17E+00 |
| KLHDC4 | 1.08E-01 | 5.63E-01 | 1.14E+00 | 3.67E-02 | 1.59E-01 | 1.15E+00 | 1.06E-01 | 4.52E-01 | 1.11E+00 | 2.12E-01 | 8.09E-01 | 1.43E+00 | 7.07E-02 | 2.52E-01 | 1.07E+00 | 6.22E-02 | 1.75E-01 | 1.11E+00 | 2.68E-04 | 9.84E-04 | -1.14E+00 | 9.73E-03 | 2.67E-02 | 1.01E+00 |
| PRR22 | 1.08E-01 | 5.64E-01 | 1.05E+00 | 8.85E-02 | 3.82E-01 | 1.02E+00 | 2.36E-02 | 1.00E-01 | 1.61E+00 | 8.38E-02 | 3.20E-01 | 1.67E+00 | 2.69E-02 | 9.61E-02 | -1.59E+00 | 2.01E-01 | 5.65E-01 | 1.17E+00 | 7.11E-02 | 2.61E-01 | -1.03E+00 | 2.80E-01 | 7.69E-01 | 1.18E+00 |
| POLG | 1.08E-01 | 5.65E-01 | 1.57E+00 | 1.10E-01 | 4.76E-01 | 1.18E+00 | 1.53E-03 | 6.49E-03 | -1.20E+00 | 1.12E-04 | 4.30E-04 | -1.30E+00 | 1.42E-01 | 5.05E-01 | 1.05E+00 | 1.48E-01 | 4.17E-01 | -1.02E+00 | 1.33E-01 | 4.89E-01 | 1.10E+00 | 1.89E-01 | 5.18E-01 | 1.09E+00 |
| AC104581.4 | 1.08E-01 | 5.65E-01 | 1.11E+00 | 2.25E-01 | 9.74E-01 | 1.39E+00 | 9.00E-03 | 3.82E-02 | -1.24E+00 | 8.94E-03 | 3.42E-02 | -1.22E+00 | 2.05E-01 | 7.29E-01 | 1.31E+00 | 3.41E-01 | 9.60E-01 | 1.48E+00 | 4.73E-02 | 1.74E-01 | -1.01E+00 | 1.14E-03 | 3.14E-03 | -1.64E+00 |
| HNRNPA1P8 | 1.08E-01 | 5.65E-01 | 1.04E+00 | 1.33E-01 | 5.74E-01 | 1.11E+00 | 1.77E-01 | 7.52E-01 | 1.06E+00 | 1.57E-01 | 6.00E-01 | 1.12E+00 | 9.50E-02 | 3.39E-01 | -1.22E+00 | 2.97E-01 | 8.37E-01 | 1.19E+00 | 5.65E-06 | 2.07E-05 | -1.71E+00 | 1.35E-03 | 3.71E-03 | -1.24E+00 |
| AKAP6 | 1.08E-01 | 5.65E-01 | -1.16E+00 | 1.27E-01 | 5.49E-01 | -1.05E+00 | 8.69E-02 | 3.69E-01 | -1.23E+00 | 1.27E-01 | 4.84E-01 | -1.07E+00 | 1.74E-01 | 6.21E-01 | 1.37E+00 | 2.03E-01 | 5.72E-01 | 1.23E+00 | 1.78E-01 | 6.52E-01 | -1.05E+00 | 1.74E-01 | 4.78E-01 | -1.23E+00 |
| ZC2HC1C | 1.09E-01 | 5.66E-01 | 1.16E+00 | 1.56E-01 | 6.72E-01 | 1.29E+00 | 7.43E-02 | 3.16E-01 | -1.21E+00 | 1.60E-01 | 6.12E-01 | 1.02E+00 | 5.71E-04 | 2.04E-03 | -1.90E+00 | 7.00E-03 | 1.97E-02 | -1.37E+00 | 1.28E-02 | 4.68E-02 | -1.43E+00 | 3.77E-02 | 1.03E-01 | -1.22E+00 |
| SLC25A10 | 1.09E-01 | 5.66E-01 | 1.45E+00 | 2.31E-01 | 9.98E-01 | 1.13E+00 | 2.04E-01 | 8.66E-01 | 1.23E+00 | 2.33E-01 | 8.90E-01 | 1.41E+00 | 4.77E-02 | 1.70E-01 | -2.33E+00 | 1.40E-01 | 3.95E-01 | -1.48E+00 | 1.23E-01 | 4.50E-01 | 1.39E+00 | 2.30E-01 | 6.31E-01 | 1.42E+00 |
| FAM72A | 1.09E-01 | 5.66E-01 | 1.49E+00 | 1.13E-01 | 4.89E-01 | 1.66E+00 | 9.02E-02 | 3.83E-01 | 1.38E+00 | 1.10E-01 | 4.19E-01 | 1.51E+00 | 2.69E-01 | 9.58E-01 | 1.18E+00 | 2.43E-01 | 6.84E-01 | -1.12E+00 | 3.36E-03 | 1.23E-02 | -1.16E+00 | 8.89E-02 | 2.44E-01 | 1.17E+00 |
| AC093663.1 | 1.09E-01 | 5.66E-01 | 1.42E+00 | 1.60E-01 | 6.90E-01 | 1.51E+00 | 1.29E-01 | 5.47E-01 | 1.00E+00 | 3.88E-02 | 1.48E-01 | -1.10E+00 | 2.34E-01 | 8.36E-01 | 1.24E+00 | 2.65E-01 | 7.46E-01 | 1.17E+00 | 2.65E-02 | 9.72E-02 | 1.73E+00 | 3.56E-01 | 9.77E-01 | 1.24E+00 |
| AC048341.2 | 1.09E-01 | 5.67E-01 | 1.30E+00 | 1.98E-02 | 8.56E-02 | 2.24E+00 | 8.95E-04 | 3.80E-03 | -1.42E+00 | 5.51E-02 | 2.10E-01 | 1.20E+00 | 1.92E-01 | 6.86E-01 | 1.62E+00 | 1.30E-01 | 3.65E-01 | 1.71E+00 | 9.80E-02 | 3.60E-01 | 1.15E+00 | 2.09E-01 | 5.73E-01 | 1.15E+00 |
| BET1L | 1.09E-01 | 5.68E-01 | 1.28E+00 | 1.51E-01 | 6.52E-01 | 1.43E+00 | 9.22E-02 | 3.92E-01 | 1.13E+00 | 1.74E-01 | 6.63E-01 | 1.31E+00 | 5.25E-02 | 1.87E-01 | 1.87E+00 | 3.17E-02 | 8.93E-02 | 1.88E+00 | 4.81E-03 | 1.76E-02 | 1.63E+00 | 1.42E-01 | 3.91E-01 | 1.38E+00 |
| SEC61G | 1.09E-01 | 5.68E-01 | 1.12E+00 | 3.09E-16 | 1.33E-15 | -1.17E+00 | 3.96E-02 | 1.68E-01 | 1.21E+00 | 1.21E-04 | 4.64E-04 | -1.16E+00 | 1.36E-05 | 4.84E-05 | -1.20E+00 | 1.22E-10 | 3.44E-10 | -1.48E+00 | 2.04E-01 | 7.49E-01 | 1.12E+00 | 1.00E-01 | 2.76E-01 | -1.10E+00 |
| AC064801.2 | 1.09E-01 | 5.68E-01 | 1.13E+00 | 2.24E-01 | 9.66E-01 | 1.39E+00 | 7.16E-02 | 3.04E-01 | -1.08E+00 | 2.51E-01 | 9.61E-01 | 1.34E+00 | 2.51E-02 | 8.94E-02 | -1.41E+00 | 1.51E-01 | 4.25E-01 | 1.13E+00 | 3.28E-02 | 1.20E-01 | -1.37E+00 | 2.09E-02 | 5.73E-02 | -1.67E+00 |
| PASK | 1.09E-01 | 5.69E-01 | -1.01E+00 | 1.13E-01 | 4.89E-01 | 1.04E+00 | 2.13E-03 | 9.04E-03 | -1.18E+00 | 1.33E-03 | 5.08E-03 | -1.11E+00 | 1.61E-01 | 5.74E-01 | -1.03E+00 | 3.32E-01 | 9.36E-01 | 1.28E+00 | 7.59E-03 | 2.79E-02 | -1.31E+00 | 2.15E-01 | 5.90E-01 | 1.34E+00 |
| ATP5F1P1 | 1.09E-01 | 5.69E-01 | 1.03E+00 | 1.74E-02 | 7.51E-02 | -1.38E+00 | 5.84E-02 | 2.48E-01 | 1.57E+00 | 3.05E-02 | 1.16E-01 | -1.40E+00 | 8.38E-04 | 2.99E-03 | 2.75E+00 | 3.02E-01 | 8.53E-01 | 1.33E+00 | 2.50E-03 | 9.17E-03 | 1.95E+00 | 3.27E-01 | 9.00E-01 | 1.16E+00 |
| RFTN2 | 1.09E-01 | 5.69E-01 | 1.50E+00 | 1.37E-02 | 5.91E-02 | -1.52E+00 | 7.45E-03 | 3.16E-02 | -1.41E+00 | 6.86E-03 | 2.62E-02 | -1.50E+00 | 2.71E-01 | 9.66E-01 | 1.25E+00 | 5.16E-02 | 1.45E-01 | -1.51E+00 | 2.47E-01 | 9.06E-01 | 1.04E+00 | 6.00E-02 | 1.65E-01 | -1.43E+00 |
| ABCA6 | 1.09E-01 | 5.69E-01 | 1.07E+00 | 2.22E-01 | 9.61E-01 | 1.45E+00 | 1.85E-01 | 7.86E-01 | -1.00E+00 | 2.02E-01 | 7.72E-01 | 1.07E+00 | 2.39E-01 | 8.54E-01 | 2.08E+00 | 2.74E-01 | 7.72E-01 | 1.09E+00 | 2.06E-01 | 7.56E-01 | 1.11E+00 | 3.54E-01 | 9.74E-01 | 1.12E+00 |
| GTF2IP13 | 1.09E-01 | 5.69E-01 | 1.00E+00 | 5.25E-02 | 2.27E-01 | -1.08E+00 | 1.64E-04 | 6.97E-04 | -1.63E+00 | 6.57E-04 | 2.51E-03 | -1.34E+00 | 6.25E-02 | 2.23E-01 | -1.06E+00 | 6.88E-02 | 1.94E-01 | -1.02E+00 | 2.68E-01 | 9.82E-01 | 1.25E+00 | 2.88E-01 | 7.91E-01 | 1.10E+00 |
| EEF1A1P24 | 1.09E-01 | 5.69E-01 | 1.16E+00 | 2.18E-02 | 9.42E-02 | -1.11E+00 | 2.27E-01 | 9.67E-01 | 1.26E+00 | 5.70E-02 | 2.18E-01 | -1.04E+00 | 1.20E-01 | 4.29E-01 | 1.07E+00 | 4.82E-02 | 1.36E-01 | -1.15E+00 | 2.42E-01 | 8.88E-01 | 1.29E+00 | 4.58E-02 | 1.26E-01 | -1.38E+00 |
| AC010168.2 | 1.09E-01 | 5.69E-01 | 1.02E+00 | 9.10E-02 | 3.93E-01 | 1.59E+00 | 1.02E-02 | 4.32E-02 | -1.31E+00 | 1.39E-01 | 5.31E-01 | 1.16E+00 | 1.73E-02 | 6.18E-02 | -1.67E+00 | 3.02E-02 | 8.52E-02 | -1.56E+00 | 7.56E-03 | 2.77E-02 | -1.37E+00 | 6.42E-02 | 1.76E-01 | -1.19E+00 |
| CCDC181 | 1.09E-01 | 5.70E-01 | 1.05E+00 | 2.01E-01 | 8.67E-01 | 1.29E+00 | 6.81E-02 | 2.89E-01 | -1.05E+00 | 5.46E-02 | 2.08E-01 | -1.03E+00 | 4.02E-02 | 1.43E-01 | -1.03E+00 | 6.16E-02 | 1.74E-01 | -1.03E+00 | 9.99E-02 | 3.67E-01 | 1.12E+00 | 1.90E-01 | 5.22E-01 | 1.06E+00 |
| HMGB1P6 | 1.09E-01 | 5.70E-01 | 1.15E+00 | 1.07E-03 | 4.62E-03 | 1.05E+00 | 7.65E-02 | 3.25E-01 | 1.59E+00 | 4.03E-03 | 1.54E-02 | 1.26E+00 | 1.20E-04 | 4.26E-04 | -1.04E+00 | 4.00E-04 | 1.13E-03 | 1.02E+00 | 2.41E-01 | 8.86E-01 | 1.28E+00 | 1.38E-01 | 3.79E-01 | 1.33E+00 |
| HIST2H2AB | 1.09E-01 | 5.70E-01 | 1.38E+00 | 1.32E-01 | 5.70E-01 | 1.59E+00 | 1.10E-01 | 4.66E-01 | 1.45E+00 | 1.26E-01 | 4.81E-01 | 1.59E+00 | 2.00E-01 | 7.14E-01 | 1.51E+00 | 1.49E-01 | 4.19E-01 | 1.89E+00 | 2.20E-01 | 8.06E-01 | 1.35E+00 | 2.78E-01 | 7.63E-01 | 1.39E+00 |
| MANEAL | 1.09E-01 | 5.71E-01 | 1.03E+00 | 6.35E-02 | 2.74E-01 | -1.07E+00 | 7.76E-02 | 3.30E-01 | 1.54E+00 | 1.88E-01 | 7.20E-01 | 1.60E+00 | 3.09E-02 | 1.10E-01 | 2.62E+00 | 4.21E-02 | 1.19E-01 | 2.64E+00 | 1.04E-01 | 3.82E-01 | 1.58E+00 | 5.32E-02 | 1.46E-01 | 1.80E+00 |
| ELANE | 1.10E-01 | 5.72E-01 | -1.55E+00 | 8.38E-02 | 3.62E-01 | -2.19E+00 | 4.14E-08 | 1.76E-07 | -2.75E+00 | 7.32E-13 | 2.80E-12 | -5.82E+00 | 2.80E-01 | 1.00E+00 | 1.00E+00 | 3.55E-01 | 1.00E+00 | -1.00E+00 | 2.45E-01 | 8.98E-01 | -1.10E+00 | 3.12E-01 | 8.56E-01 | -1.15E+00 |
| LINC01545 | 1.10E-01 | 5.73E-01 | 1.07E+00 | 5.32E-02 | 2.30E-01 | -1.05E+00 | 1.50E-03 | 6.39E-03 | -1.43E+00 | 3.83E-04 | 1.46E-03 | -1.70E+00 | 7.19E-02 | 2.56E-01 | -1.20E+00 | 1.16E-01 | 3.28E-01 | -1.06E+00 | 6.27E-03 | 2.30E-02 | -1.54E+00 | 3.39E-02 | 9.32E-02 | -1.37E+00 |
| ABCC3 | 1.10E-01 | 5.73E-01 | -1.08E+00 | 1.90E-01 | 8.23E-01 | 1.10E+00 | 4.73E-02 | 2.01E-01 | -1.32E+00 | 9.30E-02 | 3.55E-01 | -1.17E+00 | 2.42E-01 | 8.62E-01 | 1.14E+00 | 2.85E-01 | 8.04E-01 | 1.20E+00 | 5.19E-02 | 1.91E-01 | 1.07E+00 | 3.67E-03 | 1.01E-02 | -1.60E+00 |
| TBC1D22A-AS1 | 1.10E-01 | 5.73E-01 | 1.12E+00 | 2.10E-01 | 9.06E-01 | 1.54E+00 | 1.68E-01 | 7.12E-01 | 1.36E+00 | 1.90E-01 | 7.27E-01 | 1.54E+00 | 2.15E-01 | 7.66E-01 | 1.22E+00 | 2.01E-01 | 5.68E-01 | 1.65E+00 | 6.47E-02 | 2.37E-01 | -1.15E+00 | 3.13E-01 | 8.59E-01 | 1.30E+00 |
| AL391684.1 | 1.10E-01 | 5.73E-01 | 1.02E+00 | 1.85E-01 | 7.99E-01 | 1.55E+00 | 1.88E-01 | 8.00E-01 | 1.09E+00 | 2.62E-01 | 1.00E+00 | 1.24E+00 | 2.36E-01 | 8.43E-01 | 1.29E+00 | 3.27E-01 | 9.20E-01 | 1.38E+00 | 8.86E-02 | 3.25E-01 | 1.74E+00 | 2.22E-01 | 6.11E-01 | 1.36E+00 |
| MYLK-AS1 | 1.10E-01 | 5.73E-01 | 1.25E+00 | 2.38E-02 | 1.03E-01 | 1.19E+00 | 6.35E-02 | 2.70E-01 | -1.08E+00 | 1.88E-02 | 7.19E-02 | -1.09E+00 | 1.71E-01 | 6.09E-01 | 1.23E+00 | 3.36E-01 | 9.48E-01 | 1.39E+00 | 1.34E-01 | 4.92E-01 | 1.42E+00 | 2.81E-01 | 7.72E-01 | 1.42E+00 |
| AC026412.1 | 1.10E-01 | 5.73E-01 | 1.34E+00 | 1.03E-01 | 4.47E-01 | 1.55E+00 | 5.39E-02 | 2.29E-01 | -1.37E+00 | 7.53E-02 | 2.88E-01 | -1.16E+00 | 2.41E-01 | 8.58E-01 | 1.28E+00 | 3.48E-01 | 9.82E-01 | 1.27E+00 | 2.13E-01 | 7.83E-01 | 1.25E+00 | 3.21E-01 | 8.81E-01 | 1.22E+00 |
| H1FX | 1.10E-01 | 5.73E-01 | 1.03E+00 | 2.03E-01 | 8.78E-01 | 1.11E+00 | 1.54E-01 | 6.54E-01 | 1.08E+00 | 1.75E-01 | 6.67E-01 | 1.21E+00 | 1.28E-02 | 4.57E-02 | 1.30E+00 | 9.54E-04 | 2.69E-03 | 1.49E+00 | 1.33E-01 | 4.87E-01 | 1.35E+00 | 1.67E-01 | 4.58E-01 | 1.33E+00 |
| AC024293.1 | 1.10E-01 | 5.73E-01 | -1.01E+00 | 1.37E-21 | 5.94E-21 | -1.25E+00 | 2.04E-01 | 8.66E-01 | 1.01E+00 | 8.76E-13 | 3.34E-12 | -1.23E+00 | 2.79E-01 | 9.94E-01 | 1.01E+00 | 5.87E-09 | 1.65E-08 | -1.21E+00 | 2.03E-02 | 7.43E-02 | 1.02E+00 | 6.42E-11 | 1.76E-10 | -1.19E+00 |
| CACNA2D3 | 1.10E-01 | 5.73E-01 | -1.16E+00 | 1.36E-01 | 5.89E-01 | -1.16E+00 | 6.00E-02 | 2.55E-01 | -1.23E+00 | 6.98E-02 | 2.66E-01 | -1.23E+00 | 2.39E-01 | 8.52E-01 | 1.07E+00 | 1.70E-01 | 4.78E-01 | 1.42E+00 | 1.23E-28 | 4.50E-28 | -4.16E+00 | 2.51E-27 | 6.89E-27 | -3.88E+00 |
| ATP5H | 1.10E-01 | 5.74E-01 | 1.24E+00 | 1.66E-02 | 7.15E-02 | 1.23E+00 | 6.12E-02 | 2.60E-01 | 1.14E+00 | 1.13E-01 | 4.32E-01 | 1.23E+00 | 1.01E-02 | 3.58E-02 | 1.14E+00 | 9.21E-02 | 2.60E-01 | 1.32E+00 | 1.50E-01 | 5.50E-01 | 1.28E+00 | 3.06E-01 | 8.41E-01 | 1.26E+00 |
| EEF1A1P6 | 1.10E-01 | 5.74E-01 | 1.29E+00 | 5.40E-03 | 2.33E-02 | 1.23E+00 | 6.82E-02 | 2.90E-01 | 1.49E+00 | 1.73E-01 | 6.61E-01 | 1.53E+00 | 1.17E-01 | 4.18E-01 | 1.62E+00 | 3.06E-01 | 8.63E-01 | 1.40E+00 | 6.00E-02 | 2.20E-01 | 1.59E+00 | 3.58E-01 | 9.83E-01 | 1.37E+00 |
| CXADR | 1.10E-01 | 5.74E-01 | 1.64E+00 | 5.41E-02 | 2.34E-01 | 2.55E+00 | 1.97E-01 | 8.35E-01 | 1.27E+00 | 2.01E-01 | 7.68E-01 | -1.12E+00 | 1.65E-01 | 5.89E-01 | 2.06E+00 | 1.59E-02 | 4.49E-02 | 2.96E+00 | 2.44E-02 | 8.95E-02 | 1.01E+00 | 5.84E-09 | 1.60E-08 | -1.37E+00 |
| MAPK7 | 1.10E-01 | 5.75E-01 | 1.21E+00 | 6.23E-02 | 2.69E-01 | 1.12E+00 | 1.52E-04 | 6.47E-04 | -1.55E+00 | 2.49E-03 | 9.51E-03 | -1.28E+00 | 2.77E-03 | 9.87E-03 | -1.60E+00 | 4.50E-02 | 1.27E-01 | -1.08E+00 | 1.84E-03 | 6.77E-03 | -1.34E+00 | 1.04E-03 | 2.85E-03 | -1.61E+00 |
| ZNF98 | 1.10E-01 | 5.75E-01 | 1.43E+00 | 2.18E-03 | 9.44E-03 | -1.07E+00 | 4.74E-02 | 2.02E-01 | 1.11E+00 | 7.95E-04 | 3.04E-03 | -1.00E+00 | 9.19E-02 | 3.28E-01 | 1.06E+00 | 1.31E-03 | 3.68E-03 | -1.41E+00 | 2.03E-01 | 7.45E-01 | 1.45E+00 | 1.67E-01 | 4.60E-01 | 1.17E+00 |
| ITPK1 | 1.10E-01 | 5.76E-01 | 1.25E+00 | 2.73E-02 | 1.18E-01 | 1.23E+00 | 1.03E-01 | 4.38E-01 | 1.08E+00 | 8.56E-02 | 3.27E-01 | 1.20E+00 | 1.91E-02 | 6.80E-02 | -1.06E+00 | 8.83E-02 | 2.49E-01 | 1.14E+00 | 1.92E-01 | 7.05E-01 | 1.16E+00 | 2.67E-01 | 7.33E-01 | 1.44E+00 |
| CCDC88C | 1.11E-01 | 5.76E-01 | 1.39E+00 | 5.46E-02 | 2.36E-01 | 1.63E+00 | 9.17E-02 | 3.90E-01 | 1.31E+00 | 5.60E-02 | 2.14E-01 | 1.57E+00 | 2.16E-03 | 7.71E-03 | -1.07E+00 | 4.49E-02 | 1.26E-01 | 1.18E+00 | 1.73E-01 | 6.36E-01 | 1.25E+00 | 3.25E-01 | 8.92E-01 | 1.33E+00 |
| AC097634.1 | 1.11E-01 | 5.76E-01 | 1.43E+00 | 2.89E-04 | 1.25E-03 | 2.85E+00 | 1.76E-01 | 7.49E-01 | 1.23E+00 | 1.01E-01 | 3.84E-01 | 1.13E+00 | 9.99E-02 | 3.56E-01 | 1.70E+00 | 2.23E-01 | 6.30E-01 | 1.46E+00 | 1.06E-01 | 3.89E-01 | 1.58E+00 | 8.77E-02 | 2.41E-01 | 1.91E+00 |
| HMGN2P4 | 1.11E-01 | 5.76E-01 | 1.45E+00 | 2.22E-01 | 9.59E-01 | 1.23E+00 | 6.10E-03 | 2.59E-02 | 1.88E+00 | 1.48E-01 | 5.64E-01 | 1.44E+00 | 2.62E-01 | 9.33E-01 | 1.41E+00 | 2.66E-01 | 7.50E-01 | 1.08E+00 | 4.41E-02 | 1.62E-01 | 1.52E+00 | 1.78E-01 | 4.89E-01 | 1.10E+00 |
| SMDT1 | 1.11E-01 | 5.77E-01 | 1.19E+00 | 1.47E-03 | 6.37E-03 | 1.09E+00 | 2.29E-01 | 9.72E-01 | 1.07E+00 | 1.63E-02 | 6.24E-02 | -1.02E+00 | 1.10E-02 | 3.91E-02 | 1.08E+00 | 1.47E-02 | 4.15E-02 | 1.09E+00 | 8.87E-02 | 3.26E-01 | 1.31E+00 | 3.06E-01 | 8.41E-01 | 1.11E+00 |
| HMGA1 | 1.11E-01 | 5.77E-01 | 1.11E+00 | 1.30E-01 | 5.62E-01 | 1.16E+00 | 3.97E-02 | 1.69E-01 | 1.19E+00 | 1.32E-01 | 5.05E-01 | 1.18E+00 | 4.61E-07 | 1.64E-06 | -1.14E+00 | 4.81E-03 | 1.36E-02 | -1.01E+00 | 3.52E-02 | 1.29E-01 | 1.03E+00 | 6.74E-02 | 1.85E-01 | 1.28E+00 |
| LIMK2 | 1.11E-01 | 5.77E-01 | 1.55E+00 | 1.54E-01 | 6.67E-01 | 1.66E+00 | 1.87E-01 | 7.94E-01 | 1.25E+00 | 1.09E-01 | 4.15E-01 | 1.23E+00 | 6.91E-02 | 2.46E-01 | 1.15E+00 | 8.07E-02 | 2.28E-01 | 1.11E+00 | 2.58E-01 | 9.47E-01 | 1.16E+00 | 3.21E-01 | 8.81E-01 | 1.27E+00 |
| AC104651.1 | 1.11E-01 | 5.78E-01 | 1.14E+00 | 1.90E-03 | 8.19E-03 | 2.61E+00 | 2.09E-01 | 8.90E-01 | 1.24E+00 | 6.08E-03 | 2.32E-02 | 2.26E+00 | 2.16E-01 | 7.69E-01 | 1.39E+00 | 5.18E-03 | 1.46E-02 | 2.94E+00 | 1.93E-02 | 7.10E-02 | -1.07E+00 | 1.86E-02 | 5.12E-02 | 1.85E+00 |
| AC144521.1 | 1.11E-01 | 5.78E-01 | 1.03E+00 | 4.79E-06 | 2.07E-05 | 3.48E+00 | 3.27E-03 | 1.39E-02 | -1.52E+00 | 3.08E-06 | 1.18E-05 | 2.81E+00 | 1.99E-01 | 7.09E-01 | -1.02E+00 | 2.76E-01 | 7.77E-01 | 1.67E+00 | 1.99E-01 | 7.31E-01 | 1.39E+00 | 3.28E-02 | 9.01E-02 | 2.26E+00 |
| AC007250.1 | 1.11E-01 | 5.78E-01 | 1.09E+00 | 6.44E-02 | 2.78E-01 | -1.01E+00 | 1.42E-01 | 6.05E-01 | 1.12E+00 | 1.00E-02 | 3.82E-02 | -1.53E+00 | 2.24E-01 | 8.00E-01 | 1.24E+00 | 2.42E-01 | 6.83E-01 | 1.50E+00 | 1.75E-01 | 6.42E-01 | 1.31E+00 | 3.35E-01 | 9.21E-01 | 1.10E+00 |
| STRIP2 | 1.11E-01 | 5.79E-01 | -1.18E+00 | 1.33E-01 | 5.73E-01 | -1.08E+00 | 1.77E-01 | 7.53E-01 | -1.11E+00 | 1.18E-01 | 4.49E-01 | -1.24E+00 | 8.30E-02 | 2.96E-01 | -1.72E+00 | 1.54E-01 | 4.33E-01 | -1.51E+00 | 4.16E-02 | 1.53E-01 | 1.08E+00 | 3.29E-01 | 9.03E-01 | 1.33E+00 |
| LINC00641 | 1.11E-01 | 5.79E-01 | 1.26E+00 | 1.83E-01 | 7.90E-01 | 1.51E+00 | 1.50E-02 | 6.39E-02 | -1.13E+00 | 7.41E-03 | 2.83E-02 | 1.00E+00 | 2.25E-01 | 8.03E-01 | 1.24E+00 | 8.45E-02 | 2.38E-01 | 1.09E+00 | 2.13E-01 | 7.83E-01 | 1.13E+00 | 2.23E-01 | 6.13E-01 | 1.41E+00 |
| TADA2B | 1.11E-01 | 5.79E-01 | -1.06E+00 | 1.38E-01 | 5.96E-01 | -1.02E+00 | 6.74E-02 | 2.86E-01 | -1.22E+00 | 6.93E-02 | 2.65E-01 | -1.13E+00 | 1.37E-02 | 4.88E-02 | -1.06E+00 | 7.22E-03 | 2.03E-02 | -1.11E+00 | 4.30E-05 | 1.58E-04 | -1.52E+00 | 6.22E-05 | 1.71E-04 | -1.58E+00 |
| AC110769.2 | 1.11E-01 | 5.80E-01 | 1.45E+00 | 1.33E-01 | 5.76E-01 | 1.33E+00 | 2.04E-01 | 8.68E-01 | 1.22E+00 | 9.93E-02 | 3.79E-01 | 1.14E+00 | 2.00E-02 | 7.14E-02 | -1.30E+00 | 7.27E-02 | 2.05E-01 | 1.00E+00 | 2.18E-01 | 7.98E-01 | 1.10E+00 | 2.93E-01 | 8.05E-01 | 1.54E+00 |
| RTN1 | 1.11E-01 | 5.80E-01 | -1.05E+00 | 2.10E-01 | 9.06E-01 | 1.21E+00 | 1.52E-02 | 6.48E-02 | -2.09E+00 | 4.92E-02 | 1.88E-01 | -1.48E+00 | 2.55E-01 | 9.10E-01 | 1.18E+00 | 2.28E-01 | 6.44E-01 | 1.57E+00 | 1.39E-03 | 5.10E-03 | 2.06E+00 | 2.27E-02 | 6.24E-02 | 1.78E+00 |
| TSC22D1-AS1 | 1.11E-01 | 5.80E-01 | 1.36E+00 | 5.78E-02 | 2.50E-01 | 1.77E+00 | 1.35E-01 | 5.74E-01 | -1.10E+00 | 8.24E-02 | 3.15E-01 | 1.77E+00 | 3.86E-02 | 1.38E-01 | -1.39E+00 | 2.94E-01 | 8.29E-01 | 1.42E+00 | 1.69E-01 | 6.21E-01 | 1.06E+00 | 3.52E-01 | 9.67E-01 | 1.29E+00 |
| AC002553.4 | 1.12E-01 | 5.82E-01 | 1.08E+00 | 1.66E-01 | 7.15E-01 | 1.61E+00 | 2.61E-02 | 1.11E-01 | -1.14E+00 | 1.02E-01 | 3.91E-01 | 1.13E+00 | 1.35E-01 | 4.82E-01 | -1.09E+00 | 3.26E-01 | 9.20E-01 | 1.50E+00 | 1.70E-02 | 6.22E-02 | -1.14E+00 | 9.97E-03 | 2.74E-02 | -1.25E+00 |
| WTH3DI | 1.12E-01 | 5.82E-01 | 1.54E+00 | 1.06E-01 | 4.59E-01 | 1.83E+00 | 9.13E-02 | 3.88E-01 | 1.00E+00 | 2.54E-01 | 9.72E-01 | 1.38E+00 | 1.53E-01 | 5.44E-01 | 1.11E+00 | 3.20E-01 | 9.03E-01 | 1.42E+00 | 1.97E-01 | 7.23E-01 | 1.14E+00 | 2.07E-01 | 5.68E-01 | 1.36E+00 |
| HPGD | 1.12E-01 | 5.82E-01 | 1.25E+00 | 2.21E-01 | 9.56E-01 | 1.27E+00 | 2.13E-03 | 9.07E-03 | 1.78E+00 | 2.51E-01 | 9.58E-01 | 1.37E+00 | 5.11E-04 | 1.82E-03 | -2.29E+00 | 1.07E-05 | 3.01E-05 | -2.78E+00 | 1.50E-01 | 5.50E-01 | 1.11E+00 | 2.30E-01 | 6.32E-01 | 1.09E+00 |
| EEF1B2P7 | 1.12E-01 | 5.82E-01 | 1.13E+00 | 2.21E-01 | 9.53E-01 | 1.37E+00 | 1.12E-01 | 4.75E-01 | 1.07E+00 | 1.07E-03 | 4.09E-03 | -1.43E+00 | 1.10E-01 | 3.91E-01 | 1.72E+00 | 2.87E-01 | 8.09E-01 | 1.35E+00 | 1.19E-02 | 4.38E-02 | -1.31E+00 | 2.45E-01 | 6.72E-01 | 1.10E+00 |
| PADI2 | 1.12E-01 | 5.82E-01 | -1.19E+00 | 5.46E-02 | 2.36E-01 | -1.47E+00 | 1.36E-02 | 5.78E-02 | -1.35E+00 | 7.63E-04 | 2.91E-03 | -1.70E+00 | 2.33E-01 | 8.33E-01 | 1.13E+00 | 3.55E-01 | 1.00E+00 | 1.00E+00 | 1.66E-01 | 6.08E-01 | -1.16E+00 | 3.03E-01 | 8.32E-01 | -1.04E+00 |
| PRDM1 | 1.12E-01 | 5.83E-01 | -1.14E+00 | 1.50E-01 | 6.50E-01 | -1.05E+00 | 1.75E-03 | 7.42E-03 | -1.53E+00 | 1.48E-04 | 5.64E-04 | -1.95E+00 | 1.32E-01 | 4.69E-01 | 1.57E+00 | 3.16E-01 | 8.91E-01 | 1.22E+00 | 2.31E-02 | 8.47E-02 | -1.41E+00 | 3.24E-02 | 8.89E-02 | -1.45E+00 |
| SIK2 | 1.12E-01 | 5.84E-01 | 1.25E+00 | 1.32E-01 | 5.71E-01 | 1.42E+00 | 1.62E-02 | 6.88E-02 | -1.08E+00 | 1.23E-01 | 4.70E-01 | 1.23E+00 | 1.12E-01 | 3.99E-01 | 1.03E+00 | 2.85E-01 | 8.04E-01 | 1.36E+00 | 2.52E-02 | 9.23E-02 | 1.02E+00 | 1.19E-01 | 3.28E-01 | 1.09E+00 |
| SNHG26 | 1.12E-01 | 5.85E-01 | 1.16E+00 | 4.34E-02 | 1.88E-01 | 1.12E+00 | 9.90E-02 | 4.21E-01 | 1.42E+00 | 4.75E-02 | 1.82E-01 | 1.20E+00 | 6.12E-03 | 2.18E-02 | -1.18E+00 | 1.78E-01 | 5.02E-01 | 1.20E+00 | 1.13E-01 | 4.14E-01 | 1.29E+00 | 5.84E-02 | 1.61E-01 | 1.49E+00 |
| FAM171A1 | 1.12E-01 | 5.86E-01 | -1.23E+00 | 1.45E-01 | 6.26E-01 | -1.15E+00 | 2.07E-01 | 8.79E-01 | 1.10E+00 | 2.06E-01 | 7.87E-01 | 1.25E+00 | 4.78E-02 | 1.71E-01 | -1.16E+00 | 1.46E-01 | 4.11E-01 | -1.29E+00 | 3.46E-07 | 1.27E-06 | -1.48E+00 | 1.52E-11 | 4.17E-11 | -1.85E+00 |
| EEF1D | 1.12E-01 | 5.86E-01 | 1.36E+00 | 4.71E-04 | 2.03E-03 | -1.30E+00 | 2.08E-01 | 8.84E-01 | 1.18E+00 | 3.79E-04 | 1.45E-03 | -1.28E+00 | 7.94E-02 | 2.83E-01 | 1.20E+00 | 3.04E-03 | 8.57E-03 | -1.17E+00 | 2.65E-01 | 9.71E-01 | 1.21E+00 | 6.04E-04 | 1.66E-03 | -1.33E+00 |
| AC055874.1 | 1.13E-01 | 5.86E-01 | -1.66E+00 | 1.40E-01 | 6.07E-01 | -1.59E+00 | 2.23E-01 | 9.46E-01 | 1.07E+00 | 1.93E-01 | 7.38E-01 | 1.50E+00 | 6.45E-02 | 2.30E-01 | 1.09E+00 | 1.14E-01 | 3.21E-01 | 1.57E+00 | 9.05E-07 | 3.32E-06 | -1.08E+00 | 1.14E-13 | 3.13E-13 | -1.37E+00 |
| AL031777.3 | 1.13E-01 | 5.87E-01 | -1.39E+00 | 1.07E-01 | 4.63E-01 | 1.39E+00 | 1.91E-01 | 8.10E-01 | 1.17E+00 | 6.43E-02 | 2.45E-01 | 1.93E+00 | 5.87E-02 | 2.09E-01 | -1.34E+00 | 1.48E-03 | 4.17E-03 | 2.36E+00 | 8.37E-02 | 3.07E-01 | -1.07E+00 | 1.86E-01 | 5.11E-01 | 1.55E+00 |
| TMEM116 | 1.13E-01 | 5.87E-01 | 1.11E+00 | 1.17E-01 | 5.07E-01 | 1.63E+00 | 1.25E-01 | 5.32E-01 | 1.23E+00 | 2.38E-01 | 9.11E-01 | 1.19E+00 | 1.12E-02 | 4.00E-02 | -1.31E+00 | 1.74E-01 | 4.92E-01 | 1.20E+00 | 1.66E-03 | 6.09E-03 | -1.27E+00 | 6.35E-02 | 1.75E-01 | 1.02E+00 |
| AC109587.1 | 1.13E-01 | 5.88E-01 | 1.02E+00 | 2.08E-01 | 9.00E-01 | 1.26E+00 | 8.19E-02 | 3.48E-01 | -1.08E+00 | 8.49E-02 | 3.24E-01 | -1.01E+00 | 2.11E-02 | 7.51E-02 | -1.17E+00 | 2.63E-02 | 7.41E-02 | -1.33E+00 | 5.61E-02 | 2.06E-01 | 1.04E+00 | 1.55E-01 | 4.27E-01 | 1.05E+00 |
| LINC02482 | 1.13E-01 | 5.89E-01 | 1.26E+00 | 1.62E-05 | 7.02E-05 | -1.37E+00 | 4.13E-02 | 1.76E-01 | -1.09E+00 | 1.08E-03 | 4.14E-03 | -1.27E+00 | 8.50E-02 | 3.03E-01 | 1.05E+00 | 2.55E-02 | 7.19E-02 | -1.17E+00 | 9.44E-02 | 3.46E-01 | 1.02E+00 | 1.38E-01 | 3.79E-01 | 1.13E+00 |
| KRT10 | 1.13E-01 | 5.89E-01 | 1.20E+00 | 5.46E-05 | 2.36E-04 | 1.15E+00 | 1.33E-03 | 5.65E-03 | 1.50E+00 | 1.13E-01 | 4.32E-01 | 1.29E+00 | 3.88E-02 | 1.38E-01 | 1.56E+00 | 2.74E-01 | 7.71E-01 | 1.30E+00 | 9.00E-02 | 3.30E-01 | 1.35E+00 | 2.45E-01 | 6.73E-01 | 1.31E+00 |
| AC007240.1 | 1.13E-01 | 5.89E-01 | -1.29E+00 | 1.55E-01 | 6.68E-01 | -1.19E+00 | 2.32E-02 | 9.88E-02 | 1.73E+00 | 5.71E-03 | 2.18E-02 | 2.22E+00 | 2.35E-01 | 8.37E-01 | 1.20E+00 | 3.55E-01 | 1.00E+00 | -1.00E+00 | 2.27E-01 | 8.32E-01 | 1.12E+00 | 3.83E-02 | 1.05E-01 | 1.69E+00 |
| MTBP | 1.13E-01 | 5.89E-01 | 1.14E+00 | 1.55E-01 | 6.70E-01 | 1.19E+00 | 1.80E-01 | 7.64E-01 | 1.32E+00 | 1.52E-01 | 5.83E-01 | 1.23E+00 | 1.67E-01 | 5.94E-01 | 1.67E+00 | 2.63E-01 | 7.42E-01 | 1.18E+00 | 3.70E-02 | 1.36E-01 | 1.04E+00 | 2.89E-01 | 7.94E-01 | 1.27E+00 |
| COL8A2 | 1.13E-01 | 5.89E-01 | 1.07E+00 | 1.94E-01 | 8.40E-01 | 1.51E+00 | 9.43E-02 | 4.01E-01 | 1.38E+00 | 8.15E-02 | 3.11E-01 | 1.63E+00 | 2.65E-01 | 9.44E-01 | 1.02E+00 | 3.04E-01 | 8.57E-01 | -1.10E+00 | 2.16E-01 | 7.93E-01 | 1.17E+00 | 1.94E-01 | 5.32E-01 | 1.58E+00 |
| CSTA | 1.13E-01 | 5.89E-01 | -1.26E+00 | 1.30E-01 | 5.62E-01 | -1.13E+00 | 2.28E-01 | 9.69E-01 | -1.47E+00 | 2.75E-02 | 1.05E-01 | -2.50E+00 | 2.80E-01 | 1.00E+00 | 1.00E+00 | 3.55E-01 | 1.00E+00 | -1.00E+00 | 1.06E-01 | 3.89E-01 | -1.45E+00 | 1.16E-01 | 3.19E-01 | -1.83E+00 |
| AC010186.2 | 1.13E-01 | 5.89E-01 | 1.32E+00 | 4.57E-02 | 1.98E-01 | 1.83E+00 | 2.74E-03 | 1.16E-02 | -1.21E+00 | 6.70E-03 | 2.56E-02 | 1.04E+00 | 1.96E-01 | 6.98E-01 | -1.07E+00 | 2.50E-01 | 7.05E-01 | 1.63E+00 | 3.60E-02 | 1.32E-01 | -1.27E+00 | 2.70E-02 | 7.41E-02 | -1.43E+00 |
| SNHG3 | 1.13E-01 | 5.89E-01 | 1.24E+00 | 1.70E-01 | 7.37E-01 | 1.38E+00 | 7.85E-02 | 3.33E-01 | 1.18E+00 | 6.39E-02 | 2.44E-01 | 1.30E+00 | 1.26E-01 | 4.50E-01 | 1.17E+00 | 1.10E-02 | 3.10E-02 | 1.67E+00 | 1.50E-01 | 5.52E-01 | 1.23E+00 | 7.46E-04 | 2.05E-03 | 1.88E+00 |
| AL355312.2 | 1.13E-01 | 5.90E-01 | 1.35E+00 | 7.00E-02 | 3.03E-01 | 1.76E+00 | 7.16E-03 | 3.04E-02 | -1.49E+00 | 5.16E-03 | 1.97E-02 | -1.45E+00 | 5.20E-02 | 1.85E-01 | -1.07E+00 | 3.95E-03 | 1.11E-02 | -1.84E+00 | 2.10E-02 | 7.72E-02 | -1.66E+00 | 1.51E-01 | 4.13E-01 | -1.17E+00 |
| LYG2 | 1.13E-01 | 5.90E-01 | -1.06E+00 | 1.72E-01 | 7.43E-01 | 1.44E+00 | 5.08E-03 | 2.16E-02 | -1.64E+00 | 3.48E-02 | 1.33E-01 | -1.18E+00 | 6.23E-03 | 2.22E-02 | -1.82E+00 | 2.78E-01 | 7.85E-01 | 1.14E+00 | 3.70E-02 | 1.36E-01 | -1.39E+00 | 8.14E-02 | 2.24E-01 | -1.35E+00 |
| SBF2-AS1 | 1.13E-01 | 5.90E-01 | 1.07E+00 | 9.68E-02 | 4.18E-01 | 1.16E+00 | 1.12E-01 | 4.75E-01 | -1.20E+00 | 2.03E-01 | 7.75E-01 | 1.08E+00 | 1.40E-01 | 4.98E-01 | -1.43E+00 | 3.13E-01 | 8.81E-01 | 1.07E+00 | 1.00E-02 | 3.68E-02 | -1.40E+00 | 2.34E-02 | 6.44E-02 | -1.30E+00 |
| SIRPB2 | 1.13E-01 | 5.90E-01 | 1.31E+00 | 1.71E-01 | 7.37E-01 | 1.24E+00 | 4.36E-02 | 1.85E-01 | -1.03E+00 | 1.02E-01 | 3.90E-01 | 1.08E+00 | 2.15E-01 | 7.65E-01 | -1.15E+00 | 2.33E-01 | 6.56E-01 | -1.30E+00 | 2.04E-01 | 7.48E-01 | -1.05E+00 | 2.49E-01 | 6.84E-01 | -1.13E+00 |
| AC091045.1 | 1.13E-01 | 5.90E-01 | 1.06E+00 | 2.23E-01 | 9.65E-01 | 1.37E+00 | 2.76E-02 | 1.17E-01 | -1.28E+00 | 1.15E-01 | 4.40E-01 | 1.10E+00 | 1.89E-03 | 6.72E-03 | -1.31E+00 | 2.60E-02 | 7.34E-02 | -1.05E+00 | 1.28E-02 | 4.68E-02 | -1.29E+00 | 3.33E-01 | 9.14E-01 | 1.24E+00 |
| STK32C | 1.13E-01 | 5.91E-01 | 1.42E+00 | 8.40E-02 | 3.63E-01 | 1.21E+00 | 1.27E-01 | 5.40E-01 | 1.37E+00 | 9.48E-02 | 3.62E-01 | 1.24E+00 | 2.13E-01 | 7.61E-01 | 1.32E+00 | 2.13E-01 | 6.01E-01 | 1.24E+00 | 2.54E-01 | 9.33E-01 | 1.34E+00 | 2.66E-01 | 7.29E-01 | 1.35E+00 |
| AL391416.1 | 1.13E-01 | 5.91E-01 | 1.15E+00 | 8.14E-02 | 3.52E-01 | 1.15E+00 | 1.71E-01 | 7.27E-01 | 1.25E+00 | 9.84E-03 | 3.76E-02 | -1.03E+00 | 2.35E-01 | 8.38E-01 | 1.25E+00 | 1.51E-01 | 4.26E-01 | -1.01E+00 | 2.55E-01 | 9.38E-01 | 1.24E+00 | 1.88E-01 | 5.16E-01 | 1.05E+00 |
| AP002360.2 | 1.13E-01 | 5.91E-01 | 1.14E+00 | 1.38E-02 | 5.95E-02 | -1.00E+00 | 1.71E-01 | 7.27E-01 | 1.18E+00 | 7.67E-03 | 2.93E-02 | -1.03E+00 | 1.00E-01 | 3.57E-01 | 1.07E+00 | 8.90E-02 | 2.51E-01 | 1.05E+00 | 1.39E-02 | 5.09E-02 | 1.78E+00 | 3.03E-01 | 8.32E-01 | 1.45E+00 |
| ZBTB6 | 1.13E-01 | 5.91E-01 | 1.55E+00 | 1.37E-01 | 5.94E-01 | 1.67E+00 | 1.58E-01 | 6.70E-01 | 1.22E+00 | 2.21E-01 | 8.43E-01 | 1.31E+00 | 4.62E-02 | 1.65E-01 | -1.01E+00 | 3.49E-02 | 9.83E-02 | -1.09E+00 | 1.32E-01 | 4.84E-01 | 1.56E+00 | 2.61E-01 | 7.18E-01 | 1.47E+00 |
| ARTN | 1.13E-01 | 5.92E-01 | -1.04E+00 | 2.29E-01 | 9.88E-01 | 1.30E+00 | 1.19E-01 | 5.04E-01 | 1.53E+00 | 1.87E-01 | 7.15E-01 | 1.26E+00 | 1.39E-02 | 4.94E-02 | 2.72E+00 | 1.28E-01 | 3.60E-01 | 1.89E+00 | 3.16E-06 | 1.16E-05 | 2.34E+00 | 3.43E-01 | 9.42E-01 | 1.24E+00 |
| AC012146.1 | 1.14E-01 | 5.92E-01 | 1.62E+00 | 7.37E-03 | 3.18E-02 | 1.08E+00 | 1.11E-01 | 4.72E-01 | 1.38E+00 | 1.10E-02 | 4.18E-02 | 1.06E+00 | 2.56E-01 | 9.14E-01 | 1.47E+00 | 2.23E-01 | 6.29E-01 | 1.23E+00 | 1.70E-01 | 6.25E-01 | 1.25E+00 | 3.13E-02 | 8.61E-02 | 1.05E+00 |
| CDK2 | 1.14E-01 | 5.92E-01 | 1.14E+00 | 1.25E-01 | 5.38E-01 | 1.23E+00 | 1.88E-01 | 7.99E-01 | 1.04E+00 | 2.51E-01 | 9.58E-01 | 1.22E+00 | 7.28E-02 | 2.60E-01 | -1.01E+00 | 1.26E-01 | 3.55E-01 | 1.11E+00 | 8.30E-04 | 3.05E-03 | -1.00E+00 | 1.04E-01 | 2.84E-01 | 1.16E+00 |
| TRUB1 | 1.14E-01 | 5.92E-01 | 1.10E+00 | 9.45E-02 | 4.08E-01 | 1.08E+00 | 1.18E-01 | 5.02E-01 | 1.12E+00 | 5.92E-02 | 2.26E-01 | 1.15E+00 | 1.37E-02 | 4.87E-02 | 1.03E+00 | 1.91E-02 | 5.37E-02 | 1.03E+00 | 6.08E-06 | 2.23E-05 | -1.20E+00 | 1.60E-04 | 4.39E-04 | -1.07E+00 |

| | | | | | | | | | | | | | | | | | | | | | | | | |
|---|---|---|---|---|---|---|---|---|---|---|---|---|---|---|---|---|---|---|---|---|---|---|---|---|
| HBM | 1.14E-01 | 5.92E-01 | 3.25E+00 | 2.00E-01 | 8.66E-01 | 1.88E+00 | 1.77E-01 | 7.53E-01 | 1.17E+00 | 2.62E-01 | 1.00E+00 | -1.00E+00 | 1.50E-01 | 5.36E-01 | 7.78E+00 | 3.19E-01 | 8.99E-01 | 1.28E+00 | 4.00E-02 | 1.47E-01 | 1.66E+00 | 1.83E-01 | 5.03E-01 | 1.44E+00 |
| AC009005.1 | 1.14E-01 | 5.93E-01 | 1.07E+00 | 1.39E-01 | 6.00E-01 | 1.12E+00 | 4.38E-02 | 1.86E-01 | -1.11E+00 | 6.53E-04 | 2.49E-03 | -1.42E+00 | 2.70E-02 | 9.61E-02 | -1.74E+00 | 1.87E-02 | 5.28E-02 | -2.14E+00 | 9.21E-03 | 3.38E-02 | -1.41E+00 | 1.09E-03 | 3.00E-03 | -1.67E+00 |
| USP54 | 1.14E-01 | 5.93E-01 | 1.14E+00 | 2.11E-01 | 9.11E-01 | 1.58E+00 | 1.12E-01 | 4.74E-01 | 1.45E+00 | 1.61E-01 | 6.14E-01 | 1.29E+00 | 1.97E-01 | 7.02E-01 | 1.09E+00 | 2.64E-01 | 7.45E-01 | 1.67E+00 | 7.87E-02 | 2.89E-01 | -1.09E+00 | 3.58E-01 | 9.83E-01 | 1.20E+00 |
| CCR1 | 1.14E-01 | 5.94E-01 | -1.13E+00 | 1.41E-01 | 6.09E-01 | -1.13E+00 | 1.51E-07 | 6.42E-07 | -2.11E+00 | 2.66E-07 | 1.01E-06 | -2.14E+00 | 2.80E-01 | 1.00E+00 | 1.00E+00 | 3.10E-01 | 8.74E-01 | 1.11E+00 | 1.39E-01 | 5.11E-01 | 1.19E+00 | 3.64E-01 | 1.00E+00 | -1.00E+00 |
| CENPB | 1.14E-01 | 5.94E-01 | 1.33E+00 | 2.02E-01 | 8.71E-01 | 1.26E+00 | 9.86E-02 | 4.19E-01 | 1.12E+00 | 2.52E-01 | 9.63E-01 | 1.43E+00 | 1.96E-01 | 6.99E-01 | 1.19E+00 | 2.80E-01 | 7.88E-01 | 1.46E+00 | 2.22E-01 | 8.14E-01 | 1.22E+00 | 2.94E-01 | 8.07E-01 | 1.34E+00 |
| ZNF492 | 1.14E-01 | 5.96E-01 | 1.21E+00 | 1.62E-02 | 7.00E-02 | -1.13E+00 | 5.74E-02 | 2.44E-01 | -1.02E+00 | 5.27E-04 | 2.01E-03 | -1.38E+00 | 1.31E-01 | 4.66E-01 | 1.93E+00 | 3.24E-01 | 9.14E-01 | 1.34E+00 | 2.71E-04 | 9.94E-04 | 1.87E+00 | 2.68E-01 | 7.36E-01 | 1.37E+00 |
| MATN2 | 1.14E-01 | 5.96E-01 | -1.05E+00 | 1.34E-01 | 5.79E-01 | 1.64E+00 | 1.26E-01 | 5.35E-01 | -1.10E+00 | 1.81E-01 | 6.92E-01 | 1.09E+00 | 1.45E-03 | 5.17E-03 | -2.34E+00 | 3.68E-02 | 1.04E-01 | -1.38E+00 | 1.07E-02 | 3.92E-02 | -1.23E+00 | 2.88E-01 | 7.92E-01 | 1.23E+00 |
| ZWILCH | 1.14E-01 | 5.96E-01 | 1.26E+00 | 1.50E-01 | 6.47E-01 | 1.41E+00 | 1.07E-01 | 4.54E-01 | 1.11E+00 | 6.89E-02 | 2.63E-01 | 1.30E+00 | 4.09E-02 | 1.46E-01 | -1.26E+00 | 1.81E-01 | 5.10E-01 | 1.06E+00 | 4.53E-03 | 1.66E-02 | -1.03E+00 | 3.48E-01 | 9.55E-01 | 1.31E+00 |
| AL669983.1 | 1.15E-01 | 5.97E-01 | 1.23E+00 | 6.05E-19 | 2.61E-18 | -1.30E+00 | 1.48E-02 | 6.30E-02 | 1.41E+00 | 1.34E-09 | 5.12E-09 | -1.25E+00 | 2.50E-01 | 8.91E-01 | 1.31E+00 | 6.97E-08 | 1.96E-07 | -1.32E+00 | 2.68E-01 | 9.84E-01 | 1.27E+00 | 6.76E-07 | 1.86E-06 | -1.28E+00 |
| ABHD5 | 1.15E-01 | 5.97E-01 | 1.24E+00 | 3.86E-04 | 1.67E-03 | -1.29E+00 | 9.32E-02 | 3.96E-01 | 1.38E+00 | 4.48E-03 | 1.71E-02 | -1.15E+00 | 9.75E-02 | 3.48E-01 | 1.19E+00 | 8.06E-02 | 2.27E-01 | 1.12E+00 | 2.96E-02 | 1.09E-01 | 1.13E+00 | 7.49E-05 | 2.06E-04 | -1.22E+00 |
| PROSER3 | 1.15E-01 | 5.97E-01 | 1.21E+00 | 2.00E-01 | 8.65E-01 | 1.46E+00 | 1.61E-01 | 6.85E-01 | 1.10E+00 | 2.16E-01 | 8.25E-01 | 1.56E+00 | 2.50E-01 | 8.90E-01 | 1.28E+00 | 2.02E-01 | 5.70E-01 | 1.11E+00 | 2.56E-01 | 9.39E-01 | 1.42E+00 | 3.07E-01 | 8.44E-01 | 1.45E+00 |
| RGS6 | 1.15E-01 | 5.97E-01 | -1.14E+00 | 1.41E-01 | 6.11E-01 | -1.14E+00 | 1.36E-01 | 5.76E-01 | -1.13E+00 | 1.54E-01 | 5.89E-01 | -1.13E+00 | 6.19E-02 | 2.21E-01 | -1.45E+00 | 4.51E-02 | 1.27E-01 | -1.83E+00 | 1.86E-02 | 6.84E-02 | 2.39E+00 | 6.16E-02 | 1.69E-01 | 1.05E+00 |
| NBEAL2 | 1.15E-01 | 5.97E-01 | 1.29E+00 | 1.02E-01 | 4.40E-01 | 1.40E+00 | 2.10E-01 | 8.90E-01 | 1.34E+00 | 1.42E-01 | 5.41E-01 | 1.66E+00 | 2.30E-01 | 8.20E-01 | 1.43E+00 | 8.09E-02 | 2.28E-01 | 1.08E+00 | 1.93E-02 | 7.10E-02 | 1.86E+00 | 1.42E-01 | 3.91E-01 | 1.25E+00 |
| AL031056.1 | 1.15E-01 | 5.97E-01 | 2.10E+00 | 2.28E-01 | 9.86E-01 | 1.32E+00 | 1.31E-01 | 5.58E-01 | 1.15E+00 | 1.03E-01 | 3.92E-01 | 1.48E+00 | 1.25E-01 | 4.45E-01 | -3.04E+00 | 3.55E-01 | 1.00E+00 | -1.91E+00 | 2.63E-01 | 9.66E-01 | 1.12E+00 | 1.91E-01 | 5.25E-01 | -1.19E+00 |
| FOXK2 | 1.15E-01 | 5.98E-01 | 1.26E+00 | 3.20E-02 | 1.38E-01 | 1.15E+00 | 1.35E-03 | 5.75E-03 | -1.24E+00 | 1.65E-04 | 6.31E-04 | -1.21E+00 | 1.58E-01 | 5.62E-01 | 1.22E+00 | 2.93E-01 | 8.27E-01 | 1.45E+00 | 2.08E-04 | 7.64E-04 | -1.24E+00 | 2.37E-01 | 6.52E-01 | 1.24E+00 |
| KLF12 | 1.15E-01 | 5.98E-01 | 1.26E+00 | 4.69E-02 | 2.03E-01 | 2.02E+00 | 3.40E-03 | 1.45E-02 | -1.14E+00 | 4.49E-02 | 1.72E-01 | 1.18E+00 | 2.76E-01 | 9.83E-01 | 1.34E+00 | 1.96E-01 | 5.52E-01 | 1.64E+00 | 1.15E-05 | 4.21E-05 | -1.65E+00 | 3.83E-04 | 1.05E-03 | -1.45E+00 |
| SEMA6C | 1.15E-01 | 5.98E-01 | 1.53E+00 | 6.22E-02 | 2.69E-01 | 1.84E+00 | 1.09E-01 | 4.63E-01 | -1.03E+00 | 2.19E-01 | 8.38E-01 | 1.26E+00 | 1.31E-01 | 4.66E-01 | -1.00E+00 | 2.02E-01 | 5.69E-01 | 1.64E+00 | 6.40E-03 | 2.35E-02 | -1.13E+00 | 8.12E-02 | 2.23E-01 | 1.01E+00 |
| PFN2 | 1.15E-01 | 5.99E-01 | 1.35E+00 | 2.04E-02 | 8.84E-02 | 2.03E+00 | 2.34E-01 | 9.96E-01 | 1.13E+00 | 1.81E-01 | 6.90E-01 | 1.46E+00 | 1.33E-01 | 4.76E-01 | -1.14E+00 | 5.79E-02 | 1.63E-01 | -1.65E+00 | 1.47E-01 | 5.38E-01 | 1.55E+00 | 2.40E-01 | 6.59E-01 | 1.15E+00 |
| C6orf163 | 1.15E-01 | 5.99E-01 | 1.06E+00 | 1.83E-01 | 7.90E-01 | 1.27E+00 | 1.12E-01 | 4.76E-01 | 1.42E+00 | 7.85E-02 | 3.00E-01 | 1.71E+00 | 7.30E-02 | 2.60E-01 | -1.27E+00 | 1.14E-01 | 3.21E-01 | -1.15E+00 | 9.65E-04 | 3.54E-03 | -1.57E+00 | 8.43E-02 | 2.32E-01 | -1.07E+00 |
| SLC9A3-AS1 | 1.15E-01 | 5.99E-01 | 1.54E+00 | 2.00E-01 | 8.63E-01 | 1.37E+00 | 9.48E-02 | 4.03E-01 | -1.04E+00 | 1.90E-01 | 7.26E-01 | 1.24E+00 | 1.96E-01 | 6.97E-01 | 1.12E+00 | 1.59E-01 | 4.48E-01 | 1.83E+00 | 1.27E-01 | 4.65E-01 | 1.45E+00 | 2.85E-01 | 7.83E-01 | 1.26E+00 |
| AC023090.2 | 1.15E-01 | 6.00E-01 | 1.10E+00 | 5.23E-02 | 2.26E-01 | -1.07E+00 | 2.40E-04 | 1.02E-03 | -1.60E+00 | 1.39E-03 | 5.32E-03 | -1.26E+00 | 2.00E-01 | 7.14E-01 | 1.14E+00 | 1.84E-01 | 5.20E-01 | 1.01E+00 | 9.16E-04 | 3.36E-03 | -1.89E+00 | 7.74E-03 | 2.13E-02 | -1.54E+00 |
| TRAF1 | 1.15E-01 | 6.00E-01 | -1.00E+00 | 1.33E-01 | 5.74E-01 | 1.12E+00 | 6.32E-02 | 2.69E-01 | -1.24E+00 | 1.68E-01 | 6.42E-01 | 1.03E+00 | 1.88E-01 | 6.70E-01 | 1.65E+00 | 2.68E-01 | 7.56E-01 | 1.36E+00 | 2.30E-01 | 8.46E-01 | 1.22E+00 | 3.43E-01 | 9.41E-01 | 1.05E+00 |
| NEK11 | 1.15E-01 | 6.00E-01 | 1.22E+00 | 2.31E-02 | 9.98E-02 | -1.01E+00 | 2.28E-01 | 9.69E-01 | 1.23E+00 | 1.31E-01 | 5.00E-01 | 1.12E+00 | 8.48E-02 | 3.03E-01 | -1.53E+00 | 8.78E-02 | 2.48E-01 | -1.61E+00 | 1.54E-01 | 5.64E-01 | -1.14E+00 | 3.14E-01 | 8.62E-01 | 1.05E+00 |
| FOXN3-AS1 | 1.15E-01 | 6.00E-01 | 1.29E+00 | 1.63E-01 | 7.03E-01 | -1.03E+00 | 2.18E-01 | 9.28E-01 | 1.18E+00 | 3.30E-02 | 1.26E-01 | -1.29E+00 | 2.41E-01 | 8.58E-01 | 1.07E+00 | 3.40E-01 | 9.58E-01 | 1.34E+00 | 1.46E-01 | 5.34E-01 | 1.33E+00 | 1.13E-01 | 3.10E-01 | -1.08E+00 |
| EIF3K | 1.15E-01 | 6.00E-01 | 1.02E+00 | 1.38E-01 | 5.95E-01 | 1.04E+00 | 2.14E-01 | 9.10E-01 | 1.02E+00 | 2.31E-01 | 8.82E-01 | 1.04E+00 | 1.27E-01 | 4.54E-01 | -1.01E+00 | 3.55E-01 | 1.00E+00 | 1.03E+00 | 3.35E-02 | 1.23E-01 | -1.00E+00 | 2.49E-01 | 6.85E-01 | 1.07E+00 |
| LPAR5 | 1.15E-01 | 6.00E-01 | 1.77E+00 | 4.23E-02 | 1.83E-01 | 2.54E+00 | 2.03E-02 | 8.62E-02 | -1.35E+00 | 2.51E-01 | 9.58E-01 | 1.41E+00 | 5.42E-02 | 1.93E-01 | 1.14E+00 | 3.00E-01 | 8.47E-01 | 1.52E+00 | 6.75E-03 | 2.48E-02 | 2.09E+00 | 2.98E-02 | 8.19E-02 | 1.84E+00 |
| GRK3 | 1.15E-01 | 6.00E-01 | 1.28E+00 | 1.25E-01 | 5.40E-01 | 1.40E+00 | 3.03E-02 | 1.29E-01 | 1.06E+00 | 1.69E-01 | 6.44E-01 | 1.43E+00 | 6.08E-02 | 2.17E-01 | 1.03E+00 | 4.98E-02 | 1.40E-01 | -1.04E+00 | 1.74E-02 | 6.39E-02 | -1.19E+00 | 1.02E-01 | 2.79E-01 | -1.06E+00 |
| RPS3AP49 | 1.15E-01 | 6.01E-01 | 1.57E+00 | 5.29E-02 | 2.29E-01 | 1.01E+00 | 1.82E-01 | 7.74E-01 | 1.19E+00 | 5.13E-02 | 1.96E-01 | 1.04E+00 | 2.74E-01 | 9.76E-01 | 1.34E+00 | 2.46E-01 | 6.94E-01 | 1.26E+00 | 2.66E-01 | 9.76E-01 | 1.15E+00 | 3.56E-01 | 9.77E-01 | 1.16E+00 |
| AL135791.1 | 1.15E-01 | 6.01E-01 | 1.15E+00 | 2.04E-01 | 8.83E-01 | 1.26E+00 | 4.22E-02 | 1.79E-01 | -1.27E+00 | 2.15E-01 | 8.22E-01 | 1.21E+00 | 2.71E-01 | 9.68E-01 | 1.22E+00 | 2.12E-01 | 5.97E-01 | 1.73E+00 | 8.54E-02 | 3.13E-01 | 1.55E+00 | 1.42E-01 | 3.90E-01 | 1.59E+00 |
| AC092620.3 | 1.15E-01 | 6.01E-01 | 1.42E+00 | 1.53E-01 | 6.60E-01 | -1.01E+00 | 1.61E-01 | 6.83E-01 | -1.03E+00 | 1.13E-01 | 4.31E-01 | -1.18E+00 | 8.87E-02 | 3.16E-01 | -1.01E+00 | 2.39E-01 | 6.75E-01 | 1.66E+00 | 2.51E-01 | 9.23E-01 | 1.13E+00 | 4.56E-02 | 1.25E-01 | 1.94E+00 |
| HIST1H2BJ | 1.15E-01 | 6.01E-01 | 1.04E+00 | 8.13E-02 | 3.51E-01 | -1.06E+00 | 1.55E-01 | 6.58E-01 | 1.45E+00 | 2.17E-01 | 8.27E-01 | 1.22E+00 | 2.14E-02 | 7.62E-02 | -1.30E+00 | 8.25E-02 | 2.33E-01 | -1.06E+00 | 9.00E-03 | 3.30E-02 | 1.42E+00 | 3.10E-04 | 8.50E-04 | -1.04E+00 |
| ATP9A | 1.15E-01 | 6.01E-01 | 1.18E+00 | 1.70E-01 | 7.32E-01 | 1.61E+00 | 6.80E-02 | 2.89E-01 | 1.50E+00 | 2.13E-01 | 8.13E-01 | 1.38E+00 | 2.69E-01 | 9.61E-01 | 1.34E+00 | 3.30E-01 | 9.31E-01 | 1.39E+00 | 2.60E-01 | 9.54E-01 | 1.09E+00 | 2.41E-01 | 6.62E-01 | 1.40E+00 |
| RNF43 | 1.15E-01 | 6.02E-01 | 1.23E+00 | 1.82E-01 | 7.86E-01 | 1.43E+00 | 4.98E-02 | 2.12E-01 | -1.14E+00 | 1.15E-01 | 4.41E-01 | 1.62E+00 | 1.54E-01 | 5.48E-01 | 1.04E+00 | 3.16E-01 | 8.91E-01 | 1.48E+00 | 2.40E-01 | 8.82E-01 | 1.19E+00 | 6.77E-02 | 1.86E-01 | 1.72E+00 |
| PURG | 1.16E-01 | 6.02E-01 | 1.18E+00 | 7.70E-02 | 3.33E-01 | 1.69E+00 | 1.57E-04 | 6.67E-04 | -1.86E+00 | 6.94E-03 | 2.65E-02 | -1.19E+00 | 3.94E-02 | 1.40E-01 | -1.30E+00 | 5.40E-02 | 1.52E-01 | -1.22E+00 | 1.61E-02 | 5.90E-02 | -1.65E+00 | 9.09E-02 | 2.50E-01 | -1.27E+00 |
| HASPIN | 1.16E-01 | 6.02E-01 | -1.19E+00 | 1.34E-01 | 5.77E-01 | -1.18E+00 | 1.95E-01 | 8.28E-01 | 1.12E+00 | 2.28E-01 | 8.70E-01 | 1.18E+00 | 1.33E-01 | 4.74E-01 | -2.30E+00 | 2.59E-01 | 7.30E-01 | -1.88E+00 | 1.47E-01 | 5.41E-01 | 1.15E+00 | 1.70E-02 | 4.66E-02 | -1.03E+00 |
| GAS5 | 1.16E-01 | 6.02E-01 | -1.03E+00 | 2.06E-01 | 8.92E-01 | 1.00E+00 | 2.54E-02 | 1.08E-01 | -1.08E+00 | 2.04E-01 | 7.81E-01 | 1.04E+00 | 3.81E-02 | 1.36E-01 | 1.13E+00 | 1.13E-01 | 3.18E-01 | 1.08E+00 | 2.59E-01 | 9.49E-01 | 1.16E+00 | 3.04E-01 | 8.35E-01 | 1.04E+00 |
| AC092979.1 | 1.16E-01 | 6.03E-01 | -1.15E+00 | 1.90E-01 | 8.23E-01 | 1.08E+00 | 2.16E-01 | 9.20E-01 | 1.10E+00 | 2.25E-01 | 8.61E-01 | 1.26E+00 | 2.07E-01 | 7.39E-01 | -1.32E+00 | 5.71E-02 | 1.61E-01 | -1.63E+00 | 5.51E-02 | 2.02E-01 | -1.06E+00 | 7.89E-02 | 2.17E-01 | -1.01E+00 |
| C19orf57 | 1.16E-01 | 6.03E-01 | 1.57E+00 | 1.79E-01 | 7.72E-01 | 1.54E+00 | 2.12E-01 | 8.99E-01 | 1.15E+00 | 1.92E-01 | 7.33E-01 | 1.39E+00 | 2.24E-01 | 7.99E-01 | 1.01E+00 | 2.41E-01 | 6.81E-01 | -1.16E+00 | 1.99E-01 | 7.29E-01 | 1.30E+00 | 1.26E-01 | 3.45E-01 | 1.52E+00 |
| EPAS1 | 1.16E-01 | 6.03E-01 | -1.31E+00 | 1.71E-01 | 7.38E-01 | -1.07E+00 | 2.14E-01 | 9.10E-01 | 1.02E+00 | 2.45E-01 | 9.37E-01 | 1.05E+00 | 5.82E-05 | 2.07E-04 | -1.76E+00 | 1.02E-02 | 2.89E-02 | -1.13E+00 | 3.03E-11 | 1.11E-10 | -1.38E+00 | 1.33E-10 | 3.65E-10 | -1.57E+00 |
| ASTE1 | 1.16E-01 | 6.03E-01 | 1.29E+00 | 1.71E-02 | 7.39E-02 | 1.25E+00 | 9.85E-02 | 4.19E-01 | 1.10E+00 | 3.71E-02 | 1.42E-01 | 1.13E+00 | 2.48E-01 | 8.86E-01 | 1.14E+00 | 2.38E-01 | 6.70E-01 | 1.44E+00 | 1.67E-01 | 6.14E-01 | 1.29E+00 | 3.24E-01 | 8.89E-01 | 1.13E+00 |
| CD1D | 1.16E-01 | 6.04E-01 | -1.22E+00 | 1.98E-01 | 8.54E-01 | -1.04E+00 | 3.91E-03 | 1.66E-02 | -1.45E+00 | 1.01E-03 | 3.86E-03 | -1.64E+00 | 2.80E-01 | 1.00E+00 | 1.00E+00 | 3.19E-01 | 8.99E-01 | 1.09E+00 | 1.27E-03 | 4.66E-03 | -1.50E+00 | 2.74E-02 | 7.52E-02 | -1.19E+00 |
| RGPD2 | 1.16E-01 | 6.04E-01 | 1.40E+00 | 7.33E-03 | 3.17E-02 | 1.11E+00 | 4.57E-02 | 1.94E-01 | 1.43E+00 | 1.65E-03 | 6.32E-03 | -1.07E+00 | 6.96E-02 | 2.48E-01 | 1.70E+00 | 8.42E-02 | 2.37E-01 | 1.04E+00 | 2.62E-01 | 9.62E-01 | 1.37E+00 | 2.64E-03 | 7.24E-03 | -1.37E+00 |
| ATP13A2 | 1.16E-01 | 6.04E-01 | 1.32E+00 | 1.33E-02 | 5.73E-02 | 2.41E+00 | 1.42E-02 | 6.04E-02 | -1.18E+00 | 1.51E-01 | 5.76E-01 | 1.22E+00 | 7.04E-03 | 2.51E-02 | -1.35E+00 | 1.52E-01 | 4.28E-01 | 1.09E+00 | 4.70E-04 | 1.73E-03 | -1.13E+00 | 2.54E-02 | 6.98E-02 | 1.08E+00 |
| LCAT | 1.16E-01 | 6.05E-01 | 1.08E+00 | 1.41E-01 | 6.09E-01 | 1.23E+00 | 7.68E-02 | 3.26E-01 | -1.09E+00 | 1.58E-01 | 6.02E-01 | 1.12E+00 | 2.57E-01 | 9.18E-01 | 1.27E+00 | 3.30E-01 | 9.29E-01 | 1.24E+00 | 1.29E-01 | 4.75E-01 | 1.58E+00 | 2.53E-01 | 6.94E-01 | 1.42E+00 |
| STKLD1 | 1.16E-01 | 6.06E-01 | 1.14E+00 | 5.45E-02 | 2.35E-01 | -1.03E+00 | 7.38E-02 | 3.13E-01 | -1.11E+00 | 4.88E-03 | 1.87E-02 | -1.40E+00 | 1.09E-02 | 3.87E-02 | -1.43E+00 | 1.21E-02 | 3.41E-02 | -1.49E+00 | 5.19E-06 | 1.91E-05 | -2.07E+00 | 2.87E-03 | 7.89E-03 | -1.30E+00 |
| SEC31B | 1.16E-01 | 6.06E-01 | 1.13E+00 | 4.07E-02 | 1.76E-01 | 1.86E+00 | 1.04E-04 | 4.40E-04 | -1.33E+00 | 8.23E-03 | 3.14E-02 | 1.08E+00 | 3.27E-02 | 1.17E-01 | -1.05E+00 | 1.71E-01 | 4.81E-01 | 1.24E+00 | 2.44E-01 | 8.97E-01 | 1.20E+00 | 9.54E-02 | 2.62E-01 | 1.65E+00 |
| CLC | 1.16E-01 | 6.06E-01 | -5.25E+01 | 1.83E-01 | 7.90E-01 | -4.53E+01 | 1.71E-01 | 7.27E-01 | 2.81E+00 | 1.14E-01 | 4.35E-01 | 4.13E+01 | 6.31E-02 | 2.25E-01 | 1.79E+00 | 1.96E-04 | 5.52E-04 | -2.64E+00 | 2.24E-01 | 8.21E-01 | 1.12E+00 | 2.54E-01 | 6.99E-01 | 2.63E+00 |
| AC092745.1 | 1.16E-01 | 6.06E-01 | 1.02E+00 | 8.94E-02 | 3.86E-01 | -1.07E+00 | 1.38E-01 | 5.86E-01 | -1.08E+00 | 1.12E-01 | 4.29E-01 | -1.20E+00 | 1.78E-01 | 6.35E-01 | 1.42E+00 | 1.96E-01 | 5.51E-01 | 1.01E+00 | 9.22E-02 | 3.38E-01 | 1.21E+00 | 3.38E-05 | 9.29E-05 | -1.22E+00 |
| AL121944.1 | 1.16E-01 | 6.06E-01 | 1.33E+00 | 1.61E-04 | 6.95E-04 | -1.23E+00 | 3.89E-03 | 1.65E-02 | 1.81E+00 | 1.48E-01 | 5.65E-01 | 1.29E+00 | 3.79E-02 | 1.35E-01 | 1.02E+00 | 1.22E-01 | 3.43E-01 | 1.16E+00 | 9.71E-03 | 3.56E-02 | -1.21E+00 | 1.83E-03 | 5.03E-03 | -1.41E+00 |
| FBXL19 | 1.16E-01 | 6.06E-01 | 1.19E+00 | 6.87E-02 | 2.97E-01 | 1.07E+00 | 1.76E-01 | 7.47E-01 | 1.19E+00 | 1.56E-01 | 5.97E-01 | 1.46E+00 | 5.93E-02 | 2.11E-01 | -1.04E+00 | 6.60E-02 | 1.86E-01 | -1.16E+00 | 3.09E-02 | 1.14E-01 | -1.18E+00 | 1.99E-02 | 5.46E-02 | -1.22E+00 |
| AL161891.1 | 1.16E-01 | 6.07E-01 | 1.00E+00 | 1.65E-01 | 7.11E-01 | 1.17E+00 | 2.63E-02 | 1.12E-01 | -1.08E+00 | 8.18E-02 | 3.13E-01 | 1.13E+00 | 9.68E-02 | 3.45E-01 | -1.30E+00 | 2.87E-01 | 8.10E-01 | 1.53E+00 | 1.16E-01 | 4.25E-01 | -1.11E+00 | 8.86E-02 | 2.43E-01 | 1.32E+00 |
| LINC01094 | 1.17E-01 | 6.08E-01 | -1.38E+00 | 1.60E-01 | 6.91E-01 | -1.23E+00 | 2.22E-01 | 9.44E-01 | 1.03E+00 | 2.32E-01 | 8.87E-01 | 1.28E+00 | 2.68E-06 | 9.55E-06 | -1.54E+00 | 3.76E-02 | 1.06E-01 | 1.06E+00 | 3.82E-05 | 1.40E-04 | -1.51E+00 | 2.56E-02 | 7.04E-02 | -1.07E+00 |
| FAM69B | 1.17E-01 | 6.08E-01 | 1.32E+00 | 5.11E-02 | 2.21E-01 | 1.57E+00 | 5.91E-02 | 2.51E-01 | 1.37E+00 | 2.41E-03 | 9.20E-03 | 1.74E+00 | 5.61E-02 | 2.00E-01 | 1.66E+00 | 4.02E-04 | 1.13E-03 | 2.03E+00 | 5.92E-07 | 2.17E-06 | 2.02E+00 | 8.91E-06 | 2.45E-05 | 1.84E+00 |
| RABGGTB | 1.17E-01 | 6.08E-01 | 1.20E+00 | 4.32E-02 | 1.87E-01 | 1.25E+00 | 2.11E-01 | 8.96E-01 | 1.23E+00 | 6.24E-02 | 2.38E-01 | 1.19E+00 | 9.61E-03 | 3.43E-02 | 1.00E+00 | 3.61E-02 | 1.02E-01 | 1.09E+00 | 9.06E-03 | 3.33E-02 | -1.07E+00 | 4.58E-02 | 1.26E-01 | 1.05E+00 |
| AC245060.4 | 1.17E-01 | 6.08E-01 | 1.10E+00 | 1.47E-01 | 6.36E-01 | 1.23E+00 | 2.70E-02 | 1.15E-01 | -1.18E+00 | 1.51E-02 | 5.76E-02 | -1.18E+00 | 1.46E-02 | 5.20E-02 | -1.49E+00 | 1.21E-01 | 3.41E-01 | 1.04E+00 | 1.98E-01 | 7.25E-01 | 1.41E+00 | 3.32E-01 | 9.13E-01 | 1.28E+00 |
| CHAF1B | 1.17E-01 | 6.09E-01 | 1.14E+00 | 1.52E-01 | 6.55E-01 | 1.25E+00 | 1.75E-01 | 7.44E-01 | 1.20E+00 | 3.17E-02 | 1.21E-01 | 1.19E+00 | 2.08E-01 | 7.40E-01 | 1.14E+00 | 1.57E-01 | 4.42E-01 | -1.10E+00 | 1.86E-01 | 6.84E-01 | 1.20E+00 | 1.86E-01 | 5.11E-01 | 1.34E+00 |
| TTC7A | 1.17E-01 | 6.09E-01 | 1.37E+00 | 1.85E-01 | 7.99E-01 | 1.47E+00 | 5.09E-03 | 2.16E-02 | -1.02E+00 | 2.07E-03 | 7.91E-03 | 1.06E+00 | 3.59E-02 | 1.28E-01 | -1.37E+00 | 4.93E-02 | 1.39E-01 | -1.41E+00 | 2.02E-01 | 7.41E-01 | -1.04E+00 | 2.41E-01 | 6.61E-01 | 1.46E+00 |
| PLEKHG4B | 1.17E-01 | 6.09E-01 | 1.19E+00 | 8.07E-02 | 3.49E-01 | 1.76E+00 | 6.22E-02 | 2.64E-01 | 1.35E+00 | 1.61E-01 | 6.14E-01 | 1.40E+00 | 8.34E-03 | 2.97E-02 | -1.13E+00 | 2.73E-02 | 7.70E-02 | -1.04E+00 | 2.44E-01 | 8.97E-01 | -1.04E+00 | 3.47E-01 | 9.52E-01 | -1.03E+00 |
| AL359764.1 | 1.17E-01 | 6.09E-01 | 1.27E+00 | 1.66E-01 | 7.16E-01 | 1.43E+00 | 2.17E-01 | 9.24E-01 | 1.20E+00 | 1.19E-01 | 4.53E-01 | 1.19E+00 | 1.88E-01 | 6.72E-01 | 1.16E+00 | 1.09E-01 | 3.07E-01 | 1.00E+00 | 2.47E-01 | 9.06E-01 | 1.00E+00 | 1.90E-01 | 5.22E-01 | 1.48E+00 |
| AC005076.1 | 1.17E-01 | 6.10E-01 | 1.16E+00 | 1.14E-01 | 4.93E-01 | 1.25E+00 | 7.98E-02 | 3.39E-01 | 1.01E+00 | 2.50E-03 | 9.56E-03 | -1.22E+00 | 4.76E-03 | 1.70E-02 | -1.42E+00 | 1.45E-02 | 4.09E-02 | -1.27E+00 | 7.13E-04 | 2.62E-03 | -1.13E+00 | 8.98E-04 | 2.47E-03 | -1.27E+00 |
| ABCC10 | 1.17E-01 | 6.10E-01 | -1.12E+00 | 1.57E-01 | 6.78E-01 | 1.29E+00 | 3.89E-02 | 1.65E-01 | -1.10E+00 | 2.09E-01 | 7.97E-01 | 1.29E+00 | 5.15E-03 | 1.83E-02 | -1.52E+00 | 2.59E-01 | 7.29E-01 | 1.29E+00 | 7.34E-03 | 2.70E-02 | -1.33E+00 | 2.40E-01 | 6.60E-01 | 1.07E+00 |
| AP001122.1 | 1.17E-01 | 6.10E-01 | 1.09E+00 | 9.89E-03 | 4.27E-02 | 1.95E+00 | 2.14E-01 | 9.10E-01 | 1.11E+00 | 1.94E-01 | 7.39E-01 | 1.44E+00 | 1.42E-01 | 5.06E-01 | 1.37E+00 | 2.22E-01 | 6.27E-01 | 1.47E+00 | 1.34E-01 | 4.90E-01 | 2.00E+00 | 1.15E-01 | 3.15E-01 | 1.89E+00 |
| AC139530.1 | 1.17E-01 | 6.10E-01 | 1.54E+00 | 1.18E-01 | 5.12E-01 | 1.34E+00 | 9.59E-02 | 4.07E-01 | 1.37E+00 | 2.20E-01 | 8.42E-01 | 1.38E+00 | 1.13E-01 | 4.03E-01 | 1.89E+00 | 3.22E-01 | 9.07E-01 | 1.54E+00 | 2.11E-01 | 7.73E-01 | 1.26E+00 | 1.63E-01 | 4.49E-01 | 1.16E+00 |
| TBX19 | 1.17E-01 | 6.10E-01 | -1.02E+00 | 3.42E-02 | 1.48E-01 | 1.85E+00 | 1.51E-02 | 6.42E-02 | -1.44E+00 | 2.20E-01 | 8.39E-01 | 1.47E+00 | 6.52E-02 | 2.32E-01 | -1.51E+00 | 3.06E-01 | 8.62E-01 | 1.31E+00 | 3.11E-02 | 1.14E-01 | -1.49E+00 | 2.95E-01 | 8.09E-01 | 1.05E+00 |
| MASTL | 1.17E-01 | 6.10E-01 | 1.10E+00 | 2.64E-03 | 1.14E-02 | -1.66E+00 | 1.95E-02 | 8.28E-02 | 1.04E+00 | 8.01E-04 | 3.06E-03 | -1.27E+00 | 2.52E-01 | 8.98E-01 | 1.29E+00 | 3.00E-01 | 8.47E-01 | 1.18E+00 | 3.50E-02 | 1.28E-01 | 1.20E+00 | 1.38E-03 | 3.78E-03 | -1.15E+00 |
| TMEM56 | 1.17E-01 | 6.10E-01 | 1.33E+00 | 2.13E-01 | 9.21E-01 | 1.08E+00 | 2.23E-01 | 9.47E-01 | 1.00E+00 | 1.89E-01 | 7.22E-01 | -1.17E+00 | 2.17E-01 | 7.74E-01 | 1.30E+00 | 1.90E-01 | 5.35E-01 | 2.31E+00 | 1.62E-01 | 5.94E-01 | -1.16E+00 | 8.09E-02 | 2.22E-01 | 1.40E+00 |
| SAMD14 | 1.17E-01 | 6.10E-01 | -1.15E+00 | 1.51E-01 | 6.53E-01 | -1.10E+00 | 1.12E-01 | 4.75E-01 | -1.23E+00 | 1.08E-01 | 4.13E-01 | -1.29E+00 | 1.81E-01 | 6.44E-01 | -1.17E+00 | 2.32E-01 | 6.55E-01 | -1.17E+00 | 1.89E-01 | 6.93E-01 | 1.60E+00 | 5.34E-04 | 1.47E-03 | -1.23E+00 |
| TRGC2 | 1.17E-01 | 6.10E-01 | 1.20E+00 | 1.05E-01 | 4.52E-01 | 1.32E+00 | 1.30E-08 | 5.51E-08 | -1.28E+00 | 1.11E-01 | 4.23E-01 | 1.38E+00 | 1.58E-02 | 5.64E-02 | -1.04E+00 | 5.50E-02 | 1.55E-01 | 1.00E+00 | 8.12E-02 | 2.98E-01 | 1.02E+00 | 1.94E-01 | 5.34E-01 | 1.75E+00 |
| TOB2 | 1.17E-01 | 6.11E-01 | 1.52E+00 | 1.01E-03 | 4.37E-03 | -1.02E+00 | 1.85E-01 | 7.85E-01 | 1.28E+00 | 1.76E-04 | 6.73E-04 | -1.13E+00 | 2.24E-02 | 8.00E-02 | -1.02E+00 | 3.00E-07 | 8.46E-07 | -1.88E+00 | 1.62E-02 | 5.94E-02 | 1.07E+00 | 2.13E-04 | 5.86E-04 | -1.26E+00 |
| [illegible] | [illegible] | [illegible] | [illegible] | [illegible] | [illegible] | [illegible] | [illegible] | [illegible] | [illegible] | [illegible] | [illegible] | [illegible] | [illegible] | [illegible] | [illegible] | [illegible] | [illegible] | [illegible] | [illegible] | [illegible] | [illegible] | [illegible] | [illegible] | [illegible] |

| | | | | | | | | | | | | | | | | | | | | | | | | |
|---|---|---|---|---|---|---|---|---|---|---|---|---|---|---|---|---|---|---|---|---|---|---|---|---|
| STK11 | 1.26E-01 | 6.59E-01 | 1.33E+00 | 1.31E-01 | 5.66E-01 | 1.48E+00 | 2.66E-02 | 1.13E-01 | 1.05E+00 | 2.27E-01 | 8.66E-01 | 1.40E+00 | 2.07E-01 | 7.40E-01 | 1.47E+00 | 9.06E-02 | 2.55E-01 | 1.71E+00 | 1.24E-01 | 4.54E-01 | 1.15E+00 | 8.03E-02 | 2.21E-01 | 1.10E+00 |
| TSGA10 | 1.27E-01 | 6.60E-01 | 1.24E+00 | 1.55E-01 | 6.68E-01 | 1.42E+00 | 5.21E-02 | 2.21E-01 | 1.02E+00 | 7.65E-02 | 2.92E-01 | 1.14E+00 | 6.32E-03 | 2.25E-02 | -1.24E+00 | 4.37E-02 | 1.23E-01 | -1.00E+00 | 9.69E-03 | 3.55E-02 | -1.17E+00 | 3.70E-02 | 1.02E-01 | -1.05E+00 |
| TTC34 | 1.27E-01 | 6.60E-01 | 1.20E+00 | 1.28E-01 | 5.54E-01 | 1.46E+00 | 2.22E-01 | 9.45E-01 | 1.05E+00 | 1.81E-01 | 6.90E-01 | 1.03E+00 | 1.38E-01 | 4.91E-01 | -1.10E+00 | 2.07E-01 | 5.83E-01 | 1.53E+00 | 2.46E-01 | 9.04E-01 | 1.14E+00 | 3.68E-02 | 1.01E-01 | 1.74E+00 |
| CCDC68 | 1.27E-01 | 6.60E-01 | -1.27E+00 | 1.56E-01 | 6.73E-01 | -1.27E+00 | 2.00E-01 | 8.48E-01 | -1.06E+00 | 1.79E-01 | 6.84E-01 | -1.17E+00 | 1.81E-02 | 6.44E-02 | -1.75E+00 | 1.30E-02 | 3.66E-02 | -2.00E+00 | 1.14E-02 | 4.19E-02 | 1.90E+00 | 1.94E-04 | 5.34E-04 | 1.14E+00 |
| HNRNPA1P40 | 1.27E-01 | 6.60E-01 | 1.30E+00 | 1.43E-01 | 6.19E-01 | 1.43E+00 | 4.20E-02 | 1.78E-01 | -1.01E+00 | 3.46E-03 | 1.32E-02 | -1.30E+00 | 1.16E-02 | 4.13E-02 | -1.20E+00 | 8.20E-03 | 2.31E-02 | -1.35E+00 | 1.98E-02 | 7.25E-02 | -1.23E+00 | 4.44E-03 | 1.22E-02 | -1.54E+00 |
| E2F1 | 1.27E-01 | 6.60E-01 | 1.47E+00 | 2.29E-01 | 9.89E-01 | 1.22E+00 | 2.44E-03 | 1.04E-02 | 1.56E+00 | 1.29E-01 | 4.91E-01 | 1.47E+00 | 2.33E-01 | 8.31E-01 | 1.04E+00 | 2.84E-01 | 8.00E-01 | 1.02E+00 | 1.32E-02 | 4.86E-02 | 1.66E+00 | 7.57E-06 | 2.08E-05 | 2.05E+00 |
| PLBD1 | 1.27E-01 | 6.60E-01 | -1.06E+00 | 9.51E-02 | 4.11E-01 | 1.81E+00 | 2.30E-02 | 9.77E-02 | -1.46E+00 | 7.14E-02 | 2.73E-01 | -1.11E+00 | 2.51E-01 | 8.95E-01 | 1.19E+00 | 2.98E-01 | 8.40E-01 | 1.33E+00 | 2.20E-01 | 8.07E-01 | 1.12E+00 | 3.01E-01 | 8.27E-01 | 1.07E+00 |
| SMARCE1P3 | 1.27E-01 | 6.61E-01 | 1.08E+00 | 9.62E-03 | 4.16E-02 | -1.47E+00 | 3.88E-02 | 1.65E-01 | -1.06E+00 | 4.34E-04 | 1.66E-03 | -1.42E+00 | 5.65E-03 | 2.01E-02 | -1.34E+00 | 3.12E-04 | 8.78E-04 | -1.84E+00 | 2.22E-02 | 8.14E-02 | -1.21E+00 | 1.92E-04 | 5.27E-04 | -1.81E+00 |
| AC110792.3 | 1.27E-01 | 6.61E-01 | 1.14E+00 | 1.91E-01 | 8.24E-01 | 1.24E+00 | 1.16E-01 | 4.92E-01 | -1.02E+00 | 4.25E-02 | 1.62E-01 | -1.13E+00 | 1.55E-01 | 5.54E-01 | 1.03E+00 | 1.59E-01 | 4.48E-01 | -1.11E+00 | 1.71E-01 | 6.29E-01 | -1.06E+00 | 7.60E-02 | 2.09E-01 | -1.27E+00 |
| CD3E | 1.27E-01 | 6.61E-01 | 1.15E+00 | 2.31E-01 | 1.00E+00 | -1.00E+00 | 1.12E-02 | 4.75E-02 | -2.52E+00 | 1.90E-03 | 7.27E-03 | -3.44E+00 | 1.36E-01 | 4.86E-01 | 1.43E+00 | 3.55E-01 | 1.00E+00 | -1.00E+00 | 2.17E-01 | 7.98E-01 | 1.21E+00 | 3.64E-01 | 1.00E+00 | 1.00E+00 |
| AC108047.1 | 1.27E-01 | 6.62E-01 | 1.09E+00 | 2.29E-02 | 9.88E-02 | -1.16E+00 | 1.31E-01 | 5.58E-01 | 1.09E+00 | 9.42E-03 | 3.60E-02 | -1.22E+00 | 2.46E-01 | 8.77E-01 | 1.22E+00 | 2.22E-01 | 6.25E-01 | 1.66E+00 | 1.31E-02 | 4.81E-02 | -1.30E+00 | 8.13E-02 | 2.23E-01 | 1.05E+00 |
| HIST1H3B | 1.27E-01 | 6.62E-01 | -1.09E+00 | 9.80E-02 | 4.24E-01 | -1.36E+00 | 1.94E-01 | 8.23E-01 | 1.65E+00 | 1.88E-03 | 7.17E-03 | -1.28E+00 | 5.37E-02 | 1.91E-01 | -1.66E+00 | 5.74E-02 | 1.62E-01 | -1.93E+00 | 1.48E-02 | 5.44E-02 | 1.52E+00 | 8.16E-02 | 2.24E-01 | -1.11E+00 |
| SETP4 | 1.27E-01 | 6.63E-01 | 1.48E+00 | 1.90E-01 | 8.22E-01 | 1.43E+00 | 5.78E-02 | 2.46E-01 | 1.49E+00 | 4.77E-02 | 1.82E-01 | 1.82E+00 | 1.81E-01 | 6.46E-01 | -1.00E+00 | 1.09E-01 | 3.08E-01 | -1.12E+00 | 3.19E-02 | 1.17E-01 | 1.69E+00 | 1.55E-01 | 4.27E-01 | 1.49E+00 |
| AMMECR1 | 1.27E-01 | 6.63E-01 | 1.18E+00 | 1.98E-01 | 8.55E-01 | 1.48E+00 | 2.37E-03 | 1.01E-02 | -1.13E+00 | 1.36E-03 | 5.18E-03 | -1.05E+00 | 1.92E-02 | 6.84E-02 | -1.04E+00 | 1.63E-01 | 4.60E-01 | 1.24E+00 | 2.44E-17 | 8.97E-17 | -1.95E+00 | 1.19E-03 | 3.27E-03 | -1.14E+00 |
| AC139769.1 | 1.27E-01 | 6.64E-01 | 1.02E+00 | 1.42E-01 | 6.13E-01 | 1.12E+00 | 7.06E-02 | 3.00E-01 | 1.38E+00 | 1.43E-01 | 5.46E-01 | 1.40E+00 | 1.04E-01 | 3.70E-01 | 1.00E+00 | 1.04E-01 | 2.94E-01 | -1.09E+00 | 2.09E-01 | 7.67E-01 | 1.37E+00 | 1.92E-01 | 5.28E-01 | 1.52E+00 |
| PLEKHA7 | 1.27E-01 | 6.64E-01 | 1.38E+00 | 6.34E-02 | 2.74E-01 | 1.87E+00 | 6.61E-02 | 2.81E-01 | 1.42E+00 | 1.82E-01 | 6.94E-01 | 1.39E+00 | 1.44E-02 | 5.12E-02 | -1.60E+00 | 1.48E-02 | 4.18E-02 | -1.69E+00 | 1.46E-01 | 5.34E-01 | -1.21E+00 | 3.49E-01 | 9.59E-01 | 1.10E+00 |
| ZNF254 | 1.27E-01 | 6.64E-01 | 1.25E+00 | 5.18E-02 | 2.24E-01 | 1.20E+00 | 1.18E-02 | 5.01E-02 | -1.01E+00 | 4.91E-04 | 1.87E-03 | -1.02E+00 | 4.48E-06 | 1.60E-05 | -1.23E+00 | 1.45E-04 | 4.08E-04 | -1.03E+00 | 1.22E-01 | 4.46E-01 | 1.12E+00 | 8.18E-03 | 2.25E-02 | -1.11E+00 |
| CCNB2P1 | 1.28E-01 | 6.65E-01 | -1.22E+00 | 1.26E-01 | 5.46E-01 | -1.33E+00 | 2.32E-01 | 9.84E-01 | 1.16E+00 | 7.26E-02 | 2.77E-01 | 1.57E+00 | 2.55E-01 | 9.09E-01 | 1.00E+00 | 3.05E-01 | 8.60E-01 | -1.05E+00 | 2.31E-01 | 8.48E-01 | 1.19E+00 | 5.71E-03 | 1.57E-02 | 1.75E+00 |
| LINC00471 | 1.28E-01 | 6.65E-01 | 1.06E+00 | 9.36E-02 | 4.04E-01 | 1.64E+00 | 3.42E-03 | 1.45E-02 | -1.46E+00 | 1.05E-02 | 4.01E-02 | -1.28E+00 | 3.22E-02 | 1.15E-01 | -1.37E+00 | 1.38E-01 | 3.88E-01 | -1.07E+00 | 3.24E-02 | 1.19E-01 | -1.42E+00 | 3.59E-01 | 9.85E-01 | 1.20E+00 |
| SUB1P1 | 1.28E-01 | 6.65E-01 | 1.41E+00 | 2.56E-02 | 1.11E-01 | 2.17E+00 | 1.20E-01 | 5.09E-01 | 1.14E+00 | 1.47E-01 | 5.60E-01 | 1.66E+00 | 8.03E-02 | 2.86E-01 | 1.12E+00 | 3.40E-01 | 9.58E-01 | 1.56E+00 | 2.56E-01 | 9.39E-01 | 1.43E+00 | 7.97E-02 | 2.19E-01 | 1.70E+00 |
| CBLB | 1.28E-01 | 6.66E-01 | 1.30E+00 | 1.64E-01 | 7.11E-01 | 1.57E+00 | 5.60E-02 | 2.38E-01 | 1.03E+00 | 6.20E-02 | 2.37E-01 | 1.20E+00 | 8.66E-02 | 3.09E-01 | 1.13E+00 | 7.37E-02 | 2.08E-01 | 1.07E+00 | 7.12E-02 | 2.61E-01 | -1.02E+00 | 2.62E-02 | 7.21E-02 | -1.10E+00 |
| POLQ | 1.28E-01 | 6.66E-01 | -1.05E+00 | 7.04E-02 | 3.04E-01 | -1.55E+00 | 4.29E-02 | 1.82E-01 | 1.48E+00 | 1.23E-01 | 4.70E-01 | 1.44E+00 | 2.80E-01 | 1.00E+00 | 1.16E+00 | 3.41E-01 | 9.61E-01 | 1.08E+00 | 6.33E-02 | 2.32E-01 | 1.05E+00 | 3.38E-01 | 9.29E-01 | 1.27E+00 |
| MRPS31P5 | 1.28E-01 | 6.66E-01 | 1.38E+00 | 1.04E-01 | 4.50E-01 | 1.71E+00 | 5.44E-03 | 2.31E-02 | -1.37E+00 | 8.27E-03 | 3.16E-02 | -1.24E+00 | 2.49E-01 | 8.88E-01 | 1.46E+00 | 1.52E-01 | 4.27E-01 | 1.06E+00 | 4.00E-02 | 1.47E-01 | -1.08E+00 | 3.35E-02 | 9.21E-02 | -1.05E+00 |
| GRAMD1C | 1.28E-01 | 6.66E-01 | 1.11E+00 | 2.34E-02 | 1.01E-01 | 1.90E+00 | 1.09E-02 | 4.62E-02 | -1.23E+00 | 5.03E-02 | 1.92E-01 | -1.00E+00 | 2.29E-01 | 8.17E-01 | -1.05E+00 | 2.92E-01 | 8.23E-01 | -1.05E+00 | 2.39E-01 | 8.77E-01 | 1.15E+00 | 3.21E-01 | 8.83E-01 | 1.04E+00 |
| AC011825.1 | 1.28E-01 | 6.66E-01 | 1.31E+00 | 2.72E-09 | 1.18E-08 | -1.05E+00 | 8.04E-02 | 3.42E-01 | 1.29E+00 | 9.98E-07 | 3.81E-06 | -1.15E+00 | 2.58E-02 | 9.20E-02 | 1.19E+00 | 2.12E-04 | 5.99E-04 | -1.05E+00 | 1.41E-01 | 5.16E-01 | 1.24E+00 | 1.15E-04 | 3.16E-04 | -1.16E+00 |
| FOXC2 | 1.28E-01 | 6.67E-01 | -1.01E+00 | 1.71E-02 | 7.39E-02 | 1.75E+00 | 2.11E-01 | 8.98E-01 | 1.17E+00 | 6.79E-02 | 2.59E-01 | 1.77E+00 | 3.97E-04 | 1.42E-03 | 5.83E+00 | 1.33E-01 | 3.76E-01 | 2.63E+00 | 2.16E-01 | 7.94E-01 | 1.12E+00 | 3.13E-01 | 8.60E-01 | 1.29E+00 |
| GYPE | 1.28E-01 | 6.69E-01 | -1.10E+00 | 1.41E-01 | 6.09E-01 | -1.13E+00 | 2.11E-01 | 8.94E-01 | 1.16E+00 | 2.23E-01 | 8.51E-01 | 1.25E+00 | 3.23E-03 | 1.15E-02 | -1.64E+00 | 2.47E-01 | 6.96E-01 | 1.42E+00 | 7.78E-40 | 2.85E-39 | -2.07E+00 | 1.27E-28 | 3.48E-28 | -1.74E+00 |
| TSPAN18 | 1.28E-01 | 6.69E-01 | -1.17E+00 | 1.76E-01 | 7.61E-01 | -1.06E+00 | 3.57E-02 | 1.52E-01 | -1.61E+00 | 3.38E-02 | 1.29E-01 | -1.62E+00 | 2.59E-03 | 9.25E-03 | -1.69E+00 | 1.70E-01 | 4.79E-01 | -1.02E+00 | 1.86E-09 | 6.83E-09 | -1.32E+00 | 1.62E-08 | 4.46E-08 | -1.25E+00 |
| AC246787.1 | 1.28E-01 | 6.69E-01 | 1.20E+00 | 4.10E-05 | 1.77E-04 | 1.46E+00 | 1.21E-02 | 5.16E-02 | 1.27E+00 | 1.83E-05 | 6.98E-05 | 1.60E+00 | 2.06E-01 | 7.34E-01 | 1.21E+00 | 6.56E-03 | 1.85E-02 | 1.49E+00 | 5.49E-02 | 2.02E-01 | 1.29E+00 | 3.04E-07 | 8.35E-07 | 1.60E+00 |
| SLCO4A1 | 1.28E-01 | 6.70E-01 | 1.50E+00 | 2.23E-01 | 9.62E-01 | 1.47E+00 | 2.22E-01 | 9.43E-01 | 1.22E+00 | 2.21E-01 | 8.45E-01 | 1.43E+00 | 1.63E-01 | 5.80E-01 | -1.05E+00 | 3.53E-01 | 9.96E-01 | 1.29E+00 | 1.07E-01 | 3.91E-01 | -1.02E+00 | 2.76E-01 | 7.57E-01 | 1.25E+00 |
| TRIM52-AS1 | 1.29E-01 | 6.70E-01 | 1.37E+00 | 5.05E-05 | 2.18E-04 | -1.12E+00 | 3.37E-02 | 1.43E-01 | 1.50E+00 | 3.68E-02 | 1.41E-01 | 1.18E+00 | 4.96E-02 | 1.77E-01 | 1.98E+00 | 3.49E-01 | 9.83E-01 | 1.56E+00 | 1.59E-01 | 5.82E-01 | 1.49E+00 | 4.82E-02 | 1.32E-01 | 1.09E+00 |
| LGALS2 | 1.29E-01 | 6.70E-01 | -1.15E+00 | 1.90E-01 | 8.22E-01 | 1.16E+00 | 1.53E-01 | 6.50E-01 | -2.05E+00 | 1.19E-01 | 4.55E-01 | -2.47E+00 | 1.71E-01 | 6.09E-01 | -1.23E+00 | 2.20E-01 | 6.19E-01 | -1.23E+00 | 4.79E-02 | 1.76E-01 | -2.19E+00 | 9.59E-02 | 2.63E-01 | -1.56E+00 |
| AC244502.3 | 1.29E-01 | 6.71E-01 | 1.46E+00 | 5.13E-03 | 2.22E-02 | 2.52E+00 | 3.74E-03 | 1.59E-02 | -1.40E+00 | 2.22E-01 | 8.49E-01 | 1.35E+00 | 1.18E-04 | 4.21E-04 | -2.86E+00 | 8.03E-03 | 2.26E-02 | -1.46E+00 | 4.31E-02 | 1.58E-01 | -1.63E+00 | 2.72E-01 | 7.47E-01 | 1.04E+00 |
| PIK3CD-AS2 | 1.29E-01 | 6.71E-01 | 1.08E+00 | 1.38E-01 | 5.95E-01 | 1.14E+00 | 8.30E-02 | 3.53E-01 | 1.94E+00 | 1.70E-01 | 6.48E-01 | 1.69E+00 | 1.92E-03 | 6.86E-03 | 1.97E+00 | 1.37E-02 | 3.86E-02 | 1.78E+00 | 2.68E-02 | 9.84E-02 | 1.52E+00 | 1.76E-01 | 4.83E-01 | 1.23E+00 |
| AC008608.2 | 1.29E-01 | 6.72E-01 | 1.29E+00 | 1.20E-01 | 5.17E-01 | 1.33E+00 | 2.96E-02 | 1.26E-01 | 1.68E+00 | 2.34E-01 | 8.93E-01 | 1.43E+00 | 2.43E-01 | 8.66E-01 | 1.56E+00 | 2.66E-01 | 7.49E-01 | 1.58E+00 | 1.75E-01 | 6.42E-01 | -1.04E+00 | 3.55E-01 | 9.76E-01 | 1.17E+00 |
| AC021016.2 | 1.29E-01 | 6.72E-01 | 1.52E+00 | 1.80E-01 | 7.79E-01 | 1.40E+00 | 1.74E-01 | 7.41E-01 | 1.11E+00 | 8.59E-02 | 3.28E-01 | 1.02E+00 | 2.15E-02 | 7.66E-02 | -1.31E+00 | 2.91E-02 | 8.20E-02 | -1.28E+00 | 2.16E-01 | 7.92E-01 | 1.33E+00 | 2.30E-01 | 6.33E-01 | -1.06E+00 |
| AC006460.2 | 1.29E-01 | 6.73E-01 | 1.12E+00 | 2.06E-01 | 8.91E-01 | 1.38E+00 | 2.13E-01 | 9.06E-01 | 1.09E+00 | 1.60E-01 | 6.11E-01 | 1.46E+00 | 2.65E-01 | 9.46E-01 | 1.43E+00 | 2.55E-01 | 7.19E-01 | 1.75E+00 | 1.90E-02 | 6.97E-02 | -1.63E+00 | 2.41E-01 | 6.63E-01 | 1.02E+00 |
| ACTR5 | 1.29E-01 | 6.74E-01 | 1.19E+00 | 2.10E-01 | 9.07E-01 | 1.52E+00 | 1.20E-03 | 5.08E-03 | -1.25E+00 | 7.74E-03 | 2.96E-02 | -1.01E+00 | 5.74E-04 | 2.05E-03 | -1.34E+00 | 1.22E-02 | 3.45E-02 | -1.08E+00 | 5.68E-03 | 2.09E-02 | -1.10E+00 | 6.03E-02 | 1.66E-01 | -1.02E+00 |
| CR1L | 1.29E-01 | 6.74E-01 | -1.22E+00 | 1.56E-01 | 6.72E-01 | -1.14E+00 | 1.92E-01 | 8.17E-01 | -1.15E+00 | 2.46E-01 | 9.38E-01 | -1.00E+00 | 1.11E-01 | 3.97E-01 | -1.90E+00 | 1.42E-01 | 4.00E-01 | -1.72E+00 | 1.47E-01 | 5.40E-01 | 1.37E+00 | 7.05E-04 | 1.94E-03 | -1.21E+00 |
| MINDY4 | 1.29E-01 | 6.74E-01 | 1.13E+00 | 3.04E-02 | 1.31E-01 | -1.38E+00 | 5.51E-02 | 2.34E-01 | -1.13E+00 | 8.31E-03 | 3.17E-02 | -1.53E+00 | 1.72E-01 | 6.12E-01 | 1.66E+00 | 2.86E-01 | 8.05E-01 | 1.65E+00 | 7.39E-02 | 2.71E-01 | 1.07E+00 | 1.21E-01 | 3.34E-01 | 1.04E+00 |
| HMGB1P41 | 1.29E-01 | 6.75E-01 | 1.23E+00 | 7.80E-05 | 3.37E-04 | -1.84E+00 | 2.10E-01 | 8.94E-01 | 1.32E+00 | 1.60E-05 | 6.11E-05 | -2.03E+00 | 1.12E-01 | 4.01E-01 | 1.70E+00 | 4.05E-02 | 1.14E-01 | -1.38E+00 | 8.51E-02 | 3.12E-01 | -1.22E+00 | 1.23E-02 | 3.38E-02 | -1.69E+00 |
| AL353678.1 | 1.29E-01 | 6.75E-01 | 1.39E+00 | 9.91E-04 | 4.28E-03 | 1.99E+00 | 5.65E-04 | 2.40E-03 | 1.66E+00 | 8.89E-10 | 3.40E-09 | 2.54E+00 | 8.06E-02 | 2.87E-01 | 1.64E+00 | 2.12E-06 | 5.99E-06 | 2.74E+00 | 7.02E-02 | 2.58E-01 | 1.52E+00 | 4.91E-07 | 1.35E-06 | 2.33E+00 |
| PITPNM2 | 1.30E-01 | 6.75E-01 | 1.14E+00 | 2.10E-01 | 9.07E-01 | 1.45E+00 | 1.81E-01 | 7.68E-01 | 1.16E+00 | 1.03E-01 | 3.93E-01 | 1.07E+00 | 1.78E-03 | 6.34E-03 | -1.42E+00 | 2.32E-02 | 6.53E-02 | -1.11E+00 | 1.06E-07 | 3.89E-07 | -1.29E+00 | 3.34E-05 | 9.18E-05 | -1.13E+00 |
| TOMM20P1 | 1.30E-01 | 6.75E-01 | 1.60E+00 | 1.41E-01 | 6.09E-01 | 1.47E+00 | 1.22E-01 | 5.19E-01 | 1.09E+00 | 2.35E-02 | 8.97E-02 | 1.06E+00 | 7.07E-02 | 2.52E-01 | 1.13E+00 | 2.51E-02 | 7.07E-02 | 1.01E+00 | 1.86E-01 | 6.81E-01 | 1.22E+00 | 3.25E-01 | 8.93E-01 | 1.44E+00 |
| ETFDH | 1.30E-01 | 6.75E-01 | 1.39E+00 | 1.04E-02 | 4.49E-02 | 1.19E+00 | 1.42E-03 | 6.02E-03 | -1.15E+00 | 1.45E-05 | 5.53E-05 | -1.20E+00 | 2.69E-01 | 9.60E-01 | 1.61E+00 | 3.06E-01 | 8.62E-01 | 1.53E+00 | 1.81E-02 | 6.64E-02 | 1.07E+00 | 3.55E-03 | 9.74E-03 | -1.08E+00 |
| HIST2H2AC | 1.30E-01 | 6.75E-01 | 1.38E+00 | 4.32E-10 | 1.87E-09 | -1.35E+00 | 8.32E-02 | 3.53E-01 | 1.75E+00 | 1.74E-04 | 6.64E-04 | -1.05E+00 | 2.22E-01 | 7.93E-01 | 1.53E+00 | 4.56E-06 | 1.29E-05 | -1.45E+00 | 2.24E-01 | 8.21E-01 | 1.59E+00 | 5.60E-06 | 1.54E-05 | -1.33E+00 |
| MED17 | 1.30E-01 | 6.77E-01 | 1.22E+00 | 1.54E-01 | 6.67E-01 | 1.35E+00 | 2.29E-02 | 9.75E-02 | 1.00E+00 | 4.27E-02 | 1.63E-01 | 1.13E+00 | 2.19E-02 | 7.80E-02 | 1.76E+00 | 2.38E-01 | 6.72E-01 | 1.33E+00 | 1.42E-01 | 5.21E-01 | 1.35E+00 | 1.60E-01 | 4.39E-01 | 1.06E+00 |
| HGF | 1.30E-01 | 6.77E-01 | 1.13E+00 | 7.45E-02 | 3.22E-01 | 1.79E+00 | 1.31E-02 | 5.55E-02 | 1.06E+00 | 2.32E-01 | 8.84E-01 | 1.47E+00 | 2.42E-05 | 8.62E-05 | -2.20E+00 | 7.21E-03 | 2.03E-02 | -1.13E+00 | 1.72E-01 | 6.30E-01 | -1.25E+00 | 2.78E-01 | 7.65E-01 | 1.30E+00 |
| LINC00853 | 1.30E-01 | 6.77E-01 | 1.12E+00 | 6.03E-02 | 2.60E-01 | 1.01E+00 | 2.69E-03 | 1.15E-02 | -1.71E+00 | 5.72E-04 | 2.18E-03 | -1.92E+00 | 1.97E-04 | 7.02E-04 | -1.39E+00 | 3.11E-03 | 8.76E-03 | -1.15E+00 | 1.93E-01 | 7.09E-01 | 1.27E+00 | 5.74E-02 | 1.58E-01 | -1.15E+00 |
| ARMCX6 | 1.30E-01 | 6.78E-01 | 1.21E+00 | 3.49E-02 | 1.51E-01 | 1.08E+00 | 2.29E-02 | 9.75E-02 | -1.18E+00 | 1.44E-01 | 5.49E-01 | 1.20E+00 | 1.33E-01 | 4.75E-01 | 2.00E+00 | 1.45E-01 | 4.09E-01 | 1.97E+00 | 4.69E-02 | 1.72E-01 | 1.67E+00 | 3.64E-01 | 9.99E-01 | 1.19E+00 |
| TIMM13 | 1.30E-01 | 6.78E-01 | 1.21E+00 | 2.43E-02 | 1.05E-01 | 1.18E+00 | 2.13E-01 | 9.04E-01 | 1.10E+00 | 2.22E-01 | 8.46E-01 | 1.14E+00 | 2.75E-01 | 9.81E-01 | 1.21E+00 | 1.61E-01 | 4.54E-01 | 1.25E+00 | 3.28E-02 | 1.20E-01 | 1.29E+00 | 2.63E-03 | 7.23E-03 | 1.37E+00 |
| SELENOO | 1.30E-01 | 6.78E-01 | 1.24E+00 | 4.04E-02 | 1.74E-01 | 1.15E+00 | 1.15E-01 | 4.87E-01 | 1.44E+00 | 1.55E-01 | 5.90E-01 | 1.55E+00 | 3.99E-02 | 1.42E-01 | 1.00E+00 | 1.38E-01 | 3.89E-01 | 1.26E+00 | 3.04E-02 | 1.12E-01 | 1.61E+00 | 7.22E-02 | 1.98E-01 | 1.61E+00 |
| AC132216.1 | 1.30E-01 | 6.78E-01 | 1.08E+00 | 7.54E-02 | 3.26E-01 | -1.02E+00 | 1.53E-01 | 6.49E-01 | 1.43E+00 | 8.33E-02 | 3.18E-01 | 1.80E+00 | 2.58E-01 | 9.21E-01 | 1.13E+00 | 3.44E-02 | 9.71E-02 | 1.89E+00 | 8.21E-03 | 3.01E-02 | 2.07E+00 | 1.65E-02 | 4.53E-02 | 2.18E+00 |
| GPC1 | 1.30E-01 | 6.79E-01 | 1.05E+00 | 2.21E-01 | 9.54E-01 | 1.19E+00 | 1.75E-01 | 7.42E-01 | 1.25E+00 | 4.33E-03 | 1.66E-02 | 2.05E+00 | 1.33E-01 | 4.75E-01 | 1.78E+00 | 3.15E-01 | 8.88E-01 | 1.29E+00 | 4.15E-02 | 1.52E-01 | 2.05E+00 | 1.46E-03 | 4.02E-03 | 3.32E+00 |
| NTRK1 | 1.30E-01 | 6.79E-01 | -1.42E+00 | 1.61E-01 | 6.94E-01 | -1.44E+00 | 7.92E-02 | 3.36E-01 | -1.60E+00 | 1.88E-01 | 7.20E-01 | 1.14E+00 | 4.02E-18 | 1.43E-17 | -1.38E+00 | 8.67E-12 | 2.44E-11 | -1.23E+00 | 3.00E-10 | 1.10E-09 | -1.52E+00 | 4.82E-06 | 1.32E-05 | -1.23E+00 |
| KLHL11 | 1.30E-01 | 6.80E-01 | -1.06E+00 | 7.91E-02 | 3.42E-01 | -1.14E+00 | 8.57E-05 | 3.64E-04 | -1.69E+00 | 2.81E-04 | 1.07E-03 | -1.52E+00 | 1.81E-02 | 6.45E-02 | -1.25E+00 | 9.05E-03 | 2.55E-02 | -1.43E+00 | 1.62E-01 | 5.96E-01 | -1.10E+00 | 3.35E-01 | 9.20E-01 | 1.20E+00 |
| USP53 | 1.30E-01 | 6.80E-01 | 1.26E+00 | 9.30E-04 | 4.02E-03 | -1.06E+00 | 1.74E-01 | 7.41E-01 | 1.25E+00 | 1.97E-02 | 7.53E-02 | 1.12E+00 | 1.93E-01 | 6.88E-01 | 1.23E+00 | 1.24E-01 | 3.49E-01 | 1.12E+00 | 2.70E-02 | 9.89E-02 | 1.03E+00 | 1.01E-01 | 2.79E-01 | 1.12E+00 |
| CASK | 1.30E-01 | 6.80E-01 | -1.01E+00 | 1.85E-01 | 7.99E-01 | 1.31E+00 | 1.83E-01 | 7.77E-01 | 1.35E+00 | 2.36E-01 | 9.02E-01 | 1.47E+00 | 2.76E-01 | 9.83E-01 | 1.30E+00 | 2.29E-01 | 6.45E-01 | 1.08E+00 | 2.56E-01 | 9.40E-01 | 1.25E+00 | 3.24E-01 | 8.89E-01 | 1.34E+00 |
| ZNF852 | 1.31E-01 | 6.81E-01 | 1.50E+00 | 9.72E-02 | 4.20E-01 | 1.06E+00 | 1.59E-01 | 6.76E-01 | 1.12E+00 | 1.65E-01 | 6.30E-01 | 1.18E+00 | 6.02E-02 | 2.15E-01 | -1.04E+00 | 4.07E-02 | 1.15E-01 | -1.23E+00 | 5.38E-02 | 1.98E-01 | -1.31E+00 | 1.67E-01 | 4.59E-01 | -1.02E+00 |
| AL136295.2 | 1.31E-01 | 6.81E-01 | 1.15E+00 | 1.43E-01 | 6.20E-01 | 1.61E+00 | 2.13E-01 | 9.03E-01 | 1.23E+00 | 1.58E-01 | 6.05E-01 | 1.54E+00 | 1.58E-01 | 5.62E-01 | 1.14E+00 | 1.96E-01 | 5.52E-01 | 1.17E+00 | 2.27E-01 | 8.32E-01 | 1.26E+00 | 2.27E-01 | 6.22E-01 | 1.37E+00 |
| AF201337.1 | 1.31E-01 | 6.81E-01 | 1.50E+00 | 2.18E-01 | 9.42E-01 | 1.50E+00 | 7.78E-02 | 3.30E-01 | 1.08E+00 | 3.06E-02 | 1.17E-01 | 1.08E+00 | 2.19E-01 | 7.82E-01 | 1.69E+00 | 2.29E-01 | 6.45E-01 | 1.79E+00 | 5.29E-05 | 1.94E-04 | 2.33E+00 | 2.85E-03 | 7.84E-03 | 1.91E+00 |
| WDR24 | 1.31E-01 | 6.81E-01 | 1.06E+00 | 1.18E-01 | 5.11E-01 | 1.16E+00 | 3.20E-02 | 1.36E-01 | -1.09E+00 | 9.98E-02 | 3.81E-01 | 1.10E+00 | 2.92E-02 | 1.04E-01 | -1.10E+00 | 2.55E-01 | 7.18E-01 | 1.32E+00 | 5.46E-03 | 2.00E-02 | -1.23E+00 | 2.67E-01 | 7.33E-01 | 1.23E+00 |
| SMIM3 | 1.31E-01 | 6.82E-01 | 1.33E+00 | 8.12E-02 | 3.51E-01 | 1.39E+00 | 1.71E-01 | 7.28E-01 | 1.19E+00 | 2.45E-01 | 9.37E-01 | 1.32E+00 | 2.52E-01 | 8.97E-01 | 1.38E+00 | 6.87E-04 | 1.94E-03 | 1.02E+00 | 3.02E-04 | 1.11E-03 | 2.10E+00 | 2.03E-01 | 5.59E-01 | 1.48E+00 |
| A4GALT | 1.31E-01 | 6.82E-01 | -1.17E+00 | 1.61E-01 | 6.94E-01 | -1.17E+00 | 2.13E-01 | 9.04E-01 | 1.07E+00 | 2.62E-01 | 1.00E+00 | 1.00E+00 | 2.80E-01 | 1.00E+00 | 1.00E+00 | 2.88E-01 | 8.13E-01 | 1.24E+00 | 1.73E-01 | 6.36E-01 | 1.04E+00 | 3.12E-01 | 8.57E-01 | 1.03E+00 |
| AC018463.1 | 1.31E-01 | 6.82E-01 | 1.40E+00 | 1.26E-01 | 5.46E-01 | 1.13E+00 | 3.57E-02 | 1.52E-01 | 1.70E+00 | 2.50E-01 | 9.55E-01 | 1.20E+00 | 1.39E-01 | 4.95E-01 | 1.07E+00 | 1.86E-01 | 5.23E-01 | 1.11E+00 | 2.32E-02 | 8.53E-02 | 1.77E+00 | 1.23E-02 | 3.39E-02 | 2.07E+00 |
| NPHP1 | 1.31E-01 | 6.82E-01 | 1.17E+00 | 1.03E-01 | 4.46E-01 | 1.17E+00 | 6.92E-02 | 2.94E-01 | 1.00E+00 | 3.66E-02 | 1.40E-01 | -1.03E+00 | 5.35E-02 | 1.91E-01 | -1.29E+00 | 7.21E-02 | 2.03E-01 | -1.25E+00 | 9.61E-03 | 3.53E-02 | -1.60E+00 | 2.75E-02 | 7.55E-02 | -1.38E+00 |
| NQO1 | 1.31E-01 | 6.82E-01 | 1.25E+00 | 1.73E-01 | 7.48E-01 | 1.33E+00 | 2.08E-05 | 8.83E-05 | -1.35E+00 | 4.40E-08 | 1.68E-07 | -1.48E+00 | 1.37E-02 | 4.87E-02 | -1.19E+00 | 2.96E-02 | 8.34E-02 | -1.13E+00 | 8.54E-02 | 3.13E-01 | -1.06E+00 | 1.19E-01 | 3.26E-01 | -1.06E+00 |
| MINK1 | 1.31E-01 | 6.83E-01 | -1.02E+00 | 1.86E-01 | 8.03E-01 | 1.41E+00 | 3.83E-04 | 1.63E-03 | -1.77E+00 | 5.07E-03 | 1.93E-02 | -1.26E+00 | 1.83E-04 | 6.54E-04 | -1.94E+00 | 6.64E-02 | 1.87E-01 | 1.13E+00 | 6.79E-04 | 2.49E-03 | -1.19E+00 | 8.17E-03 | 2.25E-02 | -1.07E+00 |
| SIDT2 | 1.31E-01 | 6.83E-01 | 1.26E+00 | 2.07E-03 | 8.94E-03 | -1.07E+00 | 1.54E-01 | 6.53E-01 | 1.20E+00 | 1.45E-04 | 5.53E-04 | -1.53E+00 | 4.23E-02 | 1.51E-01 | 1.00E+00 | 3.60E-02 | 1.01E-01 | -1.05E+00 | 4.58E-02 | 1.68E-01 | -1.05E+00 | 4.05E-02 | 1.11E-01 | -1.10E+00 |
| EPB41L5 | 1.31E-01 | 6.83E-01 | 1.24E+00 | 6.64E-02 | 2.87E-01 | 1.26E+00 | 1.22E-02 | 5.18E-02 | -1.20E+00 | 1.79E-02 | 6.85E-02 | -1.02E+00 | 2.70E-01 | 9.63E-01 | 1.21E+00 | 1.70E-01 | 4.78E-01 | -1.01E+00 | 2.28E-02 | 8.37E-02 | -1.10E+00 | 2.07E-02 | 5.69E-02 | -1.20E+00 |
| VPS37B | 1.31E-01 | 6.83E-01 | 1.30E+00 | 2.46E-02 | 1.06E-01 | 1.29E+00 | 5.45E-02 | 2.32E-01 | 1.50E+00 | 2.43E-01 | 9.27E-01 | 1.44E+00 | 2.04E-02 | 7.26E-02 | 1.75E+00 | 3.91E-03 | 1.10E-02 | 1.99E+00 | 1.22E-03 | 4.47E-03 | -1.09E+00 | 9.46E-03 | 2.60E-02 | 1.04E+00 |
| DDX3Y | 1.31E-01 | 6.83E-01 | 1.35E+00 | 6.10E-13 | 2.64E-12 | -1.14E+00 | 2.19E-01 | 9.29E-01 | 1.27E+00 | 3.27E-05 | 1.25E-04 | 1.01E+00 | 7.94E-02 | 2.83E-01 | 1.52E+00 | 1.26E-02 | 3.56E-02 | 1.11E+00 | 2.13E-01 | 7.82E-01 | 1.22E+00 | 1.79E-03 | 4.93E-03 | -1.02E+00 |
| AL137077.2 | 1.31E-01 | 6.83E-01 | 1.44E+00 | 9.35E-02 | 4.04E-01 | 1.09E+00 | 1.35E-01 | 5.73E-01 | 1.36E+00 | 2.44E-01 | 9.31E-01 | 1.18E+00 | 1.48E-01 | 5.26E-01 | 1.77E+00 | 2.61E-01 | 7.36E-01 | -1.09E+00 | 1.45E-01 | 5.31E-01 | 1.50E+00 | 2.83E-01 | 7.78E-01 | 1.31E+00 |
| PPP1R15B | 1.31E-01 | 6.83E-01 | 1.27E+00 | 7.85E-13 | 3.39E-12 | -1.57E+00 | 2.20E-01 | 9.33E-01 | 1.28E+00 | 7.28E-07 | 2.78E-06 | -1.30E+00 | 2.52E-01 | 8.98E-01 | 1.34E+00 | 3.84E-06 | 1.08E-05 | -1.41E+00 | 5.31E-02 | 1.95E-01 | 1.10E+00 | 2.89E-06 | 7.93E-06 | -1.44E+00 |
| AP001505.2 | 1.31E-01 | 6.83E-01 | -1.06E+00 | 2.00E-01 | 8.63E-01 | 1.25E+00 | 5.96E-02 | 2.53E-01 | -1.09E+00 | 2.16E-01 | 8.26E-01 | 1.41E+00 | 1.21E-01 | 4.31E-01 | -1.37E+00 | 1.41E-01 | 3.97E-01 | -1.32E+00 | 2.52E-01 | 9.23E-01 | 1.09E+00 | 2.27E-01 | 6.24E-01 | 1.24E+00 |
| MIR222HG | 1.31E-01 | 6.84E-01 | 1.25E+00 | 1.69E-02 | 7.31E-02 | 2.03E+00 | 1.02E-01 | 4.34E-01 | 1.08E+00 | 1.15E-03 | 4.40E-03 | 2.10E+00 | 2.71E-01 | 9.68E-01 | 1.40E+00 | 2.08E-02 | 5.88E-02 | 2.00E+00 | 1.91E-02 | 7.02E-02 | 1.95E+00 | 1.83E-02 | 5.03E-02 | 2.06E+00 |
| AC116366.3 | 1.31E-01 | 6.84E-01 | 1.08E+00 | 2.23E-01 | 9.62E-01 | 1.33E+00 | 3.35E-02 | 1.42E-01 | -1.21E+00 | 1.34E-01 | 5.11E-01 | 1.13E+00 | 2.67E-01 | 9.52E-01 | 1.19E+00 | 2.38E-01 | 6.70E-01 | 1.46E+00 | 6.70E-02 | 2.46E-01 | -1.20E+00 | 1.34E-01 | 3.69E-01 | 1.49E+00 |
| SLC24A1 | 1.31E-01 | 6.84E-01 | 1.17E+00 | 1.13E-01 | 4.90E-01 | 1.22E+00 | 1.19E-01 | 5.06E-01 | 1.07E+00 | 8.59E-02 | 3.28E-01 | 1.18E+00 | 2.59E-02 | 9.23E-02 | -1.07E+00 | 1.36E-01 | 3.83E-01 | 1.07E+00 | 1.56E-01 | 5.72E-01 | 1.43E+00 | 3.59E-01 | 9.87E-01 | 1.36E+00 |
| YTHDC2 | 1.31E-01 | 6.84E-01 | 1.14E+00 | 1.40E-02 | 6.04E-02 | -1.06E+00 | 6.10E-02 | 2.59E-01 | 1.13E+00 | 2.16E-04 | 8.23E-04 | -1.11E+00 | 1.92E-03 | 6.85E-03 | -1.19E+00 | 3.61E-04 | 1.02E-03 | -1.27E+00 | 2.33E-02 | 8.56E-02 | 1.02E+00 | 8.95E-03 | 2.46E-02 | -1.06E+00 |
| SLITRK5 | 1.32E-01 | 6.86E-01 | 1.13E+00 | 2.32E-07 | 1.00E-06 | 2.26E+00 | 7.93E-02 | 3.37E-01 | -1.02E+00 | 1.60E-03 | 6.12E-03 | 2.15E+00 | 1.55E-01 | 5.53E-01 | 1.85E+00 | 2.99E-02 | 8.43E-02 | 2.59E+00 | 5.05E-02 | 1.85E-01 | 2.18E+00 | 2.47E-04 | 6.79E-04 | 3.59E+00 |
| PIP4P1 | 1.32E-01 | 6.86E-01 | 1.17E+00 | 2.56E-03 | 1.10E-02 | -1.04E+00 | 4.81E-03 | 2.04E-02 | -1.15E+00 | 2.46E-04 | 9.41E-04 | -1.18E+00 | 2.01E-01 | 7.18E-01 | 1.25E+00 | 7.12E-02 | 2.01E-01 | 1.10E+00 | 3.53E-04 | 1.30E-03 | 1.01E+00 | 3.69E-03 | 1.01E-02 | -1.05E+00 |
| RPS2P28 | 1.32E-01 | 6.86E-01 | 1.39E+00 | 2.31E-01 | 9.98E-01 | 1.38E+00 | 1.86E-03 | 7.90E-03 | 2.27E+00 | 6.88E-03 | 2.63E-02 | 2.36E+00 | 5.28E-02 | 1.88E-01 | 2.08E+00 | 3.13E-01 | 8.82E-01 | 1.41E+00 | 3.15E-03 | 1.16E-02 | 2.41E+00 | 1.71E-01 | 4.71E-01 | 1.49E+00 |
| ASB3 | 1.32E-01 | 6.87E-01 | 1.23E+00 | 1.37E-02 | 5.92E-02 | 1.01E+00 | 9.08E-03 | 3.86E-02 | -1.12E+00 | 2.73E-04 | 1.04E-03 | -1.29E+00 | 4.32E-03 | 1.54E-02 | -1.13E+00 | 4.35E-03 | 1.23E-02 | -1.15E+00 | 5.87E-03 | 2.16E-02 | -1.33E+00 | 5.70E-03 | 1.57E-02 | -1.36E+00 |
| RAB12 | 1.32E-01 | 6.87E-01 | 1.07E+00 | 5.89E-02 | 2.55E-01 | -1.07E+00 | 1.94E-01 | 8.25E-01 | 1.16E+00 | 2.85E-03 | 1.09E-02 | -1.35E+00 | 2.14E-01 | 7.64E-01 | 1.13E+00 | 1.92E-02 | 5.40E-02 | -1.59E+00 | 5.14E-02 | 1.89E-01 | -1.11E+00 | 8.51E-03 | 2.34E-02 | -1.53E+00 |
| AC138904.3 | 1.32E-01 | 6.87E-01 | 1.25E+00 | 5.42E-06 | 2.34E-05 | -2.37E+00 | 4.73E-04 | 2.01E-03 | -1.28E+00 | 1.48E-11 | 5.67E-11 | -3.61E+00 | 1.24E-01 | 4.44E-01 | 1.04E+00 | 2.93E-02 | 8.26E-02 | -1.58E+00 | 1.26E-01 | 4.61E-01 | -1.00E+00 | 8.24E-02 | 2.26E-01 | -1.25E+00 |
| ZNF530 | 1.32E-01 | 6.89E-01 | 1.18E+00 | 1.16E-01 | 5.02E-01 | 1.14E+00 | 1.83E-01 | 7.78E-01 | 1.10E+00 | 6.33E-02 | 2.42E-01 | 1.06E+00 | 2.00E-01 | 7.14E-01 | 1.24E+00 | 3.27E-02 | 9.22E-02 | -1.42E+00 | 5.94E-02 | 2.18E-01 | 1.06E+00 | 3.94E-02 | 1.08E-01 | -1.06E+00 |
| KDM6A | 1.32E-01 | 6.89E-01 | 1.24E+00 | 1.18E-02 | 5.12E-02 | 1.05E+00 | 1.44E-01 | 6.10E-01 | 1.32E+00 | 1.99E-02 | 7.62E-02 | 1.03E+00 | 2.13E-01 | 7.61E-01 | 1.42E+00 | 2.98E-02 | 8.40E-02 | 1.09E+00 | 1.32E-01 | 4.85E-01 | 1.22E+00 | 1.39E-02 | 3.82E-02 | -1.02E+00 |
| SFMBT1 | 1.32E-01 | 6.89E-01 | 1.28E+00 | 1.54E-01 | 6.65E-01 | 1.38E+00 | 1.73E-01 | 7.36E-01 | 1.23E+00 | 1.70E-01 | 6.49E-01 | 1.02E+00 | 2.54E-01 | 9.06E-01 | 1.18E+00 | 3.24E-01 | 9.13E-01 | 1.30E+00 | 4.56E-03 | 1.67E-02 | -1.28E+00 | 2.63E-01 | 7.22E-01 | 1.25E+00 |
| FAM53C | 1.32E-01 | 6.90E-01 | 1.36E+00 | 1.69E-04 | 7.30E-04 | 1.04E+00 | 1.58E-03 | 6.71E-03 | -1.02E+00 | 2.07E-04 | 7.90E-04 | 1.00E+00 | 2.52E-01 | 8.97E-01 | 1.52E+00 | 1.52E-03 | 4.29E-03 | 1.01E+00 | 2.11E-01 | 7.74E-01 | 1.41E+00 | 1.13E-03 | 3.11E-03 | -1.10E+00 |
| ADAM17 | 1.32E-01 | 6.90E-01 | 1.25E+00 | 3.60E-05 | 1.55E-04 | -1.11E+00 | 2.01E-01 | 8.53E-01 | 1.10E+00 | 1.26E-03 | 4.83E-03 | -1.06E+00 | 7.76E-03 | 2.77E-02 | -1.04E+00 | 1.46E-04 | 4.11E-04 | -1.38E+00 | 1.05E-01 | 3.84E-01 | 1.04E+00 | 2.78E-04 | 7.63E-04 | -1.56E+00 |
| RWDD4P2 | 1.33E-01 | 6.91E-01 | 1.25E+00 | 1.01E-01 | 4.36E-01 | 1.28E+00 | 2.12E-01 | 9.03E-01 | 1.24E+00 | 1.80E-01 | 6.88E-01 | 1.29E+00 | 9.21E-02 | 3.28E-01 | 1.12E+00 | 2.80E-01 | 7.89E-01 | 1.35E+00 | 2.01E-01 | 7.39E-01 | 1.30E+00 | 1.54E-01 | 4.24E-01 | 1.50E+00 |
| AC008725.1 | 1.33E-01 | 6.91E-01 | 1.49E+00 | 6.18E-02 | 2.67E-01 | 2.03E+00 | 1.20E-02 | 5.11E-02 | 1.89E+00 | 8.13E-02 | 3.11E-01 | 1.78E+00 | 1.18E-01 | 4.20E-01 | 2.04E+00 | 2.44E-02 | 6.89E-02 | 2.83E+00 | 7.74E-02 | 2.84E-01 | 1.46E+00 | 2.10E-02 | 5.76E-02 | 1.72E+00 |
| PRR34-AS1 | 1.33E-01 | 6.91E-01 | 1.57E+00 | 1.62E-01 | 6.98E-01 | 1.34E+00 | 1.98E-01 | 8.39E-01 | 1.22E+00 | 5.80E-03 | 2.22E-02 | -1.31E+00 | 1.44E-01 | 5.13E-01 | 1.12E+00 | 8.10E-02 | 2.28E-01 | 1.01E+00 | 2.40E-01 | 8.83E-01 | 1.26E+00 | 1.02E-02 | 2.81E-02 | -1.20E+00 |
| AL031432.4 | 1.33E-01 | 6.91E-01 | -1.00E+00 | 5.43E-02 | 2.34E-01 | 1.65E+00 | 1.66E-02 | 7.04E-02 | -1.21E+00 | 1.03E-02 | 3.92E-02 | -1.09E+00 | 6.25E-02 | 2.23E-01 | -1.50E+00 | 1.58E-01 | 4.47E-01 | -1.13E+00 | 2.42E-01 | 8.89E-01 | 1.11E+00 | 3.45E-01 | 9.47E-01 | 1.28E+00 |
| AC137932.3 | 1.33E-01 | 6.91E-01 | 1.12E+00 | 6.50E-02 | 2.81E-01 | 1.12E+00 | 5.40E-03 | 2.29E-02 | -1.23E+00 | 1.73E-02 | 6.59E-02 | -1.01E+00 | 1.42E-02 | 5.08E-02 | -1.39E+00 | 3.04E-01 | 8.57E-01 | 1.28E+00 | 1.91E-02 | 7.02E-02 | -1.20E+00 | 3.52E-01 | 9.68E-01 | 1.18E+00 |
| KCNH2 | 1.33E-01 | 6.91E-01 | -1.38E+00 | 1.32E-01 | 5.72E-01 | -1.91E+00 | 1.75E-01 | 7.46E-01 | -1.20E+00 | 2.39E-01 | 9.11E-01 | 1.10E+00 | 6.25E-07 | 2.23E-06 | -1.30E+00 | 1.47E-01 | 4.14E-01 | 1.38E+00 | 5.71E-09 | 2.09E-08 | -1.25E+00 | 4.56E-02 | 1.25E-01 | 1.28E+00 |
| MAP1B | 1.33E-01 | 6.91E-01 | 1.56E+00 | 6.26E-02 | 2.71E-01 | 1.95E+00 | 2.23E-01 | 9.46E-01 | 1.16E+00 | 2.26E-02 | 8.62E-02 | 1.88E+00 | 5.14E-02 | 1.83E-01 | -1.31E+00 | 1.00E-01 | 2.83E-01 | -1.21E+00 | 1.10E-03 | 4.05E-03 | -1.50E+00 | 3.96E-02 | 1.09E-01 | -1.18E+00 |
| AC006978.1 | 1.33E-01 | 6.91E-01 | 1.13E+00 | 1.09E-01 | 4.70E-01 | 1.22E+00 | 3.94E-02 | 1.67E-01 | 1.04E+00 | 3.14E-02 | 1.20E-01 | 1.10E+00 | 2.72E-01 | 9.69E-01 | 1.21E+00 | 2.67E-01 | 7.54E-01 | 1.41E+00 | 1.23E-01 | 4.53E-01 | 1.12E+00 | 1.20E-01 | 3.29E-01 | 1.14E+00 |
| AC083873.1 | 1.33E-01 | 6.92E-01 | 1.27E+00 | 1.95E-01 | 8.41E-01 | 1.60E+00 | 1.70E-02 | 7.24E-02 | 1.57E+00 | 1.50E-02 | 5.72E-02 | 1.83E+00 | 6.40E-02 | 2.28E-01 | 1.84E+00 | 6.28E-02 | 1.77E-01 | 2.07E+00 | 2.10E-03 | 7.72E-03 | 1.82E+00 | 8.16E-04 | 2.24E-03 | 2.08E+00 |
| GINS4 | 1.33E-01 | 6.92E-01 | 1.03E+00 | 2.36E-02 | 1.02E-01 | -1.47E+00 | 1.97E-01 | 8.37E-01 | 1.12E+00 | 2.27E-01 | 8.67E-01 | 1.30E+00 | 3.02E-02 | 1.08E-01 | -1.28E+00 | 9.41E-03 | 2.65E-02 | -1.71E+00 | 2.88E-02 | 1.06E-01 | 1.12E+00 | 3.52E-01 | 9.66E-01 | 1.36E+00 |
| GINS1 | 1.33E-01 | 6.93E-01 | 1.07E+00 | 1.80E-01 | 7.76E-01 | 1.70E+00 | 1.74E-01 | 7.41E-01 | 1.21E+00 | 8.73E-02 | 3.34E-01 | 1.57E+00 | 8.45E-02 | 3.01E-01 | -1.86E+00 | 1.75E-01 | 4.94E-01 | -1.34E+00 | 2.01E-05 | 7.37E-05 | -1.34E+00 | 1.32E-01 | 3.61E-01 | 1.20E+00 |
| AC124016.1 | 1.33E-01 | 6.93E-01 | 1.46E+00 | 2.12E-01 | 9.18E-01 | 1.46E+00 | 2.34E-01 | 9.93E-01 | 1.19E+00 | 1.22E-01 | 4.66E-01 | 1.20E+00 | 1.53E-01 | 5.44E-01 | 1.42E+00 | 3.47E-01 | 9.77E-01 | 1.13E+00 | 1.46E-01 | 5.37E-01 | -1.09E+00 | 7.23E-02 | 1.99E-01 | -1.35E+00 |
| SBF1 | 1.33E-01 | 6.93E-01 | 1.26E+00 | 1.50E-01 | 6.49E-01 | 1.35E+00 | 7.42E-02 | 3.15E-01 | -1.05E+00 | 1.59E-01 | 6.08E-01 | 1.16E+00 | 1.19E-01 | 4.25E-01 | 1.05E+00 | 2.99E-01 | 8.44E-01 | 1.41E+00 | 1.48E-01 | 5.42E-01 | 1.18E+00 | 1.86E-01 | 5.11E-01 | 1.48E+00 |
| TPH1 | 1.33E-01 | 6.94E-01 | 1.45E+00 | 1.61E-01 | 6.95E-01 | 1.53E+00 | 2.25E-02 | 9.58E-02 | -1.57E+00 | 1.03E-01 | 3.94E-01 | -1.08E+00 | 8.94E-02 | 3.19E-01 | -1.01E+00 | 1.96E-01 | 5.53E-01 | 1.33E+00 | 8.89E-02 | 3.26E-01 | 1.48E+00 | 1.05E-01 | 2.88E-01 | 1.49E+00 |
| AC093297.2 | 1.33E-01 | 6.94E-01 | -1.11E+00 | 5.41E-02 | 2.34E-01 | 1.87E+00 | 1.81E-02 | 7.70E-02 | -1.41E+00 | 1.80E-01 | 6.87E-01 | 1.49E+00 | 2.09E-01 | 7.44E-01 | 1.32E+00 | 9.68E-02 | 2.73E-01 | 1.99E+00 | 1.34E-01 | 4.93E-01 | -1.10E+00 | 2.95E-01 | 8.11E-01 | 1.29E+00 |
| MTHFD2L | 1.33E-01 | 6.94E-01 | 1.28E+00 | 1.32E-02 | 5.71E-02 | 1.69E+00 | 1.74E-01 | 7.41E-01 | 1.13E+00 | 4.09E-02 | 1.56E-01 | 1.49E+00 | 2.69E-01 | 9.58E-01 | 1.38E+00 | 1.24E-01 | 3.49E-01 | 1.57E+00 | 1.65E-01 | 6.05E-01 | 1.27E+00 | 3.08E-02 | 8.47E-02 | 1.45E+00 |
| AC000123.3 | 1.33E-01 | 6.94E-01 | 1.13E+00 | 1.80E-01 | 7.76E-01 | 1.21E+00 | 1.88E-02 | 7.98E-02 | -1.26E+00 | 6.37E-02 | 2.43E-01 | -1.00E+00 | 2.20E-01 | 7.83E-01 | 1.33E+00 | 2.48E-01 | 6.99E-01 | 1.57E+00 | 2.50E-01 | 9.16E-01 | 1.21E+00 | 2.91E-01 | 7.99E-01 | 1.13E+00 |
| TPT1P9 | 1.33E-01 | 6.94E-01 | 1.41E+00 | 6.47E-20 | 2.80E-19 | 2.57E+00 | 4.51E-02 | 1.91E-01 | 1.37E+00 | 3.20E-17 | 1.22E-16 | 2.78E+00 | 2.24E-02 | 7.98E-02 | 1.65E+00 | 5.77E-13 | 1.63E-12 | 2.84E+00 | 5.13E-02 | 1.88E-01 | 1.47E+00 | 3.62E-14 | 9.95E-14 | 2.44E+00 |
| RN7SL832P | 1.33E-01 | 6.95E-01 | 1.08E+00 | 2.96E-03 | 1.28E-02 | -1.74E+00 | 2.11E-01 | 8.96E-01 | 1.17E+00 | 1.32E-01 | 5.05E-01 | 1.16E+00 | 2.42E-01 | 8.64E-01 | 1.11E+00 | 2.39E-01 | 6.73E-01 | 1.09E+00 | 2.01E-01 | 7.37E-01 | 1.28E+00 | 2.82E-01 | 7.76E-01 | 1.08E+00 |
| CAPN10-AS1 | 1.33E-01 | 6.95E-01 | 1.25E+00 | 2.04E-01 | 8.80E-01 | 1.01E+00 | 6.25E-02 | 2.66E-01 | 1.58E+00 | 2.28E-01 | 8.69E-01 | 1.06E+00 | 8.84E-03 | 3.15E-02 | 2.62E+00 | 1.83E-01 | 5.15E-01 | 1.87E+00 | 2.28E-02 | 8.38E-02 | 2.10E+00 | 3.41E-02 | 9.37E-02 | 2.08E+00 |
| RARB | 1.34E-01 | 6.96E-01 | 1.29E+00 | 8.00E-02 | 3.46E-01 | -1.15E+00 | 1.56E-01 | 6.63E-01 | 1.27E+00 | 1.79E-01 | 6.86E-01 | -1.02E+00 | 1.38E-01 | 4.90E-01 | 1.82E+00 | 9.20E-02 | 2.59E-01 | -1.02E+00 | 2.45E-01 | 8.99E-01 | 1.21E+00 | 6.78E-02 | 1.86E-01 | -1.22E+00 |
| MTMR10 | 1.34E-01 | 6.97E-01 | 1.24E+00 | 3.32E-02 | 1.43E-01 | 1.15E+00 | 2.95E-03 | 1.25E-02 | -1.05E+00 | 4.70E-05 | 1.79E-04 | -1.20E+00 | 3.14E-02 | 1.12E-01 | 2.04E+00 | 3.36E-01 | 9.48E-01 | 1.32E+00 | 1.76E-02 | 6.46E-02 | 1.45E+00 | 1.56E-04 | 4.27E-04 | -1.17E+00 |
| SLC1A3 | 1.34E-01 | 6.97E-01 | -1.01E+00 | 2.19E-01 | 9.45E-01 | 1.25E+00 | 3.22E-10 | 1.37E-09 | -2.07E+00 | 7.22E-13 | 2.76E-12 | -2.39E+00 | 2.80E-01 | 1.00E+00 | 1.00E+00 | 3.29E-01 | 9.28E-01 | 1.08E+00 | 2.64E-01 | 9.69E-01 | 1.06E+00 | 2.97E-01 | 8.15E-01 | -1.02E+00 |
| CRTC3 | 1.34E-01 | 6.97E-01 | 1.15E+00 | 3.64E-02 | 1.57E-01 | 1.11E+00 | 6.27E-04 | 2.66E-03 | -1.23E+00 | 2.01E-04 | 7.66E-04 | -1.16E+00 | 2.47E-01 | 8.82E-01 | 1.20E+00 | 1.52E-01 | 4.28E-01 | 1.16E+00 | 1.04E-02 | 3.81E-02 | -1.05E+00 | 1.86E-02 | 5.11E-02 | 1.01E+00 |
| AC098614.1 | 1.34E-01 | 6.97E-01 | 1.31E+00 | 1.69E-01 | 7.32E-01 | 1.47E+00 | 1.58E-01 | 6.71E-01 | 1.06E+00 | 3.17E-02 | 1.21E-01 | 1.02E+00 | 1.53E-01 | 5.44E-01 | 1.70E+00 | 3.54E-01 | 9.99E-01 | 1.43E+00 | 3.72E-02 | 1.37E-01 | 1.53E+00 | 2.93E-01 | 8.04E-01 | 1.21E+00 |
| IL1R2 | 1.34E-01 | 6.98E-01 | 1.11E+00 | 2.31E-01 | 1.00E+00 | -1.00E+00 | 9.90E-02 | 4.21E-01 | 1.86E+00 | 2.21E-01 | 8.43E-01 | 1.06E+00 | 1.99E-01 | 7.09E-01 | 1.54E+00 | 3.55E-01 | 1.00E+00 | -1.00E+00 | 2.06E-01 | 7.57E-01 | 1.14E+00 | 2.99E-01 | 8.21E-01 | 1.05E+00 |
| OR9H1P | 1.34E-01 | 6.98E-01 | 1.21E+00 | 1.58E-01 | 6.82E-01 | 1.13E+00 | 9.95E-03 | 4.23E-02 | -1.63E+00 | 1.39E-01 | 5.31E-01 | 1.09E+00 | 6.70E-02 | 2.39E-01 | -1.14E+00 | 1.47E-01 | 4.15E-01 | 1.02E+00 | 2.38E-01 | 8.74E-01 | 1.15E+00 | 2.91E-01 | 8.01E-01 | 1.24E+00 |
| AC092718.4 | 1.34E-01 | 6.98E-01 | -1.19E+00 | 1.50E-01 | 6.49E-01 | -1.21E+00 | 3.22E-04 | 1.37E-03 | -1.63E+00 | 1.40E-02 | 5.36E-02 | -1.07E+00 | 3.03E-02 | 1.08E-01 | -1.42E+00 | 3.11E-02 | 8.78E-02 | -1.55E+00 | 5.09E-04 | 1.87E-03 | -1.34E+00 | 4.53E-03 | 1.24E-02 | -1.31E+00 |
| LIG1 | 1.34E-01 | 6.99E-01 | 1.17E+00 | 9.27E-02 | 4.01E-01 | 1.17E+00 | 1.06E-03 | 4.50E-03 | 1.63E+00 | 1.71E-01 | 6.51E-01 | 1.37E+00 | 5.34E-02 | 1.90E-01 | 1.01E+00 | 3.11E-02 | 8.76E-02 | -1.04E+00 | 1.05E-01 | 3.86E-01 | 1.15E+00 | 8.16E-02 | 2.24E-01 | 1.40E+00 |
| AC087752.1 | 1.34E-01 | 6.99E-01 | 1.48E+00 | 2.10E-01 | 9.08E-01 | 1.44E+00 | 1.80E-02 | 7.66E-02 | -1.12E+00 | 3.15E-02 | 1.20E-01 | 1.04E+00 | 9.02E-02 | 3.22E-01 | 1.95E+00 | 3.15E-01 | 8.87E-01 | 1.54E+00 | 1.03E-01 | 3.77E-01 | 1.52E+00 | 3.25E-01 | 8.94E-01 | 1.27E+00 |
| PRR16 | 1.34E-01 | 7.00E-01 | 1.35E+00 | 1.82E-02 | 7.86E-02 | 1.87E+00 | 1.29E-01 | 5.49E-01 | 1.26E+00 | 1.79E-01 | 6.86E-01 | 1.61E+00 | 2.50E-01 | 8.90E-01 | 1.18E+00 | 2.30E-01 | 6.48E-01 | 1.08E+00 | 4.30E-02 | 1.58E-01 | -1.60E+00 | 1.66E-01 | 4.55E-01 | -1.19E+00 |
| LILRA5 | 1.34E-01 | 7.00E-01 | 1.22E+00 | 1.75E-01 | 7.55E-01 | 1.51E+00 | 5.01E-02 | 2.13E-01 | -1.36E+00 | 4.56E-02 | 1.74E-01 | -1.45E+00 | 2.80E-01 | 1.00E+00 | 1.00E+00 | 3.55E-01 | 1.00E+00 | -1.00E+00 | 2.14E-01 | 7.85E-01 | -1.58E+00 | 1.88E-01 | 5.18E-01 | -1.69E+00 |
| MYO15B | 1.34E-01 | 7.00E-01 | 1.24E+00 | 2.14E-01 | 9.24E-01 | 1.53E+00 | 2.20E-01 | 9.34E-01 | 1.27E+00 | 2.57E-01 | 9.83E-01 | 1.37E+00 | 1.14E-01 | 4.08E-01 | 1.28E+00 | 5.96E-02 | 1.68E-01 | 1.15E+00 | 2.43E-01 | 8.92E-01 | 1.46E+00 | 1.74E-01 | 4.77E-01 | 1.62E+00 |
| ZNF865 | 1.34E-01 | 7.01E-01 | 1.09E+00 | 1.05E-01 | 4.53E-01 | 1.64E+00 | 7.44E-04 | 3.16E-03 | -1.55E+00 | 4.91E-03 | 1.88E-02 | -1.23E+00 | 5.85E-02 | 2.09E-01 | -1.33E+00 | 9.84E-02 | 2.77E-01 | 1.96E+00 | 6.53E-02 | 2.40E-01 | -1.13E+00 | 3.27E-01 | 8.99E-01 | 1.30E+00 |
| LINC02573 | 1.34E-01 | 7.01E-01 | 1.37E+00 | 4.05E-03 | 1.75E-02 | 1.18E+00 | 1.65E-03 | 6.99E-03 | 1.01E+00 | 3.43E-07 | 1.31E-06 | -1.24E+00 | 1.07E-03 | 3.80E-03 | 2.53E+00 | 1.63E-01 | 4.61E-01 | 1.65E+00 | 1.85E-01 | 6.79E-01 | 1.14E+00 | 3.39E-03 | 9.31E-03 | -1.44E+00 |
| AC093249.6 | 1.34E-01 | 7.01E-01 | 1.52E+00 | 1.25E-01 | 5.41E-01 | 1.29E+00 | 1.55E-01 | 6.60E-01 | 1.10E+00 | 5.71E-03 | 2.18E-02 | -1.22E+00 | 2.73E-01 | 9.74E-01 | 1.37E+00 | 3.47E-01 | 9.78E-01 | 1.35E+00 | 5.98E-02 | 2.20E-01 | 1.59E+00 | 3.45E-01 | 9.48E-01 | 1.06E+00 |
| RPL37 | 1.34E-01 | 7.01E-01 | 1.02E+00 | 8.47E-79 | 3.66E-78 | -1.60E+00 | 5.79E-02 | 2.46E-01 | 1.04E+00 | 9.30E-43 | 3.55E-42 | -1.52E+00 | 2.51E-01 | 8.95E-01 | 1.02E+00 | 7.74E-37 | 2.18E-36 | -1.55E+00 | 7.28E-05 | 2.67E-04 | -1.04E+00 | 6.27E-62 | 1.72E-61 | -1.62E+00 |
| RGS16 | 1.35E-01 | 7.01E-01 | 1.25E+00 | 2.14E-01 | 9.27E-01 | 1.02E+00 | 2.13E-01 | 9.04E-01 | 1.07E+00 | 2.26E-01 | 8.62E-01 | 1.04E+00 | 1.30E-01 | 4.62E-01 | -1.12E+00 | 2.73E-01 | 7.70E-01 | 1.02E+00 | 1.38E-01 | 5.08E-01 | 1.10E+00 | 8.85E-02 | 2.43E-01 | 1.15E+00 |
| AL513327.1 | 1.35E-01 | 7.01E-01 | 1.24E+00 | 3.66E-02 | 1.58E-01 | -1.01E+00 | 7.36E-04 | 3.13E-03 | -1.50E+00 | 7.25E-05 | 2.77E-04 | -1.63E+00 | 1.83E-01 | 6.51E-01 | 1.07E+00 | 2.55E-01 | 7.20E-01 | 1.17E+00 | 2.57E-01 | 9.42E-01 | 1.24E+00 | 3.35E-01 | 9.20E-01 | 1.20E+00 |
| PCDHGB2 | 1.35E-01 | 7.02E-01 | 1.30E+00 | 5.76E-02 | 2.49E-01 | 1.23E+00 | 1.33E-03 | 5.65E-03 | -1.47E+00 | 1.20E-02 | 4.57E-02 | -1.03E+00 | 1.42E-01 | 5.05E-01 | 1.20E+00 | 9.77E-02 | 2.75E-01 | 1.06E+00 | 2.29E-01 | 8.40E-01 | -1.01E+00 | 3.41E-01 | 9.37E-01 | 1.11E+00 |
| AL078621.1 | 1.35E-01 | 7.02E-01 | 1.44E+00 | 1.66E-01 | 7.17E-01 | 1.55E+00 | 7.34E-03 | 3.12E-02 | 1.90E+00 | 5.28E-02 | 2.02E-01 | 1.90E+00 | 1.71E-01 | 6.09E-01 | 1.75E+00 | 2.81E-01 | 7.93E-01 | 1.63E+00 | 1.21E-01 | 4.43E-01 | 1.54E+00 | 2.65E-01 | 7.27E-01 | 1.14E+00 |
| NEIL1 | 1.35E-01 | 7.02E-01 | 1.22E+00 | 5.58E-02 | 2.41E-01 | 1.81E+00 | 1.84E-01 | 7.82E-01 | 1.09E+00 | 2.40E-01 | 9.16E-01 | 1.20E+00 | 1.83E-01 | 6.53E-01 | -1.05E+00 | 3.08E-01 | 8.68E-01 | 1.46E+00 | 3.82E-02 | 1.40E-01 | -1.55E+00 | 1.36E-01 | 3.73E-01 | -1.12E+00 |
| AC012213.2 | 1.35E-01 | 7.04E-01 | -1.15E+00 | 1.08E-01 | 4.65E-01 | 1.51E+00 | 1.05E-02 | 4.44E-02 | -1.86E+00 | 2.61E-01 | 9.97E-01 | 1.05E+00 | 2.41E-01 | 8.58E-01 | 1.23E+00 | 2.48E-01 | 6.99E-01 | 1.45E+00 | 8.40E-02 | 3.08E-01 | -1.40E+00 | 3.60E-01 | 9.88E-01 | 1.10E+00 |
| AC010683.1 | 1.35E-01 | 7.05E-01 | 1.28E+00 | 8.64E-02 | 3.73E-01 | 1.25E+00 | 1.83E-01 | 7.77E-01 | 1.24E+00 | 8.26E-02 | 3.16E-01 | 1.22E+00 | 1.64E-01 | 5.85E-01 | 1.19E+00 | 2.60E-01 | 7.33E-01 | 1.21E+00 | 8.81E-02 | 3.23E-01 | 1.58E+00 | 1.23E-02 | 3.39E-02 | 1.99E+00 |
| HOXB-AS1 | 1.35E-01 | 7.05E-01 | 1.50E+00 | 1.76E-01 | 7.61E-01 | 1.53E+00 | 2.04E-01 | 8.66E-01 | 1.00E+00 | 5.68E-02 | 2.17E-01 | 1.68E+00 | 9.00E-03 | 3.21E-02 | -1.09E+00 | 6.63E-02 | 1.87E-01 | 1.73E+00 | 5.58E-04 | 2.05E-03 | -1.11E+00 | 9.55E-03 | 2.62E-02 | 1.02E+00 |
| NCKAP1 | 1.35E-01 | 7.05E-01 | -1.14E+00 | 1.09E-01 | 4.72E-01 | -1.29E+00 | 2.08E-01 | 8.82E-01 | -1.04E+00 | 2.51E-01 | 9.59E-01 | 1.06E+00 | 5.10E-02 | 1.82E-01 | -1.53E+00 | 1.61E-01 | 4.53E-01 | -1.18E+00 | 7.59E-03 | 2.79E-02 | 1.43E+00 | 1.09E-06 | 3.00E-06 | -1.55E+00 |
| Z93241.1 | 1.35E-01 | 7.06E-01 | 1.85E+00 | 7.00E-21 | 3.02E-20 | -3.68E+00 | 2.12E-02 | 9.02E-02 | 2.39E+00 | 6.46E-07 | 2.47E-06 | -1.99E+00 | 4.21E-02 | 1.50E-01 | 1.08E+00 | 2.08E-03 | 5.85E-03 | -1.70E+00 | 2.99E-02 | 1.10E-01 | -1.01E+00 | 1.59E-01 | 4.37E-01 | -1.07E+00 |
| BIN3 | 1.36E-01 | 7.06E-01 | 1.06E+00 | 2.22E-01 | 9.60E-01 | 1.26E+00 | 5.63E-04 | 2.39E-03 | -1.47E+00 | 3.17E-04 | 1.21E-03 | -1.29E+00 | 2.48E-01 | 8.85E-01 | 1.34E+00 | 2.65E-01 | 7.46E-01 | 1.50E+00 | 4.09E-02 | 1.50E-01 | -1.15E+00 | 2.28E-02 | 6.26E-02 | -1.24E+00 |
| RNVU1-4 | 1.36E-01 | 7.06E-01 | 1.46E+00 | 3.97E-03 | 1.72E-02 | -1.48E+00 | 1.08E-01 | 4.60E-01 | 1.10E+00 | 3.68E-04 | 1.41E-03 | -1.70E+00 | 2.04E-01 | 7.27E-01 | 1.08E+00 | 5.81E-02 | 1.64E-01 | -1.63E+00 | 1.53E-01 | 5.61E-01 | -1.11E+00 | 1.59E-02 | 4.37E-02 | -1.74E+00 |
| AGGF1P3 | 1.36E-01 | 7.07E-01 | 1.34E+00 | 2.80E-02 | 1.21E-01 | 1.98E+00 | 2.32E-01 | 9.84E-01 | 1.11E+00 | 1.67E-02 | 6.39E-02 | 2.17E+00 | 1.68E-01 | 5.99E-01 | -1.09E+00 | 2.52E-01 | 7.11E-01 | 1.52E+00 | 8.17E-02 | 3.00E-01 | 1.65E+00 | 4.82E-02 | 1.32E-01 | 2.00E+00 |
| RPS2P32 | 1.36E-01 | 7.07E-01 | 1.47E+00 | 6.01E-02 | 2.60E-01 | 1.14E+00 | 1.00E-01 | 4.26E-01 | 1.19E+00 | 8.61E-04 | 3.29E-03 | -1.14E+00 | 2.13E-01 | 7.58E-01 | 1.41E+00 | 1.02E-01 | 2.87E-01 | 1.02E+00 | 1.60E-01 | 5.87E-01 | 1.08E+00 | 6.00E-02 | 1.65E-01 | -1.13E+00 |
| CPEB3 | 1.36E-01 | 7.08E-01 | 1.15E+00 | 2.31E-01 | 9.99E-01 | 1.43E+00 | 1.87E-02 | 7.94E-02 | -1.46E+00 | 3.54E-02 | 1.35E-01 | -1.27E+00 | 1.63E-03 | 5.80E-03 | -1.85E+00 | 3.56E-03 | 1.00E-02 | -1.65E+00 | 1.40E-05 | 5.13E-05 | -1.73E+00 | 1.34E-05 | 3.68E-05 | -1.58E+00 |
| AC122718.2 | 1.36E-01 | 7.08E-01 | 1.16E+00 | 3.34E-02 | 1.44E-01 | -1.10E+00 | 1.37E-01 | 5.83E-01 | 1.38E+00 | 1.04E-01 | 3.99E-01 | 1.03E+00 | 2.56E-01 | 9.13E-01 | 1.47E+00 | 2.65E-01 | 7.46E-01 | 1.10E+00 | 8.58E-02 | 3.15E-01 | 1.35E+00 | 3.17E-01 | 8.71E-01 | -1.01E+00 |
| SCN3A | 1.36E-01 | 7.08E-01 | -1.01E+00 | 1.56E-01 | 6.73E-01 | -1.02E+00 | 6.17E-02 | 2.62E-01 | -1.29E+00 | 3.88E-02 | 1.48E-01 | -1.53E+00 | 1.24E-01 | 4.41E-01 | 1.86E+00 | 4.36E-02 | 1.23E-01 | 2.21E+00 | 9.67E-03 | 3.55E-02 | 2.04E+00 | 2.78E-01 | 7.63E-01 | 1.35E+00 |
| RAI1 | 1.36E-01 | 7.08E-01 | 1.43E+00 | 4.07E-03 | 1.76E-02 | -1.37E+00 | 3.71E-02 | 1.58E-01 | 1.48E+00 | 1.64E-01 | 6.27E-01 | 1.13E+00 | 1.01E-01 | 3.60E-01 | 1.15E+00 | 1.95E-02 | 5.50E-02 | -1.16E+00 | 2.42E-02 | 8.87E-02 | -1.08E+00 | 5.17E-04 | 1.42E-03 | -1.75E+00 |
| RBM17P4 | 1.36E-01 | 7.08E-01 | 1.19E+00 | 1.34E-01 | 5.78E-01 | 1.75E+00 | 1.88E-01 | 8.00E-01 | 1.30E+00 | 1.77E-01 | 6.76E-01 | 1.59E+00 | 2.32E-01 | 8.27E-01 | 1.33E+00 | 2.78E-01 | 7.84E-01 | 1.66E+00 | 6.54E-02 | 2.40E-01 | 1.50E+00 | 1.70E-01 | 4.68E-01 | 1.43E+00 |
| ZNF433 | 1.36E-01 | 7.09E-01 | 1.19E+00 | 1.13E-02 | 4.89E-02 | -1.10E+00 | 2.13E-02 | 9.04E-02 | -1.18E+00 | 9.53E-05 | 3.64E-04 | -1.63E+00 | 6.27E-02 | 2.23E-01 | 1.05E+00 | 1.72E-01 | 4.85E-01 | 1.20E+00 | 2.69E-01 | 9.88E-01 | 1.26E+00 | 1.00E-01 | 2.76E-01 | -1.02E+00 |
| RPL9P7 | 1.36E-01 | 7.09E-01 | 1.22E+00 | 1.33E-01 | 5.76E-01 | 1.65E+00 | 1.46E-01 | 6.21E-01 | 1.20E+00 | 2.81E-02 | 1.07E-01 | 1.22E+00 | 4.04E-02 | 1.44E-01 | 1.12E+00 | 3.30E-01 | 9.31E-01 | 1.52E+00 | 1.67E-01 | 6.12E-01 | 1.06E+00 | 1.72E-01 | 4.72E-01 | 1.44E+00 |
| LINC00920 | 1.36E-01 | 7.09E-01 | 1.33E+00 | 2.17E-01 | 9.38E-01 | 1.24E+00 | 1.39E-01 | 5.89E-01 | 1.05E+00 | 3.11E-02 | 1.19E-01 | -1.10E+00 | 2.78E-01 | 9.90E-01 | 1.09E+00 | 3.26E-01 | 9.19E-01 | 1.22E+00 | 5.33E-02 | 1.96E-01 | 1.80E+00 | 1.35E-01 | 3.70E-01 | 1.56E+00 |
| SLC24A4 | 1.36E-01 | 7.09E-01 | 1.14E+00 | 1.76E-01 | 7.58E-01 | 1.54E+00 | 6.81E-02 | 2.89E-01 | -1.33E+00 | 2.43E-01 | 9.28E-01 | 1.24E+00 | 3.13E-02 | 1.12E-01 | -1.17E+00 | 1.71E-01 | 4.81E-01 | 1.64E+00 | 1.84E-01 | 6.76E-01 | 1.03E+00 | 3.52E-01 | 9.67E-01 | 1.23E+00 |
| CBX3P2 | 1.36E-01 | 7.09E-01 | 1.44E+00 | 1.18E-01 | 5.09E-01 | 1.13E+00 | 1.37E-01 | 5.82E-01 | 1.05E+00 | 2.41E-01 | 9.21E-01 | 1.33E+00 | 2.23E-01 | 7.95E-01 | 1.19E+00 | 3.31E-01 | 9.34E-01 | 1.25E+00 | 2.71E-03 | 9.94E-03 | -1.47E+00 | 2.95E-03 | 8.10E-03 | -1.53E+00 |
| RPL27AP5 | 1.36E-01 | 7.09E-01 | 1.45E+00 | 8.62E-02 | 3.72E-01 | 1.82E+00 | 2.48E-02 | 1.05E-01 | 1.57E+00 | 3.56E-02 | 1.36E-01 | 1.87E+00 | 2.09E-01 | 7.44E-01 | 1.55E+00 | 4.45E-02 | 1.25E-01 | 2.16E+00 | 2.47E-01 | 9.05E-01 | 1.36E+00 | 3.57E-02 | 9.81E-02 | 1.72E+00 |
| DDX12P | 1.36E-01 | 7.09E-01 | -1.08E+00 | 9.63E-02 | 4.16E-01 | -1.16E+00 | 5.89E-02 | 2.50E-01 | 1.05E+00 | 2.21E-02 | 8.45E-02 | 1.01E+00 | 1.54E-01 | 5.48E-01 | 1.84E+00 | 3.17E-01 | 8.93E-01 | 1.32E+00 | 8.42E-02 | 3.09E-01 | 1.02E+00 | 9.65E-02 | 2.65E-01 | 1.03E+00 |
| RPL7P24 | 1.36E-01 | 7.09E-01 | 1.29E+00 | 1.44E-02 | 6.24E-02 | 1.76E+00 | 8.84E-02 | 3.75E-01 | 1.32E+00 | 4.28E-04 | 1.63E-03 | 1.96E+00 | 2.52E-02 | 8.98E-02 | 1.06E+00 | 6.68E-02 | 1.88E-01 | 1.60E+00 | 2.07E-01 | 7.61E-01 | 1.17E+00 | 3.37E-06 | 9.26E-06 | 2.07E+00 |
| PYROXD1 | 1.36E-01 | 7.09E-01 | 1.33E+00 | 1.97E-01 | 8.50E-01 | 1.21E+00 | 1.06E-01 | 4.51E-01 | 1.06E+00 | 2.49E-02 | 9.51E-02 | -1.18E+00 | 1.34E-01 | 4.77E-01 | 1.17E+00 | 9.25E-02 | 2.61E-01 | 1.08E+00 | 2.11E-02 | 7.73E-02 | -1.24E+00 | 1.88E-01 | 5.17E-01 | -1.03E+00 |
| CCNYL1 | 1.37E-01 | 7.12E-01 | 1.17E+00 | 1.45E-01 | 6.26E-01 | 1.26E+00 | 4.13E-06 | 1.75E-05 | -1.33E+00 | 2.32E-07 | 8.86E-07 | -1.37E+00 | 5.50E-03 | 1.96E-02 | -1.55E+00 | 8.60E-03 | 2.42E-02 | -1.39E+00 | 7.96E-03 | 2.92E-02 | -1.27E+00 | 5.90E-03 | 1.62E-02 | -1.37E+00 |
| PKLR | 1.37E-01 | 7.12E-01 | 1.17E+00 | 2.09E-01 | 9.01E-01 | 1.08E+00 | 2.35E-01 | 1.00E+00 | -1.00E+00 | 2.62E-01 | 1.00E+00 | 1.00E+00 | 2.46E-01 | 8.78E-01 | 1.09E+00 | 3.21E-01 | 9.05E-01 | 1.10E+00 | 2.69E-02 | 9.86E-02 | -1.08E+00 | 2.89E-02 | 7.94E-02 | -1.01E+00 |
| OR2I1P | 1.37E-01 | 7.12E-01 | -1.05E+00 | 2.07E-01 | 8.93E-01 | 1.07E+00 | 1.23E-01 | 5.24E-01 | -1.15E+00 | 1.40E-01 | 5.37E-01 | -1.15E+00 | 4.38E-05 | 1.56E-04 | -2.45E+00 | 8.84E-04 | 2.49E-03 | -1.72E+00 | 1.63E-08 | 5.97E-08 | -2.30E+00 | 5.37E-04 | 1.47E-03 | -1.35E+00 |
| GNLY | 1.37E-01 | 7.12E-01 | -1.12E+00 | 1.68E-01 | 7.24E-01 | -1.12E+00 | 1.65E-01 | 7.00E-01 | -1.08E+00 | 1.20E-01 | 4.57E-01 | -1.25E+00 | 2.45E-01 | 8.73E-01 | 1.14E+00 | 2.83E-01 | 7.97E-01 | 1.38E+00 | 1.16E-12 | 4.27E-12 | 3.81E+00 | 9.80E-03 | 2.69E-02 | 2.30E+00 |
| UNC13B | 1.37E-01 | 7.13E-01 | 1.09E+00 | 1.49E-01 | 6.45E-01 | 1.48E+00 | 8.43E-02 | 3.58E-01 | 1.07E+00 | 1.91E-01 | 7.28E-01 | 1.32E+00 | 8.11E-02 | 2.89E-01 | -1.14E+00 | 1.27E-01 | 3.59E-01 | -1.04E+00 | 1.26E-01 | 4.64E-01 | 1.29E+00 | 2.59E-01 | 7.12E-01 | 1.35E+00 |
| AC004233.3 | 1.37E-01 | 7.13E-01 | 1.31E+00 | 1.26E-01 | 5.46E-01 | 1.63E+00 | 5.09E-02 | 2.16E-01 | -1.30E+00 | 2.01E-01 | 7.68E-01 | 1.14E+00 | 2.78E-01 | 9.91E-01 | 1.24E+00 | 3.11E-01 | 8.78E-01 | 1.18E+00 | 1.61E-01 | 5.90E-01 | 1.13E+00 | 2.50E-01 | 6.87E-01 | 1.19E+00 |
| MT2P1 | 1.37E-01 | 7.13E-01 | 1.21E+00 | 7.58E-02 | 3.27E-01 | 1.04E+00 | 1.78E-01 | 7.58E-01 | 1.04E+00 | 5.05E-03 | 1.93E-02 | -1.22E+00 | 2.38E-01 | 8.48E-01 | 1.92E+00 | 2.44E-02 | 6.87E-02 | 1.16E+00 | 1.38E-01 | 5.07E-01 | 1.50E+00 | 2.10E-02 | 5.78E-02 | 1.07E+00 |
| EPX | 1.37E-01 | 7.13E-01 | -1.10E+00 | 1.43E-01 | 6.18E-01 | -1.16E+00 | 1.19E-01 | 5.06E-01 | -1.30E+00 | 2.42E-01 | 9.24E-01 | -1.01E+00 | 5.34E-20 | 1.90E-19 | -3.65E+00 | 1.17E-25 | 3.31E-25 | -3.81E+00 | 1.37E-01 | 5.02E-01 | -1.54E+00 | 1.98E-01 | 5.44E-01 | -1.54E+00 |
| AC079416.2 | 1.37E-01 | 7.14E-01 | 1.08E+00 | 1.62E-01 | 7.00E-01 | 1.19E+00 | 9.55E-02 | 4.06E-01 | -1.01E+00 | 1.49E-01 | 5.70E-01 | 1.12E+00 | 1.90E-02 | 6.77E-02 | -1.45E+00 | 1.26E-01 | 3.57E-01 | 1.03E+00 | 2.87E-03 | 1.05E-02 | -1.63E+00 | 3.04E-02 | 8.36E-02 | -1.29E+00 |
| PLK2 | 1.37E-01 | 7.14E-01 | 1.41E+00 | 1.07E-05 | 4.63E-05 | -1.13E+00 | 4.83E-02 | 2.05E-01 | 1.10E+00 | 1.51E-04 | 5.75E-04 | -1.35E+00 | 2.42E-01 | 8.64E-01 | 1.56E+00 | 1.33E-01 | 3.75E-01 | -1.61E+00 | 1.18E-01 | 4.35E-01 | -1.06E+00 | 9.97E-02 | 2.74E-01 | -1.43E+00 |
| C4orf45 | 1.37E-01 | 7.14E-01 | 1.34E+00 | 1.89E-01 | 8.17E-01 | 1.34E+00 | 1.83E-01 | 7.79E-01 | -1.03E+00 | 2.59E-01 | 9.89E-01 | 1.19E+00 | 3.67E-02 | 1.31E-01 | 2.09E+00 | 8.52E-02 | 2.40E-01 | 1.89E+00 | 1.56E-01 | 5.74E-01 | 1.42E+00 | 6.26E-02 | 1.72E-01 | 1.97E+00 |
| IFT140 | 1.37E-01 | 7.15E-01 | 1.02E+00 | 1.44E-01 | 6.24E-01 | 1.09E+00 | 2.12E-03 | 9.00E-03 | -1.32E+00 | 4.26E-02 | 1.63E-01 | 1.06E+00 | 4.11E-02 | 1.46E-01 | -1.15E+00 | 1.57E-01 | 4.41E-01 | 1.02E+00 | 1.29E-01 | 4.74E-01 | -1.18E+00 | 1.19E-01 | 3.28E-01 | -1.24E+00 |
| GNG8 | 1.37E-01 | 7.15E-01 | -1.11E+00 | 1.43E-01 | 6.18E-01 | -1.18E+00 | 2.12E-01 | 8.99E-01 | 1.04E+00 | 2.62E-01 | 1.00E+00 | 1.00E+00 | 2.80E-01 | 1.00E+00 | 1.00E+00 | 3.55E-01 | 1.00E+00 | 1.00E+00 | 2.10E-04 | 7.69E-04 | 1.02E+01 | 2.68E-01 | 7.36E-01 | 2.06E+00 |
| BX293535.1 | 1.37E-01 | 7.15E-01 | 1.29E+00 | 2.10E-01 | 9.08E-01 | 1.33E+00 | 3.01E-02 | 1.28E-01 | -1.28E+00 | 2.76E-03 | 1.06E-02 | -1.82E+00 | 2.02E-01 | 7.21E-01 | -1.19E+00 | 2.65E-01 | 7.47E-01 | -1.09E+00 | 1.77E-03 | 6.51E-03 | -1.19E+00 | 4.20E-04 | 1.15E-03 | -1.53E+00 |
| AL136320.1 | 1.37E-01 | 7.15E-01 | 1.51E+00 | 2.42E-02 | 1.05E-01 | 2.23E+00 | 1.40E-02 | 5.96E-02 | 1.64E+00 | 2.85E-02 | 1.09E-01 | 1.79E+00 | 4.66E-03 | 1.66E-02 | 2.33E+00 | 3.08E-01 | 8.69E-01 | 1.07E+00 | 2.61E-02 | 9.58E-02 | 1.97E+00 | 3.26E-01 | 8.97E-01 | 1.21E+00 |
| CLCN1 | 1.37E-01 | 7.16E-01 | 1.22E+00 | 2.01E-01 | 8.68E-01 | 1.25E+00 | 2.05E-02 | 8.70E-02 | 1.72E+00 | 2.39E-01 | 9.14E-01 | 1.25E+00 | 2.74E-01 | 9.78E-01 | -1.01E+00 | 3.42E-01 | 9.65E-01 | 1.06E+00 | 1.70E-01 | 6.25E-01 | 1.33E+00 | 3.08E-01 | 8.45E-01 | 1.10E+00 |
| NUCKS1P1 | 1.37E-01 | 7.17E-01 | 1.06E+00 | 1.73E-01 | 7.49E-01 | 1.15E+00 | 2.05E-02 | 8.70E-02 | 1.87E+00 | 3.50E-02 | 1.34E-01 | 1.97E+00 | 6.98E-02 | 2.49E-01 | -1.25E+00 | 1.36E-01 | 3.84E-01 | -1.06E+00 | 3.54E-03 | 1.30E-02 | 2.02E+00 | 6.95E-05 | 1.91E-04 | 2.30E+00 |
| ZNF460 | 1.38E-01 | 7.18E-01 | 1.22E+00 | 1.33E-04 | 5.76E-04 | -1.36E+00 | 9.76E-02 | 4.15E-01 | 1.06E+00 | 1.06E-01 | 4.06E-01 | 1.19E+00 | 1.08E-01 | 3.84E-01 | 1.24E+00 | 6.29E-02 | 1.77E-01 | -1.06E+00 | 1.95E-01 | 7.16E-01 | 1.22E+00 | 2.48E-02 | 6.82E-02 | -1.22E+00 |
| PAOX | 1.38E-01 | 7.19E-01 | 1.31E+00 | 2.16E-01 | 9.32E-01 | 1.56E+00 | 4.66E-02 | 1.98E-01 | -1.08E+00 | 5.25E-02 | 2.01E-01 | 1.08E+00 | 2.86E-02 | 1.02E-01 | 1.05E+00 | 3.01E-02 | 8.47E-02 | 1.01E+00 | 1.17E-01 | 4.29E-01 | 1.60E+00 | 2.39E-01 | 6.57E-01 | 1.23E+00 |
| CCDC17 | 1.38E-01 | 7.19E-01 | 1.43E+00 | 3.81E-02 | 1.64E-01 | -1.31E+00 | 2.20E-01 | 9.37E-01 | 1.15E+00 | 1.16E-01 | 4.42E-01 | 1.05E+00 | 1.61E-01 | 5.73E-01 | 1.45E+00 | 3.06E-01 | 8.63E-01 | 1.14E+00 | 2.09E-01 | 7.68E-01 | 1.13E+00 | 2.31E-01 | 6.35E-01 | 1.08E+00 |
| AL118558.3 | 1.38E-01 | 7.19E-01 | 1.08E+00 | 4.86E-02 | 2.10E-01 | -1.07E+00 | 1.65E-01 | 7.00E-01 | 1.32E+00 | 1.09E-01 | 4.15E-01 | 1.67E+00 | 2.03E-01 | 7.24E-01 | 1.16E+00 | 3.38E-01 | 9.53E-01 | 1.36E+00 | 1.20E-03 | 4.40E-03 | -1.21E+00 | 3.45E-02 | 9.49E-02 | -1.08E+00 |
| RPL22 | 1.38E-01 | 7.19E-01 | -1.01E+00 | 6.44E-12 | 2.78E-11 | -1.20E+00 | 1.05E-01 | 4.47E-01 | -1.03E+00 | 5.52E-07 | 2.11E-06 | -1.17E+00 | 1.03E-01 | 3.67E-01 | -1.03E+00 | 1.32E-06 | 3.73E-06 | -1.19E+00 | 1.70E-09 | 6.23E-09 | -1.11E+00 | 6.95E-14 | 1.91E-13 | -1.25E+00 |
| RPF2 | 1.38E-01 | 7.19E-01 | 1.33E+00 | 9.99E-02 | 4.32E-01 | 1.59E+00 | 1.58E-01 | 6.70E-01 | 1.14E+00 | 1.90E-01 | 7.27E-01 | 1.34E+00 | 2.48E-03 | 8.84E-03 | -1.33E+00 | 7.14E-02 | 2.01E-01 | 1.06E+00 | 2.84E-05 | 1.04E-04 | -1.16E+00 | 1.12E-01 | 3.07E-01 | 1.21E+00 |
| LRRTM3 | 1.38E-01 | 7.20E-01 | 1.18E+00 | 4.37E-02 | 1.89E-01 | 1.82E+00 | 3.95E-02 | 1.68E-01 | -1.27E+00 | 5.66E-02 | 2.16E-01 | -1.12E+00 | 9.18E-02 | 3.27E-01 | 2.06E+00 | 1.78E-01 | 5.01E-01 | 1.75E+00 | 7.35E-02 | 2.70E-01 | 1.98E+00 | 1.10E-01 | 3.02E-01 | 2.13E+00 |
| AC007952.4 | 1.38E-01 | 7.20E-01 | 1.37E+00 | 1.27E-18 | 5.51E-18 | -1.89E+00 | 3.88E-02 | 1.65E-01 | 1.38E+00 | 3.35E-08 | 1.28E-07 | -1.56E+00 | 5.98E-06 | 2.13E-05 | -1.24E+00 | 6.11E-07 | 1.72E-06 | -1.49E+00 | 5.08E-04 | 1.86E-03 | -1.23E+00 | 1.34E-03 | 3.68E-03 | -1.28E+00 |
| INTS6-AS1 | 1.38E-01 | 7.20E-01 | 1.32E+00 | 1.64E-01 | 7.10E-01 | 1.39E+00 | 1.12E-01 | 4.75E-01 | 1.15E+00 | 9.95E-02 | 3.80E-01 | 1.25E+00 | 7.81E-02 | 2.78E-01 | 1.72E+00 | 1.90E-01 | 5.35E-01 | 1.71E+00 | 3.11E-02 | 1.14E-01 | 1.81E+00 | 6.03E-02 | 1.66E-01 | 1.67E+00 |
| RPL5P23 | 1.38E-01 | 7.21E-01 | 1.28E+00 | 4.40E-03 | 1.90E-02 | 2.28E+00 | 8.95E-02 | 3.80E-01 | -1.06E+00 | 6.58E-03 | 2.51E-02 | 2.03E+00 | 4.93E-02 | 1.76E-01 | -1.22E+00 | 2.27E-01 | 6.41E-01 | 1.58E+00 | 2.08E-01 | 7.63E-01 | 1.39E+00 | 2.79E-02 | 7.66E-02 | 2.00E+00 |
| IL1RAPL1 | 1.38E-01 | 7.21E-01 | -1.10E+00 | 1.21E-01 | 5.24E-01 | -1.28E+00 | 2.18E-01 | 9.27E-01 | 1.05E+00 | 2.62E-01 | 1.00E+00 | 1.00E+00 | 4.20E-04 | 1.50E-03 | 3.96E+00 | 3.27E-02 | 9.21E-02 | 2.72E+00 | 5.09E-22 | 1.87E-21 | 4.93E+00 | 1.61E-01 | 4.43E-01 | 1.48E+00 |
| FAHD2B | 1.38E-01 | 7.21E-01 | 1.28E+00 | 1.87E-01 | 8.09E-01 | 1.59E+00 | 5.46E-02 | 2.32E-01 | 1.43E+00 | 2.39E-01 | 9.14E-01 | 1.49E+00 | 9.89E-02 | 3.53E-01 | 1.09E+00 | 3.10E-01 | 8.75E-01 | 1.27E+00 | 2.16E-01 | 7.92E-01 | 1.26E+00 | 6.68E-02 | 1.83E-01 | 1.50E+00 |
| FLJ13224 | 1.39E-01 | 7.22E-01 | 1.68E+00 | 2.06E-01 | 8.90E-01 | 1.61E+00 | 1.42E-01 | 6.05E-01 | 1.09E+00 | 1.01E-01 | 3.84E-01 | 1.71E+00 | 1.18E-01 | 4.22E-01 | 2.04E+00 | 2.31E-01 | 6.51E-01 | 1.79E+00 | 4.56E-02 | 1.67E-01 | 1.88E+00 | 4.24E-02 | 1.17E-01 | 1.86E+00 |
| LRRC29 | 1.39E-01 | 7.22E-01 | 1.15E+00 | 9.18E-02 | 3.97E-01 | 1.06E+00 | 6.97E-04 | 2.96E-03 | -1.51E+00 | 9.05E-03 | 3.46E-02 | -1.09E+00 | 2.29E-02 | 8.18E-02 | -1.10E+00 | 6.37E-02 | 1.80E-01 | 1.04E+00 | 1.99E-01 | 7.32E-01 | 1.23E+00 | 8.31E-02 | 2.28E-01 | -1.09E+00 |
| AC008802.1 | 1.39E-01 | 7.23E-01 | 1.31E+00 | 2.04E-01 | 8.80E-01 | 1.45E+00 | 2.04E-01 | 8.66E-01 | 1.28E+00 | 1.22E-01 | 4.67E-01 | 1.65E+00 | 2.67E-01 | 9.53E-01 | 1.50E+00 | 3.10E-01 | 8.73E-01 | 1.45E+00 | 2.16E-01 | 7.92E-01 | 1.27E+00 | 4.62E-02 | 1.27E-01 | 1.81E+00 |
| PRKG1-AS1 | 1.39E-01 | 7.23E-01 | 1.05E+00 | 1.46E-01 | 6.31E-01 | 1.50E+00 | 3.70E-02 | 1.57E-01 | -1.23E+00 | 1.14E-01 | 4.36E-01 | 1.06E+00 | 1.40E-01 | 5.01E-01 | 1.72E+00 | 2.49E-01 | 7.01E-01 | 1.63E+00 | 1.56E-01 | 5.71E-01 | -1.00E+00 | 3.32E-01 | 9.11E-01 | 1.10E+00 |
| TICRR | 1.39E-01 | 7.23E-01 | 1.01E+00 | 1.39E-01 | 5.99E-01 | -1.05E+00 | 2.22E-01 | 9.42E-01 | 1.19E+00 | 2.19E-01 | 8.36E-01 | 1.27E+00 | 2.60E-01 | 9.29E-01 | -1.01E+00 | 3.18E-01 | 8.96E-01 | -1.05E+00 | 8.02E-02 | 2.94E-01 | -1.08E+00 | 4.74E-04 | 1.30E-03 | 2.04E+00 |
| RPL18AP15 | 1.39E-01 | 7.23E-01 | 1.29E+00 | 1.05E-02 | 4.54E-02 | 1.16E+00 | 1.67E-01 | 7.10E-01 | 1.40E+00 | 3.49E-02 | 1.33E-01 | 1.26E+00 | 4.85E-02 | 1.73E-01 | 1.05E+00 | 1.48E-02 | 4.16E-02 | -1.04E+00 | 1.82E-01 | 6.67E-01 | 1.39E+00 | 1.07E-01 | 2.95E-01 | 1.15E+00 |
| LGALS1 | 1.39E-01 | 7.23E-01 | 1.23E+00 | 1.07E-07 | 4.60E-07 | -1.41E+00 | 4.77E-04 | 2.03E-03 | -1.10E+00 | 3.89E-09 | 1.48E-08 | -1.31E+00 | 6.71E-15 | 2.39E-14 | -1.65E+00 | 1.42E-27 | 4.01E-27 | -3.07E+00 | 3.96E-03 | 1.45E-02 | -1.45E+00 | 3.59E-04 | 9.86E-04 | -2.76E+00 |
| AZIN1-AS1 | 1.39E-01 | 7.23E-01 | 1.09E+00 | 1.26E-01 | 5.43E-01 | 1.17E+00 | 1.32E-02 | 5.59E-02 | -1.10E+00 | 7.18E-05 | 2.74E-04 | -1.42E+00 | 2.12E-01 | 7.54E-01 | 1.25E+00 | 2.76E-01 | 7.79E-01 | 1.31E+00 | 5.85E-04 | 2.15E-03 | 2.23E+00 | 1.81E-01 | 4.97E-01 | 1.51E+00 |
| NUS1P1 | 1.39E-01 | 7.24E-01 | 1.32E+00 | 1.49E-01 | 6.43E-01 | 1.08E+00 | 1.88E-01 | 7.99E-01 | 1.24E+00 | 1.73E-01 | 6.63E-01 | 1.10E+00 | 1.84E-01 | 6.54E-01 | 1.06E+00 | 2.73E-01 | 7.70E-01 | 1.16E+00 | 2.31E-01 | 8.48E-01 | 1.29E+00 | 2.73E-01 | 7.49E-01 | 1.50E+00 |
| AP003774.4 | 1.39E-01 | 7.24E-01 | 1.25E+00 | 2.10E-01 | 9.08E-01 | 1.25E+00 | 2.05E-01 | 8.71E-01 | 1.25E+00 | 2.51E-01 | 9.58E-01 | 1.03E+00 | 1.26E-02 | 4.50E-02 | -1.08E+00 | 6.28E-05 | 1.77E-04 | -1.42E+00 | 6.18E-02 | 2.27E-01 | 1.60E+00 | 2.57E-01 | 7.07E-01 | -1.10E+00 |
| AC009963.3 | 1.39E-01 | 7.25E-01 | 1.21E+00 | 7.43E-03 | 3.21E-02 | -1.33E+00 | 4.40E-02 | 1.87E-01 | -1.17E+00 | 1.47E-03 | 5.61E-03 | -1.63E+00 | 2.64E-02 | 9.42E-02 | -1.12E+00 | 1.60E-03 | 4.51E-03 | -1.80E+00 | 5.14E-03 | 1.89E-02 | -1.19E+00 | 5.84E-06 | 1.60E-05 | -1.61E+00 |
| HNRNPA1P51 | 1.39E-01 | 7.25E-01 | 1.34E+00 | 5.86E-02 | 2.53E-01 | -1.21E+00 | 1.88E-02 | 7.97E-02 | 1.69E+00 | 2.96E-02 | 1.13E-01 | 1.88E+00 | 2.15E-01 | 7.68E-01 | 1.12E+00 | 3.07E-01 | 8.65E-01 | 1.28E+00 | 1.54E-01 | 5.67E-01 | 1.35E+00 | 8.43E-02 | 2.32E-01 | 1.55E+00 |
| HIST3H2BB | 1.39E-01 | 7.25E-01 | 1.21E+00 | 9.51E-02 | 4.11E-01 | 1.13E+00 | 2.28E-01 | 9.67E-01 | 1.17E+00 | 1.14E-01 | 4.35E-01 | -1.08E+00 | 7.08E-02 | 2.53E-01 | -1.35E+00 | 3.40E-01 | 9.59E-01 | 1.29E+00 | 8.78E-02 | 3.22E-01 | 1.40E+00 | 6.82E-02 | 1.87E-01 | 1.88E+00 |
| PTGER3 | 1.39E-01 | 7.25E-01 | -1.54E+00 | 1.87E-01 | 8.07E-01 | -1.40E+00 | 2.26E-01 | 9.59E-01 | -1.00E+00 | 2.06E-01 | 7.88E-01 | -1.15E+00 | 1.19E-01 | 4.24E-01 | -1.86E+00 | 2.63E-01 | 7.42E-01 | -1.36E+00 | 1.23E-23 | 4.50E-23 | -1.96E+00 | 4.78E-14 | 1.31E-13 | -1.42E+00 |
| ZNF675 | 1.39E-01 | 7.26E-01 | 1.18E+00 | 9.38E-05 | 4.06E-04 | -1.28E+00 | 1.05E-02 | 4.46E-02 | -1.01E+00 | 1.15E-07 | 4.38E-07 | -1.41E+00 | 9.51E-04 | 3.39E-03 | -1.12E+00 | 4.09E-04 | 1.15E-03 | -1.22E+00 | 2.13E-03 | 7.82E-03 | -1.07E+00 | 2.13E-05 | 5.86E-05 | -1.26E+00 |
| CEP72 | 1.39E-01 | 7.26E-01 | -1.09E+00 | 2.31E-01 | 1.00E+00 | 1.11E+00 | 4.62E-02 | 1.96E-01 | -1.14E+00 | 1.40E-02 | 5.33E-02 | -1.18E+00 | 1.08E-03 | 3.84E-03 | -2.37E+00 | 7.32E-03 | 2.06E-02 | -1.73E+00 | 1.01E-03 | 3.72E-03 | -1.35E+00 | 2.56E-03 | 7.02E-03 | -1.25E+00 |
| AL023806.1 | 1.40E-01 | 7.27E-01 | 1.13E+00 | 1.19E-01 | 5.13E-01 | 1.09E+00 | 5.72E-02 | 2.43E-01 | 1.62E+00 | 1.72E-01 | 6.56E-01 | 1.41E+00 | 1.98E-01 | 7.08E-01 | 1.33E+00 | 2.89E-01 | 8.14E-01 | 1.11E+00 | 6.11E-02 | 2.24E-01 | 1.64E+00 | 3.56E-01 | 9.78E-01 | 1.17E+00 |
| STX12 | 1.40E-01 | 7.27E-01 | 1.26E+00 | 4.84E-04 | 2.09E-03 | 1.00E+00 | 8.86E-02 | 3.76E-01 | 1.19E+00 | 5.28E-07 | 2.02E-06 | -1.28E+00 | 2.50E-03 | 8.93E-03 | -1.11E+00 | 7.05E-04 | 1.99E-03 | -1.22E+00 | 5.17E-04 | 1.90E-03 | -1.19E+00 | 3.40E-02 | 9.34E-02 | 1.04E+00 |
| ATF7IP2 | 1.40E-01 | 7.27E-01 | 1.36E+00 | 6.87E-02 | 2.97E-01 | 1.42E+00 | 1.05E-02 | 4.46E-02 | 1.42E+00 | 2.01E-01 | 7.68E-01 | 1.37E+00 | 8.95E-10 | 3.19E-09 | 2.64E+00 | 2.33E-06 | 6.56E-06 | 2.24E+00 | 7.21E-02 | 2.65E-01 | 1.61E+00 | 2.64E-01 | 7.26E-01 | 1.31E+00 |
| AC092910.3 | 1.40E-01 | 7.27E-01 | 1.14E+00 | 6.90E-02 | 2.98E-01 | 1.02E+00 | 2.18E-01 | 9.24E-01 | 1.18E+00 | 6.17E-02 | 2.36E-01 | -1.03E+00 | 1.03E-01 | 3.68E-01 | 1.78E+00 | 1.56E-01 | 4.40E-01 | 1.90E+00 | 6.10E-02 | 2.24E-01 | -1.23E+00 | 2.00E-01 | 5.48E-01 | -1.07E+00 |
| EMC3-AS1 | 1.40E-01 | 7.28E-01 | 1.28E+00 | 1.54E-01 | 6.63E-01 | 1.48E+00 | 1.35E-01 | 5.76E-01 | 1.07E+00 | 5.14E-02 | 1.96E-01 | 1.09E+00 | 1.80E-01 | 6.42E-01 | -1.07E+00 | 2.71E-01 | 7.64E-01 | 1.09E+00 | 1.91E-01 | 7.01E-01 | 1.38E+00 | 1.65E-01 | 4.54E-01 | 1.62E+00 |
| ADSSL1 | 1.40E-01 | 7.28E-01 | 1.12E+00 | 3.19E-02 | 1.38E-01 | -1.11E+00 | 3.19E-02 | 1.35E-01 | -1.18E+00 | 1.77E-02 | 6.76E-02 | -1.16E+00 | 2.63E-01 | 9.39E-01 | 1.12E+00 | 3.55E-01 | 1.00E+00 | 1.21E+00 | 1.18E-01 | 4.33E-01 | -1.03E+00 | 2.86E-01 | 7.85E-01 | 1.33E+00 |
| AC007036.2 | 1.40E-01 | 7.29E-01 | 1.31E+00 | 1.88E-01 | 8.11E-01 | 1.63E+00 | 1.22E-01 | 5.18E-01 | 1.40E+00 | 1.84E-01 | 7.03E-01 | 1.26E+00 | 1.56E-02 | 5.55E-02 | -1.20E+00 | 8.44E-02 | 2.38E-01 | 1.01E+00 | 1.53E-01 | 5.60E-01 | 1.11E+00 | 2.34E-01 | 6.43E-01 | 1.14E+00 |
| ARG1 | 1.40E-01 | 7.30E-01 | 1.21E+00 | 2.31E-01 | 9.98E-01 | 1.06E+00 | 6.46E-02 | 2.74E-01 | 1.54E+00 | 2.03E-01 | 7.76E-01 | 1.25E+00 | 1.86E-01 | 6.64E-01 | 1.49E+00 | 1.14E-01 | 3.22E-01 | -1.18E+00 | 2.71E-01 | 9.95E-01 | 1.11E+00 | 1.78E-01 | 4.90E-01 | -1.06E+00 |
| AL031772.1 | 1.40E-01 | 7.31E-01 | 1.16E+00 | 2.07E-01 | 8.95E-01 | 1.29E+00 | 1.59E-01 | 6.77E-01 | 1.35E+00 | 5.53E-02 | 2.11E-01 | 1.96E+00 | 1.52E-01 | 5.43E-01 | 1.56E+00 | 1.08E-01 | 3.04E-01 | 2.02E+00 | 1.03E-01 | 3.79E-01 | -1.08E+00 | 1.07E-01 | 2.93E-01 | -1.08E+00 |
| MTCO1P40 | 1.40E-01 | 7.31E-01 | 1.32E+00 | 8.20E-13 | 3.54E-12 | -1.49E+00 | 1.79E-01 | 7.61E-01 | 1.25E+00 | 6.46E-18 | 2.47E-17 | -1.74E+00 | 9.67E-02 | 3.45E-01 | 1.25E+00 | 5.27E-12 | 1.49E-11 | -1.65E+00 | 1.14E-01 | 4.17E-01 | 1.25E+00 | 1.11E-17 | 3.04E-17 | -1.63E+00 |
| PRAME | 1.40E-01 | 7.31E-01 | 1.42E+00 | 1.76E-02 | 7.60E-02 | 2.15E+00 | 4.70E-02 | 2.00E-01 | 1.04E+00 | 2.37E-01 | 9.06E-01 | 1.45E+00 | 1.09E-05 | 3.88E-05 | -1.87E+00 | 9.50E-05 | 2.68E-04 | -1.53E+00 | 3.33E-03 | 1.22E-02 | -1.22E+00 | 1.54E-02 | 4.22E-02 | -1.12E+00 |
| POLE2 | 1.40E-01 | 7.31E-01 | 1.06E+00 | 1.22E-01 | 5.28E-01 | -1.01E+00 | 7.67E-02 | 3.26E-01 | 1.02E+00 | 1.40E-01 | 5.35E-01 | 1.19E+00 | 9.84E-02 | 3.51E-01 | -1.21E+00 | 3.22E-01 | 9.07E-01 | 1.13E+00 | 4.18E-08 | 1.53E-07 | -1.27E+00 | 1.96E-01 | 5.38E-01 | 1.12E+00 |
| CCDC168 | 1.40E-01 | 7.31E-01 | 1.34E+00 | 1.22E-01 | 5.29E-01 | 2.09E+00 | 4.78E-02 | 2.03E-01 | -1.29E+00 | 2.93E-02 | 1.12E-01 | -1.42E+00 | 2.69E-01 | 9.61E-01 | 1.33E+00 | 2.53E-01 | 7.14E-01 | 1.12E+00 | 1.05E-02 | 3.87E-02 | -1.83E+00 | 5.29E-02 | 1.45E-01 | -1.57E+00 |
| EPM2A | 1.40E-01 | 7.32E-01 | 1.17E+00 | 1.29E-01 | 5.56E-01 | 1.26E+00 | 6.65E-02 | 2.83E-01 | 1.13E+00 | 5.12E-02 | 1.96E-01 | 1.22E+00 | 1.74E-03 | 6.21E-03 | -1.35E+00 | 6.06E-02 | 1.71E-01 | 1.01E+00 | 1.97E-02 | 7.24E-02 | 1.00E+00 | 1.94E-02 | 5.32E-02 | -1.07E+00 |
| TNKS | 1.40E-01 | 7.32E-01 | 1.28E+00 | 1.08E-01 | 4.65E-01 | 1.43E+00 | 9.82E-04 | 4.17E-03 | -1.21E+00 | 9.62E-05 | 3.67E-04 | -1.13E+00 | 3.81E-03 | 1.36E-02 | -1.04E+00 | 5.98E-03 | 1.69E-02 | 1.03E+00 | 2.38E-03 | 8.72E-03 | -1.15E+00 | 4.09E-02 | 1.12E-01 | -1.01E+00 |
| DUS2 | 1.41E-01 | 7.32E-01 | 1.17E+00 | 1.02E-01 | 4.42E-01 | 1.83E+00 | 1.38E-01 | 5.87E-01 | -1.01E+00 | 2.52E-01 | 9.64E-01 | 1.28E+00 | 1.36E-01 | 4.86E-01 | 1.17E+00 | 2.34E-01 | 6.59E-01 | 1.25E+00 | 6.54E-03 | 2.40E-02 | -1.22E+00 | 6.80E-02 | 1.87E-01 | -1.05E+00 |
| SUMO2P20 | 1.41E-01 | 7.33E-01 | 1.42E+00 | 6.01E-02 | 2.59E-01 | 1.95E+00 | 9.81E-02 | 4.17E-01 | 1.03E+00 | 2.05E-01 | 7.83E-01 | 1.30E+00 | 5.03E-02 | 1.79E-01 | -1.04E+00 | 2.79E-01 | 7.88E-01 | 1.31E+00 | 2.59E-01 | 9.50E-01 | 1.24E+00 | 8.05E-02 | 2.21E-01 | 1.69E+00 |
| AC023481.1 | 1.41E-01 | 7.33E-01 | -1.11E+00 | 8.98E-02 | 3.88E-01 | -1.32E+00 | 6.81E-02 | 2.90E-01 | 1.51E+00 | 2.13E-01 | 8.13E-01 | 1.15E+00 | 7.12E-02 | 2.54E-01 | -1.39E+00 | 1.15E-01 | 3.24E-01 | -1.58E+00 | 1.56E-01 | 5.72E-01 | 1.24E+00 | 1.76E-01 | 4.84E-01 | -1.38E+00 |
| HAVCR2 | 1.41E-01 | 7.34E-01 | 1.22E+00 | 4.79E-03 | 2.07E-02 | -1.40E+00 | 7.52E-08 | 3.19E-07 | -1.58E+00 | 7.29E-10 | 2.78E-09 | -1.73E+00 | 2.73E-02 | 9.75E-02 | -1.42E+00 | 7.09E-04 | 2.00E-03 | -2.34E+00 | 2.64E-01 | 9.69E-01 | 1.23E+00 | 1.89E-01 | 5.19E-01 | -1.11E+00 |
| ATP5A1P2 | 1.41E-01 | 7.34E-01 | 1.21E+00 | 2.52E-02 | 1.09E-01 | 1.91E+00 | 2.00E-01 | 8.49E-01 | 1.17E+00 | 7.11E-02 | 2.72E-01 | 1.68E+00 | 2.16E-01 | 7.69E-01 | 1.36E+00 | 2.80E-02 | 7.88E-02 | 2.27E+00 | 2.50E-01 | 9.17E-01 | 1.38E+00 | 6.49E-03 | 1.78E-02 | 1.95E+00 |
| FAM161A | 1.41E-01 | 7.35E-01 | 1.17E+00 | 7.39E-02 | 3.19E-01 | 1.74E+00 | 7.56E-02 | 3.21E-01 | 1.00E+00 | 1.19E-01 | 4.53E-01 | 1.11E+00 | 2.49E-01 | 8.87E-01 | 1.08E+00 | 2.29E-01 | 6.44E-01 | 1.42E+00 | 3.75E-04 | 1.38E-03 | -1.54E+00 | 1.77E-03 | 4.87E-03 | -1.30E+00 |
| C17orf99 | 1.41E-01 | 7.35E-01 | -1.12E+00 | 2.10E-01 | 9.09E-01 | 1.34E+00 | 1.71E-07 | 7.26E-07 | -2.95E+00 | 4.30E-06 | 1.64E-05 | -2.03E+00 | 3.45E-18 | 1.23E-17 | -2.25E+00 | 1.61E-17 | 4.54E-17 | -1.95E+00 | 2.50E-03 | 9.19E-03 | -1.36E+00 | 8.87E-03 | 2.44E-02 | -1.18E+00 |
| ATP11A-AS1 | 1.41E-01 | 7.35E-01 | 1.04E+00 | 7.97E-02 | 3.44E-01 | -1.15E+00 | 9.47E-02 | 4.02E-01 | -1.23E+00 | 1.59E-02 | 6.09E-02 | -1.33E+00 | 2.61E-02 | 9.31E-02 | -1.55E+00 | 6.52E-03 | 1.84E-02 | -2.29E+00 | 7.31E-03 | 2.68E-02 | -1.39E+00 | 8.54E-03 | 2.35E-02 | -1.54E+00 |
| AC010883.1 | 1.41E-01 | 7.35E-01 | 1.45E+00 | 1.83E-01 | 7.90E-01 | 1.25E+00 | 3.01E-02 | 1.28E-01 | -1.18E+00 | 1.00E-03 | 3.83E-03 | -1.29E+00 | 2.50E-01 | 8.90E-01 | 1.22E+00 | 2.24E-01 | 6.32E-01 | 1.10E+00 | 9.41E-02 | 3.45E-01 | -1.35E+00 | 1.76E-01 | 4.84E-01 | -1.28E+00 |
| AL158211.5 | 1.41E-01 | 7.35E-01 | 1.34E+00 | 2.74E-02 | 1.19E-01 | 1.90E+00 | 2.02E-01 | 8.57E-01 | 1.29E+00 | 2.02E-01 | 7.73E-01 | 1.31E+00 | 1.10E-01 | 3.93E-01 | -1.12E+00 | 1.91E-01 | 5.37E-01 | -1.04E+00 | 1.49E-01 | 5.48E-01 | -1.15E+00 | 3.39E-01 | 9.31E-01 | 1.23E+00 |
| AC008741.2 | 1.41E-01 | 7.35E-01 | 1.47E+00 | 7.66E-04 | 3.31E-03 | 2.69E+00 | 1.60E-01 | 6.82E-01 | -1.03E+00 | 2.32E-01 | 8.87E-01 | 1.37E+00 | 1.29E-01 | 4.60E-01 | -1.01E+00 | 1.29E-01 | 3.62E-01 | 1.61E+00 | 2.64E-01 | 9.69E-01 | 1.25E+00 | 3.55E-01 | 9.76E-01 | 1.16E+00 |
| KIF4B | 1.41E-01 | 7.36E-01 | -1.07E+00 | 1.43E-01 | 6.18E-01 | -1.14E+00 | 2.90E-02 | 1.23E-01 | 1.74E+00 | 9.27E-02 | 3.54E-01 | 1.61E+00 | 2.80E-01 | 1.00E+00 | 1.00E+00 | 3.02E-01 | 8.50E-01 | 1.25E+00 | 5.27E-02 | 1.93E-01 | -1.09E+00 | 2.04E-01 | 5.60E-01 | -1.00E+00 |
| THAP12P7 | 1.41E-01 | 7.36E-01 | 1.28E+00 | 1.05E-02 | 4.55E-02 | -1.05E+00 | 1.07E-01 | 4.55E-01 | 1.41E+00 | 2.13E-01 | 8.15E-01 | 1.36E+00 | 1.65E-01 | 5.89E-01 | 1.20E+00 | 2.60E-01 | 7.33E-01 | 1.27E+00 | 6.44E-02 | 2.36E-01 | 1.47E+00 | 1.44E-01 | 3.95E-01 | 1.34E+00 |
| AC002543.1 | 1.41E-01 | 7.37E-01 | 1.24E+00 | 4.50E-02 | 1.95E-01 | 1.01E+00 | 1.47E-02 | 6.23E-02 | 2.00E+00 | 2.50E-01 | 9.54E-01 | 1.43E+00 | 2.30E-01 | 8.20E-01 | 1.25E+00 | 9.57E-02 | 2.70E-01 | -1.18E+00 | 1.12E-01 | 4.12E-01 | 1.63E+00 | 2.81E-01 | 7.72E-01 | 1.43E+00 |
| AC078778.1 | 1.41E-01 | 7.37E-01 | 1.14E+00 | 1.23E-01 | 5.30E-01 | 1.64E+00 | 2.13E-01 | 9.03E-01 | 1.23E+00 | 2.06E-01 | 7.85E-01 | 1.50E+00 | 7.44E-02 | 2.65E-01 | -1.26E+00 | 2.92E-01 | 8.24E-01 | 1.19E+00 | 1.64E-01 | 6.03E-01 | 1.41E+00 | 7.24E-03 | 1.99E-02 | 2.57E+00 |
| AC007608.4 | 1.41E-01 | 7.37E-01 | 1.11E+00 | 1.13E-01 | 4.88E-01 | 1.08E+00 | 1.56E-01 | 6.62E-01 | -1.07E+00 | 1.42E-01 | 5.41E-01 | -1.12E+00 | 1.02E-02 | 3.63E-02 | -1.31E+00 | 1.21E-03 | 3.42E-03 | -1.67E+00 | 1.09E-01 | 3.99E-01 | -1.04E+00 | 9.10E-02 | 2.50E-01 | -1.25E+00 |
| LRP5L | 1.41E-01 | 7.37E-01 | 1.21E+00 | 1.88E-01 | 8.13E-01 | 1.36E+00 | 1.64E-01 | 6.96E-01 | 1.25E+00 | 2.43E-01 | 9.28E-01 | 1.33E+00 | 6.06E-03 | 2.16E-02 | -1.12E+00 | 8.63E-05 | 2.43E-04 | -1.85E+00 | 2.91E-02 | 1.07E-01 | -1.28E+00 | 8.71E-02 | 2.39E-01 | -1.14E+00 |

| | | | | | | | | | | | | | | | | | | | | | | | | |
|---|---|---|---|---|---|---|---|---|---|---|---|---|---|---|---|---|---|---|---|---|---|---|---|---|
| SCRG1 | 1.41E-01 | 7.37E-01 | 1.11E+00 | 6.92E-05 | 2.99E-04 | 2.49E+00 | 1.06E-01 | 4.52E-01 | 1.56E+00 | 5.11E-04 | 1.95E-03 | 3.25E+00 | 4.59E-03 | 1.64E-02 | -1.81E+00 | 4.65E-02 | 1.31E-01 | -1.18E+00 | 2.22E-01 | 8.15E-01 | -1.09E+00 | 2.74E-02 | 7.52E-02 | 2.07E+00 |
| SKA1 | 1.41E-01 | 7.37E-01 | -1.06E+00 | 1.25E-01 | 5.39E-01 | -1.21E+00 | 2.31E-01 | 9.83E-01 | 1.18E+00 | 8.70E-02 | 3.32E-01 | 1.03E+00 | 2.29E-02 | 8.16E-02 | -1.98E+00 | 5.76E-02 | 1.62E-01 | -1.43E+00 | 1.96E-02 | 7.19E-02 | -1.19E+00 | 2.10E-02 | 5.78E-02 | -1.21E+00 |
| KIF18B | 1.41E-01 | 7.37E-01 | -1.11E+00 | 1.93E-01 | 8.36E-01 | -1.04E+00 | 1.34E-01 | 5.71E-01 | -1.03E+00 | 1.77E-01 | 6.77E-01 | 1.16E+00 | 1.88E-01 | 6.70E-01 | -1.22E+00 | 2.72E-01 | 7.68E-01 | -1.12E+00 | 3.06E-02 | 1.12E-01 | -1.28E+00 | 2.73E-01 | 7.49E-01 | -1.00E+00 |
| RPL35P5 | 1.41E-01 | 7.37E-01 | 1.33E+00 | 6.06E-07 | 2.62E-06 | -1.18E+00 | 1.71E-02 | 7.26E-02 | 1.47E+00 | 1.01E-03 | 3.85E-03 | -1.06E+00 | 1.52E-01 | 5.40E-01 | 1.31E+00 | 4.71E-04 | 1.33E-03 | -1.17E+00 | 1.48E-01 | 5.42E-01 | 1.38E+00 | 9.82E-04 | 2.70E-03 | -1.15E+00 |
| RAP1GDS1 | 1.41E-01 | 7.37E-01 | 1.26E+00 | 1.40E-02 | 6.05E-02 | 1.78E+00 | 1.75E-01 | 7.46E-01 | 1.12E+00 | 1.82E-01 | 6.93E-01 | 1.50E+00 | 5.55E-03 | 1.98E-02 | 1.05E+00 | 1.44E-01 | 4.07E-01 | 1.28E+00 | 1.67E-01 | 6.11E-01 | 1.12E+00 | 2.49E-01 | 6.84E-01 | 1.30E+00 |
| TCL1A | 1.41E-01 | 7.37E-01 | 1.12E+00 | 1.56E-01 | 6.75E-01 | 1.57E+00 | 9.19E-02 | 3.91E-01 | 1.42E+00 | 4.70E-03 | 1.79E-02 | 2.25E+00 | 1.97E-01 | 7.02E-01 | 1.54E+00 | 2.03E-01 | 5.71E-01 | 1.80E+00 | 2.12E-01 | 7.77E-01 | 1.32E+00 | 3.72E-02 | 1.02E-01 | 2.25E+00 |
| EXOC3L4 | 1.42E-01 | 7.38E-01 | 1.19E+00 | 1.26E-01 | 5.43E-01 | 1.40E+00 | 4.11E-02 | 1.75E-01 | -1.37E+00 | 8.37E-02 | 3.20E-01 | -1.17E+00 | 4.69E-02 | 1.67E-01 | 1.95E+00 | 2.33E-01 | 6.56E-01 | 1.34E+00 | 1.28E-04 | 4.69E-04 | 3.10E+00 | 2.56E-02 | 7.03E-02 | 1.94E+00 |
| SPAG5 | 1.42E-01 | 7.39E-01 | 1.06E+00 | 1.63E-01 | 7.04E-01 | 1.05E+00 | 1.05E-01 | 4.44E-01 | 1.34E+00 | 2.53E-01 | 9.65E-01 | 1.31E+00 | 1.73E-01 | 6.18E-01 | -1.36E+00 | 1.12E-01 | 3.15E-01 | -1.90E+00 | 5.40E-03 | 1.98E-02 | -1.24E+00 | 1.87E-01 | 5.13E-01 | -1.07E+00 |
| AC231533.2 | 1.42E-01 | 7.39E-01 | 1.08E+00 | 1.15E-01 | 4.96E-01 | 1.77E+00 | 1.97E-02 | 8.36E-02 | -1.14E+00 | 3.23E-02 | 1.23E-01 | -1.06E+00 | 1.66E-07 | 5.92E-07 | -1.99E+00 | 4.23E-02 | 1.19E-01 | 1.04E+00 | 4.08E-02 | 1.50E-01 | -1.18E+00 | 1.79E-01 | 4.92E-01 | 1.02E+00 |
| PCAT18 | 1.42E-01 | 7.39E-01 | -1.02E+00 | 1.43E-01 | 6.19E-01 | -1.11E+00 | 1.48E-01 | 6.30E-01 | 1.15E+00 | 2.62E-01 | 1.00E+00 | 1.00E+00 | 2.21E-02 | 7.87E-02 | -2.04E+00 | 6.58E-02 | 1.86E-01 | -1.70E+00 | 6.92E-04 | 2.54E-03 | -1.31E+00 | 1.17E-13 | 3.21E-13 | -2.95E+00 |
| PIN4 | 1.42E-01 | 7.40E-01 | 1.45E+00 | 6.38E-02 | 2.76E-01 | 1.31E+00 | 1.04E-01 | 4.41E-01 | 1.20E+00 | 5.34E-06 | 2.04E-05 | -1.14E+00 | 1.51E-02 | 5.40E-02 | 1.01E+00 | 1.34E-04 | 3.77E-04 | -1.34E+00 | 1.38E-01 | 5.06E-01 | 1.45E+00 | 5.72E-02 | 1.57E-01 | 1.09E+00 |
| AL035071.1 | 1.42E-01 | 7.40E-01 | 1.39E+00 | 1.71E-01 | 7.37E-01 | 1.24E+00 | 5.09E-02 | 2.16E-01 | -1.27E+00 | 2.24E-01 | 8.54E-01 | 1.19E+00 | 3.65E-02 | 1.30E-01 | 1.54E+00 | 1.51E-01 | 4.26E-01 | 1.39E+00 | 2.44E-01 | 8.94E-01 | 1.17E+00 | 3.46E-01 | 9.50E-01 | 1.15E+00 |
| PSMC1P1 | 1.42E-01 | 7.41E-01 | 1.24E+00 | 2.00E-02 | 8.63E-02 | 1.09E+00 | 1.37E-01 | 5.80E-01 | 1.31E+00 | 7.75E-02 | 2.96E-01 | 1.12E+00 | 2.59E-01 | 9.24E-01 | 1.41E+00 | 1.49E-01 | 4.20E-01 | 1.25E+00 | 1.36E-01 | 5.00E-01 | 1.24E+00 | 3.95E-03 | 1.08E-02 | 1.01E+00 |
| CD180 | 1.42E-01 | 7.41E-01 | -1.19E+00 | 1.57E-01 | 6.77E-01 | -1.22E+00 | 3.25E-02 | 1.38E-01 | -1.44E+00 | 2.75E-02 | 1.05E-01 | -1.28E+00 | 1.51E-01 | 5.38E-01 | 1.43E+00 | 3.17E-01 | 8.93E-01 | 1.12E+00 | 2.00E-01 | 7.33E-01 | 1.11E+00 | 3.14E-01 | 8.63E-01 | 1.08E+00 |
| MYO10 | 1.42E-01 | 7.41E-01 | 1.01E+00 | 1.05E-01 | 4.52E-01 | -1.19E+00 | 1.76E-01 | 7.46E-01 | -1.05E+00 | 7.94E-02 | 3.03E-01 | -1.24E+00 | 3.07E-02 | 1.09E-01 | -1.67E+00 | 2.58E-01 | 7.28E-01 | 1.03E+00 | 2.13E-03 | 7.82E-03 | -1.23E+00 | 8.07E-03 | 2.22E-02 | -1.13E+00 |
| CLCN5 | 1.42E-01 | 7.42E-01 | -1.00E+00 | 1.87E-01 | 8.06E-01 | 1.18E+00 | 1.75E-01 | 7.42E-01 | 1.10E+00 | 2.28E-01 | 8.73E-01 | 1.41E+00 | 2.37E-01 | 8.46E-01 | 1.38E+00 | 1.95E-01 | 5.49E-01 | 1.71E+00 | 8.64E-03 | 3.17E-02 | -1.10E+00 | 3.26E-01 | 8.96E-01 | 1.34E+00 |
| RPL5P5 | 1.42E-01 | 7.42E-01 | 1.38E+00 | 1.64E-01 | 7.11E-01 | 1.61E+00 | 2.27E-02 | 9.64E-02 | 1.59E+00 | 8.03E-02 | 3.07E-01 | 1.72E+00 | 1.74E-01 | 6.20E-01 | 1.25E+00 | 2.34E-01 | 6.59E-01 | 1.30E+00 | 1.35E-01 | 4.94E-01 | 1.20E+00 | 2.71E-01 | 7.44E-01 | 1.44E+00 |
| RBM11 | 1.42E-01 | 7.42E-01 | 1.30E+00 | 2.08E-01 | 8.97E-01 | 1.48E+00 | 3.42E-03 | 1.45E-02 | -1.44E+00 | 7.85E-02 | 3.00E-01 | 1.03E+00 | 2.79E-01 | 9.94E-01 | 1.14E+00 | 3.25E-01 | 9.17E-01 | 1.35E+00 | 2.25E-01 | 8.26E-01 | 1.14E+00 | 1.38E-01 | 3.79E-01 | 1.57E+00 |
| EXOC3L2 | 1.43E-01 | 7.43E-01 | -1.14E+00 | 2.14E-01 | 9.25E-01 | 1.03E+00 | 1.22E-01 | 5.19E-01 | -1.25E+00 | 1.47E-01 | 5.60E-01 | -1.18E+00 | 2.74E-01 | 9.77E-01 | 1.20E+00 | 3.49E-01 | 9.85E-01 | 1.17E+00 | 5.70E-02 | 2.09E-01 | 2.08E+00 | 1.89E-01 | 5.18E-01 | 1.01E+00 |
| AC011815.1 | 1.43E-01 | 7.44E-01 | -1.00E+00 | 2.17E-01 | 9.39E-01 | 1.21E+00 | 1.49E-01 | 6.33E-01 | -1.21E+00 | 8.38E-02 | 3.20E-01 | -1.41E+00 | 1.66E-01 | 5.91E-01 | 1.81E+00 | 3.05E-02 | 8.61E-02 | 2.89E+00 | 2.23E-01 | 8.17E-01 | 1.17E+00 | 1.52E-01 | 4.18E-01 | -1.07E+00 |
| PLIN2 | 1.43E-01 | 7.45E-01 | 1.30E+00 | 6.92E-08 | 2.99E-07 | -1.24E+00 | 9.81E-02 | 4.17E-01 | 1.11E+00 | 1.80E-08 | 6.89E-08 | -1.35E+00 | 4.48E-04 | 1.60E-03 | -1.20E+00 | 2.23E-06 | 6.28E-06 | -1.52E+00 | 8.94E-06 | 3.28E-05 | -1.35E+00 | 1.20E-09 | 3.29E-09 | -1.81E+00 |
| VAV3 | 1.43E-01 | 7.45E-01 | 1.41E+00 | 1.58E-01 | 6.84E-01 | 1.51E+00 | 6.57E-02 | 2.79E-01 | 1.42E+00 | 2.21E-01 | 8.45E-01 | 1.44E+00 | 1.06E-02 | 3.78E-02 | 1.80E+00 | 1.74E-02 | 4.91E-02 | 1.76E+00 | 1.30E-01 | 4.77E-01 | 1.36E+00 | 2.00E-01 | 5.50E-01 | 1.39E+00 |
| CALD1 | 1.43E-01 | 7.45E-01 | -1.20E+00 | 1.58E-01 | 6.85E-01 | 1.12E+00 | 9.69E-02 | 4.12E-01 | -1.29E+00 | 2.19E-01 | 8.35E-01 | 1.43E+00 | 2.63E-01 | 9.37E-01 | -1.29E+00 | 2.84E-01 | 8.01E-01 | 1.04E+00 | 1.76E-01 | 6.44E-01 | 2.06E+00 | 6.85E-02 | 1.88E-01 | -1.72E+00 |
| AC007384.1 | 1.43E-01 | 7.45E-01 | 1.15E+00 | 5.17E-02 | 2.23E-01 | -1.13E+00 | 2.15E-01 | 9.14E-01 | 1.18E+00 | 2.54E-01 | 9.70E-01 | 1.24E+00 | 2.61E-01 | 9.29E-01 | 1.05E+00 | 3.48E-01 | 9.81E-01 | 1.20E+00 | 2.13E-01 | 7.83E-01 | -1.10E+00 | 3.06E-01 | 8.39E-01 | -1.05E+00 |
| gen-01 | 1.43E-01 | 7.45E-01 | 1.30E+00 | 1.64E-01 | 7.09E-01 | 1.31E+00 | 1.18E-03 | 5.03E-03 | -1.40E+00 | 5.11E-03 | 1.95E-02 | -1.08E+00 | 7.21E-07 | 2.57E-06 | -2.00E+00 | 1.55E-05 | 4.36E-05 | -1.59E+00 | 1.91E-12 | 7.02E-12 | -1.82E+00 | 1.96E-06 | 5.38E-06 | -1.31E+00 |
| AP000873.4 | 1.43E-01 | 7.45E-01 | 1.45E+00 | 1.08E-01 | 4.66E-01 | 1.13E+00 | 6.72E-03 | 2.85E-02 | -1.28E+00 | 5.35E-05 | 2.04E-04 | -1.97E+00 | 1.21E-01 | 4.31E-01 | 1.79E+00 | 1.31E-01 | 3.69E-01 | -1.17E+00 | 2.06E-01 | 7.57E-01 | 1.17E+00 | 8.29E-03 | 2.28E-02 | -1.50E+00 |
| AL589863.1 | 1.43E-01 | 7.45E-01 | -1.05E+00 | 8.23E-02 | 3.56E-01 | -1.19E+00 | 1.14E-04 | 4.83E-04 | -1.61E+00 | 8.33E-05 | 3.18E-04 | -1.50E+00 | 4.14E-02 | 1.48E-01 | 1.09E+00 | 3.70E-03 | 1.04E-02 | -1.39E+00 | 1.86E-01 | 6.81E-01 | -1.02E+00 | 3.00E-01 | 8.23E-01 | 1.03E+00 |
| PIF1 | 1.43E-01 | 7.45E-01 | 1.25E+00 | 1.87E-01 | 8.07E-01 | 1.23E+00 | 1.31E-01 | 5.55E-01 | 1.48E+00 | 5.21E-02 | 1.99E-01 | 2.56E+00 | 9.98E-02 | 3.56E-01 | -1.15E+00 | 1.26E-01 | 3.54E-01 | -1.19E+00 | 7.91E-02 | 2.90E-01 | -1.05E+00 | 1.31E-01 | 3.60E-01 | 1.51E+00 |
| DNM1P47 | 1.43E-01 | 7.45E-01 | 1.07E+00 | 3.06E-02 | 1.32E-01 | 1.65E+00 | 2.52E-03 | 1.07E-02 | -1.33E+00 | 1.81E-02 | 6.90E-02 | -1.11E+00 | 7.55E-02 | 2.69E-01 | 1.03E+00 | 1.92E-01 | 5.42E-01 | 1.69E+00 | 7.37E-02 | 2.71E-01 | -1.49E+00 | 3.29E-01 | 9.02E-01 | 1.05E+00 |
| STX18-AS1 | 1.43E-01 | 7.46E-01 | 1.36E+00 | 8.68E-02 | 3.75E-01 | 1.70E+00 | 3.62E-03 | 1.54E-02 | -1.40E+00 | 3.71E-02 | 1.42E-01 | -1.00E+00 | 2.05E-01 | 7.30E-01 | 1.08E+00 | 1.61E-01 | 4.55E-01 | 1.87E+00 | 2.52E-02 | 9.24E-02 | -1.20E+00 | 7.96E-02 | 2.19E-01 | -1.13E+00 |
| SLC2A4 | 1.43E-01 | 7.47E-01 | 1.37E+00 | 1.53E-01 | 6.59E-01 | 1.48E+00 | 9.15E-02 | 3.89E-01 | -1.21E+00 | 9.59E-02 | 3.66E-01 | 1.63E+00 | 2.55E-01 | 9.09E-01 | -1.04E+00 | 3.08E-01 | 8.69E-01 | 1.24E+00 | 1.87E-01 | 6.86E-01 | 1.12E+00 | 3.51E-01 | 9.65E-01 | 1.47E+00 |
| AC015849.6 | 1.43E-01 | 7.47E-01 | 1.26E+00 | 1.66E-04 | 7.16E-04 | -1.26E+00 | 1.78E-01 | 7.55E-01 | 1.19E+00 | 4.05E-04 | 1.55E-03 | -1.21E+00 | 2.27E-01 | 8.08E-01 | 1.47E+00 | 1.15E-01 | 3.24E-01 | -1.08E+00 | 1.79E-01 | 6.58E-01 | 1.20E+00 | 1.13E-02 | 3.09E-02 | -1.13E+00 |
| SREK1IP1 | 1.43E-01 | 7.48E-01 | 1.44E+00 | 4.19E-05 | 1.81E-04 | 1.01E+00 | 3.91E-02 | 1.66E-01 | 1.05E+00 | 1.77E-07 | 6.76E-07 | -1.29E+00 | 2.11E-01 | 7.52E-01 | 1.30E+00 | 3.68E-05 | 1.04E-04 | -1.27E+00 | 8.73E-02 | 3.20E-01 | 1.14E+00 | 1.51E-11 | 4.16E-11 | -1.54E+00 |
| TMEM86B | 1.43E-01 | 7.48E-01 | 1.02E+00 | 1.62E-01 | 7.01E-01 | 1.44E+00 | 1.18E-01 | 5.03E-01 | 1.05E+00 | 9.65E-02 | 3.68E-01 | 1.04E+00 | 9.82E-02 | 3.50E-01 | 1.07E+00 | 8.70E-02 | 2.45E-01 | -1.10E+00 | 2.66E-01 | 9.76E-01 | 1.09E+00 | 3.52E-01 | 9.66E-01 | 1.13E+00 |
| CDKN2D | 1.43E-01 | 7.48E-01 | 1.43E+00 | 2.34E-03 | 1.01E-02 | 1.13E+00 | 2.05E-01 | 8.69E-01 | 1.36E+00 | 1.79E-02 | 6.82E-02 | 1.16E+00 | 1.56E-01 | 5.56E-01 | 1.83E+00 | 2.53E-01 | 7.14E-01 | 1.34E+00 | 3.72E-04 | 1.36E-03 | 1.79E+00 | 1.92E-02 | 5.26E-02 | -1.07E+00 |
| ITGA9 | 1.43E-01 | 7.48E-01 | 1.33E+00 | 3.38E-03 | 1.46E-02 | 1.97E+00 | 1.60E-01 | 6.81E-01 | 1.20E+00 | 2.64E-02 | 1.01E-01 | 1.86E+00 | 7.07E-02 | 2.52E-01 | 1.67E+00 | 3.63E-02 | 1.02E-01 | 1.77E+00 | 7.49E-02 | 2.75E-01 | 1.58E+00 | 1.45E-02 | 3.97E-02 | 1.88E+00 |
| AC022893.1 | 1.43E-01 | 7.48E-01 | 1.30E+00 | 1.06E-01 | 4.56E-01 | -1.06E+00 | 2.09E-01 | 8.86E-01 | 1.22E+00 | 2.46E-01 | 9.40E-01 | 1.14E+00 | 2.00E-01 | 7.13E-01 | 1.21E+00 | 1.46E-01 | 4.12E-01 | -1.12E+00 | 1.70E-01 | 6.23E-01 | 1.36E+00 | 2.41E-01 | 6.62E-01 | 1.04E+00 |
| RBKS | 1.44E-01 | 7.48E-01 | 1.45E+00 | 5.57E-03 | 2.41E-02 | -1.60E+00 | 6.44E-02 | 2.74E-01 | -1.08E+00 | 1.76E-03 | 6.73E-03 | -1.60E+00 | 1.25E-01 | 4.46E-01 | 1.57E+00 | 3.55E-01 | 1.00E+00 | 1.21E+00 | 4.55E-03 | 1.67E-02 | -1.33E+00 | 2.38E-03 | 6.53E-03 | -1.44E+00 |
| AL022322.2 | 1.44E-01 | 7.48E-01 | 1.43E+00 | 4.14E-02 | 1.79E-01 | 1.04E+00 | 9.46E-02 | 4.02E-01 | 1.14E+00 | 9.93E-02 | 3.79E-01 | 1.19E+00 | 2.13E-01 | 7.58E-01 | 1.69E+00 | 2.76E-01 | 7.78E-01 | 1.74E+00 | 2.62E-02 | 9.62E-02 | 1.52E+00 | 2.23E-02 | 6.13E-02 | -1.15E+00 |
| CLPTM1L | 1.44E-01 | 7.48E-01 | 1.40E+00 | 1.92E-01 | 8.30E-01 | 1.61E+00 | 4.09E-02 | 1.74E-01 | 1.35E+00 | 1.38E-01 | 5.28E-01 | 1.45E+00 | 8.84E-02 | 3.15E-01 | 1.22E+00 | 1.50E-01 | 4.23E-01 | 1.28E+00 | 1.16E-01 | 4.26E-01 | 1.44E+00 | 2.10E-02 | 5.77E-02 | 1.66E+00 |
| GAL | 1.44E-01 | 7.49E-01 | 1.16E+00 | 2.11E-01 | 9.10E-01 | 1.08E+00 | 9.13E-02 | 3.88E-01 | -1.11E+00 | 3.59E-02 | 1.37E-01 | -1.20E+00 | 2.80E-01 | 1.00E+00 | 1.00E+00 | 3.24E-01 | 9.13E-01 | 1.12E+00 | 8.37E-28 | 3.07E-27 | -2.30E+00 | 2.78E-32 | 7.65E-32 | -3.09E+00 |
| KDM3A | 1.44E-01 | 7.49E-01 | 1.26E+00 | 1.55E-01 | 6.69E-01 | 1.44E+00 | 1.50E-01 | 6.39E-01 | 1.14E+00 | 4.40E-02 | 1.68E-01 | 1.12E+00 | 1.32E-01 | 4.70E-01 | 1.25E+00 | 1.17E-01 | 3.30E-01 | 1.20E+00 | 2.66E-02 | 9.75E-02 | 1.20E+00 | 1.23E-02 | 3.38E-02 | 1.09E+00 |
| NPM1P12 | 1.44E-01 | 7.49E-01 | 1.14E+00 | 1.33E-01 | 5.75E-01 | 1.18E+00 | 1.85E-01 | 7.86E-01 | 1.16E+00 | 9.05E-02 | 3.46E-01 | 1.71E+00 | 5.32E-02 | 1.90E-01 | 2.08E+00 | 7.27E-02 | 2.05E-01 | 2.12E+00 | 2.45E-01 | 8.98E-01 | 1.18E+00 | 2.81E-03 | 7.72E-03 | 2.05E+00 |
| PDK4 | 1.44E-01 | 7.49E-01 | 1.08E+00 | 6.93E-02 | 3.00E-01 | 1.56E+00 | 1.63E-01 | 6.91E-01 | 1.37E+00 | 1.71E-01 | 6.51E-01 | 1.51E+00 | 1.78E-01 | 6.36E-01 | 1.52E+00 | 1.70E-01 | 4.81E-01 | 1.74E+00 | 5.75E-02 | 2.11E-01 | 1.96E+00 | 2.72E-01 | 7.46E-01 | 1.23E+00 |
| HIST1H1E | 1.44E-01 | 7.50E-01 | 1.24E+00 | 1.20E-01 | 5.16E-01 | 1.48E+00 | 9.04E-02 | 3.84E-01 | 1.39E+00 | 1.44E-02 | 5.51E-02 | 1.64E+00 | 1.69E-01 | 6.03E-01 | 1.59E+00 | 1.45E-03 | 4.09E-03 | 1.91E+00 | 1.96E-01 | 7.19E-01 | 1.41E+00 | 2.49E-01 | 6.83E-01 | 1.40E+00 |
| DYSF | 1.44E-01 | 7.50E-01 | 1.15E+00 | 2.31E-01 | 1.00E+00 | 1.00E+00 | 7.98E-02 | 3.39E-01 | 1.28E+00 | 1.80E-01 | 6.88E-01 | -1.09E+00 | 2.38E-01 | 8.49E-01 | 1.08E+00 | 3.55E-01 | 1.00E+00 | 1.00E+00 | 5.51E-02 | 2.02E-01 | -1.26E+00 | 1.83E-01 | 5.02E-01 | -1.13E+00 |
| PPP1R14BP3 | 1.44E-01 | 7.50E-01 | 1.27E+00 | 1.85E-01 | 7.98E-01 | 1.57E+00 | 1.26E-01 | 5.37E-01 | 1.40E+00 | 1.46E-03 | 5.57E-03 | 2.21E+00 | 6.28E-02 | 2.24E-01 | 1.04E+00 | 3.29E-01 | 9.26E-01 | 1.54E+00 | 2.80E-03 | 1.03E-02 | -1.06E+00 | 5.17E-02 | 1.42E-01 | 1.75E+00 |
| FAM178B | 1.44E-01 | 7.50E-01 | -1.36E+00 | 1.83E-01 | 7.89E-01 | -1.24E+00 | 1.50E-01 | 6.36E-01 | -1.17E+00 | 1.61E-01 | 6.17E-01 | -1.15E+00 | 1.06E-01 | 3.77E-01 | -1.01E+00 | 4.54E-02 | 1.28E-01 | 2.48E+00 | 3.69E-14 | 1.35E-13 | -1.44E+00 | 1.49E-02 | 4.11E-02 | 1.21E+00 |
| GPR65 | 1.44E-01 | 7.51E-01 | -1.13E+00 | 1.55E-01 | 6.71E-01 | -1.19E+00 | 1.85E-01 | 7.87E-01 | -1.11E+00 | 1.65E-02 | 6.29E-02 | -1.86E+00 | 5.14E-04 | 1.83E-03 | -2.64E+00 | 1.95E-04 | 5.51E-04 | -3.13E+00 | 1.04E-01 | 3.82E-01 | -1.09E+00 | 1.69E-02 | 4.64E-02 | -1.40E+00 |
| AC132942.1 | 1.44E-01 | 7.51E-01 | 1.12E+00 | 4.77E-02 | 2.06E-01 | -1.03E+00 | 2.06E-01 | 8.74E-01 | 1.17E+00 | 9.42E-02 | 3.60E-01 | 1.11E+00 | 1.95E-01 | 6.95E-01 | 1.13E+00 | 2.91E-01 | 8.19E-01 | 1.29E+00 | 4.62E-02 | 1.70E-01 | 1.60E+00 | 2.79E-01 | 7.65E-01 | 1.44E+00 |
| AC232271.1 | 1.44E-01 | 7.52E-01 | 1.37E+00 | 1.99E-02 | 8.59E-02 | 2.17E+00 | 8.51E-02 | 3.62E-01 | -1.10E+00 | 1.95E-01 | 7.46E-01 | 1.46E+00 | 1.37E-01 | 4.89E-01 | 1.73E+00 | 5.14E-02 | 1.45E-01 | 2.19E+00 | 1.07E-01 | 3.91E-01 | 1.08E+00 | 3.59E-01 | 9.86E-01 | 1.34E+00 |
| TIAM1 | 1.44E-01 | 7.53E-01 | 1.34E+00 | 1.93E-01 | 8.34E-01 | 1.46E+00 | 3.99E-02 | 1.70E-01 | -1.47E+00 | 1.01E-01 | 3.85E-01 | -1.19E+00 | 1.92E-01 | 6.85E-01 | 1.09E+00 | 2.28E-01 | 6.42E-01 | 1.06E+00 | 1.66E-42 | 6.08E-42 | -3.69E+00 | 4.56E-33 | 1.25E-32 | -2.60E+00 |
| LTBP3 | 1.45E-01 | 7.53E-01 | 1.37E+00 | 1.56E-01 | 6.75E-01 | 1.62E+00 | 1.81E-02 | 7.70E-02 | 1.05E+00 | 5.33E-02 | 2.04E-01 | 1.28E+00 | 1.36E-02 | 4.84E-02 | 1.19E+00 | 1.59E-01 | 4.48E-01 | 1.45E+00 | 1.41E-01 | 5.17E-01 | 1.41E+00 | 3.05E-03 | 8.39E-03 | 2.06E+00 |
| NTAN1P2 | 1.45E-01 | 7.53E-01 | 1.22E+00 | 1.63E-01 | 7.06E-01 | 1.63E+00 | 6.29E-02 | 2.67E-01 | -1.12E+00 | 7.83E-02 | 2.99E-01 | -1.02E+00 | 2.74E-01 | 9.77E-01 | 1.28E+00 | 1.61E-01 | 4.55E-01 | 1.82E+00 | 8.91E-03 | 3.27E-02 | -1.09E+00 | 6.22E-02 | 1.71E-01 | 1.02E+00 |
| AL591895.1 | 1.45E-01 | 7.54E-01 | 1.35E+00 | 1.87E-01 | 8.10E-01 | 1.58E+00 | 7.06E-02 | 3.00E-01 | -1.03E+00 | 2.05E-01 | 7.82E-01 | 1.24E+00 | 4.01E-02 | 1.43E-01 | -1.13E+00 | 6.37E-02 | 1.79E-01 | -1.10E+00 | 2.63E-02 | 9.67E-02 | -1.21E+00 | 1.03E-01 | 2.82E-01 | -1.05E+00 |
| ARL5B | 1.45E-01 | 7.54E-01 | 1.49E+00 | 3.33E-08 | 1.44E-07 | -1.20E+00 | 1.30E-01 | 5.50E-01 | 1.20E+00 | 2.30E-07 | 8.80E-07 | -1.35E+00 | 4.73E-03 | 1.69E-02 | 1.12E+00 | 2.06E-11 | 5.81E-11 | -1.79E+00 | 1.16E-04 | 4.27E-04 | -1.06E+00 | 8.74E-10 | 2.40E-09 | -1.49E+00 |
| LINC01560 | 1.45E-01 | 7.54E-01 | 1.10E+00 | 1.45E-01 | 6.27E-01 | 1.44E+00 | 3.18E-02 | 1.35E-01 | -1.29E+00 | 2.30E-01 | 8.79E-01 | 1.28E+00 | 8.42E-04 | 3.00E-03 | -1.85E+00 | 3.47E-04 | 9.79E-04 | -2.21E+00 | 2.38E-02 | 8.72E-02 | -1.19E+00 | 2.06E-01 | 5.65E-01 | 1.14E+00 |
| SRP9P1 | 1.45E-01 | 7.55E-01 | 1.08E+00 | 8.05E-02 | 3.48E-01 | -1.07E+00 | 1.07E-01 | 4.53E-01 | -1.07E+00 | 1.87E-01 | 7.15E-01 | 1.16E+00 | 1.66E-03 | 5.94E-03 | -1.75E+00 | 4.45E-03 | 1.25E-02 | -1.53E+00 | 2.57E-01 | 9.44E-01 | 1.20E+00 | 5.56E-02 | 1.53E-01 | -1.23E+00 |
| MOBP | 1.45E-01 | 7.55E-01 | 1.24E+00 | 7.58E-03 | 3.27E-02 | -1.40E+00 | 1.61E-01 | 6.83E-01 | 1.15E+00 | 2.53E-01 | 9.68E-01 | 1.22E+00 | 2.36E-01 | 8.43E-01 | 1.45E+00 | 3.46E-01 | 9.77E-01 | 1.17E+00 | 9.52E-02 | 3.49E-01 | 1.46E+00 | 3.51E-01 | 9.65E-01 | 1.06E+00 |
| C2orf48 | 1.45E-01 | 7.55E-01 | -1.25E+00 | 2.11E-01 | 9.10E-01 | -1.05E+00 | 3.04E-03 | 1.29E-02 | 1.93E+00 | 1.60E-02 | 6.12E-02 | 2.03E+00 | 2.54E-01 | 9.07E-01 | 1.17E+00 | 3.13E-01 | 8.82E-01 | 1.16E+00 | 1.62E-01 | 5.94E-01 | 1.44E+00 | 1.66E-01 | 4.56E-01 | 1.48E+00 |
| ARRDC3-AS1 | 1.45E-01 | 7.55E-01 | 1.08E+00 | 1.14E-01 | 4.91E-01 | 1.03E+00 | 1.43E-01 | 6.09E-01 | 1.04E+00 | 8.98E-02 | 3.43E-01 | -1.06E+00 | 4.35E-03 | 1.55E-02 | -1.49E+00 | 4.07E-02 | 1.15E-01 | -1.08E+00 | 6.65E-02 | 2.44E-01 | -1.11E+00 | 2.03E-01 | 5.56E-01 | 1.04E+00 |
| KCNH3 | 1.45E-01 | 7.55E-01 | 1.07E+00 | 1.78E-01 | 7.69E-01 | 1.41E+00 | 4.75E-03 | 2.02E-02 | -1.34E+00 | 1.10E-01 | 4.21E-01 | 1.09E+00 | 1.88E-01 | 6.72E-01 | 1.02E+00 | 3.29E-01 | 9.26E-01 | 1.18E+00 | 2.50E-01 | 9.19E-01 | 1.29E+00 | 2.31E-01 | 6.35E-01 | 1.50E+00 |
| U3 | 1.45E-01 | 7.56E-01 | 1.45E+00 | 9.30E-03 | 4.02E-02 | -1.04E+00 | 3.00E-02 | 1.27E-01 | -1.06E+00 | 1.23E-03 | 4.69E-03 | -1.19E+00 | 1.31E-09 | 4.67E-09 | -2.24E+00 | 8.55E-07 | 2.41E-06 | -1.57E+00 | 1.59E-13 | 5.84E-13 | -2.24E+00 | 1.11E-09 | 3.06E-09 | -1.78E+00 |
| AL049840.4 | 1.45E-01 | 7.56E-01 | 1.19E+00 | 1.70E-01 | 7.34E-01 | 1.32E+00 | 1.89E-01 | 8.02E-01 | 1.24E+00 | 1.19E-01 | 4.53E-01 | 1.58E+00 | 1.87E-01 | 6.66E-01 | 1.30E+00 | 2.45E-01 | 6.91E-01 | 1.15E+00 | 2.95E-02 | 1.08E-01 | -1.17E+00 | 1.46E-01 | 4.02E-01 | 1.43E+00 |
| ABCA3 | 1.45E-01 | 7.56E-01 | -1.01E+00 | 2.83E-02 | 1.22E-01 | 2.02E+00 | 2.22E-01 | 9.42E-01 | 1.23E+00 | 1.72E-01 | 6.56E-01 | 1.61E+00 | 1.04E-01 | 3.69E-01 | -1.26E+00 | 2.60E-01 | 7.32E-01 | 1.46E+00 | 5.51E-02 | 2.02E-01 | -1.22E+00 | 2.75E-01 | 7.55E-01 | 1.16E+00 |
| AC027307.3 | 1.45E-01 | 7.56E-01 | 1.44E+00 | 1.95E-01 | 8.41E-01 | 1.33E+00 | 1.14E-02 | 4.85E-02 | -1.18E+00 | 1.74E-02 | 6.66E-02 | 1.02E+00 | 1.86E-01 | 6.62E-01 | 1.08E+00 | 1.46E-01 | 4.12E-01 | -1.10E+00 | 1.57E-01 | 5.78E-01 | -1.06E+00 | 1.42E-01 | 3.90E-01 | -1.18E+00 |
| LINC01943 | 1.45E-01 | 7.56E-01 | 1.56E+00 | 8.38E-02 | 3.62E-01 | -1.11E+00 | 1.02E-02 | 4.33E-02 | -1.27E+00 | 7.12E-03 | 2.72E-02 | -1.28E+00 | 1.30E-01 | 4.63E-01 | 1.75E+00 | 3.42E-01 | 9.63E-01 | 1.60E+00 | 2.29E-01 | 8.39E-01 | 1.32E+00 | 6.99E-04 | 1.92E-03 | -1.07E+00 |
| TBCC | 1.45E-01 | 7.56E-01 | 1.51E+00 | 2.66E-02 | 1.15E-01 | 1.29E+00 | 9.08E-02 | 3.86E-01 | 1.18E+00 | 1.66E-04 | 6.35E-04 | -1.03E+00 | 1.48E-03 | 5.28E-03 | 1.07E+00 | 1.15E-04 | 3.25E-04 | -1.05E+00 | 2.95E-04 | 1.08E-03 | -1.13E+00 | 7.41E-03 | 2.04E-02 | 1.02E+00 |
| HIF1A-AS2 | 1.45E-01 | 7.56E-01 | 1.30E+00 | 7.80E-02 | 3.37E-01 | 1.71E+00 | 3.57E-02 | 1.52E-01 | -1.26E+00 | 2.30E-01 | 8.79E-01 | 1.47E+00 | 2.74E-01 | 9.78E-01 | 1.38E+00 | 3.07E-01 | 8.64E-01 | 1.29E+00 | 1.62E-01 | 5.96E-01 | 1.40E+00 | 2.65E-01 | 7.29E-01 | 1.28E+00 |
| HMGN2P3 | 1.45E-01 | 7.57E-01 | 1.33E+00 | 1.47E-01 | 6.33E-01 | 1.12E+00 | 2.38E-03 | 1.01E-02 | 1.93E+00 | 8.89E-02 | 3.40E-01 | -1.06E+00 | 2.55E-01 | 9.09E-01 | 1.32E+00 | 1.36E-01 | 3.83E-01 | -1.31E+00 | 1.22E-01 | 4.48E-01 | 1.34E+00 | 1.13E-02 | 3.11E-02 | -1.20E+00 |
| GATS | 1.45E-01 | 7.58E-01 | 1.15E+00 | 2.27E-01 | 9.79E-01 | 1.36E+00 | 6.46E-02 | 2.74E-01 | -1.10E+00 | 8.49E-02 | 3.24E-01 | 1.04E+00 | 3.60E-03 | 1.28E-02 | -1.63E+00 | 2.07E-01 | 5.83E-01 | 1.12E+00 | 2.71E-01 | 9.95E-01 | 1.23E+00 | 2.72E-01 | 7.46E-01 | 1.30E+00 |
| ABCB1 | 1.45E-01 | 7.58E-01 | 1.30E+00 | 6.27E-02 | 2.71E-01 | 1.80E+00 | 2.18E-01 | 9.26E-01 | 1.33E+00 | 2.09E-01 | 7.97E-01 | 1.48E+00 | 4.66E-02 | 1.66E-01 | 2.32E+00 | 2.92E-01 | 8.25E-01 | 1.57E+00 | 1.76E-03 | 6.46E-03 | 3.20E+00 | 5.89E-02 | 1.62E-01 | 2.33E+00 |
| RECQL4 | 1.45E-01 | 7.58E-01 | 1.31E+00 | 2.14E-01 | 9.26E-01 | 1.04E+00 | 1.86E-01 | 7.90E-01 | -1.00E+00 | 6.92E-02 | 2.64E-01 | -1.11E+00 | 1.62E-01 | 5.78E-01 | -1.30E+00 | 3.26E-01 | 9.19E-01 | 1.11E+00 | 1.42E-01 | 5.22E-01 | -1.01E+00 | 2.92E-02 | 8.02E-02 | 1.40E+00 |
| SERP2 | 1.45E-01 | 7.58E-01 | -1.31E+00 | 1.86E-01 | 8.02E-01 | -1.21E+00 | 1.57E-01 | 6.68E-01 | -1.21E+00 | 1.23E-01 | 4.70E-01 | -1.41E+00 | 1.15E-02 | 4.11E-02 | -1.04E+00 | 8.41E-03 | 2.37E-02 | -1.09E+00 | 8.96E-04 | 3.29E-03 | -1.33E+00 | 9.38E-02 | 2.58E-01 | -1.02E+00 |
| KIAA1671 | 1.45E-01 | 7.58E-01 | 1.03E+00 | 2.05E-01 | 8.85E-01 | 1.03E+00 | 3.83E-02 | 1.63E-01 | -1.28E+00 | 3.80E-02 | 1.45E-01 | -1.17E+00 | 1.24E-01 | 4.41E-01 | -1.49E+00 | 2.86E-01 | 8.05E-01 | -1.19E+00 | 6.86E-05 | 2.52E-04 | -1.57E+00 | 1.14E-02 | 3.12E-02 | -1.19E+00 |
| AC022182.3 | 1.45E-01 | 7.58E-01 | 1.23E+00 | 2.12E-01 | 9.16E-01 | 1.46E+00 | 3.55E-02 | 1.51E-01 | 1.64E+00 | 6.74E-02 | 2.57E-01 | 1.82E+00 | 5.23E-02 | 1.86E-01 | -1.14E+00 | 2.05E-01 | 5.78E-01 | 1.27E+00 | 1.23E-01 | 4.53E-01 | 1.50E+00 | 1.47E-03 | 4.03E-03 | 2.02E+00 |
| F2RL3 | 1.45E-01 | 7.58E-01 | -1.12E+00 | 1.84E-01 | 7.97E-01 | -1.04E+00 | 1.88E-01 | 8.01E-01 | 1.22E+00 | 2.46E-01 | 9.39E-01 | 1.06E+00 | 4.92E-02 | 1.76E-01 | -1.15E+00 | 1.67E-01 | 4.70E-01 | -1.02E+00 | 7.19E-03 | 2.64E-02 | 1.19E+00 | 1.56E-03 | 4.28E-03 | -1.15E+00 |
| EDF1 | 1.45E-01 | 7.58E-01 | 1.09E+00 | 7.50E-02 | 3.24E-01 | 1.11E+00 | 1.60E-02 | 6.81E-02 | 1.18E+00 | 1.01E-01 | 3.85E-01 | 1.17E+00 | 3.46E-03 | 1.23E-02 | -1.05E+00 | 7.26E-02 | 2.05E-01 | -1.01E+00 | 1.67E-01 | 6.14E-01 | 1.06E+00 | 3.63E-01 | 9.97E-01 | 1.09E+00 |
| AC012640.2 | 1.46E-01 | 7.59E-01 | 1.27E+00 | 4.08E-02 | 1.76E-01 | 1.04E+00 | 1.59E-01 | 6.75E-01 | 1.05E+00 | 1.21E-01 | 4.63E-01 | 1.16E+00 | 2.59E-01 | 9.25E-01 | 1.55E+00 | 2.03E-01 | 5.73E-01 | 1.69E+00 | 1.61E-01 | 5.90E-01 | 1.24E+00 | 3.59E-01 | 9.85E-01 | 1.36E+00 |
| EIF1AD | 1.46E-01 | 7.59E-01 | 1.22E+00 | 1.20E-01 | 5.18E-01 | 1.28E+00 | 8.89E-02 | 3.78E-01 | 1.06E+00 | 4.97E-02 | 1.90E-01 | 1.11E+00 | 1.67E-02 | 5.96E-02 | -1.09E+00 | 1.33E-02 | 3.74E-02 | -1.03E+00 | 5.08E-02 | 1.86E-01 | 1.07E+00 | 9.21E-02 | 2.53E-01 | 1.07E+00 |
| FUK | 1.46E-01 | 7.59E-01 | 1.05E+00 | 1.51E-01 | 6.53E-01 | 1.53E+00 | 5.60E-03 | 2.38E-02 | -1.27E+00 | 2.88E-03 | 1.10E-02 | -1.21E+00 | 1.72E-01 | 6.15E-01 | 1.62E+00 | 1.48E-01 | 4.18E-01 | 1.87E+00 | 1.09E-01 | 4.02E-01 | -1.19E+00 | 1.76E-01 | 4.83E-01 | -1.07E+00 |
| AC092279.2 | 1.46E-01 | 7.59E-01 | 1.33E+00 | 2.70E-02 | 1.17E-01 | 2.01E+00 | 1.69E-01 | 7.19E-01 | 1.38E+00 | 2.23E-01 | 8.52E-01 | 1.32E+00 | 2.60E-01 | 9.28E-01 | 1.26E+00 | 3.04E-01 | 8.57E-01 | 1.47E+00 | 4.83E-02 | 1.77E-01 | -1.07E+00 | 5.39E-02 | 1.48E-01 | -1.07E+00 |
| TSPOAP1 | 1.46E-01 | 7.60E-01 | 1.14E+00 | 2.46E-02 | 1.06E-01 | -2.16E+00 | 2.30E-05 | 9.76E-05 | -1.06E+00 | 3.94E-18 | 1.50E-17 | -1.46E+00 | 1.75E-01 | 6.25E-01 | 1.78E+00 | 3.13E-01 | 8.82E-01 | -1.26E+00 | 1.54E-01 | 5.66E-01 | -1.19E+00 | 8.71E-02 | 2.39E-01 | -1.74E+00 |
| AXIN1 | 1.46E-01 | 7.60E-01 | 1.49E+00 | 2.01E-01 | 8.68E-01 | 1.64E+00 | 1.45E-01 | 6.17E-01 | 1.10E+00 | 1.13E-01 | 4.30E-01 | 1.50E+00 | 1.38E-02 | 4.93E-02 | 1.00E+00 | 4.54E-02 | 1.28E-01 | 1.07E+00 | 2.25E-01 | 8.26E-01 | 1.41E+00 | 3.30E-01 | 9.07E-01 | 1.39E+00 |
| AC006077.2 | 1.46E-01 | 7.60E-01 | 1.28E+00 | 1.46E-01 | 6.30E-01 | 1.68E+00 | 8.54E-02 | 3.63E-01 | 1.40E+00 | 1.33E-01 | 5.07E-01 | 1.59E+00 | 2.50E-01 | 8.93E-01 | 1.42E+00 | 2.93E-01 | 8.27E-01 | 1.51E+00 | 1.22E-01 | 4.49E-01 | 1.50E+00 | 3.53E-02 | 9.70E-02 | 1.67E+00 |
| IBA57 | 1.46E-01 | 7.60E-01 | 1.32E+00 | 4.45E-04 | 1.92E-03 | -1.00E+00 | 2.21E-01 | 9.40E-01 | 1.17E+00 | 2.48E-02 | 9.48E-02 | 1.12E+00 | 8.14E-02 | 2.90E-01 | 1.81E+00 | 1.82E-01 | 5.13E-01 | 1.19E+00 | 1.86E-02 | 6.84E-02 | -1.20E+00 | 1.25E-03 | 3.43E-03 | -1.48E+00 |
| AC009299.3 | 1.46E-01 | 7.61E-01 | 1.49E+00 | 2.71E-02 | 1.17E-01 | 2.00E+00 | 1.52E-01 | 6.47E-01 | 1.15E+00 | 1.00E-01 | 3.82E-01 | 1.78E+00 | 1.75E-01 | 6.23E-01 | 1.74E+00 | 1.96E-01 | 5.52E-01 | 1.89E+00 | 2.41E-01 | 8.83E-01 | 1.12E+00 | 2.62E-01 | 7.19E-01 | 1.44E+00 |
| LAMTOR5-AS1 | 1.46E-01 | 7.61E-01 | 1.22E+00 | 6.10E-02 | 2.64E-01 | 1.14E+00 | 9.80E-03 | 4.16E-02 | -1.19E+00 | 6.92E-03 | 2.64E-02 | -1.14E+00 | 2.33E-01 | 8.30E-01 | 1.50E+00 | 3.21E-01 | 9.04E-01 | 1.44E+00 | 2.30E-01 | 8.44E-01 | 1.16E+00 | 3.22E-01 | 8.84E-01 | 1.31E+00 |
| LINC01122 | 1.46E-01 | 7.61E-01 | 1.35E+00 | 4.37E-02 | 1.89E-01 | 1.52E+00 | 1.03E-01 | 4.37E-01 | 1.04E+00 | 3.62E-02 | 1.38E-01 | 1.08E+00 | 1.99E-01 | 7.11E-01 | 1.28E+00 | 2.12E-01 | 5.98E-01 | 1.34E+00 | 1.72E-02 | 6.30E-02 | -1.42E+00 | 1.28E-01 | 3.50E-01 | -1.04E+00 |
| ZNF432 | 1.46E-01 | 7.61E-01 | 1.29E+00 | 2.72E-02 | 1.17E-01 | 1.02E+00 | 3.41E-02 | 1.45E-01 | 1.09E+00 | 6.37E-05 | 2.43E-04 | -1.27E+00 | 1.05E-01 | 3.74E-01 | 1.87E+00 | 2.07E-01 | 5.84E-01 | 1.23E+00 | 8.47E-02 | 3.11E-01 | 1.48E+00 | 2.14E-01 | 5.87E-01 | 1.08E+00 |
| SLC22A23 | 1.46E-01 | 7.61E-01 | -1.14E+00 | 1.36E-01 | 5.86E-01 | 1.41E+00 | 8.84E-02 | 3.76E-01 | -1.19E+00 | 1.25E-01 | 4.78E-01 | 1.56E+00 | 2.67E-01 | 9.51E-01 | -1.08E+00 | 2.06E-01 | 5.81E-01 | 1.42E+00 | 6.10E-02 | 2.24E-01 | -1.05E+00 | 1.02E-01 | 2.80E-01 | -1.13E+00 |
| AC004884.1 | 1.46E-01 | 7.61E-01 | 1.27E+00 | 1.36E-04 | 5.88E-04 | -1.18E+00 | 9.44E-02 | 4.01E-01 | 1.36E+00 | 3.38E-03 | 1.29E-02 | -1.09E+00 | 2.70E-01 | 9.64E-01 | 1.35E+00 | 2.74E-02 | 7.73E-02 | -1.25E+00 | 1.63E-01 | 5.97E-01 | 1.21E+00 | 1.16E-02 | 3.18E-02 | -1.14E+00 |
| PABPC4L | 1.46E-01 | 7.61E-01 | 1.26E+00 | 2.13E-01 | 9.21E-01 | 1.15E+00 | 1.50E-01 | 6.35E-01 | 1.03E+00 | 1.81E-01 | 6.93E-01 | 1.12E+00 | 1.93E-01 | 6.87E-01 | 1.85E+00 | 2.11E-01 | 5.95E-01 | 1.58E+00 | 9.43E-02 | 3.46E-01 | 1.00E+00 | 9.78E-02 | 2.69E-01 | 1.02E+00 |
| SPC25 | 1.46E-01 | 7.62E-01 | -1.24E+00 | 1.88E-01 | 8.13E-01 | -1.23E+00 | 3.04E-03 | 1.29E-02 | 1.86E+00 | 1.75E-01 | 6.69E-01 | 1.44E+00 | 5.11E-05 | 1.82E-04 | -2.21E+00 | 1.19E-03 | 3.36E-03 | -1.71E+00 | 8.84E-05 | 3.25E-04 | -1.42E+00 | 7.06E-02 | 1.94E-01 | 1.02E+00 |
| XRCC6P2 | 1.46E-01 | 7.62E-01 | 1.17E+00 | 1.51E-01 | 6.51E-01 | 1.67E+00 | 5.91E-02 | 2.51E-01 | 1.58E+00 | 1.24E-03 | 4.74E-03 | 2.34E+00 | 1.73E-01 | 6.18E-01 | 1.24E+00 | 2.20E-01 | 6.20E-01 | 1.72E+00 | 1.85E-02 | 6.80E-02 | 1.75E+00 | 4.18E-03 | 1.15E-02 | 1.90E+00 |
| FABP5P3 | 1.46E-01 | 7.62E-01 | 1.57E+00 | 6.68E-02 | 2.89E-01 | 2.23E+00 | 2.57E-02 | 1.09E-01 | 1.59E+00 | 2.90E-02 | 1.11E-01 | 1.91E+00 | 7.39E-02 | 2.64E-01 | -1.21E+00 | 3.26E-01 | 9.20E-01 | 1.53E+00 | 4.78E-02 | 1.76E-01 | 1.59E+00 | 3.32E-07 | 9.13E-07 | 2.64E+00 |
| AC243960.1 | 1.46E-01 | 7.62E-01 | 1.17E+00 | 1.07E-01 | 4.62E-01 | 1.64E+00 | 1.37E-01 | 5.81E-01 | 1.04E+00 | 1.13E-01 | 4.31E-01 | 1.11E+00 | 2.22E-01 | 7.91E-01 | 1.68E+00 | 3.41E-01 | 9.61E-01 | 1.48E+00 | 1.95E-01 | 7.17E-01 | 1.67E+00 | 3.25E-01 | 8.92E-01 | 1.42E+00 |
| USPL1 | 1.46E-01 | 7.62E-01 | 1.30E+00 | 4.52E-04 | 1.95E-03 | -1.30E+00 | 9.43E-02 | 4.01E-01 | 1.09E+00 | 2.98E-02 | 1.14E-01 | 1.08E+00 | 5.69E-02 | 2.03E-01 | 1.02E+00 | 1.02E-03 | 2.88E-03 | -1.44E+00 | 1.58E-03 | 5.79E-03 | -1.20E+00 | 2.82E-06 | 7.74E-06 | -1.59E+00 |
| NSDHL | 1.46E-01 | 7.62E-01 | 1.29E+00 | 1.00E-01 | 4.34E-01 | 1.80E+00 | 3.71E-03 | 1.58E-02 | -1.14E+00 | 2.44E-01 | 9.32E-01 | 1.46E+00 | 8.13E-02 | 2.90E-01 | -1.04E+00 | 2.72E-01 | 7.67E-01 | 1.40E+00 | 3.10E-02 | 1.14E-01 | 1.07E+00 | 1.69E-01 | 4.65E-01 | 1.46E+00 |
| CTIF | 1.46E-01 | 7.63E-01 | 1.08E+00 | 1.52E-01 | 6.57E-01 | 1.42E+00 | 8.92E-02 | 3.79E-01 | -1.16E+00 | 2.33E-01 | 8.89E-01 | 1.17E+00 | 1.03E-01 | 3.66E-01 | 1.02E+00 | 3.41E-01 | 9.62E-01 | 1.25E+00 | 1.16E-01 | 4.26E-01 | -1.13E+00 | 3.00E-01 | 8.24E-01 | 1.16E+00 |
| AP001350.1 | 1.46E-01 | 7.63E-01 | 1.23E+00 | 1.28E-01 | 5.55E-01 | 1.26E+00 | 1.56E-02 | 6.64E-02 | -1.10E+00 | 1.45E-04 | 5.54E-04 | -1.38E+00 | 2.17E-01 | 7.72E-01 | 1.56E+00 | 2.59E-01 | 7.31E-01 | 1.61E+00 | 1.83E-03 | 6.73E-03 | -1.47E+00 | 3.20E-03 | 8.79E-03 | -1.37E+00 |
| ESS2 | 1.46E-01 | 7.63E-01 | 1.31E+00 | 1.59E-01 | 6.88E-01 | 1.46E+00 | 1.17E-01 | 4.96E-01 | 1.06E+00 | 4.13E-02 | 1.58E-01 | 1.08E+00 | 6.02E-03 | 2.15E-02 | -1.02E+00 | 4.11E-04 | 1.16E-03 | -1.39E+00 | 4.09E-06 | 1.50E-05 | -1.43E+00 | 1.51E-05 | 4.15E-05 | -1.38E+00 |
| MTND1P23 | 1.47E-01 | 7.64E-01 | 1.02E+00 | 1.56E-02 | 6.72E-02 | 1.07E+00 | 6.65E-02 | 2.83E-01 | -1.03E+00 | 2.41E-01 | 9.22E-01 | 1.02E+00 | 2.09E-01 | 7.46E-01 | 1.04E+00 | 1.60E-03 | 4.52E-03 | 1.15E+00 | 2.92E-03 | 1.07E-02 | -1.02E+00 | 2.36E-03 | 6.48E-03 | 1.12E+00 |
| GNAZ | 1.47E-01 | 7.64E-01 | 1.14E+00 | 2.12E-01 | 9.14E-01 | 1.08E+00 | 1.70E-01 | 7.22E-01 | -1.08E+00 | 1.78E-01 | 6.79E-01 | -1.09E+00 | 2.80E-01 | 1.00E+00 | 1.00E+00 | 2.35E-01 | 6.63E-01 | 1.36E+00 | 3.69E-02 | 1.35E-01 | 2.28E+00 | 1.27E-01 | 3.48E-01 | 1.78E+00 |
| URB1-AS1 | 1.47E-01 | 7.64E-01 | 1.30E+00 | 1.42E-02 | 6.13E-02 | 1.10E+00 | 1.90E-01 | 8.09E-01 | 1.34E+00 | 7.08E-02 | 2.71E-01 | 1.23E+00 | 3.31E-02 | 1.18E-01 | 1.05E+00 | 4.82E-02 | 1.36E-01 | 1.06E+00 | 2.25E-01 | 8.27E-01 | 1.32E+00 | 3.45E-03 | 9.47E-03 | -1.33E+00 |
| AC068657.1 | 1.47E-01 | 7.64E-01 | 1.33E+00 | 1.32E-01 | 5.71E-01 | 1.62E+00 | 1.19E-01 | 5.05E-01 | 1.36E+00 | 3.54E-02 | 1.35E-01 | 1.82E+00 | 2.36E-01 | 8.43E-01 | 1.31E+00 | 1.24E-01 | 3.51E-01 | 1.90E+00 | 2.08E-01 | 7.62E-01 | 1.11E+00 | 1.39E-01 | 3.81E-01 | 1.52E+00 |
| PCDHB11 | 1.47E-01 | 7.64E-01 | 1.24E+00 | 1.00E-01 | 4.34E-01 | 1.76E+00 | 1.47E-01 | 6.24E-01 | 1.33E+00 | 1.69E-01 | 6.46E-01 | 1.34E+00 | 7.17E-02 | 2.56E-01 | -1.30E+00 | 1.44E-01 | 4.06E-01 | -1.17E+00 | 9.44E-02 | 3.47E-01 | 1.54E+00 | 6.13E-02 | 1.68E-01 | 1.78E+00 |
| RPF1 | 1.47E-01 | 7.64E-01 | 1.35E+00 | 1.21E-01 | 5.22E-01 | 1.46E+00 | 1.17E-01 | 4.96E-01 | 1.35E+00 | 2.11E-01 | 8.05E-01 | 1.35E+00 | 4.24E-02 | 1.51E-01 | 1.20E+00 | 1.89E-01 | 5.32E-01 | 1.37E+00 | 3.03E-02 | 1.11E-01 | 1.08E+00 | 2.28E-01 | 6.27E-01 | 1.18E+00 |
| RNF208 | 1.47E-01 | 7.64E-01 | 1.24E+00 | 1.71E-01 | 7.38E-01 | 1.16E+00 | 1.01E-02 | 4.28E-02 | -1.30E+00 | 1.98E-02 | 7.55E-02 | -1.09E+00 | 1.09E-01 | 3.88E-01 | -1.23E+00 | 2.32E-01 | 6.54E-01 | 1.10E+00 | 2.63E-01 | 9.66E-01 | 1.33E+00 | 8.93E-02 | 2.45E-01 | 1.81E+00 |
| CPAMD8 | 1.47E-01 | 7.65E-01 | 1.37E+00 | 1.06E-01 | 4.58E-01 | 1.58E+00 | 4.05E-02 | 1.72E-01 | 1.78E+00 | 1.16E-01 | 4.44E-01 | 1.68E+00 | 5.22E-02 | 1.86E-01 | -1.79E+00 | 1.44E-01 | 4.06E-01 | -1.22E+00 | 1.38E-01 | 5.05E-01 | 1.48E+00 | 2.49E-01 | 6.85E-01 | -1.08E+00 |
| YBEY | 1.47E-01 | 7.65E-01 | 1.28E+00 | 1.29E-02 | 5.59E-02 | 1.08E+00 | 1.41E-01 | 5.99E-01 | 1.21E+00 | 2.48E-02 | 9.49E-02 | 1.10E+00 | 4.08E-02 | 1.46E-01 | 1.14E+00 | 3.15E-02 | 8.88E-02 | 1.10E+00 | 9.13E-02 | 3.35E-01 | 1.14E+00 | 2.17E-02 | 5.96E-02 | 1.01E+00 |
| AL731571.1 | 1.47E-01 | 7.65E-01 | 1.47E+00 | 3.06E-02 | 1.32E-01 | 2.14E+00 | 2.29E-01 | 9.72E-01 | 1.21E+00 | 1.36E-01 | 5.19E-01 | 1.72E+00 | 1.47E-01 | 5.23E-01 | 1.04E+00 | 2.72E-01 | 7.67E-01 | 1.58E+00 | 2.20E-01 | 8.07E-01 | 1.01E+00 | 1.85E-02 | 5.07E-02 | 1.69E+00 |
| AL603766.1 | 1.47E-01 | 7.65E-01 | 1.30E+00 | 2.00E-01 | 8.62E-01 | 1.57E+00 | 1.40E-01 | 5.93E-01 | 1.32E+00 | 1.48E-01 | 5.66E-01 | 1.32E+00 | 1.61E-01 | 5.75E-01 | 1.35E+00 | 3.49E-01 | 9.83E-01 | 1.40E+00 | 1.80E-01 | 6.62E-01 | 1.23E+00 | 4.94E-02 | 1.36E-01 | 1.65E+00 |
| OR1N2 | 1.47E-01 | 7.65E-01 | 1.47E+00 | 1.29E-01 | 5.58E-01 | 1.26E+00 | 1.56E-01 | 6.61E-01 | 1.34E+00 | 1.18E-01 | 4.51E-01 | 1.54E+00 | 5.69E-02 | 2.03E-01 | 2.14E+00 | 3.88E-02 | 1.09E-01 | 2.48E+00 | 1.22E-01 | 4.50E-01 | 1.51E+00 | 5.03E-02 | 1.38E-01 | 2.08E+00 |
| SHCBP1 | 1.47E-01 | 7.65E-01 | -1.06E+00 | 1.43E-01 | 6.18E-01 | -1.32E+00 | 1.82E-01 | 7.75E-01 | -1.01E+00 | 4.81E-02 | 1.84E-01 | -1.15E+00 | 1.70E-01 | 6.05E-01 | -1.18E+00 | 2.00E-01 | 5.65E-01 | -1.37E+00 | 1.30E-03 | 4.75E-03 | -1.39E+00 | 1.71E-01 | 4.70E-01 | -1.02E+00 |
| TIMM8BP2 | 1.47E-01 | 7.65E-01 | 1.18E+00 | 1.25E-01 | 5.42E-01 | 1.21E+00 | 6.73E-02 | 2.86E-01 | 1.51E+00 | 1.71E-01 | 6.54E-01 | 1.59E+00 | 1.09E-02 | 3.90E-02 | -1.23E+00 | 2.67E-01 | 7.54E-01 | 1.38E+00 | 9.77E-02 | 3.59E-01 | 1.42E+00 | 4.65E-02 | 1.28E-01 | 1.61E+00 |
| NCAPH | 1.47E-01 | 7.65E-01 | -1.14E+00 | 1.64E-01 | 7.11E-01 | -1.16E+00 | 1.29E-01 | 5.47E-01 | 1.29E+00 | 1.50E-01 | 5.74E-01 | 1.06E+00 | 5.36E-02 | 1.91E-01 | -2.33E+00 | 1.39E-01 | 3.91E-01 | -1.43E+00 | 2.40E-01 | 8.80E-01 | -1.00E+00 | 9.80E-03 | 2.69E-02 | 1.50E+00 |
| AL731537.1 | 1.47E-01 | 7.65E-01 | 1.37E+00 | 2.23E-01 | 9.62E-01 | 1.39E+00 | 8.24E-02 | 3.50E-01 | 1.64E+00 | 2.29E-01 | 8.73E-01 | 1.31E+00 | 1.12E-02 | 4.00E-02 | 3.32E+00 | 2.12E-01 | 5.97E-01 | 1.73E+00 | 6.99E-04 | 2.57E-03 | 2.69E+00 | 3.11E-01 | 8.53E-01 | 1.37E+00 |
| FAM103A2P | 1.47E-01 | 7.66E-01 | 1.50E+00 | 1.90E-02 | 8.20E-02 | 1.04E+00 | 7.54E-02 | 3.21E-01 | 1.36E+00 | 8.09E-02 | 3.09E-01 | 1.08E+00 | 2.24E-01 | 8.00E-01 | 1.63E+00 | 1.23E-01 | 3.48E-01 | 1.24E+00 | 9.21E-03 | 3.38E-02 | 1.69E+00 | 1.87E-01 | 5.13E-01 | 1.16E+00 |
| WDR62 | 1.47E-01 | 7.66E-01 | 1.22E+00 | 7.62E-02 | 3.29E-01 | -1.15E+00 | 1.06E-02 | 4.52E-02 | 1.68E+00 | 1.72E-01 | 6.58E-01 | 1.33E+00 | 1.74E-01 | 6.22E-01 | -1.13E+00 | 2.66E-01 | 7.51E-01 | -1.06E+00 | 2.82E-02 | 1.03E-01 | -1.01E+00 | 2.14E-01 | 5.88E-01 | 1.34E+00 |
| SEPT7P6 | 1.47E-01 | 7.67E-01 | 1.33E+00 | 6.75E-02 | 2.92E-01 | 1.80E+00 | 1.98E-02 | 8.42E-02 | 1.71E+00 | 6.85E-03 | 2.62E-02 | 2.20E+00 | 2.70E-01 | 9.63E-01 | 1.33E+00 | 3.37E-01 | 9.51E-01 | 1.42E+00 | 8.57E-03 | 3.14E-02 | 1.90E+00 | 2.49E-01 | 6.85E-01 | 1.40E+00 |
| CLEC4G | 1.47E-01 | 7.68E-01 | -1.06E+00 | 1.83E-01 | 7.92E-01 | 1.07E+00 | 1.09E-01 | 4.64E-01 | -1.15E+00 | 1.88E-02 | 7.19E-02 | 1.72E+00 | 1.99E-01 | 7.11E-01 | -1.21E+00 | 3.10E-01 | 8.75E-01 | -1.03E+00 | 2.33E-01 | 8.57E-01 | -1.09E+00 | 3.14E-01 | 8.63E-01 | -1.14E+00 |
| GRPEL2 | 1.47E-01 | 7.69E-01 | 1.07E+00 | 2.04E-01 | 8.82E-01 | 1.37E+00 | 1.98E-01 | 8.40E-01 | 1.14E+00 | 2.18E-01 | 8.33E-01 | 1.24E+00 | 7.28E-02 | 2.60E-01 | -1.26E+00 | 9.11E-02 | 2.57E-01 | -1.12E+00 | 3.72E-03 | 1.36E-02 | -1.12E+00 | 1.19E-01 | 3.27E-01 | 1.10E+00 |
| AC009961.3 | 1.47E-01 | 7.69E-01 | 1.18E+00 | 7.37E-02 | 3.19E-01 | 1.05E+00 | 6.44E-02 | 2.73E-01 | 1.46E+00 | 6.12E-02 | 2.34E-01 | 1.75E+00 | 2.39E-01 | 8.53E-01 | 1.50E+00 | 2.93E-01 | 8.27E-01 | 1.44E+00 | 1.86E-01 | 6.84E-01 | 1.01E+00 | 2.65E-01 | 7.28E-01 | -1.02E+00 |
| LINC01971 | 1.48E-01 | 7.69E-01 | 1.29E+00 | 1.56E-01 | 6.75E-01 | 1.07E+00 | 7.91E-02 | 3.36E-01 | -1.15E+00 | 8.49E-02 | 3.24E-01 | 1.03E+00 | 2.52E-01 | 8.98E-01 | -1.02E+00 | 2.72E-01 | 7.67E-01 | -1.15E+00 | 2.33E-01 | 8.56E-01 | 1.04E+00 | 3.46E-01 | 9.50E-01 | -1.01E+00 |
| PEG3 | 1.48E-01 | 7.70E-01 | 1.38E+00 | 1.17E-01 | 5.03E-01 | 1.02E+00 | 5.64E-03 | 2.40E-02 | -1.63E+00 | 4.91E-02 | 1.87E-01 | -1.14E+00 | 1.28E-01 | 4.56E-01 | -1.30E+00 | 1.86E-01 | 5.24E-01 | -1.13E+00 | 2.17E-01 | 7.98E-01 | 1.10E+00 | 3.13E-01 | 8.59E-01 | 1.14E+00 |
| HCST | 1.48E-01 | 7.70E-01 | 1.24E+00 | 1.80E-15 | 7.80E-15 | -1.63E+00 | 1.26E-01 | 5.33E-01 | 1.08E+00 | 7.89E-14 | 3.01E-13 | -1.75E+00 | 2.79E-01 | 9.95E-01 | 1.50E+00 | 8.63E-08 | 2.43E-07 | -1.82E+00 | 2.04E-02 | 7.50E-02 | 1.25E+00 | 1.06E-02 | 2.92E-02 | -1.62E+00 |
| ACPP | 1.48E-01 | 7.70E-01 | 1.12E+00 | 2.31E-01 | 1.00E+00 | 1.00E+00 | 3.33E-02 | 1.42E-01 | -1.17E+00 | 7.32E-03 | 2.79E-02 | -1.35E+00 | 2.37E-01 | 8.47E-01 | 1.09E+00 | 1.77E-01 | 4.98E-01 | -1.34E+00 | 1.94E-01 | 7.12E-01 | -1.12E+00 | 2.98E-01 | 8.17E-01 | -1.06E+00 |
| ARIH2OS | 1.48E-01 | 7.71E-01 | 1.55E+00 | 5.28E-04 | 2.28E-03 | -1.02E+00 | 1.73E-01 | 7.35E-01 | 1.22E+00 | 2.97E-03 | 1.13E-02 | -1.14E+00 | 1.02E-01 | 3.64E-01 | 1.11E+00 | 1.47E-01 | 4.15E-01 | 1.18E+00 | 1.30E-01 | 4.77E-01 | 1.23E+00 | 2.06E-02 | 5.65E-02 | -1.09E+00 |
| RPS7P14 | 1.48E-01 | 7.71E-01 | 1.37E+00 | 1.79E-01 | 7.74E-01 | 1.62E+00 | 1.96E-01 | 8.32E-01 | 1.30E+00 | 5.01E-02 | 1.91E-01 | 1.68E+00 | 2.21E-01 | 7.88E-01 | 1.33E+00 | 1.47E-01 | 4.14E-01 | 1.33E+00 | 1.73E-01 | 6.34E-01 | 1.13E+00 | 2.47E-01 | 6.77E-01 | 1.32E+00 |
| ERBB2 | 1.48E-01 | 7.71E-01 | 1.07E+00 | 1.30E-01 | 5.62E-01 | 1.06E+00 | 7.53E-02 | 3.20E-01 | 1.74E+00 | 8.16E-02 | 3.12E-01 | 1.80E+00 | 2.30E-01 | 8.19E-01 | 1.08E+00 | 1.16E-01 | 3.28E-01 | -1.26E+00 | 6.44E-02 | 2.36E-01 | -1.04E+00 | 1.40E-01 | 3.83E-01 | 1.01E+00 |
| APOLD1 | 1.48E-01 | 7.71E-01 | 1.44E+00 | 1.85E-01 | 8.00E-01 | 1.70E+00 | 1.36E-02 | 5.79E-02 | -1.34E+00 | 2.44E-01 | 9.33E-01 | 1.39E+00 | 2.01E-01 | 7.15E-01 | 1.56E+00 | 3.04E-01 | 8.57E-01 | 1.41E+00 | 2.61E-02 | 9.57E-02 | -1.56E+00 | 2.79E-01 | 7.68E-01 | 1.09E+00 |
| ERVK13-1 | 1.48E-01 | 7.72E-01 | 1.18E+00 | 2.63E-02 | 1.14E-01 | 2.03E+00 | 2.53E-03 | 1.08E-02 | -1.23E+00 | 2.48E-02 | 9.47E-02 | 1.08E+00 | 2.33E-01 | 8.32E-01 | 1.61E+00 | 2.34E-01 | 6.59E-01 | 1.56E+00 | 1.12E-01 | 4.10E-01 | 1.08E+00 | 2.47E-01 | 6.78E-01 | 1.39E+00 |
| PDE1B | 1.48E-01 | 7.72E-01 | 1.14E+00 | 2.55E-04 | 1.10E-03 | -1.99E+00 | 7.03E-02 | 2.99E-01 | 1.51E+00 | 3.00E-03 | 1.15E-02 | -1.37E+00 | 8.21E-04 | 2.93E-03 | -1.45E+00 | 3.70E-06 | 1.04E-05 | -2.73E+00 | 1.54E-01 | 5.64E-01 | -1.06E+00 | 1.53E-02 | 4.20E-02 | -1.78E+00 |
| SKIV2L | 1.48E-01 | 7.72E-01 | 1.36E+00 | 3.23E-02 | 1.40E-01 | 1.17E+00 | 1.55E-01 | 6.60E-01 | 1.09E+00 | 1.92E-01 | 7.33E-01 | 1.23E+00 | 1.36E-01 | 4.85E-01 | 1.23E+00 | 4.46E-02 | 1.26E-01 | 1.06E+00 | 1.06E-02 | 3.89E-02 | -1.12E+00 | 2.94E-02 | 8.06E-02 | -1.09E+00 |
| AC008498.2 | 1.48E-01 | 7.72E-01 | 1.18E+00 | 2.00E-01 | 8.66E-01 | 1.19E+00 | 7.80E-02 | 3.32E-01 | 1.51E+00 | 5.15E-02 | 1.97E-01 | 1.90E+00 | 2.35E-01 | 8.37E-01 | -1.03E+00 | 3.44E-01 | 9.68E-01 | 1.15E+00 | 4.81E-02 | 1.77E-01 | -1.16E+00 | 2.37E-01 | 6.51E-01 | 1.06E+00 |
| HIST1H2BH | 1.48E-01 | 7.73E-01 | 1.36E+00 | 2.03E-01 | 8.75E-01 | 1.32E+00 | 1.04E-01 | 4.43E-01 | 1.16E+00 | 9.73E-03 | 3.72E-02 | -1.15E+00 | 1.77E-01 | 6.29E-01 | 1.58E+00 | 3.10E-01 | 8.74E-01 | 1.36E+00 | 7.53E-02 | 2.76E-01 | 1.03E+00 | 1.51E-02 | 4.16E-02 | -1.11E+00 |
| PSMB1 | 1.48E-01 | 7.73E-01 | 1.10E+00 | 2.01E-02 | 8.68E-02 | 1.07E+00 | 1.98E-01 | 8.42E-01 | 1.09E+00 | 1.02E-01 | 3.91E-01 | 1.02E+00 | 2.35E-02 | 8.39E-02 | 1.01E+00 | 1.85E-01 | 5.20E-01 | 1.07E+00 | 8.85E-02 | 3.25E-01 | 1.15E+00 | 1.21E-01 | 3.33E-01 | 1.14E+00 |
| CLEC16A | 1.48E-01 | 7.74E-01 | 1.30E+00 | 7.17E-02 | 3.10E-01 | 1.28E+00 | 4.17E-02 | 1.77E-01 | -1.12E+00 | 1.29E-01 | 4.94E-01 | 1.17E+00 | 4.75E-03 | 1.69E-02 | -1.18E+00 | 3.16E-02 | 8.91E-02 | -1.01E+00 | 4.43E-02 | 1.63E-01 | -1.07E+00 | 2.00E-01 | 5.49E-01 | 1.07E+00 |
| THBS1 | 1.48E-01 | 7.74E-01 | 1.09E+00 | 1.16E-01 | 5.01E-01 | -1.07E+00 | 2.07E-01 | 8.81E-01 | 1.42E+00 | 1.91E-01 | 7.28E-01 | 1.03E+00 | 1.09E-01 | 3.90E-01 | 1.45E+00 | 1.13E-01 | 3.20E-01 | -1.40E+00 | 2.29E-01 | 8.39E-01 | 1.46E+00 | 8.14E-03 | 2.24E-02 | -1.15E+00 |
| AL359697.1 | 1.49E-01 | 7.75E-01 | 1.27E+00 | 2.67E-02 | 1.15E-01 | -1.16E+00 | 4.29E-04 | 1.82E-03 | 1.82E+00 | 2.83E-02 | 1.08E-01 | -1.48E+00 | 1.54E-01 | 5.48E-01 | 1.44E+00 | 3.34E-01 | 9.43E-01 | 1.15E+00 | 2.08E-01 | 7.64E-01 | 1.17E+00 | 3.60E-01 | 9.88E-01 | 1.18E+00 |
| HIST1H3F | 1.49E-01 | 7.75E-01 | -1.09E+00 | 1.91E-01 | 8.27E-01 | 1.00E+00 | 4.21E-04 | 1.79E-03 | 2.03E+00 | 2.01E-01 | 7.68E-01 | 1.19E+00 | 2.06E-01 | 7.35E-01 | -1.03E+00 | 3.15E-01 | 8.89E-01 | 1.08E+00 | 1.33E-03 | 4.88E-03 | -1.42E+00 | 7.39E-04 | 2.03E-03 | -1.44E+00 |
| PLK1 | 1.49E-01 | 7.76E-01 | 1.30E+00 | 1.02E-01 | 4.42E-01 | 1.94E+00 | 9.98E-02 | 4.24E-01 | 1.64E+00 | 1.76E-01 | 6.73E-01 | 1.87E+00 | 2.55E-01 | 9.08E-01 | 1.09E+00 | 3.55E-01 | 1.00E+00 | 1.36E+00 | 5.67E-02 | 2.08E-01 | -1.43E+00 | 8.02E-02 | 2.20E-01 | -1.03E+00 |
| AP000873.3 | 1.49E-01 | 7.76E-01 | 1.56E+00 | 2.37E-02 | 1.02E-01 | -1.31E+00 | 1.19E-01 | 5.07E-01 | 1.51E+00 | 5.07E-03 | 1.93E-02 | -1.29E+00 | 2.69E-01 | 9.58E-01 | 1.17E+00 | 1.04E-01 | 2.94E-01 | -1.65E+00 | 5.15E-02 | 1.89E-01 | 1.59E+00 | 3.26E-01 | 8.95E-01 | 1.22E+00 |
| AC103591.3 | 1.49E-01 | 7.76E-01 | 1.54E+00 | 2.72E-07 | 1.18E-06 | -1.76E+00 | 7.50E-02 | 3.19E-01 | 1.46E+00 | 5.76E-03 | 2.20E-02 | -1.25E+00 | 2.33E-01 | 8.32E-01 | 1.47E+00 | 1.50E-01 | 4.23E-01 | 1.03E+00 | 3.29E-02 | 1.21E-01 | -1.18E+00 | 6.02E-03 | 1.65E-02 | -1.41E+00 |
| NGRN | 1.49E-01 | 7.76E-01 | 1.09E+00 | 1.53E-01 | 6.63E-01 | 1.16E+00 | 6.05E-02 | 2.57E-01 | -1.10E+00 | 1.89E-01 | 7.21E-01 | 1.18E+00 | 2.21E-03 | 7.88E-03 | -1.83E+00 | 4.60E-02 | 1.30E-01 | -1.16E+00 | 1.15E-03 | 4.22E-03 | -1.54E+00 | 3.95E-03 | 1.09E-02 | -1.40E+00 |
| MYL4 | 1.49E-01 | 7.76E-01 | 1.12E+00 | 1.47E-01 | 6.37E-01 | -1.01E+00 | 9.54E-02 | 4.06E-01 | 1.91E+00 | 1.56E-01 | 5.97E-01 | 2.18E+00 | 5.23E-08 | 1.86E-07 | 2.28E+00 | 5.75E-13 | 1.62E-12 | 2.11E+00 | 1.17E-01 | 4.29E-01 | 1.44E+00 | 3.27E-02 | 8.99E-02 | 1.11E+00 |
| WASIR1 | 1.49E-01 | 7.77E-01 | 1.32E+00 | 2.16E-01 | 9.32E-01 | 1.57E+00 | 6.13E-02 | 2.61E-01 | -1.02E+00 | 2.53E-01 | 9.68E-01 | 1.36E+00 | 7.19E-04 | 2.56E-03 | -1.26E+00 | 4.45E-02 | 1.25E-01 | 1.10E+00 | 1.94E-01 | 7.11E-01 | 1.12E+00 | 2.79E-01 | 7.68E-01 | 1.27E+00 |
| ZDHHC20 | 1.49E-01 | 7.78E-01 | 1.24E+00 | 1.08E-01 | 4.68E-01 | 1.37E+00 | 1.65E-01 | 7.02E-01 | 1.20E+00 | 1.63E-01 | 6.21E-01 | 1.14E+00 | 2.15E-03 | 7.67E-03 | -1.17E+00 | 4.68E-04 | 1.32E-03 | -1.24E+00 | 1.44E-01 | 5.30E-01 | 1.14E+00 | 4.34E-02 | 1.19E-01 | 1.01E+00 |
| AC099850.1 | 1.49E-01 | 7.78E-01 | 1.00E+00 | 2.23E-01 | 9.65E-01 | 1.21E+00 | 1.54E-01 | 6.53E-01 | -1.03E+00 | 1.30E-01 | 4.97E-01 | 1.55E+00 | 1.18E-01 | 4.20E-01 | 1.04E+00 | 2.51E-01 | 7.08E-01 | 1.18E+00 | 2.19E-01 | 8.03E-01 | 1.11E+00 | 2.26E-02 | 6.21E-02 | 1.75E+00 |
| HES6 | 1.49E-01 | 7.78E-01 | 1.51E+00 | 2.19E-01 | 9.48E-01 | 1.26E+00 | 2.17E-01 | 9.22E-01 | 1.25E+00 | 1.93E-01 | 7.38E-01 | 1.46E+00 | 6.52E-03 | 2.32E-02 | 2.52E+00 | 2.15E-08 | 6.05E-08 | 4.60E+00 | 5.04E-03 | 1.85E-02 | 1.27E+00 | 2.14E-05 | 5.89E-05 | 1.39E+00 |
| RNF187 | 1.49E-01 | 7.78E-01 | 1.35E+00 | 3.35E-02 | 1.45E-01 | 1.32E+00 | 1.38E-04 | 5.84E-04 | 1.72E+00 | 2.99E-02 | 1.14E-01 | 1.61E+00 | 1.87E-01 | 6.65E-01 | 1.52E+00 | 1.90E-01 | 5.35E-01 | 1.61E+00 | 6.51E-04 | 2.39E-03 | 1.75E+00 | 2.87E-01 | 7.88E-01 | 1.39E+00 |
| NR1I2 | 1.49E-01 | 7.78E-01 | -1.10E+00 | 1.93E-01 | 8.33E-01 | 1.03E+00 | 1.48E-02 | 6.29E-02 | -1.28E+00 | 6.23E-02 | 2.38E-01 | 1.02E+00 | 1.20E-02 | 4.30E-02 | -1.39E+00 | 5.55E-03 | 1.56E-02 | -1.77E+00 | 2.47E-01 | 9.06E-01 | -1.03E+00 | 3.45E-01 | 9.49E-01 | 1.02E+00 |
| TLR1 | 1.49E-01 | 7.78E-01 | 1.41E+00 | 1.04E-01 | 4.47E-01 | 1.21E+00 | 2.68E-03 | 1.14E-02 | -1.21E+00 | 3.59E-05 | 1.37E-04 | -1.48E+00 | 2.64E-01 | 9.41E-01 | 1.36E+00 | 2.30E-01 | 6.48E-01 | 1.50E+00 | 2.35E-01 | 8.64E-01 | 1.17E+00 | 3.56E-01 | 9.78E-01 | 1.18E+00 |
| AC025048.6 | 1.49E-01 | 7.78E-01 | 1.14E+00 | 1.49E-01 | 6.44E-01 | 1.13E+00 | 3.05E-03 | 1.30E-02 | -1.27E+00 | 1.30E-04 | 4.98E-04 | -1.56E+00 | 2.22E-02 | 7.92E-02 | -1.23E+00 | 4.77E-04 | 1.35E-03 | -2.12E+00 | 1.29E-02 | 4.73E-02 | -1.52E+00 | 5.98E-02 | 1.64E-01 | -1.24E+00 |
| ANKRD55 | 1.50E-01 | 7.79E-01 | 1.20E+00 | 2.31E-01 | 1.00E+00 | 1.00E+00 | 2.19E-01 | 9.32E-01 | 1.05E+00 | 2.44E-01 | 9.34E-01 | 1.05E+00 | 2.70E-01 | 9.62E-01 | 1.02E+00 | 2.60E-01 | 7.32E-01 | -1.23E+00 | 1.07E-02 | 3.94E-02 | 2.61E+00 | 2.85E-01 | 7.83E-01 | 1.77E+00 |
| ZNF304 | 1.50E-01 | 7.80E-01 | 1.29E+00 | 1.32E-01 | 5.72E-01 | 1.05E+00 | 2.51E-02 | 1.07E-01 | -1.20E+00 | 9.77E-03 | 3.73E-02 | -1.20E+00 | 3.14E-02 | 1.12E-01 | -1.14E+00 | 6.69E-02 | 1.89E-01 | -1.05E+00 | 6.44E-02 | 2.37E-01 | -1.06E+00 | 1.15E-01 | 3.17E-01 | -1.04E+00 |
| RPL7AP6 | 1.50E-01 | 7.80E-01 | -1.01E+00 | 2.22E-18 | 9.57E-18 | -1.23E+00 | 3.39E-02 | 1.44E-01 | 1.05E+00 | 7.88E-09 | 3.01E-08 | -1.19E+00 | 1.15E-01 | 4.09E-01 | 1.04E+00 | 7.93E-05 | 2.23E-04 | -1.14E+00 | 2.36E-01 | 8.65E-01 | 1.08E+00 | 5.23E-06 | 1.44E-05 | -1.14E+00 |
| CYB561A3 | 1.50E-01 | 7.80E-01 | 1.32E+00 | 7.51E-02 | 3.25E-01 | 1.28E+00 | 8.98E-04 | 3.82E-03 | -1.30E+00 | 8.30E-04 | 3.17E-03 | -1.20E+00 | 2.80E-01 | 1.00E+00 | 1.43E+00 | 2.12E-01 | 5.99E-01 | 1.79E+00 | 1.06E-01 | 3.88E-01 | 1.17E+00 | 3.38E-01 | 9.28E-01 | 1.32E+00 |
| HADHAP1 | 1.50E-01 | 7.80E-01 | 1.29E+00 | 8.01E-03 | 3.46E-02 | 2.40E+00 | 2.10E-01 | 8.94E-01 | 1.14E+00 | 1.27E-02 | 4.87E-02 | 2.07E+00 | 1.58E-01 | 5.62E-01 | 1.08E+00 | 2.86E-01 | 8.06E-01 | 1.54E+00 | 2.25E-02 | 8.25E-02 | 1.68E+00 | 9.23E-03 | 2.54E-02 | 2.14E+00 |
| AC026436.1 | 1.50E-01 | 7.80E-01 | 1.27E+00 | 2.02E-05 | 8.75E-05 | -1.22E+00 | 1.60E-01 | 6.80E-01 | 1.21E+00 | 9.99E-04 | 3.82E-03 | 1.03E+00 | 1.89E-01 | 6.74E-01 | 1.32E+00 | 5.55E-03 | 1.56E-02 | -1.22E+00 | 1.66E-01 | 6.11E-01 | 1.29E+00 | 8.91E-03 | 2.45E-02 | -1.09E+00 |
| CD36 | 1.50E-01 | 7.80E-01 | 1.60E+00 | 1.76E-01 | 7.59E-01 | 1.70E+00 | 1.70E-01 | 7.22E-01 | 1.22E+00 | 2.14E-01 | 8.18E-01 | 1.19E+00 | 2.08E-04 | 7.41E-04 | 6.30E+00 | 1.72E-20 | 4.84E-20 | 1.71E+01 | 1.50E-92 | 5.50E-92 | 4.01E+00 | 5.51E-45 | 1.51E-44 | 2.95E+00 |
| EXO1 | 1.50E-01 | 7.80E-01 | -1.14E+00 | 1.98E-01 | 8.55E-01 | -1.03E+00 | 1.18E-01 | 5.00E-01 | 1.31E+00 | 2.27E-01 | 8.67E-01 | 1.20E+00 | 1.79E-01 | 6.38E-01 | -1.37E+00 | 2.86E-01 | 8.05E-01 | -1.05E+00 | 1.43E-04 | 5.24E-04 | -1.36E+00 | 9.46E-02 | 2.60E-01 | 1.03E+00 |
| GYPB | 1.50E-01 | 7.81E-01 | -1.15E+00 | 1.70E-01 | 7.36E-01 | -1.39E+00 | 2.06E-01 | 8.75E-01 | -1.14E+00 | 2.34E-01 | 8.94E-01 | -1.00E+00 | 2.29E-01 | 8.17E-01 | 1.52E+00 | 5.62E-07 | 1.58E-06 | 4.59E+00 | 8.75E-03 | 3.21E-02 | -1.13E+00 | 1.19E-04 | 3.27E-04 | -1.45E+00 |
| AC103703.1 | 1.50E-01 | 7.81E-01 | 1.39E+00 | 4.12E-02 | 1.78E-01 | 1.92E+00 | 2.29E-01 | 9.75E-01 | 1.27E+00 | 1.42E-01 | 5.43E-01 | 1.22E+00 | 2.32E-01 | 8.28E-01 | 1.34E+00 | 2.97E-01 | 8.38E-01 | 1.40E+00 | 6.80E-02 | 2.49E-01 | -1.10E+00 | 1.32E-01 | 3.62E-01 | -1.03E+00 |
| ZC3H10 | 1.50E-01 | 7.82E-01 | 1.35E+00 | 1.06E-01 | 4.57E-01 | 1.23E+00 | 2.61E-04 | 1.11E-03 | -1.22E+00 | 1.68E-03 | 6.42E-03 | -1.10E+00 | 1.81E-01 | 6.44E-01 | 1.18E+00 | 2.39E-01 | 6.72E-01 | 1.21E+00 | 4.54E-02 | 1.67E-01 | -1.09E+00 | 3.12E-02 | 8.56E-02 | -1.09E+00 |
| ASAP2 | 1.50E-01 | 7.82E-01 | 1.24E+00 | 2.17E-01 | 9.37E-01 | 1.38E+00 | 8.10E-02 | 3.44E-01 | 1.54E+00 | 1.58E-01 | 6.04E-01 | 1.17E+00 | 2.12E-01 | 7.57E-01 | 1.36E+00 | 3.55E-01 | 1.00E+00 | 1.17E+00 | 2.09E-01 | 7.69E-01 | 1.74E+00 | 2.04E-01 | 5.60E-01 | 1.37E+00 |
| OGFRL1 | 1.50E-01 | 7.83E-01 | 1.25E+00 | 1.74E-02 | 7.51E-02 | 1.19E+00 | 8.85E-02 | 3.76E-01 | 1.04E+00 | 2.87E-04 | 1.10E-03 | -1.12E+00 | 5.27E-02 | 1.88E-01 | 1.09E+00 | 1.50E-03 | 4.24E-03 | -1.15E+00 | 1.61E-02 | 5.92E-02 | -1.23E+00 | 1.28E-02 | 3.51E-02 | -1.18E+00 |
| MEPCE | 1.50E-01 | 7.83E-01 | 1.23E+00 | 7.80E-04 | 3.37E-03 | -1.08E+00 | 6.55E-02 | 2.78E-01 | 1.09E+00 | 6.83E-04 | 2.61E-03 | -1.04E+00 | 4.11E-04 | 1.47E-03 | -1.16E+00 | 6.63E-06 | 1.87E-05 | -1.45E+00 | 5.17E-04 | 1.90E-03 | -1.12E+00 | 5.25E-03 | 1.44E-02 | 1.01E+00 |
| ZBTB5 | 1.50E-01 | 7.83E-01 | 1.29E+00 | 1.59E-03 | 6.89E-03 | -1.18E+00 | 2.10E-03 | 8.91E-03 | -1.11E+00 | 1.17E-04 | 4.47E-04 | -1.16E+00 | 4.29E-02 | 1.53E-01 | -1.06E+00 | 4.26E-02 | 1.20E-01 | -1.12E+00 | 9.64E-05 | 3.54E-04 | -1.38E+00 | 2.13E-05 | 5.85E-05 | -1.55E+00 |
| TMSB4Y | 1.50E-01 | 7.84E-01 | 1.39E+00 | 4.37E-02 | 1.89E-01 | 1.29E+00 | 2.13E-01 | 9.07E-01 | 1.20E+00 | 3.76E-02 | 1.44E-01 | 1.07E+00 | 9.75E-02 | 3.48E-01 | 1.16E+00 | 2.76E-01 | 7.79E-01 | 1.30E+00 | 5.32E-02 | 1.95E-01 | 1.02E+00 | 1.49E-03 | 4.10E-03 | -1.12E+00 |
| PDE1C | 1.50E-01 | 7.84E-01 | 1.04E+00 | 1.75E-01 | 7.54E-01 | 1.13E+00 | 1.49E-02 | 6.31E-02 | 1.93E+00 | 8.71E-02 | 3.33E-01 | 1.68E+00 | 2.70E-01 | 9.62E-01 | -1.07E+00 | 2.87E-01 | 8.09E-01 | -1.20E+00 | 2.17E-01 | 7.96E-01 | 1.14E+00 | 3.32E-01 | 9.13E-01 | 1.05E+00 |
| EEF1GP5 | 1.50E-01 | 7.84E-01 | 1.46E+00 | 2.11E-01 | 9.12E-01 | 1.50E+00 | 2.15E-03 | 9.15E-03 | 2.28E+00 | 2.85E-02 | 1.09E-01 | 2.16E+00 | 7.66E-02 | 2.73E-01 | 2.10E+00 | 8.91E-02 | 2.51E-01 | 2.17E+00 | 4.45E-03 | 1.63E-02 | 2.31E+00 | 1.79E-01 | 4.92E-01 | 1.56E+00 |
| CD274 | 1.50E-01 | 7.84E-01 | 1.34E+00 | 2.20E-01 | 9.51E-01 | 1.23E+00 | 1.02E-01 | 4.35E-01 | 1.63E+00 | 7.30E-02 | 2.79E-01 | 2.00E+00 | 3.89E-08 | 1.39E-07 | 7.37E+00 | 4.54E-03 | 1.28E-02 | 4.78E+00 | 3.82E-06 | 1.40E-05 | 2.06E+00 | 2.03E-02 | 5.59E-02 | 1.60E+00 |
| DNAH10 | 1.50E-01 | 7.84E-01 | 1.34E+00 | 1.20E-01 | 5.18E-01 | 1.37E+00 | 1.93E-01 | 8.18E-01 | 1.12E+00 | 1.56E-01 | 5.95E-01 | 1.15E+00 | 2.63E-01 | 9.39E-01 | 1.30E+00 | 2.16E-01 | 6.08E-01 | -1.13E+00 | 1.28E-01 | 4.69E-01 | -1.33E+00 | 2.56E-01 | 7.04E-01 | -1.04E+00 |
| HIST1H1C | 1.51E-01 | 7.85E-01 | 1.26E+00 | 3.56E-02 | 1.54E-01 | 1.18E+00 | 7.38E-02 | 3.14E-01 | 1.21E+00 | 4.58E-04 | 1.75E-03 | 1.04E+00 | 1.98E-01 | 7.07E-01 | 1.25E+00 | 1.71E-01 | 4.83E-01 | 1.21E+00 | 1.81E-01 | 6.64E-01 | 1.35E+00 | 4.37E-03 | 1.20E-02 | 1.16E+00 |
| PRKD1 | 1.51E-01 | 7.89E-01 | 1.11E+00 | 1.80E-01 | 7.77E-01 | 1.48E+00 | 9.47E-02 | 4.02E-01 | -1.50E+00 | 1.95E-01 | 7.44E-01 | -1.08E+00 | 2.62E-02 | 9.34E-02 | 2.00E+00 | 1.30E-03 | 3.67E-03 | 3.04E+00 | 2.59E-01 | 9.50E-01 | 1.12E+00 | 3.39E-01 | 9.32E-01 | 1.18E+00 |
| C4orf36 | 1.52E-01 | 7.90E-01 | 1.34E+00 | 1.28E-01 | 5.52E-01 | 1.72E+00 | 3.56E-03 | 1.51E-02 | -1.19E+00 | 8.51E-03 | 3.25E-02 | 1.04E+00 | 1.18E-02 | 4.21E-02 | -1.13E+00 | 6.49E-02 | 1.83E-01 | 1.12E+00 | 1.55E-01 | 5.67E-01 | 1.17E+00 | 7.87E-02 | 2.16E-01 | 1.04E+00 |
| DECR2 | 1.52E-01 | 7.91E-01 | 1.52E+00 | 1.99E-01 | 8.62E-01 | 1.52E+00 | 4.32E-02 | 1.84E-01 | 1.00E+00 | 4.89E-02 | 1.87E-01 | 1.16E+00 | 1.35E-02 | 4.80E-02 | -1.16E+00 | 8.22E-02 | 2.32E-01 | 1.14E+00 | 1.84E-01 | 6.77E-01 | 1.32E+00 | 3.15E-01 | 8.66E-01 | 1.49E+00 |
| AL627171.1 | 1.52E-01 | 7.91E-01 | 1.07E+00 | 1.83E-03 | 7.93E-03 | 1.97E+00 | 4.41E-02 | 1.87E-01 | -1.20E+00 | 1.34E-01 | 5.11E-01 | 1.53E+00 | 3.35E-02 | 1.20E-01 | -1.37E+00 | 2.38E-01 | 6.72E-01 | 1.63E+00 | 5.97E-02 | 2.19E-01 | -1.16E+00 | 3.39E-01 | 9.32E-01 | 1.35E+00 |
| PDK1 | 1.52E-01 | 7.91E-01 | 1.13E+00 | 9.54E-05 | 4.12E-04 | -1.43E+00 | 3.56E-04 | 1.51E-03 | -1.10E+00 | 3.21E-10 | 1.22E-09 | -1.49E+00 | 2.45E-01 | 8.73E-01 | 1.45E+00 | 2.73E-01 | 7.70E-01 | 1.14E+00 | 1.12E-08 | 4.11E-08 | -1.54E+00 | 2.94E-11 | 8.08E-11 | -1.73E+00 |
| ZNF555 | 1.52E-01 | 7.92E-01 | 1.48E+00 | 1.42E-01 | 6.14E-01 | 1.71E+00 | 1.58E-01 | 6.71E-01 | 1.24E+00 | 2.00E-01 | 7.63E-01 | 1.34E+00 | 6.57E-02 | 2.34E-01 | 1.83E+00 | 1.38E-06 | 3.89E-06 | 2.93E+00 | 4.34E-02 | 1.59E-01 | 1.11E+00 | 2.87E-01 | 7.87E-01 | 1.27E+00 |
| AC004477.1 | 1.52E-01 | 7.92E-01 | 1.43E+00 | 1.74E-01 | 7.52E-01 | 1.23E+00 | 1.96E-01 | 8.34E-01 | 1.28E+00 | 2.55E-01 | 9.74E-01 | 1.31E+00 | 1.94E-02 | 6.91E-02 | 2.83E+00 | 4.70E-02 | 1.32E-01 | 2.77E+00 | 3.83E-02 | 1.40E-01 | 1.78E+00 | 5.42E-02 | 1.49E-01 | 1.87E+00 |
| MIDN | 1.52E-01 | 7.92E-01 | 1.38E+00 | 1.95E-15 | 8.43E-15 | -1.67E+00 | 2.75E-03 | 1.17E-02 | -1.03E+00 | 7.73E-16 | 2.95E-15 | -1.76E+00 | 2.12E-06 | 7.55E-06 | -1.26E+00 | 3.81E-09 | 1.07E-08 | -1.60E+00 | 7.68E-07 | 2.82E-06 | -1.13E+00 | 8.36E-14 | 2.30E-13 | -2.17E+00 |
| FAM122C | 1.52E-01 | 7.92E-01 | 1.50E+00 | 5.18E-02 | 2.24E-01 | 1.26E+00 | 1.31E-01 | 5.58E-01 | 1.09E+00 | 1.27E-03 | 4.86E-03 | -1.36E+00 | 6.08E-02 | 2.17E-01 | 1.94E+00 | 2.67E-01 | 7.52E-01 | 1.33E+00 | 1.10E-01 | 4.05E-01 | 1.47E+00 | 1.84E-02 | 5.05E-02 | -1.10E+00 |
| DMWD | 1.52E-01 | 7.92E-01 | 1.08E+00 | 1.62E-01 | 7.00E-01 | -1.03E+00 | 1.85E-01 | 7.85E-01 | -1.01E+00 | 1.54E-01 | 5.90E-01 | -1.15E+00 | 1.82E-01 | 6.48E-01 | 1.25E+00 | 2.41E-02 | 6.80E-02 | 2.02E+00 | 3.31E-03 | 1.21E-02 | 1.01E+00 | 8.62E-03 | 2.37E-02 | -1.01E+00 |
| ZBTB46 | 1.52E-01 | 7.92E-01 | 1.26E+00 | 2.24E-01 | 9.68E-01 | 1.51E+00 | 9.62E-02 | 4.09E-01 | 1.40E+00 | 2.07E-01 | 7.92E-01 | 1.35E+00 | 1.48E-01 | 5.29E-01 | -1.03E+00 | 1.55E-01 | 4.38E-01 | -1.07E+00 | 5.62E-02 | 2.06E-01 | -1.02E+00 | 5.86E-02 | 1.61E-01 | -1.15E+00 |
| ITGA9-AS1 | 1.52E-01 | 7.92E-01 | 1.27E+00 | 2.14E-02 | 9.24E-02 | 1.06E+00 | 5.38E-02 | 2.29E-01 | 1.51E+00 | 2.08E-01 | 7.96E-01 | 1.51E+00 | 4.36E-02 | 1.56E-01 | 1.88E+00 | 3.49E-01 | 9.84E-01 | 1.35E+00 | 2.12E-01 | 7.79E-01 | 1.32E+00 | 9.28E-02 | 2.55E-01 | 1.18E+00 |
| AP000281.1 | 1.52E-01 | 7.93E-01 | 1.27E+00 | 1.72E-01 | 7.42E-01 | 1.38E+00 | 3.46E-02 | 1.47E-01 | 1.34E+00 | 2.74E-02 | 1.05E-01 | 1.45E+00 | 7.93E-02 | 2.83E-01 | 1.31E+00 | 2.93E-02 | 8.26E-02 | 1.24E+00 | 1.44E-01 | 5.28E-01 | 1.19E+00 | 1.68E-01 | 4.61E-01 | 1.24E+00 |
| DNAH1 | 1.52E-01 | 7.93E-01 | 1.29E+00 | 1.32E-01 | 5.72E-01 | 1.32E+00 | 1.76E-01 | 7.47E-01 | 1.32E+00 | 8.01E-02 | 3.06E-01 | 1.84E+00 | 2.33E-01 | 8.31E-01 | 1.62E+00 | 2.65E-01 | 7.46E-01 | 1.20E+00 | 1.86E-01 | 6.82E-01 | 1.04E+00 | 2.30E-01 | 6.33E-01 | -1.08E+00 |
| MYEOV | 1.52E-01 | 7.93E-01 | 1.15E+00 | 2.31E-01 | 1.00E+00 | 1.00E+00 | 2.35E-01 | 1.00E+00 | -1.00E+00 | 2.62E-01 | 1.00E+00 | 1.00E+00 | 2.40E-01 | 8.54E-01 | 1.16E+00 | 3.55E-01 | 1.00E+00 | 1.00E+00 | 1.56E-04 | 5.71E-04 | -1.01E+00 | 5.90E-05 | 1.62E-04 | -1.24E+00 |
| TTC25 | 1.52E-01 | 7.94E-01 | 1.02E+00 | 9.18E-02 | 3.97E-01 | -1.24E+00 | 3.64E-02 | 1.55E-01 | -1.19E+00 | 9.23E-03 | 3.53E-02 | -1.18E+00 | 1.15E-04 | 4.09E-04 | -1.82E+00 | 1.60E-03 | 4.52E-03 | -1.40E+00 | 1.28E-02 | 4.68E-02 | -1.62E+00 | 3.99E-02 | 1.10E-01 | -1.40E+00 |
| PPP1R14A | 1.52E-01 | 7.94E-01 | 1.15E+00 | 1.67E-01 | 7.22E-01 | 1.22E+00 | 2.03E-02 | 8.63E-02 | -1.32E+00 | 4.47E-02 | 1.71E-01 | -1.04E+00 | 1.28E-01 | 4.56E-01 | 1.01E+00 | 3.39E-02 | 9.55E-02 | 2.03E+00 | 2.75E-05 | 1.01E-04 | 1.02E+00 | 3.18E-15 | 8.73E-15 | -1.27E+00 |
| E2F2 | 1.52E-01 | 7.94E-01 | -1.15E+00 | 1.86E-01 | 8.04E-01 | -1.15E+00 | 1.83E-01 | 7.79E-01 | -1.13E+00 | 1.18E-01 | 4.51E-01 | -1.38E+00 | 1.76E-01 | 6.27E-01 | -1.18E+00 | 2.32E-01 | 6.55E-01 | -1.25E+00 | 1.59E-01 | 5.85E-01 | 1.26E+00 | 2.48E-01 | 6.81E-01 | 1.21E+00 |
| FAM78A | 1.52E-01 | 7.95E-01 | 1.37E+00 | 9.36E-02 | 4.04E-01 | 1.70E+00 | 1.96E-03 | 8.32E-03 | -1.18E+00 | 3.26E-02 | 1.24E-01 | 1.17E+00 | 4.38E-05 | 1.56E-04 | -1.67E+00 | 9.78E-03 | 2.76E-02 | -1.18E+00 | 1.12E-03 | 4.12E-03 | -1.09E+00 | 4.13E-03 | 1.13E-02 | -1.11E+00 |
| TNFRSF25 | 1.53E-01 | 7.95E-01 | 1.18E+00 | 1.96E-01 | 8.45E-01 | 1.37E+00 | 1.03E-01 | 4.38E-01 | -1.21E+00 | 1.72E-02 | 6.58E-02 | -1.52E+00 | 2.51E-01 | 8.93E-01 | 1.32E+00 | 4.32E-02 | 1.22E-01 | 2.83E+00 | 3.20E-12 | 1.17E-11 | -1.33E+00 | 7.79E-08 | 2.14E-07 | -1.20E+00 |
| VDAC1P6 | 1.53E-01 | 7.96E-01 | 1.15E+00 | 5.32E-02 | 2.30E-01 | -1.12E+00 | 9.54E-02 | 4.05E-01 | 1.46E+00 | 2.79E-02 | 1.06E-01 | 1.04E+00 | 2.41E-01 | 8.59E-01 | 1.26E+00 | 1.13E-01 | 3.19E-01 | -1.12E+00 | 2.52E-02 | 9.24E-02 | 1.49E+00 | 2.90E-01 | 7.96E-01 | 1.08E+00 |
| AL512625.1 | 1.53E-01 | 7.96E-01 | 1.13E+00 | 7.89E-02 | 3.41E-01 | 1.73E+00 | 1.70E-02 | 7.22E-02 | -1.17E+00 | 1.25E-01 | 4.77E-01 | 1.22E+00 | 1.72E-01 | 6.15E-01 | 1.15E+00 | 2.13E-01 | 6.00E-01 | 1.72E+00 | 4.10E-02 | 1.50E-01 | -1.02E+00 | 3.52E-01 | 9.66E-01 | 1.34E+00 |
| RAC3 | 1.53E-01 | 7.96E-01 | 1.20E+00 | 7.88E-02 | 3.40E-01 | 1.22E+00 | 1.22E-01 | 5.20E-01 | 1.24E+00 | 2.61E-02 | 9.96E-02 | 1.59E+00 | 1.31E-01 | 4.67E-01 | 1.07E+00 | 2.72E-01 | 7.68E-01 | 1.35E+00 | 2.58E-01 | 9.46E-01 | 1.29E+00 | 1.79E-02 | 4.92E-02 | 1.65E+00 |
| NDUFB8P2 | 1.53E-01 | 7.96E-01 | 1.35E+00 | 7.64E-05 | 3.30E-04 | -1.08E+00 | 6.25E-02 | 2.66E-01 | 1.37E+00 | 1.22E-02 | 4.65E-02 | 1.04E+00 | 1.06E-02 | 3.78E-02 | 1.19E+00 | 4.26E-03 | 1.20E-02 | 1.07E+00 | 1.39E-01 | 5.11E-01 | 1.33E+00 | 5.26E-03 | 1.45E-02 | -1.04E+00 |
| DPYSL3 | 1.53E-01 | 7.97E-01 | 1.36E+00 | 1.11E-01 | 4.79E-01 | 1.74E+00 | 5.03E-03 | 2.14E-02 | 1.52E+00 | 3.21E-02 | 1.22E-01 | 1.59E+00 | 2.44E-01 | 8.68E-01 | 1.59E+00 | 3.24E-01 | 9.12E-01 | 1.39E+00 | 1.14E-01 | 4.20E-01 | 1.88E+00 | 2.66E-01 | 7.30E-01 | 1.50E+00 |
| LILRB3 | 1.53E-01 | 7.98E-01 | 1.08E+00 | 7.03E-02 | 3.04E-01 | -1.12E+00 | 7.61E-04 | 3.23E-03 | -1.60E+00 | 4.32E-06 | 1.65E-05 | -2.42E+00 | 1.60E-01 | 5.72E-01 | -1.06E+00 | 2.05E-01 | 5.78E-01 | -1.27E+00 | 1.06E-01 | 3.89E-01 | -1.09E+00 | 4.42E-02 | 1.21E-01 | -1.38E+00 |
| AL158212.3 | 1.53E-01 | 7.99E-01 | 1.26E+00 | 3.78E-02 | 1.63E-01 | 1.94E+00 | 1.22E-01 | 5.19E-01 | -1.17E+00 | 1.96E-01 | 7.48E-01 | 1.40E+00 | 2.30E-01 | 8.19E-01 | 1.59E+00 | 2.52E-01 | 7.10E-01 | 1.54E+00 | 1.24E-01 | 4.54E-01 | 1.44E+00 | 6.41E-02 | 1.76E-01 | 1.65E+00 |
| HNRNPA1P10 | 1.53E-01 | 7.99E-01 | 1.29E+00 | 1.35E-01 | 5.82E-01 | 1.42E+00 | 8.41E-02 | 3.58E-01 | 1.09E+00 | 2.23E-01 | 8.51E-01 | 1.41E+00 | 6.98E-03 | 2.49E-02 | -1.21E+00 | 6.93E-02 | 1.95E-01 | 1.09E+00 | 4.84E-02 | 1.78E-01 | -1.09E+00 | 2.36E-01 | 6.48E-01 | 1.15E+00 |
| NCEH1 | 1.53E-01 | 7.99E-01 | 1.01E+00 | 1.51E-01 | 6.52E-01 | -1.01E+00 | 2.93E-02 | 1.25E-01 | -1.27E+00 | 1.90E-01 | 7.26E-01 | 1.14E+00 | 5.72E-02 | 2.04E-01 | -1.06E+00 | 1.24E-01 | 3.49E-01 | 1.11E+00 | 3.10E-02 | 1.14E-01 | -1.11E+00 | 2.05E-01 | 5.64E-01 | 1.21E+00 |
| STXBP6 | 1.53E-01 | 7.99E-01 | 2.23E+00 | 8.98E-03 | 3.88E-02 | 3.57E+00 | 1.20E-01 | 5.11E-01 | -1.01E+00 | 4.33E-02 | 1.66E-01 | 2.17E+00 | 1.68E-03 | 5.99E-03 | 1.17E+00 | 1.13E-02 | 3.18E-02 | 1.72E+00 | 5.54E-02 | 2.03E-01 | 1.02E+00 | 2.34E-01 | 6.42E-01 | 1.13E+00 |
| HDAC11 | 1.53E-01 | 7.99E-01 | 1.35E+00 | 6.36E-02 | 2.75E-01 | 1.25E+00 | 2.18E-01 | 9.27E-01 | 1.20E+00 | 2.22E-01 | 8.48E-01 | 1.25E+00 | 5.84E-02 | 2.08E-01 | -1.10E+00 | 1.79E-01 | 5.05E-01 | 1.07E+00 | 3.52E-02 | 1.29E-01 | -1.11E+00 | 5.03E-02 | 1.38E-01 | -1.16E+00 |
| GSPT2 | 1.53E-01 | 7.99E-01 | 1.29E+00 | 1.83E-01 | 7.92E-01 | 1.58E+00 | 7.39E-03 | 3.14E-02 | -1.12E+00 | 1.14E-01 | 4.34E-01 | 1.28E+00 | 4.55E-02 | 1.62E-01 | -1.12E+00 | 1.74E-01 | 4.91E-01 | 1.10E+00 | 1.46E-01 | 5.36E-01 | 1.09E+00 | 2.45E-01 | 6.72E-01 | 1.19E+00 |
| LOX | 1.53E-01 | 7.99E-01 | 1.46E+00 | 6.74E-02 | 2.91E-01 | 1.78E+00 | 1.96E-01 | 8.33E-01 | 1.43E+00 | 1.95E-01 | 7.44E-01 | 1.55E+00 | 2.30E-01 | 8.19E-01 | 1.64E+00 | 2.33E-01 | 6.56E-01 | 1.81E+00 | 4.62E-03 | 1.70E-02 | 2.03E+00 | 1.44E-02 | 3.95E-02 | 1.87E+00 |
| IFI30 | 1.53E-01 | 7.99E-01 | -1.04E+00 | 1.23E-01 | 5.32E-01 | -1.09E+00 | 1.15E-01 | 4.88E-01 | -1.04E+00 | 1.32E-01 | 5.03E-01 | -1.12E+00 | 1.05E-01 | 3.73E-01 | -1.23E+00 | 1.41E-01 | 3.96E-01 | -1.15E+00 | 1.02E-01 | 3.73E-01 | -1.05E+00 | 2.37E-01 | 6.52E-01 | -1.02E+00 |
| MCM2 | 1.53E-01 | 7.99E-01 | 1.28E+00 | 2.26E-01 | 9.78E-01 | 1.45E+00 | 7.11E-03 | 3.02E-02 | 1.33E+00 | 1.07E-01 | 4.07E-01 | 1.47E+00 | 1.01E-01 | 3.60E-01 | -1.03E+00 | 2.81E-01 | 7.93E-01 | 1.21E+00 | 6.73E-04 | 2.47E-03 | -1.09E+00 | 2.95E-02 | 8.10E-02 | 1.40E+00 |
| TEX22 | 1.53E-01 | 8.00E-01 | 1.30E+00 | 2.15E-01 | 9.31E-01 | 1.35E+00 | 1.47E-02 | 6.27E-02 | -1.26E+00 | 4.72E-02 | 1.80E-01 | -1.03E+00 | 2.71E-01 | 9.66E-01 | 1.33E+00 | 2.17E-01 | 6.13E-01 | 1.13E+00 | 9.37E-03 | 3.44E-02 | -1.45E+00 | 1.92E-01 | 5.28E-01 | 1.02E+00 |
| ZNF14 | 1.53E-01 | 8.00E-01 | 1.20E+00 | 3.42E-02 | 1.48E-01 | -1.09E+00 | 2.27E-01 | 9.64E-01 | 1.15E+00 | 5.43E-02 | 2.07E-01 | -1.03E+00 | 2.05E-01 | 7.31E-01 | 1.58E+00 | 1.13E-01 | 3.20E-01 | -1.03E+00 | 1.54E-01 | 5.67E-01 | 1.40E+00 | 3.15E-01 | 8.64E-01 | 1.23E+00 |
| EEF1B2P2 | 1.54E-01 | 8.00E-01 | 1.08E+00 | 2.34E-04 | 1.01E-03 | -1.05E+00 | 6.23E-05 | 2.65E-04 | 1.27E+00 | 7.29E-02 | 2.79E-01 | 1.12E+00 | 5.43E-03 | 1.93E-02 | 1.23E+00 | 1.42E-01 | 3.99E-01 | 1.11E+00 | 7.87E-02 | 2.89E-01 | 1.19E+00 | 8.28E-02 | 2.27E-01 | 1.13E+00 |
| OXTR | 1.54E-01 | 8.00E-01 | -1.07E+00 | 1.33E-01 | 5.76E-01 | -1.14E+00 | 1.45E-04 | 6.17E-04 | -1.46E+00 | 3.30E-03 | 1.26E-02 | -1.09E+00 | 1.72E-01 | 6.13E-01 | 1.36E+00 | 1.22E-01 | 3.44E-01 | 1.75E+00 | 1.94E-01 | 7.11E-01 | -1.01E+00 | 3.15E-01 | 8.65E-01 | -1.14E+00 |
| DISC1FP1 | 1.54E-01 | 8.00E-01 | 1.31E+00 | 2.04E-01 | 8.81E-01 | 1.29E+00 | 1.24E-01 | 5.28E-01 | 1.56E+00 | 2.09E-01 | 7.97E-01 | 1.37E+00 | 3.27E-02 | 1.17E-01 | 2.32E+00 | 2.03E-01 | 5.72E-01 | 1.89E+00 | 1.56E-01 | 5.72E-01 | 1.52E+00 | 1.90E-01 | 5.23E-01 | 1.52E+00 |
| MICALL2 | 1.54E-01 | 8.00E-01 | 1.02E+00 | 2.01E-01 | 8.68E-01 | 1.22E+00 | 2.64E-03 | 1.12E-02 | -1.36E+00 | 5.66E-05 | 2.16E-04 | -1.90E+00 | 4.69E-02 | 1.67E-01 | -1.24E+00 | 4.36E-02 | 1.23E-01 | -1.38E+00 | 5.42E-06 | 1.99E-05 | -1.34E+00 | 1.28E-07 | 3.52E-07 | -1.80E+00 |
| AL356488.3 | 1.54E-01 | 8.01E-01 | 1.26E+00 | 1.09E-01 | 4.71E-01 | 1.03E+00 | 2.15E-01 | 9.14E-01 | 1.09E+00 | 1.39E-01 | 5.32E-01 | -1.04E+00 | 1.89E-01 | 6.73E-01 | 1.63E+00 | 1.89E-01 | 5.33E-01 | 1.06E+00 | 1.57E-01 | 5.76E-01 | 1.08E+00 | 3.35E-01 | 9.21E-01 | 1.26E+00 |
| DYNLRB2 | 1.54E-01 | 8.02E-01 | 1.30E+00 | 1.26E-01 | 5.46E-01 | 1.17E+00 | 2.87E-03 | 1.22E-02 | -1.62E+00 | 1.23E-03 | 4.69E-03 | -1.48E+00 | 2.78E-01 | 9.92E-01 | 1.32E+00 | 1.59E-01 | 4.47E-01 | -1.19E+00 | 2.12E-01 | 7.77E-01 | 1.03E+00 | 2.47E-01 | 6.78E-01 | -1.04E+00 |
| PLXNB1 | 1.54E-01 | 8.02E-01 | 1.22E+00 | 1.51E-01 | 6.51E-01 | 1.12E+00 | 6.28E-02 | 2.67E-01 | -1.11E+00 | 5.99E-03 | 2.29E-02 | -1.32E+00 | 2.64E-01 | 9.42E-01 | 1.18E+00 | 2.59E-01 | 7.30E-01 | -1.06E+00 | 1.46E-01 | 5.35E-01 | -1.14E+00 | 1.80E-01 | 4.94E-01 | -1.17E+00 |
| IFIT1B | 1.54E-01 | 8.02E-01 | 1.23E+00 | 1.89E-01 | 8.17E-01 | 1.15E+00 | 2.35E-01 | 1.00E+00 | 1.00E+00 | 2.21E-01 | 8.45E-01 | 1.06E+00 | 2.05E-01 | 7.29E-01 | 1.37E+00 | 2.83E-01 | 7.98E-01 | 1.40E+00 | 1.17E-01 | 4.29E-01 | 1.09E+00 | 2.34E-02 | 6.44E-02 | -1.26E+00 |
| NMU | 1.54E-01 | 8.02E-01 | 1.01E+00 | 1.16E-01 | 5.00E-01 | -1.16E+00 | 9.87E-02 | 4.19E-01 | 1.20E+00 | 1.22E-01 | 4.66E-01 | 1.31E+00 | 2.80E-01 | 1.00E+00 | 1.00E+00 | 3.01E-01 | 8.47E-01 | 1.10E+00 | 8.44E-02 | 3.10E-01 | -1.06E+00 | 1.55E-01 | 4.25E-01 | -1.11E+00 |
| TPRXB | 1.54E-01 | 8.03E-01 | 1.35E+00 | 2.30E-02 | 9.95E-02 | 1.31E+00 | 2.97E-02 | 1.26E-01 | 1.30E+00 | 2.40E-01 | 9.16E-01 | 1.32E+00 | 1.13E-01 | 4.03E-01 | 1.22E+00 | 3.04E-01 | 8.57E-01 | 1.31E+00 | 6.88E-02 | 2.53E-01 | 1.30E+00 | 1.86E-01 | 5.10E-01 | 1.22E+00 |
| MT1P3 | 1.54E-01 | 8.03E-01 | 1.09E+00 | 1.33E-01 | 5.75E-01 | 1.00E+00 | 1.89E-02 | 8.03E-02 | -1.33E+00 | 8.67E-03 | 3.31E-02 | -1.30E+00 | 2.09E-01 | 7.47E-01 | 1.75E+00 | 2.77E-01 | 7.81E-01 | 1.19E+00 | 1.14E-02 | 4.20E-02 | 1.95E+00 | 6.30E-02 | 1.73E-01 | 1.99E+00 |
| LINC01535 | 1.54E-01 | 8.04E-01 | 1.48E+00 | 4.96E-02 | 2.14E-01 | 1.95E+00 | 5.24E-03 | 2.22E-02 | -1.66E+00 | 2.00E-01 | 7.65E-01 | 1.24E+00 | 9.87E-02 | 3.52E-01 | 1.86E+00 | 7.98E-02 | 2.25E-01 | 2.07E+00 | 2.03E-01 | 7.46E-01 | 1.08E+00 | 2.43E-01 | 6.68E-01 | 1.20E+00 |
| TMEM272 | 1.54E-01 | 8.04E-01 | 1.22E+00 | 5.67E-02 | 2.45E-01 | 1.06E+00 | 3.66E-03 | 1.56E-02 | 2.01E+00 | 1.89E-01 | 7.21E-01 | 1.44E+00 | 2.11E-01 | 7.53E-01 | 1.31E+00 | 2.46E-01 | 6.93E-01 | 1.45E+00 | 9.75E-02 | 3.58E-01 | 1.66E+00 | 1.87E-01 | 5.15E-01 | 1.44E+00 |
| SRSF12 | 1.54E-01 | 8.04E-01 | 1.24E+00 | 4.40E-02 | 1.90E-01 | -1.01E+00 | 3.73E-05 | 1.59E-04 | -1.32E+00 | 2.41E-06 | 9.22E-06 | -1.34E+00 | 3.44E-04 | 1.23E-03 | -1.67E+00 | 1.47E-04 | 4.13E-04 | -1.67E+00 | 3.88E-02 | 1.42E-01 | -1.50E+00 | 3.42E-01 | 9.40E-01 | 1.30E+00 |
| PTP4A1P2 | 1.54E-01 | 8.04E-01 | 1.22E+00 | 2.14E-01 | 9.23E-01 | 1.42E+00 | 2.15E-01 | 9.13E-01 | 1.11E+00 | 7.33E-02 | 2.80E-01 | 1.73E+00 | 1.01E-01 | 3.59E-01 | -1.21E+00 | 2.46E-01 | 6.93E-01 | 1.53E+00 | 2.08E-01 | 7.64E-01 | 1.27E+00 | 2.29E-02 | 6.29E-02 | 1.61E+00 |
| HMGB1P40 | 1.54E-01 | 8.04E-01 | 1.21E+00 | 7.76E-02 | 3.35E-01 | -1.05E+00 | 1.80E-01 | 7.66E-01 | 1.18E+00 | 3.46E-03 | 1.32E-02 | -1.28E+00 | 1.97E-01 | 7.03E-01 | 1.08E+00 | 1.20E-01 | 3.37E-01 | -1.11E+00 | 2.03E-04 | 7.44E-04 | -1.29E+00 | 2.15E-09 | 5.90E-09 | -1.88E+00 |
| PRTN3 | 1.54E-01 | 8.04E-01 | 1.83E+00 | 2.31E-01 | 1.00E+00 | 1.00E+00 | 4.58E-02 | 1.94E-01 | 1.57E+00 | 4.73E-04 | 1.81E-03 | -1.72E+00 | 2.23E-01 | 7.94E-01 | 1.28E+00 | 3.55E-01 | 1.00E+00 | 1.00E+00 | 2.33E-01 | 8.53E-01 | 1.18E+00 | 3.19E-01 | 8.76E-01 | 1.13E+00 |
| CRYBA4 | 1.54E-01 | 8.04E-01 | 1.21E+00 | 1.98E-01 | 8.56E-01 | 1.15E+00 | 1.94E-01 | 8.25E-01 | -1.05E+00 | 2.23E-01 | 8.52E-01 | -1.02E+00 | 1.32E-02 | 4.71E-02 | 4.99E+00 | 2.57E-01 | 7.25E-01 | 1.41E+00 | 4.78E-06 | 1.75E-05 | 2.59E+00 | 1.73E-01 | 4.75E-01 | 1.48E+00 |
| FAM72C | 1.54E-01 | 8.05E-01 | -1.08E+00 | 1.80E-01 | 7.77E-01 | -1.05E+00 | 6.12E-02 | 2.60E-01 | -1.34E+00 | 2.37E-01 | 9.06E-01 | 1.16E+00 | 4.04E-02 | 1.44E-01 | -1.98E+00 | 4.93E-02 | 1.39E-01 | -2.03E+00 | 1.06E-04 | 3.91E-04 | -1.72E+00 | 3.59E-01 | 9.86E-01 | 1.17E+00 |

| | | | | | | | | | | | | | | | | | | | | | | | |
|---|---|---|---|---|---|---|---|---|---|---|---|---|---|---|---|---|---|---|---|---|---|---|---|
| TRAPPC10 | 1.54E-01 | 8.05E-01 | 1.26E+00 | 2.81E-03 | 1.21E-02 | 1.02E+00 | 1.99E-01 | 8.45E-01 | 1.21E+00 | 6.12E-03 | 2.34E-02 | 1.03E+00 | 1.46E-01 | 5.21E-01 | 1.21E+00 | 2.50E-02 | 7.05E-02 | 1.02E+00 | 8.24E-03 | 3.02E-02 | -1.08E+00 | 2.34E-03 | [illegible] |
| MUC1 | 1.54E-01 | 8.05E-01 | 1.27E+00 | 5.95E-02 | 2.57E-01 | 1.12E+00 | 1.45E-01 | 6.15E-01 | 1.33E+00 | 1.84E-01 | 7.01E-01 | 1.40E+00 | 2.48E-01 | 8.83E-01 | 1.37E+00 | 3.47E-01 | 9.77E-01 | 1.38E+00 | 1.27E-01 | 4.65E-01 | 1.16E+00 | 2.41E-03 | [illegible] |
| CD14 | 1.54E-01 | 8.05E-01 | -1.01E+00 | 1.64E-01 | 7.10E-01 | 1.13E+00 | 1.05E-01 | 4.45E-01 | -4.50E+00 | 7.22E-02 | 2.76E-01 | -5.46E+00 | 2.18E-01 | 7.78E-01 | 1.28E+00 | 3.55E-01 | 1.00E+00 | -1.00E+00 | 6.62E-02 | 2.43E-01 | -1.83E+00 | 6.62E-02 | [illegible] |
| NAP1L1P3 | 1.55E-01 | 8.06E-01 | 1.31E+00 | 5.23E-02 | 2.26E-01 | 1.89E+00 | 8.03E-04 | 3.41E-03 | 1.75E+00 | 1.88E-05 | 7.17E-05 | 2.23E+00 | 2.25E-01 | 8.02E-01 | 1.33E+00 | 2.87E-01 | 8.09E-01 | 1.61E+00 | 2.18E-01 | 8.00E-01 | 1.32E+00 | 7.56E-04 | [illegible] |
| NXF3 | 1.55E-01 | 8.06E-01 | 1.42E+00 | 2.31E-01 | 1.00E+00 | 1.00E+00 | 1.98E-01 | 8.40E-01 | 1.11E+00 | 1.94E-01 | 7.39E-01 | 1.20E+00 | 1.77E-01 | 6.29E-01 | 1.52E+00 | 1.43E-01 | 4.03E-01 | 1.81E+00 | 1.99E-10 | 7.30E-10 | 6.59E+00 | 2.30E-05 | [illegible] |
| ASPSCR1 | 1.55E-01 | 8.06E-01 | 1.27E+00 | 8.65E-03 | 3.74E-02 | 1.02E+00 | 2.08E-04 | 8.85E-04 | -1.18E+00 | 1.46E-03 | 5.57E-03 | 1.03E+00 | 2.49E-03 | 8.87E-03 | -1.01E+00 | 5.01E-03 | 1.41E-02 | 1.04E+00 | 1.97E-04 | 7.21E-04 | -1.03E+00 | 3.66E-03 | [illegible] |
| GMEB2 | 1.55E-01 | 8.07E-01 | 1.28E+00 | 1.85E-02 | 7.98E-02 | -1.23E+00 | 6.51E-02 | 2.76E-01 | 1.03E+00 | 1.09E-02 | 4.17E-02 | -1.01E+00 | 1.87E-02 | 6.66E-02 | -1.20E+00 | 4.65E-02 | 1.31E-01 | -1.10E+00 | 6.52E-04 | 2.39E-03 | -1.17E+00 | 7.12E-03 | [illegible] |
| PFDN5 | 1.55E-01 | 8.07E-01 | 1.01E+00 | 2.29E-01 | 9.89E-01 | 1.01E+00 | 9.29E-02 | 3.95E-01 | 1.04E+00 | 1.57E-01 | 6.01E-01 | 1.04E+00 | 3.28E-02 | 1.17E-01 | 1.09E+00 | 3.10E-01 | 8.75E-01 | 1.02E+00 | 4.59E-02 | 1.68E-01 | 1.13E+00 | 2.45E-01 | [illegible] |
| RGS9 | 1.55E-01 | 8.07E-01 | 1.17E+00 | 1.81E-01 | 7.82E-01 | 1.21E+00 | 1.13E-01 | 4.79E-01 | -1.13E+00 | 7.85E-02 | 3.00E-01 | -1.25E+00 | 2.17E-01 | 7.74E-01 | 1.29E+00 | 2.71E-01 | 7.65E-01 | 1.30E+00 | 2.51E-01 | 9.21E-01 | 1.73E+00 | 8.21E-03 | [illegible] |
| WNT5B | 1.55E-01 | 8.07E-01 | -1.05E+00 | 1.92E-01 | 8.31E-01 | -1.01E+00 | 1.08E-01 | 4.58E-01 | -1.24E+00 | 1.29E-01 | 4.93E-01 | -1.20E+00 | 8.84E-32 | 3.15E-31 | -5.86E+00 | 2.85E-26 | 8.02E-26 | -3.05E+00 | 1.84E-34 | 6.76E-34 | -3.96E+00 | 1.61E-24 | [illegible] |
| MT-ND3 | 1.55E-01 | 8.08E-01 | 1.01E+00 | 9.48E-78 | 4.09E-77 | -1.66E+00 | 9.61E-02 | 4.09E-01 | 1.04E+00 | 1.94E-39 | 7.42E-39 | -1.54E+00 | 1.76E-03 | 6.27E-03 | 1.12E+00 | 2.41E-27 | 6.80E-27 | -1.50E+00 | 5.00E-02 | 1.83E-01 | 1.09E+00 | 1.76E-33 | [illegible] |
| EEF1A1P8 | 1.55E-01 | 8.08E-01 | 1.27E+00 | 2.34E-02 | 1.01E-01 | 1.08E+00 | 1.63E-01 | 6.91E-01 | 1.31E+00 | 3.01E-02 | 1.15E-01 | 1.02E+00 | 2.48E-01 | 8.84E-01 | 1.51E+00 | 1.79E-01 | 5.06E-01 | 1.17E+00 | 1.59E-01 | 5.82E-01 | 1.57E+00 | 2.01E-01 | [illegible] |
| KLHL29 | 1.55E-01 | 8.09E-01 | 1.13E+00 | 1.16E-01 | 5.01E-01 | 1.04E+00 | 2.06E-02 | 8.77E-02 | -1.27E+00 | 2.22E-02 | 8.48E-02 | -1.16E+00 | 1.78E-02 | 6.36E-02 | -1.37E+00 | 1.90E-01 | 5.36E-01 | 1.19E+00 | 1.22E-01 | 4.47E-01 | 1.26E+00 | 3.58E-01 | [illegible] |
| AC211486.1 | 1.55E-01 | 8.09E-01 | 1.32E+00 | 1.14E-01 | 4.92E-01 | 1.74E+00 | 1.52E-01 | 6.47E-01 | 1.04E+00 | 1.49E-01 | 5.69E-01 | 1.08E+00 | 1.03E-01 | 3.69E-01 | -1.09E+00 | 2.30E-01 | 6.50E-01 | 1.06E+00 | 5.99E-02 | 2.20E-01 | 1.66E+00 | 3.57E-01 | [illegible] |
| MCM8 | 1.55E-01 | 8.09E-01 | -1.05E+00 | 2.19E-01 | 9.47E-01 | 1.38E+00 | 2.22E-01 | 9.42E-01 | 1.35E+00 | 1.12E-01 | 4.29E-01 | 1.84E+00 | 1.40E-01 | 4.99E-01 | -1.18E+00 | 2.30E-01 | 6.48E-01 | 1.29E+00 | 2.19E-03 | 8.05E-03 | -1.16E+00 | 1.93E-01 | [illegible] |
| FNDC10 | 1.55E-01 | 8.10E-01 | 1.02E+00 | 1.08E-01 | 4.66E-01 | 1.05E+00 | 1.43E-01 | 6.09E-01 | 1.44E+00 | 1.75E-01 | 6.68E-01 | 1.51E+00 | 1.69E-01 | 6.01E-01 | 1.08E+00 | 1.56E-01 | 4.40E-01 | -1.06E+00 | 6.94E-02 | 2.55E-01 | -1.08E+00 | 2.84E-01 | [illegible] |
| MRPL57 | 1.56E-01 | 8.11E-01 | 1.19E+00 | 1.30E-01 | 5.62E-01 | 1.18E+00 | 1.03E-01 | 4.36E-01 | -1.00E+00 | 6.58E-02 | 2.51E-01 | -1.01E+00 | 4.39E-02 | 1.56E-01 | 1.14E+00 | 9.04E-03 | 2.55E-02 | 1.08E+00 | 1.57E-01 | 5.75E-01 | 1.07E+00 | 3.08E-01 | [illegible] |
| FAM196A | 1.56E-01 | 8.11E-01 | 1.27E+00 | 3.04E-05 | 1.32E-04 | 2.71E+00 | 1.81E-01 | 7.70E-01 | 1.38E+00 | 8.02E-03 | 3.06E-02 | 2.31E+00 | 2.74E-01 | 9.78E-01 | 1.23E+00 | 1.99E-01 | 5.61E-01 | 1.90E+00 | 1.91E-01 | 7.02E-01 | -1.07E+00 | 2.42E-01 | [illegible] |
| TCF19 | 1.56E-01 | 8.12E-01 | 1.35E+00 | 2.20E-01 | 9.49E-01 | 1.32E+00 | 9.53E-05 | 4.05E-04 | 1.97E+00 | 8.97E-04 | 3.43E-03 | 1.97E+00 | 1.45E-01 | 5.18E-01 | -1.78E+00 | 1.67E-01 | 4.71E-01 | -1.97E+00 | 1.45E-01 | 5.33E-01 | 1.01E+00 | 8.49E-03 | [illegible] |
| AC093484.5 | 1.56E-01 | 8.12E-01 | 1.47E+00 | 2.39E-02 | 1.03E-01 | 1.00E+00 | 1.80E-01 | 7.65E-01 | 1.16E+00 | 1.53E-01 | 5.83E-01 | 1.64E+00 | 8.10E-02 | 2.89E-01 | 1.61E+00 | 2.47E-01 | 6.96E-01 | 1.21E+00 | 7.42E-02 | 2.72E-01 | 1.49E+00 | 2.00E-01 | [illegible] |
| AFF1 | 1.56E-01 | 8.12E-01 | 1.41E+00 | 1.93E-01 | 8.34E-01 | 1.61E+00 | 5.15E-02 | 2.19E-01 | 1.45E+00 | 1.60E-01 | 6.09E-01 | 1.27E+00 | 1.22E-03 | 4.35E-03 | 1.08E+00 | 3.41E-05 | 9.60E-05 | -1.07E+00 | 8.95E-03 | 3.28E-02 | -1.00E+00 | 1.55E-03 | [illegible] |
| ZNF597 | 1.56E-01 | 8.12E-01 | 1.37E+00 | 2.19E-02 | 9.46E-02 | -1.32E+00 | 1.78E-01 | 7.56E-01 | -1.01E+00 | 5.07E-02 | 1.94E-01 | -1.27E+00 | 2.58E-02 | 9.19E-02 | -1.18E+00 | 1.32E-03 | 3.73E-03 | -1.96E+00 | 2.68E-01 | 9.83E-01 | 1.28E+00 | 1.29E-02 | [illegible] |
| [illegible] | | | | | | | | | | | | | | | | | | | | | | | |

| | | | | | | | | | | | | | | | | | | | | | | | | |
|---|---|---|---|---|---|---|---|---|---|---|---|---|---|---|---|---|---|---|---|---|---|---|---|---|
| ZFYVE26 | 1.69E-01 | 8.82E-01 | 1.27E+00 | 2.25E-01 | 9.73E-01 | 1.45E+00 | 2.02E-01 | 8.60E-01 | 1.24E+00 | 8.73E-02 | 3.33E-01 | 1.12E+00 | 1.30E-01 | 4.62E-01 | 1.03E+00 | 2.14E-01 | 6.04E-01 | 1.13E+00 | 2.63E-01 | 9.65E-01 | 1.05E+00 | 1.99E-01 | 5.47E-01 | -1.03E+00 |
| RAMP3 | 1.69E-01 | 8.83E-01 | 1.10E+00 | 2.31E-01 | 1.00E+00 | 1.00E+00 | 2.35E-01 | 1.00E+00 | 1.00E+00 | 2.62E-01 | 1.00E+00 | -1.00E+00 | 1.83E-01 | 6.51E-01 | 1.53E+00 | 2.31E-01 | 6.50E-01 | 1.39E+00 | 5.50E-07 | 2.02E-06 | 6.34E+00 | 3.60E-01 | 9.88E-01 | 2.36E+00 |
| PDE8B | 1.69E-01 | 8.83E-01 | 1.23E+00 | 1.16E-01 | 5.01E-01 | 1.76E+00 | 2.14E-03 | 9.10E-03 | -1.32E+00 | 7.57E-04 | 2.89E-03 | -1.22E+00 | 1.77E-01 | 6.30E-01 | 1.07E+00 | 2.35E-01 | 6.62E-01 | 1.15E+00 | 8.87E-02 | 3.26E-01 | -1.17E+00 | 5.33E-02 | 1.46E-01 | -1.30E+00 |
| AC073465.1 | 1.69E-01 | 8.83E-01 | 1.21E+00 | 7.70E-02 | 3.33E-01 | 1.86E+00 | 1.48E-02 | 6.27E-02 | 2.08E+00 | 1.41E-02 | 5.37E-02 | 2.27E+00 | 1.90E-02 | 6.76E-02 | -1.59E+00 | 2.56E-01 | 7.21E-01 | 1.19E+00 | 1.82E-01 | 6.68E-01 | 1.44E+00 | 3.38E-02 | 9.28E-02 | 1.74E+00 |
| EZR | 1.69E-01 | 8.83E-01 | 1.26E+00 | 1.04E-01 | 4.50E-01 | 1.24E+00 | 2.28E-01 | 9.69E-01 | 1.18E+00 | 2.13E-01 | 8.13E-01 | 1.24E+00 | 2.14E-01 | 7.64E-01 | 1.64E+00 | 2.43E-01 | 6.84E-01 | 1.38E+00 | 7.13E-03 | 2.62E-02 | -1.02E+00 | 9.10E-02 | 2.50E-01 | 1.05E+00 |
| ITLN1 | 1.70E-01 | 8.84E-01 | -1.07E+00 | 1.95E-01 | 8.42E-01 | -1.11E+00 | 2.35E-01 | 1.00E+00 | 1.00E+00 | 2.51E-01 | 9.58E-01 | 1.03E+00 | 2.17E-01 | 7.74E-01 | -1.03E+00 | 5.71E-02 | 1.61E-01 | 2.28E+00 | 2.20E-03 | 8.06E-03 | 1.62E+00 | 3.34E-01 | 9.18E-01 | 1.19E+00 |
| ESPL1 | 1.70E-01 | 8.84E-01 | -1.03E+00 | 2.31E-01 | 9.99E-01 | 1.11E+00 | 8.74E-02 | 3.71E-01 | 1.35E+00 | 1.01E-01 | 3.87E-01 | 1.56E+00 | 2.46E-01 | 8.77E-01 | 1.16E+00 | 2.94E-01 | 8.29E-01 | 1.23E+00 | 3.82E-02 | 1.40E-01 | -1.24E+00 | 2.17E-01 | 5.97E-01 | 1.25E+00 |
| ERCC6L | 1.70E-01 | 8.85E-01 | 1.18E+00 | 1.91E-01 | 8.27E-01 | 1.13E+00 | 6.84E-02 | 2.91E-01 | -1.15E+00 | 1.69E-01 | 6.45E-01 | 1.05E+00 | 2.51E-01 | 8.95E-01 | 1.31E+00 | 2.84E-01 | 8.01E-01 | 1.61E+00 | 1.25E-02 | 4.60E-02 | 1.02E+00 | 3.47E-01 | 9.54E-01 | 1.35E+00 |
| NT5C | 1.70E-01 | 8.86E-01 | 1.37E+00 | 9.40E-03 | 4.06E-02 | 1.30E+00 | 1.52E-01 | 6.46E-01 | 1.15E+00 | 4.25E-02 | 1.62E-01 | 1.19E+00 | 1.17E-01 | 4.19E-01 | 1.20E+00 | 3.06E-01 | 8.62E-01 | 1.29E+00 | 1.75E-01 | 6.42E-01 | 1.35E+00 | 2.21E-01 | 6.07E-01 | 1.17E+00 |
| RILPL1 | 1.70E-01 | 8.86E-01 | 1.20E+00 | 1.87E-01 | 8.07E-01 | 1.33E+00 | 1.87E-02 | 7.95E-02 | -1.37E+00 | 1.81E-01 | 6.92E-01 | 1.14E+00 | 8.11E-02 | 2.89E-01 | -1.04E+00 | 1.74E-01 | 4.91E-01 | 1.17E+00 | 1.66E-01 | 6.09E-01 | 1.39E+00 | 9.10E-02 | 2.50E-01 | 1.63E+00 |
| MYH11 | 1.70E-01 | 8.86E-01 | 1.19E+00 | 2.36E-02 | 1.02E-01 | -1.04E+00 | 1.15E-01 | 4.90E-01 | 1.19E+00 | 1.41E-02 | 5.38E-02 | -1.06E+00 | 6.57E-02 | 2.34E-01 | -1.20E+00 | 1.63E-01 | 4.59E-01 | 1.01E+00 | 1.47E-01 | 5.40E-01 | 1.00E+00 | 1.11E-01 | 3.06E-01 | -1.09E+00 |
| H2AFY | 1.70E-01 | 8.87E-01 | 1.03E+00 | 2.35E-02 | 1.02E-01 | 1.17E+00 | 9.33E-02 | 3.96E-01 | 1.08E+00 | 7.62E-02 | 2.91E-01 | 1.13E+00 | 5.22E-02 | 1.86E-01 | 1.02E+00 | 1.52E-01 | 4.29E-01 | 1.06E+00 | 1.80E-01 | 6.61E-01 | 1.06E+00 | 1.32E-01 | 3.62E-01 | 1.16E+00 |
| CFLAR-AS1 | 1.70E-01 | 8.87E-01 | 1.13E+00 | 2.18E-02 | 9.41E-02 | 2.15E+00 | 1.33E-01 | 5.65E-01 | -1.02E+00 | 2.22E-01 | 8.50E-01 | 1.22E+00 | 1.11E-01 | 3.96E-01 | -1.33E+00 | 3.13E-01 | 8.83E-01 | 1.21E+00 | 5.91E-02 | 2.17E-01 | -1.36E+00 | 3.38E-01 | 9.28E-01 | 1.15E+00 |
| RAB30-AS1 | 1.70E-01 | 8.88E-01 | 1.33E+00 | 2.70E-02 | 1.16E-01 | 1.26E+00 | 7.43E-02 | 3.16E-01 | 1.15E+00 | 2.53E-03 | 9.65E-03 | 1.03E+00 | 9.99E-02 | 3.56E-01 | 1.17E+00 | 2.92E-01 | 8.23E-01 | 1.35E+00 | 1.88E-01 | 6.91E-01 | 1.21E+00 | 2.05E-01 | 5.64E-01 | 1.09E+00 |
| AL136984.1 | 1.70E-01 | 8.88E-01 | 1.29E+00 | 2.21E-01 | 9.53E-01 | 1.36E+00 | 2.05E-03 | 8.70E-03 | -1.52E+00 | 1.47E-02 | 5.60E-02 | -1.14E+00 | 1.39E-01 | 4.97E-01 | 1.01E+00 | 3.14E-01 | 8.85E-01 | 1.41E+00 | 1.54E-01 | 5.64E-01 | 1.02E+00 | 2.92E-01 | 8.01E-01 | 1.11E+00 |
| CAVIN4 | 1.70E-01 | 8.88E-01 | 1.11E+00 | 2.09E-01 | 9.05E-01 | 1.16E+00 | 3.99E-02 | 1.70E-01 | 1.58E+00 | 2.61E-03 | 9.97E-03 | 2.25E+00 | 5.20E-02 | 1.85E-01 | -1.18E+00 | 1.29E-01 | 3.63E-01 | 1.02E+00 | 2.57E-02 | 9.45E-02 | -1.51E+00 | 1.35E-01 | 3.71E-01 | -1.08E+00 |
| FRG1JP | 1.70E-01 | 8.88E-01 | 1.40E+00 | 1.65E-02 | 7.14E-02 | 2.24E+00 | 1.05E-01 | 4.45E-01 | 1.43E+00 | 7.72E-03 | 2.95E-02 | 2.07E+00 | 1.21E-01 | 4.30E-01 | 1.17E+00 | 1.59E-02 | 4.49E-02 | 2.28E+00 | 6.81E-02 | 2.50E-01 | 1.54E+00 | 2.24E-03 | 6.15E-03 | 2.06E+00 |
| SLC29A4 | 1.70E-01 | 8.88E-01 | 1.21E+00 | 1.08E-01 | 4.67E-01 | 1.66E+00 | 1.84E-01 | 7.83E-01 | 1.14E+00 | 2.46E-01 | 9.41E-01 | 1.27E+00 | 2.08E-01 | 7.43E-01 | -1.59E+00 | 3.08E-01 | 8.68E-01 | -1.24E+00 | 1.76E-01 | 6.47E-01 | -1.15E+00 | 2.62E-01 | 7.19E-01 | 1.16E+00 |
| ACKR1 | 1.71E-01 | 8.89E-01 | -1.03E+00 | 1.73E-01 | 7.48E-01 | -1.11E+00 | 2.00E-01 | 8.52E-01 | -1.06E+00 | 1.81E-01 | 6.93E-01 | -1.17E+00 | 1.17E-01 | 4.18E-01 | -1.25E+00 | 2.12E-01 | 5.96E-01 | -1.02E+00 | 9.02E-04 | 3.31E-03 | -1.15E+00 | 1.20E-04 | 3.30E-04 | -1.31E+00 |
| ZNF367 | 1.71E-01 | 8.89E-01 | -1.05E+00 | 1.16E-01 | 5.00E-01 | -1.43E+00 | 9.46E-02 | 4.02E-01 | 1.36E+00 | 7.80E-02 | 2.98E-01 | 1.03E+00 | 1.98E-01 | 7.08E-01 | -1.26E+00 | 2.10E-01 | 5.91E-01 | -1.35E+00 | 2.16E-07 | 7.93E-07 | -1.41E+00 | 1.16E-03 | 3.19E-03 | -1.01E+00 |
| CDC6 | 1.71E-01 | 8.89E-01 | -1.01E+00 | 2.12E-01 | 9.18E-01 | 1.15E+00 | 3.22E-02 | 1.37E-01 | 1.37E+00 | 2.41E-01 | 9.21E-01 | 1.29E+00 | 7.00E-02 | 2.50E-01 | -2.73E+00 | 6.88E-02 | 1.94E-01 | -3.22E+00 | 6.48E-04 | 2.38E-03 | -1.22E+00 | 2.56E-01 | 7.04E-01 | 1.24E+00 |
| GRAP2 | 1.71E-01 | 8.89E-01 | -1.03E+00 | 1.93E-01 | 8.32E-01 | -1.18E+00 | 1.97E-01 | 8.39E-01 | 1.13E+00 | 2.46E-01 | 9.38E-01 | 1.35E+00 | 6.16E-24 | 2.19E-23 | -2.35E+00 | 4.77E-21 | 1.35E-20 | -1.74E+00 | 5.72E-05 | 2.10E-04 | 1.19E+00 | 2.99E-04 | 8.21E-04 | -1.09E+00 |
| LEF1 | 1.71E-01 | 8.90E-01 | -1.09E+00 | 1.78E-01 | 7.70E-01 | -1.21E+00 | 3.76E-02 | 1.60E-01 | -1.81E+00 | 5.38E-02 | 2.05E-01 | -1.68E+00 | 6.50E-02 | 2.32E-01 | -1.53E+00 | 2.70E-01 | 7.61E-01 | 1.54E+00 | 4.04E-19 | 1.48E-18 | -1.89E+00 | 2.09E-21 | 5.74E-21 | -1.88E+00 |
| DENND2C | 1.71E-01 | 8.90E-01 | 1.34E+00 | 1.81E-01 | 7.81E-01 | 1.29E+00 | 2.24E-02 | 9.53E-02 | -1.60E+00 | 5.29E-02 | 2.02E-01 | -1.22E+00 | 1.64E-04 | 5.83E-04 | -2.87E+00 | 1.62E-04 | 4.56E-04 | -2.69E+00 | 2.15E-02 | 7.90E-02 | 4.74E+00 | 3.60E-01 | 9.89E-01 | 2.02E+00 |
| AL928654.2 | 1.71E-01 | 8.91E-01 | 1.29E+00 | 1.97E-01 | 8.51E-01 | 1.31E+00 | 2.72E-02 | 1.15E-01 | -1.06E+00 | 7.78E-02 | 2.97E-01 | 1.16E+00 | 1.62E-01 | 5.77E-01 | 1.13E+00 | 3.05E-01 | 8.59E-01 | 1.46E+00 | 2.58E-03 | 9.47E-03 | -1.37E+00 | 2.46E-01 | 6.76E-01 | 1.22E+00 |
| C4orf19 | 1.71E-01 | 8.92E-01 | 1.27E+00 | 1.47E-01 | 6.36E-01 | 1.31E+00 | 1.22E-02 | 5.18E-02 | -1.08E+00 | 7.97E-04 | 3.05E-03 | -1.04E+00 | 1.84E-01 | 6.55E-01 | 1.73E+00 | 3.22E-01 | 9.07E-01 | 1.42E+00 | 1.44E-01 | 5.30E-01 | -1.20E+00 | 2.30E-01 | 6.31E-01 | -1.08E+00 |
| KDELR3 | 1.71E-01 | 8.92E-01 | 1.24E+00 | 2.01E-01 | 8.70E-01 | 1.29E+00 | 8.22E-02 | 3.49E-01 | -1.07E+00 | 2.32E-01 | 8.86E-01 | 1.32E+00 | 1.82E-01 | 6.48E-01 | -1.11E+00 | 3.44E-01 | 9.71E-01 | 1.29E+00 | 2.51E-01 | 9.23E-01 | 1.21E+00 | 2.52E-01 | 6.92E-01 | 1.35E+00 |
| AC016925.1 | 1.71E-01 | 8.92E-01 | 1.41E+00 | 6.34E-02 | 2.74E-01 | 1.09E+00 | 1.22E-01 | 5.18E-01 | 1.24E+00 | 4.63E-06 | 1.77E-05 | -1.26E+00 | 1.45E-01 | 5.18E-01 | 1.13E+00 | 1.25E-02 | 3.52E-02 | -1.43E+00 | 1.74E-02 | 6.40E-02 | 1.43E+00 | 1.91E-01 | 5.24E-01 | 1.33E+00 |
| FAM72D | 1.71E-01 | 8.92E-01 | 1.28E+00 | 2.02E-01 | 8.73E-01 | 1.34E+00 | 1.04E-01 | 4.42E-01 | -1.15E+00 | 1.86E-01 | 7.10E-01 | 1.12E+00 | 2.70E-01 | 9.63E-01 | 1.18E+00 | 2.66E-01 | 7.49E-01 | -1.11E+00 | 5.28E-04 | 1.94E-03 | -1.28E+00 | 2.77E-01 | 7.60E-01 | 1.11E+00 |
| RNASE3 | 1.71E-01 | 8.93E-01 | 1.24E+00 | 2.02E-01 | 8.74E-01 | -1.05E+00 | 3.59E-02 | 1.52E-01 | -1.10E+00 | 3.38E-03 | 1.29E-02 | -1.15E+00 | 2.54E-01 | 9.06E-01 | 1.23E+00 | 3.55E-01 | 1.00E+00 | -1.00E+00 | 2.40E-01 | 8.81E-01 | 1.10E+00 | 3.06E-01 | 8.41E-01 | -1.13E+00 |
| QPCT | 1.71E-01 | 8.93E-01 | 1.37E+00 | 1.60E-01 | 6.90E-01 | 1.65E+00 | 2.22E-01 | 9.44E-01 | 1.20E+00 | 1.66E-01 | 6.34E-01 | 1.72E+00 | 2.46E-01 | 8.77E-01 | 1.53E+00 | 1.45E-01 | 4.08E-01 | 1.54E+00 | 6.54E-02 | 2.40E-01 | 1.37E+00 | 7.83E-02 | 2.15E-01 | 1.09E+00 |
| EVA1C | 1.71E-01 | 8.93E-01 | 1.36E+00 | 1.12E-01 | 4.84E-01 | 1.72E+00 | 9.22E-02 | 3.92E-01 | 1.52E+00 | 2.83E-02 | 1.08E-01 | 1.94E+00 | 3.73E-02 | 1.33E-01 | 2.24E+00 | 1.64E-02 | 4.63E-02 | 2.67E+00 | 1.30E-04 | 4.76E-04 | 2.00E+00 | 1.21E-03 | 3.31E-03 | 1.89E+00 |
| KIFC1 | 1.71E-01 | 8.93E-01 | -1.04E+00 | 1.78E-01 | 7.69E-01 | -1.11E+00 | 1.69E-01 | 7.18E-01 | 1.45E+00 | 2.61E-01 | 9.96E-01 | 1.36E+00 | 1.25E-02 | 4.47E-02 | -3.02E+00 | 2.00E-02 | 5.64E-02 | -2.74E+00 | 3.53E-04 | 1.30E-03 | -1.72E+00 | 2.06E-01 | 5.66E-01 | -1.16E+00 |
| POLA2 | 1.71E-01 | 8.93E-01 | 1.30E+00 | 1.69E-01 | 7.30E-01 | 1.75E+00 | 2.27E-01 | 9.66E-01 | 1.25E+00 | 2.36E-01 | 9.00E-01 | 1.40E+00 | 7.94E-03 | 2.83E-02 | -1.69E+00 | 8.38E-02 | 2.36E-01 | -1.03E+00 | 4.76E-03 | 1.75E-02 | -1.00E+00 | 2.11E-01 | 5.80E-01 | 1.40E+00 |
| FCGR3A | 1.71E-01 | 8.93E-01 | 1.11E+00 | 1.91E-01 | 8.25E-01 | 3.26E+00 | 1.85E-01 | 7.85E-01 | 1.23E+00 | 2.62E-01 | 1.00E+00 | 1.00E+00 | 9.33E-02 | 3.32E-01 | 1.70E+00 | 3.24E-01 | 9.12E-01 | 1.08E+00 | 2.99E-02 | 1.10E-01 | 1.62E+00 | 3.57E-01 | 9.80E-01 | 1.01E+00 |
| KRT1 | 1.72E-01 | 8.94E-01 | 1.09E+00 | 2.31E-01 | 1.00E+00 | 1.00E+00 | 1.90E-01 | 8.07E-01 | 1.23E+00 | 2.30E-01 | 8.78E-01 | 1.12E+00 | 1.96E-43 | 6.99E-43 | -5.86E+00 | 3.43E-46 | 9.66E-46 | -4.80E+00 | 2.63E-07 | 9.66E-07 | -3.14E+00 | 8.42E-07 | 2.31E-06 | -2.50E+00 |
| ALAS2 | 1.72E-01 | 8.95E-01 | 1.41E+00 | 1.84E-01 | 7.93E-01 | 1.67E+00 | 1.89E-01 | 8.03E-01 | -1.16E+00 | 1.89E-01 | 7.23E-01 | -1.27E+00 | 2.59E-01 | 9.25E-01 | 1.82E+00 | 2.67E-01 | 7.51E-01 | 2.79E+00 | 9.30E-11 | 3.41E-10 | -1.08E+00 | 2.45E-07 | 6.73E-07 | -1.00E+00 |
| ATG9B | 1.72E-01 | 8.95E-01 | 1.24E+00 | 8.24E-02 | 3.56E-01 | -1.02E+00 | 3.03E-03 | 1.29E-02 | 1.97E+00 | 2.37E-01 | 9.06E-01 | 1.22E+00 | 1.83E-01 | 6.51E-01 | -1.26E+00 | 2.35E-01 | 6.63E-01 | -1.33E+00 | 2.57E-01 | 9.43E-01 | 1.18E+00 | 3.57E-01 | 9.80E-01 | 1.16E+00 |
| RPL39 | 1.72E-01 | 8.95E-01 | 1.01E+00 | 9.74E-16 | 4.21E-15 | -1.27E+00 | 8.04E-02 | 3.42E-01 | 1.05E+00 | 4.16E-07 | 1.59E-06 | -1.20E+00 | 1.03E-01 | 3.67E-01 | 1.05E+00 | 1.38E-06 | 3.89E-06 | -1.22E+00 | 4.93E-03 | 1.81E-02 | -1.01E+00 | 1.79E-07 | 4.90E-07 | -1.23E+00 |
| SERPINF2 | 1.72E-01 | 8.95E-01 | 1.05E+00 | 1.84E-01 | 7.95E-01 | 1.36E+00 | 2.23E-01 | 9.47E-01 | 1.20E+00 | 2.11E-01 | 8.07E-01 | 1.25E+00 | 2.20E-02 | 7.85E-02 | -1.40E+00 | 3.49E-02 | 9.83E-02 | -1.28E+00 | 2.98E-02 | 1.10E-01 | -1.73E+00 | 1.06E-01 | 2.90E-01 | -1.28E+00 |
| AMDHD2 | 1.72E-01 | 8.95E-01 | 1.27E+00 | 1.23E-01 | 5.34E-01 | -1.01E+00 | 1.41E-01 | 5.98E-01 | 1.37E+00 | 1.32E-01 | 5.03E-01 | 1.14E+00 | 4.77E-02 | 1.70E-01 | -1.14E+00 | 3.02E-02 | 8.52E-02 | -1.37E+00 | 1.20E-03 | 4.41E-03 | -1.17E+00 | 3.86E-02 | 1.06E-01 | 1.09E+00 |
| SGK494 | 1.72E-01 | 8.95E-01 | 1.18E+00 | 4.18E-02 | 1.80E-01 | 1.95E+00 | 1.10E-05 | 4.65E-05 | -1.61E+00 | 2.57E-05 | 9.82E-05 | -1.39E+00 | 1.27E-01 | 4.52E-01 | 1.04E+00 | 2.77E-01 | 7.81E-01 | 1.22E+00 | 7.43E-04 | 2.73E-03 | -1.72E+00 | 9.99E-03 | 2.74E-02 | -1.41E+00 |
| MN1 | 1.72E-01 | 8.96E-01 | 1.18E+00 | 1.01E-01 | 4.38E-01 | 1.58E+00 | 4.55E-05 | 1.93E-04 | -1.39E+00 | 3.20E-03 | 1.22E-02 | 1.03E+00 | 2.60E-01 | 9.28E-01 | 1.22E+00 | 3.37E-01 | 9.49E-01 | 1.39E+00 | 1.94E-01 | 7.12E-01 | 1.36E+00 | 1.53E-01 | 4.19E-01 | 2.08E+00 |
| CEP55 | 1.72E-01 | 8.96E-01 | 1.14E+00 | 1.93E-01 | 8.36E-01 | 1.24E+00 | 1.95E-01 | 8.29E-01 | 1.09E+00 | 1.36E-01 | 5.21E-01 | 1.08E+00 | 2.46E-01 | 8.78E-01 | -1.07E+00 | 3.10E-01 | 8.75E-01 | -1.01E+00 | 3.58E-02 | 1.32E-01 | -1.19E+00 | 3.58E-01 | 9.84E-01 | -1.04E+00 |
| ASPN | 1.72E-01 | 8.96E-01 | 1.14E+00 | 2.00E-01 | 8.62E-01 | 1.26E+00 | 5.71E-02 | 2.42E-01 | 1.58E+00 | 4.94E-02 | 1.89E-01 | 1.88E+00 | 1.58E-01 | 5.63E-01 | 1.05E+00 | 2.73E-01 | 7.70E-01 | 1.04E+00 | 9.34E-02 | 3.43E-01 | -1.01E+00 | 1.33E-01 | 3.66E-01 | 1.57E+00 |
| ABCB8 | 1.72E-01 | 8.96E-01 | 1.22E+00 | 1.85E-01 | 7.99E-01 | 1.32E+00 | 1.93E-01 | 8.19E-01 | 1.18E+00 | 5.33E-02 | 2.03E-01 | 1.08E+00 | 1.36E-01 | 4.87E-01 | 1.58E+00 | 2.89E-01 | 8.14E-01 | 1.52E+00 | 9.72E-03 | 3.57E-02 | 1.02E+00 | 1.29E-01 | 3.56E-01 | 1.21E+00 |
| FBRSL1 | 1.72E-01 | 8.97E-01 | 1.25E+00 | 1.20E-01 | 5.19E-01 | 1.33E+00 | 8.95E-02 | 3.80E-01 | 1.37E+00 | 2.21E-02 | 8.43E-02 | 1.74E+00 | 1.96E-01 | 7.00E-01 | 1.37E+00 | 2.39E-01 | 6.73E-01 | 1.38E+00 | 2.33E-02 | 8.54E-02 | 1.65E+00 | 1.44E-03 | 3.96E-03 | 1.87E+00 |
| XRCC3 | 1.72E-01 | 8.97E-01 | 1.12E+00 | 2.25E-01 | 9.72E-01 | 1.30E+00 | 1.72E-01 | 7.31E-01 | 1.35E+00 | 2.01E-01 | 7.69E-01 | 1.31E+00 | 1.40E-01 | 4.98E-01 | 1.04E+00 | 1.44E-01 | 4.07E-01 | -1.05E+00 | 1.78E-01 | 6.51E-01 | 1.48E+00 | 1.52E-01 | 4.18E-01 | 1.51E+00 |
| LINC01011 | 1.72E-01 | 8.97E-01 | 1.27E+00 | 9.53E-03 | 4.12E-02 | -1.08E+00 | 2.03E-01 | 8.61E-01 | 1.28E+00 | 5.06E-02 | 1.93E-01 | -1.16E+00 | 7.82E-03 | 2.79E-02 | -1.30E+00 | 1.31E-02 | 3.69E-02 | -1.29E+00 | 3.60E-03 | 1.32E-02 | -1.15E+00 | 1.11E-03 | 3.04E-03 | -1.38E+00 |
| PPCDC | 1.72E-01 | 8.97E-01 | 1.26E+00 | 1.93E-01 | 8.33E-01 | 1.41E+00 | 1.52E-01 | 6.45E-01 | 1.32E+00 | 3.79E-02 | 1.45E-01 | 1.09E+00 | 8.72E-02 | 3.11E-01 | 1.08E+00 | 3.96E-03 | 1.12E-02 | -1.33E+00 | 5.74E-02 | 2.11E-01 | 1.18E+00 | 4.18E-02 | 1.15E-01 | 1.01E+00 |
| TEDC1 | 1.72E-01 | 8.97E-01 | 1.36E+00 | 1.23E-01 | 5.34E-01 | 1.17E+00 | 5.89E-02 | 2.50E-01 | 1.37E+00 | 2.50E-01 | 9.54E-01 | 1.35E+00 | 5.80E-02 | 2.07E-01 | -1.26E+00 | 2.47E-01 | 6.95E-01 | 1.19E+00 | 1.67E-01 | 6.12E-01 | 1.09E+00 | 2.50E-01 | 6.88E-01 | 1.15E+00 |
| ITGAM | 1.72E-01 | 8.98E-01 | 1.22E+00 | 1.98E-01 | 8.56E-01 | 1.05E+00 | 7.95E-03 | 3.38E-02 | -1.59E+00 | 2.49E-02 | 9.53E-02 | -1.24E+00 | 1.00E-02 | 3.58E-02 | -1.78E+00 | 6.14E-02 | 1.73E-01 | -1.32E+00 | 1.03E-18 | 3.79E-18 | -1.92E+00 | 2.64E-24 | 7.26E-24 | -2.33E+00 |
| ZNF805 | 1.72E-01 | 8.99E-01 | 1.41E+00 | 2.74E-03 | 1.18E-02 | -1.14E+00 | 2.03E-01 | 8.63E-01 | 1.19E+00 | 3.15E-03 | 1.20E-02 | -1.43E+00 | 2.27E-01 | 8.10E-01 | 1.58E+00 | 1.81E-01 | 5.11E-01 | -1.02E+00 | 1.67E-02 | 6.15E-02 | 1.98E+00 | 3.25E-01 | 8.94E-01 | 1.23E+00 |
| RPL38P1 | 1.72E-01 | 8.99E-01 | -1.06E+00 | 1.99E-01 | 8.60E-01 | -1.12E+00 | 2.05E-01 | 8.71E-01 | 1.12E+00 | 2.62E-01 | 1.00E+00 | 1.00E+00 | 7.70E-02 | 2.74E-01 | 1.78E+00 | 6.38E-06 | 1.80E-05 | 4.00E+00 | 1.24E-01 | 4.55E-01 | 1.17E+00 | 1.21E-06 | 3.31E-06 | 1.95E+00 |
| C20orf203 | 1.72E-01 | 8.99E-01 | 1.24E+00 | 1.76E-01 | 7.59E-01 | 1.05E+00 | 2.89E-02 | 1.23E-01 | -1.58E+00 | 5.09E-02 | 1.95E-01 | -1.38E+00 | 2.26E-02 | 8.04E-02 | -1.29E+00 | 2.34E-01 | 6.60E-01 | 1.14E+00 | 1.58E-01 | 5.78E-01 | 1.07E+00 | 3.10E-01 | 8.52E-01 | 1.16E+00 |
| AP000845.1 | 1.73E-01 | 9.00E-01 | 1.41E+00 | 1.13E-03 | 4.87E-03 | -1.17E+00 | 1.82E-01 | 7.73E-01 | 1.16E+00 | 8.35E-03 | 3.19E-02 | -1.20E+00 | 7.90E-02 | 2.82E-01 | -1.12E+00 | 8.14E-02 | 2.29E-01 | -1.11E+00 | 1.91E-01 | 7.01E-01 | -1.04E+00 | 3.34E-01 | 9.18E-01 | 1.08E+00 |
| HTR7P1 | 1.73E-01 | 9.00E-01 | 1.16E+00 | 6.60E-02 | 2.85E-01 | -1.05E+00 | 6.45E-03 | 2.74E-02 | -1.23E+00 | 3.37E-06 | 1.29E-05 | -1.96E+00 | 2.78E-01 | 9.91E-01 | 1.26E+00 | 3.03E-01 | 8.55E-01 | 1.50E+00 | 2.17E-01 | 7.96E-01 | 1.01E+00 | 3.58E-01 | 9.83E-01 | 1.06E+00 |
| CYP51A1P1 | 1.73E-01 | 9.00E-01 | 1.39E+00 | 2.14E-01 | 9.25E-01 | 1.49E+00 | 1.29E-01 | 5.49E-01 | 1.10E+00 | 5.64E-02 | 2.15E-01 | 1.94E+00 | 1.24E-01 | 4.43E-01 | 1.48E+00 | 9.26E-02 | 2.61E-01 | 1.73E+00 | 1.52E-01 | 5.58E-01 | 1.45E+00 | 1.72E-01 | 4.71E-01 | 1.54E+00 |
| ANAPC16 | 1.73E-01 | 9.01E-01 | 1.16E+00 | 8.95E-02 | 3.87E-01 | 1.39E+00 | 1.94E-01 | 8.25E-01 | 1.20E+00 | 2.11E-01 | 8.08E-01 | 1.38E+00 | 1.12E-01 | 4.01E-01 | 1.32E+00 | 2.43E-02 | 6.84E-02 | 1.43E+00 | 1.56E-01 | 5.72E-01 | 1.29E+00 | 3.25E-01 | 8.92E-01 | 1.29E+00 |
| SERBP1P1 | 1.73E-01 | 9.01E-01 | 1.27E+00 | 8.16E-04 | 3.53E-03 | 2.15E+00 | 5.00E-04 | 2.12E-03 | 1.72E+00 | 5.43E-11 | 2.07E-10 | 2.69E+00 | 4.15E-02 | 1.48E-01 | -1.08E+00 | 1.15E-03 | 3.24E-03 | 1.86E+00 | 1.13E-01 | 4.14E-01 | 1.36E+00 | 2.11E-11 | 5.80E-11 | 2.35E+00 |
| RPL21P135 | 1.73E-01 | 9.02E-01 | 1.29E+00 | 1.96E-01 | 8.46E-01 | 1.59E+00 | 1.87E-01 | 7.93E-01 | 1.30E+00 | 1.01E-01 | 3.84E-01 | 1.73E+00 | 1.36E-01 | 4.86E-01 | 1.07E+00 | 2.86E-01 | 8.07E-01 | 1.45E+00 | 7.51E-02 | 2.76E-01 | -1.17E+00 | 3.61E-01 | 9.92E-01 | 1.36E+00 |
| NOSTRIN | 1.73E-01 | 9.02E-01 | -1.04E+00 | 2.05E-01 | 8.85E-01 | 1.13E+00 | 9.29E-02 | 3.95E-01 | 1.46E+00 | 6.85E-02 | 2.62E-01 | 1.68E+00 | 8.11E-05 | 2.89E-04 | -1.70E+00 | 2.06E-05 | 5.81E-05 | -1.86E+00 | 3.19E-03 | 1.17E-02 | -1.40E+00 | 1.15E-02 | 3.17E-02 | -1.25E+00 |
| RPL14P3 | 1.73E-01 | 9.03E-01 | 1.45E+00 | 1.38E-04 | 5.96E-04 | 2.16E+00 | 9.54E-02 | 4.06E-01 | 1.18E+00 | 1.63E-03 | 6.23E-03 | 1.94E+00 | 1.84E-01 | 6.55E-01 | 1.33E+00 | 8.68E-04 | 2.45E-03 | 2.23E+00 | 1.80E-01 | 6.61E-01 | 1.31E+00 | 3.36E-06 | 9.23E-06 | 2.00E+00 |
| SEC14L2 | 1.73E-01 | 9.03E-01 | 1.01E+00 | 1.90E-01 | 8.20E-01 | 1.14E+00 | 2.27E-01 | 9.67E-01 | -1.03E+00 | 2.41E-01 | 9.19E-01 | -1.11E+00 | 2.74E-01 | 9.77E-01 | 1.22E+00 | 1.38E-01 | 3.89E-01 | 1.85E+00 | 2.71E-02 | 9.95E-02 | 1.56E+00 | 8.68E-03 | 2.39E-02 | 1.60E+00 |
| RCAN3 | 1.73E-01 | 9.03E-01 | 1.38E+00 | 1.49E-02 | 6.42E-02 | 1.13E+00 | 4.22E-03 | 1.79E-02 | -1.12E+00 | 2.86E-04 | 1.09E-03 | -1.11E+00 | 2.71E-01 | 9.65E-01 | 1.60E+00 | 6.24E-02 | 1.76E-01 | 1.11E+00 | 1.75E-01 | 6.41E-01 | 1.60E+00 | 3.07E-01 | 8.43E-01 | 1.32E+00 |
| GPD2 | 1.73E-01 | 9.03E-01 | 1.45E+00 | 1.11E-02 | 4.81E-02 | -1.05E+00 | 2.71E-05 | 1.15E-04 | -1.53E+00 | 3.43E-05 | 1.31E-04 | -1.31E+00 | 1.30E-01 | 4.64E-01 | 1.20E+00 | 7.61E-02 | 2.15E-01 | 1.12E+00 | 1.17E-04 | 4.29E-04 | -1.13E+00 | 9.14E-04 | 2.51E-03 | -1.10E+00 |
| CTSV | 1.74E-01 | 9.04E-01 | 1.06E+00 | 1.85E-01 | 7.97E-01 | 1.13E+00 | 1.79E-01 | 7.61E-01 | 1.29E+00 | 2.10E-01 | 8.01E-01 | 1.11E+00 | 2.80E-01 | 1.00E+00 | 1.00E+00 | 2.81E-01 | 7.93E-01 | 1.23E+00 | 5.99E-02 | 2.20E-01 | 1.34E+00 | 1.02E-01 | 2.81E-01 | 1.55E+00 |
| NMB | 1.74E-01 | 9.04E-01 | 1.18E+00 | 1.53E-01 | 6.61E-01 | 1.12E+00 | 2.27E-01 | 9.65E-01 | 1.24E+00 | 9.88E-02 | 3.78E-01 | 1.22E+00 | 5.22E-04 | 1.86E-03 | -1.22E+00 | 4.23E-04 | 1.19E-03 | -1.19E+00 | 2.59E-01 | 9.52E-01 | 1.09E+00 | 2.15E-01 | 5.91E-01 | 1.29E+00 |
| AL096869.2 | 1.74E-01 | 9.04E-01 | 1.19E+00 | 6.20E-02 | 2.68E-01 | 1.90E+00 | 4.40E-03 | 1.87E-02 | -1.49E+00 | 2.21E-02 | 8.44E-02 | -1.15E+00 | 1.33E-01 | 4.76E-01 | 1.12E+00 | 2.54E-01 | 7.17E-01 | 1.27E+00 | 2.24E-01 | 8.20E-01 | 1.32E+00 | 2.67E-01 | 7.34E-01 | 1.32E+00 |
| AC012368.1 | 1.74E-01 | 9.04E-01 | 1.20E+00 | 9.17E-02 | 3.96E-01 | -1.03E+00 | 8.88E-02 | 3.77E-01 | -1.02E+00 | 2.72E-02 | 1.04E-01 | -1.06E+00 | 1.88E-01 | 6.69E-01 | 1.55E+00 | 2.81E-01 | 7.93E-01 | 1.34E+00 | 2.57E-01 | 9.44E-01 | 1.18E+00 | 8.61E-02 | 2.37E-01 | -1.25E+00 |
| POC1B-AS1 | 1.74E-01 | 9.04E-01 | 1.16E+00 | 2.03E-01 | 8.78E-01 | 1.32E+00 | 3.63E-02 | 1.54E-01 | 1.61E+00 | 2.24E-01 | 8.55E-01 | 1.34E+00 | 2.11E-01 | 7.51E-01 | 1.20E+00 | 9.16E-02 | 2.58E-01 | -1.02E+00 | 6.55E-02 | 2.40E-01 | 1.54E+00 | 3.49E-01 | 9.58E-01 | 1.16E+00 |
| GNPDA1 | 1.74E-01 | 9.05E-01 | 1.18E+00 | 1.83E-01 | 7.89E-01 | 1.23E+00 | 4.53E-04 | 1.92E-03 | -1.60E+00 | 1.02E-02 | 3.90E-02 | -1.05E+00 | 2.31E-01 | 8.23E-01 | -1.15E+00 | 3.31E-01 | 9.34E-01 | 1.19E+00 | 4.06E-02 | 1.49E-01 | -1.51E+00 | 5.88E-02 | 1.61E-01 | -1.50E+00 |
| HSPA2 | 1.74E-01 | 9.06E-01 | 1.40E+00 | 9.38E-02 | 4.05E-01 | 1.16E+00 | 1.19E-02 | 5.07E-02 | -1.26E+00 | 1.02E-01 | 3.91E-01 | 1.14E+00 | 2.31E-01 | 8.22E-01 | 1.45E+00 | 3.08E-01 | 8.68E-01 | 1.38E+00 | 2.62E-01 | 9.61E-01 | 1.29E+00 | 4.89E-02 | 1.34E-01 | 1.55E+00 |
| AC009309.1 | 1.74E-01 | 9.06E-01 | 1.26E+00 | 5.86E-02 | 2.53E-01 | -1.03E+00 | 1.51E-01 | 6.40E-01 | -1.07E+00 | 1.56E-01 | 5.97E-01 | 1.01E+00 | 3.68E-02 | 1.31E-01 | -1.16E+00 | 3.03E-02 | 8.54E-02 | -1.24E+00 | 2.30E-01 | 8.42E-01 | 1.37E+00 | 3.49E-01 | 9.58E-01 | 1.25E+00 |
| ABI3 | 1.74E-01 | 9.07E-01 | 1.06E+00 | 1.58E-01 | 6.83E-01 | 1.24E+00 | 1.94E-01 | 8.26E-01 | -1.03E+00 | 8.96E-02 | 3.42E-01 | -1.38E+00 | 2.44E-01 | 8.71E-01 | 1.10E+00 | 3.55E-01 | 1.00E+00 | 1.00E+00 | 1.35E-01 | 4.95E-01 | -1.25E+00 | 3.35E-01 | 9.21E-01 | 1.04E+00 |
| Z99129.4 | 1.74E-01 | 9.07E-01 | 1.34E+00 | 7.65E-02 | 3.30E-01 | 1.79E+00 | 2.34E-01 | 9.96E-01 | 1.17E+00 | 4.77E-02 | 1.82E-01 | 1.73E+00 | 1.65E-01 | 5.88E-01 | 1.35E+00 | 1.61E-01 | 4.54E-01 | 1.13E+00 | 2.52E-01 | 9.23E-01 | 1.34E+00 | 1.62E-01 | 4.45E-01 | 1.58E+00 |
| HMGN3 | 1.74E-01 | 9.07E-01 | 1.18E+00 | 4.60E-02 | 1.99E-01 | 1.37E+00 | 5.58E-02 | 2.37E-01 | 1.21E+00 | 3.85E-02 | 1.47E-01 | 1.28E+00 | 3.66E-04 | 1.30E-03 | -1.01E+00 | 5.40E-02 | 1.52E-01 | 1.15E+00 | 1.47E-01 | 5.40E-01 | 1.05E+00 | 2.47E-01 | 6.79E-01 | 1.13E+00 |
| GZMA | 1.74E-01 | 9.07E-01 | 1.06E+00 | 2.10E-01 | 9.08E-01 | 1.06E+00 | 2.35E-01 | 9.99E-01 | 1.23E+00 | 1.74E-01 | 6.66E-01 | -1.14E+00 | 2.43E-01 | 8.66E-01 | 1.13E+00 | 3.55E-01 | 1.00E+00 | 1.00E+00 | 9.66E-03 | 3.55E-02 | 6.99E+00 | 1.03E-02 | 2.83E-02 | 8.21E+00 |
| UPF3AP3 | 1.74E-01 | 9.08E-01 | 1.27E+00 | 1.10E-01 | 4.76E-01 | 1.19E+00 | 1.94E-02 | 8.22E-02 | -1.08E+00 | 5.07E-07 | 1.94E-06 | -2.26E+00 | 1.14E-01 | 4.05E-01 | -1.07E+00 | 6.86E-03 | 1.93E-02 | -1.78E+00 | 6.31E-02 | 2.32E-01 | 1.72E+00 | 1.99E-01 | 5.47E-01 | -1.04E+00 |
| LINC01355 | 1.74E-01 | 9.08E-01 | 1.20E+00 | 9.10E-08 | 3.93E-07 | 4.29E+00 | 1.77E-01 | 7.53E-01 | -1.10E+00 | 6.88E-03 | 2.63E-02 | 2.27E+00 | 2.57E-01 | 9.18E-01 | 1.25E+00 | 4.22E-03 | 1.19E-02 | 4.07E+00 | 2.06E-01 | 7.56E-01 | -1.10E+00 | 6.59E-03 | 1.81E-02 | 2.14E+00 |
| GRAMD4 | 1.74E-01 | 9.08E-01 | 1.46E+00 | 5.43E-03 | 2.35E-02 | 1.08E+00 | 1.81E-01 | 7.67E-01 | 1.25E+00 | 1.79E-01 | 6.82E-01 | 1.29E+00 | 2.45E-01 | 8.73E-01 | 1.50E+00 | 3.32E-01 | 9.37E-01 | 1.49E+00 | 6.52E-03 | 2.39E-02 | -1.03E+00 | 1.62E-01 | 4.46E-01 | 1.21E+00 |
| AC006033.1 | 1.74E-01 | 9.08E-01 | 1.34E+00 | 2.21E-01 | 9.53E-01 | 1.40E+00 | 2.77E-02 | 1.18E-01 | -1.14E+00 | 2.41E-02 | 9.19E-02 | -1.11E+00 | 5.97E-02 | 2.13E-01 | -1.08E+00 | 3.53E-02 | 9.94E-02 | -1.22E+00 | 9.26E-02 | 3.40E-01 | 1.62E+00 | 3.54E-01 | 9.73E-01 | 1.18E+00 |
| UBA52 | 1.74E-01 | 9.08E-01 | 1.01E+00 | 2.10E-16 | 9.06E-16 | -1.18E+00 | 2.21E-01 | 9.38E-01 | 1.01E+00 | 3.12E-09 | 1.19E-08 | -1.17E+00 | 9.27E-02 | 3.31E-01 | -1.01E+00 | 2.32E-07 | 6.54E-07 | -1.16E+00 | 4.85E-08 | 1.78E-07 | -1.09E+00 | 6.66E-14 | 1.83E-13 | -1.22E+00 |
| PPP1R12C | 1.74E-01 | 9.08E-01 | 1.39E+00 | 1.94E-01 | 8.39E-01 | 1.62E+00 | 1.14E-01 | 4.86E-01 | 1.08E+00 | 1.37E-01 | 5.22E-01 | 1.27E+00 | 2.37E-01 | 8.46E-01 | 1.32E+00 | 7.93E-02 | 2.24E-01 | 1.77E+00 | 1.67E-02 | 6.11E-02 | 1.98E+00 | 7.19E-02 | 1.97E-01 | 1.90E+00 |
| NT5DC2 | 1.74E-01 | 9.08E-01 | 1.26E+00 | 2.20E-01 | 9.50E-01 | 1.37E+00 | 4.07E-04 | 1.73E-03 | 1.66E+00 | 1.77E-01 | 6.76E-01 | 1.34E+00 | 1.60E-02 | 5.71E-02 | -1.22E+00 | 3.96E-02 | 1.12E-01 | -1.05E+00 | 3.85E-02 | 1.41E-01 | 1.53E+00 | 2.43E-03 | 6.68E-03 | 1.75E+00 |
| AF274858.3 | 1.74E-01 | 9.08E-01 | 1.27E+00 | 8.68E-02 | 3.75E-01 | 1.21E+00 | 1.91E-01 | 8.11E-01 | 1.26E+00 | 4.09E-02 | 1.56E-01 | -1.08E+00 | 2.22E-01 | 7.90E-01 | 1.22E+00 | 1.59E-01 | 4.47E-01 | 1.00E+00 | 8.22E-02 | 3.02E-01 | 1.65E+00 | 1.21E-01 | 3.33E-01 | -1.11E+00 |
| SLC35G2 | 1.74E-01 | 9.09E-01 | 1.44E+00 | 7.11E-02 | 3.07E-01 | 1.24E+00 | 1.47E-01 | 6.23E-01 | 1.01E+00 | 2.33E-02 | 8.89E-02 | -1.14E+00 | 4.19E-02 | 1.49E-01 | 1.05E+00 | 3.55E-02 | 1.00E-01 | 1.00E+00 | 1.71E-01 | 6.27E-01 | 1.41E+00 | 2.69E-01 | 7.39E-01 | 1.50E+00 |
| LINC00863 | 1.74E-01 | 9.09E-01 | 1.42E+00 | 1.60E-01 | 6.89E-01 | 1.69E+00 | 1.61E-01 | 6.85E-01 | 1.40E+00 | 8.96E-02 | 3.42E-01 | 1.73E+00 | 6.52E-02 | 2.32E-01 | 1.29E+00 | 6.28E-02 | 1.77E-01 | 1.26E+00 | 6.88E-02 | 2.52E-01 | 1.49E+00 | 1.08E-02 | 2.96E-02 | 1.64E+00 |
| INPP5A | 1.75E-01 | 9.10E-01 | 1.18E+00 | 1.26E-01 | 5.43E-01 | 1.10E+00 | 6.95E-02 | 2.95E-01 | -1.07E+00 | 5.33E-02 | 2.04E-01 | 1.02E+00 | 5.08E-04 | 1.81E-03 | -1.61E+00 | 4.78E-04 | 1.35E-03 | -1.53E+00 | 1.74E-01 | 6.40E-01 | 1.21E+00 | 1.80E-01 | 4.94E-01 | 1.18E+00 |
| LINC00893 | 1.75E-01 | 9.10E-01 | 1.25E+00 | 1.30E-01 | 5.62E-01 | 1.69E+00 | 1.52E-01 | 6.47E-01 | 1.15E+00 | 1.28E-01 | 4.89E-01 | 1.14E+00 | 2.10E-01 | 7.50E-01 | 1.55E+00 | 1.50E-01 | 4.22E-01 | 1.93E+00 | 2.01E-01 | 7.39E-01 | 1.31E+00 | 3.25E-01 | 8.93E-01 | 1.09E+00 |
| CDC20P1 | 1.75E-01 | 9.11E-01 | 1.05E+00 | 2.08E-01 | 8.97E-01 | 1.09E+00 | 4.35E-02 | 1.85E-01 | 1.52E+00 | 1.29E-01 | 4.92E-01 | 1.30E+00 | 2.42E-01 | 8.64E-01 | 1.14E+00 | 2.81E-01 | 7.93E-01 | 1.27E+00 | 2.56E-01 | 9.41E-01 | 1.13E+00 | 1.92E-01 | 5.28E-01 | 1.31E+00 |
| CDR2 | 1.75E-01 | 9.11E-01 | 1.17E+00 | 2.26E-01 | 9.76E-01 | 1.49E+00 | 7.24E-02 | 3.08E-01 | -1.06E+00 | 5.76E-02 | 2.20E-01 | 1.06E+00 | 1.82E-01 | 6.48E-01 | 1.33E+00 | 6.30E-02 | 1.78E-01 | 1.09E+00 | 5.47E-02 | 2.01E-01 | 1.03E+00 | 7.22E-02 | 1.98E-01 | 1.03E+00 |
| CDA | 1.75E-01 | 9.12E-01 | 1.06E+00 | 2.31E-01 | 1.00E+00 | -1.00E+00 | 4.92E-02 | 2.09E-01 | 1.89E+00 | 1.56E-01 | 5.96E-01 | 1.34E+00 | 2.80E-01 | 1.00E+00 | 1.00E+00 | 3.55E-01 | 1.00E+00 | -1.00E+00 | 2.46E-01 | 9.03E-01 | -1.10E+00 | 3.64E-01 | 9.99E-01 | 1.05E+00 |
| SLC16A5 | 1.75E-01 | 9.12E-01 | 1.50E+00 | 1.87E-01 | 8.08E-01 | 1.63E+00 | 1.07E-01 | 4.54E-01 | 1.02E+00 | 2.34E-01 | 8.94E-01 | 1.43E+00 | 1.33E-01 | 4.75E-01 | 1.22E+00 | 4.50E-02 | 1.27E-01 | -1.10E+00 | 2.93E-02 | 1.07E-01 | 2.17E+00 | 2.09E-01 | 5.74E-01 | 1.94E+00 |
| HMGB1P8 | 1.75E-01 | 9.13E-01 | 1.36E+00 | 5.56E-03 | 2.40E-02 | -1.12E+00 | 1.92E-01 | 8.17E-01 | 1.24E+00 | 2.76E-04 | 1.05E-03 | -1.24E+00 | 2.43E-01 | 8.68E-01 | 1.62E+00 | 2.94E-02 | 8.29E-02 | -1.02E+00 | 8.59E-02 | 3.15E-01 | 1.14E+00 | 4.27E-03 | 1.17E-02 | -1.05E+00 |
| MBLAC1 | 1.75E-01 | 9.13E-01 | 1.25E+00 | 3.26E-02 | 1.41E-01 | -1.02E+00 | 7.72E-02 | 3.28E-01 | -1.07E+00 | 2.43E-01 | 9.27E-01 | 1.32E+00 | 1.53E-02 | 5.45E-02 | -1.32E+00 | 4.05E-03 | 1.14E-02 | -1.69E+00 | 2.35E-01 | 8.64E-01 | 1.25E+00 | 2.08E-01 | 5.70E-01 | 1.05E+00 |
| EIF5AP2 | 1.75E-01 | 9.13E-01 | 1.27E+00 | 1.89E-01 | 8.15E-01 | 1.61E+00 | 9.65E-02 | 4.10E-01 | -1.06E+00 | 1.11E-01 | 4.23E-01 | 1.56E+00 | 5.68E-02 | 2.02E-01 | -1.33E+00 | 1.71E-01 | 4.82E-01 | 1.74E+00 | 2.41E-01 | 8.85E-01 | 1.17E+00 | 4.86E-03 | 1.33E-02 | 1.74E+00 |
| CLUHP3 | 1.75E-01 | 9.13E-01 | 1.40E+00 | 2.00E-01 | 8.66E-01 | 1.46E+00 | 2.28E-01 | 9.68E-01 | 1.09E+00 | 2.26E-01 | 8.65E-01 | 1.43E+00 | 9.18E-05 | 3.27E-04 | -1.55E+00 | 1.99E-03 | 5.61E-03 | -1.20E+00 | 1.91E-12 | 7.00E-12 | -2.04E+00 | 5.08E-07 | 1.39E-06 | -1.47E+00 |
| SYNGR1 | 1.75E-01 | 9.13E-01 | 1.36E+00 | 3.07E-03 | 1.33E-02 | 1.22E+00 | 1.86E-02 | 7.89E-02 | 1.34E+00 | 1.15E-01 | 4.38E-01 | 1.27E+00 | 2.00E-02 | 7.13E-02 | 1.15E+00 | 1.37E-03 | 3.86E-03 | 1.06E+00 | 1.72E-01 | 6.32E-01 | 1.42E+00 | 9.64E-03 | 2.65E-02 | 1.65E+00 |
| LINC01481 | 1.75E-01 | 9.14E-01 | 1.37E+00 | 2.31E-01 | 9.97E-01 | 1.50E+00 | 1.95E-01 | 8.27E-01 | 1.19E+00 | 1.21E-02 | 4.62E-02 | 1.11E+00 | 2.06E-01 | 7.35E-01 | 1.77E+00 | 3.37E-01 | 9.51E-01 | 1.52E+00 | 2.26E-02 | 8.31E-02 | 1.56E+00 | 5.45E-02 | 1.50E-01 | 1.56E+00 |
| LINC00884 | 1.75E-01 | 9.14E-01 | 1.30E+00 | 2.14E-01 | 9.27E-01 | 1.45E+00 | 1.34E-01 | 5.69E-01 | 1.05E+00 | 9.53E-02 | 3.64E-01 | 1.06E+00 | 7.92E-02 | 2.82E-01 | -1.55E+00 | 2.31E-01 | 6.52E-01 | 1.10E+00 | 1.61E-02 | 5.91E-02 | -1.22E+00 | 1.16E-01 | 3.20E-01 | 1.04E+00 |
| DCAF4L1 | 1.75E-01 | 9.14E-01 | 1.26E+00 | 1.89E-01 | 8.16E-01 | 1.23E+00 | 5.40E-02 | 2.30E-01 | 1.53E+00 | 7.55E-02 | 2.88E-01 | 1.55E+00 | 2.27E-01 | 8.11E-01 | 1.04E+00 | 2.81E-01 | 7.92E-01 | 1.03E+00 | 1.23E-01 | 4.53E-01 | 1.64E+00 | 8.74E-02 | 2.40E-01 | 1.83E+00 |
| PROB1 | 1.75E-01 | 9.14E-01 | 1.15E+00 | 2.09E-01 | 9.01E-01 | 1.31E+00 | 7.60E-04 | 3.23E-03 | -1.52E+00 | 2.10E-02 | 8.01E-02 | -1.11E+00 | 2.71E-01 | 9.67E-01 | 1.23E+00 | 1.74E-01 | 4.91E-01 | 1.80E+00 | 2.39E-01 | 8.75E-01 | -1.03E+00 | 3.10E-01 | 8.52E-01 | -1.01E+00 |
| PAICSP4 | 1.76E-01 | 9.15E-01 | 1.30E+00 | 9.73E-02 | 4.20E-01 | 1.90E+00 | 3.34E-02 | 1.42E-01 | 1.62E+00 | 4.42E-02 | 1.69E-01 | 1.91E+00 | 2.29E-01 | 8.18E-01 | 1.45E+00 | 2.10E-01 | 5.92E-01 | 1.80E+00 | 9.76E-02 | 3.58E-01 | 1.61E+00 | 2.13E-02 | 5.86E-02 | 1.92E+00 |
| DIAPH3 | 1.76E-01 | 9.15E-01 | 1.01E+00 | 2.15E-01 | 9.29E-01 | 1.18E+00 | 1.62E-03 | 6.87E-03 | 1.62E+00 | 1.11E-01 | 4.24E-01 | 1.32E+00 | 2.80E-01 | 1.00E+00 | 1.08E+00 | 3.15E-01 | 8.89E-01 | -1.14E+00 | 6.63E-02 | 2.43E-01 | -1.03E+00 | 1.90E-01 | 5.22E-01 | 1.27E+00 |
| VEGFC | 1.76E-01 | 9.15E-01 | 1.06E+00 | 2.31E-01 | 1.00E+00 | 1.00E+00 | 2.14E-01 | 9.08E-01 | 1.07E+00 | 2.36E-01 | 9.00E-01 | 1.10E+00 | 2.45E-01 | 8.72E-01 | 1.15E+00 | 3.27E-01 | 9.23E-01 | 1.07E+00 | 1.62E-01 | 5.96E-01 | 1.48E+00 | 7.19E-02 | 1.98E-01 | -1.09E+00 |
| RPL18AP3 | 1.76E-01 | 9.16E-01 | 1.01E+00 | 1.36E-03 | 5.86E-03 | -1.07E+00 | 9.85E-02 | 4.19E-01 | 1.03E+00 | 8.16E-02 | 3.12E-01 | -1.03E+00 | 1.89E-01 | 6.75E-01 | 1.02E+00 | 3.29E-01 | 9.29E-01 | 1.00E+00 | 1.80E-02 | 6.60E-02 | 1.01E+00 | 2.34E-01 | 6.43E-01 | -1.01E+00 |
| AC016712.1 | 1.76E-01 | 9.16E-01 | 1.42E+00 | 2.32E-05 | 1.00E-04 | 1.04E+00 | 6.59E-02 | 2.80E-01 | 1.19E+00 | 1.37E-04 | 5.24E-04 | 1.00E+00 | 2.04E-01 | 7.27E-01 | 1.35E+00 | 8.11E-02 | 2.29E-01 | 1.09E+00 | 1.73E-01 | 6.36E-01 | 1.21E+00 | 2.32E-01 | 6.36E-01 | 1.21E+00 |
| PDCD5 | 1.76E-01 | 9.16E-01 | 1.34E+00 | 1.03E-01 | 4.47E-01 | 1.57E+00 | 4.80E-02 | 2.04E-01 | 1.41E+00 | 6.42E-02 | 2.45E-01 | 1.54E+00 | 2.20E-03 | 7.84E-03 | 1.05E+00 | 3.43E-01 | 9.67E-01 | 1.53E+00 | 2.79E-02 | 1.02E-01 | 1.15E+00 | 1.66E-01 | 4.56E-01 | 1.49E+00 |
| AC073335.1 | 1.76E-01 | 9.16E-01 | 1.36E+00 | 2.01E-01 | 8.67E-01 | 1.33E+00 | 5.28E-03 | 2.24E-02 | -1.32E+00 | 2.86E-02 | 1.09E-01 | -1.00E+00 | 2.63E-01 | 9.37E-01 | 1.38E+00 | 3.23E-01 | 9.09E-01 | 1.34E+00 | 1.47E-01 | 5.39E-01 | -1.02E+00 | 1.19E-01 | 3.27E-01 | -1.15E+00 |
| AL139811.2 | 1.76E-01 | 9.16E-01 | 1.12E+00 | 1.23E-01 | 5.31E-01 | -1.18E+00 | 1.10E-01 | 4.69E-01 | 1.44E+00 | 2.02E-01 | 7.72E-01 | 1.38E+00 | 3.30E-02 | 1.18E-01 | 1.98E+00 | 1.39E-01 | 3.91E-01 | 1.72E+00 | 3.72E-04 | 1.36E-03 | 2.08E+00 | 1.45E-03 | 3.98E-03 | 2.14E+00 |
| RGS17P1 | 1.76E-01 | 9.16E-01 | 1.45E+00 | 2.07E-01 | 8.96E-01 | 1.57E+00 | 1.88E-01 | 7.99E-01 | 1.17E+00 | 1.58E-01 | 6.03E-01 | 1.27E+00 | 9.65E-02 | 3.44E-01 | 1.64E+00 | 6.98E-02 | 1.97E-01 | 1.88E+00 | 1.71E-01 | 6.29E-01 | 1.46E+00 | 2.23E-01 | 6.13E-01 | 1.50E+00 |
| BDNF | 1.76E-01 | 9.17E-01 | 1.04E+00 | 1.73E-01 | 7.50E-01 | -1.06E+00 | 1.04E-02 | 4.41E-02 | -1.29E+00 | 4.45E-03 | 1.70E-02 | -1.51E+00 | 2.37E-01 | 8.46E-01 | -1.09E+00 | 2.95E-01 | 8.32E-01 | 1.17E+00 | 1.35E-02 | 4.96E-02 | -1.04E+00 | 9.09E-03 | 2.50E-02 | -1.50E+00 |
| AL139095.1 | 1.76E-01 | 9.17E-01 | 1.26E+00 | 2.10E-01 | 9.08E-01 | 1.45E+00 | 6.44E-02 | 2.73E-01 | 1.52E+00 | 2.22E-02 | 8.48E-02 | 2.03E+00 | 7.92E-02 | 2.82E-01 | 2.08E+00 | 3.49E-02 | 9.84E-02 | 2.50E+00 | 1.47E-01 | 5.40E-01 | 1.49E+00 | 6.54E-03 | 1.80E-02 | 2.11E+00 |
| AC015912.3 | 1.76E-01 | 9.18E-01 | 1.36E+00 | 1.54E-01 | 6.66E-01 | 1.30E+00 | 2.23E-01 | 9.49E-01 | 1.08E+00 | 5.01E-02 | 1.91E-01 | -1.17E+00 | 1.77E-02 | 6.30E-02 | 2.58E+00 | 2.52E-01 | 7.12E-01 | 1.55E+00 | 1.72E-01 | 6.32E-01 | 1.25E+00 | 3.64E-01 | 9.99E-01 | 1.19E+00 |
| LY6G6F | 1.76E-01 | 9.18E-01 | 1.06E+00 | 2.31E-01 | 1.00E+00 | -1.00E+00 | 2.17E-01 | 9.21E-01 | 1.04E+00 | 2.62E-01 | 1.00E+00 | 1.00E+00 | 2.47E-01 | 8.81E-01 | 1.12E+00 | 3.55E-01 | 1.00E+00 | -1.00E+00 | 6.30E-04 | 2.31E-03 | -1.11E+00 | 5.38E-06 | 1.48E-05 | -1.98E+00 |
| AC099791.2 | 1.76E-01 | 9.18E-01 | 1.25E+00 | 7.60E-18 | 3.29E-17 | 4.88E+00 | 1.59E-01 | 6.75E-01 | 1.04E+00 | 5.80E-05 | 2.21E-04 | 3.39E+00 | 1.72E-01 | 6.13E-01 | 1.97E+00 | 1.33E-05 | 3.74E-05 | 6.18E+00 | 1.24E-01 | 4.54E-01 | 1.93E+00 | 7.58E-03 | 2.08E-02 | 3.02E+00 |
| MTND4P15 | 1.76E-01 | 9.18E-01 | 1.09E+00 | 1.35E-01 | 5.84E-01 | 1.38E+00 | 1.60E-01 | 6.80E-01 | 1.36E+00 | 1.42E-01 | 5.43E-01 | 1.52E+00 | 1.18E-01 | 4.20E-01 | 1.08E+00 | 3.07E-01 | 8.64E-01 | 1.30E+00 | 5.02E-02 | 1.84E-01 | 1.03E+00 | 2.30E-01 | 6.32E-01 | 1.11E+00 |
| VN1R110P | 1.76E-01 | 9.19E-01 | 1.20E+00 | 2.23E-01 | 9.65E-01 | 1.06E+00 | 1.88E-01 | 8.01E-01 | -1.15E+00 | 1.98E-01 | 7.56E-01 | -1.16E+00 | 3.38E-03 | 1.20E-02 | 1.17E+00 | 2.93E-01 | 8.26E-01 | 1.62E+00 | 9.26E-03 | 3.40E-02 | -1.26E+00 | 2.95E-01 | 8.10E-01 | 1.13E+00 |
| AL359513.1 | 1.76E-01 | 9.19E-01 | 1.31E+00 | 4.11E-02 | 1.78E-01 | 1.14E+00 | 1.95E-01 | 8.31E-01 | 1.07E+00 | 1.89E-02 | 7.23E-02 | -1.23E+00 | 1.75E-01 | 6.22E-01 | 1.73E+00 | 2.42E-01 | 6.83E-01 | 1.21E+00 | 2.37E-01 | 8.68E-01 | 1.29E+00 | 2.74E-01 | 7.52E-01 | 1.06E+00 |
| S100A12 | 1.76E-01 | 9.19E-01 | 1.07E+00 | 2.09E-01 | 9.03E-01 | 1.17E+00 | 2.25E-01 | 9.58E-01 | -1.05E+00 | 1.58E-01 | 6.05E-01 | -1.59E+00 | 1.58E-01 | 5.64E-01 | 3.33E+00 | 3.55E-01 | 1.00E+00 | -1.00E+00 | 5.47E-02 | 2.01E-01 | -1.45E+00 | 1.77E-01 | 4.85E-01 | 1.19E+00 |
| EYS | 1.76E-01 | 9.20E-01 | 1.33E+00 | 3.84E-02 | 1.66E-01 | 1.95E+00 | 4.62E-02 | 1.97E-01 | -1.26E+00 | 1.83E-01 | 6.98E-01 | 1.41E+00 | 2.74E-01 | 9.77E-01 | 1.23E+00 | 2.19E-01 | 6.19E-01 | 1.47E+00 | 2.18E-01 | 8.01E-01 | 1.37E+00 | 1.30E-01 | 3.57E-01 | 1.77E+00 |
| VDAC1P2 | 1.77E-01 | 9.20E-01 | 1.21E+00 | 1.53E-01 | 6.60E-01 | 1.17E+00 | 1.37E-01 | 5.83E-01 | -1.01E+00 | 1.06E-02 | 4.06E-02 | -1.16E+00 | 6.95E-02 | 2.48E-01 | -1.18E+00 | 4.19E-02 | 1.18E-01 | -1.23E+00 | 8.89E-04 | 3.26E-03 | -1.24E+00 | 4.81E-03 | 1.32E-02 | -1.20E+00 |
| AC090229.1 | 1.77E-01 | 9.22E-01 | 1.25E+00 | 2.04E-01 | 8.81E-01 | 1.41E+00 | 6.12E-02 | 2.60E-01 | -1.15E+00 | 2.42E-01 | 9.23E-01 | 1.24E+00 | 2.66E-02 | 9.49E-02 | -1.08E+00 | 2.29E-02 | 6.45E-02 | -1.20E+00 | 1.06E-04 | 3.89E-04 | -1.66E+00 | 3.70E-02 | 1.02E-01 | 1.05E+00 |
| AMN | 1.77E-01 | 9.23E-01 | 1.39E+00 | 1.87E-01 | 8.07E-01 | 1.61E+00 | 1.88E-01 | 7.97E-01 | 1.36E+00 | 7.96E-02 | 3.04E-01 | 1.72E+00 | 2.12E-01 | 7.56E-01 | 1.55E+00 | 1.56E-01 | 4.39E-01 | 1.89E+00 | 1.37E-01 | 5.01E-01 | 1.16E+00 | 3.63E-01 | 9.98E-01 | 1.32E+00 |
| ARSG | 1.77E-01 | 9.23E-01 | 1.09E+00 | 1.73E-01 | 7.48E-01 | 1.04E+00 | 8.27E-02 | 3.51E-01 | -1.20E+00 | 3.46E-02 | 1.32E-01 | -1.43E+00 | 1.42E-03 | 5.05E-03 | -1.59E+00 | 2.81E-04 | 7.93E-04 | -1.99E+00 | 2.90E-02 | 1.06E-01 | -1.03E+00 | 3.54E-02 | 9.72E-02 | -1.32E+00 |
| CTNNAL1 | 1.77E-01 | 9.23E-01 | 1.25E+00 | 7.76E-02 | 3.35E-01 | -1.17E+00 | 2.12E-01 | 8.99E-01 | 1.11E+00 | 1.65E-04 | 6.28E-04 | -1.18E+00 | 8.45E-02 | 3.01E-01 | -1.09E+00 | 2.46E-01 | 6.92E-01 | 1.37E+00 | 1.10E-03 | 4.03E-03 | 1.08E+00 | 5.08E-03 | 1.39E-02 | 1.08E+00 |
| AL132639.2 | 1.77E-01 | 9.23E-01 | 1.36E+00 | 8.08E-02 | 3.49E-01 | 1.19E+00 | 1.04E-01 | 4.42E-01 | -1.18E+00 | 8.11E-02 | 3.10E-01 | -1.19E+00 | 1.06E-01 | 3.76E-01 | -1.08E+00 | 1.99E-01 | 5.62E-01 | -1.02E+00 | 1.35E-01 | 4.94E-01 | -1.01E+00 | 3.55E-01 | 9.75E-01 | 1.27E+00 |
| EXOC3-AS1 | 1.77E-01 | 9.23E-01 | 1.38E+00 | 4.32E-02 | 1.87E-01 | 1.11E+00 | 1.68E-01 | 7.15E-01 | 1.22E+00 | 6.88E-03 | 2.63E-02 | -1.04E+00 | 2.80E-01 | 1.00E+00 | 1.47E+00 | 2.59E-01 | 7.29E-01 | 1.35E+00 | 2.14E-01 | 7.84E-01 | 1.34E+00 | 2.60E-01 | 7.15E-01 | 1.31E+00 |
| AL512770.1 | 1.77E-01 | 9.23E-01 | 1.26E+00 | 2.03E-03 | 8.79E-03 | -1.39E+00 | 9.80E-03 | 4.16E-02 | -1.28E+00 | 4.56E-04 | 1.74E-03 | -1.47E+00 | 1.42E-01 | 5.08E-01 | 1.07E+00 | 2.29E-02 | 6.46E-02 | -1.24E+00 | 3.30E-03 | 1.21E-02 | -1.36E+00 | 3.35E-02 | 9.21E-02 | -1.15E+00 |
| ALKBH1 | 1.77E-01 | 9.23E-01 | 1.33E+00 | 2.16E-01 | 9.33E-01 | 1.48E+00 | 5.29E-02 | 2.25E-01 | -1.11E+00 | 1.16E-01 | 4.44E-01 | 1.16E+00 | 7.18E-03 | 2.56E-02 | -1.71E+00 | 9.84E-02 | 2.77E-01 | -1.06E+00 | 1.29E-06 | 4.72E-06 | -1.62E+00 | 1.35E-05 | 3.70E-05 | -1.58E+00 |
| SOX4 | 1.77E-01 | 9.23E-01 | 1.05E+00 | 1.15E-01 | 4.97E-01 | 1.09E+00 | 1.03E-02 | 4.39E-02 | -1.25E+00 | 2.18E-02 | 8.31E-02 | -1.18E+00 | 1.10E-06 | 3.93E-06 | -1.42E+00 | 1.46E-02 | 4.12E-02 | -1.15E+00 | 2.35E-03 | 8.62E-03 | -1.09E+00 | 1.40E-01 | 3.83E-01 | -1.08E+00 |
| LEF1-AS1 | 1.77E-01 | 9.23E-01 | 1.05E+00 | 2.12E-01 | 9.15E-01 | 1.08E+00 | 1.03E-01 | 4.40E-01 | -1.27E+00 | 1.19E-01 | 4.54E-01 | -1.27E+00 | 2.12E-01 | 7.56E-01 | 1.14E+00 | 3.29E-01 | 9.26E-01 | 1.35E+00 | 1.03E-02 | 3.77E-02 | 1.02E+00 | 2.42E-01 | 6.66E-01 | 1.25E+00 |
| TMEM40 | 1.77E-01 | 9.23E-01 | 1.05E+00 | 2.31E-01 | 1.00E+00 | 1.00E+00 | 2.34E-01 | 9.95E-01 | 1.01E+00 | 2.07E-01 | 7.89E-01 | -1.10E+00 | 2.80E-01 | 1.00E+00 | 1.00E+00 | 3.28E-01 | 9.25E-01 | 1.08E+00 | 7.13E-07 | 2.62E-06 | 7.94E+00 | 1.31E-01 | 3.60E-01 | 4.43E+00 |
| VIM-AS1 | 1.77E-01 | 9.24E-01 | 1.26E+00 | 8.25E-03 | 3.57E-02 | -1.14E+00 | 1.47E-01 | 6.27E-01 | 1.21E+00 | 7.25E-05 | 2.77E-04 | -1.24E+00 | 1.56E-01 | 5.56E-01 | 1.50E+00 | 4.28E-02 | 1.21E-01 | -1.02E+00 | 1.69E-01 | 6.21E-01 | 1.78E+00 | 2.44E-01 | 6.71E-01 | 1.19E+00 |
| APOBEC3B | 1.77E-01 | 9.24E-01 | 1.04E+00 | 1.44E-01 | 6.22E-01 | 1.34E+00 | 1.85E-01 | 7.86E-01 | 1.16E+00 | 1.18E-01 | 4.49E-01 | 1.44E+00 | 1.12E-01 | 3.99E-01 | 1.59E+00 | 3.24E-01 | 9.15E-01 | 1.11E+00 | 9.13E-11 | 3.35E-10 | -1.93E+00 | 3.56E-02 | 9.78E-02 | 1.07E+00 |
| RPL5P1 | 1.77E-01 | 9.24E-01 | 1.38E+00 | 2.61E-05 | 1.13E-04 | 2.05E+00 | 1.21E-01 | 5.13E-01 | 1.42E+00 | 3.81E-05 | 1.45E-04 | 2.08E+00 | 1.33E-01 | 4.76E-01 | 1.20E+00 | 1.78E-02 | 5.03E-02 | 1.74E+00 | 1.19E-01 | 4.36E-01 | 1.45E+00 | 1.71E-05 | 4.71E-05 | 2.06E+00 |
| ARL6IP1P3 | 1.77E-01 | 9.25E-01 | 1.27E+00 | 7.41E-02 | 3.20E-01 | 1.17E+00 | 1.42E-01 | 6.05E-01 | 1.06E+00 | 2.02E-02 | 7.71E-02 | -1.01E+00 | 2.53E-01 | 9.00E-01 | 1.40E+00 | 1.21E-02 | 3.42E-02 | -1.42E+00 | 2.52E-01 | 9.26E-01 | 1.28E+00 | 1.91E-01 | 5.24E-01 | 1.09E+00 |
| MYO16 | 1.78E-01 | 9.25E-01 | 1.05E+00 | 2.12E-01 | 9.17E-01 | 1.08E+00 | 2.21E-01 | 9.37E-01 | 1.03E+00 | 2.62E-01 | 1.00E+00 | 1.00E+00 | 8.47E-02 | 3.02E-01 | 2.05E+00 | 1.61E-01 | 4.53E-01 | 1.90E+00 | 1.87E-12 | 6.87E-12 | -2.12E+00 | 9.73E-14 | 2.67E-13 | -2.37E+00 |
| AC130469.2 | 1.78E-01 | 9.25E-01 | 1.27E+00 | 2.14E-02 | 9.26E-02 | -1.34E+00 | 2.05E-01 | 8.73E-01 | 1.30E+00 | 1.91E-03 | 7.29E-03 | -1.41E+00 | 1.88E-01 | 6.69E-01 | 1.13E+00 | 1.01E-01 | 2.85E-01 | -1.54E+00 | 4.49E-02 | 1.65E-01 | -1.55E+00 | 6.23E-02 | 1.71E-01 | -1.62E+00 |
| PODXL | 1.78E-01 | 9.27E-01 | 1.21E+00 | 1.05E-01 | 4.53E-01 | 1.82E+00 | 1.65E-02 | 7.03E-02 | -1.29E+00 | 1.57E-01 | 5.98E-01 | 1.51E+00 | 7.84E-03 | 2.80E-02 | -1.44E+00 | 1.64E-02 | 4.61E-02 | -1.23E+00 | 4.95E-02 | 1.82E-01 | -1.13E+00 | 2.91E-01 | 7.98E-01 | 1.40E+00 |
| DNAJB7 | 1.78E-01 | 9.27E-01 | 1.24E+00 | 1.90E-01 | 8.19E-01 | 1.41E+00 | 1.75E-01 | 7.45E-01 | 1.02E+00 | 2.21E-01 | 8.43E-01 | 1.48E+00 | 2.13E-01 | 7.59E-01 | 1.61E+00 | 3.15E-01 | 8.89E-01 | 1.44E+00 | 1.54E-01 | 5.65E-01 | 1.21E+00 | 1.95E-01 | 5.37E-01 | 1.32E+00 |
| UBE2H | 1.78E-01 | 9.27E-01 | 1.31E+00 | 3.74E-03 | 1.62E-02 | 1.18E+00 | 6.52E-02 | 2.77E-01 | 1.29E+00 | 2.10E-05 | 8.01E-05 | -1.22E+00 | 6.76E-02 | 2.41E-01 | 1.29E+00 | 7.26E-05 | 2.05E-04 | -1.18E+00 | 6.40E-02 | 2.35E-01 | 1.22E+00 | 2.75E-03 | 7.55E-03 | -1.08E+00 |
| C3orf67 | 1.78E-01 | 9.27E-01 | 1.23E+00 | 1.39E-01 | 6.02E-01 | 1.20E+00 | 2.16E-01 | 9.19E-01 | 1.31E+00 | 1.51E-01 | 5.76E-01 | 1.20E+00 | 1.49E-01 | 5.32E-01 | 1.73E+00 | 2.48E-01 | 6.99E-01 | 1.64E+00 | 1.09E-02 | 4.01E-02 | -1.39E+00 | 1.56E-01 | 4.29E-01 | 1.03E+00 |
| TBC1D4 | 1.78E-01 | 9.27E-01 | 1.36E+00 | 9.30E-02 | 4.02E-01 | 1.89E+00 | 1.00E-04 | 4.26E-04 | -1.43E+00 | 4.47E-02 | 1.71E-01 | 1.20E+00 | 3.81E-05 | 1.36E-04 | -2.51E+00 | 1.95E-02 | 5.49E-02 | -1.14E+00 | 2.84E-12 | 1.04E-11 | -2.07E+00 | 7.46E-03 | 2.05E-02 | -1.03E+00 |
| RAPGEF6 | 1.78E-01 | 9.28E-01 | 1.24E+00 | 2.10E-01 | 9.06E-01 | 1.33E+00 | 8.86E-05 | 3.77E-04 | -1.38E+00 | 7.54E-05 | 2.88E-04 | -1.23E+00 | 1.97E-01 | 7.02E-01 | 1.60E+00 | 2.33E-01 | 6.57E-01 | 1.65E+00 | 3.44E-02 | 1.26E-01 | -1.11E+00 | 9.26E-02 | 2.54E-01 | 1.02E+00 |
| AC034195.1 | 1.78E-01 | 9.28E-01 | 1.37E+00 | 6.65E-02 | 2.87E-01 | 2.02E+00 | 1.50E-01 | 6.39E-01 | 1.08E+00 | 1.30E-01 | 4.98E-01 | 1.69E+00 | 2.39E-01 | 8.53E-01 | 1.10E+00 | 2.08E-01 | 5.87E-01 | 1.46E+00 | 2.23E-01 | 8.19E-01 | 1.16E+00 | 2.33E-01 | 6.40E-01 | 1.32E+00 |
| RRP12 | 1.78E-01 | 9.28E-01 | 1.28E+00 | 2.94E-02 | 1.27E-01 | -1.00E+00 | 2.98E-02 | 1.26E-01 | 1.60E+00 | 1.69E-01 | 6.45E-01 | 1.30E+00 | 1.30E-01 | 4.63E-01 | 1.16E+00 | 2.99E-01 | 8.42E-01 | 1.28E+00 | 2.59E-01 | 9.51E-01 | 1.26E+00 | 1.05E-01 | 2.89E-01 | 1.47E+00 |
| LANCL3 | 1.78E-01 | 9.29E-01 | 1.14E+00 | 2.04E-01 | 8.82E-01 | 1.27E+00 | 1.90E-01 | 8.07E-01 | -1.19E+00 | 2.58E-01 | 9.86E-01 | 1.07E+00 | 1.54E-01 | 5.49E-01 | 1.06E+00 | 2.38E-01 | 6.72E-01 | 1.51E+00 | 4.09E-05 | 1.50E-04 | -1.18E+00 | 5.23E-03 | 1.44E-02 | -1.01E+00 |
| HSPE1P9 | 1.78E-01 | 9.29E-01 | 1.21E+00 | 6.79E-02 | 2.93E-01 | 1.93E+00 | 2.30E-01 | 9.76E-01 | 1.03E+00 | 1.08E-03 | 4.12E-03 | 2.23E+00 | 1.83E-01 | 6.54E-01 | -1.17E+00 | 1.23E-01 | 3.47E-01 | 2.04E+00 | 2.00E-01 | 7.35E-01 | 1.27E+00 | 1.47E-07 | 4.04E-07 | 3.36E+00 |
| RNF168 | 1.78E-01 | 9.29E-01 | 1.45E+00 | 1.20E-01 | 5.20E-01 | 1.45E+00 | 9.84E-02 | 4.18E-01 | 1.16E+00 | 1.56E-01 | 5.97E-01 | 1.58E+00 | 5.15E-02 | 1.84E-01 | 1.44E+00 | 5.88E-03 | 1.66E-02 | 1.27E+00 | 1.75E-04 | 6.42E-04 | -1.34E+00 | 2.73E-03 | 7.50E-03 | -1.18E+00 |
| AL590705.1 | 1.78E-01 | 9.29E-01 | 1.33E+00 | 1.56E-01 | 6.76E-01 | 1.22E+00 | 1.34E-01 | 5.71E-01 | 1.12E+00 | 2.64E-03 | 1.01E-02 | -1.32E+00 | 1.23E-01 | 4.38E-01 | -2.59E+00 | 3.35E-02 | 9.43E-02 | -3.98E+00 | 2.13E-01 | 7.81E-01 | 1.27E+00 | 2.95E-01 | 8.11E-01 | 1.09E+00 |
| AC084024.2 | 1.78E-01 | 9.29E-01 | 1.13E+00 | 5.69E-02 | 2.46E-01 | 1.79E+00 | 1.72E-03 | 7.32E-03 | -1.38E+00 | 1.84E-03 | 7.02E-03 | -1.33E+00 | 1.51E-01 | 5.37E-01 | 1.06E+00 | 1.49E-01 | 4.21E-01 | 1.00E+00 | 2.98E-02 | 1.09E-01 | -1.30E+00 | 2.56E-01 | 7.04E-01 | 1.02E+00 |
| AC036214.2 | 1.78E-01 | 9.30E-01 | 1.38E+00 | 1.64E-02 | 7.08E-02 | -1.11E+00 | 2.85E-02 | 1.21E-01 | -1.38E+00 | 3.79E-02 | 1.45E-01 | -1.14E+00 | 1.13E-01 | 4.03E-01 | -1.11E+00 | 2.71E-01 | 7.65E-01 | 1.14E+00 | 4.50E-03 | 1.65E-02 | -1.56E+00 | 5.58E-04 | 1.53E-03 | -2.03E+00 |
| DRG1 | 1.78E-01 | 9.30E-01 | 1.37E+00 | 1.61E-01 | 6.97E-01 | 1.45E+00 | 1.56E-01 | 6.62E-01 | 1.21E+00 | 1.07E-01 | 4.07E-01 | 1.29E+00 | 1.45E-01 | 5.17E-01 | 1.32E+00 | 2.69E-01 | 7.58E-01 | 1.52E+00 | 1.44E-01 | 5.27E-01 | 1.19E+00 | 2.54E-01 | 6.97E-01 | 1.34E+00 |
| EIF4HP1 | 1.79E-01 | 9.31E-01 | 1.30E+00 | 6.52E-02 | 2.82E-01 | 1.15E+00 | 6.77E-02 | 2.88E-01 | 1.04E+00 | 1.73E-02 | 6.59E-02 | 1.05E+00 | 4.38E-02 | 1.56E-01 | -1.03E+00 | 1.55E-02 | 4.36E-02 | -1.17E+00 | 2.53E-01 | 9.27E-01 | 1.31E+00 | 8.46E-03 | 2.32E-02 | -1.08E+00 |
| AC015813.6 | 1.79E-01 | 9.32E-01 | 1.20E+00 | 1.75E-02 | 7.58E-02 | -1.34E+00 | 2.17E-01 | 9.22E-01 | -1.06E+00 | 2.35E-01 | 8.99E-01 | 1.05E+00 | 2.33E-01 | 8.31E-01 | 1.28E+00 | 3.38E-01 | 9.53E-01 | 1.27E+00 | 2.53E-01 | 9.29E-01 | 1.15E+00 | 2.55E-01 | 7.01E-01 | 1.31E+00 |
| AC020917.4 | 1.79E-01 | 9.32E-01 | 1.18E+00 | 5.76E-02 | 2.49E-01 | 2.03E+00 | 2.08E-01 | 8.84E-01 | 1.24E+00 | 1.61E-01 | 6.16E-01 | 1.60E+00 | 2.41E-03 | 8.59E-03 | -1.54E+00 | 2.88E-02 | 8.11E-02 | -1.11E+00 | 2.25E-01 | 8.25E-01 | 1.12E+00 | 1.94E-01 | 5.33E-01 | 1.62E+00 |
| EPB42 | 1.79E-01 | 9.32E-01 | 1.08E+00 | 1.62E-01 | 7.02E-01 | -1.04E+00 | 2.13E-01 | 9.04E-01 | -1.00E+00 | 2.62E-01 | 1.00E+00 | 1.20E+00 | 1.92E-01 | 6.84E-01 | 1.08E+00 | 3.47E-01 | 9.78E-01 | 1.21E+00 | 6.63E-02 | 2.43E-01 | 1.20E+00 | 1.76E-02 | 4.84E-02 | 1.08E+00 |
| AC092597.1 | 1.79E-01 | 9.32E-01 | 1.36E+00 | 4.25E-02 | 1.84E-01 | 1.87E+00 | 2.20E-01 | 9.34E-01 | 1.31E+00 | 7.22E-02 | 2.76E-01 | 1.66E+00 | 1.13E-01 | 4.01E-01 | 1.27E+00 | 2.67E-02 | 7.54E-02 | 1.93E+00 | 9.27E-02 | 3.40E-01 | 1.46E+00 | 7.08E-04 | 1.95E-03 | 1.99E+00 |
| HIST2H3DP1 | 1.79E-01 | 9.33E-01 | 1.06E+00 | 2.15E-01 | 9.28E-01 | 1.08E+00 | 3.66E-03 | 1.56E-02 | 2.10E+00 | 2.42E-01 | 9.25E-01 | 1.16E+00 | 1.91E-01 | 6.82E-01 | -1.36E+00 | 2.45E-01 | 6.92E-01 | -1.36E+00 | 3.95E-02 | 1.45E-01 | 1.50E+00 | 1.68E-01 | 4.60E-01 | 1.23E+00 |
| EXO3 | 1.79E-01 | 9.34E-01 | 1.34E+00 | 5.72E-02 | 2.47E-01 | 1.24E+00 | 1.88E-01 | 8.01E-01 | 1.12E+00 | 5.65E-02 | 2.16E-01 | 1.21E+00 | 5.74E-08 | 2.04E-07 | -1.45E+00 | 4.08E-09 | 1.15E-08 | -1.51E+00 | 2.49E-01 | 9.13E-01 | 1.22E+00 | 3.56E-01 | 9.78E-01 | 1.34E+00 |
| C20orf196 | 1.79E-01 | 9.34E-01 | 1.39E+00 | 2.73E-02 | 1.18E-01 | 1.05E+00 | 4.50E-02 | 1.91E-01 | 1.40E+00 | 6.49E-02 | 2.48E-01 | 1.02E+00 | 1.10E-02 | 3.92E-02 | -1.06E+00 | 1.58E-02 | 4.45E-02 | -1.02E+00 | 9.15E-05 | 3.36E-04 | -1.10E+00 | 1.72E-06 | 4.73E-06 | -1.14E+00 |
| NMRK1 | 1.79E-01 | 9.35E-01 | 1.30E+00 | 1.43E-01 | 6.17E-01 | 1.31E+00 | 8.47E-02 | 3.60E-01 | -1.02E+00 | 2.16E-02 | 8.24E-02 | -1.01E+00 | 3.22E-02 | 1.15E-01 | 1.16E+00 | 6.48E-02 | 1.83E-01 | 1.11E+00 | 9.24E-02 | 3.39E-01 | 1.23E+00 | 1.80E-02 | 4.95E-02 | 1.02E+00 |
| WDR73 | 1.80E-01 | 9.36E-01 | 1.33E+00 | 9.38E-02 | 4.05E-01 | 1.20E+00 | 5.26E-02 | 2.24E-01 | 1.58E+00 | 2.01E-01 | 7.70E-01 | 1.52E+00 | 2.57E-01 | 9.18E-01 | 1.51E+00 | 6.51E-02 | 1.84E-01 | 1.03E+00 | 2.54E-01 | 9.33E-01 | 1.25E+00 | 3.32E-01 | 9.13E-01 | 1.31E+00 |
| FCGR2C | 1.80E-01 | 9.36E-01 | 1.07E+00 | 1.82E-01 | 7.85E-01 | 1.30E+00 | 1.35E-01 | 5.74E-01 | -1.28E+00 | 1.75E-01 | 6.67E-01 | -1.17E+00 | 5.13E-02 | 1.83E-01 | 2.39E+00 | 1.27E-01 | 3.58E-01 | 1.89E+00 | 5.50E-08 | 2.02E-07 | 2.54E+00 | 1.48E-02 | 4.07E-02 | 1.87E+00 |
| C1orf141 | 1.80E-01 | 9.36E-01 | 1.07E+00 | 2.15E-01 | 9.31E-01 | 1.09E+00 | 2.35E-01 | 1.00E+00 | 1.00E+00 | 2.43E-01 | 9.29E-01 | 1.08E+00 | 2.35E-01 | 8.39E-01 | 1.09E+00 | 1.30E-01 | 3.67E-01 | 1.59E+00 | 3.72E-06 | 1.37E-05 | 2.30E+00 | 4.82E-04 | 1.33E-03 | 2.27E+00 |
| AC104109.4 | 1.80E-01 | 9.36E-01 | 1.31E+00 | 1.19E-03 | 5.13E-03 | -1.02E+00 | 1.74E-02 | 7.41E-02 | 1.62E+00 | 1.23E-01 | 4.69E-01 | 1.21E+00 | 1.85E-03 | 6.59E-03 | 2.30E+00 | 2.96E-01 | 8.35E-01 | 1.27E+00 | 2.40E-01 | 8.80E-01 | 1.19E+00 | 5.19E-03 | 1.43E-02 | -1.24E+00 |
| LINC01311 | 1.80E-01 | 9.37E-01 | 1.25E+00 | 5.59E-03 | 2.41E-02 | 2.57E+00 | 1.89E-01 | 8.02E-01 | 1.13E+00 | 3.27E-02 | 1.25E-01 | 1.65E+00 | 2.55E-02 | 9.08E-02 | -1.86E+00 | 2.39E-01 | 6.73E-01 | 1.12E+00 | 1.03E-01 | 3.78E-01 | 1.50E+00 | 2.91E-03 | 7.98E-03 | 2.60E+00 |
| AC037459.2 | 1.80E-01 | 9.37E-01 | 1.21E+00 | 1.62E-01 | 6.99E-01 | 1.48E+00 | 1.55E-01 | 6.60E-01 | -1.09E+00 | 2.06E-01 | 7.87E-01 | 1.22E+00 | 2.47E-01 | 8.82E-01 | 1.16E+00 | 1.46E-01 | 4.12E-01 | 1.56E+00 | 7.38E-02 | 2.71E-01 | -1.22E+00 | 1.96E-01 | 5.39E-01 | 1.04E+00 |
| AQP1 | 1.80E-01 | 9.37E-01 | 1.04E+00 | 2.01E-01 | 8.67E-01 | -1.01E+00 | 2.06E-01 | 8.75E-01 | 1.09E+00 | 2.29E-01 | 8.77E-01 | -1.01E+00 | 1.74E-01 | 6.19E-01 | 1.56E+00 | 7.28E-03 | 2.05E-02 | 2.53E+00 | 9.95E-05 | 3.65E-04 | 1.83E+00 | 2.31E-04 | 6.35E-04 | 1.94E+00 |
| SIGLEC12 | 1.80E-01 | 9.37E-01 | 1.05E+00 | 1.99E-01 | 8.61E-01 | 1.12E+00 | 2.33E-01 | 9.90E-01 | 1.20E+00 | 4.36E-02 | 1.67E-01 | 1.11E+00 | 4.75E-07 | 1.69E-06 | -2.66E+00 | 7.86E-07 | 2.21E-06 | -2.48E+00 | 6.10E-02 | 2.24E-01 | -1.20E+00 | 1.88E-01 | 5.17E-01 | -1.11E+00 |
| NSG1 | 1.80E-01 | 9.38E-01 | -1.04E+00 | 2.20E-01 | 9.51E-01 | 1.01E+00 | 2.12E-01 | 9.00E-01 | -1.05E+00 | 2.61E-01 | 9.96E-01 | 1.05E+00 | 2.58E-02 | 9.21E-02 | -1.67E+00 | 2.56E-02 | 7.22E-02 | 1.07E+00 | 1.23E-01 | 4.52E-01 | 1.05E+00 | 6.36E-02 | 1.75E-01 | 1.25E+00 |
| HIST1H4H | 1.80E-01 | 9.38E-01 | 1.27E+00 | 9.35E-02 | 4.04E-01 | 1.06E+00 | 5.23E-02 | 2.22E-01 | -1.11E+00 | 5.12E-03 | 1.95E-02 | -1.48E+00 | 1.94E-01 | 6.92E-01 | 1.53E+00 | 3.31E-01 | 9.33E-01 | 1.07E+00 | 4.12E-02 | 1.51E-01 | 1.69E+00 | 3.04E-01 | 8.35E-01 | 1.03E+00 |
| FAM168A | 1.80E-01 | 9.38E-01 | 1.19E+00 | 1.86E-01 | 8.03E-01 | 1.31E+00 | 1.08E-01 | 4.60E-01 | 1.19E+00 | 7.95E-02 | 3.04E-01 | 1.26E+00 | 8.32E-03 | 2.97E-02 | -1.16E+00 | 9.44E-03 | 2.66E-02 | -1.10E+00 | 1.13E-02 | 4.15E-02 | -1.22E+00 | 3.52E-02 | 9.66E-02 | -1.15E+00 |
| MELK | 1.80E-01 | 9.38E-01 | 1.11E+00 | 1.91E-01 | 8.26E-01 | 1.28E+00 | 3.73E-03 | 1.59E-02 | 1.62E+00 | 6.66E-02 | 2.55E-01 | 1.61E+00 | 9.90E-02 | 3.53E-01 | -1.96E+00 | 1.33E-01 | 3.76E-01 | -1.86E+00 | 2.62E-02 | 9.62E-02 | 1.06E+00 | 9.66E-02 | 2.65E-01 | 1.53E+00 |
| BTF3P12 | 1.80E-01 | 9.38E-01 | 1.43E+00 | 5.00E-04 | 2.16E-03 | 2.46E+00 | 1.43E-01 | 6.09E-01 | 1.34E+00 | 1.10E-05 | 4.19E-05 | 2.39E+00 | 5.40E-02 | 1.93E-01 | 1.79E+00 | 6.15E-04 | 1.73E-03 | 2.57E+00 | 1.22E-01 | 4.48E-01 | 1.39E+00 | 1.90E-04 | 5.21E-04 | 2.04E+00 |
| LONRF3 | 1.80E-01 | 9.38E-01 | 1.15E+00 | 5.51E-02 | 2.38E-01 | -1.44E+00 | 2.25E-03 | 9.57E-03 | -1.34E+00 | 3.46E-05 | 1.32E-04 | -1.69E+00 | 2.40E-01 | 8.57E-01 | 1.19E+00 | 3.55E-01 | 1.00E+00 | 1.00E+00 | 2.68E-01 | 9.84E-01 | 1.03E+00 | 3.23E-01 | 8.88E-01 | -1.03E+00 |
| TEK | 1.80E-01 | 9.38E-01 | 1.10E+00 | 7.27E-02 | 3.14E-01 | 1.74E+00 | 1.04E-01 | 4.41E-01 | -1.20E+00 | 1.35E-01 | 5.17E-01 | 1.00E+00 | 1.10E-04 | 3.93E-04 | -1.90E+00 | 1.00E-03 | 2.83E-03 | -1.62E+00 | 9.93E-17 | 3.64E-16 | -2.01E+00 | 6.63E-15 | 1.82E-14 | -2.01E+00 |
| AHSP | 1.80E-01 | 9.39E-01 | 2.40E+00 | 1.62E-01 | 6.98E-01 | 1.29E+00 | 2.16E-01 | 9.19E-01 | -1.03E+00 | 1.39E-01 | 5.31E-01 | -1.48E+00 | 2.21E-01 | 7.87E-01 | 6.66E+00 | 2.39E-01 | 6.72E-01 | 3.21E+00 | 6.07E-02 | 2.23E-01 | 1.52E+00 | 7.48E-02 | 2.05E-01 | 1.10E+00 |
| AP000942.1 | 1.80E-01 | 9.39E-01 | 1.23E+00 | 7.23E-02 | 3.12E-01 | 1.01E+00 | 1.03E-01 | 4.36E-01 | 1.02E+00 | 1.23E-03 | 4.69E-03 | -1.42E+00 | 2.90E-02 | 1.03E-01 | -1.14E+00 | 7.45E-03 | 2.10E-02 | -1.47E+00 | 1.61E-05 | 5.93E-05 | -1.45E+00 | 1.39E-05 | 3.82E-05 | -1.53E+00 |
| FAM66BP | 1.80E-01 | 9.39E-01 | 1.39E+00 | 1.97E-01 | 8.53E-01 | 1.46E+00 | 8.69E-02 | 3.69E-01 | 1.02E+00 | 2.27E-01 | 8.67E-01 | 1.39E+00 | 7.75E-03 | 2.76E-02 | -1.10E+00 | 1.98E-02 | 5.59E-02 | -1.02E+00 | 4.81E-03 | 1.77E-02 | -1.30E+00 | 1.62E-01 | 4.45E-01 | 1.10E+00 |
| ITGB5 | 1.80E-01 | 9.39E-01 | 1.09E+00 | 1.78E-01 | 7.68E-01 | -1.06E+00 | 1.88E-01 | 7.99E-01 | 1.09E+00 | 2.05E-01 | 7.83E-01 | 1.12E+00 | 1.34E-01 | 4.78E-01 | -1.14E+00 | 1.52E-01 | 4.29E-01 | -1.15E+00 | 1.36E-02 | 4.99E-02 | 2.49E+00 | 3.21E-01 | 8.81E-01 | 1.82E+00 |
| TANC1 | 1.80E-01 | 9.39E-01 | 1.06E+00 | 2.03E-01 | 8.79E-01 | 1.13E+00 | 1.52E-01 | 6.46E-01 | 1.03E+00 | 1.16E-01 | 4.42E-01 | 1.02E+00 | 2.50E-01 | 8.92E-01 | 1.12E+00 | 4.41E-03 | 1.24E-02 | 2.32E+00 | 8.46E-20 | 3.11E-19 | -1.97E+00 | 2.02E-18 | 5.55E-18 | -1.61E+00 |
| ATP1B2 | 1.80E-01 | 9.40E-01 | 1.08E+00 | 1.72E-01 | 7.43E-01 | -1.09E+00 | 1.83E-01 | 7.79E-01 | 1.11E+00 | 2.62E-01 | 1.00E+00 | 1.00E+00 | 1.92E-01 | 6.83E-01 | 1.26E+00 | 3.17E-01 | 8.95E-01 | 1.09E+00 | 2.15E-02 | 7.88E-02 | -1.22E+00 | 3.79E-03 | 1.04E-02 | -1.35E+00 |
| DDX18P4 | 1.80E-01 | 9.40E-01 | 1.27E+00 | 1.60E-01 | 6.92E-01 | 1.29E+00 | 1.14E-01 | 4.85E-01 | 1.03E+00 | 4.85E-02 | 1.85E-01 | -1.07E+00 | 2.09E-02 | 7.47E-02 | -1.37E+00 | 5.51E-02 | 1.55E-01 | -1.14E+00 | 1.67E-02 | 6.14E-02 | -1.29E+00 | 3.46E-01 | 9.50E-01 | 1.16E+00 |
| EPCAM | 1.80E-01 | 9.40E-01 | 1.03E+00 | 1.95E-01 | 8.41E-01 | -1.03E+00 | 8.69E-03 | 3.69E-02 | -1.60E+00 | 2.69E-02 | 1.03E-01 | -1.32E+00 | 2.53E-01 | 9.01E-01 | -1.01E+00 | 3.22E-01 | 9.06E-01 | -1.06E+00 | 5.36E-17 | 1.97E-16 | -2.64E+00 | 4.23E-08 | 1.16E-07 | -1.72E+00 |
| ENPP4 | 1.80E-01 | 9.40E-01 | 1.39E+00 | 1.35E-01 | 5.82E-01 | 1.70E+00 | 1.92E-01 | 8.16E-01 | 1.22E+00 | 5.36E-02 | 2.05E-01 | 1.64E+00 | 2.60E-01 | 9.27E-01 | 1.61E+00 | 6.99E-02 | 1.97E-01 | 1.84E+00 | 2.28E-01 | 8.36E-01 | 1.29E+00 | 1.34E-01 | 3.67E-01 | 1.09E+00 |
| RABAC1 | 1.81E-01 | 9.41E-01 | 1.23E+00 | 2.15E-01 | 9.31E-01 | 1.34E+00 | 1.11E-01 | 4.70E-01 | 1.21E+00 | 1.27E-01 | 4.87E-01 | 1.31E+00 | 4.31E-04 | 1.54E-03 | -1.08E+00 | 3.09E-03 | 8.70E-03 | 1.01E+00 | 4.31E-02 | 1.58E-01 | 1.38E+00 | 3.22E-01 | 8.84E-01 | 1.30E+00 |
| ANK1 | 1.81E-01 | 9.41E-01 | 1.17E+00 | 2.03E-01 | 8.76E-01 | 1.07E+00 | 2.29E-01 | 9.72E-01 | 1.19E+00 | 2.15E-01 | 8.20E-01 | 1.50E+00 | 1.67E-02 | 5.97E-02 | 1.78E+00 | 1.38E-21 | 3.90E-21 | 4.17E+00 | 8.52E-06 | 3.13E-05 | -1.27E+00 | 8.54E-04 | 2.35E-03 | -1.07E+00 |
| TRPC6 | 1.81E-01 | 9.41E-01 | 1.06E+00 | 2.31E-01 | 1.00E+00 | 1.00E+00 | 2.35E-01 | 1.00E+00 | -1.00E+00 | 2.25E-01 | 8.58E-01 | 1.18E+00 | 8.95E-02 | 3.19E-01 | -1.60E+00 | 2.85E-01 | 8.02E-01 | 1.03E+00 | 2.50E-07 | 9.18E-07 | -1.23E+00 | 7.73E-08 | 2.12E-07 | -1.30E+00 |
| TNFAIP6 | 1.81E-01 | 9.41E-01 | 1.14E+00 | 1.93E-01 | 8.34E-01 | -1.07E+00 | 1.16E-01 | 4.94E-01 | -1.17E+00 | 1.20E-01 | 4.57E-01 | -1.20E+00 | 1.04E-01 | 3.72E-01 | -1.61E+00 | 1.30E-01 | 3.68E-01 | 2.16E+00 | 3.45E-03 | 1.27E-02 | -1.41E+00 | 1.12E-01 | 3.08E-01 | -1.04E+00 |
| CPB1 | 1.81E-01 | 9.41E-01 | 1.14E+00 | 2.14E-01 | 9.26E-01 | -1.06E+00 | 1.53E-01 | 6.50E-01 | -1.38E+00 | 2.38E-01 | 9.10E-01 | 1.06E+00 | 2.38E-01 | 8.49E-01 | 1.30E+00 | 1.64E-01 | 4.63E-01 | 1.71E+00 | 1.33E-01 | 4.87E-01 | 1.21E+00 | 3.39E-01 | 9.30E-01 | 1.24E+00 |
| ZEB2-AS1 | 1.81E-01 | 9.42E-01 | 1.20E+00 | 1.81E-01 | 7.84E-01 | 1.53E+00 | 1.47E-04 | 6.25E-04 | -1.67E+00 | 3.32E-02 | 1.27E-01 | 1.08E+00 | 3.91E-03 | 1.40E-02 | -1.23E+00 | 1.34E-02 | 3.78E-02 | -1.05E+00 | 2.44E-03 | 8.97E-03 | -1.08E+00 | 7.32E-04 | 2.01E-03 | -1.21E+00 |
| AC141424.1 | 1.81E-01 | 9.42E-01 | 1.31E+00 | 1.97E-02 | 8.52E-02 | -1.07E+00 | 4.61E-02 | 1.96E-01 | 1.70E+00 | 4.54E-02 | 1.73E-01 | -1.10E+00 | 7.49E-02 | 2.67E-01 | 1.63E+00 | 2.52E-01 | 7.10E-01 | 1.03E+00 | 1.72E-01 | 6.30E-01 | 1.26E+00 | 4.35E-02 | 1.19E-01 | -1.33E+00 |
| GNA13 | 1.81E-01 | 9.42E-01 | 1.35E+00 | 1.89E-01 | 8.16E-01 | 1.56E+00 | 3.37E-02 | 1.43E-01 | 1.07E+00 | 3.38E-03 | 1.29E-02 | 1.05E+00 | 7.42E-02 | 2.65E-01 | 1.01E+00 | 5.89E-03 | 1.66E-02 | -1.25E+00 | 2.00E-04 | 7.36E-04 | -1.13E+00 | 4.88E-04 | 1.34E-03 | -1.13E+00 |
| AL137060.1 | 1.81E-01 | 9.43E-01 | 1.08E+00 | 6.34E-02 | 2.74E-01 | -1.21E+00 | 1.23E-02 | 5.21E-02 | -1.25E+00 | 2.30E-02 | 8.80E-02 | -1.05E+00 | 7.59E-02 | 2.71E-01 | -1.24E+00 | 1.36E-01 | 3.82E-01 | -1.14E+00 | 1.86E-01 | 6.84E-01 | 1.39E+00 | 3.08E-01 | 8.47E-01 | 1.08E+00 |
| SPTBN2 | 1.81E-01 | 9.43E-01 | 1.17E+00 | 2.12E-01 | 9.14E-01 | 1.01E+00 | 1.91E-01 | 8.10E-01 | 1.36E+00 | 2.54E-01 | 9.70E-01 | 1.10E+00 | 2.41E-08 | 8.59E-08 | -1.63E+00 | 1.51E-02 | 4.26E-02 | 1.16E+00 | 2.66E-22 | 9.78E-22 | -2.06E+00 | 4.61E-15 | 1.27E-14 | -1.62E+00 |
| COL23A1 | 1.81E-01 | 9.44E-01 | 1.05E+00 | 1.70E-01 | 7.34E-01 | 1.34E+00 | 2.27E-01 | 9.64E-01 | 1.09E+00 | 2.09E-01 | 7.98E-01 | 1.31E+00 | 2.17E-06 | 7.72E-06 | -1.92E+00 | 2.58E-06 | 7.27E-06 | -1.91E+00 | 2.05E-03 | 7.53E-03 | -1.30E+00 | 2.33E-03 | 6.40E-03 | -1.31E+00 |
| AC008764.8 | 1.81E-01 | 9.44E-01 | 1.29E+00 | 1.95E-01 | 8.42E-01 | 1.52E+00 | 2.34E-01 | 9.95E-01 | 1.24E+00 | 2.23E-01 | 8.51E-01 | 1.26E+00 | 1.69E-01 | 6.04E-01 | 1.59E+00 | 2.22E-01 | 6.27E-01 | 1.19E+00 | 8.60E-02 | 3.16E-01 | 1.60E+00 | 9.21E-02 | 2.53E-01 | 1.72E+00 |
| POC1A | 1.81E-01 | 9.44E-01 | 1.24E+00 | 2.19E-01 | 9.47E-01 | 1.48E+00 | 1.79E-02 | 7.59E-02 | 1.53E+00 | 1.87E-01 | 7.15E-01 | 1.30E+00 | 1.12E-01 | 4.00E-01 | -1.07E+00 | 2.29E-01 | 6.46E-01 | 1.02E+00 | 2.97E-02 | 1.09E-01 | 1.52E+00 | 6.60E-02 | 1.81E-01 | 1.53E+00 |
| ZSCAN1 | 1.81E-01 | 9.44E-01 | 1.28E+00 | 1.71E-01 | 7.38E-01 | 1.20E+00 | 9.78E-02 | 4.16E-01 | 1.52E+00 | 1.87E-01 | 7.15E-01 | 1.38E+00 | 2.54E-01 | 9.05E-01 | 1.23E+00 | 9.69E-02 | 2.73E-01 | -1.28E+00 | 5.66E-02 | 2.08E-01 | 1.73E+00 | 1.42E-01 | 3.89E-01 | 1.56E+00 |
| LIN7B | 1.81E-01 | 9.44E-01 | 1.25E+00 | 1.34E-01 | 5.79E-01 | 1.72E+00 | 2.16E-01 | 9.16E-01 | 1.11E+00 | 1.87E-01 | 7.15E-01 | 1.17E+00 | 1.16E-01 | 4.12E-01 | 1.08E+00 | 1.86E-01 | 5.25E-01 | 1.00E+00 | 6.77E-02 | 2.48E-01 | -1.07E+00 | 4.41E-02 | 1.21E-01 | -1.24E+00 |
| DHCR7 | 1.81E-01 | 9.45E-01 | 1.35E+00 | 2.07E-02 | 8.92E-02 | 1.97E+00 | 1.51E-01 | 6.43E-01 | 1.06E+00 | 4.19E-03 | 1.60E-02 | 1.86E+00 | 2.02E-01 | 7.18E-01 | 1.18E+00 | 1.82E-02 | 5.13E-02 | 1.93E+00 | 1.73E-01 | 6.34E-01 | 1.33E+00 | 2.75E-04 | 7.54E-04 | 2.20E+00 |
| ALKAL2 | 1.81E-01 | 9.46E-01 | 1.41E+00 | 4.74E-02 | 2.05E-01 | 2.50E+00 | 1.69E-01 | 7.16E-01 | 1.31E+00 | 1.78E-01 | 6.79E-01 | 1.47E+00 | 6.32E-02 | 2.25E-01 | 2.92E+00 | 1.90E-02 | 5.35E-02 | 3.56E+00 | 5.84E-15 | 2.14E-14 | 3.98E+00 | 3.59E-03 | 9.86E-03 | 1.94E+00 |
| PSMD6-AS2 | 1.81E-01 | 9.46E-01 | 1.25E+00 | 1.08E-01 | 4.66E-01 | 1.84E+00 | 8.39E-02 | 3.57E-01 | -1.10E+00 | 1.42E-01 | 5.43E-01 | 1.10E+00 | 1.44E-01 | 5.13E-01 | -1.15E+00 | 2.98E-01 | 8.40E-01 | 1.12E+00 | 2.31E-01 | 8.49E-01 | 1.28E+00 | 2.51E-01 | 6.89E-01 | 1.51E+00 |
| DDX10P1 | 1.81E-01 | 9.46E-01 | 1.13E+00 | 5.19E-02 | 2.24E-01 | -1.26E+00 | 5.32E-05 | 2.26E-04 | -1.44E+00 | 1.50E-05 | 5.74E-05 | -1.44E+00 | 1.60E-01 | 5.69E-01 | 1.70E+00 | 1.64E-01 | 4.63E-01 | 1.90E+00 | 2.61E-03 | 9.58E-03 | -1.56E+00 | 7.09E-04 | 1.95E-03 | -2.12E+00 |
| AC254633.1 | 1.81E-01 | 9.46E-01 | 1.05E+00 | 2.31E-01 | 1.00E+00 | -1.00E+00 | 2.09E-01 | 8.89E-01 | 1.09E+00 | 2.62E-01 | 1.00E+00 | 1.00E+00 | 1.85E-02 | 6.59E-02 | 2.68E+00 | 2.67E-02 | 7.52E-02 | 2.46E+00 | 1.89E-09 | 6.94E-09 | 3.00E+00 | 2.21E-01 | 6.08E-01 | 1.55E+00 |
| F12 | 1.82E-01 | 9.47E-01 | 1.11E+00 | 1.15E-01 | 4.98E-01 | 1.13E+00 | 1.53E-02 | 6.50E-02 | 1.61E+00 | 9.39E-02 | 3.59E-01 | 1.21E+00 | 5.92E-02 | 2.11E-01 | -1.38E+00 | 3.48E-02 | 9.81E-02 | -1.68E+00 | 2.31E-01 | 8.48E-01 | 1.42E+00 | 3.00E-01 | 8.23E-01 | 1.46E+00 |
| AC097534.2 | 1.82E-01 | 9.48E-01 | 1.53E+00 | 1.92E-01 | 8.31E-01 | 1.58E+00 | 1.87E-01 | 7.96E-01 | 1.30E+00 | 7.60E-02 | 2.90E-01 | 1.07E+00 | 3.24E-02 | 1.16E-01 | -1.12E+00 | 8.45E-02 | 2.38E-01 | 1.01E+00 | 6.59E-05 | 2.42E-04 | -1.48E+00 | 6.47E-06 | 1.78E-05 | -1.75E+00 |
| SNED1 | 1.82E-01 | 9.49E-01 | 1.25E+00 | 5.81E-02 | 2.51E-01 | 1.66E+00 | 1.53E-01 | 6.51E-01 | 1.22E+00 | 1.49E-01 | 5.70E-01 | 1.33E+00 | 1.25E-01 | 4.46E-01 | 2.00E+00 | 1.22E-01 | 3.44E-01 | 2.40E+00 | 2.56E-01 | 9.41E-01 | 1.14E+00 | 3.12E-01 | 8.56E-01 | -1.01E+00 |
| C12orf57 | 1.82E-01 | 9.49E-01 | 1.19E+00 | 2.05E-11 | 8.87E-11 | -1.07E+00 | 1.57E-01 | 6.68E-01 | 1.07E+00 | 1.49E-03 | 5.70E-03 | -1.13E+00 | 2.28E-01 | 8.14E-01 | -1.01E+00 | 2.39E-04 | 6.73E-04 | -1.29E+00 | 1.54E-01 | 5.63E-01 | 1.10E+00 | 1.18E-03 | 3.23E-03 | -1.17E+00 |
| AC135068.6 | 1.82E-01 | 9.49E-01 | 1.22E+00 | 2.29E-01 | 9.91E-01 | 1.35E+00 | 9.13E-04 | 3.88E-03 | 2.10E+00 | 1.63E-03 | 6.22E-03 | 2.47E+00 | 1.04E-01 | 3.72E-01 | 1.00E+00 | 1.96E-01 | 5.53E-01 | 1.87E+00 | 1.83E-01 | 6.73E-01 | 1.12E+00 | 1.09E-01 | 2.98E-01 | 1.58E+00 |
| RBM12 | 1.82E-01 | 9.49E-01 | 1.28E+00 | 3.24E-02 | 1.40E-01 | 1.19E+00 | 2.02E-02 | 8.57E-02 | -1.05E+00 | 4.33E-03 | 1.66E-02 | -1.01E+00 | 1.93E-02 | 6.90E-02 | -1.02E+00 | 8.94E-02 | 2.52E-01 | 1.08E+00 | 1.37E-04 | 5.04E-04 | -1.22E+00 | 2.32E-01 | 6.36E-01 | 1.20E+00 |
| NPM1P34 | 1.82E-01 | 9.50E-01 | 1.27E+00 | 7.58E-02 | 3.27E-01 | 1.70E+00 | 1.00E-01 | 4.27E-01 | 1.35E+00 | 2.28E-01 | 8.69E-01 | 1.44E+00 | 1.19E-01 | 4.25E-01 | 1.81E+00 | 1.47E-02 | 4.14E-02 | 2.41E+00 | 7.28E-03 | 2.67E-02 | 1.96E+00 | 5.56E-03 | 1.53E-02 | 2.18E+00 |
| MADCAM1 | 1.82E-01 | 9.51E-01 | 1.33E+00 | 1.59E-01 | 6.86E-01 | 1.60E+00 | 2.12E-01 | 9.00E-01 | 1.28E+00 | 2.23E-01 | 8.52E-01 | 1.16E+00 | 6.93E-02 | 2.47E-01 | -1.34E+00 | 1.87E-01 | 5.27E-01 | -1.10E+00 | 2.09E-01 | 7.67E-01 | -1.02E+00 | 3.50E-01 | 9.63E-01 | 1.26E+00 |
| ATP5HP2 | 1.82E-01 | 9.51E-01 | 1.22E+00 | 1.66E-01 | 7.18E-01 | 1.28E+00 | 1.67E-02 | 7.10E-02 | 1.84E+00 | 2.12E-02 | 8.09E-02 | 2.05E+00 | 1.10E-01 | 3.90E-01 | 1.93E+00 | 3.30E-01 | 9.29E-01 | 1.27E+00 | 1.20E-01 | 4.39E-01 | 1.56E+00 | 1.14E-03 | 3.12E-03 | 2.26E+00 |
| CCND1 | 1.82E-01 | 9.51E-01 | 1.05E+00 | 2.18E-01 | 9.42E-01 | 1.08E+00 | 2.29E-01 | 9.72E-01 | 1.02E+00 | 2.30E-01 | 8.78E-01 | -1.05E+00 | 9.96E-03 | 3.55E-02 | -3.66E+00 | 1.74E-02 | 4.90E-02 | -3.24E+00 | 6.88E-13 | 2.53E-12 | -1.62E+00 | 9.77E-12 | 2.68E-11 | -1.93E+00 |
| NPM1P18 | 1.82E-01 | 9.51E-01 | 1.21E+00 | 1.42E-01 | 6.15E-01 | 1.68E+00 | 8.09E-02 | 3.44E-01 | -1.02E+00 | 1.66E-01 | 6.36E-01 | 1.26E+00 | 1.07E-01 | 3.82E-01 | 1.80E+00 | 3.35E-02 | 9.45E-02 | 2.26E+00 | 2.60E-02 | 9.55E-02 | 1.72E+00 | 9.94E-02 | 2.73E-01 | 1.71E+00 |
| FBLN2 | 1.82E-01 | 9.51E-01 | 1.05E+00 | 2.05E-01 | 8.84E-01 | 1.17E+00 | 1.95E-01 | 8.27E-01 | -1.06E+00 | 2.12E-01 | 8.09E-01 | -1.04E+00 | 4.16E-04 | 1.48E-03 | -1.87E+00 | 1.80E-01 | 5.07E-01 | 1.07E+00 | 3.58E-06 | 1.31E-05 | -1.30E+00 | 1.72E-04 | 4.74E-04 | -1.08E+00 |
| ZNF266 | 1.82E-01 | 9.51E-01 | 1.28E+00 | 3.21E-02 | 1.39E-01 | 1.15E+00 | 5.00E-03 | 2.12E-02 | -1.17E+00 | 1.14E-03 | 4.36E-03 | -1.15E+00 | 1.29E-01 | 4.59E-01 | 1.18E+00 | 1.28E-02 | 3.60E-02 | -1.05E+00 | 5.50E-03 | 2.02E-02 | 1.00E+00 | 2.13E-05 | 5.84E-05 | -1.36E+00 |
| PHKA1 | 1.82E-01 | 9.51E-01 | 1.25E+00 | 8.10E-02 | 3.50E-01 | 1.86E+00 | 2.17E-02 | 9.21E-02 | -1.13E+00 | 4.96E-02 | 1.89E-01 | 1.15E+00 | 2.80E-01 | 9.98E-01 | 1.12E+00 | 1.96E-01 | 5.51E-01 | 1.65E+00 | 4.43E-03 | 1.62E-02 | -1.57E+00 | 9.96E-03 | 2.73E-02 | -1.55E+00 |
| C9orf16 | 1.83E-01 | 9.52E-01 | 1.19E+00 | 2.44E-02 | 1.06E-01 | 1.19E+00 | 8.66E-02 | 3.68E-01 | 1.22E+00 | 2.19E-01 | 8.38E-01 | 1.16E+00 | 5.71E-02 | 2.04E-01 | 1.05E+00 | 1.47E-02 | 4.14E-02 | -1.04E+00 | 1.82E-01 | 6.70E-01 | 1.26E+00 | 3.46E-01 | 9.49E-01 | 1.07E+00 |
| SVIL-AS1 | 1.83E-01 | 9.52E-01 | 1.22E+00 | 3.02E-02 | 1.30E-01 | 1.13E+00 | 8.75E-02 | 3.72E-01 | 1.03E+00 | 3.46E-03 | 1.32E-02 | -1.07E+00 | 4.52E-03 | 1.61E-02 | -1.08E+00 | 1.30E-05 | 3.66E-05 | -1.45E+00 | 9.52E-04 | 3.50E-03 | -1.18E+00 | 4.68E-03 | 1.28E-02 | -1.13E+00 |
| AC115223.1 | 1.83E-01 | 9.52E-01 | 1.45E+00 | 3.17E-02 | 1.37E-01 | 2.01E+00 | 2.20E-01 | 9.36E-01 | 1.26E+00 | 4.17E-02 | 1.59E-01 | 1.78E+00 | 2.05E-01 | 7.31E-01 | 1.27E+00 | 1.61E-02 | 4.55E-02 | 2.02E+00 | 1.86E-01 | 6.84E-01 | 1.19E+00 | 6.14E-03 | 1.69E-02 | 1.89E+00 |
| CCDC7 | 1.83E-01 | 9.52E-01 | 1.39E+00 | 9.04E-02 | 3.90E-01 | 1.32E+00 | 1.70E-04 | 7.22E-04 | -1.49E+00 | 5.48E-04 | 2.09E-03 | -1.20E+00 | 1.57E-01 | 5.59E-01 | 1.04E+00 | 3.00E-01 | 8.46E-01 | 1.18E+00 | 8.24E-03 | 3.03E-02 | -1.29E+00 | 1.39E-02 | 3.81E-02 | -1.27E+00 |
| ERFE | 1.83E-01 | 9.53E-01 | 1.03E+00 | 2.31E-01 | 1.00E+00 | -1.00E+00 | 2.07E-01 | 8.79E-01 | 1.10E+00 | 2.62E-01 | 1.00E+00 | 1.00E+00 | 2.38E-01 | 8.48E-01 | 1.00E+00 | 9.76E-02 | 2.75E-01 | 2.12E+00 | 1.37E-03 | 5.03E-03 | -1.08E+00 | 2.53E-04 | 6.94E-04 | -1.04E+00 |
| THSD7A | 1.83E-01 | 9.53E-01 | -1.09E+00 | 8.17E-03 | 3.53E-02 | -1.81E+00 | 3.41E-02 | 1.45E-01 | -1.20E+00 | 6.71E-02 | 2.56E-01 | 1.04E+00 | 2.17E-01 | 7.73E-01 | -1.14E+00 | 3.22E-01 | 9.09E-01 | 1.22E+00 | 1.40E-01 | 5.15E-01 | -1.24E+00 | 2.44E-01 | 6.70E-01 | -1.16E+00 |
| AL353771.1 | 1.83E-01 | 9.53E-01 | 1.29E+00 | 2.11E-01 | 9.13E-01 | 1.45E+00 | 1.17E-01 | 4.97E-01 | 1.43E+00 | 2.24E-01 | 8.55E-01 | 1.31E+00 | 2.61E-01 | 9.30E-01 | 1.55E+00 | 2.03E-01 | 5.71E-01 | 1.25E+00 | 1.43E-01 | 5.26E-01 | 1.22E+00 | 3.00E-01 | 8.23E-01 | 1.02E+00 |
| AL138752.1 | 1.83E-01 | 9.53E-01 | 1.26E+00 | 1.85E-01 | 8.01E-01 | 1.30E+00 | 1.86E-01 | 7.90E-01 | 1.26E+00 | 1.41E-01 | 5.39E-01 | 1.12E+00 | 1.05E-01 | 3.74E-01 | -1.07E+00 | 1.29E-01 | 3.63E-01 | -1.03E+00 | 1.54E-01 | 5.67E-01 | 1.35E+00 | 3.19E-01 | 8.76E-01 | 1.14E+00 |
| GAB2 | 1.83E-01 | 9.53E-01 | 1.33E+00 | 2.03E-02 | 8.77E-02 | 1.12E+00 | 1.25E-03 | 5.30E-03 | 1.86E+00 | 1.06E-01 | 4.05E-01 | 1.80E+00 | 5.76E-02 | 2.05E-01 | 1.02E+00 | 3.09E-02 | 8.72E-02 | -1.11E+00 | 1.59E-02 | 5.85E-02 | 1.02E+00 | 4.25E-03 | 1.17E-02 | -1.16E+00 |
| PRRT1 | 1.83E-01 | 9.54E-01 | 1.37E+00 | 9.66E-02 | 4.17E-01 | 1.18E+00 | 5.05E-02 | 2.14E-01 | -1.10E+00 | 8.01E-03 | 3.06E-02 | -1.38E+00 | 1.34E-01 | 4.77E-01 | -1.03E+00 | 2.88E-01 | 8.13E-01 | 1.17E+00 | 7.73E-02 | 2.84E-01 | 1.04E+00 | 3.83E-02 | 1.05E-01 | -1.12E+00 |

| | | | | | | | | | | | | | | | | | | | | | | | | |
|---|---|---|---|---|---|---|---|---|---|---|---|---|---|---|---|---|---|---|---|---|---|---|---|---|
| CCDC141 | 1.83E-01 | 9.54E-01 | 1.11E+00 | 2.04E-01 | 8.82E-01 | 1.02E+00 | 8.07E-02 | 3.43E-01 | 1.49E+00 | 1.49E-01 | 5.69E-01 | 1.41E+00 | 2.14E-04 | 7.62E-04 | 2.63E+00 | 2.58E-01 | 7.28E-01 | 1.40E+00 | 1.03E-01 | 3.77E-01 | 1.36E+00 | 2.45E-01 | 6.74E-01 | 1.23E+00 |
| CENPH | 1.83E-01 | 9.55E-01 | 1.24E+00 | 3.55E-02 | 1.53E-01 | 1.11E+00 | 2.02E-02 | 8.58E-02 | 1.42E+00 | 1.42E-02 | 5.44E-02 | 1.14E+00 | 9.22E-03 | 3.29E-02 | -1.20E+00 | 8.81E-03 | 2.48E-02 | -1.21E+00 | 3.62E-02 | 1.33E-01 | 1.00E+00 | 3.57E-01 | 9.80E-01 | 1.09E+00 |
| SFRP5 | 1.83E-01 | 9.55E-01 | 1.10E+00 | 2.22E-01 | 9.59E-01 | 1.06E+00 | 1.97E-01 | 8.38E-01 | -1.17E+00 | 2.44E-01 | 9.33E-01 | -1.06E+00 | 1.60E-07 | 5.72E-07 | -2.21E+00 | 5.74E-03 | 1.62E-02 | -1.03E+00 | 9.91E-40 | 3.64E-39 | -2.24E+00 | 2.02E-39 | 5.56E-39 | -2.19E+00 |
| AC015849.3 | 1.83E-01 | 9.55E-01 | 1.10E+00 | 1.35E-01 | 5.83E-01 | 1.40E+00 | 1.69E-01 | 7.16E-01 | 1.22E+00 | 1.97E-01 | 7.52E-01 | 1.29E+00 | 1.91E-02 | 6.81E-02 | -1.32E+00 | 4.28E-02 | 1.21E-01 | -1.13E+00 | 8.35E-04 | 3.06E-03 | -1.53E+00 | 1.69E-01 | 4.64E-01 | 1.08E+00 |
| SUDS3P1 | 1.83E-01 | 9.56E-01 | 1.34E+00 | 1.34E-01 | 5.78E-01 | 1.69E+00 | 8.61E-02 | 3.66E-01 | 1.49E+00 | 2.42E-01 | 9.23E-01 | 1.28E+00 | 2.80E-01 | 9.98E-01 | 1.17E+00 | 4.61E-02 | 1.30E-01 | 1.93E+00 | 6.40E-02 | 2.35E-01 | 1.61E+00 | 3.15E-02 | 8.65E-02 | 1.86E+00 |
| PTPN14 | 1.83E-01 | 9.56E-01 | 1.20E+00 | 3.68E-04 | 1.59E-03 | 2.01E+00 | 1.28E-01 | 5.42E-01 | -1.03E+00 | 1.31E-02 | 4.99E-02 | 1.66E+00 | 2.77E-01 | 9.88E-01 | 1.45E+00 | 1.79E-01 | 5.06E-01 | 2.11E+00 | 2.10E-01 | 7.72E-01 | -1.08E+00 | 1.85E-01 | 5.09E-01 | 1.82E+00 |
| ZBTB37 | 1.83E-01 | 9.56E-01 | 1.36E+00 | 2.06E-04 | 8.92E-04 | 2.09E+00 | 1.27E-03 | 5.41E-03 | -1.14E+00 | 1.65E-01 | 6.31E-01 | 1.35E+00 | 2.13E-01 | 7.59E-01 | 1.59E+00 | 2.01E-01 | 5.66E-01 | 1.67E+00 | 1.86E-03 | 6.84E-03 | -1.19E+00 | 1.51E-01 | 4.14E-01 | 1.19E+00 |
| SUMO4 | 1.83E-01 | 9.56E-01 | 1.41E+00 | 1.63E-01 | 7.04E-01 | 1.63E+00 | 2.20E-01 | 9.36E-01 | 1.25E+00 | 1.20E-01 | 4.56E-01 | 1.59E+00 | 4.13E-02 | 1.47E-01 | 2.18E+00 | 3.32E-01 | 9.35E-01 | 1.66E+00 | 8.97E-02 | 3.29E-01 | 1.62E+00 | 2.11E-01 | 5.79E-01 | 1.18E+00 |
| GYPA | 1.83E-01 | 9.56E-01 | 1.13E+00 | 2.23E-01 | 9.65E-01 | 1.12E+00 | 2.11E-01 | 8.95E-01 | 1.11E+00 | 2.15E-01 | 8.22E-01 | 1.23E+00 | 2.73E-01 | 9.73E-01 | 1.36E+00 | 1.04E-01 | 2.93E-01 | 2.69E+00 | 1.01E-04 | 3.71E-04 | -1.51E+00 | 2.20E-02 | 6.04E-02 | -1.07E+00 |
| SNX24 | 1.83E-01 | 9.56E-01 | 1.12E+00 | 2.16E-01 | 9.32E-01 | 1.19E+00 | 5.58E-02 | 2.37E-01 | -1.16E+00 | 2.12E-02 | 8.11E-02 | -1.25E+00 | 1.77E-01 | 6.29E-01 | -1.14E+00 | 1.30E-01 | 3.66E-01 | -1.34E+00 | 2.17E-01 | 7.95E-01 | 1.33E+00 | 3.50E-01 | 9.62E-01 | 1.22E+00 |
| PMP22 | 1.84E-01 | 9.57E-01 | -1.13E+00 | 2.31E-01 | 9.96E-01 | -1.04E+00 | 1.86E-01 | 7.89E-01 | 1.30E+00 | 2.37E-01 | 9.06E-01 | 1.00E+00 | 9.42E-05 | 3.36E-04 | -2.11E+00 | 2.52E-01 | 7.09E-01 | 1.36E+00 | 3.08E-37 | 1.13E-36 | -1.88E+00 | 4.10E-21 | 1.13E-20 | -1.72E+00 |
| AP001453.2 | 1.84E-01 | 9.57E-01 | -1.02E+00 | 9.27E-02 | 4.01E-01 | -1.24E+00 | 1.88E-01 | 7.99E-01 | 1.26E+00 | 1.30E-01 | 4.95E-01 | 1.02E+00 | 2.49E-01 | 8.88E-01 | 1.22E+00 | 1.72E-01 | 4.86E-01 | -1.13E+00 | 2.55E-01 | 9.37E-01 | 1.11E+00 | 2.21E-01 | 6.08E-01 | -1.10E+00 |
| FAM214A | 1.84E-01 | 9.57E-01 | 1.37E+00 | 2.96E-04 | 1.28E-03 | 1.06E+00 | 6.76E-02 | 2.87E-01 | 1.29E+00 | 1.02E-04 | 3.88E-04 | -1.22E+00 | 3.57E-02 | 1.27E-01 | 1.68E+00 | 2.91E-01 | 8.20E-01 | 1.26E+00 | 1.73E-01 | 6.36E-01 | 1.37E+00 | 7.28E-05 | 2.00E-04 | -1.33E+00 |
| ATP5J | 1.84E-01 | 9.57E-01 | 1.15E+00 | 3.73E-04 | 1.61E-03 | 1.07E+00 | 1.69E-02 | 7.19E-02 | 1.23E+00 | 2.33E-01 | 8.91E-01 | 1.07E+00 | 1.84E-02 | 6.58E-02 | 1.03E+00 | 1.08E-02 | 3.03E-02 | -1.01E+00 | 7.46E-03 | 2.74E-02 | 1.26E+00 | 3.06E-01 | 8.39E-01 | 1.06E+00 |
| ITGB3 | 1.84E-01 | 9.57E-01 | 1.05E+00 | 2.08E-01 | 8.98E-01 | 1.19E+00 | 2.22E-01 | 9.43E-01 | 1.00E+00 | 2.50E-01 | 9.55E-01 | 1.01E+00 | 8.55E-03 | 3.05E-02 | -2.08E+00 | 1.91E-02 | 5.37E-02 | -2.41E+00 | 2.33E-01 | 8.56E-01 | 1.35E+00 | 1.96E-02 | 5.39E-02 | 1.06E+00 |
| HAUS6 | 1.84E-01 | 9.57E-01 | 1.27E+00 | 2.12E-01 | 9.16E-01 | 1.47E+00 | 2.58E-02 | 1.10E-01 | 1.34E+00 | 1.17E-01 | 4.48E-01 | 1.11E+00 | 8.35E-02 | 2.98E-01 | 1.30E+00 | 1.67E-03 | 4.72E-03 | -1.21E+00 | 5.47E-05 | 2.01E-04 | -1.16E+00 | 4.43E-03 | 1.22E-02 | -1.04E+00 |
| INAFM1 | 1.84E-01 | 9.57E-01 | 1.35E+00 | 2.20E-01 | 9.50E-01 | 1.59E+00 | 2.15E-01 | 9.14E-01 | 1.20E+00 | 1.76E-01 | 6.72E-01 | 1.33E+00 | 1.86E-01 | 6.64E-01 | 1.50E+00 | 2.26E-01 | 6.37E-01 | 1.61E+00 | 3.23E-02 | 1.19E-01 | 1.69E+00 | 2.66E-01 | 7.30E-01 | 1.49E+00 |
| TRAM2-AS1 | 1.84E-01 | 9.58E-01 | 1.38E+00 | 2.00E-01 | 8.62E-01 | 1.57E+00 | 2.61E-02 | 1.11E-01 | -1.14E+00 | 1.05E-01 | 4.01E-01 | 1.13E+00 | 2.25E-01 | 8.01E-01 | 1.58E+00 | 2.20E-01 | 6.21E-01 | 1.23E+00 | 1.34E-01 | 4.92E-01 | 1.04E+00 | 1.30E-01 | 3.57E-01 | -1.02E+00 |
| TRIM16L | 1.84E-01 | 9.58E-01 | 1.39E+00 | 1.67E-01 | 7.20E-01 | 1.59E+00 | 1.02E-02 | 4.34E-02 | -1.13E+00 | 1.60E-02 | 6.13E-02 | 1.05E+00 | 2.94E-02 | 1.05E-01 | -1.12E+00 | 1.30E-01 | 3.68E-01 | 1.16E+00 | 5.78E-02 | 2.12E-01 | 1.06E+00 | 3.24E-01 | 8.89E-01 | 1.16E+00 |
| LINC01881 | 1.84E-01 | 9.58E-01 | 1.18E+00 | 1.05E-01 | 4.53E-01 | -1.01E+00 | 1.01E-02 | 4.31E-02 | -1.21E+00 | 9.70E-04 | 3.71E-03 | -1.48E+00 | 1.79E-01 | 6.40E-01 | 1.18E+00 | 6.45E-02 | 1.82E-01 | -1.23E+00 | 1.54E-02 | 5.66E-02 | -1.39E+00 | 5.24E-02 | 1.44E-01 | -1.26E+00 |
| RLN2 | 1.84E-01 | 9.59E-01 | 1.32E+00 | 2.12E-01 | 9.16E-01 | 1.43E+00 | 2.25E-01 | 9.57E-01 | 1.14E+00 | 2.62E-01 | 1.00E+00 | 1.32E+00 | 1.21E-03 | 4.30E-03 | -1.21E+00 | 1.10E-02 | 3.10E-02 | -1.00E+00 | 1.12E-01 | 4.10E-01 | 1.54E+00 | 8.54E-02 | 2.35E-01 | 1.52E+00 |
| SLITRK1 | 1.84E-01 | 9.59E-01 | 1.06E+00 | 5.69E-04 | 2.46E-03 | 2.15E+00 | 2.32E-01 | 9.85E-01 | 1.35E+00 | 2.43E-01 | 9.29E-01 | 1.14E+00 | 2.50E-01 | 8.90E-01 | 1.15E+00 | 3.29E-01 | 9.26E-01 | 1.10E+00 | 2.49E-01 | 9.13E-01 | 1.04E+00 | 2.09E-01 | 5.75E-01 | 1.33E+00 |
| NEK10 | 1.84E-01 | 9.59E-01 | 1.07E+00 | 2.21E-01 | 9.55E-01 | 1.01E+00 | 1.51E-01 | 6.42E-01 | -1.25E+00 | 1.16E-01 | 4.42E-01 | -1.26E+00 | 5.78E-02 | 2.06E-01 | -1.09E+00 | 4.74E-02 | 1.34E-01 | -1.34E+00 | 2.64E-01 | 9.69E-01 | 1.14E+00 | 1.53E-01 | 4.20E-01 | -1.12E+00 |
| PLEKHM1 | 1.84E-01 | 9.59E-01 | 1.35E+00 | 2.64E-02 | 1.14E-01 | 1.14E+00 | 2.16E-02 | 9.19E-02 | -1.10E+00 | 4.55E-03 | 1.74E-02 | -1.11E+00 | 3.90E-02 | 1.39E-01 | 1.05E+00 | 9.93E-03 | 2.80E-02 | -1.08E+00 | 6.34E-03 | 2.33E-02 | -1.17E+00 | 1.30E-02 | 3.58E-02 | -1.13E+00 |
| RIPOR2 | 1.84E-01 | 9.59E-01 | 1.42E+00 | 3.62E-03 | 1.56E-02 | 1.96E+00 | 7.91E-03 | 3.36E-02 | 1.09E+00 | 4.38E-02 | 1.67E-01 | 1.31E+00 | 5.56E-06 | 1.98E-05 | -1.12E+00 | 1.84E-03 | 5.18E-03 | -1.01E+00 | 7.23E-05 | 2.65E-04 | -1.30E+00 | 1.05E-04 | 2.89E-04 | -1.21E+00 |
| ADGRG3 | 1.84E-01 | 9.59E-01 | 1.44E+00 | 2.98E-02 | 1.29E-01 | 1.23E+00 | 3.92E-09 | 1.66E-08 | -1.45E+00 | 6.03E-09 | 2.30E-08 | -1.23E+00 | 1.84E-01 | 6.54E-01 | 1.87E+00 | 3.36E-01 | 9.47E-01 | 1.26E+00 | 2.54E-01 | 9.32E-01 | 1.40E+00 | 3.33E-01 | 9.14E-01 | 1.28E+00 |
| AC133134.1 | 1.84E-01 | 9.59E-01 | 1.18E+00 | 1.15E-08 | 4.95E-08 | -1.10E+00 | 2.28E-01 | 9.69E-01 | 1.15E+00 | 1.58E-03 | 6.03E-03 | -1.04E+00 | 3.76E-03 | 1.34E-02 | 1.10E+00 | 8.10E-07 | 2.28E-06 | -1.15E+00 | 1.85E-02 | 6.78E-02 | 1.06E+00 | 1.41E-04 | 3.86E-04 | -1.10E+00 |
| TMOD1 | 1.84E-01 | 9.60E-01 | -1.05E+00 | 2.11E-01 | 9.12E-01 | 1.10E+00 | 1.87E-01 | 7.96E-01 | 1.26E+00 | 1.91E-01 | 7.28E-01 | 1.43E+00 | 1.25E-01 | 4.44E-01 | 1.75E+00 | 4.41E-02 | 1.24E-01 | 2.52E+00 | 1.99E-01 | 7.29E-01 | 1.13E+00 | 1.55E-03 | 4.26E-03 | -1.02E+00 |
| LINC01089 | 1.84E-01 | 9.60E-01 | 1.38E+00 | 3.66E-07 | 1.58E-06 | 2.45E+00 | 2.23E-01 | 9.46E-01 | 1.24E+00 | 3.57E-04 | 1.36E-03 | 2.11E+00 | 2.54E-01 | 9.06E-01 | 1.43E+00 | 8.53E-02 | 2.41E-01 | 1.90E+00 | 7.48E-02 | 2.75E-01 | 1.34E+00 | 2.18E-02 | 6.00E-02 | 1.48E+00 |
| YTHDF3-AS1 | 1.84E-01 | 9.60E-01 | 1.33E+00 | 1.81E-02 | 7.81E-02 | 1.16E+00 | 1.73E-01 | 7.34E-01 | 1.48E+00 | 4.71E-02 | 1.80E-01 | 1.25E+00 | 5.97E-02 | 2.13E-01 | 2.05E+00 | 1.49E-01 | 4.21E-01 | 1.90E+00 | 1.71E-01 | 6.29E-01 | 1.39E+00 | 2.69E-01 | 7.39E-01 | 1.16E+00 |
| AC099522.2 | 1.84E-01 | 9.60E-01 | 1.36E+00 | 1.12E-01 | 4.83E-01 | 1.32E+00 | 4.97E-03 | 2.11E-02 | -1.27E+00 | 7.25E-03 | 2.77E-02 | -1.13E+00 | 1.29E-01 | 4.61E-01 | 1.89E+00 | 2.02E-01 | 5.69E-01 | 1.95E+00 | 1.82E-01 | 6.67E-01 | 1.30E+00 | 2.74E-01 | 7.52E-01 | 1.05E+00 |
| OTUD7B | 1.84E-01 | 9.61E-01 | 1.09E+00 | 2.17E-01 | 9.38E-01 | 1.14E+00 | 1.72E-01 | 7.31E-01 | -1.09E+00 | 1.57E-01 | 5.99E-01 | -1.13E+00 | 2.00E-01 | 7.14E-01 | 1.42E+00 | 7.39E-02 | 2.08E-01 | 2.55E+00 | 7.77E-08 | 2.85E-07 | -1.48E+00 | 9.25E-06 | 2.54E-05 | -1.24E+00 |
| ALG9 | 1.84E-01 | 9.61E-01 | 1.24E+00 | 1.95E-01 | 8.45E-01 | 1.42E+00 | 1.74E-01 | 7.38E-01 | 1.23E+00 | 1.10E-02 | 4.22E-02 | -1.05E+00 | 4.41E-02 | 1.57E-01 | -1.08E+00 | 4.07E-02 | 1.15E-01 | -1.21E+00 | 9.82E-03 | 3.61E-02 | 1.01E+00 | 7.96E-03 | 2.19E-02 | -1.01E+00 |
| AC026471.6 | 1.85E-01 | 9.62E-01 | 1.11E+00 | 1.63E-01 | 7.04E-01 | 1.10E+00 | 2.41E-02 | 1.03E-01 | 1.94E+00 | 1.23E-01 | 4.69E-01 | 1.60E+00 | 1.36E-01 | 4.83E-01 | -1.50E+00 | 3.18E-01 | 8.95E-01 | -1.01E+00 | 1.01E-01 | 3.72E-01 | 1.66E+00 | 2.98E-01 | 8.18E-01 | -1.08E+00 |
| AL683813.1 | 1.85E-01 | 9.62E-01 | 1.20E+00 | 1.14E-01 | 4.93E-01 | 1.66E+00 | 2.15E-01 | 9.14E-01 | 1.04E+00 | 1.20E-01 | 4.58E-01 | 1.39E+00 | 6.01E-02 | 2.14E-01 | -1.12E+00 | 3.38E-01 | 9.53E-01 | 1.51E+00 | 4.67E-03 | 1.72E-02 | 2.19E+00 | 3.41E-03 | 9.37E-03 | 2.72E+00 |
| CHRNA5 | 1.85E-01 | 9.62E-01 | 1.25E+00 | 1.29E-01 | 5.57E-01 | 1.68E+00 | 2.04E-05 | 8.68E-05 | -1.57E+00 | 1.73E-05 | 6.60E-05 | -1.38E+00 | 7.82E-02 | 2.79E-01 | -1.55E+00 | 1.41E-01 | 3.96E-01 | -1.31E+00 | 2.63E-01 | 9.64E-01 | 1.02E+00 | 2.19E-02 | 6.01E-02 | 1.64E+00 |
| FBXO43 | 1.85E-01 | 9.62E-01 | 1.18E+00 | 9.40E-02 | 4.06E-01 | -1.25E+00 | 2.02E-01 | 8.58E-01 | 1.12E+00 | 8.61E-02 | 3.29E-01 | 1.49E+00 | 8.49E-02 | 3.03E-01 | -1.51E+00 | 1.06E-01 | 2.99E-01 | -1.47E+00 | 2.32E-01 | 8.53E-01 | 1.20E+00 | 2.55E-01 | 6.99E-01 | 1.35E+00 |
| AGTR1 | 1.85E-01 | 9.63E-01 | 1.08E+00 | 2.23E-01 | 9.65E-01 | 1.07E+00 | 2.35E-01 | 1.00E+00 | 1.00E+00 | 2.54E-01 | 9.69E-01 | 1.03E+00 | 1.03E-02 | 3.67E-02 | 2.50E+00 | 1.58E-07 | 4.45E-07 | 5.10E+00 | 6.02E-03 | 2.21E-02 | 1.15E+00 | 1.83E-04 | 5.04E-04 | 1.48E+00 |
| AC114728.1 | 1.85E-01 | 9.63E-01 | 1.29E+00 | 1.63E-04 | 7.06E-04 | -1.33E+00 | 2.87E-02 | 1.22E-01 | 1.57E+00 | 7.88E-04 | 3.01E-03 | -1.33E+00 | 5.58E-02 | 1.99E-01 | 1.08E+00 | 4.25E-05 | 1.20E-04 | -2.02E+00 | 1.20E-01 | 4.40E-01 | 1.16E+00 | 1.88E-04 | 5.17E-04 | -1.37E+00 |
| AUNIP | 1.85E-01 | 9.64E-01 | 1.12E+00 | 1.89E-01 | 8.17E-01 | 1.10E+00 | 2.31E-01 | 9.81E-01 | 1.07E+00 | 2.53E-01 | 9.67E-01 | 1.16E+00 | 6.12E-02 | 2.18E-01 | -1.92E+00 | 9.81E-02 | 2.76E-01 | -1.71E+00 | 1.21E-07 | 4.46E-07 | -1.34E+00 | 1.32E-03 | 3.62E-03 | -1.10E+00 |
| GSTM5 | 1.85E-01 | 9.64E-01 | 1.53E+00 | 2.00E-02 | 8.64E-02 | 1.36E+00 | 1.76E-01 | 7.49E-01 | 1.42E+00 | 1.57E-01 | 6.00E-01 | 1.33E+00 | 6.79E-10 | 2.42E-09 | 4.28E+00 | 4.18E-07 | 1.18E-06 | 3.13E+00 | 8.52E-06 | 3.13E-05 | 2.11E+00 | 3.03E-07 | 8.32E-07 | 2.41E+00 |
| ADNP-AS1 | 1.85E-01 | 9.66E-01 | 1.24E+00 | 2.11E-01 | 9.12E-01 | 1.23E+00 | 1.46E-01 | 6.22E-01 | 1.07E+00 | 1.01E-01 | 3.86E-01 | -1.17E+00 | 1.54E-01 | 5.49E-01 | 1.62E+00 | 1.41E-01 | 3.97E-01 | 2.03E+00 | 3.90E-02 | 1.43E-01 | -1.21E+00 | 7.78E-02 | 2.14E-01 | -1.13E+00 |
| SF3A3P2 | 1.85E-01 | 9.66E-01 | 1.09E+00 | 1.65E-01 | 7.12E-01 | 1.49E+00 | 1.97E-01 | 8.35E-01 | 1.12E+00 | 6.23E-02 | 2.38E-01 | 1.85E+00 | 4.33E-02 | 1.54E-01 | -1.57E+00 | 2.70E-01 | 7.62E-01 | 1.26E+00 | 8.14E-02 | 2.99E-01 | 1.38E+00 | 3.30E-05 | 9.06E-05 | 2.30E+00 |
| ALOX5AP | 1.85E-01 | 9.66E-01 | 1.09E+00 | 1.83E-01 | 7.91E-01 | -1.14E+00 | 7.26E-02 | 3.08E-01 | 1.45E+00 | 1.36E-01 | 5.20E-01 | 2.08E+00 | 1.73E-07 | 6.18E-07 | -1.45E+00 | 2.18E-07 | 6.16E-07 | -1.50E+00 | 5.03E-04 | 1.85E-03 | 2.50E+00 | 3.95E-03 | 1.08E-02 | 2.73E+00 |
| SLAMF9 | 1.85E-01 | 9.66E-01 | 1.05E+00 | 2.31E-01 | 1.00E+00 | -1.00E+00 | 1.81E-01 | 7.71E-01 | 1.29E+00 | 2.20E-01 | 8.40E-01 | 1.25E+00 | 1.05E-02 | 3.75E-02 | 1.68E+00 | 1.44E-01 | 4.06E-01 | 1.50E+00 | 1.41E-02 | 5.17E-02 | 1.75E+00 | 3.55E-01 | 9.76E-01 | 1.18E+00 |
| C19orf70 | 1.85E-01 | 9.66E-01 | 1.22E+00 | 5.15E-04 | 2.23E-03 | 1.17E+00 | 1.28E-01 | 5.43E-01 | 1.27E+00 | 7.14E-02 | 2.73E-01 | 1.20E+00 | 1.29E-02 | 4.61E-02 | 1.22E+00 | 7.64E-02 | 2.15E-01 | 1.22E+00 | 2.54E-01 | 9.33E-01 | 1.22E+00 | 2.33E-01 | 6.39E-01 | 1.20E+00 |
| FAM133DP | 1.86E-01 | 9.67E-01 | 1.41E+00 | 1.31E-10 | 5.64E-10 | -1.25E+00 | 3.19E-02 | 1.35E-01 | 1.44E+00 | 8.24E-05 | 3.15E-04 | -1.22E+00 | 8.30E-02 | 2.96E-01 | 1.24E+00 | 4.98E-06 | 1.40E-05 | -1.25E+00 | 6.56E-02 | 2.41E-01 | 1.17E+00 | 1.10E-04 | 3.01E-04 | -1.18E+00 |
| LMOD3 | 1.86E-01 | 9.67E-01 | 1.01E+00 | 5.75E-02 | 2.49E-01 | 1.71E+00 | 1.55E-01 | 6.57E-01 | -1.04E+00 | 2.55E-01 | 9.72E-01 | 1.30E+00 | 1.04E-02 | 3.70E-02 | 2.93E+00 | 8.43E-02 | 2.38E-01 | 2.58E+00 | 1.93E-01 | 7.09E-01 | 1.14E+00 | 2.66E-01 | 7.31E-01 | 1.47E+00 |
| BEND3 | 1.86E-01 | 9.67E-01 | 1.09E+00 | 5.77E-02 | 2.49E-01 | -1.36E+00 | 1.90E-01 | 8.09E-01 | 1.01E+00 | 1.20E-01 | 4.57E-01 | 1.00E+00 | 1.94E-01 | 6.91E-01 | 1.43E+00 | 2.93E-01 | 8.25E-01 | 1.44E+00 | 9.14E-03 | 3.35E-02 | -1.50E+00 | 9.10E-02 | 2.50E-01 | -1.08E+00 |
| TRIP13 | 1.86E-01 | 9.68E-01 | 1.22E+00 | 2.17E-01 | 9.38E-01 | 1.30E+00 | 5.44E-02 | 2.31E-01 | 1.65E+00 | 2.71E-03 | 1.04E-02 | 2.37E+00 | 4.24E-02 | 1.51E-01 | -1.69E+00 | 7.57E-02 | 2.14E-01 | -1.39E+00 | 2.89E-02 | 1.06E-01 | -1.05E+00 | 1.86E-01 | 5.12E-01 | 1.25E+00 |
| HPS6 | 1.86E-01 | 9.68E-01 | 1.34E+00 | 7.70E-02 | 3.33E-01 | 1.82E+00 | 1.80E-01 | 7.64E-01 | 1.12E+00 | 1.33E-01 | 5.07E-01 | 1.53E+00 | 1.32E-01 | 4.69E-01 | 1.06E+00 | 2.20E-01 | 6.21E-01 | 1.13E+00 | 9.96E-03 | 3.65E-02 | -1.05E+00 | 2.98E-01 | 8.20E-01 | 1.37E+00 |
| LMCD1 | 1.86E-01 | 9.69E-01 | 1.01E+00 | 1.96E-01 | 8.49E-01 | 1.28E+00 | 2.22E-01 | 9.42E-01 | 1.00E+00 | 2.33E-01 | 8.90E-01 | 1.12E+00 | 1.82E-01 | 6.50E-01 | -1.06E+00 | 2.97E-01 | 8.37E-01 | 1.10E+00 | 5.68E-02 | 2.09E-01 | -1.09E+00 | 2.50E-01 | 6.86E-01 | 1.20E+00 |
| CENPI | 1.86E-01 | 9.69E-01 | 1.02E+00 | 2.19E-01 | 9.48E-01 | 1.07E+00 | 1.97E-01 | 8.39E-01 | 1.06E+00 | 2.41E-01 | 9.20E-01 | 1.20E+00 | 2.32E-01 | 8.26E-01 | -1.05E+00 | 3.41E-01 | 9.62E-01 | 1.08E+00 | 1.08E-02 | 3.96E-02 | -1.39E+00 | 2.66E-01 | 7.29E-01 | 1.11E+00 |
| HIST2H2BF | 1.86E-01 | 9.69E-01 | 1.41E+00 | 8.28E-02 | 3.58E-01 | 1.30E+00 | 8.22E-03 | 3.49E-02 | -1.15E+00 | 4.56E-04 | 1.74E-03 | -1.29E+00 | 1.63E-01 | 5.82E-01 | 1.75E+00 | 1.47E-01 | 4.14E-01 | 1.86E+00 | 2.43E-01 | 8.93E-01 | 1.47E+00 | 2.18E-01 | 5.98E-01 | 1.12E+00 |
| LINC01792 | 1.86E-01 | 9.69E-01 | 1.32E+00 | 9.07E-03 | 3.92E-02 | 2.34E+00 | 1.67E-01 | 7.08E-01 | 1.17E+00 | 1.32E-02 | 5.03E-02 | 1.86E+00 | 2.68E-01 | 9.56E-01 | 1.14E+00 | 3.08E-01 | 8.69E-01 | 1.49E+00 | 2.71E-01 | 9.96E-01 | -1.01E+00 | 3.00E-01 | 8.23E-01 | 1.20E+00 |
| LATS2-AS1 | 1.86E-01 | 9.69E-01 | 1.43E+00 | 2.61E-06 | 1.13E-05 | -1.35E+00 | 1.87E-01 | 7.93E-01 | 1.24E+00 | 4.41E-05 | 1.68E-04 | -1.52E+00 | 8.27E-02 | 2.95E-01 | -1.06E+00 | 1.61E-03 | 4.53E-03 | -2.07E+00 | 1.01E-03 | 3.71E-03 | -1.51E+00 | 2.22E-05 | 6.10E-05 | -2.14E+00 |
| AC008014.1 | 1.86E-01 | 9.69E-01 | 1.25E+00 | 1.01E-01 | 4.37E-01 | 1.14E+00 | 2.02E-01 | 8.59E-01 | 1.26E+00 | 1.77E-01 | 6.78E-01 | 1.18E+00 | 2.02E-01 | 7.19E-01 | 1.08E+00 | 3.46E-01 | 9.75E-01 | 1.32E+00 | 4.05E-02 | 1.49E-01 | 1.74E+00 | 3.18E-01 | 8.74E-01 | 1.30E+00 |
| MT1E | 1.86E-01 | 9.70E-01 | 1.65E+00 | 1.99E-01 | 8.61E-01 | 1.15E+00 | 6.76E-02 | 2.87E-01 | -1.27E+00 | 1.07E-02 | 4.10E-02 | -1.67E+00 | 2.49E-01 | 8.89E-01 | 2.97E+00 | 2.72E-01 | 7.66E-01 | 1.95E+00 | 8.16E-03 | 2.99E-02 | 1.52E+00 | 1.12E-02 | 3.08E-02 | 1.09E+00 |
| C5AR1 | 1.86E-01 | 9.70E-01 | 1.02E+00 | 2.05E-01 | 8.87E-01 | 1.37E+00 | 1.08E-01 | 4.57E-01 | -1.02E+00 | 1.53E-01 | 5.83E-01 | -1.03E+00 | 2.63E-01 | 9.40E-01 | 1.11E+00 | 3.40E-01 | 9.59E-01 | 1.09E+00 | 1.93E-01 | 7.08E-01 | 1.04E+00 | 1.96E-01 | 5.39E-01 | -1.15E+00 |
| ANKRD20A1 | 1.86E-01 | 9.70E-01 | 1.21E+00 | 3.17E-02 | 1.37E-01 | -1.21E+00 | 5.20E-02 | 2.21E-01 | 1.76E+00 | 1.76E-01 | 6.74E-01 | 1.49E+00 | 3.30E-04 | 1.18E-03 | 4.08E+00 | 4.99E-02 | 1.41E-01 | 2.76E+00 | 2.48E-05 | 9.09E-05 | 3.37E+00 | 8.59E-03 | 2.36E-02 | 2.45E+00 |
| AC025459.1 | 1.86E-01 | 9.70E-01 | 1.42E+00 | 1.26E-02 | 5.44E-02 | 1.35E+00 | 6.95E-03 | 2.95E-02 | 1.63E+00 | 1.05E-01 | 4.02E-01 | 1.35E+00 | 2.12E-01 | 7.57E-01 | 1.38E+00 | 1.83E-01 | 5.15E-01 | 1.35E+00 | 2.38E-01 | 8.74E-01 | 1.35E+00 | 2.45E-01 | 6.74E-01 | 1.26E+00 |
| CENPA | 1.86E-01 | 9.70E-01 | 1.08E+00 | 2.16E-01 | 9.33E-01 | 1.35E+00 | 1.60E-01 | 6.78E-01 | -1.18E+00 | 1.82E-01 | 6.94E-01 | 1.47E+00 | 2.80E-01 | 1.00E+00 | 1.00E+00 | 3.35E-01 | 9.45E-01 | 1.14E+00 | 1.20E-01 | 4.42E-01 | -1.23E+00 | 2.76E-01 | 7.58E-01 | 1.00E+00 |
| AC058823.1 | 1.86E-01 | 9.70E-01 | 1.22E+00 | 1.02E-01 | 4.42E-01 | 1.14E+00 | 7.91E-02 | 3.36E-01 | 1.02E+00 | 8.03E-02 | 3.07E-01 | 1.02E+00 | 2.35E-01 | 8.38E-01 | 1.31E+00 | 2.50E-01 | 7.06E-01 | 1.51E+00 | 1.21E-01 | 4.42E-01 | 1.41E+00 | 1.49E-01 | 4.08E-01 | -1.03E+00 |
| AC099568.2 | 1.86E-01 | 9.71E-01 | 1.32E+00 | 1.65E-01 | 7.11E-01 | 1.62E+00 | 1.24E-01 | 5.29E-01 | 1.45E+00 | 1.18E-01 | 4.53E-01 | 1.64E+00 | 2.72E-01 | 9.71E-01 | 1.33E+00 | 1.70E-01 | 4.80E-01 | 1.86E+00 | 7.22E-02 | 2.65E-01 | -1.12E+00 | 1.44E-01 | 3.96E-01 | 1.05E+00 |
| AL035458.1 | 1.86E-01 | 9.71E-01 | 1.35E+00 | 7.92E-02 | 3.42E-01 | 1.32E+00 | 1.02E-01 | 4.32E-01 | 1.08E+00 | 1.79E-02 | 6.84E-02 | 1.03E+00 | 2.36E-01 | 8.43E-01 | 1.35E+00 | 6.88E-02 | 1.94E-01 | -1.03E+00 | 2.34E-01 | 8.57E-01 | 1.24E+00 | 5.12E-02 | 1.41E-01 | 1.01E+00 |
| NID2 | 1.86E-01 | 9.71E-01 | 1.30E+00 | 2.04E-01 | 8.80E-01 | 1.33E+00 | 9.87E-02 | 4.19E-01 | 1.52E+00 | 1.53E-01 | 5.84E-01 | 1.11E+00 | 2.63E-02 | 9.38E-02 | 2.46E+00 | 8.09E-02 | 2.28E-01 | 2.01E+00 | 1.70E-01 | 6.25E-01 | 1.49E+00 | 1.08E-01 | 2.97E-01 | -1.06E+00 |
| FPR3 | 1.86E-01 | 9.71E-01 | 1.06E+00 | 2.28E-01 | 9.85E-01 | 1.04E+00 | 1.89E-01 | 8.01E-01 | -1.13E+00 | 2.07E-01 | 7.91E-01 | 1.23E+00 | 2.72E-01 | 9.71E-01 | 1.19E+00 | 1.92E-02 | 5.41E-02 | 2.81E+00 | 8.46E-02 | 3.10E-01 | 1.03E+00 | 1.18E-03 | 3.25E-03 | 1.95E+00 |
| AC010300.1 | 1.86E-01 | 9.71E-01 | 1.31E+00 | 2.27E-01 | 9.79E-01 | 1.45E+00 | 1.64E-02 | 6.96E-02 | 1.96E+00 | 9.09E-02 | 3.47E-01 | 1.70E+00 | 1.31E-01 | 4.66E-01 | 2.05E+00 | 2.79E-01 | 7.87E-01 | 1.65E+00 | 7.82E-02 | 2.87E-01 | 1.86E+00 | 1.44E-02 | 3.97E-02 | 3.02E+00 |
| EME1 | 1.87E-01 | 9.72E-01 | 1.08E+00 | 1.91E-01 | 8.25E-01 | 1.31E+00 | 3.62E-03 | 1.54E-02 | 2.23E+00 | 8.34E-03 | 3.19E-02 | 2.40E+00 | 2.15E-01 | 7.65E-01 | 1.35E+00 | 2.29E-01 | 6.45E-01 | 1.60E+00 | 2.55E-03 | 9.37E-03 | -1.08E+00 | 1.06E-01 | 2.90E-01 | 1.00E+00 |
| CMTM2 | 1.87E-01 | 9.72E-01 | 1.05E+00 | 1.28E-01 | 5.54E-01 | 1.40E+00 | 1.20E-01 | 5.12E-01 | -1.06E+00 | 8.35E-03 | 3.19E-02 | -1.66E+00 | 2.80E-01 | 1.00E+00 | 1.00E+00 | 2.77E-01 | 7.82E-01 | 1.26E+00 | 1.35E-02 | 4.94E-02 | 3.28E+00 | 2.78E-01 | 7.64E-01 | 1.26E+00 |
| AC116347.1 | 1.87E-01 | 9.73E-01 | 1.36E+00 | 1.35E-03 | 5.83E-03 | 2.71E+00 | 4.62E-02 | 1.96E-01 | 1.54E+00 | 1.07E-02 | 4.09E-02 | 2.08E+00 | 1.18E-01 | 4.22E-01 | -1.03E+00 | 2.93E-01 | 8.27E-01 | 1.56E+00 | 1.25E-01 | 4.59E-01 | 1.13E+00 | 1.45E-01 | 3.98E-01 | 1.54E+00 |
| HMGB1P1 | 1.87E-01 | 9.73E-01 | 1.38E+00 | 1.71E-02 | 7.37E-02 | 1.16E+00 | 1.63E-01 | 6.91E-01 | 1.36E+00 | 2.29E-03 | 8.74E-03 | 1.08E+00 | 1.32E-03 | 4.69E-03 | -1.10E+00 | 2.41E-02 | 6.79E-02 | 1.08E+00 | 9.94E-02 | 3.65E-01 | 1.20E+00 | 6.21E-02 | 1.70E-01 | 1.09E+00 |
| RPL13 | 1.87E-01 | 9.73E-01 | 1.00E+00 | 1.76E-04 | 7.60E-04 | -1.08E+00 | 6.51E-02 | 2.77E-01 | 1.03E+00 | 1.28E-01 | 4.90E-01 | -1.02E+00 | 5.58E-02 | 1.99E-01 | 1.05E+00 | 2.04E-01 | 5.74E-01 | -1.02E+00 | 5.91E-02 | 2.17E-01 | 1.01E+00 | 3.79E-03 | 1.04E-02 | -1.07E+00 |
| FAM86B3P | 1.87E-01 | 9.73E-01 | 1.25E+00 | 1.51E-01 | 6.52E-01 | 1.53E+00 | 9.09E-03 | 3.86E-02 | -1.24E+00 | 9.98E-02 | 3.81E-01 | 1.12E+00 | 4.70E-03 | 1.68E-02 | -2.00E+00 | 6.71E-02 | 1.89E-01 | -1.18E+00 | 3.34E-02 | 1.22E-01 | -1.48E+00 | 3.04E-01 | 8.34E-01 | -1.00E+00 |
| MYBL2 | 1.87E-01 | 9.74E-01 | -1.08E+00 | 2.11E-01 | 9.12E-01 | -1.23E+00 | 2.95E-05 | 1.25E-04 | 1.59E+00 | 1.63E-01 | 6.21E-01 | 1.15E+00 | 1.17E-01 | 4.18E-01 | -4.30E+00 | 1.80E-01 | 5.08E-01 | -3.55E+00 | 5.98E-02 | 2.19E-01 | 1.16E+00 | 2.06E-03 | 5.66E-03 | 1.36E+00 |
| STAP1 | 1.87E-01 | 9.74E-01 | 1.21E+00 | 1.81E-01 | 7.82E-01 | 1.07E+00 | 1.80E-01 | 7.64E-01 | -1.02E+00 | 3.25E-02 | 1.24E-01 | -1.83E+00 | 1.30E-03 | 4.63E-03 | -1.25E+00 | 1.80E-06 | 5.08E-06 | -1.77E+00 | 1.14E-06 | 4.17E-06 | -1.28E+00 | 6.29E-11 | 1.73E-10 | -1.54E+00 |
| PTENP1 | 1.87E-01 | 9.74E-01 | 1.33E+00 | 1.60E-01 | 6.92E-01 | 1.40E+00 | 2.24E-01 | 9.50E-01 | 1.31E+00 | 4.48E-02 | 1.71E-01 | 1.25E+00 | 1.21E-01 | 4.32E-01 | 1.17E+00 | 3.26E-01 | 9.19E-01 | 1.39E+00 | 2.48E-01 | 9.09E-01 | 1.21E+00 | 3.14E-01 | 8.63E-01 | 1.28E+00 |
| AL161421.1 | 1.87E-01 | 9.74E-01 | 1.36E+00 | 1.20E-01 | 5.20E-01 | 1.31E+00 | 1.34E-01 | 5.69E-01 | 1.16E+00 | 1.53E-02 | 5.85E-02 | -1.02E+00 | 1.44E-01 | 5.12E-01 | 1.13E+00 | 6.77E-02 | 1.91E-01 | -1.03E+00 | 1.12E-01 | 4.13E-01 | 1.42E+00 | 2.79E-01 | 7.68E-01 | 1.12E+00 |
| P2RX5-TAX1BP3 | 1.87E-01 | 9.74E-01 | 1.29E+00 | 8.51E-02 | 3.68E-01 | 1.09E+00 | 2.39E-04 | 1.02E-03 | -1.41E+00 | 1.54E-05 | 5.88E-05 | -1.56E+00 | 3.22E-02 | 1.15E-01 | 1.02E+00 | 4.02E-03 | 1.13E-02 | -1.25E+00 | 2.11E-01 | 7.73E-01 | 1.11E+00 | 2.66E-01 | 7.30E-01 | 1.11E+00 |
| ATP5O | 1.87E-01 | 9.74E-01 | 1.07E+00 | 1.86E-02 | 8.02E-02 | 1.06E+00 | 9.34E-02 | 3.97E-01 | 1.08E+00 | 1.16E-01 | 4.43E-01 | -1.01E+00 | 3.78E-03 | 1.35E-02 | -1.01E+00 | 3.19E-02 | 8.98E-02 | 1.02E+00 | 1.65E-01 | 6.05E-01 | 1.18E+00 | 2.88E-01 | 7.91E-01 | 1.09E+00 |
| TNFRSF13C | 1.87E-01 | 9.74E-01 | 1.29E+00 | 1.61E-01 | 6.96E-01 | 1.07E+00 | 1.64E-02 | 6.97E-02 | 1.81E+00 | 1.32E-01 | 5.04E-01 | 1.60E+00 | 1.61E-02 | 5.76E-02 | -1.70E+00 | 7.81E-03 | 2.20E-02 | -2.26E+00 | 6.14E-02 | 2.25E-01 | 1.04E+00 | 3.33E-03 | 9.13E-03 | -1.15E+00 |
| FAM83F | 1.87E-01 | 9.74E-01 | 1.12E+00 | 1.14E-01 | 4.94E-01 | -1.26E+00 | 1.86E-01 | 7.90E-01 | 1.02E+00 | 1.31E-01 | 4.99E-01 | -1.14E+00 | 2.10E-03 | 7.50E-03 | -1.24E+00 | 2.16E-03 | 6.10E-03 | -1.08E+00 | 2.06E-01 | 7.58E-01 | 1.06E+00 | 2.36E-01 | 6.49E-01 | -1.01E+00 |
| SLC2A14 | 1.87E-01 | 9.75E-01 | -1.20E+00 | 1.85E-01 | 7.97E-01 | -1.46E+00 | 1.71E-01 | 7.27E-01 | -1.41E+00 | 7.91E-02 | 3.02E-01 | -1.97E+00 | 1.79E-01 | 6.40E-01 | -1.16E+00 | 2.09E-01 | 5.89E-01 | -1.26E+00 | 2.34E-03 | 8.59E-03 | 1.12E+00 | 6.11E-03 | 1.68E-02 | -1.02E+00 |
| SLC30A1 | 1.87E-01 | 9.75E-01 | 1.23E+00 | 2.03E-01 | 8.79E-01 | 1.19E+00 | 1.91E-01 | 8.13E-01 | 1.19E+00 | 8.82E-03 | 3.37E-02 | 1.03E+00 | 2.66E-01 | 9.49E-01 | 1.18E+00 | 2.48E-01 | 6.98E-01 | -1.04E+00 | 5.14E-02 | 1.89E-01 | 1.08E+00 | 1.58E-02 | 4.35E-02 | -1.07E+00 |
| ENY2 | 1.87E-01 | 9.75E-01 | 1.30E+00 | 5.01E-02 | 2.17E-01 | 1.27E+00 | 2.46E-04 | 1.05E-03 | 1.45E+00 | 6.65E-03 | 2.54E-02 | 1.40E+00 | 1.57E-03 | 5.61E-03 | 1.67E+00 | 1.54E-02 | 4.34E-02 | 1.59E+00 | 7.56E-03 | 2.77E-02 | 1.51E+00 | 3.67E-02 | 1.01E-01 | 1.43E+00 |
| GTDC1 | 1.87E-01 | 9.75E-01 | 1.26E+00 | 5.04E-02 | 2.18E-01 | 1.90E+00 | 1.64E-01 | 6.98E-01 | 1.03E+00 | 2.61E-01 | 9.97E-01 | 1.33E+00 | 6.81E-03 | 2.43E-02 | 1.04E+00 | 1.54E-02 | 4.34E-02 | 1.13E+00 | 1.30E-01 | 4.76E-01 | 1.17E+00 | 1.88E-01 | 5.17E-01 | 1.16E+00 |
| RPS12P23 | 1.87E-01 | 9.76E-01 | 1.13E+00 | 5.95E-20 | 2.57E-19 | -1.29E+00 | 6.34E-02 | 2.70E-01 | 1.20E+00 | 3.55E-06 | 1.35E-05 | -1.25E+00 | 9.40E-02 | 3.35E-01 | 1.12E+00 | 4.44E-08 | 1.25E-07 | -1.26E+00 | 1.67E-01 | 6.14E-01 | 1.19E+00 | 1.91E-04 | 5.26E-04 | -1.21E+00 |
| BAMBI | 1.87E-01 | 9.76E-01 | 1.07E+00 | 2.06E-01 | 8.89E-01 | -1.07E+00 | 1.90E-01 | 8.09E-01 | 1.18E+00 | 1.80E-01 | 6.87E-01 | 1.32E+00 | 1.18E-01 | 4.21E-01 | 1.31E+00 | 2.37E-03 | 6.69E-03 | 1.92E+00 | 2.33E-01 | 8.53E-01 | 1.58E+00 | 1.23E-01 | 3.38E-01 | 1.68E+00 |
| AC113404.3 | 1.87E-01 | 9.76E-01 | 1.30E+00 | 2.11E-01 | 9.11E-01 | 1.49E+00 | 1.63E-02 | 6.91E-02 | 1.56E+00 | 1.28E-02 | 4.88E-02 | 1.86E+00 | 2.05E-01 | 7.29E-01 | 1.46E+00 | 1.71E-01 | 4.82E-01 | 1.58E+00 | 8.22E-02 | 3.02E-01 | 1.76E+00 | 1.53E-01 | 4.20E-01 | 1.35E+00 |
| PSPN | 1.87E-01 | 9.77E-01 | 1.11E+00 | 1.94E-01 | 8.37E-01 | 1.25E+00 | 6.21E-03 | 2.64E-02 | -1.34E+00 | 9.14E-03 | 3.49E-02 | -1.12E+00 | 2.00E-01 | 7.14E-01 | 1.56E+00 | 2.85E-01 | 8.05E-01 | 1.06E+00 | 4.47E-02 | 1.64E-01 | -1.12E+00 | 3.25E-02 | 8.92E-02 | -1.19E+00 |
| NCF1C | 1.87E-01 | 9.77E-01 | 1.19E+00 | 2.10E-01 | 9.06E-01 | 1.13E+00 | 1.68E-02 | 7.13E-02 | 2.57E+00 | 2.13E-01 | 8.12E-01 | -1.10E+00 | 2.49E-01 | 8.88E-01 | 1.44E+00 | 6.17E-02 | 1.74E-01 | -1.17E+00 | 7.36E-05 | 2.70E-04 | -2.38E+00 | 1.62E-04 | 4.45E-04 | -2.46E+00 |
| SND1-IT1 | 1.87E-01 | 9.77E-01 | 1.37E+00 | 1.36E-01 | 5.88E-01 | 1.47E+00 | 6.50E-03 | 2.76E-02 | 1.55E+00 | 5.57E-02 | 2.13E-01 | 1.59E+00 | 2.83E-02 | 1.01E-01 | -1.03E+00 | 1.81E-01 | 5.10E-01 | 1.24E+00 | 1.83E-01 | 6.73E-01 | 1.43E+00 | 6.64E-02 | 1.82E-01 | 1.64E+00 |
| AC006157.1 | 1.88E-01 | 9.77E-01 | 1.30E+00 | 2.00E-01 | 8.64E-01 | 1.61E+00 | 1.59E-02 | 6.76E-02 | 1.92E+00 | 1.90E-01 | 7.24E-01 | 1.54E+00 | 2.07E-01 | 7.37E-01 | 1.43E+00 | 2.42E-01 | 6.81E-01 | 1.54E+00 | 3.01E-02 | 1.10E-01 | 1.82E+00 | 2.27E-01 | 6.23E-01 | 1.36E+00 |
| AC133552.6 | 1.88E-01 | 9.77E-01 | 1.38E+00 | 1.95E-01 | 8.41E-01 | 1.47E+00 | 2.33E-02 | 9.91E-02 | 1.69E+00 | 2.02E-01 | 7.72E-01 | 1.40E+00 | 6.41E-02 | 2.29E-01 | 1.85E+00 | 1.73E-01 | 4.89E-01 | 1.56E+00 | 2.14E-01 | 7.86E-01 | 1.21E+00 | 1.60E-01 | 4.41E-01 | -1.04E+00 |
| GP6 | 1.88E-01 | 9.77E-01 | 1.04E+00 | 2.10E-01 | 9.09E-01 | -1.00E+00 | 2.04E-01 | 8.66E-01 | -1.04E+00 | 2.10E-01 | 8.01E-01 | -1.07E+00 | 2.36E-01 | 8.42E-01 | -1.18E+00 | 2.49E-01 | 7.01E-01 | -1.35E+00 | 3.32E-02 | 1.22E-01 | 1.48E+00 | 5.90E-03 | 1.62E-02 | 1.07E+00 |
| B3GALT2 | 1.88E-01 | 9.77E-01 | 1.20E+00 | 3.13E-02 | 1.35E-01 | -1.18E+00 | 1.78E-01 | 7.57E-01 | 1.24E+00 | 1.66E-01 | 6.34E-01 | 1.09E+00 | 2.85E-02 | 1.02E-01 | -1.29E+00 | 5.81E-02 | 1.64E-01 | -1.22E+00 | 4.03E-03 | 1.48E-02 | -1.38E+00 | 6.41E-04 | 1.76E-03 | -1.83E+00 |
| TK1 | 1.88E-01 | 9.78E-01 | 1.22E+00 | 2.01E-01 | 8.71E-01 | -1.01E+00 | 4.20E-05 | 1.78E-04 | 2.03E+00 | 4.47E-02 | 1.71E-01 | 1.72E+00 | 5.45E-02 | 1.94E-01 | -4.12E+00 | 7.82E-02 | 2.20E-01 | -3.67E+00 | 2.19E-05 | 8.06E-05 | 1.58E+00 | 4.18E-07 | 1.15E-06 | 1.74E+00 |
| GSTA4 | 1.88E-01 | 9.79E-01 | 1.22E+00 | 1.91E-01 | 8.25E-01 | 1.34E+00 | 1.95E-01 | 8.30E-01 | 1.30E+00 | 2.24E-01 | 8.55E-01 | 1.43E+00 | 5.54E-02 | 1.98E-01 | 1.79E+00 | 3.51E-01 | 9.90E-01 | 1.21E+00 | 1.26E-01 | 4.64E-01 | 1.50E+00 | 2.39E-01 | 6.57E-01 | 1.39E+00 |
| GGNBP2 | 1.88E-01 | 9.79E-01 | 1.23E+00 | 7.17E-02 | 3.10E-01 | 1.30E+00 | 4.08E-02 | 1.74E-01 | 1.08E+00 | 1.75E-02 | 6.68E-02 | 1.08E+00 | 4.68E-02 | 1.67E-01 | 1.12E+00 | 3.91E-05 | 1.10E-04 | -1.06E+00 | 2.25E-04 | 8.26E-04 | -1.09E+00 | 4.35E-04 | 1.20E-03 | -1.09E+00 |
| SCGB3A1 | 1.88E-01 | 9.80E-01 | 1.29E+00 | 5.41E-02 | 2.34E-01 | -1.04E+00 | 8.15E-02 | 3.46E-01 | -1.09E+00 | 5.67E-02 | 2.17E-01 | -1.02E+00 | 2.48E-01 | 8.86E-01 | 1.27E+00 | 3.12E-01 | 8.78E-01 | -1.00E+00 | 2.30E-01 | 8.45E-01 | 1.16E+00 | 2.36E-01 | 6.48E-01 | 1.32E+00 |
| LINC00612 | 1.88E-01 | 9.80E-01 | 1.28E+00 | 1.29E-01 | 5.58E-01 | 1.19E+00 | 1.05E-01 | 4.47E-01 | 1.02E+00 | 1.48E-01 | 5.64E-01 | 1.16E+00 | 5.01E-02 | 1.79E-01 | -1.09E+00 | 2.98E-02 | 8.39E-02 | -1.30E+00 | 2.69E-01 | 9.89E-01 | 1.17E+00 | 2.51E-01 | 6.88E-01 | -1.01E+00 |
| AL024508.2 | 1.88E-01 | 9.81E-01 | 1.41E+00 | 7.48E-02 | 3.23E-01 | 1.04E+00 | 1.03E-01 | 4.39E-01 | 1.01E+00 | 3.00E-02 | 1.15E-01 | -1.16E+00 | 9.05E-03 | 3.23E-02 | 2.35E+00 | 3.30E-02 | 9.30E-02 | 2.15E+00 | 1.77E-01 | 6.49E-01 | 1.28E+00 | 3.50E-01 | 9.60E-01 | 1.21E+00 |
| IL32 | 1.88E-01 | 9.81E-01 | 1.05E+00 | 1.79E-01 | 7.73E-01 | 1.30E+00 | 1.84E-02 | 7.80E-02 | -1.63E+01 | 1.38E-02 | 5.28E-02 | -2.60E+01 | 1.16E-01 | 4.12E-01 | -1.21E+00 | 1.71E-01 | 4.81E-01 | -1.14E+00 | 1.58E-01 | 5.78E-01 | -1.07E+00 | 2.75E-01 | 7.55E-01 | -1.03E+00 |
| FOXRED2 | 1.88E-01 | 9.81E-01 | 1.26E+00 | 2.30E-01 | 9.95E-01 | 1.42E+00 | 4.57E-02 | 1.94E-01 | -1.03E+00 | 5.01E-02 | 1.92E-01 | 1.12E+00 | 4.43E-02 | 1.58E-01 | -1.56E+00 | 2.35E-01 | 6.63E-01 | 1.13E+00 | 1.42E-03 | 5.21E-03 | -1.11E+00 | 2.56E-01 | 7.04E-01 | 1.23E+00 |
| AL353743.3 | 1.88E-01 | 9.81E-01 | 1.24E+00 | 1.84E-01 | 7.93E-01 | 1.48E+00 | 1.73E-01 | 7.35E-01 | 1.11E+00 | 2.10E-01 | 8.02E-01 | 1.26E+00 | 1.97E-01 | 7.03E-01 | 1.46E+00 | 3.03E-01 | 8.53E-01 | 1.30E+00 | 3.76E-02 | 1.38E-01 | 1.74E+00 | 1.08E-01 | 2.96E-01 | 1.68E+00 |
| FRMD5 | 1.88E-01 | 9.81E-01 | 1.17E+00 | 1.41E-01 | 6.10E-01 | -1.06E+00 | 1.90E-01 | 8.08E-01 | 1.29E+00 | 2.11E-01 | 8.07E-01 | 1.37E+00 | 2.59E-01 | 9.23E-01 | 1.24E+00 | 2.60E-01 | 7.32E-01 | 1.69E+00 | 2.25E-02 | 8.27E-02 | -1.09E+00 | 3.64E-01 | 1.00E+00 | 1.25E+00 |
| AC008753.1 | 1.88E-01 | 9.81E-01 | 1.39E+00 | 1.50E-03 | 6.48E-03 | -1.01E+00 | 2.34E-01 | 9.96E-01 | 1.26E+00 | 1.36E-03 | 5.19E-03 | -1.03E+00 | 1.89E-01 | 6.72E-01 | 1.61E+00 | 6.64E-02 | 1.87E-01 | 1.13E+00 | 4.36E-02 | 1.60E-01 | 1.52E+00 | 1.09E-01 | 2.99E-01 | 1.06E+00 |
| ZBTB18 | 1.88E-01 | 9.82E-01 | 1.52E+00 | 1.71E-01 | 7.39E-01 | 1.52E+00 | 2.33E-01 | 9.90E-01 | 1.25E+00 | 8.70E-02 | 3.32E-01 | 1.16E+00 | 1.20E-01 | 4.28E-01 | -1.05E+00 | 1.06E-01 | 2.99E-01 | -1.16E+00 | 4.15E-02 | 1.52E-01 | -1.34E+00 | 1.10E-01 | 3.01E-01 | -1.08E+00 |
| YBX1P2 | 1.88E-01 | 9.82E-01 | 1.30E+00 | 1.16E-01 | 5.03E-01 | 1.65E+00 | 8.97E-04 | 3.81E-03 | 1.74E+00 | 1.02E-02 | 3.90E-02 | 1.89E+00 | 1.72E-01 | 6.15E-01 | 1.60E+00 | 1.58E-01 | 4.47E-01 | 1.72E+00 | 1.83E-01 | 6.73E-01 | 1.33E+00 | 1.92E-03 | 5.28E-03 | 1.70E+00 |
| DPF1 | 1.88E-01 | 9.82E-01 | 1.13E+00 | 1.58E-03 | 6.81E-03 | -1.47E+00 | 5.85E-07 | 2.48E-06 | -1.65E+00 | 1.15E-09 | 4.40E-09 | -1.83E+00 | 1.11E-02 | 3.97E-02 | 2.25E+00 | 1.35E-01 | 3.80E-01 | -1.18E+00 | 4.40E-03 | 1.62E-02 | 2.21E+00 | 2.65E-01 | 7.29E-01 | 1.04E+00 |
| SPACA6 | 1.89E-01 | 9.83E-01 | 1.33E+00 | 2.20E-01 | 9.52E-01 | 1.47E+00 | 1.60E-01 | 6.81E-01 | 1.12E+00 | 8.75E-02 | 3.34E-01 | 1.07E+00 | 1.82E-01 | 6.50E-01 | 1.31E+00 | 1.27E-01 | 3.59E-01 | -1.00E+00 | 1.96E-01 | 7.21E-01 | -1.05E+00 | 3.18E-01 | 8.73E-01 | 1.04E+00 |
| HRASLS | 1.89E-01 | 9.85E-01 | 1.12E+00 | 1.83E-01 | 7.92E-01 | 1.10E+00 | 8.52E-03 | 3.62E-02 | 2.20E+00 | 8.24E-02 | 3.15E-01 | 1.70E+00 | 2.43E-01 | 8.65E-01 | -1.04E+00 | 2.25E-01 | 6.34E-01 | 1.62E+00 | 1.30E-03 | 4.77E-03 | 1.83E+00 | 2.20E-01 | 6.05E-01 | 1.30E+00 |
| AC093700.1 | 1.89E-01 | 9.86E-01 | 1.40E+00 | 1.76E-01 | 7.59E-01 | 1.72E+00 | 2.01E-02 | 8.55E-02 | 1.68E+00 | 1.05E-02 | 4.01E-02 | 2.04E+00 | 2.41E-01 | 8.60E-01 | 1.06E+00 | 2.31E-01 | 6.52E-01 | 1.43E+00 | 2.47E-01 | 9.05E-01 | 1.26E+00 | 7.53E-02 | 2.07E-01 | 1.62E+00 |
| CENPP | 1.89E-01 | 9.86E-01 | 1.26E+00 | 1.89E-01 | 8.17E-01 | 1.28E+00 | 2.30E-01 | 9.75E-01 | 1.19E+00 | 1.20E-03 | 4.58E-03 | -1.14E+00 | 2.17E-01 | 7.73E-01 | 1.06E+00 | 3.34E-01 | 9.42E-01 | 1.21E+00 | 6.21E-02 | 2.28E-01 | 1.63E+00 | 1.60E-01 | 4.39E-01 | 1.55E+00 |
| IL7R | 1.90E-01 | 9.88E-01 | 1.05E+00 | 1.80E-01 | 7.77E-01 | -1.11E+00 | 1.72E-02 | 7.32E-02 | -2.62E+00 | 3.15E-02 | 1.20E-01 | -2.37E+00 | 6.31E-03 | 2.25E-02 | 3.63E+00 | 1.98E-01 | 5.58E-01 | 1.83E+00 | 2.97E-03 | 1.09E-02 | 3.06E+00 | 2.52E-01 | 6.91E-01 | 1.31E+00 |
| PDE6A | 1.90E-01 | 9.88E-01 | 1.18E+00 | 1.82E-01 | 7.87E-01 | 1.38E+00 | 1.17E-01 | 4.96E-01 | -1.30E+00 | 1.86E-01 | 7.12E-01 | -1.11E+00 | 1.80E-01 | 6.41E-01 | 1.04E+00 | 1.48E-01 | 4.17E-01 | 1.50E+00 | 2.42E-01 | 8.87E-01 | 1.19E+00 | 1.94E-01 | 5.33E-01 | -1.00E+00 |
| C1orf122 | 1.90E-01 | 9.88E-01 | 1.29E+00 | 6.86E-02 | 2.96E-01 | 1.31E+00 | 9.50E-02 | 4.04E-01 | 1.31E+00 | 2.20E-01 | 8.41E-01 | 1.32E+00 | 2.46E-01 | 8.76E-01 | 1.37E+00 | 1.99E-01 | 5.61E-01 | 1.26E+00 | 9.06E-02 | 3.32E-01 | 1.46E+00 | 1.40E-01 | 3.85E-01 | 1.45E+00 |
| HIST1H2BG | 1.90E-01 | 9.88E-01 | 1.19E+00 | 3.49E-03 | 1.51E-02 | 2.07E+00 | 1.53E-01 | 6.49E-01 | -1.02E+00 | 5.70E-02 | 2.18E-01 | 1.67E+00 | 2.72E-01 | 9.69E-01 | 1.33E+00 | 2.19E-07 | 6.17E-07 | 4.18E+00 | 2.59E-02 | 9.51E-02 | 1.04E+00 | 1.80E-01 | 4.95E-01 | 2.19E+00 |
| AC106730.1 | 1.90E-01 | 9.88E-01 | 1.34E+00 | 2.24E-01 | 9.69E-01 | 1.48E+00 | 5.13E-02 | 2.18E-01 | 1.56E+00 | 6.89E-02 | 2.63E-01 | 1.69E+00 | 6.30E-02 | 2.25E-01 | -1.19E+00 | 1.79E-01 | 5.05E-01 | 1.14E+00 | 2.38E-01 | 8.74E-01 | 1.20E+00 | 1.22E-01 | 3.34E-01 | 1.61E+00 |
| PRELID2 | 1.90E-01 | 9.88E-01 | 1.37E+00 | 1.73E-01 | 7.47E-01 | 1.52E+00 | 1.01E-01 | 4.28E-01 | 1.41E+00 | 1.72E-01 | 6.56E-01 | 1.57E+00 | 1.31E-02 | 4.67E-02 | 1.03E+00 | 2.76E-01 | 7.77E-01 | 1.31E+00 | 1.12E-04 | 4.09E-04 | -1.07E+00 | 5.46E-03 | 1.50E-02 | -1.02E+00 |
| APOA1 | 1.90E-01 | 9.88E-01 | 1.31E+00 | 1.12E-02 | 4.83E-02 | -1.15E+00 | 1.63E-01 | 6.93E-01 | 1.30E+00 | 2.47E-02 | 9.43E-02 | -1.05E+00 | 1.30E-01 | 4.65E-01 | -1.28E+00 | 1.79E-01 | 5.04E-01 | -1.21E+00 | 1.40E-01 | 5.13E-01 | 1.43E+00 | 2.58E-01 | 7.08E-01 | -1.12E+00 |
| GMDS | 1.90E-01 | 9.88E-01 | 1.37E+00 | 9.03E-02 | 3.90E-01 | 1.39E+00 | 5.85E-02 | 2.48E-01 | 1.10E+00 | 2.67E-03 | 1.02E-02 | 1.09E+00 | 1.89E-01 | 6.73E-01 | 1.38E+00 | 7.94E-02 | 2.24E-01 | 1.27E+00 | 7.14E-03 | 2.62E-02 | 1.02E+00 | 1.48E-01 | 4.07E-01 | 1.25E+00 |
| TSLP | 1.90E-01 | 9.89E-01 | 1.32E+00 | 2.20E-02 | 9.50E-02 | 2.13E+00 | 1.11E-01 | 4.71E-01 | 1.07E+00 | 1.51E-01 | 5.78E-01 | 1.13E+00 | 1.87E-01 | 6.66E-01 | 1.00E+00 | 2.79E-01 | 7.85E-01 | 1.53E+00 | 8.14E-02 | 2.99E-01 | -1.00E+00 | 2.16E-01 | 5.93E-01 | 1.11E+00 |
| NEK2 | 1.90E-01 | 9.89E-01 | 1.14E+00 | 1.80E-01 | 7.76E-01 | -1.06E+00 | 1.62E-01 | 6.90E-01 | 1.23E+00 | 2.12E-01 | 8.08E-01 | 1.42E+00 | 2.14E-01 | 7.63E-01 | -1.26E+00 | 3.12E-01 | 8.80E-01 | 1.31E+00 | 2.62E-01 | 9.61E-01 | -1.03E+00 | 4.94E-03 | 1.36E-02 | 1.52E+00 |
| AL117336.3 | 1.90E-01 | 9.89E-01 | 1.29E+00 | 1.70E-02 | 7.33E-02 | 2.35E+00 | 8.07E-02 | 3.43E-01 | -1.13E+00 | 1.68E-01 | 6.41E-01 | 1.23E+00 | 8.10E-02 | 2.89E-01 | -1.03E+00 | 1.71E-01 | 4.81E-01 | 1.15E+00 | 1.46E-01 | 5.38E-01 | 1.55E+00 | 2.76E-01 | 7.59E-01 | 1.22E+00 |
| ZNF213-AS1 | 1.90E-01 | 9.89E-01 | 1.26E+00 | 3.39E-02 | 1.47E-01 | 1.71E+00 | 1.59E-01 | 6.76E-01 | 1.08E+00 | 1.54E-01 | 5.89E-01 | 1.26E+00 | 1.89E-01 | 6.74E-01 | 1.35E+00 | 2.81E-01 | 7.91E-01 | 1.69E+00 | 6.36E-02 | 2.34E-01 | 1.54E+00 | 2.58E-01 | 7.09E-01 | 1.38E+00 |
| RPL18AP7 | 1.90E-01 | 9.90E-01 | 1.24E+00 | 1.27E-01 | 5.48E-01 | 1.63E+00 | 1.97E-01 | 8.38E-01 | 1.08E+00 | 2.28E-01 | 8.72E-01 | 1.38E+00 | 2.28E-01 | 8.15E-01 | 1.17E+00 | 3.23E-01 | 9.11E-01 | 1.38E+00 | 1.61E-01 | 5.93E-01 | 1.05E+00 | 2.44E-01 | 6.71E-01 | 1.09E+00 |
| AL355353.1 | 1.90E-01 | 9.90E-01 | 1.21E+00 | 3.07E-03 | 1.33E-02 | -1.51E+00 | 2.16E-01 | 9.17E-01 | 1.22E+00 | 2.01E-02 | 7.69E-02 | -1.15E+00 | 1.05E-01 | 3.73E-01 | 1.03E+00 | 3.42E-02 | 9.64E-02 | -1.27E+00 | 1.22E-01 | 4.49E-01 | 1.37E+00 | 2.79E-02 | 7.67E-02 | -1.29E+00 |
| AC007336.1 | 1.90E-01 | 9.90E-01 | 1.12E+00 | 1.66E-02 | 7.19E-02 | 1.71E+00 | 4.22E-04 | 1.79E-03 | -1.38E+00 | 1.87E-01 | 7.15E-01 | 1.33E+00 | 5.00E-02 | 1.78E-01 | -1.04E+00 | 1.22E-01 | 3.45E-01 | 1.09E+00 | 2.71E-02 | 9.96E-02 | -1.78E+00 | 3.32E-01 | 9.11E-01 | 1.10E+00 |
| CD3G | 1.90E-01 | 9.90E-01 | 1.07E+00 | 2.11E-01 | 9.12E-01 | 1.05E+00 | 1.12E-03 | 4.78E-03 | -3.06E+00 | 9.61E-05 | 3.67E-04 | -4.33E+00 | 1.03E-01 | 3.66E-01 | 1.75E+00 | 3.20E-01 | 9.02E-01 | 1.08E+00 | 1.84E-01 | 6.74E-01 | 1.28E+00 | 3.58E-01 | 9.84E-01 | 1.05E+00 |
| DISP1 | 1.90E-01 | 9.91E-01 | 1.29E+00 | 1.06E-01 | 4.58E-01 | 1.84E+00 | 3.59E-04 | 1.53E-03 | -1.81E+00 | 3.18E-02 | 1.22E-01 | -1.11E+00 | 1.14E-02 | 4.05E-02 | -1.67E+00 | 1.13E-01 | 3.18E-01 | -1.02E+00 | 1.48E-01 | 5.43E-01 | -1.10E+00 | 1.97E-01 | 5.42E-01 | 1.39E+00 |
| SHB | 1.90E-01 | 9.91E-01 | 1.24E+00 | 2.30E-01 | 9.96E-01 | 1.37E+00 | 2.05E-01 | 8.72E-01 | 1.05E+00 | 1.78E-01 | 6.79E-01 | 1.08E+00 | 2.54E-01 | 9.06E-01 | 1.26E+00 | 2.31E-01 | 6.52E-01 | 1.57E+00 | 2.72E-01 | 9.97E-01 | 1.28E+00 | 2.70E-01 | 7.41E-01 | 1.50E+00 |
| AC005096.1 | 1.90E-01 | 9.92E-01 | 1.25E+00 | 1.30E-01 | 5.62E-01 | 1.13E+00 | 4.72E-05 | 2.01E-04 | -1.64E+00 | 7.51E-06 | 2.87E-05 | -1.76E+00 | 5.67E-02 | 2.02E-01 | -1.20E+00 | 1.18E-02 | 3.33E-02 | -1.55E+00 | 2.52E-02 | 9.25E-02 | -1.19E+00 | 4.45E-02 | 1.22E-01 | -1.15E+00 |
| SLX4 | 1.90E-01 | 9.92E-01 | 1.11E+00 | 2.31E-01 | 1.00E+00 | 1.39E+00 | 5.49E-02 | 2.33E-01 | -1.11E+00 | 5.46E-03 | 2.08E-02 | -1.52E+00 | 2.48E-01 | 8.83E-01 | 1.36E+00 | 2.30E-01 | 6.50E-01 | 1.61E+00 | 3.02E-06 | 1.11E-05 | -1.66E+00 | 4.95E-05 | 1.36E-04 | -1.53E+00 |
| MS4A2 | 1.90E-01 | 9.92E-01 | 1.37E+00 | 1.61E-01 | 6.97E-01 | 2.15E+00 | 1.17E-01 | 4.96E-01 | 4.66E+00 | 4.11E-02 | 1.57E-01 | 9.28E+00 | 8.66E-23 | 3.09E-22 | 1.67E+00 | 3.20E-28 | 9.02E-28 | 2.40E+00 | 1.04E-15 | 3.83E-15 | 4.35E+00 | 7.62E-31 | 2.09E-30 | 1.10E+01 |
| ST20-AS1 | 1.90E-01 | 9.92E-01 | 1.30E+00 | 1.01E-01 | 4.36E-01 | 1.10E+00 | 1.33E-04 | 5.64E-04 | -1.39E+00 | 1.96E-04 | 7.48E-04 | -1.28E+00 | 1.72E-01 | 6.13E-01 | 1.54E+00 | 2.51E-01 | 7.08E-01 | 1.54E+00 | 2.16E-01 | 7.95E-01 | 1.45E+00 | 2.81E-01 | 7.73E-01 | 1.45E+00 |
| C9orf66 | 1.90E-01 | 9.92E-01 | 1.32E+00 | 5.88E-02 | 2.54E-01 | 1.26E+00 | 5.95E-03 | 2.53E-02 | -1.11E+00 | 5.77E-04 | 2.20E-03 | -1.18E+00 | 2.67E-02 | 9.53E-02 | 2.09E+00 | 2.56E-01 | 7.20E-01 | 1.48E+00 | 3.05E-02 | 1.12E-01 | 1.88E+00 | 2.49E-01 | 6.83E-01 | 1.33E+00 |
| ENDOV | 1.90E-01 | 9.92E-01 | 1.38E+00 | 1.11E-01 | 4.79E-01 | 1.39E+00 | 1.55E-01 | 6.59E-01 | 1.14E+00 | 5.74E-02 | 2.19E-01 | 1.13E+00 | 2.52E-01 | 9.00E-01 | 1.39E+00 | 2.65E-01 | 7.48E-01 | 1.58E+00 | 1.42E-01 | 5.23E-01 | 1.37E+00 | 3.64E-01 | 1.00E+00 | 1.19E+00 |
| SNX5P1 | 1.91E-01 | 9.94E-01 | 1.18E+00 | 2.10E-01 | 9.07E-01 | 1.27E+00 | 5.45E-05 | 2.31E-04 | -1.94E+00 | 3.73E-04 | 1.42E-03 | -1.56E+00 | 2.30E-01 | 8.19E-01 | 1.19E+00 | 3.25E-01 | 9.17E-01 | 1.47E+00 | 1.12E-01 | 4.11E-01 | 1.10E+00 | 3.45E-01 | 9.46E-01 | 1.33E+00 |
| TTC39B | 1.91E-01 | 9.94E-01 | 1.36E+00 | 1.62E-01 | 6.99E-01 | 1.48E+00 | 1.35E-02 | 5.73E-02 | -1.19E+00 | 3.11E-03 | 1.19E-02 | -1.32E+00 | 1.10E-01 | 3.92E-01 | 1.17E+00 | 2.52E-01 | 7.11E-01 | 1.34E+00 | 2.11E-01 | 7.75E-01 | 1.25E+00 | 2.36E-01 | 6.49E-01 | 1.13E+00 |
| Z82206.1 | 1.91E-01 | 9.94E-01 | 1.26E+00 | 6.11E-02 | 2.64E-01 | -1.20E+00 | 4.70E-03 | 2.00E-02 | -1.56E+00 | 2.20E-03 | 8.38E-03 | -1.58E+00 | 8.77E-23 | 3.13E-22 | -2.94E+00 | 2.14E-16 | 6.04E-16 | -1.84E+00 | 2.27E-01 | 8.34E-01 | 1.49E+00 | 1.02E-01 | 2.80E-01 | 1.22E+00 |
| XK | 1.91E-01 | 9.94E-01 | 1.13E+00 | 1.57E-01 | 6.79E-01 | 1.81E+00 | 2.07E-01 | 8.81E-01 | -1.01E+00 | 2.58E-01 | 9.86E-01 | 1.21E+00 | 2.09E-01 | 7.46E-01 | 1.07E+00 | 6.33E-03 | 1.78E-02 | 2.09E+00 | 6.76E-05 | 2.48E-04 | -1.12E+00 | 5.59E-03 | 1.53E-02 | 1.01E+00 |
| RPL5P3 | 1.91E-01 | 9.94E-01 | 1.21E+00 | 1.87E-01 | 8.10E-01 | 1.46E+00 | 1.94E-02 | 8.25E-02 | 1.82E+00 | 9.02E-02 | 3.45E-01 | 1.69E+00 | 1.66E-01 | 5.93E-01 | 1.56E+00 | 7.58E-02 | 2.14E-01 | 2.20E+00 | 2.11E-01 | 7.76E-01 | 1.37E+00 | 9.61E-03 | 2.64E-02 | 2.13E+00 |
| CNTN4 | 1.91E-01 | 9.94E-01 | 1.22E+00 | 1.63E-01 | 7.04E-01 | 1.56E+00 | 1.41E-01 | 6.01E-01 | 1.05E+00 | 7.77E-02 | 2.97E-01 | 1.83E+00 | 2.71E-01 | 9.66E-01 | 1.30E+00 | 2.47E-01 | 6.95E-01 | 1.74E+00 | 6.25E-03 | 2.30E-02 | 2.73E+00 | 7.35E-02 | 2.02E-01 | 1.94E+00 |
| MISP3 | 1.91E-01 | 9.94E-01 | 1.26E+00 | 1.32E-01 | 5.72E-01 | 1.61E+00 | 2.02E-03 | 8.57E-03 | -1.63E+00 | 6.72E-03 | 2.57E-02 | -1.39E+00 | 2.67E-01 | 9.53E-01 | 1.38E+00 | 2.80E-01 | 7.91E-01 | 1.42E+00 | 2.27E-02 | 8.33E-02 | 2.66E+00 | 3.15E-01 | 8.67E-01 | 1.07E+00 |
| LIF | 1.91E-01 | 9.95E-01 | 1.08E+00 | 1.95E-01 | 8.41E-01 | -1.13E+00 | 1.59E-01 | 6.74E-01 | 1.41E+00 | 2.30E-01 | 8.79E-01 | -1.08E+00 | 1.08E-01 | 3.86E-01 | 1.16E+00 | 2.96E-08 | 8.35E-08 | -1.87E+00 | 8.54E-02 | 3.13E-01 | 1.88E+00 | 1.93E-01 | 5.29E-01 | 1.11E+00 |
| RPLP0P6 | 1.91E-01 | 9.95E-01 | 1.06E+00 | 7.36E-04 | 3.18E-03 | 1.29E+00 | 1.87E-01 | 7.96E-01 | 1.03E+00 | 2.29E-03 | 8.76E-03 | 1.32E+00 | 2.70E-03 | 9.64E-03 | -1.04E+00 | 5.98E-02 | 1.69E-01 | 1.25E+00 | 2.72E-02 | 9.98E-02 | -1.03E+00 | 1.07E-03 | 2.94E-03 | 1.30E+00 |
| ASAP1-IT2 | 1.91E-01 | 9.95E-01 | 1.11E+00 | 1.77E-01 | 7.63E-01 | 1.46E+00 | 1.34E-01 | 5.71E-01 | -1.06E+00 | 2.78E-02 | 1.06E-01 | -1.28E+00 | 2.78E-01 | 9.93E-01 | 1.11E+00 | 1.17E-01 | 3.29E-01 | 1.82E+00 | 8.80E-05 | 3.23E-04 | -1.56E+00 | 3.21E-03 | 8.81E-03 | -1.28E+00 |
| AP003068.2 | 1.91E-01 | 9.95E-01 | 1.27E+00 | 4.18E-02 | 1.80E-01 | 1.89E+00 | 3.50E-02 | 1.49E-01 | -1.04E+00 | 1.62E-01 | 6.20E-01 | 1.34E+00 | 6.52E-02 | 2.33E-01 | 1.93E+00 | 7.41E-02 | 2.09E-01 | 1.93E+00 | 2.56E-01 | 9.39E-01 | 1.57E+00 | 9.76E-02 | 2.68E-01 | 1.34E+00 |
| RFX3-AS1 | 1.91E-01 | 9.95E-01 | 1.29E+00 | 8.82E-02 | 3.81E-01 | -1.02E+00 | 2.58E-02 | 1.10E-01 | -1.20E+00 | 4.72E-02 | 1.80E-01 | -1.08E+00 | 1.37E-01 | 4.89E-01 | 1.01E+00 | 3.51E-01 | 9.89E-01 | 1.38E+00 | 2.22E-01 | 8.15E-01 | 1.30E+00 | 3.23E-01 | 8.86E-01 | 1.36E+00 |
| MCM7 | 1.91E-01 | 9.95E-01 | 1.29E+00 | 2.16E-02 | 9.35E-02 | 1.20E+00 | 2.13E-03 | 9.04E-03 | 1.41E+00 | 8.26E-02 | 3.16E-01 | 1.47E+00 | 1.77E-06 | 6.33E-06 | -1.08E+00 | 2.60E-06 | 7.33E-06 | -1.14E+00 | 9.80E-03 | 3.60E-02 | -1.05E+00 | 9.99E-03 | 2.75E-02 | 1.37E+00 |
| RNPEPL1 | 1.91E-01 | 9.96E-01 | 1.18E+00 | 2.14E-01 | 9.26E-01 | 1.42E+00 | 1.48E-01 | 6.31E-01 | 1.16E+00 | 2.01E-01 | 7.68E-01 | 1.37E+00 | 2.20E-01 | 7.83E-01 | 1.54E+00 | 2.00E-01 | 5.65E-01 | 1.68E+00 | 2.51E-02 | 9.21E-02 | 1.58E+00 | 2.32E-02 | 6.37E-02 | 1.52E+00 |
| RPS4XP8 | 1.91E-01 | 9.96E-01 | 1.32E+00 | 1.66E-01 | 7.16E-01 | 1.36E+00 | 4.74E-02 | 2.01E-01 | -1.01E+00 | 8.68E-03 | 3.32E-02 | -1.14E+00 | 1.18E-01 | 4.20E-01 | 1.17E+00 | 4.22E-02 | 1.19E-01 | -1.06E+00 | 5.74E-02 | 2.11E-01 | 1.66E+00 | 3.18E-01 | 8.75E-01 | 1.18E+00 |
| AL592293.1 | 1.91E-01 | 9.96E-01 | 1.23E+00 | 2.26E-01 | 9.77E-01 | 1.43E+00 | 2.34E-01 | 9.93E-01 | 1.19E+00 | 5.30E-02 | 2.02E-01 | 1.90E+00 | 2.55E-01 | 9.11E-01 | 1.49E+00 | 2.81E-01 | 7.92E-01 | 1.67E+00 | 7.25E-02 | 2.66E-01 | 1.06E+00 | 2.57E-01 | 7.06E-01 | 1.37E+00 |
| EPB41L4A | 1.91E-01 | 9.97E-01 | 1.14E+00 | 1.35E-01 | 5.81E-01 | 1.53E+00 | 5.74E-03 | 2.44E-02 | -1.28E+00 | 5.62E-02 | 2.15E-01 | 1.13E+00 | 1.61E-01 | 5.73E-01 | 1.60E+00 | 2.79E-01 | 7.85E-01 | 1.08E+00 | 1.01E-01 | 3.70E-01 | 1.56E+00 | 6.86E-02 | 1.88E-01 | 1.78E+00 |
| RPS27P29 | 1.91E-01 | 9.97E-01 | 1.34E+00 | 1.31E-03 | 5.68E-03 | -1.27E+00 | 3.66E-03 | 1.56E-02 | 1.83E+00 | 1.02E-02 | 3.91E-02 | -1.39E+00 | 7.42E-02 | 2.64E-01 | 1.65E+00 | 4.84E-02 | 1.37E-01 | -1.40E+00 | 6.10E-03 | 2.24E-02 | 1.82E+00 | 5.35E-02 | 1.47E-01 | -1.23E+00 |
| HBA2 | 1.91E-01 | 9.98E-01 | 6.65E+00 | 3.21E-11 | 1.38E-10 | 4.48E+00 | 8.04E-02 | 3.42E-01 | 1.46E+00 | 6.24E-08 | 2.38E-07 | 9.59E+00 | 1.19E-02 | 4.23E-02 | 3.25E+01 | 2.12E-03 | 5.99E-03 | 1.98E+01 | 2.30E-01 | 8.46E-01 | -1.46E+00 | 1.04E-13 | 2.86E-13 | -2.13E+00 |
| FLJ37453 | 1.92E-01 | 9.98E-01 | 1.30E+00 | 3.15E-03 | 1.36E-02 | 1.11E+00 | 1.04E-01 | 4.42E-01 | 1.09E+00 | 3.58E-03 | 1.37E-02 | 1.02E+00 | 1.12E-01 | 4.01E-01 | 1.25E+00 | 1.93E-02 | 5.44E-02 | 1.07E+00 | 7.67E-02 | 2.82E-01 | 1.05E+00 | 1.47E-01 | 4.04E-01 | 1.05E+00 |
| RN7SL4P | 1.92E-01 | 9.98E-01 | 1.29E+00 | 2.95E-04 | 1.28E-03 | -1.90E+00 | 7.96E-02 | 3.38E-01 | -1.11E+00 | 2.53E-03 | 9.66E-03 | -1.71E+00 | 4.73E-03 | 1.69E-02 | -1.24E+00 | 2.39E-05 | 6.74E-05 | -2.85E+00 | 2.08E-01 | 7.62E-01 | 1.24E+00 | 2.75E-02 | 7.54E-02 | -1.35E+00 |
| ZNF699 | 1.92E-01 | 9.98E-01 | 1.27E+00 | 2.31E-01 | 1.00E+00 | 1.40E+00 | 2.01E-01 | 8.53E-01 | 1.24E+00 | 2.55E-01 | 9.74E-01 | 1.36E+00 | 2.06E-01 | 7.36E-01 | 1.60E+00 | 2.67E-01 | 7.54E-01 | 1.20E+00 | 1.00E-01 | 3.68E-01 | -1.06E+00 | 2.99E-02 | 8.22E-02 | -1.31E+00 |
| LRRC25 | 1.92E-01 | 9.98E-01 | -1.15E+00 | 1.51E-01 | 6.53E-01 | 1.26E+00 | 1.55E-01 | 6.58E-01 | 1.03E+00 | 2.21E-01 | 8.43E-01 | -1.00E+00 | 1.93E-01 | 6.87E-01 | -1.51E+00 | 2.67E-01 | 7.53E-01 | -1.44E+00 | 2.54E-01 | 9.33E-01 | -1.01E+00 | 3.07E-01 | 8.44E-01 | 1.20E+00 |
| AC141586.5 | 1.92E-01 | 9.99E-01 | 1.25E+00 | 9.22E-02 | 3.98E-01 | 1.05E+00 | 1.44E-01 | 6.14E-01 | 1.30E+00 | 1.27E-01 | 4.85E-01 | 1.36E+00 | 7.44E-02 | 2.65E-01 | -1.48E+00 | 1.54E-01 | 4.35E-01 | -1.19E+00 | 1.54E-01 | 5.66E-01 | -1.25E+00 | 2.67E-01 | 7.34E-01 | -1.11E+00 |
| CHST2 | 1.92E-01 | 9.99E-01 | 1.27E+00 | 1.43E-01 | 6.19E-01 | 1.88E+00 | 6.44E-03 | 2.74E-02 | -1.85E+00 | 5.26E-02 | 2.01E-01 | -1.18E+00 | 8.05E-08 | 2.87E-07 | -2.43E+00 | 1.12E-02 | 3.16E-02 | -1.07E+00 | 2.98E-38 | 1.09E-37 | -2.64E+00 | 2.66E-20 | 7.29E-20 | -1.64E+00 |
| E2F3-IT1 | 1.92E-01 | 9.99E-01 | 1.30E+00 | 1.04E-01 | 4.48E-01 | 1.06E+00 | 4.75E-02 | 2.02E-01 | -1.13E+00 | 2.92E-02 | 1.11E-01 | -1.11E+00 | 2.09E-01 | 7.45E-01 | 1.45E+00 | 3.39E-01 | 9.57E-01 | 1.45E+00 | 3.83E-03 | 1.40E-02 | -1.33E+00 | 1.05E-02 | 2.88E-02 | -1.26E+00 |
| RHCE | 1.92E-01 | 9.99E-01 | 1.13E+00 | 2.27E-01 | 9.80E-01 | 1.14E+00 | 1.46E-01 | 6.20E-01 | 1.25E+00 | 9.21E-02 | 3.52E-01 | 1.67E+00 | 2.67E-01 | 9.54E-01 | 1.20E+00 | 2.82E-01 | 7.95E-01 | 1.31E+00 | 1.25E-01 | 4.58E-01 | 1.12E+00 | 7.51E-03 | 2.06E-02 | -1.18E+00 |
| LMO3 | 1.92E-01 | 1.00E+00 | 1.16E+00 | 1.80E-01 | 7.79E-01 | 1.45E+00 | 2.74E-02 | 1.17E-01 | -1.27E+00 | 5.39E-02 | 2.06E-01 | -1.16E+00 | 2.18E-01 | 7.76E-01 | 1.28E+00 | 1.21E-01 | 3.41E-01 | 2.07E+00 | 2.91E-02 | 1.07E-01 | -1.68E+00 | 3.10E-01 | 8.52E-01 | -1.01E+00 |
| NEIL3 | 1.92E-01 | 1.00E+00 | 1.11E+00 | 1.30E-01 | 5.62E-01 | -1.19E+00 | 1.38E-01 | 5.85E-01 | 1.27E+00 | 2.59E-01 | 9.89E-01 | 1.24E+00 | 2.18E-01 | 7.78E-01 | 1.43E+00 | 3.11E-01 | 8.76E-01 | -1.05E+00 | 1.23E-01 | 4.52E-01 | -1.18E+00 | 6.84E-02 | 1.88E-01 | 1.31E+00 |
| SNX30 | 1.92E-01 | 1.00E+00 | 1.42E+00 | 1.64E-01 | 7.10E-01 | 1.36E+00 | 4.73E-02 | 2.01E-01 | 1.03E+00 | 9.40E-02 | 3.59E-01 | 1.22E+00 | 1.21E-01 | 4.31E-01 | 1.63E+00 | 1.10E-01 | 3.11E-01 | -1.02E+00 | 1.50E-01 | 5.52E-01 | 1.50E+00 | 5.60E-02 | 1.54E-01 | -1.06E+00 |
| ANGPTL1 | 1.92E-01 | 1.00E+00 | 1.25E+00 | 1.25E-01 | 5.42E-01 | 1.70E+00 | 8.81E-03 | 3.74E-02 | -1.45E+00 | 2.46E-01 | 9.38E-01 | 1.33E+00 | 1.16E-01 | 4.15E-01 | 1.02E+00 | 2.44E-01 | 6.89E-01 | 1.24E+00 | 1.08E-01 | 3.95E-01 | -1.15E+00 | 2.66E-01 | 7.30E-01 | 1.14E+00 |
| TTYH1 | 1.92E-01 | 1.00E+00 | 1.27E+00 | 1.44E-01 | 6.23E-01 | 1.59E+00 | 2.08E-01 | 8.82E-01 | 1.12E+00 | 2.36E-01 | 9.01E-01 | 1.28E+00 | 9.52E-02 | 3.40E-01 | -1.63E+00 | 2.58E-01 | 7.28E-01 | -1.04E+00 | 2.72E-01 | 1.00E+00 | 1.06E+00 | 3.20E-01 | 8.80E-01 | 1.24E+00 |
| TSPAN5 | 1.92E-01 | 1.00E+00 | 1.34E+00 | 5.71E-03 | 2.47E-02 | 2.11E+00 | 1.15E-01 | 4.89E-01 | 1.20E+00 | 1.62E-01 | 6.20E-01 | 1.58E+00 | 1.23E-01 | 4.39E-01 | 1.09E+00 | 2.11E-01 | 5.95E-01 | 1.25E+00 | 2.08E-01 | 7.64E-01 | 1.22E+00 | 1.76E-01 | 4.82E-01 | 1.43E+00 |
| LINC00664 | 1.92E-01 | 1.00E+00 | 1.10E+00 | 2.29E-01 | 9.89E-01 | 1.12E+00 | 7.06E-02 | 3.00E-01 | 1.03E+00 | 1.68E-01 | 6.43E-01 | 1.12E+00 | 2.33E-01 | 8.32E-01 | 1.03E+00 | 1.70E-01 | 4.79E-01 | -1.49E+00 | 8.78E-02 | 3.22E-01 | -1.17E+00 | 4.90E-03 | 1.34E-02 | 1.56E+00 |
| B4GALT2 | 1.92E-01 | 1.00E+00 | 1.34E+00 | 1.59E-01 | 6.88E-01 | 1.35E+00 | 1.08E-01 | 4.58E-01 | 1.38E+00 | 1.71E-02 | 6.55E-02 | 1.92E+00 | 1.41E-01 | 5.02E-01 | 1.14E+00 | 1.81E-01 | 5.11E-01 | 1.88E+00 | 5.25E-02 | 1.93E-01 | 1.47E+00 | 5.09E-02 | 1.40E-01 | 1.52E+00 |
| KEL | 1.92E-01 | 1.00E+00 | 1.00E+00 | 2.09E-01 | 9.04E-01 | 1.20E+00 | 1.93E-01 | 8.21E-01 | 1.24E+00 | 2.24E-01 | 8.55E-01 | 1.22E+00 | 2.61E-01 | 9.31E-01 | -1.11E+00 | 1.62E-01 | 4.58E-01 | 1.36E+00 | 1.76E-02 | 6.47E-02 | 1.12E+00 | 1.15E-03 | 3.16E-03 | 1.03E+00 |
| ST6GALNAC1 | 1.92E-01 | 1.00E+00 | 1.00E+00 | 2.31E-01 | 1.00E+00 | 1.00E+00 | 2.35E-01 | 1.00E+00 | 1.00E+00 | 2.62E-01 | 1.00E+00 | -1.00E+00 | 2.24E-01 | 7.98E-01 | -1.23E+00 | 1.94E-01 | 5.46E-01 | 1.60E+00 | 6.32E-26 | 2.32E-25 | -1.82E+00 | 1.71E-27 | 4.69E-27 | -1.77E+00 |
| RNASE1 | 1.92E-01 | 1.00E+00 | -1.00E+00 | 2.31E-01 | 1.00E+00 | 1.00E+00 | 2.35E-01 | 1.00E+00 | -1.00E+00 | 2.62E-01 | 1.00E+00 | 1.00E+00 | 1.23E-01 | 4.37E-01 | -1.67E+00 | 1.96E-01 | 5.52E-01 | 1.59E+00 | 2.15E-139 | 7.90E-139 | -5.83E+00 | 1.68E-106 | 4.62E-106 | -4.43E+00 |
| UTS2 | 1.92E-01 | 1.00E+00 | 1.00E+00 | 2.31E-01 | 1.00E+00 | -1.00E+00 | 2.05E-01 | 8.72E-01 | -1.09E+00 | 2.40E-01 | 9.18E-01 | 1.65E+00 | 2.12E-02 | 7.56E-02 | 3.86E+00 | 1.32E-02 | 3.73E-02 | 4.69E+00 | 2.90E-16 | 1.07E-15 | 9.12E+00 | 1.57E-17 | 4.31E-17 | 1.48E+01 |
| FGF2 | 1.92E-01 | 1.00E+00 | 1.00E+00 | 2.13E-01 | 9.22E-01 | 1.10E+00 | 2.20E-01 | 9.34E-01 | 1.04E+00 | 2.41E-01 | 9.20E-01 | 1.09E+00 | 2.51E-01 | 8.94E-01 | 1.16E+00 | 1.44E-01 | 4.07E-01 | 1.84E+00 | 1.35E-02 | 4.94E-02 | 1.10E+00 | 5.20E-03 | 1.43E-02 | -1.03E+00 |
| PVALB | 1.92E-01 | 1.00E+00 | 1.00E+00 | 2.31E-01 | 1.00E+00 | -1.00E+00 | 2.19E-01 | 9.33E-01 | 1.04E+00 | 2.19E-01 | 8.38E-01 | 1.14E+00 | 1.49E-03 | 5.32E-03 | 2.80E+00 | 1.57E-01 | 4.44E-01 | 1.91E+00 | 7.78E-02 | 2.85E-01 | 1.21E+00 | 1.13E-01 | 3.10E-01 | 1.29E+00 |
| PBK | 1.92E-01 | 1.00E+00 | 1.00E+00 | 2.31E-01 | 1.00E+00 | 1.00E+00 | 2.29E-01 | 9.73E-01 | -1.06E+00 | 2.33E-01 | 8.92E-01 | -1.17E+00 | 2.80E-01 | 1.00E+00 | 1.00E+00 | 3.55E-01 | 1.00E+00 | 1.00E+00 | 3.83E-05 | 1.40E-04 | -1.59E+00 | 8.88E-02 | 2.44E-01 | -1.46E+00 |
| OOSP1 | 1.92E-01 | 1.00E+00 | 1.00E+00 | 1.91E-01 | 8.27E-01 | 1.21E+00 | 2.19E-01 | 9.32E-01 | 1.04E+00 | 1.55E-01 | 5.92E-01 | 1.34E+00 | 1.51E-01 | 5.39E-01 | 1.58E+00 | 3.14E-04 | 8.86E-04 | 3.57E+00 | 4.89E-06 | 1.80E-05 | 2.73E+00 | 2.53E-06 | 6.96E-06 | 2.91E+00 |
| SLC10A4 | 1.92E-01 | 1.00E+00 | 1.00E+00 | 2.15E-01 | 9.28E-01 | 1.07E+00 | 2.35E-01 | 1.00E+00 | 1.00E+00 | 2.40E-01 | 9.18E-01 | 1.10E+00 | 2.06E-01 | 7.35E-01 | -1.04E+00 | 8.54E-02 | 2.41E-01 | 2.04E+00 | 3.44E-31 | 1.26E-30 | -2.46E+00 | 4.63E-10 | 1.27E-09 | -1.30E+00 |
| HMGB1P30 | 1.92E-01 | 1.00E+00 | 1.00E+00 | 2.19E-01 | 9.46E-01 | 1.06E+00 | 2.13E-01 | 9.04E-01 | 1.06E+00 | 2.33E-01 | 8.91E-01 | 1.10E+00 | 7.57E-02 | 2.70E-01 | 2.35E+00 | 2.04E-04 | 5.75E-04 | 4.35E+00 | 2.68E-01 | 9.83E-01 | 1.30E+00 | 3.12E-01 | 8.58E-01 | 1.29E+00 |
| AC090409.1 | 1.92E-01 | 1.00E+00 | 1.00E+00 | 2.31E-01 | 1.00E+00 | 1.00E+00 | 2.35E-01 | 1.00E+00 | 1.00E+00 | 2.62E-01 | 1.00E+00 | 1.00E+00 | 2.80E-01 | 1.00E+00 | -1.00E+00 | 3.55E-01 | 1.00E+00 | 1.00E+00 | 6.00E-02 | 2.20E-01 | 1.28E+00 | 3.26E-02 | 8.96E-02 | 2.21E+00 |
| AL662860.1 | 1.92E-01 | 1.00E+00 | 1.00E+00 | 2.31E-01 | 1.00E+00 | -1.00E+00 | 2.35E-01 | 1.00E+00 | 1.00E+00 | 2.62E-01 | 1.00E+00 | 1.00E+00 | 1.54E-85 | 5.50E-85 | -1.27E+01 | 6.97E-78 | 1.97E-77 | -7.56E+00 | 4.24E-08 | 1.56E-07 | -2.06E+00 | 2.94E-06 | 8.07E-06 | -1.78E+00 |
| CD300E | 1.92E-01 | 1.00E+00 | 1.00E+00 | 2.31E-01 | 1.00E+00 | -1.00E+00 | 6.72E-02 | 2.86E-01 | -1.55E+00 | 5.79E-02 | 2.21E-01 | -1.74E+00 | 2.80E-01 | 1.00E+00 | -1.00E+00 | 3.55E-01 | 1.00E+00 | 1.00E+00 | 7.31E-02 | 2.68E-01 | -1.46E+00 | 8.85E-02 | 2.43E-01 | -1.38E+00 |
| CLEC4E | 1.92E-01 | 1.00E+00 | 1.00E+00 | 2.00E-01 | 8.65E-01 | 1.25E+00 | 1.85E-01 | 7.84E-01 | 1.01E+00 | 1.35E-01 | 5.17E-01 | -1.49E+00 | 1.78E-01 | 6.33E-01 | 1.75E+00 | 3.55E-01 | 1.00E+00 | 1.00E+00 | 2.61E-01 | 9.60E-01 | 1.13E+00 | 2.99E-01 | 8.23E-01 | -1.10E+00 |
| E2F7 | 1.92E-01 | 1.00E+00 | -1.00E+00 | 2.11E-01 | 9.10E-01 | 1.08E+00 | 8.36E-02 | 3.55E-01 | 1.45E+00 | 8.68E-02 | 3.32E-01 | 1.45E+00 | 2.13E-01 | 7.58E-01 | 1.24E+00 | 3.55E-01 | 1.00E+00 | 1.00E+00 | 2.21E-02 | 8.12E-02 | -1.13E+00 | 1.87E-01 | 5.12E-01 | 1.06E+00 |
| CYP27A1 | 1.92E-01 | 1.00E+00 | 1.00E+00 | 1.99E-01 | 8.59E-01 | 1.10E+00 | 2.22E-01 | 9.45E-01 | 1.10E+00 | 2.49E-01 | 9.51E-01 | 1.04E+00 | 2.80E-01 | 1.00E+00 | 1.00E+00 | 3.55E-01 | 1.00E+00 | 1.00E+00 | 1.42E-01 | 5.20E-01 | -1.27E+00 | 2.75E-01 | 7.57E-01 | -1.15E+00 |
| PTCRA | 1.92E-01 | 1.00E+00 | 1.00E+00 | 2.31E-01 | 1.00E+00 | 1.00E+00 | 1.74E-01 | 7.38E-01 | -1.02E+00 | 1.61E-01 | 6.16E-01 | -1.15E+00 | 2.41E-01 | 8.61E-01 | 1.09E+00 | 3.55E-01 | 1.00E+00 | 1.00E+00 | 6.88E-04 | 2.52E-03 | 9.48E+00 | 2.21E-01 | 6.07E-01 | 3.39E+00 |
| GAD1 | 1.92E-01 | 1.00E+00 | 1.00E+00 | 2.31E-01 | 1.00E+00 | -1.00E+00 | 2.11E-01 | 8.95E-01 | -1.02E+00 | 2.53E-01 | 9.66E-01 | 1.06E+00 | 3.88E-04 | 1.38E-03 | 5.75E+00 | 1.16E-03 | 3.28E-03 | 4.59E+00 | 8.17E-04 | 3.00E-03 | -1.14E+00 | 7.87E-06 | 2.16E-05 | -1.55E+00 |
| STUM | 1.92E-01 | 1.00E+00 | 1.00E+00 | 2.09E-01 | 9.01E-01 | 1.11E+00 | 2.14E-01 | 9.08E-01 | 1.07E+00 | 2.10E-01 | 8.03E-01 | 1.15E+00 | 1.41E-01 | 5.02E-01 | -1.12E+00 | 3.46E-01 | 9.75E-01 | 1.25E+00 | 7.83E-07 | 2.87E-06 | -1.61E+00 | 3.34E-02 | 9.17E-02 | -1.02E+00 |
| SPTB | 1.92E-01 | 1.00E+00 | 1.00E+00 | 2.31E-01 | 1.00E+00 | 1.00E+00 | 2.35E-01 | 1.00E+00 | -1.00E+00 | 2.62E-01 | 1.00E+00 | -1.00E+00 | 2.59E-01 | 9.22E-01 | 1.05E+00 | 6.39E-02 | 1.80E-01 | 2.19E+00 | 1.49E-01 | 5.45E-01 | 1.60E+00 | 4.51E-02 | 1.24E-01 | 1.76E+00 |
| TEX15 | 1.92E-01 | 1.00E+00 | -1.00E+00 | 2.04E-01 | 8.81E-01 | 1.18E+00 | 3.53E-02 | 1.50E-01 | -2.07E+00 | 1.14E-01 | 4.37E-01 | -1.36E+00 | 1.17E-13 | 4.17E-13 | -3.19E+00 | 1.28E-07 | 3.61E-07 | -1.70E+00 | 4.77E-16 | 1.75E-15 | -2.19E+00 | 1.96E-08 | 5.40E-08 | -1.49E+00 |
| CDK15 | 1.92E-01 | 1.00E+00 | -1.00E+00 | 2.10E-01 | 9.05E-01 | 1.12E+00 | 2.19E-01 | 9.29E-01 | 1.03E+00 | 1.66E-01 | 6.32E-01 | 1.27E+00 | 1.70E-08 | 6.07E-08 | -1.88E+00 | 2.58E-11 | 7.26E-11 | -1.90E+00 | 9.87E-02 | 3.62E-01 | -1.15E+00 | 1.55E-01 | 4.26E-01 | -1.18E+00 |
| DKK1 | 1.92E-01 | 1.00E+00 | 1.00E+00 | 2.31E-01 | 1.00E+00 | -1.00E+00 | 2.35E-01 | 1.00E+00 | 1.00E+00 | 2.62E-01 | 1.00E+00 | -1.00E+00 | 1.64E-01 | 5.84E-01 | -1.21E+00 | 2.62E-01 | 7.40E-01 | 1.37E+00 | 7.10E-33 | 2.61E-32 | -2.24E+00 | 1.72E-23 | 4.73E-23 | -1.62E+00 |
| UBASH3A | 1.92E-01 | 1.00E+00 | 1.00E+00 | 2.31E-01 | 1.00E+00 | 1.00E+00 | 2.35E-01 | 1.00E+00 | 1.00E+00 | 2.46E-01 | 9.41E-01 | 1.04E+00 | 1.30E-02 | 4.65E-02 | -1.79E+00 | 2.60E-01 | 7.34E-01 | 1.11E+00 | 1.49E-04 | 5.46E-04 | -1.08E+00 | 5.19E-02 | 1.42E-01 | 1.12E+00 |
| CD163 | 1.92E-01 | 1.00E+00 | 1.00E+00 | 1.44E-01 | 6.21E-01 | 1.50E+00 | 2.00E-01 | 8.48E-01 | 1.39E+00 | 2.11E-01 | 8.06E-01 | 1.02E+00 | 2.39E-01 | 8.53E-01 | 1.12E+00 | 3.55E-01 | 1.00E+00 | -1.00E+00 | 1.85E-01 | 6.81E-01 | 1.36E+00 | 3.10E-01 | 8.52E-01 | 1.10E+00 |
| HOTAIRM1 | 1.92E-01 | 1.00E+00 | 1.00E+00 | 2.31E-01 | 1.00E+00 | 1.00E+00 | 1.89E-01 | 8.04E-01 | -1.03E+00 | 1.88E-01 | 7.18E-01 | -1.10E+00 | 2.14E-14 | 7.61E-14 | -3.84E+00 | 7.74E-14 | 2.18E-13 | -3.80E+00 | 1.65E-08 | 6.07E-08 | -1.86E+00 | 1.65E-08 | 4.53E-08 | -1.74E+00 |
| TRIM10 | 1.92E-01 | 1.00E+00 | 1.00E+00 | 2.31E-01 | 1.00E+00 | -1.00E+00 | 2.05E-01 | 8.71E-01 | 1.04E+00 | 2.62E-01 | 1.00E+00 | -1.00E+00 | 2.20E-01 | 7.83E-01 | 1.18E+00 | 3.02E-01 | 8.52E-01 | 1.13E+00 | 6.73E-02 | 2.47E-01 | -1.31E+00 | 5.22E-02 | 1.44E-01 | -1.45E+00 |
| STX1A | 1.92E-01 | 1.00E+00 | 1.00E+00 | 2.31E-01 | 1.00E+00 | 1.00E+00 | 2.35E-01 | 1.00E+00 | 1.00E+00 | 2.22E-01 | 8.49E-01 | 1.09E+00 | 2.31E-01 | 8.24E-01 | 1.10E+00 | 3.09E-01 | 8.72E-01 | 1.09E+00 | 2.02E-01 | 7.41E-01 | 1.48E+00 | 2.45E-03 | 6.73E-03 | -1.66E+00 |
| SERPINA1 | 1.92E-01 | 1.00E+00 | 1.00E+00 | 1.70E-01 | 7.32E-01 | 2.48E+00 | 8.94E-02 | 3.80E-01 | -1.71E+00 | 8.18E-02 | 3.12E-01 | -2.02E+00 | 2.45E-01 | 8.74E-01 | 1.18E+00 | 3.55E-01 | 1.00E+00 | -1.00E+00 | 1.87E-01 | 6.85E-01 | -1.45E+00 | 1.67E-01 | 4.59E-01 | -1.99E+00 |
| CDKN2A | 1.92E-01 | 1.00E+00 | -1.00E+00 | 1.98E-01 | 8.57E-01 | 1.12E+00 | 2.35E-01 | 1.00E+00 | 1.00E+00 | 2.28E-01 | 8.70E-01 | 1.06E+00 | 1.14E-01 | 4.05E-01 | -1.48E+00 | 1.48E-01 | 4.16E-01 | -1.48E+00 | 4.85E-12 | 1.78E-11 | -1.70E+00 | 3.43E-15 | 9.41E-15 | -2.13E+00 |
| ADCYAP1 | 1.92E-01 | 1.00E+00 | 1.00E+00 | 2.31E-01 | 1.00E+00 | -1.00E+00 | 1.91E-01 | 8.13E-01 | -1.08E+00 | 2.08E-01 | 7.94E-01 | 1.46E+00 | 6.92E-03 | 2.47E-02 | 3.23E+00 | 1.72E-03 | 4.86E-03 | 4.01E+00 | 1.34E-01 | 4.90E-01 | 1.85E+00 | 7.48E-02 | 2.05E-01 | 1.99E+00 |
| AQP9 | 1.92E-01 | 1.00E+00 | 1.00E+00 | 2.31E-01 | 1.00E+00 | -1.00E+00 | 2.35E-01 | 1.00E+00 | -1.00E+00 | 2.62E-01 | 1.00E+00 | -1.00E+00 | 2.17E-01 | 7.74E-01 | 1.18E+00 | 3.55E-01 | 1.00E+00 | 1.00E+00 | 2.05E-01 | 7.52E-01 | -1.06E+00 | 2.28E-01 | 6.27E-01 | -1.09E+00 |
| CLEC4D | 1.92E-01 | 1.00E+00 | -1.00E+00 | 2.31E-01 | 1.00E+00 | -1.00E+00 | 1.79E-01 | 7.59E-01 | -1.03E+00 | 1.69E-01 | 6.47E-01 | -1.10E+00 | 1.47E-01 | 5.25E-01 | 1.11E+00 | 9.80E-03 | 2.76E-02 | -1.42E+00 | 4.50E-02 | 1.65E-01 | -1.33E+00 | 1.01E-01 | 2.77E-01 | 1.34E+00 |
| LRRC32 | 1.92E-01 | 1.00E+00 | 1.00E+00 | 2.11E-01 | 9.11E-01 | 1.08E+00 | 1.80E-01 | 7.66E-01 | -1.10E+00 | 1.81E-01 | 6.90E-01 | -1.15E+00 | 2.80E-01 | 1.00E+00 | 1.00E+00 | 3.55E-01 | 1.00E+00 | -1.00E+00 | 2.59E-01 | 9.52E-01 | 2.56E+00 | 1.99E-01 | 5.47E-01 | 1.58E+00 |
| HMOX1 | 1.92E-01 | 1.00E+00 | 1.00E+00 | 2.02E-01 | 8.71E-01 | 1.11E+00 | 1.48E-01 | 6.27E-01 | -1.10E+00 | 1.67E-01 | 6.38E-01 | -1.10E+00 | 4.43E-02 | 1.58E-01 | -1.56E+00 | 1.21E-01 | 3.41E-01 | -1.07E+00 | 9.54E-02 | 3.50E-01 | -1.38E+00 | 7.22E-02 | 1.98E-01 | -1.57E+00 |
| PF4V1 | 1.92E-01 | 1.00E+00 | 1.00E+00 | 2.31E-01 | 1.00E+00 | 1.00E+00 | 1.70E-01 | 7.22E-01 | 1.21E+00 | 2.62E-01 | 1.00E+00 | 1.00E+00 | 2.80E-01 | 1.00E+00 | 1.00E+00 | 3.55E-01 | 1.00E+00 | 1.00E+00 | 3.70E-03 | 1.36E-02 | 3.68E+00 | 1.23E-01 | 3.39E-01 | 1.85E+00 |
| HGD | 1.92E-01 | 1.00E+00 | 1.00E+00 | 2.31E-01 | 1.00E+00 | -1.00E+00 | 1.97E-01 | 8.37E-01 | 1.08E+00 | 2.37E-01 | 9.04E-01 | 1.08E+00 | 1.01E-01 | 3.62E-01 | 1.15E+00 | 3.16E-01 | 8.91E-01 | -1.42E+00 | 2.19E-01 | 8.02E-01 | 1.96E+00 | 2.20E-02 | 6.03E-02 | 3.49E+00 |
| CTTN | 1.92E-01 | 1.00E+00 | 1.00E+00 | 2.15E-01 | 9.30E-01 | 1.08E+00 | 1.90E-01 | 8.06E-01 | 1.13E+00 | 2.62E-01 | 1.00E+00 | -1.00E+00 | 2.10E-01 | 7.48E-01 | -1.43E+00 | 3.12E-01 | 8.79E-01 | -1.25E+00 | 1.13E-01 | 4.16E-01 | 1.15E+00 | 2.09E-02 | 5.74E-02 | -1.57E+00 |
| CD3D | 1.92E-01 | 1.00E+00 | 1.00E+00 | 2.31E-01 | 1.00E+00 | -1.00E+00 | 5.68E-02 | 2.41E-01 | -2.54E+00 | 2.75E-02 | 1.05E-01 | -3.40E+00 | 1.28E-01 | 4.55E-01 | 1.40E+00 | 3.55E-01 | 1.00E+00 | 1.00E+00 | 2.18E-01 | 8.00E-01 | 1.14E+00 | 3.64E-01 | 1.00E+00 | 1.00E+00 |
| SMIM25 | 1.92E-01 | 1.00E+00 | -1.00E+00 | 1.90E-01 | 8.19E-01 | 1.86E+00 | 1.81E-01 | 7.70E-01 | 1.02E+00 | 2.04E-01 | 7.80E-01 | -1.01E+00 | 2.37E-01 | 8.47E-01 | 1.11E+00 | 3.55E-01 | 1.00E+00 | 1.00E+00 | 1.49E-01 | 5.48E-01 | -1.27E+00 | 1.71E-01 | 4.70E-01 | -1.33E+00 |
| PDGFB | 1.92E-01 | 1.00E+00 | 1.00E+00 | 2.31E-01 | 1.00E+00 | -1.00E+00 | 2.35E-01 | 1.00E+00 | 1.00E+00 | 2.62E-01 | 1.00E+00 | -1.00E+00 | 2.80E-01 | 1.00E+00 | -1.00E+00 | 3.55E-01 | 1.00E+00 | 1.00E+00 | 4.25E-07 | 1.56E-06 | 3.80E+00 | 1.73E-01 | 4.75E-01 | 1.87E+00 |
| PF4 | 1.92E-01 | 1.00E+00 | 1.00E+00 | 2.31E-01 | 1.00E+00 | -1.00E+00 | 1.62E-01 | 6.87E-01 | 1.46E+00 | 2.62E-01 | 1.00E+00 | -1.00E+00 | 2.80E-01 | 1.00E+00 | -1.00E+00 | 3.55E-01 | 1.00E+00 | -1.00E+00 | 3.59E-03 | 1.32E-02 | 5.16E+00 | 3.04E-01 | 8.35E-01 | 1.77E+00 |
| MEFV | 1.92E-01 | 1.00E+00 | -1.00E+00 | 1.97E-01 | 8.52E-01 | 1.09E+00 | 1.42E-01 | 6.03E-01 | -1.13E+00 | 1.26E-01 | 4.82E-01 | -1.20E+00 | 2.80E-01 | 1.00E+00 | 1.00E+00 | 3.55E-01 | 1.00E+00 | -1.00E+00 | 1.77E-01 | 6.49E-01 | -1.22E+00 | 2.67E-01 | 7.34E-01 | -1.16E+00 |
| SLC35D3 | 1.92E-01 | 1.00E+00 | 1.00E+00 | 2.31E-01 | 1.00E+00 | 1.00E+00 | 2.35E-01 | 1.00E+00 | 1.00E+00 | 2.62E-01 | 1.00E+00 | 1.00E+00 | 1.18E-01 | 4.20E-01 | -1.49E+00 | 1.52E-01 | 4.30E-01 | -1.49E+00 | 8.86E-07 | 3.25E-06 | 1.03E+00 | 2.39E-09 | 6.57E-09 | -1.52E+00 |
| GIMAP4 | 1.92E-01 | 1.00E+00 | 1.00E+00 | 1.79E-01 | 7.75E-01 | 1.16E+00 | 1.89E-02 | 8.03E-02 | -2.66E+00 | 1.66E-02 | 6.35E-02 | -2.84E+00 | 2.80E-01 | 1.00E+00 | 1.00E+00 | 3.55E-01 | 1.00E+00 | 1.00E+00 | 1.44E-01 | 5.30E-01 | -1.29E+00 | 1.14E-01 | 3.14E-01 | -1.56E+00 |
