## Supplementary Data File 4 for "Mutated clones driving leukemic transformation are already detectable at the single cell level in CD34-positive cells in the chronic phase of primary myelofibrosis"

**T3 vs. T1 HSC_MPP**

| Ingenuity Canonical Pathways | -log(p-valu | Ratio | z-score | Molecules |
|---|---|---|---|---|
| TNFR1 Signaling | 1.63 | 0.1 | 2 | FOS,JUN,NFKBID,TNF,TNFAIP3 |
| Pyrimidine Ribonucleotides Interconversion | 1.36 | 0.0976 | -2 | AK4,CMPK2,ENTPD1,ENTPD6 |
| Fcgamma Receptor-mediated Phagocytosis in Macrophages and Monocytes | 1.93 | 0.0851 | -2.828 | ACTA2,CRK,FCGR1A,FYB1,HCK,PLD4,PLD6,PRKCD |
| T Cell Exhaustion Signaling Pathway | 3.19 | 0.0857 | -3 | BATF,FOS,HLA-C,HLA-DRB5,HLA-F,HLA-G,IL6R,JUN,LAG3,MAPK12,PPP2R2B,PPP2R5B,STAT1,STAT2,ZAP70 |
| Interferon Signaling | 7.92 | 0.306 | -3.317 | IFI6,IFIT1,IFIT3,IFITM1,IFITM3,ISG15,MX1,OAS1,SOCS1,STAT1,STAT2 |

**T3 vs. T1 MPP_GMP**

| Ingenuity Canonical Pathways | -log(p-value) | Ratio | z-score | Molecules |
|---|---|---|---|---|
| Chemokine Signaling | 1.3 | 0.0625 | 2.236 | CCL5,CXCR4,FOS,GNAI1,JUN |
| IL-9 Signaling | 2.05 | 0.121 | -2 | JAK3,PIK3R3,SOCS3,STAT1 |
| Sperm Motility | 1.6 | 0.0493 | -2 | AXL,CSF1R,HCK,JAK3,LMTK2,PLA2G4A,PLA2R1,PLAAT1,PRKCD,PRKG2,SLC12A2 |
| Myc Mediated Apoptosis Signaling | 1.44 | 0.08 | -2 | FAS,MCL1,MYC,TNFRSF1A |
| Role of Pattern Recognition Receptors in Recognition of Bacteria and Viruses | 5.11 | 0.0974 | -2.121 | C3AR1,CCL5,CXCL8,DDX58,EIF2AK2,IFIH1,IRF7,NOD2,OAS1,OAS2,OAS3,PIK3R3,PRKCD,TNFSF10,TNFSF13B |
| Fc-gamma Receptor-mediated Phagocytosis in Macrophages and Monocytes | 1.51 | 0.0638 | -2.449 | ACTA2,FYB1,HCK,PIK3R3,PLD6,PRKCD |
| Tec Kinase Signaling | 5.41 | 0.0976 | -2.496 | ACTA2,DIRAS3,FAS,FOS,GNAI1,GNG2,HCK,ITGA2,JAK3,PIK3R3,PRKCD,RHOB,RHOU,STAT1,STAT2,TNFSF10 |
| Systemic Lupus Erythematosus In B Cell Signaling Pathway | 4.15 | 0.0691 | -2.524 | CXCL8,FOS,HCK,IFIH1,IFIT2,IFIT3,IRF7,ISG15,JUN,MCL1,MYC,PIK3AP1,PIK3R3,PRKCD,RASGRP3,STAT1,STAT2,TNFSF10,TNFSF13B |
| Type I Diabetes Mellitus Signaling | 3.97 | 0.0991 | -2.646 | CD3E,CD3G,FAS,HLA-C,HLA-DRB5,HLA-F,HLA-G,SOCS1,SOCS3,STAT1,TNFRSF1A |
| p38 MAPK Signaling | 1.54 | 0.0593 | -2.646 | DUSP1,FAS,HSPB1,MYC,PLA2G4A,STAT1,TNFRSF1A |
| Systemic Lupus Erythematosus In T Cell Signaling Pathway | 2 | 0.048 | -2.673 | CD3E,CD3G,DIRAS3,FAS,FOS,GNAI1,HLA-C,HLA-DRB5,HLA-F,HLA-G,JUN,PIK3R3,PPP2R1B,PPP2R5B,RHOB,RHOU |
| T Cell Exhaustion Signaling Pathway | 5.05 | 0.0914 | -2.714 | BATF,FOS,HLA-C,HLA-DRB5,HLA-F,HLA-G,IL10RA,JAK3,JUN,LAG3,PIK3R3,PPP2R1B,PPP2R5B,STAT1,STAT2,TCF7 |
| Th1 Pathway | 2.47 | 0.0744 | -2.828 | CD3E,CD3G,HLA-DRB5,IL10RA,JAK3,PIK3R3,SOCS1,SOCS3,STAT1 |
| Dendritic Cell Maturation | 1.39 | 0.0492 | -2.828 | COL2A1,FCGR1B,HLA-C,HLA-DRB5,HLA-F,PIK3R3,STAT1,STAT2,TNFRSF1A |
| Necroptosis Signaling Pathway | 1.36 | 0.051 | -2.828 | AXL,EIF2AK2,FAS,PLA2G4A,STAT1,STAT2,TNFRSF1A,TNFSF10 |
| Interferon Signaling | 11.7 | 0.361 | -3.606 | BAK1,IFI6,IFIT1,IFIT3,IFITM1,IFITM3,ISG15,MX1,OAS1,SOCS1,STAT1,STAT2,TAP1 |

**T3 vs. T1 MEP_1**

| Ingenuity Canonical Pathways | -log(p-value) | Ratio | z-score | Molecules |
|---|---|---|---|---|
| PD-1, PD-L1 cancer immunotherapy pathway | 2.65 | 0.104 | 3.162 | CD274,HLA-B,HLA-C,HLA-DMB,HLA-DPB1,HLA-DRB5,HLA-F,HLA-G,MR1,PIK3C2B,PIK3R3 |
| LPS/IL-1 Mediated Inhibition of RXR Function | 2.65 | 0.0804 | 2 | ACSL4,ALAS1,ALDH1B1,ALDH4A1,ALDH6A1,ALDH9A1,APOC1,APOE,CHST2,FMO4,GSTM2,GSTM3,GSTM5,IL1RAP,IL1RAPL1,JUN,SMOX,SULT1A2 |
| Complement System | 2.61 | 0.162 | -2 | C1R,C2,C3AR1,C5,CR1,ITGAX |
| Ethanol Degradation II | 1.5 | 0.125 | -2 | ALDH1B1,ALDH4A1,ALDH9A1,DHRS9 |
| Type I Diabetes Mellitus Signaling | 4.73 | 0.135 | -2.121 | CASP9,GAD1,HLA-B,HLA-C,HLA-DMB,HLA-DRB5,HLA-F,HLA-G,IL1RAP,IRF1,NFKBIE,SOCS1,SOCS2,SOCS3,STAT1 |
| Growth Hormone Signaling | 2.28 | 0.113 | -2.121 | FOS,PIK3C2B,PIK3R3,PRKCD,SOCS1,SOCS2,SOCS3,STAT1 |
| Neuroinflammation Signaling Pathway | 2.61 | 0.0733 | -2.236 | ACVR1,BACE2,BMPR2,CASP1,CXCL8,FOS,GAD1,HLA-B,HLA-C,HLA-DMB,HLA-DRB5,HLA-F,IRAK3,IRF7,JUN,PIK3C2B,PIK3R3,PTGS2,SOD2,STAT1,TICAM1,TYROBP |
| Systemic Lupus Erythematosus In B Cell Signaling Pathway | 3.47 | 0.0836 | -2.294 | CCND1,CCND3,CXCL8,FOS,HCK,IFIH1,IFIT2,IFIT3,IRF7,ISG15,JUN,MCL1,PAG1,PIK3C2B,PIK3R3,PRKCD,RASGRP3,STAT1,STAT2,SYNJ2,TICAM1,TNFSF10,TNFSF13B |
| Th1 Pathway | 3.71 | 0.116 | -2.309 | CD274,CD4,HLA-B,HLA-DMB,HLA-DPB1,HLA-DRB5,IL10RB,IRF1,NOTCH2,PIK3C2B,PIK3R3,SOCS1,SOCS3,STAT1 |
| GM-CSF Signaling | 1.31 | 0.0857 | -2.449 | CCND1,CSF2RB,HCK,PIK3C2B,PIK3R3,STAT1 |
| NF-kB Activation by Viruses | 1.45 | 0.0854 | -2.646 | CD4,EIF2AK2,ITGB3,NFKBIE,PIK3C2B,PIK3R3,PRKCD |
| Role of Pattern Recognition Receptors in Recognition of Bacteria and Viruses | 4.66 | 0.117 | -3.162 | C3AR1,C5,CASP1,CXCL8,DDX58,EIF2AK2,IFIH1,IRF7,OAS1,OAS2,OAS3,PIK3C2B,PIK3R3,PRKCD,RNASEL,TICAM1,TNFSF10,TNFSF13B |
| Interferon Signaling | 10.7 | 0.389 | -3.742 | IFI35,IFI6,IFIT1,IFIT3,IFITM1,IFITM3,IRF1,ISG15,MX1,OAS1,SOCS1,STAT1,STAT2,TAP1 |

**T3 vs. T1 MEP_2**

| Ingenuity Canonical Pathways | -log(p-value) | Ratio | z-score | Molecules |
|---|---|---|---|---|
| PD-1, PD-L1 cancer immunotherapy pathway | 8.11 | 0.151 | 3.742 | B2M,CD274,HLA-A,HLA-B,HLA-C,HLA-DMB,HLA-DOA,HLA-DPA1,HLA-DPB1,HLA-DQA1,HLA-DQB1,HLA-DRA,HLA-DRB5,HLA-F,HLA-G,PIK3CG |
| IL-9 Signaling | 2.91 | 0.152 | -2 | CISH,NFKB2,PIK3CG,SOCS3,STAT1 |
| PEDF Signaling | 1.79 | 0.0732 | -2 | NFKB2,NFKBIA,NFKBIE,PIK3CG,SOD2,TCF7L2 |
| Role of Wnt/GSK-3Beta Signaling in the Pathogenesis of Influenza | 1.35 | 0.0641 | -2 | CTNNB1,FZD2,FZD7,TCF7L2,WNT5B |
| Factors Promoting Cardiogenesis in Vertebrates | 1.9 | 0.06 | -2.121 | ACVR1,CTNNB1,DKK1,FZD2,FZD7,LRP6,MYC,TCF7L2,WNT5B |
| Role of RIG1-like Receptors in Antiviral Innate Immunity | 4.02 | 0.159 | -2.236 | DDX58,IFIH1,IRF7,NFKB2,NFKBIA,NFKBIE,TRIM25 |
| Toll-like Receptor Signaling | 3.23 | 0.105 | -2.236 | FOS,LY96,MAP2K6,NFKB2,NFKBIA,PPARA,TICAM1,TLR2 |
| Basal Cell Carcinoma Signaling | 1.48 | 0.0694 | -2.236 | CTNNB1,FZD2,FZD7,TCF7L2,WNT5B |
| Systemic Lupus Erythematosus In B Cell Signaling Pathway | 3.72 | 0.0655 | -2.357 | CTNNB1,CXCL8,FCGR2C,FOS,IFIH1,IFIT3,IRF7,ISG15,ISG20,MYC,NFKB2,PAG1,PIK3CG,STAT1,STAT2,TICAM1,TNFSF10,TNFSF13B |
| Retinoic acid Mediated Apoptosis Signaling | 5.7 | 0.167 | -2.53 | IRF1,PARP12,PARP14,PARP3,PARP9,RARG,TIPARP,TNFRSF10A,TNFRSF10B,TNFSF10 |
| Calcium-induced T Lymphocyte Apoptosis | 5.31 | 0.152 | -2.53 | CD4,HLA-A,HLA-B,HLA-DMB,HLA-DOA,HLA-DQA1,HLA-DQB1,HLA-DRA,HLA-DRB5,NR4A1 |
| IL-15 Production | 1.51 | 0.0579 | -2.646 | EPHB4,IRF1,MAP2K6,NFKB2,STAT1,TEK,TIE1 |
| Production of Nitric Oxide and Reactive Oxygen Species in Macrophages | 2.13 | 0.0585 | -2.714 | APOE,FOS,IRF1,NCF1,NFKB2,NFKBIA,NFKBIE,PIK3CG,PPARA,STAT1,TLR2 |
| Type I Diabetes Mellitus Signaling | 10.5 | 0.171 | -2.828 | HLA-A,HLA-B,HLA-C,HLA-DMB,HLA-DOA,HLA-DQA1,HLA-DQB1,HLA-DRA,HLA-DRB5,HLA-F,HLA-G,IRF1,MAP2K6,NFKB2,NFKBIA,NFKBIE,SOCS1,SOCS3,STAT1 |
| Role of Pattern Recognition Receptors in Recognition of Bacteria and Viruses | 4.51 | 0.0909 | -2.828 | CASP1,CXCL8,DDX58,IFIH1,IRF7,NFKB2,OAS1,OAS2,OAS3,PIK3CG,TICAM1,TLR2,TNFSF10,TNFSF13B |
| Systemic Lupus Erythematosus In T Cell Signaling Pathway | 3.15 | 0.0571 | -2.982 | B2M,CASP1,FOS,GNAI1,HLA-A,HLA-B,HLA-C,HLA-DMB,HLA-DOA,HLA-DPA1,HLA-DPB1,HLA-DQA1,HLA-DQB1,HLA-DRA,HLA-DRB5,HLA-F,HLA-G,MAP2K6,PIK3CG |
| Th1 Pathway | 8.94 | 0.149 | -3 | CD274,CD4,HLA-A,HLA-B,HLA-DMB,HLA-DOA,HLA-DPA1,HLA-DPB1,HLA-DQA1,HLA-DQB1,HLA-DRA,HLA-DRB5,IRF1,MAP2K6,PIK3CG,SOCS1,SOCS3,STAT1 |
| PKCteta Signaling in T Lymphocytes | 5.79 | 0.103 | -3.207 | CACNA1C,CACNA2D3,CD4,FOS,HLA-A,HLA-B,HLA-DMB,HLA-DOA,HLA-DQA1,HLA-DQB1,HLA-DRA,HLA-DRB5,NFKB2,NFKBIA,NFKBIE,PIK3CG |
| iCOS-iCOSL Signaling in T Helper Cells | 5.42 | 0.117 | -3.317 | CD4,HLA-A,HLA-B,HLA-DMB,HLA-DOA,HLA-DQA1,HLA-DQB1,HLA-DRA,HLA-DRB5,NFKB2,NFKBIA,NFKBIE,PIK3CG |
| Neuroinflammation Signaling Pathway | 7.43 | 0.0867 | -3.545 | ACVR1,B2M,CASP1,CTNNB1,CXCL8,FOS,HLA-A,HLA-B,HLA-C,HLA-DMB,HLA-DOA,HLA-DQA1,HLA-DQB1,HLA-DRA,HLA-DRB5,HLA-F,IRF7,NCF1,NFKB2,PIK3CG,PTGS2,SOD2,STAT1,TICAM1,TLR2,TYROBP |
| Interferon Signaling | 13.2 | 0.389 | -3.742 | IFI35,IFI6,IFIT1,IFIT3,IFITM1,IFITM3,IRF1,ISG15,MX1,OAS1,SOCS1,STAT1,STAT2,TAP1 |
| Dendritic Cell Maturation | 8.22 | 0.115 | -4.025 | B2M,FCGR2C,HLA-A,HLA-B,HLA-C,HLA-DMB,HLA-DOA,HLA-DQA1,HLA-DQB1,HLA-DRA,HLA-DRB5,HLA-F,NFKB2,NFKBIA,NFKBIE,PIK3CG,RELB,STAT1,STAT2,TLR2,TYROBP |

**T2 vs. T1 HSC_MPP**

| Ingenuity Canonical Pathways | -log(p-value) | Ratio | z-score | Molecules |
|---|---|---|---|---|
| EIF2 Signaling | 1.81 | 0.0357 | 2.646 | ATF3,DDIT3,MYCN,RPL10,RPL3,RPL36A,RPL7A,RPSA |
| PD-1, PD-L1 cancer immunotherapy pathway | 1.74 | 0.0472 | 2.236 | HLA-C,HLA-DRB5,HLA-F,HLA-G,SMAD3 |
| Cyclins and Cell Cycle Regulation | 1.54 | 0.0494 | 2 | CCND3,CDKN2C,PPP2R2B,PPP2R5B |
| Type I Diabetes Mellitus Signaling | 3.71 | 0.0721 | -2 | HLA-C,HLA-DRB5,HLA-F,HLA-G,IRF1,NFKB2,NFKBIE,STAT1 |
| Ceramide Signaling | 1.43 | 0.0455 | -2 | NFKB2,PPP2R2B,PPP2R5B,S1PR2 |
| Role of PKR in Interferon Induction and Antiviral Response | 3.55 | 0.0684 | -2.121 | ATF3,FCGR1A,IRF1,IRF9,NFKB2,NFKBIE,STAT1,STAT2 |
| Dendritic Cell Maturation | 6.39 | 0.0765 | -2.887 | CD83,COL2A1,CREB5,FCGR1A,FCGR1B,HLA-C,HLA-DRB5,HLA-F,IRF8,LTB,NFKB2,NFKBIE,STAT1,STAT2 |
| Interferon Signaling | 10.3 | 0.278 | -3 | IFI6,IFIT3,IFITM1,IRF1,IRF9,ISG15,MX1,OAS1,STAT1,STAT2 |

**T2 vs. T1 MPP_GMP**

| Ingenuity Canonical Pathways | -log(p-value) | Ratio | z-score | Molecules |
|---|---|---|---|---|
| Role of Pattern Recognition Receptors in Recognition of Bacteria and Viruses | 2.83 | 0.0455 | -2 | IFIH1,IRF7,NFKB2,NOD2,OAS1,OAS2,TNFSF13B |
| Osteoarthritis Pathway | 2.08 | 0.0332 | -2 | C1QTNF4,COL2A1,DDIT4,NFKB2,S100A8,S100A9,S1PR2 |
| TREM1 Signaling | 2.05 | 0.0533 | -2 | MPO,NFKB2,NLRC5,NOD2 |
| IL-15 Production | 1.37 | 0.0331 | -2 | CSF1R,IRF1,NFKB2,STAT1 |
| Activation of IRF by Cytosolic Pattern Recognition Receptors | 6.41 | 0.127 | -2.121 | IFIH1,IFIT2,IRF7,IRF9,ISG15,NFKB2,STAT1,STAT2 |
| Type I Diabetes Mellitus Signaling | 4.54 | 0.0721 | -2.236 | CD3E,CD3G,HLA-C,HLA-F,HLA-G,IRF1,NFKB2,STAT1 |
| T Cell Exhaustion Signaling Pathway | 3.18 | 0.0457 | -2.236 | HLA-C,HLA-F,HLA-G,IL10RA,IRF9,PPP2R5B,STAT1,STAT2 |
| Systemic Lupus Erythematosus In B Cell Signaling Pathway | 4.3 | 0.0436 | -2.309 | CD72,IFIH1,IFIT2,IFIT3,IRF7,IRF9,ISG15,LILRB3,NFKB2,STAT1,STAT2,TNFSF13B |
| Neuroinflammation Signaling Pathway | 1.77 | 0.0267 | -2.449 | CSF1R,HLA-C,HLA-F,IRF7,NCF2,NFKB2,SLC1A3,STAT1 |
| Dendritic Cell Maturation | 3.74 | 0.0492 | -2.646 | COL2A1,FCGR1B,HLA-C,HLA-F,IRF8,NFKB2,RELB,STAT1,STAT2 |
| Role of PKR in Interferon Induction and Antiviral Response | 3.54 | 0.0598 | -2.646 | ATF3,IFIH1,IRF1,IRF9,NFKB2,STAT1,STAT2 |
| Interferon Signaling | 16.4 | 0.361 | -3.464 | IFI6,IFIT1,IFIT3,IFITM1,IFITM3,IRF1,IRF9,ISG15,MX1,OAS1,STAT1,STAT2,TAP1 |

**T2 vs. T1 MEP_1**

| Ingenuity Canonical Pathways | -log(p-value) | Ratio | z-score | Molecules |
|---|---|---|---|---|
| PD-1, PD-L1 cancer immunotherapy pathway | 1.94 | 0.0566 | 2.449 | CD274,HLA-B,HLA-C,HLA-DRB5,HLA-F,HLA-G |
| CD40 Signaling | 2.99 | 0.0923 | 2.236 | CD40,CD40LG,NFKBIA,NFKBIE,PTGS2,TNFAIP3 |
| Xenobiotic Metabolism PXR Signaling Pathway | 1.67 | 0.0417 | 2.121 | CITED2,GSTM2,GSTM3,GSTM5,PRKAR2B,PRKCD,PRKD1,SMOX |
| Glioma Invasiveness Signaling | 1.37 | 0.0541 | -2 | PLAU,PLAUR,PTK2,RHOU |
| Dendritic Cell Maturation | 4.61 | 0.071 | -2.111 | CD40,CD40LG,CD83,FCER1G,HLA-B,HLA-C,HLA-DRB5,HLA-F,NFKBIA,NFKBIE,RELB,STAT1,TYROBP |
| Systemic Lupus Erythematosus In T Cell Signaling Pathway | 1.44 | 0.033 | -2.111 | CASP3,CD40LG,FCER1G,GNAI1,HLA-B,HLA-C,HLA-DRB5,HLA-F,HLA-G,PTK2,RHOU |
| Interferon Signaling | 9.34 | 0.278 | -3 | IFI6,IFIT1,IFITM1,IFITM3,IRF1,IRF9,ISG15,MX1,OAS1,STAT1 |

**T2 vs. T1 MEP_2**

| Ingenuity Canonical Pathways | -log(p-value) | Ratio | z-score | Molecules |
|---|---|---|---|---|
| PD-1, PD-L1 cancer immunotherapy pathway | 5.02E+00 | 8.49E-02 | 3 | B2M,HLA-A,HLA-B,HLA-C,HLA-DMB,HLA-DRB5,HLA-F,HLA-G,PIK3CG |
| LXR/RXR Activation | 1.69E+00 | 4.13E-02 | 2.236 | ABCG1,CD36,LY96,NFKB2,S100A8 |
| EIF2 Signaling | 2.05E+00 | 3.57E-02 | 2.121 | ATF3,DDIT3,PIK3CG,RPL10,RPL3,RPL36A,RPL7,RPL7A |
| Type I Diabetes Mellitus Signaling | 6.69E+00 | 9.91E-02 | -2 | HLA-A,HLA-B,HLA-C,HLA-DMB,HLA-DRB5,HLA-F,HLA-G,IRF1,NFKB2,NFKBIA,STAT1 |
| Crosstalk between Dendritic Cells and Natural Killer Cells | 6.64E+00 | 1.12E-01 | -2 | CD83,HLA-A,HLA-B,HLA-C,HLA-DRB5,HLA-F,HLA-G,IL15RA,NFKB2,TYROBP |
| iNOS Signaling | 3.56E+00 | 1.11E-01 | -2 | IRF1,LY96,NFKB2,NFKBIA,STAT1 |
| Role of Pattern Recognition Receptors in Recognition of Bacteria and Viruses | 3.05E+00 | 5.19E-02 | -2 | IFIH1,IRF7,NFKB2,OAS1,OAS2,OAS3,PIK3CG,TNFSF13B |
| Retinoic acid Mediated Apoptosis Signaling | 2.12E+00 | 6.67E-02 | -2 | IRF1,PARP12,PARP9,TNFRSF10B |
| Calcium-induced T Lymphocyte Apoptosis | 1.98E+00 | 6.06E-02 | -2 | HLA-A,HLA-B,HLA-DMB,HLA-DRB5 |
| Pancreatic Adenocarcinoma Signaling | 1.87E+00 | 4.59E-02 | -2 | CDKN2A,E2F1,NFKB2,PIK3CG,STAT1 |
| Tec Kinase Signaling | 2.87E+00 | 4.88E-02 | -2.121 | MS4A2,NFKB2,PIK3CG,STAT1,STAT2,TNFRSF10B,TNFRSF21,YES1 |
| TREM1 Signaling | 3.38E+00 | 8.00E-02 | -2.236 | CD83,CIITA,CXCL2,NFKB2,NLRC5,TYROBP |
| IL-8 Signaling | 1.81E+00 | 3.50E-02 | -2.236 | ANGPT2,CDH1,CXCL1,ITGAM,NCF1,PIK3CG,TEK |
| IL-15 Production | 1.69E+00 | 4.13E-02 | -2.236 | IRF1,NFKB2,STAT1,TEK,YES1 |
| iCOS-iCOSL Signaling in T Helper Cells | 3.22E+00 | 6.31E-02 | -2.449 | HLA-A,HLA-B,HLA-DMB,HLA-DRB5,NFKB2,NFKBIA,PIK3CG |
| Necroptosis Signaling Pathway | 1.79E+00 | 3.82E-02 | -2.449 | BIRC3,CYLD,IRF9,STAT1,STAT2,TNFRSF10B |
| PKCteta Signaling in T Lymphocytes | 3.03E+00 | 5.16E-02 | -2.646 | CACNA2D3,HLA-A,HLA-B,HLA-DMB,HLA-DRB5,NFKB2,NFKBIA,PIK3CG |
| Production of Nitric Oxide and Reactive Oxygen Species in Macrophages | 1.95E+00 | 3.72E-02 | -2.646 | IRF1,NCF1,NFKB2,NFKBIA,PIK3CG,S100A8,STAT1 |
| Neuroinflammation Signaling Pathway | 5.04E+00 | 5.00E-02 | -2.887 | B2M,BIRC3,HLA-A,HLA-B,HLA-C,HLA-DMB,HLA-DRB5,HLA-F,IRF7,NCF1,NFKB2,PIK3CG,SOD2,STAT1,TYROBP |
| Systemic Lupus Erythematosus In T Cell Signaling Pathway | 1.93E+00 | 3.00E-02 | -3.162 | B2M,HLA-A,HLA-B,HLA-C,HLA-DMB,HLA-DRB5,HLA-F,HLA-G,PIK3CG,YES1 |
| Interferon Signaling | 1.54E+01 | 3.61E-01 | -3.464 | IFI6,IFIT1,IFIT3,IFITM1,IFITM3,IRF1,IRF9,ISG15,MX1,OAS1,STAT1,STAT2,TAP1 |
| Systemic Lupus Erythematosus In B Cell Signaling Pathway | 3.59E+00 | 4.36E-02 | -3.464 | IFIH1,IFIT3,IRF7,IRF9,ISG15,NFKB2,PIK3CG,PIM2,STAT1,STAT2,TNFSF13B,YES1 |
| Dendritic Cell Maturation | 7.77E+00 | 8.20E-02 | -3.742 | B2M,CD83,HLA-A,HLA-B,HLA-C,HLA-DMB,HLA-DRB5,HLA-F,NFKB2,NFKBIA,PIK3CG,RELB,STAT1,STAT2,TYROBP |
