## Supplementary Data File 5 for "Mutated clones driving leukemic transformation are already detectable at the single cell level in CD34-positive cells in the chronic phase of primary myelofibrosis"

**T3 vs. T1 HSC_MPP**

© 2000-2020 QIAGEN. All rights reserved.

| Upstream Regulator | Expr Fold Change | Molecule Type | Predicted Activation State | Activation z-score | Flags | p-value of overlap | Target Molecules in Dataset |
|---|---|---|---|---|---|---|---|
| MAPK1 | -1.39 | kinase | Activated | 6.042 | | 2.41E-26 | BST2,BTN3A3,CAV1,CCL3L3,CCL4,CD69,CDH1,CDKN1A,CXCL8,DUSP1,DUSP2,EGR1,EGR2,EGR4,EIF2AK2,ENDOD1,FOS,FOSB,GBP1,GBP5,HERC5,HLA-C,IFI44,IFI6,IFIH1,IFIT1,IFIT2,IFIT3,IFITM1,IFITM3,IRF7,ISG15,ISG20,JUN,JUNB,JUND,LAG3,LGALS3BP,LMNA,MCL1,MMP2,MYC,NR4A1,OAS1,OAS2,OAS3,OASL,PLSCR1,PTGS2,RGS2,SAMHD1,SERPINE1,SLC27A2,SP110,SPRY2,STAT1,STAT2,TNF,TNFSF10,TP53,TTC27,UBE2L6,USP18,VDR |
| PDGF BB | | complex | Activated | 4.83 | bias | 1.69E-17 | ACTA2,AREG,ARF5,BHLHE40,CAV2,CDKN1A,CEBPD,CP,CXCL8,DUSP1,DUSP6,EGR1,EGR2,EGR3,EREG,FBLN5,FOS,FOSB,GLUL,HES1,HOMER2,ID3,IER2,IER3,JUN,JUNB,KLF6,LMNA,MCL1,MMP2,MYC,NEXN,NFIL3,NR4A1,NR4A2,NR4A3,PHLDA1,PTGS2,RGS1,RGS2,RHOB,S1PR2,SERPINE1,SGK1,SOCS3,TNFAIP3,TP53,ZFP36 |
| EGF | | growth factor | Activated | 4.651 | bias | 1.29E-11 | AREG,B4GALT1,BTG2,CAV1,CDH1,CDKN1A,CEBPD,CLCN3,CLDN7,CLU,CXCL8,CXCR4,DUSP1,DUSP6,EGR1,EGR2,EGR3,EREG,FOS,FOSB,HES1,ID1,ID2,IDS,IER2,IER3,JUN,JUNB,JUND,MCL1,MMP2,MYC,NEXN,NR4A1,NR4A2,NR4A3,NR5A1,PER1,PLSCR1,PTGS2,REN,RHOB,S100A10,SERPINE1,SOCS3,SPRY2,STAT1,TIMP3,TNFAIP3,TP53,VDR,ZFP36 |
| TRIM24 | -1.52 | transcription regulator | Activated | 4.463 | bias | 2.62E-14 | CCL4,CMPK2,DDX60,DHX58,EPSTI1,GBP4,GLUL,IFI44,IFIH1,IFIT2,IFIT3,IRF7,ISG15,LGALS3BP,OAS1,OASL,SAMD9L,SAMHD1,SERPINE1,SOCS1,STAT1,STAT2,TLR2,TP53,USP18 |
| PNPT1 | -1.67 | enzyme | Activated | 4.325 | bias | 5.19E-17 | CDKN1A,CMPK2,CXCL8,EIF2AK2,GBP4,IFI44,IFIH1,IFIT3,IRF7,ISG15,LGALS3BP,MYC,OAS1,PARP9,SAMD9L,STAT1,STAT2,UBE2L6,USP18,XAF1 |
| leukotriene D4 | | chemical - endogenous mammalia | Activated | 4.016 | bias | 6.81E-20 | ARID5B,CCL3,CCL3L3,CCL4,CDH1,CXCL8,CYSLTR1,DUSP1,EGR1,EGR2,EGR3,KLF2,KLF4,MAP3K8,MCL1,NR4A1,NR4A2,NR4A3,PTGS2,RGS2,SIK1/SIK1B,TNF,ZFP36 |
| IGF1 | | growth factor | Activated | 3.909 | | 2.5E-12 | ACTA2,AKR1C1/AKR1C2,BHLHE40,BTG2,CCL3L3,CCL4,CDH1,CDKN1A,CEBPD,CITED2,CLU,COL2A1,CXCL8,DDIT4,DUSP1,DUSP6,EGR1,EGR2,EIF2AK2,ELN,FOS,FOSB,ID1,ID2,IER2,IER3,IFITM3,JUN,JUNB,KLF6,LARP6,MCL1,MMP2,MYC,NFE2,NOG,NR4A1,NR5A1,PHLDA1,PPP1R1B,PTGS2,SERPINE1,SGK1,SOCS3,TAP2,TNF,TP53,TXNIP,UGCG,ZFP36 |
| CREB1 | -1.53 | transcription regulator | Activated | 3.877 | | 3.24E-10 | AK4,ATF1,BHLHE40,BTG2,CCL3,CCL4,CD4,CDKN1A,CEBPD,CXCL8,CXCR4,DUSP1,EGR1,EGR2,EGR4,ENTPD1,ETV3,FGL2,FOS,FOSB,GPR12,HLA-G,ID1,IER2,IER3,IRF7,JUN,JUNB,KLF4,KLF7,MCAM,MCL1,MYC,NFIL3,NFKBID,NR4A1,NR4A2,NR4A3,PER1,PNMT,PTGS2,RASGEF1B,REN,RGS2,SERTAD1,SGK1,SHTN1,SLC16A1,TIPARP,TNF,ZFP36 |
| Pkc(s) | | group | Activated | 3.782 | bias | 0.000000135 | ACTA2,CD4,CDKN1A,CXCL8,CXCR4,DUSP1,EGR1,EGR2,FOS,ID2,JUN,JUNB,KLF6,MMP2,NOTCH4,NR4A1,NR4A2,NR4A3,PER1,PHLDA1,PTGS2,RGS2,SERPINE1,TNF,VDR |
| MAP3K8 | 2.08 | kinase | Activated | 3.744 | | 0.0000143 | ARRDC3,CCL4,CEBPD,CXCL8,DUSP6,FOS,IER3,IER5,JUN,JUNB,KDM4A,PTGS2,SESN1,SOCS3,STK17B,TNF,TXNIP |
| FGF2 | 1 | growth factor | Activated | 3.495 | bias | 4.79E-08 | ACTA2,AGAP3,AREG,BTG2,CAV1,CDH1,CDKN1A,CXCR4,DUSP6,EGR1,ELN,EREG,FOS,GBP1,HES1,ID2,ID3,JUN,JUNB,LMNA,LRP6,MCAM,MMP2,MYC,NOG,NOTCH4,NR4A1,NR4A2,PRKCD,PTGS2,S100A10,SERPINE1,SOCS3,SPRY2,TIMP3,TNF,TNFSF10,TP53 |
| GPER1 | | G-protein coupled receptor | Activated | 3.458 | bias | 5.14E-11 | ACTA2,BHLHE40,CDKN1A,CEBPD,DDIT4,DUSP1,EGR1,FOS,FOSB,IER2,JUN,MYC,NR4A2,TNF,TNFAIP3,ZFP36 |
| IRF8 | -10.9 | transcription regulator | Activated | 3.258 | | 6.13E-11 | ARID5B,CD4,CDKN1A,EGR1,EGR2,ETV3,FOS,GBP1,ID3,IFI44L,IFI6,IFIT2,IFIT3,IRF8,ISG15,KLF4,KLF6,MYC,OAS1,STAT1,STAT2,TNF,TNFSF13B,TP53 |
| EGR1 | 8.45 | transcription regulator | Activated | 3.257 | bias | 7.91E-12 | ACTA2,AREG,CAV1,CCL3L3,CCL4,CDH1,CDKN1A,CLU,COL2A1,CXCL8,DLG4,EGR1,EGR2,EREG,IL6R,JUN,JUNB,JUND,MYC,NR4A1,PNMT,PTGS2,SERPINE1,SGK1,SLC12A2,SOCS1,TNF,TNFSF10,TP53 |
| INS | | other | Activated | 3.255 | | 9.79E-08 | ACADL,CDH1,CDKN1A,CITED2,DUSP1,EGR1,EGR2,FOS,FOSB,IER2,IER3,IGKC,JUN,JUNB,JUND,NR4A1,PTGS2,SELP,TNF,TP53,UGCG |
| MAPK7 | 1.21 | kinase | Activated | 3.252 | bias | 0.000000543 | CXCL8,DUSP1,ID1,JUN,KLF2,KLF4,MCL1,NR4A1,PTGS2,TNF,TP53 |
| FOXO3 | 1.05 | transcription regulator | Activated | 3.249 | | 5.61E-08 | ACTA2,BTG1,CAV1,CDH1,CDKN1A,CDKN3,CXCL8,CXCR4,DDIT4,EGR1,EGR2,EGR4,FOS,FOSB,GLUL,IER3,IFIH1,JUNB,MCAM,MYC,PIK3IP1,POMC,PRKCD,SERPINE1,SESN1,SGK1,SLC40A1,TIE1,TNF,TNFSF10,TP53,TP53INP1,TXNIP |
| SIRT1 | -1.13 | transcription regulator | Activated | 3.206 | | 5.64E-08 | CAV1,CD72,CDH1,CDKN1A,CDKN2C,CMPK2,DDX60,DHX58,EPDR1,GSTM3,HCK,HLA-DRB5,ID1,IFI44,IFIT3,IFITM3,IRF7,JCHAIN,KLF2,LGALS3BP,MCL1,MMP2,MYC,NLGN1,NLRC5,OAS1,OAS2,PER1,RSAD2,SERPINE1,SOCS3,SP110,STAT1,TIMP3,TNF,TP53,USP18 |
| ERK | | group | Activated | 3.195 | bias | 6.19E-13 | ACTA2,ADGRG6,AREG,BTG2,CAV1,CCL4,CDH1,CDKN1A,CEBPD,CLU,CXCL8,DUSP1,EGR1,EGR2,EIF2AK2,EREG,FOS,IER3,JUN,JUNB,KYNU,MAFF,MAL,MCAM,MCL1,MMP2,MYC,OSBPL10,PTGS2,SERPINE1,STAT1,TNF,TP53,UGCG,ZFP36 |
| GLI3 | | transcription regulator | Activated | 3.162 | | 0.00013 | CD69,DUSP1,DUSP2,EGR1,FOSB,JUNB,KLF13,KLF2,NFKBID,TNFAIP3 |
| TREM1 | 1.32 | transmembrane receptor | Activated | 3.151 | bias | 0.00000102 | AREG,CCL3,CXCL8,CXCR4,DTL,EGR1,EGR2,EGR3,HES1,IFIT2,IL6R,ISG15,MAFF,MAP3K8,NR4A2,OASL,PHLDA1,PTGS2,RGS1,SPRY2,TLR2,TNF |
| CG | | complex | Activated | 3.128 | bias | 1.08E-12 | ACTA2,AKR1C3,AREG,BHLHE40,BTG1,BTG2,CCDC86,CDH1,CDKN1A,CEBPD,CLU,CXCL8,CXCR4,DTL,DUSP1,DUSP6,EGR1,EREG,FGL2,FOS,GASK1B,IER3,IFIT3,JUN,JUNB,KLF4,LGALS3BP,MCAM,MCL1,MMP2,NR4A1,PHLDA1,PRKG2,PRSS2,PTGS2,PTX3,RGS2,S100A10,SPRY2,TM4SF1,TNF,TP53,XKR8 |
| dopamine | | chemical - endogenous mammalia | Activated | 3.119 | | 0.00000036 | BTG2,CCDC146,CLCN3,CXCL8,DUSP2,EGR1,EGR3,EGR4,FOS,FOSB,GPC3,GPR12,JUN,KLF4,NR4A1,NR4A3,PER1,POMC,PPP1R1B |
| levodopa | | chemical - endogenous mammalia | Activated | 3.117 | | 0.000226 | ACTA2,AIFM2,ARAP1,ARMH4,BANF1,C1GALT1C1,CLDN5,DCAF4,DGAT2,DIXDC1,DUSP1,DUSP6,EGR2,EGR4,FAM81A,FBLN1,FGFR1OP,FKBP5,FOS,FOSB,GPC3,IER3,JUNB,KLF6,KLF9,LAG3,LMNA,MSRB1,PER1,PIEZO2,PLD4,PPP1R1B,PRKG2,RIMS1,S100A10,S1PR2,SLC29A3,SLC66A1,SMIM14,TCF4,TP53 |
| MAPK3 | -1.36 | kinase | Activated | 3.116 | bias | 7.02E-11 | B4GALT1,CCL3,CD69,CDKN1A,CXCL8,EGR1,EGR2,EGR4,FOS,FOSB,JUN,JUNB,JUND,MCL1,MYC,NR4A1,PTGS2,RGS2,TNF |
| STAT3 | 1.33 | transcription regulator | Activated | 3.108 | | 4.09E-24 | ACTA2,AREG,BATF,BST2,CCL3L3,CCL4,CDH1,CDKN1A,CEBPD,CMPK2,CXCL8,CXCR4,EGR1,EGR2,EGR3,EIF2AK2,FCGR1A,FGL2,FOS,GBP5,H2AC18/H2AC19,HERC5,HLA-DRB5,ID1,ID2,IFI44,IFI6,IFIH1,IFIT1,IFIT2,IFIT3,IFITM1,IFITM3,IKZF3,IL6R,IL7,IRF7,ISG15,ISG20,JUNB,KLF4,MCL1,MMP2,MMP7,MT- |
| NUPR1 | | transcription regulator | Activated | 3.084 | | 0.0000039 | AREG,ATP6V0A1,BTG1,CITED2,CXCL8,CXCR4,EGLN1,EREG,FUCA1,HILPDA,HLTF,IFIT2,IL6R,KLF4,KLF6,LMAN2L,LMNB1,MYC,NFIB,NFIL3,NUP50,OSBPL6,PARP9,PCTP,PHLDA1,PLK3,RAB38,RASAL2,RMI1,SAMHD1,SERPINE1,SIK1/SIK1B,TANGO6,TP53,UBIAD1,VANGL1,WDCP,ZDHHC13 |
| FOXO1 | 1.53 | transcription regulator | Activated | 3.081 | | 4.59E-08 | AIFM2,ATP6V0A1,CAV1,CCL4,CDKN1A,CDKN2C,CENPF,CITED2,CLDN5,COL2A1,CXCL8,DDIT4,EGR1,EGR2,EGR4,FOS,FOSB,IER3,JUN,JUNB,KLF2,KLF4,KLF7,MRPS18C,MYC,NKX3-1,PIK3IP1,PLSCR1,POMC,PRKCD,SERPINE1,SESN1,SGK1,SPRY2,STAT2,TIE1,TNF,TNFSF10,TP53,TXNIP |
| Insulin | | group | Activated | 3.081 | | 0.0000551 | CAV2,CCL4,CDKN1A,DDIT4,DGAT2,DUSP1,EGR1,EGR2,FOS,FOSB,HES1,HIF3A,IFIT2,IFIT3,IRF7,JUN,KLF4,LGALS3BP,MMP2,MT-ND1,MYC,NFIL3,NR0B1,NR4A1,NR4A2,NR4A3,POMC,PRKCD,SERPINE1,SGK1,STAT1,STAT2,TLR2,TNF |
| ELK1 | -1.04 | transcription regulator | Activated | 3.057 | bias | 9.77E-08 | CDKN1A,EGR1,EGR2,EGR4,FOS,FOSB,JUN,JUNB,MCL1,PTGS2,TIPARP,TNF,ZFP36 |
| ERK1/2 | | group | Activated | 3.046 | bias | 1.78E-08 | AKR1C3,AREG,CAV1,CCL3,CCL3L3,CCL4,CDKN1A,COL2A1,CXCL8,CXCR4,DUSP1,EGR1,FKBP5,FOS,FOSB,ID1,ID3,IFIT1,JUN,JUNB,JUND,KLF13,MMP2,MYC,NR4A3,POMC,PTGS2,SERPINE1,SGK1,SOCS3,TAP2,TNF |
| IL18 | 1.03 | cytokine | Activated | 3.044 | bias | 0.0036 | CCL3,CCL3L3,CCL4,CD69,CDH1,CXCL8,DDIT4,DLG4,JUN,NCR3,NFIL3,PTGS2,SAMHD1,TNF |
| MAP2K1/2 | | group | Activated | 3.028 | bias | 0.000000359 | CCL3L3,CCL4,CDH1,DUSP1,EGR1,EGR2,ELN,FOS,HSPB1,JUN,MCL1,MYC,PHLDA1,TNF |
| Growth hormone | | group | Activated | 3.026 | | 0.000374 | ARID5B,CDH1,CEBPD,CITED2,CLU,EGR1,FKBP5,FOS,FZD7,ID1,IER3,JUN,JUNB,MYC,SGK1,SOCS1,SOCS3,TNFSF10,TXNIP |
| ACKR2 | | G-protein coupled receptor | Activated | 2.982 | bias | 3.25E-17 | CCL3L3,DDX60,DHX58,EIF2AK2,IFI44,IFIT2,IFIT3,IRF7,ISG15,ISG20,OAS1,OAS2,OAS3,OASL,RSAD2,STAT1,STAT2,TNF,USP18 |
| GNRH1 | | other | Activated | 2.956 | | 0.000114 | EGR1,EGR2,FOS,FOSB,JUN,JUNB,NR5A1,PRKCD,PTGS2 |
| FOXO4 | -1.26 | transcription regulator | Activated | 2.941 | | 0.0122 | CAV1,CDKN1A,IER3,PRKCD,SERPINE1,SESN1,SGK1,TIE1,TXNIP |
| MEF2D | -1.18 | transcription regulator | Activated | 2.926 | bias | 0.000146 | CDH1,EGR2,FOS,FOSB,JUN,JUNB,JUND,KLF2,NR4A1,ZFP36 |
| AREG | 21.8 | growth factor | Activated | 2.92 | | 3.62E-08 | AREG,CENPF,CXCR4,EGR1,EREG,FOS,GBP1,H2AC18/H2AC19,IFI6,IFIT2,IFIT3,ITGB8,JUN,PTAFR,PTGS2,PTX3 |
| RAF1 | -1.11 | kinase | Activated | 2.92 | bias | 0.0000047 | AREG,BTG1,CDKN1A,DUSP2,DUSP6,EGR1,EGR3,FOS,HSPB1,ID3,IER2,IER3,JUN,MCF2L,MYC,PHLDA1,PTGS2,RGS1,SPRY2,TNF,TUBA4A,TXNIP |
| formaldehyde | | chemical - endogenous mammalia | Activated | 2.905 | bias | 0.000000154 | EGR1,EGR2,FOS,FOSB,JUN,JUNB,JUND,PTGS2,SGK1,TNF |
| NR3C2 | | ligand-dependent nuclear receptor | Activated | 2.874 | | 6.81E-09 | CCL3L3,CXCL8,CXCR4,CYP1B1,DDIT4,EGR1,FKBP5,JUNB,KLF9,MSRB1,NR4A2,PPAT,PTGS2,PTX3,RGS2,SERPINE1,SGK1,TNF,TP53,TSC22D3,TUBA4A |
| GAPDH | -1.62 | enzyme | Activated | 2.868 | | 2.1E-13 | C2,CCL3,CCL4,DUSP1,HLA-C,IFI6,IFIT2,IFITM1,OAS1,OAS2,OAS3,STAT1,TXNIP,UBE2L6,WARS1 |
| IKZF1 | 1.8 | transcription regulator | Activated | 2.854 | bias | 0.000000469 | CAV1,CAV2,CCL3L3,CCL4,CDKN1A,CITED2,CYP7B1,DOCK4,DTX3L,EPSTI1,FKBP9,GP1BA,HES1,IFI6,IFIT3,KLF4,LIG4,MYC,NFE2,PGAM1,POMC,SELENOM,SPN,TNF,TXNIP,WDFY3 |
| SMAD3 | 1.53 | transcription regulator | Activated | 2.831 | bias | 1.43E-08 | ACTA2,AREG,ARID3A,CCL3L3,CCL4,CDH1,CDKN1A,COL2A1,DOCK4,EGR1,ELN,EREG,FOS,ID1,ID2,ID3,JUN,JUNB,JUND,MYC,NR5A1,PNMT,PTGS2,PTX3,RHOB,SERPINE1,TIMP3,TNF,ZFP36 |
| Lh | | complex | Activated | 2.823 | | 0.0000049 | ACTA2,AREG,ARL4C,CDC42EP4,CXCL8,CXCR4,DUSP1,EREG,FKBP5,FOS,GBP1,MCL1,MMP2,NR5A1,PER1,PPP2R5B,PTGS2,PTX3,RASAL2,RHOB,SGK1,STAT1,TNFAIP3,TP53 |
| TGFA | | growth factor | Activated | 2.818 | bias | 0.0000949 | AREG,CDKN1A,CXCL8,EREG,FOS,JUNB,NR4A2,PTGS2,S100A10,SERPINE1,TP53 |
| FCER1G | -1.62 | transmembrane receptor | Activated | 2.772 | | 0.00000578 | CCL3,CCL4,CD69,CXCL8,IFIT2,MX1,PTGS2,RSAD2,TNF |
| PGF | | growth factor | Activated | 2.771 | bias | 0.00000719 | CCL4,CXCL8,CXCR4,DNM3OS,EGR1,EREG,FOSB,SERPINE1,TNF |
| Mek | | group | Activated | 2.759 | | 4.81E-11 | CDH1,CDKN1A,CLU,CXCL8,DUSP6,EGR1,EGR2,EGR3,EREG,FOS,HSPB1,ID1,ID2,IER2,IER3,JUN,JUNB,MAFF,MCL1,MMP2,MYC,NR4A1,PHLDA1,PPAT,PTGS2,SERPINE1,SPRY2,TNF,TNFAIP3,TP53,XAF1 |
| fatty acid | | chemical - endogenous mammalia | Activated | 2.745 | bias | 0.0299 | ACADL,CXCL8,NR4A1,NR4A2,NR4A3,PTGS2,SERPINE1,TNF |
| APEX1 | -1.22 | enzyme | Activated | 2.72 | bias | 2.74E-09 | EGR1,EGR2,EGR3,EGR4,FOS,FOSB,JUNB,NR4A1,PTGS2,RGS2,TP53 |
| KRAS | 1.16 | enzyme | Activated | 2.69 | | 7.07E-14 | AREG,ASCL2,CAV1,CDH1,CDKN1A,CDKN2C,CLU,CSF2RB,CXCL8,DUSP1,DUSP6,EFNA1,EGR1,EGR2,EIF2AK2,ELN,EREG,FOS,FOSB,GLUL,GPC3,GSTM3,HES1,HOPX,HSPB1,ID1,IER3,IFI6,IFIT1,IFITM1,IFITM3,IRF2,ISG15,JUN,JUNB,KLF6,LMNB1,MAN2A1,MAPK12,MCAM,MMP2,MMP7,MX1,MYC,NR0B1,NR4A1,OAS1,PHLDB2,PRKCD,PTGS2,RHEBL1,RHOB,SAMHD1,SPRY2,STAT1,STAT2,TIMP3,TLE3,TLR2,TMEM98,TNF,TNFSF10,TP53,TUBA4A |
| KITLG | | growth factor | Activated | 2.677 | bias | 0.000000636 | BTG1,CCL3L3,CCL4,CDKN1A,CLEC1B,CLU,CXCL8,CXCR4,EGR1,FOS,GP1BA,HCK,ID2,IRF2,KPNA2,MMP2,MRVI1,PIK3IP1,PLSCR1,PTGS2,RPS23,SOCS1,TNF,TSC22D3,TXNIP,UGCG |
| FOXL2 | | transcription regulator | Activated | 2.671 | bias | 0.0000102 | CCL3,CCL3L3,FOS,HRH2,IER3,MAFF,NR4A3,NR5A1,PTGS2,RGS2,TNFAIP3 |
| hydrogen peroxide | | chemical - endogenous mammalia | Activated | 2.643 | bias | 2.04E-10 | ACTA2,AKR1C1/AKR1C2,BTG2,CAV1,CCL3L3,CCL4,CDH1,CDKN1A,CDKN2C,CLDN7,CLU,CXCL8,CXCR4,DDIT4,DUSP1,DUSP2,EGR1,ELN,FOS,GSTM3,ID3,IER5,IFI6,IFIT2,JUN,KLF2,KLF4,MCAM,MMP2,MYC,NFIL3,NR4A1,PRDX1,PTGS2,PTX3,RCN1,SERPINE1,SESN1,SGK1,TNF,TNFAIP3,TP53,TP53INP1,TUBA4A,TXNIP,VDR,XAF1,ZBTB7A |
| EPHB1 | | kinase | Activated | 2.621 | bias | 1.42E-08 | EGR1,EGR2,FOS,JUN,JUNB,PTGS2,SERPINE1 |
| BMP15 | | growth factor | Activated | 2.616 | | 0.0000074 | AREG,EREG,ID1,ID2,ID3,PTGS2,PTX3 |
| IL33 | | cytokine | Activated | 2.605 | bias | 0.00182 | AREG,CCL3,CCL3L3,CCL4,CD69,CD72,CXCL8,DGAT2,DUSP2,EGR2,HCK,JUNB,NFKBID,SOCS3,TLR2,TNF,TNFAIP3,TNFSF10 |
| PF4 | 1 | cytokine | Activated | 2.604 | bias | 0.000172 | ACTA2,CCL3,CCL4,CDKN1A,CXCL8,KLF4,TNF |
| 10E,12Z-octadecadienoic acid | | chemical - endogenous mammalia | Activated | 2.6 | | 0.00819 | CAV1,CDKN3,CXCL8,DGAT2,EGR2,NOTCH4,NR4A1,NR4A2,NR4A3,PTGS2,RGS1,TNF |
| TRH | 1.65 | other | Activated | 2.574 | | 0.000206 | DUSP1,FOS,JUN,JUNB,NR4A1,POMC,UBOX5 |

| | | | | | | | |
|---|---|---|---|---|---|---|---|
| bisphenol A | | chemical - endogenous mammalia | Activated | 2.551 | bias | 0.000849 | CDH1,CDKN1A,CITED2,CLU,DLG4,EGR1,EREG,FOS,NR4A1,NR4A2,NR4A3,POMC,PTGS2,STK17B,TP53 |
| RELA | -1.13 | transcription regulator | Activated | 2.544 | bias | 3.04E-12 | ACTA2,ASCL2,BHLHE40,BTG2,CAV1,CCL3,CD69,CDH1,CDKN1A,CITED2,COL2A1,CXCL8,CXCR4,DUSP1,EGLN1,EGR1,ELN,FOS,FOSB,GBP1,HES1,ID1,IER2,IER3,IGKC,IRF7,ISG15,JUN,JUNB,MMP7,MYC,NFKBID,NR4A1,NR4A2,NR4A3,OAS2,PLK3,PRKCD,PTGS2,PTX3,SELP,TAP2,TLR2,TNF,TNFAIP3,TP53,WASF3 |
| TNFRSF8 | | transmembrane receptor | Activated | 2.541 | | 0.00000539 | CCL3,CCL4,CDKN1A,CXCL8,CXCR4,GBP1,JUNB,MAP4K3,MYC,TNF,TNFSF10 |
| collagenase | | group | Activated | 2.538 | bias | 5.93E-08 | CCL3L3,CCL4,CEBPD,EGR1,IER3,JUN,JUNB,PTX3,SELP,TNF |
| GATA2 | -1.23 | transcription regulator | Activated | 2.535 | | 0.0147 | ADGRE5,CCL3L3,CD69,CDKN1A,CXCR4,F13A1,HHEX,KLF2,KLF4,MMP2,MYC,NFE2,NFIL3,NRXN2,PHLDA3,PRSS2,RGS18,SELP,SPN,TNF,TP53,ZMAT3,ZNF462 |
| Ca2+ | | chemical - endogenous mammalia | Activated | 2.528 | bias | 5.64E-10 | ACTA2,CDH1,CDKN1A,CEBPD,COL2A1,CXCL8,CXCR4,CYP1B1,DUSP1,EGR1,FOS,ID1,ID3,IL6R,JUN,JUNB,JUND,KLF4,MCAM,NR4A1,NR4A3,NR5A1,PER1,PLS3,PTGS2,SELP,SERPINE1,SGK1,TCF4,TNF,UGCG,VDR |
| miR-124-3p (and other miRNAs w/seed AAGGCA | | mature microRNA | Activated | 2.474 | bias | 0.000811 | ALDH9A1,ATP6V0E1,B4GALT1,CAV1,CYP1B1,EGR1,ENDOD1,HES1,IFRD2,KLF15,LMNB1,MAN2A1,OAF,QSER1,SLC16A1,SMCO4,SUCLG2,TLCD3A |
| TLR2 | -2.67 | transmembrane receptor | Activated | 2.456 | bias | 3.22E-08 | ACTA2,CCL3L3,CCL4,CD69,CDKN1A,CEBPD,CXCL8,DUSP1,GP1BA,HLA-DRB5,KLF2,MX1,POMC,PTGS2,PTX3,SELP,SLC40A1,SOCS1,SOCS3,TLR2,TNF,TNFSF10,TP53,VDR |
| BDNF | 1.04 | growth factor | Activated | 2.451 | | 0.000000575 | BHLHE40,CAV2,CDKN1A,CP,CXCR4,DUSP1,DUSP6,EGR1,EGR2,EGR3,FOS,FOSB,GLUL,GNAI1,HSPA4L,JUN,KIF5A,MAN2A1,NR4A1,NR4A3,POMC,PPP1R1B,RPS23,S100A10,SPRY2,TIPARP,TM9SF2,TMEM45A,TXNIP,YWHAG |
| L-glutamic acid | | chemical - endogenous mammalia | Activated | 2.446 | bias | 0.00000127 | ACTA2,CDKN1A,CLDN5,EGR1,FOS,FOSB,JUN,JUNB,MMP2,NFIL3,PER1,POMC,PTGS2,TAP2,TNF,TP53 |
| Il3 | | cytokine | Activated | 2.436 | | 0.00000143 | BHLHE40,CDKN1A,COL2A1,EGR1,FOS,MCL1,MYC,NFIL3,PLA2R1,SOCS1,TP53,YWHAG,ZNF346 |
| HBEGF | 2.1 | growth factor | Activated | 2.433 | bias | 0.000625 | BHLHE40,CDH1,CXCR4,EGR1,ELN,EREG,PTGS2 |
| leukotriene C4 | | chemical - endogenous mammalia | Activated | 2.432 | bias | 1.03E-09 | CCL4,CXCL8,EGR1,EGR2,EGR3,FOS,PTGS2,SELP,TNF |
| FCGR2A | -2.23 | transmembrane receptor | Activated | 2.426 | bias | 0.000354 | CCL3,CCL4,CXCL8,PTGS2,SERPINE1,TNF |
| NKX2-3 | -1.31 | transcription regulator | Activated | 2.419 | | 5.88E-16 | ACTA2,AGAP3,BTG1,CDKN1A,CDKN3,CMPK2,CXCL8,DDX60,DHX58,EIF2AK2,FBXO6,GASK1B,GBP1,GMPR,H2AC18/H2AC19,HCP5,HLA-C,HLA-F,KLF2,KLF4,MMP7,PARP9,PLSCR1,PTGS2,RPS23,SAMD9,SAMD9L,SLC40A1,SP110,STAT1,STAT2,TAP2,TIPARP,UBE2L6,USP18,XAF1,ZMAT3 |
| CHRM1 | | G-protein coupled receptor | Activated | 2.415 | bias | 0.00000203 | EGR1,EGR2,EGR3,EGR4,FOS,JUN |
| CD244 | -1.01 | transmembrane receptor | Activated | 2.412 | bias | 0.000000111 | CCL3,CCL3L3,CCL4,EGR1,FOS,TNF |
| STAT6 | -1.27 | transcription regulator | Activated | 2.404 | | 6.85E-14 | AREG,CCDC86,CCL3L3,CD69,CD72,CDKN1A,CKAP4,CMPK2,CTNS,DHX58,EGR1,EGR2,EIF2AK2,EPHX1,FCGR1A,FGFR1OP,FKBP5,GATM,GBP5,GBP7,HCK,IER2,IFI44,IFI44L,IFIH1,IFIT3,IFITM3,IRF7,ISG15,ISG20,LDLRAD3,LGALS3BP,MMP2,NFIL3,OAS3,OASL,PLSCR1,PTGS2,RAB30,RASGRP3,S100A10,SELP,SERPINE1,SOCS1,STAT2,TNF,USP2,ZFP36,ZNF608 |
| NCR2 | | transmembrane receptor | Activated | 2.401 | bias | 0.000019 | CCL4,NR4A1,NR4A2,NR4A3,PTGS2,TNF |
| Raf | | group | Activated | 2.397 | | 6.09E-08 | AREG,CDKN1A,DUSP6,EGR1,FOS,ID2,IER2,IER3,JUN,MAFF,MCL1,MYC,SPRY2,TNFAIP3 |
| STAT5A | -1.46 | transcription regulator | Activated | 2.397 | | 1.65E-08 | AKR1C3,ASCL2,CCL3L3,CDH1,CDKN1A,CYP1B1,FOS,FOSB,GASK1B,H2AC18/H2AC19,ID1,ID2,IL6R,MCL1,MMP7,MYC,NR4A2,OAS1,RAB27B,SLC2A10,SNTB1,SOCS1,SOCS3,TLE4,TM4SF1,TNF,TP53,TXNIP,WARS1,YWHAG,ZFP36 |
| MEF2C | -1.88 | transcription regulator | Activated | 2.392 | bias | 0.00000254 | CCL3L3,CCL4,CDKN1A,COL2A1,CXCR4,EGR2,FOS,FOSB,JUN,JUNB,JUND,KLF2,NR4A1,PTGS2,SIK1/SIK1B,TNNI2,ZFP36 |
| PI3K (complex) | | complex | Activated | 2.39 | bias | 2.55E-09 | CCL3,CCL4,CDH1,CDKN1A,CXCL8,CXCR4,DDIT4,DUSP1,FBLN5,FOS,IFIT1,IRF2,JUN,KLF2,MCAM,MCL1,MMP2,MYC,NR4A2,PPP1R1B,PTGS2,RGS2,RHOB,SAMHD1,SERPINE1,SGK1,SOCS3,TNF,TNFAIP3,TNFSF10,TP53,TXNIP,UGCG |
| STAT5a/b | | group | Activated | 2.382 | | 4.65E-08 | CD69,CDKN1A,CDKN2C,CLEC1B,CLU,DUSP1,DUSP2,FCGR1A,FOS,GP1BA,ID2,MCL1,MRVI1,MYC,PTGS2,SOCS1,SOCS3,STK17B,TLR2 |
| Jnk | | group | Activated | 2.381 | bias | 1.14E-09 | ACTA2,BTG1,CCL3,CCL4,CD69,CDH1,CRIP2,CXCL8,CXCR4,DUSP1,EGR1,FOS,FOSB,HES1,JUN,JUND,KLF6,MCL1,MMP2,MYC,NR4A1,PLSCR1,PTGS2,SERPINE1,SOCS3,TNF,TP53,ZFP36 |
| CLEC7A | 1.05 | transmembrane receptor | Activated | 2.379 | bias | 0.000518 | CCL3L3,CCL4,CXCL8,PTGS2,SOCS1,TNF |
| platelet activating factor | | chemical - endogenous mammalia | Activated | 2.379 | bias | 2.04E-09 | CD69,CDH1,CDKN1A,EGR1,EGR2,FOS,MMP2,NR4A1,PTAFR,PTGS2,SELP,SERPINE1,SOCS3,TNF |
| Tlr | | group | Activated | 2.367 | bias | 0.00000942 | CCL3,CCL4,CXCL8,DUSP1,ID2,IRF7,PTGS2,RGS1,RGS18,SOCS1,SOCS3,STAT1,TNF,TNFAIP3,TNFSF13B |
| Creb | | group | Activated | 2.352 | bias | 0.0000184 | ACTA2,AREG,CDKN1A,CEBPD,CXCL8,DUSP1,EGR1,EGR3,EGR4,FOS,FOSB,GSTM3,HES1,HLA-G,IER3,JUN,JUNB,MCAM,MCL1,MED31,MMP2,NR4A1,NR4A3,PER1,PTGS2,RGS2,TNF |
| NGLY1 | -1.14 | enzyme | Activated | 2.349 | | 0.000000154 | IFI44,IFI44L,IFIT1,IFIT2,IFIT3,KEAP1,OAS1,OAS3,RSAD2,USP18 |
| IL1RN | | cytokine | Activated | 2.345 | bias | 1.47E-12 | ACTA2,BTN3A1,CXCL8,GBP1,IFI44,IFI44L,IFI6,IFIH1,IFIT3,IRF7,ISG20,KLF6,MX1,OAS1,OAS2,OAS3,OASL,PTGS2,RSAD2,SAMD9,SERPINE1,STAT2,TNF,TNFSF10,USP18 |
| NEDD9 | -1.96 | other | Activated | 2.345 | | 0.00287 | BHLHE40,BMPR1B,CDH1,DDIT4,FOS,LIN7A,MMP2,SERPINE1,TXNIP |
| TLR7/8 | | group | Activated | 2.345 | bias | 0.00000719 | ATP2B1-AS1,EREG,MAP3K8,PTGS2,RGS1,RRP7BP,TNF,TNFAIP3,VDR |
| F3 | | transmembrane receptor | Activated | 2.333 | bias | 0.0000159 | AREG,CXCL8,DOCK4,DUSP2,DUSP6,EGR1,FOS,MMP2,MMP7,SERPINE1,TNF |
| Pka | | complex | Activated | 2.31 | bias | 2.78E-09 | AREG,CAV1,CDKN1A,DUSP1,EGR1,FOS,IFI6,JUN,JUNB,KLF6,NR4A1,NR4A2,NR4A3,PNMT,POMC,PPP1R1B,PTGS2,REN,RGS2,SGK1,TNF,XAF1 |
| NOD2 | -1.47 | other | Activated | 2.295 | bias | 0.000184 | AREG,CCL3,CXCL8,IER3,KLF2,MAP3K8,RASGRP3,SOCS3,TNF,TNFAIP3 |
| MMP1 | | peptidase | Activated | 2.29 | bias | 0.0000737 | CDKN1A,CXCL8,EGR1,FOS,JUN,MYC,PTGS2,SERPINE1,TNF |
| Ap1 | | complex | Activated | 2.29 | bias | 0.0000125 | CCL3,CCL3L3,CCL4,CLU,CXCL8,DUSP6,FOS,JUN,KLF9,MMP2,MMP7,MYC,NOG,NOTCH4,PTGS2,SERPINE1,TNF,VDR |
| TNFSF12 | -1.64 | cytokine | Activated | 2.288 | | 0.144 | CCL3,CCL3L3,CXCL8,KLF15,PTX3,TNF |
| NRAS | -1.19 | enzyme | Activated | 2.288 | | 0.00000444 | CCL4,CDKN1A,CDKN3,CFH,DUSP2,DUSP6,EGR1,EPHX1,HOMER3,IFIH1,IFIT1,ISG15,JCHAIN,MCL1,PHLDA3,PTGS2,PTX3,RHOB,SESN1,STAT1,USP18 |
| VEGFA | -1.16 | growth factor | Activated | 2.278 | bias | 0.000000013 | ACTA2,AK4,BATF,CAV1,CCL3L3,CDKN1A,CITED2,CLDN7,CXCL8,CXCR4,EGR1,FOS,FOSB,GBP1,HES1,ID1,ID3,JUNB,LAG3,LMNA,MCL1,MMP2,NOTCH4,NR4A1,NR4A2,PTGS2,RASGRP3,SERPINE1,SLC25A17,TMSB10/TMSB4X,TNF,TP53 |
| IL3 | | cytokine | Activated | 2.269 | bias | 0.000016 | ADGRE5,AREG,BTG2,CD69,CDKN1A,CITED2,CXCL8,CXCR4,EGR1,EGR2,EGR3,EIF2AK2,FOS,GP1BA,JCHAIN,JUN,JUNB,KLF13,KLF9,MCL1,MYC,SOCS1,SOCS3,TLR2,TNF,TNFSF10,YWHAG,ZBTB7A |
| MIF | | cytokine | Activated | 2.239 | bias | 0.0000709 | ACTA2,CCL3L3,CCL4,CXCL8,CXCR4,DGAT2,DUSP1,FOS,IL7,JUN,MMP2,PTGS2,TNF,TP53 |
| MAP2K7 | -1.33 | kinase | Activated | 2.236 | | 0.00296 | CDKN1A,CXCL8,JUNB,MYC,RHOB,TNF,VDR |
| hydrochloric acid | | chemical - endogenous mammalia | Activated | 2.236 | bias | 0.000289 | CCL3L3,CCL4,CXCL8,FOS,JUN,TNF |
| acetic acid | | chemical - endogenous mammalia | Activated | 2.234 | bias | 0.00000945 | EGR1,FOS,JUN,JUNB,TNF |
| CCL5 | 1.58 | cytokine | Activated | 2.234 | bias | 4.41E-08 | ADGRE5,CCL3,CCL3L3,CCL4,CXCL8,CYP1B1,DUSP1,DUSP6,FOS,IER2,PPIF,SGK1,STAT1,TNF,ZFP36 |
| CSF2 | | cytokine | Activated | 2.233 | bias | 1.38E-09 | ADGRE5,CCL3,CCL3L3,CCL4,CD69,CDH1,CDKN1A,CDKN2C,CFH,CSF2RB,CXCL8,CXCR4,DUSP6,EGR1,EGR2,EGR3,ELN,FCGR1A,FOS,ID2,ID3,IER3,IFITM3,IRF2,JUN,JUNB,MCL1,MMP2,MYC,NFE2,NR4A1,NR4A2,PPIF,PTGS2,SGK1,SNTB1,SOCS1,SOCS3,TCF4,TLR2,TNF,TNFAIP3,TP53,VDR,ZFP36 |
| GNRH | | group | Activated | 2.233 | bias | 0.00138 | DUSP1,EGR1,FOS,JUN,PTGS2,SGK1 |
| TGFB1 | -1.28 | growth factor | Activated | 2.231 | | 3.09E-14 | ACTA2,AKR1C1/AKR1C2,AREG,ARID5B,ARL4A,B4GAT1,BHLHE40,BTG1,C2,CAV1,CAV2,CCL3,CCL3L3,CCL4,CD4,CD69,CD72,CDC42EP4,CDH1,CDKN1A,CDKN2C,CDKN3,CENPF,CFH,CITED2,CLU,COL2A1,CRIP2,CXCL8,CXCR4,CYB561,DDIT4,DLL3,DOCK4,DUSP1,EGLN1,EGR1,EGR2,EGR3,ELN,EREG,F13A1,FBLN5,FCGR1A,FERMT1,FOS,FOSB,GATM,GBP1,GMPR,GPR12,HAPLN3,HES1,HHEX,HLA-DRB5,HLTF,HRH2,HSPB1,ID1,ID2,ID3,IER2,IER3,IFIH1,IFIT3,IL6R,JUN,JUNB,JUND,KLF15,KLF2,KLF4,KLF9,MMP2,MMP7,MYC,NCR3,NFIB,NOC3L,NOG,NR4A1,NR4A2,NR4A3,PARP3,PDZK1IP1,PLS3,PLSCR1,PNMT,POMC,PORCN,PPP1R1B,PTAFR,PTGS2,PTX3,PWWP2A,RASGRP3,RGCC,RHOB,RSAD2,S100A10,S1PR2,SAMHD1,SELP,SERPINE1,SERTAD1,SGK1,SLC16A9,SLC39A14,SLC7A5,SOCS1,SOCS3,STAT1,TCF4,TIMP3,TLE4,TLR2,TNF,TNFAIP3,TNFSF13B,TP53,TSC22D3,TXNIP,VDR,WDFY3,ZFP36 |
| GNAI3 | 1.31 | enzyme | Activated | 2.219 | bias | 0.00056 | CCL3,CCL3L3,CCL4,CXCL8,TNF |
| ACVR1C | | kinase | Activated | 2.219 | bias | 0.000705 | CDKN1A,FOS,JUNB,KLF4,SERPINE1 |
| IL7 | -2.17 | cytokine | Activated | 2.21 | bias | 0.000139 | CCL3,CCL4,CD69,CDKN1A,CXCR4,IL7,KLF2,MCL1,MX1,MYC,PIK3IP1,SOCS1,SOCS3,TNF,TNFSF10 |
| BMP6 | | growth factor | Activated | 2.21 | | 0.00000461 | ACTA2,CDKN1A,CDKN2C,CYP1B1,EPHX1,ID1,ID2,ID3,KLF4,NOG,POMC,PTGS2,SAP30,SERPINE1 |
| Collagen type II | | complex | Activated | 2.2 | | 0.00342 | CCL3L3,MAP3K8,MMP7,NOG,TNF,TNFSF10 |
| TXN | 1.55 | enzyme | Activated | 2.2 | | 0.00342 | CDKN1A,CXCL8,CYP1B1,FOS,TNF,TP53 |
| MAPK10 | | kinase | Activated | 2.197 | | 0.000138 | CXCL8,FOS,JUN,PTGS2,TNF |
| NAMPT | 1.17 | cytokine | Activated | 2.197 | bias | 0.0158 | CXCL8,ID1,JUN,TNF,TP53 |
| TREX1 | | enzyme | Activated | 2.19 | bias | 0.0000978 | IFI44,IFIT2,ISG15,OASL,USP18 |
| FGFR3 | | kinase | Activated | 2.186 | | 0.0000262 | CCL3,CCL4,CDH1,CDKN1A,DUSP6,FOS,HES1,NOG |
| TAC1 | | other | Activated | 2.184 | bias | 0.00287 | CCL4,CXCL8,CXCR4,FOS,FOSB,PTGS2,SELP,SERPINE1,TNF |
| GAB2 | 1.33 | other | Activated | 2.184 | bias | 0.0000135 | CCL3L3,CCL4,CD69,CDH1,FOS,MCL1,SERPINE1,TNF |
| PTK2B | 1.02 | kinase | Activated | 2.183 | | 0.0000269 | CDH1,CXCL8,FOS,JUN,SOCS1,TP53 |
| MAC | | complex | Activated | 2.178 | bias | 0.00000203 | AKR1C3,AREG,EGR1,EGR2,PTGS2,RGCC,TNF,ZFP36 |
| GDF9 | | growth factor | Activated | 2.176 | | 0.00862 | ID1,ID2,ID3,PTGS2,PTX3 |
| Mapk | | group | Activated | 2.173 | bias | 0.0000164 | CDKN1A,CXCL8,DLG4,EGR1,FOS,JUN,JUNB,MMP7,MYC,NR4A3,PER1,PTGS2,STAT1,TNF |
| corticosterone | | chemical - endogenous mammalia | Activated | 2.169 | | 0.00000543 | CAV1,CDKN1A,CLDN5,DDIT4,FKBP5,FOS,GLUL,JUN,KLF9,MYC,POMC,PTGS2,SGK1,TLR2,TNF,TSC22D3 |
| PRDM16 | -1.51 | transcription regulator | Activated | 2.149 | | 0.000000207 | CDKN1A,GBP4,IFI44,IFIT2,IRF7,OAS2,OAS3,SERPINE1,STAT1,STAT2 |

| | | | | | | | |
|---|---|---|---|---|---|---|---|
| PTH | | other | Activated | 2.146 | bias | 1.71E-08 | AREG,BHLHE40,CDKN1A,COL2A1,CXCR4,DUSP1,EGR2,FOS,FOSB,IL6R,JUN,LMNA,MMP2,NFIL3,NR4A1,NR4A2,PHLDB2,PPP1R1B,PTGS2,RGS1,RGS2,SOCS3,VDR |
| MAPK8 | 1.09 | kinase | Activated | 2.138 | | 1.91E-08 | B4GALT1,CDH1,CDKN1A,CXCL8,DUSP1,DUSP6,EGR1,FOS,JUN,JUND,KLF6,MMP2,MMP7,MYC,PTGS2,PTX3,TLR2,TNF,TNFAIP3,TP53,TSC22D3,VDR,ZFP36 |
| norepinephrine | | chemical - endogenous mammalia | Activated | 2.117 | bias | 0.00000551 | CXCL8,CXCR4,DUSP1,EGR1,FOS,HILPDA,ID1,MAN2A1,MCAM,MMP2,NR4A1,NR4A3,PER1,POMC,PTGS2,SERPINE1,SGK1,TNF,WHRN |
| XIAP | -1.28 | enzyme | Activated | 2.117 | | 0.00131 | CDKN1A,CXCL8,PTGS2,SERPINE1,TNF |
| EDN1 | | cytokine | Activated | 2.107 | bias | 7.83E-08 | ACTA2,AREG,CDH1,CXCL8,CXCR4,EGR1,EREG,FOS,FOSB,JUN,JUNB,MCAM,MMP2,MMP7,MYC,POMC,PRKCD,PTGS2,REN,SERPINE1,SOCS1,SOCS3,TIMP3,TP53 |
| MAP3K1 | -1.23 | kinase | Activated | 2.1 | bias | 0.00000153 | CXCL8,DUSP1,EGR1,FOS,HSPB1,JUN,PRDX1,PTGS2,SERPINE1,TNF,TP53 |
| IgG | | complex | Activated | 2.091 | | 2.28E-12 | CCL3,CCL4,CD69,CDKN1A,CEBPD,CLDN5,CXCL8,DUSP1,EGR1,EGR2,EGR3,FCGR1A,FOSB,HSPB1,IER2,IFIH1,IFITM1,IFITM3,ISG15,ITGB8,JUND,MYC,PLS3,PPIF,PTGS2,SERPINE1,SLC40A1,TNF,TNFAIP3,UGCG,ZFP36,ZMPSTE24 |
| Vegf | | group | Activated | 2.09 | bias | 4.32E-10 | ACTA2,ARHGAP23,BMP2K,CDH1,CDKN2C,CDKN3,CENPF,CSGALNACT1,CXCL8,CXCR4,CYYR1,DUSP6,EFNA1,EGR1,EGR3,ELN,ENTPD1,FKBP5,FOSB,GASK1B,HES1,HOPX,ID1,ISG20,ITGB8,JUN,KIF20B,MALL,MAP3K8,MCL1,MMP2,MYC,NOTCH4,NR4A1,NR4A2,NR4A3,OSBPL10,PHLDA1,PPP1R<br>1B,PTGS2,RGCC,RGS2,SCML4,SGK1,SOCS3,TNF,UGCG |
| IL6R | -2.18 | transmembrane receptor | Activated | 2.089 | bias | 0.000147 | ACTA2,CCL3,CDKN1A,COL2A1,CXCL8,IL6R,MCL1,MMP2,PTGS2,SOCS3,TNF |
| hyaluronic acid | | chemical - endogenous mammalia | Activated | 2.084 | | 0.000000278 | CCL3L3,CCL4,COL2A1,CXCL8,CXCR4,FOS,JUN,MMP2,NOG,NR4A1,NR4A2,PTGS2,SERPINE1,SOCS3,TLR2,TNF,TNFAIP3 |
| EP300 | -1.17 | transcription regulator | Activated | 2.077 | bias | 1.36E-09 | ACACB,ACADL,CDKN1A,CHKA,COL2A1,CXCL8,CXCR4,DUSP1,EGR1,EGR2,EPSTI1,FOS,HLA-G,ID1,ID2,ID3,JUN,LGALS3BP,LIG4,LMNA,MAP3K8,MCF2L,MMP7,MYC,NKX3-<br>1,NR4A1,OAS3,PLEKHF1,PRKCD,PTGS2,REN,RNASEL,RSAD2,SH3GL3,SOCS3,TIPARP,TLR2,TNF,TNFAIP3,TP53,TXNIP,USP18 |
| FGF1 | | growth factor | Activated | 2.077 | | 0.0000923 | CCL3,CDH1,CDKN1A,DUSP6,EGR1,FOS,FZD7,JUN,MYC,NR4A2,PTGS2,TCF4,TIMP3,TNF |
| CGA | | other | Activated | 2.073 | | 0.0000105 | C1GALT1C1,CLDN5,CLGN,CXCL8,EGR2,HOMER2,ID1,KLF9,MMP2,NR5A1,PTGS2,SGK1,TIMP3,TP53INP1 |
| CREBBP | -1.6 | transcription regulator | Activated | 2.056 | bias | 3.68E-12 | CDH1,CDKN1A,COL2A1,CXCL8,CXCR4,DUSP1,EGR1,EGR2,EGR3,EPSTI1,FOS,FOSB,HLA-<br>G,ISG15,JUN,JUND,KLF2,LGALS3BP,LIG4,MAP3K8,MCF2L,MYC,MYCBP,NR4A1,NR4A2,NR4A3,OAS3,PLEKHF1,PRKCD,PTGS2,REN,RGS2,RNASEL,RSAD2,SELP,SH3GL3,SOCS3,TAP2,TIPARP,TLR2,TNF,TNFAIP3,TNFSF10,USP18 |
| IL1A | | cytokine | Activated | 2.055 | bias | 0.00000848 | BTN3A3,CCL3,CCL4,CDKN1A,CXCL8,FOS,FOSB,GBP1,IER3,JUN,JUNB,MCAM,MMP2,MYC,PDZK1IP1,POMC,PTGS2,PTX3,S100A10,SERPINE1,TLR2,TNF,TNFAIP3 |
| Nfat (family) | | group | Activated | 2.053 | bias | 0.000714 | B4GALT1,CCL3,CCL3L3,CXCL8,DLG4,EGR2,KLF6,LAG3,PTGS2,SERPINE1,TNF |
| PIN1 | 1.08 | enzyme | Activated | 2.053 | | 0.00554 | CDKN1A,CXCL8,HES1,MCL1,MMP2,MYC,PTGS2,TNF |
| RUNX3 | -1.13 | transcription regulator | Activated | 2.047 | bias | 0.0271 | CD4,CDKN1A,CXCL8,MMP2,MYC,RASAL2,SGPP1,TNF,UGCG |
| IL6 | | cytokine | Activated | 2.023 | bias | 3.99E-12 | ACTA2,ADGRE5,AREG,ARL4C,BATF,BST2,BTG2,CCL3L3,CCL4,CDH1,CDKN1A,CDKN3,CEBPD,CFH,CLU,COL2A1,CP,CSF2RB,CXCL8,CXCR4,CYP1B1,DUSP1,DUSP6,EGR1,EGR2,EREG,FCGR1A,FOS,GP1BA,HLA-<br>DRB5,HOMER3,ID1,ID2,IFIT1,IFIT2,IFITM3,IL6R,IL7,JCHAIN,JUN,JUNB,JUND,LAG3,LRP6,MCL1,MMP2,MMP7,MRVI1,MYC,POMC,PTGS2,SERPINE1,SGK1,SLC12A2,SLC39A14,SLC40A1,SMOX,SOCS1,SOCS3,SP110,STAT1,TLR2,TNF,TNFSF10,TP53,WARS1,WASF3 |
| MET | | kinase | Activated | 2.02 | bias | 0.00000205 | AREG,CCL3L3,CDH1,CDKN1A,CITED2,CXCL8,FGL2,FOS,HSPB1,JUN,KLF4,MAL,MMP2,MYC,NOTCH4,PTGS2,SOCS3 |
| BCR (complex) | | complex | Activated | 2.013 | bias | 0.0000132 | B4GALT1,CCL3,CCL4,CD69,CXCR4,EGR1,FOS,FOSB,JUNB,KLF4,KLF9,MCL1,MYC,PTAFR,PTGS2,S100A10 |
| Fcer1 | | complex | Activated | 2.011 | bias | 0.00000746 | CCL3,CCL3L3,CCL4,CD4,CXCL8,DUSP6,FOS,IFIT2,JUN,S1PR2,TNF |
| miR-155-5p (miRNAs w/seed UAAUGCU) | | mature microRNA | Inhibited | -2.003 | bias | 0.0103 | CCL3,CCL4,CD69,CXCL8,MARC1,MOSPD2,PTGS2,RAB27B,SERPINE1,SOCS1,TNF,TP53INP1 |
| ZMPSTE24 | -2.03 | peptidase | Inhibited | -2.005 | bias | 0.000115 | BTG2,CDKN1A,DDIT4,LMNA,SESN1,SOCS1,SOCS3,TNF |
| Hdac | | group | Inhibited | -2.037 | | 0.0000147 | AREG,CDKN1A,CXCL8,CXCR4,EGR1,EGR3,FOS,IL6R,JUN,KLF6,KLF9,MYC,NOTCH4,NR4A1,SPRY2,TNF,TNFSF10,TXNIP |
| BRCA1 | -1.18 | transcription regulator | Inhibited | -2.112 | | 0.000000196 | AREG,ASCL2,CDKN1A,CYP1B1,DDIT4,EGR1,FEN1,FKBP5,HSPB1,IFI6,IFIT1,IFIT2,IFIT3,IFITM1,IRF7,MX1,MYC,PHLDA1,PLSCR1,STAT1,TNF,TP53 |
| E2F3 | -1.01 | transcription regulator | Inhibited | -2.157 | | 0.0394 | AKR1C1/AKR1C2,AKR1C3,CAV2,CDKN1A,FBLN5,ID3,MYC,NEAT1,SERPINE1,TIMP3,TM9SF2 |
| IFNA4 | | cytokine | Inhibited | -2.159 | bias | 2.71E-08 | CD69,GBP5,IFIH1,IFIT1,IFIT2,ISG15,MAP3K8,MX1,MYC,OASL,RSAD2,USP18 |
| SPI1 | -1.6 | transcription regulator | Inhibited | -2.16 | | 7.5E-17 | ACTA2,ARID3A,CCL3,CCL4,CD72,CDKN1A,CMPK2,CSF2RB,DUSP6,EGR2,FOS,HES1,ID2,ID3,IFI44,IFI44L,IFI6,IFIT1,IFIT2,IFIT3,IFITM1,IFITM3,IRF7,ISG15,ISG20,JDP2,JUN,KLF13,KLF4,MCL1,MMP2,MX1,MYC,NIP7,OASL,PTGS2,RSAD2,SP110,TLR2,TNF,TNFSF10,USP18 |
| amino acids | | chemical - endogenous mammalia | Inhibited | -2.164 | | 0.00157 | FOS,FOSB,JUN,JUNB,MYC |
| inosine | | chemical - endogenous mammalia | Inhibited | -2.183 | bias | 0.0304 | C2,CCL3L3,IFITM3,LGALS3BP,TNF |
| EP400 | -1.12 | other | Inhibited | -2.2 | bias | 0.0131 | CDKN1A,CENPF,MAPK12,PSRC1,TP53 |
| ISGF3 | | complex | Inhibited | -2.2 | bias | 1.42E-08 | EIF2AK2,IFIH1,IFIT2,IRF7,ISG15,RSAD2,TNFSF10 |
| SGPL1 | 1.09 | enzyme | Inhibited | -2.2 | | 0.000876 | IFIT1,ISG15,MX1,OAS1,PTGS2 |
| FZD9 | | G-protein coupled receptor | Inhibited | -2.213 | bias | 0.000191 | CDH1,IFI44,IRF7,ISG15,STAT1 |
| CCN5 | | growth factor | Inhibited | -2.213 | | 0.0304 | ACTA2,CDH1,JUN,KLF4,SERPINE1 |
| CBX7 | -1 | other | Inhibited | -2.219 | bias | 0.000439 | AREG,CDH1,CXCL8,CYP1B1,KYNU |
| IFNK | | cytokine | Inhibited | -2.236 | bias | 0.000876 | EIF2AK2,IFIH1,MX1,OAS1,STAT1 |
| PML | 1.11 | transcription regulator | Inhibited | -2.259 | | 4.8E-14 | ACACB,ACADL,BST2,CDKN1A,CXCR4,DDX60,EPSTI1,HSPB1,ID2,IFI44,IFI44L,IFIH1,IFIT1,IFIT3,IFITM1,IRF7,ISG15,ISG20,MAFF,MX1,OAS1,OAS2,OAS3,PLSCR1,PRDX1,STAT1,TAP2,TNFAIP3,TP53,YWHAG |
| ZFP36 | 6.41 | transcription regulator | Inhibited | -2.286 | bias | 0.000905 | CCL3L3,CDKN1A,CLCN3,FOS,IER3,JUN,PTGS2,SOCS1,TNF,ZFP36 |
| LYN | -1.49 | kinase | Inhibited | -2.391 | | 0.00851 | CCL3L3,CCL4,CDKN1A,EGR1,MYC,SOCS1,SOCS3,TNF |
| EBI3 | | cytokine | Inhibited | -2.395 | bias | 0.0000989 | ENTPD1,FOS,HLA-C,LAG3,MX1,MYC,STAT1,STAT2,TNFSF10 |
| NR1H2 | -1.1 | ligand-dependent nuclear receptor | Inhibited | -2.425 | | 0.0233 | CCL4,CDH1,DDX60,GBP4,IFIT2,PTGS2,REN,TNF |
| CLEC12A | -1.31 | other | Inhibited | -2.425 | bias | 9.85E-09 | IFIT2,IFIT3,IRF7,ISG15,RSAD2,USP18 |
| EBF1 | | transcription regulator | Inhibited | -2.425 | bias | 0.038 | ACACB,CRK,IGKC,IRF7,NFIL3,SOCS1,SOCS3,STAT1,TLR2 |
| IRF9 | 1.05 | transcription regulator | Inhibited | -2.443 | | 3.22E-17 | CD69,CDH1,CXCL8,GBP1,IFIT1,IFIT2,IFIT3,IFITM1,IFITM3,IRF7,IRF8,ISG15,MX1,OAS2,SOCS1,SOCS3,STAT1,STAT2,TNF,TNFSF10 |
| DUSP5 | 1.01 | phosphatase | Inhibited | -2.449 | bias | 0.0000374 | DUSP2,EGR1,EGR3,EGR4,ID2,ZFP36 |
| IFN alpha/beta | | group | Inhibited | -2.59 | bias | 0.000000494 | CD69,FGL2,IFIT2,IFIT3,IL7,IRF7,IRF8,RSAD2,SOCS1,STAT1,STAT2,TLR2,TNFSF10,TNFSF13B |
| NLRP12 | | other | Inhibited | -2.607 | | 0.000172 | CXCL8,CXCR4,HLA-C,HLA-F,HLA-G,JUN,TNF |
| IFNL4 | | cytokine | Inhibited | -2.623 | bias | 5.32E-09 | DHX58,IFIH1,ISG15,MX1,OAS1,OAS2,STAT1 |
| BTNL2 | | transmembrane receptor | Inhibited | -2.63 | | 0.0132 | CDKN2C,ENTPD1,FGL2,GBP4,IFITM3,TNF,TNFSF10 |
| VCAN | 1.54 | other | Inhibited | -2.692 | | 6.76E-09 | CDKN1A,CEACAM4,CLU,CXCL8,ELN,FBLN5,IFI44,IFI44L,IFI6,IFIT1,IFIT2,IFITM1,IRF8,MX1,OAS2,OAS3,PCSK1N,RGS2,STAT1,TNF,TP53,TTC27,XAF1 |
| DACH1 | -1.32 | transcription regulator | Inhibited | -2.75 | | 5.68E-08 | CDKN1A,CXCL8,EGR1,FOS,ID1,IER2,JUN,KLF4,PTGS2,SERPINE1,TNFAIP3 |
| FASN | -1.03 | enzyme | Inhibited | -2.795 | bias | 0.000501 | ACTA2,CAV1,CDH1,CXCL8,DDIT4,DGAT2,LRRK2,PTGS2,TNF |
| FOSL1 | -1.05 | transcription regulator | Inhibited | -2.819 | | 0.00000143 | CCL3L3,CCL4,CXCL8,EGR1,EGR2,ELN,FOS,FOSB,JUN,JUNB,MMP2,SERPINE1,TNF |
| IRF5 | -1.54 | transcription regulator | Inhibited | -2.972 | bias | 4.05E-20 | CCL3,CCL4,CDKN1A,CMPK2,CXCR4,DHX58,IFI44,IFIH1,IFIT1,IFIT2,IFIT3,IFITM3,IRF7,ISG15,ISG20,MYC,OAS1,OAS2,OASL,PLSCR1,PTGS2,RSAD2,SP110,STAT1,STAT2,TNF,TNFSF10,UBE2L6 |
| MAVS | -1.54 | other | Inhibited | -2.988 | bias | 7.62E-15 | CMPK2,CXCL8,DHX58,IFIT1,IFIT2,IFIT3,IFITM3,IRF7,ISG15,ISG20,OAS1,OAS2,OASL,RSAD2,SOCS1,SOCS3,STAT1,STAT2,TNF,UBE2L6,USP18 |
| SFTPA1 | | transporter | Inhibited | -3.017 | | 0.00000224 | AKR1C1/AKR1C2,CCL3,CLEC1B,CYP1B1,EGR1,FOS,FOSB,GSTO2,KLF2,MYC,PHLDA1,SERPINE1,TNF |
| PARP9 | -3.2 | enzyme | Inhibited | -3.121 | bias | 6.18E-11 | IFI44,IFIT1,IFIT2,IFIT3,IRF2,IRF7,ISG15,OAS2,SP110,STAT1 |
| JAK | | group | Inhibited | -3.162 | bias | 9.53E-10 | EIF2AK2,IFI6,IFIH1,IFIT1,IFIT2,IFIT3,IFITM3,ISG15,PTGS2,RSAD2,SOCS1,SOCS3,STAT1 |
| DUSP1 | 2.64 | phosphatase | Inhibited | -3.242 | | 1.16E-08 | CCL3L3,CCL4,CMPK2,CXCL8,DUSP1,DUSP6,IER3,IFIT1,IFIT3,ISG20,JUN,MX1,PAQR7,PTGS2,SERPINE1,SOCS1,TLR2,TNF,ZFP36 |
| IFN type 1 | | group | Inhibited | -3.389 | bias | 8.94E-10 | BST2,CD69,CXCL8,DHX58,EIF2AK2,IFIH1,IFIT1,IFIT2,ISG15,OASL,STAT1,STAT2,TNFSF10,TNFSF13B |
| Ifn | | group | Inhibited | -3.415 | bias | 3.93E-15 | CD69,CFH,DHX58,EIF2AK2,FOS,IFIH1,IFIT1,IFITM1,IFITM3,IL7,IRF7,IRF8,ISG15,ISG20,MX1,OAS2,OAS3,OASL,PTAFR,RNASEL,RSAD2,SERPINE1,SOCS1,STAT1,TLR2,TNFSF13B,TP53 |
| STAT1 | -4.62 | transcription regulator | Inhibited | -3.468 | bias | 4.19E-31 | APOL4,BST2,BTG1,CCL3,CCL3L3,CCL4,CDKN1A,CEBPD,CMPK2,CXCL8,DDX60,EGR1,EIF2AK2,EPSTI1,FCGR1A,FGL2,FOS,GBP1,GBP1P1,GBP4,GBP5,HAPLN3,HES1,HLA-<br>DRB5,IFI44,IFI44L,IFI6,IFIH1,IFIT1,IFIT2,IFIT3,IFITM1,IFITM3,IRF2,IRF7,IRF8,ISG15,JUN,KLF4,MX1,MYC,NFE2,NLRC5,OAS1,OAS2,OAS3,OASL,PARP9,PLSCR1,POMC,PTGS2,RSAD2,S100A10,SAMD9L,SAMHD1,SLFN5,SOCS1,SOCS3,SP110,STAT1,STAT2,TNF,TNFSF10,TNFSF13B,TP53,USP18,W<br>[illegible] |
| IFNA1/IFNA13 | | cytokine | Inhibited | -3.477 | bias | 9.68E-17 | CD69,DHX58,EIF2AK2,IFI6,IFIH1,IFIT1,IFIT2,IFITM1,IRF7,ISG15,MX1,MYC,OAS1,OAS2,PLSCR1,PTGS2,RSAD2,SOCS1,STAT1,STAT2,UBE2L6 |
| IFNG | | cytokine | Inhibited | -3.487 | | 1.13E-31 | ACTA2,AREG,BST2,BTG1,BTN3A1,BTN3A2,C2,CAV1,CCL3,CCL3L3,CCL4,CD4,CD72,CDH1,CDKN1A,CEBPD,CMPK2,CP,CSF2RB,CXCL8,CXCR4,CYB561,DDX60,DLG4,DTX3L,DUSP1,EGR1,EGR2,EGR3,EIF2AK2,ELN,FBLN1,FBP1,FCGR1A,FGL2,FKBP5,FOS,FOSB,GBP1,GBP4,GBP5,GBP7,GLUL,G<br>MPR,GPR12,HCK,HCP5,HLA-C,HLA-DRB5,HLA-F,HLA-<br>G,HSPB1,ID1,IER2,IER3,IFI44,IFI44L,IFI6,IFIH1,IFIT1,IFIT2,IFIT3,IFITM1,IFITM3,IL6R,IL7,IRF2,IRF7,IRF8,ISG15,ISG20,JUN,JUNB,JUND,KLF2,KLF4,KLF6,KYNU,LAG3,LGALS3BP,MAFF,MAN2A1,MAP3K8,MMP2,MX1,MYC,NCR3,NEAT1,NLRC5,OAS1,OAS2,OAS3,OASL,PANX1,PARP9,PCTP,PHLDA1,P<br>LSCR1,POMC,PPP1R1B,PRKCD,PRKG2,PTAFR,PTGS2,PTX3,RAB38,RGCC,RHOB,RRP7BP,RSAD2,S100A10,SAMD9,SAMHD1,SCLY,SELP,SERPINE1,SLC12A2,SLC16A9,SLC40A1,SLC7A5,SLFN5,SOCS1,SOCS3,SP110,STAT1,STAT2,TAP2,TIMP3,TLR2,TMEM50B,TNF,TNFSF10,TNFSF13B,TP53,TS<br>C22D3,TXNIP,UBE2L6,USP18,VDR,WARS1,XAF1,ZFP36 |

| | | | | | | | |
|---|---|---|---|---|---|---|---|
| IFNB1 | | cytokine | Inhibited | -3.524 | | 1.7E-17 | APOL2,BHLHE40,BST2,BTN3A3,CCL3,CCL4,CDKN1A,CMPK2,CXCL8,DHX58,EIF2AK2,FKBP9,FOS,GBP4,GBP5,GBP7,HERC5,IFI6,IFIH1,IFIT1,IFIT2,IFIT3,IFITM1,IRF7,ISG15,ISG20,MMP2,MX1,MYC,OAS1,OAS2,PHLDB2,PTGS2,RNASEL,RSAD2,SLC39A14,SOCS1,SPRY2,STAT1,STAT2,TNF,TNFSF10,USP18,VDR,XAF1 |
| PRL | | cytokine | Inhibited | -3.554 | bias | 2.45E-26 | AKR1C3,AKR1C4,BST2,CAV1,CD69,CDH1,CDKN1A,CEBPD,CLDN5,CLU,CMPK2,DHX58,DTX3L,EGR1,EIF2AK2,EPSTI1,FOS,GMPR,HERC5,ID1,ID2,ID3,IER3,IFI44,IFI44L,IFI6,IFIH1,IFIT1,IFIT3,IFITM1,IRF7,ISG15,JUN,MYC,OAS1,OAS2,OAS3,PAQR7,PLSCR1,RSAD2,SAMD9,SAMD9L,SAMHD1,SOCS1,SOCS3,SP110,STAT1,STAT2,TNF,TNFSF13B,TP53,USP18,VDR,XAF1,YWHAG |
| Interferon alpha | | group | Inhibited | -3.802 | bias | 3.03E-28 | APOL2,BST2,BTG2,CCL3,CD69,CDH1,CDKN1A,CSF2RB,CXCL8,CYB561,DDIT4,DHX58,DUSP6,EIF2AK2,EPSTI1,FBXO6,FCGR1A,FOS,GBP1,GBP5,HERC5,HLA-C,HLA-F,HLA-G,IFI44,IFI44L,IFI6,IFIH1,IFIT1,IFIT2,IFIT3,IFITM1,IFITM3,IL6R,IL7,IRF7,IRF8,ISG15,ISG20,MCL1,MX1,MYC,NCR3,NFIL3,OAS1,OAS2,OAS3,PARP9,PLSCR1,POMC,PTGS2,RNASEL,RSAD2,SAMD9,SAMD9L,SLFN5,SOCS1,SOCS3,SP110,SPRY2,STAT1,STAT2,TAP2,TENT4A,TLR2,TNF,TNFSF10,TNFSF13B,UBE2L6,USP18,WARS1,ZNRF3 |
| IFN Beta | | group | Inhibited | -3.858 | bias | 1.96E-19 | BST2,CCL3,CD69,CDKN1A,CXCL8,DUSP1,EIF2AK2,HERC5,HLA-G,IFI44,IFI6,IFIH1,IFIT1,IFIT2,IFIT3,IFITM1,IRF7,ISG15,MX1,MYC,OAS1,OAS2,OAS3,OASL,RSAD2,SOCS1,SPRY2,STAT1,STAT2,TNF,TNFSF10,TP53,USP18,XAF1 |
| IRF1 | -1.39 | transcription regulator | Inhibited | -3.931 | bias | 3.05E-20 | CDKN1A,CMPK2,CXCL8,EIF2AK2,FGL2,HLA-G,IFI44,IFI44L,IFI6,IFIH1,IFIT1,IFIT2,IFIT3,IFITM1,IFITM3,IL7,IRF2,IRF7,ISG15,LIG4,MX1,MYC,NFE2,OAS1,OAS2,OAS3,OASL,PTGS2,RSAD2,SOCS1,SP110,STAT1,STAT2,TAP2,TNF,TNFSF10,TNFSF13B,TP53,XAF1 |
| RNY3 | | other | Inhibited | -4 | bias | 2.23E-17 | DDX60,EPSTI1,HERC5,IFI44,IFI44L,IFIT1,IFIT3,IFITM3,ISG15,MX1,OAS1,OAS2,OAS3,OASL,RSAD2,XAF1 |
| Ifnar | | group | Inhibited | -4.16 | bias | 5.2E-15 | EIF2AK2,HLA-G,IFIH1,IFIT2,IFIT3,IFITM3,IRF7,IRF8,ISG15,ISG20,NLRC5,OAS1,OAS2,OASL,RSAD2,STAT1,STAT2,TAP2,TLR2,TNF,TNFSF10,UBE2L6,USP18,XAF1 |
| IFNA2 | | cytokine | Inhibited | -4.277 | bias | 1.43E-22 | BST2,CAV1,CCL3,CD69,CDKN1A,CMPK2,DDX60,EIF2AK2,GBP1,GBP4,HERC5,HLA-C,IFI44,IFI44L,IFI6,IFIH1,IFIT1,IFIT2,IFIT3,IFITM1,IFITM3,IRF7,ISG15,ISG20,LGALS3BP,MAL,MX1,MYC,OAS1,OAS2,OAS3,PARP9,PLSCR1,PTAFR,RSAD2,SAMD9,SAMHD1,SOCS1,SOCS3,SP110,STAT1,TNF,TNFSF10,TP53,UBE2L6,USP18,XAF1 |
| IRF3 | -1.04 | transcription regulator | Inhibited | -4.3 | bias | 9.6E-21 | CCL3,CCL3L3,CCL4,CD69,CMPK2,CXCL8,DDX60,DHX58,EIF2AK2,FCGR1A,GBP1,GBP5,HLA-F,IFI44,IFI6,IFIH1,IFIT1,IFIT2,IFIT3,IFITM3,IRF7,ISG15,ISG20,JUNB,LAG3,NLRC5,OAS1,OAS2,OAS3,OASL,RSAD2,SAMD9L,SAP30,STAT1,STAT2,TIMP3,TNF,TNFAIP3,TNFSF10,UBE2L6,USP18 |
| IRF7 | -3.64 | transcription regulator | Inhibited | -4.858 | bias | 1.42E-23 | CD69,CMPK2,DHX58,FCGR1A,GBP1,GBP4,GBP5,HERC5,IFI44,IFI44L,IFI6,IFIH1,IFIT1,IFIT2,IFIT3,IFITM1,IFITM3,IRF7,IRF8,ISG15,ISG20,MAP3K8,MCL1,MX1,OAS1,OAS2,OAS3,OASL,PLSCR1,RSAD2,SAMD9L,SAP30,SOCS1,STAT1,STAT2,TAP2,TNFSF10,TNFSF13B,UBE2L6,USP18,XAF1 |
| IFNL1 | | cytokine | Inhibited | -4.993 | bias | 1.97E-30 | BST2,CMPK2,CXCL8,DDX60,EIF2AK2,GBP1,GBP5,HERC5,HLA-C,IFI44,IFI44L,IFI6,IFIH1,IFIT1,IFIT2,IFIT3,IFITM1,IFITM3,ISG15,ISG20,LGALS3BP,MX1,OAS1,OAS2,OAS3,OASL,PLSCR1,RSAD2,SAMD9,SP110,STAT1,STAT2,TLR2,UBE2L6,USP18,XAF1 |

**T3 vs. T1 MPP_GMP**

© 2000-2020 QIAGEN. All rights reserved.

| Upstream Regulator | Expr Fold Change | Molecule Type | Predicted Activation State | Activation z-score | Flags | p-value of overlap | Target Molecules in Dataset |
|---|---|---|---|---|---|---|---|
| MAPK1 | -1.57 | kinase | Activated | 5.534 | | 5.48E-32 | BTN3A3,CCL3L3,CCL5,CD69,CXCL8,DDX58,DUSP1,DUSP2,EGR1,EGR2,EGR4,EIF2AK2,EVI5,FN1,FOS,GBP1,GBP5,GNG2,HERC5,HLA-C,IFI44,IFI6,IFIH1,IFIT1,IFIT2,IFIT3,IFITM1,IFITM3,IRF7,ISG15,JUN,JUNB,JUND,LAG3,LGALS3BP,LMNA,MCL1,MMP2,MX2,MYC,NR4A1,OAS1,OAS2,OAS3,OASL,PLA2G4A,PLSCR1,POU2F2,PSAT1,PTGS2,RGS2,SAMHD1,SP110,SPRY2,STAT1,STAT2,TAP1,TNFSF10,TRANK1,TRIM22,TRIM25,UBE2L6,USP18 |
| TRIM24 | -1.65 | transcription regulator | Activated | 4.571 | bias | 6.28E-17 | CCL5,CMPK2,DDX58,DDX60,DHX58,EPSTI1,GBP4,HERC6,IFI44,IFIH1,IFIT2,IFIT3,IRF7,ISG15,LGALS3BP,OAS1,OASL,RTP4,SAMD9L,SAMHD1,SOCS1,STAT1,STAT2,TAP1,USP18 |
| PNPT1 | -1.45 | enzyme | Activated | 4.364 | bias | 3.52E-25 | CCL5,CMPK2,CXCL8,DDX58,EIF2AK2,GBP4,IFI44,IFIH1,IFIT3,IRF7,ISG15,LGALS3BP,MYC,OAS1,PARP14,PARP9,RTP4,SAMD9L,STAT1,STAT2,TNFRSF1A,UBE2L6,USP18,XAF1 |
| PDGF BB | | complex | Activated | 4.278 | bias | 2.05E-14 | ACTA2,ADM,AREG,CAV2,CP,CSF1R,CXCL8,DUSP1,DUSP6,EGR1,EGR2,EGR3,EREG,FN1,FOS,ID3,IER2,IER3,INHBA,JUN,JUNB,KLF6,LMNA,MCL1,MMP2,MYC,NAMPT,NEXN,NR4A1,NR4A3,PHLDA1,PTGS2,RGS1,RGS2,RHOB,SGK1,SOCS3,ZFP36 |
| TLR2 | -1.19 | transmembrane receptor | Activated | 3.779 | bias | 0.00000268 | ACTA2,CCL3L3,CCL5,CCR1,CD69,CXCL3,CXCL8,DUSP1,FN1,HLA-DRB5,KLF2,MX1,PTGS2,SLC40A1,SOCS1,SOCS3,TNFAIP8L2,TNFSF10 |
| Ca2+ | | chemical - endogenous mammalian | Activated | 3.639 | bias | 0.00000207 | ACTA2,COL2A1,CXCL8,CXCR4,DUSP1,EGR1,FN1,FOS,ID1,ID3,JUN,JUNB,JUND,KLF4,MCAM,NR4A1,NR4A3,PER1,PER2,PTGS2,SCN9A,SGK1 |
| leukotriene D4 | | chemical - endogenous mammalian | Activated | 3.638 | bias | 2.38E-14 | ARID5B,CCL3,CCL3L3,CXCL8,DUSP1,EGR1,EGR2,EGR3,KLF2,KLF4,MCL1,NR4A1,NR4A3,PTGS2,RGS2,SIK1/SIK1B,ZFP36 |
| TREM1 | 1.33 | transmembrane receptor | Activated | 3.564 | bias | 5.36E-08 | AREG,CCL3,CCL5,CXCL3,CXCL8,CXCR4,EGR1,EGR2,EGR3,IFIT2,INHBA,ISG15,MAFF,NOD2,OASL,PHLDA1,PTGS2,RGS1,RHOU,SLC1A3,SPRY2 |
| AREG | 32.8 | growth factor | Activated | 3.533 | | 0.000000429 | AREG,CXCR4,EGR1,EREG,FOS,GBP1,IFI6,IFIT2,IFIT3,ITGB8,JUN,PTAFR,PTGS2 |
| STAT3 | 1.04 | transcription regulator | Activated | 3.483 | bias | 1.68E-28 | ACTA2,ADM,AREG,BAK1,BATF,CCL3L3,CCL5,CCR1,CMPK2,CXCL3,CXCL8,CXCR4,EGR1,EGR2,EGR3,EIF2AK2,EME1,ENPP2,FAS,FN1,FOS,GBP5,HERC5,HERC6,HLA-DRB5,ID1,IFI44,IFI6,IFIH1,IFIT1,IFIT2,IFIT3,IFITM1,IFITM3,IRF7,ISG15,JUNB,KLF4,MCL1,MMP2,MMP7,MT-ND1,MX1,MX2,MYC,NAMPT,NR0B1,OAS1,OAS2,OAS3,OASL,PHLDA1,PLA2G4A,PLSCR1,PROCR,PTAFR,PTGS2,REN,RSAD2,SGK1,SLC1A3,SLFN5,SOCS1,SOCS3,SP110,STAT1,STAT2,TAP1,TNFSF10,TRIM22,USP18,XAF1,ZFP36 |
| ACKR2 | | G-protein coupled receptor | Activated | 3.441 | bias | 2.48E-19 | CCL3L3,CCL5,DDX58,DDX60,DHX58,EIF2AK2,IFI44,IFIT2,IFIT3,IRF7,ISG15,OAS1,OAS2,OAS3,OASL,RSAD2,STAT1,STAT2,USP18 |
| IL1RN | | cytokine | Activated | 3.208 | bias | 3.45E-19 | ACTA2,BTN3A1,BTRC,CXCL3,CXCL8,DDX58,GBP1,HERC6,IFI44,IFI44L,IFI6,IFIH1,IFIT3,IRF7,KLF6,MX1,MX2,OAS1,OAS2,OAS3,OASL,PTGS2,RSAD2,RTP4,SAMD9,STAT2,TNFSF10,TRIM22,USP18 |
| NUPR1 | | transcription regulator | Activated | 3.157 | | 0.0000633 | ABHD15,ADAM22,ADM,AREG,ARHGEF26,ATP6V0A1,BTG1,CARMIL1,CXCL3,CXCL8,CXCR4,EME1,EREG,GCH1,H3C7,IFIT2,KLF4,KLF6,MRS2,MX2,MYC,PARP9,PHLDA1,RASAL2,SAMD4A,SAMHD1,SIK1/SIK1B,WDCP,ZNF296 |
| dopamine | | chemical - endogenous mammalian | Activated | 3.119 | | 0.000000193 | BTG2,CLCN3,CXCL8,DUSP2,EGR1,EGR3,EGR4,FOS,GPR12,JUN,KCNK6,KLF4,NR4A1,NR4A3,PER1,SCN9A,SLC1A3 |
| SMAD7 | | transcription regulator | Activated | 3.104 | | 0.000629 | ACTA2,AREG,BMPR1B,CCL5,CDKN2C,COL2A1,FAS,FN1,ID1,ID3,MMP2,MYC |
| MAPK7 | -1.55 | kinase | Activated | 3.087 | bias | 0.000000424 | CXCL3,CXCL8,DUSP1,ID1,JUN,KLF2,KLF4,MCL1,NR4A1,PTGS2 |
| NKX2-3 | -1.19 | transcription regulator | Activated | 3.086 | | 2.04E-27 | ACTA2,ADM,BTG1,C1orf74,CAVIN2,CMPK2,CXCL8,DDX58,DDX60,DHX58,EIF2AK2,FBXO6,GASK1B,GBP1,GCA,GCH1,GMPR,HCP5,HELZ2,HLA-C,HLA-F,KLF2,KLF4,MMP7,NETO2,PARP14,PARP9,PLSCR1,PSAT1,PTGS2,RPS23,RTP4,SAMD9,SAMD9L,SLC40A1,SP110,STAT1,STAT2,TAP1,TIPARP,TRIM22,UBE2L6,USP18,XAF1,ZMAT3 |
| Mek | | group | Activated | 3.075 | | 6.08E-11 | AXL,CLU,CSF1R,CXCL8,DDX58,DUSP6,EGR1,EGR2,EGR3,EREG,FOS,HMGA2,HSPB1,ID1,IER2,IER3,JUN,JUNB,MAFF,MCL1,MMP2,MYC,NR4A1,PHLDA1,PTGS2,SPRY2,XAF1 |
| FOXO1 | 1.83 | transcription regulator | Activated | 3.029 | bias | 0.0000106 | ATP6V0A1,CDKN2C,COL2A1,CXCL8,EGR1,EGR2,EGR4,FAS,FN1,FOS,IER3,JUN,JUNB,KLF2,KLF4,KLF7,MYC,NAMPT,NKX3-1,PIK3IP1,PLSCR1,PRKAA2,PRKCD,SESN1,SGK1,SPRY2,STAT2,TNFSF10,TXNIP |
| APEX1 | -1.32 | enzyme | Activated | 2.985 | bias | 6.08E-08 | EGR1,EGR2,EGR3,EGR4,FOS,JUNB,NR4A1,PTGS2,RGS2 |
| MAPK3 | 1.04 | kinase | Activated | 2.967 | bias | 8E-11 | CCL3,CD69,CXCL8,EGR1,EGR2,EGR4,FN1,FOS,JUN,JUNB,JUND,MCL1,MYC,NR4A1,PLA2G4A,PTGS2,RGS2 |
| NGLY1 | -1.25 | enzyme | Activated | 2.91 | | 0.000000194 | IFI44,IFI44L,IFIT1,IFIT2,IFIT3,OAS1,OAS3,RSAD2,USP18 |
| GAST | | other | Activated | 2.905 | bias | 0.000249 | AREG,EGR1,FOS,HMGB2,JUN,MMP7,PTGS2,RPL7,SIK1/SIK1B |
| KRAS | 1.02 | enzyme | Activated | 2.885 | | 5.05E-14 | AREG,ASCL2,AXL,BAK1,CAVIN2,CDKN2C,CLU,CSF2RB,CXCL8,DUSP1,DUSP6,EGR1,EGR2,EIF2AK2,EREG,FAS,FN1,FOS,GSTM3,HOPX,HSPB1,ID1,IER3,IFI6,IFIT1,IFITM1,IFITM3,IRF2,ISG15,JUN,JUNB,KCNN4,KLF6,MCAM,MMP2,MMP7,MX1,MX2,MYC,NETO2,NR0B1,NR4A1,OAS1,PRKCD,PTGS2,RHOB,SAMHD1,SDC4,SPRY2,STAT1,STAT2,TAP1,TK1,TLE3,TNFSF10 |
| AHR | 1.53 | ligand-dependent nuclear receptor | Activated | 2.828 | | 0.0000922 | ACTA2,ADAM19,ADM,AFMID,AREG,BTG2,CCL5,CDKN2C,DUSP6,EGR2,FAS,FN1,FOS,JUN,JUNB,KLF2,LTBP1,MMP2,MYC,PTGS2,SOCS3,TIPARP,TK1 |
| histamine | | chemical - endogenous mammalian | Activated | 2.751 | | 0.000358 | AREG,CCL3,CCL5,CXCL8,CXCR4,FOS,JUN,NR4A1,PTGS2 |
| MAP3K8 | 1.99 | kinase | Activated | 2.745 | | 0.00000208 | CCR1,CXCL3,CXCL8,DUSP6,FOS,GCA,IER3,INHBA,JUN,JUNB,KDM4A,PTGS2,SESN1,SOCS3,STK17B,TXNIP |
| FOXO3 | 1.04 | transcription regulator | Activated | 2.726 | | 8.15E-08 | ACTA2,BTG1,CCNA1,CXCL8,CXCR4,EGR1,EGR2,EGR4,FN1,FOS,IER3,IFIH1,INHBA,JUNB,MCAM,MYC,NAMPT,PIK3IP1,PRKAA2,PRKCD,SESN1,SGK1,SLC40A1,TAL1,TNFRSF1A,TNFSF10,TP53INP1,TXNIP |
| ELK1 | -1.09 | transcription regulator | Activated | 2.725 | bias | 0.00000352 | EGR1,EGR2,EGR4,FOS,JUN,JUNB,MCL1,PTGS2,TIPARP,ZFP36 |
| STAT6 | -1.23 | transcription regulator | Activated | 2.697 | | 1.35E-13 | ADAM19,AREG,CCDC86,CCL3L3,CCL5,CD69,CMPK2,CXCL3,DHX58,EGR1,EGR2,EIF2AK2,FKBP5,GBP5,GBP7,HCK,IER2,IFI44,IFI44L,IFIH1,IFIT3,IFITM3,IL10RA,IRF7,ISG15,LDLRAD3,LGALS3BP,MMP2,OAS3,OASL,PLSCR1,POU2F2,PTGS2,RAB30,RASGRP3,SAMD4A,SCN9A,SOCS1,STAT2,TCF7,TMEM176B,ZFP36 |
| SIRT1 | -1.4 | transcription regulator | Activated | 2.687 | | 2.72E-12 | ABCG1,BTRC,CCL5,CDKN2C,CMPK2,DDX58,DDX60,DHX58,FAS,FN1,GSTM3,HCK,HELZ2,HLA-DRB5,ID1,IFI44,IFIT3,IFITM3,IRF7,KLF2,LGALS3BP,LMTK2,MCL1,MMP2,MYC,OAS1,OAS2,PARP14,PER1,PRDM16,RSAD2,RTP4,SOCS3,SP110,STAT1,TAL1,TAP1,USP18,ZNF296 |
| levodopa | | chemical - endogenous mammalian | Activated | 2.653 | | 0.00166 | ACTA2,AGPAT1,APBA2,AXL,CITED4,DCAF4,DIXDC1,DUSP1,DUSP6,EGR2,EGR4,FKBP5,FOS,GDF10,IER3,JUNB,KCNAB1,KLF6,KLF9,LAG3,LMNA,LMTK2,LTBP1,NAV1,PER1,PER2,PIEZO2,PRKG2,RIMS1,SLC29A3,TRIM25 |
| EGR1 | 9.64 | transcription regulator | Activated | 2.652 | bias | 1.36E-10 | ACTA2,AREG,CCL3L3,CCR1,CLU,COL2A1,CXCL3,CXCL8,EGR1,EGR2,EREG,FAS,FN1,JUN,JUNB,JUND,MYC,NR4A1,PNMT,PTGS2,SGK1,SLC12A2,SOCS1,TNFSF10 |
| collagenase | | group | Activated | 2.646 | bias | 0.0000146 | ADM,CCL3L3,CXCL3,EGR1,IER3,JUN,JUNB |
| Lh | | complex | Activated | 2.644 | | 0.000000187 | ACTA2,AREG,ARL4C,AXL,CXCL8,CXCR4,DUSP1,DUSP3,EREG,FKBP5,FOS,GBP1,MCL1,MMP2,PER1,PPP2R5B,PTGS2,RAB2A,RASAL2,RHOB,SGK1,STAT1,TK1 |
| BTK | -1.42 | kinase | Activated | 2.63 | | 8.24E-09 | CD69,CXCL3,CXCL8,CXCR4,ELANE,IFI44L,IFIT1,IFIT3,IFITM1,IL10RA,ISG15,JUN,MX1,MX2,OAS2,OAS3,STAT1 |
| FCER1G | -1.03 | transmembrane receptor | Activated | 2.613 | | 0.00000725 | CCL3,CCL5,CD69,CXCL8,IFIT2,MX1,PTGS2,RSAD2 |
| CCL5 | 2.94 | cytokine | Activated | 2.611 | bias | 1.32E-09 | CCL3,CCL3L3,CCL5,CCR1,CXCL3,CXCL8,DUSP1,DUSP6,FOS,IER2,NAMPT,SGK1,STAT1,TUBB4B,ZFP36 |
| DNASE2 | -1.26 | enzyme | Activated | 2.606 | bias | 5.45E-10 | CP,DHX58,IFIT3,IRF7,ISG15,OAS1,OAS3,RSAD2,RTP4,TNFSF10,USP18 |
| CG | | complex | Activated | 2.595 | bias | 7.94E-14 | ACTA2,ADM,AREG,BTG1,BTG2,CCDC86,CLU,CXCL3,CXCL8,CXCR4,DUSP1,DUSP6,EGR1,ENPP2,EREG,FAS,FOS,GASK1B,GDF10,HMGA2,IER3,IFIT3,INHBA,JUN,JUNB,KLF4,LGALS3BP,MCAM,MCL1,MMP2,NR4A1,PHLDA1,PRKG2,PRSS2,PTGS2,RGS2,SDC4,SPRY2,TM4SF1 |
| CRH | | cytokine | Activated | 2.564 | bias | 0.00158 | CXCL8,FOS,JUNB,NR4A1,PNMT,SGK1,SOCS3 |
| formaldehyde | | chemical - endogenous mammalian | Activated | 2.548 | bias | 0.00000249 | EGR1,EGR2,FOS,JUN,JUNB,JUND,PTGS2,SGK1 |
| MEF2C | -1.61 | transcription regulator | Activated | 2.537 | bias | 0.000044 | CCL3L3,COL2A1,CXCR4,EGR2,FOS,JUN,JUNB,JUND,KLF2,NR4A1,PTGS2,SIK1/SIK1B,ZFP36 |
| SOCS1 | 3.69 | other | Activated | 2.523 | bias | 4.56E-17 | BAK1,CCL3,CD69,CXCL8,DDX58,DUSP1,FAS,FKBP5,FOS,GBP5,IFI44,IFIH1,IFIT1,IFIT2,IFIT3,IRF7,ISG15,JUN,MX1,OAS1,OAS2,PTGS2,RSAD2,SOCS1,SOCS3,STAT1,TRIM22,USP18 |
| IKZF3 | 2.45 | transcription regulator | Activated | 2.52 | bias | 0.0000143 | CAV2,DDX58,FN1,IFI6,IFIT3,IRF7,KLF4,MYC,RTP4,SCN9A |
| MAP2K1/2 | | group | Activated | 2.519 | bias | 0.00000599 | CCL3L3,CXCL3,DUSP1,EGR1,EGR2,FOS,HSPB1,JUN,MCL1,MYC,PHLDA1 |
| GAPDH | -1.53 | enzyme | Activated | 2.517 | | 2.83E-12 | C2,CCL3,DUSP1,HLA-C,IFI6,IFIT2,IFITM1,OAS1,OAS2,OAS3,STAT1,TXNIP,UBE2L6 |
| IKZF1 | 1.44 | transcription regulator | Activated | 2.512 | bias | 0.000000201 | ADAM19,AXL,CAV2,CCL3L3,CCR1,DNAJC6,DTX3L,EPSTI1,FN1,FUT10,IFI6,IFIT3,KLF1,KLF4,LIG4,MYC,NFE2,PGAM1,POU2F2,RTP4,SCN9A,TNFAIP8L2,TXNIP |
| Insulin | | group | Activated | 2.493 | | 0.000192 | CAV2,DUSP1,EGR1,EGR2,ENPP2,FN1,FOS,HIF3A,IFIT2,IFIT3,INHBA,IRF7,JUN,KLF4,LGALS3BP,MMP2,MT-ND1,MYC,NR0B1,NR4A1,NR4A3,PER2,PRKAA2,PRKCD,SGK1,STAT1,STAT2 |
| leukotriene C4 | | chemical - endogenous mammalian | Activated | 2.446 | bias | 0.00000289 | CXCL8,EGR1,EGR2,EGR3,FOS,PTGS2 |
| EGF | | growth factor | Activated | 2.435 | bias | 2.5E-11 | AQP3,AREG,BAK1,BTG2,CAVIN2,CLCN3,CLDN7,CLU,CSF1R,CXCL8,CXCR4,DUSP1,DUSP6,EGR1,EGR2,EGR3,ENAH,EREG,FN1,FOS,ID1,IDS,IER2,IER3,INHBA,ITGA2,JUN,JUNB,JUND,MCL1,MMP2,MYC,NEXN,NR4A1,NR4A3,PER1,PLSCR1,PTGS2,REN,RHOB,SOCS3,SPRY2,STAT1,ZFP36 |
| EPHB1 | | kinase | Activated | 2.429 | bias | 0.000000123 | EGR1,EGR2,FOS,JUN,JUNB,PTGS2 |
| CNOT7 | -1.63 | transcription regulator | Activated | 2.425 | bias | 2.79E-16 | CMPK2,HERC6,IFI44L,IFI6,IFITM1,ISG15,LGALS3BP,OAS1,OAS2,OAS3,PLSCR1,SP110,STAT1,TAP1,UBE2L6 |
| CHRM1 | | G-protein coupled receptor | Activated | 2.415 | bias | 0.000000438 | EGR1,EGR2,EGR3,EGR4,FOS,JUN |
| Il3 | | cytokine | Activated | 2.404 | | 7.24E-08 | ADAM19,COL2A1,CSF1R,EGR1,ELANE,FAS,FOS,MCL1,MYC,PLA2R1,SOCS1,TNFRSF1A,ZNF346 |
| CD14 | -4.5 | transmembrane receptor | Activated | 2.391 | bias | 0.0000172 | CCL3,CCL3L3,CCL5,CXCL8,IFIT1,IL10RA,PTGS2,SOCS1,SOCS3 |
| chenodeoxycholic acid | | chemical - endogenous mammalian | Activated | 2.387 | | 0.027 | ABCG1,CXCL8,EGR1,FBP1,PTGS2,TK1 |
| SMAD3 | 1.13 | transcription regulator | Activated | 2.383 | bias | 0.000014 | ACTA2,AREG,ARID3A,CCL3L3,COL2A1,CXCL3,EGR1,EREG,FN1,FOS,ID1,ID3,JUN,JUNB,JUND,MYC,PNMT,PTGS2,RHOB,ZFP36 |
| deoxycholate | | chemical - endogenous mammalian | Activated | 2.382 | | 0.00199 | CXCL8,EGR1,MCL1,NR4A1,PTGS2,SLC1A3 |
| IRF8 | -1.51 | transcription regulator | Activated | 2.37 | | 2.47E-13 | ARID5B,CCL5,CSF1R,DDX58,EGR1,EGR2,FAS,FOS,GBP1,GFI1B,ID3,IFI44L,IFI6,IFIT2,IFIT3,ISG15,KLF4,KLF6,MYC,OAS1,STAT1,STAT2,TK1,TNFSF13B |
| Growth hormone | | group | Activated | 2.359 | | 0.000134 | ARID5B,CLU,EGR1,FKBP5,FN1,FOS,ID1,IER3,JUN,JUNB,MYC,NAMPT,SGK1,SOCS1,SOCS3,TNFSF10,TXNIP |
| CREB1 | -1.6 | transcription regulator | Activated | 2.358 | | 1.35E-11 | ADM,AK4,BAG3,BTG2,CCL3,CCNA1,CXCL8,CXCR4,DUSP1,EGR1,EGR2,EGR4,ENTPD1,FN1,FOS,FUT7,GPR12,HECA,HLA-G,ID1,IER2,IER3,INHBA,IRF7,JUN,JUNB,KLF4,KLF7,MCAM,MCL1,MYC,NEURL1B,NR4A1,NR4A3,PER1,PER2,PLA2G4A,PNMT,PTGS2,RASGEF1B,REN,RGS2,SERTAD1,SGK1,TIPARP,ZFP36 |

| | | | | | | | |
|---|---|---|---|---|---|---|---|
| RAF1 | -1.3 | kinase | Activated | 2.349 | bias | 1.26E-08 | AREG,BTG1,CXCL3,DUSP2,DUSP6,EGR1,EGR3,FAS,FOS,HMGA2,HSPB1,ID3,IER2,IER3,ITGA2,JUN,MYC,NINJ1,PHLDA1,PTGS2,RGS1,SPRY2,TXNIP |
| PRDM16 | -2.03 | transcription regulator | Activated | 2.321 | | 1.78E-08 | CCL5,GBP4,IFI44,IFIT2,IRF7,MX2,OAS2,OAS3,STAT1,STAT2 |
| IGF1 | | growth factor | Activated | 2.316 | bias | 1.37E-10 | ACTA2,ADM,AKR1C1/AKR1C2,BAK1,BTG2,CAVIN2,CCL3L3,CCL5,CLU,COL2A1,CXCL8,DUSP1,DUSP6,EGR1,EGR2,EIF2AK2,FN1,FOS,ID1,IER2,IER3,IFITM3,JUN,JUNB,KLF6,MCL1,MGA,MMP2,MYC,NFE2,NOG,NR4A1,PHLDA1,PTGS2,SGK1,SOCS3,TAP1,TK1,TXNIP,ZFP36 |
| NRAS | -1.13 | enzyme | Activated | 2.307 | | 1.33E-08 | CCL5,CD48,CFH,DUSP2,DUSP6,EGR1,FN1,HOMER3,IFIH1,IFIT1,IRX3,ISG15,ITGA2,LY86,MCL1,PHLDA3,PTGS2,RHOB,SESN1,STAT1,TAP1,USP18 |
| CREBBP | -1.34 | transcription regulator | Activated | 2.286 | bias | 2.03E-13 | CCL5,COL2A1,CXCL8,CXCR4,DUSP1,EDARADD,EGR1,EGR2,EGR3,ELANE,EPSTI1,FOS,FUT7,GFI1B,GNG2,HLA-G,IL10RA,ISG15,JUN,JUND,KLF2,LGALS3BP,LIG4,MGA,MYC,NR4A1,NR4A3,OAS3,PRKCD,PTGS2,REN,RGS2,RSAD2,RTP4,SDC4,SOCS3,TIPARP,TNFAIP8L2,TNFSF10,USP18 |
| ERK | | group | Activated | 2.275 | bias | 1.18E-11 | ACTA2,AREG,AXL,BTG2,CCR1,CLU,CXCL8,DUSP1,EGR1,EGR2,EIF2AK2,EREG,ESRRA,FAS,FN1,FOS,IER3,JUN,JUNB,MAFF,MAL,MCAM,MCL1,MMP2,MYC,OSBPL10,PTGS2,STAT1,ZFP36 |
| ERK1/2 | | group | Activated | 2.274 | bias | 9.72E-09 | AREG,CCL3,CCL3L3,CCL5,COL2A1,CXCL3,CXCL8,CXCR4,DUSP1,EGR1,FKBP5,FN1,FOS,ID1,ID3,IFIT1,ITGA2,JUN,JUNB,JUND,MMP2,MYC,NAMPT,NR4A3,PTGS2,SGK1,SOCS3,TAP1 |
| GNRH1 | | other | Activated | 2.271 | | 0.000106 | EGR1,EGR2,FOS,INHBA,JUN,JUNB,PRKCD,PTGS2 |
| STAT5A | -1.47 | transcription regulator | Activated | 2.269 | | 0.000619 | ABCG1,ASCL2,CCL3L3,FAS,FOS,GASK1B,ID1,MCL1,MMP7,MYC,OAS1,SOCS1,SOCS3,TLE4,TM4SF1,TNFRSF1A,TXNIP,ZFP36 |
| NORAD | -1.25 | other | Activated | 2.236 | bias | 0.00254 | ID3,JUN,JUNB,RHOB,SGK1 |
| CD244 | -1.57 | transmembrane receptor | Activated | 2.236 | bias | 0.00000134 | CCL3,CCL3L3,CCL5,EGR1,FOS |
| miR-204-5p (and other miRNAs w/seed UCCCUUU | | mature microRNA | Activated | 2.236 | bias | 0.0223 | BMPR1B,HMGA2,LTBP1,MYC,NOG |
| GNAI3 | 1.25 | enzyme | Activated | 2.236 | bias | 0.000167 | CCL3,CCL3L3,CCL5,CXCL3,CXCL8 |
| GNRH | | group | Activated | 2.233 | bias | 0.000349 | DUSP1,EGR1,FOS,JUN,PTGS2,SGK1 |
| norepinephrine | | chemical - endogenous mammalian | Activated | 2.225 | bias | 0.0000419 | CITED4,CXCL8,CXCR4,DUSP1,EGR1,FN1,FOS,ID1,MCAM,MMP2,NR4A1,NR4A3,PER1,PTGS2,SGK1 |
| ISG15 | -2.48 | other | Activated | 2.222 | bias | 0.000000233 | DDX58,IFI6,IFITM3,MX1,OAS1 |
| PAX5 | | transcription regulator | Activated | 2.213 | | 0.016 | CSF1R,FN1,MMP2,PGAM1,TXNIP |
| BMP15 | | growth factor | Activated | 2.207 | | 0.000264 | AREG,EREG,ID1,ID3,PTGS2 |
| Pkc(s) | | group | Activated | 2.205 | bias | 1.79E-08 | ACTA2,ADM,AQP3,CXCL8,CXCR4,DUSP1,EGR1,EGR2,FAS,FN1,FOS,JUN,JUNB,KLF6,MMP2,NR4A1,NR4A3,PER1,PER2,PHLDA1,POU2F2,PTGS2,RGS2 |
| PF4 | 1.46 | cytokine | Activated | 2.196 | bias | 0.00224 | ACTA2,CCL3,CXCL3,CXCL8,KLF4 |
| Raf | | group | Activated | 2.194 | | 0.000000191 | AREG,DUSP6,EGR1,FOS,HMGA2,IER2,IER3,JUN,MAFF,MCL1,MYC,SPRY2 |
| TREX1 | | enzyme | Activated | 2.19 | bias | 0.000028 | IFI44,IFIT2,ISG15,OASL,USP18 |
| VDR | 1.2 | transcription regulator | Activated | 2.183 | | 0.000173 | ACTA2,CXCL8,EGR1,ID1,IER3,IFI44L,ISG15,JUN,MYC,NOD2,PTGS2,REN,RGCC,SOCS1,SOCS3,TK1,TXNIP |
| ENG | -1.48 | transmembrane receptor | Activated | 2.18 | | 0.0036 | ACTA2,CXCR4,FN1,ID1,MYC |
| TRH | -1.29 | other | Activated | 2.169 | | 0.00254 | DUSP1,FOS,JUN,JUNB,NR4A1 |
| CGA | | other | Activated | 2.158 | | 0.00572 | CXCL8,EGR2,ID1,KLF9,MMP2,PTGS2,SGK1,TP53INP1 |
| ADRB | | group | Activated | 2.143 | | 0.000000914 | AXL,BAK1,CAVIN2,FAS,FOS,IFIT3,JUN,JUNB,KLF4,LTBP1,NR4A1,NR4A3,OASL,PTGS2,RSAD2,SLC40A1 |
| MIF | | cytokine | Activated | 2.14 | bias | 0.0000192 | ACTA2,C3AR1,CCL3L3,CCR1,CXCL3,CXCL8,CXCR4,DUSP1,FOS,JUN,MMP2,PTGS2,TRIM22 |
| IL33 | | cytokine | Activated | 2.133 | bias | 0.000255 | ABCG1,AREG,AXL,CCL3,CCL3L3,CCL5,CD69,CXCL3,CXCL8,DUSP2,EGR2,FAS,HCK,JUNB,POU2F2,SOCS3,TNFSF10 |
| SP110 | -2.6 | transcription regulator | Activated | 2.132 | | 1.49E-11 | AQP3,CD3E,CLU,EGR3,IFI6,IFIH1,IFIT1,IFIT3,IFITM1,IFITM3,LMNA,MCL1,MX1,MYC,OAS1,OAS3,PLSCR1,PTAFR,STAT1,TCF7,TNFRSF1A,TXNIP |
| LEP | | growth factor | Activated | 2.071 | | 0.00000154 | ACADL,ACTA2,ADAM8,AKR1C1/AKR1C2,AQP3,BAK1,CCL3L3,CCL5,COL2A1,DUSP6,EGR1,EGR2,FAS,FOS,JUN,JUNB,JUND,MCL1,MMP2,MMP7,MT-ND1,MYC,NAMPT,PER2,PRDM16,PRKAA2,PTGS2,SOCS3,TNFRSF1A,TNFSF10,ZFP36 |
| Jnk | | group | Activated | 2.07 | bias | 1.56E-11 | ACTA2,BTG1,BTRC,CCL3,CCL5,CD69,CRIP2,CXCL3,CXCL8,CXCR4,DUSP1,EGR1,FAS,FN1,FOS,JUN,JUND,KLF6,MCL1,MMP2,MYC,NR4A1,PLSCR1,PTGS2,RARG,SOCS3,ZFP36 |
| LDL | | complex | Activated | 2.063 | bias | 3.43E-12 | ABCG1,AXL,CCL3,CCL5,CXCL3,CXCL8,DUSP1,DUSP2,EGR1,EGR3,FAS,FOS,GNAI1,IFIT2,IFITM1,IL10RA,IRF2,IRF7,JUN,MCL1,MMP2,MX1,NAMPT,NR4A1,NR4A3,PTAFR,PTGS2,SOCS1,SOCS3,TNFAIP8L2,TNFSF10 |
| PTGER4 | 1.26 | G-protein coupled receptor | Activated | 2.058 | | 1.29E-15 | CCL3L3,CD69,CMPK2,CP,CXCL8,CXCR4,DDX58,EGR1,GBP4,HERC6,IFIH1,IFIT2,IRF7,MAFF,MMP2,PARP14,PTGS2,RNF24,RSAD2,RTP4,SAP30,SLFN5,STXBP4,TNFSF10,TP53INP1,USP18,XAF1 |
| MAPK8 | -1.05 | kinase | Activated | 2.043 | | 0.0000112 | BTRC,CCL5,CXCL8,DUSP1,DUSP6,EGR1,FOS,JUN,JUND,KLF6,MMP2,MMP7,MYC,PTGS2,TSC22D3,ZFP36 |
| IL3 | | cytokine | Activated | 2.037 | bias | 2.69E-08 | APBA2,AREG,BTG2,CD3G,CD48,CD69,CSF1R,CXCL8,CXCR4,EGR1,EGR2,EGR3,EIF2AK2,FAS,FOS,JUN,JUNB,KLF1,KLF9,MCL1,MYC,RPL7,RPS7,SOCS1,SOCS3,TAL1,TK1,TNFSF10,ZBTB7A |
| FOXL2 | | transcription regulator | Activated | 2.03 | bias | 9.58E-08 | CCL3,CCL3L3,CXCL3,FAS,FOS,IER3,INHBA,MAFF,NR4A3,PTGS2,RGS2,TNFRSF1A |
| MAP2K1 | -1.11 | kinase | Activated | 2.029 | bias | 0.00000334 | ACTA2,AREG,CCL3L3,CCL5,CXCL3,CXCL8,DUSP1,DUSP6,EGR2,FOS,JUN,JUND,MMP2,MYC,PLA2G4A,PTGS2,TK1,TNFSF10 |
| PDX1 | | transcription regulator | Activated | 2.021 | | 0.000439 | CXCR4,DUSP6,EGR1,GCH1,ID1,ID3,IER3,JUN,JUNB,KLF6,MT-ND1,MYC,RSAD2,TXNIP |
| TCR | | complex | Activated | 2.009 | bias | 2.04E-22 | ADM,BATF,CCL3,CCL3L3,CCL5,CCR1,CD69,CXCL8,CXCR4,DDX58,DHX58,EGR1,EGR2,FAS,FOS,GBP4,HERC5,ID3,IFI44,IFI44L,IFI6,IFIH1,IFIT1,IFIT2,IFIT3,IRF7,ISG15,JUN,JUNB,KLF2,MCL1,MX1,MX2,MYC,NHP2,NR4A1,OAS1,OASL,PGAM1,PIK3R3,RPL7,RPL9,RPS23,RSAD2,SOCS1,SOCS3,STAT1,TLE3,<br>TNFRSF1A,TNFSF13B,TSC22D3 |
| BRD4 | -1.51 | kinase | Inhibited | -2.043 | bias | 0.00034 | ACTA2,BTN3A2,CCR1,CXCR4,FN1,FOS,MAL,MCL1,MORC1,MYC,NAV1,ZNF485 |
| MSC | | transcription regulator | Inhibited | -2.111 | | 0.000000205 | DDX60,EPSTI1,FUT7,GPAT3,IFI44,IFI44L,IFIT1,IRF7,PIEZO2,POU2F2,XAF1 |
| JAK1/2 | | group | Inhibited | -2.138 | bias | 0.000496 | CD69,EIF2AK2,GBP5,HLA-DRB5,IRF7,ISG15,MX1,RSAD2 |
| FADD | | other | Inhibited | -2.152 | bias | 1.04E-13 | CXCL8,CXCR4,DDX58,DHX58,EGR1,EIF2AK2,FAS,FOS,IFIH1,IFIT2,IRF7,JUN,JUNB,KLF6,MYC,PARP11,SOCS1,STAT1,STAT2,TIPARP |
| EPCAM | -1.6 | other | Inhibited | -2.177 | | 0.000211 | EGR1,FOS,ID1,JUN,MYC |
| ISGF3 | | complex | Inhibited | -2.2 | bias | 2.31E-09 | EIF2AK2,IFIH1,IFIT2,IRF7,ISG15,RSAD2,TNFSF10 |
| NLRC5 | -1.9 | transcription regulator | Inhibited | -2.207 | bias | 0.000167 | DDX58,HLA-C,HLA-F,HLA-G,TAP1 |
| IFNAR1 | -1.44 | transmembrane receptor | Inhibited | -2.232 | bias | 2.88E-21 | AXL,CCL3L3,CCL5,CMPK2,CXCL3,DUSP6,EIF2AK2,GBP4,IFI44,IFI6,IFIH1,IFIT2,IFIT3,IFITM1,IRF7,ISG15,MX2,MYC,OAS1,OAS2,OAS3,OASL,PTGS2,RSAD2,RTP4,SOCS1,SOCS3,STAT1,TNFSF10,TNFSF13B,TRIM22,USP18,XAF1 |
| IFNK | | cytokine | Inhibited | -2.236 | bias | 0.000264 | EIF2AK2,IFIH1,MX1,OAS1,STAT1 |
| TNK1 | | kinase | Inhibited | -2.236 | bias | 0.0000553 | IFIH1,IFIT2,IRF7,OAS2,TNFSF10 |
| MEOX2 | | transcription regulator | Inhibited | -2.236 | bias | 0.00321 | CD69,CXCL3,CXCL8,ID1,ID3 |
| DUSP5 | -1.15 | phosphatase | Inhibited | -2.236 | bias | 0.00013 | DUSP2,EGR1,EGR3,EGR4,ZFP36 |
| SASH1 | | other | Inhibited | -2.324 | bias | 3.26E-10 | CMPK2,HELZ2,IFIT2,IFIT3,INHBA,IRF7,ISG15,LMO4,OASL,PTGS2,RSAD2,SAP30,SOCS1,STAT1,STAT2 |
| BRCA1 | 1.12 | transcription regulator | Inhibited | -2.337 | | 5.4E-11 | AREG,ASCL2,BAK1,CCL5,DDX58,DUSP3,EGR1,ENPP2,FAS,FKBP5,HMGA2,HSPB1,IFI6,IFIT1,IFIT2,IFIT3,IFITM1,IRF7,MX1,MYC,PHLDA1,PLSCR1,STAT1,TAP1 |
| IFNE | | cytokine | Inhibited | -2.343 | bias | 0.00000129 | HERC5,IFIH1,IFIT2,IFITM3,ISG15,MX2,PTGS2,STAT1,USP18 |
| DACH1 | -1.38 | transcription regulator | Inhibited | -2.377 | | 0.00000067 | CXCL3,CXCL8,EGR1,FOS,ID1,IER2,JUN,KLF4,PTGS2 |
| MUC1 | 1.33 | other | Inhibited | -2.381 | bias | 0.00674 | CXCL8,IFIT3,IFITM1,MYC,OASL,STAT1 |
| SGPL1 | -1.15 | enzyme | Inhibited | -2.401 | | 0.0000206 | DDX58,IFIT1,ISG15,MX1,OAS1,PTGS2 |
| NLRP12 | | other | Inhibited | -2.408 | | 0.000299 | CXCL8,CXCR4,HLA-C,HLA-F,HLA-G,JUN |
| LYN | -1.35 | kinase | Inhibited | -2.408 | | 0.00176 | BAK1,CCL3L3,CCL5,EGR1,KLF1,MYC,SOCS1,SOCS3 |
| CLEC12A | -1.44 | other | Inhibited | -2.425 | bias | 2.04E-09 | IFIT2,IFIT3,IRF7,ISG15,RSAD2,USP18 |
| mir-181 | | microRNA | Inhibited | -2.433 | bias | 0.000898 | CD69,CXCL8,DUSP6,KLF6,MCL1,PRKCD,PTGS2 |
| IFNAR2 | -1.17 | transmembrane receptor | Inhibited | -2.449 | bias | 1.66E-14 | DDX58,GBP4,HERC5,IFI44,IFI6,IFIH1,IFITM1,ISG15,MX2,OAS1,OAS2,TNFSF10,TRIM22,UBE2L6,USP18,XAF1 |
| TGM2 | 1.11 | enzyme | Inhibited | -2.522 | | 2.58E-17 | BTG2,CCDC169,CCL3,CXCL8,CYP2R1,DDX60,FERMT1,FFAR3,FN1,GCA,IFI6,IFIT1,IFIT2,IFIT3,IL10RA,MMP2,MYC,NLN,OAS1,OAS2,OAS3,OASL,PARP14,PARP9,PHLDA1,PLSCR1,SAMD9L,SAMHD1,SLFN5,SP110,STAT1,TAP1,TRIM22,XAF1,ZNF146 |
| EBI3 | | cytokine | Inhibited | -2.576 | bias | 0.00000156 | ENTPD1,FOS,HLA-C,LAG3,MX1,MYC,STAT1,STAT2,TAP1,TNFSF10 |
| IRF9 | 1.01 | transcription regulator | Inhibited | -2.592 | | 1.2E-16 | CD69,CXCL8,GBP1,IFIT1,IFIT2,IFIT3,IFITM1,IFITM3,IRF7,ISG15,MX1,OAS2,RTP4,SOCS1,SOCS3,STAT1,STAT2,TNFSF10 |
| POU2AF1 | | transcription regulator | Inhibited | -2.596 | bias | 0.00012 | CD69,ID3,IFI44,IFI44L,IFIT3,IFITM1,KCNN4,MX1 |
| PTX3 | 1.88 | other | Inhibited | -2.621 | bias | 0.00000926 | CCL3L3,CCL5,CXCL8,EGR2,EGR3,FOS,JUN |
| BTNL2 | | transmembrane receptor | Inhibited | -2.63 | | 0.00331 | CDKN2C,ENAH,ENTPD1,GBP4,IFITM3,KCNK6,TNFSF10 |

| | | | | | | | |
|---|---|---|---|---|---|---|---|
| SPI1 | -1.56 | transcription regulator | Inhibited | -2.679 | | 3.71E-20 | ACTA2,ARID3A,C3AR1,CCL3,CD3E,CMPK2,CSF1R,CSF2RB,DUSP6,EGR2,ELANE,EVI5,FOS,FUT7,HELZ2,ID3,IFI44,IFI44L,IFI6,IFIT1,IFIT2,IFIT3,IFITM1,IFITM3,IRF7,ISG15,JDP2,JUN,KLF4,MCL1,MMP2,MX1,MYC,OASL,PTGS2,RSAD2,SP110,TK1,TNFSF10,TRIM22,USP18 |
| SHC1 | 1.12 | other | Inhibited | -2.688 | | 0.000333 | BAK1,CD69,EGR1,FOS,HMGB2,KLF2,PHLDA1,SDC4 |
| IFNA4 | | cytokine | Inhibited | -2.799 | bias | 9.84E-11 | AP1S3,CD69,GBP5,IFIH1,IFIT1,IFIT2,ISG15,MX1,MYC,OASL,PIK3AP1,RSAD2,USP18 |
| IFNL4 | | cytokine | Inhibited | -2.807 | bias | 8.44E-12 | DDX58,DHX58,IFIH1,ISG15,MX1,OAS1,OAS2,STAT1 |
| VCAN | -1.97 | other | Inhibited | -2.824 | | 6.32E-12 | ADM,CBX6,CEACAM4,CLU,CXCL3,CXCL8,ENPP2,FAS,FN1,IFI44,IFI44L,IFI6,IFIT1,IFIT2,IFITM1,MX1,MX2,OAS2,OAS3,PARP14,RGS2,STAT1,TRIM22,XAF1 |
| IFN type 1 | | group | Inhibited | -2.864 | bias | 1.84E-12 | CCL5,CD69,CXCL8,DDX58,DHX58,EIF2AK2,IFIH1,IFIT1,IFIT2,ISG15,OASL,STAT1,STAT2,TNFSF10,TNFSF13B |
| EIF2AK2 | -2.17 | kinase | Inhibited | -2.943 | bias | 6.88E-14 | CXCL8,DDX58,EGR1,EIF2AK2,FAS,FOS,IFI6,IFIT1,IFITM1,ISG15,LGALS3BP,MYC,OAS1,OAS3,PARP9,PLSCR1,SAMHD1,SOCS3,STAT1,UBE2L6,USP18 |
| DUSP1 | 3.05 | phosphatase | Inhibited | -2.952 | | 5.45E-08 | CCL3L3,CMPK2,CXCL3,CXCL8,DUSP1,DUSP6,HELZ2,IER3,IFIT1,IFIT3,JUN,MX1,PAQR7,PTGS2,SOCS1,ZFP36 |
| FOSL1 | -1.42 | transcription regulator | Inhibited | -3.018 | | 0.0000263 | ADM,AXL,CCL3L3,CXCL8,EGR1,EGR2,FOS,JUN,JUNB,MMP2 |
| MAVS | -1.64 | other | Inhibited | -3.087 | bias | 4.1E-17 | CCL5,CMPK2,CXCL8,DDX58,DHX58,IFIT1,IFIT2,IFIT3,IFITM3,IRF7,ISG15,OAS1,OAS2,OASL,RSAD2,SOCS1,SOCS3,STAT1,STAT2,UBE2L6,USP18 |
| PARP9 | -4.17 | enzyme | Inhibited | -3.121 | bias | 4.66E-12 | IFI44,IFIT1,IFIT2,IFIT3,IRF2,IRF7,ISG15,OAS2,SP110,STAT1 |
| JAK | | group | Inhibited | -3.317 | bias | 2.11E-12 | DDX58,EIF2AK2,IFI6,IFIH1,IFIT1,IFIT2,IFIT3,IFITM3,ISG15,PTGS2,RSAD2,SOCS1,SOCS3,STAT1 |
| IFNA1/IFNA13 | | cytokine | Inhibited | -3.477 | bias | 4.76E-19 | CD69,DHX58,EIF2AK2,IFI6,IFIH1,IFIT1,IFIT2,IFITM1,IRF7,ISG15,MX1,MYC,OAS1,OAS2,PLSCR1,PTGS2,RSAD2,SOCS1,STAT1,STAT2,UBE2L6 |
| IRF5 | -1.64 | transcription regulator | Inhibited | -3.638 | bias | 3.54E-23 | BAK1,CCL3,CCL5,CMPK2,CXCR4,DDX58,DHX58,IFI44,IFIH1,IFIT1,IFIT2,IFIT3,IFITM3,IRF7,ISG15,MYC,NAMPT,OAS1,OAS2,OASL,PLSCR1,PTGS2,RSAD2,SP110,STAT1,STAT2,TNFSF10,UBE2L6 |
| PML | -1.19 | transcription regulator | Inhibited | -3.658 | | 9.41E-12 | ACADL,CCNA1,CXCR4,DDX60,EPSTI1,FAS,HERC6,HSPB1,IFI44,IFI44L,IFIH1,IFIT1,IFIT3,IFITM1,IRF7,ISG15,MAFF,MX1,OAS1,OAS2,OAS3,PLSCR1,STAT1,TAP1 |
| STAT1 | -4.89 | transcription regulator | Inhibited | -3.829 | bias | 1.66E-32 | APOL4,AXL,BTG1,CCL3,CCL3L3,CCL5,CMPK2,CXCL3,CXCL8,DDX60,EGR1,EIF2AK2,EPSTI1,FAS,FOS,GBP1,GBP1P1,GBP4,GBP5,HAPLN3,HERC6,HLA-DRB5,IFI44,IFI44L,IFI6,IFIH1,IFIT1,IFIT2,IFIT3,IFITM1,IFITM3,IRF2,IRF7,ISG15,JUN,KLF4,MX1,MYC,NFE2,OAS1,OAS2,OAS3,OASL,PARP9,PLSCR1,PTGS2,RSAD2,RTP4,SAMD9L,SAMHD1,SLFN5,SOCS1,SOCS3,SP110,STAT1,STAT2,TAP1,TNFSF10,TNFSF13B,TRIM22,USP18,XAF1 |
| Ifn | | group | Inhibited | -3.934 | bias | 6.97E-16 | CD69,CFH,DDX58,DHX58,EIF2AK2,FAS,FOS,IFIH1,IFIT1,IFITM1,IFITM3,IL10RA,IRF7,ISG15,MX1,OAS2,OAS3,OASL,PTAFR,RSAD2,SOCS1,STAT1,TAP1,TNFSF13B,TRIM22 |
| RNY3 | | other | Inhibited | -4.123 | bias | 5.78E-21 | DDX60,EPSTI1,HERC5,IFI44,IFI44L,IFIT1,IFIT3,IFITM3,ISG15,MX1,OAS1,OAS2,OAS3,OASL,RSAD2,RTP4,XAF1 |
| IFN Beta | | group | Inhibited | -4.147 | bias | 2.98E-25 | BAK1,C3AR1,CCL3,CCL5,CD69,CXCL8,DDX58,DUSP1,EIF2AK2,FN1,HERC5,HLA-G,IFI44,IFI6,IFIH1,IFIT1,IFIT2,IFIT3,IFITM1,IRF7,ISG15,MX1,MX2,MYC,OAS1,OAS2,OAS3,OASL,RSAD2,SOCS1,SPRY2,STAT1,STAT2,TNFSF10,USP18,XAF1 |
| Ifnar | | group | Inhibited | -4.247 | bias | 2.41E-15 | AXL,CCL5,DDX58,EIF2AK2,HLA-G,IFIH1,IFIT2,IFIT3,IFITM3,IRF7,ISG15,OAS1,OAS2,OASL,RSAD2,STAT1,STAT2,TAP1,TNFSF10,UBE2L6,USP18,XAF1 |
| PRL | | cytokine | Inhibited | -4.291 | bias | 3.1E-27 | CD69,CLU,CMPK2,DDX58,DHX58,DTX3L,EGR1,EIF2AK2,EPSTI1,FN1,FOS,GMPR,HELZ2,HERC5,HERC6,ID1,ID3,IER3,IFI44,IFI44L,IFI6,IFIH1,IFIT1,IFIT3,IFITM1,IRF7,ISG15,JUN,MX2,MYC,OAS1,OAS2,OAS3,PAQR7,PAQR8,PARP14,PLSCR1,RSAD2,SAMD9,SAMD9L,SAMHD1,SOCS1,SOCS3,SP110,STAT1,STAT2,TNFSF13B,TRIM25,USP18,XAF1 |
| IFNB1 | | cytokine | Inhibited | -4.299 | | 2.06E-19 | APOL2,AXL,BAK1,BTN3A3,CCL3,CCL5,CMPK2,CXCL3,CXCL8,DDX58,DHX58,EIF2AK2,FOS,GBP4,GBP5,GBP7,HERC5,HMGB2,IFI6,IFIH1,IFIT1,IFIT2,IFIT3,IFITM1,IRF7,ISG15,MMP2,MX1,MYC,NOD2,OAS1,OAS2,PARP14,PTGS2,RSAD2,SOCS1,SPRY2,STAT1,STAT2,TNFSF10,USP18,XAF1 |
| IRF3 | -1.22 | transcription regulator | Inhibited | -4.366 | bias | 4.78E-26 | CCL3,CCL3L3,CCL5,CD69,CMPK2,CXCL8,DDX58,DDX60,DHX58,EIF2AK2,FAS,FN1,GBP1,GBP5,HELZ2,HLA-F,IFI44,IFI6,IFIH1,IFIT1,IFIT2,IFIT3,IFITM3,IRF7,ISG15,JUNB,LAG3,OAS1,OAS2,OAS3,OASL,PARP14,RPS7,RSAD2,SAMD9L,SAP30,STAT1,STAT2,TAP1,TNFSF10,UBE2L6,USP18 |
| IFNG | | cytokine | Inhibited | -4.404 | | 3.92E-35 | ACTA2,ADM,AGPAT1,AREG,BAK1,BTG1,BTN3A1,BTN3A2,C2,CCL3,CCL3L3,CCL5,CCNA1,CCR1,CMPK2,CP,CSF1R,CSF2RB,CXCL3,CXCL8,CXCR4,DDX58,DDX60,DTX3L,DUSP1,EGR1,EGR2,EGR3,EIF2AK2,FAS,FBP1,FCGR1B,FKBP5,FN1,FOS,GBP1,GBP4,GBP5,GBP7,GCH1,GMPR,GNG2,GPR12,HCK,HCP5,HERC6,HLA-C,HLA-DRB5,HLA-F,HLA-G,HSPA1A/HSPA1B,HSPB1,ID1,IER2,IER3,IFI44,IFI44L,IFI6,IFIH1,IFIT1,IFIT2,IFIT3,IFITM1,IFITM3,IL10RA,INHBA,IRF2,IRF7,ISG15,JAK3,JUN,JUNB,JUND,KLF1,KLF2,KLF4,KLF6,LAG3,LGALS3BP,MAFF,MMP2,MX1,MX2,MYC,NAMPT,NEAT1,NOD2,OAS1,OAS2,OAS3,OASL,PARP14,PARP9,PHLDA1,PLSCR1,PRKCD,PRKG2,PTAFR,PTGS2,RGCC,RHOB,RSAD2,RTP4,SAMD9,SAMHD1,SDC4,SLC12A2,SLC1A3,SLC40A1,SLFN5,SOCS1,SOCS3,SP110,STAT1,STAT2,TAP1,TNFRSF1A,TNFSF10,TNFSF13B,TRIM22,TSC22D3,TXNIP,UBE2L6,USP18,XAF1,ZFP36,ZKSCAN1 |
| IRF1 | -1.21 | transcription regulator | Inhibited | -4.631 | bias | 1.76E-27 | ADAM8,BAK1,CCL5,CMPK2,CXCL3,CXCL8,DDX58,EIF2AK2,HELZ2,HLA-G,IFI44,IFI44L,IFI6,IFIH1,IFIT1,IFIT2,IFIT3,IFITM1,IFITM3,IRF2,IRF7,ISG15,LIG4,MX1,MYC,NFE2,OAS1,OAS2,OAS3,OASL,PLAAT1,PTGS2,RSAD2,SOCS1,SP110,STAT1,STAT2,TAP1,TNFSF10,TNFSF13B,TRIM22,XAF1 |
| Interferon alpha | | group | Inhibited | -4.729 | bias | 2.83E-39 | ADAM19,APOL2,AQP3,AXL,BAK1,BTG2,C3AR1,CCL3,CCL5,CCNA1,CCR1,CD69,CSF2RB,CXCL3,CXCL8,DDX58,DHX58,DUSP6,EIF2AK2,ENPP2,EPSTI1,FAS,FBXO6,FOS,GBP1,GBP5,HELZ2,HERC5,HERC6,HLA-C,HLA-F,HLA-G,HSPA1A/HSPA1B,IFI44,IFI44L,IFI6,IFIH1,IFIT1,IFIT2,IFIT3,IFITM1,IFITM3,IL10RA,IRF7,ISG15,MCL1,MOB3C,MX1,MX2,MYC,OAS1,OAS2,OAS3,PARP14,PARP9,PLSCR1,PTGS2,RSAD2,RTP4,SAMD9,SAMD9L,SLFN5,SOCS1,SOCS3,SP110,SPRY2,STAT1,STAT2,TAP1,TNFSF10,TNFSF13B,TRANK1,TRIM22,UBE2L6,USP18,ZNRF3 |
| IRF7 | -4.33 | transcription regulator | Inhibited | -5.104 | bias | 1.14E-33 | CCL5,CCNA1,CD69,CMPK2,DDX58,DHX58,GBP1,GBP4,GBP5,HELZ2,HERC5,IFI44,IFI44L,IFI6,IFIH1,IFIT1,IFIT2,IFIT3,IFITM1,IFITM3,IRF7,ISG15,MCL1,MX1,MX2,NAMPT,OAS1,OAS2,OAS3,OASL,PARP14,PLSCR1,RSAD2,RTP4,SAMD9L,SAP30,SOCS1,STAT1,STAT2,TAP1,TNFSF10,TNFSF13B,TRIM22,UBE2L6,USP18,XAF1 |
| IFNA2 | | cytokine | Inhibited | -5.11 | bias | 1.74E-30 | BAG3,CCL3,CCL5,CD69,CMPK2,DDX58,DDX60,EIF2AK2,FAS,GBP1,GBP4,HERC5,HERC6,HLA-C,IFI44,IFI44L,IFI6,IFIH1,IFIT1,IFIT2,IFIT3,IFITM1,IFITM3,IRF7,ISG15,LGALS3BP,MAL,MX1,MX2,MYC,OAS1,OAS2,OAS3,PARP9,PLSCR1,PTAFR,RSAD2,SAMD9,SAMHD1,SOCS1,SOCS3,SP110,STAT1,TAP1,TNFRSF1A,TNFSF10,TRIM22,UBE2L6,USP18,XAF1 |
| IFNL1 | | cytokine | Inhibited | -5.521 | bias | 5.95E-36 | CMPK2,CXCL8,DDX58,DDX60,EIF2AK2,GBP1,GBP5,HERC5,HERC6,HLA-C,IFI44,IFI44L,IFI6,IFIH1,IFIT1,IFIT2,IFIT3,IFITM1,IFITM3,ISG15,LGALS3BP,MX1,OAS1,OAS2,OAS3,OASL,PLSCR1,RSAD2,RTP4,SAMD9,SP110,STAT1,STAT2,TRIM22,UBE2L6,USP18,XAF1 |

**T3 vs. T1 MEP_1**

© 2000-2020 QIAGEN. All rights reserved.

| Upstream Regulator | Expr Fold Change | Molecule Type | Predicted Activation State | Activation z-score | Flags | p-value of overlap | Target Molecules in Dataset |
|---|---|---|---|---|---|---|---|
| MAPK1 | -1.63 | kinase | Activated | 5.602 | | 2.73E-27 | APOE,APOL6,BST2,BTN3A3,CCND1,CD69,CHST2,CXCL8,DDX58,DEPTOR,DSE,DUSP1,EGR1,EGR2,EIF2AK2,EMP1,ENDOD1,ENO3,FOS,GMNN,HBA1/HBA2,HDC,HERC5,HLA-B,HLA-C,IFI35,IFI44,IFI6,IFIH1,IFIT1,IFIT2,IFIT3,IFITM1,IFITM3,IRF7,ISG15,ITGB3,JUN,JUNB,JUND,LGALS3BP,LMNA,MCL1,MITF,MX2,NR4A1,OAS1,OAS2,OAS3,OASL,PLSCR1,PTGS2,RGS2,SAMHD1,SERPINE1,SOCS2,SP110,SPRY2,STAT1,STAT2,TAP1,TNFSF10,TP53,TRANK1,TRIM14,TRIM21,TRIM5,TTC27,UBE2L6,USP18 |
| TRIM24 | -1.44 | transcription regulator | Activated | 4.63 | bias | 1.55E-17 | CMPK2,DDX58,DDX60,DHX58,EPSTI1,HERC6,IFI35,IFI44,IFIH1,IFIT2,IFIT3,IRF1,IRF7,ISG15,LGALS3BP,MOV10,OAS1,OASL,PCLAF,RTP4,SAMD9L,SAMHD1,SERPINE1,SOCS1,SOCS2,STAT1,STAT2,TAP1,TP53,USP18 |
| PDGF BB | | complex | Activated | 4.535 | bias | 2.42E-11 | AREG,BCL3,BHLHE40,CAV2,CCND1,CCND3,CD44,CSF1,CSRP2,CXCL8,DUSP1,DUSP6,EGR1,EGR2,EGR3,EREG,FOS,FOSL1,FOXC2,FZD2,GPR183,HOMER2,HSD3B7,JUN,JUNB,KLF6,LMNA,MCL1,NR4A1,NR4A2,NR4A3,PTGS2,RGS1,RGS2,RHOB,SERPINE1,SGK1,SLC29A1,SOCS3,SOD2,TP53,ZFP36 |
| PNPT1 | -1.78 | enzyme | Activated | 4.447 | bias | 8.89E-16 | CMPK2,CXCL8,DDX58,EIF2AK2,IFI44,IFIH1,IFIT3,IRF7,ISG15,LGALS3BP,OAS1,PARP14,PARP9,RTP4,SAMD9L,STAT1,STAT2,UBE2L6,USP18,XAF1 |
| SIRT1 | 1.06 | transcription regulator | Activated | 4.35 | | 8.89E-13 | CCND1,CCND3,CD36,CD74,CDKN2C,CMIP,CMPK2,DBP,DDX58,DDX60,DHX58,EPDR1,FOXC2,GSTM3,HCK,HELZ2,HGF,HK2,HLA-DRB5,ID1,IFI44,IFIT3,IFITM3,IRF7,KLF2,LGALS3BP,LY6E,MCL1,NEURL1,NFKBIE,NLRC5,OAS1,OAS2,PARP14,PER1,RSAD2,RTP4,SERPINE1,SFRP5,SLC29A1,SLC43A3,SOCS3,SOD2,SP110,STAT1,TAL1,TAP1,TFPI,TP53,USP18 |
| ACKR2 | | G-protein coupled receptor | Activated | 4.123 | bias | 1.64E-13 | DDX58,DDX60,DHX58,EIF2AK2,IFI44,IFIT2,IFIT3,IRF7,ISG15,OAS1,OAS2,OAS3,OASL,RSAD2,STAT1,STAT2,USP18 |
| NUPR1 | | transcription regulator | Activated | 3.763 | | 0.000000685 | AREG,ARHGEF26,ARMC7,BTG1,CTU1,CXCL8,CXCR4,DEPTOR,ENO2,EREG,FUCA1,GRAMD2B,HILPDA,HK2,HLTF,IFIT2,KIFC1,KLF4,KLF6,LBR,LMNB1,MX2,NUP50,PARP9,PMP22,PRNP,RAB20,RAB38,RASAL2,RMI1,SAMHD1,SERPINE1,SIK1/SIK1B,SPC25,SPDL1,SPTSSA,SYNE2,SYTL2,TCEANC2,TCTN2,TP53,TRERF1,UBIAD1,VANGL1 |
| Pkc(s) | | group | Activated | 3.729 | bias | 0.00000204 | AHR,CCND3,CD36,CD4,CXCL8,CXCR4,DBP,DUSP1,EGR1,EGR2,FOS,FOSL1,HPGD,JUN,JUNB,KLF6,NR4A1,NR4A2,NR4A3,PDIA4,PER1,PER2,PTGS2,RGS2,SERPINE1 |
| MAP3K8 | 1.62 | kinase | Activated | 3.649 | | 0.0000265 | APOE,ARRDC3,CIITA,CREBL2,CXCL8,DUSP6,FOS,IER5,JUN,JUNB,PTGS2,SESN1,SLA,SOCS3,STK17B,TICAM1,TXNIP,ZC3H6 |
| leukotriene D4 | | chemical - endogenous mammalian | Activated | 3.638 | bias | 1.71E-11 | CXCL8,CYSLTR1,DUSP1,EGR1,EGR2,EGR3,HLX,KLF2,KLF4,MCL1,NR4A1,NR4A2,NR4A3,PTGS2,RGS2,SIK1/SIK1B,ZFP36 |
| CG | | complex | Activated | 3.596 | bias | 0.000000392 | ADCYAP1,AREG,BHLHE40,BTG1,BTG2,CCDC86,CCND3,CLU,CXCL8,CXCR4,DUSP1,DUSP6,EGR1,EMP1,ENPP2,EREG,FOS,GDF10,HK2,HPGD,IFIT3,ITGB3,JUN,JUNB,KLF4,LGALS3BP,MCL1,NR4A1,PRKG2,PTGS2,RGS2,SCUBE1,SPRY2,TM4SF1,TP53,UNG |
| IL1RN | | cytokine | Activated | 3.559 | bias | 1.74E-14 | BTN3A1,CD44,CIITA,CXCL8,DDX58,HDAC9,HERC6,IFI44,IFI44L,IFI6,IFIH1,IFIT3,IRF1,IRF7,KLF6,MX1,MX2,OAS1,OAS2,OAS3,OASL,PTGS2,RSAD2,RTP4,SAMD9,SERPINE1,STAT2,TNFSF10,USP18 |
| NKX2-3 | 1.9 | transcription regulator | Activated | 3.396 | | 3.87E-20 | ANGPT2,BTG1,CASP1,CAVIN2,CD36,CDKN3,CMPK2,CXCL8,DDX58,DDX60,DHX58,EIF2AK2,FBXO6,GMPR,H2AC18/H2AC19,HCP5,HELZ2,HLA-B,HLA-C,HLA-F,IL7R,KLF2,KLF4,LY6E,PARP14,PARP9,PIM3,PLSCR1,PTGS2,RPS23,RTP4,SAMD9,SAMD9L,SP110,STAT1,STAT2,TAP1,TIPARP,TRIM21,TRIM5,TYMP,UBE2L6,UNC93B1,USP18,XAF1 |
| Lh | | complex | Activated | 3.336 | | 0.0000544 | ACTN1,AREG,ARL4C,CASP9,CDC42EP4,CREBL2,CXCL8,CXCR4,DHRS3,DUSP1,EREG,FKBP5,FOS,HK2,MCL1,PER1,PTGS2,RASAL2,REXO5,RHOB,SGK1,SOD2,STAT1,TP53 |
| CREB1 | -1.55 | transcription regulator | Activated | 3.242 | | 0.00000028 | ALAS1,APOE,ARL15,ATF1,BHLHE40,BTG2,CARS1,CCND1,CCND3,CD4,CIITA,CRHBP,CSRP2,CXCL8,CXCR4,DUSP1,EGR1,EGR2,FAM3B,FOS,GAD1,HLA-G,HPGD,ID1,IRF7,JUN,JUNB,KLF4,MCL1,NR4A1,NR4A2,NR4A3,PDE3B,PER1,PER2,PIM3,PLXNC1,PNMT,PTGS2,RASGEF1B,REN,RGS2,SGK1,SOCS2,SOD2,SYNJ2,TIPARP,TRPS1,ZFP36 |
| EGF | | growth factor | Activated | 3.205 | bias | 1.85E-08 | ACSL4,ACTN1,AREG,ARTN,BET1,BTG2,CASP1,CAVIN2,CCND1,CCND3,CD44,CLU,CXCL8,CXCR4,DUSP1,DUSP6,EGR1,EGR2,EGR3,ENAH,EREG,FOS,FOSL1,HGF,HPGD,ID1,IDS,IL7R,ITGB3,JUN,JUNB,JUND,MCL1,NR4A1,NR4A2,NR4A3,PER1,PLSCR1,PTGS2,PTK2,REN,RHOB,SERPINE1,SMAD1,SOCS3,SPRY2,STAT1,TP53,ZFP36,ZFP36L2 |
| STAT3 | 1.26 | transcription regulator | Activated | 3.181 | bias | 7.65E-24 | AHR,AMER1,ANGPT2,AREG,BATF,BCL3,BST2,C5,CASP1,CASP7,CCND1,CD274,CD74,CIITA,CMPK2,COL5A1,CXCL8,CXCR4,EGR1,EGR2,EGR3,EIF2AK2,ENPP2,FOS,GPR65,H2AC18/H2AC19,HERC5,HERC6,HGF,HK2,HLA-DRB5,ID1,IFI35,IFI44,IFI6,IFIH1,IFIT1,IFIT2,IFIT3,IFITM1,IFITM3,IKZF3,IL7,IRF1,IRF7,ISG15,JUNB,KLF4,LGR4,MCL1,MITF,MT-ND1,MX1,MX2,NR4A2,OAS1,OAS2,OAS3,OASL,PDIA4,PLSCR1,PTAFR,PTGS2,PTK2,REN,RSAD2,SERPINE1,SGK1,SLFN5,SOCS1,SOCS3,SOD2,SP110,STAT1,STAT2,TAP1,TNFSF10,TP53,TRIM14,TRIM5,USP18,XAF1,ZFP36 |
| IGF1 | | growth factor | Activated | 3.157 | bias | 0.00000349 | AKR1C1/AKR1C2,ANGPT2,BHLHE40,BTG2,CAVIN2,CCND1,CCND3,CD44,CD74,CLU,CSF1,CXCL8,DUSP1,DUSP6,EGR1,EGR2,EIF2AK2,FOS,FOXC2,HDC,HGF,ID1,IFITM3,JUN,JUNB,KLF6,MCL1,MGA,NFE2,NR4A1,PPP1R1B,PRNP,PTGS2,SERPINE1,SGK1,SOCS3,TAP1,TP53,TXNIP,ZFP36,ZFP36L2 |
| SMAD4 | -1.22 | transcription regulator | Activated | 3.11 | bias | 0.0031 | ANGPT2,AREG,CCND1,CELF2,CSF1,CYB561,DOCK4,EREG,FOS,GCNT1,HHEX,ID1,IRAK3,PTGS2,RGCC,SERPINE1,SGK1,SMO,ZFP36 |
| FOXO3 | 1.15 | transcription regulator | Activated | 3.055 | | 0.00000019 | ACVR1,AHR,ANGPT2,BTG1,CCND1,CCND3,CDKN3,CXCL8,CXCR4,EGR1,EGR2,ENO2,FOS,FOXM1,GMNN,GSTM5,IFIH1,IL10RB,JUNB,PDE3B,PIK3IP1,PRDX3,PRKCD,PRNP,SAC3D1,SELL,SERPINE1,SESN1,SGK1,SOD2,TAL1,TNFSF10,TP53,TP53INP1,TXNIP |
| TREM1 | 1 | transmembrane receptor | Activated | 2.985 | bias | 0.000011 | AREG,CD274,CSF1,CXCL8,CXCR4,EGR1,EGR2,EGR3,FOSL1,HSD3B7,IFIT2,IRF1,ISG15,ITGAX,MAFF,NR4A2,OASL,PTGS2,RGS1,RIN2,SPRY2,SYNJ2 |
| STAT6 | 1.07 | transcription regulator | Activated | 2.956 | | 1.67E-09 | APOE,AREG,BCL3,CCDC86,CD69,CMPK2,CNR2,CTNS,DHX58,EGR1,EGR2,EIF2AK2,FAM118B,FKBP5,GALK2,HCK,HLA-DMB,HLX,IFI44,IFI44L,IFIH1,IFIT3,IFITM3,IRAK3,IRF1,IRF7,ISG15,LGALS3BP,MOV10,OAS3,OASL,PCYOX1L,PLSCR1,PRNP,PTGS2,RASGRP3,RNPEP,SERPINE1,SNX20,SOCS1,SOCS2,STAT2,TFEC,ZFP36,ZNF608 |
| FOXO4 | -1.09 | transcription regulator | Activated | 2.949 | | 0.0287 | CCND1,IL10RB,PRKCD,SELL,SERPINE1,SESN1,SGK1,SOD2,TXNIP |
| SMAD3 | 1.95 | transcription regulator | Activated | 2.942 | bias | 0.00125 | AREG,ARID3A,CCND1,CIITA,DEPTOR,DOCK4,EGR1,EREG,FOS,ID1,JUN,JUNB,JUND,PMEPA1,PNMT,PTGS2,RHOB,SERPINE1,SMAD1,TPSAB1/TPSB2,ZFP36 |
| dopamine | | chemical - endogenous mammalian | Activated | 2.832 | | 0.00411 | BTG2,CXCL8,EGR1,EGR3,FOS,GAD1,JUN,KCNK6,KLF4,NR4A1,NR4A3,PER1,PPP1R1B |
| Insulin | | group | Activated | 2.781 | | 0.000205 | AGTR1,APOE,BMPR2,CAV2,CCND1,CCND3,CD36,COL5A1,DUSP1,EGR1,EGR2,ENPP2,FOS,FOXC2,HIF3A,IFIT2,IFIT3,IRF7,ITGB3,JUN,KLF4,LGALS3BP,MT-ND1,NR4A1,NR4A2,NR4A3,PER2,PRKCD,PTK2,SERPINE1,SGK1,SLC29A1,SORBS1,SORT1,STAT1,STAT2 |
| NR3C2 | | ligand-dependent nuclear receptor | Activated | 2.772 | | 0.0000935 | APOE,CXCL8,CXCR4,CYP1B1,ECHDC2,EGR1,FKBP5,JUNB,KLF9,NR4A2,PTGS2,RGS2,SERPINE1,SGK1,TP53,TSC22D3 |
| KRAS | 1.14 | enzyme | Activated | 2.688 | | 7.03E-17 | AHR,AMPD3,AREG,ASCL2,CASP1,CAVIN2,CCND1,CCND3,CD274,CD44,CD74,CDKN2C,CLU,CSF2RB,CXCL8,DHFR,DUSP1,DUSP6,EGR1,EGR2,EIF2AK2,EREG,EZH2,FOS,FOSL1,FOXM1,GMNN,GSTM3,HK2,HLA-DMB,HOPX,HSPB1,ID1,IFI6,IFIT1,IFITM1,IFITM3,IQGAP2,IRF1,IRF2,ISG15,ITGB3,IVNS1ABP,JUN,JUNB,KLF6,LMNB1,LYAR,MAN2A1,MX1,MX2,NR4A1,OAS1,P2RY14,PLEKHO2,PMP22,PRKCD,PTGS2,PTK2,RAB20,RHEBL1,RHOB,SAMHD1,SLC29A1,SLC43A3,SORT1,SPRY2,STAT1,STAT2,TAP1,TESC,TLE3,TMEM98,TNFSF10,TP53,VMP1 |
| IRF8 | 1 | transcription regulator | Activated | 2.671 | bias | 3.07E-08 | CD4,CD74,CIITA,DDX58,EGR1,EGR2,FOS,GFI1B,IFI44L,IFI6,IFIT2,IFIT3,ISG15,KLF4,KLF6,OAS1,STAT1,STAT2,TNFSF13B,TP53,TRIM21,TYROBP |
| PTGER4 | 2.17 | G-protein coupled receptor | Activated | 2.647 | | 9.26E-14 | ARRDC3,CD44,CD69,CMPK2,CXCL8,CXCR4,DDX58,EGR1,HERC6,HGF,IFI35,IFIH1,IFIT2,IRF1,IRF7,MAFF,PARP14,PTGS2,RNASEL,RNF24,RSAD2,RTP4,SLFN5,TAGAP,TBC1D4,TNFSF10,TP53INP1,TRIM21,USP18,XAF1 |
| Irgm1 | | other | Activated | 2.646 | bias | 0.0017 | IFIT2,IFIT3,IRF7,OAS2,OASL,RSAD2,USP18 |
| Il3 | | cytokine | Activated | 2.623 | | 0.00961 | BHLHE40,CCND3,CSF1,EGR1,FOS,MCL1,SOCS1,TP53 |
| BTK | -1.36 | kinase | Activated | 2.622 | | 1.15E-10 | CCND3,CD274,CD69,CXCL8,CXCR4,EMP1,IFI35,IFI44L,IFIT1,IFIT3,IFITM1,IGLL1/IGLL5,IRF1,ISG15,ITGAX,ITGB3,JUN,MX1,MX2,OAS2,OAS3,STAT1,TP53 |
| EPHB1 | | kinase | Activated | 2.621 | bias | 3.95E-08 | EGR1,EGR2,FOS,JUN,JUNB,PTGS2,SERPINE1 |
| beta-estradiol | | chemical - endogenous mammalian | Activated | 2.618 | | 1.06E-13 | ACAA1,ACTL6A,AGTR1,AHR,ALDH4A1,ALDH6A1,APOE,AREG,ARL4A,BACE2,BATF,BHLHE40,BMPR2,BTG1,BTG2,C17orf53,CALR,CASP9,CAV2,CBX6,CCND1,CCND3,CD274,CD44,CD69,CD74,CDC42EP4,CDKN3,CELF2,CLU,CRYZ,CTSH,CXCL8,CXCR4,CXXC5,CYP1B1,DHRS9,DUSP1,DUSP6,DYNLL1,ECHDC2,EGR1,EGR2,EGR3,EMP1,ENPP2,EREG,ESRRA,EZH2,FOS,FOSL1,FOXC2,FOXM1,FUCA1,FZD2,GART,GSTM3,H2AC18/H2AC19,HBA1/HBA2,HERC6,HES6,HGF,HLA-DRB5,HPGD,HSPB1,ID1,IFI35,IFRD2,IGSF6,IL7,IL7R,IRF1,IRF7,ISG15,ITGB3,ITPKA,JUN,JUNB,JUND,KLF2,KLF4,KLF6,KLF9,KPNA2,KYNU,LGALS3BP,LIN7A,LMNA,LTBP1,MAL,MATN2,MDK,MEIS2,MX1,NFE2L3,NKX3-1,NR4A1,NR4A2,NR4A3,OAS1,ODF3B,PAQR7,PARD6B,PDE3B,PDIA4,PIK3C2B,PIK3R3,PMP22,PPP1R1B,PREP,PRKCD,PRMT5,PTGS2,PTK2,PVALB,PYGL,RBBP8,REN,RGS1,RGS2,RTP4,SAMHD1,SERPINE1,SFRP5,SGK1,SIK1/SIK1B,SLA,SLC26A2,SLC39A6,SLC9A3R1,SOCS1,SOCS2,SOCS3,SPDL1,SPRED1,SPRY2,TFPI,TM4SF1,TMEM150C,TMEM44,TNFSF13B,TP53,TP53INP1,TREML2,TRIB2,TSC22D3,TXNIP,UBE2T,UBIAD1,WNT5B,ZFP36,ZFP36L2 |
| STAT5A | -1.4 | transcription regulator | Activated | 2.616 | | 0.00008 | APOE,ASCL2,CASP9,CCND1,CCND3,CYP1B1,EZH2,FOS,H2AC18/H2AC19,ID1,IL7R,IRF1,MCL1,NR4A2,OAS1,RAB27B,RIN2,SOCS1,SOCS2,SOCS3,SYTL2,TLE4,TM4SF1,TP53,TXNIP,ZFP36 |
| GNRH1 | | other | Activated | 2.613 | bias | 0.00661 | EGR1,EGR2,FOS,JUN,JUNB,PRKCD,PTGS2 |
| 5-hydroxytryptamine | | chemical - endogenous mammalian | Activated | 2.6 | bias | 0.0403 | CASP9,CXCL8,CYP1B1,EGR1,FOS,GAD1,NR4A1,PTGS2,SERPINE1 |
| BDNF | -1.09 | growth factor | Activated | 2.596 | bias | 0.00018 | ADCYAP1,BHLHE40,CAV2,CD36,CXCR4,DUSP1,DUSP6,EGR1,EGR2,EGR3,FOS,GNAI1,JUN,MAN2A1,MYL4,NR4A1,NR4A3,PER2,PPP1R1B,PREP,RPS23,SOD2,SOSTDC1,SPRY2,TIPARP,TRIB2,TXNIP |
| Hsp27 | | group | Activated | 2.586 | | 0.000432 | CASP9,CCND1,CD274,CSF1,CXCL8,FOS,PPP1R1B,PTGS2 |
| PTH | | other | Activated | 2.582 | bias | 0.0000429 | AREG,BHLHE40,CCND1,CGNL1,CSF1,CXCR4,DUSP1,EGR2,FOS,JUN,LMNA,NR4A1,NR4A2,PPP1R1B,PTGS2,RGS1,RGS2,SOCS3,SORT1 |
| TLR2 | -1.82 | transmembrane receptor | Activated | 2.573 | bias | 0.0184 | CD69,CXCL8,DUSP1,GP1BA,HLA-DRB5,IRAK3,IRF1,KLF2,MX1,PTGS2,SOCS1,SOCS3,TNFSF10,TP53 |
| prostaglandin E2 | | chemical - endogenous mammalian | Activated | 2.491 | | 5.87E-11 | AHR,ANGPT2,AREG,BTG1,CASP1,CASP7,CCND1,CD274,CD44,CD69,CSF1,CXCL8,CXCR4,DUSP1,DYNLL1,EGR1,EGR2,EREG,ESRRA,FOS,FOSL1,GCNT1,HDC,IL7R,IRF1,JUN,KLF4,LMNA,MCL1,MITF,NR4A1,NR4A2,PTGS2,PTK2,RGS2,SELL,SOCS2,SOCS3,TP53,TSC22D3 |
| LDL | | complex | Activated | 2.477 | bias | 0.00000002 | APOE,CASP1,CASP7,CCND1,CD36,CXCL8,DUSP1,EGR1,EGR3,FOS,GNAI1,GPR183,IFIT2,IFITM1,IRAK3,IRF2,IRF7,ITGB3,JUN,MCL1,MX1,NR4A1,NR4A2,NR4A3,PTAFR,PTGS2,SERPINE1,SOCS1,SOCS3,SOD2,TNFSF10,TP53 |
| PPP2CA | -1.33 | phosphatase | Activated | 2.449 | | 0.0017 | CASP1,CCND1,CD36,IRF1,IRF7,MYL4,PTGS2 |
| FOXL2 | | transcription regulator | Activated | 2.449 | bias | 0.0414 | FOS,MAFF,NR4A3,PTGS2,RGS2,SOD2 |
| leukotriene C4 | | chemical - endogenous mammalian | Activated | 2.446 | bias | 0.0000302 | CXCL8,EGR1,EGR2,EGR3,FOS,PTGS2 |
| IL3 | | cytokine | Activated | 2.444 | bias | 7.83E-08 | AREG,BTG2,CALR,CASP9,CCND1,CCND3,CD69,CSF1,CXCL8,CXCR4,EGR1,EGR2,EGR3,EIF2AK2,FOS,GART,GP1BA,HBA1/HBA2,HGF,HK2,ITGAX,ITGB3,JUN,JUNB,KLF1,KLF13,KLF9,MAN2B1,MCL1,SELL,SOCS1,SOCS3,STT3A,TAL1,TNFSF10,TPSAB1/TPSB2 |
| CNOT7 | -1.24 | transcription regulator | Activated | 2.425 | bias | 4.98E-18 | BST2,C12orf75,CMPK2,HERC6,IFI35,IFI44L,IFI6,IFITM1,ISG15,LGALS3BP,OAS1,OAS2,OAS3,PLSCR1,SP110,STAT1,TAP1,UBE2L6 |
| DNASE2 | -1.66 | enzyme | Activated | 2.416 | bias | 0.000000446 | DHX58,IFIT3,IRF7,ISG15,OAS1,OAS3,RSAD2,RTP4,TNFSF10,USP18 |
| GNRH | | group | Activated | 2.411 | bias | 0.000506 | ADCYAP1,DUSP1,EGR1,FOS,JUN,PTGS2,SGK1 |
| MMP1 | | peptidase | Activated | 2.407 | bias | 0.0185 | CXCL8,EGR1,FOS,JUN,PTGS2,SERPINE1 |
| MAPK10 | | kinase | Activated | 2.401 | | 0.0000202 | CASP9,CD44,CXCL8,FOS,JUN,PTGS2 |
| MEF2C | -1.77 | transcription regulator | Activated | 2.392 | bias | 0.000226 | CALR,CXCR4,ECM2,EGR2,FOS,HDAC9,JUN,JUNB,JUND,KLF2,MYL4,NR4A1,PTGS2,SIK1/SIK1B,ZFP36 |
| PGF | | growth factor | Activated | 2.352 | bias | 0.000964 | CASP1,CASP7,CXCL8,CXCR4,EGR1,EREG,SERPINE1 |
| STAT5a/b | | group | Activated | 2.35 | | 0.000000457 | AHR,CAVIN2,CCND1,CD36,CD69,CDKN2C,CLU,DUSP1,EMP1,FOS,GFI1B,GP1BA,IRF1,MCL1,PARVB,PTGS2,SOCS1,SOCS3,STK17B |
| NGLY1 | 1 | enzyme | Activated | 2.349 | | 0.000000603 | IFI44,IFI44L,IFIT1,IFIT2,IFIT3,KEAP1,OAS1,OAS3,RSAD2,USP18 |
| APEX1 | -1.28 | enzyme | Activated | 2.345 | bias | 0.00000192 | EGR1,EGR2,EGR3,FOS,JUNB,NR4A1,PTGS2,RGS2,TP53 |

| | | | | | | | |
|---|---|---|---|---|---|---|---|
| GAPDH | -1.91 | enzyme | Activated | 2.335 | bias | 6.29E-09 | C2,DUSP1,HLA-C,IFI6,IFIT2,IFITM1,OAS1,OAS2,OAS3,STAT1,TXNIP,UBE2L6 |
| PIK3CG | -1.77 | kinase | Activated | 2.333 | | 0.00865 | C2,CD74,CXCR4,IRF1,ITGB3,NLRC5,OAS2,STAT1,TAP1,TNFSF10 |
| platelet activating factor | | chemical - endogenous mammalian | Activated | 2.313 | bias | 0.00000634 | ANGPT2,CD69,EGR1,EGR2,FOS,NR4A1,PTAFR,PTGS2,SELL,SERPINE1,SOCS3 |
| AHR | 3.64 | ligand-dependent nuclear receptor | Activated | 2.297 | | 0.00000109 | ACTL6A,ACTN1,ALDH1B1,AREG,BTG2,CCND1,CCND3,CD36,CDKN2C,CHD7,CHST2,COL5A1,CYP1B1,DHFR,DUSP6,EGR2,FOS,FOSL1,GSTM5,HDAC9,HK2,IKZF3,IRF1,JUN,JUNB,KLF2,LTBP1,MATN2,PMP22,PTGS2,SERPINE1,SOCS3,SOD2,TIPARP,TP53 |
| SOCS1 | 4.1 | other | Activated | 2.274 | bias | 1.27E-12 | CCND1,CCND3,CD44,CD69,CIITA,CXCL8,DDX58,DUSP1,FKBP5,FOS,IFI44,IFIH1,IFIT1,IFIT2,IFIT3,IRF1,IRF7,ISG15,JUN,MX1,OAS1,OAS2,PTGS2,RSAD2,SOCS1,SOCS3,STAT1,USP18 |
| arachidonic acid | | chemical - endogenous mammalian | Activated | 2.263 | | 0.00271 | ACSL4,CD36,CLU,EGR1,FOS,FOSL1,ITGB3,JUN,KLF6,PTGS2,RGS2 |
| TLR7/8 | | group | Activated | 2.236 | bias | 0.0212 | ATP2B1-AS1,EREG,LINC-PINT,PTGS2,RGS1 |
| hyaluronic acid | | chemical - endogenous mammalian | Activated | 2.229 | bias | 0.0017 | BCL3,CD44,CXCL8,CXCR4,FOS,IRAK3,JUN,NR4A1,NR4A2,PTGS2,SERPINE1,SOCS3 |
| ISG15 | -4.19 | other | Activated | 2.222 | bias | 0.00000178 | DDX58,IFI6,IFITM3,MX1,OAS1 |
| Pkg | | group | Activated | 2.219 | | 0.00109 | EGR1,FOS,HK2,JUN,SOD2 |
| LYL1 | -1.45 | transcription regulator | Activated | 2.219 | | 0.0000437 | ANGPT2,BTG2,CCND3,EGR1,ID1,TAL1 |
| ACVR1C | | kinase | Activated | 2.219 | bias | 0.00137 | CCND1,FOS,JUNB,KLF4,SERPINE1 |
| TFPI2 | | other | Activated | 2.219 | bias | 0.0017 | ADCYAP1,AREG,BTG2,CCND1,JUNB |
| MAPK7 | -1.6 | kinase | Activated | 2.215 | bias | 0.00000029 | CCND1,CXCL8,DLC1,DUSP1,ID1,JUN,KLF2,KLF4,MCL1,NR4A1,PTGS2,TP53 |
| PRKCB | 1.09 | kinase | Activated | 2.207 | bias | 0.00202 | CCND1,EGR1,FOS,JUN,PRKCD,PTGS2,SERPINE1,SOD2 |
| CHRM1 | | G-protein coupled receptor | Activated | 2.207 | bias | 0.0000898 | EGR1,EGR2,EGR3,FOS,JUN |
| USP18 | -5.12 | peptidase | Activated | 2.201 | bias | 8.55E-09 | IFI6,IFIH1,IFITM3,IRF1,IRF7,ISG15,MX1,OAS1,SOCS1,SOCS3,TNFSF10 |
| FCER1G | -1.85 | transmembrane receptor | Activated | 2.2 | | 0.00435 | CD69,CXCL8,IFIT2,MX1,PTGS2,RSAD2 |
| IL17A | | cytokine | Activated | 2.196 | bias | 0.00644 | AREG,BCL3,CCND1,CD274,CXCL8,CXCR4,EGR2,EREG,FOS,FOSL1,ISG15,JUN,KLF2,MCL1,PTGS2,SGK1,SOCS1,SOCS3,TYMP |
| MET | | kinase | Activated | 2.194 | bias | 0.000000844 | AREG,CCND1,CD274,CD44,CXCL8,FOS,HDC,HGF,HLX,HSPB1,JUN,KLF4,MAL,PIM3,PTGS2,SLA,SOCS3,SOD2,TRIM21 |
| TREX1 | | enzyme | Activated | 2.19 | bias | 0.000196 | IFI44,IFIT2,ISG15,OASL,USP18 |
| NTRK1 | -1.38 | kinase | Activated | 2.164 | bias | 0.00207 | EGR1,FOS,NR4A1,SOD2,TP53 |
| cytokine | | group | Activated | 2.163 | bias | 0.00000258 | BCL3,CCND1,CD4,CD69,CLU,CSF1,CXCL8,DUSP1,EGR1,FOS,IRF1,JUN,PTGS2,SERPINE1,SGK1,SOCS1,SOCS2,SOCS3,SOD2,TYMP |
| TOB1 | -1.83 | transcription regulator | Activated | 2.159 | | 0.0388 | CCND1,GMNN,SPDL1,TP53,UBE2T |
| MEF2D | 1.07 | transcription regulator | Activated | 2.137 | bias | 0.0000211 | CCND1,CCND3,EGR2,FOS,HDAC9,JUN,JUNB,JUND,KLF2,NR4A1,SOD2,ZFP36 |
| MAPK8 | -1.08 | kinase | Activated | 2.135 | | 0.000147 | ACSL4,APOE,CASP9,CCND1,CXCL8,DUSP1,DUSP6,EGR1,FOS,FOSL1,JUN,JUND,KLF6,PTGS2,SOD2,TP53,TSC22D3,ZFP36 |
| RUNX3 | -1.73 | transcription regulator | Activated | 2.135 | bias | 0.0106 | CCND1,CD4,CHST14,CIITA,CXCL8,FZD2,ITGAX,RASAL2,RHOBTB1,SPRED1,TPSAB1/TPSB2 |
| LEP | | growth factor | Activated | 2.127 | | 0.00256 | AGTR1,AKR1C1/AKR1C2,ANGPT2,CASP7,CASP9,CCND1,CD36,DLK1,DUSP6,EGR1,EGR2,FOS,GOT2,ITGAX,JUN,JUNB,JUND,KIF3A,KIF3B,MCL1,MT-ND1,PER2,PRDX1,PTGS2,SERPINE1,SOCS3,SOD2,TNFSF10,TP53,TPRA1,ZFP36 |
| GLI3 | | transcription regulator | Activated | 2.121 | | 0.00175 | CCND1,CD69,DUSP1,EGR1,FOXC2,IRF1,JUNB,KLF13,KLF2 |
| F3 | | transmembrane receptor | Activated | 2.121 | bias | 0.000297 | AREG,CASP7,CXCL8,DOCK4,DUSP6,EGR1,FOS,FOXC2,MDK,SERPINE1 |
| formaldehyde | | chemical - endogenous mammalian | Activated | 2.118 | bias | 0.00000583 | EGR1,EGR2,FOS,FOSL1,JUN,JUNB,JUND,PTGS2,SGK1 |
| EGR1 | 5.66 | transcription regulator | Activated | 2.115 | bias | 2.31E-08 | AREG,CASP7,CASP9,CCND1,CD44,CLU,CSF1,CXCL8,EGR1,EGR2,EREG,FOSL1,GAD1,HPGD,JUN,JUNB,JUND,NR4A1,PNMT,PTGS2,SERPINE1,SGK1,SOCS1,SOD2,TNFSF10,TP53 |
| FGF2 | 1.16 | growth factor | Activated | 2.114 | bias | 0.00263 | ANGPT2,AREG,BTG2,CCND1,CCND3,CSF1,CXCR4,DUSP6,EGR1,ENPP2,EREG,FOS,FOSL1,HGF,HK2,JUN,JUNB,LMNA,MITF,NR4A1,NR4A2,PRKCD,PTGS2,SERPINE1,SOCS3,SPRY2,TFPI,TNFSF10,TP53 |
| IRF4 | | transcription regulator | Activated | 2.109 | | 0.00374 | CIITA,CXCR4,IL7R,IRF1,IRF7,ISG15,OAS1,PLSCR1,RHOB,STAT1,STAT2,TNFSF10,TNFSF13B,TRIM21 |
| PTK2B | 1.02 | kinase | Activated | 2.105 | | 0.0000618 | CXCL8,FOS,JUN,PTK2,SOCS1,TP53 |
| PRKCE | 1.99 | kinase | Activated | 2.094 | bias | 0.00336 | CCND1,CCND3,CXCL8,EGR1,FOS,JUN,JUNB,PTGS2,RGS2 |
| sphingosine-1-phosphate | | chemical - endogenous mammalian | Activated | 2.071 | bias | 0.00318 | CCND1,CD36,CD44,CXCL8,EGR1,ENPP2,FOS,JUN,PTGS2,SERPINE1 |
| IL2 | | cytokine | Activated | 2.07 | bias | 2.85E-10 | AHR,AREG,BHLHE40,CARD6,CASP1,CCND1,CCND3,CD274,CD44,CD69,CD74,CDKN2C,CSF1,CSF2RB,CXCL8,CXCR4,CYSLTR1,DUSP6,EMP1,ENPP2,FOS,GART,GCNT1,HCK,HHEX,HK2,HSPA1A/HSPA1B,IKZF4,IL7R,IRF1,ITGB8,JUN,JUND,KLF1,KLF13,KLF6,LY6E,MX1,NAV1,NEAT1,NRGN,PDE3B,PIK3R3,PTGS2,RCN1,RGCC,RGS2,RHOB,SELL,SESN1,SOCS1,SOCS2,SOCS3,SPRED1,STK17B,TNFRSF21,TNFSF10,TP53,TP53INP1,TRIB2 |
| Ins1 | | other | Activated | 2.029 | | 0.0116 | ALAS1,CCND1,CCND3,DHFR,DLK1,DYNLL1,EGR1,EGR2,FOS,FOXC2,HK2,JUN,JUNB,LGALS3BP,PER1,PTGS2,REN,SERPINE1,SOCS2,SOCS3,SOD2 |
| AREG | 76.4 | growth factor | Activated | 2.027 | | 4.57E-08 | AREG,CCND1,CXCR4,EGR1,EREG,FOS,FOXM1,H2AC18/H2AC19,H2BC5,IFI6,IFIT2,IFIT3,ITGB8,JUN,KIF14,PTAFR,PTGS2 |
| NAMPT | 2.11 | cytokine | Activated | 2.019 | bias | 0.00148 | CXCL8,ID1,JUN,MITF,SOCS2,TFPI,TP53 |
| PDGFB | -1 | growth factor | Activated | 2.016 | bias | 0.000925 | CXCL8,EGR1,FOS,HDAC7,KLF2,NOTCH2,NR4A1,PTGS2,SMAD1 |
| CREM | 1.26 | transcription regulator | Activated | 2.01 | | 0.0000107 | APOE,BHLHE40,BTG2,CCND1,CXCL8,DUSP1,EGR1,EGR2,FOS,JUNB,MCL1,NR4A1,NR4A2,PDP2,PER1,REN,RHOB,SIK1/SIK1B,TIPARP |
| Jnk | | group | Activated | 2.009 | bias | 0.0000141 | BTG1,CCND1,CD44,CD69,CELF2,CXCL8,CXCR4,DUSP1,EGR1,FOS,JUN,JUND,KLF6,MCL1,NR4A1,PLSCR1,PTGS2,PTK2,SERPINE1,SOCS3,SOD2,TP53,ZFP36 |
| MITF | -3.83 | transcription regulator | Inhibited | -2.111 | bias | 0.00115 | APOE,CD44,CXCL8,FOS,GMPR,HPGD,IVNS1ABP,KAZN,KIFC1,MITF,MYL4,ORC6,RMI1,SERPINE1,SORT1,SPC25,TMEM251,TP53,TPSAB1/TPSB2,TXNIP |
| TIMP3 | -1.03 | other | Inhibited | -2.121 | bias | 0.000762 | EGR2,FOS,JUN,KLF4,LTBP1,SERPINE1,SMAD1,STAT1 |
| IFNA4 | | cytokine | Inhibited | -2.159 | bias | 0.000000136 | CD274,CD69,IFIH1,IFIT1,IFIT2,ISG15,MX1,OASL,PMEPA1,RSAD2,TREML2,USP18 |
| OGT | 1.04 | enzyme | Inhibited | -2.2 | | 0.168 | ARTN,CCND1,FOS,FOXM1,JUN,TP53 |
| CFTR | | ion channel | Inhibited | -2.2 | | 0.111 | ACAA1,CXCL8,HPGD,PTGS2,SOCS3,STAT1 |
| DACH1 | -1.34 | transcription regulator | Inhibited | -2.204 | bias | 0.000134 | CXCL8,EGR1,FOS,ID1,JUN,KLF4,PTGS2,SERPINE1 |
| CGAS | -1.23 | enzyme | Inhibited | -2.207 | bias | 0.000000103 | CXCL8,IFI44,IFIT2,IFIT3,IRF7,ISG15,OAS1,RSAD2,USP18 |
| carbon monoxide | | chemical - endogenous mammalian | Inhibited | -2.219 | | 0.0949 | EGR1,HGF,PRDX1,PTGS2,SERPINE1 |
| histone deacetylase | | complex | Inhibited | -2.219 | bias | 0.0192 | CCND1,CXCR4,JUNB,KEAP1,PER1 |
| PTX3 | -1.41 | other | Inhibited | -2.224 | bias | 0.00573 | CXCL8,EGR2,EGR3,FOS,JUN |
| TNK1 | | kinase | Inhibited | -2.236 | bias | 0.000379 | IFIH1,IFIT2,IRF7,OAS2,TNFSF10 |
| TMEM173 | -1.48 | other | Inhibited | -2.316 | bias | 0.000000532 | CXCL8,HDC,HOPX,IFI44,IFIT2,IFIT3,IFITM3,IRF7,ISG15,OAS1,OASL,RSAD2,USP18 |
| DDX58 | -2.18 | enzyme | Inhibited | -2.376 | bias | 1.69E-14 | CD4,CXCL8,DDX58,EIF2AK2,IFI35,IFI44,IFIH1,IFIT1,IFIT2,IFIT3,IRF1,IRF7,ISG15,KLF4,OAS1,PTGS2,RSAD2,SOCS1,SOCS3,STAT1,STAT2,TNFSF10 |
| EBF1 | | transcription regulator | Inhibited | -2.382 | bias | 0.0379 | GFI1B,HK2,IGLL1/IGLL5,IRF1,IRF7,PDE3B,PIK3R3,SOCS1,SOCS3,STAT1 |
| NLRC5 | -3.1 | transcription regulator | Inhibited | -2.387 | bias | 0.000116 | DDX58,HLA-B,HLA-C,HLA-F,HLA-G,TAP1 |
| SGPL1 | -1.49 | enzyme | Inhibited | -2.401 | | 0.000201 | DDX58,IFIT1,ISG15,MX1,OAS1,PTGS2 |
| ISGF3 | | complex | Inhibited | -2.412 | bias | 7.92E-10 | EIF2AK2,IFIH1,IFIT2,IRF1,IRF7,ISG15,RSAD2,TNFSF10 |
| CLEC12A | -1.81 | other | Inhibited | -2.425 | bias | 2.39E-08 | IFIT2,IFIT3,IRF7,ISG15,RSAD2,USP18 |
| FADD | | other | Inhibited | -2.433 | bias | 1.76E-10 | CASP7,CXCL8,CXCR4,DDX58,DHX58,EGR1,EIF2AK2,FOS,FOXM1,IFIH1,IFIT2,IRF7,JUN,JUNB,KLF6,LY6E,SOCS1,STAT1,STAT2,TIPARP |
| KLF11 | -1.36 | transcription regulator | Inhibited | -2.433 | | 0.0668 | APOL6,CXCR4,HGF,ITGB3,ITGB8,LTBP1,SERPINE1,SOD2,STAT1 |
| IFNK | | cytokine | Inhibited | -2.449 | bias | 0.000201 | EIF2AK2,IFIH1,IRF1,MX1,OAS1,STAT1 |
| IFNAR2 | -1.12 | transmembrane receptor | Inhibited | -2.449 | bias | 1.33E-09 | DDX58,HERC5,IFI44,IFI6,IFIH1,IFITM1,ISG15,MX2,OAS1,OAS2,TNFSF10,UBE2L6,USP18,XAF1 |
| CD247 | | transmembrane receptor | Inhibited | -2.449 | bias | 0.0388 | ALAS1,MTHFS,NFE2,TFEC,UBE2L6,XAF1 |
| FOSL1 | -2.58 | transcription regulator | Inhibited | -2.458 | | 0.000173 | CCND1,CD44,CXCL8,EGR1,EGR2,FOS,FOSL1,HPGD,JUN,JUNB,SERPINE1 |
| DUSP1 | 3.32 | phosphatase | Inhibited | -2.461 | | 0.000000592 | APOL6,CMPK2,CXCL8,DUSP1,DUSP6,FOSL1,HELZ2,IFIT1,IFIT3,IRF1,JUN,MX1,PAQR7,PTGS2,SERPINE1,SOCS1,SOD2,ZFP36 |
| DOCK8 | -1.24 | other | Inhibited | -2.5 | bias | 7.22E-09 | CMPK2,HDC,HELZ2,IFIT2,IFIT3,IRF1,IRF7,ISG15,LMO4,PTGS2,RGS1,RSAD2,SOCS1,STAT1,STAT2,TRIM21 |
| mir-181 | | microRNA | Inhibited | -2.621 | bias | 0.00225 | CD69,CXCL8,DUSP6,EZH2,KLF6,MCL1,PRKCD,PTGS2 |
| IFN alpha/beta | | group | Inhibited | -2.622 | bias | 0.00000283 | CD69,IFIT2,IFIT3,IL7,IRF1,IRF7,LY6E,RSAD2,SOCS1,STAT1,STAT2,TNFSF10,TNFSF13B,TRIM21 |

| | | | | | | | |
|---|---|---|---|---|---|---|---|
| SFTPA1 | | transporter | Inhibited | -2.646 | | 0.0349 | AKR1C1/AKR1C2,CYP1B1,EGR1,FOS,HBA1/HBA2,KLF2,SERPINE1 |
| MSC | | transcription regulator | Inhibited | -2.673 | | 2.86E-08 | AMPD3,DDX60,EPSTI1,HGF,IFI44,IFI44L,IFIT1,IRF7,PAG1,PDP1,PIEZO2,SELL,SYNE2,XAF1 |
| SAMSN1 | -1.5 | other | Inhibited | -2.683 | bias | 2.7E-10 | BATF,CMPK2,HDC,HELZ2,IFIT2,IFIT3,IRF1,IRF2,IRF7,ISG15,LMO4,OASL,PTGS2,RGCC,RSAD2,SOCS1,SOCS3,STAT1,STAT2,TRIM21 |
| tretinoin | | chemical - endogenous mammalian | Inhibited | -2.689 | | 5.79E-13 | AGTR1,AHR,ALAS1,ALDH9A1,APOE,AREG,BCL3,BHLHE40,BMPR2,BTG1,BTG2,C12orf29,C3AR1,CALR,CASP1,CASP7,CASP9,CCDC169,CCND1,CCND3,CD274,CD36,CD44,CD74,CDKN2C,CDKN3,CXCL8,CXCR4,DDX58,DDX60,DHRS3,DHRS9,DHX33,DHX58,DLC1,DUSP1,DYNLL1,EGR1,EIF2AK2,ENPP2,FOS,FOSL1,GMNN,GRASP,HLA-B,HLA-C,HLA-DMB,HNRNPF,HOTAIRM1,HSPB1,ID1,IFI35,IFI44,IFI44L,IFI6,IFIH1,IFIT1,IFIT2,IFIT3,IFITM1,IL1RAPL1,IRF1,ISG15,ITGAX,ITGB3,JUN,KLF1,KLF13,KLF4,KLF9,KRT1,LGALS3BP,LY6E,MCL1,MDK,MEIS2,NFE2,NLN,NRGN,OAS1,OAS2,OAS3,OASL,PARP14,PARP9,PLEKHO2,PLSCR1,PPP1R1B,PRKCD,PRNP,PTAFR,PTGS2,RASAL2,REN,RTP4,SAMD9L,SAMHD1,SELL,SERPINE1,SFRP5,SIGLEC12,SLA,SLFN5,SMAD1,SMOX,SOCS1,SP110,STAT1,STAT2,STT3A,TAL1,TAP1,TNFSF10,TOR1B,TP53,TSC22D3,TTC27,TYROBP,UBE2L6,USP18,VMP1,XAF1,ZNF146 |
| EIF2AK2 | -2.03 | kinase | Inhibited | -2.692 | bias | 4.86E-16 | BHLHE40,CASP9,CCND1,CXCL8,DDX58,EGR1,EIF2AK2,FOS,H2BC5,IFI35,IFI6,IFIT1,IFITM1,IRF1,ISG15,LGALS3BP,OAS1,OAS3,PARP9,PLSCR1,SAMHD1,SOCS3,SOD2,STAT1,TP53,UBE2L6,USP18 |
| IFNE | | cytokine | Inhibited | -2.714 | bias | 0.000000565 | BST2,HERC5,IFIH1,IFIT2,IFITM3,ISG15,MX2,PTGS2,STAT1,TRIM5,USP18 |
| PARP9 | -3.94 | enzyme | Inhibited | -2.744 | bias | 1.79E-13 | CD74,IFI44,IFIT1,IFIT2,IFIT3,IRF1,IRF2,IRF7,ISG15,OAS2,SP110,STAT1 |
| PML | 1.01 | transcription regulator | Inhibited | -2.793 | | 2.13E-12 | ACSL4,APOE,BST2,CCND1,CIITA,CXCR4,DDX60,EPSTI1,HERC6,HLA-DMB,HSPB1,IFI35,IFI44,IFI44L,IFIH1,IFIT1,IFIT3,IFITM1,IRF7,ISG15,MAFF,MX1,OAS1,OAS2,OAS3,PLSCR1,PRDX1,STAT1,TAP1,TP53 |
| VCAN | 4.99 | other | Inhibited | -2.802 | | 1.07E-09 | APOE,C1R,CASP1,CBX6,CLU,CXCL8,ENPP2,IFI44,IFI44L,IFI6,IFIT1,IFIT2,IFITM1,MX1,MX2,OAS2,OAS3,PARP14,PTK2,RGS2,SOD2,SORT1,STAT1,TP53,TTC27,XAF1 |
| IFNL4 | | cytokine | Inhibited | -2.807 | bias | 2.24E-10 | DDX58,DHX58,IFIH1,ISG15,MX1,OAS1,OAS2,STAT1 |
| JAK | | group | Inhibited | -2.887 | bias | 2.6E-12 | DDX58,EIF2AK2,HES6,IFI6,IFIH1,IFIT1,IFIT2,IFIT3,IFITM3,ISG15,PTGS2,RSAD2,SOCS1,SOCS2,SOCS3,STAT1 |
| L-triiodothyronine | | chemical - endogenous mammalian | Inhibited | -2.923 | | 0.000329 | ACOT11,APOC1,BCL3,CCND1,CD36,CD44,CDC42EP4,CLU,CTSH,DBP,DLK1,DUSP1,EGR1,EGR2,ENPP2,FKBP10,FOS,HBA1/HBA2,ID1,ITGB3,KLF9,LMNA,NR4A1,NRGN,PER1,PRDX3,PTAFR,PTGS2,PYGL,REN,RGS2,SERPINE1,SORBS1,TP53 |
| SPI1 | -1.93 | transcription regulator | Inhibited | -3.016 | | 2.89E-13 | ARID3A,C3AR1,CCND1,CIITA,CMPK2,CSF1,CSF2RB,DUSP6,EGR2,EPX,FOS,HDAC7,HELZ2,HOTAIRM1,IFI44,IFI44L,IFI6,IFIT1,IFIT2,IFIT3,IFITM1,IFITM3,IL7R,IRAK3,IRF7,ISG15,JUN,KLF13,KLF4,LBR,LY6E,MCL1,MX1,OASL,PTGS2,RSAD2,SP110,TFEC,TNFSF10,USP18 |
| JAK1/2 | | group | Inhibited | -3.053 | bias | 0.00000328 | CD69,CD74,CIITA,DGKG,EIF2AK2,HLA-DRB5,IRF1,IRF7,ISG15,ITPKA,MX1,RGS14,RSAD2 |
| TGM2 | -1.03 | enzyme | Inhibited | -3.109 | bias | 2.68E-16 | BTG2,C12orf29,CCDC169,CCND1,CD36,CD74,CXCL8,DDX60,DHX33,HLA-B,IFI35,IFI6,IFIT1,IFIT2,IFIT3,ITGAX,ITGB3,LY6E,NLN,OAS1,OAS2,OAS3,OASL,PARP14,PARP9,PLEKHO2,PLSCR1,SAMD9L,SAMHD1,SELL,SIGLEC12,SLFN5,SP110,STAT1,TAP1,TOR1B,TP53,TTC27,TYROBP,XAF1,ZNF146 |
| Hdac | | group | Inhibited | -3.142 | | 0.00000247 | AGTR1,AHR,AREG,CCND1,CXCL8,CXCR4,EGR1,EGR3,EZH2,FOS,FOSL1,GAD1,HPGD,JUN,KLF6,KLF9,NR4A1,SMO,SPRY2,TNFSF10,TXNIP |
| SASH1 | | other | Inhibited | -3.153 | bias | 1.44E-09 | CMPK2,HDC,HELZ2,HHEX,IFIT2,IFIT3,IRF1,IRF7,ISG15,LMO4,OASL,PTGS2,RSAD2,SOCS1,STAT1,STAT2,TRIM21 |
| BRCA1 | -1.06 | transcription regulator | Inhibited | -3.256 | | 0.0000273 | AREG,ASCL2,CCND1,CYP1B1,DDX58,EGR1,ENPP2,FKBP5,HSPB1,IFI6,IFIT1,IFIT2,IFIT3,IFITM1,IRF7,MX1,PLSCR1,STAT1,TAP1,TP53 |
| IFNA1/IFNA13 | | cytokine | Inhibited | -3.41 | bias | 1.24E-16 | CD274,CD69,DHX58,EIF2AK2,IFI6,IFIH1,IFIT1,IFIT2,IFITM1,IRF1,IRF7,ISG15,MX1,OAS1,OAS2,PLSCR1,PTGS2,RSAD2,SOCS1,STAT1,STAT2,UBE2L6 |
| MAVS | -1.31 | other | Inhibited | -3.478 | bias | 1.46E-12 | CMPK2,CXCL8,DDX58,DHX58,IFIT1,IFIT2,IFIT3,IFITM3,IRF7,ISG15,OAS1,OAS2,OASL,RSAD2,SOCS1,SOCS3,STAT1,STAT2,UBE2L6,USP18 |
| IFN type 1 | | group | Inhibited | -3.662 | bias | 5.24E-11 | BST2,CD69,CXCL8,DDX58,DHX58,EIF2AK2,IFIH1,IFIT1,IFIT2,IRF1,ISG15,OASL,STAT1,STAT2,TNFSF10,TNFSF13B |
| Ifn | | group | Inhibited | -3.966 | bias | 3.2E-17 | APOBEC3A,CD44,CD69,DDX58,DHX58,EIF2AK2,FOS,HGF,IFIH1,IFIT1,IFITM1,IFITM3,IL7,IRF1,IRF7,ISG15,MX1,OAS2,OAS3,OASL,PTAFR,RNASEL,RSAD2,SERPINE1,SOCS1,STAT1,TAP1,TNFSF13B,TP53,TRIM21,TYMP |
| IRF5 | -1.36 | transcription regulator | Inhibited | -4.23 | bias | 2.49E-13 | CMPK2,CXCR4,DDX58,DHX58,IFI44,IFIH1,IFIT1,IFIT2,IFIT3,IFITM3,IRF7,ISG15,OAS1,OAS2,OASL,PLSCR1,PTGS2,RSAD2,SP110,STAT1,STAT2,TNFSF10,UBE2L6 |
| RNY3 | | other | Inhibited | -4.359 | bias | 2.47E-21 | DDX60,EPSTI1,HERC5,IFI44,IFI44L,IFIT1,IFIT3,IFITM3,ISG15,LY6E,MX1,OAS1,OAS2,OAS3,OASL,RSAD2,RTP4,TRIM6,XAF1 |
| IFNB1 | | cytokine | Inhibited | -4.367 | bias | 4.14E-15 | APOL1,APOL2,APOL3,APOL6,BHLHE40,BST2,BTN3A3,CARD6,CASP1,CD274,CMPK2,CXCL8,DDX58,DHX58,EIF2AK2,FOS,HERC5,IFI6,IFIH1,IFIT1,IFIT2,IFIT3,IFITM1,IRF1,IRF7,ISG15,ITGAX,ITGB3,MX1,NFKBIE,OAS1,OAS2,PARP14,PTGS2,RNASEL,RSAD2,SOCS1,SPRY2,STAT1,STAT2,TNFSF10,TRIM14,TRIM21,USP18,XAF1 |
| STAT1 | -4.82 | transcription regulator | Inhibited | -4.551 | bias | 4.91E-28 | ANGPT2,APOE,APOL4,APOL6,BST2,BTG1,C1R,CASP1,CCND1,CCND3,CD274,CIITA,CMPK2,CXCL8,DDX60,EGR1,EIF2AK2,EPSTI1,FOS,GBP1P1,HERC6,HLA-DRB5,IFI35,IFI44,IFI44L,IFI6,IFIH1,IFIT1,IFIT2,IFIT3,IFITM1,IFITM3,IRF1,IRF2,IRF7,ISG15,ITGAX,JUN,KLF4,LY6E,MX1,NFE2,NLRC5,OAS1,OAS2,OAS3,OASL,PARP9,PLSCR1,PTGS2,RSAD2,RTP4,SAMD9L,SAMHD1,SLFN5,SOCS1,SOCS3,SORT1,SP110,STAT1,STAT2,TAP1,TNFSF10,TNFSF13B,TP53,TRIM21,TYMP,USP18,XAF1 |
| IFN Beta | | group | Inhibited | -4.604 | bias | 3.72E-22 | BST2,C3AR1,CD274,CD69,CD74,CXCL8,DDX58,DUSP1,EIF2AK2,HERC5,HLA-B,HLA-G,IFI35,IFI44,IFI6,IFIH1,IFIT1,IFIT2,IFIT3,IFITM1,IRF1,IRF7,ISG15,MX1,MX2,OAS1,OAS2,OAS3,OASL,RSAD2,SOCS1,SPRY2,STAT1,STAT2,TNFSF10,TP53,TRIM14,USP18,XAF1 |
| IRF3 | -1.37 | transcription regulator | Inhibited | -4.747 | bias | 1.77E-14 | CD274,CD69,CMPK2,CXCL8,DDX58,DDX60,DHX58,EIF2AK2,HELZ2,HLA-F,IFI44,IFI6,IFIH1,IFIT1,IFIT2,IFIT3,IFITM3,IRF1,IRF7,ISG15,JUNB,NLRC5,OAS1,OAS2,OAS3,OASL,PARP14,PRNP,RSAD2,SAMD9L,STAT1,STAT2,TAP1,TNFSF10,UBE2L6,USP18 |
| Ifnar | | group | Inhibited | -4.772 | bias | 1.43E-17 | CASP1,CD274,CD74,DDX58,EIF2AK2,HLA-G,IFI35,IFIH1,IFIT2,IFIT3,IFITM3,IRF1,IRF7,ISG15,NLRC5,OAS1,OAS2,OASL,RSAD2,STAT1,STAT2,TAP1,TNFSF10,TRIM21,UBE2L6,UNC93B1,USP18,XAF1 |
| PRL | | cytokine | Inhibited | -4.963 | bias | 1.68E-26 | AKR1C4,BST2,CCND1,CCND3,CD69,CLU,CMPK2,COL5A1,CTSH,DDX58,DHX58,DTX3L,EGR1,EIF2AK2,EPSTI1,FOS,GMPR,HELZ2,HERC5,HERC6,ID1,IFI35,IFI44,IFI44L,IFI6,IFIH1,IFIT1,IFIT3,IFITM1,IRF1,IRF7,ISG15,JUN,LY6E,MX2,OAS1,OAS2,OAS3,PAQR7,PAQR8,PARP14,PDIA4,PIM3,PLSCR1,RSAD2,SAMD9,SAMD9L,SAMHD1,SOCS1,SOCS2,SOCS3,SP110,STAT1,STAT2,TNFSF13B,TP53,TRIM14,USP18,XAF1 |
| IRF1 | -2.31 | transcription regulator | Inhibited | -5.038 | bias | 4.16E-26 | APOL6,C1R,CASP1,CASP7,CCND1,CD274,CIITA,CMPK2,CXCL8,DDX58,EIF2AK2,HELZ2,HLA-G,IFI35,IFI44,IFI44L,IFI6,IFIH1,IFIT1,IFIT2,IFIT3,IFITM1,IFITM3,IL7,IRF1,IRF2,IRF7,ISG15,LIG4,MX1,NFE2,OAS1,OAS2,OAS3,OASL,PTGS2,RSAD2,SELL,SOCS1,SP110,STAT1,STAT2,TAP1,TNFSF10,TNFSF13B,TP53,TRIM21,XAF1 |
| IFNG | | cytokine | Inhibited | -5.123 | | 7.4E-30 | AGTR1,AHR,APOL1,APOL6,AREG,BCL3,BST1,BST2,BTG1,BTN3A1,BTN3A2,C1R,C2,CARD6,CASP1,CASP7,CASP9,CCND1,CCND3,CD274,CD36,CD4,CD44,CD74,CIITA,CMPK2,CSF1,CSF2RB,CTSH,CXCL8,CXCR4,CYB561,DBP,DDX58,DDX60,DTX3L,DUSP1,EGR1,EGR2,EGR3,EIF2AK2,FKBP5,FOS,FZD2,GAD1,GART,GMPR,HCK,HCP5,HDAC9,HERC6,HK2,HLA-B,HLA-C,HLA-DMB,HLA-DRB5,HLA-F,HLA-G,HSPA1A/HSPA1B,HSPB1,ID1,IFI35,IFI44,IFI44L,IFI6,IFIH1,IFIT1,IFIT2,IFIT3,IFITM1,IFITM3,IL7,IL7R,IRF1,IRF2,IRF7,ISG15,ITGAX,ITGB3,JUN,JUNB,JUND,KLF1,KLF2,KLF4,KLF6,KYNU,LGALS3BP,LY6E,MAFF,MAN2A1,MITF,MX1,MX2,NEAT1,NFE2L3,NLRC5,OAS1,OAS2,OAS3,OASL,P2RY14,PARP14,PARP9,PARVG,PIM3,PLSCR1,PPP1R1B,PRKCD,PRKG2,PRNP,PTAFR,PTGS2,RAB20,RAB38,RGCC,RHOB,RSAD2,RTP4,SAMD9,SAMHD1,SCUBE1,SELL,SERPINE1,SLC29A1,SLFN5,SMAD1,SMTN,SOCS1,SOCS2,SOCS3,SOD2,SORT1,SP110,STAT1,STAT2,TAP1,TICAM1,TNFSF10,TNFSF13B,TP53,TRIB2,TRIM21,TRIP11,TSC22D3,TXNIP,TYMP,TYROBP,UBE2L6,USP18,XAF1,ZFP36 |
| Interferon alpha | | group | Inhibited | -5.401 | bias | 8.6E-35 | APOBEC3A,APOL1,APOL2,APOL3,BCL3,BST2,BTG2,C3AR1,CASP1,CCND1,CCND3,CD274,CD69,CIITA,CSF2RB,CXCL8,CYB561,DDX58,DHX58,DUSP6,EIF2AK2,ENPP2,EPSTI1,FBXO6,FOS,HELZ2,HERC5,HERC6,HLA-B,HLA-C,HLA-F,HLA-G,HSPA1A/HSPA1B,IFI35,IFI44,IFI44L,IFI6,IFIH1,IFIT1,IFIT2,IFIT3,IFITM1,IFITM3,IGLL1/IGLL5,IL7,IL7R,IRF1,IRF7,ISG15,MCL1,MX1,MX2,NFE2L3,OAS1,OAS2,OAS3,PARP14,PARP9,PARVG,PLSCR1,PTGS2,RNASEL,RSAD2,RTP4,SAMD9,SAMD9L,SLFN5,SOCS1,SOCS2,SOCS3,SP110,SPRY2,STAT1,STAT2,TAP1,TICAM1,TNFSF10,TNFSF13B,TRANK1,TRIB2,TRIM21,TRIM5,TYMP,UBE2L6,UNC93B1,USP18 |
| IFNA2 | | cytokine | Inhibited | -5.601 | bias | 9.02E-26 | APOL6,BST2,C1R,CD274,CD69,CMPK2,DDX58,DDX60,EIF2AK2,HERC5,HERC6,HLA-B,HLA-C,IFI35,IFI44,IFI44L,IFI6,IFIH1,IFIT1,IFIT2,IFIT3,IFITM1,IFITM3,IL10RB,IRF1,IRF7,ISG15,LGALS3BP,LY6E,MAL,MX1,MX2,OAS1,OAS2,OAS3,PARP9,PLSCR1,PTAFR,RSAD2,SAMD9,SAMHD1,SOCS1,SOCS2,SOCS3,SP110,STAT1,TAP1,TNFSF10,TP53,TRIM14,TRIM21,UBE2L6,USP18,XAF1 |
| IFNL1 | | cytokine | Inhibited | -5.688 | bias | 4.46E-32 | APOL6,BST2,CMPK2,CXCL8,DDX58,DDX60,EIF2AK2,HERC5,HERC6,HLA-B,HLA-C,IFI35,IFI44,IFI44L,IFI6,IFIH1,IFIT1,IFIT2,IFIT3,IFITM1,IFITM3,ISG15,LGALS3BP,MX1,OAS1,OAS2,OAS3,OASL,PLSCR1,RSAD2,RTP4,SAMD9,SP110,STAT1,STAT2,TRIM14,UBE2L6,USP18,XAF1 |
| IRF7 | -3.88 | transcription regulator | Inhibited | -5.835 | bias | 6.16E-25 | C5,CD69,CMPK2,DDX58,DHX58,HELZ2,HERC5,IFI35,IFI44,IFI44L,IFI6,IFIH1,IFIT1,IFIT2,IFIT3,IFITM1,IFITM3,IRF1,IRF7,ISG15,ITGAX,MCL1,MX1,MX2,OAS1,OAS2,OAS3,OASL,PARP14,PLSCR1,RSAD2,RTP4,SAMD9L,SOCS1,STAT1,STAT2,TAP1,TNFSF10,TNFSF13B,TOR1B,TRIM21,TRIM5,UBE2L6,USP18,XAF1 |

**T3 vs. T1 MEP_2**

© 2000-2020 QIAGEN. All rights reserved.

| Upstream Regulator | Expr Fold Change | Molecule Type | Predicted Activation State | Activation z-score | Flags | p-value of overlap | Target Molecules in Dataset |
|---|---|---|---|---|---|---|---|
| MAPK1 | -1.25 | kinase | Activated | 5.825 | | 3.96E-33 | APOE,APOL6,BST2,BTN3A3,CD69,CDH1,CHST2,CISH,CTNNB1,CXCL8,DDX58,DEPTOR,DUSP1,EGR1,EGR2,EPCAM,FOS,GBP1,HDC,HERC5,HLA-B,HLA-C,HOMER1,IFI27,IFI35,IFI44,IFI6,IFIH1,IFIT1,IFIT3,IFITM1,IFITM3,IRF7,ISG15,ISG20,JUNB,JUND,LGALS3BP,LOX,MYC,NR4A1,OAS1,OAS2,OAS3,OASL,PARP12,PDGFB,PLSCR1,PTGER2,PTGS2,SERPINE1,SP110,STAT1,STAT2,TAP1,TNFRSF10B,TNFSF10,TRANK1,TRIM14,TRIM21,TRIM22,TRIM25,TRIM5,USP18 |
| SIRT1 | -1.25 | transcription regulator | Activated | 4.675 | | 1.96E-16 | ABCB1,ABCG1,CD36,CD74,CDH1,CDKN2C,CMPK2,CTNNB1,DBP,DDX58,DDX60,DKK1,GABARAPL1,GSTM3,HLA-A,HLA-DQA1,HLA-DQB1,HLA-DRB5,ID1,IFI44,IFIT3,IFITM3,IRF7,KLF2,LGALS3BP,MYC,NFKBIE,NLRC5,OAS1,OAS2,PARP12,PARP14,PER1,RELB,RSAD2,RTP4,SERPINE1,SFRP5,SOCS3,SOD2,SP110,STAT1,TAP1,TNFRSF10B,USP18 |
| PNPT1 | -1.78 | enzyme | Activated | 4.55 | bias | 9.7E-21 | CMPK2,CXCL8,DDX58,GBP4,IFI44,IFIH1,IFIT3,IRF7,ISG15,LGALS3BP,MYC,OAS1,PARP12,PARP14,PARP9,RTP4,SAMD9L,STAT1,STAT2,USP18,XAF1 |
| PDGF BB | | complex | Activated | 4.478 | | 4.66E-08 | AREG,BHLHE40,CXCL8,DUSP1,EGR1,EGR2,EGR3,EREG,FOS,FZD2,GPR183,ID3,JUNB,KLF6,MICOS10-NBL1/NBL1,MYC,NR4A1,NR4A2,NR4A3,PTGS2,RGS1,SERPINE1,SGK1,SOCS3,SOD2,TGM2,VCAN,ZFP36 |
| TRIM24 | -1.35 | transcription regulator | Activated | 4.419 | | 2.47E-20 | CA2,CMPK2,DDX58,DDX60,EPSTI1,GBP4,HERC6,IFI35,IFI44,IFIH1,IFIT3,IRF1,IRF7,ISG15,LGALS3BP,OAS1,OASL,PARP12,RTP4,SAMD9L,SERPINE1,SOCS1,STAT1,STAT2,TAP1,TLR2,TRIM26,USP18 |
| IL1RN | | cytokine | Activated | 3.895 | bias | 1.85E-21 | BTN3A1,CIITA,CXCL8,DDX58,GBP1,HERC6,HLA-DQB1,IFI27,IFI44,IFI44L,IFI6,IFIH1,IFIT3,IRF1,IRF7,ISG20,KLF6,MX1,OAS1,OAS2,OAS3,OASL,PTGS2,RSAD2,RTP4,SERPINE1,STAT2,TCF7L2,TNFSF10,TRIM22,USP18 |
| STAT3 | 1.17 | transcription regulator | Activated | 3.557 | | 1.08E-32 | AHR,AREG,BST2,CASP1,CD274,CD74,CDH1,CIITA,CMPK2,CTSL,CXCL8,CXCR4,DKK1,EGR1,EGR2,EGR3,FGL2,FOS,FST,HERC5,HERC6,HLA-A,HLA-DQA1,HLA-DRB5,ID1,ID2,IFI27,IFI35,IFI44,IFI6,IFIH1,IFIT1,IFIT3,IFITM1,IFITM3,IKZF3,IL7,IRF1,IRF7,ISG15,ISG20,JUNB,KLF4,LGR4,LOX,MT-ND1,MX1,MYC,NR4A2,OAS1,OAS2,OAS3,OASL,PDGFB,PLSCR1,PTAFR,PTGS2,RSAD2,S1PR1,SERPINE1,SGK1,SLFN5,SOCS1,SOCS3,SOD2,SP110,STAT1,STAT2,TAP1,TNFRSF10B,TNFSF10,TRIM14,TRIM22,TRIM5,USP18,VCAN,XAF1,ZFP36 |
| dopamine | | chemical - endogenous mammalian | Activated | 3.385 | | 0.000335 | BTG2,CACNA2D3,CXCL8,EGR1,EGR3,FOS,FST,KLF4,NCF1,NR4A1,NR4A3,PER1 |
| VEGFA | -1.02 | growth factor | Activated | 3.38 | bias | 0.00000423 | CD36,CITED2,CTNNB1,CXCL8,CXCR4,EGR1,EPHB4,FOS,GBP1,HLA-DQB1,HLX,ID1,ID3,JUNB,NR4A1,NR4A2,PDGFB,PIK3CG,PTGS2,SCO2,SERPINE1,SOD2,TEK |
| leukotriene D4 | | chemical - endogenous mammalian | Activated | 3.378 | bias | 5.5E-12 | CDH1,CXCL8,CYSLTR1,DUSP1,EGR1,EGR2,EGR3,HLX,KLF2,KLF4,NR4A1,NR4A2,NR4A3,PTGS2,ZFP36 |
| ACKR2 | | G-protein coupled receptor | Activated | 3.357 | bias | 9.31E-14 | DDX58,DDX60,IFI44,IFIT3,IRF7,ISG15,ISG20,OAS1,OAS2,OAS3,OASL,RSAD2,STAT1,STAT2,USP18 |
| SP110 | -2.52 | transcription regulator | Activated | 3.273 | | 9.06E-11 | BST2,CD74,EGR3,IFI27,IFI6,IFIH1,IFIT1,IFIT3,IFITM1,IFITM3,LY96,MX1,MYC,OAS1,OAS3,PLSCR1,PTAFR,STAT1,TNFRSF21,TRIM5,TXNIP |
| NKX2-3 | -1.4 | transcription regulator | Activated | 3.258 | | 1E-20 | ANKRD55,BTG1,CASP1,CD36,CMPK2,CXCL8,DDX58,DDX60,FBXO6,GBP1,GIMAP7,HCP5,HLA-B,HLA-C,HLA-F,IL7R,KLF2,KLF4,LGMN,PARP12,PARP14,PARP9,PLSCR1,PTGS2,RPS23,RTP4,SAMD9L,SP110,STAT1,STAT2,TAP1,TIPARP,TRIM21,TRIM22,TRIM5,UACA,USP18,XAF1 |
| EGF | | growth factor | Activated | 3.211 | | 7.83E-08 | ACSL4,AREG,ARTN,BTG2,CASP1,CDH1,CISH,CXCL8,CXCR4,DUSP1,EGR1,EGR2,EGR3,EPCAM,EREG,FOS,FST,ID1,ID2,IDS,IL7R,JUNB,JUND,MYC,NCF1,NR4A1,NR4A2,NR4A3,PER1,PLSCR1,PTGS2,SERPINE1,SOCS3,STAT1,TGM2,VCAN,ZFP36 |
| NUPR1 | | transcription regulator | Activated | 3.128 | | 0.00624 | AREG,ARHGEF26,B3GNT5,BCL9,BTG1,CITED2,CXCL8,CXCR4,DEPTOR,EREG,KLF4,KLF6,MYC,NUP50,OSBPL6,PARP9,PCTP,PLEKHM3,RELB,SERPINE1,SLC16A10,ST3GAL2,VANGL1 |
| EGR1 | 7.47 | transcription regulator | Activated | 3.111 | bias | 1.86E-08 | ABCB1,ALOX5AP,AREG,CDH1,CXCL8,EGR1,EGR2,EREG,JUNB,JUND,MYC,NR4A1,PF4,PNMT,PTGS2,SERPINE1,SGK1,SOCS1,SOD2,TIAM1,TNFSF10 |
| AREG | 30.9 | growth factor | Activated | 3.094 | | 0.0000876 | AREG,CXCR4,EGR1,EREG,FOS,GBP1,IFI6,IFIT3,PTAFR,PTGS2 |
| Pkc(s) | | group | Activated | 3.087 | | 6.95E-08 | AHR,CACNA1C,CD36,CD4,CTNNB1,CXCL8,CXCR4,DBP,DUSP1,EGR1,EGR2,FOS,ID2,JUNB,KLF6,NR4A1,NR4A2,NR4A3,PER1,PPARA,PTGS2,SERPINE1 |
| TGFB1 | 1.1 | growth factor | Activated | 3.052 | | 6.45E-15 | ABCG1,ACVR1,ADORA2B,AHR,AKR1C1/AKR1C2,ALOX5AP,ALPP,APOE,AREG,ARL4A,B2M,B4GAT1,BHLHE40,BTG1,CASP1,CD36,CD4,CD69,CDC42EP4,CDH1,CDKN2C,CIITA,CITED2,CNN2,CTNNB1,CXCL8,CXCR4,CYB561,DBP,DEPTOR,DUSP1,EGR1,EGR2,EGR3,EPCAM,EREG,FOS,FZD2,GAL,GBP1,GCLC,GZMA,HLA-DQA1,HLA-DQB1,HLA-DRB5,HMGA2,HRH2,HSPG2,ID1,ID2,ID3,IFI27,IFIH1,IFIT3,IRF1,ITGB5,JUNB,JUND,KLF15,KLF2,KLF4,KLF9,LOX,MS4A2,MYC,NCF1,NFKBIA,NR4A1,NR4A2,NR4A3,P2RY14,PARP3,PDGFB,PLSCR1,PLXNC1,PNMT,PORCN,PTAFR,PTGER2,PTGS2,RARG,RGCC,RSAD2,S1PR1,SELENBP1,SERPINE1,SGK1,SLC12A7,SOCS1,SOCS3,SOD2,STAT1,TFAP4,TGM2,TLR2,TMEM17,TNFRSF10A,TNFSF13B,TPSAB1/TPSB2,TSC22D3,TXNIP,VCAN,WDFY3,WNT5B,ZFP36 |
| STAT6 | -1.02 | transcription regulator | Activated | 3.037 | | 4.43E-13 | ANKRD55,ANXA1,APOE,AREG,CA2,CCDC86,CD69,CISH,CMPK2,CPOX,EGR1,EGR2,FKBP5,GBP7,HLA-DMB,HLX,IFI44,IFI44L,IFIH1,IFIT3,IFITM3,IRF1,IRF7,ISG15,ISG20,KCTD12,LGALS3BP,NFKBIA,OAS3,OASL,PLSCR1,PPARA,PTGS2,SCO2,SELENBP1,SERPINE1,SNX20,SOCS1,STAT2,TFEC,ZFP36 |
| FGF2 | 1.1 | growth factor | Activated | 3.018 | | 0.000043 | AREG,BTG2,CDH1,CTNNB1,CTSL,CXCR4,DKK1,EGR1,EPCAM,EREG,FOS,GBP1,ID2,ID3,JUNB,LOX,LRP6,MYC,NR4A1,NR4A2,PDGFB,PTGS2,SERPINE1,SMIM3,SOCS3,TNFSF10 |
| IRF8 | 1.08 | transcription regulator | Activated | 3.013 | | 1.64E-12 | B2M,CD4,CD74,CIITA,DDX58,EGR1,EGR2,FOS,GBP1,ID3,IFI44L,IFI6,IFIT3,ISG15,KLF4,KLF6,MYC,OAS1,STAT1,STAT2,TNFSF13B,TRIM21,TYROBP |
| MAPK14 | 1.26 | kinase | Activated | 3.004 | | 0.000000547 | ALOX5AP,AREG,CDH1,CXCL8,DKK1,DUSP1,EREG,FOS,JUNB,JUND,LRP6,MAP2K6,PPARGC1B,PTGS2,SGK1,SOCS3,SOD2,TGM2,ZFP36 |
| platelet activating factor | | chemical - endogenous mammalian | Activated | 2.943 | | 9.6E-09 | CD69,CDH1,EGR1,EGR2,FOS,NFKBIA,NR4A1,PTAFR,PTGS2,SELL,SERPINE1,SOCS3 |
| IL10RA | -1.79 | transmembrane receptor | Activated | 2.911 | | 0.000000168 | ABCB1,ABCG1,CA2,CD36,CD69,COL14A1,GPR171,GSTM2,GSTM5,HLA-A,IL15RA,IRF1,IRF7,LMO4,NLRC5,PF4,RSAD2,S1PR1,SELENBP1,SELL,SOCS3,STAT1,TAP1,TFEC,TGM2,TLR2 |
| NGLY1 | -1.1 | enzyme | Activated | 2.91 | | 0.00000018 | IFI27,IFI44,IFI44L,IFIT1,IFIT3,OAS1,OAS3,RSAD2,USP18 |
| D-glucose | | chemical - endogenous mammalian | Activated | 2.907 | | 0.0000551 | ABCB1,ABCG1,ANXA1,CACNA1C,CASP1,CD36,CTNNB1,CXCL8,CXCR4,DUSP1,EGR1,FOS,FST,GABARAPL1,ID1,JUNB,LY96,MT-ND1,MYC,NCF1,NR4A1,NR4A2,NR4A3,PGAM1,PPARA,PTGS2,RELB,SERPINE1,SGK1,SLC39A6,SOD2,SORBS1,TGM2,TLR2,TXNIP |
| NRAS | -1.08 | enzyme | Activated | 2.905 | | 0.000706 | B2M,CTNNB1,EGR1,HLA-A,IFI35,IFIH1,IFIT1,ISG15,KCTD12,PTGS2,RNASE6,STAT1,TAP1,USP18 |
| KRAS | 1.07 | enzyme | Activated | 2.882 | | 3.51E-14 | AHR,AREG,B2M,CASP1,CD274,CD74,CDH1,CDKN2C,CTNNB1,CXCL8,DUSP1,EGR1,EGR2,EPCAM,EPHB4,EREG,FOS,GCLC,GSTM3,HLA-DMB,HOPX,ID1,IFI6,IFIT1,IFITM1,IFITM3,IRF1,ISG15,JUNB,KCNN4,KLF6,LOX,LTBP4,MICOS10-NBL1/NBL1,MX1,MYC,NCF1,NFKB2,NFKBIA,NR4A1,OAS1,P2RY14,PTGS2,ST3GAL2,STAT1,STAT2,TAP1,TLE3,TLR2,TMEM98,TNFRSF10A,TNFRSF10B,TNFSF10,VCAN,VMP1 |
| EDN1 | | cytokine | Activated | 2.865 | bias | 2.22E-09 | ANXA1,AREG,CDH1,CTNNB1,CXCL8,CXCR4,EGR1,EREG,FOS,FST,JUNB,MYC,NCF1,PPARA,PTGER2,PTGS2,RAMP3,SELL,SERPINE1,SOCS1,SOCS3,TGM2,VCAN |
| SOCS1 | 2.68 | other | Activated | 2.855 | bias | 3.63E-17 | CD69,CIITA,CISH,CXCL8,DDX58,DUSP1,FKBP5,FOS,IFI27,IFI44,IFIH1,IFIT1,IFIT3,IRF1,IRF7,ISG15,ISG20,MX1,OAS1,OAS2,PTGS2,RSAD2,SOCS1,SOCS3,STAT1,TGM2,TRIM22,USP18 |
| APEX1 | -1.27 | enzyme | Activated | 2.813 | bias | 0.000000855 | ABCB1,EGR1,EGR2,EGR3,FOS,JUNB,NR4A1,PTGS2 |
| NEDD9 | -1.05 | other | Activated | 2.813 | | 0.00193 | ARMC4,BHLHE40,CDH1,FOS,LIN7A,SERPINE1,SLC14A1,TXNIP |
| 10E,12Z-octadecadienoic acid | | chemical - endogenous mammalian | Activated | 2.811 | | 0.0086 | CD36,CXCL8,EGR2,NR4A1,NR4A2,NR4A3,PANK1,PTGS2,RGS1,SORBS1 |
| INS | | other | Activated | 2.773 | | 0.000248 | CD36,CDH1,CITED2,DEPTOR,DUSP1,EGR1,EGR2,FOS,GABARAPL1,JUNB,JUND,NR4A1,PTGS2 |
| mir-21 | | microRNA | Activated | 2.772 | | 0.0000015 | BTG2,CD274,CDH1,DBP,EGR2,EIF4E3,FGL2,GBP7,HMGA2,IRF1,MYC,NLRC5,OAS2,OAS3,PGAM1,PPARA,SLC16A10,SOD2,STAT1,STAT2,TAP1,TIAM1,TLR2 |
| CEBPA | | transcription regulator | Activated | 2.765 | | 3.99E-14 | ABCB1,AKR1C1/AKR1C2,ALOX5AP,ANXA1,ARL4C,BTG1,BTG2,CA2,CDH1,CIITA,CXCL8,CXCR4,EGR2,FOS,GAL,GBP1,GP9,HLA-B,ID1,ID2,IFI27,IFI6,IKZF3,ISG15,ITGB5,JUNB,KLF4,MSI2,MYC,NR4A2,OAS2,PPARA,PTAFR,PTGS2,SERPINE1,SOCS3,SOD2,THRB,TIAM1,TNFRSF10A,TNFRSF19,TNFSF10,TSC22D3,VCAN |
| IGF1 | | growth factor | Activated | 2.757 | | 0.000000448 | ADORA2B,AKR1C1/AKR1C2,BHLHE40,BTG2,CD74,CDH1,CISH,CITED2,CTNNB1,CXCL8,DUSP1,EGR1,EGR2,FOS,HDC,ID1,ID2,IFITM3,ITGB5,JUNB,KLF6,MYC,NFE2,NFKBIA,NR4A1,PTGS2,SERPINE1,SGK1,SLC12A7,SOCS3,TAP1,TXNIP,ZFP36 |
| arachidonic acid | | chemical - endogenous mammalian | Activated | 2.753 | | 0.00575 | ABCG1,ACSL4,CD36,EGR1,FOS,KLF6,NFKBIA,PTGS2 |
| cyclic AMP | | chemical - endogenous mammalian | Activated | 2.665 | | 0.00000601 | CA2,EGR1,EGR2,EPCAM,FOS,JUNB,LGALS3BP,MYC,NR4A1,NR4A2,PER1,PNMT,PTGER2,PTGS2,SERPINE1,SOCS3,SOD2,STAT1,TNFRSF10B,TNFSF10,TXNIP |
| Tgf beta | | group | Activated | 2.65 | | 2.86E-09 | BHLHE40,CASP1,CD69,CDH1,CIITA,CTNNB1,CXCL8,CXCR4,EGR2,FOS,FST,GCLC,HSPG2,ID1,ID3,JUNB,KRT1,LOX,MYC,NR4A2,PDGFB,PTGER2,PTGS2,SERPINE1,SGK1,SOCS3,TLR2,TNFSF10 |
| CNOT7 | -1.33 | transcription regulator | Activated | 2.621 | bias | 1.01E-26 | B2M,BST2,CMPK2,HERC6,IFI27,IFI35,IFI44L,IFI6,IFITM1,ISG15,LGALS3BP,OAS1,OAS2,OAS3,PARP12,PLSCR1,PPM1K,SLC14A1,SP110,STAT1,TAP1 |
| prostaglandin E2 | | chemical - endogenous mammalian | Activated | 2.62 | | 4.31E-11 | ABCG1,ADORA2B,AHR,AREG,BTG1,CASP1,CD274,CD69,CDH1,CXCL8,CXCR4,DUSP1,EGR1,EGR2,EREG,FOS,FST,HDC,IL15RA,IL7R,IRF1,KLF4,MYC,NFKBIA,NR4A1,NR4A2,PIK3CG,PTGER2,PTGS2,SELL,SOCS3,TSC22D3 |
| IL17A | | cytokine | Activated | 2.57 | | 0.0000432 | AREG,CD274,CDH1,CXCL8,CXCR4,CYTH3,EGR2,EREG,FOS,ISG15,KLF2,LOX,MYC,PPARA,PTGS2,SGK1,SOCS1,SOCS3,TLR2 |
| YBX1 | -1.07 | transcription regulator | Activated | 2.565 | | 0.0000329 | ABCB1,ALKAL2,APOL2,FOS,HLA-DQB1,HLA-DRA,ID1,KLF4,PDGFB,PTGS2,TRIM14 |
| NGF | | growth factor | Activated | 2.563 | | 0.00708 | BTG2,CDH1,DUSP1,EGR1,EGR2,FOS,GAL,ID1,JUNB,MYC,NR4A1,PNMT,SMIM3 |
| MAPK8 | -1.29 | kinase | Activated | 2.476 | | 0.00000235 | ABCB1,ACSL4,APOE,CDH1,CXCL8,DUSP1,EGR1,FOS,JUND,KLF6,MYC,PPARGC1B,PTGS2,SOD2,TLR2,TSC22D3,ZFP36 |
| leukotriene C4 | | chemical - endogenous mammalian | Activated | 2.446 | bias | 0.00000275 | CXCL8,EGR1,EGR2,EGR3,FOS,PTGS2 |
| GSK3B | -1.45 | kinase | Activated | 2.442 | | 0.000295 | CTNNB1,CXCL8,FOS,KIF3A,MYC,NFE2L3,PTGS2,SERPINE1,TCF7L2,TNFRSF10B,TNFSF10 |
| EPHB1 | | kinase | Activated | 2.429 | bias | 0.000000117 | EGR1,EGR2,FOS,JUNB,PTGS2,SERPINE1 |
| SAMHD1 | -1.4 | enzyme | Activated | 2.425 | bias | 0.000000416 | DDX58,IFI27,IFI6,MX1,NFKB2,NFKBIA |
| BMP15 | | growth factor | Activated | 2.423 | | 0.0000196 | AREG,EREG,ID1,ID2,ID3,PTGS2 |
| Ngf | | group | Activated | 2.388 | | 0.00696 | DUSP1,EGR1,FOS,NR4A1,NR4A2,SERPINE1 |
| ERK | | group | Activated | 2.385 | | 5.86E-09 | AREG,BTG2,CDH1,CTNNB1,CXCL8,DUSP1,EGR1,EGR2,EREG,FOS,FST,GCLC,HOMER1,JUNB,KYNU,MAFF,MYC,PTGS2,SELL,SERPINE1,STAT1,TGM2,TNFRSF10B,VCAN,ZFP36 |
| IL3 | | cytokine | Activated | 2.38 | | 0.000000981 | ALOX5AP,AREG,BTG2,CD69,CISH,CITED2,CXCL8,CXCR4,EGR1,EGR2,EGR3,FOS,GZMA,HLA-A,JUNB,KLF1,KLF9,MS4A2,MYC,NCF1,SELL,SOCS1,SOCS3,TLR2,TNFSF10,TPSAB1/TPSB2 |
| MAPK7 | -1.34 | kinase | Activated | 2.373 | | 0.00000381 | CXCL8,DUSP1,ID1,KLF2,KLF4,NR4A1,PTGER2,PTGS2,TEK |
| PGF | | growth factor | Activated | 2.368 | | 0.00000826 | ALOX5AP,CASP1,CXCL8,CXCR4,EGR1,EREG,FST,SERPINE1 |
| SMAD4 | 1.06 | transcription regulator | Activated | 2.353 | | 7.65E-08 | AREG,CDH1,CITED2,CTNNB1,CYB561,EREG,FOS,FST,HMGA2,HRH2,ID1,ID2,ITGB5,KLF15,MYC,NFKBIA,PDGFB,PTGS2,RGCC,SERPINE1,SGK1,TNFRSF10A,ZFP36 |
| formaldehyde | | chemical - endogenous mammalian | Activated | 2.351 | bias | 0.0000259 | EGR1,EGR2,FOS,JUNB,JUND,PTGS2,SGK1 |
| TAB1 | -1.28 | enzyme | Activated | 2.351 | | 3.2E-09 | CXCL8,GBP1,IFIH1,IFIT1,IRF7,PTGS2,SERPINE1,TNFSF10,TNFSF13B,XAF1 |
| HMGA1 | 1.03 | transcription regulator | Activated | 2.331 | | 0.0151 | ARL4A,B2M,EGR1,FOS,ID3,JUNB,KLF2,MYC,PTGS2 |
| Insulin | | group | Activated | 2.291 | | 0.000167 | APOE,CD36,CTNNB1,DUSP1,EGR1,EGR2,FOS,GCLC,IFIT3,IRF7,KLF4,LGALS3BP,MT-ND1,MYC,NCF1,NFKBIA,NR4A1,NR4A2,NR4A3,PPARA,PPARGC1B,SERPINE1,SGK1,SORBS1,STAT1,STAT2,TLR2 |
| PRDM16 | 1.44 | transcription regulator | Activated | 2.28 | | 0.00000293 | GBP4,IFI44,IRF7,OAS2,OAS3,SERPINE1,STAT1,STAT2 |
| CREM | -1.07 | transcription regulator | Activated | 2.264 | | 0.0000495 | APOE,BHLHE40,BTG2,CXCL8,DUSP1,EGR1,EGR2,FOS,HLA-DRA,JUNB,NR4A1,NR4A2,PER1,TIPARP |

| | | | | | | | |
|---|---|---|---|---|---|---|---|
| PPP2CA | -1.58 | phosphatase | Activated | 2.236 | | 0.00014 | CASP1,CD36,GZMA,IRF1,IRF7,MYC,PTGS2 |
| TLR7/8 | | group | Activated | 2.236 | bias | 0.000591 | ATP2B1-AS1,CA2,EREG,NFKBIA,PTGS2,RGS1 |
| BMP6 | | growth factor | Activated | 2.23 | | 0.00356 | CDKN2C,CYP1B1,ID1,ID2,ID3,KLF4,PTGS2,SERPINE1 |
| ISG15 | -3.91 | other | Activated | 2.222 | | 0.000000223 | DDX58,IFI6,IFITM3,MX1,OAS1 |
| RNF31 | | enzyme | Activated | 2.219 | | 0.0419 | AREG,CXCL8,CXCR4,EGR3,IL15RA |
| KAT5 | 1.18 | transcription regulator | Activated | 2.219 | bias | 0.0188 | CXCL8,CXCR4,EREG,MYC,PTCRA,RGS1,SOD2 |
| HBEGF | 1.39 | growth factor | Activated | 2.219 | | 0.000869 | BHLHE40,CDH1,CXCR4,EGR1,EREG,PTGS2 |
| DNASE2 | -1.43 | enzyme | Activated | 2.213 | bias | 0.000000137 | IFIT3,IRF7,ISG15,OAS1,OAS3,RSAD2,RTP4,TNFSF10,USP18 |
| PSEN1 | 1.12 | peptidase | Activated | 2.213 | | 0.0283 | ABCC6,ABCG1,APOE,CD74,CTNNB1,DBP,DUSP1,EGR1,FOS,HOMER1,IRF7,MYC,PGAM1,PTGS2,SOCS1,TIPARP,TNFRSF10B |
| KL | | enzyme | Activated | 2.207 | | 0.00476 | CTNNB1,EGR1,IRF7,MYC,SOD2 |
| N4BP1 | -1.39 | other | Activated | 2.207 | | 0.00000251 | HLA-A,HLA-B,JUNB,NFKB2,RELB |
| Lh | | complex | Activated | 2.203 | | 0.00000868 | AREG,ARL4C,CDC42EP4,CTNNB1,CXCL8,CXCR4,DUSP1,EREG,FKBP5,FOS,FST,GBP1,GNLY,ITGB5,PER1,PTGS2,RAB2A,SGK1,SOD2,STAT1 |
| USP18 | -2.97 | peptidase | Activated | 2.201 | bias | 1.04E-10 | IFI6,IFIH1,IFITM3,IRF1,IRF7,ISG15,MX1,OAS1,SOCS1,SOCS3,TNFSF10 |
| TGFA | | growth factor | Activated | 2.197 | | 0.0000476 | AREG,CASP1,CTNNB1,CXCL8,EREG,FOS,JUNB,NR4A2,PTGS2,SERPINE1 |
| ethanol | | chemical - endogenous mammalian | Activated | 2.181 | | 0.0000152 | ABCG1,CACNA1C,CD36,CXCL8,CXCR4,DKK1,DUSP1,EGR1,FOS,GNAI1,HLA-A,ID1,IRF7,KLF6,MYC,NCF1,PPARA,PTGS2,RGCC,SERPINE1,SGK1,SOD2,STAT1,TLR2,TP53INP1,TSC22D3 |
| histamine | | chemical - endogenous mammalian | Activated | 2.179 | | 0.0063 | AREG,CTNNB1,CXCL8,CXCR4,FOS,NR4A1,PTGS2 |
| TWIST1 | | transcription regulator | Activated | 2.17 | | 0.0273 | CDH1,CXCL8,EPCAM,FOS,ITGB5,KCNN4,RELB,TNFRSF21,VCAN |
| SMAD7 | | transcription regulator | Activated | 2.165 | | 0.000582 | ACVR1,AREG,CDKN2C,CITED2,DEPTOR,FST,ID1,ID2,ID3,MYC,NFKBIA,SERPINE1 |
| SMAD3 | 1.28 | transcription regulator | Activated | 2.165 | | 6.04E-08 | AREG,CDH1,CIITA,CTNNB1,DEPTOR,EGR1,EREG,FOS,FST,GZMA,ID1,ID2,ID3,ITGB5,JUNB,JUND,MYC,PDGFB,PNMT,PTGS2,S1PR1,SERPINE1,TPSAB1/TPSB2,ZFP36 |
| Smad | | complex | Activated | 2.164 | bias | 0.000384 | ID1,ID2,ID3,MYC,SERPINE1 |
| NLRX1 | -1.48 | other | Activated | 2.16 | | 0.0000379 | IRF1,NFKB2,NFKBIA,NFKBIE,OAS1,RELB,STAT1,STAT2 |
| CGA | | other | Activated | 2.158 | | 0.00543 | CXCL8,EGR2,ID1,KLF9,PTGS2,SFRP5,SGK1,TP53INP1 |
| ETV4 | | transcription regulator | Activated | 2.143 | | 0.00216 | CDH1,CTNNB1,CXCL8,CXCR4,PTGS2 |
| PTH | | other | Activated | 2.141 | | 0.0000379 | AREG,BHLHE40,CGNL1,CTNNB1,CXCR4,DUSP1,EGR2,FOS,NR4A1,NR4A2,PTGS2,RAMP3,RGS1,SLC43A2,SOCS3 |
| CREB1 | -1.53 | transcription regulator | Activated | 2.121 | | 1.31E-08 | ALKAL2,APOE,ATF1,BHLHE40,BTG2,CACNA1C,CD4,CIITA,CXCL8,CXCR4,DUSP1,EGR1,EGR2,FGL2,FOS,GAL,GPR63,HLA-A,HLA-DQB1,HLA-DRA,HLA-G,HOMER1,ID1,IRF7,JUNB,KLF4,MYC,NR4A1,NR4A2,NR4A3,PDE3B,PER1,PLXNC1,PNMT,PTGS2,RASGEF1B,SGK1,SOD2,TIPARP,ZFP36 |
| IgG | | complex | Activated | 2.086 | | 0.000000936 | APOE,B2M,CD69,CXCL8,DDX58,DUSP1,EGR1,EGR2,EGR3,FST,IFIH1,IFITM1,IFITM3,ISG15,JUND,MYC,PTGS2,S1PR1,SERPINE1,ZFP36 |
| norepinephrine | | chemical - endogenous mammalian | Activated | 2.068 | | 0.00000217 | CHST2,CXCL8,CXCR4,DUSP1,EGR1,FOS,FST,ID1,NFKBIA,NR4A1,NR4A3,PADI4,PER1,PTGS2,SERPINE1,SGK1,WHRN |
| SPP1 | | cytokine | Activated | 2.05 | | 0.049 | CA2,CDH1,CXCL8,FOS,ITGB5,LOX,MYC,PTGS2,SERPINE1 |
| Ca2+ | | chemical - endogenous mammalian | Activated | 2.043 | | 3.46E-10 | ABCB1,AHR,CDH1,CXCL8,CXCR4,CYP1B1,DBP,DUSP1,EGR1,FOS,HDC,HOMER1,ID1,ID3,IRF1,JUNB,JUND,KLF4,KRT1,NFKBIA,NR4A1,NR4A3,PER1,PPARA,PTGS2,SERPINE1,SGK1,TNFRSF10B |
| TLR2 | -2.95 | transmembrane receptor | Activated | 2.035 | | 0.0000371 | CD69,CISH,CXCL8,DUSP1,HLA-DQA1,HLA-DRB5,IL15RA,IRF1,KLF2,MX1,PTGS2,SOCS1,SOCS3,TLR2,TNFAIP8L2,TNFSF10 |
| CRH | | cytokine | Activated | 2.021 | | 0.000283 | CXCL8,FOS,JUNB,KRT1,NR4A1,PNMT,SGK1,SOCS3 |
| CLDN7 | 1.04 | other | Activated | 2.008 | | 0.00034 | APOE,CXCL8,DENND5A,DKK1,FZD2,HLA-B,IFI44,IFI6,LGR4,MX1,SLC25A22 |
| 5-hydroxytryptamine | | chemical - endogenous mammalian | Activated | 2.007 | | 0.0108 | CXCL8,CYP1B1,EGR1,FOS,MYC,NR4A1,PTGS2,SERPINE1 |
| Nfat (family) | | group | Activated | 2.007 | bias | 0.0000698 | CXCL8,DHRS9,EGR2,FGF18,KLF6,PDE3B,PDGFB,PTGS2,RELB,SERPINE1,SOD2 |
| ERK1/2 | | group | Activated | 2.002 | | 8.05E-09 | ABCC6,AREG,B2M,CACNA1C,CXCL8,CXCR4,DKK1,DUSP1,EGR1,FKBP5,FOS,HLA-A,ID1,ID3,IFIT1,JUNB,JUND,MYC,NR4A3,PTGER2,PTGS2,SERPINE1,SGK1,SLC12A7,SOCS3,TAP1,TNFRSF10B,VCAN |
| DDX58 | -2.28 | enzyme | Inhibited | -2.011 | bias | 2.75E-18 | CD4,CXCL8,DDX58,IFI27,IFI35,IFI44,IFIH1,IFIT1,IFIT3,IRF1,IRF7,ISG15,ISG20,KLF4,OAS1,PTGS2,RSAD2,SOCS1,SOCS3,STAT1,STAT2,TNFSF10 |
| FOSL1 | -1.47 | transcription regulator | Inhibited | -2.066 | | 0.0034 | CXCL8,EGR1,EGR2,FOS,GCLC,JUNB,SERPINE1 |
| MSC | | transcription regulator | Inhibited | -2.111 | | 0.000000187 | DDX60,EPSTI1,GPAT3,IFI27,IFI44,IFI44L,IFIT1,IRF7,PAG1,SELL,XAF1 |
| VCAN | 5.91 | other | Inhibited | -2.111 | | 8.12E-09 | APOE,CASP1,CXCL8,FST,IFI44,IFI44L,IFI6,IFIT1,IFITM1,MX1,NFKBIA,OAS2,OAS3,PARP14,SOD2,STAT1,TCF7L2,TRIM22,VCAN,XAF1 |
| SFTPA1 | | transporter | Inhibited | -2.121 | | 0.00096 | AKR1C1/AKR1C2,CYP1B1,EGR1,FOS,KLF2,LOX,MYC,SERPINE1 |
| MEOX2 | | transcription regulator | Inhibited | -2.177 | | 0.0031 | CD69,CXCL8,ID1,ID3,ITGB5 |
| EPCAM | -2.64 | other | Inhibited | -2.177 | | 0.0000148 | CDH1,EGR1,EPCAM,FOS,ID1,MYC |
| IFIH1 | -3.59 | enzyme | Inhibited | -2.208 | bias | 5.35E-12 | CD4,CISH,CXCL8,EGR1,IFI27,IFI44L,IFIT1,IRF7,ISG15,NFKBIA,OAS1,OAS2,RSAD2,USP18 |
| CLEC12A | -1.27 | other | Inhibited | -2.213 | bias | 0.000000223 | IFIT3,IRF7,ISG15,RSAD2,USP18 |
| FZD9 | | G-protein coupled receptor | Inhibited | -2.213 | bias | 0.000053 | CDH1,IFI44,IRF7,ISG15,STAT1 |
| Hsp90 | | group | Inhibited | -2.215 | | 0.0766 | AHR,CD4,CD69,CIITA,MYC |
| CBX7 | -1.4 | other | Inhibited | -2.219 | bias | 0.000125 | AREG,CDH1,CXCL8,CYP1B1,KYNU |
| AICAR | | chemical - endogenous mammalian | Inhibited | -2.224 | | 0.0578 | ABCG1,CTNNB1,PPARA,SERPINE1,SOCS3 |
| RXRA | -1.32 | ligand-dependent nuclear receptor | Inhibited | -2.229 | | 0.00218 | ABCB1,ABCG1,AKR1C1/AKR1C2,APOE,ARL4C,CD36,DLK1,FOS,GCLC,KLF9,PDGFB,PNMT,PPARA,RARG,SORBS1,TNFRSF10B,VCAN |
| FMR1 | -1.24 | translation regulator | Inhibited | -2.236 | | 0.183 | CLSTN3,CTNNB1,CYP1B1,EGR3,SLC38A1,SYT5 |
| IFNK | | cytokine | Inhibited | -2.236 | bias | 0.000254 | IFIH1,IRF1,MX1,OAS1,STAT1 |
| LRPAP1 | 1.16 | other | Inhibited | -2.236 | | 0.000384 | CDH1,CXCL8,PTGS2,SERPINE1,SORL1 |
| IFNAR2 | -1.09 | transmembrane receptor | Inhibited | -2.236 | bias | 2.93E-13 | DDX58,GBP4,HERC5,IFI44,IFI6,IFIH1,IFITM1,ISG15,OAS1,OAS2,TGM2,TNFSF10,TRIM22,USP18,XAF1 |
| DUSP5 | 1.47 | phosphatase | Inhibited | -2.236 | bias | 0.000125 | CA2,EGR1,EGR3,ID2,ZFP36 |
| TMEM173 | -1.03 | other | Inhibited | -2.287 | bias | 0.000000004 | CGAS,CXCL8,HDC,HOPX,IFI44,IFIT3,IFITM3,IRF7,ISG15,OAS1,OASL,RSAD2,USP18 |
| SMARCB1 | 1.02 | transcription regulator | Inhibited | -2.318 | | 0.0000405 | ABCB1,BTG1,CA8,CDH1,CDKN2C,COL14A1,CXCR4,FOS,FZD7,IFITM1,IL15RA,MX1,MYC,OAS1,OAS3 |
| BTNL2 | | transmembrane receptor | Inhibited | -2.333 | | 0.000129 | CD74,CDKN2C,CHD7,EPCAM,FGL2,GBP4,IFITM3,SELL,TNFSF10 |
| IFNAR1 | -1.39 | transmembrane receptor | Inhibited | -2.344 | | 1.3E-24 | B2M,CD274,CGAS,CIITA,CMPK2,GBP4,HLA-A,IFI44,IFI6,IFIH1,IFIT3,IFITM1,IRF1,IRF7,ISG15,MYC,OAS1,OAS2,OAS3,OASL,PARP12,PTGS2,RSAD2,RTP4,SERPINE1,SOCS1,SOCS3,SOD2,STAT1,TGM2,TLR2,TNFSF10,TNFSF13B,TRIM22,USP18,XAF1 |
| PRKCA | 1.24 | kinase | Inhibited | -2.349 | | 0.000000648 | ARL4C,CIITA,EGR1,FOS,ID1,ID2,ID3,IFIT1,IFIT3,MYC,NFKBIA,OASL,PTGS2,RSAD2,SERPINE1 |
| SASH1 | | other | Inhibited | -2.357 | bias | 1.64E-13 | CITED2,CMPK2,FGL2,HDC,IFIT3,IL15RA,IRF1,IRF7,ISG15,ISG20,LMO4,OASL,PTGS2,RSAD2,SOCS1,STAT1,STAT2,TRIM21 |
| EBF1 | | transcription regulator | Inhibited | -2.382 | | 0.00791 | CRK,IGLL1/IGLL5,IRF1,IRF7,PDE3B,SOCS1,SOCS3,STAT1,TLR2 |
| SOX4 | -1.09 | transcription regulator | Inhibited | -2.387 | | 0.0000286 | B3GNT5,CASP1,CD74,CGAS,CTNNB1,EIF4E3,FOS,GPR171,GPR183,HLA-DQB1,KYNU,MEX3A,SELL,TIAM1,TMEM98,TNFRSF19 |
| FGF7 | -1.5 | growth factor | Inhibited | -2.395 | | 0.0105 | IL7,IRF1,NFE2L3,PTAFR,SERPINE1,STAT1 |
| SGPL1 | -1.53 | enzyme | Inhibited | -2.592 | | 0.00000123 | ABCB1,DDX58,IFIT1,ISG15,MX1,OAS1,PTGS2 |
| IFNL4 | | cytokine | Inhibited | -2.623 | bias | 8.07E-10 | DDX58,IFIH1,ISG15,MX1,OAS1,OAS2,STAT1 |
| IL27 | | cytokine | Inhibited | -2.67 | | 4.7E-23 | AHR,B2M,BST2,CD274,CD69,CD74,CIITA,CXCL8,EGR1,EGR2,FOS,HLA-A,HLA-B,HLA-C,HLA-DMB,HLA-DOA,HLA-DPA1,HLA-DQA1,HLA-DQB1,HLA-DRA,ID2,IL15RA,IRF1,MX1,MYC,OAS1,PTGS2,SOCS1,SOCS3,STAT1,STAT2,TAP1,TNFSF10,TNFSF13B |
| PARP9 | -4.61 | enzyme | Inhibited | -2.729 | bias | 1.21E-15 | CD74,HLA-DQA1,HLA-DQB1,IFI44,IFIT1,IFIT3,IRF1,IRF7,ISG15,OAS2,SP110,STAT1 |
| IFNA1/IFNA13 | | cytokine | Inhibited | -2.751 | | 8.64E-18 | CD274,CD69,IFI27,IFI6,IFIH1,IFIT1,IFITM1,IRF1,IRF7,ISG15,MX1,MYC,OAS1,OAS2,PLSCR1,PTGS2,RSAD2,SOCS1,STAT1,STAT2 |
| NLRC5 | -2.82 | transcription regulator | Inhibited | -2.756 | | 2.77E-08 | B2M,DDX58,HLA-A,HLA-B,HLA-C,HLA-F,HLA-G,TAP1 |
| POU2AF1 | | transcription regulator | Inhibited | -2.78 | bias | 0.0000161 | CD69,HLA-DRA,ID3,IFI44,IFI44L,IFIT3,IFITM1,KCNN4,MX1 |
| IFN alpha/beta | | group | Inhibited | -2.881 | bias | 2.14E-10 | CD69,FGL2,HLA-A,IFIT3,IL15RA,IL7,IRF1,IRF7,RSAD2,SOCS1,STAT1,STAT2,TLR2,TNFSF10,TNFSF13B,TRIM21 |
| JAK1/2 | | group | Inhibited | -2.887 | bias | 0.000000241 | CD69,CD74,CIITA,HLA-DQA1,HLA-DRA,HLA-DRB5,IRF1,IRF7,ISG15,MX1,RSAD2,WIPF3 |
| DUSP1 | 2.96 | phosphatase | Inhibited | -2.97 | | 1.01E-09 | APOL6,CMPK2,CXCL8,DKK1,DUSP1,IFIT1,IFIT3,IRF1,ISG20,MX1,PAQR7,PDGFB,PTGS2,SERPINE1,SOCS1,SOD2,TLR2,ZFP36 |

| | | | | | | | |
|---|---|---|---|---|---|---|---|
| JAK | | group | Inhibited | -3 | | 3.41E-11 | DDX58,IFI6,IFIH1,IFIT1,IFIT3,IFITM3,ISG15,PTGER2,PTGS2,RSAD2,SOCS1,SOCS3,STAT1 |
| MAVS | -1.33 | other | Inhibited | -3.064 | bias | 5.68E-16 | CGAS,CMPK2,CXCL8,DDX58,IFIT1,IFIT3,IFITM3,IRF7,ISG15,ISG20,OAS1,OAS2,OASL,PARP12,RSAD2,SOCS1,SOCS3,STAT1,STAT2,USP18 |
| SPI1 | -1.36 | transcription regulator | Inhibited | -3.14 | | 4.12E-16 | CIITA,CMPK2,EGR2,FOS,HDAC7,ID2,ID3,IFI27,IFI44,IFI44L,IFI6,IFIT1,IFIT3,IFITM1,IFITM3,IL7R,IRF7,ISG15,ISG20,KLF4,MT1G,MX1,MYC,NCF1,OASL,PARP12,PIK3CG,PTCRA,PTGS2,RSAD2,SP110,TFEC,TLR2,TNFSF10,TRIM22,USP18 |
| PML | -1.57 | transcription regulator | Inhibited | -3.244 | | 3.78E-18 | ACSL4,APOE,BST2,CIITA,CXCR4,DDX60,EPSTI1,HERC6,HLA-DMB,HLA-DRA,ID2,IFI27,IFI35,IFI44,IFI44L,IFIH1,IFIT1,IFIT3,IFITM1,IRF7,ISG15,ISG20,MAFF,MX1,OAS1,OAS2,OAS3,PLSCR1,STAT1,TAP1,UBE2M |
| IFN type 1 | | group | Inhibited | -3.258 | | 2.56E-11 | BST2,CD69,CGAS,CXCL8,DDX58,IFIH1,IFIT1,IRF1,ISG15,OASL,STAT1,STAT2,TNFSF10,TNFSF13B |
| tretinoin | | chemical - endogenous mammalian | Inhibited | -3.263 | | 6.76E-20 | ABCG1,AHR,ALDH9A1,ANXA1,APOE,AREG,BHLHE40,BTG1,BTG2,CA2,CA8,CACNA1C,CASP1,CD274,CD36,CD74,CDH1,CDKN2C,CITED2,CTNNB1,CTSL,CXCL8,CXCR4,DDX58,DDX60,DHRS9,DKK1,DUSP1,EGR1,EPCAM,FOS,FST,GBP4,GZMA,HLA-A,HLA-B,HLA-C,HLA-DMB,HOXA3,ID1,ID2,ID3,IFI27,IFI35,IFI44,IFI44L,IFI6,IFIH1,IFIT1,IFIT3,IFITM1,IL1RAPL1,IRF1,ISG15,KLF1,KLF15,KLF4,KLF9,KRT1,LGALS3BP,LRP6,MS4A3,MYC,NCF1,NFE2,OAS1,OAS2,OAS3,OASL,PADI4,PARP14,PARP9,PDGFB,PIK3CG,PLSCR1,PPARA,PTAFR,PTGS2,RARG,RNASE1,RTP4,SAMD9L,SELL,SERPINE1,SFRP5,SLA,SLFN5,SMOX,SOCS1,SP110,STAT1,STAT2,SWT1,TAP1,TGM2,TLR2,TNFRSF10A,TNFRSF10B,TNFRSF19,TNFSF10,TRIM22,TSC22D3,TYROBP,USP18,VMP1,XAF1,ZBTB16 |
| CIITA | -3.88 | transcription regulator | Inhibited | -3.3 | | 3.49E-09 | B2M,CD74,HLA-A,HLA-B,HLA-DMB,HLA-DOA,HLA-DPA1,HLA-DQA1,HLA-DQB1,HLA-DRA,HLA-DRB5 |
| EBI3 | | cytokine | Inhibited | -3.359 | | 1.6E-15 | B2M,CD274,CIITA,FOS,HLA-A,HLA-B,HLA-C,HLA-DMB,HLA-DPA1,HLA-DQA1,HLA-DRA,IRF1,MX1,MYC,STAT1,STAT2,TAP1,TNFSF10 |
| Ifn | | group | Inhibited | -3.73 | | 3.2E-20 | B2M,CD69,DDX58,FOS,HLA-A,IFIH1,IFIT1,IFITM1,IFITM3,IL15RA,IL7,IRF1,IRF7,ISG15,ISG20,MX1,OAS2,OAS3,OASL,PTAFR,RSAD2,SERPINE1,SOCS1,STAT1,TAP1,TLR2,TNFSF13B,TRIM21,TRIM22 |
| IRF5 | -1.47 | transcription regulator | Inhibited | -3.85 | bias | 3.25E-17 | CMPK2,CXCR4,DDX58,IFI44,IFIH1,IFIT1,IFIT3,IFITM3,IRF7,ISG15,ISG20,MYC,OAS1,OAS2,OASL,PARP12,PLSCR1,PTGS2,RSAD2,SP110,STAT1,STAT2,TNFSF10 |
| TGM2 | -3.24 | enzyme | Inhibited | -4.041 | | 4.25E-14 | BTG2,CA2,CD36,CD74,CXCL8,DDX60,HLA-B,IFI35,IFI6,IFIT1,IFIT3,MS4A3,MYC,OAS1,OAS2,OAS3,OASL,PADI4,PARP14,PARP9,PLSCR1,SAMD9L,SELL,SLFN5,SP110,STAT1,SWT1,TAP1,TRIM22,TYROBP,XAF1 |
| STAT1 | -4.71 | transcription regulator | Inhibited | -4.229 | | 1.7E-40 | APOE,APOL4,APOL6,B2M,BST2,BTG1,CASP1,CD274,CIITA,CMPK2,CXCL8,DDX60,EGR1,EPSTI1,FGL2,FOS,GBP1,GBP1P1,GBP4,HERC6,HLA-DQA1,HLA-DRB5,IFI27,IFI35,IFI44,IFI44L,IFI6,IFIH1,IFIT1,IFIT3,IFITM1,IFITM3,IL15RA,IRF1,IRF7,ISG15,KCTD12,KLF4,LY96,MS4A3,MX1,MYC,NFE2,NLRC5,OAS1,OAS2,OAS3,OASL,PARP9,PF4,PLSCR1,PPARGC1B,PTGS2,RSAD2,RTP4,SAMD9L,SLFN5,SOCS1,SOCS3,SP110,STAT1,STAT2,TAP1,TNFRSF10B,TNFSF10,TNFSF13B,TRIM21,TRIM22,USP18,XAF1 |
| RNY3 | | other | Inhibited | -4.243 | bias | 7.07E-23 | DDX60,EPSTI1,HERC5,IFI44,IFI44L,IFIT1,IFIT3,IFITM3,ISG15,MX1,OAS1,OAS2,OAS3,OASL,RSAD2,RTP4,TRIM6,XAF1 |
| IFNB1 | | cytokine | Inhibited | -4.349 | | 3.02E-21 | APOL2,APOL3,APOL6,BHLHE40,BST2,BTN3A3,CASP1,CD274,CMPK2,CXCL8,DDX58,FOS,GBP4,GBP7,HERC5,HLA-A,IFI27,IFI6,IFIH1,IFIT1,IFIT3,IFITM1,IRF1,IRF7,ISG15,ISG20,MX1,MYC,NFKBIE,OAS1,OAS2,PARP12,PARP14,PTGS2,RSAD2,SOCS1,STAT1,STAT2,TNFRSF10A,TNFSF10,TRIM14,TRIM21,USP18,XAF1 |
| Ifnar | | group | Inhibited | -4.424 | bias | 2.16E-22 | B2M,CASP1,CD274,CD74,DDX58,HLA-A,HLA-G,IFI35,IFIH1,IFIT3,IFITM3,IRF1,IRF7,ISG15,ISG20,NLRC5,OAS1,OAS2,OASL,RSAD2,STAT1,STAT2,TAP1,TLR2,TNFSF10,TRIM21,USP18,XAF1 |
| IRF3 | -1.14 | transcription regulator | Inhibited | -4.427 | bias | 3.61E-22 | B2M,CD274,CD69,CMPK2,CXCL8,DDX58,DDX60,FST,GBP1,HLA-F,IFI27,IFI44,IFI6,IFIH1,IFIT1,IFIT3,IFITM3,IRF1,IRF7,ISG15,ISG20,JUNB,NLRC5,OAS1,OAS2,OAS3,OASL,PARP12,PARP14,RSAD2,SAMD9L,SORL1,STAT1,STAT2,TAP1,TNFSF10,TPST1,USP18 |
| PRL | | cytokine | Inhibited | -4.523 | | 2.29E-28 | AKR1C4,B2M,BST2,CD69,CDH1,CISH,CMPK2,DDX58,DTX3L,EGR1,EPSTI1,FOS,HERC5,HERC6,ID1,ID2,ID3,IFI35,IFI44,IFI44L,IFI6,IFIH1,IFIT1,IFIT3,IFITM1,IRF1,IRF7,ISG15,MICOS10-NBL1/NBL1,MYC,OAS1,OAS2,OAS3,PAQR7,PAQR8,PARP12,PARP14,PLSCR1,PPM1K,RSAD2,SAMD9L,SOCS1,SOCS3,SP110,STAT1,STAT2,TNFSF13B,TRIM14,TRIM25,USP18,XAF1 |
| IFN Beta | | group | Inhibited | -4.645 | bias | 1.61E-26 | BST2,CD274,CD69,CD74,CXCL8,DDX58,DUSP1,HERC5,HLA-A,HLA-B,HLA-G,IFI27,IFI35,IFI44,IFI6,IFIH1,IFIT1,IFIT3,IFITM1,IL15RA,IRF1,IRF7,ISG15,MX1,MYC,OAS1,OAS2,OAS3,OASL,RSAD2,SOCS1,STAT1,STAT2,TNFSF10,TRIM14,USP18,XAF1 |
| IRF1 | -3.08 | transcription regulator | Inhibited | -5.154 | bias | 4.69E-32 | APOL6,B2M,CASP1,CD274,CIITA,CMPK2,CXCL8,DDX58,FGL2,HLA-G,IFI27,IFI35,IFI44,IFI44L,IFI6,IFIH1,IFIT1,IFIT3,IFITM1,IFITM3,IL7,IRF1,IRF7,ISG15,LIG4,MX1,MYC,NFE2,OAS1,OAS2,OAS3,OASL,PF4,PTGS2,RSAD2,SELL,SOCS1,SP110,STAT1,STAT2,TAP1,TNFSF10,TNFSF13B,TRIM21,TRIM22,XAF1 |
| IRF7 | -4.57 | transcription regulator | Inhibited | -5.309 | bias | 2.35E-30 | CD69,CMPK2,DDX58,GBP1,GBP4,HERC5,IFI35,IFI44,IFI44L,IFI6,IFIH1,IFIT1,IFIT3,IFITM1,IFITM3,IL15RA,IRF1,IRF7,ISG15,ISG20,MX1,OAS1,OAS2,OAS3,OASL,PARP12,PARP14,PLSCR1,RSAD2,RTP4,SAMD9L,SOCS1,STAT1,STAT2,TAP1,TNFSF10,TNFSF13B,TPST1,TRIM21,TRIM22,TRIM5,USP18,XAF1 |
| IFNL1 | | cytokine | Inhibited | -5.354 | bias | 1.2E-40 | APOL6,BST2,CMPK2,CXCL8,DDX58,DDX60,GBP1,HERC5,HERC6,HLA-B,HLA-C,IFI27,IFI35,IFI44,IFI44L,IFI6,IFIH1,IFIT1,IFIT3,IFITM1,IFITM3,ISG15,ISG20,LGALS3BP,MX1,OAS1,OAS2,OAS3,OASL,PLSCR1,RSAD2,RTP4,SP110,STAT1,STAT2,TLR2,TRIM14,TRIM22,USP18,XAF1 |
| Interferon alpha | | group | Inhibited | -5.381 | | 1.96E-41 | APOL2,APOL3,B2M,BST2,BTG2,CASP1,CD274,CD69,CD81,CDH1,CGAS,CIITA,CISH,CXCL8,CYB561,DDX58,EPSTI1,FBXO6,FOS,GBP1,HERC5,HERC6,HLA-A,HLA-B,HLA-C,HLA-F,HLA-G,IFI27,IFI35,IFI44,IFI44L,IFI6,IFIH1,IFIT1,IFIT3,IFITM1,IFITM3,IGLL1/IGLL5,IL15RA,IL7,IL7R,IRF1,IRF7,ISG15,ISG20,MX1,MYC,NFE2L3,NFKBIA,OAS1,OAS2,OAS3,PARP12,PARP14,PARP9,PLSCR1,PPM1K,PTGS2,RSAD2,RTP4,SAMD9L,SLC1A5,SLFN5,SOCS1,SOCS3,SP110,STAT1,STAT2,TAP1,TICAM1,TLR2,TNFSF10,TNFSF13B,TRANK1,TRIM21,TRIM22,TRIM5,USP18 |
| IFNA2 | | cytokine | Inhibited | -5.466 | bias | 2.79E-39 | ANXA1,APOL6,B2M,BST2,CD274,CD69,CISH,CMPK2,CORO2A,DDX58,DDX60,GBP1,GBP4,HERC5,HERC6,HLA-A,HLA-B,HLA-C,IFI27,IFI35,IFI44,IFI44L,IFI6,IFIH1,IFIT1,IFIT3,IFITM1,IFITM3,IRF1,IRF7,ISG15,ISG20,LGALS3BP,MT1G,MX1,MYC,OAS1,OAS2,OAS3,PARP12,PARP9,PLSCR1,PTAFR,RSAD2,SOCS1,SOCS3,SP110,STAT1,TAP1,TGM2,TNFRSF10B,TNFSF10,TRIM14,TRIM21,TRIM22,USP18,XAF1,ZBTB11 |
| IFNG | | cytokine | Inhibited | -5.538 | | 2.46E-44 | ABCB1,ADORA2B,AHR,ALKAL2,ALOX5AP,APOL6,AREG,B2M,BST2,BTG1,BTN3A1,BTN3A2,CASP1,CD274,CD36,CD4,CD74,CDH1,CGAS,CIITA,CISH,CMPK2,CTNNB1,CXCL8,CXCR4,CYB561,DBP,DDX58,DDX60,DKK1,DTX3L,DUSP1,EGR1,EGR2,EGR3,FGL2,FKBP5,FOS,FZD2,GBP1,GBP4,GBP7,HCP5,HERC6,HLA-A,HLA-B,HLA-C,HLA-DMB,HLA-DOA,HLA-DQA1,HLA-DQB1,HLA-DRA,HLA-DRB5,HLA-F,HLA-G,HSPG2,ID1,IFI27,IFI35,IFI44,IFI44L,IFI6,IFIH1,IFIT1,IFIT3,IFITM1,IFITM3,IL15RA,IL7,IL7R,IRF1,IRF7,ISG15,ISG20,JUNB,JUND,KCTD12,KLF1,KLF2,KLF4,KLF6,KYNU,LGALS3BP,LOX,LY96,MAFF,MT1G,MX1,MYC,NEAT1,NFE2L3,NFKB2,NFKBIA,NLRC5,OAS1,OAS2,OAS3,OASL,P2RY14,PARP14,PARP9,PCTP,PDGFB,PF4,PLSCR1,PPARA,PPARGC1B,PTAFR,PTGS2,RELB,RGCC,RSAD2,RTP4,SELL,SERPINE1,SLC14A1,SLFN5,SOCS1,SOCS3,SOD2,SP110,STAT1,STAT2,TAP1,TCF7L2,TICAM1,TLR2,TNFRSF10A,TNFRSF10B,TNFSF10,TNFSF13B,TRIM21,TRIM22,TSC22D3,TXNIP,TYROBP,USP18,XAF1,ZBTB16,ZFP36 |

**T2 vs. T1 HSC_MPP**

© 2000-2020 QIAGEN. All rights reserved.

| Upstream Regulator | Expr Fold Change | Molecule Type | Predicted Activation State | Activation z-score | p-value of overlap | Target Molecules in Dataset |
|---|---|---|---|---|---|---|
| TRIM24 | -1.06 | transcription regulator | Activated | 3.199 | 9.11E-10 | EPSTI1,GLUL,IFIT3,IRF1,IRF7,IRF9,ISG15,OAS1,OASL,PCLAF,PLEC,SAMHD1,STAT1,STAT2 |
| MYC | -1.18 | transcription regulator | Activated | 3.117 | 1.18E-08 | CCND3,CD69,CLDN7,CLU,COL2A1,COX6A1,DDIT3,DDIT4,DUSP1,DUSP5,FBLN5,FKBP5,FOSL1,GADD45B,GLUL,IER3,IFI44L,IFI6,IFIT3,IGLL1/IGLL5,IRF7,IRF9,ISG20,KLF10,KLF6,MAN2A1,MX1,MYCN,OAS1,OASL,PMAIP1,REN,RHOB,RPL10,RPL3,RPL7A,RPSA,RSAD2,SLC22A4,SLC38A1,ST3GAL1,STAT1<br>,TNFSF13B,TOB1,TXNIP |
| IL1RN | | cytokine | Activated | 2.998 | 6.68E-14 | ATF3,CXCL1,GBP1,IFI44L,IFI6,IFIT3,IRF1,IRF7,IRF9,ISG20,KLF6,MX1,OAS1,OAS3,OASL,PMAIP1,RSAD2,S100A9,STAT2 |
| NKX2-3 | 2.24 | transcription regulator | Activated | 2.84 | 0.00000106 | AGAP3,BTG1,GBP1,GIMAP7,HCP5,HLA-C,HLA-F,KLF2,PARP9,PMAIP1,PTPRE,STAT1,STAT2,TYMP,XAF1 |
| PNPT1 | -1.18 | enzyme | Activated | 2.828 | 0.000000649 | IFIT3,IRF7,ISG15,OAS1,PARP9,STAT1,STAT2,XAF1 |
| IRF4 | | transcription regulator | Activated | 2.639 | 0.000000679 | CXCR4,DDIT3,GALNT2,GBP1,IRF1,IRF7,IRF9,ISG15,OAS1,RHOB,STAT1,STAT2,TNFSF13B |
| corticosterone | | chemical - endogenous mammalian | Activated | 2.6 | 0.00271 | DDIT3,DDIT4,FKBP5,GLUL,KLF9,POMC,TSC22D3 |
| Pkc(s) | | group | Activated | 2.552 | 0.000471 | ATF3,CCND3,CD4,CXCR4,DDIT3,DUSP1,FOSL1,GADD45B,KLF6,PER1,PTPRE |
| Irgm1 | | other | Activated | 2.449 | 0.0000496 | DTL,IFIT3,IRF7,OASL,RSAD2,SELP |
| AREG | 6.2 | growth factor | Activated | 2.359 | 0.00000614 | AREG,CXCR4,EREG,GBP1,H2BC8,IFI6,IFIT3,ITGB8,PTAFR |
| SP110 | -1.65 | transcription regulator | Activated | 2.324 | 2.87E-09 | ATF3,BCL3,CLU,IFI6,IFIT3,IFITM1,IRF9,MX1,MYCN,OAS1,OAS3,PANX1,PTAFR,STAT1,TXNIP |
| IKZF1 | 1.79 | transcription regulator | Activated | 2.287 | 0.0000251 | CCL3L3,CCND3,CITED2,DUSP5,EPSTI1,FKBP9,FLT3,IFI6,IFIT3,IGLL1/IGLL5,POMC,SMAD3,TXNIP,WDFY3 |
| DNMT3A | 1.05 | enzyme | Activated | 2.236 | 0.108 | CXCR4,IRF1,IRF8,JCHAIN,SKIL |
| FOXA2 | | transcription regulator | Activated | 2.236 | 0.126 | CXCR4,DDIT4,GSTM3,NFKB2,NFKBIE,RNASE2 |
| MLXIPL | | transcription regulator | Activated | 2.236 | 0.0339 | RPL10,RPL3,RPL7A,RPSA,TXNIP |
| ANXA1 | 1.11 | enzyme | Activated | 2.2 | 0.0000288 | ANXA2,CD69,CXCR4,POMC,TSC22D3,TYMP |
| NUPR1 | | transcription regulator | Activated | 2.183 | 0.00147 | AREG,ATF3,ATP6V0A1,BTG1,CITED2,CREB5,CXCR4,DDIT3,DUSP5,EPGN,EREG,IGF2BP3,KLF6,NFIB,PARP9,SAMHD1,TOB1 |
| NGLY1 | 1.06 | enzyme | Activated | 2.169 | 0.000154 | IFI44L,IFIT3,OAS1,OAS3,RSAD2 |
| USP18 | -1.7 | peptidase | Activated | 2.157 | 9.15E-09 | CCL3L3,IFI6,IRF1,IRF7,IRF9,ISG15,MX1,OAS1 |
| Lh | | complex | Activated | 2.149 | 0.000024 | AREG,CDC42EP4,CXCR4,DUSP1,EREG,FKBP5,GBP1,NR5A1,PER1,PMAIP1,PPP2R5B,PTPRE,RHOB,STAT1 |
| ACKR2 | | G-protein coupled receptor | Activated | 2.111 | 2.32E-11 | CCL3L3,IFIT3,IRF7,ISG15,ISG20,OAS1,OAS3,OASL,RSAD2,STAT1,STAT2 |
| PGR | | ligand-dependent nuclear receptor | Activated | 2.102 | 2.7E-10 | AK4,AKR1C3,AREG,BTG1,CCND3,CLDN7,FKBP5,GLUL,GSTM3,HBA1/HBA2,IER3,ITGB4,KLF11,KLF9,MYCN,PNMT,RAB27B,SERPINB8,ST3GAL1,STAT1,TSC22D3,UGCG |
| SIRT1 | -1.73 | transcription regulator | Activated | 2.034 | 0.000132 | CCND3,CD83,CDKN2C,FCMR,GSTM3,HLA-DRB5,IFIT3,IGHM,IRF7,JCHAIN,KLF2,NFKBIE,NLGN1,OAS1,PER1,RSAD2,STAT1 |
| PTGS2 | 2.08 | enzyme | Activated | 2.029 | 0.000465 | ANXA2,AREG,CCL3L3,CDKN2C,CLU,CXCR4,DUSP1,ITGB4,PTGER2,ST3GAL1 |
| STAT6 | 1.2 | transcription regulator | Activated | 2.018 | 5.62E-09 | AREG,BCL3,CCL3L3,CD69,CISH,FCGR1A,FKBP5,GADD45B,GBP5,GBP7,IFI44L,IFIT3,IRF1,IRF7,IRF9,ISG15,ISG20,LTB,OAS3,OASL,RASGRP3,SELP,STAT2,USP2,ZNF608 |
| BRCA1 | -1.05 | transcription regulator | Inhibited | -2.11 | 0.000000502 | AREG,DDIT3,DDIT4,ENPP2,FKBP5,GADD45B,HBB,IFI6,IFIT3,IFITM1,IRF7,MX1,SMAD3,STAT1 |
| cytokine | | group | Inhibited | -2.111 | 0.00000323 | BCL3,CD4,CD69,CISH,CLU,CXCL1,DDIT3,DUSP1,IRF1,RPSA,SELP,TYMP |
| JAK1/2 | | group | Inhibited | -2.121 | 0.00000984 | CD69,GBP5,HLA-DRB5,IRF1,IRF7,ISG15,MX1,RSAD2 |
| tretinoin | | chemical - endogenous mammalian | Inhibited | -2.144 | 8.22E-11 | ANXA2,AREG,BCL3,BMPR1B,BTG1,CCND3,CD53,CDKN2C,CITED2,COL2A1,CXCL1,CXCR4,DDIT3,DTL,DUSP1,ENPP2,FOSL1,GAS2,GIMAP2,GIMAP6,HLA-C,IER3,IFI44L,IFI6,IFIT3,IFITM1,IGHM,IRF1,IRF9,ISG15,JCHAIN,KLF15,KLF9,METTL7A,MT2A,MYCN,NKX2-<br>3,OAS1,OAS3,OASL,PARP9,PMAIP1,POMC,PPP2R5B,PTAFR,REN,RPL10,RPL3,S100A9,SAMHD1,SELP,SERPINB8,SLFN5,SMAD3,STAT1,STAT2,TOB1,TSC22D3,XAF1 |
| IL1B | 1.5 | cytokine | Inhibited | -2.145 | 8.63E-14 | ATF3,BCL3,CCL3L3,CD4,CD69,CD83,CISH,COL2A1,CXCL1,CXCR4,DDIT3,DDIT4,DUSP1,DUSP5,ENPP2,F13A1,FKBP5,FOSL1,GADD45B,GBP1,IER3,IFIT3,IRF1,IRF7,ISG15,ISG20,ITGB8,KLF10,MT2A,MX1,NEAT1,NFKB2,POMC,REN,RHOB,RPSA,RSAD2,S100A9,SESN1,SLC22A4,SLC2A9,STAT1,TNFSF13<br>B,TOB1,TSC22D3,TXNIP,TYMP,UGCG |
| CGAS | -2.37 | enzyme | Inhibited | -2.186 | 0.0000184 | IFIT3,IRF7,ISG15,OAS1,RSAD2 |
| IFIH1 | -1.63 | enzyme | Inhibited | -2.195 | 0.00000469 | CD4,CISH,IFI44L,IRF7,ISG15,OAS1,RSAD2 |
| OSM | -1.23 | cytokine | Inhibited | -2.201 | 5.82E-11 | AKR1C1/AKR1C2,AKR1C3,ANXA2,ATF3,BCL3,CDC42EP4,CISH,COL2A1,CXCL1,DENND5A,FOSL1,GBP1,GLUL,HLA-C,HLA-F,IER3,IRF1,IRF7,IRF9,ISG20,KLF10,MARCKS,MT2A,MX1,OAS1,PFKFB2,POMC,S100A9,SELP,SERPINB8,STAT1,TYMP |
| IFNK | | cytokine | Inhibited | -2.236 | 0.0000184 | CD83,IRF1,MX1,OAS1,STAT1 |
| IL1A | | cytokine | Inhibited | -2.257 | 0.000273 | ATF3,BCL3,CD83,CXCL1,FOSL1,GBP1,IER3,IRF1,MCAM,MT2A,POMC,S100A9 |
| STAT2 | -2.07 | transcription regulator | Inhibited | -2.269 | 6.52E-11 | GBP1,IFI6,IFIT3,IFITM1,IRF1,IRF7,IRF9,ISG15,MX1,OAS1,RSAD2,STAT1 |
| TICAM1 | -1.8 | other | Inhibited | -2.337 | 3.11E-08 | BCL3,CCL3L3,DUSP1,IFIT3,IRF1,IRF7,ISG15,ISG20,MAFF,NFKB2,OASL,RHOB,RSAD2,SAMHD1 |
| TLR4 | -1.39 | transmembrane receptor | Inhibited | -2.337 | 4.34E-09 | AREG,ATF3,CCL3L3,CISH,CITED2,DDIT3,EREG,IFIT3,IRF1,IRF7,IRF8,ISG15,ISG20,MX1,NFKB2,OASL,RGS18,RSAD2,SAP30,SELP,ST3GAL1,STAT1,STAT2,UGCG |
| MYD88 | -1.16 | other | Inhibited | -2.34 | 0.00000337 | BCL3,CCL3L3,CD83,CISH,CXCL1,DUSP1,IRF1,IRF7,IRF8,ISG15,MAFF,NFKB2,OASL,RSAD2,SAMHD1,TNFSF13B |
| JAK1 | 1.01 | kinase | Inhibited | -2.359 | 6.13E-09 | BCL3,CCL3L3,ENPP2,HLA-C,HLA-F,IRF1,IRF7,IRF9,MX1,STAT1,STAT2 |
| ECSIT | 1.04 | transcription regulator | Inhibited | -2.372 | 0.00000226 | BCL3,CD83,IER3,IRF7,NFKB2,NFKBIE |
| TMEM173 | -1.01 | other | Inhibited | -2.377 | 0.000000264 | CGAS,GBP5,HOPX,IFIT3,IRF7,ISG15,OAS1,OASL,RSAD2 |
| IFN type 1 | | group | Inhibited | -2.4 | 0.00000076 | CD69,CGAS,IRF1,ISG15,OASL,STAT1,STAT2,TNFSF13B |
| Ifn gamma | | complex | Inhibited | -2.423 | 0.00351 | GBP1,IFI44L,IRF8,PMAIP1,STAT1,XAF1 |
| SFTPA1 | | transporter | Inhibited | -2.449 | 0.00111 | AKR1C1/AKR1C2,ANKRD35,CLEC1B,HBA1/HBA2,KLF2,ST3GAL1 |
| SPI1 | -1.18 | transcription regulator | Inhibited | -2.53 | 0.000657 | IFI44L,IFI6,IFIT3,IFITM1,IRF7,IRF9,ISG15,ISG20,MT2A,MX1,OASL,RSAD2 |
| IFN alpha/beta | | group | Inhibited | -2.538 | 0.000000319 | CD69,CD83,IFIT3,IRF1,IRF7,IRF8,RSAD2,STAT1,STAT2,TNFSF13B |
| PML | 1.36 | transcription regulator | Inhibited | -2.546 | 3.62E-09 | ACADL,CXCL1,CXCR4,EPSTI1,HBB,IFI44L,IFIT3,IFITM1,IRF7,ISG15,ISG20,MAFF,MX1,OAS1,OAS3,STAT1 |
| MAVS | 1.1 | other | Inhibited | -2.578 | 7.78E-08 | CGAS,IFIT3,IRF7,ISG15,ISG20,OAS1,OASL,RSAD2,STAT1,STAT2 |
| CSF3 | | cytokine | Inhibited | -2.584 | 0.00475 | CCL3L3,CCND3,CISH,CXCR4,ENPP2,FCGR1A,GADD45B,LTB |
| IRF3 | 1.15 | transcription regulator | Inhibited | -2.612 | 2.64E-12 | CCL3L3,CD69,CXCL1,FCGR1A,GBP1,GBP5,HLA-F,IFI6,IFIT3,IRF1,IRF7,ISG15,ISG20,OAS1,OAS3,OASL,PMAIP1,RSAD2,SAP30,STAT1,STAT2 |
| IRF5 | -1.07 | transcription regulator | Inhibited | -2.741 | 7.16E-08 | CXCR4,IFIT3,IRF7,ISG15,ISG20,OAS1,OASL,PMAIP1,RSAD2,STAT1,STAT2 |
| TLR9 | | transmembrane receptor | Inhibited | -2.753 | 3.23E-14 | ATF3,CCL3L3,CD69,CD83,CISH,DUSP1,FCMR,GADD45B,IER3,IFI44L,IFIT3,IFITM1,IRF1,IRF7,IRF9,ISG15,ISG20,MX1,OAS3,PTGER2,RSAD2,STAT1,STAT2,TNFSF13B,UGCG |
| DDX58 | -1.21 | enzyme | Inhibited | -2.758 | 7.16E-09 | CD4,IFIT3,IRF1,IRF7,IRF8,ISG15,ISG20,OAS1,RSAD2,STAT1,STAT2 |
| Ifn | | group | Inhibited | -2.854 | 3.13E-11 | CD69,CD83,IFITM1,IRF1,IRF7,IRF8,ISG15,ISG20,MX1,OAS3,OASL,PTAFR,RSAD2,STAT1,TNFSF13B,TYMP |
| EIF2AK2 | -1.55 | kinase | Inhibited | -2.904 | 2.18E-09 | ATF3,DDIT3,IFI6,IFITM1,IRF1,ISG15,ISG20,NFKB2,OAS1,OAS3,PARP9,SAMHD1,STAT1 |
| IFNAR1 | -1.05 | transmembrane receptor | Inhibited | -2.935 | 1.13E-10 | ATF3,CCL3L3,CGAS,IFI6,IFIT3,IFITM1,IRF1,IRF7,ISG15,OAS1,OAS3,OASL,RNASE2,RSAD2,STAT1,TNFSF13B,XAF1 |
| IFNA1/IFNA13 | | cytokine | Inhibited | -2.949 | 4.27E-11 | CD69,CD83,IFI6,IFITM1,IRF1,IRF7,ISG15,MX1,OAS1,RSAD2,STAT1,STAT2 |
| TNFSF10 | -1.23 | cytokine | Inhibited | -2.985 | 0.0000023 | CD69,CXCR4,HLA-C,HLA-F,IFI6,IFITM1,IRF9,ISG15,PMAIP1,STAT1 |
| TGM2 | -1.49 | enzyme | Inhibited | -3.011 | 0.000000333 | GIMAP2,GIMAP6,IFI6,IFIT3,IRF9,ITGB4,MT2A,OAS1,OAS3,OASL,PARP9,SAMHD1,SELP,SLFN5,STAT1,XAF1 |
| PRL | | cytokine | Inhibited | -3.123 | 9.4E-13 | AKR1C3,ANXA2,CCND3,CD69,CISH,CLU,EPSTI1,IER3,IFI44L,IFI6,IFIT3,IFITM1,IRF1,IRF7,IRF9,ISG15,OAS1,OAS3,RPSA,RSAD2,SAMHD1,STAT1,STAT2,TNFSF13B,XAF1 |
| TLR7 | | transmembrane receptor | Inhibited | -3.144 | 1.45E-12 | ATF3,CCND3,CD69,CD83,CREB5,CXCL1,FCMR,FOSL1,IER3,IFI44L,IFIT3,IFITM1,IRF1,IRF7,IRF9,ISG15,ISG20,MX1,OAS3,RSAD2,STAT1,STAT2 |
| RNY3 | | other | Inhibited | -3.162 | 2.9E-12 | EPSTI1,IFI44L,IFIT3,ISG15,MX1,OAS1,OAS3,OASL,RSAD2,XAF1 |
| TNF | -1.19 | cytokine | Inhibited | -3.258 | 1.57E-18 | ARRDC3,ATF3,AZU1,BCL3,BTG1,CCDC80,CCL3L3,CCND3,CD4,CD69,CD83,CDC42EP4,CDKN2C,CISH,CITED2,CLDN7,CLU,COL2A1,CSGALNACT1,CXCL1,CXCR4,DDIT3,DUSP1,DUSP5,EFNB2,ENPP2,EREG,FOSL1,GADD45B,GBP1,GBP1P1,GNAI1,HLA-<br>F,IER3,IFI6,IFIT3,IFITM1,IRF1,IRF7,IRF8,ISG15,ITGB8,KLF10,KLF2,KLF6,KLF7,KYNU,LTB,MAFF,MCAM,MT2A,MX1,NEAT1,NFKB2,NFKBIE,NKX3-<br>1,NR5A1,OAS1,OAS3,OASL,PMAIP1,POMC,RHOB,RNASE2,RPSA,S100A9,SELP,SERPINB8,SLC22A4,SMAD3,STAT1,TNFSF13B,TP53INP1,TPTEP1,TSC22D3,TXNIP,TYMP,UGCG,USP2,ZBTB11 |
| Ifnar | | group | Inhibited | -3.364 | 8.15E-12 | DDIT3,HLA-G,IFIT3,IRF1,IRF7,IRF8,IRF9,ISG15,ISG20,OAS1,OASL,RSAD2,STAT1,STAT2,XAF1 |
| IFN Beta | | group | Inhibited | -3.473 | 3.77E-15 | ATF3,CD69,CD83,CXCL1,DUSP1,HLA-G,IFI6,IFIT3,IFITM1,IRF1,IRF7,IRF9,ISG15,MX1,OAS1,OAS3,OASL,RSAD2,STAT1,STAT2,XAF1 |
| IRF1 | -3.68 | transcription regulator | Inhibited | -3.679 | 4.4E-11 | HLA-G,IFI44L,IFI6,IFIT3,IFITM1,IRF1,IRF7,IRF9,ISG15,LTB,MX1,OAS1,OAS3,OASL,RSAD2,STAT1,STAT2,TNFSF13B,XAF1 |

| | | | | | | |
|---|---|---|---|---|---|---|
| IFNB1 | | cytokine | Inhibited | -3.686 | 3.19E-09 | CD83,FKBP9,GBP5,GBP7,IFI6,IFIT3,IFITM1,IRF1,IRF7,IRF9,ISG15,ISG20,MX1,NFKBIE,OAS1,PHLDB2,PMAIP1,RSAD2,STAT1,STAT2,XAF1 |
| IFNL1 | | cytokine | Inhibited | -3.77 | 1.03E-18 | ATF3,CD83,GBP1,GBP5,HLA-C,IFI44L,IFI6,IFIT3,IFITM1,IRF9,ISG15,ISG20,MX1,OAS1,OAS3,OASL,RSAD2,STAT1,STAT2,XAF1 |
| IRF7 | -2.48 | transcription regulator | Inhibited | -3.81 | 6.63E-18 | CD69,FCGR1A,GBP1,GBP5,IFI44L,IFI6,IFIT3,IFITM1,IRF1,IRF7,IRF8,IRF9,ISG15,ISG20,MX1,OAS1,OAS3,OASL,PMAIP1,RSAD2,SAP30,STAT1,STAT2,TNFSF13B,XAF1 |
| IFNA2 | | cytokine | Inhibited | -3.922 | 4.06E-17 | BAG3,CD69,CISH,GBP1,HLA-C,IFI44L,IFI6,IFIT3,IFITM1,IRF1,IRF7,IRF9,ISG15,ISG20,MAL,MT-ATP6,MT2A,MX1,OAS1,OAS3,PARP9,PMAIP1,PTAFR,RSAD2,SAMHD1,STAT1,XAF1,ZBTB11 |
| Interferon alpha | | group | Inhibited | -4.18 | 2.07E-21 | ATF3,BCL3,CCND3,CD69,CD83,CGAS,CISH,CXCL1,DDIT4,ENPP2,EPSTI1,FCGR1A,GBP1,GBP5,HLA-C,HLA-F,HLA-G,IFI44L,IFI6,IFIT3,IFITM1,IGLL1/IGLL5,IRF1,IRF7,IRF8,IRF9,ISG15,ISG20,MX1,OAS1,OAS3,PARP9,PMAIP1,POMC,RSAD2,S100A9,SLFN5,STAT1,STAT2,TBC1D10A,TNFSF13B,TYMP |
| STAT1 | -4.7 | transcription regulator | Inhibited | -4.265 | 1.68E-18 | APOL4,BTG1,CCL3L3,CCND3,EPSTI1,FCGR1A,GBP1,GBP1P1,GBP5,HLA-DRB5,IFI44L,IFI6,IFIT3,IFITM1,IRF1,IRF7,IRF8,IRF9,ISG15,MX1,OAS1,OAS3,OASL,PARP9,PMAIP1,POMC,RSAD2,SAMHD1,SLFN5,SMAD3,STAT1,STAT2,TNFSF13B,TYMP,XAF1 |
| IFNG | | cytokine | Inhibited | -5.306 | 8.33E-23 | AREG,ATF3,BCL3,BTG1,CCL3L3,CCND3,CD4,CD83,CGAS,CISH,CXCL1,CXCR4,DDIT3,DUSP1,DUSP5,EFNB2,FBLN1,FBP1,FCGR1A,FCGR1B,FKBP5,GBP1,GBP5,GBP7,GLUL,H2BC8,HCP5,HLA-C,HLA-DRB5,HLA-F,HLA-G,IER3,IFI44L,IFI6,IFIT3,IFITM1,IRF1,IRF7,IRF8,IRF9,ISG15,ISG20,KLF10,KLF2,KLF6,KYNU,LTB,MAFF,MAN2A1,MX1,MYCN,NEAT1,NFKB2,OAS1,OAS3,OASL,PANX1,PARP9,PMAIP1,POMC,PTAFR,RHOB,RSAD2,S100A9,SAMHD1,SELP,SLFN5,SMAD3,STAT1,STAT2,TBC1D10A,TNFSF13B,TSC22D3,TXNIP,TYMP,XAF1 |

**T2 vs. T1 MPP_GMP**

© 2000-2020 QIAGEN. All rights reserved.

| Upstream Regulator | Expr Fold Change | Molecule Type | Predicted Activation State | Activation z-score | Flags | p-value of overlap | Target Molecules in Dataset |
|---|---|---|---|---|---|---|---|
| TRIM24 | -1.12 | transcription regulator | Activated | 3.878 | | 1.52E-17 | CA2,EPSTI1,IFI44,IFIH1,IFIT2,IFIT3,IRF1,IRF7,IRF9,ISG15,LGALS3,OAS1,OASL,SAMD9L,SAMHD1,STAT1,STAT2,TAP1,USP18 |
| IL1RN | | cytokine | Activated | 3.748 | | 3.91E-16 | ACTA2,ATF3,GBP1,IFI44,IFI44L,IFI6,IFIH1,IFIT3,IRF1,IRF7,IRF9,MX1,OAS1,OAS2,OASL,RSAD2,S100A9,STAT2,USP18 |
| MAPK1 | 1.08 | kinase | Activated | 3.71 | | 8.81E-19 | ATF3,CD69,CISH,DUSP1,EGR1,GBP1,GBP5,HBA1/HBA2,HBB,HERC5,HIC2,HLA-C,IFI44,IFI6,IFIH1,IFIT1,IFIT2,IFIT3,IFITM1,IFITM3,IRF7,IRF9,ISG15,LGALS3,OAS1,OAS2,OASL,SAMHD1,STAT1,STAT2,TAP1,USP18 |
| STAT6 | 1.29 | transcription regulator | Activated | 3.562 | | 1.48E-11 | AREG,BCL3,CA2,CD69,CD72,CISH,EGR1,FKBP5,GADD45B,GBP5,GBP7,IFI44,IFI44L,IFIH1,IFIT3,IFITM3,IL10RA,IRF1,IRF7,IRF9,ISG15,OASL,SAMD4A,STAT2,TMEM176B |
| ACKR2 | | G-protein coupled receptor | Activated | 3.464 | | 3.16E-14 | IFI44,IFIT2,IFIT3,IRF7,ISG15,OAS1,OAS2,OASL,RSAD2,STAT1,STAT2,USP18 |
| PNPT1 | -1.12 | enzyme | Activated | 3.464 | | 2.34E-13 | IFI44,IFIH1,IFIT3,IRF7,ISG15,OAS1,PARP9,SAMD9L,STAT1,STAT2,USP18,XAF1 |
| AREG | 10 | growth factor | Activated | 3.124 | | 5.18E-08 | AREG,CXCR4,EGR1,EREG,GBP1,IFI6,IFIT2,IFIT3,ITGB8,KIF20A |
| SOCS1 | 2.31 | other | Activated | 3.116 | | 2.1E-15 | CD69,CISH,DUSP1,FKBP5,GBP5,IFI44,IFIH1,IFIT1,IFIT2,IFIT3,IRF1,IRF7,ISG15,MX1,OAS1,OAS2,RSAD2,STAT1,USP18 |
| AR | -1.17 | ligand-dependent nuclear receptor | Activated | 3.01 | | 0.0000249 | DDIT4,DHCR24,EGR1,FKBP5,HOPX,KIF15,NR0B1,NR5A1,PER1,PFKFB2,PNMT,POMC,REN,RHOB,STAT1,TMEM176A,TMEM176B,WEE1 |
| USP18 | -2.89 | peptidase | Activated | 2.975 | bias | 2.67E-11 | IFI6,IFIH1,IFITM3,IRF1,IRF7,IRF9,ISG15,MX1,OAS1 |
| SIRT1 | -1.4 | transcription regulator | Activated | 2.867 | | 0.00000018 | CD72,CDKN2C,GSTM3,HELZ2,IFI44,IFIT3,IFITM3,IRF7,KLF2,LGALS3,NLRC5,OAS1,OAS2,PER1,RELB,RSAD2,STAT1,TAP1,USP18 |
| NR3C1 | 1.13 | ligand-dependent nuclear receptor | Activated | 2.821 | | 5.7E-09 | BAG3,BTG1,C1QTNF4,CD69,CXCR4,DDIT4,DUSP1,EGR1,FKBP5,GADD45B,IFI6,IFIH1,IFIT2,INHBA,IRF1,IRF8,ISG15,MAL,MT2A,NFKB2,NOD2,NR0B1,OASL,PER1,PNMT,POMC,RHOB,TSC22D3 |
| NGLY1 | -1.24 | enzyme | Activated | 2.744 | | 3.85E-09 | IFI44,IFI44L,IFIT1,IFIT2,IFIT3,OAS1,RSAD2,USP18 |
| SOX2 | | transcription regulator | Activated | 2.681 | | 0.00000703 | ATF3,CITED2,CSF1R,CXCR4,DUSP1,DUSP5,GADD45B,HOPX,ID2,INHBA,IRX3,MYCN,NFKB2,NOG,NR0B1,SAPCD2,TCL1A,USP18 |
| levodopa | | chemical - endogenous mammalian | Activated | 2.496 | | 0.04 | ACTA2,DLK1,DUSP1,DUSP6,FKBP5,FOSL1,GADD45B,GDF10,PER1,PIEZO2,RIMS1,S1PR2,ST3GAL5 |
| TAB1 | -1.15 | enzyme | Activated | 2.433 | | 0.00000105 | GBP1,IFIH1,IFIT1,IRF7,TNFSF13B,XAF1 |
| PRDM16 | -1.02 | transcription regulator | Activated | 2.418 | | 0.00000265 | IFI44,IFIT2,IRF7,OAS2,STAT1,STAT2 |
| NLRX1 | 1.15 | other | Activated | 2.401 | | 0.000019 | IRF1,NFKB2,OAS1,RELB,STAT1,STAT2 |
| SOCS3 | -1.47 | phosphatase | Activated | 2.395 | | 0.00000107 | ATF3,EGR1,IFIT1,IFIT2,IRF1,MX1,OAS1,OAS2,POMC |
| NKX2-3 | 1.67 | transcription regulator | Activated | 2.357 | | 5.68E-11 | ACTA2,BTG1,GBP1,HCP5,HELZ2,HLA-C,HLA-F,KLF2,PARP9,PTPRE,SAMD9L,SNCAIP,STAT1,STAT2,TAP1,TNFRSF10D,USP18,XAF1 |
| Lh | | complex | Activated | 2.354 | | 0.00000525 | ACTA2,AREG,CXCR4,DUSP1,EREG,FKBP5,GBP1,NR5A1,PER1,PPP2R5B,PTPRE,RHOB,STAT1 |
| PTGER4 | -1.38 | G-protein coupled receptor | Activated | 2.35 | | 1.03E-11 | ARRDC3,CD69,CITED2,CXCR4,EGR1,IFIH1,IFIT2,IRF1,IRF7,MAFF,NCF2,RSAD2,SAP30,TP53INP1,USP18,XAF1 |
| Irgm1 | | other | Activated | 2.333 | bias | 1.43E-09 | ID2,IFIT2,IFIT3,IRF7,KIF20A,OAS2,OASL,RSAD2,USP18 |
| STAT3 | -1.04 | transcription regulator | Activated | 2.255 | | 1.27E-21 | ACTA2,AREG,BCL3,CCR1,CDK1,CXCR4,EGR1,EME1,GBP5,HBB,HERC5,ID2,IFI44,IFI6,IFIH1,IFIT1,IFIT2,IFIT3,IFITM1,IFITM3,IKZF3,IRF1,IRF7,IRF9,ISG15,MT-ATP6,MX1,NR0B1,OAS1,OAS2,OASL,POMC,REN,RSAD2,S100A9,SLC1A3,STAT1,STAT2,TAP1,TOB1,USP18,XAF1 |
| RPSA | 1.99 | translation regulator | Activated | 2.236 | | 0.0000187 | DUSP1,ISG15,RELB,STAT1,TAP1 |
| TCF7L2 | -1.57 | transcription regulator | Activated | 2.236 | | 0.308 | DHCR24,FBP1,ID2,MAL,TPRN |
| PTGER2 | 1.22 | G-protein coupled receptor | Activated | 2.207 | | 0.000994 | AREG,CSF1R,CXCR4,EGR1,KIF15,KIF20A |
| TREX1 | | enzyme | Activated | 2.19 | | 0.000000452 | IFI44,IFIT2,ISG15,OASL,USP18 |
| PIK3CG | -1.16 | kinase | Activated | 2.121 | | 0.000025 | CXCR4,GBP5,GBP7,IRF1,NLRC5,OAS2,STAT1,TAP1 |
| IL4 | | cytokine | Activated | 2.038 | | 1.48E-10 | ACTA2,BCL3,C17orf99,CA2,CD69,CD72,CDK1,CISH,CITED2,CSF1R,CXCR4,EGLN1,FKBP5,GBP5,GBP7,IFI44,IFI44L,IFIH1,IFIT3,IFITM3,IL10RA,IRF1,IRF7,IRF8,IRF9,ISG15,LGALS3,MAL,NFKB2,OASL,S100A8,S100A9,SAMD4A,STAT1,STAT2,TINAGL1,TMEM176B |
| PGR | | ligand-dependent nuclear receptor | Activated | 2.029 | | 0.0000854 | AREG,BTG1,CDK1,FKBP5,GSTM3,HBA1/HBA2,MYCN,PNMT,RAB27B,S100A8,STAT1,TSC22D3 |
| SMARCA4 | 1.09 | transcription regulator | Inhibited | -2.01 | | 4.91E-08 | ACTA2,AREG,ARRDC4,CCR1,CXCR4,DUSP6,EGR1,EREG,FBP1,FKBP5,GADD45B,GBP1,HBB,HCP5,HLA-C,HLA-F,IFIT1,IFITM1,IFITM3,INHBA,IRF1,LGALS3,MAFF,NFKB2,TAP1 |
| FADD | | other | Inhibited | -2.135 | | 0.00000067 | CDK1,CXCR4,EGR1,GADD45B,IFIH1,IFIT2,IRF7,STAT1,STAT2 |
| SNCA | 1.06 | enzyme | Inhibited | -2.138 | | 0.011 | DLK1,GBP5,GNG4,IRF1,IRF7,ISG15,MX1,NCF2,RSAD2 |
| ID2 | -3.53 | transcription regulator | Inhibited | -2.157 | | 0.000000137 | ACTA2,BCL3,CCR1,CDK1,CDKN2C,CXCR4,DUSP1,GADD45B,IKZF3,IL10RA,IRF8,JDP2,PIK3IP1,RELB |
| IFIH1 | -2.17 | enzyme | Inhibited | -2.159 | | 7.05E-11 | CISH,EGR1,IFI44L,IFIT1,IRF7,ISG15,OAS1,OAS2,RSAD2,USP18 |
| IL1B | 1.11 | cytokine | Inhibited | -2.161 | | 2.59E-20 | ACTA2,ATF3,BCL3,CCR1,CD69,CISH,COL2A1,CXCR4,DDIT4,DUSP1,DUSP5,EGLN1,EGR1,FKBP5,FOSL1,GADD45B,GBP1,HELZ2,HERC5,IFIT1,IFIT3,IL10RA,INHBA,IRF1,IRF7,ISG15,ITGB8,KIF15,MT2A,MX1,NEAT1,NFKB2,OAS2,POMC,RELB,REN,RHOB,RSAD2,S100A8,S100A9,SESN1,SLC1A3,SLC3A1,S TAT1,TMEM176B,TNFSF13B,TOB1,TSC22D3,TXNIP,USP18 |
| PARP9 | -2.9 | enzyme | Inhibited | -2.178 | | 1.02E-13 | IFI44,IFIT1,IFIT2,IFIT3,IRF1,IRF7,ISG15,OAS2,STAT1 |
| ISGF3 | | complex | Inhibited | -2.2 | bias | 7.94E-10 | IFIH1,IFIT2,IRF1,IRF7,ISG15,RSAD2 |
| JUNB | -1.47 | transcription regulator | Inhibited | -2.2 | | 0.00000306 | ATF3,BCL3,CLU,DUSP1,FOSL1,ID2,INHBA,NCF2,POMC,RELB |
| Ifn gamma | | complex | Inhibited | -2.207 | | 0.00505 | GBP1,IFI44L,IRF8,STAT1,XAF1 |
| STAT4 | -1.86 | transcription regulator | Inhibited | -2.208 | bias | 0.000988 | BCL3,FYB1,IFIH1,IFIT2,IL10RA,IRF1,ISG15,PER1,STAT1 |
| Tnf (family) | | group | Inhibited | -2.236 | | 0.0259 | ATF3,CD72,INHBA,POMC,TNFSF13B |
| IFNK | | cytokine | Inhibited | -2.236 | | 0.00000471 | IFIH1,IRF1,MX1,OAS1,STAT1 |
| IFNA4 | | cytokine | Inhibited | -2.295 | bias | 7.05E-11 | CD69,GBP5,IFIH1,IFIT1,IFIT2,ISG15,MX1,OASL,RSAD2,USP18 |
| DOCK8 | -1.1 | other | Inhibited | -2.309 | bias | 1.41E-11 | CITED2,HELZ2,IFIT2,IFIT3,INHBA,IRF1,IRF7,ISG15,RSAD2,SAP30,STAT1,STAT2 |
| NFkB (complex) | | complex | Inhibited | -2.354 | | 0.00000102 | ATF3,BCL3,CD69,CLU,COL2A1,CXCR4,DUSP5,EGR1,GADD45B,HERC5,HIF3A,HLA-F,IL10RA,IRF1,IRF7,ISG15,ITGB8,NCF2,NFKB2,RELB,RSAD2,SLC3A1,TAP1 |
| IL27 | | cytokine | Inhibited | -2.356 | | 0.000000767 | CD69,EGR1,HLA-C,ID2,IRF1,MX1,OAS1,STAT1,STAT2,TAP1,TNFSF13B |
| IL12B | | cytokine | Inhibited | -2.39 | | 0.0000495 | DUSP5,IRF9,S100A8,S100A9,STAT1,STAT2 |
| CEBPA | | transcription regulator | Inhibited | -2.401 | | 1.62E-09 | AKR1C1/AKR1C2,BTG1,CA2,CCR1,CD3G,CSF1R,CXCR4,ELANE,GBP1,ID2,IFI6,IKZF3,ISG15,MNDA,MPO,MT2A,MYCN,OAS2,PTPRE,S100A8,S100A9,TSC22D3,TUBB2A |
| EBI3 | | cytokine | Inhibited | -2.416 | | 0.0000367 | HLA-C,IRF1,MX1,STAT1,STAT2,TAP1 |
| IFNL4 | | cytokine | Inhibited | -2.425 | bias | 3.64E-10 | IFIH1,ISG15,MX1,OAS1,OAS2,STAT1 |
| CLEC12A | -1.15 | other | Inhibited | -2.425 | bias | 1.25E-11 | IFIT2,IFIT3,IRF7,ISG15,RSAD2,USP18 |
| BRCA1 | 1.44 | transcription regulator | Inhibited | -2.483 | | 2E-10 | AREG,DDIT4,EGR1,FKBP5,GADD45B,HBB,IFI6,IFIT1,IFIT2,IFIT3,IFITM1,IRF7,MX1,STAT1,TAP1,WEE1 |
| SAMSN1 | -1.19 | other | Inhibited | -2.496 | | 2.64E-11 | CITED2,HELZ2,IFIT2,IFIT3,INHBA,IRF1,IRF7,ISG15,OASL,RSAD2,SAP30,STAT1,STAT2 |
| SASH1 | | other | Inhibited | -2.496 | bias | 9.74E-13 | CITED2,HELZ2,IFIT2,IFIT3,INHBA,IRF1,IRF7,ISG15,OASL,RSAD2,SAP30,STAT1,STAT2 |
| PAF1 | -1.12 | other | Inhibited | -2.53 | bias | 1.38E-10 | CITED2,HELZ2,HERC5,IFI44,IFI44L,IFIT3,IFITM3,ISG15,OAS2,OASL |
| IFN alpha/beta | | group | Inhibited | -2.547 | | 2.31E-08 | CD69,IFIT2,IFIT3,IRF1,IRF7,IRF8,RSAD2,STAT1,STAT2,TNFSF13B |
| STAT2 | -2.32 | transcription regulator | Inhibited | -2.628 | | 3.34E-18 | GBP1,IFI6,IFIT1,IFIT2,IFIT3,IFITM1,IRF1,IRF7,IRF9,ISG15,MX1,OAS1,OAS2,RSAD2,STAT1,USP18 |
| IFNE | | cytokine | Inhibited | -2.646 | | 0.000000448 | HERC5,IFIH1,IFIT2,IFITM3,ISG15,STAT1,USP18 |
| MSC | | transcription regulator | Inhibited | -2.646 | | 0.00000322 | EPSTI1,IFI44,IFI44L,IFIT1,IRF7,PIEZO2,XAF1 |
| IL21 | | cytokine | Inhibited | -2.706 | | 1.45E-09 | CD69,EGR1,GBP5,ID2,IFIT1,IFIT2,IFIT3,IRF7,ISG15,OAS2,OASL,RSAD2,STAT2,TAP1,USP18 |
| tretinoin | | chemical - endogenous mammalian | Inhibited | -2.715 | | 9.23E-16 | AREG,BCL3,BTG1,CA2,CCR1,CDKN2C,CITED2,COL2A1,CSF1R,CXCR4,DUSP1,EGR1,FOSL1,HLA-C,ID2,IFI44,IFI44L,IFI6,IFIH1,IFIT1,IFIT2,IFIT3,IFITM1,IL10RA,INHBA,IRF1,IRF9,ISG15,KLF15,MNDA,MPO,MT2A,MYCN,NCF2,OAS1,OAS2,OASL,PARP9,POMC,PPP2R5B,REN,RPL10,RPL3,RPL7,S100A8,S100A9,SAMD9L,SAMHD1,SPINK4,STAT1,STAT2,TAP1,TCL1A,TOB1,TSC22D3,USP18,VWF,XAF1 |
| TMEM173 | -1.12 | other | Inhibited | -2.764 | bias | 3.01E-12 | GBP5,HOPX,IFI44,IFIT2,IFIT3,IFITM3,IRF7,ISG15,OAS1,OASL,RSAD2,USP18 |
| APP | -1.01 | other | Inhibited | -2.766 | | 2.43E-08 | ACTA2,ARRDC3,CD69,CDK1,CITED2,CLU,CPLX1,CSF1R,CXCR4,EGR1,GPC1,GSTM3,HBA1/HBA2,IFIH1,IFIT2,INHBA,IRF1,IRF7,LGALS3,MAFF,MPO,MT-ATP6,NOG,RSAD2,S100A8,SAP30,ST3GAL5,TP53INP1,TUBA1B,TUBB2A,USP18,XAF1 |
| CGAS | -1.81 | enzyme | Inhibited | -2.784 | bias | 9E-11 | IFI44,IFIT2,IFIT3,IRF7,ISG15,OAS1,RSAD2,USP18 |
| TICAM1 | -1.52 | other | Inhibited | -2.858 | | 6.71E-12 | BCL3,DUSP1,EGR1,IFIT1,IFIT2,IFIT3,IRF1,IRF7,ISG15,MAFF,NFKB2,OASL,RELB,RHOB,RSAD2,SAMHD1 |
| IFN type 1 | | group | Inhibited | -2.933 | | 2.1E-10 | CD69,IFIH1,IFIT1,IFIT2,IRF1,ISG15,OASL,STAT1,STAT2,TNFSF13B |
| TNFSF10 | -1.89 | cytokine | Inhibited | -2.985 | | 0.000000177 | CD69,CXCR4,HLA-C,HLA-F,IFI6,IFIT1,IFITM1,IRF9,ISG15,STAT1 |

| | | | | | | | |
|---|---|---|---|---|---|---|---|
| JAK | | group | Inhibited | -3 | | 9.27E-10 | IFI6,IFIH1,IFIT1,IFIT2,IFIT3,IFITM3,ISG15,RSAD2,STAT1 |
| VCAN | -2.37 | other | Inhibited | -3.053 | | 3.82E-08 | CLU,IFI44,IFI44L,IFI6,IFIT1,IFIT2,IFITM1,IRF8,LGALS3,MX1,OAS2,STAT1,XAF1 |
| OSM | -1.21 | cytokine | Inhibited | -3.074 | | 1.54E-10 | ACTA2,AKR1C1/AKR1C2,ATF3,BCL3,CISH,COL2A1,DENND5A,DHCR24,EGR1,FOSL1,GBP1,HLA-C,HLA-F,ID2,IRF1,IRF7,IRF9,MARCKS,MT2A,MX1,OAS1,PFKFB2,POMC,S100A8,S100A9,STAT1,TAP1 |
| IFNAR1 | -1.46 | transmembrane receptor | Inhibited | -3.094 | | 7.56E-15 | ATF3,DUSP6,IFI44,IFI6,IFIH1,IFIT2,IFIT3,IFITM1,IRF1,IRF7,ISG15,OAS1,OAS2,OASL,RSAD2,STAT1,TNFSF13B,USP18,XAF1 |
| PML | 1.14 | transcription regulator | Inhibited | -3.184 | | 4.78E-13 | CXCR4,EPSTI1,HBB,ID2,IFI44,IFI44L,IFIH1,IFIT1,IFIT3,IFITM1,IRF7,ISG15,MAFF,MX1,OAS1,OAS2,STAT1,TAP1 |
| TLR7 | | transmembrane receptor | Inhibited | -3.201 | | 4.53E-11 | ATF3,CD69,FOSL1,IFI44,IFI44L,IFIT1,IFIT3,IFITM1,IL10RA,IRF1,IRF7,IRF9,ISG15,MX1,OAS2,RSAD2,STAT1,STAT2 |
| EIF2AK2 | -1.73 | kinase | Inhibited | -3.24 | | 6.9E-11 | ATF3,EGR1,IFI6,IFIT1,IFITM1,IRF1,ISG15,NFKB2,OAS1,PARP9,SAMHD1,STAT1,USP18 |
| IL1A | | cytokine | Inhibited | -3.255 | | 0.0000912 | ATF3,BCL3,CA2,FOSL1,GBP1,INHBA,IRF1,MT2A,POMC,S100A8,S100A9 |
| TLR3 | 1.48 | transmembrane receptor | Inhibited | -3.261 | | 2.48E-17 | ARRDC4,ATF3,CD69,CLU,DUSP1,GADD45B,HERC5,IFI44,IFI44L,IFI6,IFIH1,IFIT1,IFIT2,IFIT3,IRF1,IRF7,ISG15,MX1,OAS1,OASL,RHOB,RSAD2,S100A8,STAT1,TNFSF13B,USP18 |
| JAK1 | 1.03 | kinase | Inhibited | -3.274 | | 8.03E-13 | BCL3,HLA-C,HLA-F,IFIT2,IRF1,IRF7,IRF9,MX1,SLC1A3,STAT1,STAT2,TAP1,USP18 |
| MYD88 | -1.01 | other | Inhibited | -3.28 | | 3.52E-11 | BCL3,CISH,DUSP1,EGR1,IFIT2,INHBA,IRF1,IRF7,IRF8,ISG15,MAFF,NFKB2,NOD2,OASL,RELB,RSAD2,S100A8,SAMHD1,TNFSF13B,USP18 |
| IFNA1/IFNA13 | | cytokine | Inhibited | -3.4 | | 6.41E-17 | CD69,IFI6,IFIH1,IFIT1,IFIT2,IFITM1,IRF1,IRF7,ISG15,MX1,OAS1,OAS2,RSAD2,STAT1,STAT2 |
| DDX58 | -1.52 | enzyme | Inhibited | -3.495 | | 9.74E-13 | IFI44,IFIH1,IFIT1,IFIT2,IFIT3,IRF1,IRF7,IRF8,ISG15,OAS1,RSAD2,STAT1,STAT2 |
| MAVS | 1.02 | other | Inhibited | -3.554 | | 8.03E-13 | IFIT1,IFIT2,IFIT3,IFITM3,IRF7,ISG15,OAS1,OAS2,OASL,RSAD2,STAT1,STAT2,USP18 |
| SPI1 | -1.24 | transcription regulator | Inhibited | -3.571 | | 2.34E-20 | ACTA2,CD3E,CD72,CDK1,CSF1R,DUSP6,ELANE,HELZ2,ID2,IFI44,IFI44L,IFI6,IFIT1,IFIT2,IFIT3,IFITM1,IFITM3,IRF7,IRF9,ISG15,JDP2,LILRB3,MT1F,MT2A,MX1,NCF2,OASL,RSAD2,USP18 |
| RNY3 | | other | Inhibited | -3.742 | bias | 5.53E-21 | EPSTI1,HERC5,IFI44,IFI44L,IFIT1,IFIT3,IFITM3,ISG15,MX1,OAS1,OAS2,OASL,RSAD2,XAF1 |
| IRF5 | -1.11 | transcription regulator | Inhibited | -3.786 | | 5.09E-14 | CXCR4,IFI44,IFIH1,IFIT1,IFIT2,IFIT3,IFITM3,IRF7,ISG15,OAS1,OAS2,OASL,RSAD2,STAT1,STAT2 |
| TLR4 | 1.03 | transmembrane receptor | Inhibited | -3.794 | | 2.04E-12 | AREG,ATF3,CISH,CITED2,DLK1,EREG,HELZ2,HPSE,IFIT2,IFIT3,IFITM3,INHBA,IRF1,IRF7,IRF8,ISG15,MX1,NFKB2,NOD2,OASL,RELB,RSAD2,SAP30,STAT1,STAT2 |
| TNF | -1.01 | cytokine | Inhibited | -3.87 | | 6.47E-25 | ACTA2,ARHGAP18,ARRDC3,ATF3,AZU1,BCL3,BTG1,CA2,CCR1,CD3E,CD69,CDKN2C,CISH,CITED2,CLU,COL2A1,CSF1R,CSGALNACT1,CXCR4,DUSP1,DUSP5,DUSP6,EGLN1,EGR1,EREG,FOSL1,GADD45B,GBP1,GBP1P1,GNAI1,HERC5,HLA-F,IFI6,IFIH1,IFIT1,IFIT3,IFITM1,IL10RA,INHBA,IRF1,IRF7,IRF8,ISG15,ITGB8,KIF20A,KLF2,LGALS3,MAFF,MPO,MT2A,MX1,NCF2,NEAT1,NFKB2,NOD2,NR5A1,OAS1,OAS2,OASL,POMC,RELB,RHOB,S100A8,S100A9,SAMD4A,SLC1A3,SLC43A2,ST3GAL5,STAT1,TAP1,TINAGL1,TMEM176B,TNFRSF10D,TNFSF13B,TP53INP1,TSC22D3,TXNIP |
| TGM2 | 1.12 | enzyme | Inhibited | -3.897 | | 1.28E-11 | CA2,IFI6,IFIT1,IFIT2,IFIT3,IL10RA,IRF9,MT2A,NCF2,OAS1,OAS2,OASL,PARP9,S100A8,SAMD9L,SAMHD1,STAT1,TAP1,XAF1 |
| Ifn | | group | Inhibited | -3.898 | | 1.73E-15 | CD69,IFIH1,IFIT1,IFITM1,IFITM3,IL10RA,IRF1,IRF7,IRF8,ISG15,MX1,OAS2,OASL,RSAD2,S100A8,STAT1,TAP1,TNFSF13B |
| TLR9 | | transmembrane receptor | Inhibited | -4.05 | | 3.83E-18 | ARRDC4,ATF3,CD69,CISH,DUSP1,EGR1,GADD45B,IFI44L,IFIT1,IFIT2,IFIT3,IFITM1,IL10RA,IRF1,IRF7,IRF9,ISG15,MX1,OAS2,RSAD2,S100A8,ST3GAL5,STAT1,STAT2,TNFSF13B,USP18 |
| PRL | | cytokine | Inhibited | -4.17 | | 8.23E-22 | CD69,CISH,CLU,DTX3L,EGR1,EPSTI1,HELZ2,HERC5,ID2,IFI44,IFI44L,IFI6,IFIH1,IFIT1,IFIT3,IFITM1,IRF1,IRF7,IRF9,ISG15,OAS1,OAS2,RSAD2,SAMD9L,SAMHD1,STAT1,STAT2,TNFSF13B,TUBA1B,USP18,XAF1 |
| IFN Beta | | group | Inhibited | -4.315 | | 1.04E-22 | ATF3,CD69,DUSP1,HERC5,HLA-G,IFI44,IFI6,IFIH1,IFIT1,IFIT2,IFIT3,IFITM1,IRF1,IRF7,IRF9,ISG15,MX1,OAS1,OAS2,OASL,RSAD2,STAT1,STAT2,USP18,XAF1 |
| IFNB1 | | cytokine | Inhibited | -4.363 | | 2.31E-12 | GBP5,GBP7,HERC5,IFI6,IFIH1,IFIT1,IFIT2,IFIT3,IFITM1,IRF1,IRF7,IRF9,ISG15,MX1,NOD2,OAS1,OAS2,RSAD2,STAT1,STAT2,USP18,XAF1 |
| IRF3 | -1.01 | transcription regulator | Inhibited | -4.385 | | 2.76E-20 | CD69,GBP1,GBP5,HELZ2,HLA-F,IFI44,IFI6,IFIH1,IFIT1,IFIT2,IFIT3,IFITM3,IRF1,IRF7,ISG15,NLRC5,OAS1,OAS2,OASL,RSAD2,SAMD9L,SAP30,STAT1,STAT2,TAP1,USP18 |
| Ifnar | | group | Inhibited | -4.407 | | 2.6E-20 | HLA-G,IFIH1,IFIT2,IFIT3,IFITM3,IRF1,IRF7,IRF8,IRF9,ISG15,NLRC5,OAS1,OAS2,OASL,RSAD2,STAT1,STAT2,TAP1,USP18,XAF1 |
| IRF1 | -3.2 | transcription regulator | Inhibited | -4.674 | | 5.86E-20 | HELZ2,HLA-G,IFI44,IFI44L,IFI6,IFIH1,IFIT1,IFIT2,IFIT3,IFITM1,IFITM3,IRF1,IRF7,IRF9,ISG15,MX1,OAS1,OAS2,OASL,RSAD2,STAT1,STAT2,TAP1,TNFSF13B,XAF1 |
| STAT1 | -5.56 | transcription regulator | Inhibited | -4.873 | | 1.36E-23 | BTG1,EGR1,EPSTI1,GBP1,GBP1P1,GBP5,IFI44,IFI44L,IFI6,IFIH1,IFIT1,IFIT2,IFIT3,IFITM1,IFITM3,IRF1,IRF7,IRF8,IRF9,ISG15,MX1,NLRC5,OAS1,OAS2,OASL,PARP9,POMC,RSAD2,SAMD9L,SAMHD1,STAT1,STAT2,TAP1,TNFSF13B,USP18,XAF1 |
| IFNL1 | | cytokine | Inhibited | -4.89 | | 7.31E-29 | ATF3,GBP1,GBP5,HERC5,HLA-C,IFI44,IFI44L,IFI6,IFIH1,IFIT1,IFIT2,IFIT3,IFITM1,IFITM3,IRF9,ISG15,MX1,OAS1,OAS2,OASL,RSAD2,STAT1,STAT2,USP18,XAF1 |
| Interferon alpha | | group | Inhibited | -4.896 | | 2.1E-28 | ATF3,BCL3,CCR1,CD69,CISH,DDIT4,DUSP6,EPSTI1,GBP1,GBP5,HELZ2,HERC5,HLA-C,HLA-F,HLA-G,IFI44,IFI44L,IFI6,IFIH1,IFIT1,IFIT2,IFIT3,IFITM1,IFITM3,IL10RA,IRF1,IRF7,IRF8,IRF9,ISG15,MNDA,MX1,OAS1,OAS2,PARP9,POMC,RSAD2,S100A9,SAMD9L,STAT1,STAT2,TAP1,TNFSF13B,USP18 |
| IFNA2 | | cytokine | Inhibited | -5.058 | | 2.01E-27 | BAG3,CD69,CISH,GBP1,HERC5,HLA-C,IFI44,IFI44L,IFI6,IFIH1,IFIT1,IFIT2,IFIT3,IFITM1,IFITM3,IRF1,IRF7,IRF9,ISG15,MAL,MT-ATP6,MT1F,MT2A,MX1,OAS1,OAS2,PARP9,RSAD2,SAMHD1,STAT1,TAP1,TNFRSF10D,USP18,XAF1 |
| IRF7 | -2.85 | transcription regulator | Inhibited | -5.062 | bias | 2.16E-31 | CD69,GBP1,GBP5,HELZ2,HERC5,IFI44,IFI44L,IFI6,IFIH1,IFIT1,IFIT2,IFIT3,IFITM1,IFITM3,IRF1,IRF7,IRF8,IRF9,ISG15,MX1,OAS1,OAS2,OASL,RSAD2,S100A8,SAMD9L,SAP30,STAT1,STAT2,TAP1,TNFSF13B,USP18,XAF1 |
| IFNG | | cytokine | Inhibited | -5.758 | | 9.35E-30 | ACTA2,AREG,ATF3,BCL3,BTG1,CCR1,CD72,CDK1,CISH,CSF1R,CXCR4,DTX3L,DUSP1,DUSP5,EGR1,FBP1,FCGR1B,FKBP5,GBP1,GBP5,GBP7,HCP5,HLA-C,HLA-F,HLA-G,IFI44,IFI44L,IFI6,IFIH1,IFIT1,IFIT2,IFIT3,IFITM1,IFITM3,IL10RA,INHBA,IRF1,IRF7,IRF8,IRF9,ISG15,KLF2,LGALS3,MAFF,MNDA,MT1F,MX1,MYCN,NCF2,NEAT1,NFKB2,NLRC5,NOD2,OAS1,OAS2,OASL,PARP9,POMC,RELB,RHOB,RSAD2,S100A8,S100A9,SAMHD1,SLC1A3,SLC3A1,STAT1,STAT2,TAP1,TNFSF13B,TSC22D3,TXNIP,USP18,XAF1 |

**T2 vs. T1 MEP_1**

© 2000-2020 QIAGEN. All rights reserved.

| Upstream Regulator | Expr Fold Change | Molecule Type | Predicted Activation State | Activation z-score | Flags | p-value of overlap | Target Molecules in Dataset |
|---|---|---|---|---|---|---|---|
| IL1RN | | cytokine | Activated | 4.022 | bias | 3.54E-13 | ATF3,CXCL1,CXCL3,IFI44,IFI44L,IFI6,IFIH1,IRF1,IRF7,IRF9,MX1,MX2,OAS1,OAS2,OAS3,OASL,PMAIP1,PTGS2,RSAD2,USP18 |
| PNPT1 | -1.07 | enzyme | Activated | 3.606 | bias | 4.63E-12 | CMPK2,IFI44,IFIH1,IRF7,ISG15,LGALS3BP,OAS1,PARP14,PARP9,SAMD9L,STAT1,USP18,XAF1 |
| SIRT1 | -1.08 | transcription regulator | Activated | 3.437 | | 7.09E-10 | ABCG1,CCND1,CCND3,CD36,CD83,CDKN2C,CMPK2,DHX58,GABARAPL1,GSTM3,HCK,HELZ2,HLA-DRB5,IFI44,IFITM3,IRF7,KLF2,LGALS3BP,NFKBIE,NLRC5,OAS1,OAS2,PARP14,PER1,RELB,RSAD2,SOD2,STAT1,USP18 |
| MAPK1 | -1.06 | kinase | Activated | 3.332 | | 1.06E-18 | ANPEP,APOL6,ATF3,BST2,CCND1,CD69,CISH,EGR1,ENO3,FOSL2,GBP5,HBA1/HBA2,HBB,HERC5,HLA-B,HLA-C,IFI44,IFI6,IFIH1,IFIT1,IFITM1,IFITM3,IL1RL1,IRF7,IRF9,ISG15,LGALS1,LGALS3BP,MITF,MX2,OAS1,OAS2,OAS3,OASL,PLAU,PTGS2,STAT1,TTC27,USP18,ZNF184 |
| ACKR2 | | G-protein coupled receptor | Activated | 3.317 | bias | 2.54E-10 | DHX58,IFI44,IRF7,ISG15,OAS1,OAS2,OAS3,OASL,RSAD2,STAT1,USP18 |
| STAT6 | 1.4 | transcription regulator | Activated | 3.228 | | 9.17E-13 | ADAP1,AREG,BCL3,CD40,CD69,CISH,CMPK2,CXCL2,CXCL3,DHX58,EGR1,FCER1G,FKBP5,GADD45B,GBP5,HCK,IFI44,IFI44L,IFIH1,IFITM3,IL18R1,IRF1,IRF7,IRF9,ISG15,LGALS3BP,NFKBIA,OAS3,OASL,PTGS2,RASGRP3,TFEC,USP2,ZNF608 |
| TRIM24 | -1.08 | transcription regulator | Activated | 3.193 | bias | 1.97E-11 | CMPK2,DHX58,EPSTI1,FBN1,IFI44,IFIH1,IRF1,IRF7,IRF9,ISG15,LGALS3BP,OAS1,OASL,PLAC8,SAMD9L,STAT1,USP18 |
| IRF4 | | transcription regulator | Activated | 3.116 | | 0.000736 | CXCL3,DDIT3,IRF1,IRF7,IRF9,ISG15,OAS1,PLAUR,STAT1,TNFSF13B |
| SP110 | -1.94 | transcription regulator | Activated | 3 | | 8.11E-09 | ATF3,BCL3,BST2,IFI6,IFIH1,IFIT1,IFITM1,IFITM3,IL6R,IRF9,MX1,MYCN,OAS1,OAS3,STAT1,TXNIP |
| USP18 | -6.39 | peptidase | Activated | 2.975 | bias | 2.34E-09 | IFI6,IFIH1,IFITM3,IRF1,IRF7,IRF9,ISG15,MX1,OAS1 |
| SOCS1 | -1.14 | other | Activated | 2.872 | bias | 1.98E-14 | CCND1,CCND3,CD40,CD69,CISH,CXCL2,FCER1G,FKBP5,GBP5,IFI44,IFIH1,IFIT1,IRF1,IRF7,ISG15,MX1,OAS1,OAS2,PTGS2,RSAD2,STAT1,USP18 |
| NKX2-3 | 2.67 | transcription regulator | Activated | 2.684 | | 6.27E-14 | ADAP1,ANGPT2,BTG1,CD36,CMPK2,CXADR,DHX58,FBXO6,H2AC18/H2AC19,HCP5,HELZ2,HLA-B,HLA-C,HLA-F,KLF2,PARP14,PARP9,PIM3,PMAIP1,PTGS2,SAMD9L,STAT1,TNFRSF10D,TYMP,USP18,XAF1 |
| NGLY1 | 1.05 | enzyme | Activated | 2.57 | | 0.00000301 | IFI44,IFI44L,IFIT1,OAS1,OAS3,RSAD2,USP18 |
| NEDD9 | -1.26 | other | Activated | 2.449 | | 0.00583 | BHLHE40,DDIT4,LIN7A,PLAC8,RPGR,TXNIP |
| Irgm1 | | other | Activated | 2.449 | bias | 0.00017 | ID2,IRF7,OAS2,OASL,RSAD2,USP18 |
| PKD1 | 1.26 | ion channel | Activated | 2.433 | | 0.0836 | CXCL2,DDIT3,ENPP5,IL1RL1,LTBP1,ODF3B,SLC16A4 |
| DNASE2 | -1.54 | enzyme | Activated | 2.429 | bias | 0.000000151 | CXCL2,DHX58,IRF7,ISG15,OAS1,OAS3,RSAD2,USP18 |
| AGER | 1.22 | transmembrane receptor | Activated | 2.406 | | 0.0000598 | CCND1,CD40LG,CXCL2,CXCL3,EGR1,IFIT1,PTGS2,TXNIP |
| ESR1 | 1.05 | ligand-dependent nuclear receptor | Activated | 2.401 | | 2.5E-13 | AQP1,AREG,ARRDC3,ATF3,CAV2,CCND1,CCND3,CD69,CD83,CEBPG,CENPE,CXADR,CXCL1,CXCL3,DDIT4,DUSP5,EGR1,EREG,FBLN1,FOSL1,FOSL2,GADD45B,H4C3,HBA1/HBA2,HCP5,HLA-B,HLA-C,HLA-G,IFI44,IFI44L,IFI6,IFITM1,IFITM3,IRF1,KCNK6,LGALS1,LGALS3BP,LTBP1,MARCKS,MX1,NFKBIA,NFKBIE,ODF3B,P2RY2,PLAU,PLAUR,PMAIP1,PRKCD,PTGS2,PTK2,PTX3,RELB,REN,RPL41,RSAD2,SELL,SEMA3C,SLC26A2,SLC44A1,STAT1,THRB,TNFAIP3,TNFRSF10A,TOB1,TSC22D3,TXNIP,XK,ZMAT5 |
| SERPINE1 | -1.5 | other | Activated | 2.4 | | 0.000055 | CALR,CASP3,CXCL2,DLK1,PLAU,PLAUR |
| TSC2 | 1.05 | other | Activated | 2.4 | | 0.000749 | ATF3,CCND1,CCND3,DDIT3,EGR1,IFITM3,PLAU,SOD2,STAT1 |
| KCNK9 | | ion channel | Activated | 2.236 | | 0.000912 | CA1,GADD45B,REN,RHAG,SPTA1 |
| PRDM16 | -1.84 | transcription regulator | Activated | 2.229 | | 0.0000468 | IFI44,IRF7,MX2,OAS2,OAS3,STAT1 |
| ITGB2 | -1.24 | transmembrane receptor | Activated | 2.219 | | 0.00187 | CD69,CXCL2,CXCL3,GADD45B,PTGS2 |
| Pkc(s) | | group | Activated | 2.103 | | 0.000255 | AHR,ATF3,CCND3,CD36,DDIT3,EGR1,FOSL1,GADD45B,HPGD,ID2,NR4A3,PER1,PTGS2 |
| SOCS3 | 1.36 | phosphatase | Activated | 2.069 | | 0.00000945 | ATF3,CCND1,CD40,EGR1,IFIT1,IRF1,MX1,OAS1,OAS2,POMC |
| IL1A | | cytokine | Inhibited | -2.02 | bias | 0.00000294 | ATF3,BCL3,CD274,CD40,CD83,CXCL1,CXCL2,CXCL3,FOSL1,IRF1,NFKBIA,PLAU,POMC,PTGS2,PTX3,SOD2,TNFAIP3 |
| CSF1 | 1.17 | cytokine | Inhibited | -2.043 | bias | 0.000786 | CCND1,CCND3,CXCL2,CXCL3,DUSP5,EGR1,FCER1G,IL1RL1,IRF7,PLAU,STAT1,TNFAIP3 |
| TLR4 | -1.57 | transmembrane receptor | Inhibited | -2.075 | bias | 6.81E-14 | AREG,ATF3,CD274,CD40,CISH,CITED2,CMPK2,CXCL2,CXCL3,DDIT3,DLK1,EREG,HELZ2,IFITM3,IRF1,IRF7,ISG15,LMO4,LTC4S,MX1,NFKBIA,NR4A3,OASL,PTGS2,PTX3,RELB,RSAD2,SLA,SLCO3A1,SOD2,STAT1,TNFAIP3,TNFRSF10A,TSC22D1 |
| MAP2K1 | 1.23 | kinase | Inhibited | -2.099 | bias | 0.0000162 | AHR,AREG,ATF3,CCND1,CD274,CXCL3,DUSP5,FOSL1,MITF,MYCN,NFKBIA,PLAUR,PTGS2,TNFRSF10A |
| IL21 | | cytokine | Inhibited | -2.127 | bias | 1.17E-10 | CASP3,CD40LG,CD69,CMPK2,EGR1,FCER1G,GBP5,HAVCR2,HLA-DRB5,ID2,IFIT1,IL18R1,IRF7,ISG15,OAS2,OASL,RSAD2,SELL,TYROBP,USP18 |
| HGF | -1.13 | growth factor | Inhibited | -2.131 | bias | 0.000064 | AHR,ANGPT2,ATF3,CCND1,CCNE2,CD40,CDKN2C,CITED2,CXCL2,DDIT3,EGR1,FKBP5,FOSL1,ISG15,ITGB8,NR4A3,PLAU,PLAUR,PTGS2,SKIL,SOD2,TNFAIP3,TRIB2 |
| SOX4 | -1.15 | transcription regulator | Inhibited | -2.138 | | 0.0233 | B3GNT8,EIF4E3,FCER1G,KYNU,MEX3A,PTK2,SELL,TMEM98 |
| DNMT3B | -1.61 | enzyme | Inhibited | -2.138 | | 0.0169 | AGTR1,CD36,DDIT3,IFIT1,IFITM1,MYL4,S1PR1,STAT1 |
| IFNE | | cytokine | Inhibited | -2.143 | bias | 2.54E-10 | BST2,CD40,CD83,HERC5,IFIH1,IFITM3,ISG15,MX2,PTGS2,STAT1,USP18 |
| CTNNB1 | -1.16 | transcription regulator | Inhibited | -2.151 | | 0.000000322 | AHR,ARL4A,ARMH4,BHLHE40,CCNA1,CCND1,CCNE2,CD36,CXCL1,CXCL2,DDIT3,DLK1,EGR1,ENO3,EREG,FCER1G,FOSL1,GAD1,GADD45B,ID2,IFIT1,IFITM1,KRT1,LMO2,MITF,MYCN,MYL4,PLAU,PLAUR,PTGS2,RPL10,RPL3,RPL41,RPL7A,RPSA,SEMA3C,SLC26A2,TNFAIP3,TSC22D1,WNT5B,XAF1 |
| IFNA4 | | cytokine | Inhibited | -2.159 | bias | 2.98E-11 | CD274,CD69,GBP5,IFIH1,IFIT1,ISG15,MX1,OASL,PMEPA1,RSAD2,SLCO3A1,USP18 |
| NLRC5 | -2.42 | transcription regulator | Inhibited | -2.168 | bias | 0.0000335 | CD40,HLA-B,HLA-C,HLA-F,HLA-G |
| TMEM173 | -1.14 | other | Inhibited | -2.171 | bias | 1.04E-09 | CXCL2,GBP5,GJA4,HOPX,IFI44,IFITM3,IRF7,ISG15,OAS1,OASL,RSAD2,USP18 |
| LGALS3 | 1.01 | other | Inhibited | -2.183 | | 0.0354 | CASP3,CCND1,CXCL2,CXCL3,LGALS12 |
| ETS1 | 1.16 | transcription regulator | Inhibited | -2.209 | bias | 0.00293 | ANPEP,CCND1,CD69,CITED2,EGR1,HPGD,ID2,PLAU,PTPRN2,REN,SELL |
| MAP2K3 | -1.07 | kinase | Inhibited | -2.213 | bias | 0.002 | DDIT3,IRF9,ISG15,PLAU,PTGS2,STAT1 |
| IL32 | -1.14 | cytokine | Inhibited | -2.229 | | 0.004 | CD36,CXCL1,CXCL3,MX1,PTGS2 |
| SELPLG | -1.02 | other | Inhibited | -2.236 | bias | 0.00114 | CXCL2,HAVCR2,HCK,PLAUR,PRKCD |
| ELAVL1 | 1.12 | other | Inhibited | -2.241 | | 2.54E-10 | ATF3,CD83,CENPE,HBB,HLA-DRB5,IFI44,IFIH1,IFITM3,IRF1,IRF9,LGALS3BP,NFKBIA,OAS1,OAS2,PTGS2,PTPRN2,REN,STAT1,TP53INP1,TSC22D3,USP18 |
| cholesterol | | chemical - endogenous mammalian | Inhibited | -2.25 | | 0.00000209 | ABCG1,AREG,ATF3,CAV2,CETP,CR1,CXCL2,CXCL3,DDIT3,GNAI1,GP1BA,HES6,HLA-DRB5,ITGAX,MITF,NR4A3,PTGS2 |
| IL33 | | cytokine | Inhibited | -2.296 | bias | 3.68E-11 | ABCG1,AREG,BCL3,CCND1,CD36,CD40,CD69,CISH,CXCL2,CXCL3,EPX,GADD45B,HCK,IL1RL1,MDK,NFKBIA,NFKBIE,RELB,SKIL,SOD2,TFEC,TNFAIP3,TPSAB1/TPSB2,VMP1 |
| MSC | | transcription regulator | Inhibited | -2.309 | | 4.51E-10 | EPSTI1,HAVCR2,IFI44,IFI44L,IFIT1,IRF7,LGALS12,PAG1,PDP1,PIEZO2,SELL,XAF1 |
| JAK1 | 1.03 | kinase | Inhibited | -2.343 | bias | 0.000000615 | BCL3,CD40,HLA-C,HLA-F,IRF1,IRF7,IRF9,MX1,STAT1,USP18 |
| NFkB (complex) | | complex | Inhibited | -2.388 | bias | 9.07E-14 | ABCG1,AHR,AQP1,ATF3,BCL3,CCND1,CD274,CD36,CD40,CD40LG,CD69,CD83,CETP,CXCL1,CXCL2,CXCL3,DDIT3,DUSP5,EGR1,GAD1,GADD45B,HERC5,HLA-F,IGLL1/IGLL5,IRF1,IRF7,ISG15,ITGB8,KYNU,LTC4S,NFKBIA,NFKBIE,P2RY2,PIM3,PLAU,PMAIP1,PRKCD,PTGS2,PTX3,RELB,RSAD2,SOD2,TFEC,TNFAIP3 |
| DDX58 | -1.36 | enzyme | Inhibited | -2.396 | bias | 0.000000704 | IFI44,IFIH1,IFIT1,IRF1,IRF7,ISG15,OAS1,PTGS2,RSAD2,STAT1 |
| FOSL1 | -2.82 | transcription regulator | Inhibited | -2.412 | | 0.00283 | CCND1,EGR1,FOSL1,HPGD,PLAU,PLAUR |
| OSM | -1.26 | cytokine | Inhibited | -2.413 | | 1.86E-11 | ABCG1,AHR,AKR1C1/AKR1C2,ANGPT2,ARL4A,ATF3,BCL3,BHLHE40,CCND1,CDC42EP4,CISH,CTSL,CXADR,CXCL1,CXCL2,CXCL3,DENND5A,DHRS3,EGR1,FOSL1,HLA-B,HLA-C,HLA-F,ID2,IL6R,IRF1,IRF7,IRF9,KLF10,MARCKS,MX1,OAS1,PLAU,PLLP,POMC,STAT1,TYMP |
| TNFSF12 | -1.55 | cytokine | Inhibited | -2.416 | | 0.0134 | CCND1,CXCL2,CXCL3,HES6,NFKBIA,PTX3 |
| CGAS | -1.76 | enzyme | Inhibited | -2.418 | bias | 0.00000297 | IFI44,IRF7,ISG15,OAS1,RSAD2,USP18 |
| BRCA1 | 1 | transcription regulator | Inhibited | -2.453 | | 0.0000069 | AREG,CCND1,DDIT3,DDIT4,EGR1,FKBP5,GADD45B,HBB,IFI6,IFIT1,IFITM1,IRF7,MX1,STAT1 |
| TICAM1 | -1.04 | other | Inhibited | -2.474 | bias | 2.26E-11 | BCL3,CD40,CMPK2,CXCL2,CXCL3,EGR1,IFIT1,IRF1,IRF7,ISG15,MAFF,NFKBIA,OASL,PTGS2,RELB,RSAD2,TFEC,TNFAIP3,TSC22D1 |
| IFNAR1 | -1.29 | transmembrane receptor | Inhibited | -2.508 | | 5.24E-19 | ATF3,CD274,CD40,CMPK2,CXCL2,CXCL3,IFI44,IFI6,IFIH1,IFITM1,IRF1,IRF7,ISG15,MX2,OAS1,OAS2,OAS3,OASL,PTGS2,RNASE2,RSAD2,SOD2,STAT1,TNFAIP3,TNFSF13B,USP18,XAF1 |
| C5 | -1.49 | cytokine | Inhibited | -2.522 | | 0.000112 | ATF3,CCND1,CXCL1,CXCL2,CXCL3,DAG1,EGR1,IL18R1,NFKBIA,TNFAIP3 |
| TLR3 | -1.33 | transmembrane receptor | Inhibited | -2.534 | bias | 1.35E-14 | ATF3,CD274,CD40,CD69,CMPK2,CXCL2,CXCL3,DHX58,GADD45B,HERC5,IFI44,IFI44L,IFI6,IFIH1,IFIT1,IRF1,IRF7,ISG15,MX1,MX2,OAS1,OASL,PMAIP1,PTGS2,PTX3,RSAD2,STAT1,TNFSF13B,USP18 |
| NFKBIA | -2.09 | transcription regulator | Inhibited | -2.57 | | 1.07E-09 | AGTR1,BCL3,CCND1,CCNE2,CD40,CD69,CXCL1,CXCL2,CXCL3,DAG1,DDIT3,EREG,GADD45B,GSTM5,IFI6,IRF1,ISG15,NFKBIA,NFKBIE,NINJ1,PLAU,PMEPA1,PTGS2,PTX3,RELB,RPSA,SEMA3C,SOD2,TNFAIP3,TNFRSF10A |
| JUNB | -1.27 | transcription regulator | Inhibited | -2.586 | bias | 3.51E-08 | AGTR1,ATF3,BCL3,CCND1,CD274,CXCL2,CXCL3,FOSL1,HPGD,ID2,IL1RL1,PLAU,PLAUR,POMC,RELB |
| IFNL4 | | cytokine | Inhibited | -2.623 | bias | 8E-11 | DHX58,IFIH1,ISG15,MX1,OAS1,OAS2,STAT1 |
| tretinoin | | chemical - endogenous mammalian | Inhibited | -2.642 | | 3.34E-22 | ABCG1,AGTR1,AHR,ANPEP,AQP1,AREG,BCL3,BHLHE40,BTG1,CALR,CASP3,CCNA1,CCND1,CCND3,CD274,CD36,CDKN2C,CEBPG,CFP,CITED2,CLC,CTSL,CXCL1,CXCL2,DDIT3,DHRS3,DHX58,EGR1,FCER1G,FOSL1,FOSL2,HLA-B,HLA-C,HOTAIRM1,ID2,IFI44,IFI44L,IFI6,IFIH1,IFIT1,IFITM1,IL6R,IRF1,IRF9,ISG15,ITGAX,KRT1,LGALS1,LGALS3BP,LTC4S,MDK,MS4A3,MYCN,NRGN,OAS1,OAS2,OAS3,OASL,P2RY2,PARP14,PARP9,PDE1B,PLAU,PLAUR,PMAIP1,POMC,PRKCD,PTGS2,PTX3,RASAL2,REN,RPL10,RPL3,SAMD9L,SELL,SEMA7A,SIGLEC12,SLA,SLFN5,SMOX,STAT1,TNFAIP3,TNFRSF10A,TOB1,TSC22D3,TTC27,TYROBP,USP18,VMP1,XAF1,XK |
| TRADD | 1.14 | other | Inhibited | -2.646 | bias | 0.000000592 | CXCL1,CXCL2,CXCL3,IL18R1,IRF1,NFKBIA,TNFAIP3 |
| IFNA1/IFNA13 | | cytokine | Inhibited | -2.743 | bias | 9.97E-18 | CD274,CD40,CD69,CD83,DHX58,IFI6,IFIH1,IFIT1,IFITM1,IRF1,IRF7,ISG15,MX1,OAS1,OAS2,PTGS2,RSAD2,STAT1 |
| JUN | 1.21 | transcription regulator | Inhibited | -2.769 | | 4.57E-10 | ATF3,BCL3,BTG1,CCND1,CCND3,CD274,CFP,CTSL,CXCL1,CXCL3,DUSP5,EHD4,EREG,FOSL1,HLA-B,HLA-G,HPGD,ID2,IL1RL1,ITGB8,LTBP1,MYLPF,NFKBIA,PARD6B,PLAU,PLAUR,PTGS2,PTX3,RELB,SOD2,STAT1,TNFRSF10A |

| | | | | | | | |
|---|---|---|---|---|---|---|---|
| EIF2AK2 | -1.77 | kinase | Inhibited | -2.794 | bias | 1.24E-15 | ATF3,BHLHE40,BRIP1,CCND1,DDIT3,EGR1,IFI6,IFIT1,IFITM1,IRF1,ISG15,LGALS3BP,NFKBIA,OAS1,OAS3,PARP9,SOD2,STAT1,TNFAIP3,USP18 |
| KMT2D | -1.71 | transcription regulator | Inhibited | -2.804 | bias | 0.0338 | ECHDC2,EFHC2,ENO3,GLYCTK,HOPX,IRF1,MFSD9,ZNF184 |
| PML | 1.29 | transcription regulator | Inhibited | -2.849 | | 4.51E-12 | BST2,CCNA1,CCND1,CXCL1,EPSTI1,HBB,ID2,IFI44,IFI44L,IFIH1,IFIT1,IFITM1,IRF7,ISG15,MAFF,MX1,OAS1,OAS2,OAS3,STAT1,TNFAIP3 |
| IFN type 1 | | group | Inhibited | -2.942 | | 2.71E-08 | BST2,CD69,DHX58,IFIH1,IFIT1,IRF1,ISG15,OASL,STAT1,TNFSF13B |
| MYD88 | -1.2 | other | Inhibited | -2.956 | bias | 3.57E-14 | BCL3,CD274,CD40,CD83,CISH,CMPK2,CXCL1,CXCL2,CXCL3,EGR1,FANCB,FCER1G,HCK,IRF1,IRF7,ISG15,ITGAX,MAFF,NFKBIA,OASL,PTGS2,RELB,RSAD2,SLCO3A1,TFEC,TNFAIP3,TNFSF13B,TSC22D1,USP18 |
| IRF5 | -1.2 | transcription regulator | Inhibited | -2.981 | bias | 4.54E-13 | CASP3,CMPK2,CXCL2,DHX58,IFI44,IFIH1,IFIT1,IFITM3,IRF7,ISG15,OAS1,OAS2,OASL,PMAIP1,PTGS2,RSAD2,STAT1 |
| FN1 | 1.52 | enzyme | Inhibited | -3.003 | | 0.00628 | CCND1,CD83,CXCL1,CXCL2,CXCL3,HLA-B,MARCKS,NFKBIA,PLAU,PLAUR |
| TLR9 | | transmembrane receptor | Inhibited | -3.009 | bias | 1.03E-13 | ATF3,CCND1,CD274,CD40,CD69,CD83,CISH,CXCL2,CXCL3,EGR1,GADD45B,IFI44L,IFIT1,IFITM1,IRF1,IRF7,IRF9,ISG15,MX1,MX2,OAS2,OAS3,PTGS2,RSAD2,STAT1,TNFSF13B,USP18 |
| MAVS | 1.19 | other | Inhibited | -3.074 | bias | 4.44E-10 | CMPK2,CXCL2,DHX58,IFIT1,IFITM3,IRF7,ISG15,OAS1,OAS2,OASL,RSAD2,STAT1,USP18 |
| IFNB1 | | cytokine | Inhibited | -3.077 | | 2.4E-19 | APOL6,BHLHE40,BST2,CASP3,CD274,CD40,CD83,CMPK2,CXCL2,CXCL3,DHX58,FBN1,GBP5,HERC5,IFI6,IFIH1,IFIT1,IFITM1,IRF1,IRF7,IRF9,ISG15,ITGAX,MX1,NFKBIE,OAS1,OAS2,PARP14,PMAIP1,PTGS2,RSAD2,STAT1,TNFRSF10A,UBE2E2,USP18,XAF1 |
| TLR7 | | transmembrane receptor | Inhibited | -3.109 | bias | 9.15E-20 | ATF3,CCND1,CCND3,CD274,CD40,CD69,CD83,CXCL1,CXCL2,CXCL3,FCER1G,FOSL1,IFI44,IFI44L,IFIT1,IFITM1,IRF1,IRF7,IRF9,ISG15,ITGAX,MX1,MX2,NFKBIA,OAS2,OAS3,PLAU,PTX3,RSAD2,SOD2,STAT1,TNFAIP3 |
| IRF1 | -5.15 | transcription regulator | Inhibited | -3.209 | bias | 5.88E-23 | APOL6,BRIP1,CASP3,CCND1,CD274,CD40,CMPK2,CXCL2,CXCL3,HELZ2,HLA-G,IFI44,IFI44L,IFI6,IFIH1,IFIT1,IFITM1,IFITM3,IRF1,IRF7,IRF9,ISG15,MX1,OAS1,OAS2,OAS3,OASL,PTGS2,RSAD2,SELL,STAT1,TNFSF13B,XAF1 |
| Ifnar | | group | Inhibited | -3.225 | bias | 7.81E-15 | CD274,CD40,DDIT3,FCER1G,HLA-G,IFIH1,IFITM3,IRF1,IRF7,IRF9,ISG15,NLRC5,OAS1,OAS2,OASL,RSAD2,STAT1,USP18,XAF1 |
| IL1B | 1.97 | cytokine | Inhibited | -3.321 | | 3.89E-18 | AGTR1,APOL6,ATF3,BCL3,CASP3,CD274,CD40,CD40LG,CD69,CD83,CEBPG,CISH,CMPK2,CXCL1,CXCL2,CXCL3,DDIT3,DDIT4,DUSP5,EGLN1,EGR1,FKBP5,FOSL1,GAD1,GADD45B,HELZ2,HERC5,IFIT1,IL18R1,IL1RL1,IL6R,IRF1,IRF7,ISG15,ITGB8,KLF10,MX1,NEAT1,NFKBIA,NR4A3,OAS2,PIM3,PLAU,POMC,PRKCD,PTGS2,PTX3,RELB,REN,RPSA,RSAD2,SCUBE1,SOD2,STAT1,TNFAIP3,TNFRSF10A,TNFSF13B,TOB1,TSC22D3,TXNIP,TYMP,USP18 |
| IRF3 | 1.1 | transcription regulator | Inhibited | -3.531 | bias | 9.4E-19 | CD274,CD40,CD69,CMPK2,CXCL1,DHX58,GBP5,HELZ2,HLA-F,IFI44,IFI6,IFIH1,IFIT1,IFITM3,IRF1,IRF7,ISG15,NLRC5,OAS1,OAS2,OAS3,OASL,PARP14,PLAC8,PMAIP1,RSAD2,SAMD9L,STAT1,TNFAIP3,USP18 |
| Ifn | | group | Inhibited | -3.546 | bias | 8.69E-14 | APOBEC3A,CD40,CD69,CD83,DHX58,IFIH1,IFIT1,IFITM1,IFITM3,IRF1,IRF7,ISG15,MX1,OAS2,OAS3,OASL,RSAD2,STAT1,TNFSF13B,TYMP |
| VCAN | 3.17 | other | Inhibited | -3.577 | | 3.15E-12 | CASP3,CXCL2,CXCL3,DAG1,IFI44,IFI44L,IFI6,IFIT1,IFITM1,MX1,MX2,NFKBIA,OAS2,OAS3,PARP14,PLAU,PTK2,SOD2,STAT1,TTC27,XAF1 |
| SPI1 | -1.5 | transcription regulator | Inhibited | -3.601 | | 3.49E-11 | CCND1,CCNE2,CD40LG,CMPK2,EPX,FCER1G,HELZ2,HOTAIRM1,ID2,IFI44,IFI44L,IFI6,IFIT1,IFITM1,IFITM3,IRF7,IRF9,ISG15,LMO2,MX1,OASL,PTGS2,RSAD2,TFEC,USP18 |
| STAT1 | -4.26 | transcription regulator | Inhibited | -3.749 | bias | 9.22E-30 | ANGPT2,APOL6,BST2,BTG1,CASP3,CCND1,CCND3,CD274,CD40,CMPK2,CXCL2,CXCL3,EGR1,EPSTI1,FCER1G,GBP1P1,GBP5,HAVCR2,HLA-DRB5,IFI44,IFI44L,IFI6,IFIH1,IFIT1,IFITM1,IFITM3,IRF1,IRF7,IRF9,ISG15,ITGAX,MS4A3,MX1,NLRC5,OAS1,OAS2,OAS3,OASL,PARP9,PMAIP1,POMC,PTGS2,RSAD2,SAMD9L,SLFN5,STAT1,TNFSF13B,TYMP,USP18,USP30-AS1,XAF1 |
| PRL | | cytokine | Inhibited | -3.787 | bias | 7.7E-21 | BST2,CCND1,CCND3,CD40,CD69,CISH,CMPK2,COL5A1,DHX58,EGR1,EPSTI1,HELZ2,HERC5,ID2,IFI44,IFI44L,IFI6,IFIH1,IFIT1,IFITM1,IRF1,IRF7,IRF9,ISG15,MX2,OAS1,OAS2,OAS3,PARP14,PIM3,RPSA,RSAD2,SAMD9L,STAT1,TNFSF13B,USP18,XAF1 |
| IFN Beta | | group | Inhibited | -3.8 | bias | 7.65E-22 | ATF3,BST2,CD274,CD40,CD69,CD83,CXCL1,HERC5,HLA-B,HLA-G,IFI44,IFI6,IFIH1,IFIT1,IFITM1,IRF1,IRF7,IRF9,ISG15,MX1,MX2,OAS1,OAS2,OAS3,OASL,RSAD2,STAT1,USP18,XAF1 |
| RNY3 | | other | Inhibited | -3.873 | bias | 1.1E-19 | EPSTI1,HERC5,IFI44,IFI44L,IFIT1,IFITM3,ISG15,MX1,OAS1,OAS2,OAS3,OASL,RSAD2,TRIM6,XAF1 |
| TGM2 | -1.19 | enzyme | Inhibited | -4.198 | | 1.63E-13 | CCND1,CD36,CFP,CLC,FCER1G,HLA-B,IFI6,IFIT1,IRF9,ITGAX,MS4A3,OAS1,OAS2,OAS3,OASL,PARP14,PARP9,SAMD9L,SELL,SEMA7A,SIGLEC12,SLFN5,STAT1,TTC27,TYROBP,XAF1 |
| TNF | -1.73 | cytokine | Inhibited | -4.343 | | 2.59E-26 | AGTR1,ALAD,ANGPT2,ANPEP,AQP1,ARRDC3,ATF3,BCL3,BHLHE40,BST2,BTG1,CALR,CASP3,CCND1,CCND3,CD274,CD36,CD40,CD40LG,CD69,CD83,CDC42EP4,CDKN2C,CEBPG,CISH,CITED2,CXCL1,CXCL2,CXCL3,DAG1,DDIT3,DHRS3,DUSP5,EGLN1,EGR1,EREG,FCER1G,FOSL1,FOSL2,GADD45B,GBP1P1,GNAI1,GP1BA,GSTM2,HERC5,HLA-B,HLA-F,HPGD,IFI6,IFIH1,IFIT1,IFITM1,IL18R1,IL1RL1,IRF1,IRF7,ISG15,ITGAX,ITGB8,KLF10,KLF2,KYNU,LGALS12,LTC4S,MAFF,MITF,MX1,NEAT1,NFKBIA,NFKBIE,NINJ1,NR4A3,OAS1,OAS2,OAS3,OASL,PARP14,PIM3,PLAU,PLAUR,PMAIP1,POMC,PRKCD,PRRG4,PTGS2,PTX3,RELB,RNASE2,RPSA,SCUBE1,SELL,SEMA3C,SOD2,SORBS1,STAT1,TNFAIP3,TNFRSF10A,TNFRSF10D,TNFSF13B,TP53INP1,TSC22D3,TST,TXNIP,TYMP,USP2,VMP1 |
| Interferon alpha | | group | Inhibited | -4.422 | | 1.72E-37 | APOBEC3A,ATF3,BCL3,BST2,CASP3,CCNA1,CCND1,CCND3,CD274,CD40,CD69,CD83,CISH,CXCL1,CXCL2,CXCL3,CYB561,DAG1,DDIT4,DHX58,EPSTI1,FBXO6,FOSL2,GBP5,HELZ2,HERC5,HLA-B,HLA-C,HLA-F,HLA-G,IFI44,IFI44L,IFI6,IFIH1,IFIT1,IFITM1,IFITM3,IGLL1/IGLL5,IL18R1,IL6R,IRF1,IRF7,IRF9,ISG15,MX1,MX2,NFKBIA,OAS1,OAS2,OAS3,PARP14,PARP9,PMAIP1,POMC,PTGS2,RSAD2,SAMD9L,SLC2A3,SLFN5,STAT1,TNFSF13B,TRIB2,TYMP,USP18 |
| IFNL1 | | cytokine | Inhibited | -4.873 | bias | 5.19E-29 | APOL6,ATF3,BST2,CD40,CD83,CMPK2,GBP5,HERC5,HLA-B,HLA-C,IFI44,IFI44L,IFI6,IFIH1,IFIT1,IFITM1,IFITM3,IRF9,ISG15,LGALS3BP,MX1,OAS1,OAS2,OAS3,OASL,RSAD2,STAT1,USP18,XAF1 |
| IFNA2 | | cytokine | Inhibited | -4.933 | bias | 3.6E-21 | APOL6,BST2,CD274,CD69,CISH,CMPK2,HERC5,HLA-B,HLA-C,IFI44,IFI44L,IFI6,IFIH1,IFIT1,IFITM1,IFITM3,IRF1,IRF7,IRF9,ISG15,LGALS3BP,MAL,MT-ATP6,MX1,MX2,OAS1,OAS2,OAS3,PARP9,PMAIP1,RSAD2,STAT1,TNFRSF10D,USP18,XAF1 |
| IRF7 | -2.75 | transcription regulator | Inhibited | -4.961 | bias | 1.7E-26 | CCNA1,CD40,CD69,CMPK2,DHX58,GBP5,HELZ2,HERC5,IFI44,IFI44L,IFI6,IFIH1,IFIT1,IFITM1,IFITM3,IRF1,IRF7,IRF9,ISG15,ITGAX,MX1,MX2,OAS1,OAS2,OAS3,OASL,PARP14,PLAC8,PMAIP1,RSAD2,SAMD9L,STAT1,TNFSF13B,USP18,XAF1 |
| IFNG | | cytokine | Inhibited | -5.63 | | 2.1E-34 | AGTR1,AHR,APOL6,AQP1,AREG,ATF3,BCL3,BRIP1,BST2,BTG1,CASP3,CCNA1,CCND1,CCND3,CD274,CD36,CD40,CD40LG,CD83,CFP,CISH,CMPK2,CXADR,CXCL1,CXCL2,CXCL3,CYB561,DDIT3,DUSP5,EGR1,FBLN1,FBP1,FCER1G,FKBP5,GAD1,GBP5,H2BC8,HCK,HCP5,HLA-B,HLA-C,HLA-DRB5,HLA-F,HLA-G,IFI44,IFI44L,IFI6,IFIH1,IFIT1,IFITM1,IFITM3,IL18R1,IL1RL1,IL6R,IRF1,IRF7,IRF9,ISG15,ITGAX,KLF10,KLF2,KYNU,LGALS3BP,MAFF,MITF,MX1,MX2,MYCN,NEAT1,NFKBIA,NLRC5,OAS1,OAS2,OAS3,OASL,PARP14,PARP9,PIM3,PLAU,PLAUR,PMAIP1,POMC,PRKCD,PRKG2,PTGS2,PTX3,RAB20,RELB,RSAD2,SCUBE1,SELL,SEMA7A,SLFN5,SOD2,STAT1,TNFRSF10A,TNFSF13B,TRIB2,TSC22D3,TXNIP,TYMP,TYROBP,USP18,XAF1 |

**T2 vs. T1 MEP_2**

© 2000-2020 QIAGEN. All rights reserved.

| Upstream Regulator | Expr Fold Change | Molecule Type | Predicted Activation State | Activation z-score | Flags | p-value of overlap | Target Molecules in Dataset |
|---|---|---|---|---|---|---|---|
| IL1RN | | cytokine | Activated | 4.905 | bias | 4.97E-26 | ATF3,BTN3A1,CIITA,CXCL1,CXCL3,CYLD,GBP1,HERC6,IFI27,IFI44,IFI44L,IFI6,IFIH1,IFIT3,IRF1,IRF7,IRF9,KLF6,MX1,OAS1,OAS2,OAS3,OASL,PMAIP1,RSAD2,STAT2,TRIM22,USP18 |
| MAPK1 | 1.06 | kinase | Activated | 4.811 | | 4.12E-28 | APOL6,ATF3,BST2,BTN3A3,CDH1,CISH,E2F1,EGR1,GBP1,GBP5,HBA1/HBA2,HDC,HERC5,HIC2,HLA-B,HLA-C,IFI27,IFI44,IFI6,IFIH1,IFIT1,IFIT3,IFITM1,IFITM3,IRF7,IRF9,ISG15,LGALS1,LGALS3BP,OAS1,OAS2,OAS3,OASL,PAQR6,PARP12,PTGER2,SP110,STAT1,STAT2,TAP1,TDRD7,TNFRSF10B,TRIM22,USP18 |
| TRIM24 | 1.03 | transcription regulator | Activated | 4.404 | bias | 2.26E-17 | DDX60,EPSTI1,GBP4,HERC6,IFI44,IFIH1,IFIT3,IRF1,IRF7,IRF9,ISG15,LGALS3BP,OAS1,OASL,PARP12,SAMD9L,STAT1,STAT2,TAP1,USP18 |
| NKX2-3 | 1.4 | transcription regulator | Activated | 4.379 | | 2.41E-14 | ANGPT2,BTG1,CD36,CSRNP1,DDX60,GBP1,HCP5,HELZ2,HLA-B,HLA-C,HLA-F,PARP12,PARP9,PMAIP1,RNF213,SAMD9L,SP110,STAT1,STAT2,TAP1,TRIM22,USP18,XAF1 |
| SIRT1 | -1.36 | transcription regulator | Activated | 4.274 | | 1.72E-18 | ABCG1,BBC3,BIRC3,CD36,CD83,CDH1,CDKN2A,CDKN2C,DDX60,GABARAPL1,GSTM3,HELZ2,HLA-A,HLA-DRB5,ID1,IFI44,IFIT3,IFITM3,IRF7,LGALS3BP,NLRC5,OAS1,OAS2,PARP12,PER1,RELB,RNF213,RSAD2,SFRP5,SOD2,SP110,STAT1,TAP1,TNFRSF10B,USP18 |
| PNPT1 | -1.25 | enzyme | Activated | 4 | bias | 2.02E-18 | GBP4,IFI44,IFIH1,IFIT3,IRF7,ISG15,LGALS3BP,OAS1,PARP12,PARP9,RNF213,SAMD9L,STAT1,STAT2,USP18,XAF1 |
| SOCS1 | -1.64 | other | Activated | 3.915 | bias | 2.87E-16 | CIITA,CISH,CXCL2,FKBP5,GBP5,IFI27,IFI44,IFIH1,IFIT1,IFIT3,IRF1,IRF7,ISG15,MX1,OAS1,OAS2,RSAD2,STAT1,TGM2,TRIM22,USP18 |
| ACKR2 | | G-protein coupled receptor | Activated | 3.606 | bias | 7.78E-15 | DDX60,IFI44,IFIT3,IRF7,ISG15,OAS1,OAS2,OAS3,OASL,RSAD2,STAT1,STAT2,USP18 |
| SP110 | -2.27 | transcription regulator | Activated | 3.578 | | 3.42E-15 | ATF3,BST2,CYLD,E2F1,IFI27,IFI6,IFIH1,IFIT1,IFIT3,IFITM1,IFITM3,IRF9,LY96,MX1,OAS1,OAS3,STAT1,TNFRSF21,TXNIP,YES1 |
| IL10RA | -2.53 | transmembrane receptor | Activated | 3.448 | | 4.25E-08 | ABCG1,CD36,DDIT4,GBP5,GSTM5,HLA-A,IL15RA,IRF1,IRF7,ITGAM,LMO4,NLRC5,NPL,RNF213,RSAD2,S1PR1,STAT1,TAP1,TGM2 |
| USP18 | -2.57 | peptidase | Activated | 3.138 | bias | 3.46E-12 | BBC3,IFI6,IFIH1,IFITM3,IRF1,IRF7,IRF9,ISG15,MX1,OAS1 |
| IRF4 | | transcription regulator | Activated | 3.113 | | 3.73E-10 | B2M,CDKN2A,CIITA,CXCL3,DDIT3,GBP1,IL15RA,IRF1,IRF7,IRF9,ISG15,OAS1,PIM2,STAT1,STAT2,TNFSF13B |
| STAT6 | 1.17 | transcription regulator | Activated | 3.048 | | 6.52E-13 | AREG,CISH,CXCL2,CXCL3,EGR1,FKBP5,GADD45B,GBP5,GBP7,HLA-DMB,IFI44,IFI44L,IFIH1,IFIT3,IFITM3,IL10RA,IRF1,IRF7,IRF9,ISG15,KDM6B,LGALS3BP,NFKBIA,OAS3,OASL,PIM2,RNF19B,RNF213,STAT2 |
| NLRX1 | -1 | other | Activated | 2.949 | | 1.76E-08 | BIRC3,IRF1,NFKB2,NFKBIA,OAS1,RELB,STAT1,STAT2,TNFAIP3 |
| NGLY1 | -1.02 | enzyme | Activated | 2.91 | | 5.9E-10 | IFI27,IFI44,IFI44L,IFIT1,IFIT3,OAS1,OAS3,RSAD2,USP18 |
| PTGER4 | -1.29 | G-protein coupled receptor | Activated | 2.853 | | 1.4E-10 | ARRDC3,CITED2,EGR1,GBP4,HERC6,IFIH1,IRF1,IRF7,MAFF,NCF1,RNF213,RSAD2,S1PR1,SLFN5,USP18,XAF1 |
| mir-21 | | microRNA | Activated | 2.725 | | 0.00000691 | CDH1,CDKN2A,CXCL2,EIF4E3,GBP5,GBP7,IRF1,NLRC5,OAS2,OAS3,SOD2,STAT1,STAT2,TAP1,TIAM1 |
| MYC | -1.01 | transcription regulator | Activated | 2.663 | | 1.31E-14 | ANGPT2,BBC3,CDH1,CDKN2A,DDIT3,DDIT4,E2F1,EGLN1,EGR1,FKBP5,GADD45B,GBP4,GPC1,HERC5,HES1,HLA-A,HLA-B,ID1,IFI27,IFI44,IFI44L,IFI6,IFIH1,IFIT1,IFIT3,IRF7,IRF9,ITGAM,KLF10,KLF6,LGALS1,MX1,NFKBIA,OAS1,OASL,PMAIP1,RPL10,RPL3,RPL7,RPL7A,RSAD2,SLC38A1,SOD2,ST3GAL1,STAT1,TIAM1,TNFRSF10B,TNFRSF19,TNFSF13B,TNS3,TOB1,TXNIP,USP18 |
| PSMB11 | | peptidase | Activated | 2.63 | bias | 0.000787 | CXCL3,GADD45B,ITGAM,NCF1,PIK3CG,S1PR1,TNFRSF21 |
| TAB1 | 1.05 | enzyme | Activated | 2.63 | | 0.000000135 | BIRC3,GBP1,IFIH1,IFIT1,IRF7,TNFSF13B,XAF1 |
| PRDM16 | 1.79 | transcription regulator | Activated | 2.615 | | 0.000000405 | GBP4,IFI44,IRF7,OAS2,OAS3,STAT1,STAT2 |
| Pkc(s) | | group | Activated | 2.557 | | 0.00961 | ATF3,CD36,DDIT3,EGR1,GADD45B,KLF6,PER1,S100A8 |
| NRAS | -1.03 | enzyme | Activated | 2.55 | | 0.000000501 | B2M,CDKN2A,CXCL2,EGR1,HLA-A,IFIH1,IFIT1,ISG15,LGALS1,PHLDA3,RNASE6,STAT1,TAP1,USP18 |
| SATB1 | -1.36 | transcription regulator | Activated | 2.45 | | 0.000000054 | CDH1,EPSTI1,GADD45B,HBG1,HELZ2,HLA-DMB,IL10RA,IRF7,RELB,S1PR1,TNFSF13B,TRIM22,TSC22D3,XAF1 |
| RPSA | 1.98 | translation regulator | Activated | 2.449 | | 0.00000236 | CXCL3,GBP4,ISG15,RELB,STAT1,TAP1 |
| CLDN7 | 1.02 | other | Activated | 2.449 | | 0.00605 | DENND5A,HELZ2,HLA-B,IFI44,IFI6,MX1 |
| Irgm1 | | other | Activated | 2.449 | bias | 0.0000274 | IFIT3,IRF7,OAS2,OASL,RSAD2,USP18 |
| PIK3CG | -2.07 | kinase | Activated | 2.447 | | 0.00000017 | B2M,GBP4,GBP5,GBP7,HLA-A,IRF1,NLRC5,OAS2,PIK3CG,STAT1,TAP1 |
| CNOT7 | 1.08 | transcription regulator | Activated | 2.425 | bias | 1.54E-22 | B2M,BST2,HERC6,IFI27,IFI44L,IFI6,IFITM1,ISG15,LGALS3BP,OAS1,OAS2,OAS3,PARP12,SP110,STAT1,TAP1 |
| AREG | 8.24 | growth factor | Activated | 2.412 | | 0.00113 | AREG,EGR1,EREG,GBP1,IFI6,IFIT3 |
| TNIP1 | -1.06 | other | Activated | 2.406 | | 0.0000158 | CXCL2,CXCL3,HLA-A,HLA-B,RELB,S100A8 |
| mir-155 | | microRNA | Activated | 2.401 | | 0.00516 | IFIT3,IRF7,IRF9,MX1,S1PR1,STAT1 |
| arachidonic acid | | chemical - endogenous mammalian | Activated | 2.383 | | 0.000507 | ABCG1,CD36,CD83,DDIT3,EGR1,KLF6,NFKBIA |
| KRAS | 1.06 | enzyme | Activated | 2.356 | | 4.74E-14 | AREG,ATF3,B2M,BIRC3,CDH1,CDKN2A,CDKN2C,DDIT3,E2F1,EGR1,EREG,GSTM3,HES1,HLA-DMB,HOPX,ID1,IFI6,IFIT1,IFITM1,IFITM3,IRF1,IRF9,ISG15,ITGAM,KLF6,LBH,LTBP4,MX1,NCF1,NFKB2,NFKBIA,OAS1,PMAIP1,STAT1,STAT2,TAP1,TNFRSF10B,VMP1 |
| TAL1 | -1.1 | transcription regulator | Activated | 2.324 | | 8.93E-09 | CDKN2A,CYLD,HLA-A,HLA-B,HLA-C,HLA-F,IKZF3,IL10RA,MARCKS,MS4A2,MVD,RAPGEF5,RNASE6,TNFAIP3,TOB1,TPSAB1/TPSB2,ZBTB21 |
| ITGB1 | -1.65 | transmembrane receptor | Activated | 2.236 | | 0.0107 | CDH1,CDKN2A,HSPG2,LTBP3,TGM2,TNFAIP3 |
| KCNK9 | | ion channel | Activated | 2.236 | | 0.000202 | CA1,GADD45B,GAL,HEMGN,SSTR2 |
| ADORA2A | | G-protein coupled receptor | Activated | 2.236 | | 0.0362 | CXCL2,HBA1/HBA2,IFITM3,NFKBIA,TSC22D3 |
| MEOX2 | | transcription regulator | Activated | 2.236 | | 0.000158 | CXCL1,CXCL2,CXCL3,ID1,TNFAIP3 |
| MLXIPL | | transcription regulator | Activated | 2.236 | bias | 0.0228 | RPL10,RPL3,RPL7,RPL7A,TXNIP |
| DNASE2 | -1.1 | enzyme | Activated | 2.229 | bias | 1.18E-08 | CXCL2,IFIT3,IRF7,ISG15,OAS1,OAS3,RSAD2,USP18 |
| MIR17HG | 1.56 | other | Activated | 2.219 | bias | 0.0235 | CYLD,HOPX,HSPG2,RNF11,TNFAIP3 |
| SAFB | -1.16 | other | Activated | 2.219 | | 0.0000785 | B2M,BBC3,CDH1,CDKN2A,HLA-C,S100A8 |
| SAMHD1 | -1.66 | enzyme | Activated | 2.213 | bias | 0.000000469 | IFI27,IFI6,MX1,NFKB2,NFKBIA |
| NR3C1 | -1.2 | ligand-dependent nuclear receptor | Activated | 2.196 | | 1.81E-11 | ALOX5AP,APOL3,B2M,BIRC3,BTG1,CACNA2D3,CD83,CORO2A,CSRNP1,CXCL2,CXCL3,DDIT4,EGR1,FKBP5,GADD45B,GSTP1,GZMA,IFI6,IFIH1,IL15RA,IRF1,ISG15,LMO4,NFKB2,NFKBIA,OASL,PER1,PNMT,RNF11,SLC38A1,TNFAIP3,TNFRSF10B,TNFRSF21,TSC22D3,ZFYVE28 |
| PPARGC1A | | transcription regulator | Activated | 2.166 | | 0.509 | BBC3,CD36,EGLN1,PMAIP1,SOD2 |
| ANXA1 | 1.34 | enzyme | Activated | 2.111 | | 0.000202 | ANGPT2,CDH1,CXCL2,ITGAM,TSC22D3 |
| ESR1 | -1.37 | ligand-dependent nuclear receptor | Activated | 2.075 | | 1.96E-15 | ALOX5AP,AREG,ARL4D,ARRDC3,ATF3,BIRC3,CD83,CDH1,CDKN2A,CXCL1,CXCL3,DDIT4,DHCR24,DHCR7,E2F1,EGR1,EREG,FST,GADD45B,GAL,HBA1/HBA2,HCP5,HES1,HLA-B,HLA-C,HLA-G,ID1,IFI27,IFI44,IFI44L,IFI6,IFITM1,IFITM3,IRF1,LGALS1,LGALS3BP,LTBP4,MARCKS,MX1,NFIB,NFKB2,NFKBIA,ODF3B,PIM2,PMAIP1,RELB,RENBP,RSAD2,SP110,STAT1,STX3,TEK,TGM2,THRB,TNFAIP3,TNS3,TOB1,TSC22D3,TXNIP |
| IKZF1 | 1.01 | transcription regulator | Activated | 2.034 | | 0.000000289 | B2M,CITED2,CXCL2,DTX3L,EIF4E3,EPSTI1,FLT3,HES1,IFI27,IFI6,IFIT3,RNF213,TNS3,TXNIP,TYROBP,WDFY3 |
| Alpha catenin | | group | Activated | 2.01 | bias | 0.000000654 | AREG,BIRC3,CXCL2,EREG,IRF1,ITGAM,NFKBIA,RELB,S100A8,TGM2,TNFAIP3 |
| MYCN | 1.96 | transcription regulator | Activated | 2.008 | | 0.000122 | B2M,CIITA,CITED2,E2F1,HLA-A,LGALS1,PMAIP1,RNF11,RPL10,RPL3,RPL7,TGM2,TNFRSF10B |
| NFKBIA | -3.22 | transcription regulator | Inhibited | -2.022 | | 2.95E-08 | BIRC3,CDH1,CITED4,CXCL1,CXCL2,CXCL3,DDIT3,EREG,GADD45B,GSTM5,HES1,HLA-A,IFI6,IL15RA,IRF1,ISG15,NFKB2,NFKBIA,RELB,S100A8,SOD2,TNFAIP3,TNFRSF10B |
| IKBKB | 1.15 | kinase | Inhibited | -2.028 | | 6.92E-08 | ATF3,BIRC3,CDH1,CXCL1,CXCL2,CXCL3,EGLN1,EGR1,EREG,HLA-A,IRF1,ISG15,MX1,NFKB2,NFKBIA,RELB,SOD2,TNFAIP3 |
| TNFSF12 | -1.12 | cytokine | Inhibited | -2.05 | | 0.0000801 | CXCL2,CXCL3,ITGAM,KLF15,NFKB2,NFKBIA,S100A8,TCAP |
| STAT2 | -2.08 | transcription regulator | Inhibited | -2.065 | | 1.79E-18 | CIITA,GBP1,IFI27,IFI6,IFIT1,IFIT3,IFITM1,IRF1,IRF7,IRF9,ISG15,MX1,OAS1,OAS2,RSAD2,STAT1,USP18 |
| CREB1 | -1.08 | transcription regulator | Inhibited | -2.066 | | 0.000256 | ATF3,CIITA,CSRNP1,CXCL2,EGR1,GADD45B,GAL,HLA-A,HLA-G,ID1,IRF7,MIDN,MVD,PDE3B,PER1,PNMT,SERTAD1,SOD2,TP53INP2 |
| SMAD4 | 1.02 | transcription regulator | Inhibited | -2.082 | | 0.000000158 | ANGPT2,AREG,BBC3,CDH1,CDKN2A,CITED2,EREG,FST,GADD45B,HRH2,ID1,KLF15,NFKBIA,SERTAD1,SKIL,SSTR2 |
| Map3k7 | | kinase | Inhibited | -2.1 | bias | 0.0021 | CXCL3,IL15RA,ISG15,NFKBIA,RSAD2 |
| VCAN | 2.28 | other | Inhibited | -2.101 | | 6.69E-12 | CDKN2A,CXCL2,CXCL3,FST,IFI44,IFI44L,IFI6,IFIT1,IFITM1,MX1,NFKBIA,OAS2,OAS3,PCSK1N,SOD2,STAT1,TRIM22,XAF1 |
| SYVN1 | 1.02 | transporter | Inhibited | -2.121 | bias | 0.000633 | HERC5,HLA-A,HLA-C,IFI44,LGALS3BP,NFKB2,RPL10,SLC43A2 |
| CSF3 | | cytokine | Inhibited | -2.125 | | 0.0000246 | BIRC3,CISH,CXCL2,CXCL3,EGR1,GADD45B,GNLY,HDC,ITGAM,LY96,TNFAIP3 |
| IFNE | | cytokine | Inhibited | -2.132 | bias | 4.29E-09 | BST2,CD83,HERC5,IFIH1,IFITM3,ISG15,PTGER2,STAT1,USP18 |
| palmitic acid | | chemical - endogenous mammalian | Inhibited | -2.139 | | 0.000014 | ATF3,BBC3,BIRC3,CD36,CGAS,CXCL2,CXCL3,DDIT3,MVD,NCF1,NFKBIA,TNFRSF10B,TXNIP |
| CD40LG | -1.26 | cytokine | Inhibited | -2.141 | bias | 2.95E-15 | ATF3,B2M,BIRC3,BTG1,CD83,CIITA,CXCL1,CXCL2,EGR1,IFI44,IFI6,IFIT1,IFIT3,IFITM1,IL10RA,IRF1,ISG15,ITGAM,KDM6B,MARCKS,MX1,NFKB2,NFKBIA,PMAIP1,RELB,SOD2,STAT1,TAP1,TGM2,TNFAIP3,TNFSF13B |
| C5 | 1.01 | cytokine | Inhibited | -2.155 | | 0.000269 | ATF3,CXCL1,CXCL2,CXCL3,EGR1,ITGAM,NFKBIA,TNFAIP3 |
| CXCL8 | 1.62 | cytokine | Inhibited | -2.159 | | 0.000732 | CXCL2,CXCL3,HSPG2,ITGAM,NFKBIA,TNFRSF10B |
| IL6 | | cytokine | Inhibited | -2.167 | | 2.8E-11 | AREG,ATF3,BBC3,BST2,CD36,CD83,CDH1,CDKN2A,CIITA,CISH,CXCL1,CXCL2,CXCL3,E2F1,EGR1,EREG,GADD45B,GAL,HLA-A,HLA-DRB5,ID1,IFIT1,IFITM3,IRF1,IRF9,ITGAM,LGALS1,LY96,NFKBIA,RNASE6,SLC7A7,SOD2,SP110,STAT1,TAP1,TGM2,TNFRSF10B |
| CD40 | 1.05 | transmembrane receptor | Inhibited | -2.177 | | 0.00000916 | BIRC3,CD83,CYSLTR1,IL10RA,IL15RA,IRF1,ITGAM,NFKB2,NFKBIA,PTGER2,TAP1,TNFAIP3,TNFSF13B |

| | | | | | | | |
|---|---|---|---|---|---|---|---|
| PARP9 | -3.25 | enzyme | Inhibited | -2.178 | bias | 4.95E-13 | IFI44,IFIT1,IFIT3,IRF1,IRF7,ISG15,OAS2,SP110,STAT1 |
| CD28 | | transmembrane receptor | Inhibited | -2.212 | | 0.000314 | BTG1,CISH,CXCL2,GBP1,GZMA,HBG1,IFITM1,IL10RA,IL15RA,IRF1,NFKB2,NFKBIA,TNFAIP3,TOB1 |
| CLEC12A | 1.12 | other | Inhibited | -2.213 | bias | 8.18E-09 | IFIT3,IRF7,ISG15,RSAD2,USP18 |
| FZD9 | | G-protein coupled receptor | Inhibited | -2.213 | bias | 0.00000216 | CDH1,IFI44,IRF7,ISG15,STAT1 |
| CCL5 | 2.32 | cytokine | Inhibited | -2.218 | bias | 0.00382 | CXCL2,CXCL3,HDC,ITGAM,STAT1 |
| IKBKG | 1.04 | kinase | Inhibited | -2.229 | bias | 0.000000328 | BBC3,BIRC3,CXCL2,CXCL3,EREG,HLA-A,IRF7,ISG15,NFKB2,NFKBIA,RELB,TNFAIP3 |
| CIITA | -2.15 | transcription regulator | Inhibited | -2.23 | | 0.000179 | B2M,HLA-A,HLA-B,HLA-DMB,HLA-DRB5 |
| OSCAR | 1.13 | other | Inhibited | -2.236 | | 0.0000808 | CD83,CXCL1,CXCL2,CXCL3,ITGAM |
| TRAF2 | 1.04 | enzyme | Inhibited | -2.236 | | 0.00497 | BIRC3,CDKN2A,CXCL2,CYLD,TNFAIP3 |
| SMARCA4 | 1.2 | transcription regulator | Inhibited | -2.272 | bias | 2.31E-11 | APOL3,AREG,CDH1,CDKN2A,CIITA,E2F1,EGR1,EREG,FKBP5,GADD45B,GBP1,GSTP1,HBG1,HCP5,HLA-B,HLA-C,HLA-F,IFI27,IFIT1,IFITM1,IFITM3,IL15RA,IRF1,KDM6B,KIFAP3,LGALS1,MAFF,NFKB2,RAMP1,SERTAD1,ST3GAL1,TAP1,ZNF503 |
| PAF1 | -1.18 | other | Inhibited | -2.324 | bias | 5.12E-17 | ARL4A,BIRC3,CITED2,HELZ2,HERC5,IFI44,IFI44L,IFIT3,IFITM3,ISG15,OAS2,OAS3,OASL,SERTAD1,SOD2 |
| JAK | | group | Inhibited | -2.333 | | 1.89E-10 | CD83,IFI6,IFIH1,IFIT1,IFIT3,IFITM3,ISG15,PTGER2,RSAD2,STAT1 |
| IL21 | | cytokine | Inhibited | -2.36 | | 0.000000002 | EGR1,GBP5,GZMA,HERC6,HLA-DRB5,IFIT1,IFIT3,IRF7,ISG15,OAS2,OASL,RSAD2,STAT2,TAP1,TYROBP,USP18 |
| RUNX1 | -1.24 | transcription regulator | Inhibited | -2.377 | | 2.8E-09 | BAALC,BBC3,CDKN2A,HBA1/HBA2,HBG2,HEMGN,IFITM3,IRF7,ISG15,KRT1,LMO4,OAS3,PIM2,PMAIP1,RSAD2,STAT1,STX3,TPSAB1/TPSB2 |
| Tnf (family) | | group | Inhibited | -2.404 | | 0.000174 | ATF3,BBC3,BIRC3,CXCL1,CXCL3,FST,NFKBIA,TNFAIP3,TNFSF13B |
| IFNL4 | | cytokine | Inhibited | -2.425 | bias | 1.04E-09 | IFIH1,ISG15,MX1,OAS1,OAS2,STAT1 |
| WNT1 | | cytokine | Inhibited | -2.433 | | 0.00845 | CDH1,CDKN2A,EGR1,HES1,LBH,RNASE1 |
| SOX4 | -1.08 | transcription regulator | Inhibited | -2.449 | | 0.039 | BBC3,CGAS,EIF4E3,KYNU,TIAM1,TNFRSF19 |
| IFNK | | cytokine | Inhibited | -2.449 | bias | 0.000000433 | CD83,IFIH1,IRF1,MX1,OAS1,STAT1 |
| MARK2 | -1.23 | kinase | Inhibited | -2.449 | bias | 0.00000425 | CXCL2,CXCL3,HDC,IRF1,LMO4,RSAD2 |
| TRADD | 1.3 | other | Inhibited | -2.449 | bias | 0.00000153 | CXCL1,CXCL2,CXCL3,IRF1,NFKBIA,TNFAIP3 |
| TNFSF11 | | cytokine | Inhibited | -2.481 | bias | 0.000000727 | CD83,CXCL2,CXCL3,GADD45B,HRH2,IL15RA,ITGAM,NFKB2,NFKBIA,PHLDA3,PTGER2,RELB,SLC7A7,SOD2,STAT1,USP18 |
| DDIT3 | -2.26 | transcription regulator | Inhibited | -2.521 | | 0.00000372 | ATF3,BBC3,DDIT3,ITGAM,LTBP3,PER1,TGM2,TNFRSF10B,TSC22D3,TXNIP |
| HGF | 1.3 | growth factor | Inhibited | -2.553 | | 0.0000133 | ANGPT2,ATF3,BIRC3,CDH1,CDKN2A,CDKN2C,CITED2,CXCL2,DDIT3,EGR1,FKBP5,HES1,ID1,IL15RA,ISG15,PIM2,SKIL,SOD2,TNFAIP3,TNFRSF10B |
| NLRC5 | -2.97 | transcription regulator | Inhibited | -2.572 | bias | 6.7E-09 | B2M,HLA-A,HLA-B,HLA-C,HLA-F,HLA-G,TAP1 |
| CGAS | -2.44 | enzyme | Inhibited | -2.599 | bias | 1.38E-08 | IFI44,IFIT3,IRF7,ISG15,OAS1,RSAD2,USP18 |
| IL7 | -1.71 | cytokine | Inhibited | -2.607 | | 0.000157 | CISH,CXCL1,CXCL2,CXCL3,E2F1,MX1,PIM2,PMAIP1,S1PR1 |
| Ifn gamma | | complex | Inhibited | -2.608 | | 0.000355 | CDKN2A,GBP1,IFI44L,LY96,PMAIP1,STAT1,XAF1 |
| CD3 | | complex | Inhibited | -2.717 | bias | 0.000138 | BIRC3,BST2,BTG1,CD83,CISH,GBP1,GZMA,HBG1,IFIT1,IFITM1,IL15RA,IRF1,IRF9,NFKBIA,RELB,SPINK4,STAT1,TGM2,TNFAIP3,XAF1,YES1 |
| IL33 | | cytokine | Inhibited | -2.726 | bias | 1.25E-10 | ABCG1,AREG,BIRC3,CD36,CISH,CXCL2,CXCL3,GADD45B,HDC,ITGAM,NFKB2,NFKBIA,PIM2,RELB,S100A8,SKIL,SOD2,TNFAIP3,TPSAB1/TPSB2,VMP1 |
| IFNA4 | | cytokine | Inhibited | -2.728 | bias | 0.000000147 | GBP5,IFIH1,IFIT1,ISG15,MX1,OASL,RSAD2,USP18 |
| HMGB1 | -1.27 | transcription regulator | Inhibited | -2.77 | bias | 0.0000174 | CD83,CDKN2A,CXCL2,CXCL3,HLA-G,IFIT1,MX1,RELB |
| CSF2 | | cytokine | Inhibited | -2.814 | bias | 1.03E-10 | ABCG1,ALOX5AP,BBC3,BIRC3,CD33,CD83,CDH1,CDKN2C,CIITA,CISH,CXCL1,CXCL2,DDIT3,EGR1,HDC,IFITM3,ITGAM,LY96,NFKB2,NFKBIA,PIK3CG,PIM2,PTGER2,RELB,SOD2,TGM2,TICRR,TNFAIP3 |
| APP | -1.14 | other | Inhibited | -2.821 | | 1.3E-09 | ABCG1,ARRDC3,BBC3,CDH1,CITED2,CXCL1,CXCL2,CXCL3,EGR1,GBP4,GPC1,GSTM3,GSTP1,HBA1/HBA2,HBG2,HERC6,HES1,HSPG2,IFIH1,IRF1,IRF7,ITGAM,KYNU,LMO4,MAFF,MT-ATP6,NCF1,PMAIP1,RAMP1,RNF213,RSAD2,S100A8,S1PR1,SLFN5,SOD2,TNFRSF10B,USP18,XAF1 |
| JAK1/2 | | group | Inhibited | -2.828 | bias | 0.00000451 | CIITA,GBP5,HLA-DRB5,IRF1,IRF7,ISG15,MX1,RSAD2 |
| MSC | | transcription regulator | Inhibited | -2.828 | | 0.000000823 | DDX60,EPSTI1,IFI27,IFI44,IFI44L,IFIT1,IRF7,XAF1 |
| DOCK8 | 1.02 | other | Inhibited | -2.887 | bias | 1.07E-10 | CITED2,HDC,HELZ2,IFIT3,IL15RA,IRF1,IRF7,ISG15,LMO4,RSAD2,STAT1,STAT2 |
| PRKCD | -1.49 | kinase | Inhibited | -2.909 | | 0.00000948 | BIRC3,CDKN2A,CIITA,DDIT3,DENND3,ITGAM,KLF6,LBH,NFKB2,NFKBIA,OAS1,RELB |
| SMARCB1 | -1.02 | transcription regulator | Inhibited | -2.914 | | 0.00000342 | BTG1,CDH1,CDKN2A,CDKN2C,E2F1,HBG1,HES1,IFITM1,IL15RA,MX1,OAS1,OAS3 |
| FN1 | 1.52 | enzyme | Inhibited | -2.932 | | 0.00000557 | BIRC3,CD83,CDH1,CXCL1,CXCL2,CXCL3,DHCR24,HLA-B,ITGAM,MARCKS,NFKBIA,TGM2,TNFRSF10B |
| BRCA1 | 1.32 | transcription regulator | Inhibited | -2.937 | | 1.94E-08 | AREG,DDIT3,DDIT4,EGR1,FKBP5,GADD45B,IFI27,IFI6,IFIT1,IFIT3,IFITM1,IRF7,MX1,STAT1,TAP1 |
| TRAF6 | -1.03 | enzyme | Inhibited | -2.942 | bias | 0.000000319 | BIRC3,CXCL1,CXCL2,CYLD,GABARAPL1,NCF1,RELB,SOD2,TNFAIP3 |
| IFN type 1 | | group | Inhibited | -2.942 | | 1.15E-09 | BST2,CGAS,IFIH1,IFIT1,IRF1,ISG15,OASL,STAT1,STAT2,TNFSF13B |
| TMEM173 | -1.04 | other | Inhibited | -3.008 | bias | 4.97E-14 | CGAS,CXCL2,GBP5,HDC,HOPX,IFI44,IFIT3,IFITM3,IRF7,ISG15,OAS1,OASL,RSAD2,USP18 |
| SASH1 | | other | Inhibited | -3.051 | bias | 8.87E-12 | CITED2,HDC,HELZ2,IFIT3,IL15RA,IRF1,IRF7,ISG15,LMO4,OASL,RSAD2,STAT1,STAT2 |
| Tlr | | group | Inhibited | -3.111 | bias | 0.00000372 | CD83,CISH,CXCL3,HLA-A,IRF1,IRF7,NFKBIA,STAT1,TNFAIP3,TNFSF13B |
| IFN alpha/beta | | group | Inhibited | -3.148 | bias | 0.000000119 | CD83,HLA-A,IFIT3,IL15RA,IRF1,IRF7,RSAD2,STAT1,STAT2,TNFSF13B |
| TNFSF10 | -1.82 | cytokine | Inhibited | -3.181 | | 1.91E-14 | ANGPT2,BIRC3,HLA-A,HLA-C,HLA-F,IFI27,IFI6,IFIT1,IFITM1,IRF9,ISG15,ITGAM,NFKBIA,PMAIP1,STAT1,TNFAIP3,TNFRSF10B |
| EIF2AK2 | -1.74 | kinase | Inhibited | -3.182 | bias | 2.03E-18 | ATF3,DDIT3,EGR1,IFI27,IFI6,IFIT1,IFITM1,IRF1,ISG15,LGALS3BP,NFKB2,NFKBIA,OAS1,OAS3,PARP12,PARP9,SOD2,STAT1,TNFAIP3,USP18 |
| P38 MAPK | | group | Inhibited | -3.193 | bias | 0.00000319 | ATF3,BBC3,CD36,CD83,CDKN2A,CXCL1,CXCL2,CXCL3,DDIT3,EGR1,FST,GBP1,IRF7,ITGAM,KLF15,PMAIP1,STAT1,TNFRSF10B |
| TLR4 | -1.42 | transmembrane receptor | Inhibited | -3.199 | bias | 1.86E-16 | AREG,ATF3,CDH1,CISH,CITED2,CXCL2,CXCL3,DDIT3,DLK1,EREG,HDC,HELZ2,IFIT3,IFITM3,IL15RA,IRF1,IRF7,ISG15,ITGAM,LMO4,MX1,NFKB2,NFKBIA,OASL,RELB,RSAD2,SLA,SOD2,ST3GAL1,STAT1,STAT2,TNFAIP3 |
| SAMSN1 | -1.14 | other | Inhibited | -3.207 | bias | 1.65E-11 | CITED2,CXCL2,HDC,HELZ2,IFIT3,IL15RA,IRF1,IRF7,ISG15,LMO4,OASL,RSAD2,STAT1,STAT2 |
| ELAVL1 | 1.06 | other | Inhibited | -3.212 | | 3.84E-11 | ATF3,B2M,CD83,CDH1,HLA-A,HLA-DRB5,IFI44,IFIH1,IFITM3,IRF1,IRF9,LGALS3BP,NFKBIA,OAS1,OAS2,SLC7A7,STAT1,TSC22D3,USP18 |
| EBI3 | | cytokine | Inhibited | -3.267 | | 1.64E-10 | B2M,CIITA,HLA-A,HLA-B,HLA-C,HLA-DMB,IRF1,MX1,STAT1,STAT2,TAP1 |
| JAK1 | -1.04 | kinase | Inhibited | -3.274 | bias | 7.34E-12 | HLA-A,HLA-C,HLA-F,ID1,IRF1,IRF7,IRF9,MX1,STAT1,STAT2,TAP1,TNFRSF21,USP18 |
| IL1A | | cytokine | Inhibited | -3.296 | | 0.000000167 | ATF3,BIRC3,BTN3A3,CD83,CXCL1,CXCL2,CXCL3,GBP1,HDC,HSPG2,IRF1,NFKBIA,S100A8,SOD2,SSTR2,TNFAIP3 |
| MAVS | -1.03 | other | Inhibited | -3.366 | bias | 1.83E-14 | CGAS,CXCL2,IFIT1,IFIT3,IFITM3,IRF7,ISG15,OAS1,OAS2,OASL,PARP12,RSAD2,STAT1,STAT2,USP18 |
| TICAM1 | -1.95 | other | Inhibited | -3.458 | bias | 9.22E-11 | CXCL2,CXCL3,EGR1,IFIT1,IFIT3,IL15RA,IRF1,IRF7,ISG15,MAFF,NFKB2,NFKBIA,OASL,RELB,RSAD2,TNFAIP3 |
| DDX58 | -1.67 | enzyme | Inhibited | -3.495 | bias | 8.87E-12 | IFI27,IFI44,IFIH1,IFIT1,IFIT3,IRF1,IRF7,ISG15,LTBP3,OAS1,RSAD2,STAT1,STAT2 |
| TLR3 | -1.21 | transmembrane receptor | Inhibited | -3.653 | bias | 1.79E-15 | ATF3,CXCL2,CXCL3,GADD45B,GBP4,HERC5,ID1,IFI44,IFI44L,IFI6,IFIH1,IFIT1,IFIT3,IRF1,IRF7,ISG15,MX1,OAS1,OASL,PMAIP1,PTGER2,RSAD2,S100A8,STAT1,TNFSF13B,USP18 |
| MYD88 | -1.22 | other | Inhibited | -3.674 | bias | 1.19E-10 | CD83,CISH,CXCL1,CXCL2,CXCL3,EGR1,HDC,HLA-A,IRF1,IRF7,ISG15,MAFF,NFKB2,NFKBIA,OASL,RELB,RSAD2,S100A8,TNFAIP3,TNFSF13B,USP18 |
| IL27 | | cytokine | Inhibited | -3.69 | | 1.1E-11 | B2M,BST2,CIITA,EGR1,HLA-A,HLA-B,HLA-C,HLA-DMB,IL15RA,IRF1,ITGAM,MX1,OAS1,STAT1,STAT2,TAP1,TNFSF13B |
| IFNA1/IFNA13 | | cytokine | Inhibited | -3.774 | | 8.7E-16 | CD83,IFI27,IFI6,IFIH1,IFIT1,IFITM1,IRF1,IRF7,ISG15,MX1,OAS1,OAS2,RSAD2,STAT1,STAT2 |
| IFNAR1 | -1.71 | transmembrane receptor | Inhibited | -3.92 | | 1.29E-26 | ATF3,B2M,CGAS,CIITA,CXCL2,CXCL3,GBP4,HLA-A,IFI44,IFI6,IFIH1,IFIT3,IFITM1,IRF1,IRF7,ISG15,OAS1,OAS2,OAS3,OASL,PARP12,RSAD2,SOD2,STAT1,TGM2,TNFAIP3,TNFSF13B,TRIM22,USP18,XAF1 |
| IRF5 | -1.18 | transcription regulator | Inhibited | -4.041 | bias | 2.03E-15 | CXCL2,IFI44,IFIH1,IFIT1,IFIT3,IFITM3,IRF7,ISG15,OAS1,OAS2,OASL,PARP12,PMAIP1,RSAD2,SP110,STAT1,STAT2 |
| RNY3 | | other | Inhibited | -4.123 | bias | 5.77E-26 | DDX60,EPSTI1,HERC5,IFI44,IFI44L,IFIT1,IFIT3,IFITM3,ISG15,MX1,OAS1,OAS2,OAS3,OASL,RSAD2,TRIM6,XAF1 |
| TLR7 | | transmembrane receptor | Inhibited | -4.127 | bias | 1.8E-15 | ATF3,CD83,CXCL1,CXCL2,CXCL3,IFI44,IFI44L,IFIT1,IFIT3,IFITM1,IL10RA,IRF1,IRF7,IRF9,ISG15,MX1,NFKBIA,OAS2,OAS3,RSAD2,SOD2,STAT1,STAT2,TNFAIP3 |
| OSM | -1.25 | cytokine | Inhibited | -4.13 | | 6.65E-13 | ABCG1,AKR1C1/AKR1C2,ANGPT2,ARL4A,ATF3,B2M,CDH1,CISH,CTSL,CXCL1,CXCL2,CXCL3,DENND5A,DHCR24,EGR1,GBP1,HLA-A,HLA-B,HLA-C,HLA-F,ID1,IL15RA,IRF1,IRF7,IRF9,KLF10,MARCKS,MX1,OAS1,S100A8,STAT1,TAP1,TRIM22 |
| IL1B | 1.18 | cytokine | Inhibited | -4.145 | | 2.96E-21 | APOL6,ATF3,B2M,BIRC3,CD83,CIITA,CISH,CSRNP1,CXCL1,CXCL2,CXCL3,CYSLTR1,DDIT3,DDIT4,E2F1,EGLN1,EGR1,FGF18,FKBP5,FST,GADD45B,GBP1,HELZ2,HERC5,HES1,HLA-A,HSPG2,IFIT1,IFIT3,IL10RA,IL15RA,IRF1,IRF7,ISG15,ITGAM,KLF10,LY96,MX1,NEAT1,NFKB2,NFKBIA,OAS2,RELB,RNASE1,RSAD2,S100A8,SOD2,SSTR2,STAT1,TGM2,TNFAIP3,TNFSF13B,TOB1,TSC22D3,TXNIP,USP18 |
| SPI1 | -1.16 | transcription regulator | Inhibited | -4.342 | | 2.24E-14 | CIITA,E2F1,HELZ2,HES1,IFI27,IFI44,IFI44L,IFI6,IFIT1,IFIT3,IFITM1,IFITM3,IRF7,IRF9,ISG15,ITGAM,MX1,NCF1,OASL,PARP12,PIK3CG,RSAD2,SP110,TRIM22,USP18 |
| Ifn | | group | Inhibited | -4.349 | | 5.45E-19 | B2M,CD83,HLA-A,IFIH1,IFIT1,IFITM1,IFITM3,IL10RA,IL15RA,IRF1,IRF7,ISG15,MX1,OAS2,OAS3,OASL,RSAD2,S100A8,STAT1,TAP1,TNFSF13B,TRIM22 |
| PML | -1.31 | transcription regulator | Inhibited | -4.353 | | 2.43E-19 | BST2,CDKN2A,CIITA,CXCL1,DDX60,EPSTI1,HERC6,HLA-DMB,IFI27,IFI44,IFI44L,IFIH1,IFIT1,IFIT3,IFITM1,IRF7,ISG15,MAFF,MX1,OAS1,OAS2,OAS3,STAT1,TAP1,TNFAIP3 |

| | | | | | | | |
|---|---|---|---|---|---|---|---|
| tretinoin | | chemical - endogenous mammalian | Inhibited | -4.469 | | 2.29E-26 | ABCG1,AREG,BIRC3,BTG1,CD36,CDH1,CDKN2A,CDKN2C,CITED2,CSRNP1,CTSL,CXCL1,CXCL2,DDIT3,DDX60,EGR1,FST,GBP4,GSTP1,GZMA,HES1,HLA-A,HLA-B,HLA-C,HLA-DMB,ID1,IFI27,IFI44,IFI44L,IFI6,IFIH1,IFIT1,IFIT3,IFITM1,IL10RA,IRF1,IRF9,ISG15,ITGAM,KLF15,KRT1,LGALS1,LGALS3BP,MS4A3,NCF1,OAS1,OAS2,OAS3,OASL,PARP9,PIK3CG,PMAIP1,RNASE1,RNF213,RPL10,RPL3,RPL7,S100A8,SAMD9L,SFRP5,SLA,SLC17A9,SLFN5,SMAD9,SP110,SPINK4,STAT1,STAT2,TAP1,TGM2,TNFAIP3,TNFRSF10B,TNFRSF19,TOB1,TRIM22,TSC22D3,TYROBP,USP18,USP3,VMP1,XAF1 |
| IFNB1 | | cytokine | Inhibited | -4.58 | | 4.46E-21 | APOL3,APOL6,BST2,BTN3A3,CD83,CXCL2,CXCL3,E2F1,GBP4,GBP5,GBP7,HERC5,HLA-A,IFI27,IFI6,IFIH1,IFIT1,IFIT3,IFITM1,IRF1,IRF7,IRF9,ISG15,MX1,OAS1,OAS2,PARP12,PMAIP1,RSAD2,STAT1,STAT2,USP18,XAF1 |
| TLR9 | | transmembrane receptor | Inhibited | -4.596 | bias | 2.88E-16 | ATF3,CD83,CISH,CXCL2,CXCL3,EGR1,GADD45B,IFI44L,IFIT1,IFIT3,IFITM1,IL10RA,IRF1,IRF7,IRF9,ISG15,MX1,OAS2,OAS3,PTGER2,RSAD2,S100A8,STAT1,STAT2,TNFSF13B,USP18 |
| Ifnar | | group | Inhibited | -4.617 | bias | 1.54E-21 | B2M,DDIT3,HLA-A,HLA-G,IFIH1,IFIT3,IFITM3,IRF1,IRF7,IRF9,ISG15,NLRC5,OAS1,OAS2,OASL,RNF213,RSAD2,STAT1,STAT2,TAP1,USP18,XAF1 |
| NFkB (complex) | | complex | Inhibited | -4.628 | bias | 8.05E-17 | ABCG1,ATF3,BBC3,BIRC3,CD36,CD83,CDH1,CIITA,CITED4,CXCL1,CXCL2,CXCL3,CYLD,DDIT3,EGR1,FST,GADD45B,HERC5,HLA-A,HLA-DMB,HLA-F,IL10RA,IL15RA,IRF1,IRF7,ISG15,ITGAM,KDM6B,KYNU,NFKB2,NFKBIA,PIM2,PMAIP1,RELB,RNF19B,RSAD2,SOD2,TAP1,TGM2,TNFAIP3,TNFRSF10B |
| TGM2 | -3.14 | enzyme | Inhibited | -4.677 | | 5.57E-18 | BIRC3,CD36,CSRNP1,DDX60,HLA-B,IFI6,IFIT1,IFIT3,IL10RA,IRF9,ITGAM,MS4A3,OAS1,OAS2,OAS3,OASL,PARP9,RNF213,S100A8,SAMD9L,SLFN5,SP110,STAT1,TAP1,TRIM22,TYROBP,XAF1 |
| IRF3 | 1.08 | transcription regulator | Inhibited | -4.713 | bias | 1.1E-27 | B2M,BIRC3,CXCL1,DDX60,FST,GBP1,GBP5,HELZ2,HLA-F,IFI27,IFI44,IFI6,IFIH1,IFIT1,IFIT3,IFITM3,IRF1,IRF7,ISG15,NLRC5,OAS1,OAS2,OAS3,OASL,PARP12,PMAIP1,RSAD2,SAMD9L,STAT1,STAT2,TAP1,TDRD7,TNFAIP3,USP18 |
| TNF | 1.54 | cytokine | Inhibited | -4.774 | | 1.18E-29 | ALOX5AP,ANGPT2,ARRDC3,ATF3,B2M,BBC3,BIRC3,BST2,BTG1,BTN3A3,CD36,CD83,CDH1,CDKN2A,CDKN2C,CIITA,CISH,CITED2,CXCL1,CXCL2,CXCL3,CYLD,DDIT3,E2F1,EGLN1,EGR1,EREG,FGF18,FST,GADD45B,GBP1,GBP1P1,GBP4,GSTP1,HDC,HERC5,HES1,HLA-A,HLA-B,HLA-F,HSPG2,ID1,IFI27,IFI6,IFIH1,IFIT1,IFIT3,IFITM1,IL10RA,IL15RA,IRF1,IRF7,ISG15,ITGAM,KLF10,KLF6,KYNU,LY96,MAFF,MX1,NCF1,NEAT1,NFKB2,NFKBIA,OAS1,OAS2,OAS3,OASL,PIK3CG,PIM2,PMAIP1,RAPGEF5,RELB,RNASE1,S100A8,SLC43A2,SOD2,SORBS1,STAT1,TAP1,TDRD7,TEK,TGM2,TNFAIP3,TNFRSF10B,TNFRSF21,TNFSF13B,TNS3,TSC22D3,TXNIP,VMP1,ZBTB11 |
| IRF1 | -6.27 | transcription regulator | Inhibited | -4.902 | bias | 7.28E-29 | APOL6,B2M,CIITA,CXCL2,CXCL3,E2F1,HELZ2,HLA-G,IFI27,IFI44,IFI44L,IFI6,IFIH1,IFIT1,IFIT3,IFITM1,IFITM3,IRF1,IRF7,IRF9,ISG15,MX1,OAS1,OAS2,OAS3,OASL,RSAD2,SP110,STAT1,STAT2,TAP1,TNFSF13B,TRIM22,XAF1 |
| IFN Beta | | group | Inhibited | -4.962 | bias | 2.91E-27 | ATF3,BST2,CD83,CXCL1,HERC5,HLA-A,HLA-B,HLA-G,IFI27,IFI44,IFI6,IFIH1,IFIT1,IFIT3,IFITM1,IL15RA,IRF1,IRF7,IRF9,ISG15,MX1,OAS1,OAS2,OAS3,OASL,RSAD2,STAT1,STAT2,USP18,XAF1 |
| PRL | | cytokine | Inhibited | -5.352 | | 1.01E-23 | B2M,BST2,CDH1,CISH,DTX3L,EGR1,EPSTI1,HELZ2,HERC5,HERC6,ID1,IFI44,IFI44L,IFI6,IFIH1,IFIT1,IFIT3,IFITM1,IRF1,IRF7,IRF9,ISG15,OAS1,OAS2,OAS3,PARP12,RSAD2,SAMD9L,SP110,STAT1,STAT2,TDRD7,TNFSF13B,USP18,XAF1 |
| IFNL1 | | cytokine | Inhibited | -5.798 | bias | 3.52E-45 | APOL6,ATF3,BST2,CD83,DDX60,GBP1,GBP5,HERC5,HERC6,HLA-B,HLA-C,IFI27,IFI44,IFI44L,IFI6,IFIH1,IFIT1,IFIT3,IFITM1,IFITM3,IRF9,ISG15,LGALS3BP,MX1,OAS1,OAS2,OAS3,OASL,RSAD2,SP110,STAT1,STAT2,TDRD7,TRIM22,USP18,XAF1 |
| STAT1 | -4.03 | transcription regulator | Inhibited | -5.82 | | 3.85E-44 | ANGPT2,APOL4,APOL6,B2M,BST2,BTG1,CIITA,CXCL2,CXCL3,DDX60,EGR1,EPSTI1,GBP1,GBP1P1,GBP4,GBP5,HBG1,HERC6,HES1,HLA-DRB5,IFI27,IFI44,IFI44L,IFI6,IFIH1,IFIT1,IFIT3,IFITM1,IFITM3,IL15RA,IRF1,IRF7,IRF9,ISG15,LY96,MS4A3,MX1,NLRC5,OAS1,OAS2,OAS3,OASL,PARP9,PMAIP1,RNF213,RSAD2,SAMD9L,SLFN5,SP110,STAT1,STAT2,TAP1,TNFRSF10B,TNFSF13B,TRIM22,USP18,XAF1 |
| IRF7 | -5.02 | transcription regulator | Inhibited | -5.94 | bias | 3.33E-34 | GBP1,GBP4,GBP5,HELZ2,HERC5,IFI44,IFI44L,IFI6,IFIH1,IFIT1,IFIT3,IFITM1,IFITM3,IL15RA,IRF1,IRF7,IRF9,ISG15,ITGAM,MX1,OAS1,OAS2,OAS3,OASL,PARP12,PMAIP1,RSAD2,S100A8,SAMD9L,STAT1,STAT2,TAP1,TDRD7,TNFSF13B,TRIM22,USP18,XAF1 |
| Interferon alpha | | group | Inhibited | -6.377 | | 1.39E-45 | APOL3,ATF3,B2M,BBC3,BST2,CD83,CDH1,CGAS,CIITA,CISH,CXCL1,CXCL2,CXCL3,DDIT4,E2F1,EPSTI1,GBP1,GBP5,HELZ2,HERC5,HERC6,HLA-A,HLA-B,HLA-C,HLA-F,HLA-G,IFI27,IFI44,IFI44L,IFI6,IFIH1,IFIT1,IFIT3,IFITM1,IFITM3,IL10RA,IL15RA,IRF1,IRF7,IRF9,ISG15,ITGAM,MX1,NFKBIA,OAS1,OAS2,OAS3,PARP12,PARP9,PIM2,PMAIP1,RNF213,RSAD2,SAMD9L,SLFN5,SP110,STAT1,STAT2,TAP1,TDRD7,TNFSF13B,TRIM22,USP18 |
| IFNA2 | | cytokine | Inhibited | -6.563 | bias | 6.41E-41 | APOL6,B2M,BST2,CDKN2A,CISH,CORO2A,DDX60,GBP1,GBP4,HERC5,HERC6,HLA-A,HLA-B,HLA-C,IFI27,IFI44,IFI44L,IFI6,IFIH1,IFIT1,IFIT3,IFITM1,IFITM3,IRF1,IRF7,IRF9,ISG15,LGALS3BP,MT-ATP6,MX1,OAS1,OAS2,OAS3,PARP12,PARP9,PMAIP1,RSAD2,SP110,STAT1,TAP1,TDRD7,TGM2,TNFRSF10B,TRIM22,USP18,XAF1,ZBTB11 |
| IFNG | | cytokine | Inhibited | -7.338 | | 5.41E-41 | ALOX5AP,APOL6,AREG,ATF3,B2M,BBC3,BIRC3,BST2,BTG1,BTN3A1,BTN3A2,CD36,CD83,CDH1,CDKN2A,CGAS,CIITA,CISH,CXCL1,CXCL2,CXCL3,CYLD,DDIT3,DDX60,DTX3L,E2F1,EGR1,FKBP5,GBP1,GBP4,GBP5,GBP7,GSTP1,HCP5,HERC6,HLA-A,HLA-B,HLA-C,HLA-DMB,HLA-DRB5,HLA-F,HLA-G,HSPG2,ID1,IFI27,IFI44,IFI44L,IFI6,IFIH1,IFIT1,IFIT3,IFITM1,IFITM3,IL10RA,IL15RA,IRF1,IRF7,IRF9,ISG15,ITGAM,KLF10,KLF6,KYNU,LGALS3BP,LY96,MAFF,MX1,NEAT1,NFKB2,NFKBIA,NLRC5,OAS1,OAS2,OAS3,OASL,PARP9,PIM2,PMAIP1,RELB,RSAD2,S100A8,SLFN5,SOD2,SP110,STAT1,STAT2,TAP1,TNFRSF10B,TNFSF13B,TRIM22,TSC22D3,TXNIP,TYROBP,USP18,XAF1 |
