## Supplementary Data File 6 for "Mutated clones driving leukemic transformation are already detectable at the single cell level in CD34-positive cells in the chronic phase of primary myelofibrosis"

**T3 vs. T1 HSC_MPP**

© 2000-2020 QIAGEN. All rights reserved.

| Categories | Diseases or Functions Annotation | p-value | Predicted Activation State | Activation z-score | Molecules |
|---|---|---|---|---|---|
| Organismal Survival | Morbidity or mortality | 2.17E-13 | Increased | 4.288 | ACADL,ACTA2,ADGRE5,ADGRG6,AFMID,AKR1C3,ALKBH4,ANKLE1,APBA3,AREG,ARF5,ARID3A,ARID5B,ATF1,B4GALT1,BEST1,BHLHE40,C1GALT1C1,CAV1,CBY1,CCDC86,CCL3L3,CD4,CDH1,CDKN1A,CDKN2C,CDKN3,CEBPD,CFH,CHKA,CITED2,CLCN3,CLDN5,CLDN7,CLEC1B,COL2A1,COQ9,CPLX1,CRK,CSGALNACT1,CXCL8,CXCR4,CYP7B1,DCLRE1A,DGAT2,DHX58,DLL3,DTL,DUSP1,DYRK1A,ECE2,EEF2K,EGLN1,EGR2,EGR3,EIF2AK2,ELN,F13A1,FBLN1,FBXO45,FEN1,FERMT1,FITM2,FOS,GAS2L3,GDF7,GNAI1,GPC3,H2AC18/H2AC19,HCK,HES1,HHEX,HLA-G,HLTF,HOPX,HOXA3,HRH2,HSPA4L,HSPB1,HYAL2,ICMT,ID1,ID2,ID3,IDS,IFIH1,IFIT2,IKZF3,IL6R,IRF2,IRF7,IRF8,ISG15,ITGB8,JCHAIN,JUN,JUNB,KDM4A,KEAP1,KIF5A,KLF2,KLF4,KLF6,KLF7,KLF9,LIG4,LIN7A,LMNA,LMNB1,LRP6,LRRK2,MACROH2A2,MAFF,MAL,MAN2A1,MAP3K8,MCAM,MCL1,MCOLN1,MDFI,MED24,MMACHC,MMP2,MMP7,MPG,MRVI1,MT-ND1,MYC,MYZAP,NEAT1,NEXN,NFE2,NFIB,NFIL3,NLGN1,NOC3L,NOG,NOTCH4,NR4A2,NR4A3,NR5A1,NRXN2,OASL,PARP3,PIEZO2,PLA2R1,PLK3,PLS3,POMC,POMGNT2,PORCN,PPIF,PRDX1,PRKCD,PTAFR,PTGIR,PTGS2,PTX3,PYGO1,REN,RHEBL1,RIMS1,RNASEL,S100A10,S1PR2,SALL4,SCLY,SELP,SERPINE1,SERTAD1,SH3GL3,SLC12A2,SLC40A1,SLC7A5,SNAI3,SOCS1,SOCS3,SPR,SPRY2,SRGAP3,STAT1,STAT2,SV2A,SYT5,TCF4,TIE1,TIMP3,TIPARP,TLE4,TLR2,TMEM107,TNF,TNFAIP3,TNFSF10,TNFSF13B,TP53,TP53INP1,TWNK,TXNIP,UBIAD1,UGCG,USP18,VDR,VTA1,ZAP70,ZBTB7A,ZFP36,ZMPSTE24,ZNRF3 |
| Organismal Survival | Organismal death | 4.02E-13 | Increased | 4.256 | ACADL,ACTA2,ADGRG6,AFMID,AKR1C3,ALKBH4,ANKLE1,APBA3,AREG,ARF5,ARID3A,ARID5B,ATF1,B4GALT1,BEST1,BHLHE40,C1GALT1C1,CAV1,CBY1,CCDC86,CCL3L3,CD4,CDH1,CDKN1A,CDKN2C,CDKN3,CEBPD,CFH,CHKA,CITED2,CLCN3,CLDN5,CLDN7,CLEC1B,COL2A1,COQ9,CPLX1,CRK,CSGALNACT1,CXCL8,CXCR4,CYP7B1,DCLRE1A,DGAT2,DHX58,DLL3,DTL,DUSP1,DYRK1A,ECE2,EEF2K,EGLN1,EGR2,EGR3,EIF2AK2,ELN,F13A1,FBLN1,FBXO45,FEN1,FERMT1,FITM2,FOS,GAS2L3,GDF7,GNAI1,GPC3,H2AC18/H2AC19,HCK,HES1,HHEX,HLA-G,HLTF,HOPX,HOXA3,HRH2,HSPA4L,HSPB1,HYAL2,ICMT,ID1,ID2,ID3,IDS,IFIH1,IFIT2,IKZF3,IL6R,IRF2,IRF7,IRF8,ISG15,ITGB8,JUN,JUNB,KDM4A,KEAP1,KIF5A,KLF2,KLF4,KLF6,KLF7,KLF9,LIG4,LIN7A,LMNA,LMNB1,LRP6,LRRK2,MACROH2A2,MAFF,MAL,MAN2A1,MAP3K8,MCAM,MCL1,MCOLN1,MDFI,MED24,MMACHC,MMP2,MMP7,MPG,MRVI1,MT-ND1,MYC,MYZAP,NEAT1,NEXN,NFE2,NFIB,NFIL3,NLGN1,NOC3L,NOG,NOTCH4,NR4A2,NR4A3,NR5A1,NRXN2,PARP3,PIEZO2,PLA2R1,PLK3,PLS3,POMC,POMGNT2,PORCN,PPIF,PRDX1,PRKCD,PTAFR,PTGIR,PTGS2,PTX3,PYGO1,REN,RHEBL1,RIMS1,RNASEL,S100A10,S1PR2,SALL4,SCLY,SELP,SERPINE1,SERTAD1,SH3GL3,SLC12A2,SLC40A1,SLC7A5,SNAI3,SOCS1,SOCS3,SPR,SPRY2,SRGAP3,STAT1,STAT2,SV2A,SYT5,TCF4,TIE1,TIMP3,TIPARP,TLE4,TLR2,TMEM107,TNF,TNFAIP3,TNFSF10,TNFSF13B,TP53,TP53INP1,TWNK,TXNIP,UBIAD1,UGCG,USP18,VDR,VTA1,ZAP70,ZBTB7A,ZFP36,ZMPSTE24,ZNRF3 |
| Cell Signaling | Viral life cycle | 1.25E-08 | Increased | 3.206 | BANF1,BST2,CXCR4,EIF2AK2,IFIT1,IFITM1,IFITM3,ISG15,ISG20,MX1,NUP50,OAS1,OAS3,OASL,PLSCR1,PTGS2,RNASEL,RSAD2,TNF,TP53,VTA1 |
| Cell Signaling | Replication of viral replicon | 6.44E-12 | Increased | 3.204 | BANF1,BST2,EIF2AK2,IFIT1,IFITM1,IFITM3,ISG15,ISG20,MX1,OAS1,OAS3,OASL,PLSCR1,PTGS2,RNASEL,RSAD2,TNF |
| Infectious Diseases | Infection of mammalia | 1.16E-10 | Increased | 3.057 | ADGRE5,BATF3,CAV1,CBY1,CCL3L3,CD4,CSF2RB,CXCR4,DCLRE1A,DHX58,DUSP1,EIF2AK2,FCGR1A,FOS,GPC3,HCK,HLA-G,IFIH1,IFIT2,IL6R,IRF7,IRF8,ISG15,JCHAIN,JDP2,MMP2,MMP7,NFE2,OASL,PRKCD,PTAFR,PTX3,RNASEL,SAMHD1,SELP,SERPINE1,SPON2,STAT1,STAT2,TLR2,TNF,TNFSF10,USP18 |
| Cancer,Organismal Injury and Abnormalities | Non-hematological solid tumor | 4.17E-30 | Increased | 3.036 | ABHD12B,ABHD6,ACACB,ACADL,ACTA2,ADAP2,ADGRE5,ADGRG6,AFMID,AGAP3,AGPS,AIFM2,AK4,AKR1C1/AKR1C2,AKR1C3,AKR1C4,ALDH1B1,ALDH9A1,ALKBH4,ALKBH8,ANKLE1,ANKRD20A2 (includes others),ANKRD46,ANXA2R,APBA3,APOL2,APOL4,ARAP1,AREG,ARF5,ARID3A,ARID5B,ARL4A,ARL4C,ARL4D,ARMH4,ARMT1,ARRDC3,ASCL2,ATF1,ATG101,ATP6V0A1,ATP6V0E1,AZU1,B4GALT1,B4GAT1,BATF,BATF3,BCAM,BEST1,BHLHE40,BMP2K,BMPR1B,BRD3,BST2,BTG1,BTG2,BTN3A1,BTN3A2,BTN3A3,C12orf49,C14orf119,C14orf39,C19orf54,C1GALT1C1,C1orf216,C1orf50,C2,C22orf15,C2orf69,C5orf30,CACNB4,CALHM2,CARNMT1,CAV1,CBY1,CCDC126,CCDC146,CCDC175,CCDC184,CCDC50,CCDC86,CCL3,CCL4,CD4,CD69,CD72,CDC42EP4,CDH1,CDH7,CDKN1A,CDKN2C,CDKN3,CEACAM4,CEBPD,CENPF,CFH,CHCHD10,CHKA,CITED2,CKAP4,CLCN3,CLCN4,CLDN5,CLDN7,CLEC1B,CLGN,CLU,CMPK2,CNKSR1,CNOT9,COL2A1,COLGALT1,COQ2,COQ9,CP,CPED1,CRIP2,CRK,CSF2RB,CSGALNACT1,CTNS,CTSO,CXCL8,CXCR4,CYB561,CYP1B1,CYP7B1,CYSLTR1,CYYR1,DCAF12,DCAF4,DCLRE1A,DDI2,DDIT4,DDX60,DEGS2,DENND10,DEPDC1B,DGAT2,DHX33,DHX58,DIRAS3,DIXDC1,DLAT,DLG4,DLL3,DNAJC16,DNAJC28,DNM3OS,DOCK4,DOLPP1,DSEL,DTL,DTX3L,DUSP1,DUSP2,DUSP6,DYRK1A,DYTN,DZIP1,ECE2,EDA2R,EEF2K,EEF2KMT,EFNA1,EGLN1,EGR1,EGR2,EGR3,EGR4,EIF2AK2,ELN,EME2,EML2,ENDOD1,ENPP5,ENTPD1,ENTPD6,EPDR1,EPHX1,EPSTI1,EREG,ETV3,EXOSC4,F13A1,FAM117A,FAM124B,FAM133A,FAM181B,FAM210A,FAM217B,FAM81A,FAM83D,FAM89A,FAM98A,FAXDC2,FBLN1,FBLN5,FBP1,FBXO45,FBXO48,FBXO6,FCGR1A,FCRLB,FEN1,FERMT1,FFAR3,FGFR1OP,FGL2,FITM2,FKBP5,FKBP9,FLOT1,FOS,FOSB,FREM1,FUCA1,FYB1,FZD7,GALNT2,GAPT,GAS2,GAS2L3,GASK1B,GATM,GBP1,GBP4,GBP5,GBP7,GCSAML,GDF7,GEMIN7,GGA2,GIMAP6,GLRB,GLUL,GMPR,GNAI1,GOT2,GPC3,GPR12,GPR157,GPR27,GPR35,GPT2,GRASP,GRK4,GRWD1,GSTM3,GSTO2,GTF2A1,GULP1,H2AC18/H2AC19,HAPLN3,HAUS3,HCK,HERC5,HES1,HHEX,HIF3A,HILPDA,HLA-C,HLA-DRB5,HLA-F,HLA-G,HLF,HLTF,HOMER2,HOMER3,HOPX,HOXA3,HRH2,HSPA4L,HSPB1,HYAL2,HYAL3,HYLS1,ICMT,ID1,ID2,ID3,IDS,IER3,IER5,IFI44,IFI44L,IFI6,IFIH1,IFIT1,IFIT2,IFIT3,IFITM1,IFITM3,IFRD2,IGSF6,IKZF3,IL6R,IL7,INTS14,IPO9,IQCN,IRF2,IRF2BP1,IRF7,IRF8,ISG15,ISG20,ITGB8,ITPKC,ITPRIPL2,JCHAIN,JDP2,JMJD8,JUN,JUNB,JUND,KCNA3,KDM4A,KEAP1,KIF20B,KIF21A,KIF5A,KLF11,KLF13,KLF15,KLF2,KLF4,KLF6,KLF7,KLF9,KLHDC7B,KPNA2,KYNU,L3HYPDH,LAG3,LARP6,LDLRAD3,LETM1,LETM2,LGALS3BP,LIG4,LIN52,LIN7A,LINS1,LMAN2L,LMNA,LMNB1,LONRF2,LRIF1,LRP11,LRP6,LRRC14,LRRC2,LRRC61,LRRC8D,LRRK2,LRRN1,LYSMD1,MAFF,MAGEF1,MAL,MALL,MAN2A1,MAP3K8,MAP4K3,MAPK12,MAPRE3,MARC1,MARCKS,MCAM,MCF2L,MCL1,MCOLN1,MDFI,MED24,MED31,MEIOC,METTL2A,MEX3A,MINPP1,MLEC,MMACHC,MMP2,MMP7,MOB3B,MORC1,MOSPD2,MPG,MRM2,MRPS18C,MRVI1,MT-ND1,MTIF3,MUC12,MX1,MYADM,MYC,MYCBP,MYOZ3,MYRIP,NAT8,NCR3,NEU1,NEXN,NFE2,NFIB,NFIL3,NFKBID,NIP7,NKX3-1,NLGN1,NLRC5,NOC3L,NOG,NOP16,NOS1AP,NOTCH4,NR0B1,NR4A1,NR4A2,NR4A3,NR5A1,NRXN2,NUDT22,NUP50,OAF,OAS1,OAS2,OAS3,OASL,ORC6,OSBPL10,OSBPL6,OSTM1,OTULINL,PANK4,PANX1,PAQR7,PARP3,PARP9,PCDH9,PCSK1N,PCTP,PDZK1IP1,PDZRN4,PER1,PFKFB2,PGAM1,PGS1,PHF23,PHLDA1,PHLDA3,PHLDB2,PIEZO2,PIGM,PIK3IP1,PLA2R1,PLD4,PLD6,PLEKHF1,PLK3,PLS3,PLSCR1,PNMT,POGLUT1,POLR2D,POLR3A,POMC,POMGNT2,PORCN,PPAT,PPIAL4G (includes others),PPIF,PPP1R1B,PPP2R2B,PPP2R5B,PRDM5,PRDX1,PREX2,PRKCD,PRKG2,PRX,PSRC1,PTAFR,PTGIR,PTGR1,PTGS2,PTPRD,PTX3,PWWP2A,PYGO1,QSER1,RAB1B,RAB27B,RAB30,RAB38,RAB6B,RASAL2,RASGEF1B,RASGRP3,RASSF3,RASSF6,RCN1,RDH13,REEP4,REG4,REN,RFXAP,RGCC,RGS1,RGS18,RGS2,RHEBL1,RHOB,RIMS1,RMDN2,RMI1,RNASEL,RNF175,RNF24,RPL9,RSAD2,RTN4R,RXYLT1,S100A10,S1PR2,SALL4,SAMD9,SAMD9L,SAMHD1,SAP30,SCLY,SCML4,SCPEP1,SELENOM,SELP,SEMA4B,SERPINB8,SERPINE1,SERTAD1,SESN1,SGK1,SGPP1,SH3GL3,SHE,SHQ1,SHTN1,SIK1/SIK1B,SLC12A2,SLC16A1,SLC16A9,SLC25A17,SLC26A6,SLC27A2,SLC29A3,SLC2A10,SLC35B2,SLC38A1,SLC39A14,SLC39A6,SLC40A1,SLC45A3,SLC52A3,SLC7A5,SLCO5A1,SLF2,SLFN5,SMAP2,SMCO4,SMIM14,SMOX,SMYD3,SNAI3,SNAPC3,SNTB1,SNX18,SOCS1,SOCS3,SOWAHD,SP110,SPIRE1,SPN,SPOCK2,SPON2,SPR,SPRY2,SRGAP3,SSPN,ST6GAL2,STAC3,STAT1,STAT2,STK17B,STON2,SUCLG2,SV2A,SYT5,TAF5,TANGO6,TAP2,TAS2R14,TBC1D24,TCEAL2,TCF4,TEKT4,TENT4A,TIE1,TIGD7,TIMP3,TIPARP,TLCD3A,TLE3,TLE4,TLR2,TM4SF1,TM9SF2,TMEM117,TMEM231,TMEM246,TMEM45A,TMEM50B,TMEM51,TMEM98,TMEM9B,TMSB10/TMSB4X,TNF,TNFAIP3,TNFSF10,TNFSF13B,TNNI2,TNRC6C,TP53,TP53INP1,TRAF3IP3,TRIM69,TRMT1L,TRMT9B,TSC22D3,TTC13,TTC27,TTC8,TUB,TUBA4A,TUBD1,TVP23A,TWNK,TXNIP,UBE2L6,UBE2T,UBIAD1,UBOX5,UGCG,USP18,USP2,USP28,USP37,VANGL1,VDR,VSTM4,VTA1,WASF3,WBP1L,WDCP,WDFY3,WDR7,WFDC1,WHRN,XAF1,XIRP2,XKR8,YWHAG,ZAP70,ZBTB7A,ZDHHC13,ZFP36,ZMAT3,ZMPSTE24,ZNF280B,ZNF324B,ZNF346,ZNF354B,ZNF385D,ZNF420,ZNF462,ZNF605,ZNF608,ZNF74,ZNF780A,ZNF781,ZNF785,ZNF799,ZNF835,ZNRF3,ZSWIM5 |
| Neurological Disease | Movement Disorders | 0.00000241 | Increased | 2.849 | ABHD6,ARF5,B4GAT1,BHLHE40,BTN3A1,CACNB4,CAV1,CCL4,CDKN2C,CLCN3,CLU,COQ2,COQ9,CP,CPLX1,DIXDC1,EGLN1,EGR1,EGR2,EGR3,EGR4,ENPP5,F8A1 (includes others),FAM117A,FBXO45,FCGR1A,FITM2,FLOT1,FOS,FOSB,FUCA1,GBP1,GLRB,HSPB1,ID3,IDS,IER3,IER5,IFIT2,JMJD8,JUN,JUNB,KCNA3,KIF5A,LMNA,LRRK2,LRRN1,MAFF,MCOLN1,MT-ND1,MYC,NR4A1,NR4A2,NRXN2,PCTP,PPP1R1B,PPP2R2B,PRX,PTGS2,PTPRD,RGS2,RTN4R,S100A10,S1PR2,SAMD9L,SCLY,SELP,SEPTIN5,SERPINE1,SGK1,SLC12A2,SLC16A9,SLC39A14,SMYD3,SPR,SRGAP3,SV2A,TBC1D24,TLR2,TNF,TNNI2,TP53,TSC22D3,TUBA4A,TUBD1,TWNK,UGCG,USP18,USP2,USP37,VDR,ZMPSTE24 |
| Hematological Disease | Hemorrhagic disease | 0.000000757 | Increased | 2.828 | AZU1,C1GALT1C1,CCL3,CCL3L3,CCL4,CLEC1B,CXCL8,EGR1,ENTPD1,F13A1,FCGR1A,FYB1,GP1BA,GSTM3,HYAL2,ID1,ID2,IRF7,ISG15,NFE2,OAS2,PTAFR,PTGIR,PTGS2,RGS18,SERPINE1,TNF,TNFSF10,TNFSF13B,TP53,VDR,ZBTB7A |
| Cancer,Organismal Injury and Abnormalities | Epithelial neoplasm | 1.11E-27 | Increased | 2.696 | ABHD12B,ABHD6,ACACB,ACADL,ACTA2,ADAP2,ADGRE5,ADGRG6,AFMID,AGAP3,AGPS,AIFM2,AK4,AKR1C1/AKR1C2,AKR1C3,AKR1C4,ALDH1B1,ALDH9A1,ALKBH4,ALKBH8,ANKLE1,ANKRD20A2 (includes others),ANXA2R,APBA3,APOL2,APOL4,ARAP1,AREG,ARF5,ARID3A,ARID5B,ARL4A,ARL4C,ARL4D,ARMH4,ARMT1,ARRDC3,ASCL2,ATF1,ATG101,ATP6V0A1,ATP6V0E1,AZU1,B4GALT1,B4GAT1,BATF,BATF3,BCAM,BEST1,BHLHE40,BMP2K,BMPR1B,BRD3,BST2,BTG1,BTG2,BTN3A1,BTN3A2,BTN3A3,C12orf49,C14orf119,C14orf39,C19orf54,C1GALT1C1,C1orf216,C1orf50,C2,C22orf15,C2orf69,C5orf30,CACNB4,CALHM2,CARNMT1,CAV1,CBY1,CCDC126,CCDC146,CCDC175,CCDC184,CCDC50,CCDC86,CCL4,CD4,CD69,CD72,CDC42EP4,CDH1,CDH7,CDKN1A,CDKN2C,CDKN3,CEACAM4,CEBPD,CENPF,CFH,CHCHD10,CHKA,CITED2,CKAP4,CLCN3,CLCN4,CLDN5,CLDN7,CLEC1B,CLGN,CLU,CMPK2,CNKSR1,CNOT9,COL2A1,COLGALT1,COQ2,COQ9,CP,CPED1,CRIP2,CRK,CSF2RB,CSGALNACT1,CTNS,CTSO,CXCL8,CXCR4,CYB561,CYP1B1,CYP7B1,CYSLTR1,CYYR1,DCAF12,DCAF4,DCLRE1A,DDI2,DDIT4,DDX60,DEGS2,DENND10,DEPDC1B,DGAT2,DHX33,DHX58,DIRAS3,DIXDC1,DLAT,DLG4,DLL3,DNAJC16,DNAJC28,DNM3OS,DOCK4,DOLPP1,DSEL,DTL,DTX3L,DUSP1,DUSP2,DUSP6,DYRK1A,DYTN,DZIP1,ECE2,EDA2R,EEF2K,EEF2KMT,EFNA1,EGLN1,EGR1,EGR2,EGR3,EGR4,EIF2AK2,ELN,EME2,EML2,ENDOD1,ENPP5,ENTPD1,ENTPD6,EPDR1,EPHX1,EPSTI1,EREG,ETV3,EXOSC4,F13A1,FAM117A,FAM124B,FAM133A,FAM181B,FAM210A,FAM217B,FAM81A,FAM83D,FAM89A,FAM98A,FAXDC2,FBLN1,FBLN5,FBP1,FBXO45,FBXO6,FCGR1A,FCRLB,FEN1,FERMT1,FFAR3,FGFR1OP,FGL2,FITM2,FKBP5,FKBP9,FLOT1,FOS,FOSB,FREM1,FUCA1,FYB1,FZD7,GALNT2,GAPT,GAS2,GAS2L3,GASK1B,GATM,GBP1,GBP4,GBP5,GBP7,GCSAML,GDF7,GGA2,GIMAP6,GLRB,GLUL,GMPR,GNAI1,GOT2,GPC3,GPR12,GPR157,GPR27,GPR35,GPT2,GRASP,GRK4,GRWD1,GSTO2,GTF2A1,GULP1,H2AC18/H2AC19,HAPLN3,HAUS3,HCK,HERC5,HES1,HHEX,HIF3A,HILPDA,HLA-C,HLA-DRB5,HLA-F,HLA-G,HLF,HLTF,HOMER2,HOMER3,HOPX,HOXA3,HRH2,HSPA4L,HSPB1,HYAL2,HYAL3,HYLS1,ICMT,ID1,ID2,ID3,IDS,IER3,IER5,IFI44,IFI44L,IFI6,IFIH1,IFIT1,IFIT2,IFIT3,IFITM1,IFITM3,IFRD2,IGSF6,IKZF3,IL6R,IL7,INTS14,IPO9,IQCN,IRF2,IRF2BP1,IRF7,IRF8,ISG15,ISG20,ITGB8,ITPKC,ITPRIPL2,JCHAIN,JDP2,JMJD8,JUN,JUNB,JUND,KCNA3,KDM4A,KEAP1,KIF20B,KIF21A,KIF5A,KLF11,KLF13,KLF15,KLF2,KLF4,KLF6,KLF7,KLF9,KLHDC7B,KPNA2,KYNU,L3HYPDH,LAG3,LARP6,LDLRAD3,LETM1,LETM2,LGALS3BP,LIG4,LIN52,LIN7A,LINS1,LMAN2L,LMNA,LMNB1,LONRF2,LRIF1,LRP11,LRP6,LRRC14,LRRC2,LRRC61,LRRC8D,LRRK2,LRRN1,LYSMD1,MAFF,MAGEF1,MAL,MALL,MAN2A1,MAP3K8,MAP4K3,MAPK12,MAPRE3,MARC1,MCAM,MCF2L,MCL1,MCOLN1,MDFI,MED24,MED31,MEIOC,METTL2A,MEX3A,MINPP1,MLEC,MMACHC,MMP2,MMP7,MOB3B,MORC1,MOSPD2,MPG,MRM2,MRPS18C,MRVI1,MT-ND1,MTIF3,MUC12,MX1,MYADM,MYC,MYCBP,MYOZ3,MYRIP,NAT8,NCR3,NEU1,NEXN,NFE2,NFIB,NFIL3,NFKBID,NIP7,NKX3-1,NLGN1,NLRC5,NOC3L,NOG,NOP16,NOS1AP,NOTCH4,NR0B1,NR4A1,NR4A2,NR4A3,NR5A1,NRXN2,NUDT22,NUP50,OAF,OAS1,OAS2,OAS3,OASL,ORC6,OSBPL10,OSBPL6,OSTM1,OTULINL,PANK4,PANX1,PAQR7,PARP3,PARP9,PCDH9,PCSK1N,PCTP,PDZK1IP1,PDZRN4,PER1,PFKFB2,PGAM1,PGS1,PHF23,PHLDA1,PHLDA3,PHLDB2,PIEZO2,PIK3IP1,PLA2R1,PLD4,PLD6,PLEKHF1,PLK3,PLSCR1,PNMT,POGLUT1,POLR2D,POLR3A,POMC,POMGNT2,PORCN,PPAT,PPIAL4G (includes others),PPIF,PPP1R1B,PPP2R2B,PPP2R5B,PRDM5,PRDX1,PREX2,PRKCD,PRKG2,PRX,PSRC1,PTAFR,PTGIR,PTGR1,PTGS2,PTPRD,PTX3,PWWP2A,PYGO1,QSER1,RAB1B,RAB27B,RAB30,RAB38,RAB6B,RASAL2,RASGEF1B,RASGRP3,RASSF3,RASSF6,RCN1,RDH13,REEP4,REG4,REN,RFXAP,RGCC,RGS1,RGS18,RGS2,RHEBL1,RHOB,RIMS1,RMDN2,RMI1,RNASEL,RNF175,RNF24,RPL9,RSAD2,RTN4R,RXYLT1,S100A10,S1PR2,SALL4,SAMD9,SAMD9L,SAMHD1,SAP30,SCLY,SCML4,SCPEP1,SELENOM,SELP,SEMA4B,SERPINB8,SERPINE1,SERTAD1,SESN1,SGK1,SGPP1,SH3GL3,SHE,SHQ1,SHTN1,SIK1/SIK1B,SLC12A2,SLC16A1,SLC16A9,SLC25A17,SLC26A6,SLC27A2,SLC29A3,SLC2A10,SLC35B2,SLC38A1,SLC39A14,SLC39A6,SLC40A1,SLC45A3,SLC52A3,SLC7A5,SLCO5A1,SLF2,SLFN5,SMAP2,SMIM14,SMOX,SMYD3,SNAI3,SNAPC3,SNTB1,SNX18,SOCS1,SOCS3,SOWAHD,SP110,SPIRE1,SPN,SPOCK2,SPON2,SPR,SPRY2,SRGAP3,SSPN,ST6GAL2,STAC3,STAT1,STAT2,STK17B,STON2,SUCLG2,SV2A,SYT5,TAF5,TANGO6,TAP2,TAS2R14,TBC1D24,TCEAL2,TCF4,TEKT4,TENT4A,TIE1,TIGD7,TIMP3,TIPARP,TLCD3A,TLE3,TLE4,TLR2,TM4SF1,TM9SF2,TMEM117,TMEM231,TMEM246,TMEM45A,TMEM50B,TMEM51,TMEM98,TMEM9B,TMSB10/TMSB4X,TNF,TNFAIP3,TNFSF10,TNFSF13B,TNNI2,TNRC6C,TP53,TP53INP1,TRAF3IP3,TRIM69,TRMT1L,TRMT9B,TSC22D3,TTC13,TTC27,TTC8,TUB,TUBA4A,TUBD1,TWNK,TXNIP,UBE2T,UBIAD1,UBOX5,UGCG,USP18,USP2,USP28,USP37,VANGL1,VDR,VSTM4,VTA1,WASF3,WBP1L,WDCP,WDFY3,WDR7,WFDC1,WHRN,XAF1,XIRP2,XKR8,YWHAG,ZAP70,ZBTB7A,ZDHHC13,ZFP36,ZMAT3,ZMPSTE24,ZNF280B,ZNF324B,ZNF346,ZNF354B,ZNF385D,ZNF420,ZNF462,ZNF605,ZNF608,ZNF74,ZNF780A,ZNF781,ZNF785,ZNF799,ZNF835,ZNRF3,ZSWIM5 |
| Connective Tissue Disorders,Hematological Disease,Organismal Injury and Abnormalities | Thrombocytopenia | 0.000000979 | Increased | 2.673 | AZU1,C1GALT1C1,CCL3L3,CLEC1B,CXCL8,EGR1,ENTPD1,FCGR1A,FYB1,GP1BA,GSTM3,HYAL2,ID1,ID2,IRF7,NFE2,OAS2,PTAFR,PTGIR,PTGS2,RGS18,SERPINE1,TNF,TNFSF10,TNFSF13B,TP53,VDR,ZBTB7A |
| Organismal Survival | Perinatal death | 0.000000184 | Increased | 2.57 | AFMID,APBA3,ARID5B,B4GALT1,CDKN1A,CEBPD,CITED2,CLDN5,CLEC1B,COL2A1,CPLX1,CRK,CSGALNACT1,CXCR4,DCLRE1A,DGAT2,EGR2,FBLN1,FBXO45,FOS,GPC3,HCK,HES1,HLTF,HOXA3,ID1,ID2,KIF5A,KLF4,LIG4,LIN7A,LMNB1,LRP6,MACROH2A2,MAN2A1,MMP2,NFE2,NFIB,NLGN1,NR4A2,NR5A1,NRXN2,POMC,POMGNT2,PTGS2,PYGO1,RIMS1,S1PR2,SH3GL3,SPRY2,SV2A,TCF4,TIE1,TNF,TP53,ZNRF3 |
| Cancer,Organismal Injury and Abnormalities | Tumorigenesis of epithelial neoplasm | 0.000000473 | Increased | 2.553 | ABHD12B,ABHD6,ACACB,ACTA2,ADAP2,ADGRE5,ADGRG6,AGAP3,AGPS,AKR1C1/AKR1C2,ALDH1B1,ALKBH8,APOL2,ARAP1,AREG,ARID3A,ARID5B,ARL4C,ARMT1,ARRDC3,B4GAT1,BATF,BCAM,BHLHE40,BMP2K,BMPR1B,BRD3,BTN3A1,BTN3A2,C14orf119,C14orf39,C2,C5orf30,CACNB4,CALHM2,CARNMT1,CAV1,CBY1,CCDC146,CCL4,CD4,CD69,CD72,CDH1,CDH7,CDKN1A,CDKN2C,CEBPD,CENPF,CFH,CHKA,CKAP4,CLCN3,CLCN4,CLDN7,CLEC1B,CLGN,CLU,CNOT9,COL2A1,COLGALT1,COQ2,CP,CPED1,CSF2RB,CSGALNACT1,CXCL8,CXCR4,CYP1B1,CYP7B1,CYSLTR1,CYYR1,DCAF4,DCLRE1A,DDI2,DDX60,DEGS2,DEPDC1B,DGAT2,DHX33,DIRAS3,DIXDC1,DLAT,DLL3,DNAJC28,DNM3OS,DOCK4,DOLPP1,DSEL,DTX3L,DUSP1,DUSP6,DYRK1A,DYTN,DZIP1,ECE2,EEF2K,EEF2KMT,EGR1,EGR2,EGR3,EIF2AK2,EML2,ENPP5,ENTPD6,EPDR1,EPHX1,EREG,ETV3,EXOSC4,F13A1,FAM124B,FAM133A,FAM181B,FAM217B,FAM83D,FAXDC2,FBLN1,FBLN5,FCGR1A,FCRLB,FEN1,FERMT1,FFAR3,FGFR1OP,FKBP5,FKBP9,FLOT1,FOS,FOSB,FREM1,FYB1,FZD7,GAS2L3,GASK1B,GATM,GBP1,GBP4,GBP7,GCSAML,GDF7,GGA2,GLRB,GLUL,GMPR,GOT2,GPR27,GPR35,GRK4,GULP1,H2AC18/H2AC19,HAUS3,HCK,HERC5,HILPDA,HLA-C,HLA-F,HLA-G,HRH2,HSPA4L,HSPB1,HYAL2,HYAL3,HYLS1,ICMT,ID1,IFI44,IFI44L,IFIH1,IFIT3,IFITM1,IFITM3,IGSF6,IKZF3,IL6R,IL7,IPO9,IRF2,IRF2BP1,IRF7,IRF8,ITGB8,ITPRIPL2,JCHAIN,JMJD8,JUN,JUNB,KCNA3,KDM4A,KEAP1,KIF20B,KIF21A,KIF5A,KLF4,KLF6,KLHDC7B,KPNA2,KYNU,L3HYPDH,LAG3,LARP6,LETM1,LIN7A,LINS1,LMAN2L,LONRF2,LRP6,LRRC14,LRRC2,LRRC61,LRRC8D,LRRK2,LRRN1,LYSMD1,MAGEF1,MAL,MAN2A1,MAP3K8,MAP4K3,MAPRE3,MCAM,MCL1,MED24,MINPP1,MLEC,MMP2,MMP7,MORC1,MOSPD2,MPG,MRPS18C,MRVI1,MUC12,MX1,MYC,MYOZ3,NCR3,NEXN,NFE2,NFIB,NFIL3,NLGN1,NLRC5,NOC3L,NOG,NOS1AP,NOTCH4,NR0B1,NR4A1,NR4A2,NR4A3,NR5A1,NRXN2,NUDT22,NUP50,OAS1,OAS2,OASL,OSBPL10,OSBPL6,OSTM1,OTULINL,PANK4,PAQR7,PARP3,PCDH9,PDZK1IP1,PDZRN4,PER1,PGS1,PHLDA1,PHLDB2,PIEZO2,PLA2R1,PLD4,PLEKHF1,PLK3,PLSCR1,POGLUT1,POLR3A,POMC,POMGNT2,PORCN,PPP1R1B,PPP2R2B,PPP2R5B,PRDX1,PREX2,PRKCD,PRKG2,PRX,PTAFR,PTGR1,PTGS2,PTPRD,PWWP2A,PYGO1,QSER1,RAB30,RASAL2,RASGRP3,RASSF3,REN,RGCC,RHOB,RIMS1,RMDN2,RMI1,RNASEL,RSAD2,S100A10,S1PR2,SALL4,SAMD9,SAMD9L,SAP30,SCLY,SCML4,SELP,SERPINE1,SESN1,SGK1,SGPP1,SH3GL3,SHQ1,SHTN1,SIK1/SIK1B,SLC12A2,SLC16A1,SLC16A9,SLC25A17,SLC2A10,SLC35B2,SLC38A1,SLC39A14,SLC40A1,SLC45A3,SLC52A3,SLC7A5,SLCO5A1,SLFN5,SMAP2,SMOX,SNTB1,SNX18,SOCS1,SOCS3,SOWAHD,SP110,SPIRE1,SPN,SPON2,SRGAP3,ST6GAL2,STAT1,STAT2,STK17B,STON2,SUCLG2,SV2A,TANGO6,TAP2,TBC1D24,TCEAL2,TCF4,TEKT4,TENT4A,TIE1,TIGD7,TIMP3,TIPARP,TLE4,TLR2,TM4SF1,TM9SF2,TMEM117,TMEM246,TMEM51,TNF,TNFAIP3,TNFSF10,TNRC6C,TP53,TP53INP1,TRAF3IP3,TRMT9B,TSC22D3,TTC13,TTC27,TUB,TUBA4A,TUBD1,TWNK,TXNIP,UBOX5,UGCG,USP18,USP2,USP28,USP37,VANGL1,VDR,WASF3,WDFY3,WDR7,WFDC1,WHRN,XIRP2,ZAP70,ZBTB7A,ZFP36,ZMAT3,ZMPSTE24,ZNF324B,ZNF354B,ZNF385D,ZNF420,ZNF462,ZNF74,ZNF780A,ZNF781,ZNF785,ZNF799,ZNF835,ZNRF3,ZSWIM5 |
| Cancer,Organismal Injury and Abnormalities | Nonhematologic malignant neoplasm | 2.21E-29 | Increased | 2.545 | ABHD12B,ABHD6,ACACB,ACADL,ACTA2,ADAP2,ADGRE5,ADGRG6,AFMID,AGAP3,AGPS,AIFM2,AK4,AKR1C1/AKR1C2,AKR1C3,AKR1C4,ALDH1B1,ALDH9A1,ALKBH4,ALKBH8,ANKLE1,ANKRD20A2 (includes others),ANKRD46,ANXA2R,APBA3,APOL2,APOL4,ARAP1,AREG,ARF5,ARID3A,ARID5B,ARL4A,ARL4C,ARL4D,ARMH4,ARMT1,ARRDC3,ASCL2,ATF1,ATG101,ATP6V0A1,ATP6V0E1,AZU1,B4GALT1,B4GAT1,BATF,BATF3,BCAM,BEST1,BHLHE40,BMP2K,BMPR1B,BRD3,BST2,BTG1,BTG2,BTN3A1,BTN3A2,BTN3A3,C12orf49,C14orf119,C14orf39,C19orf54,C1GALT1C1,C1orf216,C1orf50,C2,C22orf15,C2orf69,C5orf30,CACNB4,CALHM2,CARNMT1,CAV1,CBY1,CCDC126,CCDC146,CCDC175,CCDC184,CCDC50,CCDC86,CCL3,CCL4,CD4,CD69,CD72,CDC42EP4,CDH1,CDH7,CDKN1A,CDKN2C,CDKN3,CEACAM4,CENPF,CFH,CHCHD10,CHKA,CITED2,CKAP4,CLCN3,CLCN4,CLDN5,CLDN7,CLEC1B,CLGN,CLU,CMPK2,CNKSR1,CNOT9,COL2A1,COLGALT1,COQ2,COQ9,CP,CPED1,CRIP2,CRK,CSF2RB,CSGALNACT1,CTNS,CTSO,CXCL8,CXCR4,CYB561,CYP1B1,CYP7B1,CYSLTR1,CYYR1,DCAF12,DCAF4,DCLRE1A,DDI2,DDIT4,DDX60,DEGS2,DENND10,DEPDC1B,DGAT2,DHX33,DHX58,DIRAS3,DIXDC1,DLAT,DLG4,DLL3,DNAJC16,DNAJC28,DNM3OS,DOCK4,DOLPP1,DSEL,DTL,DTX3L,DUSP1,DUSP2,DUSP6,DYRK1A,DYTN,DZIP1,ECE2,EDA2R,EEF2K,EEF2KMT,EFNA1,EGLN1,EGR1,EGR2,EGR3,EGR4,EIF2AK2,ELN,EME2,EML2,ENDOD1,ENPP5,ENTPD1,ENTPD6,EPDR1,EPHX1,EPSTI1,EREG,ETV3,EXOSC4,F13A1,FAM117A,FAM124B,FAM133A,FAM181B,FAM210A,FAM217B,FAM81A,FAM83D,FAM89A,FAM98A,FAXDC2,FBLN1,FBLN5,FBP1,FBXO45,FBXO48,FBXO6,FCGR1A,FCRLB,FEN1,FERMT1,FFAR3,FGFR1OP,FGL2,FITM2,FKBP5,FKBP9,FLOT1,FOS,FOSB,FREM1,FUCA1,FYB1,FZD7,GALNT2,GAPT,GAS2,GAS2L3,GASK1B,GATM,GBP1,GBP4,GBP5,GBP7,GCSAML,GDF7,GEMIN7,GGA2,GIMAP6,GLRB,GLUL,GMPR,GNAI1,GOT2,GPC3,GPR12,GPR157,GPR27,GPR35,GPT2,GRASP,GRK4,GRWD1,GSTO2,GTF2A1,GULP1,H2AC18/H2AC19,HAPLN3,HAUS3,HCK,HERC5,HES1,HHEX,HIF3A,HILPDA,HLA-C,HLA-DRB5,HLA-F,HLA-G,HLF,HLTF,HOMER2,HOMER3,HOPX,HOXA3,HRH2,HSPA4L,HSPB1,HYAL2,HYAL3,HYLS1,ICMT,ID1,ID2,ID3,IDS,IER3,IER5,IFI44,IFI44L,IFI6,IFIH1,IFIT1,IFIT2,IFIT3,IFITM1,IFITM3,IFRD2,IGSF6,IKZF3,IL6R,IL7,INTS14,IPO9,IQCN,IRF2,IRF2BP1,IRF7,IRF8,ISG15,ISG20,ITGB8,ITPKC,ITPRIPL2,JCHAIN,JDP2,JMJD8,JUN,JUNB,JUND,KCNA3,KDM4A,KEAP1,KIF20B,KIF21A,KIF5A,KLF11,KLF13,KLF15,KLF2,KLF4,KLF6,KLF7,KLF9,KLHDC7B,KPNA2,KYNU,L3HYPDH,LAG3,LARP6,LDLRAD3,LETM1,LETM2,LGALS3BP,LIG4,LIN52,LIN7A,LINS1,LMAN2L,LMNA,LMNB1,LONRF2,LRIF1,LRP11,LRP6,LRRC14,LRRC2,LRRC61,LRRC8D,LRRK2,LRRN1,LYSMD1,MAFF,MAGEF1,MAL,MALL,MAN2A1,MAP3K8,MAP4K3,MAPK12,MAPRE3,MARC1,MARCKS,MCAM,MCF2L,MCL1,MCOLN1,MDFI,MED24,MED31,MEIOC,METTL2A,MEX3A,MINPP1,MLEC,MMACHC,MMP2,MMP7,MOB3B,MORC1,MOSPD2,MPG,MRM2,MRPS18C,MRVI1,MT-ND1,MTIF3,MUC12,MX1,MYADM,MYC,MYCBP,MYOZ3,MYRIP,NAT8,NCR3,NEU1,NEXN,NFE2,NFIB,NFIL3,NFKBID,NIP7,NKX3-1,NLGN1,NLRC5,NOC3L,NOG,NOP16,NOS1AP,NOTCH4,NR0B1,NR4A1,NR4A2,NR4A3,NR5A1,NRXN2,NUDT22,NUP50,OAF,OAS1,OAS2,OAS3,OASL,ORC6,OSBPL10,OSBPL6,OSTM1,OTULINL,PANK4,PANX1,PAQR7,PARP3,PARP9,PCDH9,PCSK1N,PCTP,PDZK1IP1,PDZRN4,PER1,PFKFB2,PGAM1,PGS1,PHF23,PHLDA1,PHLDA3,PHLDB2,PIEZO2,PIGM,PIK3IP1,PLA2R1,PLD4,PLD6,PLEKHF1,PLK3,PLS3,PLSCR1,PNMT,POGLUT1,POLR2D,POLR3A,POMC,POMGNT2,PORCN,PPAT,PPIAL4G (includes others),PPIF,PPP1R1B,PPP2R2B,PPP2R5B,PRDM5,PRDX1,PREX2,PRKCD,PRKG2,PRX,PSRC1,PTAFR,PTGIR,PTGR1,PTGS2,PTPRD,PTX3,PWWP2A,PYGO1,QSER1,RAB1B,RAB27B,RAB30,RAB38,RAB6B,RASAL2,RASGEF1B,RASGRP3,RASSF3,RASSF6,RCN1,RDH13,REEP4,REG4,REN,RFXAP,RGCC,RGS1,RGS18,RGS2,RHEBL1,RHOB,RIMS1,RMDN2,RMI1,RNASEL,RNF175,RNF24,RPL9,RSAD2,RTN4R,RXYLT1,S100A10,S1PR2,SALL4,SAMD9,SAMD9L,SAMHD1,SAP30,SCLY,SCML4,SCPEP1,SELENOM,SELP,SEMA4B,SERPINB8,SERPINE1,SERTAD1,SESN1,SGK1,SGPP1,SH3GL3,SHE,SHQ1,SHTN1,SIK1/SIK1B,SLC12A2,SLC16A1,SLC16A9,SLC25A17,SLC26A6,SLC27A2,SLC29A3,SLC2A10,SLC35B2,SLC38A1,SLC39A14,SLC39A6,SLC40A1,SLC45A3,SLC52A3,SLC7A5,SLCO5A1,SLF2,SLFN5,SMAP2,SMCO4,SMIM14,SMOX,SMYD3,SNAI3,SNAPC3,SNTB1,SNX18,SOCS1,SOCS3,SOWAHD,SP110,SPIRE1,SPN,SPOCK2,SPON2,SPR,SPRY2,SRGAP3,SSPN,ST6GAL2,STAC3,STAT1,STAT2,STK17B,STON2,SUCLG2,SV2A,SYT5,TAF5,TANGO6,TAP2,TAS2R14,TBC1D24,TCEAL2,TCF4,TEKT4,TENT4A,TIE1,TIGD7,TIMP3,TIPARP,TLCD3A,TLE3,TLE4,TLR2,TM4SF1,TM9SF2,TMEM117,TMEM231,TMEM246,TMEM45A,TMEM50B,TMEM51,TMEM98,TMEM9B,TMSB10/TMSB4X,TNF,TNFAIP3,TNFSF10,TNFSF13B,TNNI2,TNRC6C,TP53,TP53INP1,TRAF3IP3,TRIM69,TRMT1L,TRMT9B,TSC22D3,TTC13,TTC27,TTC8,TUB,TUBA4A,TUBD1,TVP23A,TWNK,TXNIP,UBE2L6,UBE2T,UBIAD1,UBOX5,UGCG,USP18,USP2,USP28,USP37,VANGL1,VDR,VSTM4,VTA1,WASF3,WBP1L,WDCP,WDFY3,WDR7,WFDC1,WHRN,XAF1,XIRP2,XKR8,YWHAG,ZAP70,ZBTB7A,ZDHHC13,ZFP36,ZMAT3,ZMPSTE24,ZNF280B,ZNF324B,ZNF346,ZNF354B,ZNF385D,ZNF420,ZNF462,ZNF605,ZNF608,ZNF74,ZNF780A,ZNF781,ZNF785,ZNF799,ZNF835,ZNRF3,ZSWIM5 |
| Cancer,Organismal Injury and Abnormalities | Pelvic tumor | 3.86E-08 | Increased | 2.539 | ABHD12B,ABHD6,ACACB,ADAP2,ADGRE5,ADGRG6,AGPS,AKR1C1/AKR1C2,AKR1C3,ALDH1B1,ALDH9A1,APOL2,APOL4,ARAP1,ARF5,ARID3A,ARID5B,ARL4C,ARL4D,ARMT1,ARRDC3,ASCL2,ATP6V0A1,ATP6V0E1,B4GAT1,BHLHE40,BMP2K,BMPR1B,BRD3,BTN3A1,BTN3A2,C14orf39,C2,CALHM2,CAV1,CBY1,CCDC146,CCDC184,CCDC50,CCL3,CCL4,CD69,CDC42EP4,CDH1,CDH7,CDKN1A,CDKN2C,CDKN3,CENPF,CFH,CKAP4,CLCN4,CLDN5,CLDN7,CLGN,CLU,CNKSR1,CNOT9,COL2A1,CP,CPED1,CSF2RB,CSGALNACT1,CXCL8,CXCR4,CYP1B1,CYSLTR1,DCAF4,DCLRE1A,DDI2,DDIT4,DDX60,DEGS2,DEPDC1B,DHX58,DIRAS3,DIXDC1,DLAT,DLL3,DOCK4,DOLPP1,DSEL,DTX3L,DUSP1,DUSP2,DUSP6,DYRK1A,DYTN,ECE2,EEF2K,EFNA1,EGR1,EGR2,EGR3,EIF2AK2,ELN,EME2,ENTPD1,ENTPD6,EPDR1,EPHX1,EPSTI1,ETV3,EXOSC4,F13A1,FAM117A,FAM124B,FAM181B,FAM217B,FAM83D,FAM89A,FBLN1,FBLN5,FEN1,FERMT1,FFAR3,FGFR1OP,FGL2,FKBP9,FLOT1,FOS,FOSB,FREM1,FYB1,FZD7,GALNT2,GAS2,GASK1B,GBP1,GCSAML,GGA2,GLRB,GLUL,GOT2,GRK4,GRWD1,GSTO2,GTF2A1,GULP1,H2AC18/H2AC19,HAPLN3,HAUS3,HERC5,HIF3A,HLA-C,HLA-G,HLF,HLTF,HOPX,HOXA3,HRH2,HSPA4L,HSPB1,HYAL2,HYLS1,ICMT,ID1,ID2,ID3,IDS,IER3,IFI44,IFI44L,IFIH1,IFIT1,IFIT2,IFIT3,IGKC,IGSF6,IKZF3,IL7,INTS14,IPO9,IRF2,IRF2BP1,IRF7,ISG15,ITGB8,JCHAIN,JUN,JUNB,KCNA3,KDM4A,KEAP1,KIF20B,KIF21A,KIF5A,KLF11,KLF2,KLF4,KLF6,KLF9,KPNA2,KYNU,LAG3,LARP6,LDLRAD3,LETM1,LETM2,LINS1,LMNA,LRP6,LRRC14,LRRC2,LRRK2,LRRN1,LYSMD1,MAGEF1,MAL,MAN2A1,MAP4K3,MCAM,MCL1,MED24,MMP2,MMP7,MORC1,MOSPD2,MPG,MRPS18C,MRVI1,MUC12,MX1,MYADM,MYC,NCR3,NEXN,NFIL3,NKX3-1,NLGN1,NLRC5,NOC3L,NOG,NOS1AP,NOTCH4,NR0B1,NR4A1,NR4A2,NR4A3,NR5A1,NRXN2,NUP50,OAS2,OAS3,OASL,OSBPL10,OSBPL6,OSTM1,OTULINL,PANK4,PAQR7,PARP3,PCDH9,PDZK1IP1,PDZRN4,PER1,PGS1,PHLDA1,PHLDA3,PHLDB2,PIEZO2,PLA2R1,PLD4,PLSCR1,POLR3A,POMC,PPAT,PPIF,PPP1R1B,PPP2R2B,PRDX1,PREX2,PRKCD,PRKG2,PRX,PTAFR,PTGR1,PTGS2,PTPRD,PYGO1,QSER1,RASAL2,RASGEF1B,RASGRP3,REN,RGCC,RGS1,RGS2,RHOB,RIMS1,RMDN2,RNASEL,RNF24,RPL9,RPS19P1,S1PR2,SALL4,SAMD9,SAMD9L,SAP30,SCLY,SCML4,SELP,SEMA4B,SERPINB8,SERPINE1,SGK1,SGPP1,SHTN1,SIK1/SIK1B,SLC12A2,SLC2A10,SLC38A1,SLC39A14,SLC39A6,SLC40A1,SLC45A3,SLC52A3,SLC7A5,SLCO5A1,SLF2,SLFN5,SMAP2,SMOX,SMYD3,SNX18,SOCS1,SOCS3,SP110,SPN,SPOCK2,SPON2,SPRY2,ST6GAL2,STAT1,STAT2,STK17B,SUCLG2,SV2A,TANGO6,TAS2R14,TBC1D24,TCF4,TEKT4,TENT4A,TIE1,TIMP3,TIPARP,TLR2,TM4SF1,TM9SF2,TMEM117,TMEM246,TMSB10/TMSB4X,TNF,TNFSF13B,TNRC6C,TP53,TP53INP1,TSC22D3,TTC13,TUBA4A,TUBD1,TWNK,TXNIP,UBE2T,UBOX5,UGCG,USP18,USP2,USP28,USP37,VDR,VSTM4,WASF3,WDCP,WDFY3,WDR7,WHRN,XAF1,XIRP2,ZAP70,ZBTB7A,ZFP36,ZMAT3,ZMPSTE24,ZNF324B,ZNF354B,ZNF462,ZNF608,ZNF780A,ZNF781,ZNF785,ZNF799,ZNF835,ZNRF3,ZSWIM5 |
| Cancer,Organismal Injury and Abnormalities | Frequency of tumor | 0.000000315 | Increased | 2.496 | ABHD12B,ABHD6,ACACB,ACTA2,ADAP2,ADGRE5,ADGRG6,AGAP3,AGPS,AKR1C1/AKR1C2,ALDH1B1,ALKBH8,APOL2,ARAP1,AREG,ARID3A,ARID5B,ARL4C,ARMT1,ARRDC3,B4GAT1,BATF,BCAM,BHLHE40,BMP2K,BMPR1B,BRD3,BTG2,BTN3A1,BTN3A2,C14orf119,C14orf39,C2,C5orf30,CACNB4,CALHM2,CARNMT1,CAV1,CBY1,CCDC146,CCL4,CD4,CD69,CD72,CDH1,CDH7,CDKN1A,CDKN2C,CEBPD,CENPF,CFH,CHKA,CKAP4,CLCN3,CLCN4,CLDN7,CLEC1B,CLGN,CLU,CNOT9,COL2A1,COLGALT1,COQ2,CP,CPED1,CSF2RB,CSGALNACT1,CXCL8,CXCR4,CYP1B1,CYP7B1,CYSLTR1,CYYR1,DCAF4,DCLRE1A,DDI2,DDX60,DEGS2,DEPDC1B,DGAT2,DHX33,DIRAS3,DIXDC1,DLAT,DLL3,DNAJC28,DNM3OS,DOCK4,DOLPP1,DSEL,DTX3L,DUSP1,DUSP6,DYRK1A,DYTN,DZIP1,ECE2,EEF2K,EEF2KMT,EGR1,EGR2,EGR3,EIF2AK2,EML2,ENPP5,ENTPD6,EPDR1,EPHX1,EREG,ETV3,EXOSC4,F13A1,FAM124B,FAM133A,FAM181B,FAM217B,FAM83D,FAXDC2,FBLN1,FBLN5,FCGR1A,FCRLB,FEN1,FERMT1,FFAR3,FGFR1OP,FKBP5,FKBP9,FLOT1,FOS,FOSB,FREM1,FYB1,FZD7,GAS2L3,GASK1B,GATM,GBP1,GBP4,GBP7,GCSAML,GDF7,GGA2,GLRB,GLUL,GMPR,GOT2,GPR27,GPR35,GRK4,GSTM3,GULP1,H2AC18/H2AC19,HAUS3,HCK,HERC5,HILPDA,HLA-C,HLA-F,HLA-G,HRH2,HSPA4L,HSPB1,HYAL2,HYAL3,HYLS1,ICMT,ID1,ID2,IER3,IFI44,IFI44L,IFIH1,IFIT3,IFITM1,IFITM3,IGSF6,IKZF3,IL6R,IL7,IPO9,IRF2,IRF2BP1,IRF7,IRF8,ITGB8,ITPRIPL2,JCHAIN,JMJD8,JUN,JUNB,KCNA3,KDM4A,KEAP1,KIF20B,KIF21A,KIF5A,KLF2,KLF4,KLF6,KLHDC7B,KPNA2,KYNU,L3HYPDH,LAG3,LARP6,LETM1,LGALS3BP,LIG4,LIN7A,LINS1,LMAN2L,LONRF2,LRP6,LRRC14,LRRC2,LRRC61,LRRC8D,LRRK2,LRRN1,LYSMD1,MAGEF1,MAL,MAN2A1,MAP3K8,MAP4K3,MAPRE3,MCAM,MCL1,MED24,MINPP1,MLEC,MMP2,MMP7,MORC1,MOSPD2,MPG,MRPS18C,MRVI1,MUC12,MX1,MYC,MYOZ3,NCR3,NEXN,NFE2,NFIB,NFIL3,NLGN1,NLRC5,NOC3L,NOG,NOS1AP,NOTCH4,NR0B1,NR4A1,NR4A2,NR4A3,NR5A1,NRXN2,NUDT22,NUP50,OAS1,OAS2,OASL,OSBPL10,OSBPL6,OSTM1,OTULINL,PANK4,PAQR7,PARP3,PCDH9,PDZK1IP1,PDZRN4,PER1,PGS1,PHLDA1,PHLDB2,PIEZO2,PLA2R1,PLD4,PLEKHF1,PLK3,PLSCR1,POGLUT1,POLR3A,POMC,POMGNT2,PORCN,PPP1R1B,PPP2R2B,PPP2R5B,PRDX1,PREX2,PRKCD,PRKG2,PRX,PTAFR,PTGR1,PTGS2,PTPRD,PWWP2A,PYGO1,QSER1,RAB30,RASAL2,RASGRP3,RASSF3,REN,RGCC,RHOB,RIMS1,RMDN2,RMI1,RNASEL,RSAD2,S100A10,S1PR2,SALL4,SAMD9,SAMD9L,SAP30,SCLY,SCML4,SELP,SERPINE1,SESN1,SGK1,SGPP1,SH3GL3,SHQ1,SHTN1,SIK1/SIK1B,SLC12A2,SLC16A1,SLC16A9,SLC25A17,SLC2A10,SLC35B2,SLC38A1,SLC39A14,SLC40A1,SLC45A3,SLC52A3,SLC7A5,SLCO5A1,SLFN5,SMAP2,SMOX,SNTB1,SNX18,SOCS1,SOCS3,SOWAHD,SP110,SPIRE1,SPN,SPON2,SRGAP3,ST6GAL2,STAT1,STAT2,STK17B,STON2,SUCLG2,SV2A,TANGO6,TAP2,TBC1D24,TCEAL2,TCF4,TEKT4,TENT4A,TIE1,TIGD7,TIMP3,TIPARP,TLE4,TLR2,TM4SF1,TM9SF2,TMEM117,TMEM246,TMEM51,TNF,TNFAIP3,TNFSF10,TNRC6C,TP53,TP53INP1,TRAF3IP3,TRMT9B,TSC22D3,TTC13,TTC27,TUB,TUBA4A,TUBD1,TWNK,TXNIP,UBOX5,UGCG,USP18,USP2,USP28,USP37,VANGL1,VDR,WASF3,WDFY3,WDR7,WFDC1,WHRN,XIRP2,ZAP70,ZBTB7A,ZFP36,ZMAT3,ZMPSTE24,ZNF324B,ZNF354B,ZNF385D,ZNF420,ZNF462,ZNF74,ZNF780A,ZNF781,ZNF785,ZNF799,ZNF835,ZNRF3,ZSWIM5 |
| Cancer,Organismal Injury and Abnormalities | Genitourinary tumor | 5.23E-11 | Increased | 2.448 | ABHD12B,ABHD6,ACACB,ACTA2,ADAP2,ADGRE5,ADGRG6,AGAP3,AGPS,AKR1C1/AKR1C2,AKR1C3,ALDH1B1,ALDH9A1,ALKBH8,APBA3,APOL2,APOL4,ARAP1,AREG,ARF5,ARID3A,ARID5B,ARL4C,ARL4D,ARMT1,ARRDC3,ASCL2,ATP6V0A1,ATP6V0E1,B4GALT1,BCAM,BHLHE40,BMP2K,BMPR1B,BRD3,BTG2,BTN3A1,BTN3A2,C14orf39,C1GALT1C1,C2,CALHM2,CAV1,CBY1,CCDC146,CCDC175,CCDC184,CCDC50,CCL3,CCL4,CD4,CD69,CDC42EP4,CDH1,CDH7,CDKN1A,CDKN2C,CDKN3,CEBPD,CENPF,CFH,CKAP4,CLCN4,CLDN5,CLDN7,CLEC1B,CLGN,CLU,CMPK2,CNKSR1,CNOT9,COL2A1,CP,CPED1,CSF2RB,CSGALNACT1,CXCL8,CXCR4,CYP1B1,CYSLTR1,DCAF4,DCLRE1A,DDI2,DDIT4,DDX60,DEGS2,DEPDC1B,DGAT2,DHX58,DIRAS3,DIXDC1,DLAT,DLL3,DOCK4,DOLPP1,DSEL,DTL,DTX3L,DUSP1,DUSP2,DUSP6,DYRK1A,DYTN,ECE2,EDA2R,EEF2K,EFNA1,EGLN1,EGR1,EGR2,EGR3,EGR4,EIF2AK2,ELN,EME2,EML2,ENPP5,ENTPD1,ENTPD6,EPDR1,EPHX1,EPSTI1,EREG,ETV3,EXOSC4,F13A1,FAM117A,FAM124B,FAM181B,FAM217B,FAM83D,FAM89A,FBLN1,FBLN5,FCGR1A,FEN1,FERMT1,FFAR3,FGFR1OP,FGL2,FKBP5,FKBP9,FLOT1,FOS,FOSB,FREM1,FYB1,FZD7,GALNT2,GAS2,GASK1B,GATM,GBP7,GCSAML,GGA2,GLRB,GLUL,GNAI1,GOT2,GPC3,GRK4,GRWD1,GSTM3,GSTO2,GTF2A1,GULP1,H2AC18/H2AC19,HAPLN3,HAUS3,HCK,HERC5,HES1,HIF3A,HILPDA,HLA-C,HLA-DRB5,HLA-G,HLF,HLTF,HOPX,HOXA3,HRH2,HSPA4L,HSPB1,HYAL2,HYLS1,ICMT,ID1,ID2,ID3,IDS,IER2,IER3,IFI44,IFI44L,IFIH1,IFIT1,IFIT2,IFIT3,IFITM1,IGKC,IGSF6,IKZF3,IL6R,IL7,INTS14,IPO9,IRF2,IRF2BP1,IRF7,ISG15,ITGB8,ITPKC,JCHAIN,JUN,JUNB,JUND,KCNA3,KDM4A,KEAP1,KIF20B,KIF21A,KIF5A,KLF11,KLF2,KLF4,KLF6,KLF9,KPNA2,KYNU,LAG3,LARP6,LDLRAD3,LETM1,LETM2,LGALS3BP,LINS1,LMAN2L,LMNA,LONRF2,LRP6,LRRC14,LRRC2,LRRK2,LRRN1,LYSMD1,MAFF,MAGEF1,MAL,MAN2A1,MAP3K8,MAP4K3,MAPRE3,MCAM,MCL1,MED24,MMP2,MMP7,MORC1,MOSPD2,MPG,MRVI1,MT-ND1,MUC12,MX1,MYADM,MYC,NCR3,NEAT1,NEU1,NEXN,NFE2,NFIL3,NKX3-1,NLGN1,NLRC5,NOC3L,NOG,NOP16,NOS1AP,NOTCH4,NR0B1,NR4A1,NR4A2,NR4A3,NR5A1,NRXN2,NUP50,OAF,OAS2,OAS3,OASL,OSBPL10,OSBPL6,OSTM1,OTULINL,PANK4,PAQR7,PARP3,PCDH9,PDZK1IP1,PDZRN4,PER1,PGAM1,PGS1,PHLDA1,PHLDA3,PHLDB2,PIEZO2,PLA2R1,PLD4,PLSCR1,POLR3A,POMC,PPAT,PPIF,PPP1R1B,PPP2R2B,PRDM5,PRDX1,PREX2,PRKCD,PRKG2,PRX,PTAFR,PTGR1,PTGS2,PTPRD,PYGO1,QSER1,RASAL2,RASGEF1B,RASGRP3,REN,RGCC,RGS1,RGS2,RHOB,RIMS1,RMDN2,RNASEL,RNF24,RPL9,RPS19P1,S1PR2,SALL4,SAMD9,SAMD9L,SAP30,SCLY,SCML4,SELP,SEMA4B,SERPINB8,SERPINE1,SGK1,SHQ1,SHTN1,SIK1/SIK1B,SLC12A2,SLC16A1,SLC2A10,SLC38A1,SLC39A14,SLC39A6,SLC40A1,SLC45A3,SLC52A3,SLC7A5,SLCO5A1,SLF2,SLFN5,SMAP2,SMOX,SMYD3,SNAPC3,SNTB1,SNX18,SOCS1,SOCS3,SP110,SPN,SPOCK2,SPON2,SPRY2,SSPN,ST6GAL2,STAT1,STAT2,STK17B,SUCLG2,SV2A,TANGO6,TAS2R14,TBC1D24,TCF4,TEKT4,TENT4A,TIE1,TIMP3,TIPARP,TLE3,TLR2,TM4SF1,TM9SF2,TMEM117,TMEM246,TMEM51,TMSB10/TMSB4X,TNF,TNFSF10,TNFSF13B,TNRC6C,TP53,TP53INP1,TRIM69,TRMT9B,TSC22D3,TTC13,TTC27,TUBA4A,TUBD1,TWNK,TXNIP,UBE2T,UBOX5,UGCG,USP18,USP2,USP28,USP37,VDR,VSTM4,WASF3,WDCP,WDFY3,WDR7,WHRN,XAF1,XIRP2,ZAP70,ZBTB7A,ZFP36,ZMAT3,ZMPSTE24,ZNF280B,ZNF324B,ZNF354B,ZNF420,ZNF462,ZNF608,ZNF780A,ZNF781,ZNF785,ZNF799,ZNF835,ZNRF3,ZSWIM5 |
| Cancer,Organismal Injury and Abnormalities | Anogenital cancer | 2.61E-10 | Increased | 2.422 | ABHD12B,ABHD6,ACACB,ACTA2,ADAP2,ADGRE5,ADGRG6,AGAP3,AGPS,AKR1C1/AKR1C2,AKR1C3,ALDH1B1,ALDH9A1,APOL2,APOL4,ARAP1,AREG,ARF5,ARID3A,ARID5B,ARL4C,ARMT1,ARRDC3,ASCL2,ATP6V0A1,BCAM,BHLHE40,BMP2K,BMPR1B,BRD3,BTG2,BTN3A1,BTN3A2,C14orf39,C1GALT1C1,C2,CAV1,CBY1,CCDC146,CCDC175,CCDC184,CCDC50,CCL3,CCL4,CD4,CD69,CDC42EP4,CDH1,CDH7,CDKN1A,CDKN2C,CDKN3,CEBPD,CENPF,CFH,CKAP4,CLCN4,CLDN5,CLDN7,CLEC1B,CLGN,CLU,CMPK2,CNOT9,COL2A1,CP,CPED1,CSF2RB,CSGALNACT1,CXCL8,CXCR4,CYP1B1,CYSLTR1,DCAF4,DCLRE1A,DDI2,DDIT4,DDX60,DEGS2,DEPDC1B,DHX58,DIRAS3,DIXDC1,DLAT,DOCK4,DOLPP1,DSEL,DTL,DTX3L,DUSP1,DUSP2,DUSP6,DYRK1A,DYTN,ECE2,EDA2R,EEF2K,EFNA1,EGLN1,EGR1,EGR2,EGR3,EGR4,EIF2AK2,ELN,EME2,EML2,ENTPD1,ENTPD6,EPDR1,EPHX1,EPSTI1,ETV3,EXOSC4,F13A1,FAM117A,FAM124B,FAM181B,FAM217B,FAM83D,FAM89A,FBLN1,FBLN5,FEN1,FERMT1,FFAR3,FGFR1OP,FGL2,FKBP5,FKBP9,FLOT1,FOS,FOSB,FREM1,FYB1,FZD7,GALNT2,GAS2,GASK1B,GBP7,GCSAML,GGA2,GLRB,GLUL,GOT2,GPC3,GRK4,GRWD1,GSTM3,GSTO2,GTF2A1,GULP1,H2AC18/H2AC19,HAPLN3,HAUS3,HCK,HERC5,HIF3A,HLA-C,HLA-DRB5,HLA-G,HLF,HLTF,HOXA3,HRH2,HSPA4L,HSPB1,HYAL2,HYLS1,ICMT,ID1,ID2,ID3,IDS,IER2,IER3,IFI44,IFI44L,IFIH1,IFIT1,IFIT2,IFIT3,IGKC,IGSF6,IKZF3,IL7,IPO9,IRF2,IRF7,ISG15,ITGB8,ITPKC,JCHAIN,JUN,JUNB,JUND,KCNA3,KDM4A,KEAP1,KIF20B,KIF21A,KIF5A,KLF11,KLF2,KLF4,KLF6,KLF9,KPNA2,LAG3,LARP6,LDLRAD3,LETM1,LETM2,LGALS3BP,LINS1,LMAN2L,LMNA,LRP6,LRRC14,LRRC2,LRRK2,LRRN1,LYSMD1,MAFF,MAGEF1,MAL,MAN2A1,MAP3K8,MAP4K3,MAPRE3,MCAM,MCL1,MED24,MMP2,MMP7,MORC1,MOSPD2,MPG,MT-ND1,MUC12,MX1,MYADM,MYC,NCR3,NEAT1,NEU1,NEXN,NFIL3,NKX3-1,NLGN1,NLRC5,NOC3L,NOP16,NOS1AP,NOTCH4,NR0B1,NR4A1,NR4A2,NR4A3,NR5A1,NRXN2,NUP50,OAF,OAS2,OAS3,OASL,OSBPL10,OSBPL6,OSTM1,OTULINL,PANK4,PAQR7,PARP3,PCDH9,PDZK1IP1,PDZRN4,PER1,PGAM1,PGS1,PHLDA1,PHLDA3,PHLDB2,PIEZO2,PLA2R1,PLD4,POLR3A,POMC,PPAT,PPIF,PPP1R1B,PRDM5,PRDX1,PREX2,PRKCD,PRKG2,PRX,PTAFR,PTGS2,PTPRD,PYGO1,QSER1,RASAL2,RASGEF1B,RASGRP3,REN,RGCC,RGS1,RGS2,RHOB,RIMS1,RMDN2,RNASEL,RNF24,RPS19P1,S1PR2,SALL4,SAMD9,SAMD9L,SAP30,SCML4,SELP,SEMA4B,SERPINB8,SERPINE1,SGK1,SHQ1,SHTN1,SIK1/SIK1B,SLC12A2,SLC16A1,SLC2A10,SLC38A1,SLC39A14,SLC39A6,SLC40A1,SLC45A3,SLC52A3,SLC7A5,SLCO5A1,SLF2,SLFN5,SMAP2,SMOX,SMYD3,SNAPC3,SNTB1,SNX18,SOCS1,SOCS3,SP110,SPN,SPOCK2,SPON2,SSPN,ST6GAL2,STAT1,STAT2,STK17B,SUCLG2,SV2A,TANGO6,TAS2R14,TBC1D24,TCF4,TEKT4,TIE1,TIMP3,TIPARP,TLE3,TLR2,TM4SF1,TM9SF2,TMEM117,TMEM246,TMSB10/TMSB4X,TNF,TNFSF10,TNFSF13B,TNRC6C,TP53,TP53INP1,TRIM69,TRMT9B,TSC22D3,TUBA4A,TUBD1,TXNIP,UBE2T,UGCG,USP18,USP2,USP28,USP37,VDR,VSTM4,WASF3,WDCP,WDFY3,WDR7,WHRN,XAF1,XIRP2,ZAP70,ZBTB7A,ZFP36,ZMAT3,ZMPSTE24,ZNF280B,ZNF324B,ZNF354B,ZNF420,ZNF462,ZNF608,ZNF780A,ZNF781,ZNF785,ZNF799,ZNF835,ZNRF3,ZSWIM5 |
| Cancer,Organismal Injury and Abnormalities | Development of malignant tumor | 0.000000857 | Increased | 2.33 | ABHD12B,ABHD6,ACACB,ACTA2,ADAP2,ADGRE5,ADGRG6,AGAP3,AGPS,AKR1C1/AKR1C2,ALDH1B1,ALKBH8,APOL2,ARAP1,AREG,ARID3A,ARID5B,ARL4C,ARMT1,ARRDC3,B4GAT1,BATF,BCAM,BHLHE40,BMP2K,BMPR1B,BRD3,BTG2,BTN3A1,BTN3A2,C14orf119,C14orf39,C2,CACNB4,CALHM2,CARNMT1,CAV1,CBY1,CCDC146,CCL4,CD4,CD69,CD72,CDH1,CDH7,CDKN1A,CDKN2C,CENPF,CFH,CHKA,CKAP4,CLCN3,CLCN4,CLDN7,CLEC1B,CLGN,CLU,CNOT9,COL2A1,COLGALT1,COQ2,CP,CPED1,CSF2RB,CSGALNACT1,CXCL8,CXCR4,CYP1B1,CYP7B1,CYSLTR1,CYYR1,DCAF4,DCLRE1A,DDI2,DDX60,DEGS2,DEPDC1B,DGAT2,DHX33,DIRAS3,DIXDC1,DLAT,DLL3,DNAJC28,DNM3OS,DOCK4,DOLPP1,DSEL,DTX3L,DUSP1,DUSP6,DYRK1A,DYTN,DZIP1,ECE2,EEF2K,EEF2KMT,EGR1,EGR2,EGR3,EIF2AK2,EML2,ENPP5,ENTPD6,EPDR1,EPHX1,EREG,ETV3,EXOSC4,F13A1,FAM124B,FAM133A,FAM181B,FAM217B,FAM83D,FAXDC2,FBLN1,FBLN5,FCGR1A,FCRLB,FEN1,FERMT1,FFAR3,FGFR1OP,FKBP5,FKBP9,FLOT1,FOS,FREM1,FYB1,FZD7,GAS2L3,GASK1B,GATM,GBP1,GBP4,GBP7,GCSAML,GDF7,GGA2,GLRB,GLUL,GMPR,GOT2,GPR27,GPR35,GRK4,GSTM3,GULP1,H2AC18/H2AC19,HAUS3,HCK,HERC5,HILPDA,HLA-C,HLA-F,HLA-G,HRH2,HSPA4L,HSPB1,HYAL2,HYAL3,HYLS1,ICMT,ID1,ID2,IER3,IFI44,IFI44L,IFIH1,IFIT3,IFITM1,IGSF6,IKZF3,IL6R,IL7,IPO9,IRF2,IRF2BP1,IRF7,IRF8,ITGB8,ITPRIPL2,JMJD8,JUN,JUNB,KCNA3,KDM4A,KEAP1,KIF20B,KIF21A,KIF5A,KLF4,KLF6,KLHDC7B,KPNA2,KYNU,L3HYPDH,LAG3,LARP6,LETM1,LGALS3BP,LIG4,LIN7A,LINS1,LMAN2L,LONRF2,LRP6,LRRC14,LRRC2,LRRC61,LRRC8D,LRRK2,LRRN1,LYSMD1,MAGEF1,MAL,MAN2A1,MAP3K8,MAP4K3,MAPRE3,MCAM,MCL1,MED24,MINPP1,MLEC,MMP2,MORC1,MOSPD2,MPG,MRPS18C,MRVI1,MUC12,MX1,MYC,MYOZ3,NCR3,NEXN,NFE2,NFIL3,NLGN1,NLRC5,NOC3L,NOG,NOS1AP,NOTCH4,NR0B1,NR4A1,NR4A2,NR4A3,NR5A1,NRXN2,NUDT22,NUP50,OAS1,OAS2,OASL,OSBPL10,OSBPL6,OSTM1,OTULINL,PANK4,PAQR7,PARP3,PCDH9,PDZK1IP1,PDZRN4,PER1,PGS1,PHLDA1,PHLDB2,PIEZO2,PLA2R1,PLD4,PLEKHF1,PLK3,PLSCR1,POGLUT1,POLR3A,POMC,POMGNT2,PORCN,PPP1R1B,PPP2R2B,PPP2R5B,PRDX1,PREX2,PRKCD,PRKG2,PRX,PTAFR,PTGR1,PTGS2,PTPRD,PWWP2A,PYGO1,QSER1,RAB30,RASAL2,RASGRP3,RASSF3,REN,RGCC,RHOB,RIMS1,RMDN2,RMI1,RNASEL,RSAD2,S100A10,S1PR2,SALL4,SAMD9,SAMD9L,SAP30,SCLY,SCML4,SELP,SERPINE1,SESN1,SGK1,SGPP1,SH3GL3,SHQ1,SHTN1,SIK1/SIK1B,SLC12A2,SLC16A1,SLC16A9,SLC25A17,SLC2A10,SLC35B2,SLC38A1,SLC39A14,SLC40A1,SLC45A3,SLC52A3,SLC7A5,SLCO5A1,SLFN5,SMAP2,SMOX,SNTB1,SNX18,SOCS1,SOCS3,SOWAHD,SP110,SPIRE1,SPN,SPON2,SRGAP3,ST6GAL2,STAT1,STAT2,STK17B,STON2,SUCLG2,SV2A,TANGO6,TAP2,TBC1D24,TCEAL2,TCF4,TEKT4,TENT4A,TIE1,TIGD7,TIMP3,TIPARP,TLE4,TLR2,TM4SF1,TM9SF2,TMEM117,TMEM246,TMEM51,TNF,TNFAIP3,TNFSF10,TNRC6C,TP53,TP53INP1,TRAF3IP3,TRMT9B,TSC22D3,TTC13,TTC27,TUB,TUBA4A,TUBD1,TWNK,TXNIP,UBOX5,UGCG,USP18,USP2,USP28,USP37,VANGL1,VDR,WASF3,WDFY3,WDR7,WFDC1,WHRN,XIRP2,ZAP70,ZBTB7A,ZFP36,ZMAT3,ZMPSTE24,ZNF324B,ZNF354B,ZNF385D,ZNF420,ZNF462,ZNF74,ZNF780A,ZNF781,ZNF785,ZNF799,ZNF835,ZNRF3,ZSWIM5 |
| Cancer,Organismal Injury and Abnormalities,Reproductive System Disease | Genital tumor | 1.23E-08 | Increased | 2.314 | ABHD12B,ABHD6,ACACB,ADAP2,ADGRE5,ADGRG6,AGPS,AKR1C1/AKR1C2,AKR1C3,ALDH1B1,ALDH9A1,APOL2,APOL4,ARAP1,ARF5,ARID3A,ARID5B,ARL4C,ARL4D,ARMT1,ARRDC3,ASCL2,ATP6V0A1,B4GAT1,BHLHE40,BMP2K,BMPR1B,BRD3,BTN3A1,BTN3A2,C14orf39,C2,CAV1,CBY1,CCDC146,CCDC184,CCDC50,CCL3,CCL4,CD69,CDC42EP4,CDH1,CDH7,CDKN1A,CDKN2C,CDKN3,CENPF,CFH,CKAP4,CLCN4,CLDN5,CLDN7,CLGN,CLU,CNOT9,COL2A1,CP,CPED1,CSF2RB,CSGALNACT1,CXCL8,CXCR4,CYP1B1,CYSLTR1,DCAF4,DCLRE1A,DDI2,DDIT4,DDX60,DEGS2,DEPDC1B,DHX58,DIRAS3,DIXDC1,DLAT,DOCK4,DOLPP1,DSEL,DTX3L,DUSP1,DUSP2,DUSP6,DYRK1A,DYTN,ECE2,EEF2K,EFNA1,EGR1,EGR2,EGR3,EIF2AK2,ELN,EME2,ENTPD1,ENTPD6,EPDR1,EPHX1,EPSTI1,ETV3,EXOSC4,F13A1,FAM117A,FAM124B,FAM181B,FAM217B,FAM83D,FAM89A,FBLN1,FBLN5,FEN1,FERMT1,FFAR3,FGFR1OP,FGL2,FKBP9,FLOT1,FOS,FREM1,FYB1,FZD7,GALNT2,GAS2,GASK1B,GCSAML,GGA2,GLRB,GLUL,GOT2,GRK4,GRWD1,GSTO2,GTF2A1,GULP1,H2AC18/H2AC19,HAPLN3,HAUS3,HERC5,HIF3A,HLA-C,HLA-G,HLF,HLTF,HOPX,HOXA3,HRH2,HSPA4L,HSPB1,HYAL2,HYLS1,ICMT,ID1,ID2,ID3,IDS,IER3,IFI44,IFI44L,IFIH1,IFIT1,IFIT2,IFIT3,IGKC,IGSF6,IKZF3,IL7,IPO9,IRF2,IRF7,ISG15,ITGB8,JCHAIN,JUN,JUNB,KCNA3,KDM4A,KEAP1,KIF20B,KIF21A,KIF5A,KLF11,KLF2,KLF4,KLF6,KLF9,KPNA2,LAG3,LARP6,LDLRAD3,LETM1,LETM2,LINS1,LMNA,LRP6,LRRC14,LRRC2,LRRK2,LRRN1,LYSMD1,MAGEF1,MAL,MAN2A1,MAP4K3,MCAM,MCL1,MED24,MMP2,MMP7,MORC1,MOSPD2,MPG,MRVI1,MUC12,MX1,MYADM,MYC,NCR3,NEXN,NFIL3,NKX3-1,NLGN1,NLRC5,NOC3L,NOS1AP,NOTCH4,NR0B1,NR4A1,NR4A2,NR4A3,NR5A1,NRXN2,NUP50,OAS2,OAS3,OASL,OSBPL10,OSBPL6,OSTM1,OTULINL,PANK4,PAQR7,PARP3,PCDH9,PDZK1IP1,PDZRN4,PER1,PGS1,PHLDA1,PHLDA3,PHLDB2,PIEZO2,PLA2R1,PLD4,POLR3A,POMC,PPAT,PPIF,PPP1R1B,PREX2,PRKCD,PRKG2,PRX,PTAFR,PTGR1,PTGS2,PTPRD,PYGO1,QSER1,RASAL2,RASGEF1B,RASGRP3,REN,RGCC,RGS1,RGS2,RHOB,RIMS1,RMDN2,RNASEL,RNF24,RPL9,RPS19P1,S1PR2,SALL4,SAMD9,SAMD9L,SAP30,SCML4,SELP,SEMA4B,SERPINB8,SERPINE1,SGK1,SHTN1,SIK1/SIK1B,SLC12A2,SLC38A1,SLC39A14,SLC39A6,SLC40A1,SLC45A3,SLC52A3,SLC7A5,SLCO5A1,SLF2,SLFN5,SMAP2,SMOX,SMYD3,SNX18,SOCS1,SOCS3,SP110,SPN,SPOCK2,SPON2,SPRY2,ST6GAL2,STAT1,STAT2,STK17B,SUCLG2,SV2A,TANGO6,TAS2R14,TBC1D24,TCF4,TEKT4,TIE1,TIMP3,TIPARP,TLE3,TLR2,TM4SF1,TM9SF2,TMEM117,TMEM246,TMSB10/TMSB4X,TNF,TNFSF13B,TNRC6C,TP53,TP53INP1,TSC22D3,TUBA4A,TUBD1,TXNIP,UBE2T,UGCG,USP18,USP2,USP28,USP37,VDR,VSTM4,WASF3,WDCP,WDFY3,WDR7,WHRN,XAF1,XIRP2,ZAP70,ZBTB7A,ZFP36,ZMAT3,ZMPSTE24,ZNF324B,ZNF354B,ZNF462,ZNF608,ZNF780A,ZNF781,ZNF785,ZNF799,ZNF835,ZNRF3,ZSWIM5 |
| Organismal Survival | Neonatal death | 0.000000125 | Increased | 2.257 | AFMID,APBA3,ARID5B,B4GALT1,CDKN1A,CEBPD,CLDN5,CLEC1B,COL2A1,CPLX1,CSGALNACT1,CXCR4,DGAT2,EGR2,FBXO45,FOS,HLTF,HOXA3,ID1,ID2,KIF5A,KLF4,LIG4,LIN7A,LMNB1,LRP6,MAN2A1,MMP2,NFE2,NFIB,NLGN1,NR4A2,NRXN2,POMGNT2,PTGS2,PYGO1,RIMS1,SH3GL3,SV2A,TCF4,TIE1,TNF,TP53 |
| Organismal Injury and Abnormalities | Benign lesion | 5.13E-10 | Increased | 2.222 | ACTA2,ADGRE5,ALDH1B1,ARL4D,B4GALT1,C5orf30,CAV1,CCL3,CCL3L3,CDH1,CDKN1A,CDKN2C,CEBPD,CFH,CHKA,CLU,COL2A1,CSF2RB,CXCR4,CYP1B1,DCLRE1A,DGAT2,DUSP1,EGR1,FBLN1,FOS,FOSB,FZD7,GLUL,GPC3,HAUS3,HOPX,HRH2,ID1,IFI44L,IFIH1,IFIT1,IFIT3,IFITM3,IL6R,JCHAIN,JUN,JUNB,KLF11,KLF2,KLF4,KLF7,LRP6,MAL,MINPP1,MMP2,MMP7,MRVI1,MYC,NFIB,NOG,NOTCH4,NR0B1,NR4A1,NR4A2,NR4A3,NR5A1,NUP50,PHLDA1,PLA2R1,POMC,PRDX1,PRKCD,PTGR1,PTGS2,RPL9,SERPINE1,SH3GL3,SIK1/SIK1B,SLC12A2,SLC39A14,SLC7A5,SOCS3,SPON2,TIE1,TLCD3A,TLR2,TNFAIP3,TNFSF10,TNFSF13B,TNNI2,TP53,TP53INP1,TUBA4A,TUBD1,UGCG,VDR,XAF1 |
| Cancer,Organismal Injury and Abnormalities | Adenocarcinoma | 2.01E-15 | Increased | 2.221 | ABHD12B,ABHD6,ACACB,ACTA2,ADAP2,ADGRE5,ADGRG6,AFMID,AGAP3,AGPS,AKR1C1/AKR1C2,AKR1C3,ALDH1B1,ALDH9A1,ALKBH4,ALKBH8,ANKLE1,ANXA2R,APOL2,APOL4,ARAP1,AREG,ARF5,ARID3A,ARID5B,ARL4C,ARL4D,ARMH4,ARMT1,ARRDC3,ATP6V0A1,AZU1,B4GAT1,BATF,BCAM,BEST1,BHLHE40,BMP2K,BMPR1B,BRD3,BST2,BTN3A1,BTN3A2,C14orf119,C14orf39,C19orf54,C1GALT1C1,C1orf216,C2,CACNB4,CALHM2,CARNMT1,CAV1,CBY1,CCDC146,CCDC175,CCDC184,CCDC50,CCDC86,CCL4,CD4,CD69,CD72,CDC42EP4,CDH1,CDH7,CDKN1A,CDKN2C,CDKN3,CEACAM4,CENPF,CFH,CHCHD10,CHKA,CITED2,CKAP4,CLCN3,CLCN4,CLDN5,CLDN7,CLEC1B,CLGN,CLU,CMPK2,CNKSR1,CNOT9,COL2A1,COLGALT1,COQ2,COQ9,CP,CPED1,CRIP2,CRK,CSF2RB,CSGALNACT1,CTNS,CXCL8,CXCR4,CYP1B1,CYP7B1,CYSLTR1,CYYR1,DCAF12,DCAF4,DCLRE1A,DDI2,DDIT4,DDX60,DEGS2,DEPDC1B,DGAT2,DHX33,DHX58,DIRAS3,DIXDC1,DLAT,DLG4,DLL3,DNAJC16,DNAJC28,DOCK4,DOLPP1,DSEL,DTL,DTX3L,DUSP1,DUSP6,DYRK1A,DYTN,DZIP1,ECE2,EDA2R,EEF2K,EEF2KMT,EFNA1,EGR1,EGR2,EGR3,EGR4,EIF2AK2,ELN,EME2,EML2,ENDOD1,ENPP5,ENTPD1,ENTPD6,EPDR1,EPHX1,EPSTI1,EREG,ETV3,EXOSC4,F13A1,FAM117A,FAM124B,FAM133A,FAM181B,FAM217B,FAM83D,FAM89A,FAM98A,FAXDC2,FBLN1,FBLN5,FBP1,FBXO45,FCGR1A,FCRLB,FEN1,FERMT1,FFAR3,FGFR1OP,FGL2,FKBP5,FKBP9,FLOT1,FOS,FOSB,FREM1,FUCA1,FYB1,FZD7,GALNT2,GAS2,GAS2L3,GASK1B,GATM,GBP1,GBP4,GBP5,GBP7,GCSAML,GDF7,GGA2,GLRB,GLUL,GMPR,GNAI1,GOT2,GPC3,GPR157,GPR27,GPR35,GPT2,GRASP,GRK4,GRWD1,GSTO2,GTF2A1,GULP1,H2AC18/H2AC19,HAPLN3,HAUS3,HCK,HERC5,HES1,HHEX,HIF3A,HILPDA,HLA-C,HLA-DRB5,HLA-F,HLA-G,HLF,HLTF,HOMER2,HOMER3,HOXA3,HRH2,HSPA4L,HSPB1,HYAL2,HYAL3,HYLS1,ICMT,ID1,ID2,ID3,IDS,IER3,IFI44,IFI44L,IFI6,IFIH1,IFIT1,IFIT2,IFIT3,IFITM1,IFRD2,IGSF6,IKZF3,IL6R,IL7,INTS14,IPO9,IQCN,IRF2,IRF2BP1,IRF7,IRF8,ISG15,ITGB8,ITPKC,ITPRIPL2,JCHAIN,JMJD8,JUN,JUNB,KCNA3,KDM4A,KEAP1,KIF20B,KIF21A,KIF5A,KLF11,KLF4,KLF6,KLF9,KLHDC7B,KPNA2,KYNU,L3HYPDH,LAG3,LARP6,LDLRAD3,LETM1,LETM2,LGALS3BP,LIG4,LIN7A,LINS1,LMAN2L,LMNA,LMNB1,LONRF2,LRP6,LRRC14,LRRC2,LRRC61,LRRC8D,LRRK2,LRRN1,LYSMD1,MAFF,MAL,MAN2A1,MAP3K8,MAP4K3,MAPK12,MAPRE3,MCAM,MCF2L,MCL1,MED24,MEX3A,MINPP1,MLEC,MMACHC,MMP2,MMP7,MOB3B,MORC1,MOSPD2,MPG,MRM2,MRPS18C,MRVI1,MT-ND1,MTIF3,MUC12,MX1,MYADM,MYC,MYOZ3,MYRIP,NAT8,NCR3,NEU1,NEXN,NFE2,NFIB,NFIL3,NFKBID,NKX3-1,NLGN1,NLRC5,NOC3L,NOG,NOP16,NOS1AP,NOTCH4,NR0B1,NR4A1,NR4A2,NR4A3,NR5A1,NRXN2,NUDT22,NUP50,OAF,OAS1,OAS2,OSBPL10,OSBPL6,OSTM1,OTULINL,PANK4,PANX1,PAQR7,PARP3,PCDH9,PCTP,PDZK1IP1,PDZRN4,PER1,PFKFB2,PGS1,PHLDA1,PHLDB2,PIEZO2,PLA2R1,PLD4,PLEKHF1,PLK3,PLSCR1,POGLUT1,POLR3A,POMC,POMGNT2,PORCN,PPIF,PPP1R1B,PPP2R2B,PPP2R5B,PRDM5,PRDX1,PREX2,PRKCD,PRKG2,PRX,PSRC1,PTAFR,PTGIR,PTGR1,PTGS2,PTPRD,PTX3,PWWP2A,PYGO1,QSER1,RAB30,RAB38,RAB6B,RASAL2,RASGEF1B,RASGRP3,RASSF3,RASSF6,RCN1,RDH13,REN,RFXAP,RGCC,RGS1,RGS2,RHOB,RIMS1,RMDN2,RMI1,RNASEL,RNF175,RNF24,RPL9,RSAD2,RXYLT1,S100A10,S1PR2,SALL4,SAMD9,SAMD9L,SAMHD1,SAP30,SCLY,SCML4,SELP,SEMA4B,SERPINB8,SERPINE1,SESN1,SGK1,SGPP1,SH3GL3,SHE,SHQ1,SHTN1,SIK1/SIK1B,SLC12A2,SLC16A1,SLC16A9,SLC25A17,SLC26A6,SLC27A2,SLC29A3,SLC2A10,SLC35B2,SLC38A1,SLC39A14,SLC39A6,SLC40A1,SLC45A3,SLC52A3,SLC7A5,SLCO5A1,SLF2,SLFN5,SMAP2,SMOX,SMYD3,SNAI3,SNAPC3,SNTB1,SNX18,SOCS1,SOCS3,SOWAHD,SP110,SPIRE1,SPN,SPOCK2,SPON2,SPRY2,SRGAP3,SSPN,ST6GAL2,STAT1,STAT2,STK17B,STON2,SUCLG2,SV2A,TAF5,TANGO6,TAP2,TAS2R14,TBC1D24,TCEAL2,TCF4,TEKT4,TENT4A,TIE1,TIGD7,TIMP3,TIPARP,TLE3,TLE4,TLR2,TM4SF1,TM9SF2,TMEM117,TMEM231,TMEM246,TMEM45A,TMEM51,TNF,TNFAIP3,TNFSF10,TNFSF13B,TNRC6C,TP53,TP53INP1,TRAF3IP3,TRIM69,TRMT9B,TSC22D3,TTC13,TTC27,TTC8,TUB,TUBA4A,TUBD1,TWNK,TXNIP,UBE2T,UBIAD1,UBOX5,UGCG,USP18,USP2,USP28,USP37,VANGL1,VDR,VSTM4,WASF3,WBP1L,WDCP,WDFY3,WDR7,WFDC1,WHRN,XAF1,XIRP2,XKR8,YWHAG,ZAP70,ZBTB7A,ZDHHC13,ZFP36,ZMAT3,ZMPSTE24,ZNF280B,ZNF324B,ZNF346,ZNF354B,ZNF385D,ZNF420,ZNF462,ZNF608,ZNF74,ZNF780A,ZNF781,ZNF785,ZNF799,ZNF835,ZNRF3,ZSWIM5 |
| Infectious Diseases | Replication of RNA virus | 4.14E-11 | Increased | 2.217 | APBA3,AREG,BST2,CAV1,CCL3,CCL3L3,CCL4,CD4,CXCL8,CXCR4,DYRK1A,EIF2AK2,F13A1,FCGR1A,GBP1,HCK,HERC5,IFI6,IFIH1,IFIT1,IFITM1,IFITM3,IGKC,IRF2,IRF7,ISG15,ISG20,JUN,MX1,MYC,NCR3,NLRC5,OAS1,OAS3,OASL,PANX1,PLK3,PRDX1,PRKCD,PTGS2,PTX3,RAB6B,RNASEL,RSAD2,SGK1,SOCS1,SOCS3,SP110,STAT1,STAT2,TLR2,TNF,TNFSF10,TP53,TXNIP,UBE2L6,USP2,ZAP70 |
| Cancer,Organismal Injury and Abnormalities | Tumorigenesis of tissue | 1.3E-28 | Increased | 2.211 | ABHD12B,ABHD6,ACACB,ACADL,ACTA2,ADAP2,ADGRE5,ADGRG6,AFMID,AGAP3,AGPS,AIFM2,AK4,AKR1C1/AKR1C2,AKR1C3,AKR1C4,ALDH1B1,ALDH9A1,ALKBH4,ALKBH8,ANKLE1,ANKRD20A2 (includes others),ANXA2R,APBA3,APOL2,APOL4,ARAP1,AREG,ARF5,ARID3A,ARID5B,ARL4A,ARL4C,ARL4D,ARMH4,ARMT1,ARRDC3,ASCL2,ATF1,ATG101,ATP6V0A1,ATP6V0E1,AZU1,B4GALT1,B4GAT1,BATF,BATF3,BCAM,BEST1,BHLHE40,BMP2K,BMPR1B,BRD3,BST2,BTG1,BTG2,BTN3A1,BTN3A2,BTN3A3,C12orf49,C14orf119,C14orf39,C19orf54,C1GALT1C1,C1orf216,C1orf50,C2,C22orf15,C2orf69,C5orf30,CACNB4,CALHM2,CARNMT1,CAV1,CBY1,CCDC126,CCDC146,CCDC175,CCDC184,CCDC50,CCDC86,CCL4,CD4,CD69,CD72,CDC42EP4,CDH1,CDH7,CDKN1A,CDKN2C,CDKN3,CEACAM4,CEBPD,CENPF,CFH,CHCHD10,CHKA,CITED2,CKAP4,CLCN3,CLCN4,CLDN5,CLDN7,CLEC1B,CLGN,CLU,CMPK2,CNKSR1,CNOT9,COL2A1,COLGALT1,COQ2,COQ9,CP,CPED1,CRIP2,CRK,CSF2RB,CSGALNACT1,CTNS,CTSO,CXCL8,CXCR4,CYB561,CYP1B1,CYP7B1,CYSLTR1,CYYR1,DCAF12,DCAF4,DCLRE1A,DDI2,DDIT4,DDX60,DEGS2,DENND10,DEPDC1B,DGAT2,DHX33,DHX58,DIRAS3,DIXDC1,DLAT,DLG4,DLL3,DNAJC16,DNAJC28,DNM3OS,DOCK4,DOLPP1,DSEL,DTL,DTX3L,DUSP1,DUSP2,DUSP6,DYRK1A,DYTN,DZIP1,ECE2,EDA2R,EEF2K,EEF2KMT,EFNA1,EGLN1,EGR1,EGR2,EGR3,EGR4,EIF2AK2,ELN,EME2,EML2,ENDOD1,ENPP5,ENTPD1,ENTPD6,EPDR1,EPHX1,EPSTI1,EREG,ETV3,EXOSC4,F13A1,FAM117A,FAM124B,FAM133A,FAM181B,FAM210A,FAM217B,FAM81A,FAM83D,FAM89A,FAM98A,FAXDC2,FBLN1,FBLN5,FBP1,FBXO45,FBXO6,FCGR1A,FCRLB,FEN1,FERMT1,FFAR3,FGFR1OP,FGL2,FITM2,FKBP5,FKBP9,FLOT1,FOS,FOSB,FREM1,FUCA1,FYB1,FZD7,GALNT2,GAPT,GAS2,GAS2L3,GASK1B,GATM,GBP1,GBP4,GBP5,GBP7,GCSAML,GDF7,GGA2,GIMAP6,GLRB,GLUL,GMPR,GNAI1,GOT2,GPC3,GPR12,GPR157,GPR27,GPR35,GPT2,GRASP,GRK4,GRWD1,GSTO2,GTF2A1,GULP1,H2AC18/H2AC19,HAPLN3,HAUS3,HCK,HERC5,HES1,HHEX,HIF3A,HILPDA,HLA-C,HLA-DRB5,HLA-F,HLA-G,HLF,HLTF,HOMER2,HOMER3,HOPX,HOXA3,HRH2,HSPA4L,HSPB1,HYAL2,HYAL3,HYLS1,ICMT,ID1,ID2,ID3,IDS,IER3,IER5,IFI44,IFI44L,IFI6,IFIH1,IFIT1,IFIT2,IFIT3,IFITM1,IFITM3,IFRD2,IGSF6,IKZF3,IL6R,IL7,INTS14,IPO9,IQCN,IRF2,IRF2BP1,IRF7,IRF8,ISG15,ISG20,ITGB8,ITPKC,ITPRIPL2,JCHAIN,JDP2,JMJD8,JUN,JUNB,JUND,KCNA3,KDM4A,KEAP1,KIF20B,KIF21A,KIF5A,KLF11,KLF13,KLF15,KLF2,KLF4,KLF6,KLF7,KLF9,KLHDC7B,KPNA2,KYNU,L3HYPDH,LAG3,LARP6,LDLRAD3,LETM1,LETM2,LGALS3BP,LIG4,LIN52,LIN7A,LINS1,LMAN2L,LMNA,LMNB1,LONRF2,LRIF1,LRP11,LRP6,LRRC14,LRRC2,LRRC61,LRRC8D,LRRK2,LRRN1,LYSMD1,MAFF,MAGEF1,MAL,MALL,MAN2A1,MAP3K8,MAP4K3,MAPK12,MAPRE3,MARC1,MCAM,MCF2L,MCL1,MCOLN1,MDFI,MED24,MED31,MEIOC,METTL2A,MEX3A,MINPP1,MLEC,MMACHC,MMP2,MMP7,MOB3B,MORC1,MOSPD2,MPG,MRM2,MRPS18C,MRVI1,MT-ND1,MTIF3,MUC12,MX1,MYADM,MYC,MYCBP,MYOZ3,MYRIP,NAT8,NCR3,NEU1,NEXN,NFE2,NFIB,NFIL3,NFKBID,NIP7,NKX3-1,NLGN1,NLRC5,NOC3L,NOG,NOP16,NOS1AP,NOTCH4,NR0B1,NR4A1,NR4A2,NR4A3,NR5A1,NRXN2,NUDT22,NUP50,OAF,OAS1,OAS2,OAS3,OASL,ORC6,OSBPL10,OSBPL6,OSTM1,OTULINL,PANK4,PANX1,PAQR7,PARP3,PARP9,PCDH9,PCSK1N,PCTP,PDZK1IP1,PDZRN4,PER1,PFKFB2,PGAM1,PGS1,PHF23,PHLDA1,PHLDA3,PHLDB2,PIEZO2,PIK3IP1,PLA2R1,PLD4,PLD6,PLEKHF1,PLK3,PLSCR1,PNMT,POGLUT1,POLR2D,POLR3A,POMC,POMGNT2,PORCN,PPAT,PPIAL4G (includes others),PPIF,PPP1R1B,PPP2R2B,PPP2R5B,PRDM5,PRDX1,PREX2,PRKCD,PRKG2,PRSS2,PRX,PSRC1,PTAFR,PTGIR,PTGR1,PTGS2,PTPRD,PTX3,PWWP2A,PYGO1,QSER1,RAB1B,RAB27B,RAB30,RAB38,RAB6B,RASAL2,RASGEF1B,RASGRP3,RASSF3,RASSF6,RCN1,RDH13,REEP4,REN,RFXAP,RGCC,RGS1,RGS18,RGS2,RHEBL1,RHOB,RIMS1,RMDN2,RMI1,RNASEL,RNF175,RNF24,RPL9,RPS23,RSAD2,RTN4R,RXYLT1,S100A10,S1PR2,SALL4,SAMD9,SAMD9L,SAMHD1,SAP30,SCLY,SCML4,SCPEP1,SELENOM,SELP,SEMA4B,SERPINB8,SERPINE1,SERTAD1,SESN1,SGK1,SGPP1,SH2B2,SH3GL3,SHE,SHQ1,SHTN1,SIK1/SIK1B,SLC12A2,SLC16A1,SLC16A9,SLC25A17,SLC26A6,SLC27A2,SLC29A3,SLC2A10,SLC35B2,SLC38A1,SLC39A14,SLC39A6,SLC40A1,SLC45A3,SLC52A3,SLC7A5,SLCO5A1,SLF2,SLFN5,SMAP2,SMIM14,SMOX,SMYD3,SNAI3,SNAPC3,SNTB1,SNX18,SOCS1,SOCS3,SOWAHD,SP110,SPIRE1,SPN,SPOCK2,SPON2,SPR,SPRY2,SRGAP3,SSPN,ST6GAL2,STAC3,STAT1,STAT2,STK17B,STON2,SUCLG2,SV2A,SYT5,TAF5,TANGO6,TAP2,TAS2R14,TBC1D24,TCEAL2,TCF4,TEKT4,TENT4A,TIE1,TIGD7,TIMP3,TIPARP,TLCD3A,TLE3,TLE4,TLR2,TM4SF1,TM9SF2,TMEM117,TMEM231,TMEM246,TMEM45A,TMEM50B,TMEM51,TMEM98,TMEM9B,TMSB10/TMSB4X,TNF,TNFAIP3,TNFSF10,TNFSF13B,TNNI2,TNRC6C,TP53,TP53INP1,TRAF3IP3,TRIM69,TRMT1L,TRMT9B,TSC22D3,TTC13,TTC27,TTC8,TUB,TUBA4A,TUBD1,TWNK,TXNIP,UBE2T,UBIAD1,UBOX5,UGCG,USP18,USP2,USP28,USP37,VANGL1,VDR,VSTM4,VTA1,WASF3,WBP1L,WDCP,WDFY3,WDR7,WFDC1,WHRN,XAF1,XIRP2,XKR8,YWHAG,ZAP70,ZBTB7A,ZDHHC13,ZFP36,ZMAT3,ZMPSTE24,ZNF280B,ZNF324B,ZNF346,ZNF354B,ZNF385D,ZNF420,ZNF462,ZNF605,ZNF608,ZNF74,ZNF780A,ZNF781,ZNF785,ZNF799,ZNF835,ZNRF3,ZSWIM5 |
| Cancer,Organismal Injury and Abnormalities,Reproductive System Disease | Mammary tumor | 1.53E-09 | Increased | 2.207 | ACACB,ACTA2,ADGRG6,AGAP3,AGPS,AKR1C1/AKR1C2,AKR1C3,ALDH1B1,APBA3,AREG,ARID5B,ARMT1,ARRDC3,BCAM,BHLHE40,BTG2,C14orf39,C1GALT1C1,C2,CAV1,CCDC175,CCL3,CCL4,CD4,CD69,CDH1,CDKN1A,CDKN2C,CEBPD,CENPF,CKAP4,CLGN,CLU,CMPK2,CP,CSF2RB,CXCL8,CXCR4,DDIT4,DHX58,DIXDC1,DOCK4,DTL,DUSP1,DUSP2,DYRK1A,EDA2R,EGLN1,EGR1,EGR3,EGR4,EIF2AK2,EML2,EPDR1,EPSTI1,FBLN1,FBLN5,FEN1,FKBP5,FOS,FOSB,FREM1,GASK1B,GBP7,GCSAML,GLUL,GPC3,GSTM3,GULP1,HAPLN3,HCK,HERC5,HES1,HLA-DRB5,HLA-G,HOXA3,HRH2,HSPB1,ID1,IER2,IER3,IFIT1,IRF7,ITPKC,JUN,JUNB,JUND,KEAP1,KLF11,KLF4,KLF6,KPNA2,LETM1,LETM2,LGALS3BP,LMAN2L,LMNA,LRP6,LRRK2,LYSMD1,MAFF,MAP3K8,MAP4K3,MAPRE3,MCAM,MCL1,MMP2,MMP7,MT-ND1,MUC12,MYC,NEAT1,NEU1,NFIL3,NOP16,NOTCH4,NR4A1,NR4A2,NR4A3,NR5A1,NRXN2,OAF,OSBPL6,OSTM1,PARP3,PCDH9,PDZK1IP1,PDZRN4,PGAM1,PHLDA1,PIEZO2,PPP1R1B,PRDM5,PRDX1,PREX2,PRKCD,PRX,PTGS2,PTPRD,RASAL2,RGS1,RGS2,RHOB,RMDN2,SALL4,SAMD9,SAMD9L,SERPINE1,SHQ1,SIK1/SIK1B,SLC16A1,SLC2A10,SLC39A6,SLC40A1,SMYD3,SNAPC3,SNTB1,SNX18,SSPN,STAT1,STK17B,SV2A,TCF4,TIE1,TIMP3,TLE3,TM4SF1,TMEM117,TNF,TNFSF10,TP53,TRIM69,TRMT9B,TUBA4A,TUBD1,UGCG,USP28,VDR,WDFY3,ZFP36,ZMPSTE24,ZNF280B,ZNF420,ZNF462,ZNF835 |
| Infectious Diseases | Replication of virus | 6.84E-11 | Increased | 2.181 | APBA3,AREG,BST2,CAV1,CCL3,CCL3L3,CCL4,CD4,CXCL8,CXCR4,DUSP1,DYRK1A,EGR1,EIF2AK2,F13A1,FCGR1A,GBP1,HCK,HERC5,IFI6,IFIH1,IFIT1,IFITM1,IFITM3,IGKC,IRF2,IRF7,ISG15,ISG20,JUN,MX1,MYC,NCR3,NLRC5,OAS1,OAS3,OASL,PANX1,PLK3,PRDX1,PRKCD,PTGS2,PTX3,RAB6B,RNASEL,RSAD2,SAMHD1,SGK1,SOCS1,SOCS3,SP110,SPRY2,STAT1,STAT2,TLR2,TNF,TNFSF10,TP53,TXNIP,UBE2L6,USP2,ZAP70 |

| | | | | | |
|---|---|---|---|---|---|
| Cancer,Organismal Injury and Abnormalities,Reproductive System Disease | Breast cancer | 3.63E-10 | Increased | 2.176 | ACACB,ACTA2,ADGRG6,AGAP3,AKR1C1/AKR1C2,AKR1C3,ALDH1B1,AREG,ARID5B,ARMT1,BCAM,BHLHE40,BTG2,C14orf39,C1GALT1C1,C2,CAV1,CCDC175,CCL3,CCL4,CD4,CD69,CDH1,CDKN1A,CDKN2C,CEBPD,CENPF,CKAP4,CLGN,CLU,CMPK2,CP,CSF2RB,CXCL8,CXCR4,DDIT4,DHX58,DIXDC1,DOCK4,DTL,DUSP1,DUSP2,DYRK1A,EDA2R,EGLN1,EGR1,EGR3,EGR4,EIF2AK2,EML2,EPDR1,EPSTI1,FBLN1,FBLN5,FKBP5,FOS,FOSB,FREM1,GASK1B,GBP7,GCSAML,GLUL,GPC3,GSTM3,GULP1,HAPLN3,HCK,HERC5,HLA-DRB5,HLA-G,HOXA3,HRH2,HSPB1,ID1,IER2,IER3,IFIT1,IRF7,ITPKC,JUN,JUNB,JUND,KEAP1,KLF11,KLF4,KLF6,KPNA2,LETM1,LETM2,LGALS3BP,LMAN2L,LMNA,LRP6,LRRK2,LYSMD1,MAFF,MAP3K8,MAP4K3,MAPRE3,MCAM,MCL1,MMP2,MMP7,MT-ND1,MUC12,MYC,NEAT1,NEU1,NFIL3,NOP16,NOTCH4,NR4A1,NR4A2,NR4A3,NR5A1,NRXN2,OAF,OSBPL6,OSTM1,PARP3,PCDH9,PDZK1IP1,PDZRN4,PGAM1,PHLDA1,PIEZO2,PPP1R1B,PRDM5,PRDX1,PREX2,PRKCD,PRX,PTGS2,PTPRD,RASAL2,RGS1,RGS2,RHOB,RMDN2,SALL4,SAMD9,SAMD9L,SERPINE1,SHQ1,SIK1/SIK1B,SLC16A1,SLC2A10,SLC39A6,SLC40A1,SMYD3,SNAPC3,SNTB1,SNX18,SSPN,STAT1,STK17B,SV2A,TCF4,TIE1,TIMP3,TLE3,TMEM117,TNF,TNFSF10,TP53,TRIM69,TRMT9B,TUBA4A,TUBD1,UGCG,USP28,VDR,WDFY3,ZFP36,ZMPSTE24,ZNF280B,ZNF420,ZNF462,ZNF835 |
| Cancer,Organismal Injury and Abnormalities,Reproductive System Disease | Breast or ovarian cancer | 7.22E-09 | Increased | 2.176 | ABHD12B,ABHD6,ACACB,ACTA2,ADGRG6,AGAP3,AKR1C1/AKR1C2,AKR1C3,ALDH1B1,AREG,ARID5B,ARL4C,ARMT1,ASCL2,BCAM,BHLHE40,BMP2K,BTG2,C14orf39,C1GALT1C1,C2,CAV1,CCDC175,CCL3,CCL4,CD4,CD69,CDH1,CDKN1A,CDKN2C,CDKN3,CEBPD,CENPF,CFH,CKAP4,CLDN7,CLGN,CLU,CMPK2,CP,CSF2RB,CXCL8,CXCR4,DCLRE1A,DDIT4,DHX58,DIRAS3,DIXDC1,DOCK4,DTL,DUSP1,DUSP2,DYRK1A,EDA2R,EGLN1,EGR1,EGR2,EGR3,EGR4,EIF2AK2,EML2,ENTPD6,EPDR1,EPHX1,EPSTI1,EXOSC4,F13A1,FAM181B,FBLN1,FBLN5,FGFR1OP,FKBP5,FLOT1,FOS,FOSB,FREM1,GASK1B,GBP7,GCSAML,GLUL,GOT2,GPC3,GSTM3,GTF2A1,GULP1,H2AC18/H2AC19,HAPLN3,HCK,HERC5,HLA-DRB5,HLA-G,HLTF,HOXA3,HRH2,HSPB1,HYAL2,ID1,IER2,IER3,IFI44L,IFIH1,IFIT1,IGSF6,IKZF3,IRF7,ITPKC,JUN,JUNB,JUND,KEAP1,KLF11,KLF4,KLF6,KPNA2,LARP6,LETM1,LETM2,LGALS3BP,LINS1,LMAN2L,LMNA,LRP6,LRRC14,LRRK2,LRRN1,LYSMD1,MAFF,MAP3K8,MAP4K3,MAPRE3,MCAM,MCL1,MMP2,MMP7,MPG,MT-ND1,MUC12,MYC,NCR3,NEAT1,NEU1,NFIL3,NLGN1,NOP16,NOTCH4,NR4A1,NR4A2,NR4A3,NR5A1,NRXN2,NUP50,OAF,OSBPL6,OSTM1,PANK4,PARP3,PCDH9,PDZK1IP1,PDZRN4,PGAM1,PHLDA1,PHLDB2,PIEZO2,PLD4,POMC,PPP1R1B,PRDM5,PRDX1,PREX2,PRKCD,PRX,PTAFR,PTGS2,PTPRD,RASAL2,REN,RGCC,RGS1,RGS2,RHOB,RIMS1,RMDN2,S1PR2,SALL4,SAMD9,SAMD9L,SELP,SERPINE1,SHQ1,SIK1/SIK1B,SLC16A1,SLC2A10,SLC39A6,SLC40A1,SLFN5,SMYD3,SNAPC3,SNTB1,SNX18,SPN,SSPN,STAT1,STAT2,STK17B,SUCLG2,SV2A,TANGO6,TCF4,TIE1,TIMP3,TIPARP,TLE3,TM4SF1,TMEM117,TNF,TNFSF10,TP53,TRIM69,TRMT9B,TSC22D3,TUBA4A,TUBD1,TXNIP,UGCG,USP18,USP28,USP37,VDR,WASF3,WDFY3,XIRP2,ZFP36,ZMPSTE24,ZNF280B,ZNF420,ZNF462,ZNF835,ZSWIM5 |
| Cancer,Organismal Injury and Abnormalities,Reproductive System Disease | Breast or gynecological cancer | 7.12E-11 | Increased | 2.174 | ABHD12B,ABHD6,ACACB,ACTA2,ADGRG6,AGAP3,AGPS,AKR1C1/AKR1C2,AKR1C3,ALDH1B1,APOL2,AREG,ARID3A,ARID5B,ARL4C,ARMT1,ARRDC3,ASCL2,BCAM,BHLHE40,BMP2K,BTG2,BTN3A1,BTN3A2,C14orf39,C1GALT1C1,C2,CAV1,CCDC146,CCDC175,CCL3,CCL4,CD4,CD69,CDH1,CDKN1A,CDKN2C,CDKN3,CEBPD,CENPF,CFH,CKAP4,CLDN5,CLDN7,CLGN,CLU,CMPK2,COL2A1,CP,CPED1,CSF2RB,CSGALNACT1,CXCL8,CXCR4,CYP1B1,CYSLTR1,DCAF4,DCLRE1A,DDIT4,DDX60,DEGS2,DEPDC1B,DHX58,DIRAS3,DIXDC1,DOCK4,DOLPP1,DSEL,DTL,DTX3L,DUSP1,DUSP2,DYRK1A,DYTN,ECE2,EDA2R,EFNA1,EGLN1,EGR1,EGR2,EGR3,EGR4,EIF2AK2,EML2,ENTPD1,ENTPD6,EPDR1,EPHX1,EPSTI1,EXOSC4,F13A1,FAM181B,FAM217B,FAM83D,FBLN1,FBLN5,FEN1,FFAR3,FGFR1OP,FGL2,FKBP5,FKBP9,FLOT1,FOS,FOSB,FREM1,FYB1,FZD7,GASK1B,GBP7,GCSAML,GGA2,GLRB,GLUL,GOT2,GPC3,GRK4,GSTM3,GTF2A1,GULP1,H2AC18/H2AC19,HAPLN3,HAUS3,HCK,HERC5,HIF3A,HLA-DRB5,HLA-G,HLF,HLTF,HOXA3,HRH2,HSPA4L,HSPB1,HYAL2,HYLS1,ICMT,ID1,IER2,IER3,IFI44,IFI44L,IFIH1,IFIT1,IFIT3,IGSF6,IKZF3,IPO9,IRF7,ISG15,ITGB8,ITPKC,JUN,JUNB,JUND,KCNA3,KDM4A,KEAP1,KIF20B,KIF21A,KIF5A,KLF11,KLF2,KLF4,KLF6,KLF9,KPNA2,LARP6,LETM1,LETM2,LGALS3BP,LINS1,LMAN2L,LMNA,LRP6,LRRC14,LRRK2,LRRN1,LYSMD1,MAFF,MAGEF1,MAL,MAN2A1,MAP3K8,MAP4K3,MAPRE3,MCAM,MCL1,MED24,MMP2,MMP7,MORC1,MOSPD2,MPG,MT-ND1,MUC12,MX1,MYC,NCR3,NEAT1,NEU1,NFIL3,NLGN1,NOC3L,NOP16,NOS1AP,NOTCH4,NR0B1,NR4A1,NR4A2,NR4A3,NR5A1,NRXN2,NUP50,OAF,OAS3,OASL,OSBPL10,OSBPL6,OSTM1,OTULINL,PANK4,PAQR7,PARP3,PCDH9,PDZK1IP1,PDZRN4,PGAM1,PGS1,PHLDA1,PHLDA3,PHLDB2,PIEZO2,PLA2R1,PLD4,POLR3A,POMC,PPIF,PPP1R1B,PRDM5,PRDX1,PREX2,PRKCD,PRX,PTAFR,PTGS2,PTPRD,PYGO1,RASAL2,RASGRP3,REN,RGCC,RGS1,RGS2,RHOB,RIMS1,RMDN2,S1PR2,SALL4,SAMD9,SAMD9L,SAP30,SELP,SERPINB8,SERPINE1,SGK1,SHQ1,SIK1/SIK1B,SLC12A2,SLC16A1,SLC2A10,SLC39A14,SLC39A6,SLC40A1,SLC45A3,SLC52A3,SLC7A5,SLFN5,SMAP2,SMOX,SMYD3,SNAPC3,SNTB1,SNX18,SP110,SPN,SPOCK2,SSPN,STAT1,STAT2,STK17B,SUCLG2,SV2A,TANGO6,TCF4,TEKT4,TIE1,TIMP3,TIPARP,TLE3,TM4SF1,TM9SF2,TMEM117,TMEM246,TMSB10/TMSB4X,TNF,TNFSF10,TNRC6C,TP53,TRIM69,TRMT9B,TSC22D3,TUBA4A,TUBD1,TXNIP,UGCG,USP18,USP2,USP28,USP37,VDR,WASF3,WDFY3,WDR7,XIRP2,ZAP70,ZBTB7A,ZFP36,ZMAT3,ZMPSTE24,ZNF280B,ZNF324B,ZNF354B,ZNF420,ZNF462,ZNF780A,ZNF781,ZNF785,ZNF835,ZNRF3,ZSWIM5 |
| Cancer,Organismal Injury and Abnormalities | Carcinoma | 4.97E-27 | Increased | 2.158 | ABHD12B,ABHD6,ACACB,ACADL,ACTA2,ADAP2,ADGRE5,ADGRG6,AFMID,AGAP3,AGPS,AIFM2,AK4,AKR1C1/AKR1C2,AKR1C3,AKR1C4,ALDH1B1,ALDH9A1,ALKBH4,ALKBH8,ANKLE1,ANKRD20A2 (includes others),ANXA2R,APBA3,APOL2,APOL4,ARAP1,AREG,ARF5,ARID3A,ARID5B,ARL4A,ARL4C,ARL4D,ARMH4,ARMT1,ARRDC3,ASCL2,ATF1,ATG101,ATP6V0A1,ATP6V0E1,AZU1,B4GALT1,B4GAT1,BATF,BATF3,BCAM,BEST1,BHLHE40,BMP2K,BMPR1B,BRD3,BST2,BTG1,BTG2,BTN3A1,BTN3A2,BTN3A3,C12orf49,C14orf119,C14orf39,C19orf54,C1GALT1C1,C1orf216,C1orf50,C2,C22orf15,C2orf69,C5orf30,CACNB4,CALHM2,CARNMT1,CAV1,CBY1,CCDC126,CCDC146,CCDC175,CCDC184,CCDC50,CCDC86,CCL4,CD4,CD69,CD72,CDC42EP4,CDH1,CDH7,CDKN1A,CDKN2C,CDKN3,CEACAM4,CENPF,CFH,CHCHD10,CHKA,CITED2,CKAP4,CLCN3,CLCN4,CLDN5,CLDN7,CLEC1B,CLGN,CLU,CMPK2,CNKSR1,CNOT9,COL2A1,COLGALT1,COQ2,COQ9,CP,CPED1,CRIP2,CRK,CSF2RB,CSGALNACT1,CTNS,CTSO,CXCL8,CXCR4,CYB561,CYP1B1,CYP7B1,CYSLTR1,CYYR1,DCAF12,DCAF4,DCLRE1A,DDI2,DDIT4,DDX60,DEGS2,DENND10,DEPDC1B,DGAT2,DHX33,DHX58,DIRAS3,DIXDC1,DLAT,DLG4,DLL3,DNAJC16,DNAJC28,DNM3OS,DOCK4,DOLPP1,DSEL,DTL,DTX3L,DUSP1,DUSP2,DUSP6,DYRK1A,DYTN,DZIP1,ECE2,EDA2R,EEF2K,EEF2KMT,EFNA1,EGLN1,EGR1,EGR2,EGR3,EGR4,EIF2AK2,ELN,EME2,EML2,ENDOD1,ENPP5,ENTPD1,ENTPD6,EPDR1,EPHX1,EPSTI1,EREG,ETV3,EXOSC4,F13A1,FAM117A,FAM124B,FAM133A,FAM181B,FAM210A,FAM217B,FAM81A,FAM83D,FAM89A,FAM98A,FAXDC2,FBLN1,FBLN5,FBP1,FBXO45,FBXO6,FCGR1A,FCRLB,FEN1,FERMT1,FFAR3,FGFR1OP,FGL2,FITM2,FKBP5,FKBP9,FLOT1,FOS,FOSB,FREM1,FUCA1,FYB1,FZD7,GALNT2,GAPT,GAS2,GAS2L3,GASK1B,GATM,GBP1,GBP4,GBP5,GBP7,GCSAML,GDF7,GGA2,GIMAP6,GLRB,GLUL,GMPR,GNAI1,GOT2,GPC3,GPR12,GPR157,GPR27,GPR35,GPT2,GRASP,GRK4,GRWD1,GSTO2,GTF2A1,GULP1,H2AC18/H2AC19,HAPLN3,HAUS3,HCK,HERC5,HES1,HHEX,HIF3A,HILPDA,HLA-C,HLA-DRB5,HLA-F,HLA-G,HLF,HLTF,HOMER2,HOMER3,HOPX,HOXA3,HRH2,HSPA4L,HSPB1,HYAL2,HYAL3,HYLS1,ICMT,ID1,ID2,ID3,IDS,IER3,IER5,IFI44,IFI44L,IFI6,IFIH1,IFIT1,IFIT2,IFIT3,IFITM1,IFITM3,IFRD2,IGSF6,IKZF3,IL6R,IL7,INTS14,IPO9,IQCN,IRF2,IRF2BP1,IRF7,IRF8,ISG15,ISG20,ITGB8,ITPKC,ITPRIPL2,JCHAIN,JDP2,JMJD8,JUN,JUNB,JUND,KCNA3,KDM4A,KEAP1,KIF20B,KIF21A,KIF5A,KLF11,KLF13,KLF15,KLF2,KLF4,KLF6,KLF7,KLF9,KLHDC7B,KPNA2,KYNU,L3HYPDH,LAG3,LARP6,LDLRAD3,LETM1,LETM2,LGALS3BP,LIG4,LIN52,LIN7A,LINS1,LMAN2L,LMNA,LMNB1,LONRF2,LRIF1,LRP11,LRP6,LRRC14,LRRC2,LRRC61,LRRC8D,LRRK2,LRRN1,LYSMD1,MAFF,MAGEF1,MAL,MALL,MAN2A1,MAP3K8,MAP4K3,MAPK12,MAPRE3,MARC1,MCAM,MCF2L,MCL1,MCOLN1,MDFI,MED24,MED31,MEIOC,METTL2A,MEX3A,MINPP1,MLEC,MMACHC,MMP2,MMP7,MOB3B,MORC1,MOSPD2,MPG,MRM2,MRPS18C,MRVI1,MT-ND1,MTIF3,MUC12,MX1,MYADM,MYC,MYCBP,MYOZ3,MYRIP,NAT8,NCR3,NEU1,NEXN,NFE2,NFIB,NFIL3,NFKBID,NIP7,NKX3-1,NLGN1,NLRC5,NOC3L,NOG,NOP16,NOS1AP,NOTCH4,NR0B1,NR4A1,NR4A2,NR4A3,NR5A1,NRXN2,NUDT22,NUP50,OAF,OAS1,OAS2,OASL,ORC6,OSBPL10,OSBPL6,OSTM1,OTULINL,PANK4,PANX1,PAQR7,PARP3,PARP9,PCDH9,PCSK1N,PCTP,PDZK1IP1,PDZRN4,PER1,PFKFB2,PGAM1,PGS1,PHF23,PHLDA1,PHLDA3,PHLDB2,PIEZO2,PIK3IP1,PLA2R1,PLD4,PLD6,PLEKHF1,PLK3,PLSCR1,PNMT,POGLUT1,POLR2D,POLR3A,POMC,POMGNT2,PORCN,PPAT,PPIAL4G (includes others),PPIF,PPP1R1B,PPP2R2B,PPP2R5B,PRDM5,PRDX1,PREX2,PRKCD,PRKG2,PRX,PSRC1,PTAFR,PTGIR,PTGR1,PTGS2,PTPRD,PTX3,PWWP2A,PYGO1,QSER1,RAB1B,RAB27B,RAB30,RAB38,RAB6B,RASAL2,RASGEF1B,RASGRP3,RASSF3,RASSF6,RCN1,RDH13,REEP4,REN,RFXAP,RGCC,RGS1,RGS18,RGS2,RHEBL1,RHOB,RIMS1,RMDN2,RMI1,RNASEL,RNF175,RNF24,RPL9,RSAD2,RTN4R,RXYLT1,S100A10,S1PR2,SALL4,SAMD9,SAMD9L,SAMHD1,SAP30,SCLY,SCML4,SCPEP1,SELENOM,SELP,SEMA4B,SERPINB8,SERPINE1,SERTAD1,SESN1,SGK1,SGPP1,SH3GL3,SHE,SHQ1,SHTN1,SIK1/SIK1B,SLC12A2,SLC16A1,SLC16A9,SLC25A17,SLC26A6,SLC27A2,SLC29A3,SLC2A10,SLC35B2,SLC38A1,SLC39A14,SLC39A6,SLC40A1,SLC45A3,SLC52A3,SLC7A5,SLCO5A1,SLF2,SLFN5,SMAP2,SMIM14,SMOX,SMYD3,SNAI3,SNAPC3,SNTB1,SNX18,SOCS1,SOCS3,SOWAHD,SP110,SPIRE1,SPN,SPOCK2,SPON2,SPR,SPRY2,SRGAP3,SSPN,ST6GAL2,STAC3,STAT1,STAT2,STK17B,STON2,SUCLG2,SV2A,SYT5,TAF5,TANGO6,TAP2,TAS2R14,TBC1D24,TCEAL2,TCF4,TEKT4,TENT4A,TIE1,TIGD7,TIMP3,TIPARP,TLCD3A,TLE3,TLE4,TLR2,TM4SF1,TM9SF2,TMEM117,TMEM231,TMEM246,TMEM45A,TMEM50B,TMEM51,TMEM98,TMEM9B,TMSB10/TMSB4X,TNF,TNFAIP3,TNFSF10,TNFSF13B,TNNI2,TNRC6C,TP53,TP53INP1,TRAF3IP3,TRIM69,TRMT1L,TRMT9B,TSC22D3,TTC13,TTC27,TTC8,TUB,TUBA4A,TUBD1,TWNK,TXNIP,UBE2T,UBIAD1,UBOX5,UGCG,USP18,USP2,USP28,USP37,VANGL1,VDR,VSTM4,VTA1,WASF3,WBP1L,WDCP,WDFY3,WDR7,WFDC1,WHRN,XAF1,XIRP2,XKR8,YWHAG,ZAP70,ZBTB7A,ZDHHC13,ZFP36,ZMAT3,ZMPSTE24,ZNF280B,ZNF324B,ZNF346,ZNF354B,ZNF385D,ZNF420,ZNF462,ZNF605,ZNF608,ZNF74,ZNF780A,ZNF781,ZNF785,ZNF799,ZNF835,ZNRF3,ZSWIM5 |
| Cancer,Organismal Injury and Abnormalities | Benign solid tumor | 3.08E-10 | Increased | 2.15 | ACTA2,ADGRE5,ALDH1B1,ARL4D,C5orf30,CAV1,CDH1,CDKN1A,CDKN2C,CEBPD,CFH,CHKA,CLU,COL2A1,CSF2RB,CXCR4,CYP1B1,DCLRE1A,DGAT2,DUSP1,EGR1,FBLN1,FOS,FOSB,FZD7,GLUL,HAUS3,HOPX,HRH2,ID1,IFI44L,IFIH1,IFIT1,IFIT3,IFITM3,IL6R,JCHAIN,JUN,JUNB,KLF11,KLF2,KLF4,MINPP1,MMP2,MMP7,MRVI1,MYC,NFIB,NOG,NOTCH4,NR0B1,NR4A1,NR4A2,NR4A3,NR5A1,NUP50,PHLDA1,PLA2R1,POMC,PRDX1,PRKCD,PTGR1,PTGS2,RPL9,SERPINE1,SIK1/SIK1B,SLC12A2,SLC39A14,SLC7A5,SOCS3,SPON2,TIE1,TLCD3A,TLR2,TNFAIP3,TNFSF10,TNFSF13B,TNNI2,TP53,TP53INP1,TUBA4A,TUBD1,UGCG,VDR,XAF1 |
| Cancer,Organismal Injury and Abnormalities | Development of carcinoma | 0.00000388 | Increased | 2.073 | ABHD12B,ABHD6,ACACB,ACTA2,ADAP2,ADGRE5,ADGRG6,AGAP3,AGPS,AKR1C1/AKR1C2,ALDH1B1,ALKBH8,APOL2,ARAP1,AREG,ARID3A,ARID5B,ARL4C,ARMT1,ARRDC3,B4GAT1,BATF,BCAM,BHLHE40,BMP2K,BMPR1B,BRD3,BTN3A1,BTN3A2,C14orf119,C14orf39,C2,CACNB4,CALHM2,CARNMT1,CAV1,CBY1,CCDC146,CCL4,CD4,CD69,CD72,CDH1,CDH7,CDKN1A,CDKN2C,CENPF,CFH,CHKA,CKAP4,CLCN3,CLCN4,CLDN7,CLEC1B,CLGN,CLU,CNOT9,COL2A1,COLGALT1,COQ2,CP,CPED1,CSF2RB,CSGALNACT1,CXCL8,CYP1B1,CYP7B1,CYSLTR1,CYYR1,DCAF4,DCLRE1A,DDI2,DDX60,DEGS2,DEPDC1B,DGAT2,DHX33,DIRAS3,DIXDC1,DLAT,DLL3,DNAJC28,DNM3OS,DOCK4,DOLPP1,DSEL,DTX3L,DUSP1,DUSP6,DYRK1A,DYTN,DZIP1,ECE2,EDA2R,EEF2K,EEF2KMT,EGR1,EGR2,EGR3,EIF2AK2,EML2,ENPP5,ENTPD6,EPDR1,EPHX1,EREG,ETV3,EXOSC4,F13A1,FAM124B,FAM133A,FAM181B,FAM217B,FAM83D,FAXDC2,FBLN1,FBLN5,FCGR1A,FCRLB,FEN1,FERMT1,FFAR3,FGFR1OP,FKBP5,FKBP9,FLOT1,FOS,FREM1,FYB1,FZD7,GAS2L3,GASK1B,GATM,GBP1,GBP4,GBP7,GCSAML,GDF7,GGA2,GLRB,GLUL,GMPR,GOT2,GPR27,GPR35,GRK4,GULP1,H2AC18/H2AC19,HAUS3,HCK,HERC5,HILPDA,HLA-C,HLA-F,HLA-G,HRH2,HSPA4L,HSPB1,HYAL2,HYAL3,HYLS1,ICMT,IFI44,IFI44L,IFIH1,IFIT3,IFITM1,IGSF6,IKZF3,IL6R,IL7,IPO9,IRF2,IRF2BP1,IRF7,IRF8,ITGB8,ITPRIPL2,JMJD8,JUN,JUNB,KCNA3,KDM4A,KEAP1,KIF20B,KIF21A,KIF5A,KLF4,KLF6,KLHDC7B,KPNA2,KYNU,L3HYPDH,LAG3,LARP6,LETM1,LIN7A,LINS1,LMAN2L,LONRF2,LRP6,LRRC14,LRRC2,LRRC61,LRRC8D,LRRK2,LRRN1,LYSMD1,MAGEF1,MAL,MAN2A1,MAP3K8,MAP4K3,MAPRE3,MCAM,MCL1,MED24,MLEC,MMP2,MORC1,MOSPD2,MPG,MRPS18C,MRVI1,MUC12,MX1,MYC,MYOZ3,NCR3,NEXN,NFE2,NFIL3,NLGN1,NLRC5,NOC3L,NOG,NOS1AP,NOTCH4,NR0B1,NR4A1,NR4A2,NR4A3,NR5A1,NRXN2,NUDT22,NUP50,OAS1,OAS2,OASL,OSBPL10,OSBPL6,OSTM1,OTULINL,PANK4,PAQR7,PARP3,PCDH9,PDZK1IP1,PDZRN4,PER1,PGS1,PHLDA1,PHLDB2,PIEZO2,PLA2R1,PLD4,PLEKHF1,PLK3,PLSCR1,POGLUT1,POLR3A,POMC,POMGNT2,PORCN,PPP1R1B,PPP2R2B,PPP2R5B,PRDX1,PREX2,PRKCD,PRKG2,PRX,PTAFR,PTGR1,PTGS2,PTPRD,PWWP2A,PYGO1,QSER1,RAB30,RASAL2,RASGRP3,RASSF3,REN,RGCC,RHOB,RIMS1,RMDN2,RMI1,RNASEL,RSAD2,S100A10,S1PR2,SALL4,SAMD9,SAMD9L,SAP30,SCLY,SCML4,SELP,SERPINE1,SESN1,SGK1,SGPP1,SH3GL3,SHQ1,SHTN1,SIK1/SIK1B,SLC12A2,SLC16A1,SLC16A9,SLC25A17,SLC2A10,SLC35B2,SLC38A1,SLC39A14,SLC40A1,SLC45A3,SLC52A3,SLC7A5,SLCO5A1,SLFN5,SMAP2,SMOX,SNTB1,SNX18,SOCS1,SOCS3,SOWAHD,SP110,SPIRE1,SPN,SPON2,SRGAP3,ST6GAL2,STAT1,STAT2,STK17B,STON2,SUCLG2,SV2A,TANGO6,TAP2,TBC1D24,TCEAL2,TCF4,TEKT4,TENT4A,TIE1,TIGD7,TIMP3,TIPARP,TLE4,TLR2,TM4SF1,TM9SF2,TMEM117,TMEM246,TMEM51,TNF,TNFAIP3,TNFSF10,TNRC6C,TP53,TRAF3IP3,TRMT9B,TSC22D3,TTC13,TTC27,TUB,TUBA4A,TUBD1,TWNK,TXNIP,UBOX5,UGCG,USP18,USP2,USP28,USP37,VANGL1,VDR,WASF3,WDFY3,WDR7,WFDC1,WHRN,XIRP2,ZAP70,ZBTB7A,ZFP36,ZMAT3,ZMPSTE24,ZNF324B,ZNF354B,ZNF385D,ZNF420,ZNF462,ZNF74,ZNF780A,ZNF781,ZNF785,ZNF799,ZNF835,ZNRF3,ZSWIM5 |
| Infectious Diseases | Replication of vesicular stomatitis virus | 0.00000933 | Increased | 2.062 | APBA3,FCGR1A,IFIH1,IFITM3,ISG20,OAS1,OASL,SOCS1,SP110,STAT2,TP53,USP2 |
| Cancer,Organismal Injury and Abnormalities | Breast or pancreatic cancer | 0.00000137 | Increased | 2.015 | ABHD12B,ABHD6,ACACB,ACTA2,ADGRG6,AGAP3,AKR1C1/AKR1C2,AKR1C3,ALDH1B1,AREG,ARID5B,ARL4C,ARMT1,ASCL2,ATP6V0A1,BCAM,BHLHE40,BMP2K,BTG2,C14orf39,C1GALT1C1,C2,CAV1,CBY1,CCDC146,CCDC175,CCDC50,CCL3,CCL4,CD4,CD69,CDH1,CDH7,CDKN1A,CDKN2C,CDKN3,CEBPD,CENPF,CFH,CHKA,CKAP4,CLDN7,CLGN,CLU,CMPK2,CNKSR1,COL2A1,CP,CSF2RB,CXCL8,CXCR4,DCLRE1A,DDIT4,DHX58,DIRAS3,DIXDC1,DLL3,DOCK4,DTL,DUSP1,DUSP2,DYRK1A,EDA2R,EGLN1,EGR1,EGR2,EGR3,EGR4,EIF2AK2,EML2,ENTPD6,EPDR1,EPHX1,EPSTI1,EXOSC4,F13A1,FAM114A,FAM181B,FBLN1,FBLN5,FGFR1OP,FKBP5,FLOT1,FOS,FOSB,FREM1,GALNT2,GAS2,GAS2L3,GASK1B,GBP7,GCSAML,GDF7,GLUL,GOT2,GPC3,GPR35,GSTM3,GTF2A1,GULP1,H2AC18/H2AC19,HAPLN3,HCK,HERC5,HLA-DRB5,HLA-G,HLTF,HOXA3,HRH2,HSPB1,HYAL2,HYAL3,ID1,IER2,IER3,IFI44L,IFIH1,IFIT1,IFITM1,IGSF6,IKZF3,INTS14,IRF2,IRF7,ITGB8,ITPKC,JUN,JUNB,JUND,KEAP1,KLF11,KLF4,KLF6,KLF9,KPNA2,LARP6,LETM1,LETM2,LGALS3BP,LINS1,LMAN2L,LMNA,LONRF2,LRP6,LRRC14,LRRK2,LRRN1,LYSMD1,MAFF,MAP3K8,MAP4K3,MAPRE3,MCAM,MCF2L,MCL1,MMACHC,MMP2,MMP7,MOSPD2,MPG,MRVI1,MT-ND1,MTIF3,MUC12,MYADM,MYC,NCR3,NEAT1,NEU1,NFIL3,NFKBID,NLGN1,NOC3L,NOP16,NOTCH4,NR4A1,NR4A2,NR4A3,NR5A1,NRXN2,NUP50,OAF,OSBPL6,OSTM1,PANK4,PARP3,PCDH9,PDZK1IP1,PDZRN4,PGAM1,PHLDA1,PHLDB2,PIEZO2,PLD4,POMC,PPP1R1B,PRDM5,PRDX1,PREX2,PRKCD,PRSS2,PRX,PTAFR,PTGS2,PTPRD,RASAL2,RASGRP3,REG4,REN,RGCC,RGS1,RGS2,RHOB,RIMS1,RMDN2,RNF175,S1PR2,SALL4,SAMD9,SAMD9L,SCLY,SELP,SERPINE1,SH3GL3,SHQ1,SIK1/SIK1B,SLC16A1,SLC2A10,SLC39A6,SLC40A1,SLCO5A1,SLFN5,SMAP2,SMOX,SMYD3,SNAPC3,SNTB1,SNX18,SOCS3,SPN,SRGAP3,SSPN,STAT1,STAT2,STK17B,SUCLG2,SV2A,TANGO6,TCF4,TEKT4,TIE1,TIGD7,TIMP3,TIPARP,TLE3,TM4SF1,TM9SF2,TMEM117,TMEM231,TNF,TNFSF10,TP53,TP53INP1,TRIM69,TRMT9B,TSC22D3,TUBA4A,TUBD1,TXNIP,UGCG,USP18,USP2,USP28,USP37,VDR,WASF3,WDCP,WDFY3,XIRP2,XKR8,ZFP36,ZMPSTE24,ZNF280B,ZNF420,ZNF462,ZNF74,ZNF835,ZSWIM5 |
| Cell Cycle,Gene Expression | Binding of protein binding site | 0.000000135 | Decreased | -2.009 | CD4,CEBPD,CRIP2,CXCL8,EGR1,EGR2,ETV3,FOS,HCK,ID1,ID2,ID3,IER3,IL7,JUN,KLF13,LMNA,MYC,NFKBID,NR0B1,PER1,PLK3,POMC,PRDX1,PRKCD,PTAFR,RGCC,RHOB,SPN,STAT1,TNF,TNFSF10,TP53 |
| Cell-To-Cell Signaling and Interaction | Activation of leukemia cell lines | 0.000000198 | Decreased | -2.041 | CXCL8,EGR2,EGR3,GIMAP6,IL7,MYC,PIK3IP1,SLC39A6,TNF,TNFSF10,ZAP70 |
| Cell Death and Survival | Necrosis | 1.57E-10 | Decreased | -2.051 | AGAP3,AREG,ARID5B,ATF1,AZU1,BHLHE40,BMPR1B,BTG1,BTG2,CAV1,CCDC86,CCL3,CCL3L3,CCL4,CD4,CD69,CDH1,CDKN1A,CDKN2C,CEBPD,CENPF,CFH,CHKA,CITED2,CLU,COL2A1,COQ9,CRK,CSF2RB,CXCL8,CXCR4,CYP1B1,CYSLTR1,DDIT4,DHX33,DHX58,DIRAS3,DLG4,DTL,DUSP1,DUSP6,EDA2R,EEF2K,EFNA1,EGLN1,EGR1,EGR2,EGR3,EIF2AK2,ENTPD1,EPHX1,EREG,F13A1,FBLN1,FCGR1A,FEN1,FGL2,FKBP5,FOS,FOSB,GBP1,GIMAP6,GPC3,GULP1,HCK,HES1,HLA-F,HLA-G,HLF,HOXA3,HRH2,HSPB1,HYAL2,ICMT,ID1,ID2,ID3,IER3,IFI6,IFIH1,IKZF3,IL6R,IL7,IRF2,IRF7,IRF8,ISG15,JUN,JUNB,JUND,KEAP1,KLF13,KLF2,KLF4,KLF6,KLF7,KLF9,KPNA2,LAG3,LETM1,LGALS3BP,LIG4,LMNA,LMNB1,LRP6,LRRK2,LRRN1,MAN2A1,MAP3K8,MAPK12,MCAM,MCF2L,MCL1,MCOLN1,MMP2,MMP7,MOB3B,MPG,MX1,MYC,MZB1,NFIB,NFIL3,NFKBID,NOG,NOTCH4,NR4A1,NR4A2,NR4A3,NR5A1,OAS1,OAS2,OAS3,PANX1,PCSK1N,PCTP,PER1,PHLDA1,PIK3IP1,PLA2R1,PLEKHF1,PLK3,PLSCR1,POMC,PORCN,PPAT,PPIF,PPP1R1B,PPP2R2B,PRDM5,PRDX1,PRKCD,PTAFR,PTGS2,RASSF3,RASSF6,RDH13,REN,RHOB,RNASEL,RPL9,RTN4R,S100A10,S1PR2,SALL4,SELP,SERPINE1,SGK1,SGPP1,SHQ1,SIK1/SIK1B,SLC16A1,SLC39A6,SMCO4,SMOX,SOCS1,SOCS3,SPN,SPRY2,STAT1,STAT2,STK17B,TBC1D24,TCF4,TIMP3,TLE4,TLR2,TMEM107,TMSB10/TMSB4X,TNF,TNFAIP3,TNFSF10,TNFSF13B,TP53,TP53INP1,TSC22D3,TXNIP,UGCG,USP18,USP28,USP2,VDR,XAF1,XKR8,YWHAG,ZAP70,ZBTB7A,ZFP36,ZMAT3,ZMPSTE24,ZNF280B,ZNF346 |
| Tissue Morphology | Quantity of epithelial cells | 2.89E-08 | Decreased | -2.078 | CDKN1A,CDKN2C,GDF7,HES1,HIF3A,HOXA3,HRH2,ID2,KLF4,LMNA,LRRK2,MCL1,MYC,NOG,PRKCD,RAB38,SPRY2,STAT1,TNF,TP53,VDR |
| Hematological System Development and Function,Lymphoid Tissue Structure and Development,Tissue Morphology | Quantity of T lymphocytes | 2.66E-12 | Decreased | -2.102 | ARID5B,BATF,BATF3,BHLHE40,CAV1,CCL4,CD4,CD69,CDKN1A,CDKN2C,CSF2RB,CXCR4,EGR2,EGR3,FOS,FOSB,FYB1,HLA-G,HOXA3,ID1,ID2,ID3,IL6R,IL7,IRF2,IRF8,ITGB8,KLF13,KLF4,LAG3,LIG4,MAN2A1,MAP3K8,MCF2L,MCL1,MPG,MYC,NFIL3,NFKBID,NLRC5,NOG,NOTCH4,NR4A1,PRDX1,PTAFR,PTGIR,REN,RGCC,S1PR2,SELP,SNAI3,SOCS1,SOCS3,STAT1,STK17B,TCF4,TLE4,TLR2,TNF,TNFAIP3,TNFSF13B,TP53,TXNIP,VDR,XKR8,ZAP70,ZFP36 |
| Gene Expression | Expression of RNA | 4.5E-14 | Decreased | -2.122 | ACACB,ACTA2,APBA3,ARID3A,ARID5B,ASCL2,ATF1,AZU1,BATF,BATF3,BHLHE40,BMPR1B,BRD3,BST2,BTG1,BTG2,CAV1,CBY1,CCL3,CCL3L3,CD4,CDH1,CDKN1A,CDKN2C,CEBPD,CENPF,CITED2,CLDN5,CLU,CNKSR1,COL2A1,CRK,CSF2RB,CXCR4,DHX33,DHX58,DIRAS3,DIXDC1,DTX3L,DUSP1,DYRK1A,EFNA1,EGLN1,EGR1,EGR2,EGR3,EGR4,EIF2AK2,EREG,ETV3,FAM98A,FBLN1,FBP1,FCGR1A,FERMT1,FOS,FOSB,FZD7,GDF7,GTF2A1,HES1,HHEX,HIF3A,HLA-DRB5,HLF,HLTF,HOPX,HSPB1,HYAL2,ID1,ID2,ID3,IER2,IER5,IKZF3,IL6R,IL7,IRF2,IRF2BP1,IRF7,IRF8,ISG20,ITGB8,JDP2,JUN,JUNB,JUND,KDM4A,KEAP1,KLF11,KLF13,KLF15,KLF2,KLF4,KLF6,KLF7,KLF9,KPNA2,LMNA,LRP6,LRRK2,MACROH2A2,MAFF,MAP3K8,MAPK12,MAPRE3,MCF2L,MDFI,MED24,MED31,MORC1,MPG,MYADM,MYC,MYCBP,NAT8,NFE2,NFIB,NFIL3,NFKBID,NKX3-1,NLRC5,NOG,NOS1AP,NOTCH4,NR0B1,NR4A1,NR4A2,NR4A3,NR5A1,PARP9,PER1,PLK3,PLSCR1,POLR2D,POMC,PPP1R1B,PPP2R5B,PRDM5,PRKCD,PSRC1,PTAFR,PTGIR,PYGO1,REN,RFXAP,RGCC,RGS2,RHOB,RIMS1,RNASEL,RPS23,SALL4,SAP30,SERPINE1,SERTAD1,SGK1,SLC40A1,SMYD3,SNAI3,SNAPC3,SOCS1,SOCS3,SPN,SPRY2,SSPN,STAT1,STAT2,TADA2A,TAF5,TCF4,TIMP3,TIPARP,TLE3,TLE4,TLR2,TNF,TNFAIP3,TNFSF10,TNNI2,TNRC6C,TP53,TP53INP1,TSC22D3,TUB,TWNK,TXNIP,USP2,VDR,WARS1,WHRN,ZAP70,ZBTB7A,ZFP36,ZMPSTE24,ZNF322,ZNF354B,ZNF462,ZNF608 |
| Cell Death and Survival | Cell death of connective tissue cells | 5.26E-08 | Decreased | -2.235 | BMPR1B,CDH1,CDKN1A,CDKN2C,CLU,COL2A1,CRK,CXCL8,CYP1B1,DDIT4,DUSP1,EGR1,EGR2,EIF2AK2,EPHX1,FCGR1A,HOXA3,HYAL2,ID1,ID2,IER3,IFIH1,IRF7,JUN,JUND,KLF2,KLF6,LMNA,MAP3K8,MCF2L,MCL1,MPG,MYC,NFKBID,NOG,NR4A1,NR4A2,PLEKHF1,PLK3,PRKCD,RHOB,RNASEL,SERPINE1,SGK1,SGPP1,SPN,STAT1,STK17B,TCF4,TIMP3,TMEM107,TNF,TNFAIP3,TNFSF10,TP53,TP53INP1,TXNIP,USP18,VDR,ZNF346 |
| Cell Death and Survival | Apoptosis of connective tissue cells | 0.00000783 | Decreased | -2.26 | BMPR1B,CDH1,CDKN1A,CDKN2C,CLU,COL2A1,CXCL8,DDIT4,DUSP1,EIF2AK2,FCGR1A,JUN,JUND,KLF2,MCF2L,MCL1,MYC,NFKBID,NOG,NR4A1,RHOB,RNASEL,SERPINE1,TCF4,TIMP3,TNF,TNFSF10,TP53,TXNIP,VDR,ZNF346 |
| Cell Cycle,Gene Expression | Binding of E box motif | 0.00000387 | Decreased | -2.382 | CD4,ID1,ID2,ID3,MYC,PER1,STAT1,TNF |
| Cellular Function and Maintenance | Engulfment of cells | 0.00000141 | Decreased | -2.397 | ATP6V0E1,AZU1,BTBD19,CAV1,CCL3,CCL4,CD4,CEACAM4,CFH,CLCN3,CRK,CSF2RB,DOCK4,EEF2K,EGR1,FCGR1A,FLOT1,GP1BA,GPC3,GULP1,HCK,HYAL2,IL7,IRF7,IRF8,ISG15,JUNB,KLF2,MYC,PLD4,PLEKHF1,PLSCR1,POMC,PRKCD,PRKG2,PTX3,RGCC,RHOB,S100A10,SERPINE1,STON2,TLE3,TLR2,TNF,TNFSF10,TP53,TUB,USP2,XKR8,ZBTB7A |
| Neurological Disease | Progressive neurological disorder | 0.0000019 | Decreased | -2.538 | ACTA2,APOL2,AREG,B4GAT1,BST2,CAV1,CCDC50,CCL3,CCL4,CDKN1A,CHCHD10,CLU,CNKSR1,CXCL8,CXCR4,CYP7B1,CYSLTR1,DYRK1A,ECE2,EEF2K,EGLN1,EGR1,EGR2,EIF2AK2,ELN,EPSTI1,FBLN1,FCGR1A,FITM2,FLOT1,FUCA1,GATM,GLRB,HLA-F,HLA-G,HRH2,HSPB1,IFIT1,IGKC,IL6R,IRF7,ISG15,JUN,KCNA3,KEAP1,KIF5A,LRP6,LRRK2,MCAM,MMP2,MT-ND1,MX1,MYC,NAT8,NEAT1,NFIL3,NOTCH4,NR4A2,OAS1,OAS3,OSBPL10,PIK3IP1,PRDX1,PRKCD,PRX,PTGS2,RHOB,RIMS1,RSAD2,RTN4R,S100A10,SELP,SERPINE1,SGK1,SLC52A3,SPR,SV2A,TAP2,TBC1D24,TIMP3,TLR2,TNF,TP53,TUBA4A,TUBD1,TWNK,USP37,VDR,WDFY3,WDR7,ZNF280B |
| Cell Cycle,Gene Expression | Binding of DNA | 1.39E-14 | Decreased | -2.758 | BMPR1B,BTG2,CAV1,CD4,CDKN1A,CEBPD,CLU,CRIP2,CXCL8,CXCR4,CYP1B1,DIRAS3,EGR1,EGR2,EGR3,ETV3,FOS,FOSB,FYB1,GTF2A1,HCK,HIF3A,ID1,ID2,ID3,IER3,IL6R,IL7,IRF7,JUN,JUNB,JUND,KEAP1,KLF13,LMNA,MAPK12,MPG,MYC,NFKBID,NR0B1,NR4A1,NR4A2,NR5A1,OAS1,PER1,PLK3,POMC,PPP1R1B,PRDX1,PRKCD,PTAFR,RGCC,RHOB,SOCS1,SOCS3,SPN,STAT1,TCF4,TNF,TNFAIP3,TNFSF10,TNFSF13B,TP53,TSC22D3,TWNK,VDR,YWHAG,ZBTB7A |
| Antimicrobial Response,Inflammatory Response | Antimicrobial response | 1.83E-16 | Decreased | -2.919 | ACTA2,AZU1,BANF1,BATF3,BST2,CCL4,CLU,CXCR4,DDIT4,DDX60,DHX58,DTX3L,EIF2AK2,EXOSC4,FCGR1A,HCK,HLA-G,HSPB1,HYAL2,HYAL3,IFI44,IFI44L,IFI6,IFIH1,IFIT1,IFIT2,IFIT3,IFITM1,IFITM3,IRF2,IRF7,IRF8,ISG15,ISG20,JCHAIN,MMP7,MX1,OAS1,OAS2,OAS3,OASL,PARP9,PLSCR1,PRKCD,PRSS2,RNASEL,RSAD2,SAMHD1,SOCS1,SOCS3,SPN,SPON2,SPRY2,STAT1,STAT2,TLR2,TNF,TP53 |
| Cell Cycle | Interaction of DNA | 5.26E-15 | Decreased | -3.009 | BMPR1B,BTG2,CAV1,CD4,CDKN1A,CEBPD,CLU,CRIP2,CXCL8,CXCR4,CYP1B1,DIRAS3,EGR1,EGR2,EGR3,ETV3,FOS,FOSB,FYB1,GTF2A1,HCK,HIF3A,ID1,ID2,ID3,IER3,IL6R,IL7,IRF7,JUN,JUNB,JUND,KEAP1,KLF13,LIG4,LMNA,MAPK12,MPG,MYC,NFKBID,NR0B1,NR4A1,NR4A2,NR5A1,OAS1,PARP3,PER1,PLK3,POMC,PPP1R1B,PRDX1,PRKCD,PTAFR,RGCC,RHOB,SOCS1,SOCS3,SPN,STAT1,TCF4,TNF,TNFAIP3,TNFSF10,TNFSF13B,TP53,TSC22D3,TWNK,VDR,YWHAG,ZBTB7A |
| Inflammatory Response | Immune response of cells | 8.29E-08 | Decreased | -3.246 | AZU1,BATF3,BHLHE40,BRD3,CAV1,CCL3,CD4,CD69,CDKN1A,CEACAM4,CFH,CLCN3,CRK,CSF2RB,CXCL8,CYP1B1,DHX58,EIF2AK2,FCGR1A,GP1BA,GULP1,HCK,HRH2,HSPB1,IFIH1,IGKC,IL6R,IL7,IRF2,IRF2BP1,IRF7,IRF8,ISG15,JUN,JUNB,KLF2,MAP3K8,MCL1,MPG,MYC,NCR3,NFIL3,NR4A2,PER1,PLD4,PRKCD,PTX3,RGCC,SERPINE1,SGK1,SMYD3,SOCS1,SOCS3,SPRY2,STAT1,TLE3,TLR2,TNF,TNFSF10,TP53,TUB,UBE2L6 |
| Antimicrobial Response,Inflammatory Response | Antiviral response | 1.3E-21 | Decreased | -3.386 | ACTA2,AZU1,BANF1,BATF3,BST2,CCL4,CLU,CXCR4,DDIT4,DDX60,DHX58,DTX3L,EIF2AK2,EXOSC4,FCGR1A,HSPB1,HYAL2,HYAL3,IFI44,IFI44L,IFI6,IFIH1,IFIT1,IFIT2,IFIT3,IFITM1,IFITM3,IRF2,IRF7,ISG15,ISG20,MX1,OAS1,OAS2,OAS3,OASL,PARP9,PLSCR1,RNASEL,RSAD2,SAMHD1,SOCS1,SOCS3,SPN,SPON2,SPRY2,STAT1,STAT2,TNF,TP53 |

**T3 vs. T1 MPP_GMP**

© 2000-2020 QIAGEN. All rights reserved.

| Categories | Diseases or Functions Annotation | p-value | Predicted Activation State | Activation z-score | Molecules |
|---|---|---|---|---|---|
| Organismal Survival | Morbidity or mortality | 5.23E-10 | Increased | 3.369 | ACADL,ACTA2,ADAM19,ADAM22,ADM,AFMID,ANKLE1,APBA2,AREG,ARID3A,ARID5B,AXL,BAG3,BAK1,BRPF1,C3AR1,CCDC86,CCL3L3,CCR1,CD3E,CDKN2C,CFH,CHKA,CLCN3,CLDN7,CLN3,COL2A1,COQ9,CPLX1,CSF1R,CSGALNACT1,CTR9,CXCL3,CXCL8,CXCR4,DDX58,DHX58,DNAJC6,DUSP1,EGR2,EGR3,EIF2AK2,ELANE,ENAH,ENPP2,F13A1,FAS,FBXO45,FERMT1,FN1,FOS,GCH1,GFI1B,GNAI1,HCK,HLA-G,HMGA2,HOPX,HSPB1,HYAL2,ID1,ID3,IDS,IFIH1,IFIT2,INHBA,IRF2,IRF7,ISG15,ITGB8,JUN,JUNB,KCNAB1,KDM4A,KIF5A,KLF1,KLF2,KLF4,KLF6,KLF7,KLF9,LIG4,LMNA,LMO4,LMTK2,LTBP1,MAFF,MAL,MCAM,MCL1,MCOLN1,MDFI,MMP2,MMP7,MT-ND1,MYC,MYCNOS,NAMPT,NAPB,NEAT1,NEXN,NFE2,NINJ1,NOD2,NOG,NR4A3,NUBP1,OASL,OR7C1,PCDHGB6,PER2,PIEZO2,PIGF,PLA2R1,POU2F2,PRKAA2,PRKCD,PROCR,PTAFR,PTGIR,PTGS2,PYGO1,RARG,REN,RIMS1,SCN9A,SDC4,SERTAD1,SLC12A2,SLC1A3,SLC22A5,SLC40A1,SNAI3,SOCS1,SOCS3,SOSTDC1,SPRY2,SRGAP3,STAT1,STAT2,TAL1,TAP1,TIPARP,TJP2,TK1,TLE4,TNFAIP8L2,TNFRSF1A,TNFSF10,TNFSF13B,TP53INP1,TXNIP,USP18,XRCC1,ZBTB7A,ZFP36,ZNF296,ZNRF3 |
| Organismal Survival | Organismal death | 6.9E-10 | Increased | 3.351 | ACADL,ACTA2,ADAM19,ADAM22,ADM,AFMID,ANKLE1,APBA2,AREG,ARID3A,ARID5B,AXL,BAG3,BAK1,BRPF1,C3AR1,CCDC86,CCL3L3,CCR1,CD3E,CDKN2C,CFH,CHKA,CLCN3,CLDN7,CLN3,COL2A1,COQ9,CPLX1,CSF1R,CSGALNACT1,CTR9,CXCL3,CXCL8,CXCR4,DDX58,DHX58,DNAJC6,DUSP1,EGR2,EGR3,EIF2AK2,ELANE,ENAH,ENPP2,F13A1,FAS,FBXO45,FERMT1,FN1,FOS,GCH1,GFI1B,GNAI1,HCK,HLA-G,HMGA2,HOPX,HSPB1,HYAL2,ID1,ID3,IDS,IFIH1,IFIT2,INHBA,IRF2,IRF7,ISG15,ITGB8,JUN,JUNB,KCNAB1,KDM4A,KIF5A,KLF1,KLF2,KLF4,KLF6,KLF7,KLF9,LIG4,LMNA,LMO4,LMTK2,LTBP1,MAFF,MAL,MCAM,MCL1,MCOLN1,MDFI,MMP2,MMP7,MT-ND1,MYC,MYCNOS,NAMPT,NAPB,NEAT1,NEXN,NFE2,NINJ1,NOD2,NOG,NR4A3,NUBP1,OR7C1,PCDHGB6,PER2,PIEZO2,PIGF,PLA2R1,POU2F2,PRKAA2,PRKCD,PROCR,PTAFR,PTGIR,PTGS2,PYGO1,RARG,REN,RIMS1,SCN9A,SDC4,SERTAD1,SLC12A2,SLC1A3,SLC22A5,SLC40A1,SNAI3,SOCS1,SOCS3,SOSTDC1,SPRY2,SRGAP3,STAT1,STAT2,TAL1,TIPARP,TJP2,TK1,TLE4,TNFAIP8L2,TNFRSF1A,TNFSF10,TNFSF13B,TP53INP1,TXNIP,USP18,XRCC1,ZBTB7A,ZFP36,ZNF296,ZNRF3 |
| Cell Signaling | Viral life cycle | 0.00000407 | Increased | 3.101 | CCL5,CHMP6,CXCR4,EIF2AK2,IFIT1,IFITM1,IFITM3,ISG15,MX1,OAS1,OAS3,OASL,PLSCR1,PTGS2,RSAD2 |
| Cell Signaling | Replication of viral replicon | 2.86E-09 | Increased | 3.1 | CCL5,EIF2AK2,IFIT1,IFITM1,IFITM3,ISG15,MX1,OAS1,OAS3,OASL,PLSCR1,PTGS2,RSAD2 |
| Cellular Development,Skeletal and Muscular System Development and Function,Tissue Development | Differentiation of muscle | 0.0000071 | Increased | 2.973 | AREG,BTG1,BTG2,EGR1,EGR2,EREG,FN1,FOS,HOPX,HYAL2,ID3,JAK3,JDP2,JUN,LMNA,MAFF,MYC,NOG,NR4A1,NR4A3,PRDM16,PRKG2,SAP30,SOCS1,SOCS3,SOSTDC1,SSPN,STAT1,TAL1 |
| Infectious Diseases | Replication of Flaviviridae | 0.00000841 | Increased | 2.674 | DDX58,EIF2AK2,IFI6,IFIT1,IFITM1,IFITM3,ISG15,OASL,RSAD2,SOCS3,TXNIP,UBE2L6 |
| Infectious Diseases | Replication of virus | 1.02E-11 | Increased | 2.519 | AREG,BAK1,CCL3,CCL3L3,CCL5,CCR1,CD48,CXCL3,CXCL8,CXCR4,DDX58,DUSP1,DUSP3,EGR1,EIF2AK2,F13A1,GBP1,HCK,HERC5,IFI6,IFIH1,IFIT1,IFITM1,IFITM3,IL10RA,IRF2,IRF7,ISG15,JUN,MX1,MX2,MYC,OAS1,OAS3,OASL,PARP11,PRKCD,PTGS2,RSAD2,SAMHD1,SGK1,SLFN12,SOCS1,SOCS3,SP110,SPRY2,STAT1,STAT2,TAP1,TNFRSF1A,TNFSF10,TRIM25,TXNIP,UBE2L6 |
| Infectious Diseases | Replication of vesicular stomatitis virus | 7.63E-08 | Increased | 2.483 | BAK1,DDX58,IFIH1,IFITM3,OAS1,OASL,PARP11,SLFN12,SOCS1,SP110,STAT2,TAP1,TRIM25 |
| Infectious Diseases | Replication of Hepatitis C virus | 0.0000136 | Increased | 2.477 | DDX58,EIF2AK2,IFI6,IFIT1,IFITM1,IFITM3,ISG15,RSAD2,SOCS3,TXNIP,UBE2L6 |
| Tissue Development | Accumulation of cells | 0.000000111 | Increased | 2.221 | ABCG1,CCL3,CCL3L3,CCL5,CCR1,CLU,CXCL3,CXCL8,CXCR4,DUSP1,EGR1,ELANE,ENPP2,ENTPD1,FAS,HCK,IER3,IL10RA,ITGA2,KLF2,MMP7,MYC,NR4A1,PIK3AP1,PTGS2,RHOB,SAMHD1,STAT1,TAL1,TNFAIP8L2,TNFRSF1A,TNFSF10,TSC22D3 |
| Infectious Diseases | Replication of RNA virus | 1.52E-09 | Increased | 2.209 | AREG,BAK1,CCL3,CCL3L3,CCL5,CCR1,CD48,CXCL8,CXCR4,DDX58,DUSP3,EIF2AK2,F13A1,GBP1,HCK,HERC5,IFI6,IFIH1,IFIT1,IFITM1,IFITM3,IRF2,IRF7,ISG15,JUN,MX1,MYC,OAS1,OAS3,OASL,PARP11,PRKCD,PTGS2,RSAD2,SGK1,SLFN12,SOCS1,SOCS3,SP110,STAT1,STAT2,TAP1,TNFSF10,TRIM25,TXNIP,UBE2L6 |
| Infectious Diseases | Infection of mammalia | 1.4E-09 | Increased | 2.106 | CCL3L3,CCR1,CSF2RB,CXCR4,DDX58,DHX58,DUSP1,EIF2AK2,ELANE,FOS,FUT7,HCK,HLA-G,IFIH1,IFIT2,IL10RA,IRF7,ISG15,JDP2,MMP2,MMP7,NFE2,NOD2,OASL,PRKCD,PTAFR,SAMHD1,SPON2,STAT1,STAT2,TAP1,TNFAIP8L2,TNFRSF1A,TNFSF10,USP18 |
| Hematological System Development and Function,Tissue Development | Accumulation of blood cells | 5.77E-08 | Increased | 2.069 | ABCG1,CCL3,CCL3L3,CCL5,CCR1,CXCL3,CXCL8,CXCR4,DUSP1,EGR1,ELANE,ENPP2,ENTPD1,FAS,HCK,IER3,IL10RA,ITGA2,KLF2,MMP7,MYC,NR4A1,PIK3AP1,PTGS2,STAT1,TAL1,TNFAIP8L2,TNFRSF1A,TNFSF10,TSC22D3 |
| Cancer,Organismal Injury and Abnormalities | Frequency of tumor | 0.00000179 | Increased | 2.062 | ABHD12B,ACTA2,ADAM19,ADAM22,ADAM8,ADM,AGPS,AKR1C1/AKR1C2,ANKRD26,ANKRD27,ANKRD9,APBA2,APOL2,AREG,ARHGEF26,ARHGEF37,ARID3A,ARID5B,ARL4C,ATP8B3,AXL,BAK1,BATF,BMP2K,BMPR1B,BRPF1,BTG2,BTN3A1,BTN3A2,BTRC,C14orf119,C2,C3AR1,C5orf30,CACNB4,CARF,CAVIN2,CBX6,CCDC169,CCNA1,CCR1,CD3G,CD48,CD69,CDKN2C,CETN3,CFH,CHKA,CHST12,CLCN3,CLDN7,CLU,COL2A1,CP,CSF1R,CSF2RB,CSGALNACT1,CTR9,CXCL8,CXCR4,DCAF4,DCP1B,DDI2,DDX58,DDX60,DIP2A,DIRAS3,DIXDC1,DNAJB5,DNAJC6,DSC2,DTX3L,DUSP1,DUSP6,DZIP1,EEF2KMT,EFNA4,EGR1,EGR2,EGR3,EIF2AK2,ENAH,ENPP2,ENTPD4,EREG,ESRRA,EVI5,F13A1,FAM217B,FAM234B,FAS,FCGR1B,FERMT1,FFAR3,FKBP5,FN1,FOS,FPGT,FREM1,FYB1,GASK1B,GBP1,GBP4,GBP7,GCSAML,GMPR,GOLPH3L,GPAT3,GPR155,GPR27,GSTM3,GULP1,HCK,HECA,HELZ2,HERC5,HLA-C,HLA-F,HLA-G,HMGA2,HMGB2,HSPB1,HYAL2,HYLS1,ID1,IER3,IFI44,IFI44L,IFIH1,IFIT3,IFITM1,IFITM3,IGSF6,IL10RA,INHBA,IRF2,IRF2BP1,IRF7,IRX3,ITGA2,ITGB8,ITPRIPL2,JAK3,JAKMIP2,JUN,JUNB,KBTBD6,KCNAB1,KCNK6,KCNN4,KDM4A,KIF21A,KIF5A,KLF2,KLF4,KLF6,KLHDC7B,LAG3,LGALS3BP,LIG4,LINS1,LMTK2,LRRC2,LRRC4C,LRRC8D,LRRIQ3,LTBP1,LY86,LYSMD1,MAGI3,MAL,MAP3K21,MCAM,MCL1,MGA,MINPP1,MMP2,MMP7,MORC1,MRS2,MUC12,MX1,MYC,MYCNOS,NAMPT,NAV1,NBPF1 (includes others),NETO2,NEURL1B,NEXN,NFE2,NHLRC1,NINJ1,NOD2,NOG,NR0B1,NR4A1,NR4A3,NTNG2,NUBP1,NUDT22,OAS1,OAS2,OASL,OR7C1,OSBPL10,PABPC3,PAQR7,PAQR8,PARP11,PARP14,PCDHGB6,PER1,PER2,PGBD4,PHLDA1,PIEZO2,PIK3R3,PLA2G4A,PLA2R1,PLSCR1,PPFIA1,PPP2R5B,PRDM16,PRKAA2,PRKCD,PRKG2,PROCR,PRR11,PRX,PSAT1,PTAFR,PTGS2,PTPRD,PYGO1,RAB30,RARG,RASAL2,RASGRP3,RASSF3,REN,RGCC,RHOB,RHOU,RIMS1,RNF170,RPS7,RSAD2,SAMD4A,SAMD9,SAMD9L,SAP30,SCARF1,SCML4,SCN9A,SDC4,SESN1,SGK1,SH2D3A,SH3BP5L,SIK1/SIK1B,SLC12A2,SLC1A3,SLC22A5,SLC35A5,SLC40A1,SLC45A3,SLC7A6,SLFN12,SLFN5,SMAP2,SNX33,SOCS1,SOCS3,SOGA1,SOWAHD,SP110,SPIRE1,SPON2,SRGAP3,SSH1,STAT1,STAT2,STK17B,STON2,TAL1,TAP1,TBC1D8B,THNSL1,TIPARP,TJP2,TK1,TLE4,TM4SF1,TMEM102,TMEM176B,TMEM51,TNFRSF1A,TNFSF10,TP53INP1,TPRN,TRAF3IP3,TRANK1,TRIM22,TRIM25,TRIM58,TRMT9B,TSC22D3,TUBB4B,TXNIP,USP18,XRCC1,ZBTB7A,ZC3H4,ZC3HAV1L,ZDHHC4,ZFP36,ZFP64,ZKSCAN1,ZMAT3,ZNF146,ZNF212,ZNF462,ZNF485,ZNF546,ZNF585B,ZNF594,ZNF616,ZNF623,ZNF780A,ZNF781,ZNF786,ZNF799,ZNF835,ZNRF3 |
| Hematological System Development and Function,Immune Cell Trafficking,Inflammatory Response,Tissue Development | Accumulation of leukocytes | 3.53E-08 | Increased | 2.055 | ABCG1,CCL3,CCL3L3,CCL5,CCR1,CXCL3,CXCL8,CXCR4,DUSP1,EGR1,ELANE,ENPP2,ENTPD1,FAS,HCK,IER3,IL10RA,ITGA2,KLF2,MMP7,NR4A1,PIK3AP1,PTGS2,STAT1,TAL1,TNFAIP8L2,TNFRSF1A,TNFSF10,TSC22D3 |
| Infectious Diseases | Replication of Herpesviridae | 7.53E-09 | Increased | 2.048 | BAK1,CXCL3,DDX58,EGR1,IFIH1,IL10RA,MX2,OAS1,OASL,RSAD2,SAMHD1,SPRY2,STAT1,STAT2 |
| Cellular Movement,Hematological System Development and Function,Immune Cell Trafficking | Mobilization of granulocytes | 0.00000257 | Increased | 2.007 | C3AR1,CCL3,CCL3L3,CCR1,CXCL3,CXCL8,CXCR4 |
| Cell Death and Survival,Endocrine System Disorders,Gastrointestinal Disease,Organismal Injury and Abnormalities | Cell death of beta islet cells | 0.00000758 | Decreased | -2.007 | FAS,JUNB,MCL1,MYC,PRKCD,SOCS1,SOCS3,STAT1,TCF7,TNFRSF1A,TXNIP,USP18 |
| Cell Death and Survival | Apoptosis of lymphatic system cells | 6.72E-08 | Decreased | -2.012 | ADAM8,BAK1,C3AR1,CCDC86,CCL3,CCL5,CD3E,CXCR4,DHX58,EGR1,EGR3,EIF2AK2,ELANE,FAS,FOS,HLA-G,ID3,IER3,JAK3,JUN,KLF2,MCL1,MYC,NR4A1,NR4A3,PARP14,PRKCD,SOCS1,STAT1,STK17B,TLE4,TNFAIP8L2,TNFRSF1A,TNFSF10,TNFSF13B,TSC22D3 |
| Cell Death and Survival | Cell death of myeloid cells | 1.15E-10 | Decreased | -2.021 | ABCG1,ADAM8,BAK1,BTG1,C3AR1,CCL5,CD69,CXCL8,CXCR4,EIF2AK2,ELANE,ENTPD1,FAS,FOS,GBP1,GFI1B,IRF2,ISG15,KLF2,MCL1,MYC,NAMPT,NR4A1,PIK3AP1,PLA2G4A,PRKCD,SOCS1,STAT1,TNFRSF1A,TNFSF10,TSC22D3 |
| Cell Death and Survival | Apoptosis of macrophages | 1.15E-09 | Decreased | -2.036 | ABCG1,ADAM8,BTG1,C3AR1,CCL5,EIF2AK2,FAS,FOS,GBP1,IRF2,ISG15,MCL1,NAMPT,PIK3AP1,PLA2G4A,STAT1,TNFRSF1A,TNFSF10 |
| Cell Death and Survival,Organismal Injury and Abnormalities | Cell death of endothelial cells | 9.16E-08 | Decreased | -2.043 | ADM,ARHGAP18,AXL,AZU1,CXCL8,CXCR4,DIP2A,DUSP6,FAS,ID1,ITGA2,KLF2,KLF4,MCAM,MCL1,MMP2,MYC,NOG,NR4A3,PROCR,RHOB,SOCS1,TNFRSF1A,TNFSF10 |
| Cell Death and Survival | Cell death of immune cells | 4.15E-11 | Decreased | -2.066 | ABCG1,ADAM8,ARID5B,AXL,BAK1,BTG1,C3AR1,CCDC86,CCL3,CCL5,CD3E,CD48,CD69,CLU,CSF1R,CSF2RB,CXCL8,CXCR4,DHX58,EGR1,EGR3,EIF2AK2,ELANE,ENTPD1,FAS,FN1,FOS,GBP1,HCK,HLA-G,ID3,IER3,IFIH1,INHBA,IRF2,ISG15,JAK3,JUN,KLF2,KLF4,LAG3,MCL1,MX1,MYC,NAMPT,NR4A1,NR4A3,PARP14,PIK3AP1,PLA2G4A,PRKCD,SOCS1,STAT1,STK17B,TCF7,TLE4,TNFAIP8L2,TNFRSF1A,TNFSF10,TNFSF13B,TSC22D3 |

| | | | | | |
|---|---|---|---|---|---|
| Cell Cycle,Gene Expression | Binding of DNA | 5.86E-13 | Decreased | -2.082 | BMPR1B,BTG2,BTRC,CD3E,CLU,CRIP2,CTR9,CXCL8,CXCR4,DDX58,DIRAS3,EGR1,EGR2,EGR3,ELANE,ESRRA,FAS,FN1,FOS,FYB1,HCK,HIF3A,HMGB2,HSPA1A/HSPA1B,ID1,ID3,IER3,IRF7,ITGA2,JAK3,JUN,JUNB,JUND,LMNA,MGA,MYC,NR0B1,NR4A1,OAS1,PER1,PRDM16,PRKCD,PTAFR,RARG,RGCC,RHOB,SOCS1,SOCS3,STAT1,TAL1,TNFRSF1A,TNFSF10,TNFSF13B,TSC22D3,ZBTB7A |
| Cell Death and Survival,Organismal Injury and Abnormalities | Apoptosis of endothelial cells | 0.000000213 | Decreased | -2.086 | ADM,ARHGAP18,AXL,AZU1,CXCL8,CXCR4,DUSP6,FAS,ID1,ITGA2,KLF4,MCAM,MCL1,MMP2,MYC,NOG,NR4A3,PROCR,RHOB,SOCS1,TNFRSF1A,TNFSF10 |
| Cell Death and Survival | Apoptosis of myeloid cells | 3.39E-11 | Decreased | -2.13 | ABCG1,ADAM8,BAK1,BTG1,C3AR1,CCL5,CD69,CXCL8,CXCR4,EIF2AK2,ELANE,FAS,FOS,GBP1,GFI1B,IRF2,ISG15,KLF2,MCL1,MYC,NAMPT,PIK3AP1,PLA2G4A,PRKCD,SOCS1,STAT1,TNFRSF1A,TNFSF10,TSC22D3 |
| Connective Tissue Disorders,Inflammatory Disease,Inflammatory Response,Organismal Injury and Abnormalities,Skeletal and Muscular Disorders | Polyarthritis | 1.43E-11 | Decreased | -2.166 | ADM,CCL3,CCL5,CCR1,CD69,COL2A1,CTR9,DUSP1,DUSP2,F13A1,FAS,FN1,FOS,HLA-G,HSPA1A/HSPA1B,MCL1,NR4A1,NR4A3,PLA2G4A,PTGIR,PTGS2,SOCS1,STAT1,TLE3,TNFRSF1A,TNFSF13B,ZFP36 |
| Cell Death and Survival,Organismal Injury and Abnormalities | Necrosis of epithelial tissue | 4.37E-09 | Decreased | -2.186 | ADM,ARHGAP18,AXL,AZU1,BAK1,CLU,CSF1R,CXCL3,CXCL8,CXCR4,DIP2A,DUSP6,EGR1,EIF2AK2,FAS,FOS,HSPB1,ID1,IER3,IFIH1,INHBA,ITGA2,JUN,KLF2,KLF4,MCAM,MCL1,MIAT,MMP2,MYC,NAMPT,NOG,NR4A1,NR4A3,OAS1,OAS2,OAS3,PLA2G4A,PRKAA2,PRKCD,PROCR,PTAFR,PTGS2,RHOB,SGK1,SOCS1,SOCS3,STAT1,TJP2,TNFAIP8L2,TNFRSF1A,TNFSF10,TP53INP1 |
| Cell Death and Survival,Endocrine System Disorders,Gastrointestinal Disease,Organismal Injury and Abnormalities | Apoptosis of pancreatic cells | 0.0000202 | Decreased | -2.201 | CXCR4,FAS,JUNB,MCL1,MYC,PRKCD,SOCS1,SOCS3,STAT1,TCF7,TXNIP,USP18 |
| Connective Tissue Disorders,Inflammatory Disease,Inflammatory Response,Organismal Injury and Abnormalities,Skeletal and Muscular Disorders | Experimentally-induced arthritis | 0.000000163 | Decreased | -2.207 | CCL3L3,CCR1,CD69,COL2A1,CTR9,CXCL8,DUSP1,F13A1,FAS,FN1,HCK,MMP2,NAMPT,NOD2,PLA2G4A,PTGIR,PTGS2,STAT1,TNFRSF1A,TNFSF13B |
| Cell Death and Survival | Apoptosis of T lymphoblasts | 0.000016 | Decreased | -2.213 | CCL3,CCL5,CD3E,FAS,MYC |
| Cellular Development,Cellular Growth and Proliferation,Embryonic Development | Proliferation of embryonic cell lines | 0.00000898 | Decreased | -2.214 | BATF,BRPF1,BTG1,BTG2,BTRC,CCNA1,CHKA,CNKSR1,DIRAS3,EGR1,EIF2AK2,FN1,FOS,GPR12,JUN,KLF2,KLF4,LIG4,LTBP1,MYC,PTGS2,SAMD9L,SOCS1,SOCS3,TSC22D3 |
| Cellular Movement | Invasion of carcinoma cell lines | 0.0000169 | Decreased | -2.215 | ACTA2,ADAM8,AREG,AXL,CCR1,CFH,CLDN7,CLU,DDX58,DUSP1,EGR1,ENAH,ENPP2,FN1,GNAI1,ID1,IFIT2,IFITM1,JUN,JUNB,JUND,KLF4,PRKCD,PTAFR |
| Inflammatory Response | Innate immune response | 0.0000006 | Decreased | -2.22 | ADAM8,CCL5,DDX58,EIF2AK2,EREG,FFAR3,FN1,GBP5,HLA-G,HMGB2,IFI6,IFIH1,IFIT2,IRF2BP1,IRF7,ISG15,MMP2,MX1,NOD2,PLSCR1,SPON2,STAT1,TRIM25,TRIM58 |
| Cell Cycle | Interaction of DNA | 5.35E-13 | Decreased | -2.224 | BMPR1B,BTG2,BTRC,CD3E,CLU,CRIP2,CTR9,CXCL8,CXCR4,DDX58,DIRAS3,EGR1,EGR2,EGR3,ELANE,ESRRA,FAS,FN1,FOS,FYB1,HCK,HIF3A,HMGB2,HSPA1A/HSPA1B,ID1,ID3,IER3,IRF7,ITGA2,JAK3,JUN,JUNB,JUND,LIG4,LMNA,MGA,MYC,NR0B1,NR4A1,OAS1,PER1,PRDM16,PRKCD,PTAFR,RARG,RGCC,RHOB,SOCS1,SOCS3,STAT1,TAL1,TNFRSF1A,TNFSF10,TNFSF13B,TSC22D3,ZBTB7A |
| Cell Death and Survival | Cell death of T lymphocytes | 3.98E-09 | Decreased | -2.247 | ADAM8,BAK1,C3AR1,CCDC86,CCL3,CCL5,CD3E,CXCR4,DHX58,EGR1,EGR3,ENTPD1,FAS,FOS,HLA-G,ID3,IER3,JAK3,JUN,KLF2,LAG3,MCL1,MYC,NR4A1,NR4A3,SOCS1,STAT1,STK17B,TLE4,TNFAIP8L2,TNFRSF1A,TNFSF10,TNFSF13B,TSC22D3 |
| Cell Death and Survival | Cell death of blood cells | 7.75E-11 | Decreased | -2.273 | ABCG1,ADAM8,ARID5B,AXL,BAK1,BTG1,C3AR1,CCDC86,CCL3,CCL3L3,CCL5,CD3E,CD48,CD69,CLU,CSF1R,CSF2RB,CXCL8,CXCR4,DHX58,EGR1,EGR3,EIF2AK2,ELANE,ENTPD1,FAS,FN1,FOS,GBP1,GFI1B,HCK,HLA-G,ID3,IER3,IFIH1,INHBA,IRF2,ISG15,JAK3,JUN,KLF2,KLF4,LAG3,MCL1,MX1,MYC,NAMPT,NR4A1,NR4A3,PARP14,PIK3AP1,PLA2G4A,PRKCD,SOCS1,STAT1,STK17B,TCF7,TLE4,TNFAIP8L2,TNFRSF1A,TNFSF10,TNFSF13B,TSC22D3 |
| Cell Death and Survival,Organismal Injury and Abnormalities | Necrosis of vascular endothelial cells | 0.00000407 | Decreased | -2.277 | ADM,ARHGAP18,AXL,CXCL8,DIP2A,DUSP6,FAS,ID1,KLF4,MCAM,MCL1,MMP2,NR4A3,TNFRSF1A,TNFSF10 |
| Cell Death and Survival | Necrosis | 1.47E-10 | Decreased | -2.281 | ABCG1,ADAM8,ADM,AREG,ARHGAP18,ARID5B,AXL,AZU1,BAG3,BAK1,BMPR1B,BTG1,BTG2,BTRC,C3AR1,CCDC86,CCL3,CCL3L3,CCL5,CCNA1,CD3E,CD48,CD69,CDKN2C,CFH,CHKA,CHMP6,CLN3,CLU,COL2A1,COQ9,CSF1R,CSF2RB,CXCL3,CXCL8,CXCR4,DDX58,DHX58,DIP2A,DIRAS3,DUSP1,DUSP6,EDA2R,EGR1,EGR2,EGR3,EIF2AK2,ELANE,ENPP2,ENTPD1,EREG,ESRRA,F13A1,FAS,FKBP5,FN1,FOS,GBP1,GCH1,GFI1B,GNG2,GULP1,HCK,HLA-F,HLA-G,HMGA2,HMGB2,HSPA1A/HSPA1B,HSPB1,HYAL2,ID1,ID3,IER3,IFI6,IFIH1,INHBA,IRF2,IRF7,ISG15,ITGA2,JAK3,JUN,JUNB,JUND,KLF2,KLF4,KLF6,KLF7,KLF9,LAG3,LGALS3BP,LIG4,LMNA,LMO4,LTBP1,MCAM,MCL1,MCOLN1,MIAT,MMP2,MMP7,MOB3C,MX1,MYC,NAMPT,NHLRC1,NOG,NR4A1,NR4A3,OAS1,OAS2,OAS3,PARP14,PCDHGB6,PER1,PHLDA1,PIK3AP1,PIK3IP1,PLA2G4A,PLA2R1,PLSCR1,PPP2R1B,PRDM5,PRKAA2,PRKCD,PROCR,PTAFR,PTGS2,RARG,RASSF3,REN,RHOB,RIOX1,RPL7,RPL9,RPS7,SDC4,SGK1,SIK1/SIK1B,SLC1A3,SLC39A6,SMCO4,SNX33,SOCS1,SOCS3,SPRY2,STAT1,STAT2,STK17B,TAL1,TCF7,TJP2,TK1,TLE4,TNFAIP8L2,TNFRSF1A,TNFSF10,TNFSF13B,TP53INP1,TRIM2,TRIM25,TSC22D3,TXNIP,USP18,XAF1,XRCC1,ZBTB7A,ZFP36,ZMAT3,ZNF346 |
| Cell Death and Survival | Cell death of connective tissue cells | 0.00000186 | Decreased | -2.368 | ADM,BAK1,BMPR1B,BTRC,CDKN2C,CLU,COL2A1,CSF1R,CXCL8,DDX58,DUSP1,EGR1,EGR2,EIF2AK2,FAS,FN1,GNG2,HYAL2,ID1,IER3,IFIH1,IRF7,JUN,JUND,KLF2,KLF6,LMNA,MCL1,MYC,NOG,NR4A1,PLA2G4A,PPP2R1B,PRKAA2,PRKCD,RHOB,SDC4,SGK1,STAT1,STK17B,TNFRSF1A,TNFSF10,TP53INP1,TXNIP,USP18,ZNF346 |
| Cell Death and Survival | Apoptosis of T lymphocytes | 7.83E-09 | Decreased | -2.394 | ADAM8,BAK1,C3AR1,CCDC86,CCL3,CCL5,CD3E,CXCR4,DHX58,EGR1,EGR3,FAS,FOS,HLA-G,ID3,IER3,JAK3,JUN,KLF2,MYC,NR4A1,NR4A3,SOCS1,STAT1,STK17B,TLE4,TNFAIP8L2,TNFRSF1A,TNFSF10,TNFSF13B,TSC22D3 |
| Cell Death and Survival | Apoptosis of lymphoblasts | 0.0000094 | Decreased | -2.425 | CCL3,CCL5,CD3E,EIF2AK2,FAS,MYC |
| Cell Death and Survival | Apoptosis of thymocytes | 0.0000145 | Decreased | -2.43 | ADAM8,BAK1,CD3E,EGR1,EGR3,FAS,FOS,ID3,JAK3,JUN,MYC,NR4A1,TLE4,TNFRSF1A,TNFSF10 |
| Cellular Movement | Cell movement | 1.76E-13 | Decreased | -2.475 | ABCG1,ACTA2,ADAM19,ADAM8,ADM,AQP3,AREG,ARHGAP18,ARID5B,ARL4C,AXL,AZU1,BAG3,BATF,BTG2,C2,C3AR1,C5orf30,CARMIL1,CCL3,CCL3L3,CCL5,CCR1,CD3E,CD48,CD69,CDKN2C,CEACAM4,CFH,CLCN3,CLDN7,CLU,COL2A1,CRIP2,CSF1R,CSF2RB,CXCL3,CXCL8,CXCR4,DDX58,DIP2A,DUSP1,DUSP3,DUSP6,EFNA4,EGR1,EGR2,EGR3,EIF2AK2,ELANE,ENAH,ENPP2,ENTPD1,EREG,ESRRA,F13A1,FAS,FBXO45,FERMT1,FN1,FOS,FUT10,FUT7,FYB1,GBP1,GNAI1,HCK,HLA-G,HMGA2,HMGB2,HOMER3,HSPA1A/HSPA1B,HSPB1,HYAL2,ID1,ID3,IER2,IFIT2,IFITM1,IFITM3,IL10RA,INHBA,IRF7,ITGA2,ITGB8,JAK3,JUN,JUNB,JUND,KCNE3,KCNN4,KLF2,KLF4,KLF6,LAG3,LGALS3BP,LMNA,LMO4,MAP3K21,MARCKS,MCAM,MCL1,MIAT,MMP2,MMP7,MT-ND1,MX1,MYADM,MYC,NAMPT,NAV1,NEXN,NINJ1,NKX3-1,NOD2,NOG,NR0B1,NR4A1,PARD6B,PARP9,PER1,PHLDA1,PIEZO2,PIGF,PIK3IP1,PLA2G4A,PLA2R1,PPFIA1,PRKAA2,PRKCD,PROCR,PTAFR,PTGIR,PTGS2,RARG,RASGRP3,REN,RGCC,RGS1,RHOB,RHOU,SCN9A,SDC4,SGK1,SLC12A2,SLC1A3,SLC7A6,SOCS1,SOCS3,SPON2,SPRY2,SSH1,STAT1,STAT2,STXBP4,TM4SF1,TMEM102,TNFAIP8L2,TNFRSF1A,TNFSF10,TNFSF13B,TP53INP1,TSC22D3,USP18,ZBTB7A |
| Cell Death and Survival,Organismal Injury and Abnormalities | Apoptosis of vascular endothelial cells | 0.00000407 | Decreased | -2.528 | ADM,ARHGAP18,AXL,CXCL8,DUSP6,FAS,ID1,KLF4,MCAM,MCL1,MMP2,NR4A3,TNFRSF1A,TNFSF10 |
| Cell Death and Survival,Endocrine System Disorders,Gastrointestinal Disease,Organismal Injury and Abnormalities | Cell death of pancreatic cells | 0.0000184 | Decreased | -2.561 | CXCR4,FAS,JUNB,MCL1,MYC,PRKCD,SOCS1,SOCS3,STAT1,TCF7,TNFRSF1A,TXNIP,USP18 |
| Cell Death and Survival | Cell death of thymocytes | 0.0000136 | Decreased | -2.568 | ADAM8,BAK1,CD3E,EGR1,EGR3,FAS,FOS,ID3,JAK3,JUN,MCL1,MYC,NR4A1,TLE4,TNFRSF1A,TNFSF10 |
| Cellular Movement | Migration of cells | 4.24E-14 | Decreased | -2.648 | ABCG1,ACTA2,ADAM19,ADAM8,ADM,AQP3,AREG,ARHGAP18,ARID5B,ARL4C,AXL,AZU1,BAG3,BATF,BTG2,C2,C3AR1,C5orf30,CARMIL1,CCL3,CCL3L3,CCL5,CCR1,CD3E,CD48,CD69,CDKN2C,CFH,CLCN3,CLDN7,CLU,CRIP2,CSF1R,CSF2RB,CXCL3,CXCL8,CXCR4,DDX58,DIP2A,DUSP1,DUSP3,DUSP6,EGR1,EGR2,EGR3,EIF2AK2,ELANE,ENAH,ENPP2,EREG,ESRRA,F13A1,FAS,FBXO45,FERMT1,FN1,FOS,FUT10,FUT7,FYB1,GBP1,GNAI1,HCK,HLA-G,HMGA2,HSPA1A/HSPA1B,HSPB1,HYAL2,ID1,ID3,IFIT2,IFITM1,IFITM3,IL10RA,INHBA,IRF7,ITGA2,ITGB8,JAK3,JUN,KCNE3,KCNN4,KLF2,KLF4,KLF6,LAG3,LGALS3BP,LMNA,LMO4,MAP3K21,MARCKS,MCAM,MCL1,MIAT,MMP2,MMP7,MT-ND1,MYADM,MYC,NAMPT,NAV1,NEXN,NINJ1,NKX3-1,NOD2,NOG,NR4A1,PARD6B,PARP9,PER1,PHLDA1,PIEZO2,PIGF,PIK3IP1,PLA2G4A,PLA2R1,PPFIA1,PRKAA2,PRKCD,PROCR,PTAFR,PTGIR,PTGS2,RARG,RASGRP3,REN,RGCC,RGS1,RHOB,RHOU,SCN9A,SDC4,SGK1,SLC12A2,SLC1A3,SLC7A6,SOCS1,SOCS3,SPON2,SPRY2,SSH1,STAT1,STAT2,STXBP4,TM4SF1,TMEM102,TNFAIP8L2,TNFRSF1A,TNFSF10,TNFSF13B,TP53INP1,TSC22D3,USP18,ZBTB7A |
| Cell Death and Survival | Apoptosis of hematopoietic progenitor cells | 0.0000021 | Decreased | -2.699 | ADAM8,BAK1,CCL3,CCL5,CD3E,EGR1,EGR3,EIF2AK2,ELANE,FAS,FOS,GFI1B,ID3,JAK3,JUN,KLF2,MYC,NR4A1,TLE4,TNFRSF1A,TNFSF10,TSC22D3 |

| | | | | | |
|---|---|---|---|---|---|
| Cell Death and Survival,Endocrine System Disorders,Gastrointestinal Disease,Organismal Injury and Abnormalities | Necrosis of pancreas | 0.00000903 | Decreased | -2.726 | CXCR4,FAS,JUNB,MCL1,MYC,PRKCD,PTGS2,SOCS1,SOCS3,STAT1,TCF7,TNFRSF1A,TXNIP,USP18 |
| Cell Death and Survival | Apoptosis of hematopoietic cells | 0.00000126 | Decreased | -2.746 | ADAM8,BAK1,CCL3,CCL3L3,CCL5,CD3E,EGR1,EGR3,EIF2AK2,ELANE,FAS,FOS,GFI1B,ID3,JAK3,JUN,KLF2,MYC,NR4A1,TLE4,TNFRSF1A,TNFSF10,TSC22D3 |
| Cell Death and Survival | Cell death of hematopoietic progenitor cells | 0.00000314 | Decreased | -2.848 | ADAM8,BAK1,CCL3,CCL5,CD3E,EGR1,EGR3,EIF2AK2,ELANE,FAS,FOS,GFI1B,ID3,JAK3,JUN,KLF2,MCL1,MYC,NR4A1,TLE4,TNFRSF1A,TNFSF10,TSC22D3 |
| Cell Death and Survival | Cell death of hematopoietic cells | 0.00000189 | Decreased | -2.893 | ADAM8,BAK1,CCL3,CCL3L3,CCL5,CD3E,EGR1,EGR3,EIF2AK2,ELANE,FAS,FOS,GFI1B,ID3,JAK3,JUN,KLF2,MCL1,MYC,NR4A1,TLE4,TNFRSF1A,TNFSF10,TSC22D3 |
| Cell Death and Survival | Cell death of pancreas | 0.00000374 | Decreased | -2.921 | CXCR4,FAS,JUNB,MCL1,MYC,NR4A1,PRKCD,PTGS2,SOCS1,SOCS3,STAT1,TCF7,TNFRSF1A,TXNIP,USP18 |
| Inflammatory Response | Immune response of cells | 4.27E-10 | Decreased | -2.926 | AXL,AZU1,BAK1,C3AR1,CCL3,CD48,CD69,CEACAM4,CFH,CLCN3,CSF1R,CSF2RB,CXCL3,CXCL8,DDX58,DHX58,EIF2AK2,ELANE,FAS,FN1,GULP1,HCK,HSPB1,IFIH1,IL10RA,IRF2,IRF2BP1,IRF7,ISG15,JUN,JUNB,KLF2,LTBP1,MCL1,MYC,NAMPT,NOD2,PER1,PIP4P2,PLA2G4A,PRKAA2,PRKCD,RGCC,SCARF1,SGK1,SOCS1,SOCS3,SPRY2,STAT1,TAP1,TLE3,TNFRSF1A,TNFSF10,TRIM25,TRIM58,UBE2L6,XRCC1 |
| Antimicrobial Response,Inflammatory Response | Antimicrobial response | 1.04E-17 | Decreased | -3.234 | ACTA2,ADM,AZU1,BAK1,CCL5,CLU,CXCR4,DDX58,DDX60,DHX58,DTX3L,EIF2AK2,ELANE,HCK,HLA-G,HSPB1,HYAL2,IFI44,IFI44L,IFI6,IFIH1,IFIT1,IFIT2,IFIT3,IFITM1,IFITM3,IRF2,IRF7,ISG15,MMP7,MX1,MX2,NOD2,OAS1,OAS2,OAS3,OASL,PARP9,PLSCR1,PRKCD,PRSS2,RSAD2,RTP4,SAMHD1,SOCS1,SOCS3,SPON2,SPRY2,STAT1,STAT2,TNFRSF1A,TRIM22 |
| Antimicrobial Response,Inflammatory Response | Antiviral response | 1.03E-19 | Decreased | -3.234 | ACTA2,AZU1,CCL5,CLU,CXCR4,DDX58,DDX60,DHX58,DTX3L,EIF2AK2,HSPB1,HYAL2,IFI44,IFI44L,IFI6,IFIH1,IFIT1,IFIT2,IFIT3,IFITM1,IFITM3,IRF2,IRF7,ISG15,MX1,MX2,OAS1,OAS2,OAS3,OASL,PARP9,PLSCR1,RSAD2,RTP4,SAMHD1,SOCS1,SOCS3,SPON2,SPRY2,STAT1,STAT2,TRIM22 |

**T3 vs. T1 MEP_1**

© 2000-2020 QIAGEN. All rights reserved.

| Categories | Diseases or Functions Annotation | p-value | Predicted Activation State | Activation z-score | Molecules |
|---|---|---|---|---|---|
| Organismal Survival | Organismal death | 8.46E-11 | Increased | 4.659 | ACTL6A,ACVR1,ADAM15,ADCYAP1,AGTR1,AHR,AMER1,ANGPT2,ANKLE1,APOE,AREG,ARID3A,ATF1,B3GALNT1,BACE2,BCL3,BHLHE40,BMPR2,BPNT1,C1GALT1C1,C3AR1,C5,CA5B,CALR,CASP1,CASP7,CASP9,CCDC86,CCND1,CCND3,CD274,CD36,CD4,CD44,CD74,CDCP1,CDKN2C,CDKN3,CELF2,CENPE,CLIP3,COL5A1,CRNDE,CSF1,CSGALNACT1,CXCL8,CXCR4,DDX58,DHFR,DHRS3,DHX58,DLC1,DLK1,DOLK,DPAGT1,DPYD,DUSP1,EGR2,EGR3,EIF2AK2,ENAH,ENPP2,EPX,EZH2,FLT3,FOS,FOSL1,FOXC2,FOXM1,GAD1,GAN,GFI1B,GMNN,GNAI1,GOLGA2,GPD1L,GPR161,GPR183,H2AC18/H2AC19,HCK,HDAC7,HGF,HHEX,HK2,HLA-G,HLTF,HLX,HOPX,HPGD,HSD3B7,HSPB1,HTRA2,HYAL2,ICMT,ID1,IDS,IFIH1,IFIT2,IKZF3,IQGAP2,IRAK3,IRF1,IRF2,IRF7,ISG15,ITGB3,ITGB8,IVNS1ABP,JUN,JUNB,KEAP1,KIF3A,KIF3B,KIFC1,KLF1,KLF2,KLF4,KLF6,KLF9,KRT1,LGR4,LIG4,LIN7A,LMNA,LMNB1,LMO4,LTBP1,LY6E,MACROH2A2,MAFF,MAL,MAN2A1,MCL1,MDFI,MDK,MED24,MEIS2,MITF,MPG,MT-ND1,MYZAP,NEAT1,NFE2,NFE2L3,NINJ1,NOC3L,NOTCH2,NR4A2,NR4A3,NTAN1,NUS1,NUSAP1,PBRM1,PCYT2,PER2,PEX11B,PIEZO2,PIGF,PMEPA1,PMP22,PRDX1,PRKCD,PRMT5,PRNP,PTAFR,PTGS2,PTK2,PTPRN2,PYGO2,RAB10,RBBP8,REN,RGS14,RHEBL1,RIMS1,RNASEL,SERPINE1,SFRP5,SLC9A3R1,SMAD1,SMTN,SNAI3,SOCS1,SOCS3,SOD2,SOSTDC1,SPRED1,SPRY2,SRGAP3,STAT1,STAT2,SYNE2,SYT5,TAL1,TARBP2,TBC1D4,TFAP4,TFPI,TICAM1,TIPARP,TLE4,TNFSF10,TNFSF13B,TOP1,TP53,TP53INP1,TRIM21,TRPS1,TXNIP,TYMP,UBIAD1,UHRF1,UNC93B1,UNG,USP18,VEZF1,VTA1,VTI1B,WNT5B,ZFP36,ZFP36L2,ZNF521 |
| Organismal Survival | Morbidity or mortality | 1.91E-11 | Increased | 4.475 | ACTL6A,ACVR1,ADAM15,ADCYAP1,AGTR1,AHR,AMER1,ANGPT2,ANKLE1,APOE,AREG,ARID3A,ATF1,B3GALNT1,BACE2,BCL3,BHLHE40,BMPR2,BPNT1,C1GALT1C1,C3AR1,C5,CA5B,CALR,CASP1,CASP7,CASP9,CCDC86,CCND1,CCND3,CD274,CD36,CD4,CD44,CD74,CDCP1,CDKN2C,CDKN3,CELF2,CENPE,CLIP3,COL5A1,CRNDE,CSF1,CSGALNACT1,CXCL8,CXCR4,DDX58,DHFR,DHRS3,DHX58,DLC1,DLK1,DOLK,DPAGT1,DPYD,DUSP1,EGR2,EGR3,EIF2AK2,ENAH,ENPP2,EPX,EZH2,FLT3,FOS,FOSL1,FOXC2,FOXM1,GAD1,GAN,GFI1B,GMNN,GNAI1,GOLGA2,GPD1L,GPR161,GPR183,H2AC18/H2AC19,HCK,HDAC7,HECTD3,HGF,HHEX,HK2,HLA-G,HLTF,HLX,HOPX,HPGD,HSD3B7,HSPB1,HTRA2,HYAL2,ICMT,ID1,IDS,IFIH1,IFIT2,IKZF3,IQGAP2,IRAK3,IRF1,IRF2,IRF7,ISG15,ITGB3,ITGB8,IVNS1ABP,JUN,JUNB,KEAP1,KIF3A,KIF3B,KIFC1,KLF1,KLF2,KLF4,KLF6,KLF9,KRT1,LGR4,LIG4,LIN7A,LMNA,LMNB1,LMO4,LTBP1,LY6E,MACROH2A2,MAFF,MAL,MAN2A1,MCL1,MDFI,MDK,MED24,MEIS2,MITF,MPG,MT-ND1,MYZAP,NEAT1,NFE2,NFE2L3,NINJ1,NOC3L,NOTCH2,NR4A2,NR4A3,NTAN1,NUS1,NUSAP1,OASL,PBRM1,PCYT2,PER2,PEX11B,PIEZO2,PIGF,PMEPA1,PMP22,PRDX1,PRKCD,PRMT5,PRNP,PTAFR,PTGS2,PTK2,PTPRN2,PYGO2,RAB10,RBBP8,REN,RGS14,RHEBL1,RIMS1,RNASEL,SCUBE1,SERPINE1,SFRP5,SLC9A3R1,SMAD1,SMTN,SNAI3,SOCS1,SOCS3,SOD2,SOSTDC1,SPRED1,SPRY2,SRGAP3,STAT1,STAT2,SYNE2,SYT5,TAL1,TAP1,TARBP2,TBC1D4,TFAP4,TFPI,TICAM1,TIPARP,TLE4,TNFSF10,TNFSF13B,TOP1,TP53,TP53INP1,TRIM21,TRIM27,TRPS1,TXNIP,TYMP,UBIAD1,UHRF1,UNC93B1,UNG,USP18,VEZF1,VTA1,VTI1B,WNT5B,ZFP36,ZFP36L2,ZNF521 |
| Cell Signaling | Replication of viral replicon | 6.27E-11 | Increased | 4.067 | APOBEC3A,APOBEC3C,BST2,EIF2AK2,IFIT1,IFITM1,IFITM3,ISG15,MX1,OAS1,OAS3,OASL,PLSCR1,PTGS2,RNASEL,RSAD2,TRIM6 |
| Cell Signaling | Viral life cycle | 3.16E-08 | Increased | 4.066 | APOBEC3A,APOBEC3C,BST2,CXCR4,EIF2AK2,IFIT1,IFITM1,IFITM3,ISG15,MX1,NUP50,OAS1,OAS3,OASL,PLSCR1,PTGS2,RNASEL,RSAD2,TP53,TRIM6,VPS37C,VTA1 |
| Infectious Diseases | Replication of vesicular stomatitis virus | 0.00000138 | Increased | 3.323 | APOBEC3C,DDX58,IFIH1,IFITM3,IRF1,LY6E,OAS1,OASL,SLFN12,SOCS1,SP110,STAT2,TAP1,TP53 |
| Infectious Diseases | Replication of virus | 8.03E-11 | Increased | 2.862 | AHR,APOBEC3C,AREG,ARTN,BCL3,BST2,CD4,CIITA,CXCL8,CXCR4,DCLK2,DDX58,DUSP1,EGR1,EIF2AK2,EZH2,HCK,HERC5,HPGD,IFI6,IFIH1,IFIT1,IFITM1,IFITM3,IRF1,IRF2,IRF7,ISG15,JUN,LY6E,MAN2B1,MOV10,MX1,MX2,NLRC5,OAS1,OAS3,OASL,PRDX1,PRKCD,PRNP,PTGS2,PVALB,RAB11FIP1,RBBP8,RNASEL,RSAD2,SAMHD1,SGK1,SLC9A3R1,SLFN12,SOCS1,SOCS2,SOCS3,SP110,SPRY2,STAT1,STAT2,TAP1,TICAM1,TNFSF10,TP53,TRERF1,TRIM14,TRIM21,TRIM5,TXNIP,UBE2L6 |
| Infectious Diseases | Infection of mammalia | 9.44E-09 | Increased | 2.761 | AHR,APOE,BCL3,CASP1,CD274,CD36,CD4,CD44,CD74,CIITA,CSF2RB,CXCR4,DDX58,DHX58,DUSP1,EIF2AK2,FOS,HCK,HECTD3,HLA-G,IFIH1,IFIT2,IRF1,IRF7,ISG15,ITGAX,NFE2,OASL,PRKCD,PRNP,PTAFR,RNASEL,SAMHD1,SERPINE1,SPON2,STAT1,STAT2,TAP1,TICAM1,TNFSF10,TPSAB1/TPSB2,UNC93B1,USP18 |
| Infectious Diseases | Replication of RNA virus | 1.5E-09 | Increased | 2.737 | APOBEC3C,AREG,ARTN,BCL3,BST2,CD4,CIITA,CXCL8,CXCR4,DCLK2,DDX58,EIF2AK2,EZH2,HCK,HERC5,HPGD,IFI6,IFIH1,IFIT1,IFITM1,IFITM3,IRF1,IRF2,IRF7,ISG15,JUN,LY6E,MAN2B1,MOV10,MX1,NLRC5,OAS1,OAS3,OASL,PRDX1,PRKCD,PRNP,PTGS2,PVALB,RAB11FIP1,RNASEL,RSAD2,SGK1,SLC9A3R1,SLFN12,SOCS1,SOCS2,SOCS3,SP110,STAT1,STAT2,TAP1,TNFSF10,TP53,TRERF1,TRIM14,TRIM21,TRIM5,TXNIP,UBE2L6 |
| Infectious Diseases | Replication of Herpesviridae | 0.000000191 | Increased | 2.703 | AHR,DDX58,EGR1,IFIH1,IRF1,MX2,OAS1,OASL,RNASEL,RSAD2,SAMHD1,SPRY2,STAT1,STAT2,TICAM1 |
| Cancer,Organismal Injury and Abnormalities | Development of carcinoma | 9.27E-11 | Increased | 2.409 | AASDH,AASDHPPT,ABHD12B,ACOT11,ACSL4,ACTN1,ACVR1,AGPAT5,AHR,AKR1C1/AKR1C2,ALAS1,ALDH1B1,ALDH4A1,ALDH6A1,ALKBH8,AMER1,ANGPT2,ANKRD28,APOBEC3A,APOE,APOL2,APPL2,AREG,ARHGEF26,ARID3A,ARL15,ARL4C,ARMC1,ARRDC3,ARSK,ASRGL1,ATF7IP2,ATG4D,ATP8B3,B3GALNT1,BACE2,BATF,BCAM,BCL3,BHLHE40,BMPR2,BOD1,BST1,BTN3A1,BTN3A2,C11orf24,C12orf29,C14orf119,C2,C3AR1,C5,CARD6,CASP1,CASP7,CAVIN2,CBX6,CCDC141,CCDC169,CCND1,CCND3,CD274,CD36,CD4,CD44,CD69,CDCP1,CDH26,CDKN2C,CELF2,CENPE,CEP72,CETN3,CGNL1,CHD7,CHST2,CIB2,CIITA,CLIP3,CLU,CMIP,CNR2,COA7,COG2,COL5A1,CR1,CRELD2,CRHBP,CRNDE,CRYZ,CSF1,CSF2RB,CSGALNACT1,CSTF2,CTTNBP2,CTU1,CXCL8,CXXC5,CYP1B1,CYSLTR1,DCAF4,DCAF7,DCP1B,DDX58,DDX59,DDX60,DEDD2,DENND2C,DGKG,DGKH,DHFR,DHX33,DLC1,DLK1,DOCK4,DOLPP1,DPAGT1,DPF1,DPP10,DPYD,DSE,DTX3L,DUSP1,DUSP6,DYTN,EBF3,ECM2,EDEM3,EFNA4,EGR1,EGR2,EGR3,EIF2AK2,EIF4E3,ELP4,ENAH,ENO2,ENPP2,ENPP5,EPDR1,EREG,ESRRA,EXOSC4,EXPH5,EZH2,FAM124B,FAM200A,FAM217B,FAM83D,FBXO21,FIGNL1,FKBP10,FKBP5,FLT3,FMO4,FOS,FOXC2,FOXM1,FYB1,FZD2,GAD1,GAN,GART,GAS2L1,GCNT1,GCSAML,GLOD5,GLYCTK,GMNN,GMPPB,GMPR,GNB1L,GOLGA2,GOT2,GPR161,GPR183,GPR27,GPR65,GSTM2,GSTM5,GULP1,GXYLT1,H2AC18/H2AC19,H2BC5,HACD2,HCK,HDAC9,HECTD3,HELZ2,HERC5,HGF,HILPDA,HLA-B,HLA-C,HLA-DPB1,HLA-F,HLA-G,HLX,HNRNPF,HOTAIRM1,HPDL,HPGD,HSPB1,HTRA2,HYAL2,ICMT,IFI44,IFI44L,IFIH1,IFIT3,IFITM1,IGLL1/IGLL5,IGSF6,IKZF3,IKZF4,IL1RAP,IL1RAPL1,IL7,IL7R,INPP4A,INTS2,INTS7,IQGAP2,IRAK3,IRF2,IRF2BP1,IRF7,ITGAX,ITGB3,ITGB8,ITPRIPL2,IVNS1ABP,JAKMIP2,JUN,JUNB,KCNA3,KCNK6,KEAP1,KIF14,KIF21A,KIF3A,KIF3B,KLF4,KLF6,KLHDC7B,KPNA2,KPTN,KRT1,KYNU,LBR,LGR4,LIN7A,LMBRD2,LMOD3,LRRC4,LRRC75B,LRRC8D,LTBP1,LY6E,LYAR,MAL,MAN2A1,MAN2B1,MCL1,MED24,MEIS2,MFSD14B,MFSD4A,MGA,MITF,MLEC,MPG,MR1,MSI2,MTA3,MTHFS,MTMR12,MX1,NAV1,NDUFS7,NEURL1,NFE2,NFE2L3,NHLRC1,NHSL2,NLRC5,NOC3L,NOTCH2,NR4A1,NR4A2,NR4A3,NTAN1,NUP210L,NUP50,NUSAP1,OAS1,OAS2,OASL,OLFM3,P2RY14,PAG1,PANK4,PAQR7,PAQR8,PARP14,PARVG,PBRM1,PCGF6,PCNX1,PCYT2,PDE3B,PDIA4,PDP2,PER1,PER2,PEX11G,PGBD2,PHACTR3,PIEZO2,PIK3C2B,PIK3R3,PIM3,PLEKHG1,PLSCR1,PLXNC1,PMEPA1,PMP22,POP1,PPIL1,PPP1R18,PPP1R1B,PRDX1,PREP,PRKCD,PRKG2,PRMT5,PRUNE1,PTAFR,PTGS2,PTK2,PTPRN2,PYGL,PYGO2,QSER1,RAB9B,RADX,RASA2,RASAL2,RASGRP3,RASGRP4,RASSF3,RBBP8,REN,REXO5,RGCC,RGS14,RHOB,RIMS1,RIN2,RMI1,RNASEL,RNF111,RNF34,RNPEP,RSAD2,RSPH3,RUNDC3B,RXFP1,SAMD9,SAMD9L,SASH3,SCAMP3,SCAMP4,SCARF1,SCUBE1,SELL,SERPINE1,SESN1,SGK1,SHQ1,SIGLEC12,SIK1/SIK1B,SLA,SLC10A7,SLC35B2,SLC38A1,SLC43A3,SLC44A1,SLC45A3,SLC52A3,SLC5A6,SLC7A6,SLC9A3R1,SLCO5A1,SLFN12,SLFN5,SMAP2,SMO,SMOX,SMTN,SNX18,SNX20,SOCS1,SOCS2,SOCS3,SOD2,SORBS1,SORT1,SP110,SPAG1,SPC25,SPDL1,SPIRE1,SPNS1,SPON2,SPRED1,SPTSSA,SRGAP3,ST6GAL2,STAT1,STAT2,STK17B,STT3A,SYNE2,SYNJ2,SYTL2,TACC1,TAGAP,TAL1,TANGO2,TAP1,TARBP2,TBC1D4,TESC,TEX15,TFEC,TICAM1,TIGD2,TIPARP,TLE4,TLNRD1,TM4SF1,TMEM102,TMEM246,TMEM51,TMX3,TNFSF10,TOP1,TP53,TRAF3IP3,TRANK1,TREML2,TRERF1,TRIB2,TRIM11,TRIM35,TRIM37,TRIM5,TRIM6,TRIP11,TRPS1,TSC22D3,TSPAN4,TTC27,TTLL7,TXNIP,TYMP,TYW1,UNC93B1,UNG,USP18,USP20,USP32,USP37,VANGL1,VEZF1,VMP1,VTI1B,WDFY3,WDR78,WNT5B,ZC3H4,ZC3H6,ZFP36,ZFP36L2,ZFP64,ZMAT5,ZNF146,ZNF331,ZNF420,ZNF462,ZNF470,ZNF521,ZNF610,ZNF616,ZNF623,ZNF653,ZNF681,ZNF74,ZNF787,ZNF799,ZNF813,ZNF835 |
| Cancer,Organismal Injury and Abnormalities,Reproductive System Disease | Mammary tumor | 0.000000882 | Increased | 2.37 | ACAA1,ADAM15,ADCYAP1,AKR1C1/AKR1C2,ALDH1B1,ANGPT2,APOE,AREG,ARRDC3,BCAM,BHLHE40,BMPR2,BTG2,C1GALT1C1,C2,C3AR1,CASP9,CCDC175,CCND1,CCND3,CD274,CD4,CD44,CD69,CDCP1,CDK2AP2,CDKN2C,CEP72,CHST14,CLU,CMPK2,COG2,CRNDE,CSF1,CSF2RB,CTU1,CXCL8,CXCR4,CXXC5,DEPTOR,DGKG,DHFR,DHX58,DLC1,DOCK4,DPYD,DSE,DUSP1,EGR1,EGR3,EIF2AK2,ENAH,EPDR1,EPSTI1,ESRRA,EZH2,FAM227A,FKBP5,FLT3,FOS,FOSL1,FOXC2,FOXM1,FUT10,GAS8,GCSAML,GMNN,GPR183,GPR65,GSTM2,GSTM3,GSTM5,GULP1,HBA1/HBA2,HCK,HERC5,HGF,HIBADH,HLA-B,HLA-DMB,HLA-DRB5,HLA-G,HSPB1,HTRA2,ID1,IFIT1,IL1RAPL1,IQGAP2,IRF1,IRF7,ITGAX,ITGB3,JUN,JUNB,JUND,KAZN,KEAP1,KIF14,KLF4,KLF6,KLHL12,KPNA2,KRT1,LGALS3BP,LMNA,LY6E,MAFF,MATN2,MCL1,MGA,MOV10,MT-ND1,MTMR12,NEAT1,NEK4,NEURL1,NFE2L3,NOTCH2,NR4A1,NR4A2,NR4A3,NUSAP1,OAF,PAQR8,PBRM1,PGAM1,PHACTR3,PIEZO2,PIK3R3,PLEKHG1,PPP1R1B,PRDX1,PRKCD,PTGS2,PTK2,RAB10,RAB11FIP1,RASAL2,RGS1,RGS2,RHOB,RIN2,SAMD9,SAMD9L,SELL,SERPINE1,SFRP5,SHQ1,SIK1/SIK1B,SLC39A6,SLC43A3,SLC44A1,SLC9A3R1,SMTN,SNX18,SOD2,SOSTDC1,SSPN,STAT1,STK17B,SYNJ2,SYTL2,TAL1,TCTN2,TEX15,TFPI,TLE3,TM4SF1,TNFSF10,TOP1,TOR1B,TP53,TPSAB1/TPSB2,TRERF1,TRIM37,TRIM5,TRIM69,TRPS1,TTLL7,TYMP,UHRF1,USP32,VEZF1,VTI1B,VWA5A,WDFY3,WNT5B,ZC3H4,ZFP36,ZFP36L2,ZFP64,ZNF420,ZNF425,ZNF462,ZNF835 |
| Cancer,Organismal Injury and Abnormalities | Genitourinary tumor | 1.69E-18 | Increased | 2.334 | AASDHPPT,ABHD12B,ACAA1,ACSL4,ACTN1,ACVR1,ADAM15,ADCYAP1,ADPRHL2,AGPAT5,AGTR1,AHR,AKR1C1/AKR1C2,ALAS1,ALDH1B1,ALDH4A1,ALDH9A1,ALKBH8,AMER1,AMPD3,ANGPT2,ANKRD28,APOBEC3A,APOE,APOL2,APOL3,APOL4,AREG,ARHGEF26,ARID3A,ARL15,ARL17A/ARL17B,ARL4C,ARL4D,ARRDC3,ARSK,ASCL2,ASRGL1,ATF7IP2,ATP8B3,B3GALNT1,BCAM,BEND6,BHLHE40,BMPR2,BST1,BTG2,BTN3A1,BTN3A2,C12orf29,C18orf54,C1GALT1C1,C1R,C2,C3AR1,C5,CALR,CARD6,CASP1,CASP7,CASP9,CAVIN2,CCDC141,CCDC169,CCDC175,CCDC184,CCND1,CCND3,CD274,CD36,CD4,CD44,CD69,CD74,CDC42EP4,CDCP1,CDK2AP2,CDKN2C,CDKN3,CELF2,CENPE,CEP72,CETN3,CGNL1,CHD7,CHST14,CHST2,CIB2,CIITA,CLIP3,CLU,CMIP,CMPK2,CNKSR1,CNR2,COG2,COL5A1,CR1,CRHBP,CRNDE,CSF1,CSF2RB,CSGALNACT1,CSTF2,CTSH,CTTNBP2,CTU1,CXCL8,CXCR4,CXXC5,CYP1B1,CYSLTR1,DCAF4,DCP1B,DDX58,DDX60,DEPTOR,DGKG,DGKH,DHFR,DHRS9,DHX58,DLC1,DLK1,DOCK4,DOLK,DOLPP1,DPAGT1,DPF1,DPP10,DPYD,DSE,DTX3L,DUSP1,DUSP6,DYTN,EBF3,ECM2,EDEM3,EEF1AKNMT,EFNA4,EGR1,EGR2,EGR3,EIF2AK2,EME2,ENAH,ENO2,ENPP2,ENPP5,EPDR1,EPSTI1,EREG,ESRRA,EXOSC4,EXPH5,EZH2,FAM117A,FAM124B,FAM217B,FAM227A,FAM3B,FAM83D,FAM89A,FBXO21,FIGNL1,FKBP10,FKBP5,FLT3,FMO4,FOS,FOSL1,FOXC2,FOXK1,FOXM1,FUT10,FYB1,FZD2,GALK2,GAN,GART,GAS8,GCNT1,GCSAML,GDF10,GFI1B,GLYCTK,GMNN,GMPPB,GNAI1,GNB1L,GOLGA2,GOT2,GPR161,GPR183,GPR65,GRWD1,GSTM2,GSTM3,GSTM5,GULP1,GXYLT1,H2AC18/H2AC19,H2BC5,HBA1/HBA2,HCK,HDAC7,HDAC9,HELZ2,HERC5,HERC6,HGF,HIBADH,HIF3A,HILPDA,HLA-B,HLA-C,HLA-DMB,HLA-DRB5,HLA-G,HLTF,HLX,HNRNPF,HOPX,HOTAIRM1,HPDL,HSPA1A/HSPA1B,HSPB1,HTRA2,HYAL2,ICMT,ID1,IDS,IFI44,IFI44L,IFIH1,IFIT1,IFIT2,IFIT3,IFITM1,IGLL1/IGLL5,IGSF6,IKZF3,IL1RAP,IL1RAPL1,IL7,IL7R,INPP4A,INTS2,INTS7,IQGAP2,IRF1,IRF2,IRF2BP1,IRF7,ISG15,ITGAX,ITGB3,ITGB8,ITPRIPL1,IVNS1ABP,JAKMIP2,JUN,JUNB,JUND,KAZN,KCNA3,KCNK6,KEAP1,KIF14,KIF21A,KIF3A,KIFC1,KLF2,KLF4,KLF6,KLF9,KLHL12,KPNA2,KPTN,KRT1,KYNU,LGALS3BP,LGR4,LMBRD2,LMNA,LMOD3,LRRC27,LRRC75B,LTBP1,LY6E,LYAR,MAFF,MAL,MAN2A1,MAN2B1,MARS2,MATN2,MCL1,MED24,MEIS2,MFSD14B,MGA,MIPEP,MITF,MOV10,MPG,MR1,MT-ND1,MTMR12,MX1,MX2,MYADM,MYL4,NAV1,NDUFS7,NEAT1,NEK4,NEURL1,NFE2,NFE2L3,NFKBIE,NHLRC1,NHSL2,NIBAN2,NINJ1,NKX3-1,NLRC5,NOC3L,NOTCH2,NR4A1,NR4A2,NR4A3,NUP210L,NUP50,NUSAP1,OAF,OAS2,OAS3,OASL,P2RY14,PAG1,PANK4,PAQR7,PAQR8,PARP14,PBRM1,PCLAF,PCNX1,PDE3B,PDIA4,PDP2,PDXDC2P-NPIPB14P,PER1,PER2,PEX11G,PGAM1,PGBD2,PHACTR3,PIEZO2,PIK3C2B,PIK3R3,PLEKHG1,PLSCR1,PLXNC1,PMP22,POP1,PPP1R1B,PRDX1,PREP,PRKCD,PRKG2,PTAFR,PTGS2,PTK2,PTPRN2,PYGL,QSER1,RAB10,RAB11FIP1,RAB9B,RADX,RAMP1,RASA2,RASAL2,RASGEF1B,RASGRP3,RASGRP4,RBBP8,REN,REXO5,RGCC,RGS1,RGS14,RGS2,RHOB,RIMS1,RIN2,RNASEL,RNF111,RNF24,RNF34,RNPEP,RPL9,RSPH3,RTKN,RTP4,RUNDC3B,RXFP1,SAMD9,SAMD9L,SAP130,SCAMP3,SCAMP4,SCUBE1,SELL,SERPINE1,SFRP5,SGK1,SHQ1,SIGLEC12,SIK1/SIK1B,SLA,SLC10A7,SLC29A1,SLC2A6,SLC38A1,SLC39A6,SLC43A3,SLC44A1,SLC45A3,SLC52A3,SLC5A6,SLC7A6,SLC9A3R1,SLCO5A1,SLFN12,SLFN5,SMAP2,SMO,SMOX,SMTN,SNX18,SNX20,SOCS1,SOCS2,SOCS3,SOD2,SORBS1,SORT1,SOSTDC1,SP110,SPAG1,SPC25,SPDL1,SPNS1,SPON2,SPRY2,SPRPB,SSPN,ST6GAL2,STAT1,STAT2,STK17B,SYNE2,SYNJ2,SYTL2,TACC1,TAGAP,TAL1,TAP1,TARBP2,TBC1D4,TCTN2,TESC,TEX15,TFAP4,TFPI,TIGD2,TIPARP,TLE3,TM4SF1,TMEM102,TMEM246,TMEM44,TMEM51,TMEM63B,TMX3,TNFSF10,TNFSF13B,TOP1,TOR1B,TP53,TP53INP1,TPRA1,TPSAB1/TPSB2,TRANK1,TREML2,TRERF1,TRIB2,TRIM11,TRIM21,TRIM37,TRIM5,TRIM6,TRIM69,TRIP11,TRPS1,TSC22D3,TSPAN4,TTC27,TTLL7,TXNIP,TYMP,TYW1,UBE2T,UHRF1,UNC93B1,USP18,USP20,USP32,USP37,VEZF1,VGLL4,VMP1,VTI1B,VWA5A,WDFY3,WDR78,WNT5B,XAF1,ZC3H4,ZC3H6,ZFP36,ZFP36L2,ZFP64,ZGLP1,ZMAT5,ZNF146,ZNF250,ZNF420,ZNF425,ZNF462,ZNF470,ZNF521,ZNF608,ZNF610,ZNF616,ZNF623,ZNF681,ZNF695,ZNF799,ZNF813,ZNF835 |
| Cancer,Organismal Injury and Abnormalities | Genitourinary carcinoma | 2.72E-13 | Increased | 2.218 | AASDHPPT,ABHD12B,ACSL4,ACTN1,ACVR1,ADCYAP1,ADPRHL2,AGPAT5,AHR,AKR1C1/AKR1C2,ALAS1,ALDH1B1,ALDH4A1,ALKBH8,AMER1,AMPD3,ANGPT2,ANKRD28,APOBEC3A,APOE,APOL2,APOL3,APOL4,ARHGEF26,ARID3A,ARL15,ARL4C,ARRDC3,ARSK,ASRGL1,ATF7IP2,ATP8B3,B3GALNT1,BEND6,BHLHE40,BMPR2,BST1,BTN3A1,BTN3A2,C12orf29,C18orf54,C1GALT1C1,C2,C3AR1,C5,CARD6,CASP1,CASP7,CASP9,CAVIN2,CCDC141,CCDC169,CCDC184,CCND1,CCND3,CD274,CD36,CD44,CDC42EP4,CDK2AP2,CDKN2C,CELF2,CENPE,CETN3,CGNL1,CHD7,CHST2,CIB2,CIITA,CLIP3,CLU,CMIP,CNKSR1,CNR2,COL5A1,CR1,CRHBP,CSF1,CSF2RB,CSGALNACT1,CSTF2,CTSH,CTTNBP2,CTU1,CXCL8,CXXC5,CYP1B1,CYSLTR1,DCAF4,DCP1B,DDX58,DDX60,DGKG,DGKH,DHFR,DHRS9,DHX58,DLC1,DLK1,DOCK4,DOLK,DOLPP1,DPAGT1,DPF1,DPP10,DPYD,DSE,DTX3L,DUSP1,DUSP6,DYTN,EBF3,ECM2,EDEM3,EEF1AKNMT,EFNA4,EGR2,EGR3,EIF2AK2,EME2,ENAH,ENO2,ENPP2,ENPP5,EPDR1,EPSTI1,EREG,ESRRA,EXOSC4,EXPH5,EZH2,FAM117A,FAM124B,FAM217B,FAM3B,FAM83D,FAM89A,FBXO21,FIGNL1,FKBP10,FLT3,FMO4,FOS,FOSL1,FOXK1,FOXM1,FYB1,FZD2,GAN,GART,GAS8,GCNT1,GCSAML,GDF10,GFI1B,GLYCTK,GMNN,GMPPB,GNB1L,GOLGA2,GOT2,GPR161,GPR65,GSTM2,GSTM5,GULP1,GXYLT1,H2AC18/H2AC19,H2BC5,HBA1/HBA2,HDAC7,HDAC9,HELZ2,HERC5,HERC6,HGF,HIF3A,HILPDA,HLA-B,HLA-C,HLA-DRB5,HLA-G,HLTF,HLX,HNRNPF,HPDL,HSPA1A/HSPA1B,HTRA2,HYAL2,ICMT,ID1,IFI44,IFI44L,IFIH1,IFIT2,IFIT3,IGLL1/IGLL5,IGSF6,IKZF3,IL1RAP,IL1RAPL1,IL7,INPP4A,INTS2,INTS7,IQGAP2,IRF2,IRF2BP1,IRF7,ISG15,ITGAX,ITGB3,ITGB8,IVNS1ABP,JAKMIP2,JUN,JUNB,KAZN,KCNA3,KCNK6,KEAP1,KIF14,KIF21A,KIF3A,KIFC1,KLF4,KLF6,KPNA2,KPTN,KRT1,KYNU,LGR4,LMBRD2,LMOD3,LRRC27,LRRC75B,LY6E,LYAR,MAL,MAN2A1,MAN2B1,MARS2,MATN2,MCL1,MED24,MEIS2,MFSD14B,MGA,MITF,MPG,MR1,MT-ND1,MTMR12,MX1,MX2,MYADM,MYL4,NAV1,NEK4,NFE2,NFE2L3,NFKBIE,NHLRC1,NHSL2,NIBAN2,NKX3-1,NLRC5,NOC3L,NOTCH2,NR4A2,NR4A3,NUP210L,NUP50,NUSAP1,OAS2,OASL,P2RY14,PAG1,PANK4,PAQR7,PARP14,PBRM1,PCNX1,PDE3B,PDIA4,PDP2,PER1,PER2,PEX11G,PGBD2,PIEZO2,PIK3C2B,PIK3R3,PLEKHG1,PLSCR1,PLXNC1,PMP22,POP1,PPP1R1B,PRDX1,PREP,PRKCD,PRKG2,PTAFR,PTGS2,PTK2,PTPRN2,PYGL,QSER1,RAB11FIP1,RAB9B,RADX,RAMP1,RASA2,RASAL2,RASGEF1B,RASGRP3,RASGRP4,RBBP8,REN,REXO5,RGCC,RGS1,RGS14,RGS2,RHOB,RIMS1,RNASEL,RNF111,RNF24,RNF34,RNPEP,RSPH3,RTKN,RTP4,RUNDC3B,RXFP1,SAMD9,SAMD9L,SAP130,SCAMP3,SCAMP4,SCUBE1,SELL,SERPINE1,SGK1,SHQ1,SIGLEC12,SIK1/SIK1B,SLA,SLC10A7,SLC29A1,SLC2A6,SLC38A1,SLC39A6,SLC43A3,SLC44A1,SLC45A3,SLC52A3,SLC5A6,SLC7A6,SLC9A3R1,SLCO5A1,SLFN12,SLFN5,SMAP2,SMO,SMOX,SMTN,SNX18,SNX20,SOCS1,SOCS2,SOCS3,SOD2,SORBS1,SORT1,SOSTDC1,SP110,SPAG1,SPC25,SPDL1,SPNS1,SPON2,SRPRB,ST6GAL2,STAT1,STAT2,STK17B,SYNE2,SYNJ2,SYTL2,TACC1,TAGAP,TAP1,TARBP2,TBC1D4,TESC,TEX15,TFAP4,TIGD2,TIPARP,TLE3,TM4SF1,TMEM102,TMEM246,TMEM44,TMEM51,TMEM63B,TMX3,TNFSF10,TNFSF13B,TOP1,TP53,TP53INP1,TPRA1,TPSAB1/TPSB2,TRANK1,TREML2,TRERF1,TRIB2,TRIM11,TRIM21,TRIM37,TRIM5,TRIM6,TRIP11,TRPS1,TSC22D3,TSPAN4,TTC27,TTLL7,TXNIP,TYMP,TYW1,UBE2T,UNC93B1,USP18,USP20,USP32,USP37,VEZF1,VGLL4,VMP1,VTI1B,VWA5A,WDFY3,WDR78,ZC3H4,ZC3H6,ZFP36L2,ZFP64,ZGLP1,ZMAT5,ZNF146,ZNF250,ZNF420,ZNF425,ZNF462,ZNF470,ZNF521,ZNF608,ZNF610,ZNF616,ZNF623,ZNF681,ZNF799,ZNF813,ZNF835 |
| Hematological System Development and Function,Inflammatory Response,Tissue Morphology | Quantity of plasmacytoid dendritic cells | 0.00000256 | Increased | 2.213 | BCL3,CSF1,FLT3,GPR183,IL7R,IRF1,MCL1,SELL,TYROBP |
| Hematological Disease | Hemorrhagic disease | 0.00000224 | Increased | 2.098 | C1GALT1C1,C5,CALR,CD44,CHD7,CXCL8,EGR1,FLT3,FUT10,FYB1,GFI1B,GP1BA,GSTM3,HYAL2,ID1,IRAK3,IRF7,ISG15,ITGB3,MITF,MX2,NFE2,OAS2,PRDX3,PTAFR,PTGS2,RASGRP4,SERPINE1,TAL1,TNFSF10,TNFSF13B,TP53,UNC93B1,ZFP36L2 |
| Cancer,Organismal Injury and Abnormalities | Adenocarcinoma | 1.07E-24 | Increased | 2.007 | AASDH,AASDHPPT,ABHD12B,ACOT11,ACSL4,ACTN1,ACVR1,ADAM15,ADCYAP1,ADPRHL2,AGPAT5,AGTR1,AHR,AKR1C1/AKR1C2,ALAS1,ALDH1B1,ALDH4A1,ALDH6A1,ALDH9A1,ALKBH8,AMER1,AMPD3,ANGPT2,ANKLE1,ANKRD28,APOBEC3A,APOE,APOL1,APOL2,APOL3,APOL4,APOL6,APPL2,AREG,ARHGEF26,ARID3A,ARL15,ARL4C,ARL4D,ARL6IP6,ARMC1,ARMH4,ARRDC3,ARSK,ASPHD2,ASRGL1,ATF7IP2,ATG4D,ATP8B3,B3GALNT1,BAALC,BACE2,BATF,BCAM,BCL3,BEND6,BHLHE40,BMPR2,BOD1,BST1,BST2,BTBD10,BTN3A1,BTN3A2,C11orf24,C12orf29,C14orf119,C17orf53,C18orf54,C19orf54,C1GALT1C1,C1orf216,C2,C3AR1,C5,CARD6,CASP1,CASP7,CASP9,CAVIN2,CBX6,CCDC141,CCDC169,CCDC175,CCDC184,CCDC86,CCND1,CCND3,CD274,CD36,CD4,CD44,CD69,CD74,CDC42EP4,CDCP1,CDH26,CDK2AP2,CDKN2C,CDKN3,CELF2,CENPE,CEP72,CETN3,CGNL1,CHCHD10,CHD7,CHST14,CHST2,CIB2,CIITA,CLIP3,CLU,CMIP,CMPK2,CNKSR1,CNR2,COA7,COG2,COL5A1,CR1,CRELD2,CRHBP,CRNDE,CRYZ,CSF1,CSF2RB,CSGALNACT1,CSRP2,CSTF2,CTNS,CTTNBP2,CTU1,CXCL8,CXCR4,CXXC5,CYP1B1,CYSLTR1,DALRD3,DCAF12,DCAF4,DCAF7,DCLK2,DCP1B,DDX58,DDX59,DDX60,DEDD2,DENND2C,DGKG,DGKH,DHFR,DHFR2,DHRS9,DHX33,DHX58,DLC1,DLK1,DNAJC12,DOCK4,DOLK,DOLPP1,DPAGT1,DPF1,DPP10,DPYD,DSE,DTX3L,DUSP1,DUSP6,DYTN,EBF3,ECHDC2,ECM2,EDEM3,EEF1AKNMT,EFNA4,EGR1,EGR2,EGR3,EIF2AK2,EIF4E3,ELP4,EME2,ENAH,ENDOD1,ENO2,ENO3,ENPP2,ENPP5,EPDR1,EPSTI1,EPX,EREG,ESRRA,EXOSC4,EXPH5,EZH2,FAM117A,FAM124B,FAM200A,FAM217B,FAM227A,FAM3B,FAM83D,FAM89A,FAM98A,FBXO21,FIGNL1,FKBP10,FKBP5,FKBPL,FLT3,FMO4,FOS,FOSL1,FOXC2,FOXK1,FOXM1,FUCA1,FUT10,FYB1,FZD2,GAD1,GALK2,GAN,GART,GAS2L1,GAS8,GCNT1,GCSAML,GDF10,GFI1B,GLOD5,GMNN,GMPPB,GMPR,GNAI1,GNB1L,GOLGA2,GOT2,GPR137C,GPR161,GPR162,GPR183,GPR27,GPR65,GRAP2,GRASP,GRWD1,GSTM2,GSTM5,GULP1,GXYLT1,H2AC18/H2AC19,H2BC5,HACD2,HBA1/HBA2,HCK,HDAC7,HDAC9,HECTD3,HELZ2,HERC5,HERC6,HGF,HHEX,HIBADH,HIF3A,HILPDA,HK2,HLA-B,HLA-C,HLA-DPB1,HLA-DRB5,HLA-F,HLA-G,HLTF,HLX,HNRNPCL1/HNRNPCL2,HNRNPF,HOMER2,HOTAIRM1,HPDL,HPGD,HSD3B7,HSPB1,HTRA2,HYAL2,ICMT,ID1,IDS,IFI44,IFI44L,IFI6,IFIH1,IFIT1,IFIT2,IFIT3,IFITM1,IFRD2,IGLL1/IGLL5,IGSF6,IKZF3,IKZF4,IL10RB,IL1RAP,IL1RAPL1,IL7,IL7R,INPP4A,INTS2,INTS7,IQGAP2,IRAK3,IRF2,IRF2BP1,IRF7,ISG15,ITGAX,ITGB3,ITGB8,ITPKA,ITPRIPL1,ITPRIPL2,IVNS1ABP,JAKMIP2,JUN,JUNB,KAZN,KCNA3,KCNK6,KEAP1,KIF14,KIF21A,KIF3A,KIF3B,KIFC1,KLF1,KLF4,KLF6,KLF9,KLHDC7B,KLHL12,KPNA2,KPTN,KRT1,KYNU,LBR,LGALS3BP,LGR4,LIG4,LIN7A,LMBRD2,LMNA,LMNB1,LMOD3,LPCAT1,LRRC27,LRRC4,LRRC75B,LRRC8D,LTBP1,LY6E,LYAR,MAFF,MAL,MAN2A1,MAN2B1,MARS2,MATN2,MCF2L,MCL1,MDK,MED24,MEIS2,MEX3A,MFSD14B,MFSD4A,MGA,MIPEP,MITF,MLEC,MOB3B,MOV10,MPG,MR1,MRM2,MS4A6A,MSI2,MT-ND1,MTA3,MTHFS,MTMR12,MX1,MX2,MYADM,MYL4,NAV1,NDUFS7,NEK4,NEURL1,NFE2,NFE2L3,NFKBIE,NHLRC1,NHSL2,NIBAN2,NINJ1,NKX3-1,NLN,NLRC5,NOC3L,NOTCH2,NR4A1,NR4A2,NR4A3,NTAN1,NUP210L,NUP50,NUSAP1,OAF,OAS1,OAS2,OLFM3,P2RY14,PAG1,PANK4,PAPOLG,PAQR7,PAQR8,PARD6B,PARP14,PARVB,PARVG,PBRM1,PCGF6,PCNX1,PCYOX1L,PCYT2,PDE3B,PDIA4,PDP1,PDP2,PER1,PER2,PEX11G,PGBD2,PHACTR3,PIEZO2,PIGF,PIK3C2B,PIK3R3,PLEKHG1,PLEKHO2,PLSCR1,PLXNC1,PMEPA1,PMP22,POP1,PPA2,PPIL1,PPP1R18,PPP1R1B,PRDX1,PREP,PRKCD,PRKG2,PRMT5,PRNP,PRUNE1,PTAFR,PTGS2,PTK2,PTPRN2,PVALB,PYGL,PYGO2,QSER1,RAB10,RAB11FIP1,RAB38,RAB9B,RADX,RAMP1,RASA2,RASAL2,RASGEF1B,RASGRP3,RASGRP4,RASSF3,RBBP8,RCN1,RDH13,REN,REXO5,RGCC,RGS1,RGS14,RGS2,RHOB,RIC3,RIMS1,RIN2,RMI1,RNASEL,RNF111,RNF24,RNF34,RNPEP,RPL9,RSAD2,RSPH3,RTKN,RTP4,RUNDC3B,RXFP1,SAMD9,SAMD9L,SAMHD1,SAP130,SASH3,SCAMP3,SCAMP4,SCARF1,SCUBE1,SELL,SERPINE1,SESN1,SFRP5,SGK1,SHQ1,SIGLEC12,SIK1/SIK1B,SLA,SLC10A7,SLC29A1,SLC2A6,SLC35B2,SLC38A1,SLC39A6,SLC43A3,SLC44A1,SLC45A3,SLC52A3,SLC5A6,SLC7A6,SLC9A3R1,SLCO5A1,SLFN12,SLFN5,SMAD1,SMAP2,SMO,SMOX,SMTN,SNAI3,SNX18,SNX20,SOCS1,SOCS2,SOCS3,SOD2,SORBS1,SORT1,SOSTDC1,SP110,SPAG1,SPC25,SPDL1,SPIRE1,SPNS1,SPON2,SPRED1,SPRY2,SPTSSA,SRGAP3,SRPRB,SSPN,ST6GAL2,STAT1,STAT2,STK17B,STT3A,SYNE2,SYNJ2,SYTL2,TACC1,TAF5,TAGAP,TAL1,TANGO2,TAP1,TARBP2,TBC1D4,TCTN2,TESC,TEX15,TFAP4,TFEC,TFPI,THAP8,TICAM1,TIGD2,TIPARP,TLE3,TLE4,TLNRD1,TM4SF1,TMEM102,TMEM121,TMEM150C,TMEM229B,TMEM246,TMEM44,TMEM51,TMEM63B,TMX3,TNFSF10,TNFSF13B,TOP1,TP53,TP53INP1,TPRA1,TPSAB1/TPSB2,TRAF3IP3,TRANK1,TREML2,TRERF1,TRIB2,TRIM11,TRIM14,TRIM21,TRIM27,TRIM35,TRIM37,TRIM5,TRIM6,TRIM68,TRIM69,TRIP11,TRPS1,TSC22D3,TSPAN4,TTC27,TTC8,TTLL7,TXNIP,TYMP,TYROBP,TYW1,UAP1L1,UBE2T,UBIAD1,UHRF1,UNC93B1,USP18,USP20,USP32,USP37,VANGL1,VASH2,VEZF1,VGLL4,VMP1,VPS37C,VTI1B,VWA5A,WBP1L,WDFY3,WDR78,WNT5B,XAF1,ZC3H4,ZC3H6,ZFP36,ZFP36L2,ZFP64,ZGLP1,ZMAT5,ZNF146,ZNF250,ZNF331,ZNF420,ZNF425,ZNF438,ZNF462,ZNF470,ZNF521,ZNF608,ZNF610,ZNF616,ZNF619,ZNF623,ZNF653,ZNF681,ZNF74,ZNF787,ZNF799,ZNF813,ZNF835 |
| Lymphoid Tissue Structure and Development,Tissue Morphology | Quantity of lymph follicle | 0.000000422 | Decreased | -2.044 | APOE,ARID3A,BCL3,BHLHE40,C17orf99,CD274,CD74,CNR2,CXCR4,DLK1,EGR2,EGR3,GAPT,IKZF3,IL7R,KLF2,NOTCH2,PRKCD,RGS1,SASH3,STAT1,TLE4,TNFSF13B,TYROBP |
| Cell-To-Cell Signaling and Interaction,Inflammatory Response | Immune response of leukocytes | 0.00000129 | Decreased | -2.046 | ADCYAP1,APOE,BHLHE40,C3AR1,C5,CALR,CASP1,CASP9,CD274,CD36,CD44,CD69,CD74,CSF2RB,CXCL8,DDX58,HCK,HGF,IL10RB,IL1RAP,IL7,IL7R,IRF2,ISG15,ITGAX,ITGB3,JUN,PRNP,RGCC,SCARF1,SERPINE1,SOCS1,SOCS3,TAP1,TICAM1,TP53,TREML2,TYROBP,UBE2L6 |
| Cell Death and Survival | Cell survival | 1.46E-10 | Decreased | -2.064 | ACTL6A,ADCYAP1,AHR,ALKBH8,ANGPT2,APOBEC3A,APOE,APPL2,AREG,ARTN,ATF1,BCL3,BHLHE40,BTG2,C5,CALR,CARS1,CASP1,CASP7,CASP9,CCND1,CCND3,CD33,CD4,CD44,CD74,CDKN2C,CLU,CR1,CRNDE,CSF1,CSF2RB,CXCL8,CXCR4,CYP1B1,DLK1,DPYD,DUSP1,DUSP6,EBF3,EFNA4,EGR1,EGR3,EMP1,EREG,ESRRA,EZH2,FKBP5,FLT3,FOS,FOSL1,FOXM1,FZD2,GALK2,GPR65,HBA1/HBA2,HCK,HGF,HHEX,HK2,HLA-F,HLA-G,HLX,HOPX,HSPA1A/HSPA1B,HSPB1,ICMT,IFIH1,IKZF3,IL7,IL7R,INPP4A,IRAK3,ITGB3,JUN,JUND,KAZN,KEAP1,KIFC1,KLF2,KLF4,KLF6,LGALS3BP,LIG4,LMNA,LMNB1,LMO4,MCF2L,MCL1,MDK,MITF,MPG,MTA3,MTMR12,MX1,NEAT1,NEK4,NFE2,NKX3-1,NOTCH2,NR4A1,NR4A2,NR4A3,PARP14,PBRM1,PDP1,PIK3IP1,PIM3,PLSCR1,PPP1R1B,PREP,PRKCD,PRNP,PTAFR,PTGS2,PTK2,PTPRN2,PYGO2,RNASEL,SELL,SERPINE1,SGK1,SMAD1,SMO,SOCS1,SOCS2,SOCS3,SOD2,SSPN,STAT1,STAT2,TAL1,TFPI,TICAM1,TNFSF10,TNFSF13B,TOP1,TP53,TRIB2,TRIM21,TRIM68,TRIM69,TSC22D3,TXNIP,TYMP,TYROBP,UHRF1,UNG,USP18,XAF1,ZFP36 |
| Cell-To-Cell Signaling and Interaction,Hematological System Development and Function,Immune Cell Trafficking,Inflammatory Response | Activation of lymphocytes | 0.000000003 | Decreased | -2.067 | ADCYAP1,AHR,APOE,BCL3,BHLHE40,BTN3A2,C3AR1,CALR,CD274,CD4,CD44,CD69,CD74,CSF2RB,CXCL8,CYSLTR1,DDX58,DUSP1,EGR2,EGR3,FLT3,FOS,GFI1B,HDAC7,HDAC9,HGF,HLA-DPB1,HLA-G,IL7,IRAK3,IRF1,KCNA3,KEAP1,KLF2,MR1,NOTCH2,NR4A1,NR4A2,NR4A3,PAG1,PRDX1,PRNP,REN,RHOB,SASH3,SOCS1,SOCS2,STAT1,TICAM1,TNFRSF21,TNFSF10,TNFSF13B,TP53,TREML2,TXNIP,TYROBP,VTI1B |
| Metabolic Disease,Organismal Injury and Abnormalities | Glucose metabolism disorder | 1.98E-10 | Decreased | -2.067 | ACAA1,ADCYAP1,AGTR1,APOC1,APOE,ARL6IP6,BCL3,BTG1,BTG2,BTN3A1,BTN3A2,BTN3A3,C2,C3AR1,C5,CA5B,CALR,CARS1,CASP1,CCDC169,CCND1,CD274,CD36,CD4,CD44,CD74,CIITA,CLU,CNR2,COG2,COL5A1,CR1,CRHBP,CSF2RB,CTSH,CXCL8,CXCR4,DDX60,DHFR,DLK1,DUSP1,EGR1,EIF2AK2,ENAH,EXPH5,FOS,FOXC2,GAS2L1,GPR65,HACD2,HBA1/HBA2,HCP5,HDAC7,HDC,HERC6,HGF,HHEX,HK2,HLA-B,HLA-C,HLA-DMB,HLA-DPB1,HLA-DRB5,HLA-F,HLTF,HPGD,HSPA1A/HSPA1B,HSPB1,HYAL2,ID1,IFI44,IFIH1,IFIT1,IFIT2,IFIT3,IFITM3,IKZF3,IL7R,IRAK3,IRF1,IRF7,ITGAX,ITGB3,JUN,KLF4,KYNU,LMNA,MPG,MR1,MS4A6A,NOTCH2,NR4A1,NR4A2,NR4A3,OAS1,P2RY14,PARP14,PDE3B,PDP2,PHACTR3,PIEZO2,PLSCR1,PRDX3,PREP,PRKCD,PTAFR,PTGS2,RAMP1,REN,RGS1,RHOB,RPL9,RSAD2,RTKN,RTP4,SAMD9L,SELL,SERPINE1,SLC45A3,SOCS1,SOCS2,SOCS3,SOD2,SORT1,SP110,SPRED1,STAT1,STAT2,SULT1A2,SYNJ2,TAP1,TBC1D4,TFPI,TNFSF10,TNFSF13B,TP53,TP53INP1,TRIM27,TRIM37,TSC22D3,TXNIP,TYROBP,UNG,WNT5B,XAF1,ZGLP1,ZNRD1ASP |

| | | | | | |
|---|---|---|---|---|---|
| Cell-To-Cell Signaling and Interaction | Activation of lymphatic system cells | 3.18E-10 | Decreased | -2.088 | ADCYAP1,AHR,ANGPT2,APOE,BCL3,BHLHE40,BTN3A2,C3AR1,CALR,CD274,CD4,CD44,CD69,CD74,CSF2RB,CXCL8,CYSLTR1,DDX58,DUSP1,EGR2,EGR3,FLT3,FOS,GFI1B,GMNN,HDAC7,HDAC9,HGF,HLA-DPB1,HLA-G,IL7,IRAK3,IRF1,ITGAX,KCNA3,KEAP1,KLF2,MR1,NOTCH2,NR4A1,NR4A2,NR4A3,PAG1,PRDX1,PRNP,REN,RHOB,SASH3,SOCS1,SOCS2,STAT1,TICAM1,TNFRSF21,TNFSF10,TNFSF13B,TP53,TREML2,TXNIP,TYROBP,VTI1B |
| Cell-To-Cell Signaling and Interaction,Hematological System Development and Function,Immune Cell Trafficking,Inflammatory Response | Activation of mononuclear leukocytes | 4.29E-10 | Decreased | -2.091 | ADCYAP1,AHR,APOE,BCL3,BHLHE40,BTN3A2,C3AR1,C5,CALR,CD274,CD4,CD44,CD69,CD74,CSF1,CSF2RB,CXCL8,CYSLTR1,DDX58,DUSP1,EGR2,EGR3,FLT3,FOS,GFI1B,HDAC7,HDAC9,HGF,HLA-DPB1,HLA-G,HYAL2,IL7,IRAK3,IRF1,KCNA3,KEAP1,KLF2,MR1,NOTCH2,NR4A1,NR4A2,NR4A3,PAG1,PRDX1,PRNP,REN,RHOB,SASH3,SOCS1,SOCS2,STAT1,TICAM1,TNFRSF21,TNFSF10,TNFSF13B,TP53,TPSAB1/TPSB2,TREML2,TXNIP,TYROBP,VTI1B |
| Cell-To-Cell Signaling and Interaction,Hematological System Development and Function,Immune Cell Trafficking,Inflammatory Response | Activation of leukocytes | 2.31E-10 | Decreased | -2.093 | ADCYAP1,AHR,ANGPT2,APOE,BCL3,BHLHE40,BTN3A2,C3AR1,C5,CALR,CASP1,CD274,CD36,CD4,CD44,CD69,CD74,CNR2,CR1,CSF1,CSF2RB,CTSH,CXCL8,CYSLTR1,DDX58,DUSP1,EGR2,EGR3,EPX,FLT3,FOS,GFI1B,HCK,HDAC7,HDAC9,HGF,HLA-DMB,HLA-DPB1,HLA-G,HYAL2,IL7,IRAK3,IRF1,KCNA3,KEAP1,KLF2,KLF4,LGALS3BP,LTBP1,MR1,NOTCH2,NR4A1,NR4A2,NR4A3,PAG1,PLSCR1,PRDX1,PRKCD,PRNP,PTGS2,RAB10,REN,RHOB,SASH3,SELL,SOCS1,SOCS2,SPON2,STAT1,TICAM1,TNFRSF21,TNFSF10,TNFSF13B,TP53,TPSAB1/TPSB2,TREML2,TXNIP,TYROBP,UNC93B1,USP18,VTI1B |
| Cellular Movement | Cell movement of tumor cell lines | 5.83E-12 | Decreased | -2.114 | ACSL4,ACTN1,ACVR1,ADAM15,AHR,ANGPT2,ANKRD28,AREG,ARL4C,ARRDC3,BCAM,BMPR2,C5,CALR,CCND1,CD36,CD44,CD69,CDCP1,CDKN3,CLU,CNR2,CRNDE,CSF1,CTSH,CXCL8,CXCR4,DDX58,DLC1,DOCK4,DPAGT1,DSE,DUSP6,EGR1,EIF2AK2,EMP1,ENAH,ENPP2,EREG,ESRRA,EZH2,FKBPL,FOS,FOSL1,FOXC2,FOXM1,FYB1,GNAI1,GOLGA2,GPR161,HCK,HDAC9,HES6,HGF,HHEX,HOTAIRM1,HSPA1A/HSPA1B,ICMT,ID1,IFIT2,IFITM3,INPP4A,ITGB3,IVNS1ABP,JUN,JUND,KEAP1,KIFC1,KLF4,KLF6,KPNA2,LGALS3BP,LGR4,LMNA,MARCKS,MCF2L,MDK,MITF,MX1,MYADM,NINJ1,NOTCH2,NR4A1,NR4A2,PAG1,PARVB,PARVG,PIEZO2,PIGF,PIK3C2B,PPP1R1B,PRKCD,PRMT5,PRNP,PRUNE1,PTAFR,PTGS2,PTK2,RAB11FIP1,RGS1,RHOB,RXFP1,SELL,SERPINE1,SLC9A3R1,SMAD1,SMO,SOCS3,SOD2,SPRY2,STAT1,TFAP4,TFPI,TNFRSF21,TNFSF10,TP53,TP53INP1,TRPS1,VGLL4 |
| Cellular Movement,Immune Cell Trafficking | Cell movement of lymphatic system cells | 1.71E-08 | Decreased | -2.133 | APOE,BATF,BCL3,BMPR2,C5,CALR,CD274,CD4,CD44,CD69,CD74,CDCP1,CHST2,CNR2,CSF2RB,CXCL8,CXCR4,CYSLTR1,DDX58,DEPTOR,EGR2,ENPP2,FOS,FOXC2,FYB1,GCNT1,GPR183,HDC,HGF,HLA-G,IL7,IL7R,ITGB3,JUN,KCNA3,KCNE3,KLF2,MR1,MYADM,NR4A1,PRKCD,PTGS2,PTK2,REN,RGS1,SELL,SOCS1,SOCS3,STAT1,TMEM102,TNFRSF21,TNFSF10,TNFSF13B,TP53 |
| Cell Death and Survival | Cell death of myeloid cells | 0.00000145 | Decreased | -2.147 | AHR,ANGPT2,APOE,BTG1,C3AR1,CASP1,CASP7,CD44,CD69,CSF1,CXCL8,CXCR4,EIF2AK2,FOS,GFI1B,IRAK3,IRF1,IRF2,ISG15,KLF2,MCL1,NR4A1,PRKCD,SOCS1,SOD2,STAT1,TICAM1,TNFSF10,TP53,TPSAB1/TPSB2,TSC22D3 |
| Cellular Movement,Hematological System Development and Function,Immune Cell Trafficking | Cell movement of lymphocytes | 6.68E-09 | Decreased | -2.156 | APOE,BATF,BCL3,BMPR2,C5,CALR,CD274,CD4,CD44,CD69,CD74,CDCP1,CHST2,CNR2,CSF2RB,CXCL8,CXCR4,CYSLTR1,DDX58,DEPTOR,EGR2,ENPP2,FOS,FYB1,GCNT1,GPR183,HDC,HGF,HLA-G,IL7,IL7R,ITGB3,JUN,KCNA3,KCNE3,KLF2,MR1,MYADM,NR4A1,PRKCD,PTGS2,PTK2,REN,RGS1,SELL,SOCS1,SOCS3,STAT1,TMEM102,TNFRSF21,TNFSF10,TNFSF13B,TP53 |
| Tissue Morphology | Quantity of cells | 7.96E-11 | Decreased | -2.158 | ADCYAP1,AGTR1,AHR,ANGPT2,APOE,ARID3A,ARL4A,BATF,BCL3,BHLHE40,BMPR2,BST1,C17orf99,C1GALT1C1,C3AR1,C5,CALR,CASP1,CASP9,CAV2,CCND1,CCND3,CD274,CD36,CD4,CD44,CD69,CD74,CDKN2C,CHD7,CHST2,CIITA,CLU,CNR2,CSF1,CSF2RB,CXCL8,CXCR4,CYSLTR1,DDX58,DHFR,DLK1,DUSP1,DUSP6,DYNLL1,EGR1,EGR2,EGR3,EIF2AK2,ESRRA,EZH2,FLT3,FOS,FOXC2,FOXM1,FYB1,GAD1,GAPT,GAS2L1,GCNT1,GOLGA2,GP1BA,GPR183,GRAP2,HBA1/HBA2,HCK,HDC,HGF,HIF3A,HLA-G,HSPB1,HTRA2,HYAL2,ID1,IGLL1/IGLL5,IKZF3,IKZF4,IL10RB,IL1RAP,IL1RAPL1,IL7,IL7R,IQGAP2,IRAK3,IRF1,IRF2,ISG15,ITGB3,ITGB8,JUN,JUNB,KLF13,KLF2,KLF4,LGR4,LIG4,LMNA,LMNB1,MAN2A1,MCF2L,MCL1,MDFI,MDK,MITF,MPG,MR1,NEURL1,NFE2,NKX3-1,NLRC5,NOTCH2,NR4A1,NR4A2,NR4A3,PAG1,PCLAF,PER1,PER2,PIM3,PLSCR1,PMP22,PRDX1,PREP,PRKCD,PRMT5,PRNP,PTAFR,PTGS2,PTK2,PTPRN2,RAB27B,RAB38,RASGRP3,RBBP8,REN,RGCC,RGS1,RHOB,RIMS1,SAC3D1,SASH3,SCARF1,SELL,SERPINE1,SFRP5,SGK1,SLA,SMAD1,SMO,SNAI3,SOCS1,SOCS3,SOD2,SORBS1,SOSTDC1,SPON2,SPRED1,SPRY2,STAT1,STK17B,SYTL2,TAL1,TAP1,TICAM1,TLE4,TNFRSF21,TNFSF10,TNFSF13B,TP53,TRERF1,TRIB2,TSC22D3,TXNIP,TYROBP,UHRF1,WNT5B,ZFP36,ZFP36L2,ZNF521 |
| Gene Expression | Transactivation of RNA | 1.28E-10 | Decreased | -2.164 | ACVR1,AHR,ATF1,BATF,BCL3,BMPR2,CCND1,CCND3,CD44,CDCA4,CHST14,CIB2,CIITA,CXCL8,DDX58,EGR1,EGR2,EGR3,EIF2AK2,ESRRA,FOS,FOSL1,FZD2,HERC5,HGF,HIF3A,ID1,IFIH1,IRF1,IRF2,IRF7,JUN,JUNB,JUND,KLF1,KLF13,KLF2,KLF4,KLF6,LMO4,MED24,MITF,MX1,NFE2,NFKBIE,NOTCH2,NR4A1,NR4A2,NR4A3,PARVB,PER1,PRDX1,PRKCD,PTGS2,RHOB,SAMHD1,SOCS1,SOCS3,STAT1,STAT2,TAL1,TICAM1,TNFSF10,TP53,TRPS1,TSC22D3 |
| Cell-To-Cell Signaling and Interaction | Activation of lymphoid cells | 1.2E-09 | Decreased | -2.178 | ADCYAP1,AHR,APOE,BCL3,BHLHE40,BTN3A2,C3AR1,CALR,CD274,CD4,CD44,CD69,CD74,CSF2RB,CXCL8,CYSLTR1,DDX58,DUSP1,EGR2,EGR3,FLT3,FOS,GFI1B,HDAC7,HDAC9,HGF,HLA-DPB1,HLA-G,IL7,IRAK3,IRF1,ITGAX,KCNA3,KEAP1,KLF2,MR1,NOTCH2,NR4A1,NR4A2,NR4A3,PAG1,PRDX1,PRNP,REN,RHOB,SASH3,SOCS1,SOCS2,STAT1,TICAM1,TNFRSF21,TNFSF10,TNFSF13B,TP53,TREML2,TXNIP,TYROBP,VTI1B |
| Gene Expression | Transactivation | 3.17E-10 | Decreased | -2.221 | ACVR1,AHR,APOE,ATF1,BATF,BCL3,BMPR2,CCND1,CCND3,CD44,CDCA4,CHST14,CIB2,CIITA,CXCL8,DDX58,EGR1,EGR2,EGR3,EIF2AK2,ESRRA,FOS,FOSL1,FZD2,HERC5,HGF,HIF3A,ID1,IFIH1,IRF1,IRF2,IRF7,JUN,JUNB,JUND,KLF1,KLF13,KLF2,KLF4,KLF6,LMO4,MED24,MITF,MX1,NFE2,NFKBIE,NOTCH2,NR4A1,NR4A2,NR4A3,PARVB,PER1,PRDX1,PRKCD,PTGS2,RHOB,SAMHD1,SOCS1,SOCS3,STAT1,STAT2,TAL1,TICAM1,TNFSF10,TP53,TRPS1,TSC22D3,ZFP36 |
| Hematological System Development and Function,Humoral Immune Response,Lymphoid Tissue Structure and Development,Tissue Morphology | Quantity of follicular B lymphocytes | 0.000000954 | Decreased | -2.223 | APOE,ARID3A,BCL3,C17orf99,CD274,CD74,CNR2,DLK1,EGR2,EGR3,GAPT,IKZF3,KLF2,NOTCH2,PRKCD,RGS1,SASH3,STAT1,TLE4,TNFSF13B,TYROBP |
| Cellular Movement | Migration of tumor cell lines | 7E-12 | Decreased | -2.232 | ACSL4,ACTN1,ACVR1,ADAM15,AHR,ANGPT2,ANKRD28,AREG,ARL4C,ARRDC3,BCAM,BMPR2,C5,CALR,CCND1,CD36,CD44,CDCP1,CDKN3,CLU,CNR2,CRNDE,CSF1,CTSH,CXCL8,CXCR4,DDX58,DLC1,DPAGT1,DSE,DUSP6,EGR1,EIF2AK2,EMP1,ENAH,ENPP2,EREG,ESRRA,EZH2,FKBPL,FOSL1,FOXC2,FOXM1,GNAI1,GOLGA2,GPR161,HES6,HGF,HHEX,HOTAIRM1,HSPA1A/HSPA1B,ICMT,ID1,IFIT2,IFITM3,INPP4A,ITGB3,IVNS1ABP,JUN,KEAP1,KIFC1,KLF4,KLF6,KPNA2,LGALS3BP,LGR4,LMNA,MCF2L,MDK,MITF,MYADM,NINJ1,NOTCH2,NR4A1,NR4A2,PAG1,PIEZO2,PIGF,PIK3C2B,PPP1R1B,PRKCD,PRMT5,PRNP,PTAFR,PTGS2,PTK2,RGS1,RHOB,RXFP1,SERPINE1,SLC9A3R1,SMAD1,SMO,SOCS3,SOD2,SPRY2,STAT1,TFAP4,TFPI,TNFRSF21,TNFSF10,TP53,TP53INP1,TRPS1,VGLL4 |
| Cell Death and Survival | Cell viability | 1.49E-10 | Decreased | -2.27 | ACTL6A,ADCYAP1,AHR,ALKBH8,ANGPT2,APOBEC3A,APOE,APPL2,AREG,ARTN,ATF1,BCL3,BHLHE40,BTG2,C5,CALR,CARS1,CASP1,CASP7,CASP9,CCND1,CCND3,CD33,CD4,CD44,CD74,CDKN2C,CLU,CR1,CRNDE,CSF1,CSF2RB,CXCL8,CXCR4,CYP1B1,DLK1,DPYD,DUSP1,DUSP6,EFNA4,EGR1,EGR3,EREG,ESRRA,EZH2,FKBP5,FLT3,FOS,FOSL1,FOXM1,FZD2,GALK2,GPR65,HBA1/HBA2,HCK,HGF,HHEX,HK2,HLA-F,HLA-G,HOPX,HSPA1A/HSPA1B,HSPB1,ICMT,IFIH1,IKZF3,IL7,IL7R,INPP4A,ITGB3,JUN,KAZN,KEAP1,KIFC1,KLF2,KLF4,KLF6,LGALS3BP,LIG4,LMNA,LMNB1,LMO4,MCF2L,MCL1,MDK,MITF,MPG,MTA3,MTMR12,MX1,NEAT1,NEK4,NFE2,NKX3-1,NOTCH2,NR4A1,NR4A2,NR4A3,PARP14,PBRM1,PDP1,PIK3IP1,PIM3,PLSCR1,PPP1R1B,PREP,PRKCD,PRNP,PTAFR,PTGS2,PTK2,PTPRN2,PYGO2,RNASEL,SELL,SERPINE1,SGK1,SMO,SOCS1,SOCS2,SOCS3,SOD2,SSPN,STAT1,STAT2,TAL1,TFPI,TICAM1,TNFSF10,TNFSF13B,TOP1,TP53,TRIB2,TRIM21,TRIM68,TRIM69,TSC22D3,TXNIP,TYMP,TYROBP,UHRF1,UNG,USP18,XAF1,ZFP36 |
| Cellular Movement,Hematological System Development and Function,Immune Cell Trafficking,Inflammatory Response | Cell movement of macrophages | 0.00000728 | Decreased | -2.296 | ANGPT2,APOE,BMPR2,C3AR1,C5,CASP1,CCND1,CD44,CD74,CNR2,CR1,CSF1,CXCL8,CXCR4,HCK,HGF,ITGB3,KLF6,LMNB1,MDK,NINJ1,PRKCD,PRNP,PTGS2,REN,RHOB,SELL,SERPINE1,SFRP5,SOCS1,STAT1,TICAM1,TNFRSF21,TP53,TYROBP |
| Cellular Development,Hematological System Development and Function,Lymphoid Tissue Structure and Development | Maturation of leukocytes | 0.000000528 | Decreased | -2.361 | AHR,C5,CD36,CD4,CD44,CD69,CSF1,CXCR4,DDX58,EGR1,EZH2,FOS,HSPA1A/HSPA1B,IFIH1,IGLL1/IGLL5,IL7,IL7R,IRF1,MITF,PARP14,PRNP,PTGS2,SELL,STAT1,TAL1,TICAM1,TNFSF10,TNFSF13B,TP53,TYROBP |
| Cell Death and Survival | Cell death of connective tissue cells | 0.00000639 | Decreased | -2.38 | AHR,CASP1,CASP7,CASP9,CCND1,CD36,CD44,CDKN2C,CENPE,CLU,CNR2,CREBL2,CSF1,CXCL8,CYP1B1,DDX58,DUSP1,EGR1,EGR2,EIF2AK2,GSTM5,HGF,HTRA2,HYAL2,ID1,IFIH1,IRF1,IRF7,ITGB3,IVNS1ABP,JUN,JUND,KLF2,KLF6,LMNA,MCF2L,MCL1,MPG,NR4A1,NR4A2,PRKCD,PRNP,PTK2,RHOB,RNASEL,SERPINE1,SGK1,SMO,SOD2,STAT1,STK17B,TACC1,TNFSF10,TP53,TP53INP1,TRIM37,TRPS1,TXNIP,UNG,USP18 |
| Cell Cycle,Gene Expression | Binding of DNA | 3.38E-11 | Decreased | -2.382 | ADCYAP1,AHR,ANGPT2,BCL3,BTG2,CALR,CCND1,CD4,CLU,CXCL8,CXCR4,CYP1B1,DDX58,EGR1,EGR2,EGR3,ESRRA,EZH2,FOS,FOSL1,FOXM1,FYB1,GMNN,GSTM5,HCK,HDC,HGF,HIF3A,HSPA1A/HSPA1B,ID1,IL7,IRF1,IRF7,ITGB3,JUN,JUNB,JUND,KEAP1,KLF13,LMNA,MGA,MPG,NFKBIE,NR4A1,NR4A2,OAS1,PER1,PPP1R1B,PRDX1,PRKCD,PTAFR,PTK2,RBBP8,RGCC,RHOB,SMAD1,SOCS1,SOCS2,SOCS3,SOD2,STAT1,TAL1,TFAP4,TNFSF10,TNFSF13B,TP53,TSC22D3 |
| Connective Tissue Disorders,Immunological Disease,Inflammatory Disease,Inflammatory Response,Organismal Injury and Abnormalities,Skeletal and Muscular Disorders | Rheumatoid arthritis | 2.81E-11 | Decreased | -2.407 | ACTL6A,ADAM15,APOE,AREG,C1R,C2,C3AR1,C5,CALR,CD274,CD33,CD36,CD4,CD44,CD69,CD74,CDH26,CELF2,CIITA,CLU,CR1,CSF1,CSF2RB,CSGALNACT1,CXCL8,CXCR4,DEDD2,DHFR,DPAGT1,DUSP1,DYNLL1,FKBP5,FLT3,FOS,GP1BA,HCK,HDAC7,HELZ2,HLA-B,HLA-C,HLA-DMB,HLA-DPB1,HLA-DRB5,HLA-G,HSPA1A/HSPA1B,IL7,IL7R,KAZN,KLF13,LMO4,LY6E,MCL1,MDK,MIPEP,MOB3B,MS4A6A,NR4A1,NR4A2,NR4A3,PHACTR3,PRDX1,PREP,PRNP,PTGS2,PTPRN2,RAB10,RASGRP4,RGCC,RIMS1,S100PBP,SAMD9L,SLFN5,SOCS1,SOCS3,STAT1,STK17B,TAGAP,TLE3,TMEM51,TNFRSF21,TNFSF10,TNFSF13B,TP53,UBE2L6,VPS37C,ZFP36,ZNF331,ZNRD1ASP |
| Connective Tissue Disorders,Organismal Injury and Abnormalities,Skeletal and Muscular Disorders | Non-traumatic arthropathy | 1.28E-10 | Decreased | -2.407 | ACTL6A,ADAM15,APOE,AREG,C1R,C2,C3AR1,C5,CALR,CASP1,CD274,CD33,CD36,CD4,CD44,CD69,CD74,CDH26,CELF2,CIITA,CLU,CR1,CSF1,CSF2RB,CSGALNACT1,CXCL8,CXCR4,DEDD2,DHFR,DPAGT1,DUSP1,DYNLL1,ESRRA,FKBP5,FLT3,FOS,GP1BA,HBA1/HBA2,HCK,HDAC7,HELZ2,HGF,HLA-B,HLA-C,HLA-DMB,HLA-DPB1,HLA-DRB5,HLA-G,HSPA1A/HSPA1B,HYAL2,IL7,IL7R,KAZN,KLF13,LMO4,LY6E,MCL1,MDK,MIPEP,MOB3B,MS4A6A,NR4A1,NR4A2,NR4A3,PHACTR3,PRDX1,PREP,PRNP,PTGS2,PTPRN2,RAB10,RASGRP4,REN,RGCC,RIMS1,S100PBP,SAMD9L,SLFN5,SOCS1,SOCS3,SOD2,STAT1,STK17B,TAGAP,TLE3,TMEM51,TNFRSF21,TNFSF10,TNFSF13B,TP53,UBE2L6,VPS37C,WNT5B,ZFP36,ZNF331,ZNRD1ASP |
| Cellular Movement | Homing of cells | 0.000000446 | Decreased | -2.437 | ACTN1,AGTR1,ANGPT2,APOE,ARTN,BATF,BST1,C3AR1,C5,CALR,CCND1,CD36,CD4,CD44,CD69,CD74,CDCP1,CHST2,CLU,CNR2,CSF1,CXCL8,CXCR4,CYSLTR1,DOCK4,DUSP6,EGR3,ENAH,ENPP2,FOSL1,FYB1,GCNT1,GPR183,HCK,HGF,HLA-G,IL7,ITGB3,JUN,MDK,MT-ND1,NINJ1,NR4A1,NUS1,PRKCD,PRNP,PTAFR,PTGS2,PTK2,RAB11FIP1,RASGRP4,RGS1,RHOB,RXFP1,SELL,SERPINE1,SFRP5,SOCS1,SOCS3,TMEM102,TPSAB1/TPSB2,TREML2,TYMP |
| Cellular Movement,Hematological System Development and Function,Immune Cell Trafficking | Cell movement of mononuclear leukocyte | 6.73E-09 | Decreased | -2.478 | ANGPT2,APOE,BATF,BCL3,BMPR2,C5,CALR,CD274,CD4,CD44,CD69,CD74,CDCP1,CHST2,CIITA,CNR2,CSF2RB,CXCL8,CXCR4,CYSLTR1,DDX58,DEPTOR,EGR2,ENPP2,FOS,FYB1,GCNT1,GPR183,HCK,HDC,HGF,HLA-G,IL7,IL7R,ITGAX,ITGB3,JUN,KCNA3,KCNE3,KLF2,MR1,MYADM,NINJ1,NR4A1,PRKCD,PRNP,PTGS2,PTK2,REN,RGS1,SELL,SERPINE1,SOCS1,SOCS3,STAT1,TMEM102,TNFRSF21,TNFSF10,TNFSF13B,TP53 |
| Cell-To-Cell Signaling and Interaction,Cellular Movement,Hematological System Development and Function,Immune Cell Trafficking | Recruitment of leukocytes | 3.42E-09 | Decreased | -2.478 | ADCYAP1,AHR,ANGPT2,APOE,BCL3,C3AR1,C5,CASP1,CD274,CD36,CD4,CD44,CD69,CD74,CLU,CNR2,CSF1,CSF2RB,CXCL8,CXCR4,CYP1B1,DDX58,EGR1,FOS,GCNT1,GP1BA,HCK,HDC,HSPA1A/HSPA1B,IL7,IRF7,JUN,MDK,MR1,NINJ1,RHOB,SELL,SERPINE1,SOCS1,SOD2,SPON2,TFPI,TICAM1,TPSAB1/TPSB2,TREML2 |
| Cellular Movement | Invasion of cells | 4.41E-09 | Decreased | -2.502 | ACSL4,ACVR1,ADAM15,AHR,ANGPT2,AREG,ARHGEF26,ARRDC3,ARTN,ATF1,BHLHE40,BMPR2,BTG2,CALR,CCND1,CD274,CD4,CD44,CD74,CDCP1,CDKN3,CELF2,CLU,CRNDE,CSF1,CTSH,CXCL8,CXCR4,CYP1B1,DDX58,DLC1,DOCK4,DSE,DUSP1,DUSP6,EFNA4,EGR1,EIF2AK2,EMP1,ENAH,ENPP2,EPSTI1,EREG,ESRRA,EZH2,FOS,FOSL1,FOXC2,FOXM1,FZD2,GNAI1,HGF,HHEX,HOTAIRM1,HSPA1A/HSPA1B,ICMT,ID1,IFIT2,IFITM1,IFITM3,ITGAX,ITGB3,ITGB8,IVNS1ABP,JUN,JUNB,JUND,KEAP1,KIFC1,KLF4,KLF6,LGR4,LMNA,MARCKS,MDK,MITF,MSI2,MX1,NIBAN2,NOTCH2,NR4A1,NR4A2,PAG1,PARVB,PRKCD,PRNP,PTAFR,PTGS2,PTK2,RAB11FIP1,RHOB,SERPINE1,SLC9A3R1,SLFN5,SMAD1,SMO,SOCS1,SOD2,SPRY2,STAT1,TFAP4,TFPI,TNFSF10,TP53,TRPS1,UBE2L6,VANGL1,WBP1L,WNT5B,ZFP36 |
| Cellular Movement,Hematological System Development and Function,Immune Cell Trafficking | Lymphocyte migration | 5.45E-08 | Decreased | -2.516 | APOE,BATF,BCL3,BMPR2,C5,CALR,CD274,CD4,CD44,CD69,CD74,CDCP1,CXCL8,CXCR4,CYSLTR1,DDX58,DEPTOR,EGR2,ENPP2,FOS,FYB1,GCNT1,GPR183,HDC,HGF,HLA-G,IL7,IL7R,ITGB3,JUN,KCNA3,KLF2,MR1,MYADM,NR4A1,PTGS2,PTK2,REN,RGS1,SELL,SOCS1,SOCS3,STAT1,TMEM102,TNFRSF21,TNFSF13B |
| Inflammatory Response | Innate immune response | 3.95E-08 | Decreased | -2.557 | ADAM15,APOBEC3A,APOL1,BCL3,BST2,CASP1,CD44,DDX58,EIF2AK2,EREG,HLA-G,IFI6,IFIH1,IFIT2,IRAK3,IRF2BP1,IRF7,ISG15,MX1,NLRC5,PLSCR1,SDHAF4,SPON2,STAT1,TRIM11,TRIM14,TRIM21,TRIM27,TRIM35,TRIM37,TRIM5,TRIM6,TYROBP |
| Cell Death and Survival,DNA Replication, Recombination, and Repair | Fragmentation of DNA | 6.78E-10 | Decreased | -2.561 | ADCYAP1,AGTR1,APOE,C5,CASP7,CASP9,CLU,CSF1,DUSP1,EIF2AK2,FOXC2,HSPB1,IL7,IRF1,IRF2,ITGB3,JUN,KEAP1,MCL1,PRKCD,PRNP,PTGS2,PTK2,SOD2,STAT1,TNFSF10,TOP1,TP53,TPSAB1/TPSB2 |
| Cellular Movement,Immune Cell Trafficking | Migration of lymphatic system cells | 0.000000039 | Decreased | -2.621 | APOE,BATF,BCL3,BMPR2,C5,CALR,CD274,CD4,CD44,CD69,CD74,CDCP1,CNR2,CXCL8,CXCR4,CYSLTR1,DDX58,DEPTOR,EGR2,ENPP2,FOS,FOXC2,FYB1,GCNT1,GPR183,HDC,HGF,HLA-G,IL7,IL7R,ITGB3,JUN,KCNA3,KLF2,MR1,MYADM,NR4A1,PTGS2,PTK2,REN,RGS1,SELL,SOCS1,SOCS3,STAT1,TMEM102,TNFRSF21,TNFSF13B |
| Cell Death and Survival | Cell death of fibroblasts | 0.00000347 | Decreased | -2.626 | AHR,CASP1,CASP7,CASP9,CCND1,CDKN2C,CENPE,CLU,CNR2,CYP1B1,DDX58,DUSP1,EIF2AK2,HGF,HTRA2,JUN,JUND,KLF2,LMNA,MCF2L,MCL1,NR4A1,PRKCD,RHOB,RNASEL,SERPINE1,SMO,SOD2,TNFSF10,TP53,TRIM37,UNG |
| Cell Cycle | Interaction of DNA | 1.59E-11 | Decreased | -2.652 | ADCYAP1,AHR,ANGPT2,BCL3,BTG2,CALR,CCND1,CD4,CLU,CXCL8,CXCR4,CYP1B1,DDX58,EGR1,EGR2,EGR3,ESRRA,EZH2,FOS,FOSL1,FOXM1,FYB1,GMNN,GSTM5,HCK,HDC,HGF,HIF3A,HSPA1A/HSPA1B,ID1,IL7,IRF1,IRF7,ITGB3,JUN,JUNB,JUND,KEAP1,KLF13,LIG4,LMNA,MGA,MPG,NFKBIE,NR4A1,NR4A2,OAS1,PER1,PPP1R1B,PRDX1,PRKCD,PTAFR,PTK2,RBBP8,RGCC,RHOB,SMAD1,SOCS1,SOCS2,SOCS3,SOD2,STAT1,TAL1,TFAP4,TNFSF10,TNFSF13B,TOP1,TP53,TSC22D3 |
| Cellular Movement,Hematological System Development and Function,Immune Cell Trafficking | Migration of mononuclear leukocytes | 8.08E-08 | Decreased | -2.657 | APOE,BATF,BCL3,BMPR2,C5,CALR,CD274,CD4,CD44,CD69,CD74,CDCP1,CXCL8,CXCR4,CYSLTR1,DDX58,DEPTOR,EGR2,ENPP2,FOS,FYB1,GCNT1,GPR183,HDC,HGF,HLA-G,IL7,IL7R,ITGAX,ITGB3,JUN,KCNA3,KLF2,MR1,MYADM,NINJ1,NR4A1,PTGS2,PTK2,REN,RGS1,SELL,SOCS1,SOCS3,STAT1,TMEM102,TNFRSF21,TNFSF13B |
| Connective Tissue Disorders,Inflammatory Disease,Organismal Injury and Abnormalities,Skeletal and Muscular Disorders | Rheumatic Disease | 1.15E-22 | Decreased | -2.67 | ACSL4,ACTL6A,ADAM15,AHR,APOBEC3A,APOE,AREG,BCL3,BST2,C1R,C2,C3AR1,C5,CALR,CASP1,CCND3,CD274,CD33,CD36,CD4,CD44,CD69,CD74,CDH26,CDK2AP2,CELF2,CIITA,CLU,CR1,CREBL2,CRYBA4,CSF1,CSF2RB,CSGALNACT1,CXCL8,CXCR4,DDX58,DDX60,DEDD2,DHFR,DHX58,DPAGT1,DTX3L,DUSP1,DYNLL1,ECHDC3,EGR2,EGR3,EIF2AK2,ENPP2,EPSTI1,ESRRA,FKBP5,FLT3,FOS,GP1BA,HBA1/HBA2,HCK,HDAC7,HELZ2,HERC5,HERC6,HGF,HLA-B,HLA-C,HLA-DMB,HLA-DPB1,HLA-DRB5,HLA-G,HLTF,HSPA1A/HSPA1B,HYAL2,IFI44,IFI44L,IFI6,IFIH1,IFIT1,IFIT3,IFITM1,IKZF3,IL7,IL7R,IRF1,IRF7,ISG15,ITGAX,JUN,JUNB,KAZN,KLF13,LBR,LINC-PINT,LMO4,LTBP1,LY6E,MCL1,MDK,MIPEP,MOB3B,MR1,MS4A6A,MT-ND1,MT-RNR2,MX1,NFE2L3,NLRC5,NR4A1,NR4A2,NR4A3,OAS1,OAS2,OASL,PARP9,PDP2,PER1,PHACTR3,PLSCR1,PRDX1,PREP,PRNP,PTGS2,PTPRN2,RAB10,RASGRP3,RASGRP4,REN,REXO5,RGCC,RGS1,RIMS1,RSAD2,RXFP1,S100PBP,SAMD9L,SAMHD1,SCARF1,SCUBE1,SELL,SGK1,SLFN5,SOCS1,SOCS3,SOD2,STAT1,STK17B,TAGAP,TAP1,TLE3,TMEM51,TNFRSF21,TNFSF10,TNFSF13B,TP53,TPSAB1/TPSB2,TRIM21,TYMP,TYROBP,UBE2L6,USP18,VPS37C,WNT5B,ZFP36,ZNF331,ZNRD1ASP |
| Cell-mediated Immune Response,Cellular Movement,Hematological System Development and Function,Immune Cell Trafficking | Cell movement of T lymphocytes | 0.00000404 | Decreased | -2.697 | APOE,BATF,BMPR2,C5,CALR,CD274,CD44,CD69,CD74,CDCP1,CXCL8,CXCR4,CYSLTR1,EGR2,ENPP2,FYB1,GCNT1,HGF,IL7,ITGB3,KCNA3,NR4A1,PTGS2,REN,RGS1,SELL,SOCS1,SOCS3,STAT1,TMEM102,TNFRSF21 |
| Neurological Disease,Skeletal and Muscular Disorders | Neuromuscular disease | 0.0000024 | Decreased | -2.713 | ACOT11,AGTR1,ALAS1,ALDH6A1,APOE,AREG,BCL3,BHLHE40,BST2,BTN3A1,C5,CALR,CASP1,CASP7,CASP9,CD44,CD74,CIITA,CLU,CNR2,CSRP2,CYSLTR1,DBP,DHFR,DPAGT1,EGR1,EGR2,ENO2,ENO3,ENPP5,EPSTI1,FAM117A,FOS,GAN,GMPPB,HLA-B,HLA-DMB,HLA-F,HLA-G,HSPA1A/HSPA1B,HSPB1,HTRA2,IER5,IFIT1,IL7R,IRF7,ISG15,ITGB3,IVNS1ABP,JUN,JUNB,KAZN,KCNA3,KEAP1,LMNA,LY6E,MAFF,MARS2,MEIS2,MR1,MT-ND1,MX1,NOTCH2,NR4A1,NR4A2,NRGN,OAS1,OAS3,PDP1,PMP22,PPP1R1B,PRNP,PTGS2,PVALB,RGS14,RSAD2,SGK1,SMTN,SORT1,TESC,TOP1,TP53,TRIM5,TSC22D3,TYW1,USP37 |
| Cellular Movement | Invasion of tumor cell lines | 2.28E-10 | Decreased | -2.733 | ACSL4,ACVR1,ADAM15,AHR,AREG,ARHGEF26,ARRDC3,ARTN,ATF1,BMPR2,BTG2,CALR,CCND1,CD274,CD44,CD74,CDCP1,CDKN3,CLU,CRNDE,CTSH,CXCL8,CXCR4,CYP1B1,DDX58,DLC1,DOCK4,DSE,DUSP1,DUSP6,EFNA4,EGR1,EIF2AK2,EMP1,ENAH,ENPP2,EPSTI1,EREG,ESRRA,EZH2,FOSL1,FOXC2,FOXM1,FZD2,GNAI1,HGF,HHEX,HOTAIRM1,HSPA1A/HSPA1B,ICMT,ID1,IFIT2,IFITM1,IFITM3,ITGB3,ITGB8,IVNS1ABP,JUN,JUNB,JUND,KEAP1,KIFC1,KLF4,KLF6,LGR4,LMNA,MARCKS,MDK,MITF,MSI2,MX1,NIBAN2,NR4A2,PARVB,PRKCD,PRNP,PTAFR,PTGS2,PTK2,RAB11FIP1,SLC9A3R1,SLFN5,SMAD1,SMO,SOD2,SPRY2,STAT1,TFAP4,TFPI,TNFSF10,TP53,TRPS1,UBE2L6,VANGL1,WBP1L,WNT5B,ZFP36 |
| Cell-To-Cell Signaling and Interaction,Embryonic Development | Response of embryonic cell lines | 0.00000885 | Decreased | -2.742 | DDX58,IFIH1,IRAK3,IRF7,SPRY2,TNFSF10,TP53,TRIM14,TRIM21,TRIM27,TRIM37,TRIM5,TRIM6 |
| Cellular Development,Cellular Growth and Proliferation,Connective Tissue Development and Function | Proliferation of fibroblast cell lines | 9.25E-11 | Decreased | -2.751 | AHR,ANGPT2,AREG,BATF,BTG1,BTG2,CCND1,CSF1,CSF2RB,CSTF2,DLK1,EGR1,EGR2,EIF2AK2,EIF4E3,EREG,EZH2,FOS,FOXM1,HGF,HSPA1A/HSPA1B,ICMT,ID1,INPP4A,IRF1,IRF2,IVNS1ABP,JUN,JUNB,JUND,KLF2,KLF4,KLF6,LIG4,LMNA,LMNB1,LTBP1,MDK,MITF,NOC3L,PCLAF,PIM3,PLSCR1,PPIL1,PRKCD,PRUNE1,PTK2,RASGRP4,RHOB,RNASEL,SAMHD1,SERPINE1,SOCS1,STAT1,TOP1,TP53,TSC22D3,TXNIP |
| Cellular Movement,Hematological System Development and Function,Immune Cell Trafficking | Cell movement of leukocytes | 3.07E-10 | Decreased | -2.756 | ADAM15,ADCYAP1,AHR,ANGPT2,APOE,BATF,BCL3,BMPR2,BST1,C2,C3AR1,C5,CALR,CASP1,CCND1,CD274,CD36,CD4,CD44,CD69,CD74,CDCP1,CHST2,CIITA,CNR2,CR1,CSF1,CSF2RB,CXCL8,CXCR4,CYSLTR1,DDX58,DEPTOR,DUSP1,EGR2,ENPP2,FOS,FOXM1,FYB1,GCNT1,GPR183,HCK,HDC,HGF,HLA-G,HSPB1,ID1,IL7,IL7R,ITGAX,ITGB3,JUN,KCNA3,KCNE3,KLF2,KLF6,LMNB1,MDK,MR1,MT-ND1,MYADM,NINJ1,NR4A1,PLXNC1,PRDX1,PRKCD,PRNP,PTGS2,PTK2,RAMP1,RASGRP4,REN,RGCC,RGS1,RHOB,SELL,SERPINE1,SFRP5,SGK1,SOCS1,SOCS3,SPON2,STAT1,STAT2,TICAM1,TMEM102,TNFRSF21,TNFSF10,TNFSF13B,TP53,TPSAB1/TPSB2,TREML2,TYROBP |
| Cellular Development | Maturation of blood cells | 3.29E-08 | Decreased | -2.784 | AHR,C5,CCND1,CCND3,CD36,CD4,CD44,CD69,CSF1,CXCR4,DDX58,EGR1,EZH2,FOS,GFI1B,HSPA1A/HSPA1B,IFIH1,IGLL1/IGLL5,IL7,IL7R,IRF1,KLF1,KLF2,MITF,NFE2,PARP14,PRNP,PTGS2,SELL,STAT1,TAL1,TICAM1,TNFSF10,TNFSF13B,TP53,TYROBP |
| Connective Tissue Disorders,Inflammatory Disease,Inflammatory Response,Organismal Injury and Abnormalities,Skeletal and Muscular Disorders | Inflammation of joint | 6.02E-10 | Decreased | -2.796 | ACTL6A,ADAM15,AHR,APOE,AREG,BCL3,C1R,C2,C3AR1,C5,CALR,CASP1,CCND3,CD274,CD33,CD36,CD4,CD44,CD69,CD74,CDH26,CDK2AP2,CELF2,CIITA,CLU,CR1,CSF1,CSF2RB,CSGALNACT1,CXCL8,CXCR4,DEDD2,DHFR,DPAGT1,DUSP1,DYNLL1,ENPP2,ESRRA,FKBP5,FLT3,FOS,GP1BA,HBA1/HBA2,HCK,HDAC7,HELZ2,HGF,HLA-B,HLA-C,HLA-DMB,HLA-DPB1,HLA-DRB5,HLA-G,HLTF,HSPA1A/HSPA1B,HYAL2,IL7,IL7R,IRF1,KAZN,KLF13,LMO4,LY6E,MCL1,MDK,MIPEP,MOB3B,MS4A6A,MT-RNR2,NR4A1,NR4A2,NR4A3,PHACTR3,PRDX1,PREP,PRNP,PTGS2,PTPRN2,RAB10,RASGRP4,REN,RGCC,RIMS1,S100PBP,SAMD9L,SLFN5,SOCS1,SOCS3,SOD2,STAT1,STK17B,TAGAP,TLE3,TMEM51,TNFRSF21,TNFSF10,TNFSF13B,TP53,TPSAB1/TPSB2,TYROBP,UBE2L6,VPS37C,WNT5B,ZFP36,ZNF331,ZNRD1ASP |
| Cell Death and Survival | Cell death of immune cells | 8.2E-09 | Decreased | -2.869 | ADCYAP1,AHR,ANGPT2,APOE,BCL3,BTG1,C3AR1,C5,CASP1,CASP7,CASP9,CCDC86,CD274,CD4,CD44,CD69,CD74,CIITA,CLU,CNR2,CSF1,CSF2RB,CXCL8,CXCR4,DHX58,DLK1,EGR1,EGR3,EIF2AK2,FLT3,FOS,FOSL1,GCNT1,GPR65,GRAP2,HCK,HGF,HLA-G,IFIH1,IKZF3,IL7,IL7R,IRAK3,IRF1,IRF2,ISG15,JUN,KLF13,KLF2,KLF4,MCF2L,MCL1,MX1,NR4A1,NR4A3,NRGN,PARP14,PRKCD,PTK2,RNASEL,SOCS1,SOD2,STAT1,STK17B,TICAM1,TLE4,TNFRSF21,TNFSF10,TNFSF13B,TP53,TPSAB1/TPSB2,TRIB2,TSC22D3,TYROBP |
| Cell-mediated Immune Response,Cellular Movement,Hematological System Development and Function,Immune Cell Trafficking | T cell migration | 0.00000224 | Decreased | -2.917 | APOE,BATF,BMPR2,C5,CALR,CD274,CD4,CD44,CD69,CD74,CDCP1,CXCL8,CXCR4,CYSLTR1,EGR2,ENPP2,FYB1,GCNT1,HGF,IL7,ITGB3,KCNA3,KLF2,NR4A1,PTGS2,PTK2,REN,RGS1,SELL,SOCS1,SOCS3,STAT1,TMEM102,TNFRSF21 |
| Cellular Movement | Invasion of carcinoma cell lines | 1.51E-09 | Decreased | -2.955 | AREG,ARTN,ATF1,BMPR2,CALR,CCND1,CD44,CLU,CTSH,DDX58,DLC1,DOCK4,DSE,DUSP1,EGR1,ENAH,ENPP2,EZH2,FOSL1,FOXM1,GNAI1,HGF,ID1,IFIT2,IFITM1,JUN,JUNB,JUND,KIFC1,KLF4,LGR4,MDK,MITF,MSI2,PRKCD,PTAFR,PTK2,SMO,TP53,WNT5B |
| Cell Death and Survival | Cell death of blood cells | 2.67E-08 | Decreased | -2.972 | ADCYAP1,AHR,ANGPT2,APOE,BCL3,BTG1,C3AR1,C5,CASP1,CASP7,CASP9,CCDC86,CD274,CD4,CD44,CD69,CD74,CIITA,CLU,CNR2,CSF1,CSF2RB,CXCL8,CXCR4,DHX58,DLK1,EGR1,EGR3,EIF2AK2,FLT3,FOS,FOSL1,GCNT1,GFI1B,GPR65,GRAP2,HCK,HGF,HLA-G,IFIH1,IKZF3,IL7,IL7R,IRAK3,IRF1,IRF2,ISG15,JUN,KLF13,KLF2,KLF4,MCF2L,MCL1,MX1,NR4A1,NR4A3,NRGN,PARP14,PCLAF,PRKCD,PTK2,RNASEL,SOCS1,SOD2,STAT1,STK17B,TICAM1,TLE4,TNFRSF21,TNFSF10,TNFSF13B,TP53,TPSAB1/TPSB2,TRIB2,TSC22D3,TYROBP |

| | | | | | |
|---|---|---|---|---|---|
| Antimicrobial Response,Inflammatory Response | Antiviral response of cells | 0.00000164 | Decreased | -3.124 | C5,DDX58,DHX58,EIF2AK2,IFIH1,IRF1,SOCS1,SOCS3,SPRY2,STAT1,TP53 |
| Cellular Movement | Migration of blood cells | 3.01E-10 | Decreased | -3.234 | ADAM15,ADCYAP1,AHR,ANGPT2,APOE,ARTN,BATF,BCL3,BMPR2,BST1,C2,C3AR1,C5,CALR,CASP1,CCND1,CD274,CD36,CD4,CD44,CD69,CD74,CDCP1,CHST2,CIITA,CLU,CNR2,CR1,CSF1,CSF2RB,CXCL8,CXCR4,CYP1B1,CYSLTR1,DDX58,DEPTOR,DUSP1,EGR1,EGR2,ENPP2,FOS,FOXC2,FOXM1,FYB1,GCNT1,GP1BA,GPR183,HCK,HDC,HGF,HLA-G,HSPA1A/HSPA1B,HSPB1,ID1,IGLL1/IGLL5,IL7,IL7R,IRF7,ITGAX,ITGB3,JUN,KCNA3,KCNE3,KLF2,KLF6,LMNB1,MDK,MR1,MT-ND1,MYADM,NINJ1,NR4A1,PER1,PLXNC1,PRDX1,PRKCD,PRNP,PTAFR,PTGS2,PTK2,RAMP1,RASGRP4,REN,RGCC,RGS1,RHOB,SELL,SERPINE1,SFRP5,SGK1,SLC7A6,SOCS1,SOCS3,SOD2,SPON2,STAT1,STAT2,TFPI,TICAM1,TMEM102,TNFRSF21,TNFSF10,TNFSF13B,TP53,TPSAB1/TPSB2,TREML2,TYROBP |
| Cellular Movement,Immune Cell Trafficking | Leukocyte migration | 6.02E-10 | Decreased | -3.317 | ADAM15,ADCYAP1,AHR,ANGPT2,APOE,BATF,BCL3,BMPR2,BST1,C2,C3AR1,C5,CALR,CASP1,CCND1,CD274,CD36,CD4,CD44,CD69,CD74,CDCP1,CHST2,CIITA,CLU,CNR2,CR1,CSF1,CSF2RB,CXCL8,CXCR4,CYP1B1,CYSLTR1,DDX58,DEPTOR,DUSP1,EGR1,EGR2,ENPP2,FOS,FOXC2,FOXM1,FYB1,GCNT1,GP1BA,GPR183,HCK,HDC,HGF,HLA-G,HSPA1A/HSPA1B,HSPB1,ID1,IGLL1/IGLL5,IL7,IL7R,IRF7,ITGAX,ITGB3,JUN,KCNA3,KCNE3,KLF2,KLF6,LMNB1,MDK,MR1,MT-ND1,MYADM,NINJ1,NR4A1,PER1,PLXNC1,PRDX1,PRKCD,PRNP,PTAFR,PTGS2,PTK2,RAMP1,RASGRP4,REN,RGCC,RGS1,RHOB,SELL,SERPINE1,SFRP5,SGK1,SLC7A6,SOCS1,SOCS3,SOD2,SPON2,STAT1,STAT2,TFPI,TICAM1,TMEM102,TNFRSF21,TNFSF10,TNFSF13B,TP53,TPSAB1/TPSB2,TREML2,TYROBP |
| Inflammatory Response | Immune response of cells | 3.19E-11 | Decreased | -3.894 | ADCYAP1,AHR,APOE,APPL2,BHLHE40,C3AR1,C5,CALR,CASP1,CASP9,CD274,CD36,CD4,CD44,CD69,CD74,CR1,CSF1,CSF2RB,CXCL8,CYP1B1,DDX58,DHX58,EIF2AK2,GP1BA,GULP1,HCK,HGF,HSPB1,IFIH1,IL10RB,IL1RAP,IL7,IL7R,IRF1,IRF2,IRF2BP1,IRF7,ISG15,ITGAX,ITGB3,JUN,JUNB,KLF2,LTBP1,LYAR,MCL1,MPG,NR4A2,PER1,PRKCD,PRNP,PTK2,RASA2,RGCC,SCARF1,SERPINE1,SGK1,SOCS1,SOCS3,SPRY2,SPTSSA,STAT1,STT3A,TAP1,TICAM1,TLE3,TNFSF10,TP53,TREML2,TRIM14,TRIM21,TRIM27,TRIM37,TRIM5,TRIM6,TYROBP,UBE2L6 |
| Antimicrobial Response,Inflammatory Response | Antiviral response | 6.05E-22 | Decreased | -3.907 | APOBEC3A,BCL3,BST2,C5,CLU,CXCR4,DDX58,DDX60,DHX58,DTX3L,EIF2AK2,EXOSC4,FOSL1,HSPB1,HYAL2,IFI44,IFI44L,IFI6,IFIH1,IFIT1,IFIT2,IFIT3,IFITM1,IFITM3,IRF1,IRF2,IRF7,ISG15,ITGAX,IVNS1ABP,MOV10,MX1,MX2,OAS1,OAS2,OAS3,OASL,PARP9,PLSCR1,RNASEL,RSAD2,RTP4,SAMHD1,SOCS1,SOCS3,SPON2,SPRY2,STAT1,STAT2,TAGAP,TICAM1,TP53,TRIM5,TRIM6 |
| Antimicrobial Response,Inflammatory Response | Antimicrobial response | 1.05E-13 | Decreased | -3.907 | APOBEC3A,BCL3,BST2,C5,CLU,CXCR4,DDX58,DDX60,DHX58,DTX3L,EIF2AK2,EXOSC4,FOSL1,HCK,HLA-G,HSPB1,HYAL2,IFI44,IFI44L,IFI6,IFIH1,IFIT1,IFIT2,IFIT3,IFITM1,IFITM3,IRF1,IRF2,IRF7,ISG15,ITGAX,IVNS1ABP,MOV10,MX1,MX2,NOTCH2,OAS1,OAS2,OAS3,OASL,PARP9,PLSCR1,PRKCD,RNASEL,RSAD2,RTP4,SAMHD1,SOCS1,SOCS3,SPON2,SPRY2,STAT1,STAT2,TAGAP,TICAM1,TP53,TRIM5,TRIM6 |
| Cellular Movement | Cell movement | 5.87E-13 | Decreased | -4.245 | ACSL4,ACTN1,ACVR1,ADAM15,ADCYAP1,AGTR1,AHR,ANGPT2,ANKRD28,APOE,APPL2,AREG,ARL4C,ARRDC3,ARTN,BATF,BCAM,BCL3,BHLHE40,BMPR2,BST1,BST2,BTG2,C2,C3AR1,C5,CALR,CASP1,CASP9,CCDC141,CCND1,CD274,CD36,CD4,CD44,CD69,CD74,CDC42EP4,CDCP1,CDKN2C,CDKN3,CHST2,CIITA,CLU,CNR2,CR1,CRNDE,CSF1,CSF2RB,CTSH,CXCL8,CXCR4,CXXC5,CYP1B1,CYSLTR1,DDX58,DEPTOR,DLC1,DOCK4,DPAGT1,DSE,DUSP1,DUSP6,EBF3,EFNA4,EGR1,EGR2,EGR3,EIF2AK2,EMP1,ENAH,ENPP2,EREG,ESRRA,EZH2,FKBPL,FOS,FOSL1,FOXC2,FOXM1,FUT10,FYB1,GAD1,GAS2L1,GAS8,GCNT1,GNAI1,GOLGA2,GP1BA,GPR161,GPR183,HCK,HDAC9,HDC,HES6,HGF,HHEX,HLA-G,HLX,HOTAIRM1,HSPA1A/HSPA1B,HSPB1,HYAL2,ICMT,ID1,IFIT2,IFITM1,IFITM3,IGLL1/IGLL5,IL1RAP,IL7,IL7R,INPP4A,IRF7,ITGAX,ITGB3,ITGB8,ITPKA,IVNS1ABP,JUN,JUNB,JUND,KCNA3,KCNE3,KEAP1,KIF14,KIFC1,KLF2,KLF4,KLF6,KPNA2,LGALS3BP,LGR4,LMNA,LMNB1,LMO4,MARCKS,MATN2,MCF2L,MCL1,MDK,MITF,MIXL1,MR1,MT-ND1,MX1,MYADM,NAV1,NEURL1,NINJ1,NKX3-1,NOTCH2,NR4A1,NR4A2,NUS1,PAG1,PARD6B,PARP9,PARVB,PARVG,PDE3B,PER1,PEX11B,PIEZO2,PIGF,PIK3C2B,PIK3IP1,PLXNC1,PMP22,PPP1R1B,PRDX1,PRKCD,PRMT5,PRNP,PRUNE1,PTAFR,PTGS2,PTK2,RAB11FIP1,RAMP1,RASGRP3,RASGRP4,REN,RGCC,RGS1,RHOB,RXFP1,SELL,SERPINE1,SFRP5,SGK1,SLC7A6,SLC9A3R1,SMAD1,SMO,SOCS1,SOCS2,SOCS3,SOD2,SORBS1,SORT1,SPON2,SPRY2,STAT1,STAT2,SYNE2,TFAP4,TFPI,TICAM1,TM4SF1,TMEM102,TNFRSF21,TNFSF10,TNFSF13B,TP53,TP53INP1,TPSAB1/TPSB2,TREML2,TRPS1,TSC22D3,TYMP,TYROBP,USP18,VASH2,VGLL4,WNT5B |
| Cellular Movement | Migration of cells | 3.88E-13 | Decreased | -4.627 | ACSL4,ACTN1,ACVR1,ADAM15,ADCYAP1,AGTR1,AHR,ANGPT2,ANKRD28,APOE,APPL2,AREG,ARL4C,ARRDC3,ARTN,BATF,BCAM,BCL3,BHLHE40,BMPR2,BST1,BST2,BTG2,C2,C3AR1,C5,CALR,CASP1,CASP9,CCDC141,CCND1,CD274,CD36,CD4,CD44,CD69,CD74,CDCP1,CDKN2C,CDKN3,CHST2,CIITA,CLU,CNR2,CR1,CRNDE,CSF1,CSF2RB,CTSH,CXCL8,CXCR4,CXXC5,CYP1B1,CYSLTR1,DDX58,DEPTOR,DLC1,DOCK4,DPAGT1,DSE,DUSP1,DUSP6,EBF3,EGR1,EGR2,EGR3,EIF2AK2,EMP1,ENAH,ENPP2,EREG,ESRRA,EZH2,FKBPL,FOS,FOSL1,FOXC2,FOXM1,FUT10,FYB1,GAD1,GCNT1,GNAI1,GOLGA2,GP1BA,GPR161,GPR183,HCK,HDC,HES6,HGF,HHEX,HLA-G,HLX,HOTAIRM1,HSPA1A/HSPA1B,HSPB1,HYAL2,ICMT,ID1,IFIT2,IFITM1,IFITM3,IGLL1/IGLL5,IL7,IL7R,INPP4A,IRF7,ITGAX,ITGB3,ITGB8,ITPKA,IVNS1ABP,JUN,KCNA3,KCNE3,KEAP1,KIF14,KIFC1,KLF2,KLF4,KLF6,KPNA2,LGALS3BP,LGR4,LMNA,LMNB1,LMO4,MARCKS,MATN2,MCF2L,MCL1,MDK,MITF,MIXL1,MR1,MT-ND1,MYADM,NAV1,NINJ1,NKX3-1,NOTCH2,NR4A1,NR4A2,NUS1,PAG1,PARD6B,PARP9,PER1,PEX11B,PIEZO2,PIGF,PIK3C2B,PIK3IP1,PLXNC1,PMP22,PPP1R1B,PRDX1,PRKCD,PRMT5,PRNP,PTAFR,PTGS2,PTK2,RAMP1,RASGRP3,RASGRP4,REN,RGCC,RGS1,RHOB,RXFP1,SELL,SERPINE1,SFRP5,SGK1,SLC7A6,SLC9A3R1,SMAD1,SMO,SOCS1,SOCS3,SOD2,SORT1,SPON2,SPRY2,STAT1,STAT2,SYNE2,TFAP4,TFPI,TICAM1,TM4SF1,TMEM102,TNFRSF21,TNFSF10,TNFSF13B,TP53,TP53INP1,TPSAB1/TPSB2,TREML2,TRPS1,TSC22D3,TYROBP,USP18,VASH2,VGLL4,WNT5B |

**T3 vs. T1 MEP_2**

© 2000-2020 QIAGEN. All rights reserved.

| Categories | Diseases or Functions Annotation | p-value | Predicted Activation State | Activation z-score | Molecules |
|---|---|---|---|---|---|
| Cell Signaling | Viral life cycle | 2.41E-08 | Increased | 3.372 | BANF1,BST2,CXCR4,IFIT1,IFITM1,IFITM3,ISG15,ISG20,MX1,NUP50,OAS1,OAS3,OASL,PLSCR1,PTGS2,RSAD2,TRIM6,VTA1 |
| Cell Signaling | Replication of viral replicon | 1.68E-11 | Increased | 3.372 | BANF1,BST2,IFIT1,IFITM1,IFITM3,ISG15,ISG20,MX1,OAS1,OAS3,OASL,PLSCR1,PTGS2,RSAD2,TRIM6 |
| Infectious Diseases | Replication of vesicular stomatitis virus | 8E-09 | Increased | 2.805 | DDX58,IFIH1,IFITM3,IRF1,ISG20,OAS1,OASL,PARP12,SLFN12,SOCS1,SP110,STAT2,TAP1,TRIM25 |
| Organismal Survival | Organismal death | 1.21E-09 | Increased | 2.759 | ABCB1,ACRBP,ACVR1,ADORA2B,AFMID,AHR,ANKLE1,ANXA1,APOE,AREG,ATF1,B2M,B3GNT5,BHLHE40,CA5B,CACNA1C,CASP1,CCDC86,CD274,CD36,CD4,CD74,CDCP1,CDH1,CDKN2C,CGAS,CITED2,CPOX,CRK,CSGALNACT1,CTNNB1,CXCL8,CXCR4,DDX58,DKK1,DLK1,DUSP1,EGR2,EGR3,FLT3,FOS,FST,GCLC,GNAI1,GNLY,GPR183,GZMA,HCG11,HDAC7,HLA-G,HLX,HMGA2,HOPX,HOXA3,HRH2,HS6ST1,HSPG2,HYAL2,ID1,ID2,ID3,IDS,IFIH1,IKZF3,IL15RA,IRF1,IRF7,ISG15,JUNB,KCTD12,KIF3A,KLF1,KLF2,KLF4,KLF6,KLF9,KMT5A,KRT1,LGR4,LIG4,LIN7A,LMO4,LOX,LRP6,LY96,MACROH2A2,MAFF,MDFI,MMUT,MT-ND1,MYC,NCF1,NEAT1,NFE2,NFE2L3,NFKB2,NFKBIA,NR4A2,NR4A3,OR7C1,PADI4,PANK1,PARP3,PDGFB,PIK3CG,PORCN,PPARA,PPARGC1B,PTAFR,PTGER2,PTGS2,PYGO1,PYGO2,RARG,RASSF2,RCE1,RELB,RFK,RFNG,RRN3,S1PR1,SELENBP1,SERPINE1,SFRP5,SLC14A1,SOCS1,SOCS3,SOD2,SRGAP3,STAT1,STAT2,SYT5,TBC1D4,TCF7L2,TEK,TFAP4,TGM2,THRB,TICAM1,TIE1,TIPARP,TLR2,TNFAIP8L2,TNFRSF10A,TNFSF10,TNFSF13B,TP53INP1,TRIM21,TXNIP,USP18,VCAN,VTA1,WNT5B,ZFP36 |
| Organismal Survival | Morbidity or mortality | 2.59E-10 | Increased | 2.419 | ABCB1,ACRBP,ACVR1,ADORA2B,AFMID,AHR,ANKLE1,ANXA1,APOE,AREG,ATF1,B2M,B3GNT5,BHLHE40,CA5B,CACNA1C,CASP1,CCDC86,CD274,CD36,CD4,CD74,CDCP1,CDH1,CDKN2C,CGAS,CITED2,CPOX,CRK,CSGALNACT1,CTNNB1,CXCL8,CXCR4,DDX58,DKK1,DLK1,DUSP1,EGR2,EGR3,FLT3,FOS,FST,GCLC,GNAI1,GNLY,GPR183,GZMA,HCG11,HDAC7,HLA-A,HLA-G,HLX,HMGA2,HOPX,HOXA3,HRH2,HS6ST1,HSPG2,HYAL2,ID1,ID2,ID3,IDS,IFIH1,IKZF3,IL15RA,IRF1,IRF7,ISG15,JUNB,KCTD12,KIF3A,KLF1,KLF2,KLF4,KLF6,KLF9,KMT5A,KRT1,LGR4,LIG4,LIN7A,LMO4,LOX,LRP6,LY96,MACROH2A2,MAFF,MDFI,MMUT,MT-ND1,MYC,NCF1,NEAT1,NFE2,NFE2L3,NFKB2,NFKBIA,NR4A2,NR4A3,OASL,OR7C1,PADI4,PANK1,PARP3,PDGFB,PIK3CG,PORCN,PPARA,PPARGC1B,PTAFR,PTGER2,PTGS2,PYGO1,PYGO2,RARG,RASSF2,RCE1,RELB,RFK,RFNG,RRN3,S1PR1,SELENBP1,SERPINE1,SFRP5,SLC14A1,SOCS1,SOCS3,SOD2,SRGAP3,STAT1,STAT2,SYT5,TAP1,TBC1D4,TCF7L2,TEK,TFAP4,TGM2,THRB,TICAM1,TIE1,TIPARP,TLR2,TNFAIP8L2,TNFRSF10A,TNFSF10,TNFSF13B,TP53INP1,TRIM21,TRIM27,TXNIP,USP18,VCAN,VTA1,WNT5B,ZFP36 |
| Infectious Diseases | Replication of RNA virus | 1.94E-14 | Increased | 2.396 | AREG,ARTN,B2M,BST2,CANT1,CD4,CD81,CIITA,CLSTN3,CXCL8,CXCR4,DDX58,DKK1,DTX2,GABARAPL1,GBP1,GCLC,HERC5,IFI27,IFI6,IFIH1,IFIT1,IFITM1,IFITM3,IRF1,IRF7,ISG15,ISG20,ITGB5,MX1,MYC,NFKBIA,NLRC5,OAS1,OAS3,OASL,PARP12,PTGS2,RSAD2,SGK1,SLFN12,SNX9,SOCS1,SOCS3,SP110,STAT1,STAT2,TAP1,TLR2,TNFSF10,TRIM14,TRIM21,TRIM25,TRIM5,TXNIP |
| Infectious Diseases | Replication of virus | 4.43E-15 | Increased | 2.379 | AHR,AREG,ARTN,B2M,BST2,CANT1,CD4,CD81,CIITA,CLSTN3,CXCL8,CXCR4,DDX58,DKK1,DTX2,DUSP1,EGR1,GABARAPL1,GBP1,GCLC,HERC5,IFI27,IFI6,IFIH1,IFIT1,IFITM1,IFITM3,IRF1,IRF7,ISG15,ISG20,ITGB5,MX1,MYC,NFKBIA,NLRC5,OAS1,OAS3,OASL,PARP12,PPARA,PTGS2,RSAD2,SGK1,SLFN12,SNX9,SOCS1,SOCS3,SP110,STAT1,STAT2,TAP1,TICAM1,TLR2,TNFSF10,TRIM14,TRIM21,TRIM25,TRIM5,TXNIP |
| Infectious Diseases | Replication of Herpesviridae | 5.88E-08 | Increased | 2.335 | AHR,DDX58,EGR1,IFIH1,IRF1,ISG20,OAS1,OASL,PARP12,RSAD2,STAT1,STAT2,TICAM1 |
| Cancer,Gastrointestinal Disease,Organismal Injury and Abnormalities | Gastrointestinal carcinoma | 0.00000036 | Increased | 2.219 | ABCB1,ABCC6,ABCG1,ABHD11,ACRBP,ACSL4,ACVR1,ADAMTS6,AFMID,AHR,AKR1C1/AKR1C2,ALDH9A1,ANKLE1,ANKRD28,ANKRD55,ANXA1,APOE,APOL4,APOL6,AREG,ARHGEF26,ARMC4,ATP10A,B2M,B3GNT5,BCL9,BEX5,BHLHE40,BST2,BTBD3,BTBD6,BTN3A1,C17orf80,C1orf141,CA1,CA2,CABLES1,CACNA1C,CACNA2D3,CANT1,CASP1,CCDC184,CCDC71,CCDC86,CD274,CD4,CD69,CD74,CD81,CD96,CDC42EP4,CDCP1,CDH1,CETN3,CGAS,CGNL1,CHD7,CHST2,CIITA,CITED2,CLSTN3,CMBL,CMPK2,CMTM2,CNKSR3,CNNM3,CNST,CNTN4,COL14A1,CORO2A,CPNE8,CPOX,CRK,CSGALNACT1,CTNNB1,CXCL8,CXCR4,CYP1B1,CYSLTR1,CYTH3,DDX58,DDX60,DENND5A,DHRS9,DIPK1B,DIXDC1,DLK1,DPF1,DTX2,DTX3L,DUSP1,DUSP19,ECHDC2,EDA2R,EGR1,EGR2,EGR3,EIF4E3,ENPP5,EPCAM,EPHB4,ERCC4,EREG,EXOC3L4,FAM222B,FAM83D,FAM98A,FGF18,FGL2,FKBP5,FLT3,FOS,FST,FZD2,FZD7,GABARAPL1,GALNT7,GAS2L1,GBP1,GBP4,GBP7,GCLC,GEMIN4,GIMAP7,GLOD5,GNAI1,GNG8,GP9,GPAT3,GPR171,GPR183,GPR27,GPR63,GSTM5,HDAC7,HDX,HEATR5A,HERC5,HERC6,HLA-A,HLA-B,HLA-DPB1,HLA-DQA1,HLA-DQB1,HLA-DRA,HLA-DRB5,HLA-F,HLA-G,HLX,HMGA2,HMGN5,HOMER1,HOPX,HOXA3,HRH2,HSPG2,HYAL2,HYPK,ID1,ID2,IDS,IFI44L,IFI6,IFIH1,IFIT1,IFIT3,IFITM1,IGSF6,IL1RAPL1,IL7R,IRF1,IRF7,IRGQ,ITGB5,ITPRIPL1,ITPRIPL2,JUNB,KCNN4,KCTD12,KIF21A,KIF3A,KIFC3,KLF4,KLF9,KMT5A,KRT1,KYNU,LGALS3BP,LGR4,LIG4,LIN7A,LMLN,LOX,LPAR5,LRP6,LRRC2,LRRC4,LTBP4,LY96,MACO1,MAFF,MAK,MAP3K21,MAP9,MDFI,MEX3A,MFAP3L,MMUT,MPV17L2,MRNIP,MS4A2,MSI2,MTHFS,MX1,MYC,MYO16,NFE2,NFE2L3,NFKB2,NFKBIA,NFKBIE,NHLRC1,NLRC5,NR4A1,NR4A2,NR4A3,NXF3,OAS1,OAS2,OLFM3,OR7C1,OSBPL6,P2RY14,PABPC3,PADI4,PAG1,PANK1,PAQR8,PARD6B,PARP14,PARP3,PCTP,PDE3B,PDGFB,PER1,PF4V1,PHACTR3,PIK3CG,PLEKHG1,PLEKHH3,PLEKHM3,PLSCR1,PLXNC1,POGLUT1,PORCN,PPARA,PPARGC1B,PPM1K,PRR11,PTAFR,PTGS2,PYGO1,PYGO2,RADX,RAMP3,RAPGEF5,RASGEF1B,RELB,RFK,RGS1,RHAG,RNF185,RNF34,RTN1,S1PR1,SAMD9L,SCN3A,SELENBP1,SELL,SERPINE1,SFRP5,SGK1,SH2D3A,SKIDA1,SLA,SLC10A4,SLC12A7,SLC16A10,SLC1A5,SLC38A1,SLC39A11,SLC39A6,SLC43A2,SLC45A3,SLC7A6,SLC9A3R2,SLC9A6,SLC9A9,SLFN12,SLFN5,SMOX,SNX18,SNX20,SNX9,SOCS1,SOCS3,SOD2,SORL1,SP110,SPIRE1,SPTBN2,SRGAP3,SSPN,ST3GAL2,ST6GAL2,STAT1,STAT2,SWT1,TAGAP,TAP1,TBC1D2,TBC1D4,TCF7L2,TEK,TEX14,TEX15,TFEC,TGIF2,TGM2,THRB,TIAM1,TICAM1,TIE1,TIPARP,TLE3,TLR2,TM4SF1,TMEM108,TMEM17,TMEM40,TMEM91,TNFRSF19,TNFSF10,TP53INP1,TPST1,TRANK1,TRIM14,TRIM21,TRIM22,TRIM26,TRIM27,TRIM5,TRIM6,TSC22D3,TSPYL2,TUBG2,TUT1,TYROBP,UACA,UBE2M,UTS2,VCAN,VMP1,VSTM4,WDFY3,WHRN,WIPF3,WNK3,WNT5B,XAF1,ZBTB11,ZBTB16,ZC3H4,ZFP36,ZHX2,ZNF212,ZNF420,ZNF462,ZNF485,ZNF503,ZNF592,ZNF594,ZNF670,ZNF74,ZNF799,ZNF804A,ZNF835 |
| Infectious Diseases | Infection of mammalia | 1.09E-12 | Increased | 2.067 | AHR,APOE,B2M,CASP1,CD274,CD36,CD4,CD74,CD81,CIITA,CXCR4,DDX58,DUSP1,FOS,GZMA,HLA-DQB1,HLA-G,IFIH1,IRF1,IRF7,ISG15,LY96,NCF1,NFE2,NFKB2,OASL,PADI4,PTAFR,RELB,SERPINE1,STAT1,STAT2,TAP1,TICAM1,TLR2,TNFAIP8L2,TNFRSF10A,TNFSF10,TPSAB1/TPSB2,USP18 |
| Cellular Movement,Hematological System Development and Function,Immune Cell Trafficking | Migration of mononuclear leukocytes | 1.1E-11 | Decreased | -2.034 | ANXA1,APOE,CD274,CD4,CD69,CD74,CD81,CDCP1,CTNNB1,CXCL8,CXCR4,CYSLTR1,DDX58,DEPTOR,DKK1,EGR2,FOS,GNLY,GPR183,HDC,HLA-A,HLA-G,IL15RA,IL7,IL7R,KCNN4,KLF2,MICOS10-NBL1/NBL1,MYADM,NFKB2,NR4A1,PF4V1,PIK3CG,PTGS2,RGS1,S1PR1,SELL,SOCS1,SOCS3,STAT1,TGM2,TLR2,TNFRSF21,TNFSF13B |
| Immunological Disease | Delayed hypersensitive reaction | 1.94E-08 | Decreased | -2.137 | AHR,ANXA1,APOE,B4GAT1,CD74,IFITM1,LGALS3BP,MX1,NCF1,NFKBIA,PIK3CG,PLSCR1,RELB,SELL,SOD2,TNFSF10,TSC22D3,ZFP36 |
| Cell-mediated Immune Response,Cellular Movement,Hematological System Development and Function,Immune Cell Trafficking | Cell movement of T lymphocytes | 8.13E-08 | Decreased | -2.153 | APOE,CD274,CD69,CD74,CD81,CDCP1,CTNNB1,CXCL8,CXCR4,CYSLTR1,EGR2,GNLY,HLA-A,IL15RA,IL7,NR4A1,PF4V1,PIK3CG,PTGS2,RGS1,S1PR1,SELL,SOCS1,SOCS3,STAT1,TGM2,TNFRSF21 |
| Cellular Development | Maturation of blood cells | 6.05E-14 | Decreased | -2.177 | ABCG1,AHR,CD36,CD4,CD69,CDH1,CXCR4,DDX58,EGR1,FOS,HLA-DRA,HOXA3,IFIH1,IGLL1/IGLL5,IL15RA,IL7,IL7R,IRF1,KLF1,KLF2,MAP2K6,MYC,NFE2,NFKB2,PARP14,PTCRA,PTGS2,RELB,RNASE1,SELL,STAT1,TICAM1,TLR2,TNFSF10,TNFSF13B,TYROBP,ZBTB16 |
| Cellular Development,Hematological System Development and Function,Lymphoid Tissue Structure and Development | Maturation of dendritic cells | 5.53E-07 | Decreased | -2.238 | CD36,CDH1,DDX58,EGR1,FOS,IFIH1,IRF1,MAP2K6,NFKB2,RELB,RNASE1,STAT1,TICAM1,TLR2,TNFSF10,TYROBP |
| Gene Expression | Transcription of RNA | 2.68E-11 | Decreased | -2.369 | ACVR1,AHR,APOE,APOL3,ATF1,BCL9,BHLHE40,BST2,BTG1,BTG2,CASP1,CD36,CD4,CD74,CD81,CDH1,CDKN2C,CHD7,CIITA,CISH,CITED2,CRK,CTNNB1,CXCR4,DBP,DDX58,DIXDC1,DKK1,DTX3L,DUSP1,EGR1,EGR2,EGR3,EPCAM,EREG,FOS,FST,FZD2,FZD7,GAL,GCLC,GPR183,HDAC7,HLA-DRB5,HMGA2,HOPX,HYAL2,ID1,ID2,ID3,IFI27,IKZF3,IRF1,IRF7,JUNB,JUND,KLF1,KLF15,KLF2,KLF4,KLF6,KLF9,KMT5A,LGR4,LMO4,LRP6,LY96,MACROH2A2,MAFF,MAP2K6,MDFI,MYC,NCF1,NFE2,NFE2L3,NFKB2,NFKBIA,NLRC5,NR4A1,NR4A2,NR4A3,PARP9,PDGFB,PER1,PF4,PLSCR1,PPARA,PPARGC1B,PTAFR,PTGER2,PYGO1,PYGO2,RARG,RELB,RGCC,RRN3,S1PR1,SERPINE1,SGK1,SLA,SOCS1,SOCS3,SOD2,SSPN,STAT1,STAT2,TCF7L2,TFAP4,TFEC,TGIF2,THRB,TLE3,TLR2,TNFRSF10B,TNFSF10,TP53INP1,TRIM22,TRIM27,TRIM6,TSC22D3,TXNIP,ZBTB16,ZFP36,ZHX2,ZKSCAN3,ZNF234,ZNF322,ZNF462 |
| Cellular Development,Hematological System Development and Function,Lymphoid Tissue Structure and Development | Maturation of leukocytes | 2.57E-13 | Decreased | -2.374 | ABCG1,AHR,CD36,CD4,CD69,CDH1,CXCR4,DDX58,EGR1,FOS,HLA-DRA,HOXA3,IFIH1,IGLL1/IGLL5,IL15RA,IL7,IL7R,IRF1,MAP2K6,MYC,NFKB2,PARP14,PTCRA,PTGS2,RELB,RNASE1,SELL,STAT1,TICAM1,TLR2,TNFSF10,TNFSF13B,TYROBP |
| Cell-mediated Immune Response,Cellular Movement,Hematological System Development and Function,Immune Cell Trafficking | T cell migration | 1.85E-08 | Decreased | -2.425 | APOE,CD274,CD4,CD69,CD74,CD81,CDCP1,CTNNB1,CXCL8,CXCR4,CYSLTR1,EGR2,GNLY,HLA-A,IL15RA,IL7,KCNN4,KLF2,NR4A1,PF4V1,PIK3CG,PTGS2,RGS1,S1PR1,SELL,SOCS1,SOCS3,STAT1,TGM2,TNFRSF21 |
| Inflammatory Response | Immune response of cells | 1.4E-13 | Decreased | -2.551 | ABCB1,ADORA2B,AHR,ANXA1,APOE,BHLHE40,CASP1,CD274,CD36,CD4,CD69,CD74,CNN2,CRK,CTNNB1,CXCL8,CYP1B1,DDX58,GULP1,HLA-A,HLA-DQB1,HRH2,IFIH1,IL15RA,IL7,IL7R,IRF1,IRF7,ISG15,ITGB5,JUNB,KLF2,LY96,MYC,NCF1,NR4A2,PADI4,PER1,PF4,PIK3CG,PTGER2,RELB,RGCC,SERPINE1,SGK1,SLC1A5,SOCS1,SOCS3,STAT1,TAP1,TGM2,TICAM1,TLE3,TLR2,TNFRSF10A,TNFSF10,TRIM14,TRIM21,TRIM25,TRIM26,TRIM27,TRIM5,TRIM6,TYROBP |
| Inflammatory Response | Innate immune response | 2.01E-09 | Decreased | -2.557 | B2M,BST2,CASP1,DDX58,EREG,HLA-G,IFI27,IFI6,IFIH1,IRF7,ISG15,MX1,NCF1,NLRC5,PIK3CG,PLSCR1,RNASE6,STAT1,TLR2,TNFRSF10A,TRIM14,TRIM21,TRIM25,TRIM26,TRIM27,TRIM5,TRIM6,TYROBP |
| Cellular Development,Hematological System Development and Function,Lymphoid Tissue Structure and Development | Maturation of antigen presenting cells | 3.68E-08 | Decreased | -2.593 | CD36,CD69,CDH1,DDX58,EGR1,FOS,HOXA3,IFIH1,IRF1,MAP2K6,NFKB2,RELB,RNASE1,STAT1,TICAM1,TLR2,TNFSF10,TYROBP |
| Cellular Development,Hematological System Development and Function,Lymphoid Tissue Structure and Development | Maturation of phagocytes | 0.00000103 | Decreased | -2.65 | CD36,CDH1,DDX58,EGR1,FOS,HOXA3,IFIH1,IRF1,MAP2K6,NFKB2,RELB,RNASE1,STAT1,TICAM1,TLR2,TNFSF10,TYROBP |
| Gene Expression | Expression of RNA | 5.44E-11 | Decreased | -2.682 | ACVR1,AHR,APOE,APOL3,ATF1,BCL9,BHLHE40,BST2,BTG1,BTG2,CASP1,CD36,CD4,CD74,CD81,CDH1,CDKN2C,CHD7,CIITA,CISH,CITED2,CNN2,CRK,CTNNB1,CXCR4,DBP,DDX58,DIXDC1,DKK1,DLK1,DTX3L,DUSP1,EGR1,EGR2,EGR3,EPCAM,EREG,FAM98A,FOS,FST,FZD2,FZD7,GAL,GAS2L1,GCLC,GPR183,HDAC7,HLA-DRB5,HMGA2,HMGN5,HOPX,HYAL2,ID1,ID2,ID3,IFI27,IKZF3,IL7,IL7R,IRF1,IRF7,ISG20,JUNB,JUND,KLF1,KLF15,KLF2,KLF4,KLF6,KLF9,KMT5A,LGMN,LGR4,LMO4,LOX,LRP6,LY96,MACROH2A2,MAFF,MAP2K6,MDFI,MYADM,MYC,NCF1,NFE2,NFE2L3,NFKB2,NFKBIA,NHLRC1,NLRC5,NR4A1,NR4A2,NR4A3,PARP14,PARP9,PDGFB,PER1,PF4,PLSCR1,PPARA,PPARGC1B,PTAFR,PTGER2,PYGO1,PYGO2,RAMP3,RARG,RELB,RGCC,RPS23,RRN3,S1PR1,SERPINE1,SGK1,SLA,SOCS1,SOCS3,SOD2,SSPN,STAT1,STAT2,TCF7L2,TFAP4,TFEC,TGIF2,TGM2,THRB,TICAM1,TIPARP,TLE3,TLR2,TNFRSF10B,TNFSF10,TP53INP1,TRIM22,TRIM27,TRIM6,TSC22D3,TXNIP,TYROBP,UTS2,WHRN,ZBTB16,ZFP36,ZHX2,ZKSCAN3,ZNF234,ZNF322,ZNF462,ZNF503,ZNF804A |
| Gene Expression | Transcription | 1.79E-12 | Decreased | -2.687 | ACVR1,AHR,APOE,APOL3,ATF1,BCL9,BHLHE40,BST2,BTG1,BTG2,CASP1,CD36,CD4,CD74,CD81,CDH1,CDKN2C,CHD7,CIITA,CISH,CITED2,CNN2,CRK,CTNNB1,CXCR4,DBP,DDX58,DIXDC1,DKK1,DLK1,DTX3L,DUSP1,EGR1,EGR2,EGR3,EPCAM,EREG,FAM98A,FOS,FST,FZD2,FZD7,GAL,GAS2L1,GCLC,GPR183,HDAC7,HLA-DRB5,HMGA2,HMGN5,HOPX,HYAL2,ID1,ID2,ID3,IFI27,IKZF3,IL7,IL7R,IRF1,IRF7,JUNB,JUND,KLF1,KLF15,KLF2,KLF4,KLF6,KLF9,KMT5A,LGMN,LGR4,LMO4,LOX,LRP6,LY96,MACROH2A2,MAFF,MAP2K6,MDFI,MX1,MYADM,MYC,NCF1,NFE2,NFE2L3,NFKB2,NFKBIA,NHLRC1,NLRC5,NR4A1,NR4A2,NR4A3,PARP14,PARP9,PDGFB,PER1,PF4,PLSCR1,PPARA,PPARGC1B,PTAFR,PTGER2,PYGO1,PYGO2,RAMP3,RARG,RELB,RGCC,RRN3,S1PR1,SERPINE1,SGK1,SLA,SOCS1,SOCS3,SOD2,SSPN,STAT1,STAT2,TCF7L2,TFAP4,TFEC,TGIF2,THRB,TICAM1,TIPARP,TLE3,TLR2,TNFRSF10B,TNFSF10,TNFSF13B,TP53INP1,TRIM22,TRIM27,TRIM6,TSC22D3,TXNIP,TYROBP,WHRN,ZBTB16,ZFP36,ZHX2,ZKSCAN3,ZNF234,ZNF322,ZNF462,ZNF503,ZNF804A |
| Inflammatory Disease,Neurological Disease,Skeletal and Muscular Disorders | Multiple Sclerosis | 5.68E-11 | Decreased | -2.813 | ABCB1,ADORA2B,ANXA1,AREG,BST2,CASP1,CYSLTR1,EGR1,EPSTI1,FCGR2C,GAL,HLA-DMB,HLA-DQA1,HLA-DQB1,HLA-DRA,HLA-F,HLA-G,HRH2,IFIT1,IL7R,IRF7,ISG15,MMUT,MX1,OAS1,OAS3,PTGS2,RSAD2,TCF7L2,TLR2,TRIM22,TRIM5 |
| Neurological Disease,Skeletal and Muscular Disorders | Progressive neuromuscular disease | 1.95E-09 | Decreased | -2.813 | ABCB1,ADORA2B,ANXA1,APOE,AREG,BST2,CACNA1C,CASP1,CD74,CYSLTR1,EGR1,EPSTI1,FCGR2C,GAL,HLA-B,HLA-DMB,HLA-DQA1,HLA-DQB1,HLA-DRA,HLA-F,HLA-G,HRH2,IFIT1,IL7R,IRF7,ISG15,MMUT,MT-ND1,MX1,NR4A2,OAS1,OAS3,PTGS2,RSAD2,RTN1,SCN3A,SGK1,TCF7L2,TGM2,TLR2,TRIM22,TRIM5 |
| Neurological Disease,Skeletal and Muscular Disorders | Neuromuscular disease | 2.1E-10 | Decreased | -2.935 | ABCB1,ADORA2B,ANXA1,APOE,AREG,B2M,B4GAT1,BHLHE40,BST2,BTN3A1,CA2,CACNA1C,CACNA2D3,CASP1,CD74,CIITA,CTNNB1,CYSLTR1,DBP,EGR1,EGR2,ENPP5,EPHB4,EPSTI1,FAM117A,FCGR2C,FOS,GAL,GBP1,HLA-B,HLA-DMB,HLA-DOA,HLA-DQA1,HLA-DQB1,HLA-DRA,HLA-F,HLA-G,HOMER1,HRH2,ID3,IFIT1,IL7R,IRF7,ISG15,JUNB,MAFF,MMUT,MT-ND1,MT1G,MX1,NR4A1,NR4A2,OAS1,OAS3,PCTP,PPARA,PPARGC1B,PTGS2,RELB,RSAD2,RTN1,S1PR1,SCN3A,SGK1,SLC14A1,SORL1,TCF7L2,TGM2,TLR2,TRIM22,TRIM5,TSC22D3,VCAN,ZBTB16 |
| Cell-To-Cell Signaling and Interaction,Embryonic Development | Response of embryonic cell lines | 6.23E-07 | Decreased | -3.111 | DDX58,IFIH1,IRF7,LY96,TNFSF10,TRIM14,TRIM21,TRIM25,TRIM26,TRIM27,TRIM5,TRIM6 |
| Neurological Disease | Progressive neurological disorder | 1.94E-08 | Decreased | -3.302 | ABCB1,ADORA2B,ANXA1,APOE,APOL2,AREG,B4GAT1,BST2,CACNA1C,CASP1,CD33,CD36,CD74,CIITA,CNKSR3,CXCL8,CXCR4,CYSLTR1,EGR1,EGR2,EPSTI1,FCGR2C,FLT3,GAL,HLA-B,HLA-DMB,HLA-DQA1,HLA-DQB1,HLA-DRA,HLA-F,HLA-G,HOMER1,HRH2,HSPG2,IFIT1,IL7R,IRF7,ISG15,KIAA0040,LGMN,LRP6,MICOS10-NBL1/NBL1,MMUT,MS4A4E,MT-ND1,MX1,MYC,NEAT1,NFKB2,NHLRC1,NR4A2,OAS1,OAS3,PDE3B,PDGFB,PIK3IP1,PPARA,PPARGC1B,PTGS2,RSAD2,RTN1,SCN3A,SELENBP1,SERPINE1,SGK1,SLC7A6,SLC9A6,SOD2,SORL1,TCF7L2,TEK,TGM2,TLR2,TRIM22,TRIM27,TRIM5,TUBG2,TYROBP,WDFY3,ZNF280B |
| Antimicrobial Response,Inflammatory Response | Antiviral response | 3.39E-18 | Decreased | -3.503 | BANF1,BST2,CGAS,CXCR4,DDX58,DDX60,DTX3L,HLA-A,HYAL2,IFI27,IFI44,IFI44L,IFI6,IFIH1,IFIT1,IFIT3,IFITM1,IFITM3,IRF1,IRF7,ISG15,ISG20,MX1,OAS1,OAS2,OAS3,OASL,PARP9,PLSCR1,RSAD2,RTP4,SOCS1,SOCS3,STAT1,STAT2,TAGAP,TICAM1,TRIM22,TRIM5,TRIM6 |
| Antimicrobial Response,Inflammatory Response | Antimicrobial response | 5.97E-14 | Decreased | -3.503 | B2M,BANF1,BST2,CGAS,CXCR4,DDX58,DDX60,DTX3L,GNLY,HLA-A,HLA-G,HYAL2,IFI27,IFI44,IFI44L,IFI6,IFIH1,IFIT1,IFIT3,IFITM1,IFITM3,IRF1,IRF7,ISG15,ISG20,MX1,NCF1,OAS1,OAS2,OAS3,OASL,PARP9,PLSCR1,RNASE6,RSAD2,RTP4,SOCS1,SOCS3,STAT1,STAT2,TAGAP,TICAM1,TLR2,TRIM22,TRIM5,TRIM6 |

**T2 vs. T1 HSC_MPP**

© 2000-2020 QIAGEN. All rights reserved.

| Categories | Diseases or Functions Annotation | p-value | Predicted Activation State | Activation z-score | Molecules |
|---|---|---|---|---|---|
| Cancer,Organismal Injury and Abnormalities | Non-hematological solid tumor | 2.47E-10 | Increased | 3.494 | ACADL,ADAP2,AGAP3,AGPS,AK4,AKR1C1/AKR1C2,AKR1C3,ANKRD35,ANXA2,APOL4,AREG,ARRDC3,ASRGL1,ATAD2,ATF3,ATP6V0A1,AZU1,BAALC-AS1,BAG3,BCL3,BMPR1B,BRD4,BTG1,C14orf39,C5orf30,CCDC146,CCDC80,CCND3,CD4,CD53,CD69,CD83,CDC42EP4,CDH7,CDKN2C,CEACAM4,CGAS,CISH,CITED2,CLCN4,CLDN7,CLEC1B,CLEC9A,CLU,COL2A1,COX6A1,CREB5,CRIP2,CSGALNACT1,CSNK1A1L,CXCL1,CXCR4,CYTOR,DDIT3,DDIT4,DENND3,DENND5A,DGAT2,DGCR6/LOC102724770,DSEL,DTL,DUSP1,DUSP5,EFCC1,EFNB2,EHD4,ENPP2,ENPP5,EPGN,EPSTI1,EREG,F13A1,FAM124B,FAM83D,FAM89A,FAM98A,FBLN1,FBLN5,FBP1,FBXO45,FCGR1A,FCGR1B,FCMR,FEM1C,FILIP1,FKBP5,FKBP9,FLT3,FOSL1,FYB1,GADD45B,GALNT2,GAS2,GBP1,GBP5,GBP7,GIMAP1,GIMAP2,GIMAP6,GIMAP7,GIMAP8,GLUL,GNAI1,GNG4,GPR27,GPR3,GPT2,GSTM3,GULP1,H4C12,H4C3,HBA1/HBA2,HBB,HIC2,HIF3A,HLA-C,HLA-DRB5,HLA-F,HLA-G,HOMER3,HOPX,HSPA4L,HYLS1,IER3,IFI44L,IFI6,IFIT3,IFITM1,IGF2BP3,IGLL1/IGLL5,IKZF3,IRF1,IRF7,IRF8,IRF9,IRGQ,ISG15,ISG20,ITGB4,ITGB8,ITPRIP,JCHAIN,KLF10,KLF11,KLF15,KLF2,KLF6,KLF7,KLF9,KLHDC7B,KPNA2,KYNU,LAYN,LIN7A,LONRF2,LTB,MAFF,MAL,MAN2A1,MARCKS,MCAM,MCF2L,METTL7A,MFAP3L,MNS1,MOB3B,MORC1,MRVI1,MT2A,MX1,MYCN,MYRIP,NACA2,NFIB,NFKB2,NFKBIE,NKX2-3,NKX3-1,NLGN1,NOS1AP,NR0B1,NR5A1,NRBP2,NRP2,NRXN2,OAS1,OAS3,OASL,ODF3B,ORC6,PANX1,PARP9,PCDH9,PCLAF,PCSK1N,PDZRN4,PER1,PEX11G,PFKFB2,PHLDB2,PIEZO2,PIK3IP1,PLD6,PLEC,PLK4,PMAIP1,PNMT,POMC,PPP2R2B,PPP2R5B,PRX,PSRC1,PTAFR,PTGER2,PTK2,PTPN14,PTPRE,PTS,PYGO1,RAB11FIP1,RAB27B,RASGRP3,RCSD1,REN,RGS18,RHOB,RHOU,RMDN2,RNF11,RNF175,RPL10,RPL3,RPL7A,RPSA,RSAD2,RUNDC3B,S100A13,S100A9,S1PR2,SAMD15,SAMHD1,SAP30,SBDS,SCRG1,SELP,SERPINB8,SESN1,SETBP1,SHD,SKIL,SLC22A4,SLC2A9,SLC38A1,SLFN5,SLITRK1,SLITRK5,SMAD3,SMAP2,SPATA2,SPOCK2,SPTY2D1,ST3GAL1,ST6GAL2,STAT1,STAT2,STXBP6,SUCLG2,TAS2R4,TAX1BP3,TBC1D10A,TCEAL2,TENT5C,TMEM117,TNFSF13B,TNNI2,TOB1,TP53INP1,TPSAB1/TPSB2,TRIM69,TSC22D3,TSPAN5,TUBB2A,TUFT1,TXNIP,TYMP,UGCG,USP2,VSTM4,WDFY3,WFDC1,XAF1,XIRP2,YOD1,ZBTB11,ZBTB37,ZFP36L2,ZNF608 |
| Cancer,Organismal Injury and Abnormalities | Tumorigenesis of tissue | 3.02E-08 | Increased | 3.045 | ACADL,ADAP2,AGAP3,AGPS,AK4,AKR1C1/AKR1C2,AKR1C3,ANKRD35,ANXA2,APOL4,AREG,ARRDC3,ASRGL1,ATAD2,ATF3,ATP6V0A1,AZU1,BAALC-AS1,BAG3,BCL3,BMPR1B,BRD4,BTG1,C14orf39,C5orf30,CCDC146,CCDC80,CCND3,CD4,CD53,CD69,CD83,CDC42EP4,CDH7,CDKN2C,CEACAM4,CGAS,CISH,CITED2,CLCN4,CLDN7,CLEC1B,CLEC9A,CLU,COL2A1,COX6A1,CREB5,CRIP2,CSGALNACT1,CSNK1A1L,CXCL1,CXCR4,CYTOR,DDIT3,DDIT4,DENND3,DENND5A,DGAT2,DGCR6/LOC102724770,DSEL,DTL,DUSP1,DUSP5,EFCC1,EFNB2,EHD4,ENPP2,ENPP5,EPGN,EPSTI1,EREG,F13A1,FAM124B,FAM83D,FAM89A,FAM98A,FBLN1,FBLN5,FBP1,FBXO45,FCGR1A,FCMR,FILIP1,FKBP5,FKBP9,FLT3,FOSL1,FYB1,GADD45B,GALNT2,GAS2,GBP1,GBP5,GBP7,GIMAP1,GIMAP2,GIMAP6,GIMAP7,GIMAP8,GLUL,GNAI1,GNG4,GPR27,GPR3,GPT2,GULP1,HBA1/HBA2,HBB,HIC2,HIF3A,HLA-C,HLA-DRB5,HLA-F,HLA-G,HOMER3,HOPX,HSPA4L,HYLS1,IER3,IFI44L,IFI6,IFIT3,IFITM1,IGF2BP3,IGLL1/IGLL5,IKZF3,IRF1,IRF7,IRF8,IRF9,IRGQ,ISG15,ISG20,ITGB4,ITGB8,ITPRIP,JCHAIN,KLF10,KLF11,KLF15,KLF2,KLF6,KLF7,KLF9,KLHDC7B,KPNA2,KYNU,LAYN,LIN7A,LONRF2,LTB,MAFF,MAL,MAN2A1,MCAM,MCF2L,METTL7A,MFAP3L,MNS1,MOB3B,MORC1,MRVI1,MT-ATP6,MT2A,MX1,MYCN,MYRIP,NACA2,NFIB,NFKB2,NFKBIE,NKX2-3,NKX3-1,NLGN1,NOS1AP,NR0B1,NR5A1,NRBP2,NRP2,NRXN2,OAS1,OAS3,OASL,ODF3B,ORC6,PANX1,PARP9,PCDH9,PCSK1N,PDZRN4,PER1,PEX11G,PFKFB2,PHLDB2,PIEZO2,PIK3IP1,PLD6,PLEC,PLK4,PMAIP1,PNMT,POMC,PPP2R2B,PPP2R5B,PRX,PSRC1,PTAFR,PTGER2,PTK2,PTPN14,PTPRE,PTS,PYGO1,RAB11FIP1,RAB27B,RASGRP3,RCSD1,REN,RGS18,RHOB,RHOU,RMDN2,RNF11,RNF175,RPL10,RPL3,RPL7A,RPSA,RSAD2,RUNDC3B,S100A9,S1PR2,SAMD15,SAMHD1,SAP30,SBDS,SCRG1,SELP,SERPINB8,SESN1,SETBP1,SHD,SKIL,SLC22A4,SLC2A9,SLC38A1,SLFN5,SLITRK1,SLITRK5,SMAD3,SMAP2,SPATA2,SPOCK2,SPTY2D1,ST3GAL1,ST6GAL2,STAT1,STAT2,STXBP6,SUCLG2,TAS2R4,TAX1BP3,TBC1D10A,TCEAL2,TENT5C,TMEM117,TNFSF13B,TNNI2,TOB1,TP53INP1,TPSAB1/TPSB2,TRIM69,TSC22D3,TSPAN5,TUBB2A,TUFT1,TXNIP,TYMP,UGCG,USP2,VSTM4,WDFY3,WFDC1,XAF1,XIRP2,YOD1,ZBTB11,ZFP36L2,ZNF608 |
| Cancer,Organismal Injury and Abnormalities | Epithelial neoplasm | 4.11E-08 | Increased | 2.934 | ACADL,ADAP2,AGAP3,AGPS,AK4,AKR1C1/AKR1C2,AKR1C3,ANKRD35,ANXA2,APOL4,AREG,ARRDC3,ASRGL1,ATAD2,ATF3,ATP6V0A1,AZU1,BAALC-AS1,BAG3,BCL3,BMPR1B,BRD4,BTG1,C14orf39,C5orf30,CCDC146,CCDC80,CCND3,CD4,CD53,CD69,CD83,CDC42EP4,CDH7,CDKN2C,CEACAM4,CGAS,CISH,CITED2,CLCN4,CLDN7,CLEC1B,CLEC9A,CLU,COL2A1,COX6A1,CREB5,CRIP2,CSGALNACT1,CSNK1A1L,CXCL1,CXCR4,CYTOR,DDIT3,DDIT4,DENND3,DENND5A,DGAT2,DGCR6/LOC102724770,DSEL,DTL,DUSP1,DUSP5,EFCC1,EFNB2,EHD4,ENPP2,ENPP5,EPGN,EPSTI1,EREG,F13A1,FAM124B,FAM83D,FAM89A,FAM98A,FBLN1,FBLN5,FBP1,FBXO45,FCGR1A,FCMR,FILIP1,FKBP5,FKBP9,FLT3,FOSL1,FYB1,GADD45B,GALNT2,GAS2,GBP1,GBP5,GBP7,GIMAP1,GIMAP2,GIMAP6,GIMAP7,GIMAP8,GLUL,GNAI1,GNG4,GPR27,GPR3,GPT2,GULP1,HBA1/HBA2,HBB,HIC2,HIF3A,HLA-C,HLA-DRB5,HLA-F,HLA-G,HOMER3,HOPX,HSPA4L,HYLS1,IER3,IFI44L,IFI6,IFIT3,IFITM1,IGF2BP3,IGLL1/IGLL5,IKZF3,IRF1,IRF7,IRF8,IRF9,IRGQ,ISG15,ISG20,ITGB4,ITGB8,ITPRIP,JCHAIN,KLF10,KLF11,KLF15,KLF2,KLF6,KLF7,KLF9,KLHDC7B,KPNA2,KYNU,LAYN,LIN7A,LONRF2,LTB,MAFF,MAL,MAN2A1,MCAM,MCF2L,METTL7A,MFAP3L,MNS1,MOB3B,MORC1,MRVI1,MT2A,MX1,MYCN,MYRIP,NACA2,NFIB,NFKB2,NFKBIE,NKX2-3,NKX3-1,NLGN1,NOS1AP,NR0B1,NR5A1,NRBP2,NRP2,NRXN2,OAS1,OAS3,OASL,ODF3B,ORC6,PANX1,PARP9,PCDH9,PCSK1N,PDZRN4,PER1,PEX11G,PFKFB2,PHLDB2,PIEZO2,PIK3IP1,PLD6,PLEC,PLK4,PMAIP1,PNMT,POMC,PPP2R2B,PPP2R5B,PRX,PSRC1,PTAFR,PTGER2,PTK2,PTPN14,PTPRE,PTS,PYGO1,RAB11FIP1,RAB27B,RASGRP3,RCSD1,REN,RGS18,RHOB,RHOU,RMDN2,RNF11,RNF175,RPL10,RPL3,RPL7A,RPSA,RSAD2,RUNDC3B,S100A9,S1PR2,SAMD15,SAMHD1,SAP30,SBDS,SCRG1,SELP,SERPINB8,SESN1,SETBP1,SHD,SKIL,SLC22A4,SLC2A9,SLC38A1,SLFN5,SLITRK1,SLITRK5,SMAD3,SMAP2,SPATA2,SPOCK2,SPTY2D1,ST3GAL1,ST6GAL2,STAT1,STAT2,STXBP6,SUCLG2,TAS2R4,TAX1BP3,TBC1D10A,TCEAL2,TENT5C,TMEM117,TNFSF13B,TNNI2,TOB1,TP53INP1,TPSAB1/TPSB2,TRIM69,TSC22D3,TSPAN5,TUBB2A,TUFT1,TXNIP,TYMP,UGCG,USP2,VSTM4,WDFY3,WFDC1,XAF1,XIRP2,YOD1,ZBTB11,ZFP36L2,ZNF608 |
| Cancer,Organismal Injury and Abnormalities | Incidence of tumor | 0.0000309 | Increased | 2.843 | ADAP2,AGAP3,AGPS,AKR1C1/AKR1C2,ANKRD35,ANXA2,AREG,ARRDC3,ASRGL1,ATAD2,ATF3,BCL3,BMPR1B,BRD4,C14orf39,C5orf30,CCDC146,CCDC80,CCND3,CD4,CD69,CD83,CDH7,CDKN2C,CGAS,CLCN4,CLDN7,CLEC1B,CLU,COL2A1,CREB5,CSGALNACT1,CXCL1,CXCR4,CYTOR,DDIT3,DENND3,DENND5A,DGAT2,DGCR6/LOC102724770,DSEL,DUSP1,DUSP5,EFNB2,EHD4,ENPP2,ENPP5,EREG,F13A1,FAM124B,FAM83D,FBLN1,FBLN5,FCGR1A,FCGR1B,FILIP1,FKBP5,FKBP9,FLT3,FYB1,GADD45B,GBP1,GBP7,GIMAP1,GIMAP8,GLUL,GNG4,GPR27,GPR3,GSTM3,GULP1,HBA1/HBA2,HBB,HIC2,HIF3A,HLA-C,HLA-F,HLA-G,HOPX,HSPA4L,HYLS1,IER3,IFI44L,IFIT3,IFITM1,IGF2BP3,IGLL1/IGLL5,IKZF3,IRF1,IRF7,IRF8,IRF9,ISG15,ITGB4,ITGB8,ITPRIP,JCHAIN,KLF10,KLF2,KLF6,KLF9,KLHDC7B,KPNA2,KYNU,LAYN,LIN7A,LONRF2,LTB,MAL,MAN2A1,MCAM,METTL7A,MFAP3L,MORC1,MRVI1,MX1,MYCN,NFIB,NFKB2,NKX3-1,NLGN1,NOS1AP,NR0B1,NR5A1,NRBP2,NRP2,NRXN2,OAS1,OAS3,OASL,PCDH9,PDZRN4,PER1,PEX11G,PHLDB2,PIEZO2,PLEC,PLK4,PMAIP1,POMC,PPP2R2B,PPP2R5B,PRX,PTAFR,PTGER2,PTK2,PTPN14,PTPRE,PYGO1,RASGRP3,REN,RHOB,RHOU,RMDN2,RSAD2,RUNDC3B,S100A9,S1PR2,SAMD15,SAP30,SELP,SERPINB8,SESN1,SETBP1,SKIL,SLC22A4,SLC2A9,SLC38A1,SLFN5,SLITRK1,SLITRK5,SMAD3,SMAP2,SPATA2,SPOCK2,ST6GAL2,STAT1,STAT2,STXBP6,SUCLG2,TBC1D10A,TCEAL2,TENT5C,TMEM117,TOB1,TP53INP1,TPSAB1/TPSB2,TSC22D3,TSPAN5,TUBB2A,TXNIP,TYMP,UGCG,USP2,WDFY3,WFDC1,XAF1,XIRP2,YOD1,ZBTB11,ZFP36L2 |
| Cancer,Organismal Injury and Abnormalities | Nonhematologic malignant neoplasm | 6.26E-09 | Increased | 2.801 | ACADL,ADAP2,AGAP3,AGPS,AK4,AKR1C1/AKR1C2,AKR1C3,ANKRD35,ANXA2,APOL4,AREG,ARRDC3,ASRGL1,ATAD2,ATF3,ATP6V0A1,AZU1,BAALC-AS1,BAG3,BCL3,BMPR1B,BRD4,BTG1,C14orf39,C5orf30,CCDC146,CCDC80,CCND3,CD4,CD53,CD69,CD83,CDC42EP4,CDH7,CDKN2C,CEACAM4,CGAS,CISH,CITED2,CLCN4,CLDN7,CLEC1B,CLEC9A,CLU,COL2A1,COX6A1,CREB5,CRIP2,CSGALNACT1,CSNK1A1L,CXCL1,CXCR4,CYTOR,DDIT3,DDIT4,DENND3,DENND5A,DGAT2,DGCR6/LOC102724770,DSEL,DTL,DUSP1,DUSP5,EFCC1,EFNB2,EHD4,ENPP2,ENPP5,EPGN,EPSTI1,EREG,F13A1,FAM124B,FAM83D,FAM89A,FAM98A,FBLN1,FBLN5,FBP1,FBXO45,FCGR1A,FCGR1B,FCMR,FEM1C,FILIP1,FKBP5,FKBP9,FLT3,FOSL1,FYB1,GADD45B,GALNT2,GAS2,GBP1,GBP5,GBP7,GIMAP1,GIMAP2,GIMAP6,GIMAP7,GIMAP8,GLUL,GNAI1,GNG4,GPR27,GPR3,GPT2,GULP1,HBA1/HBA2,HBB,HIC2,HIF3A,HLA-C,HLA-DRB5,HLA-F,HLA-G,HOMER3,HOPX,HSPA4L,HYLS1,IER3,IFI44L,IFI6,IFIT3,IFITM1,IGF2BP3,IGLL1/IGLL5,IKZF3,IRF1,IRF7,IRF8,IRF9,IRGQ,ISG15,ISG20,ITGB4,ITGB8,ITPRIP,JCHAIN,KLF10,KLF11,KLF15,KLF2,KLF6,KLF7,KLF9,KLHDC7B,KPNA2,KYNU,LAYN,LIN7A,LONRF2,LTB,MAFF,MAL,MAN2A1,MARCKS,MCAM,MCF2L,METTL7A,MFAP3L,MNS1,MOB3B,MORC1,MRVI1,MT2A,MX1,MYCN,MYRIP,NACA2,NFIB,NFKB2,NFKBIE,NKX2-3,NKX3-1,NLGN1,NOS1AP,NR0B1,NR5A1,NRBP2,NRP2,NRXN2,OAS1,OASL,ODF3B,ORC6,PANX1,PARP9,PCDH9,PCLAF,PCSK1N,PDZRN4,PER1,PEX11G,PFKFB2,PHLDB2,PIEZO2,PIK3IP1,PLD6,PLEC,PLK4,PMAIP1,PNMT,POMC,PPP2R2B,PPP2R5B,PRX,PSRC1,PTAFR,PTGER2,PTK2,PTPN14,PTPRE,PTS,PYGO1,RAB11FIP1,RAB27B,RASGRP3,RCSD1,REN,RGS18,RHOB,RHOU,RMDN2,RNF11,RNF175,RPL10,RPL3,RPL7A,RPSA,RSAD2,RUNDC3B,S100A13,S100A9,S1PR2,SAMD15,SAMHD1,SAP30,SBDS,SCRG1,SELP,SERPINB8,SESN1,SETBP1,SHD,SKIL,SLC22A4,SLC2A9,SLC38A1,SLFN5,SLITRK1,SLITRK5,SMAD3,SMAP2,SPATA2,SPOCK2,SPTY2D1,ST3GAL1,ST6GAL2,STAT1,STAT2,STXBP6,SUCLG2,TAS2R4,TAX1BP3,TBC1D10A,TCEAL2,TENT5C,TMEM117,TNFSF13B,TNNI2,TOB1,TP53INP1,TPSAB1/TPSB2,TRIM69,TSC22D3,TSPAN5,TUBB2A,TUFT1,TXNIP,TYMP,UGCG,USP2,VSTM4,WDFY3,WFDC1,XAF1,XIRP2,YOD1,ZBTB11,ZBTB37,ZFP36L2,ZNF608 |
| Cancer,Organismal Injury and Abnormalities | Genitourinary tumor | 1.63E-07 | Increased | 2.778 | ADAP2,AGAP3,AGPS,AKR1C1/AKR1C2,AKR1C3,ANKRD35,ANXA2,APOL4,AREG,ARRDC3,ASRGL1,ATAD2,ATF3,ATP6V0A1,BMPR1B,C14orf39,CCDC146,CCDC80,CCND3,CD4,CD53,CD69,CD83,CDC42EP4,CDH7,CDKN2C,CGAS,CLCN4,CLDN7,CLEC1B,CLEC9A,CLU,COL2A1,COX6A1,CREB5,CSGALNACT1,CXCL1,CXCR4,DDIT3,DDIT4,DENND3,DENND5A,DGAT2,DSEL,DTL,DUSP1,DUSP5,EFCC1,EFNB2,ENPP2,ENPP5,EPSTI1,EREG,F13A1,FAM124B,FAM83D,FAM89A,FBLN1,FBLN5,FCGR1A,FILIP1,FKBP5,FKBP9,FLT3,FOSL1,FYB1,GADD45B,GALNT2,GAS2,GBP7,GIMAP2,GIMAP7,GIMAP8,GLUL,GNAI1,GNG4,GPR3,GSTM3,GULP1,HBA1/HBA2,HBB,HIC2,HIF3A,HLA-C,HLA-DRB5,HLA-G,HOPX,HSPA4L,HYLS1,IER3,IFI44L,IFIT3,IFITM1,IGF2BP3,IGHM,IGLL1/IGLL5,IKZF3,IRF1,IRF7,IRGQ,ISG15,ITGB4,ITGB8,JCHAIN,KLF10,KLF11,KLF2,KLF6,KLF9,KPNA2,KYNU,LAYN,LONRF2,LTB,MAFF,MAL,MAN2A1,MCAM,MFAP3L,MORC1,MRVI1,MT2A,MX1,MYCN,NACA2,NEAT1,NFKBIE,NKX2-3,NKX3-1,NLGN1,NOS1AP,NR0B1,NR5A1,NRBP2,NRP2,NRXN2,OAS3,OASL,PCDH9,PCLAF,PDZRN4,PER1,PEX11G,PHLDB2,PIEZO2,PLEC,PMAIP1,POMC,PPP2R2B,PRX,PTAFR,PTGER2,PTK2,PTPN14,PTPRE,PYGO1,RAB11FIP1,RASGRP3,REN,RHOB,RMDN2,RNASE2,RUNDC3B,S100A9,S1PR2,SAMD15,SAP30,SCRG1,SELP,SERPINB8,SETBP1,SKIL,SLC22A4,SLC2A9,SLC38A1,SLFN5,SLITRK1,SLITRK5,SMAD3,SMAP2,SPOCK2,SPTY2D1,ST6GAL2,STAT1,STAT2,STXBP6,SUCLG2,TAX1BP3,TMEM117,TNFSF13B,TP53INP1,TPSAB1/TPSB2,TRIM69,TSC22D3,TSPAN5,TUBB2A,TXNIP,TYMP,UGCG,USP2,VSTM4,WDFY3,XAF1,XIRP2,YOD1,ZBTB11,ZFP36L2,ZNF608 |
| Cancer,Organismal Injury and Abnormalities | Carcinoma | 6.98E-08 | Increased | 2.444 | ACADL,ADAP2,AGAP3,AGPS,AK4,AKR1C1/AKR1C2,AKR1C3,ANKRD35,ANXA2,APOL4,AREG,ARRDC3,ASRGL1,ATAD2,ATF3,ATP6V0A1,AZU1,BAALC-AS1,BAG3,BCL3,BMPR1B,BRD4,BTG1,C14orf39,C5orf30,CCDC146,CCDC80,CCND3,CD4,CD53,CD69,CD83,CDC42EP4,CDH7,CDKN2C,CEACAM4,CGAS,CISH,CITED2,CLCN4,CLDN7,CLEC1B,CLEC9A,CLU,COL2A1,COX6A1,CREB5,CRIP2,CSGALNACT1,CSNK1A1L,CXCL1,CXCR4,CYTOR,DDIT3,DDIT4,DENND3,DENND5A,DGAT2,DGCR6/LOC102724770,DSEL,DTL,DUSP1,DUSP5,EFCC1,EFNB2,EHD4,ENPP2,ENPP5,EPGN,EPSTI1,EREG,F13A1,FAM124B,FAM83D,FAM89A,FAM98A,FBLN1,FBLN5,FBP1,FBXO45,FCGR1A,FCMR,FILIP1,FKBP5,FKBP9,FLT3,FOSL1,FYB1,GADD45B,GALNT2,GAS2,GBP1,GBP5,GBP7,GIMAP1,GIMAP2,GIMAP6,GIMAP7,GIMAP8,GLUL,GNAI1,GNG4,GPR27,GPR3,GPT2,GULP1,HBA1/HBA2,HBB,HIC2,HIF3A,HLA-C,HLA-DRB5,HLA-F,HLA-G,HOMER3,HOPX,HSPA4L,HYLS1,IER3,IFI44L,IFI6,IFIT3,IFITM1,IGF2BP3,IGLL1/IGLL5,IKZF3,IRF1,IRF7,IRF8,IRF9,IRGQ,ISG15,ISG20,ITGB4,ITGB8,ITPRIP,JCHAIN,KLF10,KLF11,KLF15,KLF2,KLF6,KLF7,KLF9,KLHDC7B,KPNA2,KYNU,LAYN,LIN7A,LONRF2,LTB,MAFF,MAL,MAN2A1,MCAM,MCF2L,METTL7A,MFAP3L,MNS1,MOB3B,MORC1,MRVI1,MT2A,MX1,MYCN,MYRIP,NACA2,NFIB,NFKB2,NFKBIE,NKX2-3,NKX3-1,NLGN1,NOS1AP,NR0B1,NR5A1,NRBP2,NRP2,NRXN2,OAS1,OASL,ODF3B,ORC6,PANX1,PARP9,PCDH9,PCSK1N,PDZRN4,PER1,PEX11G,PFKFB2,PHLDB2,PIEZO2,PIK3IP1,PLD6,PLEC,PLK4,PMAIP1,PNMT,POMC,PPP2R2B,PPP2R5B,PRX,PSRC1,PTAFR,PTGER2,PTK2,PTPN14,PTPRE,PTS,PYGO1,RAB11FIP1,RAB27B,RASGRP3,RCSD1,REN,RGS18,RHOB,RHOU,RMDN2,RNF11,RNF175,RPL10,RPL3,RPL7A,RPSA,RSAD2,RUNDC3B,S100A9,S1PR2,SAMD15,SAMHD1,SAP30,SBDS,SCRG1,SELP,SERPINB8,SESN1,SETBP1,SHD,SKIL,SLC22A4,SLC2A9,SLC38A1,SLFN5,SLITRK1,SLITRK5,SMAD3,SMAP2,SPATA2,SPOCK2,SPTY2D1,ST3GAL1,ST6GAL2,STAT1,STAT2,STXBP6,SUCLG2,TAS2R4,TAX1BP3,TBC1D10A,TCEAL2,TENT5C,TMEM117,TNFSF13B,TNNI2,TOB1,TP53INP1,TPSAB1/TPSB2,TRIM69,TSC22D3,TSPAN5,TUBB2A,TUFT1,TXNIP,TYMP,UGCG,USP2,VSTM4,WDFY3,WFDC1,XAF1,XIRP2,YOD1,ZBTB11,ZFP36L2,ZNF608 |
| Cancer,Organismal Injury and Abnormalities | Non-melanoma solid tumor | 1.71E-09 | Increased | 2.436 | ACADL,ADAP2,AGAP3,AGPS,AK4,AKR1C1/AKR1C2,AKR1C3,ANKRD35,ANXA2,APOL4,AREG,ARRDC3,ASRGL1,ATAD2,ATF3,ATP6V0A1,AZU1,BAALC-AS1,BAG3,BCL3,BMPR1B,BRD4,BTG1,C14orf39,C5orf30,CCDC146,CCDC80,CCND3,CD4,CD53,CD69,CD83,CDC42EP4,CDH7,CDKN2C,CEACAM4,CGAS,CISH,CITED2,CLCN4,CLDN7,CLEC1B,CLEC9A,CLU,COL2A1,COX6A1,CREB5,CRIP2,CSGALNACT1,CSNK1A1L,CXCL1,CXCR4,CYTOR,DDIT3,DDIT4,DENND3,DENND5A,DGAT2,DGCR6/LOC102724770,DSEL,DTL,DUSP1,DUSP5,EFCC1,EFNB2,EHD4,ENPP2,ENPP5,EPGN,EPSTI1,EREG,F13A1,FAM124B,FAM83D,FAM89A,FAM98A,FBLN1,FBLN5,FBP1,FBXO45,FCGR1A,FCMR,FILIP1,FKBP5,FKBP9,FLT3,FOSL1,FYB1,GADD45B,GALNT2,GAS2,GBP1,GBP5,GBP7,GIMAP1,GIMAP2,GIMAP6,GIMAP7,GIMAP8,GLUL,GNAI1,GNG4,GPR27,GPR3,GPT2,GSTM3,GULP1,HBA1/HBA2,HBB,HIC2,HIF3A,HLA-C,HLA-DRB5,HLA-F,HLA-G,HOMER3,HOPX,HSPA4L,HYLS1,IER3,IFI44L,IFI6,IFIT3,IFITM1,IGF2BP3,IGHM,IGLL1/IGLL5,IKZF3,IRF1,IRF7,IRF8,IRF9,IRGQ,ISG15,ISG20,ITGB4,ITGB8,ITPRIP,JCHAIN,KLF10,KLF11,KLF15,KLF2,KLF6,KLF7,KLF9,KLHDC7B,KPNA2,KYNU,LAYN,LIN7A,LONRF2,LTB,MAFF,MAL,MAN2A1,MARCKS,MCAM,MCF2L,METTL7A,MFAP3L,MNS1,MOB3B,MORC1,MRVI1,MT2A,MX1,MYCN,MYRIP,NACA2,NFIB,NFKB2,NFKBIE,NKX2-3,NKX3-1,NLGN1,NOS1AP,NR0B1,NR5A1,NRBP2,NRP2,NRXN2,OAS1,OASL,ODF3B,ORC6,PANX1,PARP9,PCDH9,PCLAF,PCSK1N,PDZRN4,PER1,PEX11G,PFKFB2,PHLDB2,PIEZO2,PIK3IP1,PLD6,PLEC,PLK4,PMAIP1,PNMT,POMC,PPP2R2B,PPP2R5B,PRX,PSRC1,PTAFR,PTGER2,PTK2,PTPN14,PTPRE,PTS,PYGO1,RAB11FIP1,RAB27B,RASGRP3,RCSD1,REN,RGS18,RHOB,RHOU,RMDN2,RNF11,RNF175,RPL10,RPL3,RPL7A,RPSA,RSAD2,RUNDC3B,S100A13,S100A9,S1PR2,SAMD15,SAMHD1,SAP30,SBDS,SCRG1,SELP,SERPINB8,SESN1,SETBP1,SHD,SKIL,SLC22A4,SLC2A9,SLC38A1,SLFN5,SLITRK1,SLITRK5,SMAD3,SMAP2,SPATA2,SPOCK2,SPTY2D1,ST3GAL1,ST6GAL2,STAT1,STAT2,STXBP6,SUCLG2,TAS2R4,TAX1BP3,TBC1D10A,TCEAL2,TENT5C,TMEM117,TNFSF13B,TNNI2,TOB1,TP53INP1,TPSAB1/TPSB2,TRIM69,TSC22D3,TSPAN5,TUBB2A,TUFT1,TXNIP,TYMP,UGCG,USP2,VSTM4,WDFY3,WFDC1,XAF1,XIRP2,YOD1,ZBTB11,ZBTB37,ZFP36L2,ZNF608 |
| Cancer,Organismal Injury and Abnormalities,Reproductive System Disease | Mammary tumor | 0.000435 | Increased | 2.399 | AGAP3,AGPS,AKR1C1/AKR1C2,AKR1C3,AREG,ARRDC3,ATAD2,ATF3,C14orf39,CCND3,CD4,CD69,CDKN2C,CLU,CXCL1,CXCR4,DDIT4,DTL,DUSP1,EPSTI1,FBLN1,FBLN5,FKBP5,FLT3,FOSL1,GADD45B,GBP7,GLUL,GSTM3,GULP1,HBA1/HBA2,HBB,HLA-DRB5,HLA-G,IER3,IRF1,IRF7,KLF10,KLF11,KLF6,KPNA2,MAFF,MCAM,MT2A,MYCN,NEAT1,NR5A1,NRP2,NRXN2,PCDH9,PDZRN4,PIEZO2,PLEC,PMAIP1,PRX,PTGER2,PTK2,PTPRE,RAB11FIP1,RHOB,RMDN2,S100A9,SCRG1,SETBP1,SKIL,SLC22A4,SMAD3,STAT1,TAX1BP3,TMEM117,TPSAB1/TPSB2,TRIM69,TUBB2A,TYMP,UGCG,WDFY3,ZFP36L2 |
| Infectious Diseases | Infection of mammalia | 8.76E-10 | Increased | 2.399 | BCL3,CCL3L3,CD4,CXCR4,DDIT3,DUSP1,FCGR1A,FCMR,HLA-G,IGHM,IRF1,IRF7,IRF8,IRF9,ISG15,JCHAIN,LTB,NFKB2,OASL,PTAFR,SAMHD1,SELP,SMAD3,STAT1,STAT2,TPSAB1/TPSB2 |
| Organ Development,Reproductive System Development and Function | Growth of ovarian follicle | 0.0000144 | Increased | 2.395 | AREG,BMPR1B,EREG,GAS2,NR5A1,PTGER2,SMAD3 |
| Hematological Disease | Hemorrhagic disease | 0.00000783 | Increased | 2.345 | AZU1,CCL3L3,CLEC1B,CXCL1,F13A1,FCGR1A,FCGR1B,FLT3,FYB1,GSTM3,IRF7,ISG15,PTAFR,RGS18,RPL3,S100A9,TNFSF13B,ZFP36L2 |
| Cellular Growth and Proliferation,Tissue Development | Proliferation of epithelial cells | 2.61E-08 | Increased | 2.298 | AREG,ATF3,CCND3,CD4,CDKN2C,CITED2,CLU,CXCR4,EFNB2,EPGN,EREG,FBLN5,GLUL,ITGB8,KLF10,KLF2,KLF9,LTB,NFIB,NKX2-3,NKX3-1,NR5A1,PIK3IP1,POMC,PTAFR,PTGER2,PTK2,S1PR2,SMAD3,TOB1,WFDC1,YOD1 |
| Cell Signaling | Viral life cycle | 0.000256 | Increased | 2.2 | CXCR4,IFITM1,ISG15,ISG20,MX1,OAS1,OAS3,OASL,RSAD2 |
| Cell Signaling | Replication of viral replicon | 0.0000024 | Increased | 2.197 | IFITM1,ISG15,ISG20,MX1,OAS1,OAS3,OASL,RSAD2 |
| Cancer,Organismal Injury and Abnormalities | Anogenital cancer | 0.0000165 | Increased | 2.184 | ADAP2,AGAP3,AGPS,AKR1C1/AKR1C2,AKR1C3,ANKRD35,ANXA2,APOL4,AREG,ARRDC3,ATAD2,ATF3,ATP6V0A1,BMPR1B,C14orf39,CCDC146,CCDC80,CCND3,CD4,CD69,CD83,CDC42EP4,CDH7,CDKN2C,CLCN4,CLDN7,CLEC9A,CLU,COL2A1,CREB5,CSGALNACT1,CXCL1,CXCR4,DDIT3,DDIT4,DENND3,DENND5A,DSEL,DTL,DUSP1,DUSP5,EFNB2,ENPP2,EPSTI1,F13A1,FAM124B,FAM83D,FAM89A,FBLN1,FBLN5,FILIP1,FKBP5,FKBP9,FLT3,FOSL1,FYB1,GADD45B,GALNT2,GAS2,GBP7,GIMAP2,GIMAP7,GLUL,GPR3,GSTM3,GULP1,HBA1/HBA2,HBB,HIC2,HIF3A,HLA-C,HLA-DRB5,HLA-G,HSPA4L,HYLS1,IER3,IFI44L,IFIT3,IFITM1,IGF2BP3,IGHM,IGLL1/IGLL5,IKZF3,IRF1,IRF7,IRGQ,ISG15,ITGB4,ITGB8,JCHAIN,KLF10,KLF11,KLF2,KLF6,KLF9,KPNA2,LAYN,LTB,MAFF,MAL,MAN2A1,MCAM,MORC1,MT2A,MX1,MYCN,NACA2,NEAT1,NKX2-3,NKX3-1,NLGN1,NOS1AP,NR0B1,NR5A1,NRBP2,NRP2,NRXN2,OAS3,OASL,PCDH9,PCLAF,PDZRN4,PER1,PEX11G,PHLDB2,PIEZO2,PLEC,PMAIP1,POMC,PRX,PTAFR,PTGER2,PTK2,PTPN14,PTPRE,PYGO1,RAB11FIP1,RASGRP3,REN,RHOB,RMDN2,RNASE2,RUNDC3B,S100A9,S1PR2,SAMD15,SAP30,SCRG1,SELP,SERPINB8,SETBP1,SKIL,SLC22A4,SLC38A1,SLFN5,SLITRK1,SMAD3,SMAP2,SPOCK2,SPTY2D1,ST6GAL2,STAT1,STAT2,STXBP6,SUCLG2,TAX1BP3,TMEM117,TNFSF13B,TP53INP1,TPSAB1/TPSB2,TRIM69,TSC22D3,TSPAN5,TUBB2A,TXNIP,TYMP,UGCG,USP2,VSTM4,WDFY3,XAF1,XIRP2,YOD1,ZFP36L2,ZNF608 |
| Organ Development,Reproductive System Development and Function | Growth of genital organ | 0.000219 | Increased | 2.183 | ANXA2,AREG,BMPR1B,EREG,GAS2,NR5A1,PTGER2,SMAD3,TSC22D3 |
| Tissue Development | Growth of epithelial tissue | 2.38E-08 | Increased | 2.095 | AREG,ATF3,CCND3,CD4,CDKN2C,CITED2,CLEC1B,CLU,CXCL1,CXCR4,EFNB2,EPGN,EREG,FBLN5,GBP1,GLUL,ITGB8,KLF10,KLF2,KLF9,LTB,MCAM,NFIB,NKX2-3,NKX3-1,NR0B1,NR5A1,NRP2,PIK3IP1,POMC,PTAFR,PTGER2,PTK2,S100A9,S1PR2,SMAD3,STAT1,TOB1,WFDC1,YOD1 |
| Cancer,Organismal Injury and Abnormalities | Malignant genitourinary solid tumor | 1.46E-07 | Increased | 2.081 | ADAP2,AGAP3,AGPS,AKR1C1/AKR1C2,AKR1C3,ANKRD35,ANXA2,APOL4,AREG,ARRDC3,ASRGL1,ATAD2,ATF3,ATP6V0A1,BMPR1B,C14orf39,CCDC146,CCDC80,CCND3,CD4,CD53,CD69,CD83,CDC42EP4,CDH7,CDKN2C,CGAS,CLCN4,CLDN7,CLEC1B,CLEC9A,CLU,COL2A1,COX6A1,CREB5,CSGALNACT1,CXCL1,CXCR4,DDIT3,DDIT4,DENND3,DENND5A,DGAT2,DSEL,DTL,DUSP1,DUSP5,EFCC1,EFNB2,ENPP2,ENPP5,EPSTI1,EREG,F13A1,FAM124B,FAM83D,FAM89A,FBLN1,FBLN5,FCGR1A,FILIP1,FKBP5,FKBP9,FLT3,FOSL1,FYB1,GADD45B,GALNT2,GAS2,GBP7,GIMAP2,GIMAP7,GIMAP8,GLUL,GNAI1,GNG4,GPR3,GSTM3,GULP1,HBA1/HBA2,HBB,HIC2,HIF3A,HLA-C,HLA-DRB5,HLA-G,HSPA4L,HYLS1,IER3,IFI44L,IFIT3,IFITM1,IGF2BP3,IGHM,IGLL1/IGLL5,IKZF3,IRF1,IRF7,IRGQ,ISG15,ITGB4,ITGB8,JCHAIN,KLF10,KLF11,KLF2,KLF6,KLF9,KPNA2,KYNU,LAYN,LONRF2,LTB,MAFF,MAL,MAN2A1,MCAM,MORC1,MT2A,MX1,MYCN,NACA2,NEAT1,NFKBIE,NKX2-3,NKX3-1,NLGN1,NOS1AP,NR0B1,NR5A1,NRBP2,NRP2,NRXN2,OAS3,OASL,PCDH9,PCLAF,PDZRN4,PER1,PEX11G,PHLDB2,PIEZO2,PLEC,PMAIP1,POMC,PPP2R2B,PRX,PTAFR,PTGER2,PTK2,PTPN14,PTPRE,PYGO1,RAB11FIP1,RASGRP3,REN,RHOB,RMDN2,RNASE2,RUNDC3B,S100A9,S1PR2,SAMD15,SAP30,SCRG1,SELP,SERPINB8,SETBP1,SKIL,SLC22A4,SLC2A9,SLC38A1,SLFN5,SLITRK1,SLITRK5,SMAD3,SMAP2,SPOCK2,SPTY2D1,ST6GAL2,STAT1,STAT2,STXBP6,SUCLG2,TAX1BP3,TMEM117,TNFSF13B,TP53INP1,TPSAB1/TPSB2,TRIM69,TSC22D3,TSPAN5,TUBB2A,TXNIP,TYMP,UGCG,USP2,VSTM4,WDFY3,XAF1,XIRP2,YOD1,ZBTB11,ZFP36L2,ZNF608 |
| Cancer,Organismal Injury and Abnormalities | Benign solid tumor | 3.98E-07 | Increased | 2.074 | ANXA2,BCL3,C5orf30,CCDC80,CCND3,CDKN2C,CLU,COL2A1,CXCR4,DGAT2,DUSP1,DUSP5,FBLN1,FLT3,GLUL,HOPX,IFI44L,IFIT3,JCHAIN,KLF10,KLF11,KLF2,MFAP3L,MRVI1,MYCN,NFIB,NR0B1,NR5A1,NRP2,POMC,PTGER2,PTK2,S100A9,SKIL,SMAD3,STXBP6,TNFSF13B,TNNI2,TP53INP1,TUBB2A,UGCG,XAF1 |
| Cancer,Organismal Injury and Abnormalities | Pelvic tumor | 0.0000116 | Increased | 2.041 | ADAP2,AGPS,AKR1C1/AKR1C2,AKR1C3,ANKRD35,ANXA2,APOL4,ARRDC3,ATAD2,ATF3,ATP6V0A1,BMPR1B,C14orf39,CCDC146,CCDC80,CCND3,CD69,CD83,CDC42EP4,CDH7,CDKN2C,CGAS,CLCN4,CLDN7,CLEC9A,CLU,COL2A1,COX6A1,CREB5,CSGALNACT1,CXCL1,CXCR4,DDIT3,DDIT4,DENND3,DENND5A,DSEL,DUSP1,DUSP5,EFCC1,EFNB2,ENPP2,EPSTI1,F13A1,FAM124B,FAM83D,FAM89A,FBLN1,FBLN5,FILIP1,FKBP5,FKBP9,FLT3,FOSL1,FYB1,GALNT2,GAS2,GBP1,GIMAP2,GIMAP7,GLUL,GNG4,GPR3,GSTM3,GULP1,HBA1/HBA2,HBB,HIC2,HIF3A,HLA-C,HLA-G,HOPX,HSPA4L,HYLS1,IER3,IFI44L,IFIT3,IGF2BP3,IGHM,IGLL1/IGLL5,IKZF3,IRF1,IRF7,IRGQ,ISG15,ITGB4,ITGB8,JCHAIN,KLF11,KLF2,KLF6,KLF9,KPNA2,KYNU,LAYN,LTB,MAL,MAN2A1,MCAM,MFAP3L,MORC1,MRVI1,MX1,MYCN,NACA2,NFKBIE,NKX2-3,NKX3-1,NLGN1,NOS1AP,NR0B1,NR5A1,NRBP2,NRP2,NRXN2,OAS3,OASL,PCDH9,PCLAF,PDZRN4,PER1,PEX11G,PHLDB2,PIEZO2,PLEC,PMAIP1,POMC,PPP2R2B,PRX,PTAFR,PTGER2,PTK2,PTPN14,PTPRE,PYGO1,RAB11FIP1,RASGRP3,REN,RHOB,RMDN2,RNASE2,RUNDC3B,S100A9,S1PR2,SAMD15,SAP30,SELP,SERPINB8,SETBP1,SKIL,SLC38A1,SLFN5,SLITRK1,SLITRK5,SMAD3,SMAP2,SPOCK2,SPTY2D1,ST6GAL2,STAT1,STAT2,STXBP6,SUCLG2,TMEM117,TNFSF13B,TP53INP1,TPSAB1/TPSB2,TSC22D3,TSPAN5,TUBB2A,TXNIP,TYMP,UGCG,USP2,VSTM4,WDFY3,XAF1,XIRP2,YOD1,ZBTB11,ZFP36L2,ZNF608 |
| Cancer,Organismal Injury and Abnormalities | Solid tumor | 5.52E-10 | Increased | 2.039 | ACADL,ADAP2,AGAP3,AGPS,AK4,AKR1C1/AKR1C2,AKR1C3,ANKRD35,ANXA2,APOL4,AREG,ARRDC3,ASRGL1,ATAD2,ATF3,ATP6V0A1,AZU1,BAALC-AS1,BAG3,BCL3,BMPR1B,BRD4,BTG1,C14orf39,C5orf30,CCDC146,CCDC80,CCND3,CD4,CD53,CD69,CD83,CDC42EP4,CDH7,CDKN2C,CEACAM4,CGAS,CISH,CITED2,CLCN4,CLDN7,CLEC1B,CLEC9A,CLU,COL2A1,COX6A1,CREB5,CRIP2,CSGALNACT1,CSNK1A1L,CXCL1,CXCR4,CYTOR,DDIT3,DDIT4,DENND3,DENND5A,DGAT2,DGCR6/LOC102724770,DSEL,DTL,DUSP1,DUSP5,EFCC1,EFNB2,EHD4,ENPP2,ENPP5,EPGN,EPSTI1,EREG,F13A1,FAM124B,FAM83D,FAM89A,FAM98A,FBLN1,FBLN5,FBP1,FBXO45,FCGR1A,FCGR1B,FCMR,FEM1C,FILIP1,FKBP5,FKBP9,FLT3,FOSL1,FYB1,GADD45B,GALNT2,GAS2,GBP1,GBP5,GBP7,GIMAP1,GIMAP2,GIMAP6,GIMAP7,GIMAP8,GLUL,GNAI1,GNG4,GPR27,GPR3,GPT2,GSTM3,GULP1,H4C12,H4C3,HBA1/HBA2,HBB,HIC2,HIF3A,HLA-C,HLA-DRB5,HLA-F,HLA-G,HOMER3,HOPX,HSPA4L,HYLS1,IER3,IFI44L,IFI6,IFIT3,IFITM1,IGF2BP3,IGHM,IGLL1/IGLL5,IKZF3,IRF1,IRF7,IRF8,IRF9,IRGQ,ISG15,ISG20,ITGB4,ITGB8,ITPRIP,JCHAIN,KLF10,KLF11,KLF15,KLF2,KLF6,KLF7,KLF9,KLHDC7B,KPNA2,KYNU,LAYN,LIN7A,LONRF2,LTB,MAFF,MAL,MAN2A1,MARCKS,MCAM,MCF2L,METTL7A,MFAP3L,MNS1,MOB3B,MORC1,MRVI1,MT2A,MX1,MYCN,MYRIP,NACA2,NEAT1,NFIB,NFKB2,NFKBIE,NKX2-3,NKX3-1,NLGN1,NOS1AP,NR0B1,NR5A1,NRBP2,NRP2,NRXN2,OAS1,OAS3,OASL,ODF3B,ORC6,PANX1,PARP9,PCDH9,PCLAF,PCSK1N,PDZRN4,PER1,PEX11G,PFKFB2,PHLDB2,PIEZO2,PIK3IP1,PLD6,PLEC,PLK4,PMAIP1,PNMT,POMC,PPP2R2B,PPP2R5B,PRX,PSRC1,PTAFR,PTGER2,PTK2,PTPN14,PTPRE,PTS,PYGO1,RAB11FIP1,RAB27B,RASGRP3,RCSD1,REN,RGS18,RHOB,RHOU,RMDN2,RNASE2,RNF11,RNF175,RPL10,RPL3,RPL36A,RPL7A,RPSA,RSAD2,RUNDC3B,S100A13,S100A9,S1PR2,SAMD15,SAMHD1,SAP30,SBDS,SCRG1,SELP,SERPINB8,SESN1,SETBP1,SHD,SKIL,SLC22A4,SLC2A9,SLC38A1,SLFN5,SLITRK1,SLITRK5,SMAD3,SMAP2,SPATA2,SPOCK2,SPTY2D1,ST3GAL1,ST6GAL2,STAT1,STAT2,STXBP6,SUCLG2,TAS2R4,TAX1BP3,TBC1D10A,TCEAL2,TENT5C,TMEM117,TNFSF13B,TNNI2,TOB1,TP53INP1,TPSAB1/TPSB2,TRIM69,TSC22D3,TSPAN5,TUBB2A,TUFT1,TXNIP,TYMP,UGCG,USP2,VSTM4,WDFY3,WFDC1,XAF1,XIRP2,YOD1,ZBTB11,ZBTB37,ZFP36L2,ZNF608 |
| Humoral Immune Response,Protein Synthesis | Quantity of IgG2a | 0.000041 | Decreased | -2.121 | ATF3,BCL3,CD69,CD83,IGHM,IKZF3,LTB,NFKB2,RASGRP3 |
| Gene Expression | Expression of RNA | 0.00000223 | Decreased | -2.125 | ATAD2,ATF3,AZU1,BCL3,BMPR1B,BRD4,BTG1,CCL3L3,CCND3,CD4,CDKN2C,CISH,CITED2,CLU,COL2A1,CREB5,CXCR4,DDIT3,DUSP1,DUSP5,EREG,FAM98A,FBLN1,FBP1,FCGR1A,FOSL1,HBB,HIF3A,HLA-DRB5,HOPX,IGF2BP3,IKZF3,IRF1,IRF7,IRF8,IRF9,ISG20,ITGB8,KLF10,KLF11,KLF15,KLF2,KLF6,KLF7,KLF9,KPNA2,MACROH2A2,MAFF,MCF2L,MORC1,MYCN,NFIB,NFKB2,NKX2-3,NKX3-1,NOS1AP,NR0B1,NR5A1,PARP9,PER1,POMC,PPP2R5B,PSRC1,PTAFR,PTGER2,PTK2,PYGO1,REN,RHOB,RNF11,RPL10,S100A9,SAP30,SKIL,SMAD3,SPTY2D1,STAT1,STAT2,TNNI2,TOB1,TP53INP1,TSC22D3,TXNIP,USP2,ZNF608 |
| Inflammatory Disease,Neurological Disease,Skeletal and Muscular Disorders | Multiple Sclerosis | 0.0000877 | Decreased | -2.138 | AREG,CXCL1,DDIT3,EPSTI1,FCGR1A,FCGR1B,HLA-F,HLA-G,IRF7,ISG15,MCAM,MX1,OAS1,OAS3,RSAD2 |
| Neurological Disease,Skeletal and Muscular Disorders | Neuromuscular disease | 0.000618 | Decreased | -2.138 | ANXA2,AREG,BCL3,CLU,CXCL1,DDIT3,DUSP5,ENPP5,EPSTI1,FCGR1A,FCGR1B,GBP1,HLA-F,HLA-G,IER3,IGHM,IRF7,IRF8,ISG15,ITGB4,MAFF,MCAM,MT2A,MX1,OAS1,OAS3,PTPRE,RPL3,RPSA,RSAD2,S1PR2,TNNI2,TSC22D3,UGCG,USP2 |
| Cell Death and Survival | Cell death of macrophages | 0.00048 | Decreased | -2.184 | BTG1,CGAS,DDIT3,GBP1,IRF1,ISG15,PMAIP1,SPATA2,STAT1 |
| Cell Death and Survival | Cell death of blood cells | 1.57E-13 | Decreased | -2.21 | ATF3,BCL3,BTG1,CCL3L3,CD4,CD69,CGAS,CISH,CLU,CXCL1,CXCR4,DDIT3,DDIT4,DUSP5,FCGR1A,FCMR,FLT3,FOSL1,GADD45B,GBP1,GIMAP1,HLA-G,IER3,IGHM,IKZF3,IRF1,IRF8,ISG15,KLF2,LTB,MCF2L,MX1,MZB1,NFKB2,NR5A1,PCLAF,PMAIP1,PPP2R2B,PTK2,S1PR2,SBDS,SKIL,SMAD3,SPATA2,ST3GAL1,STAT1,TNFSF13B,TPSAB1/TPSB2,TSC22D3 |
| Gene Expression | Transcription | 6.29E-07 | Decreased | -2.249 | ATAD2,ATF3,AZU1,BCL3,BMPR1B,BRD4,BTG1,CCL3L3,CCND3,CD4,CDKN2C,CISH,CITED2,CLU,COL2A1,CREB5,CXCR4,DDIT3,DUSP1,DUSP5,EREG,FAM98A,FBLN1,FBP1,FCGR1A,FOSL1,HBB,HIF3A,HLA-DRB5,HOPX,IKZF3,IRF1,IRF7,IRF8,IRF9,ITGB8,KLF10,KLF11,KLF15,KLF2,KLF6,KLF7,KLF9,KPNA2,MACROH2A2,MAFF,MCF2L,MORC1,MX1,MYCN,NFIB,NFKB2,NKX2-3,NKX3-1,NOS1AP,NR0B1,NR5A1,PARP9,PER1,POMC,PPP2R5B,PSRC1,PTAFR,PTGER2,PTK2,PYGO1,REN,RHOB,RNF11,RPL10,SAP30,SKIL,SMAD3,SPTY2D1,STAT1,STAT2,TNFSF13B,TNNI2,TP53INP1,TSC22D3,TXNIP,USP2,ZNF608 |
| Cellular Movement | Cell movement | 2.84E-10 | Decreased | -2.252 | ANXA2,AREG,ARRDC3,ATF3,AZU1,BAG3,BCL3,BRD4,C5orf30,CCL3L3,CD4,CD69,CDC42EP4,CDKN2C,CEACAM4,CITED2,CLCN4,CLDN7,CLEC1B,CLU,COL2A1,CRIP2,CXCL1,CXCR4,CYTOR,DDIT4,DTL,DUSP1,DUSP5,EFNB2,ENPP2,EREG,F13A1,FBLN5,FBXO45,FOSL1,FYB1,GALNT2,GBP1,GLUL,GNAI1,HLA-G,HOMER3,IFITM1,IGF2BP3,IGHM,IGLL1/IGLL5,IRF7,IRF8,ITGB4,ITGB8,JCHAIN,KLF2,KLF6,KPNA2,LTB,MARCKS,MCAM,MCF2L,MX1,NFKB2,NKX2-3,NKX3-1,NR0B1,NRP2,PARP9,PER1,PIEZO2,PIK3IP1,PLEC,PLK4,POMC,PTAFR,PTK2,PTPN14,RAB11FIP1,RASGRP3,REN,RGS18,RHOB,RHOU,RNASE2,RNF11,RPSA,S100A9,S1PR2,SBDS,SELP,SKIL,SMAD3,STAT1,STAT2,TNFSF13B,TOB1,TP53INP1,TPSAB1/TPSB2,TSC22D3,TSPAN5,TYMP,USP2 |
| Antimicrobial Response,Inflammatory Response | Antiviral response | 2.92E-14 | Decreased | -2.374 | AZU1,BCL3,CGAS,CLU,CXCR4,DDIT4,FCGR1A,FOSL1,IFI44L,IFI6,IFIT3,IFITM1,IRF1,IRF7,ISG15,ISG20,MX1,OAS1,OAS3,OASL,PARP9,PMAIP1,RNASE2,RSAD2,SAMHD1,STAT1,STAT2 |
| Cell Death and Survival | Cell death of immune cells | 8.77E-13 | Decreased | -2.594 | ATF3,BCL3,BTG1,CD4,CD69,CGAS,CISH,CLU,CXCL1,CXCR4,DDIT3,DDIT4,DUSP5,FCGR1A,FCMR,FLT3,FOSL1,GADD45B,GBP1,GIMAP1,HLA-G,IER3,IGHM,IKZF3,IRF1,IRF8,ISG15,KLF2,LTB,MCF2L,MX1,MZB1,NFKB2,NR5A1,PMAIP1,PPP2R2B,PTK2,S1PR2,SKIL,SMAD3,SPATA2,ST3GAL1,STAT1,TNFSF13B,TPSAB1/TPSB2,TSC22D3 |
| Inflammatory Disease,Neurological Disease,Skeletal and Muscular Disorders | Relapsed multiple sclerosis | 0.0000167 | Decreased | -2.63 | EPSTI1,FCGR1A,FCGR1B,HLA-G,IRF7,ISG15,MX1,OAS1,OAS3,RSAD2 |
| Inflammatory Disease,Neurological Disease,Skeletal and Muscular Disorders | Relapsing-remitting multiple sclerosis | 0.000234 | Decreased | -2.63 | EPSTI1,HLA-G,IRF7,ISG15,MX1,OAS1,OAS3,RSAD2 |

**T2 vs. T1 MPP_GMP**

© 2000-2020 QIAGEN. All rights reserved.

| Categories | Diseases or Functions Annotation | p-value | Predicted Activation State | Activation z-score | Molecules |
|---|---|---|---|---|---|
| Cell Signaling | Replication of viral replicon | 0.000000304 | Increased | 2.814 | IFIT1,IFITM1,IFITM3,ISG15,MX1,OAS1,OASL,RSAD2 |
| Cell Signaling | Viral life cycle | 0.0000316 | Increased | 2.813 | CXCR4,IFIT1,IFITM1,IFITM3,ISG15,MX1,OAS1,OASL,RSAD2 |
| Infectious Diseases | Infection of mammalia | 2.85E-13 | Increased | 2.678 | BCL3,CCR1,CXCR4,DUSP1,ELANE,HLA-G,IFIH1,IFIT2,IL10RA,IRF1,IRF7,IRF8,IRF9,ISG15,JDP2,LGALS3,MPO,NFKB2,NOD2,OASL,RELB,SAMHD1,STAT1,STAT2,TAP1,USP18,VWF |
| Infectious Diseases | Replication of Flaviviridae | 0.0000144 | Increased | 2.645 | IFI6,IFIT1,IFITM1,IFITM3,ISG15,OASL,RSAD2,TXNIP |
| Infectious Diseases | Viral Infection | 2.55E-10 | Increased | 2.588 | ACTA2,AREG,ARRDC3,ATP6V0A1,BCL3,CA2,CCR1,CD3E,CD48,CD69,CLU,CXCR4,DUSP1,EGR1,ELANE,FCGR1B,GBP1,GPT2,HBA1/HBA2,HCP5,HERC5,HLA-C,IFI6,IFIH1,IFIT1,IFIT2,IFIT3,IFITM1,IFITM3,IL10RA,IRF1,IRF7,IRF8,IRF9,ISG15,KLF15,KLF2,LGALS3,MNDA,MT2A,MX1,NACA2,NLRC5,OAS1,OAS2,OASL,PARP9,RELB,RHOB,RPL3,RSAD2,S100A8,S100A9,SAMHD1,ST3GAL5,STAT1,STAT2,TAP1,TUBB2A,TXNIP,VWF |
| Infectious Diseases | Replication of Hepatitis C virus | 0.000052 | Increased | 2.476 | IFI6,IFIT1,IFITM1,IFITM3,ISG15,RSAD2,TXNIP |
| Cancer,Immunological Disease,Organismal Injury and Abnormalities | Hyperplasia of spleen | 0.000152 | Increased | 2.433 | CDKN2C,CISH,EGLN1,GADD45B,IRF8,RELB |
| Infectious Diseases | Replication of Paramyxoviridae | 0.00000575 | Increased | 2.429 | CCR1,IFIH1,IRF9,MX1,OASL,STAT2 |
| Infectious Diseases | Replication of vesicular stomatitis virus | 0.00000228 | Increased | 2.407 | IFIH1,IFITM3,IRF1,IRF9,OAS1,OASL,STAT2,TAP1 |
| Cancer,Organismal Injury and Abnormalities | Frequency of tumor | 0.00000285 | Increased | 2.356 | ABHD12B,ACTA2,AGPS,AKR1C1/AKR1C2,AREG,ARHGAP44,ARRDC3,ATF3,ATP8B3,BCL3,C5orf30,CA2,CACNB4,CAVIN4,CCR1,CD3G,CD48,CD69,CD72,CDK1,CDKN2C,CLU,COL2A1,CSF1R,CSGALNACT1,CXCR4,DENND5A,DIRAS3,DLK1,DTX3L,DUSP1,DUSP5,DUSP6,EGR1,ELMSAN1,ENPP5,EREG,FAM193A,FCGR1B,FKBP1C,FKBP5,FLOT1,FLT3,FYB1,GADD45B,GALNT3,GBP1,GBP7,GIMAP1,GNG4,GPC1,GSTM3,HBB,HELZ2,HERC5,HIC2,HLA-C,HLA-F,HLA-G,HPSE,ID2,IFI44,IFI44L,IFIH1,IFIT3,IFITM1,IFITM3,IGSF6,IKZF3,IL10RA,INHBA,IRF1,IRF7,IRF8,IRF9,IRX3,ITGB8,KCNC1,KIF15,KLF2,KLHDC7B,LGALS3,LILRB3,LY86,MAL,MNDA,MPO,MX1,MYCN,NCF2,NHSL2,NLRC5,NOD2,NOG,NR0B1,NR5A1,NRXN2,NTNG2,OAS1,OAS2,OASL,OTULINL,PER1,PEX11G,PIEZO2,POMC,PPP2R5B,PTPRE,PYGO1,REN,RHOB,RIMS1,RSAD2,S100A8,S100A9,S1PR2,SAMD4A,SAMD9L,SAP30,SAPCD2,SESN1,SETBP1,SKIL,SLC12A5,SLC16A4,SLC1A3,SLITRK5,SNCAIP,SPATA2,ST3GAL5,STAT1,STAT2,TAP1,TCL1A,TENT5C,TMEM176A,TMEM176B,TMEM255B,TNFRSF10D,TOB1,TP53INP1,TP53INP2,TPRN,TRIP13,TRMT9B,TSC22D3,TUBB2A,TXNIP,U2AF1/U2AF1L5,USP18,VWF,WDR74,WEE1,ZC3H6,ZFP36L2,ZNF222,ZNF462 |
| Infectious Diseases | Replication of virus | 7.96E-11 | Increased | 2.261 | AREG,BCL3,CCR1,CD48,CXCR4,DUSP1,EGR1,GBP1,HERC5,IFI6,IFIH1,IFIT1,IFITM1,IFITM3,IL10RA,IRF1,IRF7,IRF9,ISG15,MNDA,MX1,NLRC5,OAS1,OASL,RSAD2,S100A8,S100A9,SAMHD1,STAT1,STAT2,TAP1,TXNIP |
| Cancer,Organismal Injury and Abnormalities | Development of malignant tumor | 0.0000161 | Increased | 2.251 | ABHD12B,ACTA2,AGPS,AKR1C1/AKR1C2,AREG,ARHGAP44,ARRDC3,ATF3,ATP8B3,BCL3,CA2,CACNB4,CAVIN4,CCR1,CD3G,CD48,CD69,CD72,CDK1,CDKN2C,CLU,COL2A1,CSF1R,CSGALNACT1,CXCR4,DENND5A,DIRAS3,DLK1,DTX3L,DUSP1,DUSP5,DUSP6,EGR1,ELMSAN1,ENPP5,EREG,FAM193A,FCGR1B,FKBP1C,FKBP5,FLOT1,FLT3,FYB1,GADD45B,GALNT3,GBP1,GBP7,GIMAP1,GNG4,GPC1,GSTM3,HBB,HELZ2,HERC5,HIC2,HLA-C,HLA-F,HLA-G,HPSE,ID2,IFI44,IFI44L,IFIH1,IFIT3,IFITM1,IGSF6,IKZF3,IL10RA,INHBA,IRF1,IRF7,IRF8,IRF9,IRX3,ITGB8,KCNC1,KIF15,KLHDC7B,LGALS3,LILRB3,LY86,MAL,MNDA,MPO,MX1,MYCN,NCF2,NHSL2,NLRC5,NOD2,NOG,NR0B1,NR5A1,NRXN2,NTNG2,OAS1,OAS2,OASL,OTULINL,PER1,PEX11G,PIEZO2,POMC,PPP2R5B,PTPRE,PYGO1,REN,RHOB,RIMS1,RSAD2,S100A8,S100A9,S1PR2,SAMD4A,SAMD9L,SAP30,SAPCD2,SESN1,SETBP1,SKIL,SLC12A5,SLC16A4,SLC1A3,SLITRK5,SNCAIP,SPATA2,ST3GAL5,STAT1,STAT2,TAP1,TCL1A,TENT5C,TMEM176A,TMEM176B,TMEM255B,TNFRSF10D,TOB1,TP53INP1,TPRN,TRIP13,TRMT9B,TSC22D3,TUBB2A,TXNIP,U2AF1/U2AF1L5,USP18,VWF,WDR74,ZC3H6,ZFP36L2,ZNF222,ZNF462 |
| Infectious Diseases | Replication of RNA virus | 8.27E-09 | Increased | 2.215 | AREG,BCL3,CCR1,CD48,CXCR4,GBP1,HERC5,IFI6,IFIH1,IFIT1,IFITM1,IFITM3,IRF1,IRF7,IRF9,ISG15,MX1,NLRC5,OAS1,OASL,RSAD2,S100A8,S100A9,STAT1,STAT2,TAP1,TXNIP |
| Cancer,Hematological Disease,Organismal Injury and Abnormalities | Hematologic cancer | 1.1E-12 | Increased | 2.169 | ATF3,BCL3,BTG1,CA2,CACNB4,CD3E,CD3G,CD48,CD69,CDK1,CDKN2C,CITED2,CLU,COL2A1,CSF1R,CXCR4,DHCR24,DUSP5,EGR1,EPSTI1,FCGR1B,FKBP1C,FLT3,FYB1,GADD45B,GNAI1,GNG4,HBA1/HBA2,HLA-C,HLA-G,ID2,IFI44,IFI44L,IFI6,IFIT1,IFIT2,IFIT3,IFITM1,IKZF3,INHBA,IRF1,IRF7,IRF8,ISG15,KLF2,MAL,MNDA,MPO,MT-ATP6,MX1,MYCN,NACA2,NFKB2,NR5A1,OAS1,OAS2,OASL,PER1,PFKFB2,PTPRE,RELB,RHOB,RIMS1,RPL10,RPL3,RPL41,RPL7,RSAD2,S100A9,S1PR2,SAMD9L,SAPCD2,SBDS,SETBP1,SKIL,SNCAIP,STAT1,STAT2,TCL1A,TENT5C,TMEM255B,TNFRSF10D,TNFSF13B,TOB1,TRIP13,TRMT9B,TUBA1B,TUBB2A,U2AF1/U2AF1L5,VSTM4,VWF,WDR74,WEE1,ZFP36L2 |
| Cancer,Organismal Injury and Abnormalities | Head and neck tumor | 0.00017 | Increased | 2.158 | ABHD12B,ACTA2,AGPS,AREG,ARHGAP18,ARHGAP44,ARRDC4,ATP6V0A1,ATP8B3,BAG3,BCL3,BTG1,C17orf99,C1QTNF4,CA2,CACNB4,CD3E,CD3G,CD48,CD72,CDK1,CDKN2C,CISH,CITED2,CLEC9A,CLU,COL2A1,CSF1R,CXCR4,DENND10,DENND5A,DIRAS3,DUSP1,DUSP5,DUSP6,EGLN1,EGR1,ELANE,ELMSAN1,EME1,EPGN,EPSTI1,EREG,FAM193A,FBP1,FKBP5,FLOT1,FLT3,FOSL1,FYB1,GADD45B,GALNT3,GBP1,GBP5,GBP7,GDF10,GIMAP1,GNAI1,GPC1,GPT2,HELZ2,HERC5,HIC2,HIF3A,HLA-C,HLA-F,HLA-G,HOPX,HPSE,ID2,IFI44,IFI44L,IFI6,IFIH1,IFIT2,IFITM1,IFITM3,IGSF6,IKZF3,IL10RA,INHBA,IRF1,IRF7,IRF8,IRX3,ITGB8,JDP2,KCNC1,KIF15,KIF20A,KLF15,KLF2,KLHDC7B,LGALS3,LILRB3,MAFF,MDFI,METRN,MNDA,MPO,MT1F,MX1,MYCN,NFKB2,NHSL2,NLRC5,NOD2,NOG,NR0B1,NRXN2,NTNG2,OAS2,OASL,ODF3B,PARP9,PER1,PEX11G,PFKFB2,PHLDA3,PIEZO2,PIK3IP1,PNMT,POMC,PPP2R5B,PRDM5,PTPRE,PYGO1,RAB27B,RELB,REN,RIMS1,RPL10,RPL3,RPL7,RPL7A,RSAD2,S100A8,S100A9,S1PR2,SAMD4A,SAMHD1,SAP30,SAPCD2,SBDS,SESN1,SETBP1,SKIL,SLC12A5,SLC1A3,SLC43A2,SLITRK5,SNCAIP,SPATA2,ST3GAL5,STAT1,STAT2,TAP1,TINAGL1,TMEM176A,TMEM176B,TMEM255B,TNFSF13B,TOB1,TP53INP1,TPRN,TRIM69,TRIP13,TUBB2A,TUFT1,U2AF1/U2AF1L5,USP18,USP42,VSTM4,VWF,WDR74,WEE1,XAF1,ZC3H6,ZFP36L2,ZNF462 |
| Infectious Diseases | Replication of Herpesviridae | 4.61E-11 | Increased | 2.145 | EGR1,IFIH1,IL10RA,IRF1,IRF9,MNDA,OAS1,OASL,RSAD2,SAMHD1,STAT1,STAT2 |
| Organismal Survival | Morbidity or mortality | 7.6E-09 | Increased | 2.105 | ACTA2,AREG,ATF3,BAG3,BCL3,CCR1,CD3E,CDK1,CDKN2C,CITED2,COL2A1,CPLX1,CSF1R,CSGALNACT1,CXCR4,DLK1,DUSP1,EGLN1,ELANE,FLT3,FOSL1,GNAI1,HLA-G,HOPX,ID2,IFIH1,IFIT2,IKZF3,INHBA,IRF1,IRF7,IRF8,IRF9,ISG15,ITGB8,KCNC1,KIF20A,KLF2,LGALS3,MAFF,MAL,MDFI,MPO,MYCN,MYZAP,NEAT1,NFKB2,NOD2,NOG,NR5A1,NRXN2,OASL,PIEZO2,POMC,PTS,PYGO1,RELB,REN,RIMS1,S100A8,S100A9,S1PR2,SBDS,SETBP1,SKIL,SLC12A5,SLC1A3,ST3GAL5,STAT1,STAT2,SUMO1P3,TAP1,TINAGL1,TNFSF13B,TP53INP1,TXNIP,U2AF1/U2AF1L5,USP18,VWF,ZFP36L2 |
| Organismal Survival | Organismal death | 2.23E-08 | Increased | 2.061 | ACTA2,AREG,ATF3,BAG3,BCL3,CCR1,CD3E,CDK1,CDKN2C,CITED2,COL2A1,CPLX1,CSF1R,CSGALNACT1,CXCR4,DLK1,DUSP1,EGLN1,ELANE,FLT3,FOSL1,GNAI1,HLA-G,HOPX,ID2,IFIH1,IFIT2,IKZF3,INHBA,IRF1,IRF7,IRF8,IRF9,ISG15,ITGB8,KCNC1,KIF20A,KLF2,LGALS3,MAFF,MAL,MDFI,MPO,MYCN,MYZAP,NEAT1,NFKB2,NOD2,NOG,NR5A1,NRXN2,PIEZO2,POMC,PTS,PYGO1,RELB,REN,RIMS1,S100A8,S100A9,S1PR2,SBDS,SETBP1,SKIL,SLC12A5,SLC1A3,ST3GAL5,STAT1,STAT2,SUMO1P3,TINAGL1,TNFSF13B,TP53INP1,TXNIP,U2AF1/U2AF1L5,USP18,VWF,ZFP36L2 |
| Neurological Disease | Motor dysfunction or movement disorder | 0.000034 | Increased | 2.054 | CA2,CACNB4,CD48,CDKN2C,CLU,CPLX1,CSF1R,DUSP5,EGLN1,EGR1,ENPP5,FCGR1B,FLOT1,GALNT3,GBP1,IFIT2,KCNC1,MAFF,MT-ATP6,MT2A,MYCN,NHSL2,NRXN2,PTPRE,PTS,RELB,RPL3,S1PR2,SAMD9L,SLC12A5,SLC1A3,SNCAIP,ST3GAL5,TMEM176B,TSC22D3,TUBA1B,TUBB2A,USP18 |
| Cancer,Cell Death and Survival,Organismal Injury and Abnormalities,Tumor Morphology | Cell death of cancer cells | 0.000223 | Decreased | -2.028 | ATF3,C1QTNF4,CDK1,CLU,CXCR4,GALNT3,INHBA,IRF1,MYCN,RPL10,RPL3,RPL7,RPL7A,TCL1A,TNFSF13B |
| Cellular Movement | Chemotaxis | 0.00000918 | Decreased | -2.074 | AZU1,CCR1,CD69,CD72,CLU,CSF1R,CXCR4,DUSP6,ELANE,FLOT1,FOSL1,HLA-G,HPSE,INHBA,LGALS3,LILRB3,NFKB2,NOD2,RELB,RHOB,S100A8,S100A9,S1PR2,SBDS |
| Cellular Movement | Cell movement | 5.06E-13 | Decreased | -2.099 | ACTA2,AREG,ARHGAP18,ARRDC3,ATF3,AZU1,BAG3,BCL3,C5orf30,CA2,CCR1,CD3E,CD48,CD69,CD72,CDK1,CDKN2C,CITED2,CLU,COL2A1,CSF1R,CXCR4,DDIT4,DUSP1,DUSP5,DUSP6,EGLN1,EGR1,ELANE,EREG,FLOT1,FOSL1,FYB1,GBP1,GNAI1,GPC1,HLA-G,HPSE,ID2,IFIT2,IFITM1,IFITM3,IL10RA,INHBA,IRF7,IRF8,ITGB8,KLF2,LGALS3,LILRB3,MARCKS,MPO,MT1F,MX1,NCF2,NFKB2,NOD2,NOG,NR0B1,PARP9,PER1,PIEZO2,PIK3IP1,POMC,RELB,REN,RHOB,S100A8,S100A9,S1PR2,SBDS,SKIL,SLC12A5,SLC16A4,SLC1A3,ST3GAL5,STAT1,STAT2,SUMO1P3,TCL1A,TNFRSF10D,TNFSF13B,TOB1,TP53INP1,TP53INP2,TSC22D3,USP18,VWF |
| Cellular Movement,Hematological System Development and Function,Immune Cell Trafficking | Cell movement of leukocytes | 8.48E-09 | Decreased | -2.179 | ARHGAP18,AZU1,BCL3,CCR1,CD3E,CD48,CD69,CD72,CSF1R,CXCR4,DUSP1,EGLN1,ELANE,FLOT1,FYB1,HLA-G,HPSE,IL10RA,INHBA,KLF2,LGALS3,LILRB3,MPO,NFKB2,NOD2,NOG,POMC,RELB,REN,RHOB,S100A8,S100A9,S1PR2,SBDS,STAT1,STAT2,TNFSF13B,VWF |
| Cellular Movement,Immune Cell Trafficking | Leukocyte migration | 2.03E-09 | Decreased | -2.214 | ARHGAP18,AZU1,BCL3,CCR1,CD3E,CD48,CD69,CD72,CLU,CSF1R,CXCR4,DUSP1,EGLN1,EGR1,ELANE,FLOT1,FYB1,GPC1,HLA-G,HPSE,IL10RA,INHBA,IRF7,KLF2,LGALS3,LILRB3,MPO,NFKB2,NOD2,NOG,PER1,POMC,RELB,REN,RHOB,S100A8,S100A9,S1PR2,SBDS,STAT1,STAT2,TNFRSF10D,TNFSF13B,VWF |
| Cellular Development,Cellular Growth and Proliferation,Hematological System Development and Function,Hematopoiesis,Lymphoid Tissue Structure and Development,Tissue Development | Leukopoiesis | 7.08E-11 | Decreased | -2.225 | BCL3,CD3E,CD3G,CD69,CD72,CISH,CITED2,CSF1R,CXCR4,DLK1,DUSP5,EGR1,FLT3,FOSL1,FYB1,GADD45B,GIMAP1,HLA-G,ID2,IKZF3,IL10RA,INHBA,IRF1,IRF7,IRF8,ITGB8,KLF2,LGALS3,LILRB3,MYCN,NFKB2,NOG,RELB,RSAD2,S100A9,SBDS,ST3GAL5,STAT1,STAT2,TNFSF13B,USP18,ZFP36L2 |
| Cellular Movement | Migration of cells | 4.89E-13 | Decreased | -2.235 | ACTA2,AREG,ARHGAP18,ARRDC3,ATF3,AZU1,BAG3,BCL3,C5orf30,CCR1,CD3E,CD48,CD69,CD72,CDK1,CDKN2C,CITED2,CLU,CSF1R,CXCR4,DDIT4,DUSP1,DUSP5,DUSP6,EGLN1,EGR1,ELANE,EREG,FLOT1,FOSL1,FYB1,GBP1,GNAI1,GPC1,HLA-G,HPSE,ID2,IFIT2,IFITM1,IFITM3,IL10RA,INHBA,IRF7,ITGB8,KLF2,LGALS3,LILRB3,MARCKS,MPO,MT1F,NCF2,NFKB2,NOD2,NOG,PARP9,PER1,PIEZO2,PIK3IP1,POMC,RELB,REN,RHOB,S100A8,S100A9,S1PR2,SBDS,SKIL,SLC12A5,SLC16A4,SLC1A3,STAT1,STAT2,SUMO1P3,TCL1A,TNFRSF10D,TNFSF13B,TOB1,TP53INP1,TP53INP2,TSC22D3,USP18,VWF |
| Cellular Movement | Homing of cells | 0.00000803 | Decreased | -2.241 | AZU1,CCR1,CD69,CD72,CLU,CSF1R,CXCR4,DUSP6,ELANE,FLOT1,FOSL1,FYB1,HLA-G,HPSE,INHBA,LGALS3,LILRB3,NFKB2,NOD2,RELB,RHOB,S100A8,S100A9,S1PR2,SBDS |
| Cellular Development,Cellular Growth and Proliferation,Embryonic Development,Hematological System Development and Function,Hematopoiesis,Lymphoid Tissue Structure and Development,Organ Development,Organismal Development,Tissue Development | Lymphopoiesis | 9.94E-10 | Decreased | -2.244 | BCL3,CD3E,CD3G,CD69,CD72,CISH,CXCR4,DUSP5,EGR1,FLT3,FOSL1,FYB1,GADD45B,GIMAP1,HLA-G,ID2,IKZF3,IL10RA,INHBA,IRF1,IRF8,ITGB8,KLF2,LGALS3,MYCN,NFKB2,NOG,RELB,RSAD2,ST3GAL5,STAT1,STAT2,TNFSF13B,USP18,ZFP36L2 |
| Gene Expression | Transcription | 3.23E-10 | Decreased | -2.324 | ACTA2,ATF3,AZU1,BCL3,BTG1,CAVIN4,CCR1,CD3E,CDK1,CDKN2C,CISH,CITED2,CLU,COL2A1,CXCR4,DIRAS3,DLK1,DTX3L,DUSP1,DUSP5,EGLN1,EGR1,ELANE,EREG,FBP1,FOSL1,HBB,HELZ2,HIF3A,HOPX,HPSE,ID2,IKZF3,INHBA,IRF1,IRF7,IRF8,IRF9,ITGB8,JDP2,KLF15,KLF2,MAFF,MDFI,MX1,MYCN,NFKB2,NLRC5,NOD2,NOG,NR0B1,NR5A1,PARP9,PER1,POMC,PPP2R5B,PRDM5,PYGO1,RELB,REN,RHOB,RIMS1,RPL10,SAP30,SKIL,SLC3A1,STAT1,STAT2,TCL1A,TNFSF13B,TP53INP1,TP53INP2,TRIP13,TSC22D3,TXNIP,ZNF462 |
| Cell Death and Survival | Cell death of immune cells | 1.16E-16 | Decreased | -2.337 | ATF3,BCL3,BTG1,CD3E,CD48,CD69,CDK1,CISH,CLU,CSF1R,CXCR4,DDIT4,DLK1,DUSP5,EGR1,ELANE,FLT3,FOSL1,GADD45B,GBP1,GIMAP1,HLA-G,ID2,IFIH1,IKZF3,INHBA,IRF1,IRF8,ISG15,KLF2,LGALS3,MPO,MX1,NCF2,NFKB2,NR5A1,RELB,S100A8,S1PR2,SKIL,SPATA2,STAT1,TCL1A,TNFSF13B,TSC22D3 |
| Cell Death and Survival | Cell death of blood cells | 2.63E-16 | Decreased | -2.346 | ATF3,BCL3,BTG1,CD3E,CD48,CD69,CDK1,CISH,CLU,CSF1R,CXCR4,DDIT4,DLK1,DUSP5,EGR1,ELANE,FLT3,FOSL1,GADD45B,GBP1,GIMAP1,HLA-G,ID2,IFIH1,IKZF3,INHBA,IRF1,IRF8,ISG15,KLF2,LGALS3,MPO,MX1,NCF2,NFKB2,NR5A1,RELB,S100A8,S1PR2,SBDS,SKIL,SPATA2,STAT1,TCL1A,TNFSF13B,TSC22D3 |
| Gene Expression | Expression of RNA | 9.71E-10 | Decreased | -2.35 | ACTA2,ATF3,AZU1,BCL3,BTG1,CAVIN4,CCR1,CD3E,CDK1,CDKN2C,CISH,CITED2,CLU,COL2A1,CXCR4,DIRAS3,DLK1,DTX3L,DUSP1,DUSP5,EGLN1,EGR1,ELANE,EREG,FBP1,FOSL1,HBB,HELZ2,HIF3A,HOPX,HPSE,ID2,IKZF3,INHBA,IRF1,IRF7,IRF8,IRF9,ITGB8,JDP2,KLF15,KLF2,MAFF,MDFI,MYCN,NFKB2,NLRC5,NOD2,NOG,NR0B1,NR5A1,PARP9,PER1,POMC,PPP2R5B,PRDM5,PYGO1,RELB,REN,RHOB,RIMS1,RPL10,RPL41,S100A9,SAMD4A,SAP30,SKIL,SLC3A1,STAT1,STAT2,TCL1A,TOB1,TP53INP1,TP53INP2,TRIP13,TSC22D3,TXNIP,ZNF462 |
| Antimicrobial Response,Inflammatory Response | Antiviral response | 2.59E-19 | Decreased | -2.412 | ACTA2,AZU1,BCL3,CLU,CXCR4,DDIT4,DTX3L,FOSL1,IFI44,IFI44L,IFI6,IFIH1,IFIT1,IFIT2,IFIT3,IFITM1,IFITM3,IRF1,IRF7,ISG15,MX1,OAS1,OAS2,OASL,PARP9,RSAD2,SAMHD1,STAT1,STAT2 |
| Cell Death and Survival | Necrosis | 7.66E-15 | Decreased | -2.479 | AREG,ARHGAP18,ATF3,AZU1,BAG3,BCL3,BTG1,C1QTNF4,CD3E,CD48,CD69,CDK1,CDKN2C,CISH,CITED2,CLU,COL2A1,CSF1R,CXCR4,DDIT4,DHCR24,DIRAS3,DLK1,DUSP1,DUSP5,DUSP6,EGLN1,EGR1,ELANE,EREG,FKBP5,FLT3,FOSL1,GADD45B,GALNT3,GBP1,GIMAP1,GPC1,HLA-F,HLA-G,HPSE,ID2,IFI6,IFIH1,IKZF3,INHBA,IRF1,IRF7,IRF8,IRF9,ISG15,KIF15,KLF2,LGALS3,LILRB3,MPO,MT1F,MT2A,MX1,MYCN,NCF2,NFKB2,NOG,NR5A1,OAS1,OAS2,PER1,PIK3IP1,POMC,PRDM5,PTPRE,RELB,REN,RHOB,RPL10,RPL3,RPL7,RPL7A,S100A8,S100A9,S1PR2,SBDS,SKIL,SLC1A3,SNCAIP,SPATA2,STAT1,STAT2,SUMO1P3,TCL1A,TNFRSF10D,TNFSF13B,TP53INP1,TSC22D3,TXNIP,USP18,VWF,WEE1,XAF1 |
| Cellular Development,Cellular Growth and Proliferation,Hematological System Development and Function,Hematopoiesis,Lymphoid Tissue Structure and Development,Tissue Development | Hematopoiesis of mononuclear leukocytes | 4.55E-10 | Decreased | -2.579 | BCL3,CD3E,CD3G,CD69,CD72,CISH,CSF1R,CXCR4,DUSP5,EGR1,FLT3,FOSL1,FYB1,GADD45B,GIMAP1,HLA-G,ID2,IKZF3,IL10RA,INHBA,IRF1,IRF7,IRF8,ITGB8,KLF2,LGALS3,MYCN,NFKB2,NOG,RELB,RSAD2,ST3GAL5,STAT1,STAT2,TNFSF13B,USP18,ZFP36L2 |
| Cell-mediated Immune Response,Cellular Development,Cellular Function and Maintenance,Cellular Growth and Proliferation,Embryonic Development,Hematological System Development and Function,Hematopoiesis,Lymphoid Tissue Structure and Development,Organ Development,Organismal Development,Tissue Development | T cell development | 0.000000141 | Decreased | -2.593 | BCL3,CD3E,CD3G,CD69,CISH,CXCR4,EGR1,FYB1,GADD45B,GIMAP1,HLA-G,ID2,IL10RA,INHBA,IRF1,IRF8,ITGB8,KLF2,NOG,RELB,RSAD2,ST3GAL5,STAT1,STAT2,TNFSF13B,USP18,ZFP36L2 |
| Neurological Disease | Progressive neurological disorder | 0.00000102 | Decreased | -2.63 | ACTA2,AREG,ATF3,CDK1,CLU,CSF1R,CXCR4,DHCR24,EGLN1,EGR1,EPSTI1,FCGR1B,FLOT1,FLT3,GALNT3,HBA1/HBA2,HLA-F,HLA-G,IFIT1,IRF7,ISG15,KCNC1,LGALS3,MPO,MS4A6A,MX1,NEAT1,NFKB2,OAS1,PIK3IP1,PTPRE,RHOB,RIMS1,RPL3,RSAD2,S100A9,SLC1A3,SNCAIP,TUBA1B,TUBB2A,ZFP36L2 |
| Inflammatory Disease,Neurological Disease,Skeletal and Muscular Disorders | Relapsed multiple sclerosis | 0.00000148 | Decreased | -2.63 | EPSTI1,FCGR1B,HLA-G,IFIT1,IRF7,ISG15,KCNC1,MX1,OAS1,RSAD2 |
| Inflammatory Disease,Neurological Disease,Skeletal and Muscular Disorders | Relapsing-remitting multiple sclerosis | 0.00000428 | Decreased | -2.63 | EPSTI1,HLA-G,IFIT1,IRF7,ISG15,KCNC1,MX1,OAS1,RSAD2 |
| Neurological Disease,Skeletal and Muscular Disorders | Progressive neuromuscular disease | 0.00001 | Decreased | -2.63 | AREG,DHCR24,EGLN1,EGR1,EPSTI1,FCGR1B,FLOT1,GALNT3,HLA-F,HLA-G,IFIT1,IRF7,ISG15,KCNC1,MX1,OAS1,RPL3,RSAD2,SNCAIP,TUBA1B |
| Neurological Disease | Progressive motor neuron disease | 0.0000112 | Decreased | -2.63 | AREG,ATF3,CLU,DHCR24,EGLN1,EGR1,EPSTI1,FCGR1B,FLOT1,GALNT3,HLA-F,HLA-G,IFIT1,IRF7,ISG15,KCNC1,MX1,NEAT1,NFKB2,OAS1,RHOB,RPL3,RSAD2,SLC1A3,SNCAIP,TUBA1B,ZFP36L2 |
| Inflammatory Disease,Neurological Disease,Skeletal and Muscular Disorders | Multiple Sclerosis | 0.0000161 | Decreased | -2.63 | AREG,DHCR24,EGR1,EPSTI1,FCGR1B,HLA-F,HLA-G,IFIT1,IRF7,ISG15,KCNC1,MX1,OAS1,RSAD2 |
| Neurological Disease | Progressive motor neuropathy | 0.0000293 | Decreased | -2.63 | AREG,ATF3,CLU,DHCR24,EGLN1,EGR1,EPSTI1,FCGR1B,FLOT1,GALNT3,HLA-F,HLA-G,IFIT1,IRF7,ISG15,KCNC1,MX1,NEAT1,OAS1,RHOB,RPL3,RSAD2,SLC1A3,SNCAIP,TUBA1B,ZFP36L2 |
| Cell-mediated Immune Response,Cellular Development,Cellular Function and Maintenance,Cellular Growth and Proliferation,Embryonic Development,Hematological System Development and Function,Hematopoiesis,Lymphoid Tissue Structure and Development,Organ Development,Organismal Development,Tissue Development | Differentiation of T lymphocytes | 1.02E-08 | Decreased | -2.726 | BCL3,CD3E,CD3G,CD69,CISH,CXCR4,EGR1,GADD45B,GIMAP1,HLA-G,ID2,IL10RA,IRF1,IRF8,ITGB8,NOG,RELB,RSAD2,ST3GAL5,STAT1,STAT2,TNFSF13B,USP18,ZFP36L2 |
| Cellular Function and Maintenance | Lymphocyte homeostasis | 5.27E-08 | Decreased | -2.735 | BCL3,CD3E,CD3G,CD69,CISH,CXCR4,EGR1,FYB1,GADD45B,GIMAP1,HLA-G,ID2,IL10RA,INHBA,IRF1,IRF8,ITGB8,KLF2,LILRB3,NOG,RELB,RSAD2,SKIL,ST3GAL5,STAT1,STAT2,TNFSF13B,USP18,ZFP36L2 |
| Cellular Function and Maintenance | Homeostasis of leukocytes | 2.28E-08 | Decreased | -2.745 | BCL3,CD3E,CD3G,CD69,CISH,CXCR4,EGR1,FLT3,FYB1,GADD45B,GIMAP1,HLA-G,ID2,IL10RA,INHBA,IRF1,IRF8,ITGB8,KLF2,LILRB3,NOG,RELB,RSAD2,SKIL,ST3GAL5,STAT1,STAT2,TNFSF13B,USP18,ZFP36L2 |
| Cell-mediated Immune Response,Cellular Development,Cellular Function and Maintenance,Cellular Growth and Proliferation,Embryonic Development,Hematological System Development and Function,Hematopoiesis,Lymphoid Tissue Structure and Development,Organ Development,Organismal Development,Tissue Development | Differentiation of helper T lymphocytes | 0.00000821 | Decreased | -2.876 | BCL3,CD69,CISH,GADD45B,IL10RA,IRF1,IRF8,RELB,RSAD2,ST3GAL5,STAT1,TNFSF13B,USP18 |
| Antimicrobial Response,Inflammatory Response | Antimicrobial response | 4.75E-19 | Decreased | -2.945 | ACTA2,AZU1,BCL3,CLU,CXCR4,DDIT4,DTX3L,ELANE,FOSL1,HLA-G,IFI44,IFI44L,IFI6,IFIH1,IFIT1,IFIT2,IFIT3,IFITM1,IFITM3,IRF1,IRF7,IRF8,ISG15,MPO,MX1,NOD2,OAS1,OAS2,OASL,PARP9,RSAD2,S100A8,S100A9,SAMHD1,STAT1,STAT2 |

**T2 vs. T1 MEP_1**

© 2000-2020 QIAGEN. All rights reserved.

| Categories | Diseases or Functions Annotation | p-value | Predicted Activation State | Activation z-score | Molecules |
|---|---|---|---|---|---|
| Cell Signaling | Replication of viral replicon | 4.21E-11 | Increased | 3.561 | APOBEC3A,BST2,IFIT1,IFITM1,IFITM3,ISG15,MX1,OAS1,OAS3,OASL,PTGS2,RSAD2,TRIM6 |
| Infectious Diseases | Replication of RNA virus | 2.96E-12 | Increased | 2.94 | ANPEP,AREG,BCL3,BST2,CASP3,CD40,CD40LG,DAG1,FBN1,GABARAPL1,H2BC15,HAVCR2,HCK,HERC5,HPGD,IFI6,IFIH1,IFIT1,IFITM1,IFITM3,IRF1,IRF7,IRF9,ISG15,MX1,NFKBIA,NLRC5,OAS1,OAS3,OASL,P2RY2,PLAU,PRKCD,PTGS2,PTX3,RAB6B,RPSA,RSAD2,STAT1,TXNIP,UBE2E2,USP2 |
| Infectious Diseases | Replication of virus | 3.87E-14 | Increased | 2.682 | AHR,ANPEP,AREG,BCL3,BST2,CASP3,CD40,CD40LG,CXCL2,CXCL3,DAG1,DDIT3,EGR1,FBN1,GABARAPL1,H2BC15,HAVCR2,HCK,HERC5,HPGD,IFI6,IFIH1,IFIT1,IFITM1,IFITM3,IRF1,IRF7,IRF9,ISG15,MX1,MX2,NFKBIA,NLRC5,OAS1,OAS3,OASL,P2RY2,PLAU,PRKCD,PTGS2,PTX3,RAB6B,RPSA,RSAD2,STAT1,TXNIP,UBE2E2,USP2 |
| Infectious Diseases | Infection of mammalia | 5.53E-11 | Increased | 2.547 | AHR,BCL3,CD274,CD36,CD40,CD40LG,CLEC12A,DDIT3,DHX58,FCER1G,HCK,HLA-G,IFIH1,IL1RL1,IL6R,IRF1,IRF7,IRF9,ISG15,ITGAX,OASL,PLAC8,PLAU,PLAUR,PRKCD,PTX3,RELB,STAT1,TNFRSF10A,TPSAB1/TPSB2,USP18 |
| Organismal Survival | Organismal death | 6.3E-10 | Increased | 2.211 | ADCYAP1,AGTR1,AHR,ANGPT2,AQP1,AREG,ATF3,ATP7B,BACE2,BCL3,BHLHE40,BRIP1,CALR,CASP3,CBX4,CCND1,CCND3,CCNE2,CD274,CD36,CD40,CD40LG,CDKN2C,CEBPG,CENPE,CFP,CITED2,COL5A1,CXADR,CXCL2,CXCL3,DAG1,DDIT3,DHRS3,DHX58,DLK1,EGLN1,EHD4,EPX,FAT4,FBLN1,FBN1,FCER1G,FOSL1,FOSL2,GAD1,GJA4,GNAI1,H2AC18/H2AC19,HAVCR2,HCK,HLA-G,HOPX,HPGD,ID2,IFIH1,IKZF3,IL18R1,IL1RL1,IL6R,IRF1,IRF7,IRF9,ISG15,ITGB8,KLF2,KRT1,LIN7A,LMO2,LMO4,LTBP1,MACROH2A2,MAFF,MAL,MDK,MITF,MYCN,MYLPF,NEAT1,NFKBIA,NINJ1,NR4A3,NTAN1,PIEZO2,PLAU,PLAUR,PMEPA1,POMC,PRKCD,PRKD1,PTGS2,PTK2,PTPRN2,PTS,PTX3,RELB,REN,RGPD4 (includes others),RIMS1,RPSA,S1PR1,SBDS,SEMA7A,SKIL,SLC2A3,SOD2,SRGAP3,STAT1,THRB,TNFAIP3,TNFRSF10A,TNFSF13B,TP53INP1,TRPS1,TXNIP,TYMP,USP18,WNT5B,ZFP36L2 |
| Infectious Diseases | Viral Infection | 2.97E-12 | Increased | 2.197 | ADCYAP1,AGTR1,AHR,ANPEP,AREG,ARRDC3,BCL3,BST2,CASP3,CAV2,CD274,CD36,CD40,CD40LG,CD69,CLC,COL5A1,CTSL,CXADR,CXCL1,CXCL2,CXCL3,DAG1,DDIT3,DHX58,EFHC2,EGR1,ENC1,FBN1,GABARAPL1,GPR162,H2AC18/H2AC19,H2BC15,HAVCR2,HBA1/HBA2,HCK,HCP5,HERC5,HLA-B,HLA-C,HPGD,IFI6,IFIH1,IFIT1,IFITM1,IFITM3,IL6R,IRF1,IRF7,IRF9,ISG15,KLF2,KPNA2,LGALS1,MX1,MX2,NACA2,NFKBIA,NLRC5,OAS1,OAS2,OAS3,OASL,P2RY2,PARP9,PDE3B,PLAU,PLAUR,PRKCD,PRKD1,PTGS2,PTPRN2,PTX3,RAB6B,RELB,RGPD4 (includes others),RNASE2,RPL3,RPSA,RSAD2,SELL,SLC2A3,SLCO3A1,STAT1,TNFRSF10A,TPSAB1/TPSB2,TXNIP,TYROBP,UBE2E2,USP2,XK |
| Infectious Diseases | Replication of Flaviviridae | 2.76E-08 | Increased | 2.057 | BST2,CD40,CD40LG,GABARAPL1,IFI6,IFIT1,IFITM1,IFITM3,ISG15,OASL,RSAD2,TXNIP,UBE2E2 |
| Cell Death and Survival | Cell viability of blood cells | 0.0000007 | Decreased | -2.025 | ADCYAP1,BCL3,CD40,CD40LG,CISH,DLK1,FCER1G,GADD45B,GPR65,HBB,HCK,KLF2,MX1,NFKBIA,PARP14,PTK2,RELB,SOD2,STAT1,TNFRSF13C,TNFSF13B,TSC22D3,TYROBP |
| Hematological System Development and Function,Immune Cell Trafficking,Inflammatory Response,Tissue Development | Accumulation of leukocytes | 1.24E-11 | Decreased | -2.052 | ABCG1,AHR,BCL3,CD274,CD40,CR1,CXCL1,CXCL2,CXCL3,DDIT3,EGR1,FOSL1,HAVCR2,HCK,IL1RL1,IRF1,ITGAX,KLF2,LTC4S,P2RY2,PLAU,POMC,PTGS2,PTX3,S1PR1,SELL,STAT1,TSC22D3,TYROBP |
| Cell Death and Survival | Cell viability of tumor cell lines | 2.3E-10 | Decreased | -2.062 | AHR,ANGPT2,AREG,ATF3,ATP7B,BHLHE40,BRD4,BRIP1,CALR,CASP3,CCND1,CCND3,CD33,CD40,CD40LG,CDKN2C,CXCL1,CXCL2,CXCL3,DUSP5,EGLN1,EGR1,EREG,FKBP5,FOSL1,HBA1/HBA2,HBB,HCK,HLA-G,HOPX,ID2,IKZF3,IL6R,LGALS3BP,MDK,MITF,MYCN,NEAT1,NFKBIA,PARP14,PDP1,PIM3,PLAU,PLAUR,PMAIP1,POMC,PRKCD,PTGS2,PTK2,PTPRN2,RELB,RNASE2,RPSA,SLC2A3,SOD2,TNFSF13B,TOB1,TRIB2,TRIM69,TYMP,USP18,USP2 |
| Cell-To-Cell Signaling and Interaction | Binding of myeloid cells | 0.00000757 | Decreased | -2.065 | ANGPT2,CCND1,CD40,CD40LG,CD69,CFP,CXCL1,CXCL2,CXCL3,HCK,ITGAX,LGALS1,MITF,PLAU,PLAUR,PTGS2,SELL |
| Gene Expression | Transcription | 5.79E-08 | Decreased | -2.084 | ADCYAP1,AHR,ATAD2,ATF3,BCL3,BHLHE40,BRD4,BRIP1,BST2,BTG1,CALR,CBX4,CCND1,CCND3,CD36,CD40,CD40LG,CDKN2C,CEBPG,CHD7,CISH,CITED2,CXCL2,DDIT3,DLK1,DUSP5,EGLN1,EGR1,EREG,FAM98A,FBLN1,FBP1,FOSL1,FOSL2,GAS2L1,HAVCR2,HBB,HELZ2,HES6,HLA-DRB5,HOPX,ID2,IKZF3,IL6R,IRF1,IRF7,IRF9,ITGAX,ITGB8,KLF10,KLF2,KPNA2,LGALS1,LMO2,LMO4,MACROH2A2,MAFF,MDK,MITF,MX1,MYCN,NFKBIA,NLRC5,NR4A3,PARP14,PARP9,PCBP3,PER1,PLAC8,POMC,PRKAR2B,PRKCD,PRKD1,PTK2,RELB,REN,RIMS1,RNF11,RPL10,RYBP,S1PR1,SKIL,SLA,SOD2,SPTY2D1,STAT1,TFEC,THRB,TLE3,TNFSF13B,TNNI2,TP53INP1,TRIM6,TRPS1,TSC22D1,TSC22D3,TXNIP,TYROBP,USP2,ZNF322,ZNF608,ZNF93 |
| Cellular Movement,Immune Cell Trafficking | Leukocyte migration | 8.59E-11 | Decreased | -2.1 | ABCG1,ADCYAP1,ADD2,AHR,ANGPT2,BCL3,CALR,CCND1,CD274,CD36,CD40,CD40LG,CD69,CR1,CXADR,CXCL1,CXCL2,CXCL3,EGLN1,EGR1,FCER1G,FOSL2,GP1BA,GYPB,HCK,HLA-G,IGLL1/IGLL5,IL1RL1,IL6R,IRF7,ITGAX,KCNE3,KLF2,LGALS1,LTC4S,MDK,NFKBIA,NINJ1,P2RY2,PER1,PLAU,PLAUR,POMC,PRKCD,PTGS2,PTK2,PTX3,RAMP1,RELB,REN,RNASE2,S1PR1,SBDS,SELL,SLC7A6,SOD2,STAT1,TNFAIP3,TNFRSF10A,TNFRSF10D,TNFSF13B,TPSAB1/TPSB2,TYROBP,UTS2 |
| Molecular Transport | Quantity of metal | 0.0000113 | Decreased | -2.114 | ADCYAP1,ADD2,AGTR1,ANGPT2,AQP1,ATP7B,CALR,CXCL2,CXCL3,FCER1G,HBA1/HBA2,KLF2,LGALS1,LTBP1,MS4A2,NRGN,P2RY2,PLAU,PLAUR,POMC,PTGS2,RASGRP3,REN,RHAG,S1PR1,SELL,TFEC,TPSAB1/TPSB2,TSC22D3,TXNIP |
| Tissue Morphology | Quantity of cells | 5.11E-15 | Decreased | -2.119 | ABCG1,ADCYAP1,ADD2,AGTR1,AHR,ANGPT2,ANPEP,AQP1,ARL4A,ATF3,BCL3,BHLHE40,CALR,CASP3,CAV2,CCND1,CCND3,CD274,CD36,CD40,CD40LG,CD69,CD83,CDKN2C,CETP,CFP,CHD7,CISH,CITED2,CXCL1,CXCL2,DDIT3,DLK1,DUSP5,EGLN1,EGR1,EHD4,FCER1G,FOSL2,GAD1,GADD45B,GAPT,GAS2L1,GCNT4,GP1BA,HAVCR2,HBA1/HBA2,HBB,HCK,HLA-G,ID2,IGLL1/IGLL5,IKZF3,IL1RL1,IL6R,IRF1,ISG15,ITGB8,KLF10,KLF2,LGALS1,LTC4S,MDK,MITF,MYCN,NANOS3,NFKBIA,NLRC5,NR4A3,PAG1,PER1,PIM3,PLAC8,PLAU,PLAUR,PMAIP1,POMC,PRKCD,PRKD1,PTGS2,PTK2,PTPRN2,RAB27B,RASGRP3,RELB,REN,RIMS1,RPGR,S1PR1,SELL,SLA,SLC2A3,SOD2,SORBS1,SPTA1,STAT1,SYTL2,THRB,TNFAIP3,TNFRSF10A,TNFRSF13C,TNFSF13B,TOB1,TRIB2,TRIP6,TSC22D3,TXNIP,TYROBP,USP2,WNT5B,ZFP36L2 |
| Cancer,Organismal Injury and Abnormalities | Growth of tumor | 8.75E-12 | Decreased | -2.127 | AHR,ANGPT2,ANPEP,AREG,ARRDC3,ATF3,BST2,CALR,CASP3,CCNA1,CCND1,CCND3,CD274,CD33,CD36,CD40,CD40LG,CXCL1,CXCL3,DDIT3,DLK1,EGLN1,EGR1,EPSTI1,FCER1G,FOSL1,GJA4,HAVCR2,HLA-G,HOTAIRM1,HPGD,ID2,IFITM3,IL1RL1,IL6R,IRF1,ITGB8,KLF10,LGALS1,LGALS12,LMO2,MYCN,NFKBIA,PDP1,PIM3,PLAU,PLAUR,PRKCD,PTGS2,PTK2,RAMP1,RELB,SELL,SEMA7A,SKIL,SOD2,STAT1,THRB,TNFRSF10A,TNFSF13B,TRIB2,TRIM69,TRPS1,TSC22D1,TXNIP |
| Hematological System Development and Function,Hematopoiesis,Humoral Immune Response,Lymphoid Tissue Structure and Development,Tissue Morphology | Quantity of pre-B lymphocytes | 0.00000408 | Decreased | -2.164 | BCL3,CCND3,CD36,CD40,CD69,DLK1,IGLL1/IGLL5,IKZF3,KLF2,PIM3,PRKCD,TNFRSF13C,TNFSF13B |
| Hematological System Development and Function,Hematopoiesis | Development of hematopoietic systen | 0.00000821 | Decreased | -2.215 | CALR,CASP3,CCND1,CCND3,CD40,CD40LG,CD83,CXCL2,CXCL3,DDIT3,EGR1,GP1BA,H4C3,IGLL1/IGLL5,IL1RL1,IL6R,IRF1,LMO2,NFKBIA,NR4A3,RASGRP3,RELB,SBDS,STAT1,TNFSF13B,TYROBP |
| Cell-To-Cell Signaling and Interaction,Inflammatory Response | Immune response of phagocytes | 0.00000411 | Decreased | -2.217 | ADCYAP1,ANPEP,CALR,CD36,CD40LG,CXCL1,CXCL3,FCER1G,HCK,IL1RL1,IL6R,ISG15,ITGAX,PLAU,PLAUR,PTX3,RELB,TYROBP |
| Hematological System Development and Function,Tissue Development | Accumulation of blood cells | 1.73E-11 | Decreased | -2.242 | ABCG1,AHR,ANGPT2,BCL3,CD274,CD40,CR1,CXCL1,CXCL2,CXCL3,DDIT3,EGR1,FOSL1,HAVCR2,HCK,IL1RL1,IRF1,ITGAX,KLF2,LTC4S,P2RY2,PLAU,POMC,PTGS2,PTX3,S1PR1,SELL,STAT1,TSC22D3,TYROBP |
| Cellular Movement,Hematological System Development and Function,Immune Cell Trafficking,Inflammatory Response | Cell movement of phagocytes | 4.09E-07 | Decreased | -2.316 | ADCYAP1,ANGPT2,CCND1,CD36,CD40,CD40LG,CD69,CR1,CXADR,CXCL1,CXCL2,CXCL3,EGLN1,FCER1G,HCK,IL1RL1,IL6R,ITGAX,LGALS1,MDK,NFKBIA,NINJ1,PLAU,PLAUR,PRKCD,PTGS2,PTX3,RAMP1,REN,RNASE2,S1PR1,SBDS,SELL,STAT1,TNFAIP3,TPSAB1/TPSB2,TYROBP,UTS2 |
| Cellular Movement | Migration of tumor cell lines | 1.85E-08 | Decreased | -2.329 | AHR,ANGPT2,ANKRD28,ANPEP,AREG,ARRDC3,ATF3,BCAM,BRD4,CALR,CCND1,CD36,CD40,CTSL,CXCL2,DUSP5,EGR1,EREG,FOSL1,GNAI1,HAVCR2,HES6,HOTAIRM1,ID2,IFITM3,IL6R,KPNA2,LGALS1,LGALS3BP,MDK,MITF,NFKBIA,NINJ1,P2RY2,PAG1,PIEZO2,PLAU,PLAUR,POMC,PRKCD,PRKD1,PTGS2,PTK2,RHOU,RNF11,SEMA3C,SLC16A4,SOD2,STAT1,THRB,TOB1,TP53INP1,TRIP6,TRPS1,USP2 |
| Antimicrobial Response,Inflammatory Response | Antiviral response | 2.8E-15 | Decreased | -2.426 | APOBEC3A,BCL3,BST2,CD40,CXADR,DDIT4,DHX58,FOSL1,IFI44,IFI44L,IFI6,IFIH1,IFIT1,IFITM1,IFITM3,IRF1,IRF7,ISG15,ITGAX,MX1,MX2,OAS1,OAS2,OAS3,OASL,PARP9,PMAIP1,RNASE2,RSAD2,STAT1,TRIM6 |
| Tissue Development | Accumulation of cells | 8.7E-11 | Decreased | -2.478 | ABCG1,AHR,ANGPT2,BCL3,CD274,CD40,CR1,CXCL1,CXCL2,CXCL3,DDIT3,EGR1,FOSL1,HAVCR2,HCK,IL1RL1,IL6R,IRF1,ITGAX,KLF2,LTC4S,P2RY2,PLAU,POMC,PTGS2,PTX3,S1PR1,SELL,SEMA7A,STAT1,TSC22D3,TYROBP |
| Cellular Movement,Hematological System Development and Function,Immune Cell Trafficking,Inflammatory Response | Migration of phagocytes | 0.00000749 | Decreased | -2.487 | ADCYAP1,ANGPT2,CCND1,CD40,CD40LG,CXCL1,CXCL2,CXCL3,HCK,ITGAX,LGALS1,MDK,NINJ1,PLAU,PLAUR,PTGS2,RAMP1,S1PR1,SBDS,SELL,TYROBP |
| Cell Death and Survival | Cell viability | 7.48E-15 | Decreased | -2.533 | ADCYAP1,AHR,ANGPT2,APOBEC3A,AREG,ATF3,ATP7B,BCL3,BHLHE40,BRD4,BRIP1,CALR,CASP3,CCND1,CCND3,CD33,CD40,CD40LG,CDKN2C,CISH,CR1,CXCL1,CXCL2,CXCL3,DDIT3,DLK1,DUSP5,EGLN1,EGR1,EHD4,EREG,FCER1G,FKBP5,FOSL1,GADD45B,GPR65,HBA1/HBA2,HBB,HCK,HLA-F,HLA-G,HOPX,ID2,IFIH1,IKZF3,IL6R,IRF9,KLF2,LGALS3BP,LMO4,MDK,MITF,MX1,MYCN,NEAT1,NFKBIA,NR4A3,PARP14,PDP1,PIK3IP1,PIM3,PLAC8,PLAU,PLAUR,PMAIP1,POMC,PRKCD,PRKD1,PTGS2,PTK2,PTPRN2,RELB,RNASE2,RPSA,S1PR1,SELL,SLC2A3,SOD2,STAT1,TNFAIP3,TNFRSF10A,TNFRSF13C,TNFSF13B,TOB1,TRIB2,TRIM69,TSC22D3,TXNIP,TYMP,TYROBP,USP18,USP2,XAF1 |
| Cellular Movement,Hematological System Development and Function,Immune Cell Trafficking,Inflammatory Response | Chemotaxis of leukocytes | 0.0000133 | Decreased | -2.591 | ANGPT2,CCND1,CD40,CD40LG,CD69,CXADR,CXCL1,CXCL2,CXCL3,FCER1G,HCK,HLA-G,IL6R,LGALS1,MDK,NINJ1,PLAU,PLAUR,RNASE2,S1PR1,SBDS,SELL,TPSAB1/TPSB2,UTS2 |
| Cellular Movement | Homing of tumor cell lines | 0.0000134 | Decreased | -2.595 | CALR,CD69,CXCL1,CXCL3,FBN1,HCK,MDK,P2RY2,PLAU,PLAUR,PTK2,RHOU,S1PR1,UTS2 |
| Inflammatory Response | Immune response of cells | 2.8E-08 | Decreased | -2.601 | ADCYAP1,AHR,ANPEP,BHLHE40,CALR,CD274,CD36,CD40,CD40LG,CD69,CD83,CR1,CXCL1,CXCL3,DHX58,FCER1G,GP1BA,HAVCR2,HCK,IFIH1,IL18R1,IL1RL1,IL6R,IRF1,IRF7,ISG15,ITGAX,KLF2,LGALS1,LTBP1,PER1,PLAU,PLAUR,PRKCD,PTK2,PTX3,RELB,STAT1,TLE3,TNFRSF10A,TRIM6,TYROBP |
| Cell Death and Survival | Cell survival | 2.57E-14 | Decreased | -2.63 | ADCYAP1,AHR,ANGPT2,APOBEC3A,AREG,ATF3,ATP7B,BCL3,BHLHE40,BRD4,BRIP1,CALR,CASP3,CCNA1,CCND1,CCND3,CD33,CD40,CD40LG,CDKN2C,CISH,CR1,CXCL1,CXCL2,CXCL3,DDIT3,DLK1,DUSP5,EGLN1,EGR1,EHD4,EREG,FCER1G,FKBP5,FOSL1,GADD45B,GPR65,HBA1/HBA2,HBB,HCK,HLA-F,HLA-G,HOPX,ID2,IFIH1,IKZF3,IL1RL1,IL6R,IRF9,KLF2,LGALS3BP,LMO4,MDK,MITF,MX1,MYCN,NEAT1,NFKBIA,NR4A3,PARP14,PDP1,PIK3IP1,PIM3,PLAC8,PLAU,PLAUR,PMAIP1,POMC,PRKCD,PRKD1,PTGS2,PTK2,PTPRN2,RELB,RNASE2,RPSA,S1PR1,SELL,SLC2A3,SOD2,STAT1,TNFAIP3,TNFRSF10A,TNFRSF13C,TNFSF13B,TOB1,TRIB2,TRIM69,TSC22D3,TXNIP,TYMP,TYROBP,USP18,USP2,XAF1 |
| Cellular Movement | Cell movement of tumor cell lines | 1E-09 | Decreased | -2.792 | AHR,ANGPT2,ANKRD28,ANPEP,AREG,ARRDC3,ATF3,BCAM,BRD4,CALR,CCND1,CD36,CD40,CD69,CTSL,CXCL1,CXCL2,CXCL3,DUSP5,EGR1,EREG,FBN1,FOSL1,GNAI1,HAVCR2,HCK,HES6,HOTAIRM1,ID2,IFITM3,IL6R,KPNA2,LGALS1,LGALS3BP,MARCKS,MDK,MITF,MX1,NFKBIA,NINJ1,P2RY2,PAG1,PIEZO2,PLAU,PLAUR,POMC,PRKCD,PRKD1,PTGS2,PTK2,RHOU,RNF11,S1PR1,SELL,SEMA3C,SLC16A4,SOD2,STAT1,THRB,TOB1,TP53INP1,TRIP6,TRPS1,USP2,UTS2 |
| Inflammatory Disease,Neurological Disease | Inflammatory demyelinating disease | 0.00000989 | Decreased | -2.813 | AGTR1,AREG,BST2,CXCL1,DDIT3,EGR1,EPSTI1,HAVCR2,HLA-F,HLA-G,IFIT1,IL6R,IRF7,ISG15,MX1,OAS1,OAS3,PTGS2,RSAD2 |
| Cellular Movement | Chemotaxis | 4.76E-07 | Decreased | -2.888 | AGTR1,ANGPT2,CALR,CCND1,CD36,CD40,CD40LG,CD69,CXADR,CXCL1,CXCL2,CXCL3,FBN1,FCER1G,FOSL1,HCK,HLA-G,IL6R,LGALS1,MDK,NINJ1,P2RY2,PLAU,PLAUR,PRKCD,PTGS2,PTK2,RELB,RHOU,RNASE2,S1PR1,SBDS,SELL,TPSAB1/TPSB2,TYMP,UTS2 |
| Cellular Movement | Migration of cells | 5.24E-14 | Decreased | -3.098 | ABCG1,ADCYAP1,ADD2,AGTR1,AHR,ANGPT2,ANKRD28,ANPEP,AQP1,AREG,ARRDC3,ATF3,BCAM,BCL3,BHLHE40,BRD4,BST2,CALR,CCND1,CD274,CD36,CD40,CD40LG,CD69,CDKN2C,CITED2,CR1,CTSL,CXADR,CXCL1,CXCL2,CXCL3,DAG1,DDIT4,DUSP5,EGLN1,EGR1,EREG,FCER1G,FOSL1,FOSL2,GAD1,GNAI1,GP1BA,GYPB,HAVCR2,HCK,HES6,HLA-G,HOTAIRM1,ID2,IFITM1,IFITM3,IGLL1/IGLL5,IL1RL1,IL6R,IRF7,ITGAX,ITGB8,KCNE3,KLF2,KPNA2,LGALS1,LGALS3BP,LMO4,LTC4S,MARCKS,MDK,MITF,NFKBIA,NINJ1,P2RY2,PAG1,PARD6B,PARP9,PER1,PIEZO2,PIK3IP1,PLAU,PLAUR,POMC,PRKCD,PRKD1,PTGS2,PTK2,PTX3,RAMP1,RASGRP3,RELB,REN,RHOU,RNASE2,RNF11,RPSA,S1PR1,SBDS,SELL,SEMA3C,SKIL,SLC16A4,SLC7A6,SOD2,STAT1,THRB,TNFAIP3,TNFRSF10A,TNFRSF10D,TNFSF13B,TOB1,TP53INP1,TPSAB1/TPSB2,TRIP6,TRPS1,TSC22D3,TYROBP,USP18,USP2,UTS2,WNT5B |
| Neurological Disease | Progressive neurological disorder | 0.00000975 | Decreased | -3.138 | AGTR1,AQP1,AREG,ATAD2,ATF3,BST2,CASP3,CCND1,CD33,CD36,CD40LG,CETP,CR1,CXADR,CXCL1,DDIT3,EGLN1,EGR1,ENO3,EPSTI1,FBLN1,HAVCR2,HBA1/HBA2,HLA-B,HLA-F,HLA-G,IFIT1,IL6R,IRF7,ISG15,LGALS1,MS4A4E,MS4A6A,MX1,NEAT1,NRGN,OAS1,OAS3,PDE3B,PIK3IP1,PLAU,PRKAR2B,PRKCD,PTGS2,RIMS1,RPL3,RSAD2,SLC2A3,SLC7A6,SOD2,TYROBP,USP37,WDFY3,ZFP36L2,ZNF184 |
| Cellular Movement | Cell movement | 7.69E-14 | Decreased | -3.289 | ABCG1,ADCYAP1,ADD2,AGTR1,AHR,ANGPT2,ANKRD28,ANPEP,AQP1,AREG,ARRDC3,ATF3,BCAM,BCL3,BHLHE40,BRD4,BST2,CALR,CATSPER1,CCND1,CD274,CD36,CD40,CD40LG,CD69,CDC42EP4,CDKN2C,CITED2,CR1,CTSL,CXADR,CXCL1,CXCL2,CXCL3,DAG1,DDIT4,DUSP5,EGLN1,EGR1,EREG,FBN1,FCER1G,FOSL1,FOSL2,GAD1,GAS2L1,GNAI1,GP1BA,GYPB,HAVCR2,HCK,HES6,HLA-G,HOTAIRM1,ID2,IFITM1,IFITM3,IGLL1/IGLL5,IL1RL1,IL6R,IRF7,ITGAX,ITGB8,KCNE3,KLF2,KPNA2,LGALS1,LGALS3BP,LMO4,LTC4S,MARCKS,MDK,MITF,MX1,NFKBIA,NINJ1,P2RY2,PAG1,PARD6B,PARP9,PDE3B,PER1,PIEZO2,PIK3IP1,PLAU,PLAUR,POMC,PRKCD,PRKD1,PTGS2,PTK2,PTX3,RAMP1,RASGRP3,RELB,REN,RHOU,RNASE2,RNF11,RPSA,S1PR1,SBDS,SELL,SEMA3C,SEMA7A,SKIL,SLC16A4,SLC7A6,SOD2,SORBS1,STAT1,THRB,TNFAIP3,TNFRSF10A,TNFRSF10D,TNFSF13B,TOB1,TP53INP1,TPSAB1/TPSB2,TRIP6,TRPS1,TSC22D3,TYMP,TYROBP,USP18,USP2,UTS2,WNT5B |

**T2 vs. T1 MEP_2**

© 2000-2020 QIAGEN. All rights reserved.

| Categories | Diseases or Functions Annotation | p-value | Predicted Activation State | Activation z-score | Molecules |
|---|---|---|---|---|---|
| Infectious Diseases | Replication of RNA virus | 2.08E-11 | Increased | 3.303 | AREG,B2M,BST2,CIITA,GABARAPL1,GBP1,H2BC15,HERC5,IFI27,IFI6,IFIH1,IFIT1,IFITM1,IFITM3,IRF1,IRF7,IRF9,ISG15,MX1,NFKBIA,NLRC5,OAS1,OAS3,OASL,PARP12,RAB11FIP1,RSAD2,S100A8,SNX9,SP110,STAT1,STAT2,TAP1,TXNIP |
| Cell Signaling | Replication of viral replicon | 3.23E-10 | Increased | 3.286 | BST2,IFIT1,IFITM1,IFITM3,ISG15,MX1,OAS1,OAS3,OASL,RSAD2,TRIM6 |
| Organismal Survival | Organismal death | 0.0000216 | Increased | 2.933 | ANGPT2,AREG,ATF3,B2M,BBC3,BIRC3,CD36,CDH1,CDKN2A,CDKN2C,CGAS,CITED2,CSRNP1,CXCL2,CXCL3,CYLD,DDIT3,DHCR7,DLK1,E2F1,EGLN1,FLT3,FST,GNLY,GSTP1,GZMA,HES1,HLA-G,HOPX,HRH2,HSPG2,ID1,IFIH1,IKZF3,IL15RA,IRF1,IRF7,IRF9,ISG15,ITGAM,KDM6B,KIFAP3,KLF6,KRT1,LMO4,LY96,MACROH2A2,MAFF,MTF1,NCF1,NEAT1,NFIB,NFKB2,NFKBIA,PIK3CG,PTGER2,RELB,S100A8,S1PR1,SERTAD1,SFRP5,SKIL,SLC17A9,SMAD9,SOD2,SRGAP3,STAT1,STAT2,TEK,TGM2,THRB,TNFAIP3,TNFSF13B,TUBG1,TXNIP,USP18,WNT5B,YES1,ZFP36L2 |
| Infectious Diseases | Replication of virus | 6.79E-14 | Increased | 2.905 | AREG,B2M,BIRC3,BST2,CIITA,CXCL2,CXCL3,DDIT3,EGR1,GABARAPL1,GBP1,H2BC15,HERC5,IFI27,IFI6,IFIH1,IFIT1,IFITM1,IFITM3,IL10RA,IRF1,IRF7,IRF9,ISG15,MX1,NFKBIA,NLRC5,OAS1,OAS3,OASL,PARP12,RAB11FIP1,RSAD2,S100A8,SNX9,SP110,STAT1,STAT2,TAP1,TXNIP |
| Cellular Growth and Proliferation,Tissue Development | Proliferation of epithelial cells | 0.0000342 | Increased | 2.867 | AREG,ATF3,B2M,CDH1,CDKN2A,CDKN2C,CITED2,CXCL3,E2F1,EREG,FGF18,FST,HES1,ID1,KLF10,LTBP4,NFIB,NFKBIA,PTGER2,RELB,SSTR2,TGM2,THRB,TOB1 |
| Infectious Diseases | Replication of vesicular stomatitis virus | 7.21E-08 | Increased | 2.779 | IFIH1,IFITM3,IRF1,IRF9,OAS1,OASL,PARP12,SP110,STAT2,TAP1 |
| Infectious Diseases | Replication of Flaviviridae | 7.77E-08 | Increased | 2.647 | BST2,GABARAPL1,IFI27,IFI6,IFIT1,IFITM1,IFITM3,ISG15,OASL,RSAD2,TXNIP |
| Organismal Survival | Morbidity or mortality | 0.00000556 | Increased | 2.623 | ANGPT2,AREG,ATF3,B2M,BBC3,BIRC3,CD36,CDH1,CDKN2A,CDKN2C,CGAS,CITED2,CSRNP1,CXCL2,CXCL3,CYLD,DDIT3,DHCR7,DLK1,E2F1,EGLN1,FLT3,FST,GNLY,GSTP1,GZMA,HES1,HLA-A,HLA-G,HOPX,HRH2,HSPG2,ID1,IFIH1,IKZF3,IL15RA,IRF1,IRF7,IRF9,ISG15,ITGAM,KDM6B,KIFAP3,KLF6,KRT1,LMO4,LY96,MACROH2A2,MAFF,MTF1,NCF1,NEAT1,NFIB,NFKB2,NFKBIA,OASL,PIK3CG,PTGER2,RELB,S100A8,S1PR1,SERTAD1,SFRP5,SKIL,SLC17A9,SMAD9,SOD2,SRGAP3,STAT1,STAT2,TAP1,TEK,TGM2,THRB,TNFAIP3,TNFSF13B,TUBG1,TXNIP,USP18,WNT5B,YES1,ZFP36L2 |
| Infectious Diseases | Replication of Hepatitis C virus | 0.000000227 | Increased | 2.455 | BST2,GABARAPL1,IFI27,IFI6,IFIT1,IFITM1,IFITM3,ISG15,RSAD2,TXNIP |
| Infectious Diseases | Replication of Paramyxoviridae | 0.0000156 | Increased | 2.429 | IFIH1,IRF9,MX1,OASL,RAB11FIP1,STAT2 |
| Tissue Development | Growth of epithelial tissue | 4.39E-08 | Increased | 2.134 | ANGPT2,AREG,ATF3,B2M,CD36,CDH1,CDKN2A,CDKN2C,CITED2,CXCL1,CXCL3,DLK1,E2F1,EGR1,EREG,FGF18,FST,GBP1,GPC1,HES1,HRH2,HSPG2,ID1,KLF10,LGALS1,LTBP4,NFIB,NFKBIA,PTGER2,RELB,S100A8,SSTR2,STAT1,TEK,TGM2,THRB,TOB1 |
| Infectious Diseases | Viral Infection | 4.7E-15 | Increased | 2.084 | AREG,ARRDC3,B2M,BBC3,BIRC3,BST2,C4orf33,CACNA2D3,CD36,CDH1,CDKN2A,CIITA,CTSL,CXCL1,CXCL2,CXCL3,DDIT3,EGR1,GABARAPL1,GBP1,GZMA,H2BC15,HBA1/HBA2,HBG1,HBG2,HCP5,HEMGN,HERC5,HES1,HLA-A,HLA-B,HLA-C,HRH2,IFI27,IFI6,IFIH1,IFIT1,IFIT3,IFITM1,IFITM3,IL10RA,IRF1,IRF7,IRF9,ISG15,ITGAM,KLF15,LGALS1,MX1,NACA2,NCF1,NFKBIA,NLRC5,OAS1,OAS2,OAS3,OASL,PARP12,PARP9,PDE3B,PIK3CG,RAB11FIP1,RELB,RPL3,RSAD2,S100A8,SNX9,SP110,STAT1,STAT2,TAP1,TNFRSF10B,TPSAB1/TPSB2,TRIM22,TUBG1,TUBG2,TXNIP,TYROBP,USP6 |
| Infectious Diseases | Infection of mammalia | 2.62E-09 | Increased | 2.019 | B2M,CD36,CIITA,CYLD,DDIT3,GZMA,HLA-G,IFIH1,IL10RA,IRF1,IRF7,IRF9,ISG15,ITGAM,LY96,NCF1,NFKB2,OASL,RELB,STAT1,STAT2,TAP1,TPSAB1/TPSB2,USP18 |
| Cell-To-Cell Signaling and Interaction | Adhesion of blood cells | 0.0000624 | Decreased | -2.027 | ANGPT2,CD36,CD83,CDH1,CXCL1,CXCL2,CXCL3,HSPG2,ITGAM,LGALS1,NFKB2,NFKBIA,PER1,PIK3CG,RELB,S100A8,STAT1,TGM2,TYROBP |
| Cardiovascular System Development and Function,Organismal Development | Vasculogenesis | 0.00000603 | Decreased | -2.091 | ANGPT2,ATF3,CD36,CDH1,CDKN2A,CHD7,CITED2,CXCL1,CXCL2,CYLD,DDIT3,DLK1,E2F1,EGLN1,EGR1,GBP1,GPC1,HES1,HRH2,HSPG2,ID1,ITGAM,KLF10,KRT1,LGALS1,NFKBIA,PIK3CG,PTGER2,S100A8,S1PR1,STAT1,TEK,TGM2,TNFAIP3,USP18 |
| Cellular Movement,Hematological System Development and Function,Immune Cell Trafficking | Cell movement of leukocytes | 1.36E-08 | Decreased | -2.092 | ABCG1,ALOX5AP,ANGPT2,CD36,CDH1,CIITA,CXCL1,CXCL2,CXCL3,CYSLTR1,EGLN1,GNLY,HDC,HLA-A,HLA-G,HRH2,ID1,IL10RA,IL15RA,ITGAM,KLF6,LGALS1,NCF1,NFKB2,NFKBIA,PIK3CG,RAMP1,RELB,S100A8,S1PR1,SFRP5,SLC17A9,STAT1,STAT2,TGM2,TIAM1,TNFAIP3,TNFRSF21,TNFSF13B,TPSAB1/TPSB2,TYROBP,UTS2 |
| Cell-To-Cell Signaling and Interaction,Hematological System Development and Function,Immune Cell Trafficking | Adhesion of immune cells | 0.0000712 | Decreased | -2.093 | ANGPT2,CD36,CD83,CDH1,CXCL1,CXCL2,CXCL3,ITGAM,LGALS1,NFKB2,NFKBIA,PER1,PIK3CG,RELB,S100A8,STAT1,TGM2,TYROBP |
| Cell Morphology,Cellular Function and Maintenance | Autophagy | 0.000013 | Decreased | -2.095 | APOL6,BBC3,BST2,CDKN2A,CTSL,DDIT3,DDIT4,DENND3,E2F1,GABARAPL1,HSPG2,IL10RA,NCF1,PIM2,PTGER2,S100A8,SOD2,STAT1,TGM2,TNFRSF10B,TP53INP2,TRIM22,VMP1,WDFY3 |
| Cellular Development,Cellular Growth and Proliferation,Hematological System Development and Function,Hematopoiesis,Lymphoid Tissue Structure and Development,Tissue Development | Hematopoiesis of mononuclear leukocytes | 1.09E-11 | Decreased | -2.099 | ABCG1,B2M,BBC3,CD36,CD83,CDKN2A,CHD7,CIITA,CISH,CXCL1,CYLD,E2F1,EGR1,FLT3,FST,GADD45B,GNLY,HLA-A,HLA-DRB5,HLA-G,ID1,IKZF3,IL10RA,IL15RA,IRF1,IRF7,ITGAM,KLF10,LGALS1,NFKB2,NFKBIA,PIK3CG,RELB,RSAD2,S1PR1,SFRP5,SLA,STAT1,STAT2,TNFAIP3,TNFSF13B,TYROBP,USP18,ZFP36L2 |
| Cellular Development,Cellular Growth and Proliferation,Hematological System Development and Function,Hematopoiesis,Lymphoid Tissue Structure and Development,Tissue Development | Leukopoiesis | 1.12E-12 | Decreased | -2.099 | ABCG1,B2M,BBC3,CD36,CD83,CDKN2A,CHD7,CIITA,CISH,CITED2,CXCL1,CXCL2,CXCL3,CYLD,CYSLTR1,DLK1,E2F1,EGR1,FLT3,FST,GADD45B,GNLY,HLA-A,HLA-DRB5,HLA-G,ID1,IKZF3,IL10RA,IL15RA,IRF1,IRF7,ITGAM,KLF10,LGALS1,NFKB2,NFKBIA,PIK3CG,RELB,RSAD2,S1PR1,SFRP5,SLA,STAT1,STAT2,TNFAIP3,TNFSF13B,TPSAB1/TPSB2,TYROBP,USP18,ZFP36L2 |
| Free Radical Scavenging | Production of superoxide | 0.0000247 | Decreased | -2.109 | ANGPT2,CD36,HBA1/HBA2,ITGAM,NCF1,NCF1C,PIK3CG,SOD2,TGM2,TYROBP |
| Cancer,Cell Death and Survival,Organismal Injury and Abnormalities,Tumor Morphology | Apoptosis of tumor cells | 9.2E-10 | Decreased | -2.119 | ATF3,B2M,BBC3,CD33,CDH1,CDKN2A,DDIT3,E2F1,GZMA,HES1,IRF1,IRF7,KLF10,LGALS1,NFKBIA,PMAIP1,PTGER2,SSTR2,THRB,TNFRSF10B,TNFRSF21,TNFSF13B,YES1 |
| Cellular Movement,Immune Cell Trafficking | Leukocyte migration | 3.01E-11 | Decreased | -2.124 | ABCG1,ALOX5AP,ANGPT2,CD36,CDH1,CIITA,CXCL1,CXCL2,CXCL3,CYSLTR1,EGLN1,EGR1,GNLY,GPC1,HDC,HLA-A,HLA-G,HRH2,ID1,IL10RA,IL15RA,IRF7,ITGAM,KDM6B,KLF6,LGALS1,LY96,NCF1,NFKB2,NFKBIA,PER1,PIK3CG,RAMP1,RELB,S100A8,S1PR1,SFRP5,SLC17A9,SLC7A7,SOD2,STAT1,STAT2,TEK,TGM2,TIAM1,TNFAIP3,TNFRSF10B,TNFRSF21,TNFSF13B,TPSAB1/TPSB2,TYROBP,UTS2,YES1 |
| Cell Death and Survival | Cell death of lymphoma cell lines | 0.00000094 | Decreased | -2.155 | BBC3,BIRC3,DDIT3,EGR1,LGALS1,NFKB2,NFKBIA,PMAIP1,SOD2,STAT1,STAT2,TGM2,TIAM1,TNFAIP3,TNFRSF10B,TNFSF13B |
| Immunological Disease | Hypersensitive reaction | 2.75E-12 | Decreased | -2.197 | ALOX5AP,ANGPT2,ANXA2P2,CD83,CISH,CTSL,CYSLTR1,E2F1,HCP5,HDC,HLA-A,HLA-B,HLA-C,HLA-DRB5,HRH2,IFI27,IFITM1,IKZF3,KRT1,LGALS1,LGALS3BP,MS4A2,MX1,NCF1,NFKBIA,PER1,PIK3CG,RELB,S100A8,SOD2,STAT1,TNFAIP3,TSC22D3 |
| Free Radical Scavenging | Generation of reactive oxygen species | 0.0000668 | Decreased | -2.206 | ANGPT2,APOL6,BBC3,CD36,DDIT4,DHCR24,GSTP1,HBA1/HBA2,ITGAM,NCF1,PIK3CG,SOD2,TXNIP |
| Cellular Movement,Hematological System Development and Function,Immune Cell Trafficking | Migration of antigen presenting cells | 0.000052 | Decreased | -2.346 | CDH1,CXCL1,CXCL2,IL10RA,ITGAM,KLF6,LGALS1,PIK3CG,RAMP1,S100A8,S1PR1,TYROBP |
| Cell Death and Survival | Cell viability | 1.1E-12 | Decreased | -2.363 | ANGPT2,AREG,ATF3,B2M,BBC3,CD33,CDH1,CDKN2A,CDKN2C,CISH,CXCL1,CXCL2,CXCL3,CYLD,DDIT3,DHCR24,DLK1,E2F1,EGLN1,EGR1,EREG,FGF18,FKBP5,FLT3,GADD45B,GBP1,GNLY,GPC1,GSTP1,HBA1/HBA2,HES1,HLA-A,HLA-F,HLA-G,HOPX,IFIH1,IKZF3,IRF9,ITGAM,KDM6B,KLF6,LGALS3BP,LMO4,MX1,NEAT1,NFKB2,NFKBIA,PCSK1N,PIK3CG,PIM2,PMAIP1,RELB,S100A8,S1PR1,SOD2,STAT1,STAT2,STX3,TEK,TGM2,TNFAIP3,TNFRSF10B,TNFSF13B,TOB1,TRIM69,TSC22D3,TXNIP,TYROBP,USP18,XAF1,YES1 |
| Cell Death and Survival | Cell viability of leukocytes | 0.0000531 | Decreased | -2.365 | BBC3,CISH,CYLD,DLK1,MX1,NFKBIA,PIK3CG,PIM2,RELB,SOD2,STAT1,TGM2,TNFSF13B,TSC22D3,TYROBP |
| Cell Death and Survival | Necroptosis | 0.00000607 | Decreased | -2.408 | BIRC3,CGAS,CYLD,ITGAM,LY96,RPL10,SOD2,SPATA2,STAT1,TNFRSF21 |
| Cell Death and Survival | Necrosis | 4.6E-16 | Decreased | -2.408 | ABCG1,ANGPT2,AREG,ATF3,B2M,BBC3,BIRC3,BTG1,CD33,CD36,CDH1,CDKN2A,CDKN2C,CGAS,CIITA,CISH,CITED2,CXCL1,CXCL2,CXCL3,CYLD,CYSLTR1,DDIT3,DDIT4,DHCR24,DHCR7,DLK1,E2F1,EGLN1,EGR1,EREG,FGF18,FKBP5,FLT3,FST,GADD45B,GAL,GBP1,GNLY,GPC1,GSTM5,GSTP1,GZMA,HES1,HLA-A,HLA-B,HLA-F,HLA-G,HRH2,ID1,IFI6,IFIH1,IKZF3,IL15RA,IRF1,IRF7,IRF9,ISG15,ITGAM,KIFAP3,KLF10,KLF6,LGALS1,LGALS3BP,LMO4,LY96,MS4A2,MTF1,MX1,NCF1,NFIB,NFKB2,NFKBIA,OAS1,OAS2,OAS3,PCSK1N,PDE3B,PER1,PIK3CG,PIM2,PMAIP1,PTGER2,RELB,RNASE1,RPL10,RPL3,RPL7,RPL7A,S100A8,S1PR1,SFRP5,SKIL,SOD2,SPATA2,SSTR2,ST3GAL1,STAT1,STAT2,TEK,TGM2,THRB,TIAM1,TNFAIP3,TNFRSF10B,TNFRSF19,TNFRSF21,TNFSF13B,TPSAB1/TPSB2,TSC22D3,TXNIP,TYROBP,USP18,XAF1,YES1 |
| Cancer,Cell Death and Survival,Organismal Injury and Abnormalities,Tumor Morphology | Necrosis of tumor | 5.44E-12 | Decreased | -2.408 | ANGPT2,ATF3,B2M,BBC3,BIRC3,BTG1,CD33,CDH1,CDKN2A,DDIT3,DHCR7,E2F1,FGF18,FLT3,GZMA,HES1,IRF1,IRF7,KLF10,LGALS1,NFKBIA,PMAIP1,PTGER2,RPL10,RPL3,RPL7,RPL7A,SSTR2,TEK,THRB,TNFRSF10B,TNFRSF21,TNFSF13B,YES1 |
| Cancer,Cell Death and Survival,Organismal Injury and Abnormalities,Tumor Morphology | Cell death of tumor cells | 6.27E-11 | Decreased | -2.408 | ATF3,B2M,BBC3,BIRC3,CD33,CDH1,CDKN2A,DDIT3,DHCR7,E2F1,FGF18,FLT3,GZMA,HES1,IRF1,IRF7,KLF10,LGALS1,NFKBIA,PMAIP1,PTGER2,RPL10,RPL3,RPL7,RPL7A,SSTR2,TEK,THRB,TNFRSF10B,TNFRSF21,TNFSF13B,YES1 |
| Immunological Disease | Delayed hypersensitive reaction | 0.0000028 | Decreased | -2.425 | ANGPT2,CD83,IFITM1,LGALS3BP,MX1,NCF1,NFKBIA,PIK3CG,RELB,SOD2,TSC22D3 |
| Gene Expression | Expression of RNA | 4.07E-09 | Decreased | -2.459 | APOL3,ATF3,BST2,BTG1,CD36,CDH1,CDKN2A,CDKN2C,CHD7,CIITA,CISH,CITED2,CITED4,CSRNP1,CXCL2,DDIT3,DLK1,DTX3L,E2F1,EGLN1,EGR1,EREG,FST,GAL,GSTP1,HBG2,HELZ2,HES1,HLA-DRB5,HOPX,ID1,IFI27,IKZF3,IRF1,IRF7,IRF9,ITGAM,KDM6B,KLF10,KLF15,KLF6,LBH,LGALS1,LMO4,LY96,MACROH2A2,MAFF,MEG3,MTF1,NCF1,NFIB,NFKB2,NFKBIA,NLRC5,PARP9,PER1,PIM2,PTGER2,RCOR2,RELB,RNF11,RPL10,S1PR1,SERTAD1,SKIL,SLA,SMAD9,SOD2,STAT1,STAT2,TGM2,THRB,TNFAIP3,TNFRSF10B,TOB1,TP53INP2,TRIM22,TRIM6,TSC22D3,TUBG1,TXNIP,TYROBP,USP3,UTS2,YES1,ZNF462,ZNF503 |
| Gene Expression | Activation of DNA endogenous promoter | 0.000000402 | Decreased | -2.481 | ATF3,CD36,CDKN2A,CHD7,CIITA,CITED2,CSRNP1,DDIT3,E2F1,EGLN1,EGR1,FST,GAL,HBG2,HELZ2,HES1,HOPX,ID1,IFI27,IKZF3,IRF1,IRF7,IRF9,KDM6B,KLF10,KLF15,KLF6,LMO4,MACROH2A2,MAFF,MTF1,NFIB,NFKB2,NFKBIA,NLRC5,PARP9,PER1,RELB,RPL10,S1PR1,SERTAD1,SKIL,SOD2,STAT1,STAT2,THRB,TSC22D3,TXNIP,USP3,YES1,ZNF462 |
| Neurological Disease | Progressive neurological disorder | 3.22E-08 | Decreased | -2.538 | AREG,ATF3,BST2,CD33,CD36,CDKN2A,CIITA,CXCL1,CYSLTR1,DDIT3,DHCR24,DHCR7,E2F1,EGLN1,EGR1,EPSTI1,FLT3,GAL,GSTP1,HBA1/HBA2,HBG2,HLA-B,HLA-DMB,HLA-F,HLA-G,HRH2,HSPG2,IFIT1,IRF7,ISG15,LGALS1,MS4A4E,MX1,NEAT1,NFKB2,OAS1,OAS3,PAQR6,PDE3B,RPL3,RSAD2,SOD2,TEK,TGM2,TRIM22,TUBG1,TUBG2,TYROBP,WDFY3,ZFP36L2 |
| Cell Death and Survival | Cell viability of blood cells | 0.0000167 | Decreased | -2.564 | BBC3,CISH,CYLD,DLK1,FLT3,GADD45B,MX1,NFKBIA,PIK3CG,PIM2,RELB,SOD2,STAT1,TGM2,TNFSF13B,TSC22D3,TYROBP |
| Cellular Function and Maintenance | Cellular homeostasis | 3.44E-14 | Decreased | -2.591 | ABCG1,ANGPT2,ANKRD26,APOL6,B2M,BBC3,BST2,CD36,CD83,CDKN2A,CHD7,CIITA,CISH,CTSL,CXCL1,CXCL2,CXCL3,CYLD,CYSLTR1,DDIT3,DDIT4,DENND3,E2F1,EGLN1,EGR1,FLT3,GABARAPL1,GADD45B,GNLY,GZMA,HDC,HELZ2,HLA-A,HLA-G,HSPG2,ID1,IFI6,IL10RA,IL15RA,IRF1,KLF10,KLF15,LGALS1,MS4A2,NCF1,NEAT1,NFKBIA,OAS1,PDE3B,PIK3CG,PIM2,PMAIP1,PTGER2,RELB,RNASE1,RSAD2,S100A8,S1PR1,SKIL,SLA,SNTA1,SOD2,SORBS1,STAT1,STAT2,TEK,TGM2,TNFAIP3,TNFRSF10B,TNFSF13B,TP53INP2,TRIM22,TSC22D3,TXNIP,TYROBP,USP18,UTS2,VMP1,WDFY3,ZFP36L2 |
| Gene Expression | Transcription of RNA | 4.13E-11 | Decreased | -2.65 | APOL3,ATF3,BST2,BTG1,CD36,CDH1,CDKN2A,CDKN2C,CHD7,CIITA,CISH,CITED2,CITED4,CSRNP1,CXCL2,DDIT3,DTX3L,E2F1,EGLN1,EGR1,EREG,FST,GAL,GSTP1,HBG2,HELZ2,HES1,HLA-DRB5,HOPX,ID1,IFI27,IKZF3,IRF1,IRF7,IRF9,KDM6B,KLF10,KLF15,KLF6,LBH,LGALS1,LMO4,LY96,MACROH2A2,MAFF,MEG3,MTF1,NCF1,NFIB,NFKB2,NFKBIA,NLRC5,PARP9,PER1,PIM2,PTGER2,RCOR2,RELB,RNF11,RPL10,S1PR1,SERTAD1,SKIL,SLA,SMAD9,SOD2,STAT1,STAT2,THRB,TNFRSF10B,TP53INP2,TRIM22,TRIM6,TSC22D3,TUBG1,TXNIP,USP3,YES1,ZNF462 |
| Cellular Movement,Hematological System Development and Function,Immune Cell Trafficking,Inflammatory Response | Migration of phagocytes | 0.0000539 | Decreased | -2.728 | ANGPT2,CDH1,CXCL1,CXCL2,CXCL3,HRH2,IL10RA,ITGAM,KLF6,LGALS1,PIK3CG,RAMP1,S100A8,S1PR1,SLC17A9,TYROBP |
| Gene Expression | Transcription of DNA | 0.000000012 | Decreased | -2.753 | ATF3,CD36,CDH1,CDKN2A,CHD7,CIITA,CITED2,CITED4,CSRNP1,DDIT3,DTX3L,E2F1,EGLN1,EGR1,EREG,FST,GAL,HBG2,HELZ2,HES1,HLA-DRB5,HOPX,ID1,IFI27,IKZF3,IRF1,IRF7,IRF9,KDM6B,KLF10,KLF15,KLF6,LBH,LMO4,MACROH2A2,MAFF,MTF1,NCF1,NFIB,NFKB2,NFKBIA,NLRC5,PARP9,PER1,PIM2,RCOR2,RELB,RPL10,S1PR1,SERTAD1,SKIL,SMAD9,SOD2,STAT1,STAT2,THRB,TP53INP2,TRIM22,TRIM6,TSC22D3,TXNIP,USP3,YES1,ZNF462 |
| Inflammatory Response | Immune response of cells | 0.0000111 | Decreased | -2.783 | BIRC3,CD36,CD83,CDKN2A,CXCL1,CXCL3,CYLD,HLA-A,HRH2,IFIH1,IL10RA,IL15RA,IRF1,IRF7,ISG15,ITGAM,LGALS1,LY96,NCF1,PER1,PIK3CG,PTGER2,RELB,S100A8,STAT1,TAP1,TGM2,TRIM6,TYROBP |
| Gene Expression | Transcription | 2.29E-09 | Decreased | -2.812 | APOL3,ATF3,BST2,BTG1,CD36,CDH1,CDKN2A,CDKN2C,CHD7,CIITA,CISH,CITED2,CITED4,CSRNP1,CXCL2,DDIT3,DLK1,DTX3L,E2F1,EGLN1,EGR1,EREG,FST,GAL,GSTP1,HBG2,HELZ2,HES1,HLA-DRB5,HOPX,ID1,IFI27,IKZF3,IRF1,IRF7,IRF9,KDM6B,KLF10,KLF15,KLF6,LBH,LGALS1,LMO4,LY96,MACROH2A2,MAFF,MEG3,MTF1,MX1,NCF1,NFIB,NFKB2,NFKBIA,NLRC5,PARP9,PER1,PIM2,PTGER2,RCOR2,RELB,RNF11,RPL10,S1PR1,SERTAD1,SKIL,SLA,SMAD9,SOD2,STAT1,STAT2,THRB,TNFRSF10B,TNFSF13B,TP53INP2,TRIM22,TRIM6,TSC22D3,TUBG1,TXNIP,TYROBP,USP3,YES1,ZNF462,ZNF503 |
| Inflammatory Disease,Neurological Disease,Skeletal and Muscular Disorders | Multiple Sclerosis | 3.91E-10 | Decreased | -2.813 | AREG,BST2,CXCL1,CYSLTR1,DDIT3,DHCR24,DHCR7,EGR1,EPSTI1,GAL,HLA-DMB,HLA-F,HLA-G,HRH2,IFIT1,IRF7,ISG15,MX1,OAS1,OAS3,RSAD2,TRIM22 |
| Neurological Disease,Skeletal and Muscular Disorders | Progressive neuromuscular disease | 1.32E-09 | Decreased | -2.813 | AREG,BST2,CXCL1,CYSLTR1,DDIT3,DHCR24,DHCR7,E2F1,EGLN1,EGR1,EPSTI1,GAL,GSTP1,HLA-B,HLA-DMB,HLA-F,HLA-G,HRH2,IFIT1,IRF7,ISG15,MX1,OAS1,OAS3,PAQR6,RPL3,RSAD2,TGM2,TRIM22 |
| Neurological Disease,Skeletal and Muscular Disorders | Neuromuscular disease | 0.00000114 | Decreased | -2.813 | AREG,B2M,BST2,BTN3A1,CACNA2D3,CIITA,CXCL1,CYSLTR1,DDIT3,DHCR24,DHCR7,E2F1,EGLN1,EGR1,EPSTI1,GAL,GBP1,GSTP1,HLA-B,HLA-DMB,HLA-F,HLA-G,HRH2,IFIT1,IRF7,ISG15,ITGAM,KCTD17,MAFF,MX1,OAS1,OAS3,PAQR6,RELB,RPL3,RSAD2,S1PR1,TGM2,TRIM22,TSC22D3 |
| Inflammatory Disease,Neurological Disease,Skeletal and Muscular Disorders | Relapsing-remitting multiple sclerosis | 0.00000229 | Decreased | -2.813 | EPSTI1,HLA-G,HRH2,IFIT1,IRF7,ISG15,MX1,OAS1,OAS3,RSAD2 |
| Antimicrobial Response,Inflammatory Response | Antiviral response | 3.47E-19 | Decreased | -2.937 | BIRC3,BST2,CGAS,DDIT4,DDX60,DTX3L,HLA-A,IFI27,IFI44,IFI44L,IFI6,IFIH1,IFIT1,IFIT3,IFITM1,IFITM3,IRF1,IRF7,ISG15,MX1,OAS1,OAS2,OAS3,OASL,PARP9,PMAIP1,RSAD2,STAT1,STAT2,TRIM22,TRIM6 |
| Neurological Disease | Progressive motor neuron disease | 0.000000076 | Decreased | -2.985 | AREG,ATF3,BST2,CD36,CXCL1,CYSLTR1,DDIT3,DHCR24,DHCR7,E2F1,EGLN1,EGR1,EPSTI1,GAL,GSTP1,HLA-B,HLA-DMB,HLA-F,HLA-G,HRH2,IFIT1,IRF7,ISG15,MX1,NEAT1,NFKB2,OAS1,OAS3,PAQR6,RPL3,RSAD2,SOD2,TGM2,TRIM22,ZFP36L2 |
| Neurological Disease | Progressive motor neuropathy | 0.000000213 | Decreased | -2.985 | AREG,ATF3,BST2,CD36,CXCL1,CYSLTR1,DDIT3,DHCR24,DHCR7,E2F1,EGLN1,EGR1,EPSTI1,GAL,GSTP1,HLA-B,HLA-DMB,HLA-F,HLA-G,HRH2,IFIT1,IRF7,ISG15,MX1,NEAT1,OAS1,OAS3,PAQR6,RPL3,RSAD2,SOD2,TGM2,TRIM22,ZFP36L2 |
| Antimicrobial Response,Inflammatory Response | Antimicrobial response | 3.32E-18 | Decreased | -3.091 | B2M,BIRC3,BST2,CGAS,DDIT4,DDX60,DTX3L,GNLY,H2BC15,HLA-A,HLA-G,IFI27,IFI44,IFI44L,IFI6,IFIH1,IFIT1,IFIT3,IFITM1,IFITM3,IRF1,IRF7,ISG15,MX1,NCF1,OAS1,OAS2,OAS3,OASL,PARP9,PMAIP1,RNASE6,RSAD2,S100A8,STAT1,STAT2,TRIM22,TRIM6 |
